# AGEAS: Automated Machine Learning based Genetic Regulatory Element Extraction System

**DOI:** 10.1101/2022.02.17.480852

**Authors:** Masayoshi Nakamoto, Jiawang Tao, Jack Yu

**Affiliations:** Shenzhen Mozhou Technology Co., Ltd, Shenzhen, China; Center for Health Research, Guangzhou Institutes of Biomedicine and Health, Chinese Academy of Sciences, Guangzhou 510530, China

## Abstract

As rapid progress in sequencing technology since last decade, numerous mechanisms underlying cell functions and developmental processes have been revealed as complex regulations of gene expressions. Since single-cell RNA sequencing (scRNA-seq) made high-resolution transcriptomic view increasingly accessible, precise identification of gene regulatory network (GRN) describing cell types and cell states became achievable. However, extracting key regulatory elements, including gene regulatory pathways (GRPs), transcription factors (TFs), and targetomes, that accurately and completely reflects functionality changes in biological phenomena remains challenging. Herein, we describe AGEAS, an semi-supervised automated machine learning (AutoML) based genetic regulatory element extraction system that assesses importances of GRPs in resulting biological phenomena, such as cell type differentiation, physiological and pathological development, and reconstructs GRNs with extracted important GRPs for comprehensive inference. With several case studies in divergent research areas, we show that AGEAS can indeed extract informative regulatory elements and reconstruct networks to indicate regulatory changes in biological phenomena of interest.

**Availability and implementation:** The AGEAS code is available at https://github.com/JackSSK/Ageas.

## Introduction

As high-resolution sequencing technologies become increasingly applicable and accessible, an efficient and robost analytical system capable of extracting key genetic features responsible for cell types and cell states difference with limited prior biological knowledge also become highly demanded. Several methodologies like SCENIC^1^ already demonstrated informativeness and robustness in studying cellular phenotypes with GRN analysis. Regulons, collections of a TF and corresponding targetomes, can successfully address cell identities with more comprehensive information compared with differential expressing genes (DEGs).^1,2^ Even though limited computational methods are capable of completely and precisly extracting key genetic regulatory elements in biological process of interest, Mogrify^3^ and CellNet^4^ have demonstrated GRN based methods can help to analyze cell type differentiation and to implement cell reprogramming. However, both methods may require large scale of additional background data to be applicable in other studies such as analyzing physiological or pathological development of selected cell type. The scREMOTE^5^ published in 2022 is a potential method to extract generalized regulatory elements since limited background data is required with sequencing data representing sample classes. But prior knowledge on key TFs and marker genes is indispensable. A regulation trajectory of DEGs or regulons showing significant activity difference after GRN reconstruction would be able to reveal the developmental pathways as generalized approach. Due to general gene regulatory nature of TFs, numerous noisy signals are expected. Therefore, here we present AGEAS, an algorithm with feasibility in extracting key regulatory elements for biological phenomena of interest and can potentially promote further discoveries, even with scarce prior knowledge.

Several reasons motivated us to develop semi-supervised AutoML based method. Firstly, accurate labeling of regulatory elements’ relatedness in biological phenomena would require extensive prior knowledge may be unavailable. While unsupervised clustering can potentially extract factors playing important roles, this approach may focus on specific aspects with regulatory elements showing similar patterns and result in losing comprehensiveness. As a result, we chose to apply biological phenomena associating sample classes, such as cell types and cell states which should be easily retrievable, as labels for classification models to predict with masked GRN as input and then interpret success predictions to extract key regulatory elements. Secondly, it is also hard to guarantee comprehensiveness when developing a generalized classification model in differentiating GRNs, considering that capturing few significant differences would be sufficient for the model to reach outstanding accuracy. Hence, an extraction system with multiple independent classification models implemented with various algorithms should be helpful to recover from potential comprehensiveness loss. Thirdly, the process of interpreting all success prediction made by every classification model and integrating corresponding interpretation results may be computational expensive. Thus, applying an AutoML-based model and feature selection procedure that can effectively decrease interpretation cost would be necessary to increase feasibility of the method.

To evaluate the performance of AGEAS, we primarily applied it on studying somatic cell reprogramming process and further addressed performance in analyzing 3 other biological processes in cell subtype differentiation, physiological and pathological development.

## Method

The basic principle of AGEAS is to find key regulatory elements associated with biological phenomena of interest through analyzing how well-performing classification models distinguish GRNs of sample with the phenomena from those without. To reconstruct sufficient GRNs for each class, RNA-seq based expression data is segmented into subsets while each one is analogized as a pseudo-sample having discrete expression data. The pseudo-sample GRNs (psGRNs) are reconstructed accordingly; thus, classification models can be trained, evaluated, and interpreted with GRPs as input factors. With heavily weighted GRPs and corresponding genes repeatedly obtained from interpretations of successful sample class predictions, GRNs could be formed and potentially play an important role in differentiating the studying sample classes. The overall workflow can be summarized in Figure 1. By default, four separate extractor units in workflow run in parallel after data preprocessing part in order to increase output stability considering the stochastic nature of AGEAS, and all extracted regulatory elements are used to form GRNs, which will later be combined into one atlas. The following sections describe each step of AGEAS in more depth.

**Figure (1).**
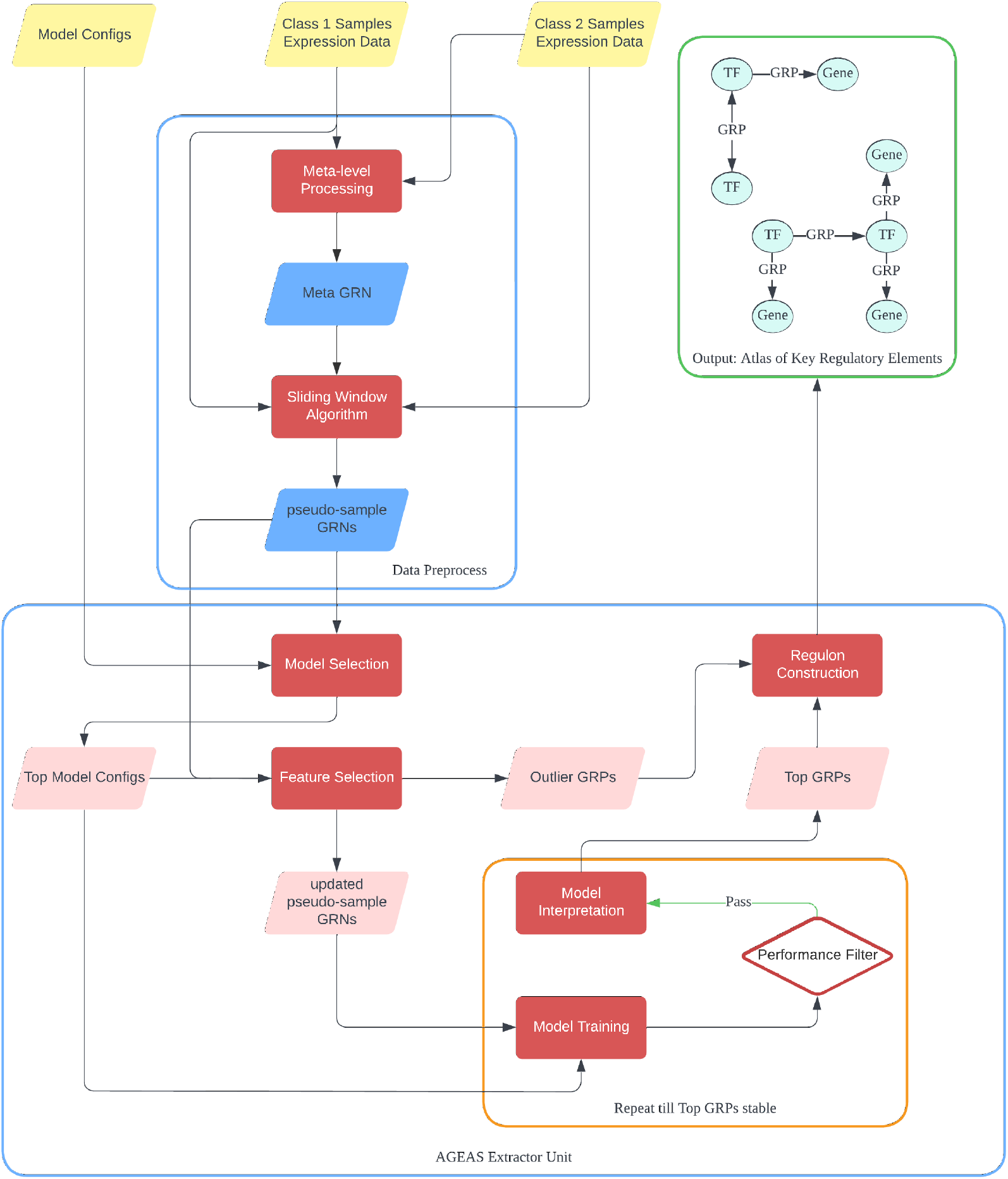
The overall workflow of AGEAS: **(*1*)** Reconstruct meta-level GRN (meta-GRN) with expression data of all samples. ***(2)*** Build pseudo-samples with sliding window algorithm and reconstruct GRNs with GRPs identified in meta-GRN accordingly. ***(3)*** Select best performing classifiers in predicting class labels of pseudo-sample GRNs (psGRNs). ***(4)*** Interpret how top models make classifications and gradually exclude GRPs with low weights or outlier-level high weights. **(*5*)** Repeatedly train classifiers with different set of psGRNs as training data to extract GRPs frequently ranked as top important features for decision. ***(6)*** Reconstruct GRNs with extracted GRPs and GRPs excluded as significant outliers.

### Step 1: Data preprocessing

The main purpose of this step is to build pseudo-samples and to reconstruct corresponding pseudo-sample GRNs (psGRNs). For each sample class, gene expression matrices (GEMs) with the same class label are concatenated as one comprehensive expression matrix. With comprehensive GEMs, a meta-level GRN (meta-GRN) is reconstructed before psGRNs to provide generic guidance on reconstruction. In general, the workflow of this step can be summarized in Figure 2.

**Figure (2).**
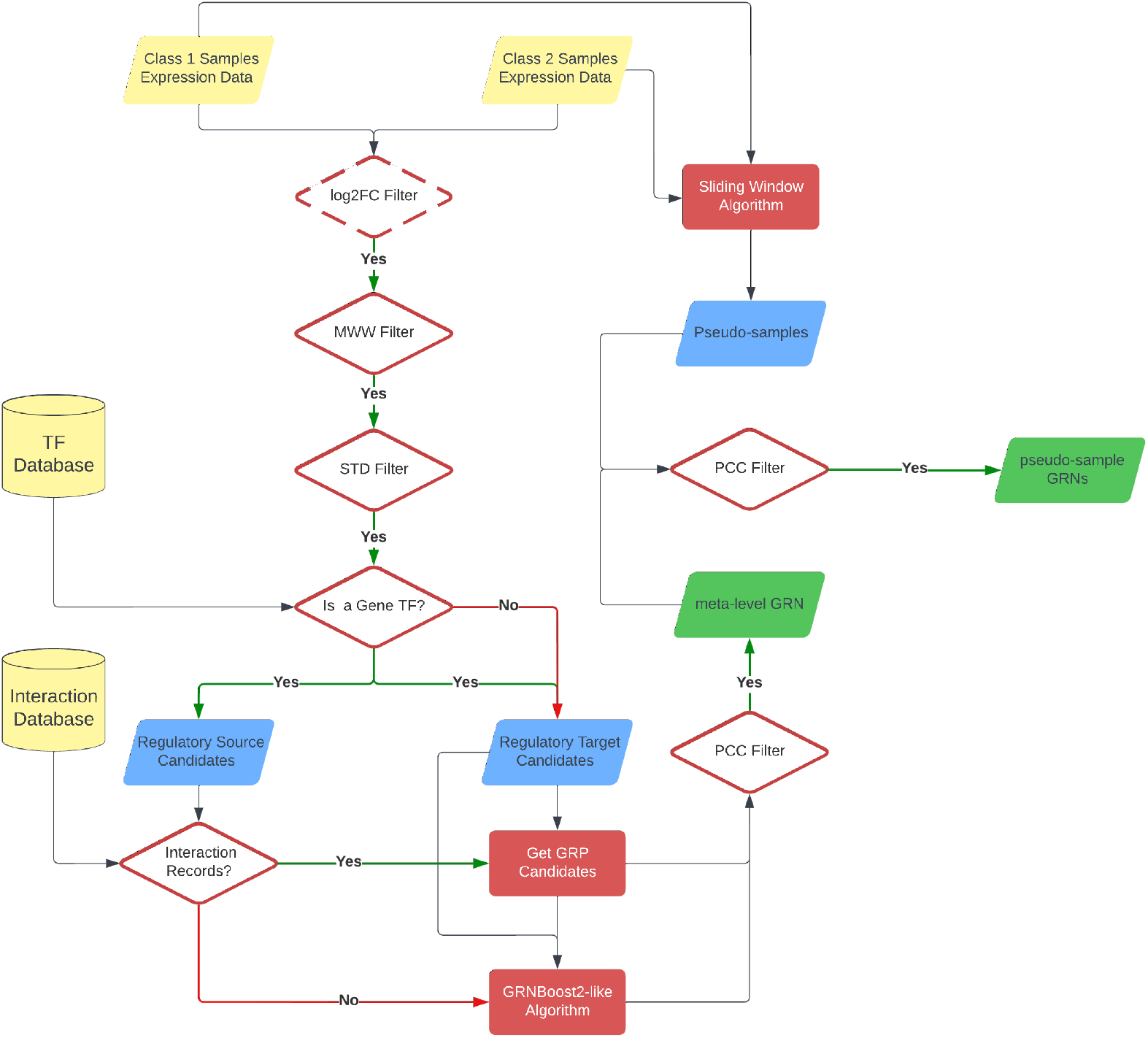
Workflow to reconstruct meta-GRN and psGRNs. ***(1)*** Filter genes from GEMs with log2FC filter(optional), MWW filter, and *σ* filter. ***(2)*** Find candidate GRP gene pairs from either interaction database or predictions made by *GRNBoost2*^10^-like algorithm. **(*3*)** Filter candidate GRP gene pairs with PCC filter and reconstruct meta-GRN with validated GRPs. **(*4*)** Generate pseudo-samples with SWA. **(*5*)** Utilize GRPs in meta-GRN as generic guidance to form candidate GRPs for pseudo-samples. ***(6)*** Filter every candidate GRP for all pseudo-samples with PCC filter and reconstruct psGRNs with validated GRPs.

#### Reconstruct meta-GRN

Firstly, genes included in the comprehensive GEMs are assessed and determined whether having potential to form informative GRPs with other genes. Commonly, DEGs are considered as important factors of studying phenomena. Here we apply the Mann-Whitney U rank test (MWW) implemented by *SciPy*^6^ to exclude genes with indistinguishable expression level distribution across GEMs of different classes. The p-value for rejecting null hypothesis, that expression profile underlying class 1 samples is the same as the expression profile underlying class 2 samples, is set to 0.05 by default. Furthermore, a log_2_ fold change (log2FC) filter is implemented in AGEAS. However, enabling the log2FC filter is not encouraged, considering that upstream TFs indirectly regulate key genes associated with phenomena of interest may not be significantly differential expressed. The log2FC filter shall mostly be used to decrease meta-GRN’s total degree in a compromising position caused by limited computational resources. After differential expression based filters, a standard deviation (σ) filter is applied to exclude genes with low expression level or merely affected by dynamic expression status of other genes. By default, the σ threshold is set to 1.0, the lowest expression level in raw gene count matrix gained from RNA-seq data. The threshold value should be adjusted based on prior knowledge of input GEMs, for example, the normalization method applied to the GEMs.

With candidate genes passing filters above, some gene pairs are formed and evaluated by the potential of representing GRPs. To reduce overall computational complexity, a gene pair shall be formed with at least one TF, which could be the regulatory source of GRP. If not further specified, a TF list will be retrieved from integrated *TRANSFAC*^7^ dataset according to the provided species information. Utilizing genetic interaction database like *GTRD*^8^ and *BioGRID*^9^, AGEAS checks whether the binding ability of gene pair is confirmed or not. By default, if the TF recorded has at least one binding site within target gene’s promoter range (−1000 to +100) by Chromatin Immunoprecipitation Sequencing (ChIP-seq) dataset retrieved from *GTRD*^8^, the potential GRP gene pair will be passed to expression correlation assessment. For TFs not covered by interaction database, *GRNBoost2*^10^-like algorithm is initiated to predict potential regulatory target genes. The prediction importance threshold can either be set manually or automatically based on recorded interactions as:

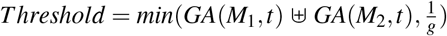

Here *GA*() denotes *GRNBoost2*^10^-like algorithm; *M*_1_ denotes concatenated class 1 samples GEM with genes passed filters; *M*_2_ denotes concatenated class 2 samples GEM with genes passed filters; *t* denotes the TF having largest amount of recorded interaction in dataset; *g* denotes total amount of unique genes in all samples passed filters.

After potential GRP gene pairs are obtained, an expression correlation filter is used to exclude gene pairs with low covariance. To assess expression correlation, AGEAS applies Pearson’s Correlation coefficient^11^ (PCC), one of the widely adopted methods.^12^ With default setting, gene pairs can reach absolute correlation coefficient of 0.2 and p-value lower than 0.05 are included in meta-GRN as validated GRP.

#### Reconstruct psGRNs

The comprehensive GEMs are divided into sample subsets, and sliding window algorithm (SWA) is used to build pseudo-samples. With SWA, we can gain GEMs of *i-th* pseudo-sample through:

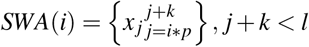

Here *l* denotes number of samples in a comprehensive GEM which can be expressed as 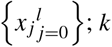 denotes window size; *p* denotes padding stride.

If sample amount is considerably low or imbalanced, customized window size and padding stride can be applied to generate sufficient pseudo-samples for later classifier training and assessment processes. Utilizing meta-GRN, psGRNs are reconstructed with GEMs of pseudo-sample. Each GRP gene pair in meta-GRN is formed with expression data in pseudo-sample and is filtered by the same PCC filter in meta-GRN reconstruction process.

After all pseudo-samples have been used to reconstruct psGRN, every psGRN can be represented as a matrix comprised of GRPs’ PCC values. The order of GRPs in each psGRN is also unified through adding GRPs included in other psGRNs with 0.0 PCC value.

### Step 2: Classification model selection

The primary goal of AGEAS is to gain insights from multiple models as divergent as possible instead of developing optimized classification model for psGRN classification. Thus, this step is designed to select model configurations capable of predicting psGRN’s masked sample class label. Considering the limited computational resources, the portion of less efficient model configurations shall be pruned at the cost of potentially losing insight in sample class difference. Therefore, we apply a simple Hyperband^13^-based algorithm 1 which performs a grid search for well-performing classification models with varying model training resources and pruning aggressiveness.

**Algorithm 1:**
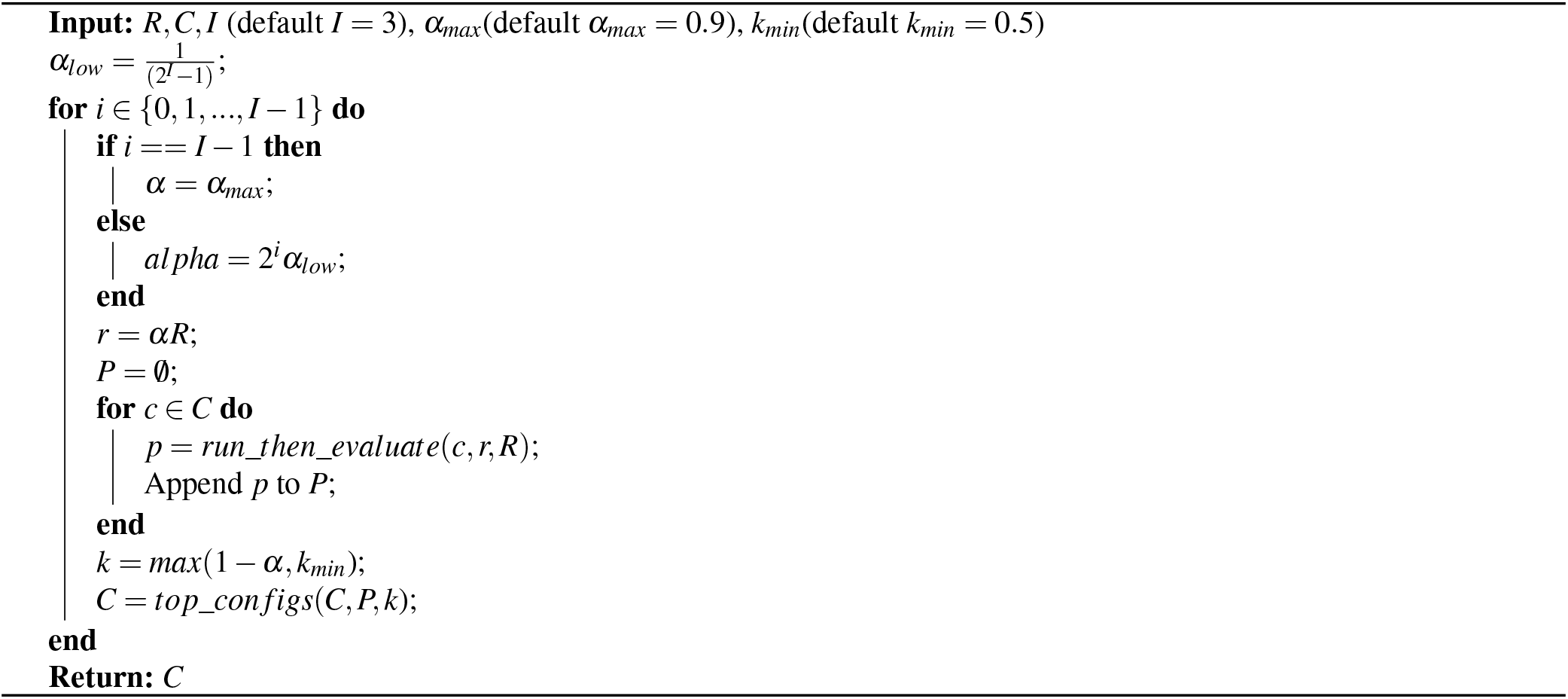
Model selection algorithm

The model selection algorithm requires five inputs (1)*R*, the maximum amount of training resource, equivalent to all available psGRNs (2)C, the total set of provided classification model configurations (3)*I*, the number of iterations for model pruning (default set as 3) (4)*α_max_*, the maximum portion of *R* can be fed to single model (default set as 0.95) (5)*k_min_*, the minimum portion of remaining model configurations will be kept by single pruning iteration (default set as 0.5). Furthermore, two required functions need to be defined based on input model configurations:

- *run_then_evaluate*(*c, r, R*): trains classification model initialized using configuration *c* for the allocated resource *r*, then returns prediction accuracy (ACC), the area under a receiver operating characteristic curve (AUROC)^14^ score, and total cross-entropy loss (*L_CE_*) calculated through predicting sample class for all psGRNs *R*.
- *top_configs*(*C, P, k*): takes a set of model configurations *C* with associated evaluation results *P* and returns configurations with ACC, AUROC score, or *L_CE_* reaching top *k* portion.

By default, AGEAS initializes with 128 model configurations utilizing 9 integrated classification algorithms listed in Table 1.

**Table (1).** Classification model algorithms integrated in AGEAS with corresponding numbers of hyperparameter and preset model configurations. Categorical hyperparameters and continuous numerical hyperparameters are clarified beside total number of hyperparameters (#λ). Conditional hyperparameters required for selected other hyperparameters are shown in brackets. # Configs indicates the default total number of configurations.

| Algorithm | #$\lambda$ | Categorical | Continuous | # Configs |
| --- | --- | --- | --- | --- |
| *Implemented with Pytorch<sup>15</sup>* |  |  |  |  |
| Transformer | 14 | 4 | 10 | 32 |
| 1D Convolutional Neural Network (1D-CNN) | 10 | 2 | 8 | 32 |
| Hybrid Convolutional Neural Network (Hybrid-CNN) | 10 | 2 | 8 | 32 |
| Gated Recurrent Unit (GRU) | 10 | 4 | 6 | 4 |
| Long Short-Term Memory (LSTM) | 11 | 4 | 7 | 4 |
| Standard Recurrent Neural Network (RNN) | 11 | 5 | 6 | 4 |
| *Implemented with XGBoost<sup>16</sup>* |  |  |  |  |
| Gradient Boosted Decision Trees (GBDT) | 18 | 5(1) | 13(4) | 16 |
| *Implemented with scikit-learn<sup>17</sup>* |  |  |  |  |
| Random Forests (RF) | 14 | 5(1) | 9(1) | 2 |
| Support Vector Machine (SVM) | 7 | 4 | 3(3) | 2 |

The general architecture designs of 1D-CNN and Hybrid-CNN are implemented based on 1D-CNN and 2D-Hybrid-CNN in a recent cancer type prediction study.^18^ However, taking one convolution layer and adjacent max-pooling layer as a layer set, we implemented both CNN models with flexibility on the number of layer set, which is fixed in the original paper. An example of 1D-CNN with two convolution layer sets is illustrated in Figure 3.

**Figure (3).**
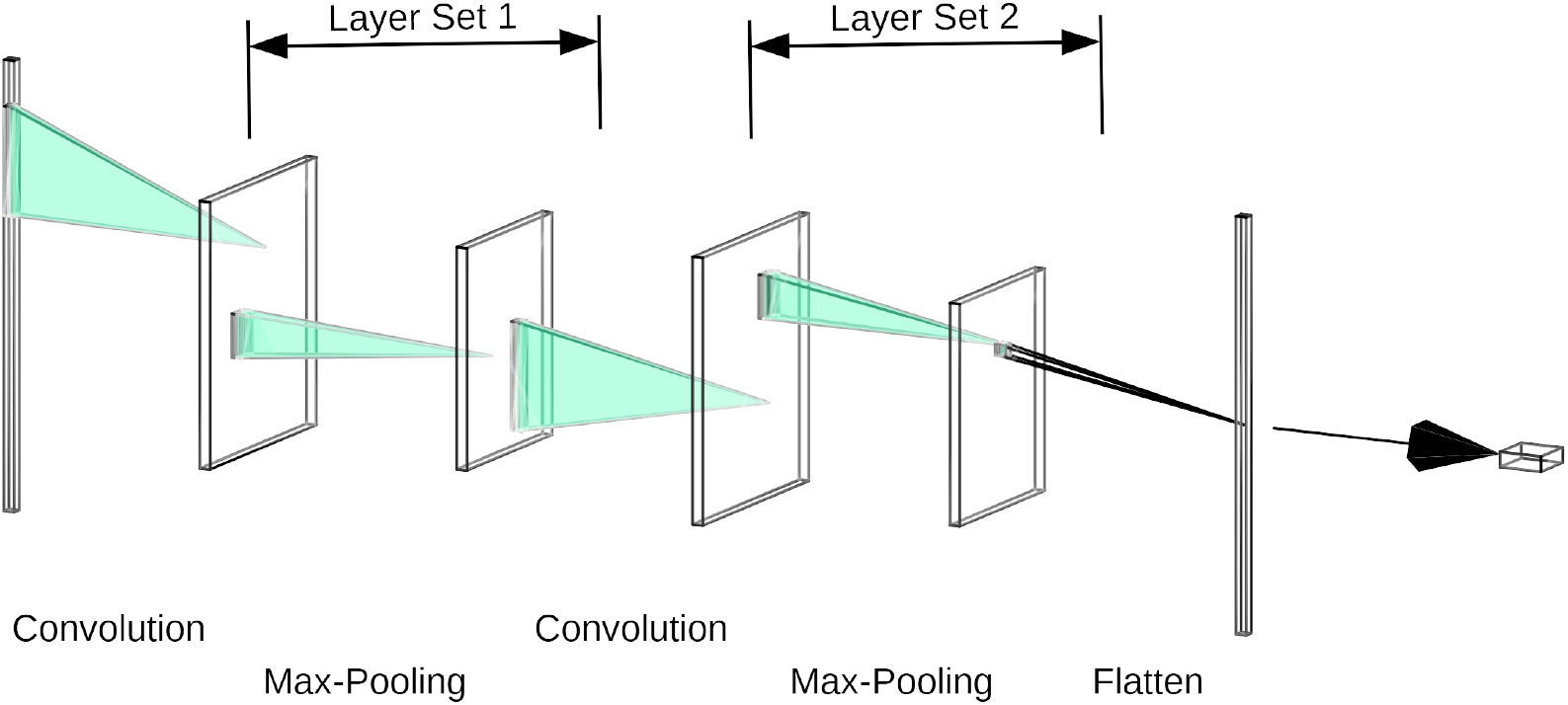
1D-CNN with 2 convolution layer set.

For transformer models, the embedding layer and positional encoding layer designed for input data tokenization in standard architecture^19^ are replaced with a single linear layer in AGEAS, considering psGRNs are already represented as numerical data matrix while GRPs should barely have positional relationships in the matrix.

### Step 3: Feature selection

With selected well-performing models, AGEAS already can start repetition of model training and interpretation in Step 4 to extract key GRPs. However, the few uninformative GRPs in training psGRNs could be pruned in advance to save computational resource. Furthermore, to prevent classification models from focusing on a small group of GRPs regardless of training psGRNs, GRPs draw excessive attention shall be separated from psGRNs to improve extraction comprehensiveness. Thus, in this step, AGEAS iteratively trains classification models with dynamic *α_max_* portion of psGRNs and obtains feature importance scores as described in subsection below to exclude GRPs either scored considerably high or relatively low.

More specifically, at each iteration, GRPs with z-scores ranked at the bottom *b* portion (default set as 0.1) are discarded. Also, an *i*-th ranked GRP will be separated from psGRNs and passed to Step 5 directly if it has a z-score fulfilling the condition:

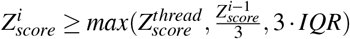

The 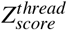 denotes input z-score threshold (default set as 3.0), and *IQR* stands for interquartile range calculated at the beginning of each iteration. With this criterion, AGEAS ensures only GRPs draw significantly more attention be selected by each iteration, despite different data distribution of z-score scaled importance values.

By default, AGEAS iterates this feature selection step for three times.

#### Feature importance estimation

AGEAS applies concept of The Shapley value^20^ for estimating the importance of each input feature, equivalent to GRP of input psGRN, in any kind of classification model when making predictions. Specific Shapley value calculation or approximation methods are implemented with *SHAP*^21^ and applied to different algorithms as shown in Table 2. We utilize *softmax* function to normalize feature importance scores and define the normalization function as:

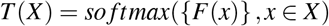

**Table (2).** Classifier algorithms with applicable Shapley value approximation methods. Algorithms marked with * have internalized feature importance estimation methods which will be applied with higher priority than Shapley value based methods. GBDT implemented with *XGBoost*^16^ can have feature importance approximated by the average weight gain at each split involving the feature. Linear SVM implemented with *scikit-learn*^17^ can have importance scores estimated with feature coefficients or using Linear Explainer. However, the feature coefficient estimation should be inappropriate for SVM with kernel function, While feature importance scores should be approximated with Kernel Explainer only.

| Algorithm | SHAP<sup>21</sup> Method |
| --- | --- |
| Transformer | Gradient Explainer |
| 1D-CNN | Deep Explainer |
| Hybrid-CNN | Deep Explainer |
| GRU | Gradient Explainer |
| LSTM | Gradient Explainer |
| RNN | Gradient Explainer |
| GBDT* | Tree Explainer |
| RF | Tree Explainer |
| SVM* | Linear Explainer / Kernel Explainer |

Here *X* denotes the total set of all features, and *F*(*x*) denotes importance score estimation method. If the feature importance can be approached with internalized method *f*(*x*), *F*(*x*) is set as:

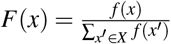

Otherwise, *F*(*x*) is defined utilizing correctly classified input samples *S*, equivalent to psGRNs, with Shapley values *φ* of feature x when predicting sample *s* as class *c*_1_ or class *c*_2_:

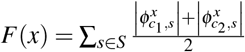

After all selected classification models *M* have been interpreted, we can integrate the feature importance matrices weighted by the corresponding models’ *L_CE_* to one matrix *A* as:

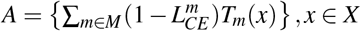

Then, generalized importance values are obtained through z-score calculation and later sorted descendingly:

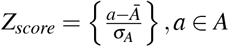

### Step 4: Top GRP extraction

To extract GRPs capable of effectively defining sample class differences, AGEAS iteratively initializes classification models with configurations kept from Step 2, trains them with randomly selected *α_max_* portion of rescaled psGRNs from Step 3, and interprets every models’ accurate predictions with methods mentioned in subsection. At each iteration, AGEAS receives a feature importance matrix A re-scaled by z-score and adds up each GRP’s score accordingly from matrix *A*′ kept from previous iteration if available. Then, top *α*, by default 100, ranked GRPs, whose scores are also greater than 0.0, are compared with GRPs extracted from *A*′ by the same setting. If changing portion of GRPs for top scored GRP sets from *A* and *A*′ is less than *d*, by default 0.05, AGEAS will consider the GRP extraction result of current iteration consistent with previous iteration. Extraction loop will terminate if AGEAS encounters *n* (default set as 3) continuous consistent result or runs out of preset iteration number (default set as 10). All feature importance scores in matrix *A* from last iteration are divided by the total iteration number processed, and top *a* ranked GRPs are considered as important GRPs for sample class differentiation.

### Step 5: Key network reconstruction

Analogizing every GRP previously extracted in Step 4 or separated as outlier in Step 3 as a directional edge connecting a TF vertex with a targetome vertex, AGEAS attempts to reconstruct a GRN graph representing regulatory differences between query sample classes. Since there is no guarantee on whether all extracted GRP edges can be connected or not, some regulatory relationships between gene vertices could be absent. Hence, AGEAS utilizes meta-GRN gained from Step 1 to find GRPs, which can further elucidate the regulatory relationships, and to use those GRPs as supportive edges.

For a limited iteration time (default set as 1), AGEAS exhaustively searches meta-GRN for TFs which can directly interact with any gene already covered in extracted GRPs and add the returned TFs as new vertices. Next, any GRP in meta-GRN capable of connecting two distinct vertices will be included as supportive GRP in extraction result. After expansion procedure above, the extracted GRN graph represents key genetic regulatory differences between input sample classes.

## Results

To assess predictive power of utilizing extraction result of AGEAS, we primarily applied AGEAS to study somatic reprogramming of induced pluripotent stem cells (iPSC) in mice,^22^ a milestone discovery of cell plasticity.^23^ With σ thread set to 2.0 for constraining total GRP amount in psGRNs, we use public scRNA-seq based GEMs of embryonic stem cells (ESCs) and mouse embryonic fibroblasts (MEFs) shown in Table 4 as inputs. The extraction result can be summarized as TF regulons, and TFs with highest regulatory degree in extracted GRN atlas are marked as top TFs (Figure 4a).

**Figure (4).**
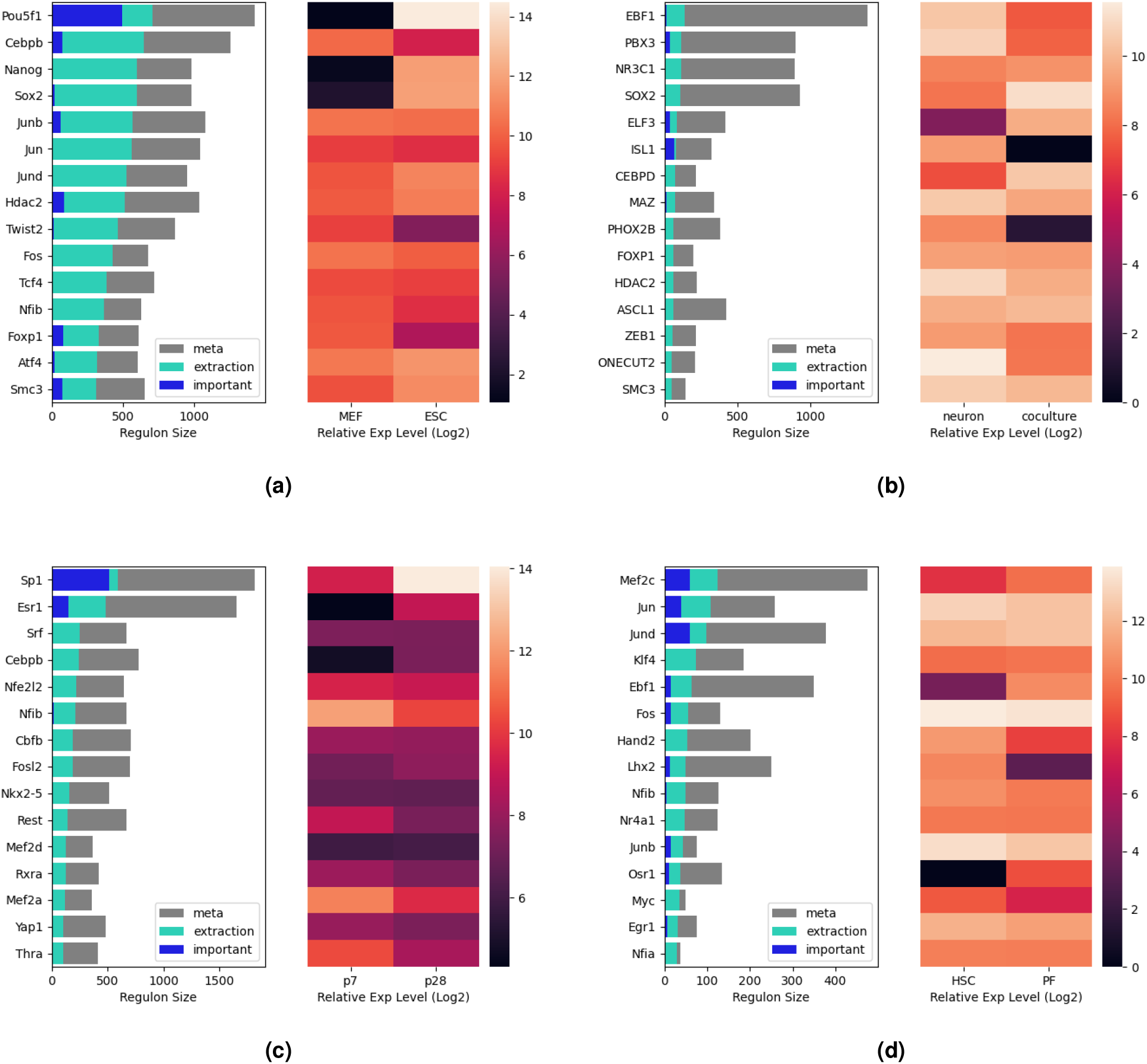
Top 15 TFs ranked by regulatory degrees in all GRPs extracted by AGEAS. ***(a)*** Mouse embryonic fibroblast vs. Embryonic stem cell ***(b)*** Purified dopaminergic neuron vs. Radial glial/neuronal co-culture ***(c)*** 7 days postnatal cardiomyocyte vs. 28 days postnatal cardiomyocyte ***(d)*** Hepatic stellate cell vs. Portal fibroblast (both after 6 weeks of CCl_4_ administration)

Among top TFs, Pou5f1 (Oct4) associates with GRPs extracted as important differential regulatory element between MEF and ESC more than any other TFs, while forming largest regulon and having expression pattern significantly favoring ESC. Thus, we infer Pou5f1 is the key regulatory element associated with ESC identity. Extensive studies already addressed Pou5f1’s important role in pluripotency maintenance.^24–26^ Moreover, to implement cell reprogramming of iPSC, Nanog and Sox2 are noteworthy, since they closely interact with Pou5f1 in extracted network, have high regulatory degrees, and are differently expressed in ESC. Multiple previous studies confirmed that all of Oct4, Nanog, and Sox2 are playing important roles to induce cell conversion from somatic cell to iPSC.^22,27–30^ Cebpb, Hdac2, and Smc3, extracted as top TFs directly affecting Pou5f1 expression, were also reported with relevant functions.^31–33^

Therefore, we infer that the investigation of TFs as common regulatory sources in important differential GRPs and corresponding regulons can effectively reflect differences between cell types or cell states. To further address AGEAS’s applicability and limitation, we applied AGEAS to three other scRNA-seq based studies in distinct research areas. With default settings described in Method section above, we evaluated AGEAS’s performance in following scenarios with public dataset (Table 4):

- **Dopaminergic neuron differentiation**: We address the applicability of AGEAS on cell subtype differentiation problems by analyzing the difference between human iPSC-derived radial glial / neuronal co-culture as neural progenitors^34^ and tyrosine hydroxylase (TH) expressing purified dopaminergic neurons (DANs). From extracted TFs (Figure 4b), we hypothesized that ISL1, ELF3, and PBX3 play important roles in DAN differentiation due to their high regulatory degree in extracted important GRPs. Previously, ISL1 was determined essential for differentiation of prethalamic DANs.^35^ ELF3 was found to be the neuronal precursor cell marker associated with neural stem cell development.^36^ This is consistent with ELF3’s high expression level in neuronal co-culture samples. Moreover, a DAN development study reported that PBX3 is required for correct differentiation of neuroblast, production of radial glial cell, into midbrain DAN and survival of midbrain DAN.^37^ Through stratifying all extracted GRPs, we only found that HDAC2 and RBPJ can potentially regulate all of ISL1, ELF3, and PBX3. Few reports demonstrated the association of HDAC2 with neurogenesis of radial glial cells.^38,39^ Nevertheless, HDAC2’s role in DAN differentiation is yet to be clarified. RBPJ was reported to be essential for DAN survival and to affect DAN development by regulating ASCL1, a essential factor in neurogenesis.^40,41^ The original research where we retrieved scRNA-seq data from also demonstrated that ASCL1 is a necessary factor to regulate DA neurotransmitter selection.^34^ Although AGEAS extracted ASCL1 as one of the top TFs, no direct regulatory relationship between RBPJ and ASCL1 was identified.
- **Postnatal cardiomyocyte maturation**: To address the performance of AGEAS to analyze cell physiological development, scRNA-seq data of cardiomyocytes (CMs) in postnatal day 7 mice (P7) and day 28 mice (P28) are used as inputs for extracting key genetic regulatory elements in postnatal CM maturation. As indicated by important differential GRPs, we first investigated Sp1 and Esr1 *(ERα*) that also have top regulatory degrees and considerably differential expression profiles (Figure 4c). Previous studies demonstrated that Sp1 promotes CM hypertrophy^42^ and maturation of electrophysiology, Ca^2+^ handling.^43,44^ Esr1 was found modulating myocardial development for postnatal cardiac growth.^45,46^ Beside the inter-regulation between Sp1 and Esr1, three other top TFs (Srf, Cebpb, Rest) and two TFs forming relatively small regulons (Gata4, Prox1), are expected to interact with both Sp1 and Esr1 based on extraction network. All five TFs were found to have CM maturation related functions. (Table 3)
- **Hepatic stellate cell activation in** *CCl*_4_ **induced liver fibrosis**: Here we address AGEAS’s applicability on cell pathological development studies through analyzing hepatic stellate cells (HSCs) and portal fibroblasts (PFs) simulating activated HSCs in mice liver administrated with chronic carbon tetrachloride (*CCl*_4_) for six weeks. Based on extracted important GRPs, we identified Mef2c, Jun, and JunD as potential representatives of key regulatory elements in HSC activation (Figure 4d). As the TF extracted with most important GRPs, Mef2c has been well-documented as the key regulator in HSC activation.^52–54^ Extensive studies also found Jun and JunD, functional components of the AP1 TF complex, are essential for fibrosis-related HSC activation and profibrogenic process.^55–58^ Analyzing TF regulons of Mef2c, Jun, and JunD, we believe that Mef2c is targetome of JunD and the inter-regulatory element of Jun. While no regulatory relationship between JunD and Jun determined, three extracted top TFs (Lhx2, Nfib, JunB) can potentially affect expressions of both JunD and Jun. JunB, another functional component of AP1, is also reported to affect HSC activation^59^. Furthermore, Lhx2 was found indispensable for quiescent HSCs.^60,61^ Although little literature had addressed Nfib’s role in HSC, Nfib was reported to be the anti-apoptotic gene for cell proliferation during *CCl*_4_ induced liver damage.^62^

**Table (3).**
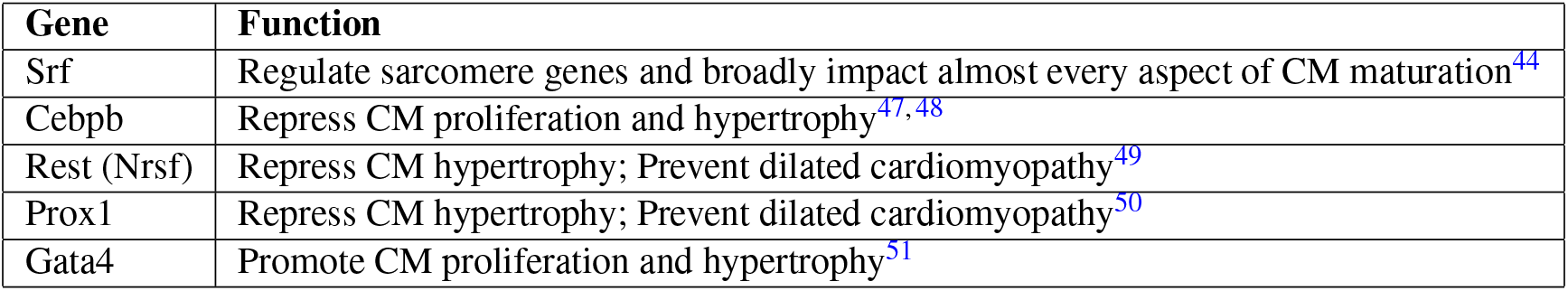
TFs interacting with both Sp1 and Esr1 order by regulatory degree in extracted GRNs.

**Table (4).**
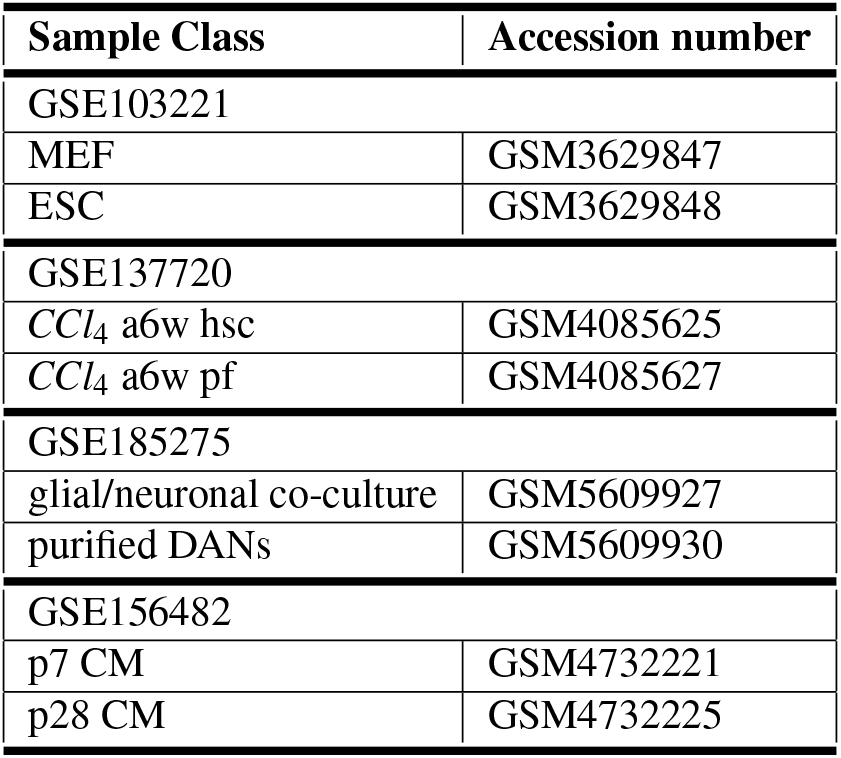

## Conclusion

In summary, AGEAS shows robustness in extracting key regulatory elements in several biological studies. Key TFs inferred by important differential GRPs and regulon analysis in all four *in silico* cases were confirmed informative to reveal main functionalities responsible for biological process of interest. Furthermore, the direct extractions of ISL1 in DAN differentiation and JunD in HSC activation demonstrated that AGEAS is neither focusing solely on the regulatory degrees in reconstructed meta-GRN nor the expression differences between sample classes. Overall, we anticipate that AGEAS is useful in providing insightful regulatory information and in promoting pioneer researches that reveal complicated mechanism behind biological phenomena.

## Discussion

Considering the rapid development of both sequencing technology and machine learning algorithms, we implemented AGEAS with modular design. In the data preprocessing part, the meta-GRN reconstruction section can be replaced with other methods to further increase fidelity of regulatory relationships or integrating more information, such as cis-regulatory elements (CREs) accessibility, which can be obtained from Assay for Transposase-Accessible Chromatin Sequencing (ATAC-seq) data. Moreover, CRE accessibility information combining with motif enrichment analysis is proved capable of identifying potential regulatory pathways to replace ChIP-seq based dataset utilized in the current study.^5^ Refining meta-GRN reconstruction method in consideration of having input data from simultaneous scRNA-seq and single cell ATAC-seq (scATAC-seq), such as SHARE-seq published in late 2020,^63^ can potentially further improve applicability of AGEAS; however, accessibility of corresponding datasets will be essential for performance assessment and method development. Similarly, initial set of classification models is also subject to change in response to future computational studies.

Also, it is important to note that AGEAS can be applied with GEMs normalized with different methods on user’s choice, as well as derived from other sequencing technologies, such as bulk RNA-seq or spatial sequencing in revealing tissue differences. Replacing transcriptiomic data with proteomic data may also be achievable in the analysis of protein interaction networks. However, larger scale of applicability tests are necessary.

Due to the stochastic nature of AGEAS, extraction results of AGEAS are not expected to stay identical for repeated applications even with exactly the same inputs. The main reason underlying the inconsistency is how AGEAS attempts to obtain top performing models for later prediction interpretations. Repeated random subsets of psGRNs used in model training and selection are expected to vary for each application, hence results in different performances of classification models and ranking results. As a result, the importance scores of GRPs might differ and slightly affect top TFs extracted later.

Two possible workarounds of the inconsistent issue would be: (i) manually set random seed for every process influencing subset selection. (ii) set multiple extractor unit and combine extraction results. In the present study, we found that n = 4 units yield relatively stable results, but this value could vary based on application scenario.

## Supporting information

Supplement 1 (MEF vs. ESC)

Supplement 2 (DAN vs. Cocul)

Supplement 4 (CCl4 a6w HSC vs. PF)

## Acknowledgement

We sincerely thank Cher Li and Jie Wang for inspiration; Augustine(Xiaoran) Yuan and Elizabeth Bania for reviewing and support.

## Author contributions statement

**J.Y.**: Methodology, Software, Writing-Original draft preparation, Project administration **M.N.**: Methodology, Software **J.T.**: Methodology, Writing-Original draft preparation, Project administration

## Additional information

All scRNA-seq datasets are retrieved from Gene Expression Omnibus(GEO) as described in the Table below.

## Notes

https://github.com/JackSSK/Ageas

### Competing Interest Statement

The authors have declared no competing interest.

### Summary of Updates

The overall method design is further improved, and more representative in silico tests are included.

