## Supplement 1 (MEF vs. ESC) for "AGEAS: Automated Machine Learning based Genetic Regulatory Element Extraction System": full_atlas.js

```
{
    "network_0": {
        "genes": {
            "Atf5": {
                "id": "Atf5",
                "type": "TF",
                "symbol": "Atf5",
                "source": [
                    "Hdac2",
                    "E2f4",
                    "Foxp1",
                    "Pou5f1",
                    "Rest",
                    "Trim28",
                    "Hmgb1",
                    "Atf4",
                    "Ctcf",
                    "Tgif1",
                    "Trp53",
                    "Hdac1",
                    "Tbx3",
                    "Ybx1",
                    "Twist2",
                    "Cebpb",
                    "Zfp57",
                    "Smarca5"
                ],
                "target": [
                    "Arf1",
                    "Cox6b1",
                    "Rnf7",
                    "Mrpl45",
                    "Mcm7",
                    "Rpl6",
                    "Bub1",
                    "Tspan17",
                    "Hspa5",
                    "Terf1",
                    "Itpa",
                    "Fosb",
                    "Spr",
                    "2810004N23Rik",
                    "Slc1a5",
                    "1110008F13Rik",
                    "Ctps",
                    "Arl6ip6",
                    "Smu1",
                    "A930018M24Rik",
                    "Eif2b2",
                    "Eif3c",
                    "Akap12",
                    "Dap3",
                    "Adrm1",
                    "Hdgfrp2",
                    "Ppfibp1",
                    "Vim",
                    "Ahnak",
                    "Tulp4",
                    "Asns",
                    "Fkbp10",
                    "Gabarap",
                    "Wars",
                    "Ddr2",
                    "Mrpl1",
                    "Rrp15",
                    "Aasdhppt",
                    "1110004F10Rik",
                    "Ubald1",
                    "Cmtm3",
                    "Calm3",
                    "Trib3",
                    "Mxd4",
                    "2810417H13Rik",
                    "Anapc15",
                    "Fam149a",
                    "Rdx",
                    "Camk1",
                    "Dmap1",
                    "1700021F05Rik",
                    "Ccdc80",
                    "Timp3",
                    "Lbh",
                    "Tmem176a",
                    "Poglut1",
                    "Uba3",
                    "1810011O10Rik",
                    "Rcc2",
                    "Ube2d2a",
                    "Tnks1bp1",
                    "Fbn2",
                    "Mapre1",
                    "Map1lc3b",
                    "Rps26-ps1",
                    "Dcp2",
                    "1110046J04Rik",
                    "Psme1",
                    "Hax1",
                    "Gm26682",
                    "Dubr",
                    "Cdc40",
                    "Atf1",
                    "Xpo1",
                    "S100a4",
                    "Kdm6b",
                    "Zyx",
                    "Nsfl1c",
                    "Cald1",
                    "Tsen34",
                    "Bcas2",
                    "Fas",
                    "0610007P14Rik",
                    "Spty2d1",
                    "H13",
                    "Marc2",
                    "Phf5a",
                    "Uchl3",
                    "Gtf2a2",
                    "2700060E02Rik",
                    "Tagln",
                    "Dnajc21",
                    "Npm1"
                ],
                "expression_sum": {
                    "class1": 10688.0,
                    "class2": 4042.0
                },
                "uniprot_ids": [
                    "O70191",
                    "Q3UJF3",
                    "A0A140LIB7",
                    "Q3TVK2"
                ]
            },
            "Asns": {
                "id": "Asns",
                "type": "Gene",
                "symbol": "Asns",
                "source": [
                    "Tcf4",
                    "Pou5f1",
                    "Atf5",
                    "Atf4",
                    "Fos",
                    "Jun",
                    "Sox2",
                    "Hes1",
                    "Nanog",
                    "Jund",
                    "Twist2",
                    "Cebpb"
                ],
                "target": [],
                "expression_sum": {
                    "class1": 9880.0,
                    "class2": 68065.0
                },
                "uniprot_ids": [
                    "Q61024",
                    "D3Z028",
                    "D3YYR0",
                    "D3Z0C2",
                    "Q3TIA9",
                    "Q3TV20"
                ]
            },
            "Tgif1": {
                "id": "Tgif1",
                "type": "TF",
                "symbol": "Tgif1",
                "source": [
                    "Tcf4",
                    "Pou5f1",
                    "Foxp1",
                    "Junb",
                    "Jun",
                    "Sox2",
                    "Hes1",
                    "Nanog",
                    "Jund",
                    "Twist2",
                    "Cebpb",
                    "Sox4"
                ],
                "target": [
                    "Mgat2",
                    "Dap",
                    "Atf5",
                    "Sar1a",
                    "Cdc42",
                    "Npc2",
                    "Gadd45g",
                    "Tcf4",
                    "Thbs1",
                    "Lims1",
                    "Tmsb4x",
                    "Cd47",
                    "Cd81",
                    "Itm2b",
                    "Pdia3",
                    "Txndc5",
                    "Tspan4",
                    "Anxa2",
                    "Copb2",
                    "Lmna",
                    "Nupr1",
                    "Copz2",
                    "Tmed2",
                    "Lgals3",
                    "Gsn",
                    "Prkar1a",
                    "Rap1b",
                    "Oat",
                    "Tagln2",
                    "Eif4g2",
                    "Surf4",
                    "Rab2a",
                    "Rexo2",
                    "Calr",
                    "Pdlim7",
                    "Tspo",
                    "Lhfp",
                    "Mbnl2",
                    "Selk",
                    "Neat1",
                    "Fam162a",
                    "Vimp",
                    "Cd63",
                    "Syncrip",
                    "Iqgap1",
                    "Sh3glb1",
                    "Fundc2",
                    "Rtn4",
                    "Ctsl",
                    "Ccnd2",
                    "Spcs1",
                    "Rpn1",
                    "Atp2b1",
                    "Tax1bp3",
                    "Nktr",
                    "Rpn2",
                    "Arpc2",
                    "Manf",
                    "Tln1",
                    "Sept11",
                    "Arhgdib",
                    "Tmed3",
                    "Timp2",
                    "Bzw1",
                    "Ubc",
                    "Sec62",
                    "Foxp1",
                    "Cdc42ep3",
                    "Cmpk1",
                    "Ptp4a2",
                    "Dnajc3",
                    "Hes1",
                    "Ap1s1",
                    "Myl12a",
                    "P4hb",
                    "Adipor1",
                    "Gnb2",
                    "Prdx5",
                    "Sri",
                    "Tm9sf3",
                    "Ociad1",
                    "Gnas",
                    "Camk2d",
                    "Sptbn1",
                    "Col1a1",
                    "Hdlbp",
                    "Ap3s1",
                    "Oxct1",
                    "Rsu1",
                    "Tmed9",
                    "Ncam1",
                    "Sepw1",
                    "Gsto1",
                    "Serp1",
                    "Fam195b",
                    "Tmem59",
                    "Vcan",
                    "Serpinb6a",
                    "Pdcl3",
                    "Junb",
                    "Csrp1",
                    "Hbegf",
                    "Ilk"
                ],
                "expression_sum": {
                    "class1": 3481.0,
                    "class2": 18140.0
                },
                "uniprot_ids": [
                    "P70284",
                    "A0A087WNP4",
                    "Q3TVD4",
                    "G3UWC5",
                    "E0CXJ4",
                    "E0CYI0",
                    "D3Z0Q5",
                    "F6TUU5"
                ]
            },
            "Tbx3": {
                "id": "Tbx3",
                "type": "TF",
                "symbol": "Tbx3",
                "source": [
                    "Pou5f1",
                    "Trim28",
                    "Sox2",
                    "Klf5",
                    "Nanog",
                    "Cebpb",
                    "Zfp57"
                ],
                "target": [
                    "Dppa5a",
                    "Dap",
                    "Cpe",
                    "Serpine2",
                    "Atf5",
                    "Tpm4",
                    "Itgb1",
                    "Ptrf",
                    "Cd44",
                    "Cdc34",
                    "Tipin",
                    "Igf2",
                    "Tmsb4x",
                    "Cd47",
                    "Efemp2",
                    "Cd81",
                    "Itm2b",
                    "Fbln2",
                    "Igfbp2",
                    "Atp5g1",
                    "Wisp1",
                    "Rps27l",
                    "Col12a1",
                    "Ngfrap1",
                    "Sod2",
                    "Trim28",
                    "Anxa2",
                    "Lmna",
                    "Cox7a2",
                    "Folr1",
                    "Prkcdbp",
                    "Vim",
                    "Lgals1",
                    "Ahnak",
                    "Fkbp10",
                    "Pam16",
                    "Phb2",
                    "Rpl9",
                    "Hmgn2",
                    "Nrp1",
                    "Sept1",
                    "Copz2",
                    "Gsn",
                    "Erh",
                    "Snrpg",
                    "Ran",
                    "Gjb3",
                    "Grb10",
                    "Jam2",
                    "Tpm1",
                    "Tuba1a",
                    "Utf1",
                    "Commd3",
                    "Gsta4",
                    "Pdlim7",
                    "Tspo",
                    "Lhfp",
                    "Mbnl2",
                    "Myl9",
                    "Spint2",
                    "Pdgfa",
                    "B2m",
                    "Uqcr10",
                    "Parva",
                    "H2afz",
                    "Gpx8",
                    "Gabarapl2",
                    "Phldb2",
                    "Rpl29",
                    "Igfbp7",
                    "Lrrc17",
                    "Cdc5l",
                    "Ung",
                    "Anxa5",
                    "S100a6",
                    "Dpy30",
                    "Odc1",
                    "Mdk",
                    "Neat1",
                    "Acta2",
                    "Ybx1",
                    "Nanog",
                    "Fstl1",
                    "Fam162a",
                    "Ifitm1",
                    "Marcks",
                    "Rhoc",
                    "Lbh",
                    "Fbn1",
                    "Nhp2",
                    "Plekhf2",
                    "Pmp22",
                    "Emp1",
                    "Rbmxl2",
                    "Tdgf1",
                    "Sh3glb1",
                    "App",
                    "Cisd1",
                    "Pole4",
                    "Impa2",
                    "Xist",
                    "Ccnd2",
                    "Tomm20",
                    "Gas1",
                    "Col6a1",
                    "Rpl21",
                    "S100a10",
                    "Mylpf",
                    "Filip1l",
                    "Col5a1",
                    "Bgn",
                    "Cav1",
                    "Fam25c",
                    "Tgfb1i1",
                    "Cdh11",
                    "Oaf",
                    "Tmed3",
                    "Tpm2",
                    "Timp2",
                    "Syngr1",
                    "Mt2",
                    "Hsd17b14",
                    "Fbln5",
                    "Morc1",
                    "Chd3",
                    "Rps12",
                    "H19",
                    "Glipr2",
                    "Dctpp1",
                    "Arpc1b",
                    "Col3a1",
                    "L1td1",
                    "Nenf",
                    "Zfp42",
                    "S100a4",
                    "Klf2",
                    "Rpp25",
                    "Zyx",
                    "Rcn3",
                    "Dut",
                    "Cald1",
                    "Sox2",
                    "Klf5",
                    "Tmem176b",
                    "Dpysl3",
                    "Mybl2",
                    "Slc2a3",
                    "Tma7",
                    "Gnas",
                    "Hcfc1r1",
                    "Camk2d",
                    "Fhl2",
                    "Calu",
                    "H3f3b",
                    "Gm7325",
                    "Rbpms2",
                    "Col1a1",
                    "Flna",
                    "Pmm1",
                    "Tpd52",
                    "Ckb",
                    "Efemp1",
                    "Gng12",
                    "Nop10",
                    "Prss23",
                    "Wls",
                    "Gdf3",
                    "Sepw1",
                    "Postn",
                    "Capn2",
                    "Laptm4a",
                    "Vcan",
                    "Dtymk",
                    "Sparc",
                    "Emp3",
                    "Ier5",
                    "Stmn2",
                    "Cebpb",
                    "Prnp",
                    "Id1",
                    "Serpinb6a",
                    "Alpl",
                    "Mkrn1",
                    "Hmga2",
                    "Nfatc2ip",
                    "Lpp",
                    "Kdelr3",
                    "Gm26917",
                    "Tagln",
                    "Kdm5b",
                    "P3h4",
                    "Anxa3",
                    "Hist1h2bc",
                    "Selm",
                    "Hells",
                    "Tfcp2l1",
                    "Axl",
                    "Cfl2",
                    "Ckap4",
                    "Col1a2"
                ],
                "expression_sum": {
                    "class1": 979.0,
                    "class2": 26585.0
                },
                "uniprot_ids": [
                    "P70324",
                    "Q8BG45",
                    "Q3ULL3",
                    "Q3U043",
                    "Q4VA43",
                    "Q8CJ82",
                    "Q8BJW3",
                    "Q8BU76"
                ]
            },
            "Zfp57": {
                "id": "Zfp57",
                "type": "TF",
                "symbol": "Zfp57",
                "source": [
                    "Pou5f1",
                    "Trim28",
                    "Sox2",
                    "Nanog",
                    "Cebpb",
                    "Sox4"
                ],
                "target": [
                    "Meg3",
                    "Lsm5",
                    "Pcbp1",
                    "Dap",
                    "Lsm7",
                    "Cpe",
                    "Serpine2",
                    "Atf5",
                    "Cdc20",
                    "Rcor2",
                    "Tpm4",
                    "Itgb1",
                    "Ptrf",
                    "Ppih",
                    "Cd44",
                    "Mat2a",
                    "Tpx2",
                    "Thbs1",
                    "G3bp2",
                    "Cd47",
                    "Efemp2",
                    "Cd81",
                    "Fbln2",
                    "Igfbp2",
                    "Atp5g1",
                    "Wisp1",
                    "Ndufs6",
                    "Ccnb1",
                    "Set",
                    "Trim28",
                    "Cebpzos",
                    "Aurka",
                    "Cks1b",
                    "Col12a1",
                    "Rpl12",
                    "Ssbp1",
                    "Anxa2",
                    "Lmna",
                    "Dnajc19",
                    "Vim",
                    "Tnfrsf12a",
                    "Ctbp2",
                    "Lgals1",
                    "Sephs2",
                    "Ahnak",
                    "Fkbp10",
                    "Gabarap",
                    "E130012A19Rik",
                    "Phb2",
                    "Rpl9",
                    "Cenpf",
                    "Copz2",
                    "Lsm3",
                    "Spag7",
                    "Arl6ip1",
                    "Tcof1",
                    "Gsn",
                    "Erh",
                    "Psmb6",
                    "Cks2",
                    "Snrpg",
                    "Hmgb2",
                    "Nhp2l1",
                    "Ran",
                    "Srsf2",
                    "Jam2",
                    "Grb10",
                    "Tpm1",
                    "Apex1",
                    "Commd3",
                    "Snrpb",
                    "Ccnb2",
                    "Fkbp4",
                    "Lhfp",
                    "Gas6",
                    "Nop56",
                    "Pdgfa",
                    "Snrpe",
                    "Btg2",
                    "Parva",
                    "Cdk1",
                    "H2afz",
                    "Lyar",
                    "Gpx8",
                    "Polr2e",
                    "Lsm2",
                    "Phldb2",
                    "Rpl29",
                    "Igfbp7",
                    "Lrrc17",
                    "Cenpa",
                    "Ranbp1",
                    "Anxa5",
                    "S100a6",
                    "Dpy30",
                    "Rbx1",
                    "H2afv",
                    "Neat1",
                    "Ybx1",
                    "Fstl1",
                    "Rhoc",
                    "Mrpl42",
                    "Timp3",
                    "Fbn1",
                    "Mrpl12",
                    "Snrpf",
                    "Exosc5",
                    "Tomm40",
                    "Rcc2",
                    "Slc29a1",
                    "Impa2",
                    "Ccnd2",
                    "Tax1bp3",
                    "Rpl21",
                    "Rangrf",
                    "Filip1l",
                    "Col5a1",
                    "Rbm8a",
                    "Bgn",
                    "Cav1",
                    "Fam25c",
                    "Rpl7a",
                    "Calm2",
                    "Cox7c",
                    "Stmn1",
                    "Ncl",
                    "Tgfb1i1",
                    "Cdh11",
                    "Ubb",
                    "Tmed3",
                    "Serpinf1",
                    "Srsf5",
                    "Rpl18",
                    "Tpm2",
                    "Timp2",
                    "Tubb4b",
                    "Parp1",
                    "Wdr74",
                    "Fbln5",
                    "Morc1",
                    "Chd3",
                    "Rps12",
                    "Psmb3",
                    "Prc1",
                    "Cdca3",
                    "Sub1",
                    "Fn1",
                    "Dctpp1",
                    "Glipr2",
                    "Arpc1b",
                    "Col3a1",
                    "Cdca8",
                    "Ppan",
                    "Ptma",
                    "Ube2s",
                    "Rpl27",
                    "Serpinh1",
                    "Klf2",
                    "Rpp25",
                    "Stra13",
                    "Rcn3",
                    "Dut",
                    "Zyx",
                    "Cald1",
                    "Fau",
                    "Mtf2",
                    "Sox2",
                    "Tmem176b",
                    "Mrpl15",
                    "Mfap2",
                    "Dpysl3",
                    "Mybl2",
                    "Tma7",
                    "Gnas",
                    "Aebp1",
                    "Fhl2",
                    "Rbpms2",
                    "Calu",
                    "Sept1",
                    "Col1a1",
                    "Gng12",
                    "Prss23",
                    "Wls",
                    "Hacd1",
                    "Eif5a",
                    "Laptm4a",
                    "Vcan",
                    "Serbp1",
                    "Tbx3",
                    "Sparc",
                    "Emp3",
                    "Hspe1",
                    "Hdgf",
                    "Cebpb",
                    "Prnp",
                    "Msn",
                    "Cryab",
                    "Ube2c",
                    "Mkrn1",
                    "Lpp",
                    "Tagln",
                    "Gm26917",
                    "Hmgb1",
                    "Hspb1",
                    "Anxa3",
                    "Birc5",
                    "Kdm5b",
                    "P3h4",
                    "Selm",
                    "Tfcp2l1",
                    "Sap30",
                    "Axl",
                    "Col1a2"
                ],
                "expression_sum": {
                    "class1": 613.0,
                    "class2": 18328.0
                },
                "uniprot_ids": [
                    "Q8C6P8",
                    "G3UYK1",
                    "G3UX77",
                    "G3UWX4",
                    "G3UYC1",
                    "G3UY05",
                    "H3BL53"
                ]
            },
            "Twist2": {
                "id": "Twist2",
                "type": "TF",
                "symbol": "Twist2",
                "source": [
                    "E2f4",
                    "Ctcf",
                    "Hdac1",
                    "Hdac2",
                    "Smarca5"
                ],
                "target": [
                    "Hsp90aa1",
                    "Lsm5",
                    "Arf1",
                    "Dap",
                    "Sar1a",
                    "Cdc20",
                    "Usmg5",
                    "Dctn3",
                    "Mcm7",
                    "Ddx24",
                    "Rps17",
                    "Rpl6",
                    "Srm",
                    "Grsf1",
                    "Rae1",
                    "Zc3h15",
                    "Cox8a",
                    "Mrps18b",
                    "Pin4",
                    "Ruvbl2",
                    "Psmb7",
                    "Spr",
                    "Txn2",
                    "Pnkd",
                    "Ppid",
                    "Ier2",
                    "2810004N23Rik",
                    "Clta",
                    "Dkc1",
                    "Marcksl1",
                    "Mrpl4",
                    "Atp5g1",
                    "1110008F13Rik",
                    "Ppp1r11",
                    "Polr2k",
                    "Sec13",
                    "Tuba1b",
                    "Gar1",
                    "Idh3g",
                    "Anapc5",
                    "Aurka",
                    "Psma6",
                    "Rpl7l1",
                    "Snrpa",
                    "St13",
                    "Ftsj3",
                    "Psmc1",
                    "Hnrnpa1",
                    "Ndufa2",
                    "Thoc7",
                    "Tpd52l2",
                    "Hnrnpf",
                    "Knop1",
                    "Mlf2",
                    "Psmd6",
                    "Rpl9",
                    "Ssrp1",
                    "Pin1",
                    "Atp5j2",
                    "Ier3ip1",
                    "Rnf187",
                    "Swi5",
                    "Eif2s1",
                    "Psmb6",
                    "Cks2",
                    "Shfm1",
                    "Med28",
                    "Hmgb2",
                    "Nubp1",
                    "Cox6c",
                    "Slbp",
                    "Ndufs3",
                    "Nedd4",
                    "Uba2",
                    "E2f4",
                    "Calr",
                    "Nup62",
                    "Cenpe",
                    "Rab18",
                    "Tubb5",
                    "Calm3",
                    "Arf6",
                    "Canx",
                    "D8Ertd738e",
                    "Hn1",
                    "Nop56",
                    "Ppp1r14b",
                    "Snrpe",
                    "Ubxn1",
                    "Mrpl14",
                    "Slc25a3",
                    "Cdk1",
                    "Phb",
                    "Lamtor1",
                    "2810417H13Rik",
                    "Psat1",
                    "Gabarapl2",
                    "Nup43",
                    "Taf7",
                    "Cdc5l",
                    "Ranbp1",
                    "Mrpl28",
                    "Psmc6",
                    "Emc6",
                    "Sms",
                    "Zwint",
                    "Rps24",
                    "Ppm1g",
                    "Eif1",
                    "Neat1",
                    "Sec61g",
                    "Ythdc1",
                    "Bola1",
                    "Ola1",
                    "Sf3b6",
                    "Snrnp27",
                    "Aurkaip1",
                    "Srsf1",
                    "Lrrc59",
                    "Uqcrc1",
                    "Exosc5",
                    "Rcc2",
                    "Atp5h",
                    "Bnip3l",
                    "Bola2",
                    "Tmem160",
                    "Kpnb1",
                    "Mapre1",
                    "C1d",
                    "Snrpd1",
                    "Rbm8a",
                    "Ppia",
                    "Hnrnpm",
                    "Capza2",
                    "Lap3",
                    "Loxl1",
                    "2310036O22Rik",
                    "Mphosph8",
                    "Supt4a",
                    "Srsf5",
                    "Aamp",
                    "Rpl35a",
                    "Chchd1",
                    "Wdr74",
                    "Gars",
                    "Psmc3",
                    "Hspa4",
                    "Luc7l2",
                    "Rpl36",
                    "Rbbp7",
                    "Psmb3",
                    "Hnrnpl",
                    "Ndufb9",
                    "Rbm39",
                    "Rps21",
                    "Arpp19",
                    "Ccdc12",
                    "Ddx21",
                    "Nedd8",
                    "Pcnp",
                    "Eef1b2",
                    "Utp3",
                    "Tmem258",
                    "Rpl24",
                    "Brix1",
                    "Fam136a",
                    "Mrpl15",
                    "Siva1",
                    "Fus",
                    "Ezh2",
                    "Mybl2",
                    "Ociad1",
                    "Pfdn1",
                    "Ddost",
                    "Rps4x",
                    "Pmm1",
                    "Fkbp8",
                    "Nip7",
                    "Ruvbl1",
                    "Clpp",
                    "Psma7",
                    "Ndufa8",
                    "Pdia4",
                    "Sept7",
                    "Gnl3",
                    "Trim27",
                    "Sssca1",
                    "Tpm3",
                    "Atp5b",
                    "Banf1",
                    "Cdc26",
                    "Dtymk",
                    "Immt",
                    "Mrps36",
                    "Emc10",
                    "Lig1",
                    "Psma1",
                    "Gpx1",
                    "Hdgf",
                    "Qdpr",
                    "Smc3",
                    "Thrap3",
                    "Tpi1",
                    "Csnk2b",
                    "Nars",
                    "Pa2g4",
                    "Pdlim1",
                    "Rpl37",
                    "Hspb1",
                    "Kdm5b",
                    "Atf4",
                    "Hist1h2bc",
                    "Ppp1cc",
                    "Eif3b",
                    "Zfp207",
                    "Pcbp1",
                    "Rpl38",
                    "Dnttip2",
                    "Sae1",
                    "Mrps24",
                    "Id2",
                    "Ndufa4",
                    "Ewsr1",
                    "Cops5",
                    "Eif4e",
                    "Ndufv1",
                    "Ndufv2",
                    "H3f3a",
                    "Nme1",
                    "Ube2k",
                    "Rfc2",
                    "Bccip",
                    "Csnk1a1",
                    "Ndufs2",
                    "Ddt",
                    "Fbxo22",
                    "Cyp51",
                    "Lamtor2",
                    "Tmsb10",
                    "Rpl3",
                    "Cebpzos",
                    "Nudt21",
                    "Trim28",
                    "Rpl12",
                    "Eif3c",
                    "Cox7a2",
                    "Cdv3",
                    "Crip2",
                    "Spc24",
                    "Ube2l3",
                    "Ctbp2",
                    "Asns",
                    "Ifitm2",
                    "Fdps",
                    "Ppp4c",
                    "Eef1d",
                    "Etf1",
                    "Ndufs7",
                    "Ube2b",
                    "Mcm2",
                    "Lsm3",
                    "Mrpl13",
                    "Gcat",
                    "Eif5",
                    "Tmed2",
                    "Tomm5",
                    "Txn1",
                    "Ube2j2",
                    "Erh",
                    "Ak2",
                    "Mrpl17",
                    "Samm50",
                    "Zmat2",
                    "Wdr77",
                    "Atp5j",
                    "Ran",
                    "Fscn1",
                    "Atp5g3",
                    "Srrm1",
                    "Atp5g2",
                    "Emc8",
                    "Srp19",
                    "Cct5",
                    "Emd",
                    "Mrpl40",
                    "Tceb1",
                    "C1qbp",
                    "Rpsa",
                    "H2afz",
                    "Bola3",
                    "Mrps21",
                    "Tardbp",
                    "Uqcr10",
                    "Cst3",
                    "Rpl36a",
                    "Eif3m",
                    "Rpl29",
                    "Iscu",
                    "Fubp1",
                    "Rpl5",
                    "Naa38",
                    "Ccdc58",
                    "Dph3",
                    "Prdx6",
                    "Odc1",
                    "Rps20",
                    "Ybx1",
                    "Dnaja2",
                    "Sys1",
                    "Tubb2b",
                    "Acat2",
                    "Tmem14c",
                    "Mrpl12",
                    "Cdc37",
                    "Ndufab1",
                    "Psmd1",
                    "Eef1a1",
                    "Mcl1",
                    "Rtn4",
                    "Cisd1",
                    "Rpl28",
                    "U2af1",
                    "Pole4",
                    "Ywhaq",
                    "Mrps17",
                    "Fbl",
                    "Pdha1",
                    "Ubald2",
                    "Dnajc8",
                    "Calm2",
                    "Rps3",
                    "Atox1",
                    "Stmn1",
                    "Srsf10",
                    "Adss",
                    "Btf3",
                    "Smarcc1",
                    "Igf2bp1",
                    "Ube2v2",
                    "Rpl18",
                    "Eno1",
                    "Hax1",
                    "Atp5d",
                    "Eef1e1",
                    "Rpl19",
                    "Taldo1",
                    "Timm13",
                    "Trnau1ap",
                    "Eif3h",
                    "Mcm3",
                    "Adprh",
                    "Rpl22l1",
                    "Vcp",
                    "Rps3a1",
                    "Mrps5",
                    "Srsf3",
                    "Hsd17b10",
                    "Smc4",
                    "Prc1",
                    "U2surp",
                    "Cmpk1",
                    "Mbd3",
                    "Rpl7",
                    "Cct7",
                    "Znrd1",
                    "Dctpp1",
                    "Ndufb2",
                    "Wdr18",
                    "Ppan",
                    "Hsp90ab1",
                    "Oaz1",
                    "Mrpl30",
                    "Uqcrq",
                    "Sec61b",
                    "BC031181",
                    "Polr2m",
                    "1810058I24Rik",
                    "Rwdd1",
                    "Snx5",
                    "Tyms",
                    "Gnai2",
                    "Rps8",
                    "Fnta",
                    "Dnaja1",
                    "Pgp",
                    "Eif4a1",
                    "Rpl8",
                    "Sumo2",
                    "Pcbp2",
                    "Magoh",
                    "Ndufc2",
                    "Polr1c",
                    "Tmed10",
                    "Snrnp25",
                    "Pmvk",
                    "H3f3b",
                    "Dnajc2",
                    "Ccdc34",
                    "Elof1",
                    "Gnb2l1",
                    "Mrps33",
                    "Ndufa12",
                    "Pold2",
                    "Nsa2",
                    "Rif1",
                    "Wdr5",
                    "Ddx46",
                    "Atxn10",
                    "Cnih4",
                    "Nudt19",
                    "Pebp1",
                    "Sgk1",
                    "Dbi",
                    "Morf4l2",
                    "Ube2d3",
                    "Aars",
                    "Rfc5",
                    "Snrpa1",
                    "Paip2",
                    "Pfdn4",
                    "Srrm2",
                    "Top1",
                    "Ywhae",
                    "Ubl5",
                    "Hmgb1",
                    "Mrps15",
                    "Psmc4",
                    "Ndufc1",
                    "Ppil1",
                    "Eif3d",
                    "Cyb5a",
                    "Rps15",
                    "Psma4",
                    "Arpc3",
                    "Scp2",
                    "Arl1",
                    "Arpc1a",
                    "Ap2s1",
                    "Rpl34",
                    "Park7",
                    "Mlec",
                    "Rab7",
                    "Sep15",
                    "Dynll1",
                    "Gtpbp4",
                    "Cenpw",
                    "Eif2s2",
                    "Hspd1",
                    "Atf5",
                    "Stoml2",
                    "Cops7a",
                    "Cdc42",
                    "Mrpl45",
                    "Rpl31",
                    "Srsf7",
                    "Smdt1",
                    "Ppih",
                    "Nap1l1",
                    "Uqcc3",
                    "Tpx2",
                    "Etfb",
                    "Dnajb6",
                    "Brd2",
                    "Actr3",
                    "Impdh2",
                    "Ccdc124",
                    "Mrps10",
                    "Gapdh",
                    "Ccnb1",
                    "Prdx2",
                    "Mpc1",
                    "Akap12",
                    "Ccnd3",
                    "Khdrbs1",
                    "Mrpl52",
                    "Adrm1",
                    "Abcf1",
                    "Psmb5",
                    "Hspa9",
                    "Chchd2",
                    "Nop58",
                    "Ube2n",
                    "Ndufa11",
                    "Smn1",
                    "Cbx5",
                    "Kif5b",
                    "Polr2c",
                    "Cct8",
                    "Hmgn2",
                    "Dohh",
                    "Ptp4a1",
                    "Ndufs5",
                    "Oat",
                    "Akr1b3",
                    "Ddx27",
                    "Ndufb8",
                    "Psmd2",
                    "Srsf2",
                    "Prdx4",
                    "Eif4g2",
                    "Jade1",
                    "Dnajc7",
                    "Uqcc2",
                    "Pak1ip1",
                    "Denr",
                    "Fxr1",
                    "Nsmce4a",
                    "Rps26",
                    "Get4",
                    "Eif3g",
                    "Nxt1",
                    "Trp53",
                    "Sra1",
                    "Tceal8",
                    "Aspscr1",
                    "Psmd11",
                    "Cnn3",
                    "Vps29",
                    "Vdac1",
                    "Rpp21",
                    "Cggbp1",
                    "Lage3",
                    "Anp32a",
                    "Rad23b",
                    "Zfp706",
                    "Polr2e",
                    "Lsm2",
                    "Nudt4",
                    "Tra2a",
                    "Pdcd5",
                    "Aprt",
                    "Ost4",
                    "Snrpc",
                    "Mrpl51",
                    "Mrpl49",
                    "Eif4g1",
                    "Tmbim6",
                    "Dynll2",
                    "Rhoa",
                    "Fcf1",
                    "Mrps18c",
                    "Atp5a1",
                    "Cnpy2",
                    "Rpl22",
                    "Pycr2",
                    "Sf3a3",
                    "Anp32e",
                    "Ssr2",
                    "Ict1",
                    "Rpl4",
                    "Glud1",
                    "March5",
                    "Cebpz",
                    "Fermt2",
                    "Hnrnpu",
                    "Ddah2",
                    "Rbmxl1",
                    "Gspt1",
                    "Arglu1",
                    "Imp4",
                    "Srsf11",
                    "Rpl18a",
                    "Hmgb3",
                    "Cbx1",
                    "Tomm20",
                    "Txndc9",
                    "Vapa",
                    "Rpl21",
                    "Map1lc3b",
                    "Eprs",
                    "Trmt112",
                    "Rpl35",
                    "Dazap1",
                    "Msmo1",
                    "Atp5l",
                    "Mrpl54",
                    "Pop5",
                    "Son",
                    "Ilf2",
                    "Cct2",
                    "Mak16",
                    "Ubb",
                    "Dtl",
                    "Ahsa1",
                    "Zfp106",
                    "Prrc1",
                    "Eif1ax",
                    "Parp1",
                    "Ak6",
                    "Rbpms",
                    "Paics",
                    "Hnrnpa3",
                    "Prmt1",
                    "Pbdc1",
                    "Rps19bp1",
                    "Cdca3",
                    "Metap2",
                    "Sub1",
                    "Asna1",
                    "Psmc5",
                    "Echdc2",
                    "Morf4l1",
                    "Mrpl43",
                    "Pafah1b1",
                    "Cdca8",
                    "Lsm6",
                    "Tecr",
                    "Sf3b1",
                    "Ube2s",
                    "Ndufs4",
                    "Mybbp1a",
                    "Snrpd3",
                    "Sf3b2",
                    "Psmd3",
                    "Suclg1",
                    "H2afy",
                    "Fam32a",
                    "Mtf2",
                    "Slc3a2",
                    "Stip1",
                    "Rpl14",
                    "Tma7",
                    "Ash2l",
                    "Hat1",
                    "Cops4",
                    "Rnps1",
                    "Lsm4",
                    "Pfdn2",
                    "Nop10",
                    "Mpdu1",
                    "Atp5f1",
                    "Eif1ad",
                    "Arid5b",
                    "Rps13",
                    "Gsto1",
                    "Tpt1",
                    "Eif5a",
                    "Rps11",
                    "Ppib",
                    "Cfl1",
                    "Rrp1b",
                    "Hspe1",
                    "Rps19",
                    "Ube2c",
                    "Exosc8",
                    "Cops6",
                    "Ptbp1",
                    "Aldoa",
                    "Mrpl20",
                    "Hnrnpdl",
                    "Srp9",
                    "Eif3a",
                    "Rtn3",
                    "Polr2h",
                    "Psmb4",
                    "Snx3",
                    "Strap",
                    "Commd1",
                    "l7Rn6",
                    "Rnaseh2a",
                    "Chtop",
                    "Srpk1",
                    "Cox6b1",
                    "Ino80e",
                    "Ppa1",
                    "Atp5e",
                    "Avpi1",
                    "Prdx1",
                    "Rsl24d1",
                    "Ssb",
                    "Tipin",
                    "Atp5o",
                    "Mrps12",
                    "Rrm2",
                    "Rpl13",
                    "Copz1",
                    "Capns1",
                    "Mrpl34",
                    "Nasp",
                    "Pdia3",
                    "Ndufb5",
                    "Rpl10a",
                    "Ndufs6",
                    "Set",
                    "Thyn1",
                    "Eif4e2",
                    "Shmt1",
                    "Alkbh1",
                    "Sod2",
                    "Ssbp1",
                    "Rps14",
                    "Tgif1",
                    "Sfpq",
                    "Mrto4",
                    "Rpl32",
                    "Parl",
                    "Lipe",
                    "Clns1a",
                    "Snrpd2",
                    "1810022K09Rik",
                    "Uqcr11",
                    "Phb2",
                    "Rps27a",
                    "Naa10",
                    "Bud31",
                    "Gtf2e2",
                    "Pdap1",
                    "Mcm5",
                    "Hsp90b1",
                    "Arl6ip1",
                    "Rpl11",
                    "Rpl15",
                    "Prkar1a",
                    "Rap1b",
                    "Tcof1",
                    "Vdac2",
                    "Pfn1",
                    "Rpl13a",
                    "Nol7",
                    "Snrpg",
                    "Tkt",
                    "Mrpl24",
                    "Snhg6",
                    "Cox5a",
                    "Minos1",
                    "Snrpb",
                    "Tmem11",
                    "Fkbp4",
                    "Zfp428",
                    "Rabggtb",
                    "Tomm22",
                    "Lyar",
                    "Ubap2l",
                    "Erp29",
                    "Snrnp70",
                    "Uchl5",
                    "Ppig",
                    "Cfdp1",
                    "Ppdpf",
                    "Dcun1d5",
                    "Eif5b",
                    "Cct4",
                    "Llph",
                    "Emg1",
                    "Lsm12",
                    "Eif4ebp1",
                    "H2afv",
                    "Nono",
                    "Csde1",
                    "Prrc2c",
                    "Gins4",
                    "Tceb2",
                    "Mrpl42",
                    "Psmg4",
                    "Rps2",
                    "Gart",
                    "Polr2g",
                    "Sumo3",
                    "Psmc2",
                    "Ndufa10",
                    "Tomm40",
                    "Fundc2",
                    "Uqcrfs1",
                    "Naa50",
                    "Ywhah",
                    "Sf3b4",
                    "Eif3i",
                    "Hdac2",
                    "Sugt1",
                    "Hnrnpc",
                    "Smc2",
                    "Fkbp3",
                    "Mtch2",
                    "Rangrf",
                    "Gins2",
                    "Bub3",
                    "Rpl7a",
                    "Uck2",
                    "Papola",
                    "Rps16",
                    "Esf1",
                    "Sin3b",
                    "Mrfap1",
                    "Tubb4b",
                    "Ccna2",
                    "Mrps16",
                    "Slc25a5",
                    "Uqcrh",
                    "Ddx18",
                    "Rrp9",
                    "Tbca",
                    "Eed",
                    "Ccne1",
                    "Srsf6",
                    "Ndufa5",
                    "Tmpo",
                    "Ndufb10",
                    "Tsn",
                    "Mrpl11",
                    "Rpl37a",
                    "Txnl1",
                    "Mesdc2",
                    "Eef1g",
                    "Rbm25",
                    "2700094K13Rik",
                    "Cycs",
                    "Myl12a",
                    "Ptma",
                    "Rac1",
                    "Rer1",
                    "Dut",
                    "Dhx16",
                    "Sdhb",
                    "Taf10",
                    "Wdr43",
                    "Ybx3",
                    "Klf5",
                    "Acin1",
                    "Rps10",
                    "Pabpc1",
                    "Eif6",
                    "Yrdc",
                    "Hspa14",
                    "Fkbp1a",
                    "Polr2j",
                    "Got2",
                    "Dad1",
                    "Slirp",
                    "0610007P14Rik",
                    "Uhrf1",
                    "Yif1b",
                    "Arpc5",
                    "Wbp11",
                    "Cox4i1",
                    "Snf8",
                    "Cwc15",
                    "Guk1",
                    "2810428I15Rik",
                    "Hspa8",
                    "Ndufs8",
                    "Psma3",
                    "Rpl39",
                    "Rplp2",
                    "Postn",
                    "Rps18",
                    "Tsfm",
                    "Mif",
                    "Ptov1",
                    "Aimp1",
                    "Phf5a",
                    "Chchd3",
                    "Gcsh",
                    "Glrx3",
                    "Gtf2a2",
                    "Mkrn1",
                    "Rrp1",
                    "Pdcl3",
                    "Rpl30",
                    "Sfr1",
                    "Myl12b",
                    "Kdm1a",
                    "Rrs1",
                    "Ncor1",
                    "Rbm17",
                    "Ly6e",
                    "Cnbp",
                    "Rsrc2"
                ],
                "expression_sum": {
                    "class1": 6849.0,
                    "class2": 547.0
                },
                "uniprot_ids": [
                    "Q9D030",
                    "A5D6P6"
                ]
            },
            "Ctcf": {
                "id": "Ctcf",
                "type": "TF",
                "symbol": "Ctcf",
                "source": [
                    "Tfcp2l1",
                    "Pou5f1",
                    "Foxp1",
                    "Junb",
                    "Id2",
                    "Jun",
                    "Sox2",
                    "Hes1",
                    "Nanog",
                    "Jund",
                    "Cebpb",
                    "Sox4"
                ],
                "target": [
                    "Arf1",
                    "Dap",
                    "Sar1a",
                    "Tpm4",
                    "Cd44",
                    "Myh9",
                    "Thbs1",
                    "Lims1",
                    "Sec13",
                    "Tuba1b",
                    "Wisp1",
                    "Slc25a4",
                    "Tspan4",
                    "Col12a1",
                    "Lmna",
                    "Cthrc1",
                    "Pck2",
                    "Rnf187",
                    "Lgals3",
                    "mt-Co1",
                    "Pgrmc1",
                    "Tagln2",
                    "Rps15a",
                    "Rexo2",
                    "Calr",
                    "Pdlim7",
                    "Tubb5",
                    "Mfge8",
                    "Mbnl2",
                    "Canx",
                    "Actn1",
                    "B2m",
                    "Slc25a3",
                    "Psat1",
                    "2810417H13Rik",
                    "Gdi2",
                    "Nr0b1",
                    "Prrx1",
                    "Neat1",
                    "Acta2",
                    "Myh10",
                    "Nanog",
                    "Ola1",
                    "Ube2r2",
                    "Tgfbi",
                    "Timp3",
                    "Gemin7",
                    "Akr1a1",
                    "Tdgf1",
                    "Sh3glb1",
                    "Atp2b1",
                    "Mapre1",
                    "Cisd2",
                    "Mylpf",
                    "Nktr",
                    "Col5a1",
                    "Tspan3",
                    "Bgn",
                    "Capza2",
                    "Klf6",
                    "Pcolce",
                    "Tgfb1i1",
                    "Loxl1",
                    "Timp2",
                    "Syngr1",
                    "Morc1",
                    "Sec62",
                    "Rps12",
                    "Maged1",
                    "Arpc1b",
                    "Col3a1",
                    "Cyr61",
                    "Kdelr1",
                    "Rab14",
                    "Serpinh1",
                    "Atp6v1g1",
                    "Gnb2",
                    "Ociad1",
                    "Dpysl3",
                    "Ddost",
                    "Hcfc1r1",
                    "Inhba",
                    "Sptbn1",
                    "Id3",
                    "Ap3s1",
                    "Gng12",
                    "Ncam1",
                    "Wls",
                    "Hacd1",
                    "Ndufa8",
                    "Pdia4",
                    "Srp14",
                    "Sept7",
                    "Fam195b",
                    "Actg1",
                    "Rcn2",
                    "Tsg101",
                    "Ier5",
                    "Prnp",
                    "Tpi1",
                    "Id1",
                    "Hmga2",
                    "Atf4",
                    "Tfcp2l1",
                    "Nrp1",
                    "Col1a2",
                    "F2r",
                    "Fkbp9",
                    "Serpine2",
                    "Igfbp4",
                    "Id2",
                    "Itgb1",
                    "Fabp5",
                    "Mettl9",
                    "Tcf4",
                    "Igf2",
                    "Cd81",
                    "Ssr1",
                    "Pgls",
                    "Lxn",
                    "Anxa2",
                    "Vmp1",
                    "Col5a2",
                    "mt-Nd4",
                    "Crip2",
                    "Cox7a2l",
                    "Vim",
                    "Basp1",
                    "Platr3",
                    "Sh3bgrl3",
                    "Wdr1",
                    "Tnc",
                    "Ube2b",
                    "Tmed2",
                    "Tpm1",
                    "Ldha",
                    "Sdcbp",
                    "Thy1",
                    "Surf4",
                    "Utf1",
                    "Phlda1",
                    "Myeov2",
                    "Vcl",
                    "Cst3",
                    "Eif3m",
                    "Lox",
                    "Higd1a",
                    "S100a6",
                    "H2-D1",
                    "Crabp1",
                    "Cd63",
                    "Capg",
                    "Lbh",
                    "S100a11",
                    "Scand1",
                    "Mtdh",
                    "Eef1a1",
                    "mt-Nd5",
                    "Maged2",
                    "Rtn4",
                    "Ctsl",
                    "Xist",
                    "Ccnd2",
                    "Tcf25",
                    "Spcs1",
                    "Cox6a1",
                    "Col6a1",
                    "Rala",
                    "Sdc2",
                    "Tax1bp3",
                    "Filip1l",
                    "Cav1",
                    "Plk2",
                    "Tln1",
                    "Cdh11",
                    "Oaf",
                    "Ostf1",
                    "Mt2",
                    "Eif3h",
                    "Fbln5",
                    "Chd3",
                    "Cmpk1",
                    "Fkbp2",
                    "Lrrc58",
                    "Fn1",
                    "Glipr2",
                    "Hes1",
                    "Ap1s1",
                    "Nenf",
                    "Rps9",
                    "Twist2",
                    "Zfp42",
                    "Rpp25",
                    "Rcn3",
                    "Malat1",
                    "Gnai2",
                    "Tmem176b",
                    "Mfap2",
                    "Tm9sf3",
                    "Tmed10",
                    "Ctsd",
                    "Reep5",
                    "Pou5f1",
                    "Fhl2",
                    "Glrx5",
                    "Col1a1",
                    "Pla2g1b",
                    "Oxct1",
                    "Peg3",
                    "Atxn10",
                    "Atp6v0e",
                    "Cnih4",
                    "Dbi",
                    "Tmem59",
                    "Eif3e",
                    "Emp3",
                    "Aspn",
                    "Cebpb",
                    "Pafah1b2",
                    "Anxa3",
                    "Psmd14",
                    "Hbegf",
                    "Arpc3",
                    "Ilk",
                    "Arl1",
                    "Mgat2",
                    "Sep15",
                    "Mlec",
                    "Lsm7",
                    "Cpe",
                    "Atf5",
                    "Arf5",
                    "Cdc42",
                    "Npc2",
                    "Eef2",
                    "Hspa5",
                    "Rnh1",
                    "Rrbp1",
                    "Actr3",
                    "Tmsb4x",
                    "Cd47",
                    "Efemp2",
                    "Fbln2",
                    "Mpc1",
                    "Copb2",
                    "Sars",
                    "Ap2m1",
                    "Nupr1",
                    "Prkcdbp",
                    "Sdpr",
                    "Lgals1",
                    "Ahnak",
                    "Col11a1",
                    "Fkbp10",
                    "Kif5b",
                    "Copz2",
                    "Ptp4a1",
                    "Lsp1",
                    "Gsn",
                    "Oat",
                    "Tubb6",
                    "Eva1b",
                    "Prdx4",
                    "Aes",
                    "Rab2a",
                    "Tspo",
                    "Lhfp",
                    "Myl9",
                    "Vps29",
                    "Btg2",
                    "Parva",
                    "Selk",
                    "Jund",
                    "Rab11a",
                    "Anxa5",
                    "Eif3f",
                    "Anxa6",
                    "Mtch1",
                    "Rhoa",
                    "Rab10",
                    "Vimp",
                    "Ccdc80",
                    "Sec61a1",
                    "Iqgap1",
                    "Kdelr2",
                    "Pmp22",
                    "Rbmxl2",
                    "Ssr3",
                    "Rpn1",
                    "Gas1",
                    "Ssr4",
                    "Txndc9",
                    "Lman1",
                    "F3",
                    "Rcn1",
                    "Arpc2",
                    "Fam25c",
                    "Ywhab",
                    "Sox4",
                    "Phlda3",
                    "Sept11",
                    "Tpm2",
                    "Ergic3",
                    "Bzw1",
                    "Sdc4",
                    "Ubc",
                    "Rock2",
                    "Cdc42ep3",
                    "Tcea3",
                    "Col8a1",
                    "L1td1",
                    "Psap",
                    "Tm4sf1",
                    "Rbms1",
                    "P4hb",
                    "Scd2",
                    "Dlk1",
                    "Zyx",
                    "Map1b",
                    "Rab34",
                    "Adipor1",
                    "Bsg",
                    "Sox2",
                    "Ppic",
                    "Camk2d",
                    "mt-Cytb",
                    "Cyba",
                    "Flna",
                    "Efemp1",
                    "H13",
                    "Rsu1",
                    "Prss23",
                    "Gdf3",
                    "Gsto1",
                    "Ppib",
                    "Vcan",
                    "H1f0",
                    "Ndufb11",
                    "Sparc",
                    "Sept2",
                    "Abracl",
                    "Msn",
                    "Aldoa",
                    "Serpinb6a",
                    "Rtn3",
                    "Lpp",
                    "Ptn",
                    "Tagln",
                    "Ctbp1",
                    "Csrp1",
                    "Selm",
                    "Ckap4",
                    "Esd",
                    "Dppa5a",
                    "Meg3",
                    "Atxn7l3b",
                    "Igfbp6",
                    "Comt",
                    "Lamp1",
                    "Rcor2",
                    "Tinagl1",
                    "Ptrf",
                    "Gadd45g",
                    "Rrm2",
                    "Sh3bgrl",
                    "Jun",
                    "Itm2b",
                    "Pdia3",
                    "Rpl10a",
                    "Txndc5",
                    "Col4a1",
                    "Hist1h1c",
                    "Cd24a",
                    "Lipe",
                    "Tnfrsf12a",
                    "Chchd10",
                    "Krtcap2",
                    "Gabarap",
                    "Phgdh",
                    "Rap1a",
                    "Hsp90b1",
                    "Mtpn",
                    "Anxa1",
                    "Prkar1a",
                    "Timp1",
                    "Ier3",
                    "Grb10",
                    "Tuba1a",
                    "Commd3",
                    "Cnn2",
                    "Gsta4",
                    "Fst",
                    "Gpx8",
                    "Gpi1",
                    "Tmem167",
                    "Phldb2",
                    "Palld",
                    "Tram1",
                    "Igfbp7",
                    "Lrrc17",
                    "Rpl23a-ps3",
                    "Cdkn1a",
                    "Fstl1",
                    "Marcks",
                    "Fam162a",
                    "Rhoc",
                    "Ifitm3",
                    "Fbn1",
                    "Srp72",
                    "Mrpl33",
                    "Emp1",
                    "App",
                    "Fundc2",
                    "Eif3i",
                    "Fbn2",
                    "Gm9843",
                    "S100a10",
                    "Bag1",
                    "Rpn2",
                    "Mfap4",
                    "Prrx2",
                    "Cetn3",
                    "Dctn2",
                    "Arhgdib",
                    "Tmed3",
                    "Serpinf1",
                    "Hsd17b14",
                    "Gap43",
                    "Tbca",
                    "Foxp1",
                    "Tmpo",
                    "Ccng1",
                    "H19",
                    "Ptp4a2",
                    "Txnl1",
                    "Cope",
                    "Dnajc3",
                    "Arhgdia",
                    "Myl12a",
                    "Rac1",
                    "Rer1",
                    "S100a4",
                    "Klf2",
                    "Cald1",
                    "Fau",
                    "Ooep",
                    "Sri",
                    "Prdx5",
                    "Clic1",
                    "Fkbp1a",
                    "Emc7",
                    "Gnas",
                    "Aebp1",
                    "Arf4",
                    "Sept1",
                    "Calu",
                    "Arpc5",
                    "Gm7325",
                    "Hdlbp",
                    "Limd2",
                    "Cox4i1",
                    "Rps18-ps3",
                    "Tmed9",
                    "Myl6",
                    "Sepw1",
                    "Serp1",
                    "Capn2",
                    "Laptm4a",
                    "Cnih1",
                    "Rheb",
                    "Carhsp1",
                    "Alpl",
                    "Pdcl3",
                    "Kdelr3",
                    "P3h4",
                    "Junb",
                    "Bcap31",
                    "Axl",
                    "Cfl2",
                    "Ogn"
                ],
                "expression_sum": {
                    "class1": 3828.0,
                    "class2": 17925.0
                },
                "uniprot_ids": [
                    "Q61164",
                    "Q3UYZ8",
                    "Q05CK6",
                    "Q3UZH8",
                    "Q3USR8"
                ]
            },
            "Cebpb": {
                "id": "Cebpb",
                "type": "TF",
                "symbol": "Cebpb",
                "source": [
                    "E2f4",
                    "Foxp1",
                    "Pou5f1",
                    "Atf4",
                    "Ctcf",
                    "Trp53",
                    "Hdac1",
                    "Tbx3",
                    "Zfp57",
                    "Hdac2",
                    "Smc3",
                    "Smarca5"
                ],
                "target": [
                    "Arf1",
                    "Dap",
                    "Srsf9",
                    "Ndufb4",
                    "Syce2",
                    "Mcm7",
                    "Grsf1",
                    "Zc3h15",
                    "Cox8a",
                    "2810004N23Rik",
                    "Sec13",
                    "Ndufa6",
                    "Polr2k",
                    "Idh3g",
                    "Rps27l",
                    "Col12a1",
                    "Lmna",
                    "Ndufa2",
                    "Hnrnpf",
                    "Lmnb1",
                    "Ddx39",
                    "Cks2",
                    "Shfm1",
                    "Med28",
                    "Hmgb2",
                    "Cox6c",
                    "Ddx3x",
                    "Slbp",
                    "Nedd4",
                    "Calr",
                    "Tubb5",
                    "Nop56",
                    "Slc25a3",
                    "Lamtor1",
                    "2810417H13Rik",
                    "Nup43",
                    "Cdc5l",
                    "Sms",
                    "Rbx1",
                    "Ctcf",
                    "Neat1",
                    "Pmf1",
                    "Ola1",
                    "Sf3b6",
                    "Lrrc59",
                    "Ube2d2a",
                    "Bnip3l",
                    "Kpnb1",
                    "Mapre1",
                    "Snrpd1",
                    "Rbm8a",
                    "Col5a1",
                    "Rpl36al",
                    "Lap3",
                    "Tgfb1i1",
                    "Supt4a",
                    "Rbbp7",
                    "Hnrnpl",
                    "Nedd8",
                    "Pcnp",
                    "Atpif1",
                    "Snrnp40",
                    "Gng12",
                    "Tpm3",
                    "Banf1",
                    "Mrps36",
                    "Qdpr",
                    "Lig1",
                    "Tpi1",
                    "Pa2g4",
                    "Gsr",
                    "Atf4",
                    "Hist1h2bc",
                    "Hprt",
                    "Nucks1",
                    "Pcbp1",
                    "Mrpl36",
                    "Itgb1",
                    "Ndufa4",
                    "Cops5",
                    "Eif4e",
                    "Ndufv1",
                    "H3f3a",
                    "Rfc2",
                    "Bccip",
                    "Cyp51",
                    "Lamtor2",
                    "Trim28",
                    "Nudt21",
                    "Crip2",
                    "Ube2l3",
                    "Eef1d",
                    "Etf1",
                    "Zmat2",
                    "Samm50",
                    "Rrp15",
                    "Nhp2l1",
                    "Ran",
                    "Atp5g3",
                    "Gjb3",
                    "Atp5g2",
                    "Chd4",
                    "Tceb1",
                    "C1qbp",
                    "Rpl36a",
                    "Eif3m",
                    "Snrpb2",
                    "Umps",
                    "Rps20",
                    "Orc6",
                    "Ybx1",
                    "Sys1",
                    "Tubb2b",
                    "Tmem14c",
                    "Rps6",
                    "U2af1",
                    "Slc29a1",
                    "Atp5c1",
                    "Mrps17",
                    "Ccnd2",
                    "Dnajc8",
                    "Cox7c",
                    "Nme4",
                    "Eno1",
                    "mt-Nd3",
                    "Rpl22l1",
                    "Lamtor4",
                    "Smc4",
                    "Mbd3",
                    "Cct7",
                    "Dctpp1",
                    "Hsp90ab1",
                    "Rpp25",
                    "Tyms",
                    "Ankrd11",
                    "Rps8",
                    "Ctsd",
                    "Magoh",
                    "Ndufc2",
                    "H3f3b",
                    "Dnajc2",
                    "Ndufa12",
                    "Cnih4",
                    "Nudt19",
                    "Dbi",
                    "Tbx3",
                    "Dstn",
                    "Cyb5r3",
                    "Ywhae",
                    "Ppil1",
                    "Rps15",
                    "Arl1",
                    "Cenpw",
                    "Eif2s2",
                    "Mrpl45",
                    "Rpl31",
                    "Smdt1",
                    "Etfb",
                    "Uqcrc2",
                    "G3bp2",
                    "Impdh2",
                    "Prps1",
                    "Akap12",
                    "Hnrnpab",
                    "Ahnak",
                    "Nop58",
                    "Ndufa11",
                    "Cbx5",
                    "Polr2c",
                    "Cct8",
                    "Dohh",
                    "Denr",
                    "Eif3g",
                    "Aspscr1",
                    "Btg2",
                    "Parva",
                    "Rpp21",
                    "Polr2e",
                    "Anapc13",
                    "Pdcd5",
                    "Tomm7",
                    "Snrpc",
                    "Mrpl49",
                    "Tmbim6",
                    "Atp5a1",
                    "Psma5",
                    "Pycr2",
                    "Sf3a3",
                    "Ccdc80",
                    "March5",
                    "Ddah2",
                    "Gspt1",
                    "Eprs",
                    "Trmt112",
                    "Rpl35",
                    "Dazap1",
                    "Ilf2",
                    "Lman2",
                    "Prmt1",
                    "Sub1",
                    "Morf4l1",
                    "Col8a1",
                    "Cdca8",
                    "L1td1",
                    "Lsm6",
                    "Sf3b1",
                    "Ndufs4",
                    "Polr1d",
                    "H2afy",
                    "Mtf2",
                    "Rpl17",
                    "mt-Nd2",
                    "Txndc17",
                    "Spty2d1",
                    "Nop10",
                    "Mpdu1",
                    "Gng5",
                    "Hspe1",
                    "Rps19",
                    "Ube2c",
                    "Msn",
                    "Exosc8",
                    "Mrpl20",
                    "Ptx3",
                    "Polr2h",
                    "Psmb4",
                    "Birc5",
                    "Commd1",
                    "Cox7b",
                    "Selm",
                    "Chtop",
                    "Dppa5a",
                    "Srpk1",
                    "Lamtor5",
                    "Avpi1",
                    "Ssb",
                    "Tinagl1",
                    "Rpa2",
                    "Mrps12",
                    "Copz1",
                    "Nfu1",
                    "Ndufb5",
                    "Set",
                    "Thyn1",
                    "Txnl4a",
                    "Ssbp1",
                    "Col4a1",
                    "Sephs2",
                    "Snrpd2",
                    "1810022K09Rik",
                    "Hdac1",
                    "Naa10",
                    "Hsp90b1",
                    "Vdac2",
                    "Mrpl24",
                    "Snhg6",
                    "Minos1",
                    "Anapc11",
                    "Gas6",
                    "Nans",
                    "Uchl5",
                    "Cxcl1",
                    "Ppig",
                    "Dcun1d5",
                    "Lsm12",
                    "Gins4",
                    "Tceb2",
                    "Rhoc",
                    "Mrpl42",
                    "Rps2",
                    "Polr2g",
                    "Sumo3",
                    "Uqcrfs1",
                    "Naa50",
                    "Ywhah",
                    "Fbn2",
                    "Hdac2",
                    "Timm17a",
                    "Rpn2",
                    "Bub3",
                    "Esf1",
                    "Plekhj1",
                    "Mrfap1",
                    "Hsd17b14",
                    "Slc25a5",
                    "Uqcrh",
                    "Foxp1",
                    "Tsn",
                    "Rpl37a",
                    "Drg1",
                    "Ptma",
                    "Ndufb6",
                    "Ybx3",
                    "Rps10",
                    "Dad1",
                    "Sept1",
                    "0610007P14Rik",
                    "Yif1b",
                    "Rpl39",
                    "Tsfm",
                    "Capn2",
                    "Ptov1",
                    "Rps27rt",
                    "Gpx4",
                    "Gcsh",
                    "Carhsp1",
                    "Gltscr2",
                    "Mkrn1",
                    "Rrp1",
                    "Pdcl3",
                    "Rpl23",
                    "Rpl30",
                    "Sfr1",
                    "Polr2f",
                    "Rrs1",
                    "Rbm17",
                    "Ly6e",
                    "Lsm5",
                    "Rnf7",
                    "Cdc20",
                    "Cd9",
                    "Ddx24",
                    "Rpl6",
                    "Srm",
                    "Thbs1",
                    "Psmb7",
                    "Spr",
                    "Pnkd",
                    "Ppid",
                    "1110008F13Rik",
                    "Anapc5",
                    "Hnrnpa1",
                    "Ssrp1",
                    "Itm2a",
                    "Rnf187",
                    "Psmb6",
                    "Ndufs3",
                    "Uba2",
                    "Mbnl2",
                    "Snrpe",
                    "B2m",
                    "Psat1",
                    "Gdi2",
                    "Taf7",
                    "Mrpl28",
                    "Psmc6",
                    "Grpel1",
                    "Sec61g",
                    "Ythdc1",
                    "Snrnp27",
                    "Srsf1",
                    "Igfbp3",
                    "Socs3",
                    "Uqcrc1",
                    "Rcc2",
                    "Impa2",
                    "Bola2",
                    "Tmem160",
                    "C1d",
                    "Cisd2",
                    "Ncl",
                    "Rpl35a",
                    "Timp2",
                    "Syngr1",
                    "Hspa4",
                    "Arpp19",
                    "Ccdc12",
                    "Ddx21",
                    "Naca",
                    "Ube2i",
                    "Rpl24",
                    "Brix1",
                    "Fam136a",
                    "Siva1",
                    "Rps4x",
                    "Fkbp8",
                    "Nip7",
                    "Ncam1",
                    "Ruvbl1",
                    "Clpp",
                    "Wls",
                    "G3bp1",
                    "Atp5b",
                    "Cdc26",
                    "Actb",
                    "Immt",
                    "Nars",
                    "Hells",
                    "Tfcp2l1",
                    "Rpl38",
                    "Glo1",
                    "Serpine2",
                    "Nme1",
                    "Crabp2",
                    "Csnk1a1",
                    "Rpl10-ps3",
                    "Rpl3",
                    "Cebpzos",
                    "Rpl12",
                    "Snw1",
                    "Ndufa7",
                    "Dnajc19",
                    "Cdv3",
                    "Hmgn1",
                    "Ctbp2",
                    "Btf3l4",
                    "Ppp4c",
                    "Imp3",
                    "Mrpl13",
                    "Ak2",
                    "Wdr77",
                    "Atp5j",
                    "Srrm1",
                    "Srp19",
                    "Emd",
                    "Pdgfa",
                    "Uqcr10",
                    "Rpsa",
                    "H2afz",
                    "Mrps21",
                    "Tardbp",
                    "Rpl5",
                    "Dph3",
                    "Dpy30",
                    "Serpine1",
                    "Fblim1",
                    "Odc1",
                    "Mrpl18",
                    "Ufc1",
                    "Ifitm1",
                    "Acat2",
                    "Mrpl12",
                    "Cdc37",
                    "Cisd1",
                    "Rpl28",
                    "Xist",
                    "Gpx3",
                    "Timm8b",
                    "Rpl26",
                    "Uqcrb",
                    "mt-Co2",
                    "Rps3",
                    "Atox1",
                    "Srsf10",
                    "Ube2v2",
                    "Rpl19",
                    "Timm13",
                    "Eif3h",
                    "Mcm3",
                    "Adprh",
                    "Rps3a1",
                    "Srsf3",
                    "Coprs",
                    "Cmpk1",
                    "Rpl7",
                    "Fn1",
                    "Znrd1",
                    "Wdr18",
                    "Polr2m",
                    "1810058I24Rik",
                    "Rpl8",
                    "Eif4a1",
                    "Pcbp2",
                    "Polr2i",
                    "Polr1c",
                    "Snrnp25",
                    "Fhl2",
                    "Ccdc34",
                    "Mrps33",
                    "Rif1",
                    "Morf4l2",
                    "Rbbp4",
                    "Eif3e",
                    "Emp3",
                    "2700060E02Rik",
                    "Park7",
                    "Dynll1",
                    "Hspd1",
                    "Stoml2",
                    "Cops7a",
                    "Sod1",
                    "Ppih",
                    "Uqcc3",
                    "Tpx2",
                    "Dnajb6",
                    "Hspa5",
                    "Gapdh",
                    "Smarca5",
                    "Idi1",
                    "Ccnb1",
                    "Ccnd3",
                    "Khdrbs1",
                    "Mrpl52",
                    "Pno1",
                    "Ecm1",
                    "Adrm1",
                    "Kif5b",
                    "Hmgn2",
                    "Hnrnpa2b1",
                    "Oat",
                    "Psmd2",
                    "Ddx27",
                    "Jade1",
                    "Dnajc7",
                    "Psmb2",
                    "Fxr1",
                    "Tceal8",
                    "Cnn3",
                    "Vps29",
                    "Vdac1",
                    "Anp32a",
                    "Tra2a",
                    "Zfp57",
                    "Dynll2",
                    "Ssu72",
                    "Cnpy2",
                    "Anp32e",
                    "Pfdn6",
                    "Ict1",
                    "Rps25",
                    "Rbmxl1",
                    "Arglu1",
                    "Imp4",
                    "Ywhab",
                    "Prelid1",
                    "Ubb",
                    "Zfp106",
                    "Prrc1",
                    "Paics",
                    "Asna1",
                    "Psmc5",
                    "Echdc2",
                    "Mrpl43",
                    "Suclg1",
                    "Hnrnpd",
                    "Ash2l",
                    "Hat1",
                    "Rnps1",
                    "H13",
                    "Tpt1",
                    "Ppib",
                    "Stmn2",
                    "Ptbp1",
                    "Aldoa",
                    "Lpp",
                    "Dnajc21",
                    "Snx3",
                    "Strap",
                    "Psmd13",
                    "Rnaseh2a",
                    "Rfc1",
                    "Cox6b1",
                    "Ptrf",
                    "Atp5o",
                    "Nasp",
                    "Ndufs6",
                    "Poldip2",
                    "Eif4e2",
                    "Sod2",
                    "Hist1h1c",
                    "Tnfrsf12a",
                    "Phgdh",
                    "Gtf2e2",
                    "Pdap1",
                    "Rpl11",
                    "Anxa1",
                    "Ube2a",
                    "Pfn1",
                    "Snrpg",
                    "Ddx39b",
                    "Grb10",
                    "Exosc4",
                    "Fkbp4",
                    "Pgam1",
                    "Snrnp70",
                    "Ppdpf",
                    "Gmnn",
                    "Rpl27a",
                    "Llph",
                    "Eif4b",
                    "Anp32b",
                    "Edf1",
                    "Fstl1",
                    "Mdm2",
                    "Eif2b5",
                    "Gart",
                    "Ndufa10",
                    "App",
                    "Tomm40",
                    "Eif3i",
                    "Smc2",
                    "Mtch2",
                    "Psmd4",
                    "Gins2",
                    "Uck2",
                    "Rps16",
                    "Bcat1",
                    "Tmed3",
                    "Mrps16",
                    "Psmd12",
                    "Ndufb10",
                    "Cth",
                    "Txnl1",
                    "Rbm25",
                    "Eef1g",
                    "Ube2e3",
                    "2700094K13Rik",
                    "Acadm",
                    "Dut",
                    "Alyref",
                    "Hspa14",
                    "Polr2j",
                    "Got2",
                    "Slirp",
                    "Ckb",
                    "Cwc15",
                    "Ndufs8",
                    "Pim1",
                    "Rplp2",
                    "Mif",
                    "Aimp1",
                    "Rheb",
                    "Chchd3",
                    "Eif4a2",
                    "Gtf2a2",
                    "Cnbp",
                    "Sdhd",
                    "Axl",
                    "Rps5",
                    "Rpl41",
                    "Cd44",
                    "Ruvbl2",
                    "Cdt1",
                    "Ier2",
                    "Dkc1",
                    "Marcksl1",
                    "Atp5g1",
                    "Tuba1b",
                    "Aurka",
                    "Psma6",
                    "Rpl7l1",
                    "Snrpa",
                    "St13",
                    "Thoc7",
                    "Mlf2",
                    "Rpl9",
                    "Vbp1",
                    "Atp5j2",
                    "Nubp1",
                    "E2f4",
                    "D8Ertd738e",
                    "Hn1",
                    "Mrpl14",
                    "Phb",
                    "Tra2b",
                    "Hmgcs1",
                    "Emc6",
                    "Rps24",
                    "Bola1",
                    "Aurkaip1",
                    "Timp3",
                    "Nhp2",
                    "Exosc5",
                    "Sh3glb1",
                    "Atp5h",
                    "Ppia",
                    "Eif3k",
                    "Mrps34",
                    "Atp5k",
                    "Mphosph8",
                    "Ppp1ca",
                    "Aamp",
                    "Chchd1",
                    "Gars",
                    "Psmc3",
                    "Luc7l2",
                    "Ndufb9",
                    "Col3a1",
                    "Utp3",
                    "Fus",
                    "Ociad1",
                    "Ddost",
                    "Rest",
                    "Cct3",
                    "Psma7",
                    "Gnl3",
                    "Trim27",
                    "Psma1",
                    "Dtymk",
                    "Emc10",
                    "2410015M20Rik",
                    "Hdgf",
                    "Prnp",
                    "Id1",
                    "Csnk2b",
                    "Rpl37",
                    "Ppp1cc",
                    "Nrp1",
                    "Dnttip2",
                    "Sae1",
                    "Hnrnpk",
                    "Mrps24",
                    "Ezr",
                    "Ewsr1",
                    "Ndufv2",
                    "Fbxo22",
                    "Fam49b",
                    "Bax",
                    "Cisd3",
                    "Gfer",
                    "Eif3c",
                    "Cox7a2",
                    "Col5a2",
                    "Ube2j2",
                    "Lsm3",
                    "Gcat",
                    "Eif5",
                    "Spag7",
                    "Tomm5",
                    "Txn1",
                    "Vapa",
                    "Erh",
                    "Fscn1",
                    "Emc8",
                    "Bclaf1",
                    "Psip1",
                    "Rpl29",
                    "Naa38",
                    "Ung",
                    "Ptges3",
                    "H2-D1",
                    "Wtap",
                    "Dnaja2",
                    "Lbh",
                    "Cenpv",
                    "Ywhaq",
                    "Pdha1",
                    "Ubald2",
                    "Filip1l",
                    "Adss",
                    "Rps7",
                    "Igf2bp1",
                    "Trnau1ap",
                    "Eef1e1",
                    "Hsd17b10",
                    "Oaz1",
                    "Stra13",
                    "Sec61b",
                    "Snx5",
                    "Got1",
                    "Bzw2",
                    "Pgp",
                    "Sumo2",
                    "Nudc",
                    "Pou5f1",
                    "Gnb2l1",
                    "Mrpl21",
                    "Pold2",
                    "Nsa2",
                    "Ddx46",
                    "Rfc5",
                    "Pfdn4",
                    "Mrps15",
                    "Psmc4",
                    "Psmd14",
                    "Npm1",
                    "Sap30",
                    "Gtpbp4",
                    "Cpe",
                    "Atf5",
                    "Srsf7",
                    "Mat2a",
                    "Csrp2",
                    "Brd2",
                    "Ccdc124",
                    "Prdx2",
                    "Abcf1",
                    "Rplp0",
                    "Psmb5",
                    "Dcn",
                    "Ube2n",
                    "Smn1",
                    "Cyb5b",
                    "Psmb1",
                    "Coa3",
                    "Akr1b3",
                    "Eif4g2",
                    "Raly",
                    "Uqcc2",
                    "Nsmce4a",
                    "Rps26",
                    "Get4",
                    "Lage3",
                    "Npm3",
                    "Lsm2",
                    "Chmp2a",
                    "Sumo1",
                    "Nudcd2",
                    "Aprt",
                    "Fcf1",
                    "Mrps18c",
                    "Cacybp",
                    "Rpl22",
                    "Nolc1",
                    "Rpl4",
                    "Glud1",
                    "Rps28",
                    "Hnrnpu",
                    "Srsf11",
                    "Hmgb3",
                    "Rpl21",
                    "Ppp1r2",
                    "Map1lc3b",
                    "Atp5l",
                    "Son",
                    "Pop5",
                    "Cct2",
                    "Acp1",
                    "Sox4",
                    "Dtl",
                    "Ahsa1",
                    "Eif1ax",
                    "Parp1",
                    "Dhfr",
                    "Mrps7",
                    "Cdca3",
                    "Metap2",
                    "Ube2s",
                    "Idh2",
                    "Snrpd3",
                    "Psmd3",
                    "Fam32a",
                    "Bcas2",
                    "Cops4",
                    "Ciapin1",
                    "Pfdn2",
                    "Atp5f1",
                    "Arid5b",
                    "Srp9",
                    "Nfatc2ip",
                    "Ino80e",
                    "Ppa1",
                    "Prdx1",
                    "Rsl24d1",
                    "Terf1",
                    "Rpl13",
                    "Mrpl34",
                    "Atrx",
                    "Nt5dc2",
                    "Tgif1",
                    "Sfpq",
                    "Mrto4",
                    "Parl",
                    "Lipe",
                    "Clns1a",
                    "Mdh2",
                    "Gabarap",
                    "Uqcr11",
                    "Phb2",
                    "Rps27a",
                    "Rpl15",
                    "Gtf2h5",
                    "Arl6ip1",
                    "Tcof1",
                    "Des",
                    "Ier3",
                    "Cuta",
                    "1500015O10Rik",
                    "Eif1a",
                    "Cox5a",
                    "Tmem11",
                    "Ubap2l",
                    "Phldb2",
                    "Cct4",
                    "Rdx",
                    "Cdkn1a",
                    "Pkm",
                    "H2afv",
                    "Prrc2c",
                    "Marcks",
                    "Psmg4",
                    "Psmc2",
                    "Sf3b4",
                    "Hnrnpc",
                    "Fkbp3",
                    "Rangrf",
                    "Rpl7a",
                    "Fam104a",
                    "Prrx2",
                    "Manf",
                    "Tpr",
                    "Ddx18",
                    "Rrp9",
                    "Hint1",
                    "Sf3a2",
                    "Ccne1",
                    "Srsf6",
                    "Ebna1bp2",
                    "Tmpo",
                    "Cycs",
                    "Klf2",
                    "Dhx16",
                    "Sdhb",
                    "Taf10",
                    "Wdr43",
                    "Klf5",
                    "Yrdc",
                    "Pabpc1",
                    "Eif6",
                    "Tsen34",
                    "Fam60a",
                    "Wbp11",
                    "2810428I15Rik",
                    "Hspa8",
                    "Postn",
                    "Ldhb",
                    "Taf1d",
                    "Serbp1",
                    "Kdm1a",
                    "Ncor1",
                    "1810037I17Rik",
                    "Rsrc2",
                    "Hsp90aa1",
                    "Usmg5",
                    "Mdh1",
                    "Ppp2ca",
                    "Rps17",
                    "Rae1",
                    "Pin4",
                    "Mrps18b",
                    "Txn2",
                    "Clta",
                    "Cox17",
                    "Ppp1r11",
                    "Mrpl4",
                    "Wisp1",
                    "Ndufb3",
                    "Gar1",
                    "Cks1b",
                    "Ftsj3",
                    "Folr1",
                    "Psmc1",
                    "Tpd52l2",
                    "Knop1",
                    "Psmd6",
                    "Pin1",
                    "Lgals3",
                    "Sfxn1",
                    "Eif2s1",
                    "Swi5",
                    "Nup62",
                    "Cenpe",
                    "Rab18",
                    "Calm3",
                    "Arf6",
                    "Ubxn1",
                    "Cdk1",
                    "Eif4h",
                    "Psmd7",
                    "Anapc15",
                    "Gabarapl2",
                    "Slc39a1",
                    "Stub1",
                    "Prrx1",
                    "Mrpl2",
                    "Ranbp1",
                    "Zwint",
                    "Ppm1g",
                    "Eif1",
                    "Tbcb",
                    "Wdr89",
                    "Pdcd6",
                    "H2afx",
                    "Hnrnpm",
                    "Capza2",
                    "Loxl1",
                    "2310036O22Rik",
                    "Fdft1",
                    "Srsf5",
                    "Hnrnph1",
                    "Wdr74",
                    "Bex4",
                    "Ywhaz",
                    "Pomp",
                    "Rpl36",
                    "Rps21",
                    "Psmb3",
                    "Ddx5",
                    "Rbm39",
                    "Pigyl",
                    "Eef1b2",
                    "Eif3l",
                    "Tmem258",
                    "Tcp1",
                    "Sarnp",
                    "Mrpl15",
                    "Ezh2",
                    "Mybl2",
                    "Pfdn1",
                    "Slc2a3",
                    "Pmm1",
                    "Actl6a",
                    "Pdia4",
                    "Sssca1",
                    "Ier5",
                    "Gpx1",
                    "Smc3",
                    "Thrap3",
                    "Hspb1",
                    "Kdm5b",
                    "Nfkbia",
                    "Eif3b",
                    "Zfp207",
                    "Ube2k",
                    "Ndufs2",
                    "Ddt",
                    "Spc24",
                    "Asns",
                    "Ifitm2",
                    "Fdps",
                    "Lsm8",
                    "Ndufs7",
                    "Cenpf",
                    "Mcm2",
                    "Gm16286",
                    "Mrpl17",
                    "Ldha",
                    "Apex1",
                    "Cct5",
                    "Ccnb2",
                    "Mrpl35",
                    "1110004F10Rik",
                    "Mrpl40",
                    "Lox",
                    "Iscu",
                    "Fubp1",
                    "Arpc5l",
                    "Rps29",
                    "Ccdc58",
                    "Cenpa",
                    "Prdx6",
                    "Mdk",
                    "Psmd1",
                    "Ndufab1",
                    "Plekhf2",
                    "Snrpf",
                    "Mcl1",
                    "Pole4",
                    "Fbl",
                    "Ndufv3",
                    "Rps26-ps1",
                    "Calm2",
                    "Stmn1",
                    "Btf3",
                    "Smarcc1",
                    "Cdh11",
                    "Rpl18",
                    "Hax1",
                    "Atp5d",
                    "Taldo1",
                    "Vcp",
                    "Mrps5",
                    "Prc1",
                    "U2surp",
                    "Fkbp2",
                    "Ppan",
                    "Rps9",
                    "Upp1",
                    "Mrpl30",
                    "Rwdd1",
                    "Cyc1",
                    "Fnta",
                    "Dnaja1",
                    "Tmem176b",
                    "Pmvk",
                    "Elof1",
                    "Lrpap1",
                    "Wdr5",
                    "Pebp1",
                    "Sgk1",
                    "Ube2d3",
                    "Aars",
                    "Marc2",
                    "Snrpa1",
                    "Paip2",
                    "Srrm2",
                    "Rnaseh2b",
                    "Top1",
                    "Ubl5",
                    "Hmgb1",
                    "Ndufc1",
                    "Eif3d",
                    "Cyb5a",
                    "Psma4",
                    "Ap2s1",
                    "Rpl34",
                    "Mlec",
                    "Rab7",
                    "Mrps14",
                    "Nap1l1",
                    "Mrps25",
                    "Actr3",
                    "Mrps10",
                    "Tbrg1",
                    "Hspa9",
                    "Fkbp10",
                    "Chchd2",
                    "Pim3",
                    "E130012A19Rik",
                    "Luc7l3",
                    "Ndufs5",
                    "Gsn",
                    "Ndufb8",
                    "Tmem97",
                    "Srsf2",
                    "Prdx4",
                    "Pak1ip1",
                    "Nxt1",
                    "Trp53",
                    "Lhfp",
                    "Rbm3",
                    "Psmd11",
                    "Cggbp1",
                    "Zfp706",
                    "Rad23b",
                    "Nudt4",
                    "Supt16",
                    "Ost4",
                    "Mrpl51",
                    "Eif4g1",
                    "Rhoa",
                    "Cdk4",
                    "Rsl1d1",
                    "Syncrip",
                    "Caprin1",
                    "Ssr2",
                    "Cebpz",
                    "Tomm20",
                    "Cbx1",
                    "Tomm70a",
                    "Msmo1",
                    "Mak16",
                    "Bzw1",
                    "Ak6",
                    "Rbpms",
                    "Hnrnpa3",
                    "Prpf40a",
                    "Pbdc1",
                    "Rps19bp1",
                    "2010107E04Rik",
                    "Tecr",
                    "Mybbp1a",
                    "Sf3b2",
                    "Atp1b1",
                    "Slc3a2",
                    "Stip1",
                    "Chchd4",
                    "Rpl14",
                    "Tma7",
                    "Camk2d",
                    "Lsm4",
                    "Tpd52",
                    "Eif1ad",
                    "Rps13",
                    "Acta1",
                    "Eif5a",
                    "Rps11",
                    "Cfl1",
                    "Rrp1b",
                    "Sparc",
                    "Uchl3",
                    "Cops6",
                    "Hnrnpdl",
                    "Eif3a",
                    "Rtn3",
                    "Adh5",
                    "l7Rn6",
                    "Atp5e",
                    "Cdc34",
                    "Gps1",
                    "Tipin",
                    "Cmtm7",
                    "Rrm2",
                    "Sh3bgrl",
                    "Itm2b",
                    "Rplp1",
                    "Rnf10",
                    "Psma2",
                    "Pfdn5",
                    "Nsun2",
                    "Ndufb7",
                    "Shmt1",
                    "Alkbh1",
                    "Slc25a39",
                    "Rps14",
                    "Rpl32",
                    "Chchd10",
                    "Bud31",
                    "Mcm5",
                    "Rpl13a",
                    "Nol7",
                    "Tkt",
                    "Apoe",
                    "Gsta4",
                    "Snrpb",
                    "Zfp428",
                    "Tomm22",
                    "Lyar",
                    "Erp29",
                    "Gpx8",
                    "Gpi1",
                    "Tmem167",
                    "Cfdp1",
                    "Eif5b",
                    "Emg1",
                    "Eif4ebp1",
                    "Rpl23a-ps3",
                    "Nono",
                    "Csde1",
                    "Mcm6",
                    "Prpf19",
                    "Sugt1",
                    "Eif4a3",
                    "Dek",
                    "Top2a",
                    "Papola",
                    "Tmem256",
                    "Cetn3",
                    "Galk1",
                    "Ccna2",
                    "Tubb4b",
                    "Eed",
                    "Ndufa5",
                    "Mrpl11",
                    "Mesdc2",
                    "Cox5b",
                    "Arhgdia",
                    "Ralbp1",
                    "Acin1",
                    "Aebp1",
                    "Rbpms2",
                    "Uhrf1",
                    "Guk1",
                    "Psma3",
                    "Skp1a",
                    "Rps18",
                    "Phf5a",
                    "Rps23",
                    "U2af2",
                    "P3h4",
                    "Myl12b"
                ],
                "expression_sum": {
                    "class1": 14867.0,
                    "class2": 3147.0
                },
                "uniprot_ids": [
                    "P28033",
                    "Q3UPN9",
                    "O70192"
                ]
            },
            "Npm1": {
                "id": "Npm1",
                "type": "Gene",
                "symbol": "Npm1",
                "source": [
                    "Tcf4",
                    "Pou5f1",
                    "Foxp1",
                    "Nfib",
                    "Atf5",
                    "Atf4",
                    "Junb",
                    "Fos",
                    "Jun",
                    "Sox2",
                    "Nanog",
                    "Jund",
                    "Cebpb",
                    "Sox4"
                ],
                "target": [],
                "expression_sum": {
                    "class1": 173806.0,
                    "class2": 794190.0
                },
                "uniprot_ids": [
                    "Q61937",
                    "Q5SQB7",
                    "Q9DAY9",
                    "Q3U536",
                    "Q5SQB0",
                    "Q5U438",
                    "Q3UDA7",
                    "Q5SQB5",
                    "Q8BSZ3",
                    "Q5BL09"
                ]
            },
            "Trim28": {
                "id": "Trim28",
                "type": "TF",
                "symbol": "Trim28",
                "source": [
                    "Pou5f1",
                    "Foxp1",
                    "Junb",
                    "Fos",
                    "Jun",
                    "Sox2",
                    "Tbx3",
                    "Nanog",
                    "Jund",
                    "Twist2",
                    "Cebpb",
                    "Zfp57",
                    "Sox4"
                ],
                "target": [
                    "Dap",
                    "Cd44",
                    "H2afj",
                    "Ddb1",
                    "Myh9",
                    "Cdt1",
                    "Thbs1",
                    "Rasgrp2",
                    "Lims1",
                    "Igfbp2",
                    "Slc25a4",
                    "Calcoco2",
                    "Lmna",
                    "Folr1",
                    "Cthrc1",
                    "Lgals3",
                    "Pgrmc1",
                    "Fos",
                    "Tagln2",
                    "Rps15a",
                    "Rexo2",
                    "Pdlim7",
                    "Rab18",
                    "Mfge8",
                    "Mbnl2",
                    "Canx",
                    "B2m",
                    "Taf7",
                    "Neat1",
                    "Myh10",
                    "Timp3",
                    "Gemin7",
                    "Akr1a1",
                    "Exosc5",
                    "Sh3glb1",
                    "Akt1s1",
                    "Impa2",
                    "Atp2b1",
                    "Cisd2",
                    "Mylpf",
                    "Nktr",
                    "Col5a1",
                    "Ctgf",
                    "Klf6",
                    "Pcolce",
                    "Crxos",
                    "Syngr1",
                    "Crip1",
                    "Morc1",
                    "Sec62",
                    "Rps12",
                    "Maged1",
                    "Cyr61",
                    "Kdelr1",
                    "Rab14",
                    "Serpinh1",
                    "Atp6v1g1",
                    "Gnb2",
                    "Dpysl3",
                    "Mybl2",
                    "Slc2a3",
                    "Ddost",
                    "Hcfc1r1",
                    "Sptbn1",
                    "Id3",
                    "Ap3s1",
                    "Gng12",
                    "Ncam1",
                    "Wls",
                    "Ndufa8",
                    "Srp14",
                    "Sept7",
                    "Actg1",
                    "Rcn2",
                    "Tsg101",
                    "Prnp",
                    "Id1",
                    "Hmga2",
                    "Hspb1",
                    "Kdm5b",
                    "Hist1h2bc",
                    "Hells",
                    "Nfkbia",
                    "Tfcp2l1",
                    "Col1a2",
                    "Atp6v0b",
                    "F2r",
                    "Fkbp9",
                    "Igfbp4",
                    "Id2",
                    "Itgb1",
                    "Fabp5",
                    "Mettl9",
                    "Chmp4b",
                    "Tcf4",
                    "Crabp2",
                    "Ssr1",
                    "Cyp51",
                    "Lxn",
                    "Pgls",
                    "Ift20",
                    "Anxa2",
                    "Vmp1",
                    "Col5a2",
                    "mt-Nd4",
                    "Crip2",
                    "Vim",
                    "Wdr1",
                    "Ube2b",
                    "Tmed2",
                    "Gjb3",
                    "Tpm1",
                    "Sdcbp",
                    "Thy1",
                    "Surf4",
                    "Utf1",
                    "Phlda1",
                    "Myeov2",
                    "Pdgfa",
                    "Cst3",
                    "Higd1a",
                    "S100a6",
                    "Dpy30",
                    "Serpine1",
                    "H2-D1",
                    "Ifitm1",
                    "Cd63",
                    "Capg",
                    "S100a11",
                    "Scand1",
                    "Mtdh",
                    "Eef1a1",
                    "Rtn4",
                    "Slc29a1",
                    "Cenpv",
                    "Ctsl",
                    "Col6a1",
                    "Rala",
                    "Sdc2",
                    "Filip1l",
                    "Plk2",
                    "Tln1",
                    "Cdh11",
                    "Snhg12",
                    "Mt2",
                    "Fbln5",
                    "Lrrc58",
                    "Btg1",
                    "Fkbp2",
                    "Fn1",
                    "Dctpp1",
                    "Glipr2",
                    "Hes1",
                    "Ap1s1",
                    "Tuba4a",
                    "Upp1",
                    "Zfp42",
                    "Malat1",
                    "Gnai2",
                    "Tmem176b",
                    "Mfap2",
                    "Tm9sf3",
                    "Ctsd",
                    "Pou5f1",
                    "Fhl2",
                    "Oxct1",
                    "Atxn10",
                    "Atp6v0e",
                    "Dbi",
                    "Tmem59",
                    "Tbx3",
                    "Emp3",
                    "Gm26917",
                    "Pafah1b2",
                    "Anxa3",
                    "Hbegf",
                    "Arpc3",
                    "Ilk",
                    "Arl1",
                    "Arpc1a",
                    "Sep15",
                    "Lsm7",
                    "Cpe",
                    "Atf5",
                    "Cdc42",
                    "Eef2",
                    "Hsbp1",
                    "Hspa5",
                    "Rnh1",
                    "Rrbp1",
                    "Actr3",
                    "Cd47",
                    "Efemp2",
                    "Fbln2",
                    "Mpc1",
                    "Copb2",
                    "Sars",
                    "Nupr1",
                    "Smim14",
                    "Ecm1",
                    "Sdpr",
                    "Lgals1",
                    "Ahnak",
                    "Kif5b",
                    "Mpc2",
                    "Copz2",
                    "Ptp4a1",
                    "Lsp1",
                    "Gsn",
                    "Oat",
                    "Cited2",
                    "Jam2",
                    "Rab2a",
                    "Tspo",
                    "Btg2",
                    "Parva",
                    "Selk",
                    "Jund",
                    "Rab11a",
                    "Zfp57",
                    "Ankrd1",
                    "Anxa5",
                    "Eif3f",
                    "Tmbim6",
                    "Mtch1",
                    "Ttc3",
                    "Sec61a1",
                    "Iqgap1",
                    "Kdelr2",
                    "Rbmxl2",
                    "Fermt2",
                    "Rpn1",
                    "Gas1",
                    "Sqstm1",
                    "Lman1",
                    "F3",
                    "Rcn1",
                    "Arpc2",
                    "Phlda3",
                    "Romo1",
                    "Tpm2",
                    "Ergic3",
                    "Sdc4",
                    "Ubc",
                    "Rock2",
                    "Cdc42ep3",
                    "Tcea3",
                    "L1td1",
                    "Rpl27",
                    "Psap",
                    "Rbms1",
                    "P4hb",
                    "Scd2",
                    "Dlk1",
                    "Zyx",
                    "Map1b",
                    "Rab34",
                    "Adipor1",
                    "Mtf2",
                    "Bsg",
                    "Sox2",
                    "Ppic",
                    "Chchd4",
                    "Trh",
                    "Sec22b",
                    "Camk2d",
                    "mt-Cytb",
                    "Cyba",
                    "Efemp1",
                    "Gdf3",
                    "Gsto1",
                    "Ppib",
                    "Vcan",
                    "Sparc",
                    "Stmn2",
                    "Sept2",
                    "Abracl",
                    "Serpinb6a",
                    "Lpp",
                    "Ctbp1",
                    "Csrp1",
                    "Selm",
                    "Ckap4",
                    "Esd",
                    "Dppa5a",
                    "Meg3",
                    "Atxn7l3b",
                    "Comt",
                    "Lamp1",
                    "Rcor2",
                    "Ndufa13",
                    "Ptrf",
                    "Gadd45g",
                    "Jun",
                    "Capns1",
                    "Itm2b",
                    "Pdia3",
                    "Rpl10a",
                    "Txndc5",
                    "Ptms",
                    "Shmt1",
                    "Hist1h1c",
                    "Cd24a",
                    "Smap1",
                    "Lipe",
                    "Tnfrsf12a",
                    "Krtcap2",
                    "Gabarap",
                    "Phgdh",
                    "Serf2",
                    "Hsp90b1",
                    "Mtpn",
                    "Ier3",
                    "Grb10",
                    "Apoe",
                    "Commd3",
                    "Gas6",
                    "Fst",
                    "Gpi1",
                    "Phldb2",
                    "Palld",
                    "Tram1",
                    "Igfbp7",
                    "Fhl1",
                    "Cdkn1a",
                    "Fstl1",
                    "Fam162a",
                    "Rhoc",
                    "Mdm2",
                    "Ifitm3",
                    "Mrpl33",
                    "Emp1",
                    "S100a10",
                    "Rpn2",
                    "Prrx2",
                    "Tmed3",
                    "Hsd17b14",
                    "Tbca",
                    "Cstb",
                    "Foxp1",
                    "H19",
                    "Ptp4a2",
                    "Fbxo15",
                    "Cope",
                    "Dnajc3",
                    "Myl12a",
                    "Rac1",
                    "Rer1",
                    "S100a4",
                    "Klf2",
                    "Cald1",
                    "Fau",
                    "Sri",
                    "Prdx5",
                    "Clic1",
                    "Emc7",
                    "Gnas",
                    "Rbpms2",
                    "Aebp1",
                    "Arf4",
                    "Calu",
                    "Arpc5",
                    "Gm7325",
                    "Sept1",
                    "Hdlbp",
                    "Limd2",
                    "Tmed9",
                    "Postn",
                    "Cnih1",
                    "Carhsp1",
                    "Alpl",
                    "Mkrn1",
                    "Pdcl3",
                    "Kdelr3",
                    "P3h4",
                    "Bcap31",
                    "Axl",
                    "Cfl2"
                ],
                "expression_sum": {
                    "class1": 7383.0,
                    "class2": 112063.0
                },
                "uniprot_ids": [
                    "Q62318",
                    "Q5EBP9",
                    "Q3UDN8"
                ]
            },
            "Smarca5": {
                "id": "Smarca5",
                "type": "TF",
                "symbol": "Smarca5",
                "source": [
                    "Tcf4",
                    "Pou5f1",
                    "Foxp1",
                    "Cebpb",
                    "Nfib",
                    "Junb",
                    "Fos",
                    "Jun",
                    "Sox2",
                    "Hes1",
                    "Nanog",
                    "Jund",
                    "Hdac2"
                ],
                "target": [
                    "Hsp90aa1",
                    "Arf1",
                    "Ppp2ca",
                    "Rps17",
                    "Fis1",
                    "Zc3h15",
                    "Cox8a",
                    "Ddb1",
                    "Myh9",
                    "Thbs1",
                    "Txn2",
                    "Lims1",
                    "Sec13",
                    "1110008F13Rik",
                    "Slc25a4",
                    "Col12a1",
                    "Lmna",
                    "Thoc7",
                    "Ndufa2",
                    "Mlf2",
                    "Psmd6",
                    "Ssrp1",
                    "Ier3ip1",
                    "Rnf187",
                    "Swi5",
                    "Cxcl12",
                    "Eif2s1",
                    "Pgrmc1",
                    "Shfm1",
                    "Fos",
                    "Tagln2",
                    "Cox6c",
                    "Slbp",
                    "Ndufs3",
                    "Nedd4",
                    "Calr",
                    "Pdlim7",
                    "Rab18",
                    "Mfge8",
                    "Mbnl2",
                    "Ppp1r14b",
                    "B2m",
                    "Mrpl14",
                    "Slc25a3",
                    "Psat1",
                    "Eif4h",
                    "Psmd7",
                    "Tra2b",
                    "Psmc6",
                    "Zwint",
                    "Rps24",
                    "Eif1",
                    "Ola1",
                    "Snrnp27",
                    "Timp3",
                    "Gemin7",
                    "Lrrc59",
                    "Uqcrc1",
                    "Akr1a1",
                    "Sh3glb1",
                    "Ube2d2a",
                    "Hras",
                    "Bnip3l",
                    "Kpnb1",
                    "Cisd2",
                    "C1d",
                    "Nktr",
                    "Col5a1",
                    "Ppia",
                    "Capza2",
                    "Klf6",
                    "Pcolce",
                    "Tgfb1i1",
                    "Loxl1",
                    "Hnrnph1",
                    "Syngr1",
                    "Ywhaz",
                    "Gars",
                    "Sec62",
                    "Rps12",
                    "Rpl36",
                    "Ddx5",
                    "Ndufb9",
                    "Rbm39",
                    "Pcnp",
                    "Ube2i",
                    "Cyr61",
                    "Kdelr1",
                    "Serpinh1",
                    "Gnb2",
                    "Fus",
                    "Ddost",
                    "Rps4x",
                    "Id3",
                    "Ap3s1",
                    "Gng12",
                    "G3bp1",
                    "Fam195b",
                    "Atp5b",
                    "Psma1",
                    "Actg1",
                    "Ier5",
                    "2410015M20Rik",
                    "Id1",
                    "Nars",
                    "Hmga2",
                    "Nrp1",
                    "Eif3b",
                    "Ube2m",
                    "Fkbp9",
                    "Rpl38",
                    "Igfbp4",
                    "Itgb1",
                    "Ndufa4",
                    "Ewsr1",
                    "Cops5",
                    "Mettl9",
                    "Nme1",
                    "Tcf4",
                    "Csnk1a1",
                    "Ssr1",
                    "Ndufs2",
                    "Pgls",
                    "Lxn",
                    "Tmsb10",
                    "Rpl3",
                    "Nudt21",
                    "Rpl12",
                    "Ube2l3",
                    "Btf3l4",
                    "Eef1d",
                    "Sh3bgrl3",
                    "Lsm8",
                    "Ddah1",
                    "Ube2b",
                    "Ube2j2",
                    "Gm16286",
                    "Eif5",
                    "Tmed2",
                    "Vapa",
                    "Ran",
                    "Fscn1",
                    "Atp5g3",
                    "Tpm1",
                    "Ldha",
                    "Surf4",
                    "Atp5g2",
                    "Srp19",
                    "Cct5",
                    "1110004F10Rik",
                    "Myeov2",
                    "Vasp",
                    "Mrps21",
                    "Bola3",
                    "Cst3",
                    "Rpl36a",
                    "Lox",
                    "Iscu",
                    "Arpc5l",
                    "Rpl5",
                    "S100a6",
                    "Rps20",
                    "Dnaja2",
                    "Sys1",
                    "Ndufab1",
                    "Mtdh",
                    "Rps6",
                    "Eef1a1",
                    "Rtn4",
                    "U2af1",
                    "Ywhaq",
                    "Tcf25",
                    "Rala",
                    "Sdc2",
                    "Rpl26",
                    "Cav1",
                    "Calm2",
                    "Srsf10",
                    "Plk2",
                    "Tln1",
                    "Oaf",
                    "Rps7",
                    "Ube2v2",
                    "Rpl18",
                    "Snhg12",
                    "Atp5d",
                    "Mt2",
                    "Chd3",
                    "Rps3a1",
                    "Lrrc58",
                    "Btg1",
                    "Cct7",
                    "Fkbp2",
                    "Fn1",
                    "Glipr2",
                    "Ap1s1",
                    "Hsp90ab1",
                    "Nenf",
                    "Rps9",
                    "Twist2",
                    "Mrpl30",
                    "Polr2m",
                    "Rwdd1",
                    "Snx5",
                    "Malat1",
                    "Gnai2",
                    "Rps8",
                    "Dnaja1",
                    "Tmem176b",
                    "Bzw2",
                    "Eif4a1",
                    "Sumo2",
                    "Tm9sf3",
                    "Pcbp2",
                    "Ctsd",
                    "Ndufc2",
                    "H3f3b",
                    "Col1a1",
                    "Mrpl21",
                    "Ndufa12",
                    "Nsa2",
                    "Oxct1",
                    "Cnih4",
                    "Dbi",
                    "Ube2d3",
                    "Tmem59",
                    "Eif3e",
                    "Pfdn4",
                    "Cebpb",
                    "Srrm2",
                    "Top1",
                    "Cyb5r3",
                    "Ubl5",
                    "Mrps15",
                    "Psmc4",
                    "Rps15",
                    "Arpc3",
                    "Scp2",
                    "Ilk",
                    "Arl1",
                    "Arpc1a",
                    "Ap2s1",
                    "Mgat2",
                    "Sep15",
                    "Zfp36l1",
                    "Dynll1",
                    "Hspd1",
                    "Cpe",
                    "Atf5",
                    "Arf5",
                    "Eef2",
                    "Hspa5",
                    "Rnh1",
                    "Rrbp1",
                    "Uqcrc2",
                    "Actr3",
                    "Impdh2",
                    "Gapdh",
                    "Prdx2",
                    "Mpc1",
                    "Fth1",
                    "Sars",
                    "Sdpr",
                    "Lgals1",
                    "Rplp0",
                    "Ahnak",
                    "Hspa9",
                    "Fkbp10",
                    "Ube2n",
                    "Kif5b",
                    "Cct8",
                    "Mpc2",
                    "Lsp1",
                    "Psmb1",
                    "Psmd2",
                    "Cited2",
                    "Eif4g2",
                    "Psmb2",
                    "Uqcc2",
                    "Denr",
                    "Lhfp",
                    "Rbm3",
                    "Psmd11",
                    "Cnn3",
                    "Btg2",
                    "Vdac1",
                    "Selk",
                    "Rad23b",
                    "Chmp2a",
                    "Pdcd5",
                    "Ankrd1",
                    "Anxa5",
                    "Eif3f",
                    "Tmbim6",
                    "Anxa6",
                    "Mtch1",
                    "Rab10",
                    "Rpl22",
                    "Rsl1d1",
                    "Anp32e",
                    "Syncrip",
                    "Sec61a1",
                    "Caprin1",
                    "Kdelr2",
                    "Ict1",
                    "Rpl4",
                    "Glud1",
                    "March5",
                    "Ssr3",
                    "Fermt2",
                    "Hnrnpu",
                    "Gspt1",
                    "Srsf11",
                    "Rpl18a",
                    "Rpn1",
                    "Gas1",
                    "Ssr4",
                    "Txndc9",
                    "Ppp1r2",
                    "Eprs",
                    "Trmt112",
                    "Lman1",
                    "Msmo1",
                    "Atp5l",
                    "Mrpl54",
                    "Son",
                    "Ywhab",
                    "Ilf2",
                    "Phlda3",
                    "Romo1",
                    "Ergic3",
                    "Bzw1",
                    "Sdc4",
                    "Ak6",
                    "Ubc",
                    "Hnrnpa3",
                    "Prpf40a",
                    "Cdc42ep3",
                    "Mrps7",
                    "Metap2",
                    "Psap",
                    "Ndufs4",
                    "Polr1d",
                    "Tm4sf1",
                    "Rbms1",
                    "Zyx",
                    "Psmd3",
                    "H2afy",
                    "Rab34",
                    "Bsg",
                    "Sec22b",
                    "Camk2d",
                    "Txndc17",
                    "Cops4",
                    "Flna",
                    "H13",
                    "Ywhag",
                    "Atp5f1",
                    "Rps13",
                    "Gsto1",
                    "Tpt1",
                    "Cfl1",
                    "Gng5",
                    "Vcan",
                    "Rps19",
                    "Sept2",
                    "Abracl",
                    "Exosc8",
                    "Cops6",
                    "Aldoa",
                    "Srp9",
                    "Hnrnpdl",
                    "Eif3a",
                    "Ctbp1",
                    "Strap",
                    "Csrp1",
                    "Commd1",
                    "Psmd13",
                    "Selm",
                    "Lamp1",
                    "Ppa1",
                    "Sat1",
                    "Lamtor5",
                    "Ssb",
                    "Ptrf",
                    "Gadd45g",
                    "Rrm2",
                    "Jun",
                    "Capns1",
                    "Pdia3",
                    "Txndc5",
                    "Pfdn5",
                    "Ndufb7",
                    "Rps14",
                    "Sfpq",
                    "Col4a1",
                    "Rpl32",
                    "Lipe",
                    "Mdh2",
                    "Snrpd2",
                    "Krtcap2",
                    "Rps27a",
                    "Bud31",
                    "Pdap1",
                    "Brk1",
                    "Gtf2h5",
                    "Rpl11",
                    "Prkar1a",
                    "Rap1b",
                    "Rpl13a",
                    "Nol7",
                    "Timp1",
                    "Tuba1a",
                    "Commd3",
                    "Fst",
                    "Tomm22",
                    "Erp29",
                    "Gpx8",
                    "Gpi1",
                    "Tmem167",
                    "Phldb2",
                    "Palld",
                    "Rpl27a",
                    "Tram1",
                    "Eif5b",
                    "Cct4",
                    "Lsm12",
                    "Pkm",
                    "Fstl1",
                    "Fam162a",
                    "Rhoc",
                    "Mdm2",
                    "Ifitm3",
                    "Mrpl33",
                    "Psmc2",
                    "Ndufa10",
                    "App",
                    "Uqcrfs1",
                    "Ywhah",
                    "Dek",
                    "Bag1",
                    "Tmem147",
                    "Rpn2",
                    "Bub3",
                    "Fam104a",
                    "Tmed3",
                    "Mrfap1",
                    "Hsd17b14",
                    "Slc25a5",
                    "Gap43",
                    "Tpr",
                    "Cstb",
                    "Foxp1",
                    "Srsf6",
                    "Ndufa5",
                    "Tmpo",
                    "Ndufb10",
                    "Ube2e3",
                    "Mesdc2",
                    "Eef1g",
                    "Rbm25",
                    "Arhgdia",
                    "Rac1",
                    "Cald1",
                    "Sri",
                    "Pabpc1",
                    "Prdx5",
                    "Tsen34",
                    "Fkbp1a",
                    "Got2",
                    "Dad1",
                    "Emc7",
                    "Gnas",
                    "Sept1",
                    "Arf4",
                    "Arpc5",
                    "Hdlbp",
                    "Cox4i1",
                    "Ccnd1",
                    "Myl6",
                    "Sepw1",
                    "Hspa8",
                    "Serp1",
                    "Rplp2",
                    "Capn2",
                    "Ptov1",
                    "Aimp1",
                    "Cnih1",
                    "Gpx4",
                    "Glrx3",
                    "Gtf2a2",
                    "Carhsp1",
                    "Rrp1",
                    "Rpl23",
                    "Sfr1",
                    "P3h4",
                    "Junb",
                    "Rbm17",
                    "Map2k2",
                    "Axl",
                    "Ogn"
                ],
                "expression_sum": {
                    "class1": 12333.0,
                    "class2": 35054.0
                },
                "uniprot_ids": [
                    "Q91ZW3",
                    "B2RT32",
                    "Q3URV3",
                    "Q3U0F8"
                ]
            },
            "Hdac2": {
                "id": "Hdac2",
                "type": "TF",
                "symbol": "Hdac2",
                "source": [
                    "Tcf4",
                    "Pou5f1",
                    "Nfib",
                    "Atf4",
                    "Junb",
                    "Fos",
                    "Jun",
                    "Sox2",
                    "Hes1",
                    "Nanog",
                    "Jund",
                    "Twist2",
                    "Cebpb"
                ],
                "target": [
                    "Rps5",
                    "Hsp90aa1",
                    "Arf1",
                    "Dap",
                    "Rpl41",
                    "Rnf7",
                    "Sar1a",
                    "Srsf9",
                    "Mdh1",
                    "Tpm4",
                    "Usmg5",
                    "Dctn3",
                    "Ppp2ca",
                    "Cd44",
                    "Ddx24",
                    "H2afj",
                    "Rps17",
                    "Fis1",
                    "Zc3h15",
                    "Cox8a",
                    "Ddb1",
                    "Myh9",
                    "Thbs1",
                    "Psmb7",
                    "Txn2",
                    "Lims1",
                    "Sec13",
                    "Clta",
                    "Ppp1r11",
                    "1110008F13Rik",
                    "Tuba1b",
                    "Wisp1",
                    "Slc25a4",
                    "Cks1b",
                    "Col12a1",
                    "Rps27l",
                    "Tspan4",
                    "Map1lc3a",
                    "Psma6",
                    "Lmna",
                    "Psmc1",
                    "Hnrnpa1",
                    "Ndufa2",
                    "Thoc7",
                    "Hnrnpf",
                    "Ddx39",
                    "Mlf2",
                    "Psmd6",
                    "Vbp1",
                    "Ssrp1",
                    "Atp5j2",
                    "Ier3ip1",
                    "Pck2",
                    "Rnf187",
                    "Lgals3",
                    "Swi5",
                    "Cxcl12",
                    "Psmb6",
                    "mt-Co1",
                    "Pgrmc1",
                    "Med28",
                    "Shfm1",
                    "Fos",
                    "Tagln2",
                    "Cox6c",
                    "Rps15a",
                    "Ddx3x",
                    "Ndufs3",
                    "Nedd4",
                    "Rexo2",
                    "Calr",
                    "Pdlim7",
                    "Rab18",
                    "Tubb5",
                    "Calm3",
                    "Mfge8",
                    "Mbnl2",
                    "Arf6",
                    "Canx",
                    "D8Ertd738e",
                    "Hn1",
                    "Ppp1r14b",
                    "Actn1",
                    "Snrpe",
                    "B2m",
                    "Mrpl14",
                    "Slc25a3",
                    "Psat1",
                    "Eif4h",
                    "Lamtor1",
                    "2810417H13Rik",
                    "Psmd7",
                    "Gdi2",
                    "Stub1",
                    "Prrx1",
                    "Tra2b",
                    "Ranbp1",
                    "Mrpl28",
                    "Psmc6",
                    "Hmgcs1",
                    "Zwint",
                    "Rbx1",
                    "Rps24",
                    "Eif1",
                    "Myh10",
                    "Neat1",
                    "Sec61g",
                    "Ythdc1",
                    "Ola1",
                    "Snrnp27",
                    "Sf3b6",
                    "Tgfbi",
                    "Ube2r2",
                    "Timp3",
                    "Srsf1",
                    "Gemin7",
                    "Wdr89",
                    "Lrrc59",
                    "Akr1a1",
                    "Tdgf1",
                    "Sh3glb1",
                    "Ssr2",
                    "Pdcd6",
                    "Akt1s1",
                    "Ube2d2a",
                    "Hras",
                    "Bnip3l",
                    "Atp2b1",
                    "Kpnb1",
                    "Mapre1",
                    "C1d",
                    "Cisd2",
                    "Mylpf",
                    "Nktr",
                    "Col5a1",
                    "Ppia",
                    "Snrpd1",
                    "Ctgf",
                    "Tspan3",
                    "Hnrnpm",
                    "Capza2",
                    "Eif3k",
                    "Klf6",
                    "Rpl36al",
                    "Pcolce",
                    "Tgfb1i1",
                    "Loxl1",
                    "2310036O22Rik",
                    "Ppp1ca",
                    "Srsf5",
                    "Hnrnph1",
                    "Rpl35a",
                    "Timp2",
                    "Syngr1",
                    "Chchd1",
                    "Crip1",
                    "Gars",
                    "Psmc3",
                    "Ywhaz",
                    "Sec62",
                    "Rps12",
                    "Pomp",
                    "Luc7l2",
                    "Rpl36",
                    "Rbbp7",
                    "Rps21",
                    "Ddx5",
                    "Ndufb9",
                    "Rbm39",
                    "Maged1",
                    "Arpp19",
                    "Nedd8",
                    "Pcnp",
                    "Eef1b2",
                    "Ube2i",
                    "Arpc1b",
                    "Col3a1",
                    "Cyr61",
                    "Eif3l",
                    "Kdelr1",
                    "Rab14",
                    "Atpif1",
                    "Serpinh1",
                    "Tmem258",
                    "Tcp1",
                    "Atp6v1g1",
                    "Rpl24",
                    "Pttg1",
                    "Gnb2",
                    "Fus",
                    "Dpysl3",
                    "Ociad1",
                    "Ddost",
                    "Hcfc1r1",
                    "Rps4x",
                    "Sptbn1",
                    "Id3",
                    "Txnl1",
                    "Ap3s1",
                    "Gng12",
                    "Cct3",
                    "Fkbp8",
                    "Ncam1",
                    "Wls",
                    "Psma7",
                    "Hacd1",
                    "G3bp1",
                    "Ndufa8",
                    "Pdia4",
                    "Sept7",
                    "Srp14",
                    "Fam195b",
                    "Tpm3",
                    "Atp5b",
                    "Banf1",
                    "Actg1",
                    "Immt",
                    "Psma1",
                    "Rcn2",
                    "Emc10",
                    "Ier5",
                    "Tsg101",
                    "2410015M20Rik",
                    "Hdgf",
                    "Prnp",
                    "Smc3",
                    "Thrap3",
                    "Id1",
                    "Csnk2b",
                    "Nars",
                    "Tpi1",
                    "Hmga2",
                    "Rpl37",
                    "Dynlrb1",
                    "Atf4",
                    "Hprt",
                    "Tmem234",
                    "Nfkbia",
                    "Nucks1",
                    "Nrp1",
                    "Col1a2",
                    "Eif3b",
                    "Ube2m",
                    "Atp6v0b",
                    "F2r",
                    "Fkbp9",
                    "Pcbp1",
                    "Rpl38",
                    "Serpine2",
                    "Hnrnpk",
                    "Igfbp4",
                    "Mrps24",
                    "Id2",
                    "Itgb1",
                    "Ndufa4",
                    "Ewsr1",
                    "Cops5",
                    "Ssr3",
                    "Eif4e",
                    "Ndufv2",
                    "Fabp5",
                    "H3f3a",
                    "Mettl9",
                    "Nme1",
                    "Chmp4b",
                    "Tcf4",
                    "Ube2k",
                    "Igf2",
                    "Csnk1a1",
                    "Cd81",
                    "Ssr1",
                    "Ndufs2",
                    "Cyp51",
                    "Lxn",
                    "Pgls",
                    "Tmsb10",
                    "Rpl3",
                    "Nudt21",
                    "Rpl12",
                    "Ift20",
                    "Anxa2",
                    "Eif3c",
                    "Vmp1",
                    "Cox7a2",
                    "Ndufa7",
                    "Col5a2",
                    "Cdv3",
                    "Crip2",
                    "Napa",
                    "Cox7a2l",
                    "mt-Nd4",
                    "Vim",
                    "Hmgn1",
                    "Ube2l3",
                    "Btf3l4",
                    "Basp1",
                    "Ppp4c",
                    "Eef1d",
                    "Sh3bgrl3",
                    "Lsm8",
                    "Wdr1",
                    "Tnc",
                    "Ube2b",
                    "Ube2j2",
                    "Mrpl13",
                    "Vapa",
                    "Eif5",
                    "Tmed2",
                    "Txn1",
                    "Erh",
                    "Mrpl17",
                    "Atp5j",
                    "Fscn1",
                    "Atp5g3",
                    "Srrm1",
                    "Tpm1",
                    "Ldha",
                    "Sdcbp",
                    "Thy1",
                    "Surf4",
                    "Atp5g2",
                    "Chd4",
                    "Srp19",
                    "Cct5",
                    "Phlda1",
                    "Emd",
                    "1110004F10Rik",
                    "Myeov2",
                    "Bclaf1",
                    "Vasp",
                    "Tceb1",
                    "Rpsa",
                    "H2afz",
                    "Bola3",
                    "Mrps21",
                    "Psip1",
                    "Vcl",
                    "Cst3",
                    "Rpl36a",
                    "Eif3m",
                    "Lox",
                    "Arpc5l",
                    "Rps29",
                    "Rpl5",
                    "Higd1a",
                    "S100a6",
                    "Prdx6",
                    "Serpine1",
                    "H2-D1",
                    "Trappc4",
                    "Rps20",
                    "Ybx1",
                    "Dnaja2",
                    "Sys1",
                    "Cd63",
                    "Capg",
                    "Lbh",
                    "Ndufa3",
                    "Tmem14c",
                    "Cdc37",
                    "Ndufab1",
                    "Mtdh",
                    "Scand1",
                    "Rps6",
                    "Snrpf",
                    "Eef1a1",
                    "Nfib",
                    "mt-Nd5",
                    "Maged2",
                    "Rtn4",
                    "Rpl28",
                    "Atp5c1",
                    "Ctsl",
                    "Ccnd2",
                    "Mrps17",
                    "Tcf25",
                    "Spcs1",
                    "Ywhaq",
                    "Cox6a1",
                    "Col6a1",
                    "Rala",
                    "Sdc2",
                    "Tax1bp3",
                    "Pdha1",
                    "Filip1l",
                    "Rpl26",
                    "Uqcrb",
                    "Cav1",
                    "Rps3",
                    "Calm2",
                    "Atox1",
                    "Cox7c",
                    "Srsf10",
                    "Adss",
                    "Btf3",
                    "Plk2",
                    "Tln1",
                    "Cdh11",
                    "Oaf",
                    "Ostf1",
                    "Rps7",
                    "Tmem126a",
                    "Ube2v2",
                    "Rpl18",
                    "Eno1",
                    "Rpl19",
                    "Atp5d",
                    "Snhg12",
                    "Taldo1",
                    "Timm13",
                    "mt-Nd3",
                    "Eif3h",
                    "Fbln5",
                    "Rpl22l1",
                    "Chd3",
                    "Rps3a1",
                    "Srsf3",
                    "Vcp",
                    "Smc4",
                    "Cmpk1",
                    "Mbd3",
                    "Rpl7",
                    "Fkbp2",
                    "Btg1",
                    "Cct7",
                    "Fn1",
                    "Glipr2",
                    "Lrrc58",
                    "Hes1",
                    "Ap1s1",
                    "Ndufb2",
                    "Hsp90ab1",
                    "Rps9",
                    "Twist2",
                    "Oaz1",
                    "Mrpl30",
                    "Rpp25",
                    "Uqcrq",
                    "Rcn3",
                    "Sec61b",
                    "BC031181",
                    "Polr2m",
                    "Rwdd1",
                    "Snx5",
                    "Malat1",
                    "Gnai2",
                    "Cyc1",
                    "Rps8",
                    "Dnaja1",
                    "Tmem176b",
                    "Eif4a1",
                    "Sumo2",
                    "Tm9sf3",
                    "Pcbp2",
                    "Ctsd",
                    "Magoh",
                    "Ndufc2",
                    "Reep5",
                    "Tmed10",
                    "Pou5f1",
                    "Fhl2",
                    "H3f3b",
                    "Glrx5",
                    "Col1a1",
                    "Elof1",
                    "Gnb2l1",
                    "Ndufa12",
                    "Ube2e3",
                    "Nsa2",
                    "Oxct1",
                    "Atxn10",
                    "Wbp5",
                    "Atp6v0e",
                    "Cnih4",
                    "Pebp1",
                    "Dbi",
                    "Morf4l2",
                    "Ube2d3",
                    "Tmem59",
                    "Rbbp4",
                    "Eif3e",
                    "Dstn",
                    "Emp3",
                    "Aspn",
                    "Pfdn4",
                    "Cebpb",
                    "Srrm2",
                    "Top1",
                    "Ywhae",
                    "Pafah1b2",
                    "Ubl5",
                    "Anxa3",
                    "Mrps15",
                    "Ndufc1",
                    "Psmc4",
                    "Eif3d",
                    "Hbegf",
                    "Psmd14",
                    "Rps15",
                    "Ghitm",
                    "Arpc3",
                    "Psma4",
                    "Scp2",
                    "Ilk",
                    "Arl1",
                    "Arpc1a",
                    "Ap2s1",
                    "Mgat2",
                    "Rpl34",
                    "Mlec",
                    "Rab7",
                    "Sep15",
                    "Dynll1",
                    "Gtpbp4",
                    "Eif2s2",
                    "Hspd1",
                    "Cpe",
                    "Atf5",
                    "Arf5",
                    "Cdc42",
                    "Npc2",
                    "Rpl31",
                    "Srsf7",
                    "Smdt1",
                    "Nap1l1",
                    "Eef2",
                    "Tpx2",
                    "Etfb",
                    "Hsbp1",
                    "Hspa5",
                    "Rnh1",
                    "Rrbp1",
                    "Uqcrc2",
                    "Actr3",
                    "Tmsb4x",
                    "Cd47",
                    "Efemp2",
                    "Ccdc124",
                    "Fbln2",
                    "Gapdh",
                    "Smarca5",
                    "Prdx2",
                    "Mpc1",
                    "Copb2",
                    "Khdrbs1",
                    "Ap2m1",
                    "Mrpl52",
                    "Nupr1",
                    "Sars",
                    "Bri3",
                    "Smim14",
                    "Prkcdbp",
                    "Hnrnpab",
                    "Sdpr",
                    "Lgals1",
                    "Rplp0",
                    "Psmb5",
                    "Ahnak",
                    "Tbrg1",
                    "Col11a1",
                    "Hspa9",
                    "Fkbp10",
                    "Nop58",
                    "Ube2n",
                    "Ndufa11",
                    "Cbx5",
                    "Kif5b",
                    "Luc7l3",
                    "Polr2c",
                    "Cct8",
                    "Mpc2",
                    "Copz2",
                    "Hnrnpa2b1",
                    "Ptp4a1",
                    "Lsp1",
                    "Ndufs5",
                    "Gsn",
                    "Psmb1",
                    "Oat",
                    "Tubb6",
                    "Coa3",
                    "Ndufb8",
                    "Cited2",
                    "Psmd2",
                    "Tmem97",
                    "Srsf2",
                    "Eva1b",
                    "Prdx4",
                    "mt-Nd1",
                    "Eif4g2",
                    "Psmb2",
                    "Rab2a",
                    "Uqcc2",
                    "Denr",
                    "Eif3g",
                    "Tspo",
                    "Sra1",
                    "Tceal8",
                    "Lhfp",
                    "Myl9",
                    "Rbm3",
                    "Psmd11",
                    "Cnn3",
                    "Vps29",
                    "Btg2",
                    "Vdac1",
                    "Selk",
                    "Cggbp1",
                    "Zfp706",
                    "Rad23b",
                    "Chmp2a",
                    "Tra2a",
                    "Nudcd2",
                    "Pdcd5",
                    "Aprt",
                    "Ost4",
                    "Rab11a",
                    "Ankrd1",
                    "Snrpc",
                    "Anxa5",
                    "Eif3f",
                    "Eif4g1",
                    "Anxa6",
                    "Mtch1",
                    "Tmbim6",
                    "Dynll2",
                    "Rhoa",
                    "Ssu72",
                    "Rab10",
                    "Atp5a1",
                    "Psma5",
                    "Ttc3",
                    "Cdk4",
                    "Cnpy2",
                    "Rpl22",
                    "Rsl1d1",
                    "Vimp",
                    "Ccdc80",
                    "Syncrip",
                    "Sec61a1",
                    "Caprin1",
                    "Iqgap1",
                    "Ict1",
                    "Kdelr2",
                    "Glud1",
                    "March5",
                    "Mrpl23",
                    "Pmp22",
                    "Rpl4",
                    "Fermt2",
                    "Hnrnpu",
                    "Rps25",
                    "Rps28",
                    "Gspt1",
                    "Srsf11",
                    "Rpl18a",
                    "Hmgb3",
                    "Rpn1",
                    "Gas1",
                    "Ssr4",
                    "Txndc9",
                    "Ppp1r2",
                    "Sqstm1",
                    "Eprs",
                    "Trmt112",
                    "Lman1",
                    "F3",
                    "Msmo1",
                    "Rcn1",
                    "Arpc2",
                    "Atp5l",
                    "Mrpl54",
                    "Rpl35",
                    "Son",
                    "Ilf2",
                    "Ywhab",
                    "Cct2",
                    "Acp1",
                    "Prelid1",
                    "Sox4",
                    "Phlda3",
                    "Sept11",
                    "Ndufa1",
                    "Romo1",
                    "Tpm2",
                    "Ergic3",
                    "Eif1ax",
                    "Bzw1",
                    "Sdc4",
                    "Ak6",
                    "Ubc",
                    "Rock2",
                    "Hnrnpa3",
                    "Prpf40a",
                    "Prmt1",
                    "Pbdc1",
                    "Cdc42ep3",
                    "Mrps7",
                    "Metap2",
                    "Mrpl57",
                    "Morf4l1",
                    "2010107E04Rik",
                    "Col8a1",
                    "Pafah1b1",
                    "Lsm6",
                    "Ube2s",
                    "Tecr",
                    "Sf3b1",
                    "Idh2",
                    "Psap",
                    "Ndufs4",
                    "Polr1d",
                    "Snrpd3",
                    "Tm4sf1",
                    "Rbms1",
                    "P4hb",
                    "Scd2",
                    "Dlk1",
                    "Sf3b2",
                    "H2afy",
                    "Map1b",
                    "Rab34",
                    "Suclg1",
                    "Zyx",
                    "Adipor1",
                    "Rpl17",
                    "Bsg",
                    "mt-Nd2",
                    "Hnrnpd",
                    "Ppic",
                    "Rpl14",
                    "Tma7",
                    "Sec22b",
                    "Camk2d",
                    "Txndc17",
                    "Cops4",
                    "Cyba",
                    "Flna",
                    "Lsm4",
                    "Efemp1",
                    "H13",
                    "Rsu1",
                    "Ywhag",
                    "Atp5f1",
                    "Prss23",
                    "Rps13",
                    "Gsto1",
                    "Tpt1",
                    "Eif5a",
                    "Rps11",
                    "Ppib",
                    "Psmd8",
                    "Cfl1",
                    "Gng5",
                    "Vcan",
                    "Sparc",
                    "Rps19",
                    "Sept2",
                    "Abracl",
                    "Exosc8",
                    "Cops6",
                    "Msn",
                    "Aldoa",
                    "Mrpl20",
                    "Hnrnpdl",
                    "Serpinb6a",
                    "Srp9",
                    "Eif3a",
                    "Rtn3",
                    "Lpp",
                    "Ptn",
                    "Tagln",
                    "Ctbp1",
                    "Psmb4",
                    "Snx3",
                    "Strap",
                    "Csrp1",
                    "Commd1",
                    "Psmd13",
                    "Selm",
                    "Ckap4",
                    "Esd",
                    "Atxn7l3b",
                    "Igfbp6",
                    "Comt",
                    "Cox6b1",
                    "Lamp1",
                    "Sat1",
                    "Lamtor5",
                    "Ndufa13",
                    "Atp5e",
                    "Prdx1",
                    "Ssb",
                    "Ptrf",
                    "Tinagl1",
                    "Gps1",
                    "Cmtm7",
                    "Gadd45g",
                    "Atp5o",
                    "Rrm2",
                    "Rpl13",
                    "Sh3bgrl",
                    "Jun",
                    "Capns1",
                    "Itm2b",
                    "Pdia3",
                    "Rplp1",
                    "Ndufb5",
                    "Rpl10a",
                    "Atrx",
                    "Psma2",
                    "Txndc5",
                    "Pfdn5",
                    "Ndufb7",
                    "Ptms",
                    "Eif4e2",
                    "Rps14",
                    "Ftl1",
                    "Sfpq",
                    "Col4a1",
                    "Rpl32",
                    "Smap1",
                    "Cd24a",
                    "Zfp36l1",
                    "Lipe",
                    "Tnfrsf12a",
                    "Mdh2",
                    "Krtcap2",
                    "Gabarap",
                    "Uqcr11",
                    "Rps27a",
                    "Phgdh",
                    "Rap1a",
                    "Serf2",
                    "Bud31",
                    "Pdap1",
                    "Brk1",
                    "Gtf2h5",
                    "Hsp90b1",
                    "Mtpn",
                    "Anxa1",
                    "Prkar1a",
                    "Rap1b",
                    "Rpl11",
                    "Vdac2",
                    "Pfn1",
                    "Rpl13a",
                    "Nol7",
                    "Timp1",
                    "Ier3",
                    "Grb10",
                    "Cuta",
                    "Apoe",
                    "Eif1a",
                    "Tuba1a",
                    "Commd3",
                    "Cnn2",
                    "Minos1",
                    "Snrpb",
                    "Pgam1",
                    "Rabggtb",
                    "Tomm22",
                    "Nans",
                    "Ubap2l",
                    "Erp29",
                    "Gpx8",
                    "Gpi1",
                    "Tmem167",
                    "Ppig",
                    "Phldb2",
                    "Palld",
                    "Rpl27a",
                    "Dcun1d5",
                    "Eif5b",
                    "Tram1",
                    "Cct4",
                    "Rdx",
                    "Lrrc17",
                    "Lsm12",
                    "Fhl1",
                    "Anp32b",
                    "Edf1",
                    "Cdkn1a",
                    "Pkm",
                    "Nono",
                    "Csde1",
                    "Fstl1",
                    "Marcks",
                    "Fam162a",
                    "Tceb2",
                    "Rhoc",
                    "Mdm2",
                    "Ifitm3",
                    "Psenen",
                    "Srp72",
                    "Mrpl33",
                    "Sumo3",
                    "Emp1",
                    "Psmc2",
                    "Ndufa10",
                    "App",
                    "Fundc2",
                    "Uqcrfs1",
                    "Naa50",
                    "Ywhah",
                    "Eif3i",
                    "Hnrnpc",
                    "Smc2",
                    "Mtch2",
                    "S100a10",
                    "Bag1",
                    "Timm17a",
                    "Psmd4",
                    "Rpn2",
                    "Tmem147",
                    "Bub3",
                    "Fam104a",
                    "Prrx2",
                    "Papola",
                    "Rps16",
                    "Manf",
                    "Tmem256",
                    "Sin3b",
                    "Cetn3",
                    "Dctn2",
                    "Arhgdib",
                    "Tmed3",
                    "Serpinf1",
                    "Mrfap1",
                    "Galk1",
                    "Hsd17b14",
                    "Slc25a5",
                    "Uqcrh",
                    "Gap43",
                    "Tpr",
                    "Hint1",
                    "Tbca",
                    "Cstb",
                    "Foxp1",
                    "Srsf6",
                    "Ndufa5",
                    "Ebna1bp2",
                    "Ccng1",
                    "Ndufb10",
                    "Tmpo",
                    "Rpl37a",
                    "Tsn",
                    "H19",
                    "Ptp4a2",
                    "Mesdc2",
                    "Eef1g",
                    "Dnajc3",
                    "Cope",
                    "Cox5b",
                    "Rbm25",
                    "Arhgdia",
                    "Cycs",
                    "Myl12a",
                    "Ndufb6",
                    "Ptma",
                    "Rac1",
                    "Rer1",
                    "S100a4",
                    "Ralbp1",
                    "Cald1",
                    "Sdhb",
                    "Fau",
                    "Ybx3",
                    "Rps10",
                    "Sri",
                    "Pabpc1",
                    "Prdx5",
                    "Alyref",
                    "Clic1",
                    "Tsen34",
                    "Fkbp1a",
                    "Got2",
                    "Dad1",
                    "Slirp",
                    "Emc7",
                    "Gnas",
                    "Aebp1",
                    "Arf4",
                    "Calu",
                    "Arpc5",
                    "Yif1b",
                    "Hdlbp",
                    "Limd2",
                    "Wbp11",
                    "Cox4i1",
                    "Ccnd1",
                    "Tmed9",
                    "Snf8",
                    "Myl6",
                    "Sepw1",
                    "2810428I15Rik",
                    "Hspa8",
                    "Ndufs8",
                    "Psma3",
                    "Rpl39",
                    "Rabac1",
                    "Rplp2",
                    "Capn2",
                    "Serp1",
                    "Mif",
                    "Laptm4a",
                    "Ptov1",
                    "Aimp1",
                    "Serbp1",
                    "Cnih1",
                    "Gpx4",
                    "Chchd3",
                    "Rheb",
                    "Rps23",
                    "Glrx3",
                    "Gtf2a2",
                    "Carhsp1",
                    "Cryab",
                    "Alpl",
                    "Rrp1",
                    "Pdcl3",
                    "Rpl23",
                    "Rpl30",
                    "Kdelr3",
                    "Sfr1",
                    "Polr2f",
                    "P3h4",
                    "Junb",
                    "Rbm17",
                    "Bcap31",
                    "Map2k2",
                    "Cnbp",
                    "Sdhd",
                    "Axl",
                    "Cfl2",
                    "Ogn",
                    "Rsrc2"
                ],
                "expression_sum": {
                    "class1": 10992.0,
                    "class2": 22123.0
                },
                "uniprot_ids": [
                    "P70288",
                    "Q8BQ10",
                    "Q3UXH8",
                    "Q3URA2",
                    "A0A0R4J008",
                    "Q3TMT1"
                ]
            },
            "E2f4": {
                "id": "E2f4",
                "type": "TF",
                "symbol": "E2f4",
                "source": [
                    "Pou5f1",
                    "Junb",
                    "Jun",
                    "Sox2",
                    "Nanog",
                    "Jund",
                    "Twist2",
                    "Cebpb"
                ],
                "target": [
                    "Mgat2",
                    "Sep15",
                    "Atxn7l3b",
                    "Dap",
                    "Lsm7",
                    "Cpe",
                    "Serpine2",
                    "Atf5",
                    "Arf5",
                    "Lamp1",
                    "Rcor2",
                    "Cdc42",
                    "Sfrp1",
                    "Itgb1",
                    "Syce2",
                    "Ptrf",
                    "Cd44",
                    "Fabp5",
                    "Gadd45g",
                    "Myh9",
                    "Tcf4",
                    "Thbs1",
                    "Rrbp1",
                    "Rasgrp2",
                    "Lims1",
                    "Tmsb4x",
                    "Cd47",
                    "Efemp2",
                    "Cd81",
                    "Itm2b",
                    "Jun",
                    "Fbln2",
                    "Pdia3",
                    "Pgls",
                    "Ssr1",
                    "Txndc5",
                    "Slc25a4",
                    "Tspan4",
                    "Col12a1",
                    "Lmna",
                    "Nupr1",
                    "Col5a2",
                    "Crip2",
                    "Prkcdbp",
                    "Vim",
                    "Lipe",
                    "Lgals1",
                    "Tnfrsf12a",
                    "Basp1",
                    "Col11a1",
                    "Fkbp10",
                    "Krtcap2",
                    "E130012A19Rik",
                    "Pam16",
                    "Kif5b",
                    "Tnc",
                    "Pck2",
                    "Tmed2",
                    "Mtpn",
                    "Lgals3",
                    "Gsn",
                    "Ilk",
                    "Tubb6",
                    "Pgrmc1",
                    "Timp1",
                    "Tagln2",
                    "Eva1b",
                    "Tpm1",
                    "Sdcbp",
                    "Surf4",
                    "Tuba1a",
                    "Rexo2",
                    "Commd3",
                    "Phlda1",
                    "Pdlim7",
                    "Lhfp",
                    "Mbnl2",
                    "Canx",
                    "Myeov2",
                    "Actn1",
                    "B2m",
                    "Btg2",
                    "Selk",
                    "Vcl",
                    "Gpi1",
                    "Lox",
                    "Palld",
                    "Tram1",
                    "Igfbp7",
                    "Lrrc17",
                    "Rab11a",
                    "Ung",
                    "Anxa5",
                    "S100a6",
                    "Anxa6",
                    "Serpine1",
                    "Cdkn1a",
                    "Neat1",
                    "Myh10",
                    "Marcks",
                    "Fam162a",
                    "Rhoc",
                    "Vimp",
                    "Cd63",
                    "Timp3",
                    "Capg",
                    "Lbh",
                    "Fbn1",
                    "Mtdh",
                    "Iqgap1",
                    "Pmp22",
                    "Emp1",
                    "Ssr3",
                    "mt-Nd5",
                    "Sh3glb1",
                    "App",
                    "Maged2",
                    "Rtn4",
                    "Impa2",
                    "Ctsl",
                    "Fbn2",
                    "Spcs1",
                    "Ssr4",
                    "Atp2b1",
                    "Gas1",
                    "Sdc2",
                    "Tax1bp3",
                    "S100a10",
                    "Nktr",
                    "Filip1l",
                    "Col5a1",
                    "Tspan3",
                    "Ctgf",
                    "Arpc2",
                    "Cav1",
                    "Klf6",
                    "Pcolce",
                    "Plk2",
                    "Loxl1",
                    "Phlda3",
                    "Oaf",
                    "Ostf1",
                    "Arhgdib",
                    "Sept11",
                    "Serpinf1",
                    "Sox4",
                    "Tgfb1i1",
                    "Tln1",
                    "Tmed3",
                    "Timp2",
                    "Syngr1",
                    "Tpm2",
                    "Sdc4",
                    "Chd3",
                    "Rock2",
                    "Rps12",
                    "Foxp1",
                    "Cdc42ep3",
                    "H19",
                    "Lrrc58",
                    "Ptp4a2",
                    "Fn1",
                    "Dnajc3",
                    "Glipr2",
                    "Arpc1b",
                    "Ap1s1",
                    "Cyr61",
                    "Hes1",
                    "Kdelr1",
                    "Nenf",
                    "Rab14",
                    "Psap",
                    "Serpinh1",
                    "Twist2",
                    "Klf2",
                    "Rbms1",
                    "P4hb",
                    "Zyx",
                    "Map1b",
                    "Rab34",
                    "Malat1",
                    "Tmem176b",
                    "Ppic",
                    "Gnb2",
                    "Sri",
                    "Mfap2",
                    "Tm9sf3",
                    "Tmed10",
                    "Ctsd",
                    "Dpysl3",
                    "Mybl2",
                    "Reep5",
                    "Emc7",
                    "Gnas",
                    "Hcfc1r1",
                    "Arf4",
                    "Camk2d",
                    "Calu",
                    "Fhl2",
                    "Gm7325",
                    "Sept1",
                    "Sptbn1",
                    "Flna",
                    "mt-Cytb",
                    "Hdlbp",
                    "Ap3s1",
                    "Gng12",
                    "Limd2",
                    "Oxct1",
                    "H13",
                    "Rsu1",
                    "Ncam1",
                    "Wls",
                    "Sepw1",
                    "Gsto1",
                    "Hacd1",
                    "Ndufa8",
                    "Sept7",
                    "Fam195b",
                    "Capn2",
                    "Laptm4a",
                    "Vcan",
                    "Actg1",
                    "Cnih1",
                    "Rcn2",
                    "Emp3",
                    "Ier5",
                    "Cebpb",
                    "Carhsp1",
                    "Abracl",
                    "Msn",
                    "Id1",
                    "Prnp",
                    "Sept2",
                    "Serpinb6a",
                    "Hmga2",
                    "Kdelr3",
                    "Pafah1b2",
                    "Ctbp1",
                    "P3h4",
                    "Junb",
                    "Csrp1",
                    "Hbegf",
                    "Selm",
                    "Axl",
                    "Cfl2",
                    "Nrp1",
                    "Col1a2",
                    "Esd"
                ],
                "expression_sum": {
                    "class1": 2638.0,
                    "class2": 24346.0
                },
                "uniprot_ids": [
                    "Q8R0K9"
                ]
            },
            "Vim": {
                "id": "Vim",
                "type": "Gene",
                "symbol": "Vim",
                "source": [
                    "E2f4",
                    "Pou5f1",
                    "Trim28",
                    "Rest",
                    "Atf5",
                    "Eno1",
                    "Ctcf",
                    "Trp53",
                    "Hdac1",
                    "Klf5",
                    "Tbx3",
                    "Zfp57",
                    "Hdac2",
                    "Smc3"
                ],
                "target": [],
                "expression_sum": {
                    "class1": 265995.0,
                    "class2": 63798.0
                },
                "uniprot_ids": [
                    "P20152",
                    "Q5FWJ3",
                    "A0A0A6YWC8",
                    "A2AKJ2",
                    "Q3UD36",
                    "Q3TWV0",
                    "Q3UAX1",
                    "Q3TFD9",
                    "Q3V2S4",
                    "A4IF59",
                    "Q3U6S1"
                ]
            },
            "Ybx1": {
                "id": "Ybx1",
                "type": "TF",
                "symbol": "Ybx1",
                "source": [
                    "Tcf4",
                    "Pou5f1",
                    "Cebpb",
                    "Nfib",
                    "Junb",
                    "Fos",
                    "Jun",
                    "Sox2",
                    "Tbx3",
                    "Zfp57",
                    "Nanog",
                    "Jund",
                    "Twist2",
                    "Hdac2",
                    "Smc3",
                    "Sox4"
                ],
                "target": [
                    "Hsp90aa1",
                    "Arf1",
                    "Dap",
                    "Hp1bp3",
                    "Clcc1",
                    "Ube2j1",
                    "Kif2a",
                    "Dpysl2",
                    "Tmem41b",
                    "Mdfic",
                    "Rbfox2",
                    "Mgat1",
                    "Cd44",
                    "Casc4",
                    "Fis1",
                    "Tead1",
                    "Colgalt1",
                    "AI314180",
                    "Igf1r",
                    "Lpcat4",
                    "Commd9",
                    "Itfg1",
                    "Npr1",
                    "Lims1",
                    "Sucla2",
                    "Ccm2",
                    "Sec13",
                    "Ryk",
                    "Tuba1b",
                    "Gm13051",
                    "Nucb1",
                    "Wisp1",
                    "Slc25a4",
                    "St7l",
                    "Ctsk",
                    "Col12a1",
                    "Calcoco2",
                    "Rpl23a",
                    "Lrrcc1",
                    "Spop",
                    "Map1lc3a",
                    "Ifngr2",
                    "Slc35a1",
                    "Tmem37",
                    "Fxyd5",
                    "Tmem70",
                    "Pik3r1",
                    "Folr1",
                    "Hsd17b12",
                    "Zfand6",
                    "Ntmt1",
                    "Lcorl",
                    "Psmd6",
                    "Cthrc1",
                    "Ctsa",
                    "Runx1",
                    "Lmbrd1",
                    "Pik3ca",
                    "Ttc14",
                    "Csf1",
                    "Hectd1",
                    "Ssrp1",
                    "Smtnl2",
                    "Serinc1",
                    "1700019D03Rik",
                    "Mkln1",
                    "Ugcg",
                    "Ier3ip1",
                    "Rnf187",
                    "mt-Co1",
                    "Pgrmc1",
                    "Anxa8",
                    "Dusp3",
                    "Tagln2",
                    "Dnajc9",
                    "Dpm3",
                    "Ccdc88a",
                    "Calr",
                    "Ptk7",
                    "Fam50a",
                    "Crlf1",
                    "Pdlim7",
                    "Rab18",
                    "Tubb5",
                    "Gm13154",
                    "Tmem120a",
                    "Actn1",
                    "Eif4h",
                    "2810417H13Rik",
                    "Rras2",
                    "Prkag1",
                    "Plod1",
                    "Tcf7l2",
                    "Nr0b1",
                    "Prrx1",
                    "Slc35e4",
                    "Fam101b",
                    "Tars",
                    "Erp44",
                    "Fam98a",
                    "Hmgcs1",
                    "Zwint",
                    "Fip1l1",
                    "Smad1",
                    "Neat1",
                    "Acta2",
                    "Aga",
                    "Ctnnd1",
                    "Ube2r2",
                    "Tgfbi",
                    "Acer3",
                    "Atp2c1",
                    "Wdr89",
                    "Tdgf1",
                    "Sh3glb1",
                    "Strn3",
                    "2310022B05Rik",
                    "Akap9",
                    "Arl6ip5",
                    "Akt1s1",
                    "Agpat4",
                    "Fech",
                    "Mospd1",
                    "Bnip3l",
                    "Atp2b1",
                    "Rit1",
                    "Apoc1",
                    "C1d",
                    "Nktr",
                    "Col5a1",
                    "Tspan3",
                    "Ctgf",
                    "Bgn",
                    "Col4a2",
                    "Klf6",
                    "Dld",
                    "Pcolce",
                    "Lysmd3",
                    "Ralb",
                    "Gnai3",
                    "Rasa1",
                    "Myadm",
                    "Timp2",
                    "Crxos",
                    "Bex4",
                    "Rab11b",
                    "Selo",
                    "Ahnak2",
                    "Psmc3",
                    "Cdc16",
                    "Sec62",
                    "Rps12",
                    "H2afy2",
                    "Gmps",
                    "Rbbp7",
                    "Gm6483",
                    "Rps27",
                    "Msrb3",
                    "Maged1",
                    "Tmem164",
                    "Tspan31",
                    "2610002M06Rik",
                    "Cfap36",
                    "Arpc1b",
                    "Col3a1",
                    "Cyr61",
                    "Eif3l",
                    "Rab14",
                    "Serpinh1",
                    "Ahsa2",
                    "Atp6v1g1",
                    "Gsk3b",
                    "Fgfr1",
                    "Arl8b",
                    "Tnfrsf23",
                    "Smim3",
                    "Acap2",
                    "Mnat1",
                    "Pitpnb",
                    "BC005537",
                    "Rbms2",
                    "Tmem165",
                    "Matr3",
                    "Ociad1",
                    "1700123O20Rik",
                    "Dpysl3",
                    "Mybl2",
                    "Mycbp",
                    "Ddost",
                    "Hcfc1r1",
                    "Sptbn1",
                    "Npdc1",
                    "Sertad2",
                    "Ap3s1",
                    "Ccdc50",
                    "Jkamp",
                    "Wwp1",
                    "Yipf5",
                    "Ncam1",
                    "Wbscr17",
                    "Dbn1",
                    "Bicd2",
                    "Hoxa5",
                    "Cdc42ep1",
                    "Pdia4",
                    "Saraf",
                    "Sept7",
                    "Srp14",
                    "Ccdc109b",
                    "Mmadhc",
                    "Atp5b",
                    "Actg1",
                    "Rcn2",
                    "Tsg101",
                    "Ier5",
                    "Mob4",
                    "Id1",
                    "Csnk2b",
                    "Pja2",
                    "Prickle1",
                    "2010111I01Rik",
                    "Dynlrb1",
                    "Hspb1",
                    "Atf4",
                    "Plekhh2",
                    "Synpo",
                    "Mcfd2",
                    "Col1a2",
                    "Tcaf1",
                    "Scd1",
                    "Stt3a",
                    "Atp6v0b",
                    "F2r",
                    "Pak2",
                    "Abhd8",
                    "Ak1",
                    "Cxx1a",
                    "Itgb1",
                    "Cd109",
                    "Boc",
                    "Fam204a",
                    "Amfr",
                    "Kmt2e",
                    "Klhdc2",
                    "Fabp5",
                    "Chmp4b",
                    "Gns",
                    "Soat1",
                    "Med13",
                    "Cd81",
                    "Vamp3",
                    "Tab2",
                    "Cyp51",
                    "Tpp2",
                    "Bdnf",
                    "Plod3",
                    "Pdgfrl",
                    "Fuca1",
                    "Tbpl1",
                    "Ift20",
                    "Anxa2",
                    "Vmp1",
                    "Ikbip",
                    "mt-Nd4",
                    "Crip2",
                    "Mycbp2",
                    "Arhgef2",
                    "Basp1",
                    "Spg20",
                    "1700020I14Rik",
                    "Wdr1",
                    "Tnc",
                    "Ube2b",
                    "Gpat4",
                    "Appl1",
                    "Ano6",
                    "Adamts5",
                    "Dnm3os",
                    "Adamts1",
                    "Slc43a3",
                    "Rbbp8",
                    "Myg1",
                    "Tpm1",
                    "Rgs4",
                    "Zfpl1",
                    "Stk17b",
                    "Ifi27",
                    "Lima1",
                    "Tank",
                    "Cct5",
                    "Uba1",
                    "Emd",
                    "1110004F10Rik",
                    "Tex261",
                    "Il6st",
                    "Myeov2",
                    "Lztr1",
                    "Unc50",
                    "Ercc1",
                    "Mcts1",
                    "Mxd4",
                    "Pdgfa",
                    "Bola3",
                    "Tmem50a",
                    "Vcl",
                    "Tmod3",
                    "Slc50a1",
                    "Eif3m",
                    "Capsl",
                    "Ppp2r1a",
                    "Bbip1",
                    "Xrn2",
                    "Smurf2",
                    "2310009B15Rik",
                    "Zfp330",
                    "Acbd3",
                    "Prdx6",
                    "Ufsp2",
                    "Far1",
                    "Lzic",
                    "Serpine1",
                    "H2-D1",
                    "Alg5",
                    "Gamt",
                    "Crabp1",
                    "Ifitm1",
                    "Afap1",
                    "Cd63",
                    "Lyrm2",
                    "Elovl5",
                    "Cdc37",
                    "Nptn",
                    "Hebp1",
                    "S100a11",
                    "Eef1a1",
                    "Sirt2",
                    "Irx3",
                    "Maged2",
                    "Gm5093",
                    "Wsb1",
                    "Ctsl",
                    "Prkca",
                    "Cox6a1",
                    "Rab5a",
                    "Slk",
                    "Ap2a2",
                    "Cops2",
                    "Tnrc18",
                    "Pold4",
                    "Pard3",
                    "Tspan5",
                    "Ndel1",
                    "Cav1",
                    "Mrps6",
                    "Prosc",
                    "Efr3a",
                    "mt-Co2",
                    "Sgcb",
                    "Uso1",
                    "Srsf10",
                    "Tln1",
                    "Cul3",
                    "Smchd1",
                    "Oaf",
                    "Ak3",
                    "Ostf1",
                    "Ndn",
                    "Ube2v2",
                    "Uba52",
                    "Eno1",
                    "Atp5d",
                    "Mt2",
                    "Eif3h",
                    "Fbln5",
                    "Acadl",
                    "Thoc2",
                    "Scoc",
                    "Cmpk1",
                    "1700061G19Rik",
                    "Figf",
                    "Btg1",
                    "Lrrc58",
                    "Fn1",
                    "Hes1",
                    "Cpsf2",
                    "Nenf",
                    "Polr3k",
                    "Rps9",
                    "Zfp42",
                    "Mrpl30",
                    "Myo1b",
                    "Rpp25",
                    "Bod1l",
                    "Malat1",
                    "mt-Nd4l",
                    "Gnai2",
                    "Tmem176b",
                    "Ap3d1",
                    "Eif4a1",
                    "Mfap2",
                    "Antxr1",
                    "Arcn1",
                    "Ctsd",
                    "Ndufc2",
                    "Borcs7",
                    "Fhl2",
                    "Elf2",
                    "Ndufa12",
                    "Krcc1",
                    "Adam19",
                    "Pla2g1b",
                    "Il1rl1",
                    "Atxn10",
                    "Ctsh",
                    "Atp6v0e",
                    "Cnih4",
                    "Medag",
                    "Adk",
                    "Ube2f",
                    "Por",
                    "Pcbp4",
                    "Prkra",
                    "Nap1l4",
                    "Tmem59",
                    "Ap1m1",
                    "Aspn",
                    "Stard4",
                    "Spcs3",
                    "Hnrnpr",
                    "Rnf166",
                    "Sav1",
                    "Gm26917",
                    "Tmbim4",
                    "Actr10",
                    "Irak1bp1",
                    "Arpc3",
                    "Zmynd11",
                    "Akr1b8",
                    "Scp2",
                    "Arl1",
                    "Arpc1a",
                    "Sep15",
                    "Arhgap28",
                    "Clptm1",
                    "Mlec",
                    "Atf5",
                    "Leprot",
                    "Cdc42",
                    "Npc2",
                    "Rnf217",
                    "Mvk",
                    "Ptbp3",
                    "Tmed5",
                    "Suv39h1",
                    "Uhrf1bp1l",
                    "Eef2",
                    "Asah1",
                    "Hspa5",
                    "Rnh1",
                    "Rrbp1",
                    "Actr3",
                    "Slit3",
                    "Tax1bp1",
                    "Pvrl2",
                    "Tmsb4x",
                    "Camk2g",
                    "Dnajc10",
                    "Gapdh",
                    "Exoc4",
                    "Adgre5",
                    "Arl3",
                    "Bex1",
                    "Mpc1",
                    "Actr1b",
                    "Gstm5",
                    "Tm2d2",
                    "Sars",
                    "Lmod1",
                    "Flnb",
                    "Yaf2",
                    "Ap2m1",
                    "Copb2",
                    "Smim14",
                    "Khdrbs1",
                    "Tmf1",
                    "B230219D22Rik",
                    "Rpl6l",
                    "Acot7",
                    "Ppp3ca",
                    "Prkcdbp",
                    "Col11a1",
                    "Itgb5",
                    "E130012A19Rik",
                    "Btbd1",
                    "Cct8",
                    "Mpc2",
                    "Copz2",
                    "Nfat5",
                    "Ptp4a1",
                    "Lsp1",
                    "Bmp1",
                    "Gsn",
                    "Tubb6",
                    "Ccny",
                    "Vma21",
                    "Cited2",
                    "N4bp3",
                    "Cap1",
                    "Hoxa9",
                    "Eva1b",
                    "Prdx4",
                    "Jam2",
                    "Eif4g2",
                    "Nudt18",
                    "Aes",
                    "mt-Nd1",
                    "Twist1",
                    "Desi2",
                    "2610305D13Rik",
                    "Trim71",
                    "Tceal8",
                    "Lhfp",
                    "Rbm3",
                    "Ankrd17",
                    "Ets1",
                    "Vps29",
                    "Parva",
                    "Tmem132a",
                    "Foxd3",
                    "Prkcsh",
                    "Rad23b",
                    "mt-Co3",
                    "Cyfip1",
                    "Coq10b",
                    "Tra2a",
                    "Cdh2",
                    "Rab11a",
                    "Sdf4",
                    "Anxa5",
                    "Aco2",
                    "Eif4g1",
                    "Anxa6",
                    "Mtch1",
                    "Rhoa",
                    "Reep3",
                    "Tusc3",
                    "Praf2",
                    "Ttc3",
                    "Shisa5",
                    "Ccdc80",
                    "Psmd9",
                    "Uba3",
                    "Caprin1",
                    "Kcnq1ot1",
                    "Kdelr2",
                    "Pmp22",
                    "Abhd5",
                    "Srsf11",
                    "Baz1a",
                    "Tmem9b",
                    "Hmgb3",
                    "Ssr4",
                    "Mapk1ip1l",
                    "Nid1",
                    "Ykt6",
                    "Fgfr1op2",
                    "Atp6ap2",
                    "Sqstm1",
                    "Lman1",
                    "Msmo1",
                    "Fyttd1",
                    "Vcam1",
                    "Arpc2",
                    "Mageh1",
                    "Fam25c",
                    "Mrpl54",
                    "Cct2",
                    "Prelid1",
                    "Ripk3",
                    "1110046J04Rik",
                    "Yae1d1",
                    "Phlda3",
                    "Trip6",
                    "Sept11",
                    "Spag9",
                    "Romo1",
                    "Ddx6",
                    "Tpm2",
                    "Eif1ax",
                    "Bzw1",
                    "Sdc4",
                    "Metrnl",
                    "Msl3",
                    "Mgst1",
                    "Ubc",
                    "Lman2",
                    "Prpf40a",
                    "0610011F06Rik",
                    "Cdc42ep3",
                    "Zfp560",
                    "Golim4",
                    "Mpp1",
                    "Fam220a",
                    "Gm5449",
                    "Serpina3g",
                    "Vasn",
                    "Pafah1b1",
                    "Bloc1s2",
                    "Ssbp3",
                    "Myc",
                    "Rpl27",
                    "Psap",
                    "Rbms1",
                    "P4hb",
                    "Scd2",
                    "Dlk1",
                    "Map1b",
                    "Rab34",
                    "Tvp23b",
                    "Bsg",
                    "Arl8a",
                    "Dync1i2",
                    "Qk",
                    "Rnase4",
                    "Fas",
                    "4933434E20Rik",
                    "Camk2d",
                    "Txndc17",
                    "Vgll3",
                    "mt-Cytb",
                    "Cyba",
                    "Sclt1",
                    "Stard3nl",
                    "Gdf3",
                    "Gsto1",
                    "Frmd6",
                    "Cers2",
                    "Rab28",
                    "H1f0",
                    "Ptprd",
                    "Vcan",
                    "Csnk1d",
                    "Sparc",
                    "Sept2",
                    "Abracl",
                    "Cops6",
                    "Serpinb6a",
                    "Aldoa",
                    "Pth1r",
                    "Tagln",
                    "Ctbp1",
                    "4921524J17Rik",
                    "Psmd13",
                    "Slc35b4",
                    "Anapc4",
                    "Selm",
                    "Mast4",
                    "Churc1",
                    "Zfyve21",
                    "Hnrnph2",
                    "Esd",
                    "Atxn7l3b",
                    "Csk",
                    "Igfbp6",
                    "Comt",
                    "En1",
                    "Vps36",
                    "Runx1t1",
                    "Lamp1",
                    "Lama4",
                    "Crim1",
                    "Actr1a",
                    "Lamtor5",
                    "Ndufa13",
                    "Sat1",
                    "Ssb",
                    "Gpr180",
                    "Zfp91",
                    "Atp5o",
                    "Mapkapk2",
                    "Gkap1",
                    "Sema3a",
                    "Tubb2a",
                    "Rrm2",
                    "Gm13242",
                    "Brcc3",
                    "Capns1",
                    "Itm2b",
                    "Pdia3",
                    "Ccdc102a",
                    "Chmp5",
                    "Smad7",
                    "Txndc5",
                    "Ptms",
                    "Mid1ip1",
                    "Osr1",
                    "Ginm1",
                    "Vezf1",
                    "Chmp2b",
                    "Ccar1",
                    "St5",
                    "Mospd2",
                    "Sfpq",
                    "Abcg2",
                    "Smap1",
                    "Cd24a",
                    "Ptpn11",
                    "Lipe",
                    "Tnfrsf12a",
                    "Chid1",
                    "Mdh2",
                    "Tulp4",
                    "Elovl1",
                    "Etfa",
                    "Phgdh",
                    "Efna5",
                    "Hsp90b1",
                    "Mtpn",
                    "Litaf",
                    "Anxa1",
                    "Herpud1",
                    "Prkar1a",
                    "Timp1",
                    "Bmpr2",
                    "Rap2a",
                    "Lrp1",
                    "Ddit4l",
                    "Vps4a",
                    "Tuba1a",
                    "Ndufa9",
                    "Scfd1",
                    "Commd3",
                    "Cnn2",
                    "Atp6v0d1",
                    "Ubqln2",
                    "Ppp1r18",
                    "Pcyt1a",
                    "Amotl1",
                    "Fst",
                    "Lasp1",
                    "Tia1",
                    "Zfp637",
                    "Tmem167",
                    "Gpr108",
                    "2410141K09Rik",
                    "Gpi1",
                    "Tcf12",
                    "Nadk",
                    "Dync1li2",
                    "Cct4",
                    "Igfbp7",
                    "Decr1",
                    "Rpl23a-ps3",
                    "Gnaq",
                    "Adm",
                    "Cdkn1a",
                    "2200002D01Rik",
                    "Aig1",
                    "Csde1",
                    "Chd9",
                    "AY036118",
                    "Pls3",
                    "Fstl1",
                    "Rhod",
                    "Zfp36l2",
                    "Prkar2b",
                    "Phc2",
                    "Ifitm3",
                    "Fbn1",
                    "Wnk1",
                    "Asap1",
                    "B3galnt1",
                    "Mef2a",
                    "Psmc2",
                    "Cd164",
                    "Has2",
                    "Ndufa10",
                    "Atp6ap1",
                    "Lztfl1",
                    "Eif3i",
                    "Anln",
                    "Gpc1",
                    "Gorasp2",
                    "Htatsf1",
                    "Epb41l4a",
                    "Bag1",
                    "Rpn2",
                    "Errfi1",
                    "Josd2",
                    "Tmem33",
                    "Manf",
                    "Dctn2",
                    "Arhgdib",
                    "Memo1",
                    "Rrm1",
                    "2610001J05Rik",
                    "Hsd17b14",
                    "Ehd1",
                    "Maoa",
                    "Lefty2",
                    "Cstb",
                    "Gm10260",
                    "Clptm1l",
                    "Fads3",
                    "Ccng1",
                    "Asph",
                    "Tsn",
                    "Ctxn1",
                    "Ptp4a2",
                    "Txnl1",
                    "Mesdc2",
                    "Emc3",
                    "Fbxo15",
                    "Arhgdia",
                    "Mxra7",
                    "Myl12a",
                    "Ndufb6",
                    "Gm9493",
                    "Rer1",
                    "Spats2l",
                    "Cdkn2b",
                    "Anapc16",
                    "Ddx3y",
                    "Acat1",
                    "Cald1",
                    "Pigp",
                    "Zdhhc3",
                    "Ppp2r5e",
                    "Fau",
                    "Ybx3",
                    "Sema3c",
                    "mt-Atp6",
                    "Akt1",
                    "Bnip2",
                    "Rbm7",
                    "Sri",
                    "Clic1",
                    "Tsen34",
                    "Got2",
                    "Bcam",
                    "Gnas",
                    "Aebp1",
                    "Emc7",
                    "Ept1",
                    "Calu",
                    "Cinp",
                    "Gm7325",
                    "Rbpms2",
                    "Rgmb",
                    "Smim15",
                    "Cox4i1",
                    "Rps18-ps3",
                    "Tmed9",
                    "Gpx7",
                    "Myl6",
                    "Cers5",
                    "Fads1",
                    "Sepw1",
                    "Pbrm1",
                    "Hspa8",
                    "Tmem214",
                    "Tnfaip1",
                    "Ccl7",
                    "Laptm4a",
                    "Pphln1",
                    "Chic2",
                    "Gpx4",
                    "Senp6",
                    "Ntan1",
                    "Bet1l",
                    "Scaf11",
                    "Fam92a",
                    "Grip2",
                    "Cryzl1",
                    "Pdcl3",
                    "Atg3",
                    "Bin1",
                    "Nek7",
                    "Cfl2",
                    "Ogn"
                ],
                "expression_sum": {
                    "class1": 176768.0,
                    "class2": 435092.0
                },
                "uniprot_ids": [
                    "P62960",
                    "Q3UBT1",
                    "Q71V06",
                    "Q8BRJ5",
                    "Q60950",
                    "Q60951",
                    "A2BGG7",
                    "Q810K5"
                ]
            },
            "Trp53": {
                "id": "Trp53",
                "type": "TF",
                "symbol": "Trp53",
                "source": [
                    "Tcf4",
                    "Pou5f1",
                    "Foxp1",
                    "Nfib",
                    "Junb",
                    "Fos",
                    "Jun",
                    "Sox2",
                    "Nanog",
                    "Jund",
                    "Twist2",
                    "Cebpb",
                    "Sox4"
                ],
                "target": [
                    "Rps5",
                    "Arf1",
                    "Dap",
                    "Rpl41",
                    "Sar1a",
                    "Tpm4",
                    "Dctn3",
                    "Cd44",
                    "H2afj",
                    "Fis1",
                    "Cox8a",
                    "Ddb1",
                    "Myh9",
                    "Thbs1",
                    "Rasgrp2",
                    "Txn2",
                    "Lims1",
                    "Sec13",
                    "Igfbp2",
                    "Wisp1",
                    "Slc25a4",
                    "Tspan4",
                    "Map1lc3a",
                    "Lmna",
                    "Folr1",
                    "Ndufa2",
                    "Itm2a",
                    "Ier3ip1",
                    "Pck2",
                    "Rnf187",
                    "Lgals3",
                    "Pgrmc1",
                    "Med28",
                    "Shfm1",
                    "Fos",
                    "Tagln2",
                    "Rps15a",
                    "Ddx3x",
                    "Nedd4",
                    "Rexo2",
                    "Calr",
                    "Pdlim7",
                    "Rab18",
                    "Mfge8",
                    "Mbnl2",
                    "Canx",
                    "Ppp1r14b",
                    "Actn1",
                    "B2m",
                    "Mrpl14",
                    "Slc25a3",
                    "Psat1",
                    "Lamtor1",
                    "2810417H13Rik",
                    "Neat1",
                    "Myh10",
                    "Ube2r2",
                    "Timp3",
                    "Gemin7",
                    "Lrrc59",
                    "Akr1a1",
                    "Sh3glb1",
                    "Akt1s1",
                    "Hras",
                    "Atp2b1",
                    "Mapre1",
                    "Cisd2",
                    "Mylpf",
                    "Nktr",
                    "Col5a1",
                    "Tspan3",
                    "Ctgf",
                    "Capza2",
                    "Klf6",
                    "Pcolce",
                    "Hnrnph1",
                    "Timp2",
                    "Syngr1",
                    "Crip1",
                    "Morc1",
                    "Sec62",
                    "Rps12",
                    "Luc7l2",
                    "Ndufb9",
                    "Maged1",
                    "Eef1b2",
                    "Arpc1b",
                    "Cyr61",
                    "Kdelr1",
                    "Rab14",
                    "Atp6v1g1",
                    "Pttg1",
                    "Gnb2",
                    "Ociad1",
                    "Dpysl3",
                    "Mybl2",
                    "Ddost",
                    "Hcfc1r1",
                    "Sptbn1",
                    "Id3",
                    "Ap3s1",
                    "Gng12",
                    "Fkbp8",
                    "Ncam1",
                    "Wls",
                    "Ndufa8",
                    "Pdia4",
                    "Srp14",
                    "Sept7",
                    "Fam195b",
                    "Actg1",
                    "Rcn2",
                    "Emc10",
                    "Ier5",
                    "Tsg101",
                    "Tpi1",
                    "Nars",
                    "Hmga2",
                    "Dynlrb1",
                    "Atf4",
                    "Hist1h2bc",
                    "Tmem234",
                    "Nfkbia",
                    "Nrp1",
                    "Ube2m",
                    "Atp6v0b",
                    "Fkbp9",
                    "Rpl38",
                    "Serpine2",
                    "Igfbp4",
                    "Id2",
                    "Itgb1",
                    "Fabp5",
                    "Mettl9",
                    "Tcf4",
                    "Chmp4b",
                    "Igf2",
                    "Csnk1a1",
                    "Cd81",
                    "Ssr1",
                    "Pgls",
                    "Lxn",
                    "Ift20",
                    "Anxa2",
                    "Eif3c",
                    "Vmp1",
                    "Napa",
                    "Crip2",
                    "mt-Nd4",
                    "Cox7a2l",
                    "Vim",
                    "Basp1",
                    "Sh3bgrl3",
                    "Wdr1",
                    "Ddah1",
                    "Ube2b",
                    "Eif5",
                    "Tmed2",
                    "Txn1",
                    "Gjb3",
                    "Tpm1",
                    "Ldha",
                    "Sdcbp",
                    "Thy1",
                    "Surf4",
                    "Utf1",
                    "Phlda1",
                    "Myeov2",
                    "Bola3",
                    "Vcl",
                    "Cst3",
                    "Higd1a",
                    "S100a6",
                    "Prdx6",
                    "Serpine1",
                    "H2-D1",
                    "Trappc4",
                    "Sys1",
                    "Cd63",
                    "Capg",
                    "Lbh",
                    "Ndufa3",
                    "S100a11",
                    "Tmem14c",
                    "Scand1",
                    "Mtdh",
                    "Nfib",
                    "Eef1a1",
                    "mt-Nd5",
                    "Maged2",
                    "Rtn4",
                    "Ctsl",
                    "Xist",
                    "Ccnd2",
                    "Tcf25",
                    "Spcs1",
                    "Cox6a1",
                    "Tax1bp3",
                    "Rala",
                    "Filip1l",
                    "mt-Co2",
                    "Atox1",
                    "Tmem126a",
                    "Plk2",
                    "Tln1",
                    "Oaf",
                    "Ostf1",
                    "Snhg12",
                    "Atp5d",
                    "Mt2",
                    "Eif3h",
                    "Rpl22l1",
                    "Chd3",
                    "Rps3a1",
                    "Cmpk1",
                    "Btg1",
                    "Fkbp2",
                    "Lrrc58",
                    "Fn1",
                    "Glipr2",
                    "Ndufb2",
                    "Hes1",
                    "Ap1s1",
                    "Rps9",
                    "Uqcrq",
                    "Mrpl30",
                    "Rcn3",
                    "Rwdd1",
                    "Malat1",
                    "Gnai2",
                    "Mfap2",
                    "Tm9sf3",
                    "Tmed10",
                    "Ctsd",
                    "Ndufc2",
                    "Reep5",
                    "Fhl2",
                    "Glrx5",
                    "Col1a1",
                    "Ndufa12",
                    "Oxct1",
                    "Atxn10",
                    "Atp6v0e",
                    "Cnih4",
                    "Pebp1",
                    "Dbi",
                    "Tmem59",
                    "Eif3e",
                    "Emp3",
                    "Cebpb",
                    "Cyb5r3",
                    "Pafah1b2",
                    "Anxa3",
                    "Hbegf",
                    "Ghitm",
                    "Arpc3",
                    "Scp2",
                    "Ilk",
                    "Arl1",
                    "Arpc1a",
                    "Ap2s1",
                    "Mgat2",
                    "Sep15",
                    "Mlec",
                    "Rab7",
                    "Lsm7",
                    "Cpe",
                    "Atf5",
                    "Cdc42",
                    "Npc2",
                    "Smdt1",
                    "Eef2",
                    "Hsbp1",
                    "Hspa5",
                    "Rnh1",
                    "Rrbp1",
                    "Actr3",
                    "Tmsb4x",
                    "Cd47",
                    "Efemp2",
                    "Fbln2",
                    "Mpc1",
                    "Fth1",
                    "Copb2",
                    "Khdrbs1",
                    "Ap2m1",
                    "Nupr1",
                    "Sars",
                    "Bri3",
                    "Ecm1",
                    "Smim14",
                    "Lgals1",
                    "Ahnak",
                    "Tbrg1",
                    "Fkbp10",
                    "E130012A19Rik",
                    "Pam16",
                    "Kif5b",
                    "Mpc2",
                    "Copz2",
                    "Ptp4a1",
                    "Lsp1",
                    "Ndufs5",
                    "Gsn",
                    "Psmb1",
                    "Oat",
                    "Tubb6",
                    "Cited2",
                    "Eva1b",
                    "Prdx4",
                    "mt-Nd1",
                    "Eif4g2",
                    "Aes",
                    "Rab2a",
                    "Uqcc2",
                    "Tspo",
                    "Sra1",
                    "Myl9",
                    "Vps29",
                    "Btg2",
                    "Parva",
                    "Selk",
                    "Rab11a",
                    "Anxa5",
                    "Eif3f",
                    "Eif4g1",
                    "Anxa6",
                    "Mtch1",
                    "Tmbim6",
                    "Dynll2",
                    "Rhoa",
                    "Rab10",
                    "Ttc3",
                    "Vimp",
                    "Syncrip",
                    "Sec61a1",
                    "Iqgap1",
                    "Kdelr2",
                    "Pmp22",
                    "Rbmxl2",
                    "Ssr3",
                    "March5",
                    "Rps28",
                    "Fermt2",
                    "Rpl18a",
                    "Hmgb3",
                    "Rpn1",
                    "Gas1",
                    "Ssr4",
                    "Sqstm1",
                    "Lman1",
                    "F3",
                    "Rcn1",
                    "Rpl35",
                    "Arpc2",
                    "Atp5l",
                    "Mrpl54",
                    "Ywhab",
                    "Prelid1",
                    "Sox4",
                    "Phlda3",
                    "Sept11",
                    "Ndufa1",
                    "Romo1",
                    "Tpm2",
                    "Ergic3",
                    "Bzw1",
                    "Sdc4",
                    "Ubc",
                    "Lman2",
                    "Rock2",
                    "Cdc42ep3",
                    "Tcea3",
                    "Pafah1b1",
                    "Psap",
                    "Rbms1",
                    "P4hb",
                    "Scd2",
                    "Zyx",
                    "H2afy",
                    "Map1b",
                    "Rab34",
                    "Adipor1",
                    "mt-Nd2",
                    "Bsg",
                    "Sec22b",
                    "Camk2d",
                    "Txndc17",
                    "mt-Cytb",
                    "Cyba",
                    "Flna",
                    "Efemp1",
                    "H13",
                    "Rsu1",
                    "Prss23",
                    "Gsto1",
                    "Tpt1",
                    "Ppib",
                    "Psmd8",
                    "Gng5",
                    "H1f0",
                    "Ndufb11",
                    "Vcan",
                    "Rps19",
                    "Sept2",
                    "Abracl",
                    "Msn",
                    "Serpinb6a",
                    "Rtn3",
                    "Lpp",
                    "Ctbp1",
                    "Csrp1",
                    "Selm",
                    "Ckap4",
                    "Esd",
                    "Dppa5a",
                    "Atxn7l3b",
                    "Comt",
                    "Cox6b1",
                    "Lamp1",
                    "Rcor2",
                    "Sat1",
                    "Ndufa13",
                    "Tinagl1",
                    "Gadd45g",
                    "Sh3bgrl",
                    "Jun",
                    "Capns1",
                    "Itm2b",
                    "Pdia3",
                    "Rplp1",
                    "Rpl10a",
                    "Txndc5",
                    "Pfdn5",
                    "Ndufb7",
                    "Ptms",
                    "Ftl1",
                    "Col4a1",
                    "Hist1h1c",
                    "Cd24a",
                    "Smap1",
                    "Lipe",
                    "Tnfrsf12a",
                    "Krtcap2",
                    "Gabarap",
                    "Phgdh",
                    "Serf2",
                    "Brk1",
                    "Hsp90b1",
                    "Mtpn",
                    "Anxa1",
                    "Prkar1a",
                    "Rap1b",
                    "Rpl13a",
                    "Ier3",
                    "Cuta",
                    "Apoe",
                    "Tuba1a",
                    "Commd3",
                    "Cnn2",
                    "Gsta4",
                    "Minos1",
                    "Erp29",
                    "Gpx8",
                    "Gpi1",
                    "Tmem167",
                    "Phldb2",
                    "Palld",
                    "Tram1",
                    "Dcun1d5",
                    "Cdkn1a",
                    "Csde1",
                    "Fstl1",
                    "Marcks",
                    "Fam162a",
                    "Rhoc",
                    "Mdm2",
                    "Ifitm3",
                    "Srp72",
                    "Mrpl33",
                    "Emp1",
                    "App",
                    "Fundc2",
                    "S100a10",
                    "Bag1",
                    "Tmem147",
                    "Rpn2",
                    "Rps16",
                    "Manf",
                    "Tmem256",
                    "Ccl2",
                    "Dctn2",
                    "Arhgdib",
                    "Tmed3",
                    "Serpinf1",
                    "Galk1",
                    "Hsd17b14",
                    "Tbca",
                    "Cstb",
                    "Foxp1",
                    "Ccng1",
                    "Ptp4a2",
                    "Cope",
                    "Dnajc3",
                    "Cox5b",
                    "Arhgdia",
                    "Myl12a",
                    "Ndufb6",
                    "Rac1",
                    "Rer1",
                    "S100a4",
                    "Klf2",
                    "Cald1",
                    "Fau",
                    "Sri",
                    "Pabpc1",
                    "Prdx5",
                    "Clic1",
                    "Tsen34",
                    "Fkbp1a",
                    "Got2",
                    "Dad1",
                    "Emc7",
                    "Gnas",
                    "Sept1",
                    "Arf4",
                    "Calu",
                    "Arpc5",
                    "Gm7325",
                    "Yif1b",
                    "Hdlbp",
                    "Ckb",
                    "Limd2",
                    "Cox4i1",
                    "Tmed9",
                    "Myl6",
                    "Sepw1",
                    "Rpl39",
                    "Ndufs8",
                    "Serp1",
                    "Rabac1",
                    "Capn2",
                    "Laptm4a",
                    "Cnih1",
                    "Rheb",
                    "Glrx3",
                    "Carhsp1",
                    "Alpl",
                    "Pdcl3",
                    "Kdelr3",
                    "Junb",
                    "Bcap31",
                    "Map2k2",
                    "Axl",
                    "Cfl2",
                    "Ogn"
                ],
                "expression_sum": {
                    "class1": 9058.0,
                    "class2": 64226.0
                },
                "uniprot_ids": [
                    "P02340",
                    "A0A158SIS7",
                    "Q549C9",
                    "Q8C526",
                    "I7HIK9",
                    "Q80ZA1",
                    "Q3UGQ1",
                    "A0A5Q0MU23",
                    "Q64451",
                    "Q9ER40",
                    "Q6TDG9",
                    "Q91XH8",
                    "W5VJU6",
                    "O70366"
                ]
            },
            "Pou5f1": {
                "id": "Pou5f1",
                "type": "TF",
                "symbol": "Pou5f1",
                "source": [
                    "Trim28",
                    "Cebpb",
                    "Klf4",
                    "Ctcf",
                    "Hdac1",
                    "Sox2",
                    "Nanog",
                    "Hdac2",
                    "Smc3"
                ],
                "target": [
                    "Arf1",
                    "Dap",
                    "Srsf9",
                    "Ndufb4",
                    "Syce2",
                    "Mcm7",
                    "Fis1",
                    "Grsf1",
                    "Zc3h15",
                    "Cox8a",
                    "2810004N23Rik",
                    "Sec13",
                    "Ndufa6",
                    "Polr2k",
                    "Idh3g",
                    "Rps27l",
                    "Col12a1",
                    "Lmna",
                    "Ndufa2",
                    "Hnrnpf",
                    "Lmnb1",
                    "Ddx39",
                    "Cks2",
                    "Shfm1",
                    "Med28",
                    "Hmgb2",
                    "Cox6c",
                    "Ddx3x",
                    "Slbp",
                    "Nedd4",
                    "Calr",
                    "Tubb5",
                    "Nop56",
                    "Sf3b5",
                    "Slc25a3",
                    "Lamtor1",
                    "2810417H13Rik",
                    "Nup43",
                    "Cdc5l",
                    "Sms",
                    "Rbx1",
                    "Ctcf",
                    "Neat1",
                    "Pmf1",
                    "Ola1",
                    "Sf3b6",
                    "Lrrc59",
                    "Sap18",
                    "Ube2d2a",
                    "Hras",
                    "Bnip3l",
                    "Enah",
                    "Kpnb1",
                    "Mapre1",
                    "Snrpd1",
                    "Rbm8a",
                    "Col5a1",
                    "Rpl36al",
                    "Lap3",
                    "Tgfb1i1",
                    "Supt4a",
                    "Crxos",
                    "Morc1",
                    "Rbbp7",
                    "Hnrnpl",
                    "Nedd8",
                    "Pcnp",
                    "Atpif1",
                    "Serpinh1",
                    "Atp6v1g1",
                    "Snrnp40",
                    "Gng12",
                    "Srp14",
                    "Tpm3",
                    "Banf1",
                    "Mrps36",
                    "Qdpr",
                    "Tsg101",
                    "Lig1",
                    "Tpi1",
                    "Pa2g4",
                    "Gsr",
                    "Atf4",
                    "Hist1h2bc",
                    "Hprt",
                    "Nucks1",
                    "Pcbp1",
                    "Mrpl36",
                    "Itgb1",
                    "Ndufa4",
                    "Rps12-ps3",
                    "Cops5",
                    "Eif4e",
                    "Ndufv1",
                    "H3f3a",
                    "Rfc2",
                    "Bccip",
                    "Lamtor2",
                    "Trim28",
                    "Nudt21",
                    "Vim",
                    "Ube2l3",
                    "Eef1d",
                    "Etf1",
                    "Zmat2",
                    "Samm50",
                    "Rrp15",
                    "Nhp2l1",
                    "Ran",
                    "Atp5g3",
                    "Gjb3",
                    "Atp5g2",
                    "Chd4",
                    "Tceb1",
                    "C1qbp",
                    "Rpl36a",
                    "Eif3m",
                    "Snrpb2",
                    "S100a6",
                    "Umps",
                    "Rps20",
                    "Orc6",
                    "Ybx1",
                    "Sys1",
                    "Tubb2b",
                    "Ndufa3",
                    "Tmem14c",
                    "Rps6",
                    "U2af1",
                    "Slc29a1",
                    "Atp5c1",
                    "Mrps17",
                    "Ccnd2",
                    "Col6a1",
                    "Rala",
                    "Dnajc8",
                    "Cox7c",
                    "Nme4",
                    "Eno1",
                    "mt-Nd3",
                    "Rpl22l1",
                    "Lamtor4",
                    "Smc4",
                    "Mbd3",
                    "Cct7",
                    "Dctpp1",
                    "Tuba4a",
                    "Hsp90ab1",
                    "Zfp42",
                    "Rpp25",
                    "Tyms",
                    "Ankrd11",
                    "Rps8",
                    "Magoh",
                    "Ndufc2",
                    "H3f3b",
                    "Dnajc2",
                    "Ndufa12",
                    "Cnih4",
                    "Nudt19",
                    "Dbi",
                    "Tbx3",
                    "Dstn",
                    "Mt1",
                    "Cyb5r3",
                    "Ywhae",
                    "Ppil1",
                    "Rps15",
                    "Arl1",
                    "Cenpw",
                    "Eif2s2",
                    "Mrpl45",
                    "Rpl31",
                    "Smdt1",
                    "Etfb",
                    "Uqcrc2",
                    "G3bp2",
                    "Cd47",
                    "Impdh2",
                    "Bex1",
                    "Prps1",
                    "Akap12",
                    "Mest",
                    "Hnrnpab",
                    "Ahnak",
                    "Nop58",
                    "Ndufa11",
                    "Cbx5",
                    "Fxyd6",
                    "Polr2c",
                    "Cct8",
                    "Mpc2",
                    "Dohh",
                    "Denr",
                    "Eif3g",
                    "Sra1",
                    "Aspscr1",
                    "Btg2",
                    "Parva",
                    "0610012G03Rik",
                    "Rpp21",
                    "Polr2e",
                    "Anapc13",
                    "Pdcd5",
                    "Tomm7",
                    "Snrpc",
                    "Mrpl49",
                    "Eif3f",
                    "Tmbim6",
                    "Atp5a1",
                    "Psma5",
                    "Pycr2",
                    "Sf3a3",
                    "Ccdc80",
                    "Rbmxl2",
                    "March5",
                    "Mrpl23",
                    "Ddah2",
                    "Gspt1",
                    "Eprs",
                    "Trmt112",
                    "Rpl35",
                    "Dazap1",
                    "Ilf2",
                    "Lman2",
                    "Prmt1",
                    "Tcea3",
                    "Sub1",
                    "Mrpl57",
                    "Morf4l1",
                    "Col8a1",
                    "Pafah1b1",
                    "Cdca8",
                    "L1td1",
                    "Lsm6",
                    "Sf3b1",
                    "Ndufs4",
                    "Polr1d",
                    "H2afy",
                    "Mtf2",
                    "Rpl17",
                    "Bsg",
                    "Sox2",
                    "mt-Nd2",
                    "Txndc17",
                    "Spty2d1",
                    "Nop10",
                    "Mpdu1",
                    "Gdf3",
                    "Gng5",
                    "Hspe1",
                    "Rps19",
                    "Ube2c",
                    "Msn",
                    "Exosc8",
                    "Mrpl20",
                    "Polr2h",
                    "Psmb4",
                    "Birc5",
                    "Commd1",
                    "Cox7b",
                    "Selm",
                    "Mien1",
                    "Chtop",
                    "Dppa5a",
                    "Srpk1",
                    "Lamtor5",
                    "Avpi1",
                    "Ssb",
                    "Rpa2",
                    "Mrps12",
                    "Copz1",
                    "Nfu1",
                    "Ly6a",
                    "Ndufb5",
                    "Rpl10a",
                    "Ptms",
                    "Set",
                    "Thyn1",
                    "Txnl4a",
                    "Ssbp1",
                    "Col4a1",
                    "Sephs2",
                    "Snrpd2",
                    "1810022K09Rik",
                    "Hdac1",
                    "Naa10",
                    "Brk1",
                    "Hsp90b1",
                    "Vdac2",
                    "Spp1",
                    "Mrpl24",
                    "Snhg6",
                    "Minos1",
                    "Anapc11",
                    "Spint2",
                    "Gas6",
                    "Rabggtb",
                    "Nans",
                    "Uchl5",
                    "Ppig",
                    "Ucp2",
                    "Dcun1d5",
                    "Lsm12",
                    "Gins4",
                    "Tceb2",
                    "Rhoc",
                    "Mrpl42",
                    "Fbn1",
                    "Rps2",
                    "Polr2g",
                    "Sumo3",
                    "Uqcrfs1",
                    "Naa50",
                    "Ywhah",
                    "Fbn2",
                    "Hdac2",
                    "Timm17a",
                    "Rpn2",
                    "Bub3",
                    "Esf1",
                    "Sin3b",
                    "Dctn2",
                    "Plekhj1",
                    "Mrfap1",
                    "Hsd17b14",
                    "Slc25a5",
                    "Uqcrh",
                    "Lefty2",
                    "Tbca",
                    "Foxp1",
                    "Tsn",
                    "Rpl37a",
                    "Drg1",
                    "Ptma",
                    "Ndufb6",
                    "Rac1",
                    "S100a4",
                    "Cald1",
                    "Ybx3",
                    "Rps10",
                    "Dad1",
                    "Sept1",
                    "0610007P14Rik",
                    "Yif1b",
                    "Rps18-ps3",
                    "Ccnd1",
                    "Rpl39",
                    "Tsfm",
                    "Capn2",
                    "Ptov1",
                    "Laptm4a",
                    "Rps27rt",
                    "Gpx4",
                    "Gcsh",
                    "Gltscr2",
                    "Mkrn1",
                    "Rrp1",
                    "Pdcl3",
                    "Rpl23",
                    "Rpl30",
                    "Sfr1",
                    "Polr2f",
                    "Rrs1",
                    "Rbm17",
                    "Ly6e",
                    "Map2k2",
                    "Lsm5",
                    "Rnf7",
                    "Cdc20",
                    "Cd9",
                    "Ddx24",
                    "Rpl6",
                    "Srm",
                    "Thbs1",
                    "Psmb7",
                    "Spr",
                    "Pnkd",
                    "Ppid",
                    "1110008F13Rik",
                    "Anapc5",
                    "Hnrnpa1",
                    "Ssrp1",
                    "Ier3ip1",
                    "Rnf187",
                    "Psmb6",
                    "Rps15a",
                    "Ndufs3",
                    "Uba2",
                    "Mbnl2",
                    "Ppp1r14b",
                    "Snrpe",
                    "B2m",
                    "Psat1",
                    "Gdi2",
                    "Taf7",
                    "Mrpl28",
                    "Psmc6",
                    "Grpel1",
                    "Sec61g",
                    "Ythdc1",
                    "Snrnp27",
                    "Srsf1",
                    "Uqcrc1",
                    "Tdgf1",
                    "Rcc2",
                    "Impa2",
                    "Bola2",
                    "Tmem160",
                    "C1d",
                    "Cisd2",
                    "Rpl10",
                    "Ncl",
                    "Rpl35a",
                    "Timp2",
                    "Syngr1",
                    "Hspa4",
                    "Rps27",
                    "Arpp19",
                    "Ccdc12",
                    "Ddx21",
                    "Naca",
                    "Ube2i",
                    "Rpl24",
                    "Brix1",
                    "Fam136a",
                    "Siva1",
                    "Rps4x",
                    "Fkbp8",
                    "Nip7",
                    "Ruvbl1",
                    "Wls",
                    "Clpp",
                    "G3bp1",
                    "Atp5b",
                    "Actg1",
                    "Cdc26",
                    "Immt",
                    "Nars",
                    "Pdlim1",
                    "Hells",
                    "Tfcp2l1",
                    "Rpl38",
                    "Glo1",
                    "Serpine2",
                    "Nme1",
                    "Csnk1a1",
                    "Rpl10-ps3",
                    "Rpl3",
                    "Cebpzos",
                    "Rpl12",
                    "Snw1",
                    "Ndufa7",
                    "Dnajc19",
                    "Cdv3",
                    "Hmgn1",
                    "Ctbp2",
                    "Btf3l4",
                    "Platr3",
                    "Ppp4c",
                    "Imp3",
                    "Mrpl13",
                    "Ak2",
                    "Wdr77",
                    "Atp5j",
                    "Srrm1",
                    "Thy1",
                    "Srp19",
                    "Emd",
                    "Vasp",
                    "Pdgfa",
                    "Uqcr10",
                    "Rpsa",
                    "H2afz",
                    "Bola3",
                    "Mrps21",
                    "Tardbp",
                    "Rpl5",
                    "Dph3",
                    "Dpy30",
                    "Serpine1",
                    "Fblim1",
                    "Odc1",
                    "Mrpl18",
                    "Ufc1",
                    "Ifitm1",
                    "Acat2",
                    "Mrpl12",
                    "Cdc37",
                    "Cisd1",
                    "Rpl28",
                    "Xist",
                    "Gpx3",
                    "Cox6a1",
                    "Timm8b",
                    "Rpl26",
                    "Uqcrb",
                    "mt-Co2",
                    "Rps3",
                    "Atox1",
                    "Srsf10",
                    "Oaf",
                    "Ube2v2",
                    "Uba52",
                    "Rpl19",
                    "Timm13",
                    "Eif3h",
                    "Mcm3",
                    "Adprh",
                    "Rps3a1",
                    "Srsf3",
                    "Coprs",
                    "Rpl7",
                    "Znrd1",
                    "Wdr18",
                    "Polr2m",
                    "1810058I24Rik",
                    "Gnai2",
                    "Rpl8",
                    "Eif4a1",
                    "Pcbp2",
                    "Polr2i",
                    "Polr1c",
                    "Snrnp25",
                    "Fhl2",
                    "Glrx5",
                    "Ccdc34",
                    "Col1a1",
                    "Mrps33",
                    "Rif1",
                    "Morf4l2",
                    "Rbbp4",
                    "Eif3e",
                    "Gm10073",
                    "Emp3",
                    "2700060E02Rik",
                    "Anxa3",
                    "Arpc3",
                    "Park7",
                    "Dynll1",
                    "Hspd1",
                    "Stoml2",
                    "Cops7a",
                    "Sod1",
                    "Ppih",
                    "Uqcc3",
                    "Eef2",
                    "Tpx2",
                    "Dnajb6",
                    "Efemp2",
                    "Gapdh",
                    "Smarca5",
                    "Idi1",
                    "Ccnb1",
                    "Ccnd3",
                    "Ap2m1",
                    "Khdrbs1",
                    "Mrpl52",
                    "Pno1",
                    "Bri3",
                    "Ecm1",
                    "Adrm1",
                    "Prkcdbp",
                    "Kif5b",
                    "Hmgn2",
                    "Hnrnpa2b1",
                    "Oat",
                    "Psmd2",
                    "Ddx27",
                    "Cbx3",
                    "Jam2",
                    "Jade1",
                    "Dnajc7",
                    "Psmb2",
                    "Fxr1",
                    "Tceal8",
                    "Myl9",
                    "Cnn3",
                    "Vps29",
                    "Vdac1",
                    "Anp32a",
                    "Tra2a",
                    "Zfp57",
                    "Dynll2",
                    "Ssu72",
                    "Cnpy2",
                    "Anp32e",
                    "Pfdn6",
                    "Pmp22",
                    "Ict1",
                    "Rps25",
                    "Rbmxl1",
                    "Arglu1",
                    "Imp4",
                    "Rpl18a",
                    "Txndc9",
                    "Mrpl54",
                    "Ywhab",
                    "Prelid1",
                    "Ubb",
                    "Tpm2",
                    "Prrc1",
                    "Zfp106",
                    "Paics",
                    "Asna1",
                    "Psmc5",
                    "Echdc2",
                    "Mrpl43",
                    "Rpl27",
                    "Suclg1",
                    "Hnrnpd",
                    "Ash2l",
                    "Hat1",
                    "Rnps1",
                    "Prss23",
                    "Tpt1",
                    "Ppib",
                    "Stmn2",
                    "Ptbp1",
                    "Aldoa",
                    "Lpp",
                    "Tagln",
                    "Dnajc21",
                    "Snx3",
                    "Strap",
                    "Psmd13",
                    "Rnaseh2a",
                    "Klf4",
                    "Rfc1",
                    "Cox6b1",
                    "Ptrf",
                    "Atp5o",
                    "Nasp",
                    "Ndufs6",
                    "Poldip2",
                    "Eif4e2",
                    "Sod2",
                    "Ftl1",
                    "Hist1h1c",
                    "Tnfrsf12a",
                    "Phgdh",
                    "Gtf2e2",
                    "Pdap1",
                    "Rpl11",
                    "Anxa1",
                    "Ube2a",
                    "Pfn1",
                    "Snrpg",
                    "Ddx39b",
                    "Grb10",
                    "Exosc4",
                    "Fkbp4",
                    "Pgam1",
                    "Pnrc1",
                    "Snrnp70",
                    "Ppdpf",
                    "Gmnn",
                    "Rpl27a",
                    "Llph",
                    "Igfbp7",
                    "Eif4b",
                    "Anp32b",
                    "Edf1",
                    "Fstl1",
                    "Mdm2",
                    "Eif2b5",
                    "Gart",
                    "Mrpl33",
                    "Ndufa10",
                    "App",
                    "Tomm40",
                    "Eif3i",
                    "Smc2",
                    "Mtch2",
                    "Psmd4",
                    "Gins2",
                    "Uck2",
                    "Rps16",
                    "Bcat1",
                    "Tmed3",
                    "Serpinf1",
                    "Mrps16",
                    "Gap43",
                    "Psmd12",
                    "Ndufb10",
                    "Cth",
                    "Txnl1",
                    "Fbxo15",
                    "Eef1g",
                    "Rbm25",
                    "Ube2e3",
                    "2700094K13Rik",
                    "Acadm",
                    "Dut",
                    "Ooep",
                    "Alyref",
                    "Hspa14",
                    "Polr2j",
                    "Got2",
                    "Slirp",
                    "Gnas",
                    "Arpc5",
                    "Ckb",
                    "Cwc15",
                    "Ndufs8",
                    "Pim1",
                    "Rabac1",
                    "Rplp2",
                    "Mif",
                    "Aimp1",
                    "Rheb",
                    "Chchd3",
                    "Eif4a2",
                    "Glrx3",
                    "Gtf2a2",
                    "Alpl",
                    "Kdelr3",
                    "Cnbp",
                    "Sdhd",
                    "Axl",
                    "Rps5",
                    "Rpl41",
                    "Dctn3",
                    "Cd44",
                    "Ruvbl2",
                    "Cdt1",
                    "Ier2",
                    "Dkc1",
                    "Igfbp2",
                    "Marcksl1",
                    "Atp5g1",
                    "Tuba1b",
                    "Aurka",
                    "Psma6",
                    "Rpl7l1",
                    "Snrpa",
                    "St13",
                    "Thoc7",
                    "Mlf2",
                    "Rpl9",
                    "Vbp1",
                    "Atp5j2",
                    "Nubp1",
                    "E2f4",
                    "D8Ertd738e",
                    "Hn1",
                    "Mrpl14",
                    "Phb",
                    "Nr0b1",
                    "Tra2b",
                    "Hmgcs1",
                    "Emc6",
                    "Rps24",
                    "Acta2",
                    "Nanog",
                    "Bola1",
                    "Aurkaip1",
                    "Timp3",
                    "Nhp2",
                    "Akr1a1",
                    "Exosc5",
                    "Akt1s1",
                    "Atp5h",
                    "Mylpf",
                    "Ppia",
                    "Eif3k",
                    "Mrps34",
                    "Atp5k",
                    "Mphosph8",
                    "Ppp1ca",
                    "Aamp",
                    "Chchd1",
                    "Gars",
                    "Psmc3",
                    "Rps12",
                    "Luc7l2",
                    "Ndufb9",
                    "Col3a1",
                    "Utp3",
                    "Fus",
                    "Ddost",
                    "Rest",
                    "Cct3",
                    "Psma7",
                    "Hacd1",
                    "Gnl3",
                    "Trim27",
                    "Psma1",
                    "Dtymk",
                    "Emc10",
                    "2410015M20Rik",
                    "Hdgf",
                    "Prnp",
                    "Id1",
                    "Csnk2b",
                    "Rpl37",
                    "Ppp1cc",
                    "Tmem234",
                    "Nrp1",
                    "Dnttip2",
                    "Sae1",
                    "Hnrnpk",
                    "Mrps24",
                    "Ezr",
                    "Ewsr1",
                    "Ndufv2",
                    "Mettl9",
                    "Fbxo22",
                    "Fam49b",
                    "Bax",
                    "Cisd3",
                    "Tmsb10",
                    "Gfer",
                    "Anxa2",
                    "Pycard",
                    "Eif3c",
                    "Cox7a2",
                    "Col5a2",
                    "Napa",
                    "Atp6v1f",
                    "Lsm3",
                    "Ube2j2",
                    "Gcat",
                    "Eif5",
                    "Spag7",
                    "Tomm5",
                    "Txn1",
                    "Vapa",
                    "Erh",
                    "Fscn1",
                    "Emc8",
                    "Utf1",
                    "Bclaf1",
                    "Psip1",
                    "Rpl29",
                    "Naa38",
                    "Ung",
                    "Ptges3",
                    "H2-D1",
                    "Wtap",
                    "Dnaja2",
                    "Lbh",
                    "Eef1a1",
                    "Cenpv",
                    "Ywhaq",
                    "Pdha1",
                    "Ubald2",
                    "Filip1l",
                    "Cav1",
                    "Adss",
                    "Rps7",
                    "Igf2bp1",
                    "Trnau1ap",
                    "Eef1e1",
                    "Fbln5",
                    "Chd3",
                    "Hsd17b10",
                    "Gadd45gip1",
                    "Nenf",
                    "Oaz1",
                    "Stra13",
                    "Uqcrq",
                    "Sec61b",
                    "Snx5",
                    "Got1",
                    "Trap1a",
                    "Bzw2",
                    "Pgp",
                    "Sumo2",
                    "Nudc",
                    "Gnb2l1",
                    "Mrpl21",
                    "Pold2",
                    "Nsa2",
                    "Ddx46",
                    "Atxn10",
                    "Rfc5",
                    "Pfdn4",
                    "Cebpb",
                    "Mrps15",
                    "Psmc4",
                    "Psmd14",
                    "Npm1",
                    "Sap30",
                    "Gtpbp4",
                    "Lsm7",
                    "Cpe",
                    "Atf5",
                    "Srsf7",
                    "Mat2a",
                    "Csrp2",
                    "Brd2",
                    "Ccdc124",
                    "Prdx2",
                    "Ngfrap1",
                    "Abcf1",
                    "Lgals1",
                    "Rplp0",
                    "Psmb5",
                    "Ube2n",
                    "Smn1",
                    "Cyb5b",
                    "Copz2",
                    "Psmb1",
                    "Coa3",
                    "Akr1b3",
                    "Eif4g2",
                    "Raly",
                    "Uqcc2",
                    "Nsmce4a",
                    "Rps26",
                    "Use1",
                    "Get4",
                    "Lage3",
                    "Npm3",
                    "Lsm2",
                    "Chmp2a",
                    "Sumo1",
                    "Nudcd2",
                    "Aprt",
                    "Anxa5",
                    "Fcf1",
                    "Mrps18c",
                    "Cacybp",
                    "Rpl22",
                    "Nolc1",
                    "Rpl4",
                    "Glud1",
                    "Rps28",
                    "Hnrnpu",
                    "Srsf11",
                    "Hmgb3",
                    "Rpl21",
                    "Ppp1r2",
                    "Map1lc3b",
                    "Atp5l",
                    "Son",
                    "Pop5",
                    "Cct2",
                    "Acp1",
                    "Dtl",
                    "Ndufa1",
                    "Ahsa1",
                    "Eif1ax",
                    "Parp1",
                    "Dhfr",
                    "Myl4",
                    "Mrps7",
                    "Cdca3",
                    "Metap2",
                    "Ube2s",
                    "Idh2",
                    "Snrpd3",
                    "Zyx",
                    "Psmd3",
                    "Fam32a",
                    "Trh",
                    "Bcas2",
                    "Cops4",
                    "Ciapin1",
                    "Pfdn2",
                    "Ywhag",
                    "Atp5f1",
                    "Arid5b",
                    "Vcan",
                    "Ndufb11",
                    "Srp9",
                    "Nfatc2ip",
                    "Ino80e",
                    "Sfrp1",
                    "Ppa1",
                    "Rcor2",
                    "Cenpm",
                    "Prdx1",
                    "Rsl24d1",
                    "Terf1",
                    "Rpl13",
                    "Mrpl34",
                    "Atrx",
                    "Tgif1",
                    "Sfpq",
                    "Mrto4",
                    "Parl",
                    "Clns1a",
                    "Mdh2",
                    "Gabarap",
                    "Uqcr11",
                    "Phb2",
                    "Rps27a",
                    "Rpl15",
                    "Gtf2h5",
                    "Arl6ip1",
                    "Tcof1",
                    "Rap1b",
                    "Ier3",
                    "Cuta",
                    "Eif1a",
                    "Cox5a",
                    "Tmem11",
                    "Ubap2l",
                    "Phldb2",
                    "Cct4",
                    "Rdx",
                    "Pkm",
                    "H2afv",
                    "Prrc2c",
                    "Marcks",
                    "Psmg4",
                    "Psmc2",
                    "Sf3b4",
                    "Hnrnpc",
                    "Fkbp3",
                    "Tmem147",
                    "Rangrf",
                    "Rpl7a",
                    "Fam104a",
                    "Prrx2",
                    "Manf",
                    "Tpr",
                    "Ddx18",
                    "Rrp9",
                    "Hint1",
                    "Sf3a2",
                    "Ccne1",
                    "Srsf6",
                    "Ebna1bp2",
                    "Tmpo",
                    "Tnnt1",
                    "Cycs",
                    "Myl12a",
                    "Klf2",
                    "Dhx16",
                    "Sdhb",
                    "Taf10",
                    "Wdr43",
                    "Klf5",
                    "Yrdc",
                    "Pabpc1",
                    "Eif6",
                    "Tsen34",
                    "Fkbp1a",
                    "Gm7325",
                    "Fam60a",
                    "Wbp11",
                    "Snf8",
                    "Myl6",
                    "2810428I15Rik",
                    "Hspa8",
                    "Postn",
                    "Ldhb",
                    "Serbp1",
                    "Taf1d",
                    "Kdm1a",
                    "Ncor1",
                    "1810037I17Rik",
                    "Rsrc2",
                    "Wbp5",
                    "Hsp90aa1",
                    "Cox14",
                    "Mdh1",
                    "Usmg5",
                    "Ppp2ca",
                    "Rps17",
                    "Rae1",
                    "Pin4",
                    "Mrps18b",
                    "Rasgrp2",
                    "Txn2",
                    "Clta",
                    "Cox17",
                    "Ppp1r11",
                    "Mrpl4",
                    "Wisp1",
                    "Ndufb3",
                    "Gar1",
                    "Cks1b",
                    "Calcoco2",
                    "Rpl23a",
                    "Gm10076",
                    "Ftsj3",
                    "Folr1",
                    "Psmc1",
                    "Tpd52l2",
                    "Knop1",
                    "Psmd6",
                    "Cthrc1",
                    "Pin1",
                    "Sfxn1",
                    "Swi5",
                    "Eif2s1",
                    "Nup62",
                    "Cenpe",
                    "Rab18",
                    "Calm3",
                    "Arf6",
                    "Ubxn1",
                    "Cdk1",
                    "Eif4h",
                    "Psmd7",
                    "Anapc15",
                    "Gabarapl2",
                    "Stub1",
                    "Prrx1",
                    "Mrpl2",
                    "Ranbp1",
                    "Zwint",
                    "Ppm1g",
                    "Eif1",
                    "Tbcb",
                    "Wdr89",
                    "Pdcd6",
                    "H2afx",
                    "Bgn",
                    "Hnrnpm",
                    "Capza2",
                    "Loxl1",
                    "2310036O22Rik",
                    "Fdft1",
                    "Srsf5",
                    "Hnrnph1",
                    "Rpl39l",
                    "Wdr74",
                    "Ywhaz",
                    "Pomp",
                    "Rpl36",
                    "Rps21",
                    "Psmb3",
                    "Ddx5",
                    "Rbm39",
                    "Pigyl",
                    "Eef1b2",
                    "Eif3l",
                    "Tmem258",
                    "Tcp1",
                    "Sarnp",
                    "Gnb2",
                    "Mrpl15",
                    "Ezh2",
                    "Mybl2",
                    "Pfdn1",
                    "Slc2a3",
                    "Pmm1",
                    "Actl6a",
                    "Sssca1",
                    "Ier5",
                    "Gpx1",
                    "Smc3",
                    "Thrap3",
                    "Dynlrb1",
                    "Hspb1",
                    "Kdm5b",
                    "Col1a2",
                    "Eif3b",
                    "Ube2m",
                    "Zfp207",
                    "Chmp4b",
                    "Ube2k",
                    "Pfkl",
                    "Ndufs2",
                    "Ddt",
                    "Cox7a2l",
                    "Spc24",
                    "Asns",
                    "Ifitm2",
                    "Fdps",
                    "Lsm8",
                    "Ndufs7",
                    "Cenpf",
                    "Mcm2",
                    "Gm16286",
                    "Mrpl17",
                    "Ldha",
                    "Apex1",
                    "Cct5",
                    "Ccnb2",
                    "Mrpl35",
                    "1110004F10Rik",
                    "Mrpl40",
                    "Lox",
                    "Iscu",
                    "Fubp1",
                    "Arpc5l",
                    "Klf9",
                    "Rps29",
                    "Ccdc58",
                    "Cenpa",
                    "Prdx6",
                    "Trappc4",
                    "Mdk",
                    "Psmd1",
                    "Ndufab1",
                    "Plekhf2",
                    "Snrpf",
                    "Mcl1",
                    "Rtn4",
                    "Pole4",
                    "Fbl",
                    "Ndufv3",
                    "Rps26-ps1",
                    "Tsc22d1",
                    "Calm2",
                    "Stmn1",
                    "Btf3",
                    "Smarcc1",
                    "Tmem126a",
                    "Cdh11",
                    "Rpl18",
                    "Hax1",
                    "Atp5d",
                    "Mt2",
                    "Taldo1",
                    "Vcp",
                    "Mrps5",
                    "Prc1",
                    "U2surp",
                    "Ndufb2",
                    "Ap1s1",
                    "Ppan",
                    "Rps9",
                    "Upp1",
                    "Mrpl30",
                    "BC031181",
                    "Rwdd1",
                    "Cyc1",
                    "Fnta",
                    "Dnaja1",
                    "Tmem176b",
                    "Mfap2",
                    "Pmvk",
                    "Elof1",
                    "Lrpap1",
                    "Wdr5",
                    "Pebp1",
                    "Sgk1",
                    "Ube2d3",
                    "Aars",
                    "Marc2",
                    "Snrpa1",
                    "Paip2",
                    "Srrm2",
                    "Rnaseh2b",
                    "Top1",
                    "Gm26917",
                    "Ubl5",
                    "Hmgb1",
                    "Ndufc1",
                    "Eif3d",
                    "Cyb5a",
                    "Ghitm",
                    "Psma4",
                    "Scp2",
                    "Arpc1a",
                    "Ap2s1",
                    "Rpl34",
                    "Mlec",
                    "Rab7",
                    "Cdc42",
                    "Mrps14",
                    "Nap1l1",
                    "Mrps25",
                    "Hsbp1",
                    "Actr3",
                    "Tmsb4x",
                    "Fbln2",
                    "Mrps10",
                    "Tbrg1",
                    "Hspa9",
                    "Fkbp10",
                    "Chchd2",
                    "Pim3",
                    "E130012A19Rik",
                    "Pam16",
                    "Luc7l3",
                    "Ndufs5",
                    "Gsn",
                    "Ndufb8",
                    "Tmem97",
                    "Srsf2",
                    "Prdx4",
                    "mt-Nd1",
                    "Pak1ip1",
                    "Higd2a",
                    "Nxt1",
                    "Trp53",
                    "Lhfp",
                    "Rbm3",
                    "Psmd11",
                    "Cggbp1",
                    "Zfp706",
                    "Rad23b",
                    "Nudt4",
                    "Supt16",
                    "Ost4",
                    "Mrpl51",
                    "Eif4g1",
                    "Rhoa",
                    "Cdk4",
                    "Rsl1d1",
                    "Syncrip",
                    "Ssr2",
                    "Caprin1",
                    "Cebpz",
                    "Tomm20",
                    "Cbx1",
                    "Tomm70a",
                    "Gas1",
                    "Msmo1",
                    "Fam25c",
                    "Mak16",
                    "Bzw1",
                    "Ak6",
                    "Rbpms",
                    "Hnrnpa3",
                    "Prpf40a",
                    "Pbdc1",
                    "Rps19bp1",
                    "2010107E04Rik",
                    "Tecr",
                    "Mybbp1a",
                    "Sf3b2",
                    "Atp1b1",
                    "Slc3a2",
                    "Stip1",
                    "Chchd4",
                    "Rpl14",
                    "Tma7",
                    "Camk2d",
                    "Lsm4",
                    "Tpd52",
                    "Eif1ad",
                    "Rps13",
                    "Eif5a",
                    "Rps11",
                    "Psmd8",
                    "Cfl1",
                    "Rrp1b",
                    "Sparc",
                    "Uchl3",
                    "Cops6",
                    "Hnrnpdl",
                    "Eif3a",
                    "Rtn3",
                    "Adh5",
                    "l7Rn6",
                    "Ndufa13",
                    "Atp5e",
                    "Cdc34",
                    "Gps1",
                    "Tipin",
                    "Cmtm7",
                    "Rrm2",
                    "Sh3bgrl",
                    "Rplp1",
                    "Capns1",
                    "Rnf10",
                    "Psma2",
                    "Pfdn5",
                    "Nsun2",
                    "Ndufb7",
                    "Shmt1",
                    "Alkbh1",
                    "Slc25a39",
                    "Rps14",
                    "Rpl32",
                    "Chchd10",
                    "Serf2",
                    "Bud31",
                    "Mcm5",
                    "Rpl13a",
                    "Nol7",
                    "Tkt",
                    "Apoe",
                    "Gsta4",
                    "Snrpb",
                    "Zfp428",
                    "Tomm22",
                    "Lyar",
                    "Erp29",
                    "Gpx8",
                    "Gpi1",
                    "Tmem167",
                    "Cfdp1",
                    "Eif5b",
                    "Emg1",
                    "Eif4ebp1",
                    "Nono",
                    "Csde1",
                    "Psenen",
                    "Emp1",
                    "Mcm6",
                    "Prpf19",
                    "Sugt1",
                    "Eif4a3",
                    "Dek",
                    "Bag1",
                    "Top2a",
                    "Papola",
                    "Tmem256",
                    "Cetn3",
                    "Galk1",
                    "Ccna2",
                    "Tubb4b",
                    "Eed",
                    "Ndufa5",
                    "Mrpl11",
                    "Mesdc2",
                    "Cox5b",
                    "Arhgdia",
                    "Rer1",
                    "Ralbp1",
                    "Fau",
                    "Acin1",
                    "Aebp1",
                    "Rbpms2",
                    "Uhrf1",
                    "Cox4i1",
                    "Guk1",
                    "Psma3",
                    "Skp1a",
                    "Rps18",
                    "Phf5a",
                    "Rps23",
                    "U2af2",
                    "P3h4",
                    "Myl12b"
                ],
                "expression_sum": {
                    "class1": 25.0,
                    "class2": 274305.0
                },
                "uniprot_ids": [
                    "P20263",
                    "E2JL30",
                    "G3UZG9",
                    "E2JL35",
                    "A0A2Z6DUK5",
                    "A0A2I6EDJ6",
                    "A0A2I6EDI9",
                    "E2JL33"
                ]
            },
            "Hdac1": {
                "id": "Hdac1",
                "type": "TF",
                "symbol": "Hdac1",
                "source": [
                    "Klf2",
                    "Tcf4",
                    "Pou5f1",
                    "Junb",
                    "Fos",
                    "Jun",
                    "Sox2",
                    "Nanog",
                    "Jund",
                    "Cebpb"
                ],
                "target": [
                    "Dap",
                    "Sar1a",
                    "Tpm4",
                    "Cd44",
                    "Fis1",
                    "Ddb1",
                    "Thbs1",
                    "Lims1",
                    "Sec13",
                    "Igfbp2",
                    "Wisp1",
                    "Slc25a4",
                    "Tspan4",
                    "Map1lc3a",
                    "Lmna",
                    "Lgals3",
                    "mt-Co1",
                    "Pgrmc1",
                    "Fos",
                    "Tagln2",
                    "Rexo2",
                    "Crlf1",
                    "Pdlim7",
                    "Mfge8",
                    "Mbnl2",
                    "Canx",
                    "Actn1",
                    "Neat1",
                    "Myh10",
                    "Ube2r2",
                    "Gemin7",
                    "Akr1a1",
                    "Sh3glb1",
                    "Akt1s1",
                    "Atp2b1",
                    "Cisd2",
                    "Nktr",
                    "Col5a1",
                    "Klf6",
                    "Pcolce",
                    "Crip1",
                    "Morc1",
                    "Rps12",
                    "Arpc1b",
                    "Rab14",
                    "Serpinh1",
                    "Gnb2",
                    "Ociad1",
                    "Dpysl3",
                    "Mybl2",
                    "Hcfc1r1",
                    "Sptbn1",
                    "Id3",
                    "Ap3s1",
                    "Gng12",
                    "Ncam1",
                    "Hacd1",
                    "Ndufa8",
                    "Pdia4",
                    "Srp14",
                    "Sept7",
                    "Fam195b",
                    "Actg1",
                    "Rcn2",
                    "Tsg101",
                    "Ier5",
                    "Prnp",
                    "Id1",
                    "Hmga2",
                    "Dynlrb1",
                    "Hist1h2bc",
                    "Nfkbia",
                    "Nrp1",
                    "Igfbp4",
                    "Id2",
                    "Itgb1",
                    "Fabp5",
                    "Mettl9",
                    "Chmp4b",
                    "Tcf4",
                    "Igf2",
                    "Cd81",
                    "Pgls",
                    "Lxn",
                    "Ift20",
                    "Anxa2",
                    "Vmp1",
                    "mt-Nd4",
                    "Crip2",
                    "Cox7a2l",
                    "Vim",
                    "Basp1",
                    "Sh3bgrl3",
                    "Wdr1",
                    "Tnc",
                    "Ube2b",
                    "Tmed2",
                    "Tpm1",
                    "Sdcbp",
                    "Surf4",
                    "Phlda1",
                    "Myeov2",
                    "Bola3",
                    "Cst3",
                    "S100a6",
                    "H2-D1",
                    "Ifitm1",
                    "Cd63",
                    "Capg",
                    "Lbh",
                    "Ndufa3",
                    "Scand1",
                    "Mtdh",
                    "Nfib",
                    "Maged2",
                    "Rtn4",
                    "Ctsl",
                    "Ccnd2",
                    "Tcf25",
                    "Spcs1",
                    "Cox6a1",
                    "Tax1bp3",
                    "Rala",
                    "Filip1l",
                    "Cav1",
                    "Tln1",
                    "Oaf",
                    "Ostf1",
                    "Snhg12",
                    "Mt2",
                    "Cmpk1",
                    "Fkbp2",
                    "Lrrc58",
                    "Fn1",
                    "Glipr2",
                    "Hes1",
                    "Ap1s1",
                    "Tuba4a",
                    "Twist2",
                    "Malat1",
                    "Gnai2",
                    "Tmem176b",
                    "Tmed10",
                    "Tm9sf3",
                    "Ctsd",
                    "Reep5",
                    "Pou5f1",
                    "Glrx5",
                    "Col1a1",
                    "Dbi",
                    "Tmem59",
                    "Cebpb",
                    "Pafah1b2",
                    "Anxa3",
                    "Hbegf",
                    "Scp2",
                    "Ilk",
                    "Mgat2",
                    "Sep15",
                    "Mlec",
                    "Lsm7",
                    "Atf5",
                    "Arf5",
                    "Cdc42",
                    "Npc2",
                    "Eef2",
                    "Hsbp1",
                    "Rnh1",
                    "Actr3",
                    "Cd47",
                    "Efemp2",
                    "Mpc1",
                    "Copb2",
                    "Ap2m1",
                    "Nupr1",
                    "Smim14",
                    "Ahnak",
                    "Fkbp10",
                    "Mpc2",
                    "Ptp4a1",
                    "Lsp1",
                    "Gsn",
                    "Oat",
                    "Tubb6",
                    "Cited2",
                    "Eva1b",
                    "mt-Nd1",
                    "Aes",
                    "Tspo",
                    "Sra1",
                    "Lhfp",
                    "Btg2",
                    "Parva",
                    "Selk",
                    "Rab11a",
                    "Anxa5",
                    "Anxa6",
                    "Mtch1",
                    "Rab10",
                    "Ttc3",
                    "Vimp",
                    "Sec61a1",
                    "Iqgap1",
                    "Kdelr2",
                    "Pmp22",
                    "March5",
                    "Fermt2",
                    "Rpl18a",
                    "Rpn1",
                    "Ssr4",
                    "F3",
                    "Rcn1",
                    "Mrpl54",
                    "Sox4",
                    "Phlda3",
                    "Sept11",
                    "Romo1",
                    "Tpm2",
                    "Ergic3",
                    "Ubc",
                    "Rock2",
                    "Psap",
                    "Tm4sf1",
                    "Rbms1",
                    "P4hb",
                    "Scd2",
                    "Dlk1",
                    "Zyx",
                    "Map1b",
                    "Rab34",
                    "Adipor1",
                    "Bsg",
                    "Sox2",
                    "Ppic",
                    "Sec22b",
                    "Camk2d",
                    "Flna",
                    "H13",
                    "Rsu1",
                    "Prss23",
                    "Gsto1",
                    "Ppib",
                    "Vcan",
                    "Rps19",
                    "Sept2",
                    "Abracl",
                    "Msn",
                    "Serpinb6a",
                    "Lpp",
                    "Ctbp1",
                    "Csrp1",
                    "Selm",
                    "Ckap4",
                    "Esd",
                    "Atxn7l3b",
                    "Comt",
                    "Lamp1",
                    "Ndufa13",
                    "Ptrf",
                    "Gadd45g",
                    "Sh3bgrl",
                    "Jun",
                    "Pdia3",
                    "Rpl10a",
                    "Txndc5",
                    "Ptms",
                    "Col4a1",
                    "Hist1h1c",
                    "Cd24a",
                    "Lipe",
                    "Tnfrsf12a",
                    "Krtcap2",
                    "Gabarap",
                    "Serf2",
                    "Mtpn",
                    "Anxa1",
                    "Prkar1a",
                    "Rap1b",
                    "Ier3",
                    "Grb10",
                    "Apoe",
                    "Commd3",
                    "Fst",
                    "Tmem167",
                    "Palld",
                    "Cdkn1a",
                    "Csde1",
                    "Marcks",
                    "Fam162a",
                    "Rhoc",
                    "Ifitm3",
                    "Srp72",
                    "Mrpl33",
                    "Emp1",
                    "App",
                    "Fundc2",
                    "Fbn2",
                    "Bag1",
                    "Rpn2",
                    "Prrx2",
                    "Arhgdib",
                    "Tmed3",
                    "Serpinf1",
                    "Hsd17b14",
                    "Tbca",
                    "Foxp1",
                    "Ccng1",
                    "Ptp4a2",
                    "Dnajc3",
                    "Arhgdia",
                    "Myl12a",
                    "Rac1",
                    "Rer1",
                    "S100a4",
                    "Cald1",
                    "Sri",
                    "Prdx5",
                    "Clic1",
                    "Fkbp1a",
                    "Gnas",
                    "Arf4",
                    "Calu",
                    "Arpc5",
                    "Gm7325",
                    "Hdlbp",
                    "Ckb",
                    "Limd2",
                    "Cox4i1",
                    "Tmed9",
                    "Myl6",
                    "Sepw1",
                    "Serp1",
                    "Capn2",
                    "Laptm4a",
                    "Carhsp1",
                    "Alpl",
                    "Kdelr3",
                    "Junb",
                    "Bcap31",
                    "Cfl2"
                ],
                "expression_sum": {
                    "class1": 3445.0,
                    "class2": 29162.0
                },
                "uniprot_ids": [
                    "O09106",
                    "Q58E49"
                ]
            },
            "Hmgb1": {
                "id": "Hmgb1",
                "type": "TF",
                "symbol": "Hmgb1",
                "source": [
                    "Klf6",
                    "Tcf4",
                    "Pou5f1",
                    "Foxp1",
                    "Nfib",
                    "Atf4",
                    "Junb",
                    "Fos",
                    "Jun",
                    "Sox2",
                    "Hes1",
                    "Nanog",
                    "Jund",
                    "Twist2",
                    "Cebpb",
                    "Zfp57",
                    "Sox4"
                ],
                "target": [
                    "Atxn7l3b",
                    "Skil",
                    "Tspan6",
                    "Amotl2",
                    "Atf5",
                    "Cuedc1",
                    "Tns1",
                    "Ak1",
                    "Svep1",
                    "Fam204a",
                    "Tmed5",
                    "Fabp5",
                    "Mettl9",
                    "Nfatc4",
                    "Hspa5",
                    "Igf2",
                    "Rasgrp2",
                    "Phtf1",
                    "Efemp2",
                    "Col12a1",
                    "Bex1",
                    "Ifngr2",
                    "Anxa2",
                    "Dcun1d1",
                    "9530068E07Rik",
                    "Ccar1",
                    "Flnb",
                    "Cox7a2l",
                    "Smap1",
                    "Acot7",
                    "Basp1",
                    "Ahnak",
                    "Elovl1",
                    "1700020I14Rik",
                    "Gabarap",
                    "Zbtb20",
                    "Metrn",
                    "Nab2",
                    "Phgdh",
                    "Rfwd2",
                    "Brk1",
                    "Hsp90b1",
                    "Pgrmc1",
                    "Adamts1",
                    "Timp1",
                    "Cited2",
                    "Celf2",
                    "Rps15a",
                    "Adam9",
                    "Setd3",
                    "Eif4g2",
                    "Sdcbp",
                    "Tuba1a",
                    "Lamb1",
                    "Emd",
                    "2610305D13Rik",
                    "Pdlim7",
                    "Micu1",
                    "Actn1",
                    "Rcbtb2",
                    "Psat1",
                    "Eif4h",
                    "Pex2",
                    "Rnf139",
                    "Trappc3",
                    "Shoc2",
                    "2310009B15Rik",
                    "Anxa5",
                    "Bad",
                    "Prdx6",
                    "Eif3f",
                    "Hadha",
                    "Serpine1",
                    "2200002D01Rik",
                    "Rab10",
                    "Alg5",
                    "AY036118",
                    "Crabp1",
                    "Ifitm3",
                    "Lix1l",
                    "Nptn",
                    "B3galnt1",
                    "Iqgap1",
                    "Casc5",
                    "Kdelr2",
                    "Tdgf1",
                    "March5",
                    "App",
                    "2310022B05Rik",
                    "Arl6ip5",
                    "Atp6ap1",
                    "Dexi",
                    "Akt1s1",
                    "Hist1h1e",
                    "Agpat4",
                    "Pde10a",
                    "P4ha1",
                    "Spcs1",
                    "Ndfip2",
                    "Ssr4",
                    "Ctdsp1",
                    "Gpc1",
                    "Ap2a2",
                    "Crebbp",
                    "Araf",
                    "Wdr26",
                    "Htatsf1",
                    "Tspan5",
                    "Dusp11",
                    "Slc25a46",
                    "Plaur",
                    "Arpc2",
                    "Mrpl54",
                    "Mageh1",
                    "Fndc4",
                    "Col4a2",
                    "Capza2",
                    "Ccl2",
                    "Chsy1",
                    "Mbnl3",
                    "Upf3a",
                    "Phlda3",
                    "Pbx1",
                    "Ralb",
                    "Trip6",
                    "Txnrd1",
                    "1600012H06Rik",
                    "Ube3a",
                    "Ergic3",
                    "Taok3",
                    "Atp5d",
                    "2610001J05Rik",
                    "Slc39a13",
                    "Maoa",
                    "H2afy2",
                    "Smim10l1",
                    "Cstb",
                    "0610011F06Rik",
                    "Gm10260",
                    "Mpp1",
                    "Btg1",
                    "Acer2",
                    "Lrrc58",
                    "Kif26b",
                    "Cope",
                    "Tspan31",
                    "L1td1",
                    "Ap1s1",
                    "Col3a1",
                    "Tuba4a",
                    "Mecr",
                    "Rab14",
                    "Serpinh1",
                    "Psap",
                    "Armcx2",
                    "H2afy",
                    "Rwdd1",
                    "4930402H24Rik",
                    "Acat1",
                    "Cald1",
                    "Arl8b",
                    "Cpox",
                    "Ext2",
                    "Sema3c",
                    "Gm17501",
                    "Rbm7",
                    "Tm9sf3",
                    "Trh",
                    "Ctsd",
                    "1700123O20Rik",
                    "Cbx7",
                    "4933434E20Rik",
                    "Vgll3",
                    "Arpc5",
                    "Elf2",
                    "Med21",
                    "Hdlbp",
                    "Cox4i1",
                    "Ephb2",
                    "Rgs16",
                    "Airn",
                    "Efnb1",
                    "Thbs2",
                    "Gdf3",
                    "Evl",
                    "Nek6",
                    "Adk",
                    "Nxn",
                    "Stard3nl",
                    "Serp1",
                    "Lrrfip1",
                    "Snx7",
                    "Laptm4a",
                    "Tmem59",
                    "Vcan",
                    "Actg1",
                    "Ptprd",
                    "Cdc27",
                    "Emp3",
                    "Fat4",
                    "Abracl",
                    "Cryab",
                    "Dnah8",
                    "Pdcl3",
                    "Map4",
                    "Cd302",
                    "Arpc3",
                    "Rassf1",
                    "Arl1",
                    "Col1a2",
                    "Sult6b1"
                ],
                "expression_sum": {
                    "class1": 34517.0,
                    "class2": 116070.0
                },
                "uniprot_ids": [
                    "P63158",
                    "Q58EV5",
                    "Q8C7C4",
                    "A0A0J9YUD8",
                    "Q497Z6",
                    "Q3UJK0",
                    "Q8BQ02",
                    "A0A0J9YUZ4",
                    "Q6P202",
                    "D3YVC6",
                    "Q3UBK2",
                    "D3YZ18",
                    "Q8BNM0"
                ]
            },
            "Cald1": {
                "id": "Cald1",
                "type": "Gene",
                "symbol": "Cald1",
                "source": [
                    "Pou5f1",
                    "Trim28",
                    "Rest",
                    "Nfib",
                    "Atf5",
                    "Hmgb1",
                    "Atf4",
                    "Ctcf",
                    "Id2",
                    "Hdac1",
                    "Klf5",
                    "Tbx3",
                    "Trp53",
                    "Zfp57",
                    "Ybx1",
                    "Hdac2",
                    "Smc3",
                    "Smarca5"
                ],
                "target": [],
                "expression_sum": {
                    "class1": 106561.0,
                    "class2": 18036.0
                },
                "uniprot_ids": [
                    "Q8VCQ8",
                    "E9Q9F3",
                    "F6QLP8",
                    "F6ZAW1",
                    "D3Z6I7",
                    "E9Q0M9",
                    "Q7TMN5",
                    "Q3TUS1",
                    "F6T2Z7",
                    "F6RGN9",
                    "E9QA16",
                    "Q8VD79",
                    "E9QA15",
                    "S4R1T7",
                    "Q3UGC3"
                ]
            },
            "Fkbp10": {
                "id": "Fkbp10",
                "type": "Gene",
                "symbol": "Fkbp10",
                "source": [
                    "E2f4",
                    "Pou5f1",
                    "Cebpb",
                    "Rest",
                    "Atf5",
                    "Ctcf",
                    "Trp53",
                    "Hdac1",
                    "Sox2",
                    "Tbx3",
                    "Nanog",
                    "Hdac2",
                    "Zfp57",
                    "Smarca5"
                ],
                "target": [],
                "expression_sum": {
                    "class1": 15737.0,
                    "class2": 1871.0
                },
                "uniprot_ids": [
                    "Q61576",
                    "Q3UNJ3",
                    "F6W360",
                    "A2A4H9"
                ]
            },
            "Foxp1": {
                "id": "Foxp1",
                "type": "TF",
                "symbol": "Foxp1",
                "source": [
                    "Hdac2",
                    "E2f4",
                    "Trim28",
                    "Pou5f1",
                    "Rest",
                    "Atf4",
                    "Ctcf",
                    "Tgif1",
                    "Trp53",
                    "Hdac1",
                    "Klf5",
                    "Cebpb",
                    "Smc3",
                    "Smarca5"
                ],
                "target": [
                    "Rps5",
                    "Hsp90aa1",
                    "Arf1",
                    "Rpl41",
                    "Rnf7",
                    "Cdc20",
                    "Sar1a",
                    "Srsf9",
                    "Usmg5",
                    "Mcm7",
                    "Ppp2ca",
                    "Ddx24",
                    "Rps17",
                    "Rae1",
                    "Zc3h15",
                    "Cox8a",
                    "Mrps18b",
                    "Cdt1",
                    "Psmb7",
                    "Txn2",
                    "Pnkd",
                    "Dkc1",
                    "Ndufa6",
                    "Ppp1r11",
                    "1110008F13Rik",
                    "Tuba1b",
                    "Polr2k",
                    "Gar1",
                    "Anapc5",
                    "St13",
                    "Thoc7",
                    "Hnrnpf",
                    "Knop1",
                    "Lmnb1",
                    "Ddx39",
                    "Mlf2",
                    "Pin1",
                    "Ier3ip1",
                    "Sfxn1",
                    "Swi5",
                    "Eif2s1",
                    "mt-Co1",
                    "Hmgb2",
                    "Slbp",
                    "Calr",
                    "Nup62",
                    "Rab18",
                    "Calm3",
                    "Arf6",
                    "Canx",
                    "Snrpe",
                    "Mrpl14",
                    "Cdk1",
                    "Phb",
                    "Lamtor1",
                    "Anapc15",
                    "Mrpl2",
                    "Tra2b",
                    "Cdc5l",
                    "Ranbp1",
                    "Psmc6",
                    "Grpel1",
                    "Rbx1",
                    "Ctcf",
                    "Ppm1g",
                    "Eif1",
                    "Neat1",
                    "Ythdc1",
                    "Bola1",
                    "Ola1",
                    "Sf3b6",
                    "Snrnp27",
                    "Aurkaip1",
                    "Srsf1",
                    "Socs3",
                    "Lrrc59",
                    "Exosc5",
                    "Rcc2",
                    "Pdcd6",
                    "Impa2",
                    "Ube2d2a",
                    "Bnip3l",
                    "Mapre1",
                    "Cisd2",
                    "Capza2",
                    "Lap3",
                    "Ncl",
                    "Fdft1",
                    "Supt4a",
                    "Ppp1ca",
                    "Srsf5",
                    "Hnrnph1",
                    "Aamp",
                    "Chchd1",
                    "Ywhaz",
                    "Psmc3",
                    "Sec62",
                    "Luc7l2",
                    "Rbbp7",
                    "Hnrnpl",
                    "Ddx5",
                    "Rbm39",
                    "Ccdc12",
                    "Nedd8",
                    "Pcnp",
                    "Ube2i",
                    "Utp3",
                    "Rab14",
                    "Tmem258",
                    "Tcp1",
                    "Brix1",
                    "Fam136a",
                    "Pttg1",
                    "Mrpl15",
                    "Siva1",
                    "Fus",
                    "Ociad1",
                    "Ddost",
                    "Pmm1",
                    "Fkbp8",
                    "Nip7",
                    "Clpp",
                    "Psma7",
                    "G3bp1",
                    "Pdia4",
                    "Sept7",
                    "Tpm3",
                    "Banf1",
                    "Cdc26",
                    "Immt",
                    "Psma1",
                    "Qdpr",
                    "Lig1",
                    "2410015M20Rik",
                    "Hdgf",
                    "Prnp",
                    "Smc3",
                    "Thrap3",
                    "Csnk2b",
                    "Pa2g4",
                    "Rpl37",
                    "Kdm5b",
                    "Ppp1cc",
                    "Atf4",
                    "Zfp207",
                    "Pcbp1",
                    "Hnrnpk",
                    "Ndufa4",
                    "Ndufv1",
                    "Ndufv2",
                    "H3f3a",
                    "Mettl9",
                    "Nme1",
                    "Pfkl",
                    "Csnk1a1",
                    "Ndufs2",
                    "Fam49b",
                    "Bax",
                    "Tmsb10",
                    "Rpl3",
                    "Cebpzos",
                    "Trim28",
                    "Snw1",
                    "Cdv3",
                    "Napa",
                    "Spc24",
                    "Btf3l4",
                    "Ppp4c",
                    "Etf1",
                    "Ube2b",
                    "Lsm3",
                    "Gm16286",
                    "Eif5",
                    "Tmed2",
                    "Txn1",
                    "Vapa",
                    "Mrpl17",
                    "Samm50",
                    "Rrp15",
                    "Wdr77",
                    "Atp5j",
                    "Ran",
                    "Fscn1",
                    "Srrm1",
                    "Emc8",
                    "Srp19",
                    "Cct5",
                    "Emd",
                    "Mrpl40",
                    "Bclaf1",
                    "Vasp",
                    "Tceb1",
                    "C1qbp",
                    "Uqcr10",
                    "H2afz",
                    "Tardbp",
                    "Psip1",
                    "Rpl36a",
                    "Rpl29",
                    "Iscu",
                    "Fubp1",
                    "Snrpb2",
                    "Arpc5l",
                    "Rps29",
                    "Ccdc58",
                    "Cenpa",
                    "Prdx6",
                    "Wtap",
                    "Odc1",
                    "Mrpl18",
                    "Rps20",
                    "Dnaja2",
                    "Sys1",
                    "Acat2",
                    "Lbh",
                    "Cdc37",
                    "Psmd1",
                    "Plekhf2",
                    "Eef1a1",
                    "Mcl1",
                    "Rtn4",
                    "U2af1",
                    "Atp5c1",
                    "Ywhaq",
                    "Spcs1",
                    "Fbl",
                    "Ndufv3",
                    "Ubald2",
                    "Tsc22d1",
                    "mt-Co2",
                    "Dnajc8",
                    "Calm2",
                    "Rps3",
                    "Atox1",
                    "Srsf10",
                    "Adss",
                    "Tmem126a",
                    "Ube2v2",
                    "Rpl18",
                    "Trnau1ap",
                    "Atp5d",
                    "Timm13",
                    "Eif3h",
                    "Rpl22l1",
                    "Adprh",
                    "Lamtor4",
                    "Rps3a1",
                    "Mrps5",
                    "U2surp",
                    "Mbd3",
                    "Cct7",
                    "Dctpp1",
                    "Ndufb2",
                    "Wdr18",
                    "Hsp90ab1",
                    "Oaz1",
                    "Mrpl30",
                    "Uqcrq",
                    "Sec61b",
                    "BC031181",
                    "Polr2m",
                    "1810058I24Rik",
                    "Rwdd1",
                    "Snx5",
                    "Got1",
                    "Fnta",
                    "Dnaja1",
                    "Eif4a1",
                    "Sumo2",
                    "Pcbp2",
                    "Snrnp25",
                    "Nudc",
                    "H3f3b",
                    "Mrps33",
                    "Rif1",
                    "Nsa2",
                    "Ddx46",
                    "Nudt19",
                    "Dbi",
                    "Morf4l2",
                    "Ube2d3",
                    "Rbbp4",
                    "Eif3e",
                    "Paip2",
                    "Cebpb",
                    "Srrm2",
                    "Rnaseh2b",
                    "Ywhae",
                    "Ubl5",
                    "Hmgb1",
                    "Ndufc1",
                    "Eif3d",
                    "Rps15",
                    "Npm1",
                    "Psma4",
                    "Sap30",
                    "Scp2",
                    "Ap2s1",
                    "Rpl34",
                    "Mlec",
                    "Dynll1",
                    "Gtpbp4",
                    "Cenpw",
                    "Hspd1",
                    "Atf5",
                    "Cops7a",
                    "Mrpl45",
                    "Mrps14",
                    "Rpl31",
                    "Srsf7",
                    "Smdt1",
                    "Sod1",
                    "Ppih",
                    "Mat2a",
                    "Nap1l1",
                    "Uqcc3",
                    "Dnajb6",
                    "Brd2",
                    "Actr3",
                    "G3bp2",
                    "Impdh2",
                    "Ccdc124",
                    "Gapdh",
                    "Smarca5",
                    "Mpc1",
                    "Prps1",
                    "Ccnd3",
                    "Ap2m1",
                    "Mrpl52",
                    "Adrm1",
                    "Abcf1",
                    "Rplp0",
                    "Psmb5",
                    "Hspa9",
                    "Pim3",
                    "Ube2n",
                    "Ndufa11",
                    "Cct8",
                    "Hmgn2",
                    "Mpc2",
                    "Dohh",
                    "Hnrnpa2b1",
                    "Ndufs5",
                    "Cited2",
                    "Srsf2",
                    "Eif4g2",
                    "Jade1",
                    "Psmb2",
                    "Raly",
                    "Denr",
                    "Fxr1",
                    "Nsmce4a",
                    "Rps26",
                    "Get4",
                    "Trp53",
                    "Sra1",
                    "Aspscr1",
                    "Psmd11",
                    "Vps29",
                    "Vdac1",
                    "Cggbp1",
                    "Zfp706",
                    "Anp32a",
                    "Polr2e",
                    "Anapc13",
                    "Sumo1",
                    "Pdcd5",
                    "Snrpc",
                    "Mrpl51",
                    "Mrpl49",
                    "Tmbim6",
                    "Dynll2",
                    "Rhoa",
                    "Fcf1",
                    "Mrps18c",
                    "Cacybp",
                    "Cdk4",
                    "Rpl22",
                    "Rsl1d1",
                    "Nolc1",
                    "Sf3a3",
                    "Sec61a1",
                    "Pfdn6",
                    "Glud1",
                    "Cebpz",
                    "Hnrnpu",
                    "Gspt1",
                    "Arglu1",
                    "Tomm20",
                    "Rpl21",
                    "Ppp1r2",
                    "Map1lc3b",
                    "Eprs",
                    "Trmt112",
                    "Dazap1",
                    "Son",
                    "Pop5",
                    "Cct2",
                    "Ubb",
                    "Dtl",
                    "Bzw1",
                    "Parp1",
                    "Dhfr",
                    "Rbpms",
                    "Paics",
                    "Hnrnpa3",
                    "Prpf40a",
                    "Pbdc1",
                    "Cdca3",
                    "Mrpl57",
                    "Morf4l1",
                    "Mrpl43",
                    "Ube2s",
                    "Tecr",
                    "Polr1d",
                    "Snrpd3",
                    "Sf3b2",
                    "Psmd3",
                    "Suclg1",
                    "Mtf2",
                    "Slc3a2",
                    "mt-Nd2",
                    "Hnrnpd",
                    "Hat1",
                    "Cops4",
                    "mt-Cytb",
                    "Rnps1",
                    "Ciapin1",
                    "Lsm4",
                    "Pfdn2",
                    "Mpdu1",
                    "Atp5f1",
                    "Eif1ad",
                    "Rps13",
                    "Tpt1",
                    "Eif5a",
                    "Ppib",
                    "Gng5",
                    "Hspe1",
                    "Rps19",
                    "Exosc8",
                    "Cops6",
                    "Ptbp1",
                    "Aldoa",
                    "Hnrnpdl",
                    "Eif3a",
                    "Rtn3",
                    "Polr2h",
                    "Snx3",
                    "Adh5",
                    "Birc5",
                    "Psmd13",
                    "Srpk1",
                    "Rfc1",
                    "Cox6b1",
                    "Ppa1",
                    "Cenpm",
                    "Prdx1",
                    "Rsl24d1",
                    "Cdc34",
                    "Rpa2",
                    "Mrpl34",
                    "Pdia3",
                    "Ndufb5",
                    "Psma2",
                    "Ndufs6",
                    "Nsun2",
                    "Set",
                    "Txnl4a",
                    "Poldip2",
                    "Eif4e2",
                    "Sod2",
                    "Ssbp1",
                    "Rps14",
                    "Tgif1",
                    "Lipe",
                    "Clns1a",
                    "Snrpd2",
                    "1810022K09Rik",
                    "Phb2",
                    "Gtf2e2",
                    "Rpl15",
                    "Brk1",
                    "Vdac2",
                    "Rap1b",
                    "Pfn1",
                    "Rpl13a",
                    "Snrpg",
                    "Ddx39b",
                    "Cuta",
                    "Snhg6",
                    "Eif1a",
                    "Cox5a",
                    "Snrpb",
                    "Tmem11",
                    "Anapc11",
                    "Fkbp4",
                    "Pgam1",
                    "Tomm22",
                    "Lyar",
                    "Ubap2l",
                    "Erp29",
                    "Snrnp70",
                    "Uchl5",
                    "Gpi1",
                    "Tmem167",
                    "Ppdpf",
                    "Llph",
                    "Emg1",
                    "Lsm12",
                    "Eif4b",
                    "Anp32b",
                    "Pkm",
                    "H2afv",
                    "Csde1",
                    "Prrc2c",
                    "Tceb2",
                    "Rps2",
                    "Eif2b5",
                    "Gart",
                    "Sumo3",
                    "Ndufa10",
                    "Uqcrfs1",
                    "Naa50",
                    "Ywhah",
                    "Sf3b4",
                    "Prpf19",
                    "Sugt1",
                    "Eif4a3",
                    "Fkbp3",
                    "Mtch2",
                    "Dek",
                    "Bag1",
                    "Psmd4",
                    "Gins2",
                    "Fam104a",
                    "Papola",
                    "Rps16",
                    "Mrfap1",
                    "Mrps16",
                    "Uqcrh",
                    "Tpr",
                    "Ddx18",
                    "Psmd12",
                    "Rrp9",
                    "Eed",
                    "Ccne1",
                    "Srsf6",
                    "Ndufa5",
                    "Tmpo",
                    "Tsn",
                    "Mrpl11",
                    "Txnl1",
                    "Mesdc2",
                    "Eef1g",
                    "Rbm25",
                    "2700094K13Rik",
                    "Cycs",
                    "Myl12a",
                    "Ptma",
                    "Rac1",
                    "Rer1",
                    "Klf2",
                    "Dut",
                    "Ralbp1",
                    "Dhx16",
                    "Sdhb",
                    "Taf10",
                    "Wdr43",
                    "Ybx3",
                    "Rps10",
                    "Yrdc",
                    "Pabpc1",
                    "Alyref",
                    "Tsen34",
                    "Hspa14",
                    "Fkbp1a",
                    "Dad1",
                    "Yif1b",
                    "Fam60a",
                    "Wbp11",
                    "Cwc15",
                    "Guk1",
                    "2810428I15Rik",
                    "Tsfm",
                    "Pim1",
                    "Postn",
                    "Rplp2",
                    "Aimp1",
                    "Serbp1",
                    "Gpx4",
                    "Gcsh",
                    "Phf5a",
                    "Eif4a2",
                    "Rheb",
                    "Gtf2a2",
                    "Mkrn1",
                    "Rrp1",
                    "U2af2",
                    "Rpl23",
                    "Sfr1",
                    "Kdm1a",
                    "Rrs1",
                    "Ly6e",
                    "Bcap31",
                    "Cnbp",
                    "Rsrc2"
                ],
                "expression_sum": {
                    "class1": 10758.0,
                    "class2": 1389.0
                },
                "uniprot_ids": [
                    "P58462",
                    "E2S038",
                    "H3BJ68",
                    "H3BJZ5",
                    "H3BLF2",
                    "Q8CCZ5",
                    "H3BJR8",
                    "D3Z6Q4",
                    "Q6P0X7",
                    "D3Z454",
                    "H3BL33",
                    "A0A0R4J1H9",
                    "H3BJ24",
                    "H3BJM1",
                    "D3Z6Q3",
                    "H3BK30",
                    "H3BKT4",
                    "A0A0R4J282",
                    "D3Z2T4",
                    "D3Z0J5",
                    "H3BK38",
                    "Q3UMG2"
                ]
            },
            "2810417H13Rik": {
                "id": "2810417H13Rik",
                "type": "Gene",
                "symbol": "2810417H13Rik",
                "source": [
                    "Klf2",
                    "Pou5f1",
                    "Hmga2",
                    "Cebpb",
                    "Atf5",
                    "Ctbp1",
                    "Ctcf",
                    "Junb",
                    "Fos",
                    "Jun",
                    "Trp53",
                    "Prrx2",
                    "Ybx1",
                    "Jund",
                    "Twist2",
                    "Hdac2"
                ],
                "target": [],
                "expression_sum": {
                    "class1": 26131.0,
                    "class2": 21217.0
                }
            },
            "Uchl3": {
                "id": "Uchl3",
                "type": "Gene",
                "symbol": "Uchl3",
                "source": [
                    "Pou5f1",
                    "Atf5",
                    "Junb",
                    "Jun",
                    "Sox2",
                    "Nanog",
                    "Jund",
                    "Cebpb"
                ],
                "target": [],
                "expression_sum": {
                    "class1": 5030.0,
                    "class2": 45415.0
                },
                "uniprot_ids": [
                    "Q9JKB1",
                    "Q8BWQ9",
                    "A0A2I3BQ39"
                ]
            },
            "Rcc2": {
                "id": "Rcc2",
                "type": "Gene",
                "symbol": "Rcc2",
                "source": [
                    "Tcf4",
                    "Pou5f1",
                    "Foxp1",
                    "Tfcp2l1",
                    "Hmga2",
                    "Atf5",
                    "Junb",
                    "Fos",
                    "Jun",
                    "Sox2",
                    "Hes1",
                    "Nanog",
                    "Jund",
                    "Twist2",
                    "Cebpb",
                    "Zfp57",
                    "Sox4"
                ],
                "target": [],
                "expression_sum": {
                    "class1": 3616.0,
                    "class2": 52274.0
                },
                "uniprot_ids": [
                    "Q8BK67",
                    "A2AWQ2"
                ]
            },
            "2700060E02Rik": {
                "id": "2700060E02Rik",
                "type": "Gene",
                "symbol": "2700060E02Rik",
                "source": [
                    "Klf2",
                    "Tcf4",
                    "Pou5f1",
                    "Hmga2",
                    "Arf4",
                    "Atf5",
                    "Atf4",
                    "Junb",
                    "Fos",
                    "Jun",
                    "Sox2",
                    "Prrx1",
                    "Nanog",
                    "Jund",
                    "Cebpb",
                    "Mybl2"
                ],
                "target": [],
                "expression_sum": {
                    "class1": 33160.0,
                    "class2": 123621.0
                },
                "uniprot_ids": [
                    "Q4VA29"
                ]
            },
            "Rest": {
                "id": "Rest",
                "type": "TF",
                "symbol": "Rest",
                "source": [
                    "Pou5f1",
                    "Junb",
                    "Fos",
                    "Jun",
                    "Sox2",
                    "Hes1",
                    "Nanog",
                    "Jund",
                    "Cebpb",
                    "Sox4"
                ],
                "target": [
                    "Dap",
                    "Sar1a",
                    "Tpm4",
                    "H2afj",
                    "Ddb1",
                    "Myh9",
                    "Thbs1",
                    "Rasgrp2",
                    "Lims1",
                    "Wisp1",
                    "Slc25a4",
                    "Tspan4",
                    "Col12a1",
                    "Map1lc3a",
                    "Lmna",
                    "Pck2",
                    "Lgals3",
                    "Cxcl12",
                    "Pgrmc1",
                    "Fos",
                    "Tagln2",
                    "Rps15a",
                    "Rexo2",
                    "Pdlim7",
                    "Mfge8",
                    "Mbnl2",
                    "Canx",
                    "Actn1",
                    "B2m",
                    "Neat1",
                    "Myh10",
                    "Nanog",
                    "Ube2r2",
                    "Timp3",
                    "Gemin7",
                    "Akr1a1",
                    "Sh3glb1",
                    "Atp2b1",
                    "Cisd2",
                    "Nktr",
                    "Col5a1",
                    "Bgn",
                    "Klf6",
                    "Tgfb1i1",
                    "Sec62",
                    "Rps12",
                    "Arpc1b",
                    "Col3a1",
                    "Cyr61",
                    "Kdelr1",
                    "Rab14",
                    "Serpinh1",
                    "Atp6v1g1",
                    "Gnb2",
                    "Ociad1",
                    "Dpysl3",
                    "Mybl2",
                    "Slc2a3",
                    "Hcfc1r1",
                    "Sptbn1",
                    "Id3",
                    "Ap3s1",
                    "Gng12",
                    "Ncam1",
                    "Wls",
                    "Hacd1",
                    "Ndufa8",
                    "Srp14",
                    "Sept7",
                    "Fam195b",
                    "Actg1",
                    "Rcn2",
                    "Ier5",
                    "Prnp",
                    "Id1",
                    "Hmga2",
                    "Hist1h2bc",
                    "Tfcp2l1",
                    "Col1a2",
                    "F2r",
                    "Serpine2",
                    "Igfbp4",
                    "Itgb1",
                    "Fabp5",
                    "Mettl9",
                    "Tcf4",
                    "Igf2",
                    "Ssr1",
                    "Pgls",
                    "Lxn",
                    "Ift20",
                    "Anxa2",
                    "Vmp1",
                    "Crip2",
                    "Cox7a2l",
                    "Vim",
                    "Basp1",
                    "Sh3bgrl3",
                    "Wdr1",
                    "Tnc",
                    "Tmed2",
                    "Gjb3",
                    "Tpm1",
                    "Sdcbp",
                    "Thy1",
                    "Surf4",
                    "Utf1",
                    "Phlda1",
                    "Myeov2",
                    "Vcl",
                    "Cst3",
                    "Lox",
                    "Ung",
                    "Higd1a",
                    "S100a6",
                    "Serpine1",
                    "Crabp1",
                    "Ifitm1",
                    "Cd63",
                    "Capg",
                    "Lbh",
                    "S100a11",
                    "Scand1",
                    "Mtdh",
                    "mt-Nd5",
                    "Maged2",
                    "Ctsl",
                    "Ccnd2",
                    "Tcf25",
                    "Spcs1",
                    "Cox6a1",
                    "Col6a1",
                    "Rala",
                    "Sdc2",
                    "Tax1bp3",
                    "Filip1l",
                    "Plk2",
                    "Tln1",
                    "Oaf",
                    "Ostf1",
                    "Snhg12",
                    "Fbln5",
                    "Chd3",
                    "Cmpk1",
                    "Lrrc58",
                    "Fn1",
                    "Glipr2",
                    "Hes1",
                    "Ap1s1",
                    "Nenf",
                    "Zfp42",
                    "Rcn3",
                    "Malat1",
                    "Tmem176b",
                    "Tmed10",
                    "Tm9sf3",
                    "Reep5",
                    "Col1a1",
                    "Oxct1",
                    "Peg3",
                    "Atxn10",
                    "Atp6v0e",
                    "Tmem59",
                    "Pafah1b2",
                    "Arpc3",
                    "Ilk",
                    "Mgat2",
                    "Sep15",
                    "Lsm7",
                    "Cpe",
                    "Atf5",
                    "Cdc42",
                    "Npc2",
                    "Eef2",
                    "Hsbp1",
                    "Rnh1",
                    "Rrbp1",
                    "Tmsb4x",
                    "Cd47",
                    "Efemp2",
                    "Mpc1",
                    "Copb2",
                    "Sars",
                    "Smim14",
                    "Lgals1",
                    "Fkbp10",
                    "E130012A19Rik",
                    "Kif5b",
                    "Ptp4a1",
                    "Lsp1",
                    "Gsn",
                    "Tubb6",
                    "Prdx4",
                    "Rab2a",
                    "Tspo",
                    "Myl9",
                    "Btg2",
                    "Selk",
                    "Rab11a",
                    "Anxa5",
                    "Anxa6",
                    "Mtch1",
                    "Rab10",
                    "Ttc3",
                    "Vimp",
                    "Sec61a1",
                    "Iqgap1",
                    "Kdelr2",
                    "Pmp22",
                    "Rbmxl2",
                    "Ssr3",
                    "Rpn1",
                    "Ssr4",
                    "Sqstm1",
                    "Lman1",
                    "Rcn1",
                    "Arpc2",
                    "Ywhab",
                    "Sox4",
                    "Phlda3",
                    "Sept11",
                    "Romo1",
                    "Tpm2",
                    "Ergic3",
                    "Sdc4",
                    "Rock2",
                    "Cdc42ep3",
                    "Col8a1",
                    "L1td1",
                    "Psap",
                    "Rbms1",
                    "P4hb",
                    "Zyx",
                    "Adipor1",
                    "Bsg",
                    "Ppic",
                    "Camk2d",
                    "mt-Cytb",
                    "Cyba",
                    "Flna",
                    "H13",
                    "Rsu1",
                    "Prss23",
                    "Gdf3",
                    "Gsto1",
                    "Ppib",
                    "Vcan",
                    "Sparc",
                    "Sept2",
                    "Abracl",
                    "Msn",
                    "Serpinb6a",
                    "Nfatc2ip",
                    "Ctbp1",
                    "Csrp1",
                    "Selm",
                    "Esd",
                    "Dppa5a",
                    "Atxn7l3b",
                    "Comt",
                    "Lamp1",
                    "Rcor2",
                    "Sat1",
                    "Ndufa13",
                    "Ptrf",
                    "Gadd45g",
                    "Sh3bgrl",
                    "Jun",
                    "Itm2b",
                    "Pdia3",
                    "Rpl10a",
                    "Txndc5",
                    "Col4a1",
                    "Hist1h1c",
                    "Cd24a",
                    "Smap1",
                    "Lipe",
                    "Tnfrsf12a",
                    "Krtcap2",
                    "Gabarap",
                    "Serf2",
                    "Mtpn",
                    "Prkar1a",
                    "Apoe",
                    "Tuba1a",
                    "Commd3",
                    "Gsta4",
                    "Fst",
                    "Gpi1",
                    "Palld",
                    "Tram1",
                    "Lrrc17",
                    "Fstl1",
                    "Marcks",
                    "Fam162a",
                    "Rhoc",
                    "Mdm2",
                    "Srp72",
                    "Mrpl33",
                    "Fundc2",
                    "S100a10",
                    "Rpn2",
                    "Prrx2",
                    "Tmed3",
                    "Serpinf1",
                    "Hsd17b14",
                    "Gap43",
                    "Foxp1",
                    "Ccng1",
                    "H19",
                    "Ptp4a2",
                    "Dnajc3",
                    "Myl12a",
                    "Rac1",
                    "Rer1",
                    "S100a4",
                    "Cald1",
                    "Sri",
                    "Prdx5",
                    "Clic1",
                    "Emc7",
                    "Gnas",
                    "Arf4",
                    "Calu",
                    "Arpc5",
                    "Gm7325",
                    "Hdlbp",
                    "Ckb",
                    "Limd2",
                    "Cox4i1",
                    "Tmed9",
                    "Myl6",
                    "Sepw1",
                    "Serp1",
                    "Capn2",
                    "Laptm4a",
                    "Carhsp1",
                    "Kdelr3",
                    "Junb",
                    "Bcap31",
                    "Axl",
                    "Cfl2"
                ],
                "expression_sum": {
                    "class1": 3691.0,
                    "class2": 33317.0
                },
                "uniprot_ids": [
                    "Q8VIG1",
                    "Q60694",
                    "F8V330"
                ]
            },
            "Atf4": {
                "id": "Atf4",
                "type": "TF",
                "symbol": "Atf4",
                "source": [
                    "Tcf4",
                    "Pou5f1",
                    "Foxp1",
                    "Cebpb",
                    "Eno1",
                    "Ctcf",
                    "Junb",
                    "Trp53",
                    "Sox2",
                    "Ybx1",
                    "Jund",
                    "Twist2",
                    "Hdac2"
                ],
                "target": [
                    "Hsp90aa1",
                    "Dap",
                    "Rnf7",
                    "Cdc20",
                    "Sar1a",
                    "Usmg5",
                    "Ppp2ca",
                    "Ddx24",
                    "Zc3h15",
                    "Myh9",
                    "Txn2",
                    "Ppid",
                    "2810004N23Rik",
                    "Clta",
                    "Igfbp2",
                    "Sec13",
                    "1110008F13Rik",
                    "Idh3g",
                    "Anapc5",
                    "Lmna",
                    "Ndufa2",
                    "Hnrnpf",
                    "Knop1",
                    "Ddx39",
                    "Mlf2",
                    "Pck2",
                    "Rnf187",
                    "Med28",
                    "Nubp1",
                    "Rps15a",
                    "Ndufs3",
                    "Nedd4",
                    "Uba2",
                    "Rexo2",
                    "Calr",
                    "Nup62",
                    "Pdlim7",
                    "Calm3",
                    "Mbnl2",
                    "Arf6",
                    "Canx",
                    "Nop56",
                    "Slc25a3",
                    "Psat1",
                    "Psmd7",
                    "Lamtor1",
                    "Tra2b",
                    "Psmc6",
                    "Zwint",
                    "Ppm1g",
                    "Eif1",
                    "Myh10",
                    "Sec61g",
                    "Ola1",
                    "Snrnp27",
                    "Aurkaip1",
                    "Gemin7",
                    "Wdr89",
                    "Lrrc59",
                    "Uqcrc1",
                    "Akt1s1",
                    "Bnip3l",
                    "H2afx",
                    "Kpnb1",
                    "Mapre1",
                    "Cisd2",
                    "Snrpd1",
                    "Nktr",
                    "Capza2",
                    "Klf6",
                    "Rpl36al",
                    "Ncl",
                    "Loxl1",
                    "Srsf5",
                    "Hnrnph1",
                    "Chchd1",
                    "Ywhaz",
                    "Gars",
                    "Psmc3",
                    "Sec62",
                    "Pomp",
                    "Hspa4",
                    "Luc7l2",
                    "Psmb3",
                    "Ddx5",
                    "Rbm39",
                    "Nedd8",
                    "Pcnp",
                    "Eef1b2",
                    "Ube2i",
                    "Arpc1b",
                    "Cyr61",
                    "Utp3",
                    "Serpinh1",
                    "Tcp1",
                    "Brix1",
                    "Gnb2",
                    "Mrpl15",
                    "Fus",
                    "Dpysl3",
                    "Ddost",
                    "Hcfc1r1",
                    "Sptbn1",
                    "Gng12",
                    "Fkbp8",
                    "Nip7",
                    "Ndufa8",
                    "Pdia4",
                    "Srp14",
                    "Sept7",
                    "Fam195b",
                    "Trim27",
                    "Mrps36",
                    "Banf1",
                    "Immt",
                    "Actg1",
                    "Dtymk",
                    "Psma1",
                    "Tsg101",
                    "Hdgf",
                    "Nars",
                    "Pa2g4",
                    "Hmga2",
                    "F2r",
                    "Pcbp1",
                    "Rpl38",
                    "Dnttip2",
                    "Sae1",
                    "Hnrnpk",
                    "Ewsr1",
                    "Eif4e",
                    "Ndufv1",
                    "Ndufv2",
                    "Fabp5",
                    "H3f3a",
                    "Nme1",
                    "Chmp4b",
                    "Tcf4",
                    "Ube2k",
                    "Ssr1",
                    "Ndufs2",
                    "Cyp51",
                    "Pgls",
                    "Tmsb10",
                    "Nudt21",
                    "Rpl12",
                    "Ift20",
                    "Anxa2",
                    "Eif3c",
                    "Snw1",
                    "Vmp1",
                    "Col5a2",
                    "Cdv3",
                    "Napa",
                    "Btf3l4",
                    "Asns",
                    "Eef1d",
                    "Etf1",
                    "Sh3bgrl3",
                    "Ndufs7",
                    "Ube2b",
                    "Lsm3",
                    "Mrpl13",
                    "Gm16286",
                    "Eif5",
                    "Tmed2",
                    "Ube2j2",
                    "Vapa",
                    "Mrpl17",
                    "Wdr77",
                    "Atp5j",
                    "Ran",
                    "Fscn1",
                    "Srrm1",
                    "Tpm1",
                    "Ldha",
                    "Sdcbp",
                    "Apex1",
                    "Surf4",
                    "Atp5g2",
                    "Emc8",
                    "Srp19",
                    "Phlda1",
                    "1110004F10Rik",
                    "Myeov2",
                    "Vasp",
                    "Tceb1",
                    "Rpsa",
                    "H2afz",
                    "Bola3",
                    "Psip1",
                    "Vcl",
                    "Iscu",
                    "Fubp1",
                    "Arpc5l",
                    "Rpl5",
                    "Dph3",
                    "Prdx6",
                    "Ptges3",
                    "Mrpl18",
                    "Rps20",
                    "Orc6",
                    "Dnaja2",
                    "Crabp1",
                    "Sys1",
                    "Mrpl12",
                    "Ndufab1",
                    "Mtdh",
                    "Eef1a1",
                    "mt-Nd5",
                    "U2af1",
                    "Atp5c1",
                    "Ywhaq",
                    "Ccnd2",
                    "Tcf25",
                    "Pdha1",
                    "Filip1l",
                    "Cav1",
                    "Dnajc8",
                    "Calm2",
                    "Stmn1",
                    "Srsf10",
                    "Btf3",
                    "Smarcc1",
                    "Plk2",
                    "Tln1",
                    "Tmem126a",
                    "Oaf",
                    "Rpl18",
                    "Atp5d",
                    "Mt2",
                    "Timm13",
                    "Rps3a1",
                    "Smc4",
                    "Cmpk1",
                    "Rpl7",
                    "Fkbp2",
                    "Lrrc58",
                    "Fn1",
                    "Glipr2",
                    "Hes1",
                    "Ap1s1",
                    "Uqcrq",
                    "Mrpl30",
                    "Sec61b",
                    "Polr2m",
                    "Rwdd1",
                    "Snx5",
                    "Gnai2",
                    "Rps8",
                    "Dnaja1",
                    "Rpl8",
                    "Eif4a1",
                    "Tm9sf3",
                    "Ndufc2",
                    "Ccdc34",
                    "Col1a1",
                    "Nsa2",
                    "Oxct1",
                    "Ddx46",
                    "Cnih4",
                    "Morf4l2",
                    "Ube2d3",
                    "Aars",
                    "Snrpa1",
                    "Dstn",
                    "Emp3",
                    "Cebpb",
                    "Srrm2",
                    "2700060E02Rik",
                    "Mt1",
                    "Ywhae",
                    "Pafah1b2",
                    "Hmgb1",
                    "Psmd14",
                    "Eif3d",
                    "Npm1",
                    "Ghitm",
                    "Scp2",
                    "Arl1",
                    "Arpc1a",
                    "Mgat2",
                    "Sep15",
                    "Rab7",
                    "Gtpbp4",
                    "Eif2s2",
                    "Hspd1",
                    "Cpe",
                    "Atf5",
                    "Cdc42",
                    "Mrpl45",
                    "Srsf7",
                    "Mrps25",
                    "Tpx2",
                    "Dnajb6",
                    "Hspa5",
                    "Uqcrc2",
                    "Actr3",
                    "G3bp2",
                    "Cd47",
                    "Impdh2",
                    "Ccdc124",
                    "Mpc1",
                    "Sars",
                    "Mrpl52",
                    "Ap2m1",
                    "Nupr1",
                    "Pno1",
                    "Hnrnpab",
                    "Rplp0",
                    "Psmb5",
                    "Ahnak",
                    "Tbrg1",
                    "Hspa9",
                    "Kif5b",
                    "Polr2c",
                    "Cct8",
                    "Hmgn2",
                    "Mpc2",
                    "Hnrnpa2b1",
                    "Ptp4a1",
                    "Lsp1",
                    "Ndufs5",
                    "Cited2",
                    "Srsf2",
                    "Prdx4",
                    "mt-Nd1",
                    "Eif4g2",
                    "Dnajc7",
                    "Psmb2",
                    "Raly",
                    "Denr",
                    "Get4",
                    "Cnn3",
                    "Parva",
                    "Vdac1",
                    "Cggbp1",
                    "Zfp706",
                    "Anp32a",
                    "Rad23b",
                    "Sumo1",
                    "Tra2a",
                    "Pdcd5",
                    "Rab11a",
                    "Ankrd1",
                    "Anxa5",
                    "Eif3f",
                    "Eif4g1",
                    "Anxa6",
                    "Mtch1",
                    "Tmbim6",
                    "Rhoa",
                    "Mrps18c",
                    "Rab10",
                    "Cacybp",
                    "Sf3a3",
                    "Ccdc80",
                    "Anp32e",
                    "Syncrip",
                    "Sec61a1",
                    "Caprin1",
                    "Kdelr2",
                    "Ict1",
                    "Pmp22",
                    "Glud1",
                    "March5",
                    "Rbmxl2",
                    "Cebpz",
                    "Ssr2",
                    "Hnrnpu",
                    "Rbmxl1",
                    "Srsf11",
                    "Arglu1",
                    "Rpl18a",
                    "Tomm20",
                    "Cbx1",
                    "Ssr4",
                    "Tomm70a",
                    "Txndc9",
                    "Ppp1r2",
                    "Eprs",
                    "Trmt112",
                    "Arpc2",
                    "Atp5l",
                    "Mrpl54",
                    "Son",
                    "Cct2",
                    "Ubb",
                    "Sept11",
                    "Romo1",
                    "Ergic3",
                    "Bzw1",
                    "Sdc4",
                    "Ubc",
                    "Lman2",
                    "Rock2",
                    "Hnrnpa3",
                    "Mrps7",
                    "Sub1",
                    "Morf4l1",
                    "2010107E04Rik",
                    "Mrpl43",
                    "Pafah1b1",
                    "Cdca8",
                    "Idh2",
                    "Polr1d",
                    "Snrpd3",
                    "Rbms1",
                    "P4hb",
                    "Sf3b2",
                    "Psmd3",
                    "Zyx",
                    "Map1b",
                    "Adipor1",
                    "mt-Nd2",
                    "Bsg",
                    "Rpl14",
                    "Camk2d",
                    "Cops4",
                    "mt-Cytb",
                    "Rnps1",
                    "Ciapin1",
                    "Flna",
                    "Pfdn2",
                    "H13",
                    "Atp5f1",
                    "Arid5b",
                    "Rps13",
                    "Tpt1",
                    "Ppib",
                    "Cfl1",
                    "Gng5",
                    "Hspe1",
                    "Rps19",
                    "Sept2",
                    "Exosc8",
                    "Cops6",
                    "Ptbp1",
                    "Mrpl20",
                    "Hnrnpdl",
                    "Eif3a",
                    "Ptx3",
                    "Ctbp1",
                    "Psmb4",
                    "Adh5",
                    "Birc5",
                    "Commd1",
                    "Csrp1",
                    "Chtop",
                    "Atxn7l3b",
                    "Ppa1",
                    "Lamtor5",
                    "Ndufa13",
                    "Gps1",
                    "Rrm2",
                    "Rpl13",
                    "Sh3bgrl",
                    "Itm2b",
                    "Pdia3",
                    "Ndufb5",
                    "Atrx",
                    "Psma2",
                    "Txndc5",
                    "Pfdn5",
                    "Nsun2",
                    "Set",
                    "Txnl4a",
                    "Poldip2",
                    "Eif4e2",
                    "Ssbp1",
                    "Rps14",
                    "Mrto4",
                    "Tnfrsf12a",
                    "Clns1a",
                    "Snrpd2",
                    "Krtcap2",
                    "Rps27a",
                    "Naa10",
                    "Bud31",
                    "Pdap1",
                    "Brk1",
                    "Hsp90b1",
                    "Vdac2",
                    "Anxa1",
                    "Rap1b",
                    "Ube2a",
                    "Rpl13a",
                    "Nol7",
                    "Eif1a",
                    "Cox5a",
                    "Commd3",
                    "Cnn2",
                    "Snrpb",
                    "Exosc4",
                    "Fkbp4",
                    "Pgam1",
                    "Rabggtb",
                    "Tomm22",
                    "Nans",
                    "Uchl5",
                    "Erp29",
                    "Ppig",
                    "Phldb2",
                    "Palld",
                    "Eif5b",
                    "Cct4",
                    "Llph",
                    "Lrrc17",
                    "Eif4b",
                    "Eif4ebp1",
                    "Anp32b",
                    "Edf1",
                    "Csde1",
                    "Marcks",
                    "Fam162a",
                    "Emp1",
                    "Psmc2",
                    "Ndufa10",
                    "Tomm40",
                    "Ywhah",
                    "Prpf19",
                    "Hdac2",
                    "Sugt1",
                    "Eif4a3",
                    "Fkbp3",
                    "Mtch2",
                    "Bag1",
                    "Tmem147",
                    "Rpn2",
                    "Bub3",
                    "Sin3b",
                    "Dctn2",
                    "Tmed3",
                    "Serpinf1",
                    "Tubb4b",
                    "Mrps16",
                    "Tpr",
                    "Hint1",
                    "Tbca",
                    "Foxp1",
                    "Srsf6",
                    "Tmpo",
                    "Tsn",
                    "Txnl1",
                    "Mesdc2",
                    "Rbm25",
                    "Ube2e3",
                    "Drg1",
                    "Arhgdia",
                    "Myl12a",
                    "Ptma",
                    "Rac1",
                    "Rer1",
                    "Ralbp1",
                    "Cald1",
                    "Fau",
                    "Yrdc",
                    "Acin1",
                    "Sri",
                    "Pabpc1",
                    "Prdx5",
                    "Alyref",
                    "Tsen34",
                    "Hspa14",
                    "Fkbp1a",
                    "Emc7",
                    "Gnas",
                    "Arf4",
                    "Calu",
                    "Yif1b",
                    "Hdlbp",
                    "Wbp11",
                    "Cox4i1",
                    "Tmed9",
                    "Snf8",
                    "Guk1",
                    "Ndufs8",
                    "Serp1",
                    "Tsfm",
                    "Capn2",
                    "Ptov1",
                    "Laptm4a",
                    "Aimp1",
                    "Serbp1",
                    "Cnih1",
                    "Gpx4",
                    "Chchd3",
                    "Gcsh",
                    "Phf5a",
                    "Rheb",
                    "Gtf2a2",
                    "Rrp1",
                    "Rpl30",
                    "Sfr1",
                    "Kdelr3",
                    "Ncor1",
                    "Rbm17",
                    "Map2k2",
                    "Cnbp",
                    "Sdhd",
                    "Ogn",
                    "Rsrc2"
                ],
                "expression_sum": {
                    "class1": 19607.0,
                    "class2": 34910.0
                },
                "uniprot_ids": [
                    "Q06507",
                    "Q8CF69",
                    "Q3U2J1",
                    "A0A2R8VI82",
                    "Q61328",
                    "Q794G7",
                    "Q3TZI8"
                ]
            },
            "Ahnak": {
                "id": "Ahnak",
                "type": "Gene",
                "symbol": "Ahnak",
                "source": [
                    "Pou5f1",
                    "Trim28",
                    "Cebpb",
                    "Atf5",
                    "Eno1",
                    "Hmgb1",
                    "Atf4",
                    "Ctcf",
                    "Trp53",
                    "Hdac1",
                    "Klf5",
                    "Tbx3",
                    "Zfp57",
                    "Hdac2",
                    "Smc3",
                    "Smarca5"
                ],
                "target": [],
                "expression_sum": {
                    "class1": 16857.0,
                    "class2": 7693.0
                },
                "uniprot_ids": [
                    "Q6UIL4",
                    "Q8BKS2",
                    "Q8CGE7",
                    "E9Q616",
                    "G5E8K8",
                    "A0PJF4",
                    "Q8VDN3",
                    "A0A494B8Y7",
                    "Q61484",
                    "A0A494BBD5",
                    "Q6UL10",
                    "A0JLR7",
                    "Q8BRB8",
                    "Q8CEX7",
                    "Q8R2L7",
                    "Q3UYW8"
                ]
            },
            "Phf5a": {
                "id": "Phf5a",
                "type": "Gene",
                "symbol": "Phf5a",
                "source": [
                    "Tcf4",
                    "Pou5f1",
                    "Foxp1",
                    "Nfib",
                    "Atf5",
                    "Atf4",
                    "Junb",
                    "Fos",
                    "Jun",
                    "Sox2",
                    "Nanog",
                    "Jund",
                    "Twist2",
                    "Cebpb",
                    "Sox4"
                ],
                "target": [],
                "expression_sum": {
                    "class1": 10206.0,
                    "class2": 56145.0
                },
                "uniprot_ids": [
                    "P83870"
                ]
            },
            "Gtf2a2": {
                "id": "Gtf2a2",
                "type": "TF",
                "symbol": "Gtf2a2",
                "source": [
                    "Tcf4",
                    "Pou5f1",
                    "Foxp1",
                    "Cebpb",
                    "Nfib",
                    "Atf5",
                    "Atf4",
                    "Junb",
                    "Fos",
                    "Jun",
                    "Sox2",
                    "Sfpq",
                    "Nanog",
                    "Jund",
                    "Twist2",
                    "Hdac2",
                    "Smc3",
                    "Smarca5"
                ],
                "target": [
                    "0610011F06Rik",
                    "Ak1",
                    "Cox8a",
                    "2010111I01Rik"
                ],
                "expression_sum": {
                    "class1": 10627.0,
                    "class2": 28035.0
                },
                "uniprot_ids": [
                    "Q80ZM7",
                    "Q05C74",
                    "Q3TE88",
                    "D3Z7S8",
                    "D3Z793",
                    "D3Z7C2",
                    "D6RIQ7",
                    "Q3UT44",
                    "Q3TN86"
                ]
            },
            "Zyx": {
                "id": "Zyx",
                "type": "Gene",
                "symbol": "Zyx",
                "source": [
                    "E2f4",
                    "Pou5f1",
                    "Trim28",
                    "Rest",
                    "Atf5",
                    "Atf4",
                    "Ctcf",
                    "Trp53",
                    "Hdac1",
                    "Klf5",
                    "Tbx3",
                    "Zfp57",
                    "Hdac2",
                    "Smc3",
                    "Smarca5"
                ],
                "target": [],
                "expression_sum": {
                    "class1": 15342.0,
                    "class2": 2787.0
                },
                "uniprot_ids": [
                    "Q62523",
                    "Q7TQE2",
                    "A0A0N4SVD2",
                    "Q8CD55",
                    "Q3TCR9",
                    "A0A0N4SVP4",
                    "Q8CBM0",
                    "A0A0N4SUX7"
                ]
            },
            "Dnajc21": {
                "id": "Dnajc21",
                "type": "Gene",
                "symbol": "Dnajc21",
                "source": [
                    "Tfcp2l1",
                    "Pou5f1",
                    "Atf5",
                    "Jun",
                    "Sox2",
                    "Hes1",
                    "Nanog",
                    "Jund",
                    "Cebpb"
                ],
                "target": [],
                "expression_sum": {
                    "class1": 2261.0,
                    "class2": 23234.0
                },
                "uniprot_ids": [
                    "E9Q8D0",
                    "Q3URP7"
                ]
            },
            "Map1lc3b": {
                "id": "Map1lc3b",
                "type": "Gene",
                "symbol": "Map1lc3b",
                "source": [
                    "Tcf4",
                    "Pou5f1",
                    "Foxp1",
                    "Atf5",
                    "Junb",
                    "Id2",
                    "Jun",
                    "Sox2",
                    "Hes1",
                    "Nanog",
                    "Twist2",
                    "Cebpb",
                    "Sox4"
                ],
                "target": [],
                "expression_sum": {
                    "class1": 22787.0,
                    "class2": 67758.0
                },
                "uniprot_ids": [
                    "Q9CQV6",
                    "M0QWC2",
                    "A0A1D5RLG5",
                    "M0QWT8",
                    "Q8K161",
                    "Q2TBF9"
                ]
            },
            "Cox6b1": {
                "id": "Cox6b1",
                "type": "Gene",
                "symbol": "Cox6b1",
                "source": [
                    "Tcf4",
                    "Pou5f1",
                    "Foxp1",
                    "Cebpb",
                    "Nfib",
                    "Atf5",
                    "Fos",
                    "Jun",
                    "Sox2",
                    "Trp53",
                    "Nanog",
                    "Jund",
                    "Twist2",
                    "Hdac2",
                    "Smc3"
                ],
                "target": [],
                "expression_sum": {
                    "class1": 57742.0,
                    "class2": 144251.0
                },
                "uniprot_ids": [
                    "P56391",
                    "A0A140LIU3"
                ]
            },
            "Gabarap": {
                "id": "Gabarap",
                "type": "Gene",
                "symbol": "Gabarap",
                "source": [
                    "Pou5f1",
                    "Trim28",
                    "Cebpb",
                    "Rest",
                    "Atf5",
                    "Hmgb1",
                    "Ctcf",
                    "Trp53",
                    "Hdac1",
                    "Klf5",
                    "Zfp57",
                    "Hdac2",
                    "Smc3"
                ],
                "target": [],
                "expression_sum": {
                    "class1": 47937.0,
                    "class2": 30478.0
                },
                "uniprot_ids": [
                    "Q9DCD6",
                    "B1AR50",
                    "A0A1S6GWH0"
                ]
            },
            "1110004F10Rik": {
                "id": "1110004F10Rik",
                "type": "Gene",
                "symbol": "1110004F10Rik",
                "source": [
                    "Atf5",
                    "Fos",
                    "Sox2",
                    "Ewsr1",
                    "Hdac2",
                    "Tcf4",
                    "Pou5f1",
                    "Arf4",
                    "Jun",
                    "Prrx2",
                    "Peg3",
                    "Smarca5",
                    "Sox4",
                    "Eno1",
                    "Prrx1",
                    "Sfpq",
                    "Jund",
                    "Cebpb",
                    "Smc3",
                    "Hmga2",
                    "Ctbp1",
                    "Nono",
                    "Atf4",
                    "Junb",
                    "Ybx1",
                    "Nanog"
                ],
                "target": [],
                "expression_sum": {
                    "class1": 13017.0,
                    "class2": 24968.0
                },
                "uniprot_ids": [
                    "D3YVS0",
                    "D3YVR9",
                    "A0A0U1RPJ7",
                    "A0A0U1RPI0",
                    "D6RI64"
                ]
            },
            "Sox2": {
                "id": "Sox2",
                "type": "TF",
                "symbol": "Sox2",
                "source": [
                    "Pou5f1",
                    "Trim28",
                    "Klf4",
                    "Ctcf",
                    "Hdac1",
                    "Tbx3",
                    "Zfp57",
                    "Nanog",
                    "Smc3"
                ],
                "target": [
                    "Rps5",
                    "Hsp90aa1",
                    "Lsm5",
                    "Arf1",
                    "Dap",
                    "Rpl41",
                    "Rnf7",
                    "Srsf9",
                    "Usmg5",
                    "Ndufb4",
                    "Syce2",
                    "Mcm7",
                    "Cd44",
                    "Rps17",
                    "Rpl6",
                    "Srm",
                    "Grsf1",
                    "Rae1",
                    "Zc3h15",
                    "Cox8a",
                    "Mrps18b",
                    "Ruvbl2",
                    "Cdt1",
                    "Psmb7",
                    "Spr",
                    "Pnkd",
                    "Ppid",
                    "Ier2",
                    "2810004N23Rik",
                    "Clta",
                    "Cox17",
                    "Dkc1",
                    "Marcksl1",
                    "Atp5g1",
                    "1110008F13Rik",
                    "Mrpl4",
                    "Ndufa6",
                    "Polr2k",
                    "Ndufb3",
                    "Gar1",
                    "Tuba1b",
                    "Anapc5",
                    "Cks1b",
                    "Rps27l",
                    "Aurka",
                    "Calcoco2",
                    "Rpl23a",
                    "Psma6",
                    "Gm10076",
                    "Rpl7l1",
                    "Lmna",
                    "Snrpa",
                    "Ftsj3",
                    "St13",
                    "Psmc1",
                    "Hnrnpa1",
                    "Ndufa2",
                    "Tpd52l2",
                    "Hnrnpf",
                    "Knop1",
                    "Lmnb1",
                    "Mlf2",
                    "Rpl9",
                    "Ssrp1",
                    "Pin1",
                    "Atp5j2",
                    "Swi5",
                    "Eif2s1",
                    "Psmb6",
                    "Cks2",
                    "Shfm1",
                    "Hmgb2",
                    "Nubp1",
                    "Cox6c",
                    "Rps15a",
                    "Uba2",
                    "E2f4",
                    "Nup62",
                    "Calm3",
                    "Arf6",
                    "D8Ertd738e",
                    "Hn1",
                    "Nop56",
                    "Snrpe",
                    "B2m",
                    "Mrpl14",
                    "Ubxn1",
                    "Cdk1",
                    "Eif4h",
                    "Phb",
                    "Psmd7",
                    "Anapc15",
                    "Gabarapl2",
                    "Stub1",
                    "Mrpl2",
                    "Taf7",
                    "Tra2b",
                    "Cdc5l",
                    "Ranbp1",
                    "Mrpl28",
                    "Emc6",
                    "Grpel1",
                    "Sms",
                    "Rbx1",
                    "Rps24",
                    "Ctcf",
                    "Ppm1g",
                    "Neat1",
                    "Sec61g",
                    "Ythdc1",
                    "Nanog",
                    "Tbcb",
                    "Sf3b6",
                    "Aurkaip1",
                    "Nhp2",
                    "Srsf1",
                    "Socs3",
                    "Lrrc59",
                    "Uqcrc1",
                    "Tdgf1",
                    "Wdr89",
                    "Exosc5",
                    "Rcc2",
                    "Pdcd6",
                    "Impa2",
                    "Ube2d2a",
                    "Atp5h",
                    "Bola2",
                    "Tmem160",
                    "H2afx",
                    "Kpnb1",
                    "Snrpd1",
                    "Mylpf",
                    "Rbm8a",
                    "Col5a1",
                    "Ppia",
                    "Hnrnpm",
                    "Rpl10",
                    "Eif3k",
                    "Rpl36al",
                    "Lap3",
                    "Ncl",
                    "Mrps34",
                    "Loxl1",
                    "Atp5k",
                    "2310036O22Rik",
                    "Supt4a",
                    "Ppp1ca",
                    "Srsf5",
                    "Rpl35a",
                    "Timp2",
                    "Syngr1",
                    "Chchd1",
                    "Wdr74",
                    "Ywhaz",
                    "Psmc3",
                    "Morc1",
                    "Rps12",
                    "Pomp",
                    "Hspa4",
                    "Rpl36",
                    "Rbbp7",
                    "Psmb3",
                    "Hnrnpl",
                    "Ddx5",
                    "Ndufb9",
                    "Rbm39",
                    "Rps21",
                    "Rps27",
                    "Arpp19",
                    "Ddx21",
                    "Naca",
                    "Nedd8",
                    "Pigyl",
                    "Eef1b2",
                    "Ube2i",
                    "Eif3l",
                    "Utp3",
                    "Atpif1",
                    "Tmem258",
                    "Tcp1",
                    "Rpl24",
                    "Snrnp40",
                    "Brix1",
                    "Fam136a",
                    "Mrpl15",
                    "Siva1",
                    "Fus",
                    "Mybl2",
                    "Pfdn1",
                    "Slc2a3",
                    "Rest",
                    "Rps4x",
                    "Pmm1",
                    "Fabp7",
                    "Actl6a",
                    "Cct3",
                    "Fkbp8",
                    "Nip7",
                    "Wls",
                    "Clpp",
                    "Psma7",
                    "G3bp1",
                    "Gnl3",
                    "Trim27",
                    "Sssca1",
                    "Tpm3",
                    "Atp5b",
                    "Banf1",
                    "Cdc26",
                    "Dtymk",
                    "Immt",
                    "Mrps36",
                    "Psma1",
                    "Lig1",
                    "2410015M20Rik",
                    "Gpx1",
                    "Smc3",
                    "Hdgf",
                    "Prnp",
                    "Id1",
                    "Csnk2b",
                    "Pa2g4",
                    "Thrap3",
                    "Rpl37",
                    "Hspb1",
                    "Kdm5b",
                    "Atf4",
                    "Hprt",
                    "Ppp1cc",
                    "Hells",
                    "Tfcp2l1",
                    "Nucks1",
                    "Pcbp1",
                    "Rpl38",
                    "Dnttip2",
                    "Sae1",
                    "Serpine2",
                    "Hnrnpk",
                    "Mrpl36",
                    "Mrps24",
                    "Ndufa4",
                    "Ewsr1",
                    "Ndufv1",
                    "Ndufv2",
                    "H3f3a",
                    "Nme1",
                    "Ube2k",
                    "Rfc2",
                    "Bccip",
                    "Ddt",
                    "Fbxo22",
                    "Lamtor2",
                    "Tmsb10",
                    "Bax",
                    "Rpl3",
                    "Gfer",
                    "Cebpzos",
                    "Nudt21",
                    "Trim28",
                    "Rpl12",
                    "Anxa2",
                    "Pycard",
                    "Eif3c",
                    "Cox7a2",
                    "Ndufa7",
                    "Dnajc19",
                    "Col5a2",
                    "Cdv3",
                    "Atp6v1f",
                    "Crip2",
                    "Spc24",
                    "Hmgn1",
                    "Ctbp2",
                    "Ube2l3",
                    "Platr3",
                    "Asns",
                    "Ifitm2",
                    "Ppp4c",
                    "Eef1d",
                    "Etf1",
                    "Imp3",
                    "Lsm8",
                    "Ndufs7",
                    "Cenpf",
                    "Mcm2",
                    "Lsm3",
                    "Mrpl13",
                    "Gcat",
                    "Eif5",
                    "Gm16286",
                    "Spag7",
                    "Tomm5",
                    "Txn1",
                    "Erh",
                    "Ak2",
                    "Mrpl17",
                    "Samm50",
                    "Zmat2",
                    "Rrp15",
                    "Atp5j",
                    "Nhp2l1",
                    "Ran",
                    "Fscn1",
                    "Atp5g3",
                    "Srrm1",
                    "Gjb3",
                    "Apex1",
                    "Atp5g2",
                    "Chd4",
                    "Emc8",
                    "Utf1",
                    "Srp19",
                    "Cct5",
                    "Ccnb2",
                    "Mrpl35",
                    "1110004F10Rik",
                    "Mrpl40",
                    "Bclaf1",
                    "Tceb1",
                    "C1qbp",
                    "Rpsa",
                    "H2afz",
                    "Mrps21",
                    "Uqcr10",
                    "Tardbp",
                    "Psip1",
                    "Rpl36a",
                    "Ppp1r11",
                    "Rpl29",
                    "Fubp1",
                    "Snrpb2",
                    "Arpc5l",
                    "Rps29",
                    "Rpl5",
                    "Naa38",
                    "Ccdc58",
                    "Cenpa",
                    "Dph3",
                    "S100a6",
                    "Ptges3",
                    "Ung",
                    "Dpy30",
                    "Wtap",
                    "Odc1",
                    "Mrpl18",
                    "Umps",
                    "Mdk",
                    "Rps20",
                    "Ufc1",
                    "Ybx1",
                    "Ifitm1",
                    "Tmem14c",
                    "Mrpl12",
                    "Cdc37",
                    "Ndufab1",
                    "Psmd1",
                    "Plekhf2",
                    "Rps6",
                    "Snrpf",
                    "Eef1a1",
                    "Cisd1",
                    "Rpl28",
                    "U2af1",
                    "Pole4",
                    "Cenpv",
                    "Atp5c1",
                    "Ywhaq",
                    "Ccnd2",
                    "Fbl",
                    "Ndufv3",
                    "Timm8b",
                    "Ubald2",
                    "Rpl26",
                    "Uqcrb",
                    "Tsc22d1",
                    "Dnajc8",
                    "Rps3",
                    "Atox1",
                    "Cox7c",
                    "Stmn1",
                    "Nme4",
                    "Adss",
                    "Btf3",
                    "Smarcc1",
                    "Srsf10",
                    "Rps7",
                    "Igf2bp1",
                    "Ube2v2",
                    "Uba52",
                    "Rpl18",
                    "Eno1",
                    "Hax1",
                    "Rpl19",
                    "Trnau1ap",
                    "Mt2",
                    "Taldo1",
                    "Timm13",
                    "Mcm3",
                    "Rpl22l1",
                    "Adprh",
                    "Rps3a1",
                    "Mrps5",
                    "Srsf3",
                    "Coprs",
                    "Prc1",
                    "Smc4",
                    "Gadd45gip1",
                    "U2surp",
                    "Mbd3",
                    "Rpl7",
                    "Cct7",
                    "Znrd1",
                    "Dctpp1",
                    "Wdr18",
                    "Ppan",
                    "Hsp90ab1",
                    "Nenf",
                    "Rps9",
                    "Zfp42",
                    "Oaz1",
                    "Mrpl30",
                    "Rpp25",
                    "Stra13",
                    "Sec61b",
                    "Uqcrq",
                    "Rwdd1",
                    "1810058I24Rik",
                    "Snx5",
                    "Ankrd11",
                    "Cyc1",
                    "Got1",
                    "Dnaja1",
                    "Rps8",
                    "Tmem176b",
                    "Trap1a",
                    "Bzw2",
                    "Rpl8",
                    "Eif4a1",
                    "Sumo2",
                    "Pcbp2",
                    "Magoh",
                    "Ndufc2",
                    "Polr1c",
                    "Polr2i",
                    "Snrnp25",
                    "Pou5f1",
                    "Nudc",
                    "H3f3b",
                    "Dnajc2",
                    "Gnb2l1",
                    "Mrps33",
                    "Mrpl21",
                    "Ndufa12",
                    "Rif1",
                    "Nsa2",
                    "Pla2g1b",
                    "Wdr5",
                    "Ddx46",
                    "Nudt19",
                    "Pebp1",
                    "Sgk1",
                    "Ube2d3",
                    "Aars",
                    "Rfc5",
                    "Rbbp4",
                    "Snrpa1",
                    "Gm10073",
                    "Dstn",
                    "Paip2",
                    "Emp3",
                    "Tbx3",
                    "Srrm2",
                    "2700060E02Rik",
                    "Rnaseh2b",
                    "Mt1",
                    "Ywhae",
                    "Gm26917",
                    "Ubl5",
                    "Hmgb1",
                    "Anxa3",
                    "Mrps15",
                    "Ndufc1",
                    "Ppil1",
                    "Eif3d",
                    "Hbegf",
                    "Cyb5a",
                    "Npm1",
                    "Psmc4",
                    "Psma4",
                    "Rps15",
                    "Rpl34",
                    "Park7",
                    "Dynll1",
                    "Gtpbp4",
                    "Cenpw",
                    "Eif2s2",
                    "Hspd1",
                    "Stoml2",
                    "Rpl31",
                    "Mrps14",
                    "Srsf7",
                    "Sod1",
                    "Ppih",
                    "Mat2a",
                    "Nap1l1",
                    "Uqcc3",
                    "Csrp2",
                    "Dnajb6",
                    "Brd2",
                    "G3bp2",
                    "Efemp2",
                    "Impdh2",
                    "Ccdc124",
                    "Gapdh",
                    "Smarca5",
                    "Ccnb1",
                    "Akap12",
                    "Ccnd3",
                    "Mrpl52",
                    "Pno1",
                    "Adrm1",
                    "Abcf1",
                    "Hnrnpab",
                    "Prkcdbp",
                    "Lgals1",
                    "Rplp0",
                    "Psmb5",
                    "Hspa9",
                    "Fkbp10",
                    "Chchd2",
                    "Pim3",
                    "Nop58",
                    "Ube2n",
                    "Ndufa11",
                    "E130012A19Rik",
                    "Pam16",
                    "Smn1",
                    "Cyb5b",
                    "Luc7l3",
                    "Polr2c",
                    "Cct8",
                    "Hmgn2",
                    "Dohh",
                    "Hnrnpa2b1",
                    "Ndufs5",
                    "Gsn",
                    "Psmb1",
                    "Coa3",
                    "Akr1b3",
                    "Ddx27",
                    "Ndufb8",
                    "Srsf2",
                    "Eif4g2",
                    "Jade1",
                    "Psmb2",
                    "Raly",
                    "Denr",
                    "Fxr1",
                    "Nsmce4a",
                    "Rps26",
                    "Get4",
                    "Eif3g",
                    "Trp53",
                    "Lhfp",
                    "Rbm3",
                    "Aspscr1",
                    "Psmd11",
                    "Vdac1",
                    "Rpp21",
                    "Cggbp1",
                    "Npm3",
                    "Anp32a",
                    "Zfp706",
                    "Polr2e",
                    "Lsm2",
                    "Anapc13",
                    "Sumo1",
                    "Tra2a",
                    "Nudcd2",
                    "Pdcd5",
                    "Aprt",
                    "Ost4",
                    "Supt16",
                    "Tomm7",
                    "Snrpc",
                    "Zfp57",
                    "Mrpl51",
                    "Mrpl49",
                    "Eif4g1",
                    "Fcf1",
                    "Ssu72",
                    "Mrps18c",
                    "Atp5a1",
                    "Cacybp",
                    "Psma5",
                    "Cdk4",
                    "Rpl22",
                    "Rsl1d1",
                    "Nolc1",
                    "Pycr2",
                    "Sf3a3",
                    "Syncrip",
                    "Caprin1",
                    "Pfdn6",
                    "Ssr2",
                    "Rbmxl2",
                    "Glud1",
                    "Rpl4",
                    "Rps28",
                    "Cebpz",
                    "Rps25",
                    "Ddah2",
                    "Rbmxl1",
                    "Gspt1",
                    "Arglu1",
                    "Imp4",
                    "Rpl18a",
                    "Tomm20",
                    "Cbx1",
                    "Tomm70a",
                    "Rpl21",
                    "Map1lc3b",
                    "Eprs",
                    "Trmt112",
                    "Rpl35",
                    "Dazap1",
                    "Atp5l",
                    "Fam25c",
                    "Pop5",
                    "Son",
                    "Ilf2",
                    "Cct2",
                    "Mak16",
                    "Acp1",
                    "Prelid1",
                    "Ubb",
                    "Ahsa1",
                    "Zfp106",
                    "Prrc1",
                    "Parp1",
                    "Dhfr",
                    "Rbpms",
                    "Paics",
                    "Hnrnpa3",
                    "Prpf40a",
                    "Prmt1",
                    "Pbdc1",
                    "Rps19bp1",
                    "Tcea3",
                    "Cdca3",
                    "Metap2",
                    "Sub1",
                    "Asna1",
                    "Mrpl57",
                    "Psmc5",
                    "Echdc2",
                    "2010107E04Rik",
                    "Morf4l1",
                    "Cdca8",
                    "L1td1",
                    "Lsm6",
                    "Sf3b1",
                    "Tecr",
                    "Ube2s",
                    "Idh2",
                    "Rpl27",
                    "Ndufs4",
                    "Polr1d",
                    "Mybbp1a",
                    "Snrpd3",
                    "Sf3b2",
                    "Psmd3",
                    "Suclg1",
                    "H2afy",
                    "Mtf2",
                    "Rpl17",
                    "Stip1",
                    "mt-Nd2",
                    "Hnrnpd",
                    "Chchd4",
                    "Bcas2",
                    "Rpl14",
                    "Ash2l",
                    "Tma7",
                    "Camk2d",
                    "Txndc17",
                    "Cops4",
                    "Rnps1",
                    "Ciapin1",
                    "Lsm4",
                    "Tpd52",
                    "Pfdn2",
                    "Nop10",
                    "Atp5f1",
                    "Eif1ad",
                    "Arid5b",
                    "Gdf3",
                    "Rps13",
                    "Tpt1",
                    "Eif5a",
                    "Rps11",
                    "Cfl1",
                    "Gng5",
                    "Rrp1b",
                    "Sparc",
                    "Uchl3",
                    "Hspe1",
                    "Rps19",
                    "Stmn2",
                    "Ube2c",
                    "Exosc8",
                    "Ptbp1",
                    "Mrpl20",
                    "Srp9",
                    "Hnrnpdl",
                    "Nfatc2ip",
                    "Eif3a",
                    "Dnajc21",
                    "Polr2h",
                    "Psmb4",
                    "Birc5",
                    "Snx3",
                    "Commd1",
                    "Psmd13",
                    "Cox7b",
                    "Strap",
                    "l7Rn6",
                    "Rnaseh2a",
                    "Dppa5a",
                    "Srpk1",
                    "Klf4",
                    "Rfc1",
                    "Cox6b1",
                    "Ino80e",
                    "Sfrp1",
                    "Ppa1",
                    "Rcor2",
                    "Lamtor5",
                    "Atp5e",
                    "Cenpm",
                    "Prdx1",
                    "Ptrf",
                    "Rsl24d1",
                    "Ssb",
                    "Cdc34",
                    "Tipin",
                    "Atp5o",
                    "Mrps12",
                    "Terf1",
                    "Rpl13",
                    "Copz1",
                    "Sh3bgrl",
                    "Nfu1",
                    "Mrpl34",
                    "Nasp",
                    "Rplp1",
                    "Rnf10",
                    "Ndufb5",
                    "Rpl10a",
                    "Atrx",
                    "Psma2",
                    "Ndufs6",
                    "Ndufb7",
                    "Nsun2",
                    "Set",
                    "Thyn1",
                    "Txnl4a",
                    "Poldip2",
                    "Eif4e2",
                    "Shmt1",
                    "Alkbh1",
                    "Sod2",
                    "Slc25a39",
                    "Ssbp1",
                    "Rps14",
                    "Tgif1",
                    "Sfpq",
                    "Rpl32",
                    "Parl",
                    "Tnfrsf12a",
                    "Sephs2",
                    "Clns1a",
                    "Mdh2",
                    "Snrpd2",
                    "1810022K09Rik",
                    "Chchd10",
                    "Uqcr11",
                    "Phb2",
                    "Hdac1",
                    "Rps27a",
                    "Naa10",
                    "Bud31",
                    "Gtf2e2",
                    "Pdap1",
                    "Mcm5",
                    "Gtf2h5",
                    "Rpl11",
                    "Rpl15",
                    "Anxa1",
                    "Tcof1",
                    "Vdac2",
                    "Ube2a",
                    "Pfn1",
                    "Rpl13a",
                    "Snrpg",
                    "Tkt",
                    "Ddx39b",
                    "Snhg6",
                    "Eif1a",
                    "Cox5a",
                    "Minos1",
                    "Gsta4",
                    "Snrpb",
                    "Tmem11",
                    "Anapc11",
                    "Exosc4",
                    "Fkbp4",
                    "Pgam1",
                    "Spint2",
                    "Tomm22",
                    "Lyar",
                    "Snrnp70",
                    "Gpx8",
                    "Ppig",
                    "Cfdp1",
                    "Ppdpf",
                    "Gmnn",
                    "Rpl27a",
                    "Dcun1d5",
                    "Eif5b",
                    "Cct4",
                    "Llph",
                    "Emg1",
                    "Igfbp7",
                    "Lsm12",
                    "Eif4ebp1",
                    "Anp32b",
                    "Edf1",
                    "Pkm",
                    "H2afv",
                    "Nono",
                    "Csde1",
                    "Prrc2c",
                    "Gins4",
                    "Fstl1",
                    "Tceb2",
                    "Rhoc",
                    "Mrpl42",
                    "Psmg4",
                    "Rps2",
                    "Eif2b5",
                    "Gart",
                    "Polr2g",
                    "Sumo3",
                    "Psmc2",
                    "Tomm40",
                    "App",
                    "Mcm6",
                    "Uqcrfs1",
                    "Naa50",
                    "Sf3b4",
                    "Eif3i",
                    "Prpf19",
                    "Hdac2",
                    "Sugt1",
                    "Hnrnpc",
                    "Smc2",
                    "Eif4a3",
                    "Fkbp3",
                    "Mtch2",
                    "Dek",
                    "Rangrf",
                    "Psmd4",
                    "Gins2",
                    "Timm17a",
                    "Top2a",
                    "Rpl7a",
                    "Fam104a",
                    "Uck2",
                    "Papola",
                    "Bcat1",
                    "Rps16",
                    "Tmed3",
                    "Mrfap1",
                    "Tubb4b",
                    "Ccna2",
                    "Mrps16",
                    "Hsd17b14",
                    "Slc25a5",
                    "Uqcrh",
                    "Tpr",
                    "Ddx18",
                    "Lefty2",
                    "Hint1",
                    "Psmd12",
                    "Eed",
                    "Rrp9",
                    "Ccne1",
                    "Sf3a2",
                    "Srsf6",
                    "Ndufa5",
                    "Ebna1bp2",
                    "Ndufb10",
                    "Tsn",
                    "Mrpl11",
                    "Rpl37a",
                    "Rbm25",
                    "Eef1g",
                    "Cox5b",
                    "2700094K13Rik",
                    "Cycs",
                    "Drg1",
                    "Ndufb6",
                    "Ptma",
                    "Klf2",
                    "Acadm",
                    "Dut",
                    "Ralbp1",
                    "Dhx16",
                    "Sdhb",
                    "Taf10",
                    "Fau",
                    "Wdr43",
                    "Ooep",
                    "Klf5",
                    "Acin1",
                    "Rps10",
                    "Pabpc1",
                    "Eif6",
                    "Alyref",
                    "Yrdc",
                    "Hspa14",
                    "Polr2j",
                    "Dad1",
                    "Slirp",
                    "Sept1",
                    "0610007P14Rik",
                    "Rbpms2",
                    "Uhrf1",
                    "Gm7325",
                    "Fam60a",
                    "Wbp11",
                    "Snf8",
                    "Cwc15",
                    "Guk1",
                    "Hspa8",
                    "Ndufs8",
                    "Psma3",
                    "Rpl39",
                    "Rplp2",
                    "Skp1a",
                    "Capn2",
                    "Rps18",
                    "Mif",
                    "Ptov1",
                    "Aimp1",
                    "Rps27rt",
                    "Serbp1",
                    "Gpx4",
                    "Gcsh",
                    "Phf5a",
                    "Rps23",
                    "Glrx3",
                    "Taf1d",
                    "Gtf2a2",
                    "Alpl",
                    "Gltscr2",
                    "Mkrn1",
                    "U2af2",
                    "Rpl23",
                    "Rpl30",
                    "Sfr1",
                    "Polr2f",
                    "Myl12b",
                    "Kdm1a",
                    "Rrs1",
                    "Ncor1",
                    "1810037I17Rik",
                    "Rbm17",
                    "Ly6e",
                    "Cnbp"
                ],
                "expression_sum": {
                    "class1": 55.0,
                    "class2": 44636.0
                },
                "uniprot_ids": [
                    "P48432",
                    "Q60I23",
                    "Q3V2B6",
                    "Q3UES8"
                ]
            },
            "Jund": {
                "id": "Jund",
                "type": "TF",
                "symbol": "Jund",
                "source": [
                    "Trim28",
                    "Ctcf"
                ],
                "target": [
                    "Hsp90aa1",
                    "Lsm5",
                    "Arf1",
                    "Rnf7",
                    "Cdc20",
                    "Srsf9",
                    "Cd9",
                    "Mdh1",
                    "Ndufb4",
                    "Usmg5",
                    "Mcm7",
                    "Ppp2ca",
                    "Ddx24",
                    "Rps17",
                    "Srm",
                    "Grsf1",
                    "Rae1",
                    "Zc3h15",
                    "Cox8a",
                    "Mrps18b",
                    "Ruvbl2",
                    "Psmb7",
                    "Spr",
                    "Ppid",
                    "2810004N23Rik",
                    "Clta",
                    "Dkc1",
                    "Marcksl1",
                    "Ppp1r11",
                    "Atp5g1",
                    "1110008F13Rik",
                    "Sec13",
                    "Polr2k",
                    "Tuba1b",
                    "Ndufb3",
                    "Gar1",
                    "Idh3g",
                    "Anapc5",
                    "Cks1b",
                    "Psma6",
                    "Rpl7l1",
                    "Snrpa",
                    "St13",
                    "Ftsj3",
                    "Psmc1",
                    "Hnrnpa1",
                    "Thoc7",
                    "Tpd52l2",
                    "Hnrnpf",
                    "Knop1",
                    "Lmnb1",
                    "Ddx39",
                    "Psmd6",
                    "Vbp1",
                    "Ssrp1",
                    "Pin1",
                    "Atp5j2",
                    "Rnf187",
                    "Sfxn1",
                    "Swi5",
                    "Eif2s1",
                    "Psmb6",
                    "Cks2",
                    "Hmgb2",
                    "Nubp1",
                    "Cox6c",
                    "Ddx3x",
                    "Slbp",
                    "Ndufs3",
                    "Nedd4",
                    "Uba2",
                    "E2f4",
                    "Calr",
                    "Nup62",
                    "Tubb5",
                    "Calm3",
                    "Canx",
                    "D8Ertd738e",
                    "Nop56",
                    "Snrpe",
                    "Ubxn1",
                    "Cdk1",
                    "Eif4h",
                    "Phb",
                    "2810417H13Rik",
                    "Psat1",
                    "Psmd7",
                    "Anapc15",
                    "Gdi2",
                    "Gabarapl2",
                    "Stub1",
                    "Mrpl2",
                    "Nup43",
                    "Tra2b",
                    "Cdc5l",
                    "Ranbp1",
                    "Mrpl28",
                    "Psmc6",
                    "Hmgcs1",
                    "Emc6",
                    "Grpel1",
                    "Sms",
                    "Zwint",
                    "Rbx1",
                    "Rps24",
                    "Ctcf",
                    "Ppm1g",
                    "Neat1",
                    "Ythdc1",
                    "Bola1",
                    "Ola1",
                    "Sf3b6",
                    "Snrnp27",
                    "Tbcb",
                    "Aurkaip1",
                    "Nhp2",
                    "Srsf1",
                    "Wdr89",
                    "Lrrc59",
                    "Uqcrc1",
                    "Exosc5",
                    "Rcc2",
                    "Pdcd6",
                    "Ube2d2a",
                    "Atp5h",
                    "H2afx",
                    "Kpnb1",
                    "Mapre1",
                    "C1d",
                    "Cisd2",
                    "Snrpd1",
                    "Rbm8a",
                    "Ppia",
                    "Hnrnpm",
                    "Capza2",
                    "Eif3k",
                    "Rpl36al",
                    "Lap3",
                    "Ncl",
                    "Atp5k",
                    "2310036O22Rik",
                    "Fdft1",
                    "Mphosph8",
                    "Ppp1ca",
                    "Srsf5",
                    "Supt4a",
                    "Hnrnph1",
                    "Aamp",
                    "Rpl35a",
                    "Chchd1",
                    "Wdr74",
                    "Ywhaz",
                    "Gars",
                    "Psmc3",
                    "Pomp",
                    "Hspa4",
                    "Luc7l2",
                    "Rpl36",
                    "Rbbp7",
                    "Psmb3",
                    "Hnrnpl",
                    "Ddx5",
                    "Rbm39",
                    "Rps21",
                    "Arpp19",
                    "Ddx21",
                    "Nedd8",
                    "Pcnp",
                    "Ube2i",
                    "Eif3l",
                    "Utp3",
                    "Atpif1",
                    "Tcp1",
                    "Snrnp40",
                    "Brix1",
                    "Sarnp",
                    "Fam136a",
                    "Pttg1",
                    "Mrpl15",
                    "Siva1",
                    "Fus",
                    "Ezh2",
                    "Ociad1",
                    "Pfdn1",
                    "Ddost",
                    "Rest",
                    "Actl6a",
                    "Cct3",
                    "Nip7",
                    "Ruvbl1",
                    "Clpp",
                    "Psma7",
                    "Ndufa8",
                    "G3bp1",
                    "Pdia4",
                    "Sept7",
                    "Gnl3",
                    "Trim27",
                    "Sssca1",
                    "Tpm3",
                    "Atp5b",
                    "Banf1",
                    "Cdc26",
                    "Dtymk",
                    "Immt",
                    "Mrps36",
                    "Psma1",
                    "Lig1",
                    "Tsg101",
                    "Gpx1",
                    "Hdgf",
                    "Smc3",
                    "Tpi1",
                    "Thrap3",
                    "Csnk2b",
                    "Nars",
                    "Pa2g4",
                    "Rpl37",
                    "Ppp1cc",
                    "Atf4",
                    "Hist1h2bc",
                    "Hprt",
                    "Nucks1",
                    "Eif3b",
                    "Zfp207",
                    "Pcbp1",
                    "Dnttip2",
                    "Glo1",
                    "Sae1",
                    "Hnrnpk",
                    "Mrpl36",
                    "Mrps24",
                    "Ndufa4",
                    "Ewsr1",
                    "Cops5",
                    "Eif4e",
                    "Ndufv1",
                    "Ndufv2",
                    "H3f3a",
                    "Nme1",
                    "Ube2k",
                    "Rfc2",
                    "Bccip",
                    "Csnk1a1",
                    "Ndufs2",
                    "Ddt",
                    "Cyp51",
                    "Lamtor2",
                    "Fam49b",
                    "Rpl3",
                    "Gfer",
                    "Cebpzos",
                    "Nudt21",
                    "Trim28",
                    "Rpl12",
                    "Eif3c",
                    "Snw1",
                    "Cox7a2",
                    "Ndufa7",
                    "Dnajc19",
                    "Cdv3",
                    "mt-Nd4",
                    "Spc24",
                    "Hmgn1",
                    "Ctbp2",
                    "Btf3l4",
                    "Ube2l3",
                    "Asns",
                    "Ifitm2",
                    "Fdps",
                    "Ppp4c",
                    "Eef1d",
                    "Etf1",
                    "Imp3",
                    "Lsm8",
                    "Ndufs7",
                    "Lsm3",
                    "Mrpl13",
                    "Gm16286",
                    "Eif5",
                    "Tomm5",
                    "Txn1",
                    "Ube2j2",
                    "Vapa",
                    "Erh",
                    "Ak2",
                    "Mrpl17",
                    "Samm50",
                    "Zmat2",
                    "Rrp15",
                    "Wdr77",
                    "Atp5j",
                    "Nhp2l1",
                    "Ran",
                    "Fscn1",
                    "Atp5g3",
                    "Srrm1",
                    "Ldha",
                    "Apex1",
                    "Atp5g2",
                    "Chd4",
                    "Emc8",
                    "Srp19",
                    "Cct5",
                    "Emd",
                    "Mrpl35",
                    "1110004F10Rik",
                    "Mrpl40",
                    "Bclaf1",
                    "Tceb1",
                    "C1qbp",
                    "Rpsa",
                    "H2afz",
                    "Uqcr10",
                    "Tardbp",
                    "Psip1",
                    "Eif3m",
                    "Rpl29",
                    "Fubp1",
                    "Snrpb2",
                    "Rps29",
                    "Rpl5",
                    "Naa38",
                    "Ccdc58",
                    "Dph3",
                    "Prdx6",
                    "Ptges3",
                    "Dpy30",
                    "Wtap",
                    "Odc1",
                    "Mrpl18",
                    "Umps",
                    "Rps20",
                    "Orc6",
                    "Dnaja2",
                    "Ybx1",
                    "Sys1",
                    "Tubb2b",
                    "Acat2",
                    "Tmem14c",
                    "Mrpl12",
                    "Cdc37",
                    "Ndufab1",
                    "Psmd1",
                    "Plekhf2",
                    "Rps6",
                    "Snrpf",
                    "mt-Nd5",
                    "Mcl1",
                    "Cisd1",
                    "U2af1",
                    "Pole4",
                    "Atp5c1",
                    "Mrps17",
                    "Ywhaq",
                    "Fbl",
                    "Timm8b",
                    "Ubald2",
                    "Rpl26",
                    "Uqcrb",
                    "Dnajc8",
                    "Calm2",
                    "Rps3",
                    "Cox7c",
                    "Stmn1",
                    "Srsf10",
                    "Adss",
                    "Btf3",
                    "Smarcc1",
                    "Tln1",
                    "Rps7",
                    "Ube2v2",
                    "Rpl18",
                    "Eno1",
                    "Hax1",
                    "Eef1e1",
                    "mt-Nd3",
                    "Taldo1",
                    "Timm13",
                    "Adprh",
                    "Vcp",
                    "Srsf3",
                    "Mrps5",
                    "Smc4",
                    "U2surp",
                    "Mbd3",
                    "Rpl7",
                    "Fkbp2",
                    "Cct7",
                    "Znrd1",
                    "Dctpp1",
                    "Wdr18",
                    "Ppan",
                    "Hsp90ab1",
                    "Oaz1",
                    "Stra13",
                    "Sec61b",
                    "Polr2m",
                    "Rwdd1",
                    "Snx5",
                    "Tyms",
                    "Cyc1",
                    "Fnta",
                    "Dnaja1",
                    "Got1",
                    "Rps8",
                    "Bzw2",
                    "Rpl8",
                    "Eif4a1",
                    "Sumo2",
                    "Pcbp2",
                    "Magoh",
                    "Ndufc2",
                    "Polr1c",
                    "Polr2i",
                    "Snrnp25",
                    "Nudc",
                    "Pmvk",
                    "H3f3b",
                    "Dnajc2",
                    "Ccdc34",
                    "Elof1",
                    "Gnb2l1",
                    "Mrpl21",
                    "Mrps33",
                    "Ndufa12",
                    "Pold2",
                    "Oxct1",
                    "Lrpap1",
                    "Ddx46",
                    "Cnih4",
                    "Nudt19",
                    "Pebp1",
                    "Sgk1",
                    "Dbi",
                    "Morf4l2",
                    "Ube2d3",
                    "Aars",
                    "Rfc5",
                    "Rbbp4",
                    "Marc2",
                    "Snrpa1",
                    "Paip2",
                    "Dstn",
                    "Pfdn4",
                    "Srrm2",
                    "2700060E02Rik",
                    "Rnaseh2b",
                    "Top1",
                    "Ywhae",
                    "Hmgb1",
                    "Mrps15",
                    "Psmc4",
                    "Ndufc1",
                    "Ppil1",
                    "Eif3d",
                    "Psmd14",
                    "Cyb5a",
                    "Npm1",
                    "Rps15",
                    "Psma4",
                    "Scp2",
                    "Arl1",
                    "Park7",
                    "Mlec",
                    "Dynll1",
                    "Gtpbp4",
                    "Cenpw",
                    "Eif2s2",
                    "Hspd1",
                    "Stoml2",
                    "Cops7a",
                    "Mrpl45",
                    "Mrps14",
                    "Rpl31",
                    "Srsf7",
                    "Smdt1",
                    "Sod1",
                    "Ppih",
                    "Mat2a",
                    "Nap1l1",
                    "Mrps25",
                    "Tpx2",
                    "Etfb",
                    "Dnajb6",
                    "Hspa5",
                    "Uqcrc2",
                    "G3bp2",
                    "Impdh2",
                    "Ccdc124",
                    "Mrps10",
                    "Gapdh",
                    "Smarca5",
                    "Idi1",
                    "Ccnb1",
                    "Mpc1",
                    "Prps1",
                    "Ccnd3",
                    "Khdrbs1",
                    "Pno1",
                    "Abcf1",
                    "Hnrnpab",
                    "Psmb5",
                    "Tbrg1",
                    "Hspa9",
                    "Chchd2",
                    "Pim3",
                    "Nop58",
                    "Ube2n",
                    "Ndufa11",
                    "Smn1",
                    "Cbx5",
                    "Cyb5b",
                    "Kif5b",
                    "Luc7l3",
                    "Polr2c",
                    "Cct8",
                    "Hmgn2",
                    "Dohh",
                    "Hnrnpa2b1",
                    "Ndufs5",
                    "Oat",
                    "Coa3",
                    "Akr1b3",
                    "Ddx27",
                    "Ndufb8",
                    "Psmd2",
                    "Tmem97",
                    "Srsf2",
                    "mt-Nd1",
                    "Eif4g2",
                    "Dnajc7",
                    "Psmb2",
                    "Raly",
                    "Uqcc2",
                    "Pak1ip1",
                    "Denr",
                    "Fxr1",
                    "Nsmce4a",
                    "Get4",
                    "Eif3g",
                    "Nxt1",
                    "Trp53",
                    "Tceal8",
                    "Rbm3",
                    "Aspscr1",
                    "Psmd11",
                    "Cnn3",
                    "Btg2",
                    "Vdac1",
                    "Rpp21",
                    "Cggbp1",
                    "Npm3",
                    "Anp32a",
                    "Rad23b",
                    "Zfp706",
                    "Polr2e",
                    "Lsm2",
                    "Anapc13",
                    "Sumo1",
                    "Tra2a",
                    "Nudcd2",
                    "Pdcd5",
                    "Aprt",
                    "Ost4",
                    "Supt16",
                    "Tomm7",
                    "Snrpc",
                    "Mrpl51",
                    "Mrpl49",
                    "Eif4g1",
                    "Tmbim6",
                    "Dynll2",
                    "Rhoa",
                    "Fcf1",
                    "Ssu72",
                    "Mrps18c",
                    "Atp5a1",
                    "Cacybp",
                    "Psma5",
                    "Ttc3",
                    "Cdk4",
                    "Rpl22",
                    "Rsl1d1",
                    "Nolc1",
                    "Pycr2",
                    "Sf3a3",
                    "Anp32e",
                    "Syncrip",
                    "Caprin1",
                    "Pfdn6",
                    "Ict1",
                    "Rpl4",
                    "Glud1",
                    "Ssr2",
                    "Cebpz",
                    "Hnrnpu",
                    "Ddah2",
                    "Rbmxl1",
                    "Gspt1",
                    "Arglu1",
                    "Imp4",
                    "Srsf11",
                    "Hmgb3",
                    "Cbx1",
                    "Tomm20",
                    "Tomm70a",
                    "Rpl21",
                    "Ppp1r2",
                    "Eprs",
                    "Trmt112",
                    "Lman1",
                    "Dazap1",
                    "Msmo1",
                    "Son",
                    "Pop5",
                    "Ilf2",
                    "Cct2",
                    "Mak16",
                    "Acp1",
                    "Ubb",
                    "Dtl",
                    "Ahsa1",
                    "Zfp106",
                    "Prrc1",
                    "Eif1ax",
                    "Bzw1",
                    "Parp1",
                    "Ak6",
                    "Dhfr",
                    "Lman2",
                    "Paics",
                    "Hnrnpa3",
                    "Prpf40a",
                    "Prmt1",
                    "Pbdc1",
                    "Rps19bp1",
                    "Mrps7",
                    "Cdca3",
                    "Metap2",
                    "Sub1",
                    "Asna1",
                    "Psmc5",
                    "Morf4l1",
                    "2010107E04Rik",
                    "Mrpl43",
                    "Pafah1b1",
                    "Cdca8",
                    "Lsm6",
                    "Tecr",
                    "Sf3b1",
                    "Ube2s",
                    "Idh2",
                    "Ndufs4",
                    "Polr1d",
                    "Mybbp1a",
                    "Snrpd3",
                    "Sf3b2",
                    "Psmd3",
                    "Suclg1",
                    "H2afy",
                    "Mtf2",
                    "Rpl17",
                    "Stip1",
                    "mt-Nd2",
                    "Hnrnpd",
                    "Bcas2",
                    "Rpl14",
                    "Ash2l",
                    "Tma7",
                    "Hat1",
                    "Cops4",
                    "mt-Cytb",
                    "Rnps1",
                    "Ciapin1",
                    "Lsm4",
                    "Pfdn2",
                    "Nop10",
                    "Mpdu1",
                    "Atp5f1",
                    "Eif1ad",
                    "Rps13",
                    "Eif5a",
                    "Rps11",
                    "Cfl1",
                    "Rrp1b",
                    "Uchl3",
                    "Hspe1",
                    "Exosc8",
                    "Cops6",
                    "Ptbp1",
                    "Aldoa",
                    "Mrpl20",
                    "Hnrnpdl",
                    "Srp9",
                    "Eif3a",
                    "Rtn3",
                    "Dnajc21",
                    "Polr2h",
                    "Psmb4",
                    "Adh5",
                    "Birc5",
                    "Commd1",
                    "Psmd13",
                    "Cox7b",
                    "Snx3",
                    "Strap",
                    "l7Rn6",
                    "Rnaseh2a",
                    "Chtop",
                    "Srpk1",
                    "Rfc1",
                    "Cox6b1",
                    "Ino80e",
                    "Ppa1",
                    "Lamtor5",
                    "Ssb",
                    "Prdx1",
                    "Rsl24d1",
                    "Cdc34",
                    "Gps1",
                    "Tipin",
                    "Atp5o",
                    "Rpa2",
                    "Mrps12",
                    "Rrm2",
                    "Terf1",
                    "Copz1",
                    "Nfu1",
                    "Mrpl34",
                    "Nasp",
                    "Pdia3",
                    "Rnf10",
                    "Ndufb5",
                    "Atrx",
                    "Psma2",
                    "Ndufs6",
                    "Ndufb7",
                    "Nsun2",
                    "Set",
                    "Thyn1",
                    "Txnl4a",
                    "Poldip2",
                    "Eif4e2",
                    "Sod2",
                    "Alkbh1",
                    "Ssbp1",
                    "Nt5dc2",
                    "Tgif1",
                    "Sfpq",
                    "Mrto4",
                    "Hist1h1c",
                    "Parl",
                    "Lipe",
                    "Clns1a",
                    "Mdh2",
                    "Snrpd2",
                    "1810022K09Rik",
                    "Uqcr11",
                    "Phb2",
                    "Hdac1",
                    "Phgdh",
                    "Naa10",
                    "Bud31",
                    "Gtf2e2",
                    "Pdap1",
                    "Mcm5",
                    "Gtf2h5",
                    "Hsp90b1",
                    "Rpl11",
                    "Tcof1",
                    "Vdac2",
                    "Ube2a",
                    "Nol7",
                    "Snrpg",
                    "Tkt",
                    "Ddx39b",
                    "Mrpl24",
                    "Eif1a",
                    "Cox5a",
                    "Snrpb",
                    "Tmem11",
                    "Anapc11",
                    "Exosc4",
                    "Fkbp4",
                    "Pgam1",
                    "Zfp428",
                    "Tomm22",
                    "Nans",
                    "Lyar",
                    "Uchl5",
                    "Erp29",
                    "Snrnp70",
                    "Ppig",
                    "Cfdp1",
                    "Gmnn",
                    "Rpl27a",
                    "Dcun1d5",
                    "Eif5b",
                    "Cct4",
                    "Llph",
                    "Rdx",
                    "Emg1",
                    "Lsm12",
                    "Eif4b",
                    "Eif4ebp1",
                    "Anp32b",
                    "Edf1",
                    "Cdkn1a",
                    "Pkm",
                    "H2afv",
                    "Nono",
                    "Csde1",
                    "Prrc2c",
                    "Gins4",
                    "Tceb2",
                    "Mrpl42",
                    "Rps2",
                    "Eif2b5",
                    "Gart",
                    "Polr2g",
                    "Sumo3",
                    "Psmc2",
                    "Ndufa10",
                    "Mcm6",
                    "Tomm40",
                    "Uqcrfs1",
                    "Naa50",
                    "Ywhah",
                    "Sf3b4",
                    "Eif3i",
                    "Prpf19",
                    "Hdac2",
                    "Sugt1",
                    "Hnrnpc",
                    "Smc2",
                    "Eif4a3",
                    "Fkbp3",
                    "Mtch2",
                    "Dek",
                    "Timm17a",
                    "Psmd4",
                    "Gins2",
                    "Rpn2",
                    "Bub3",
                    "Top2a",
                    "Rpl7a",
                    "Papola",
                    "Bcat1",
                    "Manf",
                    "Esf1",
                    "Sin3b",
                    "Cetn3",
                    "Mrfap1",
                    "Tubb4b",
                    "Ccna2",
                    "Mrps16",
                    "Slc25a5",
                    "Tpr",
                    "Ddx18",
                    "Psmd12",
                    "Hint1",
                    "Eed",
                    "Ccne1",
                    "Srsf6",
                    "Ndufa5",
                    "Ebna1bp2",
                    "Ndufb10",
                    "Tmpo",
                    "Tsn",
                    "Mrpl11",
                    "Rpl37a",
                    "Txnl1",
                    "Mesdc2",
                    "Eef1g",
                    "Rbm25",
                    "Ube2e3",
                    "Cox5b",
                    "2700094K13Rik",
                    "Arhgdia",
                    "Cycs",
                    "Drg1",
                    "Myl12a",
                    "Ndufb6",
                    "Ptma",
                    "Rer1",
                    "Acadm",
                    "Dut",
                    "Ralbp1",
                    "Sdhb",
                    "Fau",
                    "Klf5",
                    "Acin1",
                    "Rps10",
                    "Pabpc1",
                    "Eif6",
                    "Alyref",
                    "Yrdc",
                    "Tsen34",
                    "Hspa14",
                    "Polr2j",
                    "Got2",
                    "Dad1",
                    "Slirp",
                    "0610007P14Rik",
                    "Wbp11",
                    "Cwc15",
                    "Guk1",
                    "Hspa8",
                    "Ndufs8",
                    "Psma3",
                    "Tsfm",
                    "Rplp2",
                    "Skp1a",
                    "Rps18",
                    "Mif",
                    "Ptov1",
                    "Aimp1",
                    "Serbp1",
                    "Taf1d",
                    "Gpx4",
                    "Chchd3",
                    "Gcsh",
                    "Eif4a2",
                    "Phf5a",
                    "Rheb",
                    "Rps23",
                    "Gtf2a2",
                    "Gltscr2",
                    "Mkrn1",
                    "Rrp1",
                    "Pdcl3",
                    "U2af2",
                    "Rpl30",
                    "Sfr1",
                    "Polr2f",
                    "Myl12b",
                    "Kdm1a",
                    "Rrs1",
                    "Ncor1",
                    "1810037I17Rik",
                    "Rbm17",
                    "Ly6e",
                    "Bcap31",
                    "Cnbp",
                    "Sdhd",
                    "Rsrc2"
                ],
                "expression_sum": {
                    "class1": 9894.0,
                    "class2": 25816.0
                },
                "uniprot_ids": [
                    "P15066",
                    "Q3UXY7",
                    "Q52L49"
                ]
            },
            "Jun": {
                "id": "Jun",
                "type": "TF",
                "symbol": "Jun",
                "source": [
                    "E2f4",
                    "Trim28",
                    "Rest",
                    "Ctcf",
                    "Trp53",
                    "Hdac1",
                    "Hes1",
                    "Hdac2",
                    "Smc3",
                    "Smarca5"
                ],
                "target": [
                    "Hsp90aa1",
                    "Lsm5",
                    "Arf1",
                    "Rpl41",
                    "Rnf7",
                    "Cdc20",
                    "Srsf9",
                    "Cd9",
                    "Mdh1",
                    "Ndufb4",
                    "Usmg5",
                    "Mcm7",
                    "Ppp2ca",
                    "Ddx24",
                    "Rps17",
                    "Rpl6",
                    "Srm",
                    "Grsf1",
                    "Rae1",
                    "Zc3h15",
                    "Cox8a",
                    "Mrps18b",
                    "Pin4",
                    "Ruvbl2",
                    "Cdt1",
                    "Psmb7",
                    "Spr",
                    "Txn2",
                    "Pnkd",
                    "Ppid",
                    "Ier2",
                    "2810004N23Rik",
                    "Clta",
                    "Cox17",
                    "Dkc1",
                    "Marcksl1",
                    "Atp5g1",
                    "1110008F13Rik",
                    "Mrpl4",
                    "Polr2k",
                    "Ppp1r11",
                    "Ndufb3",
                    "Gar1",
                    "Idh3g",
                    "Tuba1b",
                    "Anapc5",
                    "Cks1b",
                    "Aurka",
                    "Psma6",
                    "Rpl7l1",
                    "Snrpa",
                    "St13",
                    "Ftsj3",
                    "Psmc1",
                    "Hnrnpa1",
                    "Ndufa2",
                    "Thoc7",
                    "Tpd52l2",
                    "Hnrnpf",
                    "Knop1",
                    "Lmnb1",
                    "Ddx39",
                    "Mlf2",
                    "Psmd6",
                    "Vbp1",
                    "Ssrp1",
                    "Itm2a",
                    "Atp5j2",
                    "Pin1",
                    "Rnf187",
                    "Sfxn1",
                    "Swi5",
                    "Eif2s1",
                    "Psmb6",
                    "Cks2",
                    "Shfm1",
                    "Med28",
                    "Hmgb2",
                    "Nubp1",
                    "Cox6c",
                    "Ddx3x",
                    "Slbp",
                    "Ndufs3",
                    "Uba2",
                    "E2f4",
                    "Calr",
                    "Nup62",
                    "Tubb5",
                    "Calm3",
                    "D8Ertd738e",
                    "Hn1",
                    "Nop56",
                    "Snrpe",
                    "Ubxn1",
                    "Cdk1",
                    "Eif4h",
                    "Lamtor1",
                    "2810417H13Rik",
                    "Phb",
                    "Psat1",
                    "Psmd7",
                    "Anapc15",
                    "Gdi2",
                    "Gabarapl2",
                    "Stub1",
                    "Mrpl2",
                    "Nup43",
                    "Taf7",
                    "Tra2b",
                    "Cdc5l",
                    "Ranbp1",
                    "Mrpl28",
                    "Psmc6",
                    "Hmgcs1",
                    "Emc6",
                    "Grpel1",
                    "Sms",
                    "Zwint",
                    "Rbx1",
                    "Rps24",
                    "Ctcf",
                    "Ppm1g",
                    "Eif1",
                    "Neat1",
                    "Sec61g",
                    "Pmf1",
                    "Ythdc1",
                    "Bola1",
                    "Ola1",
                    "Sf3b6",
                    "Snrnp27",
                    "Tbcb",
```
