## Supplement 1 (MEF vs. ESC) for "AGEAS: Automated Machine Learning based Genetic Regulatory Element Extraction System": key_atlas.js

```
{
    "genes": {
        "Atf5": {
            "id": "Atf5",
            "type": "TF",
            "symbol": "Atf5",
            "source": [
                "Hdac2",
                "E2f4",
                "Foxp1",
                "Pou5f1",
                "Rest",
                "Trim28",
                "Hmgb1",
                "Atf4",
                "Ctcf",
                "Tgif1",
                "Trp53",
                "Hdac1",
                "Tbx3",
                "Ybx1",
                "Twist2",
                "Cebpb",
                "Zfp57",
                "Smarca5"
            ],
            "target": [
                "Arf1",
                "Cox6b1",
                "Rnf7",
                "Mrpl45",
                "Mcm7",
                "Rpl6",
                "Bub1",
                "Tspan17",
                "Hspa5",
                "Terf1",
                "Itpa",
                "Fosb",
                "Spr",
                "2810004N23Rik",
                "Slc1a5",
                "1110008F13Rik",
                "Ctps",
                "Arl6ip6",
                "Smu1",
                "A930018M24Rik",
                "Eif2b2",
                "Eif3c",
                "Akap12",
                "Dap3",
                "Adrm1",
                "Hdgfrp2",
                "Ppfibp1",
                "Vim",
                "Ahnak",
                "Tulp4",
                "Asns",
                "Fkbp10",
                "Gabarap",
                "Wars",
                "Ddr2",
                "Mrpl1",
                "Rrp15",
                "Aasdhppt",
                "1110004F10Rik",
                "Ubald1",
                "Cmtm3",
                "Calm3",
                "Trib3",
                "Mxd4",
                "2810417H13Rik",
                "Anapc15",
                "Fam149a",
                "Rdx",
                "Camk1",
                "Dmap1",
                "1700021F05Rik",
                "Ccdc80",
                "Timp3",
                "Lbh",
                "Tmem176a",
                "Poglut1",
                "Uba3",
                "1810011O10Rik",
                "Rcc2",
                "Ube2d2a",
                "Tnks1bp1",
                "Fbn2",
                "Mapre1",
                "Map1lc3b",
                "Rps26-ps1",
                "Dcp2",
                "1110046J04Rik",
                "Psme1",
                "Hax1",
                "Gm26682",
                "Dubr",
                "Cdc40",
                "Atf1",
                "Xpo1",
                "S100a4",
                "Kdm6b",
                "Zyx",
                "Nsfl1c",
                "Cald1",
                "Tsen34",
                "Bcas2",
                "Fas",
                "0610007P14Rik",
                "Spty2d1",
                "H13",
                "Marc2",
                "Phf5a",
                "Uchl3",
                "Gtf2a2",
                "2700060E02Rik",
                "Tagln",
                "Dnajc21",
                "Npm1"
            ],
            "expression_sum": {
                "class1": 10688.0,
                "class2": 4042.0
            },
            "uniprot_ids": [
                "O70191",
                "Q3UJF3",
                "A0A140LIB7",
                "Q3TVK2"
            ],
            "net_id": 0,
            "source_num": 18,
            "target_num": 29
        },
        "Tgif1": {
            "id": "Tgif1",
            "type": "TF",
            "symbol": "Tgif1",
            "source": [
                "Tcf4",
                "Pou5f1",
                "Foxp1",
                "Junb",
                "Jun",
                "Sox2",
                "Hes1",
                "Nanog",
                "Jund",
                "Twist2",
                "Cebpb",
                "Sox4"
            ],
            "target": [
                "Mgat2",
                "Dap",
                "Atf5",
                "Sar1a",
                "Cdc42",
                "Npc2",
                "Gadd45g",
                "Tcf4",
                "Thbs1",
                "Lims1",
                "Tmsb4x",
                "Cd47",
                "Cd81",
                "Itm2b",
                "Pdia3",
                "Txndc5",
                "Tspan4",
                "Anxa2",
                "Copb2",
                "Lmna",
                "Nupr1",
                "Copz2",
                "Tmed2",
                "Lgals3",
                "Gsn",
                "Prkar1a",
                "Rap1b",
                "Oat",
                "Tagln2",
                "Eif4g2",
                "Surf4",
                "Rab2a",
                "Rexo2",
                "Calr",
                "Pdlim7",
                "Tspo",
                "Lhfp",
                "Mbnl2",
                "Selk",
                "Neat1",
                "Fam162a",
                "Vimp",
                "Cd63",
                "Syncrip",
                "Iqgap1",
                "Sh3glb1",
                "Fundc2",
                "Rtn4",
                "Ctsl",
                "Ccnd2",
                "Spcs1",
                "Rpn1",
                "Atp2b1",
                "Tax1bp3",
                "Nktr",
                "Rpn2",
                "Arpc2",
                "Manf",
                "Tln1",
                "Sept11",
                "Arhgdib",
                "Tmed3",
                "Timp2",
                "Bzw1",
                "Ubc",
                "Sec62",
                "Foxp1",
                "Cdc42ep3",
                "Cmpk1",
                "Ptp4a2",
                "Dnajc3",
                "Hes1",
                "Ap1s1",
                "Myl12a",
                "P4hb",
                "Adipor1",
                "Gnb2",
                "Prdx5",
                "Sri",
                "Tm9sf3",
                "Ociad1",
                "Gnas",
                "Camk2d",
                "Sptbn1",
                "Col1a1",
                "Hdlbp",
                "Ap3s1",
                "Oxct1",
                "Rsu1",
                "Tmed9",
                "Ncam1",
                "Sepw1",
                "Gsto1",
                "Serp1",
                "Fam195b",
                "Tmem59",
                "Vcan",
                "Serpinb6a",
                "Pdcl3",
                "Junb",
                "Csrp1",
                "Hbegf",
                "Ilk"
            ],
            "expression_sum": {
                "class1": 3481.0,
                "class2": 18140.0
            },
            "uniprot_ids": [
                "P70284",
                "A0A087WNP4",
                "Q3TVD4",
                "G3UWC5",
                "E0CXJ4",
                "E0CYI0",
                "D3Z0Q5",
                "F6TUU5"
            ],
            "net_id": 0,
            "source_num": 12,
            "target_num": 66
        },
        "Tbx3": {
            "id": "Tbx3",
            "type": "TF",
            "symbol": "Tbx3",
            "source": [
                "Pou5f1",
                "Trim28",
                "Sox2",
                "Klf5",
                "Nanog",
                "Cebpb",
                "Zfp57"
            ],
            "target": [
                "Dppa5a",
                "Dap",
                "Cpe",
                "Serpine2",
                "Atf5",
                "Tpm4",
                "Itgb1",
                "Ptrf",
                "Cd44",
                "Cdc34",
                "Tipin",
                "Igf2",
                "Tmsb4x",
                "Cd47",
                "Efemp2",
                "Cd81",
                "Itm2b",
                "Fbln2",
                "Igfbp2",
                "Atp5g1",
                "Wisp1",
                "Rps27l",
                "Col12a1",
                "Ngfrap1",
                "Sod2",
                "Trim28",
                "Anxa2",
                "Lmna",
                "Cox7a2",
                "Folr1",
                "Prkcdbp",
                "Vim",
                "Lgals1",
                "Ahnak",
                "Fkbp10",
                "Pam16",
                "Phb2",
                "Rpl9",
                "Hmgn2",
                "Nrp1",
                "Sept1",
                "Copz2",
                "Gsn",
                "Erh",
                "Snrpg",
                "Ran",
                "Gjb3",
                "Grb10",
                "Jam2",
                "Tpm1",
                "Tuba1a",
                "Utf1",
                "Commd3",
                "Gsta4",
                "Pdlim7",
                "Tspo",
                "Lhfp",
                "Mbnl2",
                "Myl9",
                "Spint2",
                "Pdgfa",
                "B2m",
                "Uqcr10",
                "Parva",
                "H2afz",
                "Gpx8",
                "Gabarapl2",
                "Phldb2",
                "Rpl29",
                "Igfbp7",
                "Lrrc17",
                "Cdc5l",
                "Ung",
                "Anxa5",
                "S100a6",
                "Dpy30",
                "Odc1",
                "Mdk",
                "Neat1",
                "Acta2",
                "Ybx1",
                "Nanog",
                "Fstl1",
                "Fam162a",
                "Ifitm1",
                "Marcks",
                "Rhoc",
                "Lbh",
                "Fbn1",
                "Nhp2",
                "Plekhf2",
                "Pmp22",
                "Emp1",
                "Rbmxl2",
                "Tdgf1",
                "Sh3glb1",
                "App",
                "Cisd1",
                "Pole4",
                "Impa2",
                "Xist",
                "Ccnd2",
                "Tomm20",
                "Gas1",
                "Col6a1",
                "Rpl21",
                "S100a10",
                "Mylpf",
                "Filip1l",
                "Col5a1",
                "Bgn",
                "Cav1",
                "Fam25c",
                "Tgfb1i1",
                "Cdh11",
                "Oaf",
                "Tmed3",
                "Tpm2",
                "Timp2",
                "Syngr1",
                "Mt2",
                "Hsd17b14",
                "Fbln5",
                "Morc1",
                "Chd3",
                "Rps12",
                "H19",
                "Glipr2",
                "Dctpp1",
                "Arpc1b",
                "Col3a1",
                "L1td1",
                "Nenf",
                "Zfp42",
                "S100a4",
                "Klf2",
                "Rpp25",
                "Zyx",
                "Rcn3",
                "Dut",
                "Cald1",
                "Sox2",
                "Klf5",
                "Tmem176b",
                "Dpysl3",
                "Mybl2",
                "Slc2a3",
                "Tma7",
                "Gnas",
                "Hcfc1r1",
                "Camk2d",
                "Fhl2",
                "Calu",
                "H3f3b",
                "Gm7325",
                "Rbpms2",
                "Col1a1",
                "Flna",
                "Pmm1",
                "Tpd52",
                "Ckb",
                "Efemp1",
                "Gng12",
                "Nop10",
                "Prss23",
                "Wls",
                "Gdf3",
                "Sepw1",
                "Postn",
                "Capn2",
                "Laptm4a",
                "Vcan",
                "Dtymk",
                "Sparc",
                "Emp3",
                "Ier5",
                "Stmn2",
                "Cebpb",
                "Prnp",
                "Id1",
                "Serpinb6a",
                "Alpl",
                "Mkrn1",
                "Hmga2",
                "Nfatc2ip",
                "Lpp",
                "Kdelr3",
                "Gm26917",
                "Tagln",
                "Kdm5b",
                "P3h4",
                "Anxa3",
                "Hist1h2bc",
                "Selm",
                "Hells",
                "Tfcp2l1",
                "Axl",
                "Cfl2",
                "Ckap4",
                "Col1a2"
            ],
            "expression_sum": {
                "class1": 979.0,
                "class2": 26585.0
            },
            "uniprot_ids": [
                "P70324",
                "Q8BG45",
                "Q3ULL3",
                "Q3U043",
                "Q4VA43",
                "Q8CJ82",
                "Q8BJW3",
                "Q8BU76"
            ],
            "net_id": 0,
            "source_num": 7,
            "target_num": 148
        },
        "Zfp57": {
            "id": "Zfp57",
            "type": "TF",
            "symbol": "Zfp57",
            "source": [
                "Pou5f1",
                "Trim28",
                "Sox2",
                "Nanog",
                "Cebpb",
                "Sox4"
            ],
            "target": [
                "Meg3",
                "Lsm5",
                "Pcbp1",
                "Dap",
                "Lsm7",
                "Cpe",
                "Serpine2",
                "Atf5",
                "Cdc20",
                "Rcor2",
                "Tpm4",
                "Itgb1",
                "Ptrf",
                "Ppih",
                "Cd44",
                "Mat2a",
                "Tpx2",
                "Thbs1",
                "G3bp2",
                "Cd47",
                "Efemp2",
                "Cd81",
                "Fbln2",
                "Igfbp2",
                "Atp5g1",
                "Wisp1",
                "Ndufs6",
                "Ccnb1",
                "Set",
                "Trim28",
                "Cebpzos",
                "Aurka",
                "Cks1b",
                "Col12a1",
                "Rpl12",
                "Ssbp1",
                "Anxa2",
                "Lmna",
                "Dnajc19",
                "Vim",
                "Tnfrsf12a",
                "Ctbp2",
                "Lgals1",
                "Sephs2",
                "Ahnak",
                "Fkbp10",
                "Gabarap",
                "E130012A19Rik",
                "Phb2",
                "Rpl9",
                "Cenpf",
                "Copz2",
                "Lsm3",
                "Spag7",
                "Arl6ip1",
                "Tcof1",
                "Gsn",
                "Erh",
                "Psmb6",
                "Cks2",
                "Snrpg",
                "Hmgb2",
                "Nhp2l1",
                "Ran",
                "Srsf2",
                "Jam2",
                "Grb10",
                "Tpm1",
                "Apex1",
                "Commd3",
                "Snrpb",
                "Ccnb2",
                "Fkbp4",
                "Lhfp",
                "Gas6",
                "Nop56",
                "Pdgfa",
                "Snrpe",
                "Btg2",
                "Parva",
                "Cdk1",
                "H2afz",
                "Lyar",
                "Gpx8",
                "Polr2e",
                "Lsm2",
                "Phldb2",
                "Rpl29",
                "Igfbp7",
                "Lrrc17",
                "Cenpa",
                "Ranbp1",
                "Anxa5",
                "S100a6",
                "Dpy30",
                "Rbx1",
                "H2afv",
                "Neat1",
                "Ybx1",
                "Fstl1",
                "Rhoc",
                "Mrpl42",
                "Timp3",
                "Fbn1",
                "Mrpl12",
                "Snrpf",
                "Exosc5",
                "Tomm40",
                "Rcc2",
                "Slc29a1",
                "Impa2",
                "Ccnd2",
                "Tax1bp3",
                "Rpl21",
                "Rangrf",
                "Filip1l",
                "Col5a1",
                "Rbm8a",
                "Bgn",
                "Cav1",
                "Fam25c",
                "Rpl7a",
                "Calm2",
                "Cox7c",
                "Stmn1",
                "Ncl",
                "Tgfb1i1",
                "Cdh11",
                "Ubb",
                "Tmed3",
                "Serpinf1",
                "Srsf5",
                "Rpl18",
                "Tpm2",
                "Timp2",
                "Tubb4b",
                "Parp1",
                "Wdr74",
                "Fbln5",
                "Morc1",
                "Chd3",
                "Rps12",
                "Psmb3",
                "Prc1",
                "Cdca3",
                "Sub1",
                "Fn1",
                "Dctpp1",
                "Glipr2",
                "Arpc1b",
                "Col3a1",
                "Cdca8",
                "Ppan",
                "Ptma",
                "Ube2s",
                "Rpl27",
                "Serpinh1",
                "Klf2",
                "Rpp25",
                "Stra13",
                "Rcn3",
                "Dut",
                "Zyx",
                "Cald1",
                "Fau",
                "Mtf2",
                "Sox2",
                "Tmem176b",
                "Mrpl15",
                "Mfap2",
                "Dpysl3",
                "Mybl2",
                "Tma7",
                "Gnas",
                "Aebp1",
                "Fhl2",
                "Rbpms2",
                "Calu",
                "Sept1",
                "Col1a1",
                "Gng12",
                "Prss23",
                "Wls",
                "Hacd1",
                "Eif5a",
                "Laptm4a",
                "Vcan",
                "Serbp1",
                "Tbx3",
                "Sparc",
                "Emp3",
                "Hspe1",
                "Hdgf",
                "Cebpb",
                "Prnp",
                "Msn",
                "Cryab",
                "Ube2c",
                "Mkrn1",
                "Lpp",
                "Tagln",
                "Gm26917",
                "Hmgb1",
                "Hspb1",
                "Anxa3",
                "Birc5",
                "Kdm5b",
                "P3h4",
                "Selm",
                "Tfcp2l1",
                "Sap30",
                "Axl",
                "Col1a2"
            ],
            "expression_sum": {
                "class1": 613.0,
                "class2": 18328.0
            },
            "uniprot_ids": [
                "Q8C6P8",
                "G3UYK1",
                "G3UX77",
                "G3UWX4",
                "G3UYC1",
                "G3UY05",
                "H3BL53"
            ],
            "net_id": 0,
            "source_num": 6,
            "target_num": 161
        },
        "Twist2": {
            "id": "Twist2",
            "type": "TF",
            "symbol": "Twist2",
            "source": [
                "E2f4",
                "Ctcf",
                "Hdac1",
                "Hdac2",
                "Smarca5"
            ],
            "target": [
                "Hsp90aa1",
                "Lsm5",
                "Arf1",
                "Dap",
                "Sar1a",
                "Cdc20",
                "Usmg5",
                "Dctn3",
                "Mcm7",
                "Ddx24",
                "Rps17",
                "Rpl6",
                "Srm",
                "Grsf1",
                "Rae1",
                "Zc3h15",
                "Cox8a",
                "Mrps18b",
                "Pin4",
                "Ruvbl2",
                "Psmb7",
                "Spr",
                "Txn2",
                "Pnkd",
                "Ppid",
                "Ier2",
                "2810004N23Rik",
                "Clta",
                "Dkc1",
                "Marcksl1",
                "Mrpl4",
                "Atp5g1",
                "1110008F13Rik",
                "Ppp1r11",
                "Polr2k",
                "Sec13",
                "Tuba1b",
                "Gar1",
                "Idh3g",
                "Anapc5",
                "Aurka",
                "Psma6",
                "Rpl7l1",
                "Snrpa",
                "St13",
                "Ftsj3",
                "Psmc1",
                "Hnrnpa1",
                "Ndufa2",
                "Thoc7",
                "Tpd52l2",
                "Hnrnpf",
                "Knop1",
                "Mlf2",
                "Psmd6",
                "Rpl9",
                "Ssrp1",
                "Pin1",
                "Atp5j2",
                "Ier3ip1",
                "Rnf187",
                "Swi5",
                "Eif2s1",
                "Psmb6",
                "Cks2",
                "Shfm1",
                "Med28",
                "Hmgb2",
                "Nubp1",
                "Cox6c",
                "Slbp",
                "Ndufs3",
                "Nedd4",
                "Uba2",
                "E2f4",
                "Calr",
                "Nup62",
                "Cenpe",
                "Rab18",
                "Tubb5",
                "Calm3",
                "Arf6",
                "Canx",
                "D8Ertd738e",
                "Hn1",
                "Nop56",
                "Ppp1r14b",
                "Snrpe",
                "Ubxn1",
                "Mrpl14",
                "Slc25a3",
                "Cdk1",
                "Phb",
                "Lamtor1",
                "2810417H13Rik",
                "Psat1",
                "Gabarapl2",
                "Nup43",
                "Taf7",
                "Cdc5l",
                "Ranbp1",
                "Mrpl28",
                "Psmc6",
                "Emc6",
                "Sms",
                "Zwint",
                "Rps24",
                "Ppm1g",
                "Eif1",
                "Neat1",
                "Sec61g",
                "Ythdc1",
                "Bola1",
                "Ola1",
                "Sf3b6",
                "Snrnp27",
                "Aurkaip1",
                "Srsf1",
                "Lrrc59",
                "Uqcrc1",
                "Exosc5",
                "Rcc2",
                "Atp5h",
                "Bnip3l",
                "Bola2",
                "Tmem160",
                "Kpnb1",
                "Mapre1",
                "C1d",
                "Snrpd1",
                "Rbm8a",
                "Ppia",
                "Hnrnpm",
                "Capza2",
                "Lap3",
                "Loxl1",
                "2310036O22Rik",
                "Mphosph8",
                "Supt4a",
                "Srsf5",
                "Aamp",
                "Rpl35a",
                "Chchd1",
                "Wdr74",
                "Gars",
                "Psmc3",
                "Hspa4",
                "Luc7l2",
                "Rpl36",
                "Rbbp7",
                "Psmb3",
                "Hnrnpl",
                "Ndufb9",
                "Rbm39",
                "Rps21",
                "Arpp19",
                "Ccdc12",
                "Ddx21",
                "Nedd8",
                "Pcnp",
                "Eef1b2",
                "Utp3",
                "Tmem258",
                "Rpl24",
                "Brix1",
                "Fam136a",
                "Mrpl15",
                "Siva1",
                "Fus",
                "Ezh2",
                "Mybl2",
                "Ociad1",
                "Pfdn1",
                "Ddost",
                "Rps4x",
                "Pmm1",
                "Fkbp8",
                "Nip7",
                "Ruvbl1",
                "Clpp",
                "Psma7",
                "Ndufa8",
                "Pdia4",
                "Sept7",
                "Gnl3",
                "Trim27",
                "Sssca1",
                "Tpm3",
                "Atp5b",
                "Banf1",
                "Cdc26",
                "Dtymk",
                "Immt",
                "Mrps36",
                "Emc10",
                "Lig1",
                "Psma1",
                "Gpx1",
                "Hdgf",
                "Qdpr",
                "Smc3",
                "Thrap3",
                "Tpi1",
                "Csnk2b",
                "Nars",
                "Pa2g4",
                "Pdlim1",
                "Rpl37",
                "Hspb1",
                "Kdm5b",
                "Atf4",
                "Hist1h2bc",
                "Ppp1cc",
                "Eif3b",
                "Zfp207",
                "Pcbp1",
                "Rpl38",
                "Dnttip2",
                "Sae1",
                "Mrps24",
                "Id2",
                "Ndufa4",
                "Ewsr1",
                "Cops5",
                "Eif4e",
                "Ndufv1",
                "Ndufv2",
                "H3f3a",
                "Nme1",
                "Ube2k",
                "Rfc2",
                "Bccip",
                "Csnk1a1",
                "Ndufs2",
                "Ddt",
                "Fbxo22",
                "Cyp51",
                "Lamtor2",
                "Tmsb10",
                "Rpl3",
                "Cebpzos",
                "Nudt21",
                "Trim28",
                "Rpl12",
                "Eif3c",
                "Cox7a2",
                "Cdv3",
                "Crip2",
                "Spc24",
                "Ube2l3",
                "Ctbp2",
                "Asns",
                "Ifitm2",
                "Fdps",
                "Ppp4c",
                "Eef1d",
                "Etf1",
                "Ndufs7",
                "Ube2b",
                "Mcm2",
                "Lsm3",
                "Mrpl13",
                "Gcat",
                "Eif5",
                "Tmed2",
                "Tomm5",
                "Txn1",
                "Ube2j2",
                "Erh",
                "Ak2",
                "Mrpl17",
                "Samm50",
                "Zmat2",
                "Wdr77",
                "Atp5j",
                "Ran",
                "Fscn1",
                "Atp5g3",
                "Srrm1",
                "Atp5g2",
                "Emc8",
                "Srp19",
                "Cct5",
                "Emd",
                "Mrpl40",
                "Tceb1",
                "C1qbp",
                "Rpsa",
                "H2afz",
                "Bola3",
                "Mrps21",
                "Tardbp",
                "Uqcr10",
                "Cst3",
                "Rpl36a",
                "Eif3m",
                "Rpl29",
                "Iscu",
                "Fubp1",
                "Rpl5",
                "Naa38",
                "Ccdc58",
                "Dph3",
                "Prdx6",
                "Odc1",
                "Rps20",
                "Ybx1",
                "Dnaja2",
                "Sys1",
                "Tubb2b",
                "Acat2",
                "Tmem14c",
                "Mrpl12",
                "Cdc37",
                "Ndufab1",
                "Psmd1",
                "Eef1a1",
                "Mcl1",
                "Rtn4",
                "Cisd1",
                "Rpl28",
                "U2af1",
                "Pole4",
                "Ywhaq",
                "Mrps17",
                "Fbl",
                "Pdha1",
                "Ubald2",
                "Dnajc8",
                "Calm2",
                "Rps3",
                "Atox1",
                "Stmn1",
                "Srsf10",
                "Adss",
                "Btf3",
                "Smarcc1",
                "Igf2bp1",
                "Ube2v2",
                "Rpl18",
                "Eno1",
                "Hax1",
                "Atp5d",
                "Eef1e1",
                "Rpl19",
                "Taldo1",
                "Timm13",
                "Trnau1ap",
                "Eif3h",
                "Mcm3",
                "Adprh",
                "Rpl22l1",
                "Vcp",
                "Rps3a1",
                "Mrps5",
                "Srsf3",
                "Hsd17b10",
                "Smc4",
                "Prc1",
                "U2surp",
                "Cmpk1",
                "Mbd3",
                "Rpl7",
                "Cct7",
                "Znrd1",
                "Dctpp1",
                "Ndufb2",
                "Wdr18",
                "Ppan",
                "Hsp90ab1",
                "Oaz1",
                "Mrpl30",
                "Uqcrq",
                "Sec61b",
                "BC031181",
                "Polr2m",
                "1810058I24Rik",
                "Rwdd1",
                "Snx5",
                "Tyms",
                "Gnai2",
                "Rps8",
                "Fnta",
                "Dnaja1",
                "Pgp",
                "Eif4a1",
                "Rpl8",
                "Sumo2",
                "Pcbp2",
                "Magoh",
                "Ndufc2",
                "Polr1c",
                "Tmed10",
                "Snrnp25",
                "Pmvk",
                "H3f3b",
                "Dnajc2",
                "Ccdc34",
                "Elof1",
                "Gnb2l1",
                "Mrps33",
                "Ndufa12",
                "Pold2",
                "Nsa2",
                "Rif1",
                "Wdr5",
                "Ddx46",
                "Atxn10",
                "Cnih4",
                "Nudt19",
                "Pebp1",
                "Sgk1",
                "Dbi",
                "Morf4l2",
                "Ube2d3",
                "Aars",
                "Rfc5",
                "Snrpa1",
                "Paip2",
                "Pfdn4",
                "Srrm2",
                "Top1",
                "Ywhae",
                "Ubl5",
                "Hmgb1",
                "Mrps15",
                "Psmc4",
                "Ndufc1",
                "Ppil1",
                "Eif3d",
                "Cyb5a",
                "Rps15",
                "Psma4",
                "Arpc3",
                "Scp2",
                "Arl1",
                "Arpc1a",
                "Ap2s1",
                "Rpl34",
                "Park7",
                "Mlec",
                "Rab7",
                "Sep15",
                "Dynll1",
                "Gtpbp4",
                "Cenpw",
                "Eif2s2",
                "Hspd1",
                "Atf5",
                "Stoml2",
                "Cops7a",
                "Cdc42",
                "Mrpl45",
                "Rpl31",
                "Srsf7",
                "Smdt1",
                "Ppih",
                "Nap1l1",
                "Uqcc3",
                "Tpx2",
                "Etfb",
                "Dnajb6",
                "Brd2",
                "Actr3",
                "Impdh2",
                "Ccdc124",
                "Mrps10",
                "Gapdh",
                "Ccnb1",
                "Prdx2",
                "Mpc1",
                "Akap12",
                "Ccnd3",
                "Khdrbs1",
                "Mrpl52",
                "Adrm1",
                "Abcf1",
                "Psmb5",
                "Hspa9",
                "Chchd2",
                "Nop58",
                "Ube2n",
                "Ndufa11",
                "Smn1",
                "Cbx5",
                "Kif5b",
                "Polr2c",
                "Cct8",
                "Hmgn2",
                "Dohh",
                "Ptp4a1",
                "Ndufs5",
                "Oat",
                "Akr1b3",
                "Ddx27",
                "Ndufb8",
                "Psmd2",
                "Srsf2",
                "Prdx4",
                "Eif4g2",
                "Jade1",
                "Dnajc7",
                "Uqcc2",
                "Pak1ip1",
                "Denr",
                "Fxr1",
                "Nsmce4a",
                "Rps26",
                "Get4",
                "Eif3g",
                "Nxt1",
                "Trp53",
                "Sra1",
                "Tceal8",
                "Aspscr1",
                "Psmd11",
                "Cnn3",
                "Vps29",
                "Vdac1",
                "Rpp21",
                "Cggbp1",
                "Lage3",
                "Anp32a",
                "Rad23b",
                "Zfp706",
                "Polr2e",
                "Lsm2",
                "Nudt4",
                "Tra2a",
                "Pdcd5",
                "Aprt",
                "Ost4",
                "Snrpc",
                "Mrpl51",
                "Mrpl49",
                "Eif4g1",
                "Tmbim6",
                "Dynll2",
                "Rhoa",
                "Fcf1",
                "Mrps18c",
                "Atp5a1",
                "Cnpy2",
                "Rpl22",
                "Pycr2",
                "Sf3a3",
                "Anp32e",
                "Ssr2",
                "Ict1",
                "Rpl4",
                "Glud1",
                "March5",
                "Cebpz",
                "Fermt2",
                "Hnrnpu",
                "Ddah2",
                "Rbmxl1",
                "Gspt1",
                "Arglu1",
                "Imp4",
                "Srsf11",
                "Rpl18a",
                "Hmgb3",
                "Cbx1",
                "Tomm20",
                "Txndc9",
                "Vapa",
                "Rpl21",
                "Map1lc3b",
                "Eprs",
                "Trmt112",
                "Rpl35",
                "Dazap1",
                "Msmo1",
                "Atp5l",
                "Mrpl54",
                "Pop5",
                "Son",
                "Ilf2",
                "Cct2",
                "Mak16",
                "Ubb",
                "Dtl",
                "Ahsa1",
                "Zfp106",
                "Prrc1",
                "Eif1ax",
                "Parp1",
                "Ak6",
                "Rbpms",
                "Paics",
                "Hnrnpa3",
                "Prmt1",
                "Pbdc1",
                "Rps19bp1",
                "Cdca3",
                "Metap2",
                "Sub1",
                "Asna1",
                "Psmc5",
                "Echdc2",
                "Morf4l1",
                "Mrpl43",
                "Pafah1b1",
                "Cdca8",
                "Lsm6",
                "Tecr",
                "Sf3b1",
                "Ube2s",
                "Ndufs4",
                "Mybbp1a",
                "Snrpd3",
                "Sf3b2",
                "Psmd3",
                "Suclg1",
                "H2afy",
                "Fam32a",
                "Mtf2",
                "Slc3a2",
                "Stip1",
                "Rpl14",
                "Tma7",
                "Ash2l",
                "Hat1",
                "Cops4",
                "Rnps1",
                "Lsm4",
                "Pfdn2",
                "Nop10",
                "Mpdu1",
                "Atp5f1",
                "Eif1ad",
                "Arid5b",
                "Rps13",
                "Gsto1",
                "Tpt1",
                "Eif5a",
                "Rps11",
                "Ppib",
                "Cfl1",
                "Rrp1b",
                "Hspe1",
                "Rps19",
                "Ube2c",
                "Exosc8",
                "Cops6",
                "Ptbp1",
                "Aldoa",
                "Mrpl20",
                "Hnrnpdl",
                "Srp9",
                "Eif3a",
                "Rtn3",
                "Polr2h",
                "Psmb4",
                "Snx3",
                "Strap",
                "Commd1",
                "l7Rn6",
                "Rnaseh2a",
                "Chtop",
                "Srpk1",
                "Cox6b1",
                "Ino80e",
                "Ppa1",
                "Atp5e",
                "Avpi1",
                "Prdx1",
                "Rsl24d1",
                "Ssb",
                "Tipin",
                "Atp5o",
                "Mrps12",
                "Rrm2",
                "Rpl13",
                "Copz1",
                "Capns1",
                "Mrpl34",
                "Nasp",
                "Pdia3",
                "Ndufb5",
                "Rpl10a",
                "Ndufs6",
                "Set",
                "Thyn1",
                "Eif4e2",
                "Shmt1",
                "Alkbh1",
                "Sod2",
                "Ssbp1",
                "Rps14",
                "Tgif1",
                "Sfpq",
                "Mrto4",
                "Rpl32",
                "Parl",
                "Lipe",
                "Clns1a",
                "Snrpd2",
                "1810022K09Rik",
                "Uqcr11",
                "Phb2",
                "Rps27a",
                "Naa10",
                "Bud31",
                "Gtf2e2",
                "Pdap1",
                "Mcm5",
                "Hsp90b1",
                "Arl6ip1",
                "Rpl11",
                "Rpl15",
                "Prkar1a",
                "Rap1b",
                "Tcof1",
                "Vdac2",
                "Pfn1",
                "Rpl13a",
                "Nol7",
                "Snrpg",
                "Tkt",
                "Mrpl24",
                "Snhg6",
                "Cox5a",
                "Minos1",
                "Snrpb",
                "Tmem11",
                "Fkbp4",
                "Zfp428",
                "Rabggtb",
                "Tomm22",
                "Lyar",
                "Ubap2l",
                "Erp29",
                "Snrnp70",
                "Uchl5",
                "Ppig",
                "Cfdp1",
                "Ppdpf",
                "Dcun1d5",
                "Eif5b",
                "Cct4",
                "Llph",
                "Emg1",
                "Lsm12",
                "Eif4ebp1",
                "H2afv",
                "Nono",
                "Csde1",
                "Prrc2c",
                "Gins4",
                "Tceb2",
                "Mrpl42",
                "Psmg4",
                "Rps2",
                "Gart",
                "Polr2g",
                "Sumo3",
                "Psmc2",
                "Ndufa10",
                "Tomm40",
                "Fundc2",
                "Uqcrfs1",
                "Naa50",
                "Ywhah",
                "Sf3b4",
                "Eif3i",
                "Hdac2",
                "Sugt1",
                "Hnrnpc",
                "Smc2",
                "Fkbp3",
                "Mtch2",
                "Rangrf",
                "Gins2",
                "Bub3",
                "Rpl7a",
                "Uck2",
                "Papola",
                "Rps16",
                "Esf1",
                "Sin3b",
                "Mrfap1",
                "Tubb4b",
                "Ccna2",
                "Mrps16",
                "Slc25a5",
                "Uqcrh",
                "Ddx18",
                "Rrp9",
                "Tbca",
                "Eed",
                "Ccne1",
                "Srsf6",
                "Ndufa5",
                "Tmpo",
                "Ndufb10",
                "Tsn",
                "Mrpl11",
                "Rpl37a",
                "Txnl1",
                "Mesdc2",
                "Eef1g",
                "Rbm25",
                "2700094K13Rik",
                "Cycs",
                "Myl12a",
                "Ptma",
                "Rac1",
                "Rer1",
                "Dut",
                "Dhx16",
                "Sdhb",
                "Taf10",
                "Wdr43",
                "Ybx3",
                "Klf5",
                "Acin1",
                "Rps10",
                "Pabpc1",
                "Eif6",
                "Yrdc",
                "Hspa14",
                "Fkbp1a",
                "Polr2j",
                "Got2",
                "Dad1",
                "Slirp",
                "0610007P14Rik",
                "Uhrf1",
                "Yif1b",
                "Arpc5",
                "Wbp11",
                "Cox4i1",
                "Snf8",
                "Cwc15",
                "Guk1",
                "2810428I15Rik",
                "Hspa8",
                "Ndufs8",
                "Psma3",
                "Rpl39",
                "Rplp2",
                "Postn",
                "Rps18",
                "Tsfm",
                "Mif",
                "Ptov1",
                "Aimp1",
                "Phf5a",
                "Chchd3",
                "Gcsh",
                "Glrx3",
                "Gtf2a2",
                "Mkrn1",
                "Rrp1",
                "Pdcl3",
                "Rpl30",
                "Sfr1",
                "Myl12b",
                "Kdm1a",
                "Rrs1",
                "Ncor1",
                "Rbm17",
                "Ly6e",
                "Cnbp",
                "Rsrc2"
            ],
            "expression_sum": {
                "class1": 6849.0,
                "class2": 547.0
            },
            "uniprot_ids": [
                "Q9D030",
                "A5D6P6"
            ],
            "net_id": 0,
            "source_num": 5,
            "target_num": 462
        },
        "Ctcf": {
            "id": "Ctcf",
            "type": "TF",
            "symbol": "Ctcf",
            "source": [
                "Tfcp2l1",
                "Pou5f1",
                "Foxp1",
                "Junb",
                "Id2",
                "Jun",
                "Sox2",
                "Hes1",
                "Nanog",
                "Jund",
                "Cebpb",
                "Sox4"
            ],
            "target": [
                "Arf1",
                "Dap",
                "Sar1a",
                "Tpm4",
                "Cd44",
                "Myh9",
                "Thbs1",
                "Lims1",
                "Sec13",
                "Tuba1b",
                "Wisp1",
                "Slc25a4",
                "Tspan4",
                "Col12a1",
                "Lmna",
                "Cthrc1",
                "Pck2",
                "Rnf187",
                "Lgals3",
                "mt-Co1",
                "Pgrmc1",
                "Tagln2",
                "Rps15a",
                "Rexo2",
                "Calr",
                "Pdlim7",
                "Tubb5",
                "Mfge8",
                "Mbnl2",
                "Canx",
                "Actn1",
                "B2m",
                "Slc25a3",
                "Psat1",
                "2810417H13Rik",
                "Gdi2",
                "Nr0b1",
                "Prrx1",
                "Neat1",
                "Acta2",
                "Myh10",
                "Nanog",
                "Ola1",
                "Ube2r2",
                "Tgfbi",
                "Timp3",
                "Gemin7",
                "Akr1a1",
                "Tdgf1",
                "Sh3glb1",
                "Atp2b1",
                "Mapre1",
                "Cisd2",
                "Mylpf",
                "Nktr",
                "Col5a1",
                "Tspan3",
                "Bgn",
                "Capza2",
                "Klf6",
                "Pcolce",
                "Tgfb1i1",
                "Loxl1",
                "Timp2",
                "Syngr1",
                "Morc1",
                "Sec62",
                "Rps12",
                "Maged1",
                "Arpc1b",
                "Col3a1",
                "Cyr61",
                "Kdelr1",
                "Rab14",
                "Serpinh1",
                "Atp6v1g1",
                "Gnb2",
                "Ociad1",
                "Dpysl3",
                "Ddost",
                "Hcfc1r1",
                "Inhba",
                "Sptbn1",
                "Id3",
                "Ap3s1",
                "Gng12",
                "Ncam1",
                "Wls",
                "Hacd1",
                "Ndufa8",
                "Pdia4",
                "Srp14",
                "Sept7",
                "Fam195b",
                "Actg1",
                "Rcn2",
                "Tsg101",
                "Ier5",
                "Prnp",
                "Tpi1",
                "Id1",
                "Hmga2",
                "Atf4",
                "Tfcp2l1",
                "Nrp1",
                "Col1a2",
                "F2r",
                "Fkbp9",
                "Serpine2",
                "Igfbp4",
                "Id2",
                "Itgb1",
                "Fabp5",
                "Mettl9",
                "Tcf4",
                "Igf2",
                "Cd81",
                "Ssr1",
                "Pgls",
                "Lxn",
                "Anxa2",
                "Vmp1",
                "Col5a2",
                "mt-Nd4",
                "Crip2",
                "Cox7a2l",
                "Vim",
                "Basp1",
                "Platr3",
                "Sh3bgrl3",
                "Wdr1",
                "Tnc",
                "Ube2b",
                "Tmed2",
                "Tpm1",
                "Ldha",
                "Sdcbp",
                "Thy1",
                "Surf4",
                "Utf1",
                "Phlda1",
                "Myeov2",
                "Vcl",
                "Cst3",
                "Eif3m",
                "Lox",
                "Higd1a",
                "S100a6",
                "H2-D1",
                "Crabp1",
                "Cd63",
                "Capg",
                "Lbh",
                "S100a11",
                "Scand1",
                "Mtdh",
                "Eef1a1",
                "mt-Nd5",
                "Maged2",
                "Rtn4",
                "Ctsl",
                "Xist",
                "Ccnd2",
                "Tcf25",
                "Spcs1",
                "Cox6a1",
                "Col6a1",
                "Rala",
                "Sdc2",
                "Tax1bp3",
                "Filip1l",
                "Cav1",
                "Plk2",
                "Tln1",
                "Cdh11",
                "Oaf",
                "Ostf1",
                "Mt2",
                "Eif3h",
                "Fbln5",
                "Chd3",
                "Cmpk1",
                "Fkbp2",
                "Lrrc58",
                "Fn1",
                "Glipr2",
                "Hes1",
                "Ap1s1",
                "Nenf",
                "Rps9",
                "Twist2",
                "Zfp42",
                "Rpp25",
                "Rcn3",
                "Malat1",
                "Gnai2",
                "Tmem176b",
                "Mfap2",
                "Tm9sf3",
                "Tmed10",
                "Ctsd",
                "Reep5",
                "Pou5f1",
                "Fhl2",
                "Glrx5",
                "Col1a1",
                "Pla2g1b",
                "Oxct1",
                "Peg3",
                "Atxn10",
                "Atp6v0e",
                "Cnih4",
                "Dbi",
                "Tmem59",
                "Eif3e",
                "Emp3",
                "Aspn",
                "Cebpb",
                "Pafah1b2",
                "Anxa3",
                "Psmd14",
                "Hbegf",
                "Arpc3",
                "Ilk",
                "Arl1",
                "Mgat2",
                "Sep15",
                "Mlec",
                "Lsm7",
                "Cpe",
                "Atf5",
                "Arf5",
                "Cdc42",
                "Npc2",
                "Eef2",
                "Hspa5",
                "Rnh1",
                "Rrbp1",
                "Actr3",
                "Tmsb4x",
                "Cd47",
                "Efemp2",
                "Fbln2",
                "Mpc1",
                "Copb2",
                "Sars",
                "Ap2m1",
                "Nupr1",
                "Prkcdbp",
                "Sdpr",
                "Lgals1",
                "Ahnak",
                "Col11a1",
                "Fkbp10",
                "Kif5b",
                "Copz2",
                "Ptp4a1",
                "Lsp1",
                "Gsn",
                "Oat",
                "Tubb6",
                "Eva1b",
                "Prdx4",
                "Aes",
                "Rab2a",
                "Tspo",
                "Lhfp",
                "Myl9",
                "Vps29",
                "Btg2",
                "Parva",
                "Selk",
                "Jund",
                "Rab11a",
                "Anxa5",
                "Eif3f",
                "Anxa6",
                "Mtch1",
                "Rhoa",
                "Rab10",
                "Vimp",
                "Ccdc80",
                "Sec61a1",
                "Iqgap1",
                "Kdelr2",
                "Pmp22",
                "Rbmxl2",
                "Ssr3",
                "Rpn1",
                "Gas1",
                "Ssr4",
                "Txndc9",
                "Lman1",
                "F3",
                "Rcn1",
                "Arpc2",
                "Fam25c",
                "Ywhab",
                "Sox4",
                "Phlda3",
                "Sept11",
                "Tpm2",
                "Ergic3",
                "Bzw1",
                "Sdc4",
                "Ubc",
                "Rock2",
                "Cdc42ep3",
                "Tcea3",
                "Col8a1",
                "L1td1",
                "Psap",
                "Tm4sf1",
                "Rbms1",
                "P4hb",
                "Scd2",
                "Dlk1",
                "Zyx",
                "Map1b",
                "Rab34",
                "Adipor1",
                "Bsg",
                "Sox2",
                "Ppic",
                "Camk2d",
                "mt-Cytb",
                "Cyba",
                "Flna",
                "Efemp1",
                "H13",
                "Rsu1",
                "Prss23",
                "Gdf3",
                "Gsto1",
                "Ppib",
                "Vcan",
                "H1f0",
                "Ndufb11",
                "Sparc",
                "Sept2",
                "Abracl",
                "Msn",
                "Aldoa",
                "Serpinb6a",
                "Rtn3",
                "Lpp",
                "Ptn",
                "Tagln",
                "Ctbp1",
                "Csrp1",
                "Selm",
                "Ckap4",
                "Esd",
                "Dppa5a",
                "Meg3",
                "Atxn7l3b",
                "Igfbp6",
                "Comt",
                "Lamp1",
                "Rcor2",
                "Tinagl1",
                "Ptrf",
                "Gadd45g",
                "Rrm2",
                "Sh3bgrl",
                "Jun",
                "Itm2b",
                "Pdia3",
                "Rpl10a",
                "Txndc5",
                "Col4a1",
                "Hist1h1c",
                "Cd24a",
                "Lipe",
                "Tnfrsf12a",
                "Chchd10",
                "Krtcap2",
                "Gabarap",
                "Phgdh",
                "Rap1a",
                "Hsp90b1",
                "Mtpn",
                "Anxa1",
                "Prkar1a",
                "Timp1",
                "Ier3",
                "Grb10",
                "Tuba1a",
                "Commd3",
                "Cnn2",
                "Gsta4",
                "Fst",
                "Gpx8",
                "Gpi1",
                "Tmem167",
                "Phldb2",
                "Palld",
                "Tram1",
                "Igfbp7",
                "Lrrc17",
                "Rpl23a-ps3",
                "Cdkn1a",
                "Fstl1",
                "Marcks",
                "Fam162a",
                "Rhoc",
                "Ifitm3",
                "Fbn1",
                "Srp72",
                "Mrpl33",
                "Emp1",
                "App",
                "Fundc2",
                "Eif3i",
                "Fbn2",
                "Gm9843",
                "S100a10",
                "Bag1",
                "Rpn2",
                "Mfap4",
                "Prrx2",
                "Cetn3",
                "Dctn2",
                "Arhgdib",
                "Tmed3",
                "Serpinf1",
                "Hsd17b14",
                "Gap43",
                "Tbca",
                "Foxp1",
                "Tmpo",
                "Ccng1",
                "H19",
                "Ptp4a2",
                "Txnl1",
                "Cope",
                "Dnajc3",
                "Arhgdia",
                "Myl12a",
                "Rac1",
                "Rer1",
                "S100a4",
                "Klf2",
                "Cald1",
                "Fau",
                "Ooep",
                "Sri",
                "Prdx5",
                "Clic1",
                "Fkbp1a",
                "Emc7",
                "Gnas",
                "Aebp1",
                "Arf4",
                "Sept1",
                "Calu",
                "Arpc5",
                "Gm7325",
                "Hdlbp",
                "Limd2",
                "Cox4i1",
                "Rps18-ps3",
                "Tmed9",
                "Myl6",
                "Sepw1",
                "Serp1",
                "Capn2",
                "Laptm4a",
                "Cnih1",
                "Rheb",
                "Carhsp1",
                "Alpl",
                "Pdcl3",
                "Kdelr3",
                "P3h4",
                "Junb",
                "Bcap31",
                "Axl",
                "Cfl2",
                "Ogn"
            ],
            "expression_sum": {
                "class1": 3828.0,
                "class2": 17925.0
            },
            "uniprot_ids": [
                "Q61164",
                "Q3UYZ8",
                "Q05CK6",
                "Q3UZH8",
                "Q3USR8"
            ],
            "net_id": 0,
            "source_num": 12,
            "target_num": 250
        },
        "Cebpb": {
            "id": "Cebpb",
            "type": "TF",
            "symbol": "Cebpb",
            "source": [
                "E2f4",
                "Foxp1",
                "Pou5f1",
                "Atf4",
                "Ctcf",
                "Trp53",
                "Hdac1",
                "Tbx3",
                "Zfp57",
                "Hdac2",
                "Smc3",
                "Smarca5"
            ],
            "target": [
                "Arf1",
                "Dap",
                "Srsf9",
                "Ndufb4",
                "Syce2",
                "Mcm7",
                "Grsf1",
                "Zc3h15",
                "Cox8a",
                "2810004N23Rik",
                "Sec13",
                "Ndufa6",
                "Polr2k",
                "Idh3g",
                "Rps27l",
                "Col12a1",
                "Lmna",
                "Ndufa2",
                "Hnrnpf",
                "Lmnb1",
                "Ddx39",
                "Cks2",
                "Shfm1",
                "Med28",
                "Hmgb2",
                "Cox6c",
                "Ddx3x",
                "Slbp",
                "Nedd4",
                "Calr",
                "Tubb5",
                "Nop56",
                "Slc25a3",
                "Lamtor1",
                "2810417H13Rik",
                "Nup43",
                "Cdc5l",
                "Sms",
                "Rbx1",
                "Ctcf",
                "Neat1",
                "Pmf1",
                "Ola1",
                "Sf3b6",
                "Lrrc59",
                "Ube2d2a",
                "Bnip3l",
                "Kpnb1",
                "Mapre1",
                "Snrpd1",
                "Rbm8a",
                "Col5a1",
                "Rpl36al",
                "Lap3",
                "Tgfb1i1",
                "Supt4a",
                "Rbbp7",
                "Hnrnpl",
                "Nedd8",
                "Pcnp",
                "Atpif1",
                "Snrnp40",
                "Gng12",
                "Tpm3",
                "Banf1",
                "Mrps36",
                "Qdpr",
                "Lig1",
                "Tpi1",
                "Pa2g4",
                "Gsr",
                "Atf4",
                "Hist1h2bc",
                "Hprt",
                "Nucks1",
                "Pcbp1",
                "Mrpl36",
                "Itgb1",
                "Ndufa4",
                "Cops5",
                "Eif4e",
                "Ndufv1",
                "H3f3a",
                "Rfc2",
                "Bccip",
                "Cyp51",
                "Lamtor2",
                "Trim28",
                "Nudt21",
                "Crip2",
                "Ube2l3",
                "Eef1d",
                "Etf1",
                "Zmat2",
                "Samm50",
                "Rrp15",
                "Nhp2l1",
                "Ran",
                "Atp5g3",
                "Gjb3",
                "Atp5g2",
                "Chd4",
                "Tceb1",
                "C1qbp",
                "Rpl36a",
                "Eif3m",
                "Snrpb2",
                "Umps",
                "Rps20",
                "Orc6",
                "Ybx1",
                "Sys1",
                "Tubb2b",
                "Tmem14c",
                "Rps6",
                "U2af1",
                "Slc29a1",
                "Atp5c1",
                "Mrps17",
                "Ccnd2",
                "Dnajc8",
                "Cox7c",
                "Nme4",
                "Eno1",
                "mt-Nd3",
                "Rpl22l1",
                "Lamtor4",
                "Smc4",
                "Mbd3",
                "Cct7",
                "Dctpp1",
                "Hsp90ab1",
                "Rpp25",
                "Tyms",
                "Ankrd11",
                "Rps8",
                "Ctsd",
                "Magoh",
                "Ndufc2",
                "H3f3b",
                "Dnajc2",
                "Ndufa12",
                "Cnih4",
                "Nudt19",
                "Dbi",
                "Tbx3",
                "Dstn",
                "Cyb5r3",
                "Ywhae",
                "Ppil1",
                "Rps15",
                "Arl1",
                "Cenpw",
                "Eif2s2",
                "Mrpl45",
                "Rpl31",
                "Smdt1",
                "Etfb",
                "Uqcrc2",
                "G3bp2",
                "Impdh2",
                "Prps1",
                "Akap12",
                "Hnrnpab",
                "Ahnak",
                "Nop58",
                "Ndufa11",
                "Cbx5",
                "Polr2c",
                "Cct8",
                "Dohh",
                "Denr",
                "Eif3g",
                "Aspscr1",
                "Btg2",
                "Parva",
                "Rpp21",
                "Polr2e",
                "Anapc13",
                "Pdcd5",
                "Tomm7",
                "Snrpc",
                "Mrpl49",
                "Tmbim6",
                "Atp5a1",
                "Psma5",
                "Pycr2",
                "Sf3a3",
                "Ccdc80",
                "March5",
                "Ddah2",
                "Gspt1",
                "Eprs",
                "Trmt112",
                "Rpl35",
                "Dazap1",
                "Ilf2",
                "Lman2",
                "Prmt1",
                "Sub1",
                "Morf4l1",
                "Col8a1",
                "Cdca8",
                "L1td1",
                "Lsm6",
                "Sf3b1",
                "Ndufs4",
                "Polr1d",
                "H2afy",
                "Mtf2",
                "Rpl17",
                "mt-Nd2",
                "Txndc17",
                "Spty2d1",
                "Nop10",
                "Mpdu1",
                "Gng5",
                "Hspe1",
                "Rps19",
                "Ube2c",
                "Msn",
                "Exosc8",
                "Mrpl20",
                "Ptx3",
                "Polr2h",
                "Psmb4",
                "Birc5",
                "Commd1",
                "Cox7b",
                "Selm",
                "Chtop",
                "Dppa5a",
                "Srpk1",
                "Lamtor5",
                "Avpi1",
                "Ssb",
                "Tinagl1",
                "Rpa2",
                "Mrps12",
                "Copz1",
                "Nfu1",
                "Ndufb5",
                "Set",
                "Thyn1",
                "Txnl4a",
                "Ssbp1",
                "Col4a1",
                "Sephs2",
                "Snrpd2",
                "1810022K09Rik",
                "Hdac1",
                "Naa10",
                "Hsp90b1",
                "Vdac2",
                "Mrpl24",
                "Snhg6",
                "Minos1",
                "Anapc11",
                "Gas6",
                "Nans",
                "Uchl5",
                "Cxcl1",
                "Ppig",
                "Dcun1d5",
                "Lsm12",
                "Gins4",
                "Tceb2",
                "Rhoc",
                "Mrpl42",
                "Rps2",
                "Polr2g",
                "Sumo3",
                "Uqcrfs1",
                "Naa50",
                "Ywhah",
                "Fbn2",
                "Hdac2",
                "Timm17a",
                "Rpn2",
                "Bub3",
                "Esf1",
                "Plekhj1",
                "Mrfap1",
                "Hsd17b14",
                "Slc25a5",
                "Uqcrh",
                "Foxp1",
                "Tsn",
                "Rpl37a",
                "Drg1",
                "Ptma",
                "Ndufb6",
                "Ybx3",
                "Rps10",
                "Dad1",
                "Sept1",
                "0610007P14Rik",
                "Yif1b",
                "Rpl39",
                "Tsfm",
                "Capn2",
                "Ptov1",
                "Rps27rt",
                "Gpx4",
                "Gcsh",
                "Carhsp1",
                "Gltscr2",
                "Mkrn1",
                "Rrp1",
                "Pdcl3",
                "Rpl23",
                "Rpl30",
                "Sfr1",
                "Polr2f",
                "Rrs1",
                "Rbm17",
                "Ly6e",
                "Lsm5",
                "Rnf7",
                "Cdc20",
                "Cd9",
                "Ddx24",
                "Rpl6",
                "Srm",
                "Thbs1",
                "Psmb7",
                "Spr",
                "Pnkd",
                "Ppid",
                "1110008F13Rik",
                "Anapc5",
                "Hnrnpa1",
                "Ssrp1",
                "Itm2a",
                "Rnf187",
                "Psmb6",
                "Ndufs3",
                "Uba2",
                "Mbnl2",
                "Snrpe",
                "B2m",
                "Psat1",
                "Gdi2",
                "Taf7",
                "Mrpl28",
                "Psmc6",
                "Grpel1",
                "Sec61g",
                "Ythdc1",
                "Snrnp27",
                "Srsf1",
                "Igfbp3",
                "Socs3",
                "Uqcrc1",
                "Rcc2",
                "Impa2",
                "Bola2",
                "Tmem160",
                "C1d",
                "Cisd2",
                "Ncl",
                "Rpl35a",
                "Timp2",
                "Syngr1",
                "Hspa4",
                "Arpp19",
                "Ccdc12",
                "Ddx21",
                "Naca",
                "Ube2i",
                "Rpl24",
                "Brix1",
                "Fam136a",
                "Siva1",
                "Rps4x",
                "Fkbp8",
                "Nip7",
                "Ncam1",
                "Ruvbl1",
                "Clpp",
                "Wls",
                "G3bp1",
                "Atp5b",
                "Cdc26",
                "Actb",
                "Immt",
                "Nars",
                "Hells",
                "Tfcp2l1",
                "Rpl38",
                "Glo1",
                "Serpine2",
                "Nme1",
                "Crabp2",
                "Csnk1a1",
                "Rpl10-ps3",
                "Rpl3",
                "Cebpzos",
                "Rpl12",
                "Snw1",
                "Ndufa7",
                "Dnajc19",
                "Cdv3",
                "Hmgn1",
                "Ctbp2",
                "Btf3l4",
                "Ppp4c",
                "Imp3",
                "Mrpl13",
                "Ak2",
                "Wdr77",
                "Atp5j",
                "Srrm1",
                "Srp19",
                "Emd",
                "Pdgfa",
                "Uqcr10",
                "Rpsa",
                "H2afz",
                "Mrps21",
                "Tardbp",
                "Rpl5",
                "Dph3",
                "Dpy30",
                "Serpine1",
                "Fblim1",
                "Odc1",
                "Mrpl18",
                "Ufc1",
                "Ifitm1",
                "Acat2",
                "Mrpl12",
                "Cdc37",
                "Cisd1",
                "Rpl28",
                "Xist",
                "Gpx3",
                "Timm8b",
                "Rpl26",
                "Uqcrb",
                "mt-Co2",
                "Rps3",
                "Atox1",
                "Srsf10",
                "Ube2v2",
                "Rpl19",
                "Timm13",
                "Eif3h",
                "Mcm3",
                "Adprh",
                "Rps3a1",
                "Srsf3",
                "Coprs",
                "Cmpk1",
                "Rpl7",
                "Fn1",
                "Znrd1",
                "Wdr18",
                "Polr2m",
                "1810058I24Rik",
                "Rpl8",
                "Eif4a1",
                "Pcbp2",
                "Polr2i",
                "Polr1c",
                "Snrnp25",
                "Fhl2",
                "Ccdc34",
                "Mrps33",
                "Rif1",
                "Morf4l2",
                "Rbbp4",
                "Eif3e",
                "Emp3",
                "2700060E02Rik",
                "Park7",
                "Dynll1",
                "Hspd1",
                "Stoml2",
                "Cops7a",
                "Sod1",
                "Ppih",
                "Uqcc3",
                "Tpx2",
                "Dnajb6",
                "Hspa5",
                "Gapdh",
                "Smarca5",
                "Idi1",
                "Ccnb1",
                "Ccnd3",
                "Khdrbs1",
                "Mrpl52",
                "Pno1",
                "Ecm1",
                "Adrm1",
                "Kif5b",
                "Hmgn2",
                "Hnrnpa2b1",
                "Oat",
                "Psmd2",
                "Ddx27",
                "Jade1",
                "Dnajc7",
                "Psmb2",
                "Fxr1",
                "Tceal8",
                "Cnn3",
                "Vps29",
                "Vdac1",
                "Anp32a",
                "Tra2a",
                "Zfp57",
                "Dynll2",
                "Ssu72",
                "Cnpy2",
                "Anp32e",
                "Pfdn6",
                "Ict1",
                "Rps25",
                "Rbmxl1",
                "Arglu1",
                "Imp4",
                "Ywhab",
                "Prelid1",
                "Ubb",
                "Zfp106",
                "Prrc1",
                "Paics",
                "Asna1",
                "Psmc5",
                "Echdc2",
                "Mrpl43",
                "Suclg1",
                "Hnrnpd",
                "Ash2l",
                "Hat1",
                "Rnps1",
                "H13",
                "Tpt1",
                "Ppib",
                "Stmn2",
                "Ptbp1",
                "Aldoa",
                "Lpp",
                "Dnajc21",
                "Snx3",
                "Strap",
                "Psmd13",
                "Rnaseh2a",
                "Rfc1",
                "Cox6b1",
                "Ptrf",
                "Atp5o",
                "Nasp",
                "Ndufs6",
                "Poldip2",
                "Eif4e2",
                "Sod2",
                "Hist1h1c",
                "Tnfrsf12a",
                "Phgdh",
                "Gtf2e2",
                "Pdap1",
                "Rpl11",
                "Anxa1",
                "Ube2a",
                "Pfn1",
                "Snrpg",
                "Ddx39b",
                "Grb10",
                "Exosc4",
                "Fkbp4",
                "Pgam1",
                "Snrnp70",
                "Ppdpf",
                "Gmnn",
                "Rpl27a",
                "Llph",
                "Eif4b",
                "Anp32b",
                "Edf1",
                "Fstl1",
                "Mdm2",
                "Eif2b5",
                "Gart",
                "Ndufa10",
                "App",
                "Tomm40",
                "Eif3i",
                "Smc2",
                "Mtch2",
                "Psmd4",
                "Gins2",
                "Uck2",
                "Rps16",
                "Bcat1",
                "Tmed3",
                "Mrps16",
                "Psmd12",
                "Ndufb10",
                "Cth",
                "Txnl1",
                "Rbm25",
                "Eef1g",
                "Ube2e3",
                "2700094K13Rik",
                "Acadm",
                "Dut",
                "Alyref",
                "Hspa14",
                "Polr2j",
                "Got2",
                "Slirp",
                "Ckb",
                "Cwc15",
                "Ndufs8",
                "Pim1",
                "Rplp2",
                "Mif",
                "Aimp1",
                "Rheb",
                "Chchd3",
                "Eif4a2",
                "Gtf2a2",
                "Cnbp",
                "Sdhd",
                "Axl",
                "Rps5",
                "Rpl41",
                "Cd44",
                "Ruvbl2",
                "Cdt1",
                "Ier2",
                "Dkc1",
                "Marcksl1",
                "Atp5g1",
                "Tuba1b",
                "Aurka",
                "Psma6",
                "Rpl7l1",
                "Snrpa",
                "St13",
                "Thoc7",
                "Mlf2",
                "Rpl9",
                "Vbp1",
                "Atp5j2",
                "Nubp1",
                "E2f4",
                "D8Ertd738e",
                "Hn1",
                "Mrpl14",
                "Phb",
                "Tra2b",
                "Hmgcs1",
                "Emc6",
                "Rps24",
                "Bola1",
                "Aurkaip1",
                "Timp3",
                "Nhp2",
                "Exosc5",
                "Sh3glb1",
                "Atp5h",
                "Ppia",
                "Eif3k",
                "Mrps34",
                "Atp5k",
                "Mphosph8",
                "Ppp1ca",
                "Aamp",
                "Chchd1",
                "Gars",
                "Psmc3",
                "Luc7l2",
                "Ndufb9",
                "Col3a1",
                "Utp3",
                "Fus",
                "Ociad1",
                "Ddost",
                "Rest",
                "Cct3",
                "Psma7",
                "Gnl3",
                "Trim27",
                "Psma1",
                "Dtymk",
                "Emc10",
                "2410015M20Rik",
                "Hdgf",
                "Prnp",
                "Id1",
                "Csnk2b",
                "Rpl37",
                "Ppp1cc",
                "Nrp1",
                "Dnttip2",
                "Sae1",
                "Hnrnpk",
                "Mrps24",
                "Ezr",
                "Ewsr1",
                "Ndufv2",
                "Fbxo22",
                "Fam49b",
                "Bax",
                "Cisd3",
                "Gfer",
                "Eif3c",
                "Cox7a2",
                "Col5a2",
                "Ube2j2",
                "Lsm3",
                "Gcat",
                "Eif5",
                "Spag7",
                "Tomm5",
                "Txn1",
                "Vapa",
                "Erh",
                "Fscn1",
                "Emc8",
                "Bclaf1",
                "Psip1",
                "Rpl29",
                "Naa38",
                "Ung",
                "Ptges3",
                "H2-D1",
                "Wtap",
                "Dnaja2",
                "Lbh",
                "Cenpv",
                "Ywhaq",
                "Pdha1",
                "Ubald2",
                "Filip1l",
                "Adss",
                "Rps7",
                "Igf2bp1",
                "Trnau1ap",
                "Eef1e1",
                "Hsd17b10",
                "Oaz1",
                "Stra13",
                "Sec61b",
                "Snx5",
                "Got1",
                "Bzw2",
                "Pgp",
                "Sumo2",
                "Nudc",
                "Pou5f1",
                "Gnb2l1",
                "Mrpl21",
                "Pold2",
                "Nsa2",
                "Ddx46",
                "Rfc5",
                "Pfdn4",
                "Mrps15",
                "Psmc4",
                "Psmd14",
                "Npm1",
                "Sap30",
                "Gtpbp4",
                "Cpe",
                "Atf5",
                "Srsf7",
                "Mat2a",
                "Csrp2",
                "Brd2",
                "Ccdc124",
                "Prdx2",
                "Abcf1",
                "Rplp0",
                "Psmb5",
                "Dcn",
                "Ube2n",
                "Smn1",
                "Cyb5b",
                "Psmb1",
                "Coa3",
                "Akr1b3",
                "Eif4g2",
                "Raly",
                "Uqcc2",
                "Nsmce4a",
                "Rps26",
                "Get4",
                "Lage3",
                "Npm3",
                "Lsm2",
                "Chmp2a",
                "Sumo1",
                "Nudcd2",
                "Aprt",
                "Fcf1",
                "Mrps18c",
                "Cacybp",
                "Rpl22",
                "Nolc1",
                "Rpl4",
                "Glud1",
                "Rps28",
                "Hnrnpu",
                "Srsf11",
                "Hmgb3",
                "Rpl21",
                "Ppp1r2",
                "Map1lc3b",
                "Atp5l",
                "Son",
                "Pop5",
                "Cct2",
                "Acp1",
                "Sox4",
                "Dtl",
                "Ahsa1",
                "Eif1ax",
                "Parp1",
                "Dhfr",
                "Mrps7",
                "Cdca3",
                "Metap2",
                "Ube2s",
                "Idh2",
                "Snrpd3",
                "Psmd3",
                "Fam32a",
                "Bcas2",
                "Cops4",
                "Ciapin1",
                "Pfdn2",
                "Atp5f1",
                "Arid5b",
                "Srp9",
                "Nfatc2ip",
                "Ino80e",
                "Ppa1",
                "Prdx1",
                "Rsl24d1",
                "Terf1",
                "Rpl13",
                "Mrpl34",
                "Atrx",
                "Nt5dc2",
                "Tgif1",
                "Sfpq",
                "Mrto4",
                "Parl",
                "Lipe",
                "Clns1a",
                "Mdh2",
                "Gabarap",
                "Uqcr11",
                "Phb2",
                "Rps27a",
                "Rpl15",
                "Gtf2h5",
                "Arl6ip1",
                "Tcof1",
                "Des",
                "Ier3",
                "Cuta",
                "1500015O10Rik",
                "Eif1a",
                "Cox5a",
                "Tmem11",
                "Ubap2l",
                "Phldb2",
                "Cct4",
                "Rdx",
                "Cdkn1a",
                "Pkm",
                "H2afv",
                "Prrc2c",
                "Marcks",
                "Psmg4",
                "Psmc2",
                "Sf3b4",
                "Hnrnpc",
                "Fkbp3",
                "Rangrf",
                "Rpl7a",
                "Fam104a",
                "Prrx2",
                "Manf",
                "Tpr",
                "Ddx18",
                "Rrp9",
                "Hint1",
                "Sf3a2",
                "Ccne1",
                "Srsf6",
                "Ebna1bp2",
                "Tmpo",
                "Cycs",
                "Klf2",
                "Dhx16",
                "Sdhb",
                "Taf10",
                "Wdr43",
                "Klf5",
                "Yrdc",
                "Pabpc1",
                "Eif6",
                "Tsen34",
                "Fam60a",
                "Wbp11",
                "2810428I15Rik",
                "Hspa8",
                "Postn",
                "Ldhb",
                "Taf1d",
                "Serbp1",
                "Kdm1a",
                "Ncor1",
                "1810037I17Rik",
                "Rsrc2",
                "Hsp90aa1",
                "Usmg5",
                "Mdh1",
                "Ppp2ca",
                "Rps17",
                "Rae1",
                "Pin4",
                "Mrps18b",
                "Txn2",
                "Clta",
                "Cox17",
                "Ppp1r11",
                "Mrpl4",
                "Wisp1",
                "Ndufb3",
                "Gar1",
                "Cks1b",
                "Ftsj3",
                "Folr1",
                "Psmc1",
                "Tpd52l2",
                "Knop1",
                "Psmd6",
                "Pin1",
                "Lgals3",
                "Sfxn1",
                "Eif2s1",
                "Swi5",
                "Nup62",
                "Cenpe",
                "Rab18",
                "Calm3",
                "Arf6",
                "Ubxn1",
                "Cdk1",
                "Eif4h",
                "Psmd7",
                "Anapc15",
                "Gabarapl2",
                "Slc39a1",
                "Stub1",
                "Prrx1",
                "Mrpl2",
                "Ranbp1",
                "Zwint",
                "Ppm1g",
                "Eif1",
                "Tbcb",
                "Wdr89",
                "Pdcd6",
                "H2afx",
                "Hnrnpm",
                "Capza2",
                "Loxl1",
                "2310036O22Rik",
                "Fdft1",
                "Srsf5",
                "Hnrnph1",
                "Wdr74",
                "Bex4",
                "Ywhaz",
                "Pomp",
                "Rpl36",
                "Rps21",
                "Psmb3",
                "Ddx5",
                "Rbm39",
                "Pigyl",
                "Eef1b2",
                "Eif3l",
                "Tmem258",
                "Tcp1",
                "Sarnp",
                "Mrpl15",
                "Ezh2",
                "Mybl2",
                "Pfdn1",
                "Slc2a3",
                "Pmm1",
                "Actl6a",
                "Pdia4",
                "Sssca1",
                "Ier5",
                "Gpx1",
                "Smc3",
                "Thrap3",
                "Hspb1",
                "Kdm5b",
                "Nfkbia",
                "Eif3b",
                "Zfp207",
                "Ube2k",
                "Ndufs2",
                "Ddt",
                "Spc24",
                "Asns",
                "Ifitm2",
                "Fdps",
                "Lsm8",
                "Ndufs7",
                "Cenpf",
                "Mcm2",
                "Gm16286",
                "Mrpl17",
                "Ldha",
                "Apex1",
                "Cct5",
                "Ccnb2",
                "Mrpl35",
                "1110004F10Rik",
                "Mrpl40",
                "Lox",
                "Iscu",
                "Fubp1",
                "Arpc5l",
                "Rps29",
                "Ccdc58",
                "Cenpa",
                "Prdx6",
                "Mdk",
                "Psmd1",
                "Ndufab1",
                "Plekhf2",
                "Snrpf",
                "Mcl1",
                "Pole4",
                "Fbl",
                "Ndufv3",
                "Rps26-ps1",
                "Calm2",
                "Stmn1",
                "Btf3",
                "Smarcc1",
                "Cdh11",
                "Rpl18",
                "Hax1",
                "Atp5d",
                "Taldo1",
                "Vcp",
                "Mrps5",
                "Prc1",
                "U2surp",
                "Fkbp2",
                "Ppan",
                "Rps9",
                "Upp1",
                "Mrpl30",
                "Rwdd1",
                "Cyc1",
                "Fnta",
                "Dnaja1",
                "Tmem176b",
                "Pmvk",
                "Elof1",
                "Lrpap1",
                "Wdr5",
                "Pebp1",
                "Sgk1",
                "Ube2d3",
                "Aars",
                "Marc2",
                "Snrpa1",
                "Paip2",
                "Srrm2",
                "Rnaseh2b",
                "Top1",
                "Ubl5",
                "Hmgb1",
                "Ndufc1",
                "Eif3d",
                "Cyb5a",
                "Psma4",
                "Ap2s1",
                "Rpl34",
                "Mlec",
                "Rab7",
                "Mrps14",
                "Nap1l1",
                "Mrps25",
                "Actr3",
                "Mrps10",
                "Tbrg1",
                "Hspa9",
                "Fkbp10",
                "Chchd2",
                "Pim3",
                "E130012A19Rik",
                "Luc7l3",
                "Ndufs5",
                "Gsn",
                "Ndufb8",
                "Tmem97",
                "Srsf2",
                "Prdx4",
                "Pak1ip1",
                "Nxt1",
                "Trp53",
                "Lhfp",
                "Rbm3",
                "Psmd11",
                "Cggbp1",
                "Zfp706",
                "Rad23b",
                "Nudt4",
                "Supt16",
                "Ost4",
                "Mrpl51",
                "Eif4g1",
                "Rhoa",
                "Cdk4",
                "Rsl1d1",
                "Syncrip",
                "Caprin1",
                "Ssr2",
                "Cebpz",
                "Tomm20",
                "Cbx1",
                "Tomm70a",
                "Msmo1",
                "Mak16",
                "Bzw1",
                "Ak6",
                "Rbpms",
                "Hnrnpa3",
                "Prpf40a",
                "Pbdc1",
                "Rps19bp1",
                "2010107E04Rik",
                "Tecr",
                "Mybbp1a",
                "Sf3b2",
                "Atp1b1",
                "Slc3a2",
                "Stip1",
                "Chchd4",
                "Rpl14",
                "Tma7",
                "Camk2d",
                "Lsm4",
                "Tpd52",
                "Eif1ad",
                "Rps13",
                "Acta1",
                "Eif5a",
                "Rps11",
                "Cfl1",
                "Rrp1b",
                "Sparc",
                "Uchl3",
                "Cops6",
                "Hnrnpdl",
                "Eif3a",
                "Rtn3",
                "Adh5",
                "l7Rn6",
                "Atp5e",
                "Cdc34",
                "Gps1",
                "Tipin",
                "Cmtm7",
                "Rrm2",
                "Sh3bgrl",
                "Itm2b",
                "Rplp1",
                "Rnf10",
                "Psma2",
                "Pfdn5",
                "Nsun2",
                "Ndufb7",
                "Shmt1",
                "Alkbh1",
                "Slc25a39",
                "Rps14",
                "Rpl32",
                "Chchd10",
                "Bud31",
                "Mcm5",
                "Rpl13a",
                "Nol7",
                "Tkt",
                "Apoe",
                "Gsta4",
                "Snrpb",
                "Zfp428",
                "Tomm22",
                "Lyar",
                "Erp29",
                "Gpx8",
                "Gpi1",
                "Tmem167",
                "Cfdp1",
                "Eif5b",
                "Emg1",
                "Eif4ebp1",
                "Rpl23a-ps3",
                "Nono",
                "Csde1",
                "Mcm6",
                "Prpf19",
                "Sugt1",
                "Eif4a3",
                "Dek",
                "Top2a",
                "Papola",
                "Tmem256",
                "Cetn3",
                "Galk1",
                "Ccna2",
                "Tubb4b",
                "Eed",
                "Ndufa5",
                "Mrpl11",
                "Mesdc2",
                "Cox5b",
                "Arhgdia",
                "Ralbp1",
                "Acin1",
                "Aebp1",
                "Rbpms2",
                "Uhrf1",
                "Guk1",
                "Psma3",
                "Skp1a",
                "Rps18",
                "Phf5a",
                "Rps23",
                "U2af2",
                "P3h4",
                "Myl12b"
            ],
            "expression_sum": {
                "class1": 14867.0,
                "class2": 3147.0
            },
            "uniprot_ids": [
                "P28033",
                "Q3UPN9",
                "O70192"
            ],
            "net_id": 0,
            "source_num": 12,
            "target_num": 647
        },
        "Trim28": {
            "id": "Trim28",
            "type": "TF",
            "symbol": "Trim28",
            "source": [
                "Pou5f1",
                "Foxp1",
                "Junb",
                "Fos",
                "Jun",
                "Sox2",
                "Tbx3",
                "Nanog",
                "Jund",
                "Twist2",
                "Cebpb",
                "Zfp57",
                "Sox4"
            ],
            "target": [
                "Dap",
                "Cd44",
                "H2afj",
                "Ddb1",
                "Myh9",
                "Cdt1",
                "Thbs1",
                "Rasgrp2",
                "Lims1",
                "Igfbp2",
                "Slc25a4",
                "Calcoco2",
                "Lmna",
                "Folr1",
                "Cthrc1",
                "Lgals3",
                "Pgrmc1",
                "Fos",
                "Tagln2",
                "Rps15a",
                "Rexo2",
                "Pdlim7",
                "Rab18",
                "Mfge8",
                "Mbnl2",
                "Canx",
                "B2m",
                "Taf7",
                "Neat1",
                "Myh10",
                "Timp3",
                "Gemin7",
                "Akr1a1",
                "Exosc5",
                "Sh3glb1",
                "Akt1s1",
                "Impa2",
                "Atp2b1",
                "Cisd2",
                "Mylpf",
                "Nktr",
                "Col5a1",
                "Ctgf",
                "Klf6",
                "Pcolce",
                "Crxos",
                "Syngr1",
                "Crip1",
                "Morc1",
                "Sec62",
                "Rps12",
                "Maged1",
                "Cyr61",
                "Kdelr1",
                "Rab14",
                "Serpinh1",
                "Atp6v1g1",
                "Gnb2",
                "Dpysl3",
                "Mybl2",
                "Slc2a3",
                "Ddost",
                "Hcfc1r1",
                "Sptbn1",
                "Id3",
                "Ap3s1",
                "Gng12",
                "Ncam1",
                "Wls",
                "Ndufa8",
                "Srp14",
                "Sept7",
                "Actg1",
                "Rcn2",
                "Tsg101",
                "Prnp",
                "Id1",
                "Hmga2",
                "Hspb1",
                "Kdm5b",
                "Hist1h2bc",
                "Hells",
                "Nfkbia",
                "Tfcp2l1",
                "Col1a2",
                "Atp6v0b",
                "F2r",
                "Fkbp9",
                "Igfbp4",
                "Id2",
                "Itgb1",
                "Fabp5",
                "Mettl9",
                "Chmp4b",
                "Tcf4",
                "Crabp2",
                "Ssr1",
                "Cyp51",
                "Lxn",
                "Pgls",
                "Ift20",
                "Anxa2",
                "Vmp1",
                "Col5a2",
                "mt-Nd4",
                "Crip2",
                "Vim",
                "Wdr1",
                "Ube2b",
                "Tmed2",
                "Gjb3",
                "Tpm1",
                "Sdcbp",
                "Thy1",
                "Surf4",
                "Utf1",
                "Phlda1",
                "Myeov2",
                "Pdgfa",
                "Cst3",
                "Higd1a",
                "S100a6",
                "Dpy30",
                "Serpine1",
                "H2-D1",
                "Ifitm1",
                "Cd63",
                "Capg",
                "S100a11",
                "Scand1",
                "Mtdh",
                "Eef1a1",
                "Rtn4",
                "Slc29a1",
                "Cenpv",
                "Ctsl",
                "Col6a1",
                "Rala",
                "Sdc2",
                "Filip1l",
                "Plk2",
                "Tln1",
                "Cdh11",
                "Snhg12",
                "Mt2",
                "Fbln5",
                "Lrrc58",
                "Btg1",
                "Fkbp2",
                "Fn1",
                "Dctpp1",
                "Glipr2",
                "Hes1",
                "Ap1s1",
                "Tuba4a",
                "Upp1",
                "Zfp42",
                "Malat1",
                "Gnai2",
                "Tmem176b",
                "Mfap2",
                "Tm9sf3",
                "Ctsd",
                "Pou5f1",
                "Fhl2",
                "Oxct1",
                "Atxn10",
                "Atp6v0e",
                "Dbi",
                "Tmem59",
                "Tbx3",
                "Emp3",
                "Gm26917",
                "Pafah1b2",
                "Anxa3",
                "Hbegf",
                "Arpc3",
                "Ilk",
                "Arl1",
                "Arpc1a",
                "Sep15",
                "Lsm7",
                "Cpe",
                "Atf5",
                "Cdc42",
                "Eef2",
                "Hsbp1",
                "Hspa5",
                "Rnh1",
                "Rrbp1",
                "Actr3",
                "Cd47",
                "Efemp2",
                "Fbln2",
                "Mpc1",
                "Copb2",
                "Sars",
                "Nupr1",
                "Smim14",
                "Ecm1",
                "Sdpr",
                "Lgals1",
                "Ahnak",
                "Kif5b",
                "Mpc2",
                "Copz2",
                "Ptp4a1",
                "Lsp1",
                "Gsn",
                "Oat",
                "Cited2",
                "Jam2",
                "Rab2a",
                "Tspo",
                "Btg2",
                "Parva",
                "Selk",
                "Jund",
                "Rab11a",
                "Zfp57",
                "Ankrd1",
                "Anxa5",
                "Eif3f",
                "Tmbim6",
                "Mtch1",
                "Ttc3",
                "Sec61a1",
                "Iqgap1",
                "Kdelr2",
                "Rbmxl2",
                "Fermt2",
                "Rpn1",
                "Gas1",
                "Sqstm1",
                "Lman1",
                "F3",
                "Rcn1",
                "Arpc2",
                "Phlda3",
                "Romo1",
                "Tpm2",
                "Ergic3",
                "Sdc4",
                "Ubc",
                "Rock2",
                "Cdc42ep3",
                "Tcea3",
                "L1td1",
                "Rpl27",
                "Psap",
                "Rbms1",
                "P4hb",
                "Scd2",
                "Dlk1",
                "Zyx",
                "Map1b",
                "Rab34",
                "Adipor1",
                "Mtf2",
                "Bsg",
                "Sox2",
                "Ppic",
                "Chchd4",
                "Trh",
                "Sec22b",
                "Camk2d",
                "mt-Cytb",
                "Cyba",
                "Efemp1",
                "Gdf3",
                "Gsto1",
                "Ppib",
                "Vcan",
                "Sparc",
                "Stmn2",
                "Sept2",
                "Abracl",
                "Serpinb6a",
                "Lpp",
                "Ctbp1",
                "Csrp1",
                "Selm",
                "Ckap4",
                "Esd",
                "Dppa5a",
                "Meg3",
                "Atxn7l3b",
                "Comt",
                "Lamp1",
                "Rcor2",
                "Ndufa13",
                "Ptrf",
                "Gadd45g",
                "Jun",
                "Capns1",
                "Itm2b",
                "Pdia3",
                "Rpl10a",
                "Txndc5",
                "Ptms",
                "Shmt1",
                "Hist1h1c",
                "Cd24a",
                "Smap1",
                "Lipe",
                "Tnfrsf12a",
                "Krtcap2",
                "Gabarap",
                "Phgdh",
                "Serf2",
                "Hsp90b1",
                "Mtpn",
                "Ier3",
                "Grb10",
                "Apoe",
                "Commd3",
                "Gas6",
                "Fst",
                "Gpi1",
                "Phldb2",
                "Palld",
                "Tram1",
                "Igfbp7",
                "Fhl1",
                "Cdkn1a",
                "Fstl1",
                "Fam162a",
                "Rhoc",
                "Mdm2",
                "Ifitm3",
                "Mrpl33",
                "Emp1",
                "S100a10",
                "Rpn2",
                "Prrx2",
                "Tmed3",
                "Hsd17b14",
                "Tbca",
                "Cstb",
                "Foxp1",
                "H19",
                "Ptp4a2",
                "Fbxo15",
                "Cope",
                "Dnajc3",
                "Myl12a",
                "Rac1",
                "Rer1",
                "S100a4",
                "Klf2",
                "Cald1",
                "Fau",
                "Sri",
                "Prdx5",
                "Clic1",
                "Emc7",
                "Gnas",
                "Rbpms2",
                "Aebp1",
                "Arf4",
                "Calu",
                "Arpc5",
                "Gm7325",
                "Sept1",
                "Hdlbp",
                "Limd2",
                "Tmed9",
                "Postn",
                "Cnih1",
                "Carhsp1",
                "Alpl",
                "Mkrn1",
                "Pdcl3",
                "Kdelr3",
                "P3h4",
                "Bcap31",
                "Axl",
                "Cfl2"
            ],
            "expression_sum": {
                "class1": 7383.0,
                "class2": 112063.0
            },
            "uniprot_ids": [
                "Q62318",
                "Q5EBP9",
                "Q3UDN8"
            ],
            "net_id": 0,
            "source_num": 13,
            "target_num": 187
        },
        "Smarca5": {
            "id": "Smarca5",
            "type": "TF",
            "symbol": "Smarca5",
            "source": [
                "Tcf4",
                "Pou5f1",
                "Foxp1",
                "Cebpb",
                "Nfib",
                "Junb",
                "Fos",
                "Jun",
                "Sox2",
                "Hes1",
                "Nanog",
                "Jund",
                "Hdac2"
            ],
            "target": [
                "Hsp90aa1",
                "Arf1",
                "Ppp2ca",
                "Rps17",
                "Fis1",
                "Zc3h15",
                "Cox8a",
                "Ddb1",
                "Myh9",
                "Thbs1",
                "Txn2",
                "Lims1",
                "Sec13",
                "1110008F13Rik",
                "Slc25a4",
                "Col12a1",
                "Lmna",
                "Thoc7",
                "Ndufa2",
                "Mlf2",
                "Psmd6",
                "Ssrp1",
                "Ier3ip1",
                "Rnf187",
                "Swi5",
                "Cxcl12",
                "Eif2s1",
                "Pgrmc1",
                "Shfm1",
                "Fos",
                "Tagln2",
                "Cox6c",
                "Slbp",
                "Ndufs3",
                "Nedd4",
                "Calr",
                "Pdlim7",
                "Rab18",
                "Mfge8",
                "Mbnl2",
                "Ppp1r14b",
                "B2m",
                "Mrpl14",
                "Slc25a3",
                "Psat1",
                "Eif4h",
                "Psmd7",
                "Tra2b",
                "Psmc6",
                "Zwint",
                "Rps24",
                "Eif1",
                "Ola1",
                "Snrnp27",
                "Timp3",
                "Gemin7",
                "Lrrc59",
                "Uqcrc1",
                "Akr1a1",
                "Sh3glb1",
                "Ube2d2a",
                "Hras",
                "Bnip3l",
                "Kpnb1",
                "Cisd2",
                "C1d",
                "Nktr",
                "Col5a1",
                "Ppia",
                "Capza2",
                "Klf6",
                "Pcolce",
                "Tgfb1i1",
                "Loxl1",
                "Hnrnph1",
                "Syngr1",
                "Ywhaz",
                "Gars",
                "Sec62",
                "Rps12",
                "Rpl36",
                "Ddx5",
                "Ndufb9",
                "Rbm39",
                "Pcnp",
                "Ube2i",
                "Cyr61",
                "Kdelr1",
                "Serpinh1",
                "Gnb2",
                "Fus",
                "Ddost",
                "Rps4x",
                "Id3",
                "Ap3s1",
                "Gng12",
                "G3bp1",
                "Fam195b",
                "Atp5b",
                "Psma1",
                "Actg1",
                "Ier5",
                "2410015M20Rik",
                "Id1",
                "Nars",
                "Hmga2",
                "Nrp1",
                "Eif3b",
                "Ube2m",
                "Fkbp9",
                "Rpl38",
                "Igfbp4",
                "Itgb1",
                "Ndufa4",
                "Ewsr1",
                "Cops5",
                "Mettl9",
                "Nme1",
                "Tcf4",
                "Csnk1a1",
                "Ssr1",
                "Ndufs2",
                "Pgls",
                "Lxn",
                "Tmsb10",
                "Rpl3",
                "Nudt21",
                "Rpl12",
                "Ube2l3",
                "Btf3l4",
                "Eef1d",
                "Sh3bgrl3",
                "Lsm8",
                "Ddah1",
                "Ube2b",
                "Ube2j2",
                "Gm16286",
                "Eif5",
                "Tmed2",
                "Vapa",
                "Ran",
                "Fscn1",
                "Atp5g3",
                "Tpm1",
                "Ldha",
                "Surf4",
                "Atp5g2",
                "Srp19",
                "Cct5",
                "1110004F10Rik",
                "Myeov2",
                "Vasp",
                "Mrps21",
                "Bola3",
                "Cst3",
                "Rpl36a",
                "Lox",
                "Iscu",
                "Arpc5l",
                "Rpl5",
                "S100a6",
                "Rps20",
                "Dnaja2",
                "Sys1",
                "Ndufab1",
                "Mtdh",
                "Rps6",
                "Eef1a1",
                "Rtn4",
                "U2af1",
                "Ywhaq",
                "Tcf25",
                "Rala",
                "Sdc2",
                "Rpl26",
                "Cav1",
                "Calm2",
                "Srsf10",
                "Plk2",
                "Tln1",
                "Oaf",
                "Rps7",
                "Ube2v2",
                "Rpl18",
                "Snhg12",
                "Atp5d",
                "Mt2",
                "Chd3",
                "Rps3a1",
                "Lrrc58",
                "Btg1",
                "Cct7",
                "Fkbp2",
                "Fn1",
                "Glipr2",
                "Ap1s1",
                "Hsp90ab1",
                "Nenf",
                "Rps9",
                "Twist2",
                "Mrpl30",
                "Polr2m",
                "Rwdd1",
                "Snx5",
                "Malat1",
                "Gnai2",
                "Rps8",
                "Dnaja1",
                "Tmem176b",
                "Bzw2",
                "Eif4a1",
                "Sumo2",
                "Tm9sf3",
                "Pcbp2",
                "Ctsd",
                "Ndufc2",
                "H3f3b",
                "Col1a1",
                "Mrpl21",
                "Ndufa12",
                "Nsa2",
                "Oxct1",
                "Cnih4",
                "Dbi",
                "Ube2d3",
                "Tmem59",
                "Eif3e",
                "Pfdn4",
                "Cebpb",
                "Srrm2",
                "Top1",
                "Cyb5r3",
                "Ubl5",
                "Mrps15",
                "Psmc4",
                "Rps15",
                "Arpc3",
                "Scp2",
                "Ilk",
                "Arl1",
                "Arpc1a",
                "Ap2s1",
                "Mgat2",
                "Sep15",
                "Zfp36l1",
                "Dynll1",
                "Hspd1",
                "Cpe",
                "Atf5",
                "Arf5",
                "Eef2",
                "Hspa5",
                "Rnh1",
                "Rrbp1",
                "Uqcrc2",
                "Actr3",
                "Impdh2",
                "Gapdh",
                "Prdx2",
                "Mpc1",
                "Fth1",
                "Sars",
                "Sdpr",
                "Lgals1",
                "Rplp0",
                "Ahnak",
                "Hspa9",
                "Fkbp10",
                "Ube2n",
                "Kif5b",
                "Cct8",
                "Mpc2",
                "Lsp1",
                "Psmb1",
                "Psmd2",
                "Cited2",
                "Eif4g2",
                "Psmb2",
                "Uqcc2",
                "Denr",
                "Lhfp",
                "Rbm3",
                "Psmd11",
                "Cnn3",
                "Btg2",
                "Vdac1",
                "Selk",
                "Rad23b",
                "Chmp2a",
                "Pdcd5",
                "Ankrd1",
                "Anxa5",
                "Eif3f",
                "Tmbim6",
                "Anxa6",
                "Mtch1",
                "Rab10",
                "Rpl22",
                "Rsl1d1",
                "Anp32e",
                "Syncrip",
                "Sec61a1",
                "Caprin1",
                "Kdelr2",
                "Ict1",
                "Rpl4",
                "Glud1",
                "March5",
                "Ssr3",
                "Fermt2",
                "Hnrnpu",
                "Gspt1",
                "Srsf11",
                "Rpl18a",
                "Rpn1",
                "Gas1",
                "Ssr4",
                "Txndc9",
                "Ppp1r2",
                "Eprs",
                "Trmt112",
                "Lman1",
                "Msmo1",
                "Atp5l",
                "Mrpl54",
                "Son",
                "Ywhab",
                "Ilf2",
                "Phlda3",
                "Romo1",
                "Ergic3",
                "Bzw1",
                "Sdc4",
                "Ak6",
                "Ubc",
                "Hnrnpa3",
                "Prpf40a",
                "Cdc42ep3",
                "Mrps7",
                "Metap2",
                "Psap",
                "Ndufs4",
                "Polr1d",
                "Tm4sf1",
                "Rbms1",
                "Zyx",
                "Psmd3",
                "H2afy",
                "Rab34",
                "Bsg",
                "Sec22b",
                "Camk2d",
                "Txndc17",
                "Cops4",
                "Flna",
                "H13",
                "Ywhag",
                "Atp5f1",
                "Rps13",
                "Gsto1",
                "Tpt1",
                "Cfl1",
                "Gng5",
                "Vcan",
                "Rps19",
                "Sept2",
                "Abracl",
                "Exosc8",
                "Cops6",
                "Aldoa",
                "Srp9",
                "Hnrnpdl",
                "Eif3a",
                "Ctbp1",
                "Strap",
                "Csrp1",
                "Commd1",
                "Psmd13",
                "Selm",
                "Lamp1",
                "Ppa1",
                "Sat1",
                "Lamtor5",
                "Ssb",
                "Ptrf",
                "Gadd45g",
                "Rrm2",
                "Jun",
                "Capns1",
                "Pdia3",
                "Txndc5",
                "Pfdn5",
                "Ndufb7",
                "Rps14",
                "Sfpq",
                "Col4a1",
                "Rpl32",
                "Lipe",
                "Mdh2",
                "Snrpd2",
                "Krtcap2",
                "Rps27a",
                "Bud31",
                "Pdap1",
                "Brk1",
                "Gtf2h5",
                "Rpl11",
                "Prkar1a",
                "Rap1b",
                "Rpl13a",
                "Nol7",
                "Timp1",
                "Tuba1a",
                "Commd3",
                "Fst",
                "Tomm22",
                "Erp29",
                "Gpx8",
                "Gpi1",
                "Tmem167",
                "Phldb2",
                "Palld",
                "Rpl27a",
                "Tram1",
                "Eif5b",
                "Cct4",
                "Lsm12",
                "Pkm",
                "Fstl1",
                "Fam162a",
                "Rhoc",
                "Mdm2",
                "Ifitm3",
                "Mrpl33",
                "Psmc2",
                "Ndufa10",
                "App",
                "Uqcrfs1",
                "Ywhah",
                "Dek",
                "Bag1",
                "Tmem147",
                "Rpn2",
                "Bub3",
                "Fam104a",
                "Tmed3",
                "Mrfap1",
                "Hsd17b14",
                "Slc25a5",
                "Gap43",
                "Tpr",
                "Cstb",
                "Foxp1",
                "Srsf6",
                "Ndufa5",
                "Tmpo",
                "Ndufb10",
                "Ube2e3",
                "Mesdc2",
                "Eef1g",
                "Rbm25",
                "Arhgdia",
                "Rac1",
                "Cald1",
                "Sri",
                "Pabpc1",
                "Prdx5",
                "Tsen34",
                "Fkbp1a",
                "Got2",
                "Dad1",
                "Emc7",
                "Gnas",
                "Sept1",
                "Arf4",
                "Arpc5",
                "Hdlbp",
                "Cox4i1",
                "Ccnd1",
                "Myl6",
                "Sepw1",
                "Hspa8",
                "Serp1",
                "Rplp2",
                "Capn2",
                "Ptov1",
                "Aimp1",
                "Cnih1",
                "Gpx4",
                "Glrx3",
                "Gtf2a2",
                "Carhsp1",
                "Rrp1",
                "Rpl23",
                "Sfr1",
                "P3h4",
                "Junb",
                "Rbm17",
                "Map2k2",
                "Axl",
                "Ogn"
            ],
            "expression_sum": {
                "class1": 12333.0,
                "class2": 35054.0
            },
            "uniprot_ids": [
                "Q91ZW3",
                "B2RT32",
                "Q3URV3",
                "Q3U0F8"
            ],
            "net_id": 0,
            "source_num": 13,
            "target_num": 250
        },
        "Hdac2": {
            "id": "Hdac2",
            "type": "TF",
            "symbol": "Hdac2",
            "source": [
                "Tcf4",
                "Pou5f1",
                "Nfib",
                "Atf4",
                "Junb",
                "Fos",
                "Jun",
                "Sox2",
                "Hes1",
                "Nanog",
                "Jund",
                "Twist2",
                "Cebpb"
            ],
            "target": [
                "Rps5",
                "Hsp90aa1",
                "Arf1",
                "Dap",
                "Rpl41",
                "Rnf7",
                "Sar1a",
                "Srsf9",
                "Mdh1",
                "Tpm4",
                "Usmg5",
                "Dctn3",
                "Ppp2ca",
                "Cd44",
                "Ddx24",
                "H2afj",
                "Rps17",
                "Fis1",
                "Zc3h15",
                "Cox8a",
                "Ddb1",
                "Myh9",
                "Thbs1",
                "Psmb7",
                "Txn2",
                "Lims1",
                "Sec13",
                "Clta",
                "Ppp1r11",
                "1110008F13Rik",
                "Tuba1b",
                "Wisp1",
                "Slc25a4",
                "Cks1b",
                "Col12a1",
                "Rps27l",
                "Tspan4",
                "Map1lc3a",
                "Psma6",
                "Lmna",
                "Psmc1",
                "Hnrnpa1",
                "Ndufa2",
                "Thoc7",
                "Hnrnpf",
                "Ddx39",
                "Mlf2",
                "Psmd6",
                "Vbp1",
                "Ssrp1",
                "Atp5j2",
                "Ier3ip1",
                "Pck2",
                "Rnf187",
                "Lgals3",
                "Swi5",
                "Cxcl12",
                "Psmb6",
                "mt-Co1",
                "Pgrmc1",
                "Med28",
                "Shfm1",
                "Fos",
                "Tagln2",
                "Cox6c",
                "Rps15a",
                "Ddx3x",
                "Ndufs3",
                "Nedd4",
                "Rexo2",
                "Calr",
                "Pdlim7",
                "Rab18",
                "Tubb5",
                "Calm3",
                "Mfge8",
                "Mbnl2",
                "Arf6",
                "Canx",
                "D8Ertd738e",
                "Hn1",
                "Ppp1r14b",
                "Actn1",
                "Snrpe",
                "B2m",
                "Mrpl14",
                "Slc25a3",
                "Psat1",
                "Eif4h",
                "Lamtor1",
                "2810417H13Rik",
                "Psmd7",
                "Gdi2",
                "Stub1",
                "Prrx1",
                "Tra2b",
                "Ranbp1",
                "Mrpl28",
                "Psmc6",
                "Hmgcs1",
                "Zwint",
                "Rbx1",
                "Rps24",
                "Eif1",
                "Myh10",
                "Neat1",
                "Sec61g",
                "Ythdc1",
                "Ola1",
                "Snrnp27",
                "Sf3b6",
                "Tgfbi",
                "Ube2r2",
                "Timp3",
                "Srsf1",
                "Gemin7",
                "Wdr89",
                "Lrrc59",
                "Akr1a1",
                "Tdgf1",
                "Sh3glb1",
                "Ssr2",
                "Pdcd6",
                "Akt1s1",
                "Ube2d2a",
                "Hras",
                "Bnip3l",
                "Atp2b1",
                "Kpnb1",
                "Mapre1",
                "C1d",
                "Cisd2",
                "Mylpf",
                "Nktr",
                "Col5a1",
                "Ppia",
                "Snrpd1",
                "Ctgf",
                "Tspan3",
                "Hnrnpm",
                "Capza2",
                "Eif3k",
                "Klf6",
                "Rpl36al",
                "Pcolce",
                "Tgfb1i1",
                "Loxl1",
                "2310036O22Rik",
                "Ppp1ca",
                "Srsf5",
                "Hnrnph1",
                "Rpl35a",
                "Timp2",
                "Syngr1",
                "Chchd1",
                "Crip1",
                "Gars",
                "Psmc3",
                "Ywhaz",
                "Sec62",
                "Rps12",
                "Pomp",
                "Luc7l2",
                "Rpl36",
                "Rbbp7",
                "Rps21",
                "Ddx5",
                "Ndufb9",
                "Rbm39",
                "Maged1",
                "Arpp19",
                "Nedd8",
                "Pcnp",
                "Eef1b2",
                "Ube2i",
                "Arpc1b",
                "Col3a1",
                "Cyr61",
                "Eif3l",
                "Kdelr1",
                "Rab14",
                "Atpif1",
                "Serpinh1",
                "Tmem258",
                "Tcp1",
                "Atp6v1g1",
                "Rpl24",
                "Pttg1",
                "Gnb2",
                "Fus",
                "Dpysl3",
                "Ociad1",
                "Ddost",
                "Hcfc1r1",
                "Rps4x",
                "Sptbn1",
                "Id3",
                "Txnl1",
                "Ap3s1",
                "Gng12",
                "Cct3",
                "Fkbp8",
                "Ncam1",
                "Wls",
                "Psma7",
                "Hacd1",
                "G3bp1",
                "Ndufa8",
                "Pdia4",
                "Sept7",
                "Srp14",
                "Fam195b",
                "Tpm3",
                "Atp5b",
                "Banf1",
                "Actg1",
                "Immt",
                "Psma1",
                "Rcn2",
                "Emc10",
                "Ier5",
                "Tsg101",
                "2410015M20Rik",
                "Hdgf",
                "Prnp",
                "Smc3",
                "Thrap3",
                "Id1",
                "Csnk2b",
                "Nars",
                "Tpi1",
                "Hmga2",
                "Rpl37",
                "Dynlrb1",
                "Atf4",
                "Hprt",
                "Tmem234",
                "Nfkbia",
                "Nucks1",
                "Nrp1",
                "Col1a2",
                "Eif3b",
                "Ube2m",
                "Atp6v0b",
                "F2r",
                "Fkbp9",
                "Pcbp1",
                "Rpl38",
                "Serpine2",
                "Hnrnpk",
                "Igfbp4",
                "Mrps24",
                "Id2",
                "Itgb1",
                "Ndufa4",
                "Ewsr1",
                "Cops5",
                "Ssr3",
                "Eif4e",
                "Ndufv2",
                "Fabp5",
                "H3f3a",
                "Mettl9",
                "Nme1",
                "Chmp4b",
                "Tcf4",
                "Ube2k",
                "Igf2",
                "Csnk1a1",
                "Cd81",
                "Ssr1",
                "Ndufs2",
                "Cyp51",
                "Lxn",
                "Pgls",
                "Tmsb10",
                "Rpl3",
                "Nudt21",
                "Rpl12",
                "Ift20",
                "Anxa2",
                "Eif3c",
                "Vmp1",
                "Cox7a2",
                "Ndufa7",
                "Col5a2",
                "Cdv3",
                "Crip2",
                "Napa",
                "Cox7a2l",
                "mt-Nd4",
                "Vim",
                "Hmgn1",
                "Ube2l3",
                "Btf3l4",
                "Basp1",
                "Ppp4c",
                "Eef1d",
                "Sh3bgrl3",
                "Lsm8",
                "Wdr1",
                "Tnc",
                "Ube2b",
                "Ube2j2",
                "Mrpl13",
                "Vapa",
                "Eif5",
                "Tmed2",
                "Txn1",
                "Erh",
                "Mrpl17",
                "Atp5j",
                "Fscn1",
                "Atp5g3",
                "Srrm1",
                "Tpm1",
                "Ldha",
                "Sdcbp",
                "Thy1",
                "Surf4",
                "Atp5g2",
                "Chd4",
                "Srp19",
                "Cct5",
                "Phlda1",
                "Emd",
                "1110004F10Rik",
                "Myeov2",
                "Bclaf1",
                "Vasp",
                "Tceb1",
                "Rpsa",
                "H2afz",
                "Bola3",
                "Mrps21",
                "Psip1",
                "Vcl",
                "Cst3",
                "Rpl36a",
                "Eif3m",
                "Lox",
                "Arpc5l",
                "Rps29",
                "Rpl5",
                "Higd1a",
                "S100a6",
                "Prdx6",
                "Serpine1",
                "H2-D1",
                "Trappc4",
                "Rps20",
                "Ybx1",
                "Dnaja2",
                "Sys1",
                "Cd63",
                "Capg",
                "Lbh",
                "Ndufa3",
                "Tmem14c",
                "Cdc37",
                "Ndufab1",
                "Mtdh",
                "Scand1",
                "Rps6",
                "Snrpf",
                "Eef1a1",
                "Nfib",
                "mt-Nd5",
                "Maged2",
                "Rtn4",
                "Rpl28",
                "Atp5c1",
                "Ctsl",
                "Ccnd2",
                "Mrps17",
                "Tcf25",
                "Spcs1",
                "Ywhaq",
                "Cox6a1",
                "Col6a1",
                "Rala",
                "Sdc2",
                "Tax1bp3",
                "Pdha1",
                "Filip1l",
                "Rpl26",
                "Uqcrb",
                "Cav1",
                "Rps3",
                "Calm2",
                "Atox1",
                "Cox7c",
                "Srsf10",
                "Adss",
                "Btf3",
                "Plk2",
                "Tln1",
                "Cdh11",
                "Oaf",
                "Ostf1",
                "Rps7",
                "Tmem126a",
                "Ube2v2",
                "Rpl18",
                "Eno1",
                "Rpl19",
                "Atp5d",
                "Snhg12",
                "Taldo1",
                "Timm13",
                "mt-Nd3",
                "Eif3h",
                "Fbln5",
                "Rpl22l1",
                "Chd3",
                "Rps3a1",
                "Srsf3",
                "Vcp",
                "Smc4",
                "Cmpk1",
                "Mbd3",
                "Rpl7",
                "Fkbp2",
                "Btg1",
                "Cct7",
                "Fn1",
                "Glipr2",
                "Lrrc58",
                "Hes1",
                "Ap1s1",
                "Ndufb2",
                "Hsp90ab1",
                "Rps9",
                "Twist2",
                "Oaz1",
                "Mrpl30",
                "Rpp25",
                "Uqcrq",
                "Rcn3",
                "Sec61b",
                "BC031181",
                "Polr2m",
                "Rwdd1",
                "Snx5",
                "Malat1",
                "Gnai2",
                "Cyc1",
                "Rps8",
                "Dnaja1",
                "Tmem176b",
                "Eif4a1",
                "Sumo2",
                "Tm9sf3",
                "Pcbp2",
                "Ctsd",
                "Magoh",
                "Ndufc2",
                "Reep5",
                "Tmed10",
                "Pou5f1",
                "Fhl2",
                "H3f3b",
                "Glrx5",
                "Col1a1",
                "Elof1",
                "Gnb2l1",
                "Ndufa12",
                "Ube2e3",
                "Nsa2",
                "Oxct1",
                "Atxn10",
                "Wbp5",
                "Atp6v0e",
                "Cnih4",
                "Pebp1",
                "Dbi",
                "Morf4l2",
                "Ube2d3",
                "Tmem59",
                "Rbbp4",
                "Eif3e",
                "Dstn",
                "Emp3",
                "Aspn",
                "Pfdn4",
                "Cebpb",
                "Srrm2",
                "Top1",
                "Ywhae",
                "Pafah1b2",
                "Ubl5",
                "Anxa3",
                "Mrps15",
                "Ndufc1",
                "Psmc4",
                "Eif3d",
                "Hbegf",
                "Psmd14",
                "Rps15",
                "Ghitm",
                "Arpc3",
                "Psma4",
                "Scp2",
                "Ilk",
                "Arl1",
                "Arpc1a",
                "Ap2s1",
                "Mgat2",
                "Rpl34",
                "Mlec",
                "Rab7",
                "Sep15",
                "Dynll1",
                "Gtpbp4",
                "Eif2s2",
                "Hspd1",
                "Cpe",
                "Atf5",
                "Arf5",
                "Cdc42",
                "Npc2",
                "Rpl31",
                "Srsf7",
                "Smdt1",
                "Nap1l1",
                "Eef2",
                "Tpx2",
                "Etfb",
                "Hsbp1",
                "Hspa5",
                "Rnh1",
                "Rrbp1",
                "Uqcrc2",
                "Actr3",
                "Tmsb4x",
                "Cd47",
                "Efemp2",
                "Ccdc124",
                "Fbln2",
                "Gapdh",
                "Smarca5",
                "Prdx2",
                "Mpc1",
                "Copb2",
                "Khdrbs1",
                "Ap2m1",
                "Mrpl52",
                "Nupr1",
                "Sars",
                "Bri3",
                "Smim14",
                "Prkcdbp",
                "Hnrnpab",
                "Sdpr",
                "Lgals1",
                "Rplp0",
                "Psmb5",
                "Ahnak",
                "Tbrg1",
                "Col11a1",
                "Hspa9",
                "Fkbp10",
                "Nop58",
                "Ube2n",
                "Ndufa11",
                "Cbx5",
                "Kif5b",
                "Luc7l3",
                "Polr2c",
                "Cct8",
                "Mpc2",
                "Copz2",
                "Hnrnpa2b1",
                "Ptp4a1",
                "Lsp1",
                "Ndufs5",
                "Gsn",
                "Psmb1",
                "Oat",
                "Tubb6",
                "Coa3",
                "Ndufb8",
                "Cited2",
                "Psmd2",
                "Tmem97",
                "Srsf2",
                "Eva1b",
                "Prdx4",
                "mt-Nd1",
                "Eif4g2",
                "Psmb2",
                "Rab2a",
                "Uqcc2",
                "Denr",
                "Eif3g",
                "Tspo",
                "Sra1",
                "Tceal8",
                "Lhfp",
                "Myl9",
                "Rbm3",
                "Psmd11",
                "Cnn3",
                "Vps29",
                "Btg2",
                "Vdac1",
                "Selk",
                "Cggbp1",
                "Zfp706",
                "Rad23b",
                "Chmp2a",
                "Tra2a",
                "Nudcd2",
                "Pdcd5",
                "Aprt",
                "Ost4",
                "Rab11a",
                "Ankrd1",
                "Snrpc",
                "Anxa5",
                "Eif3f",
                "Eif4g1",
                "Anxa6",
                "Mtch1",
                "Tmbim6",
                "Dynll2",
                "Rhoa",
                "Ssu72",
                "Rab10",
                "Atp5a1",
                "Psma5",
                "Ttc3",
                "Cdk4",
                "Cnpy2",
                "Rpl22",
                "Rsl1d1",
                "Vimp",
                "Ccdc80",
                "Syncrip",
                "Sec61a1",
                "Caprin1",
                "Iqgap1",
                "Ict1",
                "Kdelr2",
                "Glud1",
                "March5",
                "Mrpl23",
                "Pmp22",
                "Rpl4",
                "Fermt2",
                "Hnrnpu",
                "Rps25",
                "Rps28",
                "Gspt1",
                "Srsf11",
                "Rpl18a",
                "Hmgb3",
                "Rpn1",
                "Gas1",
                "Ssr4",
                "Txndc9",
                "Ppp1r2",
                "Sqstm1",
                "Eprs",
                "Trmt112",
                "Lman1",
                "F3",
                "Msmo1",
                "Rcn1",
                "Arpc2",
                "Atp5l",
                "Mrpl54",
                "Rpl35",
                "Son",
                "Ilf2",
                "Ywhab",
                "Cct2",
                "Acp1",
                "Prelid1",
                "Sox4",
                "Phlda3",
                "Sept11",
                "Ndufa1",
                "Romo1",
                "Tpm2",
                "Ergic3",
                "Eif1ax",
                "Bzw1",
                "Sdc4",
                "Ak6",
                "Ubc",
                "Rock2",
                "Hnrnpa3",
                "Prpf40a",
                "Prmt1",
                "Pbdc1",
                "Cdc42ep3",
                "Mrps7",
                "Metap2",
                "Mrpl57",
                "Morf4l1",
                "2010107E04Rik",
                "Col8a1",
                "Pafah1b1",
                "Lsm6",
                "Ube2s",
                "Tecr",
                "Sf3b1",
                "Idh2",
                "Psap",
                "Ndufs4",
                "Polr1d",
                "Snrpd3",
                "Tm4sf1",
                "Rbms1",
                "P4hb",
                "Scd2",
                "Dlk1",
                "Sf3b2",
                "H2afy",
                "Map1b",
                "Rab34",
                "Suclg1",
                "Zyx",
                "Adipor1",
                "Rpl17",
                "Bsg",
                "mt-Nd2",
                "Hnrnpd",
                "Ppic",
                "Rpl14",
                "Tma7",
                "Sec22b",
                "Camk2d",
                "Txndc17",
                "Cops4",
                "Cyba",
                "Flna",
                "Lsm4",
                "Efemp1",
                "H13",
                "Rsu1",
                "Ywhag",
                "Atp5f1",
                "Prss23",
                "Rps13",
                "Gsto1",
                "Tpt1",
                "Eif5a",
                "Rps11",
                "Ppib",
                "Psmd8",
                "Cfl1",
                "Gng5",
                "Vcan",
                "Sparc",
                "Rps19",
                "Sept2",
                "Abracl",
                "Exosc8",
                "Cops6",
                "Msn",
                "Aldoa",
                "Mrpl20",
                "Hnrnpdl",
                "Serpinb6a",
                "Srp9",
                "Eif3a",
                "Rtn3",
                "Lpp",
                "Ptn",
                "Tagln",
                "Ctbp1",
                "Psmb4",
                "Snx3",
                "Strap",
                "Csrp1",
                "Commd1",
                "Psmd13",
                "Selm",
                "Ckap4",
                "Esd",
                "Atxn7l3b",
                "Igfbp6",
                "Comt",
                "Cox6b1",
                "Lamp1",
                "Sat1",
                "Lamtor5",
                "Ndufa13",
                "Atp5e",
                "Prdx1",
                "Ssb",
                "Ptrf",
                "Tinagl1",
                "Gps1",
                "Cmtm7",
                "Gadd45g",
                "Atp5o",
                "Rrm2",
                "Rpl13",
                "Sh3bgrl",
                "Jun",
                "Capns1",
                "Itm2b",
                "Pdia3",
                "Rplp1",
                "Ndufb5",
                "Rpl10a",
                "Atrx",
                "Psma2",
                "Txndc5",
                "Pfdn5",
                "Ndufb7",
                "Ptms",
                "Eif4e2",
                "Rps14",
                "Ftl1",
                "Sfpq",
                "Col4a1",
                "Rpl32",
                "Smap1",
                "Cd24a",
                "Zfp36l1",
                "Lipe",
                "Tnfrsf12a",
                "Mdh2",
                "Krtcap2",
                "Gabarap",
                "Uqcr11",
                "Rps27a",
                "Phgdh",
                "Rap1a",
                "Serf2",
                "Bud31",
                "Pdap1",
                "Brk1",
                "Gtf2h5",
                "Hsp90b1",
                "Mtpn",
                "Anxa1",
                "Prkar1a",
                "Rap1b",
                "Rpl11",
                "Vdac2",
                "Pfn1",
                "Rpl13a",
                "Nol7",
                "Timp1",
                "Ier3",
                "Grb10",
                "Cuta",
                "Apoe",
                "Eif1a",
                "Tuba1a",
                "Commd3",
                "Cnn2",
                "Minos1",
                "Snrpb",
                "Pgam1",
                "Rabggtb",
                "Tomm22",
                "Nans",
                "Ubap2l",
                "Erp29",
                "Gpx8",
                "Gpi1",
                "Tmem167",
                "Ppig",
                "Phldb2",
                "Palld",
                "Rpl27a",
                "Dcun1d5",
                "Eif5b",
                "Tram1",
                "Cct4",
                "Rdx",
                "Lrrc17",
                "Lsm12",
                "Fhl1",
                "Anp32b",
                "Edf1",
                "Cdkn1a",
                "Pkm",
                "Nono",
                "Csde1",
                "Fstl1",
                "Marcks",
                "Fam162a",
                "Tceb2",
                "Rhoc",
                "Mdm2",
                "Ifitm3",
                "Psenen",
                "Srp72",
                "Mrpl33",
                "Sumo3",
                "Emp1",
                "Psmc2",
                "Ndufa10",
                "App",
                "Fundc2",
                "Uqcrfs1",
                "Naa50",
                "Ywhah",
                "Eif3i",
                "Hnrnpc",
                "Smc2",
                "Mtch2",
                "S100a10",
                "Bag1",
                "Timm17a",
                "Psmd4",
                "Rpn2",
                "Tmem147",
                "Bub3",
                "Fam104a",
                "Prrx2",
                "Papola",
                "Rps16",
                "Manf",
                "Tmem256",
                "Sin3b",
                "Cetn3",
                "Dctn2",
                "Arhgdib",
                "Tmed3",
                "Serpinf1",
                "Mrfap1",
                "Galk1",
                "Hsd17b14",
                "Slc25a5",
                "Uqcrh",
                "Gap43",
                "Tpr",
                "Hint1",
                "Tbca",
                "Cstb",
                "Foxp1",
                "Srsf6",
                "Ndufa5",
                "Ebna1bp2",
                "Ccng1",
                "Ndufb10",
                "Tmpo",
                "Rpl37a",
                "Tsn",
                "H19",
                "Ptp4a2",
                "Mesdc2",
                "Eef1g",
                "Dnajc3",
                "Cope",
                "Cox5b",
                "Rbm25",
                "Arhgdia",
                "Cycs",
                "Myl12a",
                "Ndufb6",
                "Ptma",
                "Rac1",
                "Rer1",
                "S100a4",
                "Ralbp1",
                "Cald1",
                "Sdhb",
                "Fau",
                "Ybx3",
                "Rps10",
                "Sri",
                "Pabpc1",
                "Prdx5",
                "Alyref",
                "Clic1",
                "Tsen34",
                "Fkbp1a",
                "Got2",
                "Dad1",
                "Slirp",
                "Emc7",
                "Gnas",
                "Aebp1",
                "Arf4",
                "Calu",
                "Arpc5",
                "Yif1b",
                "Hdlbp",
                "Limd2",
                "Wbp11",
                "Cox4i1",
                "Ccnd1",
                "Tmed9",
                "Snf8",
                "Myl6",
                "Sepw1",
                "2810428I15Rik",
                "Hspa8",
                "Ndufs8",
                "Psma3",
                "Rpl39",
                "Rabac1",
                "Rplp2",
                "Capn2",
                "Serp1",
                "Mif",
                "Laptm4a",
                "Ptov1",
                "Aimp1",
                "Serbp1",
                "Cnih1",
                "Gpx4",
                "Chchd3",
                "Rheb",
                "Rps23",
                "Glrx3",
                "Gtf2a2",
                "Carhsp1",
                "Cryab",
                "Alpl",
                "Rrp1",
                "Pdcl3",
                "Rpl23",
                "Rpl30",
                "Kdelr3",
                "Sfr1",
                "Polr2f",
                "P3h4",
                "Junb",
                "Rbm17",
                "Bcap31",
                "Map2k2",
                "Cnbp",
                "Sdhd",
                "Axl",
                "Cfl2",
                "Ogn",
                "Rsrc2"
            ],
            "expression_sum": {
                "class1": 10992.0,
                "class2": 22123.0
            },
            "uniprot_ids": [
                "P70288",
                "Q8BQ10",
                "Q3UXH8",
                "Q3URA2",
                "A0A0R4J008",
                "Q3TMT1"
            ],
            "net_id": 0,
            "source_num": 13,
            "target_num": 513
        },
        "E2f4": {
            "id": "E2f4",
            "type": "TF",
            "symbol": "E2f4",
            "source": [
                "Pou5f1",
                "Junb",
                "Jun",
                "Sox2",
                "Nanog",
                "Jund",
                "Twist2",
                "Cebpb"
            ],
            "target": [
                "Mgat2",
                "Sep15",
                "Atxn7l3b",
                "Dap",
                "Lsm7",
                "Cpe",
                "Serpine2",
                "Atf5",
                "Arf5",
                "Lamp1",
                "Rcor2",
                "Cdc42",
                "Sfrp1",
                "Itgb1",
                "Syce2",
                "Ptrf",
                "Cd44",
                "Fabp5",
                "Gadd45g",
                "Myh9",
                "Tcf4",
                "Thbs1",
                "Rrbp1",
                "Rasgrp2",
                "Lims1",
                "Tmsb4x",
                "Cd47",
                "Efemp2",
                "Cd81",
                "Itm2b",
                "Jun",
                "Fbln2",
                "Pdia3",
                "Pgls",
                "Ssr1",
                "Txndc5",
                "Slc25a4",
                "Tspan4",
                "Col12a1",
                "Lmna",
                "Nupr1",
                "Col5a2",
                "Crip2",
                "Prkcdbp",
                "Vim",
                "Lipe",
                "Lgals1",
                "Tnfrsf12a",
                "Basp1",
                "Col11a1",
                "Fkbp10",
                "Krtcap2",
                "E130012A19Rik",
                "Pam16",
                "Kif5b",
                "Tnc",
                "Pck2",
                "Tmed2",
                "Mtpn",
                "Lgals3",
                "Gsn",
                "Ilk",
                "Tubb6",
                "Pgrmc1",
                "Timp1",
                "Tagln2",
                "Eva1b",
                "Tpm1",
                "Sdcbp",
                "Surf4",
                "Tuba1a",
                "Rexo2",
                "Commd3",
                "Phlda1",
                "Pdlim7",
                "Lhfp",
                "Mbnl2",
                "Canx",
                "Myeov2",
                "Actn1",
                "B2m",
                "Btg2",
                "Selk",
                "Vcl",
                "Gpi1",
                "Lox",
                "Palld",
                "Tram1",
                "Igfbp7",
                "Lrrc17",
                "Rab11a",
                "Ung",
                "Anxa5",
                "S100a6",
                "Anxa6",
                "Serpine1",
                "Cdkn1a",
                "Neat1",
                "Myh10",
                "Marcks",
                "Fam162a",
                "Rhoc",
                "Vimp",
                "Cd63",
                "Timp3",
                "Capg",
                "Lbh",
                "Fbn1",
                "Mtdh",
                "Iqgap1",
                "Pmp22",
                "Emp1",
                "Ssr3",
                "mt-Nd5",
                "Sh3glb1",
                "App",
                "Maged2",
                "Rtn4",
                "Impa2",
                "Ctsl",
                "Fbn2",
                "Spcs1",
                "Ssr4",
                "Atp2b1",
                "Gas1",
                "Sdc2",
                "Tax1bp3",
                "S100a10",
                "Nktr",
                "Filip1l",
                "Col5a1",
                "Tspan3",
                "Ctgf",
                "Arpc2",
                "Cav1",
                "Klf6",
                "Pcolce",
                "Plk2",
                "Loxl1",
                "Phlda3",
                "Oaf",
                "Ostf1",
                "Arhgdib",
                "Sept11",
                "Serpinf1",
                "Sox4",
                "Tgfb1i1",
                "Tln1",
                "Tmed3",
                "Timp2",
                "Syngr1",
                "Tpm2",
                "Sdc4",
                "Chd3",
                "Rock2",
                "Rps12",
                "Foxp1",
                "Cdc42ep3",
                "H19",
                "Lrrc58",
                "Ptp4a2",
                "Fn1",
                "Dnajc3",
                "Glipr2",
                "Arpc1b",
                "Ap1s1",
                "Cyr61",
                "Hes1",
                "Kdelr1",
                "Nenf",
                "Rab14",
                "Psap",
                "Serpinh1",
                "Twist2",
                "Klf2",
                "Rbms1",
                "P4hb",
                "Zyx",
                "Map1b",
                "Rab34",
                "Malat1",
                "Tmem176b",
                "Ppic",
                "Gnb2",
                "Sri",
                "Mfap2",
                "Tm9sf3",
                "Tmed10",
                "Ctsd",
                "Dpysl3",
                "Mybl2",
                "Reep5",
                "Emc7",
                "Gnas",
                "Hcfc1r1",
                "Arf4",
                "Camk2d",
                "Calu",
                "Fhl2",
                "Gm7325",
                "Sept1",
                "Sptbn1",
                "Flna",
                "mt-Cytb",
                "Hdlbp",
                "Ap3s1",
                "Gng12",
                "Limd2",
                "Oxct1",
                "H13",
                "Rsu1",
                "Ncam1",
                "Wls",
                "Sepw1",
                "Gsto1",
                "Hacd1",
                "Ndufa8",
                "Sept7",
                "Fam195b",
                "Capn2",
                "Laptm4a",
                "Vcan",
                "Actg1",
                "Cnih1",
                "Rcn2",
                "Emp3",
                "Ier5",
                "Cebpb",
                "Carhsp1",
                "Abracl",
                "Msn",
                "Id1",
                "Prnp",
                "Sept2",
                "Serpinb6a",
                "Hmga2",
                "Kdelr3",
                "Pafah1b2",
                "Ctbp1",
                "P3h4",
                "Junb",
                "Csrp1",
                "Hbegf",
                "Selm",
                "Axl",
                "Cfl2",
                "Nrp1",
                "Col1a2",
                "Esd"
            ],
            "expression_sum": {
                "class1": 2638.0,
                "class2": 24346.0
            },
            "uniprot_ids": [
                "Q8R0K9"
            ],
            "net_id": 0,
            "source_num": 8,
            "target_num": 155
        },
        "Ybx1": {
            "id": "Ybx1",
            "type": "TF",
            "symbol": "Ybx1",
            "source": [
                "Tcf4",
                "Pou5f1",
                "Cebpb",
                "Nfib",
                "Junb",
                "Fos",
                "Jun",
                "Sox2",
                "Tbx3",
                "Zfp57",
                "Nanog",
                "Jund",
                "Twist2",
                "Hdac2",
                "Smc3",
                "Sox4"
            ],
            "target": [
                "Hsp90aa1",
                "Arf1",
                "Dap",
                "Hp1bp3",
                "Clcc1",
                "Ube2j1",
                "Kif2a",
                "Dpysl2",
                "Tmem41b",
                "Mdfic",
                "Rbfox2",
                "Mgat1",
                "Cd44",
                "Casc4",
                "Fis1",
                "Tead1",
                "Colgalt1",
                "AI314180",
                "Igf1r",
                "Lpcat4",
                "Commd9",
                "Itfg1",
                "Npr1",
                "Lims1",
                "Sucla2",
                "Ccm2",
                "Sec13",
                "Ryk",
                "Tuba1b",
                "Gm13051",
                "Nucb1",
                "Wisp1",
                "Slc25a4",
                "St7l",
                "Ctsk",
                "Col12a1",
                "Calcoco2",
                "Rpl23a",
                "Lrrcc1",
                "Spop",
                "Map1lc3a",
                "Ifngr2",
                "Slc35a1",
                "Tmem37",
                "Fxyd5",
                "Tmem70",
                "Pik3r1",
                "Folr1",
                "Hsd17b12",
                "Zfand6",
                "Ntmt1",
                "Lcorl",
                "Psmd6",
                "Cthrc1",
                "Ctsa",
                "Runx1",
                "Lmbrd1",
                "Pik3ca",
                "Ttc14",
                "Csf1",
                "Hectd1",
                "Ssrp1",
                "Smtnl2",
                "Serinc1",
                "1700019D03Rik",
                "Mkln1",
                "Ugcg",
                "Ier3ip1",
                "Rnf187",
                "mt-Co1",
                "Pgrmc1",
                "Anxa8",
                "Dusp3",
                "Tagln2",
                "Dnajc9",
                "Dpm3",
                "Ccdc88a",
                "Calr",
                "Ptk7",
                "Fam50a",
                "Crlf1",
                "Pdlim7",
                "Rab18",
                "Tubb5",
                "Gm13154",
                "Tmem120a",
                "Actn1",
                "Eif4h",
                "2810417H13Rik",
                "Rras2",
                "Prkag1",
                "Plod1",
                "Tcf7l2",
                "Nr0b1",
                "Prrx1",
                "Slc35e4",
                "Fam101b",
                "Tars",
                "Erp44",
                "Fam98a",
                "Hmgcs1",
                "Zwint",
                "Fip1l1",
                "Smad1",
                "Neat1",
                "Acta2",
                "Aga",
                "Ctnnd1",
                "Ube2r2",
                "Tgfbi",
                "Acer3",
                "Atp2c1",
                "Wdr89",
                "Tdgf1",
                "Sh3glb1",
                "Strn3",
                "2310022B05Rik",
                "Akap9",
                "Arl6ip5",
                "Akt1s1",
                "Agpat4",
                "Fech",
                "Mospd1",
                "Bnip3l",
                "Atp2b1",
                "Rit1",
                "Apoc1",
                "C1d",
                "Nktr",
                "Col5a1",
                "Tspan3",
                "Ctgf",
                "Bgn",
                "Col4a2",
                "Klf6",
                "Dld",
                "Pcolce",
                "Lysmd3",
                "Ralb",
                "Gnai3",
                "Rasa1",
                "Myadm",
                "Timp2",
                "Crxos",
                "Bex4",
                "Rab11b",
                "Selo",
                "Ahnak2",
                "Psmc3",
                "Cdc16",
                "Sec62",
                "Rps12",
                "H2afy2",
                "Gmps",
                "Rbbp7",
                "Gm6483",
                "Rps27",
                "Msrb3",
                "Maged1",
                "Tmem164",
                "Tspan31",
                "2610002M06Rik",
                "Cfap36",
                "Arpc1b",
                "Col3a1",
                "Cyr61",
                "Eif3l",
                "Rab14",
                "Serpinh1",
                "Ahsa2",
                "Atp6v1g1",
                "Gsk3b",
                "Fgfr1",
                "Arl8b",
                "Tnfrsf23",
                "Smim3",
                "Acap2",
                "Mnat1",
                "Pitpnb",
                "BC005537",
                "Rbms2",
                "Tmem165",
                "Matr3",
                "Ociad1",
                "1700123O20Rik",
                "Dpysl3",
                "Mybl2",
                "Mycbp",
                "Ddost",
                "Hcfc1r1",
                "Sptbn1",
                "Npdc1",
                "Sertad2",
                "Ap3s1",
                "Ccdc50",
                "Jkamp",
                "Wwp1",
                "Yipf5",
                "Ncam1",
                "Wbscr17",
                "Dbn1",
                "Bicd2",
                "Hoxa5",
                "Cdc42ep1",
                "Pdia4",
                "Saraf",
                "Sept7",
                "Srp14",
                "Ccdc109b",
                "Mmadhc",
                "Atp5b",
                "Actg1",
                "Rcn2",
                "Tsg101",
                "Ier5",
                "Mob4",
                "Id1",
                "Csnk2b",
                "Pja2",
                "Prickle1",
                "2010111I01Rik",
                "Dynlrb1",
                "Hspb1",
                "Atf4",
                "Plekhh2",
                "Synpo",
                "Mcfd2",
                "Col1a2",
                "Tcaf1",
                "Scd1",
                "Stt3a",
                "Atp6v0b",
                "F2r",
                "Pak2",
                "Abhd8",
                "Ak1",
                "Cxx1a",
                "Itgb1",
                "Cd109",
                "Boc",
                "Fam204a",
                "Amfr",
                "Kmt2e",
                "Klhdc2",
                "Fabp5",
                "Chmp4b",
                "Gns",
                "Soat1",
                "Med13",
                "Cd81",
                "Vamp3",
                "Tab2",
                "Cyp51",
                "Tpp2",
                "Bdnf",
                "Plod3",
                "Pdgfrl",
                "Fuca1",
                "Tbpl1",
                "Ift20",
                "Anxa2",
                "Vmp1",
                "Ikbip",
                "mt-Nd4",
                "Crip2",
                "Mycbp2",
                "Arhgef2",
                "Basp1",
                "Spg20",
                "1700020I14Rik",
                "Wdr1",
                "Tnc",
                "Ube2b",
                "Gpat4",
                "Appl1",
                "Ano6",
                "Adamts5",
                "Dnm3os",
                "Adamts1",
                "Slc43a3",
                "Rbbp8",
                "Myg1",
                "Tpm1",
                "Rgs4",
                "Zfpl1",
                "Stk17b",
                "Ifi27",
                "Lima1",
                "Tank",
                "Cct5",
                "Uba1",
                "Emd",
                "1110004F10Rik",
                "Tex261",
                "Il6st",
                "Myeov2",
                "Lztr1",
                "Unc50",
                "Ercc1",
                "Mcts1",
                "Mxd4",
                "Pdgfa",
                "Bola3",
                "Tmem50a",
                "Vcl",
                "Tmod3",
                "Slc50a1",
                "Eif3m",
                "Capsl",
                "Ppp2r1a",
                "Bbip1",
                "Xrn2",
                "Smurf2",
                "2310009B15Rik",
                "Zfp330",
                "Acbd3",
                "Prdx6",
                "Ufsp2",
                "Far1",
                "Lzic",
                "Serpine1",
                "H2-D1",
                "Alg5",
                "Gamt",
                "Crabp1",
                "Ifitm1",
                "Afap1",
                "Cd63",
                "Lyrm2",
                "Elovl5",
                "Cdc37",
                "Nptn",
                "Hebp1",
                "S100a11",
                "Eef1a1",
                "Sirt2",
                "Irx3",
                "Maged2",
                "Gm5093",
                "Wsb1",
                "Ctsl",
                "Prkca",
                "Cox6a1",
                "Rab5a",
                "Slk",
                "Ap2a2",
                "Cops2",
                "Tnrc18",
                "Pold4",
                "Pard3",
                "Tspan5",
                "Ndel1",
                "Cav1",
                "Mrps6",
                "Prosc",
                "Efr3a",
                "mt-Co2",
                "Sgcb",
                "Uso1",
                "Srsf10",
                "Tln1",
                "Cul3",
                "Smchd1",
                "Oaf",
                "Ak3",
                "Ostf1",
                "Ndn",
                "Ube2v2",
                "Uba52",
                "Eno1",
                "Atp5d",
                "Mt2",
                "Eif3h",
                "Fbln5",
                "Acadl",
                "Thoc2",
                "Scoc",
                "Cmpk1",
                "1700061G19Rik",
                "Figf",
                "Btg1",
                "Lrrc58",
                "Fn1",
                "Hes1",
                "Cpsf2",
                "Nenf",
                "Polr3k",
                "Rps9",
                "Zfp42",
                "Mrpl30",
                "Myo1b",
                "Rpp25",
                "Bod1l",
                "Malat1",
                "mt-Nd4l",
                "Gnai2",
                "Tmem176b",
                "Ap3d1",
                "Eif4a1",
                "Mfap2",
                "Antxr1",
                "Arcn1",
                "Ctsd",
                "Ndufc2",
                "Borcs7",
                "Fhl2",
                "Elf2",
                "Ndufa12",
                "Krcc1",
                "Adam19",
                "Pla2g1b",
                "Il1rl1",
                "Atxn10",
                "Ctsh",
                "Atp6v0e",
                "Cnih4",
                "Medag",
                "Adk",
                "Ube2f",
                "Por",
                "Pcbp4",
                "Prkra",
                "Nap1l4",
                "Tmem59",
                "Ap1m1",
                "Aspn",
                "Stard4",
                "Spcs3",
                "Hnrnpr",
                "Rnf166",
                "Sav1",
                "Gm26917",
                "Tmbim4",
                "Actr10",
                "Irak1bp1",
                "Arpc3",
                "Zmynd11",
                "Akr1b8",
                "Scp2",
                "Arl1",
                "Arpc1a",
                "Sep15",
                "Arhgap28",
                "Clptm1",
                "Mlec",
                "Atf5",
                "Leprot",
                "Cdc42",
                "Npc2",
                "Rnf217",
                "Mvk",
                "Ptbp3",
                "Tmed5",
                "Suv39h1",
                "Uhrf1bp1l",
                "Eef2",
                "Asah1",
                "Hspa5",
                "Rnh1",
                "Rrbp1",
                "Actr3",
                "Slit3",
                "Tax1bp1",
                "Pvrl2",
                "Tmsb4x",
                "Camk2g",
                "Dnajc10",
                "Gapdh",
                "Exoc4",
                "Adgre5",
                "Arl3",
                "Bex1",
                "Mpc1",
                "Actr1b",
                "Gstm5",
                "Tm2d2",
                "Sars",
                "Lmod1",
                "Flnb",
                "Yaf2",
                "Ap2m1",
                "Copb2",
                "Smim14",
                "Khdrbs1",
                "Tmf1",
                "B230219D22Rik",
                "Rpl6l",
                "Acot7",
                "Ppp3ca",
                "Prkcdbp",
                "Col11a1",
                "Itgb5",
                "E130012A19Rik",
                "Btbd1",
                "Cct8",
                "Mpc2",
                "Copz2",
                "Nfat5",
                "Ptp4a1",
                "Lsp1",
                "Bmp1",
                "Gsn",
                "Tubb6",
                "Ccny",
                "Vma21",
                "Cited2",
                "N4bp3",
                "Cap1",
                "Hoxa9",
                "Eva1b",
                "Prdx4",
                "Jam2",
                "Eif4g2",
                "Nudt18",
                "Aes",
                "mt-Nd1",
                "Twist1",
                "Desi2",
                "2610305D13Rik",
                "Trim71",
                "Tceal8",
                "Lhfp",
                "Rbm3",
                "Ankrd17",
                "Ets1",
                "Vps29",
                "Parva",
                "Tmem132a",
                "Foxd3",
                "Prkcsh",
                "Rad23b",
                "mt-Co3",
                "Cyfip1",
                "Coq10b",
                "Tra2a",
                "Cdh2",
                "Rab11a",
                "Sdf4",
                "Anxa5",
                "Aco2",
                "Eif4g1",
                "Anxa6",
                "Mtch1",
                "Rhoa",
                "Reep3",
                "Tusc3",
                "Praf2",
                "Ttc3",
                "Shisa5",
                "Ccdc80",
                "Psmd9",
                "Uba3",
                "Caprin1",
                "Kcnq1ot1",
                "Kdelr2",
                "Pmp22",
                "Abhd5",
                "Srsf11",
                "Baz1a",
                "Tmem9b",
                "Hmgb3",
                "Ssr4",
                "Mapk1ip1l",
                "Nid1",
                "Ykt6",
                "Fgfr1op2",
                "Atp6ap2",
                "Sqstm1",
                "Lman1",
                "Msmo1",
                "Fyttd1",
                "Vcam1",
                "Arpc2",
                "Mageh1",
                "Fam25c",
                "Mrpl54",
                "Cct2",
                "Prelid1",
                "Ripk3",
                "1110046J04Rik",
                "Yae1d1",
                "Phlda3",
                "Trip6",
                "Sept11",
                "Spag9",
                "Romo1",
                "Ddx6",
                "Tpm2",
                "Eif1ax",
                "Bzw1",
                "Sdc4",
                "Metrnl",
                "Msl3",
                "Mgst1",
                "Ubc",
                "Lman2",
                "Prpf40a",
                "0610011F06Rik",
                "Cdc42ep3",
                "Zfp560",
                "Golim4",
                "Mpp1",
                "Fam220a",
                "Gm5449",
                "Serpina3g",
                "Vasn",
                "Pafah1b1",
                "Bloc1s2",
                "Ssbp3",
                "Myc",
                "Rpl27",
                "Psap",
                "Rbms1",
                "P4hb",
                "Scd2",
                "Dlk1",
                "Map1b",
                "Rab34",
                "Tvp23b",
                "Bsg",
                "Arl8a",
                "Dync1i2",
                "Qk",
                "Rnase4",
                "Fas",
                "4933434E20Rik",
                "Camk2d",
                "Txndc17",
                "Vgll3",
                "mt-Cytb",
                "Cyba",
                "Sclt1",
                "Stard3nl",
                "Gdf3",
                "Gsto1",
                "Frmd6",
                "Cers2",
                "Rab28",
                "H1f0",
                "Ptprd",
                "Vcan",
                "Csnk1d",
                "Sparc",
                "Sept2",
                "Abracl",
                "Cops6",
                "Serpinb6a",
                "Aldoa",
                "Pth1r",
                "Tagln",
                "Ctbp1",
                "4921524J17Rik",
                "Psmd13",
                "Slc35b4",
                "Anapc4",
                "Selm",
                "Mast4",
                "Churc1",
                "Zfyve21",
                "Hnrnph2",
                "Esd",
                "Atxn7l3b",
                "Csk",
                "Igfbp6",
                "Comt",
                "En1",
                "Vps36",
                "Runx1t1",
                "Lamp1",
                "Lama4",
                "Crim1",
                "Actr1a",
                "Lamtor5",
                "Ndufa13",
                "Sat1",
                "Ssb",
                "Gpr180",
                "Zfp91",
                "Atp5o",
                "Mapkapk2",
                "Gkap1",
                "Sema3a",
                "Tubb2a",
                "Rrm2",
                "Gm13242",
                "Brcc3",
                "Capns1",
                "Itm2b",
                "Pdia3",
                "Ccdc102a",
                "Chmp5",
                "Smad7",
                "Txndc5",
                "Ptms",
                "Mid1ip1",
                "Osr1",
                "Ginm1",
                "Vezf1",
                "Chmp2b",
                "Ccar1",
                "St5",
                "Mospd2",
                "Sfpq",
                "Abcg2",
                "Smap1",
                "Cd24a",
                "Ptpn11",
                "Lipe",
                "Tnfrsf12a",
                "Chid1",
                "Mdh2",
                "Tulp4",
                "Elovl1",
                "Etfa",
                "Phgdh",
                "Efna5",
                "Hsp90b1",
                "Mtpn",
                "Litaf",
                "Anxa1",
                "Herpud1",
                "Prkar1a",
                "Timp1",
                "Bmpr2",
                "Rap2a",
                "Lrp1",
                "Ddit4l",
                "Vps4a",
                "Tuba1a",
                "Ndufa9",
                "Scfd1",
                "Commd3",
                "Cnn2",
                "Atp6v0d1",
                "Ubqln2",
                "Ppp1r18",
                "Pcyt1a",
                "Amotl1",
                "Fst",
                "Lasp1",
                "Tia1",
                "Zfp637",
                "Tmem167",
                "Gpr108",
                "2410141K09Rik",
                "Gpi1",
                "Tcf12",
                "Nadk",
                "Dync1li2",
                "Cct4",
                "Igfbp7",
                "Decr1",
                "Rpl23a-ps3",
                "Gnaq",
                "Adm",
                "Cdkn1a",
                "2200002D01Rik",
                "Aig1",
                "Csde1",
                "Chd9",
                "AY036118",
                "Pls3",
                "Fstl1",
                "Rhod",
                "Zfp36l2",
                "Prkar2b",
                "Phc2",
                "Ifitm3",
                "Fbn1",
                "Wnk1",
                "Asap1",
                "B3galnt1",
                "Mef2a",
                "Psmc2",
                "Cd164",
                "Has2",
                "Ndufa10",
                "Atp6ap1",
                "Lztfl1",
                "Eif3i",
                "Anln",
                "Gpc1",
                "Gorasp2",
                "Htatsf1",
                "Epb41l4a",
                "Bag1",
                "Rpn2",
                "Errfi1",
                "Josd2",
                "Tmem33",
                "Manf",
                "Dctn2",
                "Arhgdib",
                "Memo1",
                "Rrm1",
                "2610001J05Rik",
                "Hsd17b14",
                "Ehd1",
                "Maoa",
                "Lefty2",
                "Cstb",
                "Gm10260",
                "Clptm1l",
                "Fads3",
                "Ccng1",
                "Asph",
                "Tsn",
                "Ctxn1",
                "Ptp4a2",
                "Txnl1",
                "Mesdc2",
                "Emc3",
                "Fbxo15",
                "Arhgdia",
                "Mxra7",
                "Myl12a",
                "Ndufb6",
                "Gm9493",
                "Rer1",
                "Spats2l",
                "Cdkn2b",
                "Anapc16",
                "Ddx3y",
                "Acat1",
                "Cald1",
                "Pigp",
                "Zdhhc3",
                "Ppp2r5e",
                "Fau",
                "Ybx3",
                "Sema3c",
                "mt-Atp6",
                "Akt1",
                "Bnip2",
                "Rbm7",
                "Sri",
                "Clic1",
                "Tsen34",
                "Got2",
                "Bcam",
                "Gnas",
                "Aebp1",
                "Emc7",
                "Ept1",
                "Calu",
                "Cinp",
                "Gm7325",
                "Rbpms2",
                "Rgmb",
                "Smim15",
                "Cox4i1",
                "Rps18-ps3",
                "Tmed9",
                "Gpx7",
                "Myl6",
                "Cers5",
                "Fads1",
                "Sepw1",
                "Pbrm1",
                "Hspa8",
                "Tmem214",
                "Tnfaip1",
                "Ccl7",
                "Laptm4a",
                "Pphln1",
                "Chic2",
                "Gpx4",
                "Senp6",
                "Ntan1",
                "Bet1l",
                "Scaf11",
                "Fam92a",
                "Grip2",
                "Cryzl1",
                "Pdcl3",
                "Atg3",
                "Bin1",
                "Nek7",
                "Cfl2",
                "Ogn"
            ],
            "expression_sum": {
                "class1": 176768.0,
                "class2": 435092.0
            },
            "uniprot_ids": [
                "P62960",
                "Q3UBT1",
                "Q71V06",
                "Q8BRJ5",
                "Q60950",
                "Q60951",
                "A2BGG7",
                "Q810K5"
            ],
            "net_id": 0,
            "source_num": 16,
            "target_num": 206
        },
        "Trp53": {
            "id": "Trp53",
            "type": "TF",
            "symbol": "Trp53",
            "source": [
                "Tcf4",
                "Pou5f1",
                "Foxp1",
                "Nfib",
                "Junb",
                "Fos",
                "Jun",
                "Sox2",
                "Nanog",
                "Jund",
                "Twist2",
                "Cebpb",
                "Sox4"
            ],
            "target": [
                "Rps5",
                "Arf1",
                "Dap",
                "Rpl41",
                "Sar1a",
                "Tpm4",
                "Dctn3",
                "Cd44",
                "H2afj",
                "Fis1",
                "Cox8a",
                "Ddb1",
                "Myh9",
                "Thbs1",
                "Rasgrp2",
                "Txn2",
                "Lims1",
                "Sec13",
                "Igfbp2",
                "Wisp1",
                "Slc25a4",
                "Tspan4",
                "Map1lc3a",
                "Lmna",
                "Folr1",
                "Ndufa2",
                "Itm2a",
                "Ier3ip1",
                "Pck2",
                "Rnf187",
                "Lgals3",
                "Pgrmc1",
                "Med28",
                "Shfm1",
                "Fos",
                "Tagln2",
                "Rps15a",
                "Ddx3x",
                "Nedd4",
                "Rexo2",
                "Calr",
                "Pdlim7",
                "Rab18",
                "Mfge8",
                "Mbnl2",
                "Canx",
                "Ppp1r14b",
                "Actn1",
                "B2m",
                "Mrpl14",
                "Slc25a3",
                "Psat1",
                "Lamtor1",
                "2810417H13Rik",
                "Neat1",
                "Myh10",
                "Ube2r2",
                "Timp3",
                "Gemin7",
                "Lrrc59",
                "Akr1a1",
                "Sh3glb1",
                "Akt1s1",
                "Hras",
                "Atp2b1",
                "Mapre1",
                "Cisd2",
                "Mylpf",
                "Nktr",
                "Col5a1",
                "Tspan3",
                "Ctgf",
                "Capza2",
                "Klf6",
                "Pcolce",
                "Hnrnph1",
                "Timp2",
                "Syngr1",
                "Crip1",
                "Morc1",
                "Sec62",
                "Rps12",
                "Luc7l2",
                "Ndufb9",
                "Maged1",
                "Eef1b2",
                "Arpc1b",
                "Cyr61",
                "Kdelr1",
                "Rab14",
                "Atp6v1g1",
                "Pttg1",
                "Gnb2",
                "Ociad1",
                "Dpysl3",
                "Mybl2",
                "Ddost",
                "Hcfc1r1",
                "Sptbn1",
                "Id3",
                "Ap3s1",
                "Gng12",
                "Fkbp8",
                "Ncam1",
                "Wls",
                "Ndufa8",
                "Pdia4",
                "Srp14",
                "Sept7",
                "Fam195b",
                "Actg1",
                "Rcn2",
                "Emc10",
                "Ier5",
                "Tsg101",
                "Tpi1",
                "Nars",
                "Hmga2",
                "Dynlrb1",
                "Atf4",
                "Hist1h2bc",
                "Tmem234",
                "Nfkbia",
                "Nrp1",
                "Ube2m",
                "Atp6v0b",
                "Fkbp9",
                "Rpl38",
                "Serpine2",
                "Igfbp4",
                "Id2",
                "Itgb1",
                "Fabp5",
                "Mettl9",
                "Tcf4",
                "Chmp4b",
                "Igf2",
                "Csnk1a1",
                "Cd81",
                "Ssr1",
                "Pgls",
                "Lxn",
                "Ift20",
                "Anxa2",
                "Eif3c",
                "Vmp1",
                "Napa",
                "Crip2",
                "mt-Nd4",
                "Cox7a2l",
                "Vim",
                "Basp1",
                "Sh3bgrl3",
                "Wdr1",
                "Ddah1",
                "Ube2b",
                "Eif5",
                "Tmed2",
                "Txn1",
                "Gjb3",
                "Tpm1",
                "Ldha",
                "Sdcbp",
                "Thy1",
                "Surf4",
                "Utf1",
                "Phlda1",
                "Myeov2",
                "Bola3",
                "Vcl",
                "Cst3",
                "Higd1a",
                "S100a6",
                "Prdx6",
                "Serpine1",
                "H2-D1",
                "Trappc4",
                "Sys1",
                "Cd63",
                "Capg",
                "Lbh",
                "Ndufa3",
                "S100a11",
                "Tmem14c",
                "Scand1",
                "Mtdh",
                "Nfib",
                "Eef1a1",
                "mt-Nd5",
                "Maged2",
                "Rtn4",
                "Ctsl",
                "Xist",
                "Ccnd2",
                "Tcf25",
                "Spcs1",
                "Cox6a1",
                "Tax1bp3",
                "Rala",
                "Filip1l",
                "mt-Co2",
                "Atox1",
                "Tmem126a",
                "Plk2",
                "Tln1",
                "Oaf",
                "Ostf1",
                "Snhg12",
                "Atp5d",
                "Mt2",
                "Eif3h",
                "Rpl22l1",
                "Chd3",
                "Rps3a1",
                "Cmpk1",
                "Btg1",
                "Fkbp2",
                "Lrrc58",
                "Fn1",
                "Glipr2",
                "Ndufb2",
                "Hes1",
                "Ap1s1",
                "Rps9",
                "Uqcrq",
                "Mrpl30",
                "Rcn3",
                "Rwdd1",
                "Malat1",
                "Gnai2",
                "Mfap2",
                "Tm9sf3",
                "Tmed10",
                "Ctsd",
                "Ndufc2",
                "Reep5",
                "Fhl2",
                "Glrx5",
                "Col1a1",
                "Ndufa12",
                "Oxct1",
                "Atxn10",
                "Atp6v0e",
                "Cnih4",
                "Pebp1",
                "Dbi",
                "Tmem59",
                "Eif3e",
                "Emp3",
                "Cebpb",
                "Cyb5r3",
                "Pafah1b2",
                "Anxa3",
                "Hbegf",
                "Ghitm",
                "Arpc3",
                "Scp2",
                "Ilk",
                "Arl1",
                "Arpc1a",
                "Ap2s1",
                "Mgat2",
                "Sep15",
                "Mlec",
                "Rab7",
                "Lsm7",
                "Cpe",
                "Atf5",
                "Cdc42",
                "Npc2",
                "Smdt1",
                "Eef2",
                "Hsbp1",
                "Hspa5",
                "Rnh1",
                "Rrbp1",
                "Actr3",
                "Tmsb4x",
                "Cd47",
                "Efemp2",
                "Fbln2",
                "Mpc1",
                "Fth1",
                "Copb2",
                "Khdrbs1",
                "Ap2m1",
                "Nupr1",
                "Sars",
                "Bri3",
                "Ecm1",
                "Smim14",
                "Lgals1",
                "Ahnak",
                "Tbrg1",
                "Fkbp10",
                "E130012A19Rik",
                "Pam16",
                "Kif5b",
                "Mpc2",
                "Copz2",
                "Ptp4a1",
                "Lsp1",
                "Ndufs5",
                "Gsn",
                "Psmb1",
                "Oat",
                "Tubb6",
                "Cited2",
                "Eva1b",
                "Prdx4",
                "mt-Nd1",
                "Eif4g2",
                "Aes",
                "Rab2a",
                "Uqcc2",
                "Tspo",
                "Sra1",
                "Myl9",
                "Vps29",
                "Btg2",
                "Parva",
                "Selk",
                "Rab11a",
                "Anxa5",
                "Eif3f",
                "Eif4g1",
                "Anxa6",
                "Mtch1",
                "Tmbim6",
                "Dynll2",
                "Rhoa",
                "Rab10",
                "Ttc3",
                "Vimp",
                "Syncrip",
                "Sec61a1",
                "Iqgap1",
                "Kdelr2",
                "Pmp22",
                "Rbmxl2",
                "Ssr3",
                "March5",
                "Rps28",
                "Fermt2",
                "Rpl18a",
                "Hmgb3",
                "Rpn1",
                "Gas1",
                "Ssr4",
                "Sqstm1",
                "Lman1",
                "F3",
                "Rcn1",
                "Rpl35",
                "Arpc2",
                "Atp5l",
                "Mrpl54",
                "Ywhab",
                "Prelid1",
                "Sox4",
                "Phlda3",
                "Sept11",
                "Ndufa1",
                "Romo1",
                "Tpm2",
                "Ergic3",
                "Bzw1",
                "Sdc4",
                "Ubc",
                "Lman2",
                "Rock2",
                "Cdc42ep3",
                "Tcea3",
                "Pafah1b1",
                "Psap",
                "Rbms1",
                "P4hb",
                "Scd2",
                "Zyx",
                "H2afy",
                "Map1b",
                "Rab34",
                "Adipor1",
                "mt-Nd2",
                "Bsg",
                "Sec22b",
                "Camk2d",
                "Txndc17",
                "mt-Cytb",
                "Cyba",
                "Flna",
                "Efemp1",
                "H13",
                "Rsu1",
                "Prss23",
                "Gsto1",
                "Tpt1",
                "Ppib",
                "Psmd8",
                "Gng5",
                "H1f0",
                "Ndufb11",
                "Vcan",
                "Rps19",
                "Sept2",
                "Abracl",
                "Msn",
                "Serpinb6a",
                "Rtn3",
                "Lpp",
                "Ctbp1",
                "Csrp1",
                "Selm",
                "Ckap4",
                "Esd",
                "Dppa5a",
                "Atxn7l3b",
                "Comt",
                "Cox6b1",
                "Lamp1",
                "Rcor2",
                "Sat1",
                "Ndufa13",
                "Tinagl1",
                "Gadd45g",
                "Sh3bgrl",
                "Jun",
                "Capns1",
                "Itm2b",
                "Pdia3",
                "Rplp1",
                "Rpl10a",
                "Txndc5",
                "Pfdn5",
                "Ndufb7",
                "Ptms",
                "Ftl1",
                "Col4a1",
                "Hist1h1c",
                "Cd24a",
                "Smap1",
                "Lipe",
                "Tnfrsf12a",
                "Krtcap2",
                "Gabarap",
                "Phgdh",
                "Serf2",
                "Brk1",
                "Hsp90b1",
                "Mtpn",
                "Anxa1",
                "Prkar1a",
                "Rap1b",
                "Rpl13a",
                "Ier3",
                "Cuta",
                "Apoe",
                "Tuba1a",
                "Commd3",
                "Cnn2",
                "Gsta4",
                "Minos1",
                "Erp29",
                "Gpx8",
                "Gpi1",
                "Tmem167",
                "Phldb2",
                "Palld",
                "Tram1",
                "Dcun1d5",
                "Cdkn1a",
                "Csde1",
                "Fstl1",
                "Marcks",
                "Fam162a",
                "Rhoc",
                "Mdm2",
                "Ifitm3",
                "Srp72",
                "Mrpl33",
                "Emp1",
                "App",
                "Fundc2",
                "S100a10",
                "Bag1",
                "Tmem147",
                "Rpn2",
                "Rps16",
                "Manf",
                "Tmem256",
                "Ccl2",
                "Dctn2",
                "Arhgdib",
                "Tmed3",
                "Serpinf1",
                "Galk1",
                "Hsd17b14",
                "Tbca",
                "Cstb",
                "Foxp1",
                "Ccng1",
                "Ptp4a2",
                "Cope",
                "Dnajc3",
                "Cox5b",
                "Arhgdia",
                "Myl12a",
                "Ndufb6",
                "Rac1",
                "Rer1",
                "S100a4",
                "Klf2",
                "Cald1",
                "Fau",
                "Sri",
                "Pabpc1",
                "Prdx5",
                "Clic1",
                "Tsen34",
                "Fkbp1a",
                "Got2",
                "Dad1",
                "Emc7",
                "Gnas",
                "Sept1",
                "Arf4",
                "Calu",
                "Arpc5",
                "Gm7325",
                "Yif1b",
                "Hdlbp",
                "Ckb",
                "Limd2",
                "Cox4i1",
                "Tmed9",
                "Myl6",
                "Sepw1",
                "Rpl39",
                "Ndufs8",
                "Serp1",
                "Rabac1",
                "Capn2",
                "Laptm4a",
                "Cnih1",
                "Rheb",
                "Glrx3",
                "Carhsp1",
                "Alpl",
                "Pdcl3",
                "Kdelr3",
                "Junb",
                "Bcap31",
                "Map2k2",
                "Axl",
                "Cfl2",
                "Ogn"
            ],
            "expression_sum": {
                "class1": 9058.0,
                "class2": 64226.0
            },
            "uniprot_ids": [
                "P02340",
                "A0A158SIS7",
                "Q549C9",
                "Q8C526",
                "I7HIK9",
                "Q80ZA1",
                "Q3UGQ1",
                "A0A5Q0MU23",
                "Q64451",
                "Q9ER40",
                "Q6TDG9",
                "Q91XH8",
                "W5VJU6",
                "O70366"
            ],
            "net_id": 0,
            "source_num": 13,
            "target_num": 240
        },
        "Pou5f1": {
            "id": "Pou5f1",
            "type": "TF",
            "symbol": "Pou5f1",
            "source": [
                "Trim28",
                "Cebpb",
                "Klf4",
                "Ctcf",
                "Hdac1",
                "Sox2",
                "Nanog",
                "Hdac2",
                "Smc3"
            ],
            "target": [
                "Arf1",
                "Dap",
                "Srsf9",
                "Ndufb4",
                "Syce2",
                "Mcm7",
                "Fis1",
                "Grsf1",
                "Zc3h15",
                "Cox8a",
                "2810004N23Rik",
                "Sec13",
                "Ndufa6",
                "Polr2k",
                "Idh3g",
                "Rps27l",
                "Col12a1",
                "Lmna",
                "Ndufa2",
                "Hnrnpf",
                "Lmnb1",
                "Ddx39",
                "Cks2",
                "Shfm1",
                "Med28",
                "Hmgb2",
                "Cox6c",
                "Ddx3x",
                "Slbp",
                "Nedd4",
                "Calr",
                "Tubb5",
                "Nop56",
                "Sf3b5",
                "Slc25a3",
                "Lamtor1",
                "2810417H13Rik",
                "Nup43",
                "Cdc5l",
                "Sms",
                "Rbx1",
                "Ctcf",
                "Neat1",
                "Pmf1",
                "Ola1",
                "Sf3b6",
                "Lrrc59",
                "Sap18",
                "Ube2d2a",
                "Hras",
                "Bnip3l",
                "Enah",
                "Kpnb1",
                "Mapre1",
                "Snrpd1",
                "Rbm8a",
                "Col5a1",
                "Rpl36al",
                "Lap3",
                "Tgfb1i1",
                "Supt4a",
                "Crxos",
                "Morc1",
                "Rbbp7",
                "Hnrnpl",
                "Nedd8",
                "Pcnp",
                "Atpif1",
                "Serpinh1",
                "Atp6v1g1",
                "Snrnp40",
                "Gng12",
                "Srp14",
                "Tpm3",
                "Banf1",
                "Mrps36",
                "Qdpr",
                "Tsg101",
                "Lig1",
                "Tpi1",
                "Pa2g4",
                "Gsr",
                "Atf4",
                "Hist1h2bc",
                "Hprt",
                "Nucks1",
                "Pcbp1",
                "Mrpl36",
                "Itgb1",
                "Ndufa4",
                "Rps12-ps3",
                "Cops5",
                "Eif4e",
                "Ndufv1",
                "H3f3a",
                "Rfc2",
                "Bccip",
                "Lamtor2",
                "Trim28",
                "Nudt21",
                "Vim",
                "Ube2l3",
                "Eef1d",
                "Etf1",
                "Zmat2",
                "Samm50",
                "Rrp15",
                "Nhp2l1",
                "Ran",
                "Atp5g3",
                "Gjb3",
                "Atp5g2",
                "Chd4",
                "Tceb1",
                "C1qbp",
                "Rpl36a",
                "Eif3m",
                "Snrpb2",
                "S100a6",
                "Umps",
                "Rps20",
                "Orc6",
                "Ybx1",
                "Sys1",
                "Tubb2b",
                "Ndufa3",
                "Tmem14c",
                "Rps6",
                "U2af1",
                "Slc29a1",
                "Atp5c1",
                "Mrps17",
                "Ccnd2",
                "Col6a1",
                "Rala",
                "Dnajc8",
                "Cox7c",
                "Nme4",
                "Eno1",
                "mt-Nd3",
                "Rpl22l1",
                "Lamtor4",
                "Smc4",
                "Mbd3",
                "Cct7",
                "Dctpp1",
                "Tuba4a",
                "Hsp90ab1",
                "Zfp42",
                "Rpp25",
                "Tyms",
                "Ankrd11",
                "Rps8",
                "Magoh",
                "Ndufc2",
                "H3f3b",
                "Dnajc2",
                "Ndufa12",
                "Cnih4",
                "Nudt19",
                "Dbi",
                "Tbx3",
                "Dstn",
                "Mt1",
                "Cyb5r3",
                "Ywhae",
                "Ppil1",
                "Rps15",
                "Arl1",
                "Cenpw",
                "Eif2s2",
                "Mrpl45",
                "Rpl31",
                "Smdt1",
                "Etfb",
                "Uqcrc2",
                "G3bp2",
                "Cd47",
                "Impdh2",
                "Bex1",
                "Prps1",
                "Akap12",
                "Mest",
                "Hnrnpab",
                "Ahnak",
                "Nop58",
                "Ndufa11",
                "Cbx5",
                "Fxyd6",
                "Polr2c",
                "Cct8",
                "Mpc2",
                "Dohh",
                "Denr",
                "Eif3g",
                "Sra1",
                "Aspscr1",
                "Btg2",
                "Parva",
                "0610012G03Rik",
                "Rpp21",
                "Polr2e",
                "Anapc13",
                "Pdcd5",
                "Tomm7",
                "Snrpc",
                "Mrpl49",
                "Eif3f",
                "Tmbim6",
                "Atp5a1",
                "Psma5",
                "Pycr2",
                "Sf3a3",
                "Ccdc80",
                "Rbmxl2",
                "March5",
                "Mrpl23",
                "Ddah2",
                "Gspt1",
                "Eprs",
                "Trmt112",
                "Rpl35",
                "Dazap1",
                "Ilf2",
                "Lman2",
                "Prmt1",
                "Tcea3",
                "Sub1",
                "Mrpl57",
                "Morf4l1",
                "Col8a1",
                "Pafah1b1",
                "Cdca8",
                "L1td1",
                "Lsm6",
                "Sf3b1",
                "Ndufs4",
                "Polr1d",
                "H2afy",
                "Mtf2",
                "Rpl17",
                "Bsg",
                "Sox2",
                "mt-Nd2",
                "Txndc17",
                "Spty2d1",
                "Nop10",
                "Mpdu1",
                "Gdf3",
                "Gng5",
                "Hspe1",
                "Rps19",
                "Ube2c",
                "Msn",
                "Exosc8",
                "Mrpl20",
                "Polr2h",
                "Psmb4",
                "Birc5",
                "Commd1",
                "Cox7b",
                "Selm",
                "Mien1",
                "Chtop",
                "Dppa5a",
                "Srpk1",
                "Lamtor5",
                "Avpi1",
                "Ssb",
                "Rpa2",
                "Mrps12",
                "Copz1",
                "Nfu1",
                "Ly6a",
                "Ndufb5",
                "Rpl10a",
                "Ptms",
                "Set",
                "Thyn1",
                "Txnl4a",
                "Ssbp1",
                "Col4a1",
                "Sephs2",
                "Snrpd2",
                "1810022K09Rik",
                "Hdac1",
                "Naa10",
                "Brk1",
                "Hsp90b1",
                "Vdac2",
                "Spp1",
                "Mrpl24",
                "Snhg6",
                "Minos1",
                "Anapc11",
                "Spint2",
                "Gas6",
                "Rabggtb",
                "Nans",
                "Uchl5",
                "Ppig",
                "Ucp2",
                "Dcun1d5",
                "Lsm12",
                "Gins4",
                "Tceb2",
                "Rhoc",
                "Mrpl42",
                "Fbn1",
                "Rps2",
                "Polr2g",
                "Sumo3",
                "Uqcrfs1",
                "Naa50",
                "Ywhah",
                "Fbn2",
                "Hdac2",
                "Timm17a",
                "Rpn2",
                "Bub3",
                "Esf1",
                "Sin3b",
                "Dctn2",
                "Plekhj1",
                "Mrfap1",
                "Hsd17b14",
                "Slc25a5",
                "Uqcrh",
                "Lefty2",
                "Tbca",
                "Foxp1",
                "Tsn",
                "Rpl37a",
                "Drg1",
                "Ptma",
                "Ndufb6",
                "Rac1",
                "S100a4",
                "Cald1",
                "Ybx3",
                "Rps10",
                "Dad1",
                "Sept1",
                "0610007P14Rik",
                "Yif1b",
                "Rps18-ps3",
                "Ccnd1",
                "Rpl39",
                "Tsfm",
                "Capn2",
                "Ptov1",
                "Laptm4a",
                "Rps27rt",
                "Gpx4",
                "Gcsh",
                "Gltscr2",
                "Mkrn1",
                "Rrp1",
                "Pdcl3",
                "Rpl23",
                "Rpl30",
                "Sfr1",
                "Polr2f",
                "Rrs1",
                "Rbm17",
                "Ly6e",
                "Map2k2",
                "Lsm5",
                "Rnf7",
                "Cdc20",
                "Cd9",
                "Ddx24",
                "Rpl6",
                "Srm",
                "Thbs1",
                "Psmb7",
                "Spr",
                "Pnkd",
                "Ppid",
                "1110008F13Rik",
                "Anapc5",
                "Hnrnpa1",
                "Ssrp1",
                "Ier3ip1",
                "Rnf187",
                "Psmb6",
                "Rps15a",
                "Ndufs3",
                "Uba2",
                "Mbnl2",
                "Ppp1r14b",
                "Snrpe",
                "B2m",
                "Psat1",
                "Gdi2",
                "Taf7",
                "Mrpl28",
                "Psmc6",
                "Grpel1",
                "Sec61g",
                "Ythdc1",
                "Snrnp27",
                "Srsf1",
                "Uqcrc1",
                "Tdgf1",
                "Rcc2",
                "Impa2",
                "Bola2",
                "Tmem160",
                "C1d",
                "Cisd2",
                "Rpl10",
                "Ncl",
                "Rpl35a",
                "Timp2",
                "Syngr1",
                "Hspa4",
                "Rps27",
                "Arpp19",
                "Ccdc12",
                "Ddx21",
                "Naca",
                "Ube2i",
                "Rpl24",
                "Brix1",
                "Fam136a",
                "Siva1",
                "Rps4x",
                "Fkbp8",
                "Nip7",
                "Ruvbl1",
                "Wls",
                "Clpp",
                "G3bp1",
                "Atp5b",
                "Actg1",
                "Cdc26",
                "Immt",
                "Nars",
                "Pdlim1",
                "Hells",
                "Tfcp2l1",
                "Rpl38",
                "Glo1",
                "Serpine2",
                "Nme1",
                "Csnk1a1",
                "Rpl10-ps3",
                "Rpl3",
                "Cebpzos",
                "Rpl12",
                "Snw1",
                "Ndufa7",
                "Dnajc19",
                "Cdv3",
                "Hmgn1",
                "Ctbp2",
                "Btf3l4",
                "Platr3",
                "Ppp4c",
                "Imp3",
                "Mrpl13",
                "Ak2",
                "Wdr77",
                "Atp5j",
                "Srrm1",
                "Thy1",
                "Srp19",
                "Emd",
                "Vasp",
                "Pdgfa",
                "Uqcr10",
                "Rpsa",
                "H2afz",
                "Bola3",
                "Mrps21",
                "Tardbp",
                "Rpl5",
                "Dph3",
                "Dpy30",
                "Serpine1",
                "Fblim1",
                "Odc1",
                "Mrpl18",
                "Ufc1",
                "Ifitm1",
                "Acat2",
                "Mrpl12",
                "Cdc37",
                "Cisd1",
                "Rpl28",
                "Xist",
                "Gpx3",
                "Cox6a1",
                "Timm8b",
                "Rpl26",
                "Uqcrb",
                "mt-Co2",
                "Rps3",
                "Atox1",
                "Srsf10",
                "Oaf",
                "Ube2v2",
                "Uba52",
                "Rpl19",
                "Timm13",
                "Eif3h",
                "Mcm3",
                "Adprh",
                "Rps3a1",
                "Srsf3",
                "Coprs",
                "Rpl7",
                "Znrd1",
                "Wdr18",
                "Polr2m",
                "1810058I24Rik",
                "Gnai2",
                "Rpl8",
                "Eif4a1",
                "Pcbp2",
                "Polr2i",
                "Polr1c",
                "Snrnp25",
                "Fhl2",
                "Glrx5",
                "Ccdc34",
                "Col1a1",
                "Mrps33",
                "Rif1",
                "Morf4l2",
                "Rbbp4",
                "Eif3e",
                "Gm10073",
                "Emp3",
                "2700060E02Rik",
                "Anxa3",
                "Arpc3",
                "Park7",
                "Dynll1",
                "Hspd1",
                "Stoml2",
                "Cops7a",
                "Sod1",
                "Ppih",
                "Uqcc3",
                "Eef2",
                "Tpx2",
                "Dnajb6",
                "Efemp2",
                "Gapdh",
                "Smarca5",
                "Idi1",
                "Ccnb1",
                "Ccnd3",
                "Ap2m1",
                "Khdrbs1",
                "Mrpl52",
                "Pno1",
                "Bri3",
                "Ecm1",
                "Adrm1",
                "Prkcdbp",
                "Kif5b",
                "Hmgn2",
                "Hnrnpa2b1",
                "Oat",
                "Psmd2",
                "Ddx27",
                "Cbx3",
                "Jam2",
                "Jade1",
                "Dnajc7",
                "Psmb2",
                "Fxr1",
                "Tceal8",
                "Myl9",
                "Cnn3",
                "Vps29",
                "Vdac1",
                "Anp32a",
                "Tra2a",
                "Zfp57",
                "Dynll2",
                "Ssu72",
                "Cnpy2",
                "Anp32e",
                "Pfdn6",
                "Pmp22",
                "Ict1",
                "Rps25",
                "Rbmxl1",
                "Arglu1",
                "Imp4",
                "Rpl18a",
                "Txndc9",
                "Mrpl54",
                "Ywhab",
                "Prelid1",
                "Ubb",
                "Tpm2",
                "Prrc1",
                "Zfp106",
                "Paics",
                "Asna1",
                "Psmc5",
                "Echdc2",
                "Mrpl43",
                "Rpl27",
                "Suclg1",
                "Hnrnpd",
                "Ash2l",
                "Hat1",
                "Rnps1",
                "Prss23",
                "Tpt1",
                "Ppib",
                "Stmn2",
                "Ptbp1",
                "Aldoa",
                "Lpp",
                "Tagln",
                "Dnajc21",
                "Snx3",
                "Strap",
                "Psmd13",
                "Rnaseh2a",
                "Klf4",
                "Rfc1",
                "Cox6b1",
                "Ptrf",
                "Atp5o",
                "Nasp",
                "Ndufs6",
                "Poldip2",
                "Eif4e2",
                "Sod2",
                "Ftl1",
                "Hist1h1c",
                "Tnfrsf12a",
                "Phgdh",
                "Gtf2e2",
                "Pdap1",
                "Rpl11",
                "Anxa1",
                "Ube2a",
                "Pfn1",
                "Snrpg",
                "Ddx39b",
                "Grb10",
                "Exosc4",
                "Fkbp4",
                "Pgam1",
                "Pnrc1",
                "Snrnp70",
                "Ppdpf",
                "Gmnn",
                "Rpl27a",
                "Llph",
                "Igfbp7",
                "Eif4b",
                "Anp32b",
                "Edf1",
                "Fstl1",
                "Mdm2",
                "Eif2b5",
                "Gart",
                "Mrpl33",
                "Ndufa10",
                "App",
                "Tomm40",
                "Eif3i",
                "Smc2",
                "Mtch2",
                "Psmd4",
                "Gins2",
                "Uck2",
                "Rps16",
                "Bcat1",
                "Tmed3",
                "Serpinf1",
                "Mrps16",
                "Gap43",
                "Psmd12",
                "Ndufb10",
                "Cth",
                "Txnl1",
                "Fbxo15",
                "Eef1g",
                "Rbm25",
                "Ube2e3",
                "2700094K13Rik",
                "Acadm",
                "Dut",
                "Ooep",
                "Alyref",
                "Hspa14",
                "Polr2j",
                "Got2",
                "Slirp",
                "Gnas",
                "Arpc5",
                "Ckb",
                "Cwc15",
                "Ndufs8",
                "Pim1",
                "Rabac1",
                "Rplp2",
                "Mif",
                "Aimp1",
                "Rheb",
                "Chchd3",
                "Eif4a2",
                "Glrx3",
                "Gtf2a2",
                "Alpl",
                "Kdelr3",
                "Cnbp",
                "Sdhd",
                "Axl",
                "Rps5",
                "Rpl41",
                "Dctn3",
                "Cd44",
                "Ruvbl2",
                "Cdt1",
                "Ier2",
                "Dkc1",
                "Igfbp2",
                "Marcksl1",
                "Atp5g1",
                "Tuba1b",
                "Aurka",
                "Psma6",
                "Rpl7l1",
                "Snrpa",
                "St13",
                "Thoc7",
                "Mlf2",
                "Rpl9",
                "Vbp1",
                "Atp5j2",
                "Nubp1",
                "E2f4",
                "D8Ertd738e",
                "Hn1",
                "Mrpl14",
                "Phb",
                "Nr0b1",
                "Tra2b",
                "Hmgcs1",
                "Emc6",
                "Rps24",
                "Acta2",
                "Nanog",
                "Bola1",
                "Aurkaip1",
                "Timp3",
                "Nhp2",
                "Akr1a1",
                "Exosc5",
                "Akt1s1",
                "Atp5h",
                "Mylpf",
                "Ppia",
                "Eif3k",
                "Mrps34",
                "Atp5k",
                "Mphosph8",
                "Ppp1ca",
                "Aamp",
                "Chchd1",
                "Gars",
                "Psmc3",
                "Rps12",
                "Luc7l2",
                "Ndufb9",
                "Col3a1",
                "Utp3",
                "Fus",
                "Ddost",
                "Rest",
                "Cct3",
                "Psma7",
                "Hacd1",
                "Gnl3",
                "Trim27",
                "Psma1",
                "Dtymk",
                "Emc10",
                "2410015M20Rik",
                "Hdgf",
                "Prnp",
                "Id1",
                "Csnk2b",
                "Rpl37",
                "Ppp1cc",
                "Tmem234",
                "Nrp1",
                "Dnttip2",
                "Sae1",
                "Hnrnpk",
                "Mrps24",
                "Ezr",
                "Ewsr1",
                "Ndufv2",
                "Mettl9",
                "Fbxo22",
                "Fam49b",
                "Bax",
                "Cisd3",
                "Tmsb10",
                "Gfer",
                "Anxa2",
                "Pycard",
                "Eif3c",
                "Cox7a2",
                "Col5a2",
                "Napa",
                "Atp6v1f",
                "Lsm3",
                "Ube2j2",
                "Gcat",
                "Eif5",
                "Spag7",
                "Tomm5",
                "Txn1",
                "Vapa",
                "Erh",
                "Fscn1",
                "Emc8",
                "Utf1",
                "Bclaf1",
                "Psip1",
                "Rpl29",
                "Naa38",
                "Ung",
                "Ptges3",
                "H2-D1",
                "Wtap",
                "Dnaja2",
                "Lbh",
                "Eef1a1",
                "Cenpv",
                "Ywhaq",
                "Pdha1",
                "Ubald2",
                "Filip1l",
                "Cav1",
                "Adss",
                "Rps7",
                "Igf2bp1",
                "Trnau1ap",
                "Eef1e1",
                "Fbln5",
                "Chd3",
                "Hsd17b10",
                "Gadd45gip1",
                "Nenf",
                "Oaz1",
                "Stra13",
                "Uqcrq",
                "Sec61b",
                "Snx5",
                "Got1",
                "Trap1a",
                "Bzw2",
                "Pgp",
                "Sumo2",
                "Nudc",
                "Gnb2l1",
                "Mrpl21",
                "Pold2",
                "Nsa2",
                "Ddx46",
                "Atxn10",
                "Rfc5",
                "Pfdn4",
                "Cebpb",
                "Mrps15",
                "Psmc4",
                "Psmd14",
                "Npm1",
                "Sap30",
                "Gtpbp4",
                "Lsm7",
                "Cpe",
                "Atf5",
                "Srsf7",
                "Mat2a",
                "Csrp2",
                "Brd2",
                "Ccdc124",
                "Prdx2",
                "Ngfrap1",
                "Abcf1",
                "Lgals1",
                "Rplp0",
                "Psmb5",
                "Ube2n",
                "Smn1",
                "Cyb5b",
                "Copz2",
                "Psmb1",
                "Coa3",
                "Akr1b3",
                "Eif4g2",
                "Raly",
                "Uqcc2",
                "Nsmce4a",
                "Rps26",
                "Use1",
                "Get4",
                "Lage3",
                "Npm3",
                "Lsm2",
                "Chmp2a",
                "Sumo1",
                "Nudcd2",
                "Aprt",
                "Anxa5",
                "Fcf1",
                "Mrps18c",
                "Cacybp",
                "Rpl22",
                "Nolc1",
                "Rpl4",
                "Glud1",
                "Rps28",
                "Hnrnpu",
                "Srsf11",
                "Hmgb3",
                "Rpl21",
                "Ppp1r2",
                "Map1lc3b",
                "Atp5l",
                "Son",
                "Pop5",
                "Cct2",
                "Acp1",
                "Dtl",
                "Ndufa1",
                "Ahsa1",
                "Eif1ax",
                "Parp1",
                "Dhfr",
                "Myl4",
                "Mrps7",
                "Cdca3",
                "Metap2",
                "Ube2s",
                "Idh2",
                "Snrpd3",
                "Zyx",
                "Psmd3",
                "Fam32a",
                "Trh",
                "Bcas2",
                "Cops4",
                "Ciapin1",
                "Pfdn2",
                "Ywhag",
                "Atp5f1",
                "Arid5b",
                "Vcan",
                "Ndufb11",
                "Srp9",
                "Nfatc2ip",
                "Ino80e",
                "Sfrp1",
                "Ppa1",
                "Rcor2",
                "Cenpm",
                "Prdx1",
                "Rsl24d1",
                "Terf1",
                "Rpl13",
                "Mrpl34",
                "Atrx",
                "Tgif1",
                "Sfpq",
                "Mrto4",
                "Parl",
                "Clns1a",
                "Mdh2",
                "Gabarap",
                "Uqcr11",
                "Phb2",
                "Rps27a",
                "Rpl15",
                "Gtf2h5",
                "Arl6ip1",
                "Tcof1",
                "Rap1b",
                "Ier3",
                "Cuta",
                "Eif1a",
                "Cox5a",
                "Tmem11",
                "Ubap2l",
                "Phldb2",
                "Cct4",
                "Rdx",
                "Pkm",
                "H2afv",
                "Prrc2c",
                "Marcks",
                "Psmg4",
                "Psmc2",
                "Sf3b4",
                "Hnrnpc",
                "Fkbp3",
                "Tmem147",
                "Rangrf",
                "Rpl7a",
                "Fam104a",
                "Prrx2",
                "Manf",
                "Tpr",
                "Ddx18",
                "Rrp9",
                "Hint1",
                "Sf3a2",
                "Ccne1",
                "Srsf6",
                "Ebna1bp2",
                "Tmpo",
                "Tnnt1",
                "Cycs",
                "Myl12a",
                "Klf2",
                "Dhx16",
                "Sdhb",
                "Taf10",
                "Wdr43",
                "Klf5",
                "Yrdc",
                "Pabpc1",
                "Eif6",
                "Tsen34",
                "Fkbp1a",
                "Gm7325",
                "Fam60a",
                "Wbp11",
                "Snf8",
                "Myl6",
                "2810428I15Rik",
                "Hspa8",
                "Postn",
                "Ldhb",
                "Serbp1",
                "Taf1d",
                "Kdm1a",
                "Ncor1",
                "1810037I17Rik",
                "Rsrc2",
                "Wbp5",
                "Hsp90aa1",
                "Cox14",
                "Mdh1",
                "Usmg5",
                "Ppp2ca",
                "Rps17",
                "Rae1",
                "Pin4",
                "Mrps18b",
                "Rasgrp2",
                "Txn2",
                "Clta",
                "Cox17",
                "Ppp1r11",
                "Mrpl4",
                "Wisp1",
                "Ndufb3",
                "Gar1",
                "Cks1b",
                "Calcoco2",
                "Rpl23a",
                "Gm10076",
                "Ftsj3",
                "Folr1",
                "Psmc1",
                "Tpd52l2",
                "Knop1",
                "Psmd6",
                "Cthrc1",
                "Pin1",
                "Sfxn1",
                "Swi5",
                "Eif2s1",
                "Nup62",
                "Cenpe",
                "Rab18",
                "Calm3",
                "Arf6",
                "Ubxn1",
                "Cdk1",
                "Eif4h",
                "Psmd7",
                "Anapc15",
                "Gabarapl2",
                "Stub1",
                "Prrx1",
                "Mrpl2",
                "Ranbp1",
                "Zwint",
                "Ppm1g",
                "Eif1",
                "Tbcb",
                "Wdr89",
                "Pdcd6",
                "H2afx",
                "Bgn",
                "Hnrnpm",
                "Capza2",
                "Loxl1",
                "2310036O22Rik",
                "Fdft1",
                "Srsf5",
                "Hnrnph1",
                "Rpl39l",
                "Wdr74",
                "Ywhaz",
                "Pomp",
                "Rpl36",
                "Rps21",
                "Psmb3",
                "Ddx5",
                "Rbm39",
                "Pigyl",
                "Eef1b2",
                "Eif3l",
                "Tmem258",
                "Tcp1",
                "Sarnp",
                "Gnb2",
                "Mrpl15",
                "Ezh2",
                "Mybl2",
                "Pfdn1",
                "Slc2a3",
                "Pmm1",
                "Actl6a",
                "Sssca1",
                "Ier5",
                "Gpx1",
                "Smc3",
                "Thrap3",
                "Dynlrb1",
                "Hspb1",
                "Kdm5b",
                "Col1a2",
                "Eif3b",
                "Ube2m",
                "Zfp207",
                "Chmp4b",
                "Ube2k",
                "Pfkl",
                "Ndufs2",
                "Ddt",
                "Cox7a2l",
                "Spc24",
                "Asns",
                "Ifitm2",
                "Fdps",
                "Lsm8",
                "Ndufs7",
                "Cenpf",
                "Mcm2",
                "Gm16286",
                "Mrpl17",
                "Ldha",
                "Apex1",
                "Cct5",
                "Ccnb2",
                "Mrpl35",
                "1110004F10Rik",
                "Mrpl40",
                "Lox",
                "Iscu",
                "Fubp1",
                "Arpc5l",
                "Klf9",
                "Rps29",
                "Ccdc58",
                "Cenpa",
                "Prdx6",
                "Trappc4",
                "Mdk",
                "Psmd1",
                "Ndufab1",
                "Plekhf2",
                "Snrpf",
                "Mcl1",
                "Rtn4",
                "Pole4",
                "Fbl",
                "Ndufv3",
                "Rps26-ps1",
                "Tsc22d1",
                "Calm2",
                "Stmn1",
                "Btf3",
                "Smarcc1",
                "Tmem126a",
                "Cdh11",
                "Rpl18",
                "Hax1",
                "Atp5d",
                "Mt2",
                "Taldo1",
                "Vcp",
                "Mrps5",
                "Prc1",
                "U2surp",
                "Ndufb2",
                "Ap1s1",
                "Ppan",
                "Rps9",
                "Upp1",
                "Mrpl30",
                "BC031181",
                "Rwdd1",
                "Cyc1",
                "Fnta",
                "Dnaja1",
                "Tmem176b",
                "Mfap2",
                "Pmvk",
                "Elof1",
                "Lrpap1",
                "Wdr5",
                "Pebp1",
                "Sgk1",
                "Ube2d3",
                "Aars",
                "Marc2",
                "Snrpa1",
                "Paip2",
                "Srrm2",
                "Rnaseh2b",
                "Top1",
                "Gm26917",
                "Ubl5",
                "Hmgb1",
                "Ndufc1",
                "Eif3d",
                "Cyb5a",
                "Ghitm",
                "Psma4",
                "Scp2",
                "Arpc1a",
                "Ap2s1",
                "Rpl34",
                "Mlec",
                "Rab7",
                "Cdc42",
                "Mrps14",
                "Nap1l1",
                "Mrps25",
                "Hsbp1",
                "Actr3",
                "Tmsb4x",
                "Fbln2",
                "Mrps10",
                "Tbrg1",
                "Hspa9",
                "Fkbp10",
                "Chchd2",
                "Pim3",
                "E130012A19Rik",
                "Pam16",
                "Luc7l3",
                "Ndufs5",
                "Gsn",
                "Ndufb8",
                "Tmem97",
                "Srsf2",
                "Prdx4",
                "mt-Nd1",
                "Pak1ip1",
                "Higd2a",
                "Nxt1",
                "Trp53",
                "Lhfp",
                "Rbm3",
                "Psmd11",
                "Cggbp1",
                "Zfp706",
                "Rad23b",
                "Nudt4",
                "Supt16",
                "Ost4",
                "Mrpl51",
                "Eif4g1",
                "Rhoa",
                "Cdk4",
                "Rsl1d1",
                "Syncrip",
                "Ssr2",
                "Caprin1",
                "Cebpz",
                "Tomm20",
                "Cbx1",
                "Tomm70a",
                "Gas1",
                "Msmo1",
                "Fam25c",
                "Mak16",
                "Bzw1",
                "Ak6",
                "Rbpms",
                "Hnrnpa3",
                "Prpf40a",
                "Pbdc1",
                "Rps19bp1",
                "2010107E04Rik",
                "Tecr",
                "Mybbp1a",
                "Sf3b2",
                "Atp1b1",
                "Slc3a2",
                "Stip1",
                "Chchd4",
                "Rpl14",
                "Tma7",
                "Camk2d",
                "Lsm4",
                "Tpd52",
                "Eif1ad",
                "Rps13",
                "Eif5a",
                "Rps11",
                "Psmd8",
                "Cfl1",
                "Rrp1b",
                "Sparc",
                "Uchl3",
                "Cops6",
                "Hnrnpdl",
                "Eif3a",
                "Rtn3",
                "Adh5",
                "l7Rn6",
                "Ndufa13",
                "Atp5e",
                "Cdc34",
                "Gps1",
                "Tipin",
                "Cmtm7",
                "Rrm2",
                "Sh3bgrl",
                "Rplp1",
                "Capns1",
                "Rnf10",
                "Psma2",
                "Pfdn5",
                "Nsun2",
                "Ndufb7",
                "Shmt1",
                "Alkbh1",
                "Slc25a39",
                "Rps14",
                "Rpl32",
                "Chchd10",
                "Serf2",
                "Bud31",
                "Mcm5",
                "Rpl13a",
                "Nol7",
                "Tkt",
                "Apoe",
                "Gsta4",
                "Snrpb",
                "Zfp428",
                "Tomm22",
                "Lyar",
                "Erp29",
                "Gpx8",
                "Gpi1",
                "Tmem167",
                "Cfdp1",
                "Eif5b",
                "Emg1",
                "Eif4ebp1",
                "Nono",
                "Csde1",
                "Psenen",
                "Emp1",
                "Mcm6",
                "Prpf19",
                "Sugt1",
                "Eif4a3",
                "Dek",
                "Bag1",
                "Top2a",
                "Papola",
                "Tmem256",
                "Cetn3",
                "Galk1",
                "Ccna2",
                "Tubb4b",
                "Eed",
                "Ndufa5",
                "Mrpl11",
                "Mesdc2",
                "Cox5b",
                "Arhgdia",
                "Rer1",
                "Ralbp1",
                "Fau",
                "Acin1",
                "Aebp1",
                "Rbpms2",
                "Uhrf1",
                "Cox4i1",
                "Guk1",
                "Psma3",
                "Skp1a",
                "Rps18",
                "Phf5a",
                "Rps23",
                "U2af2",
                "P3h4",
                "Myl12b"
            ],
            "expression_sum": {
                "class1": 25.0,
                "class2": 274305.0
            },
            "uniprot_ids": [
                "P20263",
                "E2JL30",
                "G3UZG9",
                "E2JL35",
                "A0A2Z6DUK5",
                "A0A2I6EDJ6",
                "A0A2I6EDI9",
                "E2JL33"
            ],
            "net_id": 0,
            "source_num": 9,
            "target_num": 709
        },
        "Hdac1": {
            "id": "Hdac1",
            "type": "TF",
            "symbol": "Hdac1",
            "source": [
                "Klf2",
                "Tcf4",
                "Pou5f1",
                "Junb",
                "Fos",
                "Jun",
                "Sox2",
                "Nanog",
                "Jund",
                "Cebpb"
            ],
            "target": [
                "Dap",
                "Sar1a",
                "Tpm4",
                "Cd44",
                "Fis1",
                "Ddb1",
                "Thbs1",
                "Lims1",
                "Sec13",
                "Igfbp2",
                "Wisp1",
                "Slc25a4",
                "Tspan4",
                "Map1lc3a",
                "Lmna",
                "Lgals3",
                "mt-Co1",
                "Pgrmc1",
                "Fos",
                "Tagln2",
                "Rexo2",
                "Crlf1",
                "Pdlim7",
                "Mfge8",
                "Mbnl2",
                "Canx",
                "Actn1",
                "Neat1",
                "Myh10",
                "Ube2r2",
                "Gemin7",
                "Akr1a1",
                "Sh3glb1",
                "Akt1s1",
                "Atp2b1",
                "Cisd2",
                "Nktr",
                "Col5a1",
                "Klf6",
                "Pcolce",
                "Crip1",
                "Morc1",
                "Rps12",
                "Arpc1b",
                "Rab14",
                "Serpinh1",
                "Gnb2",
                "Ociad1",
                "Dpysl3",
                "Mybl2",
                "Hcfc1r1",
                "Sptbn1",
                "Id3",
                "Ap3s1",
                "Gng12",
                "Ncam1",
                "Hacd1",
                "Ndufa8",
                "Pdia4",
                "Srp14",
                "Sept7",
                "Fam195b",
                "Actg1",
                "Rcn2",
                "Tsg101",
                "Ier5",
                "Prnp",
                "Id1",
                "Hmga2",
                "Dynlrb1",
                "Hist1h2bc",
                "Nfkbia",
                "Nrp1",
                "Igfbp4",
                "Id2",
                "Itgb1",
                "Fabp5",
                "Mettl9",
                "Chmp4b",
                "Tcf4",
                "Igf2",
                "Cd81",
                "Pgls",
                "Lxn",
                "Ift20",
                "Anxa2",
                "Vmp1",
                "mt-Nd4",
                "Crip2",
                "Cox7a2l",
                "Vim",
                "Basp1",
                "Sh3bgrl3",
                "Wdr1",
                "Tnc",
                "Ube2b",
                "Tmed2",
                "Tpm1",
                "Sdcbp",
                "Surf4",
                "Phlda1",
                "Myeov2",
                "Bola3",
                "Cst3",
                "S100a6",
                "H2-D1",
                "Ifitm1",
                "Cd63",
                "Capg",
                "Lbh",
                "Ndufa3",
                "Scand1",
                "Mtdh",
                "Nfib",
                "Maged2",
                "Rtn4",
                "Ctsl",
                "Ccnd2",
                "Tcf25",
                "Spcs1",
                "Cox6a1",
                "Tax1bp3",
                "Rala",
                "Filip1l",
                "Cav1",
                "Tln1",
                "Oaf",
                "Ostf1",
                "Snhg12",
                "Mt2",
                "Cmpk1",
                "Fkbp2",
                "Lrrc58",
                "Fn1",
                "Glipr2",
                "Hes1",
                "Ap1s1",
                "Tuba4a",
                "Twist2",
                "Malat1",
                "Gnai2",
                "Tmem176b",
                "Tmed10",
                "Tm9sf3",
                "Ctsd",
                "Reep5",
                "Pou5f1",
                "Glrx5",
                "Col1a1",
                "Dbi",
                "Tmem59",
                "Cebpb",
                "Pafah1b2",
                "Anxa3",
                "Hbegf",
                "Scp2",
                "Ilk",
                "Mgat2",
                "Sep15",
                "Mlec",
                "Lsm7",
                "Atf5",
                "Arf5",
                "Cdc42",
                "Npc2",
                "Eef2",
                "Hsbp1",
                "Rnh1",
                "Actr3",
                "Cd47",
                "Efemp2",
                "Mpc1",
                "Copb2",
                "Ap2m1",
                "Nupr1",
                "Smim14",
                "Ahnak",
                "Fkbp10",
                "Mpc2",
                "Ptp4a1",
                "Lsp1",
                "Gsn",
                "Oat",
                "Tubb6",
                "Cited2",
                "Eva1b",
                "mt-Nd1",
                "Aes",
                "Tspo",
                "Sra1",
                "Lhfp",
                "Btg2",
                "Parva",
                "Selk",
                "Rab11a",
                "Anxa5",
                "Anxa6",
                "Mtch1",
                "Rab10",
                "Ttc3",
                "Vimp",
                "Sec61a1",
                "Iqgap1",
                "Kdelr2",
                "Pmp22",
                "March5",
                "Fermt2",
                "Rpl18a",
                "Rpn1",
                "Ssr4",
                "F3",
                "Rcn1",
                "Mrpl54",
                "Sox4",
                "Phlda3",
                "Sept11",
                "Romo1",
                "Tpm2",
                "Ergic3",
                "Ubc",
                "Rock2",
                "Psap",
                "Tm4sf1",
                "Rbms1",
                "P4hb",
                "Scd2",
                "Dlk1",
                "Zyx",
                "Map1b",
                "Rab34",
                "Adipor1",
                "Bsg",
                "Sox2",
                "Ppic",
                "Sec22b",
                "Camk2d",
                "Flna",
                "H13",
                "Rsu1",
                "Prss23",
                "Gsto1",
                "Ppib",
                "Vcan",
                "Rps19",
                "Sept2",
                "Abracl",
                "Msn",
                "Serpinb6a",
                "Lpp",
                "Ctbp1",
                "Csrp1",
                "Selm",
                "Ckap4",
                "Esd",
                "Atxn7l3b",
                "Comt",
                "Lamp1",
                "Ndufa13",
                "Ptrf",
                "Gadd45g",
                "Sh3bgrl",
                "Jun",
                "Pdia3",
                "Rpl10a",
                "Txndc5",
                "Ptms",
                "Col4a1",
                "Hist1h1c",
                "Cd24a",
                "Lipe",
                "Tnfrsf12a",
                "Krtcap2",
                "Gabarap",
                "Serf2",
                "Mtpn",
                "Anxa1",
                "Prkar1a",
                "Rap1b",
                "Ier3",
                "Grb10",
                "Apoe",
                "Commd3",
                "Fst",
                "Tmem167",
                "Palld",
                "Cdkn1a",
                "Csde1",
                "Marcks",
                "Fam162a",
                "Rhoc",
                "Ifitm3",
                "Srp72",
                "Mrpl33",
                "Emp1",
                "App",
                "Fundc2",
                "Fbn2",
                "Bag1",
                "Rpn2",
                "Prrx2",
                "Arhgdib",
                "Tmed3",
                "Serpinf1",
                "Hsd17b14",
                "Tbca",
                "Foxp1",
                "Ccng1",
                "Ptp4a2",
                "Dnajc3",
                "Arhgdia",
                "Myl12a",
                "Rac1",
                "Rer1",
                "S100a4",
                "Cald1",
                "Sri",
                "Prdx5",
                "Clic1",
                "Fkbp1a",
                "Gnas",
                "Arf4",
                "Calu",
                "Arpc5",
                "Gm7325",
                "Hdlbp",
                "Ckb",
                "Limd2",
                "Cox4i1",
                "Tmed9",
                "Myl6",
                "Sepw1",
                "Serp1",
                "Capn2",
                "Laptm4a",
                "Carhsp1",
                "Alpl",
                "Kdelr3",
                "Junb",
                "Bcap31",
                "Cfl2"
            ],
            "expression_sum": {
                "class1": 3445.0,
                "class2": 29162.0
            },
            "uniprot_ids": [
                "O09106",
                "Q58E49"
            ],
            "net_id": 0,
            "source_num": 10,
            "target_num": 175
        },
        "Hmgb1": {
            "id": "Hmgb1",
            "type": "TF",
            "symbol": "Hmgb1",
            "source": [
                "Klf6",
                "Tcf4",
                "Pou5f1",
                "Foxp1",
                "Nfib",
                "Atf4",
                "Junb",
                "Fos",
                "Jun",
                "Sox2",
                "Hes1",
                "Nanog",
                "Jund",
                "Twist2",
                "Cebpb",
                "Zfp57",
                "Sox4"
            ],
            "target": [
                "Atxn7l3b",
                "Skil",
                "Tspan6",
                "Amotl2",
                "Atf5",
                "Cuedc1",
                "Tns1",
                "Ak1",
                "Svep1",
                "Fam204a",
                "Tmed5",
                "Fabp5",
                "Mettl9",
                "Nfatc4",
                "Hspa5",
                "Igf2",
                "Rasgrp2",
                "Phtf1",
                "Efemp2",
                "Col12a1",
                "Bex1",
                "Ifngr2",
                "Anxa2",
                "Dcun1d1",
                "9530068E07Rik",
                "Ccar1",
                "Flnb",
                "Cox7a2l",
                "Smap1",
                "Acot7",
                "Basp1",
                "Ahnak",
                "Elovl1",
                "1700020I14Rik",
                "Gabarap",
                "Zbtb20",
                "Metrn",
                "Nab2",
                "Phgdh",
                "Rfwd2",
                "Brk1",
                "Hsp90b1",
                "Pgrmc1",
                "Adamts1",
                "Timp1",
                "Cited2",
                "Celf2",
                "Rps15a",
                "Adam9",
                "Setd3",
                "Eif4g2",
                "Sdcbp",
                "Tuba1a",
                "Lamb1",
                "Emd",
                "2610305D13Rik",
                "Pdlim7",
                "Micu1",
                "Actn1",
                "Rcbtb2",
                "Psat1",
                "Eif4h",
                "Pex2",
                "Rnf139",
                "Trappc3",
                "Shoc2",
                "2310009B15Rik",
                "Anxa5",
                "Bad",
                "Prdx6",
                "Eif3f",
                "Hadha",
                "Serpine1",
                "2200002D01Rik",
                "Rab10",
                "Alg5",
                "AY036118",
                "Crabp1",
                "Ifitm3",
                "Lix1l",
                "Nptn",
                "B3galnt1",
                "Iqgap1",
                "Casc5",
                "Kdelr2",
                "Tdgf1",
                "March5",
                "App",
                "2310022B05Rik",
                "Arl6ip5",
                "Atp6ap1",
                "Dexi",
                "Akt1s1",
                "Hist1h1e",
                "Agpat4",
                "Pde10a",
                "P4ha1",
                "Spcs1",
                "Ndfip2",
                "Ssr4",
                "Ctdsp1",
                "Gpc1",
                "Ap2a2",
                "Crebbp",
                "Araf",
                "Wdr26",
                "Htatsf1",
                "Tspan5",
                "Dusp11",
                "Slc25a46",
                "Plaur",
                "Arpc2",
                "Mrpl54",
                "Mageh1",
                "Fndc4",
                "Col4a2",
                "Capza2",
                "Ccl2",
                "Chsy1",
                "Mbnl3",
                "Upf3a",
                "Phlda3",
                "Pbx1",
                "Ralb",
                "Trip6",
                "Txnrd1",
                "1600012H06Rik",
                "Ube3a",
                "Ergic3",
                "Taok3",
                "Atp5d",
                "2610001J05Rik",
                "Slc39a13",
                "Maoa",
                "H2afy2",
                "Smim10l1",
                "Cstb",
                "0610011F06Rik",
                "Gm10260",
                "Mpp1",
                "Btg1",
                "Acer2",
                "Lrrc58",
                "Kif26b",
                "Cope",
                "Tspan31",
                "L1td1",
                "Ap1s1",
                "Col3a1",
                "Tuba4a",
                "Mecr",
                "Rab14",
                "Serpinh1",
                "Psap",
                "Armcx2",
                "H2afy",
                "Rwdd1",
                "4930402H24Rik",
                "Acat1",
                "Cald1",
                "Arl8b",
                "Cpox",
                "Ext2",
                "Sema3c",
                "Gm17501",
                "Rbm7",
                "Tm9sf3",
                "Trh",
                "Ctsd",
                "1700123O20Rik",
                "Cbx7",
                "4933434E20Rik",
                "Vgll3",
                "Arpc5",
                "Elf2",
                "Med21",
                "Hdlbp",
                "Cox4i1",
                "Ephb2",
                "Rgs16",
                "Airn",
                "Efnb1",
                "Thbs2",
                "Gdf3",
                "Evl",
                "Nek6",
                "Adk",
                "Nxn",
                "Stard3nl",
                "Serp1",
                "Lrrfip1",
                "Snx7",
                "Laptm4a",
                "Tmem59",
                "Vcan",
                "Actg1",
                "Ptprd",
                "Cdc27",
                "Emp3",
                "Fat4",
                "Abracl",
                "Cryab",
                "Dnah8",
                "Pdcl3",
                "Map4",
                "Cd302",
                "Arpc3",
                "Rassf1",
                "Arl1",
                "Col1a2",
                "Sult6b1"
            ],
            "expression_sum": {
                "class1": 34517.0,
                "class2": 116070.0
            },
            "uniprot_ids": [
                "P63158",
                "Q58EV5",
                "Q8C7C4",
                "A0A0J9YUD8",
                "Q497Z6",
                "Q3UJK0",
                "Q8BQ02",
                "A0A0J9YUZ4",
                "Q6P202",
                "D3YVC6",
                "Q3UBK2",
                "D3YZ18",
                "Q8BNM0"
            ],
            "net_id": 0,
            "source_num": 17,
            "target_num": 48
        },
        "Foxp1": {
            "id": "Foxp1",
            "type": "TF",
            "symbol": "Foxp1",
            "source": [
                "Hdac2",
                "E2f4",
                "Trim28",
                "Pou5f1",
                "Rest",
                "Atf4",
                "Ctcf",
                "Tgif1",
                "Trp53",
                "Hdac1",
                "Klf5",
                "Cebpb",
                "Smc3",
                "Smarca5"
            ],
            "target": [
                "Rps5",
                "Hsp90aa1",
                "Arf1",
                "Rpl41",
                "Rnf7",
                "Cdc20",
                "Sar1a",
                "Srsf9",
                "Usmg5",
                "Mcm7",
                "Ppp2ca",
                "Ddx24",
                "Rps17",
                "Rae1",
                "Zc3h15",
                "Cox8a",
                "Mrps18b",
                "Cdt1",
                "Psmb7",
                "Txn2",
                "Pnkd",
                "Dkc1",
                "Ndufa6",
                "Ppp1r11",
                "1110008F13Rik",
                "Tuba1b",
                "Polr2k",
                "Gar1",
                "Anapc5",
                "St13",
                "Thoc7",
                "Hnrnpf",
                "Knop1",
                "Lmnb1",
                "Ddx39",
                "Mlf2",
                "Pin1",
                "Ier3ip1",
                "Sfxn1",
                "Swi5",
                "Eif2s1",
                "mt-Co1",
                "Hmgb2",
                "Slbp",
                "Calr",
                "Nup62",
                "Rab18",
                "Calm3",
                "Arf6",
                "Canx",
                "Snrpe",
                "Mrpl14",
                "Cdk1",
                "Phb",
                "Lamtor1",
                "Anapc15",
                "Mrpl2",
                "Tra2b",
                "Cdc5l",
                "Ranbp1",
                "Psmc6",
                "Grpel1",
                "Rbx1",
                "Ctcf",
                "Ppm1g",
                "Eif1",
                "Neat1",
                "Ythdc1",
                "Bola1",
                "Ola1",
                "Sf3b6",
                "Snrnp27",
                "Aurkaip1",
                "Srsf1",
                "Socs3",
                "Lrrc59",
                "Exosc5",
                "Rcc2",
                "Pdcd6",
                "Impa2",
                "Ube2d2a",
                "Bnip3l",
                "Mapre1",
                "Cisd2",
                "Capza2",
                "Lap3",
                "Ncl",
                "Fdft1",
                "Supt4a",
                "Ppp1ca",
                "Srsf5",
                "Hnrnph1",
                "Aamp",
                "Chchd1",
                "Ywhaz",
                "Psmc3",
                "Sec62",
                "Luc7l2",
                "Rbbp7",
                "Hnrnpl",
                "Ddx5",
                "Rbm39",
                "Ccdc12",
                "Nedd8",
                "Pcnp",
                "Ube2i",
                "Utp3",
                "Rab14",
                "Tmem258",
                "Tcp1",
                "Brix1",
                "Fam136a",
                "Pttg1",
                "Mrpl15",
                "Siva1",
                "Fus",
                "Ociad1",
                "Ddost",
                "Pmm1",
                "Fkbp8",
                "Nip7",
                "Clpp",
                "Psma7",
                "G3bp1",
                "Pdia4",
                "Sept7",
                "Tpm3",
                "Banf1",
                "Cdc26",
                "Immt",
                "Psma1",
                "Qdpr",
                "Lig1",
                "2410015M20Rik",
                "Hdgf",
                "Prnp",
                "Smc3",
                "Thrap3",
                "Csnk2b",
                "Pa2g4",
                "Rpl37",
                "Kdm5b",
                "Ppp1cc",
                "Atf4",
                "Zfp207",
                "Pcbp1",
                "Hnrnpk",
                "Ndufa4",
                "Ndufv1",
                "Ndufv2",
                "H3f3a",
                "Mettl9",
                "Nme1",
                "Pfkl",
                "Csnk1a1",
                "Ndufs2",
                "Fam49b",
                "Bax",
                "Tmsb10",
                "Rpl3",
                "Cebpzos",
                "Trim28",
                "Snw1",
                "Cdv3",
                "Napa",
                "Spc24",
                "Btf3l4",
                "Ppp4c",
                "Etf1",
                "Ube2b",
                "Lsm3",
                "Gm16286",
                "Eif5",
                "Tmed2",
                "Txn1",
                "Vapa",
                "Mrpl17",
                "Samm50",
                "Rrp15",
                "Wdr77",
                "Atp5j",
                "Ran",
                "Fscn1",
                "Srrm1",
                "Emc8",
                "Srp19",
                "Cct5",
                "Emd",
                "Mrpl40",
                "Bclaf1",
                "Vasp",
                "Tceb1",
                "C1qbp",
                "Uqcr10",
                "H2afz",
                "Tardbp",
                "Psip1",
                "Rpl36a",
                "Rpl29",
                "Iscu",
                "Fubp1",
                "Snrpb2",
                "Arpc5l",
                "Rps29",
                "Ccdc58",
                "Cenpa",
                "Prdx6",
                "Wtap",
                "Odc1",
                "Mrpl18",
                "Rps20",
                "Dnaja2",
                "Sys1",
                "Acat2",
                "Lbh",
                "Cdc37",
                "Psmd1",
                "Plekhf2",
                "Eef1a1",
                "Mcl1",
                "Rtn4",
                "U2af1",
                "Atp5c1",
                "Ywhaq",
                "Spcs1",
                "Fbl",
                "Ndufv3",
                "Ubald2",
                "Tsc22d1",
                "mt-Co2",
                "Dnajc8",
                "Calm2",
                "Rps3",
                "Atox1",
                "Srsf10",
                "Adss",
                "Tmem126a",
                "Ube2v2",
                "Rpl18",
                "Trnau1ap",
                "Atp5d",
                "Timm13",
                "Eif3h",
                "Rpl22l1",
                "Adprh",
                "Lamtor4",
                "Rps3a1",
                "Mrps5",
                "U2surp",
                "Mbd3",
                "Cct7",
                "Dctpp1",
                "Ndufb2",
                "Wdr18",
                "Hsp90ab1",
                "Oaz1",
                "Mrpl30",
                "Uqcrq",
                "Sec61b",
                "BC031181",
                "Polr2m",
                "1810058I24Rik",
                "Rwdd1",
                "Snx5",
                "Got1",
                "Fnta",
                "Dnaja1",
                "Eif4a1",
                "Sumo2",
                "Pcbp2",
                "Snrnp25",
                "Nudc",
                "H3f3b",
                "Mrps33",
                "Rif1",
                "Nsa2",
                "Ddx46",
                "Nudt19",
                "Dbi",
                "Morf4l2",
                "Ube2d3",
                "Rbbp4",
                "Eif3e",
                "Paip2",
                "Cebpb",
                "Srrm2",
                "Rnaseh2b",
                "Ywhae",
                "Ubl5",
                "Hmgb1",
                "Ndufc1",
                "Eif3d",
                "Rps15",
                "Npm1",
                "Psma4",
                "Sap30",
                "Scp2",
                "Ap2s1",
                "Rpl34",
                "Mlec",
                "Dynll1",
                "Gtpbp4",
                "Cenpw",
                "Hspd1",
                "Atf5",
                "Cops7a",
                "Mrpl45",
                "Mrps14",
                "Rpl31",
                "Srsf7",
                "Smdt1",
                "Sod1",
                "Ppih",
                "Mat2a",
                "Nap1l1",
                "Uqcc3",
                "Dnajb6",
                "Brd2",
                "Actr3",
                "G3bp2",
                "Impdh2",
                "Ccdc124",
                "Gapdh",
                "Smarca5",
                "Mpc1",
                "Prps1",
                "Ccnd3",
                "Ap2m1",
                "Mrpl52",
                "Adrm1",
                "Abcf1",
                "Rplp0",
                "Psmb5",
                "Hspa9",
                "Pim3",
                "Ube2n",
                "Ndufa11",
                "Cct8",
                "Hmgn2",
                "Mpc2",
                "Dohh",
                "Hnrnpa2b1",
                "Ndufs5",
                "Cited2",
                "Srsf2",
                "Eif4g2",
                "Jade1",
                "Psmb2",
                "Raly",
                "Denr",
                "Fxr1",
                "Nsmce4a",
                "Rps26",
                "Get4",
                "Trp53",
                "Sra1",
                "Aspscr1",
                "Psmd11",
                "Vps29",
                "Vdac1",
                "Cggbp1",
                "Zfp706",
                "Anp32a",
                "Polr2e",
                "Anapc13",
                "Sumo1",
                "Pdcd5",
                "Snrpc",
                "Mrpl51",
                "Mrpl49",
                "Tmbim6",
                "Dynll2",
                "Rhoa",
                "Fcf1",
                "Mrps18c",
                "Cacybp",
                "Cdk4",
                "Rpl22",
                "Rsl1d1",
                "Nolc1",
                "Sf3a3",
                "Sec61a1",
                "Pfdn6",
                "Glud1",
                "Cebpz",
                "Hnrnpu",
                "Gspt1",
                "Arglu1",
                "Tomm20",
                "Rpl21",
                "Ppp1r2",
                "Map1lc3b",
                "Eprs",
                "Trmt112",
                "Dazap1",
                "Son",
                "Pop5",
                "Cct2",
                "Ubb",
                "Dtl",
                "Bzw1",
                "Parp1",
                "Dhfr",
                "Rbpms",
                "Paics",
                "Hnrnpa3",
                "Prpf40a",
                "Pbdc1",
                "Cdca3",
                "Mrpl57",
                "Morf4l1",
                "Mrpl43",
                "Ube2s",
                "Tecr",
                "Polr1d",
                "Snrpd3",
                "Sf3b2",
                "Psmd3",
                "Suclg1",
                "Mtf2",
                "Slc3a2",
                "mt-Nd2",
                "Hnrnpd",
                "Hat1",
                "Cops4",
                "mt-Cytb",
                "Rnps1",
                "Ciapin1",
                "Lsm4",
                "Pfdn2",
                "Mpdu1",
                "Atp5f1",
                "Eif1ad",
                "Rps13",
                "Tpt1",
                "Eif5a",
                "Ppib",
                "Gng5",
                "Hspe1",
                "Rps19",
                "Exosc8",
                "Cops6",
                "Ptbp1",
                "Aldoa",
                "Hnrnpdl",
                "Eif3a",
                "Rtn3",
                "Polr2h",
                "Snx3",
                "Adh5",
                "Birc5",
                "Psmd13",
                "Srpk1",
                "Rfc1",
                "Cox6b1",
                "Ppa1",
                "Cenpm",
                "Prdx1",
                "Rsl24d1",
                "Cdc34",
                "Rpa2",
                "Mrpl34",
                "Pdia3",
                "Ndufb5",
                "Psma2",
                "Ndufs6",
                "Nsun2",
                "Set",
                "Txnl4a",
                "Poldip2",
                "Eif4e2",
                "Sod2",
                "Ssbp1",
                "Rps14",
                "Tgif1",
                "Lipe",
                "Clns1a",
                "Snrpd2",
                "1810022K09Rik",
                "Phb2",
                "Gtf2e2",
                "Rpl15",
                "Brk1",
                "Vdac2",
                "Rap1b",
                "Pfn1",
                "Rpl13a",
                "Snrpg",
                "Ddx39b",
                "Cuta",
                "Snhg6",
                "Eif1a",
                "Cox5a",
                "Snrpb",
                "Tmem11",
                "Anapc11",
                "Fkbp4",
                "Pgam1",
                "Tomm22",
                "Lyar",
                "Ubap2l",
                "Erp29",
                "Snrnp70",
                "Uchl5",
                "Gpi1",
                "Tmem167",
                "Ppdpf",
                "Llph",
                "Emg1",
                "Lsm12",
                "Eif4b",
                "Anp32b",
                "Pkm",
                "H2afv",
                "Csde1",
                "Prrc2c",
                "Tceb2",
                "Rps2",
                "Eif2b5",
                "Gart",
                "Sumo3",
                "Ndufa10",
                "Uqcrfs1",
                "Naa50",
                "Ywhah",
                "Sf3b4",
                "Prpf19",
                "Sugt1",
                "Eif4a3",
                "Fkbp3",
                "Mtch2",
                "Dek",
                "Bag1",
                "Psmd4",
                "Gins2",
                "Fam104a",
                "Papola",
                "Rps16",
                "Mrfap1",
                "Mrps16",
                "Uqcrh",
                "Tpr",
                "Ddx18",
                "Psmd12",
                "Rrp9",
                "Eed",
                "Ccne1",
                "Srsf6",
                "Ndufa5",
                "Tmpo",
                "Tsn",
                "Mrpl11",
                "Txnl1",
                "Mesdc2",
                "Eef1g",
                "Rbm25",
                "2700094K13Rik",
                "Cycs",
                "Myl12a",
                "Ptma",
                "Rac1",
                "Rer1",
                "Klf2",
                "Dut",
                "Ralbp1",
                "Dhx16",
                "Sdhb",
                "Taf10",
                "Wdr43",
                "Ybx3",
                "Rps10",
                "Yrdc",
                "Pabpc1",
                "Alyref",
                "Tsen34",
                "Hspa14",
                "Fkbp1a",
                "Dad1",
                "Yif1b",
                "Fam60a",
                "Wbp11",
                "Cwc15",
                "Guk1",
                "2810428I15Rik",
                "Tsfm",
                "Pim1",
                "Postn",
                "Rplp2",
                "Aimp1",
                "Serbp1",
                "Gpx4",
                "Gcsh",
                "Phf5a",
                "Eif4a2",
                "Rheb",
                "Gtf2a2",
                "Mkrn1",
                "Rrp1",
                "U2af2",
                "Rpl23",
                "Sfr1",
                "Kdm1a",
                "Rrs1",
                "Ly6e",
                "Bcap31",
                "Cnbp",
                "Rsrc2"
            ],
            "expression_sum": {
                "class1": 10758.0,
                "class2": 1389.0
            },
            "uniprot_ids": [
                "P58462",
                "E2S038",
                "H3BJ68",
                "H3BJZ5",
                "H3BLF2",
                "Q8CCZ5",
                "H3BJR8",
                "D3Z6Q4",
                "Q6P0X7",
                "D3Z454",
                "H3BL33",
                "A0A0R4J1H9",
                "H3BJ24",
                "H3BJM1",
                "D3Z6Q3",
                "H3BK30",
                "H3BKT4",
                "A0A0R4J282",
                "D3Z2T4",
                "D3Z0J5",
                "H3BK38",
                "Q3UMG2"
            ],
            "net_id": 0,
            "source_num": 14,
            "target_num": 330
        },
        "Rest": {
            "id": "Rest",
            "type": "TF",
            "symbol": "Rest",
            "source": [
                "Pou5f1",
                "Junb",
                "Fos",
                "Jun",
                "Sox2",
                "Hes1",
                "Nanog",
                "Jund",
                "Cebpb",
                "Sox4"
            ],
            "target": [
                "Dap",
                "Sar1a",
                "Tpm4",
                "H2afj",
                "Ddb1",
                "Myh9",
                "Thbs1",
                "Rasgrp2",
                "Lims1",
                "Wisp1",
                "Slc25a4",
                "Tspan4",
                "Col12a1",
                "Map1lc3a",
                "Lmna",
                "Pck2",
                "Lgals3",
                "Cxcl12",
                "Pgrmc1",
                "Fos",
                "Tagln2",
                "Rps15a",
                "Rexo2",
                "Pdlim7",
                "Mfge8",
                "Mbnl2",
                "Canx",
                "Actn1",
                "B2m",
                "Neat1",
                "Myh10",
                "Nanog",
                "Ube2r2",
                "Timp3",
                "Gemin7",
                "Akr1a1",
                "Sh3glb1",
                "Atp2b1",
                "Cisd2",
                "Nktr",
                "Col5a1",
                "Bgn",
                "Klf6",
                "Tgfb1i1",
                "Sec62",
                "Rps12",
                "Arpc1b",
                "Col3a1",
                "Cyr61",
                "Kdelr1",
                "Rab14",
                "Serpinh1",
                "Atp6v1g1",
                "Gnb2",
                "Ociad1",
                "Dpysl3",
                "Mybl2",
                "Slc2a3",
                "Hcfc1r1",
                "Sptbn1",
                "Id3",
                "Ap3s1",
                "Gng12",
                "Ncam1",
                "Wls",
                "Hacd1",
                "Ndufa8",
                "Srp14",
                "Sept7",
                "Fam195b",
                "Actg1",
                "Rcn2",
                "Ier5",
                "Prnp",
                "Id1",
                "Hmga2",
                "Hist1h2bc",
                "Tfcp2l1",
                "Col1a2",
                "F2r",
                "Serpine2",
                "Igfbp4",
                "Itgb1",
                "Fabp5",
                "Mettl9",
                "Tcf4",
                "Igf2",
                "Ssr1",
                "Pgls",
                "Lxn",
                "Ift20",
                "Anxa2",
                "Vmp1",
                "Crip2",
                "Cox7a2l",
                "Vim",
                "Basp1",
                "Sh3bgrl3",
                "Wdr1",
                "Tnc",
                "Tmed2",
                "Gjb3",
                "Tpm1",
                "Sdcbp",
                "Thy1",
                "Surf4",
                "Utf1",
                "Phlda1",
                "Myeov2",
                "Vcl",
                "Cst3",
                "Lox",
                "Ung",
                "Higd1a",
                "S100a6",
                "Serpine1",
                "Crabp1",
                "Ifitm1",
                "Cd63",
                "Capg",
                "Lbh",
                "S100a11",
                "Scand1",
                "Mtdh",
                "mt-Nd5",
                "Maged2",
                "Ctsl",
                "Ccnd2",
                "Tcf25",
                "Spcs1",
                "Cox6a1",
                "Col6a1",
                "Rala",
                "Sdc2",
                "Tax1bp3",
                "Filip1l",
                "Plk2",
                "Tln1",
                "Oaf",
                "Ostf1",
                "Snhg12",
                "Fbln5",
                "Chd3",
                "Cmpk1",
                "Lrrc58",
                "Fn1",
                "Glipr2",
                "Hes1",
                "Ap1s1",
                "Nenf",
                "Zfp42",
                "Rcn3",
                "Malat1",
                "Tmem176b",
                "Tmed10",
                "Tm9sf3",
                "Reep5",
                "Col1a1",
                "Oxct1",
                "Peg3",
                "Atxn10",
                "Atp6v0e",
                "Tmem59",
                "Pafah1b2",
                "Arpc3",
                "Ilk",
                "Mgat2",
                "Sep15",
                "Lsm7",
                "Cpe",
                "Atf5",
                "Cdc42",
                "Npc2",
                "Eef2",
                "Hsbp1",
                "Rnh1",
                "Rrbp1",
                "Tmsb4x",
                "Cd47",
                "Efemp2",
                "Mpc1",
                "Copb2",
                "Sars",
                "Smim14",
                "Lgals1",
                "Fkbp10",
                "E130012A19Rik",
                "Kif5b",
                "Ptp4a1",
                "Lsp1",
                "Gsn",
                "Tubb6",
                "Prdx4",
                "Rab2a",
                "Tspo",
                "Myl9",
                "Btg2",
                "Selk",
                "Rab11a",
                "Anxa5",
                "Anxa6",
                "Mtch1",
                "Rab10",
                "Ttc3",
                "Vimp",
                "Sec61a1",
                "Iqgap1",
                "Kdelr2",
                "Pmp22",
                "Rbmxl2",
                "Ssr3",
                "Rpn1",
                "Ssr4",
                "Sqstm1",
                "Lman1",
                "Rcn1",
                "Arpc2",
                "Ywhab",
                "Sox4",
                "Phlda3",
                "Sept11",
                "Romo1",
                "Tpm2",
                "Ergic3",
                "Sdc4",
                "Rock2",
                "Cdc42ep3",
                "Col8a1",
                "L1td1",
                "Psap",
                "Rbms1",
                "P4hb",
                "Zyx",
                "Adipor1",
                "Bsg",
                "Ppic",
                "Camk2d",
                "mt-Cytb",
                "Cyba",
                "Flna",
                "H13",
                "Rsu1",
                "Prss23",
                "Gdf3",
                "Gsto1",
                "Ppib",
                "Vcan",
                "Sparc",
                "Sept2",
                "Abracl",
                "Msn",
                "Serpinb6a",
                "Nfatc2ip",
                "Ctbp1",
                "Csrp1",
                "Selm",
                "Esd",
                "Dppa5a",
                "Atxn7l3b",
                "Comt",
                "Lamp1",
                "Rcor2",
                "Sat1",
                "Ndufa13",
                "Ptrf",
                "Gadd45g",
                "Sh3bgrl",
                "Jun",
                "Itm2b",
                "Pdia3",
                "Rpl10a",
                "Txndc5",
                "Col4a1",
                "Hist1h1c",
                "Cd24a",
                "Smap1",
                "Lipe",
                "Tnfrsf12a",
                "Krtcap2",
                "Gabarap",
                "Serf2",
                "Mtpn",
                "Prkar1a",
                "Apoe",
                "Tuba1a",
                "Commd3",
                "Gsta4",
                "Fst",
                "Gpi1",
                "Palld",
                "Tram1",
                "Lrrc17",
                "Fstl1",
                "Marcks",
                "Fam162a",
                "Rhoc",
                "Mdm2",
                "Srp72",
                "Mrpl33",
                "Fundc2",
                "S100a10",
                "Rpn2",
                "Prrx2",
                "Tmed3",
                "Serpinf1",
                "Hsd17b14",
                "Gap43",
                "Foxp1",
                "Ccng1",
                "H19",
                "Ptp4a2",
                "Dnajc3",
                "Myl12a",
                "Rac1",
                "Rer1",
                "S100a4",
                "Cald1",
                "Sri",
                "Prdx5",
                "Clic1",
                "Emc7",
                "Gnas",
                "Arf4",
                "Calu",
                "Arpc5",
                "Gm7325",
                "Hdlbp",
                "Ckb",
                "Limd2",
                "Cox4i1",
                "Tmed9",
                "Myl6",
                "Sepw1",
                "Serp1",
                "Capn2",
                "Laptm4a",
                "Carhsp1",
                "Kdelr3",
                "Junb",
                "Bcap31",
                "Axl",
                "Cfl2"
            ],
            "expression_sum": {
                "class1": 3691.0,
                "class2": 33317.0
            },
            "uniprot_ids": [
                "Q8VIG1",
                "Q60694",
                "F8V330"
            ],
            "net_id": 0,
            "source_num": 10,
            "target_num": 185
        },
        "Atf4": {
            "id": "Atf4",
            "type": "TF",
            "symbol": "Atf4",
            "source": [
                "Tcf4",
                "Pou5f1",
                "Foxp1",
                "Cebpb",
                "Eno1",
                "Ctcf",
                "Junb",
                "Trp53",
                "Sox2",
                "Ybx1",
                "Jund",
                "Twist2",
                "Hdac2"
            ],
            "target": [
                "Hsp90aa1",
                "Dap",
                "Rnf7",
                "Cdc20",
                "Sar1a",
                "Usmg5",
                "Ppp2ca",
                "Ddx24",
                "Zc3h15",
                "Myh9",
                "Txn2",
                "Ppid",
                "2810004N23Rik",
                "Clta",
                "Igfbp2",
                "Sec13",
                "1110008F13Rik",
                "Idh3g",
                "Anapc5",
                "Lmna",
                "Ndufa2",
                "Hnrnpf",
                "Knop1",
                "Ddx39",
                "Mlf2",
                "Pck2",
                "Rnf187",
                "Med28",
                "Nubp1",
                "Rps15a",
                "Ndufs3",
                "Nedd4",
                "Uba2",
                "Rexo2",
                "Calr",
                "Nup62",
                "Pdlim7",
                "Calm3",
                "Mbnl2",
                "Arf6",
                "Canx",
                "Nop56",
                "Slc25a3",
                "Psat1",
                "Psmd7",
                "Lamtor1",
                "Tra2b",
                "Psmc6",
                "Zwint",
                "Ppm1g",
                "Eif1",
                "Myh10",
                "Sec61g",
                "Ola1",
                "Snrnp27",
                "Aurkaip1",
                "Gemin7",
                "Wdr89",
                "Lrrc59",
                "Uqcrc1",
                "Akt1s1",
                "Bnip3l",
                "H2afx",
                "Kpnb1",
                "Mapre1",
                "Cisd2",
                "Snrpd1",
                "Nktr",
                "Capza2",
                "Klf6",
                "Rpl36al",
                "Ncl",
                "Loxl1",
                "Srsf5",
                "Hnrnph1",
                "Chchd1",
                "Ywhaz",
                "Gars",
                "Psmc3",
                "Sec62",
                "Pomp",
                "Hspa4",
                "Luc7l2",
                "Psmb3",
                "Ddx5",
                "Rbm39",
                "Nedd8",
                "Pcnp",
                "Eef1b2",
                "Ube2i",
                "Arpc1b",
                "Cyr61",
                "Utp3",
                "Serpinh1",
                "Tcp1",
                "Brix1",
                "Gnb2",
                "Mrpl15",
                "Fus",
                "Dpysl3",
                "Ddost",
                "Hcfc1r1",
                "Sptbn1",
                "Gng12",
                "Fkbp8",
                "Nip7",
                "Ndufa8",
                "Pdia4",
                "Srp14",
                "Sept7",
                "Fam195b",
                "Trim27",
                "Mrps36",
                "Banf1",
                "Immt",
                "Actg1",
                "Dtymk",
                "Psma1",
                "Tsg101",
                "Hdgf",
                "Nars",
                "Pa2g4",
                "Hmga2",
                "F2r",
                "Pcbp1",
                "Rpl38",
                "Dnttip2",
                "Sae1",
                "Hnrnpk",
                "Ewsr1",
                "Eif4e",
                "Ndufv1",
                "Ndufv2",
                "Fabp5",
                "H3f3a",
                "Nme1",
                "Chmp4b",
                "Tcf4",
                "Ube2k",
                "Ssr1",
                "Ndufs2",
                "Cyp51",
                "Pgls",
                "Tmsb10",
                "Nudt21",
                "Rpl12",
                "Ift20",
                "Anxa2",
                "Eif3c",
                "Snw1",
                "Vmp1",
                "Col5a2",
                "Cdv3",
                "Napa",
                "Btf3l4",
                "Asns",
                "Eef1d",
                "Etf1",
                "Sh3bgrl3",
                "Ndufs7",
                "Ube2b",
                "Lsm3",
                "Mrpl13",
                "Gm16286",
                "Eif5",
                "Tmed2",
                "Ube2j2",
                "Vapa",
                "Mrpl17",
                "Wdr77",
                "Atp5j",
                "Ran",
                "Fscn1",
                "Srrm1",
                "Tpm1",
                "Ldha",
                "Sdcbp",
                "Apex1",
                "Surf4",
                "Atp5g2",
                "Emc8",
                "Srp19",
                "Phlda1",
                "1110004F10Rik",
                "Myeov2",
                "Vasp",
                "Tceb1",
                "Rpsa",
                "H2afz",
                "Bola3",
                "Psip1",
                "Vcl",
                "Iscu",
                "Fubp1",
                "Arpc5l",
                "Rpl5",
                "Dph3",
                "Prdx6",
                "Ptges3",
                "Mrpl18",
                "Rps20",
                "Orc6",
                "Dnaja2",
                "Crabp1",
                "Sys1",
                "Mrpl12",
                "Ndufab1",
                "Mtdh",
                "Eef1a1",
                "mt-Nd5",
                "U2af1",
                "Atp5c1",
                "Ywhaq",
                "Ccnd2",
                "Tcf25",
                "Pdha1",
                "Filip1l",
                "Cav1",
                "Dnajc8",
                "Calm2",
                "Stmn1",
                "Srsf10",
                "Btf3",
                "Smarcc1",
                "Plk2",
                "Tln1",
                "Tmem126a",
                "Oaf",
                "Rpl18",
                "Atp5d",
                "Mt2",
                "Timm13",
                "Rps3a1",
                "Smc4",
                "Cmpk1",
                "Rpl7",
                "Fkbp2",
                "Lrrc58",
                "Fn1",
                "Glipr2",
                "Hes1",
                "Ap1s1",
                "Uqcrq",
                "Mrpl30",
                "Sec61b",
                "Polr2m",
                "Rwdd1",
                "Snx5",
                "Gnai2",
                "Rps8",
                "Dnaja1",
                "Rpl8",
                "Eif4a1",
                "Tm9sf3",
                "Ndufc2",
                "Ccdc34",
                "Col1a1",
                "Nsa2",
                "Oxct1",
                "Ddx46",
                "Cnih4",
                "Morf4l2",
                "Ube2d3",
                "Aars",
                "Snrpa1",
                "Dstn",
                "Emp3",
                "Cebpb",
                "Srrm2",
                "2700060E02Rik",
                "Mt1",
                "Ywhae",
                "Pafah1b2",
                "Hmgb1",
                "Psmd14",
                "Eif3d",
                "Npm1",
                "Ghitm",
                "Scp2",
                "Arl1",
                "Arpc1a",
                "Mgat2",
                "Sep15",
                "Rab7",
                "Gtpbp4",
                "Eif2s2",
                "Hspd1",
                "Cpe",
                "Atf5",
                "Cdc42",
                "Mrpl45",
                "Srsf7",
                "Mrps25",
                "Tpx2",
                "Dnajb6",
                "Hspa5",
                "Uqcrc2",
                "Actr3",
                "G3bp2",
                "Cd47",
                "Impdh2",
                "Ccdc124",
                "Mpc1",
                "Sars",
                "Mrpl52",
                "Ap2m1",
                "Nupr1",
                "Pno1",
                "Hnrnpab",
                "Rplp0",
                "Psmb5",
                "Ahnak",
                "Tbrg1",
                "Hspa9",
                "Kif5b",
                "Polr2c",
                "Cct8",
                "Hmgn2",
                "Mpc2",
                "Hnrnpa2b1",
                "Ptp4a1",
                "Lsp1",
                "Ndufs5",
                "Cited2",
                "Srsf2",
                "Prdx4",
                "mt-Nd1",
                "Eif4g2",
                "Dnajc7",
                "Psmb2",
                "Raly",
                "Denr",
                "Get4",
                "Cnn3",
                "Parva",
                "Vdac1",
                "Cggbp1",
                "Zfp706",
                "Anp32a",
                "Rad23b",
                "Sumo1",
                "Tra2a",
                "Pdcd5",
                "Rab11a",
                "Ankrd1",
                "Anxa5",
                "Eif3f",
                "Eif4g1",
                "Anxa6",
                "Mtch1",
                "Tmbim6",
                "Rhoa",
                "Mrps18c",
                "Rab10",
                "Cacybp",
                "Sf3a3",
                "Ccdc80",
                "Anp32e",
                "Syncrip",
                "Sec61a1",
                "Caprin1",
                "Kdelr2",
                "Ict1",
                "Pmp22",
                "Glud1",
                "March5",
                "Rbmxl2",
                "Cebpz",
                "Ssr2",
                "Hnrnpu",
                "Rbmxl1",
                "Srsf11",
                "Arglu1",
                "Rpl18a",
                "Tomm20",
                "Cbx1",
                "Ssr4",
                "Tomm70a",
                "Txndc9",
                "Ppp1r2",
                "Eprs",
                "Trmt112",
                "Arpc2",
                "Atp5l",
                "Mrpl54",
                "Son",
                "Cct2",
                "Ubb",
                "Sept11",
                "Romo1",
                "Ergic3",
                "Bzw1",
                "Sdc4",
                "Ubc",
                "Lman2",
                "Rock2",
                "Hnrnpa3",
                "Mrps7",
                "Sub1",
                "Morf4l1",
                "2010107E04Rik",
                "Mrpl43",
                "Pafah1b1",
                "Cdca8",
                "Idh2",
                "Polr1d",
                "Snrpd3",
                "Rbms1",
                "P4hb",
                "Sf3b2",
                "Psmd3",
                "Zyx",
                "Map1b",
                "Adipor1",
                "mt-Nd2",
                "Bsg",
                "Rpl14",
                "Camk2d",
                "Cops4",
                "mt-Cytb",
                "Rnps1",
                "Ciapin1",
                "Flna",
                "Pfdn2",
                "H13",
                "Atp5f1",
                "Arid5b",
                "Rps13",
                "Tpt1",
                "Ppib",
                "Cfl1",
                "Gng5",
                "Hspe1",
                "Rps19",
                "Sept2",
                "Exosc8",
                "Cops6",
                "Ptbp1",
                "Mrpl20",
                "Hnrnpdl",
                "Eif3a",
                "Ptx3",
                "Ctbp1",
                "Psmb4",
                "Adh5",
                "Birc5",
                "Commd1",
                "Csrp1",
                "Chtop",
                "Atxn7l3b",
                "Ppa1",
                "Lamtor5",
                "Ndufa13",
                "Gps1",
                "Rrm2",
                "Rpl13",
                "Sh3bgrl",
                "Itm2b",
                "Pdia3",
                "Ndufb5",
                "Atrx",
                "Psma2",
                "Txndc5",
                "Pfdn5",
                "Nsun2",
                "Set",
                "Txnl4a",
                "Poldip2",
                "Eif4e2",
                "Ssbp1",
                "Rps14",
                "Mrto4",
                "Tnfrsf12a",
                "Clns1a",
                "Snrpd2",
                "Krtcap2",
                "Rps27a",
                "Naa10",
                "Bud31",
                "Pdap1",
                "Brk1",
                "Hsp90b1",
                "Vdac2",
                "Anxa1",
                "Rap1b",
                "Ube2a",
                "Rpl13a",
                "Nol7",
                "Eif1a",
                "Cox5a",
                "Commd3",
                "Cnn2",
                "Snrpb",
                "Exosc4",
                "Fkbp4",
                "Pgam1",
                "Rabggtb",
                "Tomm22",
                "Nans",
                "Uchl5",
                "Erp29",
                "Ppig",
                "Phldb2",
                "Palld",
                "Eif5b",
                "Cct4",
                "Llph",
                "Lrrc17",
                "Eif4b",
                "Eif4ebp1",
                "Anp32b",
                "Edf1",
                "Csde1",
                "Marcks",
                "Fam162a",
                "Emp1",
                "Psmc2",
                "Ndufa10",
                "Tomm40",
                "Ywhah",
                "Prpf19",
                "Hdac2",
                "Sugt1",
                "Eif4a3",
                "Fkbp3",
                "Mtch2",
                "Bag1",
                "Tmem147",
                "Rpn2",
                "Bub3",
                "Sin3b",
                "Dctn2",
                "Tmed3",
                "Serpinf1",
                "Tubb4b",
                "Mrps16",
                "Tpr",
                "Hint1",
                "Tbca",
                "Foxp1",
                "Srsf6",
                "Tmpo",
                "Tsn",
                "Txnl1",
                "Mesdc2",
                "Rbm25",
                "Ube2e3",
                "Drg1",
                "Arhgdia",
                "Myl12a",
                "Ptma",
                "Rac1",
                "Rer1",
                "Ralbp1",
                "Cald1",
                "Fau",
                "Yrdc",
                "Acin1",
                "Sri",
                "Pabpc1",
                "Prdx5",
                "Alyref",
                "Tsen34",
                "Hspa14",
                "Fkbp1a",
                "Emc7",
                "Gnas",
                "Arf4",
                "Calu",
                "Yif1b",
                "Hdlbp",
                "Wbp11",
                "Cox4i1",
                "Tmed9",
                "Snf8",
                "Guk1",
                "Ndufs8",
                "Serp1",
                "Tsfm",
                "Capn2",
                "Ptov1",
                "Laptm4a",
                "Aimp1",
                "Serbp1",
                "Cnih1",
                "Gpx4",
                "Chchd3",
                "Gcsh",
                "Phf5a",
                "Rheb",
                "Gtf2a2",
                "Rrp1",
                "Rpl30",
                "Sfr1",
                "Kdelr3",
                "Ncor1",
                "Rbm17",
                "Map2k2",
                "Cnbp",
                "Sdhd",
                "Ogn",
                "Rsrc2"
            ],
            "expression_sum": {
                "class1": 19607.0,
                "class2": 34910.0
            },
            "uniprot_ids": [
                "Q06507",
                "Q8CF69",
                "Q3U2J1",
                "A0A2R8VI82",
                "Q61328",
                "Q794G7",
                "Q3TZI8"
            ],
            "net_id": 0,
            "source_num": 13,
            "target_num": 321
        },
        "Gtf2a2": {
            "id": "Gtf2a2",
            "type": "TF",
            "symbol": "Gtf2a2",
            "source": [
                "Tcf4",
                "Pou5f1",
                "Foxp1",
                "Cebpb",
                "Nfib",
                "Atf5",
                "Atf4",
                "Junb",
                "Fos",
                "Jun",
                "Sox2",
                "Sfpq",
                "Nanog",
                "Jund",
                "Twist2",
                "Hdac2",
                "Smc3",
                "Smarca5"
            ],
            "target": [
                "0610011F06Rik",
                "Ak1",
                "Cox8a",
                "2010111I01Rik"
            ],
            "expression_sum": {
                "class1": 10627.0,
                "class2": 28035.0
            },
            "uniprot_ids": [
                "Q80ZM7",
                "Q05C74",
                "Q3TE88",
                "D3Z7S8",
                "D3Z793",
                "D3Z7C2",
                "D6RIQ7",
                "Q3UT44",
                "Q3TN86"
            ],
            "net_id": 0,
            "source_num": 18,
            "target_num": 2
        },
        "Sox2": {
            "id": "Sox2",
            "type": "TF",
            "symbol": "Sox2",
            "source": [
                "Pou5f1",
                "Trim28",
                "Klf4",
                "Ctcf",
                "Hdac1",
                "Tbx3",
                "Zfp57",
                "Nanog",
                "Smc3"
            ],
            "target": [
                "Rps5",
                "Hsp90aa1",
                "Lsm5",
                "Arf1",
                "Dap",
                "Rpl41",
                "Rnf7",
                "Srsf9",
                "Usmg5",
                "Ndufb4",
                "Syce2",
                "Mcm7",
                "Cd44",
                "Rps17",
                "Rpl6",
                "Srm",
                "Grsf1",
                "Rae1",
                "Zc3h15",
                "Cox8a",
                "Mrps18b",
                "Ruvbl2",
                "Cdt1",
                "Psmb7",
                "Spr",
                "Pnkd",
                "Ppid",
                "Ier2",
                "2810004N23Rik",
                "Clta",
                "Cox17",
                "Dkc1",
                "Marcksl1",
                "Atp5g1",
                "1110008F13Rik",
                "Mrpl4",
                "Ndufa6",
                "Polr2k",
                "Ndufb3",
                "Gar1",
                "Tuba1b",
                "Anapc5",
                "Cks1b",
                "Rps27l",
                "Aurka",
                "Calcoco2",
                "Rpl23a",
                "Psma6",
                "Gm10076",
                "Rpl7l1",
                "Lmna",
                "Snrpa",
                "Ftsj3",
                "St13",
                "Psmc1",
                "Hnrnpa1",
                "Ndufa2",
                "Tpd52l2",
                "Hnrnpf",
                "Knop1",
                "Lmnb1",
                "Mlf2",
                "Rpl9",
                "Ssrp1",
                "Pin1",
                "Atp5j2",
                "Swi5",
                "Eif2s1",
                "Psmb6",
                "Cks2",
                "Shfm1",
                "Hmgb2",
                "Nubp1",
                "Cox6c",
                "Rps15a",
                "Uba2",
                "E2f4",
                "Nup62",
                "Calm3",
                "Arf6",
                "D8Ertd738e",
                "Hn1",
                "Nop56",
                "Snrpe",
                "B2m",
                "Mrpl14",
                "Ubxn1",
                "Cdk1",
                "Eif4h",
                "Phb",
                "Psmd7",
                "Anapc15",
                "Gabarapl2",
                "Stub1",
                "Mrpl2",
                "Taf7",
                "Tra2b",
                "Cdc5l",
                "Ranbp1",
                "Mrpl28",
                "Emc6",
                "Grpel1",
                "Sms",
                "Rbx1",
                "Rps24",
                "Ctcf",
                "Ppm1g",
                "Neat1",
                "Sec61g",
                "Ythdc1",
                "Nanog",
                "Tbcb",
                "Sf3b6",
                "Aurkaip1",
                "Nhp2",
                "Srsf1",
                "Socs3",
                "Lrrc59",
                "Uqcrc1",
                "Tdgf1",
                "Wdr89",
                "Exosc5",
                "Rcc2",
                "Pdcd6",
                "Impa2",
                "Ube2d2a",
                "Atp5h",
                "Bola2",
                "Tmem160",
                "H2afx",
                "Kpnb1",
                "Snrpd1",
                "Mylpf",
                "Rbm8a",
                "Col5a1",
                "Ppia",
                "Hnrnpm",
                "Rpl10",
                "Eif3k",
                "Rpl36al",
                "Lap3",
                "Ncl",
                "Mrps34",
                "Loxl1",
                "Atp5k",
                "2310036O22Rik",
                "Supt4a",
                "Ppp1ca",
                "Srsf5",
                "Rpl35a",
                "Timp2",
                "Syngr1",
                "Chchd1",
                "Wdr74",
                "Ywhaz",
                "Psmc3",
                "Morc1",
                "Rps12",
                "Pomp",
                "Hspa4",
                "Rpl36",
                "Rbbp7",
                "Psmb3",
                "Hnrnpl",
                "Ddx5",
                "Ndufb9",
                "Rbm39",
                "Rps21",
                "Rps27",
                "Arpp19",
                "Ddx21",
                "Naca",
                "Nedd8",
                "Pigyl",
                "Eef1b2",
                "Ube2i",
                "Eif3l",
                "Utp3",
                "Atpif1",
                "Tmem258",
                "Tcp1",
                "Rpl24",
                "Snrnp40",
                "Brix1",
                "Fam136a",
                "Mrpl15",
                "Siva1",
                "Fus",
                "Mybl2",
                "Pfdn1",
                "Slc2a3",
                "Rest",
                "Rps4x",
                "Pmm1",
                "Fabp7",
                "Actl6a",
                "Cct3",
                "Fkbp8",
                "Nip7",
                "Wls",
                "Clpp",
                "Psma7",
                "G3bp1",
                "Gnl3",
                "Trim27",
                "Sssca1",
                "Tpm3",
                "Atp5b",
                "Banf1",
                "Cdc26",
                "Dtymk",
                "Immt",
                "Mrps36",
                "Psma1",
                "Lig1",
                "2410015M20Rik",
                "Gpx1",
                "Smc3",
                "Hdgf",
                "Prnp",
                "Id1",
                "Csnk2b",
                "Pa2g4",
                "Thrap3",
                "Rpl37",
                "Hspb1",
                "Kdm5b",
                "Atf4",
                "Hprt",
                "Ppp1cc",
                "Hells",
                "Tfcp2l1",
                "Nucks1",
                "Pcbp1",
                "Rpl38",
                "Dnttip2",
                "Sae1",
                "Serpine2",
                "Hnrnpk",
                "Mrpl36",
                "Mrps24",
                "Ndufa4",
                "Ewsr1",
                "Ndufv1",
                "Ndufv2",
                "H3f3a",
                "Nme1",
                "Ube2k",
                "Rfc2",
                "Bccip",
                "Ddt",
                "Fbxo22",
                "Lamtor2",
                "Tmsb10",
                "Bax",
                "Rpl3",
                "Gfer",
                "Cebpzos",
                "Nudt21",
                "Trim28",
                "Rpl12",
                "Anxa2",
                "Pycard",
                "Eif3c",
                "Cox7a2",
                "Ndufa7",
                "Dnajc19",
                "Col5a2",
                "Cdv3",
                "Atp6v1f",
                "Crip2",
                "Spc24",
                "Hmgn1",
                "Ctbp2",
                "Ube2l3",
                "Platr3",
                "Asns",
                "Ifitm2",
                "Ppp4c",
                "Eef1d",
                "Etf1",
                "Imp3",
                "Lsm8",
                "Ndufs7",
                "Cenpf",
                "Mcm2",
                "Lsm3",
                "Mrpl13",
                "Gcat",
                "Eif5",
                "Gm16286",
                "Spag7",
                "Tomm5",
                "Txn1",
                "Erh",
                "Ak2",
                "Mrpl17",
                "Samm50",
                "Zmat2",
                "Rrp15",
                "Atp5j",
                "Nhp2l1",
                "Ran",
                "Fscn1",
                "Atp5g3",
                "Srrm1",
                "Gjb3",
                "Apex1",
                "Atp5g2",
                "Chd4",
                "Emc8",
                "Utf1",
                "Srp19",
                "Cct5",
                "Ccnb2",
                "Mrpl35",
                "1110004F10Rik",
                "Mrpl40",
                "Bclaf1",
                "Tceb1",
                "C1qbp",
                "Rpsa",
                "H2afz",
                "Mrps21",
                "Uqcr10",
                "Tardbp",
                "Psip1",
                "Rpl36a",
                "Ppp1r11",
                "Rpl29",
                "Fubp1",
                "Snrpb2",
                "Arpc5l",
                "Rps29",
                "Rpl5",
                "Naa38",
                "Ccdc58",
                "Cenpa",
                "Dph3",
                "S100a6",
                "Ptges3",
                "Ung",
                "Dpy30",
                "Wtap",
                "Odc1",
                "Mrpl18",
                "Umps",
                "Mdk",
                "Rps20",
                "Ufc1",
                "Ybx1",
                "Ifitm1",
                "Tmem14c",
                "Mrpl12",
                "Cdc37",
                "Ndufab1",
                "Psmd1",
                "Plekhf2",
                "Rps6",
                "Snrpf",
                "Eef1a1",
                "Cisd1",
                "Rpl28",
                "U2af1",
                "Pole4",
                "Cenpv",
                "Atp5c1",
                "Ywhaq",
                "Ccnd2",
                "Fbl",
                "Ndufv3",
                "Timm8b",
                "Ubald2",
                "Rpl26",
                "Uqcrb",
                "Tsc22d1",
                "Dnajc8",
                "Rps3",
                "Atox1",
                "Cox7c",
                "Stmn1",
                "Nme4",
                "Adss",
                "Btf3",
                "Smarcc1",
                "Srsf10",
                "Rps7",
                "Igf2bp1",
                "Ube2v2",
                "Uba52",
                "Rpl18",
                "Eno1",
                "Hax1",
                "Rpl19",
                "Trnau1ap",
                "Mt2",
                "Taldo1",
                "Timm13",
                "Mcm3",
                "Rpl22l1",
                "Adprh",
                "Rps3a1",
                "Mrps5",
                "Srsf3",
                "Coprs",
                "Prc1",
                "Smc4",
                "Gadd45gip1",
                "U2surp",
                "Mbd3",
                "Rpl7",
                "Cct7",
                "Znrd1",
                "Dctpp1",
                "Wdr18",
                "Ppan",
                "Hsp90ab1",
                "Nenf",
                "Rps9",
                "Zfp42",
                "Oaz1",
                "Mrpl30",
                "Rpp25",
                "Stra13",
                "Sec61b",
                "Uqcrq",
                "Rwdd1",
                "1810058I24Rik",
                "Snx5",
                "Ankrd11",
                "Cyc1",
                "Got1",
                "Dnaja1",
                "Rps8",
                "Tmem176b",
                "Trap1a",
                "Bzw2",
                "Rpl8",
                "Eif4a1",
                "Sumo2",
                "Pcbp2",
                "Magoh",
                "Ndufc2",
                "Polr1c",
                "Polr2i",
                "Snrnp25",
                "Pou5f1",
                "Nudc",
                "H3f3b",
                "Dnajc2",
                "Gnb2l1",
                "Mrps33",
                "Mrpl21",
                "Ndufa12",
                "Rif1",
                "Nsa2",
                "Pla2g1b",
                "Wdr5",
                "Ddx46",
                "Nudt19",
                "Pebp1",
                "Sgk1",
                "Ube2d3",
                "Aars",
                "Rfc5",
                "Rbbp4",
                "Snrpa1",
                "Gm10073",
                "Dstn",
                "Paip2",
                "Emp3",
                "Tbx3",
                "Srrm2",
                "2700060E02Rik",
                "Rnaseh2b",
                "Mt1",
                "Ywhae",
                "Gm26917",
                "Ubl5",
                "Hmgb1",
                "Anxa3",
                "Mrps15",
                "Ndufc1",
                "Ppil1",
                "Eif3d",
                "Hbegf",
                "Cyb5a",
                "Npm1",
                "Psmc4",
                "Psma4",
                "Rps15",
                "Rpl34",
                "Park7",
                "Dynll1",
                "Gtpbp4",
                "Cenpw",
                "Eif2s2",
                "Hspd1",
                "Stoml2",
                "Rpl31",
                "Mrps14",
                "Srsf7",
                "Sod1",
                "Ppih",
                "Mat2a",
                "Nap1l1",
                "Uqcc3",
                "Csrp2",
                "Dnajb6",
                "Brd2",
                "G3bp2",
                "Efemp2",
                "Impdh2",
                "Ccdc124",
                "Gapdh",
                "Smarca5",
                "Ccnb1",
                "Akap12",
                "Ccnd3",
                "Mrpl52",
                "Pno1",
                "Adrm1",
                "Abcf1",
                "Hnrnpab",
                "Prkcdbp",
                "Lgals1",
                "Rplp0",
                "Psmb5",
                "Hspa9",
                "Fkbp10",
                "Chchd2",
                "Pim3",
                "Nop58",
                "Ube2n",
                "Ndufa11",
                "E130012A19Rik",
                "Pam16",
                "Smn1",
                "Cyb5b",
                "Luc7l3",
                "Polr2c",
                "Cct8",
                "Hmgn2",
                "Dohh",
                "Hnrnpa2b1",
                "Ndufs5",
                "Gsn",
                "Psmb1",
                "Coa3",
                "Akr1b3",
                "Ddx27",
                "Ndufb8",
                "Srsf2",
                "Eif4g2",
                "Jade1",
                "Psmb2",
                "Raly",
                "Denr",
                "Fxr1",
                "Nsmce4a",
                "Rps26",
                "Get4",
                "Eif3g",
                "Trp53",
                "Lhfp",
                "Rbm3",
                "Aspscr1",
                "Psmd11",
                "Vdac1",
                "Rpp21",
                "Cggbp1",
                "Npm3",
                "Anp32a",
                "Zfp706",
                "Polr2e",
                "Lsm2",
                "Anapc13",
                "Sumo1",
                "Tra2a",
                "Nudcd2",
                "Pdcd5",
                "Aprt",
                "Ost4",
                "Supt16",
                "Tomm7",
                "Snrpc",
                "Zfp57",
                "Mrpl51",
                "Mrpl49",
                "Eif4g1",
                "Fcf1",
                "Ssu72",
                "Mrps18c",
                "Atp5a1",
                "Cacybp",
                "Psma5",
                "Cdk4",
                "Rpl22",
                "Rsl1d1",
                "Nolc1",
                "Pycr2",
                "Sf3a3",
                "Syncrip",
                "Caprin1",
                "Pfdn6",
                "Ssr2",
                "Rbmxl2",
                "Glud1",
                "Rpl4",
                "Rps28",
                "Cebpz",
                "Rps25",
                "Ddah2",
                "Rbmxl1",
                "Gspt1",
                "Arglu1",
                "Imp4",
                "Rpl18a",
                "Tomm20",
                "Cbx1",
                "Tomm70a",
                "Rpl21",
                "Map1lc3b",
                "Eprs",
                "Trmt112",
                "Rpl35",
                "Dazap1",
                "Atp5l",
                "Fam25c",
                "Pop5",
                "Son",
                "Ilf2",
                "Cct2",
                "Mak16",
                "Acp1",
                "Prelid1",
                "Ubb",
                "Ahsa1",
                "Zfp106",
                "Prrc1",
                "Parp1",
                "Dhfr",
                "Rbpms",
                "Paics",
                "Hnrnpa3",
                "Prpf40a",
                "Prmt1",
                "Pbdc1",
                "Rps19bp1",
                "Tcea3",
                "Cdca3",
                "Metap2",
                "Sub1",
                "Asna1",
                "Mrpl57",
                "Psmc5",
                "Echdc2",
                "2010107E04Rik",
                "Morf4l1",
                "Cdca8",
                "L1td1",
                "Lsm6",
                "Sf3b1",
                "Tecr",
                "Ube2s",
                "Idh2",
                "Rpl27",
                "Ndufs4",
                "Polr1d",
                "Mybbp1a",
                "Snrpd3",
                "Sf3b2",
                "Psmd3",
                "Suclg1",
                "H2afy",
                "Mtf2",
                "Rpl17",
                "Stip1",
                "mt-Nd2",
                "Hnrnpd",
                "Chchd4",
                "Bcas2",
                "Rpl14",
                "Ash2l",
                "Tma7",
                "Camk2d",
                "Txndc17",
                "Cops4",
                "Rnps1",
                "Ciapin1",
                "Lsm4",
                "Tpd52",
                "Pfdn2",
                "Nop10",
                "Atp5f1",
                "Eif1ad",
                "Arid5b",
                "Gdf3",
                "Rps13",
                "Tpt1",
                "Eif5a",
                "Rps11",
                "Cfl1",
                "Gng5",
                "Rrp1b",
                "Sparc",
                "Uchl3",
                "Hspe1",
                "Rps19",
                "Stmn2",
                "Ube2c",
                "Exosc8",
                "Ptbp1",
                "Mrpl20",
                "Srp9",
                "Hnrnpdl",
                "Nfatc2ip",
                "Eif3a",
                "Dnajc21",
                "Polr2h",
                "Psmb4",
                "Birc5",
                "Snx3",
                "Commd1",
                "Psmd13",
                "Cox7b",
                "Strap",
                "l7Rn6",
                "Rnaseh2a",
                "Dppa5a",
                "Srpk1",
                "Klf4",
                "Rfc1",
                "Cox6b1",
                "Ino80e",
                "Sfrp1",
                "Ppa1",
                "Rcor2",
                "Lamtor5",
                "Atp5e",
                "Cenpm",
                "Prdx1",
                "Ptrf",
                "Rsl24d1",
                "Ssb",
                "Cdc34",
                "Tipin",
                "Atp5o",
                "Mrps12",
                "Terf1",
                "Rpl13",
                "Copz1",
                "Sh3bgrl",
                "Nfu1",
                "Mrpl34",
                "Nasp",
                "Rplp1",
                "Rnf10",
                "Ndufb5",
                "Rpl10a",
                "Atrx",
                "Psma2",
                "Ndufs6",
                "Ndufb7",
                "Nsun2",
                "Set",
                "Thyn1",
                "Txnl4a",
                "Poldip2",
                "Eif4e2",
                "Shmt1",
                "Alkbh1",
                "Sod2",
                "Slc25a39",
                "Ssbp1",
                "Rps14",
                "Tgif1",
                "Sfpq",
                "Rpl32",
                "Parl",
                "Tnfrsf12a",
                "Sephs2",
                "Clns1a",
                "Mdh2",
                "Snrpd2",
                "1810022K09Rik",
                "Chchd10",
                "Uqcr11",
                "Phb2",
                "Hdac1",
                "Rps27a",
                "Naa10",
                "Bud31",
                "Gtf2e2",
                "Pdap1",
                "Mcm5",
                "Gtf2h5",
                "Rpl11",
                "Rpl15",
                "Anxa1",
                "Tcof1",
                "Vdac2",
                "Ube2a",
                "Pfn1",
                "Rpl13a",
                "Snrpg",
                "Tkt",
                "Ddx39b",
                "Snhg6",
                "Eif1a",
                "Cox5a",
                "Minos1",
                "Gsta4",
                "Snrpb",
                "Tmem11",
                "Anapc11",
                "Exosc4",
                "Fkbp4",
                "Pgam1",
                "Spint2",
                "Tomm22",
                "Lyar",
                "Snrnp70",
                "Gpx8",
                "Ppig",
                "Cfdp1",
                "Ppdpf",
                "Gmnn",
                "Rpl27a",
                "Dcun1d5",
                "Eif5b",
                "Cct4",
                "Llph",
                "Emg1",
                "Igfbp7",
                "Lsm12",
                "Eif4ebp1",
                "Anp32b",
                "Edf1",
                "Pkm",
                "H2afv",
                "Nono",
                "Csde1",
                "Prrc2c",
                "Gins4",
                "Fstl1",
                "Tceb2",
                "Rhoc",
                "Mrpl42",
                "Psmg4",
                "Rps2",
                "Eif2b5",
                "Gart",
                "Polr2g",
                "Sumo3",
                "Psmc2",
                "Tomm40",
                "App",
                "Mcm6",
                "Uqcrfs1",
                "Naa50",
                "Sf3b4",
                "Eif3i",
                "Prpf19",
                "Hdac2",
                "Sugt1",
                "Hnrnpc",
                "Smc2",
                "Eif4a3",
                "Fkbp3",
                "Mtch2",
                "Dek",
                "Rangrf",
                "Psmd4",
                "Gins2",
                "Timm17a",
                "Top2a",
                "Rpl7a",
                "Fam104a",
                "Uck2",
                "Papola",
                "Bcat1",
                "Rps16",
                "Tmed3",
                "Mrfap1",
                "Tubb4b",
                "Ccna2",
                "Mrps16",
                "Hsd17b14",
                "Slc25a5",
                "Uqcrh",
                "Tpr",
                "Ddx18",
                "Lefty2",
                "Hint1",
                "Psmd12",
                "Eed",
                "Rrp9",
                "Ccne1",
                "Sf3a2",
                "Srsf6",
                "Ndufa5",
                "Ebna1bp2",
                "Ndufb10",
                "Tsn",
                "Mrpl11",
                "Rpl37a",
                "Rbm25",
                "Eef1g",
                "Cox5b",
                "2700094K13Rik",
                "Cycs",
                "Drg1",
                "Ndufb6",
                "Ptma",
                "Klf2",
                "Acadm",
                "Dut",
                "Ralbp1",
                "Dhx16",
                "Sdhb",
                "Taf10",
                "Fau",
                "Wdr43",
                "Ooep",
                "Klf5",
                "Acin1",
                "Rps10",
                "Pabpc1",
                "Eif6",
                "Alyref",
                "Yrdc",
                "Hspa14",
                "Polr2j",
                "Dad1",
                "Slirp",
                "Sept1",
                "0610007P14Rik",
                "Rbpms2",
                "Uhrf1",
                "Gm7325",
                "Fam60a",
                "Wbp11",
                "Snf8",
                "Cwc15",
                "Guk1",
                "Hspa8",
                "Ndufs8",
                "Psma3",
                "Rpl39",
                "Rplp2",
                "Skp1a",
                "Capn2",
                "Rps18",
                "Mif",
                "Ptov1",
                "Aimp1",
                "Rps27rt",
                "Serbp1",
                "Gpx4",
                "Gcsh",
                "Phf5a",
                "Rps23",
                "Glrx3",
                "Taf1d",
                "Gtf2a2",
                "Alpl",
                "Gltscr2",
                "Mkrn1",
                "U2af2",
                "Rpl23",
                "Rpl30",
                "Sfr1",
                "Polr2f",
                "Myl12b",
                "Kdm1a",
                "Rrs1",
                "Ncor1",
                "1810037I17Rik",
                "Rbm17",
                "Ly6e",
                "Cnbp"
            ],
            "expression_sum": {
                "class1": 55.0,
                "class2": 44636.0
            },
            "uniprot_ids": [
                "P48432",
                "Q60I23",
                "Q3V2B6",
                "Q3UES8"
            ],
            "net_id": 0,
            "source_num": 9,
            "target_num": 598
        },
        "Jund": {
            "id": "Jund",
            "type": "TF",
            "symbol": "Jund",
            "source": [
                "Trim28",
                "Ctcf"
            ],
            "target": [
                "Hsp90aa1",
                "Lsm5",
                "Arf1",
                "Rnf7",
                "Cdc20",
                "Srsf9",
                "Cd9",
                "Mdh1",
                "Ndufb4",
                "Usmg5",
                "Mcm7",
                "Ppp2ca",
                "Ddx24",
                "Rps17",
                "Srm",
                "Grsf1",
                "Rae1",
                "Zc3h15",
                "Cox8a",
                "Mrps18b",
                "Ruvbl2",
                "Psmb7",
                "Spr",
                "Ppid",
                "2810004N23Rik",
                "Clta",
                "Dkc1",
                "Marcksl1",
                "Ppp1r11",
                "Atp5g1",
                "1110008F13Rik",
                "Sec13",
                "Polr2k",
                "Tuba1b",
                "Ndufb3",
                "Gar1",
                "Idh3g",
                "Anapc5",
                "Cks1b",
                "Psma6",
                "Rpl7l1",
                "Snrpa",
                "St13",
                "Ftsj3",
                "Psmc1",
                "Hnrnpa1",
                "Thoc7",
                "Tpd52l2",
                "Hnrnpf",
                "Knop1",
                "Lmnb1",
                "Ddx39",
                "Psmd6",
                "Vbp1",
                "Ssrp1",
                "Pin1",
                "Atp5j2",
                "Rnf187",
                "Sfxn1",
                "Swi5",
                "Eif2s1",
                "Psmb6",
                "Cks2",
                "Hmgb2",
                "Nubp1",
                "Cox6c",
                "Ddx3x",
                "Slbp",
                "Ndufs3",
                "Nedd4",
                "Uba2",
                "E2f4",
                "Calr",
                "Nup62",
                "Tubb5",
                "Calm3",
                "Canx",
                "D8Ertd738e",
                "Nop56",
                "Snrpe",
                "Ubxn1",
                "Cdk1",
                "Eif4h",
                "Phb",
                "2810417H13Rik",
                "Psat1",
                "Psmd7",
                "Anapc15",
                "Gdi2",
                "Gabarapl2",
                "Stub1",
                "Mrpl2",
                "Nup43",
                "Tra2b",
                "Cdc5l",
                "Ranbp1",
                "Mrpl28",
                "Psmc6",
                "Hmgcs1",
                "Emc6",
                "Grpel1",
                "Sms",
                "Zwint",
                "Rbx1",
                "Rps24",
                "Ctcf",
                "Ppm1g",
                "Neat1",
                "Ythdc1",
                "Bola1",
                "Ola1",
                "Sf3b6",
                "Snrnp27",
                "Tbcb",
                "Aurkaip1",
                "Nhp2",
                "Srsf1",
                "Wdr89",
                "Lrrc59",
                "Uqcrc1",
                "Exosc5",
                "Rcc2",
                "Pdcd6",
                "Ube2d2a",
                "Atp5h",
                "H2afx",
                "Kpnb1",
                "Mapre1",
                "C1d",
                "Cisd2",
                "Snrpd1",
                "Rbm8a",
                "Ppia",
                "Hnrnpm",
                "Capza2",
                "Eif3k",
                "Rpl36al",
                "Lap3",
                "Ncl",
                "Atp5k",
                "2310036O22Rik",
                "Fdft1",
                "Mphosph8",
                "Ppp1ca",
                "Srsf5",
                "Supt4a",
                "Hnrnph1",
                "Aamp",
                "Rpl35a",
                "Chchd1",
                "Wdr74",
                "Ywhaz",
                "Gars",
                "Psmc3",
                "Pomp",
                "Hspa4",
                "Luc7l2",
                "Rpl36",
                "Rbbp7",
                "Psmb3",
                "Hnrnpl",
                "Ddx5",
                "Rbm39",
                "Rps21",
                "Arpp19",
                "Ddx21",
                "Nedd8",
                "Pcnp",
                "Ube2i",
                "Eif3l",
                "Utp3",
                "Atpif1",
                "Tcp1",
                "Snrnp40",
                "Brix1",
                "Sarnp",
                "Fam136a",
                "Pttg1",
                "Mrpl15",
                "Siva1",
                "Fus",
                "Ezh2",
                "Ociad1",
                "Pfdn1",
                "Ddost",
                "Rest",
                "Actl6a",
                "Cct3",
                "Nip7",
                "Ruvbl1",
                "Clpp",
                "Psma7",
                "Ndufa8",
                "G3bp1",
                "Pdia4",
                "Sept7",
                "Gnl3",
                "Trim27",
                "Sssca1",
                "Tpm3",
                "Atp5b",
                "Banf1",
                "Cdc26",
                "Dtymk",
                "Immt",
                "Mrps36",
                "Psma1",
                "Lig1",
                "Tsg101",
                "Gpx1",
                "Hdgf",
                "Smc3",
                "Tpi1",
                "Thrap3",
                "Csnk2b",
                "Nars",
                "Pa2g4",
                "Rpl37",
                "Ppp1cc",
                "Atf4",
                "Hist1h2bc",
                "Hprt",
                "Nucks1",
                "Eif3b",
                "Zfp207",
                "Pcbp1",
                "Dnttip2",
                "Glo1",
                "Sae1",
                "Hnrnpk",
                "Mrpl36",
                "Mrps24",
                "Ndufa4",
                "Ewsr1",
                "Cops5",
                "Eif4e",
                "Ndufv1",
                "Ndufv2",
                "H3f3a",
                "Nme1",
                "Ube2k",
                "Rfc2",
                "Bccip",
                "Csnk1a1",
                "Ndufs2",
                "Ddt",
                "Cyp51",
                "Lamtor2",
                "Fam49b",
                "Rpl3",
                "Gfer",
                "Cebpzos",
                "Nudt21",
                "Trim28",
                "Rpl12",
                "Eif3c",
                "Snw1",
                "Cox7a2",
                "Ndufa7",
                "Dnajc19",
                "Cdv3",
                "mt-Nd4",
                "Spc24",
                "Hmgn1",
                "Ctbp2",
                "Btf3l4",
                "Ube2l3",
                "Asns",
                "Ifitm2",
                "Fdps",
                "Ppp4c",
                "Eef1d",
                "Etf1",
                "Imp3",
                "Lsm8",
                "Ndufs7",
                "Lsm3",
                "Mrpl13",
                "Gm16286",
                "Eif5",
                "Tomm5",
                "Txn1",
                "Ube2j2",
                "Vapa",
                "Erh",
                "Ak2",
                "Mrpl17",
                "Samm50",
                "Zmat2",
                "Rrp15",
                "Wdr77",
                "Atp5j",
                "Nhp2l1",
                "Ran",
                "Fscn1",
                "Atp5g3",
                "Srrm1",
                "Ldha",
                "Apex1",
                "Atp5g2",
                "Chd4",
                "Emc8",
                "Srp19",
                "Cct5",
                "Emd",
                "Mrpl35",
                "1110004F10Rik",
                "Mrpl40",
                "Bclaf1",
                "Tceb1",
                "C1qbp",
                "Rpsa",
                "H2afz",
                "Uqcr10",
                "Tardbp",
                "Psip1",
                "Eif3m",
                "Rpl29",
                "Fubp1",
                "Snrpb2",
                "Rps29",
                "Rpl5",
                "Naa38",
                "Ccdc58",
                "Dph3",
                "Prdx6",
                "Ptges3",
                "Dpy30",
                "Wtap",
                "Odc1",
                "Mrpl18",
                "Umps",
                "Rps20",
                "Orc6",
                "Dnaja2",
                "Ybx1",
                "Sys1",
                "Tubb2b",
                "Acat2",
                "Tmem14c",
                "Mrpl12",
                "Cdc37",
                "Ndufab1",
                "Psmd1",
                "Plekhf2",
                "Rps6",
                "Snrpf",
                "mt-Nd5",
                "Mcl1",
                "Cisd1",
                "U2af1",
                "Pole4",
                "Atp5c1",
                "Mrps17",
                "Ywhaq",
                "Fbl",
                "Timm8b",
                "Ubald2",
                "Rpl26",
                "Uqcrb",
                "Dnajc8",
                "Calm2",
                "Rps3",
                "Cox7c",
                "Stmn1",
                "Srsf10",
                "Adss",
                "Btf3",
                "Smarcc1",
                "Tln1",
                "Rps7",
                "Ube2v2",
                "Rpl18",
                "Eno1",
                "Hax1",
                "Eef1e1",
                "mt-Nd3",
                "Taldo1",
                "Timm13",
                "Adprh",
                "Vcp",
                "Srsf3",
                "Mrps5",
                "Smc4",
                "U2surp",
                "Mbd3",
                "Rpl7",
                "Fkbp2",
                "Cct7",
                "Znrd1",
                "Dctpp1",
                "Wdr18",
                "Ppan",
                "Hsp90ab1",
                "Oaz1",
                "Stra13",
                "Sec61b",
                "Polr2m",
                "Rwdd1",
                "Snx5",
                "Tyms",
                "Cyc1",
                "Fnta",
                "Dnaja1",
                "Got1",
                "Rps8",
                "Bzw2",
                "Rpl8",
                "Eif4a1",
                "Sumo2",
                "Pcbp2",
                "Magoh",
                "Ndufc2",
                "Polr1c",
                "Polr2i",
                "Snrnp25",
                "Nudc",
                "Pmvk",
                "H3f3b",
                "Dnajc2",
                "Ccdc34",
                "Elof1",
                "Gnb2l1",
                "Mrpl21",
                "Mrps33",
                "Ndufa12",
                "Pold2",
                "Oxct1",
                "Lrpap1",
                "Ddx46",
                "Cnih4",
                "Nudt19",
                "Pebp1",
                "Sgk1",
                "Dbi",
                "Morf4l2",
                "Ube2d3",
                "Aars",
                "Rfc5",
                "Rbbp4",
                "Marc2",
                "Snrpa1",
                "Paip2",
                "Dstn",
                "Pfdn4",
                "Srrm2",
                "2700060E02Rik",
                "Rnaseh2b",
                "Top1",
                "Ywhae",
                "Hmgb1",
                "Mrps15",
                "Psmc4",
                "Ndufc1",
                "Ppil1",
                "Eif3d",
                "Psmd14",
                "Cyb5a",
                "Npm1",
                "Rps15",
                "Psma4",
                "Scp2",
                "Arl1",
                "Park7",
                "Mlec",
                "Dynll1",
                "Gtpbp4",
                "Cenpw",
                "Eif2s2",
                "Hspd1",
                "Stoml2",
                "Cops7a",
                "Mrpl45",
                "Mrps14",
                "Rpl31",
                "Srsf7",
                "Smdt1",
                "Sod1",
                "Ppih",
                "Mat2a",
                "Nap1l1",
                "Mrps25",
                "Tpx2",
                "Etfb",
                "Dnajb6",
                "Hspa5",
                "Uqcrc2",
                "G3bp2",
                "Impdh2",
                "Ccdc124",
                "Mrps10",
                "Gapdh",
                "Smarca5",
                "Idi1",
                "Ccnb1",
                "Mpc1",
                "Prps1",
                "Ccnd3",
                "Khdrbs1",
                "Pno1",
                "Abcf1",
                "Hnrnpab",
                "Psmb5",
                "Tbrg1",
                "Hspa9",
                "Chchd2",
                "Pim3",
                "Nop58",
                "Ube2n",
                "Ndufa11",
                "Smn1",
                "Cbx5",
                "Cyb5b",
                "Kif5b",
                "Luc7l3",
                "Polr2c",
                "Cct8",
                "Hmgn2",
                "Dohh",
                "Hnrnpa2b1",
                "Ndufs5",
                "Oat",
                "Coa3",
                "Akr1b3",
                "Ddx27",
                "Ndufb8",
                "Psmd2",
                "Tmem97",
                "Srsf2",
                "mt-Nd1",
                "Eif4g2",
                "Dnajc7",
                "Psmb2",
                "Raly",
                "Uqcc2",
                "Pak1ip1",
                "Denr",
                "Fxr1",
                "Nsmce4a",
                "Get4",
                "Eif3g",
                "Nxt1",
                "Trp53",
                "Tceal8",
                "Rbm3",
                "Aspscr1",
                "Psmd11",
                "Cnn3",
                "Btg2",
                "Vdac1",
                "Rpp21",
                "Cggbp1",
                "Npm3",
                "Anp32a",
                "Rad23b",
                "Zfp706",
                "Polr2e",
                "Lsm2",
                "Anapc13",
                "Sumo1",
                "Tra2a",
                "Nudcd2",
                "Pdcd5",
                "Aprt",
                "Ost4",
                "Supt16",
                "Tomm7",
                "Snrpc",
                "Mrpl51",
                "Mrpl49",
                "Eif4g1",
                "Tmbim6",
                "Dynll2",
                "Rhoa",
                "Fcf1",
                "Ssu72",
                "Mrps18c",
                "Atp5a1",
                "Cacybp",
                "Psma5",
                "Ttc3",
                "Cdk4",
                "Rpl22",
                "Rsl1d1",
                "Nolc1",
                "Pycr2",
                "Sf3a3",
                "Anp32e",
                "Syncrip",
                "Caprin1",
                "Pfdn6",
                "Ict1",
                "Rpl4",
                "Glud1",
                "Ssr2",
                "Cebpz",
                "Hnrnpu",
                "Ddah2",
                "Rbmxl1",
                "Gspt1",
                "Arglu1",
                "Imp4",
                "Srsf11",
                "Hmgb3",
                "Cbx1",
                "Tomm20",
                "Tomm70a",
                "Rpl21",
                "Ppp1r2",
                "Eprs",
                "Trmt112",
                "Lman1",
                "Dazap1",
                "Msmo1",
                "Son",
                "Pop5",
                "Ilf2",
                "Cct2",
                "Mak16",
                "Acp1",
                "Ubb",
                "Dtl",
                "Ahsa1",
                "Zfp106",
                "Prrc1",
                "Eif1ax",
                "Bzw1",
                "Parp1",
                "Ak6",
                "Dhfr",
                "Lman2",
                "Paics",
                "Hnrnpa3",
                "Prpf40a",
                "Prmt1",
                "Pbdc1",
                "Rps19bp1",
                "Mrps7",
                "Cdca3",
                "Metap2",
                "Sub1",
                "Asna1",
                "Psmc5",
                "Morf4l1",
                "2010107E04Rik",
                "Mrpl43",
                "Pafah1b1",
                "Cdca8",
                "Lsm6",
                "Tecr",
                "Sf3b1",
                "Ube2s",
                "Idh2",
                "Ndufs4",
                "Polr1d",
                "Mybbp1a",
                "Snrpd3",
                "Sf3b2",
                "Psmd3",
                "Suclg1",
                "H2afy",
                "Mtf2",
                "Rpl17",
                "Stip1",
                "mt-Nd2",
                "Hnrnpd",
                "Bcas2",
                "Rpl14",
                "Ash2l",
                "Tma7",
                "Hat1",
                "Cops4",
                "mt-Cytb",
                "Rnps1",
                "Ciapin1",
                "Lsm4",
                "Pfdn2",
                "Nop10",
                "Mpdu1",
                "Atp5f1",
                "Eif1ad",
                "Rps13",
                "Eif5a",
                "Rps11",
                "Cfl1",
                "Rrp1b",
                "Uchl3",
                "Hspe1",
                "Exosc8",
                "Cops6",
                "Ptbp1",
                "Aldoa",
                "Mrpl20",
                "Hnrnpdl",
                "Srp9",
                "Eif3a",
                "Rtn3",
                "Dnajc21",
                "Polr2h",
                "Psmb4",
                "Adh5",
                "Birc5",
                "Commd1",
                "Psmd13",
                "Cox7b",
                "Snx3",
                "Strap",
                "l7Rn6",
                "Rnaseh2a",
                "Chtop",
                "Srpk1",
                "Rfc1",
                "Cox6b1",
                "Ino80e",
                "Ppa1",
                "Lamtor5",
                "Ssb",
                "Prdx1",
                "Rsl24d1",
                "Cdc34",
                "Gps1",
                "Tipin",
                "Atp5o",
                "Rpa2",
                "Mrps12",
                "Rrm2",
                "Terf1",
                "Copz1",
                "Nfu1",
                "Mrpl34",
                "Nasp",
                "Pdia3",
                "Rnf10",
                "Ndufb5",
                "Atrx",
                "Psma2",
                "Ndufs6",
                "Ndufb7",
                "Nsun2",
                "Set",
                "Thyn1",
                "Txnl4a",
                "Poldip2",
                "Eif4e2",
                "Sod2",
                "Alkbh1",
                "Ssbp1",
                "Nt5dc2",
                "Tgif1",
                "Sfpq",
                "Mrto4",
                "Hist1h1c",
                "Parl",
                "Lipe",
                "Clns1a",
                "Mdh2",
                "Snrpd2",
                "1810022K09Rik",
                "Uqcr11",
                "Phb2",
                "Hdac1",
                "Phgdh",
                "Naa10",
                "Bud31",
                "Gtf2e2",
                "Pdap1",
                "Mcm5",
                "Gtf2h5",
                "Hsp90b1",
                "Rpl11",
                "Tcof1",
                "Vdac2",
                "Ube2a",
                "Nol7",
                "Snrpg",
                "Tkt",
                "Ddx39b",
                "Mrpl24",
                "Eif1a",
                "Cox5a",
                "Snrpb",
                "Tmem11",
                "Anapc11",
                "Exosc4",
                "Fkbp4",
                "Pgam1",
                "Zfp428",
                "Tomm22",
                "Nans",
                "Lyar",
                "Uchl5",
                "Erp29",
                "Snrnp70",
                "Ppig",
                "Cfdp1",
                "Gmnn",
                "Rpl27a",
                "Dcun1d5",
                "Eif5b",
                "Cct4",
                "Llph",
                "Rdx",
                "Emg1",
                "Lsm12",
                "Eif4b",
                "Eif4ebp1",
                "Anp32b",
                "Edf1",
                "Cdkn1a",
                "Pkm",
                "H2afv",
                "Nono",
                "Csde1",
                "Prrc2c",
                "Gins4",
                "Tceb2",
                "Mrpl42",
                "Rps2",
                "Eif2b5",
                "Gart",
                "Polr2g",
                "Sumo3",
                "Psmc2",
                "Ndufa10",
                "Mcm6",
                "Tomm40",
                "Uqcrfs1",
                "Naa50",
                "Ywhah",
                "Sf3b4",
                "Eif3i",
                "Prpf19",
                "Hdac2",
                "Sugt1",
                "Hnrnpc",
                "Smc2",
                "Eif4a3",
                "Fkbp3",
                "Mtch2",
                "Dek",
                "Timm17a",
                "Psmd4",
                "Gins2",
                "Rpn2",
                "Bub3",
                "Top2a",
                "Rpl7a",
                "Papola",
                "Bcat1",
                "Manf",
                "Esf1",
                "Sin3b",
                "Cetn3",
                "Mrfap1",
                "Tubb4b",
                "Ccna2",
                "Mrps16",
                "Slc25a5",
                "Tpr",
                "Ddx18",
                "Psmd12",
                "Hint1",
                "Eed",
                "Ccne1",
                "Srsf6",
                "Ndufa5",
                "Ebna1bp2",
                "Ndufb10",
                "Tmpo",
                "Tsn",
                "Mrpl11",
                "Rpl37a",
                "Txnl1",
                "Mesdc2",
                "Eef1g",
                "Rbm25",
                "Ube2e3",
                "Cox5b",
                "2700094K13Rik",
                "Arhgdia",
                "Cycs",
                "Drg1",
                "Myl12a",
                "Ndufb6",
                "Ptma",
                "Rer1",
                "Acadm",
                "Dut",
                "Ralbp1",
                "Sdhb",
                "Fau",
                "Klf5",
                "Acin1",
                "Rps10",
                "Pabpc1",
                "Eif6",
                "Alyref",
                "Yrdc",
                "Tsen34",
                "Hspa14",
                "Polr2j",
                "Got2",
                "Dad1",
                "Slirp",
                "0610007P14Rik",
                "Wbp11",
                "Cwc15",
                "Guk1",
                "Hspa8",
                "Ndufs8",
                "Psma3",
                "Tsfm",
                "Rplp2",
                "Skp1a",
                "Rps18",
                "Mif",
                "Ptov1",
                "Aimp1",
                "Serbp1",
                "Taf1d",
                "Gpx4",
                "Chchd3",
                "Gcsh",
                "Eif4a2",
                "Phf5a",
                "Rheb",
                "Rps23",
                "Gtf2a2",
                "Gltscr2",
                "Mkrn1",
                "Rrp1",
                "Pdcl3",
                "U2af2",
                "Rpl30",
                "Sfr1",
                "Polr2f",
                "Myl12b",
                "Kdm1a",
                "Rrs1",
                "Ncor1",
                "1810037I17Rik",
                "Rbm17",
                "Ly6e",
                "Bcap31",
                "Cnbp",
                "Sdhd",
                "Rsrc2"
            ],
            "expression_sum": {
                "class1": 9894.0,
                "class2": 25816.0
            },
            "uniprot_ids": [
                "P15066",
                "Q3UXY7",
                "Q52L49"
            ],
            "net_id": 0,
            "source_num": 2,
            "target_num": 522
        },
        "Jun": {
            "id": "Jun",
            "type": "TF",
            "symbol": "Jun",
            "source": [
                "E2f4",
                "Trim28",
                "Rest",
                "Ctcf",
                "Trp53",
                "Hdac1",
                "Hes1",
                "Hdac2",
                "Smc3",
                "Smarca5"
            ],
            "target": [
                "Hsp90aa1",
                "Lsm5",
                "Arf1",
                "Rpl41",
                "Rnf7",
                "Cdc20",
                "Srsf9",
                "Cd9",
                "Mdh1",
                "Ndufb4",
                "Usmg5",
                "Mcm7",
                "Ppp2ca",
                "Ddx24",
                "Rps17",
                "Rpl6",
                "Srm",
                "Grsf1",
                "Rae1",
                "Zc3h15",
                "Cox8a",
                "Mrps18b",
                "Pin4",
                "Ruvbl2",
                "Cdt1",
                "Psmb7",
                "Spr",
                "Txn2",
                "Pnkd",
                "Ppid",
                "Ier2",
                "2810004N23Rik",
                "Clta",
                "Cox17",
                "Dkc1",
                "Marcksl1",
                "Atp5g1",
                "1110008F13Rik",
                "Mrpl4",
                "Polr2k",
                "Ppp1r11",
                "Ndufb3",
                "Gar1",
                "Idh3g",
                "Tuba1b",
                "Anapc5",
                "Cks1b",
                "Aurka",
                "Psma6",
                "Rpl7l1",
                "Snrpa",
                "St13",
                "Ftsj3",
                "Psmc1",
                "Hnrnpa1",
                "Ndufa2",
                "Thoc7",
                "Tpd52l2",
                "Hnrnpf",
                "Knop1",
                "Lmnb1",
                "Ddx39",
                "Mlf2",
                "Psmd6",
                "Vbp1",
                "Ssrp1",
                "Itm2a",
                "Atp5j2",
                "Pin1",
                "Rnf187",
                "Sfxn1",
                "Swi5",
                "Eif2s1",
                "Psmb6",
                "Cks2",
                "Shfm1",
                "Med28",
                "Hmgb2",
                "Nubp1",
                "Cox6c",
                "Ddx3x",
                "Slbp",
                "Ndufs3",
                "Uba2",
                "E2f4",
                "Calr",
                "Nup62",
                "Tubb5",
                "Calm3",
                "D8Ertd738e",
                "Hn1",
                "Nop56",
                "Snrpe",
                "Ubxn1",
                "Cdk1",
                "Eif4h",
                "Lamtor1",
                "2810417H13Rik",
                "Phb",
                "Psat1",
                "Psmd7",
                "Anapc15",
                "Gdi2",
                "Gabarapl2",
                "Stub1",
                "Mrpl2",
                "Nup43",
                "Taf7",
                "Tra2b",
                "Cdc5l",
                "Ranbp1",
                "Mrpl28",
                "Psmc6",
                "Hmgcs1",
                "Emc6",
                "Grpel1",
                "Sms",
                "Zwint",
                "Rbx1",
                "Rps24",
                "Ctcf",
                "Ppm1g",
                "Eif1",
                "Neat1",
                "Sec61g",
                "Pmf1",
                "Ythdc1",
                "Bola1",
                "Ola1",
                "Sf3b6",
                "Snrnp27",
                "Tbcb",
                "Aurkaip1",
                "Nhp2",
                "Srsf1",
                "Wdr89",
                "Lrrc59",
                "Uqcrc1",
                "Exosc5",
                "Rcc2",
                "Pdcd6",
                "Ube2d2a",
                "Atp5h",
                "Bola2",
                "Tmem160",
                "H2afx",
                "Kpnb1",
                "Mapre1",
                "C1d",
                "Snrpd1",
                "Rbm8a",
                "Ppia",
                "Hnrnpm",
                "Capza2",
                "Eif3k",
                "Rpl36al",
                "Lap3",
                "Ncl",
                "Mrps34",
                "Atp5k",
                "2310036O22Rik",
                "Fdft1",
                "Mphosph8",
                "Ppp1ca",
                "Srsf5",
                "Supt4a",
                "Hnrnph1",
                "Aamp",
                "Rpl35a",
                "Chchd1",
                "Wdr74",
                "Ywhaz",
                "Gars",
                "Psmc3",
                "Pomp",
                "Hspa4",
                "Rpl36",
                "Rbbp7",
                "Psmb3",
                "Hnrnpl",
                "Ddx5",
                "Ndufb9",
                "Rbm39",
                "Rps21",
                "Arpp19",
                "Ccdc12",
                "Ddx21",
                "Nedd8",
                "Pcnp",
                "Ube2i",
                "Col3a1",
                "Eif3l",
                "Utp3",
                "Atpif1",
                "Tmem258",
                "Tcp1",
                "Rpl24",
                "Snrnp40",
                "Brix1",
                "Sarnp",
                "Fam136a",
                "Mrpl15",
                "Siva1",
                "Fus",
                "Ezh2",
                "Mybl2",
                "Pfdn1",
                "Rest",
                "Actl6a",
                "Cct3",
                "Nip7",
                "Ruvbl1",
                "Wls",
                "Clpp",
                "Psma7",
                "Ndufa8",
                "G3bp1",
                "Pdia4",
                "Gnl3",
                "Trim27",
                "Sssca1",
                "Tpm3",
                "Atp5b",
                "Banf1",
                "Cdc26",
                "Dtymk",
                "Immt",
                "Mrps36",
                "Psma1",
                "Lig1",
                "Qdpr",
                "Gpx1",
                "Hdgf",
                "Smc3",
                "Tsg101",
                "Thrap3",
                "Id1",
                "Csnk2b",
                "Nars",
                "Pa2g4",
                "Rpl37",
                "Gsr",
                "Hspb1",
                "Ppp1cc",
                "Hprt",
                "Hells",
                "Nucks1",
                "Eif3b",
                "Zfp207",
                "Pcbp1",
                "Rpl38",
                "Dnttip2",
                "Glo1",
                "Sae1",
                "Serpine2",
                "Hnrnpk",
                "Mrpl36",
                "Mrps24",
                "Ndufa4",
                "Ewsr1",
                "Cops5",
                "Eif4e",
                "Ndufv1",
                "Ndufv2",
                "H3f3a",
                "Nme1",
                "Ube2k",
                "Rfc2",
                "Bccip",
                "Csnk1a1",
                "Ddt",
                "Fbxo22",
                "Cyp51",
                "Lamtor2",
                "Fam49b",
                "Tmsb10",
                "Rpl3",
                "Gfer",
                "Cebpzos",
                "Nudt21",
                "Trim28",
                "Rpl12",
                "Eif3c",
                "Snw1",
                "Cox7a2",
                "Ndufa7",
                "Dnajc19",
                "Cdv3",
                "Spc24",
                "Hmgn1",
                "Ctbp2",
                "Btf3l4",
                "Ube2l3",
                "Asns",
                "Ifitm2",
                "Fdps",
                "Ppp4c",
                "Eef1d",
                "Etf1",
                "Imp3",
                "Lsm8",
                "Ndufs7",
                "Ddah1",
                "Mcm2",
                "Lsm3",
                "Mrpl13",
                "Gcat",
                "Eif5",
                "Gm16286",
                "Tomm5",
                "Txn1",
                "Ube2j2",
                "Erh",
                "Ak2",
                "Mrpl17",
                "Samm50",
                "Zmat2",
                "Rrp15",
                "Wdr77",
                "Atp5j",
                "Nhp2l1",
                "Ran",
                "Fscn1",
                "Atp5g3",
                "Srrm1",
                "Ldha",
                "Apex1",
                "Atp5g2",
                "Chd4",
                "Emc8",
                "Srp19",
                "Cct5",
                "Emd",
                "Ccnb2",
                "Mrpl35",
                "1110004F10Rik",
                "Mrpl40",
                "Bclaf1",
                "Tceb1",
                "C1qbp",
                "Rpsa",
                "H2afz",
                "Uqcr10",
                "Tardbp",
                "Psip1",
                "Rpl36a",
                "Eif3m",
                "Rpl29",
                "Iscu",
                "Fubp1",
                "Snrpb2",
                "Arpc5l",
                "Rps29",
                "Rpl5",
                "Naa38",
                "Ccdc58",
                "Cenpa",
                "Dph3",
                "Ung",
                "Prdx6",
                "Ptges3",
                "Dpy30",
                "Fblim1",
                "Wtap",
                "Odc1",
                "Mrpl18",
                "Umps",
                "Mdk",
                "Rps20",
                "Orc6",
                "Dnaja2",
                "Ufc1",
                "Ybx1",
                "Sys1",
                "Tubb2b",
                "Acat2",
                "Lbh",
                "Tmem14c",
                "Mrpl12",
                "Cdc37",
                "Ndufab1",
                "Psmd1",
                "Plekhf2",
                "Rps6",
                "Snrpf",
                "Mcl1",
                "Rtn4",
                "Cisd1",
                "Rpl28",
                "U2af1",
                "Pole4",
                "Slc29a1",
                "Atp5c1",
                "Mrps17",
                "Xist",
                "Gpx3",
                "Ywhaq",
                "Fbl",
                "Pdha1",
                "Timm8b",
                "Ubald2",
                "Rpl26",
                "Uqcrb",
                "Dnajc8",
                "Calm2",
                "Rps3",
                "Atox1",
                "Cox7c",
                "Stmn1",
                "Srsf10",
                "Adss",
                "Btf3",
                "Smarcc1",
                "Tmem126a",
                "Rps7",
                "Igf2bp1",
                "Ube2v2",
                "Rpl18",
                "Eno1",
                "Hax1",
                "Eef1e1",
                "Rpl19",
                "Taldo1",
                "Timm13",
                "Trnau1ap",
                "Mcm3",
                "Adprh",
                "Vcp",
                "Srsf3",
                "Mrps5",
                "Coprs",
                "Smc4",
                "U2surp",
                "Gadd45gip1",
                "Mbd3",
                "Rpl7",
                "Cct7",
                "Znrd1",
                "Dctpp1",
                "Wdr18",
                "Hes1",
                "Ppan",
                "Hsp90ab1",
                "Oaz1",
                "Stra13",
                "Uqcrq",
                "Sec61b",
                "Polr2m",
                "Rwdd1",
                "1810058I24Rik",
                "Snx5",
                "Tyms",
                "Cyc1",
                "Fnta",
                "Dnaja1",
                "Got1",
                "Rps8",
                "Bzw2",
                "Pgp",
                "Eif4a1",
                "Rpl8",
                "Sumo2",
                "Pcbp2",
                "Magoh",
                "Ndufc2",
                "Polr1c",
                "Polr2i",
                "Snrnp25",
                "Nudc",
                "Pmvk",
                "H3f3b",
                "Dnajc2",
                "Ccdc34",
                "Elof1",
                "Gnb2l1",
                "Mrpl21",
                "Mrps33",
                "Ndufa12",
                "Pold2",
                "Rif1",
                "Lrpap1",
                "Wdr5",
                "Ddx46",
                "Cnih4",
                "Nudt19",
                "Pebp1",
                "Sgk1",
                "Dbi",
                "Morf4l2",
                "Ube2d3",
                "Aars",
                "Rfc5",
                "Rbbp4",
                "Snrpa1",
                "Paip2",
                "Dstn",
                "Pfdn4",
                "Srrm2",
                "2700060E02Rik",
                "Rnaseh2b",
                "Top1",
                "Ywhae",
                "Hmgb1",
                "Mrps15",
                "Psmc4",
                "Ndufc1",
                "Ppil1",
                "Eif3d",
                "Psmd14",
                "Cyb5a",
                "Npm1",
                "Rps15",
                "Psma4",
                "Sap30",
                "Arl1",
                "Ap2s1",
                "Rpl34",
                "Park7",
                "Mlec",
                "Rab7",
                "Dynll1",
                "Gtpbp4",
                "Cenpw",
                "Eif2s2",
                "Cpe",
                "Hspd1",
                "Stoml2",
                "Cops7a",
                "Mrpl45",
                "Mrps14",
                "Rpl31",
                "Srsf7",
                "Smdt1",
                "Sod1",
                "Ppih",
                "Mat2a",
                "Nap1l1",
                "Mrps25",
                "Tpx2",
                "Csrp2",
                "Dnajb6",
                "Etfb",
                "Hspa5",
                "Brd2",
                "Uqcrc2",
                "Actr3",
                "G3bp2",
                "Impdh2",
                "Ccdc124",
                "Mrps10",
                "Smarca5",
                "Idi1",
                "Ccnb1",
                "Prps1",
                "Akap12",
                "Ccnd3",
                "Khdrbs1",
                "Mrpl52",
                "Eln",
                "Pno1",
                "Ecm1",
                "Adrm1",
                "Abcf1",
                "Hnrnpab",
                "Rplp0",
                "Psmb5",
                "Tbrg1",
                "Hspa9",
                "Chchd2",
                "Pim3",
                "Nop58",
                "Ube2n",
                "Ndufa11",
                "E130012A19Rik",
                "Smn1",
                "Cbx5",
                "Cyb5b",
                "Kif5b",
                "Luc7l3",
                "Polr2c",
                "Cct8",
                "Hmgn2",
                "Dohh",
                "Hnrnpa2b1",
                "Ndufs5",
                "Coa3",
                "Akr1b3",
                "Ddx27",
                "Ndufb8",
                "Psmd2",
                "Tmem97",
                "Srsf2",
                "Prdx4",
                "Eif4g2",
                "Dnajc7",
                "Psmb2",
                "Raly",
                "Uqcc2",
                "Pak1ip1",
                "Denr",
                "Fxr1",
                "Nsmce4a",
                "Rps26",
                "Get4",
                "Eif3g",
                "Nxt1",
                "Trp53",
                "Tceal8",
                "Rbm3",
                "Psmd11",
                "Cnn3",
                "Btg2",
                "Vdac1",
                "Rpp21",
                "Lage3",
                "Npm3",
                "Anp32a",
                "Rad23b",
                "Zfp706",
                "Polr2e",
                "Lsm2",
                "Anapc13",
                "Nudt4",
                "Sumo1",
                "Tra2a",
                "Nudcd2",
                "Pdcd5",
                "Aprt",
                "Ost4",
                "Supt16",
                "Tomm7",
                "Snrpc",
                "Mrpl51",
                "Mrpl49",
                "Eif4g1",
                "Tmbim6",
                "Dynll2",
                "Rhoa",
                "Fcf1",
                "Ssu72",
                "Mrps18c",
                "Atp5a1",
                "Cacybp",
                "Psma5",
                "Cdk4",
                "Cnpy2",
                "Rpl22",
                "Rsl1d1",
                "Nolc1",
                "Pycr2",
                "Sf3a3",
                "Anp32e",
                "Syncrip",
                "Caprin1",
                "Pfdn6",
                "Ict1",
                "Rpl4",
                "Glud1",
                "Rps28",
                "Ssr2",
                "Cebpz",
                "Fermt2",
                "Hnrnpu",
                "Rps25",
                "Ddah2",
                "Rbmxl1",
                "Gspt1",
                "Arglu1",
                "Imp4",
                "Srsf11",
                "Hmgb3",
                "Cbx1",
                "Tomm20",
                "Tomm70a",
                "Vapa",
                "Rpl21",
                "Ppp1r2",
                "Map1lc3b",
                "Trmt112",
                "Dazap1",
                "Son",
                "Pop5",
                "Ywhab",
                "Ilf2",
                "Cct2",
                "Mak16",
                "Acp1",
                "Prelid1",
                "Ubb",
                "Dtl",
                "Ahsa1",
                "Prrc1",
                "Eif1ax",
                "Bzw1",
                "Parp1",
                "Ak6",
                "Dhfr",
                "Lman2",
                "Paics",
                "Hnrnpa3",
                "Prpf40a",
                "Prmt1",
                "Pbdc1",
                "Rps19bp1",
                "Mrps7",
                "Cdca3",
                "Metap2",
                "Sub1",
                "Asna1",
                "Mrpl57",
                "Psmc5",
                "Echdc2",
                "2010107E04Rik",
                "Morf4l1",
                "Mrpl43",
                "Pafah1b1",
                "Cdca8",
                "Lsm6",
                "Tecr",
                "Sf3b1",
                "Ube2s",
                "Idh2",
                "Ndufs4",
                "Polr1d",
                "Mybbp1a",
                "Snrpd3",
                "Sf3b2",
                "Psmd3",
                "Suclg1",
                "H2afy",
                "Fam32a",
                "Mtf2",
                "Rpl17",
                "Stip1",
                "mt-Nd2",
                "Hnrnpd",
                "Chchd4",
                "Bcas2",
                "Rpl14",
                "Ash2l",
                "Tma7",
                "Hat1",
                "Txndc17",
                "Cops4",
                "Rnps1",
                "Ciapin1",
                "Lsm4",
                "Pfdn2",
                "Nop10",
                "Mpdu1",
                "Atp5f1",
                "Eif1ad",
                "Rps13",
                "Eif5a",
                "Rps11",
                "Cfl1",
                "Gng5",
                "Rrp1b",
                "Uchl3",
                "Hspe1",
                "Exosc8",
                "Cops6",
                "Ptbp1",
                "Aldoa",
                "Mrpl20",
                "Hnrnpdl",
                "Srp9",
                "Eif3a",
                "Rtn3",
                "Dnajc21",
                "Polr2h",
                "Psmb4",
                "Adh5",
                "Birc5",
                "Commd1",
                "Psmd13",
                "Cox7b",
                "Snx3",
                "Strap",
                "l7Rn6",
                "Rnaseh2a",
                "Chtop",
                "Srpk1",
                "Sec13",
                "Rfc1",
                "Cox6b1",
                "Ino80e",
                "Ppa1",
                "Lamtor5",
                "Atp5e",
                "Cenpm",
                "Prdx1",
                "Rsl24d1",
                "Ssb",
                "Cdc34",
                "Gps1",
                "Tipin",
                "Atp5o",
                "Rpa2",
                "Mrps12",
                "Rrm2",
                "Rpl13",
                "Terf1",
                "Copz1",
                "Nfu1",
                "Mrpl34",
                "Nasp",
                "Rplp1",
                "Ndufb5",
                "Atrx",
                "Psma2",
                "Ndufs6",
                "Ndufb7",
                "Nsun2",
                "Set",
                "Thyn1",
                "Txnl4a",
                "Poldip2",
                "Eif4e2",
                "Shmt1",
                "Alkbh1",
                "Sod2",
                "Ssbp1",
                "Nt5dc2",
                "Rps14",
                "Tgif1",
                "Sfpq",
                "Mrto4",
                "Rpl32",
                "Parl",
                "Sephs2",
                "Clns1a",
                "Mdh2",
                "Snrpd2",
                "1810022K09Rik",
                "Uqcr11",
                "Phb2",
                "Hdac1",
                "Phgdh",
                "Naa10",
                "Bud31",
                "Gtf2e2",
                "Pdap1",
                "Mcm5",
                "Gtf2h5",
                "Hsp90b1",
                "Rpl11",
                "Rpl15",
                "Tcof1",
                "Vdac2",
                "Ube2a",
                "Pfn1",
                "Nol7",
                "Snrpg",
                "Tkt",
                "Ddx39b",
                "Cuta",
                "Eif1a",
                "Cox5a",
                "Minos1",
                "Gsta4",
                "Snrpb",
                "Tmem11",
                "Anapc11",
                "Exosc4",
                "Fkbp4",
                "Pgam1",
                "Zfp428",
                "Tomm22",
                "Nans",
                "Lyar",
                "Ubap2l",
                "Erp29",
                "Snrnp70",
                "Uchl5",
                "Ppig",
                "Cfdp1",
                "Ppdpf",
                "Gmnn",
                "Rpl27a",
                "Dcun1d5",
                "Eif5b",
                "Cct4",
                "Llph",
                "Rdx",
                "Emg1",
                "Lsm12",
                "Eif4ebp1",
                "Anp32b",
                "Edf1",
                "Cdkn1a",
                "Pkm",
                "H2afv",
                "Nono",
                "Prrc2c",
                "Gins4",
                "Tceb2",
                "Mrpl42",
                "Psmg4",
                "Rps2",
                "Eif2b5",
                "Gart",
                "Polr2g",
                "Sumo3",
                "Psmc2",
                "Ndufa10",
                "Mcm6",
                "Tomm40",
                "Uqcrfs1",
                "Naa50",
                "Ywhah",
                "Sf3b4",
                "Prpf19",
                "Hdac2",
                "Sugt1",
                "Hnrnpc",
                "Smc2",
                "Eif4a3",
                "Fkbp3",
                "Mtch2",
                "Dek",
                "Timm17a",
                "Psmd4",
                "Gins2",
                "Bub3",
                "Top2a",
                "Rpl7a",
                "Fam104a",
                "Uck2",
                "Papola",
                "Bcat1",
                "Manf",
                "Rps16",
                "Esf1",
                "Sin3b",
                "Cetn3",
                "Mrfap1",
                "Galk1",
                "Ccna2",
                "Mrps16",
                "Tubb4b",
                "Slc25a5",
                "Uqcrh",
                "Tpr",
                "Ddx18",
                "Psmd12",
                "Hint1",
                "Rrp9",
                "Eed",
                "Sf3a2",
                "Ccne1",
                "Srsf6",
                "Ndufa5",
                "Ebna1bp2",
                "Ndufb10",
                "Tmpo",
                "Tsn",
                "Mrpl11",
                "Rpl37a",
                "Txnl1",
                "Mesdc2",
                "Eef1g",
                "Rbm25",
                "Ube2e3",
                "Cox5b",
                "2700094K13Rik",
                "Arhgdia",
                "Cycs",
                "Drg1",
                "Ndufb6",
                "Ptma",
                "Klf2",
                "Acadm",
                "Dut",
                "Sdhb",
                "Wdr43",
                "Ybx3",
                "Klf5",
                "Acin1",
                "Rps10",
                "Pabpc1",
                "Eif6",
                "Alyref",
                "Yrdc",
                "Tsen34",
                "Hspa14",
                "Fkbp1a",
                "Polr2j",
                "Dad1",
                "Slirp",
                "0610007P14Rik",
                "Rbpms2",
                "Uhrf1",
                "Yif1b",
                "Fam60a",
                "Ckb",
                "Wbp11",
                "Cwc15",
                "Guk1",
                "2810428I15Rik",
                "Hspa8",
                "Ndufs8",
                "Psma3",
                "Tsfm",
                "Pim1",
                "Postn",
                "Rplp2",
                "Rps18",
                "Mif",
                "Ptov1",
                "Skp1a",
                "Aimp1",
                "Serbp1",
                "Taf1d",
                "Gpx4",
                "Chchd3",
                "Gcsh",
                "Phf5a",
                "Rheb",
                "Rps23",
                "Gtf2a2",
                "Gltscr2",
                "Mkrn1",
                "Rrp1",
                "Pdcl3",
                "U2af2",
                "Rpl23",
                "Rpl30",
                "Sfr1",
                "Polr2f",
                "Kdm1a",
                "Rrs1",
                "Ncor1",
                "1810037I17Rik",
                "Rbm17",
                "Cnbp",
                "Sdhd",
                "Rsrc2"
            ],
            "expression_sum": {
                "class1": 6282.0,
                "class2": 4604.0
            },
            "uniprot_ids": [
                "P05627",
                "Q52L79",
                "Q3US19",
                "Q3V255",
                "Q6TDG5"
            ],
            "net_id": 0,
            "source_num": 10,
            "target_num": 562
        },
        "Hes1": {
            "id": "Hes1",
            "type": "TF",
            "symbol": "Hes1",
            "source": [
                "E2f4",
                "Trim28",
                "Rest",
                "Atf4",
                "Ctcf",
                "Tgif1",
                "Trp53",
                "Hdac1",
                "Jun",
                "Klf5",
                "Ybx1",
                "Hdac2",
                "Smc3"
            ],
            "target": [
                "Hsp90aa1",
                "Arf1",
                "Rnf7",
                "Srsf9",
                "Usmg5",
                "Ppp2ca",
                "Rps17",
                "Srm",
                "Grsf1",
                "Rae1",
                "Zc3h15",
                "2810004N23Rik",
                "Marcksl1",
                "1110008F13Rik",
                "Gar1",
                "Anapc5",
                "Psma6",
                "Thoc7",
                "Hnrnpf",
                "Knop1",
                "Lmnb1",
                "Mlf2",
                "Rnf187",
                "Cks2",
                "Nedd4",
                "Calm3",
                "Arf6",
                "Hn1",
                "Nop56",
                "Ubxn1",
                "Psat1",
                "Eif4h",
                "Ranbp1",
                "Sms",
                "Ctcf",
                "Ppm1g",
                "Ythdc1",
                "Bola1",
                "Ola1",
                "Srsf1",
                "Socs3",
                "Wdr89",
                "Exosc5",
                "Rcc2",
                "Kpnb1",
                "Mapre1",
                "C1d",
                "Cisd2",
                "Ppia",
                "Capza2",
                "Lap3",
                "Ncl",
                "2310036O22Rik",
                "Supt4a",
                "Srsf5",
                "Chchd1",
                "Ywhaz",
                "Gars",
                "Hspa4",
                "Rbbp7",
                "Rps21",
                "Hnrnpl",
                "Ddx5",
                "Arpp19",
                "Ube2i",
                "Utp3",
                "Mrpl15",
                "Siva1",
                "Fus",
                "Ezh2",
                "Rest",
                "Psma7",
                "G3bp1",
                "Gnl3",
                "Banf1",
                "Qdpr",
                "Actb",
                "Immt",
                "Tsg101",
                "Hdgf",
                "Pa2g4",
                "Ppp1cc",
                "Hist1h2bc",
                "Hprt",
                "Pcbp1",
                "Sae1",
                "Hnrnpk",
                "Ndufa4",
                "Ewsr1",
                "Ndufv2",
                "H3f3a",
                "Ube2k",
                "Cyp51",
                "Fam49b",
                "Cisd3",
                "Rpl3",
                "Gfer",
                "Cebpzos",
                "Cdv3",
                "Hmgn1",
                "Ctbp2",
                "Asns",
                "Eef1d",
                "Ndufs7",
                "Lsm3",
                "Gm16286",
                "Eif5",
                "Vapa",
                "Txn1",
                "Mrpl17",
                "Ran",
                "Fscn1",
                "Ldha",
                "Apex1",
                "Emc8",
                "Cct5",
                "Mrpl40",
                "Bclaf1",
                "C1qbp",
                "Rpsa",
                "H2afz",
                "Psip1",
                "Rpl29",
                "Fubp1",
                "Ccdc58",
                "Prdx6",
                "Ptges3",
                "Dpy30",
                "Wtap",
                "Odc1",
                "Rps20",
                "Dnaja2",
                "Plekhf2",
                "Rps6",
                "Cisd1",
                "U2af1",
                "Pole4",
                "Ywhaq",
                "Fbl",
                "Ubald2",
                "Calm2",
                "Cox7c",
                "Stmn1",
                "Srsf10",
                "Adss",
                "Smarcc1",
                "Ube2v2",
                "Eno1",
                "Trnau1ap",
                "Adprh",
                "Srsf3",
                "Coprs",
                "Cmpk1",
                "Rpl7",
                "Cct7",
                "Ppan",
                "Hsp90ab1",
                "Rwdd1",
                "Snx5",
                "Rps8",
                "Dnaja1",
                "Pgp",
                "Eif4a1",
                "Sumo2",
                "Pcbp2",
                "Dnajc2",
                "Rif1",
                "Pold2",
                "Wdr5",
                "Nudt19",
                "Sgk1",
                "Morf4l2",
                "Ube2d3",
                "Rbbp4",
                "Marc2",
                "Snrpa1",
                "Srrm2",
                "Top1",
                "Hmgb1",
                "Mrps15",
                "Psma4",
                "Park7",
                "Dynll1",
                "Hspd1",
                "Cpe",
                "Cops7a",
                "Mrpl45",
                "Smdt1",
                "Sod1",
                "Mat2a",
                "Nap1l1",
                "Csrp2",
                "Dnajb6",
                "Brd2",
                "Actr3",
                "G3bp2",
                "Smarca5",
                "Ccnb1",
                "Prdx2",
                "Ccnd3",
                "Khdrbs1",
                "Adrm1",
                "Abcf1",
                "Hspa9",
                "Pim3",
                "Ube2n",
                "Hmgn2",
                "Hnrnpa2b1",
                "Tmem97",
                "Srsf2",
                "Eif4g2",
                "Raly",
                "Denr",
                "Fxr1",
                "Nsmce4a",
                "Get4",
                "Vdac1",
                "Cggbp1",
                "Zfp706",
                "Anp32a",
                "Rad23b",
                "Lsm2",
                "Anapc13",
                "Nudt4",
                "Nudcd2",
                "Eif4g1",
                "Rhoa",
                "Cacybp",
                "Psma5",
                "Rpl22",
                "Nolc1",
                "Anp32e",
                "Syncrip",
                "Caprin1",
                "Glud1",
                "Hnrnpu",
                "Rbmxl1",
                "Gspt1",
                "Arglu1",
                "Imp4",
                "Srsf11",
                "Hmgb3",
                "Tomm70a",
                "Rpl21",
                "Ppp1r2",
                "Map1lc3b",
                "Eprs",
                "Dazap1",
                "Pop5",
                "Ywhab",
                "Ubb",
                "Eif1ax",
                "Parp1",
                "Ak6",
                "Hnrnpa3",
                "Prpf40a",
                "Mrps7",
                "Mrpl43",
                "Ube2s",
                "Sf3b1",
                "Idh2",
                "H2afy",
                "Hnrnpd",
                "Chchd4",
                "Bcas2",
                "Ciapin1",
                "Eif1ad",
                "Rps13",
                "Tpt1",
                "Gng5",
                "Rrp1b",
                "Hspe1",
                "Exosc8",
                "Ptbp1",
                "Aldoa",
                "Srp9",
                "Hnrnpdl",
                "Rtn3",
                "Dnajc21",
                "Polr2h",
                "Snx3",
                "Birc5",
                "Rnaseh2a",
                "Chtop",
                "Srpk1",
                "Rfc1",
                "Ppa1",
                "Lamtor5",
                "Avpi1",
                "Ssb",
                "Cdc34",
                "Gps1",
                "Jun",
                "Atrx",
                "Nsun2",
                "Set",
                "Txnl4a",
                "Eif4e2",
                "Ssbp1",
                "Tgif1",
                "Hist1h1c",
                "Clns1a",
                "Gtf2e2",
                "Rpl15",
                "Gtf2h5",
                "Hsp90b1",
                "Vdac2",
                "Pfn1",
                "Tkt",
                "Ddx39b",
                "Cuta",
                "Cox5a",
                "Anapc11",
                "Fkbp4",
                "Pgam1",
                "Nans",
                "Lyar",
                "Uchl5",
                "Snrnp70",
                "Cct4",
                "Rdx",
                "Eif4b",
                "Eif4ebp1",
                "Anp32b",
                "Edf1",
                "Psmg4",
                "Sumo3",
                "Psmc2",
                "Ndufa10",
                "Mcm6",
                "Naa50",
                "Ywhah",
                "Prpf19",
                "Hdac2",
                "Fkbp3",
                "Mtch2",
                "Dek",
                "Rpn2",
                "Rpl7a",
                "Papola",
                "Manf",
                "Cetn3",
                "Galk1",
                "Ccna2",
                "Slc25a5",
                "Tpr",
                "Hint1",
                "Eed",
                "Sf3a2",
                "Ccne1",
                "Srsf6",
                "Tmpo",
                "Tsn",
                "Txnl1",
                "Mesdc2",
                "Rbm25",
                "Drg1",
                "Cycs",
                "Arhgdia",
                "Ptma",
                "Rac1",
                "Dut",
                "Ralbp1",
                "Ybx3",
                "Klf5",
                "Yrdc",
                "Pabpc1",
                "Alyref",
                "0610007P14Rik",
                "Yif1b",
                "Guk1",
                "Tsfm",
                "Rplp2",
                "Skp1a",
                "Ptov1",
                "Gpx4",
                "Rheb",
                "Mkrn1",
                "Sfr1",
                "Ncor1"
            ],
            "expression_sum": {
                "class1": 10169.0,
                "class2": 9450.0
            },
            "uniprot_ids": [
                "P35428",
                "Q3V1C5",
                "Q3UZZ2"
            ],
            "net_id": 0,
            "source_num": 13,
            "target_num": 236
        },
        "Nanog": {
            "id": "Nanog",
            "type": "TF",
            "symbol": "Nanog",
            "source": [
                "Pou5f1",
                "Rest",
                "Ctcf",
                "Sox2",
                "Klf5",
                "Tbx3"
            ],
            "target": [
                "Rps5",
                "Hsp90aa1",
                "Lsm5",
                "Arf1",
                "Dap",
                "Rpl41",
                "Rnf7",
                "Cdc20",
                "Usmg5",
                "Cd9",
                "Mdh1",
                "Ndufb4",
                "Syce2",
                "Mcm7",
                "Rps17",
                "Rpl6",
                "Srm",
                "Grsf1",
                "Rae1",
                "Cox8a",
                "Mrps18b",
                "Ruvbl2",
                "Cdt1",
                "Psmb7",
                "Ppid",
                "2810004N23Rik",
                "Clta",
                "Cox17",
                "Dkc1",
                "Igfbp2",
                "Atp5g1",
                "1110008F13Rik",
                "Mrpl4",
                "Polr2k",
                "Ppp1r11",
                "Tuba1b",
                "Gar1",
                "Idh3g",
                "Anapc5",
                "Cks1b",
                "Rps27l",
                "Aurka",
                "Calcoco2",
                "Rpl23a",
                "Psma6",
                "Rpl7l1",
                "Lmna",
                "Snrpa",
                "Ftsj3",
                "St13",
                "Folr1",
                "Psmc1",
                "Hnrnpa1",
                "Tpd52l2",
                "Hnrnpf",
                "Knop1",
                "Lmnb1",
                "Ddx39",
                "Mlf2",
                "Rpl9",
                "Vbp1",
                "Ssrp1",
                "Pin1",
                "Atp5j2",
                "Swi5",
                "Eif2s1",
                "Psmb6",
                "Cks2",
                "Shfm1",
                "Hmgb2",
                "Nubp1",
                "Cox6c",
                "Uba2",
                "E2f4",
                "Nup62",
                "Tubb5",
                "Calm3",
                "Arf6",
                "D8Ertd738e",
                "Hn1",
                "Nop56",
                "Snrpe",
                "B2m",
                "Ubxn1",
                "Cdk1",
                "Eif4h",
                "Phb",
                "Psmd7",
                "Gdi2",
                "Gabarapl2",
                "Prrx1",
                "Mrpl2",
                "Nup43",
                "Taf7",
                "Tra2b",
                "Cdc5l",
                "Ranbp1",
                "Mrpl28",
                "Psmc6",
                "Emc6",
                "Grpel1",
                "Sms",
                "Zwint",
                "Rbx1",
                "Rps24",
                "Ctcf",
                "Ppm1g",
                "Eif1",
                "Neat1",
                "Sec61g",
                "Pmf1",
                "Ythdc1",
                "Snrnp27",
                "Sf3b6",
                "Tbcb",
                "Aurkaip1",
                "Nhp2",
                "Srsf1",
                "Lrrc59",
                "Uqcrc1",
                "Tdgf1",
                "Exosc5",
                "Rcc2",
                "Pdcd6",
                "Impa2",
                "Ube2j2",
                "Atp5h",
                "Bola2",
                "H2afx",
                "Snrpd1",
                "Mylpf",
                "Rbm8a",
                "Col5a1",
                "Ppia",
                "Bgn",
                "Hnrnpm",
                "Rpl10",
                "Eif3k",
                "Rpl36al",
                "Lap3",
                "Ncl",
                "Mrps34",
                "Atp5k",
                "2310036O22Rik",
                "Mphosph8",
                "Supt4a",
                "Ppp1ca",
                "Srsf5",
                "Aamp",
                "Crxos",
                "Rpl35a",
                "Chchd1",
                "Timp2",
                "Wdr74",
                "Ywhaz",
                "Psmc3",
                "Morc1",
                "Rps12",
                "Pomp",
                "Hspa4",
                "Rpl36",
                "Rbbp7",
                "Psmb3",
                "Hnrnpl",
                "Ddx5",
                "Ndufb9",
                "Rbm39",
                "Rps21",
                "Rps27",
                "Arpp19",
                "Ccdc12",
                "Ddx21",
                "Naca",
                "Nedd8",
                "Eef1b2",
                "Ube2i",
                "Eif3l",
                "Utp3",
                "Atpif1",
                "Serpinh1",
                "Tcp1",
                "Rpl24",
                "Snrnp40",
                "Brix1",
                "Sarnp",
                "Fam136a",
                "Mrpl15",
                "Siva1",
                "Fus",
                "Ezh2",
                "Mybl2",
                "Pfdn1",
                "Slc2a3",
                "Rest",
                "Rps4x",
                "Pmm1",
                "Actl6a",
                "Cct3",
                "Nip7",
                "Ruvbl1",
                "Wls",
                "Clpp",
                "Psma7",
                "G3bp1",
                "Gnl3",
                "Trim27",
                "Sssca1",
                "Tpm3",
                "Atp5b",
                "Banf1",
                "Cdc26",
                "Dtymk",
                "Immt",
                "Mrps36",
                "Psma1",
                "Lig1",
                "Qdpr",
                "Gpx1",
                "Hdgf",
                "Prnp",
                "Smc3",
                "Csnk2b",
                "Pa2g4",
                "Rpl37",
                "Gsr",
                "Hspb1",
                "Kdm5b",
                "Ppp1cc",
                "Hist1h2bc",
                "Hprt",
                "Hells",
                "Tfcp2l1",
                "Nucks1",
                "Eif3b",
                "Col1a2",
                "Pcbp1",
                "Rpl38",
                "Dnttip2",
                "Glo1",
                "Sae1",
                "Serpine2",
                "Hnrnpk",
                "Mrpl36",
                "Mrps24",
                "Ezr",
                "Ndufa4",
                "Ewsr1",
                "Cops5",
                "Eif4e",
                "Ndufv1",
                "Ndufv2",
                "H3f3a",
                "Nme1",
                "Ube2k",
                "Rfc2",
                "Bccip",
                "Ddt",
                "Lamtor2",
                "Fam49b",
                "Bax",
                "Tmsb10",
                "Rpl3",
                "Cebpzos",
                "Nudt21",
                "Trim28",
                "Rpl12",
                "Anxa2",
                "Pycard",
                "Snw1",
                "Cox7a2",
                "Ndufa7",
                "Dnajc19",
                "Cdv3",
                "Spc24",
                "Hmgn1",
                "Ctbp2",
                "Btf3l4",
                "Ube2l3",
                "Asns",
                "Ifitm2",
                "Ppp4c",
                "Eef1d",
                "Etf1",
                "Imp3",
                "Lsm8",
                "Ndufs7",
                "Cenpf",
                "Mcm2",
                "Lsm3",
                "Mrpl13",
                "Gcat",
                "Eif5",
                "Gm16286",
                "Spag7",
                "Tomm5",
                "Txn1",
                "Erh",
                "Ak2",
                "Mrpl17",
                "Samm50",
                "Zmat2",
                "Rrp15",
                "Wdr77",
                "Atp5j",
                "Nhp2l1",
                "Ran",
                "Fscn1",
                "Atp5g3",
                "Srrm1",
                "Gjb3",
                "Ldha",
                "Apex1",
                "Atp5g2",
                "Chd4",
                "Emc8",
                "Utf1",
                "Srp19",
                "Cct5",
                "Ccnb2",
                "Mrpl35",
                "1110004F10Rik",
                "Mrpl40",
                "Bclaf1",
                "Tceb1",
                "Pdgfa",
                "C1qbp",
                "Rpsa",
                "H2afz",
                "Uqcr10",
                "Psip1",
                "Rpl36a",
                "Eif3m",
                "Rpl29",
                "Fubp1",
                "Snrpb2",
                "Rps29",
                "Rpl5",
                "Ccdc58",
                "Cenpa",
                "Dph3",
                "S100a6",
                "Prdx6",
                "Ptges3",
                "Dpy30",
                "Ung",
                "Fblim1",
                "Wtap",
                "Odc1",
                "Mrpl18",
                "Umps",
                "Rps20",
                "Orc6",
                "Ybx1",
                "Ifitm1",
                "Tubb2b",
                "Tmem14c",
                "Mrpl12",
                "Cdc37",
                "Ndufab1",
                "Psmd1",
                "Plekhf2",
                "Rps6",
                "Snrpf",
                "Cisd1",
                "Rpl28",
                "U2af1",
                "Pole4",
                "Cenpv",
                "Atp5c1",
                "Mrps17",
                "Ccnd2",
                "Slc29a1",
                "Ywhaq",
                "Fbl",
                "Pdha1",
                "Rps26-ps1",
                "Timm8b",
                "Ubald2",
                "Rpl26",
                "Uqcrb",
                "Tsc22d1",
                "Dnajc8",
                "Calm2",
                "Rps3",
                "Atox1",
                "Cox7c",
                "Stmn1",
                "Nme4",
                "Adss",
                "Btf3",
                "Smarcc1",
                "Srsf10",
                "Rps7",
                "Igf2bp1",
                "Ube2v2",
                "Rpl18",
                "Eno1",
                "Hax1",
                "Eef1e1",
                "Rpl19",
                "Mt2",
                "Taldo1",
                "Timm13",
                "Trnau1ap",
                "Mcm3",
                "Fbln5",
                "Adprh",
                "Rps3a1",
                "Mrps5",
                "Srsf3",
                "Coprs",
                "Prc1",
                "Smc4",
                "Gadd45gip1",
                "U2surp",
                "Mbd3",
                "Rpl7",
                "Cct7",
                "Znrd1",
                "Dctpp1",
                "Wdr18",
                "Ppan",
                "Hsp90ab1",
                "Nenf",
                "Upp1",
                "Zfp42",
                "Oaz1",
                "Stra13",
                "Rpp25",
                "Sec61b",
                "Polr2m",
                "1810058I24Rik",
                "Snx5",
                "Tyms",
                "Cyc1",
                "Fnta",
                "Dnaja1",
                "Got1",
                "Rps8",
                "Tmem176b",
                "Trap1a",
                "Bzw2",
                "Pgp",
                "Eif4a1",
                "Rpl8",
                "Sumo2",
                "Pcbp2",
                "Magoh",
                "Ndufc2",
                "Polr1c",
                "Polr2i",
                "Snrnp25",
                "Nudc",
                "Pou5f1",
                "Fhl2",
                "H3f3b",
                "Dnajc2",
                "Col1a1",
                "Gnb2l1",
                "Mrpl21",
                "Mrps33",
                "Ndufa12",
                "Pold2",
                "Nsa2",
                "Rif1",
                "Wdr5",
                "Ddx46",
                "Nudt19",
                "Pebp1",
                "Sgk1",
                "Morf4l2",
                "Ube2d3",
                "Aars",
                "Rfc5",
                "Rbbp4",
                "Marc2",
                "Snrpa1",
                "Gm10073",
                "Dstn",
                "Paip2",
                "Emp3",
                "Tbx3",
                "Pfdn4",
                "Srrm2",
                "2700060E02Rik",
                "Rnaseh2b",
                "Top1",
                "Ywhae",
                "Gm26917",
                "Ubl5",
                "Hmgb1",
                "Mrps15",
                "Psmc4",
                "Ndufc1",
                "Ppil1",
                "Eif3d",
                "Psmd14",
                "Rps15",
                "Npm1",
                "Psma4",
                "Sap30",
                "Rpl34",
                "Park7",
                "Dynll1",
                "Gtpbp4",
                "Cenpw",
                "Eif2s2",
                "Cpe",
                "Hspd1",
                "Lsm7",
                "Stoml2",
                "Cops7a",
                "Mrpl45",
                "Mrps14",
                "Rpl31",
                "Srsf7",
                "Sod1",
                "Ppih",
                "Mat2a",
                "Nap1l1",
                "Uqcc3",
                "Tpx2",
                "Csrp2",
                "Dnajb6",
                "G3bp2",
                "Efemp2",
                "Impdh2",
                "Ccdc124",
                "Gapdh",
                "Smarca5",
                "Ccnb1",
                "Ngfrap1",
                "Bex1",
                "Prps1",
                "Ccnd3",
                "Mrpl52",
                "Pno1",
                "Adrm1",
                "Abcf1",
                "Hnrnpab",
                "Prkcdbp",
                "Lgals1",
                "Rplp0",
                "Psmb5",
                "Tbrg1",
                "Hspa9",
                "Fkbp10",
                "Chchd2",
                "Nop58",
                "Ube2n",
                "Ndufa11",
                "E130012A19Rik",
                "Smn1",
                "Cbx5",
                "Cyb5b",
                "Kif5b",
                "Luc7l3",
                "Polr2c",
                "Cct8",
                "Hmgn2",
                "Dohh",
                "Hnrnpa2b1",
                "Ndufs5",
                "Gsn",
                "Psmb1",
                "Coa3",
                "Akr1b3",
                "Ddx27",
                "Ndufb8",
                "Tmem97",
                "Srsf2",
                "Jam2",
                "Jade1",
                "Dnajc7",
                "Psmb2",
                "Raly",
                "Pak1ip1",
                "Denr",
                "Fxr1",
                "Nsmce4a",
                "Rps26",
                "Get4",
                "Eif3g",
                "Nxt1",
                "Trp53",
                "Lhfp",
                "Rbm3",
                "Aspscr1",
                "Psmd11",
                "Cnn3",
                "Vdac1",
                "Lage3",
                "Npm3",
                "Anp32a",
                "Rad23b",
                "Zfp706",
                "Polr2e",
                "Lsm2",
                "Nudt4",
                "Sumo1",
                "Nudcd2",
                "Pdcd5",
                "Aprt",
                "Ost4",
                "Supt16",
                "Tomm7",
                "Snrpc",
                "Zfp57",
                "Mrpl51",
                "Mrpl49",
                "Rhoa",
                "Fcf1",
                "Ssu72",
                "Mrps18c",
                "Atp5a1",
                "Cacybp",
                "Psma5",
                "Cdk4",
                "Rpl22",
                "Rsl1d1",
                "Nolc1",
                "Pycr2",
                "Sf3a3",
                "Anp32e",
                "Pfdn6",
                "Ssr2",
                "Ict1",
                "Rbmxl2",
                "Rpl4",
                "Rps28",
                "Cebpz",
                "Hnrnpu",
                "Rps25",
                "Rbmxl1",
                "Gspt1",
                "Arglu1",
                "Imp4",
                "Srsf11",
                "Tomm20",
                "Cbx1",
                "Tomm70a",
                "Rpl21",
                "Map1lc3b",
                "Trmt112",
                "Rpl35",
                "Dazap1",
                "Fam25c",
                "Pop5",
                "Ilf2",
                "Cct2",
                "Mak16",
                "Acp1",
                "Prelid1",
                "Ubb",
                "Ahsa1",
                "Eif1ax",
                "Bzw1",
                "Parp1",
                "Rbpms",
                "Paics",
                "Hnrnpa3",
                "Prpf40a",
                "Prmt1",
                "Cdca3",
                "Metap2",
                "Sub1",
                "Asna1",
                "Psmc5",
                "Echdc2",
                "2010107E04Rik",
                "Morf4l1",
                "Mrpl43",
                "Cdca8",
                "L1td1",
                "Lsm6",
                "Sf3b1",
                "Tecr",
                "Ube2s",
                "Idh2",
                "Rpl27",
                "Ndufs4",
                "Polr1d",
                "Mybbp1a",
                "Snrpd3",
                "Sf3b2",
                "Psmd3",
                "Suclg1",
                "H2afy",
                "Atp1b1",
                "Mtf2",
                "Rpl17",
                "Stip1",
                "Sox2",
                "Hnrnpd",
                "Chchd4",
                "Bcas2",
                "Rpl14",
                "Ash2l",
                "Tma7",
                "Hat1",
                "Cops4",
                "Rnps1",
                "Ciapin1",
                "Lsm4",
                "Tpd52",
                "Pfdn2",
                "Nop10",
                "Mpdu1",
                "Atp5f1",
                "Eif1ad",
                "Rps13",
                "Gdf3",
                "Tpt1",
                "Eif5a",
                "Rps11",
                "Cfl1",
                "Gng5",
                "Rrp1b",
                "Sparc",
                "Uchl3",
                "Hspe1",
                "Stmn2",
                "Ube2c",
                "Exosc8",
                "Cops6",
                "Ptbp1",
                "Aldoa",
                "Mrpl20",
                "Hnrnpdl",
                "Nfatc2ip",
                "Srp9",
                "Eif3a",
                "Dnajc21",
                "Polr2h",
                "Psmb4",
                "Adh5",
                "Birc5",
                "Psmd13",
                "Snx3",
                "Strap",
                "Rnaseh2a",
                "Chtop",
                "Dppa5a",
                "Srpk1",
                "Rfc1",
                "Cox6b1",
                "Ino80e",
                "Sfrp1",
                "Ppa1",
                "Rcor2",
                "Cenpm",
                "Prdx1",
                "Rsl24d1",
                "Ssb",
                "Cdc34",
                "Tipin",
                "Cmtm7",
                "Atp5o",
                "Rpa2",
                "Mrps12",
                "Rrm2",
                "Rpl13",
                "Terf1",
                "Sh3bgrl",
                "Mrpl34",
                "Nasp",
                "Rplp1",
                "Ndufb5",
                "Atrx",
                "Psma2",
                "Ndufs6",
                "Nsun2",
                "Set",
                "Thyn1",
                "Txnl4a",
                "Eif4e2",
                "Shmt1",
                "Sod2",
                "Ssbp1",
                "Slc25a39",
                "Rps14",
                "Tgif1",
                "Sfpq",
                "Mrto4",
                "Rpl32",
                "Parl",
                "Sephs2",
                "Clns1a",
                "Mdh2",
                "Snrpd2",
                "1810022K09Rik",
                "Uqcr11",
                "Phb2",
                "Hdac1",
                "Rps27a",
                "Naa10",
                "Bud31",
                "Rpl15",
                "Pdap1",
                "Mcm5",
                "Gtf2h5",
                "Arl6ip1",
                "Rpl11",
                "Anxa1",
                "Tcof1",
                "Vdac2",
                "Ube2a",
                "Pfn1",
                "Rpl13a",
                "Nol7",
                "Snrpg",
                "Tkt",
                "Ddx39b",
                "Mrpl24",
                "Eif1a",
                "Cox5a",
                "Gsta4",
                "Snrpb",
                "Tmem11",
                "Anapc11",
                "Exosc4",
                "Fkbp4",
                "Pgam1",
                "Zfp428",
                "Spint2",
                "Tomm22",
                "Lyar",
                "Uchl5",
                "Snrnp70",
                "Gpx8",
                "Ppig",
                "Cfdp1",
                "Gmnn",
                "Rpl27a",
                "Eif5b",
                "Cct4",
                "Llph",
                "Emg1",
                "Igfbp7",
                "Lsm12",
                "Eif4ebp1",
                "Anp32b",
                "Edf1",
                "Pkm",
                "H2afv",
                "Nono",
                "Prrc2c",
                "Gins4",
                "Fstl1",
                "Tceb2",
                "Rhoc",
                "Mrpl42",
                "Rps2",
                "Eif2b5",
                "Gart",
                "Polr2g",
                "Sumo3",
                "Psmc2",
                "Tomm40",
                "App",
                "Mcm6",
                "Uqcrfs1",
                "Naa50",
                "Ywhah",
                "Sf3b4",
                "Prpf19",
                "Hdac2",
                "Sugt1",
                "Hnrnpc",
                "Smc2",
                "Eif4a3",
                "Fkbp3",
                "Mtch2",
                "Dek",
                "Rangrf",
                "Psmd4",
                "Gins2",
                "Timm17a",
                "Bub3",
                "Top2a",
                "Rpl7a",
                "Uck2",
                "Papola",
                "Bcat1",
                "Rps16",
                "Esf1",
                "Tmed3",
                "Plekhj1",
                "Mrfap1",
                "Tubb4b",
                "Ccna2",
                "Mrps16",
                "Hsd17b14",
                "Slc25a5",
                "Uqcrh",
                "Tpr",
                "Ddx18",
                "Psmd12",
                "Hint1",
                "Rrp9",
                "Eed",
                "Sf3a2",
                "Ccne1",
                "Srsf6",
                "Ebna1bp2",
                "Ndufb10",
                "Tmpo",
                "Tsn",
                "Mrpl11",
                "Rpl37a",
                "Fbxo15",
                "Eef1g",
                "Rbm25",
                "Cox5b",
                "2700094K13Rik",
                "Cycs",
                "Drg1",
                "Ndufb6",
                "Ptma",
                "Klf2",
                "Acadm",
                "Dut",
                "Ralbp1",
                "Dhx16",
                "Sdhb",
                "Taf10",
                "Fau",
                "Wdr43",
                "Ooep",
                "Klf5",
                "Acin1",
                "Rps10",
                "Pabpc1",
                "Eif6",
                "Alyref",
                "Yrdc",
                "Hspa14",
                "Polr2j",
                "Slirp",
                "Rbpms2",
                "Uhrf1",
                "Gm7325",
                "Fam60a",
                "Ckb",
                "Wbp11",
                "Cwc15",
                "Guk1",
                "Hspa8",
                "Psma3",
                "Rpl39",
                "Tsfm",
                "Pim1",
                "Rplp2",
                "Capn2",
                "Rps18",
                "Mif",
                "Ptov1",
                "Skp1a",
                "Serbp1",
                "Taf1d",
                "Gpx4",
                "Gcsh",
                "Phf5a",
                "Rheb",
                "Glrx3",
                "Rps23",
                "Gtf2a2",
                "Alpl",
                "Gltscr2",
                "Mkrn1",
                "U2af2",
                "Rpl23",
                "Rpl30",
                "Sfr1",
                "Polr2f",
                "Myl12b",
                "Kdm1a",
                "Rrs1",
                "1810037I17Rik",
                "Rbm17",
                "Cnbp",
                "Sdhd"
            ],
            "expression_sum": {
                "class1": 30.0,
                "class2": 43776.0
            },
            "uniprot_ids": [
                "Q80Z64",
                "A2RS90",
                "B7ZN41"
            ],
            "net_id": 0,
            "source_num": 6,
            "target_num": 598
        },
        "Fos": {
            "id": "Fos",
            "type": "TF",
            "symbol": "Fos",
            "source": [
                "Trim28",
                "Rest",
                "Trp53",
                "Hdac1",
                "Hdac2",
                "Smc3",
                "Smarca5"
            ],
            "target": [
                "Hsp90aa1",
                "Lsm5",
                "Arf1",
                "Rnf7",
                "Ndufb4",
                "Ddx24",
                "Rps17",
                "Srm",
                "Zc3h15",
                "Mrps18b",
                "Psmb7",
                "Ppid",
                "Ier2",
                "Sec13",
                "Clta",
                "Marcksl1",
                "Mrpl4",
                "Atp5g1",
                "1110008F13Rik",
                "Tuba1b",
                "Polr2k",
                "Gar1",
                "Anapc5",
                "Cks1b",
                "Psma6",
                "Rpl7l1",
                "Snrpa",
                "St13",
                "Psmc1",
                "Hnrnpa1",
                "Thoc7",
                "Hnrnpf",
                "Knop1",
                "Lmnb1",
                "Psmd6",
                "Vbp1",
                "Ssrp1",
                "Pin1",
                "Atp5j2",
                "Rnf187",
                "Sfxn1",
                "Eif2s1",
                "Psmb6",
                "Hmgb2",
                "Nubp1",
                "Cox6c",
                "Slbp",
                "Ndufs3",
                "Nedd4",
                "Uba2",
                "Calr",
                "Nup62",
                "Rab18",
                "Calm3",
                "Nop56",
                "Snrpe",
                "Psat1",
                "Eif4h",
                "Psmd7",
                "2810417H13Rik",
                "Gdi2",
                "Gabarapl2",
                "Stub1",
                "Mrpl2",
                "Tra2b",
                "Cdc5l",
                "Ranbp1",
                "Mrpl28",
                "Psmc6",
                "Grpel1",
                "Sms",
                "Rbx1",
                "Ppm1g",
                "Sec61g",
                "Tbcb",
                "Sf3b6",
                "Aurkaip1",
                "Srsf1",
                "Socs3",
                "Lrrc59",
                "Uqcrc1",
                "Rcc2",
                "Pdcd6",
                "Ube2d2a",
                "Atp5h",
                "C1d",
                "Cisd2",
                "Snrpd1",
                "Rbm8a",
                "Ppia",
                "Hnrnpm",
                "Capza2",
                "Rpl36al",
                "Lap3",
                "Ncl",
                "Supt4a",
                "Ppp1ca",
                "Srsf5",
                "Hnrnph1",
                "Aamp",
                "Rpl35a",
                "Pomp",
                "Hspa4",
                "Rbbp7",
                "Psmb3",
                "Hnrnpl",
                "Ddx5",
                "Ndufb9",
                "Rbm39",
                "Rps21",
                "Arpp19",
                "Ddx21",
                "Ube2i",
                "Eif3l",
                "Utp3",
                "Atpif1",
                "Tcp1",
                "Rpl24",
                "Snrnp40",
                "Sarnp",
                "Fam136a",
                "Mrpl15",
                "Siva1",
                "Fus",
                "Ezh2",
                "Pfdn1",
                "Ddost",
                "Rest",
                "Actl6a",
                "Cct3",
                "Fkbp8",
                "Nip7",
                "Clpp",
                "Psma7",
                "G3bp1",
                "Pdia4",
                "Sssca1",
                "Tpm3",
                "Atp5b",
                "Banf1",
                "Cdc26",
                "Immt",
                "Dtymk",
                "Mrps36",
                "Emc10",
                "Lig1",
                "Psma1",
                "Hdgf",
                "Tpi1",
                "Smc3",
                "Thrap3",
                "Nars",
                "Pa2g4",
                "Ppp1cc",
                "Hprt",
                "Eif3b",
                "Zfp207",
                "Pcbp1",
                "Glo1",
                "Sae1",
                "Hnrnpk",
                "Mrpl36",
                "Mrps24",
                "Ndufa4",
                "Ewsr1",
                "Cops5",
                "Eif4e",
                "Ndufv1",
                "Ndufv2",
                "H3f3a",
                "Mettl9",
                "Nme1",
                "Ube2k",
                "Bccip",
                "Csnk1a1",
                "Lamtor2",
                "Fam49b",
                "Cebpzos",
                "Nudt21",
                "Trim28",
                "Rpl12",
                "Eif3c",
                "Snw1",
                "Cox7a2",
                "Ndufa7",
                "Dnajc19",
                "Cdv3",
                "Spc24",
                "Hmgn1",
                "Ube2l3",
                "Btf3l4",
                "Asns",
                "Ifitm2",
                "Fdps",
                "Ppp4c",
                "Eef1d",
                "Etf1",
                "Lsm8",
                "Ndufs7",
                "Mcm2",
                "Lsm3",
                "Mrpl13",
                "Tomm5",
                "Eif5",
                "Txn1",
                "Zmat2",
                "Erh",
                "Ak2",
                "Mrpl17",
                "Atp5j",
                "Nhp2l1",
                "Ran",
                "Fscn1",
                "Atp5g3",
                "Srrm1",
                "Apex1",
                "Atp5g2",
                "Chd4",
                "Emc8",
                "Cct5",
                "Mrpl35",
                "1110004F10Rik",
                "Mrpl40",
                "Bclaf1",
                "Tceb1",
                "C1qbp",
                "Rpsa",
                "H2afz",
                "Uqcr10",
                "Psip1",
                "Rpl29",
                "Fubp1",
                "Snrpb2",
                "Rps29",
                "Rpl5",
                "Dph3",
                "Prdx6",
                "Ptges3",
                "Dpy30",
                "Odc1",
                "Mrpl18",
                "Umps",
                "Rps20",
                "Orc6",
                "Dnaja2",
                "Ybx1",
                "Acat2",
                "Tmem14c",
                "Mrpl12",
                "Cdc37",
                "Ndufab1",
                "Snrpf",
                "Cisd1",
                "U2af1",
                "Pole4",
                "Atp5c1",
                "Mrps17",
                "Ywhaq",
                "Fbl",
                "Pdha1",
                "Uqcrb",
                "Dnajc8",
                "Cox7c",
                "Stmn1",
                "Srsf10",
                "Adss",
                "Btf3",
                "Smarcc1",
                "Igf2bp1",
                "Ube2v2",
                "Rpl18",
                "Hax1",
                "mt-Nd3",
                "Timm13",
                "Mcm3",
                "Rpl22l1",
                "Vcp",
                "Srsf3",
                "Mrps5",
                "U2surp",
                "Mbd3",
                "Rpl7",
                "Cct7",
                "Znrd1",
                "Dctpp1",
                "Hsp90ab1",
                "Oaz1",
                "Stra13",
                "Sec61b",
                "Polr2m",
                "Rwdd1",
                "Snx5",
                "Tyms",
                "Cyc1",
                "Rps8",
                "Dnaja1",
                "Bzw2",
                "Pgp",
                "Eif4a1",
                "Rpl8",
                "Sumo2",
                "Pcbp2",
                "Magoh",
                "Ndufc2",
                "Polr1c",
                "Polr2i",
                "Nudc",
                "H3f3b",
                "Dnajc2",
                "Gnb2l1",
                "Mrpl21",
                "Ndufa12",
                "Pold2",
                "Lrpap1",
                "Wdr5",
                "Pebp1",
                "Morf4l2",
                "Ube2d3",
                "Aars",
                "Rbbp4",
                "Marc2",
                "Snrpa1",
```
