## Supplement 1 (MEF vs. ESC) for "AGEAS: Automated Machine Learning based Genetic Regulatory Element Extraction System": meta_GRN.js

```
{
    "genes": {
        "Atf4": {
            "id": "Atf4",
            "type": "TF",
            "symbol": "Atf4",
            "source": [
                "Tcf4",
                "Pou5f1",
                "Foxp1",
                "Cebpb",
                "Eno1",
                "Ctcf",
                "Junb",
                "Trp53",
                "Sox2",
                "Ybx1",
                "Jund",
                "Twist2",
                "Hdac2"
            ],
            "target": [
                "Hsp90aa1",
                "Dap",
                "Rnf7",
                "Cdc20",
                "Sar1a",
                "Usmg5",
                "Ppp2ca",
                "Ddx24",
                "Zc3h15",
                "Myh9",
                "Txn2",
                "Ppid",
                "2810004N23Rik",
                "Clta",
                "Igfbp2",
                "Sec13",
                "1110008F13Rik",
                "Idh3g",
                "Anapc5",
                "Lmna",
                "Ndufa2",
                "Hnrnpf",
                "Knop1",
                "Ddx39",
                "Mlf2",
                "Pck2",
                "Rnf187",
                "Med28",
                "Nubp1",
                "Rps15a",
                "Ndufs3",
                "Nedd4",
                "Uba2",
                "Rexo2",
                "Calr",
                "Nup62",
                "Pdlim7",
                "Calm3",
                "Mbnl2",
                "Arf6",
                "Canx",
                "Nop56",
                "Slc25a3",
                "Psat1",
                "Psmd7",
                "Lamtor1",
                "Tra2b",
                "Psmc6",
                "Zwint",
                "Ppm1g",
                "Eif1",
                "Myh10",
                "Sec61g",
                "Ola1",
                "Snrnp27",
                "Aurkaip1",
                "Gemin7",
                "Wdr89",
                "Lrrc59",
                "Uqcrc1",
                "Akt1s1",
                "Bnip3l",
                "H2afx",
                "Kpnb1",
                "Mapre1",
                "Cisd2",
                "Snrpd1",
                "Nktr",
                "Capza2",
                "Klf6",
                "Rpl36al",
                "Ncl",
                "Loxl1",
                "Srsf5",
                "Hnrnph1",
                "Chchd1",
                "Ywhaz",
                "Gars",
                "Psmc3",
                "Sec62",
                "Pomp",
                "Hspa4",
                "Luc7l2",
                "Psmb3",
                "Ddx5",
                "Rbm39",
                "Nedd8",
                "Pcnp",
                "Eef1b2",
                "Ube2i",
                "Arpc1b",
                "Cyr61",
                "Utp3",
                "Serpinh1",
                "Tcp1",
                "Brix1",
                "Gnb2",
                "Mrpl15",
                "Fus",
                "Dpysl3",
                "Ddost",
                "Hcfc1r1",
                "Sptbn1",
                "Gng12",
                "Fkbp8",
                "Nip7",
                "Ndufa8",
                "Pdia4",
                "Srp14",
                "Sept7",
                "Fam195b",
                "Trim27",
                "Mrps36",
                "Banf1",
                "Immt",
                "Actg1",
                "Dtymk",
                "Psma1",
                "Tsg101",
                "Hdgf",
                "Nars",
                "Pa2g4",
                "Hmga2",
                "F2r",
                "Pcbp1",
                "Rpl38",
                "Dnttip2",
                "Sae1",
                "Hnrnpk",
                "Ewsr1",
                "Eif4e",
                "Ndufv1",
                "Ndufv2",
                "Fabp5",
                "H3f3a",
                "Nme1",
                "Chmp4b",
                "Tcf4",
                "Ube2k",
                "Ssr1",
                "Ndufs2",
                "Cyp51",
                "Pgls",
                "Tmsb10",
                "Nudt21",
                "Rpl12",
                "Ift20",
                "Anxa2",
                "Eif3c",
                "Snw1",
                "Vmp1",
                "Col5a2",
                "Cdv3",
                "Napa",
                "Btf3l4",
                "Asns",
                "Eef1d",
                "Etf1",
                "Sh3bgrl3",
                "Ndufs7",
                "Ube2b",
                "Lsm3",
                "Mrpl13",
                "Gm16286",
                "Eif5",
                "Tmed2",
                "Ube2j2",
                "Vapa",
                "Mrpl17",
                "Wdr77",
                "Atp5j",
                "Ran",
                "Fscn1",
                "Srrm1",
                "Tpm1",
                "Ldha",
                "Sdcbp",
                "Apex1",
                "Surf4",
                "Atp5g2",
                "Emc8",
                "Srp19",
                "Phlda1",
                "1110004F10Rik",
                "Myeov2",
                "Vasp",
                "Tceb1",
                "Rpsa",
                "H2afz",
                "Bola3",
                "Psip1",
                "Vcl",
                "Iscu",
                "Fubp1",
                "Arpc5l",
                "Rpl5",
                "Dph3",
                "Prdx6",
                "Ptges3",
                "Mrpl18",
                "Rps20",
                "Orc6",
                "Dnaja2",
                "Crabp1",
                "Sys1",
                "Mrpl12",
                "Ndufab1",
                "Mtdh",
                "Eef1a1",
                "mt-Nd5",
                "U2af1",
                "Atp5c1",
                "Ywhaq",
                "Ccnd2",
                "Tcf25",
                "Pdha1",
                "Filip1l",
                "Cav1",
                "Dnajc8",
                "Calm2",
                "Stmn1",
                "Srsf10",
                "Btf3",
                "Smarcc1",
                "Plk2",
                "Tln1",
                "Tmem126a",
                "Oaf",
                "Rpl18",
                "Atp5d",
                "Mt2",
                "Timm13",
                "Rps3a1",
                "Smc4",
                "Cmpk1",
                "Rpl7",
                "Fkbp2",
                "Lrrc58",
                "Fn1",
                "Glipr2",
                "Hes1",
                "Ap1s1",
                "Uqcrq",
                "Mrpl30",
                "Sec61b",
                "Polr2m",
                "Rwdd1",
                "Snx5",
                "Gnai2",
                "Rps8",
                "Dnaja1",
                "Rpl8",
                "Eif4a1",
                "Tm9sf3",
                "Ndufc2",
                "Ccdc34",
                "Col1a1",
                "Nsa2",
                "Oxct1",
                "Ddx46",
                "Cnih4",
                "Morf4l2",
                "Ube2d3",
                "Aars",
                "Snrpa1",
                "Dstn",
                "Emp3",
                "Cebpb",
                "Srrm2",
                "2700060E02Rik",
                "Mt1",
                "Ywhae",
                "Pafah1b2",
                "Hmgb1",
                "Psmd14",
                "Eif3d",
                "Npm1",
                "Ghitm",
                "Scp2",
                "Arl1",
                "Arpc1a",
                "Mgat2",
                "Sep15",
                "Rab7",
                "Gtpbp4",
                "Eif2s2",
                "Hspd1",
                "Cpe",
                "Atf5",
                "Cdc42",
                "Mrpl45",
                "Srsf7",
                "Mrps25",
                "Tpx2",
                "Dnajb6",
                "Hspa5",
                "Uqcrc2",
                "Actr3",
                "G3bp2",
                "Cd47",
                "Impdh2",
                "Ccdc124",
                "Mpc1",
                "Sars",
                "Mrpl52",
                "Ap2m1",
                "Nupr1",
                "Pno1",
                "Hnrnpab",
                "Rplp0",
                "Psmb5",
                "Ahnak",
                "Tbrg1",
                "Hspa9",
                "Kif5b",
                "Polr2c",
                "Cct8",
                "Hmgn2",
                "Mpc2",
                "Hnrnpa2b1",
                "Ptp4a1",
                "Lsp1",
                "Ndufs5",
                "Cited2",
                "Srsf2",
                "Prdx4",
                "mt-Nd1",
                "Eif4g2",
                "Dnajc7",
                "Psmb2",
                "Raly",
                "Denr",
                "Get4",
                "Cnn3",
                "Parva",
                "Vdac1",
                "Cggbp1",
                "Zfp706",
                "Anp32a",
                "Rad23b",
                "Sumo1",
                "Tra2a",
                "Pdcd5",
                "Rab11a",
                "Ankrd1",
                "Anxa5",
                "Eif3f",
                "Eif4g1",
                "Anxa6",
                "Mtch1",
                "Tmbim6",
                "Rhoa",
                "Mrps18c",
                "Rab10",
                "Cacybp",
                "Sf3a3",
                "Ccdc80",
                "Anp32e",
                "Syncrip",
                "Sec61a1",
                "Caprin1",
                "Kdelr2",
                "Ict1",
                "Pmp22",
                "Glud1",
                "March5",
                "Rbmxl2",
                "Cebpz",
                "Ssr2",
                "Hnrnpu",
                "Rbmxl1",
                "Srsf11",
                "Arglu1",
                "Rpl18a",
                "Tomm20",
                "Cbx1",
                "Ssr4",
                "Tomm70a",
                "Txndc9",
                "Ppp1r2",
                "Eprs",
                "Trmt112",
                "Arpc2",
                "Atp5l",
                "Mrpl54",
                "Son",
                "Cct2",
                "Ubb",
                "Sept11",
                "Romo1",
                "Ergic3",
                "Bzw1",
                "Sdc4",
                "Ubc",
                "Lman2",
                "Rock2",
                "Hnrnpa3",
                "Mrps7",
                "Sub1",
                "Morf4l1",
                "2010107E04Rik",
                "Mrpl43",
                "Pafah1b1",
                "Cdca8",
                "Idh2",
                "Polr1d",
                "Snrpd3",
                "Rbms1",
                "P4hb",
                "Sf3b2",
                "Psmd3",
                "Zyx",
                "Map1b",
                "Adipor1",
                "mt-Nd2",
                "Bsg",
                "Rpl14",
                "Camk2d",
                "Cops4",
                "mt-Cytb",
                "Rnps1",
                "Ciapin1",
                "Flna",
                "Pfdn2",
                "H13",
                "Atp5f1",
                "Arid5b",
                "Rps13",
                "Tpt1",
                "Ppib",
                "Cfl1",
                "Gng5",
                "Hspe1",
                "Rps19",
                "Sept2",
                "Exosc8",
                "Cops6",
                "Ptbp1",
                "Mrpl20",
                "Hnrnpdl",
                "Eif3a",
                "Ptx3",
                "Ctbp1",
                "Psmb4",
                "Adh5",
                "Birc5",
                "Commd1",
                "Csrp1",
                "Chtop",
                "Atxn7l3b",
                "Ppa1",
                "Lamtor5",
                "Ndufa13",
                "Gps1",
                "Rrm2",
                "Rpl13",
                "Sh3bgrl",
                "Itm2b",
                "Pdia3",
                "Ndufb5",
                "Atrx",
                "Psma2",
                "Txndc5",
                "Pfdn5",
                "Nsun2",
                "Set",
                "Txnl4a",
                "Poldip2",
                "Eif4e2",
                "Ssbp1",
                "Rps14",
                "Mrto4",
                "Tnfrsf12a",
                "Clns1a",
                "Snrpd2",
                "Krtcap2",
                "Rps27a",
                "Naa10",
                "Bud31",
                "Pdap1",
                "Brk1",
                "Hsp90b1",
                "Vdac2",
                "Anxa1",
                "Rap1b",
                "Ube2a",
                "Rpl13a",
                "Nol7",
                "Eif1a",
                "Cox5a",
                "Commd3",
                "Cnn2",
                "Snrpb",
                "Exosc4",
                "Fkbp4",
                "Pgam1",
                "Rabggtb",
                "Tomm22",
                "Nans",
                "Uchl5",
                "Erp29",
                "Ppig",
                "Phldb2",
                "Palld",
                "Eif5b",
                "Cct4",
                "Llph",
                "Lrrc17",
                "Eif4b",
                "Eif4ebp1",
                "Anp32b",
                "Edf1",
                "Csde1",
                "Marcks",
                "Fam162a",
                "Emp1",
                "Psmc2",
                "Ndufa10",
                "Tomm40",
                "Ywhah",
                "Prpf19",
                "Hdac2",
                "Sugt1",
                "Eif4a3",
                "Fkbp3",
                "Mtch2",
                "Bag1",
                "Tmem147",
                "Rpn2",
                "Bub3",
                "Sin3b",
                "Dctn2",
                "Tmed3",
                "Serpinf1",
                "Tubb4b",
                "Mrps16",
                "Tpr",
                "Hint1",
                "Tbca",
                "Foxp1",
                "Srsf6",
                "Tmpo",
                "Tsn",
                "Txnl1",
                "Mesdc2",
                "Rbm25",
                "Ube2e3",
                "Drg1",
                "Arhgdia",
                "Myl12a",
                "Ptma",
                "Rac1",
                "Rer1",
                "Ralbp1",
                "Cald1",
                "Fau",
                "Yrdc",
                "Acin1",
                "Sri",
                "Pabpc1",
                "Prdx5",
                "Alyref",
                "Tsen34",
                "Hspa14",
                "Fkbp1a",
                "Emc7",
                "Gnas",
                "Arf4",
                "Calu",
                "Yif1b",
                "Hdlbp",
                "Wbp11",
                "Cox4i1",
                "Tmed9",
                "Snf8",
                "Guk1",
                "Ndufs8",
                "Serp1",
                "Tsfm",
                "Capn2",
                "Ptov1",
                "Laptm4a",
                "Aimp1",
                "Serbp1",
                "Cnih1",
                "Gpx4",
                "Chchd3",
                "Gcsh",
                "Phf5a",
                "Rheb",
                "Gtf2a2",
                "Rrp1",
                "Rpl30",
                "Sfr1",
                "Kdelr3",
                "Ncor1",
                "Rbm17",
                "Map2k2",
                "Cnbp",
                "Sdhd",
                "Ogn",
                "Rsrc2"
            ],
            "expression_sum": {
                "class1": 19607.0,
                "class2": 34910.0
            },
            "uniprot_ids": [
                "Q06507",
                "Q8CF69",
                "Q3U2J1",
                "A0A2R8VI82",
                "Q61328",
                "Q794G7",
                "Q3TZI8"
            ]
        },
        "1110004F10Rik": {
            "id": "1110004F10Rik",
            "type": "Gene",
            "symbol": "1110004F10Rik",
            "source": [
                "Atf5",
                "Fos",
                "Sox2",
                "Ewsr1",
                "Hdac2",
                "Tcf4",
                "Pou5f1",
                "Arf4",
                "Jun",
                "Prrx2",
                "Peg3",
                "Smarca5",
                "Sox4",
                "Eno1",
                "Prrx1",
                "Sfpq",
                "Jund",
                "Cebpb",
                "Smc3",
                "Hmga2",
                "Ctbp1",
                "Nono",
                "Atf4",
                "Junb",
                "Ybx1",
                "Nanog"
            ],
            "target": [],
            "expression_sum": {
                "class1": 13017.0,
                "class2": 24968.0
            },
            "uniprot_ids": [
                "D3YVS0",
                "D3YVR9",
                "A0A0U1RPJ7",
                "A0A0U1RPI0",
                "D6RI64"
            ]
        },
        "1110008F13Rik": {
            "id": "1110008F13Rik",
            "type": "Gene",
            "symbol": "1110008F13Rik",
            "source": [
                "Klf2",
                "Atf5",
                "Fos",
                "Id2",
                "Sox2",
                "Hdac2",
                "Tcf4",
                "Pou5f1",
                "Arf4",
                "Jun",
                "Peg3",
                "Smarca5",
                "Sox4",
                "Eno1",
                "Prrx1",
                "Jund",
                "Cebpb",
                "Smc3",
                "Hmga2",
                "Foxp1",
                "Ctbp1",
                "Atf4",
                "Junb",
                "Hes1",
                "Nanog",
                "Twist2"
            ],
            "target": [],
            "expression_sum": {
                "class1": 17888.0,
                "class2": 49364.0
            }
        },
        "2010107E04Rik": {
            "id": "2010107E04Rik",
            "type": "Gene",
            "symbol": "2010107E04Rik",
            "source": [
                "Klf2",
                "Pou5f1",
                "Cebpb",
                "Arf4",
                "Eno1",
                "Ctbp1",
                "Nono",
                "Atf4",
                "Junb",
                "Id2",
                "Jun",
                "Sox2",
                "Nanog",
                "Jund",
                "Hdac2",
                "Sox4"
            ],
            "target": [],
            "expression_sum": {
                "class1": 33157.0,
                "class2": 71680.0
            }
        },
        "2700060E02Rik": {
            "id": "2700060E02Rik",
            "type": "Gene",
            "symbol": "2700060E02Rik",
            "source": [
                "Klf2",
                "Tcf4",
                "Pou5f1",
                "Hmga2",
                "Arf4",
                "Atf5",
                "Atf4",
                "Junb",
                "Fos",
                "Jun",
                "Sox2",
                "Prrx1",
                "Nanog",
                "Jund",
                "Cebpb",
                "Mybl2"
            ],
            "target": [],
            "expression_sum": {
                "class1": 33160.0,
                "class2": 123621.0
            },
            "uniprot_ids": [
                "Q4VA29"
            ]
        },
        "2810004N23Rik": {
            "id": "2810004N23Rik",
            "type": "Gene",
            "symbol": "2810004N23Rik",
            "source": [
                "Klf2",
                "Pou5f1",
                "Atf5",
                "Atf4",
                "Junb",
                "Jun",
                "Sox2",
                "Hes1",
                "Nanog",
                "Jund",
                "Twist2",
                "Cebpb",
                "Sox4"
            ],
            "target": [],
            "expression_sum": {
                "class1": 5164.0,
                "class2": 25129.0
            },
            "uniprot_ids": [
                "Q3TKJ9",
                "Q9CSU7"
            ]
        },
        "Aars": {
            "id": "Aars",
            "type": "Gene",
            "symbol": "Aars",
            "source": [
                "Klf2",
                "Tcf4",
                "Pou5f1",
                "Arf4",
                "Atf4",
                "Junb",
                "Fos",
                "Jun",
                "Sox2",
                "Nanog",
                "Tfcp2l1",
                "Jund",
                "Twist2",
                "Cebpb"
            ],
            "target": [],
            "expression_sum": {
                "class1": 8214.0,
                "class2": 46057.0
            },
            "uniprot_ids": [
                "Q8BGQ7",
                "Q3UD67",
                "A0A1Y7VE86",
                "Q6XMP4",
                "Q3TZ32",
                "Q6PDM7",
                "Q8R346"
            ]
        },
        "Acin1": {
            "id": "Acin1",
            "type": "Gene",
            "symbol": "Acin1",
            "source": [
                "Tcf4",
                "Pou5f1",
                "Hmga2",
                "Nfib",
                "Atf4",
                "Junb",
                "Jun",
                "Sox2",
                "Nanog",
                "Jund",
                "Twist2",
                "Cebpb"
            ],
            "target": [],
            "expression_sum": {
                "class1": 6145.0,
                "class2": 32225.0
            },
            "uniprot_ids": [
                "Q9JIX8",
                "Q52KR6",
                "Q80TU6",
                "B8JJ91",
                "B8JJ92",
                "Q7TSL7",
                "Q05D19",
                "Q3UZ28",
                "F6Q8C0",
                "Q3UXT9",
                "F6RJ39",
                "B8JJ89",
                "B8JJ90"
            ]
        },
        "Actg1": {
            "id": "Actg1",
            "type": "Gene",
            "symbol": "Actg1",
            "source": [
                "Tcf4",
                "E2f4",
                "Pou5f1",
                "Trim28",
                "Rest",
                "Eno1",
                "Hmgb1",
                "Atf4",
                "Ctcf",
                "Junb",
                "Id2",
                "Hdac1",
                "Trp53",
                "Ybx1",
                "Bclaf1",
                "Hdac2",
                "Smc3",
                "Smarca5"
            ],
            "target": [],
            "expression_sum": {
                "class1": 458905.0,
                "class2": 355685.0
            },
            "uniprot_ids": [
                "P63260",
                "Q4KL81",
                "Q3UD81",
                "G3UZ07",
                "B1ATY1",
                "G3UYG0",
                "F8WGM8",
                "Q9QZ83",
                "Q3TSB7"
            ]
        },
        "Actr3": {
            "id": "Actr3",
            "type": "Gene",
            "symbol": "Actr3",
            "source": [
                "Nfib",
                "Hdac2",
                "Tcf4",
                "Pou5f1",
                "Trp53",
                "Jun",
                "Prrx2",
                "Smarca5",
                "Sox4",
                "Trim28",
                "Eno1",
                "Cebpb",
                "Smc3",
                "Foxp1",
                "Atf4",
                "Ctcf",
                "Junb",
                "Hdac1",
                "Ybx1",
                "Hes1",
                "Twist2"
            ],
            "target": [],
            "expression_sum": {
                "class1": 20135.0,
                "class2": 20265.0
            },
            "uniprot_ids": [
                "Q99JY9",
                "Q3ULF7",
                "A0A087WRA1",
                "Q3TGW0",
                "Q3TGE1",
                "A0A087WQ83",
                "A0A087WQ14",
                "A0A087WPR6",
                "A0A087WS98",
                "Q3TWJ4",
                "A0A087WP86",
                "Q3UGH0",
                "Q9ERF8"
            ]
        },
        "Adh5": {
            "id": "Adh5",
            "type": "Gene",
            "symbol": "Adh5",
            "source": [
                "Tcf4",
                "Pou5f1",
                "Foxp1",
                "Hmga2",
                "Nfib",
                "Atf4",
                "Junb",
                "Fos",
                "Jun",
                "Nanog",
                "Jund",
                "Cebpb"
            ],
            "target": [],
            "expression_sum": {
                "class1": 8318.0,
                "class2": 45783.0
            },
            "uniprot_ids": [
                "P28474",
                "A0A0G2JDX2",
                "A0A0G2JGS3",
                "Q6P5I3"
            ]
        },
        "Adipor1": {
            "id": "Adipor1",
            "type": "Gene",
            "symbol": "Adipor1",
            "source": [
                "Trim28",
                "Rest",
                "Arf4",
                "Atf4",
                "Ctcf",
                "Tgif1",
                "Trp53",
                "Hdac1",
                "Hdac2",
                "Smc3"
            ],
            "target": [],
            "expression_sum": {
                "class1": 9785.0,
                "class2": 16346.0
            },
            "uniprot_ids": [
                "Q91VH1",
                "Q53YY4",
                "Q3TI02",
                "Q3UPA4"
            ]
        },
        "Ahnak": {
            "id": "Ahnak",
            "type": "Gene",
            "symbol": "Ahnak",
            "source": [
                "Pou5f1",
                "Trim28",
                "Cebpb",
                "Atf5",
                "Eno1",
                "Hmgb1",
                "Atf4",
                "Ctcf",
                "Trp53",
                "Hdac1",
                "Klf5",
                "Tbx3",
                "Zfp57",
                "Hdac2",
                "Smc3",
                "Smarca5"
            ],
            "target": [],
            "expression_sum": {
                "class1": 16857.0,
                "class2": 7693.0
            },
            "uniprot_ids": [
                "Q6UIL4",
                "Q8BKS2",
                "Q8CGE7",
                "E9Q616",
                "G5E8K8",
                "A0PJF4",
                "Q8VDN3",
                "A0A494B8Y7",
                "Q61484",
                "A0A494BBD5",
                "Q6UL10",
                "A0JLR7",
                "Q8BRB8",
                "Q8CEX7",
                "Q8R2L7",
                "Q3UYW8"
            ]
        },
        "Aimp1": {
            "id": "Aimp1",
            "type": "Gene",
            "symbol": "Aimp1",
            "source": [
                "Tcf4",
                "Pou5f1",
                "Foxp1",
                "Cebpb",
                "Nfib",
                "Atf4",
                "Junb",
                "Fos",
                "Jun",
                "Sox2",
                "Prrx1",
                "Jund",
                "Twist2",
                "Hdac2",
                "Smc3",
                "Smarca5"
            ],
            "target": [],
            "expression_sum": {
                "class1": 10808.0,
                "class2": 31695.0
            },
            "uniprot_ids": [
                "P31230",
                "Q3UZG4",
                "Q8C2U7",
                "A0A0G2JDH0",
                "A0A0G2JEU9",
                "A0A0G2JDW6",
                "Q3TUZ1"
            ]
        },
        "Akt1s1": {
            "id": "Akt1s1",
            "type": "Gene",
            "symbol": "Akt1s1",
            "source": [
                "Pou5f1",
                "Trim28",
                "Hmga2",
                "Hmgb1",
                "Atf4",
                "Trp53",
                "Hdac1",
                "Ybx1",
                "Hdac2",
                "Smc3"
            ],
            "target": [],
            "expression_sum": {
                "class1": 9585.0,
                "class2": 14154.0
            },
            "uniprot_ids": [
                "Q9D1F4",
                "E9QKI5",
                "A0A140LHA0",
                "D3Z4R0",
                "E9QKI4",
                "A0A140LJJ3",
                "F6R8S6",
                "A0A140LHN7",
                "D3Z4W3",
                "D3YWZ1"
            ]
        },
        "Alyref": {
            "id": "Alyref",
            "type": "Gene",
            "symbol": "Alyref",
            "source": [
                "Tcf4",
                "Pou5f1",
                "Foxp1",
                "Cebpb",
                "Arf4",
                "Nfib",
                "Atf4",
                "Junb",
                "Fos",
                "Id2",
                "Jun",
                "Sox2",
                "Hes1",
                "Nanog",
                "Jund",
                "Hdac2"
            ],
            "target": [],
            "expression_sum": {
                "class1": 9621.0,
                "class2": 34581.0
            },
            "uniprot_ids": [
                "O08583"
            ]
        },
        "Anapc5": {
            "id": "Anapc5",
            "type": "Gene",
            "symbol": "Anapc5",
            "source": [
                "Pou5f1",
                "Foxp1",
                "Nfib",
                "Atf4",
                "Junb",
                "Fos",
                "Jun",
                "Sox2",
                "Hes1",
                "Nanog",
                "Jund",
                "Twist2",
                "Cebpb",
                "Sox4"
            ],
            "target": [],
            "expression_sum": {
                "class1": 11364.0,
                "class2": 39999.0
            },
            "uniprot_ids": [
                "Q8BTZ4",
                "Q3U5Y4",
                "Q3TWF7",
                "A0A0G2JE03",
                "Q3TGH0",
                "A0A0G2JDM7",
                "A0A0G2JDE8",
                "A0A0G2JF95",
                "Q3U981",
                "Q3THU3",
                "A0A0G2JGE5",
                "Q3TXL9",
                "A0A0G2JF12",
                "Q3UFC2",
                "A0A0G2JG78",
                "A0A0G2JFH0"
            ]
        },
        "Ankrd1": {
            "id": "Ankrd1",
            "type": "Gene",
            "symbol": "Ankrd1",
            "source": [
                "Trim28",
                "Atf4",
                "Klf5",
                "Hdac2",
                "Smc3",
                "Smarca5"
            ],
            "target": [],
            "expression_sum": {
                "class1": 3831.0,
                "class2": 154.0
            },
            "uniprot_ids": [
                "Q9CR42",
                "A0A494BB19",
                "A0A494BA80",
                "A0A494BBH7",
                "A0A498WFQ0"
            ]
        },
        "Anp32a": {
            "id": "Anp32a",
            "type": "Gene",
            "symbol": "Anp32a",
            "source": [
                "Tcf4",
                "Pou5f1",
                "Foxp1",
                "Tfcp2l1",
                "Nfib",
                "Atf4",
                "Junb",
                "Fos",
                "Jun",
                "Sox2",
                "Hes1",
                "Nanog",
                "Jund",
                "Twist2",
                "Cebpb",
                "Sox4"
            ],
            "target": [],
            "expression_sum": {
                "class1": 11397.0,
                "class2": 76496.0
            },
            "uniprot_ids": [
                "O35381",
                "F6UFG6",
                "D3YYE1",
                "D3Z7M9"
            ]
        },
        "Anp32b": {
            "id": "Anp32b",
            "type": "Gene",
            "symbol": "Anp32b",
            "source": [
                "Tcf4",
                "Pou5f1",
                "Foxp1",
                "Cebpb",
                "Hmga2",
                "Arf4",
                "Atf4",
                "Junb",
                "Fos",
                "Jun",
                "Sox2",
                "Hes1",
                "Nanog",
                "Jund",
                "Hdac2"
            ],
            "target": [],
            "expression_sum": {
                "class1": 39874.0,
                "class2": 195117.0
            },
            "uniprot_ids": [
                "Q9EST5",
                "Q3UW32"
            ]
        },
        "Anp32e": {
            "id": "Anp32e",
            "type": "Gene",
            "symbol": "Anp32e",
            "source": [
                "Klf2",
                "Tcf4",
                "Pou5f1",
                "Nfib",
                "Atf4",
                "Junb",
                "Fos",
                "Jun",
                "Hes1",
                "Nanog",
                "Jund",
                "Twist2",
                "Cebpb",
                "Smarca5",
                "Sox4"
            ],
            "target": [],
            "expression_sum": {
                "class1": 7535.0,
                "class2": 25478.0
            },
            "uniprot_ids": [
                "P97822",
                "A0A0G2JEK9",
                "E9PZF5",
                "E9Q2G1",
                "E9Q0X5",
                "E9PVI5",
                "E9Q5H2"
            ]
        },
        "Anxa1": {
            "id": "Anxa1",
            "type": "Gene",
            "symbol": "Anxa1",
            "source": [
                "Pou5f1",
                "Cebpb",
                "Eno1",
                "Atf4",
                "Ctcf",
                "Trp53",
                "Hdac1",
                "Klf5",
                "Sox2",
                "Ybx1",
                "Nanog",
                "Hdac2",
                "Smc3"
            ],
            "target": [],
            "expression_sum": {
                "class1": 47586.0,
                "class2": 2779.0
            },
            "uniprot_ids": [
                "P10107",
                "Q4FJV4",
                "Q4FK88",
                "Q3US43",
                "A0A494BBD8",
                "Q3U5N9",
                "B7STB7"
            ]
        },
        "Anxa2": {
            "id": "Anxa2",
            "type": "Gene",
            "symbol": "Anxa2",
            "source": [
                "Klf4",
                "Sox2",
                "Klf5",
                "Hdac2",
                "Pou5f1",
                "Rest",
                "Trp53",
                "Trim28",
                "Eno1",
                "Tgif1",
                "Tbx3",
                "Sfpq",
                "Zfp57",
                "Smc3",
                "Hmgb1",
                "Atf4",
                "Ctcf",
                "Hdac1",
                "Ybx1",
                "Nanog",
                "Tfcp2l1"
            ],
            "target": [],
            "expression_sum": {
                "class1": 89791.0,
                "class2": 9262.0
            },
            "uniprot_ids": [
                "P07356",
                "Q542G9",
                "B0V2N7",
                "B0V2N5",
                "B0V2N8",
                "Q3UCD3",
                "Q9CZI7",
                "Q99KH3"
            ]
        },
        "Anxa5": {
            "id": "Anxa5",
            "type": "Gene",
            "symbol": "Anxa5",
            "source": [
                "Nfib",
                "Id2",
                "Klf5",
                "Hdac2",
                "E2f4",
                "Pou5f1",
                "Rest",
                "Arf4",
                "Trp53",
                "Smarca5",
                "Trim28",
                "Tbx3",
                "Zfp57",
                "Smc3",
                "Hmgb1",
                "Atf4",
                "Ctcf",
                "Junb",
                "Hdac1",
                "Ybx1"
            ],
            "target": [],
            "expression_sum": {
                "class1": 39422.0,
                "class2": 4580.0
            },
            "uniprot_ids": [
                "P48036",
                "A0A0G2JGQ0"
            ]
        },
        "Anxa6": {
            "id": "Anxa6",
            "type": "Gene",
            "symbol": "Anxa6",
            "source": [
                "E2f4",
                "Rest",
                "Atf4",
                "Ctcf",
                "Trp53",
                "Hdac1",
                "Klf5",
                "Ybx1",
                "Hdac2",
                "Smc3",
                "Smarca5"
            ],
            "target": [],
            "expression_sum": {
                "class1": 11023.0,
                "class2": 5301.0
            },
            "uniprot_ids": [
                "P14824",
                "F8WIT2",
                "Q8CEX0",
                "Q3UDK4",
                "Q3UI56",
                "Q99JX6",
                "Q3TT26",
                "Q3TUI1"
            ]
        },
        "Ap1s1": {
            "id": "Ap1s1",
            "type": "Gene",
            "symbol": "Ap1s1",
            "source": [
                "E2f4",
                "Pou5f1",
                "Trim28",
                "Rest",
                "Arf4",
                "Hmgb1",
                "Atf4",
                "Ctcf",
                "Junb",
                "Tgif1",
                "Hdac1",
                "Trp53",
                "Sfpq",
                "Hdac2",
                "Smc3",
                "Smarca5"
            ],
            "target": [],
            "expression_sum": {
                "class1": 13940.0,
                "class2": 15233.0
            },
            "uniprot_ids": [
                "P61967",
                "D3Z0D6",
                "D3Z268",
                "D3YXN0"
            ]
        },
        "Ap2m1": {
            "id": "Ap2m1",
            "type": "Gene",
            "symbol": "Ap2m1",
            "source": [
                "Pou5f1",
                "Foxp1",
                "Atf4",
                "Ctcf",
                "Junb",
                "Fos",
                "Hdac1",
                "Trp53",
                "Prrx1",
                "Ybx1",
                "Hdac2",
                "Smc3",
                "Sox4"
            ],
            "target": [],
            "expression_sum": {
                "class1": 23307.0,
                "class2": 30840.0
            },
            "uniprot_ids": [
                "P84091",
                "Q3TWV4",
                "Q5FWI9",
                "Q6A0C9",
                "Q3TH69",
                "A0A338P6T4",
                "A0A338P798",
                "Q3UCW9"
            ]
        },
        "Apex1": {
            "id": "Apex1",
            "type": "TF",
            "symbol": "Apex1",
            "source": [
                "Tfcp2l1",
                "Pou5f1",
                "Atf4",
                "Junb",
                "Fos",
                "Jun",
                "Sox2",
                "Hes1",
                "Nanog",
                "Jund",
                "Cebpb",
                "Zfp57",
                "Sox4"
            ],
            "target": [
                "Alpl",
                "Purb",
                "4930402H24Rik",
                "Tdgf1",
                "2610305D13Rik",
                "mt-Co1"
            ],
            "expression_sum": {
                "class1": 7723.0,
                "class2": 65512.0
            },
            "uniprot_ids": [
                "P28352",
                "D3Z124",
                "Q544Z7",
                "D3Z6R9",
                "F6QA74"
            ]
        },
        "Arf4": {
            "id": "Arf4",
            "type": "TF",
            "symbol": "Arf4",
            "source": [
                "E2f4",
                "Trim28",
                "Rest",
                "Nfib",
                "Eno1",
                "Atf4",
                "Ctcf",
                "Junb",
                "Trp53",
                "Hdac1",
                "Klf5",
                "Hdac2",
                "Smc3",
                "Smarca5"
            ],
            "target": [
                "4930453N24Rik",
                "Atxn7l3b",
                "Rbm22",
                "Dynll1",
                "Clptm1",
                "Ppa1",
                "Tmem123",
                "1700037H04Rik",
                "Lamtor5",
                "Abcf2",
                "Dctn3",
                "Mrps14",
                "1810043H04Rik",
                "Tmed5",
                "Gps1",
                "Aacs",
                "Cab39",
                "Ddb1",
                "Mrpl3",
                "Eef2",
                "Rpa2",
                "Cbfb",
                "Isoc1",
                "Pdk3",
                "Rplp1",
                "Ssr1",
                "Pgls",
                "Atp5g1",
                "1110008F13Rik",
                "Nae1",
                "Ggnbp2",
                "Prdx2",
                "Nsdhl",
                "Alkbh1",
                "Cirh1a",
                "Copb2",
                "1110059E24Rik",
                "Sars",
                "Abcg2",
                "Hdgfrp2",
                "Rpl6l",
                "Acot7",
                "Asf1b",
                "Abcf1",
                "Zfand6",
                "Ctbp2",
                "Eri1",
                "Nubp2",
                "Gm4799",
                "Mettl16",
                "1700020I14Rik",
                "Fdps",
                "Chmp1a",
                "Slc38a4",
                "Phb2",
                "2410002F23Rik",
                "0610037L13Rik",
                "Serinc1",
                "Bud31",
                "Mrpl13",
                "Pdap1",
                "Ptp4a1",
                "Ilk",
                "AK010878",
                "Psmb6",
                "Samm50",
                "Rrp15",
                "Ddx39b",
                "Rps15a",
                "Mrpl24",
                "Ccdc25",
                "Tpm1",
                "Setd3",
                "Pcif1",
                "Rab2a",
                "Gapvd1",
                "Abt1",
                "Vps35",
                "Tubb5",
                "1110004F10Rik",
                "Tceal8",
                "Mfge8",
                "Mbnl2",
                "Aspscr1",
                "Sbno1",
                "Actn1",
                "Mcts1",
                "Mrpl14",
                "Slc25a3",
                "Uqcr10",
                "Angel2",
                "Rnaseh2c",
                "Gpx8",
                "Cnpy4",
                "Pnn",
                "Cyfip1",
                "Ppp2r1a",
                "Fubp1",
                "Tars",
                "Tcf3",
                "Aprt",
                "Rpl5",
                "Rps29",
                "Anxa5",
                "Eif4ebp1",
                "Mrpl49",
                "Anp32b",
                "Ssu72",
                "Cnot2",
                "Nono",
                "Rab10",
                "Atp5a1",
                "Rsl1d1",
                "1700021F05Rik",
                "Epb41l2",
                "Tubb2b",
                "Commd6",
                "Ufd1l",
                "Syncrip",
                "Acot9",
                "Snrpf",
                "Suclg2",
                "Zfp639",
                "Ndufa10",
                "Slc38a2",
                "Tomm40",
                "Uqcrfs1",
                "Arglu1",
                "Anln",
                "Znhit1",
                "Ykt6",
                "Lmo4",
                "S100a10",
                "Snrpd1",
                "Bub3",
                "Arpc2",
                "Efr3a",
                "Ywhab",
                "Pspc1",
                "Npat",
                "Aaas",
                "1110012L19Rik",
                "2310036O22Rik",
                "Romo1",
                "Rpl35a",
                "Sdc4",
                "Ltv1",
                "Ehd1",
                "Eif3h",
                "Ndufaf4",
                "Polr2b",
                "Vcp",
                "Ada",
                "Rbbp7",
                "0610011F06Rik",
                "Gadd45gip1",
                "Cmpk1",
                "Fkbp2",
                "Ptpn2",
                "0610009B22Rik",
                "2010107E04Rik",
                "Arih1",
                "Ap1s1",
                "2700094K13Rik",
                "Gfm1",
                "Metap1",
                "Nenf",
                "Utp3",
                "Ndufs4",
                "Ptges2",
                "Vps41",
                "1810058I24Rik",
                "Farsb",
                "Nsfl1c",
                "Adipor1",
                "Mafg",
                "Cse1l",
                "Gnai2",
                "Sarnp",
                "1110038F14Rik",
                "Ppic",
                "Alyref",
                "1700123O20Rik",
                "Kansl1",
                "Fkbp1a",
                "Clspn",
                "Borcs7",
                "Ncaph2",
                "Hcfc1r1",
                "0610007P14Rik",
                "Anapc2",
                "Ccdc34",
                "Gnb2l1",
                "Sertad2",
                "Krcc1",
                "Actl6a",
                "1110004E09Rik",
                "Brd7",
                "Prss23",
                "Trappc1",
                "Carnmt1",
                "Mrps35",
                "Idh3a",
                "Rps11",
                "Kif23",
                "Aars",
                "Atp5b",
                "Dtymk",
                "Akirin2",
                "Dstn",
                "Rrp1b",
                "Glrx3",
                "2410015M20Rik",
                "2610524H06Rik",
                "Srrm2",
                "2700060E02Rik",
                "Cyb5r4",
                "1110001J03Rik",
                "Igf2bp2",
                "Nop14",
                "Atg3",
                "4921524J17Rik",
                "Cetn2",
                "Csrp1",
                "1810037I17Rik",
                "Cnbp",
                "Ncbp2",
                "9430016H08Rik",
                "Zfp36l1"
            ],
            "expression_sum": {
                "class1": 37118.0,
                "class2": 24386.0
            },
            "uniprot_ids": [
                "P61750",
                "Q14BR4",
                "E9Q2C2",
                "E9Q798",
                "F6UFB9",
                "Q9DD04"
            ]
        },
        "Arf6": {
            "id": "Arf6",
            "type": "Gene",
            "symbol": "Arf6",
            "source": [
                "Tcf4",
                "Pou5f1",
                "Foxp1",
                "Cebpb",
                "Atf4",
                "Junb",
                "Sox2",
                "Prrx2",
                "Hes1",
                "Nanog",
                "Twist2",
                "Hdac2",
                "Smc3",
                "Sox4"
            ],
            "target": [],
            "expression_sum": {
                "class1": 8595.0,
                "class2": 22159.0
            },
            "uniprot_ids": [
                "P62331",
                "Q3U0D7"
            ]
        },
        "Arglu1": {
            "id": "Arglu1",
            "type": "Gene",
            "symbol": "Arglu1",
            "source": [
                "Pou5f1",
                "Foxp1",
                "Arf4",
                "Atf4",
                "Jun",
                "Sox2",
                "Hes1",
                "Nanog",
                "Jund",
                "Twist2",
                "Cebpb",
                "Sox4"
            ],
            "target": [],
            "expression_sum": {
                "class1": 9317.0,
                "class2": 22932.0
            },
            "uniprot_ids": [
                "Q3UL36",
                "A0A140LIS5",
                "Q3TEP1",
                "Q3TDV7"
            ]
        },
        "Arhgdia": {
            "id": "Arhgdia",
            "type": "Gene",
            "symbol": "Arhgdia",
            "source": [
                "Fos",
                "Id2",
                "Hdac2",
                "Tcf4",
                "Pou5f1",
                "Trp53",
                "Jun",
                "Smarca5",
                "Sox4",
                "Eno1",
                "Jund",
                "Cebpb",
                "Hmga2",
                "Atf4",
                "Ctcf",
                "Junb",
                "Hdac1",
                "Ybx1",
                "Hes1"
            ],
            "target": [],
            "expression_sum": {
                "class1": 29275.0,
                "class2": 50135.0
            },
            "uniprot_ids": [
                "Q99PT1"
            ]
        },
        "Arid5b": {
            "id": "Arid5b",
            "type": "TF",
            "symbol": "Arid5b",
            "source": [
                "Klf2",
                "Pou5f1",
                "Eno1",
                "Atf4",
                "Sox2",
                "Twist2",
                "Cebpb"
            ],
            "target": [
                "Tcf25",
                "Ddost",
                "0610007P14Rik"
            ],
            "expression_sum": {
                "class1": 8728.0,
                "class2": 17301.0
            },
            "uniprot_ids": [
                "Q8BM75",
                "A0A1X7SB62",
                "Q3TRJ5"
            ]
        },
        "Arl1": {
            "id": "Arl1",
            "type": "Gene",
            "symbol": "Arl1",
            "source": [
                "Nfib",
                "Fos",
                "Hdac2",
                "Tcf4",
                "Pou5f1",
                "Trp53",
                "Jun",
                "Prrx2",
                "Smarca5",
                "Trim28",
                "Eno1",
                "Jund",
                "Cebpb",
                "Smc3",
                "Hmgb1",
                "Nono",
                "Atf4",
                "Ctcf",
                "Junb",
                "Ybx1",
                "Twist2"
            ],
            "target": [],
            "expression_sum": {
                "class1": 17415.0,
                "class2": 18867.0
            },
            "uniprot_ids": [
                "P61211",
                "Q14AC7",
                "F8WIB1",
                "E9Q006",
                "Q9D7H9",
                "Q3TIR2",
                "Q4V9U3"
            ]
        },
        "Arpc1a": {
            "id": "Arpc1a",
            "type": "Gene",
            "symbol": "Arpc1a",
            "source": [
                "Pou5f1",
                "Trim28",
                "Atf4",
                "Trp53",
                "Ybx1",
                "Twist2",
                "Hdac2",
                "Smarca5",
                "Sox4"
            ],
            "target": [],
            "expression_sum": {
                "class1": 11028.0,
                "class2": 18269.0
            },
            "uniprot_ids": [
                "Q9R0Q6",
                "D3YVI5",
                "A0A0G2JF52"
            ]
        },
        "Arpc1b": {
            "id": "Arpc1b",
            "type": "Gene",
            "symbol": "Arpc1b",
            "source": [
                "E2f4",
                "Klf4",
                "Rest",
                "Eno1",
                "Atf4",
                "Ctcf",
                "Trp53",
                "Hdac1",
                "Klf5",
                "Tbx3",
                "Ybx1",
                "Hdac2",
                "Zfp57"
            ],
            "target": [],
            "expression_sum": {
                "class1": 55849.0,
                "class2": 25388.0
            },
            "uniprot_ids": [
                "Q9WV32",
                "Q91Z25",
                "D3Z6S0",
                "F6THG2",
                "F6VVE6",
                "Q3U094",
                "Q3TBA2",
                "Q3TCE7"
            ]
        },
        "Arpc2": {
            "id": "Arpc2",
            "type": "Gene",
            "symbol": "Arpc2",
            "source": [
                "E2f4",
                "Trim28",
                "Rest",
                "Arf4",
                "Hmgb1",
                "Atf4",
                "Ctcf",
                "Tgif1",
                "Id2",
                "Trp53",
                "Ybx1",
                "Bclaf1",
                "Hdac2",
                "Smc3"
            ],
            "target": [],
            "expression_sum": {
                "class1": 38942.0,
                "class2": 44706.0
            },
            "uniprot_ids": [
                "Q9CVB6",
                "Q3UA52",
                "D3YXG6",
                "A0A087WRT2",
                "Q4FZG5"
            ]
        },
        "Arpc5l": {
            "id": "Arpc5l",
            "type": "Gene",
            "symbol": "Arpc5l",
            "source": [
                "Pou5f1",
                "Foxp1",
                "Cebpb",
                "Atf4",
                "Junb",
                "Jun",
                "Sox2",
                "Hdac2",
                "Smc3",
                "Smarca5"
            ],
            "target": [],
            "expression_sum": {
                "class1": 7398.0,
                "class2": 23892.0
            },
            "uniprot_ids": [
                "Q9D898",
                "A0A0R3P9C9"
            ]
        },
        "Asns": {
            "id": "Asns",
            "type": "Gene",
            "symbol": "Asns",
            "source": [
                "Tcf4",
                "Pou5f1",
                "Atf5",
                "Atf4",
                "Fos",
                "Jun",
                "Sox2",
                "Hes1",
                "Nanog",
                "Jund",
                "Twist2",
                "Cebpb"
            ],
            "target": [],
            "expression_sum": {
                "class1": 9880.0,
                "class2": 68065.0
            },
            "uniprot_ids": [
                "Q61024",
                "D3Z028",
                "D3YYR0",
                "D3Z0C2",
                "Q3TIA9",
                "Q3TV20"
            ]
        },
        "Atf5": {
            "id": "Atf5",
            "type": "TF",
            "symbol": "Atf5",
            "source": [
                "Hdac2",
                "E2f4",
                "Foxp1",
                "Pou5f1",
                "Rest",
                "Trim28",
                "Hmgb1",
                "Atf4",
                "Ctcf",
                "Tgif1",
                "Trp53",
                "Hdac1",
                "Tbx3",
                "Ybx1",
                "Twist2",
                "Cebpb",
                "Zfp57",
                "Smarca5"
            ],
            "target": [
                "Arf1",
                "Cox6b1",
                "Rnf7",
                "Mrpl45",
                "Mcm7",
                "Rpl6",
                "Bub1",
                "Tspan17",
                "Hspa5",
                "Terf1",
                "Itpa",
                "Fosb",
                "Spr",
                "2810004N23Rik",
                "Slc1a5",
                "1110008F13Rik",
                "Ctps",
                "Arl6ip6",
                "Smu1",
                "A930018M24Rik",
                "Eif2b2",
                "Eif3c",
                "Akap12",
                "Dap3",
                "Adrm1",
                "Hdgfrp2",
                "Ppfibp1",
                "Vim",
                "Ahnak",
                "Tulp4",
                "Asns",
                "Fkbp10",
                "Gabarap",
                "Wars",
                "Ddr2",
                "Mrpl1",
                "Rrp15",
                "Aasdhppt",
                "1110004F10Rik",
                "Ubald1",
                "Cmtm3",
                "Calm3",
                "Trib3",
                "Mxd4",
                "2810417H13Rik",
                "Anapc15",
                "Fam149a",
                "Rdx",
                "Camk1",
                "Dmap1",
                "1700021F05Rik",
                "Ccdc80",
                "Timp3",
                "Lbh",
                "Tmem176a",
                "Poglut1",
                "Uba3",
                "1810011O10Rik",
                "Rcc2",
                "Ube2d2a",
                "Tnks1bp1",
                "Fbn2",
                "Mapre1",
                "Map1lc3b",
                "Rps26-ps1",
                "Dcp2",
                "1110046J04Rik",
                "Psme1",
                "Hax1",
                "Gm26682",
                "Dubr",
                "Cdc40",
                "Atf1",
                "Xpo1",
                "S100a4",
                "Kdm6b",
                "Zyx",
                "Nsfl1c",
                "Cald1",
                "Tsen34",
                "Bcas2",
                "Fas",
                "0610007P14Rik",
                "Spty2d1",
                "H13",
                "Marc2",
                "Phf5a",
                "Uchl3",
                "Gtf2a2",
                "2700060E02Rik",
                "Tagln",
                "Dnajc21",
                "Npm1"
            ],
            "expression_sum": {
                "class1": 10688.0,
                "class2": 4042.0
            },
            "uniprot_ids": [
                "O70191",
                "Q3UJF3",
                "A0A140LIB7",
                "Q3TVK2"
            ]
        },
        "Atp5c1": {
            "id": "Atp5c1",
            "type": "Gene",
            "symbol": "Atp5c1",
            "source": [
                "Klf2",
                "Tcf4",
                "Pou5f1",
                "Foxp1",
                "Cebpb",
                "Atf4",
                "Junb",
                "Fos",
                "Id2",
                "Jun",
                "Sox2",
                "Nanog",
                "Tfcp2l1",
                "Jund",
                "Hdac2",
                "Smc3"
            ],
            "target": [],
            "expression_sum": {
                "class1": 37886.0,
                "class2": 104171.0
            },
            "uniprot_ids": [
                "Q91VR2",
                "Q8C2Q8",
                "Q3UD06",
                "A2AKV2",
                "A2AKV3",
                "A2AKV1",
                "A2AKV0",
                "A2AKU9",
                "Q9ERA8",
                "Q9D9D7"
            ]
        },
        "Atp5d": {
            "id": "Atp5d",
            "type": "Gene",
            "symbol": "Atp5d",
            "source": [
                "Pou5f1",
                "Foxp1",
                "Cebpb",
                "Hmga2",
                "Nfib",
                "Eno1",
                "Hmgb1",
                "Atf4",
                "Junb",
                "Id2",
                "Trp53",
                "Ybx1",
                "Twist2",
                "Hdac2",
                "Smc3",
                "Smarca5",
                "Sox4"
            ],
            "target": [],
            "expression_sum": {
                "class1": 39572.0,
                "class2": 84441.0
            },
            "uniprot_ids": [
                "Q9D3D9",
                "Q4FK74",
                "Q9D0J2",
                "D3Z7S4",
                "Q9DCZ0"
            ]
        },
        "Atp5f1": {
            "id": "Atp5f1",
            "type": "Gene",
            "symbol": "Atp5f1",
            "source": [
                "Klf2",
                "Tcf4",
                "Pou5f1",
                "Foxp1",
                "Cebpb",
                "Atf4",
                "Junb",
                "Fos",
                "Jun",
                "Sox2",
                "Nanog",
                "Jund",
                "Twist2",
                "Hdac2",
                "Smc3",
                "Smarca5",
                "Sox4"
            ],
            "target": [],
            "expression_sum": {
                "class1": 41484.0,
                "class2": 105980.0
            },
            "uniprot_ids": [
                "Q9CQQ7",
                "Q5I0W0",
                "Q64218",
                "A0A0G2JGX3",
                "Q3UF04",
                "Q3TJD4",
                "D6RGL7",
                "Q80W36"
            ]
        },
        "Atp5g2": {
            "id": "Atp5g2",
            "type": "Gene",
            "symbol": "Atp5g2",
            "source": [
                "Tcf4",
                "Pou5f1",
                "Hmga2",
                "Cebpb",
                "Nfib",
                "Atf4",
                "Junb",
                "Fos",
                "Id2",
                "Jun",
                "Sox2",
                "Nanog",
                "Jund",
                "Twist2",
                "Hdac2",
                "Smarca5",
                "Sox4"
            ],
            "target": [],
            "expression_sum": {
                "class1": 48521.0,
                "class2": 166123.0
            },
            "uniprot_ids": [
                "P56383",
                "A0A087WQS5",
                "A0A087WSE2"
            ]
        },
        "Atp5j": {
            "id": "Atp5j",
            "type": "Gene",
            "symbol": "Atp5j",
            "source": [
                "Tcf4",
                "Pou5f1",
                "Foxp1",
                "Cebpb",
                "Ctbp1",
                "Atf4",
                "Junb",
                "Fos",
                "Jun",
                "Sox2",
                "Nanog",
                "Tfcp2l1",
                "Jund",
                "Twist2",
                "Hdac2"
            ],
            "target": [],
            "expression_sum": {
                "class1": 52235.0,
                "class2": 141576.0
            },
            "uniprot_ids": [
                "P97450",
                "E9QAD6"
            ]
        },
        "Atp5l": {
            "id": "Atp5l",
            "type": "Gene",
            "symbol": "Atp5l",
            "source": [
                "Pou5f1",
                "Cebpb",
                "Nfib",
                "Atf4",
                "Junb",
                "Trp53",
                "Sox2",
                "Twist2",
                "Hdac2",
                "Smc3",
                "Smarca5"
            ],
            "target": [],
            "expression_sum": {
                "class1": 64378.0,
                "class2": 139962.0
            },
            "uniprot_ids": [
                "Q9CPQ8",
                "Q9CQY3",
                "Q9D037"
            ]
        },
        "Atrx": {
            "id": "Atrx",
            "type": "Gene",
            "symbol": "Atrx",
            "source": [
                "Pou5f1",
                "Cebpb",
                "Nfib",
                "Atf4",
                "Junb",
                "Fos",
                "Jun",
                "Sox2",
                "Hes1",
                "Nanog",
                "Jund",
                "Hdac2",
                "Sox4"
            ],
            "target": [],
            "expression_sum": {
                "class1": 7331.0,
                "class2": 23693.0
            },
            "uniprot_ids": [
                "Q61687",
                "Q3UY26",
                "Q8C2S3",
                "Q3TA30",
                "Q3TP53",
                "F6RDB7",
                "Q3UU22",
                "A0A0G2JGH3",
                "Q3TSV4",
                "A0A0G2JFA5",
                "A6PWL2",
                "A0A0G2JED2",
                "A6PWK7",
                "A6PWK9",
                "A0A0G2JFR8",
                "A0A0G2JF42",
                "A0A0G2JFG6",
                "A0A0G2JF35",
                "Q8BSH0",
                "Q8C4Z6",
                "A6PWK8",
                "A0A0G2JGR1",
                "Q3UZ76"
            ]
        },
        "Atxn7l3b": {
            "id": "Atxn7l3b",
            "type": "Gene",
            "symbol": "Atxn7l3b",
            "source": [
                "E2f4",
                "Trim28",
                "Rest",
                "Arf4",
                "Hmgb1",
                "Atf4",
                "Ctcf",
                "Trp53",
                "Hdac1",
                "Ybx1",
                "Hdac2"
            ],
            "target": [],
            "expression_sum": {
                "class1": 13373.0,
                "class2": 13154.0
            },
            "uniprot_ids": [
                "Q3UD01"
            ]
        },
        "Aurkaip1": {
            "id": "Aurkaip1",
            "type": "Gene",
            "symbol": "Aurkaip1",
            "source": [
                "Tcf4",
                "Pou5f1",
                "Foxp1",
                "Hmga2",
                "Nfib",
                "Atf4",
                "Junb",
                "Fos",
                "Jun",
                "Sox2",
                "Nanog",
                "Jund",
                "Twist2",
                "Cebpb"
            ],
            "target": [],
            "expression_sum": {
                "class1": 13527.0,
                "class2": 54785.0
            },
            "uniprot_ids": [
                "Q9DCJ7",
                "A0A0A0MQH8",
                "Q91X73",
                "Q8CCC0",
                "Q3TZ21"
            ]
        },
        "Bag1": {
            "id": "Bag1",
            "type": "Gene",
            "symbol": "Bag1",
            "source": [
                "Pou5f1",
                "Foxp1",
                "Atf4",
                "Ctcf",
                "Junb",
                "Trp53",
                "Hdac1",
                "Ybx1",
                "Hdac2",
                "Smc3",
                "Smarca5"
            ],
            "target": [],
            "expression_sum": {
                "class1": 20349.0,
                "class2": 36647.0
            },
            "uniprot_ids": [
                "Q60739",
                "F6TCF9",
                "Q9CUY1"
            ]
        },
        "Banf1": {
            "id": "Banf1",
            "type": "Gene",
            "symbol": "Banf1",
            "source": [
                "Klf2",
                "Pou5f1",
                "Foxp1",
                "Cebpb",
                "Nfib",
                "Atf4",
                "Junb",
                "Fos",
                "Jun",
                "Sox2",
                "Hes1",
                "Nanog",
                "Jund",
                "Twist2",
                "Hdac2",
                "Smc3",
                "Sox4"
            ],
            "target": [],
            "expression_sum": {
                "class1": 15993.0,
                "class2": 49281.0
            },
            "uniprot_ids": [
                "O54962"
            ]
        },
        "Birc5": {
            "id": "Birc5",
            "type": "Gene",
            "symbol": "Birc5",
            "source": [
                "Tcf4",
                "Pou5f1",
                "Foxp1",
                "Nfib",
                "Atf4",
                "Junb",
                "Fos",
                "Id2",
                "Jun",
                "Sox2",
                "Hes1",
                "Nanog",
                "Jund",
                "Cebpb",
                "Zfp57"
            ],
            "target": [],
            "expression_sum": {
                "class1": 18120.0,
                "class2": 56526.0
            },
            "uniprot_ids": [
                "O70201",
                "Q549P2"
            ]
        },
        "Bnip3l": {
            "id": "Bnip3l",
            "type": "Gene",
            "symbol": "Bnip3l",
            "source": [
                "Pou5f1",
                "Foxp1",
                "Cebpb",
                "Nfib",
                "Atf4",
                "Junb",
                "Ybx1",
                "Twist2",
                "Hdac2",
                "Smarca5"
            ],
            "target": [],
            "expression_sum": {
                "class1": 9621.0,
                "class2": 22252.0
            },
            "uniprot_ids": [
                "Q9Z2F7",
                "Q3TEW0",
                "Q91Z78"
            ]
        },
        "Bola3": {
            "id": "Bola3",
            "type": "Gene",
            "symbol": "Bola3",
            "source": [
                "Pou5f1",
                "Atf4",
                "Junb",
                "Id2",
                "Hdac1",
                "Trp53",
                "Ybx1",
                "Twist2",
                "Hdac2",
                "Smc3",
                "Smarca5",
                "Sox4"
            ],
            "target": [],
            "expression_sum": {
                "class1": 9889.0,
                "class2": 18858.0
            },
            "uniprot_ids": [
                "Q8CEI1",
                "Q14A77",
                "E9Q705",
                "E9PX89",
                "Q3TSQ5"
            ]
        },
        "Brix1": {
            "id": "Brix1",
            "type": "Gene",
            "symbol": "Brix1",
            "source": [
                "Tcf4",
                "Pou5f1",
                "Foxp1",
                "Nfib",
                "Atf4",
                "Junb",
                "Jun",
                "Sox2",
                "Nanog",
                "Jund",
                "Twist2",
                "Cebpb"
            ],
            "target": [],
            "expression_sum": {
                "class1": 5665.0,
                "class2": 35891.0
            },
            "uniprot_ids": [
                "Q9DCA5",
                "F6QAG4",
                "E9Q2V8",
                "Q8CEX3"
            ]
        },
        "Brk1": {
            "id": "Brk1",
            "type": "Gene",
            "symbol": "Brk1",
            "source": [
                "Pou5f1",
                "Foxp1",
                "Hmga2",
                "Hmgb1",
                "Atf4",
                "Trp53",
                "Hdac2",
                "Smc3",
                "Smarca5"
            ],
            "target": [],
            "expression_sum": {
                "class1": 17949.0,
                "class2": 35377.0
            },
            "uniprot_ids": [
                "Q91VR8"
            ]
        },
        "Bsg": {
            "id": "Bsg",
            "type": "Gene",
            "symbol": "Bsg",
            "source": [
                "Pou5f1",
                "Trim28",
                "Rest",
                "Atf4",
                "Junb",
                "Ctcf",
                "Trp53",
                "Hdac1",
                "Ybx1",
                "Hdac2",
                "Smc3",
                "Smarca5"
            ],
            "target": [],
            "expression_sum": {
                "class1": 39877.0,
                "class2": 106197.0
            },
            "uniprot_ids": [
                "P18572",
                "O55107",
                "J3QP71",
                "K3W4Q8",
                "O55108"
            ]
        },
        "Btf3": {
            "id": "Btf3",
            "type": "Gene",
            "symbol": "Btf3",
            "source": [
                "Klf2",
                "Pou5f1",
                "Cebpb",
                "Nfib",
                "Atf4",
                "Junb",
                "Fos",
                "Jun",
                "Sox2",
                "Nanog",
                "Tfcp2l1",
                "Jund",
                "Twist2",
                "Hdac2"
            ],
            "target": [],
            "expression_sum": {
                "class1": 97733.0,
                "class2": 297488.0
            },
            "uniprot_ids": [
                "Q64152",
                "Q3UJR8"
            ]
        },
        "Btf3l4": {
            "id": "Btf3l4",
            "type": "Gene",
            "symbol": "Btf3l4",
            "source": [
                "Tcf4",
                "Pou5f1",
                "Foxp1",
                "Cebpb",
                "Nfib",
                "Atf4",
                "Junb",
                "Fos",
                "Jun",
                "Nanog",
                "Jund",
                "Hdac2",
                "Smc3",
                "Smarca5",
                "Sox4"
            ],
            "target": [],
            "expression_sum": {
                "class1": 9975.0,
                "class2": 23061.0
            },
            "uniprot_ids": [
                "Q9CQH7",
                "A2A7Z4",
                "Q78IG7"
            ]
        },
        "Bub3": {
            "id": "Bub3",
            "type": "Gene",
            "symbol": "Bub3",
            "source": [
                "Klf2",
                "Pou5f1",
                "Cebpb",
                "Arf4",
                "Atf4",
                "Junb",
                "Jun",
                "Nanog",
                "Tfcp2l1",
                "Jund",
                "Twist2",
                "Hdac2",
                "Smarca5"
            ],
            "target": [],
            "expression_sum": {
                "class1": 10658.0,
                "class2": 30184.0
            },
            "uniprot_ids": [
                "Q9WVA3",
                "Q8BH42",
                "A0A140LHA2",
                "A0A140LJ21",
                "A0A140LI47",
                "A0A140LIM5",
                "Q9CSI6"
            ]
        },
        "Bud31": {
            "id": "Bud31",
            "type": "Gene",
            "symbol": "Bud31",
            "source": [
                "Tcf4",
                "Pou5f1",
                "Hmga2",
                "Cebpb",
                "Arf4",
                "Nfib",
                "Eno1",
                "Atf4",
                "Junb",
                "Jun",
                "Sox2",
                "Nanog",
                "Jund",
                "Twist2",
                "Hdac2",
                "Smc3",
                "Smarca5"
            ],
            "target": [],
            "expression_sum": {
                "class1": 11758.0,
                "class2": 28872.0
            },
            "uniprot_ids": [
                "Q6PGH1"
            ]
        },
        "Bzw1": {
            "id": "Bzw1",
            "type": "Gene",
            "symbol": "Bzw1",
            "source": [
                "Nfib",
                "Fos",
                "Hdac2",
                "Tcf4",
                "Pou5f1",
                "Trp53",
                "Jun",
                "Smarca5",
                "Sox4",
                "Tgif1",
                "Jund",
                "Cebpb",
                "Smc3",
                "Foxp1",
                "Atf4",
                "Ctcf",
                "Junb",
                "Ybx1",
                "Nanog"
            ],
            "target": [],
            "expression_sum": {
                "class1": 18833.0,
                "class2": 32557.0
            },
            "uniprot_ids": [
                "Q9CQC6",
                "A0A087WPF9",
                "A0A087WS48",
                "A0A087WQS2",
                "Q9CVI6"
            ]
        },
        "Cacybp": {
            "id": "Cacybp",
            "type": "Gene",
            "symbol": "Cacybp",
            "source": [
                "Tcf4",
                "Pou5f1",
                "Foxp1",
                "Nfib",
                "Atf4",
                "Junb",
                "Fos",
                "Jun",
                "Sox2",
                "Hes1",
                "Nanog",
                "Jund",
                "Cebpb",
                "Sox4"
            ],
            "target": [],
            "expression_sum": {
                "class1": 15771.0,
                "class2": 98583.0
            },
            "uniprot_ids": [
                "Q9CXW3",
                "A0A0A6YY29"
            ]
        },
        "Cald1": {
            "id": "Cald1",
            "type": "Gene",
            "symbol": "Cald1",
            "source": [
                "Pou5f1",
                "Trim28",
                "Rest",
                "Nfib",
                "Atf5",
                "Hmgb1",
                "Atf4",
                "Ctcf",
                "Id2",
                "Hdac1",
                "Klf5",
                "Tbx3",
                "Trp53",
                "Zfp57",
                "Ybx1",
                "Hdac2",
                "Smc3",
                "Smarca5"
            ],
            "target": [],
            "expression_sum": {
                "class1": 106561.0,
                "class2": 18036.0
            },
            "uniprot_ids": [
                "Q8VCQ8",
                "E9Q9F3",
                "F6QLP8",
                "F6ZAW1",
                "D3Z6I7",
                "E9Q0M9",
                "Q7TMN5",
                "Q3TUS1",
                "F6T2Z7",
                "F6RGN9",
                "E9QA16",
                "Q8VD79",
                "E9QA15",
                "S4R1T7",
                "Q3UGC3"
            ]
        },
        "Calm2": {
            "id": "Calm2",
            "type": "Gene",
            "symbol": "Calm2",
            "source": [
                "Tcf4",
                "Pou5f1",
                "Foxp1",
                "Cebpb",
                "Hmga2",
                "Nfib",
                "Atf4",
                "Junb",
                "Jun",
                "Zfp57",
                "Hes1",
                "Nanog",
                "Jund",
                "Twist2",
                "Hdac2",
                "Smc3",
                "Smarca5",
                "Sox4"
            ],
            "target": [],
            "expression_sum": {
                "class1": 53179.0,
                "class2": 90883.0
            },
            "uniprot_ids": [
                "P0DP27",
                "A0A3Q4EHJ0",
                "A0A571BE58"
            ]
        },
        "Calm3": {
            "id": "Calm3",
            "type": "Gene",
            "symbol": "Calm3",
            "source": [
                "Tcf4",
                "Pou5f1",
                "Foxp1",
                "Cebpb",
                "Nfib",
                "Atf5",
                "Atf4",
                "Junb",
                "Fos",
                "Jun",
                "Sox2",
                "Hes1",
                "Nanog",
                "Jund",
                "Twist2",
                "Hdac2",
                "Smc3",
                "Sox4"
            ],
            "target": [],
            "expression_sum": {
                "class1": 20168.0,
                "class2": 46962.0
            },
            "uniprot_ids": [
                "P0DP28",
                "G3UX57",
                "G3UZ90"
            ]
        },
        "Calr": {
            "id": "Calr",
            "type": "Gene",
            "symbol": "Calr",
            "source": [
                "Hdac2",
                "Tcf4",
                "Pou5f1",
                "Foxp1",
                "Nfib",
                "Atf4",
                "Ctcf",
                "Junb",
                "Fos",
                "Jun",
                "Tgif1",
                "Trp53",
                "Ybx1",
                "Jund",
                "Twist2",
                "Cebpb",
                "Smc3",
                "Smarca5"
            ],
            "target": [],
            "expression_sum": {
                "class1": 106640.0,
                "class2": 82779.0
            },
            "uniprot_ids": [
                "P14211",
                "Q62041",
                "B2MWM9",
                "Q3UWP8"
            ]
        },
        "Calu": {
            "id": "Calu",
            "type": "Gene",
            "symbol": "Calu",
            "source": [
                "E2f4",
                "Trim28",
                "Rest",
                "Eno1",
                "Atf4",
                "Ctcf",
                "Trp53",
                "Hdac1",
                "Klf5",
                "Tbx3",
                "Zfp57",
                "Ybx1",
                "Hdac2",
                "Smc3"
            ],
            "target": [],
            "expression_sum": {
                "class1": 14256.0,
                "class2": 8256.0
            },
            "uniprot_ids": [
                "O35887",
                "Q6XLQ8",
                "Q3TUF3",
                "G3UWR0",
                "G3UWV3",
                "G3UXA3",
                "G3UXA8",
                "G3V004",
                "G3UY49",
                "Q8CBP3",
                "Q8CCN9",
                "Q3TL33",
                "Q3TQD1",
                "Q3UG11"
            ]
        },
        "Camk2d": {
            "id": "Camk2d",
            "type": "Gene",
            "symbol": "Camk2d",
            "source": [
                "Hdac2",
                "Klf2",
                "E2f4",
                "Pou5f1",
                "Trim28",
                "Rest",
                "Atf4",
                "Ctcf",
                "Tgif1",
                "Trp53",
                "Hdac1",
                "Klf5",
                "Sox2",
                "Tbx3",
                "Ybx1",
                "Cebpb",
                "Smc3",
                "Smarca5"
            ],
            "target": [],
            "expression_sum": {
                "class1": 8252.0,
                "class2": 981.0
            },
            "uniprot_ids": [
                "P28652",
                "Q6PHZ2",
                "Q8CCM0",
                "D6RDQ8",
                "E9QAJ4",
                "E9PXV3",
                "F6RWZ9",
                "A0A0G2JGS4",
                "E9Q1T1",
                "Q8C4I3",
                "E9Q1W0",
                "E9Q1V9",
                "O70459",
                "Q3V1X9",
                "Q8C8X9"
            ]
        },
        "Canx": {
            "id": "Canx",
            "type": "Gene",
            "symbol": "Canx",
            "source": [
                "E2f4",
                "Foxp1",
                "Trim28",
                "Rest",
                "Nfib",
                "Atf4",
                "Ctcf",
                "Junb",
                "Trp53",
                "Hdac1",
                "Jund",
                "Twist2",
                "Hdac2",
                "Smc3"
            ],
            "target": [],
            "expression_sum": {
                "class1": 20380.0,
                "class2": 24724.0
            },
            "uniprot_ids": [
                "P35564",
                "Q5SUC3",
                "Q3TXE5"
            ]
        },
        "Capn2": {
            "id": "Capn2",
            "type": "Gene",
            "symbol": "Capn2",
            "source": [
                "Hdac2",
                "E2f4",
                "Pou5f1",
                "Rest",
                "Atf4",
                "Ctcf",
                "Trp53",
                "Hdac1",
                "Klf5",
                "Sox2",
                "Tbx3",
                "Nanog",
                "Cebpb",
                "Smc3",
                "Smarca5"
            ],
            "target": [],
            "expression_sum": {
                "class1": 9381.0,
                "class2": 1218.0
            },
            "uniprot_ids": [
                "O08529",
                "Q3U8S5",
                "Q3U3A8",
                "Q8BPV9"
            ]
        },
        "Caprin1": {
            "id": "Caprin1",
            "type": "Gene",
            "symbol": "Caprin1",
            "source": [
                "Tcf4",
                "Pou5f1",
                "Cebpb",
                "Nfib",
                "Atf4",
                "Junb",
                "Fos",
                "Jun",
                "Sox2",
                "Ybx1",
                "Hes1",
                "Jund",
                "Hdac2",
                "Smc3",
                "Smarca5",
                "Sox4"
            ],
            "target": [],
            "expression_sum": {
                "class1": 9849.0,
                "class2": 22876.0
            },
            "uniprot_ids": [
                "Q60865",
                "A1L0T9",
                "Q3TEY4",
                "Q3TTW8",
                "F6YP03",
                "A2BH47",
                "Q3UFZ6",
                "Q8CD82",
                "F6YLI0"
            ]
        },
        "Capza2": {
            "id": "Capza2",
            "type": "Gene",
            "symbol": "Capza2",
            "source": [
                "Hdac2",
                "Tcf4",
                "Pou5f1",
                "Foxp1",
                "Eno1",
                "Hmgb1",
                "Atf4",
                "Ctcf",
                "Junb",
                "Fos",
                "Jun",
                "Trp53",
                "Hes1",
                "Jund",
                "Twist2",
                "Cebpb",
                "Smc3",
                "Smarca5"
            ],
            "target": [],
            "expression_sum": {
                "class1": 17206.0,
                "class2": 25515.0
            },
            "uniprot_ids": [
                "P47754",
                "A0A0N4SVM0",
                "Q5DQJ3",
                "D6RCW7",
                "Q3U7G3",
                "Q3UBZ3",
                "Q3UCD2"
            ]
        },
        "Cav1": {
            "id": "Cav1",
            "type": "Gene",
            "symbol": "Cav1",
            "source": [
                "E2f4",
                "Pou5f1",
                "Atf4",
                "Ctcf",
                "Hdac1",
                "Klf5",
                "Tbx3",
                "Zfp57",
                "Prrx2",
                "Ybx1",
                "Hdac2",
                "Smc3",
                "Smarca5"
            ],
            "target": [],
            "expression_sum": {
                "class1": 19225.0,
                "class2": 1617.0
            },
            "uniprot_ids": [
                "P49817",
                "D3Z0J2",
                "D3Z148",
                "H3BLQ4",
                "H3BKG0"
            ]
        },
        "Cbx1": {
            "id": "Cbx1",
            "type": "Gene",
            "symbol": "Cbx1",
            "source": [
                "Tcf4",
                "Pou5f1",
                "Nfib",
                "Atf4",
                "Junb",
                "Nr0b1",
                "Fos",
                "Jun",
                "Id2",
                "Sox2",
                "Nanog",
                "Jund",
                "Twist2",
                "Cebpb",
                "Sox4"
            ],
            "target": [],
            "expression_sum": {
                "class1": 8817.0,
                "class2": 54367.0
            },
            "uniprot_ids": [
                "P83917",
                "Q9CYJ8",
                "Q7TPM0"
            ]
        },
        "Ccdc124": {
            "id": "Ccdc124",
            "type": "Gene",
            "symbol": "Ccdc124",
            "source": [
                "Pou5f1",
                "Foxp1",
                "Cebpb",
                "Atf4",
                "Junb",
                "Fos",
                "Jun",
                "Sox2",
                "Nanog",
                "Jund",
                "Twist2",
                "Hdac2",
                "Sox4"
            ],
            "target": [],
            "expression_sum": {
                "class1": 8448.0,
                "class2": 32236.0
            },
            "uniprot_ids": [
                "Q9D8X2",
                "A0A1D5RLD5"
            ]
        },
        "Ccdc34": {
            "id": "Ccdc34",
            "type": "Gene",
            "symbol": "Ccdc34",
            "source": [
                "Tcf4",
                "Pou5f1",
                "Arf4",
                "Nfib",
                "Atf4",
                "Jun",
                "Jund",
                "Twist2",
                "Cebpb",
                "Smc3",
                "Sox4"
            ],
            "target": [],
            "expression_sum": {
                "class1": 10348.0,
                "class2": 16837.0
            },
            "uniprot_ids": [
                "Q3UI66",
                "Q8BIA8",
                "Q9D068",
                "Q5RL17",
                "F6U3Z3",
                "F6QKZ7"
            ]
        },
        "Ccdc80": {
            "id": "Ccdc80",
            "type": "Gene",
            "symbol": "Ccdc80",
            "source": [
                "Pou5f1",
                "Cebpb",
                "Atf5",
                "Atf4",
                "Ctcf",
                "Ybx1",
                "Hdac2",
                "Smc3"
            ],
            "target": [],
            "expression_sum": {
                "class1": 5942.0,
                "class2": 2732.0
            },
            "uniprot_ids": [
                "Q8R2G6"
            ]
        },
        "Ccnd2": {
            "id": "Ccnd2",
            "type": "Gene",
            "symbol": "Ccnd2",
            "source": [
                "Pou5f1",
                "Cebpb",
                "Klf4",
                "Rest",
                "Atf4",
                "Ctcf",
                "Tgif1",
                "Trp53",
                "Hdac1",
                "Klf5",
                "Sox2",
                "Tbx3",
                "Nanog",
                "Zfp57",
                "Hdac2",
                "Smc3"
            ],
            "target": [],
            "expression_sum": {
                "class1": 27196.0,
                "class2": 6742.0
            },
            "uniprot_ids": [
                "P30280",
                "Q4FK45",
                "Q9D8L9",
                "A0A0J9YU51"
            ]
        },
        "Cct2": {
            "id": "Cct2",
            "type": "Gene",
            "symbol": "Cct2",
            "source": [
                "Klf2",
                "Pou5f1",
                "Foxp1",
                "Cebpb",
                "Hmga2",
                "Atf4",
                "Fos",
                "Jun",
                "Sox2",
                "Ybx1",
                "Nanog",
                "Jund",
                "Twist2",
                "Hdac2",
                "Sox4"
            ],
            "target": [],
            "expression_sum": {
                "class1": 21573.0,
                "class2": 76896.0
            },
            "uniprot_ids": [
                "P80314",
                "Q542X7",
                "A0A1W2P6Q3",
                "A0A1W2P828",
                "A0A1W2P8B6",
                "A0A1W2P7B7",
                "A0A1W2P871",
                "Q9JJD8"
            ]
        },
        "Cct4": {
            "id": "Cct4",
            "type": "Gene",
            "symbol": "Cct4",
            "source": [
                "Pou5f1",
                "Cebpb",
                "Nfib",
                "Atf4",
                "Junb",
                "Fos",
                "Jun",
                "Sox2",
                "Ybx1",
                "Hes1",
                "Nanog",
                "Jund",
                "Twist2",
                "Hdac2",
                "Smc3",
                "Smarca5",
                "Sox4"
            ],
            "target": [],
            "expression_sum": {
                "class1": 28926.0,
                "class2": 86606.0
            },
            "uniprot_ids": [
                "P80315",
                "Q564F4",
                "Q3UJZ8",
                "G3UXF3",
                "G3UXG2",
                "G5E839",
                "G3UYW5",
                "Q3TII0"
            ]
        },
        "Cct8": {
            "id": "Cct8",
            "type": "Gene",
            "symbol": "Cct8",
            "source": [
                "Tcf4",
                "Pou5f1",
                "Foxp1",
                "Cebpb",
                "Nfib",
                "Atf4",
                "Junb",
                "Fos",
                "Jun",
                "Sox2",
                "Ybx1",
                "Nanog",
                "Jund",
                "Twist2",
                "Hdac2",
                "Smc3",
                "Smarca5"
            ],
            "target": [],
            "expression_sum": {
                "class1": 21214.0,
                "class2": 75373.0
            },
            "uniprot_ids": [
                "P42932",
                "Q3UL22",
                "H3BJB6",
                "Q3UKQ2",
                "H3BKG2",
                "Q9CS06",
                "Q6A0F1",
                "Q8BVY8",
                "H3BLL1",
                "H3BKR8",
                "H3BL49",
                "Q9CRW7"
            ]
        },
        "Cd47": {
            "id": "Cd47",
            "type": "Gene",
            "symbol": "Cd47",
            "source": [
                "E2f4",
                "Pou5f1",
                "Trim28",
                "Rest",
                "Atf4",
                "Ctcf",
                "Tgif1",
                "Trp53",
                "Hdac1",
                "Klf5",
                "Tbx3",
                "Zfp57",
                "Hdac2",
                "Smc3"
            ],
            "target": [],
            "expression_sum": {
                "class1": 10730.0,
                "class2": 1544.0
            },
            "uniprot_ids": [
                "Q61735",
                "A0A2R8VJU9",
                "D3Z187",
                "A0A2R8VK70",
                "A0A2R8VI94",
                "A0A2R8W6P0",
                "A0A2R8VI30",
                "Q3U967",
                "Q8C7F4",
                "E5Q371"
            ]
        },
        "Cdc20": {
            "id": "Cdc20",
            "type": "Gene",
            "symbol": "Cdc20",
            "source": [
                "Klf2",
                "Tcf4",
                "Pou5f1",
                "Foxp1",
                "Atf4",
                "Junb",
                "Jun",
                "Nanog",
                "Tfcp2l1",
                "Jund",
                "Twist2",
                "Cebpb",
                "Zfp57",
                "Sox4"
            ],
            "target": [],
            "expression_sum": {
                "class1": 7604.0,
                "class2": 39758.0
            },
            "uniprot_ids": [
                "Q9JJ66",
                "V9GXW0"
            ]
        },
        "Cdc42": {
            "id": "Cdc42",
            "type": "Gene",
            "symbol": "Cdc42",
            "source": [
                "E2f4",
                "Pou5f1",
                "Trim28",
                "Rest",
                "Nfib",
                "Atf4",
                "Ctcf",
                "Junb",
                "Tgif1",
                "Hdac1",
                "Trp53",
                "Ybx1",
                "Twist2",
                "Hdac2",
                "Smc3"
            ],
            "target": [],
            "expression_sum": {
                "class1": 43679.0,
                "class2": 40418.0
            },
            "uniprot_ids": [
                "P60766",
                "Q99JI7",
                "Q3UL78"
            ]
        },
        "Cdca8": {
            "id": "Cdca8",
            "type": "Gene",
            "symbol": "Cdca8",
            "source": [
                "Pou5f1",
                "Hmga2",
                "Atf4",
                "Junb",
                "Jun",
                "Sox2",
                "Nanog",
                "Jund",
                "Twist2",
                "Cebpb",
                "Zfp57"
            ],
            "target": [],
            "expression_sum": {
                "class1": 7013.0,
                "class2": 31149.0
            },
            "uniprot_ids": [
                "Q8BHX3",
                "E9Q2C7"
            ]
        },
        "Cdv3": {
            "id": "Cdv3",
            "type": "Gene",
            "symbol": "Cdv3",
            "source": [
                "Tcf4",
                "Pou5f1",
                "Foxp1",
                "Cebpb",
                "Atf4",
                "Junb",
                "Fos",
                "Jun",
                "Sox2",
                "Hes1",
                "Nanog",
                "Jund",
                "Twist2",
                "Hdac2"
            ],
            "target": [],
            "expression_sum": {
                "class1": 13626.0,
                "class2": 50982.0
            },
            "uniprot_ids": [
                "Q4VAA2",
                "A0A087WRM0",
                "F8WGL9",
                "A0A087WNP6",
                "A0A087WS49"
            ]
        },
        "Cebpb": {
            "id": "Cebpb",
            "type": "TF",
            "symbol": "Cebpb",
            "source": [
                "E2f4",
                "Foxp1",
                "Pou5f1",
                "Atf4",
                "Ctcf",
                "Trp53",
                "Hdac1",
                "Tbx3",
                "Zfp57",
                "Hdac2",
                "Smc3",
                "Smarca5"
            ],
            "target": [
                "Arf1",
                "Dap",
                "Srsf9",
                "Ndufb4",
                "Syce2",
                "Mcm7",
                "Grsf1",
                "Zc3h15",
                "Cox8a",
                "2810004N23Rik",
                "Sec13",
                "Ndufa6",
                "Polr2k",
                "Idh3g",
                "Rps27l",
                "Col12a1",
                "Lmna",
                "Ndufa2",
                "Hnrnpf",
                "Lmnb1",
                "Ddx39",
                "Cks2",
                "Shfm1",
                "Med28",
                "Hmgb2",
                "Cox6c",
                "Ddx3x",
                "Slbp",
                "Nedd4",
                "Calr",
                "Tubb5",
                "Nop56",
                "Slc25a3",
                "Lamtor1",
                "2810417H13Rik",
                "Nup43",
                "Cdc5l",
                "Sms",
                "Rbx1",
                "Ctcf",
                "Neat1",
                "Pmf1",
                "Ola1",
                "Sf3b6",
                "Lrrc59",
                "Ube2d2a",
                "Bnip3l",
                "Kpnb1",
                "Mapre1",
                "Snrpd1",
                "Rbm8a",
                "Col5a1",
                "Rpl36al",
                "Lap3",
                "Tgfb1i1",
                "Supt4a",
                "Rbbp7",
                "Hnrnpl",
                "Nedd8",
                "Pcnp",
                "Atpif1",
                "Snrnp40",
                "Gng12",
                "Tpm3",
                "Banf1",
                "Mrps36",
                "Qdpr",
                "Lig1",
                "Tpi1",
                "Pa2g4",
                "Gsr",
                "Atf4",
                "Hist1h2bc",
                "Hprt",
                "Nucks1",
                "Pcbp1",
                "Mrpl36",
                "Itgb1",
                "Ndufa4",
                "Cops5",
                "Eif4e",
                "Ndufv1",
                "H3f3a",
                "Rfc2",
                "Bccip",
                "Cyp51",
                "Lamtor2",
                "Trim28",
                "Nudt21",
                "Crip2",
                "Ube2l3",
                "Eef1d",
                "Etf1",
                "Zmat2",
                "Samm50",
                "Rrp15",
                "Nhp2l1",
                "Ran",
                "Atp5g3",
                "Gjb3",
                "Atp5g2",
                "Chd4",
                "Tceb1",
                "C1qbp",
                "Rpl36a",
                "Eif3m",
                "Snrpb2",
                "Umps",
                "Rps20",
                "Orc6",
                "Ybx1",
                "Sys1",
                "Tubb2b",
                "Tmem14c",
                "Rps6",
                "U2af1",
                "Slc29a1",
                "Atp5c1",
                "Mrps17",
                "Ccnd2",
                "Dnajc8",
                "Cox7c",
                "Nme4",
                "Eno1",
                "mt-Nd3",
                "Rpl22l1",
                "Lamtor4",
                "Smc4",
                "Mbd3",
                "Cct7",
                "Dctpp1",
                "Hsp90ab1",
                "Rpp25",
                "Tyms",
                "Ankrd11",
                "Rps8",
                "Ctsd",
                "Magoh",
                "Ndufc2",
                "H3f3b",
                "Dnajc2",
                "Ndufa12",
                "Cnih4",
                "Nudt19",
                "Dbi",
                "Tbx3",
                "Dstn",
                "Cyb5r3",
                "Ywhae",
                "Ppil1",
                "Rps15",
                "Arl1",
                "Cenpw",
                "Eif2s2",
                "Mrpl45",
                "Rpl31",
                "Smdt1",
                "Etfb",
                "Uqcrc2",
                "G3bp2",
                "Impdh2",
                "Prps1",
                "Akap12",
                "Hnrnpab",
                "Ahnak",
                "Nop58",
                "Ndufa11",
                "Cbx5",
                "Polr2c",
                "Cct8",
                "Dohh",
                "Denr",
                "Eif3g",
                "Aspscr1",
                "Btg2",
                "Parva",
                "Rpp21",
                "Polr2e",
                "Anapc13",
                "Pdcd5",
                "Tomm7",
                "Snrpc",
                "Mrpl49",
                "Tmbim6",
                "Atp5a1",
                "Psma5",
                "Pycr2",
                "Sf3a3",
                "Ccdc80",
                "March5",
                "Ddah2",
                "Gspt1",
                "Eprs",
                "Trmt112",
                "Rpl35",
                "Dazap1",
                "Ilf2",
                "Lman2",
                "Prmt1",
                "Sub1",
                "Morf4l1",
                "Col8a1",
                "Cdca8",
                "L1td1",
                "Lsm6",
                "Sf3b1",
                "Ndufs4",
                "Polr1d",
                "H2afy",
                "Mtf2",
                "Rpl17",
                "mt-Nd2",
                "Txndc17",
                "Spty2d1",
                "Nop10",
                "Mpdu1",
                "Gng5",
                "Hspe1",
                "Rps19",
                "Ube2c",
                "Msn",
                "Exosc8",
                "Mrpl20",
                "Ptx3",
                "Polr2h",
                "Psmb4",
                "Birc5",
                "Commd1",
                "Cox7b",
                "Selm",
                "Chtop",
                "Dppa5a",
                "Srpk1",
                "Lamtor5",
                "Avpi1",
                "Ssb",
                "Tinagl1",
                "Rpa2",
                "Mrps12",
                "Copz1",
                "Nfu1",
                "Ndufb5",
                "Set",
                "Thyn1",
                "Txnl4a",
                "Ssbp1",
                "Col4a1",
                "Sephs2",
                "Snrpd2",
                "1810022K09Rik",
                "Hdac1",
                "Naa10",
                "Hsp90b1",
                "Vdac2",
                "Mrpl24",
                "Snhg6",
                "Minos1",
                "Anapc11",
                "Gas6",
                "Nans",
                "Uchl5",
                "Cxcl1",
                "Ppig",
                "Dcun1d5",
                "Lsm12",
                "Gins4",
                "Tceb2",
                "Rhoc",
                "Mrpl42",
                "Rps2",
                "Polr2g",
                "Sumo3",
                "Uqcrfs1",
                "Naa50",
                "Ywhah",
                "Fbn2",
                "Hdac2",
                "Timm17a",
                "Rpn2",
                "Bub3",
                "Esf1",
                "Plekhj1",
                "Mrfap1",
                "Hsd17b14",
                "Slc25a5",
                "Uqcrh",
                "Foxp1",
                "Tsn",
                "Rpl37a",
                "Drg1",
                "Ptma",
                "Ndufb6",
                "Ybx3",
                "Rps10",
                "Dad1",
                "Sept1",
                "0610007P14Rik",
                "Yif1b",
                "Rpl39",
                "Tsfm",
                "Capn2",
                "Ptov1",
                "Rps27rt",
                "Gpx4",
                "Gcsh",
                "Carhsp1",
                "Gltscr2",
                "Mkrn1",
                "Rrp1",
                "Pdcl3",
                "Rpl23",
                "Rpl30",
                "Sfr1",
                "Polr2f",
                "Rrs1",
                "Rbm17",
                "Ly6e",
                "Lsm5",
                "Rnf7",
                "Cdc20",
                "Cd9",
                "Ddx24",
                "Rpl6",
                "Srm",
                "Thbs1",
                "Psmb7",
                "Spr",
                "Pnkd",
                "Ppid",
                "1110008F13Rik",
                "Anapc5",
                "Hnrnpa1",
                "Ssrp1",
                "Itm2a",
                "Rnf187",
                "Psmb6",
                "Ndufs3",
                "Uba2",
                "Mbnl2",
                "Snrpe",
                "B2m",
                "Psat1",
                "Gdi2",
                "Taf7",
                "Mrpl28",
                "Psmc6",
                "Grpel1",
                "Sec61g",
                "Ythdc1",
                "Snrnp27",
                "Srsf1",
                "Igfbp3",
                "Socs3",
                "Uqcrc1",
                "Rcc2",
                "Impa2",
                "Bola2",
                "Tmem160",
                "C1d",
                "Cisd2",
                "Ncl",
                "Rpl35a",
                "Timp2",
                "Syngr1",
                "Hspa4",
                "Arpp19",
                "Ccdc12",
                "Ddx21",
                "Naca",
                "Ube2i",
                "Rpl24",
                "Brix1",
                "Fam136a",
                "Siva1",
                "Rps4x",
                "Fkbp8",
                "Nip7",
                "Ncam1",
                "Ruvbl1",
                "Clpp",
                "Wls",
                "G3bp1",
                "Atp5b",
                "Cdc26",
                "Actb",
                "Immt",
                "Nars",
                "Hells",
                "Tfcp2l1",
                "Rpl38",
                "Glo1",
                "Serpine2",
                "Nme1",
                "Crabp2",
                "Csnk1a1",
                "Rpl10-ps3",
                "Rpl3",
                "Cebpzos",
                "Rpl12",
                "Snw1",
                "Ndufa7",
                "Dnajc19",
                "Cdv3",
                "Hmgn1",
                "Ctbp2",
                "Btf3l4",
                "Ppp4c",
                "Imp3",
                "Mrpl13",
                "Ak2",
                "Wdr77",
                "Atp5j",
                "Srrm1",
                "Srp19",
                "Emd",
                "Pdgfa",
                "Uqcr10",
                "Rpsa",
                "H2afz",
                "Mrps21",
                "Tardbp",
                "Rpl5",
                "Dph3",
                "Dpy30",
                "Serpine1",
                "Fblim1",
                "Odc1",
                "Mrpl18",
                "Ufc1",
                "Ifitm1",
                "Acat2",
                "Mrpl12",
                "Cdc37",
                "Cisd1",
                "Rpl28",
                "Xist",
                "Gpx3",
                "Timm8b",
                "Rpl26",
                "Uqcrb",
                "mt-Co2",
                "Rps3",
                "Atox1",
                "Srsf10",
                "Ube2v2",
                "Rpl19",
                "Timm13",
                "Eif3h",
                "Mcm3",
                "Adprh",
                "Rps3a1",
                "Srsf3",
                "Coprs",
                "Cmpk1",
                "Rpl7",
                "Fn1",
                "Znrd1",
                "Wdr18",
                "Polr2m",
                "1810058I24Rik",
                "Rpl8",
                "Eif4a1",
                "Pcbp2",
                "Polr2i",
                "Polr1c",
                "Snrnp25",
                "Fhl2",
                "Ccdc34",
                "Mrps33",
                "Rif1",
                "Morf4l2",
                "Rbbp4",
                "Eif3e",
                "Emp3",
                "2700060E02Rik",
                "Park7",
                "Dynll1",
                "Hspd1",
                "Stoml2",
                "Cops7a",
                "Sod1",
                "Ppih",
                "Uqcc3",
                "Tpx2",
                "Dnajb6",
                "Hspa5",
                "Gapdh",
                "Smarca5",
                "Idi1",
                "Ccnb1",
                "Ccnd3",
                "Khdrbs1",
                "Mrpl52",
                "Pno1",
                "Ecm1",
                "Adrm1",
                "Kif5b",
                "Hmgn2",
                "Hnrnpa2b1",
                "Oat",
                "Psmd2",
                "Ddx27",
                "Jade1",
                "Dnajc7",
                "Psmb2",
                "Fxr1",
                "Tceal8",
                "Cnn3",
                "Vps29",
                "Vdac1",
                "Anp32a",
                "Tra2a",
                "Zfp57",
                "Dynll2",
                "Ssu72",
                "Cnpy2",
                "Anp32e",
                "Pfdn6",
                "Ict1",
                "Rps25",
                "Rbmxl1",
                "Arglu1",
                "Imp4",
                "Ywhab",
                "Prelid1",
                "Ubb",
                "Zfp106",
                "Prrc1",
                "Paics",
                "Asna1",
                "Psmc5",
                "Echdc2",
                "Mrpl43",
                "Suclg1",
                "Hnrnpd",
                "Ash2l",
                "Hat1",
                "Rnps1",
                "H13",
                "Tpt1",
                "Ppib",
                "Stmn2",
                "Ptbp1",
                "Aldoa",
                "Lpp",
                "Dnajc21",
                "Snx3",
                "Strap",
                "Psmd13",
                "Rnaseh2a",
                "Rfc1",
                "Cox6b1",
                "Ptrf",
                "Atp5o",
                "Nasp",
                "Ndufs6",
                "Poldip2",
                "Eif4e2",
                "Sod2",
                "Hist1h1c",
                "Tnfrsf12a",
                "Phgdh",
                "Gtf2e2",
                "Pdap1",
                "Rpl11",
                "Anxa1",
                "Ube2a",
                "Pfn1",
                "Snrpg",
                "Ddx39b",
                "Grb10",
                "Exosc4",
                "Fkbp4",
                "Pgam1",
                "Snrnp70",
                "Ppdpf",
                "Gmnn",
                "Rpl27a",
                "Llph",
                "Eif4b",
                "Anp32b",
                "Edf1",
                "Fstl1",
                "Mdm2",
                "Eif2b5",
                "Gart",
                "Ndufa10",
                "App",
                "Tomm40",
                "Eif3i",
                "Smc2",
                "Mtch2",
                "Psmd4",
                "Gins2",
                "Uck2",
                "Rps16",
                "Bcat1",
                "Tmed3",
                "Mrps16",
                "Psmd12",
                "Ndufb10",
                "Cth",
                "Txnl1",
                "Rbm25",
                "Eef1g",
                "Ube2e3",
                "2700094K13Rik",
                "Acadm",
                "Dut",
                "Alyref",
                "Hspa14",
                "Polr2j",
                "Got2",
                "Slirp",
                "Ckb",
                "Cwc15",
                "Ndufs8",
                "Pim1",
                "Rplp2",
                "Mif",
                "Aimp1",
                "Rheb",
                "Chchd3",
                "Eif4a2",
                "Gtf2a2",
                "Cnbp",
                "Sdhd",
                "Axl",
                "Rps5",
                "Rpl41",
                "Cd44",
                "Ruvbl2",
                "Cdt1",
                "Ier2",
                "Dkc1",
                "Marcksl1",
                "Atp5g1",
                "Tuba1b",
                "Aurka",
                "Psma6",
                "Rpl7l1",
                "Snrpa",
                "St13",
                "Thoc7",
                "Mlf2",
                "Rpl9",
                "Vbp1",
                "Atp5j2",
                "Nubp1",
                "E2f4",
                "D8Ertd738e",
                "Hn1",
                "Mrpl14",
                "Phb",
                "Tra2b",
                "Hmgcs1",
                "Emc6",
                "Rps24",
                "Bola1",
                "Aurkaip1",
                "Timp3",
                "Nhp2",
                "Exosc5",
                "Sh3glb1",
                "Atp5h",
                "Ppia",
                "Eif3k",
                "Mrps34",
                "Atp5k",
                "Mphosph8",
                "Ppp1ca",
                "Aamp",
                "Chchd1",
                "Gars",
                "Psmc3",
                "Luc7l2",
                "Ndufb9",
                "Col3a1",
                "Utp3",
                "Fus",
                "Ociad1",
                "Ddost",
                "Rest",
                "Cct3",
                "Psma7",
                "Gnl3",
                "Trim27",
                "Psma1",
                "Dtymk",
                "Emc10",
                "2410015M20Rik",
                "Hdgf",
                "Prnp",
                "Id1",
                "Csnk2b",
                "Rpl37",
                "Ppp1cc",
                "Nrp1",
                "Dnttip2",
                "Sae1",
                "Hnrnpk",
                "Mrps24",
                "Ezr",
                "Ewsr1",
                "Ndufv2",
                "Fbxo22",
                "Fam49b",
                "Bax",
                "Cisd3",
                "Gfer",
                "Eif3c",
                "Cox7a2",
                "Col5a2",
                "Ube2j2",
                "Lsm3",
                "Gcat",
                "Eif5",
                "Spag7",
                "Tomm5",
                "Txn1",
                "Vapa",
                "Erh",
                "Fscn1",
                "Emc8",
                "Bclaf1",
                "Psip1",
                "Rpl29",
                "Naa38",
                "Ung",
                "Ptges3",
                "H2-D1",
                "Wtap",
                "Dnaja2",
                "Lbh",
                "Cenpv",
                "Ywhaq",
                "Pdha1",
                "Ubald2",
                "Filip1l",
                "Adss",
                "Rps7",
                "Igf2bp1",
                "Trnau1ap",
                "Eef1e1",
                "Hsd17b10",
                "Oaz1",
                "Stra13",
                "Sec61b",
                "Snx5",
                "Got1",
                "Bzw2",
                "Pgp",
                "Sumo2",
                "Nudc",
                "Pou5f1",
                "Gnb2l1",
                "Mrpl21",
                "Pold2",
                "Nsa2",
                "Ddx46",
                "Rfc5",
                "Pfdn4",
                "Mrps15",
                "Psmc4",
                "Psmd14",
                "Npm1",
                "Sap30",
                "Gtpbp4",
                "Cpe",
                "Atf5",
                "Srsf7",
                "Mat2a",
                "Csrp2",
                "Brd2",
                "Ccdc124",
                "Prdx2",
                "Abcf1",
                "Rplp0",
                "Psmb5",
                "Dcn",
                "Ube2n",
                "Smn1",
                "Cyb5b",
                "Psmb1",
                "Coa3",
                "Akr1b3",
                "Eif4g2",
                "Raly",
                "Uqcc2",
                "Nsmce4a",
                "Rps26",
                "Get4",
                "Lage3",
                "Npm3",
                "Lsm2",
                "Chmp2a",
                "Sumo1",
                "Nudcd2",
                "Aprt",
                "Fcf1",
                "Mrps18c",
                "Cacybp",
                "Rpl22",
                "Nolc1",
                "Rpl4",
                "Glud1",
                "Rps28",
                "Hnrnpu",
                "Srsf11",
                "Hmgb3",
                "Rpl21",
                "Ppp1r2",
                "Map1lc3b",
                "Atp5l",
                "Son",
                "Pop5",
                "Cct2",
                "Acp1",
                "Sox4",
                "Dtl",
                "Ahsa1",
                "Eif1ax",
                "Parp1",
                "Dhfr",
                "Mrps7",
                "Cdca3",
                "Metap2",
                "Ube2s",
                "Idh2",
                "Snrpd3",
                "Psmd3",
                "Fam32a",
                "Bcas2",
                "Cops4",
                "Ciapin1",
                "Pfdn2",
                "Atp5f1",
                "Arid5b",
                "Srp9",
                "Nfatc2ip",
                "Ino80e",
                "Ppa1",
                "Prdx1",
                "Rsl24d1",
                "Terf1",
                "Rpl13",
                "Mrpl34",
                "Atrx",
                "Nt5dc2",
                "Tgif1",
                "Sfpq",
                "Mrto4",
                "Parl",
                "Lipe",
                "Clns1a",
                "Mdh2",
                "Gabarap",
                "Uqcr11",
                "Phb2",
                "Rps27a",
                "Rpl15",
                "Gtf2h5",
                "Arl6ip1",
                "Tcof1",
                "Des",
                "Ier3",
                "Cuta",
                "1500015O10Rik",
                "Eif1a",
                "Cox5a",
                "Tmem11",
                "Ubap2l",
                "Phldb2",
                "Cct4",
                "Rdx",
                "Cdkn1a",
                "Pkm",
                "H2afv",
                "Prrc2c",
                "Marcks",
                "Psmg4",
                "Psmc2",
                "Sf3b4",
                "Hnrnpc",
                "Fkbp3",
                "Rangrf",
                "Rpl7a",
                "Fam104a",
                "Prrx2",
                "Manf",
                "Tpr",
                "Ddx18",
                "Rrp9",
                "Hint1",
                "Sf3a2",
                "Ccne1",
                "Srsf6",
                "Ebna1bp2",
                "Tmpo",
                "Cycs",
                "Klf2",
                "Dhx16",
                "Sdhb",
                "Taf10",
                "Wdr43",
                "Klf5",
                "Yrdc",
                "Pabpc1",
                "Eif6",
                "Tsen34",
                "Fam60a",
                "Wbp11",
                "2810428I15Rik",
                "Hspa8",
                "Postn",
                "Ldhb",
                "Taf1d",
                "Serbp1",
                "Kdm1a",
                "Ncor1",
                "1810037I17Rik",
                "Rsrc2",
                "Hsp90aa1",
                "Usmg5",
                "Mdh1",
                "Ppp2ca",
                "Rps17",
                "Rae1",
                "Pin4",
                "Mrps18b",
                "Txn2",
                "Clta",
                "Cox17",
                "Ppp1r11",
                "Mrpl4",
                "Wisp1",
                "Ndufb3",
                "Gar1",
                "Cks1b",
                "Ftsj3",
                "Folr1",
                "Psmc1",
                "Tpd52l2",
                "Knop1",
                "Psmd6",
                "Pin1",
                "Lgals3",
                "Sfxn1",
                "Eif2s1",
                "Swi5",
                "Nup62",
                "Cenpe",
                "Rab18",
                "Calm3",
                "Arf6",
                "Ubxn1",
                "Cdk1",
                "Eif4h",
                "Psmd7",
                "Anapc15",
                "Gabarapl2",
                "Slc39a1",
                "Stub1",
                "Prrx1",
                "Mrpl2",
                "Ranbp1",
                "Zwint",
                "Ppm1g",
                "Eif1",
                "Tbcb",
                "Wdr89",
                "Pdcd6",
                "H2afx",
                "Hnrnpm",
                "Capza2",
                "Loxl1",
                "2310036O22Rik",
                "Fdft1",
                "Srsf5",
                "Hnrnph1",
                "Wdr74",
                "Bex4",
                "Ywhaz",
                "Pomp",
                "Rpl36",
                "Rps21",
                "Psmb3",
                "Ddx5",
                "Rbm39",
                "Pigyl",
                "Eef1b2",
                "Eif3l",
                "Tmem258",
                "Tcp1",
                "Sarnp",
                "Mrpl15",
                "Ezh2",
                "Mybl2",
                "Pfdn1",
                "Slc2a3",
                "Pmm1",
                "Actl6a",
                "Pdia4",
                "Sssca1",
                "Ier5",
                "Gpx1",
                "Smc3",
                "Thrap3",
                "Hspb1",
                "Kdm5b",
                "Nfkbia",
                "Eif3b",
                "Zfp207",
                "Ube2k",
                "Ndufs2",
                "Ddt",
                "Spc24",
                "Asns",
                "Ifitm2",
                "Fdps",
                "Lsm8",
                "Ndufs7",
                "Cenpf",
                "Mcm2",
                "Gm16286",
                "Mrpl17",
                "Ldha",
                "Apex1",
                "Cct5",
                "Ccnb2",
                "Mrpl35",
                "1110004F10Rik",
                "Mrpl40",
                "Lox",
                "Iscu",
                "Fubp1",
                "Arpc5l",
                "Rps29",
                "Ccdc58",
                "Cenpa",
                "Prdx6",
                "Mdk",
                "Psmd1",
                "Ndufab1",
                "Plekhf2",
                "Snrpf",
                "Mcl1",
                "Pole4",
                "Fbl",
                "Ndufv3",
                "Rps26-ps1",
                "Calm2",
                "Stmn1",
                "Btf3",
                "Smarcc1",
                "Cdh11",
                "Rpl18",
                "Hax1",
                "Atp5d",
                "Taldo1",
                "Vcp",
                "Mrps5",
                "Prc1",
                "U2surp",
                "Fkbp2",
                "Ppan",
                "Rps9",
                "Upp1",
                "Mrpl30",
                "Rwdd1",
                "Cyc1",
                "Fnta",
                "Dnaja1",
                "Tmem176b",
                "Pmvk",
                "Elof1",
                "Lrpap1",
                "Wdr5",
                "Pebp1",
                "Sgk1",
                "Ube2d3",
                "Aars",
                "Marc2",
                "Snrpa1",
                "Paip2",
                "Srrm2",
                "Rnaseh2b",
                "Top1",
                "Ubl5",
                "Hmgb1",
                "Ndufc1",
                "Eif3d",
                "Cyb5a",
                "Psma4",
                "Ap2s1",
                "Rpl34",
                "Mlec",
                "Rab7",
                "Mrps14",
                "Nap1l1",
                "Mrps25",
                "Actr3",
                "Mrps10",
                "Tbrg1",
                "Hspa9",
                "Fkbp10",
                "Chchd2",
                "Pim3",
                "E130012A19Rik",
                "Luc7l3",
                "Ndufs5",
                "Gsn",
                "Ndufb8",
                "Tmem97",
                "Srsf2",
                "Prdx4",
                "Pak1ip1",
                "Nxt1",
                "Trp53",
                "Lhfp",
                "Rbm3",
                "Psmd11",
                "Cggbp1",
                "Zfp706",
                "Rad23b",
                "Nudt4",
                "Supt16",
                "Ost4",
                "Mrpl51",
                "Eif4g1",
                "Rhoa",
                "Cdk4",
                "Rsl1d1",
                "Syncrip",
                "Caprin1",
                "Ssr2",
                "Cebpz",
                "Tomm20",
                "Cbx1",
                "Tomm70a",
                "Msmo1",
                "Mak16",
                "Bzw1",
                "Ak6",
                "Rbpms",
                "Hnrnpa3",
                "Prpf40a",
                "Pbdc1",
                "Rps19bp1",
                "2010107E04Rik",
                "Tecr",
                "Mybbp1a",
                "Sf3b2",
                "Atp1b1",
                "Slc3a2",
                "Stip1",
                "Chchd4",
                "Rpl14",
                "Tma7",
                "Camk2d",
                "Lsm4",
                "Tpd52",
                "Eif1ad",
                "Rps13",
                "Acta1",
                "Eif5a",
                "Rps11",
                "Cfl1",
                "Rrp1b",
                "Sparc",
                "Uchl3",
                "Cops6",
                "Hnrnpdl",
                "Eif3a",
                "Rtn3",
                "Adh5",
                "l7Rn6",
                "Atp5e",
                "Cdc34",
                "Gps1",
                "Tipin",
                "Cmtm7",
                "Rrm2",
                "Sh3bgrl",
                "Itm2b",
                "Rplp1",
                "Rnf10",
                "Psma2",
                "Pfdn5",
                "Nsun2",
                "Ndufb7",
                "Shmt1",
                "Alkbh1",
                "Slc25a39",
                "Rps14",
                "Rpl32",
                "Chchd10",
                "Bud31",
                "Mcm5",
                "Rpl13a",
                "Nol7",
                "Tkt",
                "Apoe",
                "Gsta4",
                "Snrpb",
                "Zfp428",
                "Tomm22",
                "Lyar",
                "Erp29",
                "Gpx8",
                "Gpi1",
                "Tmem167",
                "Cfdp1",
                "Eif5b",
                "Emg1",
                "Eif4ebp1",
                "Rpl23a-ps3",
                "Nono",
                "Csde1",
                "Mcm6",
                "Prpf19",
                "Sugt1",
                "Eif4a3",
                "Dek",
                "Top2a",
                "Papola",
                "Tmem256",
                "Cetn3",
                "Galk1",
                "Ccna2",
                "Tubb4b",
                "Eed",
                "Ndufa5",
                "Mrpl11",
                "Mesdc2",
                "Cox5b",
                "Arhgdia",
                "Ralbp1",
                "Acin1",
                "Aebp1",
                "Rbpms2",
                "Uhrf1",
                "Guk1",
                "Psma3",
                "Skp1a",
                "Rps18",
                "Phf5a",
                "Rps23",
                "U2af2",
                "P3h4",
                "Myl12b"
            ],
            "expression_sum": {
                "class1": 14867.0,
                "class2": 3147.0
            },
            "uniprot_ids": [
                "P28033",
                "Q3UPN9",
                "O70192"
            ]
        },
        "Cebpz": {
            "id": "Cebpz",
            "type": "TF",
            "symbol": "Cebpz",
            "source": [
                "Klf2",
                "Tcf4",
                "Pou5f1",
                "Foxp1",
                "Atf4",
                "Junb",
                "Jun",
                "Sox2",
                "Nanog",
                "Jund",
                "Twist2",
                "Cebpb",
                "Sox4"
            ],
            "target": [
                "Bbx"
            ],
            "expression_sum": {
                "class1": 3344.0,
                "class2": 23065.0
            },
            "uniprot_ids": [
                "P53569",
                "A0A3B2WCE8",
                "A0A0R4J046",
                "Q3UGK4",
                "Q3UUZ3"
            ]
        },
        "Cfl1": {
            "id": "Cfl1",
            "type": "Gene",
            "symbol": "Cfl1",
            "source": [
                "Klf2",
                "Tcf4",
                "Pou5f1",
                "Hmga2",
                "Cebpb",
                "Atf4",
                "Fos",
                "Jun",
                "Sox2",
                "Prrx1",
                "Nanog",
                "Tfcp2l1",
                "Jund",
                "Twist2",
                "Hdac2",
                "Smarca5",
                "Sox4"
            ],
            "target": [],
            "expression_sum": {
                "class1": 120744.0,
                "class2": 253376.0
            },
            "uniprot_ids": [
                "P18760",
                "Q544Y7",
                "A0A494B9A7",
                "F8WGL3",
                "Q9CX22"
            ]
        },
        "Cggbp1": {
            "id": "Cggbp1",
            "type": "Gene",
            "symbol": "Cggbp1",
            "source": [
                "Tcf4",
                "Pou5f1",
                "Foxp1",
                "Cebpb",
                "Nfib",
                "Nono",
                "Atf4",
                "Junb",
                "Sox2",
                "Hes1",
                "Jund",
                "Twist2",
                "Hdac2"
            ],
            "target": [],
            "expression_sum": {
                "class1": 7052.0,
                "class2": 17914.0
            },
            "uniprot_ids": [
                "Q8BHG9"
            ]
        },
        "Chchd1": {
            "id": "Chchd1",
            "type": "Gene",
            "symbol": "Chchd1",
            "source": [
                "Pou5f1",
                "Foxp1",
                "Cebpb",
                "Nfib",
                "Atf4",
                "Junb",
                "Jun",
                "Sox2",
                "Hes1",
                "Nanog",
                "Jund",
                "Twist2",
                "Hdac2"
            ],
            "target": [],
            "expression_sum": {
                "class1": 13867.0,
                "class2": 38913.0
            },
            "uniprot_ids": [
                "Q9CQA6",
                "Q05BF0",
                "A0A286YDP8"
            ]
        },
        "Chchd3": {
            "id": "Chchd3",
            "type": "Gene",
            "symbol": "Chchd3",
            "source": [
                "Pou5f1",
                "Cebpb",
                "Atf4",
                "Junb",
                "Fos",
                "Jun",
                "Jund",
                "Twist2",
                "Hdac2"
            ],
            "target": [],
            "expression_sum": {
                "class1": 7834.0,
                "class2": 20406.0
            },
            "uniprot_ids": [
                "Q9CRB9",
                "Q9D9P1",
                "Q91VG6",
                "D3Z0L4",
                "F6QFL0",
                "S4R238"
            ]
        },
        "Chmp4b": {
            "id": "Chmp4b",
            "type": "Gene",
            "symbol": "Chmp4b",
            "source": [
                "Pou5f1",
                "Trim28",
                "Atf4",
                "Trp53",
                "Hdac1",
                "Ybx1",
                "Hdac2"
            ],
            "target": [],
            "expression_sum": {
                "class1": 9880.0,
                "class2": 21393.0
            },
            "uniprot_ids": [
                "Q9D8B3"
            ]
        },
        "Chtop": {
            "id": "Chtop",
            "type": "Gene",
            "symbol": "Chtop",
            "source": [
                "Tcf4",
                "Pou5f1",
                "Atf4",
                "Junb",
                "Fos",
                "Jun",
                "Hes1",
                "Nanog",
                "Jund",
                "Twist2",
                "Cebpb"
            ],
            "target": [],
            "expression_sum": {
                "class1": 6104.0,
                "class2": 24610.0
            },
            "uniprot_ids": [
                "Q9CY57",
                "D3YZA1",
                "B7ZMS6",
                "E9PW20",
                "D3Z7T7",
                "Q99KL5",
                "Q2VPR3"
            ]
        },
        "Ciapin1": {
            "id": "Ciapin1",
            "type": "Gene",
            "symbol": "Ciapin1",
            "source": [
                "Klf2",
                "Tcf4",
                "Pou5f1",
                "Foxp1",
                "Nfib",
                "Atf4",
                "Junb",
                "Fos",
                "Jun",
                "Sox2",
                "Hes1",
                "Nanog",
                "Jund",
                "Cebpb",
                "Sox4"
            ],
            "target": [],
            "expression_sum": {
                "class1": 6139.0,
                "class2": 35370.0
            },
            "uniprot_ids": [
                "Q8WTY4",
                "A0A1D5RLF8",
                "E0CYY1",
                "F7BWP1"
            ]
        },
        "Cisd2": {
            "id": "Cisd2",
            "type": "Gene",
            "symbol": "Cisd2",
            "source": [
                "Hdac2",
                "Pou5f1",
                "Foxp1",
                "Trim28",
                "Rest",
                "Ctbp1",
                "Atf4",
                "Ctcf",
                "Junb",
                "Fos",
                "Hdac1",
                "Id2",
                "Trp53",
                "Hes1",
                "Jund",
                "Cebpb",
                "Smc3",
                "Smarca5"
            ],
            "target": [],
            "expression_sum": {
                "class1": 10051.0,
                "class2": 10003.0
            },
            "uniprot_ids": [
                "Q9CQB5",
                "D3Z3X4"
            ]
        },
        "Cited2": {
            "id": "Cited2",
            "type": "Gene",
            "symbol": "Cited2",
            "source": [
                "Foxp1",
                "Trim28",
                "Hmgb1",
                "Atf4",
                "Junb",
                "Trp53",
                "Hdac1",
                "Ybx1",
                "Hdac2",
                "Smc3",
                "Smarca5"
            ],
            "target": [],
            "expression_sum": {
                "class1": 12420.0,
                "class2": 7199.0
            },
            "uniprot_ids": [
                "O35740",
                "A0A1W2P7B1"
            ]
        },
        "Clns1a": {
            "id": "Clns1a",
            "type": "Gene",
            "symbol": "Clns1a",
            "source": [
                "Tfcp2l1",
                "Pou5f1",
                "Foxp1",
                "Atf4",
                "Junb",
                "Fos",
                "Jun",
                "Sox2",
                "Hes1",
                "Nanog",
                "Jund",
                "Twist2",
                "Cebpb"
            ],
            "target": [],
            "expression_sum": {
                "class1": 6543.0,
                "class2": 36940.0
            },
            "uniprot_ids": [
                "Q61189",
                "Q923F1"
            ]
        },
        "Clta": {
            "id": "Clta",
            "type": "Gene",
            "symbol": "Clta",
            "source": [
                "Klf2",
                "Tcf4",
                "Pou5f1",
                "Hmga2",
                "Cebpb",
                "Nfib",
                "Atf4",
                "Junb",
                "Fos",
                "Jun",
                "Sox2",
                "Nanog",
                "Tfcp2l1",
                "Jund",
                "Twist2",
                "Hdac2",
                "Smc3"
            ],
            "target": [],
            "expression_sum": {
                "class1": 36282.0,
                "class2": 95724.0
            },
            "uniprot_ids": [
                "O08585",
                "Q6PFA2",
                "B1AWE1",
                "B1AWD8",
                "B1AWD9",
                "B1AWE0",
                "Q3THU7"
            ]
        },
        "Cmpk1": {
            "id": "Cmpk1",
            "type": "Gene",
            "symbol": "Cmpk1",
            "source": [
                "Tcf4",
                "Cebpb",
                "Rest",
                "Arf4",
                "Nono",
                "Atf4",
                "Ctcf",
                "Junb",
                "Tgif1",
                "Hdac1",
                "Trp53",
                "Ybx1",
                "Hes1",
                "Twist2",
                "Hdac2"
            ],
            "target": [],
            "expression_sum": {
                "class1": 11454.0,
                "class2": 11250.0
            },
            "uniprot_ids": [
                "Q9DBP5",
                "F6TY56",
                "D6RIL5",
                "A0A0R4J093"
            ]
        },
        "Cnbp": {
            "id": "Cnbp",
            "type": "Gene",
            "symbol": "Cnbp",
            "source": [
                "Pou5f1",
                "Foxp1",
                "Cebpb",
                "Arf4",
                "Atf4",
                "Fos",
                "Jun",
                "Sox2",
                "Nanog",
                "Jund",
                "Twist2",
                "Hdac2",
                "Sox4"
            ],
            "target": [],
            "expression_sum": {
                "class1": 30083.0,
                "class2": 122958.0
            },
            "uniprot_ids": [
                "P53996",
                "Q3ULK8",
                "Q5QJQ9",
                "Q3U935",
                "A0A0N4SVS6",
                "Q3U5V2"
            ]
        },
        "Cnih1": {
            "id": "Cnih1",
            "type": "Gene",
            "symbol": "Cnih1",
            "source": [
                "E2f4",
                "Trim28",
                "Atf4",
                "Ctcf",
                "Trp53",
                "Hdac2",
                "Smc3",
                "Smarca5"
            ],
            "target": [],
            "expression_sum": {
                "class1": 17696.0,
                "class2": 15603.0
            },
            "uniprot_ids": [
                "O35372",
                "D6RGU4",
                "Q3U420",
                "Q3THS7"
            ]
        },
        "Cnih4": {
            "id": "Cnih4",
            "type": "Gene",
            "symbol": "Cnih4",
            "source": [
                "Tcf4",
                "Pou5f1",
                "Cebpb",
                "Nfib",
                "Eno1",
                "Atf4",
                "Ctcf",
                "Junb",
                "Trp53",
                "Jun",
                "Ybx1",
                "Jund",
                "Twist2",
                "Hdac2",
                "Smc3",
                "Smarca5",
                "Sox4"
            ],
            "target": [],
            "expression_sum": {
                "class1": 13645.0,
                "class2": 23853.0
            },
            "uniprot_ids": [
                "Q9CX13",
                "D3YZJ3",
                "A0A0A6YW87"
            ]
        },
        "Cnn2": {
            "id": "Cnn2",
            "type": "Gene",
            "symbol": "Cnn2",
            "source": [
                "Atf4",
                "Ctcf",
                "Trp53",
                "Ybx1",
                "Hdac2"
            ],
            "target": [],
            "expression_sum": {
                "class1": 55020.0,
                "class2": 8792.0
            },
            "uniprot_ids": [
                "Q08093",
                "D3Z7R6",
                "Q543F3",
                "Q922F8"
            ]
        },
        "Cnn3": {
            "id": "Cnn3",
            "type": "Gene",
            "symbol": "Cnn3",
            "source": [
                "Tcf4",
                "Pou5f1",
                "Cebpb",
                "Nfib",
                "Atf4",
                "Junb",
                "Fos",
                "Jun",
                "Nanog",
                "Jund",
                "Twist2",
                "Hdac2",
                "Smarca5",
                "Sox4"
            ],
            "target": [],
            "expression_sum": {
                "class1": 27593.0,
                "class2": 59418.0
            },
            "uniprot_ids": [
                "Q9DAW9",
                "A0A0G2JFC8",
                "A0A0G2JDV8",
                "Q99JN5",
                "A0A0G2JFG3",
                "A0A0G2JFI6"
            ]
        },
        "Col1a1": {
            "id": "Col1a1",
            "type": "Gene",
            "symbol": "Col1a1",
            "source": [
                "Pou5f1",
                "Klf4",
                "Rest",
                "Atf4",
                "Ctcf",
                "Tgif1",
                "Trp53",
                "Hdac1",
                "Klf5",
                "Tbx3",
                "Zfp57",
                "Nanog",
                "Hdac2",
                "Smc3",
                "Smarca5"
            ],
            "target": [],
            "expression_sum": {
                "class1": 139259.0,
                "class2": 28416.0
            },
            "uniprot_ids": [
                "P11087",
                "Q60785",
                "Q99LL6"
            ]
        },
        "Col5a2": {
            "id": "Col5a2",
            "type": "Gene",
            "symbol": "Col5a2",
            "source": [
                "E2f4",
                "Pou5f1",
                "Cebpb",
                "Trim28",
                "Atf4",
                "Ctcf",
                "Sox2",
                "Hdac2",
                "Smc3"
            ],
            "target": [],
            "expression_sum": {
                "class1": 16632.0,
                "class2": 3886.0
            },
            "uniprot_ids": [
                "Q3U962",
                "Q8BNA3"
            ]
        },
        "Commd1": {
            "id": "Commd1",
            "type": "Gene",
            "symbol": "Commd1",
            "source": [
                "Klf6",
                "Tcf4",
                "Pou5f1",
                "Hmga2",
                "Cebpb",
                "Nfib",
                "Atf4",
                "Junb",
                "Jun",
                "Sox2",
                "Jund",
                "Twist2",
                "Hdac2",
                "Smc3",
                "Smarca5",
                "Sox4"
            ],
            "target": [],
            "expression_sum": {
                "class1": 10469.0,
                "class2": 21679.0
            },
            "uniprot_ids": [
                "Q8K4M5",
                "Q8VI86",
                "A2RSF1",
                "Q793C9",
                "F7BZY0",
                "G8JL54",
                "F7BCN0",
                "Q3UGT3"
            ]
        },
        "Commd3": {
            "id": "Commd3",
            "type": "Gene",
            "symbol": "Commd3",
            "source": [
                "E2f4",
                "Trim28",
                "Rest",
                "Atf4",
                "Ctcf",
                "Trp53",
                "Hdac1",
                "Tbx3",
                "Zfp57",
                "Ybx1",
                "Hdac2",
                "Smc3",
                "Smarca5"
            ],
            "target": [],
            "expression_sum": {
                "class1": 12856.0,
                "class2": 6340.0
            },
            "uniprot_ids": [
                "Q63829"
            ]
        },
        "Cops4": {
            "id": "Cops4",
            "type": "Gene",
            "symbol": "Cops4",
            "source": [
                "Tcf4",
                "Pou5f1",
                "Foxp1",
                "Cebpb",
                "Nfib",
                "Atf4",
                "Junb",
                "Fos",
                "Jun",
                "Sox2",
                "Nanog",
                "Jund",
                "Twist2",
                "Hdac2",
                "Smc3",
                "Smarca5",
                "Sox4"
            ],
            "target": [],
            "expression_sum": {
                "class1": 8716.0,
                "class2": 21909.0
            },
            "uniprot_ids": [
                "O88544",
                "Q14AI7",
                "D3YV99",
                "D6RG47",
                "F6QTS1",
                "Q3TIJ1",
                "D3Z1R9"
            ]
        },
        "Cops6": {
            "id": "Cops6",
            "type": "Gene",
            "symbol": "Cops6",
            "source": [
                "Tcf4",
                "Pou5f1",
                "Foxp1",
                "Cebpb",
                "Hmga2",
                "Nfib",
                "Atf4",
                "Junb",
                "Fos",
                "Jun",
                "Prrx1",
                "Sfpq",
                "Ybx1",
                "Nanog",
                "Jund",
                "Twist2",
                "Hdac2",
                "Smarca5"
            ],
            "target": [],
            "expression_sum": {
                "class1": 15658.0,
                "class2": 43675.0
            },
            "uniprot_ids": [
                "O88545",
                "Q3UIT2",
                "F6QK86",
                "D3Z0F5"
            ]
        },
        "Cox4i1": {
            "id": "Cox4i1",
            "type": "Gene",
            "symbol": "Cox4i1",
            "source": [
                "Pou5f1",
                "Rest",
                "Hmgb1",
                "Atf4",
                "Ctcf",
                "Trp53",
                "Hdac1",
                "Ybx1",
                "Twist2",
                "Hdac2",
                "Smc3",
                "Smarca5",
                "Sox4"
            ],
            "target": [],
            "expression_sum": {
                "class1": 99007.0,
                "class2": 225403.0
            },
            "uniprot_ids": [
                "P19783",
                "A2RSV8",
                "M0QWX2",
                "M0QWX7"
            ]
        },
        "Cox5a": {
            "id": "Cox5a",
            "type": "Gene",
            "symbol": "Cox5a",
            "source": [
                "Tcf4",
                "Pou5f1",
                "Foxp1",
                "Nfib",
                "Atf4",
                "Junb",
                "Fos",
                "Jun",
                "Sox2",
                "Hes1",
                "Nanog",
                "Jund",
                "Twist2",
                "Cebpb"
            ],
            "target": [],
            "expression_sum": {
                "class1": 50696.0,
                "class2": 252822.0
            },
            "uniprot_ids": [
                "P12787"
            ]
        },
        "Cpe": {
            "id": "Cpe",
            "type": "Gene",
            "symbol": "Cpe",
            "source": [
                "E2f4",
                "Pou5f1",
                "Cebpb",
                "Rest",
                "Trim28",
                "Atf4",
                "Ctcf",
                "Trp53",
                "Jun",
                "Tbx3",
                "Zfp57",
                "Prrx1",
                "Hes1",
                "Nanog",
                "Hdac2",
                "Smc3",
                "Smarca5"
            ],
            "target": [],
            "expression_sum": {
                "class1": 9674.0,
                "class2": 1969.0
            },
            "uniprot_ids": [
                "Q00493",
                "Q543R4"
            ]
        },
        "Crabp1": {
            "id": "Crabp1",
            "type": "Gene",
            "symbol": "Crabp1",
            "source": [
                "Rest",
                "Hmgb1",
                "Atf4",
                "Ctcf",
                "Ybx1"
            ],
            "target": [],
            "expression_sum": {
                "class1": 35295.0,
                "class2": 3287.0
            },
            "uniprot_ids": [
                "P62965",
                "Q6P071"
            ]
        },
        "Csde1": {
            "id": "Csde1",
            "type": "Gene",
            "symbol": "Csde1",
            "source": [
                "Tcf4",
                "Pou5f1",
                "Foxp1",
                "Cebpb",
                "Nfib",
                "Atf4",
                "Junb",
                "Trp53",
                "Hdac1",
                "Sox2",
                "Ybx1",
                "Jund",
                "Twist2",
                "Hdac2",
                "Smc3"
            ],
            "target": [],
            "expression_sum": {
                "class1": 14985.0,
                "class2": 24969.0
            },
            "uniprot_ids": [
                "Q91W50",
                "Q3U1S6",
                "Q8JZN2",
                "Q3U672",
                "A0A0G2JF72",
                "Q05C63",
                "Q80TP8",
                "A0A0G2JDJ7",
                "A0A0G2JE62",
                "Q3TXH7",
                "Q3UJ74",
                "Q8R3R1",
                "Q3U4S7"
            ]
        },
        "Csrp1": {
            "id": "Csrp1",
            "type": "Gene",
            "symbol": "Csrp1",
            "source": [
                "E2f4",
                "Trim28",
                "Rest",
                "Arf4",
                "Nfib",
                "Atf4",
                "Ctcf",
                "Tgif1",
                "Trp53",
                "Hdac1",
                "Klf5",
                "Hdac2",
                "Smc3",
                "Smarca5"
            ],
            "target": [],
            "expression_sum": {
                "class1": 25600.0,
                "class2": 9050.0
            },
            "uniprot_ids": [
                "P97315",
                "Q4FJX4"
            ]
        },
        "Ctbp1": {
            "id": "Ctbp1",
            "type": "TF",
            "symbol": "Ctbp1",
            "source": [
                "E2f4",
                "Trim28",
                "Rest",
                "Nfib",
                "Atf4",
                "Ctcf",
                "Junb",
                "Id2",
                "Hdac1",
                "Trp53",
                "Ybx1",
                "Hdac2",
                "Smc3",
                "Smarca5"
            ],
            "target": [
                "Anapc16",
                "Actr1a",
                "Atp5j",
                "Hmgb3",
                "1700123O20Rik",
                "Fabp5",
                "Ddb1",
                "Asah1",
                "0610007P14Rik",
                "Cisd2",
                "Rpa1",
                "1110004F10Rik",
                "Hn1",
                "1110008F13Rik",
                "2310036O22Rik",
                "2810417H13Rik",
                "Mid1ip1",
                "Ube3a",
                "Asnsd1",
                "2610524H06Rik",
                "Gltscr2",
                "Cdca3",
                "1700020I14Rik",
                "0610009B22Rik",
                "2010107E04Rik",
                "Glipr2",
                "Atpif1",
                "0610037L13Rik"
            ],
            "expression_sum": {
                "class1": 9673.0,
                "class2": 8571.0
            },
            "uniprot_ids": [
                "O88712",
                "A0A0J9YUR5",
                "A0A0J9YTW3",
                "A0A0J9YUM4",
                "A0A0J9YU66",
                "A0A0J9YVI3",
                "A0A0J9YU62",
                "A0A0J9YVC3"
            ]
        },
        "Cyp51": {
            "id": "Cyp51",
            "type": "Gene",
            "symbol": "Cyp51",
            "source": [
                "Trim28",
                "Cebpb",
                "Atf4",
                "Jun",
                "Ybx1",
                "Hes1",
                "Jund",
                "Twist2",
                "Hdac2",
                "Smc3",
                "Sox4"
            ],
            "target": [],
            "expression_sum": {
                "class1": 8717.0,
                "class2": 7695.0
            },
            "uniprot_ids": [
                "Q8K0C4",
                "Q9CW87"
            ]
        },
        "Cyr61": {
            "id": "Cyr61",
            "type": "Gene",
            "symbol": "Cyr61",
            "source": [
                "E2f4",
                "Trim28",
                "Rest",
                "Eno1",
                "Atf4",
                "Ctcf",
                "Trp53",
                "Sfpq",
                "Ybx1",
                "Hdac2",
                "Smc3",
                "Smarca5"
            ],
            "target": [],
            "expression_sum": {
                "class1": 25638.0,
                "class2": 16281.0
            },
            "uniprot_ids": [
                "P18406",
                "Q3TX21"
            ]
        },
        "Dap": {
            "id": "Dap",
            "type": "Gene",
            "symbol": "Dap",
            "source": [
                "Klf4",
                "Sox2",
                "Klf5",
                "Hdac2",
                "E2f4",
                "Pou5f1",
                "Rest",
                "Trp53",
                "Trim28",
                "Tgif1",
                "Tbx3",
                "Cebpb",
                "Zfp57",
                "Atf4",
                "Ctcf",
                "Hdac1",
                "Ybx1",
                "Nanog",
                "Twist2"
            ],
            "target": [],
            "expression_sum": {
                "class1": 20440.0,
                "class2": 3224.0
            },
            "uniprot_ids": [
                "Q91XC8",
                "Q3U706",
                "A0A087WR57",
                "A0A087WRC6",
                "A0A087WSM4"
            ]
        },
        "Dctn2": {
            "id": "Dctn2",
            "type": "Gene",
            "symbol": "Dctn2",
            "source": [
                "Pou5f1",
                "Nfib",
                "Atf4",
                "Junb",
                "Ctcf",
                "Trp53",
                "Ybx1",
                "Hdac2",
                "Smc3"
            ],
            "target": [],
            "expression_sum": {
                "class1": 9024.0,
                "class2": 20051.0
            },
            "uniprot_ids": [
                "Q99KJ8",
                "Q3TPZ5",
                "Q8BW43"
            ]
        },
        "Ddost": {
            "id": "Ddost",
            "type": "Gene",
            "symbol": "Ddost",
            "source": [
                "Nfib",
                "Fos",
                "Hdac2",
                "Tcf4",
                "Pou5f1",
                "Trp53",
                "Smarca5",
                "Arid5b",
                "Sox4",
                "Trim28",
                "Eno1",
                "Jund",
                "Cebpb",
                "Smc3",
                "Foxp1",
                "Atf4",
                "Ctcf",
                "Junb",
                "Ybx1",
                "Twist2"
            ],
            "target": [],
            "expression_sum": {
                "class1": 15249.0,
                "class2": 23435.0
            },
            "uniprot_ids": [
                "O54734",
                "Q3UC51",
                "Q3UG68"
            ]
        },
        "Ddx24": {
            "id": "Ddx24",
            "type": "Gene",
            "symbol": "Ddx24",
            "source": [
                "Tcf4",
                "Pou5f1",
                "Foxp1",
                "Cebpb",
                "Atf4",
                "Junb",
                "Fos",
                "Jun",
                "Jund",
                "Twist2",
                "Hdac2",
                "Sox4"
            ],
            "target": [],
            "expression_sum": {
                "class1": 7884.0,
                "class2": 26604.0
            },
            "uniprot_ids": [
                "Q9ESV0",
                "Q3U7R4",
                "Q3UYV7",
                "A0A1Y7VK75",
                "Q9CRZ6",
                "A0A1Y7VKX7",
                "F8WJA0",
                "A0A1Y7VLU4",
                "A0A1Y7VMS6",
                "Q3TLR1"
            ]
        },
        "Ddx39": {
            "id": "Ddx39",
            "type": "Gene",
            "symbol": "Ddx39",
            "source": [
                "Tcf4",
                "Pou5f1",
                "Foxp1",
                "Cebpb",
                "Nfib",
                "Atf4",
                "Junb",
                "Jun",
                "Nanog",
                "Jund",
                "Hdac2"
            ],
            "target": [],
            "expression_sum": {
                "class1": 9542.0,
                "class2": 48475.0
            },
            "uniprot_ids": [
                "Q8VDW0",
                "Q3TPL7",
                "Q3T9L0",
                "Q3UVN0",
                "Q3TIE4",
                "Q3U837",
                "D6RHT5",
                "Q3UAF1"
            ]
        },
        "Ddx46": {
            "id": "Ddx46",
            "type": "Gene",
            "symbol": "Ddx46",
            "source": [
                "Tcf4",
                "Pou5f1",
                "Foxp1",
                "Nfib",
                "Atf4",
                "Junb",
                "Jun",
                "Sox2",
                "Nanog",
                "Jund",
                "Twist2",
                "Cebpb",
                "Sox4"
            ],
            "target": [],
            "expression_sum": {
                "class1": 5451.0,
                "class2": 24869.0
            },
            "uniprot_ids": [
                "Q569Z5",
                "Q05C73",
                "A0A2R8VHK2",
                "Q3UUW9",
                "F8WHR6",
                "Q9CVD6",
                "Q3UZ56"
            ]
        },
        "Ddx5": {
            "id": "Ddx5",
            "type": "Gene",
            "symbol": "Ddx5",
            "source": [
                "Tcf4",
                "Pou5f1",
                "Foxp1",
                "Cebpb",
                "Nfib",
                "Atf4",
                "Junb",
                "Fos",
                "Jun",
                "Sox2",
                "Hes1",
                "Nanog",
                "Jund",
                "Hdac2",
                "Smc3",
                "Smarca5",
                "Sox4"
            ],
            "target": [],
            "expression_sum": {
                "class1": 33803.0,
                "class2": 74830.0
            },
            "uniprot_ids": [
                "Q61656",
                "Q8BTS0",
                "Q3UXW3",
                "Q9CSD9",
                "B1ARC0",
                "B1ARB9",
                "A1L333",
                "S4R1I6",
                "Q3V0Z8",
                "S4R1E3",
                "Q921Y3",
                "Q5U222"
            ]
        },
        "Denr": {
            "id": "Denr",
            "type": "Gene",
            "symbol": "Denr",
            "source": [
                "Tcf4",
                "Pou5f1",
                "Foxp1",
                "Cebpb",
                "Atf4",
                "Junb",
                "Prrx1",
                "Jun",
                "Sox2",
                "Hes1",
                "Nanog",
                "Jund",
                "Twist2",
                "Hdac2",
                "Smarca5",
                "Sox4"
            ],
            "target": [],
            "expression_sum": {
                "class1": 13416.0,
                "class2": 42943.0
            },
            "uniprot_ids": [
                "Q9CQJ6",
                "E9Q0G1"
            ]
        },
        "Dnaja1": {
            "id": "Dnaja1",
            "type": "Gene",
            "symbol": "Dnaja1",
            "source": [
                "Tcf4",
                "Pou5f1",
                "Foxp1",
                "Cebpb",
                "Atf4",
                "Junb",
                "Fos",
                "Jun",
                "Sox2",
                "Hes1",
                "Nanog",
                "Jund",
                "Twist2",
                "Hdac2",
                "Smc3",
                "Smarca5",
                "Sox4"
            ],
            "target": [],
            "expression_sum": {
                "class1": 25506.0,
                "class2": 66992.0
            },
            "uniprot_ids": [
                "P63037",
                "Q5NTY0",
                "B1AXY0",
                "Q3TK61",
                "B1AXX9",
                "B1AXY1"
            ]
        },
        "Dnaja2": {
            "id": "Dnaja2",
            "type": "Gene",
            "symbol": "Dnaja2",
            "source": [
                "Pou5f1",
                "Foxp1",
                "Cebpb",
                "Hmga2",
                "Nfib",
                "Atf4",
                "Junb",
                "Fos",
                "Jun",
                "Hes1",
                "Jund",
                "Twist2",
                "Hdac2",
                "Smc3",
                "Smarca5",
                "Sox4"
            ],
            "target": [],
            "expression_sum": {
                "class1": 12302.0,
                "class2": 28544.0
            },
            "uniprot_ids": [
                "Q9QYJ0",
                "Q3TFF0",
                "Q3V472"
            ]
        },
        "Dnajb6": {
            "id": "Dnajb6",
            "type": "Gene",
            "symbol": "Dnajb6",
            "source": [
                "Klf2",
                "Tcf4",
                "Pou5f1",
                "Foxp1",
                "Tfcp2l1",
                "Atf4",
                "Junb",
                "Fos",
                "Jun",
                "Sox2",
                "Hes1",
                "Nanog",
                "Jund",
                "Twist2",
                "Cebpb"
            ],
            "target": [],
            "expression_sum": {
                "class1": 12031.0,
                "class2": 54235.0
            },
            "uniprot_ids": [
                "O54946",
                "A0A0G2JER9",
                "G3X8S5",
                "A0A0G2JGN9",
                "A0A0G2JEI3",
                "F6YRQ2",
                "Q9DAN5"
            ]
        },
        "Dnajc7": {
            "id": "Dnajc7",
            "type": "Gene",
            "symbol": "Dnajc7",
            "source": [
                "Tcf4",
                "Pou5f1",
                "Nfib",
                "Atf4",
                "Junb",
                "Jun",
                "Nanog",
                "Jund",
                "Twist2",
                "Cebpb"
            ],
            "target": [],
            "expression_sum": {
                "class1": 4439.0,
                "class2": 19419.0
            },
            "uniprot_ids": [
                "Q9QYI3",
                "A0A6I8MWZ5",
                "A0A6I8MWW2",
                "A0A6I8MWY7",
                "A0A6I8MX31",
                "A2A5E1",
                "F7BTP8",
                "A0A6I8MWW0",
                "A0A6I8MWZ9",
                "A0A6I8MWZ1",
                "A0A6I8MWX8",
                "A0A6I8MX00",
                "Q3UL32"
            ]
        },
        "Dnajc8": {
            "id": "Dnajc8",
            "type": "Gene",
            "symbol": "Dnajc8",
            "source": [
                "Klf2",
                "Tcf4",
                "Pou5f1",
                "Foxp1",
                "Nfib",
                "Atf4",
                "Junb",
                "Fos",
                "Jun",
                "Sox2",
                "Nanog",
                "Tfcp2l1",
                "Jund",
                "Twist2",
                "Cebpb"
            ],
            "target": [],
            "expression_sum": {
                "class1": 14711.0,
                "class2": 63033.0
            },
            "uniprot_ids": [
                "Q6NZB0",
                "F6TQL3",
                "E9PZT4",
                "E9Q592",
                "D3Z4I5",
                "A2ALF0",
                "A2ALF3",
                "A2ALF2",
                "F6QIL6",
                "F7CXJ2"
            ]
        },
        "Dnttip2": {
            "id": "Dnttip2",
            "type": "Gene",
            "symbol": "Dnttip2",
            "source": [
                "Pou5f1",
                "Atf4",
                "Junb",
                "Jun",
                "Sox2",
                "Nanog",
                "Jund",
                "Twist2",
                "Cebpb",
                "Sox4"
            ],
            "target": [],
            "expression_sum": {
                "class1": 4591.0,
                "class2": 21229.0
            },
            "uniprot_ids": [
                "Q8R2M2"
            ]
        },
        "Dph3": {
            "id": "Dph3",
            "type": "Gene",
            "symbol": "Dph3",
            "source": [
                "Tcf4",
                "Pou5f1",
                "Hmga2",
                "Eno1",
                "Atf4",
                "Junb",
                "Fos",
                "Jun",
                "Id2",
                "Sox2",
                "Prrx2",
                "Nanog",
                "Tfcp2l1",
                "Jund",
                "Twist2",
                "Cebpb"
            ],
            "target": [],
            "expression_sum": {
                "class1": 9680.0,
                "class2": 32550.0
            },
            "uniprot_ids": [
                "Q8K0W9",
                "D3YXV3",
                "E9PZC5"
            ]
        },
        "Dpysl3": {
            "id": "Dpysl3",
            "type": "Gene",
            "symbol": "Dpysl3",
            "source": [
                "E2f4",
                "Trim28",
                "Rest",
                "Eno1",
                "Atf4",
                "Ctcf",
                "Trp53",
                "Hdac1",
                "Klf5",
                "Tbx3",
                "Zfp57",
                "Ybx1",
                "Hdac2",
                "Smc3"
            ],
            "target": [],
            "expression_sum": {
                "class1": 8260.0,
                "class2": 1566.0
            },
            "uniprot_ids": [
                "Q62188",
                "Q3TT92",
                "A0A494BBG3",
                "E9PWE8",
                "Q71H74",
                "Q3TAS8"
            ]
        },
        "Drg1": {
            "id": "Drg1",
            "type": "Gene",
            "symbol": "Drg1",
            "source": [
                "Tcf4",
                "Pou5f1",
                "Nfib",
                "Atf4",
                "Junb",
                "Fos",
                "Jun",
                "Sox2",
                "Hes1",
                "Nanog",
                "Jund",
                "Cebpb",
                "Sox4"
            ],
            "target": [],
            "expression_sum": {
                "class1": 5243.0,
                "class2": 21979.0
            },
            "uniprot_ids": [
                "P32233",
                "Q5NBZ3",
                "Q8BNS0",
                "D6RHA5",
                "Q8CHR8",
                "F7BQ65"
            ]
        },
        "Dstn": {
            "id": "Dstn",
            "type": "Gene",
            "symbol": "Dstn",
            "source": [
                "Tcf4",
                "Pou5f1",
                "Cebpb",
                "Arf4",
                "Nfib",
                "Atf4",
                "Fos",
                "Jun",
                "Sox2",
                "Nanog",
                "Jund",
                "Hdac2"
            ],
            "target": [],
            "expression_sum": {
                "class1": 43674.0,
                "class2": 68149.0
            },
            "uniprot_ids": [
                "Q9R0P5",
                "Q4FK36"
            ]
        },
        "Dtymk": {
            "id": "Dtymk",
            "type": "Gene",
            "symbol": "Dtymk",
            "source": [
                "Tcf4",
                "Pou5f1",
                "Arf4",
                "Nfib",
                "Atf4",
                "Junb",
                "Fos",
                "Jun",
                "Sox2",
                "Tbx3",
                "Nanog",
                "Jund",
                "Twist2",
                "Cebpb"
            ],
            "target": [],
            "expression_sum": {
                "class1": 14991.0,
                "class2": 94361.0
            },
            "uniprot_ids": [
                "P97930",
                "Q6GRA7",
                "D3Z3R3"
            ]
        },
        "Edf1": {
            "id": "Edf1",
            "type": "Gene",
            "symbol": "Edf1",
            "source": [
                "Klf2",
                "Tcf4",
                "Pou5f1",
                "Cebpb",
                "Eno1",
                "Atf4",
                "Fos",
                "Jun",
                "Sox2",
                "Hes1",
                "Nanog",
                "Jund",
                "Hdac2"
            ],
            "target": [],
            "expression_sum": {
                "class1": 26934.0,
                "class2": 71835.0
            },
            "uniprot_ids": [
                "Q9JMG1"
            ]
        },
        "Eef1a1": {
            "id": "Eef1a1",
            "type": "Gene",
            "symbol": "Eef1a1",
            "source": [
                "Pou5f1",
                "Foxp1",
                "Trim28",
                "Hmga2",
                "Nfib",
                "Atf4",
                "Junb",
                "Ctcf",
                "Trp53",
                "Sox2",
                "Prrx1",
                "Ybx1",
                "Twist2",
                "Hdac2",
                "Smc3",
                "Smarca5",
                "Sox4"
            ],
            "target": [],
            "expression_sum": {
                "class1": 519359.0,
                "class2": 1081201.0
            },
            "uniprot_ids": [
                "P10126",
                "Q58E64",
                "D3YZ68",
                "D3Z3I8",
                "Q3TII3",
                "Q4FZE5",
                "Q3UZQ3",
                "Q3UA81"
            ]
        },
        "Eef1b2": {
            "id": "Eef1b2",
            "type": "Gene",
            "symbol": "Eef1b2",
            "source": [
                "Pou5f1",
                "Cebpb",
                "Atf4",
                "Junb",
                "Trp53",
                "Sox2",
                "Nanog",
                "Twist2",
                "Hdac2"
            ],
            "target": [],
            "expression_sum": {
                "class1": 103144.0,
                "class2": 301164.0
            },
            "uniprot_ids": [
                "O70251",
                "A0A087WS46",
                "M0QW60",
                "M0QWK5",
                "G3UZ47",
                "G3UX43",
                "M0QWH8"
            ]
        },
        "Eef1d": {
            "id": "Eef1d",
            "type": "Gene",
            "symbol": "Eef1d",
            "source": [
                "Tcf4",
                "Pou5f1",
                "Cebpb",
                "Nfib",
                "Atf4",
                "Junb",
                "Fos",
                "Id2",
                "Jun",
                "Sox2",
                "Hes1",
                "Nanog",
                "Jund",
                "Twist2",
                "Hdac2",
                "Smc3",
                "Smarca5"
            ],
            "target": [],
            "expression_sum": {
                "class1": 42865.0,
                "class2": 126392.0
            },
            "uniprot_ids": [
                "P57776",
                "A0A0R4J1L2",
                "Q9CVV2",
                "Q91VK2",
                "A0A0R4J1E2",
                "Q80T06",
                "D3YY68",
                "F6ZFU0",
                "D3Z7N2",
                "D3YZT9",
                "D3Z356",
                "D3YUQ9",
                "E9QN08",
                "E9PUM6",
                "Q8BW50"
            ]
        },
        "Eif1": {
            "id": "Eif1",
            "type": "Gene",
            "symbol": "Eif1",
            "source": [
                "Tcf4",
                "Pou5f1",
                "Foxp1",
                "Cebpb",
                "Nfib",
                "Atf4",
                "Junb",
                "Id2",
                "Jun",
                "Nanog",
                "Tfcp2l1",
                "Twist2",
                "Hdac2",
                "Smc3",
                "Smarca5"
            ],
            "target": [],
            "expression_sum": {
                "class1": 140733.0,
                "class2": 296333.0
            },
            "uniprot_ids": [
                "P48024",
                "Q4V9T8",
                "J3QN87",
                "J3QPZ8",
                "Q3UBT8"
            ]
        },
        "Eif1a": {
            "id": "Eif1a",
            "type": "Gene",
            "symbol": "Eif1a",
            "source": [
                "Tcf4",
                "Pou5f1",
                "Foxp1",
                "Cebpb",
                "Nfib",
                "Atf4",
                "Junb",
                "Fos",
                "Jun",
                "Sox2",
                "Nanog",
                "Tfcp2l1",
                "Jund",
                "Hdac2"
            ],
            "target": [],
            "expression_sum": {
                "class1": 13703.0,
                "class2": 33672.0
            },
            "uniprot_ids": [
                "Q60872",
                "Q4FJR7",
                "Q3UC76",
                "A0A494BB48",
                "A0A494BAI7",
                "A0A494BB95"
            ]
        },
        "Eif2s2": {
            "id": "Eif2s2",
            "type": "Gene",
            "symbol": "Eif2s2",
            "source": [
                "Pou5f1",
                "Cebpb",
                "Nfib",
                "Eno1",
                "Atf4",
                "Junb",
                "Fos",
                "Id2",
                "Jun",
                "Sox2",
                "Nanog",
                "Jund",
                "Twist2",
                "Hdac2"
            ],
            "target": [],
            "expression_sum": {
                "class1": 45839.0,
                "class2": 286345.0
            },
            "uniprot_ids": [
                "Q99L45",
                "Q3ULL5",
                "E0CXJ3",
                "Q9CS47"
            ]
        },
        "Eif3a": {
            "id": "Eif3a",
            "type": "Gene",
            "symbol": "Eif3a",
            "source": [
                "Tcf4",
                "Pou5f1",
                "Foxp1",
                "Cebpb",
                "Atf4",
                "Junb",
                "Fos",
                "Id2",
                "Jun",
                "Sox2",
                "Nanog",
                "Jund",
                "Twist2",
                "Hdac2",
                "Smarca5"
            ],
            "target": [],
            "expression_sum": {
                "class1": 20678.0,
                "class2": 52275.0
            },
            "uniprot_ids": [
                "P23116",
                "Q3TRH1",
                "A0A494B9I0",
                "Q3TKF9",
                "Q3UEK3",
                "Q6A0B9",
                "Q3UUA4",
                "Q3UL99",
                "Q3UM46"
            ]
        },
        "Eif3c": {
            "id": "Eif3c",
            "type": "Gene",
            "symbol": "Eif3c",
            "source": [
                "Pou5f1",
                "Cebpb",
                "Nfib",
                "Atf5",
                "Eno1",
                "Atf4",
                "Junb",
                "Fos",
                "Jun",
                "Sox2",
                "Trp53",
                "Prrx1",
                "Jund",
                "Twist2",
                "Hdac2"
            ],
            "target": [],
            "expression_sum": {
                "class1": 25934.0,
                "class2": 47194.0
            },
            "uniprot_ids": [
                "Q8R1B4",
                "Q3TL08",
                "Q8BVX3",
                "Q9CV26",
                "M0QWV3",
                "Q9CSD3",
                "A0A0U1RPR0",
                "Q66JU1",
                "P97398"
            ]
        },
        "Eif3d": {
            "id": "Eif3d",
            "type": "Gene",
            "symbol": "Eif3d",
            "source": [
                "Tcf4",
                "Pou5f1",
                "Foxp1",
                "Cebpb",
                "Nfib",
                "Atf4",
                "Junb",
                "Fos",
                "Jun",
                "Sox2",
                "Nanog",
                "Jund",
                "Twist2",
                "Hdac2"
            ],
            "target": [],
            "expression_sum": {
                "class1": 10206.0,
                "class2": 40858.0
            },
            "uniprot_ids": [
                "O70194",
                "A0A2R8VK20"
            ]
        },
        "Eif3f": {
            "id": "Eif3f",
            "type": "Gene",
            "symbol": "Eif3f",
            "source": [
                "Pou5f1",
                "Trim28",
                "Hmgb1",
                "Atf4",
                "Junb",
                "Ctcf",
                "Trp53",
                "Hdac2",
                "Smc3",
                "Smarca5",
                "Sox4"
            ],
            "target": [],
            "expression_sum": {
                "class1": 47201.0,
                "class2": 112596.0
            },
            "uniprot_ids": [
                "Q9DCH4",
                "Q6IRT4",
                "Q8BTW2",
                "Q3UVY8",
                "Q3U8X1",
                "Q8VH52",
                "B0LAD0"
            ]
        },
        "Eif4a1": {
            "id": "Eif4a1",
            "type": "Gene",
            "symbol": "Eif4a1",
            "source": [
                "Klf2",
                "Nfib",
                "Fos",
                "Sox2",
                "Hdac2",
                "Tcf4",
                "Pou5f1",
                "Jun",
                "Smarca5",
                "Sox4",
                "Jund",
                "Cebpb",
                "Smc3",
                "Foxp1",
                "Atf4",
                "Junb",
                "Ybx1",
                "Hes1",
                "Nanog",
                "Twist2"
            ],
            "target": [],
            "expression_sum": {
                "class1": 76800.0,
                "class2": 323196.0
            },
            "uniprot_ids": [
                "P60843",
                "Q5F2A7",
                "Q64341",
                "Q3TFG3",
                "Q3TGK7",
                "Q78WR5",
                "Q3U8I0",
                "K3W4Q6",
                "Q3UXC2",
                "Q3TLL6",
                "Q4FZL1",
                "Q3TSJ4",
                "Q99LR0"
            ]
        },
        "Eif4a3": {
            "id": "Eif4a3",
            "type": "Gene",
            "symbol": "Eif4a3",
            "source": [
                "Tcf4",
                "Pou5f1",
                "Foxp1",
                "Nfib",
                "Atf4",
                "Junb",
                "Fos",
                "Jun",
                "Sox2",
                "Nanog",
                "Jund",
                "Cebpb"
            ],
            "target": [],
            "expression_sum": {
                "class1": 6166.0,
                "class2": 25806.0
            },
            "uniprot_ids": [
                "Q91VC3",
                "A2AFK7"
            ]
        },
        "Eif4b": {
            "id": "Eif4b",
            "type": "Gene",
            "symbol": "Eif4b",
            "source": [
                "Tcf4",
                "Pou5f1",
                "Foxp1",
                "Hmga2",
                "Atf4",
                "Junb",
                "Hes1",
                "Jund",
                "Cebpb"
            ],
            "target": [],
            "expression_sum": {
                "class1": 7284.0,
                "class2": 26944.0
            },
            "uniprot_ids": [
                "Q8BGD9",
                "B2RWE8",
                "Q3TSY9",
                "Q922K6",
                "Q3TDD8",
                "Q3TLQ9",
                "Q3THB0",
                "Q3UGC0"
            ]
        },
        "Eif4e": {
            "id": "Eif4e",
            "type": "Gene",
            "symbol": "Eif4e",
            "source": [
                "Pou5f1",
                "Cebpb",
                "Nfib",
                "Atf4",
                "Junb",
                "Fos",
                "Jun",
                "Nanog",
                "Jund",
                "Twist2",
                "Hdac2",
                "Smc3"
            ],
            "target": [],
            "expression_sum": {
                "class1": 12113.0,
                "class2": 29646.0
            },
            "uniprot_ids": [
                "P63073",
                "A0A0G2JGT5",
                "Q8C470",
                "A0A0G2JG98",
                "A0A0G2JH04",
                "Q3V0N9",
                "A0A0G2JFB4",
                "Q3TK95"
            ]
        },
        "Eif4e2": {
            "id": "Eif4e2",
            "type": "Gene",
            "symbol": "Eif4e2",
            "source": [
                "Klf2",
                "Nfib",
                "Fos",
                "Sox2",
                "Hdac2",
                "Mybl2",
                "Tcf4",
                "Pou5f1",
                "Jun",
                "Sox4",
                "Eno1",
                "Jund",
                "Cebpb",
                "Foxp1",
                "Atf4",
                "Junb",
                "Hes1",
                "Nanog",
                "Twist2"
            ],
            "target": [],
            "expression_sum": {
                "class1": 14169.0,
                "class2": 57293.0
            },
            "uniprot_ids": [
                "Q8BMB3",
                "Q3UUT0",
                "Q0P688",
                "D3YUV9",
                "M0QWD1",
                "D3Z729",
                "D3Z730",
                "G3X9H1",
                "Q9D334",
                "Q80ZJ3"
            ]
        },
        "Eif4ebp1": {
            "id": "Eif4ebp1",
            "type": "Gene",
            "symbol": "Eif4ebp1",
            "source": [
                "Tcf4",
                "Pou5f1",
                "Arf4",
                "Atf4",
                "Junb",
                "Fos",
                "Jun",
                "Sox2",
                "Hes1",
                "Nanog",
                "Jund",
                "Twist2",
                "Cebpb"
            ],
            "target": [],
            "expression_sum": {
                "class1": 49232.0,
                "class2": 176913.0
            },
            "uniprot_ids": [
                "Q60876",
                "Q3TII9"
            ]
        },
        "Eif4g1": {
            "id": "Eif4g1",
            "type": "Gene",
            "symbol": "Eif4g1",
            "source": [
                "Tcf4",
                "Pou5f1",
                "Cebpb",
                "Nfib",
                "Atf4",
                "Junb",
                "Fos",
                "Jun",
                "Sox2",
                "Trp53",
                "Ybx1",
                "Hes1",
                "Tfcp2l1",
                "Jund",
                "Twist2",
                "Hdac2",
                "Smc3"
            ],
            "target": [],
            "expression_sum": {
                "class1": 9469.0,
                "class2": 24597.0
            },
            "uniprot_ids": [
                "Q6NZJ6",
                "D6RCH1",
                "Q6EVH9",
                "D3YWY7",
                "D3YWC6",
                "E9Q770",
                "A0A0J9YU44",
                "D3YWM1",
                "A0A0J9YTV0",
                "A0A0J9YUL1",
                "A0A0J9YVC1",
                "Q8R2V4",
                "Q3UYM6",
                "E9PVC6",
                "E9Q9E1",
                "Q8K0J8",
                "Q3UZD9",
                "Q6PD42",
                "D3Z439",
                "A0A338P7C5",
                "Q8BUW8",
                "A0A0J9YUS5",
                "A0A338P758",
                "Q3UQI5",
                "A0PJB5",
                "Q3UTK0",
                "E9PVC5",
                "A0A0J9YUC1"
            ]
        },
        "Eif4g2": {
            "id": "Eif4g2",
            "type": "Gene",
            "symbol": "Eif4g2",
            "source": [
                "Nfib",
                "Fos",
                "Sox2",
                "Hdac2",
                "Tcf4",
                "Pou5f1",
                "Trp53",
                "Jun",
                "Smarca5",
                "Sox4",
                "Eno1",
                "Tgif1",
                "Jund",
                "Cebpb",
                "Smc3",
                "Hmga2",
                "Foxp1",
                "Hmgb1",
                "Atf4",
                "Junb",
                "Ybx1",
                "Hes1",
                "Twist2"
            ],
            "target": [],
            "expression_sum": {
                "class1": 32841.0,
                "class2": 51455.0
            },
            "uniprot_ids": [
                "Q62448",
                "Q3U9U9",
                "A0JNY7",
                "F6UU81",
                "F6TW20",
                "F7CBP1",
                "F7DAB1",
                "G3XA17",
                "F7CHS2",
                "Q3URI0"
            ]
        },
        "Eif5": {
            "id": "Eif5",
            "type": "Gene",
            "symbol": "Eif5",
            "source": [
                "Tcf4",
                "Pou5f1",
                "Foxp1",
                "Cebpb",
                "Nfib",
                "Atf4",
                "Junb",
                "Fos",
                "Jun",
                "Sox2",
                "Trp53",
                "Hes1",
                "Nanog",
                "Jund",
                "Twist2",
                "Hdac2",
                "Smc3",
                "Smarca5"
            ],
            "target": [],
            "expression_sum": {
                "class1": 23805.0,
                "class2": 59737.0
            },
            "uniprot_ids": [
                "P59325",
                "Q8BVV6",
                "Q3TQR3",
                "Q05C65",
                "Q3TKY2",
                "A0A1Y7VLK2",
                "A0A1Y7VNP4",
                "A0A1Y7VKT5",
                "A0A1Y7VLL6",
                "A0A1Y7VJG2",
                "Q05CH7",
                "A0A1Y7VLQ0",
                "Q05CM5",
                "Q6PHA6",
                "Q3TT65",
                "Q05CY6"
            ]
        },
        "Eif5b": {
            "id": "Eif5b",
            "type": "Gene",
            "symbol": "Eif5b",
            "source": [
                "Tcf4",
                "Pou5f1",
                "Cebpb",
                "Nfib",
                "Atf4",
                "Junb",
                "Nr0b1",
                "Fos",
                "Jun",
                "Sox2",
                "Nanog",
                "Jund",
                "Twist2",
                "Hdac2",
                "Smc3",
                "Smarca5",
                "Sox4"
            ],
            "target": [],
            "expression_sum": {
                "class1": 15551.0,
                "class2": 43986.0
            },
            "uniprot_ids": [
                "Q05D44",
                "Q8BYX3",
                "Q7TPC0",
                "Q8BYY7",
                "Q6A002"
            ]
        },
        "Emc7": {
            "id": "Emc7",
            "type": "Gene",
            "symbol": "Emc7",
            "source": [
                "E2f4",
                "Trim28",
                "Rest",
                "Atf4",
                "Ctcf",
                "Trp53",
                "Ybx1",
                "Hdac2",
                "Smarca5"
            ],
            "target": [],
            "expression_sum": {
                "class1": 10321.0,
                "class2": 9995.0
            },
            "uniprot_ids": [
                "Q9EP72",
                "Q14C26",
                "Q91VV9"
            ]
        },
        "Emc8": {
            "id": "Emc8",
            "type": "Gene",
            "symbol": "Emc8",
            "source": [
                "Tcf4",
                "Pou5f1",
                "Foxp1",
                "Atf4",
                "Fos",
                "Jun",
                "Sox2",
                "Hes1",
                "Nanog",
                "Jund",
                "Twist2",
                "Cebpb",
                "Sox4"
            ],
            "target": [],
            "expression_sum": {
                "class1": 6497.0,
                "class2": 52692.0
            },
            "uniprot_ids": [
                "O70378",
                "Q3TPU5",
                "M0QWI7",
                "M0QWC9",
                "M0QWP2",
                "M0QWY0",
                "M0QWS0",
                "Q3TTC9"
            ]
        },
        "Emp1": {
            "id": "Emp1",
            "type": "Gene",
            "symbol": "Emp1",
            "source": [
                "E2f4",
                "Pou5f1",
                "Trim28",
                "Atf4",
                "Ctcf",
                "Trp53",
                "Hdac1",
                "Klf5",
                "Tbx3",
                "Hdac2",
                "Smc3"
            ],
            "target": [],
            "expression_sum": {
                "class1": 20378.0,
                "class2": 2156.0
            },
            "uniprot_ids": [
                "P47801",
                "Q4FK43",
                "S4R2U4",
                "Q05BR0",
                "Q3UXM2"
            ]
        },
        "Emp3": {
            "id": "Emp3",
            "type": "Gene",
            "symbol": "Emp3",
            "source": [
                "Klf2",
                "E2f4",
                "Pou5f1",
                "Cebpb",
                "Klf4",
                "Trim28",
                "Hmgb1",
                "Atf4",
                "Ctcf",
                "Trp53",
                "Sox2",
                "Klf5",
                "Tbx3",
                "Nanog",
                "Hdac2",
                "Zfp57"
            ],
            "target": [],
            "expression_sum": {
                "class1": 24895.0,
                "class2": 4944.0
            },
            "uniprot_ids": [
                "O35912",
                "A0A1B0GSZ1"
            ]
        },
        "Eprs": {
            "id": "Eprs",
            "type": "Gene",
            "symbol": "Eprs",
            "source": [
                "Tcf4",
                "Pou5f1",
                "Foxp1",
                "Cebpb",
                "Nfib",
                "Atf4",
                "Junb",
                "Fos",
                "Sox2",
                "Hes1",
                "Jund",
                "Twist2",
                "Hdac2",
                "Smc3",
                "Smarca5"
            ],
            "target": [],
            "expression_sum": {
                "class1": 17016.0,
                "class2": 31467.0
            },
            "uniprot_ids": [
                "Q8CGC7",
                "Q3TMX9",
                "B9EIU1",
                "Q8BTR1",
                "A0A0A6YWH3",
                "Q3UGD4",
                "Q9CRF9",
                "A0A0A6YWA4",
                "Q3TLK2"
            ]
        },
        "Ergic3": {
            "id": "Ergic3",
            "type": "Gene",
            "symbol": "Ergic3",
            "source": [
                "Trim28",
                "Rest",
                "Hmgb1",
                "Atf4",
                "Ctcf",
                "Trp53",
                "Hdac1",
                "Hdac2",
                "Smc3",
                "Smarca5"
            ],
            "target": [],
            "expression_sum": {
                "class1": 9222.0,
                "class2": 17135.0
            },
            "uniprot_ids": [
                "Q9CQE7",
                "F6UIS1",
                "F6RK81"
            ]
        },
        "Erp29": {
            "id": "Erp29",
            "type": "Gene",
            "symbol": "Erp29",
            "source": [
                "Pou5f1",
                "Foxp1",
                "Cebpb",
                "Nfib",
                "Atf4",
                "Junb",
                "Fos",
                "Id2",
                "Jun",
                "Trp53",
                "Jund",
                "Twist2",
                "Hdac2",
                "Smc3",
                "Smarca5",
                "Sox4"
            ],
            "target": [],
            "expression_sum": {
                "class1": 18925.0,
                "class2": 34274.0
            },
            "uniprot_ids": [
                "P57759",
                "F8WJI4",
                "F8WIM7",
                "D6RG87",
                "A8C1V9"
            ]
        },
        "Etf1": {
            "id": "Etf1",
            "type": "Gene",
            "symbol": "Etf1",
            "source": [
                "Klf2",
                "Tcf4",
                "Pou5f1",
                "Foxp1",
                "Atf4",
                "Junb",
                "Fos",
                "Jun",
                "Sox2",
                "Nanog",
                "Jund",
                "Twist2",
                "Cebpb"
            ],
            "target": [],
            "expression_sum": {
                "class1": 10009.0,
                "class2": 38308.0
            },
            "uniprot_ids": [
                "Q8BWY3",
                "Q3U6V5",
                "Q3TL10",
                "Q3TDF8",
                "P70199",
                "Q3TF02"
            ]
        },
        "Ewsr1": {
            "id": "Ewsr1",
            "type": "TF",
            "symbol": "Ewsr1",
            "source": [
                "Tcf4",
                "Pou5f1",
                "Cebpb",
                "Atf4",
                "Junb",
                "Fos",
                "Jun",
                "Sox2",
                "Hes1",
                "Nanog",
                "Jund",
                "Twist2",
                "Hdac2",
                "Smc3",
                "Smarca5"
            ],
            "target": [
                "Rnase4",
                "Mid1ip1",
                "Creb3l1",
                "Ptprn",
                "Cst3",
                "Dnajc1",
                "Mff",
                "Golt1b",
                "Zwint",
                "Pja1",
                "Khdrbs1",
                "1110004F10Rik",
                "Gars",
                "2310009B15Rik",
                "Mecr",
                "Ociad1"
            ],
            "expression_sum": {
                "class1": 10924.0,
                "class2": 31547.0
            },
            "uniprot_ids": [
                "Q61545",
                "Q8BQU4",
                "Q6NVA3",
                "Q3UI30",
                "Q9CRS5",
                "Q3UGI7",
                "Q5SUS8",
                "Q5SUS9",
                "Q5SUT0"
            ]
        },
        "Exosc4": {
            "id": "Exosc4",
            "type": "Gene",
            "symbol": "Exosc4",
            "source": [
                "Tcf4",
                "Pou5f1",
                "Atf4",
                "Junb",
                "Id2",
                "Jun",
                "Sox2",
                "Nanog",
                "Jund",
                "Cebpb",
                "Sox4"
            ],
            "target": [],
            "expression_sum": {
                "class1": 4238.0,
                "class2": 23398.0
            },
            "uniprot_ids": [
                "Q921I9",
                "Q542B0",
                "A0A2R8VHM9"
            ]
        },
        "Exosc8": {
            "id": "Exosc8",
            "type": "Gene",
            "symbol": "Exosc8",
            "source": [
                "Tcf4",
                "Pou5f1",
                "Foxp1",
                "Cebpb",
                "Nfib",
                "Atf4",
                "Junb",
                "Id2",
                "Jun",
                "Sox2",
                "Hes1",
                "Nanog",
                "Jund",
                "Twist2",
                "Hdac2",
                "Smc3",
                "Smarca5",
                "Sox4"
            ],
            "target": [],
            "expression_sum": {
                "class1": 6695.0,
                "class2": 20748.0
            },
            "uniprot_ids": [
                "Q9D753",
                "Q546F8",
                "D3YYN3",
                "F6SGT4",
                "Q3TEB9",
                "Q6NX62",
                "Q6PCW8",
                "Q571G2"
            ]
        },
        "F2r": {
            "id": "F2r",
            "type": "Gene",
            "symbol": "F2r",
            "source": [
                "Trim28",
                "Rest",
                "Atf4",
                "Ctcf",
                "Ybx1",
                "Hdac2",
                "Smc3"
            ],
            "target": [],
            "expression_sum": {
                "class1": 8281.0,
                "class2": 1827.0
            },
            "uniprot_ids": [
                "P30558",
                "Q3UK34"
            ]
        },
        "Fabp5": {
            "id": "Fabp5",
            "type": "Gene",
            "symbol": "Fabp5",
            "source": [
                "E2f4",
                "Trim28",
                "Rest",
                "Ctbp1",
                "Hmgb1",
                "Atf4",
                "Ctcf",
                "Trp53",
                "Hdac1",
                "Klf5",
                "Prrx1",
                "Ybx1",
                "Hdac2",
                "Smc3"
            ],
            "target": [],
            "expression_sum": {
                "class1": 50769.0,
                "class2": 8692.0
            },
            "uniprot_ids": [
                "Q05816",
                "Q497I3",
                "Q3TLH6"
            ]
        },
        "Fam162a": {
            "id": "Fam162a",
            "type": "Gene",
            "symbol": "Fam162a",
            "source": [
                "E2f4",
                "Trim28",
                "Rest",
                "Atf4",
                "Ctcf",
                "Junb",
                "Tgif1",
                "Hdac1",
                "Tbx3",
                "Trp53",
                "Hdac2",
                "Smc3",
                "Smarca5"
            ],
            "target": [],
            "expression_sum": {
                "class1": 15120.0,
                "class2": 34006.0
            },
            "uniprot_ids": [
                "Q9D6U8",
                "A0A338P6X2"
            ]
        },
        "Fam195b": {
            "id": "Fam195b",
            "type": "Gene",
            "symbol": "Fam195b",
            "source": [
                "E2f4",
                "Rest",
                "Eno1",
                "Atf4",
                "Ctcf",
                "Tgif1",
                "Trp53",
                "Hdac1",
                "Hdac2",
                "Smarca5"
            ],
            "target": [],
            "expression_sum": {
                "class1": 9603.0,
                "class2": 6586.0
            },
            "uniprot_ids": [
                "Q3UGS4"
            ]
        },
        "Fau": {
            "id": "Fau",
            "type": "Gene",
            "symbol": "Fau",
            "source": [
                "Pou5f1",
                "Trim28",
                "Atf4",
                "Ctcf",
                "Trp53",
                "Sox2",
                "Ybx1",
                "Nanog",
                "Jund",
                "Hdac2",
                "Zfp57"
            ],
            "target": [],
            "expression_sum": {
                "class1": 990.0,
                "class2": 451628.0
            },
            "uniprot_ids": [
                "P62862",
                "P35545",
                "Q642K5",
                "A0A494BBA8",
                "Q3THB2",
                "A0A494B9Z0"
            ]
        },
        "Filip1l": {
            "id": "Filip1l",
            "type": "Gene",
            "symbol": "Filip1l",
            "source": [
                "Hdac2",
                "E2f4",
                "Pou5f1",
                "Trim28",
                "Rest",
                "Atf4",
                "Ctcf",
                "Trp53",
                "Hdac1",
                "Tbx3",
                "Zfp57",
                "Cebpb",
                "Smc3"
            ],
            "target": [],
            "expression_sum": {
                "class1": 9833.0,
                "class2": 581.0
            },
            "uniprot_ids": [
                "Q6P6L0",
                "E0CYM1",
                "E0CYH7"
            ]
        },
        "Fkbp1a": {
            "id": "Fkbp1a",
            "type": "Gene",
            "symbol": "Fkbp1a",
            "source": [
                "Pou5f1",
                "Foxp1",
                "Arf4",
                "Atf4",
                "Ctcf",
                "Junb",
                "Id2",
                "Hdac1",
                "Jun",
                "Trp53",
                "Prrx1",
                "Twist2",
                "Hdac2",
                "Smc3",
                "Smarca5"
            ],
            "target": [],
            "expression_sum": {
                "class1": 53186.0,
                "class2": 83794.0
            },
            "uniprot_ids": [
                "P26883",
                "F6X9I3",
                "Q1JUQ8",
                "Q1JUQ7",
                "Q3ULN5",
                "Q3UBA0",
                "Q3UKJ3",
                "Q1JUQ6"
            ]
        },
        "Fkbp2": {
            "id": "Fkbp2",
            "type": "Gene",
            "symbol": "Fkbp2",
            "source": [
                "Tcf4",
                "Trim28",
                "Cebpb",
                "Arf4",
                "Eno1",
                "Atf4",
                "Ctcf",
                "Junb",
                "Trp53",
                "Hdac1",
                "Jund",
                "Hdac2",
                "Smc3",
                "Smarca5"
            ],
            "target": [],
            "expression_sum": {
                "class1": 9384.0,
                "class2": 9613.0
            },
            "uniprot_ids": [
                "P45878",
                "A0A494B9J3",
                "Q3TND1",
                "Q3UND3",
                "A0A494BAX7"
            ]
        },
        "Fkbp3": {
            "id": "Fkbp3",
            "type": "Gene",
            "symbol": "Fkbp3",
            "source": [
                "Tcf4",
                "Pou5f1",
                "Foxp1",
                "Tfcp2l1",
                "Nfib",
                "Atf4",
                "Junb",
                "Fos",
                "Jun",
                "Sox2",
                "Hes1",
                "Nanog",
                "Jund",
                "Twist2",
                "Cebpb",
                "Sox4"
            ],
            "target": [],
            "expression_sum": {
                "class1": 22510.0,
                "class2": 153842.0
            },
            "uniprot_ids": [
                "Q62446",
                "Q3UBU9",
                "A0A1Y7VP01",
                "A0A1Y7VLK0",
                "A0A1Y7VLV3",
                "A0A1Y7VMJ9",
                "A0A1Y7VJ86"
            ]
        },
        "Fkbp4": {
            "id": "Fkbp4",
            "type": "Gene",
            "symbol": "Fkbp4",
            "source": [
                "Tcf4",
                "Pou5f1",
                "Foxp1",
                "Hmga2",
                "Atf4",
                "Junb",
                "Fos",
                "Jun",
                "Sox2",
                "Hes1",
                "Nanog",
                "Jund",
                "Twist2",
                "Cebpb",
                "Zfp57",
                "Sox4"
            ],
            "target": [],
            "expression_sum": {
                "class1": 12551.0,
                "class2": 120503.0
            },
            "uniprot_ids": [
                "P30416",
                "F6S2D5",
                "F7CAT1",
                "Q8CBS1"
            ]
        },
        "Fkbp8": {
            "id": "Fkbp8",
            "type": "Gene",
            "symbol": "Fkbp8",
            "source": [
                "Pou5f1",
                "Foxp1",
                "Cebpb",
                "Hmga2",
                "Atf4",
                "Junb",
                "Fos",
                "Sox2",
                "Trp53",
                "Tfcp2l1",
                "Twist2",
                "Hdac2",
                "Smc3"
            ],
            "target": [],
            "expression_sum": {
                "class1": 10961.0,
                "class2": 30485.0
            },
            "uniprot_ids": [
                "O35465",
                "F6WP10",
                "D3Z597",
                "C9K0Z7"
            ]
        },
        "Flna": {
            "id": "Flna",
            "type": "Gene",
            "symbol": "Flna",
            "source": [
                "E2f4",
                "Rest",
                "Atf4",
                "Ctcf",
                "Trp53",
                "Hdac1",
                "Klf5",
                "Tbx3",
                "Hdac2",
                "Smc3",
                "Smarca5"
            ],
            "target": [],
            "expression_sum": {
                "class1": 14862.0,
                "class2": 7797.0
            },
            "uniprot_ids": [
                "Q8BTM8",
                "B7FAV1",
                "J3JS91",
                "B7FAU9",
                "F6Z2C0",
                "F6XC15",
                "A0A668KLG7",
                "F7AVL7",
                "Q9JJ37",
                "I7HPX4",
                "Q6KAM8",
                "Q3U7N9",
                "B9EKP5",
                "Q4FZG4"
            ]
        },
        "Fn1": {
            "id": "Fn1",
            "type": "Gene",
            "symbol": "Fn1",
            "source": [
                "Hdac2",
                "E2f4",
                "Trim28",
                "Rest",
                "Atf4",
                "Ctcf",
                "Trp53",
                "Hdac1",
                "Zfp57",
                "Ybx1",
                "Cebpb",
                "Smc3",
                "Smarca5"
            ],
            "target": [],
            "expression_sum": {
                "class1": 29826.0,
                "class2": 27493.0
            },
            "uniprot_ids": [
                "P11276",
                "B9EHT6",
                "B7ZNJ1",
                "Q4KL80",
                "Q3UHL6",
                "A0A087WS56",
                "A0A087WQE0",
                "A0A087WSN6",
                "A0A087WSU6",
                "A0A087WS99",
                "A0A087WR50",
                "A0A087WQW8",
                "Q3UZF9",
                "Q3UHR1",
                "Q3TCL7",
                "Q8R3F3",
                "Q3UGY5",
                "Q3TCF1",
                "Q99KD0",
                "Q3TBB4",
                "Q3UH17"
            ]
        },
        "Foxp1": {
            "id": "Foxp1",
            "type": "TF",
            "symbol": "Foxp1",
            "source": [
                "Hdac2",
                "E2f4",
                "Trim28",
                "Pou5f1",
                "Rest",
                "Atf4",
                "Ctcf",
                "Tgif1",
                "Trp53",
                "Hdac1",
                "Klf5",
                "Cebpb",
                "Smc3",
                "Smarca5"
            ],
            "target": [
                "Rps5",
                "Hsp90aa1",
                "Arf1",
                "Rpl41",
                "Rnf7",
                "Cdc20",
                "Sar1a",
                "Srsf9",
                "Usmg5",
                "Mcm7",
                "Ppp2ca",
                "Ddx24",
                "Rps17",
                "Rae1",
                "Zc3h15",
                "Cox8a",
                "Mrps18b",
                "Cdt1",
                "Psmb7",
                "Txn2",
                "Pnkd",
                "Dkc1",
                "Ndufa6",
                "Ppp1r11",
                "1110008F13Rik",
                "Tuba1b",
                "Polr2k",
                "Gar1",
                "Anapc5",
                "St13",
                "Thoc7",
                "Hnrnpf",
                "Knop1",
                "Lmnb1",
                "Ddx39",
                "Mlf2",
                "Pin1",
                "Ier3ip1",
                "Sfxn1",
                "Swi5",
                "Eif2s1",
                "mt-Co1",
                "Hmgb2",
                "Slbp",
                "Calr",
                "Nup62",
                "Rab18",
                "Calm3",
                "Arf6",
                "Canx",
                "Snrpe",
                "Mrpl14",
                "Cdk1",
                "Phb",
                "Lamtor1",
                "Anapc15",
                "Mrpl2",
                "Tra2b",
                "Cdc5l",
                "Ranbp1",
                "Psmc6",
                "Grpel1",
                "Rbx1",
                "Ctcf",
                "Ppm1g",
                "Eif1",
                "Neat1",
                "Ythdc1",
                "Bola1",
                "Ola1",
                "Sf3b6",
                "Snrnp27",
                "Aurkaip1",
                "Srsf1",
                "Socs3",
                "Lrrc59",
                "Exosc5",
                "Rcc2",
                "Pdcd6",
                "Impa2",
                "Ube2d2a",
                "Bnip3l",
                "Mapre1",
                "Cisd2",
                "Capza2",
                "Lap3",
                "Ncl",
                "Fdft1",
                "Supt4a",
                "Ppp1ca",
                "Srsf5",
                "Hnrnph1",
                "Aamp",
                "Chchd1",
                "Ywhaz",
                "Psmc3",
                "Sec62",
                "Luc7l2",
                "Rbbp7",
                "Hnrnpl",
                "Ddx5",
                "Rbm39",
                "Ccdc12",
                "Nedd8",
                "Pcnp",
                "Ube2i",
                "Utp3",
                "Rab14",
                "Tmem258",
                "Tcp1",
                "Brix1",
                "Fam136a",
                "Pttg1",
                "Mrpl15",
                "Siva1",
                "Fus",
                "Ociad1",
                "Ddost",
                "Pmm1",
                "Fkbp8",
                "Nip7",
                "Clpp",
                "Psma7",
                "G3bp1",
                "Pdia4",
                "Sept7",
                "Tpm3",
                "Banf1",
                "Cdc26",
                "Immt",
                "Psma1",
                "Qdpr",
                "Lig1",
                "2410015M20Rik",
                "Hdgf",
                "Prnp",
                "Smc3",
                "Thrap3",
                "Csnk2b",
                "Pa2g4",
                "Rpl37",
                "Kdm5b",
                "Ppp1cc",
                "Atf4",
                "Zfp207",
                "Pcbp1",
                "Hnrnpk",
                "Ndufa4",
                "Ndufv1",
                "Ndufv2",
                "H3f3a",
                "Mettl9",
                "Nme1",
                "Pfkl",
                "Csnk1a1",
                "Ndufs2",
                "Fam49b",
                "Bax",
                "Tmsb10",
                "Rpl3",
                "Cebpzos",
                "Trim28",
                "Snw1",
                "Cdv3",
                "Napa",
                "Spc24",
                "Btf3l4",
                "Ppp4c",
                "Etf1",
                "Ube2b",
                "Lsm3",
                "Gm16286",
                "Eif5",
                "Tmed2",
                "Txn1",
                "Vapa",
                "Mrpl17",
                "Samm50",
                "Rrp15",
                "Wdr77",
                "Atp5j",
                "Ran",
                "Fscn1",
                "Srrm1",
                "Emc8",
                "Srp19",
                "Cct5",
                "Emd",
                "Mrpl40",
                "Bclaf1",
                "Vasp",
                "Tceb1",
                "C1qbp",
                "Uqcr10",
                "H2afz",
                "Tardbp",
                "Psip1",
                "Rpl36a",
                "Rpl29",
                "Iscu",
                "Fubp1",
                "Snrpb2",
                "Arpc5l",
                "Rps29",
                "Ccdc58",
                "Cenpa",
                "Prdx6",
                "Wtap",
                "Odc1",
                "Mrpl18",
                "Rps20",
                "Dnaja2",
                "Sys1",
                "Acat2",
                "Lbh",
                "Cdc37",
                "Psmd1",
                "Plekhf2",
                "Eef1a1",
                "Mcl1",
                "Rtn4",
                "U2af1",
                "Atp5c1",
                "Ywhaq",
                "Spcs1",
                "Fbl",
                "Ndufv3",
                "Ubald2",
                "Tsc22d1",
                "mt-Co2",
                "Dnajc8",
                "Calm2",
                "Rps3",
                "Atox1",
                "Srsf10",
                "Adss",
                "Tmem126a",
                "Ube2v2",
                "Rpl18",
                "Trnau1ap",
                "Atp5d",
                "Timm13",
                "Eif3h",
                "Rpl22l1",
                "Adprh",
                "Lamtor4",
                "Rps3a1",
                "Mrps5",
                "U2surp",
                "Mbd3",
                "Cct7",
                "Dctpp1",
                "Ndufb2",
                "Wdr18",
                "Hsp90ab1",
                "Oaz1",
                "Mrpl30",
                "Uqcrq",
                "Sec61b",
                "BC031181",
                "Polr2m",
                "1810058I24Rik",
                "Rwdd1",
                "Snx5",
                "Got1",
                "Fnta",
                "Dnaja1",
                "Eif4a1",
                "Sumo2",
                "Pcbp2",
                "Snrnp25",
                "Nudc",
                "H3f3b",
                "Mrps33",
                "Rif1",
                "Nsa2",
                "Ddx46",
                "Nudt19",
                "Dbi",
                "Morf4l2",
                "Ube2d3",
                "Rbbp4",
                "Eif3e",
                "Paip2",
                "Cebpb",
                "Srrm2",
                "Rnaseh2b",
                "Ywhae",
                "Ubl5",
                "Hmgb1",
                "Ndufc1",
                "Eif3d",
                "Rps15",
                "Npm1",
                "Psma4",
                "Sap30",
                "Scp2",
                "Ap2s1",
                "Rpl34",
                "Mlec",
                "Dynll1",
                "Gtpbp4",
                "Cenpw",
                "Hspd1",
                "Atf5",
                "Cops7a",
                "Mrpl45",
                "Mrps14",
                "Rpl31",
                "Srsf7",
                "Smdt1",
                "Sod1",
                "Ppih",
                "Mat2a",
                "Nap1l1",
                "Uqcc3",
                "Dnajb6",
                "Brd2",
                "Actr3",
                "G3bp2",
                "Impdh2",
                "Ccdc124",
                "Gapdh",
                "Smarca5",
                "Mpc1",
                "Prps1",
                "Ccnd3",
                "Ap2m1",
                "Mrpl52",
                "Adrm1",
                "Abcf1",
                "Rplp0",
                "Psmb5",
                "Hspa9",
                "Pim3",
                "Ube2n",
                "Ndufa11",
                "Cct8",
                "Hmgn2",
                "Mpc2",
                "Dohh",
                "Hnrnpa2b1",
                "Ndufs5",
                "Cited2",
                "Srsf2",
                "Eif4g2",
                "Jade1",
                "Psmb2",
                "Raly",
                "Denr",
                "Fxr1",
                "Nsmce4a",
                "Rps26",
                "Get4",
                "Trp53",
                "Sra1",
                "Aspscr1",
                "Psmd11",
                "Vps29",
                "Vdac1",
                "Cggbp1",
                "Zfp706",
                "Anp32a",
                "Polr2e",
                "Anapc13",
                "Sumo1",
                "Pdcd5",
                "Snrpc",
                "Mrpl51",
                "Mrpl49",
                "Tmbim6",
                "Dynll2",
                "Rhoa",
                "Fcf1",
                "Mrps18c",
                "Cacybp",
                "Cdk4",
                "Rpl22",
                "Rsl1d1",
                "Nolc1",
                "Sf3a3",
                "Sec61a1",
                "Pfdn6",
                "Glud1",
                "Cebpz",
                "Hnrnpu",
                "Gspt1",
                "Arglu1",
                "Tomm20",
                "Rpl21",
                "Ppp1r2",
                "Map1lc3b",
                "Eprs",
                "Trmt112",
                "Dazap1",
                "Son",
                "Pop5",
                "Cct2",
                "Ubb",
                "Dtl",
                "Bzw1",
                "Parp1",
                "Dhfr",
                "Rbpms",
                "Paics",
                "Hnrnpa3",
                "Prpf40a",
                "Pbdc1",
                "Cdca3",
                "Mrpl57",
                "Morf4l1",
                "Mrpl43",
                "Ube2s",
                "Tecr",
                "Polr1d",
                "Snrpd3",
                "Sf3b2",
                "Psmd3",
                "Suclg1",
                "Mtf2",
                "Slc3a2",
                "mt-Nd2",
                "Hnrnpd",
                "Hat1",
                "Cops4",
                "mt-Cytb",
                "Rnps1",
                "Ciapin1",
                "Lsm4",
                "Pfdn2",
                "Mpdu1",
                "Atp5f1",
                "Eif1ad",
                "Rps13",
                "Tpt1",
                "Eif5a",
                "Ppib",
                "Gng5",
                "Hspe1",
                "Rps19",
                "Exosc8",
                "Cops6",
                "Ptbp1",
                "Aldoa",
                "Hnrnpdl",
                "Eif3a",
                "Rtn3",
                "Polr2h",
                "Snx3",
                "Adh5",
                "Birc5",
                "Psmd13",
                "Srpk1",
                "Rfc1",
                "Cox6b1",
                "Ppa1",
                "Cenpm",
                "Prdx1",
                "Rsl24d1",
                "Cdc34",
                "Rpa2",
                "Mrpl34",
                "Pdia3",
                "Ndufb5",
                "Psma2",
                "Ndufs6",
                "Nsun2",
                "Set",
                "Txnl4a",
                "Poldip2",
                "Eif4e2",
                "Sod2",
                "Ssbp1",
                "Rps14",
                "Tgif1",
                "Lipe",
                "Clns1a",
                "Snrpd2",
                "1810022K09Rik",
                "Phb2",
                "Gtf2e2",
                "Rpl15",
                "Brk1",
                "Vdac2",
                "Rap1b",
                "Pfn1",
                "Rpl13a",
                "Snrpg",
                "Ddx39b",
                "Cuta",
                "Snhg6",
                "Eif1a",
                "Cox5a",
                "Snrpb",
                "Tmem11",
                "Anapc11",
                "Fkbp4",
                "Pgam1",
                "Tomm22",
                "Lyar",
                "Ubap2l",
                "Erp29",
                "Snrnp70",
                "Uchl5",
                "Gpi1",
                "Tmem167",
                "Ppdpf",
                "Llph",
                "Emg1",
                "Lsm12",
                "Eif4b",
                "Anp32b",
                "Pkm",
                "H2afv",
                "Csde1",
                "Prrc2c",
                "Tceb2",
                "Rps2",
                "Eif2b5",
                "Gart",
                "Sumo3",
                "Ndufa10",
                "Uqcrfs1",
                "Naa50",
                "Ywhah",
                "Sf3b4",
                "Prpf19",
                "Sugt1",
                "Eif4a3",
                "Fkbp3",
                "Mtch2",
                "Dek",
                "Bag1",
                "Psmd4",
                "Gins2",
                "Fam104a",
                "Papola",
                "Rps16",
                "Mrfap1",
                "Mrps16",
                "Uqcrh",
                "Tpr",
                "Ddx18",
                "Psmd12",
                "Rrp9",
                "Eed",
                "Ccne1",
                "Srsf6",
                "Ndufa5",
                "Tmpo",
                "Tsn",
                "Mrpl11",
                "Txnl1",
                "Mesdc2",
                "Eef1g",
                "Rbm25",
                "2700094K13Rik",
                "Cycs",
                "Myl12a",
                "Ptma",
                "Rac1",
                "Rer1",
                "Klf2",
                "Dut",
                "Ralbp1",
                "Dhx16",
                "Sdhb",
                "Taf10",
                "Wdr43",
                "Ybx3",
                "Rps10",
                "Yrdc",
                "Pabpc1",
                "Alyref",
                "Tsen34",
                "Hspa14",
                "Fkbp1a",
                "Dad1",
                "Yif1b",
                "Fam60a",
                "Wbp11",
                "Cwc15",
                "Guk1",
                "2810428I15Rik",
                "Tsfm",
                "Pim1",
                "Postn",
                "Rplp2",
                "Aimp1",
                "Serbp1",
                "Gpx4",
                "Gcsh",
                "Phf5a",
                "Eif4a2",
                "Rheb",
                "Gtf2a2",
                "Mkrn1",
                "Rrp1",
                "U2af2",
                "Rpl23",
                "Sfr1",
                "Kdm1a",
                "Rrs1",
                "Ly6e",
                "Bcap31",
                "Cnbp",
                "Rsrc2"
            ],
            "expression_sum": {
                "class1": 10758.0,
                "class2": 1389.0
            },
            "uniprot_ids": [
                "P58462",
                "E2S038",
                "H3BJ68",
                "H3BJZ5",
                "H3BLF2",
                "Q8CCZ5",
                "H3BJR8",
                "D3Z6Q4",
                "Q6P0X7",
                "D3Z454",
                "H3BL33",
                "A0A0R4J1H9",
                "H3BJ24",
                "H3BJM1",
                "D3Z6Q3",
                "H3BK30",
                "H3BKT4",
                "A0A0R4J282",
                "D3Z2T4",
                "D3Z0J5",
                "H3BK38",
                "Q3UMG2"
            ]
        },
        "Fscn1": {
            "id": "Fscn1",
            "type": "Gene",
            "symbol": "Fscn1",
            "source": [
                "Klf2",
                "Tcf4",
                "Pou5f1",
                "Foxp1",
                "Cebpb",
                "Atf4",
                "Junb",
                "Fos",
                "Jun",
                "Sox2",
                "Hes1",
                "Nanog",
                "Jund",
                "Twist2",
                "Hdac2",
                "Smc3",
                "Smarca5",
                "Sox4"
            ],
            "target": [],
            "expression_sum": {
                "class1": 21099.0,
                "class2": 50984.0
            },
            "uniprot_ids": [
                "Q61553",
                "A0A0G2JDU7",
                "F7BDR1"
            ]
        },
        "Fubp1": {
            "id": "Fubp1",
            "type": "Gene",
            "symbol": "Fubp1",
            "source": [
                "Tcf4",
                "Pou5f1",
                "Foxp1",
                "Tfcp2l1",
                "Arf4",
                "Nfib",
                "Atf4",
                "Junb",
                "Fos",
                "Jun",
                "Sox2",
                "Hes1",
                "Nanog",
                "Jund",
                "Twist2",
                "Cebpb",
                "Sox4"
            ],
            "target": [],
            "expression_sum": {
                "class1": 7908.0,
                "class2": 37050.0
            },
            "uniprot_ids": [
                "Q91WJ8",
                "Q3TUE1",
                "Q3UUU2",
                "A0A0G2JFK2",
                "A0A0G2JF21",
                "A0A0G2JG00",
                "A0A0G2JGV9",
                "A0A0G2JFY5",
                "A0A0G2JGW9",
                "Q3TIS5",
                "A0A0G2JE55",
                "A0A0G2JG96",
                "A0A0G2JGT1",
                "Q0P6B2"
            ]
        },
        "Fus": {
            "id": "Fus",
            "type": "Gene",
            "symbol": "Fus",
            "source": [
                "Tcf4",
                "Pou5f1",
                "Foxp1",
                "Cebpb",
                "Atf4",
                "Junb",
                "Fos",
                "Jun",
                "Sox2",
                "Hes1",
                "Nanog",
                "Jund",
                "Twist2",
                "Hdac2",
                "Smarca5",
                "Sox4"
            ],
            "target": [],
            "expression_sum": {
                "class1": 12576.0,
                "class2": 41158.0
            },
            "uniprot_ids": [
                "P56959",
                "Q8BNR3",
                "Q91VQ2",
                "Q8CFQ9",
                "G3UXT7",
                "G3UZD2"
            ]
        },
        "G3bp2": {
            "id": "G3bp2",
            "type": "Gene",
            "symbol": "G3bp2",
            "source": [
                "Klf2",
                "Tfcp2l1",
                "Pou5f1",
                "Foxp1",
                "Nfib",
                "Atf4",
                "Junb",
                "Fos",
                "Jun",
                "Sox2",
                "Hes1",
                "Nanog",
                "Jund",
                "Cebpb",
                "Zfp57"
            ],
            "target": [],
            "expression_sum": {
                "class1": 7042.0,
                "class2": 55650.0
            },
            "uniprot_ids": [
                "P97379",
                "Q542W3",
                "Q3U541",
                "Q3U6B1",
                "A0A0J9YTQ8",
                "Q3U931",
                "Q5DU21",
                "A0A0J9YUY8",
                "Q3TEL4"
            ]
        },
        "Gars": {
            "id": "Gars",
            "type": "Gene",
            "symbol": "Gars",
            "source": [
                "Tcf4",
                "Pou5f1",
                "Cebpb",
                "Atf4",
                "Junb",
                "Id2",
                "Jun",
                "Hes1",
                "Ewsr1",
                "Jund",
                "Twist2",
                "Hdac2",
                "Smarca5"
            ],
            "target": [],
            "expression_sum": {
                "class1": 13748.0,
                "class2": 30444.0
            },
            "uniprot_ids": [
                "Q9CZD3"
            ]
        },
        "Gcsh": {
            "id": "Gcsh",
            "type": "Gene",
            "symbol": "Gcsh",
            "source": [
                "Klf2",
                "Pou5f1",
                "Foxp1",
                "Hmga2",
                "Nfib",
                "Atf4",
                "Fos",
                "Jun",
                "Sox2",
                "Nanog",
                "Jund",
                "Twist2",
                "Cebpb"
            ],
            "target": [],
            "expression_sum": {
                "class1": 9542.0,
                "class2": 59737.0
            },
            "uniprot_ids": [
                "Q91WK5",
                "Q9CV53"
            ]
        },
        "Gemin7": {
            "id": "Gemin7",
            "type": "Gene",
            "symbol": "Gemin7",
            "source": [
                "Trim28",
                "Rest",
                "Atf4",
                "Ctcf",
                "Id2",
                "Hdac1",
                "Trp53",
                "Hdac2",
                "Smc3",
                "Smarca5"
            ],
            "target": [],
            "expression_sum": {
                "class1": 11712.0,
                "class2": 17448.0
            },
            "uniprot_ids": [
                "Q9CWY4",
                "D3Z684",
                "A0A140LJ80"
            ]
        },
        "Get4": {
            "id": "Get4",
            "type": "Gene",
            "symbol": "Get4",
            "source": [
                "Tcf4",
                "Pou5f1",
                "Foxp1",
                "Tfcp2l1",
                "Atf4",
                "Junb",
                "Id2",
                "Jun",
                "Sox2",
                "Hes1",
                "Nanog",
                "Jund",
                "Twist2",
                "Cebpb",
                "Sox4"
            ],
            "target": [],
            "expression_sum": {
                "class1": 5248.0,
                "class2": 22792.0
            },
            "uniprot_ids": [
                "Q9D1H7",
                "D3Z7S0",
                "D3Z4J5"
            ]
        },
        "Ghitm": {
            "id": "Ghitm",
            "type": "Gene",
            "symbol": "Ghitm",
            "source": [
                "Pou5f1",
                "Atf4",
                "Junb",
                "Trp53",
                "Hdac2"
            ],
            "target": [],
            "expression_sum": {
                "class1": 10536.0,
                "class2": 22293.0
            },
            "uniprot_ids": [
                "Q91VC9",
                "A0A286YDK2"
            ]
        },
        "Glipr2": {
            "id": "Glipr2",
            "type": "Gene",
            "symbol": "Glipr2",
            "source": [
                "E2f4",
                "Trim28",
                "Rest",
                "Ctbp1",
                "Atf4",
                "Ctcf",
                "Trp53",
                "Hdac1",
                "Klf5",
                "Tbx3",
                "Zfp57",
                "Hdac2",
                "Smc3",
                "Smarca5"
            ],
            "target": [],
            "expression_sum": {
                "class1": 17145.0,
                "class2": 5177.0
            },
            "uniprot_ids": [
                "Q9CYL5",
                "B1AWD6"
            ]
        },
        "Glud1": {
            "id": "Glud1",
            "type": "Gene",
            "symbol": "Glud1",
            "source": [
                "Tcf4",
                "Pou5f1",
                "Foxp1",
                "Cebpb",
                "Atf4",
                "Junb",
                "Jun",
                "Sox2",
                "Hes1",
                "Jund",
                "Twist2",
                "Hdac2",
                "Smarca5"
            ],
            "target": [],
            "expression_sum": {
                "class1": 8240.0,
                "class2": 27545.0
            },
            "uniprot_ids": [
                "P26443",
                "F7CFA5",
                "Q3TSQ7"
            ]
        },
        "Gm16286": {
            "id": "Gm16286",
            "type": "Gene",
            "symbol": "Gm16286",
            "source": [
                "Tcf4",
                "Pou5f1",
                "Foxp1",
                "Atf4",
                "Junb",
                "Id2",
                "Jun",
                "Sox2",
                "Hes1",
                "Nanog",
                "Jund",
                "Cebpb",
                "Smarca5",
                "Sox4"
            ],
            "target": [],
            "expression_sum": {
                "class1": 8342.0,
                "class2": 31765.0
            }
        },
        "Gnai2": {
            "id": "Gnai2",
            "type": "Gene",
            "symbol": "Gnai2",
            "source": [
                "Pou5f1",
                "Trim28",
                "Arf4",
                "Atf4",
                "Junb",
                "Ctcf",
                "Trp53",
                "Hdac1",
                "Ybx1",
                "Twist2",
                "Hdac2",
                "Smc3",
                "Smarca5",
                "Sox4"
            ],
            "target": [],
            "expression_sum": {
                "class1": 28329.0,
                "class2": 45027.0
            },
            "uniprot_ids": [
                "P08752",
                "A0A0A6YXC2",
                "A0A0A6YXN0",
                "A0A0A6YWA9",
                "Q8JZT4",
                "Q922Y6"
            ]
        },
        "Gnas": {
            "id": "Gnas",
            "type": "Gene",
            "symbol": "Gnas",
            "source": [
                "Klf2",
                "Klf4",
                "Nfib",
                "Klf5",
                "Hdac2",
                "E2f4",
                "Pou5f1",
                "Rest",
                "Trp53",
                "Smarca5",
                "Trim28",
                "Tgif1",
                "Tbx3",
                "Zfp57",
                "Smc3",
                "Atf4",
                "Ctcf",
                "Hdac1",
                "Ybx1"
            ],
            "target": [],
            "expression_sum": {
                "class1": 106090.0,
                "class2": 32508.0
            },
            "uniprot_ids": [
                "Q6R0H6",
                "Q6R0H7",
                "P63094",
                "Q9Z0F1",
                "Z4YKV1",
                "A6PW71",
                "A6PW70",
                "Q9WUC0",
                "A0A571BGH5",
                "A0A571BDL4",
                "A0A571BEM2",
                "A0A571BE25",
                "A0A571BE38",
                "A0A571BEG2",
                "A0A571BEI3",
                "A2A610",
                "Q9JJ33",
                "Q8BUB2",
                "Q8C372"
            ]
        },
        "Gnb2": {
            "id": "Gnb2",
            "type": "Gene",
            "symbol": "Gnb2",
            "source": [
                "E2f4",
                "Pou5f1",
                "Trim28",
                "Rest",
                "Atf4",
                "Ctcf",
                "Junb",
                "Tgif1",
                "Hdac1",
                "Trp53",
                "Hdac2",
                "Smc3",
                "Smarca5"
            ],
            "target": [],
            "expression_sum": {
                "class1": 35432.0,
                "class2": 38345.0
            },
            "uniprot_ids": [
                "P62880",
                "D3YZX3",
                "Q3U9V4",
                "E9PWM7",
                "D3Z1M1",
                "D3Z1T4",
                "E9QKR0",
                "Q9QWG2"
            ]
        },
        "Gng12": {
            "id": "Gng12",
            "type": "Gene",
            "symbol": "Gng12",
            "source": [
                "Hdac2",
                "E2f4",
                "Pou5f1",
                "Trim28",
                "Rest",
                "Eno1",
                "Atf4",
                "Ctcf",
                "Trp53",
                "Hdac1",
                "Klf5",
                "Tbx3",
                "Zfp57",
                "Cebpb",
                "Smc3",
                "Smarca5"
            ],
            "target": [],
            "expression_sum": {
                "class1": 13538.0,
                "class2": 5900.0
            },
            "uniprot_ids": [
                "Q9DAS9",
                "A0A0N4SW28",
                "A0A0N4SVT3"
            ]
        },
        "Gng5": {
            "id": "Gng5",
            "type": "Gene",
            "symbol": "Gng5",
            "source": [
                "Tcf4",
                "Pou5f1",
                "Foxp1",
                "Cebpb",
                "Atf4",
                "Junb",
                "Fos",
                "Jun",
                "Sox2",
                "Trp53",
                "Hes1",
                "Nanog",
                "Hdac2",
                "Smarca5",
                "Sox4"
            ],
            "target": [],
            "expression_sum": {
                "class1": 71578.0,
                "class2": 130516.0
            },
            "uniprot_ids": [
                "Q80SZ7",
                "D3Z7Q3",
                "Q3UKC8"
            ]
        },
        "Gps1": {
            "id": "Gps1",
            "type": "Gene",
            "symbol": "Gps1",
            "source": [
                "Pou5f1",
                "Cebpb",
                "Arf4",
                "Atf4",
                "Junb",
                "Jun",
                "Hes1",
                "Jund",
                "Hdac2",
                "Smc3"
            ],
            "target": [],
            "expression_sum": {
                "class1": 7196.0,
                "class2": 20585.0
            },
            "uniprot_ids": [
                "Q99LD4",
                "Q3MIA8",
                "D6RCV3",
                "B1ATU4",
                "A0A140LJB7",
                "G3UXW9"
            ]
        },
        "Gpx4": {
            "id": "Gpx4",
            "type": "Gene",
            "symbol": "Gpx4",
            "source": [
                "Nfib",
                "Fos",
                "Sox2",
                "Hdac2",
                "Tcf4",
                "Pou5f1",
                "Jun",
                "Smarca5",
                "Sox4",
                "Jund",
                "Cebpb",
                "Smc3",
                "Foxp1",
                "Atf4",
                "Junb",
                "Ybx1",
                "Hes1",
                "Tfcp2l1",
                "Nanog"
            ],
            "target": [],
            "expression_sum": {
                "class1": 66534.0,
                "class2": 213428.0
            },
            "uniprot_ids": [
                "O70325",
                "Q3TI34",
                "Q76LV0",
                "S4R1E5",
                "Q5XJZ8",
                "Q3TIF2"
            ]
        },
        "Gtf2a2": {
            "id": "Gtf2a2",
            "type": "TF",
            "symbol": "Gtf2a2",
            "source": [
                "Tcf4",
                "Pou5f1",
                "Foxp1",
                "Cebpb",
                "Nfib",
                "Atf5",
                "Atf4",
                "Junb",
                "Fos",
                "Jun",
                "Sox2",
                "Sfpq",
                "Nanog",
                "Jund",
                "Twist2",
                "Hdac2",
                "Smc3",
                "Smarca5"
            ],
            "target": [
                "0610011F06Rik",
                "Ak1",
                "Cox8a",
                "2010111I01Rik"
            ],
            "expression_sum": {
                "class1": 10627.0,
                "class2": 28035.0
            },
            "uniprot_ids": [
                "Q80ZM7",
                "Q05C74",
                "Q3TE88",
                "D3Z7S8",
                "D3Z793",
                "D3Z7C2",
                "D6RIQ7",
                "Q3UT44",
                "Q3TN86"
            ]
        },
        "Gtpbp4": {
            "id": "Gtpbp4",
            "type": "Gene",
            "symbol": "Gtpbp4",
            "source": [
                "Tcf4",
                "Pou5f1",
                "Foxp1",
                "Cebpb",
                "Nfib",
                "Atf4",
                "Junb",
                "Jun",
                "Sox2",
                "Nanog",
                "Jund",
                "Twist2",
                "Hdac2",
                "Sox4"
            ],
            "target": [],
            "expression_sum": {
                "class1": 7484.0,
                "class2": 28578.0
            },
            "uniprot_ids": [
                "Q99ME9",
                "A0A217FL53",
                "Q8CCQ5",
                "Q3UR29",
                "Q3TPJ4"
            ]
        },
        "Guk1": {
            "id": "Guk1",
            "type": "Gene",
            "symbol": "Guk1",
            "source": [
                "Klf2",
                "Tcf4",
                "Pou5f1",
                "Foxp1",
                "Tfcp2l1",
                "Atf4",
                "Junb",
                "Fos",
                "Jun",
                "Sox2",
                "Hes1",
                "Nanog",
                "Jund",
                "Twist2",
                "Cebpb",
                "Mybl2"
            ],
            "target": [],
            "expression_sum": {
                "class1": 7310.0,
                "class2": 34729.0
            },
            "uniprot_ids": [
                "Q64520",
                "G3UYV3",
                "E9Q7K1"
            ]
        },
        "H13": {
            "id": "H13",
            "type": "Gene",
            "symbol": "H13",
            "source": [
                "Hdac2",
                "E2f4",
                "Rest",
                "Atf5",
                "Atf4",
                "Ctcf",
                "Trp53",
                "Hdac1",
                "Cebpb",
                "Smc3",
                "Smarca5"
            ],
            "target": [],
            "expression_sum": {
                "class1": 10983.0,
                "class2": 5673.0
            },
            "uniprot_ids": [
                "Q9D8V0",
                "Q6PGJ8",
                "Q8HWB1",
                "A3KGR9",
                "Q861Q5"
            ]
        },
        "H2afx": {
            "id": "H2afx",
            "type": "Gene",
            "symbol": "H2afx",
            "source": [
                "Klf2",
                "Tcf4",
                "Pou5f1",
                "Atf4",
                "Junb",
                "Jun",
                "Sox2",
                "Nanog",
                "Jund",
                "Cebpb"
            ],
            "target": [],
            "expression_sum": {
                "class1": 17696.0,
                "class2": 128708.0
            },
            "uniprot_ids": [
                "P27661"
            ]
        },
        "H2afz": {
            "id": "H2afz",
            "type": "Gene",
            "symbol": "H2afz",
            "source": [
                "Klf2",
                "Tcf4",
                "Pou5f1",
                "Foxp1",
                "Cebpb",
                "Nfib",
                "Atf4",
                "Junb",
                "Fos",
                "Jun",
                "Sox2",
                "Tbx3",
                "Hes1",
                "Nanog",
                "Jund",
                "Twist2",
                "Hdac2",
                "Zfp57"
            ],
            "target": [],
            "expression_sum": {
                "class1": 129984.0,
                "class2": 886666.0
            },
            "uniprot_ids": [
                "P0C0S6",
                "Q3UA95",
                "Q3TFU6"
            ]
        },
        "H3f3a": {
            "id": "H3f3a",
            "type": "Gene",
            "symbol": "H3f3a",
            "source": [
                "Tcf4",
                "Pou5f1",
                "Foxp1",
                "Cebpb",
                "Nfib",
                "Atf4",
                "Junb",
                "Fos",
                "Jun",
                "Sox2",
                "Prrx1",
                "Hes1",
                "Nanog",
                "Jund",
                "Twist2",
                "Hdac2",
                "Sox4"
            ],
            "target": [],
            "expression_sum": {
                "class1": 56704.0,
                "class2": 193162.0
            },
            "uniprot_ids": [
                "P84244",
                "E0CYN1",
                "E0CYR7",
                "E0CZ27",
                "F8WI35"
            ]
        },
        "Hcfc1r1": {
            "id": "Hcfc1r1",
            "type": "Gene",
            "symbol": "Hcfc1r1",
            "source": [
                "E2f4",
                "Trim28",
                "Rest",
                "Arf4",
                "Atf4",
                "Ctcf",
                "Trp53",
                "Hdac1",
                "Klf5",
                "Tbx3",
                "Ybx1",
                "Hdac2",
                "Smc3"
            ],
            "target": [],
            "expression_sum": {
                "class1": 25780.0,
                "class2": 15245.0
            },
            "uniprot_ids": [
                "Q9CYQ5",
                "J3QPD4"
            ]
        },
        "Hdac2": {
            "id": "Hdac2",
            "type": "TF",
            "symbol": "Hdac2",
            "source": [
                "Tcf4",
                "Pou5f1",
                "Nfib",
                "Atf4",
                "Junb",
                "Fos",
                "Jun",
                "Sox2",
                "Hes1",
                "Nanog",
                "Jund",
                "Twist2",
                "Cebpb"
            ],
            "target": [
                "Rps5",
                "Hsp90aa1",
                "Arf1",
                "Dap",
                "Rpl41",
                "Rnf7",
                "Sar1a",
                "Srsf9",
                "Mdh1",
                "Tpm4",
                "Usmg5",
                "Dctn3",
                "Ppp2ca",
                "Cd44",
                "Ddx24",
                "H2afj",
                "Rps17",
                "Fis1",
                "Zc3h15",
                "Cox8a",
                "Ddb1",
                "Myh9",
                "Thbs1",
                "Psmb7",
                "Txn2",
                "Lims1",
                "Sec13",
                "Clta",
                "Ppp1r11",
                "1110008F13Rik",
                "Tuba1b",
                "Wisp1",
                "Slc25a4",
                "Cks1b",
                "Col12a1",
                "Rps27l",
                "Tspan4",
                "Map1lc3a",
                "Psma6",
                "Lmna",
                "Psmc1",
                "Hnrnpa1",
                "Ndufa2",
                "Thoc7",
                "Hnrnpf",
                "Ddx39",
                "Mlf2",
                "Psmd6",
                "Vbp1",
                "Ssrp1",
                "Atp5j2",
                "Ier3ip1",
                "Pck2",
                "Rnf187",
                "Lgals3",
                "Swi5",
                "Cxcl12",
                "Psmb6",
                "mt-Co1",
                "Pgrmc1",
                "Med28",
                "Shfm1",
                "Fos",
                "Tagln2",
                "Cox6c",
                "Rps15a",
                "Ddx3x",
                "Ndufs3",
                "Nedd4",
                "Rexo2",
                "Calr",
                "Pdlim7",
                "Rab18",
                "Tubb5",
                "Calm3",
                "Mfge8",
                "Mbnl2",
                "Arf6",
                "Canx",
                "D8Ertd738e",
                "Hn1",
                "Ppp1r14b",
                "Actn1",
                "Snrpe",
                "B2m",
                "Mrpl14",
                "Slc25a3",
                "Psat1",
                "Eif4h",
                "Lamtor1",
                "2810417H13Rik",
                "Psmd7",
                "Gdi2",
                "Stub1",
                "Prrx1",
                "Tra2b",
                "Ranbp1",
                "Mrpl28",
                "Psmc6",
                "Hmgcs1",
                "Zwint",
                "Rbx1",
                "Rps24",
                "Eif1",
                "Myh10",
                "Neat1",
                "Sec61g",
                "Ythdc1",
                "Ola1",
                "Snrnp27",
                "Sf3b6",
                "Tgfbi",
                "Ube2r2",
                "Timp3",
                "Srsf1",
                "Gemin7",
                "Wdr89",
                "Lrrc59",
                "Akr1a1",
                "Tdgf1",
                "Sh3glb1",
                "Ssr2",
                "Pdcd6",
                "Akt1s1",
                "Ube2d2a",
                "Hras",
                "Bnip3l",
                "Atp2b1",
                "Kpnb1",
                "Mapre1",
                "C1d",
                "Cisd2",
                "Mylpf",
                "Nktr",
                "Col5a1",
                "Ppia",
                "Snrpd1",
                "Ctgf",
                "Tspan3",
                "Hnrnpm",
                "Capza2",
                "Eif3k",
                "Klf6",
                "Rpl36al",
                "Pcolce",
                "Tgfb1i1",
                "Loxl1",
                "2310036O22Rik",
                "Ppp1ca",
                "Srsf5",
                "Hnrnph1",
                "Rpl35a",
                "Timp2",
                "Syngr1",
                "Chchd1",
                "Crip1",
                "Gars",
                "Psmc3",
                "Ywhaz",
                "Sec62",
                "Rps12",
                "Pomp",
                "Luc7l2",
                "Rpl36",
                "Rbbp7",
                "Rps21",
                "Ddx5",
                "Ndufb9",
                "Rbm39",
                "Maged1",
                "Arpp19",
                "Nedd8",
                "Pcnp",
                "Eef1b2",
                "Ube2i",
                "Arpc1b",
                "Col3a1",
                "Cyr61",
                "Eif3l",
                "Kdelr1",
                "Rab14",
                "Atpif1",
                "Serpinh1",
                "Tmem258",
                "Tcp1",
                "Atp6v1g1",
                "Rpl24",
                "Pttg1",
                "Gnb2",
                "Fus",
                "Dpysl3",
                "Ociad1",
                "Ddost",
                "Hcfc1r1",
                "Rps4x",
                "Sptbn1",
                "Id3",
                "Txnl1",
                "Ap3s1",
                "Gng12",
                "Cct3",
                "Fkbp8",
                "Ncam1",
                "Wls",
                "Psma7",
                "Hacd1",
                "G3bp1",
                "Ndufa8",
                "Pdia4",
                "Sept7",
                "Srp14",
                "Fam195b",
                "Tpm3",
                "Atp5b",
                "Banf1",
                "Actg1",
                "Immt",
                "Psma1",
                "Rcn2",
                "Emc10",
                "Ier5",
                "Tsg101",
                "2410015M20Rik",
                "Hdgf",
                "Prnp",
                "Smc3",
                "Thrap3",
                "Id1",
                "Csnk2b",
                "Nars",
                "Tpi1",
                "Hmga2",
                "Rpl37",
                "Dynlrb1",
                "Atf4",
                "Hprt",
                "Tmem234",
                "Nfkbia",
                "Nucks1",
                "Nrp1",
                "Col1a2",
                "Eif3b",
                "Ube2m",
                "Atp6v0b",
                "F2r",
                "Fkbp9",
                "Pcbp1",
                "Rpl38",
                "Serpine2",
                "Hnrnpk",
                "Igfbp4",
                "Mrps24",
                "Id2",
                "Itgb1",
                "Ndufa4",
                "Ewsr1",
                "Cops5",
                "Ssr3",
                "Eif4e",
                "Ndufv2",
                "Fabp5",
                "H3f3a",
                "Mettl9",
                "Nme1",
                "Chmp4b",
                "Tcf4",
                "Ube2k",
                "Igf2",
                "Csnk1a1",
                "Cd81",
                "Ssr1",
                "Ndufs2",
                "Cyp51",
                "Lxn",
                "Pgls",
                "Tmsb10",
                "Rpl3",
                "Nudt21",
                "Rpl12",
                "Ift20",
                "Anxa2",
                "Eif3c",
                "Vmp1",
                "Cox7a2",
                "Ndufa7",
                "Col5a2",
                "Cdv3",
                "Crip2",
                "Napa",
                "Cox7a2l",
                "mt-Nd4",
                "Vim",
                "Hmgn1",
                "Ube2l3",
                "Btf3l4",
                "Basp1",
                "Ppp4c",
                "Eef1d",
                "Sh3bgrl3",
                "Lsm8",
                "Wdr1",
                "Tnc",
                "Ube2b",
                "Ube2j2",
                "Mrpl13",
                "Vapa",
                "Eif5",
                "Tmed2",
                "Txn1",
                "Erh",
                "Mrpl17",
                "Atp5j",
                "Fscn1",
                "Atp5g3",
                "Srrm1",
                "Tpm1",
                "Ldha",
                "Sdcbp",
                "Thy1",
                "Surf4",
                "Atp5g2",
                "Chd4",
                "Srp19",
                "Cct5",
                "Phlda1",
                "Emd",
                "1110004F10Rik",
                "Myeov2",
                "Bclaf1",
                "Vasp",
                "Tceb1",
                "Rpsa",
                "H2afz",
                "Bola3",
                "Mrps21",
                "Psip1",
                "Vcl",
                "Cst3",
                "Rpl36a",
                "Eif3m",
                "Lox",
                "Arpc5l",
                "Rps29",
                "Rpl5",
                "Higd1a",
                "S100a6",
                "Prdx6",
                "Serpine1",
                "H2-D1",
                "Trappc4",
                "Rps20",
                "Ybx1",
                "Dnaja2",
                "Sys1",
                "Cd63",
                "Capg",
                "Lbh",
                "Ndufa3",
                "Tmem14c",
                "Cdc37",
                "Ndufab1",
                "Mtdh",
                "Scand1",
                "Rps6",
                "Snrpf",
                "Eef1a1",
                "Nfib",
                "mt-Nd5",
                "Maged2",
                "Rtn4",
                "Rpl28",
                "Atp5c1",
                "Ctsl",
                "Ccnd2",
                "Mrps17",
                "Tcf25",
                "Spcs1",
                "Ywhaq",
                "Cox6a1",
                "Col6a1",
                "Rala",
                "Sdc2",
                "Tax1bp3",
                "Pdha1",
                "Filip1l",
                "Rpl26",
                "Uqcrb",
                "Cav1",
                "Rps3",
                "Calm2",
                "Atox1",
                "Cox7c",
                "Srsf10",
                "Adss",
                "Btf3",
                "Plk2",
                "Tln1",
                "Cdh11",
                "Oaf",
                "Ostf1",
                "Rps7",
                "Tmem126a",
                "Ube2v2",
                "Rpl18",
                "Eno1",
                "Rpl19",
                "Atp5d",
                "Snhg12",
                "Taldo1",
                "Timm13",
                "mt-Nd3",
                "Eif3h",
                "Fbln5",
                "Rpl22l1",
                "Chd3",
                "Rps3a1",
                "Srsf3",
                "Vcp",
                "Smc4",
                "Cmpk1",
                "Mbd3",
                "Rpl7",
                "Fkbp2",
                "Btg1",
                "Cct7",
                "Fn1",
                "Glipr2",
                "Lrrc58",
                "Hes1",
                "Ap1s1",
                "Ndufb2",
                "Hsp90ab1",
                "Rps9",
                "Twist2",
                "Oaz1",
                "Mrpl30",
                "Rpp25",
                "Uqcrq",
                "Rcn3",
                "Sec61b",
                "BC031181",
                "Polr2m",
                "Rwdd1",
                "Snx5",
                "Malat1",
                "Gnai2",
                "Cyc1",
                "Rps8",
                "Dnaja1",
                "Tmem176b",
                "Eif4a1",
                "Sumo2",
                "Tm9sf3",
                "Pcbp2",
                "Ctsd",
                "Magoh",
                "Ndufc2",
                "Reep5",
                "Tmed10",
                "Pou5f1",
                "Fhl2",
                "H3f3b",
                "Glrx5",
                "Col1a1",
                "Elof1",
                "Gnb2l1",
                "Ndufa12",
                "Ube2e3",
                "Nsa2",
                "Oxct1",
                "Atxn10",
                "Wbp5",
                "Atp6v0e",
                "Cnih4",
                "Pebp1",
                "Dbi",
                "Morf4l2",
                "Ube2d3",
                "Tmem59",
                "Rbbp4",
                "Eif3e",
                "Dstn",
                "Emp3",
                "Aspn",
                "Pfdn4",
                "Cebpb",
                "Srrm2",
                "Top1",
                "Ywhae",
                "Pafah1b2",
                "Ubl5",
                "Anxa3",
                "Mrps15",
                "Ndufc1",
                "Psmc4",
                "Eif3d",
                "Hbegf",
                "Psmd14",
                "Rps15",
                "Ghitm",
                "Arpc3",
                "Psma4",
                "Scp2",
                "Ilk",
                "Arl1",
                "Arpc1a",
                "Ap2s1",
                "Mgat2",
                "Rpl34",
                "Mlec",
                "Rab7",
                "Sep15",
                "Dynll1",
                "Gtpbp4",
                "Eif2s2",
                "Hspd1",
                "Cpe",
                "Atf5",
                "Arf5",
                "Cdc42",
                "Npc2",
                "Rpl31",
                "Srsf7",
                "Smdt1",
                "Nap1l1",
                "Eef2",
                "Tpx2",
                "Etfb",
                "Hsbp1",
                "Hspa5",
                "Rnh1",
                "Rrbp1",
                "Uqcrc2",
                "Actr3",
                "Tmsb4x",
                "Cd47",
                "Efemp2",
                "Ccdc124",
                "Fbln2",
                "Gapdh",
                "Smarca5",
                "Prdx2",
                "Mpc1",
                "Copb2",
                "Khdrbs1",
                "Ap2m1",
                "Mrpl52",
                "Nupr1",
                "Sars",
                "Bri3",
                "Smim14",
                "Prkcdbp",
                "Hnrnpab",
                "Sdpr",
                "Lgals1",
                "Rplp0",
                "Psmb5",
                "Ahnak",
                "Tbrg1",
                "Col11a1",
                "Hspa9",
                "Fkbp10",
                "Nop58",
                "Ube2n",
                "Ndufa11",
                "Cbx5",
                "Kif5b",
                "Luc7l3",
                "Polr2c",
                "Cct8",
                "Mpc2",
                "Copz2",
                "Hnrnpa2b1",
                "Ptp4a1",
                "Lsp1",
                "Ndufs5",
                "Gsn",
                "Psmb1",
                "Oat",
                "Tubb6",
                "Coa3",
                "Ndufb8",
                "Cited2",
                "Psmd2",
                "Tmem97",
                "Srsf2",
                "Eva1b",
                "Prdx4",
                "mt-Nd1",
                "Eif4g2",
                "Psmb2",
                "Rab2a",
                "Uqcc2",
                "Denr",
                "Eif3g",
                "Tspo",
                "Sra1",
                "Tceal8",
                "Lhfp",
                "Myl9",
                "Rbm3",
                "Psmd11",
                "Cnn3",
                "Vps29",
                "Btg2",
                "Vdac1",
                "Selk",
                "Cggbp1",
                "Zfp706",
                "Rad23b",
                "Chmp2a",
                "Tra2a",
                "Nudcd2",
                "Pdcd5",
                "Aprt",
                "Ost4",
                "Rab11a",
                "Ankrd1",
                "Snrpc",
                "Anxa5",
                "Eif3f",
                "Eif4g1",
                "Anxa6",
                "Mtch1",
                "Tmbim6",
                "Dynll2",
                "Rhoa",
                "Ssu72",
                "Rab10",
                "Atp5a1",
                "Psma5",
                "Ttc3",
                "Cdk4",
                "Cnpy2",
                "Rpl22",
                "Rsl1d1",
                "Vimp",
                "Ccdc80",
                "Syncrip",
                "Sec61a1",
                "Caprin1",
                "Iqgap1",
                "Ict1",
                "Kdelr2",
                "Glud1",
                "March5",
                "Mrpl23",
                "Pmp22",
                "Rpl4",
                "Fermt2",
                "Hnrnpu",
                "Rps25",
                "Rps28",
                "Gspt1",
                "Srsf11",
                "Rpl18a",
                "Hmgb3",
                "Rpn1",
                "Gas1",
                "Ssr4",
                "Txndc9",
                "Ppp1r2",
                "Sqstm1",
                "Eprs",
                "Trmt112",
                "Lman1",
                "F3",
                "Msmo1",
                "Rcn1",
                "Arpc2",
                "Atp5l",
                "Mrpl54",
                "Rpl35",
                "Son",
                "Ilf2",
                "Ywhab",
                "Cct2",
                "Acp1",
                "Prelid1",
                "Sox4",
                "Phlda3",
                "Sept11",
                "Ndufa1",
                "Romo1",
                "Tpm2",
                "Ergic3",
                "Eif1ax",
                "Bzw1",
                "Sdc4",
                "Ak6",
                "Ubc",
                "Rock2",
                "Hnrnpa3",
                "Prpf40a",
                "Prmt1",
                "Pbdc1",
                "Cdc42ep3",
                "Mrps7",
                "Metap2",
                "Mrpl57",
                "Morf4l1",
                "2010107E04Rik",
                "Col8a1",
                "Pafah1b1",
                "Lsm6",
                "Ube2s",
                "Tecr",
                "Sf3b1",
                "Idh2",
                "Psap",
                "Ndufs4",
                "Polr1d",
                "Snrpd3",
                "Tm4sf1",
                "Rbms1",
                "P4hb",
                "Scd2",
                "Dlk1",
                "Sf3b2",
                "H2afy",
                "Map1b",
                "Rab34",
                "Suclg1",
                "Zyx",
                "Adipor1",
                "Rpl17",
                "Bsg",
                "mt-Nd2",
                "Hnrnpd",
                "Ppic",
                "Rpl14",
                "Tma7",
                "Sec22b",
                "Camk2d",
                "Txndc17",
                "Cops4",
                "Cyba",
                "Flna",
                "Lsm4",
                "Efemp1",
                "H13",
                "Rsu1",
                "Ywhag",
                "Atp5f1",
                "Prss23",
                "Rps13",
                "Gsto1",
                "Tpt1",
                "Eif5a",
                "Rps11",
                "Ppib",
                "Psmd8",
                "Cfl1",
                "Gng5",
                "Vcan",
                "Sparc",
                "Rps19",
                "Sept2",
                "Abracl",
                "Exosc8",
                "Cops6",
                "Msn",
                "Aldoa",
                "Mrpl20",
                "Hnrnpdl",
                "Serpinb6a",
                "Srp9",
                "Eif3a",
                "Rtn3",
                "Lpp",
                "Ptn",
                "Tagln",
                "Ctbp1",
                "Psmb4",
                "Snx3",
                "Strap",
                "Csrp1",
                "Commd1",
                "Psmd13",
                "Selm",
                "Ckap4",
                "Esd",
                "Atxn7l3b",
                "Igfbp6",
                "Comt",
                "Cox6b1",
                "Lamp1",
                "Sat1",
                "Lamtor5",
                "Ndufa13",
                "Atp5e",
                "Prdx1",
                "Ssb",
                "Ptrf",
                "Tinagl1",
                "Gps1",
                "Cmtm7",
                "Gadd45g",
                "Atp5o",
                "Rrm2",
                "Rpl13",
                "Sh3bgrl",
                "Jun",
                "Capns1",
                "Itm2b",
                "Pdia3",
                "Rplp1",
                "Ndufb5",
                "Rpl10a",
                "Atrx",
                "Psma2",
                "Txndc5",
                "Pfdn5",
                "Ndufb7",
                "Ptms",
                "Eif4e2",
                "Rps14",
                "Ftl1",
                "Sfpq",
                "Col4a1",
                "Rpl32",
                "Smap1",
                "Cd24a",
                "Zfp36l1",
                "Lipe",
                "Tnfrsf12a",
                "Mdh2",
                "Krtcap2",
                "Gabarap",
                "Uqcr11",
                "Rps27a",
                "Phgdh",
                "Rap1a",
                "Serf2",
                "Bud31",
                "Pdap1",
                "Brk1",
                "Gtf2h5",
                "Hsp90b1",
                "Mtpn",
                "Anxa1",
                "Prkar1a",
                "Rap1b",
                "Rpl11",
                "Vdac2",
                "Pfn1",
                "Rpl13a",
                "Nol7",
                "Timp1",
                "Ier3",
                "Grb10",
                "Cuta",
                "Apoe",
                "Eif1a",
                "Tuba1a",
                "Commd3",
                "Cnn2",
                "Minos1",
                "Snrpb",
                "Pgam1",
                "Rabggtb",
                "Tomm22",
                "Nans",
                "Ubap2l",
                "Erp29",
                "Gpx8",
                "Gpi1",
                "Tmem167",
                "Ppig",
                "Phldb2",
                "Palld",
                "Rpl27a",
                "Dcun1d5",
                "Eif5b",
                "Tram1",
                "Cct4",
                "Rdx",
                "Lrrc17",
                "Lsm12",
                "Fhl1",
                "Anp32b",
                "Edf1",
                "Cdkn1a",
                "Pkm",
                "Nono",
                "Csde1",
                "Fstl1",
                "Marcks",
                "Fam162a",
                "Tceb2",
                "Rhoc",
                "Mdm2",
                "Ifitm3",
                "Psenen",
                "Srp72",
                "Mrpl33",
                "Sumo3",
                "Emp1",
                "Psmc2",
                "Ndufa10",
                "App",
                "Fundc2",
                "Uqcrfs1",
                "Naa50",
                "Ywhah",
                "Eif3i",
                "Hnrnpc",
                "Smc2",
                "Mtch2",
                "S100a10",
                "Bag1",
                "Timm17a",
                "Psmd4",
                "Rpn2",
                "Tmem147",
                "Bub3",
                "Fam104a",
                "Prrx2",
                "Papola",
                "Rps16",
                "Manf",
                "Tmem256",
                "Sin3b",
                "Cetn3",
                "Dctn2",
                "Arhgdib",
                "Tmed3",
                "Serpinf1",
                "Mrfap1",
                "Galk1",
                "Hsd17b14",
                "Slc25a5",
                "Uqcrh",
                "Gap43",
                "Tpr",
                "Hint1",
                "Tbca",
                "Cstb",
                "Foxp1",
                "Srsf6",
                "Ndufa5",
                "Ebna1bp2",
                "Ccng1",
                "Ndufb10",
                "Tmpo",
                "Rpl37a",
                "Tsn",
                "H19",
                "Ptp4a2",
                "Mesdc2",
                "Eef1g",
                "Dnajc3",
                "Cope",
                "Cox5b",
                "Rbm25",
                "Arhgdia",
                "Cycs",
                "Myl12a",
                "Ndufb6",
                "Ptma",
                "Rac1",
                "Rer1",
                "S100a4",
                "Ralbp1",
                "Cald1",
                "Sdhb",
                "Fau",
                "Ybx3",
                "Rps10",
                "Sri",
                "Pabpc1",
                "Prdx5",
                "Alyref",
                "Clic1",
                "Tsen34",
                "Fkbp1a",
                "Got2",
                "Dad1",
                "Slirp",
                "Emc7",
                "Gnas",
                "Aebp1",
                "Arf4",
                "Calu",
                "Arpc5",
                "Yif1b",
                "Hdlbp",
                "Limd2",
                "Wbp11",
                "Cox4i1",
                "Ccnd1",
                "Tmed9",
                "Snf8",
                "Myl6",
                "Sepw1",
                "2810428I15Rik",
                "Hspa8",
                "Ndufs8",
                "Psma3",
                "Rpl39",
                "Rabac1",
                "Rplp2",
                "Capn2",
                "Serp1",
                "Mif",
                "Laptm4a",
                "Ptov1",
                "Aimp1",
                "Serbp1",
                "Cnih1",
                "Gpx4",
                "Chchd3",
                "Rheb",
                "Rps23",
                "Glrx3",
                "Gtf2a2",
                "Carhsp1",
                "Cryab",
                "Alpl",
                "Rrp1",
                "Pdcl3",
                "Rpl23",
                "Rpl30",
                "Kdelr3",
                "Sfr1",
                "Polr2f",
                "P3h4",
                "Junb",
                "Rbm17",
                "Bcap31",
                "Map2k2",
                "Cnbp",
                "Sdhd",
                "Axl",
                "Cfl2",
                "Ogn",
                "Rsrc2"
            ],
            "expression_sum": {
                "class1": 10992.0,
                "class2": 22123.0
            },
            "uniprot_ids": [
                "P70288",
                "Q8BQ10",
                "Q3UXH8",
                "Q3URA2",
                "A0A0R4J008",
                "Q3TMT1"
            ]
        },
        "Hdgf": {
            "id": "Hdgf",
            "type": "Gene",
            "symbol": "Hdgf",
            "source": [
                "Nfib",
                "Fos",
                "Id2",
                "Sox2",
                "Hdac2",
                "Tcf4",
                "Pou5f1",
                "Jun",
                "Sox4",
                "Jund",
                "Cebpb",
                "Smc3",
                "Zfp57",
                "Foxp1",
                "Atf4",
                "Junb",
                "Hes1",
                "Tfcp2l1",
                "Nanog",
                "Twist2"
            ],
            "target": [],
            "expression_sum": {
                "class1": 29453.0,
                "class2": 103938.0
            },
            "uniprot_ids": [
                "P51859",
                "E0CXA0",
                "E0CYW7"
            ]
        },
        "Hdlbp": {
            "id": "Hdlbp",
            "type": "Gene",
            "symbol": "Hdlbp",
            "source": [
                "E2f4",
                "Trim28",
                "Rest",
                "Hmgb1",
                "Atf4",
                "Ctcf",
                "Tgif1",
                "Id2",
                "Hdac1",
                "Klf5",
                "Trp53",
                "Hdac2",
                "Smc3",
                "Smarca5"
            ],
            "target": [],
            "expression_sum": {
                "class1": 19355.0,
                "class2": 12145.0
            },
            "uniprot_ids": [
                "Q8VDJ3",
                "A0A087WQY9",
                "Q3V1M8",
                "A0A087WPH2",
                "Q3U4Z7",
                "A0A087WNQ4",
                "A0A087WP83",
                "A0A087WPC5",
                "A0A087WS92",
                "A0PJE9",
                "Q8BX68"
            ]
        },
        "Hes1": {
            "id": "Hes1",
            "type": "TF",
            "symbol": "Hes1",
            "source": [
                "E2f4",
                "Trim28",
                "Rest",
                "Atf4",
                "Ctcf",
                "Tgif1",
                "Trp53",
                "Hdac1",
                "Jun",
                "Klf5",
                "Ybx1",
                "Hdac2",
                "Smc3"
            ],
            "target": [
                "Hsp90aa1",
                "Arf1",
                "Rnf7",
                "Srsf9",
                "Usmg5",
                "Ppp2ca",
                "Rps17",
                "Srm",
                "Grsf1",
                "Rae1",
                "Zc3h15",
                "2810004N23Rik",
                "Marcksl1",
                "1110008F13Rik",
                "Gar1",
                "Anapc5",
                "Psma6",
                "Thoc7",
                "Hnrnpf",
                "Knop1",
                "Lmnb1",
                "Mlf2",
                "Rnf187",
                "Cks2",
                "Nedd4",
                "Calm3",
                "Arf6",
                "Hn1",
                "Nop56",
                "Ubxn1",
                "Psat1",
                "Eif4h",
                "Ranbp1",
                "Sms",
                "Ctcf",
                "Ppm1g",
                "Ythdc1",
                "Bola1",
                "Ola1",
                "Srsf1",
                "Socs3",
                "Wdr89",
                "Exosc5",
                "Rcc2",
                "Kpnb1",
                "Mapre1",
                "C1d",
                "Cisd2",
                "Ppia",
                "Capza2",
                "Lap3",
                "Ncl",
                "2310036O22Rik",
                "Supt4a",
                "Srsf5",
                "Chchd1",
                "Ywhaz",
                "Gars",
                "Hspa4",
                "Rbbp7",
                "Rps21",
                "Hnrnpl",
                "Ddx5",
                "Arpp19",
                "Ube2i",
                "Utp3",
                "Mrpl15",
                "Siva1",
                "Fus",
                "Ezh2",
                "Rest",
                "Psma7",
                "G3bp1",
                "Gnl3",
                "Banf1",
                "Qdpr",
                "Actb",
                "Immt",
                "Tsg101",
                "Hdgf",
                "Pa2g4",
                "Ppp1cc",
                "Hist1h2bc",
                "Hprt",
                "Pcbp1",
                "Sae1",
                "Hnrnpk",
                "Ndufa4",
                "Ewsr1",
                "Ndufv2",
                "H3f3a",
                "Ube2k",
                "Cyp51",
                "Fam49b",
                "Cisd3",
                "Rpl3",
                "Gfer",
                "Cebpzos",
                "Cdv3",
                "Hmgn1",
                "Ctbp2",
                "Asns",
                "Eef1d",
                "Ndufs7",
                "Lsm3",
                "Gm16286",
                "Eif5",
                "Vapa",
                "Txn1",
                "Mrpl17",
                "Ran",
                "Fscn1",
                "Ldha",
                "Apex1",
                "Emc8",
                "Cct5",
                "Mrpl40",
                "Bclaf1",
                "C1qbp",
                "Rpsa",
                "H2afz",
                "Psip1",
                "Rpl29",
                "Fubp1",
                "Ccdc58",
                "Prdx6",
                "Ptges3",
                "Dpy30",
                "Wtap",
                "Odc1",
                "Rps20",
                "Dnaja2",
                "Plekhf2",
                "Rps6",
                "Cisd1",
                "U2af1",
                "Pole4",
                "Ywhaq",
                "Fbl",
                "Ubald2",
                "Calm2",
                "Cox7c",
                "Stmn1",
                "Srsf10",
                "Adss",
                "Smarcc1",
                "Ube2v2",
                "Eno1",
                "Trnau1ap",
                "Adprh",
                "Srsf3",
                "Coprs",
                "Cmpk1",
                "Rpl7",
                "Cct7",
                "Ppan",
                "Hsp90ab1",
                "Rwdd1",
                "Snx5",
                "Rps8",
                "Dnaja1",
                "Pgp",
                "Eif4a1",
                "Sumo2",
                "Pcbp2",
                "Dnajc2",
                "Rif1",
                "Pold2",
                "Wdr5",
                "Nudt19",
                "Sgk1",
                "Morf4l2",
                "Ube2d3",
                "Rbbp4",
                "Marc2",
                "Snrpa1",
                "Srrm2",
                "Top1",
                "Hmgb1",
                "Mrps15",
                "Psma4",
                "Park7",
                "Dynll1",
                "Hspd1",
                "Cpe",
                "Cops7a",
                "Mrpl45",
                "Smdt1",
                "Sod1",
                "Mat2a",
                "Nap1l1",
                "Csrp2",
                "Dnajb6",
                "Brd2",
                "Actr3",
                "G3bp2",
                "Smarca5",
                "Ccnb1",
                "Prdx2",
                "Ccnd3",
                "Khdrbs1",
                "Adrm1",
                "Abcf1",
                "Hspa9",
                "Pim3",
                "Ube2n",
                "Hmgn2",
                "Hnrnpa2b1",
                "Tmem97",
                "Srsf2",
                "Eif4g2",
                "Raly",
                "Denr",
                "Fxr1",
                "Nsmce4a",
                "Get4",
                "Vdac1",
                "Cggbp1",
                "Zfp706",
                "Anp32a",
                "Rad23b",
                "Lsm2",
                "Anapc13",
                "Nudt4",
                "Nudcd2",
                "Eif4g1",
                "Rhoa",
                "Cacybp",
                "Psma5",
                "Rpl22",
                "Nolc1",
                "Anp32e",
                "Syncrip",
                "Caprin1",
                "Glud1",
                "Hnrnpu",
                "Rbmxl1",
                "Gspt1",
                "Arglu1",
                "Imp4",
                "Srsf11",
                "Hmgb3",
                "Tomm70a",
                "Rpl21",
                "Ppp1r2",
                "Map1lc3b",
                "Eprs",
                "Dazap1",
                "Pop5",
                "Ywhab",
                "Ubb",
                "Eif1ax",
                "Parp1",
                "Ak6",
                "Hnrnpa3",
                "Prpf40a",
                "Mrps7",
                "Mrpl43",
                "Ube2s",
                "Sf3b1",
                "Idh2",
                "H2afy",
                "Hnrnpd",
                "Chchd4",
                "Bcas2",
                "Ciapin1",
                "Eif1ad",
                "Rps13",
                "Tpt1",
                "Gng5",
                "Rrp1b",
                "Hspe1",
                "Exosc8",
                "Ptbp1",
                "Aldoa",
                "Srp9",
                "Hnrnpdl",
                "Rtn3",
                "Dnajc21",
                "Polr2h",
                "Snx3",
                "Birc5",
                "Rnaseh2a",
                "Chtop",
                "Srpk1",
                "Rfc1",
                "Ppa1",
                "Lamtor5",
                "Avpi1",
                "Ssb",
                "Cdc34",
                "Gps1",
                "Jun",
                "Atrx",
                "Nsun2",
                "Set",
                "Txnl4a",
                "Eif4e2",
                "Ssbp1",
                "Tgif1",
                "Hist1h1c",
                "Clns1a",
                "Gtf2e2",
                "Rpl15",
                "Gtf2h5",
                "Hsp90b1",
                "Vdac2",
                "Pfn1",
                "Tkt",
                "Ddx39b",
                "Cuta",
                "Cox5a",
                "Anapc11",
                "Fkbp4",
                "Pgam1",
                "Nans",
                "Lyar",
                "Uchl5",
                "Snrnp70",
                "Cct4",
                "Rdx",
                "Eif4b",
                "Eif4ebp1",
                "Anp32b",
                "Edf1",
                "Psmg4",
                "Sumo3",
                "Psmc2",
                "Ndufa10",
                "Mcm6",
                "Naa50",
                "Ywhah",
                "Prpf19",
                "Hdac2",
                "Fkbp3",
                "Mtch2",
                "Dek",
                "Rpn2",
                "Rpl7a",
                "Papola",
                "Manf",
                "Cetn3",
                "Galk1",
                "Ccna2",
                "Slc25a5",
                "Tpr",
                "Hint1",
                "Eed",
                "Sf3a2",
                "Ccne1",
                "Srsf6",
                "Tmpo",
                "Tsn",
                "Txnl1",
                "Mesdc2",
                "Rbm25",
                "Drg1",
                "Cycs",
                "Arhgdia",
                "Ptma",
                "Rac1",
                "Dut",
                "Ralbp1",
                "Ybx3",
                "Klf5",
                "Yrdc",
                "Pabpc1",
                "Alyref",
                "0610007P14Rik",
                "Yif1b",
                "Guk1",
                "Tsfm",
                "Rplp2",
                "Skp1a",
                "Ptov1",
                "Gpx4",
                "Rheb",
                "Mkrn1",
                "Sfr1",
                "Ncor1"
            ],
            "expression_sum": {
                "class1": 10169.0,
                "class2": 9450.0
            },
            "uniprot_ids": [
                "P35428",
                "Q3V1C5",
                "Q3UZZ2"
            ]
        },
        "Hint1": {
            "id": "Hint1",
            "type": "Gene",
            "symbol": "Hint1",
            "source": [
                "Pou5f1",
                "Cebpb",
                "Nfib",
                "Atf4",
                "Junb",
                "Fos",
                "Jun",
                "Sox2",
                "Hes1",
                "Nanog",
                "Jund",
                "Hdac2"
            ],
            "target": [],
            "expression_sum": {
                "class1": 79387.0,
                "class2": 177974.0
            },
            "uniprot_ids": [
                "P70349",
                "B0R1E3"
            ]
        },
        "Hmga2": {
            "id": "Hmga2",
            "type": "TF",
            "symbol": "Hmga2",
            "source": [
                "E2f4",
                "Trim28",
                "Rest",
                "Atf4",
                "Ctcf",
                "Trp53",
                "Hdac1",
                "Klf5",
                "Tbx3",
                "Hdac2",
                "Smc3",
                "Smarca5"
            ],
            "target": [
                "Me1",
                "Rnf7",
                "Srsf9",
                "Usmg5",
                "1700037H04Rik",
                "1810043H04Rik",
                "Setd6",
                "Srm",
                "Cox8a",
                "Clta",
                "Marcksl1",
                "Ppp1r11",
                "Actr2",
                "1110008F13Rik",
                "Chchd6",
                "Ndufb3",
                "Aurka",
                "Rpl23a",
                "Aggf1",
                "Mrps23",
                "1110059E24Rik",
                "Mlf2",
                "Rnf26",
                "Pstk",
                "Nop16",
                "Nlk",
                "Sfxn1",
                "Shfm1",
                "Kpna3",
                "Nedd4",
                "Hn1",
                "Cdk1",
                "Phb",
                "2810417H13Rik",
                "Anapc15",
                "Ddx1",
                "Tex30",
                "Chd1",
                "Pmf1",
                "Aurkaip1",
                "Nhp2",
                "Srsf1",
                "Golga7",
                "Wdr89",
                "Rcc2",
                "Phip",
                "Me2",
                "Pdcd6",
                "Cox20",
                "Akt1s1",
                "Ube2d2a",
                "Atp5h",
                "Cuedc2",
                "Mkrn2",
                "Dhx9",
                "Kcnj15",
                "2310036O22Rik",
                "Gnai3",
                "Ube3a",
                "Psmb3",
                "Rbm39",
                "BC023719",
                "Exosc10",
                "Rpl24",
                "Tnrc6a",
                "Copb1",
                "Arl8b",
                "Dnm1l",
                "1700123O20Rik",
                "Fam216a",
                "Rce1",
                "Fkbp8",
                "Mmadhc",
                "Nosip",
                "Pcmt1",
                "Preb",
                "Hells",
                "9430016H08Rik",
                "Tonsl",
                "Pcbp1",
                "Rps12-ps3",
                "Eps8",
                "Cops5",
                "Prrc2b",
                "Vars",
                "Csnk1a1",
                "Bax",
                "Rpl10-ps3",
                "Dnajc19",
                "Bap1",
                "Coq2",
                "1700020I14Rik",
                "Gtf3a",
                "0610037L13Rik",
                "Lsm3",
                "Prpf8",
                "Asxl1",
                "Hadh",
                "Smek2",
                "Gm8730",
                "Srrm1",
                "Aatf",
                "Diablo",
                "Sdcbp",
                "Tcn2",
                "Atp5g2",
                "Srp19",
                "Mrpl35",
                "1110004F10Rik",
                "Ncapg",
                "Ercc1",
                "Uqcr10",
                "Jmjd6",
                "Ung",
                "Dph3",
                "S100a6",
                "Ptges3",
                "Mrpl18",
                "Dnaja2",
                "Alyref2",
                "Acat2",
                "Cbll1",
                "Ndufa3",
                "Mtdh",
                "Rps6",
                "Eef1a1",
                "Ldb1",
                "Hjurp",
                "Mrps6",
                "mt-Co2",
                "Calm2",
                "Taf5l",
                "0610010K14Rik",
                "Nup35",
                "Ccnl2",
                "Atp5d",
                "Ifrd1",
                "Pabpc4",
                "Lamtor4",
                "1500011K16Rik",
                "Dctpp1",
                "Bora",
                "Mrpl50",
                "Rps9",
                "Smc1a",
                "Mafg",
                "Cyc1",
                "Prdx3",
                "Sumo2",
                "Dnttip1",
                "Ndufa12",
                "Adk",
                "Ube2f",
                "Ube2d3",
                "Chordc1",
                "Arhgap42",
                "Fam133b",
                "Eefsec",
                "2700060E02Rik",
                "1110001J03Rik",
                "Rps15",
                "Arpc3",
                "Zmynd11",
                "Mgat2",
                "Srsf7",
                "Smdt1",
                "Degs1",
                "Celf1",
                "Etfb",
                "Hsbp1",
                "Hspa5",
                "Atp1a1",
                "Tmem199",
                "Tpgs2",
                "Nup153",
                "Ppif",
                "Tsr1",
                "Actr1b",
                "Wapl",
                "Abcf1",
                "Rplp0",
                "Chchd2",
                "Smarcb1",
                "2410002F23Rik",
                "Ndufs5",
                "Polr3d",
                "Ddx42",
                "Eif4g2",
                "Uqcc2",
                "Fxr1",
                "Aasdhppt",
                "Tceal8",
                "Cpne3",
                "Rbm3",
                "Psmd11",
                "Tdg",
                "0610012G03Rik",
                "Selk",
                "Npm3",
                "Rad23b",
                "Cyfip1",
                "Qars",
                "Chmp2a",
                "Aprt",
                "Fam96a",
                "Ptrh2",
                "Supt16",
                "Snrpc",
                "Pabpn1",
                "Lsg1",
                "Adprhl2",
                "Rpl22",
                "Sf3a3",
                "Gid8",
                "Pfdn6",
                "1810011O10Rik",
                "Mrpl23",
                "Fermt2",
                "Gspt1",
                "Rpl21",
                "Ccnh",
                "Cct2",
                "Eif2b3",
                "Sept11",
                "Gcn1l1",
                "Eif1ax",
                "Ak6",
                "Mrps7",
                "Pdcd10",
                "Psmc3ip",
                "Metap2",
                "0610009B22Rik",
                "Rangap1",
                "Cdca8",
                "Metap1",
                "Taf12",
                "Tm4sf1",
                "Commd2",
                "Cep83",
                "Mtf2",
                "Hnrnpd",
                "Tma7",
                "Itga6",
                "E2f6",
                "1110004E09Rik",
                "Eif5a",
                "4833439L19Rik",
                "Cfl1",
                "Stag1",
                "Abracl",
                "Aar2",
                "Cops6",
                "Aldoa",
                "Srp9",
                "Polr2h",
                "Adh5",
                "Commd1",
                "Cox7b",
                "Chtf8",
                "Srpk1",
                "Rfc1",
                "Cmc2",
                "Ddx10",
                "Cdc34",
                "Tnk2",
                "Atp5o",
                "Timm21",
                "Mrps12",
                "Ubqln4",
                "Brcc3",
                "Wdr12",
                "Nasp",
                "Ndufb5",
                "Abhd17a",
                "Set",
                "Ssbp1",
                "Cdc25a",
                "Ago2",
                "Fez2",
                "Prmt5",
                "1810022K09Rik",
                "Slc38a4",
                "Phb2",
                "Etfa",
                "Bcl10",
                "Bud31",
                "Rpl11",
                "Brk1",
                "Rspry1",
                "Rnf220",
                "2410004B18Rik",
                "Timm17b",
                "Mpzl1",
                "Hmmr",
                "Prpf6",
                "Fkbp4",
                "Pgam1",
                "Tomm22",
                "Mier1",
                "Nadk",
                "Thap11",
                "Eif4b",
                "Rpl23a-ps3",
                "Anp32b",
                "Phf14",
                "Zfp36l2",
                "Mrpl42",
                "Pum3",
                "Ipo9",
                "Dusp9",
                "Eif3i",
                "Zfp593",
                "Mtch2",
                "Gm9843",
                "Dcp2",
                "Nelfe",
                "Depdc1a",
                "Uck2",
                "Tmem33",
                "Cetn3",
                "Arhgdib",
                "Tpr",
                "Abce1",
                "Cnpy3",
                "Ccne1",
                "Ndufb10",
                "Pnkp",
                "Ptpn2",
                "Arhgdia",
                "Cycs",
                "Dut",
                "Prpf31",
                "Med11",
                "Ubap2",
                "Gga2",
                "Ppp2r5e",
                "Ybx3",
                "Gnl1",
                "Acin1",
                "Hspa14",
                "Kansl1",
                "0610007P14Rik",
                "Smim15",
                "Nom1",
                "Glrx2",
                "2810428I15Rik",
                "Serp1",
                "Rps18",
                "Serbp1",
                "Gcsh",
                "Coq5",
                "Cars",
                "Scaf11",
                "Rpl23",
                "1810037I17Rik",
                "Ddx49",
                "Arfip2",
                "Ly6e",
                "Rbm17"
            ],
            "expression_sum": {
                "class1": 24405.0,
                "class2": 4050.0
            },
            "uniprot_ids": [
                "P52927",
                "Q6NSP9"
            ]
        },
        "Hmgb1": {
            "id": "Hmgb1",
            "type": "TF",
            "symbol": "Hmgb1",
            "source": [
                "Klf6",
                "Tcf4",
                "Pou5f1",
                "Foxp1",
                "Nfib",
                "Atf4",
                "Junb",
                "Fos",
                "Jun",
                "Sox2",
                "Hes1",
                "Nanog",
                "Jund",
                "Twist2",
                "Cebpb",
                "Zfp57",
                "Sox4"
            ],
            "target": [
                "Atxn7l3b",
                "Skil",
                "Tspan6",
                "Amotl2",
                "Atf5",
                "Cuedc1",
                "Tns1",
                "Ak1",
                "Svep1",
                "Fam204a",
                "Tmed5",
                "Fabp5",
                "Mettl9",
                "Nfatc4",
                "Hspa5",
                "Igf2",
                "Rasgrp2",
                "Phtf1",
                "Efemp2",
                "Col12a1",
                "Bex1",
                "Ifngr2",
                "Anxa2",
                "Dcun1d1",
                "9530068E07Rik",
                "Ccar1",
                "Flnb",
                "Cox7a2l",
                "Smap1",
                "Acot7",
                "Basp1",
                "Ahnak",
                "Elovl1",
                "1700020I14Rik",
                "Gabarap",
                "Zbtb20",
                "Metrn",
                "Nab2",
                "Phgdh",
                "Rfwd2",
                "Brk1",
                "Hsp90b1",
                "Pgrmc1",
                "Adamts1",
                "Timp1",
                "Cited2",
                "Celf2",
                "Rps15a",
                "Adam9",
                "Setd3",
                "Eif4g2",
                "Sdcbp",
                "Tuba1a",
                "Lamb1",
                "Emd",
                "2610305D13Rik",
                "Pdlim7",
                "Micu1",
                "Actn1",
                "Rcbtb2",
                "Psat1",
                "Eif4h",
                "Pex2",
                "Rnf139",
                "Trappc3",
                "Shoc2",
                "2310009B15Rik",
                "Anxa5",
                "Bad",
                "Prdx6",
                "Eif3f",
                "Hadha",
                "Serpine1",
                "2200002D01Rik",
                "Rab10",
                "Alg5",
                "AY036118",
                "Crabp1",
                "Ifitm3",
                "Lix1l",
                "Nptn",
                "B3galnt1",
                "Iqgap1",
                "Casc5",
                "Kdelr2",
                "Tdgf1",
                "March5",
                "App",
                "2310022B05Rik",
                "Arl6ip5",
                "Atp6ap1",
                "Dexi",
                "Akt1s1",
                "Hist1h1e",
                "Agpat4",
                "Pde10a",
                "P4ha1",
                "Spcs1",
                "Ndfip2",
                "Ssr4",
                "Ctdsp1",
                "Gpc1",
                "Ap2a2",
                "Crebbp",
                "Araf",
                "Wdr26",
                "Htatsf1",
                "Tspan5",
                "Dusp11",
                "Slc25a46",
                "Plaur",
                "Arpc2",
                "Mrpl54",
                "Mageh1",
                "Fndc4",
                "Col4a2",
                "Capza2",
                "Ccl2",
                "Chsy1",
                "Mbnl3",
                "Upf3a",
                "Phlda3",
                "Pbx1",
                "Ralb",
                "Trip6",
                "Txnrd1",
                "1600012H06Rik",
                "Ube3a",
                "Ergic3",
                "Taok3",
                "Atp5d",
                "2610001J05Rik",
                "Slc39a13",
                "Maoa",
                "H2afy2",
                "Smim10l1",
                "Cstb",
                "0610011F06Rik",
                "Gm10260",
                "Mpp1",
                "Btg1",
                "Acer2",
                "Lrrc58",
                "Kif26b",
                "Cope",
                "Tspan31",
                "L1td1",
                "Ap1s1",
                "Col3a1",
                "Tuba4a",
                "Mecr",
                "Rab14",
                "Serpinh1",
                "Psap",
                "Armcx2",
                "H2afy",
                "Rwdd1",
                "4930402H24Rik",
                "Acat1",
                "Cald1",
                "Arl8b",
                "Cpox",
                "Ext2",
                "Sema3c",
                "Gm17501",
                "Rbm7",
                "Tm9sf3",
                "Trh",
                "Ctsd",
                "1700123O20Rik",
                "Cbx7",
                "4933434E20Rik",
                "Vgll3",
                "Arpc5",
                "Elf2",
                "Med21",
                "Hdlbp",
                "Cox4i1",
                "Ephb2",
                "Rgs16",
                "Airn",
                "Efnb1",
                "Thbs2",
                "Gdf3",
                "Evl",
                "Nek6",
                "Adk",
                "Nxn",
                "Stard3nl",
                "Serp1",
                "Lrrfip1",
                "Snx7",
                "Laptm4a",
                "Tmem59",
                "Vcan",
                "Actg1",
                "Ptprd",
                "Cdc27",
                "Emp3",
                "Fat4",
                "Abracl",
                "Cryab",
                "Dnah8",
                "Pdcl3",
                "Map4",
                "Cd302",
                "Arpc3",
                "Rassf1",
                "Arl1",
                "Col1a2",
                "Sult6b1"
            ],
            "expression_sum": {
                "class1": 34517.0,
                "class2": 116070.0
            },
            "uniprot_ids": [
                "P63158",
                "Q58EV5",
                "Q8C7C4",
                "A0A0J9YUD8",
                "Q497Z6",
                "Q3UJK0",
                "Q8BQ02",
                "A0A0J9YUZ4",
                "Q6P202",
                "D3YVC6",
                "Q3UBK2",
                "D3YZ18",
                "Q8BNM0"
            ]
        },
        "Hmgn2": {
            "id": "Hmgn2",
            "type": "Gene",
            "symbol": "Hmgn2",
            "source": [
                "Klf2",
                "Tfcp2l1",
                "Pou5f1",
                "Foxp1",
                "Nfib",
                "Atf4",
                "Junb",
                "Fos",
                "Jun",
                "Sox2",
                "Tbx3",
                "Hes1",
                "Nanog",
                "Jund",
                "Twist2",
                "Cebpb",
                "Sox4"
            ],
            "target": [],
            "expression_sum": {
                "class1": 11365.0,
                "class2": 51304.0
            },
            "uniprot_ids": [
                "P09602",
                "Q5BL14",
                "Q5XK38",
                "F6W687",
                "B7ZCQ3",
                "A3KGL9",
                "Q58E57"
            ]
        },
        "Hnrnpa2b1": {
            "id": "Hnrnpa2b1",
            "type": "Gene",
            "symbol": "Hnrnpa2b1",
            "source": [
                "Tcf4",
                "Pou5f1",
                "Foxp1",
                "Cebpb",
                "Nfib",
                "Atf4",
                "Junb",
                "Fos",
                "Jun",
                "Sox2",
                "Hes1",
                "Nanog",
                "Jund",
                "Hdac2",
                "Smc3",
                "Sox4"
            ],
            "target": [],
            "expression_sum": {
                "class1": 42126.0,
                "class2": 87741.0
            },
            "uniprot_ids": [
                "O88569",
                "B7ZP22",
                "A0A0N4SUM2",
                "Q3UTH2"
            ]
        },
        "Hnrnpa3": {
            "id": "Hnrnpa3",
            "type": "Gene",
            "symbol": "Hnrnpa3",
            "source": [
                "Tcf4",
                "Pou5f1",
                "Foxp1",
                "Cebpb",
                "Nfib",
                "Atf4",
                "Junb",
                "Fos",
                "Jun",
                "Sox2",
                "Sfpq",
                "Hes1",
                "Nanog",
                "Jund",
                "Twist2",
                "Hdac2",
                "Smarca5",
                "Sox4"
            ],
            "target": [],
            "expression_sum": {
                "class1": 24535.0,
                "class2": 73867.0
            },
            "uniprot_ids": [
                "Q8BG05",
                "Q5FB19",
                "B2RXM2",
                "Q0VG47",
                "Q6P6I7",
                "A2AL12",
                "A2AL13",
                "Q3UZG3",
                "Q5U3M2",
                "Q58E39",
                "Q6PFA7"
            ]
        },
        "Hnrnpab": {
            "id": "Hnrnpab",
            "type": "Gene",
            "symbol": "Hnrnpab",
            "source": [
                "Tcf4",
                "Pou5f1",
                "Cebpb",
                "Nfib",
                "Atf4",
                "Junb",
                "Fos",
                "Jun",
                "Sox2",
                "Nanog",
                "Jund",
                "Hdac2",
                "Smc3",
                "Sox4"
            ],
            "target": [],
            "expression_sum": {
                "class1": 25134.0,
                "class2": 98881.0
            },
            "uniprot_ids": [
                "Q99020",
                "Q20BD0",
                "Q544Z3",
                "Q80XR6",
                "Q3TMZ8",
                "Q9D6G1",
                "Q8CIR3"
            ]
        },
        "Hnrnpdl": {
            "id": "Hnrnpdl",
            "type": "Gene",
            "symbol": "Hnrnpdl",
            "source": [
                "Klf2",
                "Nfib",
                "Fos",
                "Sox2",
                "Hdac2",
                "Tcf4",
                "Pou5f1",
                "Jun",
                "Bclaf1",
                "Smarca5",
                "Sox4",
                "Eno1",
                "Jund",
                "Cebpb",
                "Smc3",
                "Foxp1",
                "Atf4",
                "Junb",
                "Hes1",
                "Tfcp2l1",
                "Nanog",
                "Twist2"
            ],
            "target": [],
            "expression_sum": {
                "class1": 22110.0,
                "class2": 61323.0
            },
            "uniprot_ids": [
                "Q9Z130",
                "D3YTQ3",
                "F6ZKM7",
                "F6VQH5"
            ]
        },
        "Hnrnpf": {
            "id": "Hnrnpf",
            "type": "Gene",
            "symbol": "Hnrnpf",
            "source": [
                "Klf2",
                "Tcf4",
                "Pou5f1",
                "Foxp1",
                "Cebpb",
                "Nfib",
                "Atf4",
                "Junb",
                "Fos",
                "Jun",
                "Sox2",
                "Hes1",
                "Nanog",
                "Jund",
                "Twist2",
                "Hdac2",
                "Sox4"
            ],
            "target": [],
            "expression_sum": {
                "class1": 30092.0,
                "class2": 119058.0
            },
            "uniprot_ids": [
                "Q9Z2X1",
                "J3QP45",
                "J3QMV8",
                "J3QMT0",
                "J3QM80",
                "J3QNH2",
                "Q8R582"
            ]
        },
        "Hnrnph1": {
            "id": "Hnrnph1",
            "type": "Gene",
            "symbol": "Hnrnph1",
            "source": [
                "Tcf4",
                "Pou5f1",
                "Foxp1",
                "Cebpb",
                "Nfib",
                "Atf4",
                "Junb",
                "Fos",
                "Id2",
                "Jun",
                "Trp53",
                "Jund",
                "Hdac2",
                "Smc3",
                "Smarca5",
                "Sox4"
            ],
            "target": [],
            "expression_sum": {
                "class1": 10236.0,
                "class2": 14000.0
            },
            "uniprot_ids": [
                "O35737",
                "Q811L7",
                "Q8C2Q7",
                "Q3TNG9"
            ]
        },
        "Hnrnpk": {
            "id": "Hnrnpk",
            "type": "Gene",
            "symbol": "Hnrnpk",
            "source": [
                "Tcf4",
                "Pou5f1",
                "Foxp1",
                "Cebpb",
                "Nfib",
                "Atf4",
                "Junb",
                "Fos",
                "Id2",
                "Jun",
                "Sox2",
                "Prrx1",
                "Hes1",
                "Nanog",
                "Jund",
                "Hdac2",
                "Smc3"
            ],
            "target": [],
            "expression_sum": {
                "class1": 41108.0,
                "class2": 97907.0
            },
            "uniprot_ids": [
                "P61979",
                "H3BJ43",
                "Q5FWJ5",
                "Q8BT23",
                "B2M1R6",
                "H3BLP7",
                "H3BKD0",
                "Q3TUA1",
                "A0A286YDM3",
                "A0A286YE41",
                "H3BJS9",
                "A0A286YCM2",
                "Q3U6X2",
                "A0A286YDH1",
                "Q3TL71",
                "H3BLL4",
                "H3BKI8",
                "H3BK96",
                "Q3U9Q3",
                "A0A286YEC4",
                "Q3TJ38",
                "Q3TG17"
            ]
        },
        "Hnrnpu": {
            "id": "Hnrnpu",
            "type": "Gene",
            "symbol": "Hnrnpu",
            "source": [
                "Tcf4",
                "Pou5f1",
                "Foxp1",
                "Cebpb",
                "Nfib",
                "Atf4",
                "Junb",
                "Fos",
                "Jun",
                "Hes1",
                "Nanog",
                "Jund",
                "Twist2",
                "Hdac2",
                "Smc3",
                "Smarca5",
                "Sox4"
            ],
            "target": [],
            "expression_sum": {
                "class1": 10697.0,
                "class2": 32531.0
            },
            "uniprot_ids": [
                "Q8VEK3",
                "Q8C290",
                "Q3TGN5",
                "Q3TXW2",
                "Q3TS50",
                "Q3TWN5",
                "A0A087WPF1",
                "Q3ULH5",
                "O88568",
                "A0A087WPX7",
                "Q3TVV6"
            ]
        },
        "Hsp90aa1": {
            "id": "Hsp90aa1",
            "type": "Gene",
            "symbol": "Hsp90aa1",
            "source": [
                "Nfib",
                "Fos",
                "Id2",
                "Sox2",
                "Hdac2",
                "Tcf4",
                "Pou5f1",
                "Jun",
                "Smarca5",
                "Sox4",
                "Jund",
                "Cebpb",
                "Foxp1",
                "Atf4",
                "Junb",
                "Ybx1",
                "Hes1",
                "Tfcp2l1",
                "Nanog",
                "Twist2"
            ],
            "target": [],
            "expression_sum": {
                "class1": 73242.0,
                "class2": 675930.0
            },
            "uniprot_ids": [
                "P07901",
                "Q80Y52",
                "Q3TKG0",
                "Q3TKB9",
                "Q3TKA2",
                "Q3TJU7",
                "A2A6A2",
                "A0PJ91",
                "B7ZC50",
                "Q8C2A9",
                "Q8C5U3",
                "Q3UIF3",
                "B7ZC49"
            ]
        },
        "Hsp90b1": {
            "id": "Hsp90b1",
            "type": "Gene",
            "symbol": "Hsp90b1",
            "source": [
                "Tcf4",
                "Pou5f1",
                "Trim28",
                "Cebpb",
                "Nfib",
                "Hmgb1",
                "Atf4",
                "Ctcf",
                "Junb",
                "Fos",
                "Jun",
                "Trp53",
                "Ybx1",
                "Hes1",
                "Jund",
                "Twist2",
                "Hdac2",
                "Smc3"
            ],
            "target": [],
            "expression_sum": {
                "class1": 67214.0,
                "class2": 54362.0
            },
            "uniprot_ids": [
                "P08113",
                "Q91V38",
                "Q3UAD6",
                "F7C312",
                "Q8CCY5",
                "Q8C2R2",
                "Q3TUD6",
                "Q3UBU0",
                "Q9CW12"
            ]
        },
        "Hspa14": {
            "id": "Hspa14",
            "type": "Gene",
            "symbol": "Hspa14",
            "source": [
                "Tcf4",
                "Pou5f1",
                "Foxp1",
                "Hmga2",
                "Nfib",
                "Atf4",
                "Junb",
                "Fos",
                "Jun",
                "Sox2",
                "Nanog",
                "Jund",
                "Twist2",
                "Cebpb"
            ],
            "target": [],
            "expression_sum": {
                "class1": 3663.0,
                "class2": 20404.0
            },
            "uniprot_ids": [
                "Q99M31"
            ]
        },
        "Hspa4": {
            "id": "Hspa4",
            "type": "Gene",
            "symbol": "Hspa4",
            "source": [
                "Tcf4",
                "Pou5f1",
                "Nfib",
                "Atf4",
                "Junb",
                "Fos",
                "Jun",
                "Sox2",
                "Hes1",
                "Nanog",
                "Jund",
                "Twist2",
                "Cebpb"
            ],
            "target": [],
            "expression_sum": {
                "class1": 7741.0,
                "class2": 46965.0
            },
            "uniprot_ids": [
                "Q61316",
                "Q3U2G2",
                "Q571M2",
                "Q99L75",
                "Q8C6R3",
                "Q3UZ53"
            ]
        },
        "Hspa5": {
            "id": "Hspa5",
            "type": "Gene",
            "symbol": "Hspa5",
            "source": [
                "Tcf4",
                "Hmga2",
                "Trim28",
                "Cebpb",
                "Nfib",
                "Atf5",
                "Hmgb1",
                "Atf4",
                "Ctcf",
                "Junb",
                "Fos",
                "Jun",
                "Trp53",
                "Ybx1",
                "Jund",
                "Hdac2",
                "Smc3",
                "Smarca5"
            ],
            "target": [],
            "expression_sum": {
                "class1": 57591.0,
                "class2": 42487.0
            },
            "uniprot_ids": [
                "P20029",
                "A0A0A6YXF5",
                "Q3U7T8",
                "Q3TI47",
                "Q3UEM8",
                "Q9DC41",
                "Q3TKF8",
                "Q3TUS2",
                "Q3TWF2",
                "Q3U9G2",
                "Q3U6V3"
            ]
        },
        "Hspa9": {
            "id": "Hspa9",
            "type": "Gene",
            "symbol": "Hspa9",
            "source": [
                "Tcf4",
                "Pou5f1",
                "Foxp1",
                "Cebpb",
                "Atf4",
                "Junb",
                "Fos",
                "Jun",
                "Sox2",
                "Hes1",
                "Nanog",
                "Jund",
                "Twist2",
                "Hdac2",
                "Smarca5",
                "Sox4"
            ],
            "target": [],
            "expression_sum": {
                "class1": 24367.0,
                "class2": 61861.0
            },
            "uniprot_ids": [
                "P38647"
            ]
        },
        "Hspd1": {
            "id": "Hspd1",
            "type": "Gene",
            "symbol": "Hspd1",
            "source": [
                "Klf2",
                "Tcf4",
                "Pou5f1",
                "Foxp1",
                "Cebpb",
                "Nfib",
                "Atf4",
                "Junb",
                "Fos",
                "Jun",
                "Sox2",
                "Hes1",
                "Nanog",
                "Jund",
                "Twist2",
                "Hdac2",
                "Smarca5",
                "Sox4"
            ],
            "target": [],
            "expression_sum": {
                "class1": 36829.0,
                "class2": 132713.0
            },
            "uniprot_ids": [
                "P63038",
                "D3Z2F2",
                "D3Z7J9"
            ]
        },
        "Hspe1": {
            "id": "Hspe1",
            "type": "Gene",
            "symbol": "Hspe1",
            "source": [
                "Klf2",
                "Tcf4",
                "Pou5f1",
                "Foxp1",
                "Nfib",
                "Atf4",
                "Junb",
                "Fos",
                "Jun",
                "Sox2",
                "Hes1",
                "Nanog",
                "Jund",
                "Twist2",
                "Cebpb",
                "Zfp57",
                "Sox4"
            ],
            "target": [],
            "expression_sum": {
                "class1": 78938.0,
                "class2": 315342.0
            },
            "uniprot_ids": [
                "Q64433",
                "Q4KL76"
            ]
        },
        "Ict1": {
            "id": "Ict1",
            "type": "Gene",
            "symbol": "Ict1",
            "source": [
                "Tcf4",
                "Pou5f1",
                "Cebpb",
                "Nfib",
                "Eno1",
                "Atf4",
                "Junb",
                "Fos",
                "Id2",
                "Jun",
                "Nanog",
                "Tfcp2l1",
                "Jund",
                "Twist2",
                "Hdac2",
                "Smarca5",
                "Sox4"
            ],
            "target": [],
            "expression_sum": {
                "class1": 7814.0,
                "class2": 18878.0
            },
            "uniprot_ids": [
                "Q8R035"
            ]
        },
        "Idh2": {
            "id": "Idh2",
            "type": "Gene",
            "symbol": "Idh2",
            "source": [
                "Tcf4",
                "Pou5f1",
                "Cebpb",
                "Nfib",
                "Atf4",
                "Junb",
                "Fos",
                "Jun",
                "Sox2",
                "Hes1",
                "Nanog",
                "Jund",
                "Hdac2",
                "Smc3"
            ],
            "target": [],
            "expression_sum": {
                "class1": 13585.0,
                "class2": 26600.0
            },
            "uniprot_ids": [
                "P54071",
                "A0A0U1RP68",
                "D6RIL6",
                "A0A0U1RPR1"
            ]
        },
        "Idh3g": {
            "id": "Idh3g",
            "type": "Gene",
            "symbol": "Idh3g",
            "source": [
                "Tcf4",
                "Pou5f1",
                "Atf4",
                "Junb",
                "Jun",
                "Nanog",
                "Jund",
                "Twist2",
                "Cebpb"
            ],
            "target": [],
            "expression_sum": {
                "class1": 6456.0,
                "class2": 21502.0
            },
            "uniprot_ids": [
                "P70404",
                "Q3TGZ3",
                "Q684I8",
                "Q3TKM5",
                "Q9CW65"
            ]
        },
        "Ift20": {
            "id": "Ift20",
            "type": "Gene",
            "symbol": "Ift20",
            "source": [
                "Trim28",
                "Rest",
                "Atf4",
                "Trp53",
                "Hdac1",
                "Ybx1",
                "Hdac2",
                "Smc3"
            ],
            "target": [],
            "expression_sum": {
                "class1": 11060.0,
                "class2": 12461.0
            },
            "uniprot_ids": [
                "Q61025",
                "Z4YJL9"
            ]
        },
        "Igfbp2": {
            "id": "Igfbp2",
            "type": "Gene",
            "symbol": "Igfbp2",
            "source": [
                "Pou5f1",
                "Trim28",
                "Eno1",
                "Atf4",
                "Trp53",
                "Hdac1",
                "Klf5",
                "Tbx3",
                "Zfp57",
                "Nanog",
                "Smc3"
            ],
            "target": [],
            "expression_sum": {
                "class1": 5008.0,
                "class2": 95030.0
            },
            "uniprot_ids": [
                "P47877",
                "D3YU40",
                "Q9D057"
            ]
        },
        "Immt": {
            "id": "Immt",
            "type": "Gene",
            "symbol": "Immt",
            "source": [
                "Tcf4",
                "Pou5f1",
                "Foxp1",
                "Cebpb",
                "Nfib",
                "Eno1",
                "Atf4",
                "Fos",
                "Jun",
                "Sox2",
                "Hes1",
                "Nanog",
                "Jund",
                "Twist2",
                "Hdac2"
            ],
            "target": [],
            "expression_sum": {
                "class1": 7699.0,
                "class2": 26563.0
            },
            "uniprot_ids": [
                "Q8CAQ8",
                "A6H604",
                "Q3U7N2",
                "A0A0U1RQ14",
                "E9Q800",
                "A0A0U1RP81",
                "Q3TQL4",
                "A0A0U1RNN9",
                "E9PVS5",
                "Q3TVZ5",
                "A0A1B0GX08",
                "E9QAY6",
                "Q3TZK4",
                "Q6P8Y5",
                "Q2YDW0",
                "Q3TEY5"
            ]
        },
        "Impdh2": {
            "id": "Impdh2",
            "type": "Gene",
            "symbol": "Impdh2",
            "source": [
                "Pou5f1",
                "Foxp1",
                "Nfib",
                "Atf4",
                "Junb",
                "Fos",
                "Jun",
                "Sox2",
                "Nanog",
                "Jund",
                "Twist2",
                "Cebpb",
                "Smarca5"
            ],
            "target": [],
            "expression_sum": {
                "class1": 14625.0,
                "class2": 76027.0
            },
            "uniprot_ids": [
                "P24547",
                "Q3UAT9",
                "A0A0A6YY72",
                "A0A0A6YXS4",
                "O89058",
                "Q3UPJ2",
                "Q3U9N8"
            ]
        },
        "Iscu": {
            "id": "Iscu",
            "type": "Gene",
            "symbol": "Iscu",
            "source": [
                "Pou5f1",
                "Foxp1",
                "Atf4",
                "Junb",
                "Jun",
                "Twist2",
                "Cebpb",
                "Smc3",
                "Smarca5",
                "Sox4"
            ],
            "target": [],
            "expression_sum": {
                "class1": 8502.0,
                "class2": 20061.0
            },
            "uniprot_ids": [
                "Q9D7P6",
                "D3Z7W0",
                "D3Z0K9",
                "F6ZBG7",
                "F6XUL3",
                "D6RE47"
            ]
        },
        "Itm2b": {
            "id": "Itm2b",
            "type": "Gene",
            "symbol": "Itm2b",
            "source": [
                "E2f4",
                "Trim28",
                "Cebpb",
                "Rest",
                "Atf4",
                "Ctcf",
                "Tgif1",
                "Trp53",
                "Klf5",
                "Tbx3",
                "Ybx1",
                "Hdac2",
                "Smc3"
            ],
            "target": [],
            "expression_sum": {
                "class1": 28097.0,
                "class2": 17829.0
            },
            "uniprot_ids": [
                "O89051",
                "A0A2I3BQ30",
                "A0A2I3BPW7",
                "Q9JME4",
                "Q3TVJ0",
                "Q9D1Q3",
                "Q9CW90",
                "Q3UCT9"
            ]
        },
        "Kdelr2": {
            "id": "Kdelr2",
            "type": "Gene",
            "symbol": "Kdelr2",
            "source": [
                "Trim28",
                "Rest",
                "Hmgb1",
                "Atf4",
                "Ctcf",
                "Trp53",
                "Hdac1",
                "Ybx1",
                "Hdac2",
                "Smc3",
                "Smarca5"
            ],
            "target": [],
            "expression_sum": {
                "class1": 44621.0,
                "class2": 18002.0
            },
            "uniprot_ids": [
                "Q9CQM2"
            ]
        },
        "Kdelr3": {
            "id": "Kdelr3",
            "type": "Gene",
            "symbol": "Kdelr3",
            "source": [
                "E2f4",
                "Pou5f1",
                "Trim28",
                "Rest",
                "Atf4",
                "Ctcf",
                "Trp53",
                "Hdac1",
                "Klf5",
                "Tbx3",
                "Hdac2",
                "Smc3"
            ],
            "target": [],
            "expression_sum": {
                "class1": 14786.0,
                "class2": 2346.0
            },
            "uniprot_ids": [
                "Q8R1L4"
            ]
        },
        "Kif5b": {
            "id": "Kif5b",
            "type": "Gene",
            "symbol": "Kif5b",
            "source": [
                "Nfib",
                "Fos",
                "Hdac2",
                "Tcf4",
                "E2f4",
                "Pou5f1",
                "Rest",
                "Trp53",
                "Jun",
                "Smarca5",
                "Trim28",
                "Jund",
                "Cebpb",
                "Smc3",
                "Atf4",
                "Ctcf",
                "Junb",
                "Nanog",
                "Twist2"
            ],
            "target": [],
            "expression_sum": {
                "class1": 19017.0,
                "class2": 26639.0
            },
            "uniprot_ids": [
                "Q61768",
                "Q3UWI9",
                "Q9CUT6",
                "E9QAK5",
                "Q6NSS5",
                "Q05CG8",
                "Q8CFE7"
            ]
        },
        "Klf6": {
            "id": "Klf6",
            "type": "TF",
            "symbol": "Klf6",
            "source": [
                "E2f4",
                "Trim28",
                "Rest",
                "Atf4",
                "Ctcf",
                "Trp53",
                "Hdac1",
                "Klf5",
                "Ybx1",
                "Hdac2",
                "Smc3",
                "Smarca5"
            ],
            "target": [
                "Prmt1",
                "Atp5o",
                "Rpl24",
                "Zfp207",
                "Uqcrc1",
                "Psmd6",
                "Hmgb1",
                "Commd1",
                "Rps27a",
                "Tpr",
                "Nedd4",
                "Ociad1",
                "Ezh2"
            ],
            "expression_sum": {
                "class1": 11198.0,
                "class2": 8416.0
            },
            "uniprot_ids": [
                "O08584",
                "Q8BPQ2",
                "Q3U9J1",
                "A0A1Y7VLA9",
                "Q3UGJ1",
                "Q8VDZ7"
            ]
        },
        "Knop1": {
            "id": "Knop1",
            "type": "Gene",
            "symbol": "Knop1",
            "source": [
                "Pou5f1",
                "Foxp1",
                "Atf4",
                "Junb",
                "Fos",
                "Jun",
                "Sox2",
                "Hes1",
                "Nanog",
                "Jund",
                "Twist2",
                "Peg3",
                "Cebpb",
                "Sox4"
            ],
            "target": [],
            "expression_sum": {
                "class1": 4060.0,
                "class2": 24476.0
            },
            "uniprot_ids": [
                "Q9Z2Q2",
                "H7BX94",
                "F6ZLH4",
                "E9QKQ3",
                "D3Z5V3",
                "K4DI66"
            ]
        },
        "Kpnb1": {
            "id": "Kpnb1",
            "type": "Gene",
            "symbol": "Kpnb1",
            "source": [
                "Tcf4",
                "Pou5f1",
                "Cebpb",
                "Atf4",
                "Junb",
                "Prrx1",
                "Jun",
                "Sox2",
                "Hes1",
                "Tfcp2l1",
                "Jund",
                "Twist2",
                "Hdac2",
                "Smc3",
                "Smarca5"
            ],
            "target": [],
            "expression_sum": {
                "class1": 7531.0,
                "class2": 24219.0
            },
            "uniprot_ids": [
                "P70168",
                "Q99KM9",
                "Q3UHW8",
                "Q7TSZ6",
                "Q3TFE8"
            ]
        },
        "Krtcap2": {
            "id": "Krtcap2",
            "type": "Gene",
            "symbol": "Krtcap2",
            "source": [
                "E2f4",
                "Trim28",
                "Rest",
                "Atf4",
                "Ctcf",
                "Trp53",
                "Hdac1",
                "Klf5",
                "Hdac2",
                "Smc3",
                "Smarca5"
            ],
            "target": [],
            "expression_sum": {
                "class1": 43311.0,
                "class2": 46995.0
            },
            "uniprot_ids": [
                "Q5RL79",
                "A0A0A0MQN1"
            ]
        },
        "Lamtor1": {
            "id": "Lamtor1",
            "type": "Gene",
            "symbol": "Lamtor1",
            "source": [
                "Tcf4",
                "Pou5f1",
                "Foxp1",
                "Cebpb",
                "Atf4",
                "Junb",
                "Trp53",
                "Jun",
                "Twist2",
                "Hdac2",
                "Smc3",
                "Sox4"
            ],
            "target": [],
            "expression_sum": {
                "class1": 11940.0,
                "class2": 28205.0
            },
            "uniprot_ids": [
                "Q9CQ22",
                "A0A0A6YX02"
            ]
        },
        "Lamtor5": {
            "id": "Lamtor5",
            "type": "Gene",
            "symbol": "Lamtor5",
            "source": [
                "Pou5f1",
                "Cebpb",
                "Arf4",
                "Atf4",
                "Junb",
                "Jun",
                "Sox2",
                "Ybx1",
                "Hes1",
                "Tfcp2l1",
                "Jund",
                "Hdac2",
                "Smc3",
                "Smarca5"
            ],
            "target": [],
            "expression_sum": {
                "class1": 12512.0,
                "class2": 27130.0
            },
            "uniprot_ids": [
                "Q9D1L9",
                "G3UW70"
            ]
        },
        "Laptm4a": {
            "id": "Laptm4a",
            "type": "Gene",
            "symbol": "Laptm4a",
            "source": [
                "E2f4",
                "Pou5f1",
                "Klf4",
                "Rest",
                "Nfib",
                "Eno1",
                "Hmgb1",
                "Atf4",
                "Ctcf",
                "Trp53",
                "Hdac1",
                "Tbx3",
                "Zfp57",
                "Ybx1",
                "Hdac2",
                "Smc3"
            ],
            "target": [],
            "expression_sum": {
                "class1": 37864.0,
                "class2": 24674.0
            },
            "uniprot_ids": [
                "Q60961",
                "Q8BG66",
                "Q9CRX2",
                "E9QLR3",
                "B0LAD1"
            ]
        },
        "Ldha": {
            "id": "Ldha",
            "type": "Gene",
            "symbol": "Ldha",
            "source": [
                "Tcf4",
                "Pou5f1",
                "Cebpb",
                "Nfib",
                "Nono",
                "Atf4",
                "Ctcf",
                "Junb",
                "Trp53",
                "Jun",
                "Hes1",
                "Nanog",
                "Jund",
                "Hdac2",
                "Smc3",
                "Smarca5"
            ],
            "target": [],
            "expression_sum": {
                "class1": 82476.0,
                "class2": 268670.0
            },
            "uniprot_ids": [
                "P06151",
                "D3YZQ9",
                "Q564E2",
                "Q3THB4",
                "A0A1B0GSR9",
                "Q3UIC3",
                "A0A1B0GSX0",
                "A0A1B0GT41",
                "A0A1B0GS79",
                "A0A1B0GRS2",
                "A0A1B0GQX5",
                "A0A1B0GRW9",
                "Q3UDU4",
                "Q99K20",
                "A0A1B0GSL7",
                "Q3TCI7",
                "Q3TI99",
                "A0A1B0GRC1",
                "Q6NXW3"
            ]
        },
        "Llph": {
            "id": "Llph",
            "type": "Gene",
            "symbol": "Llph",
            "source": [
                "Pou5f1",
                "Foxp1",
                "Eno1",
                "Atf4",
                "Junb",
                "Fos",
                "Jun",
                "Sox2",
                "Nanog",
                "Jund",
                "Twist2",
                "Cebpb"
            ],
            "target": [],
            "expression_sum": {
                "class1": 10337.0,
                "class2": 53734.0
            },
            "uniprot_ids": [
                "Q9D945",
                "A0A1W2P865"
            ]
        },
        "Lman2": {
            "id": "Lman2",
            "type": "Gene",
            "symbol": "Lman2",
            "source": [
                "Tcf4",
                "Pou5f1",
                "Eno1",
                "Atf4",
                "Junb",
                "Fos",
                "Jun",
                "Trp53",
                "Ybx1",
                "Jund",
                "Cebpb",
                "Smc3"
            ],
            "target": [],
            "expression_sum": {
                "class1": 9769.0,
                "class2": 24609.0
            },
            "uniprot_ids": [
                "Q9DBH5"
            ]
        },
        "Lmna": {
            "id": "Lmna",
            "type": "Gene",
            "symbol": "Lmna",
            "source": [
                "Sox2",
                "Klf5",
                "Hdac2",
                "E2f4",
                "Pou5f1",
                "Rest",
                "Trp53",
                "Smarca5",
                "Trim28",
                "Parp1",
                "Tgif1",
                "Tbx3",
                "Cebpb",
                "Zfp57",
                "Smc3",
                "Atf4",
                "Ctcf",
                "Hdac1",
                "Nanog"
            ],
            "target": [],
            "expression_sum": {
                "class1": 39879.0,
                "class2": 8516.0
            },
            "uniprot_ids": [
                "P48678",
                "D3YUF7",
                "Q9CTL2"
            ]
        },
        "Loxl1": {
            "id": "Loxl1",
            "type": "Gene",
            "symbol": "Loxl1",
            "source": [
                "E2f4",
                "Pou5f1",
                "Cebpb",
                "Atf4",
                "Ctcf",
                "Parp1",
                "Prrx1",
                "Sox2",
                "Twist2",
                "Hdac2",
                "Smc3",
                "Smarca5"
            ],
            "target": [],
            "expression_sum": {
                "class1": 11826.0,
                "class2": 1916.0
            },
            "uniprot_ids": [
                "P97873",
                "A0A0R4J0Q4",
                "Q8BP04",
                "Q4QQJ7"
            ]
        },
        "Lrrc17": {
            "id": "Lrrc17",
            "type": "Gene",
            "symbol": "Lrrc17",
            "source": [
                "E2f4",
                "Rest",
                "Atf4",
                "Ctcf",
                "Tbx3",
                "Hdac2",
                "Zfp57"
            ],
            "target": [],
            "expression_sum": {
                "class1": 11312.0,
                "class2": 372.0
            },
            "uniprot_ids": [
                "Q9CXD9"
            ]
        },
        "Lrrc58": {
            "id": "Lrrc58",
            "type": "Gene",
            "symbol": "Lrrc58",
            "source": [
                "E2f4",
                "Trim28",
                "Rest",
                "Hmgb1",
                "Atf4",
                "Ctcf",
                "Trp53",
                "Hdac1",
                "Ybx1",
                "Hdac2",
                "Smc3",
                "Smarca5"
            ],
            "target": [],
            "expression_sum": {
                "class1": 16536.0,
                "class2": 5384.0
            },
            "uniprot_ids": [
                "Q3UGP9",
                "Q8BTI3"
            ]
        },
        "Lrrc59": {
            "id": "Lrrc59",
            "type": "Gene",
            "symbol": "Lrrc59",
            "source": [
                "Fos",
                "Id2",
                "Sox2",
                "Hdac2",
                "Tcf4",
                "Pou5f1",
                "Trp53",
                "Jun",
                "Smarca5",
                "Sox4",
                "Eno1",
                "Jund",
                "Cebpb",
                "Smc3",
                "Foxp1",
                "Atf4",
                "Junb",
                "Nanog",
                "Twist2"
            ],
            "target": [],
            "expression_sum": {
                "class1": 21107.0,
                "class2": 37917.0
            },
            "uniprot_ids": [
                "Q922Q8",
                "Q3UBL0",
                "Q3TF84"
            ]
        },
        "Lsm3": {
            "id": "Lsm3",
            "type": "Gene",
            "symbol": "Lsm3",
            "source": [
                "Tcf4",
                "Pou5f1",
                "Foxp1",
                "Hmga2",
                "Nfib",
                "Atf4",
                "Junb",
                "Fos",
                "Jun",
                "Sox2",
                "Hes1",
                "Nanog",
                "Jund",
                "Twist2",
                "Cebpb",
                "Zfp57"
            ],
            "target": [],
            "expression_sum": {
                "class1": 9991.0,
                "class2": 65992.0
            },
            "uniprot_ids": [
                "P62311"
            ]
        },
        "Lsp1": {
            "id": "Lsp1",
            "type": "Gene",
            "symbol": "Lsp1",
            "source": [
                "Trim28",
                "Rest",
                "Atf4",
                "Ctcf",
                "Trp53",
                "Hdac1",
                "Klf5",
                "Ybx1",
                "Hdac2",
                "Smc3",
                "Smarca5"
            ],
            "target": [],
            "expression_sum": {
                "class1": 6490.0,
                "class2": 315.0
            },
            "uniprot_ids": [
                "P19973",
                "A0A1B0GRF5",
                "A0A1B0GSH0",
                "A2A6J7",
                "A2A6J4"
            ]
        },
        "Luc7l2": {
            "id": "Luc7l2",
            "type": "Gene",
            "symbol": "Luc7l2",
            "source": [
                "Tcf4",
                "Pou5f1",
                "Foxp1",
                "Cebpb",
                "Nfib",
                "Eno1",
                "Atf4",
                "Junb",
                "Trp53",
                "Jund",
                "Twist2",
                "Hdac2",
                "Smc3"
            ],
            "target": [],
            "expression_sum": {
                "class1": 13261.0,
                "class2": 18901.0
            },
            "uniprot_ids": [
                "Q7TNC4",
                "E0CYQ8",
                "Q05CX5",
                "E0CZB9",
                "Q3UZ17",
                "Q3UGK2",
                "Q3U5L2",
                "E9Q715",
                "Q3USM3",
                "Q3UUS9"
            ]
        },
        "Map1b": {
            "id": "Map1b",
            "type": "Gene",
            "symbol": "Map1b",
            "source": [
                "E2f4",
                "Trim28",
                "Atf4",
                "Ctcf",
                "Trp53",
                "Hdac1",
                "Ybx1",
                "Hdac2"
            ],
            "target": [],
            "expression_sum": {
                "class1": 6986.0,
                "class2": 3619.0
            },
            "uniprot_ids": [
                "P14873",
                "B2RQQ5",
                "Q3UF78",
                "Q8CFR9",
                "Q8CAI9",
                "A0PJ96",
                "Q3TZM8",
                "A0A286YDV1",
                "Q3TYV8",
                "Q3UPV4"
            ]
        },
        "Map2k2": {
            "id": "Map2k2",
            "type": "Gene",
            "symbol": "Map2k2",
            "source": [
                "Pou5f1",
                "Atf4",
                "Junb",
                "Trp53",
                "Hdac2",
                "Smc3",
                "Smarca5"
            ],
            "target": [],
            "expression_sum": {
                "class1": 13670.0,
                "class2": 30492.0
            },
            "uniprot_ids": [
                "Q63932",
                "Q91YS7",
                "A0A1W2P769",
                "D3Z446",
                "A0A1W2P7V9",
                "M0QWN2",
                "Q3UPM3",
                "Q8CB46",
                "Q3USU3"
            ]
        },
        "Mapre1": {
            "id": "Mapre1",
            "type": "Gene",
            "symbol": "Mapre1",
            "source": [
                "Tcf4",
                "Pou5f1",
                "Foxp1",
                "Cebpb",
                "Atf5",
                "Atf4",
                "Ctcf",
                "Junb",
                "Trp53",
                "Jun",
                "Hes1",
                "Jund",
                "Twist2",
                "Hdac2"
            ],
            "target": [],
            "expression_sum": {
                "class1": 14789.0,
                "class2": 22364.0
            },
            "uniprot_ids": [
                "Q61166",
                "Q3U4H0"
            ]
        },
        "March5": {
            "id": "March5",
            "type": "Gene",
            "symbol": "March5",
            "source": [
                "Tcf4",
                "Pou5f1",
                "Cebpb",
                "Hmgb1",
                "Atf4",
                "Junb",
                "Trp53",
                "Hdac1",
                "Twist2",
                "Hdac2",
                "Smarca5",
                "Sox4"
            ],
            "target": [],
            "expression_sum": {
                "class1": 10869.0,
                "class2": 19846.0
            },
            "uniprot_ids": [
                "Q3KNM2",
                "A2RTC8",
                "F6W8T3",
                "A0A0R4J1H0"
            ]
        },
        "Marcks": {
            "id": "Marcks",
            "type": "Gene",
            "symbol": "Marcks",
            "source": [
                "Hdac2",
                "E2f4",
                "Pou5f1",
                "Rest",
                "Atf4",
                "Ctcf",
                "Trp53",
                "Hdac1",
                "Klf5",
                "Tbx3",
                "Cebpb",
                "Smc3"
            ],
            "target": [],
            "expression_sum": {
                "class1": 43984.0,
                "class2": 6802.0
            },
            "uniprot_ids": [
                "P26645",
                "Q05BL6",
                "A0A0A1HAM8"
            ]
        },
        "Mbnl2": {
            "id": "Mbnl2",
            "type": "Gene",
            "symbol": "Mbnl2",
            "source": [
                "Hdac2",
                "E2f4",
                "Pou5f1",
                "Trim28",
                "Rest",
                "Arf4",
                "Atf4",
                "Ctcf",
                "Tgif1",
                "Trp53",
                "Hdac1",
                "Klf5",
                "Tbx3",
                "Cebpb",
                "Smarca5"
            ],
            "target": [],
            "expression_sum": {
                "class1": 10571.0,
                "class2": 2086.0
            },
            "uniprot_ids": [
                "Q8C181",
                "A0A2I3BRX8",
                "A0A2K6EDM5",
                "D0EX61"
            ]
        },
        "Med28": {
            "id": "Med28",
            "type": "Gene",
            "symbol": "Med28",
            "source": [
                "Pou5f1",
                "Cebpb",
                "Nfib",
                "Eno1",
                "Atf4",
                "Junb",
                "Trp53",
                "Jun",
                "Twist2",
                "Hdac2",
                "Smc3",
                "Sox4"
            ],
            "target": [],
            "expression_sum": {
                "class1": 10954.0,
                "class2": 25293.0
            },
            "uniprot_ids": [
                "Q920D3",
                "D3Z7D6",
                "D3YWT4"
            ]
        },
        "Mesdc2": {
            "id": "Mesdc2",
            "type": "Gene",
            "symbol": "Mesdc2",
            "source": [
                "Nfib",
                "Fos",
                "Id2",
                "Hdac2",
                "Tcf4",
                "Pou5f1",
                "Jun",
                "Smarca5",
                "Sox4",
                "Eno1",
                "Jund",
                "Cebpb",
                "Smc3",
                "Foxp1",
                "Atf4",
                "Junb",
                "Ybx1",
                "Hes1",
                "Twist2"
            ],
            "target": [],
            "expression_sum": {
                "class1": 7801.0,
                "class2": 19667.0
            },
            "uniprot_ids": [
                "Q9ERE7",
                "D3YVR4",
                "F6SWV4"
            ]
        },
        "Mgat2": {
            "id": "Mgat2",
            "type": "Gene",
            "symbol": "Mgat2",
            "source": [
                "E2f4",
                "Hmga2",
                "Rest",
                "Atf4",
                "Ctcf",
                "Tgif1",
                "Trp53",
                "Hdac1",
                "Hdac2",
                "Smc3",
                "Smarca5"
            ],
            "target": [],
            "expression_sum": {
                "class1": 9456.0,
                "class2": 7431.0
            },
            "uniprot_ids": [
                "Q921V5",
                "Q3U5X7",
                "Q3TBY6",
                "Q3TXN3"
            ]
        },
        "Mlf2": {
            "id": "Mlf2",
            "type": "Gene",
            "symbol": "Mlf2",
            "source": [
                "Tcf4",
                "Pou5f1",
                "Foxp1",
                "Cebpb",
                "Hmga2",
                "Atf4",
                "Junb",
                "Jun",
                "Sox2",
                "Hes1",
                "Nanog",
                "Twist2",
                "Hdac2",
                "Smc3",
                "Smarca5"
            ],
            "target": [],
            "expression_sum": {
                "class1": 22442.0,
                "class2": 58410.0
            },
            "uniprot_ids": [
                "Q99KX1",
                "A0A0N4SVX3",
                "A0A0N4SVM2",
                "Q3URS8",
                "Q3UNV7"
            ]
        },
        "Morf4l1": {
            "id": "Morf4l1",
            "type": "Gene",
            "symbol": "Morf4l1",
            "source": [
                "Tcf4",
                "Pou5f1",
                "Foxp1",
                "Cebpb",
                "Nfib",
                "Atf4",
                "Junb",
                "Fos",
                "Jun",
                "Sox2",
                "Nanog",
                "Jund",
                "Twist2",
                "Hdac2"
            ],
            "target": [],
            "expression_sum": {
                "class1": 55910.0,
                "class2": 92315.0
            },
            "uniprot_ids": [
                "P60762",
                "Q569V4",
                "A0A087WQ34",
                "Q3U5J6",
                "Q3UIS6",
                "A0A087WSL1",
                "A0A087WQB0",
                "Q5DTS9",
                "Q3UBK1"
            ]
        },
        "Morf4l2": {
            "id": "Morf4l2",
            "type": "Gene",
            "symbol": "Morf4l2",
            "source": [
                "Tcf4",
                "Pou5f1",
                "Foxp1",
                "Cebpb",
                "Nfib",
                "Eno1",
                "Atf4",
                "Junb",
                "Fos",
                "Jun",
                "Hes1",
                "Nanog",
                "Jund",
                "Twist2",
                "Hdac2",
                "Smc3"
            ],
            "target": [],
            "expression_sum": {
                "class1": 38119.0,
                "class2": 67555.0
            },
            "uniprot_ids": [
                "Q9R0Q4",
                "A2AEB6",
                "A2AEB7",
                "A2AEB5",
                "Q3TME1",
                "A2AEB8",
                "A2AEB9",
                "A2AEB3",
                "A2AEB4",
                "Q3TXN7",
                "Q3U6G8"
            ]
        },
        "Mpc1": {
            "id": "Mpc1",
            "type": "Gene",
            "symbol": "Mpc1",
            "source": [
                "Tcf4",
                "Foxp1",
                "Trim28",
                "Rest",
                "Atf4",
                "Ctcf",
                "Junb",
                "Trp53",
                "Hdac1",
                "Ybx1",
                "Jund",
                "Twist2",
                "Hdac2",
                "Smc3",
                "Smarca5"
            ],
            "target": [],
            "expression_sum": {
                "class1": 11932.0,
                "class2": 11356.0
            },
            "uniprot_ids": [
                "P63030",
                "Q3UX28",
                "D3YWY6",
                "Q9D6B6",
                "Q3TCV4",
                "E9Q0V4",
                "D3Z5S0",
                "D3Z786"
            ]
        },
        "Mpc2": {
            "id": "Mpc2",
            "type": "Gene",
            "symbol": "Mpc2",
            "source": [
                "Pou5f1",
                "Foxp1",
                "Trim28",
                "Eno1",
                "Atf4",
                "Junb",
                "Trp53",
                "Hdac1",
                "Ybx1",
                "Peg3",
                "Hdac2",
                "Smc3",
                "Smarca5"
            ],
            "target": [],
            "expression_sum": {
                "class1": 10535.0,
                "class2": 12891.0
            },
            "uniprot_ids": [
                "Q9D023",
                "A0A0A6YY89"
            ]
        },
        "Mrpl12": {
            "id": "Mrpl12",
            "type": "Gene",
            "symbol": "Mrpl12",
            "source": [
                "Pou5f1",
                "Atf4",
                "Junb",
                "Fos",
                "Jun",
                "Id2",
                "Sox2",
                "Nanog",
                "Tfcp2l1",
                "Jund",
                "Twist2",
                "Cebpb",
                "Zfp57",
                "Sox4"
            ],
            "target": [],
            "expression_sum": {
                "class1": 12214.0,
                "class2": 64957.0
            },
            "uniprot_ids": [
                "Q9DB15"
            ]
        },
        "Mrpl13": {
            "id": "Mrpl13",
            "type": "Gene",
            "symbol": "Mrpl13",
            "source": [
                "Tcf4",
                "Pou5f1",
                "Cebpb",
                "Arf4",
                "Nfib",
                "Atf4",
                "Junb",
                "Fos",
                "Jun",
                "Sox2",
                "Nanog",
                "Jund",
                "Twist2",
                "Hdac2",
                "Smc3"
            ],
            "target": [],
            "expression_sum": {
                "class1": 7722.0,
                "class2": 20048.0
            },
            "uniprot_ids": [
                "Q9D1P0",
                "G5E909",
                "F7CXG9",
                "Q05CB9",
                "Q7TMH5"
            ]
        },
        "Mrpl15": {
            "id": "Mrpl15",
            "type": "Gene",
            "symbol": "Mrpl15",
            "source": [
                "Klf2",
                "Tcf4",
                "Pou5f1",
                "Foxp1",
                "Tfcp2l1",
                "Nfib",
                "Atf4",
                "Fos",
                "Jun",
                "Sox2",
                "Hes1",
                "Nanog",
                "Jund",
                "Twist2",
                "Cebpb",
                "Zfp57",
                "Sox4"
            ],
            "target": [],
            "expression_sum": {
                "class1": 8109.0,
                "class2": 66425.0
            },
            "uniprot_ids": [
                "Q9CPR5",
                "F8WHN1",
                "A0A0A6YVP4"
            ]
        },
        "Mrpl17": {
            "id": "Mrpl17",
            "type": "Gene",
            "symbol": "Mrpl17",
            "source": [
                "Tcf4",
                "Pou5f1",
                "Foxp1",
                "Cebpb",
                "Tfcp2l1",
                "Eno1",
                "Atf4",
                "Junb",
                "Fos",
                "Jun",
                "Sox2",
                "Hes1",
                "Nanog",
                "Jund",
                "Twist2",
                "Hdac2",
                "Smc3"
            ],
            "target": [],
            "expression_sum": {
                "class1": 16031.0,
                "class2": 65213.0
            },
            "uniprot_ids": [
                "Q9D8P4",
                "H3BJT9",
                "A0A1B0GSV1",
                "H3BIZ3"
            ]
        },
        "Mrpl18": {
            "id": "Mrpl18",
            "type": "Gene",
            "symbol": "Mrpl18",
            "source": [
                "Tcf4",
                "Pou5f1",
                "Foxp1",
                "Hmga2",
                "Nfib",
                "Atf4",
                "Junb",
                "Fos",
                "Jun",
                "Sox2",
                "Nanog",
                "Tfcp2l1",
                "Jund",
                "Cebpb"
            ],
            "target": [],
            "expression_sum": {
                "class1": 13291.0,
                "class2": 73858.0
            },
            "uniprot_ids": [
                "Q9CQL5",
                "Q8BTQ6"
            ]
        },
        "Mrpl20": {
            "id": "Mrpl20",
            "type": "Gene",
            "symbol": "Mrpl20",
            "source": [
                "Tcf4",
                "Pou5f1",
                "Cebpb",
                "Nfib",
                "Eno1",
                "Atf4",
                "Junb",
                "Fos",
                "Id2",
                "Jun",
                "Sox2",
                "Nanog",
                "Jund",
                "Twist2",
                "Hdac2",
                "Smc3",
                "Sox4"
            ],
            "target": [],
            "expression_sum": {
                "class1": 18594.0,
                "class2": 47682.0
            },
            "uniprot_ids": [
                "Q9CQL4",
                "V9GXI7",
                "V9GX65",
                "Q91YJ4"
            ]
        },
        "Mrpl30": {
            "id": "Mrpl30",
            "type": "Gene",
            "symbol": "Mrpl30",
            "source": [
                "Pou5f1",
                "Foxp1",
                "Cebpb",
                "Atf4",
                "Junb",
                "Trp53",
                "Sox2",
                "Ybx1",
                "Twist2",
                "Hdac2",
                "Smc3",
                "Smarca5",
                "Sox4"
            ],
            "target": [],
            "expression_sum": {
                "class1": 18091.0,
                "class2": 38137.0
            },
            "uniprot_ids": [
                "Q9D7N6",
                "A0A0A6YXW4"
            ]
        },
        "Mrpl43": {
            "id": "Mrpl43",
            "type": "Gene",
            "symbol": "Mrpl43",
            "source": [
                "Tcf4",
                "Pou5f1",
                "Foxp1",
                "Atf4",
                "Junb",
                "Jun",
                "Hes1",
                "Nanog",
                "Jund",
                "Twist2",
                "Cebpb"
            ],
            "target": [],
            "expression_sum": {
                "class1": 6772.0,
                "class2": 26776.0
            },
            "uniprot_ids": [
                "Q99N89",
                "Q5RL20",
                "Q810B1",
                "Q6P060"
            ]
        },
        "Mrpl45": {
            "id": "Mrpl45",
            "type": "Gene",
            "symbol": "Mrpl45",
            "source": [
                "Pou5f1",
                "Foxp1",
                "Atf5",
                "Atf4",
                "Junb",
                "Jun",
                "Hes1",
                "Nanog",
                "Jund",
                "Twist2",
                "Cebpb"
            ],
            "target": [],
            "expression_sum": {
                "class1": 3878.0,
                "class2": 28679.0
            },
            "uniprot_ids": [
                "Q9D0Q7",
                "F6QAU7",
                "Q9CSZ8"
            ]
        },
        "Mrpl52": {
            "id": "Mrpl52",
            "type": "Gene",
            "symbol": "Mrpl52",
            "source": [
                "Pou5f1",
                "Foxp1",
                "Cebpb",
                "Atf4",
                "Junb",
                "Jun",
                "Sox2",
                "Nanog",
                "Twist2",
                "Hdac2",
                "Sox4"
            ],
            "target": [],
            "expression_sum": {
                "class1": 20895.0,
                "class2": 51147.0
            },
            "uniprot_ids": [
                "Q9D0Y8",
                "A0A0G2JE60"
            ]
        },
        "Mrpl54": {
            "id": "Mrpl54",
            "type": "Gene",
            "symbol": "Mrpl54",
            "source": [
                "Pou5f1",
                "Eno1",
                "Hmgb1",
                "Atf4",
                "Junb",
                "Trp53",
                "Hdac1",
                "Ybx1",
                "Twist2",
                "Hdac2",
                "Smarca5",
                "Sox4"
            ],
            "target": [],
            "expression_sum": {
                "class1": 19078.0,
                "class2": 27796.0
            },
            "uniprot_ids": [
                "Q9CPW3"
            ]
        },
        "Mrps16": {
            "id": "Mrps16",
            "type": "Gene",
            "symbol": "Mrps16",
            "source": [
                "Tcf4",
                "Pou5f1",
                "Foxp1",
                "Atf4",
                "Junb",
                "Fos",
                "Id2",
                "Jun",
                "Sox2",
                "Nanog",
                "Jund",
                "Twist2",
                "Cebpb",
                "Sox4"
            ],
            "target": [],
            "expression_sum": {
                "class1": 13487.0,
                "class2": 53852.0
            },
            "uniprot_ids": [
                "Q9CPX7"
            ]
        },
        "Mrps18c": {
            "id": "Mrps18c",
            "type": "Gene",
            "symbol": "Mrps18c",
            "source": [
                "Tcf4",
                "Pou5f1",
                "Foxp1",
                "Atf4",
                "Junb",
                "Jun",
                "Sox2",
                "Nanog",
                "Jund",
                "Twist2",
                "Cebpb"
            ],
            "target": [],
            "expression_sum": {
                "class1": 8135.0,
                "class2": 29051.0
            },
            "uniprot_ids": [
                "Q8R2L5",
                "Q8BTZ9",
                "D3Z3D3",
                "A0A0G2JFG7"
            ]
        },
        "Mrps25": {
            "id": "Mrps25",
            "type": "Gene",
            "symbol": "Mrps25",
            "source": [
                "Pou5f1",
                "Atf4",
                "Junb",
                "Jun",
                "Jund",
                "Cebpb"
            ],
            "target": [],
            "expression_sum": {
                "class1": 5273.0,
                "class2": 20454.0
            },
            "uniprot_ids": [
                "Q9D125",
                "Q6KAU3"
            ]
        },
        "Mrps36": {
            "id": "Mrps36",
            "type": "Gene",
            "symbol": "Mrps36",
            "source": [
                "Tcf4",
                "Pou5f1",
                "Nfib",
                "Eno1",
                "Atf4",
                "Junb",
                "Fos",
                "Jun",
                "Sox2",
                "Nanog",
                "Jund",
                "Twist2",
                "Cebpb"
            ],
            "target": [],
            "expression_sum": {
                "class1": 11595.0,
                "class2": 28174.0
            },
            "uniprot_ids": [
                "Q9CQX8",
                "Q9D6T9"
            ]
        },
        "Mrps7": {
            "id": "Mrps7",
            "type": "Gene",
            "symbol": "Mrps7",
            "source": [
                "Tcf4",
                "Pou5f1",
                "Hmga2",
                "Cebpb",
                "Atf4",
                "Junb",
                "Fos",
                "Id2",
                "Jun",
                "Hes1",
                "Jund",
                "Hdac2",
                "Smc3",
                "Smarca5"
            ],
            "target": [],
            "expression_sum": {
                "class1": 8813.0,
                "class2": 24217.0
            },
            "uniprot_ids": [
                "Q80X85"
            ]
        },
        "Mrto4": {
            "id": "Mrto4",
            "type": "Gene",
            "symbol": "Mrto4",
            "source": [
                "Tcf4",
                "Pou5f1",
                "Atf4",
                "Prrx1",
                "Jun",
                "Nanog",
                "Jund",
                "Twist2",
                "Cebpb",
                "Smc3",
                "Sox4"
            ],
            "target": [],
            "expression_sum": {
                "class1": 5982.0,
                "class2": 31890.0
            },
            "uniprot_ids": [
                "Q9D0I8",
                "A2AMU9",
                "D6RCI7",
                "A2AMV1"
            ]
        },
        "Mt1": {
            "id": "Mt1",
            "type": "Gene",
            "symbol": "Mt1",
            "source": [
                "Sox2",
                "Pou5f1",
                "Atf4"
            ],
            "target": [],
            "expression_sum": {
                "class1": 11823.0,
                "class2": 258126.0
            },
            "uniprot_ids": [
                "P02802",
                "A0A1D5RLN7",
                "A0A1D5RMG1",
                "Q3U340"
            ]
        },
        "Mt2": {
            "id": "Mt2",
            "type": "Gene",
            "symbol": "Mt2",
            "source": [
                "Klf2",
                "Pou5f1",
                "Trim28",
                "Atf4",
                "Ctcf",
                "Trp53",
                "Hdac1",
                "Klf5",
                "Sox2",
                "Tbx3",
                "Nanog",
                "Tfcp2l1",
                "Ybx1",
                "Smarca5"
            ],
            "target": [],
            "expression_sum": {
                "class1": 3164.0,
                "class2": 253848.0
            },
            "uniprot_ids": [
                "P02798",
                "A0A1D5RLC3"
            ]
        },
        "Mtch1": {
            "id": "Mtch1",
            "type": "Gene",
            "symbol": "Mtch1",
            "source": [
                "Trim28",
                "Rest",
                "Atf4",
                "Ctcf",
                "Trp53",
                "Hdac1",
                "Ybx1",
                "Hdac2",
                "Smc3",
                "Smarca5"
            ],
            "target": [],
            "expression_sum": {
                "class1": 12272.0,
                "class2": 12940.0
            },
            "uniprot_ids": [
                "Q791T5",
                "A0A3Q4EGL4",
                "D3YXC2",
                "A0A3Q4EC24",
                "A0A3Q4EG99",
                "A0A3Q4EC97",
                "A0A3Q4EBT8",
                "A0A3Q4EGX8",
                "A0A3Q4EIA3",
                "Q8CEY5"
            ]
        },
        "Mtch2": {
            "id": "Mtch2",
            "type": "Gene",
            "symbol": "Mtch2",
            "source": [
                "Nfib",
                "Fos",
                "Id2",
                "Sox2",
                "Hdac2",
                "Tcf4",
                "Pou5f1",
                "Jun",
                "Sox4",
                "Jund",
                "Cebpb",
                "Hmga2",
                "Foxp1",
                "Atf4",
                "Junb",
                "Hes1",
                "Tfcp2l1",
                "Nanog",
                "Twist2"
            ],
            "target": [],
            "expression_sum": {
                "class1": 16018.0,
                "class2": 48857.0
            },
            "uniprot_ids": [
                "Q791V5",
                "Q9D050",
                "A2AFW6",
                "D6RCZ1",
                "Q05C68",
                "Q9CUI0",
                "Q3UA17"
            ]
        },
        "Mtdh": {
            "id": "Mtdh",
            "type": "Gene",
            "symbol": "Mtdh",
            "source": [
                "E2f4",
                "Trim28",
                "Hmga2",
                "Rest",
                "Nfib",
                "Atf4",
                "Ctcf",
                "Junb",
                "Parp1",
                "Hdac1",
                "Trp53",
                "Hdac2",
                "Smc3",
                "Smarca5"
            ],
            "target": [],
            "expression_sum": {
                "class1": 16259.0,
                "class2": 24323.0
            },
            "uniprot_ids": [
                "Q80WJ7",
                "F6ZSG0",
                "F6QHD1",
                "F6QFT1",
                "F6ZQL0",
                "A0A2I3BRF4",
                "Q3TAB3",
                "E9PUX0"
            ]
        },
        "Myeov2": {
            "id": "Myeov2",
            "type": "Gene",
            "symbol": "Myeov2",
            "source": [
                "E2f4",
                "Trim28",
                "Rest",
                "Atf4",
                "Ctcf",
                "Trp53",
                "Hdac1",
                "Ybx1",
                "Hdac2",
                "Smc3",
                "Smarca5"
            ],
            "target": [],
            "expression_sum": {
                "class1": 16647.0,
                "class2": 23837.0
            },
            "uniprot_ids": [
                "Q3U898",
                "B2RVT1",
                "F6VAI7",
                "D3Z159"
            ]
        },
        "Myh10": {
            "id": "Myh10",
            "type": "Gene",
            "symbol": "Myh10",
            "source": [
                "E2f4",
                "Trim28",
                "Rest",
                "Atf4",
                "Ctcf",
                "Trp53",
                "Hdac1",
                "Klf5",
                "Hdac2",
                "Smc3"
            ],
            "target": [],
            "expression_sum": {
                "class1": 7269.0,
                "class2": 2656.0
            },
            "uniprot_ids": [
                "Q61879",
                "Q3UH59",
                "Q8BXF2",
                "Q80T68",
                "Q4KMN4",
                "Q5SV64",
                "Q6PB65",
                "Q8BWI3",
                "Q8BLI3",
                "Q8BLI1"
            ]
        },
        "Myh9": {
            "id": "Myh9",
            "type": "Gene",
            "symbol": "Myh9",
            "source": [
                "E2f4",
                "Trim28",
                "Rest",
                "Nono",
                "Atf4",
                "Ctcf",
                "Trp53",
                "Klf5",
                "Hdac2",
                "Smc3",
                "Smarca5"
            ],
            "target": [],
            "expression_sum": {
                "class1": 10150.0,
                "class2": 5310.0
            },
            "uniprot_ids": [
                "Q8VDD5",
                "Q14B05",
                "A0A2R8W6V7",
                "Q3UFT0",
                "Q7TQJ6",
                "Q811J9",
                "Q99JJ3",
                "F2Z494",
                "A0A2R8VKI5",
                "Q8BIM8",
                "O89055",
                "Q8BIE2",
                "A2VCK0"
            ]
        },
        "Myl12a": {
            "id": "Myl12a",
            "type": "Gene",
            "symbol": "Myl12a",
            "source": [
                "Tcf4",
                "Pou5f1",
                "Foxp1",
                "Trim28",
                "Rest",
                "Nfib",
                "Eno1",
                "Atf4",
                "Ctcf",
                "Junb",
                "Tgif1",
                "Hdac1",
                "Trp53",
                "Ybx1",
                "Jund",
                "Twist2",
                "Hdac2",
                "Smc3"
            ],
            "target": [],
            "expression_sum": {
                "class1": 91135.0,
                "class2": 80877.0
            },
            "uniprot_ids": [
                "Q6ZWQ9",
                "D3Z249",
                "D3YV37"
            ]
        },
        "Naa10": {
            "id": "Naa10",
            "type": "Gene",
            "symbol": "Naa10",
            "source": [
                "Tcf4",
                "Pou5f1",
                "Nfib",
                "Atf4",
                "Fos",
                "Jun",
                "Sox2",
                "Nanog",
                "Jund",
                "Twist2",
                "Cebpb",
                "Sox4"
            ],
            "target": [],
            "expression_sum": {
                "class1": 5493.0,
                "class2": 19669.0
            },
            "uniprot_ids": [
                "Q9QY36",
                "Q8R2U5",
                "Q3V4D5",
                "Q540H0",
                "B1AUY9",
                "Q684Q8",
                "B1AUZ1",
                "B1AUY7",
                "B1AUY8"
            ]
        },
        "Nans": {
            "id": "Nans",
            "type": "Gene",
            "symbol": "Nans",
            "source": [
                "Tcf4",
                "Pou5f1",
                "Cebpb",
                "Nfib",
                "Atf4",
                "Jun",
                "Hes1",
                "Jund",
                "Hdac2"
            ],
            "target": [],
            "expression_sum": {
                "class1": 9290.0,
                "class2": 19130.0
            },
            "uniprot_ids": [
                "Q99J77",
                "Q3TFB5"
            ]
        },
        "Napa": {
            "id": "Napa",
            "type": "Gene",
            "symbol": "Napa",
            "source": [
                "Pou5f1",
                "Foxp1",
                "Atf4",
                "Trp53",
                "Hdac2",
                "Smc3"
            ],
            "target": [],
            "expression_sum": {
                "class1": 8748.0,
                "class2": 19441.0
            },
            "uniprot_ids": [
                "Q9DB05",
                "A0A1B0GR35",
                "Q9CXX1"
            ]
        },
        "Nars": {
            "id": "Nars",
            "type": "Gene",
            "symbol": "Nars",
            "source": [
                "Tcf4",
                "Pou5f1",
                "Cebpb",
                "Atf4",
                "Junb",
                "Fos",
                "Jun",
                "Trp53",
                "Jund",
                "Twist2",
                "Hdac2",
                "Smarca5"
            ],
            "target": [],
            "expression_sum": {
                "class1": 15348.0,
                "class2": 28798.0
            },
            "uniprot_ids": [
                "Q8BP47",
                "A0A494BB89",
                "A0A494B996",
                "A0A494B9G0",
                "A0A494B927",
                "A0A494BAW6",
                "A0A494BB81",
                "A0A494BAX5",
                "A0A498WGK2",
                "A0A494BB92",
                "A0A494B949"
            ]
        },
        "Ncl": {
            "id": "Ncl",
            "type": "Gene",
            "symbol": "Ncl",
            "source": [
                "Tcf4",
                "Pou5f1",
                "Foxp1",
                "Nfib",
                "Atf4",
                "Junb",
                "Fos",
                "Jun",
                "Sox2",
                "Hes1",
                "Nanog",
                "Jund",
                "Cebpb",
                "Zfp57"
            ],
            "target": [],
            "expression_sum": {
                "class1": 45421.0,
                "class2": 325413.0
            },
            "uniprot_ids": [
                "P09405",
                "Q3TGR3",
                "Q9CT46",
                "Q8CE30",
                "Q3TT41",
                "Q8CD23",
                "Q3TL52",
                "A0A087WRM5"
            ]
        },
        "Ncor1": {
            "id": "Ncor1",
            "type": "Gene",
            "symbol": "Ncor1",
            "source": [
                "Tcf4",
                "Pou5f1",
                "Nfib",
                "Atf4",
                "Junb",
                "Fos",
                "Jun",
                "Sox2",
                "Hes1",
                "Tfcp2l1",
                "Jund",
                "Twist2",
                "Cebpb",
                "Sox4"
            ],
            "target": [],
            "expression_sum": {
                "class1": 6455.0,
                "class2": 19546.0
            },
            "uniprot_ids": [
                "Q60974",
                "Q8BK32",
                "Q8CHB6",
                "Q3UV08",
                "Q5RIM6",
                "Q5RIN0",
                "Q3UV34",
                "E9Q9Y2",
                "F6TRR6",
                "Q5PRE9",
                "Q3UWZ3",
                "E9Q8K6",
                "Q05C35",
                "Q8VDE8",
                "E0CYX5",
                "Q9CUV3",
                "F7C134",
                "E9Q2B2",
                "F6SFD2",
                "Q05CF0",
                "F6U338",
                "F6TWR1"
            ]
        },
        "Ndufa10": {
            "id": "Ndufa10",
            "type": "Gene",
            "symbol": "Ndufa10",
            "source": [
                "Tcf4",
                "Pou5f1",
                "Foxp1",
                "Cebpb",
                "Arf4",
                "Nfib",
                "Atf4",
                "Junb",
                "Jun",
                "Ybx1",
                "Hes1",
                "Jund",
                "Twist2",
                "Hdac2",
                "Smc3",
                "Smarca5",
                "Sox4"
            ],
            "target": [],
            "expression_sum": {
                "class1": 13126.0,
                "class2": 28688.0
            },
            "uniprot_ids": [
                "Q99LC3",
                "A0A087WR38",
                "A0A087WRD4"
            ]
        },
        "Ndufa13": {
            "id": "Ndufa13",
            "type": "Gene",
            "symbol": "Ndufa13",
            "source": [
                "Pou5f1",
                "Trim28",
                "Rest",
                "Nfib",
                "Atf4",
                "Trp53",
                "Hdac1",
                "Ybx1",
                "Hdac2",
                "Smc3"
            ],
            "target": [],
            "expression_sum": {
                "class1": 31209.0,
                "class2": 58898.0
            },
            "uniprot_ids": [
                "Q9ERS2"
            ]
        },
        "Ndufa2": {
            "id": "Ndufa2",
            "type": "Gene",
            "symbol": "Ndufa2",
            "source": [
                "Pou5f1",
                "Cebpb",
                "Atf4",
                "Trp53",
                "Jun",
                "Sox2",
                "Twist2",
                "Hdac2",
                "Smarca5",
                "Sox4"
            ],
            "target": [],
            "expression_sum": {
                "class1": 40893.0,
                "class2": 73128.0
            },
            "uniprot_ids": [
                "Q9CQ75"
            ]
        },
        "Ndufa8": {
            "id": "Ndufa8",
            "type": "Gene",
            "symbol": "Ndufa8",
            "source": [
                "E2f4",
                "Trim28",
                "Rest",
                "Atf4",
                "Ctcf",
                "Junb",
                "Id2",
                "Hdac1",
                "Jun",
                "Trp53",
                "Jund",
                "Twist2",
                "Hdac2",
                "Smc3"
            ],
            "target": [],
            "expression_sum": {
                "class1": 14874.0,
                "class2": 29733.0
            },
            "uniprot_ids": [
                "Q9DCJ5",
                "A0A1S6GWI0"
            ]
        },
        "Ndufab1": {
            "id": "Ndufab1",
            "type": "Gene",
            "symbol": "Ndufab1",
            "source": [
                "Tcf4",
                "Pou5f1",
                "Cebpb",
                "Atf4",
                "Junb",
                "Fos",
                "Jun",
                "Sox2",
                "Nanog",
                "Jund",
                "Twist2",
                "Hdac2",
                "Smc3",
                "Smarca5"
            ],
            "target": [],
            "expression_sum": {
                "class1": 24864.0,
                "class2": 67496.0
            },
            "uniprot_ids": [
                "Q9CR21",
                "Q569N0",
                "F8WJ64",
                "F6ZFT1",
                "Q3TJG2"
            ]
        },
        "Ndufb5": {
            "id": "Ndufb5",
            "type": "Gene",
            "symbol": "Ndufb5",
            "source": [
                "Tcf4",
                "Pou5f1",
                "Foxp1",
                "Cebpb",
                "Hmga2",
                "Nfib",
                "Atf4",
                "Junb",
                "Fos",
                "Jun",
                "Sox2",
                "Nanog",
                "Jund",
                "Twist2",
                "Hdac2",
                "Smc3"
            ],
            "target": [],
            "expression_sum": {
                "class1": 21445.0,
                "class2": 59093.0
            },
            "uniprot_ids": [
                "Q9CQH3",
                "D3Z568",
                "D3YX99",
                "F6Y6V5",
                "D3Z6W9",
                "F8WI84",
                "F6VXF9",
                "Q8BU20"
            ]
        },
        "Ndufc2": {
            "id": "Ndufc2",
            "type": "Gene",
            "symbol": "Ndufc2",
            "source": [
                "Pou5f1",
                "Cebpb",
                "Eno1",
                "Atf4",
                "Junb",
                "Fos",
                "Jun",
                "Sox2",
                "Trp53",
                "Ybx1",
                "Nanog",
                "Jund",
                "Twist2",
                "Hdac2",
                "Smc3",
                "Smarca5"
            ],
            "target": [],
            "expression_sum": {
                "class1": 20590.0,
                "class2": 48755.0
            },
            "uniprot_ids": [
                "Q9CQ54",
                "A0A0U1RPJ5",
                "Q9D846"
            ]
        },
        "Ndufs2": {
            "id": "Ndufs2",
            "type": "Gene",
            "symbol": "Ndufs2",
            "source": [
                "Pou5f1",
                "Foxp1",
                "Cebpb",
                "Atf4",
                "Junb",
                "Jund",
                "Twist2",
                "Hdac2",
                "Smc3",
                "Smarca5",
                "Sox4"
            ],
            "target": [],
            "expression_sum": {
                "class1": 13450.0,
                "class2": 34619.0
            },
            "uniprot_ids": [
                "Q91WD5",
                "A0A0A6YXD3",
                "A0A0A6YW30",
                "D3YXT0"
            ]
        },
        "Ndufs3": {
            "id": "Ndufs3",
            "type": "Gene",
            "symbol": "Ndufs3",
            "source": [
                "Pou5f1",
                "Cebpb",
                "Nfib",
                "Atf4",
                "Junb",
                "Fos",
                "Jun",
                "Jund",
                "Twist2",
                "Hdac2",
                "Smarca5",
                "Sox4"
            ],
            "target": [],
            "expression_sum": {
                "class1": 11419.0,
                "class2": 24876.0
            },
            "uniprot_ids": [
                "Q9DCT2"
            ]
        },
        "Ndufs5": {
            "id": "Ndufs5",
            "type": "Gene",
            "symbol": "Ndufs5",
            "source": [
                "Pou5f1",
                "Foxp1",
                "Cebpb",
                "Hmga2",
                "Nfib",
                "Atf4",
                "Junb",
                "Fos",
                "Jun",
                "Sox2",
                "Trp53",
                "Prrx2",
                "Nanog",
                "Jund",
                "Twist2",
                "Hdac2",
                "Smc3",
                "Sox4"
            ],
            "target": [],
            "expression_sum": {
                "class1": 25852.0,
                "class2": 59407.0
            },
            "uniprot_ids": [
                "Q99LY9",
                "B1ARW4",
                "A0A1S6GWI4"
            ]
        },
        "Ndufs7": {
            "id": "Ndufs7",
            "type": "Gene",
            "symbol": "Ndufs7",
            "source": [
                "Tcf4",
                "Pou5f1",
                "Nfib",
                "Atf4",
                "Junb",
                "Fos",
                "Id2",
                "Jun",
                "Sox2",
                "Hes1",
                "Nanog",
                "Jund",
                "Twist2",
                "Cebpb",
                "Sox4"
            ],
            "target": [],
            "expression_sum": {
                "class1": 11085.0,
                "class2": 35895.0
            },
            "uniprot_ids": [
                "Q9DC70"
            ]
        },
        "Ndufs8": {
            "id": "Ndufs8",
            "type": "Gene",
            "symbol": "Ndufs8",
            "source": [
                "Tcf4",
                "Pou5f1",
                "Cebpb",
                "Nfib",
                "Atf4",
                "Junb",
                "Trp53",
                "Jun",
                "Sox2",
                "Tfcp2l1",
                "Jund",
                "Twist2",
                "Hdac2",
                "Smc3"
            ],
            "target": [],
            "expression_sum": {
                "class1": 12302.0,
                "class2": 23207.0
            },
            "uniprot_ids": [
                "Q8K3J1",
                "Q5M9P5",
                "A0A494BAW8",
                "Q8VC72",
                "Q3UY05"
            ]
        },
        "Ndufv1": {
            "id": "Ndufv1",
            "type": "Gene",
            "symbol": "Ndufv1",
            "source": [
                "Tcf4",
                "Pou5f1",
                "Foxp1",
                "Nfib",
                "Atf4",
                "Junb",
                "Fos",
                "Id2",
                "Jun",
                "Sox2",
                "Nanog",
                "Jund",
                "Twist2",
                "Cebpb",
                "Sox4"
            ],
            "target": [],
            "expression_sum": {
                "class1": 6790.0,
                "class2": 23682.0
            },
            "uniprot_ids": [
                "Q91YT0",
                "D3Z1U9",
                "D3YUM1",
                "A0A494BAS8",
                "D3YXX5",
                "D3Z0K1",
                "A0A494BA02",
                "D6RG60"
            ]
        },
        "Ndufv2": {
            "id": "Ndufv2",
            "type": "Gene",
            "symbol": "Ndufv2",
            "source": [
                "Tcf4",
                "Pou5f1",
                "Foxp1",
                "Cebpb",
                "Nfib",
                "Atf4",
                "Junb",
                "Fos",
                "Jun",
                "Sox2",
                "Hes1",
                "Nanog",
                "Jund",
                "Twist2",
                "Hdac2",
                "Sox4"
            ],
            "target": [],
            "expression_sum": {
                "class1": 15369.0,
                "class2": 53142.0
            },
            "uniprot_ids": [
                "Q9D6J6",
                "M0QWP9",
                "D6RGN4"
            ]
        },
        "Nedd4": {
            "id": "Nedd4",
            "type": "Gene",
            "symbol": "Nedd4",
            "source": [
                "Klf6",
                "Pou5f1",
                "Hmga2",
                "Cebpb",
                "Nfib",
                "Atf4",
                "Fos",
                "Trp53",
                "Hes1",
                "Jund",
                "Twist2",
                "Hdac2",
                "Smc3",
                "Smarca5"
            ],
            "target": [],
            "expression_sum": {
                "class1": 32473.0,
                "class2": 36740.0
            },
            "uniprot_ids": [
                "P46935",
                "Q8BU52",
                "Q3V335",
                "B2RSC8",
                "Q3TUU1",
                "V9GXS6",
                "A0A571BEP7",
                "V9GWV8",
                "V9GXK3",
                "A0A571BDN7",
                "Q8BNU7",
                "A0JNZ0"
            ]
        },
        "Nedd8": {
            "id": "Nedd8",
            "type": "Gene",
            "symbol": "Nedd8",
            "source": [
                "Tcf4",
                "Pou5f1",
                "Foxp1",
                "Cebpb",
                "Atf4",
                "Junb",
                "Jun",
                "Sox2",
                "Nanog",
                "Jund",
                "Twist2",
                "Hdac2",
                "Smc3",
                "Sox4"
            ],
            "target": [],
            "expression_sum": {
                "class1": 34309.0,
                "class2": 83106.0
            },
            "uniprot_ids": [
                "P29595",
                "Q3UI46",
                "A0A2I3BRG0",
                "Q3TY05"
            ]
        },
        "Nip7": {
            "id": "Nip7",
            "type": "Gene",
            "symbol": "Nip7",
            "source": [
                "Klf2",
                "Tcf4",
                "Pou5f1",
                "Foxp1",
                "Atf4",
                "Junb",
                "Fos",
                "Jun",
                "Sox2",
                "Nanog",
                "Jund",
                "Twist2",
                "Cebpb",
                "Sox4"
            ],
            "target": [],
            "expression_sum": {
                "class1": 5329.0,
                "class2": 37438.0
            },
            "uniprot_ids": [
                "Q9CXK8",
                "E9Q109"
            ]
        },
        "Nktr": {
            "id": "Nktr",
            "type": "Gene",
            "symbol": "Nktr",
            "source": [
                "E2f4",
                "Trim28",
                "Rest",
                "Nfib",
                "Atf4",
                "Ctcf",
                "Tgif1",
                "Trp53",
                "Hdac1",
                "Ybx1",
                "Hdac2",
                "Smarca5"
            ],
            "target": [],
            "expression_sum": {
                "class1": 10219.0,
                "class2": 7691.0
            },
            "uniprot_ids": [
                "P30415",
                "Q8CBP6",
                "Q3TQ51",
                "S4R2L3",
                "S4R2C2",
                "S4R1C3",
                "S4R1L7",
                "S4R220"
            ]
        },
        "Nme1": {
            "id": "Nme1",
            "type": "Gene",
            "symbol": "Nme1",
            "source": [
                "Nfib",
                "Fos",
                "Id2",
                "Sox2",
                "Hdac2",
                "Tcf4",
                "Pou5f1",
                "Jun",
                "Smarca5",
                "Sox4",
                "Jund",
                "Cebpb",
                "Smc3",
                "Foxp1",
                "Atf4",
                "Junb",
                "Nanog",
                "Tfcp2l1",
                "Twist2"
            ],
            "target": [],
            "expression_sum": {
                "class1": 59662.0,
                "class2": 112366.0
            },
            "uniprot_ids": [
                "P15532",
                "Q5NC81",
                "Q5NC79",
                "Q5NC80"
            ]
        },
        "Nol7": {
            "id": "Nol7",
            "type": "Gene",
            "symbol": "Nol7",
            "source": [
                "Klf2",
                "Tcf4",
                "Pou5f1",
                "Cebpb",
                "Nfib",
                "Atf4",
                "Junb",
                "Fos",
                "Jun",
                "Nanog",
                "Jund",
                "Twist2",
                "Hdac2",
                "Smarca5",
                "Sox4"
            ],
            "target": [],
            "expression_sum": {
                "class1": 10992.0,
                "class2": 29467.0
            },
            "uniprot_ids": [
                "Q9D7Z3",
                "A1L3B6",
                "A0A1Y7VMS4"
            ]
        },
        "Nop56": {
            "id": "Nop56",
            "type": "Gene",
            "symbol": "Nop56",
            "source": [
                "Klf2",
                "Tcf4",
                "Pou5f1",
                "Atf4",
                "Junb",
                "Fos",
                "Jun",
                "Sox2",
                "Hes1",
                "Nanog",
                "Jund",
                "Twist2",
                "Cebpb",
                "Zfp57",
                "Sox4"
            ],
            "target": [],
            "expression_sum": {
                "class1": 9544.0,
                "class2": 64815.0
            },
            "uniprot_ids": [
                "Q9D6Z1",
                "F6USW7",
                "A2APD7",
                "F7CHQ7",
                "E0CXZ0",
                "F6V095",
                "F6TLG3",
                "Q3USZ5",
                "Q3UJZ7",
                "F6U250",
                "F7CHP9"
            ]
        },
        "Npm1": {
            "id": "Npm1",
            "type": "Gene",
            "symbol": "Npm1",
            "source": [
                "Tcf4",
                "Pou5f1",
                "Foxp1",
                "Nfib",
                "Atf5",
                "Atf4",
                "Junb",
                "Fos",
                "Jun",
                "Sox2",
                "Nanog",
                "Jund",
                "Cebpb",
                "Sox4"
            ],
            "target": [],
            "expression_sum": {
                "class1": 173806.0,
```
