## Supplement 1 (MEF vs. ESC) for "AGEAS: Automated Machine Learning based Genetic Regulatory Element Extraction System": outlier_grps.js

```
{
    "Smarca5_Gng12": 37.7299644540652,
    "Sarnp_Pou5f1": 27.18349279818091,
    "Ybx1_Cav1": 21.722016107788964,
    "Sox4_Hspd1": 20.644335272398717,
    "Ybx1_Actg1": 20.540151778619617,
    "Pou5f1_Impa2": 20.30698222730161,
    "Rpl19_Pou5f1": 16.49233378801502,
    "Trp53_Anxa3": 16.43200605566026,
    "Sumo1_Junb": 15.897515146481782,
    "Pou5f1_Eif5a": 13.751124691313363,
    "Pou5f1_Chchd4": 13.411722376369942,
    "Trp53_Rcn3": 13.334339228323088,
    "Reep5_E2f4": 13.31648028261263,
    "Uchl3_Pou5f1": 13.205721209195719,
    "Pou5f1_H3f3b": 13.172729817178606,
    "Smarca5_Serpinh1": 13.126280619626307,
    "Tnfrsf12a_Smc3": 13.122294152195447,
    "Pou5f1_Hmgn2": 13.096551253970157,
    "Snrpf_Pou5f1": 13.06982712285176,
    "Tipin_Pou5f1": 13.053899525593797,
    "Pou5f1_Cenpa": 13.03179219927653,
    "Sdcbp_Hdac1": 12.95339525698891,
    "Smc3_Marcks": 12.88880167807471,
    "Trim28_Rbms1": 12.88827580856661,
    "Smarca5_Nenf": 12.845228474560642,
    "Mbnl2_Hdac2": 12.827869208593004,
    "Smc3_Kdelr3": 12.75934902319567,
    "Pou5f1_Emc8": 12.750765634635652,
    "Sox4_H3f3a": 12.750667495647907,
    "Trp53_Itm2b": 12.748599040023969,
    "Srsf6_Foxp1": 12.695197542132158,
    "Serbp1_Foxp1": 12.638944913722055,
    "Trp53_Calu": 12.625941717547422,
    "Smarca5_Sh3glb1": 12.61272931403217,
    "Pou5f1_Hnrnpdl": 12.541885557151842,
    "Sox4_Rbm39": 12.488146270157616,
    "Sox4_Hmgn1": 12.483484434078855,
    "Npm1_Cebpb": 12.472251687031884,
    "Ube2a_Pou5f1": 12.448983351861148,
    "Cebpb_Atp5g1": 12.392069116216172,
    "Trp53_Glipr2": 12.386617549861178,
    "Trim28_Sri": 12.35199001393722,
    "Pou5f1_Ccnd3": 12.282259545680267,
    "Ubxn1_Pou5f1": 12.259689716489984,
    "Ctcf_Basp1": 12.210971877205528,
    "Ppa1_Jund": 12.139106545763415,
    "Pou5f1_Eif1a": 11.955650999342344,
    "Rpl12_Pou5f1": 9.059755101421048,
    "Rpl26_Pou5f1": 9.018127023542075,
    "Rpl14_Pou5f1": 8.975007888712234,
    "Ubb_Pou5f1": 8.873215967114389,
    "Mrpl35_Hmga2": 8.841964396244249,
    "Rcc2_Pou5f1": 8.77689698762181,
    "Rpl13a_Pou5f1": 8.732925373496046,
    "Serpinh1_Hmgb1": 8.713716024448019,
    "Ppia_Pou5f1": 8.676786342256804,
    "Cebpb_2700060E02Rik": 8.642952955711628,
    "Pou5f1_Bex1": 8.639333181585743,
    "Zfp42_Klf2": 8.637544720620218,
    "Ywhaq_Pou5f1": 8.630891619854783,
    "Rbm3_Pou5f1": 8.629166916962628,
    "Pou5f1_Pomp": 8.625668956847324,
    "Smarca5_Anxa5": 8.593730565749441,
    "Pou5f1_Mrpl36": 8.535766196673544,
    "Pou5f1_Birc5": 8.515177508794906,
    "Pou5f1_Pfdn6": 8.499721725582697,
    "Foxp1_Cct8": 8.496354466873393,
    "Ybx1_Pla2g1b": 8.474867348773778,
    "Ybx1_Bola3": 8.472387737430928,
    "Utf1_Trim28": 8.447174097436122,
    "Son_Foxp1": 8.426446379279167,
    "Rest_Lgals1": 8.425078338833623,
    "Hdac2_Cdc42": 8.424956247876356,
    "Trp53_Cald1": 8.421134219715197,
    "Psmb6_Pou5f1": 8.420208755108712,
    "Pou5f1_Gapdh": 8.40392400693104,
    "Pou5f1_Akr1b3": 8.402365138407676,
    "Tdgf1_Pou5f1": 8.400620922485118,
    "Ybx1_Txnl1": 8.397114202018864,
    "Trp53_Tmsb4x": 8.384520511179321,
    "Slc25a5_Junb": 8.373067097934287,
    "Rps3a1_Pou5f1": 8.36845520316534,
    "Pcbp1_Hmga2": 8.355909570755902,
    "Ybx1_Cct4": 8.34942112853969,
    "Hmga2_Cyc1": 8.346933036384653,
    "Ptges3_Pou5f1": 8.344112774922573,
    "Ybx1_Anxa1": 8.34183040037124,
    "Hdac2_Actg1": 8.330584578824473,
    "Trp53_S100a6": 8.328214261279264,
    "Pou5f1_Cdk1": 8.324574372234277,
    "Zfp42_Sox2": 8.322687663873698,
    "Pou5f1_Cct5": 8.311826439518091,
    "Pou5f1_Ost4": 8.306611063133765,
    "Trp53_Pou5f1": 8.294842020051686,
    "Pou5f1_Atp5j": 8.285990777070747,
    "Pou5f1_Cct8": 8.27920231093949,
    "Rbpms2_Pou5f1": 8.274117879604557,
    "Hmga2_Cetn3": 8.26454405818755,
    "Psip1_Pou5f1": 8.26241686723412,
    "Trp53_Itgb1": 8.259825593218316,
    "Stra13_Pou5f1": 8.254535393057743,
    "Pou5f1_Cfdp1": 8.25445489516856,
    "Smc3_Selk": 8.25326184464126,
    "Trim28_Anxa3": 8.235935064150205,
    "Junb_Hnrnpa2b1": 8.225901849635546,
    "Srp9_Pou5f1": 8.22351024043115,
    "Eno1_Basp1": 8.222009820730793,
    "Pou5f1_Aurkaip1": 8.220196088130676,
    "Pou5f1_Eif5": 8.21590099908131,
    "Tmed10_Hdac2": 8.20936046652538,
    "Rps16_Pou5f1": 8.205636855118305,
    "Nap1l1_Junb": 8.203720163502044,
    "Wls_Smc3": 8.19697921550924,
    "Rpl38_Pou5f1": 8.192023392467465,
    "Pou5f1_Hells": 8.184169856610753,
    "Ybx1_Laptm4a": 8.179164777725925,
    "Gng12_Eno1": 8.166513271363922,
    "Hmga2_Gcsh": 8.160322361509447,
    "Hmga2_Gspt1": 8.15803042446739,
    "Snrpd1_Pou5f1": 8.156263486860976,
    "Ybx1_Timp1": 8.155720467747752,
    "Sparc_Hdac2": 8.152551087201802,
    "Ybx1_Eva1b": 8.14586145971783,
    "Atf4_Actg1": 8.140562733172102,
    "L1td1_Hmgb1": 8.138299717553414,
    "Atf5_Asns": 8.12849421171278,
    "Tomm70a_Pou5f1": 8.11273651505171,
    "Snrpd1_Arf4": 8.112481966617606,
    "Ybx1_Cd44": 8.10886900662441,
    "Sox2_Cox7c": 8.106849899426166,
    "Sox4_Eif4e2": 8.106733622524755,
    "Pou5f1_Morc1": 8.103537794332038,
    "Pou5f1_Impdh2": 8.101647909947827,
    "Pou5f1_Mbd3": 8.085240598991575,
    "Psmc4_Pou5f1": 8.08416474079271,
    "Pou5f1_G3bp2": 8.083838570458665,
    "Ppa1_Arf4": 8.081855373596701,
    "Smarca5_Cnih1": 8.081678864523996,
    "Hsp90aa1_Cebpb": 8.077518729684453,
    "Sox4_Ptma": 8.071741171201085,
    "Ybx1_Gnas": 8.068984384450518,
    "Fkbp3_Cebpb": 8.063787217565059,
    "Hdac2_Arpc1b": 8.06039078273605,
    "Slc25a4_Hdac2": 8.058772066578511,
    "Ctcf_Cald1": 8.050702523563567,
    "Snrpa_Pou5f1": 8.039741464372256,
    "Serpinh1_Ctcf": 8.03240092306852,
    "Hdac2_Gnb2": 8.02768077177797,
    "Smarca5_Slc25a4": 8.026341275451754,
    "Trp53_S100a11": 8.022211377365938,
    "Rrp1b_Pou5f1": 8.016930058518389,
    "Pou5f1_Mrpl21": 8.015179388310349,
    "Ybx1_Cnn2": 8.013977844587977,
    "Ybx1_Gsn": 8.003163103102715,
    "Pou5f1_Dazap1": 7.99238557130544,
    "Hmgb1_Gdf3": 7.991961934417451,
    "Pou5f1_Gmnn": 7.984293152226233,
    "Ddx39b_Arf4": 7.952575846978511,
    "Pou5f1_Cox8a": 7.944979899207902,
    "Psmb7_Junb": 7.934327926699716,
    "Srsf7_Pou5f1": 7.916059543334124,
    "Ybx1_Tmed5": 7.914561884129938,
    "Nhp2_Hmga2": 7.902328448361087,
    "Smc3_Arf4": 7.890793320268882,
    "Pou5f1_Fam49b": 7.885083214234243,
    "Smc3_Ppic": 7.879369807828928,
    "Smarca5_Calr": 7.873679893070482,
    "Fus_Foxp1": 7.869365109263608,
    "Ybx1_Sept7": 7.868683013696691,
    "Ybx1_Acbd3": 7.863022071119593,
    "Uhrf1_Pou5f1": 7.855068519229861,
    "Sox2_Eif2s2": 7.849760212654985,
    "Smc3_Canx": 7.805451539374967,
    "S100a6_Hdac1": 7.7832306190166864,
    "Pcbp1_Junb": 7.775541441654179,
    "Junb_Hnrnpa1": 7.773162982043471,
    "Tomm22_Hmga2": 7.7494523184515485,
    "Smarca5_Rrbp1": 7.698763815138154,
    "l7Rn6_Pou5f1": 7.688540541767165,
    "Junb_Atp5b": 7.671532383623449,
    "Sox4_Pfdn1": 7.5745061954577935,
    "Nudc_Cebpb": 7.517255721218203,
    "Sox2_Erh": 7.484498116008014,
    "Smarca5_Fam162a": 7.484322107905873,
    "Hmgb1_Elovl1": 7.463305005860957,
    "Ybx1_Rer1": 7.362151877334733,
    "Snrpd1_Junb": 7.351439412831105,
    "Smarca5_Arpc5": 7.338998292754545,
    "Raly_Foxp1": 7.319053221595314,
    "Gtf2a2_2010111I01Rik": 7.316721448253342,
    "Trp53_Cfl2": 7.267121913585511,
    "Hsp90ab1_Cebpb": 36.798690114567634,
    "S100a6_Ctcf": 36.728697212882196,
    "Pou5f1_2700060E02Rik": 20.479259974999867,
    "Pou5f1_Ak2": 20.149184375189968,
    "Rpl29_Cebpb": 16.053704312492197,
    "Rps6_Pou5f1": 15.62009719361934,
    "Sox4_Bclaf1": 12.983617095431667,
    "Foxp1_Anapc11": 12.658178972258643,
    "U2af1_Prrx1": 12.642585252756794,
    "Trp53_Arpc1b": 12.61775845021631,
    "Pou5f1_Pa2g4": 12.5654271802602,
    "Uqcr10_Pou5f1": 12.554238600131338,
    "Pou5f1_Odc1": 12.553570546780742,
    "Ctcf_Anxa2": 12.53397592172945,
    "Pou5f1_Lsm3": 12.525376329071582,
    "Smarca5_Cav1": 12.489883325870867,
    "Sox4_Hnrnpa3": 12.44500823444734,
    "Rnps1_Pou5f1": 12.431715855605,
    "Smc3_S100a11": 12.429550223645851,
    "Twist2_Lsm5": 12.415750856042322,
    "Sox4_Atp5j2": 12.395625506694715,
    "Cyc1_Cebpb": 12.370255033241056,
    "Ppa1_Junb": 12.369298589943975,
    "Smc3_Cald1": 12.362214402102842,
    "Sox2_Snrpd2": 12.358017774365003,
    "Trp53_Tax1bp3": 12.34635152639331,
    "Ube2n_Pou5f1": 12.334906788351692,
    "Pou5f1_Hdac2": 12.329179689351637,
    "Rbm39_Pou5f1": 12.287448044659781,
    "Cebpb_Bzw1": 12.277632046040008,
    "Vim_Rest": 12.227343951571168,
    "Prrx1_Cct7": 12.209287484441704,
    "Hdac2_Col1a1": 12.108340969716473,
    "Pou5f1_Mybbp1a": 12.100137725841515,
    "Smarca5_Ahnak": 12.088975473861792,
    "U2surp_Pou5f1": 11.805756392862099,
    "Rbm8a_Klf2": 11.738262962168523,
    "Hdac2_Chd3": 11.658057434341274,
    "Trp53_Pgrmc1": 11.581822800847133,
    "Pou5f1_Kdm1a": 11.399126894434447,
    "Hdac2_Fundc2": 8.896019025621609,
    "Rhoa_Pou5f1": 8.761830364291406,
    "Smc3_Ptp4a2": 8.618220245350937,
    "Sox2_Ptges3": 8.569843201012997,
    "Snrpd2_Atf4": 8.502895564012345,
    "Srrm1_Hmga2": 8.411022902948437,
    "Smc3_Anxa5": 8.379947300091509,
    "Pou5f1_Ndufc1": 8.32734453559645,
    "Tceal8_Arf4": 8.308015998086207,
    "Pou5f1_Cox6c": 8.302757880239234,
    "Smc3_Rhoa": 8.283295114574793,
    "Pcbp1_Id2": 8.253885741447524,
    "Mrpl42_Klf2": 8.223524995959254,
    "Trim28_Esd": 8.221825679069612,
    "Timm13_Foxp1": 8.1725254608399,
    "Trim28_Pou5f1": 8.169724445866809,
    "Pou5f1_Hint1": 8.169254379698957,
    "Vim_Hdac2": 8.134671575984669,
    "Smarca5_Mgat2": 8.124431948681178,
    "Smc3_S100a6": 8.117593735569699,
    "Sub1_Pou5f1": 8.11642611421973,
    "Ybx1_Gpx4": 8.111852387965829,
    "Psmb6_Junb": 8.107660463441754,
    "Ptma_Pou5f1": 8.101526336790496,
    "Rps2_Pou5f1": 8.097209430537125,
    "Rps18_Hmga2": 8.085796034204186,
    "Pou5f1_Atp5j2": 8.083063081004216,
    "Rps3_Pou5f1": 8.082215932816284,
    "Smarca5_Bsg": 8.079217325644613,
    "Ptma_Junb": 8.05855453492486,
    "Ybx1_Emc7": 8.054290297533456,
    "Siva1_Pou5f1": 8.042267459391358,
    "Foxp1_Aurkaip1": 8.03705901065291,
    "Pou5f1_Hsp90aa1": 8.025454184348224,
    "Snrpc_Hmga2": 8.025407396609912,
    "Pou5f1_Hspd1": 8.009809620596297,
    "Trp53_Sept7": 8.009672424981463,
    "Trp53_Parva": 8.002536211062527,
    "Smarca5_Oxct1": 7.99876970026743,
    "Pou5f1_Ewsr1": 7.993458782915246,
    "Trim28_S100a6": 7.989853331050689,
    "Eno1_Anxa2": 7.986120521338453,
    "Srrm1_Pou5f1": 7.981665324560436,
    "Smc3_Abracl": 7.974824367341525,
    "Txndc5_Trp53": 7.963570079643296,
    "Pou5f1_Bcat1": 7.959003250609138,
    "Pou5f1_Gpx4": 7.957117213513914,
    "Pou5f1_Eif3k": 7.9501567098811625,
    "Ybx1_Selm": 7.946806189414246,
    "Junb_H2afz": 7.946052838385887,
    "Pou5f1_Mlf2": 7.944653689326114,
    "Pou5f1_Gsta4": 7.9362155231221765,
    "Rps17_Junb": 7.936184116343617,
    "Srsf1_Hmga2": 7.929641015971048,
    "Mrpl18_Hmga2": 7.927241041365846,
    "Trp53_Rhoc": 7.920877687182265,
    "Sod2_Pou5f1": 7.913516852608424,
    "Pou5f1_Mrpl51": 7.912624670128874,
    "Trp53_Laptm4a": 7.909683959925096,
    "Smc3_Nenf": 7.908067906743671,
    "Pou5f1_Ooep": 7.906379861933894,
    "Hdac2_Glipr2": 7.903820400063852,
    "Ppic_Hdac2": 7.9001887045769905,
    "Ybx1_Fam25c": 7.899470083975307,
    "Ptma_Foxp1": 7.898467663316851,
    "Foxp1_Btf3l4": 7.895187392279698,
    "Rpl41_Pou5f1": 7.894589008724252,
    "Pou5f1_Nucks1": 7.888201774581483,
    "Ybx1_Tagln2": 7.87933387482355,
    "Pou5f1_Gm7325": 7.877999006265764,
    "Ybx1_Cfl2": 7.875909828946483,
    "Pou5f1_Cox7c": 7.87530596084193,
    "Pou5f1_Marcksl1": 7.874807172877382,
    "Smc3_Lmna": 7.868651296381543,
    "Prelid1_Pou5f1": 7.867706775704786,
    "Sox4_Set": 7.8674056417093094,
    "Pou5f1_Cdca3": 7.8654717025140615,
    "Rangrf_Pou5f1": 7.8647855632479455,
    "Rpl31_Pou5f1": 7.864263856481698,
    "Hnrnpa3_Cebpb": 7.857034669319972,
    "Rps8_Pou5f1": 7.8525563189053464,
    "Smarca5_Selk": 7.851236612229138,
    "Lgals1_Ctcf": 7.850398783000659,
    "Rpl36_Pou5f1": 7.840749698738069,
    "Smarca5_Actg1": 7.836562614834177,
    "Ybx1_Sri": 7.836367817698857,
    "Rpl17_Pou5f1": 7.833224339250407,
    "Trp53_Timp2": 7.828286227468908,
    "Smarca5_Pdia3": 7.827970888150837,
    "Ybx1_Parva": 7.826894173613137,
    "Pou5f1_Fscn1": 7.824850349892777,
    "Rpl9_Pou5f1": 7.79233545185511,
    "Trp53_Dppa5a": 7.7922074161467405,
    "Tra2b_Atf4": 7.7901533918525105,
    "Pou5f1_Polr1c": 7.790095544493092,
    "Trp53_Pafah1b2": 7.789787621855348,
    "Pou5f1_Park7": 7.789400472552343,
    "Trp53_Emp1": 7.788485613785897,
    "Sox4_Psma3": 7.776274487067037,
    "Rps7_Pou5f1": 7.77035446306071,
    "Pou5f1_Aurka": 7.767576631737148,
    "Smc3_Rbms1": 7.765900741778497,
    "Ubald2_Pou5f1": 7.765199405138855,
    "Junb_Hnrnpf": 7.7468304096191485,
    "Twist2_Ptma": 7.745790146066578,
    "Trim28_L1td1": 7.745626808075205,
    "Tma7_Hmga2": 7.745070387184092,
    "Wdr18_Pou5f1": 7.745000972265579,
    "Eif4a1_Atf4": 7.742855334829515,
    "Smarca5_Gapdh": 7.741997776431559,
    "Npm1_Foxp1": 7.741488307830459,
    "Trp53_Tpm4": 7.738536298120599,
    "Smarca5_Pgrmc1": 7.7380929685226185,
    "Fkbp2_Arf4": 7.737080351916048,
    "Pou5f1_Mrps34": 7.7118183576576875,
    "Pou5f1_Eif3a": 7.707119837502557,
    "Pou5f1_Phf5a": 7.705781366733639,
    "Smarca5_Rpn1": 7.705284765011227,
    "Ybx1_Cdc42": 7.698936528409467,
    "Pou5f1_Atp5g2": 7.694029124309022,
    "Ybx1_Mtpn": 7.691934013172456,
    "Rab14_Hmgb1": 7.6885407988010295,
    "Hdac2_Calr": 7.686382392598309,
    "Sox2_Mrpl18": 7.6795534306968225,
    "Ybx1_Fbln5": 7.659063019968849,
    "Srsf3_Junb": 7.650762997907476,
    "Pou5f1_Pmf1": 7.638445398955234,
    "Ybx1_Spop": 7.633579416216308,
    "Ybx1_Nenf": 7.63316522919355,
    "Ybx1_Prrx1": 7.627312033409459,
    "Trp53_Cdc42": 7.624367478995845,
    "Rpl35a_Arf4": 7.622931137390778,
    "Trp53_Minos1": 7.6150087203543695,
    "Pou5f1_Paics": 7.611991372322627,
    "Ybx1_Esd": 7.599499559577334,
    "Uqcr10_Sox2": 7.585286136721712,
    "Ybx1_2810417H13Rik": 7.583366878011531,
    "Psmc6_Foxp1": 7.557877504687303,
    "Foxp1_2700094K13Rik": 7.535328327328505,
    "Ncl_Cebpb": 7.530728736869166,
    "Pou5f1_Adprh": 7.529923091231538,
    "Ppic_Eno1": 7.522855293350869,
    "Sox2_Nhp2l1": 7.486751866038561,
    "Hdgf_Foxp1": 7.477840359872436,
    "Sae1_Pou5f1": 7.472144469174464,
    "Trp53_Cnn2": 7.437384892613084,
    "Sri_Smc3": 7.436595517535624,
    "Ybx1_Khdrbs1": 7.408718381768149,
    "Junb_Eif4h": 7.341794991431949,
    "Gtf2a2_Atf4": 7.32208881150167,
    "Ybx1_Eif4g2": 7.231758630382088,
    "Psmb6_Arf4": 7.19559752809031,
    "Fdps_Arf4": 7.004637647393337,
    "Ybx1_Fabp5": 6.501775978731659,
    "Zyx_Smarca5": 34.40596565577896,
    "Ran_Pou5f1": 25.873646455082003,
    "Junb_Anp32b": 25.309900454157546,
    "Pou5f1_Hdgf": 19.406247712511192,
    "Pou5f1_Hnrnpa3": 19.189570013057253,
    "Sfpq_Abracl": 18.680835857603146,
    "Ybx1_Lims1": 18.506444076526428,
    "Ybx1_Gpi1": 15.323787220732338,
    "Raly_Junb": 15.025188868819955,
    "Ybx1_Grip2": 12.340285686102199,
    "Ctcf_Anxa1": 12.287947995341275,
    "Sumo1_Pou5f1": 11.854221794585145,
    "Nenf_Ctcf": 11.825603685767227,
    "Prmt1_Cebpb": 11.818719694666784,
    "Smarca5_Gnas": 11.799805742401348,
    "Ctcf_Bgn": 11.779660630786191,
    "Twist2_Hsp90aa1": 11.773582555816951,
    "Phf5a_Junb": 11.762975106915682,
    "Pou5f1_Ciapin1": 11.714544396981323,
    "Pou5f1_Fbl": 11.68546351576669,
    "Pou5f1_Dnajc21": 11.669922427617596,
    "Pou5f1_Asna1": 11.663775904589418,
    "Pou5f1_Gcsh": 11.648881910939705,
    "Pou5f1_Dph3": 11.629099130920096,
    "Tomm22_Pou5f1": 11.624225336253279,
    "Sumo2_Hmga2": 11.606080593462579,
    "Smarca5_Foxp1": 11.597104151225288,
    "Nap1l1_Cebpb": 11.572427175540716,
    "Snrpe_Foxp1": 11.56063247475627,
    "Foxp1_Cdk4": 11.536981232755846,
    "Pcbp1_Foxp1": 11.531651354437457,
    "Sox4_Serbp1": 11.52260138692865,
    "Snrpb_Foxp1": 11.489472047711883,
    "Trim28_Serpinb6a": 11.452276892594133,
    "Sumo3_Foxp1": 11.413231283139742,
    "Smc3_Sh3glb1": 11.378368650303207,
    "Hnrnpa2b1_Foxp1": 11.290405392462123,
    "Foxp1_Cox5a": 11.272815046022627,
    "Pou5f1_Mrpl12": 11.26158229858789,
    "Tomm7_Sox4": 11.149809502747885,
    "Foxp1_Cdv3": 11.115275560494764,
    "Ybx1_Ndn": 10.988611275499588,
    "Trp53_Tm9sf3": 10.834216280457538,
    "Eno1_Arpc4": 8.283398945238597,
    "Id2_2310036O22Rik": 8.063941553748487,
    "Tpd52l2_Pou5f1": 8.028127270700365,
    "Gnb2_Atf4": 7.951101062321447,
    "Smarca5_Cstb": 7.893879649980394,
    "Ybx1_Gm7325": 7.83414007993731,
    "Hdac2_Anxa2": 7.795982688438572,
    "Smc3_Anxa1": 7.784886756550952,
    "Trp53_Lipe": 7.764049790139759,
    "Stub1_Pou5f1": 7.729957383314949,
    "Hdac2_Gnas": 7.666406757360481,
    "Trp53_Lmna": 7.653981336387827,
    "Dnaja1_Atf4": 7.643745447787841,
    "Ndufa4_Cebpb": 7.630511431040795,
    "Pou5f1_E2f4": 7.6197788983429735,
    "Pou5f1_Mybl2": 7.596896366831975,
    "Serbp1_Pou5f1": 7.5806445353322705,
    "Ybx1_Smchd1": 7.5731345728752535,
    "Skp1a_Pou5f1": 7.56239714749396,
    "Srsf11_Junb": 7.55412401167296,
    "Trp53_Sri": 7.553767923234988,
    "Jund_Hspd1": 7.5511913653986245,
    "Pou5f1_Ifitm1": 7.536896532303901,
    "Pou5f1_Cnbp": 7.5359680534404445,
    "Rps20_Pou5f1": 7.531206981429083,
    "Pou5f1_Map1lc3b": 7.531048496049879,
    "Junb_Erh": 7.526310606592853,
    "Pcbp2_Id2": 7.524113204263519,
    "Kdelr1_Hdac2": 7.503786899055971,
    "Pou5f1_Kdm5b": 7.502315692800322,
    "Pou5f1_Pim3": 7.484901028179682,
    "Pou5f1_Cox6b1": 7.479384976381982,
    "Rps12_Pou5f1": 7.47572149560268,
    "Smarca5_Arpc3": 7.474414775045278,
    "Set_Hmga2": 7.466206627686979,
    "Tuba1a_Hmgb1": 7.464089736502987,
    "Rps6_Hmga2": 7.459714157329323,
    "Trim28_Anxa2": 7.454349330680033,
    "Tma7_Pou5f1": 7.453154590568292,
    "Trim28_Gabarap": 7.447145066143109,
    "Tubb6_Ctcf": 7.438947400425239,
    "Ppa1_Pou5f1": 7.433201682911275,
    "Hmgb1_Bex1": 7.4306266088495265,
    "Trp53_Cnih1": 7.4273282607756865,
    "Smc3_Prrx1": 7.4241813612233125,
    "Smc3_Cd63": 7.422517274876333,
    "Rpl30_Pou5f1": 7.4184478920016765,
    "Sdhb_Pou5f1": 7.4074379892927205,
    "Pou5f1_Pole4": 7.407288741802937,
    "Sox2_Marcksl1": 7.399124149961559,
    "Ybx1_Sparc": 7.396952809692504,
    "Utf1_Pou5f1": 7.385995593876762,
    "Ybx1_Tdgf1": 7.385069780384149,
    "Hdac2_Eprs": 7.380162774377818,
    "Serbp1_Cebpb": 7.376544817366318,
    "Wls_Trp53": 7.373709789864915,
    "Pou5f1_Nudc": 7.37341677417327,
    "Tcf4_Snrpb2": 7.36836235787861,
    "Smc3_Hdlbp": 7.367435008595264,
    "Pou5f1_Mif": 7.365891503834771,
    "Rps21_Pou5f1": 7.362385643569518,
    "Twist2_Rpl29": 7.359359686935936,
    "Smc3_Kdelr1": 7.354721655628736,
    "Pou5f1_Fkbp4": 7.353682889849773,
    "Pou5f1_Alpl": 7.351001071537364,
    "Sept11_Eno1": 7.3505119341647305,
    "Tcp1_Pou5f1": 7.350317186593473,
    "Prrx1_Ppp2ca": 7.345920827584141,
    "Pou5f1_Cct7": 7.345573608010695,
    "Smarca5_Arf4": 7.345386871866771,
    "Smarca5_Hmga2": 7.34155848996304,
    "Pou5f1_Pcbp1": 7.341123864331861,
    "Pou5f1_Hnrnpm": 7.340387386132713,
    "Hdac1_Capn2": 7.340331424544367,
    "H2afz_Foxp1": 7.339600922564044,
    "Pou5f1_Mylpf": 7.33119324089369,
    "Ybx1_Actn1": 7.330754751657126,
    "Trp53_Tnfrsf12a": 7.3268375545192,
    "Pou5f1_Ndufb9": 7.326474441959241,
    "Ybx1_Ccm2": 7.326228729014993,
    "Trp53_Gpx8": 7.312504595969643,
    "Trp53_Anxa5": 7.307810083699885,
    "Ptbp1_Pou5f1": 7.306564212039534,
    "Pou5f1_Nubp1": 7.303094085368425,
    "Sox2_Pou5f1": 7.300743814940146,
    "Pou5f1_Anapc5": 7.297973860383017,
    "Pou5f1_Cbx1": 7.295604397161349,
    "Smc3_Arpc2": 7.294134784229033,
    "Ranbp1_Foxp1": 7.2923654725764315,
    "Pou5f1_Copz1": 7.283338109943936,
    "Pou5f1_Ndufv3": 7.279726284458913,
    "Tubb6_Smc3": 7.276091508896408,
    "Hdac1_Anxa5": 7.2730170356860935,
    "Syncrip_Arf4": 7.270785960703658,
    "Oaz1_Foxp1": 7.269882931617511,
    "Pou5f1_Fam136a": 7.269757268323948,
    "Sox2_Set": 7.264520130947032,
    "Stmn1_Pou5f1": 7.260620419987253,
    "Tuba1a_Smc3": 7.257528025465047,
    "Trp53_B2m": 7.253385015201734,
    "Ywhae_Foxp1": 7.249510016541392,
    "Eif2s2_Cebpb": 7.249192833029712,
    "Trim28_Tpm2": 7.247573644088382,
    "Eno1_Dppa5a": 7.241530849938524,
    "Mylpf_Mybl2": 7.2355129509946305,
    "Sparc_Rest": 7.234957031884258,
    "Canx_Atf4": 7.232316740457965,
    "Sumo2_Foxp1": 7.229714900163607,
    "Pou5f1_Eif1ax": 7.228001011758799,
    "Ybx1_Acta2": 7.21646196061122,
    "Sumo3_Pou5f1": 7.209396060306393,
    "Pou5f1_Mrps16": 7.2069540637052825,
    "Pou5f1_H3f3a": 7.204152384027583,
    "Pou5f1_Mat2a": 7.20385805574739,
    "Hmgb1_Efemp2": 7.177888082408375,
    "Ybx1_Sept11": 7.155861645273011,
    "Tra2b_Pou5f1": 7.153115892902698,
    "Hdac2_Cd63": 7.14407111529306,
    "Trp53_Abracl": 7.128916143499938,
    "Rps11_Pou5f1": 7.116455332356983,
    "Smarca5_Nktr": 7.11051902207843,
    "Hmga2_Eif5a": 7.1089564222924215,
    "Txndc5_Trim28": 7.10444717938078,
    "Ybx1_Ccng1": 7.1039190401474785,
    "Psma2_Pou5f1": 7.090052358235143,
    "Trim28_Col1a2": 7.083006663430977,
    "Pou5f1_Pfdn2": 7.074498345494486,
    "Smarca5_Emc7": 7.041591306356138,
    "Spag7_Pou5f1": 7.035975629909738,
    "Mtpn_Hdac2": 7.035156728566732,
    "Pou5f1_Pdgfa": 7.030250703778039,
    "Trim28_Gdf3": 7.000661600345579,
    "Smc3_Gng12": 6.976864662017318,
    "Junb_Atpif1": 6.974569159400291,
    "Sfpq_1110004F10Rik": 6.953399920984324,
    "Tagln2_Hdac2": 6.951612033245949,
    "Pou5f1_Ndufv2": 6.937516691374128,
    "Snrpf_Junb": 6.802142055509803,
    "Sox2_H3f3b": 6.800090267791159,
    "Sox4_Snx5": 6.79873533097307,
    "Rest_Bgn": 6.7688566034906685,
    "Cnbp_Cebpb": 6.674061679124141,
    "Psmg4_Pou5f1": 6.640894335058586,
    "Ybx1_Rab14": 6.491730471737821,
    "Jund_Ahsa1": 6.425443060847365,
    "Junb_Banf1": 6.405487093602978,
    "Pgls_Hdac1": 6.164920071831532,
    "Ybx1_Tead1": 6.110366233673462
}
```
