## Supplement 1 (MEF vs. ESC) for "AGEAS: Automated Machine Learning based Genetic Regulatory Element Extraction System": outlier_grps.js

```
{
    "Bzw1_Atf4": 66.31120265167813,
    "Srsf10_Foxp1": 33.02049439569706,
    "Smc3_Ptrf": 22.76496243981318,
    "Tmed5_Hmgb1": 18.756830401163736,
    "Smc3_Cd63": 18.602193028347692,
    "Ybx1_Gnas": 18.556032338163075,
    "Rpl12_Pou5f1": 18.517441085063723,
    "Pou5f1_Dnaja1": 18.394955496081423,
    "Smc3_Ilk": 18.26231530472231,
    "Sfpq_Abracl": 17.97216312895666,
    "Ube2d3_Pou5f1": 14.322229356230551,
    "Spag7_Pou5f1": 13.767025677950269,
    "Pou5f1_Dctpp1": 11.70792370326775,
    "Lmna_Hdac1": 11.705774810004325,
    "Pou5f1_Nhp2l1": 11.62963777958375,
    "Sf3b5_Pou5f1": 11.60932219876877,
    "Pou5f1_Lsm4": 11.583786789344426,
    "Pou5f1_Dtymk": 11.465653756375831,
    "l7Rn6_Pou5f1": 11.432808820181506,
    "Twist2_Fkbp3": 11.41501040126676,
    "Pou5f1_Hnrnpa2b1": 11.375627469716315,
    "Twist2_Hsp90aa1": 11.36075422160838,
    "Pou5f1_H3f3b": 11.333854518054261,
    "Smarca5_Cald1": 11.324245319949156,
    "Tomm22_Foxp1": 11.324060833192755,
    "Ctcf_Bgn": 11.323219543410733,
    "Pou5f1_Cdc34": 11.28442569971491,
    "Trp53_Rbms1": 11.282096383678853,
    "Snrpd1_Pou5f1": 11.266232562291274,
    "Hsp90ab1_Foxp1": 11.24919738165178,
    "Sox4_Dnaja1": 11.227653904500546,
    "Rbm39_Pou5f1": 11.207231039141096,
    "Sox4_Hnrnpf": 11.188293427839131,
    "Hspa8_Cebpb": 11.173978511369597,
    "Supt4a_Pou5f1": 11.169205283408683,
    "Pou5f1_Ndufb8": 11.143372909207358,
    "Hnrnpk_Foxp1": 11.139022031012756,
    "Laptm4a_Ctcf": 11.135722883840748,
    "Trp53_Gng12": 11.131152683130093,
    "Smarca5_Ahnak": 11.113288030620172,
    "Smarca5_Nenf": 11.10266572292584,
    "Trp53_Dnajc3": 11.10190452759769,
    "Sumo2_Junb": 11.09479108282208,
    "Zfp706_Foxp1": 11.019099048582646,
    "Junb_Fkbp3": 10.968309871941354,
    "Klf2_Cks1b": 10.936185391515446,
    "Snrpe_Cebpb": 10.91568959961695,
    "Mrpl18_Hmga2": 10.832341528698157,
    "Junb_Anp32b": 10.767584376394613,
    "Twist2_Psmb3": 10.61512114962013,
    "Reep5_Ctcf": 10.577499168298896,
    "Hmga2_1810022K09Rik": 10.255032824734775,
    "Pou5f1_Fdps": 7.7904242887475466,
    "Rpsa_Junb": 7.779385177482992,
    "Nfib_Hsp90ab1": 7.717637086486333,
    "Trim28_S100a6": 7.710533717140501,
    "Pou5f1_Btf3": 7.65945864846141,
    "Junb_Gpx4": 7.650081381148753,
    "Pou5f1_2810004N23Rik": 7.611695697093077,
    "Pou5f1_H3f3a": 7.582377983493376,
    "Tma7_Pou5f1": 7.574495615901612,
    "Ybx1_Serpinh1": 7.546436774336864,
    "Pou5f1_Platr3": 7.539502537293132,
    "Vim_Smc3": 7.526920554911792,
    "Rpl37_Pou5f1": 7.489535942357265,
    "Rpl7_Pou5f1": 7.466015160858833,
    "Serpinh1_Rest": 7.456250784650731,
    "Prrx1_H3f3a": 7.448167491112057,
    "Trim28_Cald1": 7.434118737101319,
    "Hdac2_Ckap4": 7.402482256313722,
    "Ranbp1_Foxp1": 7.3998190488486415,
    "Pou5f1_Hint1": 7.396349323620627,
    "S100a6_Eno1": 7.390658779556676,
    "Tmem258_Pou5f1": 7.387028269891077,
    "Rplp0_Pou5f1": 7.384174904314967,
    "Pou5f1_Eef1g": 7.383686187628755,
    "Smc3_Nenf": 7.375711786138505,
    "Rpl17_Pou5f1": 7.372525461353981,
    "Tomm7_Pou5f1": 7.370206022327842,
    "Ybx1_Timp1": 7.367952189370861,
    "Hmgb1_Hdlbp": 7.36140523700719,
    "Pou5f1_Ooep": 7.359008634756368,
    "Pou5f1_Hsp90aa1": 7.354223387456546,
    "Trp53_S100a11": 7.3423154540535664,
    "Pou5f1_Hmgb2": 7.3297923864397845,
    "Snrpd1_Junb": 7.3276550895618024,
    "Hnrnpd_Hmga2": 7.321912821382281,
    "Wdr74_Pou5f1": 7.316912583163923,
    "Trim28_Pou5f1": 7.297323724524795,
    "Pou5f1_Eif4ebp1": 7.290066539350581,
    "Pou5f1_Eif2s2": 7.277644216906578,
    "Stra13_Pou5f1": 7.265482277311648,
    "Pou5f1_Hells": 7.259368693637417,
    "Trp53_Mtpn": 7.248883629489844,
    "Sumo2_Cebpb": 7.244864472846796,
    "Npm1_Cebpb": 7.2440960242746035,
    "Ybx1_Gsn": 7.240824640445817,
    "Trp53_Anxa2": 7.23765275849176,
    "Rpl3_Pou5f1": 7.230030262486915,
    "Sparc_Hdac2": 7.226353384911085,
    "Tuba1a_Ctcf": 7.224153913904137,
    "Srp9_Pou5f1": 7.221620301253452,
    "Psmb6_Pou5f1": 7.211505582683895,
    "Ybx1_Ndn": 7.20920698229936,
    "Ybx1_Itm2b": 7.207332119416787,
    "Smc3_Anxa1": 7.1878973781197,
    "Ybx1_Tmed9": 7.180272047806402,
    "Ybx1_Timp2": 7.179087196324512,
    "Trp53_Ssr3": 7.173227862069985,
    "Trp53_Tnfrsf12a": 7.161748703266242,
    "Set_Cebpb": 7.161621599822701,
    "Smarca5_Serpinh1": 7.1571024324216586,
    "Smarca5_Pgrmc1": 7.15499856961667,
    "Upp1_Eno1": 7.1524531809349154,
    "Ybx1_Bnip2": 7.146684994582334,
    "Ybx1_Fam25c": 7.143289865106091,
    "Trp53_Tmsb4x": 7.1353020447743445,
    "Smc3_Sep15": 7.129602201845872,
    "Wls_Trp53": 7.1285241548999,
    "Pou5f1_Ifitm1": 7.127369481612913,
    "Wls_Smc3": 7.124953628404737,
    "Rae1_Pou5f1": 7.121355842283578,
    "Txndc5_Hdac2": 7.1111293384649015,
    "Ybx1_Tmsb4x": 7.110860216768412,
    "Pou5f1_Ddx46": 7.103320168639539,
    "Smarca5_Abracl": 7.100096545296069,
    "Sf3a2_Pou5f1": 7.0896518095688075,
    "Zyx_Trp53": 7.089131865757108,
    "Tfcp2l1_Sumo2": 7.086189413133255,
    "Trp53_Emp1": 7.084620254663775,
    "Smc3_Glipr2": 7.077737199515404,
    "Rbms1_Hdac2": 7.0720294939264035,
    "Hdac2_Cnn2": 7.069652931554416,
    "Snrpg_Klf2": 7.068868897749176,
    "Rps26_Pou5f1": 7.066397128133428,
    "Lrrc58_Hmgb1": 7.066068284434404,
    "Sumo2_Foxp1": 7.0564017671920825,
    "Rpl11_Pou5f1": 7.05449773862006,
    "Smc3_Kdelr1": 7.051141458072855,
    "Trim28_Serpinh1": 7.048269995044116,
    "Txndc5_Trp53": 7.039272859562683,
    "Pou5f1_Actl6a": 7.035032047544873,
    "Sec61g_Pou5f1": 7.022512664857498,
    "Ppid_Pou5f1": 7.010782243571368,
    "Pou5f1_Ak2": 7.00890472197141,
    "Zfp706_Pou5f1": 6.997186177737368,
    "Pou5f1_Paip2": 6.994882522352271,
    "Rest_Pou5f1": 6.989238588850989,
    "Pou5f1_Aurka": 6.984685028196232,
    "Pou5f1_Ldha": 6.980912759596413,
    "Ybx1_Akt1": 6.97970143786433,
    "Sox4_Hmgn1": 6.956166677017029,
    "Mrpl42_Hmga2": 6.931756443214838,
    "Junb_Hnrnpf": 6.9296600847043175,
    "Trim28_Gsn": 6.901257150074119,
    "Pou5f1_Ndufa6": 6.8994207893488815,
    "Trim28_Mylpf": 6.8917585853884855,
    "Trim28_L1td1": 6.881157022346927,
    "Pou5f1_Pdgfa": 6.869228997005151,
    "Ybx1_Sept7": 6.862180611331056,
    "Hnrnpf_Cebpb": 6.85817263922919,
    "Hdac2_Anxa3": 6.858005686813042,
    "Lgals1_Ctcf": 6.857300615337291,
    "Prrx1_Ppp2ca": 6.847202616018767,
    "S100a11_Ctcf": 6.836631548344424,
    "Tuba1a_Hmgb1": 6.835300594723464,
    "Smarca5_Lims1": 6.820555736587823,
    "Trp53_Surf4": 6.819242385813218,
    "Pou5f1_Mat2a": 6.806473048054755,
    "Cebpb_Cct3": 6.796887288152904,
    "Cnbp_Arf4": 6.764561790096633,
    "Hmga2_Dut": 6.751257504683163,
    "Junb_Dpy30": 6.7199692974575,
    "Oaz1_Cebpb": 6.7134642717668696,
    "Rpl5_Junb": 6.683147629492487,
    "Mrpl15_Foxp1": 6.681891561813345,
    "Ybx1_Sec62": 6.662076018283711,
    "Sumo1_Cebpb": 6.653915253317032,
    "Umps_Pou5f1": 6.651091798895528,
    "Psmb6_Cebpb": 6.626615164184501,
    "H3f3b_Cebpb": 6.60728337111879,
    "Trp53_Emc7": 6.573116049151217,
    "Sf3b6_Junb": 6.528992417485569,
    "Ybx1_Bzw1": 6.501258018207801,
    "Eif4g2_Atf4": 6.430037757604938,
    "Ybx1_Tead1": 6.406266247539202,
    "Tspan4_Hdac2": 6.373415892732007,
    "Ctcf_B2m": 6.304530911604713,
    "Ybx1_Rpl23a-ps3": 6.24333368129737,
    "Ssb_Smarca5": 5.771633996801645,
    "Eif4a1_Atf4": 33.18553942966644,
    "Trp53_Itm2b": 32.946106161219255,
    "Smc3_Chd3": 32.73943356909012,
    "Pou5f1_Nop10": 25.264474705893544,
    "Pou5f1_Lsm6": 24.57804697288533,
    "Pou5f1_Polr2f": 22.40121889931078,
    "Ppia_Pou5f1": 18.62013875293697,
    "Trp53_Ier5": 18.105210668266384,
    "Hdgf_Foxp1": 17.74023556422207,
    "Trim28_Morc1": 14.758154072461346,
    "Ybx1_Pla2g1b": 14.10075751804515,
    "Pou5f1_Pam16": 13.864401658202974,
    "Twist2_Mrpl13": 12.656173991430377,
    "Sox4_Cnbp": 11.734293939334881,
    "Ran_Pou5f1": 11.63024272956374,
    "Prkcdbp_Ctcf": 11.467688870040126,
    "Ptges3_Pou5f1": 11.438772075312501,
    "Pou5f1_Mrpl42": 11.392695227665786,
    "Pou5f1_Lsm3": 11.38967070278267,
    "Fubp1_Foxp1": 11.295281485488468,
    "Ppa1_Pou5f1": 11.294480374086879,
    "Rnps1_Pou5f1": 11.228531431582962,
    "Klf2_H2afz": 11.208890588275793,
    "Pou5f1_Hdac2": 11.199770358043484,
    "Ssrp1_Junb": 11.189162284564224,
    "Sox4_Erh": 11.17227237823908,
    "P4ha1_Hmgb1": 11.17088193140938,
    "Sumo1_Foxp1": 11.14054553786464,
    "Sox4_Serbp1": 11.087824332688347,
    "Smarca5_Cav1": 11.059162847444876,
    "Tspo_Trp53": 11.046884646615414,
    "Sox4_Rpl29": 11.040505062135075,
    "Trp53_Tax1bp3": 11.03540938889895,
    "Smarca5_Fstl1": 11.032583364353094,
    "Ube2i_Pou5f1": 11.020649645981889,
    "Sf3b6_Cebpb": 10.98624957346076,
    "Mrps16_Foxp1": 10.976496292473014,
    "Trp53_Hcfc1r1": 10.954742217043547,
    "Pou5f1_Gtf2a2": 10.929155313728717,
    "Rpl7l1_Junb": 10.879909644379824,
    "Smarca5_Itgb1": 10.852126091107307,
    "Trim28_Calu": 10.814244731210586,
    "Ywhae_Sox4": 10.803572176475472,
    "Sox4_Psma6": 10.764427227430357,
    "Hdac2_Camk2d": 10.74974294160202,
    "Pou5f1_E2f4": 10.723937349350052,
    "Pou5f1_Cenpm": 10.687831661850135,
    "Npm1_Foxp1": 10.649273882090096,
    "Foxp1_Aurkaip1": 10.57446697418847,
    "Vdac2_Foxp1": 10.50994729875069,
    "Tnfrsf12a_Hdac2": 10.504979759357608,
    "Hspe1_Cebpb": 10.502133275303791,
    "Hdac2_Cav1": 10.479946638521081,
    "Papola_Foxp1": 10.476203674087133,
    "Rbx1_Foxp1": 10.42645546785286,
    "Pin1_Klf2": 10.394443770244344,
    "Srsf11_Junb": 10.335249419733131,
    "Smarca5_Csrp1": 10.29054330263057,
    "Smc3_Ppic": 9.85421751403567,
    "Pou5f1_Cops7a": 7.9649407851630665,
    "Pou5f1_Cox5a": 7.688390013306492,
    "Lrrc58_Hdac2": 7.627736195115365,
    "Pou5f1_Fkbp4": 7.624221384409442,
    "Ptma_Pou5f1": 7.560513878594601,
    "Rpl29_Pou5f1": 7.525629351094675,
    "Hdac2_Fkbp1a": 7.513585371756197,
    "Hdac2_Copb2": 7.501768108833948,
    "Pou5f1_Cks1b": 7.458211665944051,
    "Hmgb1_Anxa2": 7.43919083640381,
    "Trp53_Tmed2": 7.431585054972004,
    "Pou5f1_Fam25c": 7.428559289969107,
    "Sox2_Rps29": 7.4152425966934725,
    "Snrpf_Arf4": 7.411930424209864,
    "Pou5f1_Atp5g2": 7.402493215996369,
    "Rps13_Pou5f1": 7.401645303430078,
    "Rpl32_Pou5f1": 7.381813406469639,
    "Pou5f1_Pfdn1": 7.374765485627259,
    "Rps18_Pou5f1": 7.36400771200645,
    "Pou5f1_Calm3": 7.355439559995739,
    "Snrpc_Pou5f1": 7.331699601338456,
    "Slc25a5_Pou5f1": 7.322853113640296,
    "Pou5f1_Polr1c": 7.318002279029912,
    "Taf10_Pou5f1": 7.314945585703537,
    "Srrm2_Pou5f1": 7.290982195901747,
    "Rcn1_Hdac2": 7.28127790280328,
    "Srm_Pou5f1": 7.256696138913458,
    "Pou5f1_Park7": 7.245588972088135,
    "Trp53_Rbmxl2": 7.23694168663833,
    "Rps10_Pou5f1": 7.2169872734317595,
    "Srsf5_Pou5f1": 7.19969217592525,
    "Sri_Smc3": 7.1874613107090095,
    "Timm13_Pou5f1": 7.186528866240998,
    "Pou5f1_Kdm5b": 7.185286400865922,
    "Tomm40_Pou5f1": 7.184749769158182,
    "Pou5f1_Mbd3": 7.181270563896534,
    "Pou5f1_Asns": 7.176990875765549,
    "Pou5f1_Anp32b": 7.172309669388445,
    "Psmc2_Pou5f1": 7.1705474652595385,
    "Trim28_Tagln2": 7.16174864202391,
    "Ybx1_Mtpn": 7.159303265488548,
    "Ybx1_Lamp1": 7.15701377125899,
    "Snrpe_Junb": 7.1556373719204105,
    "U2af1_Foxp1": 7.153340649219626,
    "Pou5f1_Nanog": 7.153297528887809,
    "Hdac2_Anxa2": 7.151146175347684,
    "Smc3_P4hb": 7.14763349817807,
    "Vim_Eno1": 7.146644304867253,
    "Tfcp2l1_Hdgf": 7.1359871512223565,
    "Pou5f1_Pcbp1": 7.13125526094331,
    "Smc3_S100a6": 7.12711364055148,
    "Rbbp7_Arf4": 7.10660304980061,
    "Ybx1_Rer1": 7.103022655560358,
    "Tomm70a_Pou5f1": 7.094820994492913,
    "Tuba1a_Smarca5": 7.093465386967202,
    "Smarca5_Calr": 7.092224634824237,
    "Uqcrfs1_Pou5f1": 7.08768119549326,
    "Ybx1_Elovl1": 7.087665563867327,
    "Pou5f1_Arl6ip1": 7.085213545013358,
    "Trp53_Itgb1": 7.079453961606307,
    "Tpm4_Ctcf": 7.071980017951999,
    "Uqcr10_Pou5f1": 7.058117168836373,
    "Ybx1_Cald1": 7.0572537728612454,
    "Smarca5_S100a6": 7.051960165182669,
    "Trp53_Cyba": 7.051366623478768,
    "Ybx1_Fstl1": 7.050633725577415,
    "Trp53_Cnih1": 7.049829885320679,
    "Ybx1_Anxa1": 7.048541740440586,
    "Pou5f1_Eif5": 7.039296807898112,
    "Smc3_Cnih1": 7.028464531436539,
    "Pou5f1_Alyref": 7.022538853268718,
    "Smarca5_Rrbp1": 7.015362352897373,
    "Trp53_Gnb2": 7.014096906292928,
    "Ybx1_Selm": 7.007125893183435,
    "Pafah1b2_Ctcf": 7.000316792410527,
    "Snrpc_Hmga2": 7.000255734027683,
    "Ybx1_Acta2": 6.997440641468998,
    "Hmgn2_Foxp1": 6.993432861675027,
    "Pou5f1_Impdh2": 6.9932840624022115,
    "Hsp90aa1_Foxp1": 6.992055929852096,
    "Pou5f1_Mrpl12": 6.989282736959438,
    "Trp53_Rab14": 6.981342263110818,
    "Ruvbl1_Pou5f1": 6.979534188968803,
    "Tmed9_Smc3": 6.967012149849111,
    "Smarca5_Cnih4": 6.964176825135997,
    "Hdac2_Cd81": 6.959748634499096,
    "Ybx1_Emc7": 6.956689811307444,
    "Pou5f1_Mrps18c": 6.9553153087861865,
    "Rps12-ps3_Pou5f1": 6.946615829147568,
    "Sumo2_Jund": 6.945898832631046,
    "Ybx1_Tln1": 6.940121405250573,
    "Ppp2r1a_Arf4": 6.938738341835898,
    "Pou5f1_H2afx": 6.937536998344459,
    "Ybx1_Sri": 6.921385079139604,
    "Ybx1_Cd81": 6.9036342639859685,
    "Jund_Erh": 6.899796139358837,
    "Pou5f1_Apex1": 6.869791956069799,
    "Ybx1_Rpp25": 6.8696182051978045,
    "Tfcp2l1_Rpl18": 6.864717006469489,
    "Junb_Hnrnpa3": 6.85727043147678,
    "Pou5f1_Mrpl36": 6.856674711745351,
    "Smc3_Sept7": 6.845526160969786,
    "Eno1_Actg1": 6.843274688759227,
    "Rnf7_Pou5f1": 6.839493188365515,
    "Ybx1_Khdrbs1": 6.839285952571578,
    "Smc3_Basp1": 6.83765194029096,
    "Nanog_Hmgb2": 6.8142347254732405,
    "Psma7_Pou5f1": 6.7955898945085,
    "Smc3_Actg1": 6.790248624226253,
    "Ybx1_Eif3i": 6.783488620435017,
    "Trp53_Cd81": 6.763982230728522,
    "Trim28_Ilk": 6.76051291974262,
    "Hmga2_Cyc1": 6.7429848177517595,
    "Rpl21_Foxp1": 6.741307262222834,
    "Trp53_Basp1": 6.736560168543498,
    "Smc3_Arf4": 6.705633630181191,
    "Ybx1_Pcolce": 6.679360687134642,
    "Sox4_Rpl18": 6.661925043715321,
    "Nono_Arf4": 6.654100878198045,
    "Sec13_Hdac2": 6.643271593385813,
    "Spc24_Pou5f1": 6.577997088736822,
    "Sox4_Hspa8": 6.548912555508536,
    "Sox4_Gspt1": 6.546383558822702,
    "Rbm39_Foxp1": 6.502657404264892,
    "Trim28_Lpp": 6.438252517994622,
    "Pou5f1_Eif1a": 6.437535122046115,
    "Psmc4_Pou5f1": 6.4036967295480185,
    "Srsf6_Junb": 6.385138743470728,
    "Smarca5_Cpe": 6.3201141007916535,
    "Mrps36_Cebpb": 6.2763258524913,
    "Junb_Bcat1": 5.72244720961959,
    "Serpinh1_Atf4": 59.65403508167303,
    "Pou5f1_Ccnb2": 31.782484316000936,
    "Siva1_Pou5f1": 21.649498663838056,
    "Hmgn1_Cebpb": 21.518056700397672,
    "Srsf2_Pou5f1": 19.580688238759237,
    "Smc3_Gnas": 19.57760313690401,
    "Smarca5_Gpx8": 19.572038564963837,
    "Pou5f1_Gadd45gip1": 16.534782396082594,
    "Sod1_Pou5f1": 15.999516069352284,
    "Ybx1_Cyba": 15.995713098869587,
    "Ndufa12_Hdac2": 15.441570035622927,
    "Ssr3_Hdac2": 12.773300040551483,
    "Ybx1_Cfl2": 12.275073763985475,
    "Pou5f1_Dnajb6": 12.115730718542363,
    "Dazap1_Cebpb": 11.428632704716408,
    "Twist2_Snrpd1": 10.360424249897411,
    "Tpm3_Junb": 10.15705921734485,
    "Sox4_Bclaf1": 10.15693558877626,
    "Hnrnpab_Cebpb": 9.952298991169359,
    "Prmt1_Cebpb": 9.95220102267964,
    "Ctcf_Anxa3": 9.949376318239267,
    "Pou5f1_Impa2": 9.759973321686187,
    "Trp53_Ckap4": 9.757796882227824,
    "Ppp4c_Pou5f1": 9.75768223036799,
    "Smc3_Itgb1": 9.755878182153305,
    "Vim_Rest": 9.754945408036349,
    "Smarca5_Gnas": 9.75488208040045,
    "Pou5f1_Nxt1": 9.753884972314964,
    "Hsp90ab1_Cebpb": 9.752641051819522,
    "Raly_Cebpb": 9.749695460821641,
    "Pou5f1_Mkrn1": 9.747031571473665,
    "Pou5f1_Cyc1": 9.74638869900607,
    "Pou5f1_Ndufb10": 9.744230925872108,
    "Sparc_Nono": 9.74313376355143,
    "Sparc_Smc3": 9.742533952846532,
    "Smarca5_Rhoc": 9.738093097342464,
    "Foxp1_Eif5a": 9.737790010304616,
    "Twist2_Hnrnpf": 9.736849334301338,
    "Smarca5_Selm": 9.733680893359173,
    "Sumo2_Hmga2": 9.56240932337706,
    "Serpinh1_Ctcf": 9.554267186119896,
    "Sfpq_Asap1": 9.553044890158853,
    "Trp53_Serpinb6a": 9.551124320921184,
    "Eif5a_Cebpb": 9.548801391269219,
    "Ran_Cebpb": 9.547910398429716,
    "Siva1_Cebpb": 9.546675260293505,
    "Pou5f1_Mrpl51": 9.544325885484554,
    "Pou5f1_Nucks1": 9.542326447258876,
    "Psma5_Pou5f1": 9.540018449835323,
    "Ctcf_Clic1": 9.537264959837447,
    "Lsm4_Cebpb": 9.53309140259671,
    "Polr2h_Hmga2": 9.53063443722393,
    "Hdac2_Cald1": 9.355611031167756,
    "Pou5f1_Pa2g4": 9.344271451382166,
    "Tmsb4x_Hdac2": 9.15590349877926,
    "Pou5f1_Magoh": 9.146759479429726,
    "Srsf7_Jund": 9.142493275713584,
    "Rps10_Klf2": 8.953015017140567,
    "Tpm1_Atf4": 6.946123047893302,
    "Sox4_Ndufa11": 6.762198056295752,
    "Rab14_Hdac2": 6.75812673621261,
    "Rpl17_Cebpb": 6.757159283812603,
    "Cct8_Atf4": 6.60166307291656,
    "Rpl36_Cebpb": 6.58802378118141,
    "Pou5f1_Pim3": 6.437150153147918,
    "Smarca5_Sec62": 6.432531176285381,
    "Id2_Hnrnpk": 6.425685695293806,
    "Ranbp1_Junb": 6.421781285474992,
    "Ssbp1_Pou5f1": 6.273766516549738,
    "Pgrmc1_Hdac2": 6.273551143995632,
    "Junb_Dut": 6.270030197467922,
    "Pou5f1_Ddx27": 6.268496780876416,
    "Nono_Cmpk1": 6.2682960975989435,
    "Rps15_Pou5f1": 6.26566584626588,
    "Rpl28_Pou5f1": 6.265342574044302,
    "Rps25_Pou5f1": 6.265068056156643,
    "Smarca5_Mgat2": 6.264768367691777,
    "Ctcf_Basp1": 6.264486439141198,
    "Ybx1_Ube2b": 6.263332897826901,
    "Ybx1_Bsg": 6.2618769442887485,
    "Pou5f1_Pfdn2": 6.260249386134029,
    "Pou5f1_Cct8": 6.258802792984576,
    "Junb_Hnrnpa1": 6.25873070502368,
    "Twist2_Tmem14c": 6.2586408379418295,
    "Trnau1ap_Pou5f1": 6.258266386492312,
    "Rheb_Atf4": 6.257383896314701,
    "Junb_Fscn1": 6.255771160531067,
    "Ubxn1_Pou5f1": 6.255764869703011,
    "Tuba1a_Eno1": 6.255365076082293,
    "Ybx1_Calr": 6.25531544173384,
    "Cebpb_Cct2": 6.255126196293524,
    "Psma7_Foxp1": 6.254489528100789,
    "Mrfap1_Cebpb": 6.254256894759894,
    "Ybx1_Cd63": 6.254225010942757,
    "Hmgb1_Cald1": 6.110408082861465,
    "Trp53_Dppa5a": 6.109701941909413,
    "Trim28_Anxa5": 6.109556205513089,
    "Rpl22_Pou5f1": 6.108866152582117,
    "Utf1_Pou5f1": 6.108549312010872,
    "Pou5f1_Clns1a": 6.107308188751961,
    "Psma2_Pou5f1": 6.10712608294679,
    "Ybx1_Cnn2": 6.106394958076024,
    "Rpl29_Cebpb": 6.106078849483605,
    "H3f3a_Foxp1": 6.105418266891083,
    "Ybx1_Tpm1": 6.105191240288746,
    "Tcp1_Pou5f1": 6.104969806734081,
    "Smarca5_Selk": 6.1048885826074635,
    "Ube2l3_Pou5f1": 6.104824519132685,
    "Tdgf1_Hmgb1": 6.104803627328443,
    "Wdr18_Pou5f1": 6.1046091890000405,
    "Rpl21_Pou5f1": 6.1046052929855925,
    "Sumo2_Pou5f1": 6.104540571373881,
    "Set_Pou5f1": 6.104437478006698,
    "Pou5f1_Gdf3": 6.104332487124387,
    "Pou5f1_Eif1ax": 6.104177933011004,
    "Ybx1_Anxa5": 6.103672297805943,
    "Hdac1_Anxa2": 6.102482193707215,
    "S100a6_Hdac2": 6.10220844216357,
    "Junb_Hnrnpa2b1": 6.101724817585425,
    "Pou5f1_Hmgn2": 6.100667448284216,
    "Eif4a1_Cebpb": 6.100607087006619,
    "Eno1_Anxa2": 6.098605578314122,
    "Hmgb1_Anxa5": 6.098327635579506,
    "Trp53_Cald1": 6.097857732997263,
    "Rpl38_Pou5f1": 6.097365865517362,
    "Erh_Cebpb": 6.095285973250049,
    "Sumo2_Sox4": 6.095251536683726,
    "Srsf7_Hmga2": 6.094824846827884,
    "Atp5a1_Arf4": 6.09468294193872,
    "Yrdc_Pou5f1": 6.09369238716226,
    "Ybx1_Tagln": 6.093475613038643,
    "Pou5f1_Nubp1": 6.093387265510873,
    "Trp53_Pcolce": 6.092779643598015,
    "Lsm4_Foxp1": 6.092454638770113,
    "Smarca5_Hmga2": 6.091037627667011,
    "Tfcp2l1_Pou5f1": 6.0906022886302384,
    "Sox2_Marcksl1": 6.087958528167333,
    "Chchd2_Cebpb": 6.087597429485261,
    "Ybx1_Dnajc10": 6.087384554675392,
    "Junb_Hnrnpk": 6.0873449175289895,
    "Hmga2_Cct2": 6.087293790588467,
    "Canx_Atf4": 6.0861717159818784,
    "Pou5f1_2700094K13Rik": 5.950103090069128,
    "Pou5f1_Pin1": 5.943758742239479,
    "Zfp428_Pou5f1": 5.943699264618543,
    "Pou5f1_Exosc5": 5.942307056598493,
    "Ybx1_Fau": 5.9422653895388,
    "Pou5f1_Fam136a": 5.941458638225265,
    "Pou5f1_Dpy30": 5.9413459889476234,
    "Pou5f1_Lmnb1": 5.94093805369325,
    "Sparc_Rest": 5.939646607470631,
    "Tnfrsf12a_Smc3": 5.939130276706014,
    "Sparc_Eno1": 5.937637691847688,
    "Ybx1_Commd3": 5.936467654833022,
    "Ppil1_Pou5f1": 5.936414393083839,
    "Trim28_Tpm2": 5.935803185977885,
    "Pou5f1_Fbl": 5.935172797887108,
    "Pou5f1_Ier2": 5.93346822806712,
    "Tecr_Pou5f1": 5.931781386340539,
    "Mrps33_Foxp1": 5.930075657468062,
    "Trp53_Gpx8": 5.929253953556287,
    "Hes1_H3f3a": 5.927998055561502,
    "Ybx1_Prdx6": 5.925308749237547,
    "Ybx1_Tmem167": 5.922087899777329,
    "Twist2_Mrfap1": 5.917211911995597,
    "Rpl23a_Pou5f1": 5.782879505627093,
    "Wdr5_Pou5f1": 5.781618446016744,
    "Pou5f1_Arpp19": 5.779318504557705,
    "Pou5f1_Cenpv": 5.777927249453346,
    "Ybx1_Sptbn1": 5.7738147308271754,
    "Trp53_Efemp2": 5.773699377570459,
    "Snrpd3_Pou5f1": 5.772798235333387,
    "Pou5f1_Phb": 5.770596466294985,
    "Pou5f1_Hn1": 5.769063234799037,
    "Pou5f1_Hspa4": 5.768434521717117,
    "Pou5f1_Bola2": 5.768346238749597,
    "Srsf6_Atf4": 5.767312560337092,
    "Tgfb1i1_Smc3": 5.7646971735883294,
    "Trim28_Kdelr3": 5.762509544161819,
    "Sep15_Hdac2": 5.7601338765594505,
    "Rps7_Pou5f1": 5.615687079448102,
    "Vdac1_Foxp1": 5.613542717501115,
    "Taf1d_Pou5f1": 5.608506355457673,
    "Pou5f1_Pold2": 5.60747656411147,
    "Id2_H3f3b": 5.605704280182338,
    "Vcl_Hdac2": 5.598557023854692
}
```
