## Supplement 1 (MEF vs. ESC) for "AGEAS: Automated Machine Learning based Genetic Regulatory Element Extraction System": outlier_grps.js

```
{
    "Hdac2_Grb10": 37.594623678144075,
    "Sox4_Ranbp1": 36.92629214645116,
    "Trim28_Anxa2": 21.141757537539014,
    "Sf3a2_Pou5f1": 20.47116771712004,
    "Rest_Gnas": 20.361425230040577,
    "Pou5f1_Ddx46": 20.34430042885119,
    "Trp53_Cnih1": 20.336200569375762,
    "Rpl6_Pou5f1": 20.262720983350345,
    "Pou5f1_Hint1": 16.132395647530643,
    "Rpl9_Pou5f1": 16.00738632837459,
    "Pou5f1_Mrpl49": 15.770008073301947,
    "Tuba1a_Hdac2": 15.541141493031294,
    "Pou5f1_Polr2f": 12.932037152149634,
    "Smarca5_Fkbp10": 12.840235535480263,
    "Pou5f1_Mrps16": 12.817393641350925,
    "Pou5f1_Emc8": 12.803864003010904,
    "Snx7_Hmgb1": 12.781057994360054,
    "Pou5f1_Odc1": 12.738385728102456,
    "Cyc1_Cebpb": 12.662090525837094,
    "Sparc_Nono": 12.661969760267832,
    "Hsp90ab1_Foxp1": 12.658716793897444,
    "Sox4_Serbp1": 12.635278502765619,
    "Sox4_H3f3a": 12.576056271849405,
    "Tpm1_Smc3": 12.569800907258058,
    "Pou5f1_Arpp19": 12.530835740862281,
    "Smarca5_P3h4": 12.520511928989935,
    "Ran_Foxp1": 12.50283532210417,
    "Hspa8_Cebpb": 12.50005260940511,
    "Pcbp1_Foxp1": 12.483175383279335,
    "Ube2n_Pou5f1": 12.405661735300447,
    "Sri_Smarca5": 12.401444893515697,
    "Srsf10_Foxp1": 12.380393650847406,
    "Hnrnpk_Foxp1": 12.36903737533626,
    "Junb_Hmgb1": 12.35914229428579,
    "Twist2_Hnrnpf": 12.346564895134449,
    "Sox4_Eif5a": 12.325531744571885,
    "Eif5b_Cebpb": 12.273442623761062,
    "Slc38a4_Arf4": 12.246213233411403,
    "Sox4_Rbm39": 12.223942217847965,
    "Siva1_Cebpb": 12.116970756150979,
    "Psmb6_Cebpb": 12.105229822440208,
    "Sh3glb1_Ctcf": 12.095222760689952,
    "Hmga2_Fkbp4": 12.064273301423585,
    "Trim28_Tagln2": 11.881139834429801,
    "Sox4_Hspa8": 11.722814032796363,
    "Pou5f1_Aspscr1": 11.344925707615655,
    "Rps20_Pou5f1": 8.983585735963958,
    "Smarca5_Hdlbp": 8.778168798680479,
    "Ybx3_Hmga2": 8.770444764402237,
    "Rpl12_Pou5f1": 8.737756268674437,
    "Ybx1_Tmed5": 8.734419780030455,
    "Ubb_Pou5f1": 8.561445855828119,
    "Ybx1_Oaf": 8.556598643511851,
    "Snrpb_Pou5f1": 8.548867629099076,
    "Pou5f1_Oaz1": 8.514712209111456,
    "Ybx1_Pou5f1": 8.490393464801317,
    "Rpl26_Pou5f1": 8.461187473111492,
    "Rpl30_Pou5f1": 8.438172025544905,
    "Snf8_Pou5f1": 8.41387582393996,
    "Rpl37a_Pou5f1": 8.401560712953149,
    "Sfpq_Anxa2": 8.394132119964894,
    "Rpl37_Pou5f1": 8.393933852012488,
    "Ssbp1_Pou5f1": 8.370882195482126,
    "Selk_Hdac2": 8.339863769791915,
    "Trp53_Anxa5": 8.335038783666802,
    "Npm1_Junb": 8.320886297067997,
    "Hmgb1_Basp1": 8.319652868213176,
    "Pou5f1_Cdk4": 8.308321637615359,
    "Pou5f1_Mybl2": 8.306690831512247,
    "Sms_Pou5f1": 8.268704035305126,
    "Atf4_Arf4": 8.266649593522002,
    "Pou5f1_Dctpp1": 8.258940975822318,
    "Rpl11_Pou5f1": 8.256740963153131,
    "Psma2_Pou5f1": 8.247907473788686,
    "Pou5f1_Hnrnpa1": 8.229010634132214,
    "Pou5f1_Eif2s2": 8.228798207162136,
    "Trp53_Anxa2": 8.219403070686605,
    "Eno1_Dppa5a": 8.213450994716727,
    "Trim28_Ckap4": 8.203680575527244,
    "Wdr77_Pou5f1": 8.201289670848121,
    "Cebpb_Anp32b": 8.196661710555677,
    "Pou5f1_Map1lc3b": 8.194755633625133,
    "Trim28_Rhoc": 8.181217486295822,
    "Ube2d3_Pou5f1": 8.177824542846807,
    "Ptges3_Hmga2": 8.176386546904796,
    "Ybx1_Cct4": 8.175532859629076,
    "Smarca5_Cnih4": 8.17055811084514,
    "Pou5f1_Chchd10": 8.160563643199968,
    "Trim28_Anxa5": 8.157522269595972,
    "Ybx1_Txndc5": 8.157099492171218,
    "Pou5f1_Ooep": 8.152458829692907,
    "Smc3_S100a6": 8.145020724176698,
    "Tubb6_Ctcf": 8.142170562064077,
    "Hmgb1_Anxa5": 8.140698872658033,
    "Pou5f1_Polr1d": 8.134543969672686,
    "Prelid1_Pou5f1": 8.130246997821612,
    "Tmsb4x_Hdac2": 8.127643038613503,
    "U2af1_Prrx1": 8.122397603851999,
    "Trim28_Itgb1": 8.115034459149694,
    "Pou5f1_D8Ertd738e": 8.108885899113078,
    "Pou5f1_Paics": 8.108810839161848,
    "Serbp1_Hmga2": 8.108310932183455,
    "Ybx1_Pla2g1b": 8.10406875199371,
    "Pdlim7_Ctcf": 8.094168334363605,
    "Hdac2_Ckap4": 8.078067378020732,
    "Pou5f1_Mrps18b": 8.075778371075524,
    "Srrm1_Pou5f1": 8.068450093524389,
    "Rest_Dppa5a": 8.065895704870961,
    "Pou5f1_Fus": 8.064493172407388,
    "Pou5f1_Hmgn2": 8.060899296185712,
    "Smc3_Cd63": 8.054137903182156,
    "Pou5f1_Cox7b": 8.052513401599226,
    "Rbmxl2_Pou5f1": 8.051336122458043,
    "Trim28_Dppa5a": 8.048935268981833,
    "Mrpl42_Hmga2": 8.047528856676237,
    "Smn1_Pou5f1": 8.047211125825278,
    "Snrpf_Pou5f1": 8.047001173650388,
    "Tpr_Foxp1": 8.037267016861454,
    "Rcn2_Hdac2": 8.028488026791559,
    "Tagln2_Smc3": 8.025604640979722,
    "Smc3_Hdlbp": 8.021270131863139,
    "Tuba1a_Smc3": 7.998741123461736,
    "Surf4_Hdac2": 7.992054039648782,
    "Pou5f1_Hells": 7.985197616811698,
    "Rpl22_Pou5f1": 7.982272557904036,
    "Tdgf1_Hmgb1": 7.981242183560656,
    "Hmgb1_Emp3": 7.975643895080575,
    "Sumo3_Pou5f1": 7.966248220749877,
    "Pou5f1_Dazap1": 7.964991727753537,
    "Tmed2_Hdac2": 7.957677447677514,
    "Lipe_Hdac2": 7.952257958875727,
    "Ybx1_Cd81": 7.95187869927038,
    "Prrx1_Aimp1": 7.945630739556695,
    "Vim_Eno1": 7.934352493924062,
    "Hdac2_Foxp1": 7.932625744692035,
    "Hdac1_Anxa5": 7.932516269571497,
    "Ubald2_Pou5f1": 7.932178726701428,
    "Hmga2_2810417H13Rik": 7.921753941084115,
    "Pou5f1_2310036O22Rik": 7.9209127386572264,
    "Rbm39_Hmga2": 7.90974397440619,
    "Smc3_Prrx1": 7.892185466381723,
    "Ybx1_Cyba": 7.891807129226232,
    "Pou5f1_Mrpl4": 7.886094752285878,
    "Sox2_Set": 7.883296986126034,
    "Bzw1_Atf4": 7.876675330265334,
    "Ppa1_Arf4": 7.871263201964912,
    "Junb_Idh2": 7.85722528872228,
    "Trim28_Mylpf": 7.857130284376587,
    "Txndc5_Smc3": 7.852062315812061,
    "Pou5f1_Asns": 7.8507061684750346,
    "Trp53_Gpx8": 7.850127843235097,
    "Sumo1_Junb": 7.849084131607317,
    "Pou5f1_Nudt21": 7.825476679312412,
    "Yrdc_Pou5f1": 7.813840466721097,
    "Pou5f1_Pam16": 7.811614660048371,
    "Ybx1_Cfl2": 7.806174188473251,
    "Hdac2_Copb2": 7.804149106152574,
    "Eif4a1_Atf4": 7.8013700075209504,
    "Sox2_Ptma": 7.786126311000462,
    "Laptm4a_Hdac2": 7.784982976316776,
    "Nap1l1_Foxp1": 7.775857256854935,
    "Junb_Eif5a": 7.7755149510645305,
    "Snrpa1_Pou5f1": 7.764671184639007,
    "Ube2i_Pou5f1": 7.753720288889134,
    "Sox4_Ddx5": 7.751723678504857,
    "Ybx1_Gnas": 7.746049333290206,
    "Rfc1_Pou5f1": 7.739163601767069,
    "Ybx1_Pdlim7": 7.71472415673071,
    "Pou5f1_Nup62": 7.698434292576773,
    "Twist2_Ranbp1": 7.687204847035765,
    "Hdac2_Gnb2": 7.663434492710257,
    "Sfpq_Pou5f1": 7.660507270758876,
    "Trp53_Rab14": 7.654155914547484,
    "Pou5f1_Nolc1": 7.6481693032793014,
    "Smc3_Cdc42": 7.606775782086077,
    "Ctcf_Clic1": 7.605109345791694,
    "Txnl1_Hdac2": 7.507002193559392,
    "Trim28_Esd": 7.46741499964228,
    "Hdac2_Cnih1": 7.442957078458812,
    "Smc3_Rac1": 7.3907596103504165,
    "Rest_Bgn": 7.349477082477719,
    "Rbpms2_Pou5f1": 7.3261065389728035,
    "Sox4_Gnb2l1": 7.2392021004393765,
    "Smarca5_Pcolce": 7.202456133052206,
    "Ybx1_Bag1": 7.166314674282441,
    "Cebpb_Atp5a1": 7.080360450343734,
    "Rnf7_Jund": 6.751063863874501,
    "Mrpl51_Cebpb": 6.226409466360937,
    "Pou5f1_Fbl": 48.09193007584724,
    "Pou5f1_Dnajc19": 24.183738487398443,
    "Tubb6_Hdac2": 20.892326853919556,
    "Ssr3_Smarca5": 20.498317072558624,
    "Ybx1_Serpinh1": 20.42305485724099,
    "Tuba1a_Hmgb1": 19.85267119172265,
    "Rpl31_Pou5f1": 15.43226663460093,
    "Hmgb1_Col1a2": 15.206531852055091,
    "Pou5f1_Ndufv2": 14.870235162639935,
    "Spag7_Pou5f1": 14.759718244750957,
    "Prkcdbp_Ctcf": 12.632324117294194,
    "Pou5f1_Npm1": 12.489844107714971,
    "Smarca5_Mbnl2": 12.396460684449588,
    "Ubb_Cebpb": 12.384666591658696,
    "Trp53_Rcn3": 12.338627564425034,
    "Fus_Cebpb": 12.313327487348376,
    "Zfp706_Sox4": 12.304827034138098,
    "Hdac2_Fbln5": 12.288519135852876,
    "Smarca5_Ptrf": 12.276705562959576,
    "Smc3_Cald1": 12.271419019877474,
    "Smc3_Marcks": 12.243916711121312,
    "Tipin_Pou5f1": 12.23181404097828,
    "Snrpb_Foxp1": 12.191390042316062,
    "Slc25a5_Cebpb": 12.187590590258466,
    "Nptn_Hmgb1": 12.177244677706735,
    "Smc3_Lmna": 12.144638167105132,
    "Pou5f1_Cacybp": 12.14169387654389,
    "Sox4_Lipe": 12.134210937217157,
    "Ranbp1_Pou5f1": 12.123370581650065,
    "Pou5f1_Mbd3": 12.115533667626547,
    "Sox4_Erh": 12.11481578746794,
    "Hsp90ab1_Cebpb": 12.099976278474456,
    "Smc3_Nenf": 12.043293808326052,
    "Sumo1_Foxp1": 12.000799738077228,
    "Hdac2_Anxa5": 11.945330158147893,
    "Tspo_Trp53": 11.925043130632385,
    "Snrpd2_Foxp1": 11.869245207345221,
    "Pou5f1_Hnrnpm": 11.743298174084988,
    "Smc3_Basp1": 11.638765088968269,
    "Pou5f1_Hdac2": 11.601852920738658,
    "Vim_Ctcf": 11.583246569836309,
    "Rpl5_Cebpb": 11.571762354019544,
    "Pou5f1_Polr2k": 11.053096179201763,
    "Rplp0_Hmga2": 8.520635397011146,
    "Ybx1_Akt1s1": 8.307495645314484,
    "Hdac2_Basp1": 8.214242762063371,
    "Pou5f1_Ndufv1": 8.209918414576022,
    "Ybx1_Gm7325": 8.2016613018232,
    "Psmd7_Pou5f1": 8.038941524311083,
    "Ybx1_Cav1": 8.037391506946625,
    "Pou5f1_Mrpl45": 7.971899492161101,
    "Vapa_Pou5f1": 7.970363157631878,
    "Hmga2_Anp32b": 7.969330037433467,
    "Pou5f1_Edf1": 7.962837540923042,
    "Smc3_Gpx8": 7.958139096989598,
    "Smc3_Actg1": 7.9446270285102365,
    "Smc3_Sep15": 7.940191551091825,
    "Ssr2_Pou5f1": 7.918722296266354,
    "Pou5f1_Lsm2": 7.8982271588965896,
    "Ybx1_Cnn2": 7.886334824283217,
    "Id2_H3f3b": 7.886075782117029,
    "Pou5f1_Ckb": 7.885589618670069,
    "Rpl35a_Pou5f1": 7.8795291336451,
    "Ybx1_Prrx1": 7.87772298889521,
    "Ppia_Pou5f1": 7.876050386758272,
    "Rpl23a_Pou5f1": 7.8734080859676325,
    "Rpl29_Cebpb": 7.871924835427137,
    "Pou5f1_Ifitm1": 7.871186873847418,
    "Lrrc58_Hdac2": 7.860711013037078,
    "Ube2s_Foxp1": 7.853022805601892,
    "Pou5f1_Cnbp": 7.841774121219587,
    "Tagln2_Smarca5": 7.830935366985883,
    "Pou5f1_Erh": 7.8307365754807945,
    "Rps14_Pou5f1": 7.823920409785158,
    "Ybx1_Klhdc2": 7.823361369169405,
    "Pou5f1_Fam25c": 7.8225467808889935,
    "Pou5f1_Cox6b1": 7.820969535759345,
    "Pou5f1_Cdca3": 7.81958557805929,
    "Pou5f1_Atpif1": 7.813266048757851,
    "Set_Nanog": 7.811608611364548,
    "Tdgf1_Pou5f1": 7.810855887312952,
    "Suclg1_Pou5f1": 7.805244766688437,
    "Nap1l1_Junb": 7.7922880801024865,
    "Ybx1_Acta2": 7.791898739026563,
    "Slc25a5_Pou5f1": 7.78597200435751,
    "Sox4_Cnn3": 7.784300158902567,
    "Pou5f1_H2afz": 7.773531055663131,
    "Tceb2_Pou5f1": 7.769207029398145,
    "Pou5f1_Anapc5": 7.762002623847131,
    "Pou5f1_Nudc": 7.749872816332353,
    "Eno1_Anxa2": 7.742114352692196,
    "Hmgb1_Efemp2": 7.731037640047531,
    "Pdap1_Junb": 7.723739235874299,
    "Lrrc58_Hmgb1": 7.723059396850982,
    "Raly_Pou5f1": 7.717439109004415,
    "Zmat2_Pou5f1": 7.7139024513965095,
    "Pou5f1_Mrpl51": 7.710081735209855,
    "Pou5f1_Nucks1": 7.70467060771294,
    "Smarca5_Kdelr1": 7.702857210268319,
    "Rpl36_Pou5f1": 7.702534705407953,
    "Ybx1_Clic1": 7.69690901285223,
    "Zfp706_Pou5f1": 7.693592845077142,
    "Trim28_Clic1": 7.685036539637969,
    "Pou5f1_Eif1": 7.672733669739255,
    "Pou5f1_Hnrnpd": 7.665426961486758,
    "Dnaja1_Atf4": 7.662198192535496,
    "Pou5f1_Nhp2": 7.661145919930383,
    "Lmna_Hdac2": 7.65926526305634,
    "Hmgb1_Bex1": 7.657897381903912,
    "Pou5f1_Cox7c": 7.653352619685818,
    "Ybx1_Bsg": 7.650197743354391,
    "Hnrnpa2b1_Foxp1": 7.648025577698981,
    "Hsp90aa1_Cebpb": 7.627004399146117,
    "Ybx1_Ndn": 7.623223255186151,
    "Taf7_Pou5f1": 7.618702124740334,
    "Reep5_Ctcf": 7.617321604944567,
    "Ybx1_Timp1": 7.614046613209798,
    "Trim28_Tpm2": 7.612167996240321,
    "Pin1_Klf2": 7.610802703184945,
    "Eno1_Basp1": 7.606501881174325,
    "Utf1_Trp53": 7.601860761781343,
    "Trp53_Pgrmc1": 7.6014996712076375,
    "Srsf7_Hmga2": 7.599332701551708,
    "Sumo1_Pou5f1": 7.59730265383685,
    "Sumo2_Foxp1": 7.596625093053725,
    "Pou5f1_Klf5": 7.594863287260972,
    "Lmna_Ctcf": 7.582642834824443,
    "Timm13_Pou5f1": 7.574978840274039,
    "Hmga2_Eif5a": 7.573921105870584,
    "Utf1_Pou5f1": 7.5683683917841975,
    "Wdr18_Pou5f1": 7.56610796153829,
    "Trp53_Ier5": 7.563390007312729,
    "Pou5f1_Eif5": 7.560890451837366,
    "Ybx1_Rbms1": 7.560322708871809,
    "Hdac2_Ddost": 7.559288761624095,
    "Ybx1_Dap": 7.556744639765084,
    "Rae1_Pou5f1": 7.550345154856982,
    "Zyx_Hdac2": 7.548456157007176,
    "Pou5f1_Fkbp8": 7.545983048819995,
    "Rfc5_Pou5f1": 7.544086087243428,
    "Smc3_Canx": 7.5340560653311925,
    "Trp53_Emp1": 7.532946998457311,
    "Hdlbp_Hdac2": 7.531354202293919,
    "Rpl17_Pou5f1": 7.5310726623450295,
    "Rps15_Pou5f1": 7.524955341538155,
    "Mtdh_Hdac2": 7.524289912724723,
    "Ssr1_Arf4": 7.523776148031538,
    "Ybx1_Cmpk1": 7.5054772190267,
    "Sumo2_Hmga2": 7.499629714344411,
    "Tubb6_Trp53": 7.498646618453704,
    "Ybx1_Tpm1": 7.486188497790359,
    "Rpl3_Pou5f1": 7.470314386106906,
    "Pou5f1_Mybbp1a": 7.444351541884104,
    "Sox4_Hmgn1": 7.439882820945719,
    "Pou5f1_Aars": 7.437477275866369,
    "Smarca5_B2m": 7.421442106398741,
    "Rest_Cald1": 7.409749165637664,
    "Uqcr10_Sox2": 7.406928103054238,
    "Phlda3_Nono": 7.395110010251462,
    "Dnaja2_Atf4": 7.394930382381105,
    "Smarca5_Cnih1": 7.387296691321644,
    "Smc3_Arf4": 7.329408782800594,
    "Snrpd1_Cebpb": 7.326725201862956,
    "Hdac2_Anxa3": 7.323012844297243,
    "Smarca5_Pdia3": 7.31424283502787,
    "Slc29a1_Pou5f1": 7.313515490280597,
    "Timp1_Eno1": 7.310417725664986,
    "Pou5f1_Fam136a": 7.303954410423199,
    "Slc25a4_Hdac1": 7.3037991681243275,
    "Pou5f1_Cox7a2": 7.30287068666683,
    "Uhrf1_Pou5f1": 7.2822487156025,
    "Srp19_Pou5f1": 7.280777008339711,
    "Ran_Junb": 7.278977640398071,
    "Ssr3_Smc3": 7.27849802229843,
    "Psmc6_Atf4": 7.274994154773006,
    "Uqcrh_Pou5f1": 7.271558880640726,
    "Pou5f1_Immt": 7.233606678683131,
    "Pou5f1_Dnaja1": 7.226200177546665,
    "Tomm20_Pou5f1": 7.219040898177584,
    "Pou5f1_E2f4": 7.218703101845778,
    "Smarca5_Fam162a": 7.1546116193481595,
    "Ranbp1_Junb": 7.144568775759927,
    "Ybx1_Tuba1b": 7.10606994548866,
    "Immt_Atf4": 7.073629152605266,
    "Ranbp1_Cebpb": 7.047513044926714,
    "Hdac2_H13": 7.001303642466611,
    "Pou5f1_Arglu1": 6.9973747438423874,
    "Ybx1_Serpinb6a": 6.957443153637906,
    "Rnps1_Junb": 6.7113972195032074,
    "Smc3_Rhoc": 27.61372704290027,
    "Ybx1_Spop": 26.58830375687052,
    "S100a11_Rest": 17.25210385844196,
    "Pmp22_Hdac2": 17.199410144912054,
    "Set_Pou5f1": 17.074955671592853,
    "Timp2_Ctcf": 17.03448028826798,
    "Pou5f1_Eif5a": 16.87862205696514,
    "Smc3_S100a11": 16.84431539795891,
    "Tomm22_Pou5f1": 16.839702005964025,
    "Pou5f1_Cdc34": 16.830812258801977,
    "Twist2_Tecr": 16.811086381989504,
    "Smarca5_Gnas": 16.799189920801005,
    "Stmn1_Pou5f1": 16.74417729459101,
    "Smarca5_Cald1": 16.727048186951073,
    "Hmgb1_Ahnak": 16.69927822679962,
    "Foxp1_Eif4a1": 16.68733165861012,
    "Tcf4_Sfpq": 16.676095677883687,
    "Pou5f1_Park7": 16.629886544114694,
    "Trp53_Basp1": 16.592727867070646,
    "Pou5f1_Chchd2": 16.59066716427501,
    "Uqcr10_Pou5f1": 16.522595133703653,
    "Twist2_Ssb": 16.496919991262516,
    "Pou5f1_Mrpl12": 16.46326730456741,
    "Smarca5_Itgb1": 16.395419261936333,
    "Hdac2_Emp3": 16.338713866543703,
    "Tnfrsf12a_Hdac2": 16.055165199956374,
    "Pou5f1_Gcat": 15.94603666373953,
    "Junb_Dnaja1": 15.656871704548983,
    "Rpl29_Mybl2": 15.416368890057107,
    "Junb_Cnn3": 15.050525093435867,
    "Hdlbp_Ctcf": 12.069571974671538,
    "Hmgb1_Anxa2": 11.429342967672802,
    "Cebpb_2700060E02Rik": 11.421774111389471,
    "Sdhb_Pou5f1": 11.153703781946291,
    "Ube2d3_Foxp1": 11.142271456360463,
    "Srsf11_Junb": 11.092030766542516,
    "Ndufa5_Junb": 11.087116476790655,
    "Pou5f1_Drg1": 11.077674361262195,
    "Rbm8a_Pou5f1": 11.006645372481506,
    "Ybx1_Tex261": 10.98058701219447,
    "Trp53_Esd": 10.959641364655376,
    "Hdac2_Gnas": 10.947211301501074,
    "Rpl24_Pou5f1": 10.896932063736735,
    "Rpl4_Pou5f1": 10.85581644810812,
    "Pou5f1_Grsf1": 10.84169243410234,
    "Ybx1_Camk2d": 10.833272062554359,
    "Rps17_Pou5f1": 10.826340643365732,
    "Pou5f1_Mrps17": 10.813036825622707,
    "Rbm25_Foxp1": 10.770652329344758,
    "Pou5f1_Pole4": 10.764316039704303,
    "Smarca5_Lrrc58": 10.756911358379046,
    "Stub1_Pou5f1": 10.745566099650928,
    "Ppp1ca_Pou5f1": 10.738851903153193,
    "Hmgb1_Gdf3": 10.734252593743614,
    "Hdac2_Cdc42": 10.728387533110656,
    "Gnas_Ctcf": 10.712638642003027,
    "Ybx1_Ube2j1": 10.706378753756121,
    "Trim28_Lgals1": 10.70089699920882,
    "Pou5f1_Gdf3": 10.697959406349522,
    "Smarca5_Axl": 10.688402935469195,
    "Ybx1_Timp2": 10.687541616833705,
    "Junb_Fkbp3": 10.669758800858126,
    "Swi5_Pou5f1": 10.668004867024736,
    "Mrfap1_Hdac2": 10.66431136601044,
    "Ybx1_Anxa1": 10.64986491036422,
    "Ybx1_Ptp4a2": 10.642215150362937,
    "Pou5f1_Ndufb4": 10.639084533812586,
    "Smc3_Kdelr1": 10.631986615337532,
    "Serpinh1_Rest": 10.626183756977966,
    "Trp53_Gng12": 10.616486484810716,
    "Prss23_Hdac2": 10.594866406823035,
    "Smarca5_Slc25a4": 10.593794158112557,
    "Pou5f1_Gapdh": 10.587113634309611,
    "Wls_Smc3": 10.56896957989176,
    "Hmga2_Cdc34": 10.563466987560938,
    "Trim28_Anxa3": 10.55756459552698,
    "Ybx1_Nenf": 10.55739187964348,
    "Smc3_Ppic": 10.536601294910815,
    "Pou5f1_Alyref": 10.514769056583757,
    "Smc3_Pdia3": 10.470134465532174,
    "Rpl35a_Cebpb": 10.466328415841833,
    "Trp53_Tpm1": 10.445274002877177,
    "Ybx1_Scoc": 10.437661341640247,
    "Pou5f1_Bcat1": 10.424711385888108,
    "Rcn3_Hdac2": 10.41189727971257,
    "Ybx1_Sept7": 10.389166590979048,
    "Pou5f1_Nfu1": 10.355857022757334,
    "Pou5f1_Adprh": 10.355281494237662,
    "Pou5f1_Polr2j": 10.338120953362148,
    "Trim28_Gnas": 10.311477636904288,
    "Klf2_H2afx": 10.298760960065428,
    "Smc3_Mgat2": 10.236125765479121,
    "Hmga2_Dctpp1": 10.235932729204867,
    "Son_Junb": 9.95234137619696,
    "Ybx1_Hcfc1r1": 9.908522807828726,
    "Pou5f1_Ost4": 9.890933498576551,
    "Ybx1_Tm2d2": 9.840038261432943,
    "Sumo2_Sox2": 9.691314975996976
}
```
