## Supplement 1 (MEF vs. ESC) for "AGEAS: Automated Machine Learning based Genetic Regulatory Element Extraction System": outlier_grps.js

```
{
    "Pou5f1_Pole4": 23.038942812891747,
    "Ybx1_Lipe": 22.46861742238946,
    "Swi5_Pou5f1": 21.074091851573147,
    "Pou5f1_Ndufs6": 17.12029252963813,
    "Rest_Rcn3": 14.460549689993732,
    "Twist2_Eif3g": 14.43823834061034,
    "Twist2_Erh": 14.32513518689268,
    "Lmna_E2f4": 14.24183749316934,
    "Snrpe_Pou5f1": 14.14322831833552,
    "Smarca5_Capn2": 14.117816291377578,
    "Sox4_Lsm4": 14.051120365372965,
    "Tm4sf1_Smarca5": 14.050066829341247,
    "Klf2_Eif5a": 14.036114066517719,
    "Hdac2_Fbln5": 13.926210852550222,
    "Sox4_Hnrnpdl": 13.894549196830491,
    "Tipin_Pou5f1": 13.894480566700134,
    "Tpm1_E2f4": 13.882747309078232,
    "Smarca5_Cald1": 13.871816383030245,
    "P4ha1_Hmgb1": 13.862407095204397,
    "Timp2_Ctcf": 13.813835340929185,
    "Pou5f1_H3f3b": 13.8085003439208,
    "Ybx1_Rps12": 13.779583516445285,
    "Vim_E2f4": 13.776454293416087,
    "Sox4_Hnrnpf": 13.774894445369586,
    "Marcks_Ctcf": 13.752776838449885,
    "Smarca5_Itgb1": 13.721212671479732,
    "Pou5f1_Lsm6": 13.70934535610806,
    "Pou5f1_Atp5j": 13.701646893436545,
    "Rnps1_Pou5f1": 13.691983505759822,
    "Smarca5_H13": 13.687796126222187,
    "Smarca5_Psap": 13.628921357156303,
    "Pou5f1_Adrm1": 13.615359775596067,
    "Trp53_Gng12": 13.597452432501132,
    "Hnrnpa3_Cebpb": 13.569128977921391,
    "Sox4_Dnaja1": 13.564677790451103,
    "Hdac2_Dap": 13.536172706360247,
    "Trp53_Gabarap": 13.460434345068506,
    "Rbx1_Pou5f1": 13.427696254186777,
    "Srsf5_Cebpb": 13.318468118959478,
    "Sox2_Eif2s2": 13.308278916705515,
    "Pou5f1_E2f4": 13.185743567895095,
    "Ctcf_Ckap4": 13.135131699612396,
    "Tomm20_Cebpb": 13.039757554730064,
    "Rbbp7_Junb": 12.935090000011703,
    "Rpl21_Junb": 9.354638530592677,
    "Rpl4_Pou5f1": 9.346361753895476,
    "Tubb4b_Klf2": 9.337744101356042,
    "Pou5f1_Gltscr2": 9.276836268528443,
    "Ybx1_Prrx1": 9.226329679969751,
    "Hdac2_Cfl2": 9.205305712338356,
    "Pou5f1_Ndufb7": 9.138610309397917,
    "Rps17_Pou5f1": 9.136964860833743,
    "Rps4x_Pou5f1": 9.126383101705633,
    "Dnaja1_Atf4": 9.123749049466099,
    "Pou5f1_Mrpl36": 9.116552139057447,
    "Cebpb_Atpif1": 9.085724513007033,
    "Ybx1_Fbln5": 9.075033477729487,
    "Trp53_Abracl": 9.049929729619762,
    "Naa50_Junb": 9.036759553433297,
    "Pou5f1_Atp5j2": 9.029445738394426,
    "Pou5f1_Clpp": 9.025362773461518,
    "Tuba1b_Pou5f1": 9.019950498042261,
    "Pou5f1_Polr1c": 8.999636394936024,
    "Pou5f1_Nap1l1": 8.988952252109735,
    "Trim28_Rhoc": 8.98152738726052,
    "Rps15_Pou5f1": 8.977547468341026,
    "Pcbp2_Foxp1": 8.964326217169127,
    "Ssr3_Smarca5": 8.964311484517276,
    "Trp53_B2m": 8.963571132671978,
    "Rplp0_Pou5f1": 8.957512351264835,
    "Rpl22_Pou5f1": 8.951561479659816,
    "S100a6_Hdac2": 8.940153087730808,
    "Rbms1_Hdac2": 8.918126071093761,
    "Sod1_Pou5f1": 8.917775232270309,
    "Tagln2_Smc3": 8.912352761753843,
    "Nap1l1_Junb": 8.90518259635033,
    "Smarca5_Lrrc58": 8.904628783077971,
    "Snrnp70_Pou5f1": 8.901121103849942,
    "Timm13_Pou5f1": 8.899355755089992,
    "Wdr77_Pou5f1": 8.885574516325061,
    "Junb_Erh": 8.883499219742413,
    "Sec61g_Pou5f1": 8.882739606271015,
    "Rpl6_Pou5f1": 8.880144353231804,
    "Foxp1_Cdv3": 8.873248111582106,
    "Trp53_Cyba": 8.871279573113739,
    "Ybx1_Col1a2": 8.871234700341681,
    "Pou5f1_Cdca8": 8.864865699555697,
    "Pou5f1_Anp32a": 8.862643544511872,
    "Top2a_Pou5f1": 8.858237878871867,
    "Pou5f1_Cfdp1": 8.851427961967543,
    "Rps24_Pou5f1": 8.847617350644633,
    "Trp53_Ssr3": 8.830244793726976,
    "Pou5f1_Eif3k": 8.828218713570433,
    "Pou5f1_Mrps34": 8.825838850496462,
    "Ybx1_S100a11": 8.81587728261113,
    "Ybx1_Emc7": 8.814382725822247,
    "Ybx1_Cav1": 8.812883257954613,
    "Rpl31_Pou5f1": 8.81075867449897,
    "Hdac1_Anxa5": 8.80865695410755,
    "Pou5f1_Cks1b": 8.807310438985363,
    "Nap1l1_Nanog": 8.806650449846295,
    "Zfp42_Pou5f1": 8.804191927756582,
    "Rpl7_Pou5f1": 8.801921917897484,
    "Hmgb1_Bex1": 8.799307890536053,
    "Pou5f1_Cyc1": 8.79024118039474,
    "Trp53_Fstl1": 8.790200056282014,
    "Smc3_Sept2": 8.786100791641369,
    "Pou5f1_G3bp2": 8.782823997568928,
    "Snrpc_Pou5f1": 8.780689542020482,
    "Rps7_Pou5f1": 8.780035798359886,
    "Utf1_Pou5f1": 8.77391417021314,
    "Trim28_Ckap4": 8.763312978283134,
    "Rps18_Pou5f1": 8.756216707525965,
    "Smarca5_Abracl": 8.74790931396448,
    "Trp53_Mettl9": 8.747714870952892,
    "Pou5f1_Eif2s2": 8.742583357785843,
    "Tomm40_Pou5f1": 8.731466551804981,
    "Smc3_Clic1": 8.730145170096389,
    "Id2_H3f3b": 8.72953449834817,
    "Pou5f1_Fnta": 8.72755276884381,
    "Smc3_Lgals1": 8.717039253452086,
    "Tagln2_Smarca5": 8.707999240977106,
    "Ppp1ca_Pou5f1": 8.707754293826923,
    "Pou5f1_Fscn1": 8.686345865076904,
    "Pou5f1_Dtymk": 8.680792684300423,
    "Junb_Fkbp3": 8.67711055544749,
    "Tuba1a_Smarca5": 8.674754631786172,
    "Trp53_Tpm4": 8.672325419387432,
    "Gnas_Ctcf": 8.667878440833837,
    "Atf4_Actg1": 8.65855226912995,
    "Trp53_S100a11": 8.658081672000657,
    "Sox4_Psma2": 8.657246954228418,
    "Tdgf1_Pou5f1": 8.656724599617574,
    "Pou5f1_Anapc5": 8.656256640038121,
    "Pcbp1_Hmga2": 8.655502518207504,
    "Ybx1_Tpm2": 8.653649408167377,
    "Pou5f1_Chchd2": 8.651683974328156,
    "Eno1_Anxa1": 8.648984230356985,
    "Pou5f1_Hdgf": 8.64557404329622,
    "Ube2c_Pou5f1": 8.640668862771442,
    "Fkbp1a_Atf4": 8.640619803055104,
    "Ybx1_Tln1": 8.63871170812894,
    "Snrpg_Klf2": 8.623681089146663,
    "Uqcrfs1_Pou5f1": 8.617394801473719,
    "Slc25a4_Hdac2": 8.594110028209556,
    "Ybx1_Cfl2": 8.580999389897608,
    "Zfp42_Sox2": 8.572819242027583,
    "Syncrip_Arf4": 8.559709214590619,
    "Smarca5_Pou5f1": 8.556352792552408,
    "Pou5f1_Pdap1": 8.554953326142215,
    "Pou5f1_Gsta4": 8.54921812649952,
    "Pou5f1_Akr1b3": 8.541018605322055,
    "Zfp42_Trim28": 8.533384956412418,
    "Ywhae_Foxp1": 8.528498402520553,
    "Pou5f1_Eif3c": 8.520584924063538,
    "Trp53_Pafah1b2": 8.508905179271673,
    "Smc3_Ppic": 8.500223177586868,
    "Tspo_Trp53": 8.499327956102324,
    "Smc3_Ilk": 8.498606573459913,
    "Serpinh1_Atf4": 8.48955832180354,
    "Srp9_Pou5f1": 8.487168082011033,
    "Trp53_Pdia3": 8.482204871880969,
    "Trp53_Arf4": 8.475675504790853,
    "Gtf2a2_Foxp1": 8.468532152620554,
    "Ybx1_Ccng1": 8.457630468553736,
    "Hdgf_Foxp1": 8.440884978972402,
    "Eif4a1_Atf4": 8.440282498021979,
    "Pou5f1_Eif3a": 8.437321043644562,
    "Txndc5_Smc3": 8.401133058236033,
    "Pou5f1_Nsmce4a": 8.399843542898271,
    "Ybx1_Tex261": 8.39407398367559,
    "Ctcf_Arpc1b": 8.370136546784003,
    "Srsf3_Junb": 8.361088172029332,
    "Raly_Pou5f1": 8.315601191412213,
    "Pou5f1_Cisd1": 8.306077691168879,
    "Ndufb9_Eno1": 8.301364985408362,
    "Ranbp1_Cebpb": 8.2995243625857,
    "Ppic_Hdac2": 8.279638819209834,
    "Ppp2r1a_Arf4": 8.276974798358102,
    "Trp53_Col1a1": 8.270944191642945,
    "Pou5f1_Dazap1": 8.267937799411666,
    "Ybx1_Spop": 8.244920153688428,
    "H3f3b_Foxp1": 8.243750500843973,
    "Smc3_Rac1": 8.172707793980965,
    "Sox4_Rbm39": 8.171032585478246,
    "Twist2_Mrfap1": 8.169934531348318,
    "Ybx1_Hsd17b12": 8.140270459878467,
    "Sox2_Dpy30": 8.085615319860317,
    "Rsl1d1_Foxp1": 8.079930563518852,
    "Rrs1_Pou5f1": 8.07913798826479,
    "Pou5f1_Mrpl17": 8.075608302682085,
    "Hdac2_Arpc1b": 8.06271283220666,
    "Smc3_Lamp1": 7.936441585227518,
    "Trim28_Ssr1": 7.9019693953050005,
    "Ybx1_Pafah1b1": 7.759601006407063,
    "Smc3_Ap2m1": 7.758046351139837,
    "Ybx1_Bnip2": 28.922280305988817,
    "Atf5_Asns": 28.227260298819733,
    "Ctcf_Anxa5": 25.441653979592957,
    "Hdac2_Gnas": 21.535201896946273,
    "Smc3_Slc25a4": 20.13571728758273,
    "Stmn2_Pou5f1": 16.219435154352144,
    "Rps6_Hmga2": 16.205044349630427,
    "Tgfb1i1_Smarca5": 13.478732561347465,
    "Zyx_E2f4": 13.304424881567753,
    "Sf3b5_Pou5f1": 13.130306716242954,
    "Pou5f1_E130012A19Rik": 13.110391279960766,
    "Hes1_Fus": 12.954605953100291,
    "Ppa1_Pou5f1": 12.954292332151223,
    "Zfp706_Sox4": 12.866805652230685,
    "Sox4_Hsp90ab1": 12.807383525969048,
    "Smarca5_Rhoc": 12.803939651503962,
    "Trp53_Ckap4": 12.765656283631566,
    "Ppp4c_Pou5f1": 12.76377042190127,
    "Hnrnpk_Foxp1": 12.757402792973842,
    "Foxp1_Eif5a": 12.723419048802107,
    "Smc3_Emp1": 12.72177422081052,
    "Sumo3_Cebpb": 12.7126591096412,
    "Smc3_Marcks": 12.711495006682688,
    "Trp53_Commd3": 12.710820579312363,
    "Pou5f1_Pa2g4": 12.710806735858919,
    "Pou5f1_Ak2": 12.707231150269031,
    "Stoml2_Prrx1": 12.695934753585817,
    "Pou5f1_Gabarapl2": 12.587168595650683,
    "Pou5f1_Polr2j": 12.564355339246612,
    "l7Rn6_Pou5f1": 12.511059857454306,
    "Foxp1_Dek": 12.473010354939206,
    "Itm2b_Hdac2": 12.460294182202805,
    "Tmed3_Hdac2": 12.454785556454034,
    "Twist2_Ssb": 12.447509629583147,
    "Rpl8_Klf2": 12.432240383140108,
    "Twist2_Ran": 12.398324484788445,
    "Rbm39_Cebpb": 12.261967613077754,
    "Raly_Foxp1": 12.189928091983312,
    "Cebpb_Acp1": 12.182819060704944,
    "Serpinh1_Hdac2": 11.959551410559087,
    "Smc3_Rbms1": 11.911271484101064,
    "Rps3a1_Pou5f1": 9.143010527880753,
    "Ybx1_Eif4a1": 8.661420856760385,
    "Rpl26_Pou5f1": 8.645854652383457,
    "Hdac2_Arf4": 8.620843428059747,
    "Pou5f1_Ccdc12": 8.594785556458264,
    "Foxp1_Atp5j": 8.57678426346441,
    "Pou5f1_Ndufab1": 8.531399674636274,
    "Rwdd1_Pou5f1": 8.49439084828239,
    "Sox4_Rpl31": 8.432840268918483,
    "Hmgb1_Anxa2": 8.404626547628684,
    "Ybx1_Calu": 8.378139444107356,
    "Rpl28_Pou5f1": 8.339835632093132,
    "Trim28_Itgb1": 8.301555379349848,
    "Rpl21_Foxp1": 8.293196459869748,
    "Smc3_Sep15": 8.28981434582822,
    "Serbp1_Hmga2": 8.288777958778983,
    "Trp53_Anxa1": 8.287790939651563,
    "Pou5f1_Cox5a": 8.277566385471555,
    "Nanog_Anp32b": 8.275277043833022,
    "Tagln2_Hdac2": 8.255559561985997,
    "Ybx1_Tulp4": 8.251447246180259,
    "Sri_Smc3": 8.243029267916903,
    "Ube2n_Junb": 8.23681391365277,
    "Ybx1_Basp1": 8.235449411344803,
    "Junb_Fscn1": 8.230483273915095,
    "Ybx1_Wdr1": 8.220242963145127,
    "Tomm7_Sox2": 8.220102681535153,
    "Rps10_Pou5f1": 8.21057964585357,
    "Ybx1_Itm2b": 8.206602600233676,
    "Hdac2_Cnih1": 8.198294791846925,
    "Foxp1_Cct2": 8.191968673449823,
    "Surf4_Smarca5": 8.185681050243785,
    "Pou5f1_Cdca3": 8.179029462042331,
    "Ybx1_Lamp1": 8.178715839805433,
    "Rpl36_Pou5f1": 8.176697267025832,
    "Ybx1_Serpinb6a": 8.176433247215936,
    "Ywhaq_Sox4": 8.170834160091115,
    "Trim28_Fstl1": 8.16392313328393,
    "Pou5f1_Eif5a": 8.157368238189514,
    "Smc3_Efemp2": 8.150809474935773,
    "Smarca5_Pdia3": 8.147138608699315,
    "Prrx1_2700060E02Rik": 8.141035558271662,
    "Sep15_Hdac2": 8.13907127480487,
    "Hmga2_Fkbp4": 8.135636850318505,
    "Rpl36al_Cebpb": 8.134122185468867,
    "Ybx1_Tagln": 8.133950888959848,
    "Pou5f1_Gadd45gip1": 8.128056945533036,
    "Rpl8_Pou5f1": 8.11848129475058,
    "Ybx1_Klhdc2": 8.11593492833811,
    "Hdac1_Anxa2": 8.115809207963954,
    "Pou5f1_Atp5g1": 8.102147188499863,
    "Sumo2_Foxp1": 8.097676318602357,
    "Pou5f1_Ccnb1": 8.090707188555117,
    "Hmgb1_Efemp2": 8.088759597262376,
    "Pou5f1_Hspe1": 8.086921105628726,
    "Trp53_Rhoc": 8.084953848416106,
    "Srsf10_Atf4": 8.07837958446058,
    "Pou5f1_Hspa8": 8.067373575125776,
    "Rcc2_Pou5f1": 8.066821286243325,
    "Pou5f1_H2afz": 8.064211545005028,
    "Smc3_Lrrc58": 8.06240005014507,
    "Rps13_Pou5f1": 8.056041805349071,
    "Trp53_Fam162a": 8.04928146320219,
    "Ybx1_Cd63": 8.044373825236475,
    "Junb_Hnrnpf": 8.043725409196126,
    "Rbmxl2_Pou5f1": 8.039227424940531,
    "Pou5f1_Hspd1": 8.03829026777111,
    "Pou5f1_Dctpp1": 8.037936261379587,
    "Pou5f1_Pdha1": 8.037353761016696,
    "Oaz1_Foxp1": 8.031334735838861,
    "Smarca5_Pdlim7": 8.029300284879076,
    "Pou5f1_Ooep": 8.02667675685167,
    "Ybx1_Anxa6": 8.022041924370583,
    "Smarca5_Prkar1a": 8.021238424218403,
    "Psip1_Pou5f1": 8.021075516889757,
    "Ybx1_Tuba1a": 8.019003868017004,
    "Smc3_Rhoc": 8.018961203752102,
    "Trp53_Cnih1": 8.017498726490027,
    "Ptma_Foxp1": 8.01471696900548,
    "Ybx1_Clic1": 8.012894388220499,
    "Trp53_Clic1": 8.006524570751763,
    "Klf2_H2afz": 8.004895891583068,
    "Hdac2_Cd81": 8.002491242801147,
    "Hdac1_Capn2": 7.990286466028446,
    "Trp53_Esd": 7.986962438791473,
    "Tomm7_Pou5f1": 7.9864340299559196,
    "Pou5f1_Fam136a": 7.986309907346996,
    "Tdgf1_Hmgb1": 7.985003288557837,
    "Yrdc_Pou5f1": 7.975167093113656,
    "Serpinh1_Rest": 7.974003491961844,
    "Ybx1_Timp2": 7.968426525392111,
    "Smarca5_P3h4": 7.960984377747724,
    "Hmgb1_Gdf3": 7.960234721818291,
    "Txndc5_Ctcf": 7.95458546193521,
    "Lmna_Hdac2": 7.954339688352124,
    "Txndc5_Hdac2": 7.946608410022731,
    "Ybx1_Lrrc58": 7.945779858822081,
    "Nfib_Hsp90ab1": 7.941339169309771,
    "U2af1_Nanog": 7.93782737501627,
    "Ybx1_Nenf": 7.935443037553993,
    "Sox4_Set": 7.934089991755331,
    "Pou5f1_Pam16": 7.92859696059872,
    "Txndc5_Trp53": 7.927903067397324,
    "Rbm39_Hmga2": 7.926645896283386,
    "Zmat2_Pou5f1": 7.925954919838205,
    "Ybx1_Hspa8": 7.909236606465143,
    "Pou5f1_Copz1": 7.902610591309627,
    "Snrpa1_Pou5f1": 7.8975257063860065,
    "Trim28_Slc25a4": 7.896699261517133,
    "Sox2_Dppa5a": 7.895629215033346,
    "Pou5f1_Hnrnpa3": 7.8842293363355225,
    "Tra2b_Atf4": 7.875230805362106,
    "Slc25a5_Pou5f1": 7.868032202989599,
    "Smc3_Mtdh": 7.866087766995155,
    "Tpm1_Hdac2": 7.858073363042033,
    "Srrm2_Pou5f1": 7.857147531469137,
    "Tcof1_Pou5f1": 7.854496303841087,
    "Rps16_Pou5f1": 7.849201141353808,
    "Psma7_Pou5f1": 7.842689866490863,
    "Trp53_Dppa5a": 7.839684467511706,
    "Dnaja2_Atf4": 7.839021053560303,
    "Smc3_Cnih1": 7.82958050998919,
    "Tcp1_Pou5f1": 7.810202285413804,
    "Hmga2_Bud31": 7.797065604627053,
    "Trim28_Commd3": 7.774386838030645,
    "Mtpn_Hdac2": 7.766579575039774,
    "Trim28_Tmed3": 7.758483434950713,
    "Tcea3_Pou5f1": 7.745975813846254,
    "Sox4_Hmgn1": 7.7401359121294915,
    "Sept7_Atf4": 7.728368125875935,
    "Ube2n_Pou5f1": 7.718437449882575,
    "Hdac2_Abracl": 7.711030186247881,
    "Pou5f1_Cenpf": 7.705029118826781,
    "Pou5f1_Dnajc19": 7.689561766719666,
    "Ybx1_Gsn": 7.682778644108742,
    "Serbp1_Junb": 7.676488305866028,
    "Nfib_Gspt1": 7.674418945245849,
    "Phlda3_Hmgb1": 7.606178850220682,
    "Uchl3_Pou5f1": 7.588406888323434,
    "Pou5f1_Mbd3": 7.484869196441488,
    "Ybx1_Ptp4a2": 7.484395405116937,
    "Psmb6_Cebpb": 7.463428760365689,
    "Eif5a_Cebpb": 7.437259509151418,
    "Sox2_Ran": 7.430129404475255,
    "Ybx1_Rbm3": 7.429908465320014,
    "mt-Cytb_Eno1": 7.330733216543228,
    "Hdac2_Ddost": 7.278191628626089,
    "Ncl_Junb": 7.266824110476998,
    "Sox4_Rpl7a": 7.223684710885694,
    "Marcksl1_Hmga2": 7.1893327952439465,
    "Hdac2_Fstl1": 6.692063363492072,
    "Smc3_Kdelr3": 34.81109416198286,
    "Rbm39_Pou5f1": 25.872187563771938,
    "Ybx1_Lhfp": 14.844695360528892,
    "Trp53_Itgb1": 14.795040254211361,
    "Eno1_Actg1": 14.72350508061639,
    "Pfn1_Eno1": 14.710398495768246,
    "Etf1_Cebpb": 12.271649110978105,
    "Sox4_Cct3": 12.17655499131264,
    "Trp53_Rcn3": 12.066885780422545,
    "Hnrnpdl_Foxp1": 12.057363627952682,
    "Smarca5_Cebpb": 11.99766987524247,
    "Pou5f1_Cox7c": 11.976006377810487,
    "Hnrnpab_Cebpb": 11.967181513877351,
    "Hdac2_Grb10": 11.955335724428712,
    "Pou5f1_Emc8": 11.886192375386935,
    "Pou5f1_2700094K13Rik": 11.851838795881667,
    "Sox4_Rpl29": 11.842520632874038,
    "Tpm4_Rest": 11.833878195845267,
    "Pou5f1_Odc1": 11.800154815363822,
    "Ssr2_Pou5f1": 11.79266536424897,
    "Hsp90ab1_Cebpb": 11.761450611806376,
    "Pou5f1_Dnajc2": 11.75900281882254,
    "Pou5f1_Nhp2": 11.75644431273674,
    "Rnf7_Cebpb": 11.742345130702072,
    "Prrx1_0610007P14Rik": 11.73788765578071,
    "Smarca5_Phldb2": 11.730316611299777,
    "Gspt1_Cebpb": 11.706302333850692,
    "Hsp90aa1_Foxp1": 11.704570798693707,
    "U2af1_Pou5f1": 11.701792021419452,
    "Ctcf_Anxa2": 11.695573772346592,
    "Smarca5_Cav1": 11.68999450571626,
    "Nono_Cebpb": 11.684972545217644,
    "Smarca5_Nenf": 11.671695627413605,
    "Sumo1_Foxp1": 11.666308529973637,
    "Pou5f1_Ciapin1": 11.662507457811598,
    "Hdac2_Anxa5": 11.6465461855825,
    "Tomm5_Pou5f1": 11.62851129329749,
    "Taf1d_Pou5f1": 11.593750714437027,
    "Pou5f1_Lig1": 11.59299999759014,
    "Smarca5_Glipr2": 11.59260878268577,
    "Pou5f1_Gpx1": 11.568024760843475,
    "Smc3_Laptm4a": 11.567064356281184,
    "Sumo2_Junb": 11.540912378983773,
    "Ywhae_Sox4": 11.528219714782985,
    "Sumo3_Foxp1": 11.513236574708355,
    "Sfpq_Asap1": 11.469463892388736,
    "Ybx1_Bex4": 11.461278045466159,
    "Trp53_Cd81": 11.454234875556034,
    "Snrpd2_Foxp1": 11.434284628971806,
    "Trim28_Sh3glb1": 11.249950937138951,
    "Foxp1_C1qbp": 11.245558471805904,
    "Junb_Hnrnpa2b1": 11.243811170896565,
    "Sox4_Ndufs7": 11.225579919586151,
    "Psma2_Cebpb": 11.1314570209143,
    "Nanog_G3bp1": 11.031282723660212,
    "Sox4_Pfdn1": 11.01173000957538,
    "Junb_Atp5j2": 7.925312538866642,
    "Smarca5_Rpn1": 7.898700847208733,
    "Ybx1_Rab11a": 7.889042494221577,
    "Srsf11_Junb": 7.8301275853204135,
    "Hdac1_Cd81": 7.744562146528723,
    "Trp53_Pgls": 7.7356007985635875,
    "Timm13_Foxp1": 7.732781356542149,
    "Ube2d3_Foxp1": 7.732213222608711,
    "Tpm1_Ctcf": 7.717409062257594,
    "Tgif1_Pou5f1": 7.7152722507838485,
    "Smc3_Cisd2": 7.687135115185578,
    "Sparc_Eno1": 7.682822157958691,
    "Rpl15_Pou5f1": 7.673544699347143,
    "Smc3_Cdc42": 7.6385393199764655,
    "Selk_Hdac2": 7.627749789802108,
    "Trp53_Lgals1": 7.614282349818423,
    "Sox4_Rpl21": 7.592269894745049,
    "Rab14_Hmgb1": 7.59012726111152,
    "Snrnp25_Pou5f1": 7.589222453081652,
    "Rcn2_Hdac1": 7.563934154975626,
    "Pou5f1_Arl6ip1": 7.561114273488015,
    "Ybx1_Tspan3": 7.55608347996439,
    "Sub1_Pou5f1": 7.551409531245017,
    "Tomm22_Junb": 7.523496603032332,
    "Junb_Dnaja1": 7.522817355678441,
    "Atp5a1_Arf4": 7.521633764066558,
    "Fkbp3_Cebpb": 7.5206581997384365,
    "Pou5f1_Gnb2l1": 7.516066770736136,
    "Snrpe_Cebpb": 7.5035606453384736,
    "Rps15_Hmga2": 7.502060641403186,
    "Samm50_Pou5f1": 7.498175609653353,
    "Pou5f1_Cebpzos": 7.491972716294742,
    "Rpl35a_Pou5f1": 7.490258002740686,
    "Rpn1_Eno1": 7.489243292248278,
    "Trim28_Anxa5": 7.483444442884175,
    "Pou5f1_Marcksl1": 7.479382721869804,
    "Smarca5_Eef1a1": 7.478848045057612,
    "Hmgb1_Anxa5": 7.475000964122088,
    "Smc3_Anxa5": 7.473459900601973,
    "Vim_Trp53": 7.469764753304324,
    "Npm1_Foxp1": 7.447339973201931,
    "Trim28_Mylpf": 7.44653066270804,
    "Trp53_Anxa2": 7.445449896208696,
    "Pou5f1_Naca": 7.445253125029697,
    "Tubb6_Trp53": 7.439033281151756,
    "Pou5f1_Aurka": 7.434584586509376,
    "Ybx1_Arpc1b": 7.434499979287466,
    "Wls_Trp53": 7.430732168127693,
    "Ybx1_Wisp1": 7.406917152501107,
    "Pou5f1_Hnrnpab": 7.398386262766458,
    "Pou5f1_Mrpl51": 7.397996837674519,
    "Serpinh1_Hmgb1": 7.397065192888276,
    "Rbm8a_Pou5f1": 7.3919018081883285,
    "Rpl5_Cebpb": 7.388910422695582,
    "Rps12-ps3_Pou5f1": 7.384649375753681,
    "Vim_Hdac2": 7.383458553811974,
    "Smarca5_Anxa5": 7.383051755439206,
    "Ybx1_Tpm1": 7.379396391847999,
    "Srsf7_Hmga2": 7.375485088701745,
    "Ybx1_Gstm5": 7.3749436985564145,
    "Ybx1_Sept7": 7.374795784550163,
    "Pou5f1_Nop10": 7.3708293085799,
    "Pou5f1_Plekhf2": 7.361441262618922,
    "Ctcf_Cnn2": 7.360121131275503,
    "Ranbp1_Foxp1": 7.356842166519488,
    "Pou5f1_Gpx4": 7.356746914531341,
    "Ybx1_Laptm4a": 7.356563376688822,
    "Smarca5_Sh3glb1": 7.355560041727109,
    "H3f3a_Foxp1": 7.352317918311624,
    "Utf1_Trp53": 7.351693816565187,
    "Foxp1_Eif3d": 7.349120863982253,
    "Nhp2l1_Junb": 7.3442900317982875,
    "Hmgb1_Cald1": 7.323109757708681,
    "Klf2_Bex4": 7.3228182867893326,
    "Smarca5_Cnih1": 7.31852605535402,
    "Srm_Pou5f1": 7.30900054791578,
    "Ctcf_Cald1": 7.308197064850735,
    "Smc3_Arpc2": 7.307390298641931,
    "Tuba1a_Smc3": 7.304044182494492,
    "Junb_Hnrnpa3": 7.302032460249938,
    "Psma2_Pou5f1": 7.300437680337739,
    "Smarca5_S100a6": 7.299397213985105,
    "Rps27a_Pou5f1": 7.296797501576221,
    "Trp53_Tmsb4x": 7.296446741695307,
    "Hnrnpa3_Foxp1": 7.292776746932766,
    "Ybx1_Cdc42": 7.2924775923767795,
    "Hmga2_Cyc1": 7.289569262744642,
    "Pou5f1_Eif1ax": 7.288134557415096,
    "Myl12a_Eno1": 7.285772452262864,
    "Smarca5_C1d": 7.28422168564803,
    "Sf3a2_Pou5f1": 7.284203656067825,
    "Smarca5_Hmga2": 7.283255513764141,
    "Tmem59_Smarca5": 7.279484036381894,
    "Pou5f1_Ict1": 7.278866260227395,
    "Pdia3_Eno1": 7.256485136644741,
    "Pou5f1_Anapc11": 7.229305112595494,
    "Ybx1_Sec62": 7.224935678157735,
    "Pou5f1_Hnrnpd": 7.198041425925974,
    "Hmgb1_Emp3": 7.187505044849835,
    "Mrpl42_Hmga2": 7.182428571108091,
    "Pou5f1_Mrpl20": 7.18186101638084,
    "Pou5f1_Ost4": 7.17920651859206,
    "Srsf6_Atf4": 7.175718689114274,
    "Trim28_Col1a2": 7.173719182504851,
    "Snrpb2_Foxp1": 7.16278865376368,
    "Oaz1_Cebpb": 7.162006707321434,
    "Pou5f1_Impa2": 7.161410647667457,
    "Trp53_Cd63": 7.155548217298501,
    "Pou5f1_Lyar": 7.145527780766957,
    "Rbm25_Pou5f1": 7.1429665539750395,
    "Tkt_Pou5f1": 7.129202174214466,
    "Ybx1_Sept11": 7.115108931154156,
    "Sec13_Hdac2": 7.112146013929053,
    "Ppp1cc_Junb": 7.108515098338174,
    "Eno1_Basp1": 7.098057264773355,
    "Pou5f1_Lsm2": 7.094434947337842,
    "Sox2_Hmgn1": 7.087841373117927,
    "Smarca5_Arf4": 7.086561399444098,
    "Trp53_Igf2": 7.071794732784169,
    "Nap1l1_Cebpb": 6.959844231358052,
    "Pou5f1_Mrpl40": 6.9522828807845345,
    "Slc25a4_Hdac1": 6.931827071627381,
    "Trp53_Emp3": 6.924959925585035,
    "Ybx1_Cmpk1": 6.923497282667253,
    "Sox4_Hspa8": 6.914415778382018,
    "Smc3_Gpx8": 6.910855626084021,
    "Mtdh_Atf4": 6.90673925923259,
    "Sox2_Hmgb1": 6.889358720838972,
    "Tm9sf3_Smc3": 6.792886896029971,
    "Rps20_Foxp1": 6.776504553872396,
    "Smarca5_Lipe": 6.772483304984658,
    "Prss23_Hdac2": 6.770223475912246,
    "Junb_Fbl": 6.762846382509894,
    "Rnf7_Junb": 6.7344454483048315,
    "Smarca5_Sepw1": 6.733248866194454,
    "Ybx1_Cct8": 6.704001747850834,
    "Smc3_Phldb2": 6.698419968308312,
    "Trp53_Myl12a": 6.551779299445632
}
```
