## Supplementary figures and images for "AGEAS: Automated Machine Learning based Genetic Regulatory Element Extraction System"

### CMp7d_CMp28d.png

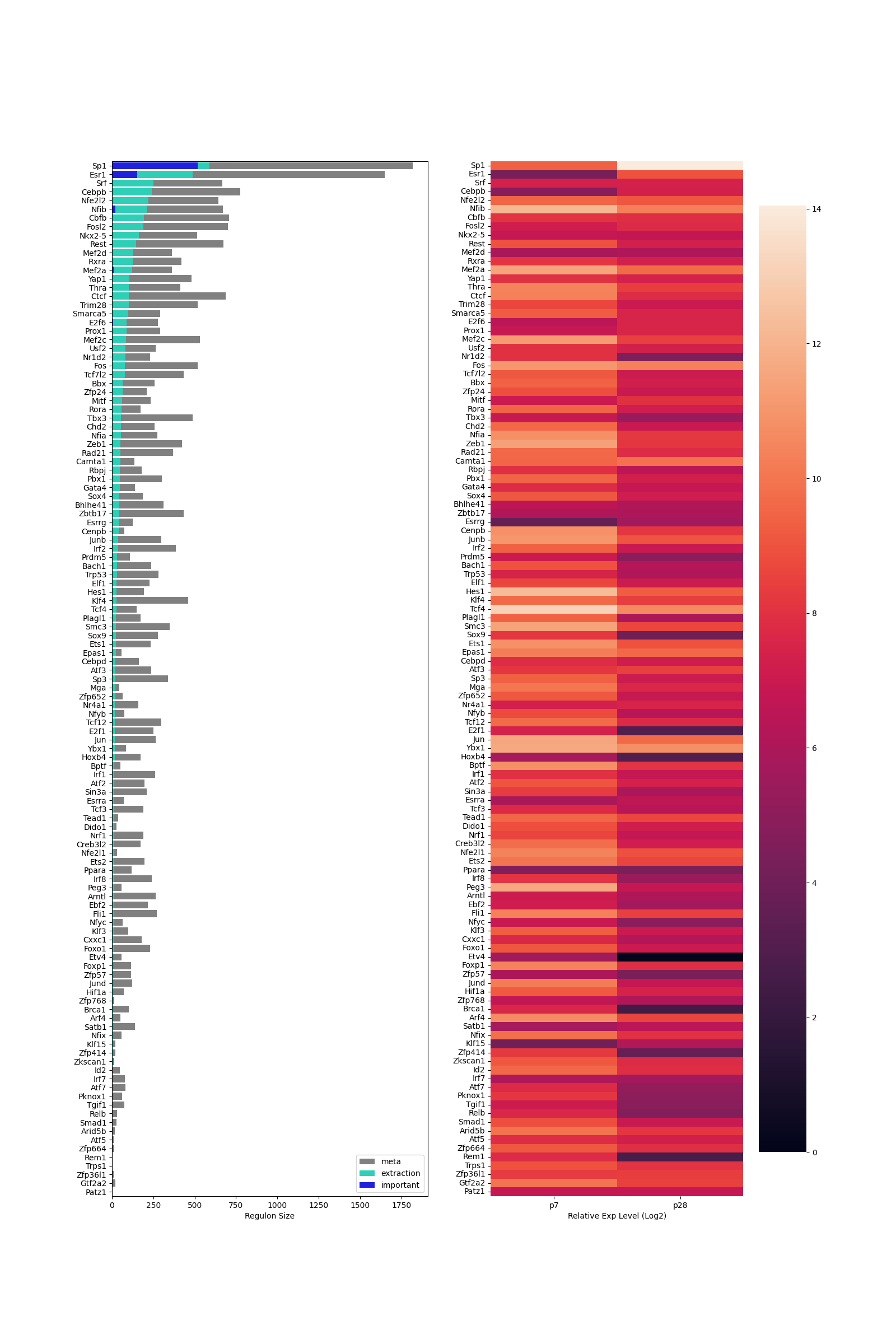

### HSCp6w_PFp6w.png

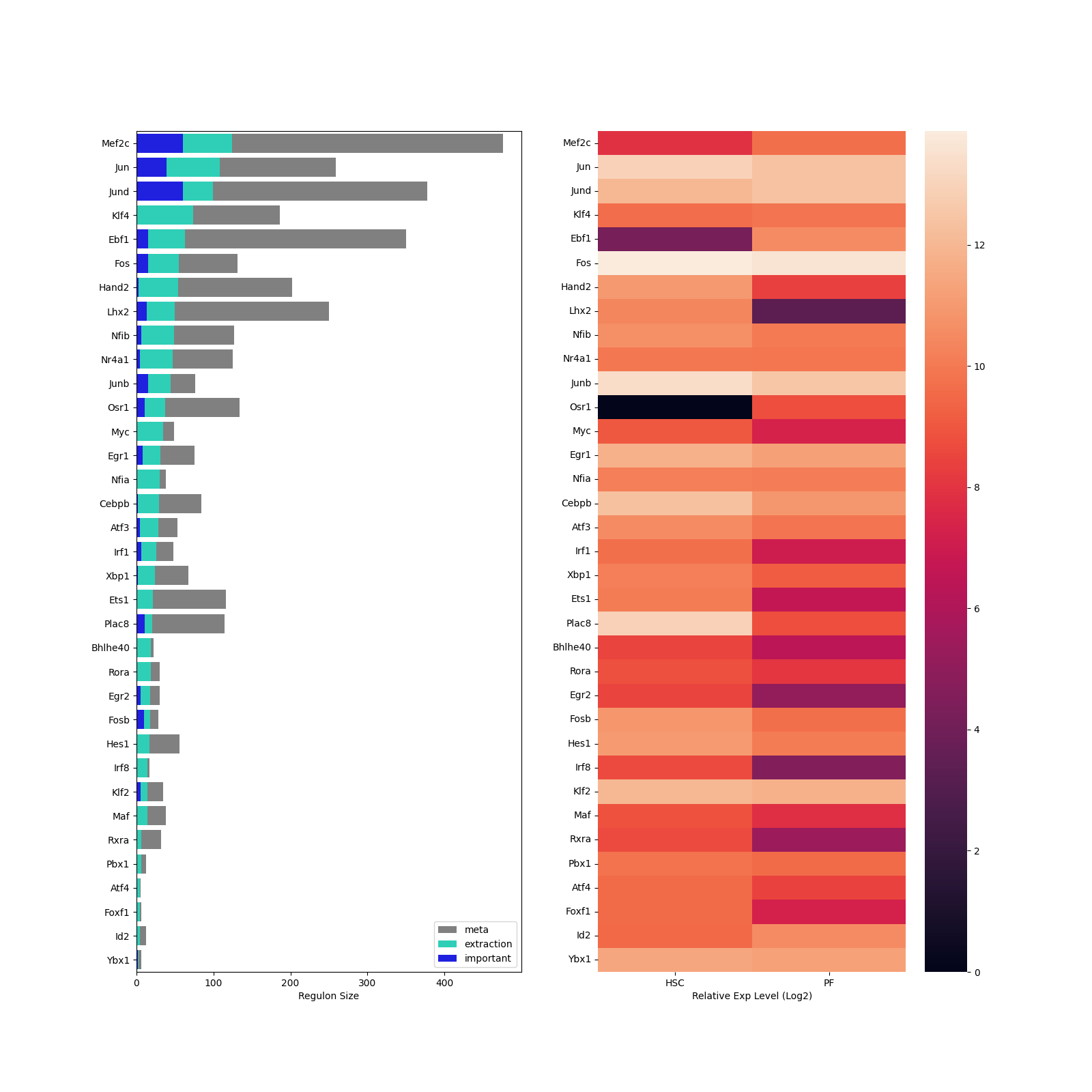

### MEF_ESC.png

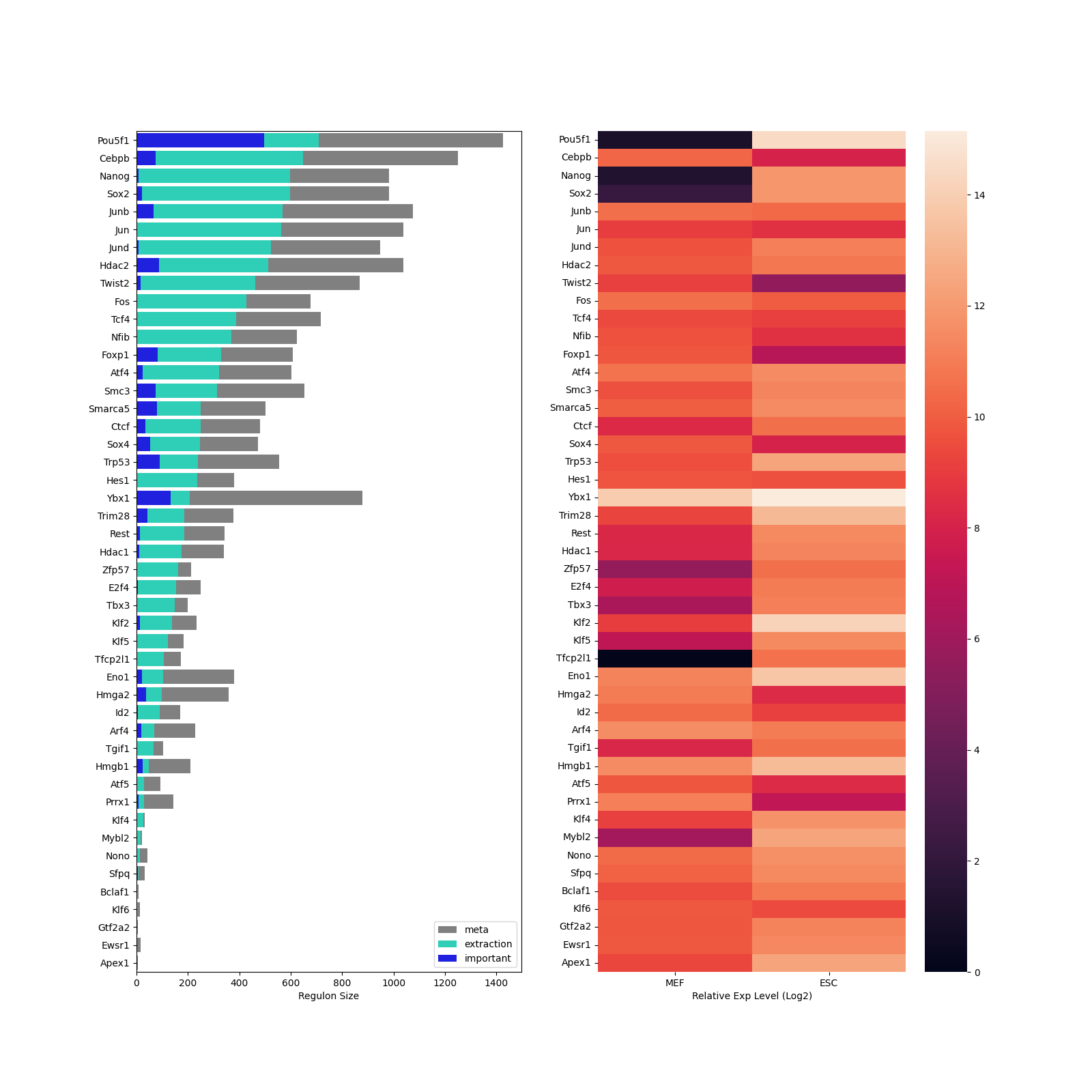

### N_cocul.png

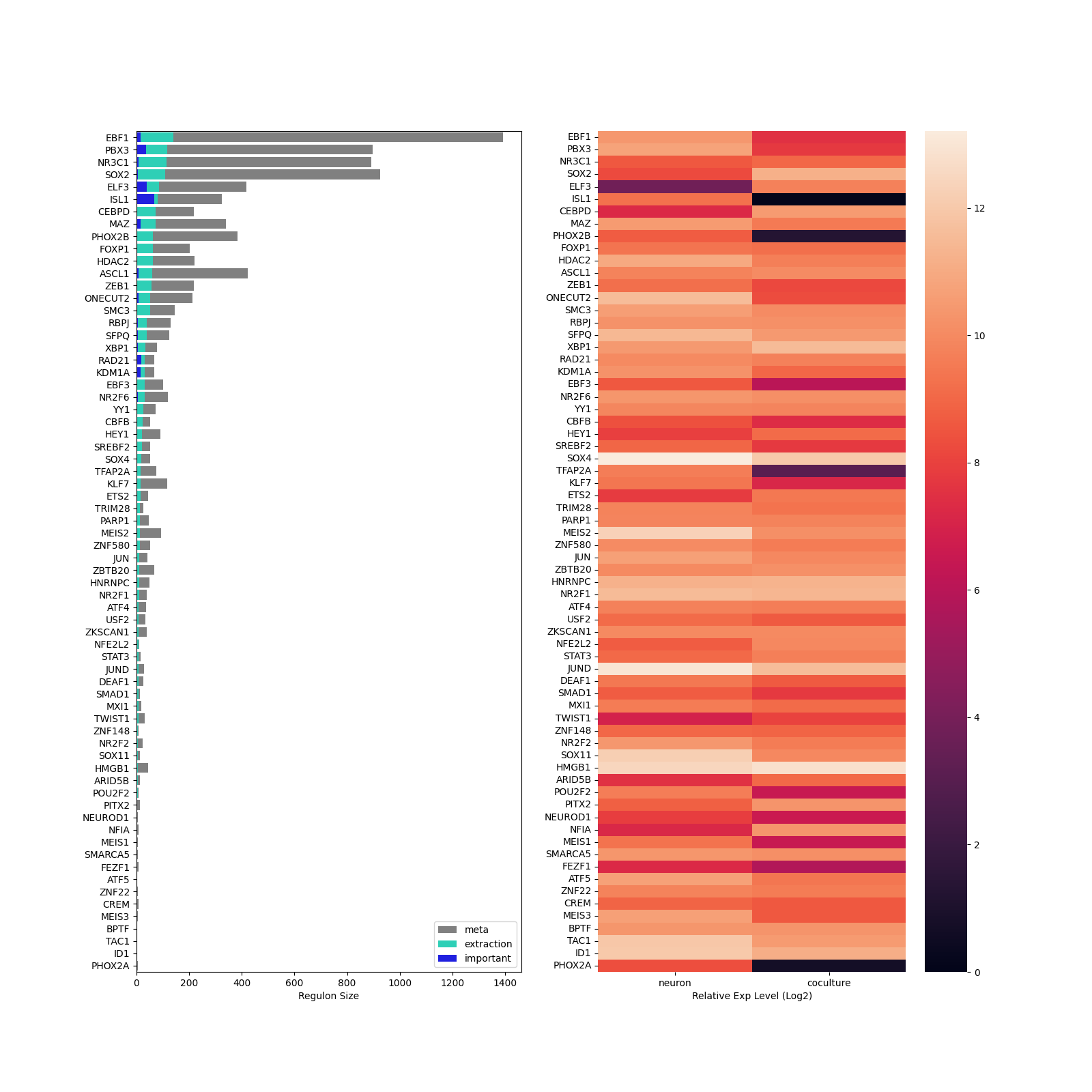
