## Supplement 3 (P7 CM vs. P28 CM) for "AGEAS: Automated Machine Learning based Genetic Regulatory Element Extraction System": full_atlas.js

```
{
    "network_0": {
        "genes": {
            "ENSMUSG00000001280": {
                "id": "ENSMUSG00000001280",
                "type": "TF",
                "symbol": "Sp1",
                "source": [
                    "ENSMUSG00000029135",
                    "ENSMUSG00000021944",
                    "ENSMUSG00000008575",
                    "ENSMUSG00000015605",
                    "ENSMUSG00000022528",
                    "ENSMUSG00000057469",
                    "ENSMUSG00000010175",
                    "ENSMUSG00000058239",
                    "ENSMUSG00000015579",
                    "ENSMUSG00000051469",
                    "ENSMUSG00000019768",
                    "ENSMUSG00000031885",
                    "ENSMUSG00000053110",
                    "ENSMUSG00000031715",
                    "ENSMUSG00000015846",
                    "ENSMUSG00000058440",
                    "ENSMUSG00000015839",
                    "ENSMUSG00000032238",
                    "ENSMUSG00000053477",
                    "ENSMUSG00000058756",
                    "ENSMUSG00000056501",
                    "ENSMUSG00000029249",
                    "ENSMUSG00000032035"
                ],
                "target": [
                    "ENSMUSG00000000538",
                    "ENSMUSG00000029860",
                    "ENSMUSG00000029681",
                    "ENSMUSG00000054455",
                    "ENSMUSG00000032737",
                    "ENSMUSG00000073702",
                    "ENSMUSG00000031403",
                    "ENSMUSG00000028081",
                    "ENSMUSG00000091478",
                    "ENSMUSG00000024228",
                    "ENSMUSG00000020694",
                    "ENSMUSG00000006676",
                    "ENSMUSG00000020307",
                    "ENSMUSG00000043683",
                    "ENSMUSG00000021248",
                    "ENSMUSG00000031885",
                    "ENSMUSG00000026176",
                    "ENSMUSG00000092074",
                    "ENSMUSG00000021235",
                    "ENSMUSG00000070283",
                    "ENSMUSG00000039640",
                    "ENSMUSG00000034903",
                    "ENSMUSG00000030739",
                    "ENSMUSG00000061461",
                    "ENSMUSG00000004044",
                    "ENSMUSG00000008036",
                    "ENSMUSG00000028086",
                    "ENSMUSG00000044221",
                    "ENSMUSG00000035202",
                    "ENSMUSG00000026087",
                    "ENSMUSG00000026626",
                    "ENSMUSG00000031790",
                    "ENSMUSG00000031543",
                    "ENSMUSG00000007891",
                    "ENSMUSG00000069744",
                    "ENSMUSG00000004789",
                    "ENSMUSG00000083853",
                    "ENSMUSG00000018363",
                    "ENSMUSG00000018286",
                    "ENSMUSG00000064349",
                    "ENSMUSG00000080242",
                    "ENSMUSG00000007656",
                    "ENSMUSG00000027288",
                    "ENSMUSG00000032314",
                    "ENSMUSG00000008892",
                    "ENSMUSG00000030557",
                    "ENSMUSG00000020149",
                    "ENSMUSG00000026858",
                    "ENSMUSG00000027698",
                    "ENSMUSG00000081344",
                    "ENSMUSG00000028229",
                    "ENSMUSG00000030852",
                    "ENSMUSG00000010048",
                    "ENSMUSG00000053329",
                    "ENSMUSG00000031197",
                    "ENSMUSG00000034981",
                    "ENSMUSG00000029499",
                    "ENSMUSG00000032216",
                    "ENSMUSG00000090066",
                    "ENSMUSG00000026755",
                    "ENSMUSG00000018446",
                    "ENSMUSG00000064302",
                    "ENSMUSG00000026775",
                    "ENSMUSG00000029392",
                    "ENSMUSG00000044365",
                    "ENSMUSG00000001552",
                    "ENSMUSG00000020155",
                    "ENSMUSG00000000056",
                    "ENSMUSG00000024962",
                    "ENSMUSG00000057133",
                    "ENSMUSG00000020949",
                    "ENSMUSG00000039103",
                    "ENSMUSG00000073471",
                    "ENSMUSG00000009628",
                    "ENSMUSG00000029148",
                    "ENSMUSG00000026158",
                    "ENSMUSG00000020349",
                    "ENSMUSG00000038708",
                    "ENSMUSG00000032171",
                    "ENSMUSG00000056501",
                    "ENSMUSG00000029505",
                    "ENSMUSG00000026615",
                    "ENSMUSG00000055320",
                    "ENSMUSG00000031698",
                    "ENSMUSG00000022174",
                    "ENSMUSG00000021024",
                    "ENSMUSG00000027602",
                    "ENSMUSG00000025745",
                    "ENSMUSG00000049265",
                    "ENSMUSG00000024304",
                    "ENSMUSG00000006219",
                    "ENSMUSG00000030057",
                    "ENSMUSG00000025369",
                    "ENSMUSG00000078139",
                    "ENSMUSG00000055322",
                    "ENSMUSG00000039016",
                    "ENSMUSG00000006057",
                    "ENSMUSG00000018196",
                    "ENSMUSG00000068923",
                    "ENSMUSG00000047284",
                    "ENSMUSG00000087943",
                    "ENSMUSG00000052253",
                    "ENSMUSG00000057841",
                    "ENSMUSG00000044533",
                    "ENSMUSG00000064345",
                    "ENSMUSG00000021666",
                    "ENSMUSG00000019471",
                    "ENSMUSG00000028884",
                    "ENSMUSG00000018585",
                    "ENSMUSG00000032648",
                    "ENSMUSG00000052962",
                    "ENSMUSG00000026374",
                    "ENSMUSG00000024871",
                    "ENSMUSG00000053929",
                    "ENSMUSG00000096842",
                    "ENSMUSG00000020315",
                    "ENSMUSG00000026123",
                    "ENSMUSG00000059981",
                    "ENSMUSG00000059796",
                    "ENSMUSG00000005882",
                    "ENSMUSG00000020455",
                    "ENSMUSG00000025208",
                    "ENSMUSG00000060002",
                    "ENSMUSG00000026672",
                    "ENSMUSG00000029221",
                    "ENSMUSG00000047419",
                    "ENSMUSG00000025234",
                    "ENSMUSG00000006998",
                    "ENSMUSG00000025616",
                    "ENSMUSG00000020459",
                    "ENSMUSG00000079184",
                    "ENSMUSG00000001334",
                    "ENSMUSG00000001415",
                    "ENSMUSG00000058076",
                    "ENSMUSG00000007815",
                    "ENSMUSG00000028165",
                    "ENSMUSG00000018765",
                    "ENSMUSG00000073557",
                    "ENSMUSG00000066637",
                    "ENSMUSG00000033060",
                    "ENSMUSG00000025939",
                    "ENSMUSG00000005610",
                    "ENSMUSG00000021282",
                    "ENSMUSG00000031812",
                    "ENSMUSG00000064367",
                    "ENSMUSG00000021361",
                    "ENSMUSG00000024695",
                    "ENSMUSG00000028955",
                    "ENSMUSG00000037706",
                    "ENSMUSG00000026701",
                    "ENSMUSG00000037905",
                    "ENSMUSG00000015806",
                    "ENSMUSG00000025271",
                    "ENSMUSG00000022185",
                    "ENSMUSG00000084168",
                    "ENSMUSG00000014313",
                    "ENSMUSG00000016253",
                    "ENSMUSG00000035936",
                    "ENSMUSG00000029422",
                    "ENSMUSG00000027668",
                    "ENSMUSG00000002015",
                    "ENSMUSG00000026107",
                    "ENSMUSG00000024845",
                    "ENSMUSG00000008429",
                    "ENSMUSG00000064339",
                    "ENSMUSG00000060261",
                    "ENSMUSG00000028245",
                    "ENSMUSG00000035529",
                    "ENSMUSG00000042978",
                    "ENSMUSG00000066839",
                    "ENSMUSG00000028849",
                    "ENSMUSG00000034951",
                    "ENSMUSG00000015149",
                    "ENSMUSG00000022108",
                    "ENSMUSG00000040033",
                    "ENSMUSG00000024712",
                    "ENSMUSG00000037197",
                    "ENSMUSG00000006024",
                    "ENSMUSG00000022682",
                    "ENSMUSG00000025348",
                    "ENSMUSG00000053617",
                    "ENSMUSG00000030603",
                    "ENSMUSG00000052512",
                    "ENSMUSG00000019578",
                    "ENSMUSG00000020647",
                    "ENSMUSG00000067925",
                    "ENSMUSG00000019791",
                    "ENSMUSG00000027332",
                    "ENSMUSG00000015850",
                    "ENSMUSG00000025733",
                    "ENSMUSG00000020864",
                    "ENSMUSG00000022214",
                    "ENSMUSG00000022110",
                    "ENSMUSG00000025404",
                    "ENSMUSG00000004356",
                    "ENSMUSG00000037916",
                    "ENSMUSG00000030779",
                    "ENSMUSG00000029156",
                    "ENSMUSG00000078572",
                    "ENSMUSG00000040321",
                    "ENSMUSG00000052681",
                    "ENSMUSG00000045316",
                    "ENSMUSG00000000628",
                    "ENSMUSG00000040771",
                    "ENSMUSG00000021870",
                    "ENSMUSG00000032724",
                    "ENSMUSG00000033161",
                    "ENSMUSG00000056342",
                    "ENSMUSG00000062006",
                    "ENSMUSG00000003072",
                    "ENSMUSG00000037349",
                    "ENSMUSG00000031534",
                    "ENSMUSG00000027133",
                    "ENSMUSG00000027253",
                    "ENSMUSG00000008730",
                    "ENSMUSG00000009293",
                    "ENSMUSG00000027488",
                    "ENSMUSG00000019809",
                    "ENSMUSG00000000168",
                    "ENSMUSG00000026554",
                    "ENSMUSG00000029249",
                    "ENSMUSG00000028902",
                    "ENSMUSG00000097487",
                    "ENSMUSG00000030061",
                    "ENSMUSG00000038459",
                    "ENSMUSG00000021906",
                    "ENSMUSG00000021313",
                    "ENSMUSG00000026496",
                    "ENSMUSG00000025241",
                    "ENSMUSG00000015112",
                    "ENSMUSG00000075054",
                    "ENSMUSG00000021453",
                    "ENSMUSG00000032299",
                    "ENSMUSG00000028936",
                    "ENSMUSG00000062981",
                    "ENSMUSG00000028639",
                    "ENSMUSG00000037601",
                    "ENSMUSG00000002496",
                    "ENSMUSG00000064342",
                    "ENSMUSG00000027227",
                    "ENSMUSG00000074884",
                    "ENSMUSG00000031983",
                    "ENSMUSG00000038028",
                    "ENSMUSG00000003531",
                    "ENSMUSG00000003923",
                    "ENSMUSG00000020523",
                    "ENSMUSG00000028792",
                    "ENSMUSG00000001445",
                    "ENSMUSG00000035325",
                    "ENSMUSG00000055239",
                    "ENSMUSG00000035203",
                    "ENSMUSG00000029580",
                    "ENSMUSG00000028247",
                    "ENSMUSG00000027823",
                    "ENSMUSG00000008822",
                    "ENSMUSG00000029430",
                    "ENSMUSG00000004069",
                    "ENSMUSG00000040785",
                    "ENSMUSG00000021178",
                    "ENSMUSG00000022336",
                    "ENSMUSG00000027708",
                    "ENSMUSG00000019797",
                    "ENSMUSG00000058600",
                    "ENSMUSG00000064351",
                    "ENSMUSG00000040850",
                    "ENSMUSG00000046756",
                    "ENSMUSG00000029020",
                    "ENSMUSG00000028173",
                    "ENSMUSG00000027706",
                    "ENSMUSG00000049643",
                    "ENSMUSG00000038375",
                    "ENSMUSG00000022452",
                    "ENSMUSG00000039509",
                    "ENSMUSG00000023106",
                    "ENSMUSG00000036206",
                    "ENSMUSG00000034898",
                    "ENSMUSG00000024527",
                    "ENSMUSG00000031166",
                    "ENSMUSG00000038393",
                    "ENSMUSG00000075467",
                    "ENSMUSG00000025374",
                    "ENSMUSG00000019577",
                    "ENSMUSG00000033429",
                    "ENSMUSG00000027894",
                    "ENSMUSG00000002010",
                    "ENSMUSG00000022415",
                    "ENSMUSG00000020593",
                    "ENSMUSG00000010064",
                    "ENSMUSG00000058997",
                    "ENSMUSG00000059923",
                    "ENSMUSG00000032328",
                    "ENSMUSG00000060904",
                    "ENSMUSG00000025508",
                    "ENSMUSG00000028273",
                    "ENSMUSG00000019158",
                    "ENSMUSG00000008604",
                    "ENSMUSG00000058594",
                    "ENSMUSG00000021591",
                    "ENSMUSG00000032786",
                    "ENSMUSG00000022865",
                    "ENSMUSG00000050856",
                    "ENSMUSG00000025856",
                    "ENSMUSG00000038762",
                    "ENSMUSG00000098973",
                    "ENSMUSG00000053768",
                    "ENSMUSG00000025781",
                    "ENSMUSG00000020340",
                    "ENSMUSG00000033781",
                    "ENSMUSG00000018796",
                    "ENSMUSG00000008690",
                    "ENSMUSG00000032518",
                    "ENSMUSG00000021311",
                    "ENSMUSG00000022403",
                    "ENSMUSG00000059734",
                    "ENSMUSG00000022223",
                    "ENSMUSG00000022969",
                    "ENSMUSG00000091537",
                    "ENSMUSG00000029227",
                    "ENSMUSG00000078670",
                    "ENSMUSG00000062070",
                    "ENSMUSG00000047905",
                    "ENSMUSG00000039630",
                    "ENSMUSG00000024608",
                    "ENSMUSG00000038764",
                    "ENSMUSG00000026489",
                    "ENSMUSG00000045948",
                    "ENSMUSG00000022553",
                    "ENSMUSG00000009079",
                    "ENSMUSG00000007739",
                    "ENSMUSG00000027573",
                    "ENSMUSG00000028633",
                    "ENSMUSG00000032035",
                    "ENSMUSG00000028394",
                    "ENSMUSG00000057914",
                    "ENSMUSG00000025968",
                    "ENSMUSG00000002844",
                    "ENSMUSG00000064370",
                    "ENSMUSG00000020133",
                    "ENSMUSG00000030137",
                    "ENSMUSG00000020821",
                    "ENSMUSG00000018293",
                    "ENSMUSG00000044033",
                    "ENSMUSG00000024140",
                    "ENSMUSG00000028691",
                    "ENSMUSG00000022321",
                    "ENSMUSG00000061477",
                    "ENSMUSG00000037805",
                    "ENSMUSG00000024346",
                    "ENSMUSG00000017119",
                    "ENSMUSG00000031701",
                    "ENSMUSG00000020738",
                    "ENSMUSG00000029467",
                    "ENSMUSG00000039219",
                    "ENSMUSG00000022193",
                    "ENSMUSG00000011148",
                    "ENSMUSG00000062580",
                    "ENSMUSG00000026817",
                    "ENSMUSG00000035891",
                    "ENSMUSG00000022031",
                    "ENSMUSG00000040242",
                    "ENSMUSG00000078453",
                    "ENSMUSG00000020572",
                    "ENSMUSG00000021039",
                    "ENSMUSG00000021218",
                    "ENSMUSG00000062270",
                    "ENSMUSG00000062397",
                    "ENSMUSG00000025060",
                    "ENSMUSG00000028937",
                    "ENSMUSG00000015357",
                    "ENSMUSG00000034566",
                    "ENSMUSG00000024985",
                    "ENSMUSG00000021243",
                    "ENSMUSG00000013593",
                    "ENSMUSG00000068267",
                    "ENSMUSG00000021417",
                    "ENSMUSG00000062373",
                    "ENSMUSG00000024053",
                    "ENSMUSG00000032578",
                    "ENSMUSG00000004951",
                    "ENSMUSG00000031916",
                    "ENSMUSG00000071533",
                    "ENSMUSG00000029328",
                    "ENSMUSG00000029432",
                    "ENSMUSG00000038520",
                    "ENSMUSG00000038058",
                    "ENSMUSG00000024963",
                    "ENSMUSG00000058569",
                    "ENSMUSG00000036781",
                    "ENSMUSG00000006418",
                    "ENSMUSG00000074754",
                    "ENSMUSG00000040964",
                    "ENSMUSG00000020988",
                    "ENSMUSG00000024012",
                    "ENSMUSG00000020530",
                    "ENSMUSG00000039670",
                    "ENSMUSG00000018340",
                    "ENSMUSG00000022111",
                    "ENSMUSG00000046364",
                    "ENSMUSG00000037236",
                    "ENSMUSG00000023307",
                    "ENSMUSG00000038650",
                    "ENSMUSG00000055053",
                    "ENSMUSG00000051391",
                    "ENSMUSG00000015120",
                    "ENSMUSG00000058056",
                    "ENSMUSG00000064369",
                    "ENSMUSG00000026154",
                    "ENSMUSG00000022265",
                    "ENSMUSG00000028041",
                    "ENSMUSG00000047746",
                    "ENSMUSG00000025509",
                    "ENSMUSG00000027245",
                    "ENSMUSG00000083863",
                    "ENSMUSG00000036880",
                    "ENSMUSG00000053819",
                    "ENSMUSG00000022551",
                    "ENSMUSG00000037110",
                    "ENSMUSG00000034353",
                    "ENSMUSG00000020440",
                    "ENSMUSG00000024158",
                    "ENSMUSG00000042607",
                    "ENSMUSG00000023094",
                    "ENSMUSG00000021236",
                    "ENSMUSG00000030847",
                    "ENSMUSG00000008575",
                    "ENSMUSG00000022400",
                    "ENSMUSG00000035173",
                    "ENSMUSG00000004558",
                    "ENSMUSG00000037062",
                    "ENSMUSG00000067722",
                    "ENSMUSG00000090258",
                    "ENSMUSG00000019787",
                    "ENSMUSG00000026003",
                    "ENSMUSG00000067288",
                    "ENSMUSG00000021156",
                    "ENSMUSG00000040209",
                    "ENSMUSG00000038462",
                    "ENSMUSG00000022797",
                    "ENSMUSG00000029864",
                    "ENSMUSG00000032966",
                    "ENSMUSG00000034936",
                    "ENSMUSG00000082195",
                    "ENSMUSG00000043483",
                    "ENSMUSG00000074994",
                    "ENSMUSG00000084105",
                    "ENSMUSG00000024099",
                    "ENSMUSG00000028977",
                    "ENSMUSG00000037656",
                    "ENSMUSG00000028295",
                    "ENSMUSG00000096942",
                    "ENSMUSG00000032563",
                    "ENSMUSG00000028053",
                    "ENSMUSG00000014294",
                    "ENSMUSG00000034875",
                    "ENSMUSG00000038975",
                    "ENSMUSG00000021939",
                    "ENSMUSG00000028104",
                    "ENSMUSG00000090935",
                    "ENSMUSG00000021096",
                    "ENSMUSG00000027777",
                    "ENSMUSG00000022020",
                    "ENSMUSG00000040481",
                    "ENSMUSG00000002661",
                    "ENSMUSG00000040269",
                    "ENSMUSG00000046434",
                    "ENSMUSG00000018411",
                    "ENSMUSG00000003444",
                    "ENSMUSG00000015837",
                    "ENSMUSG00000029823",
                    "ENSMUSG00000031382",
                    "ENSMUSG00000091086",
                    "ENSMUSG00000028690",
                    "ENSMUSG00000002028",
                    "ENSMUSG00000071540",
                    "ENSMUSG00000097414",
                    "ENSMUSG00000081179",
                    "ENSMUSG00000034751",
                    "ENSMUSG00000022707",
                    "ENSMUSG00000029629",
                    "ENSMUSG00000027984",
                    "ENSMUSG00000041193",
                    "ENSMUSG00000074698",
                    "ENSMUSG00000031353",
                    "ENSMUSG00000031960",
                    "ENSMUSG00000024097",
                    "ENSMUSG00000044477",
                    "ENSMUSG00000022354",
                    "ENSMUSG00000023951",
                    "ENSMUSG00000028063",
                    "ENSMUSG00000032355",
                    "ENSMUSG00000005354",
                    "ENSMUSG00000020477",
                    "ENSMUSG00000026623",
                    "ENSMUSG00000035158",
                    "ENSMUSG00000029198",
                    "ENSMUSG00000030747",
                    "ENSMUSG00000020496",
                    "ENSMUSG00000023087",
                    "ENSMUSG00000057469",
                    "ENSMUSG00000005469",
                    "ENSMUSG00000097867",
                    "ENSMUSG00000026803",
                    "ENSMUSG00000005299",
                    "ENSMUSG00000052539",
                    "ENSMUSG00000029152",
                    "ENSMUSG00000040653",
                    "ENSMUSG00000022519",
                    "ENSMUSG00000051331",
                    "ENSMUSG00000025260",
                    "ENSMUSG00000016528",
                    "ENSMUSG00000024038",
                    "ENSMUSG00000025190",
                    "ENSMUSG00000033065",
                    "ENSMUSG00000057738",
                    "ENSMUSG00000042010",
                    "ENSMUSG00000030850",
                    "ENSMUSG00000085684",
                    "ENSMUSG00000014606",
                    "ENSMUSG00000026260",
                    "ENSMUSG00000055839",
                    "ENSMUSG00000020612",
                    "ENSMUSG00000039891",
                    "ENSMUSG00000048490",
                    "ENSMUSG00000031059",
                    "ENSMUSG00000024359",
                    "ENSMUSG00000062461",
                    "ENSMUSG00000039063",
                    "ENSMUSG00000032435",
                    "ENSMUSG00000038690",
                    "ENSMUSG00000044700",
                    "ENSMUSG00000028954",
                    "ENSMUSG00000021111",
                    "ENSMUSG00000039067",
                    "ENSMUSG00000003865",
                    "ENSMUSG00000024539",
                    "ENSMUSG00000020460",
                    "ENSMUSG00000031871",
                    "ENSMUSG00000071054",
                    "ENSMUSG00000041028",
                    "ENSMUSG00000028700",
                    "ENSMUSG00000021079",
                    "ENSMUSG00000026112",
                    "ENSMUSG00000021772",
                    "ENSMUSG00000002007",
                    "ENSMUSG00000018566",
                    "ENSMUSG00000020321",
                    "ENSMUSG00000025141",
                    "ENSMUSG00000059447",
                    "ENSMUSG00000024277",
                    "ENSMUSG00000024120",
                    "ENSMUSG00000029614",
                    "ENSMUSG00000053199",
                    "ENSMUSG00000026473",
                    "ENSMUSG00000044086",
                    "ENSMUSG00000001508",
                    "ENSMUSG00000047675",
                    "ENSMUSG00000013419",
                    "ENSMUSG00000031880",
                    "ENSMUSG00000038615",
                    "ENSMUSG00000029064",
                    "ENSMUSG00000045210",
                    "ENSMUSG00000079677",
                    "ENSMUSG00000025130",
                    "ENSMUSG00000030795",
                    "ENSMUSG00000023990",
                    "ENSMUSG00000015757",
                    "ENSMUSG00000025290",
                    "ENSMUSG00000021917",
                    "ENSMUSG00000027879",
                    "ENSMUSG00000033918",
                    "ENSMUSG00000028443",
                    "ENSMUSG00000022517",
                    "ENSMUSG00000016344",
                    "ENSMUSG00000024236",
                    "ENSMUSG00000039001",
                    "ENSMUSG00000063904",
                    "ENSMUSG00000061518",
                    "ENSMUSG00000033400",
                    "ENSMUSG00000036570",
                    "ENSMUSG00000001056",
                    "ENSMUSG00000025935",
                    "ENSMUSG00000003299",
                    "ENSMUSG00000040351",
                    "ENSMUSG00000050812",
                    "ENSMUSG00000027452",
                    "ENSMUSG00000025869",
                    "ENSMUSG00000007617",
                    "ENSMUSG00000021147",
                    "ENSMUSG00000024083",
                    "ENSMUSG00000051469",
                    "ENSMUSG00000019768",
                    "ENSMUSG00000029723",
                    "ENSMUSG00000036371",
                    "ENSMUSG00000070493",
                    "ENSMUSG00000030335",
                    "ENSMUSG00000021748",
                    "ENSMUSG00000036580",
                    "ENSMUSG00000071369",
                    "ENSMUSG00000030357",
                    "ENSMUSG00000038695",
                    "ENSMUSG00000026456",
                    "ENSMUSG00000028932",
                    "ENSMUSG00000027184",
                    "ENSMUSG00000029576",
                    "ENSMUSG00000068921",
                    "ENSMUSG00000001441",
                    "ENSMUSG00000059714",
                    "ENSMUSG00000030122",
                    "ENSMUSG00000034220",
                    "ENSMUSG00000016534",
                    "ENSMUSG00000051319",
                    "ENSMUSG00000031231",
                    "ENSMUSG00000071653",
                    "ENSMUSG00000026895",
                    "ENSMUSG00000026179",
                    "ENSMUSG00000033059",
                    "ENSMUSG00000002342",
                    "ENSMUSG00000033813",
                    "ENSMUSG00000000708",
                    "ENSMUSG00000000088",
                    "ENSMUSG00000032038",
                    "ENSMUSG00000038204",
                    "ENSMUSG00000042462",
                    "ENSMUSG00000022257",
                    "ENSMUSG00000025393",
                    "ENSMUSG00000048279",
                    "ENSMUSG00000029319",
                    "ENSMUSG00000026930",
                    "ENSMUSG00000031641",
                    "ENSMUSG00000038967",
                    "ENSMUSG00000025410",
                    "ENSMUSG00000036112",
                    "ENSMUSG00000058070",
                    "ENSMUSG00000024588",
                    "ENSMUSG00000006299",
                    "ENSMUSG00000028495",
                    "ENSMUSG00000002416",
                    "ENSMUSG00000008668",
                    "ENSMUSG00000003526",
                    "ENSMUSG00000025791",
                    "ENSMUSG00000078154",
                    "ENSMUSG00000066043",
                    "ENSMUSG00000021460",
                    "ENSMUSG00000022477",
                    "ENSMUSG00000022337",
                    "ENSMUSG00000036698",
                    "ENSMUSG00000042298",
                    "ENSMUSG00000052146",
                    "ENSMUSG00000012405",
                    "ENSMUSG00000015290",
                    "ENSMUSG00000074800",
                    "ENSMUSG00000030695",
                    "ENSMUSG00000064341",
                    "ENSMUSG00000028618",
                    "ENSMUSG00000040740",
                    "ENSMUSG00000013539",
                    "ENSMUSG00000001366",
                    "ENSMUSG00000025940",
                    "ENSMUSG00000010376",
                    "ENSMUSG00000054640",
                    "ENSMUSG00000035242",
                    "ENSMUSG00000043716",
                    "ENSMUSG00000039768",
                    "ENSMUSG00000028398",
                    "ENSMUSG00000050608",
                    "ENSMUSG00000021706",
                    "ENSMUSG00000004891",
                    "ENSMUSG00000036160",
                    "ENSMUSG00000027206",
                    "ENSMUSG00000019851",
                    "ENSMUSG00000059208",
                    "ENSMUSG00000022016",
                    "ENSMUSG00000026150",
                    "ENSMUSG00000038412",
                    "ENSMUSG00000020390",
                    "ENSMUSG00000036323",
                    "ENSMUSG00000039474",
                    "ENSMUSG00000038370",
                    "ENSMUSG00000044475",
                    "ENSMUSG00000020453",
                    "ENSMUSG00000029128",
                    "ENSMUSG00000029474",
                    "ENSMUSG00000099250",
                    "ENSMUSG00000013736",
                    "ENSMUSG00000082884",
                    "ENSMUSG00000055737",
                    "ENSMUSG00000063870",
                    "ENSMUSG00000020982",
                    "ENSMUSG00000032902",
                    "ENSMUSG00000031633",
                    "ENSMUSG00000063694",
                    "ENSMUSG00000068487",
                    "ENSMUSG00000020368",
                    "ENSMUSG00000039473",
                    "ENSMUSG00000092014",
                    "ENSMUSG00000019795",
                    "ENSMUSG00000040385",
                    "ENSMUSG00000038086",
                    "ENSMUSG00000016252",
                    "ENSMUSG00000041571",
                    "ENSMUSG00000084178",
                    "ENSMUSG00000082874",
                    "ENSMUSG00000022889",
                    "ENSMUSG00000019528",
                    "ENSMUSG00000021843",
                    "ENSMUSG00000039611",
                    "ENSMUSG00000067274",
                    "ENSMUSG00000063787",
                    "ENSMUSG00000063882",
                    "ENSMUSG00000028607",
                    "ENSMUSG00000039105",
                    "ENSMUSG00000026028",
                    "ENSMUSG00000026078",
                    "ENSMUSG00000039910",
                    "ENSMUSG00000021557",
                    "ENSMUSG00000028745",
                    "ENSMUSG00000038886",
                    "ENSMUSG00000053702",
                    "ENSMUSG00000020267",
                    "ENSMUSG00000022051",
                    "ENSMUSG00000006818",
                    "ENSMUSG00000026860",
                    "ENSMUSG00000076431",
                    "ENSMUSG00000042541",
                    "ENSMUSG00000094344",
                    "ENSMUSG00000069601",
                    "ENSMUSG00000028692",
                    "ENSMUSG00000031928",
                    "ENSMUSG00000021375",
                    "ENSMUSG00000060739",
                    "ENSMUSG00000033039",
                    "ENSMUSG00000006931",
                    "ENSMUSG00000025499",
                    "ENSMUSG00000032399",
                    "ENSMUSG00000059586",
                    "ENSMUSG00000078566",
                    "ENSMUSG00000002102",
                    "ENSMUSG00000021390",
                    "ENSMUSG00000002343",
                    "ENSMUSG00000039652",
                    "ENSMUSG00000024613",
                    "ENSMUSG00000027472",
                    "ENSMUSG00000054889",
                    "ENSMUSG00000070544",
                    "ENSMUSG00000046351",
                    "ENSMUSG00000020166",
                    "ENSMUSG00000024953",
                    "ENSMUSG00000027774",
                    "ENSMUSG00000022982",
                    "ENSMUSG00000058558",
                    "ENSMUSG00000026020",
                    "ENSMUSG00000050323",
                    "ENSMUSG00000026032",
                    "ENSMUSG00000068479",
                    "ENSMUSG00000021868",
                    "ENSMUSG00000037376",
                    "ENSMUSG00000028452",
                    "ENSMUSG00000028156",
                    "ENSMUSG00000032407",
                    "ENSMUSG00000039987",
                    "ENSMUSG00000041548",
                    "ENSMUSG00000044502",
                    "ENSMUSG00000070802",
                    "ENSMUSG00000024576",
                    "ENSMUSG00000035478",
                    "ENSMUSG00000028333",
                    "ENSMUSG00000041058",
                    "ENSMUSG00000036202",
                    "ENSMUSG00000032112",
                    "ENSMUSG00000027257",
                    "ENSMUSG00000020372",
                    "ENSMUSG00000032523",
                    "ENSMUSG00000028834",
                    "ENSMUSG00000031352",
                    "ENSMUSG00000073838",
                    "ENSMUSG00000041870",
                    "ENSMUSG00000039536",
                    "ENSMUSG00000000740",
                    "ENSMUSG00000040752",
                    "ENSMUSG00000023723",
                    "ENSMUSG00000003934",
                    "ENSMUSG00000030884",
                    "ENSMUSG00000025466",
                    "ENSMUSG00000018567",
                    "ENSMUSG00000051518",
                    "ENSMUSG00000022000",
                    "ENSMUSG00000023025",
                    "ENSMUSG00000021476",
                    "ENSMUSG00000032051",
                    "ENSMUSG00000034040",
                    "ENSMUSG00000023460",
                    "ENSMUSG00000002767",
                    "ENSMUSG00000034994",
                    "ENSMUSG00000020108",
                    "ENSMUSG00000030541",
                    "ENSMUSG00000031438",
                    "ENSMUSG00000026208",
                    "ENSMUSG00000031696",
                    "ENSMUSG00000090266",
                    "ENSMUSG00000036052",
                    "ENSMUSG00000015217",
                    "ENSMUSG00000024213",
                    "ENSMUSG00000030652",
                    "ENSMUSG00000089917",
                    "ENSMUSG00000027238",
                    "ENSMUSG00000004364",
                    "ENSMUSG00000047388",
                    "ENSMUSG00000021109",
                    "ENSMUSG00000074412",
                    "ENSMUSG00000061306",
                    "ENSMUSG00000024869",
                    "ENSMUSG00000017686",
                    "ENSMUSG00000033307",
                    "ENSMUSG00000057406",
                    "ENSMUSG00000046580",
                    "ENSMUSG00000069089",
                    "ENSMUSG00000033543",
                    "ENSMUSG00000020415",
                    "ENSMUSG00000024146",
                    "ENSMUSG00000006800",
                    "ENSMUSG00000002957",
                    "ENSMUSG00000041957",
                    "ENSMUSG00000024073",
                    "ENSMUSG00000028282",
                    "ENSMUSG00000062997",
                    "ENSMUSG00000081431",
                    "ENSMUSG00000020430",
                    "ENSMUSG00000009112",
                    "ENSMUSG00000031311",
                    "ENSMUSG00000026603",
                    "ENSMUSG00000027180",
                    "ENSMUSG00000016308",
                    "ENSMUSG00000027422",
                    "ENSMUSG00000050043",
                    "ENSMUSG00000018882",
                    "ENSMUSG00000028470",
                    "ENSMUSG00000000787",
                    "ENSMUSG00000015214",
                    "ENSMUSG00000037710",
                    "ENSMUSG00000024187",
                    "ENSMUSG00000019362",
                    "ENSMUSG00000016427",
                    "ENSMUSG00000038240",
                    "ENSMUSG00000027293",
                    "ENSMUSG00000003948",
                    "ENSMUSG00000019055",
                    "ENSMUSG00000003559",
                    "ENSMUSG00000001127",
                    "ENSMUSG00000015656",
                    "ENSMUSG00000025868",
                    "ENSMUSG00000040520",
                    "ENSMUSG00000058756",
                    "ENSMUSG00000025651",
                    "ENSMUSG00000028603",
                    "ENSMUSG00000071037",
                    "ENSMUSG00000037887",
                    "ENSMUSG00000028820",
                    "ENSMUSG00000004268",
                    "ENSMUSG00000025085",
                    "ENSMUSG00000035329",
                    "ENSMUSG00000065979",
                    "ENSMUSG00000028261",
                    "ENSMUSG00000024259",
                    "ENSMUSG00000025068",
                    "ENSMUSG00000045103",
                    "ENSMUSG00000022205",
                    "ENSMUSG00000034312",
                    "ENSMUSG00000032602",
                    "ENSMUSG00000032042",
                    "ENSMUSG00000061032",
                    "ENSMUSG00000023089",
                    "ENSMUSG00000019843",
                    "ENSMUSG00000020225",
                    "ENSMUSG00000073676",
                    "ENSMUSG00000032060",
                    "ENSMUSG00000023456",
                    "ENSMUSG00000022010",
                    "ENSMUSG00000027673",
                    "ENSMUSG00000083878",
                    "ENSMUSG00000059811",
                    "ENSMUSG00000013662",
                    "ENSMUSG00000026914",
                    "ENSMUSG00000083757",
                    "ENSMUSG00000024587",
                    "ENSMUSG00000024121",
                    "ENSMUSG00000029616",
                    "ENSMUSG00000020003",
                    "ENSMUSG00000019659",
                    "ENSMUSG00000061904",
                    "ENSMUSG00000027305",
                    "ENSMUSG00000029454",
                    "ENSMUSG00000083240",
                    "ENSMUSG00000004937",
                    "ENSMUSG00000015605",
                    "ENSMUSG00000020485",
                    "ENSMUSG00000006740",
                    "ENSMUSG00000024197",
                    "ENSMUSG00000007777",
                    "ENSMUSG00000019432",
                    "ENSMUSG00000022914",
                    "ENSMUSG00000063457",
                    "ENSMUSG00000008859",
                    "ENSMUSG00000020088",
                    "ENSMUSG00000039148",
                    "ENSMUSG00000058440",
                    "ENSMUSG00000019689",
                    "ENSMUSG00000020741",
                    "ENSMUSG00000037266",
                    "ENSMUSG00000038095",
                    "ENSMUSG00000037339",
                    "ENSMUSG00000030982",
                    "ENSMUSG00000027679",
                    "ENSMUSG00000041841",
                    "ENSMUSG00000026576",
                    "ENSMUSG00000041881",
                    "ENSMUSG00000021764",
                    "ENSMUSG00000038763",
                    "ENSMUSG00000043445",
                    "ENSMUSG00000039648",
                    "ENSMUSG00000020163",
                    "ENSMUSG00000001767",
                    "ENSMUSG00000009863",
                    "ENSMUSG00000090553",
                    "ENSMUSG00000031770",
                    "ENSMUSG00000038400",
                    "ENSMUSG00000030591",
                    "ENSMUSG00000036309",
                    "ENSMUSG00000031715",
                    "ENSMUSG00000028945",
                    "ENSMUSG00000056900",
                    "ENSMUSG00000032238",
                    "ENSMUSG00000062210",
                    "ENSMUSG00000060376",
                    "ENSMUSG00000014592",
                    "ENSMUSG00000090150",
                    "ENSMUSG00000017428",
                    "ENSMUSG00000079508",
                    "ENSMUSG00000081051",
                    "ENSMUSG00000062694",
                    "ENSMUSG00000020153",
                    "ENSMUSG00000093674",
                    "ENSMUSG00000083563",
                    "ENSMUSG00000022312",
                    "ENSMUSG00000040760",
                    "ENSMUSG00000069729",
                    "ENSMUSG00000036452",
                    "ENSMUSG00000031818",
                    "ENSMUSG00000031782",
                    "ENSMUSG00000048486",
                    "ENSMUSG00000055681",
                    "ENSMUSG00000032511",
                    "ENSMUSG00000038526",
                    "ENSMUSG00000024683",
                    "ENSMUSG00000096449",
                    "ENSMUSG00000044894",
                    "ENSMUSG00000007097",
                    "ENSMUSG00000029404",
                    "ENSMUSG00000024937",
                    "ENSMUSG00000056032",
                    "ENSMUSG00000022820",
                    "ENSMUSG00000037234",
                    "ENSMUSG00000042148",
                    "ENSMUSG00000034295",
                    "ENSMUSG00000031783",
                    "ENSMUSG00000022594",
                    "ENSMUSG00000063229",
                    "ENSMUSG00000001604",
                    "ENSMUSG00000005417",
                    "ENSMUSG00000021577",
                    "ENSMUSG00000038936",
                    "ENSMUSG00000029776",
                    "ENSMUSG00000029135",
                    "ENSMUSG00000025898",
                    "ENSMUSG00000036427",
                    "ENSMUSG00000070867",
                    "ENSMUSG00000038417",
                    "ENSMUSG00000045160",
                    "ENSMUSG00000024942",
                    "ENSMUSG00000022698",
                    "ENSMUSG00000024335",
                    "ENSMUSG00000027371",
                    "ENSMUSG00000028743",
                    "ENSMUSG00000037075",
                    "ENSMUSG00000040407",
                    "ENSMUSG00000030647",
                    "ENSMUSG00000073664",
                    "ENSMUSG00000025545",
                    "ENSMUSG00000045503",
                    "ENSMUSG00000091831",
                    "ENSMUSG00000026888",
                    "ENSMUSG00000022094",
                    "ENSMUSG00000052712",
                    "ENSMUSG00000030087",
                    "ENSMUSG00000031776",
                    "ENSMUSG00000006333",
                    "ENSMUSG00000026568",
                    "ENSMUSG00000028328",
                    "ENSMUSG00000068328",
                    "ENSMUSG00000005683",
                    "ENSMUSG00000029106",
                    "ENSMUSG00000031700",
                    "ENSMUSG00000046516",
                    "ENSMUSG00000022332",
                    "ENSMUSG00000040112",
                    "ENSMUSG00000037960",
                    "ENSMUSG00000058655",
                    "ENSMUSG00000022961",
                    "ENSMUSG00000020085",
                    "ENSMUSG00000031536",
                    "ENSMUSG00000032348",
                    "ENSMUSG00000039218",
                    "ENSMUSG00000021133",
                    "ENSMUSG00000029594",
                    "ENSMUSG00000023286",
                    "ENSMUSG00000031516",
                    "ENSMUSG00000052214",
                    "ENSMUSG00000033228",
                    "ENSMUSG00000046032",
                    "ENSMUSG00000032418",
                    "ENSMUSG00000032826",
                    "ENSMUSG00000028266",
                    "ENSMUSG00000081382",
                    "ENSMUSG00000026064",
                    "ENSMUSG00000047371",
                    "ENSMUSG00000029048",
                    "ENSMUSG00000022848",
                    "ENSMUSG00000020737",
                    "ENSMUSG00000031232",
                    "ENSMUSG00000001419",
                    "ENSMUSG00000051671",
                    "ENSMUSG00000022765",
                    "ENSMUSG00000026360",
                    "ENSMUSG00000047260",
                    "ENSMUSG00000004460",
                    "ENSMUSG00000042198",
                    "ENSMUSG00000035674",
                    "ENSMUSG00000009927",
                    "ENSMUSG00000083832",
                    "ENSMUSG00000026571",
                    "ENSMUSG00000025465",
                    "ENSMUSG00000042605",
                    "ENSMUSG00000028919",
                    "ENSMUSG00000040990",
                    "ENSMUSG00000027423",
                    "ENSMUSG00000001833",
                    "ENSMUSG00000025204",
                    "ENSMUSG00000023055",
                    "ENSMUSG00000098178",
                    "ENSMUSG00000031672",
                    "ENSMUSG00000075704",
                    "ENSMUSG00000031660",
                    "ENSMUSG00000040713",
                    "ENSMUSG00000026977",
                    "ENSMUSG00000030172",
                    "ENSMUSG00000059895",
                    "ENSMUSG00000016319",
                    "ENSMUSG00000030706",
                    "ENSMUSG00000024780",
                    "ENSMUSG00000021488",
                    "ENSMUSG00000002948",
                    "ENSMUSG00000015575",
                    "ENSMUSG00000034290",
                    "ENSMUSG00000005981",
                    "ENSMUSG00000023150",
                    "ENSMUSG00000022216",
                    "ENSMUSG00000063800",
                    "ENSMUSG00000030298",
                    "ENSMUSG00000025795",
                    "ENSMUSG00000036181",
                    "ENSMUSG00000028291",
                    "ENSMUSG00000017404",
                    "ENSMUSG00000020190",
                    "ENSMUSG00000021944",
                    "ENSMUSG00000028648",
                    "ENSMUSG00000026991",
                    "ENSMUSG00000029623",
                    "ENSMUSG00000018848",
                    "ENSMUSG00000004535",
                    "ENSMUSG00000014633",
                    "ENSMUSG00000007892",
                    "ENSMUSG00000003380",
                    "ENSMUSG00000074064",
                    "ENSMUSG00000060938",
                    "ENSMUSG00000010406",
                    "ENSMUSG00000030607",
                    "ENSMUSG00000025428",
                    "ENSMUSG00000024538",
                    "ENSMUSG00000021585",
                    "ENSMUSG00000029017",
                    "ENSMUSG00000030016",
                    "ENSMUSG00000041426",
                    "ENSMUSG00000031634",
                    "ENSMUSG00000020849",
                    "ENSMUSG00000068706",
                    "ENSMUSG00000002748",
                    "ENSMUSG00000041297",
                    "ENSMUSG00000026766",
                    "ENSMUSG00000024924",
                    "ENSMUSG00000042744",
                    "ENSMUSG00000031987",
                    "ENSMUSG00000040952",
                    "ENSMUSG00000043311",
                    "ENSMUSG00000038970",
                    "ENSMUSG00000028367",
                    "ENSMUSG00000055302",
                    "ENSMUSG00000028013",
                    "ENSMUSG00000000346",
                    "ENSMUSG00000039089",
                    "ENSMUSG00000019782",
                    "ENSMUSG00000026687",
                    "ENSMUSG00000053898",
                    "ENSMUSG00000060038",
                    "ENSMUSG00000030409",
                    "ENSMUSG00000062691",
                    "ENSMUSG00000083820",
                    "ENSMUSG00000001105",
                    "ENSMUSG00000025221",
                    "ENSMUSG00000029632",
                    "ENSMUSG00000026207",
                    "ENSMUSG00000036620",
                    "ENSMUSG00000022641",
                    "ENSMUSG00000054766",
                    "ENSMUSG00000015846",
                    "ENSMUSG00000052337",
                    "ENSMUSG00000028223",
                    "ENSMUSG00000031262",
                    "ENSMUSG00000026342",
                    "ENSMUSG00000025920",
                    "ENSMUSG00000074466",
                    "ENSMUSG00000001794",
                    "ENSMUSG00000031791",
                    "ENSMUSG00000048264",
                    "ENSMUSG00000037820",
                    "ENSMUSG00000071644",
                    "ENSMUSG00000090000",
                    "ENSMUSG00000026027",
                    "ENSMUSG00000020785",
                    "ENSMUSG00000030615",
                    "ENSMUSG00000053012",
                    "ENSMUSG00000020483",
                    "ENSMUSG00000009090",
                    "ENSMUSG00000004748",
                    "ENSMUSG00000072692",
                    "ENSMUSG00000032679",
                    "ENSMUSG00000026878",
                    "ENSMUSG00000024055",
                    "ENSMUSG00000029518",
                    "ENSMUSG00000006304",
                    "ENSMUSG00000031320",
                    "ENSMUSG00000032959",
                    "ENSMUSG00000028964",
                    "ENSMUSG00000025893",
                    "ENSMUSG00000042032",
                    "ENSMUSG00000060802",
                    "ENSMUSG00000002728",
                    "ENSMUSG00000027406",
                    "ENSMUSG00000060032",
                    "ENSMUSG00000025487",
                    "ENSMUSG00000078784",
                    "ENSMUSG00000028756",
                    "ENSMUSG00000063273",
                    "ENSMUSG00000060675",
                    "ENSMUSG00000051154",
                    "ENSMUSG00000020029",
                    "ENSMUSG00000037712",
                    "ENSMUSG00000034708",
                    "ENSMUSG00000036932",
                    "ENSMUSG00000003955",
                    "ENSMUSG00000028234",
                    "ENSMUSG00000061838",
                    "ENSMUSG00000019179",
                    "ENSMUSG00000051495",
                    "ENSMUSG00000079477",
                    "ENSMUSG00000022841",
                    "ENSMUSG00000000253",
                    "ENSMUSG00000071415",
                    "ENSMUSG00000037152",
                    "ENSMUSG00000019494",
                    "ENSMUSG00000020708",
                    "ENSMUSG00000023010",
                    "ENSMUSG00000036278",
                    "ENSMUSG00000022391",
                    "ENSMUSG00000028788",
                    "ENSMUSG00000062169",
                    "ENSMUSG00000053907",
                    "ENSMUSG00000067369",
                    "ENSMUSG00000003316",
                    "ENSMUSG00000022629",
                    "ENSMUSG00000002980",
                    "ENSMUSG00000023806",
                    "ENSMUSG00000005510",
                    "ENSMUSG00000020547",
                    "ENSMUSG00000024208",
                    "ENSMUSG00000020232",
                    "ENSMUSG00000024507",
                    "ENSMUSG00000049832",
                    "ENSMUSG00000029469",
                    "ENSMUSG00000087260",
                    "ENSMUSG00000038374",
                    "ENSMUSG00000019428",
                    "ENSMUSG00000024803",
                    "ENSMUSG00000022255",
                    "ENSMUSG00000025967",
                    "ENSMUSG00000005034",
                    "ENSMUSG00000031683",
                    "ENSMUSG00000036822",
                    "ENSMUSG00000022442",
                    "ENSMUSG00000028248",
                    "ENSMUSG00000078812",
                    "ENSMUSG00000082474",
                    "ENSMUSG00000001158",
                    "ENSMUSG00000090733",
                    "ENSMUSG00000036352",
                    "ENSMUSG00000059534",
                    "ENSMUSG00000038467",
                    "ENSMUSG00000027367",
                    "ENSMUSG00000029108",
                    "ENSMUSG00000007338",
                    "ENSMUSG00000080836",
                    "ENSMUSG00000044308",
                    "ENSMUSG00000053536",
                    "ENSMUSG00000022300",
                    "ENSMUSG00000003429",
                    "ENSMUSG00000021185",
                    "ENSMUSG00000018599",
                    "ENSMUSG00000037822",
                    "ENSMUSG00000084319",
                    "ENSMUSG00000064344",
                    "ENSMUSG00000029095",
                    "ENSMUSG00000015839",
                    "ENSMUSG00000027499",
                    "ENSMUSG00000027784",
                    "ENSMUSG00000065990",
                    "ENSMUSG00000060438",
                    "ENSMUSG00000003541",
                    "ENSMUSG00000029068",
                    "ENSMUSG00000064340",
                    "ENSMUSG00000032047",
                    "ENSMUSG00000042133",
                    "ENSMUSG00000044393",
                    "ENSMUSG00000014470",
                    "ENSMUSG00000031586",
                    "ENSMUSG00000001472",
                    "ENSMUSG00000048076",
                    "ENSMUSG00000036002",
                    "ENSMUSG00000072640",
                    "ENSMUSG00000038582",
                    "ENSMUSG00000026864",
                    "ENSMUSG00000023939",
                    "ENSMUSG00000035885",
                    "ENSMUSG00000059325",
                    "ENSMUSG00000031078",
                    "ENSMUSG00000026341",
                    "ENSMUSG00000038900",
                    "ENSMUSG00000032431",
                    "ENSMUSG00000032330",
                    "ENSMUSG00000022023",
                    "ENSMUSG00000038170",
                    "ENSMUSG00000006315",
                    "ENSMUSG00000078713",
                    "ENSMUSG00000051695",
                    "ENSMUSG00000049090",
                    "ENSMUSG00000041361",
                    "ENSMUSG00000049327",
                    "ENSMUSG00000006932",
                    "ENSMUSG00000091449",
                    "ENSMUSG00000022748",
                    "ENSMUSG00000060981",
                    "ENSMUSG00000031865",
                    "ENSMUSG00000030089",
                    "ENSMUSG00000000149",
                    "ENSMUSG00000023206",
                    "ENSMUSG00000028518",
                    "ENSMUSG00000074129",
                    "ENSMUSG00000031839",
                    "ENSMUSG00000001175",
                    "ENSMUSG00000038084",
                    "ENSMUSG00000008136",
                    "ENSMUSG00000049881",
                    "ENSMUSG00000005873",
                    "ENSMUSG00000054405",
                    "ENSMUSG00000030744",
                    "ENSMUSG00000029019",
                    "ENSMUSG00000023944",
                    "ENSMUSG00000015932",
                    "ENSMUSG00000029780",
                    "ENSMUSG00000020358",
                    "ENSMUSG00000022681",
                    "ENSMUSG00000068699",
                    "ENSMUSG00000003970",
                    "ENSMUSG00000030662",
                    "ENSMUSG00000072946",
                    "ENSMUSG00000010911",
                    "ENSMUSG00000050953",
                    "ENSMUSG00000027765",
                    "ENSMUSG00000020308",
                    "ENSMUSG00000021546",
                    "ENSMUSG00000064372",
                    "ENSMUSG00000030002",
                    "ENSMUSG00000032495",
                    "ENSMUSG00000028772",
                    "ENSMUSG00000025034",
                    "ENSMUSG00000025794",
                    "ENSMUSG00000046997",
                    "ENSMUSG00000020022",
                    "ENSMUSG00000034088",
                    "ENSMUSG00000071655",
                    "ENSMUSG00000001911",
                    "ENSMUSG00000034832",
                    "ENSMUSG00000020664",
                    "ENSMUSG00000022389",
                    "ENSMUSG00000075595",
                    "ENSMUSG00000033722",
                    "ENSMUSG00000040118",
                    "ENSMUSG00000029203",
                    "ENSMUSG00000022635",
                    "ENSMUSG00000067851",
                    "ENSMUSG00000018770",
                    "ENSMUSG00000052911",
                    "ENSMUSG00000039782",
                    "ENSMUSG00000097088",
                    "ENSMUSG00000035781",
                    "ENSMUSG00000087380",
                    "ENSMUSG00000067736",
                    "ENSMUSG00000028737",
                    "ENSMUSG00000031068",
                    "ENSMUSG00000031954",
                    "ENSMUSG00000030621",
                    "ENSMUSG00000056602",
                    "ENSMUSG00000062908",
                    "ENSMUSG00000022253",
                    "ENSMUSG00000021057",
                    "ENSMUSG00000068264",
                    "ENSMUSG00000025613",
                    "ENSMUSG00000022462",
                    "ENSMUSG00000027447",
                    "ENSMUSG00000021993",
                    "ENSMUSG00000031533",
                    "ENSMUSG00000029390",
                    "ENSMUSG00000000355",
                    "ENSMUSG00000020290",
                    "ENSMUSG00000037032",
                    "ENSMUSG00000024283",
                    "ENSMUSG00000036585",
                    "ENSMUSG00000029394",
                    "ENSMUSG00000030319",
                    "ENSMUSG00000059070",
                    "ENSMUSG00000062647",
                    "ENSMUSG00000025492",
                    "ENSMUSG00000062083",
                    "ENSMUSG00000099021",
                    "ENSMUSG00000042709",
                    "ENSMUSG00000033916",
                    "ENSMUSG00000032525",
                    "ENSMUSG00000020402",
                    "ENSMUSG00000060073",
                    "ENSMUSG00000022217",
                    "ENSMUSG00000026853",
                    "ENSMUSG00000034880",
                    "ENSMUSG00000054428",
                    "ENSMUSG00000066900",
                    "ENSMUSG00000028959",
                    "ENSMUSG00000008035",
                    "ENSMUSG00000026021",
                    "ENSMUSG00000019920",
                    "ENSMUSG00000014551",
                    "ENSMUSG00000021270",
                    "ENSMUSG00000011257",
                    "ENSMUSG00000021930",
                    "ENSMUSG00000031840",
                    "ENSMUSG00000039943",
                    "ENSMUSG00000006576",
                    "ENSMUSG00000038914",
                    "ENSMUSG00000064037",
                    "ENSMUSG00000004085",
                    "ENSMUSG00000056851",
                    "ENSMUSG00000028062",
                    "ENSMUSG00000014426",
                    "ENSMUSG00000020048",
                    "ENSMUSG00000063316",
                    "ENSMUSG00000022867",
                    "ENSMUSG00000028447",
                    "ENSMUSG00000048277",
                    "ENSMUSG00000020705",
                    "ENSMUSG00000038538",
                    "ENSMUSG00000084786",
                    "ENSMUSG00000020634",
                    "ENSMUSG00000036678",
                    "ENSMUSG00000030861",
                    "ENSMUSG00000028161",
                    "ENSMUSG00000018102",
                    "ENSMUSG00000029056",
                    "ENSMUSG00000025809",
                    "ENSMUSG00000026688",
                    "ENSMUSG00000015092",
                    "ENSMUSG00000029833",
                    "ENSMUSG00000034574",
                    "ENSMUSG00000040666",
                    "ENSMUSG00000025184",
                    "ENSMUSG00000003809",
                    "ENSMUSG00000010175",
                    "ENSMUSG00000000399",
                    "ENSMUSG00000004931",
                    "ENSMUSG00000044792",
                    "ENSMUSG00000039976",
                    "ENSMUSG00000022437",
                    "ENSMUSG00000051343",
                    "ENSMUSG00000019872",
                    "ENSMUSG00000031708",
                    "ENSMUSG00000026353",
                    "ENSMUSG00000014504",
                    "ENSMUSG00000022037",
                    "ENSMUSG00000022035",
                    "ENSMUSG00000033938",
                    "ENSMUSG00000024997",
                    "ENSMUSG00000057657",
                    "ENSMUSG00000017778",
                    "ENSMUSG00000020952",
                    "ENSMUSG00000048546",
                    "ENSMUSG00000035105",
                    "ENSMUSG00000030753",
                    "ENSMUSG00000073639",
                    "ENSMUSG00000045776",
                    "ENSMUSG00000020580",
                    "ENSMUSG00000020671",
                    "ENSMUSG00000028207",
                    "ENSMUSG00000040414",
                    "ENSMUSG00000001289",
                    "ENSMUSG00000021537",
                    "ENSMUSG00000060798",
                    "ENSMUSG00000036564",
                    "ENSMUSG00000074093",
                    "ENSMUSG00000037049",
                    "ENSMUSG00000020079",
                    "ENSMUSG00000054452",
                    "ENSMUSG00000002981",
                    "ENSMUSG00000024150",
                    "ENSMUSG00000020585",
                    "ENSMUSG00000040694",
                    "ENSMUSG00000012848",
                    "ENSMUSG00000032480",
                    "ENSMUSG00000022800",
                    "ENSMUSG00000000184",
                    "ENSMUSG00000091383",
                    "ENSMUSG00000010122",
                    "ENSMUSG00000050541",
                    "ENSMUSG00000038679",
                    "ENSMUSG00000031447",
                    "ENSMUSG00000020621",
                    "ENSMUSG00000008682",
                    "ENSMUSG00000034157",
                    "ENSMUSG00000020180",
                    "ENSMUSG00000031765",
                    "ENSMUSG00000023020",
                    "ENSMUSG00000024293",
                    "ENSMUSG00000045201",
                    "ENSMUSG00000052934",
                    "ENSMUSG00000030421",
                    "ENSMUSG00000033382",
                    "ENSMUSG00000034424",
                    "ENSMUSG00000071793",
                    "ENSMUSG00000031521",
                    "ENSMUSG00000064366",
                    "ENSMUSG00000019558",
                    "ENSMUSG00000031029",
                    "ENSMUSG00000031441",
                    "ENSMUSG00000029145",
                    "ENSMUSG00000006494",
                    "ENSMUSG00000021102",
                    "ENSMUSG00000023832",
                    "ENSMUSG00000005220",
                    "ENSMUSG00000004980",
                    "ENSMUSG00000005625",
                    "ENSMUSG00000032580",
                    "ENSMUSG00000071014",
                    "ENSMUSG00000044864",
                    "ENSMUSG00000049751",
                    "ENSMUSG00000025666",
                    "ENSMUSG00000021373",
                    "ENSMUSG00000031201",
                    "ENSMUSG00000000171",
                    "ENSMUSG00000000441",
                    "ENSMUSG00000041685",
                    "ENSMUSG00000034620",
                    "ENSMUSG00000038784",
                    "ENSMUSG00000024661",
                    "ENSMUSG00000036748",
                    "ENSMUSG00000053483",
                    "ENSMUSG00000025156",
                    "ENSMUSG00000002608",
                    "ENSMUSG00000025103",
                    "ENSMUSG00000039601",
                    "ENSMUSG00000057322",
                    "ENSMUSG00000073889",
                    "ENSMUSG00000022270",
                    "ENSMUSG00000026234",
                    "ENSMUSG00000029059",
                    "ENSMUSG00000035783",
                    "ENSMUSG00000045690",
                    "ENSMUSG00000032526",
                    "ENSMUSG00000038132",
                    "ENSMUSG00000000340",
                    "ENSMUSG00000037949",
                    "ENSMUSG00000055762",
                    "ENSMUSG00000034868",
                    "ENSMUSG00000064343",
                    "ENSMUSG00000021709",
                    "ENSMUSG00000043323",
                    "ENSMUSG00000022186",
                    "ENSMUSG00000021597",
                    "ENSMUSG00000056091",
                    "ENSMUSG00000045624",
                    "ENSMUSG00000086567",
                    "ENSMUSG00000064337",
                    "ENSMUSG00000020116",
                    "ENSMUSG00000071528",
                    "ENSMUSG00000031913",
                    "ENSMUSG00000071654",
                    "ENSMUSG00000030846",
                    "ENSMUSG00000021905",
                    "ENSMUSG00000031841",
                    "ENSMUSG00000061130",
                    "ENSMUSG00000036854",
                    "ENSMUSG00000026131",
                    "ENSMUSG00000026202",
                    "ENSMUSG00000020721",
                    "ENSMUSG00000027566",
                    "ENSMUSG00000049960",
                    "ENSMUSG00000048175",
                    "ENSMUSG00000020326",
                    "ENSMUSG00000031751",
                    "ENSMUSG00000032010",
                    "ENSMUSG00000041278",
                    "ENSMUSG00000021838",
                    "ENSMUSG00000062279",
                    "ENSMUSG00000031996",
                    "ENSMUSG00000067279",
                    "ENSMUSG00000064347",
                    "ENSMUSG00000023277",
                    "ENSMUSG00000034892",
                    "ENSMUSG00000017756",
                    "ENSMUSG00000023092",
                    "ENSMUSG00000059291",
                    "ENSMUSG00000032177",
                    "ENSMUSG00000003420",
                    "ENSMUSG00000002379",
                    "ENSMUSG00000027177",
                    "ENSMUSG00000039087",
                    "ENSMUSG00000029701",
                    "ENSMUSG00000052912",
                    "ENSMUSG00000029455",
                    "ENSMUSG00000046727",
                    "ENSMUSG00000064346",
                    "ENSMUSG00000058690",
                    "ENSMUSG00000027010",
                    "ENSMUSG00000049606",
                    "ENSMUSG00000030876",
                    "ENSMUSG00000060147",
                    "ENSMUSG00000059852",
                    "ENSMUSG00000051285",
                    "ENSMUSG00000024063",
                    "ENSMUSG00000037270",
                    "ENSMUSG00000040370",
                    "ENSMUSG00000074218",
                    "ENSMUSG00000021371",
                    "ENSMUSG00000042323",
                    "ENSMUSG00000030417",
                    "ENSMUSG00000026674",
                    "ENSMUSG00000034345",
                    "ENSMUSG00000037818",
                    "ENSMUSG00000037826",
                    "ENSMUSG00000029651",
                    "ENSMUSG00000023921",
                    "ENSMUSG00000032126",
                    "ENSMUSG00000057649",
                    "ENSMUSG00000021000",
                    "ENSMUSG00000042650",
                    "ENSMUSG00000000563",
                    "ENSMUSG00000065037",
                    "ENSMUSG00000037815",
                    "ENSMUSG00000027809",
                    "ENSMUSG00000047888",
                    "ENSMUSG00000034342",
                    "ENSMUSG00000038949",
                    "ENSMUSG00000047187",
                    "ENSMUSG00000061462",
                    "ENSMUSG00000037958",
                    "ENSMUSG00000031024",
                    "ENSMUSG00000020456",
                    "ENSMUSG00000021061",
                    "ENSMUSG00000045983",
                    "ENSMUSG00000032312",
                    "ENSMUSG00000050627",
                    "ENSMUSG00000051747",
                    "ENSMUSG00000025911",
                    "ENSMUSG00000035458",
                    "ENSMUSG00000032030",
                    "ENSMUSG00000032481",
                    "ENSMUSG00000022040",
                    "ENSMUSG00000028180",
                    "ENSMUSG00000026824",
                    "ENSMUSG00000022013",
                    "ENSMUSG00000081400",
                    "ENSMUSG00000081331",
                    "ENSMUSG00000011305",
                    "ENSMUSG00000014226",
                    "ENSMUSG00000006356",
                    "ENSMUSG00000039197",
                    "ENSMUSG00000022890",
                    "ENSMUSG00000053062",
                    "ENSMUSG00000019810",
                    "ENSMUSG00000024993",
                    "ENSMUSG00000022684",
                    "ENSMUSG00000032199",
                    "ENSMUSG00000001211",
                    "ENSMUSG00000056692",
                    "ENSMUSG00000070436",
                    "ENSMUSG00000052738",
                    "ENSMUSG00000031985",
                    "ENSMUSG00000029918",
                    "ENSMUSG00000029359",
                    "ENSMUSG00000023175",
                    "ENSMUSG00000052374",
                    "ENSMUSG00000044627",
                    "ENSMUSG00000015176",
                    "ENSMUSG00000024991",
                    "ENSMUSG00000037523",
                    "ENSMUSG00000024687",
                    "ENSMUSG00000022201",
                    "ENSMUSG00000027030",
                    "ENSMUSG00000080893",
                    "ENSMUSG00000021226",
                    "ENSMUSG00000022940",
                    "ENSMUSG00000030934",
                    "ENSMUSG00000032590",
                    "ENSMUSG00000026385",
                    "ENSMUSG00000041444",
                    "ENSMUSG00000020859",
                    "ENSMUSG00000024309",
                    "ENSMUSG00000050199",
                    "ENSMUSG00000025362",
                    "ENSMUSG00000058239",
                    "ENSMUSG00000071604",
                    "ENSMUSG00000022964",
                    "ENSMUSG00000082687",
                    "ENSMUSG00000029836",
                    "ENSMUSG00000025871",
                    "ENSMUSG00000025364",
                    "ENSMUSG00000038612",
                    "ENSMUSG00000042670",
                    "ENSMUSG00000021807",
                    "ENSMUSG00000022803",
                    "ENSMUSG00000064338",
                    "ENSMUSG00000061104",
                    "ENSMUSG00000000738",
                    "ENSMUSG00000039100",
                    "ENSMUSG00000009621",
                    "ENSMUSG00000044080",
                    "ENSMUSG00000046671",
                    "ENSMUSG00000024442",
                    "ENSMUSG00000004096",
                    "ENSMUSG00000028035",
                    "ENSMUSG00000047459",
                    "ENSMUSG00000029649",
                    "ENSMUSG00000022528",
                    "ENSMUSG00000060679",
                    "ENSMUSG00000038844",
                    "ENSMUSG00000052698",
                    "ENSMUSG00000005686",
                    "ENSMUSG00000048416",
                    "ENSMUSG00000053110",
                    "ENSMUSG00000010608",
                    "ENSMUSG00000027742",
                    "ENSMUSG00000032279",
                    "ENSMUSG00000033788",
                    "ENSMUSG00000086324",
                    "ENSMUSG00000023861",
                    "ENSMUSG00000067847",
                    "ENSMUSG00000026664",
                    "ENSMUSG00000068523",
                    "ENSMUSG00000043822",
                    "ENSMUSG00000021520",
                    "ENSMUSG00000006717",
                    "ENSMUSG00000001666",
                    "ENSMUSG00000090862",
                    "ENSMUSG00000036810",
                    "ENSMUSG00000008683",
                    "ENSMUSG00000003438",
                    "ENSMUSG00000042099",
                    "ENSMUSG00000083219",
                    "ENSMUSG00000005204",
                    "ENSMUSG00000097277",
                    "ENSMUSG00000028538",
                    "ENSMUSG00000007721",
                    "ENSMUSG00000020777",
                    "ENSMUSG00000034842",
                    "ENSMUSG00000041733",
                    "ENSMUSG00000062328",
                    "ENSMUSG00000029190",
                    "ENSMUSG00000039873",
                    "ENSMUSG00000004610",
                    "ENSMUSG00000015944",
                    "ENSMUSG00000021265",
                    "ENSMUSG00000021290",
                    "ENSMUSG00000037573",
                    "ENSMUSG00000049470",
                    "ENSMUSG00000069049",
                    "ENSMUSG00000046330",
                    "ENSMUSG00000029401",
                    "ENSMUSG00000074247",
                    "ENSMUSG00000003534",
                    "ENSMUSG00000025453",
                    "ENSMUSG00000025860",
                    "ENSMUSG00000030612",
                    "ENSMUSG00000004035",
                    "ENSMUSG00000074918",
                    "ENSMUSG00000024725",
                    "ENSMUSG00000021606",
                    "ENSMUSG00000019877",
                    "ENSMUSG00000033416",
                    "ENSMUSG00000023068",
                    "ENSMUSG00000021192",
                    "ENSMUSG00000036860",
                    "ENSMUSG00000079641",
                    "ENSMUSG00000075705",
                    "ENSMUSG00000045636",
                    "ENSMUSG00000020462",
                    "ENSMUSG00000062300",
                    "ENSMUSG00000063077",
                    "ENSMUSG00000022200",
                    "ENSMUSG00000031299",
                    "ENSMUSG00000053477",
                    "ENSMUSG00000021771",
                    "ENSMUSG00000032267",
                    "ENSMUSG00000035247",
                    "ENSMUSG00000040048",
                    "ENSMUSG00000028461",
                    "ENSMUSG00000064365",
                    "ENSMUSG00000024976",
                    "ENSMUSG00000043639",
                    "ENSMUSG00000043866",
                    "ENSMUSG00000071662",
                    "ENSMUSG00000028557",
                    "ENSMUSG00000023938",
                    "ENSMUSG00000027274",
                    "ENSMUSG00000036707",
                    "ENSMUSG00000032527",
                    "ENSMUSG00000035268",
                    "ENSMUSG00000085779",
                    "ENSMUSG00000009549",
                    "ENSMUSG00000068798",
                    "ENSMUSG00000020608",
                    "ENSMUSG00000022450",
                    "ENSMUSG00000015579",
                    "ENSMUSG00000073563",
                    "ENSMUSG00000024805",
                    "ENSMUSG00000078622",
                    "ENSMUSG00000082877",
                    "ENSMUSG00000032366",
                    "ENSMUSG00000032370",
                    "ENSMUSG00000098343",
                    "ENSMUSG00000028383",
                    "ENSMUSG00000041115",
                    "ENSMUSG00000025066",
                    "ENSMUSG00000017548",
                    "ENSMUSG00000020309",
                    "ENSMUSG00000064348",
                    "ENSMUSG00000094483",
                    "ENSMUSG00000027709",
                    "ENSMUSG00000024498",
                    "ENSMUSG00000022965",
                    "ENSMUSG00000022884",
                    "ENSMUSG00000015733",
                    "ENSMUSG00000027099",
                    "ENSMUSG00000025220",
                    "ENSMUSG00000035027",
                    "ENSMUSG00000034042",
                    "ENSMUSG00000052812",
                    "ENSMUSG00000037685",
                    "ENSMUSG00000024290",
                    "ENSMUSG00000026526",
                    "ENSMUSG00000022951",
                    "ENSMUSG00000024132",
                    "ENSMUSG00000029338",
                    "ENSMUSG00000021621",
                    "ENSMUSG00000006005",
                    "ENSMUSG00000034525",
                    "ENSMUSG00000046312",
                    "ENSMUSG00000081929",
                    "ENSMUSG00000073002",
                    "ENSMUSG00000055204",
                    "ENSMUSG00000042115",
                    "ENSMUSG00000023904",
                    "ENSMUSG00000054312",
                    "ENSMUSG00000033295",
                    "ENSMUSG00000060803",
                    "ENSMUSG00000030246",
                    "ENSMUSG00000061474",
                    "ENSMUSG00000096232",
                    "ENSMUSG00000010914",
                    "ENSMUSG00000042699",
                    "ENSMUSG00000017760",
                    "ENSMUSG00000055128",
                    "ENSMUSG00000030869",
                    "ENSMUSG00000031767",
                    "ENSMUSG00000064371",
                    "ENSMUSG00000084314",
                    "ENSMUSG00000027620",
                    "ENSMUSG00000017817",
                    "ENSMUSG00000018395",
                    "ENSMUSG00000002908",
                    "ENSMUSG00000031229",
                    "ENSMUSG00000074656",
                    "ENSMUSG00000039126",
                    "ENSMUSG00000045639",
                    "ENSMUSG00000028847",
                    "ENSMUSG00000033047",
                    "ENSMUSG00000024913"
                ],
                "expression_sum": {
                    "class1": 1988.0,
                    "class2": 50963.0
                },
                "uniprot_ids": [
                    "O89090",
                    "E9Q0S7",
                    "E9Q113",
                    "E9Q2V3",
                    "G3X8Q0"
                ]
            },
            "ENSMUSG00000026003": {
                "id": "ENSMUSG00000026003",
                "type": "Gene",
                "symbol": "Acadl",
                "source": [
                    "ENSMUSG00000015839",
                    "ENSMUSG00000056501",
                    "ENSMUSG00000001280",
                    "ENSMUSG00000019768",
                    "ENSMUSG00000005698",
                    "ENSMUSG00000031885",
                    "ENSMUSG00000015846"
                ],
                "target": [],
                "expression_sum": {
                    "class1": 5318.0,
                    "class2": 6090.0
                },
                "uniprot_ids": [
                    "A0A0R4J083"
                ]
            },
            "ENSMUSG00000007656": {
                "id": "ENSMUSG00000007656",
                "type": "Gene",
                "symbol": "Arpp19",
                "source": [
                    "ENSMUSG00000021250",
                    "ENSMUSG00000031627",
                    "ENSMUSG00000018604",
                    "ENSMUSG00000015579",
                    "ENSMUSG00000019768",
                    "ENSMUSG00000005698",
                    "ENSMUSG00000022641",
                    "ENSMUSG00000029178",
                    "ENSMUSG00000053110",
                    "ENSMUSG00000035632",
                    "ENSMUSG00000022811",
                    "ENSMUSG00000015839",
                    "ENSMUSG00000036461",
                    "ENSMUSG00000005583",
                    "ENSMUSG00000056501",
                    "ENSMUSG00000029249",
                    "ENSMUSG00000001280",
                    "ENSMUSG00000009079",
                    "ENSMUSG00000032035"
                ],
                "target": [],
                "expression_sum": {
                    "class1": 13814.0,
                    "class2": 2924.0
                },
                "uniprot_ids": [
                    "P56212",
                    "E9Q4B9",
                    "E9Q827",
                    "E9PXD7"
                ]
            },
            "ENSMUSG00000000563": {
                "id": "ENSMUSG00000000563",
                "type": "Gene",
                "symbol": "Atp5pb",
                "source": [
                    "ENSMUSG00000037174",
                    "ENSMUSG00000024695",
                    "ENSMUSG00000015605",
                    "ENSMUSG00000021775",
                    "ENSMUSG00000040321",
                    "ENSMUSG00000057469",
                    "ENSMUSG00000015579",
                    "ENSMUSG00000090258",
                    "ENSMUSG00000019768",
                    "ENSMUSG00000031885",
                    "ENSMUSG00000036721",
                    "ENSMUSG00000039191",
                    "ENSMUSG00000031715",
                    "ENSMUSG00000015839",
                    "ENSMUSG00000033943",
                    "ENSMUSG00000068267",
                    "ENSMUSG00000014592",
                    "ENSMUSG00000003923",
                    "ENSMUSG00000001280",
                    "ENSMUSG00000024498"
                ],
                "target": [],
                "expression_sum": {
                    "class1": 18039.0,
                    "class2": 18055.0
                },
                "uniprot_ids": [
                    "Q9CQQ7",
                    "A0A0G2JGX3",
                    "D6RGL7"
                ]
            },
            "ENSMUSG00000020821": {
                "id": "ENSMUSG00000020821",
                "type": "Gene",
                "symbol": "Kif1c",
                "source": [
                    "ENSMUSG00000017724",
                    "ENSMUSG00000021944",
                    "ENSMUSG00000015839",
                    "ENSMUSG00000015579",
                    "ENSMUSG00000001280",
                    "ENSMUSG00000005566",
                    "ENSMUSG00000005583",
                    "ENSMUSG00000019768",
                    "ENSMUSG00000024140",
                    "ENSMUSG00000024498",
                    "ENSMUSG00000031885",
                    "ENSMUSG00000058756",
                    "ENSMUSG00000015605",
                    "ENSMUSG00000029913"
                ],
                "target": [],
                "expression_sum": {
                    "class1": 1817.0,
                    "class2": 1453.0
                },
                "uniprot_ids": [
                    "O35071",
                    "Q3V3Y9",
                    "E9Q9B0"
                ]
            },
            "ENSMUSG00000024913": {
                "id": "ENSMUSG00000024913",
                "type": "Gene",
                "symbol": "Lrp5",
                "source": [
                    "ENSMUSG00000057469",
                    "ENSMUSG00000027490",
                    "ENSMUSG00000001280",
                    "ENSMUSG00000019768",
                    "ENSMUSG00000009079",
                    "ENSMUSG00000068267",
                    "ENSMUSG00000058756"
                ],
                "target": [],
                "expression_sum": {
                    "class1": 5088.0,
                    "class2": 1277.0
                },
                "uniprot_ids": [
                    "Q91VN0",
                    "H3BJH0",
                    "H3BL52",
                    "H3BIY3"
                ]
            },
            "ENSMUSG00000062908": {
                "id": "ENSMUSG00000062908",
                "type": "Gene",
                "symbol": "Acadm",
                "source": [
                    "ENSMUSG00000010175",
                    "ENSMUSG00000015839",
                    "ENSMUSG00000015605",
                    "ENSMUSG00000058239",
                    "ENSMUSG00000056501",
                    "ENSMUSG00000015579",
                    "ENSMUSG00000029249",
                    "ENSMUSG00000001280",
                    "ENSMUSG00000005566",
                    "ENSMUSG00000019768",
                    "ENSMUSG00000005698",
                    "ENSMUSG00000046351",
                    "ENSMUSG00000053110",
                    "ENSMUSG00000058756",
                    "ENSMUSG00000008575",
                    "ENSMUSG00000014592",
                    "ENSMUSG00000021775"
                ],
                "target": [],
                "expression_sum": {
                    "class1": 4130.0,
                    "class2": 4788.0
                },
                "uniprot_ids": [
                    "P45952",
                    "D6RFD7",
                    "D3Z2A5"
                ]
            },
            "ENSMUSG00000062981": {
                "id": "ENSMUSG00000062981",
                "type": "Gene",
                "symbol": "Mrpl42",
                "source": [
                    "ENSMUSG00000024238",
                    "ENSMUSG00000001419",
                    "ENSMUSG00000029135",
                    "ENSMUSG00000010175",
                    "ENSMUSG00000015605",
                    "ENSMUSG00000056501",
                    "ENSMUSG00000015579",
                    "ENSMUSG00000049878",
                    "ENSMUSG00000001280",
                    "ENSMUSG00000019768",
                    "ENSMUSG00000031885",
                    "ENSMUSG00000008575",
                    "ENSMUSG00000039191",
                    "ENSMUSG00000042557",
                    "ENSMUSG00000031715"
                ],
                "target": [],
                "expression_sum": {
                    "class1": 18496.0,
                    "class2": 12399.0
                },
                "uniprot_ids": [
                    "Q9CPV3",
                    "A0A1W2P6M3",
                    "A0A1W2P7E4"
                ]
            },
            "ENSMUSG00000016427": {
                "id": "ENSMUSG00000016427",
                "type": "Gene",
                "symbol": "Ndufa1",
                "source": [
                    "ENSMUSG00000015839",
                    "ENSMUSG00000030256",
                    "ENSMUSG00000027490",
                    "ENSMUSG00000001280",
                    "ENSMUSG00000019768",
                    "ENSMUSG00000031885",
                    "ENSMUSG00000015605",
                    "ENSMUSG00000021775"
                ],
                "target": [],
                "expression_sum": {
                    "class1": 10490.0,
                    "class2": 16150.0
                },
                "uniprot_ids": [
                    "O35683"
                ]
            },
            "ENSMUSG00000019768": {
                "id": "ENSMUSG00000019768",
                "type": "TF",
                "symbol": "Esr1",
                "source": [
                    "ENSMUSG00000021944",
                    "ENSMUSG00000010175",
                    "ENSMUSG00000056501",
                    "ENSMUSG00000030557",
                    "ENSMUSG00000029249",
                    "ENSMUSG00000001280",
                    "ENSMUSG00000005566",
                    "ENSMUSG00000028565",
                    "ENSMUSG00000005698",
                    "ENSMUSG00000015605"
                ],
                "target": [
                    "ENSMUSG00000000538",
                    "ENSMUSG00000054455",
                    "ENSMUSG00000001741",
                    "ENSMUSG00000031403",
                    "ENSMUSG00000064363",
                    "ENSMUSG00000091478",
                    "ENSMUSG00000024228",
                    "ENSMUSG00000020694",
                    "ENSMUSG00000030314",
                    "ENSMUSG00000020307",
                    "ENSMUSG00000043683",
                    "ENSMUSG00000021248",
                    "ENSMUSG00000031885",
                    "ENSMUSG00000021235",
                    "ENSMUSG00000070283",
                    "ENSMUSG00000039640",
                    "ENSMUSG00000042302",
                    "ENSMUSG00000081792",
                    "ENSMUSG00000034903",
                    "ENSMUSG00000030739",
                    "ENSMUSG00000014402",
                    "ENSMUSG00000004044",
                    "ENSMUSG00000081773",
                    "ENSMUSG00000028086",
                    "ENSMUSG00000044221",
                    "ENSMUSG00000035202",
                    "ENSMUSG00000026087",
                    "ENSMUSG00000026626",
                    "ENSMUSG00000031543",
                    "ENSMUSG00000007891",
                    "ENSMUSG00000004789",
                    "ENSMUSG00000080746",
                    "ENSMUSG00000018286",
                    "ENSMUSG00000064349",
                    "ENSMUSG00000080242",
                    "ENSMUSG00000020743",
                    "ENSMUSG00000007656",
                    "ENSMUSG00000027288",
                    "ENSMUSG00000041459",
                    "ENSMUSG00000032314",
                    "ENSMUSG00000008892",
                    "ENSMUSG00000030557",
                    "ENSMUSG00000036097",
                    "ENSMUSG00000026858",
                    "ENSMUSG00000027698",
                    "ENSMUSG00000028229",
                    "ENSMUSG00000030852",
                    "ENSMUSG00000010048",
                    "ENSMUSG00000053329",
                    "ENSMUSG00000081485",
                    "ENSMUSG00000031197",
                    "ENSMUSG00000034981",
                    "ENSMUSG00000032216",
                    "ENSMUSG00000071796",
                    "ENSMUSG00000090066",
                    "ENSMUSG00000026755",
                    "ENSMUSG00000018446",
                    "ENSMUSG00000064302",
                    "ENSMUSG00000029158",
                    "ENSMUSG00000026775",
                    "ENSMUSG00000029392",
                    "ENSMUSG00000044365",
                    "ENSMUSG00000000056",
                    "ENSMUSG00000024962",
                    "ENSMUSG00000010392",
                    "ENSMUSG00000057133",
                    "ENSMUSG00000020949",
                    "ENSMUSG00000039103",
                    "ENSMUSG00000029148",
                    "ENSMUSG00000026158",
                    "ENSMUSG00000020349",
                    "ENSMUSG00000038708",
                    "ENSMUSG00000039515",
                    "ENSMUSG00000031728",
                    "ENSMUSG00000056501",
                    "ENSMUSG00000039741",
                    "ENSMUSG00000026615",
                    "ENSMUSG00000055320",
                    "ENSMUSG00000031698",
                    "ENSMUSG00000021024",
                    "ENSMUSG00000027602",
                    "ENSMUSG00000029167",
                    "ENSMUSG00000061286",
                    "ENSMUSG00000025745",
                    "ENSMUSG00000026414",
                    "ENSMUSG00000049265",
                    "ENSMUSG00000082043",
                    "ENSMUSG00000024304",
                    "ENSMUSG00000006219",
                    "ENSMUSG00000030057",
                    "ENSMUSG00000094249",
                    "ENSMUSG00000025369",
                    "ENSMUSG00000041130",
                    "ENSMUSG00000055322",
                    "ENSMUSG00000006057",
                    "ENSMUSG00000068923",
                    "ENSMUSG00000047284",
                    "ENSMUSG00000032536",
                    "ENSMUSG00000044533",
                    "ENSMUSG00000064345",
                    "ENSMUSG00000019471",
                    "ENSMUSG00000028884",
                    "ENSMUSG00000032648",
                    "ENSMUSG00000026374",
                    "ENSMUSG00000024871",
                    "ENSMUSG00000053929",
                    "ENSMUSG00000096842",
                    "ENSMUSG00000026123",
                    "ENSMUSG00000034673",
                    "ENSMUSG00000024177",
                    "ENSMUSG00000059981",
                    "ENSMUSG00000097652",
                    "ENSMUSG00000005882",
                    "ENSMUSG00000020455",
                    "ENSMUSG00000025208",
                    "ENSMUSG00000060002",
                    "ENSMUSG00000029221",
                    "ENSMUSG00000047419",
                    "ENSMUSG00000025234",
                    "ENSMUSG00000006998",
                    "ENSMUSG00000025616",
                    "ENSMUSG00000020459",
                    "ENSMUSG00000001334",
                    "ENSMUSG00000001415",
                    "ENSMUSG00000058076",
                    "ENSMUSG00000028165",
                    "ENSMUSG00000018765",
                    "ENSMUSG00000015843",
                    "ENSMUSG00000073557",
                    "ENSMUSG00000066637",
                    "ENSMUSG00000060882",
                    "ENSMUSG00000033060",
                    "ENSMUSG00000005610",
                    "ENSMUSG00000021282",
                    "ENSMUSG00000031812",
                    "ENSMUSG00000034520",
                    "ENSMUSG00000064367",
                    "ENSMUSG00000024695",
                    "ENSMUSG00000068206",
                    "ENSMUSG00000028955",
                    "ENSMUSG00000037706",
                    "ENSMUSG00000026701",
                    "ENSMUSG00000037905",
                    "ENSMUSG00000015806",
                    "ENSMUSG00000025271",
                    "ENSMUSG00000022185",
                    "ENSMUSG00000014313",
                    "ENSMUSG00000027668",
                    "ENSMUSG00000002015",
                    "ENSMUSG00000024750",
                    "ENSMUSG00000024845",
                    "ENSMUSG00000008429",
                    "ENSMUSG00000064339",
                    "ENSMUSG00000024298",
                    "ENSMUSG00000053093",
                    "ENSMUSG00000042215",
                    "ENSMUSG00000002500",
                    "ENSMUSG00000028849",
                    "ENSMUSG00000034951",
                    "ENSMUSG00000015149",
                    "ENSMUSG00000040033",
                    "ENSMUSG00000024940",
                    "ENSMUSG00000037197",
                    "ENSMUSG00000006024",
                    "ENSMUSG00000022682",
                    "ENSMUSG00000027552",
                    "ENSMUSG00000025348",
                    "ENSMUSG00000053617",
                    "ENSMUSG00000052512",
                    "ENSMUSG00000020647",
                    "ENSMUSG00000040506",
                    "ENSMUSG00000067925",
                    "ENSMUSG00000019791",
                    "ENSMUSG00000027332",
                    "ENSMUSG00000015850",
                    "ENSMUSG00000025733",
                    "ENSMUSG00000020864",
                    "ENSMUSG00000022214",
                    "ENSMUSG00000022110",
                    "ENSMUSG00000004356",
                    "ENSMUSG00000037916",
                    "ENSMUSG00000082641",
                    "ENSMUSG00000030779",
                    "ENSMUSG00000029708",
                    "ENSMUSG00000029156",
                    "ENSMUSG00000078572",
                    "ENSMUSG00000040321",
                    "ENSMUSG00000025437",
                    "ENSMUSG00000045316",
                    "ENSMUSG00000000628",
                    "ENSMUSG00000027805",
                    "ENSMUSG00000028565",
                    "ENSMUSG00000040771",
                    "ENSMUSG00000021870",
                    "ENSMUSG00000032724",
                    "ENSMUSG00000033161",
                    "ENSMUSG00000003072",
                    "ENSMUSG00000037349",
                    "ENSMUSG00000033943",
                    "ENSMUSG00000008730",
                    "ENSMUSG00000000168",
                    "ENSMUSG00000048661",
                    "ENSMUSG00000029249",
                    "ENSMUSG00000026113",
                    "ENSMUSG00000027861",
                    "ENSMUSG00000030061",
                    "ENSMUSG00000021906",
                    "ENSMUSG00000081278",
                    "ENSMUSG00000021313",
                    "ENSMUSG00000025241",
                    "ENSMUSG00000059430",
                    "ENSMUSG00000015112",
                    "ENSMUSG00000075054",
                    "ENSMUSG00000021453",
                    "ENSMUSG00000018433",
                    "ENSMUSG00000062981",
                    "ENSMUSG00000030754",
                    "ENSMUSG00000002496",
                    "ENSMUSG00000064342",
                    "ENSMUSG00000002233",
                    "ENSMUSG00000074884",
                    "ENSMUSG00000031983",
                    "ENSMUSG00000038028",
                    "ENSMUSG00000003531",
                    "ENSMUSG00000030835",
                    "ENSMUSG00000020523",
                    "ENSMUSG00000030470",
                    "ENSMUSG00000035325",
                    "ENSMUSG00000055239",
                    "ENSMUSG00000027381",
                    "ENSMUSG00000035203",
                    "ENSMUSG00000028247",
                    "ENSMUSG00000027823",
                    "ENSMUSG00000029094",
                    "ENSMUSG00000008822",
                    "ENSMUSG00000002228",
                    "ENSMUSG00000045886",
                    "ENSMUSG00000004069",
                    "ENSMUSG00000040785",
                    "ENSMUSG00000022336",
                    "ENSMUSG00000019797",
                    "ENSMUSG00000058600",
                    "ENSMUSG00000064351",
                    "ENSMUSG00000040850",
                    "ENSMUSG00000046756",
                    "ENSMUSG00000029020",
                    "ENSMUSG00000028173",
                    "ENSMUSG00000027706",
                    "ENSMUSG00000022131",
                    "ENSMUSG00000049643",
                    "ENSMUSG00000042529",
                    "ENSMUSG00000038375",
                    "ENSMUSG00000022452",
                    "ENSMUSG00000039509",
                    "ENSMUSG00000060600",
                    "ENSMUSG00000026283",
                    "ENSMUSG00000034898",
                    "ENSMUSG00000022228",
                    "ENSMUSG00000038393",
                    "ENSMUSG00000024143",
                    "ENSMUSG00000009647",
                    "ENSMUSG00000025374",
                    "ENSMUSG00000019577",
                    "ENSMUSG00000033429",
                    "ENSMUSG00000002010",
                    "ENSMUSG00000020593",
                    "ENSMUSG00000010064",
                    "ENSMUSG00000058997",
                    "ENSMUSG00000060904",
                    "ENSMUSG00000028273",
                    "ENSMUSG00000019158",
                    "ENSMUSG00000021200",
                    "ENSMUSG00000008604",
                    "ENSMUSG00000058594",
                    "ENSMUSG00000021591",
                    "ENSMUSG00000048732",
                    "ENSMUSG00000032786",
                    "ENSMUSG00000050856",
                    "ENSMUSG00000025856",
                    "ENSMUSG00000038762",
                    "ENSMUSG00000053768",
                    "ENSMUSG00000082978",
                    "ENSMUSG00000025781",
                    "ENSMUSG00000033781",
                    "ENSMUSG00000018796",
                    "ENSMUSG00000008690",
                    "ENSMUSG00000032518",
                    "ENSMUSG00000014195",
                    "ENSMUSG00000031461",
                    "ENSMUSG00000049422",
                    "ENSMUSG00000087679",
                    "ENSMUSG00000022403",
                    "ENSMUSG00000026409",
                    "ENSMUSG00000059734",
                    "ENSMUSG00000022223",
                    "ENSMUSG00000022969",
                    "ENSMUSG00000095687",
                    "ENSMUSG00000091537",
                    "ENSMUSG00000078670",
                    "ENSMUSG00000062070",
                    "ENSMUSG00000059878",
                    "ENSMUSG00000047905",
                    "ENSMUSG00000053964",
                    "ENSMUSG00000037904",
                    "ENSMUSG00000038764",
                    "ENSMUSG00000019173",
                    "ENSMUSG00000022553",
                    "ENSMUSG00000091898",
                    "ENSMUSG00000009079",
                    "ENSMUSG00000007739",
                    "ENSMUSG00000028633",
                    "ENSMUSG00000028394",
                    "ENSMUSG00000025968",
                    "ENSMUSG00000002844",
                    "ENSMUSG00000064370",
                    "ENSMUSG00000020133",
                    "ENSMUSG00000020821",
                    "ENSMUSG00000097626",
                    "ENSMUSG00000044033",
                    "ENSMUSG00000024140",
                    "ENSMUSG00000028691",
                    "ENSMUSG00000028399",
                    "ENSMUSG00000061477",
                    "ENSMUSG00000024346",
                    "ENSMUSG00000037805",
                    "ENSMUSG00000046201",
                    "ENSMUSG00000017119",
                    "ENSMUSG00000028911",
                    "ENSMUSG00000031701",
                    "ENSMUSG00000020738",
                    "ENSMUSG00000075701",
                    "ENSMUSG00000029467",
                    "ENSMUSG00000022545",
                    "ENSMUSG00000038022",
                    "ENSMUSG00000044751",
                    "ENSMUSG00000022193",
                    "ENSMUSG00000011148",
                    "ENSMUSG00000062580",
                    "ENSMUSG00000026817",
                    "ENSMUSG00000022031",
                    "ENSMUSG00000026187",
                    "ENSMUSG00000021039",
                    "ENSMUSG00000021218",
                    "ENSMUSG00000062270",
                    "ENSMUSG00000062397",
                    "ENSMUSG00000028937",
                    "ENSMUSG00000015357",
                    "ENSMUSG00000034566",
                    "ENSMUSG00000024985",
                    "ENSMUSG00000021243",
                    "ENSMUSG00000013593",
                    "ENSMUSG00000068267",
                    "ENSMUSG00000021417",
                    "ENSMUSG00000042293",
                    "ENSMUSG00000062373",
                    "ENSMUSG00000029432",
                    "ENSMUSG00000038520",
                    "ENSMUSG00000038058",
                    "ENSMUSG00000024963",
                    "ENSMUSG00000020034",
                    "ENSMUSG00000058569",
                    "ENSMUSG00000022215",
                    "ENSMUSG00000009545",
                    "ENSMUSG00000040964",
                    "ENSMUSG00000020988",
                    "ENSMUSG00000020530",
                    "ENSMUSG00000039670",
                    "ENSMUSG00000032897",
                    "ENSMUSG00000018340",
                    "ENSMUSG00000022111",
                    "ENSMUSG00000037236",
                    "ENSMUSG00000038650",
                    "ENSMUSG00000055053",
                    "ENSMUSG00000051391",
                    "ENSMUSG00000058056",
                    "ENSMUSG00000064369",
                    "ENSMUSG00000047746",
                    "ENSMUSG00000025509",
                    "ENSMUSG00000029655",
                    "ENSMUSG00000036880",
                    "ENSMUSG00000053819",
                    "ENSMUSG00000022551",
                    "ENSMUSG00000037110",
                    "ENSMUSG00000029571",
                    "ENSMUSG00000005779",
                    "ENSMUSG00000082768",
                    "ENSMUSG00000020440",
                    "ENSMUSG00000023094",
                    "ENSMUSG00000002455",
                    "ENSMUSG00000030847",
                    "ENSMUSG00000008575",
                    "ENSMUSG00000022400",
                    "ENSMUSG00000035173",
                    "ENSMUSG00000004558",
                    "ENSMUSG00000067722",
                    "ENSMUSG00000090258",
                    "ENSMUSG00000019787",
                    "ENSMUSG00000026003",
                    "ENSMUSG00000040209",
                    "ENSMUSG00000038462",
                    "ENSMUSG00000022797",
                    "ENSMUSG00000067121",
                    "ENSMUSG00000034936",
                    "ENSMUSG00000082195",
                    "ENSMUSG00000074994",
                    "ENSMUSG00000026610",
                    "ENSMUSG00000024099",
                    "ENSMUSG00000061311",
                    "ENSMUSG00000037656",
                    "ENSMUSG00000028295",
                    "ENSMUSG00000045694",
                    "ENSMUSG00000032563",
                    "ENSMUSG00000028053",
                    "ENSMUSG00000014294",
                    "ENSMUSG00000034875",
                    "ENSMUSG00000021939",
                    "ENSMUSG00000028104",
                    "ENSMUSG00000045838",
                    "ENSMUSG00000047547",
                    "ENSMUSG00000090935",
                    "ENSMUSG00000027777",
                    "ENSMUSG00000022020",
                    "ENSMUSG00000040481",
                    "ENSMUSG00000045302",
                    "ENSMUSG00000002661",
                    "ENSMUSG00000018501",
                    "ENSMUSG00000040269",
                    "ENSMUSG00000003444",
                    "ENSMUSG00000044938",
                    "ENSMUSG00000015837",
                    "ENSMUSG00000029823",
                    "ENSMUSG00000031382",
                    "ENSMUSG00000002028",
                    "ENSMUSG00000097414",
                    "ENSMUSG00000081179",
                    "ENSMUSG00000034751",
                    "ENSMUSG00000022707",
                    "ENSMUSG00000029629",
                    "ENSMUSG00000027984",
                    "ENSMUSG00000031353",
                    "ENSMUSG00000024097",
                    "ENSMUSG00000022354",
                    "ENSMUSG00000023951",
                    "ENSMUSG00000005354",
                    "ENSMUSG00000020477",
                    "ENSMUSG00000026623",
                    "ENSMUSG00000024811",
                    "ENSMUSG00000035158",
                    "ENSMUSG00000080848",
                    "ENSMUSG00000029198",
                    "ENSMUSG00000030747",
                    "ENSMUSG00000020496",
                    "ENSMUSG00000023087",
                    "ENSMUSG00000057469",
                    "ENSMUSG00000004347",
                    "ENSMUSG00000005469",
                    "ENSMUSG00000026803",
                    "ENSMUSG00000005299",
                    "ENSMUSG00000052539",
                    "ENSMUSG00000073684",
                    "ENSMUSG00000029152",
                    "ENSMUSG00000051048",
                    "ENSMUSG00000022519",
                    "ENSMUSG00000051331",
                    "ENSMUSG00000025260",
                    "ENSMUSG00000016528",
                    "ENSMUSG00000024038",
                    "ENSMUSG00000025190",
                    "ENSMUSG00000033065",
                    "ENSMUSG00000057738",
                    "ENSMUSG00000042010",
                    "ENSMUSG00000083186",
                    "ENSMUSG00000030850",
                    "ENSMUSG00000014606",
                    "ENSMUSG00000036893",
                    "ENSMUSG00000026260",
                    "ENSMUSG00000055839",
                    "ENSMUSG00000041921",
                    "ENSMUSG00000020612",
                    "ENSMUSG00000039891",
                    "ENSMUSG00000048490",
                    "ENSMUSG00000031059",
                    "ENSMUSG00000024359",
                    "ENSMUSG00000019943",
                    "ENSMUSG00000039063",
                    "ENSMUSG00000032883",
                    "ENSMUSG00000032263",
                    "ENSMUSG00000032435",
                    "ENSMUSG00000038690",
                    "ENSMUSG00000028954",
                    "ENSMUSG00000039067",
                    "ENSMUSG00000003865",
                    "ENSMUSG00000014956",
                    "ENSMUSG00000024539",
                    "ENSMUSG00000020460",
                    "ENSMUSG00000023004",
                    "ENSMUSG00000083932",
                    "ENSMUSG00000041028",
                    "ENSMUSG00000028700",
                    "ENSMUSG00000026112",
                    "ENSMUSG00000021772",
                    "ENSMUSG00000018566",
                    "ENSMUSG00000020321",
                    "ENSMUSG00000059447",
                    "ENSMUSG00000024277",
                    "ENSMUSG00000024120",
                    "ENSMUSG00000029614",
                    "ENSMUSG00000035296",
                    "ENSMUSG00000053199",
                    "ENSMUSG00000001508",
                    "ENSMUSG00000036902",
                    "ENSMUSG00000013419",
                    "ENSMUSG00000031880",
                    "ENSMUSG00000038615",
                    "ENSMUSG00000029064",
                    "ENSMUSG00000045210",
                    "ENSMUSG00000025130",
                    "ENSMUSG00000023990",
                    "ENSMUSG00000025290",
                    "ENSMUSG00000021917",
                    "ENSMUSG00000009563",
                    "ENSMUSG00000033918",
                    "ENSMUSG00000028443",
                    "ENSMUSG00000024236",
                    "ENSMUSG00000063904",
                    "ENSMUSG00000061518",
                    "ENSMUSG00000034252",
                    "ENSMUSG00000033400",
                    "ENSMUSG00000036570",
                    "ENSMUSG00000001056",
                    "ENSMUSG00000020220",
                    "ENSMUSG00000025935",
                    "ENSMUSG00000003299",
                    "ENSMUSG00000039911",
                    "ENSMUSG00000040351",
                    "ENSMUSG00000021549",
                    "ENSMUSG00000051223",
                    "ENSMUSG00000027091",
                    "ENSMUSG00000050812",
                    "ENSMUSG00000027452",
                    "ENSMUSG00000038827",
                    "ENSMUSG00000025869",
                    "ENSMUSG00000051469",
                    "ENSMUSG00000029723",
                    "ENSMUSG00000036371",
                    "ENSMUSG00000058135",
                    "ENSMUSG00000070493",
                    "ENSMUSG00000030335",
                    "ENSMUSG00000021748",
                    "ENSMUSG00000071369",
                    "ENSMUSG00000025006",
                    "ENSMUSG00000038695",
                    "ENSMUSG00000026456",
                    "ENSMUSG00000027184",
                    "ENSMUSG00000028116",
                    "ENSMUSG00000068921",
                    "ENSMUSG00000001441",
                    "ENSMUSG00000059714",
                    "ENSMUSG00000034220",
                    "ENSMUSG00000070713",
                    "ENSMUSG00000031231",
                    "ENSMUSG00000084111",
                    "ENSMUSG00000026895",
                    "ENSMUSG00000092341",
                    "ENSMUSG00000026179",
                    "ENSMUSG00000033059",
                    "ENSMUSG00000002342",
                    "ENSMUSG00000000088",
                    "ENSMUSG00000031626",
                    "ENSMUSG00000048154",
                    "ENSMUSG00000042462",
                    "ENSMUSG00000026393",
                    "ENSMUSG00000009995",
                    "ENSMUSG00000025393",
                    "ENSMUSG00000048279",
                    "ENSMUSG00000026930",
                    "ENSMUSG00000026229",
                    "ENSMUSG00000031641",
                    "ENSMUSG00000001435",
                    "ENSMUSG00000038967",
                    "ENSMUSG00000025410",
                    "ENSMUSG00000036112",
                    "ENSMUSG00000017485",
                    "ENSMUSG00000024588",
                    "ENSMUSG00000023249",
                    "ENSMUSG00000068874",
                    "ENSMUSG00000006299",
                    "ENSMUSG00000032340",
                    "ENSMUSG00000002416",
                    "ENSMUSG00000036197",
                    "ENSMUSG00000025791",
                    "ENSMUSG00000066043",
                    "ENSMUSG00000022404",
                    "ENSMUSG00000067058",
                    "ENSMUSG00000021460",
                    "ENSMUSG00000036867",
                    "ENSMUSG00000022477",
                    "ENSMUSG00000022337",
                    "ENSMUSG00000036698",
                    "ENSMUSG00000019988",
                    "ENSMUSG00000041144",
                    "ENSMUSG00000042298",
                    "ENSMUSG00000028649",
                    "ENSMUSG00000052146",
                    "ENSMUSG00000012405",
                    "ENSMUSG00000030695",
                    "ENSMUSG00000064341",
                    "ENSMUSG00000024049",
                    "ENSMUSG00000013539",
                    "ENSMUSG00000001366",
                    "ENSMUSG00000035242",
                    "ENSMUSG00000043716",
                    "ENSMUSG00000050608",
                    "ENSMUSG00000021706",
                    "ENSMUSG00000036160",
                    "ENSMUSG00000027206",
                    "ENSMUSG00000074182",
                    "ENSMUSG00000019851",
                    "ENSMUSG00000059208",
                    "ENSMUSG00000022016",
                    "ENSMUSG00000022503",
                    "ENSMUSG00000026150",
                    "ENSMUSG00000038412",
                    "ENSMUSG00000020230",
                    "ENSMUSG00000020390",
                    "ENSMUSG00000021687",
                    "ENSMUSG00000036323",
                    "ENSMUSG00000039474",
                    "ENSMUSG00000038370",
                    "ENSMUSG00000044475",
                    "ENSMUSG00000029128",
                    "ENSMUSG00000099250",
                    "ENSMUSG00000013736",
                    "ENSMUSG00000055737",
                    "ENSMUSG00000062604",
                    "ENSMUSG00000071748",
                    "ENSMUSG00000025607",
                    "ENSMUSG00000063870",
                    "ENSMUSG00000032902",
                    "ENSMUSG00000031633",
                    "ENSMUSG00000063694",
                    "ENSMUSG00000041235",
                    "ENSMUSG00000057580",
                    "ENSMUSG00000020368",
                    "ENSMUSG00000039473",
                    "ENSMUSG00000034610",
                    "ENSMUSG00000042772",
                    "ENSMUSG00000019795",
                    "ENSMUSG00000040385",
                    "ENSMUSG00000016252",
                    "ENSMUSG00000030793",
                    "ENSMUSG00000026586",
                    "ENSMUSG00000035776",
                    "ENSMUSG00000022889",
                    "ENSMUSG00000021281",
                    "ENSMUSG00000021843",
                    "ENSMUSG00000039611",
                    "ENSMUSG00000038239",
                    "ENSMUSG00000067274",
                    "ENSMUSG00000063787",
                    "ENSMUSG00000063882",
                    "ENSMUSG00000019773",
                    "ENSMUSG00000040928",
                    "ENSMUSG00000031010",
                    "ENSMUSG00000026028",
                    "ENSMUSG00000039910",
                    "ENSMUSG00000028745",
                    "ENSMUSG00000038886",
                    "ENSMUSG00000035152",
                    "ENSMUSG00000053702",
                    "ENSMUSG00000020267",
                    "ENSMUSG00000036550",
                    "ENSMUSG00000022051",
                    "ENSMUSG00000006818",
                    "ENSMUSG00000026860",
                    "ENSMUSG00000034621",
                    "ENSMUSG00000001017",
                    "ENSMUSG00000042541",
                    "ENSMUSG00000021375",
                    "ENSMUSG00000029545",
                    "ENSMUSG00000058793",
                    "ENSMUSG00000033039",
                    "ENSMUSG00000032519",
                    "ENSMUSG00000056608",
                    "ENSMUSG00000032399",
                    "ENSMUSG00000059586",
                    "ENSMUSG00000078566",
                    "ENSMUSG00000002102",
                    "ENSMUSG00000002343",
                    "ENSMUSG00000039652",
                    "ENSMUSG00000024613",
                    "ENSMUSG00000070544",
                    "ENSMUSG00000054889",
                    "ENSMUSG00000046351",
                    "ENSMUSG00000053838",
                    "ENSMUSG00000024953",
                    "ENSMUSG00000083793",
                    "ENSMUSG00000022982",
                    "ENSMUSG00000050323",
                    "ENSMUSG00000026032",
                    "ENSMUSG00000068479",
                    "ENSMUSG00000021868",
                    "ENSMUSG00000028312",
                    "ENSMUSG00000037376",
                    "ENSMUSG00000028452",
                    "ENSMUSG00000028156",
                    "ENSMUSG00000064360",
                    "ENSMUSG00000021144",
                    "ENSMUSG00000039987",
                    "ENSMUSG00000041548",
                    "ENSMUSG00000070802",
                    "ENSMUSG00000035478",
                    "ENSMUSG00000028333",
                    "ENSMUSG00000041058",
                    "ENSMUSG00000036202",
                    "ENSMUSG00000027257",
                    "ENSMUSG00000020372",
                    "ENSMUSG00000044573",
                    "ENSMUSG00000032523",
                    "ENSMUSG00000028834",
                    "ENSMUSG00000031352",
                    "ENSMUSG00000073838",
                    "ENSMUSG00000041870",
                    "ENSMUSG00000005102",
                    "ENSMUSG00000039536",
                    "ENSMUSG00000000740",
                    "ENSMUSG00000040752",
                    "ENSMUSG00000023723",
                    "ENSMUSG00000003934",
                    "ENSMUSG00000030884",
                    "ENSMUSG00000025466",
                    "ENSMUSG00000042997",
                    "ENSMUSG00000092274",
                    "ENSMUSG00000022000",
                    "ENSMUSG00000023025",
                    "ENSMUSG00000028419",
                    "ENSMUSG00000032051",
                    "ENSMUSG00000034040",
                    "ENSMUSG00000038143",
                    "ENSMUSG00000023460",
                    "ENSMUSG00000002767",
                    "ENSMUSG00000034994",
                    "ENSMUSG00000023971",
                    "ENSMUSG00000030541",
                    "ENSMUSG00000070003",
                    "ENSMUSG00000031438",
                    "ENSMUSG00000081100",
                    "ENSMUSG00000026208",
                    "ENSMUSG00000036052",
                    "ENSMUSG00000015217",
                    "ENSMUSG00000024213",
                    "ENSMUSG00000030652",
                    "ENSMUSG00000046314",
                    "ENSMUSG00000027238",
                    "ENSMUSG00000047388",
                    "ENSMUSG00000060923",
                    "ENSMUSG00000061306",
                    "ENSMUSG00000017686",
                    "ENSMUSG00000013936",
                    "ENSMUSG00000057406",
                    "ENSMUSG00000069089",
                    "ENSMUSG00000020415",
                    "ENSMUSG00000026811",
                    "ENSMUSG00000006800",
                    "ENSMUSG00000059741",
                    "ENSMUSG00000030255",
                    "ENSMUSG00000002957",
                    "ENSMUSG00000024073",
                    "ENSMUSG00000028282",
                    "ENSMUSG00000020430",
                    "ENSMUSG00000009112",
                    "ENSMUSG00000039130",
                    "ENSMUSG00000078486",
                    "ENSMUSG00000033565",
                    "ENSMUSG00000027180",
                    "ENSMUSG00000016308",
                    "ENSMUSG00000027422",
                    "ENSMUSG00000050875",
                    "ENSMUSG00000050043",
                    "ENSMUSG00000031843",
                    "ENSMUSG00000018882",
                    "ENSMUSG00000028470",
                    "ENSMUSG00000027671",
                    "ENSMUSG00000000787",
                    "ENSMUSG00000070319",
                    "ENSMUSG00000031808",
                    "ENSMUSG00000015214",
                    "ENSMUSG00000037710",
                    "ENSMUSG00000024457",
                    "ENSMUSG00000024187",
                    "ENSMUSG00000019362",
                    "ENSMUSG00000026308",
                    "ENSMUSG00000016427",
                    "ENSMUSG00000024205",
                    "ENSMUSG00000027293",
                    "ENSMUSG00000001127",
                    "ENSMUSG00000015656",
                    "ENSMUSG00000025868",
                    "ENSMUSG00000027832",
                    "ENSMUSG00000040520",
                    "ENSMUSG00000058756",
                    "ENSMUSG00000025651",
                    "ENSMUSG00000033102",
                    "ENSMUSG00000028603",
                    "ENSMUSG00000004268",
                    "ENSMUSG00000025085",
                    "ENSMUSG00000065979",
                    "ENSMUSG00000028261",
                    "ENSMUSG00000024077",
                    "ENSMUSG00000025068",
                    "ENSMUSG00000045103",
                    "ENSMUSG00000028675",
                    "ENSMUSG00000032042",
                    "ENSMUSG00000061032",
                    "ENSMUSG00000023089",
                    "ENSMUSG00000058927",
                    "ENSMUSG00000030996",
                    "ENSMUSG00000078851",
                    "ENSMUSG00000073676",
                    "ENSMUSG00000032060",
                    "ENSMUSG00000023456",
                    "ENSMUSG00000046573",
                    "ENSMUSG00000048731",
                    "ENSMUSG00000022010",
                    "ENSMUSG00000025810",
                    "ENSMUSG00000027673",
                    "ENSMUSG00000059811",
                    "ENSMUSG00000024587",
                    "ENSMUSG00000024121",
                    "ENSMUSG00000083911",
                    "ENSMUSG00000015568",
                    "ENSMUSG00000031504",
                    "ENSMUSG00000031448",
                    "ENSMUSG00000061904",
                    "ENSMUSG00000074582",
                    "ENSMUSG00000029454",
                    "ENSMUSG00000015605",
                    "ENSMUSG00000020485",
                    "ENSMUSG00000006740",
                    "ENSMUSG00000024197",
                    "ENSMUSG00000007777",
                    "ENSMUSG00000019432",
                    "ENSMUSG00000022914",
                    "ENSMUSG00000026491",
                    "ENSMUSG00000018845",
                    "ENSMUSG00000039148",
                    "ENSMUSG00000089838",
                    "ENSMUSG00000044791",
                    "ENSMUSG00000020741",
                    "ENSMUSG00000037266",
                    "ENSMUSG00000037339",
                    "ENSMUSG00000030982",
                    "ENSMUSG00000028676",
                    "ENSMUSG00000027679",
                    "ENSMUSG00000022521",
                    "ENSMUSG00000026576",
                    "ENSMUSG00000041881",
                    "ENSMUSG00000020691",
                    "ENSMUSG00000021764",
                    "ENSMUSG00000038763",
                    "ENSMUSG00000021209",
                    "ENSMUSG00000038535",
                    "ENSMUSG00000043445",
                    "ENSMUSG00000039648",
                    "ENSMUSG00000020163",
                    "ENSMUSG00000001767",
                    "ENSMUSG00000039956",
                    "ENSMUSG00000009863",
                    "ENSMUSG00000033411",
                    "ENSMUSG00000031770",
                    "ENSMUSG00000038400",
                    "ENSMUSG00000031715",
                    "ENSMUSG00000056900",
                    "ENSMUSG00000032238",
                    "ENSMUSG00000062210",
                    "ENSMUSG00000060376",
                    "ENSMUSG00000014592",
                    "ENSMUSG00000090150",
                    "ENSMUSG00000017428",
                    "ENSMUSG00000062694",
                    "ENSMUSG00000040661",
                    "ENSMUSG00000020153",
                    "ENSMUSG00000093674",
                    "ENSMUSG00000083563",
                    "ENSMUSG00000040760",
                    "ENSMUSG00000036452",
                    "ENSMUSG00000031818",
                    "ENSMUSG00000031782",
                    "ENSMUSG00000048486",
                    "ENSMUSG00000055148",
                    "ENSMUSG00000032511",
                    "ENSMUSG00000044894",
                    "ENSMUSG00000007097",
                    "ENSMUSG00000029404",
                    "ENSMUSG00000048578",
                    "ENSMUSG00000021134",
                    "ENSMUSG00000022820",
                    "ENSMUSG00000037234",
                    "ENSMUSG00000002129",
                    "ENSMUSG00000042148",
                    "ENSMUSG00000034295",
                    "ENSMUSG00000031783",
                    "ENSMUSG00000022594",
                    "ENSMUSG00000063229",
                    "ENSMUSG00000027076",
                    "ENSMUSG00000005417",
                    "ENSMUSG00000021577",
                    "ENSMUSG00000022994",
                    "ENSMUSG00000029209",
                    "ENSMUSG00000038936",
                    "ENSMUSG00000029776",
                    "ENSMUSG00000029135",
                    "ENSMUSG00000025898",
                    "ENSMUSG00000036427",
                    "ENSMUSG00000026849",
                    "ENSMUSG00000070867",
                    "ENSMUSG00000038417",
                    "ENSMUSG00000046329",
                    "ENSMUSG00000045160",
                    "ENSMUSG00000018040",
                    "ENSMUSG00000024942",
                    "ENSMUSG00000022698",
                    "ENSMUSG00000027371",
                    "ENSMUSG00000037075",
                    "ENSMUSG00000029036",
                    "ENSMUSG00000040407",
                    "ENSMUSG00000030647",
                    "ENSMUSG00000001424",
                    "ENSMUSG00000073664",
                    "ENSMUSG00000025545",
                    "ENSMUSG00000026888",
                    "ENSMUSG00000022094",
                    "ENSMUSG00000052712",
                    "ENSMUSG00000030087",
                    "ENSMUSG00000019979",
                    "ENSMUSG00000006333",
                    "ENSMUSG00000026568",
                    "ENSMUSG00000028328",
                    "ENSMUSG00000005683",
                    "ENSMUSG00000046516",
                    "ENSMUSG00000037960",
                    "ENSMUSG00000058655",
                    "ENSMUSG00000039239",
                    "ENSMUSG00000022961",
                    "ENSMUSG00000020085",
                    "ENSMUSG00000037443",
                    "ENSMUSG00000031536",
                    "ENSMUSG00000032348",
                    "ENSMUSG00000029913",
                    "ENSMUSG00000039218",
                    "ENSMUSG00000029594",
                    "ENSMUSG00000042834",
                    "ENSMUSG00000032352",
                    "ENSMUSG00000003033",
                    "ENSMUSG00000052214",
                    "ENSMUSG00000033228",
                    "ENSMUSG00000046032",
                    "ENSMUSG00000064181",
                    "ENSMUSG00000032826",
                    "ENSMUSG00000028266",
                    "ENSMUSG00000033594",
                    "ENSMUSG00000047371",
                    "ENSMUSG00000020737",
                    "ENSMUSG00000031232",
                    "ENSMUSG00000001419",
                    "ENSMUSG00000038583",
                    "ENSMUSG00000022765",
                    "ENSMUSG00000047260",
                    "ENSMUSG00000004460",
                    "ENSMUSG00000042198",
                    "ENSMUSG00000035674",
                    "ENSMUSG00000022261",
                    "ENSMUSG00000009927",
                    "ENSMUSG00000026571",
                    "ENSMUSG00000025465",
                    "ENSMUSG00000042605",
                    "ENSMUSG00000028919",
                    "ENSMUSG00000027423",
                    "ENSMUSG00000001833",
                    "ENSMUSG00000025204",
                    "ENSMUSG00000023055",
                    "ENSMUSG00000098178",
                    "ENSMUSG00000031672",
                    "ENSMUSG00000030172",
                    "ENSMUSG00000059895",
                    "ENSMUSG00000016319",
                    "ENSMUSG00000024780",
                    "ENSMUSG00000021488",
                    "ENSMUSG00000034290",
                    "ENSMUSG00000023150",
                    "ENSMUSG00000038718",
                    "ENSMUSG00000063800",
                    "ENSMUSG00000030298",
                    "ENSMUSG00000025795",
                    "ENSMUSG00000097729",
                    "ENSMUSG00000017404",
                    "ENSMUSG00000020190",
                    "ENSMUSG00000021944",
                    "ENSMUSG00000028648",
                    "ENSMUSG00000029623",
                    "ENSMUSG00000004535",
                    "ENSMUSG00000002949",
                    "ENSMUSG00000014633",
                    "ENSMUSG00000007892",
                    "ENSMUSG00000003380",
                    "ENSMUSG00000074064",
                    "ENSMUSG00000060938",
                    "ENSMUSG00000010406",
                    "ENSMUSG00000025428",
                    "ENSMUSG00000036214",
                    "ENSMUSG00000024538",
                    "ENSMUSG00000057335",
                    "ENSMUSG00000021585",
                    "ENSMUSG00000029017",
                    "ENSMUSG00000030016",
                    "ENSMUSG00000013921",
                    "ENSMUSG00000041426",
                    "ENSMUSG00000031634",
                    "ENSMUSG00000020849",
                    "ENSMUSG00000028343",
                    "ENSMUSG00000081233",
                    "ENSMUSG00000068706",
                    "ENSMUSG00000002748",
                    "ENSMUSG00000021238",
                    "ENSMUSG00000041297",
                    "ENSMUSG00000026766",
                    "ENSMUSG00000024924",
                    "ENSMUSG00000039989",
                    "ENSMUSG00000042744",
                    "ENSMUSG00000031987",
                    "ENSMUSG00000038005",
                    "ENSMUSG00000043311",
                    "ENSMUSG00000038970",
                    "ENSMUSG00000020171",
                    "ENSMUSG00000005483",
                    "ENSMUSG00000055302",
                    "ENSMUSG00000028013",
                    "ENSMUSG00000039089",
                    "ENSMUSG00000053898",
                    "ENSMUSG00000060038",
                    "ENSMUSG00000002831",
                    "ENSMUSG00000030409",
                    "ENSMUSG00000062691",
                    "ENSMUSG00000040648",
                    "ENSMUSG00000001105",
                    "ENSMUSG00000083820",
                    "ENSMUSG00000025221",
                    "ENSMUSG00000029632",
                    "ENSMUSG00000026207",
                    "ENSMUSG00000036620",
                    "ENSMUSG00000022641",
                    "ENSMUSG00000054766",
                    "ENSMUSG00000052337",
                    "ENSMUSG00000083307",
                    "ENSMUSG00000031367",
                    "ENSMUSG00000028223",
                    "ENSMUSG00000027087",
                    "ENSMUSG00000026342",
                    "ENSMUSG00000074466",
                    "ENSMUSG00000001794",
                    "ENSMUSG00000031791",
                    "ENSMUSG00000048264",
                    "ENSMUSG00000027478",
                    "ENSMUSG00000030399",
                    "ENSMUSG00000071644",
                    "ENSMUSG00000026027",
                    "ENSMUSG00000053012",
                    "ENSMUSG00000020483",
                    "ENSMUSG00000009090",
                    "ENSMUSG00000081974",
                    "ENSMUSG00000032679",
                    "ENSMUSG00000026878",
                    "ENSMUSG00000098274",
                    "ENSMUSG00000024055",
                    "ENSMUSG00000029518",
                    "ENSMUSG00000006304",
                    "ENSMUSG00000031320",
                    "ENSMUSG00000032959",
                    "ENSMUSG00000028964",
                    "ENSMUSG00000042032",
                    "ENSMUSG00000002728",
                    "ENSMUSG00000018893",
                    "ENSMUSG00000027406",
                    "ENSMUSG00000019863",
                    "ENSMUSG00000060032",
                    "ENSMUSG00000028756",
                    "ENSMUSG00000063273",
                    "ENSMUSG00000000811",
                    "ENSMUSG00000059974",
                    "ENSMUSG00000026927",
                    "ENSMUSG00000037712",
                    "ENSMUSG00000003955",
                    "ENSMUSG00000028234",
                    "ENSMUSG00000019179",
                    "ENSMUSG00000018474",
                    "ENSMUSG00000051495",
                    "ENSMUSG00000079477",
                    "ENSMUSG00000022841",
                    "ENSMUSG00000042682",
                    "ENSMUSG00000066613",
                    "ENSMUSG00000025732",
                    "ENSMUSG00000031575",
                    "ENSMUSG00000037152",
                    "ENSMUSG00000023010",
                    "ENSMUSG00000036278",
                    "ENSMUSG00000022391",
                    "ENSMUSG00000028788",
                    "ENSMUSG00000062169",
                    "ENSMUSG00000067369",
                    "ENSMUSG00000003316",
                    "ENSMUSG00000030058",
                    "ENSMUSG00000002980",
                    "ENSMUSG00000005510",
                    "ENSMUSG00000020547",
                    "ENSMUSG00000024208",
                    "ENSMUSG00000020232",
                    "ENSMUSG00000049246",
                    "ENSMUSG00000029686",
                    "ENSMUSG00000004591",
                    "ENSMUSG00000026098",
                    "ENSMUSG00000006221",
                    "ENSMUSG00000087260",
                    "ENSMUSG00000062588",
                    "ENSMUSG00000019428",
                    "ENSMUSG00000024803",
                    "ENSMUSG00000049134",
                    "ENSMUSG00000028541",
                    "ENSMUSG00000031683",
                    "ENSMUSG00000054178",
                    "ENSMUSG00000021395",
                    "ENSMUSG00000078812",
                    "ENSMUSG00000022235",
                    "ENSMUSG00000054619",
                    "ENSMUSG00000030714",
                    "ENSMUSG00000020542",
                    "ENSMUSG00000059534",
                    "ENSMUSG00000027367",
                    "ENSMUSG00000029447",
                    "ENSMUSG00000029108",
                    "ENSMUSG00000037300",
                    "ENSMUSG00000044308",
                    "ENSMUSG00000053536",
                    "ENSMUSG00000021631",
                    "ENSMUSG00000022300",
                    "ENSMUSG00000003429",
                    "ENSMUSG00000018599",
                    "ENSMUSG00000005698",
                    "ENSMUSG00000064344",
                    "ENSMUSG00000029095",
                    "ENSMUSG00000015839",
                    "ENSMUSG00000027499",
                    "ENSMUSG00000027784",
                    "ENSMUSG00000049878",
                    "ENSMUSG00000003541",
                    "ENSMUSG00000029068",
                    "ENSMUSG00000007613",
                    "ENSMUSG00000027455",
                    "ENSMUSG00000064340",
                    "ENSMUSG00000032047",
                    "ENSMUSG00000042133",
                    "ENSMUSG00000044393",
                    "ENSMUSG00000015377",
                    "ENSMUSG00000014470",
                    "ENSMUSG00000048076",
                    "ENSMUSG00000001472",
                    "ENSMUSG00000067818",
                    "ENSMUSG00000036002",
                    "ENSMUSG00000026864",
                    "ENSMUSG00000023939",
                    "ENSMUSG00000035885",
                    "ENSMUSG00000059325",
                    "ENSMUSG00000028134",
                    "ENSMUSG00000031078",
                    "ENSMUSG00000031333",
                    "ENSMUSG00000026341",
                    "ENSMUSG00000032330",
                    "ENSMUSG00000038170",
                    "ENSMUSG00000006315",
                    "ENSMUSG00000078713",
                    "ENSMUSG00000051695",
                    "ENSMUSG00000041361",
                    "ENSMUSG00000049327",
                    "ENSMUSG00000086863",
                    "ENSMUSG00000048280",
                    "ENSMUSG00000022748",
                    "ENSMUSG00000033392",
                    "ENSMUSG00000031865",
                    "ENSMUSG00000019087",
                    "ENSMUSG00000029004",
                    "ENSMUSG00000000149",
                    "ENSMUSG00000023206",
                    "ENSMUSG00000028518",
                    "ENSMUSG00000056973",
                    "ENSMUSG00000074129",
                    "ENSMUSG00000039234",
                    "ENSMUSG00000008136",
                    "ENSMUSG00000020516",
                    "ENSMUSG00000049881",
                    "ENSMUSG00000005873",
                    "ENSMUSG00000023944",
                    "ENSMUSG00000015932",
                    "ENSMUSG00000091989",
                    "ENSMUSG00000020358",
                    "ENSMUSG00000022681",
                    "ENSMUSG00000068699",
                    "ENSMUSG00000029201",
                    "ENSMUSG00000003970",
                    "ENSMUSG00000020475",
                    "ENSMUSG00000030662",
                    "ENSMUSG00000078671",
                    "ENSMUSG00000036941",
                    "ENSMUSG00000055670",
                    "ENSMUSG00000050953",
                    "ENSMUSG00000027765",
                    "ENSMUSG00000020308",
                    "ENSMUSG00000021546",
                    "ENSMUSG00000064372",
                    "ENSMUSG00000030002",
                    "ENSMUSG00000032495",
                    "ENSMUSG00000028772",
                    "ENSMUSG00000024639",
                    "ENSMUSG00000028145",
                    "ENSMUSG00000025794",
                    "ENSMUSG00000046997",
                    "ENSMUSG00000020022",
                    "ENSMUSG00000034088",
                    "ENSMUSG00000001911",
                    "ENSMUSG00000034832",
                    "ENSMUSG00000020664",
                    "ENSMUSG00000022389",
                    "ENSMUSG00000086938",
                    "ENSMUSG00000075595",
                    "ENSMUSG00000033722",
                    "ENSMUSG00000037095",
                    "ENSMUSG00000007415",
                    "ENSMUSG00000040118",
                    "ENSMUSG00000022635",
                    "ENSMUSG00000018770",
                    "ENSMUSG00000055775",
                    "ENSMUSG00000052911",
                    "ENSMUSG00000039782",
                    "ENSMUSG00000009569",
                    "ENSMUSG00000066406",
                    "ENSMUSG00000005566",
                    "ENSMUSG00000026564",
                    "ENSMUSG00000067736",
                    "ENSMUSG00000027996",
                    "ENSMUSG00000028737",
                    "ENSMUSG00000031954",
                    "ENSMUSG00000062908",
                    "ENSMUSG00000068264",
                    "ENSMUSG00000022462",
                    "ENSMUSG00000038070",
                    "ENSMUSG00000027447",
                    "ENSMUSG00000021993",
                    "ENSMUSG00000037032",
                    "ENSMUSG00000024283",
                    "ENSMUSG00000032423",
                    "ENSMUSG00000036585",
                    "ENSMUSG00000030319",
                    "ENSMUSG00000062647",
                    "ENSMUSG00000021681",
                    "ENSMUSG00000020393",
                    "ENSMUSG00000042709",
                    "ENSMUSG00000031540",
                    "ENSMUSG00000032525",
                    "ENSMUSG00000020402",
                    "ENSMUSG00000022217",
                    "ENSMUSG00000026853",
                    "ENSMUSG00000034880",
                    "ENSMUSG00000054428",
                    "ENSMUSG00000028953",
                    "ENSMUSG00000028959",
                    "ENSMUSG00000022119",
                    "ENSMUSG00000019920",
                    "ENSMUSG00000021270",
                    "ENSMUSG00000041729",
                    "ENSMUSG00000011257",
                    "ENSMUSG00000039943",
                    "ENSMUSG00000006576",
                    "ENSMUSG00000064037",
                    "ENSMUSG00000004085",
                    "ENSMUSG00000056851",
                    "ENSMUSG00000020048",
                    "ENSMUSG00000022867",
                    "ENSMUSG00000028878",
                    "ENSMUSG00000038538",
                    "ENSMUSG00000036678",
                    "ENSMUSG00000029056",
                    "ENSMUSG00000026688",
                    "ENSMUSG00000056679",
                    "ENSMUSG00000058498",
                    "ENSMUSG00000029833",
                    "ENSMUSG00000030213",
                    "ENSMUSG00000034574",
                    "ENSMUSG00000040666",
                    "ENSMUSG00000041731",
                    "ENSMUSG00000025184",
                    "ENSMUSG00000003809",
                    "ENSMUSG00000010175",
                    "ENSMUSG00000020176",
                    "ENSMUSG00000020863",
                    "ENSMUSG00000022095",
                    "ENSMUSG00000000399",
                    "ENSMUSG00000004931",
                    "ENSMUSG00000044792",
                    "ENSMUSG00000039976",
                    "ENSMUSG00000022437",
                    "ENSMUSG00000019872",
                    "ENSMUSG00000067653",
                    "ENSMUSG00000026353",
                    "ENSMUSG00000014504",
                    "ENSMUSG00000022037",
                    "ENSMUSG00000026276",
                    "ENSMUSG00000033938",
                    "ENSMUSG00000024997",
                    "ENSMUSG00000038663",
                    "ENSMUSG00000017778",
                    "ENSMUSG00000020952",
                    "ENSMUSG00000048546",
                    "ENSMUSG00000019132",
                    "ENSMUSG00000073639",
                    "ENSMUSG00000045776",
                    "ENSMUSG00000028207",
                    "ENSMUSG00000034973",
                    "ENSMUSG00000040414",
                    "ENSMUSG00000021537",
                    "ENSMUSG00000026434",
                    "ENSMUSG00000063801",
                    "ENSMUSG00000036564",
                    "ENSMUSG00000074093",
                    "ENSMUSG00000037049",
                    "ENSMUSG00000020079",
                    "ENSMUSG00000054452",
                    "ENSMUSG00000024150",
                    "ENSMUSG00000062456",
                    "ENSMUSG00000032480",
                    "ENSMUSG00000000184",
                    "ENSMUSG00000091383",
                    "ENSMUSG00000010122",
                    "ENSMUSG00000050541",
                    "ENSMUSG00000015937",
                    "ENSMUSG00000031447",
                    "ENSMUSG00000066279",
                    "ENSMUSG00000008682",
                    "ENSMUSG00000031765",
                    "ENSMUSG00000023020",
                    "ENSMUSG00000018339",
                    "ENSMUSG00000024392",
                    "ENSMUSG00000052934",
                    "ENSMUSG00000030421",
                    "ENSMUSG00000033382",
                    "ENSMUSG00000034424",
                    "ENSMUSG00000071793",
                    "ENSMUSG00000064366",
                    "ENSMUSG00000019558",
                    "ENSMUSG00000031029",
                    "ENSMUSG00000021149",
                    "ENSMUSG00000006494",
                    "ENSMUSG00000023832",
                    "ENSMUSG00000004980",
                    "ENSMUSG00000062963",
                    "ENSMUSG00000032580",
                    "ENSMUSG00000055027",
                    "ENSMUSG00000071014",
                    "ENSMUSG00000044864",
                    "ENSMUSG00000083326",
                    "ENSMUSG00000025666",
                    "ENSMUSG00000000171",
                    "ENSMUSG00000000441",
                    "ENSMUSG00000041685",
                    "ENSMUSG00000038784",
                    "ENSMUSG00000024661",
                    "ENSMUSG00000036748",
                    "ENSMUSG00000053483",
                    "ENSMUSG00000024526",
                    "ENSMUSG00000025156",
                    "ENSMUSG00000002608",
                    "ENSMUSG00000025103",
                    "ENSMUSG00000039601",
                    "ENSMUSG00000057322",
                    "ENSMUSG00000022270",
                    "ENSMUSG00000026234",
                    "ENSMUSG00000022136",
                    "ENSMUSG00000029191",
                    "ENSMUSG00000035783",
                    "ENSMUSG00000045690",
                    "ENSMUSG00000089909",
                    "ENSMUSG00000032526",
                    "ENSMUSG00000081228",
                    "ENSMUSG00000038132",
                    "ENSMUSG00000037949",
                    "ENSMUSG00000055762",
                    "ENSMUSG00000064343",
                    "ENSMUSG00000022186",
                    "ENSMUSG00000048814",
                    "ENSMUSG00000021597",
                    "ENSMUSG00000056091",
                    "ENSMUSG00000045624",
                    "ENSMUSG00000064337",
                    "ENSMUSG00000020116",
                    "ENSMUSG00000071528",
                    "ENSMUSG00000026754",
                    "ENSMUSG00000063382",
                    "ENSMUSG00000047710",
                    "ENSMUSG00000031841",
                    "ENSMUSG00000036854",
                    "ENSMUSG00000026131",
                    "ENSMUSG00000026202",
                    "ENSMUSG00000020721",
                    "ENSMUSG00000027566",
                    "ENSMUSG00000049960",
                    "ENSMUSG00000048175",
                    "ENSMUSG00000021639",
                    "ENSMUSG00000020326",
                    "ENSMUSG00000033538",
                    "ENSMUSG00000028124",
                    "ENSMUSG00000031751",
                    "ENSMUSG00000032010",
                    "ENSMUSG00000041278",
                    "ENSMUSG00000021838",
                    "ENSMUSG00000031996",
                    "ENSMUSG00000067279",
                    "ENSMUSG00000064347",
                    "ENSMUSG00000034892",
                    "ENSMUSG00000017756",
                    "ENSMUSG00000023092",
                    "ENSMUSG00000059291",
                    "ENSMUSG00000028300",
                    "ENSMUSG00000026577",
                    "ENSMUSG00000027618",
                    "ENSMUSG00000032177",
                    "ENSMUSG00000048758",
                    "ENSMUSG00000002379",
                    "ENSMUSG00000022570",
                    "ENSMUSG00000027177",
                    "ENSMUSG00000020994",
                    "ENSMUSG00000029455",
                    "ENSMUSG00000064346",
                    "ENSMUSG00000058690",
                    "ENSMUSG00000027010",
                    "ENSMUSG00000065947",
                    "ENSMUSG00000059852",
                    "ENSMUSG00000024063",
                    "ENSMUSG00000037270",
                    "ENSMUSG00000040370",
                    "ENSMUSG00000074218",
                    "ENSMUSG00000021371",
                    "ENSMUSG00000021798",
                    "ENSMUSG00000042323",
                    "ENSMUSG00000001280",
                    "ENSMUSG00000030417",
                    "ENSMUSG00000037826",
                    "ENSMUSG00000029651",
                    "ENSMUSG00000057649",
                    "ENSMUSG00000000751",
                    "ENSMUSG00000021000",
                    "ENSMUSG00000000563",
                    "ENSMUSG00000065037",
                    "ENSMUSG00000037815",
                    "ENSMUSG00000067344",
                    "ENSMUSG00000027809",
                    "ENSMUSG00000051169",
                    "ENSMUSG00000034342",
                    "ENSMUSG00000028759",
                    "ENSMUSG00000038949",
                    "ENSMUSG00000047187",
                    "ENSMUSG00000025488",
                    "ENSMUSG00000018417",
                    "ENSMUSG00000037958",
                    "ENSMUSG00000022840",
                    "ENSMUSG00000020456",
                    "ENSMUSG00000031024",
                    "ENSMUSG00000045983",
                    "ENSMUSG00000051747",
                    "ENSMUSG00000081967",
                    "ENSMUSG00000027304",
                    "ENSMUSG00000035458",
                    "ENSMUSG00000032481",
                    "ENSMUSG00000022040",
                    "ENSMUSG00000033004",
                    "ENSMUSG00000022013",
                    "ENSMUSG00000011305",
                    "ENSMUSG00000006356",
                    "ENSMUSG00000022890",
                    "ENSMUSG00000030505",
                    "ENSMUSG00000040723",
                    "ENSMUSG00000045482",
                    "ENSMUSG00000019810",
                    "ENSMUSG00000032199",
                    "ENSMUSG00000025982",
                    "ENSMUSG00000032224",
                    "ENSMUSG00000001211",
                    "ENSMUSG00000007877",
                    "ENSMUSG00000056692",
                    "ENSMUSG00000052738",
                    "ENSMUSG00000031985",
                    "ENSMUSG00000070436",
                    "ENSMUSG00000029918",
                    "ENSMUSG00000029359",
                    "ENSMUSG00000023175",
                    "ENSMUSG00000027598",
                    "ENSMUSG00000052374",
                    "ENSMUSG00000024991",
                    "ENSMUSG00000024687",
                    "ENSMUSG00000022201",
                    "ENSMUSG00000027030",
                    "ENSMUSG00000080893",
                    "ENSMUSG00000021226",
                    "ENSMUSG00000022940",
                    "ENSMUSG00000030934",
                    "ENSMUSG00000055093",
                    "ENSMUSG00000032590",
                    "ENSMUSG00000019998",
                    "ENSMUSG00000024309",
                    "ENSMUSG00000050199",
                    "ENSMUSG00000025362",
                    "ENSMUSG00000071604",
                    "ENSMUSG00000027881",
                    "ENSMUSG00000022964",
                    "ENSMUSG00000025364",
                    "ENSMUSG00000021807",
                    "ENSMUSG00000022803",
                    "ENSMUSG00000064338",
                    "ENSMUSG00000039100",
                    "ENSMUSG00000040473",
                    "ENSMUSG00000044080",
                    "ENSMUSG00000046671",
                    "ENSMUSG00000047459",
                    "ENSMUSG00000029649",
                    "ENSMUSG00000060679",
                    "ENSMUSG00000038844",
                    "ENSMUSG00000020074",
                    "ENSMUSG00000021877",
                    "ENSMUSG00000048416",
                    "ENSMUSG00000064193",
                    "ENSMUSG00000031490",
                    "ENSMUSG00000010608",
                    "ENSMUSG00000032279",
                    "ENSMUSG00000023861",
                    "ENSMUSG00000067847",
                    "ENSMUSG00000026664",
                    "ENSMUSG00000021520",
                    "ENSMUSG00000006717",
                    "ENSMUSG00000008683",
                    "ENSMUSG00000003438",
                    "ENSMUSG00000034235",
                    "ENSMUSG00000097277",
                    "ENSMUSG00000028538",
                    "ENSMUSG00000061666",
                    "ENSMUSG00000065701",
                    "ENSMUSG00000034842",
                    "ENSMUSG00000029190",
                    "ENSMUSG00000004610",
                    "ENSMUSG00000015944",
                    "ENSMUSG00000021265",
                    "ENSMUSG00000021290",
                    "ENSMUSG00000049470",
                    "ENSMUSG00000078695",
                    "ENSMUSG00000029401",
                    "ENSMUSG00000029673",
                    "ENSMUSG00000074247",
                    "ENSMUSG00000003534",
                    "ENSMUSG00000025453",
                    "ENSMUSG00000004035",
                    "ENSMUSG00000090110",
                    "ENSMUSG00000072720",
                    "ENSMUSG00000074918",
                    "ENSMUSG00000021606",
                    "ENSMUSG00000032621",
                    "ENSMUSG00000039753",
                    "ENSMUSG00000019877",
                    "ENSMUSG00000033416",
                    "ENSMUSG00000023068",
                    "ENSMUSG00000036860",
                    "ENSMUSG00000079641",
                    "ENSMUSG00000045636",
                    "ENSMUSG00000020462",
                    "ENSMUSG00000026074",
                    "ENSMUSG00000062300",
                    "ENSMUSG00000031299",
                    "ENSMUSG00000021771",
                    "ENSMUSG00000032267",
                    "ENSMUSG00000035247",
                    "ENSMUSG00000040048",
                    "ENSMUSG00000028461",
                    "ENSMUSG00000045664",
                    "ENSMUSG00000064365",
                    "ENSMUSG00000048478",
                    "ENSMUSG00000043639",
                    "ENSMUSG00000043866",
                    "ENSMUSG00000023938",
                    "ENSMUSG00000024188",
                    "ENSMUSG00000035268",
                    "ENSMUSG00000085779",
                    "ENSMUSG00000022450",
                    "ENSMUSG00000073563",
                    "ENSMUSG00000024805",
                    "ENSMUSG00000032366",
                    "ENSMUSG00000050503",
                    "ENSMUSG00000032370",
                    "ENSMUSG00000098343",
                    "ENSMUSG00000021910",
                    "ENSMUSG00000028383",
                    "ENSMUSG00000041115",
                    "ENSMUSG00000020309",
                    "ENSMUSG00000064348",
                    "ENSMUSG00000094483",
                    "ENSMUSG00000024498",
                    "ENSMUSG00000022965",
                    "ENSMUSG00000022884",
                    "ENSMUSG00000027099",
                    "ENSMUSG00000035027",
                    "ENSMUSG00000052812",
                    "ENSMUSG00000044067",
                    "ENSMUSG00000037685",
                    "ENSMUSG00000024290",
                    "ENSMUSG00000026526",
                    "ENSMUSG00000024132",
                    "ENSMUSG00000029338",
                    "ENSMUSG00000021621",
                    "ENSMUSG00000006005",
                    "ENSMUSG00000034525",
                    "ENSMUSG00000046312",
                    "ENSMUSG00000055204",
                    "ENSMUSG00000042115",
                    "ENSMUSG00000071151",
                    "ENSMUSG00000054312",
                    "ENSMUSG00000060803",
                    "ENSMUSG00000030246",
                    "ENSMUSG00000061474",
                    "ENSMUSG00000096232",
                    "ENSMUSG00000010914",
                    "ENSMUSG00000017760",
                    "ENSMUSG00000039914",
                    "ENSMUSG00000030869",
                    "ENSMUSG00000031767",
                    "ENSMUSG00000064371",
                    "ENSMUSG00000084314",
                    "ENSMUSG00000027620",
                    "ENSMUSG00000017817",
                    "ENSMUSG00000018395",
                    "ENSMUSG00000002908",
                    "ENSMUSG00000031229",
                    "ENSMUSG00000074656",
                    "ENSMUSG00000039126",
                    "ENSMUSG00000030095",
                    "ENSMUSG00000026489",
                    "ENSMUSG00000024913"
                ],
                "expression_sum": {
                    "class1": 60.0,
                    "class2": 1510.0
                },
                "uniprot_ids": [
                    "P19785",
                    "D3Z6V3"
                ]
            },
            "ENSMUSG00000064369": {
                "id": "ENSMUSG00000064369",
                "type": "Gene",
                "symbol": "TrnE",
                "source": [
                    "ENSMUSG00000001419",
                    "ENSMUSG00000021109",
                    "ENSMUSG00000056501",
                    "ENSMUSG00000028639",
                    "ENSMUSG00000030557",
                    "ENSMUSG00000001280",
                    "ENSMUSG00000019768",
                    "ENSMUSG00000022383",
                    "ENSMUSG00000015605",
                    "ENSMUSG00000014592"
                ],
                "target": [],
                "expression_sum": {
                    "class1": 7230.0,
                    "class2": 24642.0
                }
            },
            "ENSMUSG00000025393": {
                "id": "ENSMUSG00000025393",
                "type": "Gene",
                "symbol": "Atp5b",
                "source": [
                    "ENSMUSG00000006215",
                    "ENSMUSG00000058239",
                    "ENSMUSG00000015605",
                    "ENSMUSG00000015579",
                    "ENSMUSG00000049878",
                    "ENSMUSG00000051469",
                    "ENSMUSG00000001280",
                    "ENSMUSG00000019768",
                    "ENSMUSG00000014592",
                    "ENSMUSG00000031885",
                    "ENSMUSG00000076431",
                    "ENSMUSG00000008575",
                    "ENSMUSG00000039191",
                    "ENSMUSG00000021775",
                    "ENSMUSG00000031715"
                ],
                "target": [],
                "expression_sum": {
                    "class1": 11185.0,
                    "class2": 29636.0
                },
                "uniprot_ids": [
                    "P56480"
                ]
            },
            "ENSMUSG00000070493": {
                "id": "ENSMUSG00000070493",
                "type": "Gene",
                "symbol": "Chchd2",
                "source": [
                    "ENSMUSG00000020248",
                    "ENSMUSG00000056501",
                    "ENSMUSG00000015579",
                    "ENSMUSG00000018899",
                    "ENSMUSG00000030557",
                    "ENSMUSG00000001280",
                    "ENSMUSG00000019768",
                    "ENSMUSG00000027551",
                    "ENSMUSG00000009079",
                    "ENSMUSG00000031627",
                    "ENSMUSG00000031885",
                    "ENSMUSG00000008575",
                    "ENSMUSG00000035158",
                    "ENSMUSG00000031715"
                ],
                "target": [],
                "expression_sum": {
                    "class1": 16823.0,
                    "class2": 7536.0
                },
                "uniprot_ids": [
                    "Q9D1L0"
                ]
            },
            "ENSMUSG00000031231": {
                "id": "ENSMUSG00000031231",
                "type": "Gene",
                "symbol": "Cox7b",
                "source": [
                    "ENSMUSG00000015839",
                    "ENSMUSG00000056501",
                    "ENSMUSG00000001280",
                    "ENSMUSG00000019768",
                    "ENSMUSG00000033943",
                    "ENSMUSG00000025958",
                    "ENSMUSG00000038914",
                    "ENSMUSG00000053110",
                    "ENSMUSG00000068267",
                    "ENSMUSG00000015605",
                    "ENSMUSG00000021775",
                    "ENSMUSG00000015846"
                ],
                "target": [],
                "expression_sum": {
                    "class1": 12993.0,
                    "class2": 18769.0
                },
                "uniprot_ids": [
                    "P56393"
                ]
            },
            "ENSMUSG00000024049": {
                "id": "ENSMUSG00000024049",
                "type": "Gene",
                "symbol": "Myom1",
                "source": [
                    "ENSMUSG00000019768",
                    "ENSMUSG00000001419",
                    "ENSMUSG00000068267",
                    "ENSMUSG00000015846"
                ],
                "target": [],
                "expression_sum": {
                    "class1": 2502.0,
                    "class2": 4834.0
                },
                "uniprot_ids": [
                    "Q62234",
                    "A0A3B2W4I4",
                    "Z4YJF5"
                ]
            },
            "ENSMUSG00000026568": {
                "id": "ENSMUSG00000026568",
                "type": "Gene",
                "symbol": "Mpc2",
                "source": [
                    "ENSMUSG00000029135",
                    "ENSMUSG00000020248",
                    "ENSMUSG00000005566",
                    "ENSMUSG00000015605",
                    "ENSMUSG00000008575",
                    "ENSMUSG00000021775",
                    "ENSMUSG00000010175",
                    "ENSMUSG00000058239",
                    "ENSMUSG00000030256",
                    "ENSMUSG00000015579",
                    "ENSMUSG00000051469",
                    "ENSMUSG00000019768",
                    "ENSMUSG00000028565",
                    "ENSMUSG00000053110",
                    "ENSMUSG00000031715",
                    "ENSMUSG00000015846",
                    "ENSMUSG00000015839",
                    "ENSMUSG00000030557",
                    "ENSMUSG00000075595",
                    "ENSMUSG00000056501",
                    "ENSMUSG00000029249",
                    "ENSMUSG00000001280",
                    "ENSMUSG00000024498",
                    "ENSMUSG00000030087"
                ],
                "target": [],
                "expression_sum": {
                    "class1": 9768.0,
                    "class2": 9176.0
                },
                "uniprot_ids": [
                    "Q9D023",
                    "A0A0A6YY89"
                ]
            },
            "ENSMUSG00000025428": {
                "id": "ENSMUSG00000025428",
                "type": "Gene",
                "symbol": "Atp5a1",
                "source": [
                    "ENSMUSG00000001419",
                    "ENSMUSG00000015839",
                    "ENSMUSG00000058239",
                    "ENSMUSG00000015605",
                    "ENSMUSG00000026610",
                    "ENSMUSG00000056501",
                    "ENSMUSG00000078671",
                    "ENSMUSG00000030557",
                    "ENSMUSG00000029249",
                    "ENSMUSG00000001280",
                    "ENSMUSG00000005566",
                    "ENSMUSG00000019768",
                    "ENSMUSG00000032238",
                    "ENSMUSG00000053110",
                    "ENSMUSG00000031885",
                    "ENSMUSG00000058756",
                    "ENSMUSG00000008575",
                    "ENSMUSG00000031715"
                ],
                "target": [],
                "expression_sum": {
                    "class1": 22288.0,
                    "class2": 25058.0
                },
                "uniprot_ids": [
                    "Q03265",
                    "D6RJ16",
                    "D3Z6F5"
                ]
            },
            "ENSMUSG00000061904": {
                "id": "ENSMUSG00000061904",
                "type": "Gene",
                "symbol": "Slc25a3",
                "source": [
                    "ENSMUSG00000029135",
                    "ENSMUSG00000025612",
                    "ENSMUSG00000010175",
                    "ENSMUSG00000038615",
                    "ENSMUSG00000015605",
                    "ENSMUSG00000058239",
                    "ENSMUSG00000056501",
                    "ENSMUSG00000029249",
                    "ENSMUSG00000001280",
                    "ENSMUSG00000005566",
                    "ENSMUSG00000019768",
                    "ENSMUSG00000038914",
                    "ENSMUSG00000053110",
                    "ENSMUSG00000031885",
                    "ENSMUSG00000058756",
                    "ENSMUSG00000008575",
                    "ENSMUSG00000031715",
                    "ENSMUSG00000024985"
                ],
                "target": [],
                "expression_sum": {
                    "class1": 18939.0,
                    "class2": 16886.0
                },
                "uniprot_ids": [
                    "Q8VEM8",
                    "G5E902",
                    "E9PX16"
                ]
            },
            "ENSMUSG00000006818": {
                "id": "ENSMUSG00000006818",
                "type": "Gene",
                "symbol": "Sod2",
                "source": [
                    "ENSMUSG00000029135",
                    "ENSMUSG00000052534",
                    "ENSMUSG00000022811",
                    "ENSMUSG00000056501",
                    "ENSMUSG00000024955",
                    "ENSMUSG00000001280",
                    "ENSMUSG00000005566",
                    "ENSMUSG00000019768",
                    "ENSMUSG00000014592",
                    "ENSMUSG00000024498",
                    "ENSMUSG00000053110",
                    "ENSMUSG00000031885",
                    "ENSMUSG00000008575",
                    "ENSMUSG00000035158",
                    "ENSMUSG00000021775",
                    "ENSMUSG00000015846"
                ],
                "target": [],
                "expression_sum": {
                    "class1": 3751.0,
                    "class2": 5759.0
                },
                "uniprot_ids": [
                    "P09671",
                    "A0A3B2WBF0",
                    "A0A3B2W479"
                ]
            },
            "ENSMUSG00000029632": {
                "id": "ENSMUSG00000029632",
                "type": "Gene",
                "symbol": "Ndufa4",
                "source": [
                    "ENSMUSG00000001419",
                    "ENSMUSG00000022528",
                    "ENSMUSG00000015839",
                    "ENSMUSG00000010175",
                    "ENSMUSG00000039191",
                    "ENSMUSG00000056501",
                    "ENSMUSG00000003923",
                    "ENSMUSG00000001280",
                    "ENSMUSG00000019768",
                    "ENSMUSG00000032238",
                    "ENSMUSG00000053110",
                    "ENSMUSG00000031885",
                    "ENSMUSG00000058756",
                    "ENSMUSG00000008575",
                    "ENSMUSG00000031715",
                    "ENSMUSG00000021775",
                    "ENSMUSG00000015846"
                ],
                "target": [],
                "expression_sum": {
                    "class1": 11538.0,
                    "class2": 29630.0
                },
                "uniprot_ids": [
                    "Q62425",
                    "A0A0N4SVQ1"
                ]
            },
            "ENSMUSG00000051747": {
                "id": "ENSMUSG00000051747",
                "type": "Gene",
                "symbol": "Ttn",
                "source": [
                    "ENSMUSG00000001419",
                    "ENSMUSG00000056501",
                    "ENSMUSG00000015579",
                    "ENSMUSG00000024955",
                    "ENSMUSG00000030557",
                    "ENSMUSG00000029249",
                    "ENSMUSG00000001280",
                    "ENSMUSG00000019768",
                    "ENSMUSG00000005698",
                    "ENSMUSG00000022641",
                    "ENSMUSG00000015605"
                ],
                "target": [],
                "expression_sum": {
                    "class1": 48594.0,
                    "class2": 76727.0
                },
                "uniprot_ids": [
                    "A2ASS6",
                    "Q8BUJ6",
                    "A2AT62",
                    "A2AT70",
                    "A0A087WRP0",
                    "F6Q6Z0",
                    "A0A5K1VVQ9",
                    "A2AT63",
                    "F6RSJ3",
                    "A2AT65",
                    "A2AT61",
                    "A2AT66",
                    "F6ZRX7",
                    "F7CR78",
                    "E9Q8N1",
                    "F6ZZ75",
                    "A2AT68",
                    "A2AT64",
                    "A0A5K1VVQ1",
                    "E9Q8K5",
                    "A2AT67"
                ]
            },
            "ENSMUSG00000026526": {
                "id": "ENSMUSG00000026526",
                "type": "Gene",
                "symbol": "Fh1",
                "source": [
                    "ENSMUSG00000029135",
                    "ENSMUSG00000015839",
                    "ENSMUSG00000010175",
                    "ENSMUSG00000030557",
                    "ENSMUSG00000001280",
                    "ENSMUSG00000019768",
                    "ENSMUSG00000053110",
                    "ENSMUSG00000031885",
                    "ENSMUSG00000008575"
                ],
                "target": [],
                "expression_sum": {
                    "class1": 756.0,
                    "class2": 2289.0
                },
                "uniprot_ids": [
                    "P97807",
                    "H3BKG7",
                    "H3BJV5"
                ]
            },
            "ENSMUSG00000060002": {
                "id": "ENSMUSG00000060002",
                "type": "Gene",
                "symbol": "Chpt1",
                "source": [
                    "ENSMUSG00000015839",
                    "ENSMUSG00000056501",
                    "ENSMUSG00000029249",
                    "ENSMUSG00000001280",
                    "ENSMUSG00000005566",
                    "ENSMUSG00000005583",
                    "ENSMUSG00000005698",
                    "ENSMUSG00000019768",
                    "ENSMUSG00000032238",
                    "ENSMUSG00000031885",
                    "ENSMUSG00000058756",
                    "ENSMUSG00000015605",
                    "ENSMUSG00000031715"
                ],
                "target": [],
                "expression_sum": {
                    "class1": 1779.0,
                    "class2": 3030.0
                },
                "uniprot_ids": [
                    "Q8C025",
                    "D6RCW3",
                    "F6X1K1",
                    "F7DAN4",
                    "D3YU39",
                    "F6ZDL8",
                    "D3Z6W4"
                ]
            },
            "ENSMUSG00000078812": {
                "id": "ENSMUSG00000078812",
                "type": "Gene",
                "symbol": "Eif5a",
                "source": [
                    "ENSMUSG00000059552",
                    "ENSMUSG00000029135",
                    "ENSMUSG00000055116",
                    "ENSMUSG00000001280",
                    "ENSMUSG00000019768",
                    "ENSMUSG00000002265",
                    "ENSMUSG00000076431"
                ],
                "target": [],
                "expression_sum": {
                    "class1": 8125.0,
                    "class2": 1084.0
                },
                "uniprot_ids": [
                    "P63242",
                    "J3QPS8",
                    "A0A0A0MQM0"
                ]
            },
            "ENSMUSG00000024099": {
                "id": "ENSMUSG00000024099",
                "type": "Gene",
                "symbol": "Ndufv2",
                "source": [
                    "ENSMUSG00000029135",
                    "ENSMUSG00000019564",
                    "ENSMUSG00000021540",
                    "ENSMUSG00000015605",
                    "ENSMUSG00000021775",
                    "ENSMUSG00000006215",
                    "ENSMUSG00000010175",
                    "ENSMUSG00000022346",
                    "ENSMUSG00000058239",
                    "ENSMUSG00000051469",
                    "ENSMUSG00000019768",
                    "ENSMUSG00000031885",
                    "ENSMUSG00000031715",
                    "ENSMUSG00000015846",
                    "ENSMUSG00000015839",
                    "ENSMUSG00000075595",
                    "ENSMUSG00000014592",
                    "ENSMUSG00000056501",
                    "ENSMUSG00000024955",
                    "ENSMUSG00000003923",
                    "ENSMUSG00000027490",
                    "ENSMUSG00000001280"
                ],
                "target": [],
                "expression_sum": {
                    "class1": 8390.0,
                    "class2": 7406.0
                },
                "uniprot_ids": [
                    "Q9D6J6",
                    "M0QWP9",
                    "D6RGN4"
                ]
            },
            "ENSMUSG00000083820": {
                "id": "ENSMUSG00000083820",
                "type": "Gene",
                "symbol": "Ndufs6b",
                "source": [
                    "ENSMUSG00000015605",
                    "ENSMUSG00000052837",
                    "ENSMUSG00000056501",
                    "ENSMUSG00000030557",
                    "ENSMUSG00000001280",
                    "ENSMUSG00000019768",
                    "ENSMUSG00000025958",
                    "ENSMUSG00000008575",
                    "ENSMUSG00000015846"
                ],
                "target": [],
                "expression_sum": {
                    "class1": 1852.0,
                    "class2": 3990.0
                }
            },
            "ENSMUSG00000030695": {
                "id": "ENSMUSG00000030695",
                "type": "Gene",
                "symbol": "Aldoa",
                "source": [
                    "ENSMUSG00000029135",
                    "ENSMUSG00000001419",
                    "ENSMUSG00000003032",
                    "ENSMUSG00000015839",
                    "ENSMUSG00000015605",
                    "ENSMUSG00000026610",
                    "ENSMUSG00000052837",
                    "ENSMUSG00000015579",
                    "ENSMUSG00000056501",
                    "ENSMUSG00000014303",
                    "ENSMUSG00000029249",
                    "ENSMUSG00000001280",
                    "ENSMUSG00000019768",
                    "ENSMUSG00000058756",
                    "ENSMUSG00000008575",
                    "ENSMUSG00000026803",
                    "ENSMUSG00000021775",
                    "ENSMUSG00000031715"
                ],
                "target": [],
                "expression_sum": {
                    "class1": 19841.0,
                    "class2": 16711.0
                },
                "uniprot_ids": [
                    "P05064",
                    "A6ZI44",
                    "D3YWI1",
                    "A0A0U1RPT5",
                    "D3Z510",
                    "A0A0U1RPN8",
                    "D3YV98"
                ]
            },
            "ENSMUSG00000064348": {
                "id": "ENSMUSG00000064348",
                "type": "Gene",
                "symbol": "TrnN",
                "source": [
                    "ENSMUSG00000056501",
                    "ENSMUSG00000090258",
                    "ENSMUSG00000001280",
                    "ENSMUSG00000019768",
                    "ENSMUSG00000024695",
                    "ENSMUSG00000075595",
                    "ENSMUSG00000031885",
                    "ENSMUSG00000038679",
                    "ENSMUSG00000015605"
                ],
                "target": [],
                "expression_sum": {
                    "class1": 7274.0,
                    "class2": 12846.0
                }
            },
            "ENSMUSG00000040752": {
                "id": "ENSMUSG00000040752",
                "type": "Gene",
                "symbol": "Myh6",
                "source": [
                    "ENSMUSG00000026610",
                    "ENSMUSG00000015579",
                    "ENSMUSG00000001280",
                    "ENSMUSG00000019768",
                    "ENSMUSG00000005698",
                    "ENSMUSG00000053110",
                    "ENSMUSG00000058756",
                    "ENSMUSG00000015605"
                ],
                "target": [],
                "expression_sum": {
                    "class1": 57016.0,
                    "class2": 83323.0
                },
                "uniprot_ids": [
                    "Q02566",
                    "Q1WNP4",
                    "B8JJH3",
                    "A0A2I3BPY4"
                ]
            },
            "ENSMUSG00000024661": {
                "id": "ENSMUSG00000024661",
                "type": "Gene",
                "symbol": "Fth1",
                "source": [
                    "ENSMUSG00000021944",
                    "ENSMUSG00000015839",
                    "ENSMUSG00000015579",
                    "ENSMUSG00000001280",
                    "ENSMUSG00000019768",
                    "ENSMUSG00000068267",
                    "ENSMUSG00000031627",
                    "ENSMUSG00000031885",
                    "ENSMUSG00000008575",
                    "ENSMUSG00000015846"
                ],
                "target": [],
                "expression_sum": {
                    "class1": 26778.0,
                    "class2": 23997.0
                },
                "uniprot_ids": [
                    "P09528",
                    "A0A494B9D4",
                    "A0A494BAP3",
                    "A0A494BA92"
                ]
            },
            "ENSMUSG00000035458": {
                "id": "ENSMUSG00000035458",
                "type": "Gene",
                "symbol": "Tnni3",
                "source": [
                    "ENSMUSG00000006215",
                    "ENSMUSG00000058239",
                    "ENSMUSG00000056501",
                    "ENSMUSG00000015579",
                    "ENSMUSG00000051469",
                    "ENSMUSG00000001280",
                    "ENSMUSG00000019768",
                    "ENSMUSG00000025958",
                    "ENSMUSG00000053110",
                    "ENSMUSG00000031885",
                    "ENSMUSG00000015605",
                    "ENSMUSG00000015846"
                ],
                "target": [],
                "expression_sum": {
                    "class1": 62934.0,
                    "class2": 75488.0
                },
                "uniprot_ids": [
                    "P48787",
                    "D3YXP5",
                    "D3Z160",
                    "A0A140LIV0"
                ]
            },
            "ENSMUSG00000031818": {
                "id": "ENSMUSG00000031818",
                "type": "Gene",
                "symbol": "Cox4i1",
                "source": [
                    "ENSMUSG00000024238",
                    "ENSMUSG00000015839",
                    "ENSMUSG00000015605",
                    "ENSMUSG00000056501",
                    "ENSMUSG00000015579",
                    "ENSMUSG00000030557",
                    "ENSMUSG00000090258",
                    "ENSMUSG00000001280",
                    "ENSMUSG00000019768",
                    "ENSMUSG00000033943",
                    "ENSMUSG00000058756",
                    "ENSMUSG00000008575",
                    "ENSMUSG00000039191",
                    "ENSMUSG00000021775",
                    "ENSMUSG00000015846"
                ],
                "target": [],
                "expression_sum": {
                    "class1": 53822.0,
                    "class2": 41890.0
                },
                "uniprot_ids": [
                    "P19783",
                    "M0QWX2",
                    "M0QWX7"
                ]
            },
            "ENSMUSG00000036880": {
                "id": "ENSMUSG00000036880",
                "type": "Gene",
                "symbol": "Acaa2",
                "source": [
                    "ENSMUSG00000057469",
                    "ENSMUSG00000008575",
                    "ENSMUSG00000040481",
                    "ENSMUSG00000015839",
                    "ENSMUSG00000078671",
                    "ENSMUSG00000026610",
                    "ENSMUSG00000015579",
                    "ENSMUSG00000029249",
                    "ENSMUSG00000001280",
                    "ENSMUSG00000005566",
                    "ENSMUSG00000019768",
                    "ENSMUSG00000022641",
                    "ENSMUSG00000024695",
                    "ENSMUSG00000031885",
                    "ENSMUSG00000058756",
                    "ENSMUSG00000015605",
                    "ENSMUSG00000031715"
                ],
                "target": [],
                "expression_sum": {
                    "class1": 9273.0,
                    "class2": 6375.0
                },
                "uniprot_ids": [
                    "Q8BWT1",
                    "A0A494B9J2"
                ]
            },
            "ENSMUSG00000059534": {
                "id": "ENSMUSG00000059534",
                "type": "Gene",
                "symbol": "Uqcr10",
                "source": [
                    "ENSMUSG00000015839",
                    "ENSMUSG00000026610",
                    "ENSMUSG00000052837",
                    "ENSMUSG00000056501",
                    "ENSMUSG00000001280",
                    "ENSMUSG00000019768",
                    "ENSMUSG00000033943",
                    "ENSMUSG00000075595",
                    "ENSMUSG00000031885",
                    "ENSMUSG00000015605",
                    "ENSMUSG00000015846"
                ],
                "target": [],
                "expression_sum": {
                    "class1": 31182.0,
                    "class2": 32494.0
                },
                "uniprot_ids": [
                    "Q8R1I1"
                ]
            },
            "ENSMUSG00000064337": {
                "id": "ENSMUSG00000064337",
                "type": "Gene",
                "symbol": "mt-Rnr1",
                "source": [
                    "ENSMUSG00000001280",
                    "ENSMUSG00000019768",
                    "ENSMUSG00000052837"
                ],
                "target": [],
                "expression_sum": {
                    "class1": 4181321.0,
                    "class2": 3023384.0
                }
            },
            "ENSMUSG00000023944": {
                "id": "ENSMUSG00000023944",
                "type": "Gene",
                "symbol": "Hsp90ab1",
                "source": [
                    "ENSMUSG00000029135",
                    "ENSMUSG00000001419",
                    "ENSMUSG00000030256",
                    "ENSMUSG00000056501",
                    "ENSMUSG00000015579",
                    "ENSMUSG00000001280",
                    "ENSMUSG00000019768",
                    "ENSMUSG00000068267",
                    "ENSMUSG00000020919",
                    "ENSMUSG00000076431",
                    "ENSMUSG00000015605",
                    "ENSMUSG00000015846"
                ],
                "target": [],
                "expression_sum": {
                    "class1": 51426.0,
                    "class2": 13787.0
                },
                "uniprot_ids": [
                    "P11499",
                    "E9PX27",
                    "E9Q0C3",
                    "E9Q3D6",
                    "D3Z1R1"
                ]
            },
            "ENSMUSG00000083863": {
                "id": "ENSMUSG00000083863",
                "type": "Gene",
                "symbol": "Gm13341",
                "source": [
                    "ENSMUSG00000001280",
                    "ENSMUSG00000052837"
                ],
                "target": [],
                "expression_sum": {
                    "class1": 332083.0,
                    "class2": 544689.0
                }
            },
            "ENSMUSG00000005683": {
                "id": "ENSMUSG00000005683",
                "type": "Gene",
                "symbol": "Cs",
                "source": [
                    "ENSMUSG00000021944",
                    "ENSMUSG00000015605",
                    "ENSMUSG00000021775",
                    "ENSMUSG00000010175",
                    "ENSMUSG00000015579",
                    "ENSMUSG00000019768",
                    "ENSMUSG00000005698",
                    "ENSMUSG00000053110",
                    "ENSMUSG00000031715",
                    "ENSMUSG00000024985",
                    "ENSMUSG00000015839",
                    "ENSMUSG00000023994",
                    "ENSMUSG00000020453",
                    "ENSMUSG00000058756",
                    "ENSMUSG00000029913",
                    "ENSMUSG00000026610",
                    "ENSMUSG00000056501",
                    "ENSMUSG00000029249",
                    "ENSMUSG00000001280"
                ],
                "target": [],
                "expression_sum": {
                    "class1": 2326.0,
                    "class2": 5470.0
                },
                "uniprot_ids": [
                    "Q9CZU6"
                ]
            },
            "ENSMUSG00000007892": {
                "id": "ENSMUSG00000007892",
                "type": "Gene",
                "symbol": "Rplp1",
                "source": [
                    "ENSMUSG00000001280",
                    "ENSMUSG00000019768",
                    "ENSMUSG00000015579"
                ],
                "target": [],
                "expression_sum": {
                    "class1": 14503.0,
                    "class2": 4682.0
                },
                "uniprot_ids": [
                    "P47955"
                ]
            },
            "ENSMUSG00000033060": {
                "id": "ENSMUSG00000033060",
                "type": "Gene",
                "symbol": "Lmo7",
                "source": [
                    "ENSMUSG00000001419",
                    "ENSMUSG00000057469",
                    "ENSMUSG00000010175",
                    "ENSMUSG00000015839",
                    "ENSMUSG00000026610",
                    "ENSMUSG00000056501",
                    "ENSMUSG00000029249",
                    "ENSMUSG00000001280",
                    "ENSMUSG00000005566",
                    "ENSMUSG00000019768",
                    "ENSMUSG00000032238",
                    "ENSMUSG00000058756",
                    "ENSMUSG00000015605",
                    "ENSMUSG00000031715",
                    "ENSMUSG00000024985"
                ],
                "target": [],
                "expression_sum": {
                    "class1": 3093.0,
                    "class2": 2348.0
                },
                "uniprot_ids": [
                    "Q3UPW3",
                    "E0CZD8",
                    "A0A5F8MPW2",
                    "F6VG99",
                    "E9PYI7",
                    "F6TFN2",
                    "E9PYF4"
                ]
            },
            "ENSMUSG00000064341": {
                "id": "ENSMUSG00000064341",
                "type": "Gene",
                "symbol": "ND1",
                "source": [
                    "ENSMUSG00000001419",
                    "ENSMUSG00000021944",
                    "ENSMUSG00000022346",
                    "ENSMUSG00000052837",
                    "ENSMUSG00000001280",
                    "ENSMUSG00000019768"
                ],
                "target": [],
                "expression_sum": {
                    "class1": 2775363.0,
                    "class2": 2282892.0
                },
                "uniprot_ids": [
                    "P03888"
                ]
            },
            "ENSMUSG00000023020": {
                "id": "ENSMUSG00000023020",
                "type": "Gene",
                "symbol": "Cox14",
                "source": [
                    "ENSMUSG00000029135",
                    "ENSMUSG00000001419",
                    "ENSMUSG00000015605",
                    "ENSMUSG00000030557",
                    "ENSMUSG00000001280",
                    "ENSMUSG00000019768",
                    "ENSMUSG00000053110",
                    "ENSMUSG00000068267",
                    "ENSMUSG00000022053",
                    "ENSMUSG00000031885",
                    "ENSMUSG00000008575",
                    "ENSMUSG00000031715"
                ],
                "target": [],
                "expression_sum": {
                    "class1": 5155.0,
                    "class2": 3481.0
                },
                "uniprot_ids": [
                    "Q8BH51"
                ]
            },
            "ENSMUSG00000020402": {
                "id": "ENSMUSG00000020402",
                "type": "Gene",
                "symbol": "Vdac1",
                "source": [
                    "ENSMUSG00000019564",
                    "ENSMUSG00000005566",
                    "ENSMUSG00000038615",
                    "ENSMUSG00000015605",
                    "ENSMUSG00000008575",
                    "ENSMUSG00000035158",
                    "ENSMUSG00000021775",
                    "ENSMUSG00000010175",
                    "ENSMUSG00000058239",
                    "ENSMUSG00000090258",
                    "ENSMUSG00000019768",
                    "ENSMUSG00000031885",
                    "ENSMUSG00000015846",
                    "ENSMUSG00000020167",
                    "ENSMUSG00000015839",
                    "ENSMUSG00000028639",
                    "ENSMUSG00000001419",
                    "ENSMUSG00000078671",
                    "ENSMUSG00000003923",
                    "ENSMUSG00000001280"
                ],
                "target": [],
                "expression_sum": {
                    "class1": 4293.0,
                    "class2": 5335.0
                },
                "uniprot_ids": [
                    "Q60932",
                    "F2Z471"
                ]
            },
            "ENSMUSG00000020163": {
                "id": "ENSMUSG00000020163",
                "type": "Gene",
                "symbol": "Uqcr11",
                "source": [
                    "ENSMUSG00000015605",
                    "ENSMUSG00000001280",
                    "ENSMUSG00000019768",
                    "ENSMUSG00000031885",
                    "ENSMUSG00000008575",
                    "ENSMUSG00000021775"
                ],
                "target": [],
                "expression_sum": {
                    "class1": 30681.0,
                    "class2": 28881.0
                },
                "uniprot_ids": [
                    "Q9CPX8"
                ]
            },
            "ENSMUSG00000096449": {
                "id": "ENSMUSG00000096449",
                "type": "Gene",
                "symbol": "LOC118568041",
                "source": [
                    "ENSMUSG00000001280",
                    "ENSMUSG00000015846"
                ],
                "target": [],
                "expression_sum": {
                    "class1": 63886.0,
                    "class2": 40321.0
                }
            },
            "ENSMUSG00000064365": {
                "id": "ENSMUSG00000064365",
                "type": "Gene",
                "symbol": "TrnS2",
                "source": [
                    "ENSMUSG00000001419",
                    "ENSMUSG00000030557",
                    "ENSMUSG00000001280",
                    "ENSMUSG00000019768",
                    "ENSMUSG00000053110",
                    "ENSMUSG00000015605"
                ],
                "target": [],
                "expression_sum": {
                    "class1": 1230.0,
                    "class2": 1958.0
                }
            },
            "ENSMUSG00000016252": {
                "id": "ENSMUSG00000016252",
                "type": "Gene",
                "symbol": "Atp5e",
                "source": [
                    "ENSMUSG00000015605",
                    "ENSMUSG00000056501",
                    "ENSMUSG00000003923",
                    "ENSMUSG00000001280",
                    "ENSMUSG00000019768",
                    "ENSMUSG00000033943",
                    "ENSMUSG00000008575",
                    "ENSMUSG00000015846"
                ],
                "target": [],
                "expression_sum": {
                    "class1": 14112.0,
                    "class2": 20607.0
                },
                "uniprot_ids": [
                    "P56382"
                ]
            },
            "ENSMUSG00000023089": {
                "id": "ENSMUSG00000023089",
                "type": "Gene",
                "symbol": "Ndufa5",
                "source": [
                    "ENSMUSG00000029135",
                    "ENSMUSG00000057469",
                    "ENSMUSG00000038615",
                    "ENSMUSG00000015839",
                    "ENSMUSG00000010175",
                    "ENSMUSG00000015605",
                    "ENSMUSG00000058239",
                    "ENSMUSG00000056501",
                    "ENSMUSG00000030557",
                    "ENSMUSG00000001280",
                    "ENSMUSG00000019768",
                    "ENSMUSG00000024498",
                    "ENSMUSG00000055320",
                    "ENSMUSG00000068267",
                    "ENSMUSG00000031885",
                    "ENSMUSG00000008575",
                    "ENSMUSG00000031715"
                ],
                "target": [],
                "expression_sum": {
                    "class1": 22015.0,
                    "class2": 21494.0
                },
                "uniprot_ids": [
                    "Q9CPP6",
                    "D3YTQ8"
                ]
            },
            "ENSMUSG00000038462": {
                "id": "ENSMUSG00000038462",
                "type": "Gene",
                "symbol": "Uqcrfs1",
                "source": [
                    "ENSMUSG00000029135",
                    "ENSMUSG00000025612",
                    "ENSMUSG00000015839",
                    "ENSMUSG00000015605",
                    "ENSMUSG00000056501",
                    "ENSMUSG00000015579",
                    "ENSMUSG00000029249",
                    "ENSMUSG00000001280",
                    "ENSMUSG00000019768",
                    "ENSMUSG00000038914",
                    "ENSMUSG00000022641",
                    "ENSMUSG00000008575"
                ],
                "target": [],
                "expression_sum": {
                    "class1": 9844.0,
                    "class2": 14923.0
                },
                "uniprot_ids": [
                    "Q9CR68"
                ]
            },
            "ENSMUSG00000045983": {
                "id": "ENSMUSG00000045983",
                "type": "Gene",
                "symbol": "Eif4g1",
                "source": [
                    "ENSMUSG00000029135",
                    "ENSMUSG00000026890",
                    "ENSMUSG00000005566",
                    "ENSMUSG00000031627",
                    "ENSMUSG00000015605",
                    "ENSMUSG00000008575",
                    "ENSMUSG00000035158",
                    "ENSMUSG00000057469",
                    "ENSMUSG00000058239",
                    "ENSMUSG00000030256",
                    "ENSMUSG00000019768",
                    "ENSMUSG00000022641",
                    "ENSMUSG00000053110",
                    "ENSMUSG00000024985",
                    "ENSMUSG00000030557",
                    "ENSMUSG00000058756",
                    "ENSMUSG00000056501",
                    "ENSMUSG00000029249",
                    "ENSMUSG00000001280"
                ],
                "target": [],
                "expression_sum": {
                    "class1": 6166.0,
                    "class2": 1232.0
                },
                "uniprot_ids": [
                    "Q6NZJ6",
                    "D6RCH1",
                    "D3YWY7",
                    "D3YWC6",
                    "E9Q770",
                    "A0A0J9YU44",
                    "D3YWM1",
                    "A0A0J9YTV0",
                    "A0A0J9YUL1",
                    "A0A0J9YVC1",
                    "E9PVC6",
                    "E9Q9E1",
                    "D3Z439",
                    "A0A338P7C5",
                    "A0A0J9YUS5",
                    "A0A338P758",
                    "E9PVC5",
                    "A0A0J9YUC1"
                ]
            },
            "ENSMUSG00000064372": {
                "id": "ENSMUSG00000064372",
                "type": "Gene",
                "symbol": "TrnP",
                "source": [
                    "ENSMUSG00000001419",
                    "ENSMUSG00000021944",
                    "ENSMUSG00000001280",
                    "ENSMUSG00000019768",
                    "ENSMUSG00000015846"
                ],
                "target": [],
                "expression_sum": {
                    "class1": 17830.0,
                    "class2": 35531.0
                }
            },
            "ENSMUSG00000021606": {
                "id": "ENSMUSG00000021606",
                "type": "Gene",
                "symbol": "Ndufs6",
                "source": [
                    "ENSMUSG00000029135",
                    "ENSMUSG00000015839",
                    "ENSMUSG00000078671",
                    "ENSMUSG00000015605",
                    "ENSMUSG00000052837",
                    "ENSMUSG00000056501",
                    "ENSMUSG00000030557",
                    "ENSMUSG00000029249",
                    "ENSMUSG00000001280",
                    "ENSMUSG00000019768",
                    "ENSMUSG00000028565",
                    "ENSMUSG00000031885",
                    "ENSMUSG00000008575",
                    "ENSMUSG00000015846"
                ],
                "target": [],
                "expression_sum": {
                    "class1": 15587.0,
                    "class2": 12789.0
                },
                "uniprot_ids": [
                    "P52503",
                    "D6RJ54",
                    "A0A1Y7VM38",
                    "A0A1Y7VKP8"
                ]
            },
            "ENSMUSG00000044080": {
                "id": "ENSMUSG00000044080",
                "type": "Gene",
                "symbol": "S100a1",
                "source": [
                    "ENSMUSG00000029135",
                    "ENSMUSG00000010175",
                    "ENSMUSG00000015839",
                    "ENSMUSG00000015605",
                    "ENSMUSG00000058239",
                    "ENSMUSG00000056501",
                    "ENSMUSG00000015579",
                    "ENSMUSG00000029249",
                    "ENSMUSG00000001280",
                    "ENSMUSG00000019768",
                    "ENSMUSG00000028565",
                    "ENSMUSG00000032238",
                    "ENSMUSG00000022641",
                    "ENSMUSG00000008575",
                    "ENSMUSG00000021775"
                ],
                "target": [],
                "expression_sum": {
                    "class1": 8499.0,
                    "class2": 13758.0
                },
                "uniprot_ids": [
                    "Q91V77",
                    "D3YUT6"
                ]
            },
            "ENSMUSG00000037916": {
                "id": "ENSMUSG00000037916",
                "type": "Gene",
                "symbol": "Ndufv1",
                "source": [
                    "ENSMUSG00000015839",
                    "ENSMUSG00000026628",
                    "ENSMUSG00000015605",
                    "ENSMUSG00000026610",
                    "ENSMUSG00000056501",
                    "ENSMUSG00000015579",
                    "ENSMUSG00000078671",
                    "ENSMUSG00000029249",
                    "ENSMUSG00000001280",
                    "ENSMUSG00000005566",
                    "ENSMUSG00000019768",
                    "ENSMUSG00000005698",
                    "ENSMUSG00000022641",
                    "ENSMUSG00000031885",
                    "ENSMUSG00000008575",
                    "ENSMUSG00000014592",
                    "ENSMUSG00000029913",
                    "ENSMUSG00000024985"
                ],
                "target": [],
                "expression_sum": {
                    "class1": 4135.0,
                    "class2": 3683.0
                },
                "uniprot_ids": [
                    "Q91YT0",
                    "D3Z1U9",
                    "D3YUM1",
                    "A0A494BAS8",
                    "D3YXX5",
                    "D3Z0K1",
                    "A0A494BA02",
                    "D6RG60"
                ]
            },
            "ENSMUSG00000030869": {
                "id": "ENSMUSG00000030869",
                "type": "Gene",
                "symbol": "Ndufab1",
                "source": [
                    "ENSMUSG00000001419",
                    "ENSMUSG00000021944",
                    "ENSMUSG00000015839",
                    "ENSMUSG00000058239",
                    "ENSMUSG00000056501",
                    "ENSMUSG00000001280",
                    "ENSMUSG00000019768",
                    "ENSMUSG00000033943",
                    "ENSMUSG00000025958",
                    "ENSMUSG00000031627",
                    "ENSMUSG00000075595",
                    "ENSMUSG00000031885",
                    "ENSMUSG00000015605",
                    "ENSMUSG00000021775",
                    "ENSMUSG00000015846"
                ],
                "target": [],
                "expression_sum": {
                    "class1": 7704.0,
                    "class2": 7737.0
                },
                "uniprot_ids": [
                    "Q9CR21",
                    "F8WJ64",
                    "F6ZFT1"
                ]
            },
            "ENSMUSG00000011257": {
                "id": "ENSMUSG00000011257",
                "type": "Gene",
                "symbol": "Pabpc4",
                "source": [
                    "ENSMUSG00000029135",
                    "ENSMUSG00000015839",
                    "ENSMUSG00000026610",
                    "ENSMUSG00000056501",
                    "ENSMUSG00000015579",
                    "ENSMUSG00000030557",
                    "ENSMUSG00000001280",
                    "ENSMUSG00000019768",
                    "ENSMUSG00000014592",
                    "ENSMUSG00000024695",
                    "ENSMUSG00000015605",
                    "ENSMUSG00000035158"
                ],
                "target": [],
                "expression_sum": {
                    "class1": 1049.0,
                    "class2": 1139.0
                },
                "uniprot_ids": [
                    "Q6PHQ9",
                    "Q91YZ8",
                    "A3KFU8",
                    "V9GXG3",
                    "A3KFU5"
                ]
            },
            "ENSMUSG00000024608": {
                "id": "ENSMUSG00000024608",
                "type": "Gene",
                "symbol": "Rps14",
                "source": [
                    "ENSMUSG00000001419",
                    "ENSMUSG00000036461",
                    "ENSMUSG00000001280",
                    "ENSMUSG00000005583",
                    "ENSMUSG00000040732",
                    "ENSMUSG00000024140",
                    "ENSMUSG00000027439"
                ],
                "target": [],
                "expression_sum": {
                    "class1": 35608.0,
                    "class2": 9929.0
                },
                "uniprot_ids": [
                    "P62264",
                    "A0A494BAS4",
                    "D3Z7I1",
                    "D3YVF4"
                ]
            },
            "ENSMUSG00000041028": {
                "id": "ENSMUSG00000041028",
                "type": "Gene",
                "symbol": "Ghitm",
                "source": [
                    "ENSMUSG00000006215",
                    "ENSMUSG00000015839",
                    "ENSMUSG00000015605",
                    "ENSMUSG00000056501",
                    "ENSMUSG00000029249",
                    "ENSMUSG00000001280",
                    "ENSMUSG00000005566",
                    "ENSMUSG00000019768",
                    "ENSMUSG00000032238",
                    "ENSMUSG00000068267",
                    "ENSMUSG00000031627",
                    "ENSMUSG00000031885",
                    "ENSMUSG00000008575"
                ],
                "target": [],
                "expression_sum": {
                    "class1": 8154.0,
                    "class2": 5059.0
                },
                "uniprot_ids": [
                    "Q91VC9",
                    "A0A286YDK2"
                ]
            },
            "ENSMUSG00000003072": {
                "id": "ENSMUSG00000003072",
                "type": "Gene",
                "symbol": "Atp5d",
                "source": [
                    "ENSMUSG00000029135",
                    "ENSMUSG00000044452",
                    "ENSMUSG00000031627",
                    "ENSMUSG00000015605",
                    "ENSMUSG00000008575",
                    "ENSMUSG00000021775",
                    "ENSMUSG00000035158",
                    "ENSMUSG00000057469",
                    "ENSMUSG00000006215",
                    "ENSMUSG00000058239",
                    "ENSMUSG00000030256",
                    "ENSMUSG00000019768",
                    "ENSMUSG00000038914",
                    "ENSMUSG00000028565",
                    "ENSMUSG00000031885",
                    "ENSMUSG00000024985",
                    "ENSMUSG00000031715",
                    "ENSMUSG00000015846",
                    "ENSMUSG00000015839",
                    "ENSMUSG00000030557",
                    "ENSMUSG00000014592",
                    "ENSMUSG00000029913",
                    "ENSMUSG00000001419",
                    "ENSMUSG00000078671",
                    "ENSMUSG00000052837",
                    "ENSMUSG00000056501",
                    "ENSMUSG00000029249",
                    "ENSMUSG00000001280",
                    "ENSMUSG00000024498"
                ],
                "target": [],
                "expression_sum": {
                    "class1": 9390.0,
                    "class2": 7454.0
                },
                "uniprot_ids": [
                    "Q9D3D9",
                    "D3Z7S4"
                ]
            },
            "ENSMUSG00000020321": {
                "id": "ENSMUSG00000020321",
                "type": "Gene",
                "symbol": "Mdh1",
                "source": [
                    "ENSMUSG00000057469",
                    "ENSMUSG00000010175",
                    "ENSMUSG00000058239",
                    "ENSMUSG00000056501",
                    "ENSMUSG00000001280",
                    "ENSMUSG00000005566",
                    "ENSMUSG00000019768",
                    "ENSMUSG00000031885",
                    "ENSMUSG00000015605",
                    "ENSMUSG00000026803"
                ],
                "target": [],
                "expression_sum": {
                    "class1": 11753.0,
                    "class2": 16362.0
                },
                "uniprot_ids": [
                    "P14152",
                    "A0A5F8MPN8",
                    "B1ATQ3"
                ]
            },
            "ENSMUSG00000038615": {
                "id": "ENSMUSG00000038615",
                "type": "TF",
                "symbol": "Nfe2l1",
                "source": [
                    "ENSMUSG00000001419",
                    "ENSMUSG00000015579",
                    "ENSMUSG00000024955",
                    "ENSMUSG00000001280",
                    "ENSMUSG00000019768",
                    "ENSMUSG00000044167",
                    "ENSMUSG00000053110",
                    "ENSMUSG00000031885",
                    "ENSMUSG00000008575",
                    "ENSMUSG00000031715",
                    "ENSMUSG00000015846"
                ],
                "target": [
                    "ENSMUSG00000054619",
                    "ENSMUSG00000020483",
                    "ENSMUSG00000020190",
                    "ENSMUSG00000061904",
                    "ENSMUSG00000009927",
                    "ENSMUSG00000026817",
                    "ENSMUSG00000040451",
                    "ENSMUSG00000044308",
                    "ENSMUSG00000027206",
                    "ENSMUSG00000027805",
                    "ENSMUSG00000040269",
                    "ENSMUSG00000040850",
                    "ENSMUSG00000042632",
                    "ENSMUSG00000018819",
                    "ENSMUSG00000023089",
                    "ENSMUSG00000018217",
                    "ENSMUSG00000015672",
                    "ENSMUSG00000038121",
                    "ENSMUSG00000073676",
                    "ENSMUSG00000032348",
                    "ENSMUSG00000048731",
                    "ENSMUSG00000034160",
                    "ENSMUSG00000069844",
                    "ENSMUSG00000087270",
                    "ENSMUSG00000026766",
                    "ENSMUSG00000008429",
                    "ENSMUSG00000067925",
                    "ENSMUSG00000020402",
                    "ENSMUSG00000059714",
                    "ENSMUSG00000035202"
                ],
                "expression_sum": {
                    "class1": 4094.0,
                    "class2": 1452.0
                },
                "uniprot_ids": [
                    "Q61985",
                    "Q3V3M1",
                    "A2A6D1",
                    "A2A6D0",
                    "A2A6D2"
                ]
            },
            "ENSMUSG00000018770": {
                "id": "ENSMUSG00000018770",
                "type": "Gene",
                "symbol": "Atp5g3",
                "source": [
                    "ENSMUSG00000029135",
                    "ENSMUSG00000015839",
                    "ENSMUSG00000015605",
                    "ENSMUSG00000056501",
                    "ENSMUSG00000001280",
                    "ENSMUSG00000019768",
                    "ENSMUSG00000030087",
                    "ENSMUSG00000031885",
                    "ENSMUSG00000008575",
                    "ENSMUSG00000021775",
                    "ENSMUSG00000031715"
                ],
                "target": [],
                "expression_sum": {
                    "class1": 21391.0,
                    "class2": 36740.0
                },
                "uniprot_ids": [
                    "P56384"
                ]
            },
            "ENSMUSG00000031059": {
                "id": "ENSMUSG00000031059",
                "type": "Gene",
                "symbol": "Ndufb11",
                "source": [
                    "ENSMUSG00000022346",
                    "ENSMUSG00000001280",
                    "ENSMUSG00000019768",
                    "ENSMUSG00000053110",
                    "ENSMUSG00000031885",
                    "ENSMUSG00000015605",
                    "ENSMUSG00000015846"
                ],
                "target": [],
                "expression_sum": {
                    "class1": 7502.0,
                    "class2": 8518.0
                },
                "uniprot_ids": [
                    "O09111"
                ]
            },
            "ENSMUSG00000068487": {
                "id": "ENSMUSG00000068487",
                "type": "Gene",
                "symbol": "Ndufa11b",
                "source": [
                    "ENSMUSG00000001280",
                    "ENSMUSG00000003032",
                    "ENSMUSG00000038914",
                    "ENSMUSG00000015846"
                ],
                "target": [],
                "expression_sum": {
                    "class1": 2712.0,
                    "class2": 2506.0
                }
            },
            "ENSMUSG00000022450": {
                "id": "ENSMUSG00000022450",
                "type": "Gene",
                "symbol": "Ndufa6",
                "source": [
                    "ENSMUSG00000029135",
                    "ENSMUSG00000021944",
                    "ENSMUSG00000030256",
                    "ENSMUSG00000052837",
                    "ENSMUSG00000024985",
                    "ENSMUSG00000030557",
                    "ENSMUSG00000001280",
                    "ENSMUSG00000019768",
                    "ENSMUSG00000028565",
                    "ENSMUSG00000035158",
                    "ENSMUSG00000024140",
                    "ENSMUSG00000075595",
                    "ENSMUSG00000031885",
                    "ENSMUSG00000008575",
                    "ENSMUSG00000031715",
                    "ENSMUSG00000021775",
                    "ENSMUSG00000015846"
                ],
                "target": [],
                "expression_sum": {
                    "class1": 15920.0,
                    "class2": 12759.0
                },
                "uniprot_ids": [
                    "Q9CQZ5",
                    "A0A2R8W6V1",
                    "A0A2R8VHX1",
                    "A0A2R8VI90"
                ]
            },
            "ENSMUSG00000022890": {
                "id": "ENSMUSG00000022890",
                "type": "Gene",
                "symbol": "Atp5j",
                "source": [
                    "ENSMUSG00000057469",
                    "ENSMUSG00000015839",
                    "ENSMUSG00000010175",
                    "ENSMUSG00000056501",
                    "ENSMUSG00000015579",
                    "ENSMUSG00000001280",
                    "ENSMUSG00000019768",
                    "ENSMUSG00000031885",
                    "ENSMUSG00000015605",
                    "ENSMUSG00000021775",
                    "ENSMUSG00000031715"
                ],
                "target": [],
                "expression_sum": {
                    "class1": 12832.0,
                    "class2": 37118.0
                },
                "uniprot_ids": [
                    "P97450",
                    "E9QAD6"
                ]
            },
            "ENSMUSG00000080836": {
                "id": "ENSMUSG00000080836",
                "type": "Gene",
                "symbol": "Gm6444",
                "source": [
                    "ENSMUSG00000001280",
                    "ENSMUSG00000056501",
                    "ENSMUSG00000015605"
                ],
                "target": [],
                "expression_sum": {
                    "class1": 413.0,
                    "class2": 1889.0
                }
            },
            "ENSMUSG00000096842": {
                "id": "ENSMUSG00000096842",
                "type": "Gene",
                "symbol": "Gm10736",
                "source": [
                    "ENSMUSG00000001280",
                    "ENSMUSG00000019768"
                ],
                "target": [],
                "expression_sum": {
                    "class1": 206.0,
                    "class2": 593.0
                }
            },
            "ENSMUSG00000004610": {
                "id": "ENSMUSG00000004610",
                "type": "Gene",
                "symbol": "Etfb",
                "source": [
                    "ENSMUSG00000015839",
                    "ENSMUSG00000078671",
                    "ENSMUSG00000056501",
                    "ENSMUSG00000015579",
                    "ENSMUSG00000051469",
                    "ENSMUSG00000047371",
                    "ENSMUSG00000001280",
                    "ENSMUSG00000019768",
                    "ENSMUSG00000038914",
                    "ENSMUSG00000024498",
                    "ENSMUSG00000055320",
                    "ENSMUSG00000053110",
                    "ENSMUSG00000024695",
                    "ENSMUSG00000068267",
                    "ENSMUSG00000031885",
                    "ENSMUSG00000015605",
                    "ENSMUSG00000015846"
                ],
                "target": [],
                "expression_sum": {
                    "class1": 16226.0,
                    "class2": 9774.0
                },
                "uniprot_ids": [
                    "Q9DCW4",
                    "A0A0U1RNR3",
                    "A0A0U1RNP5",
                    "A0A0U1RQB4",
                    "A0A0U1RNK9"
                ]
            },
            "ENSMUSG00000022354": {
                "id": "ENSMUSG00000022354",
                "type": "Gene",
                "symbol": "Ndufb9",
                "source": [
                    "ENSMUSG00000029135",
                    "ENSMUSG00000021944",
                    "ENSMUSG00000031627",
                    "ENSMUSG00000015605",
                    "ENSMUSG00000008575",
                    "ENSMUSG00000022346",
                    "ENSMUSG00000058239",
                    "ENSMUSG00000015579",
                    "ENSMUSG00000019768",
                    "ENSMUSG00000022641",
                    "ENSMUSG00000039191",
                    "ENSMUSG00000031715",
                    "ENSMUSG00000015839",
                    "ENSMUSG00000030557",
                    "ENSMUSG00000033943",
                    "ENSMUSG00000008958",
                    "ENSMUSG00000075595",
                    "ENSMUSG00000056501",
                    "ENSMUSG00000001280",
                    "ENSMUSG00000025958"
                ],
                "target": [],
                "expression_sum": {
                    "class1": 22185.0,
                    "class2": 17371.0
                },
                "uniprot_ids": [
                    "Q9CQJ8"
                ]
            },
            "ENSMUSG00000026817": {
                "id": "ENSMUSG00000026817",
                "type": "Gene",
                "symbol": "Ak1",
                "source": [
                    "ENSMUSG00000029135",
                    "ENSMUSG00000001419",
                    "ENSMUSG00000015605",
                    "ENSMUSG00000026610",
                    "ENSMUSG00000056501",
                    "ENSMUSG00000015579",
                    "ENSMUSG00000030557",
                    "ENSMUSG00000001280",
                    "ENSMUSG00000005566",
                    "ENSMUSG00000015843",
                    "ENSMUSG00000005698",
                    "ENSMUSG00000019768",
                    "ENSMUSG00000068267",
                    "ENSMUSG00000038615",
                    "ENSMUSG00000058756",
                    "ENSMUSG00000008575",
                    "ENSMUSG00000031715"
                ],
                "target": [],
                "expression_sum": {
                    "class1": 3795.0,
                    "class2": 2402.0
                },
                "uniprot_ids": [
                    "Q9R0Y5",
                    "Z4YN97",
                    "A0A0A6YXW8"
                ]
            },
            "ENSMUSG00000015733": {
                "id": "ENSMUSG00000015733",
                "type": "Gene",
                "symbol": "Capza2",
                "source": [
                    "ENSMUSG00000024238",
                    "ENSMUSG00000021250",
                    "ENSMUSG00000019817",
                    "ENSMUSG00000052534",
                    "ENSMUSG00000027109",
                    "ENSMUSG00000001280",
                    "ENSMUSG00000035158",
                    "ENSMUSG00000005583",
                    "ENSMUSG00000044167",
                    "ENSMUSG00000022314"
                ],
                "target": [],
                "expression_sum": {
                    "class1": 11290.0,
                    "class2": 3116.0
                },
                "uniprot_ids": [
                    "P47754",
                    "A0A0N4SVM0",
                    "D6RCW7"
                ]
            },
            "ENSMUSG00000032399": {
                "id": "ENSMUSG00000032399",
                "type": "Gene",
                "symbol": "Rpl4",
                "source": [
                    "ENSMUSG00000001280",
                    "ENSMUSG00000019768",
                    "ENSMUSG00000015579",
                    "ENSMUSG00000029135"
                ],
                "target": [],
                "expression_sum": {
                    "class1": 40612.0,
                    "class2": 4982.0
                },
                "uniprot_ids": [
                    "Q9D8E6"
                ]
            },
            "ENSMUSG00000083563": {
                "id": "ENSMUSG00000083563",
                "type": "Gene",
                "symbol": "Gm13340",
                "source": [
                    "ENSMUSG00000001280",
                    "ENSMUSG00000019768"
                ],
                "target": [],
                "expression_sum": {
                    "class1": 513886.0,
                    "class2": 1573504.0
                }
            },
            "ENSMUSG00000064338": {
                "id": "ENSMUSG00000064338",
                "type": "Gene",
                "symbol": "TrnV",
                "source": [
                    "ENSMUSG00000001419",
                    "ENSMUSG00000028639",
                    "ENSMUSG00000030557",
                    "ENSMUSG00000029249",
                    "ENSMUSG00000001280",
                    "ENSMUSG00000019768",
                    "ENSMUSG00000015843",
                    "ENSMUSG00000031885",
                    "ENSMUSG00000058756"
                ],
                "target": [],
                "expression_sum": {
                    "class1": 1115.0,
                    "class2": 1859.0
                }
            },
            "ENSMUSG00000064367": {
                "id": "ENSMUSG00000064367",
                "type": "Gene",
                "symbol": "ND5",
                "source": [
                    "ENSMUSG00000001419",
                    "ENSMUSG00000021944",
                    "ENSMUSG00000052837",
                    "ENSMUSG00000001280",
                    "ENSMUSG00000019768",
                    "ENSMUSG00000015846"
                ],
                "target": [],
                "expression_sum": {
                    "class1": 781750.0,
                    "class2": 893789.0
                },
                "uniprot_ids": [
                    "P03921"
                ]
            },
            "ENSMUSG00000027288": {
                "id": "ENSMUSG00000027288",
                "type": "Gene",
                "symbol": "Zfp106",
                "source": [
                    "ENSMUSG00000029135",
                    "ENSMUSG00000001419",
                    "ENSMUSG00000015839",
                    "ENSMUSG00000015605",
                    "ENSMUSG00000056501",
                    "ENSMUSG00000015579",
                    "ENSMUSG00000030557",
                    "ENSMUSG00000029249",
                    "ENSMUSG00000001280",
                    "ENSMUSG00000005566",
                    "ENSMUSG00000019768",
                    "ENSMUSG00000005698",
                    "ENSMUSG00000022641",
                    "ENSMUSG00000032238",
                    "ENSMUSG00000031885",
                    "ENSMUSG00000008575",
                    "ENSMUSG00000035158",
                    "ENSMUSG00000031715"
                ],
                "target": [],
                "expression_sum": {
                    "class1": 2134.0,
                    "class2": 2114.0
                },
                "uniprot_ids": [
                    "F6QFS1",
                    "E9PVH6",
                    "F6SID8",
                    "E9Q7S1",
                    "R4GML0",
                    "F6UL00"
                ]
            },
            "ENSMUSG00000035242": {
                "id": "ENSMUSG00000035242",
                "type": "Gene",
                "symbol": "Oaz1",
                "source": [
                    "ENSMUSG00000032897",
                    "ENSMUSG00000001419",
                    "ENSMUSG00000022346",
                    "ENSMUSG00000015839",
                    "ENSMUSG00000056501",
                    "ENSMUSG00000051469",
                    "ENSMUSG00000001280",
                    "ENSMUSG00000019768",
                    "ENSMUSG00000068267"
                ],
                "target": [],
                "expression_sum": {
                    "class1": 33627.0,
                    "class2": 3141.0
                },
                "uniprot_ids": [
                    "P54369",
                    "J3QM96",
                    "F8WI29",
                    "J3QMV2",
                    "J3QM60",
                    "J3QME8"
                ]
            },
            "ENSMUSG00000091537": {
                "id": "ENSMUSG00000091537",
                "type": "Gene",
                "symbol": "Tma7",
                "source": [
                    "ENSMUSG00000001419",
                    "ENSMUSG00000021250",
                    "ENSMUSG00000015579",
                    "ENSMUSG00000001280",
                    "ENSMUSG00000019768",
                    "ENSMUSG00000015605",
                    "ENSMUSG00000035158",
                    "ENSMUSG00000015846"
                ],
                "target": [],
                "expression_sum": {
                    "class1": 3915.0,
                    "class2": 1802.0
                },
                "uniprot_ids": [
                    "Q8K003",
                    "A0A0A6YXZ5"
                ]
            },
            "ENSMUSG00000064343": {
                "id": "ENSMUSG00000064343",
                "type": "Gene",
                "symbol": "TrnQ",
                "source": [
                    "ENSMUSG00000001419",
                    "ENSMUSG00000056501",
                    "ENSMUSG00000001280",
                    "ENSMUSG00000019768",
                    "ENSMUSG00000015605"
                ],
                "target": [],
                "expression_sum": {
                    "class1": 106516.0,
                    "class2": 451310.0
                }
            },
            "ENSMUSG00000002416": {
                "id": "ENSMUSG00000002416",
                "type": "Gene",
                "symbol": "Ndufb2",
                "source": [
                    "ENSMUSG00000029135",
                    "ENSMUSG00000015839",
                    "ENSMUSG00000052534",
                    "ENSMUSG00000015605",
                    "ENSMUSG00000055116",
                    "ENSMUSG00000056501",
                    "ENSMUSG00000015579",
                    "ENSMUSG00000028639",
                    "ENSMUSG00000029249",
                    "ENSMUSG00000001280",
                    "ENSMUSG00000019768",
                    "ENSMUSG00000047371",
                    "ENSMUSG00000068267",
                    "ENSMUSG00000031885",
                    "ENSMUSG00000008575",
                    "ENSMUSG00000014592",
                    "ENSMUSG00000015846"
                ],
                "target": [],
                "expression_sum": {
                    "class1": 12451.0,
                    "class2": 9814.0
                },
                "uniprot_ids": [
                    "Q9CPU2"
                ]
            },
            "ENSMUSG00000022186": {
                "id": "ENSMUSG00000022186",
                "type": "Gene",
                "symbol": "Oxct1",
                "source": [
                    "ENSMUSG00000015839",
                    "ENSMUSG00000019817",
                    "ENSMUSG00000058239",
                    "ENSMUSG00000056501",
                    "ENSMUSG00000030557",
                    "ENSMUSG00000029249",
                    "ENSMUSG00000001280",
                    "ENSMUSG00000019768",
                    "ENSMUSG00000005583",
                    "ENSMUSG00000005698",
                    "ENSMUSG00000024498",
                    "ENSMUSG00000033943",
                    "ENSMUSG00000058756",
                    "ENSMUSG00000008575",
                    "ENSMUSG00000039191",
                    "ENSMUSG00000031715"
                ],
                "target": [],
                "expression_sum": {
                    "class1": 4321.0,
                    "class2": 8337.0
                },
                "uniprot_ids": [
                    "Q9D0K2",
                    "Q3UJQ9"
                ]
            },
            "ENSMUSG00000032060": {
                "id": "ENSMUSG00000032060",
                "type": "Gene",
                "symbol": "Cryab",
                "source": [
                    "ENSMUSG00000024955",
                    "ENSMUSG00000001280",
                    "ENSMUSG00000019768",
                    "ENSMUSG00000025958",
                    "ENSMUSG00000024140",
                    "ENSMUSG00000008575",
                    "ENSMUSG00000015846"
                ],
                "target": [],
                "expression_sum": {
                    "class1": 10916.0,
                    "class2": 16314.0
                },
                "uniprot_ids": [
                    "P23927",
                    "A0A1L1SRG5"
                ]
            },
            "ENSMUSG00000023861": {
                "id": "ENSMUSG00000023861",
                "type": "Gene",
                "symbol": "Mpc1",
                "source": [
                    "ENSMUSG00000029135",
                    "ENSMUSG00000015605",
                    "ENSMUSG00000008575",
                    "ENSMUSG00000021775",
                    "ENSMUSG00000058239",
                    "ENSMUSG00000015579",
                    "ENSMUSG00000051469",
                    "ENSMUSG00000019768",
                    "ENSMUSG00000028565",
                    "ENSMUSG00000031885",
                    "ENSMUSG00000053110",
                    "ENSMUSG00000031715",
                    "ENSMUSG00000015846",
                    "ENSMUSG00000015839",
                    "ENSMUSG00000028639",
                    "ENSMUSG00000030557",
                    "ENSMUSG00000060601",
                    "ENSMUSG00000001419",
                    "ENSMUSG00000078671",
                    "ENSMUSG00000001280",
                    "ENSMUSG00000025958"
                ],
                "target": [],
                "expression_sum": {
                    "class1": 127.0,
                    "class2": 3511.0
                },
                "uniprot_ids": [
                    "P63030",
                    "D3YWY6",
                    "E9Q0V4",
                    "D3Z5S0",
                    "D3Z786"
                ]
            },
            "ENSMUSG00000028367": {
                "id": "ENSMUSG00000028367",
                "type": "Gene",
                "symbol": "Txn1",
                "source": [
                    "ENSMUSG00000003032",
                    "ENSMUSG00000021250",
                    "ENSMUSG00000018604",
                    "ENSMUSG00000023034",
                    "ENSMUSG00000056501",
                    "ENSMUSG00000032228",
                    "ENSMUSG00000018899",
                    "ENSMUSG00000001280",
                    "ENSMUSG00000040732",
                    "ENSMUSG00000052684",
                    "ENSMUSG00000024140",
                    "ENSMUSG00000020919",
                    "ENSMUSG00000016087",
                    "ENSMUSG00000015605"
                ],
                "target": [],
                "expression_sum": {
                    "class1": 8901.0,
                    "class2": 3884.0
                },
                "uniprot_ids": [
                    "P10639"
                ]
            },
            "ENSMUSG00000004980": {
                "id": "ENSMUSG00000004980",
                "type": "Gene",
                "symbol": "Hnrnpa2b1",
                "source": [
                    "ENSMUSG00000029135",
                    "ENSMUSG00000010175",
                    "ENSMUSG00000015839",
                    "ENSMUSG00000056501",
                    "ENSMUSG00000015579",
                    "ENSMUSG00000001280",
                    "ENSMUSG00000019768",
                    "ENSMUSG00000028565",
                    "ENSMUSG00000014592",
                    "ENSMUSG00000008958",
                    "ENSMUSG00000053110",
                    "ENSMUSG00000022389",
                    "ENSMUSG00000068267",
                    "ENSMUSG00000076431",
                    "ENSMUSG00000008575",
                    "ENSMUSG00000039191",
                    "ENSMUSG00000029913",
                    "ENSMUSG00000015846"
                ],
                "target": [],
                "expression_sum": {
                    "class1": 29892.0,
                    "class2": 7972.0
                },
                "uniprot_ids": [
                    "O88569",
                    "A0A0N4SUM2"
                ]
            },
            "ENSMUSG00000026260": {
                "id": "ENSMUSG00000026260",
                "type": "Gene",
                "symbol": "Ndufa10",
                "source": [
                    "ENSMUSG00000029135",
                    "ENSMUSG00000057469",
                    "ENSMUSG00000015605",
                    "ENSMUSG00000052837",
                    "ENSMUSG00000056501",
                    "ENSMUSG00000030557",
                    "ENSMUSG00000029249",
                    "ENSMUSG00000001280",
                    "ENSMUSG00000005566",
                    "ENSMUSG00000019768",
                    "ENSMUSG00000004151",
                    "ENSMUSG00000022641",
                    "ENSMUSG00000031885",
                    "ENSMUSG00000008575",
                    "ENSMUSG00000039191",
                    "ENSMUSG00000021775",
                    "ENSMUSG00000031715"
                ],
                "target": [],
                "expression_sum": {
                    "class1": 8426.0,
                    "class2": 5351.0
                },
                "uniprot_ids": [
                    "Q99LC3",
                    "A0A087WR38",
                    "A0A087WRD4"
                ]
            },
            "ENSMUSG00000061518": {
                "id": "ENSMUSG00000061518",
                "type": "Gene",
                "symbol": "Cox5b",
                "source": [
                    "ENSMUSG00000029135",
                    "ENSMUSG00000001419",
                    "ENSMUSG00000057469",
                    "ENSMUSG00000015605",
                    "ENSMUSG00000056501",
                    "ENSMUSG00000001280",
                    "ENSMUSG00000019768",
                    "ENSMUSG00000031885",
                    "ENSMUSG00000008575",
                    "ENSMUSG00000022314",
                    "ENSMUSG00000015846"
                ],
                "target": [],
                "expression_sum": {
                    "class1": 8730.0,
                    "class2": 13479.0
                },
                "uniprot_ids": [
                    "Q9D881",
                    "A0A0A6YVR0",
                    "F7C106"
                ]
            },
            "ENSMUSG00000040666": {
                "id": "ENSMUSG00000040666",
                "type": "Gene",
                "symbol": "Sh3bgr",
                "source": [
                    "ENSMUSG00000015605",
                    "ENSMUSG00000001280",
                    "ENSMUSG00000019768",
                    "ENSMUSG00000005698",
                    "ENSMUSG00000025958",
                    "ENSMUSG00000008575",
                    "ENSMUSG00000015846"
                ],
                "target": [],
                "expression_sum": {
                    "class1": 11244.0,
                    "class2": 4704.0
                },
                "uniprot_ids": [
                    "Q9WUZ7",
                    "D3Z3Z9",
                    "A0A3B2W7J5",
                    "F7CKA6",
                    "F6QLU8",
                    "A0A3B2WCC8",
                    "A0A3B2WB97",
                    "E9Q2Q9",
                    "F6X737"
                ]
            },
            "ENSMUSG00000064345": {
                "id": "ENSMUSG00000064345",
                "type": "Gene",
                "symbol": "ND2",
                "source": [
                    "ENSMUSG00000001419",
                    "ENSMUSG00000021944",
                    "ENSMUSG00000052837",
                    "ENSMUSG00000001280",
                    "ENSMUSG00000019768"
                ],
                "target": [],
                "expression_sum": {
                    "class1": 695799.0,
                    "class2": 2075792.0
                },
                "uniprot_ids": [
                    "P03893"
                ]
            },
            "ENSMUSG00000033161": {
                "id": "ENSMUSG00000033161",
                "type": "Gene",
                "symbol": "Atp1a1",
                "source": [
                    "ENSMUSG00000029135",
                    "ENSMUSG00000001419",
                    "ENSMUSG00000057469",
                    "ENSMUSG00000059552",
                    "ENSMUSG00000015839",
                    "ENSMUSG00000015605",
                    "ENSMUSG00000026610",
                    "ENSMUSG00000056501",
                    "ENSMUSG00000029249",
                    "ENSMUSG00000001280",
                    "ENSMUSG00000019768",
                    "ENSMUSG00000005583",
                    "ENSMUSG00000005698",
                    "ENSMUSG00000058756",
                    "ENSMUSG00000008575"
                ],
                "target": [],
                "expression_sum": {
                    "class1": 3091.0,
                    "class2": 7460.0
                },
                "uniprot_ids": [
                    "Q8VDN2"
                ]
            },
            "ENSMUSG00000028820": {
                "id": "ENSMUSG00000028820",
                "type": "Gene",
                "symbol": "Sfpq",
                "source": [
                    "ENSMUSG00000079509",
                    "ENSMUSG00000071076",
                    "ENSMUSG00000018604",
                    "ENSMUSG00000001280",
                    "ENSMUSG00000005698",
                    "ENSMUSG00000017146",
                    "ENSMUSG00000076431",
                    "ENSMUSG00000031715"
                ],
                "target": [],
                "expression_sum": {
                    "class1": 6742.0,
                    "class2": 1572.0
                },
                "uniprot_ids": [
                    "Q8VIJ6"
                ]
            },
            "ENSMUSG00000022110": {
                "id": "ENSMUSG00000022110",
                "type": "Gene",
                "symbol": "Sucla2",
                "source": [
                    "ENSMUSG00000029135",
                    "ENSMUSG00000076431",
                    "ENSMUSG00000015605",
                    "ENSMUSG00000008575",
                    "ENSMUSG00000021775",
                    "ENSMUSG00000006215",
                    "ENSMUSG00000010175",
                    "ENSMUSG00000019768",
                    "ENSMUSG00000028565",
                    "ENSMUSG00000005698",
                    "ENSMUSG00000031885",
                    "ENSMUSG00000039191",
                    "ENSMUSG00000031715",
                    "ENSMUSG00000015839",
                    "ENSMUSG00000030557",
                    "ENSMUSG00000022389",
                    "ENSMUSG00000001419",
                    "ENSMUSG00000078671",
                    "ENSMUSG00000026610",
                    "ENSMUSG00000056501",
                    "ENSMUSG00000001280"
                ],
                "target": [],
                "expression_sum": {
                    "class1": 5648.0,
                    "class2": 6979.0
                },
                "uniprot_ids": [
                    "Q9Z2I9"
                ]
            },
            "ENSMUSG00000021843": {
                "id": "ENSMUSG00000021843",
                "type": "Gene",
                "symbol": "Ktn1",
                "source": [
                    "ENSMUSG00000015579",
                    "ENSMUSG00000001280",
                    "ENSMUSG00000019768",
                    "ENSMUSG00000017146",
                    "ENSMUSG00000031715",
                    "ENSMUSG00000022314",
                    "ENSMUSG00000015846"
                ],
                "target": [],
                "expression_sum": {
                    "class1": 13474.0,
                    "class2": 6036.0
                },
                "uniprot_ids": [
                    "A0A087WP48",
                    "F8VQC7",
                    "A0A087WQQ5",
                    "A0A087WNW3",
                    "A0A087WQI3",
                    "A0A087WP85",
                    "A0A087WRS1",
                    "A0A087WQG4",
                    "A0A087WPX0",
                    "A0A087WQF8",
                    "A0A087WS23",
                    "A0A087WQ25",
                    "A0A087WS29",
                    "A0A087WP14",
                    "A0A087WPY8",
                    "A0A087WPW5",
                    "A0A087WQD0",
                    "A0A087WR91",
                    "A0A087WS04"
                ]
            },
            "ENSMUSG00000006219": {
                "id": "ENSMUSG00000006219",
                "type": "Gene",
                "symbol": "Fblim1",
                "source": [
                    "ENSMUSG00000029135",
                    "ENSMUSG00000019789",
                    "ENSMUSG00000015839",
                    "ENSMUSG00000049577",
                    "ENSMUSG00000056501",
                    "ENSMUSG00000018899",
                    "ENSMUSG00000030557",
                    "ENSMUSG00000001280",
                    "ENSMUSG00000005566",
                    "ENSMUSG00000019768",
                    "ENSMUSG00000005698",
                    "ENSMUSG00000058756",
                    "ENSMUSG00000015605"
                ],
                "target": [],
                "expression_sum": {
                    "class1": 1043.0,
                    "class2": 1448.0
                },
                "uniprot_ids": [
                    "Q71FD7",
                    "A2ADE8",
                    "A2ADE6",
                    "A2ADE7",
                    "A2ADE9",
                    "A2ADF3",
                    "A2ADF1",
                    "A2ADF2",
                    "A2ADE5",
                    "A2ADF4"
                ]
            },
            "ENSMUSG00000035674": {
                "id": "ENSMUSG00000035674",
                "type": "Gene",
                "symbol": "Ndufa3",
                "source": [
                    "ENSMUSG00000015839",
                    "ENSMUSG00000026610",
                    "ENSMUSG00000056501",
                    "ENSMUSG00000001280",
                    "ENSMUSG00000019768",
                    "ENSMUSG00000055320",
                    "ENSMUSG00000031885",
                    "ENSMUSG00000015605",
                    "ENSMUSG00000021775",
                    "ENSMUSG00000015846"
                ],
                "target": [],
                "expression_sum": {
                    "class1": 26340.0,
                    "class2": 25769.0
                },
                "uniprot_ids": [
                    "Q9CQ91",
                    "D3Z670",
                    "F7BLS8",
                    "D6RGT3"
                ]
            },
            "ENSMUSG00000037152": {
                "id": "ENSMUSG00000037152",
                "type": "Gene",
                "symbol": "Ndufc1",
                "source": [
                    "ENSMUSG00000001419",
                    "ENSMUSG00000015839",
                    "ENSMUSG00000052534",
                    "ENSMUSG00000056501",
                    "ENSMUSG00000051469",
                    "ENSMUSG00000001280",
                    "ENSMUSG00000019768",
                    "ENSMUSG00000031885",
                    "ENSMUSG00000031715",
                    "ENSMUSG00000021775",
                    "ENSMUSG00000015846"
                ],
                "target": [],
                "expression_sum": {
                    "class1": 27708.0,
                    "class2": 24125.0
                },
                "uniprot_ids": [
                    "Q9CQY9"
                ]
            },
            "ENSMUSG00000023456": {
                "id": "ENSMUSG00000023456",
                "type": "Gene",
                "symbol": "Tpi1",
                "source": [
                    "ENSMUSG00000029135",
                    "ENSMUSG00000001419",
                    "ENSMUSG00000001911",
                    "ENSMUSG00000015605",
                    "ENSMUSG00000056501",
                    "ENSMUSG00000015579",
                    "ENSMUSG00000051469",
                    "ENSMUSG00000001280",
                    "ENSMUSG00000019768",
                    "ENSMUSG00000008575",
                    "ENSMUSG00000039191",
                    "ENSMUSG00000000567"
                ],
                "target": [],
                "expression_sum": {
                    "class1": 4685.0,
                    "class2": 2207.0
                },
                "uniprot_ids": [
                    "P17751",
                    "H7BXC3"
                ]
            },
            "ENSMUSG00000025290": {
                "id": "ENSMUSG00000025290",
                "type": "Gene",
                "symbol": "Rps24",
                "source": [
                    "ENSMUSG00000018604",
                    "ENSMUSG00000036461",
                    "ENSMUSG00000001280",
                    "ENSMUSG00000019768",
                    "ENSMUSG00000005583",
                    "ENSMUSG00000028565",
                    "ENSMUSG00000040732",
                    "ENSMUSG00000053110",
                    "ENSMUSG00000008575"
                ],
                "target": [],
                "expression_sum": {
                    "class1": 9707.0,
                    "class2": 2805.0
                },
                "uniprot_ids": [
                    "P62849",
                    "A0A286YEB7"
                ]
            },
            "ENSMUSG00000071528": {
                "id": "ENSMUSG00000071528",
                "type": "Gene",
                "symbol": "Atp5md",
                "source": [
                    "ENSMUSG00000015839",
                    "ENSMUSG00000006215",
                    "ENSMUSG00000015605",
                    "ENSMUSG00000021250",
                    "ENSMUSG00000023927",
                    "ENSMUSG00000056501",
                    "ENSMUSG00000058239",
                    "ENSMUSG00000001280",
                    "ENSMUSG00000019768",
                    "ENSMUSG00000031885",
                    "ENSMUSG00000008575",
                    "ENSMUSG00000021775",
                    "ENSMUSG00000015846"
                ],
                "target": [],
                "expression_sum": {
                    "class1": 432.0,
                    "class2": 3467.0
                },
                "uniprot_ids": [
                    "Q78IK2"
                ]
            },
            "ENSMUSG00000053329": {
                "id": "ENSMUSG00000053329",
                "type": "Gene",
                "symbol": "Gatd3a",
                "source": [
                    "ENSMUSG00000021250",
                    "ENSMUSG00000052534",
                    "ENSMUSG00000024985",
                    "ENSMUSG00000001280",
                    "ENSMUSG00000019768",
                    "ENSMUSG00000008575",
                    "ENSMUSG00000015846",
                    "ENSMUSG00000000567"
                ],
                "target": [],
                "expression_sum": {
                    "class1": 1568.0,
                    "class2": 1607.0
                },
                "uniprot_ids": [
                    "Q9D172",
                    "A0A1W2P870",
                    "A0A1W2P7B6"
                ]
            },
            "ENSMUSG00000038690": {
                "id": "ENSMUSG00000038690",
                "type": "Gene",
                "symbol": "Atp5j2",
                "source": [
                    "ENSMUSG00000057469",
                    "ENSMUSG00000030341",
                    "ENSMUSG00000015839",
                    "ENSMUSG00000052837",
                    "ENSMUSG00000015579",
                    "ENSMUSG00000056501",
                    "ENSMUSG00000001280",
                    "ENSMUSG00000019768",
                    "ENSMUSG00000015605",
                    "ENSMUSG00000014592",
                    "ENSMUSG00000021775",
                    "ENSMUSG00000031715"
                ],
                "target": [],
                "expression_sum": {
                    "class1": 21333.0,
                    "class2": 20451.0
                },
                "uniprot_ids": [
                    "P56135",
                    "F6USF4",
                    "M0QWW1",
                    "F8WHP8"
                ]
            },
            "ENSMUSG00000021000": {
                "id": "ENSMUSG00000021000",
                "type": "Gene",
                "symbol": "Mia2",
                "source": [
                    "ENSMUSG00000058440",
                    "ENSMUSG00000024238",
                    "ENSMUSG00000057469",
                    "ENSMUSG00000025612",
                    "ENSMUSG00000015839",
                    "ENSMUSG00000010175",
                    "ENSMUSG00000019564",
                    "ENSMUSG00000022346",
                    "ENSMUSG00000030256",
                    "ENSMUSG00000056749",
                    "ENSMUSG00000051469",
                    "ENSMUSG00000030557",
                    "ENSMUSG00000001280",
                    "ENSMUSG00000019768",
                    "ENSMUSG00000031885",
                    "ENSMUSG00000035158"
                ],
                "target": [],
                "expression_sum": {
                    "class1": 2604.0,
                    "class2": 2197.0
                },
                "uniprot_ids": [
                    "H3BL86",
                    "H3BJ71",
                    "H3BKL6",
                    "H3BJS5",
                    "H3BJ84",
                    "H3BKH2",
                    "H3BL19",
                    "H3BJ35",
                    "H3BJY7",
                    "H3BJS0",
                    "E9Q3S2",
                    "H3BK44",
                    "A0A1W2P711",
                    "H3BJI6",
                    "E9PY90",
                    "H3BK48"
                ]
            },
            "ENSMUSG00000001604": {
                "id": "ENSMUSG00000001604",
                "type": "Gene",
                "symbol": "Tcea3",
                "source": [
                    "ENSMUSG00000001280",
                    "ENSMUSG00000017299"
                ],
                "target": [],
                "expression_sum": {
                    "class1": 942.0,
                    "class2": 1838.0
                },
                "uniprot_ids": [
                    "P23881",
                    "A2AW45"
                ]
            },
            "ENSMUSG00000032199": {
                "id": "ENSMUSG00000032199",
                "type": "Gene",
                "symbol": "Polr2m",
                "source": [
                    "ENSMUSG00000029135",
                    "ENSMUSG00000031627",
                    "ENSMUSG00000024695",
                    "ENSMUSG00000057469",
                    "ENSMUSG00000022346",
                    "ENSMUSG00000030256",
                    "ENSMUSG00000015579",
                    "ENSMUSG00000051469",
                    "ENSMUSG00000019768",
                    "ENSMUSG00000024974",
                    "ENSMUSG00000058440",
                    "ENSMUSG00000015839",
                    "ENSMUSG00000030557",
                    "ENSMUSG00000068267",
                    "ENSMUSG00000026610",
                    "ENSMUSG00000055116",
                    "ENSMUSG00000056501",
                    "ENSMUSG00000029249",
                    "ENSMUSG00000001280",
                    "ENSMUSG00000099083"
                ],
                "target": [],
                "expression_sum": {
                    "class1": 3426.0,
                    "class2": 2081.0
                },
                "uniprot_ids": [
                    "Q6P6I6",
                    "A0A3B2WBC6",
                    "E9PZ03"
                ]
            },
            "ENSMUSG00000064370": {
                "id": "ENSMUSG00000064370",
                "type": "Gene",
                "symbol": "CYTB",
                "source": [
                    "ENSMUSG00000001280",
                    "ENSMUSG00000019768",
                    "ENSMUSG00000021944",
                    "ENSMUSG00000052837"
                ],
                "target": [],
                "expression_sum": {
                    "class1": 1943163.0,
                    "class2": 2762249.0
                },
                "uniprot_ids": [
                    "P00158"
                ]
            },
            "ENSMUSG00000038370": {
                "id": "ENSMUSG00000038370",
                "type": "Gene",
                "symbol": "Pcp4l1",
                "source": [
                    "ENSMUSG00000015839",
                    "ENSMUSG00000010175",
                    "ENSMUSG00000078671",
                    "ENSMUSG00000056501",
                    "ENSMUSG00000001280",
                    "ENSMUSG00000019768",
                    "ENSMUSG00000058756",
                    "ENSMUSG00000015846"
                ],
                "target": [],
                "expression_sum": {
                    "class1": 3251.0,
                    "class2": 2245.0
                },
                "uniprot_ids": [
                    "Q6W8Q3"
                ]
            },
            "ENSMUSG00000093674": {
                "id": "ENSMUSG00000093674",
                "type": "Gene",
                "symbol": "Rpl41",
                "source": [
                    "ENSMUSG00000048387",
                    "ENSMUSG00000001419",
                    "ENSMUSG00000001280",
                    "ENSMUSG00000019768",
                    "ENSMUSG00000038692",
                    "ENSMUSG00000015846"
                ],
                "target": [],
                "expression_sum": {
                    "class1": 14482.0,
                    "class2": 4102.0
                },
                "uniprot_ids": [
                    "P62947",
                    "H3BK91"
                ]
            },
            "ENSMUSG00000000088": {
                "id": "ENSMUSG00000000088",
                "type": "Gene",
                "symbol": "Cox5a",
                "source": [
                    "ENSMUSG00000029135",
                    "ENSMUSG00000034673",
                    "ENSMUSG00000037174",
                    "ENSMUSG00000024695",
                    "ENSMUSG00000044452",
                    "ENSMUSG00000015605",
                    "ENSMUSG00000008575",
                    "ENSMUSG00000057469",
                    "ENSMUSG00000010175",
                    "ENSMUSG00000001156",
                    "ENSMUSG00000015579",
                    "ENSMUSG00000090258",
                    "ENSMUSG00000019768",
                    "ENSMUSG00000028565",
                    "ENSMUSG00000031885",
                    "ENSMUSG00000015846",
                    "ENSMUSG00000015839",
                    "ENSMUSG00000001911",
                    "ENSMUSG00000049878",
                    "ENSMUSG00000028639",
                    "ENSMUSG00000030557",
                    "ENSMUSG00000031734",
                    "ENSMUSG00000033943",
                    "ENSMUSG00000068267",
                    "ENSMUSG00000059878",
                    "ENSMUSG00000014592",
                    "ENSMUSG00000056501",
                    "ENSMUSG00000001280",
                    "ENSMUSG00000025958",
                    "ENSMUSG00000024498",
                    "ENSMUSG00000040841",
                    "ENSMUSG00000055320"
                ],
                "target": [],
                "expression_sum": {
                    "class1": 264.0,
                    "class2": 3902.0
                },
                "uniprot_ids": [
                    "P12787"
                ]
            },
            "ENSMUSG00000025794": {
                "id": "ENSMUSG00000025794",
                "type": "Gene",
                "symbol": "Rpl14",
                "source": [
                    "ENSMUSG00000055148",
                    "ENSMUSG00000018604",
                    "ENSMUSG00000001280",
                    "ENSMUSG00000019768",
                    "ENSMUSG00000040732",
                    "ENSMUSG00000022641",
                    "ENSMUSG00000022895",
                    "ENSMUSG00000000567"
                ],
                "target": [],
                "expression_sum": {
                    "class1": 17351.0,
                    "class2": 2678.0
                },
                "uniprot_ids": [
                    "Q9CR57",
                    "A0A1L1SUF6"
                ]
            },
            "ENSMUSG00000017404": {
                "id": "ENSMUSG00000017404",
                "type": "Gene",
                "symbol": "Rpl19",
                "source": [
                    "ENSMUSG00000024238",
                    "ENSMUSG00000057469",
                    "ENSMUSG00000025612",
                    "ENSMUSG00000052534",
                    "ENSMUSG00000056501",
                    "ENSMUSG00000015579",
                    "ENSMUSG00000001280",
                    "ENSMUSG00000019768",
                    "ENSMUSG00000040732",
                    "ENSMUSG00000025025",
                    "ENSMUSG00000035158",
                    "ENSMUSG00000020644"
                ],
                "target": [],
                "expression_sum": {
                    "class1": 4028.0,
                    "class2": 1165.0
                },
                "uniprot_ids": [
                    "P84099",
                    "A2A547"
                ]
            },
            "ENSMUSG00000033938": {
                "id": "ENSMUSG00000033938",
                "type": "Gene",
                "symbol": "Ndufb7",
                "source": [
                    "ENSMUSG00000026398",
                    "ENSMUSG00000015839",
                    "ENSMUSG00000078671",
                    "ENSMUSG00000015605",
                    "ENSMUSG00000026610",
                    "ENSMUSG00000052837",
                    "ENSMUSG00000056501",
                    "ENSMUSG00000030557",
                    "ENSMUSG00000029249",
                    "ENSMUSG00000001280",
                    "ENSMUSG00000019768",
                    "ENSMUSG00000025958",
                    "ENSMUSG00000014592",
                    "ENSMUSG00000031885",
                    "ENSMUSG00000008575",
                    "ENSMUSG00000039191",
                    "ENSMUSG00000015846"
                ],
                "target": [],
                "expression_sum": {
                    "class1": 15274.0,
                    "class2": 9998.0
                },
                "uniprot_ids": [
                    "Q9CR61"
                ]
            },
            "ENSMUSG00000082884": {
                "id": "ENSMUSG00000082884",
                "type": "Gene",
                "symbol": "Gm13339",
                "source": [
                    "ENSMUSG00000001280"
                ],
                "target": [],
                "expression_sum": {
                    "class1": 35272.0,
                    "class2": 234847.0
                }
            },
            "ENSMUSG00000063694": {
                "id": "ENSMUSG00000063694",
                "type": "Gene",
                "symbol": "Cycs",
                "source": [
                    "ENSMUSG00000029135",
                    "ENSMUSG00000015839",
                    "ENSMUSG00000010175",
                    "ENSMUSG00000015605",
                    "ENSMUSG00000058239",
                    "ENSMUSG00000052837",
                    "ENSMUSG00000056501",
                    "ENSMUSG00000001280",
                    "ENSMUSG00000019768",
                    "ENSMUSG00000032238",
                    "ENSMUSG00000031885",
                    "ENSMUSG00000008575",
                    "ENSMUSG00000039191",
                    "ENSMUSG00000021775",
                    "ENSMUSG00000015846"
                ],
                "target": [],
                "expression_sum": {
                    "class1": 13499.0,
                    "class2": 15508.0
                },
                "uniprot_ids": [
                    "P62897"
                ]
            },
            "ENSMUSG00000091478": {
                "id": "ENSMUSG00000091478",
                "type": "Gene",
                "symbol": "Gm10039",
                "source": [
                    "ENSMUSG00000015839",
                    "ENSMUSG00000058239",
                    "ENSMUSG00000056501",
                    "ENSMUSG00000001280",
                    "ENSMUSG00000019768",
                    "ENSMUSG00000015846",
                    "ENSMUSG00000031885",
                    "ENSMUSG00000039191",
                    "ENSMUSG00000031715"
                ],
                "target": [],
                "expression_sum": {
                    "class1": 8331.0,
                    "class2": 22931.0
                }
            },
            "ENSMUSG00000087943": {
                "id": "ENSMUSG00000087943",
                "type": "Gene",
                "symbol": null,
                "source": [
                    "ENSMUSG00000001280"
                ],
                "target": [],
                "expression_sum": {
                    "class1": 370.0,
                    "class2": 865.0
                }
            },
            "ENSMUSG00000014313": {
                "id": "ENSMUSG00000014313",
                "type": "Gene",
                "symbol": "Cox6c",
                "source": [
                    "ENSMUSG00000015605",
                    "ENSMUSG00000056501",
                    "ENSMUSG00000074733",
                    "ENSMUSG00000001280",
                    "ENSMUSG00000019768",
                    "ENSMUSG00000031885",
                    "ENSMUSG00000008575",
                    "ENSMUSG00000031715",
                    "ENSMUSG00000021775",
                    "ENSMUSG00000015846"
                ],
                "target": [],
                "expression_sum": {
                    "class1": 36766.0,
                    "class2": 38457.0
                },
                "uniprot_ids": [
                    "Q9CPQ1",
                    "A0A0G2JDA6"
                ]
            },
            "ENSMUSG00000084319": {
                "id": "ENSMUSG00000084319",
                "type": "Gene",
                "symbol": "Tpt1-ps3",
                "source": [
                    "ENSMUSG00000001280",
                    "ENSMUSG00000018604"
                ],
                "target": [],
                "expression_sum": {
                    "class1": 7086.0,
                    "class2": 4401.0
                }
            },
            "ENSMUSG00000027133": {
                "id": "ENSMUSG00000027133",
                "type": "Gene",
                "symbol": "Nop10",
                "source": [
                    "ENSMUSG00000022346",
                    "ENSMUSG00000030256",
                    "ENSMUSG00000015579",
                    "ENSMUSG00000001280",
                    "ENSMUSG00000028565",
                    "ENSMUSG00000020919",
                    "ENSMUSG00000014592",
                    "ENSMUSG00000031715"
                ],
                "target": [],
                "expression_sum": {
                    "class1": 9465.0,
                    "class2": 3417.0
                },
                "uniprot_ids": [
                    "Q9CQS2"
                ]
            },
            "ENSMUSG00000025508": {
                "id": "ENSMUSG00000025508",
                "type": "Gene",
                "symbol": "Rplp2",
                "source": [
                    "ENSMUSG00000015839",
                    "ENSMUSG00000018604",
                    "ENSMUSG00000030256",
                    "ENSMUSG00000001280",
                    "ENSMUSG00000032238",
                    "ENSMUSG00000035158",
                    "ENSMUSG00000031715",
                    "ENSMUSG00000015846"
                ],
                "target": [],
                "expression_sum": {
                    "class1": 788.0,
                    "class2": 1417.0
                },
                "uniprot_ids": [
                    "P99027"
                ]
            },
            "ENSMUSG00000059734": {
                "id": "ENSMUSG00000059734",
                "type": "Gene",
                "symbol": "Ndufs8",
                "source": [
                    "ENSMUSG00000024238",
                    "ENSMUSG00000020248",
                    "ENSMUSG00000010175",
                    "ENSMUSG00000022346",
                    "ENSMUSG00000015605",
                    "ENSMUSG00000052534",
                    "ENSMUSG00000015579",
                    "ENSMUSG00000001280",
                    "ENSMUSG00000019768",
                    "ENSMUSG00000005583",
                    "ENSMUSG00000053477",
                    "ENSMUSG00000024974",
                    "ENSMUSG00000008575",
                    "ENSMUSG00000022314"
                ],
                "target": [],
                "expression_sum": {
                    "class1": 12482.0,
                    "class2": 6822.0
                },
                "uniprot_ids": [
                    "Q8K3J1",
                    "A0A494BAW8"
                ]
            },
            "ENSMUSG00000023277": {
                "id": "ENSMUSG00000023277",
                "type": "Gene",
                "symbol": "Twf2",
                "source": [
                    "ENSMUSG00000059552",
                    "ENSMUSG00000018604",
                    "ENSMUSG00000027109",
                    "ENSMUSG00000001280",
                    "ENSMUSG00000031885"
                ],
                "target": [],
                "expression_sum": {
                    "class1": 479.0,
                    "class2": 539.0
                },
                "uniprot_ids": [
                    "Q9Z0P5",
                    "A0A1L1SU53",
                    "A0A1L1STC8",
                    "A0A087WPT3",
                    "A0A087WRG4"
                ]
            },
            "ENSMUSG00000031767": {
                "id": "ENSMUSG00000031767",
                "type": "Gene",
                "symbol": "Nudt7",
                "source": [
                    "ENSMUSG00000026398",
                    "ENSMUSG00000071076",
                    "ENSMUSG00000021944",
                    "ENSMUSG00000001280",
                    "ENSMUSG00000005566",
                    "ENSMUSG00000019768"
                ],
                "target": [],
                "expression_sum": {
                    "class1": 268.0,
                    "class2": 335.0
                },
                "uniprot_ids": [
                    "Q99P30"
                ]
            },
            "ENSMUSG00000056501": {
                "id": "ENSMUSG00000056501",
                "type": "TF",
                "symbol": "Cebpb",
                "source": [
                    "ENSMUSG00000059552",
                    "ENSMUSG00000015839",
                    "ENSMUSG00000041515",
                    "ENSMUSG00000018899",
                    "ENSMUSG00000001280",
                    "ENSMUSG00000019768",
                    "ENSMUSG00000005698"
                ],
                "target": [
                    "ENSMUSG00000015568",
                    "ENSMUSG00000061603",
                    "ENSMUSG00000055775",
                    "ENSMUSG00000061904",
                    "ENSMUSG00000068566",
                    "ENSMUSG00000097088",
                    "ENSMUSG00000035781",
                    "ENSMUSG00000038366",
                    "ENSMUSG00000044330",
                    "ENSMUSG00000074582",
                    "ENSMUSG00000036835",
                    "ENSMUSG00000031403",
                    "ENSMUSG00000034674",
                    "ENSMUSG00000045665",
                    "ENSMUSG00000062908",
                    "ENSMUSG00000007777",
                    "ENSMUSG00000091478",
                    "ENSMUSG00000063457",
                    "ENSMUSG00000002661",
                    "ENSMUSG00000068264",
                    "ENSMUSG00000027022",
                    "ENSMUSG00000042828",
                    "ENSMUSG00000021248",
                    "ENSMUSG00000027447",
                    "ENSMUSG00000015837",
                    "ENSMUSG00000043644",
                    "ENSMUSG00000068040",
                    "ENSMUSG00000044791",
                    "ENSMUSG00000020741",
                    "ENSMUSG00000033991",
                    "ENSMUSG00000021235",
                    "ENSMUSG00000070283",
                    "ENSMUSG00000027984",
                    "ENSMUSG00000041193",
                    "ENSMUSG00000021629",
                    "ENSMUSG00000001962",
                    "ENSMUSG00000004044",
                    "ENSMUSG00000041779",
                    "ENSMUSG00000003363",
                    "ENSMUSG00000024097",
                    "ENSMUSG00000031540",
                    "ENSMUSG00000041881",
                    "ENSMUSG00000030214",
                    "ENSMUSG00000060073",
                    "ENSMUSG00000081773",
                    "ENSMUSG00000022354",
                    "ENSMUSG00000062593",
                    "ENSMUSG00000054428",
                    "ENSMUSG00000020070",
                    "ENSMUSG00000031543",
                    "ENSMUSG00000007891",
                    "ENSMUSG00000001674",
                    "ENSMUSG00000038763",
                    "ENSMUSG00000005354",
                    "ENSMUSG00000027247",
                    "ENSMUSG00000009863",
                    "ENSMUSG00000064349",
                    "ENSMUSG00000011257",
                    "ENSMUSG00000019579",
                    "ENSMUSG00000032397",
                    "ENSMUSG00000006576",
                    "ENSMUSG00000038914",
                    "ENSMUSG00000022433",
                    "ENSMUSG00000005469",
                    "ENSMUSG00000020300",
                    "ENSMUSG00000039298",
                    "ENSMUSG00000007656",
                    "ENSMUSG00000026803",
                    "ENSMUSG00000027288",
                    "ENSMUSG00000052539",
                    "ENSMUSG00000032314",
                    "ENSMUSG00000081153",
                    "ENSMUSG00000008892",
                    "ENSMUSG00000051331",
                    "ENSMUSG00000036097",
                    "ENSMUSG00000029103",
                    "ENSMUSG00000020048",
                    "ENSMUSG00000024038",
                    "ENSMUSG00000098772",
                    "ENSMUSG00000027698",
                    "ENSMUSG00000057738",
                    "ENSMUSG00000016510",
                    "ENSMUSG00000014592",
                    "ENSMUSG00000081344",
                    "ENSMUSG00000030850",
                    "ENSMUSG00000045211",
                    "ENSMUSG00000042225",
                    "ENSMUSG00000014606",
                    "ENSMUSG00000020153",
                    "ENSMUSG00000025439",
                    "ENSMUSG00000026260",
                    "ENSMUSG00000081485",
                    "ENSMUSG00000056698",
                    "ENSMUSG00000017737",
                    "ENSMUSG00000032216",
                    "ENSMUSG00000020612",
                    "ENSMUSG00000094664",
                    "ENSMUSG00000024359",
                    "ENSMUSG00000003847",
                    "ENSMUSG00000031818",
                    "ENSMUSG00000029833",
                    "ENSMUSG00000031782",
                    "ENSMUSG00000032511",
                    "ENSMUSG00000029392",
                    "ENSMUSG00000036825",
                    "ENSMUSG00000001552",
                    "ENSMUSG00000044365",
                    "ENSMUSG00000000056",
                    "ENSMUSG00000044894",
                    "ENSMUSG00000038690",
                    "ENSMUSG00000034006",
                    "ENSMUSG00000039068",
                    "ENSMUSG00000054414",
                    "ENSMUSG00000024962",
                    "ENSMUSG00000068758",
                    "ENSMUSG00000045251",
                    "ENSMUSG00000020949",
                    "ENSMUSG00000021134",
                    "ENSMUSG00000039067",
                    "ENSMUSG00000004931",
                    "ENSMUSG00000039103",
                    "ENSMUSG00000037070",
                    "ENSMUSG00000029148",
                    "ENSMUSG00000020329",
                    "ENSMUSG00000022820",
                    "ENSMUSG00000044792",
                    "ENSMUSG00000021244",
                    "ENSMUSG00000061111",
                    "ENSMUSG00000083932",
                    "ENSMUSG00000041028",
                    "ENSMUSG00000026504",
                    "ENSMUSG00000022594",
                    "ENSMUSG00000063229",
                    "ENSMUSG00000026249",
                    "ENSMUSG00000035168",
                    "ENSMUSG00000021577",
                    "ENSMUSG00000022994",
                    "ENSMUSG00000082283",
                    "ENSMUSG00000020321",
                    "ENSMUSG00000059447",
                    "ENSMUSG00000033938",
                    "ENSMUSG00000077450",
                    "ENSMUSG00000029135",
                    "ENSMUSG00000027593",
                    "ENSMUSG00000022324",
                    "ENSMUSG00000025745",
                    "ENSMUSG00000017778",
                    "ENSMUSG00000026414",
                    "ENSMUSG00000047675",
                    "ENSMUSG00000024304",
                    "ENSMUSG00000038417",
                    "ENSMUSG00000030753",
                    "ENSMUSG00000045210",
                    "ENSMUSG00000006219",
                    "ENSMUSG00000019132",
                    "ENSMUSG00000030057",
                    "ENSMUSG00000094249",
                    "ENSMUSG00000078139",
                    "ENSMUSG00000055322",
                    "ENSMUSG00000039016",
                    "ENSMUSG00000006057",
                    "ENSMUSG00000020580",
                    "ENSMUSG00000034329",
                    "ENSMUSG00000040414",
                    "ENSMUSG00000018196",
                    "ENSMUSG00000082258",
                    "ENSMUSG00000030647",
                    "ENSMUSG00000032911",
                    "ENSMUSG00000026434",
                    "ENSMUSG00000060798",
                    "ENSMUSG00000028443",
                    "ENSMUSG00000051502",
                    "ENSMUSG00000036564",
                    "ENSMUSG00000025545",
                    "ENSMUSG00000081359",
                    "ENSMUSG00000038371",
                    "ENSMUSG00000081413",
                    "ENSMUSG00000054452",
                    "ENSMUSG00000061518",
                    "ENSMUSG00000033400",
                    "ENSMUSG00000036570",
                    "ENSMUSG00000012848",
                    "ENSMUSG00000062456",
                    "ENSMUSG00000049130",
                    "ENSMUSG00000022800",
                    "ENSMUSG00000043885",
                    "ENSMUSG00000029186",
                    "ENSMUSG00000029452",
                    "ENSMUSG00000044533",
                    "ENSMUSG00000032648",
                    "ENSMUSG00000003299",
                    "ENSMUSG00000022538",
                    "ENSMUSG00000026568",
                    "ENSMUSG00000005683",
                    "ENSMUSG00000032562",
                    "ENSMUSG00000059981",
                    "ENSMUSG00000031765",
                    "ENSMUSG00000057858",
                    "ENSMUSG00000060730",
                    "ENSMUSG00000005882",
                    "ENSMUSG00000046516",
                    "ENSMUSG00000007617",
                    "ENSMUSG00000083831",
                    "ENSMUSG00000028641",
                    "ENSMUSG00000068220",
                    "ENSMUSG00000037960",
                    "ENSMUSG00000058655",
                    "ENSMUSG00000052934",
                    "ENSMUSG00000060002",
                    "ENSMUSG00000019768",
                    "ENSMUSG00000024331",
                    "ENSMUSG00000028573",
                    "ENSMUSG00000063406",
                    "ENSMUSG00000036371",
                    "ENSMUSG00000034424",
                    "ENSMUSG00000058135",
                    "ENSMUSG00000070493",
                    "ENSMUSG00000040761",
                    "ENSMUSG00000047419",
                    "ENSMUSG00000022197",
                    "ENSMUSG00000003778",
                    "ENSMUSG00000020085",
                    "ENSMUSG00000032497",
                    "ENSMUSG00000021374",
                    "ENSMUSG00000030357",
                    "ENSMUSG00000052748",
                    "ENSMUSG00000063663",
                    "ENSMUSG00000081824",
                    "ENSMUSG00000030309",
                    "ENSMUSG00000004980",
                    "ENSMUSG00000030785",
                    "ENSMUSG00000032116",
                    "ENSMUSG00000028049",
                    "ENSMUSG00000045098",
                    "ENSMUSG00000055027",
                    "ENSMUSG00000028932",
                    "ENSMUSG00000010054",
                    "ENSMUSG00000071014",
                    "ENSMUSG00000027184",
                    "ENSMUSG00000049804",
                    "ENSMUSG00000023919",
                    "ENSMUSG00000049751",
                    "ENSMUSG00000066637",
                    "ENSMUSG00000025666",
                    "ENSMUSG00000059714",
                    "ENSMUSG00000000171",
                    "ENSMUSG00000034220",
                    "ENSMUSG00000010830",
                    "ENSMUSG00000041187",
                    "ENSMUSG00000033060",
                    "ENSMUSG00000089862",
                    "ENSMUSG00000005610",
                    "ENSMUSG00000038594",
                    "ENSMUSG00000039000",
                    "ENSMUSG00000032418",
                    "ENSMUSG00000017176",
                    "ENSMUSG00000031627",
                    "ENSMUSG00000031231",
                    "ENSMUSG00000032826",
                    "ENSMUSG00000002608",
                    "ENSMUSG00000039601",
                    "ENSMUSG00000026701",
                    "ENSMUSG00000022270",
                    "ENSMUSG00000030536",
                    "ENSMUSG00000089909",
                    "ENSMUSG00000014313",
                    "ENSMUSG00000041453",
                    "ENSMUSG00000033554",
                    "ENSMUSG00000000088",
                    "ENSMUSG00000081228",
                    "ENSMUSG00000084830",
                    "ENSMUSG00000055762",
                    "ENSMUSG00000064343",
                    "ENSMUSG00000017291",
                    "ENSMUSG00000074971",
                    "ENSMUSG00000031232",
                    "ENSMUSG00000036632",
                    "ENSMUSG00000022186",
                    "ENSMUSG00000048814",
                    "ENSMUSG00000026229",
                    "ENSMUSG00000031641",
                    "ENSMUSG00000038967",
                    "ENSMUSG00000056091",
                    "ENSMUSG00000045624",
                    "ENSMUSG00000024870",
                    "ENSMUSG00000043153",
                    "ENSMUSG00000071528",
                    "ENSMUSG00000046311",
                    "ENSMUSG00000053093",
                    "ENSMUSG00000083856",
                    "ENSMUSG00000059820",
                    "ENSMUSG00000017264",
                    "ENSMUSG00000035674",
                    "ENSMUSG00000033720",
                    "ENSMUSG00000023886",
                    "ENSMUSG00000002416",
                    "ENSMUSG00000031841",
                    "ENSMUSG00000002500",
                    "ENSMUSG00000055963",
                    "ENSMUSG00000026131",
                    "ENSMUSG00000026202",
                    "ENSMUSG00000054823",
                    "ENSMUSG00000071657",
                    "ENSMUSG00000034714",
                    "ENSMUSG00000034951",
                    "ENSMUSG00000049960",
                    "ENSMUSG00000066491",
                    "ENSMUSG00000028729",
                    "ENSMUSG00000027423",
                    "ENSMUSG00000082051",
                    "ENSMUSG00000025204",
                    "ENSMUSG00000043024",
                    "ENSMUSG00000001440",
                    "ENSMUSG00000022477",
                    "ENSMUSG00000068614",
                    "ENSMUSG00000031672",
                    "ENSMUSG00000041278",
                    "ENSMUSG00000016319",
                    "ENSMUSG00000064347",
                    "ENSMUSG00000034892",
                    "ENSMUSG00000062038",
                    "ENSMUSG00000028649",
                    "ENSMUSG00000027018",
                    "ENSMUSG00000023150",
                    "ENSMUSG00000041515",
                    "ENSMUSG00000012405",
                    "ENSMUSG00000038718",
                    "ENSMUSG00000074622",
                    "ENSMUSG00000030695",
                    "ENSMUSG00000067148",
                    "ENSMUSG00000017404",
                    "ENSMUSG00000022110",
                    "ENSMUSG00000028648",
                    "ENSMUSG00000036478",
                    "ENSMUSG00000004356",
                    "ENSMUSG00000037916",
                    "ENSMUSG00000021831",
                    "ENSMUSG00000035242",
                    "ENSMUSG00000052783",
                    "ENSMUSG00000038503",
                    "ENSMUSG00000050608",
                    "ENSMUSG00000029156",
                    "ENSMUSG00000034601",
                    "ENSMUSG00000033282",
                    "ENSMUSG00000081352",
                    "ENSMUSG00000052681",
                    "ENSMUSG00000019851",
                    "ENSMUSG00000059208",
                    "ENSMUSG00000015289",
                    "ENSMUSG00000015133",
                    "ENSMUSG00000024054",
                    "ENSMUSG00000010110",
                    "ENSMUSG00000025428",
                    "ENSMUSG00000033161",
                    "ENSMUSG00000058690",
                    "ENSMUSG00000038412",
                    "ENSMUSG00000081214",
                    "ENSMUSG00000003072",
                    "ENSMUSG00000020390",
                    "ENSMUSG00000024063",
                    "ENSMUSG00000038370",
                    "ENSMUSG00000046982",
                    "ENSMUSG00000082487",
                    "ENSMUSG00000013921",
                    "ENSMUSG00000029128",
                    "ENSMUSG00000000743",
                    "ENSMUSG00000020849",
                    "ENSMUSG00000074218",
                    "ENSMUSG00000081233",
                    "ENSMUSG00000030894",
                    "ENSMUSG00000041297",
                    "ENSMUSG00000068706",
                    "ENSMUSG00000034686",
                    "ENSMUSG00000000168",
                    "ENSMUSG00000024924",
                    "ENSMUSG00000048661",
                    "ENSMUSG00000039989",
                    "ENSMUSG00000001280",
                    "ENSMUSG00000031987",
                    "ENSMUSG00000038005",
                    "ENSMUSG00000062014",
                    "ENSMUSG00000062604",
                    "ENSMUSG00000080866",
                    "ENSMUSG00000034345",
                    "ENSMUSG00000027861",
                    "ENSMUSG00000022390",
                    "ENSMUSG00000031633",
                    "ENSMUSG00000063694",
                    "ENSMUSG00000028367",
                    "ENSMUSG00000059552",
                    "ENSMUSG00000051817",
                    "ENSMUSG00000041319",
                    "ENSMUSG00000059743",
                    "ENSMUSG00000049119",
                    "ENSMUSG00000058624",
                    "ENSMUSG00000060038",
                    "ENSMUSG00000081278",
                    "ENSMUSG00000042650",
                    "ENSMUSG00000005410",
                    "ENSMUSG00000083820",
                    "ENSMUSG00000061533",
                    "ENSMUSG00000021313",
                    "ENSMUSG00000026662",
                    "ENSMUSG00000029632",
                    "ENSMUSG00000016252",
                    "ENSMUSG00000020740",
                    "ENSMUSG00000025241",
                    "ENSMUSG00000038172",
                    "ENSMUSG00000059430",
                    "ENSMUSG00000028843",
                    "ENSMUSG00000061462",
                    "ENSMUSG00000080831",
                    "ENSMUSG00000035776",
                    "ENSMUSG00000052337",
                    "ENSMUSG00000031367",
                    "ENSMUSG00000018417",
                    "ENSMUSG00000022889",
                    "ENSMUSG00000062981",
                    "ENSMUSG00000025962",
                    "ENSMUSG00000020456",
                    "ENSMUSG00000045983",
                    "ENSMUSG00000051747",
                    "ENSMUSG00000060198",
                    "ENSMUSG00000021281",
                    "ENSMUSG00000035458",
                    "ENSMUSG00000001794",
                    "ENSMUSG00000038239",
                    "ENSMUSG00000086298",
                    "ENSMUSG00000021767",
                    "ENSMUSG00000028007",
                    "ENSMUSG00000031791",
                    "ENSMUSG00000022536",
                    "ENSMUSG00000022040",
                    "ENSMUSG00000048264",
                    "ENSMUSG00000003923",
                    "ENSMUSG00000030399",
                    "ENSMUSG00000055612",
                    "ENSMUSG00000021557",
                    "ENSMUSG00000022678",
                    "ENSMUSG00000036167",
                    "ENSMUSG00000040325",
                    "ENSMUSG00000028745",
                    "ENSMUSG00000020483",
                    "ENSMUSG00000028228",
                    "ENSMUSG00000053702",
                    "ENSMUSG00000014226",
                    "ENSMUSG00000020267",
                    "ENSMUSG00000009575",
                    "ENSMUSG00000058427",
                    "ENSMUSG00000022890",
                    "ENSMUSG00000018481",
                    "ENSMUSG00000024767",
                    "ENSMUSG00000021835",
                    "ENSMUSG00000040723",
                    "ENSMUSG00000006818",
                    "ENSMUSG00000020733",
                    "ENSMUSG00000019810",
                    "ENSMUSG00000021112",
                    "ENSMUSG00000027708",
                    "ENSMUSG00000095588",
                    "ENSMUSG00000032199",
                    "ENSMUSG00000042541",
                    "ENSMUSG00000049517",
                    "ENSMUSG00000058600",
                    "ENSMUSG00000028692",
                    "ENSMUSG00000018068",
                    "ENSMUSG00000040850",
                    "ENSMUSG00000058793",
                    "ENSMUSG00000034795",
                    "ENSMUSG00000001211",
                    "ENSMUSG00000033039",
                    "ENSMUSG00000031985",
                    "ENSMUSG00000038375",
                    "ENSMUSG00000052738",
                    "ENSMUSG00000082715",
                    "ENSMUSG00000005225",
                    "ENSMUSG00000025135",
                    "ENSMUSG00000029359",
                    "ENSMUSG00000005897",
                    "ENSMUSG00000029918",
                    "ENSMUSG00000039968",
                    "ENSMUSG00000023175",
                    "ENSMUSG00000039509",
                    "ENSMUSG00000018893",
                    "ENSMUSG00000018899",
                    "ENSMUSG00000060600",
                    "ENSMUSG00000091742",
                    "ENSMUSG00000027406",
                    "ENSMUSG00000062421",
                    "ENSMUSG00000060032",
                    "ENSMUSG00000031667",
                    "ENSMUSG00000056071",
                    "ENSMUSG00000025792",
                    "ENSMUSG00000038393",
                    "ENSMUSG00000078784",
                    "ENSMUSG00000028756",
                    "ENSMUSG00000080893",
                    "ENSMUSG00000002102",
                    "ENSMUSG00000056054",
                    "ENSMUSG00000030934",
                    "ENSMUSG00000040254",
                    "ENSMUSG00000086229",
                    "ENSMUSG00000002010",
                    "ENSMUSG00000054889",
                    "ENSMUSG00000004018",
                    "ENSMUSG00000024792",
                    "ENSMUSG00000047022",
                    "ENSMUSG00000020409",
                    "ENSMUSG00000024953",
                    "ENSMUSG00000030060",
                    "ENSMUSG00000032328",
                    "ENSMUSG00000073982",
                    "ENSMUSG00000011831",
                    "ENSMUSG00000019179",
                    "ENSMUSG00000028273",
                    "ENSMUSG00000019158",
                    "ENSMUSG00000018474",
                    "ENSMUSG00000028811",
                    "ENSMUSG00000050199",
                    "ENSMUSG00000021200",
                    "ENSMUSG00000056498",
                    "ENSMUSG00000001156",
                    "ENSMUSG00000022964",
                    "ENSMUSG00000079477",
                    "ENSMUSG00000083804",
                    "ENSMUSG00000039680",
                    "ENSMUSG00000050856",
                    "ENSMUSG00000074513",
                    "ENSMUSG00000071415",
                    "ENSMUSG00000037152",
                    "ENSMUSG00000020027",
                    "ENSMUSG00000031917",
                    "ENSMUSG00000026032",
                    "ENSMUSG00000008658",
                    "ENSMUSG00000021868",
                    "ENSMUSG00000053768",
                    "ENSMUSG00000032231",
                    "ENSMUSG00000043091",
                    "ENSMUSG00000022120",
                    "ENSMUSG00000018648",
                    "ENSMUSG00000025781",
                    "ENSMUSG00000028452",
                    "ENSMUSG00000039987",
                    "ENSMUSG00000026429",
                    "ENSMUSG00000062169",
                    "ENSMUSG00000000295",
                    "ENSMUSG00000020053",
                    "ENSMUSG00000018796",
                    "ENSMUSG00000002980",
                    "ENSMUSG00000044080",
                    "ENSMUSG00000024208",
                    "ENSMUSG00000031461",
                    "ENSMUSG00000029924",
                    "ENSMUSG00000005958",
                    "ENSMUSG00000049246",
                    "ENSMUSG00000022403",
                    "ENSMUSG00000083076",
                    "ENSMUSG00000027257",
                    "ENSMUSG00000020372",
                    "ENSMUSG00000028834",
                    "ENSMUSG00000029649",
                    "ENSMUSG00000000416",
                    "ENSMUSG00000027852",
                    "ENSMUSG00000073838",
                    "ENSMUSG00000035370",
                    "ENSMUSG00000000740",
                    "ENSMUSG00000010608",
                    "ENSMUSG00000033788",
                    "ENSMUSG00000083483",
                    "ENSMUSG00000034593",
                    "ENSMUSG00000022516",
                    "ENSMUSG00000037795",
                    "ENSMUSG00000062070",
                    "ENSMUSG00000021520",
                    "ENSMUSG00000023460",
                    "ENSMUSG00000031458",
                    "ENSMUSG00000006717",
                    "ENSMUSG00000020680",
                    "ENSMUSG00000091898",
                    "ENSMUSG00000026208",
                    "ENSMUSG00000031696",
                    "ENSMUSG00000081661",
                    "ENSMUSG00000070425",
                    "ENSMUSG00000085566",
                    "ENSMUSG00000059534",
                    "ENSMUSG00000046314",
                    "ENSMUSG00000030652",
                    "ENSMUSG00000028394",
                    "ENSMUSG00000018548",
                    "ENSMUSG00000025968",
                    "ENSMUSG00000067780",
                    "ENSMUSG00000080836",
                    "ENSMUSG00000060187",
                    "ENSMUSG00000053536",
                    "ENSMUSG00000022300",
                    "ENSMUSG00000053841",
                    "ENSMUSG00000065701",
                    "ENSMUSG00000005698",
                    "ENSMUSG00000026278",
                    "ENSMUSG00000028691",
                    "ENSMUSG00000060923",
                    "ENSMUSG00000081700",
                    "ENSMUSG00000020077",
                    "ENSMUSG00000028869",
                    "ENSMUSG00000029373",
                    "ENSMUSG00000029190",
                    "ENSMUSG00000089648",
                    "ENSMUSG00000015839",
                    "ENSMUSG00000037805",
                    "ENSMUSG00000024070",
                    "ENSMUSG00000024869",
                    "ENSMUSG00000004610",
                    "ENSMUSG00000021290",
                    "ENSMUSG00000020546",
                    "ENSMUSG00000019303",
                    "ENSMUSG00000049470",
                    "ENSMUSG00000028911",
                    "ENSMUSG00000013936",
                    "ENSMUSG00000031701",
                    "ENSMUSG00000069049",
                    "ENSMUSG00000022836",
                    "ENSMUSG00000064340",
                    "ENSMUSG00000055632",
                    "ENSMUSG00000029401",
                    "ENSMUSG00000075701",
                    "ENSMUSG00000032047",
                    "ENSMUSG00000024151",
                    "ENSMUSG00000069089",
                    "ENSMUSG00000085442",
                    "ENSMUSG00000020415",
                    "ENSMUSG00000072720",
                    "ENSMUSG00000080806",
                    "ENSMUSG00000054027",
                    "ENSMUSG00000044393",
                    "ENSMUSG00000062190",
                    "ENSMUSG00000059741",
                    "ENSMUSG00000030255",
                    "ENSMUSG00000021606",
                    "ENSMUSG00000034675",
                    "ENSMUSG00000089672",
                    "ENSMUSG00000033083",
                    "ENSMUSG00000027303",
                    "ENSMUSG00000038539",
                    "ENSMUSG00000036002",
                    "ENSMUSG00000026817",
                    "ENSMUSG00000038582",
                    "ENSMUSG00000048126",
                    "ENSMUSG00000025959",
                    "ENSMUSG00000023883",
                    "ENSMUSG00000019877",
                    "ENSMUSG00000035885",
                    "ENSMUSG00000096210",
                    "ENSMUSG00000020649",
                    "ENSMUSG00000083341",
                    "ENSMUSG00000023068",
                    "ENSMUSG00000062270",
                    "ENSMUSG00000050592",
                    "ENSMUSG00000026341",
                    "ENSMUSG00000087635",
                    "ENSMUSG00000034566",
                    "ENSMUSG00000032330",
                    "ENSMUSG00000067336",
                    "ENSMUSG00000045636",
                    "ENSMUSG00000026074",
                    "ENSMUSG00000050555",
                    "ENSMUSG00000038170",
                    "ENSMUSG00000031299",
                    "ENSMUSG00000021771",
                    "ENSMUSG00000040048",
                    "ENSMUSG00000020122",
                    "ENSMUSG00000034781",
                    "ENSMUSG00000038403",
                    "ENSMUSG00000097790",
                    "ENSMUSG00000092674",
                    "ENSMUSG00000049090",
                    "ENSMUSG00000037710",
                    "ENSMUSG00000083116",
                    "ENSMUSG00000041476",
                    "ENSMUSG00000046722",
                    "ENSMUSG00000049327",
                    "ENSMUSG00000027712",
                    "ENSMUSG00000038668",
                    "ENSMUSG00000021737",
                    "ENSMUSG00000038058",
                    "ENSMUSG00000097282",
                    "ENSMUSG00000033632",
                    "ENSMUSG00000007050",
                    "ENSMUSG00000025133",
                    "ENSMUSG00000034729",
                    "ENSMUSG00000082634",
                    "ENSMUSG00000036781",
                    "ENSMUSG00000024205",
                    "ENSMUSG00000022215",
                    "ENSMUSG00000043140",
                    "ENSMUSG00000034083",
                    "ENSMUSG00000020988",
                    "ENSMUSG00000028518",
                    "ENSMUSG00000009207",
                    "ENSMUSG00000001783",
                    "ENSMUSG00000074129",
                    "ENSMUSG00000003948",
                    "ENSMUSG00000038084",
                    "ENSMUSG00000039670",
                    "ENSMUSG00000070343",
                    "ENSMUSG00000016554",
                    "ENSMUSG00000022111",
                    "ENSMUSG00000024805",
                    "ENSMUSG00000046364",
                    "ENSMUSG00000028793",
                    "ENSMUSG00000032366",
                    "ENSMUSG00000037236",
                    "ENSMUSG00000023307",
                    "ENSMUSG00000051391",
                    "ENSMUSG00000064369",
                    "ENSMUSG00000023944",
                    "ENSMUSG00000015932",
                    "ENSMUSG00000091989",
                    "ENSMUSG00000000823",
                    "ENSMUSG00000019837",
                    "ENSMUSG00000020358",
                    "ENSMUSG00000022681",
                    "ENSMUSG00000040536",
                    "ENSMUSG00000028364",
                    "ENSMUSG00000029922",
                    "ENSMUSG00000043895",
                    "ENSMUSG00000041115",
                    "ENSMUSG00000053819",
                    "ENSMUSG00000003970",
                    "ENSMUSG00000020475",
                    "ENSMUSG00000022551",
                    "ENSMUSG00000037110",
                    "ENSMUSG00000064348",
                    "ENSMUSG00000068699",
                    "ENSMUSG00000029571",
                    "ENSMUSG00000072946",
                    "ENSMUSG00000025651",
                    "ENSMUSG00000028603",
                    "ENSMUSG00000010911",
                    "ENSMUSG00000050953",
                    "ENSMUSG00000040086",
                    "ENSMUSG00000022965",
                    "ENSMUSG00000027765",
                    "ENSMUSG00000032508",
                    "ENSMUSG00000068882",
                    "ENSMUSG00000080823",
                    "ENSMUSG00000042607",
                    "ENSMUSG00000001300",
                    "ENSMUSG00000039697",
                    "ENSMUSG00000035329",
                    "ENSMUSG00000046312",
                    "ENSMUSG00000081043",
                    "ENSMUSG00000041650",
                    "ENSMUSG00000004558",
                    "ENSMUSG00000037062",
                    "ENSMUSG00000026003",
                    "ENSMUSG00000054312",
                    "ENSMUSG00000029683",
                    "ENSMUSG00000023089",
                    "ENSMUSG00000020022",
                    "ENSMUSG00000046997",
                    "ENSMUSG00000078851",
                    "ENSMUSG00000073676",
                    "ENSMUSG00000023456",
                    "ENSMUSG00000030970",
                    "ENSMUSG00000038462",
                    "ENSMUSG00000020664",
                    "ENSMUSG00000048410",
                    "ENSMUSG00000030869",
                    "ENSMUSG00000033722",
                    "ENSMUSG00000083567",
                    "ENSMUSG00000040699",
                    "ENSMUSG00000026610",
                    "ENSMUSG00000024099",
                    "ENSMUSG00000028949",
                    "ENSMUSG00000018770",
                    "ENSMUSG00000044408"
                ],
                "expression_sum": {
                    "class1": 83.0,
                    "class2": 485.0
                },
                "uniprot_ids": [
                    "P28033"
                ]
            },
            "ENSMUSG00000020453": {
                "id": "ENSMUSG00000020453",
                "type": "TF",
                "symbol": "Patz1",
                "source": [
                    "ENSMUSG00000001280",
                    "ENSMUSG00000005698",
                    "ENSMUSG00000024955"
                ],
                "target": [
                    "ENSMUSG00000005683"
                ],
                "expression_sum": {
                    "class1": 336.0,
                    "class2": 334.0
                },
                "uniprot_ids": [
                    "Q5NBZ2",
                    "Q7TNQ3",
                    "Q9JMG9",
                    "Q5NBY8",
                    "Q5NBZ1",
                    "Q5NBY9"
                ]
            },
            "ENSMUSG00000083240": {
                "id": "ENSMUSG00000083240",
                "type": "Gene",
                "symbol": "Gm13453",
                "source": [
                    "ENSMUSG00000001280"
                ],
                "target": [],
                "expression_sum": {
                    "class1": 198.0,
                    "class2": 330.0
                }
            },
            "ENSMUSG00000030470": {
                "id": "ENSMUSG00000030470",
                "type": "Gene",
                "symbol": "Csrp3",
                "source": [
                    "ENSMUSG00000001419",
                    "ENSMUSG00000015839",
                    "ENSMUSG00000010175",
                    "ENSMUSG00000015579",
                    "ENSMUSG00000019768",
                    "ENSMUSG00000053110",
                    "ENSMUSG00000021775",
                    "ENSMUSG00000015846"
                ],
                "target": [],
                "expression_sum": {
                    "class1": 12429.0,
                    "class2": 14512.0
                },
                "uniprot_ids": [
                    "P50462",
                    "A0A140LHC6"
                ]
            },
            "ENSMUSG00000063882": {
                "id": "ENSMUSG00000063882",
                "type": "Gene",
                "symbol": "Uqcrh",
                "source": [
                    "ENSMUSG00000006215",
                    "ENSMUSG00000058239",
                    "ENSMUSG00000015605",
                    "ENSMUSG00000029249",
                    "ENSMUSG00000001280",
                    "ENSMUSG00000019768",
                    "ENSMUSG00000014592",
                    "ENSMUSG00000031885",
                    "ENSMUSG00000008575",
                    "ENSMUSG00000039191",
                    "ENSMUSG00000021775",
                    "ENSMUSG00000015846"
                ],
                "target": [],
                "expression_sum": {
                    "class1": 18321.0,
                    "class2": 14132.0
                },
                "uniprot_ids": [
                    "P99028",
                    "Q9D2Q7"
                ]
            },
            "ENSMUSG00000020849": {
                "id": "ENSMUSG00000020849",
                "type": "Gene",
                "symbol": "Ywhae",
                "source": [
                    "ENSMUSG00000029135",
                    "ENSMUSG00000001419",
                    "ENSMUSG00000031681",
                    "ENSMUSG00000057469",
                    "ENSMUSG00000015605",
                    "ENSMUSG00000026610",
                    "ENSMUSG00000056501",
                    "ENSMUSG00000001280",
                    "ENSMUSG00000019768",
                    "ENSMUSG00000030067",
                    "ENSMUSG00000068267",
                    "ENSMUSG00000058756",
                    "ENSMUSG00000008575",
                    "ENSMUSG00000035632",
                    "ENSMUSG00000029913",
                    "ENSMUSG00000024985"
                ],
                "target": [],
                "expression_sum": {
                    "class1": 14344.0,
                    "class2": 3338.0
                },
                "uniprot_ids": [
                    "P62259",
                    "F6WA09",
                    "D6REF3"
                ]
            },
            "ENSMUSG00000005610": {
                "id": "ENSMUSG00000005610",
                "type": "Gene",
                "symbol": "Eif4g2",
                "source": [
                    "ENSMUSG00000029135",
                    "ENSMUSG00000076431",
                    "ENSMUSG00000015605",
                    "ENSMUSG00000008575",
                    "ENSMUSG00000040481",
                    "ENSMUSG00000036144",
                    "ENSMUSG00000015579",
                    "ENSMUSG00000040033",
                    "ENSMUSG00000051469",
                    "ENSMUSG00000019768",
                    "ENSMUSG00000022556",
                    "ENSMUSG00000028565",
                    "ENSMUSG00000031885",
                    "ENSMUSG00000053110",
                    "ENSMUSG00000024985",
                    "ENSMUSG00000015839",
                    "ENSMUSG00000030557",
                    "ENSMUSG00000047371",
                    "ENSMUSG00000056501",
                    "ENSMUSG00000029249",
                    "ENSMUSG00000001280",
                    "ENSMUSG00000049755",
                    "ENSMUSG00000055320"
                ],
                "target": [],
                "expression_sum": {
                    "class1": 6285.0,
                    "class2": 4554.0
                },
                "uniprot_ids": [
                    "F6UU81",
                    "F6TW20",
                    "F7CBP1",
                    "F7DAB1",
                    "G3XA17",
                    "F7CHS2"
                ]
            },
            "ENSMUSG00000022477": {
                "id": "ENSMUSG00000022477",
                "type": "Gene",
                "symbol": "Aco2",
                "source": [
                    "ENSMUSG00000029135",
                    "ENSMUSG00000055799",
                    "ENSMUSG00000005566",
                    "ENSMUSG00000031627",
                    "ENSMUSG00000076431",
                    "ENSMUSG00000015605",
                    "ENSMUSG00000008575",
                    "ENSMUSG00000006215",
                    "ENSMUSG00000010175",
                    "ENSMUSG00000058239",
                    "ENSMUSG00000019768",
                    "ENSMUSG00000005698",
                    "ENSMUSG00000053110",
                    "ENSMUSG00000031715",
                    "ENSMUSG00000015839",
                    "ENSMUSG00000032238",
                    "ENSMUSG00000005583",
                    "ENSMUSG00000058756",
                    "ENSMUSG00000056501",
                    "ENSMUSG00000029249",
                    "ENSMUSG00000001280"
                ],
                "target": [],
                "expression_sum": {
                    "class1": 1088.0,
                    "class2": 3823.0
                },
                "uniprot_ids": [
                    "Q99KI0",
                    "A0A2R8VHM8",
                    "A0A2R8W744",
                    "A0A2R8VJM1",
                    "A0A2R8VJW0"
                ]
            },
            "ENSMUSG00000029455": {
                "id": "ENSMUSG00000029455",
                "type": "Gene",
                "symbol": "Aldh2",
                "source": [
                    "ENSMUSG00000015839",
                    "ENSMUSG00000010175",
                    "ENSMUSG00000001280",
                    "ENSMUSG00000019768",
                    "ENSMUSG00000031885",
                    "ENSMUSG00000035158"
                ],
                "target": [],
                "expression_sum": {
                    "class1": 941.0,
                    "class2": 695.0
                },
                "uniprot_ids": [
                    "P47738",
                    "A0A0G2JF60",
                    "D3YYF3",
                    "A0A0G2JEU1",
                    "A0A0G2JFQ0"
                ]
            },
            "ENSMUSG00000026385": {
                "id": "ENSMUSG00000026385",
                "type": "Gene",
                "symbol": "Dbi",
                "source": [
                    "ENSMUSG00000024238",
                    "ENSMUSG00000020248",
                    "ENSMUSG00000021944",
                    "ENSMUSG00000029135",
                    "ENSMUSG00000019817",
                    "ENSMUSG00000015839",
                    "ENSMUSG00000019564",
                    "ENSMUSG00000021250",
                    "ENSMUSG00000030256",
                    "ENSMUSG00000039191",
                    "ENSMUSG00000052534",
                    "ENSMUSG00000025959",
                    "ENSMUSG00000001280",
                    "ENSMUSG00000005583",
                    "ENSMUSG00000071637",
                    "ENSMUSG00000022314",
                    "ENSMUSG00000031715"
                ],
                "target": [],
                "expression_sum": {
                    "class1": 15549.0,
                    "class2": 4575.0
                },
                "uniprot_ids": [
                    "P31786",
                    "Q4VWZ5",
                    "D3Z563",
                    "M0QWU8"
                ]
            },
            "ENSMUSG00000089838": {
                "id": "ENSMUSG00000089838",
                "type": "Gene",
                "symbol": "Gm2962",
                "source": [
                    "ENSMUSG00000019768",
                    "ENSMUSG00000031885",
                    "ENSMUSG00000015846"
                ],
                "target": [],
                "expression_sum": {
                    "class1": 1117.0,
                    "class2": 3445.0
                }
            },
            "ENSMUSG00000021577": {
                "id": "ENSMUSG00000021577",
                "type": "Gene",
                "symbol": "Sdha",
                "source": [
                    "ENSMUSG00000006215",
                    "ENSMUSG00000015839",
                    "ENSMUSG00000015605",
                    "ENSMUSG00000056501",
                    "ENSMUSG00000015579",
                    "ENSMUSG00000001280",
                    "ENSMUSG00000019768",
                    "ENSMUSG00000031885",
                    "ENSMUSG00000008575",
                    "ENSMUSG00000014592",
                    "ENSMUSG00000021775",
                    "ENSMUSG00000024985"
                ],
                "target": [],
                "expression_sum": {
                    "class1": 4491.0,
                    "class2": 6644.0
                },
                "uniprot_ids": [
                    "Q8K2B3",
                    "A0A1Y7VJ55"
                ]
            },
            "ENSMUSG00000067736": {
                "id": "ENSMUSG00000067736",
                "type": "Gene",
                "symbol": "Gm10222",
                "source": [
                    "ENSMUSG00000021944",
                    "ENSMUSG00000030557",
                    "ENSMUSG00000001280",
                    "ENSMUSG00000019768",
                    "ENSMUSG00000015846"
                ],
                "target": [],
                "expression_sum": {
                    "class1": 172.0,
                    "class2": 701.0
                }
            },
            "ENSMUSG00000032047": {
                "id": "ENSMUSG00000032047",
                "type": "Gene",
                "symbol": "Acat1",
                "source": [
                    "ENSMUSG00000040321",
                    "ENSMUSG00000057469",
                    "ENSMUSG00000003032",
                    "ENSMUSG00000015839",
                    "ENSMUSG00000058239",
                    "ENSMUSG00000015605",
                    "ENSMUSG00000026610",
                    "ENSMUSG00000056501",
                    "ENSMUSG00000029249",
                    "ENSMUSG00000001280",
                    "ENSMUSG00000005566",
                    "ENSMUSG00000019768",
                    "ENSMUSG00000005698",
                    "ENSMUSG00000030087",
                    "ENSMUSG00000008575",
                    "ENSMUSG00000035158"
                ],
                "target": [],
                "expression_sum": {
                    "class1": 4452.0,
                    "class2": 3198.0
                },
                "uniprot_ids": [
                    "Q8QZT1"
                ]
            },
            "ENSMUSG00000029017": {
                "id": "ENSMUSG00000029017",
                "type": "Gene",
                "symbol": "Pmpcb",
                "source": [
                    "ENSMUSG00000001280",
                    "ENSMUSG00000019768",
                    "ENSMUSG00000015605"
                ],
                "target": [],
                "expression_sum": {
                    "class1": 716.0,
                    "class2": 1642.0
                },
                "uniprot_ids": [
                    "Q9CXT8",
                    "D3Z1T9",
                    "F7BPA5"
                ]
            },
            "ENSMUSG00000039891": {
                "id": "ENSMUSG00000039891",
                "type": "Gene",
                "symbol": "Txlnb",
                "source": [
                    "ENSMUSG00000015839",
                    "ENSMUSG00000058239",
                    "ENSMUSG00000078671",
                    "ENSMUSG00000030557",
                    "ENSMUSG00000001280",
                    "ENSMUSG00000005566",
                    "ENSMUSG00000019768",
                    "ENSMUSG00000031885",
                    "ENSMUSG00000015605",
                    "ENSMUSG00000031715"
                ],
                "target": [],
                "expression_sum": {
                    "class1": 2872.0,
                    "class2": 4957.0
                },
                "uniprot_ids": [
                    "Q8VBT1"
                ]
            },
            "ENSMUSG00000004096": {
                "id": "ENSMUSG00000004096",
                "type": "Gene",
                "symbol": "Cwc15",
                "source": [
                    "ENSMUSG00000001280",
                    "ENSMUSG00000032897",
                    "ENSMUSG00000005698",
                    "ENSMUSG00000020248"
                ],
                "target": [],
                "expression_sum": {
                    "class1": 3268.0,
                    "class2": 939.0
                },
                "uniprot_ids": [
                    "Q9JHS9",
                    "A0A1L1SVD8"
                ]
            },
            "ENSMUSG00000019797": {
                "id": "ENSMUSG00000019797",
                "type": "Gene",
                "symbol": "Mtres1",
                "source": [
                    "ENSMUSG00000006215",
                    "ENSMUSG00000015579",
                    "ENSMUSG00000001280",
                    "ENSMUSG00000019768",
                    "ENSMUSG00000038692",
                    "ENSMUSG00000031885"
                ],
                "target": [],
                "expression_sum": {
                    "class1": 2788.0,
                    "class2": 1536.0
                },
                "uniprot_ids": [
                    "Q9CQF4"
                ]
            },
            "ENSMUSG00000008668": {
                "id": "ENSMUSG00000008668",
                "type": "Gene",
                "symbol": "Rps18",
                "source": [
                    "ENSMUSG00000024238",
                    "ENSMUSG00000021250",
                    "ENSMUSG00000019817",
                    "ENSMUSG00000052534",
                    "ENSMUSG00000001280",
                    "ENSMUSG00000005583",
                    "ENSMUSG00000040732",
                    "ENSMUSG00000022314"
                ],
                "target": [],
                "expression_sum": {
                    "class1": 3997.0,
                    "class2": 2150.0
                },
                "uniprot_ids": [
                    "P62270",
                    "A0A3Q4EGP3",
                    "S4R1N6"
                ]
            },
            "ENSMUSG00000064346": {
                "id": "ENSMUSG00000064346",
                "type": "Gene",
                "symbol": "TrnW",
                "source": [
                    "ENSMUSG00000001280",
                    "ENSMUSG00000019768",
                    "ENSMUSG00000001419"
                ],
                "target": [],
                "expression_sum": {
                    "class1": 1283.0,
                    "class2": 1333.0
                }
            },
            "ENSMUSG00000064339": {
                "id": "ENSMUSG00000064339",
                "type": "Gene",
                "symbol": "mt-Rnr2",
                "source": [
                    "ENSMUSG00000001419",
                    "ENSMUSG00000021944",
                    "ENSMUSG00000022346",
                    "ENSMUSG00000052837",
                    "ENSMUSG00000001280",
                    "ENSMUSG00000019768"
                ],
                "target": [],
                "expression_sum": {
                    "class1": 16265703.0,
                    "class2": 10674107.0
                }
            },
            "ENSMUSG00000019877": {
                "id": "ENSMUSG00000019877",
                "type": "Gene",
                "symbol": "Serinc1",
                "source": [
                    "ENSMUSG00000029135",
                    "ENSMUSG00000006705",
                    "ENSMUSG00000015579",
                    "ENSMUSG00000056501",
                    "ENSMUSG00000051469",
                    "ENSMUSG00000001280",
                    "ENSMUSG00000019768",
                    "ENSMUSG00000015605",
                    "ENSMUSG00000035158"
                ],
                "target": [],
                "expression_sum": {
                    "class1": 3523.0,
                    "class2": 1599.0
                },
                "uniprot_ids": [
                    "Q9QZI8",
                    "E9PY81",
                    "E9Q553"
                ]
            },
            "ENSMUSG00000027447": {
                "id": "ENSMUSG00000027447",
                "type": "Gene",
                "symbol": "Cst3",
                "source": [
                    "ENSMUSG00000029135",
                    "ENSMUSG00000021250",
                    "ENSMUSG00000058239",
                    "ENSMUSG00000030256",
                    "ENSMUSG00000073423",
                    "ENSMUSG00000023034",
                    "ENSMUSG00000056501",
                    "ENSMUSG00000041515",
                    "ENSMUSG00000001280",
                    "ENSMUSG00000019768",
                    "ENSMUSG00000052684",
                    "ENSMUSG00000035158"
                ],
                "target": [],
                "expression_sum": {
                    "class1": 20310.0,
                    "class2": 14063.0
                },
                "uniprot_ids": [
                    "P21460",
                    "A2APX3"
                ]
            },
            "ENSMUSG00000073676": {
                "id": "ENSMUSG00000073676",
                "type": "Gene",
                "symbol": "Hspe1",
                "source": [
                    "ENSMUSG00000015605",
                    "ENSMUSG00000030256",
                    "ENSMUSG00000056501",
                    "ENSMUSG00000074622",
                    "ENSMUSG00000001280",
                    "ENSMUSG00000019768",
                    "ENSMUSG00000035158",
                    "ENSMUSG00000055320",
                    "ENSMUSG00000038615",
                    "ENSMUSG00000008575",
                    "ENSMUSG00000031715",
                    "ENSMUSG00000015846"
                ],
                "target": [],
                "expression_sum": {
                    "class1": 3594.0,
                    "class2": 2183.0
                },
                "uniprot_ids": [
                    "Q64433"
                ]
            },
            "ENSMUSG00000030647": {
                "id": "ENSMUSG00000030647",
                "type": "Gene",
                "symbol": "Ndufc2",
                "source": [
                    "ENSMUSG00000029135",
                    "ENSMUSG00000001419",
                    "ENSMUSG00000056501",
                    "ENSMUSG00000030557",
                    "ENSMUSG00000001280",
                    "ENSMUSG00000019768",
                    "ENSMUSG00000035158",
                    "ENSMUSG00000068267",
                    "ENSMUSG00000031885",
                    "ENSMUSG00000027104",
                    "ENSMUSG00000031715",
                    "ENSMUSG00000015846"
                ],
                "target": [],
                "expression_sum": {
                    "class1": 6911.0,
                    "class2": 4839.0
                },
                "uniprot_ids": [
                    "Q9CQ54",
                    "A0A0U1RPJ5"
                ]
            },
            "ENSMUSG00000003955": {
                "id": "ENSMUSG00000003955",
                "type": "Gene",
                "symbol": "Fam162a",
                "source": [
                    "ENSMUSG00000017724",
                    "ENSMUSG00000029135",
                    "ENSMUSG00000001419",
                    "ENSMUSG00000020248",
                    "ENSMUSG00000006215",
                    "ENSMUSG00000015839",
                    "ENSMUSG00000058239",
                    "ENSMUSG00000015579",
                    "ENSMUSG00000051469",
                    "ENSMUSG00000001280",
                    "ENSMUSG00000019768",
                    "ENSMUSG00000005583",
                    "ENSMUSG00000005698",
                    "ENSMUSG00000053477",
                    "ENSMUSG00000031885",
                    "ENSMUSG00000008575",
                    "ENSMUSG00000022314"
                ],
                "target": [],
                "expression_sum": {
                    "class1": 4987.0,
                    "class2": 2717.0
                },
                "uniprot_ids": [
                    "Q9D6U8",
                    "A0A338P6X2"
                ]
            },
            "ENSMUSG00000024871": {
                "id": "ENSMUSG00000024871",
                "type": "Gene",
                "symbol": "Doc2g",
                "source": [
                    "ENSMUSG00000001419",
                    "ENSMUSG00000001280",
                    "ENSMUSG00000005566",
                    "ENSMUSG00000019768",
                    "ENSMUSG00000005698",
                    "ENSMUSG00000022641",
                    "ENSMUSG00000058756"
                ],
                "target": [],
                "expression_sum": {
                    "class1": 693.0,
                    "class2": 850.0
                },
                "uniprot_ids": [
                    "A0A494BAY3",
                    "A0A0R4J053"
                ]
            },
            "ENSMUSG00000060938": {
                "id": "ENSMUSG00000060938",
                "type": "Gene",
                "symbol": "Rpl26",
                "source": [
                    "ENSMUSG00000001280",
                    "ENSMUSG00000019768",
                    "ENSMUSG00000076431",
                    "ENSMUSG00000035158",
                    "ENSMUSG00000015846"
                ],
                "target": [],
                "expression_sum": {
                    "class1": 10370.0,
                    "class2": 1892.0
                },
                "uniprot_ids": [
                    "P61255",
                    "B1ARA3",
                    "B1ARA5"
                ]
            },
            "ENSMUSG00000048416": {
                "id": "ENSMUSG00000048416",
                "type": "Gene",
                "symbol": "Mlf1",
                "source": [
                    "ENSMUSG00000058239",
                    "ENSMUSG00000001280",
                    "ENSMUSG00000019768",
                    "ENSMUSG00000075595",
                    "ENSMUSG00000031885",
                    "ENSMUSG00000000567"
                ],
                "target": [],
                "expression_sum": {
                    "class1": 5501.0,
                    "class2": 4146.0
                },
                "uniprot_ids": [
                    "Q9QWV4",
                    "Q8JZS2",
                    "A0A0A6YVP7"
                ]
            },
            "ENSMUSG00000028639": {
                "id": "ENSMUSG00000028639",
                "type": "TF",
                "symbol": "Ybx1",
                "source": [
                    "ENSMUSG00000032897",
                    "ENSMUSG00000021250",
                    "ENSMUSG00000015839",
                    "ENSMUSG00000018604",
                    "ENSMUSG00000001280",
                    "ENSMUSG00000035158"
                ],
                "target": [
                    "ENSMUSG00000014599",
                    "ENSMUSG00000003037",
                    "ENSMUSG00000040025",
                    "ENSMUSG00000004127",
                    "ENSMUSG00000056941",
                    "ENSMUSG00000026617",
                    "ENSMUSG00000043683",
                    "ENSMUSG00000063684",
                    "ENSMUSG00000018819",
                    "ENSMUSG00000000031",
                    "ENSMUSG00000007080",
                    "ENSMUSG00000052384",
                    "ENSMUSG00000081792",
                    "ENSMUSG00000025077",
                    "ENSMUSG00000027750",
                    "ENSMUSG00000028165",
                    "ENSMUSG00000095280",
                    "ENSMUSG00000000568",
                    "ENSMUSG00000020402",
                    "ENSMUSG00000025395",
                    "ENSMUSG00000021645",
                    "ENSMUSG00000024066",
                    "ENSMUSG00000008035",
                    "ENSMUSG00000063019",
                    "ENSMUSG00000004789",
                    "ENSMUSG00000020477",
                    "ENSMUSG00000040242",
                    "ENSMUSG00000082684",
                    "ENSMUSG00000000127",
                    "ENSMUSG00000029729",
                    "ENSMUSG00000004085",
                    "ENSMUSG00000030536",
                    "ENSMUSG00000025175",
                    "ENSMUSG00000031821",
                    "ENSMUSG00000000088",
                    "ENSMUSG00000040048",
                    "ENSMUSG00000003423",
                    "ENSMUSG00000075700",
                    "ENSMUSG00000027160",
                    "ENSMUSG00000049658",
                    "ENSMUSG00000006519",
                    "ENSMUSG00000081620",
                    "ENSMUSG00000027439",
                    "ENSMUSG00000061559",
                    "ENSMUSG00000021190",
                    "ENSMUSG00000002416",
                    "ENSMUSG00000083832",
                    "ENSMUSG00000029502",
                    "ENSMUSG00000022744",
                    "ENSMUSG00000008136",
                    "ENSMUSG00000025647",
                    "ENSMUSG00000016256",
                    "ENSMUSG00000064369",
                    "ENSMUSG00000024400",
                    "ENSMUSG00000038612",
                    "ENSMUSG00000068749",
                    "ENSMUSG00000097991",
                    "ENSMUSG00000064338",
                    "ENSMUSG00000028452",
                    "ENSMUSG00000083932",
                    "ENSMUSG00000052146",
                    "ENSMUSG00000021772",
                    "ENSMUSG00000022216",
                    "ENSMUSG00000019791",
                    "ENSMUSG00000001525",
                    "ENSMUSG00000063698",
                    "ENSMUSG00000045103",
                    "ENSMUSG00000025130",
                    "ENSMUSG00000000628",
                    "ENSMUSG00000005686",
                    "ENSMUSG00000021622",
                    "ENSMUSG00000022122",
                    "ENSMUSG00000001627",
                    "ENSMUSG00000032279",
                    "ENSMUSG00000001424",
                    "ENSMUSG00000023861",
                    "ENSMUSG00000066861",
                    "ENSMUSG00000029864",
                    "ENSMUSG00000081359",
                    "ENSMUSG00000028832",
                    "ENSMUSG00000020067",
                    "ENSMUSG00000027861",
                    "ENSMUSG00000081049"
                ],
                "expression_sum": {
                    "class1": 9831.0,
                    "class2": 5203.0
                },
                "uniprot_ids": [
                    "P62960",
                    "A2BGG7"
                ]
            },
            "ENSMUSG00000050608": {
                "id": "ENSMUSG00000050608",
                "type": "Gene",
                "symbol": "Micos10",
                "source": [
                    "ENSMUSG00000010175",
                    "ENSMUSG00000015839",
                    "ENSMUSG00000015605",
                    "ENSMUSG00000056501",
                    "ENSMUSG00000015579",
                    "ENSMUSG00000029249",
                    "ENSMUSG00000001280",
                    "ENSMUSG00000019768",
                    "ENSMUSG00000032238",
                    "ENSMUSG00000022641",
                    "ENSMUSG00000008575",
                    "ENSMUSG00000039191",
                    "ENSMUSG00000021775",
                    "ENSMUSG00000015846"
                ],
                "target": [],
                "expression_sum": {
                    "class1": 23119.0,
                    "class2": 18454.0
                },
                "uniprot_ids": [
                    "Q7TNS2"
                ]
            },
            "ENSMUSG00000022205": {
                "id": "ENSMUSG00000022205",
                "type": "Gene",
                "symbol": "Sub1",
                "source": [
                    "ENSMUSG00000029135",
                    "ENSMUSG00000024515",
                    "ENSMUSG00000059552",
                    "ENSMUSG00000036461",
                    "ENSMUSG00000027490",
                    "ENSMUSG00000001280",
                    "ENSMUSG00000028565",
                    "ENSMUSG00000040732",
                    "ENSMUSG00000020919",
                    "ENSMUSG00000024974",
                    "ENSMUSG00000022314"
                ],
                "target": [],
                "expression_sum": {
                    "class1": 24823.0,
                    "class2": 4551.0
                },
                "uniprot_ids": [
                    "P11031"
                ]
            },
            "ENSMUSG00000038170": {
                "id": "ENSMUSG00000038170",
                "type": "Gene",
                "symbol": "Pde4dip",
                "source": [
                    "ENSMUSG00000005566",
                    "ENSMUSG00000015605",
                    "ENSMUSG00000008575",
                    "ENSMUSG00000010175",
                    "ENSMUSG00000015579",
                    "ENSMUSG00000019768",
                    "ENSMUSG00000028565",
                    "ENSMUSG00000005698",
                    "ENSMUSG00000022641",
                    "ENSMUSG00000024985",
                    "ENSMUSG00000030557",
                    "ENSMUSG00000022389",
                    "ENSMUSG00000005583",
                    "ENSMUSG00000058756",
                    "ENSMUSG00000029913",
                    "ENSMUSG00000001419",
                    "ENSMUSG00000056501",
                    "ENSMUSG00000029249",
                    "ENSMUSG00000001280"
                ],
                "target": [],
                "expression_sum": {
                    "class1": 2456.0,
                    "class2": 4810.0
                },
                "uniprot_ids": [
                    "Q80YT7",
                    "Q3UR03",
                    "A0A0G2JEX6",
                    "H3BJ38",
                    "E9Q1A0",
                    "G3X9L9"
                ]
            },
            "ENSMUSG00000020267": {
                "id": "ENSMUSG00000020267",
                "type": "Gene",
                "symbol": "Hint1",
                "source": [
                    "ENSMUSG00000029135",
                    "ENSMUSG00000057469",
                    "ENSMUSG00000056501",
                    "ENSMUSG00000015579",
                    "ENSMUSG00000027109",
                    "ENSMUSG00000029249",
                    "ENSMUSG00000001280",
                    "ENSMUSG00000019768",
                    "ENSMUSG00000028565",
                    "ENSMUSG00000044167",
                    "ENSMUSG00000015846"
                ],
                "target": [],
                "expression_sum": {
                    "class1": 22029.0,
                    "class2": 7562.0
                },
                "uniprot_ids": [
                    "P70349",
                    "B0R1E3"
                ]
            },
            "ENSMUSG00000064349": {
                "id": "ENSMUSG00000064349",
                "type": "Gene",
                "symbol": "TrnC",
                "source": [
                    "ENSMUSG00000056501",
                    "ENSMUSG00000030557",
                    "ENSMUSG00000001280",
                    "ENSMUSG00000019768",
                    "ENSMUSG00000032238",
                    "ENSMUSG00000053110",
                    "ENSMUSG00000015605",
                    "ENSMUSG00000015846"
                ],
                "target": [],
                "expression_sum": {
                    "class1": 1257.0,
                    "class2": 4446.0
                }
            },
            "ENSMUSG00000021313": {
                "id": "ENSMUSG00000021313",
                "type": "Gene",
                "symbol": "Ryr2",
                "source": [
                    "ENSMUSG00000040481",
                    "ENSMUSG00000056501",
                    "ENSMUSG00000038718",
                    "ENSMUSG00000001280",
                    "ENSMUSG00000005566",
                    "ENSMUSG00000019768",
                    "ENSMUSG00000033943",
                    "ENSMUSG00000053110",
                    "ENSMUSG00000059878",
                    "ENSMUSG00000015605"
                ],
                "target": [],
                "expression_sum": {
                    "class1": 3087.0,
                    "class2": 5871.0
                },
                "uniprot_ids": [
                    "E9Q401",
                    "Q80ZZ5",
                    "F6U7V1",
                    "A0A1Y7VJP1",
                    "A0A1Y7VK09"
                ]
            },
            "ENSMUSG00000026208": {
                "id": "ENSMUSG00000026208",
                "type": "Gene",
                "symbol": "Des",
                "source": [
                    "ENSMUSG00000001419",
                    "ENSMUSG00000040481",
                    "ENSMUSG00000022346",
                    "ENSMUSG00000058239",
                    "ENSMUSG00000056501",
                    "ENSMUSG00000030557",
                    "ENSMUSG00000029249",
                    "ENSMUSG00000001280",
                    "ENSMUSG00000005566",
                    "ENSMUSG00000019768",
                    "ENSMUSG00000005698",
                    "ENSMUSG00000075595",
                    "ENSMUSG00000031885",
                    "ENSMUSG00000058756",
                    "ENSMUSG00000015605"
                ],
                "target": [],
                "expression_sum": {
                    "class1": 3427.0,
                    "class2": 3604.0
                },
                "uniprot_ids": [
                    "P31001"
                ]
            },
            "ENSMUSG00000091449": {
                "id": "ENSMUSG00000091449",
                "type": "Gene",
                "symbol": "Gm10269",
                "source": [
                    "ENSMUSG00000001280",
                    "ENSMUSG00000018604"
                ],
                "target": [],
                "expression_sum": {
                    "class1": 34702.0,
                    "class2": 6531.0
                }
            },
            "ENSMUSG00000040850": {
                "id": "ENSMUSG00000040850",
                "type": "Gene",
                "symbol": "Psme4",
                "source": [
                    "ENSMUSG00000029135",
                    "ENSMUSG00000005566",
                    "ENSMUSG00000038615",
                    "ENSMUSG00000076431",
                    "ENSMUSG00000015605",
                    "ENSMUSG00000008575",
                    "ENSMUSG00000035158",
                    "ENSMUSG00000040481",
                    "ENSMUSG00000052534",
                    "ENSMUSG00000015579",
                    "ENSMUSG00000051469",
                    "ENSMUSG00000019768",
                    "ENSMUSG00000005698",
                    "ENSMUSG00000031885",
                    "ENSMUSG00000053110",
                    "ENSMUSG00000031715",
                    "ENSMUSG00000030557",
                    "ENSMUSG00000005583",
                    "ENSMUSG00000014592",
                    "ENSMUSG00000056501",
                    "ENSMUSG00000029249",
                    "ENSMUSG00000001280"
                ],
                "target": [],
                "expression_sum": {
                    "class1": 2105.0,
                    "class2": 2061.0
                },
                "uniprot_ids": [
                    "Q5SSW2",
                    "Q5SSW0",
                    "F6R7K9"
                ]
            },
            "ENSMUSG00000017778": {
                "id": "ENSMUSG00000017778",
                "type": "Gene",
                "symbol": "Cox7c",
                "source": [
                    "ENSMUSG00000057469",
                    "ENSMUSG00000015839",
                    "ENSMUSG00000058239",
                    "ENSMUSG00000056501",
                    "ENSMUSG00000030557",
                    "ENSMUSG00000001280",
                    "ENSMUSG00000019768",
                    "ENSMUSG00000025958",
                    "ENSMUSG00000014592",
                    "ENSMUSG00000031885",
                    "ENSMUSG00000058756",
                    "ENSMUSG00000015605",
                    "ENSMUSG00000039191"
                ],
                "target": [],
                "expression_sum": {
                    "class1": 3123.0,
                    "class2": 12206.0
                },
                "uniprot_ids": [
                    "P17665",
                    "B8JJA9",
                    "B8JJB0"
                ]
            },
            "ENSMUSG00000020326": {
                "id": "ENSMUSG00000020326",
                "type": "Gene",
                "symbol": "Ccng1",
                "source": [
                    "ENSMUSG00000030557",
                    "ENSMUSG00000001280",
                    "ENSMUSG00000019768",
                    "ENSMUSG00000031885",
                    "ENSMUSG00000022228",
                    "ENSMUSG00000015605",
                    "ENSMUSG00000015846"
                ],
                "target": [],
                "expression_sum": {
                    "class1": 3003.0,
                    "class2": 3442.0
                },
                "uniprot_ids": [
                    "P51945"
                ]
            },
            "ENSMUSG00000041841": {
                "id": "ENSMUSG00000041841",
                "type": "Gene",
                "symbol": "Rpl37",
                "source": [
                    "ENSMUSG00000059552",
                    "ENSMUSG00000022346",
                    "ENSMUSG00000018604",
                    "ENSMUSG00000019564",
                    "ENSMUSG00000001280",
                    "ENSMUSG00000028565",
                    "ENSMUSG00000040732"
                ],
                "target": [],
                "expression_sum": {
                    "class1": 6200.0,
                    "class2": 906.0
                },
                "uniprot_ids": [
                    "Q9D823"
                ]
            },
            "ENSMUSG00000081331": {
                "id": "ENSMUSG00000081331",
                "type": "Gene",
                "symbol": "Gm15151",
                "source": [
                    "ENSMUSG00000001280",
                    "ENSMUSG00000033943"
                ],
                "target": [],
                "expression_sum": {
                    "class1": 1363.0,
                    "class2": 4179.0
                }
            },
            "ENSMUSG00000041881": {
                "id": "ENSMUSG00000041881",
                "type": "Gene",
                "symbol": "Ndufa7",
                "source": [
                    "ENSMUSG00000029135",
                    "ENSMUSG00000058239",
                    "ENSMUSG00000015605",
                    "ENSMUSG00000030256",
                    "ENSMUSG00000056501",
                    "ENSMUSG00000030557",
                    "ENSMUSG00000014303",
                    "ENSMUSG00000001280",
                    "ENSMUSG00000019768",
                    "ENSMUSG00000053110",
                    "ENSMUSG00000008575"
                ],
                "target": [],
                "expression_sum": {
                    "class1": 21994.0,
                    "class2": 10717.0
                },
                "uniprot_ids": [
                    "Q9Z1P6",
                    "G3UXZ4"
                ]
            },
            "ENSMUSG00000030246": {
                "id": "ENSMUSG00000030246",
                "type": "Gene",
                "symbol": "Ldhb",
                "source": [
                    "ENSMUSG00000057469",
                    "ENSMUSG00000021944",
                    "ENSMUSG00000015839",
                    "ENSMUSG00000058239",
                    "ENSMUSG00000026610",
                    "ENSMUSG00000028466",
                    "ENSMUSG00000029249",
                    "ENSMUSG00000001280",
                    "ENSMUSG00000005566",
                    "ENSMUSG00000019768",
                    "ENSMUSG00000031885",
                    "ENSMUSG00000015605",
                    "ENSMUSG00000014592"
                ],
                "target": [],
                "expression_sum": {
                    "class1": 9981.0,
                    "class2": 12296.0
                },
                "uniprot_ids": [
                    "P16125",
                    "A0A0N4SVV8",
                    "D3Z7F0",
                    "A0A0N4SWC9",
                    "A0A6I8MX27"
                ]
            },
            "ENSMUSG00000064340": {
                "id": "ENSMUSG00000064340",
                "type": "Gene",
                "symbol": "TrnL1",
                "source": [
                    "ENSMUSG00000001419",
                    "ENSMUSG00000021944",
                    "ENSMUSG00000056501",
                    "ENSMUSG00000029249",
                    "ENSMUSG00000001280",
                    "ENSMUSG00000005566",
                    "ENSMUSG00000015843",
                    "ENSMUSG00000005698",
                    "ENSMUSG00000019768",
                    "ENSMUSG00000028565",
                    "ENSMUSG00000032238",
                    "ENSMUSG00000053477",
                    "ENSMUSG00000058756",
                    "ENSMUSG00000015605",
                    "ENSMUSG00000035158",
                    "ENSMUSG00000029913"
                ],
                "target": [],
                "expression_sum": {
                    "class1": 528.0,
                    "class2": 1066.0
                }
            },
            "ENSMUSG00000006057": {
                "id": "ENSMUSG00000006057",
                "type": "Gene",
                "symbol": "Atp5g1",
                "source": [
                    "ENSMUSG00000029135",
                    "ENSMUSG00000057469",
                    "ENSMUSG00000006215",
                    "ENSMUSG00000015839",
                    "ENSMUSG00000015605",
                    "ENSMUSG00000026610",
                    "ENSMUSG00000056501",
                    "ENSMUSG00000015579",
                    "ENSMUSG00000001280",
                    "ENSMUSG00000019768",
                    "ENSMUSG00000014592",
                    "ENSMUSG00000068267",
                    "ENSMUSG00000031885",
                    "ENSMUSG00000008575",
                    "ENSMUSG00000039191",
                    "ENSMUSG00000021775",
                    "ENSMUSG00000015846"
                ],
                "target": [],
                "expression_sum": {
                    "class1": 25985.0,
                    "class2": 16923.0
                },
                "uniprot_ids": [
                    "Q9CR84"
                ]
            },
            "ENSMUSG00000081051": {
                "id": "ENSMUSG00000081051",
                "type": "Gene",
                "symbol": "Gm15427",
                "source": [
                    "ENSMUSG00000001280"
                ],
                "target": [],
                "expression_sum": {
                    "class1": 112800.0,
                    "class2": 12039.0
                }
            },
            "ENSMUSG00000025241": {
                "id": "ENSMUSG00000025241",
                "type": "Gene",
                "symbol": "Fyco1",
                "source": [
                    "ENSMUSG00000015839",
                    "ENSMUSG00000058239",
                    "ENSMUSG00000078671",
                    "ENSMUSG00000056501",
                    "ENSMUSG00000001280",
                    "ENSMUSG00000005566",
                    "ENSMUSG00000019768",
                    "ENSMUSG00000075595",
                    "ENSMUSG00000031885",
                    "ENSMUSG00000008575",
                    "ENSMUSG00000024985"
                ],
                "target": [],
                "expression_sum": {
                    "class1": 1435.0,
                    "class2": 1371.0
                },
                "uniprot_ids": [
                    "Q8VDC1",
                    "A0A140T8V9"
                ]
            },
            "ENSMUSG00000074800": {
                "id": "ENSMUSG00000074800",
                "type": "Gene",
                "symbol": "Gm4149",
                "source": [
                    "ENSMUSG00000001280",
                    "ENSMUSG00000019564",
                    "ENSMUSG00000015846"
                ],
                "target": [],
                "expression_sum": {
                    "class1": 16870.0,
                    "class2": 1931.0
                }
            },
            "ENSMUSG00000027602": {
                "id": "ENSMUSG00000027602",
                "type": "Gene",
                "symbol": "Map1lc3a",
                "source": [
                    "ENSMUSG00000057469",
                    "ENSMUSG00000022346",
                    "ENSMUSG00000015605",
                    "ENSMUSG00000029249",
                    "ENSMUSG00000001280",
                    "ENSMUSG00000019768",
                    "ENSMUSG00000038914",
                    "ENSMUSG00000076431",
                    "ENSMUSG00000008575",
                    "ENSMUSG00000015846"
                ],
                "target": [],
                "expression_sum": {
                    "class1": 8210.0,
                    "class2": 4798.0
                },
                "uniprot_ids": [
                    "Q91VR7"
                ]
            },
            "ENSMUSG00000068706": {
                "id": "ENSMUSG00000068706",
                "type": "Gene",
                "symbol": "Gm10250",
                "source": [
                    "ENSMUSG00000056501",
                    "ENSMUSG00000001280",
                    "ENSMUSG00000033943",
                    "ENSMUSG00000019768",
                    "ENSMUSG00000014592"
                ],
                "target": [],
                "expression_sum": {
                    "class1": 23884.0,
                    "class2": 30612.0
                }
            },
            "ENSMUSG00000063229": {
                "id": "ENSMUSG00000063229",
                "type": "Gene",
                "symbol": "Ldha",
                "source": [
                    "ENSMUSG00000029135",
                    "ENSMUSG00000001419",
                    "ENSMUSG00000057469",
                    "ENSMUSG00000010175",
                    "ENSMUSG00000015839",
                    "ENSMUSG00000030256",
                    "ENSMUSG00000052837",
                    "ENSMUSG00000015579",
                    "ENSMUSG00000056501",
                    "ENSMUSG00000001280",
                    "ENSMUSG00000005566",
                    "ENSMUSG00000019768",
                    "ENSMUSG00000008575",
                    "ENSMUSG00000039191"
                ],
                "target": [],
                "expression_sum": {
                    "class1": 10566.0,
                    "class2": 6624.0
                },
                "uniprot_ids": [
                    "P06151",
                    "D3YZQ9",
                    "A0A1B0GSR9",
                    "A0A1B0GSX0",
                    "A0A1B0GT41",
                    "A0A1B0GS79",
                    "A0A1B0GRS2",
                    "A0A1B0GQX5",
                    "A0A1B0GRW9",
                    "A0A1B0GSL7",
                    "A0A1B0GRC1"
                ]
            },
            "ENSMUSG00000032366": {
                "id": "ENSMUSG00000032366",
                "type": "Gene",
                "symbol": "Tpm1",
                "source": [
                    "ENSMUSG00000029135",
                    "ENSMUSG00000001419",
                    "ENSMUSG00000015839",
                    "ENSMUSG00000058239",
                    "ENSMUSG00000015605",
                    "ENSMUSG00000056501",
                    "ENSMUSG00000051469",
                    "ENSMUSG00000030557",
                    "ENSMUSG00000001280",
                    "ENSMUSG00000019768",
                    "ENSMUSG00000053110",
                    "ENSMUSG00000031885",
                    "ENSMUSG00000008575",
                    "ENSMUSG00000031715"
                ],
                "target": [],
                "expression_sum": {
                    "class1": 121065.0,
                    "class2": 110163.0
                },
                "uniprot_ids": [
                    "P58771",
                    "Q8BP43",
                    "Q8BSH3",
                    "B7ZNL3",
                    "E9Q448",
                    "F8WID5",
                    "E9Q450",
                    "E9Q456",
                    "G5E8R0",
                    "S4R2U0",
                    "E9Q455",
                    "S4R261",
                    "E9Q452",
                    "E9Q453",
                    "E9Q454",
                    "G5E8R1",
                    "G5E8R2"
                ]
            },
            "ENSMUSG00000055320": {
                "id": "ENSMUSG00000055320",
                "type": "TF",
                "symbol": "Tead1",
                "source": [
                    "ENSMUSG00000029135",
                    "ENSMUSG00000001419",
                    "ENSMUSG00000026610",
                    "ENSMUSG00000015579",
                    "ENSMUSG00000029249",
                    "ENSMUSG00000001280",
                    "ENSMUSG00000019768",
                    "ENSMUSG00000031715"
                ],
                "target": [
                    "ENSMUSG00000035674",
                    "ENSMUSG00000005958",
                    "ENSMUSG00000002010",
                    "ENSMUSG00000005610",
                    "ENSMUSG00000001767",
                    "ENSMUSG00000032119",
                    "ENSMUSG00000014633",
                    "ENSMUSG00000024197",
                    "ENSMUSG00000020530",
                    "ENSMUSG00000029158",
                    "ENSMUSG00000001105",
                    "ENSMUSG00000046516",
                    "ENSMUSG00000025130",
                    "ENSMUSG00000000056",
                    "ENSMUSG00000039239",
                    "ENSMUSG00000071793",
                    "ENSMUSG00000000031",
                    "ENSMUSG00000000753",
                    "ENSMUSG00000023089",
                    "ENSMUSG00000000088",
                    "ENSMUSG00000082165",
                    "ENSMUSG00000050222",
                    "ENSMUSG00000073676",
                    "ENSMUSG00000039509",
                    "ENSMUSG00000004610",
                    "ENSMUSG00000020664",
                    "ENSMUSG00000069049",
                    "ENSMUSG00000013936",
                    "ENSMUSG00000017817",
                    "ENSMUSG00000001962",
                    "ENSMUSG00000000581",
                    "ENSMUSG00000028007",
                    "ENSMUSG00000023460",
                    "ENSMUSG00000001240",
                    "ENSMUSG00000003531",
                    "ENSMUSG00000028633",
                    "ENSMUSG00000029661"
                ],
                "expression_sum": {
                    "class1": 2182.0,
                    "class2": 1186.0
                },
                "uniprot_ids": [
                    "Q3USK5",
                    "Q3UFP5",
                    "Q6PAQ8",
                    "E9Q387",
                    "E9PZ73",
                    "F6QIG2",
                    "E9PVS2",
                    "F6QH05",
                    "F7B3M8",
                    "F6ZIM6",
                    "F6TBI7",
                    "E9PVJ5",
                    "E9QAK6"
                ]
            },
            "ENSMUSG00000084105": {
                "id": "ENSMUSG00000084105",
                "type": "Gene",
                "symbol": "Gm13252",
                "source": [
                    "ENSMUSG00000001280",
                    "ENSMUSG00000015605",
                    "ENSMUSG00000015846"
                ],
                "target": [],
                "expression_sum": {
                    "class1": 615.0,
                    "class2": 3356.0
                }
            },
            "ENSMUSG00000053898": {
                "id": "ENSMUSG00000053898",
                "type": "Gene",
                "symbol": "Ech1",
                "source": [
                    "ENSMUSG00000010175",
                    "ENSMUSG00000015839",
                    "ENSMUSG00000015605",
                    "ENSMUSG00000052534",
                    "ENSMUSG00000052837",
                    "ENSMUSG00000015579",
                    "ENSMUSG00000024985",
                    "ENSMUSG00000051469",
                    "ENSMUSG00000030557",
                    "ENSMUSG00000058239",
                    "ENSMUSG00000001280",
                    "ENSMUSG00000019768",
                    "ENSMUSG00000031885",
                    "ENSMUSG00000008575",
                    "ENSMUSG00000031715",
                    "ENSMUSG00000021775",
                    "ENSMUSG00000015846"
                ],
                "target": [],
                "expression_sum": {
                    "class1": 8744.0,
                    "class2": 6690.0
                },
                "uniprot_ids": [
                    "O35459",
                    "F7B227"
                ]
            },
            "ENSMUSG00000030541": {
                "id": "ENSMUSG00000030541",
                "type": "Gene",
                "symbol": "Idh2",
                "source": [
                    "ENSMUSG00000057469",
                    "ENSMUSG00000015839",
                    "ENSMUSG00000015605",
                    "ENSMUSG00000074622",
                    "ENSMUSG00000029249",
                    "ENSMUSG00000001280",
                    "ENSMUSG00000005566",
                    "ENSMUSG00000019768",
                    "ENSMUSG00000022641",
                    "ENSMUSG00000068267",
                    "ENSMUSG00000008575",
                    "ENSMUSG00000039191",
                    "ENSMUSG00000024985"
                ],
                "target": [],
                "expression_sum": {
                    "class1": 9027.0,
                    "class2": 9998.0
                },
                "uniprot_ids": [
                    "P54071",
                    "A0A0U1RP68",
                    "D6RIL6",
                    "A0A0U1RPR1"
                ]
            },
            "ENSMUSG00000080893": {
                "id": "ENSMUSG00000080893",
                "type": "Gene",
                "symbol": "Ndufa12-ps",
                "source": [
                    "ENSMUSG00000029135",
                    "ENSMUSG00000018604",
                    "ENSMUSG00000056501",
                    "ENSMUSG00000001280",
                    "ENSMUSG00000019768",
                    "ENSMUSG00000015846"
                ],
                "target": [],
                "expression_sum": {
                    "class1": 16164.0,
                    "class2": 11861.0
                }
            },
            "ENSMUSG00000081344": {
                "id": "ENSMUSG00000081344",
                "type": "Gene",
                "symbol": "Gm14303",
                "source": [
                    "ENSMUSG00000001280",
                    "ENSMUSG00000056501",
                    "ENSMUSG00000090258"
                ],
                "target": [],
                "expression_sum": {
                    "class1": 27056.0,
                    "class2": 8022.0
                }
            },
            "ENSMUSG00000064363": {
                "id": "ENSMUSG00000064363",
                "type": "Gene",
                "symbol": "ND4",
                "source": [
                    "ENSMUSG00000019768",
                    "ENSMUSG00000052837",
                    "ENSMUSG00000021944",
                    "ENSMUSG00000022346"
                ],
                "target": [],
                "expression_sum": {
                    "class1": 2285058.0,
                    "class2": 4070488.0
                },
                "uniprot_ids": [
                    "P03911"
                ]
            },
            "ENSMUSG00000058076": {
                "id": "ENSMUSG00000058076",
                "type": "Gene",
                "symbol": "Sdhc",
                "source": [
                    "ENSMUSG00000015839",
                    "ENSMUSG00000019817",
                    "ENSMUSG00000015605",
                    "ENSMUSG00000052534",
                    "ENSMUSG00000029249",
                    "ENSMUSG00000001280",
                    "ENSMUSG00000005566",
                    "ENSMUSG00000005583",
                    "ENSMUSG00000019768",
                    "ENSMUSG00000032238",
                    "ENSMUSG00000008575",
                    "ENSMUSG00000035158"
                ],
                "target": [],
                "expression_sum": {
                    "class1": 3154.0,
                    "class2": 3722.0
                },
                "uniprot_ids": [
                    "Q9CZB0",
                    "D3Z1A8",
                    "F8WGB3"
                ]
            },
            "ENSMUSG00000027673": {
                "id": "ENSMUSG00000027673",
                "type": "Gene",
                "symbol": "Ndufb5",
                "source": [
                    "ENSMUSG00000015839",
                    "ENSMUSG00000058239",
                    "ENSMUSG00000026610",
                    "ENSMUSG00000024985",
                    "ENSMUSG00000001280",
                    "ENSMUSG00000005566",
                    "ENSMUSG00000019768",
                    "ENSMUSG00000031885",
                    "ENSMUSG00000008575",
                    "ENSMUSG00000014592",
                    "ENSMUSG00000015846"
                ],
                "target": [],
                "expression_sum": {
                    "class1": 10246.0,
                    "class2": 8846.0
                },
                "uniprot_ids": [
                    "Q9CQH3",
                    "D3Z568",
                    "D3YX99",
                    "F6Y6V5",
                    "D3Z6W9",
                    "F8WI84",
                    "F6VXF9"
                ]
            },
            "ENSMUSG00000023951": {
                "id": "ENSMUSG00000023951",
                "type": "Gene",
                "symbol": "Vegfa",
                "source": [
                    "ENSMUSG00000015839",
                    "ENSMUSG00000015579",
                    "ENSMUSG00000024955",
                    "ENSMUSG00000001280",
                    "ENSMUSG00000019768",
                    "ENSMUSG00000060601",
                    "ENSMUSG00000020919",
                    "ENSMUSG00000038692",
                    "ENSMUSG00000031885",
                    "ENSMUSG00000008575",
                    "ENSMUSG00000031715"
                ],
                "target": [],
                "expression_sum": {
                    "class1": 783.0,
                    "class2": 1944.0
                },
                "uniprot_ids": [
                    "Q00731",
                    "F8WH81",
                    "E9Q4N8",
                    "E9Q4N9",
                    "F8WH10"
                ]
            },
            "ENSMUSG00000098178": {
                "id": "ENSMUSG00000098178",
                "type": "Gene",
                "symbol": null,
                "source": [
                    "ENSMUSG00000022346",
                    "ENSMUSG00000021250",
                    "ENSMUSG00000018604",
                    "ENSMUSG00000052534",
                    "ENSMUSG00000001280",
                    "ENSMUSG00000019768",
                    "ENSMUSG00000021877",
                    "ENSMUSG00000035158"
                ],
                "target": [],
                "expression_sum": {
                    "class1": 1897553.0,
                    "class2": 328598.0
                }
            },
            "ENSMUSG00000040048": {
                "id": "ENSMUSG00000040048",
                "type": "Gene",
                "symbol": "Ndufb10",
                "source": [
                    "ENSMUSG00000058239",
                    "ENSMUSG00000056501",
                    "ENSMUSG00000028639",
                    "ENSMUSG00000001280",
                    "ENSMUSG00000019768",
                    "ENSMUSG00000015846",
                    "ENSMUSG00000076431",
                    "ENSMUSG00000021775",
                    "ENSMUSG00000031715"
                ],
                "target": [],
                "expression_sum": {
                    "class1": 27373.0,
                    "class2": 19705.0
                },
                "uniprot_ids": [
                    "Q9DCS9",
                    "D3YUK4"
                ]
            },
            "ENSMUSG00000044894": {
                "id": "ENSMUSG00000044894",
                "type": "Gene",
                "symbol": "Uqcrq",
                "source": [
                    "ENSMUSG00000029135",
                    "ENSMUSG00000015839",
                    "ENSMUSG00000010175",
                    "ENSMUSG00000015605",
                    "ENSMUSG00000056501",
                    "ENSMUSG00000001280",
                    "ENSMUSG00000019768",
                    "ENSMUSG00000008575",
                    "ENSMUSG00000039191",
                    "ENSMUSG00000021775",
                    "ENSMUSG00000015846"
                ],
                "target": [],
                "expression_sum": {
                    "class1": 38662.0,
                    "class2": 36514.0
                },
                "uniprot_ids": [
                    "Q9CQ69",
                    "I7HPX6"
                ]
            },
            "ENSMUSG00000053477": {
                "id": "ENSMUSG00000053477",
                "type": "TF",
                "symbol": "Tcf4",
                "source": [
                    "ENSMUSG00000019947",
                    "ENSMUSG00000024238",
                    "ENSMUSG00000029135",
                    "ENSMUSG00000019817",
                    "ENSMUSG00000018604",
                    "ENSMUSG00000052534",
                    "ENSMUSG00000015579",
                    "ENSMUSG00000036461",
                    "ENSMUSG00000001280",
                    "ENSMUSG00000005583",
                    "ENSMUSG00000030067",
                    "ENSMUSG00000022641",
                    "ENSMUSG00000071637",
                    "ENSMUSG00000053110",
                    "ENSMUSG00000022314",
                    "ENSMUSG00000022528"
                ],
                "target": [
                    "ENSMUSG00000051615",
                    "ENSMUSG00000063320",
                    "ENSMUSG00000002844",
                    "ENSMUSG00000032737",
                    "ENSMUSG00000029454",
                    "ENSMUSG00000020841",
                    "ENSMUSG00000015766",
                    "ENSMUSG00000028643",
                    "ENSMUSG00000001016",
                    "ENSMUSG00000040147",
                    "ENSMUSG00000047888",
                    "ENSMUSG00000044702",
                    "ENSMUSG00000022641",
                    "ENSMUSG00000042901",
                    "ENSMUSG00000054766",
                    "ENSMUSG00000018819",
                    "ENSMUSG00000021484",
                    "ENSMUSG00000020077",
                    "ENSMUSG00000029823",
                    "ENSMUSG00000021453",
                    "ENSMUSG00000026176",
                    "ENSMUSG00000078247",
                    "ENSMUSG00000042029",
                    "ENSMUSG00000020493",
                    "ENSMUSG00000043415",
                    "ENSMUSG00000032423",
                    "ENSMUSG00000060771",
                    "ENSMUSG00000038628",
                    "ENSMUSG00000021619",
                    "ENSMUSG00000021400",
                    "ENSMUSG00000027800",
                    "ENSMUSG00000079184",
                    "ENSMUSG00000025492",
                    "ENSMUSG00000045098",
                    "ENSMUSG00000064340",
                    "ENSMUSG00000025077",
                    "ENSMUSG00000052926",
                    "ENSMUSG00000039219",
                    "ENSMUSG00000092116",
                    "ENSMUSG00000029576",
                    "ENSMUSG00000017493",
                    "ENSMUSG00000024422",
                    "ENSMUSG00000024528",
                    "ENSMUSG00000041685",
                    "ENSMUSG00000008822",
                    "ENSMUSG00000020430",
                    "ENSMUSG00000029290",
                    "ENSMUSG00000025959",
                    "ENSMUSG00000042389",
                    "ENSMUSG00000025534",
                    "ENSMUSG00000090553",
                    "ENSMUSG00000019951",
                    "ENSMUSG00000036860",
                    "ENSMUSG00000033813",
                    "ENSMUSG00000022905",
                    "ENSMUSG00000060802",
                    "ENSMUSG00000038803",
                    "ENSMUSG00000038671",
                    "ENSMUSG00000028461",
                    "ENSMUSG00000063316",
                    "ENSMUSG00000004951",
                    "ENSMUSG00000053317",
                    "ENSMUSG00000034154",
                    "ENSMUSG00000020076",
                    "ENSMUSG00000071533",
                    "ENSMUSG00000038967",
                    "ENSMUSG00000042426",
                    "ENSMUSG00000008429",
                    "ENSMUSG00000018102",
                    "ENSMUSG00000096472",
                    "ENSMUSG00000020009",
                    "ENSMUSG00000026755",
                    "ENSMUSG00000018171",
                    "ENSMUSG00000020248",
                    "ENSMUSG00000021500",
                    "ENSMUSG00000001930",
                    "ENSMUSG00000070544",
                    "ENSMUSG00000026365",
                    "ENSMUSG00000003099",
                    "ENSMUSG00000042506",
                    "ENSMUSG00000045954",
                    "ENSMUSG00000030067",
                    "ENSMUSG00000003955",
                    "ENSMUSG00000053838",
                    "ENSMUSG00000014329",
                    "ENSMUSG00000026202",
                    "ENSMUSG00000025184",
                    "ENSMUSG00000028107",
                    "ENSMUSG00000022108",
                    "ENSMUSG00000024309",
                    "ENSMUSG00000020773",
                    "ENSMUSG00000040990",
                    "ENSMUSG00000026219",
                    "ENSMUSG00000040033",
                    "ENSMUSG00000019970",
                    "ENSMUSG00000024878",
                    "ENSMUSG00000033538",
                    "ENSMUSG00000003200",
                    "ENSMUSG00000025574",
                    "ENSMUSG00000038780",
                    "ENSMUSG00000030172",
                    "ENSMUSG00000019494",
                    "ENSMUSG00000036461",
                    "ENSMUSG00000023010",
                    "ENSMUSG00000009555",
                    "ENSMUSG00000015575",
                    "ENSMUSG00000005981",
                    "ENSMUSG00000034109",
                    "ENSMUSG00000003420",
                    "ENSMUSG00000026615",
                    "ENSMUSG00000030682",
                    "ENSMUSG00000029135",
                    "ENSMUSG00000022174",
                    "ENSMUSG00000040124",
                    "ENSMUSG00000022528",
                    "ENSMUSG00000035266",
                    "ENSMUSG00000073639",
                    "ENSMUSG00000052534",
                    "ENSMUSG00000023990",
                    "ENSMUSG00000020074",
                    "ENSMUSG00000027326",
                    "ENSMUSG00000021917",
                    "ENSMUSG00000059734",
                    "ENSMUSG00000001289",
                    "ENSMUSG00000032932",
                    "ENSMUSG00000053110",
                    "ENSMUSG00000022314",
                    "ENSMUSG00000042524",
                    "ENSMUSG00000001627",
                    "ENSMUSG00000002031",
                    "ENSMUSG00000022206",
                    "ENSMUSG00000016344",
                    "ENSMUSG00000024491",
                    "ENSMUSG00000044475",
                    "ENSMUSG00000053565",
                    "ENSMUSG00000019868",
                    "ENSMUSG00000022094",
                    "ENSMUSG00000028248",
                    "ENSMUSG00000020585",
                    "ENSMUSG00000032265",
                    "ENSMUSG00000002748",
                    "ENSMUSG00000026766",
                    "ENSMUSG00000001280",
                    "ENSMUSG00000090266",
                    "ENSMUSG00000026674",
                    "ENSMUSG00000005078",
                    "ENSMUSG00000028949",
                    "ENSMUSG00000039660"
                ],
                "expression_sum": {
                    "class1": 26199.0,
                    "class2": 4655.0
                },
                "uniprot_ids": [
                    "Q60722",
                    "A0A0J9YTU7",
                    "A0A0J9YUZ0",
                    "Q91YV0",
                    "A0A0J9YUG4",
                    "Q8BPX7",
                    "A0A0J9YUP9",
                    "A0A0J9YVG8",
                    "A0A0J9YTX3",
                    "A0A1B0GXC8",
                    "A0A140LJA2",
                    "A0A0J9YUK1",
                    "A0A0J9YUS2",
                    "A0A0J9YVG2",
                    "A0A0J9YU75",
                    "A0A0J9YV39",
                    "A0A140LIL4",
                    "E9Q8G4",
                    "A0A0J9YTR9",
                    "A0A0J9YV54",
                    "A0A0J9YU37",
                    "A0A0J9YUT7",
                    "A0A0J9YUX6",
                    "A0A0J9YVC6",
                    "A0A0J9YUF5",
                    "A0A0J9YU89",
                    "A0A0J9YUY3",
                    "A0A0J9YVI1",
                    "A0A0J9YU91",
                    "A0A0J9YUT3",
                    "A0A0J9YUW9",
                    "A0A0J9YUR0",
                    "A0A0J9YU99",
                    "A0A0J9YV50",
                    "A0A0J9YTW1",
                    "A0A140LJC0",
                    "A0A0J9YV88",
                    "A0A0J9YVB3"
                ]
            },
            "ENSMUSG00000014294": {
                "id": "ENSMUSG00000014294",
                "type": "Gene",
                "symbol": "Ndufa2",
                "source": [
                    "ENSMUSG00000058239",
                    "ENSMUSG00000015579",
                    "ENSMUSG00000090258",
                    "ENSMUSG00000001280",
                    "ENSMUSG00000019768",
                    "ENSMUSG00000031885",
                    "ENSMUSG00000076431",
                    "ENSMUSG00000015605",
                    "ENSMUSG00000039191",
                    "ENSMUSG00000021775"
                ],
                "target": [],
                "expression_sum": {
                    "class1": 27962.0,
                    "class2": 13811.0
                },
                "uniprot_ids": [
                    "Q9CQ75"
                ]
            },
            "ENSMUSG00000074218": {
                "id": "ENSMUSG00000074218",
                "type": "Gene",
                "symbol": "Cox7a1",
                "source": [
                    "ENSMUSG00000029135",
                    "ENSMUSG00000001419",
                    "ENSMUSG00000015839",
                    "ENSMUSG00000058239",
                    "ENSMUSG00000056501",
                    "ENSMUSG00000015579",
                    "ENSMUSG00000001280",
                    "ENSMUSG00000005566",
                    "ENSMUSG00000019768",
                    "ENSMUSG00000028565",
                    "ENSMUSG00000032238",
                    "ENSMUSG00000024140",
                    "ENSMUSG00000031885",
                    "ENSMUSG00000015605",
                    "ENSMUSG00000015846"
                ],
                "target": [],
                "expression_sum": {
                    "class1": 13581.0,
                    "class2": 45730.0
                },
                "uniprot_ids": [
                    "P56392",
                    "A0A140LIU4"
                ]
            },
            "ENSMUSG00000054889": {
                "id": "ENSMUSG00000054889",
                "type": "Gene",
                "symbol": "Dsp",
                "source": [
                    "ENSMUSG00000029135",
                    "ENSMUSG00000057469",
                    "ENSMUSG00000025612",
                    "ENSMUSG00000010175",
                    "ENSMUSG00000015839",
                    "ENSMUSG00000015605",
                    "ENSMUSG00000056501",
                    "ENSMUSG00000001280",
                    "ENSMUSG00000019768",
                    "ENSMUSG00000005583",
                    "ENSMUSG00000005698",
                    "ENSMUSG00000022641",
                    "ENSMUSG00000032238",
                    "ENSMUSG00000053110",
                    "ENSMUSG00000031885",
                    "ENSMUSG00000008575",
                    "ENSMUSG00000021775",
                    "ENSMUSG00000024985"
                ],
                "target": [],
                "expression_sum": {
                    "class1": 3045.0,
                    "class2": 4006.0
                },
                "uniprot_ids": [
                    "E9Q557",
                    "E9PZW0"
                ]
            },
            "ENSMUSG00000009863": {
                "id": "ENSMUSG00000009863",
                "type": "Gene",
                "symbol": "Sdhb",
                "source": [
                    "ENSMUSG00000015839",
                    "ENSMUSG00000058239",
                    "ENSMUSG00000015605",
                    "ENSMUSG00000001911",
                    "ENSMUSG00000056501",
                    "ENSMUSG00000001156",
                    "ENSMUSG00000040033",
                    "ENSMUSG00000029249",
                    "ENSMUSG00000001280",
                    "ENSMUSG00000019768",
                    "ENSMUSG00000028565",
                    "ENSMUSG00000025958",
                    "ENSMUSG00000026803",
                    "ENSMUSG00000033943",
                    "ENSMUSG00000008575",
                    "ENSMUSG00000039191"
                ],
                "target": [],
                "expression_sum": {
                    "class1": 11335.0,
                    "class2": 10601.0
                },
                "uniprot_ids": [
                    "Q9CQA3"
                ]
            },
            "ENSMUSG00000025190": {
                "id": "ENSMUSG00000025190",
                "type": "Gene",
                "symbol": "Got1",
                "source": [
                    "ENSMUSG00000001419",
                    "ENSMUSG00000010175",
                    "ENSMUSG00000015605",
                    "ENSMUSG00000030557",
                    "ENSMUSG00000001280",
                    "ENSMUSG00000019768",
                    "ENSMUSG00000032238",
                    "ENSMUSG00000008575",
                    "ENSMUSG00000015846"
                ],
                "target": [],
                "expression_sum": {
                    "class1": 800.0,
                    "class2": 3149.0
                },
                "uniprot_ids": [
                    "P05201",
                    "F7ALS6"
                ]
            },
            "ENSMUSG00000022452": {
                "id": "ENSMUSG00000022452",
                "type": "Gene",
                "symbol": "Smdt1",
                "source": [
                    "ENSMUSG00000029135",
                    "ENSMUSG00000057469",
                    "ENSMUSG00000021944",
                    "ENSMUSG00000026890",
                    "ENSMUSG00000018604",
                    "ENSMUSG00000026610",
                    "ENSMUSG00000015579",
                    "ENSMUSG00000029249",
                    "ENSMUSG00000001280",
                    "ENSMUSG00000005566",
                    "ENSMUSG00000019768",
                    "ENSMUSG00000001493",
                    "ENSMUSG00000022641",
                    "ENSMUSG00000031885",
                    "ENSMUSG00000076431",
                    "ENSMUSG00000008575"
                ],
                "target": [],
                "expression_sum": {
                    "class1": 17534.0,
                    "class2": 4783.0
                },
                "uniprot_ids": [
                    "Q9DB10",
                    "E0CZF0",
                    "A0A2R8VHK6",
                    "E0CXP9"
                ]
            },
            "ENSMUSG00000053702": {
                "id": "ENSMUSG00000053702",
                "type": "Gene",
                "symbol": "Nebl",
                "source": [
                    "ENSMUSG00000029135",
                    "ENSMUSG00000056501",
                    "ENSMUSG00000051469",
                    "ENSMUSG00000030557",
                    "ENSMUSG00000001280",
                    "ENSMUSG00000019768",
                    "ENSMUSG00000024974",
                    "ENSMUSG00000031885"
                ],
                "target": [],
                "expression_sum": {
                    "class1": 4810.0,
                    "class2": 4096.0
                },
                "uniprot_ids": [
                    "Q9DC07",
                    "Q0II04",
                    "Q3TR73",
                    "B7ZCI2",
                    "A0A1B0GXL1",
                    "B7ZCI3",
                    "F6RM14"
                ]
            },
            "ENSMUSG00000031299": {
                "id": "ENSMUSG00000031299",
                "type": "Gene",
                "symbol": "Pdha1",
                "source": [
                    "ENSMUSG00000010175",
                    "ENSMUSG00000015839",
                    "ENSMUSG00000021250",
                    "ENSMUSG00000056501",
                    "ENSMUSG00000015579",
                    "ENSMUSG00000001280",
                    "ENSMUSG00000019768",
                    "ENSMUSG00000005583",
                    "ENSMUSG00000005698",
                    "ENSMUSG00000025958",
                    "ENSMUSG00000031885",
                    "ENSMUSG00000058756",
                    "ENSMUSG00000024985"
                ],
                "target": [],
                "expression_sum": {
                    "class1": 3919.0,
                    "class2": 5152.0
                },
                "uniprot_ids": [
                    "P35486"
                ]
            },
            "ENSMUSG00000031701": {
                "id": "ENSMUSG00000031701",
                "type": "Gene",
                "symbol": "Dnaja2",
                "source": [
                    "ENSMUSG00000001419",
                    "ENSMUSG00000015839",
                    "ENSMUSG00000049577",
                    "ENSMUSG00000078671",
                    "ENSMUSG00000056501",
                    "ENSMUSG00000030557",
                    "ENSMUSG00000029249",
                    "ENSMUSG00000001280",
                    "ENSMUSG00000005566",
                    "ENSMUSG00000019768",
                    "ENSMUSG00000003184",
                    "ENSMUSG00000008575"
                ],
                "target": [],
                "expression_sum": {
                    "class1": 2672.0,
                    "class2": 1612.0
                },
                "uniprot_ids": [
                    "Q9QYJ0"
                ]
            },
            "ENSMUSG00000062997": {
                "id": "ENSMUSG00000062997",
                "type": "Gene",
                "symbol": "Rpl35",
                "source": [
                    "ENSMUSG00000021250",
                    "ENSMUSG00000018604",
                    "ENSMUSG00000036461",
                    "ENSMUSG00000001280",
                    "ENSMUSG00000005583",
                    "ENSMUSG00000040732",
                    "ENSMUSG00000022641",
                    "ENSMUSG00000020919"
                ],
                "target": [],
                "expression_sum": {
                    "class1": 4281.0,
                    "class2": 1045.0
                },
                "uniprot_ids": [
                    "Q6ZWV7"
                ]
            },
            "ENSMUSG00000057580": {
                "id": "ENSMUSG00000057580",
                "type": "Gene",
                "symbol": "Cox7c-ps1",
                "source": [
                    "ENSMUSG00000019768",
                    "ENSMUSG00000024081",
                    "ENSMUSG00000051469"
                ],
                "target": [],
                "expression_sum": {
                    "class1": 634.0,
                    "class2": 7269.0
                }
            },
            "ENSMUSG00000074129": {
                "id": "ENSMUSG00000074129",
                "type": "Gene",
                "symbol": "Rpl13a",
                "source": [
                    "ENSMUSG00000024238",
                    "ENSMUSG00000052837",
                    "ENSMUSG00000056501",
                    "ENSMUSG00000001280",
                    "ENSMUSG00000019768",
                    "ENSMUSG00000042557"
                ],
                "target": [],
                "expression_sum": {
                    "class1": 12609.0,
                    "class2": 1752.0
                },
                "uniprot_ids": [
                    "P19253",
                    "A0A1B0GSQ6",
                    "A0A1B0GTA1",
                    "A0A1B0GSC2",
                    "A0A1B0GRH1",
                    "A0A1B0GT00",
                    "A0A1B0GSL5",
                    "A0A1B0GQW6",
                    "A0A1B0GSF0",
                    "A0A1B0GSJ2",
                    "A0A1B0GSB2"
                ]
            },
            "ENSMUSG00000025488": {
                "id": "ENSMUSG00000025488",
                "type": "Gene",
                "symbol": "Cox8b",
                "source": [
                    "ENSMUSG00000019768",
                    "ENSMUSG00000021250",
                    "ENSMUSG00000015846"
                ],
                "target": [],
                "expression_sum": {
                    "class1": 22153.0,
                    "class2": 54979.0
                },
                "uniprot_ids": [
                    "P48772"
                ]
            },
            "ENSMUSG00000030747": {
                "id": "ENSMUSG00000030747",
                "type": "Gene",
                "symbol": "Dgat2",
                "source": [
                    "ENSMUSG00000008575",
                    "ENSMUSG00000019789",
                    "ENSMUSG00000015579",
                    "ENSMUSG00000029249",
                    "ENSMUSG00000001280",
                    "ENSMUSG00000005566",
                    "ENSMUSG00000019768",
                    "ENSMUSG00000015605",
                    "ENSMUSG00000024985"
                ],
                "target": [],
                "expression_sum": {
                    "class1": 186.0,
                    "class2": 719.0
                },
                "uniprot_ids": [
                    "Q9DCV3",
                    "A0A140LHA9",
                    "A0A140LJ48",
                    "A0A140LHF3"
                ]
            },
            "ENSMUSG00000086567": {
                "id": "ENSMUSG00000086567",
                "type": "Gene",
                "symbol": "Gm2830",
                "source": [
                    "ENSMUSG00000001280"
                ],
                "target": [],
                "expression_sum": {
                    "class1": 1528.0,
                    "class2": 704.0
                }
            },
            "ENSMUSG00000049832": {
                "id": "ENSMUSG00000049832",
                "type": "Gene",
                "symbol": "Rbx1-ps",
                "source": [
                    "ENSMUSG00000001280",
                    "ENSMUSG00000014592"
                ],
                "target": [],
                "expression_sum": {
                    "class1": 6440.0,
                    "class2": 3030.0
                }
            },
            "ENSMUSG00000027984": {
                "id": "ENSMUSG00000027984",
                "type": "Gene",
                "symbol": "Hadh",
                "source": [
                    "ENSMUSG00000015839",
                    "ENSMUSG00000010175",
                    "ENSMUSG00000021109",
                    "ENSMUSG00000056501",
                    "ENSMUSG00000029249",
                    "ENSMUSG00000001280",
                    "ENSMUSG00000019768",
                    "ENSMUSG00000005698",
                    "ENSMUSG00000031885",
                    "ENSMUSG00000008575",
                    "ENSMUSG00000021775"
                ],
                "target": [],
                "expression_sum": {
                    "class1": 1649.0,
                    "class2": 2325.0
                },
                "uniprot_ids": [
                    "Q61425"
                ]
            },
            "ENSMUSG00000049265": {
                "id": "ENSMUSG00000049265",
                "type": "Gene",
                "symbol": "Kcnk3",
                "source": [
                    "ENSMUSG00000021250",
                    "ENSMUSG00000001280",
                    "ENSMUSG00000019768",
                    "ENSMUSG00000025958",
                    "ENSMUSG00000058756",
                    "ENSMUSG00000015846"
                ],
                "target": [],
                "expression_sum": {
                    "class1": 233.0,
                    "class2": 658.0
                },
                "uniprot_ids": [
                    "O35111"
                ]
            },
            "ENSMUSG00000025204": {
                "id": "ENSMUSG00000025204",
                "type": "Gene",
                "symbol": "Ndufb8",
                "source": [
                    "ENSMUSG00000015839",
                    "ENSMUSG00000006215",
                    "ENSMUSG00000026610",
                    "ENSMUSG00000056501",
                    "ENSMUSG00000051469",
                    "ENSMUSG00000001280",
                    "ENSMUSG00000019768",
                    "ENSMUSG00000025958",
                    "ENSMUSG00000031885",
                    "ENSMUSG00000008575",
                    "ENSMUSG00000039191",
                    "ENSMUSG00000024985"
                ],
                "target": [],
                "expression_sum": {
                    "class1": 18334.0,
                    "class2": 10697.0
                },
                "uniprot_ids": [
                    "Q9D6J5",
                    "Q3V406",
                    "E9Q9J5",
                    "E9PY23",
                    "F6VCY0"
                ]
            },
            "ENSMUSG00000027809": {
                "id": "ENSMUSG00000027809",
                "type": "Gene",
                "symbol": "Etfdh",
                "source": [
                    "ENSMUSG00000029135",
                    "ENSMUSG00000015839",
                    "ENSMUSG00000058239",
                    "ENSMUSG00000015605",
                    "ENSMUSG00000078671",
                    "ENSMUSG00000015579",
                    "ENSMUSG00000024955",
                    "ENSMUSG00000051469",
                    "ENSMUSG00000001280",
                    "ENSMUSG00000019768",
                    "ENSMUSG00000028565",
                    "ENSMUSG00000025958",
                    "ENSMUSG00000053110",
                    "ENSMUSG00000031627",
                    "ENSMUSG00000075595",
                    "ENSMUSG00000031885",
                    "ENSMUSG00000008575",
                    "ENSMUSG00000024985"
                ],
                "target": [],
                "expression_sum": {
                    "class1": 1620.0,
                    "class2": 2035.0
                },
                "uniprot_ids": [
                    "Q921G7",
                    "Q6PF96"
                ]
            },
            "ENSMUSG00000030417": {
                "id": "ENSMUSG00000030417",
                "type": "Gene",
                "symbol": "Pdcd5",
                "source": [
                    "ENSMUSG00000040481",
                    "ENSMUSG00000019817",
                    "ENSMUSG00000015605",
                    "ENSMUSG00000051469",
                    "ENSMUSG00000030557",
                    "ENSMUSG00000029249",
                    "ENSMUSG00000001280",
                    "ENSMUSG00000019768",
                    "ENSMUSG00000005583",
                    "ENSMUSG00000024140",
                    "ENSMUSG00000068267",
                    "ENSMUSG00000008575",
                    "ENSMUSG00000022314"
                ],
                "target": [],
                "expression_sum": {
                    "class1": 3505.0,
                    "class2": 1163.0
                },
                "uniprot_ids": [
                    "P56812",
                    "D3Z7Q5"
                ]
            },
            "ENSMUSG00000058927": {
                "id": "ENSMUSG00000058927",
                "type": "Gene",
                "symbol": "Gm10053",
                "source": [
                    "ENSMUSG00000019768",
                    "ENSMUSG00000014592"
                ],
                "target": [],
                "expression_sum": {
                    "class1": 15278.0,
                    "class2": 25592.0
                },
                "uniprot_ids": [
                    "P62897"
                ]
            },
            "ENSMUSG00000032279": {
                "id": "ENSMUSG00000032279",
                "type": "Gene",
                "symbol": "Idh3a",
                "source": [
                    "ENSMUSG00000029135",
                    "ENSMUSG00000015605",
                    "ENSMUSG00000015579",
                    "ENSMUSG00000028639",
                    "ENSMUSG00000001280",
                    "ENSMUSG00000019768",
                    "ENSMUSG00000033943",
                    "ENSMUSG00000053110",
                    "ENSMUSG00000037174",
                    "ENSMUSG00000008575",
                    "ENSMUSG00000039191"
                ],
                "target": [],
                "expression_sum": {
                    "class1": 2398.0,
                    "class2": 4195.0
                },
                "uniprot_ids": [
                    "Q9D6R2",
                    "A0A1L1STE6",
                    "A0A1L1SQF9"
                ]
            },
            "ENSMUSG00000073702": {
                "id": "ENSMUSG00000073702",
                "type": "Gene",
                "symbol": "Rpl31",
                "source": [
                    "ENSMUSG00000001280",
                    "ENSMUSG00000005583",
                    "ENSMUSG00000024238",
                    "ENSMUSG00000022314"
                ],
                "target": [],
                "expression_sum": {
                    "class1": 2510.0,
                    "class2": 553.0
                },
                "uniprot_ids": [
                    "P62900",
                    "A0A0A6YXL3",
                    "A0A0A6YX26"
                ]
            },
            "ENSMUSG00000019689": {
                "id": "ENSMUSG00000019689",
                "type": "Gene",
                "symbol": "Fmc1",
                "source": [
                    "ENSMUSG00000001419",
                    "ENSMUSG00000001280",
                    "ENSMUSG00000038914",
                    "ENSMUSG00000076431",
                    "ENSMUSG00000021775",
                    "ENSMUSG00000031715",
                    "ENSMUSG00000000567"
                ],
                "target": [],
                "expression_sum": {
                    "class1": 4516.0,
                    "class2": 4608.0
                },
                "uniprot_ids": [
                    "Q9CR13"
                ]
            },
            "ENSMUSG00000064360": {
                "id": "ENSMUSG00000064360",
                "type": "Gene",
                "symbol": "ND3",
                "source": [
                    "ENSMUSG00000019768",
                    "ENSMUSG00000052837",
                    "ENSMUSG00000022346"
                ],
                "target": [],
                "expression_sum": {
                    "class1": 65910.0,
                    "class2": 329907.0
                },
                "uniprot_ids": [
                    "P03899"
                ]
            },
            "ENSMUSG00000025781": {
                "id": "ENSMUSG00000025781",
                "type": "Gene",
                "symbol": "Atp5c1",
                "source": [
                    "ENSMUSG00000015839",
                    "ENSMUSG00000078671",
                    "ENSMUSG00000026610",
                    "ENSMUSG00000056501",
                    "ENSMUSG00000015579",
                    "ENSMUSG00000030557",
                    "ENSMUSG00000029249",
                    "ENSMUSG00000001280",
                    "ENSMUSG00000005566",
                    "ENSMUSG00000019768",
                    "ENSMUSG00000014592",
                    "ENSMUSG00000022641",
                    "ENSMUSG00000031885",
                    "ENSMUSG00000015605",
                    "ENSMUSG00000035158",
                    "ENSMUSG00000015846"
                ],
                "target": [],
                "expression_sum": {
                    "class1": 10481.0,
                    "class2": 9354.0
                },
                "uniprot_ids": [
                    "Q91VR2",
                    "Q8C2Q8",
                    "A2AKV2",
                    "A2AKV3",
                    "A2AKV1",
                    "A2AKV0",
                    "A2AKU9"
                ]
            },
            "ENSMUSG00000049751": {
                "id": "ENSMUSG00000049751",
                "type": "Gene",
                "symbol": "Rpl36al",
                "source": [
                    "ENSMUSG00000059552",
                    "ENSMUSG00000024515",
                    "ENSMUSG00000056501",
                    "ENSMUSG00000001280",
                    "ENSMUSG00000028565",
                    "ENSMUSG00000040732"
                ],
                "target": [],
                "expression_sum": {
                    "class1": 3604.0,
                    "class2": 753.0
                },
                "uniprot_ids": [
                    "P83882"
                ]
            },
            "ENSMUSG00000062397": {
                "id": "ENSMUSG00000062397",
                "type": "Gene",
                "symbol": "Zfp706",
                "source": [
                    "ENSMUSG00000029135",
                    "ENSMUSG00000057469",
                    "ENSMUSG00000025612",
                    "ENSMUSG00000015839",
                    "ENSMUSG00000019564",
                    "ENSMUSG00000024955",
                    "ENSMUSG00000001280",
                    "ENSMUSG00000019768",
                    "ENSMUSG00000031885"
                ],
                "target": [],
                "expression_sum": {
                    "class1": 5617.0,
                    "class2": 2159.0
                },
                "uniprot_ids": [
                    "Q9D115"
                ]
            },
            "ENSMUSG00000028452": {
                "id": "ENSMUSG00000028452",
                "type": "Gene",
                "symbol": "Vcp",
                "source": [
                    "ENSMUSG00000015839",
                    "ENSMUSG00000078671",
                    "ENSMUSG00000015605",
                    "ENSMUSG00000030256",
                    "ENSMUSG00000056501",
                    "ENSMUSG00000028639",
                    "ENSMUSG00000029249",
```
