## Supplement 3 (P7 CM vs. P28 CM) for "AGEAS: Automated Machine Learning based Genetic Regulatory Element Extraction System": key_atlas.js

```
{
    "genes": {
        "ENSMUSG00000001280": {
            "id": "ENSMUSG00000001280",
            "type": "TF",
            "symbol": "Sp1",
            "source": [
                "ENSMUSG00000029135",
                "ENSMUSG00000021944",
                "ENSMUSG00000008575",
                "ENSMUSG00000015605",
                "ENSMUSG00000022528",
                "ENSMUSG00000057469",
                "ENSMUSG00000010175",
                "ENSMUSG00000058239",
                "ENSMUSG00000015579",
                "ENSMUSG00000051469",
                "ENSMUSG00000019768",
                "ENSMUSG00000031885",
                "ENSMUSG00000053110",
                "ENSMUSG00000031715",
                "ENSMUSG00000015846",
                "ENSMUSG00000058440",
                "ENSMUSG00000015839",
                "ENSMUSG00000032238",
                "ENSMUSG00000053477",
                "ENSMUSG00000058756",
                "ENSMUSG00000056501",
                "ENSMUSG00000029249",
                "ENSMUSG00000032035"
            ],
            "target": [
                "ENSMUSG00000000538",
                "ENSMUSG00000029860",
                "ENSMUSG00000029681",
                "ENSMUSG00000054455",
                "ENSMUSG00000032737",
                "ENSMUSG00000073702",
                "ENSMUSG00000031403",
                "ENSMUSG00000028081",
                "ENSMUSG00000091478",
                "ENSMUSG00000024228",
                "ENSMUSG00000020694",
                "ENSMUSG00000006676",
                "ENSMUSG00000020307",
                "ENSMUSG00000043683",
                "ENSMUSG00000021248",
                "ENSMUSG00000031885",
                "ENSMUSG00000026176",
                "ENSMUSG00000092074",
                "ENSMUSG00000021235",
                "ENSMUSG00000070283",
                "ENSMUSG00000039640",
                "ENSMUSG00000034903",
                "ENSMUSG00000030739",
                "ENSMUSG00000061461",
                "ENSMUSG00000004044",
                "ENSMUSG00000008036",
                "ENSMUSG00000028086",
                "ENSMUSG00000044221",
                "ENSMUSG00000035202",
                "ENSMUSG00000026087",
                "ENSMUSG00000026626",
                "ENSMUSG00000031790",
                "ENSMUSG00000031543",
                "ENSMUSG00000007891",
                "ENSMUSG00000069744",
                "ENSMUSG00000004789",
                "ENSMUSG00000083853",
                "ENSMUSG00000018363",
                "ENSMUSG00000018286",
                "ENSMUSG00000064349",
                "ENSMUSG00000080242",
                "ENSMUSG00000007656",
                "ENSMUSG00000027288",
                "ENSMUSG00000032314",
                "ENSMUSG00000008892",
                "ENSMUSG00000030557",
                "ENSMUSG00000020149",
                "ENSMUSG00000026858",
                "ENSMUSG00000027698",
                "ENSMUSG00000081344",
                "ENSMUSG00000028229",
                "ENSMUSG00000030852",
                "ENSMUSG00000010048",
                "ENSMUSG00000053329",
                "ENSMUSG00000031197",
                "ENSMUSG00000034981",
                "ENSMUSG00000029499",
                "ENSMUSG00000032216",
                "ENSMUSG00000090066",
                "ENSMUSG00000026755",
                "ENSMUSG00000018446",
                "ENSMUSG00000064302",
                "ENSMUSG00000026775",
                "ENSMUSG00000029392",
                "ENSMUSG00000044365",
                "ENSMUSG00000001552",
                "ENSMUSG00000020155",
                "ENSMUSG00000000056",
                "ENSMUSG00000024962",
                "ENSMUSG00000057133",
                "ENSMUSG00000020949",
                "ENSMUSG00000039103",
                "ENSMUSG00000073471",
                "ENSMUSG00000009628",
                "ENSMUSG00000029148",
                "ENSMUSG00000026158",
                "ENSMUSG00000020349",
                "ENSMUSG00000038708",
                "ENSMUSG00000032171",
                "ENSMUSG00000056501",
                "ENSMUSG00000029505",
                "ENSMUSG00000026615",
                "ENSMUSG00000055320",
                "ENSMUSG00000031698",
                "ENSMUSG00000022174",
                "ENSMUSG00000021024",
                "ENSMUSG00000027602",
                "ENSMUSG00000025745",
                "ENSMUSG00000049265",
                "ENSMUSG00000024304",
                "ENSMUSG00000006219",
                "ENSMUSG00000030057",
                "ENSMUSG00000025369",
                "ENSMUSG00000078139",
                "ENSMUSG00000055322",
                "ENSMUSG00000039016",
                "ENSMUSG00000006057",
                "ENSMUSG00000018196",
                "ENSMUSG00000068923",
                "ENSMUSG00000047284",
                "ENSMUSG00000087943",
                "ENSMUSG00000052253",
                "ENSMUSG00000057841",
                "ENSMUSG00000044533",
                "ENSMUSG00000064345",
                "ENSMUSG00000021666",
                "ENSMUSG00000019471",
                "ENSMUSG00000028884",
                "ENSMUSG00000018585",
                "ENSMUSG00000032648",
                "ENSMUSG00000052962",
                "ENSMUSG00000026374",
                "ENSMUSG00000024871",
                "ENSMUSG00000053929",
                "ENSMUSG00000096842",
                "ENSMUSG00000020315",
                "ENSMUSG00000026123",
                "ENSMUSG00000059981",
                "ENSMUSG00000059796",
                "ENSMUSG00000005882",
                "ENSMUSG00000020455",
                "ENSMUSG00000025208",
                "ENSMUSG00000060002",
                "ENSMUSG00000026672",
                "ENSMUSG00000029221",
                "ENSMUSG00000047419",
                "ENSMUSG00000025234",
                "ENSMUSG00000006998",
                "ENSMUSG00000025616",
                "ENSMUSG00000020459",
                "ENSMUSG00000079184",
                "ENSMUSG00000001334",
                "ENSMUSG00000001415",
                "ENSMUSG00000058076",
                "ENSMUSG00000007815",
                "ENSMUSG00000028165",
                "ENSMUSG00000018765",
                "ENSMUSG00000073557",
                "ENSMUSG00000066637",
                "ENSMUSG00000033060",
                "ENSMUSG00000025939",
                "ENSMUSG00000005610",
                "ENSMUSG00000021282",
                "ENSMUSG00000031812",
                "ENSMUSG00000064367",
                "ENSMUSG00000021361",
                "ENSMUSG00000024695",
                "ENSMUSG00000028955",
                "ENSMUSG00000037706",
                "ENSMUSG00000026701",
                "ENSMUSG00000037905",
                "ENSMUSG00000015806",
                "ENSMUSG00000025271",
                "ENSMUSG00000022185",
                "ENSMUSG00000084168",
                "ENSMUSG00000014313",
                "ENSMUSG00000016253",
                "ENSMUSG00000035936",
                "ENSMUSG00000029422",
                "ENSMUSG00000027668",
                "ENSMUSG00000002015",
                "ENSMUSG00000026107",
                "ENSMUSG00000024845",
                "ENSMUSG00000008429",
                "ENSMUSG00000064339",
                "ENSMUSG00000060261",
                "ENSMUSG00000028245",
                "ENSMUSG00000035529",
                "ENSMUSG00000042978",
                "ENSMUSG00000066839",
                "ENSMUSG00000028849",
                "ENSMUSG00000034951",
                "ENSMUSG00000015149",
                "ENSMUSG00000022108",
                "ENSMUSG00000040033",
                "ENSMUSG00000024712",
                "ENSMUSG00000037197",
                "ENSMUSG00000006024",
                "ENSMUSG00000022682",
                "ENSMUSG00000025348",
                "ENSMUSG00000053617",
                "ENSMUSG00000030603",
                "ENSMUSG00000052512",
                "ENSMUSG00000019578",
                "ENSMUSG00000020647",
                "ENSMUSG00000067925",
                "ENSMUSG00000019791",
                "ENSMUSG00000027332",
                "ENSMUSG00000015850",
                "ENSMUSG00000025733",
                "ENSMUSG00000020864",
                "ENSMUSG00000022214",
                "ENSMUSG00000022110",
                "ENSMUSG00000025404",
                "ENSMUSG00000004356",
                "ENSMUSG00000037916",
                "ENSMUSG00000030779",
                "ENSMUSG00000029156",
                "ENSMUSG00000078572",
                "ENSMUSG00000040321",
                "ENSMUSG00000052681",
                "ENSMUSG00000045316",
                "ENSMUSG00000000628",
                "ENSMUSG00000040771",
                "ENSMUSG00000021870",
                "ENSMUSG00000032724",
                "ENSMUSG00000033161",
                "ENSMUSG00000056342",
                "ENSMUSG00000062006",
                "ENSMUSG00000003072",
                "ENSMUSG00000037349",
                "ENSMUSG00000031534",
                "ENSMUSG00000027133",
                "ENSMUSG00000027253",
                "ENSMUSG00000008730",
                "ENSMUSG00000009293",
                "ENSMUSG00000027488",
                "ENSMUSG00000019809",
                "ENSMUSG00000000168",
                "ENSMUSG00000026554",
                "ENSMUSG00000029249",
                "ENSMUSG00000028902",
                "ENSMUSG00000097487",
                "ENSMUSG00000030061",
                "ENSMUSG00000038459",
                "ENSMUSG00000021906",
                "ENSMUSG00000021313",
                "ENSMUSG00000026496",
                "ENSMUSG00000025241",
                "ENSMUSG00000015112",
                "ENSMUSG00000075054",
                "ENSMUSG00000021453",
                "ENSMUSG00000032299",
                "ENSMUSG00000028936",
                "ENSMUSG00000062981",
                "ENSMUSG00000028639",
                "ENSMUSG00000037601",
                "ENSMUSG00000002496",
                "ENSMUSG00000064342",
                "ENSMUSG00000027227",
                "ENSMUSG00000074884",
                "ENSMUSG00000031983",
                "ENSMUSG00000038028",
                "ENSMUSG00000003531",
                "ENSMUSG00000003923",
                "ENSMUSG00000020523",
                "ENSMUSG00000028792",
                "ENSMUSG00000001445",
                "ENSMUSG00000035325",
                "ENSMUSG00000055239",
                "ENSMUSG00000035203",
                "ENSMUSG00000029580",
                "ENSMUSG00000028247",
                "ENSMUSG00000027823",
                "ENSMUSG00000008822",
                "ENSMUSG00000029430",
                "ENSMUSG00000004069",
                "ENSMUSG00000040785",
                "ENSMUSG00000021178",
                "ENSMUSG00000022336",
                "ENSMUSG00000027708",
                "ENSMUSG00000019797",
                "ENSMUSG00000058600",
                "ENSMUSG00000064351",
                "ENSMUSG00000040850",
                "ENSMUSG00000046756",
                "ENSMUSG00000029020",
                "ENSMUSG00000028173",
                "ENSMUSG00000027706",
                "ENSMUSG00000049643",
                "ENSMUSG00000038375",
                "ENSMUSG00000022452",
                "ENSMUSG00000039509",
                "ENSMUSG00000023106",
                "ENSMUSG00000036206",
                "ENSMUSG00000034898",
                "ENSMUSG00000024527",
                "ENSMUSG00000031166",
                "ENSMUSG00000038393",
                "ENSMUSG00000075467",
                "ENSMUSG00000025374",
                "ENSMUSG00000019577",
                "ENSMUSG00000033429",
                "ENSMUSG00000027894",
                "ENSMUSG00000002010",
                "ENSMUSG00000022415",
                "ENSMUSG00000020593",
                "ENSMUSG00000010064",
                "ENSMUSG00000058997",
                "ENSMUSG00000059923",
                "ENSMUSG00000032328",
                "ENSMUSG00000060904",
                "ENSMUSG00000025508",
                "ENSMUSG00000028273",
                "ENSMUSG00000019158",
                "ENSMUSG00000008604",
                "ENSMUSG00000058594",
                "ENSMUSG00000021591",
                "ENSMUSG00000032786",
                "ENSMUSG00000022865",
                "ENSMUSG00000050856",
                "ENSMUSG00000025856",
                "ENSMUSG00000038762",
                "ENSMUSG00000098973",
                "ENSMUSG00000053768",
                "ENSMUSG00000025781",
                "ENSMUSG00000020340",
                "ENSMUSG00000033781",
                "ENSMUSG00000018796",
                "ENSMUSG00000008690",
                "ENSMUSG00000032518",
                "ENSMUSG00000021311",
                "ENSMUSG00000022403",
                "ENSMUSG00000059734",
                "ENSMUSG00000022223",
                "ENSMUSG00000022969",
                "ENSMUSG00000091537",
                "ENSMUSG00000029227",
                "ENSMUSG00000078670",
                "ENSMUSG00000062070",
                "ENSMUSG00000047905",
                "ENSMUSG00000039630",
                "ENSMUSG00000024608",
                "ENSMUSG00000038764",
                "ENSMUSG00000026489",
                "ENSMUSG00000045948",
                "ENSMUSG00000022553",
                "ENSMUSG00000009079",
                "ENSMUSG00000007739",
                "ENSMUSG00000027573",
                "ENSMUSG00000028633",
                "ENSMUSG00000032035",
                "ENSMUSG00000028394",
                "ENSMUSG00000057914",
                "ENSMUSG00000025968",
                "ENSMUSG00000002844",
                "ENSMUSG00000064370",
                "ENSMUSG00000020133",
                "ENSMUSG00000030137",
                "ENSMUSG00000020821",
                "ENSMUSG00000018293",
                "ENSMUSG00000044033",
                "ENSMUSG00000024140",
                "ENSMUSG00000028691",
                "ENSMUSG00000022321",
                "ENSMUSG00000061477",
                "ENSMUSG00000037805",
                "ENSMUSG00000024346",
                "ENSMUSG00000017119",
                "ENSMUSG00000031701",
                "ENSMUSG00000020738",
                "ENSMUSG00000029467",
                "ENSMUSG00000039219",
                "ENSMUSG00000022193",
                "ENSMUSG00000011148",
                "ENSMUSG00000062580",
                "ENSMUSG00000026817",
                "ENSMUSG00000035891",
                "ENSMUSG00000022031",
                "ENSMUSG00000040242",
                "ENSMUSG00000078453",
                "ENSMUSG00000020572",
                "ENSMUSG00000021039",
                "ENSMUSG00000021218",
                "ENSMUSG00000062270",
                "ENSMUSG00000062397",
                "ENSMUSG00000025060",
                "ENSMUSG00000028937",
                "ENSMUSG00000015357",
                "ENSMUSG00000034566",
                "ENSMUSG00000024985",
                "ENSMUSG00000021243",
                "ENSMUSG00000013593",
                "ENSMUSG00000068267",
                "ENSMUSG00000021417",
                "ENSMUSG00000062373",
                "ENSMUSG00000024053",
                "ENSMUSG00000032578",
                "ENSMUSG00000004951",
                "ENSMUSG00000031916",
                "ENSMUSG00000071533",
                "ENSMUSG00000029328",
                "ENSMUSG00000029432",
                "ENSMUSG00000038520",
                "ENSMUSG00000038058",
                "ENSMUSG00000024963",
                "ENSMUSG00000058569",
                "ENSMUSG00000036781",
                "ENSMUSG00000006418",
                "ENSMUSG00000074754",
                "ENSMUSG00000040964",
                "ENSMUSG00000020988",
                "ENSMUSG00000024012",
                "ENSMUSG00000020530",
                "ENSMUSG00000039670",
                "ENSMUSG00000018340",
                "ENSMUSG00000022111",
                "ENSMUSG00000046364",
                "ENSMUSG00000037236",
                "ENSMUSG00000023307",
                "ENSMUSG00000038650",
                "ENSMUSG00000055053",
                "ENSMUSG00000051391",
                "ENSMUSG00000015120",
                "ENSMUSG00000058056",
                "ENSMUSG00000064369",
                "ENSMUSG00000026154",
                "ENSMUSG00000022265",
                "ENSMUSG00000028041",
                "ENSMUSG00000047746",
                "ENSMUSG00000025509",
                "ENSMUSG00000027245",
                "ENSMUSG00000083863",
                "ENSMUSG00000036880",
                "ENSMUSG00000053819",
                "ENSMUSG00000022551",
                "ENSMUSG00000037110",
                "ENSMUSG00000034353",
                "ENSMUSG00000020440",
                "ENSMUSG00000024158",
                "ENSMUSG00000042607",
                "ENSMUSG00000023094",
                "ENSMUSG00000021236",
                "ENSMUSG00000030847",
                "ENSMUSG00000008575",
                "ENSMUSG00000022400",
                "ENSMUSG00000035173",
                "ENSMUSG00000004558",
                "ENSMUSG00000037062",
                "ENSMUSG00000067722",
                "ENSMUSG00000090258",
                "ENSMUSG00000019787",
                "ENSMUSG00000026003",
                "ENSMUSG00000067288",
                "ENSMUSG00000021156",
                "ENSMUSG00000040209",
                "ENSMUSG00000038462",
                "ENSMUSG00000022797",
                "ENSMUSG00000029864",
                "ENSMUSG00000032966",
                "ENSMUSG00000034936",
                "ENSMUSG00000082195",
                "ENSMUSG00000043483",
                "ENSMUSG00000074994",
                "ENSMUSG00000084105",
                "ENSMUSG00000024099",
                "ENSMUSG00000028977",
                "ENSMUSG00000037656",
                "ENSMUSG00000028295",
                "ENSMUSG00000096942",
                "ENSMUSG00000032563",
                "ENSMUSG00000028053",
                "ENSMUSG00000014294",
                "ENSMUSG00000034875",
                "ENSMUSG00000038975",
                "ENSMUSG00000021939",
                "ENSMUSG00000028104",
                "ENSMUSG00000090935",
                "ENSMUSG00000021096",
                "ENSMUSG00000027777",
                "ENSMUSG00000022020",
                "ENSMUSG00000040481",
                "ENSMUSG00000002661",
                "ENSMUSG00000040269",
                "ENSMUSG00000046434",
                "ENSMUSG00000018411",
                "ENSMUSG00000003444",
                "ENSMUSG00000015837",
                "ENSMUSG00000029823",
                "ENSMUSG00000031382",
                "ENSMUSG00000091086",
                "ENSMUSG00000028690",
                "ENSMUSG00000002028",
                "ENSMUSG00000071540",
                "ENSMUSG00000097414",
                "ENSMUSG00000081179",
                "ENSMUSG00000034751",
                "ENSMUSG00000022707",
                "ENSMUSG00000029629",
                "ENSMUSG00000027984",
                "ENSMUSG00000041193",
                "ENSMUSG00000074698",
                "ENSMUSG00000031353",
                "ENSMUSG00000031960",
                "ENSMUSG00000024097",
                "ENSMUSG00000044477",
                "ENSMUSG00000022354",
                "ENSMUSG00000023951",
                "ENSMUSG00000028063",
                "ENSMUSG00000032355",
                "ENSMUSG00000005354",
                "ENSMUSG00000020477",
                "ENSMUSG00000026623",
                "ENSMUSG00000035158",
                "ENSMUSG00000029198",
                "ENSMUSG00000030747",
                "ENSMUSG00000020496",
                "ENSMUSG00000023087",
                "ENSMUSG00000057469",
                "ENSMUSG00000005469",
                "ENSMUSG00000097867",
                "ENSMUSG00000026803",
                "ENSMUSG00000005299",
                "ENSMUSG00000052539",
                "ENSMUSG00000029152",
                "ENSMUSG00000040653",
                "ENSMUSG00000022519",
                "ENSMUSG00000051331",
                "ENSMUSG00000025260",
                "ENSMUSG00000016528",
                "ENSMUSG00000024038",
                "ENSMUSG00000025190",
                "ENSMUSG00000033065",
                "ENSMUSG00000057738",
                "ENSMUSG00000042010",
                "ENSMUSG00000030850",
                "ENSMUSG00000085684",
                "ENSMUSG00000014606",
                "ENSMUSG00000026260",
                "ENSMUSG00000055839",
                "ENSMUSG00000020612",
                "ENSMUSG00000039891",
                "ENSMUSG00000048490",
                "ENSMUSG00000031059",
                "ENSMUSG00000024359",
                "ENSMUSG00000062461",
                "ENSMUSG00000039063",
                "ENSMUSG00000032435",
                "ENSMUSG00000038690",
                "ENSMUSG00000044700",
                "ENSMUSG00000028954",
                "ENSMUSG00000021111",
                "ENSMUSG00000039067",
                "ENSMUSG00000003865",
                "ENSMUSG00000024539",
                "ENSMUSG00000020460",
                "ENSMUSG00000031871",
                "ENSMUSG00000071054",
                "ENSMUSG00000041028",
                "ENSMUSG00000028700",
                "ENSMUSG00000021079",
                "ENSMUSG00000026112",
                "ENSMUSG00000021772",
                "ENSMUSG00000002007",
                "ENSMUSG00000018566",
                "ENSMUSG00000020321",
                "ENSMUSG00000025141",
                "ENSMUSG00000059447",
                "ENSMUSG00000024277",
                "ENSMUSG00000024120",
                "ENSMUSG00000029614",
                "ENSMUSG00000053199",
                "ENSMUSG00000026473",
                "ENSMUSG00000044086",
                "ENSMUSG00000001508",
                "ENSMUSG00000047675",
                "ENSMUSG00000013419",
                "ENSMUSG00000031880",
                "ENSMUSG00000038615",
                "ENSMUSG00000029064",
                "ENSMUSG00000045210",
                "ENSMUSG00000079677",
                "ENSMUSG00000025130",
                "ENSMUSG00000030795",
                "ENSMUSG00000023990",
                "ENSMUSG00000015757",
                "ENSMUSG00000025290",
                "ENSMUSG00000021917",
                "ENSMUSG00000027879",
                "ENSMUSG00000033918",
                "ENSMUSG00000028443",
                "ENSMUSG00000022517",
                "ENSMUSG00000016344",
                "ENSMUSG00000024236",
                "ENSMUSG00000039001",
                "ENSMUSG00000063904",
                "ENSMUSG00000061518",
                "ENSMUSG00000033400",
                "ENSMUSG00000036570",
                "ENSMUSG00000001056",
                "ENSMUSG00000025935",
                "ENSMUSG00000003299",
                "ENSMUSG00000040351",
                "ENSMUSG00000050812",
                "ENSMUSG00000027452",
                "ENSMUSG00000025869",
                "ENSMUSG00000007617",
                "ENSMUSG00000021147",
                "ENSMUSG00000024083",
                "ENSMUSG00000051469",
                "ENSMUSG00000019768",
                "ENSMUSG00000029723",
                "ENSMUSG00000036371",
                "ENSMUSG00000070493",
                "ENSMUSG00000030335",
                "ENSMUSG00000021748",
                "ENSMUSG00000036580",
                "ENSMUSG00000071369",
                "ENSMUSG00000030357",
                "ENSMUSG00000038695",
                "ENSMUSG00000026456",
                "ENSMUSG00000028932",
                "ENSMUSG00000027184",
                "ENSMUSG00000029576",
                "ENSMUSG00000068921",
                "ENSMUSG00000001441",
                "ENSMUSG00000059714",
                "ENSMUSG00000030122",
                "ENSMUSG00000034220",
                "ENSMUSG00000016534",
                "ENSMUSG00000051319",
                "ENSMUSG00000031231",
                "ENSMUSG00000071653",
                "ENSMUSG00000026895",
                "ENSMUSG00000026179",
                "ENSMUSG00000033059",
                "ENSMUSG00000002342",
                "ENSMUSG00000033813",
                "ENSMUSG00000000708",
                "ENSMUSG00000000088",
                "ENSMUSG00000032038",
                "ENSMUSG00000038204",
                "ENSMUSG00000042462",
                "ENSMUSG00000022257",
                "ENSMUSG00000025393",
                "ENSMUSG00000048279",
                "ENSMUSG00000029319",
                "ENSMUSG00000026930",
                "ENSMUSG00000031641",
                "ENSMUSG00000038967",
                "ENSMUSG00000025410",
                "ENSMUSG00000036112",
                "ENSMUSG00000058070",
                "ENSMUSG00000024588",
                "ENSMUSG00000006299",
                "ENSMUSG00000028495",
                "ENSMUSG00000002416",
                "ENSMUSG00000008668",
                "ENSMUSG00000003526",
                "ENSMUSG00000025791",
                "ENSMUSG00000078154",
                "ENSMUSG00000066043",
                "ENSMUSG00000021460",
                "ENSMUSG00000022477",
                "ENSMUSG00000022337",
                "ENSMUSG00000036698",
                "ENSMUSG00000042298",
                "ENSMUSG00000052146",
                "ENSMUSG00000012405",
                "ENSMUSG00000015290",
                "ENSMUSG00000074800",
                "ENSMUSG00000030695",
                "ENSMUSG00000064341",
                "ENSMUSG00000028618",
                "ENSMUSG00000040740",
                "ENSMUSG00000013539",
                "ENSMUSG00000001366",
                "ENSMUSG00000025940",
                "ENSMUSG00000010376",
                "ENSMUSG00000054640",
                "ENSMUSG00000035242",
                "ENSMUSG00000043716",
                "ENSMUSG00000039768",
                "ENSMUSG00000028398",
                "ENSMUSG00000050608",
                "ENSMUSG00000021706",
                "ENSMUSG00000004891",
                "ENSMUSG00000036160",
                "ENSMUSG00000027206",
                "ENSMUSG00000019851",
                "ENSMUSG00000059208",
                "ENSMUSG00000022016",
                "ENSMUSG00000026150",
                "ENSMUSG00000038412",
                "ENSMUSG00000020390",
                "ENSMUSG00000036323",
                "ENSMUSG00000039474",
                "ENSMUSG00000038370",
                "ENSMUSG00000044475",
                "ENSMUSG00000020453",
                "ENSMUSG00000029128",
                "ENSMUSG00000029474",
                "ENSMUSG00000099250",
                "ENSMUSG00000013736",
                "ENSMUSG00000082884",
                "ENSMUSG00000055737",
                "ENSMUSG00000063870",
                "ENSMUSG00000020982",
                "ENSMUSG00000032902",
                "ENSMUSG00000031633",
                "ENSMUSG00000063694",
                "ENSMUSG00000068487",
                "ENSMUSG00000020368",
                "ENSMUSG00000039473",
                "ENSMUSG00000092014",
                "ENSMUSG00000019795",
                "ENSMUSG00000040385",
                "ENSMUSG00000038086",
                "ENSMUSG00000016252",
                "ENSMUSG00000041571",
                "ENSMUSG00000084178",
                "ENSMUSG00000082874",
                "ENSMUSG00000022889",
                "ENSMUSG00000019528",
                "ENSMUSG00000021843",
                "ENSMUSG00000039611",
                "ENSMUSG00000067274",
                "ENSMUSG00000063787",
                "ENSMUSG00000063882",
                "ENSMUSG00000028607",
                "ENSMUSG00000039105",
                "ENSMUSG00000026028",
                "ENSMUSG00000026078",
                "ENSMUSG00000039910",
                "ENSMUSG00000021557",
                "ENSMUSG00000028745",
                "ENSMUSG00000038886",
                "ENSMUSG00000053702",
                "ENSMUSG00000020267",
                "ENSMUSG00000022051",
                "ENSMUSG00000006818",
                "ENSMUSG00000026860",
                "ENSMUSG00000076431",
                "ENSMUSG00000042541",
                "ENSMUSG00000094344",
                "ENSMUSG00000069601",
                "ENSMUSG00000028692",
                "ENSMUSG00000031928",
                "ENSMUSG00000021375",
                "ENSMUSG00000060739",
                "ENSMUSG00000033039",
                "ENSMUSG00000006931",
                "ENSMUSG00000025499",
                "ENSMUSG00000032399",
                "ENSMUSG00000059586",
                "ENSMUSG00000078566",
                "ENSMUSG00000002102",
                "ENSMUSG00000021390",
                "ENSMUSG00000002343",
                "ENSMUSG00000039652",
                "ENSMUSG00000024613",
                "ENSMUSG00000027472",
                "ENSMUSG00000054889",
                "ENSMUSG00000070544",
                "ENSMUSG00000046351",
                "ENSMUSG00000020166",
                "ENSMUSG00000024953",
                "ENSMUSG00000027774",
                "ENSMUSG00000022982",
                "ENSMUSG00000058558",
                "ENSMUSG00000026020",
                "ENSMUSG00000050323",
                "ENSMUSG00000026032",
                "ENSMUSG00000068479",
                "ENSMUSG00000021868",
                "ENSMUSG00000037376",
                "ENSMUSG00000028452",
                "ENSMUSG00000028156",
                "ENSMUSG00000032407",
                "ENSMUSG00000039987",
                "ENSMUSG00000041548",
                "ENSMUSG00000044502",
                "ENSMUSG00000070802",
                "ENSMUSG00000024576",
                "ENSMUSG00000035478",
                "ENSMUSG00000028333",
                "ENSMUSG00000041058",
                "ENSMUSG00000036202",
                "ENSMUSG00000032112",
                "ENSMUSG00000027257",
                "ENSMUSG00000020372",
                "ENSMUSG00000032523",
                "ENSMUSG00000028834",
                "ENSMUSG00000031352",
                "ENSMUSG00000073838",
                "ENSMUSG00000041870",
                "ENSMUSG00000039536",
                "ENSMUSG00000000740",
                "ENSMUSG00000040752",
                "ENSMUSG00000023723",
                "ENSMUSG00000003934",
                "ENSMUSG00000030884",
                "ENSMUSG00000025466",
                "ENSMUSG00000018567",
                "ENSMUSG00000051518",
                "ENSMUSG00000022000",
                "ENSMUSG00000023025",
                "ENSMUSG00000021476",
                "ENSMUSG00000032051",
                "ENSMUSG00000034040",
                "ENSMUSG00000023460",
                "ENSMUSG00000002767",
                "ENSMUSG00000034994",
                "ENSMUSG00000020108",
                "ENSMUSG00000030541",
                "ENSMUSG00000031438",
                "ENSMUSG00000026208",
                "ENSMUSG00000031696",
                "ENSMUSG00000090266",
                "ENSMUSG00000036052",
                "ENSMUSG00000015217",
                "ENSMUSG00000024213",
                "ENSMUSG00000030652",
                "ENSMUSG00000089917",
                "ENSMUSG00000027238",
                "ENSMUSG00000004364",
                "ENSMUSG00000047388",
                "ENSMUSG00000021109",
                "ENSMUSG00000074412",
                "ENSMUSG00000061306",
                "ENSMUSG00000024869",
                "ENSMUSG00000017686",
                "ENSMUSG00000033307",
                "ENSMUSG00000057406",
                "ENSMUSG00000046580",
                "ENSMUSG00000069089",
                "ENSMUSG00000033543",
                "ENSMUSG00000020415",
                "ENSMUSG00000024146",
                "ENSMUSG00000006800",
                "ENSMUSG00000002957",
                "ENSMUSG00000041957",
                "ENSMUSG00000024073",
                "ENSMUSG00000028282",
                "ENSMUSG00000062997",
                "ENSMUSG00000081431",
                "ENSMUSG00000020430",
                "ENSMUSG00000009112",
                "ENSMUSG00000031311",
                "ENSMUSG00000026603",
                "ENSMUSG00000027180",
                "ENSMUSG00000016308",
                "ENSMUSG00000027422",
                "ENSMUSG00000050043",
                "ENSMUSG00000018882",
                "ENSMUSG00000028470",
                "ENSMUSG00000000787",
                "ENSMUSG00000015214",
                "ENSMUSG00000037710",
                "ENSMUSG00000024187",
                "ENSMUSG00000019362",
                "ENSMUSG00000016427",
                "ENSMUSG00000038240",
                "ENSMUSG00000027293",
                "ENSMUSG00000003948",
                "ENSMUSG00000019055",
                "ENSMUSG00000003559",
                "ENSMUSG00000001127",
                "ENSMUSG00000015656",
                "ENSMUSG00000025868",
                "ENSMUSG00000040520",
                "ENSMUSG00000058756",
                "ENSMUSG00000025651",
                "ENSMUSG00000028603",
                "ENSMUSG00000071037",
                "ENSMUSG00000037887",
                "ENSMUSG00000028820",
                "ENSMUSG00000004268",
                "ENSMUSG00000025085",
                "ENSMUSG00000035329",
                "ENSMUSG00000065979",
                "ENSMUSG00000028261",
                "ENSMUSG00000024259",
                "ENSMUSG00000025068",
                "ENSMUSG00000045103",
                "ENSMUSG00000022205",
                "ENSMUSG00000034312",
                "ENSMUSG00000032602",
                "ENSMUSG00000032042",
                "ENSMUSG00000061032",
                "ENSMUSG00000023089",
                "ENSMUSG00000019843",
                "ENSMUSG00000020225",
                "ENSMUSG00000073676",
                "ENSMUSG00000032060",
                "ENSMUSG00000023456",
                "ENSMUSG00000022010",
                "ENSMUSG00000027673",
                "ENSMUSG00000083878",
                "ENSMUSG00000059811",
                "ENSMUSG00000013662",
                "ENSMUSG00000026914",
                "ENSMUSG00000083757",
                "ENSMUSG00000024587",
                "ENSMUSG00000024121",
                "ENSMUSG00000029616",
                "ENSMUSG00000020003",
                "ENSMUSG00000019659",
                "ENSMUSG00000061904",
                "ENSMUSG00000027305",
                "ENSMUSG00000029454",
                "ENSMUSG00000083240",
                "ENSMUSG00000004937",
                "ENSMUSG00000015605",
                "ENSMUSG00000020485",
                "ENSMUSG00000006740",
                "ENSMUSG00000024197",
                "ENSMUSG00000007777",
                "ENSMUSG00000019432",
                "ENSMUSG00000022914",
                "ENSMUSG00000063457",
                "ENSMUSG00000008859",
                "ENSMUSG00000020088",
                "ENSMUSG00000039148",
                "ENSMUSG00000058440",
                "ENSMUSG00000019689",
                "ENSMUSG00000020741",
                "ENSMUSG00000037266",
                "ENSMUSG00000038095",
                "ENSMUSG00000037339",
                "ENSMUSG00000030982",
                "ENSMUSG00000027679",
                "ENSMUSG00000041841",
                "ENSMUSG00000026576",
                "ENSMUSG00000041881",
                "ENSMUSG00000021764",
                "ENSMUSG00000038763",
                "ENSMUSG00000043445",
                "ENSMUSG00000039648",
                "ENSMUSG00000020163",
                "ENSMUSG00000001767",
                "ENSMUSG00000009863",
                "ENSMUSG00000090553",
                "ENSMUSG00000031770",
                "ENSMUSG00000038400",
                "ENSMUSG00000030591",
                "ENSMUSG00000036309",
                "ENSMUSG00000031715",
                "ENSMUSG00000028945",
                "ENSMUSG00000056900",
                "ENSMUSG00000032238",
                "ENSMUSG00000062210",
                "ENSMUSG00000060376",
                "ENSMUSG00000014592",
                "ENSMUSG00000090150",
                "ENSMUSG00000017428",
                "ENSMUSG00000079508",
                "ENSMUSG00000081051",
                "ENSMUSG00000062694",
                "ENSMUSG00000020153",
                "ENSMUSG00000093674",
                "ENSMUSG00000083563",
                "ENSMUSG00000022312",
                "ENSMUSG00000040760",
                "ENSMUSG00000069729",
                "ENSMUSG00000036452",
                "ENSMUSG00000031818",
                "ENSMUSG00000031782",
                "ENSMUSG00000048486",
                "ENSMUSG00000055681",
                "ENSMUSG00000032511",
                "ENSMUSG00000038526",
                "ENSMUSG00000024683",
                "ENSMUSG00000096449",
                "ENSMUSG00000044894",
                "ENSMUSG00000007097",
                "ENSMUSG00000029404",
                "ENSMUSG00000024937",
                "ENSMUSG00000056032",
                "ENSMUSG00000022820",
                "ENSMUSG00000037234",
                "ENSMUSG00000042148",
                "ENSMUSG00000034295",
                "ENSMUSG00000031783",
                "ENSMUSG00000022594",
                "ENSMUSG00000063229",
                "ENSMUSG00000001604",
                "ENSMUSG00000005417",
                "ENSMUSG00000021577",
                "ENSMUSG00000038936",
                "ENSMUSG00000029776",
                "ENSMUSG00000029135",
                "ENSMUSG00000025898",
                "ENSMUSG00000036427",
                "ENSMUSG00000070867",
                "ENSMUSG00000038417",
                "ENSMUSG00000045160",
                "ENSMUSG00000024942",
                "ENSMUSG00000022698",
                "ENSMUSG00000024335",
                "ENSMUSG00000027371",
                "ENSMUSG00000028743",
                "ENSMUSG00000037075",
                "ENSMUSG00000040407",
                "ENSMUSG00000030647",
                "ENSMUSG00000073664",
                "ENSMUSG00000025545",
                "ENSMUSG00000045503",
                "ENSMUSG00000091831",
                "ENSMUSG00000026888",
                "ENSMUSG00000022094",
                "ENSMUSG00000052712",
                "ENSMUSG00000030087",
                "ENSMUSG00000031776",
                "ENSMUSG00000006333",
                "ENSMUSG00000026568",
                "ENSMUSG00000028328",
                "ENSMUSG00000068328",
                "ENSMUSG00000005683",
                "ENSMUSG00000029106",
                "ENSMUSG00000031700",
                "ENSMUSG00000046516",
                "ENSMUSG00000022332",
                "ENSMUSG00000040112",
                "ENSMUSG00000037960",
                "ENSMUSG00000058655",
                "ENSMUSG00000022961",
                "ENSMUSG00000020085",
                "ENSMUSG00000031536",
                "ENSMUSG00000032348",
                "ENSMUSG00000039218",
                "ENSMUSG00000021133",
                "ENSMUSG00000029594",
                "ENSMUSG00000023286",
                "ENSMUSG00000031516",
                "ENSMUSG00000052214",
                "ENSMUSG00000033228",
                "ENSMUSG00000046032",
                "ENSMUSG00000032418",
                "ENSMUSG00000032826",
                "ENSMUSG00000028266",
                "ENSMUSG00000081382",
                "ENSMUSG00000026064",
                "ENSMUSG00000047371",
                "ENSMUSG00000029048",
                "ENSMUSG00000022848",
                "ENSMUSG00000020737",
                "ENSMUSG00000031232",
                "ENSMUSG00000001419",
                "ENSMUSG00000051671",
                "ENSMUSG00000022765",
                "ENSMUSG00000026360",
                "ENSMUSG00000047260",
                "ENSMUSG00000004460",
                "ENSMUSG00000042198",
                "ENSMUSG00000035674",
                "ENSMUSG00000009927",
                "ENSMUSG00000083832",
                "ENSMUSG00000026571",
                "ENSMUSG00000025465",
                "ENSMUSG00000042605",
                "ENSMUSG00000028919",
                "ENSMUSG00000040990",
                "ENSMUSG00000027423",
                "ENSMUSG00000001833",
                "ENSMUSG00000025204",
                "ENSMUSG00000023055",
                "ENSMUSG00000098178",
                "ENSMUSG00000031672",
                "ENSMUSG00000075704",
                "ENSMUSG00000031660",
                "ENSMUSG00000040713",
                "ENSMUSG00000026977",
                "ENSMUSG00000030172",
                "ENSMUSG00000059895",
                "ENSMUSG00000016319",
                "ENSMUSG00000030706",
                "ENSMUSG00000024780",
                "ENSMUSG00000021488",
                "ENSMUSG00000002948",
                "ENSMUSG00000015575",
                "ENSMUSG00000034290",
                "ENSMUSG00000005981",
                "ENSMUSG00000023150",
                "ENSMUSG00000022216",
                "ENSMUSG00000063800",
                "ENSMUSG00000030298",
                "ENSMUSG00000025795",
                "ENSMUSG00000036181",
                "ENSMUSG00000028291",
                "ENSMUSG00000017404",
                "ENSMUSG00000020190",
                "ENSMUSG00000021944",
                "ENSMUSG00000028648",
                "ENSMUSG00000026991",
                "ENSMUSG00000029623",
                "ENSMUSG00000018848",
                "ENSMUSG00000004535",
                "ENSMUSG00000014633",
                "ENSMUSG00000007892",
                "ENSMUSG00000003380",
                "ENSMUSG00000074064",
                "ENSMUSG00000060938",
                "ENSMUSG00000010406",
                "ENSMUSG00000030607",
                "ENSMUSG00000025428",
                "ENSMUSG00000024538",
                "ENSMUSG00000021585",
                "ENSMUSG00000029017",
                "ENSMUSG00000030016",
                "ENSMUSG00000041426",
                "ENSMUSG00000031634",
                "ENSMUSG00000020849",
                "ENSMUSG00000068706",
                "ENSMUSG00000002748",
                "ENSMUSG00000041297",
                "ENSMUSG00000026766",
                "ENSMUSG00000024924",
                "ENSMUSG00000042744",
                "ENSMUSG00000031987",
                "ENSMUSG00000040952",
                "ENSMUSG00000043311",
                "ENSMUSG00000038970",
                "ENSMUSG00000028367",
                "ENSMUSG00000055302",
                "ENSMUSG00000028013",
                "ENSMUSG00000000346",
                "ENSMUSG00000039089",
                "ENSMUSG00000019782",
                "ENSMUSG00000026687",
                "ENSMUSG00000053898",
                "ENSMUSG00000060038",
                "ENSMUSG00000030409",
                "ENSMUSG00000062691",
                "ENSMUSG00000083820",
                "ENSMUSG00000001105",
                "ENSMUSG00000025221",
                "ENSMUSG00000029632",
                "ENSMUSG00000026207",
                "ENSMUSG00000036620",
                "ENSMUSG00000022641",
                "ENSMUSG00000054766",
                "ENSMUSG00000015846",
                "ENSMUSG00000052337",
                "ENSMUSG00000028223",
                "ENSMUSG00000031262",
                "ENSMUSG00000026342",
                "ENSMUSG00000025920",
                "ENSMUSG00000074466",
                "ENSMUSG00000001794",
                "ENSMUSG00000031791",
                "ENSMUSG00000048264",
                "ENSMUSG00000037820",
                "ENSMUSG00000071644",
                "ENSMUSG00000090000",
                "ENSMUSG00000026027",
                "ENSMUSG00000020785",
                "ENSMUSG00000030615",
                "ENSMUSG00000053012",
                "ENSMUSG00000020483",
                "ENSMUSG00000009090",
                "ENSMUSG00000004748",
                "ENSMUSG00000072692",
                "ENSMUSG00000032679",
                "ENSMUSG00000026878",
                "ENSMUSG00000024055",
                "ENSMUSG00000029518",
                "ENSMUSG00000006304",
                "ENSMUSG00000031320",
                "ENSMUSG00000032959",
                "ENSMUSG00000028964",
                "ENSMUSG00000025893",
                "ENSMUSG00000042032",
                "ENSMUSG00000060802",
                "ENSMUSG00000002728",
                "ENSMUSG00000027406",
                "ENSMUSG00000060032",
                "ENSMUSG00000025487",
                "ENSMUSG00000078784",
                "ENSMUSG00000028756",
                "ENSMUSG00000063273",
                "ENSMUSG00000060675",
                "ENSMUSG00000051154",
                "ENSMUSG00000020029",
                "ENSMUSG00000037712",
                "ENSMUSG00000034708",
                "ENSMUSG00000036932",
                "ENSMUSG00000003955",
                "ENSMUSG00000028234",
                "ENSMUSG00000061838",
                "ENSMUSG00000019179",
                "ENSMUSG00000051495",
                "ENSMUSG00000079477",
                "ENSMUSG00000022841",
                "ENSMUSG00000000253",
                "ENSMUSG00000071415",
                "ENSMUSG00000037152",
                "ENSMUSG00000019494",
                "ENSMUSG00000020708",
                "ENSMUSG00000023010",
                "ENSMUSG00000036278",
                "ENSMUSG00000022391",
                "ENSMUSG00000028788",
                "ENSMUSG00000062169",
                "ENSMUSG00000053907",
                "ENSMUSG00000067369",
                "ENSMUSG00000003316",
                "ENSMUSG00000022629",
                "ENSMUSG00000002980",
                "ENSMUSG00000023806",
                "ENSMUSG00000005510",
                "ENSMUSG00000020547",
                "ENSMUSG00000024208",
                "ENSMUSG00000020232",
                "ENSMUSG00000024507",
                "ENSMUSG00000049832",
                "ENSMUSG00000029469",
                "ENSMUSG00000087260",
                "ENSMUSG00000038374",
                "ENSMUSG00000019428",
                "ENSMUSG00000024803",
                "ENSMUSG00000022255",
                "ENSMUSG00000025967",
                "ENSMUSG00000005034",
                "ENSMUSG00000031683",
                "ENSMUSG00000036822",
                "ENSMUSG00000022442",
                "ENSMUSG00000028248",
                "ENSMUSG00000078812",
                "ENSMUSG00000082474",
                "ENSMUSG00000001158",
                "ENSMUSG00000090733",
                "ENSMUSG00000036352",
                "ENSMUSG00000059534",
                "ENSMUSG00000038467",
                "ENSMUSG00000027367",
                "ENSMUSG00000029108",
                "ENSMUSG00000007338",
                "ENSMUSG00000080836",
                "ENSMUSG00000044308",
                "ENSMUSG00000053536",
                "ENSMUSG00000022300",
                "ENSMUSG00000003429",
                "ENSMUSG00000021185",
                "ENSMUSG00000018599",
                "ENSMUSG00000037822",
                "ENSMUSG00000084319",
                "ENSMUSG00000064344",
                "ENSMUSG00000029095",
                "ENSMUSG00000015839",
                "ENSMUSG00000027499",
                "ENSMUSG00000027784",
                "ENSMUSG00000065990",
                "ENSMUSG00000060438",
                "ENSMUSG00000003541",
                "ENSMUSG00000029068",
                "ENSMUSG00000064340",
                "ENSMUSG00000032047",
                "ENSMUSG00000042133",
                "ENSMUSG00000044393",
                "ENSMUSG00000014470",
                "ENSMUSG00000031586",
                "ENSMUSG00000001472",
                "ENSMUSG00000048076",
                "ENSMUSG00000036002",
                "ENSMUSG00000072640",
                "ENSMUSG00000038582",
                "ENSMUSG00000026864",
                "ENSMUSG00000023939",
                "ENSMUSG00000035885",
                "ENSMUSG00000059325",
                "ENSMUSG00000031078",
                "ENSMUSG00000026341",
                "ENSMUSG00000038900",
                "ENSMUSG00000032431",
                "ENSMUSG00000032330",
                "ENSMUSG00000022023",
                "ENSMUSG00000038170",
                "ENSMUSG00000006315",
                "ENSMUSG00000078713",
                "ENSMUSG00000051695",
                "ENSMUSG00000049090",
                "ENSMUSG00000041361",
                "ENSMUSG00000049327",
                "ENSMUSG00000006932",
                "ENSMUSG00000091449",
                "ENSMUSG00000022748",
                "ENSMUSG00000060981",
                "ENSMUSG00000031865",
                "ENSMUSG00000030089",
                "ENSMUSG00000000149",
                "ENSMUSG00000023206",
                "ENSMUSG00000028518",
                "ENSMUSG00000074129",
                "ENSMUSG00000031839",
                "ENSMUSG00000001175",
                "ENSMUSG00000038084",
                "ENSMUSG00000008136",
                "ENSMUSG00000049881",
                "ENSMUSG00000005873",
                "ENSMUSG00000054405",
                "ENSMUSG00000030744",
                "ENSMUSG00000029019",
                "ENSMUSG00000023944",
                "ENSMUSG00000015932",
                "ENSMUSG00000029780",
                "ENSMUSG00000020358",
                "ENSMUSG00000022681",
                "ENSMUSG00000068699",
                "ENSMUSG00000003970",
                "ENSMUSG00000030662",
                "ENSMUSG00000072946",
                "ENSMUSG00000010911",
                "ENSMUSG00000050953",
                "ENSMUSG00000027765",
                "ENSMUSG00000020308",
                "ENSMUSG00000021546",
                "ENSMUSG00000064372",
                "ENSMUSG00000030002",
                "ENSMUSG00000032495",
                "ENSMUSG00000028772",
                "ENSMUSG00000025034",
                "ENSMUSG00000025794",
                "ENSMUSG00000046997",
                "ENSMUSG00000020022",
                "ENSMUSG00000034088",
                "ENSMUSG00000071655",
                "ENSMUSG00000001911",
                "ENSMUSG00000034832",
                "ENSMUSG00000020664",
                "ENSMUSG00000022389",
                "ENSMUSG00000075595",
                "ENSMUSG00000033722",
                "ENSMUSG00000040118",
                "ENSMUSG00000029203",
                "ENSMUSG00000022635",
                "ENSMUSG00000067851",
                "ENSMUSG00000018770",
                "ENSMUSG00000052911",
                "ENSMUSG00000039782",
                "ENSMUSG00000097088",
                "ENSMUSG00000035781",
                "ENSMUSG00000087380",
                "ENSMUSG00000067736",
                "ENSMUSG00000028737",
                "ENSMUSG00000031068",
                "ENSMUSG00000031954",
                "ENSMUSG00000030621",
                "ENSMUSG00000056602",
                "ENSMUSG00000062908",
                "ENSMUSG00000022253",
                "ENSMUSG00000021057",
                "ENSMUSG00000068264",
                "ENSMUSG00000025613",
                "ENSMUSG00000022462",
                "ENSMUSG00000027447",
                "ENSMUSG00000021993",
                "ENSMUSG00000031533",
                "ENSMUSG00000029390",
                "ENSMUSG00000000355",
                "ENSMUSG00000020290",
                "ENSMUSG00000037032",
                "ENSMUSG00000024283",
                "ENSMUSG00000036585",
                "ENSMUSG00000029394",
                "ENSMUSG00000030319",
                "ENSMUSG00000059070",
                "ENSMUSG00000062647",
                "ENSMUSG00000025492",
                "ENSMUSG00000062083",
                "ENSMUSG00000099021",
                "ENSMUSG00000042709",
                "ENSMUSG00000033916",
                "ENSMUSG00000032525",
                "ENSMUSG00000020402",
                "ENSMUSG00000060073",
                "ENSMUSG00000022217",
                "ENSMUSG00000026853",
                "ENSMUSG00000034880",
                "ENSMUSG00000054428",
                "ENSMUSG00000066900",
                "ENSMUSG00000028959",
                "ENSMUSG00000008035",
                "ENSMUSG00000026021",
                "ENSMUSG00000019920",
                "ENSMUSG00000014551",
                "ENSMUSG00000021270",
                "ENSMUSG00000011257",
                "ENSMUSG00000021930",
                "ENSMUSG00000031840",
                "ENSMUSG00000039943",
                "ENSMUSG00000006576",
                "ENSMUSG00000038914",
                "ENSMUSG00000064037",
                "ENSMUSG00000004085",
                "ENSMUSG00000056851",
                "ENSMUSG00000028062",
                "ENSMUSG00000014426",
                "ENSMUSG00000020048",
                "ENSMUSG00000063316",
                "ENSMUSG00000022867",
                "ENSMUSG00000028447",
                "ENSMUSG00000048277",
                "ENSMUSG00000020705",
                "ENSMUSG00000038538",
                "ENSMUSG00000084786",
                "ENSMUSG00000020634",
                "ENSMUSG00000036678",
                "ENSMUSG00000030861",
                "ENSMUSG00000028161",
                "ENSMUSG00000018102",
                "ENSMUSG00000029056",
                "ENSMUSG00000025809",
                "ENSMUSG00000026688",
                "ENSMUSG00000015092",
                "ENSMUSG00000029833",
                "ENSMUSG00000034574",
                "ENSMUSG00000040666",
                "ENSMUSG00000025184",
                "ENSMUSG00000003809",
                "ENSMUSG00000010175",
                "ENSMUSG00000000399",
                "ENSMUSG00000004931",
                "ENSMUSG00000044792",
                "ENSMUSG00000039976",
                "ENSMUSG00000022437",
                "ENSMUSG00000051343",
                "ENSMUSG00000019872",
                "ENSMUSG00000031708",
                "ENSMUSG00000026353",
                "ENSMUSG00000014504",
                "ENSMUSG00000022037",
                "ENSMUSG00000022035",
                "ENSMUSG00000033938",
                "ENSMUSG00000024997",
                "ENSMUSG00000057657",
                "ENSMUSG00000017778",
                "ENSMUSG00000020952",
                "ENSMUSG00000048546",
                "ENSMUSG00000035105",
                "ENSMUSG00000030753",
                "ENSMUSG00000073639",
                "ENSMUSG00000045776",
                "ENSMUSG00000020580",
                "ENSMUSG00000020671",
                "ENSMUSG00000028207",
                "ENSMUSG00000040414",
                "ENSMUSG00000001289",
                "ENSMUSG00000021537",
                "ENSMUSG00000060798",
                "ENSMUSG00000036564",
                "ENSMUSG00000074093",
                "ENSMUSG00000037049",
                "ENSMUSG00000020079",
                "ENSMUSG00000054452",
                "ENSMUSG00000002981",
                "ENSMUSG00000024150",
                "ENSMUSG00000020585",
                "ENSMUSG00000040694",
                "ENSMUSG00000012848",
                "ENSMUSG00000032480",
                "ENSMUSG00000022800",
                "ENSMUSG00000000184",
                "ENSMUSG00000091383",
                "ENSMUSG00000010122",
                "ENSMUSG00000050541",
                "ENSMUSG00000038679",
                "ENSMUSG00000031447",
                "ENSMUSG00000020621",
                "ENSMUSG00000008682",
                "ENSMUSG00000034157",
                "ENSMUSG00000020180",
                "ENSMUSG00000031765",
                "ENSMUSG00000023020",
                "ENSMUSG00000024293",
                "ENSMUSG00000045201",
                "ENSMUSG00000052934",
                "ENSMUSG00000030421",
                "ENSMUSG00000033382",
                "ENSMUSG00000034424",
                "ENSMUSG00000071793",
                "ENSMUSG00000031521",
                "ENSMUSG00000064366",
                "ENSMUSG00000019558",
                "ENSMUSG00000031029",
                "ENSMUSG00000031441",
                "ENSMUSG00000029145",
                "ENSMUSG00000006494",
                "ENSMUSG00000021102",
                "ENSMUSG00000023832",
                "ENSMUSG00000005220",
                "ENSMUSG00000004980",
                "ENSMUSG00000005625",
                "ENSMUSG00000032580",
                "ENSMUSG00000071014",
                "ENSMUSG00000044864",
                "ENSMUSG00000049751",
                "ENSMUSG00000025666",
                "ENSMUSG00000021373",
                "ENSMUSG00000031201",
                "ENSMUSG00000000171",
                "ENSMUSG00000000441",
                "ENSMUSG00000041685",
                "ENSMUSG00000034620",
                "ENSMUSG00000038784",
                "ENSMUSG00000024661",
                "ENSMUSG00000036748",
                "ENSMUSG00000053483",
                "ENSMUSG00000025156",
                "ENSMUSG00000002608",
                "ENSMUSG00000025103",
                "ENSMUSG00000039601",
                "ENSMUSG00000057322",
                "ENSMUSG00000073889",
                "ENSMUSG00000022270",
                "ENSMUSG00000026234",
                "ENSMUSG00000029059",
                "ENSMUSG00000035783",
                "ENSMUSG00000045690",
                "ENSMUSG00000032526",
                "ENSMUSG00000038132",
                "ENSMUSG00000000340",
                "ENSMUSG00000037949",
                "ENSMUSG00000055762",
                "ENSMUSG00000034868",
                "ENSMUSG00000064343",
                "ENSMUSG00000021709",
                "ENSMUSG00000043323",
                "ENSMUSG00000022186",
                "ENSMUSG00000021597",
                "ENSMUSG00000056091",
                "ENSMUSG00000045624",
                "ENSMUSG00000086567",
                "ENSMUSG00000064337",
                "ENSMUSG00000020116",
                "ENSMUSG00000071528",
                "ENSMUSG00000031913",
                "ENSMUSG00000071654",
                "ENSMUSG00000030846",
                "ENSMUSG00000021905",
                "ENSMUSG00000031841",
                "ENSMUSG00000061130",
                "ENSMUSG00000036854",
                "ENSMUSG00000026131",
                "ENSMUSG00000026202",
                "ENSMUSG00000020721",
                "ENSMUSG00000027566",
                "ENSMUSG00000049960",
                "ENSMUSG00000048175",
                "ENSMUSG00000020326",
                "ENSMUSG00000031751",
                "ENSMUSG00000032010",
                "ENSMUSG00000041278",
                "ENSMUSG00000021838",
                "ENSMUSG00000062279",
                "ENSMUSG00000031996",
                "ENSMUSG00000067279",
                "ENSMUSG00000064347",
                "ENSMUSG00000023277",
                "ENSMUSG00000034892",
                "ENSMUSG00000017756",
                "ENSMUSG00000023092",
                "ENSMUSG00000059291",
                "ENSMUSG00000032177",
                "ENSMUSG00000003420",
                "ENSMUSG00000002379",
                "ENSMUSG00000027177",
                "ENSMUSG00000039087",
                "ENSMUSG00000029701",
                "ENSMUSG00000052912",
                "ENSMUSG00000029455",
                "ENSMUSG00000046727",
                "ENSMUSG00000064346",
                "ENSMUSG00000058690",
                "ENSMUSG00000027010",
                "ENSMUSG00000049606",
                "ENSMUSG00000030876",
                "ENSMUSG00000060147",
                "ENSMUSG00000059852",
                "ENSMUSG00000051285",
                "ENSMUSG00000024063",
                "ENSMUSG00000037270",
                "ENSMUSG00000040370",
                "ENSMUSG00000074218",
                "ENSMUSG00000021371",
                "ENSMUSG00000042323",
                "ENSMUSG00000030417",
                "ENSMUSG00000026674",
                "ENSMUSG00000034345",
                "ENSMUSG00000037818",
                "ENSMUSG00000037826",
                "ENSMUSG00000029651",
                "ENSMUSG00000023921",
                "ENSMUSG00000032126",
                "ENSMUSG00000057649",
                "ENSMUSG00000021000",
                "ENSMUSG00000042650",
                "ENSMUSG00000000563",
                "ENSMUSG00000065037",
                "ENSMUSG00000037815",
                "ENSMUSG00000027809",
                "ENSMUSG00000047888",
                "ENSMUSG00000034342",
                "ENSMUSG00000038949",
                "ENSMUSG00000047187",
                "ENSMUSG00000061462",
                "ENSMUSG00000037958",
                "ENSMUSG00000031024",
                "ENSMUSG00000020456",
                "ENSMUSG00000021061",
                "ENSMUSG00000045983",
                "ENSMUSG00000032312",
                "ENSMUSG00000050627",
                "ENSMUSG00000051747",
                "ENSMUSG00000025911",
                "ENSMUSG00000035458",
                "ENSMUSG00000032030",
                "ENSMUSG00000032481",
                "ENSMUSG00000022040",
                "ENSMUSG00000028180",
                "ENSMUSG00000026824",
                "ENSMUSG00000022013",
                "ENSMUSG00000081400",
                "ENSMUSG00000081331",
                "ENSMUSG00000011305",
                "ENSMUSG00000014226",
                "ENSMUSG00000006356",
                "ENSMUSG00000039197",
                "ENSMUSG00000022890",
                "ENSMUSG00000053062",
                "ENSMUSG00000019810",
                "ENSMUSG00000024993",
                "ENSMUSG00000022684",
                "ENSMUSG00000032199",
                "ENSMUSG00000001211",
                "ENSMUSG00000056692",
                "ENSMUSG00000070436",
                "ENSMUSG00000052738",
                "ENSMUSG00000031985",
                "ENSMUSG00000029918",
                "ENSMUSG00000029359",
                "ENSMUSG00000023175",
                "ENSMUSG00000052374",
                "ENSMUSG00000044627",
                "ENSMUSG00000015176",
                "ENSMUSG00000024991",
                "ENSMUSG00000037523",
                "ENSMUSG00000024687",
                "ENSMUSG00000022201",
                "ENSMUSG00000027030",
                "ENSMUSG00000080893",
                "ENSMUSG00000021226",
                "ENSMUSG00000022940",
                "ENSMUSG00000030934",
                "ENSMUSG00000032590",
                "ENSMUSG00000026385",
                "ENSMUSG00000041444",
                "ENSMUSG00000020859",
                "ENSMUSG00000024309",
                "ENSMUSG00000050199",
                "ENSMUSG00000025362",
                "ENSMUSG00000058239",
                "ENSMUSG00000071604",
                "ENSMUSG00000022964",
                "ENSMUSG00000082687",
                "ENSMUSG00000029836",
                "ENSMUSG00000025871",
                "ENSMUSG00000025364",
                "ENSMUSG00000038612",
                "ENSMUSG00000042670",
                "ENSMUSG00000021807",
                "ENSMUSG00000022803",
                "ENSMUSG00000064338",
                "ENSMUSG00000061104",
                "ENSMUSG00000000738",
                "ENSMUSG00000039100",
                "ENSMUSG00000009621",
                "ENSMUSG00000044080",
                "ENSMUSG00000046671",
                "ENSMUSG00000024442",
                "ENSMUSG00000004096",
                "ENSMUSG00000028035",
                "ENSMUSG00000047459",
                "ENSMUSG00000029649",
                "ENSMUSG00000022528",
                "ENSMUSG00000060679",
                "ENSMUSG00000038844",
                "ENSMUSG00000052698",
                "ENSMUSG00000005686",
                "ENSMUSG00000048416",
                "ENSMUSG00000053110",
                "ENSMUSG00000010608",
                "ENSMUSG00000027742",
                "ENSMUSG00000032279",
                "ENSMUSG00000033788",
                "ENSMUSG00000086324",
                "ENSMUSG00000023861",
                "ENSMUSG00000067847",
                "ENSMUSG00000026664",
                "ENSMUSG00000068523",
                "ENSMUSG00000043822",
                "ENSMUSG00000021520",
                "ENSMUSG00000006717",
                "ENSMUSG00000001666",
                "ENSMUSG00000090862",
                "ENSMUSG00000036810",
                "ENSMUSG00000008683",
                "ENSMUSG00000003438",
                "ENSMUSG00000042099",
                "ENSMUSG00000083219",
                "ENSMUSG00000005204",
                "ENSMUSG00000097277",
                "ENSMUSG00000028538",
                "ENSMUSG00000007721",
                "ENSMUSG00000020777",
                "ENSMUSG00000034842",
                "ENSMUSG00000041733",
                "ENSMUSG00000062328",
                "ENSMUSG00000029190",
                "ENSMUSG00000039873",
                "ENSMUSG00000004610",
                "ENSMUSG00000015944",
                "ENSMUSG00000021265",
                "ENSMUSG00000021290",
                "ENSMUSG00000037573",
                "ENSMUSG00000049470",
                "ENSMUSG00000069049",
                "ENSMUSG00000046330",
                "ENSMUSG00000029401",
                "ENSMUSG00000074247",
                "ENSMUSG00000003534",
                "ENSMUSG00000025453",
                "ENSMUSG00000025860",
                "ENSMUSG00000030612",
                "ENSMUSG00000004035",
                "ENSMUSG00000074918",
                "ENSMUSG00000024725",
                "ENSMUSG00000021606",
                "ENSMUSG00000019877",
                "ENSMUSG00000033416",
                "ENSMUSG00000023068",
                "ENSMUSG00000021192",
                "ENSMUSG00000036860",
                "ENSMUSG00000079641",
                "ENSMUSG00000075705",
                "ENSMUSG00000045636",
                "ENSMUSG00000020462",
                "ENSMUSG00000062300",
                "ENSMUSG00000063077",
                "ENSMUSG00000022200",
                "ENSMUSG00000031299",
                "ENSMUSG00000053477",
                "ENSMUSG00000021771",
                "ENSMUSG00000032267",
                "ENSMUSG00000035247",
                "ENSMUSG00000040048",
                "ENSMUSG00000028461",
                "ENSMUSG00000064365",
                "ENSMUSG00000024976",
                "ENSMUSG00000043639",
                "ENSMUSG00000043866",
                "ENSMUSG00000071662",
                "ENSMUSG00000028557",
                "ENSMUSG00000023938",
                "ENSMUSG00000027274",
                "ENSMUSG00000036707",
                "ENSMUSG00000032527",
                "ENSMUSG00000035268",
                "ENSMUSG00000085779",
                "ENSMUSG00000009549",
                "ENSMUSG00000068798",
                "ENSMUSG00000020608",
                "ENSMUSG00000022450",
                "ENSMUSG00000015579",
                "ENSMUSG00000073563",
                "ENSMUSG00000024805",
                "ENSMUSG00000078622",
                "ENSMUSG00000082877",
                "ENSMUSG00000032366",
                "ENSMUSG00000032370",
                "ENSMUSG00000098343",
                "ENSMUSG00000028383",
                "ENSMUSG00000041115",
                "ENSMUSG00000025066",
                "ENSMUSG00000017548",
                "ENSMUSG00000020309",
                "ENSMUSG00000064348",
                "ENSMUSG00000094483",
                "ENSMUSG00000027709",
                "ENSMUSG00000024498",
                "ENSMUSG00000022965",
                "ENSMUSG00000022884",
                "ENSMUSG00000015733",
                "ENSMUSG00000027099",
                "ENSMUSG00000025220",
                "ENSMUSG00000035027",
                "ENSMUSG00000034042",
                "ENSMUSG00000052812",
                "ENSMUSG00000037685",
                "ENSMUSG00000024290",
                "ENSMUSG00000026526",
                "ENSMUSG00000022951",
                "ENSMUSG00000024132",
                "ENSMUSG00000029338",
                "ENSMUSG00000021621",
                "ENSMUSG00000006005",
                "ENSMUSG00000034525",
                "ENSMUSG00000046312",
                "ENSMUSG00000081929",
                "ENSMUSG00000073002",
                "ENSMUSG00000055204",
                "ENSMUSG00000042115",
                "ENSMUSG00000023904",
                "ENSMUSG00000054312",
                "ENSMUSG00000033295",
                "ENSMUSG00000060803",
                "ENSMUSG00000030246",
                "ENSMUSG00000061474",
                "ENSMUSG00000096232",
                "ENSMUSG00000010914",
                "ENSMUSG00000042699",
                "ENSMUSG00000017760",
                "ENSMUSG00000055128",
                "ENSMUSG00000030869",
                "ENSMUSG00000031767",
                "ENSMUSG00000064371",
                "ENSMUSG00000084314",
                "ENSMUSG00000027620",
                "ENSMUSG00000017817",
                "ENSMUSG00000018395",
                "ENSMUSG00000002908",
                "ENSMUSG00000031229",
                "ENSMUSG00000074656",
                "ENSMUSG00000039126",
                "ENSMUSG00000045639",
                "ENSMUSG00000028847",
                "ENSMUSG00000033047",
                "ENSMUSG00000024913"
            ],
            "expression_sum": {
                "class1": 1988.0,
                "class2": 50963.0
            },
            "uniprot_ids": [
                "O89090",
                "E9Q0S7",
                "E9Q113",
                "E9Q2V3",
                "G3X8Q0"
            ],
            "net_id": 0,
            "source_num": 23,
            "target_num": 589
        },
        "ENSMUSG00000019768": {
            "id": "ENSMUSG00000019768",
            "type": "TF",
            "symbol": "Esr1",
            "source": [
                "ENSMUSG00000021944",
                "ENSMUSG00000010175",
                "ENSMUSG00000056501",
                "ENSMUSG00000030557",
                "ENSMUSG00000029249",
                "ENSMUSG00000001280",
                "ENSMUSG00000005566",
                "ENSMUSG00000028565",
                "ENSMUSG00000005698",
                "ENSMUSG00000015605"
            ],
            "target": [
                "ENSMUSG00000000538",
                "ENSMUSG00000054455",
                "ENSMUSG00000001741",
                "ENSMUSG00000031403",
                "ENSMUSG00000064363",
                "ENSMUSG00000091478",
                "ENSMUSG00000024228",
                "ENSMUSG00000020694",
                "ENSMUSG00000030314",
                "ENSMUSG00000020307",
                "ENSMUSG00000043683",
                "ENSMUSG00000021248",
                "ENSMUSG00000031885",
                "ENSMUSG00000021235",
                "ENSMUSG00000070283",
                "ENSMUSG00000039640",
                "ENSMUSG00000042302",
                "ENSMUSG00000081792",
                "ENSMUSG00000034903",
                "ENSMUSG00000030739",
                "ENSMUSG00000014402",
                "ENSMUSG00000004044",
                "ENSMUSG00000081773",
                "ENSMUSG00000028086",
                "ENSMUSG00000044221",
                "ENSMUSG00000035202",
                "ENSMUSG00000026087",
                "ENSMUSG00000026626",
                "ENSMUSG00000031543",
                "ENSMUSG00000007891",
                "ENSMUSG00000004789",
                "ENSMUSG00000080746",
                "ENSMUSG00000018286",
                "ENSMUSG00000064349",
                "ENSMUSG00000080242",
                "ENSMUSG00000020743",
                "ENSMUSG00000007656",
                "ENSMUSG00000027288",
                "ENSMUSG00000041459",
                "ENSMUSG00000032314",
                "ENSMUSG00000008892",
                "ENSMUSG00000030557",
                "ENSMUSG00000036097",
                "ENSMUSG00000026858",
                "ENSMUSG00000027698",
                "ENSMUSG00000028229",
                "ENSMUSG00000030852",
                "ENSMUSG00000010048",
                "ENSMUSG00000053329",
                "ENSMUSG00000081485",
                "ENSMUSG00000031197",
                "ENSMUSG00000034981",
                "ENSMUSG00000032216",
                "ENSMUSG00000071796",
                "ENSMUSG00000090066",
                "ENSMUSG00000026755",
                "ENSMUSG00000018446",
                "ENSMUSG00000064302",
                "ENSMUSG00000029158",
                "ENSMUSG00000026775",
                "ENSMUSG00000029392",
                "ENSMUSG00000044365",
                "ENSMUSG00000000056",
                "ENSMUSG00000024962",
                "ENSMUSG00000010392",
                "ENSMUSG00000057133",
                "ENSMUSG00000020949",
                "ENSMUSG00000039103",
                "ENSMUSG00000029148",
                "ENSMUSG00000026158",
                "ENSMUSG00000020349",
                "ENSMUSG00000038708",
                "ENSMUSG00000039515",
                "ENSMUSG00000031728",
                "ENSMUSG00000056501",
                "ENSMUSG00000039741",
                "ENSMUSG00000026615",
                "ENSMUSG00000055320",
                "ENSMUSG00000031698",
                "ENSMUSG00000021024",
                "ENSMUSG00000027602",
                "ENSMUSG00000029167",
                "ENSMUSG00000061286",
                "ENSMUSG00000025745",
                "ENSMUSG00000026414",
                "ENSMUSG00000049265",
                "ENSMUSG00000082043",
                "ENSMUSG00000024304",
                "ENSMUSG00000006219",
                "ENSMUSG00000030057",
                "ENSMUSG00000094249",
                "ENSMUSG00000025369",
                "ENSMUSG00000041130",
                "ENSMUSG00000055322",
                "ENSMUSG00000006057",
                "ENSMUSG00000068923",
                "ENSMUSG00000047284",
                "ENSMUSG00000032536",
                "ENSMUSG00000044533",
                "ENSMUSG00000064345",
                "ENSMUSG00000019471",
                "ENSMUSG00000028884",
                "ENSMUSG00000032648",
                "ENSMUSG00000026374",
                "ENSMUSG00000024871",
                "ENSMUSG00000053929",
                "ENSMUSG00000096842",
                "ENSMUSG00000026123",
                "ENSMUSG00000034673",
                "ENSMUSG00000024177",
                "ENSMUSG00000059981",
                "ENSMUSG00000097652",
                "ENSMUSG00000005882",
                "ENSMUSG00000020455",
                "ENSMUSG00000025208",
                "ENSMUSG00000060002",
                "ENSMUSG00000029221",
                "ENSMUSG00000047419",
                "ENSMUSG00000025234",
                "ENSMUSG00000006998",
                "ENSMUSG00000025616",
                "ENSMUSG00000020459",
                "ENSMUSG00000001334",
                "ENSMUSG00000001415",
                "ENSMUSG00000058076",
                "ENSMUSG00000028165",
                "ENSMUSG00000018765",
                "ENSMUSG00000015843",
                "ENSMUSG00000073557",
                "ENSMUSG00000066637",
                "ENSMUSG00000060882",
                "ENSMUSG00000033060",
                "ENSMUSG00000005610",
                "ENSMUSG00000021282",
                "ENSMUSG00000031812",
                "ENSMUSG00000034520",
                "ENSMUSG00000064367",
                "ENSMUSG00000024695",
                "ENSMUSG00000068206",
                "ENSMUSG00000028955",
                "ENSMUSG00000037706",
                "ENSMUSG00000026701",
                "ENSMUSG00000037905",
                "ENSMUSG00000015806",
                "ENSMUSG00000025271",
                "ENSMUSG00000022185",
                "ENSMUSG00000014313",
                "ENSMUSG00000027668",
                "ENSMUSG00000002015",
                "ENSMUSG00000024750",
                "ENSMUSG00000024845",
                "ENSMUSG00000008429",
                "ENSMUSG00000064339",
                "ENSMUSG00000024298",
                "ENSMUSG00000053093",
                "ENSMUSG00000042215",
                "ENSMUSG00000002500",
                "ENSMUSG00000028849",
                "ENSMUSG00000034951",
                "ENSMUSG00000015149",
                "ENSMUSG00000040033",
                "ENSMUSG00000024940",
                "ENSMUSG00000037197",
                "ENSMUSG00000006024",
                "ENSMUSG00000022682",
                "ENSMUSG00000027552",
                "ENSMUSG00000025348",
                "ENSMUSG00000053617",
                "ENSMUSG00000052512",
                "ENSMUSG00000020647",
                "ENSMUSG00000040506",
                "ENSMUSG00000067925",
                "ENSMUSG00000019791",
                "ENSMUSG00000027332",
                "ENSMUSG00000015850",
                "ENSMUSG00000025733",
                "ENSMUSG00000020864",
                "ENSMUSG00000022214",
                "ENSMUSG00000022110",
                "ENSMUSG00000004356",
                "ENSMUSG00000037916",
                "ENSMUSG00000082641",
                "ENSMUSG00000030779",
                "ENSMUSG00000029708",
                "ENSMUSG00000029156",
                "ENSMUSG00000078572",
                "ENSMUSG00000040321",
                "ENSMUSG00000025437",
                "ENSMUSG00000045316",
                "ENSMUSG00000000628",
                "ENSMUSG00000027805",
                "ENSMUSG00000028565",
                "ENSMUSG00000040771",
                "ENSMUSG00000021870",
                "ENSMUSG00000032724",
                "ENSMUSG00000033161",
                "ENSMUSG00000003072",
                "ENSMUSG00000037349",
                "ENSMUSG00000033943",
                "ENSMUSG00000008730",
                "ENSMUSG00000000168",
                "ENSMUSG00000048661",
                "ENSMUSG00000029249",
                "ENSMUSG00000026113",
                "ENSMUSG00000027861",
                "ENSMUSG00000030061",
                "ENSMUSG00000021906",
                "ENSMUSG00000081278",
                "ENSMUSG00000021313",
                "ENSMUSG00000025241",
                "ENSMUSG00000059430",
                "ENSMUSG00000015112",
                "ENSMUSG00000075054",
                "ENSMUSG00000021453",
                "ENSMUSG00000018433",
                "ENSMUSG00000062981",
                "ENSMUSG00000030754",
                "ENSMUSG00000002496",
                "ENSMUSG00000064342",
                "ENSMUSG00000002233",
                "ENSMUSG00000074884",
                "ENSMUSG00000031983",
                "ENSMUSG00000038028",
                "ENSMUSG00000003531",
                "ENSMUSG00000030835",
                "ENSMUSG00000020523",
                "ENSMUSG00000030470",
                "ENSMUSG00000035325",
                "ENSMUSG00000055239",
                "ENSMUSG00000027381",
                "ENSMUSG00000035203",
                "ENSMUSG00000028247",
                "ENSMUSG00000027823",
                "ENSMUSG00000029094",
                "ENSMUSG00000008822",
                "ENSMUSG00000002228",
                "ENSMUSG00000045886",
                "ENSMUSG00000004069",
                "ENSMUSG00000040785",
                "ENSMUSG00000022336",
                "ENSMUSG00000019797",
                "ENSMUSG00000058600",
                "ENSMUSG00000064351",
                "ENSMUSG00000040850",
                "ENSMUSG00000046756",
                "ENSMUSG00000029020",
                "ENSMUSG00000028173",
                "ENSMUSG00000027706",
                "ENSMUSG00000022131",
                "ENSMUSG00000049643",
                "ENSMUSG00000042529",
                "ENSMUSG00000038375",
                "ENSMUSG00000022452",
                "ENSMUSG00000039509",
                "ENSMUSG00000060600",
                "ENSMUSG00000026283",
                "ENSMUSG00000034898",
                "ENSMUSG00000022228",
                "ENSMUSG00000038393",
                "ENSMUSG00000024143",
                "ENSMUSG00000009647",
                "ENSMUSG00000025374",
                "ENSMUSG00000019577",
                "ENSMUSG00000033429",
                "ENSMUSG00000002010",
                "ENSMUSG00000020593",
                "ENSMUSG00000010064",
                "ENSMUSG00000058997",
                "ENSMUSG00000060904",
                "ENSMUSG00000028273",
                "ENSMUSG00000019158",
                "ENSMUSG00000021200",
                "ENSMUSG00000008604",
                "ENSMUSG00000058594",
                "ENSMUSG00000021591",
                "ENSMUSG00000048732",
                "ENSMUSG00000032786",
                "ENSMUSG00000050856",
                "ENSMUSG00000025856",
                "ENSMUSG00000038762",
                "ENSMUSG00000053768",
                "ENSMUSG00000082978",
                "ENSMUSG00000025781",
                "ENSMUSG00000033781",
                "ENSMUSG00000018796",
                "ENSMUSG00000008690",
                "ENSMUSG00000032518",
                "ENSMUSG00000014195",
                "ENSMUSG00000031461",
                "ENSMUSG00000049422",
                "ENSMUSG00000087679",
                "ENSMUSG00000022403",
                "ENSMUSG00000026409",
                "ENSMUSG00000059734",
                "ENSMUSG00000022223",
                "ENSMUSG00000022969",
                "ENSMUSG00000095687",
                "ENSMUSG00000091537",
                "ENSMUSG00000078670",
                "ENSMUSG00000062070",
                "ENSMUSG00000059878",
                "ENSMUSG00000047905",
                "ENSMUSG00000053964",
                "ENSMUSG00000037904",
                "ENSMUSG00000038764",
                "ENSMUSG00000019173",
                "ENSMUSG00000022553",
                "ENSMUSG00000091898",
                "ENSMUSG00000009079",
                "ENSMUSG00000007739",
                "ENSMUSG00000028633",
                "ENSMUSG00000028394",
                "ENSMUSG00000025968",
                "ENSMUSG00000002844",
                "ENSMUSG00000064370",
                "ENSMUSG00000020133",
                "ENSMUSG00000020821",
                "ENSMUSG00000097626",
                "ENSMUSG00000044033",
                "ENSMUSG00000024140",
                "ENSMUSG00000028691",
                "ENSMUSG00000028399",
                "ENSMUSG00000061477",
                "ENSMUSG00000024346",
                "ENSMUSG00000037805",
                "ENSMUSG00000046201",
                "ENSMUSG00000017119",
                "ENSMUSG00000028911",
                "ENSMUSG00000031701",
                "ENSMUSG00000020738",
                "ENSMUSG00000075701",
                "ENSMUSG00000029467",
                "ENSMUSG00000022545",
                "ENSMUSG00000038022",
                "ENSMUSG00000044751",
                "ENSMUSG00000022193",
                "ENSMUSG00000011148",
                "ENSMUSG00000062580",
                "ENSMUSG00000026817",
                "ENSMUSG00000022031",
                "ENSMUSG00000026187",
                "ENSMUSG00000021039",
                "ENSMUSG00000021218",
                "ENSMUSG00000062270",
                "ENSMUSG00000062397",
                "ENSMUSG00000028937",
                "ENSMUSG00000015357",
                "ENSMUSG00000034566",
                "ENSMUSG00000024985",
                "ENSMUSG00000021243",
                "ENSMUSG00000013593",
                "ENSMUSG00000068267",
                "ENSMUSG00000021417",
                "ENSMUSG00000042293",
                "ENSMUSG00000062373",
                "ENSMUSG00000029432",
                "ENSMUSG00000038520",
                "ENSMUSG00000038058",
                "ENSMUSG00000024963",
                "ENSMUSG00000020034",
                "ENSMUSG00000058569",
                "ENSMUSG00000022215",
                "ENSMUSG00000009545",
                "ENSMUSG00000040964",
                "ENSMUSG00000020988",
                "ENSMUSG00000020530",
                "ENSMUSG00000039670",
                "ENSMUSG00000032897",
                "ENSMUSG00000018340",
                "ENSMUSG00000022111",
                "ENSMUSG00000037236",
                "ENSMUSG00000038650",
                "ENSMUSG00000055053",
                "ENSMUSG00000051391",
                "ENSMUSG00000058056",
                "ENSMUSG00000064369",
                "ENSMUSG00000047746",
                "ENSMUSG00000025509",
                "ENSMUSG00000029655",
                "ENSMUSG00000036880",
                "ENSMUSG00000053819",
                "ENSMUSG00000022551",
                "ENSMUSG00000037110",
                "ENSMUSG00000029571",
                "ENSMUSG00000005779",
                "ENSMUSG00000082768",
                "ENSMUSG00000020440",
                "ENSMUSG00000023094",
                "ENSMUSG00000002455",
                "ENSMUSG00000030847",
                "ENSMUSG00000008575",
                "ENSMUSG00000022400",
                "ENSMUSG00000035173",
                "ENSMUSG00000004558",
                "ENSMUSG00000067722",
                "ENSMUSG00000090258",
                "ENSMUSG00000019787",
                "ENSMUSG00000026003",
                "ENSMUSG00000040209",
                "ENSMUSG00000038462",
                "ENSMUSG00000022797",
                "ENSMUSG00000067121",
                "ENSMUSG00000034936",
                "ENSMUSG00000082195",
                "ENSMUSG00000074994",
                "ENSMUSG00000026610",
                "ENSMUSG00000024099",
                "ENSMUSG00000061311",
                "ENSMUSG00000037656",
                "ENSMUSG00000028295",
                "ENSMUSG00000045694",
                "ENSMUSG00000032563",
                "ENSMUSG00000028053",
                "ENSMUSG00000014294",
                "ENSMUSG00000034875",
                "ENSMUSG00000021939",
                "ENSMUSG00000028104",
                "ENSMUSG00000045838",
                "ENSMUSG00000047547",
                "ENSMUSG00000090935",
                "ENSMUSG00000027777",
                "ENSMUSG00000022020",
                "ENSMUSG00000040481",
                "ENSMUSG00000045302",
                "ENSMUSG00000002661",
                "ENSMUSG00000018501",
                "ENSMUSG00000040269",
                "ENSMUSG00000003444",
                "ENSMUSG00000044938",
                "ENSMUSG00000015837",
                "ENSMUSG00000029823",
                "ENSMUSG00000031382",
                "ENSMUSG00000002028",
                "ENSMUSG00000097414",
                "ENSMUSG00000081179",
                "ENSMUSG00000034751",
                "ENSMUSG00000022707",
                "ENSMUSG00000029629",
                "ENSMUSG00000027984",
                "ENSMUSG00000031353",
                "ENSMUSG00000024097",
                "ENSMUSG00000022354",
                "ENSMUSG00000023951",
                "ENSMUSG00000005354",
                "ENSMUSG00000020477",
                "ENSMUSG00000026623",
                "ENSMUSG00000024811",
                "ENSMUSG00000035158",
                "ENSMUSG00000080848",
                "ENSMUSG00000029198",
                "ENSMUSG00000030747",
                "ENSMUSG00000020496",
                "ENSMUSG00000023087",
                "ENSMUSG00000057469",
                "ENSMUSG00000004347",
                "ENSMUSG00000005469",
                "ENSMUSG00000026803",
                "ENSMUSG00000005299",
                "ENSMUSG00000052539",
                "ENSMUSG00000073684",
                "ENSMUSG00000029152",
                "ENSMUSG00000051048",
                "ENSMUSG00000022519",
                "ENSMUSG00000051331",
                "ENSMUSG00000025260",
                "ENSMUSG00000016528",
                "ENSMUSG00000024038",
                "ENSMUSG00000025190",
                "ENSMUSG00000033065",
                "ENSMUSG00000057738",
                "ENSMUSG00000042010",
                "ENSMUSG00000083186",
                "ENSMUSG00000030850",
                "ENSMUSG00000014606",
                "ENSMUSG00000036893",
                "ENSMUSG00000026260",
                "ENSMUSG00000055839",
                "ENSMUSG00000041921",
                "ENSMUSG00000020612",
                "ENSMUSG00000039891",
                "ENSMUSG00000048490",
                "ENSMUSG00000031059",
                "ENSMUSG00000024359",
                "ENSMUSG00000019943",
                "ENSMUSG00000039063",
                "ENSMUSG00000032883",
                "ENSMUSG00000032263",
                "ENSMUSG00000032435",
                "ENSMUSG00000038690",
                "ENSMUSG00000028954",
                "ENSMUSG00000039067",
                "ENSMUSG00000003865",
                "ENSMUSG00000014956",
                "ENSMUSG00000024539",
                "ENSMUSG00000020460",
                "ENSMUSG00000023004",
                "ENSMUSG00000083932",
                "ENSMUSG00000041028",
                "ENSMUSG00000028700",
                "ENSMUSG00000026112",
                "ENSMUSG00000021772",
                "ENSMUSG00000018566",
                "ENSMUSG00000020321",
                "ENSMUSG00000059447",
                "ENSMUSG00000024277",
                "ENSMUSG00000024120",
                "ENSMUSG00000029614",
                "ENSMUSG00000035296",
                "ENSMUSG00000053199",
                "ENSMUSG00000001508",
                "ENSMUSG00000036902",
                "ENSMUSG00000013419",
                "ENSMUSG00000031880",
                "ENSMUSG00000038615",
                "ENSMUSG00000029064",
                "ENSMUSG00000045210",
                "ENSMUSG00000025130",
                "ENSMUSG00000023990",
                "ENSMUSG00000025290",
                "ENSMUSG00000021917",
                "ENSMUSG00000009563",
                "ENSMUSG00000033918",
                "ENSMUSG00000028443",
                "ENSMUSG00000024236",
                "ENSMUSG00000063904",
                "ENSMUSG00000061518",
                "ENSMUSG00000034252",
                "ENSMUSG00000033400",
                "ENSMUSG00000036570",
                "ENSMUSG00000001056",
                "ENSMUSG00000020220",
                "ENSMUSG00000025935",
                "ENSMUSG00000003299",
                "ENSMUSG00000039911",
                "ENSMUSG00000040351",
                "ENSMUSG00000021549",
                "ENSMUSG00000051223",
                "ENSMUSG00000027091",
                "ENSMUSG00000050812",
                "ENSMUSG00000027452",
                "ENSMUSG00000038827",
                "ENSMUSG00000025869",
                "ENSMUSG00000051469",
                "ENSMUSG00000029723",
                "ENSMUSG00000036371",
                "ENSMUSG00000058135",
                "ENSMUSG00000070493",
                "ENSMUSG00000030335",
                "ENSMUSG00000021748",
                "ENSMUSG00000071369",
                "ENSMUSG00000025006",
                "ENSMUSG00000038695",
                "ENSMUSG00000026456",
                "ENSMUSG00000027184",
                "ENSMUSG00000028116",
                "ENSMUSG00000068921",
                "ENSMUSG00000001441",
                "ENSMUSG00000059714",
                "ENSMUSG00000034220",
                "ENSMUSG00000070713",
                "ENSMUSG00000031231",
                "ENSMUSG00000084111",
                "ENSMUSG00000026895",
                "ENSMUSG00000092341",
                "ENSMUSG00000026179",
                "ENSMUSG00000033059",
                "ENSMUSG00000002342",
                "ENSMUSG00000000088",
                "ENSMUSG00000031626",
                "ENSMUSG00000048154",
                "ENSMUSG00000042462",
                "ENSMUSG00000026393",
                "ENSMUSG00000009995",
                "ENSMUSG00000025393",
                "ENSMUSG00000048279",
                "ENSMUSG00000026930",
                "ENSMUSG00000026229",
                "ENSMUSG00000031641",
                "ENSMUSG00000001435",
                "ENSMUSG00000038967",
                "ENSMUSG00000025410",
                "ENSMUSG00000036112",
                "ENSMUSG00000017485",
                "ENSMUSG00000024588",
                "ENSMUSG00000023249",
                "ENSMUSG00000068874",
                "ENSMUSG00000006299",
                "ENSMUSG00000032340",
                "ENSMUSG00000002416",
                "ENSMUSG00000036197",
                "ENSMUSG00000025791",
                "ENSMUSG00000066043",
                "ENSMUSG00000022404",
                "ENSMUSG00000067058",
                "ENSMUSG00000021460",
                "ENSMUSG00000036867",
                "ENSMUSG00000022477",
                "ENSMUSG00000022337",
                "ENSMUSG00000036698",
                "ENSMUSG00000019988",
                "ENSMUSG00000041144",
                "ENSMUSG00000042298",
                "ENSMUSG00000028649",
                "ENSMUSG00000052146",
                "ENSMUSG00000012405",
                "ENSMUSG00000030695",
                "ENSMUSG00000064341",
                "ENSMUSG00000024049",
                "ENSMUSG00000013539",
                "ENSMUSG00000001366",
                "ENSMUSG00000035242",
                "ENSMUSG00000043716",
                "ENSMUSG00000050608",
                "ENSMUSG00000021706",
                "ENSMUSG00000036160",
                "ENSMUSG00000027206",
                "ENSMUSG00000074182",
                "ENSMUSG00000019851",
                "ENSMUSG00000059208",
                "ENSMUSG00000022016",
                "ENSMUSG00000022503",
                "ENSMUSG00000026150",
                "ENSMUSG00000038412",
                "ENSMUSG00000020230",
                "ENSMUSG00000020390",
                "ENSMUSG00000021687",
                "ENSMUSG00000036323",
                "ENSMUSG00000039474",
                "ENSMUSG00000038370",
                "ENSMUSG00000044475",
                "ENSMUSG00000029128",
                "ENSMUSG00000099250",
                "ENSMUSG00000013736",
                "ENSMUSG00000055737",
                "ENSMUSG00000062604",
                "ENSMUSG00000071748",
                "ENSMUSG00000025607",
                "ENSMUSG00000063870",
                "ENSMUSG00000032902",
                "ENSMUSG00000031633",
                "ENSMUSG00000063694",
                "ENSMUSG00000041235",
                "ENSMUSG00000057580",
                "ENSMUSG00000020368",
                "ENSMUSG00000039473",
                "ENSMUSG00000034610",
                "ENSMUSG00000042772",
                "ENSMUSG00000019795",
                "ENSMUSG00000040385",
                "ENSMUSG00000016252",
                "ENSMUSG00000030793",
                "ENSMUSG00000026586",
                "ENSMUSG00000035776",
                "ENSMUSG00000022889",
                "ENSMUSG00000021281",
                "ENSMUSG00000021843",
                "ENSMUSG00000039611",
                "ENSMUSG00000038239",
                "ENSMUSG00000067274",
                "ENSMUSG00000063787",
                "ENSMUSG00000063882",
                "ENSMUSG00000019773",
                "ENSMUSG00000040928",
                "ENSMUSG00000031010",
                "ENSMUSG00000026028",
                "ENSMUSG00000039910",
                "ENSMUSG00000028745",
                "ENSMUSG00000038886",
                "ENSMUSG00000035152",
                "ENSMUSG00000053702",
                "ENSMUSG00000020267",
                "ENSMUSG00000036550",
                "ENSMUSG00000022051",
                "ENSMUSG00000006818",
                "ENSMUSG00000026860",
                "ENSMUSG00000034621",
                "ENSMUSG00000001017",
                "ENSMUSG00000042541",
                "ENSMUSG00000021375",
                "ENSMUSG00000029545",
                "ENSMUSG00000058793",
                "ENSMUSG00000033039",
                "ENSMUSG00000032519",
                "ENSMUSG00000056608",
                "ENSMUSG00000032399",
                "ENSMUSG00000059586",
                "ENSMUSG00000078566",
                "ENSMUSG00000002102",
                "ENSMUSG00000002343",
                "ENSMUSG00000039652",
                "ENSMUSG00000024613",
                "ENSMUSG00000070544",
                "ENSMUSG00000054889",
                "ENSMUSG00000046351",
                "ENSMUSG00000053838",
                "ENSMUSG00000024953",
                "ENSMUSG00000083793",
                "ENSMUSG00000022982",
                "ENSMUSG00000050323",
                "ENSMUSG00000026032",
                "ENSMUSG00000068479",
                "ENSMUSG00000021868",
                "ENSMUSG00000028312",
                "ENSMUSG00000037376",
                "ENSMUSG00000028452",
                "ENSMUSG00000028156",
                "ENSMUSG00000064360",
                "ENSMUSG00000021144",
                "ENSMUSG00000039987",
                "ENSMUSG00000041548",
                "ENSMUSG00000070802",
                "ENSMUSG00000035478",
                "ENSMUSG00000028333",
                "ENSMUSG00000041058",
                "ENSMUSG00000036202",
                "ENSMUSG00000027257",
                "ENSMUSG00000020372",
                "ENSMUSG00000044573",
                "ENSMUSG00000032523",
                "ENSMUSG00000028834",
                "ENSMUSG00000031352",
                "ENSMUSG00000073838",
                "ENSMUSG00000041870",
                "ENSMUSG00000005102",
                "ENSMUSG00000039536",
                "ENSMUSG00000000740",
                "ENSMUSG00000040752",
                "ENSMUSG00000023723",
                "ENSMUSG00000003934",
                "ENSMUSG00000030884",
                "ENSMUSG00000025466",
                "ENSMUSG00000042997",
                "ENSMUSG00000092274",
                "ENSMUSG00000022000",
                "ENSMUSG00000023025",
                "ENSMUSG00000028419",
                "ENSMUSG00000032051",
                "ENSMUSG00000034040",
                "ENSMUSG00000038143",
                "ENSMUSG00000023460",
                "ENSMUSG00000002767",
                "ENSMUSG00000034994",
                "ENSMUSG00000023971",
                "ENSMUSG00000030541",
                "ENSMUSG00000070003",
                "ENSMUSG00000031438",
                "ENSMUSG00000081100",
                "ENSMUSG00000026208",
                "ENSMUSG00000036052",
                "ENSMUSG00000015217",
                "ENSMUSG00000024213",
                "ENSMUSG00000030652",
                "ENSMUSG00000046314",
                "ENSMUSG00000027238",
                "ENSMUSG00000047388",
                "ENSMUSG00000060923",
                "ENSMUSG00000061306",
                "ENSMUSG00000017686",
                "ENSMUSG00000013936",
                "ENSMUSG00000057406",
                "ENSMUSG00000069089",
                "ENSMUSG00000020415",
                "ENSMUSG00000026811",
                "ENSMUSG00000006800",
                "ENSMUSG00000059741",
                "ENSMUSG00000030255",
                "ENSMUSG00000002957",
                "ENSMUSG00000024073",
                "ENSMUSG00000028282",
                "ENSMUSG00000020430",
                "ENSMUSG00000009112",
                "ENSMUSG00000039130",
                "ENSMUSG00000078486",
                "ENSMUSG00000033565",
                "ENSMUSG00000027180",
                "ENSMUSG00000016308",
                "ENSMUSG00000027422",
                "ENSMUSG00000050875",
                "ENSMUSG00000050043",
                "ENSMUSG00000031843",
                "ENSMUSG00000018882",
                "ENSMUSG00000028470",
                "ENSMUSG00000027671",
                "ENSMUSG00000000787",
                "ENSMUSG00000070319",
                "ENSMUSG00000031808",
                "ENSMUSG00000015214",
                "ENSMUSG00000037710",
                "ENSMUSG00000024457",
                "ENSMUSG00000024187",
                "ENSMUSG00000019362",
                "ENSMUSG00000026308",
                "ENSMUSG00000016427",
                "ENSMUSG00000024205",
                "ENSMUSG00000027293",
                "ENSMUSG00000001127",
                "ENSMUSG00000015656",
                "ENSMUSG00000025868",
                "ENSMUSG00000027832",
                "ENSMUSG00000040520",
                "ENSMUSG00000058756",
                "ENSMUSG00000025651",
                "ENSMUSG00000033102",
                "ENSMUSG00000028603",
                "ENSMUSG00000004268",
                "ENSMUSG00000025085",
                "ENSMUSG00000065979",
                "ENSMUSG00000028261",
                "ENSMUSG00000024077",
                "ENSMUSG00000025068",
                "ENSMUSG00000045103",
                "ENSMUSG00000028675",
                "ENSMUSG00000032042",
                "ENSMUSG00000061032",
                "ENSMUSG00000023089",
                "ENSMUSG00000058927",
                "ENSMUSG00000030996",
                "ENSMUSG00000078851",
                "ENSMUSG00000073676",
                "ENSMUSG00000032060",
                "ENSMUSG00000023456",
                "ENSMUSG00000046573",
                "ENSMUSG00000048731",
                "ENSMUSG00000022010",
                "ENSMUSG00000025810",
                "ENSMUSG00000027673",
                "ENSMUSG00000059811",
                "ENSMUSG00000024587",
                "ENSMUSG00000024121",
                "ENSMUSG00000083911",
                "ENSMUSG00000015568",
                "ENSMUSG00000031504",
                "ENSMUSG00000031448",
                "ENSMUSG00000061904",
                "ENSMUSG00000074582",
                "ENSMUSG00000029454",
                "ENSMUSG00000015605",
                "ENSMUSG00000020485",
                "ENSMUSG00000006740",
                "ENSMUSG00000024197",
                "ENSMUSG00000007777",
                "ENSMUSG00000019432",
                "ENSMUSG00000022914",
                "ENSMUSG00000026491",
                "ENSMUSG00000018845",
                "ENSMUSG00000039148",
                "ENSMUSG00000089838",
                "ENSMUSG00000044791",
                "ENSMUSG00000020741",
                "ENSMUSG00000037266",
                "ENSMUSG00000037339",
                "ENSMUSG00000030982",
                "ENSMUSG00000028676",
                "ENSMUSG00000027679",
                "ENSMUSG00000022521",
                "ENSMUSG00000026576",
                "ENSMUSG00000041881",
                "ENSMUSG00000020691",
                "ENSMUSG00000021764",
                "ENSMUSG00000038763",
                "ENSMUSG00000021209",
                "ENSMUSG00000038535",
                "ENSMUSG00000043445",
                "ENSMUSG00000039648",
                "ENSMUSG00000020163",
                "ENSMUSG00000001767",
                "ENSMUSG00000039956",
                "ENSMUSG00000009863",
                "ENSMUSG00000033411",
                "ENSMUSG00000031770",
                "ENSMUSG00000038400",
                "ENSMUSG00000031715",
                "ENSMUSG00000056900",
                "ENSMUSG00000032238",
                "ENSMUSG00000062210",
                "ENSMUSG00000060376",
                "ENSMUSG00000014592",
                "ENSMUSG00000090150",
                "ENSMUSG00000017428",
                "ENSMUSG00000062694",
                "ENSMUSG00000040661",
                "ENSMUSG00000020153",
                "ENSMUSG00000093674",
                "ENSMUSG00000083563",
                "ENSMUSG00000040760",
                "ENSMUSG00000036452",
                "ENSMUSG00000031818",
                "ENSMUSG00000031782",
                "ENSMUSG00000048486",
                "ENSMUSG00000055148",
                "ENSMUSG00000032511",
                "ENSMUSG00000044894",
                "ENSMUSG00000007097",
                "ENSMUSG00000029404",
                "ENSMUSG00000048578",
                "ENSMUSG00000021134",
                "ENSMUSG00000022820",
                "ENSMUSG00000037234",
                "ENSMUSG00000002129",
                "ENSMUSG00000042148",
                "ENSMUSG00000034295",
                "ENSMUSG00000031783",
                "ENSMUSG00000022594",
                "ENSMUSG00000063229",
                "ENSMUSG00000027076",
                "ENSMUSG00000005417",
                "ENSMUSG00000021577",
                "ENSMUSG00000022994",
                "ENSMUSG00000029209",
                "ENSMUSG00000038936",
                "ENSMUSG00000029776",
                "ENSMUSG00000029135",
                "ENSMUSG00000025898",
                "ENSMUSG00000036427",
                "ENSMUSG00000026849",
                "ENSMUSG00000070867",
                "ENSMUSG00000038417",
                "ENSMUSG00000046329",
                "ENSMUSG00000045160",
                "ENSMUSG00000018040",
                "ENSMUSG00000024942",
                "ENSMUSG00000022698",
                "ENSMUSG00000027371",
                "ENSMUSG00000037075",
                "ENSMUSG00000029036",
                "ENSMUSG00000040407",
                "ENSMUSG00000030647",
                "ENSMUSG00000001424",
                "ENSMUSG00000073664",
                "ENSMUSG00000025545",
                "ENSMUSG00000026888",
                "ENSMUSG00000022094",
                "ENSMUSG00000052712",
                "ENSMUSG00000030087",
                "ENSMUSG00000019979",
                "ENSMUSG00000006333",
                "ENSMUSG00000026568",
                "ENSMUSG00000028328",
                "ENSMUSG00000005683",
                "ENSMUSG00000046516",
                "ENSMUSG00000037960",
                "ENSMUSG00000058655",
                "ENSMUSG00000039239",
                "ENSMUSG00000022961",
                "ENSMUSG00000020085",
                "ENSMUSG00000037443",
                "ENSMUSG00000031536",
                "ENSMUSG00000032348",
                "ENSMUSG00000029913",
                "ENSMUSG00000039218",
                "ENSMUSG00000029594",
                "ENSMUSG00000042834",
                "ENSMUSG00000032352",
                "ENSMUSG00000003033",
                "ENSMUSG00000052214",
                "ENSMUSG00000033228",
                "ENSMUSG00000046032",
                "ENSMUSG00000064181",
                "ENSMUSG00000032826",
                "ENSMUSG00000028266",
                "ENSMUSG00000033594",
                "ENSMUSG00000047371",
                "ENSMUSG00000020737",
                "ENSMUSG00000031232",
                "ENSMUSG00000001419",
                "ENSMUSG00000038583",
                "ENSMUSG00000022765",
                "ENSMUSG00000047260",
                "ENSMUSG00000004460",
                "ENSMUSG00000042198",
                "ENSMUSG00000035674",
                "ENSMUSG00000022261",
                "ENSMUSG00000009927",
                "ENSMUSG00000026571",
                "ENSMUSG00000025465",
                "ENSMUSG00000042605",
                "ENSMUSG00000028919",
                "ENSMUSG00000027423",
                "ENSMUSG00000001833",
                "ENSMUSG00000025204",
                "ENSMUSG00000023055",
                "ENSMUSG00000098178",
                "ENSMUSG00000031672",
                "ENSMUSG00000030172",
                "ENSMUSG00000059895",
                "ENSMUSG00000016319",
                "ENSMUSG00000024780",
                "ENSMUSG00000021488",
                "ENSMUSG00000034290",
                "ENSMUSG00000023150",
                "ENSMUSG00000038718",
                "ENSMUSG00000063800",
                "ENSMUSG00000030298",
                "ENSMUSG00000025795",
                "ENSMUSG00000097729",
                "ENSMUSG00000017404",
                "ENSMUSG00000020190",
                "ENSMUSG00000021944",
                "ENSMUSG00000028648",
                "ENSMUSG00000029623",
                "ENSMUSG00000004535",
                "ENSMUSG00000002949",
                "ENSMUSG00000014633",
                "ENSMUSG00000007892",
                "ENSMUSG00000003380",
                "ENSMUSG00000074064",
                "ENSMUSG00000060938",
                "ENSMUSG00000010406",
                "ENSMUSG00000025428",
                "ENSMUSG00000036214",
                "ENSMUSG00000024538",
                "ENSMUSG00000057335",
                "ENSMUSG00000021585",
                "ENSMUSG00000029017",
                "ENSMUSG00000030016",
                "ENSMUSG00000013921",
                "ENSMUSG00000041426",
                "ENSMUSG00000031634",
                "ENSMUSG00000020849",
                "ENSMUSG00000028343",
                "ENSMUSG00000081233",
                "ENSMUSG00000068706",
                "ENSMUSG00000002748",
                "ENSMUSG00000021238",
                "ENSMUSG00000041297",
                "ENSMUSG00000026766",
                "ENSMUSG00000024924",
                "ENSMUSG00000039989",
                "ENSMUSG00000042744",
                "ENSMUSG00000031987",
                "ENSMUSG00000038005",
                "ENSMUSG00000043311",
                "ENSMUSG00000038970",
                "ENSMUSG00000020171",
                "ENSMUSG00000005483",
                "ENSMUSG00000055302",
                "ENSMUSG00000028013",
                "ENSMUSG00000039089",
                "ENSMUSG00000053898",
                "ENSMUSG00000060038",
                "ENSMUSG00000002831",
                "ENSMUSG00000030409",
                "ENSMUSG00000062691",
                "ENSMUSG00000040648",
                "ENSMUSG00000001105",
                "ENSMUSG00000083820",
                "ENSMUSG00000025221",
                "ENSMUSG00000029632",
                "ENSMUSG00000026207",
                "ENSMUSG00000036620",
                "ENSMUSG00000022641",
                "ENSMUSG00000054766",
                "ENSMUSG00000052337",
                "ENSMUSG00000083307",
                "ENSMUSG00000031367",
                "ENSMUSG00000028223",
                "ENSMUSG00000027087",
                "ENSMUSG00000026342",
                "ENSMUSG00000074466",
                "ENSMUSG00000001794",
                "ENSMUSG00000031791",
                "ENSMUSG00000048264",
                "ENSMUSG00000027478",
                "ENSMUSG00000030399",
                "ENSMUSG00000071644",
                "ENSMUSG00000026027",
                "ENSMUSG00000053012",
                "ENSMUSG00000020483",
                "ENSMUSG00000009090",
                "ENSMUSG00000081974",
                "ENSMUSG00000032679",
                "ENSMUSG00000026878",
                "ENSMUSG00000098274",
                "ENSMUSG00000024055",
                "ENSMUSG00000029518",
                "ENSMUSG00000006304",
                "ENSMUSG00000031320",
                "ENSMUSG00000032959",
                "ENSMUSG00000028964",
                "ENSMUSG00000042032",
                "ENSMUSG00000002728",
                "ENSMUSG00000018893",
                "ENSMUSG00000027406",
                "ENSMUSG00000019863",
                "ENSMUSG00000060032",
                "ENSMUSG00000028756",
                "ENSMUSG00000063273",
                "ENSMUSG00000000811",
                "ENSMUSG00000059974",
                "ENSMUSG00000026927",
                "ENSMUSG00000037712",
                "ENSMUSG00000003955",
                "ENSMUSG00000028234",
                "ENSMUSG00000019179",
                "ENSMUSG00000018474",
                "ENSMUSG00000051495",
                "ENSMUSG00000079477",
                "ENSMUSG00000022841",
                "ENSMUSG00000042682",
                "ENSMUSG00000066613",
                "ENSMUSG00000025732",
                "ENSMUSG00000031575",
                "ENSMUSG00000037152",
                "ENSMUSG00000023010",
                "ENSMUSG00000036278",
                "ENSMUSG00000022391",
                "ENSMUSG00000028788",
                "ENSMUSG00000062169",
                "ENSMUSG00000067369",
                "ENSMUSG00000003316",
                "ENSMUSG00000030058",
                "ENSMUSG00000002980",
                "ENSMUSG00000005510",
                "ENSMUSG00000020547",
                "ENSMUSG00000024208",
                "ENSMUSG00000020232",
                "ENSMUSG00000049246",
                "ENSMUSG00000029686",
                "ENSMUSG00000004591",
                "ENSMUSG00000026098",
                "ENSMUSG00000006221",
                "ENSMUSG00000087260",
                "ENSMUSG00000062588",
                "ENSMUSG00000019428",
                "ENSMUSG00000024803",
                "ENSMUSG00000049134",
                "ENSMUSG00000028541",
                "ENSMUSG00000031683",
                "ENSMUSG00000054178",
                "ENSMUSG00000021395",
                "ENSMUSG00000078812",
                "ENSMUSG00000022235",
                "ENSMUSG00000054619",
                "ENSMUSG00000030714",
                "ENSMUSG00000020542",
                "ENSMUSG00000059534",
                "ENSMUSG00000027367",
                "ENSMUSG00000029447",
                "ENSMUSG00000029108",
                "ENSMUSG00000037300",
                "ENSMUSG00000044308",
                "ENSMUSG00000053536",
                "ENSMUSG00000021631",
                "ENSMUSG00000022300",
                "ENSMUSG00000003429",
                "ENSMUSG00000018599",
                "ENSMUSG00000005698",
                "ENSMUSG00000064344",
                "ENSMUSG00000029095",
                "ENSMUSG00000015839",
                "ENSMUSG00000027499",
                "ENSMUSG00000027784",
                "ENSMUSG00000049878",
                "ENSMUSG00000003541",
                "ENSMUSG00000029068",
                "ENSMUSG00000007613",
                "ENSMUSG00000027455",
                "ENSMUSG00000064340",
                "ENSMUSG00000032047",
                "ENSMUSG00000042133",
                "ENSMUSG00000044393",
                "ENSMUSG00000015377",
                "ENSMUSG00000014470",
                "ENSMUSG00000048076",
                "ENSMUSG00000001472",
                "ENSMUSG00000067818",
                "ENSMUSG00000036002",
                "ENSMUSG00000026864",
                "ENSMUSG00000023939",
                "ENSMUSG00000035885",
                "ENSMUSG00000059325",
                "ENSMUSG00000028134",
                "ENSMUSG00000031078",
                "ENSMUSG00000031333",
                "ENSMUSG00000026341",
                "ENSMUSG00000032330",
                "ENSMUSG00000038170",
                "ENSMUSG00000006315",
                "ENSMUSG00000078713",
                "ENSMUSG00000051695",
                "ENSMUSG00000041361",
                "ENSMUSG00000049327",
                "ENSMUSG00000086863",
                "ENSMUSG00000048280",
                "ENSMUSG00000022748",
                "ENSMUSG00000033392",
                "ENSMUSG00000031865",
                "ENSMUSG00000019087",
                "ENSMUSG00000029004",
                "ENSMUSG00000000149",
                "ENSMUSG00000023206",
                "ENSMUSG00000028518",
                "ENSMUSG00000056973",
                "ENSMUSG00000074129",
                "ENSMUSG00000039234",
                "ENSMUSG00000008136",
                "ENSMUSG00000020516",
                "ENSMUSG00000049881",
                "ENSMUSG00000005873",
                "ENSMUSG00000023944",
                "ENSMUSG00000015932",
                "ENSMUSG00000091989",
                "ENSMUSG00000020358",
                "ENSMUSG00000022681",
                "ENSMUSG00000068699",
                "ENSMUSG00000029201",
                "ENSMUSG00000003970",
                "ENSMUSG00000020475",
                "ENSMUSG00000030662",
                "ENSMUSG00000078671",
                "ENSMUSG00000036941",
                "ENSMUSG00000055670",
                "ENSMUSG00000050953",
                "ENSMUSG00000027765",
                "ENSMUSG00000020308",
                "ENSMUSG00000021546",
                "ENSMUSG00000064372",
                "ENSMUSG00000030002",
                "ENSMUSG00000032495",
                "ENSMUSG00000028772",
                "ENSMUSG00000024639",
                "ENSMUSG00000028145",
                "ENSMUSG00000025794",
                "ENSMUSG00000046997",
                "ENSMUSG00000020022",
                "ENSMUSG00000034088",
                "ENSMUSG00000001911",
                "ENSMUSG00000034832",
                "ENSMUSG00000020664",
                "ENSMUSG00000022389",
                "ENSMUSG00000086938",
                "ENSMUSG00000075595",
                "ENSMUSG00000033722",
                "ENSMUSG00000037095",
                "ENSMUSG00000007415",
                "ENSMUSG00000040118",
                "ENSMUSG00000022635",
                "ENSMUSG00000018770",
                "ENSMUSG00000055775",
                "ENSMUSG00000052911",
                "ENSMUSG00000039782",
                "ENSMUSG00000009569",
                "ENSMUSG00000066406",
                "ENSMUSG00000005566",
                "ENSMUSG00000026564",
                "ENSMUSG00000067736",
                "ENSMUSG00000027996",
                "ENSMUSG00000028737",
                "ENSMUSG00000031954",
                "ENSMUSG00000062908",
                "ENSMUSG00000068264",
                "ENSMUSG00000022462",
                "ENSMUSG00000038070",
                "ENSMUSG00000027447",
                "ENSMUSG00000021993",
                "ENSMUSG00000037032",
                "ENSMUSG00000024283",
                "ENSMUSG00000032423",
                "ENSMUSG00000036585",
                "ENSMUSG00000030319",
                "ENSMUSG00000062647",
                "ENSMUSG00000021681",
                "ENSMUSG00000020393",
                "ENSMUSG00000042709",
                "ENSMUSG00000031540",
                "ENSMUSG00000032525",
                "ENSMUSG00000020402",
                "ENSMUSG00000022217",
                "ENSMUSG00000026853",
                "ENSMUSG00000034880",
                "ENSMUSG00000054428",
                "ENSMUSG00000028953",
                "ENSMUSG00000028959",
                "ENSMUSG00000022119",
                "ENSMUSG00000019920",
                "ENSMUSG00000021270",
                "ENSMUSG00000041729",
                "ENSMUSG00000011257",
                "ENSMUSG00000039943",
                "ENSMUSG00000006576",
                "ENSMUSG00000064037",
                "ENSMUSG00000004085",
                "ENSMUSG00000056851",
                "ENSMUSG00000020048",
                "ENSMUSG00000022867",
                "ENSMUSG00000028878",
                "ENSMUSG00000038538",
                "ENSMUSG00000036678",
                "ENSMUSG00000029056",
                "ENSMUSG00000026688",
                "ENSMUSG00000056679",
                "ENSMUSG00000058498",
                "ENSMUSG00000029833",
                "ENSMUSG00000030213",
                "ENSMUSG00000034574",
                "ENSMUSG00000040666",
                "ENSMUSG00000041731",
                "ENSMUSG00000025184",
                "ENSMUSG00000003809",
                "ENSMUSG00000010175",
                "ENSMUSG00000020176",
                "ENSMUSG00000020863",
                "ENSMUSG00000022095",
                "ENSMUSG00000000399",
                "ENSMUSG00000004931",
                "ENSMUSG00000044792",
                "ENSMUSG00000039976",
                "ENSMUSG00000022437",
                "ENSMUSG00000019872",
                "ENSMUSG00000067653",
                "ENSMUSG00000026353",
                "ENSMUSG00000014504",
                "ENSMUSG00000022037",
                "ENSMUSG00000026276",
                "ENSMUSG00000033938",
                "ENSMUSG00000024997",
                "ENSMUSG00000038663",
                "ENSMUSG00000017778",
                "ENSMUSG00000020952",
                "ENSMUSG00000048546",
                "ENSMUSG00000019132",
                "ENSMUSG00000073639",
                "ENSMUSG00000045776",
                "ENSMUSG00000028207",
                "ENSMUSG00000034973",
                "ENSMUSG00000040414",
                "ENSMUSG00000021537",
                "ENSMUSG00000026434",
                "ENSMUSG00000063801",
                "ENSMUSG00000036564",
                "ENSMUSG00000074093",
                "ENSMUSG00000037049",
                "ENSMUSG00000020079",
                "ENSMUSG00000054452",
                "ENSMUSG00000024150",
                "ENSMUSG00000062456",
                "ENSMUSG00000032480",
                "ENSMUSG00000000184",
                "ENSMUSG00000091383",
                "ENSMUSG00000010122",
                "ENSMUSG00000050541",
                "ENSMUSG00000015937",
                "ENSMUSG00000031447",
                "ENSMUSG00000066279",
                "ENSMUSG00000008682",
                "ENSMUSG00000031765",
                "ENSMUSG00000023020",
                "ENSMUSG00000018339",
                "ENSMUSG00000024392",
                "ENSMUSG00000052934",
                "ENSMUSG00000030421",
                "ENSMUSG00000033382",
                "ENSMUSG00000034424",
                "ENSMUSG00000071793",
                "ENSMUSG00000064366",
                "ENSMUSG00000019558",
                "ENSMUSG00000031029",
                "ENSMUSG00000021149",
                "ENSMUSG00000006494",
                "ENSMUSG00000023832",
                "ENSMUSG00000004980",
                "ENSMUSG00000062963",
                "ENSMUSG00000032580",
                "ENSMUSG00000055027",
                "ENSMUSG00000071014",
                "ENSMUSG00000044864",
                "ENSMUSG00000083326",
                "ENSMUSG00000025666",
                "ENSMUSG00000000171",
                "ENSMUSG00000000441",
                "ENSMUSG00000041685",
                "ENSMUSG00000038784",
                "ENSMUSG00000024661",
                "ENSMUSG00000036748",
                "ENSMUSG00000053483",
                "ENSMUSG00000024526",
                "ENSMUSG00000025156",
                "ENSMUSG00000002608",
                "ENSMUSG00000025103",
                "ENSMUSG00000039601",
                "ENSMUSG00000057322",
                "ENSMUSG00000022270",
                "ENSMUSG00000026234",
                "ENSMUSG00000022136",
                "ENSMUSG00000029191",
                "ENSMUSG00000035783",
                "ENSMUSG00000045690",
                "ENSMUSG00000089909",
                "ENSMUSG00000032526",
                "ENSMUSG00000081228",
                "ENSMUSG00000038132",
                "ENSMUSG00000037949",
                "ENSMUSG00000055762",
                "ENSMUSG00000064343",
                "ENSMUSG00000022186",
                "ENSMUSG00000048814",
                "ENSMUSG00000021597",
                "ENSMUSG00000056091",
                "ENSMUSG00000045624",
                "ENSMUSG00000064337",
                "ENSMUSG00000020116",
                "ENSMUSG00000071528",
                "ENSMUSG00000026754",
                "ENSMUSG00000063382",
                "ENSMUSG00000047710",
                "ENSMUSG00000031841",
                "ENSMUSG00000036854",
                "ENSMUSG00000026131",
                "ENSMUSG00000026202",
                "ENSMUSG00000020721",
                "ENSMUSG00000027566",
                "ENSMUSG00000049960",
                "ENSMUSG00000048175",
                "ENSMUSG00000021639",
                "ENSMUSG00000020326",
                "ENSMUSG00000033538",
                "ENSMUSG00000028124",
                "ENSMUSG00000031751",
                "ENSMUSG00000032010",
                "ENSMUSG00000041278",
                "ENSMUSG00000021838",
                "ENSMUSG00000031996",
                "ENSMUSG00000067279",
                "ENSMUSG00000064347",
                "ENSMUSG00000034892",
                "ENSMUSG00000017756",
                "ENSMUSG00000023092",
                "ENSMUSG00000059291",
                "ENSMUSG00000028300",
                "ENSMUSG00000026577",
                "ENSMUSG00000027618",
                "ENSMUSG00000032177",
                "ENSMUSG00000048758",
                "ENSMUSG00000002379",
                "ENSMUSG00000022570",
                "ENSMUSG00000027177",
                "ENSMUSG00000020994",
                "ENSMUSG00000029455",
                "ENSMUSG00000064346",
                "ENSMUSG00000058690",
                "ENSMUSG00000027010",
                "ENSMUSG00000065947",
                "ENSMUSG00000059852",
                "ENSMUSG00000024063",
                "ENSMUSG00000037270",
                "ENSMUSG00000040370",
                "ENSMUSG00000074218",
                "ENSMUSG00000021371",
                "ENSMUSG00000021798",
                "ENSMUSG00000042323",
                "ENSMUSG00000001280",
                "ENSMUSG00000030417",
                "ENSMUSG00000037826",
                "ENSMUSG00000029651",
                "ENSMUSG00000057649",
                "ENSMUSG00000000751",
                "ENSMUSG00000021000",
                "ENSMUSG00000000563",
                "ENSMUSG00000065037",
                "ENSMUSG00000037815",
                "ENSMUSG00000067344",
                "ENSMUSG00000027809",
                "ENSMUSG00000051169",
                "ENSMUSG00000034342",
                "ENSMUSG00000028759",
                "ENSMUSG00000038949",
                "ENSMUSG00000047187",
                "ENSMUSG00000025488",
                "ENSMUSG00000018417",
                "ENSMUSG00000037958",
                "ENSMUSG00000022840",
                "ENSMUSG00000020456",
                "ENSMUSG00000031024",
                "ENSMUSG00000045983",
                "ENSMUSG00000051747",
                "ENSMUSG00000081967",
                "ENSMUSG00000027304",
                "ENSMUSG00000035458",
                "ENSMUSG00000032481",
                "ENSMUSG00000022040",
                "ENSMUSG00000033004",
                "ENSMUSG00000022013",
                "ENSMUSG00000011305",
                "ENSMUSG00000006356",
                "ENSMUSG00000022890",
                "ENSMUSG00000030505",
                "ENSMUSG00000040723",
                "ENSMUSG00000045482",
                "ENSMUSG00000019810",
                "ENSMUSG00000032199",
                "ENSMUSG00000025982",
                "ENSMUSG00000032224",
                "ENSMUSG00000001211",
                "ENSMUSG00000007877",
                "ENSMUSG00000056692",
                "ENSMUSG00000052738",
                "ENSMUSG00000031985",
                "ENSMUSG00000070436",
                "ENSMUSG00000029918",
                "ENSMUSG00000029359",
                "ENSMUSG00000023175",
                "ENSMUSG00000027598",
                "ENSMUSG00000052374",
                "ENSMUSG00000024991",
                "ENSMUSG00000024687",
                "ENSMUSG00000022201",
                "ENSMUSG00000027030",
                "ENSMUSG00000080893",
                "ENSMUSG00000021226",
                "ENSMUSG00000022940",
                "ENSMUSG00000030934",
                "ENSMUSG00000055093",
                "ENSMUSG00000032590",
                "ENSMUSG00000019998",
                "ENSMUSG00000024309",
                "ENSMUSG00000050199",
                "ENSMUSG00000025362",
                "ENSMUSG00000071604",
                "ENSMUSG00000027881",
                "ENSMUSG00000022964",
                "ENSMUSG00000025364",
                "ENSMUSG00000021807",
                "ENSMUSG00000022803",
                "ENSMUSG00000064338",
                "ENSMUSG00000039100",
                "ENSMUSG00000040473",
                "ENSMUSG00000044080",
                "ENSMUSG00000046671",
                "ENSMUSG00000047459",
                "ENSMUSG00000029649",
                "ENSMUSG00000060679",
                "ENSMUSG00000038844",
                "ENSMUSG00000020074",
                "ENSMUSG00000021877",
                "ENSMUSG00000048416",
                "ENSMUSG00000064193",
                "ENSMUSG00000031490",
                "ENSMUSG00000010608",
                "ENSMUSG00000032279",
                "ENSMUSG00000023861",
                "ENSMUSG00000067847",
                "ENSMUSG00000026664",
                "ENSMUSG00000021520",
                "ENSMUSG00000006717",
                "ENSMUSG00000008683",
                "ENSMUSG00000003438",
                "ENSMUSG00000034235",
                "ENSMUSG00000097277",
                "ENSMUSG00000028538",
                "ENSMUSG00000061666",
                "ENSMUSG00000065701",
                "ENSMUSG00000034842",
                "ENSMUSG00000029190",
                "ENSMUSG00000004610",
                "ENSMUSG00000015944",
                "ENSMUSG00000021265",
                "ENSMUSG00000021290",
                "ENSMUSG00000049470",
                "ENSMUSG00000078695",
                "ENSMUSG00000029401",
                "ENSMUSG00000029673",
                "ENSMUSG00000074247",
                "ENSMUSG00000003534",
                "ENSMUSG00000025453",
                "ENSMUSG00000004035",
                "ENSMUSG00000090110",
                "ENSMUSG00000072720",
                "ENSMUSG00000074918",
                "ENSMUSG00000021606",
                "ENSMUSG00000032621",
                "ENSMUSG00000039753",
                "ENSMUSG00000019877",
                "ENSMUSG00000033416",
                "ENSMUSG00000023068",
                "ENSMUSG00000036860",
                "ENSMUSG00000079641",
                "ENSMUSG00000045636",
                "ENSMUSG00000020462",
                "ENSMUSG00000026074",
                "ENSMUSG00000062300",
                "ENSMUSG00000031299",
                "ENSMUSG00000021771",
                "ENSMUSG00000032267",
                "ENSMUSG00000035247",
                "ENSMUSG00000040048",
                "ENSMUSG00000028461",
                "ENSMUSG00000045664",
                "ENSMUSG00000064365",
                "ENSMUSG00000048478",
                "ENSMUSG00000043639",
                "ENSMUSG00000043866",
                "ENSMUSG00000023938",
                "ENSMUSG00000024188",
                "ENSMUSG00000035268",
                "ENSMUSG00000085779",
                "ENSMUSG00000022450",
                "ENSMUSG00000073563",
                "ENSMUSG00000024805",
                "ENSMUSG00000032366",
                "ENSMUSG00000050503",
                "ENSMUSG00000032370",
                "ENSMUSG00000098343",
                "ENSMUSG00000021910",
                "ENSMUSG00000028383",
                "ENSMUSG00000041115",
                "ENSMUSG00000020309",
                "ENSMUSG00000064348",
                "ENSMUSG00000094483",
                "ENSMUSG00000024498",
                "ENSMUSG00000022965",
                "ENSMUSG00000022884",
                "ENSMUSG00000027099",
                "ENSMUSG00000035027",
                "ENSMUSG00000052812",
                "ENSMUSG00000044067",
                "ENSMUSG00000037685",
                "ENSMUSG00000024290",
                "ENSMUSG00000026526",
                "ENSMUSG00000024132",
                "ENSMUSG00000029338",
                "ENSMUSG00000021621",
                "ENSMUSG00000006005",
                "ENSMUSG00000034525",
                "ENSMUSG00000046312",
                "ENSMUSG00000055204",
                "ENSMUSG00000042115",
                "ENSMUSG00000071151",
                "ENSMUSG00000054312",
                "ENSMUSG00000060803",
                "ENSMUSG00000030246",
                "ENSMUSG00000061474",
                "ENSMUSG00000096232",
                "ENSMUSG00000010914",
                "ENSMUSG00000017760",
                "ENSMUSG00000039914",
                "ENSMUSG00000030869",
                "ENSMUSG00000031767",
                "ENSMUSG00000064371",
                "ENSMUSG00000084314",
                "ENSMUSG00000027620",
                "ENSMUSG00000017817",
                "ENSMUSG00000018395",
                "ENSMUSG00000002908",
                "ENSMUSG00000031229",
                "ENSMUSG00000074656",
                "ENSMUSG00000039126",
                "ENSMUSG00000030095",
                "ENSMUSG00000026489",
                "ENSMUSG00000024913"
            ],
            "expression_sum": {
                "class1": 60.0,
                "class2": 1510.0
            },
            "uniprot_ids": [
                "P19785",
                "D3Z6V3"
            ],
            "net_id": 0,
            "source_num": 10,
            "target_num": 487
        },
        "ENSMUSG00000038615": {
            "id": "ENSMUSG00000038615",
            "type": "TF",
            "symbol": "Nfe2l1",
            "source": [
                "ENSMUSG00000001419",
                "ENSMUSG00000015579",
                "ENSMUSG00000024955",
                "ENSMUSG00000001280",
                "ENSMUSG00000019768",
                "ENSMUSG00000044167",
                "ENSMUSG00000053110",
                "ENSMUSG00000031885",
                "ENSMUSG00000008575",
                "ENSMUSG00000031715",
                "ENSMUSG00000015846"
            ],
            "target": [
                "ENSMUSG00000054619",
                "ENSMUSG00000020483",
                "ENSMUSG00000020190",
                "ENSMUSG00000061904",
                "ENSMUSG00000009927",
                "ENSMUSG00000026817",
                "ENSMUSG00000040451",
                "ENSMUSG00000044308",
                "ENSMUSG00000027206",
                "ENSMUSG00000027805",
                "ENSMUSG00000040269",
                "ENSMUSG00000040850",
                "ENSMUSG00000042632",
                "ENSMUSG00000018819",
                "ENSMUSG00000023089",
                "ENSMUSG00000018217",
                "ENSMUSG00000015672",
                "ENSMUSG00000038121",
                "ENSMUSG00000073676",
                "ENSMUSG00000032348",
                "ENSMUSG00000048731",
                "ENSMUSG00000034160",
                "ENSMUSG00000069844",
                "ENSMUSG00000087270",
                "ENSMUSG00000026766",
                "ENSMUSG00000008429",
                "ENSMUSG00000067925",
                "ENSMUSG00000020402",
                "ENSMUSG00000059714",
                "ENSMUSG00000035202"
            ],
            "expression_sum": {
                "class1": 4094.0,
                "class2": 1452.0
            },
            "uniprot_ids": [
                "Q61985",
                "Q3V3M1",
                "A2A6D1",
                "A2A6D0",
                "A2A6D2"
            ],
            "net_id": 0,
            "source_num": 11,
            "target_num": 12
        },
        "ENSMUSG00000056501": {
            "id": "ENSMUSG00000056501",
            "type": "TF",
            "symbol": "Cebpb",
            "source": [
                "ENSMUSG00000059552",
                "ENSMUSG00000015839",
                "ENSMUSG00000041515",
                "ENSMUSG00000018899",
                "ENSMUSG00000001280",
                "ENSMUSG00000019768",
                "ENSMUSG00000005698"
            ],
            "target": [
                "ENSMUSG00000015568",
                "ENSMUSG00000061603",
                "ENSMUSG00000055775",
                "ENSMUSG00000061904",
                "ENSMUSG00000068566",
                "ENSMUSG00000097088",
                "ENSMUSG00000035781",
                "ENSMUSG00000038366",
                "ENSMUSG00000044330",
                "ENSMUSG00000074582",
                "ENSMUSG00000036835",
                "ENSMUSG00000031403",
                "ENSMUSG00000034674",
                "ENSMUSG00000045665",
                "ENSMUSG00000062908",
                "ENSMUSG00000007777",
                "ENSMUSG00000091478",
                "ENSMUSG00000063457",
                "ENSMUSG00000002661",
                "ENSMUSG00000068264",
                "ENSMUSG00000027022",
                "ENSMUSG00000042828",
                "ENSMUSG00000021248",
                "ENSMUSG00000027447",
                "ENSMUSG00000015837",
                "ENSMUSG00000043644",
                "ENSMUSG00000068040",
                "ENSMUSG00000044791",
                "ENSMUSG00000020741",
                "ENSMUSG00000033991",
                "ENSMUSG00000021235",
                "ENSMUSG00000070283",
                "ENSMUSG00000027984",
                "ENSMUSG00000041193",
                "ENSMUSG00000021629",
                "ENSMUSG00000001962",
                "ENSMUSG00000004044",
                "ENSMUSG00000041779",
                "ENSMUSG00000003363",
                "ENSMUSG00000024097",
                "ENSMUSG00000031540",
                "ENSMUSG00000041881",
                "ENSMUSG00000030214",
                "ENSMUSG00000060073",
                "ENSMUSG00000081773",
                "ENSMUSG00000022354",
                "ENSMUSG00000062593",
                "ENSMUSG00000054428",
                "ENSMUSG00000020070",
                "ENSMUSG00000031543",
                "ENSMUSG00000007891",
                "ENSMUSG00000001674",
                "ENSMUSG00000038763",
                "ENSMUSG00000005354",
                "ENSMUSG00000027247",
                "ENSMUSG00000009863",
                "ENSMUSG00000064349",
                "ENSMUSG00000011257",
                "ENSMUSG00000019579",
                "ENSMUSG00000032397",
                "ENSMUSG00000006576",
                "ENSMUSG00000038914",
                "ENSMUSG00000022433",
                "ENSMUSG00000005469",
                "ENSMUSG00000020300",
                "ENSMUSG00000039298",
                "ENSMUSG00000007656",
                "ENSMUSG00000026803",
                "ENSMUSG00000027288",
                "ENSMUSG00000052539",
                "ENSMUSG00000032314",
                "ENSMUSG00000081153",
                "ENSMUSG00000008892",
                "ENSMUSG00000051331",
                "ENSMUSG00000036097",
                "ENSMUSG00000029103",
                "ENSMUSG00000020048",
                "ENSMUSG00000024038",
                "ENSMUSG00000098772",
                "ENSMUSG00000027698",
                "ENSMUSG00000057738",
                "ENSMUSG00000016510",
                "ENSMUSG00000014592",
                "ENSMUSG00000081344",
                "ENSMUSG00000030850",
                "ENSMUSG00000045211",
                "ENSMUSG00000042225",
                "ENSMUSG00000014606",
                "ENSMUSG00000020153",
                "ENSMUSG00000025439",
                "ENSMUSG00000026260",
                "ENSMUSG00000081485",
                "ENSMUSG00000056698",
                "ENSMUSG00000017737",
                "ENSMUSG00000032216",
                "ENSMUSG00000020612",
                "ENSMUSG00000094664",
                "ENSMUSG00000024359",
                "ENSMUSG00000003847",
                "ENSMUSG00000031818",
                "ENSMUSG00000029833",
                "ENSMUSG00000031782",
                "ENSMUSG00000032511",
                "ENSMUSG00000029392",
                "ENSMUSG00000036825",
                "ENSMUSG00000001552",
                "ENSMUSG00000044365",
                "ENSMUSG00000000056",
                "ENSMUSG00000044894",
                "ENSMUSG00000038690",
                "ENSMUSG00000034006",
                "ENSMUSG00000039068",
                "ENSMUSG00000054414",
                "ENSMUSG00000024962",
                "ENSMUSG00000068758",
                "ENSMUSG00000045251",
                "ENSMUSG00000020949",
                "ENSMUSG00000021134",
                "ENSMUSG00000039067",
                "ENSMUSG00000004931",
                "ENSMUSG00000039103",
                "ENSMUSG00000037070",
                "ENSMUSG00000029148",
                "ENSMUSG00000020329",
                "ENSMUSG00000022820",
                "ENSMUSG00000044792",
                "ENSMUSG00000021244",
                "ENSMUSG00000061111",
                "ENSMUSG00000083932",
                "ENSMUSG00000041028",
                "ENSMUSG00000026504",
                "ENSMUSG00000022594",
                "ENSMUSG00000063229",
                "ENSMUSG00000026249",
                "ENSMUSG00000035168",
                "ENSMUSG00000021577",
                "ENSMUSG00000022994",
                "ENSMUSG00000082283",
                "ENSMUSG00000020321",
                "ENSMUSG00000059447",
                "ENSMUSG00000033938",
                "ENSMUSG00000077450",
                "ENSMUSG00000029135",
                "ENSMUSG00000027593",
                "ENSMUSG00000022324",
                "ENSMUSG00000025745",
                "ENSMUSG00000017778",
                "ENSMUSG00000026414",
                "ENSMUSG00000047675",
                "ENSMUSG00000024304",
                "ENSMUSG00000038417",
                "ENSMUSG00000030753",
                "ENSMUSG00000045210",
                "ENSMUSG00000006219",
                "ENSMUSG00000019132",
                "ENSMUSG00000030057",
                "ENSMUSG00000094249",
                "ENSMUSG00000078139",
                "ENSMUSG00000055322",
                "ENSMUSG00000039016",
                "ENSMUSG00000006057",
                "ENSMUSG00000020580",
                "ENSMUSG00000034329",
                "ENSMUSG00000040414",
                "ENSMUSG00000018196",
                "ENSMUSG00000082258",
                "ENSMUSG00000030647",
                "ENSMUSG00000032911",
                "ENSMUSG00000026434",
                "ENSMUSG00000060798",
                "ENSMUSG00000028443",
                "ENSMUSG00000051502",
                "ENSMUSG00000036564",
                "ENSMUSG00000025545",
                "ENSMUSG00000081359",
                "ENSMUSG00000038371",
                "ENSMUSG00000081413",
                "ENSMUSG00000054452",
                "ENSMUSG00000061518",
                "ENSMUSG00000033400",
                "ENSMUSG00000036570",
                "ENSMUSG00000012848",
                "ENSMUSG00000062456",
                "ENSMUSG00000049130",
                "ENSMUSG00000022800",
                "ENSMUSG00000043885",
                "ENSMUSG00000029186",
                "ENSMUSG00000029452",
                "ENSMUSG00000044533",
                "ENSMUSG00000032648",
                "ENSMUSG00000003299",
                "ENSMUSG00000022538",
                "ENSMUSG00000026568",
                "ENSMUSG00000005683",
                "ENSMUSG00000032562",
                "ENSMUSG00000059981",
                "ENSMUSG00000031765",
                "ENSMUSG00000057858",
                "ENSMUSG00000060730",
                "ENSMUSG00000005882",
                "ENSMUSG00000046516",
                "ENSMUSG00000007617",
                "ENSMUSG00000083831",
                "ENSMUSG00000028641",
                "ENSMUSG00000068220",
                "ENSMUSG00000037960",
                "ENSMUSG00000058655",
                "ENSMUSG00000052934",
                "ENSMUSG00000060002",
                "ENSMUSG00000019768",
                "ENSMUSG00000024331",
                "ENSMUSG00000028573",
                "ENSMUSG00000063406",
                "ENSMUSG00000036371",
                "ENSMUSG00000034424",
                "ENSMUSG00000058135",
                "ENSMUSG00000070493",
                "ENSMUSG00000040761",
                "ENSMUSG00000047419",
                "ENSMUSG00000022197",
                "ENSMUSG00000003778",
                "ENSMUSG00000020085",
                "ENSMUSG00000032497",
                "ENSMUSG00000021374",
                "ENSMUSG00000030357",
                "ENSMUSG00000052748",
                "ENSMUSG00000063663",
                "ENSMUSG00000081824",
                "ENSMUSG00000030309",
                "ENSMUSG00000004980",
                "ENSMUSG00000030785",
                "ENSMUSG00000032116",
                "ENSMUSG00000028049",
                "ENSMUSG00000045098",
                "ENSMUSG00000055027",
                "ENSMUSG00000028932",
                "ENSMUSG00000010054",
                "ENSMUSG00000071014",
                "ENSMUSG00000027184",
                "ENSMUSG00000049804",
                "ENSMUSG00000023919",
                "ENSMUSG00000049751",
                "ENSMUSG00000066637",
                "ENSMUSG00000025666",
                "ENSMUSG00000059714",
                "ENSMUSG00000000171",
                "ENSMUSG00000034220",
                "ENSMUSG00000010830",
                "ENSMUSG00000041187",
                "ENSMUSG00000033060",
                "ENSMUSG00000089862",
                "ENSMUSG00000005610",
                "ENSMUSG00000038594",
                "ENSMUSG00000039000",
                "ENSMUSG00000032418",
                "ENSMUSG00000017176",
                "ENSMUSG00000031627",
                "ENSMUSG00000031231",
                "ENSMUSG00000032826",
                "ENSMUSG00000002608",
                "ENSMUSG00000039601",
                "ENSMUSG00000026701",
                "ENSMUSG00000022270",
                "ENSMUSG00000030536",
                "ENSMUSG00000089909",
                "ENSMUSG00000014313",
                "ENSMUSG00000041453",
                "ENSMUSG00000033554",
                "ENSMUSG00000000088",
                "ENSMUSG00000081228",
                "ENSMUSG00000084830",
                "ENSMUSG00000055762",
                "ENSMUSG00000064343",
                "ENSMUSG00000017291",
                "ENSMUSG00000074971",
                "ENSMUSG00000031232",
                "ENSMUSG00000036632",
                "ENSMUSG00000022186",
                "ENSMUSG00000048814",
                "ENSMUSG00000026229",
                "ENSMUSG00000031641",
                "ENSMUSG00000038967",
                "ENSMUSG00000056091",
                "ENSMUSG00000045624",
                "ENSMUSG00000024870",
                "ENSMUSG00000043153",
                "ENSMUSG00000071528",
                "ENSMUSG00000046311",
                "ENSMUSG00000053093",
                "ENSMUSG00000083856",
                "ENSMUSG00000059820",
                "ENSMUSG00000017264",
                "ENSMUSG00000035674",
                "ENSMUSG00000033720",
                "ENSMUSG00000023886",
                "ENSMUSG00000002416",
                "ENSMUSG00000031841",
                "ENSMUSG00000002500",
                "ENSMUSG00000055963",
                "ENSMUSG00000026131",
                "ENSMUSG00000026202",
                "ENSMUSG00000054823",
                "ENSMUSG00000071657",
                "ENSMUSG00000034714",
                "ENSMUSG00000034951",
                "ENSMUSG00000049960",
                "ENSMUSG00000066491",
                "ENSMUSG00000028729",
                "ENSMUSG00000027423",
                "ENSMUSG00000082051",
                "ENSMUSG00000025204",
                "ENSMUSG00000043024",
                "ENSMUSG00000001440",
                "ENSMUSG00000022477",
                "ENSMUSG00000068614",
                "ENSMUSG00000031672",
                "ENSMUSG00000041278",
                "ENSMUSG00000016319",
                "ENSMUSG00000064347",
                "ENSMUSG00000034892",
                "ENSMUSG00000062038",
                "ENSMUSG00000028649",
                "ENSMUSG00000027018",
                "ENSMUSG00000023150",
                "ENSMUSG00000041515",
                "ENSMUSG00000012405",
                "ENSMUSG00000038718",
                "ENSMUSG00000074622",
                "ENSMUSG00000030695",
                "ENSMUSG00000067148",
                "ENSMUSG00000017404",
                "ENSMUSG00000022110",
                "ENSMUSG00000028648",
                "ENSMUSG00000036478",
                "ENSMUSG00000004356",
                "ENSMUSG00000037916",
                "ENSMUSG00000021831",
                "ENSMUSG00000035242",
                "ENSMUSG00000052783",
                "ENSMUSG00000038503",
                "ENSMUSG00000050608",
                "ENSMUSG00000029156",
                "ENSMUSG00000034601",
                "ENSMUSG00000033282",
                "ENSMUSG00000081352",
                "ENSMUSG00000052681",
                "ENSMUSG00000019851",
                "ENSMUSG00000059208",
                "ENSMUSG00000015289",
                "ENSMUSG00000015133",
                "ENSMUSG00000024054",
                "ENSMUSG00000010110",
                "ENSMUSG00000025428",
                "ENSMUSG00000033161",
                "ENSMUSG00000058690",
                "ENSMUSG00000038412",
                "ENSMUSG00000081214",
                "ENSMUSG00000003072",
                "ENSMUSG00000020390",
                "ENSMUSG00000024063",
                "ENSMUSG00000038370",
                "ENSMUSG00000046982",
                "ENSMUSG00000082487",
                "ENSMUSG00000013921",
                "ENSMUSG00000029128",
                "ENSMUSG00000000743",
                "ENSMUSG00000020849",
                "ENSMUSG00000074218",
                "ENSMUSG00000081233",
                "ENSMUSG00000030894",
                "ENSMUSG00000041297",
                "ENSMUSG00000068706",
                "ENSMUSG00000034686",
                "ENSMUSG00000000168",
                "ENSMUSG00000024924",
                "ENSMUSG00000048661",
                "ENSMUSG00000039989",
                "ENSMUSG00000001280",
                "ENSMUSG00000031987",
                "ENSMUSG00000038005",
                "ENSMUSG00000062014",
                "ENSMUSG00000062604",
                "ENSMUSG00000080866",
                "ENSMUSG00000034345",
                "ENSMUSG00000027861",
                "ENSMUSG00000022390",
                "ENSMUSG00000031633",
                "ENSMUSG00000063694",
                "ENSMUSG00000028367",
                "ENSMUSG00000059552",
                "ENSMUSG00000051817",
                "ENSMUSG00000041319",
                "ENSMUSG00000059743",
                "ENSMUSG00000049119",
                "ENSMUSG00000058624",
                "ENSMUSG00000060038",
                "ENSMUSG00000081278",
                "ENSMUSG00000042650",
                "ENSMUSG00000005410",
                "ENSMUSG00000083820",
                "ENSMUSG00000061533",
                "ENSMUSG00000021313",
                "ENSMUSG00000026662",
                "ENSMUSG00000029632",
                "ENSMUSG00000016252",
                "ENSMUSG00000020740",
                "ENSMUSG00000025241",
                "ENSMUSG00000038172",
                "ENSMUSG00000059430",
                "ENSMUSG00000028843",
                "ENSMUSG00000061462",
                "ENSMUSG00000080831",
                "ENSMUSG00000035776",
                "ENSMUSG00000052337",
                "ENSMUSG00000031367",
                "ENSMUSG00000018417",
                "ENSMUSG00000022889",
                "ENSMUSG00000062981",
                "ENSMUSG00000025962",
                "ENSMUSG00000020456",
                "ENSMUSG00000045983",
                "ENSMUSG00000051747",
                "ENSMUSG00000060198",
                "ENSMUSG00000021281",
                "ENSMUSG00000035458",
                "ENSMUSG00000001794",
                "ENSMUSG00000038239",
                "ENSMUSG00000086298",
                "ENSMUSG00000021767",
                "ENSMUSG00000028007",
                "ENSMUSG00000031791",
                "ENSMUSG00000022536",
                "ENSMUSG00000022040",
                "ENSMUSG00000048264",
                "ENSMUSG00000003923",
                "ENSMUSG00000030399",
                "ENSMUSG00000055612",
                "ENSMUSG00000021557",
                "ENSMUSG00000022678",
                "ENSMUSG00000036167",
                "ENSMUSG00000040325",
                "ENSMUSG00000028745",
                "ENSMUSG00000020483",
                "ENSMUSG00000028228",
                "ENSMUSG00000053702",
                "ENSMUSG00000014226",
                "ENSMUSG00000020267",
                "ENSMUSG00000009575",
                "ENSMUSG00000058427",
                "ENSMUSG00000022890",
                "ENSMUSG00000018481",
                "ENSMUSG00000024767",
                "ENSMUSG00000021835",
                "ENSMUSG00000040723",
                "ENSMUSG00000006818",
                "ENSMUSG00000020733",
                "ENSMUSG00000019810",
                "ENSMUSG00000021112",
                "ENSMUSG00000027708",
                "ENSMUSG00000095588",
                "ENSMUSG00000032199",
                "ENSMUSG00000042541",
                "ENSMUSG00000049517",
                "ENSMUSG00000058600",
                "ENSMUSG00000028692",
                "ENSMUSG00000018068",
                "ENSMUSG00000040850",
                "ENSMUSG00000058793",
                "ENSMUSG00000034795",
                "ENSMUSG00000001211",
                "ENSMUSG00000033039",
                "ENSMUSG00000031985",
                "ENSMUSG00000038375",
                "ENSMUSG00000052738",
                "ENSMUSG00000082715",
                "ENSMUSG00000005225",
                "ENSMUSG00000025135",
                "ENSMUSG00000029359",
                "ENSMUSG00000005897",
                "ENSMUSG00000029918",
                "ENSMUSG00000039968",
                "ENSMUSG00000023175",
                "ENSMUSG00000039509",
                "ENSMUSG00000018893",
                "ENSMUSG00000018899",
                "ENSMUSG00000060600",
                "ENSMUSG00000091742",
                "ENSMUSG00000027406",
                "ENSMUSG00000062421",
                "ENSMUSG00000060032",
                "ENSMUSG00000031667",
                "ENSMUSG00000056071",
                "ENSMUSG00000025792",
                "ENSMUSG00000038393",
                "ENSMUSG00000078784",
                "ENSMUSG00000028756",
                "ENSMUSG00000080893",
                "ENSMUSG00000002102",
                "ENSMUSG00000056054",
                "ENSMUSG00000030934",
                "ENSMUSG00000040254",
                "ENSMUSG00000086229",
                "ENSMUSG00000002010",
                "ENSMUSG00000054889",
                "ENSMUSG00000004018",
                "ENSMUSG00000024792",
                "ENSMUSG00000047022",
                "ENSMUSG00000020409",
                "ENSMUSG00000024953",
                "ENSMUSG00000030060",
                "ENSMUSG00000032328",
                "ENSMUSG00000073982",
                "ENSMUSG00000011831",
                "ENSMUSG00000019179",
                "ENSMUSG00000028273",
                "ENSMUSG00000019158",
                "ENSMUSG00000018474",
                "ENSMUSG00000028811",
                "ENSMUSG00000050199",
                "ENSMUSG00000021200",
                "ENSMUSG00000056498",
                "ENSMUSG00000001156",
                "ENSMUSG00000022964",
                "ENSMUSG00000079477",
                "ENSMUSG00000083804",
                "ENSMUSG00000039680",
                "ENSMUSG00000050856",
                "ENSMUSG00000074513",
                "ENSMUSG00000071415",
                "ENSMUSG00000037152",
                "ENSMUSG00000020027",
                "ENSMUSG00000031917",
                "ENSMUSG00000026032",
                "ENSMUSG00000008658",
                "ENSMUSG00000021868",
                "ENSMUSG00000053768",
                "ENSMUSG00000032231",
                "ENSMUSG00000043091",
                "ENSMUSG00000022120",
                "ENSMUSG00000018648",
                "ENSMUSG00000025781",
                "ENSMUSG00000028452",
                "ENSMUSG00000039987",
                "ENSMUSG00000026429",
                "ENSMUSG00000062169",
                "ENSMUSG00000000295",
                "ENSMUSG00000020053",
                "ENSMUSG00000018796",
                "ENSMUSG00000002980",
                "ENSMUSG00000044080",
                "ENSMUSG00000024208",
                "ENSMUSG00000031461",
                "ENSMUSG00000029924",
                "ENSMUSG00000005958",
                "ENSMUSG00000049246",
                "ENSMUSG00000022403",
                "ENSMUSG00000083076",
                "ENSMUSG00000027257",
                "ENSMUSG00000020372",
                "ENSMUSG00000028834",
                "ENSMUSG00000029649",
                "ENSMUSG00000000416",
                "ENSMUSG00000027852",
                "ENSMUSG00000073838",
                "ENSMUSG00000035370",
                "ENSMUSG00000000740",
                "ENSMUSG00000010608",
                "ENSMUSG00000033788",
                "ENSMUSG00000083483",
                "ENSMUSG00000034593",
                "ENSMUSG00000022516",
                "ENSMUSG00000037795",
                "ENSMUSG00000062070",
                "ENSMUSG00000021520",
                "ENSMUSG00000023460",
                "ENSMUSG00000031458",
                "ENSMUSG00000006717",
                "ENSMUSG00000020680",
                "ENSMUSG00000091898",
                "ENSMUSG00000026208",
                "ENSMUSG00000031696",
                "ENSMUSG00000081661",
                "ENSMUSG00000070425",
                "ENSMUSG00000085566",
                "ENSMUSG00000059534",
                "ENSMUSG00000046314",
                "ENSMUSG00000030652",
                "ENSMUSG00000028394",
                "ENSMUSG00000018548",
                "ENSMUSG00000025968",
                "ENSMUSG00000067780",
                "ENSMUSG00000080836",
                "ENSMUSG00000060187",
                "ENSMUSG00000053536",
                "ENSMUSG00000022300",
                "ENSMUSG00000053841",
                "ENSMUSG00000065701",
                "ENSMUSG00000005698",
                "ENSMUSG00000026278",
                "ENSMUSG00000028691",
                "ENSMUSG00000060923",
                "ENSMUSG00000081700",
                "ENSMUSG00000020077",
                "ENSMUSG00000028869",
                "ENSMUSG00000029373",
                "ENSMUSG00000029190",
                "ENSMUSG00000089648",
                "ENSMUSG00000015839",
                "ENSMUSG00000037805",
                "ENSMUSG00000024070",
                "ENSMUSG00000024869",
                "ENSMUSG00000004610",
                "ENSMUSG00000021290",
                "ENSMUSG00000020546",
                "ENSMUSG00000019303",
                "ENSMUSG00000049470",
                "ENSMUSG00000028911",
                "ENSMUSG00000013936",
                "ENSMUSG00000031701",
                "ENSMUSG00000069049",
                "ENSMUSG00000022836",
                "ENSMUSG00000064340",
                "ENSMUSG00000055632",
                "ENSMUSG00000029401",
                "ENSMUSG00000075701",
                "ENSMUSG00000032047",
                "ENSMUSG00000024151",
                "ENSMUSG00000069089",
                "ENSMUSG00000085442",
                "ENSMUSG00000020415",
                "ENSMUSG00000072720",
                "ENSMUSG00000080806",
                "ENSMUSG00000054027",
                "ENSMUSG00000044393",
                "ENSMUSG00000062190",
                "ENSMUSG00000059741",
                "ENSMUSG00000030255",
                "ENSMUSG00000021606",
                "ENSMUSG00000034675",
                "ENSMUSG00000089672",
                "ENSMUSG00000033083",
                "ENSMUSG00000027303",
                "ENSMUSG00000038539",
                "ENSMUSG00000036002",
                "ENSMUSG00000026817",
                "ENSMUSG00000038582",
                "ENSMUSG00000048126",
                "ENSMUSG00000025959",
                "ENSMUSG00000023883",
                "ENSMUSG00000019877",
                "ENSMUSG00000035885",
                "ENSMUSG00000096210",
                "ENSMUSG00000020649",
                "ENSMUSG00000083341",
                "ENSMUSG00000023068",
                "ENSMUSG00000062270",
                "ENSMUSG00000050592",
                "ENSMUSG00000026341",
                "ENSMUSG00000087635",
                "ENSMUSG00000034566",
                "ENSMUSG00000032330",
                "ENSMUSG00000067336",
                "ENSMUSG00000045636",
                "ENSMUSG00000026074",
                "ENSMUSG00000050555",
                "ENSMUSG00000038170",
                "ENSMUSG00000031299",
                "ENSMUSG00000021771",
                "ENSMUSG00000040048",
                "ENSMUSG00000020122",
                "ENSMUSG00000034781",
                "ENSMUSG00000038403",
                "ENSMUSG00000097790",
                "ENSMUSG00000092674",
                "ENSMUSG00000049090",
                "ENSMUSG00000037710",
                "ENSMUSG00000083116",
                "ENSMUSG00000041476",
                "ENSMUSG00000046722",
                "ENSMUSG00000049327",
                "ENSMUSG00000027712",
                "ENSMUSG00000038668",
                "ENSMUSG00000021737",
                "ENSMUSG00000038058",
                "ENSMUSG00000097282",
                "ENSMUSG00000033632",
                "ENSMUSG00000007050",
                "ENSMUSG00000025133",
                "ENSMUSG00000034729",
                "ENSMUSG00000082634",
                "ENSMUSG00000036781",
                "ENSMUSG00000024205",
                "ENSMUSG00000022215",
                "ENSMUSG00000043140",
                "ENSMUSG00000034083",
                "ENSMUSG00000020988",
                "ENSMUSG00000028518",
                "ENSMUSG00000009207",
                "ENSMUSG00000001783",
                "ENSMUSG00000074129",
                "ENSMUSG00000003948",
                "ENSMUSG00000038084",
                "ENSMUSG00000039670",
                "ENSMUSG00000070343",
                "ENSMUSG00000016554",
                "ENSMUSG00000022111",
                "ENSMUSG00000024805",
                "ENSMUSG00000046364",
                "ENSMUSG00000028793",
                "ENSMUSG00000032366",
                "ENSMUSG00000037236",
                "ENSMUSG00000023307",
                "ENSMUSG00000051391",
                "ENSMUSG00000064369",
                "ENSMUSG00000023944",
                "ENSMUSG00000015932",
                "ENSMUSG00000091989",
                "ENSMUSG00000000823",
                "ENSMUSG00000019837",
                "ENSMUSG00000020358",
                "ENSMUSG00000022681",
                "ENSMUSG00000040536",
                "ENSMUSG00000028364",
                "ENSMUSG00000029922",
                "ENSMUSG00000043895",
                "ENSMUSG00000041115",
                "ENSMUSG00000053819",
                "ENSMUSG00000003970",
                "ENSMUSG00000020475",
                "ENSMUSG00000022551",
                "ENSMUSG00000037110",
                "ENSMUSG00000064348",
                "ENSMUSG00000068699",
                "ENSMUSG00000029571",
                "ENSMUSG00000072946",
                "ENSMUSG00000025651",
                "ENSMUSG00000028603",
                "ENSMUSG00000010911",
                "ENSMUSG00000050953",
                "ENSMUSG00000040086",
                "ENSMUSG00000022965",
                "ENSMUSG00000027765",
                "ENSMUSG00000032508",
                "ENSMUSG00000068882",
                "ENSMUSG00000080823",
                "ENSMUSG00000042607",
                "ENSMUSG00000001300",
                "ENSMUSG00000039697",
                "ENSMUSG00000035329",
                "ENSMUSG00000046312",
                "ENSMUSG00000081043",
                "ENSMUSG00000041650",
                "ENSMUSG00000004558",
                "ENSMUSG00000037062",
                "ENSMUSG00000026003",
                "ENSMUSG00000054312",
                "ENSMUSG00000029683",
                "ENSMUSG00000023089",
                "ENSMUSG00000020022",
                "ENSMUSG00000046997",
                "ENSMUSG00000078851",
                "ENSMUSG00000073676",
                "ENSMUSG00000023456",
                "ENSMUSG00000030970",
                "ENSMUSG00000038462",
                "ENSMUSG00000020664",
                "ENSMUSG00000048410",
                "ENSMUSG00000030869",
                "ENSMUSG00000033722",
                "ENSMUSG00000083567",
                "ENSMUSG00000040699",
                "ENSMUSG00000026610",
                "ENSMUSG00000024099",
                "ENSMUSG00000028949",
                "ENSMUSG00000018770",
                "ENSMUSG00000044408"
            ],
            "expression_sum": {
                "class1": 83.0,
                "class2": 485.0
            },
            "uniprot_ids": [
                "P28033"
            ],
            "net_id": 0,
            "source_num": 7,
            "target_num": 241
        },
        "ENSMUSG00000020453": {
            "id": "ENSMUSG00000020453",
            "type": "TF",
            "symbol": "Patz1",
            "source": [
                "ENSMUSG00000001280",
                "ENSMUSG00000005698",
                "ENSMUSG00000024955"
            ],
            "target": [
                "ENSMUSG00000005683"
            ],
            "expression_sum": {
                "class1": 336.0,
                "class2": 334.0
            },
            "uniprot_ids": [
                "Q5NBZ2",
                "Q7TNQ3",
                "Q9JMG9",
                "Q5NBY8",
                "Q5NBZ1",
                "Q5NBY9"
            ],
            "net_id": 0,
            "source_num": 3,
            "target_num": 1
        },
        "ENSMUSG00000028639": {
            "id": "ENSMUSG00000028639",
            "type": "TF",
            "symbol": "Ybx1",
            "source": [
                "ENSMUSG00000032897",
                "ENSMUSG00000021250",
                "ENSMUSG00000015839",
                "ENSMUSG00000018604",
                "ENSMUSG00000001280",
                "ENSMUSG00000035158"
            ],
            "target": [
                "ENSMUSG00000014599",
                "ENSMUSG00000003037",
                "ENSMUSG00000040025",
                "ENSMUSG00000004127",
                "ENSMUSG00000056941",
                "ENSMUSG00000026617",
                "ENSMUSG00000043683",
                "ENSMUSG00000063684",
                "ENSMUSG00000018819",
                "ENSMUSG00000000031",
                "ENSMUSG00000007080",
                "ENSMUSG00000052384",
                "ENSMUSG00000081792",
                "ENSMUSG00000025077",
                "ENSMUSG00000027750",
                "ENSMUSG00000028165",
                "ENSMUSG00000095280",
                "ENSMUSG00000000568",
                "ENSMUSG00000020402",
                "ENSMUSG00000025395",
                "ENSMUSG00000021645",
                "ENSMUSG00000024066",
                "ENSMUSG00000008035",
                "ENSMUSG00000063019",
                "ENSMUSG00000004789",
                "ENSMUSG00000020477",
                "ENSMUSG00000040242",
                "ENSMUSG00000082684",
                "ENSMUSG00000000127",
                "ENSMUSG00000029729",
                "ENSMUSG00000004085",
                "ENSMUSG00000030536",
                "ENSMUSG00000025175",
                "ENSMUSG00000031821",
                "ENSMUSG00000000088",
                "ENSMUSG00000040048",
                "ENSMUSG00000003423",
                "ENSMUSG00000075700",
                "ENSMUSG00000027160",
                "ENSMUSG00000049658",
                "ENSMUSG00000006519",
                "ENSMUSG00000081620",
                "ENSMUSG00000027439",
                "ENSMUSG00000061559",
                "ENSMUSG00000021190",
                "ENSMUSG00000002416",
                "ENSMUSG00000083832",
                "ENSMUSG00000029502",
                "ENSMUSG00000022744",
                "ENSMUSG00000008136",
                "ENSMUSG00000025647",
                "ENSMUSG00000016256",
                "ENSMUSG00000064369",
                "ENSMUSG00000024400",
                "ENSMUSG00000038612",
                "ENSMUSG00000068749",
                "ENSMUSG00000097991",
                "ENSMUSG00000064338",
                "ENSMUSG00000028452",
                "ENSMUSG00000083932",
                "ENSMUSG00000052146",
                "ENSMUSG00000021772",
                "ENSMUSG00000022216",
                "ENSMUSG00000019791",
                "ENSMUSG00000001525",
                "ENSMUSG00000063698",
                "ENSMUSG00000045103",
                "ENSMUSG00000025130",
                "ENSMUSG00000000628",
                "ENSMUSG00000005686",
                "ENSMUSG00000021622",
                "ENSMUSG00000022122",
                "ENSMUSG00000001627",
                "ENSMUSG00000032279",
                "ENSMUSG00000001424",
                "ENSMUSG00000023861",
                "ENSMUSG00000066861",
                "ENSMUSG00000029864",
                "ENSMUSG00000081359",
                "ENSMUSG00000028832",
                "ENSMUSG00000020067",
                "ENSMUSG00000027861",
                "ENSMUSG00000081049"
            ],
            "expression_sum": {
                "class1": 9831.0,
                "class2": 5203.0
            },
            "uniprot_ids": [
                "P62960",
                "A2BGG7"
            ],
            "net_id": 0,
            "source_num": 6,
            "target_num": 17
        },
        "ENSMUSG00000055320": {
            "id": "ENSMUSG00000055320",
            "type": "TF",
            "symbol": "Tead1",
            "source": [
                "ENSMUSG00000029135",
                "ENSMUSG00000001419",
                "ENSMUSG00000026610",
                "ENSMUSG00000015579",
                "ENSMUSG00000029249",
                "ENSMUSG00000001280",
                "ENSMUSG00000019768",
                "ENSMUSG00000031715"
            ],
            "target": [
                "ENSMUSG00000035674",
                "ENSMUSG00000005958",
                "ENSMUSG00000002010",
                "ENSMUSG00000005610",
                "ENSMUSG00000001767",
                "ENSMUSG00000032119",
                "ENSMUSG00000014633",
                "ENSMUSG00000024197",
                "ENSMUSG00000020530",
                "ENSMUSG00000029158",
                "ENSMUSG00000001105",
                "ENSMUSG00000046516",
                "ENSMUSG00000025130",
                "ENSMUSG00000000056",
                "ENSMUSG00000039239",
                "ENSMUSG00000071793",
                "ENSMUSG00000000031",
                "ENSMUSG00000000753",
                "ENSMUSG00000023089",
                "ENSMUSG00000000088",
                "ENSMUSG00000082165",
                "ENSMUSG00000050222",
                "ENSMUSG00000073676",
                "ENSMUSG00000039509",
                "ENSMUSG00000004610",
                "ENSMUSG00000020664",
                "ENSMUSG00000069049",
                "ENSMUSG00000013936",
                "ENSMUSG00000017817",
                "ENSMUSG00000001962",
                "ENSMUSG00000000581",
                "ENSMUSG00000028007",
                "ENSMUSG00000023460",
                "ENSMUSG00000001240",
                "ENSMUSG00000003531",
                "ENSMUSG00000028633",
                "ENSMUSG00000029661"
            ],
            "expression_sum": {
                "class1": 2182.0,
                "class2": 1186.0
            },
            "uniprot_ids": [
                "Q3USK5",
                "Q3UFP5",
                "Q6PAQ8",
                "E9Q387",
                "E9PZ73",
                "F6QIG2",
                "E9PVS2",
                "F6QH05",
                "F7B3M8",
                "F6ZIM6",
                "F6TBI7",
                "E9PVJ5",
                "E9QAK6"
            ],
            "net_id": 0,
            "source_num": 8,
            "target_num": 13
        },
        "ENSMUSG00000053477": {
            "id": "ENSMUSG00000053477",
            "type": "TF",
            "symbol": "Tcf4",
            "source": [
                "ENSMUSG00000019947",
                "ENSMUSG00000024238",
                "ENSMUSG00000029135",
                "ENSMUSG00000019817",
                "ENSMUSG00000018604",
                "ENSMUSG00000052534",
                "ENSMUSG00000015579",
                "ENSMUSG00000036461",
                "ENSMUSG00000001280",
                "ENSMUSG00000005583",
                "ENSMUSG00000030067",
                "ENSMUSG00000022641",
                "ENSMUSG00000071637",
                "ENSMUSG00000053110",
                "ENSMUSG00000022314",
                "ENSMUSG00000022528"
            ],
            "target": [
                "ENSMUSG00000051615",
                "ENSMUSG00000063320",
                "ENSMUSG00000002844",
                "ENSMUSG00000032737",
                "ENSMUSG00000029454",
                "ENSMUSG00000020841",
                "ENSMUSG00000015766",
                "ENSMUSG00000028643",
                "ENSMUSG00000001016",
                "ENSMUSG00000040147",
                "ENSMUSG00000047888",
                "ENSMUSG00000044702",
                "ENSMUSG00000022641",
                "ENSMUSG00000042901",
                "ENSMUSG00000054766",
                "ENSMUSG00000018819",
                "ENSMUSG00000021484",
                "ENSMUSG00000020077",
                "ENSMUSG00000029823",
                "ENSMUSG00000021453",
                "ENSMUSG00000026176",
                "ENSMUSG00000078247",
                "ENSMUSG00000042029",
                "ENSMUSG00000020493",
                "ENSMUSG00000043415",
                "ENSMUSG00000032423",
                "ENSMUSG00000060771",
                "ENSMUSG00000038628",
                "ENSMUSG00000021619",
                "ENSMUSG00000021400",
                "ENSMUSG00000027800",
                "ENSMUSG00000079184",
                "ENSMUSG00000025492",
                "ENSMUSG00000045098",
                "ENSMUSG00000064340",
                "ENSMUSG00000025077",
                "ENSMUSG00000052926",
                "ENSMUSG00000039219",
                "ENSMUSG00000092116",
                "ENSMUSG00000029576",
                "ENSMUSG00000017493",
                "ENSMUSG00000024422",
                "ENSMUSG00000024528",
                "ENSMUSG00000041685",
                "ENSMUSG00000008822",
                "ENSMUSG00000020430",
                "ENSMUSG00000029290",
                "ENSMUSG00000025959",
                "ENSMUSG00000042389",
                "ENSMUSG00000025534",
                "ENSMUSG00000090553",
                "ENSMUSG00000019951",
                "ENSMUSG00000036860",
                "ENSMUSG00000033813",
                "ENSMUSG00000022905",
                "ENSMUSG00000060802",
                "ENSMUSG00000038803",
                "ENSMUSG00000038671",
                "ENSMUSG00000028461",
                "ENSMUSG00000063316",
                "ENSMUSG00000004951",
                "ENSMUSG00000053317",
                "ENSMUSG00000034154",
                "ENSMUSG00000020076",
                "ENSMUSG00000071533",
                "ENSMUSG00000038967",
                "ENSMUSG00000042426",
                "ENSMUSG00000008429",
                "ENSMUSG00000018102",
                "ENSMUSG00000096472",
                "ENSMUSG00000020009",
                "ENSMUSG00000026755",
                "ENSMUSG00000018171",
                "ENSMUSG00000020248",
                "ENSMUSG00000021500",
                "ENSMUSG00000001930",
                "ENSMUSG00000070544",
                "ENSMUSG00000026365",
                "ENSMUSG00000003099",
                "ENSMUSG00000042506",
                "ENSMUSG00000045954",
                "ENSMUSG00000030067",
                "ENSMUSG00000003955",
                "ENSMUSG00000053838",
                "ENSMUSG00000014329",
                "ENSMUSG00000026202",
                "ENSMUSG00000025184",
                "ENSMUSG00000028107",
                "ENSMUSG00000022108",
                "ENSMUSG00000024309",
                "ENSMUSG00000020773",
                "ENSMUSG00000040990",
                "ENSMUSG00000026219",
                "ENSMUSG00000040033",
                "ENSMUSG00000019970",
                "ENSMUSG00000024878",
                "ENSMUSG00000033538",
                "ENSMUSG00000003200",
                "ENSMUSG00000025574",
                "ENSMUSG00000038780",
                "ENSMUSG00000030172",
                "ENSMUSG00000019494",
                "ENSMUSG00000036461",
                "ENSMUSG00000023010",
                "ENSMUSG00000009555",
                "ENSMUSG00000015575",
                "ENSMUSG00000005981",
                "ENSMUSG00000034109",
                "ENSMUSG00000003420",
                "ENSMUSG00000026615",
                "ENSMUSG00000030682",
                "ENSMUSG00000029135",
                "ENSMUSG00000022174",
                "ENSMUSG00000040124",
                "ENSMUSG00000022528",
                "ENSMUSG00000035266",
                "ENSMUSG00000073639",
                "ENSMUSG00000052534",
                "ENSMUSG00000023990",
                "ENSMUSG00000020074",
                "ENSMUSG00000027326",
                "ENSMUSG00000021917",
                "ENSMUSG00000059734",
                "ENSMUSG00000001289",
                "ENSMUSG00000032932",
                "ENSMUSG00000053110",
                "ENSMUSG00000022314",
                "ENSMUSG00000042524",
                "ENSMUSG00000001627",
                "ENSMUSG00000002031",
                "ENSMUSG00000022206",
                "ENSMUSG00000016344",
                "ENSMUSG00000024491",
                "ENSMUSG00000044475",
                "ENSMUSG00000053565",
                "ENSMUSG00000019868",
                "ENSMUSG00000022094",
                "ENSMUSG00000028248",
                "ENSMUSG00000020585",
                "ENSMUSG00000032265",
                "ENSMUSG00000002748",
                "ENSMUSG00000026766",
                "ENSMUSG00000001280",
                "ENSMUSG00000090266",
                "ENSMUSG00000026674",
                "ENSMUSG00000005078",
                "ENSMUSG00000028949",
                "ENSMUSG00000039660"
            ],
            "expression_sum": {
                "class1": 26199.0,
                "class2": 4655.0
            },
            "uniprot_ids": [
                "Q60722",
                "A0A0J9YTU7",
                "A0A0J9YUZ0",
                "Q91YV0",
                "A0A0J9YUG4",
                "Q8BPX7",
                "A0A0J9YUP9",
                "A0A0J9YVG8",
                "A0A0J9YTX3",
                "A0A1B0GXC8",
                "A0A140LJA2",
                "A0A0J9YUK1",
                "A0A0J9YUS2",
                "A0A0J9YVG2",
                "A0A0J9YU75",
                "A0A0J9YV39",
                "A0A140LIL4",
                "E9Q8G4",
                "A0A0J9YTR9",
                "A0A0J9YV54",
                "A0A0J9YU37",
                "A0A0J9YUT7",
                "A0A0J9YUX6",
                "A0A0J9YVC6",
                "A0A0J9YUF5",
                "A0A0J9YU89",
                "A0A0J9YUY3",
                "A0A0J9YVI1",
                "A0A0J9YU91",
                "A0A0J9YUT3",
                "A0A0J9YUW9",
                "A0A0J9YUR0",
                "A0A0J9YU99",
                "A0A0J9YV50",
                "A0A0J9YTW1",
                "A0A140LJC0",
                "A0A0J9YV88",
                "A0A0J9YVB3"
            ],
            "net_id": 0,
            "source_num": 16,
            "target_num": 26
        },
        "ENSMUSG00000008575": {
            "id": "ENSMUSG00000008575",
            "type": "TF",
            "symbol": "Nfib",
            "source": [
                "ENSMUSG00000057469",
                "ENSMUSG00000010175",
                "ENSMUSG00000001280",
                "ENSMUSG00000032238",
                "ENSMUSG00000005583",
                "ENSMUSG00000019768",
                "ENSMUSG00000022641",
                "ENSMUSG00000053110",
                "ENSMUSG00000036036"
            ],
            "target": [
                "ENSMUSG00000000538",
                "ENSMUSG00000039782",
                "ENSMUSG00000061904",
                "ENSMUSG00000009569",
                "ENSMUSG00000025747",
                "ENSMUSG00000029454",
                "ENSMUSG00000024948",
                "ENSMUSG00000034875",
                "ENSMUSG00000021939",
                "ENSMUSG00000031403",
                "ENSMUSG00000045838",
                "ENSMUSG00000024197",
                "ENSMUSG00000027777",
                "ENSMUSG00000022020",
                "ENSMUSG00000062908",
                "ENSMUSG00000007777",
                "ENSMUSG00000026491",
                "ENSMUSG00000068264",
                "ENSMUSG00000083985",
                "ENSMUSG00000020307",
                "ENSMUSG00000022462",
                "ENSMUSG00000043683",
                "ENSMUSG00000031885",
                "ENSMUSG00000029823",
                "ENSMUSG00000068040",
                "ENSMUSG00000070047",
                "ENSMUSG00000020741",
                "ENSMUSG00000076490",
                "ENSMUSG00000037266",
                "ENSMUSG00000064138",
                "ENSMUSG00000021235",
                "ENSMUSG00000034751",
                "ENSMUSG00000036585",
                "ENSMUSG00000070283",
                "ENSMUSG00000037339",
                "ENSMUSG00000039640",
                "ENSMUSG00000030982",
                "ENSMUSG00000027984",
                "ENSMUSG00000034903",
                "ENSMUSG00000027679",
                "ENSMUSG00000021681",
                "ENSMUSG00000030739",
                "ENSMUSG00000052397",
                "ENSMUSG00000042709",
                "ENSMUSG00000026576",
                "ENSMUSG00000041881",
                "ENSMUSG00000020402",
                "ENSMUSG00000028086",
                "ENSMUSG00000022217",
                "ENSMUSG00000026853",
                "ENSMUSG00000022354",
                "ENSMUSG00000044221",
                "ENSMUSG00000028959",
                "ENSMUSG00000021764",
                "ENSMUSG00000007891",
                "ENSMUSG00000023951",
                "ENSMUSG00000004789",
                "ENSMUSG00000005354",
                "ENSMUSG00000020163",
                "ENSMUSG00000013465",
                "ENSMUSG00000009863",
                "ENSMUSG00000024811",
                "ENSMUSG00000035158",
                "ENSMUSG00000030747",
                "ENSMUSG00000090553",
                "ENSMUSG00000020496",
                "ENSMUSG00000023087",
                "ENSMUSG00000086763",
                "ENSMUSG00000006576",
                "ENSMUSG00000064037",
                "ENSMUSG00000004085",
                "ENSMUSG00000004347",
                "ENSMUSG00000026803",
                "ENSMUSG00000027288",
                "ENSMUSG00000052539",
                "ENSMUSG00000056851",
                "ENSMUSG00000032314",
                "ENSMUSG00000073684",
                "ENSMUSG00000028062",
                "ENSMUSG00000029152",
                "ENSMUSG00000022519",
                "ENSMUSG00000016528",
                "ENSMUSG00000032238",
                "ENSMUSG00000038803",
                "ENSMUSG00000024038",
                "ENSMUSG00000025190",
                "ENSMUSG00000033065",
                "ENSMUSG00000062210",
                "ENSMUSG00000027160",
                "ENSMUSG00000042010",
                "ENSMUSG00000090150",
                "ENSMUSG00000038538",
                "ENSMUSG00000030850",
                "ENSMUSG00000079508",
                "ENSMUSG00000033184",
                "ENSMUSG00000014606",
                "ENSMUSG00000028760",
                "ENSMUSG00000030861",
                "ENSMUSG00000040661",
                "ENSMUSG00000020153",
                "ENSMUSG00000026260",
                "ENSMUSG00000053329",
                "ENSMUSG00000018102",
                "ENSMUSG00000034981",
                "ENSMUSG00000029499",
                "ENSMUSG00000020612",
                "ENSMUSG00000090066",
                "ENSMUSG00000048490",
                "ENSMUSG00000094664",
                "ENSMUSG00000024359",
                "ENSMUSG00000036452",
                "ENSMUSG00000031818",
                "ENSMUSG00000040883",
                "ENSMUSG00000064302",
                "ENSMUSG00000040666",
                "ENSMUSG00000025184",
                "ENSMUSG00000031782",
                "ENSMUSG00000048486",
                "ENSMUSG00000026775",
                "ENSMUSG00000026558",
                "ENSMUSG00000044365",
                "ENSMUSG00000000056",
                "ENSMUSG00000032435",
                "ENSMUSG00000044894",
                "ENSMUSG00000034006",
                "ENSMUSG00000048578",
                "ENSMUSG00000028954",
                "ENSMUSG00000020949",
                "ENSMUSG00000000399",
                "ENSMUSG00000039103",
                "ENSMUSG00000003865",
                "ENSMUSG00000014956",
                "ENSMUSG00000024539",
                "ENSMUSG00000022820",
                "ENSMUSG00000037234",
                "ENSMUSG00000044792",
                "ENSMUSG00000039976",
                "ENSMUSG00000002129",
                "ENSMUSG00000022437",
                "ENSMUSG00000042148",
                "ENSMUSG00000038708",
                "ENSMUSG00000041028",
                "ENSMUSG00000031783",
                "ENSMUSG00000039515",
                "ENSMUSG00000026112",
                "ENSMUSG00000021772",
                "ENSMUSG00000063229",
                "ENSMUSG00000014504",
                "ENSMUSG00000021577",
                "ENSMUSG00000022994",
                "ENSMUSG00000022037",
                "ENSMUSG00000059447",
                "ENSMUSG00000026615",
                "ENSMUSG00000033938",
                "ENSMUSG00000029776",
                "ENSMUSG00000031698",
                "ENSMUSG00000029135",
                "ENSMUSG00000022174",
                "ENSMUSG00000027602",
                "ENSMUSG00000029167",
                "ENSMUSG00000061286",
                "ENSMUSG00000025745",
                "ENSMUSG00000026849",
                "ENSMUSG00000039210",
                "ENSMUSG00000048546",
                "ENSMUSG00000070867",
                "ENSMUSG00000013419",
                "ENSMUSG00000038615",
                "ENSMUSG00000038417",
                "ENSMUSG00000046329",
                "ENSMUSG00000073639",
                "ENSMUSG00000023990",
                "ENSMUSG00000055322",
                "ENSMUSG00000006057",
                "ENSMUSG00000025290",
                "ENSMUSG00000021917",
                "ENSMUSG00000024942",
                "ENSMUSG00000040414",
                "ENSMUSG00000028443",
                "ENSMUSG00000016344",
                "ENSMUSG00000037049",
                "ENSMUSG00000025545",
                "ENSMUSG00000020079",
                "ENSMUSG00000061518",
                "ENSMUSG00000032536",
                "ENSMUSG00000033400",
                "ENSMUSG00000032480",
                "ENSMUSG00000033885",
                "ENSMUSG00000040351",
                "ENSMUSG00000019979",
                "ENSMUSG00000021549",
                "ENSMUSG00000031447",
                "ENSMUSG00000053929",
                "ENSMUSG00000031776",
                "ENSMUSG00000026568",
                "ENSMUSG00000026123",
                "ENSMUSG00000027452",
                "ENSMUSG00000059981",
                "ENSMUSG00000023020",
                "ENSMUSG00000031700",
                "ENSMUSG00000024033",
                "ENSMUSG00000046516",
                "ENSMUSG00000063236",
                "ENSMUSG00000058655",
                "ENSMUSG00000052934",
                "ENSMUSG00000022961",
                "ENSMUSG00000034424",
                "ENSMUSG00000029221",
                "ENSMUSG00000070493",
                "ENSMUSG00000030335",
                "ENSMUSG00000021748",
                "ENSMUSG00000047419",
                "ENSMUSG00000071369",
                "ENSMUSG00000029145",
                "ENSMUSG00000043415",
                "ENSMUSG00000006494",
                "ENSMUSG00000026456",
                "ENSMUSG00000004980",
                "ENSMUSG00000058076",
                "ENSMUSG00000025077",
                "ENSMUSG00000029594",
                "ENSMUSG00000071014",
                "ENSMUSG00000044864",
                "ENSMUSG00000018765",
                "ENSMUSG00000035992",
                "ENSMUSG00000073557",
                "ENSMUSG00000066637",
                "ENSMUSG00000068921",
                "ENSMUSG00000005973",
                "ENSMUSG00000000441",
                "ENSMUSG00000034220",
                "ENSMUSG00000024066",
                "ENSMUSG00000041685",
                "ENSMUSG00000025939",
                "ENSMUSG00000005610",
                "ENSMUSG00000033228",
                "ENSMUSG00000024661",
                "ENSMUSG00000053483",
                "ENSMUSG00000024695",
                "ENSMUSG00000031627",
                "ENSMUSG00000032826",
                "ENSMUSG00000057322",
                "ENSMUSG00000033059",
                "ENSMUSG00000073700",
                "ENSMUSG00000029059",
                "ENSMUSG00000033813",
                "ENSMUSG00000024513",
                "ENSMUSG00000025271",
                "ENSMUSG00000014313",
                "ENSMUSG00000000088",
                "ENSMUSG00000033594",
                "ENSMUSG00000025393",
                "ENSMUSG00000048279",
                "ENSMUSG00000047371",
                "ENSMUSG00000027665",
                "ENSMUSG00000031232",
                "ENSMUSG00000001419",
                "ENSMUSG00000022186",
                "ENSMUSG00000048814",
                "ENSMUSG00000021597",
                "ENSMUSG00000022765",
                "ENSMUSG00000038967",
                "ENSMUSG00000042426",
                "ENSMUSG00000008429",
                "ENSMUSG00000045624",
                "ENSMUSG00000024298",
                "ENSMUSG00000056091",
                "ENSMUSG00000071528",
                "ENSMUSG00000024588",
                "ENSMUSG00000068874",
                "ENSMUSG00000002416",
                "ENSMUSG00000031841",
                "ENSMUSG00000026571",
                "ENSMUSG00000036854",
                "ENSMUSG00000026131",
                "ENSMUSG00000026202",
                "ENSMUSG00000042605",
                "ENSMUSG00000028849",
                "ENSMUSG00000028107",
                "ENSMUSG00000034951",
                "ENSMUSG00000022114",
                "ENSMUSG00000028919",
                "ENSMUSG00000040033",
                "ENSMUSG00000033365",
                "ENSMUSG00000028124",
                "ENSMUSG00000025204",
                "ENSMUSG00000036867",
                "ENSMUSG00000023055",
                "ENSMUSG00000022477",
                "ENSMUSG00000032010",
                "ENSMUSG00000041278",
                "ENSMUSG00000006024",
                "ENSMUSG00000021838",
                "ENSMUSG00000030172",
                "ENSMUSG00000067279",
                "ENSMUSG00000036698",
                "ENSMUSG00000031774",
                "ENSMUSG00000034290",
                "ENSMUSG00000023150",
                "ENSMUSG00000027618",
                "ENSMUSG00000038718",
                "ENSMUSG00000031429",
                "ENSMUSG00000040506",
                "ENSMUSG00000019791",
                "ENSMUSG00000030695",
                "ENSMUSG00000063800",
                "ENSMUSG00000067925",
                "ENSMUSG00000025795",
                "ENSMUSG00000027332",
                "ENSMUSG00000015850",
                "ENSMUSG00000001366",
                "ENSMUSG00000025733",
                "ENSMUSG00000025813",
                "ENSMUSG00000002379",
                "ENSMUSG00000022110",
                "ENSMUSG00000020190",
                "ENSMUSG00000028648",
                "ENSMUSG00000097729",
                "ENSMUSG00000004356",
                "ENSMUSG00000037916",
                "ENSMUSG00000027177",
                "ENSMUSG00000039768",
                "ENSMUSG00000037306",
                "ENSMUSG00000050608",
                "ENSMUSG00000029156",
                "ENSMUSG00000078572",
                "ENSMUSG00000021706",
                "ENSMUSG00000036160",
                "ENSMUSG00000027206",
                "ENSMUSG00000019851",
                "ENSMUSG00000022016",
                "ENSMUSG00000074064",
                "ENSMUSG00000027805",
                "ENSMUSG00000040771",
                "ENSMUSG00000025428",
                "ENSMUSG00000027010",
                "ENSMUSG00000033161",
                "ENSMUSG00000058690",
                "ENSMUSG00000001627",
                "ENSMUSG00000003072",
                "ENSMUSG00000021687",
                "ENSMUSG00000024063",
                "ENSMUSG00000037349",
                "ENSMUSG00000039474",
                "ENSMUSG00000044475",
                "ENSMUSG00000021585",
                "ENSMUSG00000041426",
                "ENSMUSG00000020849",
                "ENSMUSG00000002748",
                "ENSMUSG00000021238",
                "ENSMUSG00000042744",
                "ENSMUSG00000001280",
                "ENSMUSG00000031987",
                "ENSMUSG00000030417",
                "ENSMUSG00000038005",
                "ENSMUSG00000038970",
                "ENSMUSG00000055737",
                "ENSMUSG00000063870",
                "ENSMUSG00000054400",
                "ENSMUSG00000031633",
                "ENSMUSG00000063694",
                "ENSMUSG00000037826",
                "ENSMUSG00000028013",
                "ENSMUSG00000023921",
                "ENSMUSG00000063320",
                "ENSMUSG00000097487",
                "ENSMUSG00000020368",
                "ENSMUSG00000053898",
                "ENSMUSG00000021906",
                "ENSMUSG00000060038",
                "ENSMUSG00000000751",
                "ENSMUSG00000083820",
                "ENSMUSG00000025221",
                "ENSMUSG00000029632",
                "ENSMUSG00000037815",
                "ENSMUSG00000027809",
                "ENSMUSG00000016252",
                "ENSMUSG00000025241",
                "ENSMUSG00000022641",
                "ENSMUSG00000054766",
                "ENSMUSG00000034342",
                "ENSMUSG00000038949",
                "ENSMUSG00000075054",
                "ENSMUSG00000018433",
                "ENSMUSG00000062981",
                "ENSMUSG00000031024",
                "ENSMUSG00000020456",
                "ENSMUSG00000045983",
                "ENSMUSG00000026342",
                "ENSMUSG00000038028",
                "ENSMUSG00000032481",
                "ENSMUSG00000063882",
                "ENSMUSG00000026027",
                "ENSMUSG00000027381",
                "ENSMUSG00000026028",
                "ENSMUSG00000039910",
                "ENSMUSG00000032187",
                "ENSMUSG00000028745",
                "ENSMUSG00000038886",
                "ENSMUSG00000020483",
                "ENSMUSG00000008822",
                "ENSMUSG00000006356",
                "ENSMUSG00000004069",
                "ENSMUSG00000032679",
                "ENSMUSG00000022051",
                "ENSMUSG00000006818",
                "ENSMUSG00000026860",
                "ENSMUSG00000024055",
                "ENSMUSG00000032959",
                "ENSMUSG00000031320",
                "ENSMUSG00000001506",
                "ENSMUSG00000058600",
                "ENSMUSG00000028692",
                "ENSMUSG00000040850",
                "ENSMUSG00000032224",
                "ENSMUSG00000021375",
                "ENSMUSG00000022131",
                "ENSMUSG00000001211",
                "ENSMUSG00000049643",
                "ENSMUSG00000031985",
                "ENSMUSG00000038375",
                "ENSMUSG00000022452",
                "ENSMUSG00000023175",
                "ENSMUSG00000039509",
                "ENSMUSG00000001228",
                "ENSMUSG00000027406",
                "ENSMUSG00000006931",
                "ENSMUSG00000075700",
                "ENSMUSG00000006728",
                "ENSMUSG00000059586",
                "ENSMUSG00000024687",
                "ENSMUSG00000027030",
                "ENSMUSG00000024143",
                "ENSMUSG00000078566",
                "ENSMUSG00000002102",
                "ENSMUSG00000098090",
                "ENSMUSG00000025374",
                "ENSMUSG00000021226",
                "ENSMUSG00000026927",
                "ENSMUSG00000020029",
                "ENSMUSG00000030934",
                "ENSMUSG00000002343",
                "ENSMUSG00000019577",
                "ENSMUSG00000021500",
                "ENSMUSG00000039652",
                "ENSMUSG00000002010",
                "ENSMUSG00000032590",
                "ENSMUSG00000070544",
                "ENSMUSG00000054889",
                "ENSMUSG00000046351",
                "ENSMUSG00000003955",
                "ENSMUSG00000053838",
                "ENSMUSG00000024953",
                "ENSMUSG00000058997",
                "ENSMUSG00000019179",
                "ENSMUSG00000028273",
                "ENSMUSG00000050199",
                "ENSMUSG00000051495",
                "ENSMUSG00000022964",
                "ENSMUSG00000008604",
                "ENSMUSG00000058594",
                "ENSMUSG00000032786",
                "ENSMUSG00000025856",
                "ENSMUSG00000025364",
                "ENSMUSG00000038612",
                "ENSMUSG00000068479",
                "ENSMUSG00000023010",
                "ENSMUSG00000028452",
                "ENSMUSG00000036278",
                "ENSMUSG00000022391",
                "ENSMUSG00000039987",
                "ENSMUSG00000041548",
                "ENSMUSG00000039100",
                "ENSMUSG00000062169",
                "ENSMUSG00000003316",
                "ENSMUSG00000018796",
                "ENSMUSG00000008690",
                "ENSMUSG00000044080",
                "ENSMUSG00000005510",
                "ENSMUSG00000046671",
                "ENSMUSG00000087679",
                "ENSMUSG00000026098",
                "ENSMUSG00000047459",
                "ENSMUSG00000032523",
                "ENSMUSG00000028834",
                "ENSMUSG00000038844",
                "ENSMUSG00000020074",
                "ENSMUSG00000059734",
                "ENSMUSG00000039536",
                "ENSMUSG00000053110",
                "ENSMUSG00000010608",
                "ENSMUSG00000027742",
                "ENSMUSG00000032279",
                "ENSMUSG00000023861",
                "ENSMUSG00000026664",
                "ENSMUSG00000025466",
                "ENSMUSG00000009630",
                "ENSMUSG00000022000",
                "ENSMUSG00000062070",
                "ENSMUSG00000021395",
                "ENSMUSG00000028419",
                "ENSMUSG00000032051",
                "ENSMUSG00000021520",
                "ENSMUSG00000038764",
                "ENSMUSG00000006717",
                "ENSMUSG00000034994",
                "ENSMUSG00000023971",
                "ENSMUSG00000020108",
                "ENSMUSG00000030541",
                "ENSMUSG00000031438",
                "ENSMUSG00000090266",
                "ENSMUSG00000022235",
                "ENSMUSG00000028633",
                "ENSMUSG00000030652",
                "ENSMUSG00000003438",
                "ENSMUSG00000027367",
                "ENSMUSG00000002844",
                "ENSMUSG00000003345",
                "ENSMUSG00000097277",
                "ENSMUSG00000028538",
                "ENSMUSG00000061666",
                "ENSMUSG00000020133",
                "ENSMUSG00000047388",
                "ENSMUSG00000026483",
                "ENSMUSG00000022300",
                "ENSMUSG00000041236",
                "ENSMUSG00000018599",
                "ENSMUSG00000024140",
                "ENSMUSG00000066324",
                "ENSMUSG00000034842",
                "ENSMUSG00000029190",
                "ENSMUSG00000027499",
                "ENSMUSG00000061306",
                "ENSMUSG00000037805",
                "ENSMUSG00000021265",
                "ENSMUSG00000007080",
                "ENSMUSG00000017686",
                "ENSMUSG00000049470",
                "ENSMUSG00000017119",
                "ENSMUSG00000078695",
                "ENSMUSG00000028911",
                "ENSMUSG00000021400",
                "ENSMUSG00000031701",
                "ENSMUSG00000020738",
                "ENSMUSG00000038807",
                "ENSMUSG00000029401",
                "ENSMUSG00000032047",
                "ENSMUSG00000029467",
                "ENSMUSG00000003534",
                "ENSMUSG00000025453",
                "ENSMUSG00000042133",
                "ENSMUSG00000020415",
                "ENSMUSG00000038022",
                "ENSMUSG00000044393",
                "ENSMUSG00000059741",
                "ENSMUSG00000014470",
                "ENSMUSG00000021606",
                "ENSMUSG00000002957",
                "ENSMUSG00000022193",
                "ENSMUSG00000011148",
                "ENSMUSG00000036002",
                "ENSMUSG00000062580",
                "ENSMUSG00000026817",
                "ENSMUSG00000009112",
                "ENSMUSG00000026864",
                "ENSMUSG00000059325",
                "ENSMUSG00000033416",
                "ENSMUSG00000023068",
                "ENSMUSG00000026341",
                "ENSMUSG00000028937",
                "ENSMUSG00000034566",
                "ENSMUSG00000057561",
                "ENSMUSG00000024059",
                "ENSMUSG00000045636",
                "ENSMUSG00000024985",
                "ENSMUSG00000026074",
                "ENSMUSG00000069763",
                "ENSMUSG00000022023",
                "ENSMUSG00000038170",
                "ENSMUSG00000021771",
                "ENSMUSG00000006315",
                "ENSMUSG00000031808",
                "ENSMUSG00000028461",
                "ENSMUSG00000042293",
                "ENSMUSG00000004951",
                "ENSMUSG00000049327",
                "ENSMUSG00000071533",
                "ENSMUSG00000029333",
                "ENSMUSG00000029432",
                "ENSMUSG00000043866",
                "ENSMUSG00000038520",
                "ENSMUSG00000022748",
                "ENSMUSG00000038058",
                "ENSMUSG00000019362",
                "ENSMUSG00000021975",
                "ENSMUSG00000019087",
                "ENSMUSG00000042506",
                "ENSMUSG00000027293",
                "ENSMUSG00000020988",
                "ENSMUSG00000023206",
                "ENSMUSG00000028518",
                "ENSMUSG00000019467",
                "ENSMUSG00000002768",
                "ENSMUSG00000054263",
                "ENSMUSG00000020530",
                "ENSMUSG00000008136",
                "ENSMUSG00000022364",
                "ENSMUSG00000022450",
                "ENSMUSG00000022111",
                "ENSMUSG00000073563",
                "ENSMUSG00000024805",
                "ENSMUSG00000032366",
                "ENSMUSG00000038102",
                "ENSMUSG00000037236",
                "ENSMUSG00000051391",
                "ENSMUSG00000032370",
                "ENSMUSG00000035310",
                "ENSMUSG00000028383",
                "ENSMUSG00000029655",
                "ENSMUSG00000022681",
                "ENSMUSG00000029433",
                "ENSMUSG00000036880",
                "ENSMUSG00000040520",
                "ENSMUSG00000053819",
                "ENSMUSG00000058756",
                "ENSMUSG00000041115",
                "ENSMUSG00000068699",
                "ENSMUSG00000030662",
                "ENSMUSG00000029571",
                "ENSMUSG00000025651",
                "ENSMUSG00000055670",
                "ENSMUSG00000028603",
                "ENSMUSG00000027765",
                "ENSMUSG00000022884",
                "ENSMUSG00000052812",
                "ENSMUSG00000004268",
                "ENSMUSG00000037685",
                "ENSMUSG00000026526",
                "ENSMUSG00000028772",
                "ENSMUSG00000024639",
                "ENSMUSG00000030847",
                "ENSMUSG00000024259",
                "ENSMUSG00000055204",
                "ENSMUSG00000058492",
                "ENSMUSG00000028145",
                "ENSMUSG00000054312",
                "ENSMUSG00000023089",
                "ENSMUSG00000046997",
                "ENSMUSG00000034088",
                "ENSMUSG00000061474",
                "ENSMUSG00000001911",
                "ENSMUSG00000034832",
                "ENSMUSG00000073676",
                "ENSMUSG00000032060",
                "ENSMUSG00000023456",
                "ENSMUSG00000024491",
                "ENSMUSG00000038462",
                "ENSMUSG00000022389",
                "ENSMUSG00000080921",
                "ENSMUSG00000033722",
                "ENSMUSG00000027673",
                "ENSMUSG00000018395",
                "ENSMUSG00000059811",
                "ENSMUSG00000013662",
                "ENSMUSG00000026914",
                "ENSMUSG00000030095",
                "ENSMUSG00000018770",
                "ENSMUSG00000026489",
                "ENSMUSG00000003068"
            ],
            "expression_sum": {
                "class1": 13763.0,
                "class2": 3754.0
            },
            "uniprot_ids": [
                "P97863",
                "I7HIP8",
                "A2BG75",
                "A2BG76",
                "A2BG77",
                "A2ADI3"
            ],
            "net_id": 0,
            "source_num": 9,
            "target_num": 211
        },
        "ENSMUSG00000035158": {
            "id": "ENSMUSG00000035158",
            "type": "TF",
            "symbol": "Mitf",
            "source": [
                "ENSMUSG00000058440",
                "ENSMUSG00000001419",
                "ENSMUSG00000015839",
                "ENSMUSG00000001280",
                "ENSMUSG00000019768",
                "ENSMUSG00000044167",
                "ENSMUSG00000075595",
                "ENSMUSG00000008575",
                "ENSMUSG00000039191"
            ],
            "target": [
                "ENSMUSG00000051223",
                "ENSMUSG00000027091",
                "ENSMUSG00000008682",
                "ENSMUSG00000029860",
                "ENSMUSG00000039782",
                "ENSMUSG00000025968",
                "ENSMUSG00000027367",
                "ENSMUSG00000020368",
                "ENSMUSG00000039473",
                "ENSMUSG00000009013",
                "ENSMUSG00000029454",
                "ENSMUSG00000020330",
                "ENSMUSG00000034875",
                "ENSMUSG00000000751",
                "ENSMUSG00000004937",
                "ENSMUSG00000037572",
                "ENSMUSG00000021000",
                "ENSMUSG00000042650",
                "ENSMUSG00000028081",
                "ENSMUSG00000033423",
                "ENSMUSG00000044308",
                "ENSMUSG00000020777",
                "ENSMUSG00000029050",
                "ENSMUSG00000068264",
                "ENSMUSG00000030793",
                "ENSMUSG00000070493",
                "ENSMUSG00000054766",
                "ENSMUSG00000014177",
                "ENSMUSG00000022247",
                "ENSMUSG00000038949",
                "ENSMUSG00000027447",
                "ENSMUSG00000029823",
                "ENSMUSG00000031533",
                "ENSMUSG00000018433",
                "ENSMUSG00000091086",
                "ENSMUSG00000015839",
                "ENSMUSG00000037958",
                "ENSMUSG00000039873",
                "ENSMUSG00000061306",
                "ENSMUSG00000061477",
                "ENSMUSG00000028639",
                "ENSMUSG00000065990",
                "ENSMUSG00000045983",
                "ENSMUSG00000020456",
                "ENSMUSG00000021102",
                "ENSMUSG00000049470",
                "ENSMUSG00000064342",
                "ENSMUSG00000048000",
                "ENSMUSG00000074466",
                "ENSMUSG00000001415",
                "ENSMUSG00000007613",
                "ENSMUSG00000058076",
                "ENSMUSG00000064340",
                "ENSMUSG00000027679",
                "ENSMUSG00000032047",
                "ENSMUSG00000032481",
                "ENSMUSG00000044864",
                "ENSMUSG00000035992",
                "ENSMUSG00000042133",
                "ENSMUSG00000032525",
                "ENSMUSG00000020818",
                "ENSMUSG00000021373",
                "ENSMUSG00000020402",
                "ENSMUSG00000028086",
                "ENSMUSG00000039910",
                "ENSMUSG00000029920",
                "ENSMUSG00000032352",
                "ENSMUSG00000028745",
                "ENSMUSG00000035152",
                "ENSMUSG00000001054",
                "ENSMUSG00000020691",
                "ENSMUSG00000024073",
                "ENSMUSG00000033228",
                "ENSMUSG00000036748",
                "ENSMUSG00000013465",
                "ENSMUSG00000026864",
                "ENSMUSG00000006818",
                "ENSMUSG00000019877",
                "ENSMUSG00000020572",
                "ENSMUSG00000039130",
                "ENSMUSG00000011257",
                "ENSMUSG00000031840",
                "ENSMUSG00000020496",
                "ENSMUSG00000032437",
                "ENSMUSG00000036777",
                "ENSMUSG00000022136",
                "ENSMUSG00000040850",
                "ENSMUSG00000046756",
                "ENSMUSG00000062270",
                "ENSMUSG00000015806",
                "ENSMUSG00000028937",
                "ENSMUSG00000025175",
                "ENSMUSG00000049643",
                "ENSMUSG00000027288",
                "ENSMUSG00000005299",
                "ENSMUSG00000031447",
                "ENSMUSG00000056851",
                "ENSMUSG00000029918",
                "ENSMUSG00000031681",
                "ENSMUSG00000018882",
                "ENSMUSG00000038121",
                "ENSMUSG00000030095",
                "ENSMUSG00000024038",
                "ENSMUSG00000027698",
                "ENSMUSG00000032615",
                "ENSMUSG00000021709",
                "ENSMUSG00000014592",
                "ENSMUSG00000042293",
                "ENSMUSG00000051232",
                "ENSMUSG00000060376",
                "ENSMUSG00000024991",
                "ENSMUSG00000001419",
                "ENSMUSG00000028651",
                "ENSMUSG00000036678",
                "ENSMUSG00000014856",
                "ENSMUSG00000029432",
                "ENSMUSG00000028161",
                "ENSMUSG00000038393",
                "ENSMUSG00000038967",
                "ENSMUSG00000008429",
                "ENSMUSG00000040661",
                "ENSMUSG00000047710",
                "ENSMUSG00000033429",
                "ENSMUSG00000030089",
                "ENSMUSG00000034708",
                "ENSMUSG00000070544",
                "ENSMUSG00000029486",
                "ENSMUSG00000053553",
                "ENSMUSG00000024953",
                "ENSMUSG00000028278",
                "ENSMUSG00000025465",
                "ENSMUSG00000042605",
                "ENSMUSG00000031782",
                "ENSMUSG00000025508",
                "ENSMUSG00000035765",
                "ENSMUSG00000020522",
                "ENSMUSG00000015149",
                "ENSMUSG00000022450",
                "ENSMUSG00000024683",
                "ENSMUSG00000073563",
                "ENSMUSG00000026885",
                "ENSMUSG00000027423",
                "ENSMUSG00000028124",
                "ENSMUSG00000024962",
                "ENSMUSG00000007097",
                "ENSMUSG00000029404",
                "ENSMUSG00000037236",
                "ENSMUSG00000000374",
                "ENSMUSG00000098178",
                "ENSMUSG00000031575",
                "ENSMUSG00000032370",
                "ENSMUSG00000022265",
                "ENSMUSG00000034059",
                "ENSMUSG00000021807",
                "ENSMUSG00000019988",
                "ENSMUSG00000025781",
                "ENSMUSG00000028156",
                "ENSMUSG00000053819",
                "ENSMUSG00000005981",
                "ENSMUSG00000023150",
                "ENSMUSG00000027618",
                "ENSMUSG00000018796",
                "ENSMUSG00000040506",
                "ENSMUSG00000019791",
                "ENSMUSG00000054843",
                "ENSMUSG00000022884",
                "ENSMUSG00000015733",
                "ENSMUSG00000020328",
                "ENSMUSG00000028479",
                "ENSMUSG00000024277",
                "ENSMUSG00000017404",
                "ENSMUSG00000024120",
                "ENSMUSG00000003680",
                "ENSMUSG00000053199",
                "ENSMUSG00000027177",
                "ENSMUSG00000048546",
                "ENSMUSG00000028035",
                "ENSMUSG00000026857",
                "ENSMUSG00000026098",
                "ENSMUSG00000030779",
                "ENSMUSG00000002949",
                "ENSMUSG00000025591",
                "ENSMUSG00000014418",
                "ENSMUSG00000028683",
                "ENSMUSG00000030814",
                "ENSMUSG00000027206",
                "ENSMUSG00000055204",
                "ENSMUSG00000031352",
                "ENSMUSG00000073838",
                "ENSMUSG00000020074",
                "ENSMUSG00000029455",
                "ENSMUSG00000074064",
                "ENSMUSG00000037062",
                "ENSMUSG00000060938",
                "ENSMUSG00000028207",
                "ENSMUSG00000046727",
                "ENSMUSG00000018196",
                "ENSMUSG00000042505",
                "ENSMUSG00000054312",
                "ENSMUSG00000029146",
                "ENSMUSG00000022369",
                "ENSMUSG00000036721",
                "ENSMUSG00000010608",
                "ENSMUSG00000030647",
                "ENSMUSG00000026150",
                "ENSMUSG00000034088",
                "ENSMUSG00000096232",
                "ENSMUSG00000003072",
                "ENSMUSG00000020225",
                "ENSMUSG00000051502",
                "ENSMUSG00000073676",
                "ENSMUSG00000091537",
                "ENSMUSG00000022797",
                "ENSMUSG00000033943",
                "ENSMUSG00000075595",
                "ENSMUSG00000020079",
                "ENSMUSG00000039630",
                "ENSMUSG00000009293",
                "ENSMUSG00000002748",
                "ENSMUSG00000042323",
                "ENSMUSG00000032480",
                "ENSMUSG00000044533",
                "ENSMUSG00000029452",
                "ENSMUSG00000021666",
                "ENSMUSG00000038005",
                "ENSMUSG00000031438",
                "ENSMUSG00000031696",
                "ENSMUSG00000038970",
                "ENSMUSG00000028847",
                "ENSMUSG00000007739",
                "ENSMUSG00000032902",
                "ENSMUSG00000021549"
            ],
            "expression_sum": {
                "class1": 377.0,
                "class2": 773.0
            },
            "uniprot_ids": [
                "Q08874",
                "E9PZ28",
                "D6RCW2",
                "A0A0N4SV79",
                "A0A5H1ZRM6"
            ],
            "net_id": 0,
            "source_num": 9,
            "target_num": 62
        },
        "ENSMUSG00000021944": {
            "id": "ENSMUSG00000021944",
            "type": "TF",
            "symbol": "Gata4",
            "source": [
                "ENSMUSG00000001280",
                "ENSMUSG00000019768",
                "ENSMUSG00000010175"
            ],
            "target": [
                "ENSMUSG00000028902",
                "ENSMUSG00000028013",
                "ENSMUSG00000005683",
                "ENSMUSG00000046709",
                "ENSMUSG00000067736",
                "ENSMUSG00000038696",
                "ENSMUSG00000064363",
                "ENSMUSG00000024197",
                "ENSMUSG00000027777",
                "ENSMUSG00000064370",
                "ENSMUSG00000033285",
                "ENSMUSG00000034684",
                "ENSMUSG00000035770",
                "ENSMUSG00000038384",
                "ENSMUSG00000071317",
                "ENSMUSG00000042686",
                "ENSMUSG00000019768",
                "ENSMUSG00000020821",
                "ENSMUSG00000027496",
                "ENSMUSG00000029221",
                "ENSMUSG00000022247",
                "ENSMUSG00000024816",
                "ENSMUSG00000020077",
                "ENSMUSG00000097583",
                "ENSMUSG00000091086",
                "ENSMUSG00000070283",
                "ENSMUSG00000063594",
                "ENSMUSG00000034614",
                "ENSMUSG00000030231",
                "ENSMUSG00000038239",
                "ENSMUSG00000064340",
                "ENSMUSG00000007815",
                "ENSMUSG00000031791",
                "ENSMUSG00000021681",
                "ENSMUSG00000073557",
                "ENSMUSG00000066637",
                "ENSMUSG00000072720",
                "ENSMUSG00000022354",
                "ENSMUSG00000023286",
                "ENSMUSG00000030255",
                "ENSMUSG00000020070",
                "ENSMUSG00000031543",
                "ENSMUSG00000024302",
                "ENSMUSG00000024661",
                "ENSMUSG00000039648",
                "ENSMUSG00000031137",
                "ENSMUSG00000064367",
                "ENSMUSG00000005533",
                "ENSMUSG00000006699",
                "ENSMUSG00000021930",
                "ENSMUSG00000023087",
                "ENSMUSG00000025144",
                "ENSMUSG00000075307",
                "ENSMUSG00000039943",
                "ENSMUSG00000064351",
                "ENSMUSG00000025271",
                "ENSMUSG00000054206",
                "ENSMUSG00000032431",
                "ENSMUSG00000050043",
                "ENSMUSG00000040007",
                "ENSMUSG00000025135",
                "ENSMUSG00000038204",
                "ENSMUSG00000022452",
                "ENSMUSG00000004994",
                "ENSMUSG00000034898",
                "ENSMUSG00000086863",
                "ENSMUSG00000031641",
                "ENSMUSG00000064339",
                "ENSMUSG00000024963",
                "ENSMUSG00000028121",
                "ENSMUSG00000063382",
                "ENSMUSG00000045273",
                "ENSMUSG00000028322",
                "ENSMUSG00000029486",
                "ENSMUSG00000026385",
                "ENSMUSG00000053931",
                "ENSMUSG00000020747",
                "ENSMUSG00000070469",
                "ENSMUSG00000026131",
                "ENSMUSG00000023467",
                "ENSMUSG00000028262",
                "ENSMUSG00000010175",
                "ENSMUSG00000022450",
                "ENSMUSG00000044365",
                "ENSMUSG00000020186",
                "ENSMUSG00000032786",
                "ENSMUSG00000007097",
                "ENSMUSG00000031672",
                "ENSMUSG00000041057",
                "ENSMUSG00000026154",
                "ENSMUSG00000005846",
                "ENSMUSG00000022853",
                "ENSMUSG00000026781",
                "ENSMUSG00000020592",
                "ENSMUSG00000022803",
                "ENSMUSG00000021910",
                "ENSMUSG00000067279",
                "ENSMUSG00000053819",
                "ENSMUSG00000052512",
                "ENSMUSG00000020053",
                "ENSMUSG00000026353",
                "ENSMUSG00000007850",
                "ENSMUSG00000064341",
                "ENSMUSG00000022037",
                "ENSMUSG00000029614",
                "ENSMUSG00000064372",
                "ENSMUSG00000032495",
                "ENSMUSG00000037685",
                "ENSMUSG00000035274",
                "ENSMUSG00000024077",
                "ENSMUSG00000028035",
                "ENSMUSG00000036186",
                "ENSMUSG00000020917",
                "ENSMUSG00000079677",
                "ENSMUSG00000046010",
                "ENSMUSG00000019132",
                "ENSMUSG00000000305",
                "ENSMUSG00000021622",
                "ENSMUSG00000019787",
                "ENSMUSG00000024335",
                "ENSMUSG00000028207",
                "ENSMUSG00000041671",
                "ENSMUSG00000049606",
                "ENSMUSG00000046997",
                "ENSMUSG00000039607",
                "ENSMUSG00000030246",
                "ENSMUSG00000065947",
                "ENSMUSG00000030884",
                "ENSMUSG00000051518",
                "ENSMUSG00000020664",
                "ENSMUSG00000030869",
                "ENSMUSG00000031767",
                "ENSMUSG00000060427",
                "ENSMUSG00000064371",
                "ENSMUSG00000020400",
                "ENSMUSG00000044857",
                "ENSMUSG00000021371",
                "ENSMUSG00000037904",
                "ENSMUSG00000064345",
                "ENSMUSG00000001280"
            ],
            "expression_sum": {
                "class1": 651.0,
                "class2": 333.0
            },
            "uniprot_ids": [
                "Q08369",
                "Q3UYJ1",
                "A4K4Z6",
                "E9PXW9",
                "D3YTQ2"
            ],
            "net_id": 0,
            "source_num": 3,
            "target_num": 46
        },
        "ENSMUSG00000010175": {
            "id": "ENSMUSG00000010175",
            "type": "TF",
            "symbol": "Prox1",
            "source": [
                "ENSMUSG00000021944",
                "ENSMUSG00000025612",
                "ENSMUSG00000015839",
                "ENSMUSG00000052534",
                "ENSMUSG00000051469",
                "ENSMUSG00000001280",
                "ENSMUSG00000019768",
                "ENSMUSG00000047371",
                "ENSMUSG00000053110"
            ],
            "target": [
                "ENSMUSG00000002985",
                "ENSMUSG00000037826",
                "ENSMUSG00000026568",
                "ENSMUSG00000029860",
                "ENSMUSG00000031448",
                "ENSMUSG00000061904",
                "ENSMUSG00000005683",
                "ENSMUSG00000087380",
                "ENSMUSG00000029447",
                "ENSMUSG00000020368",
                "ENSMUSG00000053898",
                "ENSMUSG00000034875",
                "ENSMUSG00000028737",
                "ENSMUSG00000060038",
                "ENSMUSG00000038696",
                "ENSMUSG00000021000",
                "ENSMUSG00000031700",
                "ENSMUSG00000040648",
                "ENSMUSG00000062908",
                "ENSMUSG00000020777",
                "ENSMUSG00000022332",
                "ENSMUSG00000029632",
                "ENSMUSG00000021147",
                "ENSMUSG00000002661",
                "ENSMUSG00000058655",
                "ENSMUSG00000017144",
                "ENSMUSG00000022300",
                "ENSMUSG00000019768",
                "ENSMUSG00000024331",
                "ENSMUSG00000029723",
                "ENSMUSG00000020307",
                "ENSMUSG00000042632",
                "ENSMUSG00000034424",
                "ENSMUSG00000015112",
                "ENSMUSG00000038949",
                "ENSMUSG00000029823",
                "ENSMUSG00000029095",
                "ENSMUSG00000017221",
                "ENSMUSG00000027433",
                "ENSMUSG00000029190",
                "ENSMUSG00000015839",
                "ENSMUSG00000020741",
                "ENSMUSG00000022889",
                "ENSMUSG00000037805",
                "ENSMUSG00000046201",
                "ENSMUSG00000060591",
                "ENSMUSG00000028223",
                "ENSMUSG00000062981",
                "ENSMUSG00000006494",
                "ENSMUSG00000032348",
                "ENSMUSG00000036585",
                "ENSMUSG00000042302",
                "ENSMUSG00000026342",
                "ENSMUSG00000074466",
                "ENSMUSG00000027984",
                "ENSMUSG00000025911",
                "ENSMUSG00000052384",
                "ENSMUSG00000069844",
                "ENSMUSG00000004980",
                "ENSMUSG00000027455",
                "ENSMUSG00000034245",
                "ENSMUSG00000038807",
                "ENSMUSG00000022040",
                "ENSMUSG00000029076",
                "ENSMUSG00000030470",
                "ENSMUSG00000063888",
                "ENSMUSG00000020402",
                "ENSMUSG00000001441",
                "ENSMUSG00000017493",
                "ENSMUSG00000026078",
                "ENSMUSG00000029580",
                "ENSMUSG00000021557",
                "ENSMUSG00000034880",
                "ENSMUSG00000000441",
                "ENSMUSG00000039910",
                "ENSMUSG00000041012",
                "ENSMUSG00000044393",
                "ENSMUSG00000056211",
                "ENSMUSG00000020483",
                "ENSMUSG00000003033",
                "ENSMUSG00000047067",
                "ENSMUSG00000014226",
                "ENSMUSG00000063450",
                "ENSMUSG00000033060",
                "ENSMUSG00000062580",
                "ENSMUSG00000030451",
                "ENSMUSG00000006356",
                "ENSMUSG00000022890",
                "ENSMUSG00000004069",
                "ENSMUSG00000031592",
                "ENSMUSG00000032554",
                "ENSMUSG00000032679",
                "ENSMUSG00000020572",
                "ENSMUSG00000039703",
                "ENSMUSG00000020733",
                "ENSMUSG00000023087",
                "ENSMUSG00000032959",
                "ENSMUSG00000033059",
                "ENSMUSG00000022270",
                "ENSMUSG00000069601",
                "ENSMUSG00000028270",
                "ENSMUSG00000031613",
                "ENSMUSG00000036860",
                "ENSMUSG00000004085",
                "ENSMUSG00000015806",
                "ENSMUSG00000021375",
                "ENSMUSG00000025271",
                "ENSMUSG00000027287",
                "ENSMUSG00000006476",
                "ENSMUSG00000029020",
                "ENSMUSG00000032330",
                "ENSMUSG00000056692",
                "ENSMUSG00000000088",
                "ENSMUSG00000045636",
                "ENSMUSG00000052539",
                "ENSMUSG00000056851",
                "ENSMUSG00000027875",
                "ENSMUSG00000038170",
                "ENSMUSG00000022519",
                "ENSMUSG00000029311",
                "ENSMUSG00000016528",
                "ENSMUSG00000031299",
                "ENSMUSG00000013593",
                "ENSMUSG00000030095",
                "ENSMUSG00000025190",
                "ENSMUSG00000021417",
                "ENSMUSG00000029048",
                "ENSMUSG00000030108",
                "ENSMUSG00000033065",
                "ENSMUSG00000001663",
                "ENSMUSG00000060376",
                "ENSMUSG00000001419",
                "ENSMUSG00000032578",
                "ENSMUSG00000049327",
                "ENSMUSG00000019122",
                "ENSMUSG00000030852",
                "ENSMUSG00000030861",
                "ENSMUSG00000038967",
                "ENSMUSG00000028756",
                "ENSMUSG00000056091",
                "ENSMUSG00000008429",
                "ENSMUSG00000078566",
                "ENSMUSG00000017485",
                "ENSMUSG00000019362",
                "ENSMUSG00000024963",
                "ENSMUSG00000028121",
                "ENSMUSG00000063382",
                "ENSMUSG00000086922",
                "ENSMUSG00000004460",
                "ENSMUSG00000060981",
                "ENSMUSG00000041351",
                "ENSMUSG00000019577",
                "ENSMUSG00000033429",
                "ENSMUSG00000039652",
                "ENSMUSG00000054889",
                "ENSMUSG00000020593",
                "ENSMUSG00000046351",
                "ENSMUSG00000039063",
                "ENSMUSG00000019960",
                "ENSMUSG00000062980",
                "ENSMUSG00000023206",
                "ENSMUSG00000024953",
                "ENSMUSG00000034574",
                "ENSMUSG00000025465",
                "ENSMUSG00000042605",
                "ENSMUSG00000028849",
                "ENSMUSG00000052406",
                "ENSMUSG00000019179",
                "ENSMUSG00000048486",
                "ENSMUSG00000003809",
                "ENSMUSG00000028273",
                "ENSMUSG00000051495",
                "ENSMUSG00000022964",
                "ENSMUSG00000026219",
                "ENSMUSG00000044894",
                "ENSMUSG00000022477",
                "ENSMUSG00000055053",
                "ENSMUSG00000033610",
                "ENSMUSG00000021134",
                "ENSMUSG00000075704",
                "ENSMUSG00000026029",
                "ENSMUSG00000022248",
                "ENSMUSG00000030172",
                "ENSMUSG00000060678",
                "ENSMUSG00000029162",
                "ENSMUSG00000067279",
                "ENSMUSG00000025509",
                "ENSMUSG00000041828",
                "ENSMUSG00000019988",
                "ENSMUSG00000028156",
                "ENSMUSG00000017756",
                "ENSMUSG00000041548",
                "ENSMUSG00000037110",
                "ENSMUSG00000050107",
                "ENSMUSG00000039515",
                "ENSMUSG00000025651",
                "ENSMUSG00000052146",
                "ENSMUSG00000023150",
                "ENSMUSG00000028603",
                "ENSMUSG00000018796",
                "ENSMUSG00000063229",
                "ENSMUSG00000024498",
                "ENSMUSG00000003420",
                "ENSMUSG00000044080",
                "ENSMUSG00000040740",
                "ENSMUSG00000018566",
                "ENSMUSG00000020321",
                "ENSMUSG00000027332",
                "ENSMUSG00000001366",
                "ENSMUSG00000005470",
                "ENSMUSG00000059447",
                "ENSMUSG00000029776",
                "ENSMUSG00000025813",
                "ENSMUSG00000022214",
                "ENSMUSG00000021236",
                "ENSMUSG00000021944",
                "ENSMUSG00000022110",
                "ENSMUSG00000024120",
                "ENSMUSG00000029135",
                "ENSMUSG00000032491",
                "ENSMUSG00000029167",
                "ENSMUSG00000022403",
                "ENSMUSG00000030002",
                "ENSMUSG00000029528",
                "ENSMUSG00000025085",
                "ENSMUSG00000026526",
                "ENSMUSG00000025745",
                "ENSMUSG00000036427",
                "ENSMUSG00000048546",
                "ENSMUSG00000004535",
                "ENSMUSG00000030847",
                "ENSMUSG00000024132",
                "ENSMUSG00000028398",
                "ENSMUSG00000028673",
                "ENSMUSG00000058709",
                "ENSMUSG00000070867",
                "ENSMUSG00000050608",
                "ENSMUSG00000090673",
                "ENSMUSG00000008575",
                "ENSMUSG00000031930",
                "ENSMUSG00000019851",
                "ENSMUSG00000055204",
                "ENSMUSG00000052534",
                "ENSMUSG00000023990",
                "ENSMUSG00000055322",
                "ENSMUSG00000029455",
                "ENSMUSG00000045776",
                "ENSMUSG00000020150",
                "ENSMUSG00000028207",
                "ENSMUSG00000059734",
                "ENSMUSG00000039536",
                "ENSMUSG00000049422",
                "ENSMUSG00000032724",
                "ENSMUSG00000058690",
                "ENSMUSG00000023089",
                "ENSMUSG00000038793",
                "ENSMUSG00000028906",
                "ENSMUSG00000010608",
                "ENSMUSG00000011382",
                "ENSMUSG00000047284",
                "ENSMUSG00000034088",
                "ENSMUSG00000040209",
                "ENSMUSG00000026664",
                "ENSMUSG00000008540",
                "ENSMUSG00000024236",
                "ENSMUSG00000039474",
                "ENSMUSG00000038370",
                "ENSMUSG00000022797",
                "ENSMUSG00000063856",
                "ENSMUSG00000092274",
                "ENSMUSG00000026888",
                "ENSMUSG00000038371",
                "ENSMUSG00000040524",
                "ENSMUSG00000032051",
                "ENSMUSG00000036570",
                "ENSMUSG00000026489",
                "ENSMUSG00000000168",
                "ENSMUSG00000034994",
                "ENSMUSG00000020108",
                "ENSMUSG00000052962",
                "ENSMUSG00000001280",
                "ENSMUSG00000026113",
                "ENSMUSG00000050541",
                "ENSMUSG00000024099",
                "ENSMUSG00000036611",
                "ENSMUSG00000022241",
                "ENSMUSG00000045639",
                "ENSMUSG00000028847",
                "ENSMUSG00000055737",
                "ENSMUSG00000025232",
                "ENSMUSG00000063694",
                "ENSMUSG00000028082"
            ],
            "expression_sum": {
                "class1": 359.0,
                "class2": 558.0
            },
            "uniprot_ids": [
                "P48437"
            ],
            "net_id": 0,
            "source_num": 9,
            "target_num": 87
        },
        "ENSMUSG00000029249": {
            "id": "ENSMUSG00000029249",
            "type": "TF",
            "symbol": "Rest",
            "source": [
                "ENSMUSG00000003032",
                "ENSMUSG00000015839",
                "ENSMUSG00000021250",
                "ENSMUSG00000026628",
                "ENSMUSG00000015579",
                "ENSMUSG00000027109",
                "ENSMUSG00000001280",
                "ENSMUSG00000005566",
                "ENSMUSG00000019768",
                "ENSMUSG00000052684",
                "ENSMUSG00000030087",
                "ENSMUSG00000024974",
                "ENSMUSG00000058756",
                "ENSMUSG00000042557",
                "ENSMUSG00000020644"
            ],
            "target": [
                "ENSMUSG00000015568",
                "ENSMUSG00000000538",
                "ENSMUSG00000023104",
                "ENSMUSG00000061904",
                "ENSMUSG00000009569",
                "ENSMUSG00000005566",
                "ENSMUSG00000027996",
                "ENSMUSG00000017009",
                "ENSMUSG00000021939",
                "ENSMUSG00000028104",
                "ENSMUSG00000031403",
                "ENSMUSG00000020485",
                "ENSMUSG00000024197",
                "ENSMUSG00000045838",
                "ENSMUSG00000059005",
                "ENSMUSG00000022020",
                "ENSMUSG00000062908",
                "ENSMUSG00000007777",
                "ENSMUSG00000040481",
                "ENSMUSG00000019432",
                "ENSMUSG00000020694",
                "ENSMUSG00000026491",
                "ENSMUSG00000068264",
                "ENSMUSG00000045679",
                "ENSMUSG00000027109",
                "ENSMUSG00000003444",
                "ENSMUSG00000022462",
                "ENSMUSG00000043683",
                "ENSMUSG00000038070",
                "ENSMUSG00000015837",
                "ENSMUSG00000021993",
                "ENSMUSG00000002028",
                "ENSMUSG00000021235",
                "ENSMUSG00000024283",
                "ENSMUSG00000037339",
                "ENSMUSG00000030598",
                "ENSMUSG00000030982",
                "ENSMUSG00000039640",
                "ENSMUSG00000027984",
                "ENSMUSG00000074698",
                "ENSMUSG00000034903",
                "ENSMUSG00000024487",
                "ENSMUSG00000021681",
                "ENSMUSG00000030739",
                "ENSMUSG00000042709",
                "ENSMUSG00000034297",
                "ENSMUSG00000024097",
                "ENSMUSG00000026576",
                "ENSMUSG00000031540",
                "ENSMUSG00000044477",
                "ENSMUSG00000026853",
                "ENSMUSG00000026626",
                "ENSMUSG00000024165",
                "ENSMUSG00000031543",
                "ENSMUSG00000097039",
                "ENSMUSG00000038763",
                "ENSMUSG00000004789",
                "ENSMUSG00000025337",
                "ENSMUSG00000005354",
                "ENSMUSG00000009863",
                "ENSMUSG00000024811",
                "ENSMUSG00000029198",
                "ENSMUSG00000030747",
                "ENSMUSG00000020496",
                "ENSMUSG00000023087",
                "ENSMUSG00000006395",
                "ENSMUSG00000031770",
                "ENSMUSG00000064037",
                "ENSMUSG00000004347",
                "ENSMUSG00000005469",
                "ENSMUSG00000063931",
                "ENSMUSG00000007656",
                "ENSMUSG00000026803",
                "ENSMUSG00000027288",
                "ENSMUSG00000073684",
                "ENSMUSG00000022519",
                "ENSMUSG00000024529",
                "ENSMUSG00000041084",
                "ENSMUSG00000036097",
                "ENSMUSG00000020661",
                "ENSMUSG00000057738",
                "ENSMUSG00000090150",
                "ENSMUSG00000042010",
                "ENSMUSG00000017428",
                "ENSMUSG00000038538",
                "ENSMUSG00000030850",
                "ENSMUSG00000023966",
                "ENSMUSG00000014606",
                "ENSMUSG00000010048",
                "ENSMUSG00000020153",
                "ENSMUSG00000026260",
                "ENSMUSG00000034981",
                "ENSMUSG00000032216",
                "ENSMUSG00000042557",
                "ENSMUSG00000020612",
                "ENSMUSG00000020280",
                "ENSMUSG00000029833",
                "ENSMUSG00000041731",
                "ENSMUSG00000034574",
                "ENSMUSG00000025184",
                "ENSMUSG00000031782",
                "ENSMUSG00000048486",
                "ENSMUSG00000041390",
                "ENSMUSG00000000056",
                "ENSMUSG00000034006",
                "ENSMUSG00000029404",
                "ENSMUSG00000048578",
                "ENSMUSG00000057133",
                "ENSMUSG00000028954",
                "ENSMUSG00000022095",
                "ENSMUSG00000000399",
                "ENSMUSG00000004931",
                "ENSMUSG00000017288",
                "ENSMUSG00000003865",
                "ENSMUSG00000024217",
                "ENSMUSG00000037234",
                "ENSMUSG00000044792",
                "ENSMUSG00000023004",
                "ENSMUSG00000002129",
                "ENSMUSG00000022437",
                "ENSMUSG00000042148",
                "ENSMUSG00000038708",
                "ENSMUSG00000041028",
                "ENSMUSG00000031783",
                "ENSMUSG00000022594",
                "ENSMUSG00000039741",
                "ENSMUSG00000022037",
                "ENSMUSG00000018566",
                "ENSMUSG00000034998",
                "ENSMUSG00000033938",
                "ENSMUSG00000055320",
                "ENSMUSG00000059447",
                "ENSMUSG00000029776",
                "ENSMUSG00000027602",
                "ENSMUSG00000025898",
                "ENSMUSG00000026473",
                "ENSMUSG00000061286",
                "ENSMUSG00000025745",
                "ENSMUSG00000026849",
                "ENSMUSG00000048546",
                "ENSMUSG00000020952",
                "ENSMUSG00000038417",
                "ENSMUSG00000046329",
                "ENSMUSG00000045210",
                "ENSMUSG00000019132",
                "ENSMUSG00000073639",
                "ENSMUSG00000025130",
                "ENSMUSG00000023990",
                "ENSMUSG00000055322",
                "ENSMUSG00000024942",
                "ENSMUSG00000028207",
                "ENSMUSG00000022698",
                "ENSMUSG00000040414",
                "ENSMUSG00000038525",
                "ENSMUSG00000001424",
                "ENSMUSG00000031527",
                "ENSMUSG00000028443",
                "ENSMUSG00000055720",
                "ENSMUSG00000036564",
                "ENSMUSG00000063904",
                "ENSMUSG00000020079",
                "ENSMUSG00000024150",
                "ENSMUSG00000052712",
                "ENSMUSG00000032480",
                "ENSMUSG00000029186",
                "ENSMUSG00000028884",
                "ENSMUSG00000032648",
                "ENSMUSG00000032383",
                "ENSMUSG00000030560",
                "ENSMUSG00000040351",
                "ENSMUSG00000031447",
                "ENSMUSG00000053929",
                "ENSMUSG00000031776",
                "ENSMUSG00000026568",
                "ENSMUSG00000026123",
                "ENSMUSG00000005683",
                "ENSMUSG00000028960",
                "ENSMUSG00000059981",
                "ENSMUSG00000021127",
                "ENSMUSG00000046516",
                "ENSMUSG00000027509",
                "ENSMUSG00000058655",
                "ENSMUSG00000052934",
                "ENSMUSG00000039239",
                "ENSMUSG00000019768",
                "ENSMUSG00000060002",
                "ENSMUSG00000071793",
                "ENSMUSG00000058135",
                "ENSMUSG00000029221",
                "ENSMUSG00000047419",
                "ENSMUSG00000021748",
                "ENSMUSG00000064366",
                "ENSMUSG00000019558",
                "ENSMUSG00000041773",
                "ENSMUSG00000032348",
                "ENSMUSG00000023832",
                "ENSMUSG00000026456",
                "ENSMUSG00000029913",
                "ENSMUSG00000058076",
                "ENSMUSG00000028165",
                "ENSMUSG00000029594",
                "ENSMUSG00000044864",
                "ENSMUSG00000027184",
                "ENSMUSG00000066637",
                "ENSMUSG00000025666",
                "ENSMUSG00000000441",
                "ENSMUSG00000031516",
                "ENSMUSG00000037465",
                "ENSMUSG00000033060",
                "ENSMUSG00000038784",
                "ENSMUSG00000005610",
                "ENSMUSG00000036748",
                "ENSMUSG00000053483",
                "ENSMUSG00000024695",
                "ENSMUSG00000025156",
                "ENSMUSG00000002608",
                "ENSMUSG00000020709",
                "ENSMUSG00000026701",
                "ENSMUSG00000022270",
                "ENSMUSG00000033059",
                "ENSMUSG00000028266",
                "ENSMUSG00000022185",
                "ENSMUSG00000033594",
                "ENSMUSG00000048154",
                "ENSMUSG00000027668",
                "ENSMUSG00000009995",
                "ENSMUSG00000055762",
                "ENSMUSG00000048279",
                "ENSMUSG00000037568",
                "ENSMUSG00000017291",
                "ENSMUSG00000001663",
                "ENSMUSG00000031232",
                "ENSMUSG00000022186",
                "ENSMUSG00000026930",
                "ENSMUSG00000048814",
                "ENSMUSG00000031641",
                "ENSMUSG00000021597",
                "ENSMUSG00000022765",
                "ENSMUSG00000038967",
                "ENSMUSG00000024845",
                "ENSMUSG00000042520",
                "ENSMUSG00000045624",
                "ENSMUSG00000056091",
                "ENSMUSG00000047260",
                "ENSMUSG00000024588",
                "ENSMUSG00000042198",
                "ENSMUSG00000009927",
                "ENSMUSG00000002416",
                "ENSMUSG00000026571",
                "ENSMUSG00000036854",
                "ENSMUSG00000026131",
                "ENSMUSG00000026202",
                "ENSMUSG00000027993",
                "ENSMUSG00000034951",
                "ENSMUSG00000025791",
                "ENSMUSG00000022114",
                "ENSMUSG00000028919",
                "ENSMUSG00000025010",
                "ENSMUSG00000040033",
                "ENSMUSG00000027423",
                "ENSMUSG00000031751",
                "ENSMUSG00000038497",
                "ENSMUSG00000021460",
                "ENSMUSG00000023055",
                "ENSMUSG00000022477",
                "ENSMUSG00000036867",
                "ENSMUSG00000032010",
                "ENSMUSG00000031207",
                "ENSMUSG00000068614",
                "ENSMUSG00000031672",
                "ENSMUSG00000041278",
                "ENSMUSG00000044786",
                "ENSMUSG00000006024",
                "ENSMUSG00000030172",
                "ENSMUSG00000022682",
                "ENSMUSG00000025348",
                "ENSMUSG00000036698",
                "ENSMUSG00000024780",
                "ENSMUSG00000017756",
                "ENSMUSG00000059291",
                "ENSMUSG00000034290",
                "ENSMUSG00000023150",
                "ENSMUSG00000020647",
                "ENSMUSG00000012405",
                "ENSMUSG00000029110",
                "ENSMUSG00000038718",
                "ENSMUSG00000040506",
                "ENSMUSG00000063800",
                "ENSMUSG00000030695",
                "ENSMUSG00000030298",
                "ENSMUSG00000013539",
                "ENSMUSG00000001366",
                "ENSMUSG00000015850",
                "ENSMUSG00000027332",
                "ENSMUSG00000060681",
                "ENSMUSG00000097729",
                "ENSMUSG00000022214",
                "ENSMUSG00000020190",
                "ENSMUSG00000053286",
                "ENSMUSG00000004356",
                "ENSMUSG00000037916",
                "ENSMUSG00000039046",
                "ENSMUSG00000050608",
                "ENSMUSG00000078572",
                "ENSMUSG00000027006",
                "ENSMUSG00000036160",
                "ENSMUSG00000040321",
                "ENSMUSG00000025437",
                "ENSMUSG00000059208",
                "ENSMUSG00000015289",
                "ENSMUSG00000074064",
                "ENSMUSG00000040771",
                "ENSMUSG00000025428",
                "ENSMUSG00000027010",
                "ENSMUSG00000032724",
                "ENSMUSG00000033161",
                "ENSMUSG00000059852",
                "ENSMUSG00000003072",
                "ENSMUSG00000024063",
                "ENSMUSG00000037349",
                "ENSMUSG00000039474",
                "ENSMUSG00000044475",
                "ENSMUSG00000021585",
                "ENSMUSG00000030016",
                "ENSMUSG00000013921",
                "ENSMUSG00000041426",
                "ENSMUSG00000029128",
                "ENSMUSG00000028343",
                "ENSMUSG00000031634",
                "ENSMUSG00000031548",
                "ENSMUSG00000045594",
                "ENSMUSG00000000168",
                "ENSMUSG00000024924",
                "ENSMUSG00000042744",
                "ENSMUSG00000001280",
                "ENSMUSG00000038005",
                "ENSMUSG00000030417",
                "ENSMUSG00000052684",
                "ENSMUSG00000018442",
                "ENSMUSG00000063870",
                "ENSMUSG00000032902",
                "ENSMUSG00000031633",
                "ENSMUSG00000041235",
                "ENSMUSG00000055302",
                "ENSMUSG00000037826",
                "ENSMUSG00000029651",
                "ENSMUSG00000039089",
                "ENSMUSG00000020368",
                "ENSMUSG00000060038",
                "ENSMUSG00000021906",
                "ENSMUSG00000042772",
                "ENSMUSG00000040648",
                "ENSMUSG00000059456",
                "ENSMUSG00000068735",
                "ENSMUSG00000039220",
                "ENSMUSG00000038172",
                "ENSMUSG00000021614",
                "ENSMUSG00000015112",
                "ENSMUSG00000034342",
                "ENSMUSG00000029176",
                "ENSMUSG00000038949",
                "ENSMUSG00000075054",
                "ENSMUSG00000021453",
                "ENSMUSG00000031367",
                "ENSMUSG00000031024",
                "ENSMUSG00000020456",
                "ENSMUSG00000045983",
                "ENSMUSG00000051747",
                "ENSMUSG00000027087",
                "ENSMUSG00000001794",
                "ENSMUSG00000030316",
                "ENSMUSG00000038028",
                "ENSMUSG00000031791",
                "ENSMUSG00000032481",
                "ENSMUSG00000022040",
                "ENSMUSG00000063882",
                "ENSMUSG00000003531",
                "ENSMUSG00000019773",
                "ENSMUSG00000028871",
                "ENSMUSG00000071644",
                "ENSMUSG00000026027",
                "ENSMUSG00000055239",
                "ENSMUSG00000035203",
                "ENSMUSG00000027210",
                "ENSMUSG00000026028",
                "ENSMUSG00000028745",
                "ENSMUSG00000038886",
                "ENSMUSG00000020483",
                "ENSMUSG00000029094",
                "ENSMUSG00000031360",
                "ENSMUSG00000020267",
                "ENSMUSG00000035960",
                "ENSMUSG00000040723",
                "ENSMUSG00000019810",
                "ENSMUSG00000024055",
                "ENSMUSG00000029518",
                "ENSMUSG00000031320",
                "ENSMUSG00000032199",
                "ENSMUSG00000058600",
                "ENSMUSG00000040850",
                "ENSMUSG00000046756",
                "ENSMUSG00000032224",
                "ENSMUSG00000028173",
                "ENSMUSG00000039221",
                "ENSMUSG00000058793",
                "ENSMUSG00000015961",
                "ENSMUSG00000049643",
                "ENSMUSG00000033039",
                "ENSMUSG00000031985",
                "ENSMUSG00000038375",
                "ENSMUSG00000022452",
                "ENSMUSG00000023175",
                "ENSMUSG00000039509",
                "ENSMUSG00000009907",
                "ENSMUSG00000032519",
                "ENSMUSG00000060600",
                "ENSMUSG00000026283",
                "ENSMUSG00000027406",
                "ENSMUSG00000056608",
                "ENSMUSG00000015176",
                "ENSMUSG00000024164",
                "ENSMUSG00000060032",
                "ENSMUSG00000026628",
                "ENSMUSG00000059586",
                "ENSMUSG00000028756",
                "ENSMUSG00000022201",
                "ENSMUSG00000027030",
                "ENSMUSG00000078566",
                "ENSMUSG00000000811",
                "ENSMUSG00000025374",
                "ENSMUSG00000059974",
                "ENSMUSG00000030934",
                "ENSMUSG00000040254",
                "ENSMUSG00000002343",
                "ENSMUSG00000019577",
                "ENSMUSG00000002010",
                "ENSMUSG00000024613",
                "ENSMUSG00000098107",
                "ENSMUSG00000032590",
                "ENSMUSG00000046351",
                "ENSMUSG00000004317",
                "ENSMUSG00000036432",
                "ENSMUSG00000058997",
                "ENSMUSG00000019998",
                "ENSMUSG00000028273",
                "ENSMUSG00000018474",
                "ENSMUSG00000050199",
                "ENSMUSG00000025362",
                "ENSMUSG00000021200",
                "ENSMUSG00000051495",
                "ENSMUSG00000079477",
                "ENSMUSG00000014245",
                "ENSMUSG00000022964",
                "ENSMUSG00000058594",
                "ENSMUSG00000042682",
                "ENSMUSG00000032786",
                "ENSMUSG00000082687",
                "ENSMUSG00000025856",
                "ENSMUSG00000040028",
                "ENSMUSG00000020027",
                "ENSMUSG00000050323",
                "ENSMUSG00000049728",
                "ENSMUSG00000038612",
                "ENSMUSG00000021868",
                "ENSMUSG00000068479",
                "ENSMUSG00000022803",
                "ENSMUSG00000068732",
                "ENSMUSG00000055723",
                "ENSMUSG00000064338",
                "ENSMUSG00000025781",
                "ENSMUSG00000028452",
                "ENSMUSG00000039987",
                "ENSMUSG00000041548",
                "ENSMUSG00000028788",
                "ENSMUSG00000070802",
                "ENSMUSG00000033781",
                "ENSMUSG00000003316",
                "ENSMUSG00000033488",
                "ENSMUSG00000018796",
                "ENSMUSG00000044080",
                "ENSMUSG00000021186",
                "ENSMUSG00000036202",
                "ENSMUSG00000020232",
                "ENSMUSG00000046671",
                "ENSMUSG00000087679",
                "ENSMUSG00000005803",
                "ENSMUSG00000032523",
                "ENSMUSG00000044224",
                "ENSMUSG00000060679",
                "ENSMUSG00000032178",
                "ENSMUSG00000073838",
                "ENSMUSG00000038844",
                "ENSMUSG00000041870",
                "ENSMUSG00000005102",
                "ENSMUSG00000034320",
                "ENSMUSG00000025381",
                "ENSMUSG00000010608",
                "ENSMUSG00000019428",
                "ENSMUSG00000030216",
                "ENSMUSG00000003934",
                "ENSMUSG00000030884",
                "ENSMUSG00000033760",
                "ENSMUSG00000023025",
                "ENSMUSG00000062070",
                "ENSMUSG00000021395",
                "ENSMUSG00000032051",
                "ENSMUSG00000034994",
                "ENSMUSG00000022553",
                "ENSMUSG00000030541",
                "ENSMUSG00000091898",
                "ENSMUSG00000026208",
                "ENSMUSG00000009079",
                "ENSMUSG00000022235",
                "ENSMUSG00000028633",
                "ENSMUSG00000030714",
                "ENSMUSG00000046314",
                "ENSMUSG00000030652",
                "ENSMUSG00000028394",
                "ENSMUSG00000003438",
                "ENSMUSG00000032388",
                "ENSMUSG00000015981",
                "ENSMUSG00000030787",
                "ENSMUSG00000037300",
                "ENSMUSG00000097277",
                "ENSMUSG00000028538",
                "ENSMUSG00000027238",
                "ENSMUSG00000020133",
                "ENSMUSG00000061666",
                "ENSMUSG00000020777",
                "ENSMUSG00000053536",
                "ENSMUSG00000022300",
                "ENSMUSG00000018599",
                "ENSMUSG00000024974",
                "ENSMUSG00000085636",
                "ENSMUSG00000029190",
                "ENSMUSG00000015839",
                "ENSMUSG00000027499",
                "ENSMUSG00000027784",
                "ENSMUSG00000021265",
                "ENSMUSG00000015944",
                "ENSMUSG00000037573",
                "ENSMUSG00000000827",
                "ENSMUSG00000026000",
                "ENSMUSG00000028911",
                "ENSMUSG00000031701",
                "ENSMUSG00000007613",
                "ENSMUSG00000027455",
                "ENSMUSG00000064340",
                "ENSMUSG00000038807",
                "ENSMUSG00000029401",
                "ENSMUSG00000032047",
                "ENSMUSG00000029467",
                "ENSMUSG00000025453",
                "ENSMUSG00000074247",
                "ENSMUSG00000086841",
                "ENSMUSG00000069089",
                "ENSMUSG00000020415",
                "ENSMUSG00000038022",
                "ENSMUSG00000072720",
                "ENSMUSG00000006800",
                "ENSMUSG00000014470",
                "ENSMUSG00000074918",
                "ENSMUSG00000021606",
                "ENSMUSG00000002957",
                "ENSMUSG00000021250",
                "ENSMUSG00000028282",
                "ENSMUSG00000036002",
                "ENSMUSG00000030451",
                "ENSMUSG00000022031",
                "ENSMUSG00000009112",
                "ENSMUSG00000026187",
                "ENSMUSG00000033416",
                "ENSMUSG00000021039",
                "ENSMUSG00000021218",
                "ENSMUSG00000016308",
                "ENSMUSG00000027180",
                "ENSMUSG00000028134",
                "ENSMUSG00000031078",
                "ENSMUSG00000028937",
                "ENSMUSG00000015357",
                "ENSMUSG00000026739",
                "ENSMUSG00000020644",
                "ENSMUSG00000024985",
                "ENSMUSG00000026074",
                "ENSMUSG00000045636",
                "ENSMUSG00000031843",
                "ENSMUSG00000007041",
                "ENSMUSG00000018882",
                "ENSMUSG00000038170",
                "ENSMUSG00000000787",
                "ENSMUSG00000021771",
                "ENSMUSG00000068267",
                "ENSMUSG00000028461",
                "ENSMUSG00000042293",
                "ENSMUSG00000015214",
                "ENSMUSG00000041361",
                "ENSMUSG00000022463",
                "ENSMUSG00000049327",
                "ENSMUSG00000095403",
                "ENSMUSG00000024187",
                "ENSMUSG00000043866",
                "ENSMUSG00000037475",
                "ENSMUSG00000038520",
                "ENSMUSG00000019362",
                "ENSMUSG00000029004",
                "ENSMUSG00000027293",
                "ENSMUSG00000020988",
                "ENSMUSG00000005893",
                "ENSMUSG00000039234",
                "ENSMUSG00000020530",
                "ENSMUSG00000039670",
                "ENSMUSG00000022111",
                "ENSMUSG00000073563",
                "ENSMUSG00000024805",
                "ENSMUSG00000051391",
                "ENSMUSG00000015932",
                "ENSMUSG00000021910",
                "ENSMUSG00000048612",
                "ENSMUSG00000029655",
                "ENSMUSG00000020358",
                "ENSMUSG00000022681",
                "ENSMUSG00000036880",
                "ENSMUSG00000053819",
                "ENSMUSG00000058756",
                "ENSMUSG00000029201",
                "ENSMUSG00000003970",
                "ENSMUSG00000022551",
                "ENSMUSG00000030662",
                "ENSMUSG00000037110",
                "ENSMUSG00000005779",
                "ENSMUSG00000025651",
                "ENSMUSG00000033102",
                "ENSMUSG00000078671",
                "ENSMUSG00000055670",
                "ENSMUSG00000028603",
                "ENSMUSG00000007850",
                "ENSMUSG00000037486",
                "ENSMUSG00000027765",
                "ENSMUSG00000020308",
                "ENSMUSG00000035027",
                "ENSMUSG00000052812",
                "ENSMUSG00000064141",
                "ENSMUSG00000065979",
                "ENSMUSG00000024639",
                "ENSMUSG00000030847",
                "ENSMUSG00000021621",
                "ENSMUSG00000042115",
                "ENSMUSG00000032042",
                "ENSMUSG00000021622",
                "ENSMUSG00000032244",
                "ENSMUSG00000046997",
                "ENSMUSG00000030246",
                "ENSMUSG00000034088",
                "ENSMUSG00000061474",
                "ENSMUSG00000078851",
                "ENSMUSG00000001911",
                "ENSMUSG00000066861",
                "ENSMUSG00000010914",
                "ENSMUSG00000038462",
                "ENSMUSG00000020664",
                "ENSMUSG00000022389",
                "ENSMUSG00000064371",
                "ENSMUSG00000033722",
                "ENSMUSG00000017817",
                "ENSMUSG00000034936",
                "ENSMUSG00000040118",
                "ENSMUSG00000018395",
                "ENSMUSG00000074994",
                "ENSMUSG00000059811",
                "ENSMUSG00000022241",
                "ENSMUSG00000024587",
                "ENSMUSG00000024121",
                "ENSMUSG00000026489"
            ],
            "expression_sum": {
                "class1": 1514.0,
                "class2": 476.0
            },
            "uniprot_ids": [
                "Q8VIG1"
            ],
            "net_id": 0,
            "source_num": 15,
            "target_num": 147
        },
        "ENSMUSG00000005566": {
            "id": "ENSMUSG00000005566",
            "type": "TF",
            "symbol": "Trim28",
            "source": [
                "ENSMUSG00000057469",
                "ENSMUSG00000015839",
                "ENSMUSG00000029249",
                "ENSMUSG00000019768",
                "ENSMUSG00000031885",
                "ENSMUSG00000029913"
            ],
            "target": [
                "ENSMUSG00000061904",
                "ENSMUSG00000025747",
                "ENSMUSG00000020841",
                "ENSMUSG00000021939",
                "ENSMUSG00000028104",
                "ENSMUSG00000031403",
                "ENSMUSG00000059005",
                "ENSMUSG00000022020",
                "ENSMUSG00000062908",
                "ENSMUSG00000007777",
                "ENSMUSG00000040481",
                "ENSMUSG00000020694",
                "ENSMUSG00000002661",
                "ENSMUSG00000031539",
                "ENSMUSG00000045679",
                "ENSMUSG00000068264",
                "ENSMUSG00000020099",
                "ENSMUSG00000022462",
                "ENSMUSG00000031885",
                "ENSMUSG00000047146",
                "ENSMUSG00000019066",
                "ENSMUSG00000020741",
                "ENSMUSG00000097414",
                "ENSMUSG00000045414",
                "ENSMUSG00000036585",
                "ENSMUSG00000070283",
                "ENSMUSG00000059647",
                "ENSMUSG00000030982",
                "ENSMUSG00000034903",
                "ENSMUSG00000021681",
                "ENSMUSG00000030739",
                "ENSMUSG00000042709",
                "ENSMUSG00000041779",
                "ENSMUSG00000026576",
                "ENSMUSG00000020402",
                "ENSMUSG00000026853",
                "ENSMUSG00000015749",
                "ENSMUSG00000026626",
                "ENSMUSG00000028959",
                "ENSMUSG00000031543",
                "ENSMUSG00000007891",
                "ENSMUSG00000038763",
                "ENSMUSG00000006301",
                "ENSMUSG00000043445",
                "ENSMUSG00000005354",
                "ENSMUSG00000027394",
                "ENSMUSG00000030747",
                "ENSMUSG00000020496",
                "ENSMUSG00000031770",
                "ENSMUSG00000006576",
                "ENSMUSG00000026803",
                "ENSMUSG00000027288",
                "ENSMUSG00000073684",
                "ENSMUSG00000032314",
                "ENSMUSG00000056900",
                "ENSMUSG00000078773",
                "ENSMUSG00000038872",
                "ENSMUSG00000033065",
                "ENSMUSG00000057738",
                "ENSMUSG00000090150",
                "ENSMUSG00000027165",
                "ENSMUSG00000030850",
                "ENSMUSG00000038116",
                "ENSMUSG00000010048",
                "ENSMUSG00000026260",
                "ENSMUSG00000039891",
                "ENSMUSG00000018446",
                "ENSMUSG00000040760",
                "ENSMUSG00000024359",
                "ENSMUSG00000036452",
                "ENSMUSG00000015714",
                "ENSMUSG00000029833",
                "ENSMUSG00000064302",
                "ENSMUSG00000034574",
                "ENSMUSG00000025184",
                "ENSMUSG00000029122",
                "ENSMUSG00000031782",
                "ENSMUSG00000048486",
                "ENSMUSG00000036825",
                "ENSMUSG00000026558",
                "ENSMUSG00000044365",
                "ENSMUSG00000000056",
                "ENSMUSG00000029174",
                "ENSMUSG00000037353",
                "ENSMUSG00000029404",
                "ENSMUSG00000000374",
                "ENSMUSG00000028954",
                "ENSMUSG00000046679",
                "ENSMUSG00000004931",
                "ENSMUSG00000039103",
                "ENSMUSG00000003865",
                "ENSMUSG00000037234",
                "ENSMUSG00000022437",
                "ENSMUSG00000041028",
                "ENSMUSG00000032221",
                "ENSMUSG00000063229",
                "ENSMUSG00000039741",
                "ENSMUSG00000022994",
                "ENSMUSG00000038936",
                "ENSMUSG00000022037",
                "ENSMUSG00000020321",
                "ENSMUSG00000029217",
                "ENSMUSG00000029776",
                "ENSMUSG00000039828",
                "ENSMUSG00000061286",
                "ENSMUSG00000025745",
                "ENSMUSG00000048546",
                "ENSMUSG00000020952",
                "ENSMUSG00000093846",
                "ENSMUSG00000024304",
                "ENSMUSG00000026944",
                "ENSMUSG00000006219",
                "ENSMUSG00000023990",
                "ENSMUSG00000055322",
                "ENSMUSG00000022698",
                "ENSMUSG00000025104",
                "ENSMUSG00000040414",
                "ENSMUSG00000022724",
                "ENSMUSG00000010663",
                "ENSMUSG00000055720",
                "ENSMUSG00000036564",
                "ENSMUSG00000026888",
                "ENSMUSG00000054452",
                "ENSMUSG00000033400",
                "ENSMUSG00000052712",
                "ENSMUSG00000032480",
                "ENSMUSG00000028884",
                "ENSMUSG00000097574",
                "ENSMUSG00000032383",
                "ENSMUSG00000003299",
                "ENSMUSG00000024871",
                "ENSMUSG00000031447",
                "ENSMUSG00000022142",
                "ENSMUSG00000066279",
                "ENSMUSG00000026568",
                "ENSMUSG00000034673",
                "ENSMUSG00000027363",
                "ENSMUSG00000003810",
                "ENSMUSG00000027342",
                "ENSMUSG00000045201",
                "ENSMUSG00000058655",
                "ENSMUSG00000052934",
                "ENSMUSG00000035024",
                "ENSMUSG00000060002",
                "ENSMUSG00000019768",
                "ENSMUSG00000071793",
                "ENSMUSG00000033382",
                "ENSMUSG00000029221",
                "ENSMUSG00000033386",
                "ENSMUSG00000058135",
                "ENSMUSG00000085741",
                "ENSMUSG00000039886",
                "ENSMUSG00000027860",
                "ENSMUSG00000016349",
                "ENSMUSG00000041773",
                "ENSMUSG00000021149",
                "ENSMUSG00000026456",
                "ENSMUSG00000029913",
                "ENSMUSG00000058076",
                "ENSMUSG00000028165",
                "ENSMUSG00000024640",
                "ENSMUSG00000029594",
                "ENSMUSG00000047721",
                "ENSMUSG00000098990",
                "ENSMUSG00000066637",
                "ENSMUSG00000025666",
                "ENSMUSG00000059714",
                "ENSMUSG00000030587",
                "ENSMUSG00000033060",
                "ENSMUSG00000033228",
                "ENSMUSG00000053483",
                "ENSMUSG00000025156",
                "ENSMUSG00000048668",
                "ENSMUSG00000032826",
                "ENSMUSG00000002608",
                "ENSMUSG00000026701",
                "ENSMUSG00000028266",
                "ENSMUSG00000021377",
                "ENSMUSG00000032038",
                "ENSMUSG00000033594",
                "ENSMUSG00000048154",
                "ENSMUSG00000055762",
                "ENSMUSG00000048279",
                "ENSMUSG00000047371",
                "ENSMUSG00000031232",
                "ENSMUSG00000026930",
                "ENSMUSG00000048814",
                "ENSMUSG00000031641",
                "ENSMUSG00000021597",
                "ENSMUSG00000038967",
                "ENSMUSG00000023249",
                "ENSMUSG00000030846",
                "ENSMUSG00000031841",
                "ENSMUSG00000020807",
                "ENSMUSG00000026571",
                "ENSMUSG00000036854",
                "ENSMUSG00000026202",
                "ENSMUSG00000039842",
                "ENSMUSG00000027993",
                "ENSMUSG00000034951",
                "ENSMUSG00000020392",
                "ENSMUSG00000028919",
                "ENSMUSG00000034906",
                "ENSMUSG00000066043",
                "ENSMUSG00000031751",
                "ENSMUSG00000021460",
                "ENSMUSG00000023055",
                "ENSMUSG00000022477",
                "ENSMUSG00000036867",
                "ENSMUSG00000032010",
                "ENSMUSG00000030172",
                "ENSMUSG00000022682",
                "ENSMUSG00000032462",
                "ENSMUSG00000064347",
                "ENSMUSG00000053617",
                "ENSMUSG00000020277",
                "ENSMUSG00000059291",
                "ENSMUSG00000029781",
                "ENSMUSG00000034290",
                "ENSMUSG00000023150",
                "ENSMUSG00000020647",
                "ENSMUSG00000012405",
                "ENSMUSG00000040506",
                "ENSMUSG00000043207",
                "ENSMUSG00000030298",
                "ENSMUSG00000072082",
                "ENSMUSG00000019054",
                "ENSMUSG00000001366",
                "ENSMUSG00000022214",
                "ENSMUSG00000020190",
                "ENSMUSG00000037916",
                "ENSMUSG00000054640",
                "ENSMUSG00000020716",
                "ENSMUSG00000064080",
                "ENSMUSG00000027006",
                "ENSMUSG00000036160",
                "ENSMUSG00000040321",
                "ENSMUSG00000019851",
                "ENSMUSG00000045316",
                "ENSMUSG00000052912",
                "ENSMUSG00000030607",
                "ENSMUSG00000040771",
                "ENSMUSG00000020473",
                "ENSMUSG00000025428",
                "ENSMUSG00000032724",
                "ENSMUSG00000026655",
                "ENSMUSG00000020390",
                "ENSMUSG00000039474",
                "ENSMUSG00000028832",
                "ENSMUSG00000030016",
                "ENSMUSG00000041426",
                "ENSMUSG00000031634",
                "ENSMUSG00000008730",
                "ENSMUSG00000074218",
                "ENSMUSG00000026766",
                "ENSMUSG00000000168",
                "ENSMUSG00000024924",
                "ENSMUSG00000029249",
                "ENSMUSG00000083435",
                "ENSMUSG00000028522",
                "ENSMUSG00000063870",
                "ENSMUSG00000032902",
                "ENSMUSG00000031633",
                "ENSMUSG00000022390",
                "ENSMUSG00000062329",
                "ENSMUSG00000041415",
                "ENSMUSG00000037826",
                "ENSMUSG00000052957",
                "ENSMUSG00000026708",
                "ENSMUSG00000029651",
                "ENSMUSG00000039089",
                "ENSMUSG00000060038",
                "ENSMUSG00000021906",
                "ENSMUSG00000038644",
                "ENSMUSG00000042942",
                "ENSMUSG00000040648",
                "ENSMUSG00000025221",
                "ENSMUSG00000021313",
                "ENSMUSG00000061533",
                "ENSMUSG00000026207",
                "ENSMUSG00000025241",
                "ENSMUSG00000021614",
                "ENSMUSG00000034342",
                "ENSMUSG00000027883",
                "ENSMUSG00000079111",
                "ENSMUSG00000031024",
                "ENSMUSG00000020456",
                "ENSMUSG00000045983",
                "ENSMUSG00000064342",
                "ENSMUSG00000027087",
                "ENSMUSG00000001794",
                "ENSMUSG00000031983",
                "ENSMUSG00000050174",
                "ENSMUSG00000032481",
                "ENSMUSG00000003531",
                "ENSMUSG00000026027",
                "ENSMUSG00000035203",
                "ENSMUSG00000034437",
                "ENSMUSG00000028745",
                "ENSMUSG00000038886",
                "ENSMUSG00000020483",
                "ENSMUSG00000027823",
                "ENSMUSG00000031906",
                "ENSMUSG00000022051",
                "ENSMUSG00000006818",
                "ENSMUSG00000024055",
                "ENSMUSG00000013584",
                "ENSMUSG00000032959",
                "ENSMUSG00000095588",
                "ENSMUSG00000021485",
                "ENSMUSG00000040850",
                "ENSMUSG00000042032",
                "ENSMUSG00000032224",
                "ENSMUSG00000021375",
                "ENSMUSG00000028173",
                "ENSMUSG00000058793",
                "ENSMUSG00000049643",
                "ENSMUSG00000031985",
                "ENSMUSG00000029359",
                "ENSMUSG00000022452",
                "ENSMUSG00000026283",
                "ENSMUSG00000001228",
                "ENSMUSG00000027406",
                "ENSMUSG00000091742",
                "ENSMUSG00000056608",
                "ENSMUSG00000015176",
                "ENSMUSG00000024164",
                "ENSMUSG00000001240",
                "ENSMUSG00000059586",
                "ENSMUSG00000026707",
                "ENSMUSG00000028756",
                "ENSMUSG00000024143",
                "ENSMUSG00000078566",
                "ENSMUSG00000025374",
                "ENSMUSG00000059974",
                "ENSMUSG00000063052",
                "ENSMUSG00000002343",
                "ENSMUSG00000027894",
                "ENSMUSG00000037643",
                "ENSMUSG00000024613",
                "ENSMUSG00000044042",
                "ENSMUSG00000058997",
                "ENSMUSG00000007827",
                "ENSMUSG00000050199",
                "ENSMUSG00000051495",
                "ENSMUSG00000022964",
                "ENSMUSG00000008604",
                "ENSMUSG00000022841",
                "ENSMUSG00000021591",
                "ENSMUSG00000024590",
                "ENSMUSG00000058594",
                "ENSMUSG00000032786",
                "ENSMUSG00000021215",
                "ENSMUSG00000074671",
                "ENSMUSG00000042700",
                "ENSMUSG00000025856",
                "ENSMUSG00000040028",
                "ENSMUSG00000041926",
                "ENSMUSG00000026281",
                "ENSMUSG00000053768",
                "ENSMUSG00000031756",
                "ENSMUSG00000023010",
                "ENSMUSG00000028312",
                "ENSMUSG00000055723",
                "ENSMUSG00000061104",
                "ENSMUSG00000025781",
                "ENSMUSG00000036278",
                "ENSMUSG00000038683",
                "ENSMUSG00000039987",
                "ENSMUSG00000041548",
                "ENSMUSG00000094410",
                "ENSMUSG00000067369",
                "ENSMUSG00000033781",
                "ENSMUSG00000003316",
                "ENSMUSG00000008690",
                "ENSMUSG00000021186",
                "ENSMUSG00000005510",
                "ENSMUSG00000036202",
                "ENSMUSG00000059689",
                "ENSMUSG00000005687",
                "ENSMUSG00000044337",
                "ENSMUSG00000036572",
                "ENSMUSG00000037148",
                "ENSMUSG00000041870",
                "ENSMUSG00000025466",
                "ENSMUSG00000023025",
                "ENSMUSG00000062070",
                "ENSMUSG00000048911",
                "ENSMUSG00000021395",
                "ENSMUSG00000038764",
                "ENSMUSG00000022553",
                "ENSMUSG00000020108",
                "ENSMUSG00000030541",
                "ENSMUSG00000070003",
                "ENSMUSG00000026208",
                "ENSMUSG00000031787",
                "ENSMUSG00000056394",
                "ENSMUSG00000027238",
                "ENSMUSG00000020133",
                "ENSMUSG00000053536",
                "ENSMUSG00000022300",
                "ENSMUSG00000020821",
                "ENSMUSG00000026278",
                "ENSMUSG00000063684",
                "ENSMUSG00000029190",
                "ENSMUSG00000036223",
                "ENSMUSG00000015839",
                "ENSMUSG00000027499",
                "ENSMUSG00000045752",
                "ENSMUSG00000021265",
                "ENSMUSG00000015944",
                "ENSMUSG00000017686",
                "ENSMUSG00000017119",
                "ENSMUSG00000078695",
                "ENSMUSG00000028911",
                "ENSMUSG00000031701",
                "ENSMUSG00000069049",
                "ENSMUSG00000007613",
                "ENSMUSG00000027455",
                "ENSMUSG00000020738",
                "ENSMUSG00000038807",
                "ENSMUSG00000064340",
                "ENSMUSG00000032047",
                "ENSMUSG00000029467",
                "ENSMUSG00000086841",
                "ENSMUSG00000069089",
                "ENSMUSG00000020415",
                "ENSMUSG00000018189",
                "ENSMUSG00000020818",
                "ENSMUSG00000044393",
                "ENSMUSG00000006800",
                "ENSMUSG00000014470",
                "ENSMUSG00000031762",
                "ENSMUSG00000028403",
                "ENSMUSG00000030255",
                "ENSMUSG00000036002",
                "ENSMUSG00000026817",
                "ENSMUSG00000022031",
                "ENSMUSG00000092486",
                "ENSMUSG00000033416",
                "ENSMUSG00000021039",
                "ENSMUSG00000027180",
                "ENSMUSG00000028134",
                "ENSMUSG00000024212",
                "ENSMUSG00000028937",
                "ENSMUSG00000015357",
                "ENSMUSG00000045636",
                "ENSMUSG00000024985",
                "ENSMUSG00000027420",
                "ENSMUSG00000062300",
                "ENSMUSG00000038170",
                "ENSMUSG00000024043",
                "ENSMUSG00000042293",
                "ENSMUSG00000027223",
                "ENSMUSG00000049327",
                "ENSMUSG00000006932",
                "ENSMUSG00000024187",
                "ENSMUSG00000038520",
                "ENSMUSG00000038058",
                "ENSMUSG00000007050",
                "ENSMUSG00000001930",
                "ENSMUSG00000009545",
                "ENSMUSG00000040964",
                "ENSMUSG00000020988",
                "ENSMUSG00000061086",
                "ENSMUSG00000039670",
                "ENSMUSG00000006191",
                "ENSMUSG00000027605",
                "ENSMUSG00000020974",
                "ENSMUSG00000040998",
                "ENSMUSG00000073563",
                "ENSMUSG00000024805",
                "ENSMUSG00000051391",
                "ENSMUSG00000015932",
                "ENSMUSG00000029655",
                "ENSMUSG00000036880",
                "ENSMUSG00000040520",
                "ENSMUSG00000053819",
                "ENSMUSG00000021280",
                "ENSMUSG00000022551",
                "ENSMUSG00000029571",
                "ENSMUSG00000025651",
                "ENSMUSG00000078671",
                "ENSMUSG00000050953",
                "ENSMUSG00000037887",
                "ENSMUSG00000021236",
                "ENSMUSG00000052812",
                "ENSMUSG00000035093",
                "ENSMUSG00000030847",
                "ENSMUSG00000046312",
                "ENSMUSG00000042115",
                "ENSMUSG00000004558",
                "ENSMUSG00000020467",
                "ENSMUSG00000046997",
                "ENSMUSG00000030246",
                "ENSMUSG00000034088",
                "ENSMUSG00000019843",
                "ENSMUSG00000010914",
                "ENSMUSG00000052139",
                "ENSMUSG00000020664",
                "ENSMUSG00000022389",
                "ENSMUSG00000048731",
                "ENSMUSG00000061755",
                "ENSMUSG00000031767",
                "ENSMUSG00000064371",
                "ENSMUSG00000022837",
                "ENSMUSG00000027673",
                "ENSMUSG00000036402",
                "ENSMUSG00000040118",
                "ENSMUSG00000018395",
                "ENSMUSG00000059811",
                "ENSMUSG00000032376",
                "ENSMUSG00000028977",
                "ENSMUSG00000002059",
                "ENSMUSG00000022322",
                "ENSMUSG00000046179",
                "ENSMUSG00000067851",
                "ENSMUSG00000026489"
            ],
            "expression_sum": {
                "class1": 1187.0,
                "class2": 381.0
            },
            "uniprot_ids": [
                "Q62318"
            ],
            "net_id": 0,
            "source_num": 6,
            "target_num": 100
        },
        "ENSMUSG00000028565": {
            "id": "ENSMUSG00000028565",
            "type": "TF",
            "symbol": "Nfia",
            "source": [
                "ENSMUSG00000059552",
                "ENSMUSG00000024515",
                "ENSMUSG00000019768",
                "ENSMUSG00000040732",
                "ENSMUSG00000042557",
                "ENSMUSG00000022528"
            ],
            "target": [
                "ENSMUSG00000022617",
                "ENSMUSG00000027642",
                "ENSMUSG00000054619",
                "ENSMUSG00000059552",
                "ENSMUSG00000002020",
                "ENSMUSG00000026568",
                "ENSMUSG00000026123",
                "ENSMUSG00000030652",
                "ENSMUSG00000004771",
                "ENSMUSG00000030352",
                "ENSMUSG00000038366",
                "ENSMUSG00000025869",
                "ENSMUSG00000028053",
                "ENSMUSG00000004105",
                "ENSMUSG00000096255",
                "ENSMUSG00000050043",
                "ENSMUSG00000036555",
                "ENSMUSG00000048100",
                "ENSMUSG00000034684",
                "ENSMUSG00000053536",
                "ENSMUSG00000021592",
                "ENSMUSG00000027809",
                "ENSMUSG00000017747",
                "ENSMUSG00000017144",
                "ENSMUSG00000019768",
                "ENSMUSG00000019774",
                "ENSMUSG00000053841",
                "ENSMUSG00000006676",
                "ENSMUSG00000025613",
                "ENSMUSG00000021701",
                "ENSMUSG00000029710",
                "ENSMUSG00000043683",
                "ENSMUSG00000006134",
                "ENSMUSG00000018819",
                "ENSMUSG00000034342",
                "ENSMUSG00000047419",
                "ENSMUSG00000056763",
                "ENSMUSG00000035776",
                "ENSMUSG00000055371",
                "ENSMUSG00000020389",
                "ENSMUSG00000004642",
                "ENSMUSG00000034263",
                "ENSMUSG00000020423",
                "ENSMUSG00000026404",
                "ENSMUSG00000037032",
                "ENSMUSG00000090516",
                "ENSMUSG00000028223",
                "ENSMUSG00000024283",
                "ENSMUSG00000058013",
                "ENSMUSG00000002496",
                "ENSMUSG00000020456",
                "ENSMUSG00000021033",
                "ENSMUSG00000059070",
                "ENSMUSG00000026342",
                "ENSMUSG00000011427",
                "ENSMUSG00000062647",
                "ENSMUSG00000033454",
                "ENSMUSG00000020319",
                "ENSMUSG00000078695",
                "ENSMUSG00000027193",
                "ENSMUSG00000004980",
                "ENSMUSG00000022816",
                "ENSMUSG00000035878",
                "ENSMUSG00000020738",
                "ENSMUSG00000031353",
                "ENSMUSG00000050737",
                "ENSMUSG00000027750",
                "ENSMUSG00000029718",
                "ENSMUSG00000035934",
                "ENSMUSG00000041841",
                "ENSMUSG00000064340",
                "ENSMUSG00000003923",
                "ENSMUSG00000031434",
                "ENSMUSG00000049751",
                "ENSMUSG00000022722",
                "ENSMUSG00000029276",
                "ENSMUSG00000002881",
                "ENSMUSG00000027997",
                "ENSMUSG00000029576",
                "ENSMUSG00000030659",
                "ENSMUSG00000017493",
                "ENSMUSG00000026853",
                "ENSMUSG00000041354",
                "ENSMUSG00000002052",
                "ENSMUSG00000026087",
                "ENSMUSG00000024725",
                "ENSMUSG00000021606",
                "ENSMUSG00000024066",
                "ENSMUSG00000038831",
                "ENSMUSG00000024302",
                "ENSMUSG00000020267",
                "ENSMUSG00000069744",
                "ENSMUSG00000005610",
                "ENSMUSG00000074305",
                "ENSMUSG00000034520",
                "ENSMUSG00000020477",
                "ENSMUSG00000019978",
                "ENSMUSG00000021540",
                "ENSMUSG00000004069",
                "ENSMUSG00000032679",
                "ENSMUSG00000013465",
                "ENSMUSG00000040690",
                "ENSMUSG00000053062",
                "ENSMUSG00000023939",
                "ENSMUSG00000009863",
                "ENSMUSG00000078453",
                "ENSMUSG00000021693",
                "ENSMUSG00000036863",
                "ENSMUSG00000021027",
                "ENSMUSG00000024515",
                "ENSMUSG00000032353",
                "ENSMUSG00000043964",
                "ENSMUSG00000022136",
                "ENSMUSG00000017615",
                "ENSMUSG00000026341",
                "ENSMUSG00000023905",
                "ENSMUSG00000015357",
                "ENSMUSG00000022204",
                "ENSMUSG00000000088",
                "ENSMUSG00000031447",
                "ENSMUSG00000027669",
                "ENSMUSG00000026743",
                "ENSMUSG00000031715",
                "ENSMUSG00000006498",
                "ENSMUSG00000028945",
                "ENSMUSG00000029918",
                "ENSMUSG00000029163",
                "ENSMUSG00000045636",
                "ENSMUSG00000054302",
                "ENSMUSG00000061479",
                "ENSMUSG00000038170",
                "ENSMUSG00000022519",
                "ENSMUSG00000026393",
                "ENSMUSG00000041736",
                "ENSMUSG00000030516",
                "ENSMUSG00000022867",
                "ENSMUSG00000031485",
                "ENSMUSG00000031697",
                "ENSMUSG00000048277",
                "ENSMUSG00000051674",
                "ENSMUSG00000031232",
                "ENSMUSG00000015176",
                "ENSMUSG00000041577",
                "ENSMUSG00000058240",
                "ENSMUSG00000022361",
                "ENSMUSG00000037608",
                "ENSMUSG00000020076",
                "ENSMUSG00000037523",
                "ENSMUSG00000040717",
                "ENSMUSG00000024687",
                "ENSMUSG00000061136",
                "ENSMUSG00000021737",
                "ENSMUSG00000001844",
                "ENSMUSG00000000811",
                "ENSMUSG00000053774",
                "ENSMUSG00000060261",
                "ENSMUSG00000027439",
                "ENSMUSG00000061559",
                "ENSMUSG00000038387",
                "ENSMUSG00000042557",
                "ENSMUSG00000021975",
                "ENSMUSG00000026755",
                "ENSMUSG00000005262",
                "ENSMUSG00000028322",
                "ENSMUSG00000023249",
                "ENSMUSG00000009927",
                "ENSMUSG00000021905",
                "ENSMUSG00000018425",
                "ENSMUSG00000031841",
                "ENSMUSG00000028597",
                "ENSMUSG00000026202",
                "ENSMUSG00000054263",
                "ENSMUSG00000022401",
                "ENSMUSG00000060904",
                "ENSMUSG00000022450",
                "ENSMUSG00000025362",
                "ENSMUSG00000073563",
                "ENSMUSG00000032786",
                "ENSMUSG00000024182",
                "ENSMUSG00000039031",
                "ENSMUSG00000057133",
                "ENSMUSG00000036438",
                "ENSMUSG00000029836",
                "ENSMUSG00000039103",
                "ENSMUSG00000026922",
                "ENSMUSG00000015932",
                "ENSMUSG00000037286",
                "ENSMUSG00000059895",
                "ENSMUSG00000003190",
                "ENSMUSG00000028452",
                "ENSMUSG00000034295",
                "ENSMUSG00000004530",
                "ENSMUSG00000050721",
                "ENSMUSG00000043895",
                "ENSMUSG00000058756",
                "ENSMUSG00000028788",
                "ENSMUSG00000002733",
                "ENSMUSG00000032625",
                "ENSMUSG00000005779",
                "ENSMUSG00000025503",
                "ENSMUSG00000019841",
                "ENSMUSG00000024052",
                "ENSMUSG00000037370",
                "ENSMUSG00000029110",
                "ENSMUSG00000018796",
                "ENSMUSG00000037098",
                "ENSMUSG00000039982",
                "ENSMUSG00000044080",
                "ENSMUSG00000028618",
                "ENSMUSG00000059447",
                "ENSMUSG00000022808",
                "ENSMUSG00000041959",
                "ENSMUSG00000065126",
                "ENSMUSG00000022110",
                "ENSMUSG00000024997",
                "ENSMUSG00000052812",
                "ENSMUSG00000037685",
                "ENSMUSG00000025085",
                "ENSMUSG00000052727",
                "ENSMUSG00000024304",
                "ENSMUSG00000029790",
                "ENSMUSG00000035686",
                "ENSMUSG00000022528",
                "ENSMUSG00000022205",
                "ENSMUSG00000042104",
                "ENSMUSG00000042115",
                "ENSMUSG00000070372",
                "ENSMUSG00000031451",
                "ENSMUSG00000025246",
                "ENSMUSG00000025290",
                "ENSMUSG00000021917",
                "ENSMUSG00000031483",
                "ENSMUSG00000036968",
                "ENSMUSG00000066440",
                "ENSMUSG00000034724",
                "ENSMUSG00000046603",
                "ENSMUSG00000071042",
                "ENSMUSG00000023367",
                "ENSMUSG00000022858",
                "ENSMUSG00000023723",
                "ENSMUSG00000023861",
                "ENSMUSG00000003072",
                "ENSMUSG00000040034",
                "ENSMUSG00000032667",
                "ENSMUSG00000034462",
                "ENSMUSG00000097325",
                "ENSMUSG00000020044",
                "ENSMUSG00000020451",
                "ENSMUSG00000073664",
                "ENSMUSG00000052253",
                "ENSMUSG00000027752",
                "ENSMUSG00000023025",
                "ENSMUSG00000042688",
                "ENSMUSG00000027133",
                "ENSMUSG00000047881",
                "ENSMUSG00000046062",
                "ENSMUSG00000032051",
                "ENSMUSG00000021520",
                "ENSMUSG00000074218",
                "ENSMUSG00000024975",
                "ENSMUSG00000029033",
                "ENSMUSG00000032480",
                "ENSMUSG00000001056",
                "ENSMUSG00000022800",
                "ENSMUSG00000024924",
                "ENSMUSG00000026639",
                "ENSMUSG00000036019",
                "ENSMUSG00000026574",
                "ENSMUSG00000038970",
                "ENSMUSG00000014496",
                "ENSMUSG00000022090",
                "ENSMUSG00000019979",
                "ENSMUSG00000071337",
                "ENSMUSG00000021549",
                "ENSMUSG00000071252",
                "ENSMUSG00000053929"
            ],
            "expression_sum": {
                "class1": 5350.0,
                "class2": 901.0
            },
            "uniprot_ids": [
                "Q02780",
                "B1AUB9",
                "B1AUC0",
                "F7CDR2",
                "B1AUB8",
                "B1AUB6"
            ],
            "net_id": 0,
            "source_num": 4,
            "target_num": 53
        },
        "ENSMUSG00000005698": {
            "id": "ENSMUSG00000005698",
            "type": "TF",
            "symbol": "Ctcf",
            "source": [
                "ENSMUSG00000021109",
                "ENSMUSG00000018604",
                "ENSMUSG00000056501",
                "ENSMUSG00000019768",
                "ENSMUSG00000022641",
                "ENSMUSG00000076431",
                "ENSMUSG00000022528"
            ],
            "target": [
                "ENSMUSG00000059182",
                "ENSMUSG00000061603",
                "ENSMUSG00000055775",
                "ENSMUSG00000052911",
                "ENSMUSG00000031448",
                "ENSMUSG00000009569",
                "ENSMUSG00000032563",
                "ENSMUSG00000020841",
                "ENSMUSG00000024197",
                "ENSMUSG00000022020",
                "ENSMUSG00000044548",
                "ENSMUSG00000024228",
                "ENSMUSG00000062908",
                "ENSMUSG00000021057",
                "ENSMUSG00000018476",
                "ENSMUSG00000040269",
                "ENSMUSG00000058388",
                "ENSMUSG00000027531",
                "ENSMUSG00000022462",
                "ENSMUSG00000038070",
                "ENSMUSG00000015837",
                "ENSMUSG00000043644",
                "ENSMUSG00000020741",
                "ENSMUSG00000097414",
                "ENSMUSG00000022339",
                "ENSMUSG00000036585",
                "ENSMUSG00000070283",
                "ENSMUSG00000037339",
                "ENSMUSG00000030982",
                "ENSMUSG00000011427",
                "ENSMUSG00000027984",
                "ENSMUSG00000021629",
                "ENSMUSG00000024487",
                "ENSMUSG00000030739",
                "ENSMUSG00000021339",
                "ENSMUSG00000042709",
                "ENSMUSG00000094320",
                "ENSMUSG00000022636",
                "ENSMUSG00000026576",
                "ENSMUSG00000029998",
                "ENSMUSG00000031540",
                "ENSMUSG00000026853",
                "ENSMUSG00000030403",
                "ENSMUSG00000044221",
                "ENSMUSG00000031790",
                "ENSMUSG00000066900",
                "ENSMUSG00000020070",
                "ENSMUSG00000027660",
                "ENSMUSG00000038763",
                "ENSMUSG00000019920",
                "ENSMUSG00000043445",
                "ENSMUSG00000019978",
                "ENSMUSG00000026623",
                "ENSMUSG00000018363",
                "ENSMUSG00000090553",
                "ENSMUSG00000039943",
                "ENSMUSG00000053754",
                "ENSMUSG00000032397",
                "ENSMUSG00000030562",
                "ENSMUSG00000006576",
                "ENSMUSG00000064037",
                "ENSMUSG00000022433",
                "ENSMUSG00000026313",
                "ENSMUSG00000005469",
                "ENSMUSG00000000753",
                "ENSMUSG00000007656",
                "ENSMUSG00000026803",
                "ENSMUSG00000027288",
                "ENSMUSG00000045411",
                "ENSMUSG00000066026",
                "ENSMUSG00000026201",
                "ENSMUSG00000090812",
                "ENSMUSG00000022519",
                "ENSMUSG00000051331",
                "ENSMUSG00000030921",
                "ENSMUSG00000030516",
                "ENSMUSG00000036097",
                "ENSMUSG00000029103",
                "ENSMUSG00000016510",
                "ENSMUSG00000042010",
                "ENSMUSG00000019838",
                "ENSMUSG00000090150",
                "ENSMUSG00000030850",
                "ENSMUSG00000058240",
                "ENSMUSG00000033184",
                "ENSMUSG00000020282",
                "ENSMUSG00000042225",
                "ENSMUSG00000025439",
                "ENSMUSG00000029578",
                "ENSMUSG00000029499",
                "ENSMUSG00000027884",
                "ENSMUSG00000094664",
                "ENSMUSG00000036452",
                "ENSMUSG00000064302",
                "ENSMUSG00000040666",
                "ENSMUSG00000025184",
                "ENSMUSG00000031922",
                "ENSMUSG00000055148",
                "ENSMUSG00000036825",
                "ENSMUSG00000032883",
                "ENSMUSG00000001552",
                "ENSMUSG00000042197",
                "ENSMUSG00000000056",
                "ENSMUSG00000046916",
                "ENSMUSG00000039068",
                "ENSMUSG00000022095",
                "ENSMUSG00000017288",
                "ENSMUSG00000005481",
                "ENSMUSG00000003865",
                "ENSMUSG00000039103",
                "ENSMUSG00000044792",
                "ENSMUSG00000021244",
                "ENSMUSG00000026504",
                "ENSMUSG00000067653",
                "ENSMUSG00000056501",
                "ENSMUSG00000026249",
                "ENSMUSG00000022052",
                "ENSMUSG00000082283",
                "ENSMUSG00000077450",
                "ENSMUSG00000031698",
                "ENSMUSG00000029135",
                "ENSMUSG00000025898",
                "ENSMUSG00000061286",
                "ENSMUSG00000039233",
                "ENSMUSG00000031785",
                "ENSMUSG00000024304",
                "ENSMUSG00000037321",
                "ENSMUSG00000030753",
                "ENSMUSG00000006219",
                "ENSMUSG00000019132",
                "ENSMUSG00000094886",
                "ENSMUSG00000024381",
                "ENSMUSG00000078139",
                "ENSMUSG00000023990",
                "ENSMUSG00000079225",
                "ENSMUSG00000018040",
                "ENSMUSG00000024942",
                "ENSMUSG00000028497",
                "ENSMUSG00000041303",
                "ENSMUSG00000086167",
                "ENSMUSG00000037049",
                "ENSMUSG00000084416",
                "ENSMUSG00000081359",
                "ENSMUSG00000063904",
                "ENSMUSG00000080896",
                "ENSMUSG00000054452",
                "ENSMUSG00000036570",
                "ENSMUSG00000032480",
                "ENSMUSG00000029186",
                "ENSMUSG00000029452",
                "ENSMUSG00000033885",
                "ENSMUSG00000010122",
                "ENSMUSG00000026374",
                "ENSMUSG00000003299",
                "ENSMUSG00000005078",
                "ENSMUSG00000024871",
                "ENSMUSG00000030087",
                "ENSMUSG00000040351",
                "ENSMUSG00000051223",
                "ENSMUSG00000022538",
                "ENSMUSG00000005683",
                "ENSMUSG00000034673",
                "ENSMUSG00000049588",
                "ENSMUSG00000034274",
                "ENSMUSG00000097137",
                "ENSMUSG00000020846",
                "ENSMUSG00000056313",
                "ENSMUSG00000026197",
                "ENSMUSG00000040112",
                "ENSMUSG00000045201",
                "ENSMUSG00000029177",
                "ENSMUSG00000023868",
                "ENSMUSG00000058655",
                "ENSMUSG00000039239",
                "ENSMUSG00000019768",
                "ENSMUSG00000028573",
                "ENSMUSG00000060002",
                "ENSMUSG00000082088",
                "ENSMUSG00000075284",
                "ENSMUSG00000058135",
                "ENSMUSG00000029221",
                "ENSMUSG00000038013",
                "ENSMUSG00000020085",
                "ENSMUSG00000041482",
                "ENSMUSG00000032497",
                "ENSMUSG00000041797",
                "ENSMUSG00000031029",
                "ENSMUSG00000035104",
                "ENSMUSG00000031536",
                "ENSMUSG00000030309",
                "ENSMUSG00000026456",
                "ENSMUSG00000001334",
                "ENSMUSG00000064061",
                "ENSMUSG00000055027",
                "ENSMUSG00000039108",
                "ENSMUSG00000049804",
                "ENSMUSG00000015843",
                "ENSMUSG00000032750",
                "ENSMUSG00000079481",
                "ENSMUSG00000026411",
                "ENSMUSG00000039849",
                "ENSMUSG00000084349",
                "ENSMUSG00000021645",
                "ENSMUSG00000028973",
                "ENSMUSG00000000441",
                "ENSMUSG00000010830",
                "ENSMUSG00000016534",
                "ENSMUSG00000041685",
                "ENSMUSG00000083899",
                "ENSMUSG00000089862",
                "ENSMUSG00000038594",
                "ENSMUSG00000017176",
                "ENSMUSG00000053483",
                "ENSMUSG00000022941",
                "ENSMUSG00000002608",
                "ENSMUSG00000039601",
                "ENSMUSG00000083396",
                "ENSMUSG00000026701",
                "ENSMUSG00000025271",
                "ENSMUSG00000041453",
                "ENSMUSG00000081228",
                "ENSMUSG00000067430",
                "ENSMUSG00000033594",
                "ENSMUSG00000027668",
                "ENSMUSG00000037949",
                "ENSMUSG00000055762",
                "ENSMUSG00000074971",
                "ENSMUSG00000002396",
                "ENSMUSG00000027665",
                "ENSMUSG00000031232",
                "ENSMUSG00000020205",
                "ENSMUSG00000027306",
                "ENSMUSG00000043323",
                "ENSMUSG00000022186",
                "ENSMUSG00000048814",
                "ENSMUSG00000037204",
                "ENSMUSG00000022765",
                "ENSMUSG00000038967",
                "ENSMUSG00000024544",
                "ENSMUSG00000061136",
                "ENSMUSG00000045624",
                "ENSMUSG00000036112",
                "ENSMUSG00000083856",
                "ENSMUSG00000053093",
                "ENSMUSG00000047260",
                "ENSMUSG00000017264",
                "ENSMUSG00000089715",
                "ENSMUSG00000025059",
                "ENSMUSG00000045534",
                "ENSMUSG00000083892",
                "ENSMUSG00000030846",
                "ENSMUSG00000023886",
                "ENSMUSG00000006920",
                "ENSMUSG00000095306",
                "ENSMUSG00000002500",
                "ENSMUSG00000055963",
                "ENSMUSG00000026131",
                "ENSMUSG00000054823",
                "ENSMUSG00000030315",
                "ENSMUSG00000034951",
                "ENSMUSG00000015149",
                "ENSMUSG00000028919",
                "ENSMUSG00000078154",
                "ENSMUSG00000005233",
                "ENSMUSG00000028729",
                "ENSMUSG00000027423",
                "ENSMUSG00000033538",
                "ENSMUSG00000043024",
                "ENSMUSG00000022089",
                "ENSMUSG00000022477",
                "ENSMUSG00000023055",
                "ENSMUSG00000068614",
                "ENSMUSG00000031672",
                "ENSMUSG00000046410",
                "ENSMUSG00000030172",
                "ENSMUSG00000059493",
                "ENSMUSG00000062279",
                "ENSMUSG00000025348",
                "ENSMUSG00000034584",
                "ENSMUSG00000022194",
                "ENSMUSG00000017756",
                "ENSMUSG00000068854",
                "ENSMUSG00000020277",
                "ENSMUSG00000023150",
                "ENSMUSG00000074622",
                "ENSMUSG00000067925",
                "ENSMUSG00000092118",
                "ENSMUSG00000013539",
                "ENSMUSG00000001366",
                "ENSMUSG00000031740",
                "ENSMUSG00000067148",
                "ENSMUSG00000022110",
                "ENSMUSG00000020190",
                "ENSMUSG00000028648",
                "ENSMUSG00000004356",
                "ENSMUSG00000037916",
                "ENSMUSG00000034601",
                "ENSMUSG00000029670",
                "ENSMUSG00000019851",
                "ENSMUSG00000059208",
                "ENSMUSG00000015289",
                "ENSMUSG00000022016",
                "ENSMUSG00000074064",
                "ENSMUSG00000024054",
                "ENSMUSG00000042207",
                "ENSMUSG00000010110",
                "ENSMUSG00000030607",
                "ENSMUSG00000040771",
                "ENSMUSG00000020473",
                "ENSMUSG00000030220",
                "ENSMUSG00000021870",
                "ENSMUSG00000033161",
                "ENSMUSG00000058690",
                "ENSMUSG00000066632",
                "ENSMUSG00000024538",
                "ENSMUSG00000081214",
                "ENSMUSG00000046691",
                "ENSMUSG00000024063",
                "ENSMUSG00000082516",
                "ENSMUSG00000047557",
                "ENSMUSG00000082487",
                "ENSMUSG00000020453",
                "ENSMUSG00000030016",
                "ENSMUSG00000013921",
                "ENSMUSG00000029804",
                "ENSMUSG00000029128",
                "ENSMUSG00000021798",
                "ENSMUSG00000030047",
                "ENSMUSG00000034686",
                "ENSMUSG00000000168",
                "ENSMUSG00000024924",
                "ENSMUSG00000021806",
                "ENSMUSG00000038039",
                "ENSMUSG00000042744",
                "ENSMUSG00000060594",
                "ENSMUSG00000036611",
                "ENSMUSG00000025268",
                "ENSMUSG00000028522",
                "ENSMUSG00000022390",
                "ENSMUSG00000041235",
                "ENSMUSG00000033082",
                "ENSMUSG00000037826",
                "ENSMUSG00000029651",
                "ENSMUSG00000023921",
                "ENSMUSG00000039089",
                "ENSMUSG00000092837",
                "ENSMUSG00000049119",
                "ENSMUSG00000005410",
                "ENSMUSG00000046324",
                "ENSMUSG00000065037",
                "ENSMUSG00000022747",
                "ENSMUSG00000026662",
                "ENSMUSG00000020740",
                "ENSMUSG00000026207",
                "ENSMUSG00000036620",
                "ENSMUSG00000038172",
                "ENSMUSG00000022641",
                "ENSMUSG00000039478",
                "ENSMUSG00000059430",
                "ENSMUSG00000075054",
                "ENSMUSG00000021959",
                "ENSMUSG00000035776",
                "ENSMUSG00000042165",
                "ENSMUSG00000031367",
                "ENSMUSG00000028936",
                "ENSMUSG00000031024",
                "ENSMUSG00000020456",
                "ENSMUSG00000051747",
                "ENSMUSG00000064342",
                "ENSMUSG00000078853",
                "ENSMUSG00000050627",
                "ENSMUSG00000025911",
                "ENSMUSG00000082180",
                "ENSMUSG00000039611",
                "ENSMUSG00000078202",
                "ENSMUSG00000038239",
                "ENSMUSG00000086298",
                "ENSMUSG00000021767",
                "ENSMUSG00000028007",
                "ENSMUSG00000031791",
                "ENSMUSG00000050174",
                "ENSMUSG00000022040",
                "ENSMUSG00000083327",
                "ENSMUSG00000003923",
                "ENSMUSG00000030399",
                "ENSMUSG00000090000",
                "ENSMUSG00000055239",
                "ENSMUSG00000035203",
                "ENSMUSG00000055612",
                "ENSMUSG00000040325",
                "ENSMUSG00000020483",
                "ENSMUSG00000026890",
                "ENSMUSG00000082935",
                "ENSMUSG00000020075",
                "ENSMUSG00000022051",
                "ENSMUSG00000024483",
                "ENSMUSG00000025332",
                "ENSMUSG00000034621",
                "ENSMUSG00000026878",
                "ENSMUSG00000076431",
                "ENSMUSG00000022336",
                "ENSMUSG00000070565",
                "ENSMUSG00000029518",
                "ENSMUSG00000042364",
                "ENSMUSG00000028755",
                "ENSMUSG00000058600",
                "ENSMUSG00000040850",
                "ENSMUSG00000021375",
                "ENSMUSG00000028173",
                "ENSMUSG00000029111",
                "ENSMUSG00000034795",
                "ENSMUSG00000058793",
                "ENSMUSG00000049643",
                "ENSMUSG00000031985",
                "ENSMUSG00000025135",
                "ENSMUSG00000029359",
                "ENSMUSG00000005897",
                "ENSMUSG00000023175",
                "ENSMUSG00000003808",
                "ENSMUSG00000038121",
                "ENSMUSG00000018893",
                "ENSMUSG00000046876",
                "ENSMUSG00000027406",
                "ENSMUSG00000069255",
                "ENSMUSG00000070034",
                "ENSMUSG00000060032",
                "ENSMUSG00000031111",
                "ENSMUSG00000028756",
                "ENSMUSG00000025374",
                "ENSMUSG00000038387",
                "ENSMUSG00000063052",
                "ENSMUSG00000049791",
                "ENSMUSG00000040254",
                "ENSMUSG00000050002",
                "ENSMUSG00000024613",
                "ENSMUSG00000032590",
                "ENSMUSG00000045954",
                "ENSMUSG00000054889",
                "ENSMUSG00000046351",
                "ENSMUSG00000047022",
                "ENSMUSG00000020409",
                "ENSMUSG00000003955",
                "ENSMUSG00000030060",
                "ENSMUSG00000032328",
                "ENSMUSG00000083793",
                "ENSMUSG00000004054",
                "ENSMUSG00000022982",
                "ENSMUSG00000063021",
                "ENSMUSG00000028273",
                "ENSMUSG00000019158",
                "ENSMUSG00000024309",
                "ENSMUSG00000021200",
                "ENSMUSG00000058594",
                "ENSMUSG00000074671",
                "ENSMUSG00000042700",
                "ENSMUSG00000074513",
                "ENSMUSG00000020027",
                "ENSMUSG00000025871",
                "ENSMUSG00000053768",
                "ENSMUSG00000022120",
                "ENSMUSG00000018648",
                "ENSMUSG00000028452",
                "ENSMUSG00000000738",
                "ENSMUSG00000039987",
                "ENSMUSG00000041548",
                "ENSMUSG00000062169",
                "ENSMUSG00000028348",
                "ENSMUSG00000033781",
                "ENSMUSG00000020053",
                "ENSMUSG00000018796",
                "ENSMUSG00000036202",
                "ENSMUSG00000032112",
                "ENSMUSG00000087679",
                "ENSMUSG00000004096",
                "ENSMUSG00000043241",
                "ENSMUSG00000081302",
                "ENSMUSG00000022781",
                "ENSMUSG00000028834",
                "ENSMUSG00000027239",
                "ENSMUSG00000022528",
                "ENSMUSG00000060679",
                "ENSMUSG00000073838",
                "ENSMUSG00000053453",
                "ENSMUSG00000035370",
                "ENSMUSG00000039536",
                "ENSMUSG00000042524",
                "ENSMUSG00000022969",
                "ENSMUSG00000040752",
                "ENSMUSG00000031749",
                "ENSMUSG00000074207",
                "ENSMUSG00000034593",
                "ENSMUSG00000022516",
                "ENSMUSG00000037795",
                "ENSMUSG00000051235",
                "ENSMUSG00000040139",
                "ENSMUSG00000033760",
                "ENSMUSG00000023025",
                "ENSMUSG00000062070",
                "ENSMUSG00000043822",
                "ENSMUSG00000028248",
                "ENSMUSG00000024975",
                "ENSMUSG00000038143",
                "ENSMUSG00000059901",
                "ENSMUSG00000045948",
                "ENSMUSG00000020680",
                "ENSMUSG00000026208",
                "ENSMUSG00000009079",
                "ENSMUSG00000031696",
                "ENSMUSG00000048583",
                "ENSMUSG00000028394",
                "ENSMUSG00000018548",
                "ENSMUSG00000025968",
                "ENSMUSG00000067780",
                "ENSMUSG00000051811",
                "ENSMUSG00000060187",
                "ENSMUSG00000097277",
                "ENSMUSG00000028538",
                "ENSMUSG00000027238",
                "ENSMUSG00000020133",
                "ENSMUSG00000053536",
                "ENSMUSG00000021109",
                "ENSMUSG00000040147",
                "ENSMUSG00000053841",
                "ENSMUSG00000065701",
                "ENSMUSG00000024140",
                "ENSMUSG00000026278",
                "ENSMUSG00000070284",
                "ENSMUSG00000028436",
                "ENSMUSG00000029190",
                "ENSMUSG00000034210",
                "ENSMUSG00000027784",
                "ENSMUSG00000028494",
                "ENSMUSG00000081629",
                "ENSMUSG00000078496",
                "ENSMUSG00000021265",
                "ENSMUSG00000015944",
                "ENSMUSG00000026879",
                "ENSMUSG00000037664",
                "ENSMUSG00000017119",
                "ENSMUSG00000078695",
                "ENSMUSG00000024691",
                "ENSMUSG00000007613",
                "ENSMUSG00000064340",
                "ENSMUSG00000009291",
                "ENSMUSG00000004655",
                "ENSMUSG00000029401",
                "ENSMUSG00000055632",
                "ENSMUSG00000032047",
                "ENSMUSG00000075701",
                "ENSMUSG00000025453",
                "ENSMUSG00000039219",
                "ENSMUSG00000033931",
                "ENSMUSG00000069089",
                "ENSMUSG00000033543",
                "ENSMUSG00000026880",
                "ENSMUSG00000038022",
                "ENSMUSG00000021929",
                "ENSMUSG00000044393",
                "ENSMUSG00000062190",
                "ENSMUSG00000048076",
                "ENSMUSG00000067818",
                "ENSMUSG00000030255",
                "ENSMUSG00000038539",
                "ENSMUSG00000036002",
                "ENSMUSG00000062580",
                "ENSMUSG00000020430",
                "ENSMUSG00000026817",
                "ENSMUSG00000009112",
                "ENSMUSG00000048126",
                "ENSMUSG00000025959",
                "ENSMUSG00000004455",
                "ENSMUSG00000022272",
                "ENSMUSG00000015536",
                "ENSMUSG00000087635",
                "ENSMUSG00000015357",
                "ENSMUSG00000037622",
                "ENSMUSG00000024059",
                "ENSMUSG00000039007",
                "ENSMUSG00000022905",
                "ENSMUSG00000024985",
                "ENSMUSG00000026074",
                "ENSMUSG00000040818",
                "ENSMUSG00000029071",
                "ENSMUSG00000050555",
                "ENSMUSG00000038170",
                "ENSMUSG00000030447",
                "ENSMUSG00000031299",
                "ENSMUSG00000006315",
                "ENSMUSG00000020122",
                "ENSMUSG00000031808",
                "ENSMUSG00000042293",
                "ENSMUSG00000038403",
                "ENSMUSG00000020380",
                "ENSMUSG00000031176",
                "ENSMUSG00000026418",
                "ENSMUSG00000092674",
                "ENSMUSG00000041361",
                "ENSMUSG00000046722",
                "ENSMUSG00000024457",
                "ENSMUSG00000038668",
                "ENSMUSG00000038520",
                "ENSMUSG00000038058",
                "ENSMUSG00000046402",
                "ENSMUSG00000026308",
                "ENSMUSG00000033632",
                "ENSMUSG00000046908",
                "ENSMUSG00000034729",
                "ENSMUSG00000063511",
                "ENSMUSG00000023959",
                "ENSMUSG00000019087",
                "ENSMUSG00000029004",
                "ENSMUSG00000000149",
                "ENSMUSG00000034083",
                "ENSMUSG00000036707",
                "ENSMUSG00000020988",
                "ENSMUSG00000023206",
                "ENSMUSG00000028518",
                "ENSMUSG00000084166",
                "ENSMUSG00000009207",
                "ENSMUSG00000063180",
                "ENSMUSG00000039670",
                "ENSMUSG00000073563",
                "ENSMUSG00000038518",
                "ENSMUSG00000024805",
                "ENSMUSG00000056671",
                "ENSMUSG00000023307",
                "ENSMUSG00000051391",
                "ENSMUSG00000023345",
                "ENSMUSG00000084304",
                "ENSMUSG00000025509",
                "ENSMUSG00000053819",
                "ENSMUSG00000040520",
                "ENSMUSG00000028364",
                "ENSMUSG00000029922",
                "ENSMUSG00000058756",
                "ENSMUSG00000068699",
                "ENSMUSG00000012609",
                "ENSMUSG00000020475",
                "ENSMUSG00000030662",
                "ENSMUSG00000096252",
                "ENSMUSG00000025651",
                "ENSMUSG00000027641",
                "ENSMUSG00000024158",
                "ENSMUSG00000040086",
                "ENSMUSG00000024498",
                "ENSMUSG00000027765",
                "ENSMUSG00000022884",
                "ENSMUSG00000032508",
                "ENSMUSG00000037486",
                "ENSMUSG00000059237",
                "ENSMUSG00000028820",
                "ENSMUSG00000003528",
                "ENSMUSG00000024639",
                "ENSMUSG00000024077",
                "ENSMUSG00000030847",
                "ENSMUSG00000059474",
                "ENSMUSG00000065808",
                "ENSMUSG00000028293",
                "ENSMUSG00000046312",
                "ENSMUSG00000018604",
                "ENSMUSG00000004558",
                "ENSMUSG00000062127",
                "ENSMUSG00000037062",
                "ENSMUSG00000071151",
                "ENSMUSG00000021622",
                "ENSMUSG00000026003",
                "ENSMUSG00000034390",
                "ENSMUSG00000085385",
                "ENSMUSG00000029683",
                "ENSMUSG00000046997",
                "ENSMUSG00000018001",
                "ENSMUSG00000061474",
                "ENSMUSG00000050222",
                "ENSMUSG00000034832",
                "ENSMUSG00000056749",
                "ENSMUSG00000030970",
                "ENSMUSG00000048410",
                "ENSMUSG00000055128",
                "ENSMUSG00000086938",
                "ENSMUSG00000033722",
                "ENSMUSG00000043483",
                "ENSMUSG00000018395",
                "ENSMUSG00000029512",
                "ENSMUSG00000002908",
                "ENSMUSG00000027204",
                "ENSMUSG00000022883",
                "ENSMUSG00000028278",
                "ENSMUSG00000044408",
                "ENSMUSG00000026489"
            ],
            "expression_sum": {
                "class1": 3975.0,
                "class2": 674.0
            },
            "uniprot_ids": [
                "Q61164"
            ],
            "net_id": 0,
            "source_num": 7,
            "target_num": 101
        },
        "ENSMUSG00000015605": {
            "id": "ENSMUSG00000015605",
            "type": "TF",
            "symbol": "Srf",
            "source": [
                "ENSMUSG00000029135",
                "ENSMUSG00000020248",
                "ENSMUSG00000001280",
                "ENSMUSG00000019768",
                "ENSMUSG00000017146"
            ],
            "target": [
                "ENSMUSG00000000538",
                "ENSMUSG00000061904",
                "ENSMUSG00000032563",
                "ENSMUSG00000029454",
                "ENSMUSG00000014294",
                "ENSMUSG00000020485",
                "ENSMUSG00000024197",
                "ENSMUSG00000006740",
                "ENSMUSG00000022020",
                "ENSMUSG00000062908",
                "ENSMUSG00000007777",
                "ENSMUSG00000024228",
                "ENSMUSG00000040481",
                "ENSMUSG00000068264",
                "ENSMUSG00000046434",
                "ENSMUSG00000020307",
                "ENSMUSG00000043683",
                "ENSMUSG00000038070",
                "ENSMUSG00000015837",
                "ENSMUSG00000020741",
                "ENSMUSG00000097414",
                "ENSMUSG00000070283",
                "ENSMUSG00000037339",
                "ENSMUSG00000030982",
                "ENSMUSG00000031353",
                "ENSMUSG00000030739",
                "ENSMUSG00000024097",
                "ENSMUSG00000026576",
                "ENSMUSG00000041881",
                "ENSMUSG00000020402",
                "ENSMUSG00000026853",
                "ENSMUSG00000044221",
                "ENSMUSG00000022354",
                "ENSMUSG00000054428",
                "ENSMUSG00000031543",
                "ENSMUSG00000007891",
                "ENSMUSG00000004789",
                "ENSMUSG00000019920",
                "ENSMUSG00000005354",
                "ENSMUSG00000039648",
                "ENSMUSG00000043445",
                "ENSMUSG00000020163",
                "ENSMUSG00000026632",
                "ENSMUSG00000009863",
                "ENSMUSG00000064349",
                "ENSMUSG00000011257",
                "ENSMUSG00000030747",
                "ENSMUSG00000020496",
                "ENSMUSG00000057469",
                "ENSMUSG00000025078",
                "ENSMUSG00000039943",
                "ENSMUSG00000006576",
                "ENSMUSG00000064037",
                "ENSMUSG00000004085",
                "ENSMUSG00000026803",
                "ENSMUSG00000027288",
                "ENSMUSG00000052539",
                "ENSMUSG00000032314",
                "ENSMUSG00000056900",
                "ENSMUSG00000021945",
                "ENSMUSG00000022519",
                "ENSMUSG00000051331",
                "ENSMUSG00000025260",
                "ENSMUSG00000038416",
                "ENSMUSG00000024038",
                "ENSMUSG00000025190",
                "ENSMUSG00000033065",
                "ENSMUSG00000090150",
                "ENSMUSG00000014592",
                "ENSMUSG00000038538",
                "ENSMUSG00000038446",
                "ENSMUSG00000033184",
                "ENSMUSG00000047446",
                "ENSMUSG00000010048",
                "ENSMUSG00000020153",
                "ENSMUSG00000026260",
                "ENSMUSG00000031197",
                "ENSMUSG00000022312",
                "ENSMUSG00000029499",
                "ENSMUSG00000039891",
                "ENSMUSG00000040760",
                "ENSMUSG00000031059",
                "ENSMUSG00000024359",
                "ENSMUSG00000001687",
                "ENSMUSG00000031818",
                "ENSMUSG00000040666",
                "ENSMUSG00000025184",
                "ENSMUSG00000031782",
                "ENSMUSG00000032511",
                "ENSMUSG00000040865",
                "ENSMUSG00000024683",
                "ENSMUSG00000000056",
                "ENSMUSG00000044894",
                "ENSMUSG00000038690",
                "ENSMUSG00000039068",
                "ENSMUSG00000024962",
                "ENSMUSG00000000374",
                "ENSMUSG00000022095",
                "ENSMUSG00000079509",
                "ENSMUSG00000000399",
                "ENSMUSG00000039103",
                "ENSMUSG00000003865",
                "ENSMUSG00000020460",
                "ENSMUSG00000037234",
                "ENSMUSG00000044792",
                "ENSMUSG00000026158",
                "ENSMUSG00000022437",
                "ENSMUSG00000042148",
                "ENSMUSG00000034295",
                "ENSMUSG00000038708",
                "ENSMUSG00000041028",
                "ENSMUSG00000067653",
                "ENSMUSG00000022438",
                "ENSMUSG00000021577",
                "ENSMUSG00000022994",
                "ENSMUSG00000038936",
                "ENSMUSG00000039205",
                "ENSMUSG00000018566",
                "ENSMUSG00000020321",
                "ENSMUSG00000033938",
                "ENSMUSG00000031698",
                "ENSMUSG00000029135",
                "ENSMUSG00000024120",
                "ENSMUSG00000024997",
                "ENSMUSG00000027602",
                "ENSMUSG00000025745",
                "ENSMUSG00000017778",
                "ENSMUSG00000026414",
                "ENSMUSG00000035105",
                "ENSMUSG00000070867",
                "ENSMUSG00000024304",
                "ENSMUSG00000006219",
                "ENSMUSG00000005611",
                "ENSMUSG00000019132",
                "ENSMUSG00000040548",
                "ENSMUSG00000039887",
                "ENSMUSG00000039016",
                "ENSMUSG00000015757",
                "ENSMUSG00000006057",
                "ENSMUSG00000045776",
                "ENSMUSG00000055322",
                "ENSMUSG00000087443",
                "ENSMUSG00000040414",
                "ENSMUSG00000037075",
                "ENSMUSG00000028443",
                "ENSMUSG00000036564",
                "ENSMUSG00000037049",
                "ENSMUSG00000024236",
                "ENSMUSG00000091831",
                "ENSMUSG00000057841",
                "ENSMUSG00000020079",
                "ENSMUSG00000061518",
                "ENSMUSG00000033400",
                "ENSMUSG00000073176",
                "ENSMUSG00000032480",
                "ENSMUSG00000032648",
                "ENSMUSG00000050541",
                "ENSMUSG00000003299",
                "ENSMUSG00000015937",
                "ENSMUSG00000021549",
                "ENSMUSG00000000001",
                "ENSMUSG00000031447",
                "ENSMUSG00000053929",
                "ENSMUSG00000026568",
                "ENSMUSG00000022706",
                "ENSMUSG00000005683",
                "ENSMUSG00000023020",
                "ENSMUSG00000039682",
                "ENSMUSG00000005882",
                "ENSMUSG00000046516",
                "ENSMUSG00000083831",
                "ENSMUSG00000040112",
                "ENSMUSG00000058655",
                "ENSMUSG00000052934",
                "ENSMUSG00000060002",
                "ENSMUSG00000019768",
                "ENSMUSG00000030335",
                "ENSMUSG00000047419",
                "ENSMUSG00000020085",
                "ENSMUSG00000016349",
                "ENSMUSG00000082165",
                "ENSMUSG00000031402",
                "ENSMUSG00000031536",
                "ENSMUSG00000063594",
                "ENSMUSG00000001334",
                "ENSMUSG00000030785",
                "ENSMUSG00000039218",
                "ENSMUSG00000058076",
                "ENSMUSG00000055027",
                "ENSMUSG00000028165",
                "ENSMUSG00000029594",
                "ENSMUSG00000044864",
                "ENSMUSG00000039461",
                "ENSMUSG00000032750",
                "ENSMUSG00000066637",
                "ENSMUSG00000059714",
                "ENSMUSG00000000171",
                "ENSMUSG00000000441",
                "ENSMUSG00000034220",
                "ENSMUSG00000030393",
                "ENSMUSG00000033060",
                "ENSMUSG00000025939",
                "ENSMUSG00000005610",
                "ENSMUSG00000038784",
                "ENSMUSG00000036748",
                "ENSMUSG00000053483",
                "ENSMUSG00000031231",
                "ENSMUSG00000032826",
                "ENSMUSG00000002608",
                "ENSMUSG00000039601",
                "ENSMUSG00000026895",
                "ENSMUSG00000033059",
                "ENSMUSG00000014313",
                "ENSMUSG00000035936",
                "ENSMUSG00000000088",
                "ENSMUSG00000081228",
                "ENSMUSG00000013275",
                "ENSMUSG00000069272",
                "ENSMUSG00000033594",
                "ENSMUSG00000027668",
                "ENSMUSG00000037949",
                "ENSMUSG00000025393",
                "ENSMUSG00000055485",
                "ENSMUSG00000064343",
                "ENSMUSG00000031232",
                "ENSMUSG00000032846",
                "ENSMUSG00000048814",
                "ENSMUSG00000038583",
                "ENSMUSG00000031641",
                "ENSMUSG00000038967",
                "ENSMUSG00000047945",
                "ENSMUSG00000045624",
                "ENSMUSG00000036112",
                "ENSMUSG00000056091",
                "ENSMUSG00000020116",
                "ENSMUSG00000071528",
                "ENSMUSG00000027333",
                "ENSMUSG00000026585",
                "ENSMUSG00000024588",
                "ENSMUSG00000040312",
                "ENSMUSG00000035674",
                "ENSMUSG00000036109",
                "ENSMUSG00000042198",
                "ENSMUSG00000009927",
                "ENSMUSG00000002416",
                "ENSMUSG00000031841",
                "ENSMUSG00000026571",
                "ENSMUSG00000036854",
                "ENSMUSG00000025465",
                "ENSMUSG00000026131",
                "ENSMUSG00000026202",
                "ENSMUSG00000066839",
                "ENSMUSG00000001383",
                "ENSMUSG00000034951",
                "ENSMUSG00000028919",
                "ENSMUSG00000040250",
                "ENSMUSG00000022404",
                "ENSMUSG00000020326",
                "ENSMUSG00000028809",
                "ENSMUSG00000017146",
                "ENSMUSG00000021460",
                "ENSMUSG00000022089",
                "ENSMUSG00000022477",
                "ENSMUSG00000023055",
                "ENSMUSG00000032010",
                "ENSMUSG00000068614",
                "ENSMUSG00000031672",
                "ENSMUSG00000006024",
                "ENSMUSG00000064347",
                "ENSMUSG00000050721",
                "ENSMUSG00000022194",
                "ENSMUSG00000017756",
                "ENSMUSG00000033706",
                "ENSMUSG00000059291",
                "ENSMUSG00000023852",
                "ENSMUSG00000023150",
                "ENSMUSG00000012405",
                "ENSMUSG00000040506",
                "ENSMUSG00000030695",
                "ENSMUSG00000092118",
                "ENSMUSG00000013539",
                "ENSMUSG00000015850",
                "ENSMUSG00000001366",
                "ENSMUSG00000054934",
                "ENSMUSG00000025813",
                "ENSMUSG00000002379",
                "ENSMUSG00000022110",
                "ENSMUSG00000022214",
                "ENSMUSG00000028648",
                "ENSMUSG00000097729",
                "ENSMUSG00000025404",
                "ENSMUSG00000037916",
                "ENSMUSG00000043336",
                "ENSMUSG00000027177",
                "ENSMUSG00000000555",
                "ENSMUSG00000050608",
                "ENSMUSG00000039763",
                "ENSMUSG00000025949",
                "ENSMUSG00000019851",
                "ENSMUSG00000059208",
                "ENSMUSG00000000628",
                "ENSMUSG00000074064",
                "ENSMUSG00000020677",
                "ENSMUSG00000046727",
                "ENSMUSG00000040771",
                "ENSMUSG00000025428",
                "ENSMUSG00000027010",
                "ENSMUSG00000032724",
                "ENSMUSG00000033161",
                "ENSMUSG00000038412",
                "ENSMUSG00000003072",
                "ENSMUSG00000051285",
                "ENSMUSG00000024063",
                "ENSMUSG00000037349",
                "ENSMUSG00000039474",
                "ENSMUSG00000044475",
                "ENSMUSG00000071454",
                "ENSMUSG00000029017",
                "ENSMUSG00000030016",
                "ENSMUSG00000013921",
                "ENSMUSG00000041426",
                "ENSMUSG00000029128",
                "ENSMUSG00000020849",
                "ENSMUSG00000074218",
                "ENSMUSG00000021798",
                "ENSMUSG00000024924",
                "ENSMUSG00000027792",
                "ENSMUSG00000021806",
                "ENSMUSG00000042744",
                "ENSMUSG00000001280",
                "ENSMUSG00000060594",
                "ENSMUSG00000030417",
                "ENSMUSG00000055737",
                "ENSMUSG00000062604",
                "ENSMUSG00000063870",
                "ENSMUSG00000032902",
                "ENSMUSG00000031633",
                "ENSMUSG00000047921",
                "ENSMUSG00000063694",
                "ENSMUSG00000028367",
                "ENSMUSG00000055302",
                "ENSMUSG00000037826",
                "ENSMUSG00000029651",
                "ENSMUSG00000024853",
                "ENSMUSG00000039089",
                "ENSMUSG00000053898",
                "ENSMUSG00000021906",
                "ENSMUSG00000000751",
                "ENSMUSG00000000563",
                "ENSMUSG00000083820",
                "ENSMUSG00000001105",
                "ENSMUSG00000040385",
                "ENSMUSG00000037815",
                "ENSMUSG00000021313",
                "ENSMUSG00000071317",
                "ENSMUSG00000027809",
                "ENSMUSG00000016252",
                "ENSMUSG00000026207",
                "ENSMUSG00000059430",
                "ENSMUSG00000054766",
                "ENSMUSG00000015112",
                "ENSMUSG00000038949",
                "ENSMUSG00000047187",
                "ENSMUSG00000075054",
                "ENSMUSG00000035776",
                "ENSMUSG00000021453",
                "ENSMUSG00000052337",
                "ENSMUSG00000062981",
                "ENSMUSG00000028223",
                "ENSMUSG00000028552",
                "ENSMUSG00000031024",
                "ENSMUSG00000020456",
                "ENSMUSG00000045983",
                "ENSMUSG00000051747",
                "ENSMUSG00000064342",
                "ENSMUSG00000035458",
                "ENSMUSG00000031791",
                "ENSMUSG00000032481",
                "ENSMUSG00000048264",
                "ENSMUSG00000063882",
                "ENSMUSG00000001436",
                "ENSMUSG00000030399",
                "ENSMUSG00000053007",
                "ENSMUSG00000028745",
                "ENSMUSG00000020483",
                "ENSMUSG00000027823",
                "ENSMUSG00000097589",
                "ENSMUSG00000022890",
                "ENSMUSG00000040723",
                "ENSMUSG00000022051",
                "ENSMUSG00000018770",
                "ENSMUSG00000026878",
                "ENSMUSG00000001017",
                "ENSMUSG00000032959",
                "ENSMUSG00000031320",
                "ENSMUSG00000040850",
                "ENSMUSG00000049957",
                "ENSMUSG00000042032",
                "ENSMUSG00000032224",
                "ENSMUSG00000029020",
                "ENSMUSG00000058793",
                "ENSMUSG00000049643",
                "ENSMUSG00000033039",
                "ENSMUSG00000031985",
                "ENSMUSG00000038375",
                "ENSMUSG00000052738",
                "ENSMUSG00000029359",
                "ENSMUSG00000023175",
                "ENSMUSG00000038121",
                "ENSMUSG00000060600",
                "ENSMUSG00000090035",
                "ENSMUSG00000027406",
                "ENSMUSG00000024527",
                "ENSMUSG00000024991",
                "ENSMUSG00000060032",
                "ENSMUSG00000027968",
                "ENSMUSG00000022201",
                "ENSMUSG00000028756",
                "ENSMUSG00000024143",
                "ENSMUSG00000026927",
                "ENSMUSG00000020248",
                "ENSMUSG00000002343",
                "ENSMUSG00000019577",
                "ENSMUSG00000002010",
                "ENSMUSG00000032590",
                "ENSMUSG00000044042",
                "ENSMUSG00000054889",
                "ENSMUSG00000046351",
                "ENSMUSG00000076437",
                "ENSMUSG00000019998",
                "ENSMUSG00000006021",
                "ENSMUSG00000022982",
                "ENSMUSG00000019179",
                "ENSMUSG00000028273",
                "ENSMUSG00000019158",
                "ENSMUSG00000024309",
                "ENSMUSG00000050199",
                "ENSMUSG00000051495",
                "ENSMUSG00000021200",
                "ENSMUSG00000022964",
                "ENSMUSG00000058594",
                "ENSMUSG00000032786",
                "ENSMUSG00000050856",
                "ENSMUSG00000025856",
                "ENSMUSG00000071415",
                "ENSMUSG00000045613",
                "ENSMUSG00000050323",
                "ENSMUSG00000026032",
                "ENSMUSG00000020708",
                "ENSMUSG00000021807",
                "ENSMUSG00000022803",
                "ENSMUSG00000053768",
                "ENSMUSG00000025781",
                "ENSMUSG00000028452",
                "ENSMUSG00000062169",
                "ENSMUSG00000070802",
                "ENSMUSG00000033781",
                "ENSMUSG00000003316",
                "ENSMUSG00000024576",
                "ENSMUSG00000018796",
                "ENSMUSG00000022673",
                "ENSMUSG00000008690",
                "ENSMUSG00000044080",
                "ENSMUSG00000031461",
                "ENSMUSG00000036202",
                "ENSMUSG00000029924",
                "ENSMUSG00000046671",
                "ENSMUSG00000028622",
                "ENSMUSG00000028834",
                "ENSMUSG00000060679",
                "ENSMUSG00000073838",
                "ENSMUSG00000038844",
                "ENSMUSG00000031312",
                "ENSMUSG00000059734",
                "ENSMUSG00000040752",
                "ENSMUSG00000023723",
                "ENSMUSG00000030884",
                "ENSMUSG00000032279",
                "ENSMUSG00000023861",
                "ENSMUSG00000066771",
                "ENSMUSG00000026664",
                "ENSMUSG00000074207",
                "ENSMUSG00000034271",
                "ENSMUSG00000018567",
                "ENSMUSG00000042558",
                "ENSMUSG00000036822",
                "ENSMUSG00000091537",
                "ENSMUSG00000068523",
                "ENSMUSG00000043822",
                "ENSMUSG00000072407",
                "ENSMUSG00000032051",
                "ENSMUSG00000021520",
                "ENSMUSG00000006717",
                "ENSMUSG00000034994",
                "ENSMUSG00000030541",
                "ENSMUSG00000026208",
                "ENSMUSG00000022235",
                "ENSMUSG00000036352",
                "ENSMUSG00000059534",
                "ENSMUSG00000020542",
                "ENSMUSG00000030059",
                "ENSMUSG00000046314",
                "ENSMUSG00000003438",
                "ENSMUSG00000025968",
                "ENSMUSG00000027367",
                "ENSMUSG00000057363",
                "ENSMUSG00000080836",
                "ENSMUSG00000061666",
                "ENSMUSG00000027238",
                "ENSMUSG00000044308",
                "ENSMUSG00000022300",
                "ENSMUSG00000020821",
                "ENSMUSG00000018599",
                "ENSMUSG00000024140",
                "ENSMUSG00000081700",
                "ENSMUSG00000064344",
                "ENSMUSG00000047649",
                "ENSMUSG00000029190",
                "ENSMUSG00000027499",
                "ENSMUSG00000037805",
                "ENSMUSG00000004610",
                "ENSMUSG00000015944",
                "ENSMUSG00000021265",
                "ENSMUSG00000021290",
                "ENSMUSG00000049470",
                "ENSMUSG00000033307",
                "ENSMUSG00000017119",
                "ENSMUSG00000069049",
                "ENSMUSG00000013936",
                "ENSMUSG00000027455",
                "ENSMUSG00000009291",
                "ENSMUSG00000064340",
                "ENSMUSG00000029401",
                "ENSMUSG00000032047",
                "ENSMUSG00000029467",
                "ENSMUSG00000025453",
                "ENSMUSG00000024151",
                "ENSMUSG00000039219",
                "ENSMUSG00000069089",
                "ENSMUSG00000020415",
                "ENSMUSG00000038022",
                "ENSMUSG00000072720",
                "ENSMUSG00000028706",
                "ENSMUSG00000044393",
                "ENSMUSG00000014470",
                "ENSMUSG00000021982",
                "ENSMUSG00000021606",
                "ENSMUSG00000022193",
                "ENSMUSG00000024073",
                "ENSMUSG00000036002",
                "ENSMUSG00000062580",
                "ENSMUSG00000020430",
                "ENSMUSG00000026817",
                "ENSMUSG00000022031",
                "ENSMUSG00000009112",
                "ENSMUSG00000026864",
                "ENSMUSG00000019877",
                "ENSMUSG00000023939",
                "ENSMUSG00000059325",
                "ENSMUSG00000033416",
                "ENSMUSG00000026135",
                "ENSMUSG00000029414",
                "ENSMUSG00000027422",
                "ENSMUSG00000062270",
                "ENSMUSG00000028937",
                "ENSMUSG00000015357",
                "ENSMUSG00000037622",
                "ENSMUSG00000032330",
                "ENSMUSG00000050043",
                "ENSMUSG00000022905",
                "ENSMUSG00000026074",
                "ENSMUSG00000062300",
                "ENSMUSG00000018882",
                "ENSMUSG00000038170",
                "ENSMUSG00000013593",
                "ENSMUSG00000021771",
                "ENSMUSG00000032267",
                "ENSMUSG00000006315",
                "ENSMUSG00000064365",
                "ENSMUSG00000068267",
                "ENSMUSG00000042293",
                "ENSMUSG00000062373",
                "ENSMUSG00000028601",
                "ENSMUSG00000037710",
                "ENSMUSG00000041361",
                "ENSMUSG00000041476",
                "ENSMUSG00000049327",
                "ENSMUSG00000043866",
                "ENSMUSG00000038520",
                "ENSMUSG00000059810",
                "ENSMUSG00000016427",
                "ENSMUSG00000031865",
                "ENSMUSG00000022215",
                "ENSMUSG00000019087",
                "ENSMUSG00000023959",
                "ENSMUSG00000027293",
                "ENSMUSG00000000149",
                "ENSMUSG00000009545",
                "ENSMUSG00000020988",
                "ENSMUSG00000023206",
                "ENSMUSG00000028518",
                "ENSMUSG00000035268",
                "ENSMUSG00000008136",
                "ENSMUSG00000039670",
                "ENSMUSG00000005873",
                "ENSMUSG00000027189",
                "ENSMUSG00000024805",
                "ENSMUSG00000073563",
                "ENSMUSG00000032366",
                "ENSMUSG00000037236",
                "ENSMUSG00000051391",
                "ENSMUSG00000064369",
                "ENSMUSG00000022265",
                "ENSMUSG00000023944",
                "ENSMUSG00000020358",
                "ENSMUSG00000022681",
                "ENSMUSG00000036880",
                "ENSMUSG00000053819",
                "ENSMUSG00000068699",
                "ENSMUSG00000032381",
                "ENSMUSG00000021280",
                "ENSMUSG00000003970",
                "ENSMUSG00000022551",
                "ENSMUSG00000030662",
                "ENSMUSG00000037110",
                "ENSMUSG00000064348",
                "ENSMUSG00000025651",
                "ENSMUSG00000010911",
                "ENSMUSG00000024158",
                "ENSMUSG00000050953",
                "ENSMUSG00000022965",
                "ENSMUSG00000027765",
                "ENSMUSG00000024639",
                "ENSMUSG00000030847",
                "ENSMUSG00000038299",
                "ENSMUSG00000029338",
                "ENSMUSG00000006005",
                "ENSMUSG00000028293",
                "ENSMUSG00000035293",
                "ENSMUSG00000037062",
                "ENSMUSG00000071151",
                "ENSMUSG00000019787",
                "ENSMUSG00000029683",
                "ENSMUSG00000023089",
                "ENSMUSG00000046997",
                "ENSMUSG00000020121",
                "ENSMUSG00000030246",
                "ENSMUSG00000034088",
                "ENSMUSG00000078851",
                "ENSMUSG00000073676",
                "ENSMUSG00000023456",
                "ENSMUSG00000038462",
                "ENSMUSG00000020664",
                "ENSMUSG00000030869",
                "ENSMUSG00000086938",
                "ENSMUSG00000022360",
                "ENSMUSG00000022191",
                "ENSMUSG00000047881",
                "ENSMUSG00000017817",
                "ENSMUSG00000034936",
                "ENSMUSG00000033684",
                "ENSMUSG00000059811",
                "ENSMUSG00000084105",
                "ENSMUSG00000024099",
                "ENSMUSG00000024587",
                "ENSMUSG00000026489"
            ],
            "expression_sum": {
                "class1": 510.0,
                "class2": 502.0
            },
            "uniprot_ids": [
                "Q9JM73",
                "A0A3B2WCW1"
            ],
            "net_id": 0,
            "source_num": 5,
            "target_num": 252
        },
        "ENSMUSG00000029135": {
            "id": "ENSMUSG00000029135",
            "type": "TF",
            "symbol": "Fosl2",
            "source": [
                "ENSMUSG00000021250",
                "ENSMUSG00000015605",
                "ENSMUSG00000008575",
                "ENSMUSG00000057469",
                "ENSMUSG00000010175",
                "ENSMUSG00000018604",
                "ENSMUSG00000019817",
                "ENSMUSG00000015579",
                "ENSMUSG00000019768",
                "ENSMUSG00000005698",
                "ENSMUSG00000022641",
                "ENSMUSG00000053110",
                "ENSMUSG00000022895",
                "ENSMUSG00000015839",
                "ENSMUSG00000032228",
                "ENSMUSG00000053477",
                "ENSMUSG00000058756",
                "ENSMUSG00000052837",
                "ENSMUSG00000056501",
                "ENSMUSG00000001280"
            ],
            "target": [
                "ENSMUSG00000029860",
                "ENSMUSG00000039782",
                "ENSMUSG00000061904",
                "ENSMUSG00000032563",
                "ENSMUSG00000029454",
                "ENSMUSG00000001741",
                "ENSMUSG00000021939",
                "ENSMUSG00000004937",
                "ENSMUSG00000015605",
                "ENSMUSG00000020485",
                "ENSMUSG00000024197",
                "ENSMUSG00000031954",
                "ENSMUSG00000027777",
                "ENSMUSG00000007777",
                "ENSMUSG00000019432",
                "ENSMUSG00000027720",
                "ENSMUSG00000002661",
                "ENSMUSG00000068264",
                "ENSMUSG00000040269",
                "ENSMUSG00000022462",
                "ENSMUSG00000021248",
                "ENSMUSG00000002486",
                "ENSMUSG00000027447",
                "ENSMUSG00000015837",
                "ENSMUSG00000002028",
                "ENSMUSG00000020741",
                "ENSMUSG00000044791",
                "ENSMUSG00000021676",
                "ENSMUSG00000037266",
                "ENSMUSG00000076490",
                "ENSMUSG00000021235",
                "ENSMUSG00000030319",
                "ENSMUSG00000062647",
                "ENSMUSG00000011427",
                "ENSMUSG00000035356",
                "ENSMUSG00000031216",
                "ENSMUSG00000003038",
                "ENSMUSG00000052397",
                "ENSMUSG00000014402",
                "ENSMUSG00000042709",
                "ENSMUSG00000024097",
                "ENSMUSG00000031540",
                "ENSMUSG00000032525",
                "ENSMUSG00000041881",
                "ENSMUSG00000019850",
                "ENSMUSG00000022354",
                "ENSMUSG00000028926",
                "ENSMUSG00000066900",
                "ENSMUSG00000058503",
                "ENSMUSG00000021764",
                "ENSMUSG00000007891",
                "ENSMUSG00000019920",
                "ENSMUSG00000020477",
                "ENSMUSG00000021270",
                "ENSMUSG00000018286",
                "ENSMUSG00000011257",
                "ENSMUSG00000020496",
                "ENSMUSG00000057469",
                "ENSMUSG00000031770",
                "ENSMUSG00000039943",
                "ENSMUSG00000002289",
                "ENSMUSG00000005469",
                "ENSMUSG00000027288",
                "ENSMUSG00000052539",
                "ENSMUSG00000029152",
                "ENSMUSG00000051048",
                "ENSMUSG00000069516",
                "ENSMUSG00000016528",
                "ENSMUSG00000036097",
                "ENSMUSG00000020661",
                "ENSMUSG00000057738",
                "ENSMUSG00000014592",
                "ENSMUSG00000038538",
                "ENSMUSG00000030850",
                "ENSMUSG00000079508",
                "ENSMUSG00000033184",
                "ENSMUSG00000036678",
                "ENSMUSG00000010067",
                "ENSMUSG00000011884",
                "ENSMUSG00000014606",
                "ENSMUSG00000030120",
                "ENSMUSG00000030861",
                "ENSMUSG00000028161",
                "ENSMUSG00000036893",
                "ENSMUSG00000040661",
                "ENSMUSG00000026260",
                "ENSMUSG00000025809",
                "ENSMUSG00000049047",
                "ENSMUSG00000032216",
                "ENSMUSG00000026688",
                "ENSMUSG00000026254",
                "ENSMUSG00000026755",
                "ENSMUSG00000035385",
                "ENSMUSG00000024359",
                "ENSMUSG00000069729",
                "ENSMUSG00000019943",
                "ENSMUSG00000064302",
                "ENSMUSG00000034574",
                "ENSMUSG00000025184",
                "ENSMUSG00000031782",
                "ENSMUSG00000048486",
                "ENSMUSG00000055148",
                "ENSMUSG00000026775",
                "ENSMUSG00000032511",
                "ENSMUSG00000019817",
                "ENSMUSG00000024683",
                "ENSMUSG00000029392",
                "ENSMUSG00000032883",
                "ENSMUSG00000042197",
                "ENSMUSG00000044894",
                "ENSMUSG00000029404",
                "ENSMUSG00000036438",
                "ENSMUSG00000048578",
                "ENSMUSG00000057133",
                "ENSMUSG00000028954",
                "ENSMUSG00000000399",
                "ENSMUSG00000017288",
                "ENSMUSG00000083022",
                "ENSMUSG00000003865",
                "ENSMUSG00000020460",
                "ENSMUSG00000044792",
                "ENSMUSG00000066037",
                "ENSMUSG00000020349",
                "ENSMUSG00000078180",
                "ENSMUSG00000039515",
                "ENSMUSG00000057596",
                "ENSMUSG00000021772",
                "ENSMUSG00000063229",
                "ENSMUSG00000026353",
                "ENSMUSG00000049755",
                "ENSMUSG00000005417",
                "ENSMUSG00000022037",
                "ENSMUSG00000022048",
                "ENSMUSG00000045427",
                "ENSMUSG00000055320",
                "ENSMUSG00000045128",
                "ENSMUSG00000024277",
                "ENSMUSG00000029776",
                "ENSMUSG00000029614",
                "ENSMUSG00000025134",
                "ENSMUSG00000025898",
                "ENSMUSG00000056220",
                "ENSMUSG00000020952",
                "ENSMUSG00000039233",
                "ENSMUSG00000031785",
                "ENSMUSG00000037321",
                "ENSMUSG00000046329",
                "ENSMUSG00000029064",
                "ENSMUSG00000045210",
                "ENSMUSG00000006219",
                "ENSMUSG00000019132",
                "ENSMUSG00000024381",
                "ENSMUSG00000025130",
                "ENSMUSG00000055322",
                "ENSMUSG00000045776",
                "ENSMUSG00000006057",
                "ENSMUSG00000020580",
                "ENSMUSG00000009687",
                "ENSMUSG00000068923",
                "ENSMUSG00000022369",
                "ENSMUSG00000030647",
                "ENSMUSG00000027879",
                "ENSMUSG00000060798",
                "ENSMUSG00000036564",
                "ENSMUSG00000061518",
                "ENSMUSG00000037072",
                "ENSMUSG00000043885",
                "ENSMUSG00000000184",
                "ENSMUSG00000030180",
                "ENSMUSG00000025935",
                "ENSMUSG00000026374",
                "ENSMUSG00000050541",
                "ENSMUSG00000015937",
                "ENSMUSG00000030560",
                "ENSMUSG00000000001",
                "ENSMUSG00000031447",
                "ENSMUSG00000053929",
                "ENSMUSG00000006333",
                "ENSMUSG00000026568",
                "ENSMUSG00000026123",
                "ENSMUSG00000020180",
                "ENSMUSG00000026096",
                "ENSMUSG00000023020",
                "ENSMUSG00000005882",
                "ENSMUSG00000020455",
                "ENSMUSG00000021127",
                "ENSMUSG00000023868",
                "ENSMUSG00000037960",
                "ENSMUSG00000052934",
                "ENSMUSG00000030421",
                "ENSMUSG00000034424",
                "ENSMUSG00000058135",
                "ENSMUSG00000038013",
                "ENSMUSG00000071369",
                "ENSMUSG00000031029",
                "ENSMUSG00000031536",
                "ENSMUSG00000006494",
                "ENSMUSG00000032348",
                "ENSMUSG00000023832",
                "ENSMUSG00000079184",
                "ENSMUSG00000004980",
                "ENSMUSG00000039218",
                "ENSMUSG00000032580",
                "ENSMUSG00000029594",
                "ENSMUSG00000055027",
                "ENSMUSG00000044864",
                "ENSMUSG00000071014",
                "ENSMUSG00000029576",
                "ENSMUSG00000032750",
                "ENSMUSG00000028116",
                "ENSMUSG00000028973",
                "ENSMUSG00000000441",
                "ENSMUSG00000010362",
                "ENSMUSG00000041685",
                "ENSMUSG00000037465",
                "ENSMUSG00000038784",
                "ENSMUSG00000005610",
                "ENSMUSG00000031812",
                "ENSMUSG00000079597",
                "ENSMUSG00000028226",
                "ENSMUSG00000032418",
                "ENSMUSG00000046032",
                "ENSMUSG00000024695",
                "ENSMUSG00000032826",
                "ENSMUSG00000028955",
                "ENSMUSG00000057322",
                "ENSMUSG00000026179",
                "ENSMUSG00000026701",
                "ENSMUSG00000022270",
                "ENSMUSG00000022136",
                "ENSMUSG00000045690",
                "ENSMUSG00000033813",
                "ENSMUSG00000079435",
                "ENSMUSG00000022185",
                "ENSMUSG00000032526",
                "ENSMUSG00000000088",
                "ENSMUSG00000031626",
                "ENSMUSG00000000340",
                "ENSMUSG00000029422",
                "ENSMUSG00000029465",
                "ENSMUSG00000042462",
                "ENSMUSG00000026393",
                "ENSMUSG00000037949",
                "ENSMUSG00000028790",
                "ENSMUSG00000041560",
                "ENSMUSG00000083743",
                "ENSMUSG00000043510",
                "ENSMUSG00000002396",
                "ENSMUSG00000027665",
                "ENSMUSG00000001419",
                "ENSMUSG00000022765",
                "ENSMUSG00000024845",
                "ENSMUSG00000008429",
                "ENSMUSG00000036112",
                "ENSMUSG00000042520",
                "ENSMUSG00000045624",
                "ENSMUSG00000027333",
                "ENSMUSG00000026754",
                "ENSMUSG00000064899",
                "ENSMUSG00000047260",
                "ENSMUSG00000069833",
                "ENSMUSG00000042198",
                "ENSMUSG00000047710",
                "ENSMUSG00000023249",
                "ENSMUSG00000006299",
                "ENSMUSG00000002416",
                "ENSMUSG00000095306",
                "ENSMUSG00000026571",
                "ENSMUSG00000036854",
                "ENSMUSG00000026131",
                "ENSMUSG00000026202",
                "ENSMUSG00000042605",
                "ENSMUSG00000028849",
                "ENSMUSG00000027566",
                "ENSMUSG00000030315",
                "ENSMUSG00000028919",
                "ENSMUSG00000033538",
                "ENSMUSG00000028124",
                "ENSMUSG00000067058",
                "ENSMUSG00000038127",
                "ENSMUSG00000036867",
                "ENSMUSG00000023055",
                "ENSMUSG00000022477",
                "ENSMUSG00000031672",
                "ENSMUSG00000019988",
                "ENSMUSG00000022194",
                "ENSMUSG00000034892",
                "ENSMUSG00000059291",
                "ENSMUSG00000028649",
                "ENSMUSG00000028300",
                "ENSMUSG00000026577",
                "ENSMUSG00000023150",
                "ENSMUSG00000020647",
                "ENSMUSG00000012405",
                "ENSMUSG00000069014",
                "ENSMUSG00000063800",
                "ENSMUSG00000029062",
                "ENSMUSG00000030695",
                "ENSMUSG00000032177",
                "ENSMUSG00000027332",
                "ENSMUSG00000015850",
                "ENSMUSG00000022110",
                "ENSMUSG00000004356",
                "ENSMUSG00000027177",
                "ENSMUSG00000004535",
                "ENSMUSG00000030779",
                "ENSMUSG00000002949",
                "ENSMUSG00000060510",
                "ENSMUSG00000029670",
                "ENSMUSG00000019851",
                "ENSMUSG00000059208",
                "ENSMUSG00000030204",
                "ENSMUSG00000010406",
                "ENSMUSG00000040771",
                "ENSMUSG00000040488",
                "ENSMUSG00000058690",
                "ENSMUSG00000027010",
                "ENSMUSG00000033161",
                "ENSMUSG00000026150",
                "ENSMUSG00000081214",
                "ENSMUSG00000003072",
                "ENSMUSG00000020390",
                "ENSMUSG00000024063",
                "ENSMUSG00000021585",
                "ENSMUSG00000031556",
                "ENSMUSG00000041426",
                "ENSMUSG00000029128",
                "ENSMUSG00000020849",
                "ENSMUSG00000028343",
                "ENSMUSG00000031634",
                "ENSMUSG00000021798",
                "ENSMUSG00000074218",
                "ENSMUSG00000002748",
                "ENSMUSG00000041297",
                "ENSMUSG00000000168",
                "ENSMUSG00000048661",
                "ENSMUSG00000001280",
                "ENSMUSG00000026113",
                "ENSMUSG00000040952",
                "ENSMUSG00000043505",
                "ENSMUSG00000026674",
                "ENSMUSG00000038970",
                "ENSMUSG00000040435",
                "ENSMUSG00000062604",
                "ENSMUSG00000034345",
                "ENSMUSG00000027861",
                "ENSMUSG00000063694",
                "ENSMUSG00000041235",
                "ENSMUSG00000039089",
                "ENSMUSG00000020368",
                "ENSMUSG00000039473",
                "ENSMUSG00000020674",
                "ENSMUSG00000021906",
                "ENSMUSG00000035352",
                "ENSMUSG00000042772",
                "ENSMUSG00000025351",
                "ENSMUSG00000002831",
                "ENSMUSG00000030409",
                "ENSMUSG00000040648",
                "ENSMUSG00000046324",
                "ENSMUSG00000040385",
                "ENSMUSG00000027809",
                "ENSMUSG00000030793",
                "ENSMUSG00000022641",
                "ENSMUSG00000039478",
                "ENSMUSG00000054766",
                "ENSMUSG00000030056",
                "ENSMUSG00000018433",
                "ENSMUSG00000062981",
                "ENSMUSG00000028223",
                "ENSMUSG00000022309",
                "ENSMUSG00000037601",
                "ENSMUSG00000002496",
                "ENSMUSG00000020456",
                "ENSMUSG00000045983",
                "ENSMUSG00000021281",
                "ENSMUSG00000001794",
                "ENSMUSG00000032030",
                "ENSMUSG00000038642",
                "ENSMUSG00000064210",
                "ENSMUSG00000031791",
                "ENSMUSG00000048264",
                "ENSMUSG00000067274",
                "ENSMUSG00000037820",
                "ENSMUSG00000019505",
                "ENSMUSG00000020523",
                "ENSMUSG00000022013",
                "ENSMUSG00000029580",
                "ENSMUSG00000028247",
                "ENSMUSG00000028745",
                "ENSMUSG00000035152",
                "ENSMUSG00000053702",
                "ENSMUSG00000035373",
                "ENSMUSG00000020267",
                "ENSMUSG00000090516",
                "ENSMUSG00000040723",
                "ENSMUSG00000022051",
                "ENSMUSG00000006818",
                "ENSMUSG00000018770",
                "ENSMUSG00000040785",
                "ENSMUSG00000022336",
                "ENSMUSG00000019810",
                "ENSMUSG00000032199",
                "ENSMUSG00000029767",
                "ENSMUSG00000034681",
                "ENSMUSG00000058600",
                "ENSMUSG00000028964",
                "ENSMUSG00000040850",
                "ENSMUSG00000027706",
                "ENSMUSG00000028410",
                "ENSMUSG00000001211",
                "ENSMUSG00000049643",
                "ENSMUSG00000033039",
                "ENSMUSG00000031985",
                "ENSMUSG00000038375",
                "ENSMUSG00000029918",
                "ENSMUSG00000029359",
                "ENSMUSG00000040152",
                "ENSMUSG00000022452",
                "ENSMUSG00000023175",
                "ENSMUSG00000038145",
                "ENSMUSG00000006931",
                "ENSMUSG00000024991",
                "ENSMUSG00000032399",
                "ENSMUSG00000024687",
                "ENSMUSG00000028756",
                "ENSMUSG00000022201",
                "ENSMUSG00000080893",
                "ENSMUSG00000002102",
                "ENSMUSG00000025374",
                "ENSMUSG00000086922",
                "ENSMUSG00000056054",
                "ENSMUSG00000026927",
                "ENSMUSG00000022940",
                "ENSMUSG00000002343",
                "ENSMUSG00000039652",
                "ENSMUSG00000050002",
                "ENSMUSG00000034708",
                "ENSMUSG00000002010",
                "ENSMUSG00000026385",
                "ENSMUSG00000045954",
                "ENSMUSG00000054889",
                "ENSMUSG00000003955",
                "ENSMUSG00000074781",
                "ENSMUSG00000024953",
                "ENSMUSG00000030060",
                "ENSMUSG00000028234",
                "ENSMUSG00000061838",
                "ENSMUSG00000034118",
                "ENSMUSG00000019179",
                "ENSMUSG00000058558",
                "ENSMUSG00000019158",
                "ENSMUSG00000025362",
                "ENSMUSG00000001156",
                "ENSMUSG00000027881",
                "ENSMUSG00000022841",
                "ENSMUSG00000058594",
                "ENSMUSG00000021591",
                "ENSMUSG00000079477",
                "ENSMUSG00000071415",
                "ENSMUSG00000068479",
                "ENSMUSG00000021807",
                "ENSMUSG00000022803",
                "ENSMUSG00000023010",
                "ENSMUSG00000032231",
                "ENSMUSG00000021607",
                "ENSMUSG00000036278",
                "ENSMUSG00000032407",
                "ENSMUSG00000096403",
                "ENSMUSG00000062169",
                "ENSMUSG00000028348",
                "ENSMUSG00000067369",
                "ENSMUSG00000033781",
                "ENSMUSG00000003316",
                "ENSMUSG00000024576",
                "ENSMUSG00000018796",
                "ENSMUSG00000038774",
                "ENSMUSG00000008690",
                "ENSMUSG00000044080",
                "ENSMUSG00000005510",
                "ENSMUSG00000024208",
                "ENSMUSG00000020720",
                "ENSMUSG00000055900",
                "ENSMUSG00000046671",
                "ENSMUSG00000087679",
                "ENSMUSG00000020372",
                "ENSMUSG00000022781",
                "ENSMUSG00000041654",
                "ENSMUSG00000031352",
                "ENSMUSG00000021877",
                "ENSMUSG00000087260",
                "ENSMUSG00000039536",
                "ENSMUSG00000027272",
                "ENSMUSG00000000740",
                "ENSMUSG00000053110",
                "ENSMUSG00000010608",
                "ENSMUSG00000031749",
                "ENSMUSG00000024803",
                "ENSMUSG00000030884",
                "ENSMUSG00000023861",
                "ENSMUSG00000032279",
                "ENSMUSG00000022255",
                "ENSMUSG00000031683",
                "ENSMUSG00000018567",
                "ENSMUSG00000022000",
                "ENSMUSG00000033760",
                "ENSMUSG00000023025",
                "ENSMUSG00000062070",
                "ENSMUSG00000068523",
                "ENSMUSG00000028419",
                "ENSMUSG00000032051",
                "ENSMUSG00000021520",
                "ENSMUSG00000038143",
                "ENSMUSG00000023460",
                "ENSMUSG00000030513",
                "ENSMUSG00000078812",
                "ENSMUSG00000034994",
                "ENSMUSG00000023971",
                "ENSMUSG00000039967",
                "ENSMUSG00000009079",
                "ENSMUSG00000028633",
                "ENSMUSG00000046314",
                "ENSMUSG00000008683",
                "ENSMUSG00000030652",
                "ENSMUSG00000034243",
                "ENSMUSG00000002844",
                "ENSMUSG00000020133",
                "ENSMUSG00000033793",
                "ENSMUSG00000044308",
                "ENSMUSG00000047388",
                "ENSMUSG00000053536",
                "ENSMUSG00000074412",
                "ENSMUSG00000022300",
                "ENSMUSG00000003429",
                "ENSMUSG00000086583",
                "ENSMUSG00000018293",
                "ENSMUSG00000044033",
                "ENSMUSG00000065701",
                "ENSMUSG00000028436",
                "ENSMUSG00000034842",
                "ENSMUSG00000047649",
                "ENSMUSG00000029190",
                "ENSMUSG00000034210",
                "ENSMUSG00000015839",
                "ENSMUSG00000061306",
                "ENSMUSG00000037805",
                "ENSMUSG00000046201",
                "ENSMUSG00000061477",
                "ENSMUSG00000049878",
                "ENSMUSG00000026879",
                "ENSMUSG00000017119",
                "ENSMUSG00000023830",
                "ENSMUSG00000013936",
                "ENSMUSG00000024691",
                "ENSMUSG00000009291",
                "ENSMUSG00000055632",
                "ENSMUSG00000029467",
                "ENSMUSG00000032612",
                "ENSMUSG00000052837",
                "ENSMUSG00000069089",
                "ENSMUSG00000033543",
                "ENSMUSG00000042133",
                "ENSMUSG00000000078",
                "ENSMUSG00000006800",
                "ENSMUSG00000014470",
                "ENSMUSG00000021606",
                "ENSMUSG00000089672",
                "ENSMUSG00000079017",
                "ENSMUSG00000011148",
                "ENSMUSG00000024073",
                "ENSMUSG00000028282",
                "ENSMUSG00000036002",
                "ENSMUSG00000030451",
                "ENSMUSG00000072640",
                "ENSMUSG00000026817",
                "ENSMUSG00000031592",
                "ENSMUSG00000039130",
                "ENSMUSG00000026864",
                "ENSMUSG00000078486",
                "ENSMUSG00000019877",
                "ENSMUSG00000020572",
                "ENSMUSG00000004455",
                "ENSMUSG00000035885",
                "ENSMUSG00000043535",
                "ENSMUSG00000033416",
                "ENSMUSG00000015536",
                "ENSMUSG00000023068",
                "ENSMUSG00000031078",
                "ENSMUSG00000062270",
                "ENSMUSG00000062397",
                "ENSMUSG00000026341",
                "ENSMUSG00000028937",
                "ENSMUSG00000015357",
                "ENSMUSG00000050043",
                "ENSMUSG00000022811",
                "ENSMUSG00000022905",
                "ENSMUSG00000026074",
                "ENSMUSG00000040818",
                "ENSMUSG00000029071",
                "ENSMUSG00000062300",
                "ENSMUSG00000030447",
                "ENSMUSG00000026656",
                "ENSMUSG00000053477",
                "ENSMUSG00000013593",
                "ENSMUSG00000032267",
                "ENSMUSG00000006315",
                "ENSMUSG00000028461",
                "ENSMUSG00000042293",
                "ENSMUSG00000037710",
                "ENSMUSG00000049327",
                "ENSMUSG00000026042",
                "ENSMUSG00000029432",
                "ENSMUSG00000043866",
                "ENSMUSG00000071662",
                "ENSMUSG00000033392",
                "ENSMUSG00000019362",
                "ENSMUSG00000060981",
                "ENSMUSG00000031865",
                "ENSMUSG00000063511",
                "ENSMUSG00000029004",
                "ENSMUSG00000027293",
                "ENSMUSG00000036707",
                "ENSMUSG00000020988",
                "ENSMUSG00000023206",
                "ENSMUSG00000001175",
                "ENSMUSG00000009549",
                "ENSMUSG00000039670",
                "ENSMUSG00000018340",
                "ENSMUSG00000022450",
                "ENSMUSG00000025014",
                "ENSMUSG00000054405",
                "ENSMUSG00000073563",
                "ENSMUSG00000024805",
                "ENSMUSG00000019970",
                "ENSMUSG00000032366",
                "ENSMUSG00000001127",
                "ENSMUSG00000024002",
                "ENSMUSG00000037236",
                "ENSMUSG00000041219",
                "ENSMUSG00000055053",
                "ENSMUSG00000051391",
                "ENSMUSG00000056234",
                "ENSMUSG00000015656",
                "ENSMUSG00000023944",
                "ENSMUSG00000015932",
                "ENSMUSG00000025509",
                "ENSMUSG00000053819",
                "ENSMUSG00000068699",
                "ENSMUSG00000003970",
                "ENSMUSG00000037110",
                "ENSMUSG00000030662",
                "ENSMUSG00000082768",
                "ENSMUSG00000094483",
                "ENSMUSG00000025651",
                "ENSMUSG00000028603",
                "ENSMUSG00000021546",
                "ENSMUSG00000032253",
                "ENSMUSG00000037887",
                "ENSMUSG00000021903",
                "ENSMUSG00000052812",
                "ENSMUSG00000004268",
                "ENSMUSG00000039814",
                "ENSMUSG00000003528",
                "ENSMUSG00000026526",
                "ENSMUSG00000035329",
                "ENSMUSG00000084284",
                "ENSMUSG00000029338",
                "ENSMUSG00000045103",
                "ENSMUSG00000022205",
                "ENSMUSG00000036931",
                "ENSMUSG00000055204",
                "ENSMUSG00000018604",
                "ENSMUSG00000032042",
                "ENSMUSG00000035173",
                "ENSMUSG00000061032",
                "ENSMUSG00000067288",
                "ENSMUSG00000085385",
                "ENSMUSG00000023089",
                "ENSMUSG00000020022",
                "ENSMUSG00000022858",
                "ENSMUSG00000029146",
                "ENSMUSG00000036721",
                "ENSMUSG00000060803",
                "ENSMUSG00000034088",
                "ENSMUSG00000001911",
                "ENSMUSG00000023456",
                "ENSMUSG00000024491",
                "ENSMUSG00000032228",
                "ENSMUSG00000038462",
                "ENSMUSG00000020664",
                "ENSMUSG00000022389",
                "ENSMUSG00000022797",
                "ENSMUSG00000022010",
                "ENSMUSG00000027620",
                "ENSMUSG00000033684",
                "ENSMUSG00000074656",
                "ENSMUSG00000024099",
                "ENSMUSG00000019929",
                "ENSMUSG00000024587",
                "ENSMUSG00000016559"
            ],
            "expression_sum": {
                "class1": 424.0,
                "class2": 660.0
            },
            "uniprot_ids": [
                "P47930"
            ],
            "net_id": 0,
            "source_num": 20,
            "target_num": 190
        },
        "ENSMUSG00000022528": {
            "id": "ENSMUSG00000022528",
            "type": "TF",
            "symbol": "Hes1",
            "source": [
                "ENSMUSG00000032897",
                "ENSMUSG00000022346",
                "ENSMUSG00000021250",
                "ENSMUSG00000030256",
                "ENSMUSG00000036461",
                "ENSMUSG00000001280",
                "ENSMUSG00000053477",
                "ENSMUSG00000005583",
                "ENSMUSG00000005698",
                "ENSMUSG00000028565",
                "ENSMUSG00000040732",
                "ENSMUSG00000031885",
                "ENSMUSG00000076431",
                "ENSMUSG00000032035",
                "ENSMUSG00000071637",
                "ENSMUSG00000024985"
            ],
            "target": [
                "ENSMUSG00000027405",
                "ENSMUSG00000009569",
                "ENSMUSG00000040025",
                "ENSMUSG00000074582",
                "ENSMUSG00000027452",
                "ENSMUSG00000028484",
                "ENSMUSG00000032737",
                "ENSMUSG00000009013",
                "ENSMUSG00000026869",
                "ENSMUSG00000021939",
                "ENSMUSG00000019795",
                "ENSMUSG00000027635",
                "ENSMUSG00000097277",
                "ENSMUSG00000020846",
                "ENSMUSG00000022253",
                "ENSMUSG00000022346",
                "ENSMUSG00000023973",
                "ENSMUSG00000029632",
                "ENSMUSG00000056941",
                "ENSMUSG00000029050",
                "ENSMUSG00000029919",
                "ENSMUSG00000020441",
                "ENSMUSG00000036620",
                "ENSMUSG00000042632",
                "ENSMUSG00000020346",
                "ENSMUSG00000018293",
                "ENSMUSG00000005698",
                "ENSMUSG00000022641",
                "ENSMUSG00000026134",
                "ENSMUSG00000028843",
                "ENSMUSG00000016018",
                "ENSMUSG00000031885",
                "ENSMUSG00000071637",
                "ENSMUSG00000032020",
                "ENSMUSG00000025969",
                "ENSMUSG00000044791",
                "ENSMUSG00000046201",
                "ENSMUSG00000030105",
                "ENSMUSG00000030243",
                "ENSMUSG00000056153",
                "ENSMUSG00000053289",
                "ENSMUSG00000041439",
                "ENSMUSG00000035992",
                "ENSMUSG00000045658",
                "ENSMUSG00000079481",
                "ENSMUSG00000037089",
                "ENSMUSG00000025198",
                "ENSMUSG00000039849",
                "ENSMUSG00000068921",
                "ENSMUSG00000015542",
                "ENSMUSG00000036023",
                "ENSMUSG00000030615",
                "ENSMUSG00000034203",
                "ENSMUSG00000032187",
                "ENSMUSG00000054027",
                "ENSMUSG00000022978",
                "ENSMUSG00000028224",
                "ENSMUSG00000009090",
                "ENSMUSG00000017132",
                "ENSMUSG00000063450",
                "ENSMUSG00000037965",
                "ENSMUSG00000029290",
                "ENSMUSG00000021540",
                "ENSMUSG00000032554",
                "ENSMUSG00000031627",
                "ENSMUSG00000037022",
                "ENSMUSG00000030287",
                "ENSMUSG00000047842",
                "ENSMUSG00000020572",
                "ENSMUSG00000026878",
                "ENSMUSG00000036890",
                "ENSMUSG00000059325",
                "ENSMUSG00000027540",
                "ENSMUSG00000035559",
                "ENSMUSG00000041598",
                "ENSMUSG00000006395",
                "ENSMUSG00000030256",
                "ENSMUSG00000028266",
                "ENSMUSG00000045690",
                "ENSMUSG00000026341",
                "ENSMUSG00000015806",
                "ENSMUSG00000026313",
                "ENSMUSG00000034795",
                "ENSMUSG00000067336",
                "ENSMUSG00000033554",
                "ENSMUSG00000024985",
                "ENSMUSG00000046876",
                "ENSMUSG00000021650",
                "ENSMUSG00000070348",
                "ENSMUSG00000053477",
                "ENSMUSG00000030516",
                "ENSMUSG00000005583",
                "ENSMUSG00000055762",
                "ENSMUSG00000020661",
                "ENSMUSG00000069255",
                "ENSMUSG00000029319",
                "ENSMUSG00000047879",
                "ENSMUSG00000060032",
                "ENSMUSG00000032846",
                "ENSMUSG00000030120",
                "ENSMUSG00000006932",
                "ENSMUSG00000028580",
                "ENSMUSG00000008429",
                "ENSMUSG00000042520",
                "ENSMUSG00000018102",
                "ENSMUSG00000025809",
                "ENSMUSG00000027479",
                "ENSMUSG00000063382",
                "ENSMUSG00000049791",
                "ENSMUSG00000020029",
                "ENSMUSG00000022940",
                "ENSMUSG00000048490",
                "ENSMUSG00000004633",
                "ENSMUSG00000019907",
                "ENSMUSG00000001687",
                "ENSMUSG00000006920",
                "ENSMUSG00000050890",
                "ENSMUSG00000061838",
                "ENSMUSG00000054263",
                "ENSMUSG00000042605",
                "ENSMUSG00000030315",
                "ENSMUSG00000068798",
                "ENSMUSG00000027351",
                "ENSMUSG00000028273",
                "ENSMUSG00000042197",
                "ENSMUSG00000005873",
                "ENSMUSG00000019970",
                "ENSMUSG00000000058",
                "ENSMUSG00000055917",
                "ENSMUSG00000019726",
                "ENSMUSG00000024357",
                "ENSMUSG00000032370",
                "ENSMUSG00000041264",
                "ENSMUSG00000037070",
                "ENSMUSG00000025364",
                "ENSMUSG00000038612",
                "ENSMUSG00000036461",
                "ENSMUSG00000044792",
                "ENSMUSG00000028684",
                "ENSMUSG00000022476",
                "ENSMUSG00000039831",
                "ENSMUSG00000042705",
                "ENSMUSG00000028059",
                "ENSMUSG00000027641",
                "ENSMUSG00000037098",
                "ENSMUSG00000018796",
                "ENSMUSG00000097162",
                "ENSMUSG00000028917",
                "ENSMUSG00000038936",
                "ENSMUSG00000028393",
                "ENSMUSG00000025795",
                "ENSMUSG00000032253",
                "ENSMUSG00000021420",
                "ENSMUSG00000026484",
                "ENSMUSG00000025404",
                "ENSMUSG00000026473",
                "ENSMUSG00000052563",
                "ENSMUSG00000029338",
                "ENSMUSG00000025584",
                "ENSMUSG00000030844",
                "ENSMUSG00000034525",
                "ENSMUSG00000024085",
                "ENSMUSG00000038160",
                "ENSMUSG00000095115",
                "ENSMUSG00000023990",
                "ENSMUSG00000027805",
                "ENSMUSG00000028565",
                "ENSMUSG00000022698",
                "ENSMUSG00000034320",
                "ENSMUSG00000042524",
                "ENSMUSG00000001424",
                "ENSMUSG00000002031",
                "ENSMUSG00000041440",
                "ENSMUSG00000015882",
                "ENSMUSG00000020390",
                "ENSMUSG00000016344",
                "ENSMUSG00000098496",
                "ENSMUSG00000038225",
                "ENSMUSG00000047557",
                "ENSMUSG00000056412",
                "ENSMUSG00000029202",
                "ENSMUSG00000041297",
                "ENSMUSG00000006717",
                "ENSMUSG00000046111",
                "ENSMUSG00000020023",
                "ENSMUSG00000001280",
                "ENSMUSG00000036611",
                "ENSMUSG00000070858",
                "ENSMUSG00000090266",
                "ENSMUSG00000026643",
                "ENSMUSG00000032035",
                "ENSMUSG00000028070"
            ],
            "expression_sum": {
                "class1": 14631.0,
                "class2": 1929.0
            },
            "uniprot_ids": [
                "P35428"
            ],
            "net_id": 0,
            "source_num": 14,
            "target_num": 27
        },
        "ENSMUSG00000057469": {
            "id": "ENSMUSG00000057469",
            "type": "TF",
            "symbol": "E2f6",
            "source": [
                "ENSMUSG00000029135",
                "ENSMUSG00000001280",
                "ENSMUSG00000019768",
                "ENSMUSG00000015605",
                "ENSMUSG00000029913",
                "ENSMUSG00000000567"
            ],
            "target": [
                "ENSMUSG00000027091",
                "ENSMUSG00000036052",
                "ENSMUSG00000037826",
                "ENSMUSG00000026123",
                "ENSMUSG00000003438",
                "ENSMUSG00000009569",
                "ENSMUSG00000034673",
                "ENSMUSG00000066406",
                "ENSMUSG00000057649",
                "ENSMUSG00000059981",
                "ENSMUSG00000002844",
                "ENSMUSG00000005566",
                "ENSMUSG00000060038",
                "ENSMUSG00000034610",
                "ENSMUSG00000034875",
                "ENSMUSG00000031403",
                "ENSMUSG00000020485",
                "ENSMUSG00000021000",
                "ENSMUSG00000006740",
                "ENSMUSG00000000563",
                "ENSMUSG00000027238",
                "ENSMUSG00000028081",
                "ENSMUSG00000037300",
                "ENSMUSG00000053536",
                "ENSMUSG00000019432",
                "ENSMUSG00000026491",
                "ENSMUSG00000018599",
                "ENSMUSG00000022641",
                "ENSMUSG00000036371",
                "ENSMUSG00000058135",
                "ENSMUSG00000015112",
                "ENSMUSG00000038070",
                "ENSMUSG00000075054",
                "ENSMUSG00000021453",
                "ENSMUSG00000052337",
                "ENSMUSG00000031367",
                "ENSMUSG00000027499",
                "ENSMUSG00000027784",
                "ENSMUSG00000032348",
                "ENSMUSG00000024283",
                "ENSMUSG00000032423",
                "ENSMUSG00000045983",
                "ENSMUSG00000029629",
                "ENSMUSG00000017119",
                "ENSMUSG00000028911",
                "ENSMUSG00000029913",
                "ENSMUSG00000081792",
                "ENSMUSG00000068267",
                "ENSMUSG00000031983",
                "ENSMUSG00000038028",
                "ENSMUSG00000032481",
                "ENSMUSG00000032047",
                "ENSMUSG00000029594",
                "ENSMUSG00000019773",
                "ENSMUSG00000025453",
                "ENSMUSG00000067274",
                "ENSMUSG00000030788",
                "ENSMUSG00000015843",
                "ENSMUSG00000026576",
                "ENSMUSG00000055239",
                "ENSMUSG00000066637",
                "ENSMUSG00000032525",
                "ENSMUSG00000038022",
                "ENSMUSG00000044393",
                "ENSMUSG00000000441",
                "ENSMUSG00000026626",
                "ENSMUSG00000074918",
                "ENSMUSG00000034220",
                "ENSMUSG00000027823",
                "ENSMUSG00000031543",
                "ENSMUSG00000007891",
                "ENSMUSG00000041685",
                "ENSMUSG00000024073",
                "ENSMUSG00000028282",
                "ENSMUSG00000020267",
                "ENSMUSG00000033060",
                "ENSMUSG00000038784",
                "ENSMUSG00000062580",
                "ENSMUSG00000021209",
                "ENSMUSG00000005354",
                "ENSMUSG00000022890",
                "ENSMUSG00000032826",
                "ENSMUSG00000023939",
                "ENSMUSG00000035885",
                "ENSMUSG00000022336",
                "ENSMUSG00000026187",
                "ENSMUSG00000029198",
                "ENSMUSG00000033411",
                "ENSMUSG00000032959",
                "ENSMUSG00000032437",
                "ENSMUSG00000026701",
                "ENSMUSG00000033416",
                "ENSMUSG00000021218",
                "ENSMUSG00000032199",
                "ENSMUSG00000028266",
                "ENSMUSG00000045690",
                "ENSMUSG00000062397",
                "ENSMUSG00000026341",
                "ENSMUSG00000025271",
                "ENSMUSG00000022185",
                "ENSMUSG00000027706",
                "ENSMUSG00000058793",
                "ENSMUSG00000022131",
                "ENSMUSG00000049643",
                "ENSMUSG00000000088",
                "ENSMUSG00000031447",
                "ENSMUSG00000024985",
                "ENSMUSG00000031985",
                "ENSMUSG00000052539",
                "ENSMUSG00000026064",
                "ENSMUSG00000032314",
                "ENSMUSG00000073684",
                "ENSMUSG00000022452",
                "ENSMUSG00000023175",
                "ENSMUSG00000022519",
                "ENSMUSG00000026283",
                "ENSMUSG00000036097",
                "ENSMUSG00000006315",
                "ENSMUSG00000020048",
                "ENSMUSG00000027406",
                "ENSMUSG00000026858",
                "ENSMUSG00000028461",
                "ENSMUSG00000037568",
                "ENSMUSG00000020737",
                "ENSMUSG00000014592",
                "ENSMUSG00000042293",
                "ENSMUSG00000052374",
                "ENSMUSG00000015214",
                "ENSMUSG00000030850",
                "ENSMUSG00000024991",
                "ENSMUSG00000037710",
                "ENSMUSG00000028229",
                "ENSMUSG00000049327",
                "ENSMUSG00000048814",
                "ENSMUSG00000041075",
                "ENSMUSG00000021597",
                "ENSMUSG00000010048",
                "ENSMUSG00000022201",
                "ENSMUSG00000025410",
                "ENSMUSG00000026260",
                "ENSMUSG00000036112",
                "ENSMUSG00000002102",
                "ENSMUSG00000022748",
                "ENSMUSG00000043866",
                "ENSMUSG00000056091",
                "ENSMUSG00000025374",
                "ENSMUSG00000018446",
                "ENSMUSG00000030934",
                "ENSMUSG00000042198",
                "ENSMUSG00000019577",
                "ENSMUSG00000023249",
                "ENSMUSG00000024613",
                "ENSMUSG00000027274",
                "ENSMUSG00000054889",
                "ENSMUSG00000027293",
                "ENSMUSG00000070544",
                "ENSMUSG00000046351",
                "ENSMUSG00000020988",
                "ENSMUSG00000028234",
                "ENSMUSG00000035268",
                "ENSMUSG00000022982",
                "ENSMUSG00000025184",
                "ENSMUSG00000048486",
                "ENSMUSG00000032897",
                "ENSMUSG00000060904",
                "ENSMUSG00000028273",
                "ENSMUSG00000034951",
                "ENSMUSG00000026775",
                "ENSMUSG00000018474",
                "ENSMUSG00000032883",
                "ENSMUSG00000044365",
                "ENSMUSG00000066043",
                "ENSMUSG00000073563",
                "ENSMUSG00000038690",
                "ENSMUSG00000020176",
                "ENSMUSG00000027423",
                "ENSMUSG00000031751",
                "ENSMUSG00000023055",
                "ENSMUSG00000037236",
                "ENSMUSG00000048578",
                "ENSMUSG00000038650",
                "ENSMUSG00000025856",
                "ENSMUSG00000018559",
                "ENSMUSG00000020027",
                "ENSMUSG00000025364",
                "ENSMUSG00000022682",
                "ENSMUSG00000054469",
                "ENSMUSG00000021910",
                "ENSMUSG00000036880",
                "ENSMUSG00000030603",
                "ENSMUSG00000017756",
                "ENSMUSG00000039987",
                "ENSMUSG00000026429",
                "ENSMUSG00000028788",
                "ENSMUSG00000031774",
                "ENSMUSG00000031783",
                "ENSMUSG00000003316",
                "ENSMUSG00000033102",
                "ENSMUSG00000078671",
                "ENSMUSG00000034290",
                "ENSMUSG00000023150",
                "ENSMUSG00000027618",
                "ENSMUSG00000018796",
                "ENSMUSG00000028603",
                "ENSMUSG00000038718",
                "ENSMUSG00000039741",
                "ENSMUSG00000063229",
                "ENSMUSG00000032518",
                "ENSMUSG00000021546",
                "ENSMUSG00000020321",
                "ENSMUSG00000018566",
                "ENSMUSG00000027332",
                "ENSMUSG00000029135",
                "ENSMUSG00000017404",
                "ENSMUSG00000022214",
                "ENSMUSG00000004268",
                "ENSMUSG00000027602",
                "ENSMUSG00000061286",
                "ENSMUSG00000017778",
                "ENSMUSG00000044573",
                "ENSMUSG00000004591",
                "ENSMUSG00000021621",
                "ENSMUSG00000024304",
                "ENSMUSG00000008575",
                "ENSMUSG00000014633",
                "ENSMUSG00000021706",
                "ENSMUSG00000060679",
                "ENSMUSG00000030057",
                "ENSMUSG00000074182",
                "ENSMUSG00000019851",
                "ENSMUSG00000059208",
                "ENSMUSG00000032042",
                "ENSMUSG00000052912",
                "ENSMUSG00000022016",
                "ENSMUSG00000055322",
                "ENSMUSG00000074064",
                "ENSMUSG00000006057",
                "ENSMUSG00000067722",
                "ENSMUSG00000028207",
                "ENSMUSG00000040414",
                "ENSMUSG00000040771",
                "ENSMUSG00000032724",
                "ENSMUSG00000033161",
                "ENSMUSG00000023089",
                "ENSMUSG00000030246",
                "ENSMUSG00000001424",
                "ENSMUSG00000026434",
                "ENSMUSG00000061474",
                "ENSMUSG00000003072",
                "ENSMUSG00000025466",
                "ENSMUSG00000036564",
                "ENSMUSG00000024236",
                "ENSMUSG00000039474",
                "ENSMUSG00000020664",
                "ENSMUSG00000021585",
                "ENSMUSG00000022797",
                "ENSMUSG00000020079",
                "ENSMUSG00000061518",
                "ENSMUSG00000021395",
                "ENSMUSG00000022010",
                "ENSMUSG00000020849",
                "ENSMUSG00000024150",
                "ENSMUSG00000028343",
                "ENSMUSG00000031634",
                "ENSMUSG00000082195",
                "ENSMUSG00000006717",
                "ENSMUSG00000032480",
                "ENSMUSG00000001056",
                "ENSMUSG00000024924",
                "ENSMUSG00000059811",
                "ENSMUSG00000028884",
                "ENSMUSG00000030541",
                "ENSMUSG00000001280",
                "ENSMUSG00000038005",
                "ENSMUSG00000070003",
                "ENSMUSG00000026489",
                "ENSMUSG00000024913"
            ],
            "expression_sum": {
                "class1": 276.0,
                "class2": 554.0
            },
            "uniprot_ids": [
                "O54917",
                "Q8K455",
                "A0A1Y7VJU3",
                "A0A1Y7VP85"
            ],
            "net_id": 0,
            "source_num": 6,
            "target_num": 88
        },
        "ENSMUSG00000058239": {
            "id": "ENSMUSG00000058239",
            "type": "TF",
            "symbol": "Usf2",
            "source": [
                "ENSMUSG00000019817",
                "ENSMUSG00000015579",
                "ENSMUSG00000047407",
                "ENSMUSG00000001280",
                "ENSMUSG00000075595",
                "ENSMUSG00000031885"
            ],
            "target": [
                "ENSMUSG00000039671",
                "ENSMUSG00000072969",
                "ENSMUSG00000026568",
                "ENSMUSG00000026918",
                "ENSMUSG00000054728",
                "ENSMUSG00000061904",
                "ENSMUSG00000025968",
                "ENSMUSG00000020700",
                "ENSMUSG00000039473",
                "ENSMUSG00000014294",
                "ENSMUSG00000053898",
                "ENSMUSG00000031883",
                "ENSMUSG00000020133",
                "ENSMUSG00000046516",
                "ENSMUSG00000062908",
                "ENSMUSG00000047388",
                "ENSMUSG00000091478",
                "ENSMUSG00000053536",
                "ENSMUSG00000027809",
                "ENSMUSG00000024392",
                "ENSMUSG00000025241",
                "ENSMUSG00000018599",
                "ENSMUSG00000031669",
                "ENSMUSG00000041329",
                "ENSMUSG00000065701",
                "ENSMUSG00000026134",
                "ENSMUSG00000031885",
                "ENSMUSG00000014177",
                "ENSMUSG00000034342",
                "ENSMUSG00000034842",
                "ENSMUSG00000027447",
                "ENSMUSG00000060923",
                "ENSMUSG00000049866",
                "ENSMUSG00000027784",
                "ENSMUSG00000049878",
                "ENSMUSG00000021265",
                "ENSMUSG00000021290",
                "ENSMUSG00000028223",
                "ENSMUSG00000045983",
                "ENSMUSG00000037339",
                "ENSMUSG00000062647",
                "ENSMUSG00000049076",
                "ENSMUSG00000041268",
                "ENSMUSG00000035458",
                "ENSMUSG00000079184",
                "ENSMUSG00000020689",
                "ENSMUSG00000020738",
                "ENSMUSG00000032047",
                "ENSMUSG00000021339",
                "ENSMUSG00000023015",
                "ENSMUSG00000029467",
                "ENSMUSG00000001436",
                "ENSMUSG00000063882",
                "ENSMUSG00000014776",
                "ENSMUSG00000042834",
                "ENSMUSG00000041881",
                "ENSMUSG00000020402",
                "ENSMUSG00000022354",
                "ENSMUSG00000026087",
                "ENSMUSG00000053012",
                "ENSMUSG00000011305",
                "ENSMUSG00000035152",
                "ENSMUSG00000021764",
                "ENSMUSG00000025212",
                "ENSMUSG00000027303",
                "ENSMUSG00000029910",
                "ENSMUSG00000084968",
                "ENSMUSG00000047407",
                "ENSMUSG00000021427",
                "ENSMUSG00000031812",
                "ENSMUSG00000009575",
                "ENSMUSG00000037514",
                "ENSMUSG00000072568",
                "ENSMUSG00000039648",
                "ENSMUSG00000037174",
                "ENSMUSG00000040242",
                "ENSMUSG00000039130",
                "ENSMUSG00000009863",
                "ENSMUSG00000059325",
                "ENSMUSG00000026135",
                "ENSMUSG00000019853",
                "ENSMUSG00000036860",
                "ENSMUSG00000001062",
                "ENSMUSG00000079179",
                "ENSMUSG00000079641",
                "ENSMUSG00000021375",
                "ENSMUSG00000029020",
                "ENSMUSG00000028410",
                "ENSMUSG00000021245",
                "ENSMUSG00000034566",
                "ENSMUSG00000089909",
                "ENSMUSG00000020462",
                "ENSMUSG00000021044",
                "ENSMUSG00000049792",
                "ENSMUSG00000022519",
                "ENSMUSG00000008892",
                "ENSMUSG00000009907",
                "ENSMUSG00000060600",
                "ENSMUSG00000021243",
                "ENSMUSG00000037949",
                "ENSMUSG00000020149",
                "ENSMUSG00000025393",
                "ENSMUSG00000037580",
                "ENSMUSG00000040048",
                "ENSMUSG00000024038",
                "ENSMUSG00000022867",
                "ENSMUSG00000027698",
                "ENSMUSG00000029048",
                "ENSMUSG00000031934",
                "ENSMUSG00000052374",
                "ENSMUSG00000001663",
                "ENSMUSG00000001467",
                "ENSMUSG00000001419",
                "ENSMUSG00000079508",
                "ENSMUSG00000001240",
                "ENSMUSG00000022186",
                "ENSMUSG00000022772",
                "ENSMUSG00000011884",
                "ENSMUSG00000024687",
                "ENSMUSG00000028760",
                "ENSMUSG00000030861",
                "ENSMUSG00000028756",
                "ENSMUSG00000024976",
                "ENSMUSG00000040661",
                "ENSMUSG00000008429",
                "ENSMUSG00000059810",
                "ENSMUSG00000071528",
                "ENSMUSG00000026754",
                "ENSMUSG00000031347",
                "ENSMUSG00000039615",
                "ENSMUSG00000031913",
                "ENSMUSG00000020601",
                "ENSMUSG00000039891",
                "ENSMUSG00000089715",
                "ENSMUSG00000047710",
                "ENSMUSG00000024359",
                "ENSMUSG00000006920",
```
