## Supplement 3 (P7 CM vs. P28 CM) for "AGEAS: Automated Machine Learning based Genetic Regulatory Element Extraction System": meta_GRN.js

```
{
    "genes": {
        "ENSMUSG00000000567": {
            "id": "ENSMUSG00000000567",
            "type": "TF",
            "symbol": "Sox9",
            "source": [
                "ENSMUSG00000005583",
                "ENSMUSG00000026586",
                "ENSMUSG00000031099"
            ],
            "target": [
                "ENSMUSG00000022621",
                "ENSMUSG00000057894",
                "ENSMUSG00000039231",
                "ENSMUSG00000006333",
                "ENSMUSG00000023104",
                "ENSMUSG00000029994",
                "ENSMUSG00000032388",
                "ENSMUSG00000051817",
                "ENSMUSG00000030352",
                "ENSMUSG00000026687",
                "ENSMUSG00000092837",
                "ENSMUSG00000039356",
                "ENSMUSG00000026657",
                "ENSMUSG00000028901",
                "ENSMUSG00000029536",
                "ENSMUSG00000020485",
                "ENSMUSG00000031253",
                "ENSMUSG00000020646",
                "ENSMUSG00000024959",
                "ENSMUSG00000040112",
                "ENSMUSG00000053460",
                "ENSMUSG00000033544",
                "ENSMUSG00000004151",
                "ENSMUSG00000055707",
                "ENSMUSG00000030782",
                "ENSMUSG00000041236",
                "ENSMUSG00000046572",
                "ENSMUSG00000022961",
                "ENSMUSG00000070284",
                "ENSMUSG00000021036",
                "ENSMUSG00000032997",
                "ENSMUSG00000028869",
                "ENSMUSG00000026843",
                "ENSMUSG00000078247",
                "ENSMUSG00000022686",
                "ENSMUSG00000029190",
                "ENSMUSG00000019689",
                "ENSMUSG00000026031",
                "ENSMUSG00000026404",
                "ENSMUSG00000028936",
                "ENSMUSG00000028466",
                "ENSMUSG00000032012",
                "ENSMUSG00000041773",
                "ENSMUSG00000052748",
                "ENSMUSG00000022159",
                "ENSMUSG00000022339",
                "ENSMUSG00000026879",
                "ENSMUSG00000028552",
                "ENSMUSG00000000827",
                "ENSMUSG00000027411",
                "ENSMUSG00000050627",
                "ENSMUSG00000053877",
                "ENSMUSG00000031422",
                "ENSMUSG00000031730",
                "ENSMUSG00000032440",
                "ENSMUSG00000071014",
                "ENSMUSG00000001436",
                "ENSMUSG00000040928",
                "ENSMUSG00000022722",
                "ENSMUSG00000029998",
                "ENSMUSG00000002881",
                "ENSMUSG00000004947",
                "ENSMUSG00000018189",
                "ENSMUSG00000017999",
                "ENSMUSG00000024146",
                "ENSMUSG00000026566",
                "ENSMUSG00000022150",
                "ENSMUSG00000026028",
                "ENSMUSG00000062190",
                "ENSMUSG00000018217",
                "ENSMUSG00000040274",
                "ENSMUSG00000028403",
                "ENSMUSG00000030522",
                "ENSMUSG00000029659",
                "ENSMUSG00000031543",
                "ENSMUSG00000022119",
                "ENSMUSG00000030393",
                "ENSMUSG00000050188",
                "ENSMUSG00000036565",
                "ENSMUSG00000069744",
                "ENSMUSG00000024921",
                "ENSMUSG00000031812",
                "ENSMUSG00000005354",
                "ENSMUSG00000053483",
                "ENSMUSG00000026864",
                "ENSMUSG00000004562",
                "ENSMUSG00000021027",
                "ENSMUSG00000033565",
                "ENSMUSG00000057469",
                "ENSMUSG00000074457",
                "ENSMUSG00000023068",
                "ENSMUSG00000009894",
                "ENSMUSG00000079435",
                "ENSMUSG00000034566",
                "ENSMUSG00000021245",
                "ENSMUSG00000057561",
                "ENSMUSG00000000753",
                "ENSMUSG00000016503",
                "ENSMUSG00000031821",
                "ENSMUSG00000003662",
                "ENSMUSG00000073684",
                "ENSMUSG00000027067",
                "ENSMUSG00000038244",
                "ENSMUSG00000037852",
                "ENSMUSG00000049792",
                "ENSMUSG00000046876",
                "ENSMUSG00000063077",
                "ENSMUSG00000090812",
                "ENSMUSG00000042462",
                "ENSMUSG00000021243",
                "ENSMUSG00000021771",
                "ENSMUSG00000052763",
                "ENSMUSG00000068267",
                "ENSMUSG00000025782",
                "ENSMUSG00000043323",
                "ENSMUSG00000020282",
                "ENSMUSG00000051671",
                "ENSMUSG00000041075",
                "ENSMUSG00000022203",
                "ENSMUSG00000037204",
                "ENSMUSG00000044201",
                "ENSMUSG00000027030",
                "ENSMUSG00000053329",
                "ENSMUSG00000021318",
                "ENSMUSG00000025140",
                "ENSMUSG00000025374",
                "ENSMUSG00000023022",
                "ENSMUSG00000001173",
                "ENSMUSG00000026749",
                "ENSMUSG00000026585",
                "ENSMUSG00000026754",
                "ENSMUSG00000026791",
                "ENSMUSG00000032212",
                "ENSMUSG00000040359",
                "ENSMUSG00000024588",
                "ENSMUSG00000004460",
                "ENSMUSG00000021130",
                "ENSMUSG00000044442",
                "ENSMUSG00000046402",
                "ENSMUSG00000027884",
                "ENSMUSG00000006299",
                "ENSMUSG00000055172",
                "ENSMUSG00000040151",
                "ENSMUSG00000054893",
                "ENSMUSG00000019467",
                "ENSMUSG00000026202",
                "ENSMUSG00000066839",
                "ENSMUSG00000020530",
                "ENSMUSG00000020522",
                "ENSMUSG00000091028",
                "ENSMUSG00000018340",
                "ENSMUSG00000026775",
                "ENSMUSG00000022103",
                "ENSMUSG00000070697",
                "ENSMUSG00000026219",
                "ENSMUSG00000026748",
                "ENSMUSG00000030082",
                "ENSMUSG00000031400",
                "ENSMUSG00000042700",
                "ENSMUSG00000042606",
                "ENSMUSG00000055053",
                "ENSMUSG00000021408",
                "ENSMUSG00000052676",
                "ENSMUSG00000054770",
                "ENSMUSG00000039976",
                "ENSMUSG00000019837",
                "ENSMUSG00000034485",
                "ENSMUSG00000034584",
                "ENSMUSG00000044629",
                "ENSMUSG00000082978",
                "ENSMUSG00000029433",
                "ENSMUSG00000022378",
                "ENSMUSG00000022754",
                "ENSMUSG00000033781",
                "ENSMUSG00000005881",
                "ENSMUSG00000006050",
                "ENSMUSG00000031217",
                "ENSMUSG00000035478",
                "ENSMUSG00000017421",
                "ENSMUSG00000028333",
                "ENSMUSG00000040658",
                "ENSMUSG00000025484",
                "ENSMUSG00000028069",
                "ENSMUSG00000045128",
                "ENSMUSG00000031740",
                "ENSMUSG00000022325",
                "ENSMUSG00000029838",
                "ENSMUSG00000029924",
                "ENSMUSG00000016262",
                "ENSMUSG00000032491",
                "ENSMUSG00000035376",
                "ENSMUSG00000037674",
                "ENSMUSG00000037211",
                "ENSMUSG00000024769",
                "ENSMUSG00000050705",
                "ENSMUSG00000036862",
                "ENSMUSG00000024639",
                "ENSMUSG00000028034",
                "ENSMUSG00000044167",
                "ENSMUSG00000051159",
                "ENSMUSG00000061477",
                "ENSMUSG00000025591",
                "ENSMUSG00000031799",
                "ENSMUSG00000026634",
                "ENSMUSG00000027239",
                "ENSMUSG00000007872",
                "ENSMUSG00000048388",
                "ENSMUSG00000021706",
                "ENSMUSG00000041801",
                "ENSMUSG00000078201",
                "ENSMUSG00000025949",
                "ENSMUSG00000018861",
                "ENSMUSG00000030057",
                "ENSMUSG00000027797",
                "ENSMUSG00000053475",
                "ENSMUSG00000000359",
                "ENSMUSG00000006585",
                "ENSMUSG00000055204",
                "ENSMUSG00000022016",
                "ENSMUSG00000091803",
                "ENSMUSG00000061878",
                "ENSMUSG00000024054",
                "ENSMUSG00000010110",
                "ENSMUSG00000048416",
                "ENSMUSG00000001964",
                "ENSMUSG00000010406",
                "ENSMUSG00000041375",
                "ENSMUSG00000030528",
                "ENSMUSG00000025794",
                "ENSMUSG00000008200",
                "ENSMUSG00000038374",
                "ENSMUSG00000039497",
                "ENSMUSG00000022466",
                "ENSMUSG00000019775",
                "ENSMUSG00000024538",
                "ENSMUSG00000026150",
                "ENSMUSG00000032334",
                "ENSMUSG00000085208",
                "ENSMUSG00000067847",
                "ENSMUSG00000073725",
                "ENSMUSG00000018567",
                "ENSMUSG00000023456",
                "ENSMUSG00000027167",
                "ENSMUSG00000026082",
                "ENSMUSG00000056749",
                "ENSMUSG00000049690",
                "ENSMUSG00000060288",
                "ENSMUSG00000040339",
                "ENSMUSG00000062309",
                "ENSMUSG00000042208",
                "ENSMUSG00000026761",
                "ENSMUSG00000026932",
                "ENSMUSG00000021714",
                "ENSMUSG00000025533",
                "ENSMUSG00000029804",
                "ENSMUSG00000002981",
                "ENSMUSG00000027620",
                "ENSMUSG00000020766",
                "ENSMUSG00000033684",
                "ENSMUSG00000024104",
                "ENSMUSG00000028048",
                "ENSMUSG00000036570",
                "ENSMUSG00000033055",
                "ENSMUSG00000066551",
                "ENSMUSG00000044533",
                "ENSMUSG00000013662",
                "ENSMUSG00000040952",
                "ENSMUSG00000031438",
                "ENSMUSG00000050541",
                "ENSMUSG00000014769",
                "ENSMUSG00000020755",
                "ENSMUSG00000022090",
                "ENSMUSG00000038970",
                "ENSMUSG00000040351",
                "ENSMUSG00000040435",
                "ENSMUSG00000054400",
                "ENSMUSG00000031979"
            ],
            "expression_sum": {
                "class1": 840.0,
                "class2": 46.0
            },
            "uniprot_ids": [
                "Q04887"
            ]
        },
        "ENSMUSG00000000359": {
            "id": "ENSMUSG00000000359",
            "type": "TF",
            "symbol": "Rem1",
            "source": [
                "ENSMUSG00000066551",
                "ENSMUSG00000001493",
                "ENSMUSG00000016503",
                "ENSMUSG00000015846",
                "ENSMUSG00000000567"
            ],
            "target": [
                "ENSMUSG00000022722",
                "ENSMUSG00000038695",
                "ENSMUSG00000002265"
            ],
            "expression_sum": {
                "class1": 644.0,
                "class2": 20.0
            },
            "uniprot_ids": [
                "O35929",
                "A3KGR6"
            ]
        },
        "ENSMUSG00000000753": {
            "id": "ENSMUSG00000000753",
            "type": "Gene",
            "symbol": "Serpinf1",
            "source": [
                "ENSMUSG00000005897",
                "ENSMUSG00000003923",
                "ENSMUSG00000074622",
                "ENSMUSG00000005698",
                "ENSMUSG00000031885",
                "ENSMUSG00000055320",
                "ENSMUSG00000071637",
                "ENSMUSG00000000567"
            ],
            "target": [],
            "expression_sum": {
                "class1": 1540.0,
                "class2": 264.0
            },
            "uniprot_ids": [
                "P97298",
                "E9PWS2",
                "Q5ND37",
                "E9PZX2",
                "E9PXR9",
                "B7ZC25",
                "F6S1M4"
            ]
        },
        "ENSMUSG00000000827": {
            "id": "ENSMUSG00000000827",
            "type": "Gene",
            "symbol": "Tpd52l2",
            "source": [
                "ENSMUSG00000059552",
                "ENSMUSG00000003032",
                "ENSMUSG00000026628",
                "ENSMUSG00000021250",
                "ENSMUSG00000001911",
                "ENSMUSG00000066551",
                "ENSMUSG00000041515",
                "ENSMUSG00000029249",
                "ENSMUSG00000040732",
                "ENSMUSG00000022383",
                "ENSMUSG00000027210",
                "ENSMUSG00000034998",
                "ENSMUSG00000032035",
                "ENSMUSG00000039191",
                "ENSMUSG00000000567"
            ],
            "target": [],
            "expression_sum": {
                "class1": 3327.0,
                "class2": 630.0
            },
            "uniprot_ids": [
                "Q9CYZ2",
                "Q3TUJ9",
                "Q8BKP1",
                "Q3TAI4",
                "F6VQ81",
                "V9GWU5",
                "V9GXN2",
                "A2AUD5"
            ]
        },
        "ENSMUSG00000001173": {
            "id": "ENSMUSG00000001173",
            "type": "Gene",
            "symbol": "Ocrl",
            "source": [
                "ENSMUSG00000013419",
                "ENSMUSG00000000567"
            ],
            "target": [],
            "expression_sum": {
                "class1": 405.0,
                "class2": 86.0
            },
            "uniprot_ids": [
                "Q6NVF0",
                "Q7TPR1",
                "F7CAV3"
            ]
        },
        "ENSMUSG00000001436": {
            "id": "ENSMUSG00000001436",
            "type": "Gene",
            "symbol": "Slc19a1",
            "source": [
                "ENSMUSG00000058440",
                "ENSMUSG00000058239",
                "ENSMUSG00000019817",
                "ENSMUSG00000044167",
                "ENSMUSG00000015605",
                "ENSMUSG00000000567"
            ],
            "target": [],
            "expression_sum": {
                "class1": 206.0,
                "class2": 40.0
            },
            "uniprot_ids": [
                "P41438",
                "D3YZE1",
                "D3Z0I6",
                "E9Q8X6",
                "D3Z4F3",
                "D3YU74",
                "F6YWA3"
            ]
        },
        "ENSMUSG00000001964": {
            "id": "ENSMUSG00000001964",
            "type": "Gene",
            "symbol": "Emd",
            "source": [
                "ENSMUSG00000015839",
                "ENSMUSG00000015579",
                "ENSMUSG00000051469",
                "ENSMUSG00000027109",
                "ENSMUSG00000000567"
            ],
            "target": [],
            "expression_sum": {
                "class1": 1043.0,
                "class2": 205.0
            },
            "uniprot_ids": [
                "O08579",
                "I7HJS1",
                "Q3THM8",
                "B7FAU5",
                "D6RFS0"
            ]
        },
        "ENSMUSG00000002881": {
            "id": "ENSMUSG00000002881",
            "type": "Gene",
            "symbol": "Nab1",
            "source": [
                "ENSMUSG00000059552",
                "ENSMUSG00000024515",
                "ENSMUSG00000028565",
                "ENSMUSG00000040732",
                "ENSMUSG00000000567"
            ],
            "target": [],
            "expression_sum": {
                "class1": 1283.0,
                "class2": 440.0
            },
            "uniprot_ids": [
                "Q61122",
                "A0A087WSU4",
                "E9PZ64"
            ]
        },
        "ENSMUSG00000002981": {
            "id": "ENSMUSG00000002981",
            "type": "Gene",
            "symbol": "Clptm1",
            "source": [
                "ENSMUSG00000030393",
                "ENSMUSG00000056749",
                "ENSMUSG00000001280",
                "ENSMUSG00000027104",
                "ENSMUSG00000000567"
            ],
            "target": [],
            "expression_sum": {
                "class1": 717.0,
                "class2": 287.0
            },
            "uniprot_ids": [
                "Q8VBZ3"
            ]
        },
        "ENSMUSG00000003662": {
            "id": "ENSMUSG00000003662",
            "type": "Gene",
            "symbol": "Ciao1",
            "source": [
                "ENSMUSG00000058440",
                "ENSMUSG00000000567"
            ],
            "target": [],
            "expression_sum": {
                "class1": 572.0,
                "class2": 345.0
            },
            "uniprot_ids": [
                "Q99KN2",
                "G3UYS4",
                "G3UYN9",
                "G3UYR5"
            ]
        },
        "ENSMUSG00000004151": {
            "id": "ENSMUSG00000004151",
            "type": "TF",
            "symbol": "Etv1",
            "source": [
                "ENSMUSG00000030256",
                "ENSMUSG00000015579",
                "ENSMUSG00000022383",
                "ENSMUSG00000027210",
                "ENSMUSG00000000567"
            ],
            "target": [
                "ENSMUSG00000078815",
                "ENSMUSG00000060288",
                "ENSMUSG00000026260",
                "ENSMUSG00000000058",
                "ENSMUSG00000001418",
                "ENSMUSG00000000374",
                "ENSMUSG00000032381",
                "ENSMUSG00000029649",
                "ENSMUSG00000028788"
            ],
            "expression_sum": {
                "class1": 859.0,
                "class2": 58.0
            },
            "uniprot_ids": [
                "P41164",
                "Q8CCR6",
                "E0CXX3",
                "E0CYN8",
                "E0CZ37",
                "E0CXD2",
                "E0CYI1",
                "E0CZ54",
                "E0CXE6"
            ]
        },
        "ENSMUSG00000004460": {
            "id": "ENSMUSG00000004460",
            "type": "Gene",
            "symbol": "Dnajb11",
            "source": [
                "ENSMUSG00000010175",
                "ENSMUSG00000015579",
                "ENSMUSG00000001280",
                "ENSMUSG00000019768",
                "ENSMUSG00000025025",
                "ENSMUSG00000000567"
            ],
            "target": [],
            "expression_sum": {
                "class1": 1919.0,
                "class2": 492.0
            },
            "uniprot_ids": [
                "Q99KV1",
                "A0A338P778"
            ]
        },
        "ENSMUSG00000004562": {
            "id": "ENSMUSG00000004562",
            "type": "Gene",
            "symbol": "Arhgef40",
            "source": [
                "ENSMUSG00000074733",
                "ENSMUSG00000001493",
                "ENSMUSG00000005583",
                "ENSMUSG00000040270",
                "ENSMUSG00000000567"
            ],
            "target": [],
            "expression_sum": {
                "class1": 1899.0,
                "class2": 182.0
            },
            "uniprot_ids": [
                "Q3UPH7",
                "S4R1Y4",
                "S4R237",
                "S4R1R5",
                "S4R1X0",
                "S4R2G8",
                "S4R2U9",
                "S4R189",
                "S4R1S6",
                "S4R2R3",
                "S4R2A6"
            ]
        },
        "ENSMUSG00000004947": {
            "id": "ENSMUSG00000004947",
            "type": "Gene",
            "symbol": "Dtx2",
            "source": [
                "ENSMUSG00000005583",
                "ENSMUSG00000037638",
                "ENSMUSG00000000567"
            ],
            "target": [],
            "expression_sum": {
                "class1": 283.0,
                "class2": 97.0
            },
            "uniprot_ids": [
                "Q8R3P2",
                "A0A0G2JDR7",
                "A0A0R4IZY1",
                "A0A1C7ZMZ6"
            ]
        },
        "ENSMUSG00000005354": {
            "id": "ENSMUSG00000005354",
            "type": "Gene",
            "symbol": "Txn2",
            "source": [
                "ENSMUSG00000057469",
                "ENSMUSG00000015605",
                "ENSMUSG00000026610",
                "ENSMUSG00000056501",
                "ENSMUSG00000015579",
                "ENSMUSG00000030557",
                "ENSMUSG00000029249",
                "ENSMUSG00000001280",
                "ENSMUSG00000005566",
                "ENSMUSG00000019768",
                "ENSMUSG00000008575",
                "ENSMUSG00000000567"
            ],
            "target": [],
            "expression_sum": {
                "class1": 3147.0,
                "class2": 1109.0
            },
            "uniprot_ids": [
                "P97493",
                "Q3TUS3",
                "G3UZY2",
                "A2A439",
                "G3UX99"
            ]
        },
        "ENSMUSG00000005881": {
            "id": "ENSMUSG00000005881",
            "type": "Gene",
            "symbol": "Ergic3",
            "source": [
                "ENSMUSG00000011427",
                "ENSMUSG00000078671",
                "ENSMUSG00000022811",
                "ENSMUSG00000000567"
            ],
            "target": [],
            "expression_sum": {
                "class1": 2442.0,
                "class2": 353.0
            },
            "uniprot_ids": [
                "Q9CQE7",
                "F6UIS1",
                "F6RK81"
            ]
        },
        "ENSMUSG00000006050": {
            "id": "ENSMUSG00000006050",
            "type": "Gene",
            "symbol": "Sra1",
            "source": [
                "ENSMUSG00000022895",
                "ENSMUSG00000006215",
                "ENSMUSG00000044167",
                "ENSMUSG00000024974",
                "ENSMUSG00000035632",
                "ENSMUSG00000015846",
                "ENSMUSG00000000567"
            ],
            "target": [],
            "expression_sum": {
                "class1": 6472.0,
                "class2": 1050.0
            },
            "uniprot_ids": [
                "Q80VJ2",
                "G3X8R2",
                "G3UZQ4",
                "A0A494BBA2"
            ]
        },
        "ENSMUSG00000006299": {
            "id": "ENSMUSG00000006299",
            "type": "Gene",
            "symbol": "Aamp",
            "source": [
                "ENSMUSG00000029135",
                "ENSMUSG00000043099",
                "ENSMUSG00000001280",
                "ENSMUSG00000033943",
                "ENSMUSG00000019768",
                "ENSMUSG00000000567"
            ],
            "target": [],
            "expression_sum": {
                "class1": 2207.0,
                "class2": 440.0
            },
            "uniprot_ids": [
                "Q3TJ22",
                "A0A087WNM5",
                "J3QN89",
                "A0A087WR11"
            ]
        },
        "ENSMUSG00000006333": {
            "id": "ENSMUSG00000006333",
            "type": "Gene",
            "symbol": "Rps9",
            "source": [
                "ENSMUSG00000029135",
                "ENSMUSG00000001419",
                "ENSMUSG00000018604",
                "ENSMUSG00000023034",
                "ENSMUSG00000001280",
                "ENSMUSG00000019768",
                "ENSMUSG00000000567"
            ],
            "target": [],
            "expression_sum": {
                "class1": 10298.0,
                "class2": 2655.0
            },
            "uniprot_ids": [
                "Q6ZWN5",
                "Q9CXW7",
                "D3YWH9",
                "D3YUV6",
                "F7CJS8",
                "D3Z673"
            ]
        },
        "ENSMUSG00000006585": {
            "id": "ENSMUSG00000006585",
            "type": "Gene",
            "symbol": "Cdt1",
            "source": [
                "ENSMUSG00000040481",
                "ENSMUSG00000000567"
            ],
            "target": [],
            "expression_sum": {
                "class1": 227.0,
                "class2": 66.0
            },
            "uniprot_ids": [
                "Q8R4E9"
            ]
        },
        "ENSMUSG00000007872": {
            "id": "ENSMUSG00000007872",
            "type": "Gene",
            "symbol": "Id3",
            "source": [
                "ENSMUSG00000003032",
                "ENSMUSG00000018604",
                "ENSMUSG00000027361",
                "ENSMUSG00000026872",
                "ENSMUSG00000031715",
                "ENSMUSG00000000567"
            ],
            "target": [],
            "expression_sum": {
                "class1": 8263.0,
                "class2": 2143.0
            },
            "uniprot_ids": [
                "P41133"
            ]
        },
        "ENSMUSG00000008200": {
            "id": "ENSMUSG00000008200",
            "type": "Gene",
            "symbol": "Fnbp4",
            "source": [
                "ENSMUSG00000024515",
                "ENSMUSG00000030341",
                "ENSMUSG00000044167",
                "ENSMUSG00000053110",
                "ENSMUSG00000057098",
                "ENSMUSG00000024985",
                "ENSMUSG00000000567"
            ],
            "target": [],
            "expression_sum": {
                "class1": 1739.0,
                "class2": 289.0
            },
            "uniprot_ids": [
                "A0A0R4IZZ6",
                "A0A5F8MPU2"
            ]
        },
        "ENSMUSG00000009894": {
            "id": "ENSMUSG00000009894",
            "type": "Gene",
            "symbol": "Snap47",
            "source": [
                "ENSMUSG00000078671",
                "ENSMUSG00000000567"
            ],
            "target": [],
            "expression_sum": {
                "class1": 790.0,
                "class2": 189.0
            },
            "uniprot_ids": [
                "Q8R570",
                "I7HPB1",
                "B2FDF7",
                "I7HFU4",
                "I7HIK2",
                "J3JS29",
                "B2FDF6",
                "A0A0G2JDA4"
            ]
        },
        "ENSMUSG00000010110": {
            "id": "ENSMUSG00000010110",
            "type": "Gene",
            "symbol": "Stx5a",
            "source": [
                "ENSMUSG00000015839",
                "ENSMUSG00000006215",
                "ENSMUSG00000021250",
                "ENSMUSG00000056501",
                "ENSMUSG00000023079",
                "ENSMUSG00000005583",
                "ENSMUSG00000005698",
                "ENSMUSG00000000567"
            ],
            "target": [],
            "expression_sum": {
                "class1": 2172.0,
                "class2": 237.0
            },
            "uniprot_ids": [
                "Q8K1E0",
                "H3BKP3",
                "H3BKJ7",
                "H3BJE3",
                "H3BL25",
                "H3BKD6",
                "H3BJ02",
                "H3BJ89",
                "I1E4X1",
                "H3BKP0"
            ]
        },
        "ENSMUSG00000010406": {
            "id": "ENSMUSG00000010406",
            "type": "Gene",
            "symbol": "Mrpl52",
            "source": [
                "ENSMUSG00000029135",
                "ENSMUSG00000024560",
                "ENSMUSG00000030557",
                "ENSMUSG00000027490",
                "ENSMUSG00000001280",
                "ENSMUSG00000019768",
                "ENSMUSG00000044167",
                "ENSMUSG00000024140",
                "ENSMUSG00000031885",
                "ENSMUSG00000076431",
                "ENSMUSG00000014592",
                "ENSMUSG00000021775",
                "ENSMUSG00000000567"
            ],
            "target": [],
            "expression_sum": {
                "class1": 4360.0,
                "class2": 1217.0
            },
            "uniprot_ids": [
                "Q9D0Y8",
                "A0A0G2JE60"
            ]
        },
        "ENSMUSG00000013662": {
            "id": "ENSMUSG00000013662",
            "type": "Gene",
            "symbol": "Atad1",
            "source": [
                "ENSMUSG00000001280",
                "ENSMUSG00000031885",
                "ENSMUSG00000008575",
                "ENSMUSG00000042557",
                "ENSMUSG00000031715",
                "ENSMUSG00000000567"
            ],
            "target": [],
            "expression_sum": {
                "class1": 585.0,
                "class2": 388.0
            },
            "uniprot_ids": [
                "Q9D5T0",
                "A0A494BAB5",
                "A0A494B9L7",
                "A0A494B9Y0"
            ]
        },
        "ENSMUSG00000014769": {
            "id": "ENSMUSG00000014769",
            "type": "Gene",
            "symbol": "Psmb1",
            "source": [
                "ENSMUSG00000001419",
                "ENSMUSG00000024560",
                "ENSMUSG00000038535",
                "ENSMUSG00000040732",
                "ENSMUSG00000046470",
                "ENSMUSG00000031715",
                "ENSMUSG00000000567"
            ],
            "target": [],
            "expression_sum": {
                "class1": 7146.0,
                "class2": 1773.0
            },
            "uniprot_ids": [
                "O09061",
                "A0A338P7C0",
                "A0A338P7F1"
            ]
        },
        "ENSMUSG00000016262": {
            "id": "ENSMUSG00000016262",
            "type": "Gene",
            "symbol": "Sertad4",
            "source": [
                "ENSMUSG00000019817",
                "ENSMUSG00000052534",
                "ENSMUSG00000015846",
                "ENSMUSG00000000567"
            ],
            "target": [],
            "expression_sum": {
                "class1": 734.0,
                "class2": 65.0
            },
            "uniprot_ids": [
                "A7DTG3",
                "A0A0A6YXT5"
            ]
        },
        "ENSMUSG00000016503": {
            "id": "ENSMUSG00000016503",
            "type": "TF",
            "symbol": "Gtf3a",
            "source": [
                "ENSMUSG00000032238",
                "ENSMUSG00000042557",
                "ENSMUSG00000000567"
            ],
            "target": [
                "ENSMUSG00000000817",
                "ENSMUSG00000000359",
                "ENSMUSG00000024052",
                "ENSMUSG00000000982",
                "ENSMUSG00000002844"
            ],
            "expression_sum": {
                "class1": 204.0,
                "class2": 44.0
            },
            "uniprot_ids": [
                "Q8VHT7",
                "F6SIJ4",
                "F6RCF6"
            ]
        },
        "ENSMUSG00000017421": {
            "id": "ENSMUSG00000017421",
            "type": "Gene",
            "symbol": "Zfp207",
            "source": [
                "ENSMUSG00000026398",
                "ENSMUSG00000027490",
                "ENSMUSG00000052684",
                "ENSMUSG00000031885",
                "ENSMUSG00000000567"
            ],
            "target": [],
            "expression_sum": {
                "class1": 3697.0,
                "class2": 846.0
            },
            "uniprot_ids": [
                "Q9JMD0",
                "Q8CCB2",
                "M0QWF0",
                "B1AQG7"
            ]
        },
        "ENSMUSG00000017999": {
            "id": "ENSMUSG00000017999",
            "type": "Gene",
            "symbol": "Ddx27",
            "source": [
                "ENSMUSG00000022346",
                "ENSMUSG00000055116",
                "ENSMUSG00000028466",
                "ENSMUSG00000047407",
                "ENSMUSG00000031627",
                "ENSMUSG00000000567"
            ],
            "target": [],
            "expression_sum": {
                "class1": 1934.0,
                "class2": 243.0
            },
            "uniprot_ids": [
                "Q921N6",
                "H3BLJ6"
            ]
        },
        "ENSMUSG00000018189": {
            "id": "ENSMUSG00000018189",
            "type": "Gene",
            "symbol": "Uchl5",
            "source": [
                "ENSMUSG00000000567",
                "ENSMUSG00000005566",
                "ENSMUSG00000042557",
                "ENSMUSG00000022895",
                "ENSMUSG00000020167"
            ],
            "target": [],
            "expression_sum": {
                "class1": 2634.0,
                "class2": 570.0
            },
            "uniprot_ids": [
                "Q9WUP7",
                "A0A087WP81",
                "A0A087WRL3"
            ]
        },
        "ENSMUSG00000018217": {
            "id": "ENSMUSG00000018217",
            "type": "Gene",
            "symbol": "Pmp22",
            "source": [
                "ENSMUSG00000026398",
                "ENSMUSG00000038615",
                "ENSMUSG00000002265",
                "ENSMUSG00000022053",
                "ENSMUSG00000027104",
                "ENSMUSG00000000567"
            ],
            "target": [],
            "expression_sum": {
                "class1": 6082.0,
                "class2": 731.0
            },
            "uniprot_ids": [
                "Q5SXS3"
            ]
        },
        "ENSMUSG00000018340": {
            "id": "ENSMUSG00000018340",
            "type": "Gene",
            "symbol": "Anxa6",
            "source": [
                "ENSMUSG00000026398",
                "ENSMUSG00000029135",
                "ENSMUSG00000006215",
                "ENSMUSG00000023034",
                "ENSMUSG00000001280",
                "ENSMUSG00000019768",
                "ENSMUSG00000024498",
                "ENSMUSG00000000567"
            ],
            "target": [],
            "expression_sum": {
                "class1": 3577.0,
                "class2": 771.0
            },
            "uniprot_ids": [
                "P14824",
                "F8WIT2"
            ]
        },
        "ENSMUSG00000018567": {
            "id": "ENSMUSG00000018567",
            "type": "Gene",
            "symbol": "Gabarap",
            "source": [
                "ENSMUSG00000029135",
                "ENSMUSG00000023034",
                "ENSMUSG00000001280",
                "ENSMUSG00000015605",
                "ENSMUSG00000000567"
            ],
            "target": [],
            "expression_sum": {
                "class1": 10539.0,
                "class2": 2900.0
            },
            "uniprot_ids": [
                "Q9DCD6",
                "B1AR50"
            ]
        },
        "ENSMUSG00000018861": {
            "id": "ENSMUSG00000018861",
            "type": "Gene",
            "symbol": "Fdxr",
            "source": [
                "ENSMUSG00000052684",
                "ENSMUSG00000005583",
                "ENSMUSG00000021250",
                "ENSMUSG00000000567"
            ],
            "target": [],
            "expression_sum": {
                "class1": 506.0,
                "class2": 64.0
            },
            "uniprot_ids": [
                "Q61578"
            ]
        },
        "ENSMUSG00000019467": {
            "id": "ENSMUSG00000019467",
            "type": "Gene",
            "symbol": "Arhgef25",
            "source": [
                "ENSMUSG00000008575",
                "ENSMUSG00000000567"
            ],
            "target": [],
            "expression_sum": {
                "class1": 895.0,
                "class2": 130.0
            },
            "uniprot_ids": [
                "A0A1Y7VNM7",
                "A0A1W2P813",
                "A0A1W2P718",
                "A0A1W2P6R9",
                "A0A1W2P7S3",
                "G5E825",
                "A0A0R4J211"
            ]
        },
        "ENSMUSG00000019689": {
            "id": "ENSMUSG00000019689",
            "type": "Gene",
            "symbol": "Fmc1",
            "source": [
                "ENSMUSG00000001419",
                "ENSMUSG00000001280",
                "ENSMUSG00000038914",
                "ENSMUSG00000076431",
                "ENSMUSG00000021775",
                "ENSMUSG00000031715",
                "ENSMUSG00000000567"
            ],
            "target": [],
            "expression_sum": {
                "class1": 4516.0,
                "class2": 4608.0
            },
            "uniprot_ids": [
                "Q9CR13"
            ]
        },
        "ENSMUSG00000019775": {
            "id": "ENSMUSG00000019775",
            "type": "Gene",
            "symbol": "Rgs17",
            "source": [
                "ENSMUSG00000058440",
                "ENSMUSG00000031099",
                "ENSMUSG00000000567"
            ],
            "target": [],
            "expression_sum": {
                "class1": 208.0,
                "class2": 140.0
            },
            "uniprot_ids": [
                "Q9QZB0",
                "G5E8E0",
                "F8WH97"
            ]
        },
        "ENSMUSG00000019837": {
            "id": "ENSMUSG00000019837",
            "type": "Gene",
            "symbol": "Gtf3c6",
            "source": [
                "ENSMUSG00000059552",
                "ENSMUSG00000024515",
                "ENSMUSG00000056501",
                "ENSMUSG00000038648",
                "ENSMUSG00000027490",
                "ENSMUSG00000005583",
                "ENSMUSG00000027104",
                "ENSMUSG00000053110",
                "ENSMUSG00000039191",
                "ENSMUSG00000000567"
            ],
            "target": [],
            "expression_sum": {
                "class1": 3640.0,
                "class2": 635.0
            },
            "uniprot_ids": [
                "Q9D8P7",
                "Q9DB36",
                "A0A1L1SQC4",
                "A0A1L1SUI7",
                "A0A1L1STD0"
            ]
        },
        "ENSMUSG00000020282": {
            "id": "ENSMUSG00000020282",
            "type": "Gene",
            "symbol": "Rhbdf1",
            "source": [
                "ENSMUSG00000021250",
                "ENSMUSG00000052684",
                "ENSMUSG00000005583",
                "ENSMUSG00000005698",
                "ENSMUSG00000000567"
            ],
            "target": [],
            "expression_sum": {
                "class1": 741.0,
                "class2": 116.0
            },
            "uniprot_ids": [
                "Q6PIX5",
                "F2Z3W3",
                "A2AVE4",
                "A7M7T5",
                "A2AVE5"
            ]
        },
        "ENSMUSG00000020485": {
            "id": "ENSMUSG00000020485",
            "type": "Gene",
            "symbol": "Supt4a",
            "source": [
                "ENSMUSG00000029135",
                "ENSMUSG00000057469",
                "ENSMUSG00000047407",
                "ENSMUSG00000029249",
                "ENSMUSG00000001280",
                "ENSMUSG00000019768",
                "ENSMUSG00000022641",
                "ENSMUSG00000068267",
                "ENSMUSG00000032035",
                "ENSMUSG00000016087",
                "ENSMUSG00000015605",
                "ENSMUSG00000000567"
            ],
            "target": [],
            "expression_sum": {
                "class1": 2517.0,
                "class2": 395.0
            },
            "uniprot_ids": [
                "P63271",
                "G3UVU8",
                "D6RHM5"
            ]
        },
        "ENSMUSG00000020522": {
            "id": "ENSMUSG00000020522",
            "type": "Gene",
            "symbol": "Mfap3",
            "source": [
                "ENSMUSG00000027490",
                "ENSMUSG00000044167",
                "ENSMUSG00000040732",
                "ENSMUSG00000035158",
                "ENSMUSG00000000567"
            ],
            "target": [],
            "expression_sum": {
                "class1": 1866.0,
                "class2": 216.0
            },
            "uniprot_ids": [
                "A0A0R4J1C7",
                "D3Z5B4"
            ]
        },
        "ENSMUSG00000020530": {
            "id": "ENSMUSG00000020530",
            "type": "Gene",
            "symbol": "Ggnbp2",
            "source": [
                "ENSMUSG00000001419",
                "ENSMUSG00000078671",
                "ENSMUSG00000024560",
                "ENSMUSG00000030557",
                "ENSMUSG00000029249",
                "ENSMUSG00000001280",
                "ENSMUSG00000019768",
                "ENSMUSG00000022383",
                "ENSMUSG00000055320",
                "ENSMUSG00000058756",
                "ENSMUSG00000008575",
                "ENSMUSG00000031715",
                "ENSMUSG00000024985",
                "ENSMUSG00000000567"
            ],
            "target": [],
            "expression_sum": {
                "class1": 4257.0,
                "class2": 1071.0
            },
            "uniprot_ids": [
                "Q5SV77",
                "H7BX83",
                "F6Y247",
                "E9Q6H9",
                "E9Q389"
            ]
        },
        "ENSMUSG00000020646": {
            "id": "ENSMUSG00000020646",
            "type": "Gene",
            "symbol": "Mboat2",
            "source": [
                "ENSMUSG00000078671",
                "ENSMUSG00000005583",
                "ENSMUSG00000025958",
                "ENSMUSG00000015846",
                "ENSMUSG00000000567"
            ],
            "target": [],
            "expression_sum": {
                "class1": 341.0,
                "class2": 74.0
            },
            "uniprot_ids": [
                "Q8R3I2",
                "A0A1Y7VKU2",
                "A0A217FL69",
                "A0A1Y7VNY8"
            ]
        },
        "ENSMUSG00000020755": {
            "id": "ENSMUSG00000020755",
            "type": "Gene",
            "symbol": "Sap30bp",
            "source": [
                "ENSMUSG00000052534",
                "ENSMUSG00000006705",
                "ENSMUSG00000005583",
                "ENSMUSG00000031627",
                "ENSMUSG00000000567"
            ],
            "target": [],
            "expression_sum": {
                "class1": 1472.0,
                "class2": 125.0
            },
            "uniprot_ids": [
                "Q02614"
            ]
        },
        "ENSMUSG00000020766": {
            "id": "ENSMUSG00000020766",
            "type": "Gene",
            "symbol": "Galk1",
            "source": [
                "ENSMUSG00000024560",
                "ENSMUSG00000000567"
            ],
            "target": [],
            "expression_sum": {
                "class1": 2465.0,
                "class2": 102.0
            },
            "uniprot_ids": [
                "Q9R0N0"
            ]
        },
        "ENSMUSG00000021027": {
            "id": "ENSMUSG00000021027",
            "type": "Gene",
            "symbol": "Ralgapa1",
            "source": [
                "ENSMUSG00000026398",
                "ENSMUSG00000024515",
                "ENSMUSG00000059552",
                "ENSMUSG00000028565",
                "ENSMUSG00000040732",
                "ENSMUSG00000060261",
                "ENSMUSG00000027439",
                "ENSMUSG00000000567"
            ],
            "target": [],
            "expression_sum": {
                "class1": 1459.0,
                "class2": 791.0
            },
            "uniprot_ids": [
                "Q6GYP7",
                "A0A1W2P7P2",
                "A0A1W2P712",
                "A0A1W2P832",
                "A0A2I3BRX9",
                "A0A1W2P6Q0",
                "A0A1W2P7Y4"
            ]
        },
        "ENSMUSG00000021036": {
            "id": "ENSMUSG00000021036",
            "type": "Gene",
            "symbol": "Sptlc2",
            "source": [
                "ENSMUSG00000027490",
                "ENSMUSG00000000567"
            ],
            "target": [],
            "expression_sum": {
                "class1": 1317.0,
                "class2": 378.0
            },
            "uniprot_ids": [
                "P97363"
            ]
        },
        "ENSMUSG00000021130": {
            "id": "ENSMUSG00000021130",
            "type": "Gene",
            "symbol": "Galnt16",
            "source": [
                "ENSMUSG00000042557",
                "ENSMUSG00000000567"
            ],
            "target": [],
            "expression_sum": {
                "class1": 892.0,
                "class2": 39.0
            },
            "uniprot_ids": [
                "Q9JJ61",
                "A0A1W2P6P9",
                "A0A1W2P7U3"
            ]
        },
        "ENSMUSG00000021243": {
            "id": "ENSMUSG00000021243",
            "type": "Gene",
            "symbol": "Fcf1",
            "source": [
                "ENSMUSG00000058239",
                "ENSMUSG00000018604",
                "ENSMUSG00000001280",
                "ENSMUSG00000019768",
                "ENSMUSG00000025958",
                "ENSMUSG00000071637",
                "ENSMUSG00000000567"
            ],
            "target": [],
            "expression_sum": {
                "class1": 2813.0,
                "class2": 908.0
            },
            "uniprot_ids": [
                "Q9CTH6",
                "E9Q186"
            ]
        },
        "ENSMUSG00000021245": {
            "id": "ENSMUSG00000021245",
            "type": "Gene",
            "symbol": "Mlh3",
            "source": [
                "ENSMUSG00000058440",
                "ENSMUSG00000058239",
                "ENSMUSG00000044167",
                "ENSMUSG00000031627",
                "ENSMUSG00000000567"
            ],
            "target": [],
            "expression_sum": {
                "class1": 588.0,
                "class2": 94.0
            },
            "uniprot_ids": [
                "Q68FG1",
                "A0A1Y7VMP7",
                "A0A1Y7VJX4"
            ]
        },
        "ENSMUSG00000021318": {
            "id": "ENSMUSG00000021318",
            "type": "TF",
            "symbol": "Gli3",
            "source": [
                "ENSMUSG00000027490",
                "ENSMUSG00000000567"
            ],
            "target": [
                "ENSMUSG00000022020",
                "ENSMUSG00000046324",
                "ENSMUSG00000028950",
                "ENSMUSG00000029190",
                "ENSMUSG00000028556",
                "ENSMUSG00000053907",
                "ENSMUSG00000063904",
                "ENSMUSG00000073838",
                "ENSMUSG00000097729",
                "ENSMUSG00000090150",
                "ENSMUSG00000025745",
                "ENSMUSG00000029655",
                "ENSMUSG00000041375",
                "ENSMUSG00000053483",
                "ENSMUSG00000019518",
                "ENSMUSG00000027304",
                "ENSMUSG00000026456",
                "ENSMUSG00000022515"
            ],
            "expression_sum": {
                "class1": 400.0,
                "class2": 14.0
            },
            "uniprot_ids": [
                "Q61602",
                "Q3UMJ8",
                "A0A1Y7VKQ8"
            ]
        },
        "ENSMUSG00000021408": {
            "id": "ENSMUSG00000021408",
            "type": "Gene",
            "symbol": "Ripk1",
            "source": [
                "ENSMUSG00000058440",
                "ENSMUSG00000044167",
                "ENSMUSG00000032228",
                "ENSMUSG00000000567"
            ],
            "target": [],
            "expression_sum": {
                "class1": 552.0,
                "class2": 132.0
            },
            "uniprot_ids": [
                "Q60855",
                "F7D1J2"
            ]
        },
        "ENSMUSG00000021706": {
            "id": "ENSMUSG00000021706",
            "type": "Gene",
            "symbol": "Zfyve16",
            "source": [
                "ENSMUSG00000057469",
                "ENSMUSG00000025612",
                "ENSMUSG00000015839",
                "ENSMUSG00000058239",
                "ENSMUSG00000015579",
                "ENSMUSG00000051469",
                "ENSMUSG00000001280",
                "ENSMUSG00000019768",
                "ENSMUSG00000025958",
                "ENSMUSG00000002265",
                "ENSMUSG00000031885",
                "ENSMUSG00000008575",
                "ENSMUSG00000000567"
            ],
            "target": [],
            "expression_sum": {
                "class1": 2557.0,
                "class2": 339.0
            },
            "uniprot_ids": [
                "Q80U44"
            ]
        },
        "ENSMUSG00000021714": {
            "id": "ENSMUSG00000021714",
            "type": "Gene",
            "symbol": "Cenpk",
            "source": [
                "ENSMUSG00000006215",
                "ENSMUSG00000052837",
                "ENSMUSG00000047407",
                "ENSMUSG00000031627",
                "ENSMUSG00000031885",
                "ENSMUSG00000024985",
                "ENSMUSG00000000567"
            ],
            "target": [],
            "expression_sum": {
                "class1": 1780.0,
                "class2": 22.0
            },
            "uniprot_ids": [
                "Q9ESN5",
                "A0A286YE26",
                "A0A0R4J037",
                "A0A286YDG4"
            ]
        },
        "ENSMUSG00000021771": {
            "id": "ENSMUSG00000021771",
            "type": "Gene",
            "symbol": "Vdac2",
            "source": [
                "ENSMUSG00000001419",
                "ENSMUSG00000015605",
                "ENSMUSG00000056501",
                "ENSMUSG00000024560",
                "ENSMUSG00000027490",
                "ENSMUSG00000001280",
                "ENSMUSG00000019768",
                "ENSMUSG00000005583",
                "ENSMUSG00000029249",
                "ENSMUSG00000024695",
                "ENSMUSG00000008575",
                "ENSMUSG00000024985",
                "ENSMUSG00000000567"
            ],
            "target": [],
            "expression_sum": {
                "class1": 5761.0,
                "class2": 2799.0
            },
            "uniprot_ids": [
                "Q60930",
                "G3UX26",
                "A0A286YCR8",
                "D3YZT5",
                "D3YUN8"
            ]
        },
        "ENSMUSG00000022016": {
            "id": "ENSMUSG00000022016",
            "type": "Gene",
            "symbol": "Akap11",
            "source": [
                "ENSMUSG00000058440",
                "ENSMUSG00000057469",
                "ENSMUSG00000031965",
                "ENSMUSG00000001280",
                "ENSMUSG00000019768",
                "ENSMUSG00000044167",
                "ENSMUSG00000005698",
                "ENSMUSG00000022383",
                "ENSMUSG00000025025",
                "ENSMUSG00000024974",
                "ENSMUSG00000008575",
                "ENSMUSG00000000567"
            ],
            "target": [],
            "expression_sum": {
                "class1": 1440.0,
                "class2": 372.0
            },
            "uniprot_ids": [
                "E9Q777"
            ]
        },
        "ENSMUSG00000022090": {
            "id": "ENSMUSG00000022090",
            "type": "Gene",
            "symbol": "Pdlim2",
            "source": [
                "ENSMUSG00000059552",
                "ENSMUSG00000024515",
                "ENSMUSG00000022346",
                "ENSMUSG00000019564",
                "ENSMUSG00000028565",
                "ENSMUSG00000040732",
                "ENSMUSG00000021775",
                "ENSMUSG00000000567"
            ],
            "target": [],
            "expression_sum": {
                "class1": 2233.0,
                "class2": 176.0
            },
            "uniprot_ids": [
                "Q8R1G6",
                "A0A087WPL1",
                "E9Q996",
                "F7C957"
            ]
        },
        "ENSMUSG00000022103": {
            "id": "ENSMUSG00000022103",
            "type": "Gene",
            "symbol": "Gfra2",
            "source": [
                "ENSMUSG00000000567"
            ],
            "target": [],
            "expression_sum": {
                "class1": 380.0,
                "class2": 50.0
            },
            "uniprot_ids": [
                "O08842",
                "A0A2I3BQ61"
            ]
        },
        "ENSMUSG00000022119": {
            "id": "ENSMUSG00000022119",
            "type": "Gene",
            "symbol": "Rbm26",
            "source": [
                "ENSMUSG00000019768",
                "ENSMUSG00000026890",
                "ENSMUSG00000013419",
                "ENSMUSG00000000567"
            ],
            "target": [],
            "expression_sum": {
                "class1": 1494.0,
                "class2": 261.0
            },
            "uniprot_ids": [
                "Q6NZN0",
                "E9PYZ7",
                "E9Q640",
                "A0A0R4J214",
                "E9PUF4"
            ]
        },
        "ENSMUSG00000022150": {
            "id": "ENSMUSG00000022150",
            "type": "TF",
            "symbol": "Dab2",
            "source": [
                "ENSMUSG00000026628",
                "ENSMUSG00000021250",
                "ENSMUSG00000023034",
                "ENSMUSG00000052837",
                "ENSMUSG00000031099",
                "ENSMUSG00000029178",
                "ENSMUSG00000032035",
                "ENSMUSG00000000567"
            ],
            "target": [
                "ENSMUSG00000025580",
                "ENSMUSG00000027679",
                "ENSMUSG00000029379",
                "ENSMUSG00000029089",
                "ENSMUSG00000011256",
                "ENSMUSG00000082016",
                "ENSMUSG00000004996",
                "ENSMUSG00000026955",
                "ENSMUSG00000018387",
                "ENSMUSG00000018196",
                "ENSMUSG00000032384",
                "ENSMUSG00000000058",
                "ENSMUSG00000051172",
                "ENSMUSG00000025815",
                "ENSMUSG00000030879",
                "ENSMUSG00000028869"
            ],
            "expression_sum": {
                "class1": 2759.0,
                "class2": 869.0
            },
            "uniprot_ids": [
                "Q3TRE6",
                "Q9DCE6",
                "E9PX84",
                "E9QL31",
                "E0CXT5",
                "F6TQN9",
                "A0A0R4J104",
                "A0A1Y7VGS2",
                "E0CZ53",
                "E0CYJ2"
            ]
        },
        "ENSMUSG00000022159": {
            "id": "ENSMUSG00000022159",
            "type": "Gene",
            "symbol": "Rab2b",
            "source": [
                "ENSMUSG00000015579",
                "ENSMUSG00000024560",
                "ENSMUSG00000023994",
                "ENSMUSG00000031715",
                "ENSMUSG00000000567"
            ],
            "target": [],
            "expression_sum": {
                "class1": 484.0,
                "class2": 57.0
            },
            "uniprot_ids": [
                "P59279",
                "Q3TEG7",
                "G3UZ24",
                "G3V022",
                "G3UXQ7"
            ]
        },
        "ENSMUSG00000022203": {
            "id": "ENSMUSG00000022203",
            "type": "Gene",
            "symbol": "Efs",
            "source": [
                "ENSMUSG00000000567"
            ],
            "target": [],
            "expression_sum": {
                "class1": 254.0,
                "class2": 52.0
            },
            "uniprot_ids": [
                "Q64355",
                "A0A2I3BQ52",
                "A0A2I3BRP5"
            ]
        },
        "ENSMUSG00000022325": {
            "id": "ENSMUSG00000022325",
            "type": "Gene",
            "symbol": "Pop1",
            "source": [
                "ENSMUSG00000044167",
                "ENSMUSG00000000567"
            ],
            "target": [],
            "expression_sum": {
                "class1": 296.0,
                "class2": 46.0
            },
            "uniprot_ids": [
                "Q8K205",
                "Q9D4G5"
            ]
        },
        "ENSMUSG00000022339": {
            "id": "ENSMUSG00000022339",
            "type": "Gene",
            "symbol": "Ebag9",
            "source": [
                "ENSMUSG00000005583",
                "ENSMUSG00000005698",
                "ENSMUSG00000000567"
            ],
            "target": [],
            "expression_sum": {
                "class1": 741.0,
                "class2": 82.0
            },
            "uniprot_ids": [
                "Q9D0V7",
                "A0A2I3BPQ0"
            ]
        },
        "ENSMUSG00000022378": {
            "id": "ENSMUSG00000022378",
            "type": "Gene",
            "symbol": "Cyrib",
            "source": [
                "ENSMUSG00000031885",
                "ENSMUSG00000058239",
                "ENSMUSG00000000567"
            ],
            "target": [],
            "expression_sum": {
                "class1": 508.0,
                "class2": 237.0
            },
            "uniprot_ids": [
                "Q921M7",
                "A0A2I3BQK1",
                "A0A2I3BPH9",
                "A0A2I3BRN5"
            ]
        },
        "ENSMUSG00000022466": {
            "id": "ENSMUSG00000022466",
            "type": "Gene",
            "symbol": "Rpap3",
            "source": [
                "ENSMUSG00000015839",
                "ENSMUSG00000044167",
                "ENSMUSG00000075595",
                "ENSMUSG00000031885",
                "ENSMUSG00000000567"
            ],
            "target": [],
            "expression_sum": {
                "class1": 1108.0,
                "class2": 228.0
            },
            "uniprot_ids": [
                "Q9D706"
            ]
        },
        "ENSMUSG00000022621": {
            "id": "ENSMUSG00000022621",
            "type": "Gene",
            "symbol": "Rabl2",
            "source": [
                "ENSMUSG00000000567"
            ],
            "target": [],
            "expression_sum": {
                "class1": 763.0,
                "class2": 46.0
            },
            "uniprot_ids": [
                "E9Q9D5",
                "E9PZU3",
                "E9Q8G6",
                "F6Y6I9",
                "E9Q0F8",
                "E9Q473"
            ]
        },
        "ENSMUSG00000022686": {
            "id": "ENSMUSG00000022686",
            "type": "Gene",
            "symbol": "B3gnt5",
            "source": [
                "ENSMUSG00000018604",
                "ENSMUSG00000025958",
                "ENSMUSG00000020919",
                "ENSMUSG00000016087",
                "ENSMUSG00000000567"
            ],
            "target": [],
            "expression_sum": {
                "class1": 191.0,
                "class2": 57.0
            },
            "uniprot_ids": [
                "Q8BGY6"
            ]
        },
        "ENSMUSG00000022722": {
            "id": "ENSMUSG00000022722",
            "type": "Gene",
            "symbol": "Arl6",
            "source": [
                "ENSMUSG00000000359",
                "ENSMUSG00000028565",
                "ENSMUSG00000018899",
                "ENSMUSG00000000567"
            ],
            "target": [],
            "expression_sum": {
                "class1": 160.0,
                "class2": 33.0
            },
            "uniprot_ids": [
                "O88848",
                "D3Z067"
            ]
        },
        "ENSMUSG00000022754": {
            "id": "ENSMUSG00000022754",
            "type": "Gene",
            "symbol": "Tmem45a",
            "source": [
                "ENSMUSG00000021250",
                "ENSMUSG00000041515",
                "ENSMUSG00000032228",
                "ENSMUSG00000052684",
                "ENSMUSG00000053110",
                "ENSMUSG00000000567"
            ],
            "target": [],
            "expression_sum": {
                "class1": 642.0,
                "class2": 26.0
            },
            "uniprot_ids": [
                "Q8BFU6",
                "A9C432",
                "A0A338P6Q2"
            ]
        },
        "ENSMUSG00000022961": {
            "id": "ENSMUSG00000022961",
            "type": "Gene",
            "symbol": "Son",
            "source": [
                "ENSMUSG00000024238",
                "ENSMUSG00000020248",
                "ENSMUSG00000019817",
                "ENSMUSG00000052534",
                "ENSMUSG00000029729",
                "ENSMUSG00000015579",
                "ENSMUSG00000036461",
                "ENSMUSG00000074733",
                "ENSMUSG00000027109",
                "ENSMUSG00000001280",
                "ENSMUSG00000019768",
                "ENSMUSG00000005583",
                "ENSMUSG00000008575",
                "ENSMUSG00000022314",
                "ENSMUSG00000000567"
            ],
            "target": [],
            "expression_sum": {
                "class1": 37006.0,
                "class2": 8182.0
            },
            "uniprot_ids": [
                "Q9QX47",
                "H9KV01",
                "H9KV02",
                "H9KV15",
                "F7CU80",
                "H9KV00"
            ]
        },
        "ENSMUSG00000023022": {
            "id": "ENSMUSG00000023022",
            "type": "Gene",
            "symbol": "Lima1",
            "source": [
                "ENSMUSG00000018604",
                "ENSMUSG00000043456",
                "ENSMUSG00000000567"
            ],
            "target": [],
            "expression_sum": {
                "class1": 3484.0,
                "class2": 484.0
            },
            "uniprot_ids": [
                "Q9ERG0",
                "A0A2R8VI67"
            ]
        },
        "ENSMUSG00000023068": {
            "id": "ENSMUSG00000023068",
            "type": "Gene",
            "symbol": "Nus1",
            "source": [
                "ENSMUSG00000029135",
                "ENSMUSG00000015839",
                "ENSMUSG00000018604",
                "ENSMUSG00000056501",
                "ENSMUSG00000032228",
                "ENSMUSG00000051469",
                "ENSMUSG00000001280",
                "ENSMUSG00000019768",
                "ENSMUSG00000021877",
                "ENSMUSG00000033943",
                "ENSMUSG00000053110",
                "ENSMUSG00000031885",
                "ENSMUSG00000016087",
                "ENSMUSG00000076431",
                "ENSMUSG00000008575",
                "ENSMUSG00000000567"
            ],
            "target": [],
            "expression_sum": {
                "class1": 1446.0,
                "class2": 1078.0
            },
            "uniprot_ids": [
                "Q99LJ8"
            ]
        },
        "ENSMUSG00000023104": {
            "id": "ENSMUSG00000023104",
            "type": "Gene",
            "symbol": "Rfc2",
            "source": [
                "ENSMUSG00000027109",
                "ENSMUSG00000029249",
                "ENSMUSG00000005583",
                "ENSMUSG00000022383",
                "ENSMUSG00000000567"
            ],
            "target": [],
            "expression_sum": {
                "class1": 1941.0,
                "class2": 139.0
            },
            "uniprot_ids": [
                "Q9WUK4",
                "A0A0J9YV68"
            ]
        },
        "ENSMUSG00000023456": {
            "id": "ENSMUSG00000023456",
            "type": "Gene",
            "symbol": "Tpi1",
            "source": [
                "ENSMUSG00000029135",
                "ENSMUSG00000001419",
                "ENSMUSG00000001911",
                "ENSMUSG00000015605",
                "ENSMUSG00000056501",
                "ENSMUSG00000015579",
                "ENSMUSG00000051469",
                "ENSMUSG00000001280",
                "ENSMUSG00000019768",
                "ENSMUSG00000008575",
                "ENSMUSG00000039191",
                "ENSMUSG00000000567"
            ],
            "target": [],
            "expression_sum": {
                "class1": 4685.0,
                "class2": 2207.0
            },
            "uniprot_ids": [
                "P17751",
                "H7BXC3"
            ]
        },
        "ENSMUSG00000024054": {
            "id": "ENSMUSG00000024054",
            "type": "Gene",
            "symbol": "Smchd1",
            "source": [
                "ENSMUSG00000021250",
                "ENSMUSG00000021109",
                "ENSMUSG00000056501",
                "ENSMUSG00000041515",
                "ENSMUSG00000024560",
                "ENSMUSG00000005698",
                "ENSMUSG00000000567"
            ],
            "target": [],
            "expression_sum": {
                "class1": 2336.0,
                "class2": 457.0
            },
            "uniprot_ids": [
                "Q6P5D8",
                "A0A3B2W494",
                "A0A3B2WD81"
            ]
        },
        "ENSMUSG00000024104": {
            "id": "ENSMUSG00000024104",
            "type": "Gene",
            "symbol": "Washc2",
            "source": [
                "ENSMUSG00000038692",
                "ENSMUSG00000059552",
                "ENSMUSG00000000567"
            ],
            "target": [],
            "expression_sum": {
                "class1": 1578.0,
                "class2": 302.0
            },
            "uniprot_ids": [
                "Q6PGL7",
                "A0A0N4SVI5",
                "A0A0N4SV74",
                "A0A0N4SUJ0",
                "A0A0N4SVK9",
                "A0A0N4SUQ0"
            ]
        },
        "ENSMUSG00000024146": {
            "id": "ENSMUSG00000024146",
            "type": "Gene",
            "symbol": "Cript",
            "source": [
                "ENSMUSG00000001280",
                "ENSMUSG00000031885",
                "ENSMUSG00000053110",
                "ENSMUSG00000000567"
            ],
            "target": [],
            "expression_sum": {
                "class1": 3091.0,
                "class2": 708.0
            },
            "uniprot_ids": [
                "O70333"
            ]
        },
        "ENSMUSG00000024538": {
            "id": "ENSMUSG00000024538",
            "type": "Gene",
            "symbol": "Ppic",
            "source": [
                "ENSMUSG00000003032",
                "ENSMUSG00000058239",
                "ENSMUSG00000027490",
                "ENSMUSG00000001280",
                "ENSMUSG00000019768",
                "ENSMUSG00000033943",
                "ENSMUSG00000005698",
                "ENSMUSG00000025958",
                "ENSMUSG00000031885",
                "ENSMUSG00000053110",
                "ENSMUSG00000000567"
            ],
            "target": [],
            "expression_sum": {
                "class1": 5705.0,
                "class2": 1061.0
            },
            "uniprot_ids": [
                "P30412"
            ]
        },
        "ENSMUSG00000024588": {
            "id": "ENSMUSG00000024588",
            "type": "Gene",
            "symbol": "Fech",
            "source": [
                "ENSMUSG00000006215",
                "ENSMUSG00000015605",
                "ENSMUSG00000029249",
                "ENSMUSG00000001280",
                "ENSMUSG00000019768",
                "ENSMUSG00000008575",
                "ENSMUSG00000024985",
                "ENSMUSG00000000567"
            ],
            "target": [],
            "expression_sum": {
                "class1": 1621.0,
                "class2": 1333.0
            },
            "uniprot_ids": [
                "Q91W09",
                "Q544X6",
                "A0A494BAI5",
                "A0A494B9Y8",
                "A0A494BBL4",
                "A0A494BAV9",
                "F6T2S0",
                "A0A494BA61",
                "A0A494BB70",
                "A0A494BAR8"
            ]
        },
        "ENSMUSG00000024639": {
            "id": "ENSMUSG00000024639",
            "type": "Gene",
            "symbol": "Gnaq",
            "source": [
                "ENSMUSG00000026398",
                "ENSMUSG00000008575",
                "ENSMUSG00000026610",
                "ENSMUSG00000029249",
                "ENSMUSG00000019768",
                "ENSMUSG00000005583",
                "ENSMUSG00000005698",
                "ENSMUSG00000015605",
                "ENSMUSG00000042557",
                "ENSMUSG00000000567"
            ],
            "target": [],
            "expression_sum": {
                "class1": 1729.0,
                "class2": 597.0
            },
            "uniprot_ids": [
                "P21279"
            ]
        },
        "ENSMUSG00000024769": {
            "id": "ENSMUSG00000024769",
            "type": "Gene",
            "symbol": "Cdc42bpg",
            "source": [
                "ENSMUSG00000026398",
                "ENSMUSG00000022383",
                "ENSMUSG00000031103",
                "ENSMUSG00000000567"
            ],
            "target": [],
            "expression_sum": {
                "class1": 210.0,
                "class2": 81.0
            },
            "uniprot_ids": [
                "Q80UW5"
            ]
        },
        "ENSMUSG00000024921": {
            "id": "ENSMUSG00000024921",
            "type": "Gene",
            "symbol": "Smarca2",
            "source": [
                "ENSMUSG00000018604",
                "ENSMUSG00000022641",
                "ENSMUSG00000000567"
            ],
            "target": [],
            "expression_sum": {
                "class1": 6338.0,
                "class2": 896.0
            },
            "uniprot_ids": [
                "H3BK47",
                "Q9D007",
                "H3BKJ3",
                "A0A140LI08",
                "A0A140LJD7",
                "A0A140LHZ8",
                "H3BLH0",
                "H3BLI4",
                "A0A140LHF4",
                "A0A140LJK1",
                "A0A140LIV3",
                "A0A140LI14",
                "A0A140LI12",
                "A0A140LHE2",
                "A0A140LIQ9",
                "A0A140LHP6",
                "A0A140LIK7",
                "E9QAB8",
                "H3BJI1",
                "A0A140LHJ8",
                "A0A140LID0",
                "H3BKN5",
                "H3BJK2",
                "H3BJM2",
                "A0A140LIM0",
                "H3BKD2",
                "A0A140LHL7",
                "F2Z4A9"
            ]
        },
        "ENSMUSG00000024959": {
            "id": "ENSMUSG00000024959",
            "type": "Gene",
            "symbol": "Bad",
            "source": [
                "ENSMUSG00000051469",
                "ENSMUSG00000038648",
                "ENSMUSG00000020167",
                "ENSMUSG00000000567"
            ],
            "target": [],
            "expression_sum": {
                "class1": 3799.0,
                "class2": 309.0
            },
            "uniprot_ids": [
                "Q61337",
                "Q3U9H3",
                "D3YZR8",
                "F6S493",
                "A0A494BBC4",
                "F7ABX5"
            ]
        },
        "ENSMUSG00000025140": {
            "id": "ENSMUSG00000025140",
            "type": "Gene",
            "symbol": "Pycr1",
            "source": [
                "ENSMUSG00000005583",
                "ENSMUSG00000021250",
                "ENSMUSG00000000567"
            ],
            "target": [],
            "expression_sum": {
                "class1": 290.0,
                "class2": 10.0
            },
            "uniprot_ids": [
                "Q922W5",
                "A2ABZ0",
                "A2ABZ2",
                "A2ABZ3"
            ]
        },
        "ENSMUSG00000025374": {
            "id": "ENSMUSG00000025374",
            "type": "Gene",
            "symbol": "Nabp2",
            "source": [
                "ENSMUSG00000029135",
                "ENSMUSG00000001419",
                "ENSMUSG00000057469",
                "ENSMUSG00000078671",
                "ENSMUSG00000029249",
                "ENSMUSG00000001280",
                "ENSMUSG00000005566",
                "ENSMUSG00000005583",
                "ENSMUSG00000005698",
                "ENSMUSG00000019768",
                "ENSMUSG00000032238",
                "ENSMUSG00000031885",
                "ENSMUSG00000008575",
                "ENSMUSG00000029913",
                "ENSMUSG00000024985",
                "ENSMUSG00000000567"
            ],
            "target": [],
            "expression_sum": {
                "class1": 541.0,
                "class2": 124.0
            },
            "uniprot_ids": [
                "Q8R2Y9",
                "E9PV68",
                "E9Q1I2",
                "E9Q199",
                "E9Q2V9",
                "E9PW70",
                "E9Q5F7"
            ]
        },
        "ENSMUSG00000025484": {
            "id": "ENSMUSG00000025484",
            "type": "Gene",
            "symbol": "Bet1l",
            "source": [
                "ENSMUSG00000017146",
                "ENSMUSG00000000567"
            ],
            "target": [],
            "expression_sum": {
                "class1": 1790.0,
                "class2": 290.0
            },
            "uniprot_ids": [
                "O35153",
                "A0A1B0GSF3",
                "A0A1B0GR53",
                "A0A1B0GR52",
                "A0A1B0GQV3",
                "A0A1B0GS34"
            ]
        },
        "ENSMUSG00000025533": {
            "id": "ENSMUSG00000025533",
            "type": "Gene",
            "symbol": "Asl",
            "source": [
                "ENSMUSG00000024974",
                "ENSMUSG00000000567"
            ],
            "target": [],
            "expression_sum": {
                "class1": 229.0,
                "class2": 14.0
            },
            "uniprot_ids": [
                "Q91YI0",
                "E0CY49",
                "F7D439",
                "E0CYV3"
            ]
        },
        "ENSMUSG00000025591": {
            "id": "ENSMUSG00000025591",
            "type": "Gene",
            "symbol": "Tma16",
            "source": [
                "ENSMUSG00000001419",
                "ENSMUSG00000024515",
                "ENSMUSG00000024955",
                "ENSMUSG00000035158",
                "ENSMUSG00000000567"
            ],
            "target": [],
            "expression_sum": {
                "class1": 558.0,
                "class2": 155.0
            },
            "uniprot_ids": [
                "Q9CR02",
                "A0A1D5RM14"
            ]
        },
        "ENSMUSG00000025782": {
            "id": "ENSMUSG00000025782",
            "type": "Gene",
            "symbol": "Taf3",
            "source": [
                "ENSMUSG00000058440",
                "ENSMUSG00000017146",
                "ENSMUSG00000022895",
                "ENSMUSG00000000567"
            ],
            "target": [],
            "expression_sum": {
                "class1": 2397.0,
                "class2": 343.0
            },
            "uniprot_ids": [
                "Q5HZG4",
                "A2ASX9",
                "A2ASY0",
                "A2ASX8"
            ]
        },
        "ENSMUSG00000025794": {
            "id": "ENSMUSG00000025794",
            "type": "Gene",
            "symbol": "Rpl14",
            "source": [
                "ENSMUSG00000055148",
                "ENSMUSG00000018604",
                "ENSMUSG00000001280",
                "ENSMUSG00000019768",
                "ENSMUSG00000040732",
                "ENSMUSG00000022641",
                "ENSMUSG00000022895",
                "ENSMUSG00000000567"
            ],
            "target": [],
            "expression_sum": {
                "class1": 17351.0,
                "class2": 2678.0
            },
            "uniprot_ids": [
                "Q9CR57",
                "A0A1L1SUF6"
            ]
        },
        "ENSMUSG00000025949": {
            "id": "ENSMUSG00000025949",
            "type": "Gene",
            "symbol": "Pikfyve",
            "source": [
                "ENSMUSG00000024560",
                "ENSMUSG00000005583",
                "ENSMUSG00000040732",
                "ENSMUSG00000015605",
                "ENSMUSG00000031715",
                "ENSMUSG00000000567"
            ],
            "target": [],
            "expression_sum": {
                "class1": 351.0,
                "class2": 55.0
            },
            "uniprot_ids": [
                "Q9Z1T6",
                "D3Z5N5",
                "A0A087WS78",
                "A0A087WQI1"
            ]
        },
        "ENSMUSG00000026028": {
            "id": "ENSMUSG00000026028",
            "type": "Gene",
            "symbol": "Trak2",
            "source": [
                "ENSMUSG00000026398",
                "ENSMUSG00000001419",
                "ENSMUSG00000051469",
                "ENSMUSG00000029249",
                "ENSMUSG00000001280",
                "ENSMUSG00000019768",
                "ENSMUSG00000005583",
                "ENSMUSG00000022383",
                "ENSMUSG00000075595",
                "ENSMUSG00000008575",
                "ENSMUSG00000015846",
                "ENSMUSG00000000567"
            ],
            "target": [],
            "expression_sum": {
                "class1": 2031.0,
                "class2": 971.0
            },
            "uniprot_ids": [
                "Q6P9N8",
                "G3UZH5",
                "G3UYW9"
            ]
        },
        "ENSMUSG00000026031": {
            "id": "ENSMUSG00000026031",
            "type": "Gene",
            "symbol": "Cflar",
            "source": [
                "ENSMUSG00000042557",
                "ENSMUSG00000022346",
                "ENSMUSG00000000567"
            ],
            "target": [],
            "expression_sum": {
                "class1": 1714.0,
                "class2": 668.0
            },
            "uniprot_ids": [
                "O35732",
                "A0A2C9F2D9"
            ]
        },
        "ENSMUSG00000026082": {
            "id": "ENSMUSG00000026082",
            "type": "Gene",
            "symbol": "Rev1",
            "source": [
                "ENSMUSG00000015839",
                "ENSMUSG00000044167",
                "ENSMUSG00000005583",
                "ENSMUSG00000017146",
                "ENSMUSG00000024826",
                "ENSMUSG00000000567"
            ],
            "target": [],
            "expression_sum": {
                "class1": 1219.0,
                "class2": 100.0
            },
            "uniprot_ids": [
                "Q920Q2",
                "A0A0A6YX32",
                "A0A0A6YX77",
                "A0A0A6YWQ5",
                "A0A0A6YW35",
                "A0A0A6YWI0"
            ]
        },
        "ENSMUSG00000026150": {
            "id": "ENSMUSG00000026150",
            "type": "Gene",
            "symbol": "Mff",
            "source": [
                "ENSMUSG00000029135",
                "ENSMUSG00000001419",
                "ENSMUSG00000027201",
                "ENSMUSG00000025612",
                "ENSMUSG00000023927",
                "ENSMUSG00000015579",
                "ENSMUSG00000030557",
                "ENSMUSG00000027490",
                "ENSMUSG00000001280",
                "ENSMUSG00000019768",
                "ENSMUSG00000022641",
                "ENSMUSG00000031627",
                "ENSMUSG00000053110",
                "ENSMUSG00000035158",
                "ENSMUSG00000000567"
            ],
            "target": [],
            "expression_sum": {
                "class1": 2527.0,
                "class2": 1078.0
            },
            "uniprot_ids": [
                "Q6PCP5",
                "F7B1A4",
                "F7CTF8",
                "F6VAL0",
                "F7BC60",
                "E0CYB9",
                "E0CYI3"
            ]
        },
        "ENSMUSG00000026202": {
            "id": "ENSMUSG00000026202",
            "type": "Gene",
            "symbol": "Tuba4a",
            "source": [
                "ENSMUSG00000029135",
                "ENSMUSG00000005566",
                "ENSMUSG00000015605",
                "ENSMUSG00000008575",
                "ENSMUSG00000021775",
                "ENSMUSG00000000567",
                "ENSMUSG00000015579",
                "ENSMUSG00000019768",
                "ENSMUSG00000028565",
                "ENSMUSG00000037868",
                "ENSMUSG00000031885",
                "ENSMUSG00000031715",
                "ENSMUSG00000030557",
                "ENSMUSG00000032238",
                "ENSMUSG00000053477",
                "ENSMUSG00000058756",
                "ENSMUSG00000014592",
                "ENSMUSG00000026628",
                "ENSMUSG00000078671",
                "ENSMUSG00000056501",
                "ENSMUSG00000029249",
                "ENSMUSG00000001280"
            ],
            "target": [],
            "expression_sum": {
                "class1": 314.0,
                "class2": 1551.0
            },
            "uniprot_ids": [
                "P68368",
                "A0A087WSL5",
                "A0A087WQS4",
                "A0A0A0MQA5",
                "A0A087WS35",
                "A0A087WSB0",
                "A0A087WRB4"
            ]
        },
        "ENSMUSG00000026219": {
            "id": "ENSMUSG00000026219",
            "type": "Gene",
            "symbol": "Trip12",
            "source": [
                "ENSMUSG00000024238",
                "ENSMUSG00000019817",
                "ENSMUSG00000010175",
                "ENSMUSG00000015579",
                "ENSMUSG00000053477",
                "ENSMUSG00000005583",
                "ENSMUSG00000030067",
                "ENSMUSG00000076431",
                "ENSMUSG00000022314",
                "ENSMUSG00000071637",
                "ENSMUSG00000000567"
            ],
            "target": [],
            "expression_sum": {
                "class1": 2975.0,
                "class2": 823.0
            },
            "uniprot_ids": [
                "G5E870",
                "Q3TP48",
                "A0A087WRV6",
                "A0A087WRK9",
                "A0A087WP92",
                "A0A087WQ02",
                "A0A087WSG4",
                "A0A087WNZ7",
                "A0A087WS65",
                "A0A087WS33",
                "A0A0B4J1N9"
            ]
        },
        "ENSMUSG00000026404": {
            "id": "ENSMUSG00000026404",
            "type": "Gene",
            "symbol": "Ddx59",
            "source": [
                "ENSMUSG00000026398",
                "ENSMUSG00000001419",
                "ENSMUSG00000024515",
                "ENSMUSG00000059552",
                "ENSMUSG00000028565",
                "ENSMUSG00000040732",
                "ENSMUSG00000000567"
            ],
            "target": [],
            "expression_sum": {
                "class1": 322.0,
                "class2": 45.0
            },
            "uniprot_ids": [
                "Q9DBN9"
            ]
        },
        "ENSMUSG00000026566": {
            "id": "ENSMUSG00000026566",
            "type": "Gene",
            "symbol": "Mpzl1",
            "source": [
                "ENSMUSG00000016477",
                "ENSMUSG00000018604",
                "ENSMUSG00000049577",
                "ENSMUSG00000076431",
                "ENSMUSG00000015846",
                "ENSMUSG00000000567"
            ],
            "target": [],
            "expression_sum": {
                "class1": 1426.0,
                "class2": 279.0
            },
            "uniprot_ids": [
                "Q3TEW6",
                "A0A0A6YWA2",
                "A0A0A6YXL8",
                "A0A0A6YXX8",
                "A0A0A6YX14",
                "A0A0A6YW22",
                "A0A0A6YXW1",
                "A0A0R4J0V2"
            ]
        },
        "ENSMUSG00000026585": {
            "id": "ENSMUSG00000026585",
            "type": "Gene",
            "symbol": "Kifap3",
            "source": [
                "ENSMUSG00000056749",
                "ENSMUSG00000032228",
                "ENSMUSG00000005583",
                "ENSMUSG00000015605",
                "ENSMUSG00000014592",
                "ENSMUSG00000000567"
            ],
            "target": [],
            "expression_sum": {
                "class1": 2309.0,
                "class2": 323.0
            },
            "uniprot_ids": [
                "P70188"
            ]
        },
        "ENSMUSG00000026634": {
            "id": "ENSMUSG00000026634",
            "type": "Gene",
            "symbol": "Angel2",
            "source": [
                "ENSMUSG00000016477",
                "ENSMUSG00000015839",
                "ENSMUSG00000044167",
                "ENSMUSG00000024974",
                "ENSMUSG00000031885",
                "ENSMUSG00000000567"
            ],
            "target": [],
            "expression_sum": {
                "class1": 1742.0,
                "class2": 404.0
            },
            "uniprot_ids": [
                "Q8K1C0",
                "F6SBQ6",
                "A0A0A6YY82"
            ]
        },
        "ENSMUSG00000026657": {
            "id": "ENSMUSG00000026657",
            "type": "Gene",
            "symbol": "Frmd4a",
            "source": [
                "ENSMUSG00000021250",
                "ENSMUSG00000005583",
                "ENSMUSG00000052684",
                "ENSMUSG00000024974",
                "ENSMUSG00000039191",
                "ENSMUSG00000000567"
            ],
            "target": [],
            "expression_sum": {
                "class1": 2729.0,
                "class2": 276.0
            },
            "uniprot_ids": [
                "Q8BIE6",
                "Q3TB04",
                "H3BJN6",
                "H3BL79",
                "H3BIZ7",
                "H3BKK8",
                "H3BK97",
                "A0A5F8MPJ8",
                "F7CDA2"
            ]
        },
        "ENSMUSG00000026687": {
            "id": "ENSMUSG00000026687",
            "type": "Gene",
            "symbol": "Aldh9a1",
            "source": [
                "ENSMUSG00000025612",
                "ENSMUSG00000015839",
                "ENSMUSG00000030256",
                "ENSMUSG00000051469",
                "ENSMUSG00000001280",
                "ENSMUSG00000021877",
                "ENSMUSG00000031885",
                "ENSMUSG00000053110",
                "ENSMUSG00000000567"
            ],
            "target": [],
            "expression_sum": {
                "class1": 795.0,
                "class2": 211.0
            },
            "uniprot_ids": [
                "Q3U367"
            ]
        },
        "ENSMUSG00000026748": {
            "id": "ENSMUSG00000026748",
            "type": "Gene",
            "symbol": "Plxdc2",
            "source": [
                "ENSMUSG00000024238",
                "ENSMUSG00000006215",
                "ENSMUSG00000000567"
            ],
            "target": [],
            "expression_sum": {
                "class1": 1057.0,
                "class2": 235.0
            },
            "uniprot_ids": [
                "Q9DC11",
                "B1AY85",
                "B1AY86"
            ]
        },
        "ENSMUSG00000026749": {
            "id": "ENSMUSG00000026749",
            "type": "Gene",
            "symbol": "Nek6",
            "source": [
                "ENSMUSG00000022346",
                "ENSMUSG00000002983",
                "ENSMUSG00000000567"
            ],
            "target": [],
            "expression_sum": {
                "class1": 903.0,
                "class2": 70.0
            },
            "uniprot_ids": [
                "Q9ES70",
                "A2ATP0",
                "A2ATN9",
                "A2ATP1"
            ]
        },
        "ENSMUSG00000026754": {
            "id": "ENSMUSG00000026754",
            "type": "Gene",
            "symbol": "Golga1",
            "source": [
                "ENSMUSG00000029135",
                "ENSMUSG00000025612",
                "ENSMUSG00000015839",
                "ENSMUSG00000058239",
                "ENSMUSG00000015579",
                "ENSMUSG00000030557",
                "ENSMUSG00000019768",
                "ENSMUSG00000031885",
                "ENSMUSG00000053110",
                "ENSMUSG00000031715",
                "ENSMUSG00000000567"
            ],
            "target": [],
            "expression_sum": {
                "class1": 1179.0,
                "class2": 211.0
            },
            "uniprot_ids": [
                "Q9CW79",
                "A0A0N4SVQ8",
                "A0A0N4SVD1",
                "A0A0N4SVV1"
            ]
        },
        "ENSMUSG00000026761": {
            "id": "ENSMUSG00000026761",
            "type": "Gene",
            "symbol": "Orc4",
            "source": [
                "ENSMUSG00000000567",
                "ENSMUSG00000020167",
                "ENSMUSG00000005583",
                "ENSMUSG00000042557",
                "ENSMUSG00000022895",
                "ENSMUSG00000036036"
            ],
            "target": [],
            "expression_sum": {
                "class1": 942.0,
                "class2": 403.0
            },
            "uniprot_ids": [
                "O88708",
                "Q9CZF7",
                "A2AI40",
                "A2AI43",
                "A2AI42"
            ]
        },
        "ENSMUSG00000026775": {
            "id": "ENSMUSG00000026775",
            "type": "Gene",
            "symbol": "Yme1l1",
            "source": [
                "ENSMUSG00000029135",
                "ENSMUSG00000057469",
                "ENSMUSG00000001280",
                "ENSMUSG00000019768",
                "ENSMUSG00000058756",
                "ENSMUSG00000008575",
                "ENSMUSG00000014592",
                "ENSMUSG00000000567"
            ],
            "target": [],
            "expression_sum": {
                "class1": 2936.0,
                "class2": 692.0
            },
            "uniprot_ids": [
                "O88967"
            ]
        },
        "ENSMUSG00000026791": {
            "id": "ENSMUSG00000026791",
            "type": "Gene",
            "symbol": "Slc2a8",
            "source": [
                "ENSMUSG00000032897",
                "ENSMUSG00000018899",
                "ENSMUSG00000000567"
            ],
            "target": [],
            "expression_sum": {
                "class1": 488.0,
                "class2": 59.0
            },
            "uniprot_ids": [
                "Q9JIF3",
                "Q2TK27",
                "Q2TK26",
                "A0A0A6YXF7",
                "A0A0A6YX81"
            ]
        },
        "ENSMUSG00000026843": {
            "id": "ENSMUSG00000026843",
            "type": "Gene",
            "symbol": "Fubp3",
            "source": [
                "ENSMUSG00000005583",
                "ENSMUSG00000000567"
            ],
            "target": [],
            "expression_sum": {
                "class1": 600.0,
                "class2": 157.0
            },
            "uniprot_ids": [
                "Q3TIX6",
                "F7AM43",
                "A0A0A6YX76",
                "A2AJ72",
                "A0A0A6YW42",
                "A0A0A6YVV5",
                "A0A0A6YY39",
                "F6W2V3",
                "A2AJ71"
            ]
        },
        "ENSMUSG00000026864": {
            "id": "ENSMUSG00000026864",
            "type": "Gene",
            "symbol": "Hspa5",
            "source": [
                "ENSMUSG00000029135",
                "ENSMUSG00000020248",
                "ENSMUSG00000046351",
                "ENSMUSG00000031627",
                "ENSMUSG00000015605",
                "ENSMUSG00000008575",
                "ENSMUSG00000035158",
                "ENSMUSG00000000567",
                "ENSMUSG00000015579",
                "ENSMUSG00000051469",
                "ENSMUSG00000019768",
                "ENSMUSG00000024140",
                "ENSMUSG00000039191",
                "ENSMUSG00000035632",
                "ENSMUSG00000031715",
                "ENSMUSG00000015839",
                "ENSMUSG00000030557",
                "ENSMUSG00000058756",
                "ENSMUSG00000014592",
                "ENSMUSG00000001419",
                "ENSMUSG00000001280"
            ],
            "target": [],
            "expression_sum": {
                "class1": 45612.0,
                "class2": 8593.0
            },
            "uniprot_ids": [
                "P20029",
                "A0A0A6YXF5"
            ]
        },
        "ENSMUSG00000026879": {
            "id": "ENSMUSG00000026879",
            "type": "Gene",
            "symbol": "Gsn",
            "source": [
                "ENSMUSG00000029135",
                "ENSMUSG00000021250",
                "ENSMUSG00000030256",
                "ENSMUSG00000005583",
                "ENSMUSG00000005698",
                "ENSMUSG00000002265",
                "ENSMUSG00000027104",
                "ENSMUSG00000053110",
                "ENSMUSG00000000567"
            ],
            "target": [],
            "expression_sum": {
                "class1": 6550.0,
                "class2": 5136.0
            },
            "uniprot_ids": [
                "P13020",
                "A6PWS5",
                "A0A0J9YUQ8",
                "A0A0J9YUJ8"
            ]
        },
        "ENSMUSG00000026932": {
            "id": "ENSMUSG00000026932",
            "type": "Gene",
            "symbol": "Nacc2",
            "source": [
                "ENSMUSG00000000567"
            ],
            "target": [],
            "expression_sum": {
                "class1": 162.0,
                "class2": 159.0
            },
            "uniprot_ids": [
                "Q9DCM7",
                "A2ARL3"
            ]
        },
        "ENSMUSG00000027030": {
            "id": "ENSMUSG00000027030",
            "type": "Gene",
            "symbol": "Stk39",
            "source": [
                "ENSMUSG00000015839",
                "ENSMUSG00000029249",
                "ENSMUSG00000001280",
                "ENSMUSG00000019768",
                "ENSMUSG00000031885",
                "ENSMUSG00000008575",
                "ENSMUSG00000057098",
                "ENSMUSG00000000567"
            ],
            "target": [],
            "expression_sum": {
                "class1": 1196.0,
                "class2": 689.0
            },
            "uniprot_ids": [
                "Q9Z1W9"
            ]
        },
        "ENSMUSG00000027067": {
            "id": "ENSMUSG00000027067",
            "type": "Gene",
            "symbol": "Ssrp1",
            "source": [
                "ENSMUSG00000018604",
                "ENSMUSG00000016087",
                "ENSMUSG00000026398",
                "ENSMUSG00000000567"
            ],
            "target": [],
            "expression_sum": {
                "class1": 3112.0,
                "class2": 346.0
            },
            "uniprot_ids": [
                "Q08943",
                "A2AW05"
            ]
        },
        "ENSMUSG00000027167": {
            "id": "ENSMUSG00000027167",
            "type": "Gene",
            "symbol": "Elp4",
            "source": [
                "ENSMUSG00000071076",
                "ENSMUSG00000015839",
                "ENSMUSG00000024974",
                "ENSMUSG00000038648",
                "ENSMUSG00000005583",
                "ENSMUSG00000022053",
                "ENSMUSG00000027104",
                "ENSMUSG00000000567"
            ],
            "target": [],
            "expression_sum": {
                "class1": 474.0,
                "class2": 105.0
            },
            "uniprot_ids": [
                "Q9ER73",
                "A2A414"
            ]
        },
        "ENSMUSG00000027239": {
            "id": "ENSMUSG00000027239",
            "type": "Gene",
            "symbol": "Mdk",
            "source": [
                "ENSMUSG00000024238",
                "ENSMUSG00000032228",
                "ENSMUSG00000027109",
                "ENSMUSG00000005698",
                "ENSMUSG00000031885",
                "ENSMUSG00000031311",
                "ENSMUSG00000000567"
            ],
            "target": [],
            "expression_sum": {
                "class1": 6358.0,
                "class2": 63.0
            },
            "uniprot_ids": [
                "P12025"
            ]
        },
        "ENSMUSG00000027411": {
            "id": "ENSMUSG00000027411",
            "type": "Gene",
            "symbol": "Vps16",
            "source": [
                "ENSMUSG00000026890",
                "ENSMUSG00000000567"
            ],
            "target": [],
            "expression_sum": {
                "class1": 492.0,
                "class2": 76.0
            },
            "uniprot_ids": [
                "G3X8X7",
                "A2BI90"
            ]
        },
        "ENSMUSG00000027620": {
            "id": "ENSMUSG00000027620",
            "type": "Gene",
            "symbol": "Rbm39",
            "source": [
                "ENSMUSG00000024238",
                "ENSMUSG00000029135",
                "ENSMUSG00000078671",
                "ENSMUSG00000001280",
                "ENSMUSG00000019768",
                "ENSMUSG00000038692",
                "ENSMUSG00000000567"
            ],
            "target": [],
            "expression_sum": {
                "class1": 21930.0,
                "class2": 4459.0
            },
            "uniprot_ids": [
                "Q8VH51",
                "D6RCZ8",
                "B7ZD63",
                "B7ZD64",
                "E9Q8F0",
                "E9Q999",
                "D6RJ85",
                "E9PZF9",
                "B7ZD61",
                "F7AA45"
            ]
        },
        "ENSMUSG00000027797": {
            "id": "ENSMUSG00000027797",
            "type": "Gene",
            "symbol": "Dclk1",
            "source": [
                "ENSMUSG00000019564",
                "ENSMUSG00000044167",
                "ENSMUSG00000022346",
                "ENSMUSG00000000567"
            ],
            "target": [],
            "expression_sum": {
                "class1": 1865.0,
                "class2": 54.0
            },
            "uniprot_ids": [
                "Q9JLM8",
                "Q8CCN4",
                "Q80VB6",
                "A8IP62",
                "Q1EDH1",
                "A0A0G2JE54",
                "A0A0G2JGR9",
                "A0A0G2JGH1",
                "H7BX36",
                "A0A0G2JEN4",
                "A0A0G2JGQ6"
            ]
        },
        "ENSMUSG00000027884": {
            "id": "ENSMUSG00000027884",
            "type": "Gene",
            "symbol": "Clcc1",
            "source": [
                "ENSMUSG00000005583",
                "ENSMUSG00000005698",
                "ENSMUSG00000000567"
            ],
            "target": [],
            "expression_sum": {
                "class1": 1338.0,
                "class2": 171.0
            },
            "uniprot_ids": [
                "Q99LI2",
                "Z4YMG1",
                "A2AEM2"
            ]
        },
        "ENSMUSG00000028034": {
            "id": "ENSMUSG00000028034",
            "type": "Gene",
            "symbol": "Fubp1",
            "source": [
                "ENSMUSG00000024974",
                "ENSMUSG00000038692",
                "ENSMUSG00000049577",
                "ENSMUSG00000000567"
            ],
            "target": [],
            "expression_sum": {
                "class1": 3917.0,
                "class2": 793.0
            },
            "uniprot_ids": [
                "Q91WJ8",
                "Q3TUE1",
                "Q3UUU2",
                "A0A0G2JFK2",
                "A0A0G2JF21",
                "A0A0G2JG00",
                "A0A0G2JGV9",
                "A0A0G2JFY5",
                "A0A0G2JGW9",
                "A0A0G2JE55",
                "A0A0G2JG96",
                "A0A0G2JGT1"
            ]
        },
        "ENSMUSG00000028048": {
            "id": "ENSMUSG00000028048",
            "type": "Gene",
            "symbol": "Gba",
            "source": [
                "ENSMUSG00000031311",
                "ENSMUSG00000000567"
            ],
            "target": [],
            "expression_sum": {
                "class1": 488.0,
                "class2": 65.0
            },
            "uniprot_ids": [
                "P17439",
                "A0A0G2JDK2"
            ]
        },
        "ENSMUSG00000028069": {
            "id": "ENSMUSG00000028069",
            "type": "Gene",
            "symbol": "Gpatch4",
            "source": [
                "ENSMUSG00000044167",
                "ENSMUSG00000015839",
                "ENSMUSG00000000567"
            ],
            "target": [],
            "expression_sum": {
                "class1": 867.0,
                "class2": 205.0
            },
            "uniprot_ids": [
                "Q3TFK5",
                "A0A0A6YWB3"
            ]
        },
        "ENSMUSG00000028333": {
            "id": "ENSMUSG00000028333",
            "type": "Gene",
            "symbol": "Anp32b",
            "source": [
                "ENSMUSG00000001280",
                "ENSMUSG00000018604",
                "ENSMUSG00000019768",
                "ENSMUSG00000000567"
            ],
            "target": [],
            "expression_sum": {
                "class1": 7078.0,
                "class2": 970.0
            },
            "uniprot_ids": [
                "Q9EST5"
            ]
        },
        "ENSMUSG00000028403": {
            "id": "ENSMUSG00000028403",
            "type": "Gene",
            "symbol": "Zdhhc21",
            "source": [
                "ENSMUSG00000005566",
                "ENSMUSG00000058440",
                "ENSMUSG00000006215",
                "ENSMUSG00000000567"
            ],
            "target": [],
            "expression_sum": {
                "class1": 939.0,
                "class2": 169.0
            },
            "uniprot_ids": [
                "Q9D270",
                "Q5SRN8",
                "G3UXC1",
                "Q5SRP0"
            ]
        },
        "ENSMUSG00000028466": {
            "id": "ENSMUSG00000028466",
            "type": "TF",
            "symbol": "Creb3",
            "source": [
                "ENSMUSG00000024238",
                "ENSMUSG00000071637",
                "ENSMUSG00000022346",
                "ENSMUSG00000052534",
                "ENSMUSG00000005583",
                "ENSMUSG00000031627",
                "ENSMUSG00000024974",
                "ENSMUSG00000022314",
                "ENSMUSG00000022895",
                "ENSMUSG00000000567"
            ],
            "target": [
                "ENSMUSG00000030246",
                "ENSMUSG00000099250",
                "ENSMUSG00000004951",
                "ENSMUSG00000006024",
                "ENSMUSG00000015932",
                "ENSMUSG00000026435",
                "ENSMUSG00000034109",
                "ENSMUSG00000085812",
                "ENSMUSG00000034868",
                "ENSMUSG00000017999",
                "ENSMUSG00000001127",
                "ENSMUSG00000081434"
            ],
            "expression_sum": {
                "class1": 1713.0,
                "class2": 302.0
            },
            "uniprot_ids": [
                "Q3UNH6",
                "E9PW77",
                "E0CYW8"
            ]
        },
        "ENSMUSG00000028552": {
            "id": "ENSMUSG00000028552",
            "type": "Gene",
            "symbol": "Eps15",
            "source": [
                "ENSMUSG00000024238",
                "ENSMUSG00000052534",
                "ENSMUSG00000005583",
                "ENSMUSG00000015605",
                "ENSMUSG00000000567"
            ],
            "target": [],
            "expression_sum": {
                "class1": 948.0,
                "class2": 401.0
            },
            "uniprot_ids": [
                "P42567",
                "Q5JC28",
                "H3BK65",
                "H3BJB8",
                "F6W2Q5",
                "H3BLE4",
                "A0A0R4J0A0"
            ]
        },
        "ENSMUSG00000028869": {
            "id": "ENSMUSG00000028869",
            "type": "Gene",
            "symbol": "Gnl2",
            "source": [
                "ENSMUSG00000022150",
                "ENSMUSG00000015839",
                "ENSMUSG00000006215",
                "ENSMUSG00000051817",
                "ENSMUSG00000056501",
                "ENSMUSG00000018899",
                "ENSMUSG00000044167",
                "ENSMUSG00000037235",
                "ENSMUSG00000027104",
                "ENSMUSG00000000567"
            ],
            "target": [],
            "expression_sum": {
                "class1": 1586.0,
                "class2": 374.0
            },
            "uniprot_ids": [
                "Q99LH1",
                "B1ASC2"
            ]
        },
        "ENSMUSG00000028901": {
            "id": "ENSMUSG00000028901",
            "type": "TF",
            "symbol": "Gmeb1",
            "source": [
                "ENSMUSG00000052837",
                "ENSMUSG00000059552",
                "ENSMUSG00000024985",
                "ENSMUSG00000000567"
            ],
            "target": [
                "ENSMUSG00000068324",
                "ENSMUSG00000000058",
                "ENSMUSG00000001440"
            ],
            "expression_sum": {
                "class1": 545.0,
                "class2": 77.0
            },
            "uniprot_ids": [
                "Q9JL60"
            ]
        },
        "ENSMUSG00000028936": {
            "id": "ENSMUSG00000028936",
            "type": "Gene",
            "symbol": "Rpl22",
            "source": [
                "ENSMUSG00000079509",
                "ENSMUSG00000024238",
                "ENSMUSG00000003032",
                "ENSMUSG00000055116",
                "ENSMUSG00000001280",
                "ENSMUSG00000040732",
                "ENSMUSG00000005698",
                "ENSMUSG00000030067",
                "ENSMUSG00000031627",
                "ENSMUSG00000005893",
                "ENSMUSG00000027104",
                "ENSMUSG00000003154",
                "ENSMUSG00000024985",
                "ENSMUSG00000000567"
            ],
            "target": [],
            "expression_sum": {
                "class1": 4172.0,
                "class2": 1373.0
            },
            "uniprot_ids": [
                "P67984"
            ]
        },
        "ENSMUSG00000029190": {
            "id": "ENSMUSG00000029190",
            "type": "Gene",
            "symbol": "D5Ertd579e",
            "source": [
                "ENSMUSG00000029135",
                "ENSMUSG00000059552",
                "ENSMUSG00000010175",
                "ENSMUSG00000078671",
                "ENSMUSG00000015605",
                "ENSMUSG00000026610",
                "ENSMUSG00000056501",
                "ENSMUSG00000029249",
                "ENSMUSG00000001280",
                "ENSMUSG00000005566",
                "ENSMUSG00000019768",
                "ENSMUSG00000005698",
                "ENSMUSG00000022383",
                "ENSMUSG00000032238",
                "ENSMUSG00000021318",
                "ENSMUSG00000008575",
                "ENSMUSG00000000567"
            ],
            "target": [],
            "expression_sum": {
                "class1": 951.0,
                "class2": 569.0
            },
            "uniprot_ids": [
                "Q80U59",
                "E9QPJ5",
                "F7D143"
            ]
        },
        "ENSMUSG00000029433": {
            "id": "ENSMUSG00000029433",
            "type": "Gene",
            "symbol": "Diablo",
            "source": [
                "ENSMUSG00000022383",
                "ENSMUSG00000031885",
                "ENSMUSG00000008575",
                "ENSMUSG00000057098",
                "ENSMUSG00000000567"
            ],
            "target": [],
            "expression_sum": {
                "class1": 981.0,
                "class2": 625.0
            },
            "uniprot_ids": [
                "Q9JIQ3",
                "Q8R1D8",
                "D3Z3K5",
                "A0A0G2JGS0",
                "D3Z2Q3"
            ]
        },
        "ENSMUSG00000029536": {
            "id": "ENSMUSG00000029536",
            "type": "Gene",
            "symbol": "Gatc",
            "source": [
                "ENSMUSG00000044167",
                "ENSMUSG00000039191",
                "ENSMUSG00000000567"
            ],
            "target": [],
            "expression_sum": {
                "class1": 864.0,
                "class2": 134.0
            },
            "uniprot_ids": [
                "Q8CBY0"
            ]
        },
        "ENSMUSG00000029659": {
            "id": "ENSMUSG00000029659",
            "type": "Gene",
            "symbol": "Medag",
            "source": [
                "ENSMUSG00000057098",
                "ENSMUSG00000000567"
            ],
            "target": [],
            "expression_sum": {
                "class1": 1276.0,
                "class2": 224.0
            },
            "uniprot_ids": [
                "Q14BA6"
            ]
        },
        "ENSMUSG00000029804": {
            "id": "ENSMUSG00000029804",
            "type": "Gene",
            "symbol": "Herc3",
            "source": [
                "ENSMUSG00000005583",
                "ENSMUSG00000005698",
                "ENSMUSG00000022811",
                "ENSMUSG00000000567"
            ],
            "target": [],
            "expression_sum": {
                "class1": 203.0,
                "class2": 269.0
            },
            "uniprot_ids": [
                "A6H6S0",
                "G5E876",
                "D3Z2S3",
                "D3Z3U0",
                "A0A0N4SVW7",
                "A0A0N4SW02",
                "D3Z6D4"
            ]
        },
        "ENSMUSG00000029838": {
            "id": "ENSMUSG00000029838",
            "type": "Gene",
            "symbol": "Ptn",
            "source": [
                "ENSMUSG00000052684",
                "ENSMUSG00000000567"
            ],
            "target": [],
            "expression_sum": {
                "class1": 7167.0,
                "class2": 15.0
            },
            "uniprot_ids": [
                "P63089",
                "Q3UZI5"
            ]
        },
        "ENSMUSG00000029924": {
            "id": "ENSMUSG00000029924",
            "type": "Gene",
            "symbol": "Slc37a3",
            "source": [
                "ENSMUSG00000056501",
                "ENSMUSG00000034041",
                "ENSMUSG00000025958",
                "ENSMUSG00000015605",
                "ENSMUSG00000000567"
            ],
            "target": [],
            "expression_sum": {
                "class1": 533.0,
                "class2": 97.0
            },
            "uniprot_ids": [
                "A0A0J9YV99",
                "A0A0J9YUP6",
                "A0A0J9YUL8",
                "A0A0J9YV97",
                "A0A0J9YVF5",
                "A0A0R4J133",
                "A0A0J9YUS3",
                "A0A0J9YU14"
            ]
        },
        "ENSMUSG00000029994": {
            "id": "ENSMUSG00000029994",
            "type": "Gene",
            "symbol": "Anxa4",
            "source": [
                "ENSMUSG00000032238",
                "ENSMUSG00000036461",
                "ENSMUSG00000042557",
                "ENSMUSG00000000567"
            ],
            "target": [],
            "expression_sum": {
                "class1": 707.0,
                "class2": 152.0
            },
            "uniprot_ids": [
                "P97429",
                "A0A0N4SV57",
                "S4R1F2",
                "A0A0N4SW89",
                "F7ANV6",
                "D3Z0S1"
            ]
        },
        "ENSMUSG00000029998": {
            "id": "ENSMUSG00000029998",
            "type": "Gene",
            "symbol": "Pcyox1",
            "source": [
                "ENSMUSG00000019789",
                "ENSMUSG00000021250",
                "ENSMUSG00000018604",
                "ENSMUSG00000015579",
                "ENSMUSG00000032228",
                "ENSMUSG00000005698",
                "ENSMUSG00000024974",
                "ENSMUSG00000076431",
                "ENSMUSG00000039191",
                "ENSMUSG00000000567"
            ],
            "target": [],
            "expression_sum": {
                "class1": 778.0,
                "class2": 316.0
            },
            "uniprot_ids": [
                "Q9CQF9",
                "F7CIP8",
                "D3Z275",
                "A0A0N4SWD3"
            ]
        },
        "ENSMUSG00000030057": {
            "id": "ENSMUSG00000030057",
            "type": "Gene",
            "symbol": "Cnbp",
            "source": [
                "ENSMUSG00000001419",
                "ENSMUSG00000057469",
                "ENSMUSG00000022346",
                "ENSMUSG00000019564",
                "ENSMUSG00000056501",
                "ENSMUSG00000001280",
                "ENSMUSG00000019768",
                "ENSMUSG00000000567"
            ],
            "target": [],
            "expression_sum": {
                "class1": 7257.0,
                "class2": 2840.0
            },
            "uniprot_ids": [
                "P53996",
                "A0A0N4SVS6"
            ]
        },
        "ENSMUSG00000030082": {
            "id": "ENSMUSG00000030082",
            "type": "Gene",
            "symbol": "Sec61a1",
            "source": [
                "ENSMUSG00000021250",
                "ENSMUSG00000000567"
            ],
            "target": [],
            "expression_sum": {
                "class1": 3170.0,
                "class2": 411.0
            },
            "uniprot_ids": [
                "P61620"
            ]
        },
        "ENSMUSG00000030352": {
            "id": "ENSMUSG00000030352",
            "type": "Gene",
            "symbol": "Tspan9",
            "source": [
                "ENSMUSG00000028565",
                "ENSMUSG00000029178",
                "ENSMUSG00000032035",
                "ENSMUSG00000042557",
                "ENSMUSG00000000567"
            ],
            "target": [],
            "expression_sum": {
                "class1": 1205.0,
                "class2": 249.0
            },
            "uniprot_ids": [
                "Q8BJU2",
                "A0A0J9YTS4",
                "A0A0J9YVG5",
                "D3YU98",
                "D3YV18",
                "D3YXN7",
                "D3Z6M6"
            ]
        },
        "ENSMUSG00000030393": {
            "id": "ENSMUSG00000030393",
            "type": "TF",
            "symbol": "Zik1",
            "source": [
                "ENSMUSG00000015605",
                "ENSMUSG00000000567"
            ],
            "target": [
                "ENSMUSG00000002981",
                "ENSMUSG00000000934"
            ],
            "expression_sum": {
                "class1": 429.0,
                "class2": 5.0
            },
            "uniprot_ids": [
                "Q80YP6"
            ]
        },
        "ENSMUSG00000030522": {
            "id": "ENSMUSG00000030522",
            "type": "Gene",
            "symbol": "Mtmr10",
            "source": [
                "ENSMUSG00000018604",
                "ENSMUSG00000024955",
                "ENSMUSG00000032228",
                "ENSMUSG00000037235",
                "ENSMUSG00000016087",
                "ENSMUSG00000076431",
                "ENSMUSG00000000567"
            ],
            "target": [],
            "expression_sum": {
                "class1": 324.0,
                "class2": 94.0
            },
            "uniprot_ids": [
                "Q7TPM9",
                "A0A0U1RNK5",
                "A0A0U1RQ60"
            ]
        },
        "ENSMUSG00000030528": {
            "id": "ENSMUSG00000030528",
            "type": "Gene",
            "symbol": "Blm",
            "source": [
                "ENSMUSG00000006215",
                "ENSMUSG00000018604",
                "ENSMUSG00000044167",
                "ENSMUSG00000025958",
                "ENSMUSG00000038692",
                "ENSMUSG00000024974",
                "ENSMUSG00000016087",
                "ENSMUSG00000031885",
                "ENSMUSG00000000567"
            ],
            "target": [],
            "expression_sum": {
                "class1": 567.0,
                "class2": 58.0
            },
            "uniprot_ids": [
                "O88700",
                "E9PZ97",
                "A0A0U1RPS3",
                "A0A0U1RNI0",
                "A0A0U1RPP0"
            ]
        },
        "ENSMUSG00000030782": {
            "id": "ENSMUSG00000030782",
            "type": "Gene",
            "symbol": "Tgfb1i1",
            "source": [
                "ENSMUSG00000040732",
                "ENSMUSG00000038648",
                "ENSMUSG00000000567"
            ],
            "target": [],
            "expression_sum": {
                "class1": 2453.0,
                "class2": 360.0
            },
            "uniprot_ids": [
                "Q62219",
                "E9PYQ1",
                "E9Q1V4",
                "E9Q1D5"
            ]
        },
        "ENSMUSG00000031217": {
            "id": "ENSMUSG00000031217",
            "type": "Gene",
            "symbol": "Efnb1",
            "source": [
                "ENSMUSG00000025958",
                "ENSMUSG00000003032",
                "ENSMUSG00000000567"
            ],
            "target": [],
            "expression_sum": {
                "class1": 960.0,
                "class2": 206.0
            },
            "uniprot_ids": [
                "P52795"
            ]
        },
        "ENSMUSG00000031253": {
            "id": "ENSMUSG00000031253",
            "type": "Gene",
            "symbol": "Srpx2",
            "source": [
                "ENSMUSG00000000567"
            ],
            "target": [],
            "expression_sum": {
                "class1": 2121.0,
                "class2": 116.0
            },
            "uniprot_ids": [
                "Q8R054"
            ]
        },
        "ENSMUSG00000031400": {
            "id": "ENSMUSG00000031400",
            "type": "Gene",
            "symbol": "G6pdx",
            "source": [
                "ENSMUSG00000055116",
                "ENSMUSG00000041515",
                "ENSMUSG00000005583",
                "ENSMUSG00000042557",
                "ENSMUSG00000000567"
            ],
            "target": [],
            "expression_sum": {
                "class1": 281.0,
                "class2": 41.0
            },
            "uniprot_ids": [
                "Q00612",
                "A3KG36"
            ]
        },
        "ENSMUSG00000031422": {
            "id": "ENSMUSG00000031422",
            "type": "Gene",
            "symbol": "Morf4l2",
            "source": [
                "ENSMUSG00000074622",
                "ENSMUSG00000002265",
                "ENSMUSG00000000567"
            ],
            "target": [],
            "expression_sum": {
                "class1": 2219.0,
                "class2": 271.0
            },
            "uniprot_ids": [
                "Q9R0Q4",
                "A2AEB6",
                "A2AEB7",
                "A2AEB5",
                "A2AEB8",
                "A2AEB9",
                "A2AEB3",
                "A2AEB4"
            ]
        },
        "ENSMUSG00000031438": {
            "id": "ENSMUSG00000031438",
            "type": "Gene",
            "symbol": "Rnf128",
            "source": [
                "ENSMUSG00000031681",
                "ENSMUSG00000025369",
                "ENSMUSG00000015839",
                "ENSMUSG00000058239",
                "ENSMUSG00000001280",
                "ENSMUSG00000019768",
                "ENSMUSG00000025958",
                "ENSMUSG00000053110",
                "ENSMUSG00000075595",
                "ENSMUSG00000008575",
                "ENSMUSG00000035158",
                "ENSMUSG00000000567"
            ],
            "target": [],
            "expression_sum": {
                "class1": 246.0,
                "class2": 662.0
            },
            "uniprot_ids": [
                "Q9D304",
                "A2AGL8"
            ]
        },
        "ENSMUSG00000031543": {
            "id": "ENSMUSG00000031543",
            "type": "Gene",
            "symbol": "Ank1",
            "source": [
                "ENSMUSG00000024515",
                "ENSMUSG00000021944",
                "ENSMUSG00000057469",
                "ENSMUSG00000022346",
                "ENSMUSG00000056501",
                "ENSMUSG00000029249",
                "ENSMUSG00000001280",
                "ENSMUSG00000005566",
                "ENSMUSG00000019768",
                "ENSMUSG00000058756",
                "ENSMUSG00000015605",
                "ENSMUSG00000039191",
                "ENSMUSG00000029913",
                "ENSMUSG00000024985",
                "ENSMUSG00000000567"
            ],
            "target": [],
            "expression_sum": {
                "class1": 733.0,
                "class2": 2034.0
            },
            "uniprot_ids": [
                "Q02357",
                "B7ZW98",
                "Q0VGY9",
                "D6RJ51",
                "G8JL84",
                "G5E8J2",
                "A0A0R4J1N7",
                "F7D1P5",
                "D3YTV8",
                "G3UXF4",
                "E9QNT8",
                "G3UY11",
                "D3Z5M4"
            ]
        },
        "ENSMUSG00000031730": {
            "id": "ENSMUSG00000031730",
            "type": "Gene",
            "symbol": "Dhodh",
            "source": [
                "ENSMUSG00000037868",
                "ENSMUSG00000000567"
            ],
            "target": [],
            "expression_sum": {
                "class1": 304.0,
                "class2": 54.0
            },
            "uniprot_ids": [
                "O35435",
                "F6ZQJ5",
                "D3YZY5",
                "E9Q1V3",
                "F6TCT7"
            ]
        },
        "ENSMUSG00000031740": {
            "id": "ENSMUSG00000031740",
            "type": "Gene",
            "symbol": "Mmp2",
            "source": [
                "ENSMUSG00000005698",
                "ENSMUSG00000053110",
                "ENSMUSG00000000567"
            ],
            "target": [],
            "expression_sum": {
                "class1": 6310.0,
                "class2": 786.0
            },
            "uniprot_ids": [
                "P33434",
                "A0A1B0GS88"
            ]
        },
        "ENSMUSG00000031799": {
            "id": "ENSMUSG00000031799",
            "type": "Gene",
            "symbol": "Tpm4",
            "source": [
                "ENSMUSG00000018604",
                "ENSMUSG00000016087",
                "ENSMUSG00000020919",
                "ENSMUSG00000000567"
            ],
            "target": [],
            "expression_sum": {
                "class1": 16707.0,
                "class2": 2318.0
            },
            "uniprot_ids": [
                "Q6IRU2",
                "A0A571BEU1"
            ]
        },
        "ENSMUSG00000031812": {
            "id": "ENSMUSG00000031812",
            "type": "Gene",
            "symbol": "Map1lc3b",
            "source": [
                "ENSMUSG00000029135",
                "ENSMUSG00000025612",
                "ENSMUSG00000015839",
                "ENSMUSG00000052534",
                "ENSMUSG00000030256",
                "ENSMUSG00000058239",
                "ENSMUSG00000030557",
                "ENSMUSG00000001280",
                "ENSMUSG00000019768",
                "ENSMUSG00000033943",
                "ENSMUSG00000031885",
                "ENSMUSG00000031715",
                "ENSMUSG00000000567"
            ],
            "target": [],
            "expression_sum": {
                "class1": 7773.0,
                "class2": 2253.0
            },
            "uniprot_ids": [
                "Q9CQV6",
                "M0QWC2",
                "A0A1D5RLG5",
                "M0QWT8"
            ]
        },
        "ENSMUSG00000031821": {
            "id": "ENSMUSG00000031821",
            "type": "Gene",
            "symbol": "Gins2",
            "source": [
                "ENSMUSG00000038692",
                "ENSMUSG00000028639",
                "ENSMUSG00000024985",
                "ENSMUSG00000000567"
            ],
            "target": [],
            "expression_sum": {
                "class1": 207.0,
                "class2": 16.0
            },
            "uniprot_ids": [
                "Q9D600"
            ]
        },
        "ENSMUSG00000031979": {
            "id": "ENSMUSG00000031979",
            "type": "Gene",
            "symbol": "Cog2",
            "source": [
                "ENSMUSG00000059552",
                "ENSMUSG00000014592",
                "ENSMUSG00000000567"
            ],
            "target": [],
            "expression_sum": {
                "class1": 720.0,
                "class2": 77.0
            },
            "uniprot_ids": [
                "Q921L5",
                "H3BL04",
                "H3BJK5"
            ]
        },
        "ENSMUSG00000032012": {
            "id": "ENSMUSG00000032012",
            "type": "Gene",
            "symbol": "Nectin1",
            "source": [
                "ENSMUSG00000057098",
                "ENSMUSG00000000567"
            ],
            "target": [],
            "expression_sum": {
                "class1": 263.0,
                "class2": 42.0
            },
            "uniprot_ids": [
                "Q9JKF6",
                "A0A1L1SQP1",
                "A0A668KLC0"
            ]
        },
        "ENSMUSG00000032212": {
            "id": "ENSMUSG00000032212",
            "type": "Gene",
            "symbol": "Sltm",
            "source": [
                "ENSMUSG00000040270",
                "ENSMUSG00000024515",
                "ENSMUSG00000000567"
            ],
            "target": [],
            "expression_sum": {
                "class1": 4309.0,
                "class2": 769.0
            },
            "uniprot_ids": [
                "Q8CH25",
                "A0A1L1STF9",
                "A0A1L1SRL0"
            ]
        },
        "ENSMUSG00000032334": {
            "id": "ENSMUSG00000032334",
            "type": "Gene",
            "symbol": "Loxl1",
            "source": [
                "ENSMUSG00000053110",
                "ENSMUSG00000002265",
                "ENSMUSG00000000567"
            ],
            "target": [],
            "expression_sum": {
                "class1": 2952.0,
                "class2": 450.0
            },
            "uniprot_ids": [
                "A0A0R4J0Q4"
            ]
        },
        "ENSMUSG00000032388": {
            "id": "ENSMUSG00000032388",
            "type": "Gene",
            "symbol": "Spg21",
            "source": [
                "ENSMUSG00000022676",
                "ENSMUSG00000029249",
                "ENSMUSG00000006215",
                "ENSMUSG00000000567"
            ],
            "target": [],
            "expression_sum": {
                "class1": 1127.0,
                "class2": 75.0
            },
            "uniprot_ids": [
                "Q9CQC8",
                "A0A1L1SST5",
                "A0A1L1SSV6"
            ]
        },
        "ENSMUSG00000032440": {
            "id": "ENSMUSG00000032440",
            "type": "Gene",
            "symbol": "Tgfbr2",
            "source": [
                "ENSMUSG00000016477",
                "ENSMUSG00000026628",
                "ENSMUSG00000021250",
                "ENSMUSG00000027109",
                "ENSMUSG00000030067",
                "ENSMUSG00000027104",
                "ENSMUSG00000000567"
            ],
            "target": [],
            "expression_sum": {
                "class1": 1364.0,
                "class2": 465.0
            },
            "uniprot_ids": [
                "Q62312"
            ]
        },
        "ENSMUSG00000032491": {
            "id": "ENSMUSG00000032491",
            "type": "Gene",
            "symbol": "Nradd",
            "source": [
                "ENSMUSG00000010175",
                "ENSMUSG00000031627",
                "ENSMUSG00000027104",
                "ENSMUSG00000024985",
                "ENSMUSG00000000567"
            ],
            "target": [],
            "expression_sum": {
                "class1": 127.0,
                "class2": 6.0
            },
            "uniprot_ids": [
                "Q8CJ26",
                "A0A0G2JH13",
                "A0A0G2JEW0"
            ]
        },
        "ENSMUSG00000032997": {
            "id": "ENSMUSG00000032997",
            "type": "Gene",
            "symbol": "Chpf",
            "source": [
                "ENSMUSG00000006705",
                "ENSMUSG00000015846",
                "ENSMUSG00000000567"
            ],
            "target": [],
            "expression_sum": {
                "class1": 897.0,
                "class2": 124.0
            },
            "uniprot_ids": [
                "Q6IQX7",
                "D6RFA4",
                "D6RGS3"
            ]
        },
        "ENSMUSG00000033055": {
            "id": "ENSMUSG00000033055",
            "type": "Gene",
            "symbol": "Ankrd54",
            "source": [
                "ENSMUSG00000036461",
                "ENSMUSG00000000567"
            ],
            "target": [],
            "expression_sum": {
                "class1": 503.0,
                "class2": 75.0
            },
            "uniprot_ids": [
                "Q91WK7",
                "B7ZBY0",
                "F6QTQ3",
                "F6UYM2",
                "D6REF6"
            ]
        },
        "ENSMUSG00000033544": {
            "id": "ENSMUSG00000033544",
            "type": "Gene",
            "symbol": "Angptl1",
            "source": [
                "ENSMUSG00000000567"
            ],
            "target": [],
            "expression_sum": {
                "class1": 359.0,
                "class2": 18.0
            },
            "uniprot_ids": [
                "Q640P2"
            ]
        },
        "ENSMUSG00000033565": {
            "id": "ENSMUSG00000033565",
            "type": "Gene",
            "symbol": "Rbfox2",
            "source": [
                "ENSMUSG00000027490",
                "ENSMUSG00000019768",
                "ENSMUSG00000022641",
                "ENSMUSG00000038692",
                "ENSMUSG00000000567"
            ],
            "target": [],
            "expression_sum": {
                "class1": 1660.0,
                "class2": 521.0
            },
            "uniprot_ids": [
                "Q8BP71",
                "A0A2I3BRA9",
                "A0A2I3BRU6",
                "A0A2I3BQD1",
                "A0A2I3BR20",
                "A0A2K6EDK7"
            ]
        },
        "ENSMUSG00000033684": {
            "id": "ENSMUSG00000033684",
            "type": "Gene",
            "symbol": "Qsox1",
            "source": [
                "ENSMUSG00000029135",
                "ENSMUSG00000018604",
                "ENSMUSG00000056749",
                "ENSMUSG00000032228",
                "ENSMUSG00000016087",
                "ENSMUSG00000015605",
                "ENSMUSG00000000567"
            ],
            "target": [],
            "expression_sum": {
                "class1": 958.0,
                "class2": 126.0
            },
            "uniprot_ids": [
                "Q8BND5"
            ]
        },
        "ENSMUSG00000033781": {
            "id": "ENSMUSG00000033781",
            "type": "Gene",
            "symbol": "Asb13",
            "source": [
                "ENSMUSG00000029135",
                "ENSMUSG00000006215",
                "ENSMUSG00000029249",
                "ENSMUSG00000001280",
                "ENSMUSG00000005566",
                "ENSMUSG00000019768",
                "ENSMUSG00000005698",
                "ENSMUSG00000015605",
                "ENSMUSG00000031715",
                "ENSMUSG00000024985",
                "ENSMUSG00000000567"
            ],
            "target": [],
            "expression_sum": {
                "class1": 176.0,
                "class2": 209.0
            },
            "uniprot_ids": [
                "Q8VBX0"
            ]
        },
        "ENSMUSG00000034485": {
            "id": "ENSMUSG00000034485",
            "type": "Gene",
            "symbol": "Uaca",
            "source": [
                "ENSMUSG00000044167",
                "ENSMUSG00000000567"
            ],
            "target": [],
            "expression_sum": {
                "class1": 4221.0,
                "class2": 607.0
            },
            "uniprot_ids": [
                "A0A1L1SVG0",
                "A0A1L1STY4",
                "A0A0R4J0S7"
            ]
        },
        "ENSMUSG00000034566": {
            "id": "ENSMUSG00000034566",
            "type": "Gene",
            "symbol": "Atp5h",
            "source": [
                "ENSMUSG00000006215",
                "ENSMUSG00000015839",
                "ENSMUSG00000058239",
                "ENSMUSG00000056501",
                "ENSMUSG00000015579",
                "ENSMUSG00000001280",
                "ENSMUSG00000019768",
                "ENSMUSG00000053110",
                "ENSMUSG00000031885",
                "ENSMUSG00000008575",
                "ENSMUSG00000021775",
                "ENSMUSG00000000567"
            ],
            "target": [],
            "expression_sum": {
                "class1": 25943.0,
                "class2": 23000.0
            },
            "uniprot_ids": [
                "Q9DCX2",
                "B1ASE2"
            ]
        },
        "ENSMUSG00000034584": {
            "id": "ENSMUSG00000034584",
            "type": "Gene",
            "symbol": "Exph5",
            "source": [
                "ENSMUSG00000021250",
                "ENSMUSG00000056749",
                "ENSMUSG00000005583",
                "ENSMUSG00000005698",
                "ENSMUSG00000000567"
            ],
            "target": [],
            "expression_sum": {
                "class1": 142.0,
                "class2": 10.0
            },
            "uniprot_ids": [
                "Q0VAV2"
            ]
        },
        "ENSMUSG00000035376": {
            "id": "ENSMUSG00000035376",
            "type": "Gene",
            "symbol": "Hacd2",
            "source": [
                "ENSMUSG00000030256",
                "ENSMUSG00000037674",
                "ENSMUSG00000047407",
                "ENSMUSG00000027490",
                "ENSMUSG00000000567"
            ],
            "target": [],
            "expression_sum": {
                "class1": 179.0,
                "class2": 129.0
            },
            "uniprot_ids": [
                "Q9D3B1"
            ]
        },
        "ENSMUSG00000035478": {
            "id": "ENSMUSG00000035478",
            "type": "Gene",
            "symbol": "Mbd3",
            "source": [
                "ENSMUSG00000032897",
                "ENSMUSG00000071076",
                "ENSMUSG00000019564",
                "ENSMUSG00000030557",
                "ENSMUSG00000001280",
                "ENSMUSG00000019768",
                "ENSMUSG00000075595",
                "ENSMUSG00000000567"
            ],
            "target": [],
            "expression_sum": {
                "class1": 4206.0,
                "class2": 421.0
            },
            "uniprot_ids": [
                "Q9Z2D8",
                "F6WZH1",
                "D3YTR4",
                "D3YTR5"
            ]
        },
        "ENSMUSG00000036565": {
            "id": "ENSMUSG00000036565",
            "type": "Gene",
            "symbol": "Ttyh3",
            "source": [
                "ENSMUSG00000016477",
                "ENSMUSG00000003032",
                "ENSMUSG00000025612",
                "ENSMUSG00000026628",
                "ENSMUSG00000015579",
                "ENSMUSG00000027490",
                "ENSMUSG00000005583",
                "ENSMUSG00000030067",
                "ENSMUSG00000000567"
            ],
            "target": [],
            "expression_sum": {
                "class1": 240.0,
                "class2": 68.0
            },
            "uniprot_ids": [
                "Q6P5F7",
                "A0A0G2JGJ4"
            ]
        },
        "ENSMUSG00000036570": {
            "id": "ENSMUSG00000036570",
            "type": "Gene",
            "symbol": "Fxyd1",
            "source": [
                "ENSMUSG00000010175",
                "ENSMUSG00000056501",
                "ENSMUSG00000001280",
                "ENSMUSG00000019768",
                "ENSMUSG00000032238",
                "ENSMUSG00000005698",
                "ENSMUSG00000024695",
                "ENSMUSG00000058756",
                "ENSMUSG00000053110",
                "ENSMUSG00000021775",
                "ENSMUSG00000015846",
                "ENSMUSG00000000567"
            ],
            "target": [],
            "expression_sum": {
                "class1": 4627.0,
                "class2": 3459.0
            },
            "uniprot_ids": [
                "Q9Z239",
                "A0A0U1RPI1",
                "A0A0U1RPU6"
            ]
        },
        "ENSMUSG00000036862": {
            "id": "ENSMUSG00000036862",
            "type": "Gene",
            "symbol": "Dchs1",
            "source": [
                "ENSMUSG00000003032",
                "ENSMUSG00000037465",
                "ENSMUSG00000027109",
                "ENSMUSG00000040732",
                "ENSMUSG00000000567"
            ],
            "target": [],
            "expression_sum": {
                "class1": 637.0,
                "class2": 130.0
            },
            "uniprot_ids": [
                "E9PVD3",
                "A0A1B0GSA3"
            ]
        },
        "ENSMUSG00000037204": {
            "id": "ENSMUSG00000037204",
            "type": "Gene",
            "symbol": "Atg101",
            "source": [
                "ENSMUSG00000018899",
                "ENSMUSG00000005583",
                "ENSMUSG00000025958",
                "ENSMUSG00000005698",
                "ENSMUSG00000052684",
                "ENSMUSG00000031627",
                "ENSMUSG00000000567"
            ],
            "target": [],
            "expression_sum": {
                "class1": 3441.0,
                "class2": 218.0
            },
            "uniprot_ids": [
                "Q9D8Z6",
                "A0A2R8VH88",
                "A0A2R8VJN5"
            ]
        },
        "ENSMUSG00000037211": {
            "id": "ENSMUSG00000037211",
            "type": "Gene",
            "symbol": "Spry1",
            "source": [
                "ENSMUSG00000000567"
            ],
            "target": [],
            "expression_sum": {
                "class1": 904.0,
                "class2": 257.0
            },
            "uniprot_ids": [
                "Q9QXV9"
            ]
        },
        "ENSMUSG00000037674": {
            "id": "ENSMUSG00000037674",
            "type": "TF",
            "symbol": "Rfx7",
            "source": [
                "ENSMUSG00000044167",
                "ENSMUSG00000015846",
                "ENSMUSG00000000567"
            ],
            "target": [
                "ENSMUSG00000000957",
                "ENSMUSG00000035376",
                "ENSMUSG00000001506",
                "ENSMUSG00000001493",
                "ENSMUSG00000012519",
                "ENSMUSG00000000838"
            ],
            "expression_sum": {
                "class1": 518.0,
                "class2": 165.0
            },
            "uniprot_ids": [
                "Q3TDE2",
                "F8VPJ6"
            ]
        },
        "ENSMUSG00000037852": {
            "id": "ENSMUSG00000037852",
            "type": "Gene",
            "symbol": "Cpe",
            "source": [
                "ENSMUSG00000024238",
                "ENSMUSG00000052534",
                "ENSMUSG00000044167",
                "ENSMUSG00000005583",
                "ENSMUSG00000022314",
                "ENSMUSG00000000567"
            ],
            "target": [],
            "expression_sum": {
                "class1": 4035.0,
                "class2": 2248.0
            },
            "uniprot_ids": [
                "Q00493"
            ]
        },
        "ENSMUSG00000038244": {
            "id": "ENSMUSG00000038244",
            "type": "Gene",
            "symbol": "Mical2",
            "source": [
                "ENSMUSG00000005583",
                "ENSMUSG00000021775",
                "ENSMUSG00000015846",
                "ENSMUSG00000000567"
            ],
            "target": [],
            "expression_sum": {
                "class1": 1128.0,
                "class2": 188.0
            },
            "uniprot_ids": [
                "Q8BML1",
                "Q9D5U9",
                "A0A1L1SVE5",
                "F6WY65",
                "A0A1L1ST75",
                "A0A1L1SQF0"
            ]
        },
        "ENSMUSG00000038374": {
            "id": "ENSMUSG00000038374",
            "type": "Gene",
            "symbol": "Rbm8a",
            "source": [
                "ENSMUSG00000018604",
                "ENSMUSG00000024560",
                "ENSMUSG00000018899",
                "ENSMUSG00000001280",
                "ENSMUSG00000000567"
            ],
            "target": [],
            "expression_sum": {
                "class1": 4285.0,
                "class2": 1098.0
            },
            "uniprot_ids": [
                "Q9CWZ3",
                "A0A0G2JFX7",
                "A0A0G2JEA9"
            ]
        },
        "ENSMUSG00000038970": {
            "id": "ENSMUSG00000038970",
            "type": "Gene",
            "symbol": "Lmtk2",
            "source": [
                "ENSMUSG00000029135",
                "ENSMUSG00000059552",
                "ENSMUSG00000025612",
                "ENSMUSG00000015839",
                "ENSMUSG00000015579",
                "ENSMUSG00000001280",
                "ENSMUSG00000019768",
                "ENSMUSG00000028565",
                "ENSMUSG00000052684",
                "ENSMUSG00000053110",
                "ENSMUSG00000031885",
                "ENSMUSG00000008575",
                "ENSMUSG00000035158",
                "ENSMUSG00000031715",
                "ENSMUSG00000000567"
            ],
            "target": [],
            "expression_sum": {
                "class1": 100.0,
                "class2": 158.0
            },
            "uniprot_ids": [
                "Q3TYD6"
            ]
        },
        "ENSMUSG00000039231": {
            "id": "ENSMUSG00000039231",
            "type": "Gene",
            "symbol": "Suv39h1",
            "source": [
                "ENSMUSG00000024515",
                "ENSMUSG00000026628",
                "ENSMUSG00000074622",
                "ENSMUSG00000005583",
                "ENSMUSG00000038692",
                "ENSMUSG00000031885",
                "ENSMUSG00000000567"
            ],
            "target": [],
            "expression_sum": {
                "class1": 635.0,
                "class2": 51.0
            },
            "uniprot_ids": [
                "O54864",
                "A2AC19"
            ]
        },
        "ENSMUSG00000039356": {
            "id": "ENSMUSG00000039356",
            "type": "Gene",
            "symbol": "Exosc2",
            "source": [
                "ENSMUSG00000000567"
            ],
            "target": [],
            "expression_sum": {
                "class1": 327.0,
                "class2": 119.0
            },
            "uniprot_ids": [
                "Q8VBV3",
                "A0A0A6YW57",
                "A2AV19"
            ]
        },
        "ENSMUSG00000039497": {
            "id": "ENSMUSG00000039497",
            "type": "Gene",
            "symbol": "Dse",
            "source": [
                "ENSMUSG00000000567"
            ],
            "target": [],
            "expression_sum": {
                "class1": 208.0,
                "class2": 94.0
            },
            "uniprot_ids": [
                "Q8BLI4",
                "A0A1L1SSJ9",
                "A0A1L1SUI1"
            ]
        },
        "ENSMUSG00000039976": {
            "id": "ENSMUSG00000039976",
            "type": "Gene",
            "symbol": "Tbc1d16",
            "source": [
                "ENSMUSG00000001280",
                "ENSMUSG00000019768",
                "ENSMUSG00000008575",
                "ENSMUSG00000015846",
                "ENSMUSG00000000567"
            ],
            "target": [],
            "expression_sum": {
                "class1": 509.0,
                "class2": 360.0
            },
            "uniprot_ids": [
                "Q6PFC0",
                "A2ABG4",
                "A2AFK9"
            ]
        },
        "ENSMUSG00000040112": {
            "id": "ENSMUSG00000040112",
            "type": "Gene",
            "symbol": "Mrps35",
            "source": [
                "ENSMUSG00000030557",
                "ENSMUSG00000001280",
                "ENSMUSG00000005698",
                "ENSMUSG00000040270",
                "ENSMUSG00000024974",
                "ENSMUSG00000015605",
                "ENSMUSG00000015846",
                "ENSMUSG00000000567"
            ],
            "target": [],
            "expression_sum": {
                "class1": 2425.0,
                "class2": 973.0
            },
            "uniprot_ids": [
                "A0A0R4J0L6",
                "A0A0N4SV23"
            ]
        },
        "ENSMUSG00000040151": {
            "id": "ENSMUSG00000040151",
            "type": "Gene",
            "symbol": "Hs2st1",
            "source": [
                "ENSMUSG00000044167",
                "ENSMUSG00000071076",
                "ENSMUSG00000053110",
                "ENSMUSG00000000567"
            ],
            "target": [],
            "expression_sum": {
                "class1": 961.0,
                "class2": 123.0
            },
            "uniprot_ids": [
                "Q8R3H7",
                "E0CYX6"
            ]
        },
        "ENSMUSG00000040274": {
            "id": "ENSMUSG00000040274",
            "type": "Gene",
            "symbol": "Cdk6",
            "source": [
                "ENSMUSG00000044167",
                "ENSMUSG00000022383",
                "ENSMUSG00000030796",
                "ENSMUSG00000000567"
            ],
            "target": [],
            "expression_sum": {
                "class1": 834.0,
                "class2": 70.0
            },
            "uniprot_ids": [
                "Q64261",
                "A0A0G2JGH2",
                "A0A0G2JEJ6",
                "A0A0G2JGA8"
            ]
        },
        "ENSMUSG00000040339": {
            "id": "ENSMUSG00000040339",
            "type": "Gene",
            "symbol": "Fam102b",
            "source": [
                "ENSMUSG00000052684",
                "ENSMUSG00000027490",
                "ENSMUSG00000000567"
            ],
            "target": [],
            "expression_sum": {
                "class1": 994.0,
                "class2": 268.0
            },
            "uniprot_ids": [
                "Q8BQS4",
                "E9Q4R1"
            ]
        },
        "ENSMUSG00000040351": {
            "id": "ENSMUSG00000040351",
            "type": "Gene",
            "symbol": "Ankib1",
            "source": [
                "ENSMUSG00000015839",
                "ENSMUSG00000078671",
                "ENSMUSG00000029249",
                "ENSMUSG00000001280",
                "ENSMUSG00000019768",
                "ENSMUSG00000005583",
                "ENSMUSG00000005698",
                "ENSMUSG00000022641",
                "ENSMUSG00000008575",
                "ENSMUSG00000000567"
            ],
            "target": [],
            "expression_sum": {
                "class1": 1059.0,
                "class2": 313.0
            },
            "uniprot_ids": [
                "Q6ZPS6",
                "A0A0G2JEA0"
            ]
        },
        "ENSMUSG00000040359": {
            "id": "ENSMUSG00000040359",
            "type": "Gene",
            "symbol": "Ufl1",
            "source": [
                "ENSMUSG00000025612",
                "ENSMUSG00000038648",
                "ENSMUSG00000025025",
                "ENSMUSG00000022053",
                "ENSMUSG00000027104",
                "ENSMUSG00000015846",
                "ENSMUSG00000000567"
            ],
            "target": [],
            "expression_sum": {
                "class1": 2204.0,
                "class2": 506.0
            },
            "uniprot_ids": [
                "Q8CCJ3"
            ]
        },
        "ENSMUSG00000040435": {
            "id": "ENSMUSG00000040435",
            "type": "Gene",
            "symbol": "Ppp1r15a",
            "source": [
                "ENSMUSG00000029135",
                "ENSMUSG00000024515",
                "ENSMUSG00000022346",
                "ENSMUSG00000026628",
                "ENSMUSG00000019564",
                "ENSMUSG00000023034",
                "ENSMUSG00000000567"
            ],
            "target": [],
            "expression_sum": {
                "class1": 6786.0,
                "class2": 615.0
            },
            "uniprot_ids": [
                "P17564",
                "A0A1B0GRX4"
            ]
        },
        "ENSMUSG00000040658": {
            "id": "ENSMUSG00000040658",
            "type": "Gene",
            "symbol": "Dnph1",
            "source": [
                "ENSMUSG00000039191",
                "ENSMUSG00000000567"
            ],
            "target": [],
            "expression_sum": {
                "class1": 171.0,
                "class2": 19.0
            },
            "uniprot_ids": [
                "Q80VJ3"
            ]
        },
        "ENSMUSG00000040928": {
            "id": "ENSMUSG00000040928",
            "type": "Gene",
            "symbol": "S100pbp",
            "source": [
                "ENSMUSG00000059552",
                "ENSMUSG00000055116",
                "ENSMUSG00000019768",
                "ENSMUSG00000005583",
                "ENSMUSG00000022641",
                "ENSMUSG00000022053",
                "ENSMUSG00000000567"
            ],
            "target": [],
            "expression_sum": {
                "class1": 471.0,
                "class2": 164.0
            },
            "uniprot_ids": [
                "Q9D5K4",
                "E9Q5M3"
            ]
        },
        "ENSMUSG00000040952": {
            "id": "ENSMUSG00000040952",
            "type": "Gene",
            "symbol": "Rps19",
            "source": [
                "ENSMUSG00000029135",
                "ENSMUSG00000021250",
                "ENSMUSG00000018604",
                "ENSMUSG00000022346",
                "ENSMUSG00000036461",
                "ENSMUSG00000001280",
                "ENSMUSG00000016087",
                "ENSMUSG00000032035",
                "ENSMUSG00000024985",
                "ENSMUSG00000000567"
            ],
            "target": [],
            "expression_sum": {
                "class1": 24279.0,
                "class2": 2718.0
            },
            "uniprot_ids": [
                "Q9CZX8",
                "D3YUG3",
                "D3YUT3",
                "S4R223",
                "D3Z722",
                "D3Z5R8"
            ]
        },
        "ENSMUSG00000041075": {
            "id": "ENSMUSG00000041075",
            "type": "Gene",
            "symbol": "Fzd7",
            "source": [
                "ENSMUSG00000057469",
                "ENSMUSG00000026565",
                "ENSMUSG00000023994",
                "ENSMUSG00000044167",
                "ENSMUSG00000036721",
                "ENSMUSG00000000567"
            ],
            "target": [],
            "expression_sum": {
                "class1": 115.0,
                "class2": 78.0
            },
            "uniprot_ids": [
                "Q61090"
            ]
        },
        "ENSMUSG00000041236": {
            "id": "ENSMUSG00000041236",
            "type": "Gene",
            "symbol": "Vps41",
            "source": [
                "ENSMUSG00000019817",
                "ENSMUSG00000018604",
                "ENSMUSG00000032228",
                "ENSMUSG00000005583",
                "ENSMUSG00000008575",
                "ENSMUSG00000022314",
                "ENSMUSG00000000567"
            ],
            "target": [],
            "expression_sum": {
                "class1": 1481.0,
                "class2": 253.0
            },
            "uniprot_ids": [
                "Q5KU39"
            ]
        },
        "ENSMUSG00000041375": {
            "id": "ENSMUSG00000041375",
            "type": "Gene",
            "symbol": "Ccdc9",
            "source": [
                "ENSMUSG00000052837",
                "ENSMUSG00000031099",
                "ENSMUSG00000021318",
                "ENSMUSG00000000567"
            ],
            "target": [],
            "expression_sum": {
                "class1": 648.0,
                "class2": 91.0
            },
            "uniprot_ids": [
                "G3UYE3",
                "D6RFU6",
                "G3X927",
                "A0A0G2JDC5",
                "G3UXR7",
                "G3UXX6",
                "D3Z1Q6",
                "D3YW42"
            ]
        },
        "ENSMUSG00000041773": {
            "id": "ENSMUSG00000041773",
            "type": "Gene",
            "symbol": "Enc1",
            "source": [
                "ENSMUSG00000003032",
                "ENSMUSG00000021250",
                "ENSMUSG00000026628",
                "ENSMUSG00000027109",
                "ENSMUSG00000029249",
                "ENSMUSG00000005566",
                "ENSMUSG00000032035",
                "ENSMUSG00000020644",
                "ENSMUSG00000000567"
            ],
            "target": [],
            "expression_sum": {
                "class1": 380.0,
                "class2": 112.0
            },
            "uniprot_ids": [
                "O35709"
            ]
        },
        "ENSMUSG00000041801": {
            "id": "ENSMUSG00000041801",
            "type": "Gene",
            "symbol": "Phlda3",
            "source": [
                "ENSMUSG00000032228",
                "ENSMUSG00000005583",
                "ENSMUSG00000058756",
                "ENSMUSG00000053110",
                "ENSMUSG00000000567"
            ],
            "target": [],
            "expression_sum": {
                "class1": 794.0,
                "class2": 38.0
            },
            "uniprot_ids": [
                "Q9WV95"
            ]
        },
        "ENSMUSG00000042208": {
            "id": "ENSMUSG00000042208",
            "type": "Gene",
            "symbol": "Sanbr",
            "source": [
                "ENSMUSG00000000567"
            ],
            "target": [],
            "expression_sum": {
                "class1": 1429.0,
                "class2": 167.0
            },
            "uniprot_ids": [
                "Q68FF0",
                "M0QWF3",
                "Z4YM60",
                "Z4YN77",
                "Z4YN22"
            ]
        },
        "ENSMUSG00000042462": {
            "id": "ENSMUSG00000042462",
            "type": "Gene",
            "symbol": "Dctpp1",
            "source": [
                "ENSMUSG00000029135",
                "ENSMUSG00000049577",
                "ENSMUSG00000015579",
                "ENSMUSG00000027109",
                "ENSMUSG00000001280",
                "ENSMUSG00000019768",
                "ENSMUSG00000053110",
                "ENSMUSG00000000567"
            ],
            "target": [],
            "expression_sum": {
                "class1": 2995.0,
                "class2": 623.0
            },
            "uniprot_ids": [
                "Q9QY93"
            ]
        },
        "ENSMUSG00000042606": {
            "id": "ENSMUSG00000042606",
            "type": "Gene",
            "symbol": "Hirip3",
            "source": [
                "ENSMUSG00000031103",
                "ENSMUSG00000017146",
                "ENSMUSG00000021779",
                "ENSMUSG00000000567"
            ],
            "target": [],
            "expression_sum": {
                "class1": 1135.0,
                "class2": 73.0
            },
            "uniprot_ids": [
                "Q8BLH7",
                "D3YWU7"
            ]
        },
        "ENSMUSG00000042700": {
            "id": "ENSMUSG00000042700",
            "type": "Gene",
            "symbol": "Sipa1l1",
            "source": [
                "ENSMUSG00000000567",
                "ENSMUSG00000005566",
                "ENSMUSG00000005583",
                "ENSMUSG00000005698",
                "ENSMUSG00000022676",
                "ENSMUSG00000042557",
                "ENSMUSG00000022895",
                "ENSMUSG00000020167"
            ],
            "target": [],
            "expression_sum": {
                "class1": 708.0,
                "class2": 87.0
            },
            "uniprot_ids": [
                "Q8C0T5",
                "Q4VBF8",
                "A0A1Y7VJM1"
            ]
        },
        "ENSMUSG00000043323": {
            "id": "ENSMUSG00000043323",
            "type": "Gene",
            "symbol": "Fbrsl1",
            "source": [
                "ENSMUSG00000001280",
                "ENSMUSG00000005583",
                "ENSMUSG00000005698",
                "ENSMUSG00000025958",
                "ENSMUSG00000075595",
                "ENSMUSG00000015846",
                "ENSMUSG00000000567"
            ],
            "target": [],
            "expression_sum": {
                "class1": 247.0,
                "class2": 347.0
            },
            "uniprot_ids": [
                "E9PUB6",
                "A0A0G2JE44",
                "A0A0G2JG68",
                "A0A571BFB2",
                "A0A0G2JDI2",
                "A0A0G2JFF2",
                "E9Q9T0"
            ]
        },
        "ENSMUSG00000044167": {
            "id": "ENSMUSG00000044167",
            "type": "TF",
            "symbol": "Foxo1",
            "source": [
                "ENSMUSG00000024974",
                "ENSMUSG00000027109",
                "ENSMUSG00000022895",
                "ENSMUSG00000000567"
            ],
            "target": [
                "ENSMUSG00000002732",
                "ENSMUSG00000027642",
                "ENSMUSG00000051223",
                "ENSMUSG00000006010",
                "ENSMUSG00000051817",
                "ENSMUSG00000055799",
                "ENSMUSG00000026708",
                "ENSMUSG00000025579",
                "ENSMUSG00000025747",
                "ENSMUSG00000029447",
                "ENSMUSG00000047731",
                "ENSMUSG00000031787",
                "ENSMUSG00000041133",
                "ENSMUSG00000038459",
                "ENSMUSG00000038991",
                "ENSMUSG00000029536",
                "ENSMUSG00000037572",
                "ENSMUSG00000027342",
                "ENSMUSG00000023224",
                "ENSMUSG00000020777",
                "ENSMUSG00000028641",
                "ENSMUSG00000041891",
                "ENSMUSG00000053460",
                "ENSMUSG00000002661",
                "ENSMUSG00000024392",
                "ENSMUSG00000047003",
                "ENSMUSG00000026496",
                "ENSMUSG00000027109",
                "ENSMUSG00000035024",
                "ENSMUSG00000039239",
                "ENSMUSG00000020346",
                "ENSMUSG00000026134",
                "ENSMUSG00000024974",
                "ENSMUSG00000015112",
                "ENSMUSG00000022247",
                "ENSMUSG00000028869",
                "ENSMUSG00000047146",
                "ENSMUSG00000018433",
                "ENSMUSG00000036223",
                "ENSMUSG00000046201",
                "ENSMUSG00000005732",
                "ENSMUSG00000031441",
                "ENSMUSG00000052748",
                "ENSMUSG00000030254",
                "ENSMUSG00000020946",
                "ENSMUSG00000020459",
                "ENSMUSG00000026923",
                "ENSMUSG00000021400",
                "ENSMUSG00000074466",
                "ENSMUSG00000053877",
                "ENSMUSG00000039218",
                "ENSMUSG00000044037",
                "ENSMUSG00000022521",
                "ENSMUSG00000003402",
                "ENSMUSG00000019773",
                "ENSMUSG00000001436",
                "ENSMUSG00000015474",
                "ENSMUSG00000024640",
                "ENSMUSG00000041779",
                "ENSMUSG00000044847",
                "ENSMUSG00000047721",
                "ENSMUSG00000022217",
                "ENSMUSG00000024422",
                "ENSMUSG00000015749",
                "ENSMUSG00000027349",
                "ENSMUSG00000040274",
                "ENSMUSG00000040605",
                "ENSMUSG00000035152",
                "ENSMUSG00000041187",
                "ENSMUSG00000029910",
                "ENSMUSG00000031906",
                "ENSMUSG00000020267",
                "ENSMUSG00000030494",
                "ENSMUSG00000021209",
                "ENSMUSG00000009112",
                "ENSMUSG00000029267",
                "ENSMUSG00000065954",
                "ENSMUSG00000035158",
                "ENSMUSG00000092486",
                "ENSMUSG00000021112",
                "ENSMUSG00000027533",
                "ENSMUSG00000028266",
                "ENSMUSG00000031834",
                "ENSMUSG00000028212",
                "ENSMUSG00000035248",
                "ENSMUSG00000021245",
                "ENSMUSG00000015961",
                "ENSMUSG00000021377",
                "ENSMUSG00000051177",
                "ENSMUSG00000026739",
                "ENSMUSG00000020462",
                "ENSMUSG00000021713",
                "ENSMUSG00000026064",
                "ENSMUSG00000078773",
                "ENSMUSG00000026192",
                "ENSMUSG00000037852",
                "ENSMUSG00000037278",
                "ENSMUSG00000039128",
                "ENSMUSG00000034868",
                "ENSMUSG00000030095",
                "ENSMUSG00000029863",
                "ENSMUSG00000016510",
                "ENSMUSG00000044768",
                "ENSMUSG00000025499",
                "ENSMUSG00000054387",
                "ENSMUSG00000020024",
                "ENSMUSG00000027429",
                "ENSMUSG00000045078",
                "ENSMUSG00000041075",
                "ENSMUSG00000048120",
                "ENSMUSG00000028757",
                "ENSMUSG00000038116",
                "ENSMUSG00000029578",
                "ENSMUSG00000027255",
                "ENSMUSG00000058799",
                "ENSMUSG00000035199",
                "ENSMUSG00000001930",
                "ENSMUSG00000019907",
                "ENSMUSG00000027353",
                "ENSMUSG00000026365",
                "ENSMUSG00000047832",
                "ENSMUSG00000032527",
                "ENSMUSG00000040151",
                "ENSMUSG00000020807",
                "ENSMUSG00000036854",
                "ENSMUSG00000042712",
                "ENSMUSG00000024754",
                "ENSMUSG00000054263",
                "ENSMUSG00000042605",
                "ENSMUSG00000006021",
                "ENSMUSG00000026289",
                "ENSMUSG00000039842",
                "ENSMUSG00000035765",
                "ENSMUSG00000020522",
                "ENSMUSG00000057113",
                "ENSMUSG00000026558",
                "ENSMUSG00000054405",
                "ENSMUSG00000024182",
                "ENSMUSG00000028124",
                "ENSMUSG00000010392",
                "ENSMUSG00000024002",
                "ENSMUSG00000055917",
                "ENSMUSG00000031207",
                "ENSMUSG00000021408",
                "ENSMUSG00000035632",
                "ENSMUSG00000022895",
                "ENSMUSG00000075704",
                "ENSMUSG00000005846",
                "ENSMUSG00000026281",
                "ENSMUSG00000031904",
                "ENSMUSG00000034059",
                "ENSMUSG00000042670",
                "ENSMUSG00000023004",
                "ENSMUSG00000034485",
                "ENSMUSG00000036180",
                "ENSMUSG00000018761",
                "ENSMUSG00000020349",
                "ENSMUSG00000030663",
                "ENSMUSG00000037110",
                "ENSMUSG00000019578",
                "ENSMUSG00000039100",
                "ENSMUSG00000032221",
                "ENSMUSG00000039515",
                "ENSMUSG00000006050",
                "ENSMUSG00000024052",
                "ENSMUSG00000052146",
                "ENSMUSG00000034109",
                "ENSMUSG00000075254",
                "ENSMUSG00000022673",
                "ENSMUSG00000027551",
                "ENSMUSG00000040506",
                "ENSMUSG00000029505",
                "ENSMUSG00000043207",
                "ENSMUSG00000015247",
                "ENSMUSG00000015733",
                "ENSMUSG00000028069",
                "ENSMUSG00000029217",
                "ENSMUSG00000022325",
                "ENSMUSG00000034042",
                "ENSMUSG00000025580",
                "ENSMUSG00000032097",
                "ENSMUSG00000037674",
                "ENSMUSG00000046185",
                "ENSMUSG00000034311",
                "ENSMUSG00000053199",
                "ENSMUSG00000058709",
                "ENSMUSG00000044573",
                "ENSMUSG00000028673",
                "ENSMUSG00000038503",
                "ENSMUSG00000005687",
                "ENSMUSG00000038615",
                "ENSMUSG00000026634",
                "ENSMUSG00000003226",
                "ENSMUSG00000026563",
                "ENSMUSG00000036572",
                "ENSMUSG00000031729",
                "ENSMUSG00000027797",
                "ENSMUSG00000042104",
                "ENSMUSG00000069309",
                "ENSMUSG00000032042",
                "ENSMUSG00000022016",
                "ENSMUSG00000042207",
                "ENSMUSG00000010406",
                "ENSMUSG00000025104",
                "ENSMUSG00000030528",
                "ENSMUSG00000008200",
                "ENSMUSG00000034317",
                "ENSMUSG00000038793",
                "ENSMUSG00000022466",
                "ENSMUSG00000022724",
                "ENSMUSG00000056342",
                "ENSMUSG00000040209",
                "ENSMUSG00000037795",
                "ENSMUSG00000026082",
                "ENSMUSG00000029227",
                "ENSMUSG00000033152",
                "ENSMUSG00000034403",
                "ENSMUSG00000041241",
                "ENSMUSG00000029538",
                "ENSMUSG00000045237",
                "ENSMUSG00000022360",
                "ENSMUSG00000063904",
                "ENSMUSG00000028248",
                "ENSMUSG00000032376",
                "ENSMUSG00000026554",
                "ENSMUSG00000022322",
                "ENSMUSG00000028522",
                "ENSMUSG00000007739",
                "ENSMUSG00000062014",
                "ENSMUSG00000056919",
                "ENSMUSG00000022142"
            ],
            "expression_sum": {
                "class1": 1606.0,
                "class2": 381.0
            },
            "uniprot_ids": [
                "Q9R1E0"
            ]
        },
        "ENSMUSG00000044201": {
            "id": "ENSMUSG00000044201",
            "type": "Gene",
            "symbol": "Cdc25c",
            "source": [
                "ENSMUSG00000000567"
            ],
            "target": [],
            "expression_sum": {
                "class1": 263.0,
                "class2": 2.0
            },
            "uniprot_ids": [
                "P48967",
                "A0A494BAB9"
            ]
        },
        "ENSMUSG00000044442": {
            "id": "ENSMUSG00000044442",
            "type": "Gene",
            "symbol": "N6amt1",
            "source": [
                "ENSMUSG00000052684",
                "ENSMUSG00000005583",
                "ENSMUSG00000000567"
            ],
            "target": [],
            "expression_sum": {
                "class1": 2753.0,
                "class2": 278.0
            },
            "uniprot_ids": [
                "Q6SKR2",
                "A0A4W7N3B2"
            ]
        },
        "ENSMUSG00000044533": {
            "id": "ENSMUSG00000044533",
            "type": "Gene",
            "symbol": "Rps2",
            "source": [
                "ENSMUSG00000031681",
                "ENSMUSG00000025612",
                "ENSMUSG00000015839",
                "ENSMUSG00000056501",
                "ENSMUSG00000015579",
                "ENSMUSG00000030557",
                "ENSMUSG00000001280",
                "ENSMUSG00000019768",
                "ENSMUSG00000022641",
                "ENSMUSG00000053110",
                "ENSMUSG00000035158",
                "ENSMUSG00000021775",
                "ENSMUSG00000000567"
            ],
            "target": [],
            "expression_sum": {
                "class1": 1468.0,
                "class2": 886.0
            },
            "uniprot_ids": [
                "P25444",
                "D3YWJ3",
                "D3YVC1"
            ]
        },
        "ENSMUSG00000044629": {
            "id": "ENSMUSG00000044629",
            "type": "Gene",
            "symbol": "Cnrip1",
            "source": [
                "ENSMUSG00000019564",
                "ENSMUSG00000034041",
                "ENSMUSG00000052837",
                "ENSMUSG00000000567"
            ],
            "target": [],
            "expression_sum": {
                "class1": 1430.0,
                "class2": 84.0
            },
            "uniprot_ids": [
                "Q5M8N0",
                "F7C0H6"
            ]
        },
        "ENSMUSG00000045128": {
            "id": "ENSMUSG00000045128",
            "type": "Gene",
            "symbol": "Rpl18a",
            "source": [
                "ENSMUSG00000029135",
                "ENSMUSG00000021250",
                "ENSMUSG00000036461",
                "ENSMUSG00000023079",
                "ENSMUSG00000017146",
                "ENSMUSG00000025025",
                "ENSMUSG00000020919",
                "ENSMUSG00000031715",
                "ENSMUSG00000000567"
            ],
            "target": [],
            "expression_sum": {
                "class1": 1491.0,
                "class2": 262.0
            },
            "uniprot_ids": [
                "P62717",
                "A0A1D5RMC7",
                "A0A1D5RM79",
                "A0A1D5RLW5",
                "A0A1D5RM85",
                "A0A1D5RLA0",
                "A0A1D5RME4"
            ]
        },
        "ENSMUSG00000046402": {
            "id": "ENSMUSG00000046402",
            "type": "Gene",
            "symbol": "Rbp1",
            "source": [
                "ENSMUSG00000005698",
                "ENSMUSG00000022314",
                "ENSMUSG00000000567"
            ],
            "target": [],
            "expression_sum": {
                "class1": 3695.0,
                "class2": 369.0
            },
            "uniprot_ids": [
                "Q00915"
            ]
        },
        "ENSMUSG00000046572": {
            "id": "ENSMUSG00000046572",
            "type": "Gene",
            "symbol": "Zfp518b",
            "source": [
                "ENSMUSG00000024974",
                "ENSMUSG00000031627",
                "ENSMUSG00000022346",
                "ENSMUSG00000000567"
            ],
            "target": [],
            "expression_sum": {
                "class1": 304.0,
                "class2": 35.0
            },
            "uniprot_ids": [
                "J3QPN0",
                "J3QNM2",
                "J3QNT5"
            ]
        },
        "ENSMUSG00000046876": {
            "id": "ENSMUSG00000046876",
            "type": "Gene",
            "symbol": "Atxn1",
            "source": [
                "ENSMUSG00000024238",
                "ENSMUSG00000021250",
                "ENSMUSG00000018604",
                "ENSMUSG00000052837",
                "ENSMUSG00000005583",
                "ENSMUSG00000005698",
                "ENSMUSG00000053110",
                "ENSMUSG00000031885",
                "ENSMUSG00000016087",
                "ENSMUSG00000032035",
                "ENSMUSG00000071637",
                "ENSMUSG00000022528",
                "ENSMUSG00000000567"
            ],
            "target": [],
            "expression_sum": {
                "class1": 1522.0,
                "class2": 455.0
            },
            "uniprot_ids": [
                "P54254",
                "J3QPR1"
            ]
        },
        "ENSMUSG00000048388": {
            "id": "ENSMUSG00000048388",
            "type": "Gene",
            "symbol": "Fam171b",
            "source": [
                "ENSMUSG00000000567"
            ],
            "target": [],
            "expression_sum": {
                "class1": 315.0,
                "class2": 49.0
            },
            "uniprot_ids": [
                "Q14CH0"
            ]
        },
        "ENSMUSG00000048416": {
            "id": "ENSMUSG00000048416",
            "type": "Gene",
            "symbol": "Mlf1",
            "source": [
                "ENSMUSG00000058239",
                "ENSMUSG00000001280",
                "ENSMUSG00000019768",
                "ENSMUSG00000075595",
                "ENSMUSG00000031885",
                "ENSMUSG00000000567"
            ],
            "target": [],
            "expression_sum": {
                "class1": 5501.0,
                "class2": 4146.0
            },
            "uniprot_ids": [
                "Q9QWV4",
                "Q8JZS2",
                "A0A0A6YVP7"
            ]
        },
        "ENSMUSG00000049690": {
            "id": "ENSMUSG00000049690",
            "type": "Gene",
            "symbol": "Nckap5",
            "source": [
                "ENSMUSG00000003032",
                "ENSMUSG00000055116",
                "ENSMUSG00000099083",
                "ENSMUSG00000053110",
                "ENSMUSG00000000567"
            ],
            "target": [],
            "expression_sum": {
                "class1": 410.0,
                "class2": 67.0
            },
            "uniprot_ids": [
                "Q8C6S8",
                "Q6NZK2",
                "Q29SA9",
                "E0CYS4",
                "E2QRQ1",
                "E9QAE1",
                "E0CYQ5",
                "E9Q3G9"
            ]
        },
        "ENSMUSG00000049792": {
            "id": "ENSMUSG00000049792",
            "type": "Gene",
            "symbol": "Bag5",
            "source": [
                "ENSMUSG00000003032",
                "ENSMUSG00000058239",
                "ENSMUSG00000018604",
                "ENSMUSG00000024560",
                "ENSMUSG00000027109",
                "ENSMUSG00000031715",
                "ENSMUSG00000000567"
            ],
            "target": [],
            "expression_sum": {
                "class1": 2179.0,
                "class2": 391.0
            },
            "uniprot_ids": [
                "Q8CI32",
                "E0CYW3",
                "E0CX76"
            ]
        },
        "ENSMUSG00000050188": {
            "id": "ENSMUSG00000050188",
            "type": "Gene",
            "symbol": "Lsm10",
            "source": [
                "ENSMUSG00000052534",
                "ENSMUSG00000026628",
                "ENSMUSG00000000567"
            ],
            "target": [],
            "expression_sum": {
                "class1": 1147.0,
                "class2": 199.0
            },
            "uniprot_ids": [
                "Q8QZX5",
                "A8Y5G9"
            ]
        },
        "ENSMUSG00000050541": {
            "id": "ENSMUSG00000050541",
            "type": "Gene",
            "symbol": "Adra1b",
            "source": [
                "ENSMUSG00000029135",
                "ENSMUSG00000010175",
                "ENSMUSG00000001280",
                "ENSMUSG00000019768",
                "ENSMUSG00000032238",
                "ENSMUSG00000015605",
                "ENSMUSG00000000567"
            ],
            "target": [],
            "expression_sum": {
                "class1": 198.0,
                "class2": 293.0
            },
            "uniprot_ids": [
                "Q9DBL0",
                "B1AU41"
            ]
        },
        "ENSMUSG00000050627": {
            "id": "ENSMUSG00000050627",
            "type": "Gene",
            "symbol": "Gpd1l",
            "source": [
                "ENSMUSG00000001280",
                "ENSMUSG00000005583",
                "ENSMUSG00000005698",
                "ENSMUSG00000000567"
            ],
            "target": [],
            "expression_sum": {
                "class1": 234.0,
                "class2": 227.0
            },
            "uniprot_ids": [
                "Q3ULJ0",
                "D3Z0L6"
            ]
        },
        "ENSMUSG00000050705": {
            "id": "ENSMUSG00000050705",
            "type": "Gene",
            "symbol": "2310061I04Rik",
            "source": [
                "ENSMUSG00000032228",
                "ENSMUSG00000027490",
                "ENSMUSG00000005583",
                "ENSMUSG00000024140",
                "ENSMUSG00000016087",
                "ENSMUSG00000053110",
                "ENSMUSG00000000567"
            ],
            "target": [],
            "expression_sum": {
                "class1": 452.0,
                "class2": 809.0
            },
            "uniprot_ids": [
                "B8JJ66",
                "B8JJ68",
                "B8JJ67",
                "F6X1P1",
                "B8JJ69",
                "F6W3U3"
            ]
        },
        "ENSMUSG00000051159": {
            "id": "ENSMUSG00000051159",
            "type": "Gene",
            "symbol": "Cited1",
            "source": [
                "ENSMUSG00000000567"
            ],
            "target": [],
            "expression_sum": {
                "class1": 279.0,
                "class2": 27.0
            },
            "uniprot_ids": [
                "P97769",
                "F6ZC52",
                "A0A0A0MQI7",
                "F6YUL1"
            ]
        },
        "ENSMUSG00000051671": {
            "id": "ENSMUSG00000051671",
            "type": "Gene",
            "symbol": "Coa6",
            "source": [
                "ENSMUSG00000015839",
                "ENSMUSG00000001280",
                "ENSMUSG00000031885",
                "ENSMUSG00000029178",
                "ENSMUSG00000000567"
            ],
            "target": [],
            "expression_sum": {
                "class1": 813.0,
                "class2": 558.0
            },
            "uniprot_ids": [
                "Q8BGD8"
            ]
        },
        "ENSMUSG00000051817": {
            "id": "ENSMUSG00000051817",
            "type": "TF",
            "symbol": "Sox12",
            "source": [
                "ENSMUSG00000058440",
                "ENSMUSG00000059552",
                "ENSMUSG00000015839",
                "ENSMUSG00000056501",
                "ENSMUSG00000018899",
                "ENSMUSG00000044167",
                "ENSMUSG00000000567"
            ],
            "target": [
                "ENSMUSG00000028869",
                "ENSMUSG00000004035",
                "ENSMUSG00000029563",
                "ENSMUSG00000000743"
            ],
            "expression_sum": {
                "class1": 409.0,
                "class2": 159.0
            },
            "uniprot_ids": [
                "Q04890"
            ]
        },
        "ENSMUSG00000052676": {
            "id": "ENSMUSG00000052676",
            "type": "Gene",
            "symbol": "Zmat1",
            "source": [
                "ENSMUSG00000000567"
            ],
            "target": [],
            "expression_sum": {
                "class1": 588.0,
                "class2": 62.0
            },
            "uniprot_ids": [
                "Z4YL92",
                "G5E8E4"
            ]
        },
        "ENSMUSG00000052748": {
            "id": "ENSMUSG00000052748",
            "type": "Gene",
            "symbol": "Swt1",
            "source": [
                "ENSMUSG00000006215",
                "ENSMUSG00000056501",
                "ENSMUSG00000044167",
                "ENSMUSG00000031885",
                "ENSMUSG00000000567"
            ],
            "target": [],
            "expression_sum": {
                "class1": 765.0,
                "class2": 190.0
            },
            "uniprot_ids": [
                "Q9DBQ9",
                "F7ABE5",
                "F6V527"
            ]
        },
        "ENSMUSG00000052763": {
            "id": "ENSMUSG00000052763",
            "type": "TF",
            "symbol": "Zfp212",
            "source": [
                "ENSMUSG00000006215",
                "ENSMUSG00000000567"
            ],
            "target": [
                "ENSMUSG00000000957",
                "ENSMUSG00000047347",
                "ENSMUSG00000024158",
                "ENSMUSG00000008734",
                "ENSMUSG00000030309",
                "ENSMUSG00000001383"
            ],
            "expression_sum": {
                "class1": 224.0,
                "class2": 18.0
            },
            "uniprot_ids": [
                "G3X8R7"
            ]
        },
        "ENSMUSG00000053329": {
            "id": "ENSMUSG00000053329",
            "type": "Gene",
            "symbol": "Gatd3a",
            "source": [
                "ENSMUSG00000021250",
                "ENSMUSG00000052534",
                "ENSMUSG00000024985",
                "ENSMUSG00000001280",
                "ENSMUSG00000019768",
                "ENSMUSG00000008575",
                "ENSMUSG00000015846",
                "ENSMUSG00000000567"
            ],
            "target": [],
            "expression_sum": {
                "class1": 1568.0,
                "class2": 1607.0
            },
            "uniprot_ids": [
                "Q9D172",
                "A0A1W2P870",
                "A0A1W2P7B6"
            ]
        },
        "ENSMUSG00000053460": {
            "id": "ENSMUSG00000053460",
            "type": "Gene",
            "symbol": "Ggcx",
            "source": [
                "ENSMUSG00000058440",
                "ENSMUSG00000032228",
                "ENSMUSG00000018899",
                "ENSMUSG00000044167",
                "ENSMUSG00000049606",
                "ENSMUSG00000024985",
                "ENSMUSG00000000567"
            ],
            "target": [],
            "expression_sum": {
                "class1": 733.0,
                "class2": 155.0
            },
            "uniprot_ids": [
                "Q9QYC7",
                "A0A0U1RNG6",
                "A0A0U1RPT2",
                "A0A0U1RPG7",
                "A0A0U1RPU0"
            ]
        },
        "ENSMUSG00000053475": {
            "id": "ENSMUSG00000053475",
            "type": "Gene",
            "symbol": "Tnfaip6",
            "source": [
                "ENSMUSG00000005583",
                "ENSMUSG00000022346",
                "ENSMUSG00000000567"
            ],
            "target": [],
            "expression_sum": {
                "class1": 292.0,
                "class2": 20.0
            },
            "uniprot_ids": [
                "O08859"
            ]
        },
        "ENSMUSG00000053483": {
            "id": "ENSMUSG00000053483",
            "type": "Gene",
            "symbol": "Usp21",
            "source": [
                "ENSMUSG00000032897",
                "ENSMUSG00000015839",
                "ENSMUSG00000015605",
                "ENSMUSG00000015579",
                "ENSMUSG00000029249",
                "ENSMUSG00000001280",
                "ENSMUSG00000005566",
                "ENSMUSG00000019768",
                "ENSMUSG00000005698",
                "ENSMUSG00000032238",
                "ENSMUSG00000021318",
                "ENSMUSG00000058756",
                "ENSMUSG00000008575",
                "ENSMUSG00000000567"
            ],
            "target": [],
            "expression_sum": {
                "class1": 390.0,
                "class2": 129.0
            },
            "uniprot_ids": [
                "E9PUE2",
                "D6RJJ7",
                "G3UWN7"
            ]
        },
        "ENSMUSG00000053877": {
            "id": "ENSMUSG00000053877",
            "type": "Gene",
            "symbol": "Srcap",
            "source": [
                "ENSMUSG00000031965",
                "ENSMUSG00000036461",
                "ENSMUSG00000027490",
                "ENSMUSG00000032238",
                "ENSMUSG00000044167",
                "ENSMUSG00000031627",
                "ENSMUSG00000042557",
                "ENSMUSG00000000567"
            ],
            "target": [],
            "expression_sum": {
                "class1": 489.0,
                "class2": 74.0
            },
            "uniprot_ids": [
                "Q8BQA6",
                "A0A087WQ44",
                "A0A087WNL7",
                "A0A087WS36",
                "E9Q9V7",
                "A0A087WNX7",
                "A0A087WQY5",
                "A0A087WSL6"
            ]
        },
        "ENSMUSG00000054400": {
            "id": "ENSMUSG00000054400",
            "type": "Gene",
            "symbol": "Cklf",
            "source": [
                "ENSMUSG00000024238",
                "ENSMUSG00000052534",
                "ENSMUSG00000022641",
                "ENSMUSG00000008575",
                "ENSMUSG00000022314",
                "ENSMUSG00000000567"
            ],
            "target": [],
            "expression_sum": {
                "class1": 681.0,
                "class2": 242.0
            },
            "uniprot_ids": [
                "Q9DAS1",
                "A0A1D5RLG8",
                "A0A1D5RMJ6",
                "A0A1D5RLH5"
            ]
        },
        "ENSMUSG00000054770": {
            "id": "ENSMUSG00000054770",
            "type": "Gene",
            "symbol": "Kctd18",
            "source": [
                "ENSMUSG00000023034",
                "ENSMUSG00000052837",
                "ENSMUSG00000005583",
                "ENSMUSG00000024974",
                "ENSMUSG00000000567"
            ],
            "target": [],
            "expression_sum": {
                "class1": 406.0,
                "class2": 65.0
            },
            "uniprot_ids": [
                "E9PYP3",
                "E0CZ26",
                "E9Q945",
                "E0CXJ5"
            ]
        },
        "ENSMUSG00000054893": {
            "id": "ENSMUSG00000054893",
            "type": "TF",
            "symbol": "Zfp667",
            "source": [
                "ENSMUSG00000058440",
                "ENSMUSG00000024238",
                "ENSMUSG00000023927",
                "ENSMUSG00000049577",
                "ENSMUSG00000000567"
            ],
            "target": [
                "ENSMUSG00000028126",
                "ENSMUSG00000008373",
                "ENSMUSG00000001986",
                "ENSMUSG00000002524"
            ],
            "expression_sum": {
                "class1": 465.0,
                "class2": 22.0
            },
            "uniprot_ids": [
                "Q2TL60",
                "A0A140LHW4"
            ]
        },
        "ENSMUSG00000055053": {
            "id": "ENSMUSG00000055053",
            "type": "TF",
            "symbol": "Nfic",
            "source": [
                "ENSMUSG00000029135",
                "ENSMUSG00000001419",
                "ENSMUSG00000010175",
                "ENSMUSG00000006215",
                "ENSMUSG00000058239",
                "ENSMUSG00000051469",
                "ENSMUSG00000001280",
                "ENSMUSG00000019768",
                "ENSMUSG00000039191",
                "ENSMUSG00000031715",
                "ENSMUSG00000000567"
            ],
            "target": [],
            "expression_sum": {
                "class1": 3135.0,
                "class2": 592.0
            },
            "uniprot_ids": [
                "P70255",
                "F6XYU7",
                "A0A1Y7VK55"
            ]
        },
        "ENSMUSG00000055172": {
            "id": "ENSMUSG00000055172",
            "type": "Gene",
            "symbol": "C1ra",
            "source": [
                "ENSMUSG00000071076",
                "ENSMUSG00000055116",
                "ENSMUSG00000052837",
                "ENSMUSG00000074622",
                "ENSMUSG00000032228",
                "ENSMUSG00000022053",
                "ENSMUSG00000016087",
                "ENSMUSG00000053110",
                "ENSMUSG00000000567"
            ],
            "target": [],
            "expression_sum": {
                "class1": 161.0,
                "class2": 159.0
            },
            "uniprot_ids": [
                "Q8CG16"
            ]
        },
        "ENSMUSG00000055204": {
            "id": "ENSMUSG00000055204",
            "type": "Gene",
            "symbol": "Ankrd17",
            "source": [
                "ENSMUSG00000029135",
                "ENSMUSG00000076431",
                "ENSMUSG00000008575",
                "ENSMUSG00000035158",
                "ENSMUSG00000000567",
                "ENSMUSG00000006215",
                "ENSMUSG00000010175",
                "ENSMUSG00000058239",
                "ENSMUSG00000015579",
                "ENSMUSG00000051469",
                "ENSMUSG00000027109",
                "ENSMUSG00000019768",
                "ENSMUSG00000031885",
                "ENSMUSG00000053110",
                "ENSMUSG00000031715",
                "ENSMUSG00000015839",
                "ENSMUSG00000030557",
                "ENSMUSG00000075595",
                "ENSMUSG00000001419",
                "ENSMUSG00000025612",
                "ENSMUSG00000001280"
            ],
            "target": [],
            "expression_sum": {
                "class1": 3084.0,
                "class2": 958.0
            },
            "uniprot_ids": [
                "Q99NH0",
                "A0A1W2P6X1",
                "E9QKG6",
                "A0A0G2JDZ9",
                "E9Q804"
            ]
        },
        "ENSMUSG00000055707": {
            "id": "ENSMUSG00000055707",
            "type": "Gene",
            "symbol": "Klhl26",
            "source": [
                "ENSMUSG00000032228",
                "ENSMUSG00000000567"
            ],
            "target": [],
            "expression_sum": {
                "class1": 192.0,
                "class2": 32.0
            },
            "uniprot_ids": [
                "Q8BGY4",
                "A0A1B0GR96",
                "E9Q4C1"
            ]
        },
        "ENSMUSG00000056749": {
            "id": "ENSMUSG00000056749",
            "type": "TF",
            "symbol": "Nfil3",
            "source": [
                "ENSMUSG00000005698",
                "ENSMUSG00000000567"
            ],
            "target": [
                "ENSMUSG00000028953",
                "ENSMUSG00000032014",
                "ENSMUSG00000031596",
                "ENSMUSG00000028398",
                "ENSMUSG00000034674",
                "ENSMUSG00000078201",
                "ENSMUSG00000021000",
                "ENSMUSG00000022300",
                "ENSMUSG00000074748",
                "ENSMUSG00000050732",
                "ENSMUSG00000015806",
                "ENSMUSG00000025613",
                "ENSMUSG00000043962",
                "ENSMUSG00000020778",
                "ENSMUSG00000039531",
                "ENSMUSG00000024357",
                "ENSMUSG00000017221",
                "ENSMUSG00000056234",
                "ENSMUSG00000057177",
                "ENSMUSG00000020098",
                "ENSMUSG00000007987",
                "ENSMUSG00000032624",
                "ENSMUSG00000034584",
                "ENSMUSG00000040536",
                "ENSMUSG00000002981",
                "ENSMUSG00000033684",
                "ENSMUSG00000059518",
                "ENSMUSG00000030861",
                "ENSMUSG00000007850",
                "ENSMUSG00000026585"
            ],
            "expression_sum": {
                "class1": 191.0,
                "class2": 195.0
            },
            "uniprot_ids": [
                "O08750"
            ]
        },
        "ENSMUSG00000057469": {
            "id": "ENSMUSG00000057469",
            "type": "TF",
            "symbol": "E2f6",
            "source": [
                "ENSMUSG00000029135",
                "ENSMUSG00000001280",
                "ENSMUSG00000019768",
                "ENSMUSG00000015605",
                "ENSMUSG00000029913",
                "ENSMUSG00000000567"
            ],
            "target": [
                "ENSMUSG00000027091",
                "ENSMUSG00000036052",
                "ENSMUSG00000037826",
                "ENSMUSG00000026123",
                "ENSMUSG00000003438",
                "ENSMUSG00000009569",
                "ENSMUSG00000034673",
                "ENSMUSG00000066406",
                "ENSMUSG00000057649",
                "ENSMUSG00000059981",
                "ENSMUSG00000002844",
                "ENSMUSG00000005566",
                "ENSMUSG00000060038",
                "ENSMUSG00000034610",
                "ENSMUSG00000034875",
                "ENSMUSG00000031403",
                "ENSMUSG00000020485",
                "ENSMUSG00000021000",
                "ENSMUSG00000006740",
                "ENSMUSG00000000563",
                "ENSMUSG00000027238",
                "ENSMUSG00000028081",
                "ENSMUSG00000037300",
                "ENSMUSG00000053536",
                "ENSMUSG00000019432",
                "ENSMUSG00000026491",
                "ENSMUSG00000018599",
                "ENSMUSG00000022641",
                "ENSMUSG00000036371",
                "ENSMUSG00000058135",
                "ENSMUSG00000015112",
                "ENSMUSG00000038070",
                "ENSMUSG00000075054",
                "ENSMUSG00000021453",
                "ENSMUSG00000052337",
                "ENSMUSG00000031367",
                "ENSMUSG00000027499",
                "ENSMUSG00000027784",
                "ENSMUSG00000032348",
                "ENSMUSG00000024283",
                "ENSMUSG00000032423",
                "ENSMUSG00000045983",
                "ENSMUSG00000029629",
                "ENSMUSG00000017119",
                "ENSMUSG00000028911",
                "ENSMUSG00000029913",
                "ENSMUSG00000081792",
                "ENSMUSG00000068267",
                "ENSMUSG00000031983",
                "ENSMUSG00000038028",
                "ENSMUSG00000032481",
                "ENSMUSG00000032047",
                "ENSMUSG00000029594",
                "ENSMUSG00000019773",
                "ENSMUSG00000025453",
                "ENSMUSG00000067274",
                "ENSMUSG00000030788",
                "ENSMUSG00000015843",
                "ENSMUSG00000026576",
                "ENSMUSG00000055239",
                "ENSMUSG00000066637",
                "ENSMUSG00000032525",
                "ENSMUSG00000038022",
                "ENSMUSG00000044393",
                "ENSMUSG00000000441",
                "ENSMUSG00000026626",
                "ENSMUSG00000074918",
                "ENSMUSG00000034220",
                "ENSMUSG00000027823",
                "ENSMUSG00000031543",
                "ENSMUSG00000007891",
                "ENSMUSG00000041685",
                "ENSMUSG00000024073",
                "ENSMUSG00000028282",
                "ENSMUSG00000020267",
                "ENSMUSG00000033060",
                "ENSMUSG00000038784",
                "ENSMUSG00000062580",
                "ENSMUSG00000021209",
                "ENSMUSG00000005354",
                "ENSMUSG00000022890",
                "ENSMUSG00000032826",
                "ENSMUSG00000023939",
                "ENSMUSG00000035885",
                "ENSMUSG00000022336",
                "ENSMUSG00000026187",
                "ENSMUSG00000029198",
                "ENSMUSG00000033411",
                "ENSMUSG00000032959",
                "ENSMUSG00000032437",
                "ENSMUSG00000026701",
                "ENSMUSG00000033416",
                "ENSMUSG00000021218",
                "ENSMUSG00000032199",
                "ENSMUSG00000028266",
                "ENSMUSG00000045690",
                "ENSMUSG00000062397",
                "ENSMUSG00000026341",
                "ENSMUSG00000025271",
                "ENSMUSG00000022185",
                "ENSMUSG00000027706",
                "ENSMUSG00000058793",
                "ENSMUSG00000022131",
                "ENSMUSG00000049643",
                "ENSMUSG00000000088",
                "ENSMUSG00000031447",
                "ENSMUSG00000024985",
                "ENSMUSG00000031985",
                "ENSMUSG00000052539",
                "ENSMUSG00000026064",
                "ENSMUSG00000032314",
                "ENSMUSG00000073684",
                "ENSMUSG00000022452",
                "ENSMUSG00000023175",
                "ENSMUSG00000022519",
                "ENSMUSG00000026283",
                "ENSMUSG00000036097",
                "ENSMUSG00000006315",
                "ENSMUSG00000020048",
                "ENSMUSG00000027406",
                "ENSMUSG00000026858",
                "ENSMUSG00000028461",
                "ENSMUSG00000037568",
                "ENSMUSG00000020737",
                "ENSMUSG00000014592",
                "ENSMUSG00000042293",
                "ENSMUSG00000052374",
                "ENSMUSG00000015214",
                "ENSMUSG00000030850",
                "ENSMUSG00000024991",
                "ENSMUSG00000037710",
                "ENSMUSG00000028229",
                "ENSMUSG00000049327",
                "ENSMUSG00000048814",
                "ENSMUSG00000041075",
                "ENSMUSG00000021597",
                "ENSMUSG00000010048",
                "ENSMUSG00000022201",
                "ENSMUSG00000025410",
                "ENSMUSG00000026260",
                "ENSMUSG00000036112",
                "ENSMUSG00000002102",
                "ENSMUSG00000022748",
                "ENSMUSG00000043866",
                "ENSMUSG00000056091",
                "ENSMUSG00000025374",
                "ENSMUSG00000018446",
                "ENSMUSG00000030934",
                "ENSMUSG00000042198",
                "ENSMUSG00000019577",
                "ENSMUSG00000023249",
                "ENSMUSG00000024613",
                "ENSMUSG00000027274",
                "ENSMUSG00000054889",
                "ENSMUSG00000027293",
                "ENSMUSG00000070544",
                "ENSMUSG00000046351",
                "ENSMUSG00000020988",
                "ENSMUSG00000028234",
                "ENSMUSG00000035268",
                "ENSMUSG00000022982",
                "ENSMUSG00000025184",
                "ENSMUSG00000048486",
                "ENSMUSG00000032897",
                "ENSMUSG00000060904",
                "ENSMUSG00000028273",
                "ENSMUSG00000034951",
                "ENSMUSG00000026775",
                "ENSMUSG00000018474",
                "ENSMUSG00000032883",
                "ENSMUSG00000044365",
                "ENSMUSG00000066043",
                "ENSMUSG00000073563",
                "ENSMUSG00000038690",
                "ENSMUSG00000020176",
                "ENSMUSG00000027423",
                "ENSMUSG00000031751",
                "ENSMUSG00000023055",
                "ENSMUSG00000037236",
                "ENSMUSG00000048578",
                "ENSMUSG00000038650",
                "ENSMUSG00000025856",
                "ENSMUSG00000018559",
                "ENSMUSG00000020027",
                "ENSMUSG00000025364",
                "ENSMUSG00000022682",
                "ENSMUSG00000054469",
                "ENSMUSG00000021910",
                "ENSMUSG00000036880",
                "ENSMUSG00000030603",
                "ENSMUSG00000017756",
                "ENSMUSG00000039987",
                "ENSMUSG00000026429",
                "ENSMUSG00000028788",
                "ENSMUSG00000031774",
                "ENSMUSG00000031783",
                "ENSMUSG00000003316",
                "ENSMUSG00000033102",
                "ENSMUSG00000078671",
                "ENSMUSG00000034290",
                "ENSMUSG00000023150",
                "ENSMUSG00000027618",
                "ENSMUSG00000018796",
                "ENSMUSG00000028603",
                "ENSMUSG00000038718",
                "ENSMUSG00000039741",
                "ENSMUSG00000063229",
                "ENSMUSG00000032518",
                "ENSMUSG00000021546",
                "ENSMUSG00000020321",
                "ENSMUSG00000018566",
                "ENSMUSG00000027332",
                "ENSMUSG00000029135",
                "ENSMUSG00000017404",
                "ENSMUSG00000022214",
                "ENSMUSG00000004268",
                "ENSMUSG00000027602",
                "ENSMUSG00000061286",
                "ENSMUSG00000017778",
                "ENSMUSG00000044573",
                "ENSMUSG00000004591",
                "ENSMUSG00000021621",
                "ENSMUSG00000024304",
                "ENSMUSG00000008575",
                "ENSMUSG00000014633",
                "ENSMUSG00000021706",
                "ENSMUSG00000060679",
                "ENSMUSG00000030057",
                "ENSMUSG00000074182",
                "ENSMUSG00000019851",
                "ENSMUSG00000059208",
                "ENSMUSG00000032042",
                "ENSMUSG00000052912",
                "ENSMUSG00000022016",
                "ENSMUSG00000055322",
                "ENSMUSG00000074064",
                "ENSMUSG00000006057",
                "ENSMUSG00000067722",
                "ENSMUSG00000028207",
                "ENSMUSG00000040414",
                "ENSMUSG00000040771",
                "ENSMUSG00000032724",
                "ENSMUSG00000033161",
                "ENSMUSG00000023089",
                "ENSMUSG00000030246",
                "ENSMUSG00000001424",
                "ENSMUSG00000026434",
                "ENSMUSG00000061474",
                "ENSMUSG00000003072",
                "ENSMUSG00000025466",
                "ENSMUSG00000036564",
                "ENSMUSG00000024236",
                "ENSMUSG00000039474",
                "ENSMUSG00000020664",
                "ENSMUSG00000021585",
                "ENSMUSG00000022797",
                "ENSMUSG00000020079",
                "ENSMUSG00000061518",
                "ENSMUSG00000021395",
                "ENSMUSG00000022010",
                "ENSMUSG00000020849",
                "ENSMUSG00000024150",
                "ENSMUSG00000028343",
                "ENSMUSG00000031634",
                "ENSMUSG00000082195",
                "ENSMUSG00000006717",
                "ENSMUSG00000032480",
                "ENSMUSG00000001056",
                "ENSMUSG00000024924",
                "ENSMUSG00000059811",
                "ENSMUSG00000028884",
                "ENSMUSG00000030541",
                "ENSMUSG00000001280",
                "ENSMUSG00000038005",
                "ENSMUSG00000070003",
                "ENSMUSG00000026489",
                "ENSMUSG00000024913"
            ],
            "expression_sum": {
                "class1": 276.0,
                "class2": 554.0
            },
            "uniprot_ids": [
                "O54917",
                "Q8K455",
                "A0A1Y7VJU3",
                "A0A1Y7VP85"
            ]
        },
        "ENSMUSG00000057561": {
            "id": "ENSMUSG00000057561",
            "type": "Gene",
            "symbol": "Eif1a",
            "source": [
                "ENSMUSG00000058440",
                "ENSMUSG00000024238",
                "ENSMUSG00000019817",
                "ENSMUSG00000055116",
                "ENSMUSG00000008575",
                "ENSMUSG00000022314",
                "ENSMUSG00000000567"
            ],
            "target": [],
            "expression_sum": {
                "class1": 2900.0,
                "class2": 472.0
            },
            "uniprot_ids": [
                "Q60872",
                "A0A494BB48",
                "A0A494BAI7",
                "A0A494BB95"
            ]
        },
        "ENSMUSG00000057894": {
            "id": "ENSMUSG00000057894",
            "type": "TF",
            "symbol": "Zfp329",
            "source": [
                "ENSMUSG00000003032",
                "ENSMUSG00000000567"
            ],
            "target": [
                "ENSMUSG00000024055",
                "ENSMUSG00000000957",
                "ENSMUSG00000024539",
                "ENSMUSG00000001029",
                "ENSMUSG00000018293",
                "ENSMUSG00000005417"
            ],
            "expression_sum": {
                "class1": 961.0,
                "class2": 117.0
            },
            "uniprot_ids": [
                "Q6GQR8",
                "D3Z6L7"
            ]
        },
        "ENSMUSG00000060288": {
            "id": "ENSMUSG00000060288",
            "type": "Gene",
            "symbol": "Ppih",
            "source": [
                "ENSMUSG00000058440",
                "ENSMUSG00000004151",
                "ENSMUSG00000031627",
                "ENSMUSG00000031885",
                "ENSMUSG00000057098",
                "ENSMUSG00000024985",
                "ENSMUSG00000000567"
            ],
            "target": [],
            "expression_sum": {
                "class1": 1006.0,
                "class2": 85.0
            },
            "uniprot_ids": [
                "Q9D868",
                "A2BGI9",
                "A2BGI8"
            ]
        },
        "ENSMUSG00000061477": {
            "id": "ENSMUSG00000061477",
            "type": "Gene",
            "symbol": "Rps7",
            "source": [
                "ENSMUSG00000029135",
                "ENSMUSG00000001419",
                "ENSMUSG00000015579",
                "ENSMUSG00000001280",
                "ENSMUSG00000019768",
                "ENSMUSG00000035158",
                "ENSMUSG00000000567"
            ],
            "target": [],
            "expression_sum": {
                "class1": 5575.0,
                "class2": 1559.0
            },
            "uniprot_ids": [
                "P62082"
            ]
        },
        "ENSMUSG00000061878": {
            "id": "ENSMUSG00000061878",
            "type": "Gene",
            "symbol": "Sphk1",
            "source": [
                "ENSMUSG00000022053",
                "ENSMUSG00000000567"
            ],
            "target": [],
            "expression_sum": {
                "class1": 313.0,
                "class2": 12.0
            },
            "uniprot_ids": [
                "Q8CI15",
                "B1AT86",
                "B1AT88",
                "B1AT87",
                "B1AT89",
                "B1AT85"
            ]
        },
        "ENSMUSG00000062190": {
            "id": "ENSMUSG00000062190",
            "type": "Gene",
            "symbol": "Lancl2",
            "source": [
                "ENSMUSG00000026398",
                "ENSMUSG00000056501",
                "ENSMUSG00000005698",
                "ENSMUSG00000025498",
                "ENSMUSG00000000567"
            ],
            "target": [],
            "expression_sum": {
                "class1": 541.0,
                "class2": 62.0
            },
            "uniprot_ids": [
                "Q9JJK2",
                "F6RJV6"
            ]
        },
        "ENSMUSG00000062309": {
            "id": "ENSMUSG00000062309",
            "type": "Gene",
            "symbol": "Rpp25",
            "source": [
                "ENSMUSG00000021250",
                "ENSMUSG00000027109",
                "ENSMUSG00000052684",
                "ENSMUSG00000005583",
                "ENSMUSG00000000567"
            ],
            "target": [],
            "expression_sum": {
                "class1": 219.0,
                "class2": 4.0
            },
            "uniprot_ids": [
                "Q91WE3"
            ]
        },
        "ENSMUSG00000063077": {
            "id": "ENSMUSG00000063077",
            "type": "Gene",
            "symbol": "Kif1b",
            "source": [
                "ENSMUSG00000024238",
                "ENSMUSG00000015839",
                "ENSMUSG00000015579",
                "ENSMUSG00000051469",
                "ENSMUSG00000030557",
                "ENSMUSG00000001280",
                "ENSMUSG00000000567"
            ],
            "target": [],
            "expression_sum": {
                "class1": 3571.0,
                "class2": 1817.0
            },
            "uniprot_ids": [
                "Q60575",
                "A0A5F8MPF0",
                "F6XNB6",
                "A2AH75"
            ]
        },
        "ENSMUSG00000066551": {
            "id": "ENSMUSG00000066551",
            "type": "TF",
            "symbol": "Hmgb1",
            "source": [
                "ENSMUSG00000026398",
                "ENSMUSG00000021250",
                "ENSMUSG00000030256",
                "ENSMUSG00000005583",
                "ENSMUSG00000022053",
                "ENSMUSG00000000567"
            ],
            "target": [
                "ENSMUSG00000000827",
                "ENSMUSG00000000787",
                "ENSMUSG00000000359"
            ],
            "expression_sum": {
                "class1": 1081.0,
                "class2": 169.0
            },
            "uniprot_ids": [
                "P63158",
                "A0A0J9YUD8",
                "A0A0J9YUZ4",
                "D3YVC6",
                "D3YZ18"
            ]
        },
        "ENSMUSG00000066839": {
            "id": "ENSMUSG00000066839",
            "type": "Gene",
            "symbol": "Ecsit",
            "source": [
                "ENSMUSG00000001280",
                "ENSMUSG00000015605",
                "ENSMUSG00000000567"
            ],
            "target": [],
            "expression_sum": {
                "class1": 243.0,
                "class2": 643.0
            },
            "uniprot_ids": [
                "A0A0R4J174",
                "J3KMM6"
            ]
        },
        "ENSMUSG00000067847": {
            "id": "ENSMUSG00000067847",
            "type": "Gene",
            "symbol": "Romo1",
            "source": [
                "ENSMUSG00000049577",
                "ENSMUSG00000001156",
                "ENSMUSG00000001280",
                "ENSMUSG00000019768",
                "ENSMUSG00000000567"
            ],
            "target": [],
            "expression_sum": {
                "class1": 8412.0,
                "class2": 4583.0
            },
            "uniprot_ids": [
                "P60603"
            ]
        },
        "ENSMUSG00000068267": {
            "id": "ENSMUSG00000068267",
            "type": "TF",
            "symbol": "Cenpb",
            "source": [
                "ENSMUSG00000031681",
                "ENSMUSG00000057469",
                "ENSMUSG00000024985",
                "ENSMUSG00000038648",
                "ENSMUSG00000029249",
                "ENSMUSG00000001280",
                "ENSMUSG00000019768",
                "ENSMUSG00000015605",
                "ENSMUSG00000015846",
                "ENSMUSG00000000567"
            ],
            "target": [
                "ENSMUSG00000036002",
                "ENSMUSG00000022215",
                "ENSMUSG00000000149",
                "ENSMUSG00000006356",
                "ENSMUSG00000024661",
                "ENSMUSG00000035242",
                "ENSMUSG00000002416",
                "ENSMUSG00000026817",
                "ENSMUSG00000082043",
                "ENSMUSG00000031231",
                "ENSMUSG00000024304",
                "ENSMUSG00000025465",
                "ENSMUSG00000020485",
                "ENSMUSG00000023020",
                "ENSMUSG00000000563",
                "ENSMUSG00000028955",
                "ENSMUSG00000038296",
                "ENSMUSG00000016308",
                "ENSMUSG00000000056",
                "ENSMUSG00000000628",
                "ENSMUSG00000082895",
                "ENSMUSG00000006057",
                "ENSMUSG00000040147",
                "ENSMUSG00000032199",
                "ENSMUSG00000023991",
                "ENSMUSG00000021917",
                "ENSMUSG00000020717",
                "ENSMUSG00000040414",
                "ENSMUSG00000021622",
                "ENSMUSG00000001211",
                "ENSMUSG00000023089",
                "ENSMUSG00000000088",
                "ENSMUSG00000020022",
                "ENSMUSG00000025856",
                "ENSMUSG00000030647",
                "ENSMUSG00000000399",
                "ENSMUSG00000023944",
                "ENSMUSG00000001911",
                "ENSMUSG00000031367",
                "ENSMUSG00000034164",
                "ENSMUSG00000037032",
                "ENSMUSG00000040097",
                "ENSMUSG00000018893",
                "ENSMUSG00000004610",
                "ENSMUSG00000031029",
                "ENSMUSG00000055762",
                "ENSMUSG00000024038",
                "ENSMUSG00000028996",
                "ENSMUSG00000041028",
                "ENSMUSG00000054452",
                "ENSMUSG00000001794",
                "ENSMUSG00000017817",
                "ENSMUSG00000004980",
                "ENSMUSG00000039218",
                "ENSMUSG00000020849",
                "ENSMUSG00000022551",
                "ENSMUSG00000041297",
                "ENSMUSG00000006717",
                "ENSMUSG00000048814",
                "ENSMUSG00000000168",
                "ENSMUSG00000030399",
                "ENSMUSG00000000184",
                "ENSMUSG00000024913",
                "ENSMUSG00000007850",
                "ENSMUSG00000030541",
                "ENSMUSG00000039684",
                "ENSMUSG00000030417",
                "ENSMUSG00000055239",
                "ENSMUSG00000038022",
                "ENSMUSG00000081661",
                "ENSMUSG00000026853",
                "ENSMUSG00000044393",
                "ENSMUSG00000024049",
                "ENSMUSG00000000171",
                "ENSMUSG00000044751"
            ],
            "expression_sum": {
                "class1": 5419.0,
                "class2": 860.0
            },
            "uniprot_ids": [
                "P27790"
            ]
        },
        "ENSMUSG00000069744": {
            "id": "ENSMUSG00000069744",
            "type": "Gene",
            "symbol": "Psmb3",
            "source": [
                "ENSMUSG00000059552",
                "ENSMUSG00000024515",
                "ENSMUSG00000038648",
                "ENSMUSG00000018899",
                "ENSMUSG00000001280",
                "ENSMUSG00000005583",
                "ENSMUSG00000028565",
                "ENSMUSG00000040732",
                "ENSMUSG00000031627",
                "ENSMUSG00000027104",
                "ENSMUSG00000024985",
                "ENSMUSG00000000567"
            ],
            "target": [],
            "expression_sum": {
                "class1": 3202.0,
                "class2": 1003.0
            },
            "uniprot_ids": [
                "Q9R1P1"
            ]
        },
        "ENSMUSG00000070284": {
            "id": "ENSMUSG00000070284",
            "type": "Gene",
            "symbol": "Gmppb",
            "source": [
                "ENSMUSG00000021250",
                "ENSMUSG00000015839",
                "ENSMUSG00000005583",
                "ENSMUSG00000005698",
                "ENSMUSG00000000567"
            ],
            "target": [],
            "expression_sum": {
                "class1": 80.0,
                "class2": 2.0
            },
            "uniprot_ids": [
                "Q8BTZ7"
            ]
        },
        "ENSMUSG00000070697": {
            "id": "ENSMUSG00000070697",
            "type": "Gene",
            "symbol": "Utp3",
            "source": [
                "ENSMUSG00000021250",
                "ENSMUSG00000006215",
                "ENSMUSG00000041515",
                "ENSMUSG00000005583",
                "ENSMUSG00000052684",
                "ENSMUSG00000031885",
                "ENSMUSG00000000567"
            ],
            "target": [],
            "expression_sum": {
                "class1": 2441.0,
                "class2": 344.0
            },
            "uniprot_ids": [
                "Q9JI13"
            ]
        },
        "ENSMUSG00000071014": {
            "id": "ENSMUSG00000071014",
            "type": "Gene",
            "symbol": "Ndufb6",
            "source": [
                "ENSMUSG00000029135",
                "ENSMUSG00000056501",
                "ENSMUSG00000003923",
                "ENSMUSG00000051469",
                "ENSMUSG00000001280",
                "ENSMUSG00000019768",
                "ENSMUSG00000031627",
                "ENSMUSG00000031885",
                "ENSMUSG00000008575",
                "ENSMUSG00000000567"
            ],
            "target": [],
            "expression_sum": {
                "class1": 1661.0,
                "class2": 3928.0
            },
            "uniprot_ids": [
                "Q3UIU2",
                "A2AP32"
            ]
        },
        "ENSMUSG00000073684": {
            "id": "ENSMUSG00000073684",
            "type": "Gene",
            "symbol": "Faap20",
            "source": [
                "ENSMUSG00000024238",
                "ENSMUSG00000057469",
                "ENSMUSG00000052534",
                "ENSMUSG00000029249",
                "ENSMUSG00000005566",
                "ENSMUSG00000019768",
                "ENSMUSG00000005583",
                "ENSMUSG00000058756",
                "ENSMUSG00000008575",
                "ENSMUSG00000022314",
                "ENSMUSG00000000567"
            ],
            "target": [],
            "expression_sum": {
                "class1": 1012.0,
                "class2": 130.0
            },
            "uniprot_ids": [
                "Q3UN58",
                "F2Z421",
                "F6QQ55"
            ]
        },
        "ENSMUSG00000073725": {
            "id": "ENSMUSG00000073725",
            "type": "Gene",
            "symbol": "Lmbrd1",
            "source": [
                "ENSMUSG00000030256",
                "ENSMUSG00000015839",
                "ENSMUSG00000006215",
                "ENSMUSG00000000567"
            ],
            "target": [],
            "expression_sum": {
                "class1": 803.0,
                "class2": 585.0
            },
            "uniprot_ids": [
                "Q8K0B2",
                "A0A087WS60",
                "A0A087WR85",
                "A0A087WPS2"
            ]
        },
        "ENSMUSG00000074457": {
            "id": "ENSMUSG00000074457",
            "type": "Gene",
            "symbol": "S100a16",
            "source": [
                "ENSMUSG00000071076",
                "ENSMUSG00000024560",
                "ENSMUSG00000000567"
            ],
            "target": [],
            "expression_sum": {
                "class1": 11863.0,
                "class2": 1261.0
            },
            "uniprot_ids": [
                "Q9D708",
                "D3Z2Y6"
            ]
        },
        "ENSMUSG00000078201": {
            "id": "ENSMUSG00000078201",
            "type": "Gene",
            "symbol": "Tmem203",
            "source": [
                "ENSMUSG00000006215",
                "ENSMUSG00000041515",
                "ENSMUSG00000056749",
                "ENSMUSG00000052684",
                "ENSMUSG00000005583",
                "ENSMUSG00000000567"
            ],
            "target": [],
            "expression_sum": {
                "class1": 385.0,
                "class2": 57.0
            },
            "uniprot_ids": [
                "Q8R235"
            ]
        },
        "ENSMUSG00000078247": {
            "id": "ENSMUSG00000078247",
            "type": "Gene",
            "symbol": "Airn",
            "source": [
                "ENSMUSG00000053477",
                "ENSMUSG00000005583",
                "ENSMUSG00000022053",
                "ENSMUSG00000022314",
                "ENSMUSG00000000567"
            ],
            "target": [],
            "expression_sum": {
                "class1": 802.0,
                "class2": 138.0
            }
        },
        "ENSMUSG00000079435": {
            "id": "ENSMUSG00000079435",
            "type": "Gene",
            "symbol": "Rpl36a",
            "source": [
                "ENSMUSG00000029135",
                "ENSMUSG00000055148",
                "ENSMUSG00000021250",
                "ENSMUSG00000052534",
                "ENSMUSG00000005583",
                "ENSMUSG00000040732",
                "ENSMUSG00000076431",
                "ENSMUSG00000022314",
                "ENSMUSG00000000567"
            ],
            "target": [],
            "expression_sum": {
                "class1": 2928.0,
                "class2": 560.0
            },
            "uniprot_ids": [
                "P83882",
                "E9Q131"
            ]
        },
        "ENSMUSG00000082978": {
            "id": "ENSMUSG00000082978",
            "type": "Gene",
            "symbol": "Rpsa-ps11",
            "source": [
                "ENSMUSG00000019768",
                "ENSMUSG00000052837",
                "ENSMUSG00000017146",
                "ENSMUSG00000000567"
            ],
            "target": [],
            "expression_sum": {
                "class1": 2036.0,
                "class2": 209.0
            }
        },
        "ENSMUSG00000085208": {
            "id": "ENSMUSG00000085208",
            "type": "Gene",
            "symbol": "Brip1os",
            "source": [
                "ENSMUSG00000053110",
                "ENSMUSG00000000567"
            ],
            "target": [],
            "expression_sum": {
                "class1": 1169.0,
                "class2": 188.0
            }
        },
        "ENSMUSG00000090812": {
            "id": "ENSMUSG00000090812",
            "type": "Gene",
            "symbol": "Samd15",
            "source": [
                "ENSMUSG00000021250",
                "ENSMUSG00000000567",
                "ENSMUSG00000005583",
                "ENSMUSG00000005698",
                "ENSMUSG00000020167"
            ],
            "target": [],
            "expression_sum": {
                "class1": 345.0,
                "class2": 27.0
            },
            "uniprot_ids": [
                "F6XZJ7",
                "H9H9T8"
            ]
        },
        "ENSMUSG00000091028": {
            "id": "ENSMUSG00000091028",
            "type": "Gene",
            "symbol": "Gm10722",
            "source": [
                "ENSMUSG00000005583",
                "ENSMUSG00000000567"
            ],
            "target": [],
            "expression_sum": {
                "class1": 341.0,
                "class2": 4.0
            },
            "uniprot_ids": [
                "F6VLK5"
            ]
        },
        "ENSMUSG00000091803": {
            "id": "ENSMUSG00000091803",
            "type": "Gene",
            "symbol": "Cox16",
            "source": [
                "ENSMUSG00000000567"
            ],
            "target": [],
            "expression_sum": {
                "class1": 250.0,
                "class2": 140.0
            },
            "uniprot_ids": [
                "Q9CR63",
                "G3UWC6",
                "E9Q3U7",
                "E9Q5H8"
            ]
        },
        "ENSMUSG00000092837": {
            "id": "ENSMUSG00000092837",
            "type": "Gene",
            "symbol": "Rpph1",
            "source": [
                "ENSMUSG00000021250",
                "ENSMUSG00000041515",
                "ENSMUSG00000032238",
                "ENSMUSG00000099083",
                "ENSMUSG00000005583",
                "ENSMUSG00000005698",
                "ENSMUSG00000052684",
                "ENSMUSG00000000567"
            ],
            "target": [],
            "expression_sum": {
                "class1": 755.0,
                "class2": 60.0
            }
        },
        "ENSMUSG00000001280": {
            "id": "ENSMUSG00000001280",
            "type": "TF",
            "symbol": "Sp1",
            "source": [
                "ENSMUSG00000029135",
                "ENSMUSG00000021944",
                "ENSMUSG00000008575",
                "ENSMUSG00000015605",
                "ENSMUSG00000022528",
                "ENSMUSG00000057469",
                "ENSMUSG00000010175",
                "ENSMUSG00000058239",
                "ENSMUSG00000015579",
                "ENSMUSG00000051469",
                "ENSMUSG00000019768",
                "ENSMUSG00000031885",
                "ENSMUSG00000053110",
                "ENSMUSG00000031715",
                "ENSMUSG00000015846",
                "ENSMUSG00000058440",
                "ENSMUSG00000015839",
                "ENSMUSG00000032238",
                "ENSMUSG00000053477",
                "ENSMUSG00000058756",
                "ENSMUSG00000056501",
                "ENSMUSG00000029249",
                "ENSMUSG00000032035"
            ],
            "target": [
                "ENSMUSG00000000538",
                "ENSMUSG00000029860",
                "ENSMUSG00000029681",
                "ENSMUSG00000054455",
                "ENSMUSG00000032737",
                "ENSMUSG00000073702",
                "ENSMUSG00000031403",
                "ENSMUSG00000028081",
                "ENSMUSG00000091478",
                "ENSMUSG00000024228",
                "ENSMUSG00000020694",
                "ENSMUSG00000006676",
                "ENSMUSG00000020307",
                "ENSMUSG00000043683",
                "ENSMUSG00000021248",
                "ENSMUSG00000031885",
                "ENSMUSG00000026176",
                "ENSMUSG00000092074",
                "ENSMUSG00000021235",
                "ENSMUSG00000070283",
                "ENSMUSG00000039640",
                "ENSMUSG00000034903",
                "ENSMUSG00000030739",
                "ENSMUSG00000061461",
                "ENSMUSG00000004044",
                "ENSMUSG00000008036",
                "ENSMUSG00000028086",
                "ENSMUSG00000044221",
                "ENSMUSG00000035202",
                "ENSMUSG00000026087",
                "ENSMUSG00000026626",
                "ENSMUSG00000031790",
                "ENSMUSG00000031543",
                "ENSMUSG00000007891",
                "ENSMUSG00000069744",
                "ENSMUSG00000004789",
                "ENSMUSG00000083853",
                "ENSMUSG00000018363",
                "ENSMUSG00000018286",
                "ENSMUSG00000064349",
                "ENSMUSG00000080242",
                "ENSMUSG00000007656",
                "ENSMUSG00000027288",
                "ENSMUSG00000032314",
                "ENSMUSG00000008892",
                "ENSMUSG00000030557",
                "ENSMUSG00000020149",
                "ENSMUSG00000026858",
                "ENSMUSG00000027698",
                "ENSMUSG00000081344",
                "ENSMUSG00000028229",
                "ENSMUSG00000030852",
                "ENSMUSG00000010048",
                "ENSMUSG00000053329",
                "ENSMUSG00000031197",
                "ENSMUSG00000034981",
                "ENSMUSG00000029499",
                "ENSMUSG00000032216",
                "ENSMUSG00000090066",
                "ENSMUSG00000026755",
                "ENSMUSG00000018446",
                "ENSMUSG00000064302",
                "ENSMUSG00000026775",
                "ENSMUSG00000029392",
                "ENSMUSG00000044365",
                "ENSMUSG00000001552",
                "ENSMUSG00000020155",
                "ENSMUSG00000000056",
                "ENSMUSG00000024962",
                "ENSMUSG00000057133",
                "ENSMUSG00000020949",
                "ENSMUSG00000039103",
                "ENSMUSG00000073471",
                "ENSMUSG00000009628",
                "ENSMUSG00000029148",
                "ENSMUSG00000026158",
                "ENSMUSG00000020349",
                "ENSMUSG00000038708",
                "ENSMUSG00000032171",
                "ENSMUSG00000056501",
                "ENSMUSG00000029505",
                "ENSMUSG00000026615",
                "ENSMUSG00000055320",
                "ENSMUSG00000031698",
                "ENSMUSG00000022174",
                "ENSMUSG00000021024",
                "ENSMUSG00000027602",
                "ENSMUSG00000025745",
                "ENSMUSG00000049265",
                "ENSMUSG00000024304",
                "ENSMUSG00000006219",
                "ENSMUSG00000030057",
                "ENSMUSG00000025369",
                "ENSMUSG00000078139",
                "ENSMUSG00000055322",
                "ENSMUSG00000039016",
                "ENSMUSG00000006057",
                "ENSMUSG00000018196",
                "ENSMUSG00000068923",
                "ENSMUSG00000047284",
                "ENSMUSG00000087943",
                "ENSMUSG00000052253",
                "ENSMUSG00000057841",
                "ENSMUSG00000044533",
                "ENSMUSG00000064345",
                "ENSMUSG00000021666",
                "ENSMUSG00000019471",
                "ENSMUSG00000028884",
                "ENSMUSG00000018585",
                "ENSMUSG00000032648",
                "ENSMUSG00000052962",
                "ENSMUSG00000026374",
                "ENSMUSG00000024871",
                "ENSMUSG00000053929",
                "ENSMUSG00000096842",
                "ENSMUSG00000020315",
                "ENSMUSG00000026123",
                "ENSMUSG00000059981",
                "ENSMUSG00000059796",
                "ENSMUSG00000005882",
                "ENSMUSG00000020455",
                "ENSMUSG00000025208",
                "ENSMUSG00000060002",
                "ENSMUSG00000026672",
                "ENSMUSG00000029221",
                "ENSMUSG00000047419",
                "ENSMUSG00000025234",
                "ENSMUSG00000006998",
                "ENSMUSG00000025616",
                "ENSMUSG00000020459",
                "ENSMUSG00000079184",
                "ENSMUSG00000001334",
                "ENSMUSG00000001415",
                "ENSMUSG00000058076",
                "ENSMUSG00000007815",
                "ENSMUSG00000028165",
                "ENSMUSG00000018765",
                "ENSMUSG00000073557",
                "ENSMUSG00000066637",
                "ENSMUSG00000033060",
                "ENSMUSG00000025939",
                "ENSMUSG00000005610",
                "ENSMUSG00000021282",
                "ENSMUSG00000031812",
                "ENSMUSG00000064367",
                "ENSMUSG00000021361",
                "ENSMUSG00000024695",
                "ENSMUSG00000028955",
                "ENSMUSG00000037706",
                "ENSMUSG00000026701",
                "ENSMUSG00000037905",
                "ENSMUSG00000015806",
                "ENSMUSG00000025271",
                "ENSMUSG00000022185",
                "ENSMUSG00000084168",
                "ENSMUSG00000014313",
                "ENSMUSG00000016253",
                "ENSMUSG00000035936",
                "ENSMUSG00000029422",
                "ENSMUSG00000027668",
                "ENSMUSG00000002015",
                "ENSMUSG00000026107",
                "ENSMUSG00000024845",
                "ENSMUSG00000008429",
                "ENSMUSG00000064339",
                "ENSMUSG00000060261",
                "ENSMUSG00000028245",
                "ENSMUSG00000035529",
                "ENSMUSG00000042978",
                "ENSMUSG00000066839",
                "ENSMUSG00000028849",
                "ENSMUSG00000034951",
                "ENSMUSG00000015149",
                "ENSMUSG00000022108",
                "ENSMUSG00000040033",
                "ENSMUSG00000024712",
                "ENSMUSG00000037197",
                "ENSMUSG00000006024",
                "ENSMUSG00000022682",
                "ENSMUSG00000025348",
                "ENSMUSG00000053617",
                "ENSMUSG00000030603",
                "ENSMUSG00000052512",
                "ENSMUSG00000019578",
                "ENSMUSG00000020647",
                "ENSMUSG00000067925",
                "ENSMUSG00000019791",
                "ENSMUSG00000027332",
                "ENSMUSG00000015850",
                "ENSMUSG00000025733",
                "ENSMUSG00000020864",
                "ENSMUSG00000022214",
                "ENSMUSG00000022110",
                "ENSMUSG00000025404",
                "ENSMUSG00000004356",
                "ENSMUSG00000037916",
                "ENSMUSG00000030779",
                "ENSMUSG00000029156",
                "ENSMUSG00000078572",
                "ENSMUSG00000040321",
                "ENSMUSG00000052681",
                "ENSMUSG00000045316",
                "ENSMUSG00000000628",
                "ENSMUSG00000040771",
                "ENSMUSG00000021870",
                "ENSMUSG00000032724",
                "ENSMUSG00000033161",
                "ENSMUSG00000056342",
                "ENSMUSG00000062006",
                "ENSMUSG00000003072",
                "ENSMUSG00000037349",
                "ENSMUSG00000031534",
                "ENSMUSG00000027133",
                "ENSMUSG00000027253",
                "ENSMUSG00000008730",
                "ENSMUSG00000009293",
                "ENSMUSG00000027488",
                "ENSMUSG00000019809",
                "ENSMUSG00000000168",
                "ENSMUSG00000026554",
                "ENSMUSG00000029249",
                "ENSMUSG00000028902",
                "ENSMUSG00000097487",
                "ENSMUSG00000030061",
                "ENSMUSG00000038459",
                "ENSMUSG00000021906",
                "ENSMUSG00000021313",
                "ENSMUSG00000026496",
                "ENSMUSG00000025241",
                "ENSMUSG00000015112",
                "ENSMUSG00000075054",
                "ENSMUSG00000021453",
                "ENSMUSG00000032299",
                "ENSMUSG00000028936",
                "ENSMUSG00000062981",
                "ENSMUSG00000028639",
                "ENSMUSG00000037601",
                "ENSMUSG00000002496",
                "ENSMUSG00000064342",
                "ENSMUSG00000027227",
                "ENSMUSG00000074884",
                "ENSMUSG00000031983",
                "ENSMUSG00000038028",
                "ENSMUSG00000003531",
                "ENSMUSG00000003923",
                "ENSMUSG00000020523",
                "ENSMUSG00000028792",
                "ENSMUSG00000001445",
                "ENSMUSG00000035325",
                "ENSMUSG00000055239",
                "ENSMUSG00000035203",
                "ENSMUSG00000029580",
                "ENSMUSG00000028247",
                "ENSMUSG00000027823",
                "ENSMUSG00000008822",
                "ENSMUSG00000029430",
                "ENSMUSG00000004069",
                "ENSMUSG00000040785",
                "ENSMUSG00000021178",
                "ENSMUSG00000022336",
                "ENSMUSG00000027708",
                "ENSMUSG00000019797",
                "ENSMUSG00000058600",
                "ENSMUSG00000064351",
                "ENSMUSG00000040850",
                "ENSMUSG00000046756",
                "ENSMUSG00000029020",
                "ENSMUSG00000028173",
                "ENSMUSG00000027706",
                "ENSMUSG00000049643",
                "ENSMUSG00000038375",
                "ENSMUSG00000022452",
                "ENSMUSG00000039509",
                "ENSMUSG00000023106",
                "ENSMUSG00000036206",
                "ENSMUSG00000034898",
                "ENSMUSG00000024527",
                "ENSMUSG00000031166",
                "ENSMUSG00000038393",
                "ENSMUSG00000075467",
                "ENSMUSG00000025374",
                "ENSMUSG00000019577",
                "ENSMUSG00000033429",
                "ENSMUSG00000027894",
                "ENSMUSG00000002010",
                "ENSMUSG00000022415",
                "ENSMUSG00000020593",
                "ENSMUSG00000010064",
                "ENSMUSG00000058997",
                "ENSMUSG00000059923",
                "ENSMUSG00000032328",
                "ENSMUSG00000060904",
                "ENSMUSG00000025508",
                "ENSMUSG00000028273",
                "ENSMUSG00000019158",
                "ENSMUSG00000008604",
                "ENSMUSG00000058594",
                "ENSMUSG00000021591",
                "ENSMUSG00000032786",
                "ENSMUSG00000022865",
                "ENSMUSG00000050856",
                "ENSMUSG00000025856",
                "ENSMUSG00000038762",
                "ENSMUSG00000098973",
                "ENSMUSG00000053768",
                "ENSMUSG00000025781",
                "ENSMUSG00000020340",
                "ENSMUSG00000033781",
                "ENSMUSG00000018796",
                "ENSMUSG00000008690",
                "ENSMUSG00000032518",
                "ENSMUSG00000021311",
                "ENSMUSG00000022403",
                "ENSMUSG00000059734",
                "ENSMUSG00000022223",
                "ENSMUSG00000022969",
                "ENSMUSG00000091537",
                "ENSMUSG00000029227",
                "ENSMUSG00000078670",
                "ENSMUSG00000062070",
                "ENSMUSG00000047905",
                "ENSMUSG00000039630",
                "ENSMUSG00000024608",
                "ENSMUSG00000038764",
                "ENSMUSG00000026489",
                "ENSMUSG00000045948",
                "ENSMUSG00000022553",
                "ENSMUSG00000009079",
                "ENSMUSG00000007739",
                "ENSMUSG00000027573",
                "ENSMUSG00000028633",
                "ENSMUSG00000032035",
                "ENSMUSG00000028394",
                "ENSMUSG00000057914",
                "ENSMUSG00000025968",
                "ENSMUSG00000002844",
                "ENSMUSG00000064370",
                "ENSMUSG00000020133",
                "ENSMUSG00000030137",
                "ENSMUSG00000020821",
                "ENSMUSG00000018293",
                "ENSMUSG00000044033",
                "ENSMUSG00000024140",
                "ENSMUSG00000028691",
                "ENSMUSG00000022321",
                "ENSMUSG00000061477",
                "ENSMUSG00000037805",
                "ENSMUSG00000024346",
                "ENSMUSG00000017119",
                "ENSMUSG00000031701",
                "ENSMUSG00000020738",
                "ENSMUSG00000029467",
                "ENSMUSG00000039219",
                "ENSMUSG00000022193",
                "ENSMUSG00000011148",
                "ENSMUSG00000062580",
                "ENSMUSG00000026817",
                "ENSMUSG00000035891",
                "ENSMUSG00000022031",
                "ENSMUSG00000040242",
                "ENSMUSG00000078453",
                "ENSMUSG00000020572",
                "ENSMUSG00000021039",
                "ENSMUSG00000021218",
                "ENSMUSG00000062270",
                "ENSMUSG00000062397",
                "ENSMUSG00000025060",
                "ENSMUSG00000028937",
                "ENSMUSG00000015357",
                "ENSMUSG00000034566",
                "ENSMUSG00000024985",
                "ENSMUSG00000021243",
                "ENSMUSG00000013593",
                "ENSMUSG00000068267",
                "ENSMUSG00000021417",
                "ENSMUSG00000062373",
                "ENSMUSG00000024053",
                "ENSMUSG00000032578",
                "ENSMUSG00000004951",
                "ENSMUSG00000031916",
                "ENSMUSG00000071533",
                "ENSMUSG00000029328",
                "ENSMUSG00000029432",
                "ENSMUSG00000038520",
                "ENSMUSG00000038058",
                "ENSMUSG00000024963",
                "ENSMUSG00000058569",
                "ENSMUSG00000036781",
                "ENSMUSG00000006418",
                "ENSMUSG00000074754",
                "ENSMUSG00000040964",
                "ENSMUSG00000020988",
                "ENSMUSG00000024012",
                "ENSMUSG00000020530",
                "ENSMUSG00000039670",
                "ENSMUSG00000018340",
                "ENSMUSG00000022111",
                "ENSMUSG00000046364",
                "ENSMUSG00000037236",
                "ENSMUSG00000023307",
                "ENSMUSG00000038650",
                "ENSMUSG00000055053",
                "ENSMUSG00000051391",
                "ENSMUSG00000015120",
                "ENSMUSG00000058056",
                "ENSMUSG00000064369",
                "ENSMUSG00000026154",
                "ENSMUSG00000022265",
                "ENSMUSG00000028041",
                "ENSMUSG00000047746",
                "ENSMUSG00000025509",
                "ENSMUSG00000027245",
                "ENSMUSG00000083863",
                "ENSMUSG00000036880",
                "ENSMUSG00000053819",
                "ENSMUSG00000022551",
                "ENSMUSG00000037110",
                "ENSMUSG00000034353",
                "ENSMUSG00000020440",
                "ENSMUSG00000024158",
                "ENSMUSG00000042607",
                "ENSMUSG00000023094",
                "ENSMUSG00000021236",
                "ENSMUSG00000030847",
                "ENSMUSG00000008575",
                "ENSMUSG00000022400",
                "ENSMUSG00000035173",
                "ENSMUSG00000004558",
                "ENSMUSG00000037062",
                "ENSMUSG00000067722",
                "ENSMUSG00000090258",
                "ENSMUSG00000019787",
                "ENSMUSG00000026003",
                "ENSMUSG00000067288",
                "ENSMUSG00000021156",
                "ENSMUSG00000040209",
                "ENSMUSG00000038462",
                "ENSMUSG00000022797",
                "ENSMUSG00000029864",
                "ENSMUSG00000032966",
                "ENSMUSG00000034936",
                "ENSMUSG00000082195",
                "ENSMUSG00000043483",
                "ENSMUSG00000074994",
                "ENSMUSG00000084105",
                "ENSMUSG00000024099",
                "ENSMUSG00000028977",
                "ENSMUSG00000037656",
                "ENSMUSG00000028295",
                "ENSMUSG00000096942",
                "ENSMUSG00000032563",
                "ENSMUSG00000028053",
                "ENSMUSG00000014294",
                "ENSMUSG00000034875",
                "ENSMUSG00000038975",
                "ENSMUSG00000021939",
                "ENSMUSG00000028104",
                "ENSMUSG00000090935",
                "ENSMUSG00000021096",
                "ENSMUSG00000027777",
                "ENSMUSG00000022020",
                "ENSMUSG00000040481",
                "ENSMUSG00000002661",
                "ENSMUSG00000040269",
                "ENSMUSG00000046434",
                "ENSMUSG00000018411",
                "ENSMUSG00000003444",
                "ENSMUSG00000015837",
                "ENSMUSG00000029823",
                "ENSMUSG00000031382",
                "ENSMUSG00000091086",
                "ENSMUSG00000028690",
                "ENSMUSG00000002028",
                "ENSMUSG00000071540",
                "ENSMUSG00000097414",
                "ENSMUSG00000081179",
                "ENSMUSG00000034751",
                "ENSMUSG00000022707",
                "ENSMUSG00000029629",
                "ENSMUSG00000027984",
                "ENSMUSG00000041193",
                "ENSMUSG00000074698",
                "ENSMUSG00000031353",
                "ENSMUSG00000031960",
                "ENSMUSG00000024097",
                "ENSMUSG00000044477",
                "ENSMUSG00000022354",
                "ENSMUSG00000023951",
                "ENSMUSG00000028063",
                "ENSMUSG00000032355",
                "ENSMUSG00000005354",
                "ENSMUSG00000020477",
                "ENSMUSG00000026623",
                "ENSMUSG00000035158",
                "ENSMUSG00000029198",
                "ENSMUSG00000030747",
                "ENSMUSG00000020496",
                "ENSMUSG00000023087",
                "ENSMUSG00000057469",
                "ENSMUSG00000005469",
                "ENSMUSG00000097867",
                "ENSMUSG00000026803",
                "ENSMUSG00000005299",
                "ENSMUSG00000052539",
                "ENSMUSG00000029152",
                "ENSMUSG00000040653",
                "ENSMUSG00000022519",
                "ENSMUSG00000051331",
                "ENSMUSG00000025260",
                "ENSMUSG00000016528",
                "ENSMUSG00000024038",
                "ENSMUSG00000025190",
                "ENSMUSG00000033065",
                "ENSMUSG00000057738",
                "ENSMUSG00000042010",
                "ENSMUSG00000030850",
                "ENSMUSG00000085684",
                "ENSMUSG00000014606",
                "ENSMUSG00000026260",
                "ENSMUSG00000055839",
                "ENSMUSG00000020612",
                "ENSMUSG00000039891",
                "ENSMUSG00000048490",
                "ENSMUSG00000031059",
                "ENSMUSG00000024359",
                "ENSMUSG00000062461",
                "ENSMUSG00000039063",
                "ENSMUSG00000032435",
                "ENSMUSG00000038690",
                "ENSMUSG00000044700",
                "ENSMUSG00000028954",
                "ENSMUSG00000021111",
                "ENSMUSG00000039067",
                "ENSMUSG00000003865",
                "ENSMUSG00000024539",
                "ENSMUSG00000020460",
                "ENSMUSG00000031871",
                "ENSMUSG00000071054",
                "ENSMUSG00000041028",
                "ENSMUSG00000028700",
                "ENSMUSG00000021079",
                "ENSMUSG00000026112",
                "ENSMUSG00000021772",
                "ENSMUSG00000002007",
                "ENSMUSG00000018566",
                "ENSMUSG00000020321",
                "ENSMUSG00000025141",
                "ENSMUSG00000059447",
                "ENSMUSG00000024277",
                "ENSMUSG00000024120",
                "ENSMUSG00000029614",
                "ENSMUSG00000053199",
                "ENSMUSG00000026473",
                "ENSMUSG00000044086",
                "ENSMUSG00000001508",
                "ENSMUSG00000047675",
                "ENSMUSG00000013419",
                "ENSMUSG00000031880",
                "ENSMUSG00000038615",
                "ENSMUSG00000029064",
                "ENSMUSG00000045210",
                "ENSMUSG00000079677",
                "ENSMUSG00000025130",
                "ENSMUSG00000030795",
                "ENSMUSG00000023990",
                "ENSMUSG00000015757",
                "ENSMUSG00000025290",
                "ENSMUSG00000021917",
                "ENSMUSG00000027879",
                "ENSMUSG00000033918",
                "ENSMUSG00000028443",
                "ENSMUSG00000022517",
                "ENSMUSG00000016344",
                "ENSMUSG00000024236",
                "ENSMUSG00000039001",
                "ENSMUSG00000063904",
                "ENSMUSG00000061518",
                "ENSMUSG00000033400",
                "ENSMUSG00000036570",
                "ENSMUSG00000001056",
                "ENSMUSG00000025935",
                "ENSMUSG00000003299",
                "ENSMUSG00000040351",
                "ENSMUSG00000050812",
                "ENSMUSG00000027452",
                "ENSMUSG00000025869",
                "ENSMUSG00000007617",
                "ENSMUSG00000021147",
                "ENSMUSG00000024083",
                "ENSMUSG00000051469",
                "ENSMUSG00000019768",
                "ENSMUSG00000029723",
                "ENSMUSG00000036371",
                "ENSMUSG00000070493",
                "ENSMUSG00000030335",
                "ENSMUSG00000021748",
                "ENSMUSG00000036580",
                "ENSMUSG00000071369",
                "ENSMUSG00000030357",
                "ENSMUSG00000038695",
                "ENSMUSG00000026456",
                "ENSMUSG00000028932",
                "ENSMUSG00000027184",
                "ENSMUSG00000029576",
                "ENSMUSG00000068921",
                "ENSMUSG00000001441",
                "ENSMUSG00000059714",
                "ENSMUSG00000030122",
                "ENSMUSG00000034220",
                "ENSMUSG00000016534",
                "ENSMUSG00000051319",
                "ENSMUSG00000031231",
                "ENSMUSG00000071653",
                "ENSMUSG00000026895",
                "ENSMUSG00000026179",
                "ENSMUSG00000033059",
                "ENSMUSG00000002342",
                "ENSMUSG00000033813",
                "ENSMUSG00000000708",
                "ENSMUSG00000000088",
                "ENSMUSG00000032038",
                "ENSMUSG00000038204",
                "ENSMUSG00000042462",
                "ENSMUSG00000022257",
                "ENSMUSG00000025393",
                "ENSMUSG00000048279",
                "ENSMUSG00000029319",
                "ENSMUSG00000026930",
                "ENSMUSG00000031641",
                "ENSMUSG00000038967",
                "ENSMUSG00000025410",
                "ENSMUSG00000036112",
                "ENSMUSG00000058070",
                "ENSMUSG00000024588",
                "ENSMUSG00000006299",
                "ENSMUSG00000028495",
                "ENSMUSG00000002416",
                "ENSMUSG00000008668",
                "ENSMUSG00000003526",
                "ENSMUSG00000025791",
                "ENSMUSG00000078154",
                "ENSMUSG00000066043",
                "ENSMUSG00000021460",
                "ENSMUSG00000022477",
                "ENSMUSG00000022337",
                "ENSMUSG00000036698",
                "ENSMUSG00000042298",
                "ENSMUSG00000052146",
                "ENSMUSG00000012405",
                "ENSMUSG00000015290",
                "ENSMUSG00000074800",
                "ENSMUSG00000030695",
                "ENSMUSG00000064341",
                "ENSMUSG00000028618",
                "ENSMUSG00000040740",
                "ENSMUSG00000013539",
                "ENSMUSG00000001366",
                "ENSMUSG00000025940",
                "ENSMUSG00000010376",
                "ENSMUSG00000054640",
                "ENSMUSG00000035242",
                "ENSMUSG00000043716",
                "ENSMUSG00000039768",
                "ENSMUSG00000028398",
                "ENSMUSG00000050608",
                "ENSMUSG00000021706",
                "ENSMUSG00000004891",
                "ENSMUSG00000036160",
                "ENSMUSG00000027206",
                "ENSMUSG00000019851",
                "ENSMUSG00000059208",
                "ENSMUSG00000022016",
                "ENSMUSG00000026150",
                "ENSMUSG00000038412",
                "ENSMUSG00000020390",
                "ENSMUSG00000036323",
                "ENSMUSG00000039474",
                "ENSMUSG00000038370",
                "ENSMUSG00000044475",
                "ENSMUSG00000020453",
                "ENSMUSG00000029128",
                "ENSMUSG00000029474",
                "ENSMUSG00000099250",
                "ENSMUSG00000013736",
                "ENSMUSG00000082884",
                "ENSMUSG00000055737",
                "ENSMUSG00000063870",
                "ENSMUSG00000020982",
                "ENSMUSG00000032902",
                "ENSMUSG00000031633",
                "ENSMUSG00000063694",
                "ENSMUSG00000068487",
                "ENSMUSG00000020368",
                "ENSMUSG00000039473",
                "ENSMUSG00000092014",
                "ENSMUSG00000019795",
                "ENSMUSG00000040385",
                "ENSMUSG00000038086",
                "ENSMUSG00000016252",
                "ENSMUSG00000041571",
                "ENSMUSG00000084178",
                "ENSMUSG00000082874",
                "ENSMUSG00000022889",
                "ENSMUSG00000019528",
                "ENSMUSG00000021843",
                "ENSMUSG00000039611",
                "ENSMUSG00000067274",
                "ENSMUSG00000063787",
                "ENSMUSG00000063882",
                "ENSMUSG00000028607",
                "ENSMUSG00000039105",
                "ENSMUSG00000026028",
                "ENSMUSG00000026078",
                "ENSMUSG00000039910",
                "ENSMUSG00000021557",
                "ENSMUSG00000028745",
                "ENSMUSG00000038886",
                "ENSMUSG00000053702",
                "ENSMUSG00000020267",
                "ENSMUSG00000022051",
                "ENSMUSG00000006818",
                "ENSMUSG00000026860",
                "ENSMUSG00000076431",
                "ENSMUSG00000042541",
                "ENSMUSG00000094344",
                "ENSMUSG00000069601",
                "ENSMUSG00000028692",
                "ENSMUSG00000031928",
                "ENSMUSG00000021375",
                "ENSMUSG00000060739",
                "ENSMUSG00000033039",
                "ENSMUSG00000006931",
                "ENSMUSG00000025499",
                "ENSMUSG00000032399",
                "ENSMUSG00000059586",
                "ENSMUSG00000078566",
                "ENSMUSG00000002102",
                "ENSMUSG00000021390",
                "ENSMUSG00000002343",
                "ENSMUSG00000039652",
                "ENSMUSG00000024613",
                "ENSMUSG00000027472",
                "ENSMUSG00000054889",
                "ENSMUSG00000070544",
                "ENSMUSG00000046351",
                "ENSMUSG00000020166",
                "ENSMUSG00000024953",
                "ENSMUSG00000027774",
                "ENSMUSG00000022982",
                "ENSMUSG00000058558",
                "ENSMUSG00000026020",
                "ENSMUSG00000050323",
                "ENSMUSG00000026032",
                "ENSMUSG00000068479",
                "ENSMUSG00000021868",
                "ENSMUSG00000037376",
                "ENSMUSG00000028452",
                "ENSMUSG00000028156",
                "ENSMUSG00000032407",
                "ENSMUSG00000039987",
                "ENSMUSG00000041548",
                "ENSMUSG00000044502",
                "ENSMUSG00000070802",
                "ENSMUSG00000024576",
                "ENSMUSG00000035478",
                "ENSMUSG00000028333",
                "ENSMUSG00000041058",
                "ENSMUSG00000036202",
                "ENSMUSG00000032112",
                "ENSMUSG00000027257",
                "ENSMUSG00000020372",
                "ENSMUSG00000032523",
                "ENSMUSG00000028834",
                "ENSMUSG00000031352",
                "ENSMUSG00000073838",
                "ENSMUSG00000041870",
                "ENSMUSG00000039536",
                "ENSMUSG00000000740",
                "ENSMUSG00000040752",
                "ENSMUSG00000023723",
                "ENSMUSG00000003934",
                "ENSMUSG00000030884",
                "ENSMUSG00000025466",
                "ENSMUSG00000018567",
                "ENSMUSG00000051518",
                "ENSMUSG00000022000",
                "ENSMUSG00000023025",
                "ENSMUSG00000021476",
                "ENSMUSG00000032051",
                "ENSMUSG00000034040",
                "ENSMUSG00000023460",
                "ENSMUSG00000002767",
                "ENSMUSG00000034994",
                "ENSMUSG00000020108",
                "ENSMUSG00000030541",
                "ENSMUSG00000031438",
                "ENSMUSG00000026208",
                "ENSMUSG00000031696",
                "ENSMUSG00000090266",
                "ENSMUSG00000036052",
                "ENSMUSG00000015217",
                "ENSMUSG00000024213",
                "ENSMUSG00000030652",
                "ENSMUSG00000089917",
                "ENSMUSG00000027238",
                "ENSMUSG00000004364",
                "ENSMUSG00000047388",
                "ENSMUSG00000021109",
                "ENSMUSG00000074412",
                "ENSMUSG00000061306",
                "ENSMUSG00000024869",
                "ENSMUSG00000017686",
                "ENSMUSG00000033307",
                "ENSMUSG00000057406",
                "ENSMUSG00000046580",
                "ENSMUSG00000069089",
                "ENSMUSG00000033543",
                "ENSMUSG00000020415",
                "ENSMUSG00000024146",
                "ENSMUSG00000006800",
                "ENSMUSG00000002957",
                "ENSMUSG00000041957",
                "ENSMUSG00000024073",
                "ENSMUSG00000028282",
                "ENSMUSG00000062997",
                "ENSMUSG00000081431",
                "ENSMUSG00000020430",
                "ENSMUSG00000009112",
                "ENSMUSG00000031311",
                "ENSMUSG00000026603",
                "ENSMUSG00000027180",
                "ENSMUSG00000016308",
                "ENSMUSG00000027422",
                "ENSMUSG00000050043",
                "ENSMUSG00000018882",
                "ENSMUSG00000028470",
                "ENSMUSG00000000787",
                "ENSMUSG00000015214",
                "ENSMUSG00000037710",
                "ENSMUSG00000024187",
                "ENSMUSG00000019362",
                "ENSMUSG00000016427",
                "ENSMUSG00000038240",
                "ENSMUSG00000027293",
                "ENSMUSG00000003948",
                "ENSMUSG00000019055",
                "ENSMUSG00000003559",
                "ENSMUSG00000001127",
                "ENSMUSG00000015656",
                "ENSMUSG00000025868",
                "ENSMUSG00000040520",
                "ENSMUSG00000058756",
                "ENSMUSG00000025651",
                "ENSMUSG00000028603",
                "ENSMUSG00000071037",
                "ENSMUSG00000037887",
                "ENSMUSG00000028820",
                "ENSMUSG00000004268",
                "ENSMUSG00000025085",
                "ENSMUSG00000035329",
                "ENSMUSG00000065979",
                "ENSMUSG00000028261",
                "ENSMUSG00000024259",
                "ENSMUSG00000025068",
                "ENSMUSG00000045103",
                "ENSMUSG00000022205",
                "ENSMUSG00000034312",
                "ENSMUSG00000032602",
                "ENSMUSG00000032042",
                "ENSMUSG00000061032",
                "ENSMUSG00000023089",
                "ENSMUSG00000019843",
                "ENSMUSG00000020225",
                "ENSMUSG00000073676",
                "ENSMUSG00000032060",
                "ENSMUSG00000023456",
                "ENSMUSG00000022010",
                "ENSMUSG00000027673",
                "ENSMUSG00000083878",
                "ENSMUSG00000059811",
                "ENSMUSG00000013662",
                "ENSMUSG00000026914",
                "ENSMUSG00000083757",
                "ENSMUSG00000024587",
                "ENSMUSG00000024121",
                "ENSMUSG00000029616",
                "ENSMUSG00000020003",
                "ENSMUSG00000019659",
                "ENSMUSG00000061904",
                "ENSMUSG00000027305",
                "ENSMUSG00000029454",
                "ENSMUSG00000083240",
                "ENSMUSG00000004937",
                "ENSMUSG00000015605",
                "ENSMUSG00000020485",
                "ENSMUSG00000006740",
                "ENSMUSG00000024197",
                "ENSMUSG00000007777",
                "ENSMUSG00000019432",
                "ENSMUSG00000022914",
                "ENSMUSG00000063457",
                "ENSMUSG00000008859",
                "ENSMUSG00000020088",
                "ENSMUSG00000039148",
                "ENSMUSG00000058440",
                "ENSMUSG00000019689",
                "ENSMUSG00000020741",
                "ENSMUSG00000037266",
                "ENSMUSG00000038095",
                "ENSMUSG00000037339",
                "ENSMUSG00000030982",
                "ENSMUSG00000027679",
                "ENSMUSG00000041841",
                "ENSMUSG00000026576",
                "ENSMUSG00000041881",
                "ENSMUSG00000021764",
                "ENSMUSG00000038763",
                "ENSMUSG00000043445",
                "ENSMUSG00000039648",
                "ENSMUSG00000020163",
                "ENSMUSG00000001767",
                "ENSMUSG00000009863",
                "ENSMUSG00000090553",
                "ENSMUSG00000031770",
                "ENSMUSG00000038400",
                "ENSMUSG00000030591",
                "ENSMUSG00000036309",
                "ENSMUSG00000031715",
                "ENSMUSG00000028945",
                "ENSMUSG00000056900",
                "ENSMUSG00000032238",
                "ENSMUSG00000062210",
                "ENSMUSG00000060376",
                "ENSMUSG00000014592",
                "ENSMUSG00000090150",
                "ENSMUSG00000017428",
                "ENSMUSG00000079508",
                "ENSMUSG00000081051",
                "ENSMUSG00000062694",
                "ENSMUSG00000020153",
                "ENSMUSG00000093674",
                "ENSMUSG00000083563",
                "ENSMUSG00000022312",
                "ENSMUSG00000040760",
                "ENSMUSG00000069729",
                "ENSMUSG00000036452",
                "ENSMUSG00000031818",
                "ENSMUSG00000031782",
                "ENSMUSG00000048486",
                "ENSMUSG00000055681",
                "ENSMUSG00000032511",
                "ENSMUSG00000038526",
                "ENSMUSG00000024683",
                "ENSMUSG00000096449",
                "ENSMUSG00000044894",
                "ENSMUSG00000007097",
                "ENSMUSG00000029404",
                "ENSMUSG00000024937",
                "ENSMUSG00000056032",
                "ENSMUSG00000022820",
                "ENSMUSG00000037234",
                "ENSMUSG00000042148",
                "ENSMUSG00000034295",
                "ENSMUSG00000031783",
                "ENSMUSG00000022594",
                "ENSMUSG00000063229",
                "ENSMUSG00000001604",
                "ENSMUSG00000005417",
                "ENSMUSG00000021577",
                "ENSMUSG00000038936",
                "ENSMUSG00000029776",
                "ENSMUSG00000029135",
                "ENSMUSG00000025898",
                "ENSMUSG00000036427",
                "ENSMUSG00000070867",
                "ENSMUSG00000038417",
                "ENSMUSG00000045160",
                "ENSMUSG00000024942",
                "ENSMUSG00000022698",
                "ENSMUSG00000024335",
                "ENSMUSG00000027371",
                "ENSMUSG00000028743",
                "ENSMUSG00000037075",
                "ENSMUSG00000040407",
                "ENSMUSG00000030647",
                "ENSMUSG00000073664",
                "ENSMUSG00000025545",
                "ENSMUSG00000045503",
                "ENSMUSG00000091831",
                "ENSMUSG00000026888",
                "ENSMUSG00000022094",
                "ENSMUSG00000052712",
                "ENSMUSG00000030087",
                "ENSMUSG00000031776",
                "ENSMUSG00000006333",
                "ENSMUSG00000026568",
                "ENSMUSG00000028328",
                "ENSMUSG00000068328",
                "ENSMUSG00000005683",
                "ENSMUSG00000029106",
                "ENSMUSG00000031700",
                "ENSMUSG00000046516",
                "ENSMUSG00000022332",
                "ENSMUSG00000040112",
                "ENSMUSG00000037960",
                "ENSMUSG00000058655",
                "ENSMUSG00000022961",
                "ENSMUSG00000020085",
                "ENSMUSG00000031536",
                "ENSMUSG00000032348",
                "ENSMUSG00000039218",
                "ENSMUSG00000021133",
                "ENSMUSG00000029594",
                "ENSMUSG00000023286",
                "ENSMUSG00000031516",
                "ENSMUSG00000052214",
                "ENSMUSG00000033228",
                "ENSMUSG00000046032",
                "ENSMUSG00000032418",
                "ENSMUSG00000032826",
                "ENSMUSG00000028266",
                "ENSMUSG00000081382",
                "ENSMUSG00000026064",
                "ENSMUSG00000047371",
                "ENSMUSG00000029048",
                "ENSMUSG00000022848",
                "ENSMUSG00000020737",
                "ENSMUSG00000031232",
                "ENSMUSG00000001419",
                "ENSMUSG00000051671",
                "ENSMUSG00000022765",
                "ENSMUSG00000026360",
                "ENSMUSG00000047260",
                "ENSMUSG00000004460",
                "ENSMUSG00000042198",
                "ENSMUSG00000035674",
                "ENSMUSG00000009927",
                "ENSMUSG00000083832",
                "ENSMUSG00000026571",
                "ENSMUSG00000025465",
                "ENSMUSG00000042605",
                "ENSMUSG00000028919",
                "ENSMUSG00000040990",
                "ENSMUSG00000027423",
                "ENSMUSG00000001833",
                "ENSMUSG00000025204",
                "ENSMUSG00000023055",
                "ENSMUSG00000098178",
                "ENSMUSG00000031672",
                "ENSMUSG00000075704",
                "ENSMUSG00000031660",
                "ENSMUSG00000040713",
                "ENSMUSG00000026977",
                "ENSMUSG00000030172",
                "ENSMUSG00000059895",
                "ENSMUSG00000016319",
                "ENSMUSG00000030706",
                "ENSMUSG00000024780",
                "ENSMUSG00000021488",
                "ENSMUSG00000002948",
                "ENSMUSG00000015575",
                "ENSMUSG00000034290",
                "ENSMUSG00000005981",
                "ENSMUSG00000023150",
                "ENSMUSG00000022216",
                "ENSMUSG00000063800",
                "ENSMUSG00000030298",
                "ENSMUSG00000025795",
                "ENSMUSG00000036181",
                "ENSMUSG00000028291",
                "ENSMUSG00000017404",
                "ENSMUSG00000020190",
                "ENSMUSG00000021944",
                "ENSMUSG00000028648",
                "ENSMUSG00000026991",
                "ENSMUSG00000029623",
                "ENSMUSG00000018848",
                "ENSMUSG00000004535",
                "ENSMUSG00000014633",
                "ENSMUSG00000007892",
                "ENSMUSG00000003380",
                "ENSMUSG00000074064",
                "ENSMUSG00000060938",
                "ENSMUSG00000010406",
                "ENSMUSG00000030607",
                "ENSMUSG00000025428",
                "ENSMUSG00000024538",
                "ENSMUSG00000021585",
                "ENSMUSG00000029017",
                "ENSMUSG00000030016",
                "ENSMUSG00000041426",
                "ENSMUSG00000031634",
                "ENSMUSG00000020849",
                "ENSMUSG00000068706",
                "ENSMUSG00000002748",
                "ENSMUSG00000041297",
                "ENSMUSG00000026766",
                "ENSMUSG00000024924",
                "ENSMUSG00000042744",
                "ENSMUSG00000031987",
                "ENSMUSG00000040952",
                "ENSMUSG00000043311",
                "ENSMUSG00000038970",
                "ENSMUSG00000028367",
                "ENSMUSG00000055302",
                "ENSMUSG00000028013",
                "ENSMUSG00000000346",
                "ENSMUSG00000039089",
                "ENSMUSG00000019782",
                "ENSMUSG00000026687",
                "ENSMUSG00000053898",
                "ENSMUSG00000060038",
                "ENSMUSG00000030409",
                "ENSMUSG00000062691",
                "ENSMUSG00000083820",
                "ENSMUSG00000001105",
                "ENSMUSG00000025221",
                "ENSMUSG00000029632",
                "ENSMUSG00000026207",
                "ENSMUSG00000036620",
                "ENSMUSG00000022641",
                "ENSMUSG00000054766",
                "ENSMUSG00000015846",
                "ENSMUSG00000052337",
                "ENSMUSG00000028223",
                "ENSMUSG00000031262",
                "ENSMUSG00000026342",
                "ENSMUSG00000025920",
                "ENSMUSG00000074466",
                "ENSMUSG00000001794",
                "ENSMUSG00000031791",
                "ENSMUSG00000048264",
                "ENSMUSG00000037820",
                "ENSMUSG00000071644",
                "ENSMUSG00000090000",
                "ENSMUSG00000026027",
                "ENSMUSG00000020785",
                "ENSMUSG00000030615",
                "ENSMUSG00000053012",
                "ENSMUSG00000020483",
                "ENSMUSG00000009090",
                "ENSMUSG00000004748",
                "ENSMUSG00000072692",
                "ENSMUSG00000032679",
                "ENSMUSG00000026878",
                "ENSMUSG00000024055",
                "ENSMUSG00000029518",
                "ENSMUSG00000006304",
                "ENSMUSG00000031320",
                "ENSMUSG00000032959",
                "ENSMUSG00000028964",
                "ENSMUSG00000025893",
                "ENSMUSG00000042032",
                "ENSMUSG00000060802",
                "ENSMUSG00000002728",
                "ENSMUSG00000027406",
                "ENSMUSG00000060032",
                "ENSMUSG00000025487",
                "ENSMUSG00000078784",
                "ENSMUSG00000028756",
                "ENSMUSG00000063273",
                "ENSMUSG00000060675",
                "ENSMUSG00000051154",
                "ENSMUSG00000020029",
                "ENSMUSG00000037712",
                "ENSMUSG00000034708",
                "ENSMUSG00000036932",
                "ENSMUSG00000003955",
                "ENSMUSG00000028234",
                "ENSMUSG00000061838",
                "ENSMUSG00000019179",
                "ENSMUSG00000051495",
                "ENSMUSG00000079477",
                "ENSMUSG00000022841",
                "ENSMUSG00000000253",
                "ENSMUSG00000071415",
                "ENSMUSG00000037152",
                "ENSMUSG00000019494",
                "ENSMUSG00000020708",
                "ENSMUSG00000023010",
                "ENSMUSG00000036278",
                "ENSMUSG00000022391",
                "ENSMUSG00000028788",
                "ENSMUSG00000062169",
                "ENSMUSG00000053907",
                "ENSMUSG00000067369",
                "ENSMUSG00000003316",
                "ENSMUSG00000022629",
                "ENSMUSG00000002980",
                "ENSMUSG00000023806",
                "ENSMUSG00000005510",
                "ENSMUSG00000020547",
                "ENSMUSG00000024208",
                "ENSMUSG00000020232",
                "ENSMUSG00000024507",
                "ENSMUSG00000049832",
                "ENSMUSG00000029469",
                "ENSMUSG00000087260",
                "ENSMUSG00000038374",
                "ENSMUSG00000019428",
                "ENSMUSG00000024803",
                "ENSMUSG00000022255",
                "ENSMUSG00000025967",
                "ENSMUSG00000005034",
                "ENSMUSG00000031683",
                "ENSMUSG00000036822",
                "ENSMUSG00000022442",
                "ENSMUSG00000028248",
                "ENSMUSG00000078812",
                "ENSMUSG00000082474",
                "ENSMUSG00000001158",
                "ENSMUSG00000090733",
                "ENSMUSG00000036352",
                "ENSMUSG00000059534",
                "ENSMUSG00000038467",
                "ENSMUSG00000027367",
                "ENSMUSG00000029108",
                "ENSMUSG00000007338",
                "ENSMUSG00000080836",
                "ENSMUSG00000044308",
                "ENSMUSG00000053536",
                "ENSMUSG00000022300",
                "ENSMUSG00000003429",
                "ENSMUSG00000021185",
                "ENSMUSG00000018599",
                "ENSMUSG00000037822",
                "ENSMUSG00000084319",
                "ENSMUSG00000064344",
                "ENSMUSG00000029095",
                "ENSMUSG00000015839",
                "ENSMUSG00000027499",
                "ENSMUSG00000027784",
                "ENSMUSG00000065990",
                "ENSMUSG00000060438",
                "ENSMUSG00000003541",
                "ENSMUSG00000029068",
                "ENSMUSG00000064340",
                "ENSMUSG00000032047",
                "ENSMUSG00000042133",
                "ENSMUSG00000044393",
                "ENSMUSG00000014470",
                "ENSMUSG00000031586",
                "ENSMUSG00000001472",
                "ENSMUSG00000048076",
                "ENSMUSG00000036002",
                "ENSMUSG00000072640",
                "ENSMUSG00000038582",
                "ENSMUSG00000026864",
                "ENSMUSG00000023939",
                "ENSMUSG00000035885",
                "ENSMUSG00000059325",
                "ENSMUSG00000031078",
                "ENSMUSG00000026341",
                "ENSMUSG00000038900",
                "ENSMUSG00000032431",
                "ENSMUSG00000032330",
                "ENSMUSG00000022023",
                "ENSMUSG00000038170",
                "ENSMUSG00000006315",
                "ENSMUSG00000078713",
                "ENSMUSG00000051695",
                "ENSMUSG00000049090",
                "ENSMUSG00000041361",
                "ENSMUSG00000049327",
                "ENSMUSG00000006932",
                "ENSMUSG00000091449",
                "ENSMUSG00000022748",
                "ENSMUSG00000060981",
                "ENSMUSG00000031865",
                "ENSMUSG00000030089",
                "ENSMUSG00000000149",
                "ENSMUSG00000023206",
                "ENSMUSG00000028518",
                "ENSMUSG00000074129",
                "ENSMUSG00000031839",
                "ENSMUSG00000001175",
                "ENSMUSG00000038084",
                "ENSMUSG00000008136",
                "ENSMUSG00000049881",
                "ENSMUSG00000005873",
                "ENSMUSG00000054405",
                "ENSMUSG00000030744",
                "ENSMUSG00000029019",
                "ENSMUSG00000023944",
                "ENSMUSG00000015932",
                "ENSMUSG00000029780",
                "ENSMUSG00000020358",
                "ENSMUSG00000022681",
                "ENSMUSG00000068699",
                "ENSMUSG00000003970",
                "ENSMUSG00000030662",
                "ENSMUSG00000072946",
                "ENSMUSG00000010911",
                "ENSMUSG00000050953",
                "ENSMUSG00000027765",
                "ENSMUSG00000020308",
                "ENSMUSG00000021546",
                "ENSMUSG00000064372",
                "ENSMUSG00000030002",
                "ENSMUSG00000032495",
                "ENSMUSG00000028772",
                "ENSMUSG00000025034",
                "ENSMUSG00000025794",
                "ENSMUSG00000046997",
                "ENSMUSG00000020022",
                "ENSMUSG00000034088",
                "ENSMUSG00000071655",
                "ENSMUSG00000001911",
                "ENSMUSG00000034832",
                "ENSMUSG00000020664",
                "ENSMUSG00000022389",
                "ENSMUSG00000075595",
                "ENSMUSG00000033722",
                "ENSMUSG00000040118",
                "ENSMUSG00000029203",
                "ENSMUSG00000022635",
                "ENSMUSG00000067851",
                "ENSMUSG00000018770",
                "ENSMUSG00000052911",
                "ENSMUSG00000039782",
                "ENSMUSG00000097088",
                "ENSMUSG00000035781",
                "ENSMUSG00000087380",
                "ENSMUSG00000067736",
                "ENSMUSG00000028737",
                "ENSMUSG00000031068",
                "ENSMUSG00000031954",
                "ENSMUSG00000030621",
                "ENSMUSG00000056602",
                "ENSMUSG00000062908",
                "ENSMUSG00000022253",
                "ENSMUSG00000021057",
                "ENSMUSG00000068264",
                "ENSMUSG00000025613",
                "ENSMUSG00000022462",
                "ENSMUSG00000027447",
                "ENSMUSG00000021993",
                "ENSMUSG00000031533",
                "ENSMUSG00000029390",
                "ENSMUSG00000000355",
                "ENSMUSG00000020290",
                "ENSMUSG00000037032",
                "ENSMUSG00000024283",
                "ENSMUSG00000036585",
                "ENSMUSG00000029394",
                "ENSMUSG00000030319",
                "ENSMUSG00000059070",
                "ENSMUSG00000062647",
                "ENSMUSG00000025492",
                "ENSMUSG00000062083",
                "ENSMUSG00000099021",
                "ENSMUSG00000042709",
                "ENSMUSG00000033916",
                "ENSMUSG00000032525",
                "ENSMUSG00000020402",
                "ENSMUSG00000060073",
                "ENSMUSG00000022217",
                "ENSMUSG00000026853",
                "ENSMUSG00000034880",
                "ENSMUSG00000054428",
                "ENSMUSG00000066900",
                "ENSMUSG00000028959",
                "ENSMUSG00000008035",
                "ENSMUSG00000026021",
                "ENSMUSG00000019920",
                "ENSMUSG00000014551",
                "ENSMUSG00000021270",
                "ENSMUSG00000011257",
                "ENSMUSG00000021930",
                "ENSMUSG00000031840",
                "ENSMUSG00000039943",
                "ENSMUSG00000006576",
                "ENSMUSG00000038914",
                "ENSMUSG00000064037",
                "ENSMUSG00000004085",
                "ENSMUSG00000056851",
                "ENSMUSG00000028062",
                "ENSMUSG00000014426",
                "ENSMUSG00000020048",
                "ENSMUSG00000063316",
                "ENSMUSG00000022867",
                "ENSMUSG00000028447",
                "ENSMUSG00000048277",
                "ENSMUSG00000020705",
                "ENSMUSG00000038538",
                "ENSMUSG00000084786",
                "ENSMUSG00000020634",
                "ENSMUSG00000036678",
                "ENSMUSG00000030861",
                "ENSMUSG00000028161",
                "ENSMUSG00000018102",
                "ENSMUSG00000029056",
                "ENSMUSG00000025809",
                "ENSMUSG00000026688",
                "ENSMUSG00000015092",
                "ENSMUSG00000029833",
                "ENSMUSG00000034574",
                "ENSMUSG00000040666",
                "ENSMUSG00000025184",
                "ENSMUSG00000003809",
                "ENSMUSG00000010175",
                "ENSMUSG00000000399",
                "ENSMUSG00000004931",
                "ENSMUSG00000044792",
                "ENSMUSG00000039976",
                "ENSMUSG00000022437",
                "ENSMUSG00000051343",
                "ENSMUSG00000019872",
                "ENSMUSG00000031708",
                "ENSMUSG00000026353",
                "ENSMUSG00000014504",
                "ENSMUSG00000022037",
                "ENSMUSG00000022035",
                "ENSMUSG00000033938",
                "ENSMUSG00000024997",
                "ENSMUSG00000057657",
                "ENSMUSG00000017778",
                "ENSMUSG00000020952",
                "ENSMUSG00000048546",
                "ENSMUSG00000035105",
                "ENSMUSG00000030753",
                "ENSMUSG00000073639",
                "ENSMUSG00000045776",
                "ENSMUSG00000020580",
                "ENSMUSG00000020671",
                "ENSMUSG00000028207",
                "ENSMUSG00000040414",
                "ENSMUSG00000001289",
                "ENSMUSG00000021537",
                "ENSMUSG00000060798",
                "ENSMUSG00000036564",
                "ENSMUSG00000074093",
                "ENSMUSG00000037049",
                "ENSMUSG00000020079",
                "ENSMUSG00000054452",
                "ENSMUSG00000002981",
                "ENSMUSG00000024150",
                "ENSMUSG00000020585",
                "ENSMUSG00000040694",
                "ENSMUSG00000012848",
                "ENSMUSG00000032480",
                "ENSMUSG00000022800",
                "ENSMUSG00000000184",
                "ENSMUSG00000091383",
                "ENSMUSG00000010122",
                "ENSMUSG00000050541",
                "ENSMUSG00000038679",
                "ENSMUSG00000031447",
                "ENSMUSG00000020621",
                "ENSMUSG00000008682",
                "ENSMUSG00000034157",
                "ENSMUSG00000020180",
                "ENSMUSG00000031765",
                "ENSMUSG00000023020",
                "ENSMUSG00000024293",
                "ENSMUSG00000045201",
                "ENSMUSG00000052934",
                "ENSMUSG00000030421",
                "ENSMUSG00000033382",
                "ENSMUSG00000034424",
                "ENSMUSG00000071793",
                "ENSMUSG00000031521",
                "ENSMUSG00000064366",
                "ENSMUSG00000019558",
                "ENSMUSG00000031029",
                "ENSMUSG00000031441",
                "ENSMUSG00000029145",
                "ENSMUSG00000006494",
                "ENSMUSG00000021102",
                "ENSMUSG00000023832",
                "ENSMUSG00000005220",
                "ENSMUSG00000004980",
                "ENSMUSG00000005625",
                "ENSMUSG00000032580",
                "ENSMUSG00000071014",
                "ENSMUSG00000044864",
                "ENSMUSG00000049751",
                "ENSMUSG00000025666",
                "ENSMUSG00000021373",
                "ENSMUSG00000031201",
                "ENSMUSG00000000171",
                "ENSMUSG00000000441",
                "ENSMUSG00000041685",
                "ENSMUSG00000034620",
                "ENSMUSG00000038784",
                "ENSMUSG00000024661",
                "ENSMUSG00000036748",
                "ENSMUSG00000053483",
                "ENSMUSG00000025156",
                "ENSMUSG00000002608",
                "ENSMUSG00000025103",
                "ENSMUSG00000039601",
                "ENSMUSG00000057322",
                "ENSMUSG00000073889",
                "ENSMUSG00000022270",
                "ENSMUSG00000026234",
                "ENSMUSG00000029059",
                "ENSMUSG00000035783",
                "ENSMUSG00000045690",
                "ENSMUSG00000032526",
                "ENSMUSG00000038132",
                "ENSMUSG00000000340",
                "ENSMUSG00000037949",
                "ENSMUSG00000055762",
                "ENSMUSG00000034868",
                "ENSMUSG00000064343",
                "ENSMUSG00000021709",
                "ENSMUSG00000043323",
                "ENSMUSG00000022186",
                "ENSMUSG00000021597",
                "ENSMUSG00000056091",
                "ENSMUSG00000045624",
                "ENSMUSG00000086567",
                "ENSMUSG00000064337",
                "ENSMUSG00000020116",
                "ENSMUSG00000071528",
                "ENSMUSG00000031913",
                "ENSMUSG00000071654",
                "ENSMUSG00000030846",
                "ENSMUSG00000021905",
                "ENSMUSG00000031841",
                "ENSMUSG00000061130",
                "ENSMUSG00000036854",
                "ENSMUSG00000026131",
                "ENSMUSG00000026202",
                "ENSMUSG00000020721",
                "ENSMUSG00000027566",
                "ENSMUSG00000049960",
                "ENSMUSG00000048175",
                "ENSMUSG00000020326",
                "ENSMUSG00000031751",
                "ENSMUSG00000032010",
                "ENSMUSG00000041278",
                "ENSMUSG00000021838",
                "ENSMUSG00000062279",
                "ENSMUSG00000031996",
                "ENSMUSG00000067279",
                "ENSMUSG00000064347",
                "ENSMUSG00000023277",
                "ENSMUSG00000034892",
                "ENSMUSG00000017756",
                "ENSMUSG00000023092",
                "ENSMUSG00000059291",
                "ENSMUSG00000032177",
                "ENSMUSG00000003420",
                "ENSMUSG00000002379",
                "ENSMUSG00000027177",
                "ENSMUSG00000039087",
                "ENSMUSG00000029701",
                "ENSMUSG00000052912",
                "ENSMUSG00000029455",
                "ENSMUSG00000046727",
                "ENSMUSG00000064346",
                "ENSMUSG00000058690",
                "ENSMUSG00000027010",
                "ENSMUSG00000049606",
                "ENSMUSG00000030876",
                "ENSMUSG00000060147",
                "ENSMUSG00000059852",
                "ENSMUSG00000051285",
                "ENSMUSG00000024063",
                "ENSMUSG00000037270",
                "ENSMUSG00000040370",
                "ENSMUSG00000074218",
                "ENSMUSG00000021371",
                "ENSMUSG00000042323",
                "ENSMUSG00000030417",
                "ENSMUSG00000026674",
                "ENSMUSG00000034345",
                "ENSMUSG00000037818",
                "ENSMUSG00000037826",
                "ENSMUSG00000029651",
                "ENSMUSG00000023921",
                "ENSMUSG00000032126",
                "ENSMUSG00000057649",
                "ENSMUSG00000021000",
                "ENSMUSG00000042650",
                "ENSMUSG00000000563",
                "ENSMUSG00000065037",
                "ENSMUSG00000037815",
                "ENSMUSG00000027809",
                "ENSMUSG00000047888",
                "ENSMUSG00000034342",
                "ENSMUSG00000038949",
                "ENSMUSG00000047187",
                "ENSMUSG00000061462",
                "ENSMUSG00000037958",
                "ENSMUSG00000031024",
                "ENSMUSG00000020456",
                "ENSMUSG00000021061",
                "ENSMUSG00000045983",
                "ENSMUSG00000032312",
                "ENSMUSG00000050627",
                "ENSMUSG00000051747",
                "ENSMUSG00000025911",
                "ENSMUSG00000035458",
                "ENSMUSG00000032030",
                "ENSMUSG00000032481",
                "ENSMUSG00000022040",
                "ENSMUSG00000028180",
                "ENSMUSG00000026824",
                "ENSMUSG00000022013",
                "ENSMUSG00000081400",
                "ENSMUSG00000081331",
                "ENSMUSG00000011305",
                "ENSMUSG00000014226",
                "ENSMUSG00000006356",
                "ENSMUSG00000039197",
                "ENSMUSG00000022890",
                "ENSMUSG00000053062",
                "ENSMUSG00000019810",
                "ENSMUSG00000024993",
                "ENSMUSG00000022684",
                "ENSMUSG00000032199",
                "ENSMUSG00000001211",
                "ENSMUSG00000056692",
                "ENSMUSG00000070436",
                "ENSMUSG00000052738",
                "ENSMUSG00000031985",
                "ENSMUSG00000029918",
                "ENSMUSG00000029359",
                "ENSMUSG00000023175",
                "ENSMUSG00000052374",
                "ENSMUSG00000044627",
                "ENSMUSG00000015176",
                "ENSMUSG00000024991",
                "ENSMUSG00000037523",
                "ENSMUSG00000024687",
                "ENSMUSG00000022201",
                "ENSMUSG00000027030",
                "ENSMUSG00000080893",
                "ENSMUSG00000021226",
                "ENSMUSG00000022940",
                "ENSMUSG00000030934",
                "ENSMUSG00000032590",
                "ENSMUSG00000026385",
                "ENSMUSG00000041444",
                "ENSMUSG00000020859",
                "ENSMUSG00000024309",
                "ENSMUSG00000050199",
                "ENSMUSG00000025362",
                "ENSMUSG00000058239",
                "ENSMUSG00000071604",
                "ENSMUSG00000022964",
                "ENSMUSG00000082687",
                "ENSMUSG00000029836",
                "ENSMUSG00000025871",
                "ENSMUSG00000025364",
                "ENSMUSG00000038612",
                "ENSMUSG00000042670",
                "ENSMUSG00000021807",
                "ENSMUSG00000022803",
                "ENSMUSG00000064338",
                "ENSMUSG00000061104",
                "ENSMUSG00000000738",
                "ENSMUSG00000039100",
                "ENSMUSG00000009621",
                "ENSMUSG00000044080",
                "ENSMUSG00000046671",
                "ENSMUSG00000024442",
                "ENSMUSG00000004096",
                "ENSMUSG00000028035",
                "ENSMUSG00000047459",
                "ENSMUSG00000029649",
                "ENSMUSG00000022528",
                "ENSMUSG00000060679",
                "ENSMUSG00000038844",
                "ENSMUSG00000052698",
                "ENSMUSG00000005686",
                "ENSMUSG00000048416",
                "ENSMUSG00000053110",
                "ENSMUSG00000010608",
                "ENSMUSG00000027742",
                "ENSMUSG00000032279",
                "ENSMUSG00000033788",
                "ENSMUSG00000086324",
                "ENSMUSG00000023861",
                "ENSMUSG00000067847",
                "ENSMUSG00000026664",
                "ENSMUSG00000068523",
                "ENSMUSG00000043822",
                "ENSMUSG00000021520",
                "ENSMUSG00000006717",
                "ENSMUSG00000001666",
                "ENSMUSG00000090862",
                "ENSMUSG00000036810",
                "ENSMUSG00000008683",
                "ENSMUSG00000003438",
                "ENSMUSG00000042099",
                "ENSMUSG00000083219",
                "ENSMUSG00000005204",
                "ENSMUSG00000097277",
                "ENSMUSG00000028538",
                "ENSMUSG00000007721",
                "ENSMUSG00000020777",
                "ENSMUSG00000034842",
                "ENSMUSG00000041733",
                "ENSMUSG00000062328",
                "ENSMUSG00000029190",
                "ENSMUSG00000039873",
                "ENSMUSG00000004610",
                "ENSMUSG00000015944",
                "ENSMUSG00000021265",
                "ENSMUSG00000021290",
                "ENSMUSG00000037573",
                "ENSMUSG00000049470",
                "ENSMUSG00000069049",
                "ENSMUSG00000046330",
                "ENSMUSG00000029401",
                "ENSMUSG00000074247",
                "ENSMUSG00000003534",
                "ENSMUSG00000025453",
                "ENSMUSG00000025860",
                "ENSMUSG00000030612",
                "ENSMUSG00000004035",
                "ENSMUSG00000074918",
                "ENSMUSG00000024725",
                "ENSMUSG00000021606",
                "ENSMUSG00000019877",
                "ENSMUSG00000033416",
                "ENSMUSG00000023068",
                "ENSMUSG00000021192",
                "ENSMUSG00000036860",
                "ENSMUSG00000079641",
                "ENSMUSG00000075705",
                "ENSMUSG00000045636",
                "ENSMUSG00000020462",
                "ENSMUSG00000062300",
                "ENSMUSG00000063077",
                "ENSMUSG00000022200",
                "ENSMUSG00000031299",
                "ENSMUSG00000053477",
                "ENSMUSG00000021771",
                "ENSMUSG00000032267",
                "ENSMUSG00000035247",
                "ENSMUSG00000040048",
                "ENSMUSG00000028461",
                "ENSMUSG00000064365",
                "ENSMUSG00000024976",
                "ENSMUSG00000043639",
                "ENSMUSG00000043866",
                "ENSMUSG00000071662",
                "ENSMUSG00000028557",
                "ENSMUSG00000023938",
                "ENSMUSG00000027274",
                "ENSMUSG00000036707",
                "ENSMUSG00000032527",
                "ENSMUSG00000035268",
                "ENSMUSG00000085779",
                "ENSMUSG00000009549",
                "ENSMUSG00000068798",
                "ENSMUSG00000020608",
                "ENSMUSG00000022450",
                "ENSMUSG00000015579",
                "ENSMUSG00000073563",
                "ENSMUSG00000024805",
                "ENSMUSG00000078622",
                "ENSMUSG00000082877",
                "ENSMUSG00000032366",
                "ENSMUSG00000032370",
                "ENSMUSG00000098343",
                "ENSMUSG00000028383",
                "ENSMUSG00000041115",
                "ENSMUSG00000025066",
                "ENSMUSG00000017548",
                "ENSMUSG00000020309",
                "ENSMUSG00000064348",
                "ENSMUSG00000094483",
                "ENSMUSG00000027709",
                "ENSMUSG00000024498",
                "ENSMUSG00000022965",
                "ENSMUSG00000022884",
                "ENSMUSG00000015733",
                "ENSMUSG00000027099",
                "ENSMUSG00000025220",
                "ENSMUSG00000035027",
                "ENSMUSG00000034042",
                "ENSMUSG00000052812",
                "ENSMUSG00000037685",
                "ENSMUSG00000024290",
                "ENSMUSG00000026526",
                "ENSMUSG00000022951",
                "ENSMUSG00000024132",
                "ENSMUSG00000029338",
                "ENSMUSG00000021621",
                "ENSMUSG00000006005",
                "ENSMUSG00000034525",
                "ENSMUSG00000046312",
                "ENSMUSG00000081929",
                "ENSMUSG00000073002",
                "ENSMUSG00000055204",
                "ENSMUSG00000042115",
                "ENSMUSG00000023904",
                "ENSMUSG00000054312",
                "ENSMUSG00000033295",
                "ENSMUSG00000060803",
                "ENSMUSG00000030246",
                "ENSMUSG00000061474",
                "ENSMUSG00000096232",
                "ENSMUSG00000010914",
                "ENSMUSG00000042699",
                "ENSMUSG00000017760",
                "ENSMUSG00000055128",
                "ENSMUSG00000030869",
                "ENSMUSG00000031767",
                "ENSMUSG00000064371",
                "ENSMUSG00000084314",
                "ENSMUSG00000027620",
                "ENSMUSG00000017817",
                "ENSMUSG00000018395",
                "ENSMUSG00000002908",
                "ENSMUSG00000031229",
                "ENSMUSG00000074656",
                "ENSMUSG00000039126",
                "ENSMUSG00000045639",
                "ENSMUSG00000028847",
                "ENSMUSG00000033047",
                "ENSMUSG00000024913"
            ],
            "expression_sum": {
                "class1": 1988.0,
                "class2": 50963.0
            },
            "uniprot_ids": [
                "O89090",
                "E9Q0S7",
                "E9Q113",
                "E9Q2V3",
                "G3X8Q0"
            ]
        },
        "ENSMUSG00000085684": {
            "id": "ENSMUSG00000085684",
            "type": "Gene",
            "symbol": "4930469K13Rik",
            "source": [
                "ENSMUSG00000001280"
            ],
            "target": [],
            "expression_sum": {
                "class1": 91.0,
                "class2": 0.0
            },
            "uniprot_ids": [
                "A0A286YDX3",
                "A0A286YDB2"
            ]
        },
        "ENSMUSG00000000056": {
            "id": "ENSMUSG00000000056",
            "type": "Gene",
            "symbol": "Narf",
            "source": [
                "ENSMUSG00000026890",
                "ENSMUSG00000034673",
                "ENSMUSG00000005566",
                "ENSMUSG00000013419",
                "ENSMUSG00000024695",
                "ENSMUSG00000015605",
                "ENSMUSG00000008575",
                "ENSMUSG00000040321",
                "ENSMUSG00000040481",
                "ENSMUSG00000055148",
                "ENSMUSG00000023990",
                "ENSMUSG00000049739",
                "ENSMUSG00000015579",
                "ENSMUSG00000040033",
                "ENSMUSG00000019768",
                "ENSMUSG00000005698",
                "ENSMUSG00000024140",
                "ENSMUSG00000053110",
                "ENSMUSG00000026803",
                "ENSMUSG00000022389",
                "ENSMUSG00000005583",
                "ENSMUSG00000068267",
                "ENSMUSG00000047371",
                "ENSMUSG00000078671",
                "ENSMUSG00000026610",
                "ENSMUSG00000052837",
                "ENSMUSG00000056501",
                "ENSMUSG00000029249",
                "ENSMUSG00000001280",
                "ENSMUSG00000009079",
                "ENSMUSG00000055320"
            ],
            "target": [],
            "expression_sum": {
                "class1": 1281.0,
                "class2": 986.0
            },
            "uniprot_ids": [
                "Q9CYQ7"
            ]
        },
        "ENSMUSG00000000088": {
            "id": "ENSMUSG00000000088",
            "type": "Gene",
            "symbol": "Cox5a",
            "source": [
                "ENSMUSG00000029135",
                "ENSMUSG00000034673",
                "ENSMUSG00000037174",
                "ENSMUSG00000024695",
                "ENSMUSG00000044452",
                "ENSMUSG00000015605",
                "ENSMUSG00000008575",
                "ENSMUSG00000057469",
                "ENSMUSG00000010175",
                "ENSMUSG00000001156",
                "ENSMUSG00000015579",
                "ENSMUSG00000090258",
                "ENSMUSG00000019768",
                "ENSMUSG00000028565",
                "ENSMUSG00000031885",
                "ENSMUSG00000015846",
                "ENSMUSG00000015839",
                "ENSMUSG00000001911",
                "ENSMUSG00000049878",
                "ENSMUSG00000028639",
                "ENSMUSG00000030557",
                "ENSMUSG00000031734",
                "ENSMUSG00000033943",
                "ENSMUSG00000068267",
                "ENSMUSG00000059878",
                "ENSMUSG00000014592",
                "ENSMUSG00000056501",
                "ENSMUSG00000001280",
                "ENSMUSG00000025958",
                "ENSMUSG00000024498",
                "ENSMUSG00000040841",
                "ENSMUSG00000055320"
            ],
            "target": [],
            "expression_sum": {
                "class1": 264.0,
                "class2": 3902.0
            },
            "uniprot_ids": [
                "P12787"
            ]
        },
        "ENSMUSG00000000149": {
            "id": "ENSMUSG00000000149",
            "type": "Gene",
            "symbol": "Gna12",
            "source": [
                "ENSMUSG00000018604",
                "ENSMUSG00000078671",
                "ENSMUSG00000024081",
                "ENSMUSG00000040033",
                "ENSMUSG00000001280",
                "ENSMUSG00000019768",
                "ENSMUSG00000022389",
                "ENSMUSG00000005698",
                "ENSMUSG00000022641",
                "ENSMUSG00000024140",
                "ENSMUSG00000033943",
                "ENSMUSG00000037235",
                "ENSMUSG00000047371",
                "ENSMUSG00000068267",
                "ENSMUSG00000068959",
                "ENSMUSG00000015605",
                "ENSMUSG00000014592",
                "ENSMUSG00000022811"
            ],
            "target": [],
            "expression_sum": {
                "class1": 357.0,
                "class2": 429.0
            },
            "uniprot_ids": [
                "P27600",
                "A0A0G2JG40"
            ]
        },
        "ENSMUSG00000000168": {
            "id": "ENSMUSG00000000168",
            "type": "Gene",
            "symbol": "Dlat",
            "source": [
                "ENSMUSG00000029135",
                "ENSMUSG00000034673",
                "ENSMUSG00000005566",
                "ENSMUSG00000013419",
                "ENSMUSG00000024695",
                "ENSMUSG00000037174",
                "ENSMUSG00000021775",
                "ENSMUSG00000040481",
                "ENSMUSG00000010175",
                "ENSMUSG00000023990",
                "ENSMUSG00000040033",
                "ENSMUSG00000090258",
                "ENSMUSG00000019768",
                "ENSMUSG00000005698",
                "ENSMUSG00000017146",
                "ENSMUSG00000039191",
                "ENSMUSG00000015846",
                "ENSMUSG00000015839",
                "ENSMUSG00000001911",
                "ENSMUSG00000030557",
                "ENSMUSG00000032238",
                "ENSMUSG00000068267",
                "ENSMUSG00000058756",
                "ENSMUSG00000014592",
                "ENSMUSG00000026610",
                "ENSMUSG00000056501",
                "ENSMUSG00000038718",
                "ENSMUSG00000029249",
                "ENSMUSG00000001280",
                "ENSMUSG00000024498",
                "ENSMUSG00000009079"
            ],
            "target": [],
            "expression_sum": {
                "class1": 2565.0,
                "class2": 2529.0
            },
            "uniprot_ids": [
                "Q8BMF4"
            ]
        },
        "ENSMUSG00000000171": {
            "id": "ENSMUSG00000000171",
            "type": "Gene",
            "symbol": "Sdhd",
            "source": [
                "ENSMUSG00000073423",
                "ENSMUSG00000056501",
                "ENSMUSG00000001156",
                "ENSMUSG00000001280",
                "ENSMUSG00000019768",
                "ENSMUSG00000022389",
                "ENSMUSG00000038914",
                "ENSMUSG00000045639",
                "ENSMUSG00000040841",
                "ENSMUSG00000036180",
                "ENSMUSG00000068267",
                "ENSMUSG00000031311",
                "ENSMUSG00000015605",
                "ENSMUSG00000015846"
            ],
            "target": [],
            "expression_sum": {
                "class1": 4757.0,
                "class2": 4659.0
            },
            "uniprot_ids": [
                "Q9CXV1"
            ]
        },
        "ENSMUSG00000000184": {
            "id": "ENSMUSG00000000184",
            "type": "Gene",
            "symbol": "Ccnd2",
            "source": [
                "ENSMUSG00000058440",
                "ENSMUSG00000029135",
                "ENSMUSG00000019947",
                "ENSMUSG00000067430",
                "ENSMUSG00000047003",
                "ENSMUSG00000027109",
                "ENSMUSG00000001280",
                "ENSMUSG00000019768",
                "ENSMUSG00000022389",
                "ENSMUSG00000030067",
                "ENSMUSG00000068267",
                "ENSMUSG00000027210",
                "ENSMUSG00000031311",
                "ENSMUSG00000035632",
                "ENSMUSG00000049421"
            ],
            "target": [],
            "expression_sum": {
                "class1": 9096.0,
                "class2": 999.0
            },
            "uniprot_ids": [
                "P30280",
                "Q9D8L9",
                "A0A0J9YU51"
            ]
        },
        "ENSMUSG00000000253": {
            "id": "ENSMUSG00000000253",
            "type": "Gene",
            "symbol": "Gmpr",
            "source": [
                "ENSMUSG00000001280",
                "ENSMUSG00000040841"
            ],
            "target": [],
            "expression_sum": {
                "class1": 1028.0,
                "class2": 649.0
            },
            "uniprot_ids": [
                "Q9DCZ1",
                "F6VY18"
            ]
        },
        "ENSMUSG00000000340": {
            "id": "ENSMUSG00000000340",
            "type": "Gene",
            "symbol": "Dbt",
            "source": [
                "ENSMUSG00000029135",
                "ENSMUSG00000076431",
                "ENSMUSG00000018604",
                "ENSMUSG00000051469",
                "ENSMUSG00000090258",
                "ENSMUSG00000021877",
                "ENSMUSG00000024974",
                "ENSMUSG00000031885",
                "ENSMUSG00000036721",
                "ENSMUSG00000015839",
                "ENSMUSG00000001911",
                "ENSMUSG00000049878",
                "ENSMUSG00000033943",
                "ENSMUSG00000016087",
                "ENSMUSG00000014592",
                "ENSMUSG00000001419",
                "ENSMUSG00000001280",
                "ENSMUSG00000025958",
                "ENSMUSG00000024498"
            ],
            "target": [],
            "expression_sum": {
                "class1": 292.0,
                "class2": 581.0
            },
            "uniprot_ids": [
                "P53395",
                "A0A0G2JGU0",
                "A0A0G2JGV4"
            ]
        },
        "ENSMUSG00000000346": {
            "id": "ENSMUSG00000000346",
            "type": "Gene",
            "symbol": "Dazap2",
            "source": [
                "ENSMUSG00000033454",
                "ENSMUSG00000030256",
                "ENSMUSG00000039087",
                "ENSMUSG00000021485",
                "ENSMUSG00000023079",
                "ENSMUSG00000073423",
                "ENSMUSG00000014704",
                "ENSMUSG00000001280",
                "ENSMUSG00000021877",
                "ENSMUSG00000031627",
                "ENSMUSG00000018697",
                "ENSMUSG00000026872"
            ],
            "target": [],
            "expression_sum": {
                "class1": 6539.0,
                "class2": 1604.0
            },
            "uniprot_ids": [
                "Q9DCP9"
            ]
        },
        "ENSMUSG00000000355": {
            "id": "ENSMUSG00000000355",
            "type": "Gene",
            "symbol": "Mcts1",
            "source": [
                "ENSMUSG00000001280",
                "ENSMUSG00000005893"
            ],
            "target": [],
            "expression_sum": {
                "class1": 1699.0,
                "class2": 711.0
            },
            "uniprot_ids": [
                "Q9DB27"
            ]
        },
        "ENSMUSG00000000399": {
            "id": "ENSMUSG00000000399",
            "type": "Gene",
            "symbol": "Ndufa9",
            "source": [
                "ENSMUSG00000029135",
                "ENSMUSG00000034673",
                "ENSMUSG00000024695",
                "ENSMUSG00000015605",
                "ENSMUSG00000008575",
                "ENSMUSG00000040321",
                "ENSMUSG00000058239",
                "ENSMUSG00000052534",
                "ENSMUSG00000015579",
                "ENSMUSG00000090258",
                "ENSMUSG00000019768",
                "ENSMUSG00000038914",
                "ENSMUSG00000031715",
                "ENSMUSG00000015839",
                "ENSMUSG00000001911",
                "ENSMUSG00000049878",
                "ENSMUSG00000030557",
                "ENSMUSG00000033943",
                "ENSMUSG00000068267",
                "ENSMUSG00000047371",
                "ENSMUSG00000014592",
                "ENSMUSG00000024955",
                "ENSMUSG00000029249",
                "ENSMUSG00000001280",
                "ENSMUSG00000009079"
            ],
            "target": [],
            "expression_sum": {
                "class1": 6844.0,
                "class2": 4677.0
            },
            "uniprot_ids": [
                "Q9DC69",
                "A0A0R3P9C8"
            ]
        },
        "ENSMUSG00000000441": {
            "id": "ENSMUSG00000000441",
            "type": "Gene",
            "symbol": "Raf1",
            "source": [
                "ENSMUSG00000029135",
                "ENSMUSG00000051550",
                "ENSMUSG00000015605",
                "ENSMUSG00000008575",
                "ENSMUSG00000057469",
                "ENSMUSG00000055148",
                "ENSMUSG00000010175",
                "ENSMUSG00000018604",
                "ENSMUSG00000019768",
                "ENSMUSG00000005698",
                "ENSMUSG00000024985",
                "ENSMUSG00000001911",
                "ENSMUSG00000023034",
                "ENSMUSG00000032238",
                "ENSMUSG00000019789",
                "ENSMUSG00000015501",
                "ENSMUSG00000038718",
                "ENSMUSG00000029249",
                "ENSMUSG00000001280",
                "ENSMUSG00000009079"
            ],
            "target": [],
            "expression_sum": {
                "class1": 296.0,
                "class2": 393.0
            },
            "uniprot_ids": [
                "Q99N57",
                "F6TUC4",
                "A0A0N4SV36",
                "A0A0N4SWH5"
            ]
        },
        "ENSMUSG00000000538": {
            "id": "ENSMUSG00000000538",
            "type": "Gene",
            "symbol": "Tom1l2",
            "source": [
                "ENSMUSG00000013419",
                "ENSMUSG00000015605",
                "ENSMUSG00000008575",
                "ENSMUSG00000032897",
                "ENSMUSG00000055148",
                "ENSMUSG00000019768",
                "ENSMUSG00000022641",
                "ENSMUSG00000031885",
                "ENSMUSG00000015839",
                "ENSMUSG00000032228",
                "ENSMUSG00000032238",
                "ENSMUSG00000005583",
                "ENSMUSG00000029913",
                "ENSMUSG00000029322",
                "ENSMUSG00000078671",
                "ENSMUSG00000003923",
                "ENSMUSG00000038718",
                "ENSMUSG00000029249",
                "ENSMUSG00000001280",
                "ENSMUSG00000099083"
            ],
            "target": [],
            "expression_sum": {
                "class1": 777.0,
                "class2": 521.0
            },
            "uniprot_ids": [
                "Q5SRX1",
                "Q5SXA5",
                "Q5SXA4",
                "F6RBX1",
                "F6ZDJ1"
            ]
        },
        "ENSMUSG00000000563": {
            "id": "ENSMUSG00000000563",
            "type": "Gene",
            "symbol": "Atp5pb",
            "source": [
                "ENSMUSG00000037174",
                "ENSMUSG00000024695",
                "ENSMUSG00000015605",
                "ENSMUSG00000021775",
                "ENSMUSG00000040321",
                "ENSMUSG00000057469",
                "ENSMUSG00000015579",
                "ENSMUSG00000090258",
                "ENSMUSG00000019768",
                "ENSMUSG00000031885",
                "ENSMUSG00000036721",
                "ENSMUSG00000039191",
                "ENSMUSG00000031715",
                "ENSMUSG00000015839",
                "ENSMUSG00000033943",
                "ENSMUSG00000068267",
                "ENSMUSG00000014592",
                "ENSMUSG00000003923",
                "ENSMUSG00000001280",
                "ENSMUSG00000024498"
            ],
            "target": [],
            "expression_sum": {
                "class1": 18039.0,
                "class2": 18055.0
            },
            "uniprot_ids": [
                "Q9CQQ7",
                "A0A0G2JGX3",
                "D6RGL7"
            ]
        },
        "ENSMUSG00000000628": {
            "id": "ENSMUSG00000000628",
            "type": "Gene",
            "symbol": "Hk2",
            "source": [
                "ENSMUSG00000024238",
                "ENSMUSG00000021250",
                "ENSMUSG00000024695",
                "ENSMUSG00000039087",
                "ENSMUSG00000015605",
                "ENSMUSG00000040481",
                "ENSMUSG00000066721",
                "ENSMUSG00000019768",
                "ENSMUSG00000038914",
                "ENSMUSG00000017146",
                "ENSMUSG00000031885",
                "ENSMUSG00000031715",
                "ENSMUSG00000020644",
                "ENSMUSG00000001911",
                "ENSMUSG00000028639",
                "ENSMUSG00000068267",
                "ENSMUSG00000014592",
                "ENSMUSG00000071076",
                "ENSMUSG00000001280"
            ],
            "target": [],
            "expression_sum": {
                "class1": 706.0,
                "class2": 1965.0
            },
            "uniprot_ids": [
                "O08528",
                "E9Q9M6",
                "E9Q5B5"
            ]
        },
        "ENSMUSG00000000708": {
            "id": "ENSMUSG00000000708",
            "type": "Gene",
            "symbol": "Kat2b",
            "source": [
                "ENSMUSG00000001280",
                "ENSMUSG00000040270",
                "ENSMUSG00000002265"
            ],
            "target": [],
            "expression_sum": {
                "class1": 430.0,
                "class2": 364.0
            },
            "uniprot_ids": [
                "Q9JHD1",
                "F6SIC5",
                "E9Q1F7",
                "F6ZCH1"
            ]
        },
        "ENSMUSG00000000738": {
            "id": "ENSMUSG00000000738",
            "type": "Gene",
            "symbol": "Spg7",
            "source": [
                "ENSMUSG00000001280",
                "ENSMUSG00000005698",
                "ENSMUSG00000041515",
                "ENSMUSG00000030557"
            ],
            "target": [],
            "expression_sum": {
                "class1": 391.0,
                "class2": 307.0
            },
            "uniprot_ids": [
                "Q3ULF4",
                "A0A1I7Q4C2",
                "D3Z342",
                "D3YZN4",
                "F6VTG4",
                "F6W695",
                "G3UX97",
                "D3YXB7"
            ]
        },
        "ENSMUSG00000000740": {
            "id": "ENSMUSG00000000740",
            "type": "Gene",
            "symbol": "Rpl13",
            "source": [
                "ENSMUSG00000029135",
                "ENSMUSG00000026890",
                "ENSMUSG00000002983",
                "ENSMUSG00000023990",
                "ENSMUSG00000056501",
                "ENSMUSG00000015579",
                "ENSMUSG00000001280",
                "ENSMUSG00000019768",
                "ENSMUSG00000040732",
                "ENSMUSG00000021877",
                "ENSMUSG00000002265",
                "ENSMUSG00000076431",
                "ENSMUSG00000042557",
                "ENSMUSG00000015846"
            ],
            "target": [],
            "expression_sum": {
                "class1": 12535.0,
                "class2": 1680.0
            },
            "uniprot_ids": [
                "P47963"
            ]
        },
        "ENSMUSG00000000787": {
            "id": "ENSMUSG00000000787",
            "type": "Gene",
            "symbol": "Ddx3x",
            "source": [
                "ENSMUSG00000038539",
                "ENSMUSG00000060510",
                "ENSMUSG00000055148",
                "ENSMUSG00000030256",
                "ENSMUSG00000031737",
                "ENSMUSG00000038648",
                "ENSMUSG00000019768",
                "ENSMUSG00000024985",
                "ENSMUSG00000049658",
                "ENSMUSG00000001419",
                "ENSMUSG00000037608",
                "ENSMUSG00000078671",
                "ENSMUSG00000015501",
                "ENSMUSG00000066551",
                "ENSMUSG00000029249",
                "ENSMUSG00000001280",
                "ENSMUSG00000009079",
                "ENSMUSG00000022053",
                "ENSMUSG00000027104"
            ],
            "target": [],
            "expression_sum": {
                "class1": 5552.0,
                "class2": 1734.0
            },
            "uniprot_ids": [
                "Q62167"
            ]
        },
        "ENSMUSG00000001056": {
            "id": "ENSMUSG00000001056",
            "type": "Gene",
            "symbol": "Nhp2",
            "source": [
                "ENSMUSG00000058440",
                "ENSMUSG00000057469",
                "ENSMUSG00000018604",
                "ENSMUSG00000001280",
                "ENSMUSG00000019768",
                "ENSMUSG00000028565",
                "ENSMUSG00000033943",
                "ENSMUSG00000017146",
                "ENSMUSG00000060601"
            ],
            "target": [],
            "expression_sum": {
                "class1": 5992.0,
                "class2": 867.0
            },
            "uniprot_ids": [
                "Q9CRB2"
            ]
        },
        "ENSMUSG00000001105": {
            "id": "ENSMUSG00000001105",
            "type": "Gene",
            "symbol": "Ift20",
            "source": [
                "ENSMUSG00000006215",
                "ENSMUSG00000052534",
                "ENSMUSG00000038872",
                "ENSMUSG00000055116",
                "ENSMUSG00000038648",
                "ENSMUSG00000024431",
                "ENSMUSG00000001280",
                "ENSMUSG00000019768",
                "ENSMUSG00000052684",
                "ENSMUSG00000021877",
                "ENSMUSG00000055320",
                "ENSMUSG00000002265",
                "ENSMUSG00000047371",
                "ENSMUSG00000027104",
                "ENSMUSG00000015605",
                "ENSMUSG00000024985",
                "ENSMUSG00000049658"
            ],
            "target": [],
            "expression_sum": {
                "class1": 4845.0,
                "class2": 890.0
            },
            "uniprot_ids": [
                "Q61025",
                "Z4YJL9"
            ]
        },
        "ENSMUSG00000001127": {
            "id": "ENSMUSG00000001127",
            "type": "Gene",
            "symbol": "Araf",
            "source": [
                "ENSMUSG00000029135",
                "ENSMUSG00000026565",
                "ENSMUSG00000034673",
                "ENSMUSG00000021775",
                "ENSMUSG00000032897",
                "ENSMUSG00000001156",
                "ENSMUSG00000015579",
                "ENSMUSG00000058886",
                "ENSMUSG00000090258",
                "ENSMUSG00000019768",
                "ENSMUSG00000021877",
                "ENSMUSG00000024140",
                "ENSMUSG00000031885",
                "ENSMUSG00000015846",
                "ENSMUSG00000001911",
                "ENSMUSG00000028466",
                "ENSMUSG00000027552",
                "ENSMUSG00000033943",
                "ENSMUSG00000078671",
                "ENSMUSG00000001280"
            ],
            "target": [],
            "expression_sum": {
                "class1": 813.0,
                "class2": 843.0
            },
            "uniprot_ids": [
                "P04627",
                "Q8CAD1",
                "B1AUP0",
                "E9Q0P8",
                "B1AUN8"
            ]
        },
        "ENSMUSG00000001158": {
            "id": "ENSMUSG00000001158",
            "type": "Gene",
            "symbol": "Snrnp27",
            "source": [
                "ENSMUSG00000025612",
                "ENSMUSG00000001280",
                "ENSMUSG00000099083",
                "ENSMUSG00000025025",
                "ENSMUSG00000039087",
                "ENSMUSG00000039191",
                "ENSMUSG00000020644"
            ],
            "target": [],
            "expression_sum": {
                "class1": 5920.0,
                "class2": 1614.0
            },
            "uniprot_ids": [
                "Q8K194"
            ]
        },
        "ENSMUSG00000001175": {
            "id": "ENSMUSG00000001175",
            "type": "Gene",
            "symbol": "Calm1",
            "source": [
                "ENSMUSG00000029135",
                "ENSMUSG00000026890",
                "ENSMUSG00000021250",
                "ENSMUSG00000018604",
                "ENSMUSG00000036144",
                "ENSMUSG00000073423",
                "ENSMUSG00000029729",
                "ENSMUSG00000036461",
                "ENSMUSG00000049739",
                "ENSMUSG00000001280",
                "ENSMUSG00000020919",
                "ENSMUSG00000031627",
                "ENSMUSG00000016087",
                "ENSMUSG00000024985"
            ],
            "target": [],
            "expression_sum": {
                "class1": 81479.0,
                "class2": 13694.0
            },
            "uniprot_ids": [
                "P0DP26",
                "P0DP27",
                "P0DP28",
                "A0A3Q4EHJ0"
            ]
        },
        "ENSMUSG00000001211": {
            "id": "ENSMUSG00000001211",
            "type": "Gene",
            "symbol": "Agpat3",
            "source": [
                "ENSMUSG00000029135",
                "ENSMUSG00000001419",
                "ENSMUSG00000015839",
                "ENSMUSG00000019817",
                "ENSMUSG00000056501",
                "ENSMUSG00000015579",
                "ENSMUSG00000032228",
                "ENSMUSG00000051469",
                "ENSMUSG00000001280",
                "ENSMUSG00000019768",
                "ENSMUSG00000068267",
                "ENSMUSG00000008575",
                "ENSMUSG00000024985"
            ],
            "target": [],
            "expression_sum": {
                "class1": 1033.0,
                "class2": 901.0
            },
            "uniprot_ids": [
                "Q9D517",
                "D3YTN7",
                "A0A1W2P764",
                "D3YUW0",
                "D3YV65",
                "D3Z1M6"
            ]
        },
        "ENSMUSG00000001289": {
            "id": "ENSMUSG00000001289",
            "type": "Gene",
            "symbol": "Pfdn5",
            "source": [
                "ENSMUSG00000024238",
                "ENSMUSG00000021250",
                "ENSMUSG00000019817",
                "ENSMUSG00000022346",
                "ENSMUSG00000052534",
                "ENSMUSG00000036461",
                "ENSMUSG00000014303",
                "ENSMUSG00000001280",
                "ENSMUSG00000053477",
                "ENSMUSG00000005583",
                "ENSMUSG00000035310",
                "ENSMUSG00000025959",
                "ENSMUSG00000022314",
                "ENSMUSG00000060510",
                "ENSMUSG00000015846"
            ],
            "target": [],
            "expression_sum": {
                "class1": 12873.0,
                "class2": 2735.0
            },
            "uniprot_ids": [
                "Q9WU28",
                "E9PVG3",
                "E9Q093",
                "H7BWX1",
                "E9PZ62",
                "E9Q5Q8"
            ]
        },
        "ENSMUSG00000001334": {
            "id": "ENSMUSG00000001334",
            "type": "Gene",
            "symbol": "Fndc5",
            "source": [
                "ENSMUSG00000001280",
                "ENSMUSG00000019768",
                "ENSMUSG00000005698",
                "ENSMUSG00000024140",
                "ENSMUSG00000031885",
                "ENSMUSG00000015605",
                "ENSMUSG00000014592"
            ],
            "target": [],
            "expression_sum": {
                "class1": 914.0,
                "class2": 1866.0
            },
            "uniprot_ids": [
                "Q8K4Z2"
            ]
        },
        "ENSMUSG00000001366": {
            "id": "ENSMUSG00000001366",
            "type": "Gene",
            "symbol": "Fbxo9",
            "source": [
                "ENSMUSG00000010175",
                "ENSMUSG00000078671",
                "ENSMUSG00000015605",
                "ENSMUSG00000026610",
                "ENSMUSG00000015579",
                "ENSMUSG00000029249",
                "ENSMUSG00000001280",
                "ENSMUSG00000005566",
                "ENSMUSG00000005583",
                "ENSMUSG00000005698",
                "ENSMUSG00000019768",
                "ENSMUSG00000008575",
                "ENSMUSG00000029913"
            ],
            "target": [],
            "expression_sum": {
                "class1": 734.0,
                "class2": 444.0
            },
            "uniprot_ids": [
                "Q8BK06",
                "E0CYX8"
            ]
        },
        "ENSMUSG00000001415": {
            "id": "ENSMUSG00000001415",
            "type": "Gene",
            "symbol": "Smg5",
            "source": [
                "ENSMUSG00000052837",
                "ENSMUSG00000001280",
                "ENSMUSG00000019768",
                "ENSMUSG00000040712",
                "ENSMUSG00000035158",
                "ENSMUSG00000024985"
            ],
            "target": [],
            "expression_sum": {
                "class1": 555.0,
                "class2": 143.0
            },
            "uniprot_ids": [
                "Q6ZPY2",
                "A0A0A6YWQ7"
            ]
        },
        "ENSMUSG00000001419": {
            "id": "ENSMUSG00000001419",
            "type": "TF",
            "symbol": "Mef2d",
            "source": [
                "ENSMUSG00000029135",
                "ENSMUSG00000010175",
                "ENSMUSG00000078671",
                "ENSMUSG00000015839",
                "ENSMUSG00000058239",
                "ENSMUSG00000015579",
                "ENSMUSG00000047407",
                "ENSMUSG00000030557",
                "ENSMUSG00000001280",
                "ENSMUSG00000019768",
                "ENSMUSG00000053110",
                "ENSMUSG00000038692",
                "ENSMUSG00000008575",
                "ENSMUSG00000035158",
                "ENSMUSG00000031715"
            ],
            "target": [
                "ENSMUSG00000034875",
                "ENSMUSG00000007777",
                "ENSMUSG00000004383",
                "ENSMUSG00000002661",
                "ENSMUSG00000018501",
                "ENSMUSG00000042828",
                "ENSMUSG00000018845",
                "ENSMUSG00000019689",
                "ENSMUSG00000037266",
                "ENSMUSG00000034751",
                "ENSMUSG00000036585",
                "ENSMUSG00000070283",
                "ENSMUSG00000029629",
                "ENSMUSG00000039640",
                "ENSMUSG00000042717",
                "ENSMUSG00000042709",
                "ENSMUSG00000033916",
                "ENSMUSG00000031540",
                "ENSMUSG00000032525",
                "ENSMUSG00000020402",
                "ENSMUSG00000026087",
                "ENSMUSG00000038535",
                "ENSMUSG00000020477",
                "ENSMUSG00000021270",
                "ENSMUSG00000037174",
                "ENSMUSG00000026623",
                "ENSMUSG00000035158",
                "ENSMUSG00000036309",
                "ENSMUSG00000027288",
                "ENSMUSG00000056851",
                "ENSMUSG00000030557",
                "ENSMUSG00000020048",
                "ENSMUSG00000025190",
                "ENSMUSG00000090150",
                "ENSMUSG00000014592",
                "ENSMUSG00000083186",
                "ENSMUSG00000038538",
                "ENSMUSG00000062694",
                "ENSMUSG00000014606",
                "ENSMUSG00000040661",
                "ENSMUSG00000020153",
                "ENSMUSG00000093674",
                "ENSMUSG00000030213",
                "ENSMUSG00000029158",
                "ENSMUSG00000024683",
                "ENSMUSG00000044365",
                "ENSMUSG00000029404",
                "ENSMUSG00000048578",
                "ENSMUSG00000020949",
                "ENSMUSG00000046679",
                "ENSMUSG00000039103",
                "ENSMUSG00000023004",
                "ENSMUSG00000067653",
                "ENSMUSG00000028700",
                "ENSMUSG00000039515",
                "ENSMUSG00000027378",
                "ENSMUSG00000063229",
                "ENSMUSG00000029505",
                "ENSMUSG00000038936",
                "ENSMUSG00000018566",
                "ENSMUSG00000055320",
                "ENSMUSG00000077450",
                "ENSMUSG00000035296",
                "ENSMUSG00000044086",
                "ENSMUSG00000038663",
                "ENSMUSG00000001508",
                "ENSMUSG00000026414",
                "ENSMUSG00000048546",
                "ENSMUSG00000031880",
                "ENSMUSG00000038615",
                "ENSMUSG00000030057",
                "ENSMUSG00000023990",
                "ENSMUSG00000055322",
                "ENSMUSG00000045776",
                "ENSMUSG00000028207",
                "ENSMUSG00000030647",
                "ENSMUSG00000027879",
                "ENSMUSG00000034931",
                "ENSMUSG00000037049",
                "ENSMUSG00000024236",
                "ENSMUSG00000039001",
                "ENSMUSG00000034160",
                "ENSMUSG00000061518",
                "ENSMUSG00000033400",
                "ENSMUSG00000040694",
                "ENSMUSG00000064345",
                "ENSMUSG00000032648",
                "ENSMUSG00000026374",
                "ENSMUSG00000014769",
                "ENSMUSG00000024871",
                "ENSMUSG00000031447",
                "ENSMUSG00000006333",
                "ENSMUSG00000004127",
                "ENSMUSG00000023020",
                "ENSMUSG00000051469",
                "ENSMUSG00000052934",
                "ENSMUSG00000029723",
                "ENSMUSG00000071793",
                "ENSMUSG00000064366",
                "ENSMUSG00000071369",
                "ENSMUSG00000037443",
                "ENSMUSG00000049872",
                "ENSMUSG00000029145",
                "ENSMUSG00000030357",
                "ENSMUSG00000026456",
                "ENSMUSG00000030785",
                "ENSMUSG00000055027",
                "ENSMUSG00000073557",
                "ENSMUSG00000042834",
                "ENSMUSG00000028116",
                "ENSMUSG00000033060",
                "ENSMUSG00000046032",
                "ENSMUSG00000064367",
                "ENSMUSG00000032826",
                "ENSMUSG00000039601",
                "ENSMUSG00000057322",
                "ENSMUSG00000092341",
                "ENSMUSG00000001062",
                "ENSMUSG00000031626",
                "ENSMUSG00000000340",
                "ENSMUSG00000055762",
                "ENSMUSG00000024750",
                "ENSMUSG00000064343",
                "ENSMUSG00000047371",
                "ENSMUSG00000028233",
                "ENSMUSG00000026930",
                "ENSMUSG00000021597",
                "ENSMUSG00000045624",
                "ENSMUSG00000064339",
                "ENSMUSG00000017485",
                "ENSMUSG00000031913",
                "ENSMUSG00000020601",
                "ENSMUSG00000051373",
                "ENSMUSG00000042198",
                "ENSMUSG00000047710",
                "ENSMUSG00000031841",
                "ENSMUSG00000036854",
                "ENSMUSG00000026131",
                "ENSMUSG00000033365",
                "ENSMUSG00000068614",
                "ENSMUSG00000032010",
                "ENSMUSG00000031672",
                "ENSMUSG00000021838",
                "ENSMUSG00000034892",
                "ENSMUSG00000023092",
                "ENSMUSG00000059291",
                "ENSMUSG00000028649",
                "ENSMUSG00000028300",
                "ENSMUSG00000052146",
                "ENSMUSG00000069014",
                "ENSMUSG00000067925",
                "ENSMUSG00000030695",
                "ENSMUSG00000030298",
                "ENSMUSG00000064341",
                "ENSMUSG00000048758",
                "ENSMUSG00000024049",
                "ENSMUSG00000015850",
                "ENSMUSG00000020864",
                "ENSMUSG00000022110",
                "ENSMUSG00000035242",
                "ENSMUSG00000030779",
                "ENSMUSG00000027206",
                "ENSMUSG00000003380",
                "ENSMUSG00000064346",
                "ENSMUSG00000021870",
                "ENSMUSG00000025428",
                "ENSMUSG00000033161",
                "ENSMUSG00000058690",
                "ENSMUSG00000026150",
                "ENSMUSG00000039376",
                "ENSMUSG00000003072",
                "ENSMUSG00000032249",
                "ENSMUSG00000057335",
                "ENSMUSG00000082487",
                "ENSMUSG00000020849",
                "ENSMUSG00000074218",
                "ENSMUSG00000037533",
                "ENSMUSG00000009293",
                "ENSMUSG00000021798",
                "ENSMUSG00000041297",
                "ENSMUSG00000026766",
                "ENSMUSG00000038039",
                "ENSMUSG00000055737",
                "ENSMUSG00000062604",
                "ENSMUSG00000032902",
                "ENSMUSG00000031633",
                "ENSMUSG00000027861",
                "ENSMUSG00000028013",
                "ENSMUSG00000000751",
                "ENSMUSG00000030409",
                "ENSMUSG00000065037",
                "ENSMUSG00000029632",
                "ENSMUSG00000026207",
                "ENSMUSG00000052337",
                "ENSMUSG00000022889",
                "ENSMUSG00000062981",
                "ENSMUSG00000064342",
                "ENSMUSG00000020456",
                "ENSMUSG00000021061",
                "ENSMUSG00000051747",
                "ENSMUSG00000026342",
                "ENSMUSG00000074466",
                "ENSMUSG00000086298",
                "ENSMUSG00000038239",
                "ENSMUSG00000063787",
                "ENSMUSG00000030399",
                "ENSMUSG00000033004",
                "ENSMUSG00000030470",
                "ENSMUSG00000026028",
                "ENSMUSG00000039910",
                "ENSMUSG00000021557",
                "ENSMUSG00000034621",
                "ENSMUSG00000027298",
                "ENSMUSG00000046756",
                "ENSMUSG00000001211",
                "ENSMUSG00000007877",
                "ENSMUSG00000049643",
                "ENSMUSG00000018893",
                "ENSMUSG00000034898",
                "ENSMUSG00000024527",
                "ENSMUSG00000038393",
                "ENSMUSG00000028756",
                "ENSMUSG00000025374",
                "ENSMUSG00000033429",
                "ENSMUSG00000003955",
                "ENSMUSG00000059923",
                "ENSMUSG00000021200",
                "ENSMUSG00000032786",
                "ENSMUSG00000071415",
                "ENSMUSG00000037152",
                "ENSMUSG00000019494",
                "ENSMUSG00000026032",
                "ENSMUSG00000022803",
                "ENSMUSG00000053768",
                "ENSMUSG00000023010",
                "ENSMUSG00000028312",
                "ENSMUSG00000064338",
                "ENSMUSG00000036278",
                "ENSMUSG00000028156",
                "ENSMUSG00000022391",
                "ENSMUSG00000041548",
                "ENSMUSG00000062169",
                "ENSMUSG00000028348",
                "ENSMUSG00000018796",
                "ENSMUSG00000005510",
                "ENSMUSG00000031461",
                "ENSMUSG00000055900",
                "ENSMUSG00000022403",
                "ENSMUSG00000027257",
                "ENSMUSG00000025591",
                "ENSMUSG00000028834",
                "ENSMUSG00000020074",
                "ENSMUSG00000036636",
                "ENSMUSG00000010608",
                "ENSMUSG00000024803",
                "ENSMUSG00000023861",
                "ENSMUSG00000031683",
                "ENSMUSG00000091537",
                "ENSMUSG00000039630",
                "ENSMUSG00000053964",
                "ENSMUSG00000024608",
                "ENSMUSG00000034994",
                "ENSMUSG00000023971",
                "ENSMUSG00000090862",
                "ENSMUSG00000026208",
                "ENSMUSG00000057914",
                "ENSMUSG00000034243",
                "ENSMUSG00000025968",
                "ENSMUSG00000029447",
                "ENSMUSG00000027238",
                "ENSMUSG00000053536",
                "ENSMUSG00000044033",
                "ENSMUSG00000065701",
                "ENSMUSG00000034842",
                "ENSMUSG00000064344",
                "ENSMUSG00000026404",
                "ENSMUSG00000027499",
                "ENSMUSG00000037805",
                "ENSMUSG00000061477",
                "ENSMUSG00000037573",
                "ENSMUSG00000033044",
                "ENSMUSG00000031701",
                "ENSMUSG00000027455",
                "ENSMUSG00000064340",
                "ENSMUSG00000029401",
                "ENSMUSG00000025453",
                "ENSMUSG00000032612",
                "ENSMUSG00000069089",
                "ENSMUSG00000072720",
                "ENSMUSG00000027303",
                "ENSMUSG00000036002",
                "ENSMUSG00000026817",
                "ENSMUSG00000039130",
                "ENSMUSG00000026864",
                "ENSMUSG00000078486",
                "ENSMUSG00000033416",
                "ENSMUSG00000075307",
                "ENSMUSG00000023942",
                "ENSMUSG00000079179",
                "ENSMUSG00000032330",
                "ENSMUSG00000045636",
                "ENSMUSG00000027671",
                "ENSMUSG00000038170",
                "ENSMUSG00000000787",
                "ENSMUSG00000013593",
                "ENSMUSG00000021771",
                "ENSMUSG00000032267",
                "ENSMUSG00000006315",
                "ENSMUSG00000064365",
                "ENSMUSG00000028461",
                "ENSMUSG00000042293",
                "ENSMUSG00000038403",
                "ENSMUSG00000026698",
                "ENSMUSG00000024457",
                "ENSMUSG00000038520",
                "ENSMUSG00000026308",
                "ENSMUSG00000038240",
                "ENSMUSG00000060981",
                "ENSMUSG00000029486",
                "ENSMUSG00000040964",
                "ENSMUSG00000027961",
                "ENSMUSG00000085779",
                "ENSMUSG00000020530",
                "ENSMUSG00000049881",
                "ENSMUSG00000032366",
                "ENSMUSG00000055053",
                "ENSMUSG00000058056",
                "ENSMUSG00000064369",
                "ENSMUSG00000015656",
                "ENSMUSG00000023944",
                "ENSMUSG00000025509",
                "ENSMUSG00000028383",
                "ENSMUSG00000029655",
                "ENSMUSG00000020358",
                "ENSMUSG00000053819",
                "ENSMUSG00000020475",
                "ENSMUSG00000022551",
                "ENSMUSG00000078671",
                "ENSMUSG00000083462",
                "ENSMUSG00000022884",
                "ENSMUSG00000021546",
                "ENSMUSG00000037887",
                "ENSMUSG00000052812",
                "ENSMUSG00000064372",
                "ENSMUSG00000032495",
                "ENSMUSG00000030847",
                "ENSMUSG00000034525",
                "ENSMUSG00000045103",
                "ENSMUSG00000055204",
                "ENSMUSG00000037062",
                "ENSMUSG00000021622",
                "ENSMUSG00000042505",
                "ENSMUSG00000029683",
                "ENSMUSG00000004698",
                "ENSMUSG00000020022",
                "ENSMUSG00000034088",
                "ENSMUSG00000023456",
                "ENSMUSG00000022389",
                "ENSMUSG00000030869",
                "ENSMUSG00000064371",
                "ENSMUSG00000022010",
                "ENSMUSG00000017817",
                "ENSMUSG00000059811",
                "ENSMUSG00000028847"
            ],
            "expression_sum": {
                "class1": 184.0,
                "class2": 204.0
            },
            "uniprot_ids": [
                "Q63943",
                "Q921S6",
                "E9QKT0",
                "E9Q5E0"
            ]
        },
        "ENSMUSG00000001441": {
            "id": "ENSMUSG00000001441",
            "type": "Gene",
            "symbol": "Npepps",
            "source": [
                "ENSMUSG00000010175",
                "ENSMUSG00000001280",
                "ENSMUSG00000019768",
                "ENSMUSG00000063524",
                "ENSMUSG00000045639",
                "ENSMUSG00000048410",
                "ENSMUSG00000016087"
            ],
            "target": [],
            "expression_sum": {
                "class1": 1201.0,
                "class2": 482.0
            },
            "uniprot_ids": [
                "Q11011",
                "F6Z7Y0",
                "F7ANF4",
                "F2Z3V5",
                "F6QYF8",
                "F6V7K3",
                "E9Q6F4",
                "E9Q039",
                "F6RAU6"
            ]
        },
        "ENSMUSG00000001445": {
            "id": "ENSMUSG00000001445",
            "type": "Gene",
            "symbol": "Mrpl10",
            "source": [
                "ENSMUSG00000001280",
                "ENSMUSG00000067430",
                "ENSMUSG00000063108"
            ],
            "target": [],
            "expression_sum": {
                "class1": 536.0,
                "class2": 239.0
            },
            "uniprot_ids": [
                "Q3TBW2",
                "A2A715"
            ]
        },
        "ENSMUSG00000001472": {
            "id": "ENSMUSG00000001472",
            "type": "Gene",
            "symbol": "Tcf25",
            "source": [
                "ENSMUSG00000015839",
                "ENSMUSG00000006215",
                "ENSMUSG00000024431",
                "ENSMUSG00000001280",
                "ENSMUSG00000019768",
                "ENSMUSG00000002265",
                "ENSMUSG00000020644"
            ],
            "target": [],
            "expression_sum": {
                "class1": 5523.0,
                "class2": 996.0
            },
            "uniprot_ids": [
                "Q8R3L2",
                "B2ZAC8",
                "A0A1D5RM90",
                "A0A1D5RLU0"
            ]
        },
        "ENSMUSG00000001508": {
            "id": "ENSMUSG00000001508",
            "type": "Gene",
            "symbol": "Sgca",
            "source": [
                "ENSMUSG00000040481",
                "ENSMUSG00000001419",
                "ENSMUSG00000001280",
                "ENSMUSG00000019768",
                "ENSMUSG00000058756"
            ],
            "target": [],
            "expression_sum": {
                "class1": 71.0,
                "class2": 518.0
            },
            "uniprot_ids": [
                "P82350",
                "Q5SWB0",
                "Q5SWB4"
            ]
        },
        "ENSMUSG00000001552": {
            "id": "ENSMUSG00000001552",
            "type": "Gene",
            "symbol": "Jup",
            "source": [
                "ENSMUSG00000003032",
                "ENSMUSG00000056501",
                "ENSMUSG00000041515",
                "ENSMUSG00000001280",
                "ENSMUSG00000005698"
            ],
            "target": [],
            "expression_sum": {
                "class1": 1716.0,
                "class2": 891.0
            },
            "uniprot_ids": [
                "Q02257"
            ]
        },
        "ENSMUSG00000001604": {
            "id": "ENSMUSG00000001604",
            "type": "Gene",
            "symbol": "Tcea3",
            "source": [
                "ENSMUSG00000001280",
                "ENSMUSG00000017299"
            ],
            "target": [],
            "expression_sum": {
                "class1": 942.0,
                "class2": 1838.0
            },
            "uniprot_ids": [
                "P23881",
                "A2AW45"
            ]
        },
        "ENSMUSG00000001666": {
            "id": "ENSMUSG00000001666",
            "type": "Gene",
            "symbol": "Ddt",
            "source": [
                "ENSMUSG00000001280",
                "ENSMUSG00000058239",
                "ENSMUSG00000023927"
            ],
            "target": [],
            "expression_sum": {
                "class1": 255.0,
                "class2": 303.0
            },
            "uniprot_ids": [
                "O35215",
                "G3UZN1"
            ]
        },
        "ENSMUSG00000001767": {
            "id": "ENSMUSG00000001767",
            "type": "Gene",
            "symbol": "Crnkl1",
            "source": [
                "ENSMUSG00000026663",
                "ENSMUSG00000044807",
                "ENSMUSG00000001280",
                "ENSMUSG00000019768",
                "ENSMUSG00000096795",
                "ENSMUSG00000055320",
                "ENSMUSG00000026803"
            ],
            "target": [],
            "expression_sum": {
                "class1": 1469.0,
                "class2": 309.0
            },
            "uniprot_ids": [
                "P63154"
            ]
        },
        "ENSMUSG00000001794": {
            "id": "ENSMUSG00000001794",
            "type": "Gene",
            "symbol": "Capns1",
            "source": [
                "ENSMUSG00000026398",
                "ENSMUSG00000029135",
                "ENSMUSG00000019817",
                "ENSMUSG00000021250",
                "ENSMUSG00000056501",
                "ENSMUSG00000018899",
                "ENSMUSG00000029249",
                "ENSMUSG00000001280",
                "ENSMUSG00000005566",
                "ENSMUSG00000005583",
                "ENSMUSG00000019768",
                "ENSMUSG00000030557",
                "ENSMUSG00000014704",
                "ENSMUSG00000024140",
                "ENSMUSG00000068267",
                "ENSMUSG00000031885",
                "ENSMUSG00000022811",
                "ENSMUSG00000015846"
            ],
            "target": [],
            "expression_sum": {
                "class1": 6084.0,
                "class2": 1890.0
            },
            "uniprot_ids": [
                "A0A0R4IZW8",
                "A0A0R4J1C2"
            ]
        },
        "ENSMUSG00000001833": {
            "id": "ENSMUSG00000001833",
            "type": "Gene",
            "symbol": "Septin7",
            "source": [
                "ENSMUSG00000017724",
                "ENSMUSG00000024515",
                "ENSMUSG00000030256",
                "ENSMUSG00000041515",
                "ENSMUSG00000001280",
                "ENSMUSG00000019768",
                "ENSMUSG00000040732"
            ],
            "target": [],
            "expression_sum": {
                "class1": 38775.0,
                "class2": 3985.0
            },
            "uniprot_ids": [
                "E9Q1G8",
                "E9Q9F5"
            ]
        },
        "ENSMUSG00000001911": {
            "id": "ENSMUSG00000001911",
            "type": "TF",
            "symbol": "Nfix",
            "source": [
                "ENSMUSG00000029135",
                "ENSMUSG00000019817",
                "ENSMUSG00000030256",
                "ENSMUSG00000029249",
                "ENSMUSG00000001280",
                "ENSMUSG00000019768",
                "ENSMUSG00000005583",
                "ENSMUSG00000017146",
                "ENSMUSG00000068267",
                "ENSMUSG00000008575",
                "ENSMUSG00000022314",
                "ENSMUSG00000029913"
            ],
            "target": [
                "ENSMUSG00000003037",
                "ENSMUSG00000016206",
                "ENSMUSG00000024290",
                "ENSMUSG00000070427",
                "ENSMUSG00000062846",
                "ENSMUSG00000000125",
                "ENSMUSG00000009863",
                "ENSMUSG00000058558",
                "ENSMUSG00000000127",
                "ENSMUSG00000005410",
                "ENSMUSG00000081352",
                "ENSMUSG00000084384",
                "ENSMUSG00000053536",
                "ENSMUSG00000021928",
                "ENSMUSG00000000628",
                "ENSMUSG00000022111",
                "ENSMUSG00000031613",
                "ENSMUSG00000045776",
                "ENSMUSG00000020441",
                "ENSMUSG00000030889",
                "ENSMUSG00000075705",
                "ENSMUSG00000000058",
                "ENSMUSG00000001127",
                "ENSMUSG00000000031",
                "ENSMUSG00000000088",
                "ENSMUSG00000020022",
                "ENSMUSG00000027669",
                "ENSMUSG00000091747",
                "ENSMUSG00000003778",
                "ENSMUSG00000000399",
                "ENSMUSG00000001424",
                "ENSMUSG00000000340",
                "ENSMUSG00000043889",
                "ENSMUSG00000038612",
                "ENSMUSG00000018893",
                "ENSMUSG00000023456",
                "ENSMUSG00000040265",
                "ENSMUSG00000043415",
                "ENSMUSG00000030243",
                "ENSMUSG00000000827",
                "ENSMUSG00000001119",
                "ENSMUSG00000005220",
                "ENSMUSG00000034484",
                "ENSMUSG00000030512",
                "ENSMUSG00000022194",
                "ENSMUSG00000029401",
                "ENSMUSG00000048355",
                "ENSMUSG00000015575",
                "ENSMUSG00000000168",
                "ENSMUSG00000057572",
                "ENSMUSG00000018585",
                "ENSMUSG00000000884",
                "ENSMUSG00000000378",
                "ENSMUSG00000022371",
                "ENSMUSG00000001525",
                "ENSMUSG00000008690",
                "ENSMUSG00000063870",
                "ENSMUSG00000034998",
                "ENSMUSG00000000441"
            ],
            "expression_sum": {
                "class1": 2637.0,
                "class2": 758.0
            },
            "uniprot_ids": [
                "P70257",
                "Q3TYK3",
                "D3YZ00",
                "E9PUH7"
            ]
        },
        "ENSMUSG00000002007": {
            "id": "ENSMUSG00000002007",
            "type": "Gene",
            "symbol": "Srpk3",
            "source": [
                "ENSMUSG00000001280",
                "ENSMUSG00000015579",
                "ENSMUSG00000038914"
            ],
            "target": [],
            "expression_sum": {
                "class1": 108.0,
                "class2": 206.0
            },
            "uniprot_ids": [
                "Q9Z0G2"
            ]
        },
        "ENSMUSG00000002010": {
            "id": "ENSMUSG00000002010",
            "type": "Gene",
            "symbol": "Idh3g",
            "source": [
                "ENSMUSG00000029135",
                "ENSMUSG00000003847",
                "ENSMUSG00000015605",
                "ENSMUSG00000008575",
                "ENSMUSG00000032897",
                "ENSMUSG00000040481",
                "ENSMUSG00000055148",
                "ENSMUSG00000040033",
                "ENSMUSG00000090258",
                "ENSMUSG00000019768",
                "ENSMUSG00000024140",
                "ENSMUSG00000026803",
                "ENSMUSG00000031715",
                "ENSMUSG00000024985",
                "ENSMUSG00000015839",
                "ENSMUSG00000058756",
                "ENSMUSG00000078671",
                "ENSMUSG00000056501",
                "ENSMUSG00000029249",
                "ENSMUSG00000001280",
                "ENSMUSG00000024498",
                "ENSMUSG00000009079",
                "ENSMUSG00000055320"
            ],
            "target": [],
            "expression_sum": {
                "class1": 2643.0,
                "class2": 3492.0
            },
            "uniprot_ids": [
                "P70404"
            ]
        },
        "ENSMUSG00000002015": {
            "id": "ENSMUSG00000002015",
            "type": "Gene",
            "symbol": "Bcap31",
            "source": [
                "ENSMUSG00000059552",
                "ENSMUSG00000022346",
                "ENSMUSG00000006215",
                "ENSMUSG00000030256",
                "ENSMUSG00000055116",
                "ENSMUSG00000001280",
                "ENSMUSG00000019768",
                "ENSMUSG00000005583",
                "ENSMUSG00000099083",
                "ENSMUSG00000038692"
            ],
            "target": [],
            "expression_sum": {
                "class1": 3913.0,
                "class2": 742.0
            },
            "uniprot_ids": [
                "Q61335"
            ]
        },
        "ENSMUSG00000002028": {
            "id": "ENSMUSG00000002028",
            "type": "Gene",
            "symbol": "Kmt2a",
            "source": [
                "ENSMUSG00000029135",
                "ENSMUSG00000003032",
                "ENSMUSG00000079215",
                "ENSMUSG00000029249",
                "ENSMUSG00000001280",
                "ENSMUSG00000027551",
                "ENSMUSG00000019768",
                "ENSMUSG00000030067",
                "ENSMUSG00000034429",
                "ENSMUSG00000029913"
            ],
            "target": [],
            "expression_sum": {
                "class1": 2542.0,
                "class2": 2082.0
            },
            "uniprot_ids": [
                "P55200",
                "S4R2S5",
                "S4R199",
                "D3Z5H5",
                "A0A1L1SQS9"
            ]
        },
        "ENSMUSG00000002102": {
            "id": "ENSMUSG00000002102",
            "type": "Gene",
            "symbol": "Psmc3",
            "source": [
                "ENSMUSG00000029135",
                "ENSMUSG00000057469",
                "ENSMUSG00000056501",
                "ENSMUSG00000001280",
                "ENSMUSG00000019768",
                "ENSMUSG00000026803",
                "ENSMUSG00000045251",
                "ENSMUSG00000031885",
                "ENSMUSG00000058756",
                "ENSMUSG00000008575",
                "ENSMUSG00000014592",
                "ENSMUSG00000015846"
            ],
            "target": [],
            "expression_sum": {
                "class1": 14347.0,
                "class2": 2496.0
            },
            "uniprot_ids": [
                "O88685",
                "F6Q2E3",
                "A0A087WPH7",
                "B7ZCF1",
                "A2AGN7"
            ]
        },
        "ENSMUSG00000002342": {
            "id": "ENSMUSG00000002342",
            "type": "Gene",
            "symbol": "Tmem161a",
            "source": [
                "ENSMUSG00000071076",
                "ENSMUSG00000078671",
                "ENSMUSG00000001280",
                "ENSMUSG00000019768",
                "ENSMUSG00000042557"
            ],
            "target": [],
            "expression_sum": {
                "class1": 445.0,
                "class2": 418.0
            },
            "uniprot_ids": [
                "Q8VCA6",
                "M0QWE7",
                "S4R1Z2",
                "F6UNW2",
                "S4R2H0",
                "M0QWV0"
            ]
        },
        "ENSMUSG00000002343": {
            "id": "ENSMUSG00000002343",
            "type": "Gene",
            "symbol": "Armc6",
            "source": [
                "ENSMUSG00000029135",
                "ENSMUSG00000059552",
                "ENSMUSG00000015605",
                "ENSMUSG00000026610",
                "ENSMUSG00000051469",
                "ENSMUSG00000029249",
                "ENSMUSG00000001280",
                "ENSMUSG00000005566",
                "ENSMUSG00000019768",
                "ENSMUSG00000022641",
                "ENSMUSG00000058756",
                "ENSMUSG00000008575",
                "ENSMUSG00000029913",
                "ENSMUSG00000024985"
            ],
            "target": [],
            "expression_sum": {
                "class1": 110.0,
                "class2": 82.0
            },
            "uniprot_ids": [
                "Q8BNU0",
                "D3Z3J0"
            ]
        },
        "ENSMUSG00000002379": {
            "id": "ENSMUSG00000002379",
            "type": "Gene",
            "symbol": "Ndufa11",
            "source": [
                "ENSMUSG00000003032",
                "ENSMUSG00000052534",
                "ENSMUSG00000015605",
                "ENSMUSG00000027490",
                "ENSMUSG00000001280",
                "ENSMUSG00000019768",
                "ENSMUSG00000040841",
                "ENSMUSG00000008575",
                "ENSMUSG00000014592",
                "ENSMUSG00000021775",
                "ENSMUSG00000015846"
            ],
            "target": [],
            "expression_sum": {
                "class1": 21470.0,
                "class2": 11435.0
            },
            "uniprot_ids": [
                "A0A3B2W4A5",
                "A0A3B2W441",
                "G5E814"
            ]
        },
        "ENSMUSG00000002416": {
            "id": "ENSMUSG00000002416",
            "type": "Gene",
            "symbol": "Ndufb2",
            "source": [
                "ENSMUSG00000029135",
                "ENSMUSG00000015839",
                "ENSMUSG00000052534",
                "ENSMUSG00000015605",
                "ENSMUSG00000055116",
                "ENSMUSG00000056501",
                "ENSMUSG00000015579",
                "ENSMUSG00000028639",
                "ENSMUSG00000029249",
                "ENSMUSG00000001280",
                "ENSMUSG00000019768",
                "ENSMUSG00000047371",
                "ENSMUSG00000068267",
                "ENSMUSG00000031885",
                "ENSMUSG00000008575",
                "ENSMUSG00000014592",
                "ENSMUSG00000015846"
            ],
            "target": [],
            "expression_sum": {
                "class1": 12451.0,
                "class2": 9814.0
            },
            "uniprot_ids": [
                "Q9CPU2"
            ]
        },
        "ENSMUSG00000002496": {
            "id": "ENSMUSG00000002496",
            "type": "Gene",
            "symbol": "Tsc2",
            "source": [
                "ENSMUSG00000029135",
                "ENSMUSG00000015839",
                "ENSMUSG00000078671",
                "ENSMUSG00000015579",
                "ENSMUSG00000001280",
                "ENSMUSG00000019768",
                "ENSMUSG00000028565",
                "ENSMUSG00000025958",
                "ENSMUSG00000022641",
                "ENSMUSG00000031885",
                "ENSMUSG00000016087",
                "ENSMUSG00000021775",
                "ENSMUSG00000031715"
            ],
            "target": [],
            "expression_sum": {
                "class1": 298.0,
                "class2": 253.0
            },
            "uniprot_ids": [
                "Q3UHB2",
                "Q3TQ10",
                "Q7TT21",
                "A0A2I3BRT5",
                "A0A2I3BRA1",
                "A0A2I3BPR7",
                "A0A2I3BPP1",
                "A0A2I3BPE9"
            ]
        },
        "ENSMUSG00000002608": {
            "id": "ENSMUSG00000002608",
            "type": "Gene",
            "symbol": "Ccdc97",
            "source": [
                "ENSMUSG00000025612",
                "ENSMUSG00000078671",
                "ENSMUSG00000056501",
                "ENSMUSG00000015579",
                "ENSMUSG00000023994",
                "ENSMUSG00000029249",
                "ENSMUSG00000001280",
                "ENSMUSG00000005566",
                "ENSMUSG00000019768",
                "ENSMUSG00000005698",
                "ENSMUSG00000032238",
                "ENSMUSG00000015605",
                "ENSMUSG00000026803",
                "ENSMUSG00000029913"
            ],
            "target": [],
            "expression_sum": {
                "class1": 195.0,
                "class2": 165.0
            },
            "uniprot_ids": [
                "Q9DBT3",
                "A0A0U1RP69"
            ]
        },
        "ENSMUSG00000002661": {
            "id": "ENSMUSG00000002661",
            "type": "Gene",
            "symbol": "Alkbh7",
            "source": [
                "ENSMUSG00000029135",
                "ENSMUSG00000001419",
                "ENSMUSG00000010175",
                "ENSMUSG00000056501",
                "ENSMUSG00000018899",
                "ENSMUSG00000027109",
                "ENSMUSG00000001280",
                "ENSMUSG00000005566",
                "ENSMUSG00000019768",
                "ENSMUSG00000030557",
                "ENSMUSG00000044167",
                "ENSMUSG00000031885",
                "ENSMUSG00000014592",
                "ENSMUSG00000015846"
            ],
            "target": [],
            "expression_sum": {
                "class1": 869.0,
                "class2": 662.0
            },
            "uniprot_ids": [
                "Q9D6Z0"
            ]
        },
        "ENSMUSG00000002728": {
            "id": "ENSMUSG00000002728",
            "type": "Gene",
            "symbol": "Naa20",
            "source": [
                "ENSMUSG00000024238",
                "ENSMUSG00000015579",
                "ENSMUSG00000030557",
                "ENSMUSG00000001280",
                "ENSMUSG00000019768",
                "ENSMUSG00000026872",
                "ENSMUSG00000036036"
            ],
            "target": [],
            "expression_sum": {
                "class1": 745.0,
                "class2": 566.0
            },
            "uniprot_ids": [
                "P61600",
                "Q9DB82",
                "G5E8X4",
                "E9PYH1",
                "E9Q023",
                "E9PXA4",
                "E9PZX8"
            ]
        },
        "ENSMUSG00000002748": {
            "id": "ENSMUSG00000002748",
            "type": "Gene",
            "symbol": "Baz1b",
            "source": [
                "ENSMUSG00000024238",
                "ENSMUSG00000020248",
                "ENSMUSG00000029135",
                "ENSMUSG00000021250",
                "ENSMUSG00000008575",
                "ENSMUSG00000035158",
                "ENSMUSG00000052534",
                "ENSMUSG00000030256",
                "ENSMUSG00000027109",
                "ENSMUSG00000090258",
                "ENSMUSG00000019768",
                "ENSMUSG00000022641",
                "ENSMUSG00000024974",
                "ENSMUSG00000031885",
                "ENSMUSG00000022314",
                "ENSMUSG00000015839",
                "ENSMUSG00000053477",
                "ENSMUSG00000005583",
                "ENSMUSG00000037608",
                "ENSMUSG00000001280"
            ],
            "target": [],
            "expression_sum": {
                "class1": 6200.0,
                "class2": 1020.0
            },
            "uniprot_ids": [
                "Q9Z277"
            ]
        },
        "ENSMUSG00000002767": {
            "id": "ENSMUSG00000002767",
            "type": "Gene",
            "symbol": "Mrpl2",
            "source": [
                "ENSMUSG00000015839",
                "ENSMUSG00000030256",
                "ENSMUSG00000055116",
                "ENSMUSG00000015579",
                "ENSMUSG00000018899",
                "ENSMUSG00000001280",
                "ENSMUSG00000019768",
                "ENSMUSG00000024985",
                "ENSMUSG00000020167"
            ],
            "target": [],
            "expression_sum": {
                "class1": 1552.0,
                "class2": 731.0
            },
            "uniprot_ids": [
                "Q9D773",
                "B1B1D7",
                "B1B1D8"
            ]
        },
        "ENSMUSG00000002844": {
            "id": "ENSMUSG00000002844",
            "type": "Gene",
            "symbol": "Adprh",
            "source": [
                "ENSMUSG00000058440",
                "ENSMUSG00000024238",
                "ENSMUSG00000029135",
                "ENSMUSG00000057469",
                "ENSMUSG00000052534",
                "ENSMUSG00000041515",
                "ENSMUSG00000001280",
                "ENSMUSG00000019768",
                "ENSMUSG00000005583",
                "ENSMUSG00000053477",
                "ENSMUSG00000016503",
                "ENSMUSG00000008575",
                "ENSMUSG00000022314"
            ],
            "target": [],
            "expression_sum": {
                "class1": 1689.0,
                "class2": 473.0
            },
            "uniprot_ids": [
                "P54923"
            ]
        },
        "ENSMUSG00000002908": {
            "id": "ENSMUSG00000002908",
            "type": "Gene",
            "symbol": "Kcnn1",
            "source": [
                "ENSMUSG00000001280",
                "ENSMUSG00000019768",
                "ENSMUSG00000005698"
            ],
            "target": [],
            "expression_sum": {
                "class1": 19.0,
                "class2": 118.0
            },
            "uniprot_ids": [
                "A0A140T8Q8",
                "A0A1D5RMF8",
                "Z4YL28",
                "A0A1D5RLS4",
                "A0A1D5RLR9",
                "A0A1D5RMB2",
                "A0A1D5RM38"
            ]
        },
        "ENSMUSG00000002948": {
            "id": "ENSMUSG00000002948",
            "type": "Gene",
            "symbol": "Map2k7",
            "source": [
                "ENSMUSG00000001280",
                "ENSMUSG00000041420",
                "ENSMUSG00000031885",
                "ENSMUSG00000032419"
            ],
            "target": [],
            "expression_sum": {
                "class1": 639.0,
                "class2": 396.0
            },
            "uniprot_ids": [
                "Q8CE90",
                "A0A140LHN8"
            ]
        },
        "ENSMUSG00000002957": {
            "id": "ENSMUSG00000002957",
            "type": "Gene",
            "symbol": "Ap2a2",
            "source": [
                "ENSMUSG00000040481",
                "ENSMUSG00000019564",
                "ENSMUSG00000030256",
                "ENSMUSG00000029249",
                "ENSMUSG00000001280",
                "ENSMUSG00000019768",
                "ENSMUSG00000031627",
                "ENSMUSG00000008575"
            ],
            "target": [],
            "expression_sum": {
                "class1": 2618.0,
                "class2": 461.0
            },
            "uniprot_ids": [
                "P17427",
                "A0A0J9YUA7"
            ]
        },
        "ENSMUSG00000002980": {
            "id": "ENSMUSG00000002980",
            "type": "Gene",
            "symbol": "Bcam",
            "source": [
                "ENSMUSG00000001280",
                "ENSMUSG00000019768",
                "ENSMUSG00000058756",
                "ENSMUSG00000056501"
            ],
            "target": [],
            "expression_sum": {
                "class1": 731.0,
                "class2": 401.0
            },
            "uniprot_ids": [
                "Q9R069",
                "D6RE44",
                "D3YTK7"
            ]
        },
        "ENSMUSG00000003072": {
            "id": "ENSMUSG00000003072",
            "type": "Gene",
            "symbol": "Atp5d",
            "source": [
                "ENSMUSG00000029135",
                "ENSMUSG00000044452",
                "ENSMUSG00000031627",
                "ENSMUSG00000015605",
                "ENSMUSG00000008575",
                "ENSMUSG00000021775",
                "ENSMUSG00000035158",
                "ENSMUSG00000057469",
                "ENSMUSG00000006215",
                "ENSMUSG00000058239",
                "ENSMUSG00000030256",
                "ENSMUSG00000019768",
                "ENSMUSG00000038914",
                "ENSMUSG00000028565",
                "ENSMUSG00000031885",
                "ENSMUSG00000024985",
                "ENSMUSG00000031715",
                "ENSMUSG00000015846",
                "ENSMUSG00000015839",
                "ENSMUSG00000030557",
                "ENSMUSG00000014592",
                "ENSMUSG00000029913",
                "ENSMUSG00000001419",
                "ENSMUSG00000078671",
                "ENSMUSG00000052837",
                "ENSMUSG00000056501",
                "ENSMUSG00000029249",
                "ENSMUSG00000001280",
                "ENSMUSG00000024498"
            ],
            "target": [],
            "expression_sum": {
                "class1": 9390.0,
                "class2": 7454.0
            },
            "uniprot_ids": [
                "Q9D3D9",
                "D3Z7S4"
            ]
        },
        "ENSMUSG00000003299": {
            "id": "ENSMUSG00000003299",
            "type": "Gene",
            "symbol": "Mrpl4",
            "source": [
                "ENSMUSG00000026610",
                "ENSMUSG00000056501",
                "ENSMUSG00000001280",
                "ENSMUSG00000005566",
                "ENSMUSG00000019768",
                "ENSMUSG00000005698",
                "ENSMUSG00000058756",
                "ENSMUSG00000015605",
                "ENSMUSG00000026803"
            ],
            "target": [],
            "expression_sum": {
                "class1": 1168.0,
                "class2": 600.0
            },
            "uniprot_ids": [
                "Q9DCU6",
                "A0A1L1SV70",
                "A0A1L1SU24",
                "A0A1L1STN5",
                "A0A1L1SSY9",
                "A0A1L1SRY8"
            ]
        },
        "ENSMUSG00000003316": {
            "id": "ENSMUSG00000003316",
            "type": "Gene",
            "symbol": "Glg1",
            "source": [
                "ENSMUSG00000029135",
                "ENSMUSG00000057469",
                "ENSMUSG00000015605",
                "ENSMUSG00000040033",
                "ENSMUSG00000029249",
                "ENSMUSG00000001280",
                "ENSMUSG00000005566",
                "ENSMUSG00000019768",
                "ENSMUSG00000032238",
                "ENSMUSG00000022641",
                "ENSMUSG00000013419",
                "ENSMUSG00000008575"
            ],
            "target": [],
            "expression_sum": {
                "class1": 3298.0,
                "class2": 745.0
            },
            "uniprot_ids": [
                "Q61543",
                "E9PWD8",
                "F8WHM5",
                "F6RSH1"
            ]
        },
        "ENSMUSG00000003380": {
            "id": "ENSMUSG00000003380",
            "type": "Gene",
            "symbol": "Rabac1",
            "source": [
                "ENSMUSG00000001419",
                "ENSMUSG00000037608",
                "ENSMUSG00000078671",
                "ENSMUSG00000023990",
                "ENSMUSG00000034673",
                "ENSMUSG00000001280",
                "ENSMUSG00000019768",
                "ENSMUSG00000031885",
                "ENSMUSG00000014592"
            ],
            "target": [],
            "expression_sum": {
                "class1": 5574.0,
                "class2": 1949.0
            },
            "uniprot_ids": [
                "Q9Z0S9",
                "A0A0U1RPM3"
            ]
        },
        "ENSMUSG00000003420": {
            "id": "ENSMUSG00000003420",
            "type": "Gene",
            "symbol": "Fcgrt",
            "source": [
                "ENSMUSG00000021250",
                "ENSMUSG00000010175",
                "ENSMUSG00000049739",
                "ENSMUSG00000001280",
                "ENSMUSG00000053477",
                "ENSMUSG00000005583",
                "ENSMUSG00000050619",
                "ENSMUSG00000016087",
                "ENSMUSG00000032035"
            ],
            "target": [],
            "expression_sum": {
                "class1": 4088.0,
                "class2": 608.0
            },
            "uniprot_ids": [
                "Q61559",
                "Q6PKB0",
                "A0A1B0GSK0",
                "A0A1B0GS17"
            ]
        },
        "ENSMUSG00000003429": {
            "id": "ENSMUSG00000003429",
            "type": "Gene",
            "symbol": "Rps11",
            "source": [
                "ENSMUSG00000029135",
                "ENSMUSG00000059552",
                "ENSMUSG00000021250",
                "ENSMUSG00000076431",
                "ENSMUSG00000024515",
                "ENSMUSG00000006215",
                "ENSMUSG00000022346",
                "ENSMUSG00000030256",
                "ENSMUSG00000015579",
                "ENSMUSG00000038648",
                "ENSMUSG00000019768",
                "ENSMUSG00000022641",
                "ENSMUSG00000031885",
                "ENSMUSG00000029178",
                "ENSMUSG00000053110",
                "ENSMUSG00000071637",
                "ENSMUSG00000015839",
                "ENSMUSG00000036461",
                "ENSMUSG00000005583",
                "ENSMUSG00000016087",
                "ENSMUSG00000001280",
                "ENSMUSG00000040732",
                "ENSMUSG00000032035"
            ],
            "target": [],
            "expression_sum": {
                "class1": 812.0,
                "class2": 1228.0
            },
            "uniprot_ids": [
                "P62281",
                "A0A1B0GSE8",
                "A0A1B0GRR3"
            ]
        },
        "ENSMUSG00000003438": {
            "id": "ENSMUSG00000003438",
            "type": "Gene",
            "symbol": "Timm50",
            "source": [
                "ENSMUSG00000057469",
                "ENSMUSG00000040481",
                "ENSMUSG00000015839",
                "ENSMUSG00000078671",
                "ENSMUSG00000015605",
                "ENSMUSG00000021485",
                "ENSMUSG00000029249",
                "ENSMUSG00000001280",
                "ENSMUSG00000019768",
                "ENSMUSG00000022641",
                "ENSMUSG00000046179",
                "ENSMUSG00000008575"
            ],
            "target": [],
            "expression_sum": {
                "class1": 2392.0,
                "class2": 369.0
            },
            "uniprot_ids": [
                "Q9D880"
            ]
        },
        "ENSMUSG00000003444": {
            "id": "ENSMUSG00000003444",
            "type": "Gene",
            "symbol": "Med29",
            "source": [
                "ENSMUSG00000001280",
                "ENSMUSG00000019768",
                "ENSMUSG00000027109",
                "ENSMUSG00000029249"
            ],
            "target": [],
            "expression_sum": {
                "class1": 810.0,
                "class2": 145.0
            },
            "uniprot_ids": [
                "Q9DB91"
            ]
        },
        "ENSMUSG00000003526": {
            "id": "ENSMUSG00000003526",
            "type": "Gene",
            "symbol": "Prodh",
            "source": [
                "ENSMUSG00000001280"
            ],
            "target": [],
            "expression_sum": {
                "class1": 127.0,
                "class2": 67.0
            },
            "uniprot_ids": [
                "Q9WU79",
                "F6YFQ5",
                "A0A338P738",
                "F6PYI8",
                "F6XMC0",
                "E9PUR5",
                "A0A0R4J1T9"
            ]
        },
        "ENSMUSG00000003531": {
            "id": "ENSMUSG00000003531",
            "type": "Gene",
            "symbol": "Dgcr6",
            "source": [
                "ENSMUSG00000026398",
                "ENSMUSG00000041515",
                "ENSMUSG00000029249",
                "ENSMUSG00000001280",
                "ENSMUSG00000005566",
                "ENSMUSG00000019768",
                "ENSMUSG00000040270",
                "ENSMUSG00000058756",
                "ENSMUSG00000055320"
            ],
            "target": [],
            "expression_sum": {
                "class1": 1538.0,
                "class2": 398.0
            },
            "uniprot_ids": [
                "Q3TV71",
                "E9QP31",
                "D3Z558",
                "F6W2M9",
                "D3YUN0"
            ]
        },
        "ENSMUSG00000003534": {
            "id": "ENSMUSG00000003534",
            "type": "Gene",
            "symbol": "Ddr1",
            "source": [
                "ENSMUSG00000078671",
                "ENSMUSG00000001280",
                "ENSMUSG00000019768",
                "ENSMUSG00000030087",
                "ENSMUSG00000058756",
                "ENSMUSG00000008575",
                "ENSMUSG00000022314"
            ],
            "target": [],
            "expression_sum": {
                "class1": 208.0,
                "class2": 127.0
            },
            "uniprot_ids": [
                "Q03146",
                "B0V2H8",
                "B0V2I0",
                "G3UXE5",
                "F7BZR8",
                "D6RII6",
                "B0V2H9",
                "E9PW79"
            ]
        },
        "ENSMUSG00000003541": {
            "id": "ENSMUSG00000003541",
            "type": "Gene",
            "symbol": "Ier3",
            "source": [
                "ENSMUSG00000071076",
                "ENSMUSG00000001280",
                "ENSMUSG00000019768",
                "ENSMUSG00000052684",
                "ENSMUSG00000031627"
            ],
            "target": [],
            "expression_sum": {
                "class1": 1309.0,
                "class2": 483.0
            },
            "uniprot_ids": [
                "Q91VZ5"
            ]
        },
        "ENSMUSG00000003559": {
            "id": "ENSMUSG00000003559",
            "type": "Gene",
            "symbol": "As3mt",
            "source": [
                "ENSMUSG00000001280",
                "ENSMUSG00000022346"
            ],
            "target": [],
            "expression_sum": {
                "class1": 532.0,
                "class2": 561.0
            },
            "uniprot_ids": [
                "Q91WU5",
                "A0A494B9D3",
                "A0A494BBJ9",
                "A0A494B985"
            ]
        },
        "ENSMUSG00000003809": {
            "id": "ENSMUSG00000003809",
            "type": "Gene",
            "symbol": "Gcdh",
            "source": [
                "ENSMUSG00000015839",
                "ENSMUSG00000010175",
                "ENSMUSG00000001280",
                "ENSMUSG00000019768",
                "ENSMUSG00000042557"
            ],
            "target": [],
            "expression_sum": {
                "class1": 350.0,
                "class2": 597.0
            },
            "uniprot_ids": [
                "Q60759",
                "A0A0A0MQ68",
                "D3Z4I2"
            ]
        },
        "ENSMUSG00000003865": {
            "id": "ENSMUSG00000003865",
            "type": "Gene",
            "symbol": "Gys1",
            "source": [
                "ENSMUSG00000029135",
                "ENSMUSG00000015839",
                "ENSMUSG00000049577",
                "ENSMUSG00000015605",
                "ENSMUSG00000078671",
                "ENSMUSG00000029249",
                "ENSMUSG00000001280",
                "ENSMUSG00000005566",
                "ENSMUSG00000019768",
                "ENSMUSG00000005698",
                "ENSMUSG00000032238",
                "ENSMUSG00000009079",
                "ENSMUSG00000031627",
                "ENSMUSG00000038692",
                "ENSMUSG00000008575"
            ],
            "target": [],
            "expression_sum": {
                "class1": 489.0,
                "class2": 393.0
            },
            "uniprot_ids": [
                "Q9Z1E4",
                "A0A1B0GR90",
                "A0A1B0GT92"
            ]
        },
        "ENSMUSG00000003923": {
            "id": "ENSMUSG00000003923",
            "type": "TF",
            "symbol": "Tfam",
            "source": [
                "ENSMUSG00000022346",
                "ENSMUSG00000021250",
                "ENSMUSG00000056501",
                "ENSMUSG00000041515",
                "ENSMUSG00000001280",
                "ENSMUSG00000028565",
                "ENSMUSG00000005698",
                "ENSMUSG00000031627",
                "ENSMUSG00000037235",
                "ENSMUSG00000031885",
                "ENSMUSG00000021775"
            ],
            "target": [
                "ENSMUSG00000028339",
                "ENSMUSG00000000538",
                "ENSMUSG00000028295",
                "ENSMUSG00000035781",
                "ENSMUSG00000000320",
                "ENSMUSG00000000563",
                "ENSMUSG00000029632",
                "ENSMUSG00000016252",
                "ENSMUSG00000001440",
                "ENSMUSG00000061462",
                "ENSMUSG00000000753",
                "ENSMUSG00000034911",
                "ENSMUSG00000062038",
                "ENSMUSG00000071014",
                "ENSMUSG00000019897",
                "ENSMUSG00000018796",
                "ENSMUSG00000030824",
                "ENSMUSG00000024099",
                "ENSMUSG00000020402",
                "ENSMUSG00000071796"
            ],
            "expression_sum": {
                "class1": 767.0,
                "class2": 391.0
            },
            "uniprot_ids": [
                "P40630",
                "A0A0R4J1N9",
                "D3Z0F3"
            ]
        },
        "ENSMUSG00000003934": {
            "id": "ENSMUSG00000003934",
            "type": "Gene",
            "symbol": "Efnb3",
            "source": [
                "ENSMUSG00000029249",
                "ENSMUSG00000001280",
                "ENSMUSG00000019768",
                "ENSMUSG00000024695",
                "ENSMUSG00000024974"
            ],
            "target": [],
            "expression_sum": {
                "class1": 104.0,
                "class2": 309.0
            },
            "uniprot_ids": [
                "O35393"
            ]
        },
        "ENSMUSG00000003948": {
            "id": "ENSMUSG00000003948",
            "type": "Gene",
            "symbol": "Mmd",
            "source": [
                "ENSMUSG00000001280",
                "ENSMUSG00000056501",
                "ENSMUSG00000027109",
                "ENSMUSG00000033543"
            ],
            "target": [],
            "expression_sum": {
                "class1": 2075.0,
                "class2": 810.0
            },
            "uniprot_ids": [
                "Q9CQY7",
                "Z4YKP7"
            ]
        },
        "ENSMUSG00000003955": {
            "id": "ENSMUSG00000003955",
            "type": "Gene",
            "symbol": "Fam162a",
            "source": [
                "ENSMUSG00000017724",
                "ENSMUSG00000029135",
                "ENSMUSG00000001419",
                "ENSMUSG00000020248",
                "ENSMUSG00000006215",
                "ENSMUSG00000015839",
                "ENSMUSG00000058239",
                "ENSMUSG00000015579",
                "ENSMUSG00000051469",
                "ENSMUSG00000001280",
                "ENSMUSG00000019768",
                "ENSMUSG00000005583",
                "ENSMUSG00000005698",
                "ENSMUSG00000053477",
                "ENSMUSG00000031885",
                "ENSMUSG00000008575",
                "ENSMUSG00000022314"
            ],
            "target": [],
            "expression_sum": {
                "class1": 4987.0,
                "class2": 2717.0
            },
            "uniprot_ids": [
                "Q9D6U8",
                "A0A338P6X2"
            ]
        },
        "ENSMUSG00000003970": {
            "id": "ENSMUSG00000003970",
            "type": "Gene",
            "symbol": "Rpl8",
            "source": [
                "ENSMUSG00000029135",
                "ENSMUSG00000040481",
                "ENSMUSG00000055148",
                "ENSMUSG00000019564",
                "ENSMUSG00000023990",
                "ENSMUSG00000056501",
                "ENSMUSG00000040033",
                "ENSMUSG00000030557",
                "ENSMUSG00000029249",
                "ENSMUSG00000001280",
                "ENSMUSG00000019768",
                "ENSMUSG00000002265",
                "ENSMUSG00000015605",
                "ENSMUSG00000026803"
            ],
            "target": [],
            "expression_sum": {
                "class1": 12731.0,
                "class2": 2987.0
            },
            "uniprot_ids": [
                "P62918"
            ]
        },
        "ENSMUSG00000004035": {
            "id": "ENSMUSG00000004035",
            "type": "Gene",
            "symbol": "Gstm7",
            "source": [
                "ENSMUSG00000051817",
                "ENSMUSG00000030557",
                "ENSMUSG00000001280",
                "ENSMUSG00000019768",
                "ENSMUSG00000022676",
                "ENSMUSG00000031715"
            ],
            "target": [],
            "expression_sum": {
                "class1": 148.0,
                "class2": 229.0
            },
            "uniprot_ids": [
                "Q80W21",
                "D3YVP6",
                "D3YVP5",
                "D3YZ29",
                "E9QAC8"
            ]
        },
        "ENSMUSG00000004044": {
            "id": "ENSMUSG00000004044",
            "type": "Gene",
            "symbol": "Cavin1",
            "source": [
                "ENSMUSG00000001280",
                "ENSMUSG00000019768",
                "ENSMUSG00000022346",
                "ENSMUSG00000056501"
            ],
            "target": [],
            "expression_sum": {
                "class1": 14144.0,
                "class2": 3680.0
            },
            "uniprot_ids": [
                "O54724"
            ]
        },
        "ENSMUSG00000004069": {
            "id": "ENSMUSG00000004069",
            "type": "Gene",
            "symbol": "Dnaja3",
            "source": [
                "ENSMUSG00000010175",
                "ENSMUSG00000022346",
                "ENSMUSG00000019564",
                "ENSMUSG00000039191",
                "ENSMUSG00000001280",
                "ENSMUSG00000019768",
                "ENSMUSG00000028565",
                "ENSMUSG00000036281",
                "ENSMUSG00000008575",
                "ENSMUSG00000031715",
                "ENSMUSG00000015846"
            ],
            "target": [],
            "expression_sum": {
                "class1": 293.0,
                "class2": 702.0
            },
            "uniprot_ids": [
                "Q99M87"
            ]
        },
        "ENSMUSG00000004085": {
            "id": "ENSMUSG00000004085",
            "type": "Gene",
            "symbol": "Map3k20",
            "source": [
                "ENSMUSG00000015839",
                "ENSMUSG00000010175",
                "ENSMUSG00000015605",
                "ENSMUSG00000022346",
                "ENSMUSG00000028639",
                "ENSMUSG00000001280",
                "ENSMUSG00000019768",
                "ENSMUSG00000031885",
                "ENSMUSG00000008575"
            ],
            "target": [],
            "expression_sum": {
                "class1": 2726.0,
                "class2": 1421.0
            },
            "uniprot_ids": [
                "Q9ESL4",
                "A2ASW6"
            ]
        },
        "ENSMUSG00000004096": {
            "id": "ENSMUSG00000004096",
            "type": "Gene",
            "symbol": "Cwc15",
            "source": [
                "ENSMUSG00000001280",
                "ENSMUSG00000032897",
                "ENSMUSG00000005698",
                "ENSMUSG00000020248"
            ],
            "target": [],
            "expression_sum": {
                "class1": 3268.0,
                "class2": 939.0
            },
            "uniprot_ids": [
                "Q9JHS9",
                "A0A1L1SVD8"
            ]
        },
        "ENSMUSG00000004268": {
            "id": "ENSMUSG00000004268",
            "type": "Gene",
            "symbol": "Emg1",
            "source": [
                "ENSMUSG00000029135",
                "ENSMUSG00000057469",
                "ENSMUSG00000015839",
                "ENSMUSG00000006215",
                "ENSMUSG00000073423",
                "ENSMUSG00000001280",
                "ENSMUSG00000019768",
                "ENSMUSG00000008575",
                "ENSMUSG00000031715"
            ],
            "target": [],
            "expression_sum": {
                "class1": 1415.0,
                "class2": 589.0
            },
            "uniprot_ids": [
                "O35130",
                "A0A0N4SUL6"
            ]
        },
        "ENSMUSG00000004356": {
            "id": "ENSMUSG00000004356",
            "type": "Gene",
            "symbol": "Utp20",
            "source": [
                "ENSMUSG00000029135",
                "ENSMUSG00000056501",
                "ENSMUSG00000029249",
                "ENSMUSG00000001280",
                "ENSMUSG00000019768",
                "ENSMUSG00000005698",
                "ENSMUSG00000008575",
                "ENSMUSG00000042557",
                "ENSMUSG00000031715"
            ],
            "target": [],
            "expression_sum": {
                "class1": 692.0,
                "class2": 113.0
            },
            "uniprot_ids": [
                "A0A1W2P8D7",
                "A0A1W2P6J9",
                "E9QK83",
                "A0A1W2P7L8"
            ]
        },
        "ENSMUSG00000004364": {
            "id": "ENSMUSG00000004364",
            "type": "Gene",
            "symbol": "Cul3",
            "source": [
                "ENSMUSG00000001280",
                "ENSMUSG00000029178",
                "ENSMUSG00000031885"
            ],
            "target": [],
            "expression_sum": {
                "class1": 1856.0,
                "class2": 1170.0
            },
            "uniprot_ids": [
                "Q9JLV5",
                "F6UY44",
                "E9Q4T8",
                "F6R0N8",
                "F6ZZK0"
            ]
        },
        "ENSMUSG00000004535": {
            "id": "ENSMUSG00000004535",
            "type": "Gene",
            "symbol": "Tax1bp1",
            "source": [
                "ENSMUSG00000029135",
                "ENSMUSG00000010175",
                "ENSMUSG00000001280",
                "ENSMUSG00000019768",
                "ENSMUSG00000040270",
                "ENSMUSG00000020919"
            ],
            "target": [],
            "expression_sum": {
                "class1": 12827.0,
                "class2": 2554.0
            },
            "uniprot_ids": [
                "Q3UKC1",
                "D3Z213",
                "D3Z152",
                "D3Z715"
            ]
        },
        "ENSMUSG00000004558": {
            "id": "ENSMUSG00000004558",
            "type": "Gene",
            "symbol": "Ndrg2",
            "source": [
                "ENSMUSG00000038539",
                "ENSMUSG00000026610",
                "ENSMUSG00000056501",
                "ENSMUSG00000001280",
                "ENSMUSG00000005566",
                "ENSMUSG00000005583",
                "ENSMUSG00000005698",
                "ENSMUSG00000019768",
                "ENSMUSG00000032238"
            ],
            "target": [],
            "expression_sum": {
                "class1": 3935.0,
                "class2": 5600.0
            },
            "uniprot_ids": [
                "Q9QYG0",
                "A0A2I3BPL1",
                "A0A2I3BQR5",
                "A0A2I3BPW0",
                "A0A2I3BS20",
                "A0A2I3BQM5"
            ]
        },
        "ENSMUSG00000004610": {
            "id": "ENSMUSG00000004610",
            "type": "Gene",
            "symbol": "Etfb",
            "source": [
                "ENSMUSG00000015839",
                "ENSMUSG00000078671",
                "ENSMUSG00000056501",
                "ENSMUSG00000015579",
                "ENSMUSG00000051469",
                "ENSMUSG00000047371",
                "ENSMUSG00000001280",
                "ENSMUSG00000019768",
                "ENSMUSG00000038914",
                "ENSMUSG00000024498",
                "ENSMUSG00000055320",
                "ENSMUSG00000053110",
                "ENSMUSG00000024695",
                "ENSMUSG00000068267",
                "ENSMUSG00000031885",
                "ENSMUSG00000015605",
                "ENSMUSG00000015846"
            ],
            "target": [],
            "expression_sum": {
                "class1": 16226.0,
                "class2": 9774.0
            },
            "uniprot_ids": [
                "Q9DCW4",
                "A0A0U1RNR3",
                "A0A0U1RNP5",
                "A0A0U1RQB4",
                "A0A0U1RNK9"
            ]
        },
        "ENSMUSG00000004748": {
            "id": "ENSMUSG00000004748",
            "type": "Gene",
            "symbol": "Mtfp1",
            "source": [
                "ENSMUSG00000001280",
                "ENSMUSG00000055116"
            ],
            "target": [],
            "expression_sum": {
                "class1": 619.0,
                "class2": 1114.0
            },
            "uniprot_ids": [
                "Q9CRB8"
            ]
        },
        "ENSMUSG00000004789": {
            "id": "ENSMUSG00000004789",
            "type": "Gene",
            "symbol": "Dlst",
            "source": [
                "ENSMUSG00000040481",
                "ENSMUSG00000019564",
                "ENSMUSG00000015605",
                "ENSMUSG00000028639",
                "ENSMUSG00000030557",
                "ENSMUSG00000029249",
                "ENSMUSG00000001280",
                "ENSMUSG00000019768",
                "ENSMUSG00000022641",
                "ENSMUSG00000031885",
                "ENSMUSG00000008575",
                "ENSMUSG00000021775",
                "ENSMUSG00000022895"
            ],
            "target": [],
            "expression_sum": {
                "class1": 1197.0,
                "class2": 2741.0
            },
            "uniprot_ids": [
                "Q9D2G2",
                "A0A1Y7VN44"
            ]
        },
        "ENSMUSG00000004891": {
            "id": "ENSMUSG00000004891",
            "type": "Gene",
            "symbol": "Nes",
            "source": [
                "ENSMUSG00000001280",
                "ENSMUSG00000036721",
                "ENSMUSG00000022314"
            ],
            "target": [],
            "expression_sum": {
                "class1": 13080.0,
                "class2": 1459.0
            },
            "uniprot_ids": [
                "Q6P5H2"
            ]
        },
        "ENSMUSG00000004931": {
            "id": "ENSMUSG00000004931",
            "type": "Gene",
            "symbol": "Apba3",
            "source": [
                "ENSMUSG00000058440",
                "ENSMUSG00000006215",
                "ENSMUSG00000056501",
                "ENSMUSG00000029249",
                "ENSMUSG00000001280",
                "ENSMUSG00000005566",
                "ENSMUSG00000019768",
                "ENSMUSG00000017146",
                "ENSMUSG00000025025",
                "ENSMUSG00000031715",
                "ENSMUSG00000029913",
                "ENSMUSG00000022895"
            ],
            "target": [],
            "expression_sum": {
                "class1": 118.0,
                "class2": 244.0
            },
            "uniprot_ids": [
                "O88888",
                "A0A1W2P7G9",
                "A0A1W2P6Z9",
                "A0A1W2P837",
                "A0A1W2P6S0"
            ]
        },
        "ENSMUSG00000004937": {
            "id": "ENSMUSG00000004937",
            "type": "Gene",
            "symbol": "Sgta",
            "source": [
                "ENSMUSG00000029135",
                "ENSMUSG00000021250",
                "ENSMUSG00000018604",
                "ENSMUSG00000029178",
                "ENSMUSG00000030557",
                "ENSMUSG00000001280",
                "ENSMUSG00000005583",
                "ENSMUSG00000016087",
                "ENSMUSG00000053110",
                "ENSMUSG00000035158",
                "ENSMUSG00000020167"
            ],
            "target": [],
            "expression_sum": {
                "class1": 2010.0,
                "class2": 411.0
            },
            "uniprot_ids": [
                "Q8BJU0",
                "A0A1W2P7I5",
                "A0A1W2P6P1"
            ]
        },
        "ENSMUSG00000004951": {
            "id": "ENSMUSG00000004951",
            "type": "Gene",
            "symbol": "Hspb1",
            "source": [
                "ENSMUSG00000019947",
                "ENSMUSG00000024238",
                "ENSMUSG00000028466",
                "ENSMUSG00000001280",
                "ENSMUSG00000053477",
                "ENSMUSG00000005583",
                "ENSMUSG00000031627",
                "ENSMUSG00000032035",
                "ENSMUSG00000008575",
                "ENSMUSG00000022314"
            ],
            "target": [],
            "expression_sum": {
                "class1": 2026.0,
                "class2": 1703.0
            },
            "uniprot_ids": [
                "P14602",
                "D3YZ06"
            ]
        },
        "ENSMUSG00000004980": {
            "id": "ENSMUSG00000004980",
            "type": "Gene",
            "symbol": "Hnrnpa2b1",
            "source": [
                "ENSMUSG00000029135",
                "ENSMUSG00000010175",
                "ENSMUSG00000015839",
                "ENSMUSG00000056501",
                "ENSMUSG00000015579",
                "ENSMUSG00000001280",
                "ENSMUSG00000019768",
                "ENSMUSG00000028565",
                "ENSMUSG00000014592",
                "ENSMUSG00000008958",
                "ENSMUSG00000053110",
                "ENSMUSG00000022389",
                "ENSMUSG00000068267",
                "ENSMUSG00000076431",
                "ENSMUSG00000008575",
                "ENSMUSG00000039191",
                "ENSMUSG00000029913",
                "ENSMUSG00000015846"
            ],
            "target": [],
            "expression_sum": {
                "class1": 29892.0,
                "class2": 7972.0
            },
            "uniprot_ids": [
                "O88569",
                "A0A0N4SUM2"
            ]
        },
        "ENSMUSG00000005034": {
            "id": "ENSMUSG00000005034",
            "type": "Gene",
            "symbol": "Prkacb",
            "source": [
                "ENSMUSG00000024238",
                "ENSMUSG00000019817",
                "ENSMUSG00000052534",
                "ENSMUSG00000001280",
                "ENSMUSG00000005583",
                "ENSMUSG00000022314"
            ],
```
