## Supplement 3 (P7 CM vs. P28 CM) for "AGEAS: Automated Machine Learning based Genetic Regulatory Element Extraction System": outlier_grps.js

```
{
    "ENSMUSG00000023456_ENSMUSG00000001280": 173.15586930040922,
    "ENSMUSG00000025290_ENSMUSG00000001280": 161.842946578805,
    "ENSMUSG00000071528_ENSMUSG00000019768": 43.266341219259154,
    "ENSMUSG00000053329_ENSMUSG00000001280": 26.47930589253777,
    "ENSMUSG00000038690_ENSMUSG00000019768": 24.800752189561333,
    "ENSMUSG00000021000_ENSMUSG00000001280": 22.914135689341947,
    "ENSMUSG00000001604_ENSMUSG00000001280": 22.87282326192546,
    "ENSMUSG00000032199_ENSMUSG00000001280": 22.053109841600637,
    "ENSMUSG00000064370_ENSMUSG00000001280": 21.96812720002869,
    "ENSMUSG00000038370_ENSMUSG00000001280": 21.73511564057947,
    "ENSMUSG00000093674_ENSMUSG00000001280": 21.68490770267772,
    "ENSMUSG00000001280_ENSMUSG00000000088": 21.00657408745297,
    "ENSMUSG00000025794_ENSMUSG00000001280": 20.97942851639826,
    "ENSMUSG00000017404_ENSMUSG00000001280": 20.976587028740816,
    "ENSMUSG00000033938_ENSMUSG00000001280": 20.932851329875138,
    "ENSMUSG00000082884_ENSMUSG00000001280": 19.907901285550196,
    "ENSMUSG00000063694_ENSMUSG00000001280": 19.84206388776648,
    "ENSMUSG00000091478_ENSMUSG00000001280": 19.66111091150793,
    "ENSMUSG00000087943_ENSMUSG00000001280": 19.62715601981537,
    "ENSMUSG00000019768_ENSMUSG00000014313": 19.459806041261007,
    "ENSMUSG00000084319_ENSMUSG00000001280": 19.124626964981037,
    "ENSMUSG00000027133_ENSMUSG00000001280": 17.863244202693892,
    "ENSMUSG00000025508_ENSMUSG00000001280": 17.271238245345483,
    "ENSMUSG00000059734_ENSMUSG00000001280": 16.092638612813168,
    "ENSMUSG00000023277_ENSMUSG00000001280": 14.467224241073021,
    "ENSMUSG00000078671_ENSMUSG00000066979": 14.155335562233743,
    "ENSMUSG00000031767_ENSMUSG00000001280": 13.655507634812759,
    "ENSMUSG00000056501_ENSMUSG00000035674": 13.619485279052382,
    "ENSMUSG00000020453_ENSMUSG00000001280": 13.10583974720821,
    "ENSMUSG00000019768_ENSMUSG00000000088": 12.869019768613287,
    "ENSMUSG00000083240_ENSMUSG00000001280": 12.819991649850511,
    "ENSMUSG00000030470_ENSMUSG00000019768": 12.379201967964663,
    "ENSMUSG00000063882_ENSMUSG00000019768": 12.28846093796364,
    "ENSMUSG00000020849_ENSMUSG00000019768": 12.283707478501782,
    "ENSMUSG00000019768_ENSMUSG00000005610": 12.195206203660145,
    "ENSMUSG00000022890_ENSMUSG00000001280": 11.996461993544312,
    "ENSMUSG00000035458_ENSMUSG00000019768": 11.75454546394766,
    "ENSMUSG00000022477_ENSMUSG00000001280": 11.698580111170946,
    "ENSMUSG00000029455_ENSMUSG00000001280": 11.670768267938689,
    "ENSMUSG00000026385_ENSMUSG00000001280": 11.596272330825647,
    "ENSMUSG00000089838_ENSMUSG00000019768": 11.574636556644887,
    "ENSMUSG00000021577_ENSMUSG00000019768": 11.534729629302037,
    "ENSMUSG00000091478_ENSMUSG00000019768": 11.527657680130927,
    "ENSMUSG00000014313_ENSMUSG00000001280": 11.460410049832365,
    "ENSMUSG00000067736_ENSMUSG00000001280": 11.413733869795692,
    "ENSMUSG00000032047_ENSMUSG00000001280": 11.390133555302192,
    "ENSMUSG00000038690_ENSMUSG00000001280": 11.389694743503208,
    "ENSMUSG00000035674_ENSMUSG00000019768": 11.335759519031372,
    "ENSMUSG00000031818_ENSMUSG00000019768": 11.306066170116308,
    "ENSMUSG00000029017_ENSMUSG00000001280": 11.28943276058486,
    "ENSMUSG00000039891_ENSMUSG00000001280": 11.28661863476335,
    "ENSMUSG00000004096_ENSMUSG00000001280": 11.279157632351096,
    "ENSMUSG00000019797_ENSMUSG00000001280": 11.193003957127688,
    "ENSMUSG00000008668_ENSMUSG00000001280": 11.191062420950498,
    "ENSMUSG00000064346_ENSMUSG00000001280": 11.159365809262281,
    "ENSMUSG00000064339_ENSMUSG00000019768": 11.153410123867864,
    "ENSMUSG00000019877_ENSMUSG00000001280": 11.080822597880783,
    "ENSMUSG00000027447_ENSMUSG00000001280": 11.076580329464191,
    "ENSMUSG00000073676_ENSMUSG00000001280": 10.961663885293905,
    "ENSMUSG00000030647_ENSMUSG00000001280": 10.926336426386156,
    "ENSMUSG00000064339_ENSMUSG00000001280": 10.89996027311148,
    "ENSMUSG00000003955_ENSMUSG00000001280": 10.8781525303297,
    "ENSMUSG00000024871_ENSMUSG00000001280": 10.854251023674186,
    "ENSMUSG00000025393_ENSMUSG00000001280": 10.797237202658938,
    "ENSMUSG00000060938_ENSMUSG00000001280": 10.792826205153634,
    "ENSMUSG00000048416_ENSMUSG00000001280": 10.777103883264376,
    "ENSMUSG00000028639_ENSMUSG00000001280": 10.75021556110606,
    "ENSMUSG00000050608_ENSMUSG00000001280": 10.723613107459027,
    "ENSMUSG00000021843_ENSMUSG00000001280": 10.709558238164163,
    "ENSMUSG00000022205_ENSMUSG00000001280": 10.684441590521109,
    "ENSMUSG00000038170_ENSMUSG00000001280": 10.669760774311866,
    "ENSMUSG00000063694_ENSMUSG00000019768": 10.66087204852009,
    "ENSMUSG00000020267_ENSMUSG00000001280": 10.641890893852832,
    "ENSMUSG00000064349_ENSMUSG00000001280": 10.609598275760904,
    "ENSMUSG00000002265_ENSMUSG00000001506": 10.606288439587463,
    "ENSMUSG00000021313_ENSMUSG00000019768": 10.602463807488325,
    "ENSMUSG00000026208_ENSMUSG00000001280": 10.596061768372742,
    "ENSMUSG00000091449_ENSMUSG00000001280": 10.594787265448133,
    "ENSMUSG00000040850_ENSMUSG00000001280": 10.579896748788604,
    "ENSMUSG00000017778_ENSMUSG00000001280": 10.503468046365155,
    "ENSMUSG00000020326_ENSMUSG00000001280": 10.489603178053224,
    "ENSMUSG00000029632_ENSMUSG00000019768": 10.463887826780002,
    "ENSMUSG00000023089_ENSMUSG00000019768": 10.430411953932081,
    "ENSMUSG00000041841_ENSMUSG00000001280": 10.406253298640715,
    "ENSMUSG00000016252_ENSMUSG00000001280": 10.384449626835629,
    "ENSMUSG00000021577_ENSMUSG00000001280": 10.379801577545656,
    "ENSMUSG00000022354_ENSMUSG00000001280": 10.264816568450989,
    "ENSMUSG00000081331_ENSMUSG00000001280": 10.247744675675746,
    "ENSMUSG00000041881_ENSMUSG00000001280": 10.24607877461018,
    "ENSMUSG00000030246_ENSMUSG00000001280": 10.210816244461244,
    "ENSMUSG00000064340_ENSMUSG00000001280": 10.186819199815288,
    "ENSMUSG00000006057_ENSMUSG00000001280": 10.164282008744795,
    "ENSMUSG00000081051_ENSMUSG00000001280": 10.16243026155263,
    "ENSMUSG00000025241_ENSMUSG00000001280": 10.155595346376202,
    "ENSMUSG00000074800_ENSMUSG00000001280": 10.141350689077484,
    "ENSMUSG00000027602_ENSMUSG00000001280": 10.136288226188547,
    "ENSMUSG00000068706_ENSMUSG00000001280": 10.09986639644704,
    "ENSMUSG00000063229_ENSMUSG00000019768": 10.099532535561067,
    "ENSMUSG00000032366_ENSMUSG00000019768": 10.09750327138814,
    "ENSMUSG00000020163_ENSMUSG00000019768": 10.074990678526264,
    "ENSMUSG00000055320_ENSMUSG00000001280": 10.057244711061255,
    "ENSMUSG00000084105_ENSMUSG00000001280": 10.056179305453329,
    "ENSMUSG00000053898_ENSMUSG00000001280": 10.03641250372757,
    "ENSMUSG00000030541_ENSMUSG00000001280": 10.029697229443514,
    "ENSMUSG00000080893_ENSMUSG00000001280": 10.012542789925979,
    "ENSMUSG00000005610_ENSMUSG00000001280": 10.00896096630181,
    "ENSMUSG00000081344_ENSMUSG00000001280": 10.004126870997704,
    "ENSMUSG00000064363_ENSMUSG00000019768": 9.988034648063982,
    "ENSMUSG00000058076_ENSMUSG00000001280": 9.973586718861347,
    "ENSMUSG00000027673_ENSMUSG00000001280": 9.966454602135272,
    "ENSMUSG00000023951_ENSMUSG00000001280": 9.949397801267734,
    "ENSMUSG00000098178_ENSMUSG00000001280": 9.923705583212223,
    "ENSMUSG00000040048_ENSMUSG00000001280": 9.906695363903106,
    "ENSMUSG00000044894_ENSMUSG00000001280": 9.903536562329652,
    "ENSMUSG00000053477_ENSMUSG00000001280": 9.836568729376868,
    "ENSMUSG00000014294_ENSMUSG00000001280": 9.828190470606632,
    "ENSMUSG00000074218_ENSMUSG00000001280": 9.816208797418229,
    "ENSMUSG00000054889_ENSMUSG00000001280": 9.815224957848926,
    "ENSMUSG00000009863_ENSMUSG00000001280": 9.789977652393665,
    "ENSMUSG00000074218_ENSMUSG00000019768": 9.772417111957967,
    "ENSMUSG00000025190_ENSMUSG00000001280": 9.74778037332574,
    "ENSMUSG00000022452_ENSMUSG00000001280": 9.746721936741396,
    "ENSMUSG00000053702_ENSMUSG00000001280": 9.693404042895446,
    "ENSMUSG00000031299_ENSMUSG00000001280": 9.678570921945898,
    "ENSMUSG00000031701_ENSMUSG00000001280": 9.674069745145362,
    "ENSMUSG00000038462_ENSMUSG00000001280": 9.671172480508238,
    "ENSMUSG00000062997_ENSMUSG00000001280": 9.663837053799014,
    "ENSMUSG00000057580_ENSMUSG00000019768": 9.64522154492427,
    "ENSMUSG00000074129_ENSMUSG00000001280": 9.636389477030223,
    "ENSMUSG00000025488_ENSMUSG00000019768": 9.620380274327921,
    "ENSMUSG00000030747_ENSMUSG00000001280": 9.603719985955369,
    "ENSMUSG00000086567_ENSMUSG00000001280": 9.545380252542534,
    "ENSMUSG00000049832_ENSMUSG00000001280": 9.450306067325586,
    "ENSMUSG00000064369_ENSMUSG00000001280": 9.367130242904237,
    "ENSMUSG00000044894_ENSMUSG00000019768": 9.284560874461333,
    "ENSMUSG00000027984_ENSMUSG00000001280": 9.268253909925093,
    "ENSMUSG00000049265_ENSMUSG00000001280": 9.26067670620855,
    "ENSMUSG00000025204_ENSMUSG00000001280": 9.209869243695888,
    "ENSMUSG00000027809_ENSMUSG00000001280": 9.206290276020798,
    "ENSMUSG00000030417_ENSMUSG00000001280": 9.199971049106002,
    "ENSMUSG00000058927_ENSMUSG00000019768": 9.15569461421031,
    "ENSMUSG00000019768_ENSMUSG00000005683": 9.054539090116807,
    "ENSMUSG00000021313_ENSMUSG00000001280": 9.026741275748558,
    "ENSMUSG00000073411_ENSMUSG00000052684": 9.003814665406995,
    "ENSMUSG00000032279_ENSMUSG00000001280": 8.996443647442572,
    "ENSMUSG00000073702_ENSMUSG00000001280": 8.99642096187753,
    "ENSMUSG00000019689_ENSMUSG00000001280": 8.991285689416884,
    "ENSMUSG00000064360_ENSMUSG00000019768": 8.977628960343143,
    "ENSMUSG00000025781_ENSMUSG00000019768": 8.975875732866031,
    "ENSMUSG00000049751_ENSMUSG00000001280": 8.845211383950755,
    "ENSMUSG00000062397_ENSMUSG00000001280": 8.810568364466716,
    "ENSMUSG00000028452_ENSMUSG00000001280": 8.787905120668688,
    "ENSMUSG00000082687_ENSMUSG00000001280": 8.768746304551938,
    "ENSMUSG00000039105_ENSMUSG00000001280": 8.490316024789069,
    "ENSMUSG00000031633_ENSMUSG00000019768": 8.424498449308759,
    "ENSMUSG00000062908_ENSMUSG00000008575": 8.394797598852138,
    "ENSMUSG00000070493_ENSMUSG00000035158": 8.056766393783697,
    "ENSMUSG00000024991_ENSMUSG00000001280": 8.055939025130233,
    "ENSMUSG00000027293_ENSMUSG00000001280": 8.025234097867845,
    "ENSMUSG00000048486_ENSMUSG00000001280": 7.990839750362555,
    "ENSMUSG00000028936_ENSMUSG00000001280": 7.990091765407867,
    "ENSMUSG00000025190_ENSMUSG00000008575": 7.979581964810497,
    "ENSMUSG00000025453_ENSMUSG00000019768": 7.64207391139255,
    "ENSMUSG00000061104_ENSMUSG00000001280": 7.570082562381152,
    "ENSMUSG00000006005_ENSMUSG00000001280": 7.361598380066219,
    "ENSMUSG00000022820_ENSMUSG00000019768": 7.306802379306191,
    "ENSMUSG00000001280_ENSMUSG00000001211": 7.175804166718762,
    "ENSMUSG00000030557_ENSMUSG00000025085": 6.7948156079054565,
    "ENSMUSG00000028207_ENSMUSG00000001280": 6.7690573817251956,
    "ENSMUSG00000030409_ENSMUSG00000001280": 6.6023044332143614,
    "ENSMUSG00000053093_ENSMUSG00000019768": 6.587412102552988,
    "ENSMUSG00000025066_ENSMUSG00000001280": 6.557357769127576,
    "ENSMUSG00000063787_ENSMUSG00000019768": 6.5228749216010655,
    "ENSMUSG00000020088_ENSMUSG00000001280": 6.439907295734913,
    "ENSMUSG00000090553_ENSMUSG00000001280": 6.428191812218424,
    "ENSMUSG00000029723_ENSMUSG00000001280": 6.398798239157052,
    "ENSMUSG00000070544_ENSMUSG00000001280": 6.056925992558757,
    "ENSMUSG00000020153_ENSMUSG00000001280": 5.846659506835455,
    "ENSMUSG00000059741_ENSMUSG00000019768": 5.845938414700699,
    "ENSMUSG00000030357_ENSMUSG00000001280": 5.773394137507462,
    "ENSMUSG00000025968_ENSMUSG00000019768": 5.493464547328771,
    "ENSMUSG00000022108_ENSMUSG00000001280": 5.372276392158416,
    "ENSMUSG00000058558_ENSMUSG00000001280": 5.128734070037296,
    "ENSMUSG00000056501_ENSMUSG00000045983": 4.897225150014417,
    "ENSMUSG00000075595_ENSMUSG00000034220": 4.1918863254300875,
    "ENSMUSG00000021760_ENSMUSG00000003032": 3.0084832208600725
}
```
