## Supplement 3 (P7 CM vs. P28 CM) for "AGEAS: Automated Machine Learning based Genetic Regulatory Element Extraction System": outlier_grps.js

```
{
    "ENSMUSG00000023456_ENSMUSG00000001280": 176.98248704303603,
    "ENSMUSG00000021313_ENSMUSG00000001280": 91.21681015990701,
    "ENSMUSG00000020326_ENSMUSG00000001280": 32.81266598881995,
    "ENSMUSG00000022354_ENSMUSG00000019768": 29.09973145664068,
    "ENSMUSG00000022890_ENSMUSG00000019768": 22.253508108799164,
    "ENSMUSG00000038615_ENSMUSG00000001280": 21.3448216861397,
    "ENSMUSG00000064371_ENSMUSG00000001280": 21.320635667720403,
    "ENSMUSG00000019179_ENSMUSG00000001280": 20.598118163774252,
    "ENSMUSG00000064367_ENSMUSG00000019768": 20.12642557015844,
    "ENSMUSG00000061104_ENSMUSG00000001280": 20.1170674094318,
    "ENSMUSG00000023861_ENSMUSG00000001280": 19.04290878428217,
    "ENSMUSG00000032959_ENSMUSG00000001280": 18.84833532023267,
    "ENSMUSG00000093674_ENSMUSG00000001280": 18.556629820315493,
    "ENSMUSG00000031059_ENSMUSG00000001280": 18.508225964619903,
    "ENSMUSG00000030884_ENSMUSG00000001280": 18.38005129999674,
    "ENSMUSG00000064351_ENSMUSG00000001280": 17.712736136781817,
    "ENSMUSG00000035242_ENSMUSG00000001280": 16.085740209135928,
    "ENSMUSG00000029661_ENSMUSG00000002265": 16.040192636247617,
    "ENSMUSG00000015733_ENSMUSG00000001280": 14.979014213106332,
    "ENSMUSG00000037685_ENSMUSG00000030557": 13.95032307085265,
    "ENSMUSG00000022217_ENSMUSG00000019768": 13.357533676877209,
    "ENSMUSG00000067653_ENSMUSG00000019768": 12.852415640739713,
    "ENSMUSG00000029614_ENSMUSG00000019768": 12.450110043990406,
    "ENSMUSG00000025453_ENSMUSG00000019768": 12.209249983808897,
    "ENSMUSG00000032238_ENSMUSG00000022477": 12.086270787970713,
    "ENSMUSG00000038170_ENSMUSG00000010175": 11.718160361318992,
    "ENSMUSG00000064347_ENSMUSG00000019768": 11.629785204474468,
    "ENSMUSG00000064337_ENSMUSG00000001280": 11.610442801395656,
    "ENSMUSG00000021877_ENSMUSG00000016559": 11.39380700663306,
    "ENSMUSG00000044033_ENSMUSG00000001280": 11.205013339271423,
    "ENSMUSG00000063882_ENSMUSG00000019768": 11.039170425357856,
    "ENSMUSG00000018585_ENSMUSG00000001280": 11.031580117194279,
    "ENSMUSG00000019768_ENSMUSG00000000563": 11.003150348224251,
    "ENSMUSG00000024049_ENSMUSG00000019768": 10.993167166446002,
    "ENSMUSG00000032786_ENSMUSG00000001280": 10.90901789581017,
    "ENSMUSG00000001280_ENSMUSG00000001211": 10.878531233337574,
    "ENSMUSG00000061904_ENSMUSG00000019768": 10.853546177640855,
    "ENSMUSG00000031812_ENSMUSG00000019768": 10.81789647651326,
    "ENSMUSG00000019768_ENSMUSG00000014294": 10.806874155137486,
    "ENSMUSG00000029729_ENSMUSG00000028603": 10.724935126513635,
    "ENSMUSG00000035158_ENSMUSG00000015733": 10.670087441626288,
    "ENSMUSG00000020741_ENSMUSG00000001280": 10.525277578048676,
    "ENSMUSG00000064372_ENSMUSG00000019768": 10.458265771876873,
    "ENSMUSG00000006717_ENSMUSG00000001280": 10.41936552767762,
    "ENSMUSG00000011257_ENSMUSG00000001280": 10.407128030854569,
    "ENSMUSG00000048486_ENSMUSG00000001280": 10.40537854202438,
    "ENSMUSG00000031818_ENSMUSG00000019768": 10.390141772844673,
    "ENSMUSG00000019768_ENSMUSG00000016427": 10.364148376141413,
    "ENSMUSG00000021290_ENSMUSG00000019768": 10.3316087448725,
    "ENSMUSG00000091478_ENSMUSG00000019768": 10.279133902532042,
    "ENSMUSG00000035674_ENSMUSG00000019768": 10.270115779646758,
    "ENSMUSG00000025651_ENSMUSG00000001280": 10.257927973623184,
    "ENSMUSG00000062908_ENSMUSG00000001280": 10.244283821658275,
    "ENSMUSG00000053093_ENSMUSG00000019768": 10.226172405201872,
    "ENSMUSG00000030335_ENSMUSG00000001280": 10.168723272040097,
    "ENSMUSG00000005610_ENSMUSG00000001280": 10.082540912064578,
    "ENSMUSG00000060036_ENSMUSG00000027109": 10.045887086879679,
    "ENSMUSG00000079508_ENSMUSG00000001280": 10.040187463925546,
    "ENSMUSG00000019797_ENSMUSG00000001280": 10.02912633849366,
    "ENSMUSG00000015837_ENSMUSG00000001280": 10.025172041970954,
    "ENSMUSG00000020153_ENSMUSG00000001280": 10.010869973571392,
    "ENSMUSG00000026260_ENSMUSG00000001280": 9.98564424217982,
    "ENSMUSG00000020477_ENSMUSG00000001280": 9.957382700506225,
    "ENSMUSG00000083563_ENSMUSG00000019768": 9.9508015385185,
    "ENSMUSG00000057322_ENSMUSG00000001280": 9.907448463793706,
    "ENSMUSG00000019768_ENSMUSG00000006356": 9.895802128801718,
    "ENSMUSG00000053819_ENSMUSG00000001280": 9.894805321196447,
    "ENSMUSG00000026526_ENSMUSG00000001280": 9.891081887397057,
    "ENSMUSG00000064348_ENSMUSG00000019768": 9.880724852130863,
    "ENSMUSG00000052684_ENSMUSG00000021127": 9.847435971815042,
    "ENSMUSG00000033065_ENSMUSG00000019768": 9.845644826833224,
    "ENSMUSG00000030246_ENSMUSG00000001280": 9.829238295629038,
    "ENSMUSG00000064367_ENSMUSG00000001280": 9.811521026219161,
    "ENSMUSG00000012848_ENSMUSG00000001280": 9.760578449191602,
    "ENSMUSG00000021520_ENSMUSG00000001280": 9.73607943551233,
    "ENSMUSG00000027245_ENSMUSG00000001280": 9.625638370836048,
    "ENSMUSG00000006818_ENSMUSG00000001280": 9.594047409378742,
    "ENSMUSG00000053898_ENSMUSG00000001280": 9.54827882892487,
    "ENSMUSG00000019768_ENSMUSG00000008136": 9.495543371327337,
    "ENSMUSG00000064346_ENSMUSG00000001280": 9.48195168701092,
    "ENSMUSG00000054428_ENSMUSG00000001280": 9.465456689966633,
    "ENSMUSG00000019768_ENSMUSG00000013936": 9.443560134720395,
    "ENSMUSG00000050608_ENSMUSG00000001280": 9.442046148737164,
    "ENSMUSG00000055762_ENSMUSG00000001280": 9.408864783717263,
    "ENSMUSG00000009863_ENSMUSG00000001280": 9.400594829908519,
    "ENSMUSG00000034566_ENSMUSG00000001280": 9.357527937089595,
    "ENSMUSG00000072946_ENSMUSG00000001280": 9.344588067670161,
    "ENSMUSG00000059534_ENSMUSG00000019768": 9.343107257711514,
    "ENSMUSG00000064341_ENSMUSG00000019768": 9.34125504392261,
    "ENSMUSG00000051747_ENSMUSG00000001280": 9.32805960324118,
    "ENSMUSG00000035458_ENSMUSG00000001280": 9.313827119598761,
    "ENSMUSG00000040666_ENSMUSG00000001280": 9.300714353378332,
    "ENSMUSG00000020483_ENSMUSG00000001280": 9.2907731872286,
    "ENSMUSG00000038370_ENSMUSG00000001280": 9.284517158637042,
    "ENSMUSG00000083219_ENSMUSG00000001280": 9.276368197874872,
    "ENSMUSG00000020163_ENSMUSG00000019768": 9.265548808027162,
    "ENSMUSG00000022110_ENSMUSG00000001280": 9.263844574450644,
    "ENSMUSG00000080836_ENSMUSG00000001280": 9.248912520588197,
    "ENSMUSG00000096449_ENSMUSG00000001280": 9.239654237770042,
    "ENSMUSG00000094249_ENSMUSG00000019768": 9.234683868465524,
    "ENSMUSG00000020372_ENSMUSG00000001280": 9.231573909842645,
    "ENSMUSG00000032330_ENSMUSG00000001280": 9.218123553308317,
    "ENSMUSG00000029918_ENSMUSG00000001280": 9.175086351435752,
    "ENSMUSG00000007097_ENSMUSG00000001280": 9.138040002255325,
    "ENSMUSG00000022450_ENSMUSG00000001280": 9.134999237128492,
    "ENSMUSG00000059534_ENSMUSG00000001280": 9.12252802845187,
    "ENSMUSG00000064369_ENSMUSG00000001280": 9.09672824490974,
    "ENSMUSG00000063316_ENSMUSG00000001280": 9.073401323704735,
    "ENSMUSG00000023944_ENSMUSG00000001280": 9.070426712937659,
    "ENSMUSG00000074218_ENSMUSG00000001280": 9.069832466106812,
    "ENSMUSG00000041881_ENSMUSG00000001280": 9.057286765553993,
    "ENSMUSG00000026208_ENSMUSG00000001280": 9.031085281364783,
    "ENSMUSG00000032399_ENSMUSG00000001280": 9.014081677062475,
    "ENSMUSG00000037152_ENSMUSG00000001280": 9.012690725166818,
    "ENSMUSG00000098178_ENSMUSG00000001280": 8.992351235323433,
    "ENSMUSG00000064343_ENSMUSG00000001280": 8.950199831422195,
    "ENSMUSG00000022803_ENSMUSG00000001280": 8.941919456342763,
    "ENSMUSG00000063229_ENSMUSG00000019768": 8.921723772968958,
    "ENSMUSG00000026087_ENSMUSG00000001280": 8.916968518285886,
    "ENSMUSG00000053702_ENSMUSG00000001280": 8.910538956446935,
    "ENSMUSG00000036880_ENSMUSG00000001280": 8.893631730962932,
    "ENSMUSG00000020022_ENSMUSG00000001280": 8.84052907939638,
    "ENSMUSG00000068487_ENSMUSG00000001280": 8.834175851416964,
    "ENSMUSG00000021606_ENSMUSG00000019768": 8.807253348251205,
    "ENSMUSG00000030541_ENSMUSG00000001280": 8.762919300058016,
    "ENSMUSG00000018770_ENSMUSG00000001280": 8.76217352036937,
    "ENSMUSG00000062397_ENSMUSG00000001280": 8.75061297779602,
    "ENSMUSG00000025794_ENSMUSG00000001280": 8.735735537702656,
    "ENSMUSG00000068706_ENSMUSG00000001280": 8.698818136091818,
    "ENSMUSG00000038170_ENSMUSG00000001280": 8.676234274329337,
    "ENSMUSG00000074218_ENSMUSG00000019768": 8.673600229856435,
    "ENSMUSG00000064342_ENSMUSG00000001280": 8.670580311142546,
    "ENSMUSG00000064340_ENSMUSG00000001280": 8.643603959192012,
    "ENSMUSG00000081051_ENSMUSG00000001280": 8.64086932126284,
    "ENSMUSG00000030869_ENSMUSG00000001280": 8.635913140644037,
    "ENSMUSG00000037815_ENSMUSG00000001280": 8.625237769815481,
    "ENSMUSG00000020456_ENSMUSG00000001280": 8.617943232040428,
    "ENSMUSG00000091478_ENSMUSG00000001280": 8.614942420464306,
    "ENSMUSG00000052337_ENSMUSG00000001280": 8.591838219612555,
    "ENSMUSG00000074129_ENSMUSG00000001280": 8.590346228561382,
    "ENSMUSG00000032366_ENSMUSG00000019768": 8.56327012489934,
    "ENSMUSG00000064372_ENSMUSG00000001280": 8.56010197433857,
    "ENSMUSG00000087943_ENSMUSG00000001280": 8.55967219306344,
    "ENSMUSG00000031320_ENSMUSG00000001280": 8.554906704599107,
    "ENSMUSG00000084319_ENSMUSG00000001280": 8.548844083009294,
    "ENSMUSG00000032314_ENSMUSG00000001280": 8.547984163396361,
    "ENSMUSG00000042541_ENSMUSG00000001280": 8.52748582149722,
    "ENSMUSG00000031633_ENSMUSG00000001280": 8.512918074273736,
    "ENSMUSG00000025428_ENSMUSG00000001280": 8.476270198739105,
    "ENSMUSG00000059734_ENSMUSG00000001280": 8.397249495720645,
    "ENSMUSG00000025465_ENSMUSG00000001280": 8.074888368355484,
    "ENSMUSG00000064360_ENSMUSG00000019768": 8.054219334266415,
    "ENSMUSG00000005683_ENSMUSG00000001280": 8.010916687023903,
    "ENSMUSG00000026817_ENSMUSG00000001280": 7.965175829023219,
    "ENSMUSG00000035202_ENSMUSG00000001280": 7.936448508504494,
    "ENSMUSG00000083832_ENSMUSG00000001280": 7.856681249965769,
    "ENSMUSG00000019768_ENSMUSG00000016252": 7.8065958822526555,
    "ENSMUSG00000049265_ENSMUSG00000001280": 7.752358146307093,
    "ENSMUSG00000061477_ENSMUSG00000001280": 7.751994945809098,
    "ENSMUSG00000058076_ENSMUSG00000001280": 7.746352383665731,
    "ENSMUSG00000081382_ENSMUSG00000001280": 7.697841868042629,
    "ENSMUSG00000030470_ENSMUSG00000019768": 7.490438138707922,
    "ENSMUSG00000023089_ENSMUSG00000019768": 7.328017990744143,
    "ENSMUSG00000024063_ENSMUSG00000001280": 7.092081048101451,
    "ENSMUSG00000037236_ENSMUSG00000001280": 6.986330108881274,
    "ENSMUSG00000022193_ENSMUSG00000001280": 6.960169122982657,
    "ENSMUSG00000045690_ENSMUSG00000019768": 6.638662962511477,
    "ENSMUSG00000059895_ENSMUSG00000001280": 6.627300176124094,
    "ENSMUSG00000038967_ENSMUSG00000008575": 6.605639849487218,
    "ENSMUSG00000025967_ENSMUSG00000001280": 6.531115069347214,
    "ENSMUSG00000071528_ENSMUSG00000001280": 6.491536327011836,
    "ENSMUSG00000055839_ENSMUSG00000001280": 6.438053175002956,
    "ENSMUSG00000058056_ENSMUSG00000001280": 6.276894525679913,
    "ENSMUSG00000047187_ENSMUSG00000001280": 5.992480876854286,
    "ENSMUSG00000091831_ENSMUSG00000001280": 5.90333913675371,
    "ENSMUSG00000068267_ENSMUSG00000044751": 5.879300165946291,
    "ENSMUSG00000026610_ENSMUSG00000025428": 5.428107749861071,
    "ENSMUSG00000030753_ENSMUSG00000001280": 5.15030203373771,
    "ENSMUSG00000028557_ENSMUSG00000001280": 4.920556203412716,
    "ENSMUSG00000019851_ENSMUSG00000001280": 4.853234634591883,
    "ENSMUSG00000026991_ENSMUSG00000001280": 4.829368242287752,
    "ENSMUSG00000070493_ENSMUSG00000035158": 4.792365132112365,
    "ENSMUSG00000057469_ENSMUSG00000036880": 4.186305488207534,
    "ENSMUSG00000051495_ENSMUSG00000008575": 3.9888880340218327,
    "ENSMUSG00000058239_ENSMUSG00000025241": 3.8743099998793395,
    "ENSMUSG00000051747_ENSMUSG00000019768": 3.8012033641335594,
    "ENSMUSG00000006219_ENSMUSG00000001280": 3.6658953369955167,
    "ENSMUSG00000028013_ENSMUSG00000008575": 3.4461879288398753,
    "ENSMUSG00000025290_ENSMUSG00000001280": 127.95981226652168
}
```
