## Supplement 3 (P7 CM vs. P28 CM) for "AGEAS: Automated Machine Learning based Genetic Regulatory Element Extraction System": outlier_grps.js

```
{
    "ENSMUSG00000023456_ENSMUSG00000001280": 176.95343511903744,
    "ENSMUSG00000025290_ENSMUSG00000001280": 168.0639612345917,
    "ENSMUSG00000007656_ENSMUSG00000001280": 86.89821706608994,
    "ENSMUSG00000006356_ENSMUSG00000001280": 37.11036693485854,
    "ENSMUSG00000031818_ENSMUSG00000019768": 30.311756371439138,
    "ENSMUSG00000074800_ENSMUSG00000001280": 27.67542789626417,
    "ENSMUSG00000023944_ENSMUSG00000001280": 27.43577008773136,
    "ENSMUSG00000021024_ENSMUSG00000001280": 23.230473620811164,
    "ENSMUSG00000064343_ENSMUSG00000019768": 20.43997708836555,
    "ENSMUSG00000026526_ENSMUSG00000001280": 20.424141383725026,
    "ENSMUSG00000033161_ENSMUSG00000019768": 18.39209909328601,
    "ENSMUSG00000020163_ENSMUSG00000001280": 18.171673506237074,
    "ENSMUSG00000096449_ENSMUSG00000001280": 17.73137236016031,
    "ENSMUSG00000064343_ENSMUSG00000001280": 17.443336522425735,
    "ENSMUSG00000006057_ENSMUSG00000001280": 17.41219785518362,
    "ENSMUSG00000038170_ENSMUSG00000001280": 17.213451135337873,
    "ENSMUSG00000044792_ENSMUSG00000001280": 17.031448208070227,
    "ENSMUSG00000014313_ENSMUSG00000001280": 16.91132955827591,
    "ENSMUSG00000045690_ENSMUSG00000001280": 16.87039839873483,
    "ENSMUSG00000060002_ENSMUSG00000019768": 16.77965073015242,
    "ENSMUSG00000044894_ENSMUSG00000001280": 16.255031434655933,
    "ENSMUSG00000028234_ENSMUSG00000001280": 16.080961242083543,
    "ENSMUSG00000064348_ENSMUSG00000019768": 15.829706017312345,
    "ENSMUSG00000067288_ENSMUSG00000001280": 14.94847786683424,
    "ENSMUSG00000046671_ENSMUSG00000001280": 12.079457290361603,
    "ENSMUSG00000056501_ENSMUSG00000039016": 11.80973452241966,
    "ENSMUSG00000056692_ENSMUSG00000019768": 11.473487730681931,
    "ENSMUSG00000024073_ENSMUSG00000001280": 11.42826776147341,
    "ENSMUSG00000042148_ENSMUSG00000008575": 11.06680422721335,
    "ENSMUSG00000053093_ENSMUSG00000024140": 11.004173231984,
    "ENSMUSG00000026021_ENSMUSG00000001280": 10.784526374128848,
    "ENSMUSG00000043716_ENSMUSG00000001280": 10.626860092730203,
    "ENSMUSG00000073557_ENSMUSG00000001280": 10.413212326857046,
    "ENSMUSG00000027668_ENSMUSG00000001280": 10.306423141683204,
    "ENSMUSG00000019768_ENSMUSG00000013593": 10.261196560324915,
    "ENSMUSG00000019158_ENSMUSG00000001280": 10.191045412762364,
    "ENSMUSG00000027422_ENSMUSG00000001280": 10.110494325345327,
    "ENSMUSG00000091537_ENSMUSG00000001280": 10.09587104908413,
    "ENSMUSG00000006333_ENSMUSG00000001280": 10.04888335795915,
    "ENSMUSG00000071528_ENSMUSG00000019768": 10.026953332206817,
    "ENSMUSG00000022186_ENSMUSG00000019768": 10.02091820170877,
    "ENSMUSG00000030869_ENSMUSG00000019768": 10.006109633854559,
    "ENSMUSG00000070493_ENSMUSG00000019768": 9.958430270962934,
    "ENSMUSG00000019768_ENSMUSG00000000563": 9.950677225847619,
    "ENSMUSG00000060376_ENSMUSG00000001280": 9.92233817945089,
    "ENSMUSG00000021000_ENSMUSG00000001280": 9.60360613211536,
    "ENSMUSG00000022477_ENSMUSG00000001280": 9.554300722421935,
    "ENSMUSG00000021313_ENSMUSG00000001280": 9.553887312678526,
    "ENSMUSG00000021290_ENSMUSG00000019768": 9.447028379370668,
    "ENSMUSG00000026568_ENSMUSG00000019768": 9.442216714209598,
    "ENSMUSG00000025795_ENSMUSG00000001280": 9.435185573545766,
    "ENSMUSG00000038412_ENSMUSG00000001280": 9.428687940596138,
    "ENSMUSG00000032279_ENSMUSG00000019768": 9.427436570432999,
    "ENSMUSG00000025745_ENSMUSG00000019768": 9.410609465793934,
    "ENSMUSG00000064369_ENSMUSG00000019768": 9.395765061371161,
    "ENSMUSG00000064370_ENSMUSG00000019768": 9.379395686677896,
    "ENSMUSG00000033065_ENSMUSG00000019768": 9.368157664318318,
    "ENSMUSG00000032495_ENSMUSG00000001280": 9.35314083658238,
    "ENSMUSG00000027472_ENSMUSG00000001280": 9.350813742330116,
    "ENSMUSG00000003072_ENSMUSG00000001280": 9.347637017856941,
    "ENSMUSG00000019768_ENSMUSG00000016427": 9.336462938671994,
    "ENSMUSG00000020477_ENSMUSG00000001280": 9.029160849400709,
    "ENSMUSG00000034088_ENSMUSG00000001280": 8.954510103325088,
    "ENSMUSG00000052837_ENSMUSG00000036896": 8.866804769153164,
    "ENSMUSG00000064366_ENSMUSG00000001280": 8.858710168484976,
    "ENSMUSG00000053898_ENSMUSG00000001280": 8.835217911208282,
    "ENSMUSG00000027452_ENSMUSG00000001280": 8.80932644380139,
    "ENSMUSG00000031791_ENSMUSG00000019768": 8.796146949732375,
    "ENSMUSG00000055762_ENSMUSG00000001280": 8.7824859929252,
    "ENSMUSG00000022450_ENSMUSG00000001280": 8.769693472847129,
    "ENSMUSG00000020326_ENSMUSG00000001280": 8.761950463546398,
    "ENSMUSG00000025241_ENSMUSG00000001280": 8.758854827187074,
    "ENSMUSG00000005683_ENSMUSG00000001280": 8.754681475615062,
    "ENSMUSG00000028820_ENSMUSG00000001280": 8.732185279603318,
    "ENSMUSG00000030695_ENSMUSG00000001280": 8.724594211629153,
    "ENSMUSG00000062908_ENSMUSG00000019768": 8.722892420867108,
    "ENSMUSG00000031543_ENSMUSG00000001280": 8.716367006795917,
    "ENSMUSG00000020153_ENSMUSG00000001280": 8.714377482796662,
    "ENSMUSG00000026895_ENSMUSG00000001280": 8.710558173789824,
    "ENSMUSG00000030744_ENSMUSG00000001280": 8.708102944734302,
    "ENSMUSG00000004558_ENSMUSG00000001280": 8.706797811308611,
    "ENSMUSG00000020475_ENSMUSG00000019768": 8.700400507169276,
    "ENSMUSG00000065947_ENSMUSG00000019768": 8.68403034456525,
    "ENSMUSG00000024099_ENSMUSG00000001280": 8.667855899227357,
    "ENSMUSG00000022354_ENSMUSG00000019768": 8.658048858378669,
    "ENSMUSG00000019689_ENSMUSG00000001280": 8.637675704405838,
    "ENSMUSG00000020163_ENSMUSG00000019768": 8.636674076131436,
    "ENSMUSG00000064367_ENSMUSG00000001280": 8.478362993370913,
    "ENSMUSG00000026003_ENSMUSG00000001280": 8.393907287929675,
    "ENSMUSG00000064349_ENSMUSG00000001280": 8.320090075140547,
    "ENSMUSG00000024063_ENSMUSG00000001280": 8.270327464078761,
    "ENSMUSG00000081344_ENSMUSG00000001280": 8.264555013056645,
    "ENSMUSG00000061904_ENSMUSG00000001280": 8.252923235937704,
    "ENSMUSG00000035458_ENSMUSG00000001280": 8.241427042916854,
    "ENSMUSG00000070493_ENSMUSG00000001280": 8.235327740299175,
    "ENSMUSG00000038690_ENSMUSG00000001280": 8.219743536264723,
    "ENSMUSG00000098178_ENSMUSG00000001280": 8.216763369810463,
    "ENSMUSG00000061518_ENSMUSG00000001280": 8.176142292232825,
    "ENSMUSG00000016427_ENSMUSG00000001280": 8.165377937789524,
    "ENSMUSG00000064370_ENSMUSG00000001280": 8.159419323214726,
    "ENSMUSG00000064337_ENSMUSG00000001280": 8.15715662259763,
    "ENSMUSG00000001280_ENSMUSG00000000563": 8.147940382418884,
    "ENSMUSG00000039103_ENSMUSG00000001280": 8.144659604947302,
    "ENSMUSG00000020483_ENSMUSG00000001280": 8.142627261863328,
    "ENSMUSG00000064348_ENSMUSG00000001280": 8.141579368784257,
    "ENSMUSG00000027984_ENSMUSG00000001280": 8.13456787872487,
    "ENSMUSG00000026087_ENSMUSG00000001280": 8.134493478156813,
    "ENSMUSG00000032314_ENSMUSG00000001280": 8.133164517207343,
    "ENSMUSG00000082884_ENSMUSG00000001280": 8.132907443326587,
    "ENSMUSG00000004610_ENSMUSG00000001280": 8.110313915747929,
    "ENSMUSG00000020022_ENSMUSG00000001280": 8.105552589796524,
    "ENSMUSG00000033938_ENSMUSG00000001280": 8.103998176615312,
    "ENSMUSG00000061474_ENSMUSG00000001280": 8.100831938524491,
    "ENSMUSG00000059534_ENSMUSG00000001280": 8.098447266469146,
    "ENSMUSG00000081485_ENSMUSG00000019768": 8.097500567192364,
    "ENSMUSG00000053477_ENSMUSG00000001280": 8.089213143857709,
    "ENSMUSG00000064369_ENSMUSG00000001280": 8.086359851287687,
    "ENSMUSG00000017404_ENSMUSG00000001280": 8.075732793829696,
    "ENSMUSG00000081331_ENSMUSG00000001280": 8.075561877199435,
    "ENSMUSG00000024038_ENSMUSG00000001280": 8.074853155861424,
    "ENSMUSG00000046330_ENSMUSG00000001280": 8.070610079950674,
    "ENSMUSG00000044080_ENSMUSG00000001280": 8.064994407611001,
    "ENSMUSG00000063229_ENSMUSG00000019768": 8.040358471997934,
    "ENSMUSG00000068706_ENSMUSG00000001280": 8.030836828041984,
    "ENSMUSG00000064365_ENSMUSG00000001280": 8.02160399683228,
    "ENSMUSG00000035242_ENSMUSG00000001280": 8.01706869970867,
    "ENSMUSG00000005610_ENSMUSG00000001280": 8.006263152640747,
    "ENSMUSG00000025794_ENSMUSG00000001280": 8.00488826697726,
    "ENSMUSG00000001280_ENSMUSG00000000740": 7.995599494648907,
    "ENSMUSG00000031818_ENSMUSG00000001280": 7.993077866207709,
    "ENSMUSG00000015092_ENSMUSG00000001280": 7.985581407468985,
    "ENSMUSG00000074884_ENSMUSG00000001280": 7.980943437845407,
    "ENSMUSG00000031672_ENSMUSG00000001280": 7.975492887480824,
    "ENSMUSG00000022013_ENSMUSG00000001280": 7.974659679567965,
    "ENSMUSG00000074218_ENSMUSG00000019768": 7.974299048876372,
    "ENSMUSG00000023861_ENSMUSG00000001280": 7.972137140146233,
    "ENSMUSG00000019494_ENSMUSG00000001280": 7.971054813242392,
    "ENSMUSG00000063229_ENSMUSG00000001280": 7.959813115367368,
    "ENSMUSG00000014592_ENSMUSG00000001280": 7.9535574027402,
    "ENSMUSG00000059741_ENSMUSG00000019768": 7.823033390074048,
    "ENSMUSG00000025204_ENSMUSG00000001280": 7.787972020412303,
    "ENSMUSG00000034295_ENSMUSG00000001280": 7.660947462885199,
    "ENSMUSG00000050856_ENSMUSG00000001280": 7.615129098548383,
    "ENSMUSG00000025651_ENSMUSG00000001280": 7.602279203679816,
    "ENSMUSG00000015837_ENSMUSG00000001280": 7.542016525173431,
    "ENSMUSG00000061462_ENSMUSG00000001280": 7.484687535585244,
    "ENSMUSG00000081051_ENSMUSG00000001280": 7.482057650415139,
    "ENSMUSG00000027566_ENSMUSG00000001280": 7.470668741901817,
    "ENSMUSG00000021748_ENSMUSG00000001280": 7.449018267801307,
    "ENSMUSG00000022205_ENSMUSG00000001280": 7.440018351420257,
    "ENSMUSG00000060938_ENSMUSG00000001280": 7.438675356958954,
    "ENSMUSG00000068487_ENSMUSG00000001280": 7.383278335116411,
    "ENSMUSG00000058076_ENSMUSG00000001280": 7.367303041564824,
    "ENSMUSG00000057580_ENSMUSG00000019768": 7.2964383384741005,
    "ENSMUSG00000019768_ENSMUSG00000006057": 7.290656191985378,
    "ENSMUSG00000022108_ENSMUSG00000001280": 7.287527357687171,
    "ENSMUSG00000031633_ENSMUSG00000008575": 7.036414983297292,
    "ENSMUSG00000002102_ENSMUSG00000001280": 6.922125384095717,
    "ENSMUSG00000025290_ENSMUSG00000019768": 6.7724027417449415,
    "ENSMUSG00000028452_ENSMUSG00000001280": 6.753721291705401,
    "ENSMUSG00000040666_ENSMUSG00000001280": 6.702898105159643,
    "ENSMUSG00000020580_ENSMUSG00000001280": 6.6569096081633194,
    "ENSMUSG00000022452_ENSMUSG00000019768": 6.277423245541707,
    "ENSMUSG00000001833_ENSMUSG00000001280": 6.13575785736343,
    "ENSMUSG00000021843_ENSMUSG00000001280": 6.115417175619965,
    "ENSMUSG00000015656_ENSMUSG00000001280": 6.086696760244999,
    "ENSMUSG00000093674_ENSMUSG00000001280": 6.062390120736068,
    "ENSMUSG00000071528_ENSMUSG00000008575": 6.0574726109441945,
    "ENSMUSG00000022193_ENSMUSG00000001280": 6.01518175338307,
    "ENSMUSG00000057469_ENSMUSG00000003072": 5.418974233331608,
    "ENSMUSG00000061904_ENSMUSG00000058756": 4.866971669218132,
    "ENSMUSG00000058239_ENSMUSG00000026977": 4.850669942515068,
    "ENSMUSG00000058239_ENSMUSG00000014956": 4.80263047577396,
    "ENSMUSG00000026864_ENSMUSG00000001419": 3.6760732754166794,
    "ENSMUSG00000003970_ENSMUSG00000001280": 3.511657688764235,
    "ENSMUSG00000024991_ENSMUSG00000001280": 3.2296443279337166
}
```
