## Supplement 3 (P7 CM vs. P28 CM) for "AGEAS: Automated Machine Learning based Genetic Regulatory Element Extraction System": outlier_grps.js

```
{
    "ENSMUSG00000046330_ENSMUSG00000001280": 42.04280430488623,
    "ENSMUSG00000021178_ENSMUSG00000001280": 34.66559452989648,
    "ENSMUSG00000064349_ENSMUSG00000001280": 33.376370350891385,
    "ENSMUSG00000062691_ENSMUSG00000001280": 28.496899288251388,
    "ENSMUSG00000064348_ENSMUSG00000019768": 26.86892681355487,
    "ENSMUSG00000038967_ENSMUSG00000001280": 24.88217895451096,
    "ENSMUSG00000064372_ENSMUSG00000019768": 24.857914700350012,
    "ENSMUSG00000025488_ENSMUSG00000019768": 24.70373416758651,
    "ENSMUSG00000080836_ENSMUSG00000001280": 24.61377138355299,
    "ENSMUSG00000084319_ENSMUSG00000001280": 23.777148230983695,
    "ENSMUSG00000064372_ENSMUSG00000001280": 23.26617369797211,
    "ENSMUSG00000022820_ENSMUSG00000001280": 23.03150036632883,
    "ENSMUSG00000040752_ENSMUSG00000001280": 22.798010164409465,
    "ENSMUSG00000074129_ENSMUSG00000001280": 22.761428081794502,
    "ENSMUSG00000025204_ENSMUSG00000001280": 22.613905624197013,
    "ENSMUSG00000019768_ENSMUSG00000005683": 22.472792365019224,
    "ENSMUSG00000035242_ENSMUSG00000001280": 22.078214193950576,
    "ENSMUSG00000038170_ENSMUSG00000001280": 21.775207557689367,
    "ENSMUSG00000007097_ENSMUSG00000001280": 21.583003191632482,
    "ENSMUSG00000030695_ENSMUSG00000001280": 21.497114319877753,
    "ENSMUSG00000022354_ENSMUSG00000001280": 21.33738528283662,
    "ENSMUSG00000050856_ENSMUSG00000001280": 21.320011524413196,
    "ENSMUSG00000050608_ENSMUSG00000001280": 21.30188667963755,
    "ENSMUSG00000091449_ENSMUSG00000001280": 21.229171898562694,
    "ENSMUSG00000025809_ENSMUSG00000001280": 18.28190241642258,
    "ENSMUSG00000019768_ENSMUSG00000006740": 16.916060321473783,
    "ENSMUSG00000022217_ENSMUSG00000019768": 15.198880192212096,
    "ENSMUSG00000030785_ENSMUSG00000001419": 14.337483605137233,
    "ENSMUSG00000020326_ENSMUSG00000019768": 14.224160843653545,
    "ENSMUSG00000027698_ENSMUSG00000001280": 13.994804654637495,
    "ENSMUSG00000038412_ENSMUSG00000019768": 13.895509768886171,
    "ENSMUSG00000053898_ENSMUSG00000019768": 13.422639019153143,
    "ENSMUSG00000022186_ENSMUSG00000019768": 13.418717344769439,
    "ENSMUSG00000064347_ENSMUSG00000019768": 13.379722702889199,
    "ENSMUSG00000030409_ENSMUSG00000001280": 13.378215528922139,
    "ENSMUSG00000028367_ENSMUSG00000001280": 13.376574653612558,
    "ENSMUSG00000019768_ENSMUSG00000000399": 13.321156544158871,
    "ENSMUSG00000019768_ENSMUSG00000005610": 13.294010098434823,
    "ENSMUSG00000032786_ENSMUSG00000001280": 13.27497395454267,
    "ENSMUSG00000058427_ENSMUSG00000026628": 13.213569873487371,
    "ENSMUSG00000098178_ENSMUSG00000019768": 13.19203499561834,
    "ENSMUSG00000064348_ENSMUSG00000001280": 13.067272744992275,
    "ENSMUSG00000017119_ENSMUSG00000001280": 13.061200825075792,
    "ENSMUSG00000062006_ENSMUSG00000001280": 13.008612442427196,
    "ENSMUSG00000064367_ENSMUSG00000001280": 12.878210172556422,
    "ENSMUSG00000020849_ENSMUSG00000019768": 12.759162908592453,
    "ENSMUSG00000018286_ENSMUSG00000001280": 12.648616981237803,
    "ENSMUSG00000003072_ENSMUSG00000001280": 12.58423536477705,
    "ENSMUSG00000020849_ENSMUSG00000001280": 12.577868883920216,
    "ENSMUSG00000060002_ENSMUSG00000001280": 12.56196958099671,
    "ENSMUSG00000064345_ENSMUSG00000019768": 12.560631880622733,
    "ENSMUSG00000091478_ENSMUSG00000019768": 12.508243674995477,
    "ENSMUSG00000019768_ENSMUSG00000017778": 12.499749817520597,
    "ENSMUSG00000064343_ENSMUSG00000019768": 12.493977288460663,
    "ENSMUSG00000004044_ENSMUSG00000001280": 12.487766749057453,
    "ENSMUSG00000038690_ENSMUSG00000019768": 12.434050914042906,
    "ENSMUSG00000019768_ENSMUSG00000016427": 12.420934439920554,
    "ENSMUSG00000008668_ENSMUSG00000001280": 12.419822489130386,
    "ENSMUSG00000019797_ENSMUSG00000001280": 12.309671310529003,
    "ENSMUSG00000027180_ENSMUSG00000001280": 12.281534511249466,
    "ENSMUSG00000029017_ENSMUSG00000001280": 12.242427832127934,
    "ENSMUSG00000035783_ENSMUSG00000001280": 12.123927936551734,
    "ENSMUSG00000021520_ENSMUSG00000001280": 11.90662453371384,
    "ENSMUSG00000024099_ENSMUSG00000001280": 11.85718540933456,
    "ENSMUSG00000031672_ENSMUSG00000001280": 11.851321645236041,
    "ENSMUSG00000063882_ENSMUSG00000001280": 11.846583822886979,
    "ENSMUSG00000024608_ENSMUSG00000001280": 11.64193923106751,
    "ENSMUSG00000030884_ENSMUSG00000001280": 11.633595663477934,
    "ENSMUSG00000063457_ENSMUSG00000001280": 11.620013569953102,
    "ENSMUSG00000064351_ENSMUSG00000019768": 11.613021258001025,
    "ENSMUSG00000026003_ENSMUSG00000019768": 11.57821049537281,
    "ENSMUSG00000065947_ENSMUSG00000019768": 11.570931949496845,
    "ENSMUSG00000038370_ENSMUSG00000001280": 11.544948479973685,
    "ENSMUSG00000020163_ENSMUSG00000019768": 11.523626827734896,
    "ENSMUSG00000031299_ENSMUSG00000019768": 11.508646890165437,
    "ENSMUSG00000062908_ENSMUSG00000019768": 11.46265982322821,
    "ENSMUSG00000038583_ENSMUSG00000019768": 11.441077942278113,
    "ENSMUSG00000026087_ENSMUSG00000001280": 11.428854365488096,
    "ENSMUSG00000040752_ENSMUSG00000019768": 11.35288168678363,
    "ENSMUSG00000064345_ENSMUSG00000001280": 11.33487270969831,
    "ENSMUSG00000019768_ENSMUSG00000003072": 11.332433155597483,
    "ENSMUSG00000064302_ENSMUSG00000001280": 11.292560484024625,
    "ENSMUSG00000064339_ENSMUSG00000001280": 11.235997813272352,
    "ENSMUSG00000032366_ENSMUSG00000001280": 11.179782772827219,
    "ENSMUSG00000064337_ENSMUSG00000001280": 11.17243483046003,
    "ENSMUSG00000083863_ENSMUSG00000001280": 11.170635111793558,
    "ENSMUSG00000064342_ENSMUSG00000001280": 11.15293860101806,
    "ENSMUSG00000025428_ENSMUSG00000001280": 11.152489568455872,
    "ENSMUSG00000035458_ENSMUSG00000001280": 11.077659878386104,
    "ENSMUSG00000018770_ENSMUSG00000001280": 11.058766148312658,
    "ENSMUSG00000025781_ENSMUSG00000001280": 10.99123244017683,
    "ENSMUSG00000029632_ENSMUSG00000001280": 10.990014261365921,
    "ENSMUSG00000064338_ENSMUSG00000001280": 10.974216525777617,
    "ENSMUSG00000091831_ENSMUSG00000001280": 10.966684373474292,
    "ENSMUSG00000072946_ENSMUSG00000001280": 10.965842888204197,
    "ENSMUSG00000021606_ENSMUSG00000001280": 10.964056443612199,
    "ENSMUSG00000027673_ENSMUSG00000001280": 10.963849994603274,
    "ENSMUSG00000063694_ENSMUSG00000001280": 10.952899509589377,
    "ENSMUSG00000014294_ENSMUSG00000001280": 10.947267246835509,
    "ENSMUSG00000044894_ENSMUSG00000001280": 10.932623912396375,
    "ENSMUSG00000017778_ENSMUSG00000001280": 10.885226404876727,
    "ENSMUSG00000044080_ENSMUSG00000001280": 10.883235342718995,
    "ENSMUSG00000001280_ENSMUSG00000000563": 10.875686819201402,
    "ENSMUSG00000035674_ENSMUSG00000001280": 10.860952184839118,
    "ENSMUSG00000070493_ENSMUSG00000001280": 10.858085869212585,
    "ENSMUSG00000016427_ENSMUSG00000001280": 10.814616003779468,
    "ENSMUSG00000031299_ENSMUSG00000001280": 10.813007948477898,
    "ENSMUSG00000074884_ENSMUSG00000001280": 10.80002804329054,
    "ENSMUSG00000019768_ENSMUSG00000013936": 10.799372749029189,
    "ENSMUSG00000091478_ENSMUSG00000001280": 10.783497406920253,
    "ENSMUSG00000026003_ENSMUSG00000001280": 10.777802568921443,
    "ENSMUSG00000067736_ENSMUSG00000001280": 10.777087760708893,
    "ENSMUSG00000098973_ENSMUSG00000001280": 10.769574219589797,
    "ENSMUSG00000052337_ENSMUSG00000001280": 10.71806077025906,
    "ENSMUSG00000068706_ENSMUSG00000001280": 10.715849725332216,
    "ENSMUSG00000020402_ENSMUSG00000001280": 10.697362524660585,
    "ENSMUSG00000020585_ENSMUSG00000001280": 10.690558617409073,
    "ENSMUSG00000052146_ENSMUSG00000001280": 10.69051420561625,
    "ENSMUSG00000074800_ENSMUSG00000001280": 10.67483012802601,
    "ENSMUSG00000034566_ENSMUSG00000001280": 10.67380221412478,
    "ENSMUSG00000045690_ENSMUSG00000001280": 10.672275087883627,
    "ENSMUSG00000061904_ENSMUSG00000001280": 10.65852288939676,
    "ENSMUSG00000031231_ENSMUSG00000001280": 10.651042050689762,
    "ENSMUSG00000037916_ENSMUSG00000001280": 10.630722951772984,
    "ENSMUSG00000041841_ENSMUSG00000001280": 10.612820199371436,
    "ENSMUSG00000054889_ENSMUSG00000001280": 10.609345162928001,
    "ENSMUSG00000078812_ENSMUSG00000001280": 10.60845983980669,
    "ENSMUSG00000024038_ENSMUSG00000001280": 10.601747270290785,
    "ENSMUSG00000032518_ENSMUSG00000001280": 10.567421204051813,
    "ENSMUSG00000025290_ENSMUSG00000001280": 10.559162948574437,
    "ENSMUSG00000040952_ENSMUSG00000001280": 10.49494506917855,
    "ENSMUSG00000034842_ENSMUSG00000001280": 10.48169724912814,
    "ENSMUSG00000028207_ENSMUSG00000001280": 10.283836526452996,
    "ENSMUSG00000071528_ENSMUSG00000021250": 10.250072465750863,
    "ENSMUSG00000019787_ENSMUSG00000001280": 10.084500589699354,
    "ENSMUSG00000039105_ENSMUSG00000001280": 10.06434240966616,
    "ENSMUSG00000051747_ENSMUSG00000001280": 9.97621003475246,
    "ENSMUSG00000022551_ENSMUSG00000001280": 9.926724156029946,
    "ENSMUSG00000071528_ENSMUSG00000001280": 9.89128103017814,
    "ENSMUSG00000068706_ENSMUSG00000019768": 9.836001928501515,
    "ENSMUSG00000052512_ENSMUSG00000001280": 9.829937800315923,
    "ENSMUSG00000081051_ENSMUSG00000001280": 9.81374548997548,
    "ENSMUSG00000007656_ENSMUSG00000001280": 9.810275052309816,
    "ENSMUSG00000026043_ENSMUSG00000002265": 9.795998853278634,
    "ENSMUSG00000071415_ENSMUSG00000001280": 9.782594218929887,
    "ENSMUSG00000020547_ENSMUSG00000001280": 9.781800491204676,
    "ENSMUSG00000023089_ENSMUSG00000019768": 9.7525564134257,
    "ENSMUSG00000068523_ENSMUSG00000001280": 9.729416992051796,
    "ENSMUSG00000036371_ENSMUSG00000001280": 9.058610915523316,
    "ENSMUSG00000029729_ENSMUSG00000028639": 8.98501056820326,
    "ENSMUSG00000090862_ENSMUSG00000001280": 8.957532549643044,
    "ENSMUSG00000033059_ENSMUSG00000001280": 8.893916454444993,
    "ENSMUSG00000061477_ENSMUSG00000001280": 8.833179552679116,
    "ENSMUSG00000022010_ENSMUSG00000001280": 8.821883340458141,
    "ENSMUSG00000047675_ENSMUSG00000005583": 8.489071613325489,
    "ENSMUSG00000037236_ENSMUSG00000001280": 7.982397903246177,
    "ENSMUSG00000030557_ENSMUSG00000026526": 7.942457067945978,
    "ENSMUSG00000022890_ENSMUSG00000010175": 7.67824102133893,
    "ENSMUSG00000082687_ENSMUSG00000001280": 7.3493448928543925,
    "ENSMUSG00000015839_ENSMUSG00000003429": 7.159600285629612,
    "ENSMUSG00000068921_ENSMUSG00000019768": 7.131149301085926,
    "ENSMUSG00000026817_ENSMUSG00000001280": 6.895677229654485,
    "ENSMUSG00000056692_ENSMUSG00000030557": 6.874767331662653,
    "ENSMUSG00000044894_ENSMUSG00000019768": 6.8629186369433075,
    "ENSMUSG00000042032_ENSMUSG00000001280": 6.822380743179227,
    "ENSMUSG00000026414_ENSMUSG00000019768": 6.56087059523302,
    "ENSMUSG00000020326_ENSMUSG00000015605": 6.196400661552758,
    "ENSMUSG00000031590_ENSMUSG00000023034": 6.0653892473939335,
    "ENSMUSG00000034525_ENSMUSG00000001280": 5.403513441608667,
    "ENSMUSG00000074412_ENSMUSG00000001280": 5.222039983712009,
    "ENSMUSG00000062981_ENSMUSG00000008575": 5.065127777026477,
    "ENSMUSG00000015656_ENSMUSG00000001419": 5.039834079630343,
    "ENSMUSG00000052374_ENSMUSG00000001280": 4.844279937310955,
    "ENSMUSG00000032348_ENSMUSG00000005583": 4.631061936065264,
    "ENSMUSG00000030846_ENSMUSG00000005698": 4.309025556962441,
    "ENSMUSG00000058756_ENSMUSG00000025465": 3.8566948908032965,
    "ENSMUSG00000029864_ENSMUSG00000001280": 3.4513901395947313,
    "ENSMUSG00000052337_ENSMUSG00000001419": 3.0258665939425105,
    "ENSMUSG00000022890_ENSMUSG00000001280": 49.55809143610088,
    "ENSMUSG00000035458_ENSMUSG00000019768": 39.402150151793144,
    "ENSMUSG00000025465_ENSMUSG00000001280": 28.83697997245992,
    "ENSMUSG00000057469_ENSMUSG00000006057": 27.110140432132226,
    "ENSMUSG00000031818_ENSMUSG00000019768": 24.252778851595256,
    "ENSMUSG00000019768_ENSMUSG00000014294": 24.100905374653067,
    "ENSMUSG00000019768_ENSMUSG00000000563": 23.608470035327173,
    "ENSMUSG00000096842_ENSMUSG00000001280": 22.494564660053296,
    "ENSMUSG00000027447_ENSMUSG00000001280": 22.33053857784716,
    "ENSMUSG00000059534_ENSMUSG00000019768": 22.238676994789216,
    "ENSMUSG00000020153_ENSMUSG00000001280": 22.22317344025309,
    "ENSMUSG00000009863_ENSMUSG00000001280": 21.655372875626213,
    "ENSMUSG00000057322_ENSMUSG00000001280": 21.580627578810958,
    "ENSMUSG00000031633_ENSMUSG00000019768": 21.397403535243384,
    "ENSMUSG00000064346_ENSMUSG00000001280": 20.969538162952823,
    "ENSMUSG00000033060_ENSMUSG00000001280": 20.87569947545673,
    "ENSMUSG00000023020_ENSMUSG00000001280": 20.830313981551015,
    "ENSMUSG00000015733_ENSMUSG00000001280": 20.681568454392156,
    "ENSMUSG00000033938_ENSMUSG00000001280": 20.519030972547636,
    "ENSMUSG00000026032_ENSMUSG00000001280": 20.434107008096902,
    "ENSMUSG00000028820_ENSMUSG00000001280": 20.02309267292917,
    "ENSMUSG00000064363_ENSMUSG00000019768": 19.63609235646859,
    "ENSMUSG00000041881_ENSMUSG00000001280": 19.480347758556864,
    "ENSMUSG00000025393_ENSMUSG00000001280": 19.34450264125693,
    "ENSMUSG00000022400_ENSMUSG00000001280": 16.07346868481367,
    "ENSMUSG00000029135_ENSMUSG00000020485": 16.069740570111918,
    "ENSMUSG00000023020_ENSMUSG00000019768": 15.173302981605199,
    "ENSMUSG00000052684_ENSMUSG00000036887": 14.390619704007094,
    "ENSMUSG00000024869_ENSMUSG00000001280": 14.11966849003534,
    "ENSMUSG00000059430_ENSMUSG00000019768": 13.955016947157548,
    "ENSMUSG00000040653_ENSMUSG00000001280": 13.799512034325845,
    "ENSMUSG00000031715_ENSMUSG00000021764": 13.653463576171475,
    "ENSMUSG00000035458_ENSMUSG00000015605": 13.620088539833233,
    "ENSMUSG00000057469_ENSMUSG00000003072": 13.530668771733827,
    "ENSMUSG00000020593_ENSMUSG00000001280": 13.166799318996464,
    "ENSMUSG00000061838_ENSMUSG00000001280": 12.836412675667049,
    "ENSMUSG00000031715_ENSMUSG00000026568": 12.689259713335227,
    "ENSMUSG00000029614_ENSMUSG00000019768": 12.62813139743414,
    "ENSMUSG00000056501_ENSMUSG00000040048": 12.576572930048352,
    "ENSMUSG00000063882_ENSMUSG00000019768": 12.220998481114869,
    "ENSMUSG00000026664_ENSMUSG00000008575": 12.14998129339845,
    "ENSMUSG00000025103_ENSMUSG00000001280": 12.082302388847888,
    "ENSMUSG00000021868_ENSMUSG00000019768": 12.005362638559715,
    "ENSMUSG00000018585_ENSMUSG00000001280": 11.958285496850024,
    "ENSMUSG00000024197_ENSMUSG00000001280": 11.545074680589451,
    "ENSMUSG00000022519_ENSMUSG00000019768": 11.537477093445348,
    "ENSMUSG00000064369_ENSMUSG00000019768": 11.51566305561367,
    "ENSMUSG00000027861_ENSMUSG00000019768": 11.512073885696992,
    "ENSMUSG00000033065_ENSMUSG00000001280": 11.486734681106928,
    "ENSMUSG00000052698_ENSMUSG00000001280": 11.447735411520513,
    "ENSMUSG00000022477_ENSMUSG00000001280": 11.389008049643666,
    "ENSMUSG00000026526_ENSMUSG00000001280": 11.35653595059177,
    "ENSMUSG00000027472_ENSMUSG00000001280": 11.350551191795477,
    "ENSMUSG00000019768_ENSMUSG00000019179": 11.319002898080159,
    "ENSMUSG00000022890_ENSMUSG00000019768": 11.307670307847506,
    "ENSMUSG00000029455_ENSMUSG00000001280": 11.298755169691693,
    "ENSMUSG00000070493_ENSMUSG00000019768": 11.29606196979797,
    "ENSMUSG00000083563_ENSMUSG00000019768": 11.269371722237285,
    "ENSMUSG00000039001_ENSMUSG00000001280": 11.171741556557862,
    "ENSMUSG00000012848_ENSMUSG00000001280": 11.171555772000152,
    "ENSMUSG00000062580_ENSMUSG00000057469": 11.169760614612875,
    "ENSMUSG00000061104_ENSMUSG00000001280": 10.963449060029507,
    "ENSMUSG00000030869_ENSMUSG00000001280": 10.868932133485222,
    "ENSMUSG00000013593_ENSMUSG00000001280": 10.860585485119579,
    "ENSMUSG00000053768_ENSMUSG00000001280": 10.707424393624922,
    "ENSMUSG00000028603_ENSMUSG00000001280": 10.664157972443311,
    "ENSMUSG00000004610_ENSMUSG00000001280": 10.657632561530491,
    "ENSMUSG00000063694_ENSMUSG00000019768": 10.644387390805653,
    "ENSMUSG00000059534_ENSMUSG00000001280": 10.64238267279395,
    "ENSMUSG00000020048_ENSMUSG00000001280": 10.611300007901116,
    "ENSMUSG00000025241_ENSMUSG00000001280": 10.60379960433411,
    "ENSMUSG00000037710_ENSMUSG00000001280": 10.600507377147418,
    "ENSMUSG00000054428_ENSMUSG00000001280": 10.52763350024288,
    "ENSMUSG00000039103_ENSMUSG00000019768": 10.502524637414075,
    "ENSMUSG00000047675_ENSMUSG00000001280": 10.50217910004817,
    "ENSMUSG00000024576_ENSMUSG00000001280": 10.500676648399063,
    "ENSMUSG00000001604_ENSMUSG00000001280": 10.49131609522672,
    "ENSMUSG00000064337_ENSMUSG00000019768": 10.464202225589188,
    "ENSMUSG00000027452_ENSMUSG00000001280": 10.445418564609108,
    "ENSMUSG00000004558_ENSMUSG00000001280": 10.44533181484054,
    "ENSMUSG00000036309_ENSMUSG00000001280": 10.42974561486315,
    "ENSMUSG00000035202_ENSMUSG00000001280": 10.409843864587627,
    "ENSMUSG00000038375_ENSMUSG00000001280": 10.396340697408133,
    "ENSMUSG00000032959_ENSMUSG00000001280": 10.32874489298712,
    "ENSMUSG00000064370_ENSMUSG00000001280": 10.263248781425805,
    "ENSMUSG00000001280_ENSMUSG00000000088": 10.232357082688225,
    "ENSMUSG00000022982_ENSMUSG00000001280": 10.116116387001634,
    "ENSMUSG00000081485_ENSMUSG00000019768": 10.104370838860532,
    "ENSMUSG00000038375_ENSMUSG00000008575": 10.081861649278146,
    "ENSMUSG00000082884_ENSMUSG00000001280": 10.076362594646186,
    "ENSMUSG00000032314_ENSMUSG00000001280": 10.076012195126996,
    "ENSMUSG00000029467_ENSMUSG00000001280": 10.00138632570227,
    "ENSMUSG00000064340_ENSMUSG00000001280": 9.976082523518132,
    "ENSMUSG00000037152_ENSMUSG00000001280": 9.973583111363654,
    "ENSMUSG00000064365_ENSMUSG00000001280": 9.960573289975152,
    "ENSMUSG00000033161_ENSMUSG00000001280": 9.949680171310044,
    "ENSMUSG00000074218_ENSMUSG00000001280": 9.940740393200919,
    "ENSMUSG00000024913_ENSMUSG00000001280": 9.9404148418187,
    "ENSMUSG00000026260_ENSMUSG00000019768": 9.936787721045013,
    "ENSMUSG00000021313_ENSMUSG00000019768": 9.919057285504055,
    "ENSMUSG00000059741_ENSMUSG00000019768": 9.915771416714124,
    "ENSMUSG00000098178_ENSMUSG00000001280": 9.897997594498564,
    "ENSMUSG00000083219_ENSMUSG00000001280": 9.87924023075642,
    "ENSMUSG00000031059_ENSMUSG00000001280": 9.876505950757307,
    "ENSMUSG00000032060_ENSMUSG00000001280": 9.836734009098018,
    "ENSMUSG00000006057_ENSMUSG00000001280": 9.832895256219217,
    "ENSMUSG00000068487_ENSMUSG00000001280": 9.82411251986486,
    "ENSMUSG00000040048_ENSMUSG00000001280": 9.822170887603722,
    "ENSMUSG00000017404_ENSMUSG00000001280": 9.820613473433509,
    "ENSMUSG00000064360_ENSMUSG00000019768": 9.814493043228074,
    "ENSMUSG00000025968_ENSMUSG00000001280": 9.787876120222656,
    "ENSMUSG00000038462_ENSMUSG00000001280": 9.785193948431898,
    "ENSMUSG00000032279_ENSMUSG00000001280": 9.775797142844771,
    "ENSMUSG00000019362_ENSMUSG00000001280": 9.772525908419675,
    "ENSMUSG00000021577_ENSMUSG00000001280": 9.765858284917776,
    "ENSMUSG00000029632_ENSMUSG00000019768": 9.765827342502481,
    "ENSMUSG00000021290_ENSMUSG00000001280": 9.718161473131985,
    "ENSMUSG00000038583_ENSMUSG00000014592": 9.71285728237253,
    "ENSMUSG00000028691_ENSMUSG00000001280": 9.692234760196555,
    "ENSMUSG00000062397_ENSMUSG00000001280": 9.6892283685008,
    "ENSMUSG00000020321_ENSMUSG00000001280": 9.689207285608482,
    "ENSMUSG00000002010_ENSMUSG00000001280": 9.658580003160798,
    "ENSMUSG00000032399_ENSMUSG00000001280": 9.652325503909747,
    "ENSMUSG00000055320_ENSMUSG00000001280": 9.561418251870661,
    "ENSMUSG00000093674_ENSMUSG00000001280": 9.553363916456151,
    "ENSMUSG00000021282_ENSMUSG00000001280": 9.548661251641752,
    "ENSMUSG00000018340_ENSMUSG00000001280": 9.51650624834309,
    "ENSMUSG00000064369_ENSMUSG00000028639": 9.276240817394902,
    "ENSMUSG00000063787_ENSMUSG00000001280": 9.226313248462356,
    "ENSMUSG00000057580_ENSMUSG00000019768": 9.184636730370219,
    "ENSMUSG00000026688_ENSMUSG00000001280": 9.107286107182418,
    "ENSMUSG00000064369_ENSMUSG00000001280": 9.07475600504669,
    "ENSMUSG00000028452_ENSMUSG00000001280": 8.93507479100597,
    "ENSMUSG00000005683_ENSMUSG00000001280": 8.919746757532078,
    "ENSMUSG00000032199_ENSMUSG00000001280": 8.8709494004476,
    "ENSMUSG00000021748_ENSMUSG00000001280": 8.852107186641975,
    "ENSMUSG00000036860_ENSMUSG00000001280": 8.837292508818189,
    "ENSMUSG00000028756_ENSMUSG00000001280": 8.803864763725459,
    "ENSMUSG00000079641_ENSMUSG00000001280": 8.766317712145986,
    "ENSMUSG00000071014_ENSMUSG00000001280": 8.504857626137424,
    "ENSMUSG00000027984_ENSMUSG00000001280": 8.30218319748121,
    "ENSMUSG00000037916_ENSMUSG00000014592": 8.29749264612638,
    "ENSMUSG00000015656_ENSMUSG00000001280": 8.235515799828358,
    "ENSMUSG00000039640_ENSMUSG00000001280": 8.231050767658676,
    "ENSMUSG00000060679_ENSMUSG00000001280": 8.1689966194986,
    "ENSMUSG00000021577_ENSMUSG00000019768": 8.165660492312178,
    "ENSMUSG00000049327_ENSMUSG00000001280": 8.164897805112156,
    "ENSMUSG00000084168_ENSMUSG00000001280": 8.106607727594541,
    "ENSMUSG00000060938_ENSMUSG00000001280": 8.07768622590113,
    "ENSMUSG00000037152_ENSMUSG00000019768": 8.04063287995397,
    "ENSMUSG00000047905_ENSMUSG00000001280": 8.004981415050715,
    "ENSMUSG00000053477_ENSMUSG00000022174": 7.9463945638278695,
    "ENSMUSG00000030557_ENSMUSG00000022110": 7.916074823197848,
    "ENSMUSG00000026817_ENSMUSG00000019768": 7.721706102747295,
    "ENSMUSG00000022519_ENSMUSG00000001280": 7.66063982411248,
    "ENSMUSG00000029392_ENSMUSG00000001280": 7.61114399068689,
    "ENSMUSG00000040666_ENSMUSG00000001280": 7.392507942488433,
    "ENSMUSG00000081100_ENSMUSG00000019768": 7.3610339490053835,
    "ENSMUSG00000061904_ENSMUSG00000058756": 7.149747952002246,
    "ENSMUSG00000061518_ENSMUSG00000057469": 6.754048086061417,
    "ENSMUSG00000031701_ENSMUSG00000001280": 6.680898473673975,
    "ENSMUSG00000083563_ENSMUSG00000001280": 6.431292484650507,
    "ENSMUSG00000020580_ENSMUSG00000001280": 6.2900086739405054,
    "ENSMUSG00000028081_ENSMUSG00000001280": 5.802763467405232,
    "ENSMUSG00000035674_ENSMUSG00000019768": 5.653122193020531,
    "ENSMUSG00000024974_ENSMUSG00000023460": 5.6023111132374845,
    "ENSMUSG00000001280_ENSMUSG00000000399": 5.390594901508978,
    "ENSMUSG00000024140_ENSMUSG00000001334": 5.384456011568297,
    "ENSMUSG00000073702_ENSMUSG00000001280": 5.378864267649398,
    "ENSMUSG00000030557_ENSMUSG00000020456": 5.33319755652402,
    "ENSMUSG00000064369_ENSMUSG00000014592": 5.148816934509415,
    "ENSMUSG00000073411_ENSMUSG00000052684": 5.0313932946534035,
    "ENSMUSG00000052684_ENSMUSG00000020423": 4.378251745660982,
    "ENSMUSG00000027305_ENSMUSG00000001280": 4.311652179952762,
    "ENSMUSG00000019179_ENSMUSG00000008575": 3.6161920828496457,
    "ENSMUSG00000076431_ENSMUSG00000020315": 3.4376583128857434,
    "ENSMUSG00000036452_ENSMUSG00000001280": 3.1879627746371475,
    "ENSMUSG00000058239_ENSMUSG00000030246": 3.1776240295169806,
    "ENSMUSG00000014313_ENSMUSG00000001280": 32.046741904689675,
    "ENSMUSG00000001280_ENSMUSG00000000538": 26.483458872481318,
    "ENSMUSG00000062908_ENSMUSG00000001280": 22.841180549103434,
    "ENSMUSG00000053898_ENSMUSG00000001280": 21.584457953002534,
    "ENSMUSG00000042541_ENSMUSG00000001280": 21.556788383467428,
    "ENSMUSG00000058076_ENSMUSG00000001280": 21.551962766510165,
    "ENSMUSG00000064351_ENSMUSG00000001280": 21.399848085342988,
    "ENSMUSG00000002265_ENSMUSG00000001506": 20.90723390569587,
    "ENSMUSG00000005610_ENSMUSG00000001280": 20.88544591021294,
    "ENSMUSG00000007892_ENSMUSG00000001280": 20.605211832006724,
    "ENSMUSG00000084105_ENSMUSG00000001280": 20.59305469636408,
    "ENSMUSG00000064371_ENSMUSG00000001280": 20.274228348494304,
    "ENSMUSG00000064341_ENSMUSG00000019768": 20.173658896371887,
    "ENSMUSG00000059447_ENSMUSG00000001280": 20.162745699340924,
    "ENSMUSG00000020949_ENSMUSG00000001280": 19.698909912990498,
    "ENSMUSG00000098343_ENSMUSG00000001280": 19.692194799737518,
    "ENSMUSG00000064344_ENSMUSG00000001280": 19.51390970468268,
    "ENSMUSG00000081344_ENSMUSG00000001280": 19.304204183051,
    "ENSMUSG00000026568_ENSMUSG00000001280": 19.21699894688383,
    "ENSMUSG00000020326_ENSMUSG00000001280": 18.58905543988568,
    "ENSMUSG00000027706_ENSMUSG00000001280": 18.113299561125196,
    "ENSMUSG00000087943_ENSMUSG00000001280": 17.845608282399084,
    "ENSMUSG00000074218_ENSMUSG00000019768": 17.792082783337847,
    "ENSMUSG00000019768_ENSMUSG00000002416": 17.418004380507522,
    "ENSMUSG00000052738_ENSMUSG00000001280": 16.811201455654384,
    "ENSMUSG00000046516_ENSMUSG00000001280": 16.784727507395058,
    "ENSMUSG00000061518_ENSMUSG00000001280": 16.700785612096062,
    "ENSMUSG00000057469_ENSMUSG00000025453": 15.429087636455497,
    "ENSMUSG00000032112_ENSMUSG00000005698": 15.318280592756132,
    "ENSMUSG00000020390_ENSMUSG00000015839": 14.708022196768644,
    "ENSMUSG00000083326_ENSMUSG00000019768": 14.488959878699239,
    "ENSMUSG00000097487_ENSMUSG00000001280": 14.383103018494028,
    "ENSMUSG00000031320_ENSMUSG00000015579": 13.975690286265033,
    "ENSMUSG00000030557_ENSMUSG00000026360": 13.959904664079648,
    "ENSMUSG00000021877_ENSMUSG00000016559": 13.825951112303244,
    "ENSMUSG00000024073_ENSMUSG00000001280": 13.788408475857423,
    "ENSMUSG00000001280_ENSMUSG00000001211": 13.776094985063594,
    "ENSMUSG00000019768_ENSMUSG00000003970": 13.631409152913038,
    "ENSMUSG00000058756_ENSMUSG00000002500": 13.297315579320658,
    "ENSMUSG00000068267_ENSMUSG00000001794": 13.233359230389766,
    "ENSMUSG00000040370_ENSMUSG00000001280": 13.171616257373739,
    "ENSMUSG00000052934_ENSMUSG00000001280": 13.117403097683233,
    "ENSMUSG00000064365_ENSMUSG00000019768": 12.98426431669067,
    "ENSMUSG00000037266_ENSMUSG00000001280": 12.95460958470767,
    "ENSMUSG00000030002_ENSMUSG00000001280": 12.947241205841307,
    "ENSMUSG00000045983_ENSMUSG00000019768": 12.902863359413764,
    "ENSMUSG00000037032_ENSMUSG00000001280": 12.844925086801872,
    "ENSMUSG00000031698_ENSMUSG00000019768": 12.790662691311269,
    "ENSMUSG00000020153_ENSMUSG00000019768": 12.681273760435506,
    "ENSMUSG00000027774_ENSMUSG00000001280": 12.165879490605262,
    "ENSMUSG00000063077_ENSMUSG00000001280": 11.987565731505251,
    "ENSMUSG00000023150_ENSMUSG00000001280": 11.972020755243978,
    "ENSMUSG00000019768_ENSMUSG00000013593": 11.676519439799765,
    "ENSMUSG00000043683_ENSMUSG00000001280": 11.659197359005793,
    "ENSMUSG00000021000_ENSMUSG00000001280": 11.60217726193083,
    "ENSMUSG00000038690_ENSMUSG00000001280": 11.53196140004053,
    "ENSMUSG00000025745_ENSMUSG00000019768": 11.529479846344614,
    "ENSMUSG00000038412_ENSMUSG00000001280": 11.524302735824325,
    "ENSMUSG00000059447_ENSMUSG00000019768": 11.515643993286119,
    "ENSMUSG00000024049_ENSMUSG00000019768": 11.49457116491082,
    "ENSMUSG00000021290_ENSMUSG00000019768": 11.42362417446954,
    "ENSMUSG00000054889_ENSMUSG00000008575": 11.404103756974402,
    "ENSMUSG00000030541_ENSMUSG00000019768": 11.330798282916964,
    "ENSMUSG00000026853_ENSMUSG00000001280": 11.327643205400635,
    "ENSMUSG00000092014_ENSMUSG00000001280": 11.30874335276894,
    "ENSMUSG00000054312_ENSMUSG00000001280": 11.27332340018842,
    "ENSMUSG00000031782_ENSMUSG00000001280": 11.25636851827584,
    "ENSMUSG00000026202_ENSMUSG00000019768": 11.25355547480333,
    "ENSMUSG00000001911_ENSMUSG00000001280": 11.24802023562183,
    "ENSMUSG00000029198_ENSMUSG00000001280": 11.22531983431544,
    "ENSMUSG00000035268_ENSMUSG00000001280": 11.212319135481403,
    "ENSMUSG00000020477_ENSMUSG00000001280": 11.199918870031489,
    "ENSMUSG00000020664_ENSMUSG00000001280": 11.14785373869165,
    "ENSMUSG00000033543_ENSMUSG00000001280": 11.085459535640146,
    "ENSMUSG00000024238_ENSMUSG00000014402": 11.05596428403179,
    "ENSMUSG00000023944_ENSMUSG00000019768": 10.99537908654845,
    "ENSMUSG00000041028_ENSMUSG00000019768": 10.886985777938971,
    "ENSMUSG00000002379_ENSMUSG00000001280": 10.860422368919565,
    "ENSMUSG00000091537_ENSMUSG00000001280": 10.843430256412349,
    "ENSMUSG00000008136_ENSMUSG00000001280": 10.809203380977811,
    "ENSMUSG00000024527_ENSMUSG00000001280": 10.808329586730276,
    "ENSMUSG00000026626_ENSMUSG00000001280": 10.79233891909301,
    "ENSMUSG00000024661_ENSMUSG00000001280": 10.757301157682935,
    "ENSMUSG00000020483_ENSMUSG00000001280": 10.709477134145715,
    "ENSMUSG00000022450_ENSMUSG00000001280": 10.704367717230728,
    "ENSMUSG00000064366_ENSMUSG00000001280": 10.680872070613894,
    "ENSMUSG00000040850_ENSMUSG00000001280": 10.640553774460848,
    "ENSMUSG00000044080_ENSMUSG00000019768": 10.596154199449476,
    "ENSMUSG00000047459_ENSMUSG00000001280": 10.524728110614523,
    "ENSMUSG00000026193_ENSMUSG00000002265": 10.51672254354766,
    "ENSMUSG00000008575_ENSMUSG00000001280": 10.511502948864546,
    "ENSMUSG00000025393_ENSMUSG00000019768": 10.510960021678537,
    "ENSMUSG00000094249_ENSMUSG00000019768": 10.503744869266662,
    "ENSMUSG00000096942_ENSMUSG00000001280": 10.459415566161773,
    "ENSMUSG00000068706_ENSMUSG00000014592": 10.390601584844731,
    "ENSMUSG00000080242_ENSMUSG00000001280": 10.371571628081218,
    "ENSMUSG00000023456_ENSMUSG00000001280": 10.371308719947848,
    "ENSMUSG00000005354_ENSMUSG00000001280": 10.35617347437722,
    "ENSMUSG00000026766_ENSMUSG00000001280": 10.355419036962482,
    "ENSMUSG00000047419_ENSMUSG00000001280": 10.33419131349003,
    "ENSMUSG00000060802_ENSMUSG00000001280": 10.33059036101056,
    "ENSMUSG00000027777_ENSMUSG00000001280": 10.157359281316955,
    "ENSMUSG00000036880_ENSMUSG00000001280": 10.120991247750043,
    "ENSMUSG00000029432_ENSMUSG00000001280": 10.069746336643997,
    "ENSMUSG00000052911_ENSMUSG00000001280": 10.06334558142719,
    "ENSMUSG00000064341_ENSMUSG00000001280": 10.041113067370125,
    "ENSMUSG00000020022_ENSMUSG00000001280": 10.011513183325793,
    "ENSMUSG00000032511_ENSMUSG00000001280": 9.989303704565605,
    "ENSMUSG00000081331_ENSMUSG00000001280": 9.94411598199988,
    "ENSMUSG00000031791_ENSMUSG00000001280": 9.885485494224222,
    "ENSMUSG00000081661_ENSMUSG00000014592": 9.87050493355843,
    "ENSMUSG00000024997_ENSMUSG00000001280": 9.839328520881942,
    "ENSMUSG00000024962_ENSMUSG00000001280": 9.8227436442332,
    "ENSMUSG00000025794_ENSMUSG00000001280": 9.790967835204212,
    "ENSMUSG00000070544_ENSMUSG00000001280": 9.773679321605362,
    "ENSMUSG00000036427_ENSMUSG00000001280": 9.770804342614595,
    "ENSMUSG00000019768_ENSMUSG00000008136": 9.764197939783065,
    "ENSMUSG00000023089_ENSMUSG00000001280": 9.759036001584578,
    "ENSMUSG00000009927_ENSMUSG00000001280": 9.732163518846368,
    "ENSMUSG00000032648_ENSMUSG00000001280": 9.725842714378196,
    "ENSMUSG00000041028_ENSMUSG00000001280": 9.714156865305641,
    "ENSMUSG00000019768_ENSMUSG00000016252": 9.700055486044134,
    "ENSMUSG00000057322_ENSMUSG00000008575": 9.689722692801963,
    "ENSMUSG00000025428_ENSMUSG00000019768": 9.658610516797854,
    "ENSMUSG00000062997_ENSMUSG00000001280": 9.639874551489445,
    "ENSMUSG00000063229_ENSMUSG00000019768": 9.636740704708895,
    "ENSMUSG00000023861_ENSMUSG00000001280": 9.624148647839162,
    "ENSMUSG00000001833_ENSMUSG00000001280": 9.610469890723088,
    "ENSMUSG00000064344_ENSMUSG00000030557": 9.462804302550976,
    "ENSMUSG00000010608_ENSMUSG00000001280": 9.439561335466344,
    "ENSMUSG00000031698_ENSMUSG00000001280": 9.41970753066159,
    "ENSMUSG00000061518_ENSMUSG00000019768": 9.35313016544013,
    "ENSMUSG00000015837_ENSMUSG00000001280": 9.016484314008204,
    "ENSMUSG00000022186_ENSMUSG00000001280": 9.015219190393283,
    "ENSMUSG00000063229_ENSMUSG00000001280": 8.979295358020819,
    "ENSMUSG00000024359_ENSMUSG00000001280": 8.958786534549633,
    "ENSMUSG00000021313_ENSMUSG00000001280": 8.928265159054767,
    "ENSMUSG00000025260_ENSMUSG00000001280": 8.839003958978084,
    "ENSMUSG00000028223_ENSMUSG00000001280": 8.832220567256483,
    "ENSMUSG00000019768_ENSMUSG00000006057": 8.82475311625106,
    "ENSMUSG00000032966_ENSMUSG00000001280": 8.80106437204736,
    "ENSMUSG00000025508_ENSMUSG00000001280": 8.632678347341917,
    "ENSMUSG00000007891_ENSMUSG00000001280": 8.545322346694077,
    "ENSMUSG00000082474_ENSMUSG00000001280": 8.534169356824139,
    "ENSMUSG00000006717_ENSMUSG00000001280": 8.397679138184044,
    "ENSMUSG00000026260_ENSMUSG00000001280": 8.318061236522759,
    "ENSMUSG00000006219_ENSMUSG00000001280": 7.856402833541235,
    "ENSMUSG00000019689_ENSMUSG00000001280": 7.846260019366253,
    "ENSMUSG00000024683_ENSMUSG00000001280": 7.320512270209525,
    "ENSMUSG00000005873_ENSMUSG00000001280": 7.287288376478506,
    "ENSMUSG00000019768_ENSMUSG00000006818": 7.278474467171327,
    "ENSMUSG00000035329_ENSMUSG00000001280": 7.004565122346684,
    "ENSMUSG00000027447_ENSMUSG00000021250": 6.890746404891511,
    "ENSMUSG00000031885_ENSMUSG00000020402": 6.643373147707506,
    "ENSMUSG00000053768_ENSMUSG00000030557": 6.614567121759081,
    "ENSMUSG00000073411_ENSMUSG00000021250": 6.574047057336408,
    "ENSMUSG00000027673_ENSMUSG00000019768": 6.5715142372303355,
    "ENSMUSG00000027010_ENSMUSG00000008575": 6.500844507537435,
    "ENSMUSG00000019768_ENSMUSG00000004069": 6.449117423434171,
    "ENSMUSG00000044533_ENSMUSG00000030557": 5.842143290243047,
    "ENSMUSG00000028234_ENSMUSG00000001280": 5.81733243971842,
    "ENSMUSG00000031660_ENSMUSG00000001280": 5.752970516083994,
    "ENSMUSG00000030852_ENSMUSG00000001280": 5.468940628524868,
    "ENSMUSG00000043483_ENSMUSG00000001280": 5.411247272653108,
    "ENSMUSG00000091086_ENSMUSG00000001280": 5.305519054762978,
    "ENSMUSG00000028518_ENSMUSG00000008575": 5.245831644372669,
    "ENSMUSG00000008575_ENSMUSG00000007891": 5.00455942922336,
    "ENSMUSG00000020372_ENSMUSG00000001280": 4.765291332131981,
    "ENSMUSG00000044894_ENSMUSG00000008575": 4.604904023887949,
    "ENSMUSG00000019055_ENSMUSG00000001280": 4.6044053689632385,
    "ENSMUSG00000092074_ENSMUSG00000001280": 4.58288708900055,
    "ENSMUSG00000067288_ENSMUSG00000001280": 4.397744528552969,
    "ENSMUSG00000005510_ENSMUSG00000001280": 4.369043120831976,
    "ENSMUSG00000052837_ENSMUSG00000018669": 4.305184425388483,
    "ENSMUSG00000040269_ENSMUSG00000001280": 4.275347494861212,
    "ENSMUSG00000021371_ENSMUSG00000001280": 4.234467086607963,
    "ENSMUSG00000058756_ENSMUSG00000018363": 4.2305241761344465,
    "ENSMUSG00000020456_ENSMUSG00000005583": 4.169615383758787,
    "ENSMUSG00000029135_ENSMUSG00000019505": 4.120734139658008,
    "ENSMUSG00000063180_ENSMUSG00000052684": 3.8398808954377714,
    "ENSMUSG00000029249_ENSMUSG00000027602": 3.8075908445489786,
    "ENSMUSG00000019179_ENSMUSG00000001280": 3.6257225282939043,
    "ENSMUSG00000036461_ENSMUSG00000025810": 3.5984263252793696,
    "ENSMUSG00000022617_ENSMUSG00000015839": 3.5146700409360916,
    "ENSMUSG00000083261_ENSMUSG00000033543": 3.089917969848105
}
```
