## Supplement 2 (DAN vs. Cocul) for "AGEAS: Automated Machine Learning based Genetic Regulatory Element Extraction System": full_atlas.js

```
{
    "network_0": {
        "genes": {
            "ENSG00000168214": {
                "id": "ENSG00000168214",
                "type": "TF",
                "symbol": "RBPJ",
                "source": [
                    "ENSG00000163435",
                    "ENSG00000157557",
                    "ENSG00000167081",
                    "ENSG00000113580",
                    "ENSG00000148516",
                    "ENSG00000164330",
                    "ENSG00000100219",
                    "ENSG00000221869"
                ],
                "target": [
                    "ENSG00000169856",
                    "ENSG00000152558",
                    "ENSG00000198840",
                    "ENSG00000132698",
                    "ENSG00000164093",
                    "ENSG00000107105",
                    "ENSG00000101443",
                    "ENSG00000166582",
                    "ENSG00000179981",
                    "ENSG00000169435",
                    "ENSG00000172183",
                    "ENSG00000165215",
                    "ENSG00000163191",
                    "ENSG00000189143",
                    "ENSG00000123416",
                    "ENSG00000198125",
                    "ENSG00000181885",
                    "ENSG00000245532",
                    "ENSG00000102466",
                    "ENSG00000265972",
                    "ENSG00000154556",
                    "ENSG00000167642",
                    "ENSG00000102007",
                    "ENSG00000163956",
                    "ENSG00000087258",
                    "ENSG00000187244",
                    "ENSG00000198899",
                    "ENSG00000111728",
                    "ENSG00000119888",
                    "ENSG00000100604",
                    "ENSG00000169862",
                    "ENSG00000180879",
                    "ENSG00000099194",
                    "ENSG00000104722",
                    "ENSG00000171345",
                    "ENSG00000176887",
                    "ENSG00000169783",
                    "ENSG00000187193",
                    "ENSG00000148516",
                    "ENSG00000102934",
                    "ENSG00000170266",
                    "ENSG00000117525",
                    "ENSG00000213190",
                    "ENSG00000060982",
                    "ENSG00000196361",
                    "ENSG00000113732",
                    "ENSG00000147889",
                    "ENSG00000130303",
                    "ENSG00000131711",
                    "ENSG00000075618",
                    "ENSG00000144218",
                    "ENSG00000121039",
                    "ENSG00000107130",
                    "ENSG00000161249",
                    "ENSG00000170091",
                    "ENSG00000138085",
                    "ENSG00000170293",
                    "ENSG00000064601",
                    "ENSG00000113580",
                    "ENSG00000068615",
                    "ENSG00000100219",
                    "ENSG00000185650",
                    "ENSG00000215845",
                    "ENSG00000111057",
                    "ENSG00000104870",
                    "ENSG00000165092",
                    "ENSG00000176788",
                    "ENSG00000149557",
                    "ENSG00000145428",
                    "ENSG00000148180",
                    "ENSG00000145335",
                    "ENSG00000163399",
                    "ENSG00000138772",
                    "ENSG00000112773",
                    "ENSG00000244187",
                    "ENSG00000174498",
                    "ENSG00000213760",
                    "ENSG00000124107",
                    "ENSG00000117691",
                    "ENSG00000092964",
                    "ENSG00000143702",
                    "ENSG00000135535",
                    "ENSG00000112972",
                    "ENSG00000122012",
                    "ENSG00000107438",
                    "ENSG00000016082",
                    "ENSG00000142156",
                    "ENSG00000152503",
                    "ENSG00000120437",
                    "ENSG00000103316",
                    "ENSG00000149294",
                    "ENSG00000124942",
                    "ENSG00000157557",
                    "ENSG00000096696",
                    "ENSG00000167552",
                    "ENSG00000104833",
                    "ENSG00000164330",
                    "ENSG00000137267",
                    "ENSG00000143819",
                    "ENSG00000172915",
                    "ENSG00000030582",
                    "ENSG00000179218",
                    "ENSG00000100345",
                    "ENSG00000214530",
                    "ENSG00000198727",
                    "ENSG00000147676",
                    "ENSG00000105784",
                    "ENSG00000137880",
                    "ENSG00000115461",
                    "ENSG00000148834",
                    "ENSG00000072832",
                    "ENSG00000168938",
                    "ENSG00000221866",
                    "ENSG00000246705",
                    "ENSG00000198712",
                    "ENSG00000117155",
                    "ENSG00000089356",
                    "ENSG00000146674",
                    "ENSG00000213614",
                    "ENSG00000137285",
                    "ENSG00000118640",
                    "ENSG00000177459",
                    "ENSG00000198938",
                    "ENSG00000139970",
                    "ENSG00000242265",
                    "ENSG00000163435",
                    "ENSG00000145730",
                    "ENSG00000109062",
                    "ENSG00000167081",
                    "ENSG00000108679"
                ],
                "expression_sum": {
                    "class1": 13084.0,
                    "class2": 12713.0
                },
                "ens_id": "ENSG00000168214",
                "uniprot_ids": [
                    "Q06330",
                    "A0A7P0T821",
                    "A0A7P0TA33",
                    "D6RCM1",
                    "D6RB37",
                    "D6REC2",
                    "D6RA45",
                    "A0A7P0T964",
                    "A0A7P0T947",
                    "A0A7P0T9W9",
                    "A0A7P0TBJ4",
                    "A0A7P0T848",
                    "D6R946",
                    "D6R9X3",
                    "D6REP3",
                    "A0A7P0Z4N7",
                    "D6R9K5",
                    "D6RIZ8",
                    "A0A7P0T9Y9",
                    "D6RAT2",
                    "D6RE93",
                    "A0A7P0T7Z3",
                    "D6RF98",
                    "A0A7P0T806",
                    "D6RIV8",
                    "D6RBQ8",
                    "H0Y8R3",
                    "D6R927"
                ]
            },
            "ENSG00000181885": {
                "id": "ENSG00000181885",
                "type": "Gene",
                "symbol": "CLDN7",
                "source": [
                    "ENSG00000123933",
                    "ENSG00000163435",
                    "ENSG00000160113",
                    "ENSG00000113580",
                    "ENSG00000153147",
                    "ENSG00000105698",
                    "ENSG00000168610",
                    "ENSG00000100219",
                    "ENSG00000221869",
                    "ENSG00000139352",
                    "ENSG00000168214",
                    "ENSG00000196591",
                    "ENSG00000124766",
                    "ENSG00000173575",
                    "ENSG00000177606"
                ],
                "target": [],
                "expression_sum": {
                    "class1": 425.0,
                    "class2": 17786.0
                },
                "ens_id": "ENSG00000181885",
                "uniprot_ids": [
                    "O95471",
                    "I3L3X1",
                    "I3L3L6",
                    "K7EL87",
                    "F5H496",
                    "K7EP40"
                ]
            },
            "ENSG00000160113": {
                "id": "ENSG00000160113",
                "type": "TF",
                "symbol": "NR2F6",
                "source": [
                    "ENSG00000163435",
                    "ENSG00000113580",
                    "ENSG00000148516",
                    "ENSG00000164330",
                    "ENSG00000100219",
                    "ENSG00000181722",
                    "ENSG00000221869"
                ],
                "target": [
                    "ENSG00000169856",
                    "ENSG00000152558",
                    "ENSG00000111674",
                    "ENSG00000166710",
                    "ENSG00000087085",
                    "ENSG00000108830",
                    "ENSG00000247095",
                    "ENSG00000166582",
                    "ENSG00000182534",
                    "ENSG00000085871",
                    "ENSG00000129355",
                    "ENSG00000105255",
                    "ENSG00000196230",
                    "ENSG00000086300",
                    "ENSG00000187134",
                    "ENSG00000172183",
                    "ENSG00000163191",
                    "ENSG00000123416",
                    "ENSG00000198125",
                    "ENSG00000181885",
                    "ENSG00000245532",
                    "ENSG00000265972",
                    "ENSG00000154556",
                    "ENSG00000167642",
                    "ENSG00000119865",
                    "ENSG00000108515",
                    "ENSG00000173267",
                    "ENSG00000181722",
                    "ENSG00000105088",
                    "ENSG00000079462",
                    "ENSG00000130066",
                    "ENSG00000119888",
                    "ENSG00000115310",
                    "ENSG00000100867",
                    "ENSG00000221869",
                    "ENSG00000159335",
                    "ENSG00000099194",
                    "ENSG00000180879",
                    "ENSG00000164929",
                    "ENSG00000109084",
                    "ENSG00000170266",
                    "ENSG00000101856",
                    "ENSG00000141934",
                    "ENSG00000084628",
                    "ENSG00000060982",
                    "ENSG00000131981",
                    "ENSG00000075618",
                    "ENSG00000138758",
                    "ENSG00000213719",
                    "ENSG00000132688",
                    "ENSG00000075043",
                    "ENSG00000175567",
                    "ENSG00000189334",
                    "ENSG00000170540",
                    "ENSG00000161249",
                    "ENSG00000129521",
                    "ENSG00000170091",
                    "ENSG00000170293",
                    "ENSG00000064601",
                    "ENSG00000113580",
                    "ENSG00000100219",
                    "ENSG00000185650",
                    "ENSG00000111275",
                    "ENSG00000104870",
                    "ENSG00000069849",
                    "ENSG00000165092",
                    "ENSG00000111907",
                    "ENSG00000148180",
                    "ENSG00000112773",
                    "ENSG00000163399",
                    "ENSG00000156675",
                    "ENSG00000244187",
                    "ENSG00000117472",
                    "ENSG00000174498",
                    "ENSG00000128564",
                    "ENSG00000186868",
                    "ENSG00000108106",
                    "ENSG00000213760",
                    "ENSG00000143702",
                    "ENSG00000135535",
                    "ENSG00000112972",
                    "ENSG00000051523",
                    "ENSG00000107438",
                    "ENSG00000144674",
                    "ENSG00000120437",
                    "ENSG00000103316",
                    "ENSG00000149294",
                    "ENSG00000124942",
                    "ENSG00000115758",
                    "ENSG00000113328",
                    "ENSG00000157557",
                    "ENSG00000096696",
                    "ENSG00000141505",
                    "ENSG00000137267",
                    "ENSG00000105048",
                    "ENSG00000163931",
                    "ENSG00000030582",
                    "ENSG00000144339",
                    "ENSG00000214530",
                    "ENSG00000161082",
                    "ENSG00000060138",
                    "ENSG00000174021",
                    "ENSG00000152952",
                    "ENSG00000070961",
                    "ENSG00000137880",
                    "ENSG00000148834",
                    "ENSG00000135480",
                    "ENSG00000182287",
                    "ENSG00000104549",
                    "ENSG00000213614",
                    "ENSG00000118640",
                    "ENSG00000170421",
                    "ENSG00000097021",
                    "ENSG00000152082",
                    "ENSG00000163435",
                    "ENSG00000213937",
                    "ENSG00000135406",
                    "ENSG00000170017",
                    "ENSG00000104419",
                    "ENSG00000108679"
                ],
                "expression_sum": {
                    "class1": 14139.0,
                    "class2": 12251.0
                },
                "ens_id": "ENSG00000160113",
                "uniprot_ids": [
                    "P10588",
                    "M0R175"
                ]
            },
            "ENSG00000119888": {
                "id": "ENSG00000119888",
                "type": "Gene",
                "symbol": "EPCAM",
                "source": [
                    "ENSG00000163435",
                    "ENSG00000160113",
                    "ENSG00000113580",
                    "ENSG00000105698",
                    "ENSG00000181722",
                    "ENSG00000139352",
                    "ENSG00000173575",
                    "ENSG00000168214",
                    "ENSG00000119547",
                    "ENSG00000177606",
                    "ENSG00000092199"
                ],
                "target": [],
                "expression_sum": {
                    "class1": 3257.0,
                    "class2": 27181.0
                },
                "ens_id": "ENSG00000119888",
                "uniprot_ids": [
                    "P16422",
                    "B5MCA4",
                    "C9JKY3"
                ]
            },
            "ENSG00000100219": {
                "id": "ENSG00000100219",
                "type": "TF",
                "symbol": "XBP1",
                "source": [
                    "ENSG00000168214",
                    "ENSG00000106261",
                    "ENSG00000177606",
                    "ENSG00000160113"
                ],
                "target": [
                    "ENSG00000106261",
                    "ENSG00000152558",
                    "ENSG00000126005",
                    "ENSG00000166710",
                    "ENSG00000198840",
                    "ENSG00000132698",
                    "ENSG00000171604",
                    "ENSG00000160113",
                    "ENSG00000177606",
                    "ENSG00000076554",
                    "ENSG00000163191",
                    "ENSG00000189143",
                    "ENSG00000184076",
                    "ENSG00000181885",
                    "ENSG00000245532",
                    "ENSG00000265972",
                    "ENSG00000154556",
                    "ENSG00000114439",
                    "ENSG00000181722",
                    "ENSG00000141279",
                    "ENSG00000111229",
                    "ENSG00000051620",
                    "ENSG00000150687",
                    "ENSG00000197930",
                    "ENSG00000135069",
                    "ENSG00000101040",
                    "ENSG00000198886",
                    "ENSG00000144560",
                    "ENSG00000134138",
                    "ENSG00000166347",
                    "ENSG00000213719",
                    "ENSG00000121039",
                    "ENSG00000143013",
                    "ENSG00000148444",
                    "ENSG00000026025",
                    "ENSG00000129521",
                    "ENSG00000263001",
                    "ENSG00000109099",
                    "ENSG00000163399",
                    "ENSG00000138772",
                    "ENSG00000156675",
                    "ENSG00000118680",
                    "ENSG00000128564",
                    "ENSG00000168214",
                    "ENSG00000124107",
                    "ENSG00000070831",
                    "ENSG00000144674",
                    "ENSG00000106537",
                    "ENSG00000114405",
                    "ENSG00000091844",
                    "ENSG00000172915",
                    "ENSG00000030582",
                    "ENSG00000002586",
                    "ENSG00000123636",
                    "ENSG00000214530",
                    "ENSG00000174938",
                    "ENSG00000198727",
                    "ENSG00000147676",
                    "ENSG00000104763",
                    "ENSG00000160613",
                    "ENSG00000060138",
                    "ENSG00000169764",
                    "ENSG00000196586",
                    "ENSG00000198786",
                    "ENSG00000146674",
                    "ENSG00000118640",
                    "ENSG00000170421",
                    "ENSG00000164096",
                    "ENSG00000049618",
                    "ENSG00000139970",
                    "ENSG00000182718",
                    "ENSG00000163435",
                    "ENSG00000133639",
                    "ENSG00000109062",
                    "ENSG00000138640",
                    "ENSG00000104419",
                    "ENSG00000198888"
                ],
                "expression_sum": {
                    "class1": 15335.0,
                    "class2": 32457.0
                },
                "ens_id": "ENSG00000100219",
                "uniprot_ids": [
                    "P17861",
                    "B1AHH2",
                    "B1AHH1"
                ]
            },
            "ENSG00000124107": {
                "id": "ENSG00000124107",
                "type": "Gene",
                "symbol": "SLPI",
                "source": [
                    "ENSG00000100219",
                    "ENSG00000163435",
                    "ENSG00000168214"
                ],
                "target": [],
                "expression_sum": {
                    "class1": 10.0,
                    "class2": 24968.0
                },
                "ens_id": "ENSG00000124107",
                "uniprot_ids": [
                    "P03973"
                ]
            },
            "ENSG00000109099": {
                "id": "ENSG00000109099",
                "type": "Gene",
                "symbol": "PMP22",
                "source": [
                    "ENSG00000175745",
                    "ENSG00000148516",
                    "ENSG00000100219"
                ],
                "target": [],
                "expression_sum": {
                    "class1": 994.0,
                    "class2": 5019.0
                },
                "ens_id": "ENSG00000109099",
                "uniprot_ids": [
                    "Q01453",
                    "J3KT36",
                    "A0A6Q8PF08",
                    "A8MU75",
                    "A0A2R8Y5L5",
                    "J3KQW0",
                    "A0A6Q8PF92",
                    "A0A6Q8PFD5",
                    "A0A6Q8PFS0"
                ]
            },
            "ENSG00000182718": {
                "id": "ENSG00000182718",
                "type": "Gene",
                "symbol": "ANXA2",
                "source": [
                    "ENSG00000163435",
                    "ENSG00000016082",
                    "ENSG00000113580",
                    "ENSG00000164683",
                    "ENSG00000167081",
                    "ENSG00000148516",
                    "ENSG00000164330",
                    "ENSG00000108001",
                    "ENSG00000100219",
                    "ENSG00000221869",
                    "ENSG00000116044"
                ],
                "target": [],
                "expression_sum": {
                    "class1": 38391.0,
                    "class2": 72712.0
                },
                "ens_id": "ENSG00000182718",
                "uniprot_ids": [
                    "P07355",
                    "H0YKS4",
                    "H0YL33",
                    "H0YKZ7",
                    "H0YKN4",
                    "H0YKV8",
                    "H0YMT9",
                    "H0YMD9",
                    "H0YKL9",
                    "H0YMW4",
                    "H0YM50",
                    "H0YNP5",
                    "H0YLE2",
                    "H0YKX9",
                    "H0YN28",
                    "H0YNA0",
                    "H0YLV6",
                    "H0YMU9",
                    "H0YMM1",
                    "H0YN42",
                    "A0A7I2V4P4",
                    "H0YMD0",
                    "H0YNB8"
                ]
            },
            "ENSG00000167081": {
                "id": "ENSG00000167081",
                "type": "TF",
                "symbol": "PBX3",
                "source": [
                    "ENSG00000100811",
                    "ENSG00000134138",
                    "ENSG00000004487",
                    "ENSG00000103495",
                    "ENSG00000105698",
                    "ENSG00000213015",
                    "ENSG00000189403",
                    "ENSG00000119950",
                    "ENSG00000196591",
                    "ENSG00000168214",
                    "ENSG00000092199"
                ],
                "target": [
                    "ENSG00000078668",
                    "ENSG00000090621",
                    "ENSG00000206573",
                    "ENSG00000112081",
                    "ENSG00000232119",
                    "ENSG00000107223",
                    "ENSG00000162704",
                    "ENSG00000163634",
                    "ENSG00000165655",
                    "ENSG00000153130",
                    "ENSG00000111674",
                    "ENSG00000111716",
                    "ENSG00000139644",
                    "ENSG00000005810",
                    "ENSG00000169612",
                    "ENSG00000099622",
                    "ENSG00000137955",
                    "ENSG00000185651",
                    "ENSG00000090060",
                    "ENSG00000005020",
                    "ENSG00000106603",
                    "ENSG00000140941",
                    "ENSG00000161016",
                    "ENSG00000134248",
                    "ENSG00000127540",
                    "ENSG00000075945",
                    "ENSG00000125743",
                    "ENSG00000099797",
                    "ENSG00000119632",
                    "ENSG00000189403",
                    "ENSG00000159377",
                    "ENSG00000136261",
                    "ENSG00000075415",
                    "ENSG00000147155",
                    "ENSG00000077380",
                    "ENSG00000154556",
                    "ENSG00000052802",
                    "ENSG00000134283",
                    "ENSG00000198931",
                    "ENSG00000173812",
                    "ENSG00000197451",
                    "ENSG00000152256",
                    "ENSG00000237190",
                    "ENSG00000173915",
                    "ENSG00000065911",
                    "ENSG00000151729",
                    "ENSG00000172292",
                    "ENSG00000170310",
                    "ENSG00000129636",
                    "ENSG00000115540",
                    "ENSG00000123240",
                    "ENSG00000041357",
                    "ENSG00000102103",
                    "ENSG00000141858",
                    "ENSG00000178913",
                    "ENSG00000163875",
                    "ENSG00000163319",
                    "ENSG00000167136",
                    "ENSG00000164924",
                    "ENSG00000074319",
                    "ENSG00000149196",
                    "ENSG00000141759",
                    "ENSG00000174574",
                    "ENSG00000146007",
                    "ENSG00000106554",
                    "ENSG00000084764",
                    "ENSG00000277791",
                    "ENSG00000125868",
                    "ENSG00000153815",
                    "ENSG00000047249",
                    "ENSG00000147548",
                    "ENSG00000171703",
                    "ENSG00000196504",
                    "ENSG00000131236",
                    "ENSG00000089220",
                    "ENSG00000055163",
                    "ENSG00000141447",
                    "ENSG00000116898",
                    "ENSG00000175581",
                    "ENSG00000196072",
                    "ENSG00000179431",
                    "ENSG00000187164",
                    "ENSG00000188243",
                    "ENSG00000067225",
                    "ENSG00000109390",
                    "ENSG00000113712",
                    "ENSG00000126698",
                    "ENSG00000136143",
                    "ENSG00000233927",
                    "ENSG00000117139",
                    "ENSG00000148444",
                    "ENSG00000140105",
                    "ENSG00000159685",
                    "ENSG00000034713",
                    "ENSG00000130811",
                    "ENSG00000144746",
                    "ENSG00000095380",
                    "ENSG00000133703",
                    "ENSG00000063854",
                    "ENSG00000103769",
                    "ENSG00000034510",
                    "ENSG00000146425",
                    "ENSG00000005194",
                    "ENSG00000138071",
                    "ENSG00000203778",
                    "ENSG00000117410",
                    "ENSG00000131100",
                    "ENSG00000075142",
                    "ENSG00000273841",
                    "ENSG00000124802",
                    "ENSG00000263001",
                    "ENSG00000142657",
                    "ENSG00000196405",
                    "ENSG00000183011",
                    "ENSG00000174437",
                    "ENSG00000114520",
                    "ENSG00000213741",
                    "ENSG00000126934",
                    "ENSG00000145592",
                    "ENSG00000187051",
                    "ENSG00000069849",
                    "ENSG00000240344",
                    "ENSG00000129515",
                    "ENSG00000100519",
                    "ENSG00000144635",
                    "ENSG00000187514",
                    "ENSG00000163041",
                    "ENSG00000155115",
                    "ENSG00000160194",
                    "ENSG00000141027",
                    "ENSG00000092199",
                    "ENSG00000110955",
                    "ENSG00000178980",
                    "ENSG00000204463",
                    "ENSG00000164346",
                    "ENSG00000150316",
                    "ENSG00000010256",
                    "ENSG00000120053",
                    "ENSG00000174953",
                    "ENSG00000169100",
                    "ENSG00000162191",
                    "ENSG00000118363",
                    "ENSG00000108654",
                    "ENSG00000135940",
                    "ENSG00000170634",
                    "ENSG00000108106",
                    "ENSG00000198836",
                    "ENSG00000107581",
                    "ENSG00000033800",
                    "ENSG00000105258",
                    "ENSG00000106052",
                    "ENSG00000147677",
                    "ENSG00000231500",
                    "ENSG00000108666",
                    "ENSG00000153037",
                    "ENSG00000175792",
                    "ENSG00000102317",
                    "ENSG00000265491",
                    "ENSG00000213465",
                    "ENSG00000109332",
                    "ENSG00000075624",
                    "ENSG00000072062",
                    "ENSG00000168275",
                    "ENSG00000075785",
                    "ENSG00000184900",
                    "ENSG00000095059",
                    "ENSG00000110696",
                    "ENSG00000111850",
                    "ENSG00000102898",
                    "ENSG00000107937",
                    "ENSG00000166974",
                    "ENSG00000063322",
                    "ENSG00000232388",
                    "ENSG00000141562",
                    "ENSG00000114391",
                    "ENSG00000163931",
                    "ENSG00000175166",
                    "ENSG00000168303",
                    "ENSG00000167088",
                    "ENSG00000100320",
                    "ENSG00000162852",
                    "ENSG00000105419",
                    "ENSG00000141030",
                    "ENSG00000075292",
                    "ENSG00000164985",
                    "ENSG00000239672",
                    "ENSG00000105723",
                    "ENSG00000153310",
                    "ENSG00000256060",
                    "ENSG00000131508",
                    "ENSG00000197223",
                    "ENSG00000005022",
                    "ENSG00000180957",
                    "ENSG00000131469",
                    "ENSG00000080824",
                    "ENSG00000160271",
                    "ENSG00000090266",
                    "ENSG00000136810",
                    "ENSG00000142937",
                    "ENSG00000048649",
                    "ENSG00000162374",
                    "ENSG00000169764",
                    "ENSG00000106682",
                    "ENSG00000116209",
                    "ENSG00000072832",
                    "ENSG00000130312",
                    "ENSG00000136950",
                    "ENSG00000079459",
                    "ENSG00000104341",
                    "ENSG00000197170",
                    "ENSG00000104388",
                    "ENSG00000083750",
                    "ENSG00000168159",
                    "ENSG00000267100",
                    "ENSG00000170144",
                    "ENSG00000097021",
                    "ENSG00000152082",
                    "ENSG00000087191",
                    "ENSG00000257704",
                    "ENSG00000168710",
                    "ENSG00000127914",
                    "ENSG00000124614",
                    "ENSG00000014641",
                    "ENSG00000126267",
                    "ENSG00000175582",
                    "ENSG00000104765",
                    "ENSG00000171130",
                    "ENSG00000198612",
                    "ENSG00000120860",
                    "ENSG00000081154",
                    "ENSG00000122566",
                    "ENSG00000131473",
                    "ENSG00000112305",
                    "ENSG00000108774",
                    "ENSG00000269893",
                    "ENSG00000047932",
                    "ENSG00000091140",
                    "ENSG00000253738",
                    "ENSG00000135250",
                    "ENSG00000165119",
                    "ENSG00000067064",
                    "ENSG00000163636",
                    "ENSG00000135486",
                    "ENSG00000115128",
                    "ENSG00000126005",
                    "ENSG00000137818",
                    "ENSG00000082153",
                    "ENSG00000241343",
                    "ENSG00000100347",
                    "ENSG00000105677",
                    "ENSG00000089737",
                    "ENSG00000114784",
                    "ENSG00000184752",
                    "ENSG00000197302",
                    "ENSG00000129355",
                    "ENSG00000188343",
                    "ENSG00000275052",
                    "ENSG00000135002",
                    "ENSG00000100575",
                    "ENSG00000127445",
                    "ENSG00000183172",
                    "ENSG00000114744",
                    "ENSG00000146282",
                    "ENSG00000170027",
                    "ENSG00000147274",
                    "ENSG00000171055",
                    "ENSG00000078674",
                    "ENSG00000076554",
                    "ENSG00000010270",
                    "ENSG00000103495",
                    "ENSG00000147604",
                    "ENSG00000204308",
                    "ENSG00000214046",
                    "ENSG00000123416",
                    "ENSG00000171314",
                    "ENSG00000168291",
                    "ENSG00000090238",
                    "ENSG00000130520",
                    "ENSG00000170315",
                    "ENSG00000167792",
                    "ENSG00000138413",
                    "ENSG00000147684",
                    "ENSG00000096746",
                    "ENSG00000103266",
                    "ENSG00000079785",
                    "ENSG00000134779",
                    "ENSG00000119950",
                    "ENSG00000099795",
                    "ENSG00000129084",
                    "ENSG00000069275",
                    "ENSG00000138777",
                    "ENSG00000072042",
                    "ENSG00000126756",
                    "ENSG00000162585",
                    "ENSG00000103145",
                    "ENSG00000163714",
                    "ENSG00000051620",
                    "ENSG00000028839",
                    "ENSG00000114200",
                    "ENSG00000189043",
                    "ENSG00000163939",
                    "ENSG00000117523",
                    "ENSG00000152154",
                    "ENSG00000137100",
                    "ENSG00000123106",
                    "ENSG00000100764",
                    "ENSG00000109475",
                    "ENSG00000196199",
                    "ENSG00000187109",
                    "ENSG00000198886",
                    "ENSG00000147533",
                    "ENSG00000155506",
                    "ENSG00000122643",
                    "ENSG00000196262",
                    "ENSG00000104904",
                    "ENSG00000100316",
                    "ENSG00000125971",
                    "ENSG00000173726",
                    "ENSG00000134287",
                    "ENSG00000155959",
                    "ENSG00000137692",
                    "ENSG00000116586",
                    "ENSG00000089693",
                    "ENSG00000105254",
                    "ENSG00000168092",
                    "ENSG00000115286",
                    "ENSG00000105698",
                    "ENSG00000120438",
                    "ENSG00000084234",
                    "ENSG00000105640",
                    "ENSG00000114023",
                    "ENSG00000125691",
                    "ENSG00000154723",
                    "ENSG00000122026",
                    "ENSG00000166347",
                    "ENSG00000149357",
                    "ENSG00000198843",
                    "ENSG00000125901",
                    "ENSG00000109654",
                    "ENSG00000168288",
                    "ENSG00000130956",
                    "ENSG00000130558",
                    "ENSG00000234745",
                    "ENSG00000113387",
                    "ENSG00000158864",
                    "ENSG00000137409",
                    "ENSG00000006125",
                    "ENSG00000156508",
                    "ENSG00000196937",
                    "ENSG00000135900",
                    "ENSG00000082701",
                    "ENSG00000110717",
                    "ENSG00000120727",
                    "ENSG00000100442",
                    "ENSG00000143799",
                    "ENSG00000106028",
                    "ENSG00000173917",
                    "ENSG00000115091",
                    "ENSG00000204628",
                    "ENSG00000115053",
                    "ENSG00000178234",
                    "ENSG00000134986",
                    "ENSG00000121022",
                    "ENSG00000164615",
                    "ENSG00000113369",
                    "ENSG00000197111",
                    "ENSG00000163682",
                    "ENSG00000175334",
                    "ENSG00000146729",
                    "ENSG00000171421",
                    "ENSG00000256537",
                    "ENSG00000124380",
                    "ENSG00000123562",
                    "ENSG00000105701",
                    "ENSG00000166260",
                    "ENSG00000111669",
                    "ENSG00000136100",
                    "ENSG00000159720",
                    "ENSG00000184007",
                    "ENSG00000140307",
                    "ENSG00000132821",
                    "ENSG00000102401",
                    "ENSG00000167863",
                    "ENSG00000105409",
                    "ENSG00000197747",
                    "ENSG00000004779",
                    "ENSG00000184277",
                    "ENSG00000106355",
                    "ENSG00000090273",
                    "ENSG00000162368",
                    "ENSG00000254682",
                    "ENSG00000004478",
                    "ENSG00000175756",
                    "ENSG00000134333",
                    "ENSG00000197345",
                    "ENSG00000116251",
                    "ENSG00000091640",
                    "ENSG00000147123",
                    "ENSG00000161960",
                    "ENSG00000143575",
                    "ENSG00000038219",
                    "ENSG00000130560",
                    "ENSG00000159140",
                    "ENSG00000160014",
                    "ENSG00000125818",
                    "ENSG00000105649",
                    "ENSG00000146731",
                    "ENSG00000103187",
                    "ENSG00000105220",
                    "ENSG00000105372",
                    "ENSG00000053254",
                    "ENSG00000111678",
                    "ENSG00000011295",
                    "ENSG00000176871",
                    "ENSG00000007168",
                    "ENSG00000169288",
                    "ENSG00000111142",
                    "ENSG00000164713",
                    "ENSG00000132507",
                    "ENSG00000176209",
                    "ENSG00000105379",
                    "ENSG00000175550",
                    "ENSG00000099901",
                    "ENSG00000167508",
                    "ENSG00000173692",
                    "ENSG00000111237",
                    "ENSG00000132824",
                    "ENSG00000112110",
                    "ENSG00000023191",
                    "ENSG00000111911",
                    "ENSG00000135070",
                    "ENSG00000151366",
                    "ENSG00000108298",
                    "ENSG00000145782",
                    "ENSG00000166226",
                    "ENSG00000112695",
                    "ENSG00000174444",
                    "ENSG00000165416",
                    "ENSG00000203667",
                    "ENSG00000133318",
                    "ENSG00000130414",
                    "ENSG00000115738",
                    "ENSG00000103429",
                    "ENSG00000138430",
                    "ENSG00000169738",
                    "ENSG00000169213",
                    "ENSG00000166780",
                    "ENSG00000176340",
                    "ENSG00000172354",
                    "ENSG00000273611",
                    "ENSG00000182670",
                    "ENSG00000177410",
                    "ENSG00000115514",
                    "ENSG00000101745",
                    "ENSG00000145337",
                    "ENSG00000113068",
                    "ENSG00000182718",
                    "ENSG00000221983",
                    "ENSG00000173436",
                    "ENSG00000175348",
                    "ENSG00000069956",
                    "ENSG00000063046",
                    "ENSG00000122565",
                    "ENSG00000163541",
                    "ENSG00000146416",
                    "ENSG00000164258",
                    "ENSG00000116560",
                    "ENSG00000106261",
                    "ENSG00000198492",
                    "ENSG00000141002",
                    "ENSG00000171604",
                    "ENSG00000179262",
                    "ENSG00000183207",
                    "ENSG00000073792",
                    "ENSG00000143355",
                    "ENSG00000178952",
                    "ENSG00000196230",
                    "ENSG00000225663",
                    "ENSG00000186106",
                    "ENSG00000198034",
                    "ENSG00000129518",
                    "ENSG00000104408",
                    "ENSG00000105373",
                    "ENSG00000129235",
                    "ENSG00000173418",
                    "ENSG00000102144",
                    "ENSG00000167526",
                    "ENSG00000130770",
                    "ENSG00000265681",
                    "ENSG00000142864",
                    "ENSG00000119013",
                    "ENSG00000161203",
                    "ENSG00000100823",
                    "ENSG00000142534",
                    "ENSG00000151746",
                    "ENSG00000101782",
                    "ENSG00000177889",
                    "ENSG00000142541",
                    "ENSG00000109971",
                    "ENSG00000109536",
                    "ENSG00000156136",
                    "ENSG00000118816",
                    "ENSG00000121851",
                    "ENSG00000151743",
                    "ENSG00000135624",
                    "ENSG00000104671",
                    "ENSG00000147669",
                    "ENSG00000004487",
                    "ENSG00000163468",
                    "ENSG00000111843",
                    "ENSG00000137312",
                    "ENSG00000091039",
                    "ENSG00000167522",
                    "ENSG00000099800",
                    "ENSG00000147403",
                    "ENSG00000134056",
                    "ENSG00000167220",
                    "ENSG00000277443",
                    "ENSG00000172115",
                    "ENSG00000119547",
                    "ENSG00000145912",
                    "ENSG00000233429",
                    "ENSG00000006451",
                    "ENSG00000177156",
                    "ENSG00000145293",
                    "ENSG00000167468",
                    "ENSG00000187634",
                    "ENSG00000111640",
                    "ENSG00000125652",
                    "ENSG00000135387",
                    "ENSG00000124596",
                    "ENSG00000084628",
                    "ENSG00000115241",
                    "ENSG00000156976",
                    "ENSG00000180964",
                    "ENSG00000111481",
                    "ENSG00000253719",
                    "ENSG00000134138",
                    "ENSG00000161970",
                    "ENSG00000160888",
                    "ENSG00000128245",
                    "ENSG00000077721",
                    "ENSG00000177879",
                    "ENSG00000205531",
                    "ENSG00000154001",
                    "ENSG00000138758",
                    "ENSG00000115541",
                    "ENSG00000069329",
                    "ENSG00000138069",
                    "ENSG00000162244",
                    "ENSG00000083099",
                    "ENSG00000144034",
                    "ENSG00000171033",
                    "ENSG00000071082",
                    "ENSG00000225733",
                    "ENSG00000071994",
                    "ENSG00000005483",
                    "ENSG00000115484",
                    "ENSG00000177030",
                    "ENSG00000138107",
                    "ENSG00000126261",
                    "ENSG00000130748",
                    "ENSG00000138326",
                    "ENSG00000204568",
                    "ENSG00000157778",
                    "ENSG00000175203",
                    "ENSG00000111642",
                    "ENSG00000154380",
                    "ENSG00000104695",
                    "ENSG00000123144",
                    "ENSG00000100216",
                    "ENSG00000101557",
                    "ENSG00000172531",
                    "ENSG00000113575",
                    "ENSG00000149084",
                    "ENSG00000127022",
                    "ENSG00000074800",
                    "ENSG00000162961",
                    "ENSG00000148303",
                    "ENSG00000104863",
                    "ENSG00000225921",
                    "ENSG00000165502",
                    "ENSG00000114302",
                    "ENSG00000084733",
                    "ENSG00000142676",
                    "ENSG00000119705",
                    "ENSG00000165283",
                    "ENSG00000075151",
                    "ENSG00000122034",
                    "ENSG00000185664",
                    "ENSG00000100528",
                    "ENSG00000154582",
                    "ENSG00000130255",
                    "ENSG00000067836",
                    "ENSG00000214113",
                    "ENSG00000096384",
                    "ENSG00000109107",
                    "ENSG00000147687",
                    "ENSG00000168214",
                    "ENSG00000197958",
                    "ENSG00000100911",
                    "ENSG00000160213",
                    "ENSG00000171497",
                    "ENSG00000188994",
                    "ENSG00000174775",
                    "ENSG00000143947",
                    "ENSG00000164978",
                    "ENSG00000112237",
                    "ENSG00000164830",
                    "ENSG00000213015",
                    "ENSG00000175416",
                    "ENSG00000128609",
                    "ENSG00000090470",
                    "ENSG00000170632",
                    "ENSG00000063177",
                    "ENSG00000155368",
                    "ENSG00000198918",
                    "ENSG00000169564",
                    "ENSG00000094916",
                    "ENSG00000103035",
                    "ENSG00000132305",
                    "ENSG00000138663",
                    "ENSG00000086061",
                    "ENSG00000204310",
                    "ENSG00000070756",
                    "ENSG00000136930",
                    "ENSG00000166922",
                    "ENSG00000149806",
                    "ENSG00000140386",
                    "ENSG00000140740",
                    "ENSG00000146386",
                    "ENSG00000204272",
                    "ENSG00000166012",
                    "ENSG00000167658",
                    "ENSG00000113161",
                    "ENSG00000197006",
                    "ENSG00000185608",
                    "ENSG00000142549",
                    "ENSG00000084070",
                    "ENSG00000174173",
                    "ENSG00000068745",
                    "ENSG00000196465",
                    "ENSG00000168028",
                    "ENSG00000108848",
                    "ENSG00000136193",
                    "ENSG00000156928",
                    "ENSG00000104427",
                    "ENSG00000124702",
                    "ENSG00000111786",
                    "ENSG00000109919",
                    "ENSG00000143933",
                    "ENSG00000166352",
                    "ENSG00000165629",
                    "ENSG00000136603",
                    "ENSG00000167815",
                    "ENSG00000177971",
                    "ENSG00000126457",
                    "ENSG00000167996",
                    "ENSG00000132963",
                    "ENSG00000042753",
                    "ENSG00000175390",
                    "ENSG00000168522",
                    "ENSG00000148154",
                    "ENSG00000119048",
                    "ENSG00000198408",
                    "ENSG00000177600",
                    "ENSG00000148450",
                    "ENSG00000134419",
                    "ENSG00000178127",
                    "ENSG00000174456",
                    "ENSG00000184117",
                    "ENSG00000167699",
                    "ENSG00000166946",
                    "ENSG00000092148",
                    "ENSG00000181610",
                    "ENSG00000054148",
                    "ENSG00000270170",
                    "ENSG00000143621",
                    "ENSG00000156411",
                    "ENSG00000108960",
                    "ENSG00000137343",
                    "ENSG00000177200",
                    "ENSG00000115944",
                    "ENSG00000080822",
                    "ENSG00000111639",
                    "ENSG00000196531",
                    "ENSG00000100138",
                    "ENSG00000174695",
                    "ENSG00000116030",
                    "ENSG00000129968",
                    "ENSG00000115268",
                    "ENSG00000182534",
                    "ENSG00000093144",
                    "ENSG00000125534",
                    "ENSG00000160075",
                    "ENSG00000170043",
                    "ENSG00000089157",
                    "ENSG00000149231",
                    "ENSG00000131435",
                    "ENSG00000130723",
                    "ENSG00000198363",
                    "ENSG00000140988",
                    "ENSG00000168090",
                    "ENSG00000122033",
                    "ENSG00000197728",
                    "ENSG00000178741",
                    "ENSG00000156515",
                    "ENSG00000115947",
                    "ENSG00000101266",
                    "ENSG00000177885",
                    "ENSG00000233016",
                    "ENSG00000143774",
                    "ENSG00000182512",
                    "ENSG00000088832",
                    "ENSG00000179271",
                    "ENSG00000122545",
                    "ENSG00000134330",
                    "ENSG00000143612",
                    "ENSG00000135720",
                    "ENSG00000007520",
                    "ENSG00000166848",
                    "ENSG00000149547",
                    "ENSG00000125870",
                    "ENSG00000159335",
                    "ENSG00000120686",
                    "ENSG00000102189",
                    "ENSG00000123352",
                    "ENSG00000108518",
                    "ENSG00000181061",
                    "ENSG00000128739",
                    "ENSG00000086666",
                    "ENSG00000278311",
                    "ENSG00000172809",
                    "ENSG00000145425",
                    "ENSG00000114902",
                    "ENSG00000157514",
                    "ENSG00000106615",
                    "ENSG00000107341",
                    "ENSG00000089818",
                    "ENSG00000099341",
                    "ENSG00000173273",
                    "ENSG00000172500",
                    "ENSG00000230989",
                    "ENSG00000087302",
                    "ENSG00000091164",
                    "ENSG00000136146",
                    "ENSG00000066557",
                    "ENSG00000090263",
                    "ENSG00000172757",
                    "ENSG00000105669",
                    "ENSG00000188290",
                    "ENSG00000149313",
                    "ENSG00000063245",
                    "ENSG00000136942",
                    "ENSG00000126858",
                    "ENSG00000185658",
                    "ENSG00000050426",
                    "ENSG00000119185",
                    "ENSG00000160752",
                    "ENSG00000143252",
                    "ENSG00000141367",
                    "ENSG00000144381",
                    "ENSG00000136521",
                    "ENSG00000100811",
                    "ENSG00000183648",
                    "ENSG00000159352",
                    "ENSG00000213246",
                    "ENSG00000155099",
                    "ENSG00000172687",
                    "ENSG00000133872",
                    "ENSG00000137106",
                    "ENSG00000179010",
                    "ENSG00000198668",
                    "ENSG00000108107",
                    "ENSG00000134308",
                    "ENSG00000111832",
                    "ENSG00000061794",
                    "ENSG00000134684",
                    "ENSG00000008324",
                    "ENSG00000166228",
                    "ENSG00000213619",
                    "ENSG00000153774",
                    "ENSG00000129197",
                    "ENSG00000138757",
                    "ENSG00000108561",
                    "ENSG00000033627",
                    "ENSG00000178982",
                    "ENSG00000108010",
                    "ENSG00000083845",
                    "ENSG00000138382",
                    "ENSG00000111726",
                    "ENSG00000125821",
                    "ENSG00000137547",
                    "ENSG00000100387",
                    "ENSG00000140319",
                    "ENSG00000132466",
                    "ENSG00000149809",
                    "ENSG00000181704",
                    "ENSG00000172428",
                    "ENSG00000118680",
                    "ENSG00000243317",
                    "ENSG00000070669",
                    "ENSG00000165233",
                    "ENSG00000117450",
                    "ENSG00000184220",
                    "ENSG00000103485",
                    "ENSG00000009307",
                    "ENSG00000143995",
                    "ENSG00000138138",
                    "ENSG00000204856",
                    "ENSG00000124172",
                    "ENSG00000182196",
                    "ENSG00000057608",
                    "ENSG00000150991",
                    "ENSG00000065548",
                    "ENSG00000152795",
                    "ENSG00000212907",
                    "ENSG00000196591",
                    "ENSG00000137210",
                    "ENSG00000166681",
                    "ENSG00000132432",
                    "ENSG00000184009",
                    "ENSG00000023734",
                    "ENSG00000120437",
                    "ENSG00000125944",
                    "ENSG00000054523",
                    "ENSG00000183726",
                    "ENSG00000148925",
                    "ENSG00000128626",
                    "ENSG00000214941",
                    "ENSG00000113328",
                    "ENSG00000186184",
                    "ENSG00000166164",
                    "ENSG00000119640",
                    "ENSG00000113648",
                    "ENSG00000170899",
                    "ENSG00000185787",
                    "ENSG00000170889",
                    "ENSG00000137947",
                    "ENSG00000100554",
                    "ENSG00000130638",
                    "ENSG00000169045",
                    "ENSG00000111775",
                    "ENSG00000123636",
                    "ENSG00000087502",
                    "ENSG00000139684",
                    "ENSG00000197102",
                    "ENSG00000114120",
                    "ENSG00000166562",
                    "ENSG00000168724",
                    "ENSG00000126602",
                    "ENSG00000180817",
                    "ENSG00000086065",
                    "ENSG00000164587",
                    "ENSG00000116288",
                    "ENSG00000181163",
                    "ENSG00000008018",
                    "ENSG00000160948",
                    "ENSG00000117569",
                    "ENSG00000198898",
                    "ENSG00000115685",
                    "ENSG00000215021",
                    "ENSG00000165487",
                    "ENSG00000103275",
                    "ENSG00000134313",
                    "ENSG00000101654",
                    "ENSG00000104549",
                    "ENSG00000044115",
                    "ENSG00000265241",
                    "ENSG00000136238",
                    "ENSG00000091527",
                    "ENSG00000123545",
                    "ENSG00000224032",
                    "ENSG00000047849",
                    "ENSG00000182154",
                    "ENSG00000221914",
                    "ENSG00000122873",
                    "ENSG00000103415",
                    "ENSG00000065978",
                    "ENSG00000087460",
                    "ENSG00000100567",
                    "ENSG00000151247",
                    "ENSG00000113558",
                    "ENSG00000143153",
                    "ENSG00000109180",
                    "ENSG00000123983",
                    "ENSG00000116489",
                    "ENSG00000143742",
                    "ENSG00000133169",
                    "ENSG00000205629",
                    "ENSG00000174886",
                    "ENSG00000164167",
                    "ENSG00000092841",
                    "ENSG00000109929",
                    "ENSG00000089009",
                    "ENSG00000173163"
                ],
                "expression_sum": {
                    "class1": 19184.0,
                    "class2": 2356.0
                },
                "ens_id": "ENSG00000167081",
                "uniprot_ids": [
                    "P40426",
                    "H3BLX0",
                    "Q5JS98",
                    "U3KQA2",
                    "H0Y5D0"
                ]
            },
            "ENSG00000113558": {
                "id": "ENSG00000113558",
                "type": "Gene",
                "symbol": "SKP1",
                "source": [
                    "ENSG00000181449",
                    "ENSG00000163435",
                    "ENSG00000109132",
                    "ENSG00000167081",
                    "ENSG00000113580",
                    "ENSG00000114861",
                    "ENSG00000221869"
                ],
                "target": [],
                "expression_sum": {
                    "class1": 107742.0,
                    "class2": 85135.0
                },
                "ens_id": "ENSG00000113558",
                "uniprot_ids": [
                    "P63208",
                    "E5RGM3",
                    "F8W8N3",
                    "E5RK33",
                    "E5RJR5",
                    "E5RGM4",
                    "E7ERH2",
                    "E5RHM3"
                ]
            },
            "ENSG00000156976": {
                "id": "ENSG00000156976",
                "type": "Gene",
                "symbol": "EIF4A2",
                "source": [
                    "ENSG00000181449",
                    "ENSG00000163435",
                    "ENSG00000157557",
                    "ENSG00000109132",
                    "ENSG00000016082",
                    "ENSG00000113580",
                    "ENSG00000167081",
                    "ENSG00000148516",
                    "ENSG00000103495",
                    "ENSG00000164330",
                    "ENSG00000114861",
                    "ENSG00000221869",
                    "ENSG00000139352",
                    "ENSG00000137203"
                ],
                "target": [],
                "expression_sum": {
                    "class1": 81149.0,
                    "class2": 40818.0
                },
                "ens_id": "ENSG00000156976",
                "uniprot_ids": [
                    "Q14240",
                    "E9PBH4",
                    "F8WE11",
                    "C9JUF0",
                    "J3KSN7",
                    "E7EMV8",
                    "E7EQG2"
                ]
            },
            "ENSG00000111716": {
                "id": "ENSG00000111716",
                "type": "Gene",
                "symbol": "LDHB",
                "source": [
                    "ENSG00000181449",
                    "ENSG00000109132",
                    "ENSG00000016082",
                    "ENSG00000113580",
                    "ENSG00000167081",
                    "ENSG00000103495",
                    "ENSG00000164330",
                    "ENSG00000108001",
                    "ENSG00000114861",
                    "ENSG00000137947",
                    "ENSG00000119547",
                    "ENSG00000196591",
                    "ENSG00000181315",
                    "ENSG00000137309"
                ],
                "target": [],
                "expression_sum": {
                    "class1": 74907.0,
                    "class2": 46249.0
                },
                "ens_id": "ENSG00000111716",
                "uniprot_ids": [
                    "P07195",
                    "F5H793",
                    "A0A5F9ZHM4",
                    "A8MW50",
                    "C9J7H8"
                ]
            },
            "ENSG00000016082": {
                "id": "ENSG00000016082",
                "type": "TF",
                "symbol": "ISL1",
                "source": [
                    "ENSG00000170365",
                    "ENSG00000100811",
                    "ENSG00000130726",
                    "ENSG00000109132",
                    "ENSG00000153147",
                    "ENSG00000004487",
                    "ENSG00000103495",
                    "ENSG00000198911",
                    "ENSG00000108055",
                    "ENSG00000164754",
                    "ENSG00000168214",
                    "ENSG00000196591"
                ],
                "target": [
                    "ENSG00000017260",
                    "ENSG00000162704",
                    "ENSG00000111716",
                    "ENSG00000005810",
                    "ENSG00000140941",
                    "ENSG00000197959",
                    "ENSG00000145817",
                    "ENSG00000075415",
                    "ENSG00000077380",
                    "ENSG00000154556",
                    "ENSG00000118733",
                    "ENSG00000052802",
                    "ENSG00000087274",
                    "ENSG00000165669",
                    "ENSG00000065911",
                    "ENSG00000106443",
                    "ENSG00000129636",
                    "ENSG00000197061",
                    "ENSG00000242173",
                    "ENSG00000139767",
                    "ENSG00000174574",
                    "ENSG00000139116",
                    "ENSG00000075914",
                    "ENSG00000047249",
                    "ENSG00000060982",
                    "ENSG00000141447",
                    "ENSG00000175581",
                    "ENSG00000067225",
                    "ENSG00000144218",
                    "ENSG00000117139",
                    "ENSG00000140105",
                    "ENSG00000159685",
                    "ENSG00000205981",
                    "ENSG00000157216",
                    "ENSG00000144746",
                    "ENSG00000157168",
                    "ENSG00000133606",
                    "ENSG00000125835",
                    "ENSG00000075142",
                    "ENSG00000150722",
                    "ENSG00000263001",
                    "ENSG00000213741",
                    "ENSG00000086598",
                    "ENSG00000129515",
                    "ENSG00000076604",
                    "ENSG00000142546",
                    "ENSG00000110955",
                    "ENSG00000175216",
                    "ENSG00000168137",
                    "ENSG00000149187",
                    "ENSG00000213760",
                    "ENSG00000107581",
                    "ENSG00000112972",
                    "ENSG00000147677",
                    "ENSG00000231500",
                    "ENSG00000204842",
                    "ENSG00000167196",
                    "ENSG00000180370",
                    "ENSG00000079246",
                    "ENSG00000168275",
                    "ENSG00000057019",
                    "ENSG00000111850",
                    "ENSG00000105185",
                    "ENSG00000166974",
                    "ENSG00000138801",
                    "ENSG00000100320",
                    "ENSG00000114391",
                    "ENSG00000164985",
                    "ENSG00000135390",
                    "ENSG00000239672",
                    "ENSG00000141867",
                    "ENSG00000153310",
                    "ENSG00000256060",
                    "ENSG00000197223",
                    "ENSG00000154277",
                    "ENSG00000131469",
                    "ENSG00000128654",
                    "ENSG00000090266",
                    "ENSG00000171634",
                    "ENSG00000162374",
                    "ENSG00000169764",
                    "ENSG00000166200",
                    "ENSG00000104388",
                    "ENSG00000204954",
                    "ENSG00000143222",
                    "ENSG00000151914",
                    "ENSG00000161057",
                    "ENSG00000177565",
                    "ENSG00000125827",
                    "ENSG00000171130",
                    "ENSG00000073578",
                    "ENSG00000081154",
                    "ENSG00000122566",
                    "ENSG00000143878",
                    "ENSG00000135486",
                    "ENSG00000109670",
                    "ENSG00000275052",
                    "ENSG00000128951",
                    "ENSG00000100575",
                    "ENSG00000172159",
                    "ENSG00000167792",
                    "ENSG00000138413",
                    "ENSG00000070761",
                    "ENSG00000163848",
                    "ENSG00000069275",
                    "ENSG00000119729",
                    "ENSG00000113742",
                    "ENSG00000170515",
                    "ENSG00000144895",
                    "ENSG00000115875",
                    "ENSG00000198886",
                    "ENSG00000164929",
                    "ENSG00000196262",
                    "ENSG00000115355",
                    "ENSG00000214078",
                    "ENSG00000173726",
                    "ENSG00000134287",
                    "ENSG00000089280",
                    "ENSG00000139117",
                    "ENSG00000101132",
                    "ENSG00000122026",
                    "ENSG00000151327",
                    "ENSG00000106400",
                    "ENSG00000140836",
                    "ENSG00000113387",
                    "ENSG00000156508",
                    "ENSG00000196937",
                    "ENSG00000082701",
                    "ENSG00000234608",
                    "ENSG00000170142",
                    "ENSG00000173141",
                    "ENSG00000084463",
                    "ENSG00000185630",
                    "ENSG00000100442",
                    "ENSG00000115091",
                    "ENSG00000204628",
                    "ENSG00000178234",
                    "ENSG00000121022",
                    "ENSG00000197111",
                    "ENSG00000163069",
                    "ENSG00000134825",
                    "ENSG00000184007",
                    "ENSG00000099250",
                    "ENSG00000197345",
                    "ENSG00000116251",
                    "ENSG00000038219",
                    "ENSG00000053254",
                    "ENSG00000164330",
                    "ENSG00000156639",
                    "ENSG00000175550",
                    "ENSG00000144339",
                    "ENSG00000156011",
                    "ENSG00000135070",
                    "ENSG00000214413",
                    "ENSG00000145782",
                    "ENSG00000153187",
                    "ENSG00000174444",
                    "ENSG00000110711",
                    "ENSG00000106829",
                    "ENSG00000138430",
                    "ENSG00000126214",
                    "ENSG00000097033",
                    "ENSG00000273611",
                    "ENSG00000164733",
                    "ENSG00000179933",
                    "ENSG00000148943",
                    "ENSG00000189159",
                    "ENSG00000177410",
                    "ENSG00000186472",
                    "ENSG00000170365",
                    "ENSG00000182718",
                    "ENSG00000089053",
                    "ENSG00000122565",
                    "ENSG00000163541",
                    "ENSG00000164258",
                    "ENSG00000147862",
                    "ENSG00000006715",
                    "ENSG00000079102",
                    "ENSG00000197756",
                    "ENSG00000065135",
                    "ENSG00000008294",
                    "ENSG00000104408",
                    "ENSG00000105373",
                    "ENSG00000265681",
                    "ENSG00000084207",
                    "ENSG00000058272",
                    "ENSG00000165434",
                    "ENSG00000122068",
                    "ENSG00000170873",
                    "ENSG00000162434",
                    "ENSG00000118816",
                    "ENSG00000182827",
                    "ENSG00000204442",
                    "ENSG00000163468",
                    "ENSG00000172115",
                    "ENSG00000118257",
                    "ENSG00000156976",
                    "ENSG00000078369",
                    "ENSG00000171033",
                    "ENSG00000138107",
                    "ENSG00000149485",
                    "ENSG00000071994",
                    "ENSG00000157778",
                    "ENSG00000154380",
                    "ENSG00000182621",
                    "ENSG00000172531",
                    "ENSG00000148303",
                    "ENSG00000054118",
                    "ENSG00000023228",
                    "ENSG00000136717",
                    "ENSG00000168175",
                    "ENSG00000136854",
                    "ENSG00000196305",
                    "ENSG00000117859",
                    "ENSG00000113845",
                    "ENSG00000197958",
                    "ENSG00000077279",
                    "ENSG00000188994",
                    "ENSG00000108946",
                    "ENSG00000071794",
                    "ENSG00000172586",
                    "ENSG00000063177",
                    "ENSG00000094916",
                    "ENSG00000103035",
                    "ENSG00000138663",
                    "ENSG00000140386",
                    "ENSG00000072041",
                    "ENSG00000113108",
                    "ENSG00000165462",
                    "ENSG00000128699",
                    "ENSG00000108848",
                    "ENSG00000108826",
                    "ENSG00000156928",
                    "ENSG00000186469",
                    "ENSG00000166352",
                    "ENSG00000126457",
                    "ENSG00000070087",
                    "ENSG00000174547",
                    "ENSG00000131876",
                    "ENSG00000148700",
                    "ENSG00000167699",
                    "ENSG00000092148",
                    "ENSG00000135406",
                    "ENSG00000134824",
                    "ENSG00000177675",
                    "ENSG00000151552",
                    "ENSG00000061676",
                    "ENSG00000080822",
                    "ENSG00000168268",
                    "ENSG00000138594",
                    "ENSG00000179915",
                    "ENSG00000159593",
                    "ENSG00000180228",
                    "ENSG00000139613",
                    "ENSG00000131116",
                    "ENSG00000197728",
                    "ENSG00000104823",
                    "ENSG00000130779",
                    "ENSG00000229117",
                    "ENSG00000166206",
                    "ENSG00000126773",
                    "ENSG00000115350",
                    "ENSG00000086666",
                    "ENSG00000154518",
                    "ENSG00000087302",
                    "ENSG00000182628",
                    "ENSG00000090263",
                    "ENSG00000105669",
                    "ENSG00000160752",
                    "ENSG00000136521",
                    "ENSG00000204220",
                    "ENSG00000164741",
                    "ENSG00000155330",
                    "ENSG00000108100",
                    "ENSG00000122958",
                    "ENSG00000111832",
                    "ENSG00000118263",
                    "ENSG00000068615",
                    "ENSG00000153774",
                    "ENSG00000165525",
                    "ENSG00000083845",
                    "ENSG00000132466",
                    "ENSG00000128923",
                    "ENSG00000184220",
                    "ENSG00000009307",
                    "ENSG00000101882",
                    "ENSG00000018189",
                    "ENSG00000107372",
                    "ENSG00000109132",
                    "ENSG00000204856",
                    "ENSG00000182117",
                    "ENSG00000150991",
                    "ENSG00000212907",
                    "ENSG00000007080",
                    "ENSG00000104679",
                    "ENSG00000142875",
                    "ENSG00000173660",
                    "ENSG00000214941",
                    "ENSG00000186184",
                    "ENSG00000170899",
                    "ENSG00000123636",
                    "ENSG00000162604",
                    "ENSG00000163964",
                    "ENSG00000102003",
                    "ENSG00000010810",
                    "ENSG00000196781",
                    "ENSG00000188846",
                    "ENSG00000078018",
                    "ENSG00000111530",
                    "ENSG00000070961",
                    "ENSG00000103275",
                    "ENSG00000167232",
                    "ENSG00000134313",
                    "ENSG00000198712",
                    "ENSG00000113448",
                    "ENSG00000186468",
                    "ENSG00000183527",
                    "ENSG00000100567",
                    "ENSG00000144567",
                    "ENSG00000123983",
                    "ENSG00000172270",
                    "ENSG00000198888",
                    "ENSG00000089009"
                ],
                "expression_sum": {
                    "class1": 6727.0,
                    "class2": 11.0
                },
                "ens_id": "ENSG00000016082",
                "uniprot_ids": [
                    "P61371",
                    "D6RBJ1"
                ]
            },
            "ENSG00000174444": {
                "id": "ENSG00000174444",
                "type": "Gene",
                "symbol": "RPL4",
                "source": [
                    "ENSG00000181449",
                    "ENSG00000016082",
                    "ENSG00000167081",
                    "ENSG00000164330",
                    "ENSG00000114861",
                    "ENSG00000221869",
                    "ENSG00000119547"
                ],
                "target": [],
                "expression_sum": {
                    "class1": 108249.0,
                    "class2": 88709.0
                },
                "ens_id": "ENSG00000174444",
                "uniprot_ids": [
                    "P36578",
                    "H3BM89",
                    "H3BU31",
                    "H3BTP7"
                ]
            },
            "ENSG00000075142": {
                "id": "ENSG00000075142",
                "type": "Gene",
                "symbol": "SRI",
                "source": [
                    "ENSG00000181449",
                    "ENSG00000109132",
                    "ENSG00000016082",
                    "ENSG00000167081",
                    "ENSG00000169136",
                    "ENSG00000164330",
                    "ENSG00000108001",
                    "ENSG00000119547"
                ],
                "target": [],
                "expression_sum": {
                    "class1": 28378.0,
                    "class2": 20792.0
                },
                "ens_id": "ENSG00000075142",
                "uniprot_ids": [
                    "P30626",
                    "B4DHQ6",
                    "E9PG82",
                    "C9J0K6"
                ]
            },
            "ENSG00000139116": {
                "id": "ENSG00000139116",
                "type": "Gene",
                "symbol": "KIF21A",
                "source": [
                    "ENSG00000181449",
                    "ENSG00000109132",
                    "ENSG00000016082",
                    "ENSG00000143799",
                    "ENSG00000164330",
                    "ENSG00000124766"
                ],
                "target": [],
                "expression_sum": {
                    "class1": 22491.0,
                    "class2": 11405.0
                },
                "ens_id": "ENSG00000139116",
                "uniprot_ids": [
                    "Q7Z4S6",
                    "H0YIM6",
                    "H0YHG9",
                    "H0YHT2",
                    "A0A1B0GV47",
                    "H0YI78",
                    "H0YIM7"
                ]
            },
            "ENSG00000128951": {
                "id": "ENSG00000128951",
                "type": "Gene",
                "symbol": "DUT",
                "source": [
                    "ENSG00000016082",
                    "ENSG00000181449",
                    "ENSG00000164330"
                ],
                "target": [],
                "expression_sum": {
                    "class1": 24345.0,
                    "class2": 20216.0
                },
                "ens_id": "ENSG00000128951",
                "uniprot_ids": [
                    "P33316",
                    "A0A0C4DGL3",
                    "H0YKI0",
                    "H0YKC5",
                    "H0YNJ9",
                    "H0YNW5",
                    "H0YMM5",
                    "H0YMP1"
                ]
            },
            "ENSG00000144339": {
                "id": "ENSG00000144339",
                "type": "Gene",
                "symbol": "TMEFF2",
                "source": [
                    "ENSG00000170365",
                    "ENSG00000016082",
                    "ENSG00000004487",
                    "ENSG00000103495",
                    "ENSG00000198911",
                    "ENSG00000196591",
                    "ENSG00000160113"
                ],
                "target": [],
                "expression_sum": {
                    "class1": 7361.0,
                    "class2": 1097.0
                },
                "ens_id": "ENSG00000144339",
                "uniprot_ids": [
                    "Q9UIK5"
                ]
            },
            "ENSG00000164741": {
                "id": "ENSG00000164741",
                "type": "Gene",
                "symbol": "DLC1",
                "source": [
                    "ENSG00000016082",
                    "ENSG00000147862",
                    "ENSG00000109132"
                ],
                "target": [],
                "expression_sum": {
                    "class1": 1303.0,
                    "class2": 583.0
                },
                "ens_id": "ENSG00000164741",
                "uniprot_ids": [
                    "Q96QB1",
                    "A0A0J9YWS8",
                    "E5RI70",
                    "R4GMP5"
                ]
            },
            "ENSG00000157168": {
                "id": "ENSG00000157168",
                "type": "TF",
                "symbol": "NRG1",
                "source": [
                    "ENSG00000109132",
                    "ENSG00000016082",
                    "ENSG00000113580",
                    "ENSG00000103495",
                    "ENSG00000139352",
                    "ENSG00000116560"
                ],
                "target": [
                    "ENSG00000070669",
                    "ENSG00000134198",
                    "ENSG00000101134"
                ],
                "expression_sum": {
                    "class1": 5138.0,
                    "class2": 954.0
                },
                "ens_id": "ENSG00000157168",
                "uniprot_ids": [
                    "Q02297",
                    "H0YBA3",
                    "A0A494C114",
                    "A0A494C0Q4",
                    "A0A494C1B5",
                    "E5RHQ1",
                    "A0A087X0X9",
                    "A0A5F9ZHA5",
                    "E5RHP6",
                    "A0A494C1F5",
                    "A0A494C0K4",
                    "A0A494BZT4",
                    "E5RIG8",
                    "A0A494C0L9",
                    "H0YCP0",
                    "A0A494C054",
                    "A0A494C1F8",
                    "H0YDC2",
                    "A0A494C0T5",
                    "A0A494C1G2",
                    "F6RJN6",
                    "A0A494C043"
                ]
            },
            "ENSG00000057019": {
                "id": "ENSG00000057019",
                "type": "Gene",
                "symbol": "DCBLD2",
                "source": [
                    "ENSG00000016082",
                    "ENSG00000164330"
                ],
                "target": [],
                "expression_sum": {
                    "class1": 6090.0,
                    "class2": 3668.0
                },
                "ens_id": "ENSG00000057019",
                "uniprot_ids": [
                    "Q96PD2",
                    "C9JIW6"
                ]
            },
            "ENSG00000072041": {
                "id": "ENSG00000072041",
                "type": "Gene",
                "symbol": "SLC6A15",
                "source": [
                    "ENSG00000016082",
                    "ENSG00000103495",
                    "ENSG00000109132"
                ],
                "target": [],
                "expression_sum": {
                    "class1": 3430.0,
                    "class2": 873.0
                },
                "ens_id": "ENSG00000072041",
                "uniprot_ids": [
                    "Q9H2J7",
                    "A0A7P0T9Z9",
                    "F8VSG1",
                    "A0A7P0TB74",
                    "A0A7P0TAT1",
                    "A0A7P0TAZ8",
                    "A0A7P0TB05",
                    "A0A7P0Z4F0",
                    "A0A7P0T8I1",
                    "A0A7P0T8B7",
                    "A0A7P0T9P8",
                    "F8WJN6",
                    "A0A7P0Z4M4",
                    "A0A7P0T9G8",
                    "A0A7P0TA72",
                    "A0A7P0TBL2",
                    "A0A7P0T826"
                ]
            },
            "ENSG00000162374": {
                "id": "ENSG00000162374",
                "type": "Gene",
                "symbol": "ELAVL4",
                "source": [
                    "ENSG00000100811",
                    "ENSG00000130726",
                    "ENSG00000109132",
                    "ENSG00000016082",
                    "ENSG00000167081",
                    "ENSG00000148516",
                    "ENSG00000004487",
                    "ENSG00000108055",
                    "ENSG00000139352",
                    "ENSG00000164754",
                    "ENSG00000116560",
                    "ENSG00000196591",
                    "ENSG00000143995"
                ],
                "target": [],
                "expression_sum": {
                    "class1": 31005.0,
                    "class2": 1862.0
                },
                "ens_id": "ENSG00000162374",
                "uniprot_ids": [
                    "P26378",
                    "B1APY9",
                    "A0A0R4J2E6",
                    "A0A494C0Q5",
                    "A0A494C015",
                    "A0A494BZW4",
                    "A0A494C173",
                    "A0A494C0M6",
                    "A0A494C011",
                    "A0A494C0F8",
                    "A0A494C147",
                    "A0A494C1D9"
                ]
            },
            "ENSG00000118257": {
                "id": "ENSG00000118257",
                "type": "Gene",
                "symbol": "NRP2",
                "source": [
                    "ENSG00000016082"
                ],
                "target": [],
                "expression_sum": {
                    "class1": 4560.0,
                    "class2": 4429.0
                },
                "ens_id": "ENSG00000118257",
                "uniprot_ids": [
                    "O60462",
                    "C9JH98"
                ]
            },
            "ENSG00000173726": {
                "id": "ENSG00000173726",
                "type": "Gene",
                "symbol": "TOMM20",
                "source": [
                    "ENSG00000181449",
                    "ENSG00000016082",
                    "ENSG00000113580",
                    "ENSG00000167081",
                    "ENSG00000164330",
                    "ENSG00000114861",
                    "ENSG00000221869",
                    "ENSG00000139352"
                ],
                "target": [],
                "expression_sum": {
                    "class1": 33673.0,
                    "class2": 25445.0
                },
                "ens_id": "ENSG00000173726",
                "uniprot_ids": [
                    "Q15388"
                ]
            },
            "ENSG00000196262": {
                "id": "ENSG00000196262",
                "type": "Gene",
                "symbol": "PPIA",
                "source": [
                    "ENSG00000181449",
                    "ENSG00000109132",
                    "ENSG00000114315",
                    "ENSG00000016082",
                    "ENSG00000113580",
                    "ENSG00000167081",
                    "ENSG00000164330",
                    "ENSG00000108001",
                    "ENSG00000221869",
                    "ENSG00000067955",
                    "ENSG00000119547"
                ],
                "target": [],
                "expression_sum": {
                    "class1": 112820.0,
                    "class2": 72384.0
                },
                "ens_id": "ENSG00000196262",
                "uniprot_ids": [
                    "P62937",
                    "A0A7I2V4V1",
                    "C9J5S7",
                    "F8WE65",
                    "E5RIZ5",
                    "A0A7I2V5J5",
                    "A0A7P0S768"
                ]
            },
            "ENSG00000131469": {
                "id": "ENSG00000131469",
                "type": "Gene",
                "symbol": "RPL27",
                "source": [
                    "ENSG00000181449",
                    "ENSG00000016082",
                    "ENSG00000167081",
                    "ENSG00000118263",
                    "ENSG00000164330",
                    "ENSG00000119547"
                ],
                "target": [],
                "expression_sum": {
                    "class1": 116994.0,
                    "class2": 98367.0
                },
                "ens_id": "ENSG00000131469",
                "uniprot_ids": [
                    "P61353",
                    "K7ERY7",
                    "K7ELC7",
                    "K7EQQ9"
                ]
            },
            "ENSG00000164330": {
                "id": "ENSG00000164330",
                "type": "TF",
                "symbol": "EBF1",
                "source": [
                    "ENSG00000100811",
                    "ENSG00000106261",
                    "ENSG00000016082",
                    "ENSG00000148516",
                    "ENSG00000143799",
                    "ENSG00000105698",
                    "ENSG00000103495",
                    "ENSG00000128272",
                    "ENSG00000108055",
                    "ENSG00000119950",
                    "ENSG00000163848",
                    "ENSG00000168214",
                    "ENSG00000116560",
                    "ENSG00000196591",
                    "ENSG00000164754",
                    "ENSG00000165512",
                    "ENSG00000067955"
                ],
                "target": [
                    "ENSG00000206573",
                    "ENSG00000005810",
                    "ENSG00000117395",
                    "ENSG00000134248",
                    "ENSG00000189403",
                    "ENSG00000022840",
                    "ENSG00000164091",
                    "ENSG00000144029",
                    "ENSG00000102172",
                    "ENSG00000150779",
                    "ENSG00000077380",
                    "ENSG00000154556",
                    "ENSG00000134283",
                    "ENSG00000173812",
                    "ENSG00000117592",
                    "ENSG00000106443",
                    "ENSG00000123240",
                    "ENSG00000102103",
                    "ENSG00000178913",
                    "ENSG00000074319",
                    "ENSG00000146007",
                    "ENSG00000106554",
                    "ENSG00000157045",
                    "ENSG00000163923",
                    "ENSG00000125356",
                    "ENSG00000255302",
                    "ENSG00000144218",
                    "ENSG00000113712",
                    "ENSG00000153317",
                    "ENSG00000026025",
                    "ENSG00000157216",
                    "ENSG00000133606",
                    "ENSG00000022267",
                    "ENSG00000103769",
                    "ENSG00000170584",
                    "ENSG00000075142",
                    "ENSG00000273841",
                    "ENSG00000124802",
                    "ENSG00000142657",
                    "ENSG00000196405",
                    "ENSG00000126934",
                    "ENSG00000151500",
                    "ENSG00000086598",
                    "ENSG00000121774",
                    "ENSG00000149136",
                    "ENSG00000127314",
                    "ENSG00000142546",
                    "ENSG00000185627",
                    "ENSG00000092199",
                    "ENSG00000107262",
                    "ENSG00000175216",
                    "ENSG00000174953",
                    "ENSG00000118363",
                    "ENSG00000122417",
                    "ENSG00000112972",
                    "ENSG00000231500",
                    "ENSG00000265491",
                    "ENSG00000113013",
                    "ENSG00000109332",
                    "ENSG00000079246",
                    "ENSG00000057019",
                    "ENSG00000095059",
                    "ENSG00000232388",
                    "ENSG00000114391",
                    "ENSG00000168303",
                    "ENSG00000112701",
                    "ENSG00000162852",
                    "ENSG00000135390",
                    "ENSG00000131508",
                    "ENSG00000154277",
                    "ENSG00000198015",
                    "ENSG00000048649",
                    "ENSG00000169764",
                    "ENSG00000120694",
                    "ENSG00000127184",
                    "ENSG00000160551",
                    "ENSG00000103363",
                    "ENSG00000064607",
                    "ENSG00000171530",
                    "ENSG00000149273",
                    "ENSG00000165943",
                    "ENSG00000137207",
                    "ENSG00000108774",
                    "ENSG00000115128",
                    "ENSG00000134152",
                    "ENSG00000138760",
                    "ENSG00000183172",
                    "ENSG00000169567",
                    "ENSG00000275052",
                    "ENSG00000114744",
                    "ENSG00000147274",
                    "ENSG00000123179",
                    "ENSG00000101752",
                    "ENSG00000108175",
                    "ENSG00000084623",
                    "ENSG00000096746",
                    "ENSG00000119950",
                    "ENSG00000138777",
                    "ENSG00000069345",
                    "ENSG00000162585",
                    "ENSG00000075089",
                    "ENSG00000028839",
                    "ENSG00000189043",
                    "ENSG00000163939",
                    "ENSG00000075790",
                    "ENSG00000099246",
                    "ENSG00000115307",
                    "ENSG00000116661",
                    "ENSG00000134287",
                    "ENSG00000105254",
                    "ENSG00000119335",
                    "ENSG00000120438",
                    "ENSG00000127774",
                    "ENSG00000172893",
                    "ENSG00000106400",
                    "ENSG00000122482",
                    "ENSG00000125901",
                    "ENSG00000113387",
                    "ENSG00000149100",
                    "ENSG00000158864",
                    "ENSG00000137409",
                    "ENSG00000006125",
                    "ENSG00000136888",
                    "ENSG00000170142",
                    "ENSG00000127884",
                    "ENSG00000121653",
                    "ENSG00000140612",
                    "ENSG00000163069",
                    "ENSG00000111669",
                    "ENSG00000124383",
                    "ENSG00000197183",
                    "ENSG00000102401",
                    "ENSG00000132463",
                    "ENSG00000124098",
                    "ENSG00000197982",
                    "ENSG00000165280",
                    "ENSG00000116251",
                    "ENSG00000091640",
                    "ENSG00000147123",
                    "ENSG00000143575",
                    "ENSG00000176485",
                    "ENSG00000184203",
                    "ENSG00000183617",
                    "ENSG00000105372",
                    "ENSG00000143353",
                    "ENSG00000111678",
                    "ENSG00000169288",
                    "ENSG00000130706",
                    "ENSG00000161547",
                    "ENSG00000135070",
                    "ENSG00000108298",
                    "ENSG00000174444",
                    "ENSG00000130414",
                    "ENSG00000169738",
                    "ENSG00000172354",
                    "ENSG00000150753",
                    "ENSG00000138385",
                    "ENSG00000164733",
                    "ENSG00000115514",
                    "ENSG00000113068",
                    "ENSG00000221983",
                    "ENSG00000069956",
                    "ENSG00000122565",
                    "ENSG00000164258",
                    "ENSG00000137815",
                    "ENSG00000141002",
                    "ENSG00000171604",
                    "ENSG00000170430",
                    "ENSG00000166582",
                    "ENSG00000225663",
                    "ENSG00000065135",
                    "ENSG00000105373",
                    "ENSG00000124733",
                    "ENSG00000272325",
                    "ENSG00000100612",
                    "ENSG00000166295",
                    "ENSG00000168259",
                    "ENSG00000119013",
                    "ENSG00000161203",
                    "ENSG00000025796",
                    "ENSG00000151746",
                    "ENSG00000101782",
                    "ENSG00000099800",
                    "ENSG00000147403",
                    "ENSG00000140905",
                    "ENSG00000101040",
                    "ENSG00000277258",
                    "ENSG00000115241",
                    "ENSG00000161970",
                    "ENSG00000100632",
                    "ENSG00000129083",
                    "ENSG00000078369",
                    "ENSG00000133112",
                    "ENSG00000138069",
                    "ENSG00000110321",
                    "ENSG00000115484",
                    "ENSG00000110651",
                    "ENSG00000157778",
                    "ENSG00000145362",
                    "ENSG00000182621",
                    "ENSG00000123144",
                    "ENSG00000100216",
                    "ENSG00000143256",
                    "ENSG00000149084",
                    "ENSG00000102931",
                    "ENSG00000165502",
                    "ENSG00000114302",
                    "ENSG00000104660",
                    "ENSG00000185664",
                    "ENSG00000067955",
                    "ENSG00000130255",
                    "ENSG00000214113",
                    "ENSG00000022277",
                    "ENSG00000147687",
                    "ENSG00000160310",
                    "ENSG00000158417",
                    "ENSG00000197958",
                    "ENSG00000171497",
                    "ENSG00000198380",
                    "ENSG00000108946",
                    "ENSG00000213015",
                    "ENSG00000175416",
                    "ENSG00000170632",
                    "ENSG00000172586",
                    "ENSG00000063177",
                    "ENSG00000160953",
                    "ENSG00000155368",
                    "ENSG00000169564",
                    "ENSG00000103035",
                    "ENSG00000128050",
                    "ENSG00000165775",
                    "ENSG00000086061",
                    "ENSG00000167658",
                    "ENSG00000113161",
                    "ENSG00000189180",
                    "ENSG00000196465",
                    "ENSG00000128699",
                    "ENSG00000143761",
                    "ENSG00000111860",
                    "ENSG00000031698",
                    "ENSG00000109919",
                    "ENSG00000167815",
                    "ENSG00000126457",
                    "ENSG00000177971",
                    "ENSG00000070087",
                    "ENSG00000177600",
                    "ENSG00000148450",
                    "ENSG00000136522",
                    "ENSG00000166946",
                    "ENSG00000136628",
                    "ENSG00000092148",
                    "ENSG00000054148",
                    "ENSG00000070010",
                    "ENSG00000181904",
                    "ENSG00000104529",
                    "ENSG00000118181",
                    "ENSG00000080822",
                    "ENSG00000186591",
                    "ENSG00000168653",
                    "ENSG00000159593",
                    "ENSG00000156261",
                    "ENSG00000198363",
                    "ENSG00000151465",
                    "ENSG00000140988",
                    "ENSG00000122033",
                    "ENSG00000163950",
                    "ENSG00000143774",
                    "ENSG00000165209",
                    "ENSG00000182512",
                    "ENSG00000110367",
                    "ENSG00000100142",
                    "ENSG00000115310",
                    "ENSG00000122545",
                    "ENSG00000175029",
                    "ENSG00000108055",
                    "ENSG00000099194",
                    "ENSG00000086666",
                    "ENSG00000188290",
                    "ENSG00000167515",
                    "ENSG00000134717",
                    "ENSG00000169599",
                    "ENSG00000141367",
                    "ENSG00000213719",
                    "ENSG00000105438",
                    "ENSG00000112335",
                    "ENSG00000204217",
                    "ENSG00000132341",
                    "ENSG00000124562",
                    "ENSG00000100300",
                    "ENSG00000145741",
                    "ENSG00000087448",
                    "ENSG00000128923",
                    "ENSG00000175826",
                    "ENSG00000009307",
                    "ENSG00000138138",
                    "ENSG00000168958",
                    "ENSG00000057608",
                    "ENSG00000165169",
                    "ENSG00000065548",
                    "ENSG00000150991",
                    "ENSG00000017797",
                    "ENSG00000107281",
                    "ENSG00000132432",
                    "ENSG00000184009",
                    "ENSG00000120437",
                    "ENSG00000054523",
                    "ENSG00000146457",
                    "ENSG00000100380",
                    "ENSG00000128626",
                    "ENSG00000166164",
                    "ENSG00000119640",
                    "ENSG00000104979",
                    "ENSG00000114062",
                    "ENSG00000128989",
                    "ENSG00000213123",
                    "ENSG00000168724",
                    "ENSG00000126602",
                    "ENSG00000036257",
                    "ENSG00000164587",
                    "ENSG00000008018",
                    "ENSG00000160948",
                    "ENSG00000112531",
                    "ENSG00000082898",
                    "ENSG00000115685",
                    "ENSG00000101654",
                    "ENSG00000167232",
                    "ENSG00000144369",
                    "ENSG00000091527",
                    "ENSG00000182154",
                    "ENSG00000181817",
                    "ENSG00000143153",
                    "ENSG00000187735",
                    "ENSG00000087365",
                    "ENSG00000174886",
                    "ENSG00000017260",
                    "ENSG00000111716",
                    "ENSG00000169612",
                    "ENSG00000099622",
                    "ENSG00000114416",
                    "ENSG00000131143",
                    "ENSG00000184983",
                    "ENSG00000127540",
                    "ENSG00000180304",
                    "ENSG00000108424",
                    "ENSG00000147155",
                    "ENSG00000051128",
                    "ENSG00000250317",
                    "ENSG00000141279",
                    "ENSG00000237190",
                    "ENSG00000135441",
                    "ENSG00000172292",
                    "ENSG00000129636",
                    "ENSG00000134453",
                    "ENSG00000182220",
                    "ENSG00000141858",
                    "ENSG00000167136",
                    "ENSG00000149196",
                    "ENSG00000204392",
                    "ENSG00000114354",
                    "ENSG00000153815",
                    "ENSG00000196504",
                    "ENSG00000055163",
                    "ENSG00000037241",
                    "ENSG00000109390",
                    "ENSG00000170540",
                    "ENSG00000144746",
                    "ENSG00000133703",
                    "ENSG00000146425",
                    "ENSG00000120963",
                    "ENSG00000131100",
                    "ENSG00000176261",
                    "ENSG00000114520",
                    "ENSG00000215845",
                    "ENSG00000069849",
                    "ENSG00000240344",
                    "ENSG00000123200",
                    "ENSG00000147649",
                    "ENSG00000144635",
                    "ENSG00000187514",
                    "ENSG00000076604",
                    "ENSG00000160194",
                    "ENSG00000178980",
                    "ENSG00000083807",
                    "ENSG00000168137",
                    "ENSG00000172239",
                    "ENSG00000129562",
                    "ENSG00000108106",
                    "ENSG00000116857",
                    "ENSG00000256053",
                    "ENSG00000105258",
                    "ENSG00000263465",
                    "ENSG00000169710",
                    "ENSG00000163170",
                    "ENSG00000184900",
                    "ENSG00000110696",
                    "ENSG00000088986",
                    "ENSG00000107937",
                    "ENSG00000131051",
                    "ENSG00000166974",
                    "ENSG00000029993",
                    "ENSG00000239672",
                    "ENSG00000005022",
                    "ENSG00000131469",
                    "ENSG00000047410",
                    "ENSG00000136810",
                    "ENSG00000142937",
                    "ENSG00000171634",
                    "ENSG00000232112",
                    "ENSG00000106682",
                    "ENSG00000079459",
                    "ENSG00000178719",
                    "ENSG00000119314",
                    "ENSG00000170144",
                    "ENSG00000143222",
                    "ENSG00000127914",
                    "ENSG00000177565",
                    "ENSG00000106803",
                    "ENSG00000175582",
                    "ENSG00000171130",
                    "ENSG00000198612",
                    "ENSG00000130725",
                    "ENSG00000122566",
                    "ENSG00000142507",
                    "ENSG00000047932",
                    "ENSG00000237765",
                    "ENSG00000165410",
                    "ENSG00000089737",
                    "ENSG00000130313",
                    "ENSG00000129355",
                    "ENSG00000188343",
                    "ENSG00000128951",
                    "ENSG00000100575",
                    "ENSG00000120656",
                    "ENSG00000241837",
                    "ENSG00000010270",
                    "ENSG00000056097",
                    "ENSG00000131495",
                    "ENSG00000173267",
                    "ENSG00000147684",
                    "ENSG00000072042",
                    "ENSG00000126756",
                    "ENSG00000103145",
                    "ENSG00000137941",
                    "ENSG00000167283",
                    "ENSG00000187109",
                    "ENSG00000115875",
                    "ENSG00000164929",
                    "ENSG00000204130",
                    "ENSG00000196262",
                    "ENSG00000125977",
                    "ENSG00000089280",
                    "ENSG00000130332",
                    "ENSG00000168092",
                    "ENSG00000115286",
                    "ENSG00000105698",
                    "ENSG00000105640",
                    "ENSG00000114023",
                    "ENSG00000166347",
                    "ENSG00000131174",
                    "ENSG00000106588",
                    "ENSG00000196937",
                    "ENSG00000188690",
                    "ENSG00000135900",
                    "ENSG00000170445",
                    "ENSG00000143799",
                    "ENSG00000204628",
                    "ENSG00000102241",
                    "ENSG00000134986",
                    "ENSG00000121022",
                    "ENSG00000113369",
                    "ENSG00000163682",
                    "ENSG00000171421",
                    "ENSG00000085231",
                    "ENSG00000124380",
                    "ENSG00000105701",
                    "ENSG00000134825",
                    "ENSG00000090013",
                    "ENSG00000167863",
                    "ENSG00000125037",
                    "ENSG00000004478",
                    "ENSG00000105058",
                    "ENSG00000175756",
                    "ENSG00000145734",
                    "ENSG00000197345",
                    "ENSG00000005448",
                    "ENSG00000250479",
                    "ENSG00000160014",
                    "ENSG00000146731",
                    "ENSG00000130724",
                    "ENSG00000164713",
                    "ENSG00000132507",
                    "ENSG00000175550",
                    "ENSG00000173692",
                    "ENSG00000132824",
                    "ENSG00000164163",
                    "ENSG00000134769",
                    "ENSG00000166226",
                    "ENSG00000103429",
                    "ENSG00000102309",
                    "ENSG00000167674",
                    "ENSG00000258289",
                    "ENSG00000164096",
                    "ENSG00000156587",
                    "ENSG00000116754",
                    "ENSG00000175348",
                    "ENSG00000141429",
                    "ENSG00000100941",
                    "ENSG00000164898",
                    "ENSG00000152944",
                    "ENSG00000141698",
                    "ENSG00000162521",
                    "ENSG00000196230",
                    "ENSG00000100028",
                    "ENSG00000029363",
                    "ENSG00000100823",
                    "ENSG00000105887",
                    "ENSG00000169136",
                    "ENSG00000160131",
                    "ENSG00000109971",
                    "ENSG00000159199",
                    "ENSG00000118816",
                    "ENSG00000172590",
                    "ENSG00000151474",
                    "ENSG00000137312",
                    "ENSG00000167522",
                    "ENSG00000111371",
                    "ENSG00000172115",
                    "ENSG00000101558",
                    "ENSG00000125970",
                    "ENSG00000006451",
                    "ENSG00000167468",
                    "ENSG00000111640",
                    "ENSG00000183978",
                    "ENSG00000125652",
                    "ENSG00000124596",
                    "ENSG00000169021",
                    "ENSG00000130294",
                    "ENSG00000077721",
                    "ENSG00000099624",
                    "ENSG00000120265",
                    "ENSG00000138758",
                    "ENSG00000150093",
                    "ENSG00000162441",
                    "ENSG00000130748",
                    "ENSG00000111642",
                    "ENSG00000137575",
                    "ENSG00000145833",
                    "ENSG00000103121",
                    "ENSG00000104412",
                    "ENSG00000113575",
                    "ENSG00000225921",
                    "ENSG00000105193",
                    "ENSG00000009954",
                    "ENSG00000084733",
                    "ENSG00000136717",
                    "ENSG00000149557",
                    "ENSG00000206503",
                    "ENSG00000100528",
                    "ENSG00000120533",
                    "ENSG00000162734",
                    "ENSG00000168214",
                    "ENSG00000071794",
                    "ENSG00000112237",
                    "ENSG00000106591",
                    "ENSG00000130311",
                    "ENSG00000130985",
                    "ENSG00000094916",
                    "ENSG00000138663",
                    "ENSG00000186063",
                    "ENSG00000140740",
                    "ENSG00000140995",
                    "ENSG00000146386",
                    "ENSG00000197006",
                    "ENSG00000185608",
                    "ENSG00000108468",
                    "ENSG00000205302",
                    "ENSG00000104427",
                    "ENSG00000115756",
                    "ENSG00000113141",
                    "ENSG00000166503",
                    "ENSG00000132963",
                    "ENSG00000175390",
                    "ENSG00000168522",
                    "ENSG00000148154",
                    "ENSG00000119048",
                    "ENSG00000172775",
                    "ENSG00000134419",
                    "ENSG00000174456",
                    "ENSG00000135406",
                    "ENSG00000143621",
                    "ENSG00000177200",
                    "ENSG00000135334",
                    "ENSG00000100138",
                    "ENSG00000168268",
                    "ENSG00000107798",
                    "ENSG00000125534",
                    "ENSG00000131435",
                    "ENSG00000126432",
                    "ENSG00000130723",
                    "ENSG00000124767",
                    "ENSG00000178741",
                    "ENSG00000156515",
                    "ENSG00000101266",
                    "ENSG00000233016",
                    "ENSG00000166595",
                    "ENSG00000134330",
                    "ENSG00000184207",
                    "ENSG00000167969",
                    "ENSG00000007520",
                    "ENSG00000159335",
                    "ENSG00000137726",
                    "ENSG00000141522",
                    "ENSG00000114902",
                    "ENSG00000099341",
                    "ENSG00000116095",
                    "ENSG00000103342",
                    "ENSG00000172757",
                    "ENSG00000105669",
                    "ENSG00000050426",
                    "ENSG00000136156",
                    "ENSG00000119185",
                    "ENSG00000169020",
                    "ENSG00000171159",
                    "ENSG00000144381",
                    "ENSG00000136521",
                    "ENSG00000213246",
                    "ENSG00000155330",
                    "ENSG00000112667",
                    "ENSG00000133872",
                    "ENSG00000198668",
                    "ENSG00000134308",
                    "ENSG00000134684",
                    "ENSG00000089063",
                    "ENSG00000204525",
                    "ENSG00000168439",
                    "ENSG00000138757",
                    "ENSG00000067900",
                    "ENSG00000214253",
                    "ENSG00000138382",
                    "ENSG00000118680",
                    "ENSG00000184220",
                    "ENSG00000205937",
                    "ENSG00000115694",
                    "ENSG00000065000",
                    "ENSG00000125445",
                    "ENSG00000165916",
                    "ENSG00000182196",
                    "ENSG00000152795",
                    "ENSG00000165716",
                    "ENSG00000072803",
                    "ENSG00000100568",
                    "ENSG00000176407",
                    "ENSG00000125944",
                    "ENSG00000183726",
                    "ENSG00000214941",
                    "ENSG00000186184",
                    "ENSG00000100804",
                    "ENSG00000179222",
                    "ENSG00000163964",
                    "ENSG00000166562",
                    "ENSG00000104763",
                    "ENSG00000181163",
                    "ENSG00000158710",
                    "ENSG00000215021",
                    "ENSG00000122873",
                    "ENSG00000113851",
                    "ENSG00000145247",
                    "ENSG00000137776",
                    "ENSG00000139637",
                    "ENSG00000106244",
                    "ENSG00000078668",
                    "ENSG00000090621",
                    "ENSG00000100410",
                    "ENSG00000116171",
                    "ENSG00000162704",
                    "ENSG00000139644",
                    "ENSG00000175745",
                    "ENSG00000182552",
                    "ENSG00000090060",
                    "ENSG00000116161",
                    "ENSG00000160209",
                    "ENSG00000161016",
                    "ENSG00000125743",
                    "ENSG00000105819",
                    "ENSG00000179958",
                    "ENSG00000075415",
                    "ENSG00000107874",
                    "ENSG00000152904",
                    "ENSG00000163466",
                    "ENSG00000198931",
                    "ENSG00000197451",
                    "ENSG00000152256",
                    "ENSG00000173915",
                    "ENSG00000111728",
                    "ENSG00000167085",
                    "ENSG00000170310",
                    "ENSG00000115540",
                    "ENSG00000041357",
                    "ENSG00000163875",
                    "ENSG00000163319",
                    "ENSG00000164924",
                    "ENSG00000141759",
                    "ENSG00000165264",
                    "ENSG00000174574",
                    "ENSG00000110619",
                    "ENSG00000139116",
                    "ENSG00000277791",
                    "ENSG00000149923",
                    "ENSG00000118482",
                    "ENSG00000116898",
                    "ENSG00000179431",
                    "ENSG00000188243",
                    "ENSG00000100591",
                    "ENSG00000140105",
                    "ENSG00000034713",
                    "ENSG00000130811",
                    "ENSG00000005194",
                    "ENSG00000153048",
                    "ENSG00000125835",
                    "ENSG00000183011",
                    "ENSG00000174437",
                    "ENSG00000187051",
                    "ENSG00000129515",
                    "ENSG00000155115",
                    "ENSG00000204463",
                    "ENSG00000010256",
                    "ENSG00000120053",
                    "ENSG00000135940",
                    "ENSG00000172336",
                    "ENSG00000131966",
                    "ENSG00000107438",
                    "ENSG00000153037",
                    "ENSG00000106245",
                    "ENSG00000075624",
                    "ENSG00000072062",
                    "ENSG00000168275",
                    "ENSG00000102898",
                    "ENSG00000152700",
                    "ENSG00000167088",
                    "ENSG00000240972",
                    "ENSG00000196655",
                    "ENSG00000141030",
                    "ENSG00000164985",
                    "ENSG00000250303",
                    "ENSG00000141867",
                    "ENSG00000165948",
                    "ENSG00000197223",
                    "ENSG00000180957",
                    "ENSG00000099817",
                    "ENSG00000143862",
                    "ENSG00000176731",
                    "ENSG00000161981",
                    "ENSG00000162910",
                    "ENSG00000104388",
                    "ENSG00000097021",
                    "ENSG00000087191",
                    "ENSG00000166734",
                    "ENSG00000257704",
                    "ENSG00000130204",
                    "ENSG00000014641",
                    "ENSG00000121067",
                    "ENSG00000176171",
                    "ENSG00000131473",
                    "ENSG00000112305",
                    "ENSG00000253738",
                    "ENSG00000165119",
                    "ENSG00000126005",
                    "ENSG00000241343",
                    "ENSG00000184752",
                    "ENSG00000198258",
                    "ENSG00000170027",
                    "ENSG00000023287",
                    "ENSG00000111707",
                    "ENSG00000176101",
                    "ENSG00000123416",
                    "ENSG00000171314",
                    "ENSG00000144713",
                    "ENSG00000067248",
                    "ENSG00000164975",
                    "ENSG00000130520",
                    "ENSG00000170315",
                    "ENSG00000167792",
                    "ENSG00000138413",
                    "ENSG00000163848",
                    "ENSG00000099795",
                    "ENSG00000079462",
                    "ENSG00000166037",
                    "ENSG00000163344",
                    "ENSG00000051620",
                    "ENSG00000119729",
                    "ENSG00000135404",
                    "ENSG00000113742",
                    "ENSG00000142192",
                    "ENSG00000131828",
                    "ENSG00000171858",
                    "ENSG00000144895",
                    "ENSG00000123106",
                    "ENSG00000147533",
                    "ENSG00000155506",
                    "ENSG00000100316",
                    "ENSG00000170348",
                    "ENSG00000140259",
                    "ENSG00000143771",
                    "ENSG00000089693",
                    "ENSG00000169432",
                    "ENSG00000144028",
                    "ENSG00000164182",
                    "ENSG00000122026",
                    "ENSG00000198843",
                    "ENSG00000109654",
                    "ENSG00000145495",
                    "ENSG00000254999",
                    "ENSG00000130741",
                    "ENSG00000177556",
                    "ENSG00000186222",
                    "ENSG00000156508",
                    "ENSG00000082701",
                    "ENSG00000120727",
                    "ENSG00000101052",
                    "ENSG00000106367",
                    "ENSG00000101126",
                    "ENSG00000197111",
                    "ENSG00000164615",
                    "ENSG00000169813",
                    "ENSG00000175334",
                    "ENSG00000099783",
                    "ENSG00000159720",
                    "ENSG00000182093",
                    "ENSG00000184007",
                    "ENSG00000140307",
                    "ENSG00000106355",
                    "ENSG00000169139",
                    "ENSG00000109736",
                    "ENSG00000203950",
                    "ENSG00000070831",
                    "ENSG00000161960",
                    "ENSG00000167004",
                    "ENSG00000249915",
                    "ENSG00000110700",
                    "ENSG00000159140",
                    "ENSG00000170892",
                    "ENSG00000067560",
                    "ENSG00000176871",
                    "ENSG00000156639",
                    "ENSG00000143727",
                    "ENSG00000167508",
                    "ENSG00000111237",
                    "ENSG00000182985",
                    "ENSG00000023191",
                    "ENSG00000151366",
                    "ENSG00000214413",
                    "ENSG00000145782",
                    "ENSG00000116747",
                    "ENSG00000153187",
                    "ENSG00000106628",
                    "ENSG00000100354",
                    "ENSG00000131149",
                    "ENSG00000183520",
                    "ENSG00000138430",
                    "ENSG00000176340",
                    "ENSG00000273611",
                    "ENSG00000166441",
                    "ENSG00000101745",
                    "ENSG00000145337",
                    "ENSG00000170365",
                    "ENSG00000182718",
                    "ENSG00000140990",
                    "ENSG00000163541",
                    "ENSG00000146416",
                    "ENSG00000028203",
                    "ENSG00000112245",
                    "ENSG00000092201",
                    "ENSG00000141543",
                    "ENSG00000173272",
                    "ENSG00000179262",
                    "ENSG00000101210",
                    "ENSG00000197756",
                    "ENSG00000178952",
                    "ENSG00000260032",
                    "ENSG00000129235",
                    "ENSG00000103005",
                    "ENSG00000117448",
                    "ENSG00000142864",
                    "ENSG00000137288",
                    "ENSG00000142534",
                    "ENSG00000211450",
                    "ENSG00000106399",
                    "ENSG00000142541",
                    "ENSG00000122068",
                    "ENSG00000181722",
                    "ENSG00000109046",
                    "ENSG00000143753",
                    "ENSG00000135624",
                    "ENSG00000163468",
                    "ENSG00000155096",
                    "ENSG00000091039",
                    "ENSG00000117713",
                    "ENSG00000177156",
                    "ENSG00000140350",
                    "ENSG00000128789",
                    "ENSG00000197746",
                    "ENSG00000135387",
                    "ENSG00000160888",
                    "ENSG00000205531",
                    "ENSG00000143549",
                    "ENSG00000187231",
                    "ENSG00000225733",
                    "ENSG00000177030",
                    "ENSG00000171490",
                    "ENSG00000154380",
                    "ENSG00000127125",
                    "ENSG00000127022",
                    "ENSG00000074800",
                    "ENSG00000162961",
                    "ENSG00000148303",
                    "ENSG00000133026",
                    "ENSG00000122034",
                    "ENSG00000138592",
                    "ENSG00000154582",
                    "ENSG00000067836",
                    "ENSG00000114209",
                    "ENSG00000186298",
                    "ENSG00000164902",
                    "ENSG00000144224",
                    "ENSG00000137996",
                    "ENSG00000188994",
                    "ENSG00000164830",
                    "ENSG00000133226",
                    "ENSG00000135241",
                    "ENSG00000108953",
                    "ENSG00000105655",
                    "ENSG00000132305",
                    "ENSG00000204310",
                    "ENSG00000136930",
                    "ENSG00000149806",
                    "ENSG00000132475",
                    "ENSG00000108848",
                    "ENSG00000156928",
                    "ENSG00000111786",
                    "ENSG00000124702",
                    "ENSG00000143933",
                    "ENSG00000167996",
                    "ENSG00000131876",
                    "ENSG00000134824",
                    "ENSG00000156411",
                    "ENSG00000099864",
                    "ENSG00000138443",
                    "ENSG00000196531",
                    "ENSG00000115268",
                    "ENSG00000182534",
                    "ENSG00000242485",
                    "ENSG00000086300",
                    "ENSG00000170043",
                    "ENSG00000149231",
                    "ENSG00000182481",
                    "ENSG00000188229",
                    "ENSG00000118292",
                    "ENSG00000180228",
                    "ENSG00000126247",
                    "ENSG00000136758",
                    "ENSG00000006015",
                    "ENSG00000173457",
                    "ENSG00000088832",
                    "ENSG00000135720",
                    "ENSG00000136485",
                    "ENSG00000013275",
                    "ENSG00000125870",
                    "ENSG00000123352",
                    "ENSG00000112514",
                    "ENSG00000137133",
                    "ENSG00000278311",
                    "ENSG00000172809",
                    "ENSG00000157514",
                    "ENSG00000107341",
                    "ENSG00000172500",
                    "ENSG00000091164",
                    "ENSG00000136146",
                    "ENSG00000155366",
                    "ENSG00000160752",
                    "ENSG00000075618",
                    "ENSG00000143252",
                    "ENSG00000134809",
                    "ENSG00000198961",
                    "ENSG00000159352",
                    "ENSG00000155099",
                    "ENSG00000137106",
                    "ENSG00000008324",
                    "ENSG00000166228",
                    "ENSG00000118263",
                    "ENSG00000213619",
                    "ENSG00000129197",
                    "ENSG00000164919",
                    "ENSG00000165525",
                    "ENSG00000033627",
                    "ENSG00000140319",
                    "ENSG00000132466",
                    "ENSG00000149809",
                    "ENSG00000243317",
                    "ENSG00000165233",
                    "ENSG00000107372",
                    "ENSG00000085719",
                    "ENSG00000137876",
                    "ENSG00000138398",
                    "ENSG00000136527",
                    "ENSG00000136819",
                    "ENSG00000110958",
                    "ENSG00000101608",
                    "ENSG00000164323",
                    "ENSG00000122705",
                    "ENSG00000178035",
                    "ENSG00000140374",
                    "ENSG00000173660",
                    "ENSG00000113648",
                    "ENSG00000104643",
                    "ENSG00000227051",
                    "ENSG00000162385",
                    "ENSG00000169045",
                    "ENSG00000165732",
                    "ENSG00000008988",
                    "ENSG00000196781",
                    "ENSG00000180817",
                    "ENSG00000086065",
                    "ENSG00000136938",
                    "ENSG00000220205",
                    "ENSG00000136238",
                    "ENSG00000164754",
                    "ENSG00000143183",
                    "ENSG00000123545",
                    "ENSG00000224032",
                    "ENSG00000105401",
                    "ENSG00000221914",
                    "ENSG00000103415",
                    "ENSG00000065978",
                    "ENSG00000087460",
                    "ENSG00000151247",
                    "ENSG00000170906",
                    "ENSG00000109180",
                    "ENSG00000123983",
                    "ENSG00000143742",
                    "ENSG00000205629",
                    "ENSG00000172270",
                    "ENSG00000177733",
                    "ENSG00000092841",
                    "ENSG00000129351",
                    "ENSG00000089009",
                    "ENSG00000141741",
                    "ENSG00000134884",
                    "ENSG00000163634",
                    "ENSG00000095787",
                    "ENSG00000169976",
                    "ENSG00000106603",
                    "ENSG00000140941",
                    "ENSG00000184840",
                    "ENSG00000075945",
                    "ENSG00000099797",
                    "ENSG00000196419",
                    "ENSG00000105993",
                    "ENSG00000213722",
                    "ENSG00000088833",
                    "ENSG00000119421",
                    "ENSG00000114439",
                    "ENSG00000087274",
                    "ENSG00000165669",
                    "ENSG00000184697",
                    "ENSG00000065911",
                    "ENSG00000151729",
                    "ENSG00000272398",
                    "ENSG00000197061",
                    "ENSG00000125995",
                    "ENSG00000135829",
                    "ENSG00000135069",
                    "ENSG00000162664",
                    "ENSG00000112308",
                    "ENSG00000148908",
                    "ENSG00000139112",
                    "ENSG00000156482",
                    "ENSG00000165637",
                    "ENSG00000241685",
                    "ENSG00000060982",
                    "ENSG00000147548",
                    "ENSG00000171703",
                    "ENSG00000131236",
                    "ENSG00000129625",
                    "ENSG00000196072",
                    "ENSG00000100077",
                    "ENSG00000067225",
                    "ENSG00000144642",
                    "ENSG00000141552",
                    "ENSG00000136143",
                    "ENSG00000205581",
                    "ENSG00000117139",
                    "ENSG00000104131",
                    "ENSG00000095380",
                    "ENSG00000034510",
                    "ENSG00000138071",
                    "ENSG00000263001",
                    "ENSG00000213741",
                    "ENSG00000145592",
                    "ENSG00000134001",
                    "ENSG00000138698",
                    "ENSG00000123143",
                    "ENSG00000110955",
                    "ENSG00000164687",
                    "ENSG00000169100",
                    "ENSG00000162191",
                    "ENSG00000147677",
                    "ENSG00000108666",
                    "ENSG00000070770",
                    "ENSG00000175792",
                    "ENSG00000167196",
                    "ENSG00000180370",
                    "ENSG00000171222",
                    "ENSG00000181222",
                    "ENSG00000105185",
                    "ENSG00000153310",
                    "ENSG00000100348",
                    "ENSG00000213585",
                    "ENSG00000145919",
                    "ENSG00000146223",
                    "ENSG00000131437",
                    "ENSG00000143106",
                    "ENSG00000127616",
                    "ENSG00000174720",
                    "ENSG00000128654",
                    "ENSG00000080824",
                    "ENSG00000160271",
                    "ENSG00000166200",
                    "ENSG00000130312",
                    "ENSG00000136950",
                    "ENSG00000197694",
                    "ENSG00000115816",
                    "ENSG00000168159",
                    "ENSG00000152082",
                    "ENSG00000115677",
                    "ENSG00000151914",
                    "ENSG00000168710",
                    "ENSG00000124614",
                    "ENSG00000092203",
                    "ENSG00000104765",
                    "ENSG00000073578",
                    "ENSG00000182199",
                    "ENSG00000269893",
                    "ENSG00000067064",
                    "ENSG00000135486",
                    "ENSG00000163636",
                    "ENSG00000109670",
                    "ENSG00000137818",
                    "ENSG00000082153",
                    "ENSG00000002330",
                    "ENSG00000129559",
                    "ENSG00000134265",
                    "ENSG00000105677",
                    "ENSG00000135002",
                    "ENSG00000171735",
                    "ENSG00000146282",
                    "ENSG00000078674",
                    "ENSG00000103495",
                    "ENSG00000126768",
                    "ENSG00000147604",
                    "ENSG00000204308",
                    "ENSG00000165512",
                    "ENSG00000086758",
                    "ENSG00000134779",
                    "ENSG00000079785",
                    "ENSG00000070761",
                    "ENSG00000185825",
                    "ENSG00000134369",
                    "ENSG00000163714",
                    "ENSG00000173473",
                    "ENSG00000196636",
                    "ENSG00000170515",
                    "ENSG00000164405",
                    "ENSG00000196199",
                    "ENSG00000104904",
                    "ENSG00000134970",
                    "ENSG00000173726",
                    "ENSG00000167460",
                    "ENSG00000155959",
                    "ENSG00000116586",
                    "ENSG00000186010",
                    "ENSG00000084234",
                    "ENSG00000125691",
                    "ENSG00000151327",
                    "ENSG00000167978",
                    "ENSG00000186187",
                    "ENSG00000089289",
                    "ENSG00000168288",
                    "ENSG00000004059",
                    "ENSG00000144848",
                    "ENSG00000095739",
                    "ENSG00000234608",
                    "ENSG00000241468",
                    "ENSG00000110717",
                    "ENSG00000115053",
                    "ENSG00000111275",
                    "ENSG00000142949",
                    "ENSG00000136100",
                    "ENSG00000099250",
                    "ENSG00000197747",
                    "ENSG00000162368",
                    "ENSG00000254682",
                    "ENSG00000100353",
                    "ENSG00000116459",
                    "ENSG00000134375",
                    "ENSG00000134333",
                    "ENSG00000038219",
                    "ENSG00000086232",
                    "ENSG00000125818",
                    "ENSG00000103187",
                    "ENSG00000115758",
                    "ENSG00000105220",
                    "ENSG00000053254",
                    "ENSG00000272047",
                    "ENSG00000011295",
                    "ENSG00000119707",
                    "ENSG00000007168",
                    "ENSG00000176209",
                    "ENSG00000112110",
                    "ENSG00000067369",
                    "ENSG00000104886",
                    "ENSG00000097033",
                    "ENSG00000029725",
                    "ENSG00000177410",
                    "ENSG00000146701",
                    "ENSG00000116560",
                    "ENSG00000114956",
                    "ENSG00000198492",
                    "ENSG00000183207",
                    "ENSG00000073792",
                    "ENSG00000182004",
                    "ENSG00000198034",
                    "ENSG00000141526",
                    "ENSG00000104408",
                    "ENSG00000173418",
                    "ENSG00000160633",
                    "ENSG00000102144",
                    "ENSG00000167526",
                    "ENSG00000084207",
                    "ENSG00000058272",
                    "ENSG00000184076",
                    "ENSG00000177889",
                    "ENSG00000166913",
                    "ENSG00000182774",
                    "ENSG00000111906",
                    "ENSG00000204435",
                    "ENSG00000109536",
                    "ENSG00000162434",
                    "ENSG00000090487",
                    "ENSG00000126524",
                    "ENSG00000081870",
                    "ENSG00000111843",
                    "ENSG00000074071",
                    "ENSG00000109255",
                    "ENSG00000134056",
                    "ENSG00000145293",
                    "ENSG00000181924",
                    "ENSG00000148516",
                    "ENSG00000074696",
                    "ENSG00000169750",
                    "ENSG00000253719",
                    "ENSG00000156976",
                    "ENSG00000169230",
                    "ENSG00000105404",
                    "ENSG00000177879",
                    "ENSG00000154001",
                    "ENSG00000115541",
                    "ENSG00000168066",
                    "ENSG00000162244",
                    "ENSG00000083099",
                    "ENSG00000198420",
                    "ENSG00000149485",
                    "ENSG00000071994",
                    "ENSG00000109270",
                    "ENSG00000104695",
                    "ENSG00000141404",
                    "ENSG00000172531",
                    "ENSG00000054118",
                    "ENSG00000179950",
                    "ENSG00000142676",
                    "ENSG00000165283",
                    "ENSG00000054116",
                    "ENSG00000196305",
                    "ENSG00000096384",
                    "ENSG00000102753",
                    "ENSG00000109107",
                    "ENSG00000113845",
                    "ENSG00000124486",
                    "ENSG00000198900",
                    "ENSG00000100911",
                    "ENSG00000214517",
                    "ENSG00000115524",
                    "ENSG00000174775",
                    "ENSG00000143947",
                    "ENSG00000128609",
                    "ENSG00000182809",
                    "ENSG00000198918",
                    "ENSG00000147140",
                    "ENSG00000133706",
                    "ENSG00000205155",
                    "ENSG00000161013",
                    "ENSG00000088930",
                    "ENSG00000168028",
                    "ENSG00000042753",
                    "ENSG00000198408",
                    "ENSG00000178127",
                    "ENSG00000184117",
                    "ENSG00000167699",
                    "ENSG00000105518",
                    "ENSG00000143198",
                    "ENSG00000170017",
                    "ENSG00000145494",
                    "ENSG00000122378",
                    "ENSG00000115944",
                    "ENSG00000111639",
                    "ENSG00000093144",
                    "ENSG00000160113",
                    "ENSG00000138594",
                    "ENSG00000160075",
                    "ENSG00000120992",
                    "ENSG00000089157",
                    "ENSG00000166165",
                    "ENSG00000131116",
                    "ENSG00000168090",
                    "ENSG00000197728",
                    "ENSG00000104823",
                    "ENSG00000177885",
                    "ENSG00000130779",
                    "ENSG00000122406",
                    "ENSG00000215712",
                    "ENSG00000179271",
                    "ENSG00000196683",
                    "ENSG00000149547",
                    "ENSG00000120686",
                    "ENSG00000102189",
                    "ENSG00000149600",
                    "ENSG00000108518",
                    "ENSG00000106615",
                    "ENSG00000087302",
                    "ENSG00000136942",
                    "ENSG00000105176",
                    "ENSG00000119318",
                    "ENSG00000166963",
                    "ENSG00000104177",
                    "ENSG00000152234",
                    "ENSG00000108107",
                    "ENSG00000143158",
                    "ENSG00000111832",
                    "ENSG00000061794",
                    "ENSG00000153774",
                    "ENSG00000108561",
                    "ENSG00000178982",
                    "ENSG00000108010",
                    "ENSG00000083845",
                    "ENSG00000155876",
                    "ENSG00000171867",
                    "ENSG00000111726",
                    "ENSG00000125821",
                    "ENSG00000070669",
                    "ENSG00000117450",
                    "ENSG00000018189",
                    "ENSG00000101220",
                    "ENSG00000182117",
                    "ENSG00000124172",
                    "ENSG00000196591",
                    "ENSG00000142669",
                    "ENSG00000100075",
                    "ENSG00000025293",
                    "ENSG00000178104",
                    "ENSG00000115306",
                    "ENSG00000114125",
                    "ENSG00000185787",
                    "ENSG00000105568",
                    "ENSG00000100554",
                    "ENSG00000111775",
                    "ENSG00000133142",
                    "ENSG00000114120",
                    "ENSG00000163938",
                    "ENSG00000188612",
                    "ENSG00000137154",
                    "ENSG00000116288",
                    "ENSG00000205542",
                    "ENSG00000103275",
                    "ENSG00000128524",
                    "ENSG00000137274",
                    "ENSG00000044115",
                    "ENSG00000047849",
                    "ENSG00000130402",
                    "ENSG00000198648",
                    "ENSG00000085733",
                    "ENSG00000114115",
                    "ENSG00000181610",
                    "ENSG00000173163"
                ],
                "expression_sum": {
                    "class1": 14988.0,
                    "class2": 1997.0
                },
                "ens_id": "ENSG00000164330",
                "uniprot_ids": [
                    "Q9UH73",
                    "E5RFQ1",
                    "A0A087WTE5"
                ]
            },
            "ENSG00000111775": {
                "id": "ENSG00000111775",
                "type": "Gene",
                "symbol": "COX6A1",
                "source": [
                    "ENSG00000181449",
                    "ENSG00000163435",
                    "ENSG00000109132",
                    "ENSG00000167081",
                    "ENSG00000113580",
                    "ENSG00000148516",
                    "ENSG00000164330",
                    "ENSG00000108001",
                    "ENSG00000114861",
                    "ENSG00000221869",
                    "ENSG00000139352"
                ],
                "target": [],
                "expression_sum": {
                    "class1": 86953.0,
                    "class2": 56143.0
                },
                "ens_id": "ENSG00000111775",
                "uniprot_ids": [
                    "P12074"
                ]
            },
            "ENSG00000134138": {
                "id": "ENSG00000134138",
                "type": "TF",
                "symbol": "MEIS2",
                "source": [
                    "ENSG00000162599",
                    "ENSG00000167081",
                    "ENSG00000113580",
                    "ENSG00000168610",
                    "ENSG00000175745",
                    "ENSG00000100219",
                    "ENSG00000114861",
                    "ENSG00000221869",
                    "ENSG00000139352",
                    "ENSG00000137203",
                    "ENSG00000119547",
                    "ENSG00000143995"
                ],
                "target": [
                    "ENSG00000049130",
                    "ENSG00000196230",
                    "ENSG00000164199",
                    "ENSG00000102265",
                    "ENSG00000187134",
                    "ENSG00000117984",
                    "ENSG00000135506",
                    "ENSG00000184486",
                    "ENSG00000265972",
                    "ENSG00000151746",
                    "ENSG00000162490",
                    "ENSG00000104805",
                    "ENSG00000183091",
                    "ENSG00000163956",
                    "ENSG00000137203",
                    "ENSG00000166250",
                    "ENSG00000150593",
                    "ENSG00000124570",
                    "ENSG00000196141",
                    "ENSG00000251562",
                    "ENSG00000135404",
                    "ENSG00000153234",
                    "ENSG00000180879",
                    "ENSG00000112514",
                    "ENSG00000119547",
                    "ENSG00000065613",
                    "ENSG00000175130",
                    "ENSG00000170266",
                    "ENSG00000213064",
                    "ENSG00000197746",
                    "ENSG00000188290",
                    "ENSG00000130005",
                    "ENSG00000169432",
                    "ENSG00000255302",
                    "ENSG00000131711",
                    "ENSG00000122884",
                    "ENSG00000153113",
                    "ENSG00000068697",
                    "ENSG00000140939",
                    "ENSG00000181744",
                    "ENSG00000143013",
                    "ENSG00000174851",
                    "ENSG00000130511",
                    "ENSG00000163382",
                    "ENSG00000205981",
                    "ENSG00000135549",
                    "ENSG00000166228",
                    "ENSG00000162496",
                    "ENSG00000119655",
                    "ENSG00000149557",
                    "ENSG00000185664",
                    "ENSG00000163399",
                    "ENSG00000123131",
                    "ENSG00000244187",
                    "ENSG00000102804",
                    "ENSG00000143995",
                    "ENSG00000196220",
                    "ENSG00000105223",
                    "ENSG00000071553",
                    "ENSG00000158604",
                    "ENSG00000131773",
                    "ENSG00000144674",
                    "ENSG00000114861",
                    "ENSG00000106537",
                    "ENSG00000103316",
                    "ENSG00000120742",
                    "ENSG00000146540",
                    "ENSG00000124155",
                    "ENSG00000185115",
                    "ENSG00000143543",
                    "ENSG00000137267",
                    "ENSG00000151150",
                    "ENSG00000146278",
                    "ENSG00000090932",
                    "ENSG00000145675",
                    "ENSG00000100345",
                    "ENSG00000005249",
                    "ENSG00000157851",
                    "ENSG00000078018",
                    "ENSG00000137880",
                    "ENSG00000132475",
                    "ENSG00000197043",
                    "ENSG00000133318",
                    "ENSG00000026297",
                    "ENSG00000213614",
                    "ENSG00000137285",
                    "ENSG00000131037",
                    "ENSG00000189159",
                    "ENSG00000112290",
                    "ENSG00000185624",
                    "ENSG00000167081",
                    "ENSG00000172270",
                    "ENSG00000138640",
                    "ENSG00000107872"
                ],
                "expression_sum": {
                    "class1": 55941.0,
                    "class2": 12368.0
                },
                "ens_id": "ENSG00000134138",
                "uniprot_ids": [
                    "O14770",
                    "H0YM65",
                    "H0YMX5",
                    "H0YKE5",
                    "U3KQI2",
                    "H0YKN2",
                    "U3KQ95"
                ]
            },
            "ENSG00000173812": {
                "id": "ENSG00000173812",
                "type": "Gene",
                "symbol": "EIF1",
                "source": [
                    "ENSG00000181449",
                    "ENSG00000163435",
                    "ENSG00000167081",
                    "ENSG00000113580",
                    "ENSG00000148516",
                    "ENSG00000118263",
                    "ENSG00000164330",
                    "ENSG00000114861",
                    "ENSG00000221869",
                    "ENSG00000139352",
                    "ENSG00000184486"
                ],
                "target": [],
                "expression_sum": {
                    "class1": 238688.0,
                    "class2": 146076.0
                },
                "ens_id": "ENSG00000173812",
                "uniprot_ids": [
                    "P41567",
                    "K7EM18",
                    "K7EQP2"
                ]
            },
            "ENSG00000162704": {
                "id": "ENSG00000162704",
                "type": "Gene",
                "symbol": "ARPC5",
                "source": [
                    "ENSG00000181449",
                    "ENSG00000163435",
                    "ENSG00000109132",
                    "ENSG00000016082",
                    "ENSG00000113580",
                    "ENSG00000167081",
                    "ENSG00000103495",
                    "ENSG00000164330",
                    "ENSG00000108001",
                    "ENSG00000114861",
                    "ENSG00000221869",
                    "ENSG00000196591",
                    "ENSG00000119547"
                ],
                "target": [],
                "expression_sum": {
                    "class1": 34566.0,
                    "class2": 18610.0
                },
                "ens_id": "ENSG00000162704",
                "uniprot_ids": [
                    "O15511",
                    "B1ALC0"
                ]
            },
            "ENSG00000143995": {
                "id": "ENSG00000143995",
                "type": "TF",
                "symbol": "MEIS1",
                "source": [
                    "ENSG00000167081",
                    "ENSG00000175745",
                    "ENSG00000119547",
                    "ENSG00000134138"
                ],
                "target": [
                    "ENSG00000134138",
                    "ENSG00000162374",
                    "ENSG00000165655",
                    "ENSG00000175745",
                    "ENSG00000151150",
                    "ENSG00000149294"
                ],
                "expression_sum": {
                    "class1": 6891.0,
                    "class2": 992.0
                },
                "ens_id": "ENSG00000143995",
                "uniprot_ids": [
                    "O00470",
                    "U3KPR8",
                    "F5GYS8",
                    "E9PDC4",
                    "H0YNY8"
                ]
            },
            "ENSG00000239672": {
                "id": "ENSG00000239672",
                "type": "Gene",
                "symbol": "NME1",
                "source": [
                    "ENSG00000181449",
                    "ENSG00000016082",
                    "ENSG00000167081",
                    "ENSG00000118263",
                    "ENSG00000103495",
                    "ENSG00000164330",
                    "ENSG00000221869",
                    "ENSG00000067955",
                    "ENSG00000119547"
                ],
                "target": [],
                "expression_sum": {
                    "class1": 25152.0,
                    "class2": 19027.0
                },
                "ens_id": "ENSG00000239672",
                "uniprot_ids": [
                    "P15531",
                    "E5RHP0",
                    "E7ERL0",
                    "C9K028"
                ]
            },
            "ENSG00000204628": {
                "id": "ENSG00000204628",
                "type": "Gene",
                "symbol": "RACK1",
                "source": [
                    "ENSG00000016082",
                    "ENSG00000181449",
                    "ENSG00000167081",
                    "ENSG00000164330"
                ],
                "target": [],
                "expression_sum": {
                    "class1": 192757.0,
                    "class2": 175312.0
                },
                "ens_id": "ENSG00000204628",
                "uniprot_ids": [
                    "P63244",
                    "E9PD14",
                    "D6RHJ5",
                    "D6REE5",
                    "D6RFZ9",
                    "D6RBD0",
                    "D6R909",
                    "H0Y8W2",
                    "D6RF23",
                    "J3KPE3",
                    "D6RAU2",
                    "H0YAF8",
                    "D6RDI0",
                    "D6RAC2",
                    "D6RHH4",
                    "D6RFX4",
                    "D6RGK8",
                    "H0Y9P0",
                    "H0YAM7",
                    "D6R9Z1",
                    "D6R9L0",
                    "H0Y8R5"
                ]
            },
            "ENSG00000135486": {
                "id": "ENSG00000135486",
                "type": "Gene",
                "symbol": "HNRNPA1",
                "source": [
                    "ENSG00000181449",
                    "ENSG00000109132",
                    "ENSG00000016082",
                    "ENSG00000167081",
                    "ENSG00000164330",
                    "ENSG00000108001",
                    "ENSG00000119547"
                ],
                "target": [],
                "expression_sum": {
                    "class1": 87578.0,
                    "class2": 59517.0
                },
                "ens_id": "ENSG00000135486",
                "uniprot_ids": [
                    "P09651",
                    "A0A7I2YQW0",
                    "A0A7I2V5W8",
                    "A0A7I2V2M7",
                    "A0A7I2YQV6",
                    "A0A7I2V4R7",
                    "H0YH80",
                    "A0A7I2V5N7",
                    "A0A7I2YQU4",
                    "A0A7I2V3W0",
                    "A0A7I2V3W4",
                    "F8VTQ5",
                    "A0A7I2V3S1",
                    "A0A7I2V4M0",
                    "A0A7I2V2F1",
                    "A0A7I2V666",
                    "A0A7I2V2Q7",
                    "A0A7I2V5U9",
                    "A0A7I2V5T1",
                    "A0A7I2YQR6",
                    "A0A7I2V2Z4",
                    "A0A7I2V3C7",
                    "A0A7I2V434",
                    "A0A7I2V5P1",
                    "A0A7I2V2X1",
                    "A0A7I2V453",
                    "A0A7I2V3K5",
                    "A0A7I2V3U8",
                    "A0A7I2YQJ2",
                    "A0A7I2V497",
                    "A0A7I2V360",
                    "A0A7I2YQ85",
                    "A0A7I2V4F8",
                    "A0A7I2V2L6",
                    "A0A7I2YQV3",
                    "A0A7I2YQY2",
                    "F8VYN5",
                    "A0A7I2V2R4",
                    "A0A7I2V2W8",
                    "A0A7I2V459",
                    "A0A7I2V4P7",
                    "A0A7I2V3F4",
                    "A0A7I2V3J5",
                    "F8W646",
                    "A0A7I2V4E2",
                    "A0A7I2V5H0",
                    "A0A7I2V3R8",
                    "F8W6I7",
                    "A0A7I2V661",
                    "A0A7I2V334",
                    "A0A7I2V3U9",
                    "A0A7I2V595",
                    "A0A7I2YQX9",
                    "F8VZ49",
                    "A0A7I2V520",
                    "A0A7I2V5L7"
                ]
            },
            "ENSG00000189043": {
                "id": "ENSG00000189043",
                "type": "Gene",
                "symbol": "NDUFA4",
                "source": [
                    "ENSG00000181449",
                    "ENSG00000114315",
                    "ENSG00000167081",
                    "ENSG00000004487",
                    "ENSG00000122691",
                    "ENSG00000164330",
                    "ENSG00000108001",
                    "ENSG00000137947",
                    "ENSG00000221869",
                    "ENSG00000196591",
                    "ENSG00000067955",
                    "ENSG00000119547"
                ],
                "target": [],
                "expression_sum": {
                    "class1": 124471.0,
                    "class2": 73711.0
                },
                "ens_id": "ENSG00000189043",
                "uniprot_ids": [
                    "O00483"
                ]
            },
            "ENSG00000188846": {
                "id": "ENSG00000188846",
                "type": "Gene",
                "symbol": "RPL14",
                "source": [
                    "ENSG00000016082",
                    "ENSG00000119547"
                ],
                "target": [],
                "expression_sum": {
                    "class1": 179167.0,
                    "class2": 175270.0
                },
                "ens_id": "ENSG00000188846",
                "uniprot_ids": [
                    "P50914",
                    "F8WDZ7",
                    "E7EPB3"
                ]
            },
            "ENSG00000009307": {
                "id": "ENSG00000009307",
                "type": "Gene",
                "symbol": "CSDE1",
                "source": [
                    "ENSG00000181449",
                    "ENSG00000163435",
                    "ENSG00000016082",
                    "ENSG00000113580",
                    "ENSG00000167081",
                    "ENSG00000148516",
                    "ENSG00000164330",
                    "ENSG00000114861",
                    "ENSG00000221869",
                    "ENSG00000139352"
                ],
                "target": [],
                "expression_sum": {
                    "class1": 33475.0,
                    "class2": 20721.0
                },
                "ens_id": "ENSG00000009307",
                "uniprot_ids": [
                    "O75534",
                    "E9PLD4",
                    "E9PJK7",
                    "E9PLT0",
                    "A0A1Y8EMC9",
                    "E9PNG3",
                    "E9PKN4"
                ]
            },
            "ENSG00000197111": {
                "id": "ENSG00000197111",
                "type": "Gene",
                "symbol": "PCBP2",
                "source": [
                    "ENSG00000181449",
                    "ENSG00000163435",
                    "ENSG00000109132",
                    "ENSG00000016082",
                    "ENSG00000113580",
                    "ENSG00000167081",
                    "ENSG00000148516",
                    "ENSG00000164330",
                    "ENSG00000108001",
                    "ENSG00000114861",
                    "ENSG00000221869",
                    "ENSG00000139352"
                ],
                "target": [],
                "expression_sum": {
                    "class1": 33932.0,
                    "class2": 21711.0
                },
                "ens_id": "ENSG00000197111",
                "uniprot_ids": [
                    "Q15366",
                    "F8W0G4",
                    "F8VZX2",
                    "H3BSP4",
                    "F8VXH9",
                    "H3BND9",
                    "H3BSS4",
                    "F8VTZ0",
                    "F8VRH0",
                    "F8W1G6",
                    "H3BRU6"
                ]
            },
            "ENSG00000089220": {
                "id": "ENSG00000089220",
                "type": "Gene",
                "symbol": "PEBP1",
                "source": [
                    "ENSG00000181449",
                    "ENSG00000167081",
                    "ENSG00000113580",
                    "ENSG00000140262",
                    "ENSG00000114861",
                    "ENSG00000221869"
                ],
                "target": [],
                "expression_sum": {
                    "class1": 86379.0,
                    "class2": 60084.0
                },
                "ens_id": "ENSG00000089220",
                "uniprot_ids": [
                    "P30086"
                ]
            },
            "ENSG00000126214": {
                "id": "ENSG00000126214",
                "type": "Gene",
                "symbol": "KLC1",
                "source": [
                    "ENSG00000016082",
                    "ENSG00000113580",
                    "ENSG00000148516",
                    "ENSG00000143799",
                    "ENSG00000118263",
                    "ENSG00000103495",
                    "ENSG00000108055",
                    "ENSG00000213015",
                    "ENSG00000139352",
                    "ENSG00000116560",
                    "ENSG00000196591"
                ],
                "target": [],
                "expression_sum": {
                    "class1": 25362.0,
                    "class2": 5579.0
                },
                "ens_id": "ENSG00000126214",
                "uniprot_ids": [
                    "Q07866",
                    "G3V2P7",
                    "G3V2E7",
                    "H0YJL0",
                    "H0YGB8",
                    "H0YG16",
                    "F8W6L3",
                    "G3V3H3",
                    "H0YJK1",
                    "H0YJT3",
                    "H0YJU9",
                    "G5E9S8",
                    "G3V5R9"
                ]
            },
            "ENSG00000105185": {
                "id": "ENSG00000105185",
                "type": "Gene",
                "symbol": "PDCD5",
                "source": [
                    "ENSG00000181449",
                    "ENSG00000016082",
                    "ENSG00000113580",
                    "ENSG00000164330",
                    "ENSG00000119547"
                ],
                "target": [],
                "expression_sum": {
                    "class1": 22760.0,
                    "class2": 16647.0
                },
                "ens_id": "ENSG00000105185",
                "uniprot_ids": [
                    "O14737",
                    "Q3HM38",
                    "K7EQA1",
                    "X6R2P6",
                    "K7ESJ4"
                ]
            },
            "ENSG00000070961": {
                "id": "ENSG00000070961",
                "type": "Gene",
                "symbol": "ATP2B1",
                "source": [
                    "ENSG00000181449",
                    "ENSG00000016082",
                    "ENSG00000122691",
                    "ENSG00000175745",
                    "ENSG00000160113"
                ],
                "target": [],
                "expression_sum": {
                    "class1": 15671.0,
                    "class2": 29562.0
                },
                "ens_id": "ENSG00000070961",
                "uniprot_ids": [
                    "P20020",
                    "H0YHH6",
                    "A0A0U1RQU3",
                    "F8W1V5",
                    "E7ERY9"
                ]
            },
            "ENSG00000163435": {
                "id": "ENSG00000163435",
                "type": "TF",
                "symbol": "ELF3",
                "source": [
                    "ENSG00000106261",
                    "ENSG00000100219",
                    "ENSG00000221869",
                    "ENSG00000196591",
                    "ENSG00000168214",
                    "ENSG00000124766",
                    "ENSG00000116044",
                    "ENSG00000160113"
                ],
                "target": [
                    "ENSG00000206573",
                    "ENSG00000141741",
                    "ENSG00000162704",
                    "ENSG00000163634",
                    "ENSG00000132698",
                    "ENSG00000139644",
                    "ENSG00000090060",
                    "ENSG00000173467",
                    "ENSG00000134248",
                    "ENSG00000131143",
                    "ENSG00000127540",
                    "ENSG00000160932",
                    "ENSG00000165215",
                    "ENSG00000146247",
                    "ENSG00000108424",
                    "ENSG00000113282",
                    "ENSG00000188483",
                    "ENSG00000265972",
                    "ENSG00000154556",
                    "ENSG00000163466",
                    "ENSG00000198931",
                    "ENSG00000114439",
                    "ENSG00000173812",
                    "ENSG00000141279",
                    "ENSG00000198899",
                    "ENSG00000123240",
                    "ENSG00000041357",
                    "ENSG00000008517",
                    "ENSG00000163875",
                    "ENSG00000167136",
                    "ENSG00000164924",
                    "ENSG00000075213",
                    "ENSG00000197930",
                    "ENSG00000180879",
                    "ENSG00000104964",
                    "ENSG00000156482",
                    "ENSG00000125868",
                    "ENSG00000165637",
                    "ENSG00000241685",
                    "ENSG00000125356",
                    "ENSG00000147548",
                    "ENSG00000196504",
                    "ENSG00000129625",
                    "ENSG00000116898",
                    "ENSG00000130513",
                    "ENSG00000147889",
                    "ENSG00000067225",
                    "ENSG00000153113",
                    "ENSG00000113712",
                    "ENSG00000233927",
                    "ENSG00000150459",
                    "ENSG00000117139",
                    "ENSG00000148444",
                    "ENSG00000205581",
                    "ENSG00000161249",
                    "ENSG00000129521",
                    "ENSG00000062716",
                    "ENSG00000095380",
                    "ENSG00000086548",
                    "ENSG00000133703",
                    "ENSG00000133606",
                    "ENSG00000116044",
                    "ENSG00000103769",
                    "ENSG00000138071",
                    "ENSG00000170293",
                    "ENSG00000153048",
                    "ENSG00000263001",
                    "ENSG00000110092",
                    "ENSG00000213741",
                    "ENSG00000215845",
                    "ENSG00000111907",
                    "ENSG00000187514",
                    "ENSG00000076604",
                    "ENSG00000163041",
                    "ENSG00000092199",
                    "ENSG00000110955",
                    "ENSG00000107262",
                    "ENSG00000164346",
                    "ENSG00000172239",
                    "ENSG00000174953",
                    "ENSG00000064651",
                    "ENSG00000169100",
                    "ENSG00000162191",
                    "ENSG00000108654",
                    "ENSG00000135940",
                    "ENSG00000124107",
                    "ENSG00000107581",
                    "ENSG00000092010",
                    "ENSG00000106052",
                    "ENSG00000147677",
                    "ENSG00000107438",
                    "ENSG00000079246",
                    "ENSG00000171222",
                    "ENSG00000135046",
                    "ENSG00000095059",
                    "ENSG00000138801",
                    "ENSG00000240972",
                    "ENSG00000054267",
                    "ENSG00000172915",
                    "ENSG00000030582",
                    "ENSG00000135390",
                    "ENSG00000136770",
                    "ENSG00000005022",
                    "ENSG00000127616",
                    "ENSG00000128654",
                    "ENSG00000153250",
                    "ENSG00000047410",
                    "ENSG00000136810",
                    "ENSG00000174021",
                    "ENSG00000169764",
                    "ENSG00000127184",
                    "ENSG00000116209",
                    "ENSG00000136950",
                    "ENSG00000104388",
                    "ENSG00000105835",
                    "ENSG00000119314",
                    "ENSG00000170144",
                    "ENSG00000171530",
                    "ENSG00000198938",
                    "ENSG00000168710",
                    "ENSG00000127914",
                    "ENSG00000126267",
                    "ENSG00000108679",
                    "ENSG00000253738",
                    "ENSG00000152558",
                    "ENSG00000154640",
                    "ENSG00000126005",
                    "ENSG00000166710",
                    "ENSG00000198840",
                    "ENSG00000082153",
                    "ENSG00000247095",
                    "ENSG00000101443",
                    "ENSG00000085871",
                    "ENSG00000184752",
                    "ENSG00000169567",
                    "ENSG00000198258",
                    "ENSG00000023287",
                    "ENSG00000076554",
                    "ENSG00000164190",
                    "ENSG00000189143",
                    "ENSG00000135821",
                    "ENSG00000131495",
                    "ENSG00000171314",
                    "ENSG00000067248",
                    "ENSG00000090238",
                    "ENSG00000167642",
                    "ENSG00000111229",
                    "ENSG00000130066",
                    "ENSG00000126756",
                    "ENSG00000006007",
                    "ENSG00000051620",
                    "ENSG00000173473",
                    "ENSG00000163939",
                    "ENSG00000059804",
                    "ENSG00000117523",
                    "ENSG00000127838",
                    "ENSG00000167283",
                    "ENSG00000198886",
                    "ENSG00000122643",
                    "ENSG00000104904",
                    "ENSG00000125977",
                    "ENSG00000170348",
                    "ENSG00000116586",
                    "ENSG00000119335",
                    "ENSG00000084234",
                    "ENSG00000154723",
                    "ENSG00000166347",
                    "ENSG00000131174",
                    "ENSG00000143013",
                    "ENSG00000168288",
                    "ENSG00000004059",
                    "ENSG00000234745",
                    "ENSG00000113387",
                    "ENSG00000196937",
                    "ENSG00000106028",
                    "ENSG00000100697",
                    "ENSG00000168118",
                    "ENSG00000106367",
                    "ENSG00000185650",
                    "ENSG00000100815",
                    "ENSG00000197111",
                    "ENSG00000101182",
                    "ENSG00000142949",
                    "ENSG00000092820",
                    "ENSG00000185043",
                    "ENSG00000184007",
                    "ENSG00000090013",
                    "ENSG00000197747",
                    "ENSG00000004779",
                    "ENSG00000004478",
                    "ENSG00000135968",
                    "ENSG00000070831",
                    "ENSG00000051523",
                    "ENSG00000250479",
                    "ENSG00000144674",
                    "ENSG00000106537",
                    "ENSG00000146731",
                    "ENSG00000053254",
                    "ENSG00000105048",
                    "ENSG00000164713",
                    "ENSG00000143819",
                    "ENSG00000111237",
                    "ENSG00000183742",
                    "ENSG00000132824",
                    "ENSG00000254505",
                    "ENSG00000112695",
                    "ENSG00000118640",
                    "ENSG00000176340",
                    "ENSG00000097033",
                    "ENSG00000117335",
                    "ENSG00000170421",
                    "ENSG00000049618",
                    "ENSG00000164733",
                    "ENSG00000140264",
                    "ENSG00000177410",
                    "ENSG00000101745",
                    "ENSG00000140990",
                    "ENSG00000182718",
                    "ENSG00000138640",
                    "ENSG00000164258",
                    "ENSG00000106261",
                    "ENSG00000198492",
                    "ENSG00000171604",
                    "ENSG00000233098",
                    "ENSG00000129235",
                    "ENSG00000102144",
                    "ENSG00000172183",
                    "ENSG00000130770",
                    "ENSG00000084207",
                    "ENSG00000186395",
                    "ENSG00000117448",
                    "ENSG00000166295",
                    "ENSG00000142864",
                    "ENSG00000119013",
                    "ENSG00000184076",
                    "ENSG00000196154",
                    "ENSG00000211450",
                    "ENSG00000166337",
                    "ENSG00000166913",
                    "ENSG00000162434",
                    "ENSG00000118816",
                    "ENSG00000119888",
                    "ENSG00000100906",
                    "ENSG00000221869",
                    "ENSG00000150687",
                    "ENSG00000134056",
                    "ENSG00000101040",
                    "ENSG00000177156",
                    "ENSG00000187193",
                    "ENSG00000197956",
                    "ENSG00000144560",
                    "ENSG00000117525",
                    "ENSG00000156976",
                    "ENSG00000129083",
                    "ENSG00000160888",
                    "ENSG00000113732",
                    "ENSG00000197448",
                    "ENSG00000133112",
                    "ENSG00000131981",
                    "ENSG00000166145",
                    "ENSG00000124762",
                    "ENSG00000187231",
                    "ENSG00000121039",
                    "ENSG00000150093",
                    "ENSG00000138069",
                    "ENSG00000110321",
                    "ENSG00000144034",
                    "ENSG00000156642",
                    "ENSG00000123144",
                    "ENSG00000127125",
                    "ENSG00000143256",
                    "ENSG00000149084",
                    "ENSG00000165502",
                    "ENSG00000084733",
                    "ENSG00000138061",
                    "ENSG00000206503",
                    "ENSG00000122034",
                    "ENSG00000138592",
                    "ENSG00000138772",
                    "ENSG00000149932",
                    "ENSG00000114209",
                    "ENSG00000168214",
                    "ENSG00000100911",
                    "ENSG00000160213",
                    "ENSG00000144224",
                    "ENSG00000115524",
                    "ENSG00000138764",
                    "ENSG00000155368",
                    "ENSG00000096696",
                    "ENSG00000169564",
                    "ENSG00000143543",
                    "ENSG00000070756",
                    "ENSG00000140740",
                    "ENSG00000146386",
                    "ENSG00000198727",
                    "ENSG00000167658",
                    "ENSG00000145632",
                    "ENSG00000147676",
                    "ENSG00000185608",
                    "ENSG00000060138",
                    "ENSG00000132475",
                    "ENSG00000135480",
                    "ENSG00000196586",
                    "ENSG00000168938",
                    "ENSG00000143933",
                    "ENSG00000146674",
                    "ENSG00000165629",
                    "ENSG00000167996",
                    "ENSG00000132963",
                    "ENSG00000100664",
                    "ENSG00000136522",
                    "ENSG00000178127",
                    "ENSG00000010244",
                    "ENSG00000105518",
                    "ENSG00000133639",
                    "ENSG00000092148",
                    "ENSG00000170017",
                    "ENSG00000124214",
                    "ENSG00000163683",
                    "ENSG00000116030",
                    "ENSG00000186591",
                    "ENSG00000160113",
                    "ENSG00000160075",
                    "ENSG00000120992",
                    "ENSG00000174748",
                    "ENSG00000148346",
                    "ENSG00000163191",
                    "ENSG00000126432",
                    "ENSG00000181885",
                    "ENSG00000126247",
                    "ENSG00000245532",
                    "ENSG00000197728",
                    "ENSG00000178741",
                    "ENSG00000085063",
                    "ENSG00000171206",
                    "ENSG00000173457",
                    "ENSG00000088832",
                    "ENSG00000108518",
                    "ENSG00000278311",
                    "ENSG00000086666",
                    "ENSG00000114902",
                    "ENSG00000230989",
                    "ENSG00000154518",
                    "ENSG00000178449",
                    "ENSG00000106541",
                    "ENSG00000134717",
                    "ENSG00000136156",
                    "ENSG00000125844",
                    "ENSG00000171159",
                    "ENSG00000134809",
                    "ENSG00000213719",
                    "ENSG00000112335",
                    "ENSG00000204525",
                    "ENSG00000164919",
                    "ENSG00000145741",
                    "ENSG00000178982",
                    "ENSG00000067900",
                    "ENSG00000111057",
                    "ENSG00000104870",
                    "ENSG00000100387",
                    "ENSG00000114573",
                    "ENSG00000140319",
                    "ENSG00000148180",
                    "ENSG00000163399",
                    "ENSG00000118680",
                    "ENSG00000117450",
                    "ENSG00000244187",
                    "ENSG00000117472",
                    "ENSG00000198763",
                    "ENSG00000009307",
                    "ENSG00000107372",
                    "ENSG00000135679",
                    "ENSG00000124172",
                    "ENSG00000150991",
                    "ENSG00000196591",
                    "ENSG00000017797",
                    "ENSG00000100568",
                    "ENSG00000101608",
                    "ENSG00000122705",
                    "ENSG00000183726",
                    "ENSG00000113648",
                    "ENSG00000111775",
                    "ENSG00000123636",
                    "ENSG00000087502",
                    "ENSG00000197102",
                    "ENSG00000163938",
                    "ENSG00000008394",
                    "ENSG00000188612",
                    "ENSG00000154080",
                    "ENSG00000104763",
                    "ENSG00000205542",
                    "ENSG00000158710",
                    "ENSG00000160948",
                    "ENSG00000137880",
                    "ENSG00000115685",
                    "ENSG00000128524",
                    "ENSG00000167232",
                    "ENSG00000165678",
                    "ENSG00000198712",
                    "ENSG00000044115",
                    "ENSG00000136238",
                    "ENSG00000186352",
                    "ENSG00000130402",
                    "ENSG00000182154",
                    "ENSG00000122873",
                    "ENSG00000087460",
                    "ENSG00000158195",
                    "ENSG00000170759",
                    "ENSG00000145247",
                    "ENSG00000113558",
                    "ENSG00000143153",
                    "ENSG00000109062",
                    "ENSG00000113407",
                    "ENSG00000143742",
                    "ENSG00000104419",
                    "ENSG00000085733",
                    "ENSG00000092841",
                    "ENSG00000173230"
                ],
                "expression_sum": {
                    "class1": 152.0,
                    "class2": 9183.0
                },
                "ens_id": "ENSG00000163435",
                "uniprot_ids": [
                    "P78545",
                    "Q5SR36"
                ]
            },
            "ENSG00000107438": {
                "id": "ENSG00000107438",
                "type": "Gene",
                "symbol": "PDLIM1",
                "source": [
                    "ENSG00000163435",
                    "ENSG00000160113",
                    "ENSG00000164683",
                    "ENSG00000113580",
                    "ENSG00000164330",
                    "ENSG00000168610",
                    "ENSG00000221869",
                    "ENSG00000168214",
                    "ENSG00000177606",
                    "ENSG00000092199"
                ],
                "target": [],
                "expression_sum": {
                    "class1": 3744.0,
                    "class2": 10802.0
                },
                "ens_id": "ENSG00000107438",
                "uniprot_ids": [
                    "O00151"
                ]
            },
            "ENSG00000154080": {
                "id": "ENSG00000154080",
                "type": "Gene",
                "symbol": "CHST9",
                "source": [
                    "ENSG00000124766",
                    "ENSG00000106261",
                    "ENSG00000163435",
                    "ENSG00000164683"
                ],
                "target": [],
                "expression_sum": {
                    "class1": 165.0,
                    "class2": 2971.0
                },
                "ens_id": "ENSG00000154080",
                "uniprot_ids": [
                    "Q7L1S5"
                ]
            },
            "ENSG00000177410": {
                "id": "ENSG00000177410",
                "type": "Gene",
                "symbol": "ZFAS1",
                "source": [
                    "ENSG00000163435",
                    "ENSG00000016082",
                    "ENSG00000167081",
                    "ENSG00000148516",
                    "ENSG00000143799",
                    "ENSG00000164330",
                    "ENSG00000108055",
                    "ENSG00000221869"
                ],
                "target": [],
                "expression_sum": {
                    "class1": 98029.0,
                    "class2": 78142.0
                },
                "ens_id": "ENSG00000177410"
            },
            "ENSG00000118640": {
                "id": "ENSG00000118640",
                "type": "Gene",
                "symbol": "VAMP8",
                "source": [
                    "ENSG00000100219",
                    "ENSG00000163435",
                    "ENSG00000168214",
                    "ENSG00000160113"
                ],
                "target": [],
                "expression_sum": {
                    "class1": 334.0,
                    "class2": 6604.0
                },
                "ens_id": "ENSG00000118640",
                "uniprot_ids": [
                    "Q9BV40",
                    "C9JXZ5",
                    "B8ZZT4"
                ]
            },
            "ENSG00000143153": {
                "id": "ENSG00000143153",
                "type": "Gene",
                "symbol": "ATP1B1",
                "source": [
                    "ENSG00000163435",
                    "ENSG00000109132",
                    "ENSG00000167081",
                    "ENSG00000113580",
                    "ENSG00000148516",
                    "ENSG00000164330",
                    "ENSG00000108055",
                    "ENSG00000114861",
                    "ENSG00000139352",
                    "ENSG00000116044"
                ],
                "target": [],
                "expression_sum": {
                    "class1": 31529.0,
                    "class2": 34067.0
                },
                "ens_id": "ENSG00000143153",
                "uniprot_ids": [
                    "P05026",
                    "V9GYR2"
                ]
            },
            "ENSG00000109062": {
                "id": "ENSG00000109062",
                "type": "Gene",
                "symbol": "SLC9A3R1",
                "source": [
                    "ENSG00000163435",
                    "ENSG00000164683",
                    "ENSG00000100219",
                    "ENSG00000168214",
                    "ENSG00000116044"
                ],
                "target": [],
                "expression_sum": {
                    "class1": 2277.0,
                    "class2": 6516.0
                },
                "ens_id": "ENSG00000109062",
                "uniprot_ids": [
                    "O14745",
                    "J3QRA3",
                    "J3QRP6"
                ]
            },
            "ENSG00000170017": {
                "id": "ENSG00000170017",
                "type": "Gene",
                "symbol": "ALCAM",
                "source": [
                    "ENSG00000163435",
                    "ENSG00000109132",
                    "ENSG00000164330",
                    "ENSG00000221869",
                    "ENSG00000160113"
                ],
                "target": [],
                "expression_sum": {
                    "class1": 5616.0,
                    "class2": 10940.0
                },
                "ens_id": "ENSG00000170017",
                "uniprot_ids": [
                    "Q13740",
                    "F5GXJ9",
                    "H7C543"
                ]
            },
            "ENSG00000197930": {
                "id": "ENSG00000197930",
                "type": "Gene",
                "symbol": "ERO1A",
                "source": [
                    "ENSG00000113580",
                    "ENSG00000163435",
                    "ENSG00000100219"
                ],
                "target": [],
                "expression_sum": {
                    "class1": 6272.0,
                    "class2": 8478.0
                },
                "ens_id": "ENSG00000197930",
                "uniprot_ids": [
                    "Q96HE7",
                    "G3V3E6",
                    "G3V503",
                    "G3V5B3",
                    "G3V2H0"
                ]
            },
            "ENSG00000148346": {
                "id": "ENSG00000148346",
                "type": "Gene",
                "symbol": "LCN2",
                "source": [
                    "ENSG00000163435"
                ],
                "target": [],
                "expression_sum": {
                    "class1": 6.0,
                    "class2": 5487.0
                },
                "ens_id": "ENSG00000148346",
                "uniprot_ids": [
                    "P80188",
                    "X6R8F3"
                ]
            },
            "ENSG00000130066": {
                "id": "ENSG00000130066",
                "type": "Gene",
                "symbol": "SAT1",
                "source": [
                    "ENSG00000164683",
                    "ENSG00000116044",
                    "ENSG00000163435",
                    "ENSG00000160113"
                ],
                "target": [],
                "expression_sum": {
                    "class1": 5582.0,
                    "class2": 109201.0
                },
                "ens_id": "ENSG00000130066",
                "uniprot_ids": [
                    "P21673",
                    "Q2TM25",
                    "E9PD37"
                ]
            },
            "ENSG00000060138": {
                "id": "ENSG00000060138",
                "type": "Gene",
                "symbol": "YBX3",
                "source": [
                    "ENSG00000164683",
                    "ENSG00000100219",
                    "ENSG00000163435",
                    "ENSG00000160113"
                ],
                "target": [],
                "expression_sum": {
                    "class1": 4287.0,
                    "class2": 23580.0
                },
                "ens_id": "ENSG00000060138",
                "uniprot_ids": [
                    "P16989",
                    "A0A0D9SEI8"
                ]
            },
            "ENSG00000168938": {
                "id": "ENSG00000168938",
                "type": "Gene",
                "symbol": "PPIC",
                "source": [
                    "ENSG00000164683",
                    "ENSG00000163435",
                    "ENSG00000168214"
                ],
                "target": [],
                "expression_sum": {
                    "class1": 821.0,
                    "class2": 4611.0
                },
                "ens_id": "ENSG00000168938",
                "uniprot_ids": [
                    "P45877"
                ]
            },
            "ENSG00000130513": {
                "id": "ENSG00000130513",
                "type": "Gene",
                "symbol": "GDF15",
                "source": [
                    "ENSG00000124766",
                    "ENSG00000163435"
                ],
                "target": [],
                "expression_sum": {
                    "class1": 2362.0,
                    "class2": 9765.0
                },
                "ens_id": "ENSG00000130513",
                "uniprot_ids": [
                    "Q99988",
                    "A0A0A0MTT8"
                ]
            },
            "ENSG00000150687": {
                "id": "ENSG00000150687",
                "type": "Gene",
                "symbol": "PRSS23",
                "source": [
                    "ENSG00000181449",
                    "ENSG00000163435",
                    "ENSG00000164683",
                    "ENSG00000100219",
                    "ENSG00000221869",
                    "ENSG00000128610"
                ],
                "target": [],
                "expression_sum": {
                    "class1": 3193.0,
                    "class2": 10284.0
                },
                "ens_id": "ENSG00000150687",
                "uniprot_ids": [
                    "O95084",
                    "E9PIB7",
                    "E9PRR2",
                    "E9PMX2"
                ]
            },
            "ENSG00000181449": {
                "id": "ENSG00000181449",
                "type": "TF",
                "symbol": "SOX2",
                "source": [
                    "ENSG00000100811",
                    "ENSG00000105698",
                    "ENSG00000103495",
                    "ENSG00000108055",
                    "ENSG00000163848",
                    "ENSG00000116560",
                    "ENSG00000164754",
                    "ENSG00000128610",
                    "ENSG00000130522"
                ],
                "target": [
                    "ENSG00000078668",
                    "ENSG00000206573",
                    "ENSG00000112081",
                    "ENSG00000107223",
                    "ENSG00000162704",
                    "ENSG00000134884",
                    "ENSG00000163634",
                    "ENSG00000165655",
                    "ENSG00000153130",
                    "ENSG00000111716",
                    "ENSG00000139644",
                    "ENSG00000099622",
                    "ENSG00000090060",
                    "ENSG00000116161",
                    "ENSG00000106603",
                    "ENSG00000117395",
                    "ENSG00000116350",
                    "ENSG00000161016",
                    "ENSG00000134248",
                    "ENSG00000131143",
                    "ENSG00000184983",
                    "ENSG00000127540",
                    "ENSG00000099797",
                    "ENSG00000119632",
                    "ENSG00000189403",
                    "ENSG00000108424",
                    "ENSG00000159377",
                    "ENSG00000196419",
                    "ENSG00000075415",
                    "ENSG00000107874",
                    "ENSG00000105993",
                    "ENSG00000102172",
                    "ENSG00000149212",
                    "ENSG00000213722",
                    "ENSG00000150779",
                    "ENSG00000165898",
                    "ENSG00000077380",
                    "ENSG00000154556",
                    "ENSG00000048740",
                    "ENSG00000163466",
                    "ENSG00000119421",
                    "ENSG00000052802",
                    "ENSG00000134283",
                    "ENSG00000173812",
                    "ENSG00000117592",
                    "ENSG00000173915",
                    "ENSG00000167085",
                    "ENSG00000151729",
                    "ENSG00000135441",
                    "ENSG00000106443",
                    "ENSG00000134453",
                    "ENSG00000041357",
                    "ENSG00000182220",
                    "ENSG00000163875",
                    "ENSG00000163319",
                    "ENSG00000164924",
                    "ENSG00000149196",
                    "ENSG00000141759",
                    "ENSG00000135069",
                    "ENSG00000162664",
                    "ENSG00000204392",
                    "ENSG00000165264",
                    "ENSG00000224877",
                    "ENSG00000139116",
                    "ENSG00000114354",
                    "ENSG00000277791",
                    "ENSG00000156482",
                    "ENSG00000125868",
                    "ENSG00000165637",
                    "ENSG00000165672",
                    "ENSG00000173674",
                    "ENSG00000147548",
                    "ENSG00000171703",
                    "ENSG00000167775",
                    "ENSG00000129625",
                    "ENSG00000089220",
                    "ENSG00000116898",
                    "ENSG00000196072",
                    "ENSG00000179431",
                    "ENSG00000255302",
                    "ENSG00000100526",
                    "ENSG00000188243",
                    "ENSG00000161542",
                    "ENSG00000067225",
                    "ENSG00000109390",
                    "ENSG00000141552",
                    "ENSG00000113712",
                    "ENSG00000126698",
                    "ENSG00000136143",
                    "ENSG00000174851",
                    "ENSG00000150459",
                    "ENSG00000170540",
                    "ENSG00000233927",
                    "ENSG00000205581",
                    "ENSG00000159685",
                    "ENSG00000026025",
                    "ENSG00000034713",
                    "ENSG00000144746",
                    "ENSG00000034510",
                    "ENSG00000146425",
                    "ENSG00000138071",
                    "ENSG00000120963",
                    "ENSG00000117410",
                    "ENSG00000170584",
                    "ENSG00000075142",
                    "ENSG00000273841",
                    "ENSG00000124802",
                    "ENSG00000263001",
                    "ENSG00000166275",
                    "ENSG00000183011",
                    "ENSG00000126934",
                    "ENSG00000213741",
                    "ENSG00000215845",
                    "ENSG00000175701",
                    "ENSG00000121774",
                    "ENSG00000069849",
                    "ENSG00000174917",
                    "ENSG00000147649",
                    "ENSG00000129515",
                    "ENSG00000100519",
                    "ENSG00000187514",
                    "ENSG00000163041",
                    "ENSG00000155115",
                    "ENSG00000141027",
                    "ENSG00000092199",
                    "ENSG00000110955",
                    "ENSG00000178980",
                    "ENSG00000175216",
                    "ENSG00000089335",
                    "ENSG00000083807",
                    "ENSG00000150316",
                    "ENSG00000010256",
                    "ENSG00000172239",
                    "ENSG00000065518",
                    "ENSG00000174953",
                    "ENSG00000152767",
                    "ENSG00000109775",
                    "ENSG00000162191",
                    "ENSG00000118363",
                    "ENSG00000129562",
                    "ENSG00000135940",
                    "ENSG00000108654",
                    "ENSG00000108106",
                    "ENSG00000162599",
                    "ENSG00000172336",
                    "ENSG00000107581",
                    "ENSG00000131966",
                    "ENSG00000105258",
                    "ENSG00000112972",
                    "ENSG00000204842",
                    "ENSG00000085224",
                    "ENSG00000153037",
                    "ENSG00000113013",
                    "ENSG00000265491",
                    "ENSG00000092621",
                    "ENSG00000106245",
                    "ENSG00000213465",
                    "ENSG00000109332",
                    "ENSG00000163170",
                    "ENSG00000079246",
                    "ENSG00000168275",
                    "ENSG00000075785",
                    "ENSG00000198937",
                    "ENSG00000075624",
                    "ENSG00000244879",
                    "ENSG00000171222",
                    "ENSG00000095059",
                    "ENSG00000105185",
                    "ENSG00000131051",
                    "ENSG00000167552",
                    "ENSG00000063322",
                    "ENSG00000205726",
                    "ENSG00000175166",
                    "ENSG00000168303",
                    "ENSG00000167088",
                    "ENSG00000240972",
                    "ENSG00000196655",
                    "ENSG00000029993",
                    "ENSG00000164985",
                    "ENSG00000239672",
                    "ENSG00000213585",
                    "ENSG00000131508",
                    "ENSG00000005022",
                    "ENSG00000127616",
                    "ENSG00000154277",
                    "ENSG00000131469",
                    "ENSG00000099817",
                    "ENSG00000171603",
                    "ENSG00000128654",
                    "ENSG00000047410",
                    "ENSG00000169714",
                    "ENSG00000090266",
                    "ENSG00000142937",
                    "ENSG00000136810",
                    "ENSG00000171634",
                    "ENSG00000125656",
                    "ENSG00000232112",
                    "ENSG00000127184",
                    "ENSG00000106682",
                    "ENSG00000130312",
                    "ENSG00000197694",
                    "ENSG00000079459",
                    "ENSG00000104388",
                    "ENSG00000110917",
                    "ENSG00000168159",
                    "ENSG00000170144",
                    "ENSG00000064607",
                    "ENSG00000171530",
                    "ENSG00000152082",
                    "ENSG00000087191",
                    "ENSG00000166734",
                    "ENSG00000123349",
                    "ENSG00000105583",
                    "ENSG00000143870",
                    "ENSG00000147655",
                    "ENSG00000014641",
                    "ENSG00000167614",
                    "ENSG00000164683",
                    "ENSG00000106803",
                    "ENSG00000126267",
                    "ENSG00000104765",
                    "ENSG00000120860",
                    "ENSG00000198612",
                    "ENSG00000137710",
                    "ENSG00000081154",
                    "ENSG00000122566",
                    "ENSG00000182199",
                    "ENSG00000130725",
                    "ENSG00000100129",
                    "ENSG00000112305",
                    "ENSG00000142507",
                    "ENSG00000108774",
                    "ENSG00000237765",
                    "ENSG00000165119",
                    "ENSG00000067064",
                    "ENSG00000135486",
                    "ENSG00000115128",
                    "ENSG00000132718",
                    "ENSG00000137818",
                    "ENSG00000129559",
                    "ENSG00000068796",
                    "ENSG00000105677",
                    "ENSG00000114784",
                    "ENSG00000184752",
                    "ENSG00000169567",
                    "ENSG00000198258",
                    "ENSG00000183172",
                    "ENSG00000127445",
                    "ENSG00000128951",
                    "ENSG00000100575",
                    "ENSG00000105810",
                    "ENSG00000188186",
                    "ENSG00000171735",
                    "ENSG00000241837",
                    "ENSG00000147274",
                    "ENSG00000103495",
                    "ENSG00000204308",
                    "ENSG00000214046",
                    "ENSG00000176101",
                    "ENSG00000123416",
                    "ENSG00000131495",
                    "ENSG00000144713",
                    "ENSG00000155957",
                    "ENSG00000130520",
                    "ENSG00000084623",
                    "ENSG00000086758",
                    "ENSG00000108671",
                    "ENSG00000100852",
                    "ENSG00000167792",
                    "ENSG00000147684",
                    "ENSG00000096746",
                    "ENSG00000170315",
                    "ENSG00000079785",
                    "ENSG00000119950",
                    "ENSG00000163848",
                    "ENSG00000099795",
                    "ENSG00000129084",
                    "ENSG00000069275",
                    "ENSG00000079462",
                    "ENSG00000138777",
                    "ENSG00000072042",
                    "ENSG00000069345",
                    "ENSG00000162585",
                    "ENSG00000163714",
                    "ENSG00000189043",
                    "ENSG00000099246",
                    "ENSG00000170515",
                    "ENSG00000142192",
                    "ENSG00000127838",
                    "ENSG00000131828",
                    "ENSG00000234545",
                    "ENSG00000137100",
                    "ENSG00000167283",
                    "ENSG00000164405",
                    "ENSG00000170035",
                    "ENSG00000196199",
                    "ENSG00000187109",
                    "ENSG00000155506",
                    "ENSG00000044574",
                    "ENSG00000196262",
                    "ENSG00000104904",
                    "ENSG00000115355",
                    "ENSG00000125977",
                    "ENSG00000170348",
                    "ENSG00000173726",
                    "ENSG00000089280",
                    "ENSG00000155959",
                    "ENSG00000137692",
                    "ENSG00000116586",
                    "ENSG00000089693",
                    "ENSG00000105254",
                    "ENSG00000117419",
                    "ENSG00000119335",
                    "ENSG00000115286",
                    "ENSG00000077152",
                    "ENSG00000144028",
                    "ENSG00000084234",
                    "ENSG00000114023",
                    "ENSG00000127774",
                    "ENSG00000101367",
                    "ENSG00000172893",
                    "ENSG00000154723",
                    "ENSG00000122026",
                    "ENSG00000166598",
                    "ENSG00000167978",
                    "ENSG00000198729",
                    "ENSG00000149357",
                    "ENSG00000131174",
                    "ENSG00000106400",
                    "ENSG00000143013",
                    "ENSG00000109654",
                    "ENSG00000179091",
                    "ENSG00000254999",
                    "ENSG00000133935",
                    "ENSG00000177556",
                    "ENSG00000113387",
                    "ENSG00000158864",
                    "ENSG00000137409",
                    "ENSG00000006125",
                    "ENSG00000188690",
                    "ENSG00000153944",
                    "ENSG00000241468",
                    "ENSG00000136888",
                    "ENSG00000110717",
                    "ENSG00000120727",
                    "ENSG00000127884",
                    "ENSG00000185630",
                    "ENSG00000100442",
                    "ENSG00000143799",
                    "ENSG00000106028",
                    "ENSG00000140612",
                    "ENSG00000115091",
                    "ENSG00000204628",
                    "ENSG00000115053",
                    "ENSG00000102241",
                    "ENSG00000106367",
                    "ENSG00000121022",
                    "ENSG00000164615",
                    "ENSG00000139168",
                    "ENSG00000197111",
                    "ENSG00000111275",
                    "ENSG00000139343",
                    "ENSG00000175334",
                    "ENSG00000146729",
                    "ENSG00000171421",
                    "ENSG00000085231",
                    "ENSG00000133134",
                    "ENSG00000123562",
                    "ENSG00000099783",
                    "ENSG00000105701",
                    "ENSG00000111669",
                    "ENSG00000136100",
                    "ENSG00000182093",
                    "ENSG00000134825",
                    "ENSG00000185745",
                    "ENSG00000140307",
                    "ENSG00000132463",
                    "ENSG00000167863",
                    "ENSG00000004779",
                    "ENSG00000106355",
                    "ENSG00000071553",
                    "ENSG00000165280",
                    "ENSG00000169139",
                    "ENSG00000254682",
                    "ENSG00000116459",
                    "ENSG00000175756",
                    "ENSG00000203950",
                    "ENSG00000174684",
                    "ENSG00000134333",
                    "ENSG00000197345",
                    "ENSG00000006625",
                    "ENSG00000091640",
                    "ENSG00000147123",
                    "ENSG00000143575",
                    "ENSG00000143977",
                    "ENSG00000038219",
                    "ENSG00000249915",
                    "ENSG00000130560",
                    "ENSG00000110700",
                    "ENSG00000167004",
                    "ENSG00000160014",
                    "ENSG00000100201",
                    "ENSG00000213639",
                    "ENSG00000159140",
                    "ENSG00000183617",
                    "ENSG00000125818",
                    "ENSG00000146540",
                    "ENSG00000105220",
                    "ENSG00000143353",
                    "ENSG00000170892",
                    "ENSG00000111678",
                    "ENSG00000067560",
                    "ENSG00000163001",
                    "ENSG00000130724",
                    "ENSG00000007168",
                    "ENSG00000156639",
                    "ENSG00000164713",
                    "ENSG00000116649",
                    "ENSG00000132507",
                    "ENSG00000176209",
                    "ENSG00000105379",
                    "ENSG00000175550",
                    "ENSG00000099901",
                    "ENSG00000143727",
                    "ENSG00000126067",
                    "ENSG00000173692",
                    "ENSG00000111237",
                    "ENSG00000132670",
                    "ENSG00000112110",
                    "ENSG00000023191",
                    "ENSG00000215301",
                    "ENSG00000104886",
                    "ENSG00000151366",
                    "ENSG00000108298",
                    "ENSG00000214413",
                    "ENSG00000166226",
                    "ENSG00000112695",
                    "ENSG00000116747",
                    "ENSG00000153187",
                    "ENSG00000106628",
                    "ENSG00000077782",
                    "ENSG00000174444",
                    "ENSG00000203667",
                    "ENSG00000100354",
                    "ENSG00000133318",
                    "ENSG00000130414",
                    "ENSG00000169738",
                    "ENSG00000185236",
                    "ENSG00000137285",
                    "ENSG00000176340",
                    "ENSG00000172354",
                    "ENSG00000150753",
                    "ENSG00000164096",
                    "ENSG00000049618",
                    "ENSG00000136824",
                    "ENSG00000138385",
                    "ENSG00000083457",
                    "ENSG00000140264",
                    "ENSG00000189159",
                    "ENSG00000116754",
                    "ENSG00000101745",
                    "ENSG00000145337",
                    "ENSG00000140990",
                    "ENSG00000221983",
                    "ENSG00000173436",
                    "ENSG00000175348",
                    "ENSG00000063046",
                    "ENSG00000122565",
                    "ENSG00000163541",
                    "ENSG00000008952",
                    "ENSG00000127483",
                    "ENSG00000146416",
                    "ENSG00000230124",
                    "ENSG00000164258",
                    "ENSG00000116560",
                    "ENSG00000100941",
                    "ENSG00000109472",
                    "ENSG00000114956",
                    "ENSG00000147862",
                    "ENSG00000140391",
                    "ENSG00000173272",
                    "ENSG00000137815",
                    "ENSG00000141002",
                    "ENSG00000171604",
                    "ENSG00000179262",
                    "ENSG00000196976",
                    "ENSG00000197756",
                    "ENSG00000178952",
                    "ENSG00000143155",
                    "ENSG00000196230",
                    "ENSG00000225663",
                    "ENSG00000182004",
                    "ENSG00000141526",
                    "ENSG00000124733",
                    "ENSG00000173418",
                    "ENSG00000102144",
                    "ENSG00000272325",
                    "ENSG00000130770",
                    "ENSG00000029363",
                    "ENSG00000156467",
                    "ENSG00000167526",
                    "ENSG00000100028",
                    "ENSG00000168259",
                    "ENSG00000142864",
                    "ENSG00000119013",
                    "ENSG00000161203",
                    "ENSG00000182944",
                    "ENSG00000184076",
                    "ENSG00000137288",
                    "ENSG00000105887",
                    "ENSG00000100836",
                    "ENSG00000266472",
                    "ENSG00000101782",
                    "ENSG00000177889",
                    "ENSG00000142541",
                    "ENSG00000109971",
                    "ENSG00000166913",
                    "ENSG00000204435",
                    "ENSG00000159199",
                    "ENSG00000118816",
                    "ENSG00000121851",
                    "ENSG00000135624",
                    "ENSG00000104671",
                    "ENSG00000147669",
                    "ENSG00000168003",
                    "ENSG00000126524",
                    "ENSG00000163468",
                    "ENSG00000111843",
                    "ENSG00000074071",
                    "ENSG00000099800",
                    "ENSG00000150687",
                    "ENSG00000140905",
                    "ENSG00000134056",
                    "ENSG00000277443",
                    "ENSG00000172115",
                    "ENSG00000101558",
                    "ENSG00000145912",
                    "ENSG00000140350",
                    "ENSG00000145293",
                    "ENSG00000181924",
                    "ENSG00000074696",
                    "ENSG00000175130",
                    "ENSG00000167468",
                    "ENSG00000111640",
                    "ENSG00000197746",
                    "ENSG00000183978",
                    "ENSG00000125652",
                    "ENSG00000115241",
                    "ENSG00000253719",
                    "ENSG00000156976",
                    "ENSG00000100632",
                    "ENSG00000161970",
                    "ENSG00000130294",
                    "ENSG00000078369",
                    "ENSG00000119396",
                    "ENSG00000105404",
                    "ENSG00000177879",
                    "ENSG00000205531",
                    "ENSG00000099624",
                    "ENSG00000120265",
                    "ENSG00000154001",
                    "ENSG00000138758",
                    "ENSG00000115541",
                    "ENSG00000168066",
                    "ENSG00000133083",
                    "ENSG00000069329",
                    "ENSG00000138069",
                    "ENSG00000083099",
                    "ENSG00000110321",
                    "ENSG00000225733",
                    "ENSG00000111875",
                    "ENSG00000071994",
                    "ENSG00000130748",
                    "ENSG00000115484",
                    "ENSG00000169689",
                    "ENSG00000168785",
                    "ENSG00000110651",
                    "ENSG00000138326",
                    "ENSG00000204568",
                    "ENSG00000175203",
                    "ENSG00000102409",
                    "ENSG00000154380",
                    "ENSG00000145833",
                    "ENSG00000104695",
                    "ENSG00000123144",
                    "ENSG00000113575",
                    "ENSG00000143256",
                    "ENSG00000127022",
                    "ENSG00000149084",
                    "ENSG00000074800",
                    "ENSG00000128610",
                    "ENSG00000054118",
                    "ENSG00000105193",
                    "ENSG00000084733",
                    "ENSG00000142676",
                    "ENSG00000138668",
                    "ENSG00000179950",
                    "ENSG00000119705",
                    "ENSG00000133026",
                    "ENSG00000149557",
                    "ENSG00000165283",
                    "ENSG00000075151",
                    "ENSG00000122034",
                    "ENSG00000154582",
                    "ENSG00000130255",
                    "ENSG00000120533",
                    "ENSG00000162734",
                    "ENSG00000196305",
                    "ENSG00000103024",
                    "ENSG00000096384",
                    "ENSG00000022277",
                    "ENSG00000105223",
                    "ENSG00000160310",
                    "ENSG00000174469",
                    "ENSG00000151665",
                    "ENSG00000164902",
                    "ENSG00000144224",
                    "ENSG00000188994",
                    "ENSG00000174775",
                    "ENSG00000143947",
                    "ENSG00000071794",
                    "ENSG00000213015",
                    "ENSG00000175416",
                    "ENSG00000128609",
                    "ENSG00000063177",
                    "ENSG00000108953",
                    "ENSG00000155368",
                    "ENSG00000050393",
                    "ENSG00000125611",
                    "ENSG00000169564",
                    "ENSG00000094916",
                    "ENSG00000103035",
                    "ENSG00000132305",
                    "ENSG00000174721",
                    "ENSG00000128050",
                    "ENSG00000242372",
                    "ENSG00000138663",
                    "ENSG00000133706",
                    "ENSG00000111011",
                    "ENSG00000245694",
                    "ENSG00000086061",
                    "ENSG00000070756",
                    "ENSG00000136930",
                    "ENSG00000140740",
                    "ENSG00000149806",
                    "ENSG00000166012",
                    "ENSG00000167658",
                    "ENSG00000142168",
                    "ENSG00000186577",
                    "ENSG00000197006",
                    "ENSG00000185608",
                    "ENSG00000108468",
                    "ENSG00000196465",
                    "ENSG00000168028",
                    "ENSG00000108826",
                    "ENSG00000108848",
                    "ENSG00000031698",
                    "ENSG00000111786",
                    "ENSG00000109919",
                    "ENSG00000130731",
                    "ENSG00000143933",
                    "ENSG00000172172",
                    "ENSG00000165629",
                    "ENSG00000166503",
                    "ENSG00000126457",
                    "ENSG00000132963",
                    "ENSG00000042753",
                    "ENSG00000100664",
                    "ENSG00000177600",
                    "ENSG00000148450",
                    "ENSG00000136522",
                    "ENSG00000178127",
                    "ENSG00000131876",
                    "ENSG00000174456",
                    "ENSG00000148700",
                    "ENSG00000167699",
                    "ENSG00000105518",
                    "ENSG00000143198",
                    "ENSG00000054148",
                    "ENSG00000143621",
                    "ENSG00000145494",
                    "ENSG00000122378",
                    "ENSG00000156411",
                    "ENSG00000061676",
                    "ENSG00000177200",
                    "ENSG00000115944",
                    "ENSG00000111639",
                    "ENSG00000138443",
                    "ENSG00000135334",
                    "ENSG00000196531",
                    "ENSG00000100138",
                    "ENSG00000116030",
                    "ENSG00000115268",
                    "ENSG00000182534",
                    "ENSG00000081138",
                    "ENSG00000160075",
                    "ENSG00000174748",
                    "ENSG00000170043",
                    "ENSG00000089157",
                    "ENSG00000168653",
                    "ENSG00000141504",
                    "ENSG00000182481",
                    "ENSG00000156261",
                    "ENSG00000180228",
                    "ENSG00000126432",
                    "ENSG00000198363",
                    "ENSG00000151465",
                    "ENSG00000131116",
                    "ENSG00000197728",
                    "ENSG00000156515",
                    "ENSG00000057757",
                    "ENSG00000006015",
                    "ENSG00000171206",
                    "ENSG00000104823",
                    "ENSG00000233016",
                    "ENSG00000122406",
                    "ENSG00000166595",
                    "ENSG00000143774",
                    "ENSG00000182512",
                    "ENSG00000179271",
                    "ENSG00000166136",
                    "ENSG00000196141",
                    "ENSG00000115310",
                    "ENSG00000122545",
                    "ENSG00000134330",
                    "ENSG00000143612",
                    "ENSG00000159884",
                    "ENSG00000167969",
                    "ENSG00000007520",
                    "ENSG00000135720",
                    "ENSG00000196683",
                    "ENSG00000149547",
                    "ENSG00000108055",
                    "ENSG00000159335",
                    "ENSG00000136485",
                    "ENSG00000149600",
                    "ENSG00000123352",
                    "ENSG00000099194",
                    "ENSG00000137133",
                    "ENSG00000115350",
                    "ENSG00000128739",
                    "ENSG00000172809",
                    "ENSG00000145425",
                    "ENSG00000114902",
                    "ENSG00000107341",
                    "ENSG00000106615",
                    "ENSG00000099341",
                    "ENSG00000172500",
                    "ENSG00000230989",
                    "ENSG00000154518",
                    "ENSG00000109084",
                    "ENSG00000087302",
                    "ENSG00000101856",
                    "ENSG00000091164",
                    "ENSG00000136146",
                    "ENSG00000103342",
                    "ENSG00000090263",
                    "ENSG00000172757",
                    "ENSG00000182628",
                    "ENSG00000105669",
                    "ENSG00000188290",
                    "ENSG00000101460",
                    "ENSG00000136942",
                    "ENSG00000138175",
                    "ENSG00000105176",
                    "ENSG00000134717",
                    "ENSG00000169599",
                    "ENSG00000119185",
                    "ENSG00000215386",
                    "ENSG00000160752",
                    "ENSG00000143252",
                    "ENSG00000198961",
                    "ENSG00000141367",
                    "ENSG00000144381",
                    "ENSG00000119318",
                    "ENSG00000136521",
                    "ENSG00000132688",
                    "ENSG00000100811",
                    "ENSG00000145354",
                    "ENSG00000104177",
                    "ENSG00000159352",
                    "ENSG00000060762",
                    "ENSG00000166794",
                    "ENSG00000204220",
                    "ENSG00000112335",
                    "ENSG00000152234",
                    "ENSG00000133872",
                    "ENSG00000113811",
                    "ENSG00000179010",
                    "ENSG00000198668",
                    "ENSG00000108107",
                    "ENSG00000134308",
                    "ENSG00000108100",
                    "ENSG00000143158",
                    "ENSG00000132341",
                    "ENSG00000111832",
                    "ENSG00000153774",
                    "ENSG00000129197",
                    "ENSG00000164919",
                    "ENSG00000145741",
                    "ENSG00000138757",
                    "ENSG00000108561",
                    "ENSG00000118418",
                    "ENSG00000178982",
                    "ENSG00000108010",
                    "ENSG00000083845",
                    "ENSG00000214253",
                    "ENSG00000138382",
                    "ENSG00000155876",
                    "ENSG00000171867",
                    "ENSG00000102316",
                    "ENSG00000125821",
                    "ENSG00000137547",
                    "ENSG00000140319",
                    "ENSG00000132466",
                    "ENSG00000172428",
                    "ENSG00000175826",
                    "ENSG00000184220",
                    "ENSG00000009307",
                    "ENSG00000018625",
                    "ENSG00000168958",
                    "ENSG00000107372",
                    "ENSG00000182117",
                    "ENSG00000124172",
                    "ENSG00000057608",
                    "ENSG00000150991",
                    "ENSG00000065548",
                    "ENSG00000152795",
                    "ENSG00000137876",
                    "ENSG00000138398",
                    "ENSG00000137210",
                    "ENSG00000136527",
                    "ENSG00000162076",
                    "ENSG00000110958",
                    "ENSG00000100568",
                    "ENSG00000107281",
                    "ENSG00000007080",
                    "ENSG00000164323",
                    "ENSG00000166681",
                    "ENSG00000122705",
                    "ENSG00000132432",
                    "ENSG00000140374",
                    "ENSG00000115306",
                    "ENSG00000184009",
                    "ENSG00000023734",
                    "ENSG00000114405",
                    "ENSG00000120437",
                    "ENSG00000125944",
                    "ENSG00000054523",
                    "ENSG00000100380",
                    "ENSG00000173660",
                    "ENSG00000186184",
                    "ENSG00000100804",
                    "ENSG00000119640",
                    "ENSG00000113648",
                    "ENSG00000114125",
                    "ENSG00000185787",
                    "ENSG00000179222",
                    "ENSG00000105568",
                    "ENSG00000160799",
                    "ENSG00000130638",
                    "ENSG00000143333",
                    "ENSG00000169045",
                    "ENSG00000111775",
                    "ENSG00000179218",
                    "ENSG00000139684",
                    "ENSG00000128989",
                    "ENSG00000162604",
                    "ENSG00000144834",
                    "ENSG00000114120",
                    "ENSG00000163964",
                    "ENSG00000126602",
                    "ENSG00000188612",
                    "ENSG00000010810",
                    "ENSG00000180817",
                    "ENSG00000086065",
                    "ENSG00000181163",
                    "ENSG00000116288",
                    "ENSG00000008018",
                    "ENSG00000112531",
                    "ENSG00000136938",
                    "ENSG00000070961",
                    "ENSG00000198898",
                    "ENSG00000082898",
                    "ENSG00000115685",
                    "ENSG00000220205",
                    "ENSG00000128524",
                    "ENSG00000101654",
                    "ENSG00000101940",
                    "ENSG00000104549",
                    "ENSG00000044115",
                    "ENSG00000167113",
                    "ENSG00000265241",
                    "ENSG00000136238",
                    "ENSG00000164754",
                    "ENSG00000143183",
                    "ENSG00000132424",
                    "ENSG00000130522",
                    "ENSG00000047849",
                    "ENSG00000182154",
                    "ENSG00000221914",
                    "ENSG00000122873",
                    "ENSG00000065978",
                    "ENSG00000151923",
                    "ENSG00000158195",
                    "ENSG00000087460",
                    "ENSG00000100567",
                    "ENSG00000151247",
                    "ENSG00000183513",
                    "ENSG00000113558",
                    "ENSG00000181817",
                    "ENSG00000115457",
                    "ENSG00000109180",
                    "ENSG00000187735",
                    "ENSG00000115233",
                    "ENSG00000156052",
                    "ENSG00000143742",
                    "ENSG00000174886",
                    "ENSG00000172270",
                    "ENSG00000139637",
                    "ENSG00000106244",
                    "ENSG00000164167",
                    "ENSG00000092841",
                    "ENSG00000129351",
                    "ENSG00000173163"
                ],
                "expression_sum": {
                    "class1": 3253.0,
                    "class2": 24950.0
                },
                "ens_id": "ENSG00000181449",
                "uniprot_ids": [
                    "P48431"
                ]
            },
            "ENSG00000138772": {
                "id": "ENSG00000138772",
                "type": "Gene",
                "symbol": "ANXA3",
                "source": [
                    "ENSG00000100219",
                    "ENSG00000163435",
                    "ENSG00000168214"
                ],
                "target": [],
                "expression_sum": {
                    "class1": 187.0,
                    "class2": 3446.0
                },
                "ens_id": "ENSG00000138772",
                "uniprot_ids": [
                    "P12429",
                    "D6RA82",
                    "D6RFG5",
                    "D6RCA8",
                    "D6RAZ8",
                    "D6RFJ9"
                ]
            },
            "ENSG00000166710": {
                "id": "ENSG00000166710",
                "type": "Gene",
                "symbol": "B2M",
                "source": [
                    "ENSG00000163435",
                    "ENSG00000164683",
                    "ENSG00000100219",
                    "ENSG00000043355",
                    "ENSG00000160113"
                ],
                "target": [],
                "expression_sum": {
                    "class1": 26051.0,
                    "class2": 58091.0
                },
                "ens_id": "ENSG00000166710",
                "uniprot_ids": [
                    "P61769",
                    "H0YLF3",
                    "Q16446",
                    "F5H6I0",
                    "J3KNU0"
                ]
            },
            "ENSG00000136238": {
                "id": "ENSG00000136238",
                "type": "Gene",
                "symbol": "RAC1",
                "source": [
                    "ENSG00000181449",
                    "ENSG00000163435",
                    "ENSG00000167081",
                    "ENSG00000113580",
                    "ENSG00000148516",
                    "ENSG00000118263",
                    "ENSG00000164330",
                    "ENSG00000177030",
                    "ENSG00000114861",
                    "ENSG00000221869",
                    "ENSG00000119547"
                ],
                "target": [],
                "expression_sum": {
                    "class1": 48391.0,
                    "class2": 42293.0
                },
                "ens_id": "ENSG00000136238",
                "uniprot_ids": [
                    "P63000"
                ]
            },
            "ENSG00000084207": {
                "id": "ENSG00000084207",
                "type": "Gene",
                "symbol": "GSTP1",
                "source": [
                    "ENSG00000016082",
                    "ENSG00000164683",
                    "ENSG00000163435",
                    "ENSG00000164330"
                ],
                "target": [],
                "expression_sum": {
                    "class1": 70861.0,
                    "class2": 126160.0
                },
                "ens_id": "ENSG00000084207",
                "uniprot_ids": [
                    "P09211",
                    "A0A087X2E9",
                    "A8MX94",
                    "A0A2R8Y5E5",
                    "A0A087X243"
                ]
            },
            "ENSG00000147676": {
                "id": "ENSG00000147676",
                "type": "Gene",
                "symbol": "MAL2",
                "source": [
                    "ENSG00000100219",
                    "ENSG00000163435",
                    "ENSG00000173575",
                    "ENSG00000168214"
                ],
                "target": [],
                "expression_sum": {
                    "class1": 949.0,
                    "class2": 4134.0
                },
                "ens_id": "ENSG00000147676",
                "uniprot_ids": [
                    "Q969L2",
                    "A0A087WZL1"
                ]
            },
            "ENSG00000121039": {
                "id": "ENSG00000121039",
                "type": "Gene",
                "symbol": "RDH10",
                "source": [
                    "ENSG00000100219",
                    "ENSG00000163435",
                    "ENSG00000168214"
                ],
                "target": [],
                "expression_sum": {
                    "class1": 393.0,
                    "class2": 2648.0
                },
                "ens_id": "ENSG00000121039",
                "uniprot_ids": [
                    "Q8IZV5",
                    "E5RK48",
                    "E5RHJ9"
                ]
            },
            "ENSG00000076554": {
                "id": "ENSG00000076554",
                "type": "Gene",
                "symbol": "TPD52",
                "source": [
                    "ENSG00000163435",
                    "ENSG00000109132",
                    "ENSG00000167081",
                    "ENSG00000113580",
                    "ENSG00000148516",
                    "ENSG00000103495",
                    "ENSG00000100219",
                    "ENSG00000114861",
                    "ENSG00000139352"
                ],
                "target": [],
                "expression_sum": {
                    "class1": 9166.0,
                    "class2": 5233.0
                },
                "ens_id": "ENSG00000076554",
                "uniprot_ids": [
                    "P55327",
                    "E5RGI1",
                    "E5RFR7",
                    "H0YC44",
                    "E5RK35",
                    "E5RJ27"
                ]
            },
            "ENSG00000131981": {
                "id": "ENSG00000131981",
                "type": "Gene",
                "symbol": "LGALS3",
                "source": [
                    "ENSG00000164683",
                    "ENSG00000116044",
                    "ENSG00000163435",
                    "ENSG00000160113"
                ],
                "target": [],
                "expression_sum": {
                    "class1": 661.0,
                    "class2": 13809.0
                },
                "ens_id": "ENSG00000131981",
                "uniprot_ids": [
                    "P17931",
                    "G3V3R6",
                    "G3V407"
                ]
            },
            "ENSG00000178741": {
                "id": "ENSG00000178741",
                "type": "Gene",
                "symbol": "COX5A",
                "source": [
                    "ENSG00000163435",
                    "ENSG00000157557",
                    "ENSG00000167081",
                    "ENSG00000113580",
                    "ENSG00000148516",
                    "ENSG00000164330",
                    "ENSG00000114861",
                    "ENSG00000221869",
                    "ENSG00000139352"
                ],
                "target": [],
                "expression_sum": {
                    "class1": 50131.0,
                    "class2": 31266.0
                },
                "ens_id": "ENSG00000178741",
                "uniprot_ids": [
                    "P20674",
                    "H3BRM5",
                    "H3BRI0",
                    "H3BNX8",
                    "H3BV69"
                ]
            },
            "ENSG00000167658": {
                "id": "ENSG00000167658",
                "type": "Gene",
                "symbol": "EEF2",
                "source": [
                    "ENSG00000181449",
                    "ENSG00000163435",
                    "ENSG00000109132",
                    "ENSG00000167081",
                    "ENSG00000113580",
                    "ENSG00000164330",
                    "ENSG00000114861",
                    "ENSG00000221869",
                    "ENSG00000067955",
                    "ENSG00000137203"
                ],
                "target": [],
                "expression_sum": {
                    "class1": 84901.0,
                    "class2": 64690.0
                },
                "ens_id": "ENSG00000167658",
                "uniprot_ids": [
                    "P13639",
                    "M0R0I6"
                ]
            },
            "ENSG00000213760": {
                "id": "ENSG00000213760",
                "type": "Gene",
                "symbol": "ATP6V1G2",
                "source": [
                    "ENSG00000170365",
                    "ENSG00000100811",
                    "ENSG00000109132",
                    "ENSG00000175197",
                    "ENSG00000016082",
                    "ENSG00000004487",
                    "ENSG00000103495",
                    "ENSG00000128272",
                    "ENSG00000108055",
                    "ENSG00000213015",
                    "ENSG00000168214",
                    "ENSG00000116560",
                    "ENSG00000196591",
                    "ENSG00000147140",
                    "ENSG00000160113"
                ],
                "target": [],
                "expression_sum": {
                    "class1": 21218.0,
                    "class2": 3011.0
                },
                "ens_id": "ENSG00000213760",
                "uniprot_ids": [
                    "O95670",
                    "H0Y474"
                ]
            },
            "ENSG00000135940": {
                "id": "ENSG00000135940",
                "type": "Gene",
                "symbol": "COX5B",
                "source": [
                    "ENSG00000181449",
                    "ENSG00000163435",
                    "ENSG00000167081",
                    "ENSG00000113580",
                    "ENSG00000148516",
                    "ENSG00000164330",
                    "ENSG00000114861"
                ],
                "target": [],
                "expression_sum": {
                    "class1": 55957.0,
                    "class2": 46762.0
                },
                "ens_id": "ENSG00000135940",
                "uniprot_ids": [
                    "P10606"
                ]
            },
            "ENSG00000099622": {
                "id": "ENSG00000099622",
                "type": "Gene",
                "symbol": "CIRBP",
                "source": [
                    "ENSG00000181449",
                    "ENSG00000157557",
                    "ENSG00000109132",
                    "ENSG00000167081",
                    "ENSG00000113580",
                    "ENSG00000164330",
                    "ENSG00000181722",
                    "ENSG00000221869",
                    "ENSG00000139352",
                    "ENSG00000137203"
                ],
                "target": [],
                "expression_sum": {
                    "class1": 105777.0,
                    "class2": 50275.0
                },
                "ens_id": "ENSG00000099622",
                "uniprot_ids": [
                    "Q14011",
                    "K7EIF7",
                    "D6W5Y5",
                    "K7ELT6",
                    "K7EQL0",
                    "K7EPM4",
                    "K7ENX8",
                    "K7ELL4",
                    "K7EJV1",
                    "K7EQX4",
                    "K7ELV6",
                    "K7ER40",
                    "K7ENN6",
                    "K7EMY9",
                    "K7EJV5",
                    "F6WMK9",
                    "K7EQR7"
                ]
            },
            "ENSG00000118363": {
                "id": "ENSG00000118363",
                "type": "Gene",
                "symbol": "SPCS2",
                "source": [
                    "ENSG00000181449",
                    "ENSG00000167081",
                    "ENSG00000113580",
                    "ENSG00000164330",
                    "ENSG00000114861",
                    "ENSG00000221869",
                    "ENSG00000119547"
                ],
                "target": [],
                "expression_sum": {
                    "class1": 30945.0,
                    "class2": 26055.0
                },
                "ens_id": "ENSG00000118363",
                "uniprot_ids": [
                    "Q15005",
                    "E9PL01",
                    "A0A087WUC6",
                    "E9PRB9",
                    "H0YE04",
                    "E9PI68"
                ]
            },
            "ENSG00000122545": {
                "id": "ENSG00000122545",
                "type": "Gene",
                "symbol": "SEPT7",
                "source": [
                    "ENSG00000181449",
                    "ENSG00000167081",
                    "ENSG00000113580",
                    "ENSG00000103495",
                    "ENSG00000164330",
                    "ENSG00000108001",
                    "ENSG00000221869",
                    "ENSG00000196591",
                    "ENSG00000124766",
                    "ENSG00000119547",
                    "ENSG00000162599"
                ],
                "target": [],
                "expression_sum": {
                    "class1": 47839.0,
                    "class2": 32964.0
                },
                "ens_id": "ENSG00000122545",
                "uniprot_ids": [
                    "Q16181",
                    "Q5JXL7",
                    "A0A0U1RQW0",
                    "A0A0U1RRD1",
                    "A0A0U1RRM2",
                    "G3V1Q4",
                    "A0A0U1RRE1",
                    "A0A5F9ZGZ7",
                    "A0A0U1RRH9",
                    "E7EPK1",
                    "E7ES33"
                ]
            },
            "ENSG00000143198": {
                "id": "ENSG00000143198",
                "type": "Gene",
                "symbol": "MGST3",
                "source": [
                    "ENSG00000181449",
                    "ENSG00000109132",
                    "ENSG00000113580",
                    "ENSG00000103495",
                    "ENSG00000164330",
                    "ENSG00000108001",
                    "ENSG00000114861",
                    "ENSG00000067955",
                    "ENSG00000196591",
                    "ENSG00000124766",
                    "ENSG00000119547"
                ],
                "target": [],
                "expression_sum": {
                    "class1": 47478.0,
                    "class2": 36395.0
                },
                "ens_id": "ENSG00000143198",
                "uniprot_ids": [
                    "O14880",
                    "Q5VV89",
                    "Q5VV87"
                ]
            },
            "ENSG00000166974": {
                "id": "ENSG00000166974",
                "type": "Gene",
                "symbol": "MAPRE2",
                "source": [
                    "ENSG00000109132",
                    "ENSG00000016082",
                    "ENSG00000113580",
                    "ENSG00000167081",
                    "ENSG00000143799",
                    "ENSG00000148516",
                    "ENSG00000103495",
                    "ENSG00000164330",
                    "ENSG00000108055",
                    "ENSG00000139352",
                    "ENSG00000116560",
                    "ENSG00000196591"
                ],
                "target": [],
                "expression_sum": {
                    "class1": 8544.0,
                    "class2": 2412.0
                },
                "ens_id": "ENSG00000166974",
                "uniprot_ids": [
                    "Q15555",
                    "M0QX52",
                    "K7ERD8",
                    "K7EL66",
                    "K7ENB3"
                ]
            },
            "ENSG00000172159": {
                "id": "ENSG00000172159",
                "type": "Gene",
                "symbol": "FRMD3",
                "source": [
                    "ENSG00000109132",
                    "ENSG00000016082",
                    "ENSG00000113580",
                    "ENSG00000148516",
                    "ENSG00000103495",
                    "ENSG00000139352",
                    "ENSG00000116560",
                    "ENSG00000124766",
                    "ENSG00000137309"
                ],
                "target": [],
                "expression_sum": {
                    "class1": 3118.0,
                    "class2": 2014.0
                },
                "ens_id": "ENSG00000172159",
                "uniprot_ids": [
                    "A2A2Y4",
                    "Q5JV61"
                ]
            },
            "ENSG00000170365": {
                "id": "ENSG00000170365",
                "type": "TF",
                "symbol": "SMAD1",
                "source": [
                    "ENSG00000016082",
                    "ENSG00000164330"
                ],
                "target": [
                    "ENSG00000131016",
                    "ENSG00000143702",
                    "ENSG00000176788",
                    "ENSG00000144339",
                    "ENSG00000016082",
                    "ENSG00000135406",
                    "ENSG00000221866",
                    "ENSG00000182481",
                    "ENSG00000112773",
                    "ENSG00000139970",
                    "ENSG00000087258",
                    "ENSG00000213760",
                    "ENSG00000144642"
                ],
                "expression_sum": {
                    "class1": 4497.0,
                    "class2": 2361.0
                },
                "ens_id": "ENSG00000170365",
                "uniprot_ids": [
                    "Q15797",
                    "D6RDN7",
                    "D6RAY1",
                    "D6RD62",
                    "D6REQ3",
                    "A0A1D5RMR4",
                    "D6RBH9"
                ]
            },
            "ENSG00000100811": {
                "id": "ENSG00000100811",
                "type": "TF",
                "symbol": "YY1",
                "source": [
                    "ENSG00000167081",
                    "ENSG00000181449"
                ],
                "target": [
                    "ENSG00000104722",
                    "ENSG00000181449",
                    "ENSG00000103034",
                    "ENSG00000176788",
                    "ENSG00000172020",
                    "ENSG00000169856",
                    "ENSG00000152558",
                    "ENSG00000169783",
                    "ENSG00000145428",
                    "ENSG00000161082",
                    "ENSG00000132698",
                    "ENSG00000087085",
                    "ENSG00000167654",
                    "ENSG00000164093",
                    "ENSG00000107105",
                    "ENSG00000122756",
                    "ENSG00000148798",
                    "ENSG00000129951",
                    "ENSG00000130540",
                    "ENSG00000105784",
                    "ENSG00000198910",
                    "ENSG00000084628",
                    "ENSG00000166257",
                    "ENSG00000174021",
                    "ENSG00000128564",
                    "ENSG00000100167",
                    "ENSG00000197959",
                    "ENSG00000162374",
                    "ENSG00000196361",
                    "ENSG00000105409",
                    "ENSG00000072832",
                    "ENSG00000221866",
                    "ENSG00000155980",
                    "ENSG00000151320",
                    "ENSG00000186868",
                    "ENSG00000168280",
                    "ENSG00000005108",
                    "ENSG00000198125",
                    "ENSG00000213760",
                    "ENSG00000137285",
                    "ENSG00000092964",
                    "ENSG00000143702",
                    "ENSG00000144218",
                    "ENSG00000104435",
                    "ENSG00000122012",
                    "ENSG00000103154",
                    "ENSG00000016082",
                    "ENSG00000171033",
                    "ENSG00000124194",
                    "ENSG00000173267",
                    "ENSG00000087258",
                    "ENSG00000111728",
                    "ENSG00000149294",
                    "ENSG00000170091",
                    "ENSG00000139970",
                    "ENSG00000205922",
                    "ENSG00000084731",
                    "ENSG00000185624",
                    "ENSG00000167081",
                    "ENSG00000135406",
                    "ENSG00000168824",
                    "ENSG00000159409",
                    "ENSG00000118263",
                    "ENSG00000068615",
                    "ENSG00000164330",
                    "ENSG00000137267",
                    "ENSG00000104833",
                    "ENSG00000177675",
                    "ENSG00000174521",
                    "ENSG00000185650",
                    "ENSG00000135447",
                    "ENSG00000277586",
                    "ENSG00000104870"
                ],
                "expression_sum": {
                    "class1": 10338.0,
                    "class2": 9993.0
                },
                "ens_id": "ENSG00000100811",
                "uniprot_ids": [
                    "P25490",
                    "G3V3M8",
                    "A0A494C032",
                    "H0YJU4",
                    "H0YJV7"
                ]
            },
            "ENSG00000130726": {
                "id": "ENSG00000130726",
                "type": "TF",
                "symbol": "TRIM28",
                "source": [],
                "target": [
                    "ENSG00000104722",
                    "ENSG00000176788",
                    "ENSG00000172020",
                    "ENSG00000087085",
                    "ENSG00000107105",
                    "ENSG00000148798",
                    "ENSG00000213190",
                    "ENSG00000171551",
                    "ENSG00000142549",
                    "ENSG00000198910",
                    "ENSG00000162188",
                    "ENSG00000196361",
                    "ENSG00000162374",
                    "ENSG00000132872",
                    "ENSG00000105409",
                    "ENSG00000072832",
                    "ENSG00000221866",
                    "ENSG00000155980",
                    "ENSG00000186868",
                    "ENSG00000016082",
                    "ENSG00000124194",
                    "ENSG00000139970",
                    "ENSG00000135406",
                    "ENSG00000168824",
                    "ENSG00000135447",
                    "ENSG00000277586"
                ],
                "expression_sum": {
                    "class1": 9647.0,
                    "class2": 6926.0
                },
                "ens_id": "ENSG00000130726",
                "uniprot_ids": [
                    "Q13263",
                    "M0R0K9",
                    "M0QZE6",
                    "M0R3C0",
                    "M0R2I3"
                ]
            },
            "ENSG00000109132": {
                "id": "ENSG00000109132",
                "type": "TF",
                "symbol": "PHOX2B",
                "source": [
                    "ENSG00000016082",
                    "ENSG00000196591"
                ],
                "target": [
                    "ENSG00000078668",
                    "ENSG00000017260",
                    "ENSG00000116171",
                    "ENSG00000162704",
                    "ENSG00000111716",
                    "ENSG00000099622",
                    "ENSG00000115226",
                    "ENSG00000116161",
                    "ENSG00000140941",
                    "ENSG00000197959",
                    "ENSG00000179958",
                    "ENSG00000189403",
                    "ENSG00000159377",
                    "ENSG00000075415",
                    "ENSG00000107874",
                    "ENSG00000102172",
                    "ENSG00000077380",
                    "ENSG00000154556",
                    "ENSG00000163466",
                    "ENSG00000108515",
                    "ENSG00000134283",
                    "ENSG00000167085",
                    "ENSG00000272398",
                    "ENSG00000106443",
                    "ENSG00000115540",
                    "ENSG00000123240",
                    "ENSG00000163319",
                    "ENSG00000164924",
                    "ENSG00000139116",
                    "ENSG00000156482",
                    "ENSG00000047249",
                    "ENSG00000060982",
                    "ENSG00000147548",
                    "ENSG00000171703",
                    "ENSG00000196504",
                    "ENSG00000188243",
                    "ENSG00000067225",
                    "ENSG00000144642",
                    "ENSG00000144218",
                    "ENSG00000113712",
                    "ENSG00000140105",
                    "ENSG00000104131",
                    "ENSG00000159685",
                    "ENSG00000026025",
                    "ENSG00000095380",
                    "ENSG00000157168",
                    "ENSG00000103769",
                    "ENSG00000075142",
                    "ENSG00000150722",
                    "ENSG00000124802",
                    "ENSG00000187051",
                    "ENSG00000213741",
                    "ENSG00000151500",
                    "ENSG00000069849",
                    "ENSG00000123200",
                    "ENSG00000129515",
                    "ENSG00000100519",
                    "ENSG00000142546",
                    "ENSG00000163041",
                    "ENSG00000155115",
                    "ENSG00000110955",
                    "ENSG00000164346",
                    "ENSG00000164687",
                    "ENSG00000065518",
                    "ENSG00000174953",
                    "ENSG00000213760",
                    "ENSG00000137309",
                    "ENSG00000198836",
                    "ENSG00000143702",
                    "ENSG00000131966",
                    "ENSG00000231500",
                    "ENSG00000113013",
                    "ENSG00000109332",
                    "ENSG00000075624",
                    "ENSG00000079246",
                    "ENSG00000171222",
                    "ENSG00000244879",
                    "ENSG00000166974",
                    "ENSG00000164438",
                    "ENSG00000167552",
                    "ENSG00000167088",
                    "ENSG00000078618",
                    "ENSG00000136783",
                    "ENSG00000135390",
                    "ENSG00000153310",
                    "ENSG00000197223",
                    "ENSG00000154277",
                    "ENSG00000082397",
                    "ENSG00000105784",
                    "ENSG00000090266",
                    "ENSG00000198015",
                    "ENSG00000162374",
                    "ENSG00000171204",
                    "ENSG00000127184",
                    "ENSG00000103363",
                    "ENSG00000104388",
                    "ENSG00000134982",
                    "ENSG00000143222",
                    "ENSG00000151914",
                    "ENSG00000170903",
                    "ENSG00000165943",
                    "ENSG00000014641",
                    "ENSG00000125827",
                    "ENSG00000104723",
                    "ENSG00000175582",
                    "ENSG00000198612",
                    "ENSG00000081154",
                    "ENSG00000122566",
                    "ENSG00000139163",
                    "ENSG00000135486",
                    "ENSG00000109670",
                    "ENSG00000068796",
                    "ENSG00000138760",
                    "ENSG00000197302",
                    "ENSG00000170027",
                    "ENSG00000076554",
                    "ENSG00000103495",
                    "ENSG00000010270",
                    "ENSG00000147604",
                    "ENSG00000204308",
                    "ENSG00000108175",
                    "ENSG00000144713",
                    "ENSG00000172159",
                    "ENSG00000170315",
                    "ENSG00000096746",
                    "ENSG00000079785",
                    "ENSG00000067715",
                    "ENSG00000129084",
                    "ENSG00000069345",
                    "ENSG00000113742",
                    "ENSG00000117523",
                    "ENSG00000142192",
                    "ENSG00000137100",
                    "ENSG00000170035",
                    "ENSG00000187109",
                    "ENSG00000196262",
                    "ENSG00000125977",
                    "ENSG00000139117",
                    "ENSG00000137692",
                    "ENSG00000116586",
                    "ENSG00000089693",
                    "ENSG00000120438",
                    "ENSG00000077152",
                    "ENSG00000101132",
                    "ENSG00000154723",
                    "ENSG00000149357",
                    "ENSG00000131174",
                    "ENSG00000089289",
                    "ENSG00000106588",
                    "ENSG00000113387",
                    "ENSG00000137409",
                    "ENSG00000006125",
                    "ENSG00000196937",
                    "ENSG00000188613",
                    "ENSG00000170445",
                    "ENSG00000100442",
                    "ENSG00000106028",
                    "ENSG00000100697",
                    "ENSG00000134986",
                    "ENSG00000121022",
                    "ENSG00000197111",
                    "ENSG00000163682",
                    "ENSG00000101182",
                    "ENSG00000172020",
                    "ENSG00000124380",
                    "ENSG00000166260",
                    "ENSG00000051108",
                    "ENSG00000182093",
                    "ENSG00000134825",
                    "ENSG00000184007",
                    "ENSG00000096063",
                    "ENSG00000186868",
                    "ENSG00000134333",
                    "ENSG00000122012",
                    "ENSG00000116251",
                    "ENSG00000091640",
                    "ENSG00000143977",
                    "ENSG00000038219",
                    "ENSG00000110700",
                    "ENSG00000146731",
                    "ENSG00000143353",
                    "ENSG00000053254",
                    "ENSG00000011295",
                    "ENSG00000130706",
                    "ENSG00000173692",
                    "ENSG00000182985",
                    "ENSG00000161547",
                    "ENSG00000134769",
                    "ENSG00000145782",
                    "ENSG00000100354",
                    "ENSG00000106829",
                    "ENSG00000103429",
                    "ENSG00000138430",
                    "ENSG00000137285",
                    "ENSG00000104435",
                    "ENSG00000273611",
                    "ENSG00000186472",
                    "ENSG00000069956",
                    "ENSG00000146416",
                    "ENSG00000164258",
                    "ENSG00000116560",
                    "ENSG00000109472",
                    "ENSG00000114956",
                    "ENSG00000145725",
                    "ENSG00000092201",
                    "ENSG00000147862",
                    "ENSG00000079102",
                    "ENSG00000182004",
                    "ENSG00000008294",
                    "ENSG00000104408",
                    "ENSG00000156467",
                    "ENSG00000168259",
                    "ENSG00000161203",
                    "ENSG00000100823",
                    "ENSG00000101782",
                    "ENSG00000142541",
                    "ENSG00000111906",
                    "ENSG00000143753",
                    "ENSG00000118816",
                    "ENSG00000106105",
                    "ENSG00000147669",
                    "ENSG00000151474",
                    "ENSG00000126524",
                    "ENSG00000163468",
                    "ENSG00000091039",
                    "ENSG00000167522",
                    "ENSG00000134056",
                    "ENSG00000277443",
                    "ENSG00000135387",
                    "ENSG00000156976",
                    "ENSG00000169021",
                    "ENSG00000078369",
                    "ENSG00000138758",
                    "ENSG00000115541",
                    "ENSG00000150093",
                    "ENSG00000138107",
                    "ENSG00000149485",
                    "ENSG00000130748",
                    "ENSG00000116641",
                    "ENSG00000138326",
                    "ENSG00000154380",
                    "ENSG00000145362",
                    "ENSG00000104695",
                    "ENSG00000182621",
                    "ENSG00000123144",
                    "ENSG00000113575",
                    "ENSG00000149084",
                    "ENSG00000127022",
                    "ENSG00000114302",
                    "ENSG00000142676",
                    "ENSG00000119705",
                    "ENSG00000165283",
                    "ENSG00000075151",
                    "ENSG00000100528",
                    "ENSG00000154582",
                    "ENSG00000151320",
                    "ENSG00000077279",
                    "ENSG00000137996",
                    "ENSG00000016082",
                    "ENSG00000108946",
                    "ENSG00000164830",
                    "ENSG00000108953",
                    "ENSG00000146267",
                    "ENSG00000094916",
                    "ENSG00000177875",
                    "ENSG00000186063",
                    "ENSG00000204310",
                    "ENSG00000010404",
                    "ENSG00000070756",
                    "ENSG00000136930",
                    "ENSG00000140740",
                    "ENSG00000167658",
                    "ENSG00000113161",
                    "ENSG00000072041",
                    "ENSG00000197006",
                    "ENSG00000159176",
                    "ENSG00000196465",
                    "ENSG00000108848",
                    "ENSG00000143933",
                    "ENSG00000132963",
                    "ENSG00000042753",
                    "ENSG00000175390",
                    "ENSG00000168522",
                    "ENSG00000119048",
                    "ENSG00000198408",
                    "ENSG00000111897",
                    "ENSG00000166946",
                    "ENSG00000143198",
                    "ENSG00000135406",
                    "ENSG00000170017",
                    "ENSG00000177675",
                    "ENSG00000115944",
                    "ENSG00000111639",
                    "ENSG00000138443",
                    "ENSG00000168268",
                    "ENSG00000107105",
                    "ENSG00000086300",
                    "ENSG00000112186",
                    "ENSG00000203930",
                    "ENSG00000182481",
                    "ENSG00000159593",
                    "ENSG00000156261",
                    "ENSG00000198363",
                    "ENSG00000197728",
                    "ENSG00000170606",
                    "ENSG00000182512",
                    "ENSG00000179271",
                    "ENSG00000184207",
                    "ENSG00000135720",
                    "ENSG00000166848",
                    "ENSG00000196683",
                    "ENSG00000166206",
                    "ENSG00000099194",
                    "ENSG00000172809",
                    "ENSG00000089818",
                    "ENSG00000099341",
                    "ENSG00000087302",
                    "ENSG00000100220",
                    "ENSG00000105176",
                    "ENSG00000169599",
                    "ENSG00000160752",
                    "ENSG00000143252",
                    "ENSG00000144381",
                    "ENSG00000136521",
                    "ENSG00000104177",
                    "ENSG00000164741",
                    "ENSG00000152234",
                    "ENSG00000172687",
                    "ENSG00000108100",
                    "ENSG00000061794",
                    "ENSG00000135549",
                    "ENSG00000153774",
                    "ENSG00000145741",
                    "ENSG00000138757",
                    "ENSG00000033627",
                    "ENSG00000155876",
                    "ENSG00000171867",
                    "ENSG00000137547",
                    "ENSG00000185565",
                    "ENSG00000184220",
                    "ENSG00000018189",
                    "ENSG00000174498",
                    "ENSG00000137876",
                    "ENSG00000196591",
                    "ENSG00000138398",
                    "ENSG00000122705",
                    "ENSG00000115306",
                    "ENSG00000125944",
                    "ENSG00000054523",
                    "ENSG00000100380",
                    "ENSG00000100804",
                    "ENSG00000119640",
                    "ENSG00000170899",
                    "ENSG00000114125",
                    "ENSG00000185787",
                    "ENSG00000141424",
                    "ENSG00000130638",
                    "ENSG00000111775",
                    "ENSG00000123636",
                    "ENSG00000139684",
                    "ENSG00000114120",
                    "ENSG00000164587",
                    "ENSG00000181163",
                    "ENSG00000008018",
                    "ENSG00000078018",
                    "ENSG00000204843",
                    "ENSG00000215021",
                    "ENSG00000137274",
                    "ENSG00000044115",
                    "ENSG00000265241",
                    "ENSG00000134590",
                    "ENSG00000103415",
                    "ENSG00000065978",
                    "ENSG00000113558",
                    "ENSG00000139970",
                    "ENSG00000143153",
                    "ENSG00000123983",
                    "ENSG00000116489",
                    "ENSG00000143742",
                    "ENSG00000133169",
                    "ENSG00000205629",
                    "ENSG00000164167",
                    "ENSG00000089335",
                    "ENSG00000181610",
                    "ENSG00000089009"
                ],
                "expression_sum": {
                    "class1": 4511.0,
                    "class2": 26.0
                },
                "ens_id": "ENSG00000109132",
                "uniprot_ids": [
                    "Q99453"
                ]
            },
            "ENSG00000153147": {
                "id": "ENSG00000153147",
                "type": "TF",
                "symbol": "SMARCA5",
                "source": [],
                "target": [
                    "ENSG00000016082",
                    "ENSG00000167654",
                    "ENSG00000107105",
                    "ENSG00000198125",
                    "ENSG00000181885",
                    "ENSG00000104870"
                ],
                "expression_sum": {
                    "class1": 14120.0,
                    "class2": 12380.0
                },
                "ens_id": "ENSG00000153147",
                "uniprot_ids": [
                    "O60264"
                ]
            },
            "ENSG00000198911": {
                "id": "ENSG00000198911",
                "type": "TF",
                "symbol": "SREBF2",
                "source": [],
                "target": [
                    "ENSG00000104722",
                    "ENSG00000176788",
                    "ENSG00000144339",
                    "ENSG00000169783",
                    "ENSG00000113657",
                    "ENSG00000113161",
                    "ENSG00000107105",
                    "ENSG00000129951",
                    "ENSG00000080824",
                    "ENSG00000129355",
                    "ENSG00000166226",
                    "ENSG00000198910",
                    "ENSG00000060982",
                    "ENSG00000149313",
                    "ENSG00000105409",
                    "ENSG00000197747",
                    "ENSG00000164687",
                    "ENSG00000072832",
                    "ENSG00000104427",
                    "ENSG00000101132",
                    "ENSG00000182481",
                    "ENSG00000197694",
                    "ENSG00000168280",
                    "ENSG00000186868",
                    "ENSG00000047849",
                    "ENSG00000144642",
                    "ENSG00000092964",
                    "ENSG00000143702",
                    "ENSG00000144218",
                    "ENSG00000163539",
                    "ENSG00000122012",
                    "ENSG00000125354",
                    "ENSG00000077380",
                    "ENSG00000162490",
                    "ENSG00000016082",
                    "ENSG00000097021",
                    "ENSG00000124194",
                    "ENSG00000120437",
                    "ENSG00000160014",
                    "ENSG00000117632",
                    "ENSG00000111728",
                    "ENSG00000154917",
                    "ENSG00000034510",
                    "ENSG00000131016",
                    "ENSG00000146425",
                    "ENSG00000139970",
                    "ENSG00000079462",
                    "ENSG00000084731",
                    "ENSG00000135406",
                    "ENSG00000115665",
                    "ENSG00000068615",
                    "ENSG00000277586"
                ],
                "expression_sum": {
                    "class1": 5454.0,
                    "class2": 2413.0
                },
                "ens_id": "ENSG00000198911",
                "uniprot_ids": [
                    "Q12772",
                    "G3V0I8",
                    "H0Y7E5",
                    "A0A087X1T2"
                ]
            },
            "ENSG00000196591": {
                "id": "ENSG00000196591",
                "type": "TF",
                "symbol": "HDAC2",
                "source": [
                    "ENSG00000163435",
                    "ENSG00000109132",
                    "ENSG00000167081",
                    "ENSG00000148516",
                    "ENSG00000164330"
                ],
                "target": [
                    "ENSG00000099864",
                    "ENSG00000103034",
                    "ENSG00000169856",
                    "ENSG00000162704",
                    "ENSG00000111716",
                    "ENSG00000087085",
                    "ENSG00000108830",
                    "ENSG00000113657",
                    "ENSG00000161682",
                    "ENSG00000141698",
                    "ENSG00000101210",
                    "ENSG00000164093",
                    "ENSG00000107105",
                    "ENSG00000122756",
                    "ENSG00000115226",
                    "ENSG00000166582",
                    "ENSG00000182534",
                    "ENSG00000129355",
                    "ENSG00000173467",
                    "ENSG00000196230",
                    "ENSG00000086300",
                    "ENSG00000107742",
                    "ENSG00000105255",
                    "ENSG00000100167",
                    "ENSG00000197959",
                    "ENSG00000170027",
                    "ENSG00000134594",
                    "ENSG00000075945",
                    "ENSG00000010270",
                    "ENSG00000182481",
                    "ENSG00000168280",
                    "ENSG00000123416",
                    "ENSG00000181885",
                    "ENSG00000177432",
                    "ENSG00000126950",
                    "ENSG00000165434",
                    "ENSG00000108175",
                    "ENSG00000077380",
                    "ENSG00000162490",
                    "ENSG00000173267",
                    "ENSG00000138413",
                    "ENSG00000071054",
                    "ENSG00000184697",
                    "ENSG00000087258",
                    "ENSG00000105088",
                    "ENSG00000117632",
                    "ENSG00000111728",
                    "ENSG00000154917",
                    "ENSG00000272398",
                    "ENSG00000015592",
                    "ENSG00000079462",
                    "ENSG00000134369",
                    "ENSG00000103145",
                    "ENSG00000115665",
                    "ENSG00000115310",
                    "ENSG00000122545",
                    "ENSG00000137312",
                    "ENSG00000105270",
                    "ENSG00000018236",
                    "ENSG00000189043",
                    "ENSG00000099194",
                    "ENSG00000277443",
                    "ENSG00000104722",
                    "ENSG00000162545",
                    "ENSG00000169783",
                    "ENSG00000157514",
                    "ENSG00000148516",
                    "ENSG00000221946",
                    "ENSG00000167654",
                    "ENSG00000066557",
                    "ENSG00000105254",
                    "ENSG00000084628",
                    "ENSG00000188290",
                    "ENSG00000166257",
                    "ENSG00000196361",
                    "ENSG00000108828",
                    "ENSG00000196072",
                    "ENSG00000104112",
                    "ENSG00000131711",
                    "ENSG00000155980",
                    "ENSG00000075618",
                    "ENSG00000101132",
                    "ENSG00000120265",
                    "ENSG00000144642",
                    "ENSG00000138758",
                    "ENSG00000144218",
                    "ENSG00000125354",
                    "ENSG00000133083",
                    "ENSG00000213420",
                    "ENSG00000171033",
                    "ENSG00000155099",
                    "ENSG00000107130",
                    "ENSG00000034713",
                    "ENSG00000175203",
                    "ENSG00000034510",
                    "ENSG00000170091",
                    "ENSG00000146425",
                    "ENSG00000158164",
                    "ENSG00000206579",
                    "ENSG00000141404",
                    "ENSG00000118263",
                    "ENSG00000068615",
                    "ENSG00000243449",
                    "ENSG00000185650",
                    "ENSG00000168118",
                    "ENSG00000134986",
                    "ENSG00000277586",
                    "ENSG00000111275",
                    "ENSG00000176788",
                    "ENSG00000172020",
                    "ENSG00000125821",
                    "ENSG00000149557",
                    "ENSG00000151640",
                    "ENSG00000144635",
                    "ENSG00000185664",
                    "ENSG00000148798",
                    "ENSG00000145335",
                    "ENSG00000130540",
                    "ENSG00000171551",
                    "ENSG00000198910",
                    "ENSG00000018189",
                    "ENSG00000134198",
                    "ENSG00000138138",
                    "ENSG00000128564",
                    "ENSG00000101220",
                    "ENSG00000109132",
                    "ENSG00000132821",
                    "ENSG00000105409",
                    "ENSG00000197747",
                    "ENSG00000164687",
                    "ENSG00000151320",
                    "ENSG00000186868",
                    "ENSG00000077279",
                    "ENSG00000213760",
                    "ENSG00000092964",
                    "ENSG00000143702",
                    "ENSG00000112972",
                    "ENSG00000122012",
                    "ENSG00000016082",
                    "ENSG00000142156",
                    "ENSG00000152503",
                    "ENSG00000164221",
                    "ENSG00000166736",
                    "ENSG00000120437",
                    "ENSG00000160014",
                    "ENSG00000054523",
                    "ENSG00000149294",
                    "ENSG00000171533",
                    "ENSG00000103187",
                    "ENSG00000084731",
                    "ENSG00000166974",
                    "ENSG00000167552",
                    "ENSG00000174521",
                    "ENSG00000164330",
                    "ENSG00000141505",
                    "ENSG00000137267",
                    "ENSG00000177875",
                    "ENSG00000227051",
                    "ENSG00000163931",
                    "ENSG00000203879",
                    "ENSG00000135447",
                    "ENSG00000176973",
                    "ENSG00000105419",
                    "ENSG00000144339",
                    "ENSG00000250303",
                    "ENSG00000197457",
                    "ENSG00000131437",
                    "ENSG00000113161",
                    "ENSG00000154277",
                    "ENSG00000171724",
                    "ENSG00000050165",
                    "ENSG00000129951",
                    "ENSG00000137942",
                    "ENSG00000080824",
                    "ENSG00000105784",
                    "ENSG00000166226",
                    "ENSG00000107643",
                    "ENSG00000162188",
                    "ENSG00000143862",
                    "ENSG00000157851",
                    "ENSG00000078018",
                    "ENSG00000117569",
                    "ENSG00000162374",
                    "ENSG00000182287",
                    "ENSG00000072832",
                    "ENSG00000104427",
                    "ENSG00000221866",
                    "ENSG00000133318",
                    "ENSG00000197694",
                    "ENSG00000186469",
                    "ENSG00000134590",
                    "ENSG00000047849",
                    "ENSG00000137285",
                    "ENSG00000103154",
                    "ENSG00000104435",
                    "ENSG00000126214",
                    "ENSG00000167996",
                    "ENSG00000097021",
                    "ENSG00000124194",
                    "ENSG00000148154",
                    "ENSG00000152082",
                    "ENSG00000131037",
                    "ENSG00000189159",
                    "ENSG00000111897",
                    "ENSG00000131016",
                    "ENSG00000139970",
                    "ENSG00000090097",
                    "ENSG00000112290",
                    "ENSG00000163435",
                    "ENSG00000143198",
                    "ENSG00000167081",
                    "ENSG00000135406",
                    "ENSG00000168824",
                    "ENSG00000159409",
                    "ENSG00000175582",
                    "ENSG00000177675",
                    "ENSG00000131473",
                    "ENSG00000122378",
                    "ENSG00000108960",
                    "ENSG00000137343"
                ],
                "expression_sum": {
                    "class1": 21723.0,
                    "class2": 9054.0
                },
                "ens_id": "ENSG00000196591",
                "uniprot_ids": [
                    "Q92769",
                    "E5RGV4",
                    "E5RK19",
                    "E5RG37",
                    "H3BM24",
                    "E5RFP9",
                    "E5RH52",
                    "E5RFI6",
                    "E5RJ04",
                    "E5RHE7"
                ]
            },
            "ENSG00000105698": {
                "id": "ENSG00000105698",
                "type": "TF",
                "symbol": "USF2",
                "source": [
                    "ENSG00000167081",
                    "ENSG00000113580",
                    "ENSG00000139352",
                    "ENSG00000164330"
                ],
                "target": [
                    "ENSG00000104722",
                    "ENSG00000181449",
                    "ENSG00000176788",
                    "ENSG00000149557",
                    "ENSG00000198840",
                    "ENSG00000221946",
                    "ENSG00000107105",
                    "ENSG00000129951",
                    "ENSG00000105784",
                    "ENSG00000198910",
                    "ENSG00000060982",
                    "ENSG00000102265",
                    "ENSG00000174021",
                    "ENSG00000113732",
                    "ENSG00000148834",
                    "ENSG00000072832",
                    "ENSG00000182481",
                    "ENSG00000198125",
                    "ENSG00000181885",
                    "ENSG00000117691",
                    "ENSG00000092964",
                    "ENSG00000124194",
                    "ENSG00000111728",
                    "ENSG00000158164",
                    "ENSG00000167081",
                    "ENSG00000113580",
                    "ENSG00000119888",
                    "ENSG00000143545",
                    "ENSG00000104833",
                    "ENSG00000174521",
                    "ENSG00000164330",
                    "ENSG00000137267",
                    "ENSG00000118418"
                ],
                "expression_sum": {
                    "class1": 6116.0,
                    "class2": 4359.0
                },
                "ens_id": "ENSG00000105698",
                "uniprot_ids": [
                    "Q15853",
                    "B4DLJ1",
                    "M0R1C6",
                    "M0QY46",
                    "M0QZ39",
                    "V9GYR5",
                    "M0QXT0",
                    "M0QYU7"
                ]
            },
            "ENSG00000213015": {
                "id": "ENSG00000213015",
                "type": "TF",
                "symbol": "ZNF580",
                "source": [
                    "ENSG00000167081",
                    "ENSG00000181449",
                    "ENSG00000164330"
                ],
                "target": [
                    "ENSG00000176788",
                    "ENSG00000169856",
                    "ENSG00000107105",
                    "ENSG00000080824",
                    "ENSG00000129355",
                    "ENSG00000084628",
                    "ENSG00000176749",
                    "ENSG00000128564",
                    "ENSG00000187134",
                    "ENSG00000197959",
                    "ENSG00000197747",
                    "ENSG00000010270",
                    "ENSG00000104427",
                    "ENSG00000090273",
                    "ENSG00000133318",
                    "ENSG00000198513",
                    "ENSG00000186868",
                    "ENSG00000168280",
                    "ENSG00000123416",
                    "ENSG00000115756",
                    "ENSG00000182481",
                    "ENSG00000108106",
                    "ENSG00000213760",
                    "ENSG00000137285",
                    "ENSG00000165434",
                    "ENSG00000143702",
                    "ENSG00000126214",
                    "ENSG00000213420",
                    "ENSG00000097021",
                    "ENSG00000152503",
                    "ENSG00000164221",
                    "ENSG00000131037",
                    "ENSG00000168243",
                    "ENSG00000189159",
                    "ENSG00000120437",
                    "ENSG00000160014",
                    "ENSG00000117632",
                    "ENSG00000272398",
                    "ENSG00000131016",
                    "ENSG00000170091",
                    "ENSG00000146425",
                    "ENSG00000149294",
                    "ENSG00000105649",
                    "ENSG00000079462",
                    "ENSG00000167081",
                    "ENSG00000135406",
                    "ENSG00000143545",
                    "ENSG00000164032",
                    "ENSG00000137267",
                    "ENSG00000185650",
                    "ENSG00000149926"
                ],
                "expression_sum": {
                    "class1": 11545.0,
                    "class2": 8422.0
                },
                "ens_id": "ENSG00000213015",
                "uniprot_ids": [
                    "Q9UK33",
                    "K7EJ53",
                    "K7ERU0",
                    "K7ENY2"
                ]
            },
            "ENSG00000119950": {
                "id": "ENSG00000119950",
                "type": "TF",
                "symbol": "MXI1",
                "source": [
                    "ENSG00000181449",
                    "ENSG00000167081",
                    "ENSG00000113580",
                    "ENSG00000164330",
                    "ENSG00000164093",
                    "ENSG00000139352"
                ],
                "target": [
                    "ENSG00000104722",
                    "ENSG00000169783",
                    "ENSG00000112773",
                    "ENSG00000105784",
                    "ENSG00000196230",
                    "ENSG00000084628",
                    "ENSG00000105409",
                    "ENSG00000072832",
                    "ENSG00000182481",
                    "ENSG00000117155",
                    "ENSG00000117691",
                    "ENSG00000139970",
                    "ENSG00000170091",
                    "ENSG00000157557",
                    "ENSG00000167081",
                    "ENSG00000113580",
                    "ENSG00000100604",
                    "ENSG00000164330",
                    "ENSG00000137267"
                ],
                "expression_sum": {
                    "class1": 8238.0,
                    "class2": 6134.0
                },
                "ens_id": "ENSG00000119950",
                "uniprot_ids": [
                    "P50539",
                    "A0A494BZS3",
                    "A0A494C0W2",
                    "A0A494C1P8",
                    "A0A0S2Z429",
                    "B1ANN8",
                    "F6U3F6",
                    "A0A494C1C1",
                    "F6WCI9",
                    "A0A494C0I8"
                ]
            },
            "ENSG00000092199": {
                "id": "ENSG00000092199",
                "type": "TF",
                "symbol": "HNRNPC",
                "source": [
                    "ENSG00000181449",
                    "ENSG00000163435",
                    "ENSG00000167081",
                    "ENSG00000113580",
                    "ENSG00000164330"
                ],
                "target": [
                    "ENSG00000162496",
                    "ENSG00000130643",
                    "ENSG00000103034",
                    "ENSG00000087085",
                    "ENSG00000185664",
                    "ENSG00000148798",
                    "ENSG00000105784",
                    "ENSG00000171551",
                    "ENSG00000244187",
                    "ENSG00000141934",
                    "ENSG00000084628",
                    "ENSG00000166257",
                    "ENSG00000174021",
                    "ENSG00000128564",
                    "ENSG00000060982",
                    "ENSG00000197959",
                    "ENSG00000174498",
                    "ENSG00000176749",
                    "ENSG00000196361",
                    "ENSG00000105409",
                    "ENSG00000182287",
                    "ENSG00000221866",
                    "ENSG00000131711",
                    "ENSG00000166145",
                    "ENSG00000005108",
                    "ENSG00000213719",
                    "ENSG00000092964",
                    "ENSG00000143702",
                    "ENSG00000103154",
                    "ENSG00000213420",
                    "ENSG00000107438",
                    "ENSG00000162490",
                    "ENSG00000171033",
                    "ENSG00000152503",
                    "ENSG00000128228",
                    "ENSG00000087258",
                    "ENSG00000100167",
                    "ENSG00000139970",
                    "ENSG00000171533",
                    "ENSG00000124942",
                    "ENSG00000167081",
                    "ENSG00000113580",
                    "ENSG00000119888",
                    "ENSG00000135406",
                    "ENSG00000174521",
                    "ENSG00000151150",
                    "ENSG00000118418",
                    "ENSG00000185650",
                    "ENSG00000104870"
                ],
                "expression_sum": {
                    "class1": 26156.0,
                    "class2": 27085.0
                },
                "ens_id": "ENSG00000092199",
                "uniprot_ids": [
                    "P07910",
                    "G3V576",
                    "B2R5W2",
                    "B4DY08",
                    "B4DSU6",
                    "G3V4C1",
                    "G3V4M8",
                    "G3V2H6",
                    "G3V4W0",
                    "G3V5X6",
                    "G3V2Q1",
                    "G3V251",
                    "G3V555",
                    "G3V575",
                    "G3V3K6",
                    "G3V2D6",
                    "G3V5V7"
                ]
            },
            "ENSG00000106261": {
                "id": "ENSG00000106261",
                "type": "TF",
                "symbol": "ZKSCAN1",
                "source": [
                    "ENSG00000163435",
                    "ENSG00000167081",
                    "ENSG00000113580",
                    "ENSG00000148516",
                    "ENSG00000100219",
                    "ENSG00000221869"
                ],
                "target": [
                    "ENSG00000103034",
                    "ENSG00000148516",
                    "ENSG00000167654",
                    "ENSG00000101443",
                    "ENSG00000163399",
                    "ENSG00000154080",
                    "ENSG00000196230",
                    "ENSG00000060982",
                    "ENSG00000152952",
                    "ENSG00000130303",
                    "ENSG00000163191",
                    "ENSG00000155980",
                    "ENSG00000123416",
                    "ENSG00000198125",
                    "ENSG00000138758",
                    "ENSG00000092964",
                    "ENSG00000132688",
                    "ENSG00000143702",
                    "ENSG00000135535",
                    "ENSG00000245532",
                    "ENSG00000075043",
                    "ENSG00000167642",
                    "ENSG00000102007",
                    "ENSG00000107130",
                    "ENSG00000149485",
                    "ENSG00000129521",
                    "ENSG00000189159",
                    "ENSG00000117632",
                    "ENSG00000170091",
                    "ENSG00000163435",
                    "ENSG00000096696",
                    "ENSG00000167552",
                    "ENSG00000118263",
                    "ENSG00000164330",
                    "ENSG00000137267",
                    "ENSG00000100219",
                    "ENSG00000185650",
                    "ENSG00000104870"
                ],
                "expression_sum": {
                    "class1": 11131.0,
                    "class2": 11221.0
                },
                "ens_id": "ENSG00000106261",
                "uniprot_ids": [
                    "P17029",
                    "B3KRF7",
                    "E9PC66",
                    "C9JRM9"
                ]
            },
            "ENSG00000148516": {
                "id": "ENSG00000148516",
                "type": "TF",
                "symbol": "ZEB1",
                "source": [
                    "ENSG00000106261",
                    "ENSG00000164330",
                    "ENSG00000168214",
                    "ENSG00000196591",
                    "ENSG00000124766",
                    "ENSG00000119547"
                ],
                "target": [
                    "ENSG00000103034",
                    "ENSG00000106261",
                    "ENSG00000163634",
                    "ENSG00000111674",
                    "ENSG00000087085",
                    "ENSG00000005810",
                    "ENSG00000073792",
                    "ENSG00000107105",
                    "ENSG00000114784",
                    "ENSG00000197302",
                    "ENSG00000160113",
                    "ENSG00000179915",
                    "ENSG00000120992",
                    "ENSG00000186106",
                    "ENSG00000128872",
                    "ENSG00000170027",
                    "ENSG00000075945",
                    "ENSG00000078674",
                    "ENSG00000023287",
                    "ENSG00000076554",
                    "ENSG00000156467",
                    "ENSG00000166295",
                    "ENSG00000168280",
                    "ENSG00000119013",
                    "ENSG00000164692",
                    "ENSG00000184076",
                    "ENSG00000165434",
                    "ENSG00000108175",
                    "ENSG00000172159",
                    "ENSG00000077380",
                    "ENSG00000154556",
                    "ENSG00000178741",
                    "ENSG00000173267",
                    "ENSG00000138413",
                    "ENSG00000173812",
                    "ENSG00000067715",
                    "ENSG00000184697",
                    "ENSG00000154917",
                    "ENSG00000272398",
                    "ENSG00000172292",
                    "ENSG00000123240",
                    "ENSG00000051620",
                    "ENSG00000166136",
                    "ENSG00000251562",
                    "ENSG00000115310",
                    "ENSG00000166848",
                    "ENSG00000137312",
                    "ENSG00000074319",
                    "ENSG00000113742",
                    "ENSG00000164924",
                    "ENSG00000091039",
                    "ENSG00000163939",
                    "ENSG00000111371",
                    "ENSG00000137941",
                    "ENSG00000119547",
                    "ENSG00000104722",
                    "ENSG00000101040",
                    "ENSG00000159023",
                    "ENSG00000198886",
                    "ENSG00000204130",
                    "ENSG00000108309",
                    "ENSG00000125868",
                    "ENSG00000104904",
                    "ENSG00000125971",
                    "ENSG00000153815",
                    "ENSG00000106565",
                    "ENSG00000135387",
                    "ENSG00000166257",
                    "ENSG00000147548",
                    "ENSG00000156976",
                    "ENSG00000168092",
                    "ENSG00000253719",
                    "ENSG00000196361",
                    "ENSG00000119335",
                    "ENSG00000130294",
                    "ENSG00000078369",
                    "ENSG00000100077",
                    "ENSG00000169432",
                    "ENSG00000104112",
                    "ENSG00000155980",
                    "ENSG00000185201",
                    "ENSG00000187231",
                    "ENSG00000154723",
                    "ENSG00000166347",
                    "ENSG00000163539",
                    "ENSG00000131174",
                    "ENSG00000186187",
                    "ENSG00000114739",
                    "ENSG00000110321",
                    "ENSG00000117139",
                    "ENSG00000109654",
                    "ENSG00000198420",
                    "ENSG00000234745",
                    "ENSG00000034713",
                    "ENSG00000205403",
                    "ENSG00000198668",
                    "ENSG00000063854",
                    "ENSG00000103769",
                    "ENSG00000146425",
                    "ENSG00000170091",
                    "ENSG00000111642",
                    "ENSG00000117410",
                    "ENSG00000145362",
                    "ENSG00000141404",
                    "ENSG00000118263",
                    "ENSG00000273841",
                    "ENSG00000068615",
                    "ENSG00000243449",
                    "ENSG00000204525",
                    "ENSG00000263001",
                    "ENSG00000149257",
                    "ENSG00000196405",
                    "ENSG00000129197",
                    "ENSG00000033627",
                    "ENSG00000100697",
                    "ENSG00000162961",
                    "ENSG00000197111",
                    "ENSG00000225783",
                    "ENSG00000084733",
                    "ENSG00000111726",
                    "ENSG00000133134",
                    "ENSG00000109099",
                    "ENSG00000142949",
                    "ENSG00000114573",
                    "ENSG00000144635",
                    "ENSG00000196233",
                    "ENSG00000185664",
                    "ENSG00000163041",
                    "ENSG00000148798",
                    "ENSG00000118680",
                    "ENSG00000197183",
                    "ENSG00000009307",
                    "ENSG00000136854",
                    "ENSG00000018189",
                    "ENSG00000138138",
                    "ENSG00000110955",
                    "ENSG00000096384",
                    "ENSG00000175274",
                    "ENSG00000197747",
                    "ENSG00000196591",
                    "ENSG00000168214",
                    "ENSG00000108654",
                    "ENSG00000122417",
                    "ENSG00000135940",
                    "ENSG00000143702",
                    "ENSG00000131966",
                    "ENSG00000038219",
                    "ENSG00000178104",
                    "ENSG00000128609",
                    "ENSG00000114405",
                    "ENSG00000149294",
                    "ENSG00000244879",
                    "ENSG00000105220",
                    "ENSG00000166974",
                    "ENSG00000104643",
                    "ENSG00000164330",
                    "ENSG00000007168",
                    "ENSG00000151150",
                    "ENSG00000100320",
                    "ENSG00000111775",
                    "ENSG00000123636",
                    "ENSG00000162852",
                    "ENSG00000139146",
                    "ENSG00000166562",
                    "ENSG00000132824",
                    "ENSG00000131437",
                    "ENSG00000102003",
                    "ENSG00000127616",
                    "ENSG00000196781",
                    "ENSG00000082397",
                    "ENSG00000137942",
                    "ENSG00000080824",
                    "ENSG00000107643",
                    "ENSG00000112695",
                    "ENSG00000160271",
                    "ENSG00000162188",
                    "ENSG00000162374",
                    "ENSG00000120694",
                    "ENSG00000220205",
                    "ENSG00000196586",
                    "ENSG00000104427",
                    "ENSG00000184545",
                    "ENSG00000131149",
                    "ENSG00000136238",
                    "ENSG00000198513",
                    "ENSG00000143933",
                    "ENSG00000169213",
                    "ENSG00000113141",
                    "ENSG00000165629",
                    "ENSG00000166503",
                    "ENSG00000103154",
                    "ENSG00000104435",
                    "ENSG00000126214",
                    "ENSG00000167815",
                    "ENSG00000097033",
                    "ENSG00000119314",
                    "ENSG00000156587",
                    "ENSG00000124194",
                    "ENSG00000170759",
                    "ENSG00000131037",
                    "ENSG00000177410",
                    "ENSG00000145247",
                    "ENSG00000178127",
                    "ENSG00000165943",
                    "ENSG00000104381",
                    "ENSG00000186472",
                    "ENSG00000139970",
                    "ENSG00000127914",
                    "ENSG00000090097",
                    "ENSG00000182718",
                    "ENSG00000143153",
                    "ENSG00000168824",
                    "ENSG00000133169",
                    "ENSG00000175582",
                    "ENSG00000169439",
                    "ENSG00000018408",
                    "ENSG00000149926"
                ],
                "expression_sum": {
                    "class1": 6453.0,
                    "class2": 3224.0
                },
                "ens_id": "ENSG00000148516",
                "uniprot_ids": [
                    "P37275",
                    "F6TDF5",
                    "H0YND9",
                    "F6U0D0",
                    "H0YMD3"
                ]
            },
            "ENSG00000143799": {
                "id": "ENSG00000143799",
                "type": "TF",
                "symbol": "PARP1",
                "source": [
                    "ENSG00000124766",
                    "ENSG00000181449",
                    "ENSG00000167081",
                    "ENSG00000164330"
                ],
                "target": [
                    "ENSG00000169783",
                    "ENSG00000139116",
                    "ENSG00000087085",
                    "ENSG00000131437",
                    "ENSG00000185664",
                    "ENSG00000166803",
                    "ENSG00000099250",
                    "ENSG00000018189",
                    "ENSG00000166257",
                    "ENSG00000186106",
                    "ENSG00000197959",
                    "ENSG00000105409",
                    "ENSG00000104427",
                    "ENSG00000221866",
                    "ENSG00000155980",
                    "ENSG00000151320",
                    "ENSG00000197694",
                    "ENSG00000129682",
                    "ENSG00000198513",
                    "ENSG00000186469",
                    "ENSG00000144642",
                    "ENSG00000143702",
                    "ENSG00000126214",
                    "ENSG00000152503",
                    "ENSG00000107130",
                    "ENSG00000173267",
                    "ENSG00000177410",
                    "ENSG00000111728",
                    "ENSG00000149294",
                    "ENSG00000131016",
                    "ENSG00000170091",
                    "ENSG00000116729",
                    "ENSG00000139970",
                    "ENSG00000090097",
                    "ENSG00000156298",
                    "ENSG00000123240",
                    "ENSG00000145362",
                    "ENSG00000135406",
                    "ENSG00000166974",
                    "ENSG00000150722",
                    "ENSG00000184144",
                    "ENSG00000164330",
                    "ENSG00000151150",
                    "ENSG00000033627",
                    "ENSG00000018236",
                    "ENSG00000185650",
                    "ENSG00000149926"
                ],
                "expression_sum": {
                    "class1": 10029.0,
                    "class2": 9650.0
                },
                "ens_id": "ENSG00000143799",
                "uniprot_ids": [
                    "P09874",
                    "A0A7I2V625",
                    "A0A7I2V5E9",
                    "A0A7I2V384",
                    "A0A7I2V3E1"
                ]
            },
            "ENSG00000128272": {
                "id": "ENSG00000128272",
                "type": "TF",
                "symbol": "ATF4",
                "source": [],
                "target": [
                    "ENSG00000166922",
                    "ENSG00000172020",
                    "ENSG00000079102",
                    "ENSG00000182985",
                    "ENSG00000005810",
                    "ENSG00000113161",
                    "ENSG00000154277",
                    "ENSG00000182093",
                    "ENSG00000082397",
                    "ENSG00000086300",
                    "ENSG00000166257",
                    "ENSG00000060982",
                    "ENSG00000130294",
                    "ENSG00000111860",
                    "ENSG00000131711",
                    "ENSG00000151320",
                    "ENSG00000198513",
                    "ENSG00000168280",
                    "ENSG00000213760",
                    "ENSG00000137285",
                    "ENSG00000156515",
                    "ENSG00000152503",
                    "ENSG00000173267",
                    "ENSG00000138413",
                    "ENSG00000176046",
                    "ENSG00000087258",
                    "ENSG00000090097",
                    "ENSG00000145362",
                    "ENSG00000108821",
                    "ENSG00000115310",
                    "ENSG00000118263",
                    "ENSG00000081181",
                    "ENSG00000164330",
                    "ENSG00000033627",
                    "ENSG00000185650",
                    "ENSG00000149926"
                ],
                "expression_sum": {
                    "class1": 9228.0,
                    "class2": 8636.0
                },
                "ens_id": "ENSG00000128272",
                "uniprot_ids": [
                    "P18848",
                    "B4DJD4",
                    "A0A6Q8PGY1",
                    "A0A6Q8PHB3",
                    "A0A6Q8PFH6",
                    "A0A6Q8PG17",
                    "A0A7P0Z4J5",
                    "A0A6Q8PF56"
                ]
            },
            "ENSG00000163848": {
                "id": "ENSG00000163848",
                "type": "TF",
                "symbol": "ZNF148",
                "source": [
                    "ENSG00000016082",
                    "ENSG00000181449",
                    "ENSG00000164330"
                ],
                "target": [
                    "ENSG00000181449",
                    "ENSG00000143702",
                    "ENSG00000060982",
                    "ENSG00000115461",
                    "ENSG00000176788",
                    "ENSG00000149557",
                    "ENSG00000135406",
                    "ENSG00000164330",
                    "ENSG00000198910"
                ],
                "expression_sum": {
                    "class1": 5593.0,
                    "class2": 5287.0
                },
                "ens_id": "ENSG00000163848",
                "uniprot_ids": [
                    "Q9UQR1",
                    "G5E9X2",
                    "C9K0U4",
                    "C9J6Y6",
                    "C9JRX0"
                ]
            },
            "ENSG00000165512": {
                "id": "ENSG00000165512",
                "type": "TF",
                "symbol": "ZNF22",
                "source": [
                    "ENSG00000164330"
                ],
                "target": [
                    "ENSG00000084628",
                    "ENSG00000131016",
                    "ENSG00000164330",
                    "ENSG00000161082"
                ],
                "expression_sum": {
                    "class1": 9759.0,
                    "class2": 8278.0
                },
                "ens_id": "ENSG00000165512",
                "uniprot_ids": [
                    "P17026"
                ]
            },
            "ENSG00000067955": {
                "id": "ENSG00000067955",
                "type": "TF",
                "symbol": "CBFB",
                "source": [
                    "ENSG00000113580",
                    "ENSG00000164330"
                ],
                "target": [
                    "ENSG00000067064",
                    "ENSG00000138698",
                    "ENSG00000197102",
                    "ENSG00000136717",
                    "ENSG00000173272",
                    "ENSG00000239672",
                    "ENSG00000109670",
                    "ENSG00000157514",
                    "ENSG00000106615",
                    "ENSG00000005810",
                    "ENSG00000167658",
                    "ENSG00000109084",
                    "ENSG00000196262",
                    "ENSG00000137955",
                    "ENSG00000166226",
                    "ENSG00000196230",
                    "ENSG00000178980",
                    "ENSG00000136942",
                    "ENSG00000089157",
                    "ENSG00000101132",
                    "ENSG00000075618",
                    "ENSG00000197694",
                    "ENSG00000123416",
                    "ENSG00000198961",
                    "ENSG00000047849",
                    "ENSG00000108175",
                    "ENSG00000134982",
                    "ENSG00000144713",
                    "ENSG00000143702",
                    "ENSG00000144218",
                    "ENSG00000265972",
                    "ENSG00000065978",
                    "ENSG00000231500",
                    "ENSG00000138413",
                    "ENSG00000140374",
                    "ENSG00000169710",
                    "ENSG00000118200",
                    "ENSG00000189159",
                    "ENSG00000006125",
                    "ENSG00000117632",
                    "ENSG00000065911",
                    "ENSG00000168275",
                    "ENSG00000034510",
                    "ENSG00000143198",
                    "ENSG00000113580",
                    "ENSG00000115310",
                    "ENSG00000118263",
                    "ENSG00000164330",
                    "ENSG00000033627",
                    "ENSG00000164924",
                    "ENSG00000189043",
                    "ENSG00000108010"
                ],
                "expression_sum": {
                    "class1": 3646.0,
                    "class2": 1787.0
                },
                "ens_id": "ENSG00000067955",
                "uniprot_ids": [
                    "Q13951",
                    "H3BSC0",
                    "J3KS23",
                    "A0A494C0A9",
                    "J3KTD8",
                    "J3KRT0"
                ]
            },
            "ENSG00000162599": {
                "id": "ENSG00000162599",
                "type": "TF",
                "symbol": "NFIA",
                "source": [
                    "ENSG00000175745",
                    "ENSG00000181449"
                ],
                "target": [
                    "ENSG00000134138",
                    "ENSG00000134245",
                    "ENSG00000122545",
                    "ENSG00000175745",
                    "ENSG00000215386",
                    "ENSG00000120885",
                    "ENSG00000134308"
                ],
                "expression_sum": {
                    "class1": 1610.0,
                    "class2": 14292.0
                },
                "ens_id": "ENSG00000162599",
                "uniprot_ids": [
                    "Q12857",
                    "A0A590UJU2",
                    "A0A590UK38",
                    "B1AKN8",
                    "A0A590UJ67",
                    "S4R3W2",
                    "S4R3K5",
                    "S4R3K4",
                    "S4R3W6",
                    "A0A590UJA3",
                    "A0A590UK09",
                    "A0A590UJ82",
                    "B1AKN6",
                    "S4R308",
                    "B1AKN5",
                    "B1AKN7",
                    "A0A590UK73"
                ]
            },
            "ENSG00000168610": {
                "id": "ENSG00000168610",
                "type": "TF",
                "symbol": "STAT3",
                "source": [],
                "target": [
                    "ENSG00000128564",
                    "ENSG00000122012",
                    "ENSG00000197959",
                    "ENSG00000265972",
                    "ENSG00000154556",
                    "ENSG00000183742",
                    "ENSG00000107438",
                    "ENSG00000135406",
                    "ENSG00000132698",
                    "ENSG00000010278",
                    "ENSG00000134138",
                    "ENSG00000221866",
                    "ENSG00000139970",
                    "ENSG00000089356",
                    "ENSG00000181885",
                    "ENSG00000118816"
                ],
                "expression_sum": {
                    "class1": 5863.0,
                    "class2": 8873.0
                },
                "ens_id": "ENSG00000168610",
                "uniprot_ids": [
                    "P40763",
                    "A0A7I2V5N9",
                    "A0A7I2V2T1",
                    "A0A7I2V4C8",
                    "A0A7I2V4R3",
                    "A0A7I2V395",
                    "A0A7I2V2G1",
                    "A0A7I2V552",
                    "G8JLH9",
                    "A0A7I2YQD2",
                    "A0A7I2V3V0",
                    "A0A7I2YQR5",
                    "A0A7I2V444",
                    "A0A7I2V4F6",
                    "A0A7I2YQI1",
                    "A0A7I2V4R2"
                ]
            },
            "ENSG00000114861": {
                "id": "ENSG00000114861",
                "type": "TF",
                "symbol": "FOXP1",
                "source": [
                    "ENSG00000119547",
                    "ENSG00000134138"
                ],
                "target": [
                    "ENSG00000078668",
                    "ENSG00000232119",
                    "ENSG00000107223",
                    "ENSG00000162704",
                    "ENSG00000196531",
                    "ENSG00000163634",
                    "ENSG00000111716",
                    "ENSG00000082153",
                    "ENSG00000114784",
                    "ENSG00000184752",
                    "ENSG00000169567",
                    "ENSG00000129355",
                    "ENSG00000242485",
                    "ENSG00000140941",
                    "ENSG00000134248",
                    "ENSG00000131143",
                    "ENSG00000168653",
                    "ENSG00000102144",
                    "ENSG00000076554",
                    "ENSG00000130770",
                    "ENSG00000156467",
                    "ENSG00000131507",
                    "ENSG00000075415",
                    "ENSG00000184076",
                    "ENSG00000171314",
                    "ENSG00000154556",
                    "ENSG00000178741",
                    "ENSG00000057757",
                    "ENSG00000177889",
                    "ENSG00000173267",
                    "ENSG00000147684",
                    "ENSG00000166913",
                    "ENSG00000173812",
                    "ENSG00000152256",
                    "ENSG00000173915",
                    "ENSG00000065911",
                    "ENSG00000111229",
                    "ENSG00000118816",
                    "ENSG00000124570",
                    "ENSG00000041357",
                    "ENSG00000115310",
                    "ENSG00000163875",
                    "ENSG00000167136",
                    "ENSG00000149547",
                    "ENSG00000164924",
                    "ENSG00000096092",
                    "ENSG00000147403",
                    "ENSG00000172115",
                    "ENSG00000119547",
                    "ENSG00000101558",
                    "ENSG00000162545",
                    "ENSG00000167283",
                    "ENSG00000170035",
                    "ENSG00000137726",
                    "ENSG00000106615",
                    "ENSG00000099341",
                    "ENSG00000198886",
                    "ENSG00000230989",
                    "ENSG00000154518",
                    "ENSG00000111640",
                    "ENSG00000125868",
                    "ENSG00000104904",
                    "ENSG00000165637",
                    "ENSG00000125977",
                    "ENSG00000241685",
                    "ENSG00000173726",
                    "ENSG00000125971",
                    "ENSG00000089693",
                    "ENSG00000173674",
                    "ENSG00000161970",
                    "ENSG00000156976",
                    "ENSG00000128245",
                    "ENSG00000129625",
                    "ENSG00000089220",
                    "ENSG00000134138",
                    "ENSG00000101460",
                    "ENSG00000117758",
                    "ENSG00000134717",
                    "ENSG00000099624",
                    "ENSG00000114023",
                    "ENSG00000120265",
                    "ENSG00000067225",
                    "ENSG00000154723",
                    "ENSG00000113712",
                    "ENSG00000105438",
                    "ENSG00000138069",
                    "ENSG00000110321",
                    "ENSG00000109654",
                    "ENSG00000145495",
                    "ENSG00000254999",
                    "ENSG00000112335",
                    "ENSG00000133872",
                    "ENSG00000130558",
                    "ENSG00000150459",
                    "ENSG00000034713",
                    "ENSG00000152234",
                    "ENSG00000113387",
                    "ENSG00000095380",
                    "ENSG00000198668",
                    "ENSG00000108107",
                    "ENSG00000134308",
                    "ENSG00000156508",
                    "ENSG00000034510",
                    "ENSG00000134684",
                    "ENSG00000117410",
                    "ENSG00000145362",
                    "ENSG00000123144",
                    "ENSG00000204525",
                    "ENSG00000263001",
                    "ENSG00000145741",
                    "ENSG00000143256",
                    "ENSG00000149084",
                    "ENSG00000174437",
                    "ENSG00000166619",
                    "ENSG00000074800",
                    "ENSG00000197111",
                    "ENSG00000165502",
                    "ENSG00000123562",
                    "ENSG00000122034",
                    "ENSG00000051108",
                    "ENSG00000111669",
                    "ENSG00000163041",
                    "ENSG00000134825",
                    "ENSG00000009307",
                    "ENSG00000004779",
                    "ENSG00000150991",
                    "ENSG00000152795",
                    "ENSG00000169139",
                    "ENSG00000118363",
                    "ENSG00000142669",
                    "ENSG00000135940",
                    "ENSG00000175756",
                    "ENSG00000070831",
                    "ENSG00000134333",
                    "ENSG00000100568",
                    "ENSG00000122705",
                    "ENSG00000132432",
                    "ENSG00000102317",
                    "ENSG00000110700",
                    "ENSG00000175416",
                    "ENSG00000128609",
                    "ENSG00000160014",
                    "ENSG00000063177",
                    "ENSG00000105649",
                    "ENSG00000173660",
                    "ENSG00000108953",
                    "ENSG00000105220",
                    "ENSG00000186184",
                    "ENSG00000100804",
                    "ENSG00000155368",
                    "ENSG00000169564",
                    "ENSG00000114125",
                    "ENSG00000176871",
                    "ENSG00000240972",
                    "ENSG00000111775",
                    "ENSG00000105379",
                    "ENSG00000197102",
                    "ENSG00000135390",
                    "ENSG00000149806",
                    "ENSG00000132824",
                    "ENSG00000213585",
                    "ENSG00000131508",
                    "ENSG00000131437",
                    "ENSG00000167658",
                    "ENSG00000188612",
                    "ENSG00000180817",
                    "ENSG00000112695",
                    "ENSG00000116288",
                    "ENSG00000136810",
                    "ENSG00000174444",
                    "ENSG00000220205",
                    "ENSG00000127184",
                    "ENSG00000128524",
                    "ENSG00000165678",
                    "ENSG00000133318",
                    "ENSG00000265241",
                    "ENSG00000136238",
                    "ENSG00000143933",
                    "ENSG00000165629",
                    "ENSG00000132963",
                    "ENSG00000164096",
                    "ENSG00000087460",
                    "ENSG00000182670",
                    "ENSG00000171530",
                    "ENSG00000148450",
                    "ENSG00000123349",
                    "ENSG00000114115",
                    "ENSG00000178127",
                    "ENSG00000113558",
                    "ENSG00000140990",
                    "ENSG00000143153",
                    "ENSG00000143198",
                    "ENSG00000136628",
                    "ENSG00000014641",
                    "ENSG00000109180",
                    "ENSG00000123983",
                    "ENSG00000133639",
                    "ENSG00000126267",
                    "ENSG00000143742",
                    "ENSG00000100129",
                    "ENSG00000109472"
                ],
                "expression_sum": {
                    "class1": 7147.0,
                    "class2": 6582.0
                },
                "ens_id": "ENSG00000114861",
                "uniprot_ids": [
                    "Q9H334",
                    "G5E965",
                    "A0A3B3IRW5",
                    "A0A3B3ITZ4",
                    "A0A3B3IT66",
                    "A0A0B4J2F3",
                    "A0A3B3IRS5",
                    "A0A3B3IU08",
                    "A0A3B3IST6",
                    "A0A3B3IS87",
                    "A0A3B3IT14",
                    "A0A087WVT2",
                    "A0A3B3IT26",
                    "A0A3B3IRL7",
                    "A0A3B3ISQ6",
                    "A0A087X2G3",
                    "A0A3B3IST0",
                    "A0A3B3ISQ3",
                    "C9IYY1",
                    "A0A3B3IUB6",
                    "C9J0F0",
                    "C9J5T4",
                    "A0A3B3ISQ7",
                    "A0A3B3IRY1",
                    "A0A3B3ISA4",
                    "H0Y882",
                    "A0A3B3IU15"
                ]
            },
            "ENSG00000221869": {
                "id": "ENSG00000221869",
                "type": "TF",
                "symbol": "CEBPD",
                "source": [
                    "ENSG00000163435",
                    "ENSG00000160113",
                    "ENSG00000164683",
                    "ENSG00000130522",
                    "ENSG00000177606"
                ],
                "target": [
                    "ENSG00000206573",
                    "ENSG00000106261",
                    "ENSG00000115944",
                    "ENSG00000162704",
                    "ENSG00000196531",
                    "ENSG00000126005",
                    "ENSG00000171604",
                    "ENSG00000116030",
                    "ENSG00000099622",
                    "ENSG00000180011",
                    "ENSG00000129559",
                    "ENSG00000184752",
                    "ENSG00000169567",
                    "ENSG00000125534",
                    "ENSG00000177606",
                    "ENSG00000160113",
                    "ENSG00000174748",
                    "ENSG00000131143",
                    "ENSG00000168653",
                    "ENSG00000241837",
                    "ENSG00000156467",
                    "ENSG00000142864",
                    "ENSG00000161203",
                    "ENSG00000075415",
                    "ENSG00000166165",
                    "ENSG00000141753",
                    "ENSG00000181885",
                    "ENSG00000184076",
                    "ENSG00000171314",
                    "ENSG00000266472",
                    "ENSG00000150779",
                    "ENSG00000154556",
                    "ENSG00000178741",
                    "ENSG00000163466",
                    "ENSG00000198931",
                    "ENSG00000166913",
                    "ENSG00000173812",
                    "ENSG00000141279",
                    "ENSG00000099795",
                    "ENSG00000111229",
                    "ENSG00000151729",
                    "ENSG00000118816",
                    "ENSG00000088832",
                    "ENSG00000162585",
                    "ENSG00000051620",
                    "ENSG00000122545",
                    "ENSG00000149547",
                    "ENSG00000159335",
                    "ENSG00000164924",
                    "ENSG00000189043",
                    "ENSG00000150687",
                    "ENSG00000142192",
                    "ENSG00000277443",
                    "ENSG00000101558",
                    "ENSG00000162545",
                    "ENSG00000101040",
                    "ENSG00000128739",
                    "ENSG00000109475",
                    "ENSG00000154518",
                    "ENSG00000196262",
                    "ENSG00000111640",
                    "ENSG00000104904",
                    "ENSG00000165637",
                    "ENSG00000183978",
                    "ENSG00000241685",
                    "ENSG00000173726",
                    "ENSG00000091164",
                    "ENSG00000172757",
                    "ENSG00000125971",
                    "ENSG00000130332",
                    "ENSG00000116586",
                    "ENSG00000188290",
                    "ENSG00000156976",
                    "ENSG00000173674",
                    "ENSG00000147548",
                    "ENSG00000089220",
                    "ENSG00000134138",
                    "ENSG00000119335",
                    "ENSG00000115286",
                    "ENSG00000134717",
                    "ENSG00000119396",
                    "ENSG00000136156",
                    "ENSG00000177879",
                    "ENSG00000105640",
                    "ENSG00000099624",
                    "ENSG00000171159",
                    "ENSG00000067225",
                    "ENSG00000154723",
                    "ENSG00000166347",
                    "ENSG00000213719",
                    "ENSG00000113712",
                    "ENSG00000110321",
                    "ENSG00000233927",
                    "ENSG00000254999",
                    "ENSG00000004059",
                    "ENSG00000152234",
                    "ENSG00000137106",
                    "ENSG00000177556",
                    "ENSG00000034713",
                    "ENSG00000113387",
                    "ENSG00000100592",
                    "ENSG00000198668",
                    "ENSG00000134308",
                    "ENSG00000034510",
                    "ENSG00000123144",
                    "ENSG00000115091",
                    "ENSG00000149257",
                    "ENSG00000143256",
                    "ENSG00000149084",
                    "ENSG00000126934",
                    "ENSG00000154639",
                    "ENSG00000168118",
                    "ENSG00000106367",
                    "ENSG00000197111",
                    "ENSG00000214253",
                    "ENSG00000101182",
                    "ENSG00000162496",
                    "ENSG00000142676",
                    "ENSG00000171421",
                    "ENSG00000129515",
                    "ENSG00000100387",
                    "ENSG00000140319",
                    "ENSG00000206503",
                    "ENSG00000122034",
                    "ENSG00000187514",
                    "ENSG00000111669",
                    "ENSG00000163041",
                    "ENSG00000118680",
                    "ENSG00000243317",
                    "ENSG00000175826",
                    "ENSG00000184007",
                    "ENSG00000009307",
                    "ENSG00000178980",
                    "ENSG00000107262",
                    "ENSG00000004779",
                    "ENSG00000182117",
                    "ENSG00000124172",
                    "ENSG00000182196",
                    "ENSG00000169139",
                    "ENSG00000168214",
                    "ENSG00000118363",
                    "ENSG00000162191",
                    "ENSG00000100911",
                    "ENSG00000070831",
                    "ENSG00000162076",
                    "ENSG00000107438",
                    "ENSG00000122705",
                    "ENSG00000166482",
                    "ENSG00000110700",
                    "ENSG00000184009",
                    "ENSG00000075624",
                    "ENSG00000095059",
                    "ENSG00000173660",
                    "ENSG00000103187",
                    "ENSG00000108953",
                    "ENSG00000186184",
                    "ENSG00000100804",
                    "ENSG00000155368",
                    "ENSG00000272047",
                    "ENSG00000114125",
                    "ENSG00000185787",
                    "ENSG00000163931",
                    "ENSG00000164713",
                    "ENSG00000130638",
                    "ENSG00000240972",
                    "ENSG00000111775",
                    "ENSG00000105379",
                    "ENSG00000123636",
                    "ENSG00000135390",
                    "ENSG00000149806",
                    "ENSG00000239672",
                    "ENSG00000213585",
                    "ENSG00000167658",
                    "ENSG00000005022",
                    "ENSG00000188612",
                    "ENSG00000128654",
                    "ENSG00000112715",
                    "ENSG00000180817",
                    "ENSG00000112695",
                    "ENSG00000116288",
                    "ENSG00000090266",
                    "ENSG00000158710",
                    "ENSG00000205542",
                    "ENSG00000160948",
                    "ENSG00000136810",
                    "ENSG00000174444",
                    "ENSG00000169764",
                    "ENSG00000168028",
                    "ENSG00000127184",
                    "ENSG00000136238",
                    "ENSG00000104341",
                    "ENSG00000130522",
                    "ENSG00000224032",
                    "ENSG00000167815",
                    "ENSG00000122873",
                    "ENSG00000126457",
                    "ENSG00000176340",
                    "ENSG00000167996",
                    "ENSG00000132963",
                    "ENSG00000065978",
                    "ENSG00000087460",
                    "ENSG00000170759",
                    "ENSG00000171530",
                    "ENSG00000148450",
                    "ENSG00000177410",
                    "ENSG00000178127",
                    "ENSG00000113558",
                    "ENSG00000101745",
                    "ENSG00000140990",
                    "ENSG00000182718",
                    "ENSG00000105518",
                    "ENSG00000163435",
                    "ENSG00000133639",
                    "ENSG00000126267",
                    "ENSG00000100129",
                    "ENSG00000170017",
                    "ENSG00000092841",
                    "ENSG00000145494"
                ],
                "expression_sum": {
                    "class1": 1701.0,
                    "class2": 15658.0
                },
                "ens_id": "ENSG00000221869",
                "uniprot_ids": [
                    "P49716"
                ]
            },
            "ENSG00000157557": {
                "id": "ENSG00000157557",
                "type": "TF",
                "symbol": "ETS2",
                "source": [
                    "ENSG00000160113",
                    "ENSG00000119950",
                    "ENSG00000168214",
                    "ENSG00000119547",
                    "ENSG00000177606"
                ],
                "target": [
                    "ENSG00000206573",
                    "ENSG00000181061",
                    "ENSG00000214530",
                    "ENSG00000165655",
                    "ENSG00000099622",
                    "ENSG00000111640",
                    "ENSG00000243317",
                    "ENSG00000067836",
                    "ENSG00000120071",
                    "ENSG00000110955",
                    "ENSG00000156976",
                    "ENSG00000102144",
                    "ENSG00000169432",
                    "ENSG00000136710",
                    "ENSG00000104112",
                    "ENSG00000168214",
                    "ENSG00000115738",
                    "ENSG00000143933",
                    "ENSG00000114023",
                    "ENSG00000132424",
                    "ENSG00000186352",
                    "ENSG00000130338",
                    "ENSG00000067225",
                    "ENSG00000137500",
                    "ENSG00000160714",
                    "ENSG00000122873",
                    "ENSG00000176340",
                    "ENSG00000178741",
                    "ENSG00000140416",
                    "ENSG00000055609",
                    "ENSG00000181722",
                    "ENSG00000164830",
                    "ENSG00000172115",
                    "ENSG00000181315",
                    "ENSG00000151729",
                    "ENSG00000143158",
                    "ENSG00000118816",
                    "ENSG00000272398",
                    "ENSG00000186472",
                    "ENSG00000101745",
                    "ENSG00000105649",
                    "ENSG00000163875",
                    "ENSG00000100320",
                    "ENSG00000119547"
                ],
                "expression_sum": {
                    "class1": 2470.0,
                    "class2": 7721.0
                },
                "ens_id": "ENSG00000157557",
                "uniprot_ids": [
                    "P15036",
                    "A0A590UJP9",
                    "C9JAG2",
                    "A0A590UJR2",
                    "A0A0C4DG47"
                ]
            },
            "ENSG00000181722": {
                "id": "ENSG00000181722",
                "type": "TF",
                "symbol": "ZBTB20",
                "source": [
                    "ENSG00000157557",
                    "ENSG00000113580",
                    "ENSG00000164330",
                    "ENSG00000100219",
                    "ENSG00000139352",
                    "ENSG00000160113"
                ],
                "target": [
                    "ENSG00000146278",
                    "ENSG00000181061",
                    "ENSG00000143321",
                    "ENSG00000065613",
                    "ENSG00000145675",
                    "ENSG00000257727",
                    "ENSG00000152558",
                    "ENSG00000114902",
                    "ENSG00000178467",
                    "ENSG00000099622",
                    "ENSG00000180011",
                    "ENSG00000164093",
                    "ENSG00000104964",
                    "ENSG00000163399",
                    "ENSG00000167397",
                    "ENSG00000165637",
                    "ENSG00000179981",
                    "ENSG00000244187",
                    "ENSG00000102804",
                    "ENSG00000160113",
                    "ENSG00000108176",
                    "ENSG00000077782",
                    "ENSG00000129625",
                    "ENSG00000134248",
                    "ENSG00000113732",
                    "ENSG00000220205",
                    "ENSG00000116209",
                    "ENSG00000117155",
                    "ENSG00000109472",
                    "ENSG00000132424",
                    "ENSG00000184076",
                    "ENSG00000135968",
                    "ENSG00000198125",
                    "ENSG00000184164",
                    "ENSG00000068697",
                    "ENSG00000198168",
                    "ENSG00000134333",
                    "ENSG00000140939",
                    "ENSG00000143013",
                    "ENSG00000125851",
                    "ENSG00000054356",
                    "ENSG00000102007",
                    "ENSG00000087460",
                    "ENSG00000205981",
                    "ENSG00000205922",
                    "ENSG00000143774",
                    "ENSG00000152128",
                    "ENSG00000143870",
                    "ENSG00000138085",
                    "ENSG00000120742",
                    "ENSG00000197261",
                    "ENSG00000172663",
                    "ENSG00000133639",
                    "ENSG00000185624",
                    "ENSG00000089063",
                    "ENSG00000119888",
                    "ENSG00000113580",
                    "ENSG00000166228",
                    "ENSG00000143545",
                    "ENSG00000124155",
                    "ENSG00000064601",
                    "ENSG00000143543",
                    "ENSG00000151552",
                    "ENSG00000185650",
                    "ENSG00000114115",
                    "ENSG00000115828",
                    "ENSG00000225783",
                    "ENSG00000104870"
                ],
                "expression_sum": {
                    "class1": 10960.0,
                    "class2": 13009.0
                },
                "ens_id": "ENSG00000181722",
                "uniprot_ids": [
                    "Q9HC78",
                    "C9JUH0",
                    "C9JCX0"
                ]
            },
            "ENSG00000177606": {
                "id": "ENSG00000177606",
                "type": "TF",
                "symbol": "JUN",
                "source": [
                    "ENSG00000100219",
                    "ENSG00000221869",
                    "ENSG00000139352"
                ],
                "target": [
                    "ENSG00000176887",
                    "ENSG00000198840",
                    "ENSG00000132698",
                    "ENSG00000185664",
                    "ENSG00000164093",
                    "ENSG00000122756",
                    "ENSG00000165973",
                    "ENSG00000112773",
                    "ENSG00000169435",
                    "ENSG00000115461",
                    "ENSG00000148834",
                    "ENSG00000010278",
                    "ENSG00000104870",
                    "ENSG00000182481",
                    "ENSG00000189143",
                    "ENSG00000166145",
                    "ENSG00000198125",
                    "ENSG00000181885",
                    "ENSG00000132386",
                    "ENSG00000166598",
                    "ENSG00000117691",
                    "ENSG00000135535",
                    "ENSG00000164305",
                    "ENSG00000107438",
                    "ENSG00000167642",
                    "ENSG00000161249",
                    "ENSG00000187244",
                    "ENSG00000198938",
                    "ENSG00000198899",
                    "ENSG00000167641",
                    "ENSG00000015592",
                    "ENSG00000143870",
                    "ENSG00000135549",
                    "ENSG00000213937",
                    "ENSG00000157557",
                    "ENSG00000119888",
                    "ENSG00000100219",
                    "ENSG00000221869",
                    "ENSG00000185650",
                    "ENSG00000108679",
                    "ENSG00000172915",
                    "ENSG00000179218"
                ],
                "expression_sum": {
                    "class1": 17839.0,
                    "class2": 10689.0
                },
                "ens_id": "ENSG00000177606",
                "uniprot_ids": [
                    "P05412"
                ]
            },
            "ENSG00000124766": {
                "id": "ENSG00000124766",
                "type": "TF",
                "symbol": "SOX4",
                "source": [],
                "target": [
                    "ENSG00000132718",
                    "ENSG00000149557",
                    "ENSG00000148516",
                    "ENSG00000139116",
                    "ENSG00000132698",
                    "ENSG00000109339",
                    "ENSG00000074696",
                    "ENSG00000164929",
                    "ENSG00000185664",
                    "ENSG00000050165",
                    "ENSG00000182534",
                    "ENSG00000080824",
                    "ENSG00000154080",
                    "ENSG00000166226",
                    "ENSG00000166257",
                    "ENSG00000112531",
                    "ENSG00000130513",
                    "ENSG00000147889",
                    "ENSG00000171055",
                    "ENSG00000172183",
                    "ENSG00000104112",
                    "ENSG00000090273",
                    "ENSG00000182481",
                    "ENSG00000123416",
                    "ENSG00000108106",
                    "ENSG00000181885",
                    "ENSG00000113141",
                    "ENSG00000138758",
                    "ENSG00000092964",
                    "ENSG00000166963",
                    "ENSG00000172159",
                    "ENSG00000105258",
                    "ENSG00000173267",
                    "ENSG00000152082",
                    "ENSG00000175792",
                    "ENSG00000120885",
                    "ENSG00000198668",
                    "ENSG00000054523",
                    "ENSG00000146425",
                    "ENSG00000188042",
                    "ENSG00000163435",
                    "ENSG00000143198",
                    "ENSG00000135406",
                    "ENSG00000119640",
                    "ENSG00000122545",
                    "ENSG00000143799",
                    "ENSG00000141404",
                    "ENSG00000169862",
                    "ENSG00000205726",
                    "ENSG00000185650",
                    "ENSG00000163931",
                    "ENSG00000140905"
                ],
                "expression_sum": {
                    "class1": 103965.0,
                    "class2": 44155.0
                },
                "ens_id": "ENSG00000124766",
                "uniprot_ids": [
                    "Q06945"
                ]
            },
            "ENSG00000116044": {
                "id": "ENSG00000116044",
                "type": "TF",
                "symbol": "NFE2L2",
                "source": [
                    "ENSG00000164683",
                    "ENSG00000163435"
                ],
                "target": [
                    "ENSG00000130066",
                    "ENSG00000182718",
                    "ENSG00000163435",
                    "ENSG00000143153",
                    "ENSG00000109062",
                    "ENSG00000169764",
                    "ENSG00000131981",
                    "ENSG00000164093",
                    "ENSG00000118816",
                    "ENSG00000135046"
                ],
                "expression_sum": {
                    "class1": 4576.0,
                    "class2": 10732.0
                },
                "ens_id": "ENSG00000116044",
                "uniprot_ids": [
                    "Q16236",
                    "K7ER33",
                    "K7EIW5",
                    "C9J1A8",
                    "C9J3K4",
                    "C9JBM0",
                    "H7C498",
                    "C9JFL6",
                    "C9J0Y1"
                ]
            },
            "ENSG00000128610": {
                "id": "ENSG00000128610",
                "type": "TF",
                "symbol": "FEZF1",
                "source": [
                    "ENSG00000181449"
                ],
                "target": [
                    "ENSG00000181449",
                    "ENSG00000114353",
                    "ENSG00000043355",
                    "ENSG00000173404",
                    "ENSG00000150687",
                    "ENSG00000163931",
                    "ENSG00000125462",
                    "ENSG00000245526"
                ],
                "expression_sum": {
                    "class1": 1699.0,
                    "class2": 607.0
                },
                "ens_id": "ENSG00000128610",
                "uniprot_ids": [
                    "A0PJY2",
                    "C9J0C5"
                ]
            },
            "ENSG00000130522": {
                "id": "ENSG00000130522",
                "type": "TF",
                "symbol": "JUND",
                "source": [
                    "ENSG00000181449",
                    "ENSG00000221869"
                ],
                "target": [
                    "ENSG00000181449",
                    "ENSG00000162496",
                    "ENSG00000176887",
                    "ENSG00000172020",
                    "ENSG00000087085",
                    "ENSG00000185664",
                    "ENSG00000107105",
                    "ENSG00000213190",
                    "ENSG00000162188",
                    "ENSG00000115461",
                    "ENSG00000128564",
                    "ENSG00000130303",
                    "ENSG00000182481",
                    "ENSG00000108106",
                    "ENSG00000182916",
                    "ENSG00000125354",
                    "ENSG00000104435",
                    "ENSG00000164305",
                    "ENSG00000108515",
                    "ENSG00000124194",
                    "ENSG00000015592",
                    "ENSG00000135046",
                    "ENSG00000090097",
                    "ENSG00000177675",
                    "ENSG00000221869",
                    "ENSG00000185650",
                    "ENSG00000149926",
                    "ENSG00000104870"
                ],
                "expression_sum": {
                    "class1": 87153.0,
                    "class2": 33345.0
                },
                "ens_id": "ENSG00000130522",
                "uniprot_ids": [
                    "P17535",
                    "U3KPR5"
                ]
            },
            "ENSG00000119547": {
                "id": "ENSG00000119547",
                "type": "TF",
                "symbol": "ONECUT2",
                "source": [
                    "ENSG00000185551",
                    "ENSG00000157557",
                    "ENSG00000134138",
                    "ENSG00000167081",
                    "ENSG00000113580",
                    "ENSG00000148516",
                    "ENSG00000103495",
                    "ENSG00000108055",
                    "ENSG00000114861",
                    "ENSG00000139352"
                ],
                "target": [
                    "ENSG00000090621",
                    "ENSG00000135486",
                    "ENSG00000162373",
                    "ENSG00000058668",
                    "ENSG00000173272",
                    "ENSG00000162704",
                    "ENSG00000135334",
                    "ENSG00000196531",
                    "ENSG00000156475",
                    "ENSG00000111716",
                    "ENSG00000175745",
                    "ENSG00000164093",
                    "ENSG00000184752",
                    "ENSG00000196230",
                    "ENSG00000117395",
                    "ENSG00000182004",
                    "ENSG00000179915",
                    "ENSG00000065135",
                    "ENSG00000104408",
                    "ENSG00000103495",
                    "ENSG00000171055",
                    "ENSG00000149231",
                    "ENSG00000147604",
                    "ENSG00000182481",
                    "ENSG00000123416",
                    "ENSG00000142864",
                    "ENSG00000100823",
                    "ENSG00000198125",
                    "ENSG00000108175",
                    "ENSG00000140988",
                    "ENSG00000197728",
                    "ENSG00000102466",
                    "ENSG00000028277",
                    "ENSG00000077380",
                    "ENSG00000167642",
                    "ENSG00000163466",
                    "ENSG00000096746",
                    "ENSG00000138413",
                    "ENSG00000134779",
                    "ENSG00000071054",
                    "ENSG00000109536",
                    "ENSG00000111728",
                    "ENSG00000069275",
                    "ENSG00000118816",
                    "ENSG00000041357",
                    "ENSG00000119888",
                    "ENSG00000122545",
                    "ENSG00000121766",
                    "ENSG00000153234",
                    "ENSG00000189043",
                    "ENSG00000123352",
                    "ENSG00000180879",
                    "ENSG00000277443",
                    "ENSG00000183654",
                    "ENSG00000233429",
                    "ENSG00000171858",
                    "ENSG00000115350",
                    "ENSG00000138650",
                    "ENSG00000148516",
                    "ENSG00000187109",
                    "ENSG00000106615",
                    "ENSG00000156482",
                    "ENSG00000196262",
                    "ENSG00000091164",
                    "ENSG00000182628",
                    "ENSG00000167460",
                    "ENSG00000172757",
                    "ENSG00000171951",
                    "ENSG00000060982",
                    "ENSG00000188290",
                    "ENSG00000100632",
                    "ENSG00000161970",
                    "ENSG00000141447",
                    "ENSG00000134138",
                    "ENSG00000164114",
                    "ENSG00000119335",
                    "ENSG00000168894",
                    "ENSG00000108828",
                    "ENSG00000176840",
                    "ENSG00000077264",
                    "ENSG00000177879",
                    "ENSG00000215386",
                    "ENSG00000101132",
                    "ENSG00000188243",
                    "ENSG00000125691",
                    "ENSG00000143549",
                    "ENSG00000138758",
                    "ENSG00000133083",
                    "ENSG00000143013",
                    "ENSG00000233927",
                    "ENSG00000117139",
                    "ENSG00000137106",
                    "ENSG00000157216",
                    "ENSG00000168785",
                    "ENSG00000108107",
                    "ENSG00000138326",
                    "ENSG00000134308",
                    "ENSG00000122958",
                    "ENSG00000022267",
                    "ENSG00000034510",
                    "ENSG00000133703",
                    "ENSG00000137575",
                    "ENSG00000135549",
                    "ENSG00000154380",
                    "ENSG00000151789",
                    "ENSG00000064601",
                    "ENSG00000104695",
                    "ENSG00000113580",
                    "ENSG00000075142",
                    "ENSG00000106028",
                    "ENSG00000175161",
                    "ENSG00000115091",
                    "ENSG00000115053",
                    "ENSG00000117152",
                    "ENSG00000127022",
                    "ENSG00000213741",
                    "ENSG00000074800",
                    "ENSG00000145592",
                    "ENSG00000069849",
                    "ENSG00000133026",
                    "ENSG00000147649",
                    "ENSG00000123562",
                    "ENSG00000163041",
                    "ENSG00000130255",
                    "ENSG00000184007",
                    "ENSG00000143995",
                    "ENSG00000174498",
                    "ENSG00000185551",
                    "ENSG00000197747",
                    "ENSG00000105223",
                    "ENSG00000164687",
                    "ENSG00000106355",
                    "ENSG00000057608",
                    "ENSG00000174469",
                    "ENSG00000065548",
                    "ENSG00000130396",
                    "ENSG00000118363",
                    "ENSG00000108106",
                    "ENSG00000005108",
                    "ENSG00000117691",
                    "ENSG00000112972",
                    "ENSG00000006625",
                    "ENSG00000116251",
                    "ENSG00000071794",
                    "ENSG00000122705",
                    "ENSG00000110700",
                    "ENSG00000114861",
                    "ENSG00000196776",
                    "ENSG00000125944",
                    "ENSG00000054523",
                    "ENSG00000063177",
                    "ENSG00000100380",
                    "ENSG00000115758",
                    "ENSG00000113328",
                    "ENSG00000129167",
                    "ENSG00000155368",
                    "ENSG00000117906",
                    "ENSG00000105185",
                    "ENSG00000105372",
                    "ENSG00000157557",
                    "ENSG00000094916",
                    "ENSG00000205726",
                    "ENSG00000128050",
                    "ENSG00000043355",
                    "ENSG00000163931",
                    "ENSG00000164713",
                    "ENSG00000176973",
                    "ENSG00000105379",
                    "ENSG00000186063",
                    "ENSG00000107863",
                    "ENSG00000136930",
                    "ENSG00000164985",
                    "ENSG00000239672",
                    "ENSG00000144724",
                    "ENSG00000146386",
                    "ENSG00000213585",
                    "ENSG00000131508",
                    "ENSG00000127955",
                    "ENSG00000180957",
                    "ENSG00000131469",
                    "ENSG00000188846",
                    "ENSG00000137942",
                    "ENSG00000080824",
                    "ENSG00000166226",
                    "ENSG00000108468",
                    "ENSG00000181163",
                    "ENSG00000136810",
                    "ENSG00000142937",
                    "ENSG00000078018",
                    "ENSG00000196465",
                    "ENSG00000112531",
                    "ENSG00000174444",
                    "ENSG00000104549",
                    "ENSG00000136238",
                    "ENSG00000117155",
                    "ENSG00000103363",
                    "ENSG00000079459",
                    "ENSG00000138430",
                    "ENSG00000047849",
                    "ENSG00000168916",
                    "ENSG00000132437",
                    "ENSG00000167996",
                    "ENSG00000065978",
                    "ENSG00000170144",
                    "ENSG00000152082",
                    "ENSG00000100664",
                    "ENSG00000189159",
                    "ENSG00000242265",
                    "ENSG00000143198",
                    "ENSG00000122565",
                    "ENSG00000054148",
                    "ENSG00000134824",
                    "ENSG00000165119"
                ],
                "expression_sum": {
                    "class1": 33288.0,
                    "class2": 3372.0
                },
                "ens_id": "ENSG00000119547",
                "uniprot_ids": [
                    "O95948"
                ]
            },
            "ENSG00000277443": {
                "id": "ENSG00000277443",
                "type": "Gene",
                "symbol": "MARCKS",
                "source": [
                    "ENSG00000181449",
                    "ENSG00000109132",
                    "ENSG00000167081",
                    "ENSG00000108001",
                    "ENSG00000221869",
                    "ENSG00000196591",
                    "ENSG00000119547"
                ],
                "target": [],
                "expression_sum": {
                    "class1": 82788.0,
                    "class2": 78453.0
                },
                "ens_id": "ENSG00000277443",
                "uniprot_ids": [
                    "P29966"
                ]
            },
            "ENSG00000175745": {
                "id": "ENSG00000175745",
                "type": "TF",
                "symbol": "NR2F1",
                "source": [
                    "ENSG00000162599",
                    "ENSG00000113580",
                    "ENSG00000122691",
                    "ENSG00000164330",
                    "ENSG00000108001",
                    "ENSG00000184486",
                    "ENSG00000119547",
                    "ENSG00000143995"
                ],
                "target": [
                    "ENSG00000147862",
                    "ENSG00000140937",
                    "ENSG00000109099",
                    "ENSG00000144366",
                    "ENSG00000185664",
                    "ENSG00000198910",
                    "ENSG00000143995",
                    "ENSG00000084628",
                    "ENSG00000164199",
                    "ENSG00000105810",
                    "ENSG00000070961",
                    "ENSG00000134138",
                    "ENSG00000110492",
                    "ENSG00000197043",
                    "ENSG00000185650",
                    "ENSG00000130005",
                    "ENSG00000104112",
                    "ENSG00000133302",
                    "ENSG00000122884",
                    "ENSG00000185201",
                    "ENSG00000162599",
                    "ENSG00000196562",
                    "ENSG00000181744",
                    "ENSG00000132437",
                    "ENSG00000213420",
                    "ENSG00000187957",
                    "ENSG00000026025",
                    "ENSG00000131037",
                    "ENSG00000149294",
                    "ENSG00000147655",
                    "ENSG00000123240",
                    "ENSG00000185630",
                    "ENSG00000196141",
                    "ENSG00000113580",
                    "ENSG00000159409",
                    "ENSG00000123989",
                    "ENSG00000047457",
                    "ENSG00000018408",
                    "ENSG00000164056",
                    "ENSG00000129244"
                ],
                "expression_sum": {
                    "class1": 32875.0,
                    "class2": 29213.0
                },
                "ens_id": "ENSG00000175745",
                "uniprot_ids": [
                    "P10589",
                    "F1DAL7",
                    "F1DAL9"
                ]
            },
            "ENSG00000171951": {
                "id": "ENSG00000171951",
                "type": "Gene",
                "symbol": "SCG2",
                "source": [
                    "ENSG00000119547"
                ],
                "target": [],
                "expression_sum": {
                    "class1": 26419.0,
                    "class2": 18983.0
                },
                "ens_id": "ENSG00000171951",
                "uniprot_ids": [
                    "P13521",
                    "C9JQI2",
                    "C9JDT0"
                ]
            },
            "ENSG00000156475": {
                "id": "ENSG00000156475",
                "type": "Gene",
                "symbol": "PPP2R2B",
                "source": [
                    "ENSG00000139352",
                    "ENSG00000119547"
                ],
                "target": [],
                "expression_sum": {
                    "class1": 19128.0,
                    "class2": 9807.0
                },
                "ens_id": "ENSG00000156475",
                "uniprot_ids": [
                    "Q00005",
                    "E9PCT7",
                    "D6RF04",
                    "D6RB57",
                    "H0YA16",
                    "E5RFC4",
                    "D6RGF9"
                ]
            },
            "ENSG00000185551": {
                "id": "ENSG00000185551",
                "type": "TF",
                "symbol": "NR2F2",
                "source": [
                    "ENSG00000119547"
                ],
                "target": [
                    "ENSG00000174938",
                    "ENSG00000198727",
                    "ENSG00000198886",
                    "ENSG00000104964",
                    "ENSG00000118680",
                    "ENSG00000156675",
                    "ENSG00000198763",
                    "ENSG00000245526",
                    "ENSG00000153956",
                    "ENSG00000179915",
                    "ENSG00000104112",
                    "ENSG00000198712",
                    "ENSG00000167642",
                    "ENSG00000198938",
                    "ENSG00000145247",
                    "ENSG00000198899",
                    "ENSG00000242265",
                    "ENSG00000251562",
                    "ENSG00000148053",
                    "ENSG00000127990",
                    "ENSG00000102109",
                    "ENSG00000215845",
                    "ENSG00000119547"
                ],
                "expression_sum": {
                    "class1": 14979.0,
                    "class2": 8249.0
                },
                "ens_id": "ENSG00000185551",
                "uniprot_ids": [
                    "P24468",
                    "H3BTC2"
                ]
            },
            "ENSG00000171634": {
                "id": "ENSG00000171634",
                "type": "TF",
                "symbol": "BPTF",
                "source": [
                    "ENSG00000016082",
                    "ENSG00000181449",
                    "ENSG00000113580",
                    "ENSG00000164330"
                ],
                "target": [
                    "ENSG00000149557",
                    "ENSG00000113580",
                    "ENSG00000111728"
                ],
                "expression_sum": {
                    "class1": 14482.0,
                    "class2": 13915.0
                },
                "ens_id": "ENSG00000171634",
                "uniprot_ids": [
                    "Q12830",
                    "J3QQQ8",
                    "A0A0A0MR81",
                    "F5H176",
                    "J3KSK9",
                    "E7ETD6",
                    "F5GXF5",
                    "J3QQK4",
                    "A0A2R8Y7Q1",
                    "E9PE19",
                    "J3QLW4",
                    "J3KSY0"
                ]
            },
            "ENSG00000113580": {
                "id": "ENSG00000113580",
                "type": "TF",
                "symbol": "NR3C1",
                "source": [
                    "ENSG00000171634",
                    "ENSG00000125968",
                    "ENSG00000105698",
                    "ENSG00000175745",
                    "ENSG00000181722",
                    "ENSG00000119950",
                    "ENSG00000067955",
                    "ENSG00000168214",
                    "ENSG00000119547",
                    "ENSG00000160113",
                    "ENSG00000092199"
                ],
                "target": [
                    "ENSG00000078668",
                    "ENSG00000206573",
                    "ENSG00000141741",
                    "ENSG00000232119",
                    "ENSG00000017260",
                    "ENSG00000162704",
                    "ENSG00000163634",
                    "ENSG00000165655",
                    "ENSG00000132698",
                    "ENSG00000153130",
                    "ENSG00000111674",
                    "ENSG00000111716",
                    "ENSG00000139644",
                    "ENSG00000175745",
                    "ENSG00000005810",
                    "ENSG00000169612",
                    "ENSG00000099622",
                    "ENSG00000079950",
                    "ENSG00000090060",
                    "ENSG00000005020",
                    "ENSG00000140941",
                    "ENSG00000122641",
                    "ENSG00000134248",
                    "ENSG00000124785",
                    "ENSG00000131143",
                    "ENSG00000075945",
                    "ENSG00000145817",
                    "ENSG00000146247",
                    "ENSG00000196419",
                    "ENSG00000075415",
                    "ENSG00000022840",
                    "ENSG00000105993",
                    "ENSG00000102172",
                    "ENSG00000160714",
                    "ENSG00000178567",
                    "ENSG00000102466",
                    "ENSG00000265972",
                    "ENSG00000077380",
                    "ENSG00000154556",
                    "ENSG00000163466",
                    "ENSG00000119421",
                    "ENSG00000134283",
                    "ENSG00000114439",
                    "ENSG00000173812",
                    "ENSG00000141279",
                    "ENSG00000183091",
                    "ENSG00000152256",
                    "ENSG00000237190",
                    "ENSG00000198899",
                    "ENSG00000147416",
                    "ENSG00000065911",
                    "ENSG00000173915",
                    "ENSG00000151729",
                    "ENSG00000272398",
                    "ENSG00000129636",
                    "ENSG00000123240",
                    "ENSG00000081181",
                    "ENSG00000144320",
                    "ENSG00000178913",
                    "ENSG00000163875",
                    "ENSG00000167136",
                    "ENSG00000138083",
                    "ENSG00000164924",
                    "ENSG00000074319",
                    "ENSG00000197930",
                    "ENSG00000141759",
                    "ENSG00000162545",
                    "ENSG00000165264",
                    "ENSG00000180773",
                    "ENSG00000146007",
                    "ENSG00000084764",
                    "ENSG00000277791",
                    "ENSG00000104964",
                    "ENSG00000170242",
                    "ENSG00000139910",
                    "ENSG00000125868",
                    "ENSG00000165637",
                    "ENSG00000241685",
                    "ENSG00000153815",
                    "ENSG00000047249",
                    "ENSG00000165672",
                    "ENSG00000173674",
                    "ENSG00000147548",
                    "ENSG00000125968",
                    "ENSG00000131236",
                    "ENSG00000129625",
                    "ENSG00000055163",
                    "ENSG00000089220",
                    "ENSG00000116898",
                    "ENSG00000117758",
                    "ENSG00000167733",
                    "ENSG00000196072",
                    "ENSG00000104112",
                    "ENSG00000255302",
                    "ENSG00000187164",
                    "ENSG00000188243",
                    "ENSG00000067225",
                    "ENSG00000109390",
                    "ENSG00000123933",
                    "ENSG00000072401",
                    "ENSG00000107771",
                    "ENSG00000140939",
                    "ENSG00000113712",
                    "ENSG00000136143",
                    "ENSG00000150459",
                    "ENSG00000148444",
                    "ENSG00000034713",
                    "ENSG00000182247",
                    "ENSG00000100592",
                    "ENSG00000144746",
                    "ENSG00000157168",
                    "ENSG00000133703",
                    "ENSG00000063854",
                    "ENSG00000133606",
                    "ENSG00000103769",
                    "ENSG00000146425",
                    "ENSG00000138071",
                    "ENSG00000117410",
                    "ENSG00000131100",
                    "ENSG00000273841",
                    "ENSG00000124802",
                    "ENSG00000263001",
                    "ENSG00000196405",
                    "ENSG00000174437",
                    "ENSG00000126934",
                    "ENSG00000154639",
                    "ENSG00000215845",
                    "ENSG00000185808",
                    "ENSG00000225783",
                    "ENSG00000078140",
                    "ENSG00000129515",
                    "ENSG00000100519",
                    "ENSG00000144366",
                    "ENSG00000121741",
                    "ENSG00000144635",
                    "ENSG00000163041",
                    "ENSG00000155115",
                    "ENSG00000160194",
                    "ENSG00000141027",
                    "ENSG00000092199",
                    "ENSG00000110955",
                    "ENSG00000107262",
                    "ENSG00000164346",
                    "ENSG00000168137",
                    "ENSG00000010256",
                    "ENSG00000120053",
                    "ENSG00000065518",
                    "ENSG00000174953",
                    "ENSG00000169100",
                    "ENSG00000162191",
                    "ENSG00000118363",
                    "ENSG00000122417",
                    "ENSG00000135940",
                    "ENSG00000108654",
                    "ENSG00000163888",
                    "ENSG00000198836",
                    "ENSG00000172336",
                    "ENSG00000107581",
                    "ENSG00000033800",
                    "ENSG00000131966",
                    "ENSG00000106052",
                    "ENSG00000147677",
                    "ENSG00000107438",
                    "ENSG00000244754",
                    "ENSG00000085224",
                    "ENSG00000102317",
                    "ENSG00000265491",
                    "ENSG00000180370",
                    "ENSG00000213465",
                    "ENSG00000173404",
                    "ENSG00000109332",
                    "ENSG00000163170",
                    "ENSG00000079246",
                    "ENSG00000075785",
                    "ENSG00000171222",
                    "ENSG00000095059",
                    "ENSG00000244879",
                    "ENSG00000105185",
                    "ENSG00000166974",
                    "ENSG00000189241",
                    "ENSG00000141562",
                    "ENSG00000100320",
                    "ENSG00000175166",
                    "ENSG00000168303",
                    "ENSG00000203879",
                    "ENSG00000240972",
                    "ENSG00000054267",
                    "ENSG00000136783",
                    "ENSG00000162852",
                    "ENSG00000196655",
                    "ENSG00000172915",
                    "ENSG00000141030",
                    "ENSG00000075292",
                    "ENSG00000105419",
                    "ENSG00000135390",
                    "ENSG00000153310",
                    "ENSG00000174938",
                    "ENSG00000213585",
                    "ENSG00000131508",
                    "ENSG00000197223",
                    "ENSG00000131437",
                    "ENSG00000005022",
                    "ENSG00000127616",
                    "ENSG00000076043",
                    "ENSG00000128654",
                    "ENSG00000160271",
                    "ENSG00000160613",
                    "ENSG00000090266",
                    "ENSG00000136810",
                    "ENSG00000143862",
                    "ENSG00000198015",
                    "ENSG00000171634",
                    "ENSG00000048649",
                    "ENSG00000125656",
                    "ENSG00000127184",
                    "ENSG00000176731",
                    "ENSG00000072832",
                    "ENSG00000130312",
                    "ENSG00000136950",
                    "ENSG00000160551",
                    "ENSG00000104388",
                    "ENSG00000100095",
                    "ENSG00000168159",
                    "ENSG00000119314",
                    "ENSG00000135472",
                    "ENSG00000267100",
                    "ENSG00000064607",
                    "ENSG00000171530",
                    "ENSG00000151914",
                    "ENSG00000123349",
                    "ENSG00000105583",
                    "ENSG00000165943",
                    "ENSG00000127914",
                    "ENSG00000014641",
                    "ENSG00000126267",
                    "ENSG00000175582",
                    "ENSG00000152377",
                    "ENSG00000104765",
                    "ENSG00000171130",
                    "ENSG00000198612",
                    "ENSG00000168300",
                    "ENSG00000176171",
                    "ENSG00000130725",
                    "ENSG00000100129",
                    "ENSG00000112305",
                    "ENSG00000142507",
                    "ENSG00000253738",
                    "ENSG00000047932",
                    "ENSG00000237765",
                    "ENSG00000165119",
                    "ENSG00000135250",
                    "ENSG00000139163",
                    "ENSG00000170871",
                    "ENSG00000126005",
                    "ENSG00000198840",
                    "ENSG00000082153",
                    "ENSG00000113657",
                    "ENSG00000247095",
                    "ENSG00000129559",
                    "ENSG00000089737",
                    "ENSG00000114784",
                    "ENSG00000184752",
                    "ENSG00000169567",
                    "ENSG00000197302",
                    "ENSG00000198258",
                    "ENSG00000275052",
                    "ENSG00000135002",
                    "ENSG00000127445",
                    "ENSG00000129355",
                    "ENSG00000183172",
                    "ENSG00000145545",
                    "ENSG00000146282",
                    "ENSG00000170027",
                    "ENSG00000241837",
                    "ENSG00000023287",
                    "ENSG00000076554",
                    "ENSG00000164190",
                    "ENSG00000120509",
                    "ENSG00000198125",
                    "ENSG00000131495",
                    "ENSG00000171314",
                    "ENSG00000168291",
                    "ENSG00000164975",
                    "ENSG00000172159",
                    "ENSG00000090238",
                    "ENSG00000170315",
                    "ENSG00000103266",
                    "ENSG00000147684",
                    "ENSG00000134779",
                    "ENSG00000077254",
                    "ENSG00000152932",
                    "ENSG00000067715",
                    "ENSG00000119950",
                    "ENSG00000049759",
                    "ENSG00000099795",
                    "ENSG00000129084",
                    "ENSG00000144747",
                    "ENSG00000111229",
                    "ENSG00000069345",
                    "ENSG00000189060",
                    "ENSG00000103145",
                    "ENSG00000051620",
                    "ENSG00000113742",
                    "ENSG00000163939",
                    "ENSG00000137941",
                    "ENSG00000167283",
                    "ENSG00000137100",
                    "ENSG00000123106",
                    "ENSG00000196199",
                    "ENSG00000198886",
                    "ENSG00000267313",
                    "ENSG00000102934",
                    "ENSG00000155506",
                    "ENSG00000116661",
                    "ENSG00000122643",
                    "ENSG00000196262",
                    "ENSG00000104904",
                    "ENSG00000125971",
                    "ENSG00000173726",
                    "ENSG00000130332",
                    "ENSG00000155959",
                    "ENSG00000137692",
                    "ENSG00000116586",
                    "ENSG00000089693",
                    "ENSG00000105254",
                    "ENSG00000168092",
                    "ENSG00000119335",
                    "ENSG00000115286",
                    "ENSG00000105698",
                    "ENSG00000092051",
                    "ENSG00000169432",
                    "ENSG00000101132",
                    "ENSG00000114023",
                    "ENSG00000154723",
                    "ENSG00000166347",
                    "ENSG00000151327",
                    "ENSG00000131174",
                    "ENSG00000166405",
                    "ENSG00000143013",
                    "ENSG00000198843",
                    "ENSG00000109654",
                    "ENSG00000145495",
                    "ENSG00000168288",
                    "ENSG00000254999",
                    "ENSG00000004059",
                    "ENSG00000089289",
                    "ENSG00000183779",
                    "ENSG00000177556",
                    "ENSG00000130558",
                    "ENSG00000113387",
                    "ENSG00000151893",
                    "ENSG00000158864",
                    "ENSG00000137409",
                    "ENSG00000082701",
                    "ENSG00000241468",
                    "ENSG00000136888",
                    "ENSG00000100442",
                    "ENSG00000121653",
                    "ENSG00000106028",
                    "ENSG00000173917",
                    "ENSG00000100697",
                    "ENSG00000168118",
                    "ENSG00000106367",
                    "ENSG00000121022",
                    "ENSG00000164615",
                    "ENSG00000197111",
                    "ENSG00000101182",
                    "ENSG00000146729",
                    "ENSG00000142949",
                    "ENSG00000105701",
                    "ENSG00000111669",
                    "ENSG00000136100",
                    "ENSG00000159720",
                    "ENSG00000148798",
                    "ENSG00000134825",
                    "ENSG00000184007",
                    "ENSG00000140307",
                    "ENSG00000167863",
                    "ENSG00000197747",
                    "ENSG00000004779",
                    "ENSG00000184277",
                    "ENSG00000127995",
                    "ENSG00000169139",
                    "ENSG00000117318",
                    "ENSG00000162368",
                    "ENSG00000004478",
                    "ENSG00000116459",
                    "ENSG00000186868",
                    "ENSG00000175756",
                    "ENSG00000135968",
                    "ENSG00000070831",
                    "ENSG00000137500",
                    "ENSG00000134333",
                    "ENSG00000091640",
                    "ENSG00000250479",
                    "ENSG00000147123",
                    "ENSG00000112320",
                    "ENSG00000161960",
                    "ENSG00000144674",
                    "ENSG00000143575",
                    "ENSG00000038219",
                    "ENSG00000130560",
                    "ENSG00000091844",
                    "ENSG00000182050",
                    "ENSG00000160014",
                    "ENSG00000184203",
                    "ENSG00000125818",
                    "ENSG00000183617",
                    "ENSG00000105649",
                    "ENSG00000103187",
                    "ENSG00000105220",
                    "ENSG00000053254",
                    "ENSG00000272047",
                    "ENSG00000198791",
                    "ENSG00000011295",
                    "ENSG00000176871",
                    "ENSG00000165152",
                    "ENSG00000007168",
                    "ENSG00000164713",
                    "ENSG00000134049",
                    "ENSG00000134744",
                    "ENSG00000139146",
                    "ENSG00000105379",
                    "ENSG00000126067",
                    "ENSG00000173692",
                    "ENSG00000197457",
                    "ENSG00000214530",
                    "ENSG00000111237",
                    "ENSG00000132670",
                    "ENSG00000132824",
                    "ENSG00000111911",
                    "ENSG00000254615",
                    "ENSG00000135070",
                    "ENSG00000104886",
                    "ENSG00000151366",
                    "ENSG00000214413",
                    "ENSG00000145782",
                    "ENSG00000112695",
                    "ENSG00000162188",
                    "ENSG00000108176",
                    "ENSG00000165416",
                    "ENSG00000203667",
                    "ENSG00000136710",
                    "ENSG00000100354",
                    "ENSG00000130414",
                    "ENSG00000115738",
                    "ENSG00000138430",
                    "ENSG00000169738",
                    "ENSG00000169213",
                    "ENSG00000166780",
                    "ENSG00000126214",
                    "ENSG00000176340",
                    "ENSG00000097033",
                    "ENSG00000258289",
                    "ENSG00000150753",
                    "ENSG00000049618",
                    "ENSG00000156587",
                    "ENSG00000273611",
                    "ENSG00000182670",
                    "ENSG00000177459",
                    "ENSG00000148943",
                    "ENSG00000181315",
                    "ENSG00000146701",
                    "ENSG00000115514",
                    "ENSG00000186472",
                    "ENSG00000101745",
                    "ENSG00000140990",
                    "ENSG00000182718",
                    "ENSG00000173436",
                    "ENSG00000069956",
                    "ENSG00000063046",
                    "ENSG00000163541",
                    "ENSG00000102109",
                    "ENSG00000169439",
                    "ENSG00000164258",
                    "ENSG00000169925",
                    "ENSG00000109472",
                    "ENSG00000114956",
                    "ENSG00000149926",
                    "ENSG00000106261",
                    "ENSG00000171604",
                    "ENSG00000101210",
                    "ENSG00000143155",
                    "ENSG00000146066",
                    "ENSG00000065135",
                    "ENSG00000129235",
                    "ENSG00000132639",
                    "ENSG00000102144",
                    "ENSG00000130770",
                    "ENSG00000156467",
                    "ENSG00000168280",
                    "ENSG00000119013",
                    "ENSG00000161203",
                    "ENSG00000175324",
                    "ENSG00000184076",
                    "ENSG00000105887",
                    "ENSG00000165434",
                    "ENSG00000266472",
                    "ENSG00000101782",
                    "ENSG00000177889",
                    "ENSG00000166913",
                    "ENSG00000181722",
                    "ENSG00000109046",
                    "ENSG00000159199",
                    "ENSG00000170873",
                    "ENSG00000151743",
                    "ENSG00000162434",
                    "ENSG00000121851",
                    "ENSG00000118816",
                    "ENSG00000105290",
                    "ENSG00000147669",
                    "ENSG00000107679",
                    "ENSG00000119888",
                    "ENSG00000204442",
                    "ENSG00000163468",
                    "ENSG00000137312",
                    "ENSG00000074071",
                    "ENSG00000091039",
                    "ENSG00000096092",
                    "ENSG00000111371",
                    "ENSG00000134056",
                    "ENSG00000172115",
                    "ENSG00000119547",
                    "ENSG00000101558",
                    "ENSG00000101040",
                    "ENSG00000091542",
                    "ENSG00000175130",
                    "ENSG00000128789",
                    "ENSG00000111640",
                    "ENSG00000169750",
                    "ENSG00000120071",
                    "ENSG00000124596",
                    "ENSG00000138078",
                    "ENSG00000115241",
                    "ENSG00000253719",
                    "ENSG00000156976",
                    "ENSG00000128245",
                    "ENSG00000134138",
                    "ENSG00000130294",
                    "ENSG00000078369",
                    "ENSG00000119396",
                    "ENSG00000077264",
                    "ENSG00000205531",
                    "ENSG00000099624",
                    "ENSG00000120265",
                    "ENSG00000187231",
                    "ENSG00000138495",
                    "ENSG00000154001",
                    "ENSG00000101489",
                    "ENSG00000138069",
                    "ENSG00000083099",
                    "ENSG00000187257",
                    "ENSG00000171033",
                    "ENSG00000110321",
                    "ENSG00000198420",
                    "ENSG00000138107",
                    "ENSG00000055609",
                    "ENSG00000005483",
                    "ENSG00000139726",
                    "ENSG00000204568",
                    "ENSG00000175203",
                    "ENSG00000111642",
                    "ENSG00000102409",
                    "ENSG00000145362",
                    "ENSG00000123144",
                    "ENSG00000101557",
                    "ENSG00000127125",
                    "ENSG00000113575",
                    "ENSG00000143256",
                    "ENSG00000149084",
                    "ENSG00000102931",
                    "ENSG00000166619",
                    "ENSG00000074800",
                    "ENSG00000104863",
                    "ENSG00000165502",
                    "ENSG00000114302",
                    "ENSG00000084733",
                    "ENSG00000119705",
                    "ENSG00000075151",
                    "ENSG00000122034",
                    "ENSG00000154582",
                    "ENSG00000067955",
                    "ENSG00000067836",
                    "ENSG00000120533",
                    "ENSG00000136854",
                    "ENSG00000214113",
                    "ENSG00000114209",
                    "ENSG00000109107",
                    "ENSG00000168214",
                    "ENSG00000214517",
                    "ENSG00000135535",
                    "ENSG00000137996",
                    "ENSG00000115524",
                    "ENSG00000188994",
                    "ENSG00000174775",
                    "ENSG00000108946",
                    "ENSG00000112237",
                    "ENSG00000164830",
                    "ENSG00000175416",
                    "ENSG00000128609",
                    "ENSG00000172586",
                    "ENSG00000103316",
                    "ENSG00000160953",
                    "ENSG00000108953",
                    "ENSG00000155368",
                    "ENSG00000169564",
                    "ENSG00000094916",
                    "ENSG00000132305",
                    "ENSG00000138663",
                    "ENSG00000086061",
                    "ENSG00000166922",
                    "ENSG00000145675",
                    "ENSG00000149806",
                    "ENSG00000140386",
                    "ENSG00000140740",
                    "ENSG00000198727",
                    "ENSG00000204272",
                    "ENSG00000167658",
                    "ENSG00000142168",
                    "ENSG00000189180",
                    "ENSG00000084070",
                    "ENSG00000129946",
                    "ENSG00000159176",
                    "ENSG00000068745",
                    "ENSG00000143761",
                    "ENSG00000108848",
                    "ENSG00000136193",
                    "ENSG00000196586",
                    "ENSG00000104427",
                    "ENSG00000109919",
                    "ENSG00000198786",
                    "ENSG00000143933",
                    "ENSG00000172172",
                    "ENSG00000165629",
                    "ENSG00000166503",
                    "ENSG00000167815",
                    "ENSG00000126457",
                    "ENSG00000177971",
                    "ENSG00000132963",
                    "ENSG00000131089",
                    "ENSG00000070087",
                    "ENSG00000168522",
                    "ENSG00000177311",
                    "ENSG00000119048",
                    "ENSG00000198408",
                    "ENSG00000087470",
                    "ENSG00000103254",
                    "ENSG00000131370",
                    "ENSG00000148450",
                    "ENSG00000178127",
                    "ENSG00000131876",
                    "ENSG00000104381",
                    "ENSG00000174456",
                    "ENSG00000168701",
                    "ENSG00000167699",
                    "ENSG00000010244",
                    "ENSG00000136628",
                    "ENSG00000143198",
                    "ENSG00000133639",
                    "ENSG00000092148",
                    "ENSG00000151552",
                    "ENSG00000145494",
                    "ENSG00000156411",
                    "ENSG00000061676",
                    "ENSG00000108960",
                    "ENSG00000177200",
                    "ENSG00000115944",
                    "ENSG00000174939",
                    "ENSG00000138443",
                    "ENSG00000135334",
                    "ENSG00000196531",
                    "ENSG00000100138",
                    "ENSG00000116030",
                    "ENSG00000091592",
                    "ENSG00000109133",
                    "ENSG00000169435",
                    "ENSG00000242485",
                    "ENSG00000160113",
                    "ENSG00000160075",
                    "ENSG00000179915",
                    "ENSG00000120992",
                    "ENSG00000128872",
                    "ENSG00000170043",
                    "ENSG00000168653",
                    "ENSG00000131507",
                    "ENSG00000156261",
                    "ENSG00000181885",
                    "ENSG00000130723",
                    "ENSG00000178741",
                    "ENSG00000156515",
                    "ENSG00000107758",
                    "ENSG00000170606",
                    "ENSG00000101266",
                    "ENSG00000171206",
                    "ENSG00000177885",
                    "ENSG00000166595",
                    "ENSG00000143774",
                    "ENSG00000165209",
                    "ENSG00000182512",
                    "ENSG00000088832",
                    "ENSG00000182463",
                    "ENSG00000166136",
                    "ENSG00000122545",
                    "ENSG00000134330",
                    "ENSG00000143612",
                    "ENSG00000135720",
                    "ENSG00000136485",
                    "ENSG00000166848",
                    "ENSG00000196683",
                    "ENSG00000149547",
                    "ENSG00000166206",
                    "ENSG00000125870",
                    "ENSG00000102189",
                    "ENSG00000149600",
                    "ENSG00000112514",
                    "ENSG00000181061",
                    "ENSG00000150787",
                    "ENSG00000128739",
                    "ENSG00000086666",
                    "ENSG00000137726",
                    "ENSG00000159023",
                    "ENSG00000114902",
                    "ENSG00000089818",
                    "ENSG00000099341",
                    "ENSG00000173273",
                    "ENSG00000230989",
                    "ENSG00000108309",
                    "ENSG00000154518",
                    "ENSG00000087302",
                    "ENSG00000213064",
                    "ENSG00000091164",
                    "ENSG00000172757",
                    "ENSG00000178449",
                    "ENSG00000101460",
                    "ENSG00000138175",
                    "ENSG00000126858",
                    "ENSG00000134717",
                    "ENSG00000050426",
                    "ENSG00000151778",
                    "ENSG00000119782",
                    "ENSG00000143252",
                    "ENSG00000141367",
                    "ENSG00000213719",
                    "ENSG00000198168",
                    "ENSG00000105438",
                    "ENSG00000155099",
                    "ENSG00000152234",
                    "ENSG00000133872",
                    "ENSG00000131094",
                    "ENSG00000198668",
                    "ENSG00000134308",
                    "ENSG00000143158",
                    "ENSG00000134684",
                    "ENSG00000008324",
                    "ENSG00000166228",
                    "ENSG00000204525",
                    "ENSG00000243449",
                    "ENSG00000213619",
                    "ENSG00000164919",
                    "ENSG00000153774",
                    "ENSG00000129197",
                    "ENSG00000138757",
                    "ENSG00000108561",
                    "ENSG00000033627",
                    "ENSG00000178982",
                    "ENSG00000067900",
                    "ENSG00000087448",
                    "ENSG00000214253",
                    "ENSG00000138382",
                    "ENSG00000155876",
                    "ENSG00000125821",
                    "ENSG00000137547",
                    "ENSG00000100387",
                    "ENSG00000114573",
                    "ENSG00000140319",
                    "ENSG00000132466",
                    "ENSG00000196233",
                    "ENSG00000181704",
                    "ENSG00000128923",
                    "ENSG00000163399",
                    "ENSG00000243317",
                    "ENSG00000070669",
                    "ENSG00000118680",
                    "ENSG00000009307",
                    "ENSG00000205937",
                    "ENSG00000018189",
                    "ENSG00000138138",
                    "ENSG00000168958",
                    "ENSG00000204856",
                    "ENSG00000124172",
                    "ENSG00000057608",
                    "ENSG00000165169",
                    "ENSG00000152795",
                    "ENSG00000065548",
                    "ENSG00000182195",
                    "ENSG00000137876",
                    "ENSG00000017797",
                    "ENSG00000138398",
                    "ENSG00000212907",
                    "ENSG00000142669",
                    "ENSG00000162076",
                    "ENSG00000100568",
                    "ENSG00000107281",
                    "ENSG00000164323",
                    "ENSG00000025293",
                    "ENSG00000122705",
                    "ENSG00000114631",
                    "ENSG00000132432",
                    "ENSG00000140374",
                    "ENSG00000178104",
                    "ENSG00000115306",
                    "ENSG00000023734",
                    "ENSG00000114405",
                    "ENSG00000184009",
                    "ENSG00000183726",
                    "ENSG00000173660",
                    "ENSG00000164292",
                    "ENSG00000214941",
                    "ENSG00000186184",
                    "ENSG00000100804",
                    "ENSG00000119640",
                    "ENSG00000113648",
                    "ENSG00000104643",
                    "ENSG00000114125",
                    "ENSG00000185787",
                    "ENSG00000137947",
                    "ENSG00000105568",
                    "ENSG00000100554",
                    "ENSG00000130638",
                    "ENSG00000169045",
                    "ENSG00000111775",
                    "ENSG00000123636",
                    "ENSG00000133142",
                    "ENSG00000139684",
                    "ENSG00000197102",
                    "ENSG00000128989",
                    "ENSG00000114120",
                    "ENSG00000166562",
                    "ENSG00000168724",
                    "ENSG00000237651",
                    "ENSG00000165704",
                    "ENSG00000102003",
                    "ENSG00000188612",
                    "ENSG00000196781",
                    "ENSG00000104763",
                    "ENSG00000180817",
                    "ENSG00000086065",
                    "ENSG00000116288",
                    "ENSG00000008018",
                    "ENSG00000160948",
                    "ENSG00000111530",
                    "ENSG00000188322",
                    "ENSG00000165487",
                    "ENSG00000220205",
                    "ENSG00000128524",
                    "ENSG00000134313",
                    "ENSG00000167232",
                    "ENSG00000165678",
                    "ENSG00000198712",
                    "ENSG00000133302",
                    "ENSG00000167113",
                    "ENSG00000136238",
                    "ENSG00000132424",
                    "ENSG00000130338",
                    "ENSG00000224032",
                    "ENSG00000047849",
                    "ENSG00000183527",
                    "ENSG00000221914",
                    "ENSG00000122873",
                    "ENSG00000113851",
                    "ENSG00000103415",
                    "ENSG00000151923",
                    "ENSG00000170759",
                    "ENSG00000087460",
                    "ENSG00000145247",
                    "ENSG00000113558",
                    "ENSG00000176978",
                    "ENSG00000143153",
                    "ENSG00000109180",
                    "ENSG00000213937",
                    "ENSG00000115233",
                    "ENSG00000123983",
                    "ENSG00000156052",
                    "ENSG00000143742",
                    "ENSG00000198648",
                    "ENSG00000133169",
                    "ENSG00000106244",
                    "ENSG00000104419",
                    "ENSG00000164167",
                    "ENSG00000100201",
                    "ENSG00000114115",
                    "ENSG00000198888",
                    "ENSG00000173163"
                ],
                "expression_sum": {
                    "class1": 4215.0,
                    "class2": 5563.0
                },
                "ens_id": "ENSG00000113580",
                "uniprot_ids": [
                    "P04150",
                    "Q3MSN4",
                    "A0A494C0P1",
                    "D6RDA9"
                ]
            },
            "ENSG00000171530": {
                "id": "ENSG00000171530",
                "type": "Gene",
                "symbol": "TBCA",
                "source": [
                    "ENSG00000181449",
                    "ENSG00000163435",
                    "ENSG00000113580",
                    "ENSG00000164330",
                    "ENSG00000114861",
                    "ENSG00000221869",
                    "ENSG00000184486"
                ],
                "target": [],
                "expression_sum": {
                    "class1": 60465.0,
                    "class2": 43721.0
                },
                "ens_id": "ENSG00000171530",
                "uniprot_ids": [
                    "O75347",
                    "A0A494C144",
                    "E5RHG6",
                    "E5RJD8",
                    "E5RJM1",
                    "E5RIX8",
                    "E5RIW3"
                ]
            },
            "ENSG00000220205": {
                "id": "ENSG00000220205",
                "type": "Gene",
                "symbol": "VAMP2",
                "source": [
                    "ENSG00000181449",
                    "ENSG00000113580",
                    "ENSG00000148516",
                    "ENSG00000103495",
                    "ENSG00000164330",
                    "ENSG00000140262",
                    "ENSG00000181722",
                    "ENSG00000114861",
                    "ENSG00000137203"
                ],
                "target": [],
                "expression_sum": {
                    "class1": 90691.0,
                    "class2": 33391.0
                },
                "ens_id": "ENSG00000220205",
                "uniprot_ids": [
                    "P63027",
                    "F8WCA0",
                    "J3QRU4",
                    "K7ENK9"
                ]
            },
            "ENSG00000169439": {
                "id": "ENSG00000169439",
                "type": "Gene",
                "symbol": "SDC2",
                "source": [
                    "ENSG00000108055",
                    "ENSG00000113580",
```
