## Supplement 2 (DAN vs. Cocul) for "AGEAS: Automated Machine Learning based Genetic Regulatory Element Extraction System": key_atlas.js

```
{
    "genes": {
        "ENSG00000168214": {
            "id": "ENSG00000168214",
            "type": "TF",
            "symbol": "RBPJ",
            "source": [
                "ENSG00000163435",
                "ENSG00000157557",
                "ENSG00000167081",
                "ENSG00000113580",
                "ENSG00000148516",
                "ENSG00000164330",
                "ENSG00000100219",
                "ENSG00000221869"
            ],
            "target": [
                "ENSG00000169856",
                "ENSG00000152558",
                "ENSG00000198840",
                "ENSG00000132698",
                "ENSG00000164093",
                "ENSG00000107105",
                "ENSG00000101443",
                "ENSG00000166582",
                "ENSG00000179981",
                "ENSG00000169435",
                "ENSG00000172183",
                "ENSG00000165215",
                "ENSG00000163191",
                "ENSG00000189143",
                "ENSG00000123416",
                "ENSG00000198125",
                "ENSG00000181885",
                "ENSG00000245532",
                "ENSG00000102466",
                "ENSG00000265972",
                "ENSG00000154556",
                "ENSG00000167642",
                "ENSG00000102007",
                "ENSG00000163956",
                "ENSG00000087258",
                "ENSG00000187244",
                "ENSG00000198899",
                "ENSG00000111728",
                "ENSG00000119888",
                "ENSG00000100604",
                "ENSG00000169862",
                "ENSG00000180879",
                "ENSG00000099194",
                "ENSG00000104722",
                "ENSG00000171345",
                "ENSG00000176887",
                "ENSG00000169783",
                "ENSG00000187193",
                "ENSG00000148516",
                "ENSG00000102934",
                "ENSG00000170266",
                "ENSG00000117525",
                "ENSG00000213190",
                "ENSG00000060982",
                "ENSG00000196361",
                "ENSG00000113732",
                "ENSG00000147889",
                "ENSG00000130303",
                "ENSG00000131711",
                "ENSG00000075618",
                "ENSG00000144218",
                "ENSG00000121039",
                "ENSG00000107130",
                "ENSG00000161249",
                "ENSG00000170091",
                "ENSG00000138085",
                "ENSG00000170293",
                "ENSG00000064601",
                "ENSG00000113580",
                "ENSG00000068615",
                "ENSG00000100219",
                "ENSG00000185650",
                "ENSG00000215845",
                "ENSG00000111057",
                "ENSG00000104870",
                "ENSG00000165092",
                "ENSG00000176788",
                "ENSG00000149557",
                "ENSG00000145428",
                "ENSG00000148180",
                "ENSG00000145335",
                "ENSG00000163399",
                "ENSG00000138772",
                "ENSG00000112773",
                "ENSG00000244187",
                "ENSG00000174498",
                "ENSG00000213760",
                "ENSG00000124107",
                "ENSG00000117691",
                "ENSG00000092964",
                "ENSG00000143702",
                "ENSG00000135535",
                "ENSG00000112972",
                "ENSG00000122012",
                "ENSG00000107438",
                "ENSG00000016082",
                "ENSG00000142156",
                "ENSG00000152503",
                "ENSG00000120437",
                "ENSG00000103316",
                "ENSG00000149294",
                "ENSG00000124942",
                "ENSG00000157557",
                "ENSG00000096696",
                "ENSG00000167552",
                "ENSG00000104833",
                "ENSG00000164330",
                "ENSG00000137267",
                "ENSG00000143819",
                "ENSG00000172915",
                "ENSG00000030582",
                "ENSG00000179218",
                "ENSG00000100345",
                "ENSG00000214530",
                "ENSG00000198727",
                "ENSG00000147676",
                "ENSG00000105784",
                "ENSG00000137880",
                "ENSG00000115461",
                "ENSG00000148834",
                "ENSG00000072832",
                "ENSG00000168938",
                "ENSG00000221866",
                "ENSG00000246705",
                "ENSG00000198712",
                "ENSG00000117155",
                "ENSG00000089356",
                "ENSG00000146674",
                "ENSG00000213614",
                "ENSG00000137285",
                "ENSG00000118640",
                "ENSG00000177459",
                "ENSG00000198938",
                "ENSG00000139970",
                "ENSG00000242265",
                "ENSG00000163435",
                "ENSG00000145730",
                "ENSG00000109062",
                "ENSG00000167081",
                "ENSG00000108679"
            ],
            "expression_sum": {
                "class1": 13084.0,
                "class2": 12713.0
            },
            "ens_id": "ENSG00000168214",
            "uniprot_ids": [
                "Q06330",
                "A0A7P0T821",
                "A0A7P0TA33",
                "D6RCM1",
                "D6RB37",
                "D6REC2",
                "D6RA45",
                "A0A7P0T964",
                "A0A7P0T947",
                "A0A7P0T9W9",
                "A0A7P0TBJ4",
                "A0A7P0T848",
                "D6R946",
                "D6R9X3",
                "D6REP3",
                "A0A7P0Z4N7",
                "D6R9K5",
                "D6RIZ8",
                "A0A7P0T9Y9",
                "D6RAT2",
                "D6RE93",
                "A0A7P0T7Z3",
                "D6RF98",
                "A0A7P0T806",
                "D6RIV8",
                "D6RBQ8",
                "H0Y8R3",
                "D6R927"
            ],
            "net_id": 0,
            "source_num": 8,
            "target_num": 40
        },
        "ENSG00000160113": {
            "id": "ENSG00000160113",
            "type": "TF",
            "symbol": "NR2F6",
            "source": [
                "ENSG00000163435",
                "ENSG00000113580",
                "ENSG00000148516",
                "ENSG00000164330",
                "ENSG00000100219",
                "ENSG00000181722",
                "ENSG00000221869"
            ],
            "target": [
                "ENSG00000169856",
                "ENSG00000152558",
                "ENSG00000111674",
                "ENSG00000166710",
                "ENSG00000087085",
                "ENSG00000108830",
                "ENSG00000247095",
                "ENSG00000166582",
                "ENSG00000182534",
                "ENSG00000085871",
                "ENSG00000129355",
                "ENSG00000105255",
                "ENSG00000196230",
                "ENSG00000086300",
                "ENSG00000187134",
                "ENSG00000172183",
                "ENSG00000163191",
                "ENSG00000123416",
                "ENSG00000198125",
                "ENSG00000181885",
                "ENSG00000245532",
                "ENSG00000265972",
                "ENSG00000154556",
                "ENSG00000167642",
                "ENSG00000119865",
                "ENSG00000108515",
                "ENSG00000173267",
                "ENSG00000181722",
                "ENSG00000105088",
                "ENSG00000079462",
                "ENSG00000130066",
                "ENSG00000119888",
                "ENSG00000115310",
                "ENSG00000100867",
                "ENSG00000221869",
                "ENSG00000159335",
                "ENSG00000099194",
                "ENSG00000180879",
                "ENSG00000164929",
                "ENSG00000109084",
                "ENSG00000170266",
                "ENSG00000101856",
                "ENSG00000141934",
                "ENSG00000084628",
                "ENSG00000060982",
                "ENSG00000131981",
                "ENSG00000075618",
                "ENSG00000138758",
                "ENSG00000213719",
                "ENSG00000132688",
                "ENSG00000075043",
                "ENSG00000175567",
                "ENSG00000189334",
                "ENSG00000170540",
                "ENSG00000161249",
                "ENSG00000129521",
                "ENSG00000170091",
                "ENSG00000170293",
                "ENSG00000064601",
                "ENSG00000113580",
                "ENSG00000100219",
                "ENSG00000185650",
                "ENSG00000111275",
                "ENSG00000104870",
                "ENSG00000069849",
                "ENSG00000165092",
                "ENSG00000111907",
                "ENSG00000148180",
                "ENSG00000112773",
                "ENSG00000163399",
                "ENSG00000156675",
                "ENSG00000244187",
                "ENSG00000117472",
                "ENSG00000174498",
                "ENSG00000128564",
                "ENSG00000186868",
                "ENSG00000108106",
                "ENSG00000213760",
                "ENSG00000143702",
                "ENSG00000135535",
                "ENSG00000112972",
                "ENSG00000051523",
                "ENSG00000107438",
                "ENSG00000144674",
                "ENSG00000120437",
                "ENSG00000103316",
                "ENSG00000149294",
                "ENSG00000124942",
                "ENSG00000115758",
                "ENSG00000113328",
                "ENSG00000157557",
                "ENSG00000096696",
                "ENSG00000141505",
                "ENSG00000137267",
                "ENSG00000105048",
                "ENSG00000163931",
                "ENSG00000030582",
                "ENSG00000144339",
                "ENSG00000214530",
                "ENSG00000161082",
                "ENSG00000060138",
                "ENSG00000174021",
                "ENSG00000152952",
                "ENSG00000070961",
                "ENSG00000137880",
                "ENSG00000148834",
                "ENSG00000135480",
                "ENSG00000182287",
                "ENSG00000104549",
                "ENSG00000213614",
                "ENSG00000118640",
                "ENSG00000170421",
                "ENSG00000097021",
                "ENSG00000152082",
                "ENSG00000163435",
                "ENSG00000213937",
                "ENSG00000135406",
                "ENSG00000170017",
                "ENSG00000104419",
                "ENSG00000108679"
            ],
            "expression_sum": {
                "class1": 14139.0,
                "class2": 12251.0
            },
            "ens_id": "ENSG00000160113",
            "uniprot_ids": [
                "P10588",
                "M0R175"
            ],
            "net_id": 0,
            "source_num": 7,
            "target_num": 30
        },
        "ENSG00000100219": {
            "id": "ENSG00000100219",
            "type": "TF",
            "symbol": "XBP1",
            "source": [
                "ENSG00000168214",
                "ENSG00000106261",
                "ENSG00000177606",
                "ENSG00000160113"
            ],
            "target": [
                "ENSG00000106261",
                "ENSG00000152558",
                "ENSG00000126005",
                "ENSG00000166710",
                "ENSG00000198840",
                "ENSG00000132698",
                "ENSG00000171604",
                "ENSG00000160113",
                "ENSG00000177606",
                "ENSG00000076554",
                "ENSG00000163191",
                "ENSG00000189143",
                "ENSG00000184076",
                "ENSG00000181885",
                "ENSG00000245532",
                "ENSG00000265972",
                "ENSG00000154556",
                "ENSG00000114439",
                "ENSG00000181722",
                "ENSG00000141279",
                "ENSG00000111229",
                "ENSG00000051620",
                "ENSG00000150687",
                "ENSG00000197930",
                "ENSG00000135069",
                "ENSG00000101040",
                "ENSG00000198886",
                "ENSG00000144560",
                "ENSG00000134138",
                "ENSG00000166347",
                "ENSG00000213719",
                "ENSG00000121039",
                "ENSG00000143013",
                "ENSG00000148444",
                "ENSG00000026025",
                "ENSG00000129521",
                "ENSG00000263001",
                "ENSG00000109099",
                "ENSG00000163399",
                "ENSG00000138772",
                "ENSG00000156675",
                "ENSG00000118680",
                "ENSG00000128564",
                "ENSG00000168214",
                "ENSG00000124107",
                "ENSG00000070831",
                "ENSG00000144674",
                "ENSG00000106537",
                "ENSG00000114405",
                "ENSG00000091844",
                "ENSG00000172915",
                "ENSG00000030582",
                "ENSG00000002586",
                "ENSG00000123636",
                "ENSG00000214530",
                "ENSG00000174938",
                "ENSG00000198727",
                "ENSG00000147676",
                "ENSG00000104763",
                "ENSG00000160613",
                "ENSG00000060138",
                "ENSG00000169764",
                "ENSG00000196586",
                "ENSG00000198786",
                "ENSG00000146674",
                "ENSG00000118640",
                "ENSG00000170421",
                "ENSG00000164096",
                "ENSG00000049618",
                "ENSG00000139970",
                "ENSG00000182718",
                "ENSG00000163435",
                "ENSG00000133639",
                "ENSG00000109062",
                "ENSG00000138640",
                "ENSG00000104419",
                "ENSG00000198888"
            ],
            "expression_sum": {
                "class1": 15335.0,
                "class2": 32457.0
            },
            "ens_id": "ENSG00000100219",
            "uniprot_ids": [
                "P17861",
                "B1AHH2",
                "B1AHH1"
            ],
            "net_id": 0,
            "source_num": 4,
            "target_num": 33
        },
        "ENSG00000167081": {
            "id": "ENSG00000167081",
            "type": "TF",
            "symbol": "PBX3",
            "source": [
                "ENSG00000100811",
                "ENSG00000134138",
                "ENSG00000004487",
                "ENSG00000103495",
                "ENSG00000105698",
                "ENSG00000213015",
                "ENSG00000189403",
                "ENSG00000119950",
                "ENSG00000196591",
                "ENSG00000168214",
                "ENSG00000092199"
            ],
            "target": [
                "ENSG00000078668",
                "ENSG00000090621",
                "ENSG00000206573",
                "ENSG00000112081",
                "ENSG00000232119",
                "ENSG00000107223",
                "ENSG00000162704",
                "ENSG00000163634",
                "ENSG00000165655",
                "ENSG00000153130",
                "ENSG00000111674",
                "ENSG00000111716",
                "ENSG00000139644",
                "ENSG00000005810",
                "ENSG00000169612",
                "ENSG00000099622",
                "ENSG00000137955",
                "ENSG00000185651",
                "ENSG00000090060",
                "ENSG00000005020",
                "ENSG00000106603",
                "ENSG00000140941",
                "ENSG00000161016",
                "ENSG00000134248",
                "ENSG00000127540",
                "ENSG00000075945",
                "ENSG00000125743",
                "ENSG00000099797",
                "ENSG00000119632",
                "ENSG00000189403",
                "ENSG00000159377",
                "ENSG00000136261",
                "ENSG00000075415",
                "ENSG00000147155",
                "ENSG00000077380",
                "ENSG00000154556",
                "ENSG00000052802",
                "ENSG00000134283",
                "ENSG00000198931",
                "ENSG00000173812",
                "ENSG00000197451",
                "ENSG00000152256",
                "ENSG00000237190",
                "ENSG00000173915",
                "ENSG00000065911",
                "ENSG00000151729",
                "ENSG00000172292",
                "ENSG00000170310",
                "ENSG00000129636",
                "ENSG00000115540",
                "ENSG00000123240",
                "ENSG00000041357",
                "ENSG00000102103",
                "ENSG00000141858",
                "ENSG00000178913",
                "ENSG00000163875",
                "ENSG00000163319",
                "ENSG00000167136",
                "ENSG00000164924",
                "ENSG00000074319",
                "ENSG00000149196",
                "ENSG00000141759",
                "ENSG00000174574",
                "ENSG00000146007",
                "ENSG00000106554",
                "ENSG00000084764",
                "ENSG00000277791",
                "ENSG00000125868",
                "ENSG00000153815",
                "ENSG00000047249",
                "ENSG00000147548",
                "ENSG00000171703",
                "ENSG00000196504",
                "ENSG00000131236",
                "ENSG00000089220",
                "ENSG00000055163",
                "ENSG00000141447",
                "ENSG00000116898",
                "ENSG00000175581",
                "ENSG00000196072",
                "ENSG00000179431",
                "ENSG00000187164",
                "ENSG00000188243",
                "ENSG00000067225",
                "ENSG00000109390",
                "ENSG00000113712",
                "ENSG00000126698",
                "ENSG00000136143",
                "ENSG00000233927",
                "ENSG00000117139",
                "ENSG00000148444",
                "ENSG00000140105",
                "ENSG00000159685",
                "ENSG00000034713",
                "ENSG00000130811",
                "ENSG00000144746",
                "ENSG00000095380",
                "ENSG00000133703",
                "ENSG00000063854",
                "ENSG00000103769",
                "ENSG00000034510",
                "ENSG00000146425",
                "ENSG00000005194",
                "ENSG00000138071",
                "ENSG00000203778",
                "ENSG00000117410",
                "ENSG00000131100",
                "ENSG00000075142",
                "ENSG00000273841",
                "ENSG00000124802",
                "ENSG00000263001",
                "ENSG00000142657",
                "ENSG00000196405",
                "ENSG00000183011",
                "ENSG00000174437",
                "ENSG00000114520",
                "ENSG00000213741",
                "ENSG00000126934",
                "ENSG00000145592",
                "ENSG00000187051",
                "ENSG00000069849",
                "ENSG00000240344",
                "ENSG00000129515",
                "ENSG00000100519",
                "ENSG00000144635",
                "ENSG00000187514",
                "ENSG00000163041",
                "ENSG00000155115",
                "ENSG00000160194",
                "ENSG00000141027",
                "ENSG00000092199",
                "ENSG00000110955",
                "ENSG00000178980",
                "ENSG00000204463",
                "ENSG00000164346",
                "ENSG00000150316",
                "ENSG00000010256",
                "ENSG00000120053",
                "ENSG00000174953",
                "ENSG00000169100",
                "ENSG00000162191",
                "ENSG00000118363",
                "ENSG00000108654",
                "ENSG00000135940",
                "ENSG00000170634",
                "ENSG00000108106",
                "ENSG00000198836",
                "ENSG00000107581",
                "ENSG00000033800",
                "ENSG00000105258",
                "ENSG00000106052",
                "ENSG00000147677",
                "ENSG00000231500",
                "ENSG00000108666",
                "ENSG00000153037",
                "ENSG00000175792",
                "ENSG00000102317",
                "ENSG00000265491",
                "ENSG00000213465",
                "ENSG00000109332",
                "ENSG00000075624",
                "ENSG00000072062",
                "ENSG00000168275",
                "ENSG00000075785",
                "ENSG00000184900",
                "ENSG00000095059",
                "ENSG00000110696",
                "ENSG00000111850",
                "ENSG00000102898",
                "ENSG00000107937",
                "ENSG00000166974",
                "ENSG00000063322",
                "ENSG00000232388",
                "ENSG00000141562",
                "ENSG00000114391",
                "ENSG00000163931",
                "ENSG00000175166",
                "ENSG00000168303",
                "ENSG00000167088",
                "ENSG00000100320",
                "ENSG00000162852",
                "ENSG00000105419",
                "ENSG00000141030",
                "ENSG00000075292",
                "ENSG00000164985",
                "ENSG00000239672",
                "ENSG00000105723",
                "ENSG00000153310",
                "ENSG00000256060",
                "ENSG00000131508",
                "ENSG00000197223",
                "ENSG00000005022",
                "ENSG00000180957",
                "ENSG00000131469",
                "ENSG00000080824",
                "ENSG00000160271",
                "ENSG00000090266",
                "ENSG00000136810",
                "ENSG00000142937",
                "ENSG00000048649",
                "ENSG00000162374",
                "ENSG00000169764",
                "ENSG00000106682",
                "ENSG00000116209",
                "ENSG00000072832",
                "ENSG00000130312",
                "ENSG00000136950",
                "ENSG00000079459",
                "ENSG00000104341",
                "ENSG00000197170",
                "ENSG00000104388",
                "ENSG00000083750",
                "ENSG00000168159",
                "ENSG00000267100",
                "ENSG00000170144",
                "ENSG00000097021",
                "ENSG00000152082",
                "ENSG00000087191",
                "ENSG00000257704",
                "ENSG00000168710",
                "ENSG00000127914",
                "ENSG00000124614",
                "ENSG00000014641",
                "ENSG00000126267",
                "ENSG00000175582",
                "ENSG00000104765",
                "ENSG00000171130",
                "ENSG00000198612",
                "ENSG00000120860",
                "ENSG00000081154",
                "ENSG00000122566",
                "ENSG00000131473",
                "ENSG00000112305",
                "ENSG00000108774",
                "ENSG00000269893",
                "ENSG00000047932",
                "ENSG00000091140",
                "ENSG00000253738",
                "ENSG00000135250",
                "ENSG00000165119",
                "ENSG00000067064",
                "ENSG00000163636",
                "ENSG00000135486",
                "ENSG00000115128",
                "ENSG00000126005",
                "ENSG00000137818",
                "ENSG00000082153",
                "ENSG00000241343",
                "ENSG00000100347",
                "ENSG00000105677",
                "ENSG00000089737",
                "ENSG00000114784",
                "ENSG00000184752",
                "ENSG00000197302",
                "ENSG00000129355",
                "ENSG00000188343",
                "ENSG00000275052",
                "ENSG00000135002",
                "ENSG00000100575",
                "ENSG00000127445",
                "ENSG00000183172",
                "ENSG00000114744",
                "ENSG00000146282",
                "ENSG00000170027",
                "ENSG00000147274",
                "ENSG00000171055",
                "ENSG00000078674",
                "ENSG00000076554",
                "ENSG00000010270",
                "ENSG00000103495",
                "ENSG00000147604",
                "ENSG00000204308",
                "ENSG00000214046",
                "ENSG00000123416",
                "ENSG00000171314",
                "ENSG00000168291",
                "ENSG00000090238",
                "ENSG00000130520",
                "ENSG00000170315",
                "ENSG00000167792",
                "ENSG00000138413",
                "ENSG00000147684",
                "ENSG00000096746",
                "ENSG00000103266",
                "ENSG00000079785",
                "ENSG00000134779",
                "ENSG00000119950",
                "ENSG00000099795",
                "ENSG00000129084",
                "ENSG00000069275",
                "ENSG00000138777",
                "ENSG00000072042",
                "ENSG00000126756",
                "ENSG00000162585",
                "ENSG00000103145",
                "ENSG00000163714",
                "ENSG00000051620",
                "ENSG00000028839",
                "ENSG00000114200",
                "ENSG00000189043",
                "ENSG00000163939",
                "ENSG00000117523",
                "ENSG00000152154",
                "ENSG00000137100",
                "ENSG00000123106",
                "ENSG00000100764",
                "ENSG00000109475",
                "ENSG00000196199",
                "ENSG00000187109",
                "ENSG00000198886",
                "ENSG00000147533",
                "ENSG00000155506",
                "ENSG00000122643",
                "ENSG00000196262",
                "ENSG00000104904",
                "ENSG00000100316",
                "ENSG00000125971",
                "ENSG00000173726",
                "ENSG00000134287",
                "ENSG00000155959",
                "ENSG00000137692",
                "ENSG00000116586",
                "ENSG00000089693",
                "ENSG00000105254",
                "ENSG00000168092",
                "ENSG00000115286",
                "ENSG00000105698",
                "ENSG00000120438",
                "ENSG00000084234",
                "ENSG00000105640",
                "ENSG00000114023",
                "ENSG00000125691",
                "ENSG00000154723",
                "ENSG00000122026",
                "ENSG00000166347",
                "ENSG00000149357",
                "ENSG00000198843",
                "ENSG00000125901",
                "ENSG00000109654",
                "ENSG00000168288",
                "ENSG00000130956",
                "ENSG00000130558",
                "ENSG00000234745",
                "ENSG00000113387",
                "ENSG00000158864",
                "ENSG00000137409",
                "ENSG00000006125",
                "ENSG00000156508",
                "ENSG00000196937",
                "ENSG00000135900",
                "ENSG00000082701",
                "ENSG00000110717",
                "ENSG00000120727",
                "ENSG00000100442",
                "ENSG00000143799",
                "ENSG00000106028",
                "ENSG00000173917",
                "ENSG00000115091",
                "ENSG00000204628",
                "ENSG00000115053",
                "ENSG00000178234",
                "ENSG00000134986",
                "ENSG00000121022",
                "ENSG00000164615",
                "ENSG00000113369",
                "ENSG00000197111",
                "ENSG00000163682",
                "ENSG00000175334",
                "ENSG00000146729",
                "ENSG00000171421",
                "ENSG00000256537",
                "ENSG00000124380",
                "ENSG00000123562",
                "ENSG00000105701",
                "ENSG00000166260",
                "ENSG00000111669",
                "ENSG00000136100",
                "ENSG00000159720",
                "ENSG00000184007",
                "ENSG00000140307",
                "ENSG00000132821",
                "ENSG00000102401",
                "ENSG00000167863",
                "ENSG00000105409",
                "ENSG00000197747",
                "ENSG00000004779",
                "ENSG00000184277",
                "ENSG00000106355",
                "ENSG00000090273",
                "ENSG00000162368",
                "ENSG00000254682",
                "ENSG00000004478",
                "ENSG00000175756",
                "ENSG00000134333",
                "ENSG00000197345",
                "ENSG00000116251",
                "ENSG00000091640",
                "ENSG00000147123",
                "ENSG00000161960",
                "ENSG00000143575",
                "ENSG00000038219",
                "ENSG00000130560",
                "ENSG00000159140",
                "ENSG00000160014",
                "ENSG00000125818",
                "ENSG00000105649",
                "ENSG00000146731",
                "ENSG00000103187",
                "ENSG00000105220",
                "ENSG00000105372",
                "ENSG00000053254",
                "ENSG00000111678",
                "ENSG00000011295",
                "ENSG00000176871",
                "ENSG00000007168",
                "ENSG00000169288",
                "ENSG00000111142",
                "ENSG00000164713",
                "ENSG00000132507",
                "ENSG00000176209",
                "ENSG00000105379",
                "ENSG00000175550",
                "ENSG00000099901",
                "ENSG00000167508",
                "ENSG00000173692",
                "ENSG00000111237",
                "ENSG00000132824",
                "ENSG00000112110",
                "ENSG00000023191",
                "ENSG00000111911",
                "ENSG00000135070",
                "ENSG00000151366",
                "ENSG00000108298",
                "ENSG00000145782",
                "ENSG00000166226",
                "ENSG00000112695",
                "ENSG00000174444",
                "ENSG00000165416",
                "ENSG00000203667",
                "ENSG00000133318",
                "ENSG00000130414",
                "ENSG00000115738",
                "ENSG00000103429",
                "ENSG00000138430",
                "ENSG00000169738",
                "ENSG00000169213",
                "ENSG00000166780",
                "ENSG00000176340",
                "ENSG00000172354",
                "ENSG00000273611",
                "ENSG00000182670",
                "ENSG00000177410",
                "ENSG00000115514",
                "ENSG00000101745",
                "ENSG00000145337",
                "ENSG00000113068",
                "ENSG00000182718",
                "ENSG00000221983",
                "ENSG00000173436",
                "ENSG00000175348",
                "ENSG00000069956",
                "ENSG00000063046",
                "ENSG00000122565",
                "ENSG00000163541",
                "ENSG00000146416",
                "ENSG00000164258",
                "ENSG00000116560",
                "ENSG00000106261",
                "ENSG00000198492",
                "ENSG00000141002",
                "ENSG00000171604",
                "ENSG00000179262",
                "ENSG00000183207",
                "ENSG00000073792",
                "ENSG00000143355",
                "ENSG00000178952",
                "ENSG00000196230",
                "ENSG00000225663",
                "ENSG00000186106",
                "ENSG00000198034",
                "ENSG00000129518",
                "ENSG00000104408",
                "ENSG00000105373",
                "ENSG00000129235",
                "ENSG00000173418",
                "ENSG00000102144",
                "ENSG00000167526",
                "ENSG00000130770",
                "ENSG00000265681",
                "ENSG00000142864",
                "ENSG00000119013",
                "ENSG00000161203",
                "ENSG00000100823",
                "ENSG00000142534",
                "ENSG00000151746",
                "ENSG00000101782",
                "ENSG00000177889",
                "ENSG00000142541",
                "ENSG00000109971",
                "ENSG00000109536",
                "ENSG00000156136",
                "ENSG00000118816",
                "ENSG00000121851",
                "ENSG00000151743",
                "ENSG00000135624",
                "ENSG00000104671",
                "ENSG00000147669",
                "ENSG00000004487",
                "ENSG00000163468",
                "ENSG00000111843",
                "ENSG00000137312",
                "ENSG00000091039",
                "ENSG00000167522",
                "ENSG00000099800",
                "ENSG00000147403",
                "ENSG00000134056",
                "ENSG00000167220",
                "ENSG00000277443",
                "ENSG00000172115",
                "ENSG00000119547",
                "ENSG00000145912",
                "ENSG00000233429",
                "ENSG00000006451",
                "ENSG00000177156",
                "ENSG00000145293",
                "ENSG00000167468",
                "ENSG00000187634",
                "ENSG00000111640",
                "ENSG00000125652",
                "ENSG00000135387",
                "ENSG00000124596",
                "ENSG00000084628",
                "ENSG00000115241",
                "ENSG00000156976",
                "ENSG00000180964",
                "ENSG00000111481",
                "ENSG00000253719",
                "ENSG00000134138",
                "ENSG00000161970",
                "ENSG00000160888",
                "ENSG00000128245",
                "ENSG00000077721",
                "ENSG00000177879",
                "ENSG00000205531",
                "ENSG00000154001",
                "ENSG00000138758",
                "ENSG00000115541",
                "ENSG00000069329",
                "ENSG00000138069",
                "ENSG00000162244",
                "ENSG00000083099",
                "ENSG00000144034",
                "ENSG00000171033",
                "ENSG00000071082",
                "ENSG00000225733",
                "ENSG00000071994",
                "ENSG00000005483",
                "ENSG00000115484",
                "ENSG00000177030",
                "ENSG00000138107",
                "ENSG00000126261",
                "ENSG00000130748",
                "ENSG00000138326",
                "ENSG00000204568",
                "ENSG00000157778",
                "ENSG00000175203",
                "ENSG00000111642",
                "ENSG00000154380",
                "ENSG00000104695",
                "ENSG00000123144",
                "ENSG00000100216",
                "ENSG00000101557",
                "ENSG00000172531",
                "ENSG00000113575",
                "ENSG00000149084",
                "ENSG00000127022",
                "ENSG00000074800",
                "ENSG00000162961",
                "ENSG00000148303",
                "ENSG00000104863",
                "ENSG00000225921",
                "ENSG00000165502",
                "ENSG00000114302",
                "ENSG00000084733",
                "ENSG00000142676",
                "ENSG00000119705",
                "ENSG00000165283",
                "ENSG00000075151",
                "ENSG00000122034",
                "ENSG00000185664",
                "ENSG00000100528",
                "ENSG00000154582",
                "ENSG00000130255",
                "ENSG00000067836",
                "ENSG00000214113",
                "ENSG00000096384",
                "ENSG00000109107",
                "ENSG00000147687",
                "ENSG00000168214",
                "ENSG00000197958",
                "ENSG00000100911",
                "ENSG00000160213",
                "ENSG00000171497",
                "ENSG00000188994",
                "ENSG00000174775",
                "ENSG00000143947",
                "ENSG00000164978",
                "ENSG00000112237",
                "ENSG00000164830",
                "ENSG00000213015",
                "ENSG00000175416",
                "ENSG00000128609",
                "ENSG00000090470",
                "ENSG00000170632",
                "ENSG00000063177",
                "ENSG00000155368",
                "ENSG00000198918",
                "ENSG00000169564",
                "ENSG00000094916",
                "ENSG00000103035",
                "ENSG00000132305",
                "ENSG00000138663",
                "ENSG00000086061",
                "ENSG00000204310",
                "ENSG00000070756",
                "ENSG00000136930",
                "ENSG00000166922",
                "ENSG00000149806",
                "ENSG00000140386",
                "ENSG00000140740",
                "ENSG00000146386",
                "ENSG00000204272",
                "ENSG00000166012",
                "ENSG00000167658",
                "ENSG00000113161",
                "ENSG00000197006",
                "ENSG00000185608",
                "ENSG00000142549",
                "ENSG00000084070",
                "ENSG00000174173",
                "ENSG00000068745",
                "ENSG00000196465",
                "ENSG00000168028",
                "ENSG00000108848",
                "ENSG00000136193",
                "ENSG00000156928",
                "ENSG00000104427",
                "ENSG00000124702",
                "ENSG00000111786",
                "ENSG00000109919",
                "ENSG00000143933",
                "ENSG00000166352",
                "ENSG00000165629",
                "ENSG00000136603",
                "ENSG00000167815",
                "ENSG00000177971",
                "ENSG00000126457",
                "ENSG00000167996",
                "ENSG00000132963",
                "ENSG00000042753",
                "ENSG00000175390",
                "ENSG00000168522",
                "ENSG00000148154",
                "ENSG00000119048",
                "ENSG00000198408",
                "ENSG00000177600",
                "ENSG00000148450",
                "ENSG00000134419",
                "ENSG00000178127",
                "ENSG00000174456",
                "ENSG00000184117",
                "ENSG00000167699",
                "ENSG00000166946",
                "ENSG00000092148",
                "ENSG00000181610",
                "ENSG00000054148",
                "ENSG00000270170",
                "ENSG00000143621",
                "ENSG00000156411",
                "ENSG00000108960",
                "ENSG00000137343",
                "ENSG00000177200",
                "ENSG00000115944",
                "ENSG00000080822",
                "ENSG00000111639",
                "ENSG00000196531",
                "ENSG00000100138",
                "ENSG00000174695",
                "ENSG00000116030",
                "ENSG00000129968",
                "ENSG00000115268",
                "ENSG00000182534",
                "ENSG00000093144",
                "ENSG00000125534",
                "ENSG00000160075",
                "ENSG00000170043",
                "ENSG00000089157",
                "ENSG00000149231",
                "ENSG00000131435",
                "ENSG00000130723",
                "ENSG00000198363",
                "ENSG00000140988",
                "ENSG00000168090",
                "ENSG00000122033",
                "ENSG00000197728",
                "ENSG00000178741",
                "ENSG00000156515",
                "ENSG00000115947",
                "ENSG00000101266",
                "ENSG00000177885",
                "ENSG00000233016",
                "ENSG00000143774",
                "ENSG00000182512",
                "ENSG00000088832",
                "ENSG00000179271",
                "ENSG00000122545",
                "ENSG00000134330",
                "ENSG00000143612",
                "ENSG00000135720",
                "ENSG00000007520",
                "ENSG00000166848",
                "ENSG00000149547",
                "ENSG00000125870",
                "ENSG00000159335",
                "ENSG00000120686",
                "ENSG00000102189",
                "ENSG00000123352",
                "ENSG00000108518",
                "ENSG00000181061",
                "ENSG00000128739",
                "ENSG00000086666",
                "ENSG00000278311",
                "ENSG00000172809",
                "ENSG00000145425",
                "ENSG00000114902",
                "ENSG00000157514",
                "ENSG00000106615",
                "ENSG00000107341",
                "ENSG00000089818",
                "ENSG00000099341",
                "ENSG00000173273",
                "ENSG00000172500",
                "ENSG00000230989",
                "ENSG00000087302",
                "ENSG00000091164",
                "ENSG00000136146",
                "ENSG00000066557",
                "ENSG00000090263",
                "ENSG00000172757",
                "ENSG00000105669",
                "ENSG00000188290",
                "ENSG00000149313",
                "ENSG00000063245",
                "ENSG00000136942",
                "ENSG00000126858",
                "ENSG00000185658",
                "ENSG00000050426",
                "ENSG00000119185",
                "ENSG00000160752",
                "ENSG00000143252",
                "ENSG00000141367",
                "ENSG00000144381",
                "ENSG00000136521",
                "ENSG00000100811",
                "ENSG00000183648",
                "ENSG00000159352",
                "ENSG00000213246",
                "ENSG00000155099",
                "ENSG00000172687",
                "ENSG00000133872",
                "ENSG00000137106",
                "ENSG00000179010",
                "ENSG00000198668",
                "ENSG00000108107",
                "ENSG00000134308",
                "ENSG00000111832",
                "ENSG00000061794",
                "ENSG00000134684",
                "ENSG00000008324",
                "ENSG00000166228",
                "ENSG00000213619",
                "ENSG00000153774",
                "ENSG00000129197",
                "ENSG00000138757",
                "ENSG00000108561",
                "ENSG00000033627",
                "ENSG00000178982",
                "ENSG00000108010",
                "ENSG00000083845",
                "ENSG00000138382",
                "ENSG00000111726",
                "ENSG00000125821",
                "ENSG00000137547",
                "ENSG00000100387",
                "ENSG00000140319",
                "ENSG00000132466",
                "ENSG00000149809",
                "ENSG00000181704",
                "ENSG00000172428",
                "ENSG00000118680",
                "ENSG00000243317",
                "ENSG00000070669",
                "ENSG00000165233",
                "ENSG00000117450",
                "ENSG00000184220",
                "ENSG00000103485",
                "ENSG00000009307",
                "ENSG00000143995",
                "ENSG00000138138",
                "ENSG00000204856",
                "ENSG00000124172",
                "ENSG00000182196",
                "ENSG00000057608",
                "ENSG00000150991",
                "ENSG00000065548",
                "ENSG00000152795",
                "ENSG00000212907",
                "ENSG00000196591",
                "ENSG00000137210",
                "ENSG00000166681",
                "ENSG00000132432",
                "ENSG00000184009",
                "ENSG00000023734",
                "ENSG00000120437",
                "ENSG00000125944",
                "ENSG00000054523",
                "ENSG00000183726",
                "ENSG00000148925",
                "ENSG00000128626",
                "ENSG00000214941",
                "ENSG00000113328",
                "ENSG00000186184",
                "ENSG00000166164",
                "ENSG00000119640",
                "ENSG00000113648",
                "ENSG00000170899",
                "ENSG00000185787",
                "ENSG00000170889",
                "ENSG00000137947",
                "ENSG00000100554",
                "ENSG00000130638",
                "ENSG00000169045",
                "ENSG00000111775",
                "ENSG00000123636",
                "ENSG00000087502",
                "ENSG00000139684",
                "ENSG00000197102",
                "ENSG00000114120",
                "ENSG00000166562",
                "ENSG00000168724",
                "ENSG00000126602",
                "ENSG00000180817",
                "ENSG00000086065",
                "ENSG00000164587",
                "ENSG00000116288",
                "ENSG00000181163",
                "ENSG00000008018",
                "ENSG00000160948",
                "ENSG00000117569",
                "ENSG00000198898",
                "ENSG00000115685",
                "ENSG00000215021",
                "ENSG00000165487",
                "ENSG00000103275",
                "ENSG00000134313",
                "ENSG00000101654",
                "ENSG00000104549",
                "ENSG00000044115",
                "ENSG00000265241",
                "ENSG00000136238",
                "ENSG00000091527",
                "ENSG00000123545",
                "ENSG00000224032",
                "ENSG00000047849",
                "ENSG00000182154",
                "ENSG00000221914",
                "ENSG00000122873",
                "ENSG00000103415",
                "ENSG00000065978",
                "ENSG00000087460",
                "ENSG00000100567",
                "ENSG00000151247",
                "ENSG00000113558",
                "ENSG00000143153",
                "ENSG00000109180",
                "ENSG00000123983",
                "ENSG00000116489",
                "ENSG00000143742",
                "ENSG00000133169",
                "ENSG00000205629",
                "ENSG00000174886",
                "ENSG00000164167",
                "ENSG00000092841",
                "ENSG00000109929",
                "ENSG00000089009",
                "ENSG00000173163"
            ],
            "expression_sum": {
                "class1": 19184.0,
                "class2": 2356.0
            },
            "ens_id": "ENSG00000167081",
            "uniprot_ids": [
                "P40426",
                "H3BLX0",
                "Q5JS98",
                "U3KQA2",
                "H0Y5D0"
            ],
            "net_id": 0,
            "source_num": 11,
            "target_num": 116
        },
        "ENSG00000016082": {
            "id": "ENSG00000016082",
            "type": "TF",
            "symbol": "ISL1",
            "source": [
                "ENSG00000170365",
                "ENSG00000100811",
                "ENSG00000130726",
                "ENSG00000109132",
                "ENSG00000153147",
                "ENSG00000004487",
                "ENSG00000103495",
                "ENSG00000198911",
                "ENSG00000108055",
                "ENSG00000164754",
                "ENSG00000168214",
                "ENSG00000196591"
            ],
            "target": [
                "ENSG00000017260",
                "ENSG00000162704",
                "ENSG00000111716",
                "ENSG00000005810",
                "ENSG00000140941",
                "ENSG00000197959",
                "ENSG00000145817",
                "ENSG00000075415",
                "ENSG00000077380",
                "ENSG00000154556",
                "ENSG00000118733",
                "ENSG00000052802",
                "ENSG00000087274",
                "ENSG00000165669",
                "ENSG00000065911",
                "ENSG00000106443",
                "ENSG00000129636",
                "ENSG00000197061",
                "ENSG00000242173",
                "ENSG00000139767",
                "ENSG00000174574",
                "ENSG00000139116",
                "ENSG00000075914",
                "ENSG00000047249",
                "ENSG00000060982",
                "ENSG00000141447",
                "ENSG00000175581",
                "ENSG00000067225",
                "ENSG00000144218",
                "ENSG00000117139",
                "ENSG00000140105",
                "ENSG00000159685",
                "ENSG00000205981",
                "ENSG00000157216",
                "ENSG00000144746",
                "ENSG00000157168",
                "ENSG00000133606",
                "ENSG00000125835",
                "ENSG00000075142",
                "ENSG00000150722",
                "ENSG00000263001",
                "ENSG00000213741",
                "ENSG00000086598",
                "ENSG00000129515",
                "ENSG00000076604",
                "ENSG00000142546",
                "ENSG00000110955",
                "ENSG00000175216",
                "ENSG00000168137",
                "ENSG00000149187",
                "ENSG00000213760",
                "ENSG00000107581",
                "ENSG00000112972",
                "ENSG00000147677",
                "ENSG00000231500",
                "ENSG00000204842",
                "ENSG00000167196",
                "ENSG00000180370",
                "ENSG00000079246",
                "ENSG00000168275",
                "ENSG00000057019",
                "ENSG00000111850",
                "ENSG00000105185",
                "ENSG00000166974",
                "ENSG00000138801",
                "ENSG00000100320",
                "ENSG00000114391",
                "ENSG00000164985",
                "ENSG00000135390",
                "ENSG00000239672",
                "ENSG00000141867",
                "ENSG00000153310",
                "ENSG00000256060",
                "ENSG00000197223",
                "ENSG00000154277",
                "ENSG00000131469",
                "ENSG00000128654",
                "ENSG00000090266",
                "ENSG00000171634",
                "ENSG00000162374",
                "ENSG00000169764",
                "ENSG00000166200",
                "ENSG00000104388",
                "ENSG00000204954",
                "ENSG00000143222",
                "ENSG00000151914",
                "ENSG00000161057",
                "ENSG00000177565",
                "ENSG00000125827",
                "ENSG00000171130",
                "ENSG00000073578",
                "ENSG00000081154",
                "ENSG00000122566",
                "ENSG00000143878",
                "ENSG00000135486",
                "ENSG00000109670",
                "ENSG00000275052",
                "ENSG00000128951",
                "ENSG00000100575",
                "ENSG00000172159",
                "ENSG00000167792",
                "ENSG00000138413",
                "ENSG00000070761",
                "ENSG00000163848",
                "ENSG00000069275",
                "ENSG00000119729",
                "ENSG00000113742",
                "ENSG00000170515",
                "ENSG00000144895",
                "ENSG00000115875",
                "ENSG00000198886",
                "ENSG00000164929",
                "ENSG00000196262",
                "ENSG00000115355",
                "ENSG00000214078",
                "ENSG00000173726",
                "ENSG00000134287",
                "ENSG00000089280",
                "ENSG00000139117",
                "ENSG00000101132",
                "ENSG00000122026",
                "ENSG00000151327",
                "ENSG00000106400",
                "ENSG00000140836",
                "ENSG00000113387",
                "ENSG00000156508",
                "ENSG00000196937",
                "ENSG00000082701",
                "ENSG00000234608",
                "ENSG00000170142",
                "ENSG00000173141",
                "ENSG00000084463",
                "ENSG00000185630",
                "ENSG00000100442",
                "ENSG00000115091",
                "ENSG00000204628",
                "ENSG00000178234",
                "ENSG00000121022",
                "ENSG00000197111",
                "ENSG00000163069",
                "ENSG00000134825",
                "ENSG00000184007",
                "ENSG00000099250",
                "ENSG00000197345",
                "ENSG00000116251",
                "ENSG00000038219",
                "ENSG00000053254",
                "ENSG00000164330",
                "ENSG00000156639",
                "ENSG00000175550",
                "ENSG00000144339",
                "ENSG00000156011",
                "ENSG00000135070",
                "ENSG00000214413",
                "ENSG00000145782",
                "ENSG00000153187",
                "ENSG00000174444",
                "ENSG00000110711",
                "ENSG00000106829",
                "ENSG00000138430",
                "ENSG00000126214",
                "ENSG00000097033",
                "ENSG00000273611",
                "ENSG00000164733",
                "ENSG00000179933",
                "ENSG00000148943",
                "ENSG00000189159",
                "ENSG00000177410",
                "ENSG00000186472",
                "ENSG00000170365",
                "ENSG00000182718",
                "ENSG00000089053",
                "ENSG00000122565",
                "ENSG00000163541",
                "ENSG00000164258",
                "ENSG00000147862",
                "ENSG00000006715",
                "ENSG00000079102",
                "ENSG00000197756",
                "ENSG00000065135",
                "ENSG00000008294",
                "ENSG00000104408",
                "ENSG00000105373",
                "ENSG00000265681",
                "ENSG00000084207",
                "ENSG00000058272",
                "ENSG00000165434",
                "ENSG00000122068",
                "ENSG00000170873",
                "ENSG00000162434",
                "ENSG00000118816",
                "ENSG00000182827",
                "ENSG00000204442",
                "ENSG00000163468",
                "ENSG00000172115",
                "ENSG00000118257",
                "ENSG00000156976",
                "ENSG00000078369",
                "ENSG00000171033",
                "ENSG00000138107",
                "ENSG00000149485",
                "ENSG00000071994",
                "ENSG00000157778",
                "ENSG00000154380",
                "ENSG00000182621",
                "ENSG00000172531",
                "ENSG00000148303",
                "ENSG00000054118",
                "ENSG00000023228",
                "ENSG00000136717",
                "ENSG00000168175",
                "ENSG00000136854",
                "ENSG00000196305",
                "ENSG00000117859",
                "ENSG00000113845",
                "ENSG00000197958",
                "ENSG00000077279",
                "ENSG00000188994",
                "ENSG00000108946",
                "ENSG00000071794",
                "ENSG00000172586",
                "ENSG00000063177",
                "ENSG00000094916",
                "ENSG00000103035",
                "ENSG00000138663",
                "ENSG00000140386",
                "ENSG00000072041",
                "ENSG00000113108",
                "ENSG00000165462",
                "ENSG00000128699",
                "ENSG00000108848",
                "ENSG00000108826",
                "ENSG00000156928",
                "ENSG00000186469",
                "ENSG00000166352",
                "ENSG00000126457",
                "ENSG00000070087",
                "ENSG00000174547",
                "ENSG00000131876",
                "ENSG00000148700",
                "ENSG00000167699",
                "ENSG00000092148",
                "ENSG00000135406",
                "ENSG00000134824",
                "ENSG00000177675",
                "ENSG00000151552",
                "ENSG00000061676",
                "ENSG00000080822",
                "ENSG00000168268",
                "ENSG00000138594",
                "ENSG00000179915",
                "ENSG00000159593",
                "ENSG00000180228",
                "ENSG00000139613",
                "ENSG00000131116",
                "ENSG00000197728",
                "ENSG00000104823",
                "ENSG00000130779",
                "ENSG00000229117",
                "ENSG00000166206",
                "ENSG00000126773",
                "ENSG00000115350",
                "ENSG00000086666",
                "ENSG00000154518",
                "ENSG00000087302",
                "ENSG00000182628",
                "ENSG00000090263",
                "ENSG00000105669",
                "ENSG00000160752",
                "ENSG00000136521",
                "ENSG00000204220",
                "ENSG00000164741",
                "ENSG00000155330",
                "ENSG00000108100",
                "ENSG00000122958",
                "ENSG00000111832",
                "ENSG00000118263",
                "ENSG00000068615",
                "ENSG00000153774",
                "ENSG00000165525",
                "ENSG00000083845",
                "ENSG00000132466",
                "ENSG00000128923",
                "ENSG00000184220",
                "ENSG00000009307",
                "ENSG00000101882",
                "ENSG00000018189",
                "ENSG00000107372",
                "ENSG00000109132",
                "ENSG00000204856",
                "ENSG00000182117",
                "ENSG00000150991",
                "ENSG00000212907",
                "ENSG00000007080",
                "ENSG00000104679",
                "ENSG00000142875",
                "ENSG00000173660",
                "ENSG00000214941",
                "ENSG00000186184",
                "ENSG00000170899",
                "ENSG00000123636",
                "ENSG00000162604",
                "ENSG00000163964",
                "ENSG00000102003",
                "ENSG00000010810",
                "ENSG00000196781",
                "ENSG00000188846",
                "ENSG00000078018",
                "ENSG00000111530",
                "ENSG00000070961",
                "ENSG00000103275",
                "ENSG00000167232",
                "ENSG00000134313",
                "ENSG00000198712",
                "ENSG00000113448",
                "ENSG00000186468",
                "ENSG00000183527",
                "ENSG00000100567",
                "ENSG00000144567",
                "ENSG00000123983",
                "ENSG00000172270",
                "ENSG00000198888",
                "ENSG00000089009"
            ],
            "expression_sum": {
                "class1": 6727.0,
                "class2": 11.0
            },
            "ens_id": "ENSG00000016082",
            "uniprot_ids": [
                "P61371",
                "D6RBJ1"
            ],
            "net_id": 0,
            "source_num": 12,
            "target_num": 80
        },
        "ENSG00000164330": {
            "id": "ENSG00000164330",
            "type": "TF",
            "symbol": "EBF1",
            "source": [
                "ENSG00000100811",
                "ENSG00000106261",
                "ENSG00000016082",
                "ENSG00000148516",
                "ENSG00000143799",
                "ENSG00000105698",
                "ENSG00000103495",
                "ENSG00000128272",
                "ENSG00000108055",
                "ENSG00000119950",
                "ENSG00000163848",
                "ENSG00000168214",
                "ENSG00000116560",
                "ENSG00000196591",
                "ENSG00000164754",
                "ENSG00000165512",
                "ENSG00000067955"
            ],
            "target": [
                "ENSG00000206573",
                "ENSG00000005810",
                "ENSG00000117395",
                "ENSG00000134248",
                "ENSG00000189403",
                "ENSG00000022840",
                "ENSG00000164091",
                "ENSG00000144029",
                "ENSG00000102172",
                "ENSG00000150779",
                "ENSG00000077380",
                "ENSG00000154556",
                "ENSG00000134283",
                "ENSG00000173812",
                "ENSG00000117592",
                "ENSG00000106443",
                "ENSG00000123240",
                "ENSG00000102103",
                "ENSG00000178913",
                "ENSG00000074319",
                "ENSG00000146007",
                "ENSG00000106554",
                "ENSG00000157045",
                "ENSG00000163923",
                "ENSG00000125356",
                "ENSG00000255302",
                "ENSG00000144218",
                "ENSG00000113712",
                "ENSG00000153317",
                "ENSG00000026025",
                "ENSG00000157216",
                "ENSG00000133606",
                "ENSG00000022267",
                "ENSG00000103769",
                "ENSG00000170584",
                "ENSG00000075142",
                "ENSG00000273841",
                "ENSG00000124802",
                "ENSG00000142657",
                "ENSG00000196405",
                "ENSG00000126934",
                "ENSG00000151500",
                "ENSG00000086598",
                "ENSG00000121774",
                "ENSG00000149136",
                "ENSG00000127314",
                "ENSG00000142546",
                "ENSG00000185627",
                "ENSG00000092199",
                "ENSG00000107262",
                "ENSG00000175216",
                "ENSG00000174953",
                "ENSG00000118363",
                "ENSG00000122417",
                "ENSG00000112972",
                "ENSG00000231500",
                "ENSG00000265491",
                "ENSG00000113013",
                "ENSG00000109332",
                "ENSG00000079246",
                "ENSG00000057019",
                "ENSG00000095059",
                "ENSG00000232388",
                "ENSG00000114391",
                "ENSG00000168303",
                "ENSG00000112701",
                "ENSG00000162852",
                "ENSG00000135390",
                "ENSG00000131508",
                "ENSG00000154277",
                "ENSG00000198015",
                "ENSG00000048649",
                "ENSG00000169764",
                "ENSG00000120694",
                "ENSG00000127184",
                "ENSG00000160551",
                "ENSG00000103363",
                "ENSG00000064607",
                "ENSG00000171530",
                "ENSG00000149273",
                "ENSG00000165943",
                "ENSG00000137207",
                "ENSG00000108774",
                "ENSG00000115128",
                "ENSG00000134152",
                "ENSG00000138760",
                "ENSG00000183172",
                "ENSG00000169567",
                "ENSG00000275052",
                "ENSG00000114744",
                "ENSG00000147274",
                "ENSG00000123179",
                "ENSG00000101752",
                "ENSG00000108175",
                "ENSG00000084623",
                "ENSG00000096746",
                "ENSG00000119950",
                "ENSG00000138777",
                "ENSG00000069345",
                "ENSG00000162585",
                "ENSG00000075089",
                "ENSG00000028839",
                "ENSG00000189043",
                "ENSG00000163939",
                "ENSG00000075790",
                "ENSG00000099246",
                "ENSG00000115307",
                "ENSG00000116661",
                "ENSG00000134287",
                "ENSG00000105254",
                "ENSG00000119335",
                "ENSG00000120438",
                "ENSG00000127774",
                "ENSG00000172893",
                "ENSG00000106400",
                "ENSG00000122482",
                "ENSG00000125901",
                "ENSG00000113387",
                "ENSG00000149100",
                "ENSG00000158864",
                "ENSG00000137409",
                "ENSG00000006125",
                "ENSG00000136888",
                "ENSG00000170142",
                "ENSG00000127884",
                "ENSG00000121653",
                "ENSG00000140612",
                "ENSG00000163069",
                "ENSG00000111669",
                "ENSG00000124383",
                "ENSG00000197183",
                "ENSG00000102401",
                "ENSG00000132463",
                "ENSG00000124098",
                "ENSG00000197982",
                "ENSG00000165280",
                "ENSG00000116251",
                "ENSG00000091640",
                "ENSG00000147123",
                "ENSG00000143575",
                "ENSG00000176485",
                "ENSG00000184203",
                "ENSG00000183617",
                "ENSG00000105372",
                "ENSG00000143353",
                "ENSG00000111678",
                "ENSG00000169288",
                "ENSG00000130706",
                "ENSG00000161547",
                "ENSG00000135070",
                "ENSG00000108298",
                "ENSG00000174444",
                "ENSG00000130414",
                "ENSG00000169738",
                "ENSG00000172354",
                "ENSG00000150753",
                "ENSG00000138385",
                "ENSG00000164733",
                "ENSG00000115514",
                "ENSG00000113068",
                "ENSG00000221983",
                "ENSG00000069956",
                "ENSG00000122565",
                "ENSG00000164258",
                "ENSG00000137815",
                "ENSG00000141002",
                "ENSG00000171604",
                "ENSG00000170430",
                "ENSG00000166582",
                "ENSG00000225663",
                "ENSG00000065135",
                "ENSG00000105373",
                "ENSG00000124733",
                "ENSG00000272325",
                "ENSG00000100612",
                "ENSG00000166295",
                "ENSG00000168259",
                "ENSG00000119013",
                "ENSG00000161203",
                "ENSG00000025796",
                "ENSG00000151746",
                "ENSG00000101782",
                "ENSG00000099800",
                "ENSG00000147403",
                "ENSG00000140905",
                "ENSG00000101040",
                "ENSG00000277258",
                "ENSG00000115241",
                "ENSG00000161970",
                "ENSG00000100632",
                "ENSG00000129083",
                "ENSG00000078369",
                "ENSG00000133112",
                "ENSG00000138069",
                "ENSG00000110321",
                "ENSG00000115484",
                "ENSG00000110651",
                "ENSG00000157778",
                "ENSG00000145362",
                "ENSG00000182621",
                "ENSG00000123144",
                "ENSG00000100216",
                "ENSG00000143256",
                "ENSG00000149084",
                "ENSG00000102931",
                "ENSG00000165502",
                "ENSG00000114302",
                "ENSG00000104660",
                "ENSG00000185664",
                "ENSG00000067955",
                "ENSG00000130255",
                "ENSG00000214113",
                "ENSG00000022277",
                "ENSG00000147687",
                "ENSG00000160310",
                "ENSG00000158417",
                "ENSG00000197958",
                "ENSG00000171497",
                "ENSG00000198380",
                "ENSG00000108946",
                "ENSG00000213015",
                "ENSG00000175416",
                "ENSG00000170632",
                "ENSG00000172586",
                "ENSG00000063177",
                "ENSG00000160953",
                "ENSG00000155368",
                "ENSG00000169564",
                "ENSG00000103035",
                "ENSG00000128050",
                "ENSG00000165775",
                "ENSG00000086061",
                "ENSG00000167658",
                "ENSG00000113161",
                "ENSG00000189180",
                "ENSG00000196465",
                "ENSG00000128699",
                "ENSG00000143761",
                "ENSG00000111860",
                "ENSG00000031698",
                "ENSG00000109919",
                "ENSG00000167815",
                "ENSG00000126457",
                "ENSG00000177971",
                "ENSG00000070087",
                "ENSG00000177600",
                "ENSG00000148450",
                "ENSG00000136522",
                "ENSG00000166946",
                "ENSG00000136628",
                "ENSG00000092148",
                "ENSG00000054148",
                "ENSG00000070010",
                "ENSG00000181904",
                "ENSG00000104529",
                "ENSG00000118181",
                "ENSG00000080822",
                "ENSG00000186591",
                "ENSG00000168653",
                "ENSG00000159593",
                "ENSG00000156261",
                "ENSG00000198363",
                "ENSG00000151465",
                "ENSG00000140988",
                "ENSG00000122033",
                "ENSG00000163950",
                "ENSG00000143774",
                "ENSG00000165209",
                "ENSG00000182512",
                "ENSG00000110367",
                "ENSG00000100142",
                "ENSG00000115310",
                "ENSG00000122545",
                "ENSG00000175029",
                "ENSG00000108055",
                "ENSG00000099194",
                "ENSG00000086666",
                "ENSG00000188290",
                "ENSG00000167515",
                "ENSG00000134717",
                "ENSG00000169599",
                "ENSG00000141367",
                "ENSG00000213719",
                "ENSG00000105438",
                "ENSG00000112335",
                "ENSG00000204217",
                "ENSG00000132341",
                "ENSG00000124562",
                "ENSG00000100300",
                "ENSG00000145741",
                "ENSG00000087448",
                "ENSG00000128923",
                "ENSG00000175826",
                "ENSG00000009307",
                "ENSG00000138138",
                "ENSG00000168958",
                "ENSG00000057608",
                "ENSG00000165169",
                "ENSG00000065548",
                "ENSG00000150991",
                "ENSG00000017797",
                "ENSG00000107281",
                "ENSG00000132432",
                "ENSG00000184009",
                "ENSG00000120437",
                "ENSG00000054523",
                "ENSG00000146457",
                "ENSG00000100380",
                "ENSG00000128626",
                "ENSG00000166164",
                "ENSG00000119640",
                "ENSG00000104979",
                "ENSG00000114062",
                "ENSG00000128989",
                "ENSG00000213123",
                "ENSG00000168724",
                "ENSG00000126602",
                "ENSG00000036257",
                "ENSG00000164587",
                "ENSG00000008018",
                "ENSG00000160948",
                "ENSG00000112531",
                "ENSG00000082898",
                "ENSG00000115685",
                "ENSG00000101654",
                "ENSG00000167232",
                "ENSG00000144369",
                "ENSG00000091527",
                "ENSG00000182154",
                "ENSG00000181817",
                "ENSG00000143153",
                "ENSG00000187735",
                "ENSG00000087365",
                "ENSG00000174886",
                "ENSG00000017260",
                "ENSG00000111716",
                "ENSG00000169612",
                "ENSG00000099622",
                "ENSG00000114416",
                "ENSG00000131143",
                "ENSG00000184983",
                "ENSG00000127540",
                "ENSG00000180304",
                "ENSG00000108424",
                "ENSG00000147155",
                "ENSG00000051128",
                "ENSG00000250317",
                "ENSG00000141279",
                "ENSG00000237190",
                "ENSG00000135441",
                "ENSG00000172292",
                "ENSG00000129636",
                "ENSG00000134453",
                "ENSG00000182220",
                "ENSG00000141858",
                "ENSG00000167136",
                "ENSG00000149196",
                "ENSG00000204392",
                "ENSG00000114354",
                "ENSG00000153815",
                "ENSG00000196504",
                "ENSG00000055163",
                "ENSG00000037241",
                "ENSG00000109390",
                "ENSG00000170540",
                "ENSG00000144746",
                "ENSG00000133703",
                "ENSG00000146425",
                "ENSG00000120963",
                "ENSG00000131100",
                "ENSG00000176261",
                "ENSG00000114520",
                "ENSG00000215845",
                "ENSG00000069849",
                "ENSG00000240344",
                "ENSG00000123200",
                "ENSG00000147649",
                "ENSG00000144635",
                "ENSG00000187514",
                "ENSG00000076604",
                "ENSG00000160194",
                "ENSG00000178980",
                "ENSG00000083807",
                "ENSG00000168137",
                "ENSG00000172239",
                "ENSG00000129562",
                "ENSG00000108106",
                "ENSG00000116857",
                "ENSG00000256053",
                "ENSG00000105258",
                "ENSG00000263465",
                "ENSG00000169710",
                "ENSG00000163170",
                "ENSG00000184900",
                "ENSG00000110696",
                "ENSG00000088986",
                "ENSG00000107937",
                "ENSG00000131051",
                "ENSG00000166974",
                "ENSG00000029993",
                "ENSG00000239672",
                "ENSG00000005022",
                "ENSG00000131469",
                "ENSG00000047410",
                "ENSG00000136810",
                "ENSG00000142937",
                "ENSG00000171634",
                "ENSG00000232112",
                "ENSG00000106682",
                "ENSG00000079459",
                "ENSG00000178719",
                "ENSG00000119314",
                "ENSG00000170144",
                "ENSG00000143222",
                "ENSG00000127914",
                "ENSG00000177565",
                "ENSG00000106803",
                "ENSG00000175582",
                "ENSG00000171130",
                "ENSG00000198612",
                "ENSG00000130725",
                "ENSG00000122566",
                "ENSG00000142507",
                "ENSG00000047932",
                "ENSG00000237765",
                "ENSG00000165410",
                "ENSG00000089737",
                "ENSG00000130313",
                "ENSG00000129355",
                "ENSG00000188343",
                "ENSG00000128951",
                "ENSG00000100575",
                "ENSG00000120656",
                "ENSG00000241837",
                "ENSG00000010270",
                "ENSG00000056097",
                "ENSG00000131495",
                "ENSG00000173267",
                "ENSG00000147684",
                "ENSG00000072042",
                "ENSG00000126756",
                "ENSG00000103145",
                "ENSG00000137941",
                "ENSG00000167283",
                "ENSG00000187109",
                "ENSG00000115875",
                "ENSG00000164929",
                "ENSG00000204130",
                "ENSG00000196262",
                "ENSG00000125977",
                "ENSG00000089280",
                "ENSG00000130332",
                "ENSG00000168092",
                "ENSG00000115286",
                "ENSG00000105698",
                "ENSG00000105640",
                "ENSG00000114023",
                "ENSG00000166347",
                "ENSG00000131174",
                "ENSG00000106588",
                "ENSG00000196937",
                "ENSG00000188690",
                "ENSG00000135900",
                "ENSG00000170445",
                "ENSG00000143799",
                "ENSG00000204628",
                "ENSG00000102241",
                "ENSG00000134986",
                "ENSG00000121022",
                "ENSG00000113369",
                "ENSG00000163682",
                "ENSG00000171421",
                "ENSG00000085231",
                "ENSG00000124380",
                "ENSG00000105701",
                "ENSG00000134825",
                "ENSG00000090013",
                "ENSG00000167863",
                "ENSG00000125037",
                "ENSG00000004478",
                "ENSG00000105058",
                "ENSG00000175756",
                "ENSG00000145734",
                "ENSG00000197345",
                "ENSG00000005448",
                "ENSG00000250479",
                "ENSG00000160014",
                "ENSG00000146731",
                "ENSG00000130724",
                "ENSG00000164713",
                "ENSG00000132507",
                "ENSG00000175550",
                "ENSG00000173692",
                "ENSG00000132824",
                "ENSG00000164163",
                "ENSG00000134769",
                "ENSG00000166226",
                "ENSG00000103429",
                "ENSG00000102309",
                "ENSG00000167674",
                "ENSG00000258289",
                "ENSG00000164096",
                "ENSG00000156587",
                "ENSG00000116754",
                "ENSG00000175348",
                "ENSG00000141429",
                "ENSG00000100941",
                "ENSG00000164898",
                "ENSG00000152944",
                "ENSG00000141698",
                "ENSG00000162521",
                "ENSG00000196230",
                "ENSG00000100028",
                "ENSG00000029363",
                "ENSG00000100823",
                "ENSG00000105887",
                "ENSG00000169136",
                "ENSG00000160131",
                "ENSG00000109971",
                "ENSG00000159199",
                "ENSG00000118816",
                "ENSG00000172590",
                "ENSG00000151474",
                "ENSG00000137312",
                "ENSG00000167522",
                "ENSG00000111371",
                "ENSG00000172115",
                "ENSG00000101558",
                "ENSG00000125970",
                "ENSG00000006451",
                "ENSG00000167468",
                "ENSG00000111640",
                "ENSG00000183978",
                "ENSG00000125652",
                "ENSG00000124596",
                "ENSG00000169021",
                "ENSG00000130294",
                "ENSG00000077721",
                "ENSG00000099624",
                "ENSG00000120265",
                "ENSG00000138758",
                "ENSG00000150093",
                "ENSG00000162441",
                "ENSG00000130748",
                "ENSG00000111642",
                "ENSG00000137575",
                "ENSG00000145833",
                "ENSG00000103121",
                "ENSG00000104412",
                "ENSG00000113575",
                "ENSG00000225921",
                "ENSG00000105193",
                "ENSG00000009954",
                "ENSG00000084733",
                "ENSG00000136717",
                "ENSG00000149557",
                "ENSG00000206503",
                "ENSG00000100528",
                "ENSG00000120533",
                "ENSG00000162734",
                "ENSG00000168214",
                "ENSG00000071794",
                "ENSG00000112237",
                "ENSG00000106591",
                "ENSG00000130311",
                "ENSG00000130985",
                "ENSG00000094916",
                "ENSG00000138663",
                "ENSG00000186063",
                "ENSG00000140740",
                "ENSG00000140995",
                "ENSG00000146386",
                "ENSG00000197006",
                "ENSG00000185608",
                "ENSG00000108468",
                "ENSG00000205302",
                "ENSG00000104427",
                "ENSG00000115756",
                "ENSG00000113141",
                "ENSG00000166503",
                "ENSG00000132963",
                "ENSG00000175390",
                "ENSG00000168522",
                "ENSG00000148154",
                "ENSG00000119048",
                "ENSG00000172775",
                "ENSG00000134419",
                "ENSG00000174456",
                "ENSG00000135406",
                "ENSG00000143621",
                "ENSG00000177200",
                "ENSG00000135334",
                "ENSG00000100138",
                "ENSG00000168268",
                "ENSG00000107798",
                "ENSG00000125534",
                "ENSG00000131435",
                "ENSG00000126432",
                "ENSG00000130723",
                "ENSG00000124767",
                "ENSG00000178741",
                "ENSG00000156515",
                "ENSG00000101266",
                "ENSG00000233016",
                "ENSG00000166595",
                "ENSG00000134330",
                "ENSG00000184207",
                "ENSG00000167969",
                "ENSG00000007520",
                "ENSG00000159335",
                "ENSG00000137726",
                "ENSG00000141522",
                "ENSG00000114902",
                "ENSG00000099341",
                "ENSG00000116095",
                "ENSG00000103342",
                "ENSG00000172757",
                "ENSG00000105669",
                "ENSG00000050426",
                "ENSG00000136156",
                "ENSG00000119185",
                "ENSG00000169020",
                "ENSG00000171159",
                "ENSG00000144381",
                "ENSG00000136521",
                "ENSG00000213246",
                "ENSG00000155330",
                "ENSG00000112667",
                "ENSG00000133872",
                "ENSG00000198668",
                "ENSG00000134308",
                "ENSG00000134684",
                "ENSG00000089063",
                "ENSG00000204525",
                "ENSG00000168439",
                "ENSG00000138757",
                "ENSG00000067900",
                "ENSG00000214253",
                "ENSG00000138382",
                "ENSG00000118680",
                "ENSG00000184220",
                "ENSG00000205937",
                "ENSG00000115694",
                "ENSG00000065000",
                "ENSG00000125445",
                "ENSG00000165916",
                "ENSG00000182196",
                "ENSG00000152795",
                "ENSG00000165716",
                "ENSG00000072803",
                "ENSG00000100568",
                "ENSG00000176407",
                "ENSG00000125944",
                "ENSG00000183726",
                "ENSG00000214941",
                "ENSG00000186184",
                "ENSG00000100804",
                "ENSG00000179222",
                "ENSG00000163964",
                "ENSG00000166562",
                "ENSG00000104763",
                "ENSG00000181163",
                "ENSG00000158710",
                "ENSG00000215021",
                "ENSG00000122873",
                "ENSG00000113851",
                "ENSG00000145247",
                "ENSG00000137776",
                "ENSG00000139637",
                "ENSG00000106244",
                "ENSG00000078668",
                "ENSG00000090621",
                "ENSG00000100410",
                "ENSG00000116171",
                "ENSG00000162704",
                "ENSG00000139644",
                "ENSG00000175745",
                "ENSG00000182552",
                "ENSG00000090060",
                "ENSG00000116161",
                "ENSG00000160209",
                "ENSG00000161016",
                "ENSG00000125743",
                "ENSG00000105819",
                "ENSG00000179958",
                "ENSG00000075415",
                "ENSG00000107874",
                "ENSG00000152904",
                "ENSG00000163466",
                "ENSG00000198931",
                "ENSG00000197451",
                "ENSG00000152256",
                "ENSG00000173915",
                "ENSG00000111728",
                "ENSG00000167085",
                "ENSG00000170310",
                "ENSG00000115540",
                "ENSG00000041357",
                "ENSG00000163875",
                "ENSG00000163319",
                "ENSG00000164924",
                "ENSG00000141759",
                "ENSG00000165264",
                "ENSG00000174574",
                "ENSG00000110619",
                "ENSG00000139116",
                "ENSG00000277791",
                "ENSG00000149923",
                "ENSG00000118482",
                "ENSG00000116898",
                "ENSG00000179431",
                "ENSG00000188243",
                "ENSG00000100591",
                "ENSG00000140105",
                "ENSG00000034713",
                "ENSG00000130811",
                "ENSG00000005194",
                "ENSG00000153048",
                "ENSG00000125835",
                "ENSG00000183011",
                "ENSG00000174437",
                "ENSG00000187051",
                "ENSG00000129515",
                "ENSG00000155115",
                "ENSG00000204463",
                "ENSG00000010256",
                "ENSG00000120053",
                "ENSG00000135940",
                "ENSG00000172336",
                "ENSG00000131966",
                "ENSG00000107438",
                "ENSG00000153037",
                "ENSG00000106245",
                "ENSG00000075624",
                "ENSG00000072062",
                "ENSG00000168275",
                "ENSG00000102898",
                "ENSG00000152700",
                "ENSG00000167088",
                "ENSG00000240972",
                "ENSG00000196655",
                "ENSG00000141030",
                "ENSG00000164985",
                "ENSG00000250303",
                "ENSG00000141867",
                "ENSG00000165948",
                "ENSG00000197223",
                "ENSG00000180957",
                "ENSG00000099817",
                "ENSG00000143862",
                "ENSG00000176731",
                "ENSG00000161981",
                "ENSG00000162910",
                "ENSG00000104388",
                "ENSG00000097021",
                "ENSG00000087191",
                "ENSG00000166734",
                "ENSG00000257704",
                "ENSG00000130204",
                "ENSG00000014641",
                "ENSG00000121067",
                "ENSG00000176171",
                "ENSG00000131473",
                "ENSG00000112305",
                "ENSG00000253738",
                "ENSG00000165119",
                "ENSG00000126005",
                "ENSG00000241343",
                "ENSG00000184752",
                "ENSG00000198258",
                "ENSG00000170027",
                "ENSG00000023287",
                "ENSG00000111707",
                "ENSG00000176101",
                "ENSG00000123416",
                "ENSG00000171314",
                "ENSG00000144713",
                "ENSG00000067248",
                "ENSG00000164975",
                "ENSG00000130520",
                "ENSG00000170315",
                "ENSG00000167792",
                "ENSG00000138413",
                "ENSG00000163848",
                "ENSG00000099795",
                "ENSG00000079462",
                "ENSG00000166037",
                "ENSG00000163344",
                "ENSG00000051620",
                "ENSG00000119729",
                "ENSG00000135404",
                "ENSG00000113742",
                "ENSG00000142192",
                "ENSG00000131828",
                "ENSG00000171858",
                "ENSG00000144895",
                "ENSG00000123106",
                "ENSG00000147533",
                "ENSG00000155506",
                "ENSG00000100316",
                "ENSG00000170348",
                "ENSG00000140259",
                "ENSG00000143771",
                "ENSG00000089693",
                "ENSG00000169432",
                "ENSG00000144028",
                "ENSG00000164182",
                "ENSG00000122026",
                "ENSG00000198843",
                "ENSG00000109654",
                "ENSG00000145495",
                "ENSG00000254999",
                "ENSG00000130741",
                "ENSG00000177556",
                "ENSG00000186222",
                "ENSG00000156508",
                "ENSG00000082701",
                "ENSG00000120727",
                "ENSG00000101052",
                "ENSG00000106367",
                "ENSG00000101126",
                "ENSG00000197111",
                "ENSG00000164615",
                "ENSG00000169813",
                "ENSG00000175334",
                "ENSG00000099783",
                "ENSG00000159720",
                "ENSG00000182093",
                "ENSG00000184007",
                "ENSG00000140307",
                "ENSG00000106355",
                "ENSG00000169139",
                "ENSG00000109736",
                "ENSG00000203950",
                "ENSG00000070831",
                "ENSG00000161960",
                "ENSG00000167004",
                "ENSG00000249915",
                "ENSG00000110700",
                "ENSG00000159140",
                "ENSG00000170892",
                "ENSG00000067560",
                "ENSG00000176871",
                "ENSG00000156639",
                "ENSG00000143727",
                "ENSG00000167508",
                "ENSG00000111237",
                "ENSG00000182985",
                "ENSG00000023191",
                "ENSG00000151366",
                "ENSG00000214413",
                "ENSG00000145782",
                "ENSG00000116747",
                "ENSG00000153187",
                "ENSG00000106628",
                "ENSG00000100354",
                "ENSG00000131149",
                "ENSG00000183520",
                "ENSG00000138430",
                "ENSG00000176340",
                "ENSG00000273611",
                "ENSG00000166441",
                "ENSG00000101745",
                "ENSG00000145337",
                "ENSG00000170365",
                "ENSG00000182718",
                "ENSG00000140990",
                "ENSG00000163541",
                "ENSG00000146416",
                "ENSG00000028203",
                "ENSG00000112245",
                "ENSG00000092201",
                "ENSG00000141543",
                "ENSG00000173272",
                "ENSG00000179262",
                "ENSG00000101210",
                "ENSG00000197756",
                "ENSG00000178952",
                "ENSG00000260032",
                "ENSG00000129235",
                "ENSG00000103005",
                "ENSG00000117448",
                "ENSG00000142864",
                "ENSG00000137288",
                "ENSG00000142534",
                "ENSG00000211450",
                "ENSG00000106399",
                "ENSG00000142541",
                "ENSG00000122068",
                "ENSG00000181722",
                "ENSG00000109046",
                "ENSG00000143753",
                "ENSG00000135624",
                "ENSG00000163468",
                "ENSG00000155096",
                "ENSG00000091039",
                "ENSG00000117713",
                "ENSG00000177156",
                "ENSG00000140350",
                "ENSG00000128789",
                "ENSG00000197746",
                "ENSG00000135387",
                "ENSG00000160888",
                "ENSG00000205531",
                "ENSG00000143549",
                "ENSG00000187231",
                "ENSG00000225733",
                "ENSG00000177030",
                "ENSG00000171490",
                "ENSG00000154380",
                "ENSG00000127125",
                "ENSG00000127022",
                "ENSG00000074800",
                "ENSG00000162961",
                "ENSG00000148303",
                "ENSG00000133026",
                "ENSG00000122034",
                "ENSG00000138592",
                "ENSG00000154582",
                "ENSG00000067836",
                "ENSG00000114209",
                "ENSG00000186298",
                "ENSG00000164902",
                "ENSG00000144224",
                "ENSG00000137996",
                "ENSG00000188994",
                "ENSG00000164830",
                "ENSG00000133226",
                "ENSG00000135241",
                "ENSG00000108953",
                "ENSG00000105655",
                "ENSG00000132305",
                "ENSG00000204310",
                "ENSG00000136930",
                "ENSG00000149806",
                "ENSG00000132475",
                "ENSG00000108848",
                "ENSG00000156928",
                "ENSG00000111786",
                "ENSG00000124702",
                "ENSG00000143933",
                "ENSG00000167996",
                "ENSG00000131876",
                "ENSG00000134824",
                "ENSG00000156411",
                "ENSG00000099864",
                "ENSG00000138443",
                "ENSG00000196531",
                "ENSG00000115268",
                "ENSG00000182534",
                "ENSG00000242485",
                "ENSG00000086300",
                "ENSG00000170043",
                "ENSG00000149231",
                "ENSG00000182481",
                "ENSG00000188229",
                "ENSG00000118292",
                "ENSG00000180228",
                "ENSG00000126247",
                "ENSG00000136758",
                "ENSG00000006015",
                "ENSG00000173457",
                "ENSG00000088832",
                "ENSG00000135720",
                "ENSG00000136485",
                "ENSG00000013275",
                "ENSG00000125870",
                "ENSG00000123352",
                "ENSG00000112514",
                "ENSG00000137133",
                "ENSG00000278311",
                "ENSG00000172809",
                "ENSG00000157514",
                "ENSG00000107341",
                "ENSG00000172500",
                "ENSG00000091164",
                "ENSG00000136146",
                "ENSG00000155366",
                "ENSG00000160752",
                "ENSG00000075618",
                "ENSG00000143252",
                "ENSG00000134809",
                "ENSG00000198961",
                "ENSG00000159352",
                "ENSG00000155099",
                "ENSG00000137106",
                "ENSG00000008324",
                "ENSG00000166228",
                "ENSG00000118263",
                "ENSG00000213619",
                "ENSG00000129197",
                "ENSG00000164919",
                "ENSG00000165525",
                "ENSG00000033627",
                "ENSG00000140319",
                "ENSG00000132466",
                "ENSG00000149809",
                "ENSG00000243317",
                "ENSG00000165233",
                "ENSG00000107372",
                "ENSG00000085719",
                "ENSG00000137876",
                "ENSG00000138398",
                "ENSG00000136527",
                "ENSG00000136819",
                "ENSG00000110958",
                "ENSG00000101608",
                "ENSG00000164323",
                "ENSG00000122705",
                "ENSG00000178035",
                "ENSG00000140374",
                "ENSG00000173660",
                "ENSG00000113648",
                "ENSG00000104643",
                "ENSG00000227051",
                "ENSG00000162385",
                "ENSG00000169045",
                "ENSG00000165732",
                "ENSG00000008988",
                "ENSG00000196781",
                "ENSG00000180817",
                "ENSG00000086065",
                "ENSG00000136938",
                "ENSG00000220205",
                "ENSG00000136238",
                "ENSG00000164754",
                "ENSG00000143183",
                "ENSG00000123545",
                "ENSG00000224032",
                "ENSG00000105401",
                "ENSG00000221914",
                "ENSG00000103415",
                "ENSG00000065978",
                "ENSG00000087460",
                "ENSG00000151247",
                "ENSG00000170906",
                "ENSG00000109180",
                "ENSG00000123983",
                "ENSG00000143742",
                "ENSG00000205629",
                "ENSG00000172270",
                "ENSG00000177733",
                "ENSG00000092841",
                "ENSG00000129351",
                "ENSG00000089009",
                "ENSG00000141741",
                "ENSG00000134884",
                "ENSG00000163634",
                "ENSG00000095787",
                "ENSG00000169976",
                "ENSG00000106603",
                "ENSG00000140941",
                "ENSG00000184840",
                "ENSG00000075945",
                "ENSG00000099797",
                "ENSG00000196419",
                "ENSG00000105993",
                "ENSG00000213722",
                "ENSG00000088833",
                "ENSG00000119421",
                "ENSG00000114439",
                "ENSG00000087274",
                "ENSG00000165669",
                "ENSG00000184697",
                "ENSG00000065911",
                "ENSG00000151729",
                "ENSG00000272398",
                "ENSG00000197061",
                "ENSG00000125995",
                "ENSG00000135829",
                "ENSG00000135069",
                "ENSG00000162664",
                "ENSG00000112308",
                "ENSG00000148908",
                "ENSG00000139112",
                "ENSG00000156482",
                "ENSG00000165637",
                "ENSG00000241685",
                "ENSG00000060982",
                "ENSG00000147548",
                "ENSG00000171703",
                "ENSG00000131236",
                "ENSG00000129625",
                "ENSG00000196072",
                "ENSG00000100077",
                "ENSG00000067225",
                "ENSG00000144642",
                "ENSG00000141552",
                "ENSG00000136143",
                "ENSG00000205581",
                "ENSG00000117139",
                "ENSG00000104131",
                "ENSG00000095380",
                "ENSG00000034510",
                "ENSG00000138071",
                "ENSG00000263001",
                "ENSG00000213741",
                "ENSG00000145592",
                "ENSG00000134001",
                "ENSG00000138698",
                "ENSG00000123143",
                "ENSG00000110955",
                "ENSG00000164687",
                "ENSG00000169100",
                "ENSG00000162191",
                "ENSG00000147677",
                "ENSG00000108666",
                "ENSG00000070770",
                "ENSG00000175792",
                "ENSG00000167196",
                "ENSG00000180370",
                "ENSG00000171222",
                "ENSG00000181222",
                "ENSG00000105185",
                "ENSG00000153310",
                "ENSG00000100348",
                "ENSG00000213585",
                "ENSG00000145919",
                "ENSG00000146223",
                "ENSG00000131437",
                "ENSG00000143106",
                "ENSG00000127616",
                "ENSG00000174720",
                "ENSG00000128654",
                "ENSG00000080824",
                "ENSG00000160271",
                "ENSG00000166200",
                "ENSG00000130312",
                "ENSG00000136950",
                "ENSG00000197694",
                "ENSG00000115816",
                "ENSG00000168159",
                "ENSG00000152082",
                "ENSG00000115677",
                "ENSG00000151914",
                "ENSG00000168710",
                "ENSG00000124614",
                "ENSG00000092203",
                "ENSG00000104765",
                "ENSG00000073578",
                "ENSG00000182199",
                "ENSG00000269893",
                "ENSG00000067064",
                "ENSG00000135486",
                "ENSG00000163636",
                "ENSG00000109670",
                "ENSG00000137818",
                "ENSG00000082153",
                "ENSG00000002330",
                "ENSG00000129559",
                "ENSG00000134265",
                "ENSG00000105677",
                "ENSG00000135002",
                "ENSG00000171735",
                "ENSG00000146282",
                "ENSG00000078674",
                "ENSG00000103495",
                "ENSG00000126768",
                "ENSG00000147604",
                "ENSG00000204308",
                "ENSG00000165512",
                "ENSG00000086758",
                "ENSG00000134779",
                "ENSG00000079785",
                "ENSG00000070761",
                "ENSG00000185825",
                "ENSG00000134369",
                "ENSG00000163714",
                "ENSG00000173473",
                "ENSG00000196636",
                "ENSG00000170515",
                "ENSG00000164405",
                "ENSG00000196199",
                "ENSG00000104904",
                "ENSG00000134970",
                "ENSG00000173726",
                "ENSG00000167460",
                "ENSG00000155959",
                "ENSG00000116586",
                "ENSG00000186010",
                "ENSG00000084234",
                "ENSG00000125691",
                "ENSG00000151327",
                "ENSG00000167978",
                "ENSG00000186187",
                "ENSG00000089289",
                "ENSG00000168288",
                "ENSG00000004059",
                "ENSG00000144848",
                "ENSG00000095739",
                "ENSG00000234608",
                "ENSG00000241468",
                "ENSG00000110717",
                "ENSG00000115053",
                "ENSG00000111275",
                "ENSG00000142949",
                "ENSG00000136100",
                "ENSG00000099250",
                "ENSG00000197747",
                "ENSG00000162368",
                "ENSG00000254682",
                "ENSG00000100353",
                "ENSG00000116459",
                "ENSG00000134375",
                "ENSG00000134333",
                "ENSG00000038219",
                "ENSG00000086232",
                "ENSG00000125818",
                "ENSG00000103187",
                "ENSG00000115758",
                "ENSG00000105220",
                "ENSG00000053254",
                "ENSG00000272047",
                "ENSG00000011295",
                "ENSG00000119707",
                "ENSG00000007168",
                "ENSG00000176209",
                "ENSG00000112110",
                "ENSG00000067369",
                "ENSG00000104886",
                "ENSG00000097033",
                "ENSG00000029725",
                "ENSG00000177410",
                "ENSG00000146701",
                "ENSG00000116560",
                "ENSG00000114956",
                "ENSG00000198492",
                "ENSG00000183207",
                "ENSG00000073792",
                "ENSG00000182004",
                "ENSG00000198034",
                "ENSG00000141526",
                "ENSG00000104408",
                "ENSG00000173418",
                "ENSG00000160633",
                "ENSG00000102144",
                "ENSG00000167526",
                "ENSG00000084207",
                "ENSG00000058272",
                "ENSG00000184076",
                "ENSG00000177889",
                "ENSG00000166913",
                "ENSG00000182774",
                "ENSG00000111906",
                "ENSG00000204435",
                "ENSG00000109536",
                "ENSG00000162434",
                "ENSG00000090487",
                "ENSG00000126524",
                "ENSG00000081870",
                "ENSG00000111843",
                "ENSG00000074071",
                "ENSG00000109255",
                "ENSG00000134056",
                "ENSG00000145293",
                "ENSG00000181924",
                "ENSG00000148516",
                "ENSG00000074696",
                "ENSG00000169750",
                "ENSG00000253719",
                "ENSG00000156976",
                "ENSG00000169230",
                "ENSG00000105404",
                "ENSG00000177879",
                "ENSG00000154001",
                "ENSG00000115541",
                "ENSG00000168066",
                "ENSG00000162244",
                "ENSG00000083099",
                "ENSG00000198420",
                "ENSG00000149485",
                "ENSG00000071994",
                "ENSG00000109270",
                "ENSG00000104695",
                "ENSG00000141404",
                "ENSG00000172531",
                "ENSG00000054118",
                "ENSG00000179950",
                "ENSG00000142676",
                "ENSG00000165283",
                "ENSG00000054116",
                "ENSG00000196305",
                "ENSG00000096384",
                "ENSG00000102753",
                "ENSG00000109107",
                "ENSG00000113845",
                "ENSG00000124486",
                "ENSG00000198900",
                "ENSG00000100911",
                "ENSG00000214517",
                "ENSG00000115524",
                "ENSG00000174775",
                "ENSG00000143947",
                "ENSG00000128609",
                "ENSG00000182809",
                "ENSG00000198918",
                "ENSG00000147140",
                "ENSG00000133706",
                "ENSG00000205155",
                "ENSG00000161013",
                "ENSG00000088930",
                "ENSG00000168028",
                "ENSG00000042753",
                "ENSG00000198408",
                "ENSG00000178127",
                "ENSG00000184117",
                "ENSG00000167699",
                "ENSG00000105518",
                "ENSG00000143198",
                "ENSG00000170017",
                "ENSG00000145494",
                "ENSG00000122378",
                "ENSG00000115944",
                "ENSG00000111639",
                "ENSG00000093144",
                "ENSG00000160113",
                "ENSG00000138594",
                "ENSG00000160075",
                "ENSG00000120992",
                "ENSG00000089157",
                "ENSG00000166165",
                "ENSG00000131116",
                "ENSG00000168090",
                "ENSG00000197728",
                "ENSG00000104823",
                "ENSG00000177885",
                "ENSG00000130779",
                "ENSG00000122406",
                "ENSG00000215712",
                "ENSG00000179271",
                "ENSG00000196683",
                "ENSG00000149547",
                "ENSG00000120686",
                "ENSG00000102189",
                "ENSG00000149600",
                "ENSG00000108518",
                "ENSG00000106615",
                "ENSG00000087302",
                "ENSG00000136942",
                "ENSG00000105176",
                "ENSG00000119318",
                "ENSG00000166963",
                "ENSG00000104177",
                "ENSG00000152234",
                "ENSG00000108107",
                "ENSG00000143158",
                "ENSG00000111832",
                "ENSG00000061794",
                "ENSG00000153774",
                "ENSG00000108561",
                "ENSG00000178982",
                "ENSG00000108010",
                "ENSG00000083845",
                "ENSG00000155876",
                "ENSG00000171867",
                "ENSG00000111726",
                "ENSG00000125821",
                "ENSG00000070669",
                "ENSG00000117450",
                "ENSG00000018189",
                "ENSG00000101220",
                "ENSG00000182117",
                "ENSG00000124172",
                "ENSG00000196591",
                "ENSG00000142669",
                "ENSG00000100075",
                "ENSG00000025293",
                "ENSG00000178104",
                "ENSG00000115306",
                "ENSG00000114125",
                "ENSG00000185787",
                "ENSG00000105568",
                "ENSG00000100554",
                "ENSG00000111775",
                "ENSG00000133142",
                "ENSG00000114120",
                "ENSG00000163938",
                "ENSG00000188612",
                "ENSG00000137154",
                "ENSG00000116288",
                "ENSG00000205542",
                "ENSG00000103275",
                "ENSG00000128524",
                "ENSG00000137274",
                "ENSG00000044115",
                "ENSG00000047849",
                "ENSG00000130402",
                "ENSG00000198648",
                "ENSG00000085733",
                "ENSG00000114115",
                "ENSG00000181610",
                "ENSG00000173163"
            ],
            "expression_sum": {
                "class1": 14988.0,
                "class2": 1997.0
            },
            "ens_id": "ENSG00000164330",
            "uniprot_ids": [
                "Q9UH73",
                "E5RFQ1",
                "A0A087WTE5"
            ],
            "net_id": 0,
            "source_num": 17,
            "target_num": 141
        },
        "ENSG00000134138": {
            "id": "ENSG00000134138",
            "type": "TF",
            "symbol": "MEIS2",
            "source": [
                "ENSG00000162599",
                "ENSG00000167081",
                "ENSG00000113580",
                "ENSG00000168610",
                "ENSG00000175745",
                "ENSG00000100219",
                "ENSG00000114861",
                "ENSG00000221869",
                "ENSG00000139352",
                "ENSG00000137203",
                "ENSG00000119547",
                "ENSG00000143995"
            ],
            "target": [
                "ENSG00000049130",
                "ENSG00000196230",
                "ENSG00000164199",
                "ENSG00000102265",
                "ENSG00000187134",
                "ENSG00000117984",
                "ENSG00000135506",
                "ENSG00000184486",
                "ENSG00000265972",
                "ENSG00000151746",
                "ENSG00000162490",
                "ENSG00000104805",
                "ENSG00000183091",
                "ENSG00000163956",
                "ENSG00000137203",
                "ENSG00000166250",
                "ENSG00000150593",
                "ENSG00000124570",
                "ENSG00000196141",
                "ENSG00000251562",
                "ENSG00000135404",
                "ENSG00000153234",
                "ENSG00000180879",
                "ENSG00000112514",
                "ENSG00000119547",
                "ENSG00000065613",
                "ENSG00000175130",
                "ENSG00000170266",
                "ENSG00000213064",
                "ENSG00000197746",
                "ENSG00000188290",
                "ENSG00000130005",
                "ENSG00000169432",
                "ENSG00000255302",
                "ENSG00000131711",
                "ENSG00000122884",
                "ENSG00000153113",
                "ENSG00000068697",
                "ENSG00000140939",
                "ENSG00000181744",
                "ENSG00000143013",
                "ENSG00000174851",
                "ENSG00000130511",
                "ENSG00000163382",
                "ENSG00000205981",
                "ENSG00000135549",
                "ENSG00000166228",
                "ENSG00000162496",
                "ENSG00000119655",
                "ENSG00000149557",
                "ENSG00000185664",
                "ENSG00000163399",
                "ENSG00000123131",
                "ENSG00000244187",
                "ENSG00000102804",
                "ENSG00000143995",
                "ENSG00000196220",
                "ENSG00000105223",
                "ENSG00000071553",
                "ENSG00000158604",
                "ENSG00000131773",
                "ENSG00000144674",
                "ENSG00000114861",
                "ENSG00000106537",
                "ENSG00000103316",
                "ENSG00000120742",
                "ENSG00000146540",
                "ENSG00000124155",
                "ENSG00000185115",
                "ENSG00000143543",
                "ENSG00000137267",
                "ENSG00000151150",
                "ENSG00000146278",
                "ENSG00000090932",
                "ENSG00000145675",
                "ENSG00000100345",
                "ENSG00000005249",
                "ENSG00000157851",
                "ENSG00000078018",
                "ENSG00000137880",
                "ENSG00000132475",
                "ENSG00000197043",
                "ENSG00000133318",
                "ENSG00000026297",
                "ENSG00000213614",
                "ENSG00000137285",
                "ENSG00000131037",
                "ENSG00000189159",
                "ENSG00000112290",
                "ENSG00000185624",
                "ENSG00000167081",
                "ENSG00000172270",
                "ENSG00000138640",
                "ENSG00000107872"
            ],
            "expression_sum": {
                "class1": 55941.0,
                "class2": 12368.0
            },
            "ens_id": "ENSG00000134138",
            "uniprot_ids": [
                "O14770",
                "H0YM65",
                "H0YMX5",
                "H0YKE5",
                "U3KQI2",
                "H0YKN2",
                "U3KQ95"
            ],
            "net_id": 0,
            "source_num": 12,
            "target_num": 12
        },
        "ENSG00000143995": {
            "id": "ENSG00000143995",
            "type": "TF",
            "symbol": "MEIS1",
            "source": [
                "ENSG00000167081",
                "ENSG00000175745",
                "ENSG00000119547",
                "ENSG00000134138"
            ],
            "target": [
                "ENSG00000134138",
                "ENSG00000162374",
                "ENSG00000165655",
                "ENSG00000175745",
                "ENSG00000151150",
                "ENSG00000149294"
            ],
            "expression_sum": {
                "class1": 6891.0,
                "class2": 992.0
            },
            "ens_id": "ENSG00000143995",
            "uniprot_ids": [
                "O00470",
                "U3KPR8",
                "F5GYS8",
                "E9PDC4",
                "H0YNY8"
            ],
            "net_id": 0,
            "source_num": 4,
            "target_num": 3
        },
        "ENSG00000163435": {
            "id": "ENSG00000163435",
            "type": "TF",
            "symbol": "ELF3",
            "source": [
                "ENSG00000106261",
                "ENSG00000100219",
                "ENSG00000221869",
                "ENSG00000196591",
                "ENSG00000168214",
                "ENSG00000124766",
                "ENSG00000116044",
                "ENSG00000160113"
            ],
            "target": [
                "ENSG00000206573",
                "ENSG00000141741",
                "ENSG00000162704",
                "ENSG00000163634",
                "ENSG00000132698",
                "ENSG00000139644",
                "ENSG00000090060",
                "ENSG00000173467",
                "ENSG00000134248",
                "ENSG00000131143",
                "ENSG00000127540",
                "ENSG00000160932",
                "ENSG00000165215",
                "ENSG00000146247",
                "ENSG00000108424",
                "ENSG00000113282",
                "ENSG00000188483",
                "ENSG00000265972",
                "ENSG00000154556",
                "ENSG00000163466",
                "ENSG00000198931",
                "ENSG00000114439",
                "ENSG00000173812",
                "ENSG00000141279",
                "ENSG00000198899",
                "ENSG00000123240",
                "ENSG00000041357",
                "ENSG00000008517",
                "ENSG00000163875",
                "ENSG00000167136",
                "ENSG00000164924",
                "ENSG00000075213",
                "ENSG00000197930",
                "ENSG00000180879",
                "ENSG00000104964",
                "ENSG00000156482",
                "ENSG00000125868",
                "ENSG00000165637",
                "ENSG00000241685",
                "ENSG00000125356",
                "ENSG00000147548",
                "ENSG00000196504",
                "ENSG00000129625",
                "ENSG00000116898",
                "ENSG00000130513",
                "ENSG00000147889",
                "ENSG00000067225",
                "ENSG00000153113",
                "ENSG00000113712",
                "ENSG00000233927",
                "ENSG00000150459",
                "ENSG00000117139",
                "ENSG00000148444",
                "ENSG00000205581",
                "ENSG00000161249",
                "ENSG00000129521",
                "ENSG00000062716",
                "ENSG00000095380",
                "ENSG00000086548",
                "ENSG00000133703",
                "ENSG00000133606",
                "ENSG00000116044",
                "ENSG00000103769",
                "ENSG00000138071",
                "ENSG00000170293",
                "ENSG00000153048",
                "ENSG00000263001",
                "ENSG00000110092",
                "ENSG00000213741",
                "ENSG00000215845",
                "ENSG00000111907",
                "ENSG00000187514",
                "ENSG00000076604",
                "ENSG00000163041",
                "ENSG00000092199",
                "ENSG00000110955",
                "ENSG00000107262",
                "ENSG00000164346",
                "ENSG00000172239",
                "ENSG00000174953",
                "ENSG00000064651",
                "ENSG00000169100",
                "ENSG00000162191",
                "ENSG00000108654",
                "ENSG00000135940",
                "ENSG00000124107",
                "ENSG00000107581",
                "ENSG00000092010",
                "ENSG00000106052",
                "ENSG00000147677",
                "ENSG00000107438",
                "ENSG00000079246",
                "ENSG00000171222",
                "ENSG00000135046",
                "ENSG00000095059",
                "ENSG00000138801",
                "ENSG00000240972",
                "ENSG00000054267",
                "ENSG00000172915",
                "ENSG00000030582",
                "ENSG00000135390",
                "ENSG00000136770",
                "ENSG00000005022",
                "ENSG00000127616",
                "ENSG00000128654",
                "ENSG00000153250",
                "ENSG00000047410",
                "ENSG00000136810",
                "ENSG00000174021",
                "ENSG00000169764",
                "ENSG00000127184",
                "ENSG00000116209",
                "ENSG00000136950",
                "ENSG00000104388",
                "ENSG00000105835",
                "ENSG00000119314",
                "ENSG00000170144",
                "ENSG00000171530",
                "ENSG00000198938",
                "ENSG00000168710",
                "ENSG00000127914",
                "ENSG00000126267",
                "ENSG00000108679",
                "ENSG00000253738",
                "ENSG00000152558",
                "ENSG00000154640",
                "ENSG00000126005",
                "ENSG00000166710",
                "ENSG00000198840",
                "ENSG00000082153",
                "ENSG00000247095",
                "ENSG00000101443",
                "ENSG00000085871",
                "ENSG00000184752",
                "ENSG00000169567",
                "ENSG00000198258",
                "ENSG00000023287",
                "ENSG00000076554",
                "ENSG00000164190",
                "ENSG00000189143",
                "ENSG00000135821",
                "ENSG00000131495",
                "ENSG00000171314",
                "ENSG00000067248",
                "ENSG00000090238",
                "ENSG00000167642",
                "ENSG00000111229",
                "ENSG00000130066",
                "ENSG00000126756",
                "ENSG00000006007",
                "ENSG00000051620",
                "ENSG00000173473",
                "ENSG00000163939",
                "ENSG00000059804",
                "ENSG00000117523",
                "ENSG00000127838",
                "ENSG00000167283",
                "ENSG00000198886",
                "ENSG00000122643",
                "ENSG00000104904",
                "ENSG00000125977",
                "ENSG00000170348",
                "ENSG00000116586",
                "ENSG00000119335",
                "ENSG00000084234",
                "ENSG00000154723",
                "ENSG00000166347",
                "ENSG00000131174",
                "ENSG00000143013",
                "ENSG00000168288",
                "ENSG00000004059",
                "ENSG00000234745",
                "ENSG00000113387",
                "ENSG00000196937",
                "ENSG00000106028",
                "ENSG00000100697",
                "ENSG00000168118",
                "ENSG00000106367",
                "ENSG00000185650",
                "ENSG00000100815",
                "ENSG00000197111",
                "ENSG00000101182",
                "ENSG00000142949",
                "ENSG00000092820",
                "ENSG00000185043",
                "ENSG00000184007",
                "ENSG00000090013",
                "ENSG00000197747",
                "ENSG00000004779",
                "ENSG00000004478",
                "ENSG00000135968",
                "ENSG00000070831",
                "ENSG00000051523",
                "ENSG00000250479",
                "ENSG00000144674",
                "ENSG00000106537",
                "ENSG00000146731",
                "ENSG00000053254",
                "ENSG00000105048",
                "ENSG00000164713",
                "ENSG00000143819",
                "ENSG00000111237",
                "ENSG00000183742",
                "ENSG00000132824",
                "ENSG00000254505",
                "ENSG00000112695",
                "ENSG00000118640",
                "ENSG00000176340",
                "ENSG00000097033",
                "ENSG00000117335",
                "ENSG00000170421",
                "ENSG00000049618",
                "ENSG00000164733",
                "ENSG00000140264",
                "ENSG00000177410",
                "ENSG00000101745",
                "ENSG00000140990",
                "ENSG00000182718",
                "ENSG00000138640",
                "ENSG00000164258",
                "ENSG00000106261",
                "ENSG00000198492",
                "ENSG00000171604",
                "ENSG00000233098",
                "ENSG00000129235",
                "ENSG00000102144",
                "ENSG00000172183",
                "ENSG00000130770",
                "ENSG00000084207",
                "ENSG00000186395",
                "ENSG00000117448",
                "ENSG00000166295",
                "ENSG00000142864",
                "ENSG00000119013",
                "ENSG00000184076",
                "ENSG00000196154",
                "ENSG00000211450",
                "ENSG00000166337",
                "ENSG00000166913",
                "ENSG00000162434",
                "ENSG00000118816",
                "ENSG00000119888",
                "ENSG00000100906",
                "ENSG00000221869",
                "ENSG00000150687",
                "ENSG00000134056",
                "ENSG00000101040",
                "ENSG00000177156",
                "ENSG00000187193",
                "ENSG00000197956",
                "ENSG00000144560",
                "ENSG00000117525",
                "ENSG00000156976",
                "ENSG00000129083",
                "ENSG00000160888",
                "ENSG00000113732",
                "ENSG00000197448",
                "ENSG00000133112",
                "ENSG00000131981",
                "ENSG00000166145",
                "ENSG00000124762",
                "ENSG00000187231",
                "ENSG00000121039",
                "ENSG00000150093",
                "ENSG00000138069",
                "ENSG00000110321",
                "ENSG00000144034",
                "ENSG00000156642",
                "ENSG00000123144",
                "ENSG00000127125",
                "ENSG00000143256",
                "ENSG00000149084",
                "ENSG00000165502",
                "ENSG00000084733",
                "ENSG00000138061",
                "ENSG00000206503",
                "ENSG00000122034",
                "ENSG00000138592",
                "ENSG00000138772",
                "ENSG00000149932",
                "ENSG00000114209",
                "ENSG00000168214",
                "ENSG00000100911",
                "ENSG00000160213",
                "ENSG00000144224",
                "ENSG00000115524",
                "ENSG00000138764",
                "ENSG00000155368",
                "ENSG00000096696",
                "ENSG00000169564",
                "ENSG00000143543",
                "ENSG00000070756",
                "ENSG00000140740",
                "ENSG00000146386",
                "ENSG00000198727",
                "ENSG00000167658",
                "ENSG00000145632",
                "ENSG00000147676",
                "ENSG00000185608",
                "ENSG00000060138",
                "ENSG00000132475",
                "ENSG00000135480",
                "ENSG00000196586",
                "ENSG00000168938",
                "ENSG00000143933",
                "ENSG00000146674",
                "ENSG00000165629",
                "ENSG00000167996",
                "ENSG00000132963",
                "ENSG00000100664",
                "ENSG00000136522",
                "ENSG00000178127",
                "ENSG00000010244",
                "ENSG00000105518",
                "ENSG00000133639",
                "ENSG00000092148",
                "ENSG00000170017",
                "ENSG00000124214",
                "ENSG00000163683",
                "ENSG00000116030",
                "ENSG00000186591",
                "ENSG00000160113",
                "ENSG00000160075",
                "ENSG00000120992",
                "ENSG00000174748",
                "ENSG00000148346",
                "ENSG00000163191",
                "ENSG00000126432",
                "ENSG00000181885",
                "ENSG00000126247",
                "ENSG00000245532",
                "ENSG00000197728",
                "ENSG00000178741",
                "ENSG00000085063",
                "ENSG00000171206",
                "ENSG00000173457",
                "ENSG00000088832",
                "ENSG00000108518",
                "ENSG00000278311",
                "ENSG00000086666",
                "ENSG00000114902",
                "ENSG00000230989",
                "ENSG00000154518",
                "ENSG00000178449",
                "ENSG00000106541",
                "ENSG00000134717",
                "ENSG00000136156",
                "ENSG00000125844",
                "ENSG00000171159",
                "ENSG00000134809",
                "ENSG00000213719",
                "ENSG00000112335",
                "ENSG00000204525",
                "ENSG00000164919",
                "ENSG00000145741",
                "ENSG00000178982",
                "ENSG00000067900",
                "ENSG00000111057",
                "ENSG00000104870",
                "ENSG00000100387",
                "ENSG00000114573",
                "ENSG00000140319",
                "ENSG00000148180",
                "ENSG00000163399",
                "ENSG00000118680",
                "ENSG00000117450",
                "ENSG00000244187",
                "ENSG00000117472",
                "ENSG00000198763",
                "ENSG00000009307",
                "ENSG00000107372",
                "ENSG00000135679",
                "ENSG00000124172",
                "ENSG00000150991",
                "ENSG00000196591",
                "ENSG00000017797",
                "ENSG00000100568",
                "ENSG00000101608",
                "ENSG00000122705",
                "ENSG00000183726",
                "ENSG00000113648",
                "ENSG00000111775",
                "ENSG00000123636",
                "ENSG00000087502",
                "ENSG00000197102",
                "ENSG00000163938",
                "ENSG00000008394",
                "ENSG00000188612",
                "ENSG00000154080",
                "ENSG00000104763",
                "ENSG00000205542",
                "ENSG00000158710",
                "ENSG00000160948",
                "ENSG00000137880",
                "ENSG00000115685",
                "ENSG00000128524",
                "ENSG00000167232",
                "ENSG00000165678",
                "ENSG00000198712",
                "ENSG00000044115",
                "ENSG00000136238",
                "ENSG00000186352",
                "ENSG00000130402",
                "ENSG00000182154",
                "ENSG00000122873",
                "ENSG00000087460",
                "ENSG00000158195",
                "ENSG00000170759",
                "ENSG00000145247",
                "ENSG00000113558",
                "ENSG00000143153",
                "ENSG00000109062",
                "ENSG00000113407",
                "ENSG00000143742",
                "ENSG00000104419",
                "ENSG00000085733",
                "ENSG00000092841",
                "ENSG00000173230"
            ],
            "expression_sum": {
                "class1": 152.0,
                "class2": 9183.0
            },
            "ens_id": "ENSG00000163435",
            "uniprot_ids": [
                "P78545",
                "Q5SR36"
            ],
            "net_id": 0,
            "source_num": 8,
            "target_num": 85
        },
        "ENSG00000181449": {
            "id": "ENSG00000181449",
            "type": "TF",
            "symbol": "SOX2",
            "source": [
                "ENSG00000100811",
                "ENSG00000105698",
                "ENSG00000103495",
                "ENSG00000108055",
                "ENSG00000163848",
                "ENSG00000116560",
                "ENSG00000164754",
                "ENSG00000128610",
                "ENSG00000130522"
            ],
            "target": [
                "ENSG00000078668",
                "ENSG00000206573",
                "ENSG00000112081",
                "ENSG00000107223",
                "ENSG00000162704",
                "ENSG00000134884",
                "ENSG00000163634",
                "ENSG00000165655",
                "ENSG00000153130",
                "ENSG00000111716",
                "ENSG00000139644",
                "ENSG00000099622",
                "ENSG00000090060",
                "ENSG00000116161",
                "ENSG00000106603",
                "ENSG00000117395",
                "ENSG00000116350",
                "ENSG00000161016",
                "ENSG00000134248",
                "ENSG00000131143",
                "ENSG00000184983",
                "ENSG00000127540",
                "ENSG00000099797",
                "ENSG00000119632",
                "ENSG00000189403",
                "ENSG00000108424",
                "ENSG00000159377",
                "ENSG00000196419",
                "ENSG00000075415",
                "ENSG00000107874",
                "ENSG00000105993",
                "ENSG00000102172",
                "ENSG00000149212",
                "ENSG00000213722",
                "ENSG00000150779",
                "ENSG00000165898",
                "ENSG00000077380",
                "ENSG00000154556",
                "ENSG00000048740",
                "ENSG00000163466",
                "ENSG00000119421",
                "ENSG00000052802",
                "ENSG00000134283",
                "ENSG00000173812",
                "ENSG00000117592",
                "ENSG00000173915",
                "ENSG00000167085",
                "ENSG00000151729",
                "ENSG00000135441",
                "ENSG00000106443",
                "ENSG00000134453",
                "ENSG00000041357",
                "ENSG00000182220",
                "ENSG00000163875",
                "ENSG00000163319",
                "ENSG00000164924",
                "ENSG00000149196",
                "ENSG00000141759",
                "ENSG00000135069",
                "ENSG00000162664",
                "ENSG00000204392",
                "ENSG00000165264",
                "ENSG00000224877",
                "ENSG00000139116",
                "ENSG00000114354",
                "ENSG00000277791",
                "ENSG00000156482",
                "ENSG00000125868",
                "ENSG00000165637",
                "ENSG00000165672",
                "ENSG00000173674",
                "ENSG00000147548",
                "ENSG00000171703",
                "ENSG00000167775",
                "ENSG00000129625",
                "ENSG00000089220",
                "ENSG00000116898",
                "ENSG00000196072",
                "ENSG00000179431",
                "ENSG00000255302",
                "ENSG00000100526",
                "ENSG00000188243",
                "ENSG00000161542",
                "ENSG00000067225",
                "ENSG00000109390",
                "ENSG00000141552",
                "ENSG00000113712",
                "ENSG00000126698",
                "ENSG00000136143",
                "ENSG00000174851",
                "ENSG00000150459",
                "ENSG00000170540",
                "ENSG00000233927",
                "ENSG00000205581",
                "ENSG00000159685",
                "ENSG00000026025",
                "ENSG00000034713",
                "ENSG00000144746",
                "ENSG00000034510",
                "ENSG00000146425",
                "ENSG00000138071",
                "ENSG00000120963",
                "ENSG00000117410",
                "ENSG00000170584",
                "ENSG00000075142",
                "ENSG00000273841",
                "ENSG00000124802",
                "ENSG00000263001",
                "ENSG00000166275",
                "ENSG00000183011",
                "ENSG00000126934",
                "ENSG00000213741",
                "ENSG00000215845",
                "ENSG00000175701",
                "ENSG00000121774",
                "ENSG00000069849",
                "ENSG00000174917",
                "ENSG00000147649",
                "ENSG00000129515",
                "ENSG00000100519",
                "ENSG00000187514",
                "ENSG00000163041",
                "ENSG00000155115",
                "ENSG00000141027",
                "ENSG00000092199",
                "ENSG00000110955",
                "ENSG00000178980",
                "ENSG00000175216",
                "ENSG00000089335",
                "ENSG00000083807",
                "ENSG00000150316",
                "ENSG00000010256",
                "ENSG00000172239",
                "ENSG00000065518",
                "ENSG00000174953",
                "ENSG00000152767",
                "ENSG00000109775",
                "ENSG00000162191",
                "ENSG00000118363",
                "ENSG00000129562",
                "ENSG00000135940",
                "ENSG00000108654",
                "ENSG00000108106",
                "ENSG00000162599",
                "ENSG00000172336",
                "ENSG00000107581",
                "ENSG00000131966",
                "ENSG00000105258",
                "ENSG00000112972",
                "ENSG00000204842",
                "ENSG00000085224",
                "ENSG00000153037",
                "ENSG00000113013",
                "ENSG00000265491",
                "ENSG00000092621",
                "ENSG00000106245",
                "ENSG00000213465",
                "ENSG00000109332",
                "ENSG00000163170",
                "ENSG00000079246",
                "ENSG00000168275",
                "ENSG00000075785",
                "ENSG00000198937",
                "ENSG00000075624",
                "ENSG00000244879",
                "ENSG00000171222",
                "ENSG00000095059",
                "ENSG00000105185",
                "ENSG00000131051",
                "ENSG00000167552",
                "ENSG00000063322",
                "ENSG00000205726",
                "ENSG00000175166",
                "ENSG00000168303",
                "ENSG00000167088",
                "ENSG00000240972",
                "ENSG00000196655",
                "ENSG00000029993",
                "ENSG00000164985",
                "ENSG00000239672",
                "ENSG00000213585",
                "ENSG00000131508",
                "ENSG00000005022",
                "ENSG00000127616",
                "ENSG00000154277",
                "ENSG00000131469",
                "ENSG00000099817",
                "ENSG00000171603",
                "ENSG00000128654",
                "ENSG00000047410",
                "ENSG00000169714",
                "ENSG00000090266",
                "ENSG00000142937",
                "ENSG00000136810",
                "ENSG00000171634",
                "ENSG00000125656",
                "ENSG00000232112",
                "ENSG00000127184",
                "ENSG00000106682",
                "ENSG00000130312",
                "ENSG00000197694",
                "ENSG00000079459",
                "ENSG00000104388",
                "ENSG00000110917",
                "ENSG00000168159",
                "ENSG00000170144",
                "ENSG00000064607",
                "ENSG00000171530",
                "ENSG00000152082",
                "ENSG00000087191",
                "ENSG00000166734",
                "ENSG00000123349",
                "ENSG00000105583",
                "ENSG00000143870",
                "ENSG00000147655",
                "ENSG00000014641",
                "ENSG00000167614",
                "ENSG00000164683",
                "ENSG00000106803",
                "ENSG00000126267",
                "ENSG00000104765",
                "ENSG00000120860",
                "ENSG00000198612",
                "ENSG00000137710",
                "ENSG00000081154",
                "ENSG00000122566",
                "ENSG00000182199",
                "ENSG00000130725",
                "ENSG00000100129",
                "ENSG00000112305",
                "ENSG00000142507",
                "ENSG00000108774",
                "ENSG00000237765",
                "ENSG00000165119",
                "ENSG00000067064",
                "ENSG00000135486",
                "ENSG00000115128",
                "ENSG00000132718",
                "ENSG00000137818",
                "ENSG00000129559",
                "ENSG00000068796",
                "ENSG00000105677",
                "ENSG00000114784",
                "ENSG00000184752",
                "ENSG00000169567",
                "ENSG00000198258",
                "ENSG00000183172",
                "ENSG00000127445",
                "ENSG00000128951",
                "ENSG00000100575",
                "ENSG00000105810",
                "ENSG00000188186",
                "ENSG00000171735",
                "ENSG00000241837",
                "ENSG00000147274",
                "ENSG00000103495",
                "ENSG00000204308",
                "ENSG00000214046",
                "ENSG00000176101",
                "ENSG00000123416",
                "ENSG00000131495",
                "ENSG00000144713",
                "ENSG00000155957",
                "ENSG00000130520",
                "ENSG00000084623",
                "ENSG00000086758",
                "ENSG00000108671",
                "ENSG00000100852",
                "ENSG00000167792",
                "ENSG00000147684",
                "ENSG00000096746",
                "ENSG00000170315",
                "ENSG00000079785",
                "ENSG00000119950",
                "ENSG00000163848",
                "ENSG00000099795",
                "ENSG00000129084",
                "ENSG00000069275",
                "ENSG00000079462",
                "ENSG00000138777",
                "ENSG00000072042",
                "ENSG00000069345",
                "ENSG00000162585",
                "ENSG00000163714",
                "ENSG00000189043",
                "ENSG00000099246",
                "ENSG00000170515",
                "ENSG00000142192",
                "ENSG00000127838",
                "ENSG00000131828",
                "ENSG00000234545",
                "ENSG00000137100",
                "ENSG00000167283",
                "ENSG00000164405",
                "ENSG00000170035",
                "ENSG00000196199",
                "ENSG00000187109",
                "ENSG00000155506",
                "ENSG00000044574",
                "ENSG00000196262",
                "ENSG00000104904",
                "ENSG00000115355",
                "ENSG00000125977",
                "ENSG00000170348",
                "ENSG00000173726",
                "ENSG00000089280",
                "ENSG00000155959",
                "ENSG00000137692",
                "ENSG00000116586",
                "ENSG00000089693",
                "ENSG00000105254",
                "ENSG00000117419",
                "ENSG00000119335",
                "ENSG00000115286",
                "ENSG00000077152",
                "ENSG00000144028",
                "ENSG00000084234",
                "ENSG00000114023",
                "ENSG00000127774",
                "ENSG00000101367",
                "ENSG00000172893",
                "ENSG00000154723",
                "ENSG00000122026",
                "ENSG00000166598",
                "ENSG00000167978",
                "ENSG00000198729",
                "ENSG00000149357",
                "ENSG00000131174",
                "ENSG00000106400",
                "ENSG00000143013",
                "ENSG00000109654",
                "ENSG00000179091",
                "ENSG00000254999",
                "ENSG00000133935",
                "ENSG00000177556",
                "ENSG00000113387",
                "ENSG00000158864",
                "ENSG00000137409",
                "ENSG00000006125",
                "ENSG00000188690",
                "ENSG00000153944",
                "ENSG00000241468",
                "ENSG00000136888",
                "ENSG00000110717",
                "ENSG00000120727",
                "ENSG00000127884",
                "ENSG00000185630",
                "ENSG00000100442",
                "ENSG00000143799",
                "ENSG00000106028",
                "ENSG00000140612",
                "ENSG00000115091",
                "ENSG00000204628",
                "ENSG00000115053",
                "ENSG00000102241",
                "ENSG00000106367",
                "ENSG00000121022",
                "ENSG00000164615",
                "ENSG00000139168",
                "ENSG00000197111",
                "ENSG00000111275",
                "ENSG00000139343",
                "ENSG00000175334",
                "ENSG00000146729",
                "ENSG00000171421",
                "ENSG00000085231",
                "ENSG00000133134",
                "ENSG00000123562",
                "ENSG00000099783",
                "ENSG00000105701",
                "ENSG00000111669",
                "ENSG00000136100",
                "ENSG00000182093",
                "ENSG00000134825",
                "ENSG00000185745",
                "ENSG00000140307",
                "ENSG00000132463",
                "ENSG00000167863",
                "ENSG00000004779",
                "ENSG00000106355",
                "ENSG00000071553",
                "ENSG00000165280",
                "ENSG00000169139",
                "ENSG00000254682",
                "ENSG00000116459",
                "ENSG00000175756",
                "ENSG00000203950",
                "ENSG00000174684",
                "ENSG00000134333",
                "ENSG00000197345",
                "ENSG00000006625",
                "ENSG00000091640",
                "ENSG00000147123",
                "ENSG00000143575",
                "ENSG00000143977",
                "ENSG00000038219",
                "ENSG00000249915",
                "ENSG00000130560",
                "ENSG00000110700",
                "ENSG00000167004",
                "ENSG00000160014",
                "ENSG00000100201",
                "ENSG00000213639",
                "ENSG00000159140",
                "ENSG00000183617",
                "ENSG00000125818",
                "ENSG00000146540",
                "ENSG00000105220",
                "ENSG00000143353",
                "ENSG00000170892",
                "ENSG00000111678",
                "ENSG00000067560",
                "ENSG00000163001",
                "ENSG00000130724",
                "ENSG00000007168",
                "ENSG00000156639",
                "ENSG00000164713",
                "ENSG00000116649",
                "ENSG00000132507",
                "ENSG00000176209",
                "ENSG00000105379",
                "ENSG00000175550",
                "ENSG00000099901",
                "ENSG00000143727",
                "ENSG00000126067",
                "ENSG00000173692",
                "ENSG00000111237",
                "ENSG00000132670",
                "ENSG00000112110",
                "ENSG00000023191",
                "ENSG00000215301",
                "ENSG00000104886",
                "ENSG00000151366",
                "ENSG00000108298",
                "ENSG00000214413",
                "ENSG00000166226",
                "ENSG00000112695",
                "ENSG00000116747",
                "ENSG00000153187",
                "ENSG00000106628",
                "ENSG00000077782",
                "ENSG00000174444",
                "ENSG00000203667",
                "ENSG00000100354",
                "ENSG00000133318",
                "ENSG00000130414",
                "ENSG00000169738",
                "ENSG00000185236",
                "ENSG00000137285",
                "ENSG00000176340",
                "ENSG00000172354",
                "ENSG00000150753",
                "ENSG00000164096",
                "ENSG00000049618",
                "ENSG00000136824",
                "ENSG00000138385",
                "ENSG00000083457",
                "ENSG00000140264",
                "ENSG00000189159",
                "ENSG00000116754",
                "ENSG00000101745",
                "ENSG00000145337",
                "ENSG00000140990",
                "ENSG00000221983",
                "ENSG00000173436",
                "ENSG00000175348",
                "ENSG00000063046",
                "ENSG00000122565",
                "ENSG00000163541",
                "ENSG00000008952",
                "ENSG00000127483",
                "ENSG00000146416",
                "ENSG00000230124",
                "ENSG00000164258",
                "ENSG00000116560",
                "ENSG00000100941",
                "ENSG00000109472",
                "ENSG00000114956",
                "ENSG00000147862",
                "ENSG00000140391",
                "ENSG00000173272",
                "ENSG00000137815",
                "ENSG00000141002",
                "ENSG00000171604",
                "ENSG00000179262",
                "ENSG00000196976",
                "ENSG00000197756",
                "ENSG00000178952",
                "ENSG00000143155",
                "ENSG00000196230",
                "ENSG00000225663",
                "ENSG00000182004",
                "ENSG00000141526",
                "ENSG00000124733",
                "ENSG00000173418",
                "ENSG00000102144",
                "ENSG00000272325",
                "ENSG00000130770",
                "ENSG00000029363",
                "ENSG00000156467",
                "ENSG00000167526",
                "ENSG00000100028",
                "ENSG00000168259",
                "ENSG00000142864",
                "ENSG00000119013",
                "ENSG00000161203",
                "ENSG00000182944",
                "ENSG00000184076",
                "ENSG00000137288",
                "ENSG00000105887",
                "ENSG00000100836",
                "ENSG00000266472",
                "ENSG00000101782",
                "ENSG00000177889",
                "ENSG00000142541",
                "ENSG00000109971",
                "ENSG00000166913",
                "ENSG00000204435",
                "ENSG00000159199",
                "ENSG00000118816",
                "ENSG00000121851",
                "ENSG00000135624",
                "ENSG00000104671",
                "ENSG00000147669",
                "ENSG00000168003",
                "ENSG00000126524",
                "ENSG00000163468",
                "ENSG00000111843",
                "ENSG00000074071",
                "ENSG00000099800",
                "ENSG00000150687",
                "ENSG00000140905",
                "ENSG00000134056",
                "ENSG00000277443",
                "ENSG00000172115",
                "ENSG00000101558",
                "ENSG00000145912",
                "ENSG00000140350",
                "ENSG00000145293",
                "ENSG00000181924",
                "ENSG00000074696",
                "ENSG00000175130",
                "ENSG00000167468",
                "ENSG00000111640",
                "ENSG00000197746",
                "ENSG00000183978",
                "ENSG00000125652",
                "ENSG00000115241",
                "ENSG00000253719",
                "ENSG00000156976",
                "ENSG00000100632",
                "ENSG00000161970",
                "ENSG00000130294",
                "ENSG00000078369",
                "ENSG00000119396",
                "ENSG00000105404",
                "ENSG00000177879",
                "ENSG00000205531",
                "ENSG00000099624",
                "ENSG00000120265",
                "ENSG00000154001",
                "ENSG00000138758",
                "ENSG00000115541",
                "ENSG00000168066",
                "ENSG00000133083",
                "ENSG00000069329",
                "ENSG00000138069",
                "ENSG00000083099",
                "ENSG00000110321",
                "ENSG00000225733",
                "ENSG00000111875",
                "ENSG00000071994",
                "ENSG00000130748",
                "ENSG00000115484",
                "ENSG00000169689",
                "ENSG00000168785",
                "ENSG00000110651",
                "ENSG00000138326",
                "ENSG00000204568",
                "ENSG00000175203",
                "ENSG00000102409",
                "ENSG00000154380",
                "ENSG00000145833",
                "ENSG00000104695",
                "ENSG00000123144",
                "ENSG00000113575",
                "ENSG00000143256",
                "ENSG00000127022",
                "ENSG00000149084",
                "ENSG00000074800",
                "ENSG00000128610",
                "ENSG00000054118",
                "ENSG00000105193",
                "ENSG00000084733",
                "ENSG00000142676",
                "ENSG00000138668",
                "ENSG00000179950",
                "ENSG00000119705",
                "ENSG00000133026",
                "ENSG00000149557",
                "ENSG00000165283",
                "ENSG00000075151",
                "ENSG00000122034",
                "ENSG00000154582",
                "ENSG00000130255",
                "ENSG00000120533",
                "ENSG00000162734",
                "ENSG00000196305",
                "ENSG00000103024",
                "ENSG00000096384",
                "ENSG00000022277",
                "ENSG00000105223",
                "ENSG00000160310",
                "ENSG00000174469",
                "ENSG00000151665",
                "ENSG00000164902",
                "ENSG00000144224",
                "ENSG00000188994",
                "ENSG00000174775",
                "ENSG00000143947",
                "ENSG00000071794",
                "ENSG00000213015",
                "ENSG00000175416",
                "ENSG00000128609",
                "ENSG00000063177",
                "ENSG00000108953",
                "ENSG00000155368",
                "ENSG00000050393",
                "ENSG00000125611",
                "ENSG00000169564",
                "ENSG00000094916",
                "ENSG00000103035",
                "ENSG00000132305",
                "ENSG00000174721",
                "ENSG00000128050",
                "ENSG00000242372",
                "ENSG00000138663",
                "ENSG00000133706",
                "ENSG00000111011",
                "ENSG00000245694",
                "ENSG00000086061",
                "ENSG00000070756",
                "ENSG00000136930",
                "ENSG00000140740",
                "ENSG00000149806",
                "ENSG00000166012",
                "ENSG00000167658",
                "ENSG00000142168",
                "ENSG00000186577",
                "ENSG00000197006",
                "ENSG00000185608",
                "ENSG00000108468",
                "ENSG00000196465",
                "ENSG00000168028",
                "ENSG00000108826",
                "ENSG00000108848",
                "ENSG00000031698",
                "ENSG00000111786",
                "ENSG00000109919",
                "ENSG00000130731",
                "ENSG00000143933",
                "ENSG00000172172",
                "ENSG00000165629",
                "ENSG00000166503",
                "ENSG00000126457",
                "ENSG00000132963",
                "ENSG00000042753",
                "ENSG00000100664",
                "ENSG00000177600",
                "ENSG00000148450",
                "ENSG00000136522",
                "ENSG00000178127",
                "ENSG00000131876",
                "ENSG00000174456",
                "ENSG00000148700",
                "ENSG00000167699",
                "ENSG00000105518",
                "ENSG00000143198",
                "ENSG00000054148",
                "ENSG00000143621",
                "ENSG00000145494",
                "ENSG00000122378",
                "ENSG00000156411",
                "ENSG00000061676",
                "ENSG00000177200",
                "ENSG00000115944",
                "ENSG00000111639",
                "ENSG00000138443",
                "ENSG00000135334",
                "ENSG00000196531",
                "ENSG00000100138",
                "ENSG00000116030",
                "ENSG00000115268",
                "ENSG00000182534",
                "ENSG00000081138",
                "ENSG00000160075",
                "ENSG00000174748",
                "ENSG00000170043",
                "ENSG00000089157",
                "ENSG00000168653",
                "ENSG00000141504",
                "ENSG00000182481",
                "ENSG00000156261",
                "ENSG00000180228",
                "ENSG00000126432",
                "ENSG00000198363",
                "ENSG00000151465",
                "ENSG00000131116",
                "ENSG00000197728",
                "ENSG00000156515",
                "ENSG00000057757",
                "ENSG00000006015",
                "ENSG00000171206",
                "ENSG00000104823",
                "ENSG00000233016",
                "ENSG00000122406",
                "ENSG00000166595",
                "ENSG00000143774",
                "ENSG00000182512",
                "ENSG00000179271",
                "ENSG00000166136",
                "ENSG00000196141",
                "ENSG00000115310",
                "ENSG00000122545",
                "ENSG00000134330",
                "ENSG00000143612",
                "ENSG00000159884",
                "ENSG00000167969",
                "ENSG00000007520",
                "ENSG00000135720",
                "ENSG00000196683",
                "ENSG00000149547",
                "ENSG00000108055",
                "ENSG00000159335",
                "ENSG00000136485",
                "ENSG00000149600",
                "ENSG00000123352",
                "ENSG00000099194",
                "ENSG00000137133",
                "ENSG00000115350",
                "ENSG00000128739",
                "ENSG00000172809",
                "ENSG00000145425",
                "ENSG00000114902",
                "ENSG00000107341",
                "ENSG00000106615",
                "ENSG00000099341",
                "ENSG00000172500",
                "ENSG00000230989",
                "ENSG00000154518",
                "ENSG00000109084",
                "ENSG00000087302",
                "ENSG00000101856",
                "ENSG00000091164",
                "ENSG00000136146",
                "ENSG00000103342",
                "ENSG00000090263",
                "ENSG00000172757",
                "ENSG00000182628",
                "ENSG00000105669",
                "ENSG00000188290",
                "ENSG00000101460",
                "ENSG00000136942",
                "ENSG00000138175",
                "ENSG00000105176",
                "ENSG00000134717",
                "ENSG00000169599",
                "ENSG00000119185",
                "ENSG00000215386",
                "ENSG00000160752",
                "ENSG00000143252",
                "ENSG00000198961",
                "ENSG00000141367",
                "ENSG00000144381",
                "ENSG00000119318",
                "ENSG00000136521",
                "ENSG00000132688",
                "ENSG00000100811",
                "ENSG00000145354",
                "ENSG00000104177",
                "ENSG00000159352",
                "ENSG00000060762",
                "ENSG00000166794",
                "ENSG00000204220",
                "ENSG00000112335",
                "ENSG00000152234",
                "ENSG00000133872",
                "ENSG00000113811",
                "ENSG00000179010",
                "ENSG00000198668",
                "ENSG00000108107",
                "ENSG00000134308",
                "ENSG00000108100",
                "ENSG00000143158",
                "ENSG00000132341",
                "ENSG00000111832",
                "ENSG00000153774",
                "ENSG00000129197",
                "ENSG00000164919",
                "ENSG00000145741",
                "ENSG00000138757",
                "ENSG00000108561",
                "ENSG00000118418",
                "ENSG00000178982",
                "ENSG00000108010",
                "ENSG00000083845",
                "ENSG00000214253",
                "ENSG00000138382",
                "ENSG00000155876",
                "ENSG00000171867",
                "ENSG00000102316",
                "ENSG00000125821",
                "ENSG00000137547",
                "ENSG00000140319",
                "ENSG00000132466",
                "ENSG00000172428",
                "ENSG00000175826",
                "ENSG00000184220",
                "ENSG00000009307",
                "ENSG00000018625",
                "ENSG00000168958",
                "ENSG00000107372",
                "ENSG00000182117",
                "ENSG00000124172",
                "ENSG00000057608",
                "ENSG00000150991",
                "ENSG00000065548",
                "ENSG00000152795",
                "ENSG00000137876",
                "ENSG00000138398",
                "ENSG00000137210",
                "ENSG00000136527",
                "ENSG00000162076",
                "ENSG00000110958",
                "ENSG00000100568",
                "ENSG00000107281",
                "ENSG00000007080",
                "ENSG00000164323",
                "ENSG00000166681",
                "ENSG00000122705",
                "ENSG00000132432",
                "ENSG00000140374",
                "ENSG00000115306",
                "ENSG00000184009",
                "ENSG00000023734",
                "ENSG00000114405",
                "ENSG00000120437",
                "ENSG00000125944",
                "ENSG00000054523",
                "ENSG00000100380",
                "ENSG00000173660",
                "ENSG00000186184",
                "ENSG00000100804",
                "ENSG00000119640",
                "ENSG00000113648",
                "ENSG00000114125",
                "ENSG00000185787",
                "ENSG00000179222",
                "ENSG00000105568",
                "ENSG00000160799",
                "ENSG00000130638",
                "ENSG00000143333",
                "ENSG00000169045",
                "ENSG00000111775",
                "ENSG00000179218",
                "ENSG00000139684",
                "ENSG00000128989",
                "ENSG00000162604",
                "ENSG00000144834",
                "ENSG00000114120",
                "ENSG00000163964",
                "ENSG00000126602",
                "ENSG00000188612",
                "ENSG00000010810",
                "ENSG00000180817",
                "ENSG00000086065",
                "ENSG00000181163",
                "ENSG00000116288",
                "ENSG00000008018",
                "ENSG00000112531",
                "ENSG00000136938",
                "ENSG00000070961",
                "ENSG00000198898",
                "ENSG00000082898",
                "ENSG00000115685",
                "ENSG00000220205",
                "ENSG00000128524",
                "ENSG00000101654",
                "ENSG00000101940",
                "ENSG00000104549",
                "ENSG00000044115",
                "ENSG00000167113",
                "ENSG00000265241",
                "ENSG00000136238",
                "ENSG00000164754",
                "ENSG00000143183",
                "ENSG00000132424",
                "ENSG00000130522",
                "ENSG00000047849",
                "ENSG00000182154",
                "ENSG00000221914",
                "ENSG00000122873",
                "ENSG00000065978",
                "ENSG00000151923",
                "ENSG00000158195",
                "ENSG00000087460",
                "ENSG00000100567",
                "ENSG00000151247",
                "ENSG00000183513",
                "ENSG00000113558",
                "ENSG00000181817",
                "ENSG00000115457",
                "ENSG00000109180",
                "ENSG00000187735",
                "ENSG00000115233",
                "ENSG00000156052",
                "ENSG00000143742",
                "ENSG00000174886",
                "ENSG00000172270",
                "ENSG00000139637",
                "ENSG00000106244",
                "ENSG00000164167",
                "ENSG00000092841",
                "ENSG00000129351",
                "ENSG00000173163"
            ],
            "expression_sum": {
                "class1": 3253.0,
                "class2": 24950.0
            },
            "ens_id": "ENSG00000181449",
            "uniprot_ids": [
                "P48431"
            ],
            "net_id": 0,
            "source_num": 9,
            "target_num": 108
        },
        "ENSG00000170365": {
            "id": "ENSG00000170365",
            "type": "TF",
            "symbol": "SMAD1",
            "source": [
                "ENSG00000016082",
                "ENSG00000164330"
            ],
            "target": [
                "ENSG00000131016",
                "ENSG00000143702",
                "ENSG00000176788",
                "ENSG00000144339",
                "ENSG00000016082",
                "ENSG00000135406",
                "ENSG00000221866",
                "ENSG00000182481",
                "ENSG00000112773",
                "ENSG00000139970",
                "ENSG00000087258",
                "ENSG00000213760",
                "ENSG00000144642"
            ],
            "expression_sum": {
                "class1": 4497.0,
                "class2": 2361.0
            },
            "ens_id": "ENSG00000170365",
            "uniprot_ids": [
                "Q15797",
                "D6RDN7",
                "D6RAY1",
                "D6RD62",
                "D6REQ3",
                "A0A1D5RMR4",
                "D6RBH9"
            ],
            "net_id": 0,
            "source_num": 2,
            "target_num": 7
        },
        "ENSG00000100811": {
            "id": "ENSG00000100811",
            "type": "TF",
            "symbol": "YY1",
            "source": [
                "ENSG00000167081",
                "ENSG00000181449"
            ],
            "target": [
                "ENSG00000104722",
                "ENSG00000181449",
                "ENSG00000103034",
                "ENSG00000176788",
                "ENSG00000172020",
                "ENSG00000169856",
                "ENSG00000152558",
                "ENSG00000169783",
                "ENSG00000145428",
                "ENSG00000161082",
                "ENSG00000132698",
                "ENSG00000087085",
                "ENSG00000167654",
                "ENSG00000164093",
                "ENSG00000107105",
                "ENSG00000122756",
                "ENSG00000148798",
                "ENSG00000129951",
                "ENSG00000130540",
                "ENSG00000105784",
                "ENSG00000198910",
                "ENSG00000084628",
                "ENSG00000166257",
                "ENSG00000174021",
                "ENSG00000128564",
                "ENSG00000100167",
                "ENSG00000197959",
                "ENSG00000162374",
                "ENSG00000196361",
                "ENSG00000105409",
                "ENSG00000072832",
                "ENSG00000221866",
                "ENSG00000155980",
                "ENSG00000151320",
                "ENSG00000186868",
                "ENSG00000168280",
                "ENSG00000005108",
                "ENSG00000198125",
                "ENSG00000213760",
                "ENSG00000137285",
                "ENSG00000092964",
                "ENSG00000143702",
                "ENSG00000144218",
                "ENSG00000104435",
                "ENSG00000122012",
                "ENSG00000103154",
                "ENSG00000016082",
                "ENSG00000171033",
                "ENSG00000124194",
                "ENSG00000173267",
                "ENSG00000087258",
                "ENSG00000111728",
                "ENSG00000149294",
                "ENSG00000170091",
                "ENSG00000139970",
                "ENSG00000205922",
                "ENSG00000084731",
                "ENSG00000185624",
                "ENSG00000167081",
                "ENSG00000135406",
                "ENSG00000168824",
                "ENSG00000159409",
                "ENSG00000118263",
                "ENSG00000068615",
                "ENSG00000164330",
                "ENSG00000137267",
                "ENSG00000104833",
                "ENSG00000177675",
                "ENSG00000174521",
                "ENSG00000185650",
                "ENSG00000135447",
                "ENSG00000277586",
                "ENSG00000104870"
            ],
            "expression_sum": {
                "class1": 10338.0,
                "class2": 9993.0
            },
            "ens_id": "ENSG00000100811",
            "uniprot_ids": [
                "P25490",
                "G3V3M8",
                "A0A494C032",
                "H0YJU4",
                "H0YJV7"
            ],
            "net_id": 0,
            "source_num": 2,
            "target_num": 27
        },
        "ENSG00000130726": {
            "id": "ENSG00000130726",
            "type": "TF",
            "symbol": "TRIM28",
            "source": [],
            "target": [
                "ENSG00000104722",
                "ENSG00000176788",
                "ENSG00000172020",
                "ENSG00000087085",
                "ENSG00000107105",
                "ENSG00000148798",
                "ENSG00000213190",
                "ENSG00000171551",
                "ENSG00000142549",
                "ENSG00000198910",
                "ENSG00000162188",
                "ENSG00000196361",
                "ENSG00000162374",
                "ENSG00000132872",
                "ENSG00000105409",
                "ENSG00000072832",
                "ENSG00000221866",
                "ENSG00000155980",
                "ENSG00000186868",
                "ENSG00000016082",
                "ENSG00000124194",
                "ENSG00000139970",
                "ENSG00000135406",
                "ENSG00000168824",
                "ENSG00000135447",
                "ENSG00000277586"
            ],
            "expression_sum": {
                "class1": 9647.0,
                "class2": 6926.0
            },
            "ens_id": "ENSG00000130726",
            "uniprot_ids": [
                "Q13263",
                "M0R0K9",
                "M0QZE6",
                "M0R3C0",
                "M0R2I3"
            ],
            "net_id": 0,
            "source_num": 0,
            "target_num": 13
        },
        "ENSG00000109132": {
            "id": "ENSG00000109132",
            "type": "TF",
            "symbol": "PHOX2B",
            "source": [
                "ENSG00000016082",
                "ENSG00000196591"
            ],
            "target": [
                "ENSG00000078668",
                "ENSG00000017260",
                "ENSG00000116171",
                "ENSG00000162704",
                "ENSG00000111716",
                "ENSG00000099622",
                "ENSG00000115226",
                "ENSG00000116161",
                "ENSG00000140941",
                "ENSG00000197959",
                "ENSG00000179958",
                "ENSG00000189403",
                "ENSG00000159377",
                "ENSG00000075415",
                "ENSG00000107874",
                "ENSG00000102172",
                "ENSG00000077380",
                "ENSG00000154556",
                "ENSG00000163466",
                "ENSG00000108515",
                "ENSG00000134283",
                "ENSG00000167085",
                "ENSG00000272398",
                "ENSG00000106443",
                "ENSG00000115540",
                "ENSG00000123240",
                "ENSG00000163319",
                "ENSG00000164924",
                "ENSG00000139116",
                "ENSG00000156482",
                "ENSG00000047249",
                "ENSG00000060982",
                "ENSG00000147548",
                "ENSG00000171703",
                "ENSG00000196504",
                "ENSG00000188243",
                "ENSG00000067225",
                "ENSG00000144642",
                "ENSG00000144218",
                "ENSG00000113712",
                "ENSG00000140105",
                "ENSG00000104131",
                "ENSG00000159685",
                "ENSG00000026025",
                "ENSG00000095380",
                "ENSG00000157168",
                "ENSG00000103769",
                "ENSG00000075142",
                "ENSG00000150722",
                "ENSG00000124802",
                "ENSG00000187051",
                "ENSG00000213741",
                "ENSG00000151500",
                "ENSG00000069849",
                "ENSG00000123200",
                "ENSG00000129515",
                "ENSG00000100519",
                "ENSG00000142546",
                "ENSG00000163041",
                "ENSG00000155115",
                "ENSG00000110955",
                "ENSG00000164346",
                "ENSG00000164687",
                "ENSG00000065518",
                "ENSG00000174953",
                "ENSG00000213760",
                "ENSG00000137309",
                "ENSG00000198836",
                "ENSG00000143702",
                "ENSG00000131966",
                "ENSG00000231500",
                "ENSG00000113013",
                "ENSG00000109332",
                "ENSG00000075624",
                "ENSG00000079246",
                "ENSG00000171222",
                "ENSG00000244879",
                "ENSG00000166974",
                "ENSG00000164438",
                "ENSG00000167552",
                "ENSG00000167088",
                "ENSG00000078618",
                "ENSG00000136783",
                "ENSG00000135390",
                "ENSG00000153310",
                "ENSG00000197223",
                "ENSG00000154277",
                "ENSG00000082397",
                "ENSG00000105784",
                "ENSG00000090266",
                "ENSG00000198015",
                "ENSG00000162374",
                "ENSG00000171204",
                "ENSG00000127184",
                "ENSG00000103363",
                "ENSG00000104388",
                "ENSG00000134982",
                "ENSG00000143222",
                "ENSG00000151914",
                "ENSG00000170903",
                "ENSG00000165943",
                "ENSG00000014641",
                "ENSG00000125827",
                "ENSG00000104723",
                "ENSG00000175582",
                "ENSG00000198612",
                "ENSG00000081154",
                "ENSG00000122566",
                "ENSG00000139163",
                "ENSG00000135486",
                "ENSG00000109670",
                "ENSG00000068796",
                "ENSG00000138760",
                "ENSG00000197302",
                "ENSG00000170027",
                "ENSG00000076554",
                "ENSG00000103495",
                "ENSG00000010270",
                "ENSG00000147604",
                "ENSG00000204308",
                "ENSG00000108175",
                "ENSG00000144713",
                "ENSG00000172159",
                "ENSG00000170315",
                "ENSG00000096746",
                "ENSG00000079785",
                "ENSG00000067715",
                "ENSG00000129084",
                "ENSG00000069345",
                "ENSG00000113742",
                "ENSG00000117523",
                "ENSG00000142192",
                "ENSG00000137100",
                "ENSG00000170035",
                "ENSG00000187109",
                "ENSG00000196262",
                "ENSG00000125977",
                "ENSG00000139117",
                "ENSG00000137692",
                "ENSG00000116586",
                "ENSG00000089693",
                "ENSG00000120438",
                "ENSG00000077152",
                "ENSG00000101132",
                "ENSG00000154723",
                "ENSG00000149357",
                "ENSG00000131174",
                "ENSG00000089289",
                "ENSG00000106588",
                "ENSG00000113387",
                "ENSG00000137409",
                "ENSG00000006125",
                "ENSG00000196937",
                "ENSG00000188613",
                "ENSG00000170445",
                "ENSG00000100442",
                "ENSG00000106028",
                "ENSG00000100697",
                "ENSG00000134986",
                "ENSG00000121022",
                "ENSG00000197111",
                "ENSG00000163682",
                "ENSG00000101182",
                "ENSG00000172020",
                "ENSG00000124380",
                "ENSG00000166260",
                "ENSG00000051108",
                "ENSG00000182093",
                "ENSG00000134825",
                "ENSG00000184007",
                "ENSG00000096063",
                "ENSG00000186868",
                "ENSG00000134333",
                "ENSG00000122012",
                "ENSG00000116251",
                "ENSG00000091640",
                "ENSG00000143977",
                "ENSG00000038219",
                "ENSG00000110700",
                "ENSG00000146731",
                "ENSG00000143353",
                "ENSG00000053254",
                "ENSG00000011295",
                "ENSG00000130706",
                "ENSG00000173692",
                "ENSG00000182985",
                "ENSG00000161547",
                "ENSG00000134769",
                "ENSG00000145782",
                "ENSG00000100354",
                "ENSG00000106829",
                "ENSG00000103429",
                "ENSG00000138430",
                "ENSG00000137285",
                "ENSG00000104435",
                "ENSG00000273611",
                "ENSG00000186472",
                "ENSG00000069956",
                "ENSG00000146416",
                "ENSG00000164258",
                "ENSG00000116560",
                "ENSG00000109472",
                "ENSG00000114956",
                "ENSG00000145725",
                "ENSG00000092201",
                "ENSG00000147862",
                "ENSG00000079102",
                "ENSG00000182004",
                "ENSG00000008294",
                "ENSG00000104408",
                "ENSG00000156467",
                "ENSG00000168259",
                "ENSG00000161203",
                "ENSG00000100823",
                "ENSG00000101782",
                "ENSG00000142541",
                "ENSG00000111906",
                "ENSG00000143753",
                "ENSG00000118816",
                "ENSG00000106105",
                "ENSG00000147669",
                "ENSG00000151474",
                "ENSG00000126524",
                "ENSG00000163468",
                "ENSG00000091039",
                "ENSG00000167522",
                "ENSG00000134056",
                "ENSG00000277443",
                "ENSG00000135387",
                "ENSG00000156976",
                "ENSG00000169021",
                "ENSG00000078369",
                "ENSG00000138758",
                "ENSG00000115541",
                "ENSG00000150093",
                "ENSG00000138107",
                "ENSG00000149485",
                "ENSG00000130748",
                "ENSG00000116641",
                "ENSG00000138326",
                "ENSG00000154380",
                "ENSG00000145362",
                "ENSG00000104695",
                "ENSG00000182621",
                "ENSG00000123144",
                "ENSG00000113575",
                "ENSG00000149084",
                "ENSG00000127022",
                "ENSG00000114302",
                "ENSG00000142676",
                "ENSG00000119705",
                "ENSG00000165283",
                "ENSG00000075151",
                "ENSG00000100528",
                "ENSG00000154582",
                "ENSG00000151320",
                "ENSG00000077279",
                "ENSG00000137996",
                "ENSG00000016082",
                "ENSG00000108946",
                "ENSG00000164830",
                "ENSG00000108953",
                "ENSG00000146267",
                "ENSG00000094916",
                "ENSG00000177875",
                "ENSG00000186063",
                "ENSG00000204310",
                "ENSG00000010404",
                "ENSG00000070756",
                "ENSG00000136930",
                "ENSG00000140740",
                "ENSG00000167658",
                "ENSG00000113161",
                "ENSG00000072041",
                "ENSG00000197006",
                "ENSG00000159176",
                "ENSG00000196465",
                "ENSG00000108848",
                "ENSG00000143933",
                "ENSG00000132963",
                "ENSG00000042753",
                "ENSG00000175390",
                "ENSG00000168522",
                "ENSG00000119048",
                "ENSG00000198408",
                "ENSG00000111897",
                "ENSG00000166946",
                "ENSG00000143198",
                "ENSG00000135406",
                "ENSG00000170017",
                "ENSG00000177675",
                "ENSG00000115944",
                "ENSG00000111639",
                "ENSG00000138443",
                "ENSG00000168268",
                "ENSG00000107105",
                "ENSG00000086300",
                "ENSG00000112186",
                "ENSG00000203930",
                "ENSG00000182481",
                "ENSG00000159593",
                "ENSG00000156261",
                "ENSG00000198363",
                "ENSG00000197728",
                "ENSG00000170606",
                "ENSG00000182512",
                "ENSG00000179271",
                "ENSG00000184207",
                "ENSG00000135720",
                "ENSG00000166848",
                "ENSG00000196683",
                "ENSG00000166206",
                "ENSG00000099194",
                "ENSG00000172809",
                "ENSG00000089818",
                "ENSG00000099341",
                "ENSG00000087302",
                "ENSG00000100220",
                "ENSG00000105176",
                "ENSG00000169599",
                "ENSG00000160752",
                "ENSG00000143252",
                "ENSG00000144381",
                "ENSG00000136521",
                "ENSG00000104177",
                "ENSG00000164741",
                "ENSG00000152234",
                "ENSG00000172687",
                "ENSG00000108100",
                "ENSG00000061794",
                "ENSG00000135549",
                "ENSG00000153774",
                "ENSG00000145741",
                "ENSG00000138757",
                "ENSG00000033627",
                "ENSG00000155876",
                "ENSG00000171867",
                "ENSG00000137547",
                "ENSG00000185565",
                "ENSG00000184220",
                "ENSG00000018189",
                "ENSG00000174498",
                "ENSG00000137876",
                "ENSG00000196591",
                "ENSG00000138398",
                "ENSG00000122705",
                "ENSG00000115306",
                "ENSG00000125944",
                "ENSG00000054523",
                "ENSG00000100380",
                "ENSG00000100804",
                "ENSG00000119640",
                "ENSG00000170899",
                "ENSG00000114125",
                "ENSG00000185787",
                "ENSG00000141424",
                "ENSG00000130638",
                "ENSG00000111775",
                "ENSG00000123636",
                "ENSG00000139684",
                "ENSG00000114120",
                "ENSG00000164587",
                "ENSG00000181163",
                "ENSG00000008018",
                "ENSG00000078018",
                "ENSG00000204843",
                "ENSG00000215021",
                "ENSG00000137274",
                "ENSG00000044115",
                "ENSG00000265241",
                "ENSG00000134590",
                "ENSG00000103415",
                "ENSG00000065978",
                "ENSG00000113558",
                "ENSG00000139970",
                "ENSG00000143153",
                "ENSG00000123983",
                "ENSG00000116489",
                "ENSG00000143742",
                "ENSG00000133169",
                "ENSG00000205629",
                "ENSG00000164167",
                "ENSG00000089335",
                "ENSG00000181610",
                "ENSG00000089009"
            ],
            "expression_sum": {
                "class1": 4511.0,
                "class2": 26.0
            },
            "ens_id": "ENSG00000109132",
            "uniprot_ids": [
                "Q99453"
            ],
            "net_id": 0,
            "source_num": 2,
            "target_num": 63
        },
        "ENSG00000153147": {
            "id": "ENSG00000153147",
            "type": "TF",
            "symbol": "SMARCA5",
            "source": [],
            "target": [
                "ENSG00000016082",
                "ENSG00000167654",
                "ENSG00000107105",
                "ENSG00000198125",
                "ENSG00000181885",
                "ENSG00000104870"
            ],
            "expression_sum": {
                "class1": 14120.0,
                "class2": 12380.0
            },
            "ens_id": "ENSG00000153147",
            "uniprot_ids": [
                "O60264"
            ],
            "net_id": 0,
            "source_num": 0,
            "target_num": 3
        },
        "ENSG00000198911": {
            "id": "ENSG00000198911",
            "type": "TF",
            "symbol": "SREBF2",
            "source": [],
            "target": [
                "ENSG00000104722",
                "ENSG00000176788",
                "ENSG00000144339",
                "ENSG00000169783",
                "ENSG00000113657",
                "ENSG00000113161",
                "ENSG00000107105",
                "ENSG00000129951",
                "ENSG00000080824",
                "ENSG00000129355",
                "ENSG00000166226",
                "ENSG00000198910",
                "ENSG00000060982",
                "ENSG00000149313",
                "ENSG00000105409",
                "ENSG00000197747",
                "ENSG00000164687",
                "ENSG00000072832",
                "ENSG00000104427",
                "ENSG00000101132",
                "ENSG00000182481",
                "ENSG00000197694",
                "ENSG00000168280",
                "ENSG00000186868",
                "ENSG00000047849",
                "ENSG00000144642",
                "ENSG00000092964",
                "ENSG00000143702",
                "ENSG00000144218",
                "ENSG00000163539",
                "ENSG00000122012",
                "ENSG00000125354",
                "ENSG00000077380",
                "ENSG00000162490",
                "ENSG00000016082",
                "ENSG00000097021",
                "ENSG00000124194",
                "ENSG00000120437",
                "ENSG00000160014",
                "ENSG00000117632",
                "ENSG00000111728",
                "ENSG00000154917",
                "ENSG00000034510",
                "ENSG00000131016",
                "ENSG00000146425",
                "ENSG00000139970",
                "ENSG00000079462",
                "ENSG00000084731",
                "ENSG00000135406",
                "ENSG00000115665",
                "ENSG00000068615",
                "ENSG00000277586"
            ],
            "expression_sum": {
                "class1": 5454.0,
                "class2": 2413.0
            },
            "ens_id": "ENSG00000198911",
            "uniprot_ids": [
                "Q12772",
                "G3V0I8",
                "H0Y7E5",
                "A0A087X1T2"
            ],
            "net_id": 0,
            "source_num": 0,
            "target_num": 20
        },
        "ENSG00000196591": {
            "id": "ENSG00000196591",
            "type": "TF",
            "symbol": "HDAC2",
            "source": [
                "ENSG00000163435",
                "ENSG00000109132",
                "ENSG00000167081",
                "ENSG00000148516",
                "ENSG00000164330"
            ],
            "target": [
                "ENSG00000099864",
                "ENSG00000103034",
                "ENSG00000169856",
                "ENSG00000162704",
                "ENSG00000111716",
                "ENSG00000087085",
                "ENSG00000108830",
                "ENSG00000113657",
                "ENSG00000161682",
                "ENSG00000141698",
                "ENSG00000101210",
                "ENSG00000164093",
                "ENSG00000107105",
                "ENSG00000122756",
                "ENSG00000115226",
                "ENSG00000166582",
                "ENSG00000182534",
                "ENSG00000129355",
                "ENSG00000173467",
                "ENSG00000196230",
                "ENSG00000086300",
                "ENSG00000107742",
                "ENSG00000105255",
                "ENSG00000100167",
                "ENSG00000197959",
                "ENSG00000170027",
                "ENSG00000134594",
                "ENSG00000075945",
                "ENSG00000010270",
                "ENSG00000182481",
                "ENSG00000168280",
                "ENSG00000123416",
                "ENSG00000181885",
                "ENSG00000177432",
                "ENSG00000126950",
                "ENSG00000165434",
                "ENSG00000108175",
                "ENSG00000077380",
                "ENSG00000162490",
                "ENSG00000173267",
                "ENSG00000138413",
                "ENSG00000071054",
                "ENSG00000184697",
                "ENSG00000087258",
                "ENSG00000105088",
                "ENSG00000117632",
                "ENSG00000111728",
                "ENSG00000154917",
                "ENSG00000272398",
                "ENSG00000015592",
                "ENSG00000079462",
                "ENSG00000134369",
                "ENSG00000103145",
                "ENSG00000115665",
                "ENSG00000115310",
                "ENSG00000122545",
                "ENSG00000137312",
                "ENSG00000105270",
                "ENSG00000018236",
                "ENSG00000189043",
                "ENSG00000099194",
                "ENSG00000277443",
                "ENSG00000104722",
                "ENSG00000162545",
                "ENSG00000169783",
                "ENSG00000157514",
                "ENSG00000148516",
                "ENSG00000221946",
                "ENSG00000167654",
                "ENSG00000066557",
                "ENSG00000105254",
                "ENSG00000084628",
                "ENSG00000188290",
                "ENSG00000166257",
                "ENSG00000196361",
                "ENSG00000108828",
                "ENSG00000196072",
                "ENSG00000104112",
                "ENSG00000131711",
                "ENSG00000155980",
                "ENSG00000075618",
                "ENSG00000101132",
                "ENSG00000120265",
                "ENSG00000144642",
                "ENSG00000138758",
                "ENSG00000144218",
                "ENSG00000125354",
                "ENSG00000133083",
                "ENSG00000213420",
                "ENSG00000171033",
                "ENSG00000155099",
                "ENSG00000107130",
                "ENSG00000034713",
                "ENSG00000175203",
                "ENSG00000034510",
                "ENSG00000170091",
                "ENSG00000146425",
                "ENSG00000158164",
                "ENSG00000206579",
                "ENSG00000141404",
                "ENSG00000118263",
                "ENSG00000068615",
                "ENSG00000243449",
                "ENSG00000185650",
                "ENSG00000168118",
                "ENSG00000134986",
                "ENSG00000277586",
                "ENSG00000111275",
                "ENSG00000176788",
                "ENSG00000172020",
                "ENSG00000125821",
                "ENSG00000149557",
                "ENSG00000151640",
                "ENSG00000144635",
                "ENSG00000185664",
                "ENSG00000148798",
                "ENSG00000145335",
                "ENSG00000130540",
                "ENSG00000171551",
                "ENSG00000198910",
                "ENSG00000018189",
                "ENSG00000134198",
                "ENSG00000138138",
                "ENSG00000128564",
                "ENSG00000101220",
                "ENSG00000109132",
                "ENSG00000132821",
                "ENSG00000105409",
                "ENSG00000197747",
                "ENSG00000164687",
                "ENSG00000151320",
                "ENSG00000186868",
                "ENSG00000077279",
                "ENSG00000213760",
                "ENSG00000092964",
                "ENSG00000143702",
                "ENSG00000112972",
                "ENSG00000122012",
                "ENSG00000016082",
                "ENSG00000142156",
                "ENSG00000152503",
                "ENSG00000164221",
                "ENSG00000166736",
                "ENSG00000120437",
                "ENSG00000160014",
                "ENSG00000054523",
                "ENSG00000149294",
                "ENSG00000171533",
                "ENSG00000103187",
                "ENSG00000084731",
                "ENSG00000166974",
                "ENSG00000167552",
                "ENSG00000174521",
                "ENSG00000164330",
                "ENSG00000141505",
                "ENSG00000137267",
                "ENSG00000177875",
                "ENSG00000227051",
                "ENSG00000163931",
                "ENSG00000203879",
                "ENSG00000135447",
                "ENSG00000176973",
                "ENSG00000105419",
                "ENSG00000144339",
                "ENSG00000250303",
                "ENSG00000197457",
                "ENSG00000131437",
                "ENSG00000113161",
                "ENSG00000154277",
                "ENSG00000171724",
                "ENSG00000050165",
                "ENSG00000129951",
                "ENSG00000137942",
                "ENSG00000080824",
                "ENSG00000105784",
                "ENSG00000166226",
                "ENSG00000107643",
                "ENSG00000162188",
                "ENSG00000143862",
                "ENSG00000157851",
                "ENSG00000078018",
                "ENSG00000117569",
                "ENSG00000162374",
                "ENSG00000182287",
                "ENSG00000072832",
                "ENSG00000104427",
                "ENSG00000221866",
                "ENSG00000133318",
                "ENSG00000197694",
                "ENSG00000186469",
                "ENSG00000134590",
                "ENSG00000047849",
                "ENSG00000137285",
                "ENSG00000103154",
                "ENSG00000104435",
                "ENSG00000126214",
                "ENSG00000167996",
                "ENSG00000097021",
                "ENSG00000124194",
                "ENSG00000148154",
                "ENSG00000152082",
                "ENSG00000131037",
                "ENSG00000189159",
                "ENSG00000111897",
                "ENSG00000131016",
                "ENSG00000139970",
                "ENSG00000090097",
                "ENSG00000112290",
                "ENSG00000163435",
                "ENSG00000143198",
                "ENSG00000167081",
                "ENSG00000135406",
                "ENSG00000168824",
                "ENSG00000159409",
                "ENSG00000175582",
                "ENSG00000177675",
                "ENSG00000131473",
                "ENSG00000122378",
                "ENSG00000108960",
                "ENSG00000137343"
            ],
            "expression_sum": {
                "class1": 21723.0,
                "class2": 9054.0
            },
            "ens_id": "ENSG00000196591",
            "uniprot_ids": [
                "Q92769",
                "E5RGV4",
                "E5RK19",
                "E5RG37",
                "H3BM24",
                "E5RFP9",
                "E5RH52",
                "E5RFI6",
                "E5RJ04",
                "E5RHE7"
            ],
            "net_id": 0,
            "source_num": 5,
            "target_num": 61
        },
        "ENSG00000105698": {
            "id": "ENSG00000105698",
            "type": "TF",
            "symbol": "USF2",
            "source": [
                "ENSG00000167081",
                "ENSG00000113580",
                "ENSG00000139352",
                "ENSG00000164330"
            ],
            "target": [
                "ENSG00000104722",
                "ENSG00000181449",
                "ENSG00000176788",
                "ENSG00000149557",
                "ENSG00000198840",
                "ENSG00000221946",
                "ENSG00000107105",
                "ENSG00000129951",
                "ENSG00000105784",
                "ENSG00000198910",
                "ENSG00000060982",
                "ENSG00000102265",
                "ENSG00000174021",
                "ENSG00000113732",
                "ENSG00000148834",
                "ENSG00000072832",
                "ENSG00000182481",
                "ENSG00000198125",
                "ENSG00000181885",
                "ENSG00000117691",
                "ENSG00000092964",
                "ENSG00000124194",
                "ENSG00000111728",
                "ENSG00000158164",
                "ENSG00000167081",
                "ENSG00000113580",
                "ENSG00000119888",
                "ENSG00000143545",
                "ENSG00000104833",
                "ENSG00000174521",
                "ENSG00000164330",
                "ENSG00000137267",
                "ENSG00000118418"
            ],
            "expression_sum": {
                "class1": 6116.0,
                "class2": 4359.0
            },
            "ens_id": "ENSG00000105698",
            "uniprot_ids": [
                "Q15853",
                "B4DLJ1",
                "M0R1C6",
                "M0QY46",
                "M0QZ39",
                "V9GYR5",
                "M0QXT0",
                "M0QYU7"
            ],
            "net_id": 0,
            "source_num": 4,
            "target_num": 9
        },
        "ENSG00000213015": {
            "id": "ENSG00000213015",
            "type": "TF",
            "symbol": "ZNF580",
            "source": [
                "ENSG00000167081",
                "ENSG00000181449",
                "ENSG00000164330"
            ],
            "target": [
                "ENSG00000176788",
                "ENSG00000169856",
                "ENSG00000107105",
                "ENSG00000080824",
                "ENSG00000129355",
                "ENSG00000084628",
                "ENSG00000176749",
                "ENSG00000128564",
                "ENSG00000187134",
                "ENSG00000197959",
                "ENSG00000197747",
                "ENSG00000010270",
                "ENSG00000104427",
                "ENSG00000090273",
                "ENSG00000133318",
                "ENSG00000198513",
                "ENSG00000186868",
                "ENSG00000168280",
                "ENSG00000123416",
                "ENSG00000115756",
                "ENSG00000182481",
                "ENSG00000108106",
                "ENSG00000213760",
                "ENSG00000137285",
                "ENSG00000165434",
                "ENSG00000143702",
                "ENSG00000126214",
                "ENSG00000213420",
                "ENSG00000097021",
                "ENSG00000152503",
                "ENSG00000164221",
                "ENSG00000131037",
                "ENSG00000168243",
                "ENSG00000189159",
                "ENSG00000120437",
                "ENSG00000160014",
                "ENSG00000117632",
                "ENSG00000272398",
                "ENSG00000131016",
                "ENSG00000170091",
                "ENSG00000146425",
                "ENSG00000149294",
                "ENSG00000105649",
                "ENSG00000079462",
                "ENSG00000167081",
                "ENSG00000135406",
                "ENSG00000143545",
                "ENSG00000164032",
                "ENSG00000137267",
                "ENSG00000185650",
                "ENSG00000149926"
            ],
            "expression_sum": {
                "class1": 11545.0,
                "class2": 8422.0
            },
            "ens_id": "ENSG00000213015",
            "uniprot_ids": [
                "Q9UK33",
                "K7EJ53",
                "K7ERU0",
                "K7ENY2"
            ],
            "net_id": 0,
            "source_num": 3,
            "target_num": 12
        },
        "ENSG00000119950": {
            "id": "ENSG00000119950",
            "type": "TF",
            "symbol": "MXI1",
            "source": [
                "ENSG00000181449",
                "ENSG00000167081",
                "ENSG00000113580",
                "ENSG00000164330",
                "ENSG00000164093",
                "ENSG00000139352"
            ],
            "target": [
                "ENSG00000104722",
                "ENSG00000169783",
                "ENSG00000112773",
                "ENSG00000105784",
                "ENSG00000196230",
                "ENSG00000084628",
                "ENSG00000105409",
                "ENSG00000072832",
                "ENSG00000182481",
                "ENSG00000117155",
                "ENSG00000117691",
                "ENSG00000139970",
                "ENSG00000170091",
                "ENSG00000157557",
                "ENSG00000167081",
                "ENSG00000113580",
                "ENSG00000100604",
                "ENSG00000164330",
                "ENSG00000137267"
            ],
            "expression_sum": {
                "class1": 8238.0,
                "class2": 6134.0
            },
            "ens_id": "ENSG00000119950",
            "uniprot_ids": [
                "P50539",
                "A0A494BZS3",
                "A0A494C0W2",
                "A0A494C1P8",
                "A0A0S2Z429",
                "B1ANN8",
                "F6U3F6",
                "A0A494C1C1",
                "F6WCI9",
                "A0A494C0I8"
            ],
            "net_id": 0,
            "source_num": 6,
            "target_num": 7
        },
        "ENSG00000092199": {
            "id": "ENSG00000092199",
            "type": "TF",
            "symbol": "HNRNPC",
            "source": [
                "ENSG00000181449",
                "ENSG00000163435",
                "ENSG00000167081",
                "ENSG00000113580",
                "ENSG00000164330"
            ],
            "target": [
                "ENSG00000162496",
                "ENSG00000130643",
                "ENSG00000103034",
                "ENSG00000087085",
                "ENSG00000185664",
                "ENSG00000148798",
                "ENSG00000105784",
                "ENSG00000171551",
                "ENSG00000244187",
                "ENSG00000141934",
                "ENSG00000084628",
                "ENSG00000166257",
                "ENSG00000174021",
                "ENSG00000128564",
                "ENSG00000060982",
                "ENSG00000197959",
                "ENSG00000174498",
                "ENSG00000176749",
                "ENSG00000196361",
                "ENSG00000105409",
                "ENSG00000182287",
                "ENSG00000221866",
                "ENSG00000131711",
                "ENSG00000166145",
                "ENSG00000005108",
                "ENSG00000213719",
                "ENSG00000092964",
                "ENSG00000143702",
                "ENSG00000103154",
                "ENSG00000213420",
                "ENSG00000107438",
                "ENSG00000162490",
                "ENSG00000171033",
                "ENSG00000152503",
                "ENSG00000128228",
                "ENSG00000087258",
                "ENSG00000100167",
                "ENSG00000139970",
                "ENSG00000171533",
                "ENSG00000124942",
                "ENSG00000167081",
                "ENSG00000113580",
                "ENSG00000119888",
                "ENSG00000135406",
                "ENSG00000174521",
                "ENSG00000151150",
                "ENSG00000118418",
                "ENSG00000185650",
                "ENSG00000104870"
            ],
            "expression_sum": {
                "class1": 26156.0,
                "class2": 27085.0
            },
            "ens_id": "ENSG00000092199",
            "uniprot_ids": [
                "P07910",
                "G3V576",
                "B2R5W2",
                "B4DY08",
                "B4DSU6",
                "G3V4C1",
                "G3V4M8",
                "G3V2H6",
                "G3V4W0",
                "G3V5X6",
                "G3V2Q1",
                "G3V251",
                "G3V555",
                "G3V575",
                "G3V3K6",
                "G3V2D6",
                "G3V5V7"
            ],
            "net_id": 0,
            "source_num": 5,
            "target_num": 10
        },
        "ENSG00000106261": {
            "id": "ENSG00000106261",
            "type": "TF",
            "symbol": "ZKSCAN1",
            "source": [
                "ENSG00000163435",
                "ENSG00000167081",
                "ENSG00000113580",
                "ENSG00000148516",
                "ENSG00000100219",
                "ENSG00000221869"
            ],
            "target": [
                "ENSG00000103034",
                "ENSG00000148516",
                "ENSG00000167654",
                "ENSG00000101443",
                "ENSG00000163399",
                "ENSG00000154080",
                "ENSG00000196230",
                "ENSG00000060982",
                "ENSG00000152952",
                "ENSG00000130303",
                "ENSG00000163191",
                "ENSG00000155980",
                "ENSG00000123416",
                "ENSG00000198125",
                "ENSG00000138758",
                "ENSG00000092964",
                "ENSG00000132688",
                "ENSG00000143702",
                "ENSG00000135535",
                "ENSG00000245532",
                "ENSG00000075043",
                "ENSG00000167642",
                "ENSG00000102007",
                "ENSG00000107130",
                "ENSG00000149485",
                "ENSG00000129521",
                "ENSG00000189159",
                "ENSG00000117632",
                "ENSG00000170091",
                "ENSG00000163435",
                "ENSG00000096696",
                "ENSG00000167552",
                "ENSG00000118263",
                "ENSG00000164330",
                "ENSG00000137267",
                "ENSG00000100219",
                "ENSG00000185650",
                "ENSG00000104870"
            ],
            "expression_sum": {
                "class1": 11131.0,
                "class2": 11221.0
            },
            "ens_id": "ENSG00000106261",
            "uniprot_ids": [
                "P17029",
                "B3KRF7",
                "E9PC66",
                "C9JRM9"
            ],
            "net_id": 0,
            "source_num": 6,
            "target_num": 9
        },
        "ENSG00000148516": {
            "id": "ENSG00000148516",
            "type": "TF",
            "symbol": "ZEB1",
            "source": [
                "ENSG00000106261",
                "ENSG00000164330",
                "ENSG00000168214",
                "ENSG00000196591",
                "ENSG00000124766",
                "ENSG00000119547"
            ],
            "target": [
                "ENSG00000103034",
                "ENSG00000106261",
                "ENSG00000163634",
                "ENSG00000111674",
                "ENSG00000087085",
                "ENSG00000005810",
                "ENSG00000073792",
                "ENSG00000107105",
                "ENSG00000114784",
                "ENSG00000197302",
                "ENSG00000160113",
                "ENSG00000179915",
                "ENSG00000120992",
                "ENSG00000186106",
                "ENSG00000128872",
                "ENSG00000170027",
                "ENSG00000075945",
                "ENSG00000078674",
                "ENSG00000023287",
                "ENSG00000076554",
                "ENSG00000156467",
                "ENSG00000166295",
                "ENSG00000168280",
                "ENSG00000119013",
                "ENSG00000164692",
                "ENSG00000184076",
                "ENSG00000165434",
                "ENSG00000108175",
                "ENSG00000172159",
                "ENSG00000077380",
                "ENSG00000154556",
                "ENSG00000178741",
                "ENSG00000173267",
                "ENSG00000138413",
                "ENSG00000173812",
                "ENSG00000067715",
                "ENSG00000184697",
                "ENSG00000154917",
                "ENSG00000272398",
                "ENSG00000172292",
                "ENSG00000123240",
                "ENSG00000051620",
                "ENSG00000166136",
                "ENSG00000251562",
                "ENSG00000115310",
                "ENSG00000166848",
                "ENSG00000137312",
                "ENSG00000074319",
                "ENSG00000113742",
                "ENSG00000164924",
                "ENSG00000091039",
                "ENSG00000163939",
                "ENSG00000111371",
                "ENSG00000137941",
                "ENSG00000119547",
                "ENSG00000104722",
                "ENSG00000101040",
                "ENSG00000159023",
                "ENSG00000198886",
                "ENSG00000204130",
                "ENSG00000108309",
                "ENSG00000125868",
                "ENSG00000104904",
                "ENSG00000125971",
                "ENSG00000153815",
                "ENSG00000106565",
                "ENSG00000135387",
                "ENSG00000166257",
                "ENSG00000147548",
                "ENSG00000156976",
                "ENSG00000168092",
                "ENSG00000253719",
                "ENSG00000196361",
                "ENSG00000119335",
                "ENSG00000130294",
                "ENSG00000078369",
                "ENSG00000100077",
                "ENSG00000169432",
                "ENSG00000104112",
                "ENSG00000155980",
                "ENSG00000185201",
                "ENSG00000187231",
                "ENSG00000154723",
                "ENSG00000166347",
                "ENSG00000163539",
                "ENSG00000131174",
                "ENSG00000186187",
                "ENSG00000114739",
                "ENSG00000110321",
                "ENSG00000117139",
                "ENSG00000109654",
                "ENSG00000198420",
                "ENSG00000234745",
                "ENSG00000034713",
                "ENSG00000205403",
                "ENSG00000198668",
                "ENSG00000063854",
                "ENSG00000103769",
                "ENSG00000146425",
                "ENSG00000170091",
                "ENSG00000111642",
                "ENSG00000117410",
                "ENSG00000145362",
                "ENSG00000141404",
                "ENSG00000118263",
                "ENSG00000273841",
                "ENSG00000068615",
                "ENSG00000243449",
                "ENSG00000204525",
                "ENSG00000263001",
                "ENSG00000149257",
                "ENSG00000196405",
                "ENSG00000129197",
                "ENSG00000033627",
                "ENSG00000100697",
                "ENSG00000162961",
                "ENSG00000197111",
                "ENSG00000225783",
                "ENSG00000084733",
                "ENSG00000111726",
                "ENSG00000133134",
                "ENSG00000109099",
                "ENSG00000142949",
                "ENSG00000114573",
                "ENSG00000144635",
                "ENSG00000196233",
                "ENSG00000185664",
                "ENSG00000163041",
                "ENSG00000148798",
                "ENSG00000118680",
                "ENSG00000197183",
                "ENSG00000009307",
                "ENSG00000136854",
                "ENSG00000018189",
                "ENSG00000138138",
                "ENSG00000110955",
                "ENSG00000096384",
                "ENSG00000175274",
                "ENSG00000197747",
                "ENSG00000196591",
                "ENSG00000168214",
                "ENSG00000108654",
                "ENSG00000122417",
                "ENSG00000135940",
                "ENSG00000143702",
                "ENSG00000131966",
                "ENSG00000038219",
                "ENSG00000178104",
                "ENSG00000128609",
                "ENSG00000114405",
                "ENSG00000149294",
                "ENSG00000244879",
                "ENSG00000105220",
                "ENSG00000166974",
                "ENSG00000104643",
                "ENSG00000164330",
                "ENSG00000007168",
                "ENSG00000151150",
                "ENSG00000100320",
                "ENSG00000111775",
                "ENSG00000123636",
                "ENSG00000162852",
                "ENSG00000139146",
                "ENSG00000166562",
                "ENSG00000132824",
                "ENSG00000131437",
                "ENSG00000102003",
                "ENSG00000127616",
                "ENSG00000196781",
                "ENSG00000082397",
                "ENSG00000137942",
                "ENSG00000080824",
                "ENSG00000107643",
                "ENSG00000112695",
                "ENSG00000160271",
                "ENSG00000162188",
                "ENSG00000162374",
                "ENSG00000120694",
                "ENSG00000220205",
                "ENSG00000196586",
                "ENSG00000104427",
                "ENSG00000184545",
                "ENSG00000131149",
                "ENSG00000136238",
                "ENSG00000198513",
                "ENSG00000143933",
                "ENSG00000169213",
                "ENSG00000113141",
                "ENSG00000165629",
                "ENSG00000166503",
                "ENSG00000103154",
                "ENSG00000104435",
                "ENSG00000126214",
                "ENSG00000167815",
                "ENSG00000097033",
                "ENSG00000119314",
                "ENSG00000156587",
                "ENSG00000124194",
                "ENSG00000170759",
                "ENSG00000131037",
                "ENSG00000177410",
                "ENSG00000145247",
                "ENSG00000178127",
                "ENSG00000165943",
                "ENSG00000104381",
                "ENSG00000186472",
                "ENSG00000139970",
                "ENSG00000127914",
                "ENSG00000090097",
                "ENSG00000182718",
                "ENSG00000143153",
                "ENSG00000168824",
                "ENSG00000133169",
                "ENSG00000175582",
                "ENSG00000169439",
                "ENSG00000018408",
                "ENSG00000149926"
            ],
            "expression_sum": {
                "class1": 6453.0,
                "class2": 3224.0
            },
            "ens_id": "ENSG00000148516",
            "uniprot_ids": [
                "P37275",
                "F6TDF5",
                "H0YND9",
                "F6U0D0",
                "H0YMD3"
            ],
            "net_id": 0,
            "source_num": 6,
            "target_num": 56
        },
        "ENSG00000143799": {
            "id": "ENSG00000143799",
            "type": "TF",
            "symbol": "PARP1",
            "source": [
                "ENSG00000124766",
                "ENSG00000181449",
                "ENSG00000167081",
                "ENSG00000164330"
            ],
            "target": [
                "ENSG00000169783",
                "ENSG00000139116",
                "ENSG00000087085",
                "ENSG00000131437",
                "ENSG00000185664",
                "ENSG00000166803",
                "ENSG00000099250",
                "ENSG00000018189",
                "ENSG00000166257",
                "ENSG00000186106",
                "ENSG00000197959",
                "ENSG00000105409",
                "ENSG00000104427",
                "ENSG00000221866",
                "ENSG00000155980",
                "ENSG00000151320",
                "ENSG00000197694",
                "ENSG00000129682",
                "ENSG00000198513",
                "ENSG00000186469",
                "ENSG00000144642",
                "ENSG00000143702",
                "ENSG00000126214",
                "ENSG00000152503",
                "ENSG00000107130",
                "ENSG00000173267",
                "ENSG00000177410",
                "ENSG00000111728",
                "ENSG00000149294",
                "ENSG00000131016",
                "ENSG00000170091",
                "ENSG00000116729",
                "ENSG00000139970",
                "ENSG00000090097",
                "ENSG00000156298",
                "ENSG00000123240",
                "ENSG00000145362",
                "ENSG00000135406",
                "ENSG00000166974",
                "ENSG00000150722",
                "ENSG00000184144",
                "ENSG00000164330",
                "ENSG00000151150",
                "ENSG00000033627",
                "ENSG00000018236",
                "ENSG00000185650",
                "ENSG00000149926"
            ],
            "expression_sum": {
                "class1": 10029.0,
                "class2": 9650.0
            },
            "ens_id": "ENSG00000143799",
            "uniprot_ids": [
                "P09874",
                "A0A7I2V625",
                "A0A7I2V5E9",
                "A0A7I2V384",
                "A0A7I2V3E1"
            ],
            "net_id": 0,
            "source_num": 4,
            "target_num": 12
        },
        "ENSG00000128272": {
            "id": "ENSG00000128272",
            "type": "TF",
            "symbol": "ATF4",
            "source": [],
            "target": [
                "ENSG00000166922",
                "ENSG00000172020",
                "ENSG00000079102",
                "ENSG00000182985",
                "ENSG00000005810",
                "ENSG00000113161",
                "ENSG00000154277",
                "ENSG00000182093",
                "ENSG00000082397",
                "ENSG00000086300",
                "ENSG00000166257",
                "ENSG00000060982",
                "ENSG00000130294",
                "ENSG00000111860",
                "ENSG00000131711",
                "ENSG00000151320",
                "ENSG00000198513",
                "ENSG00000168280",
                "ENSG00000213760",
                "ENSG00000137285",
                "ENSG00000156515",
                "ENSG00000152503",
                "ENSG00000173267",
                "ENSG00000138413",
                "ENSG00000176046",
                "ENSG00000087258",
                "ENSG00000090097",
                "ENSG00000145362",
                "ENSG00000108821",
                "ENSG00000115310",
                "ENSG00000118263",
                "ENSG00000081181",
                "ENSG00000164330",
                "ENSG00000033627",
                "ENSG00000185650",
                "ENSG00000149926"
            ],
            "expression_sum": {
                "class1": 9228.0,
                "class2": 8636.0
            },
            "ens_id": "ENSG00000128272",
            "uniprot_ids": [
                "P18848",
                "B4DJD4",
                "A0A6Q8PGY1",
                "A0A6Q8PHB3",
                "A0A6Q8PFH6",
                "A0A6Q8PG17",
                "A0A7P0Z4J5",
                "A0A6Q8PF56"
            ],
            "net_id": 0,
            "source_num": 0,
            "target_num": 9
        },
        "ENSG00000163848": {
            "id": "ENSG00000163848",
            "type": "TF",
            "symbol": "ZNF148",
            "source": [
                "ENSG00000016082",
                "ENSG00000181449",
                "ENSG00000164330"
            ],
            "target": [
                "ENSG00000181449",
                "ENSG00000143702",
                "ENSG00000060982",
                "ENSG00000115461",
                "ENSG00000176788",
                "ENSG00000149557",
                "ENSG00000135406",
                "ENSG00000164330",
                "ENSG00000198910"
            ],
            "expression_sum": {
                "class1": 5593.0,
                "class2": 5287.0
            },
            "ens_id": "ENSG00000163848",
            "uniprot_ids": [
                "Q9UQR1",
                "G5E9X2",
                "C9K0U4",
                "C9J6Y6",
                "C9JRX0"
            ],
            "net_id": 0,
            "source_num": 3,
            "target_num": 4
        },
        "ENSG00000165512": {
            "id": "ENSG00000165512",
            "type": "TF",
            "symbol": "ZNF22",
            "source": [
                "ENSG00000164330"
            ],
            "target": [
                "ENSG00000084628",
                "ENSG00000131016",
                "ENSG00000164330",
                "ENSG00000161082"
            ],
            "expression_sum": {
                "class1": 9759.0,
                "class2": 8278.0
            },
            "ens_id": "ENSG00000165512",
            "uniprot_ids": [
                "P17026"
            ],
            "net_id": 0,
            "source_num": 1,
            "target_num": 2
        },
        "ENSG00000067955": {
            "id": "ENSG00000067955",
            "type": "TF",
            "symbol": "CBFB",
            "source": [
                "ENSG00000113580",
                "ENSG00000164330"
            ],
            "target": [
                "ENSG00000067064",
                "ENSG00000138698",
                "ENSG00000197102",
                "ENSG00000136717",
                "ENSG00000173272",
                "ENSG00000239672",
                "ENSG00000109670",
                "ENSG00000157514",
                "ENSG00000106615",
                "ENSG00000005810",
                "ENSG00000167658",
                "ENSG00000109084",
                "ENSG00000196262",
                "ENSG00000137955",
                "ENSG00000166226",
                "ENSG00000196230",
                "ENSG00000178980",
                "ENSG00000136942",
                "ENSG00000089157",
                "ENSG00000101132",
                "ENSG00000075618",
                "ENSG00000197694",
                "ENSG00000123416",
                "ENSG00000198961",
                "ENSG00000047849",
                "ENSG00000108175",
                "ENSG00000134982",
                "ENSG00000144713",
                "ENSG00000143702",
                "ENSG00000144218",
                "ENSG00000265972",
                "ENSG00000065978",
                "ENSG00000231500",
                "ENSG00000138413",
                "ENSG00000140374",
                "ENSG00000169710",
                "ENSG00000118200",
                "ENSG00000189159",
                "ENSG00000006125",
                "ENSG00000117632",
                "ENSG00000065911",
                "ENSG00000168275",
                "ENSG00000034510",
                "ENSG00000143198",
                "ENSG00000113580",
                "ENSG00000115310",
                "ENSG00000118263",
                "ENSG00000164330",
                "ENSG00000033627",
                "ENSG00000164924",
                "ENSG00000189043",
                "ENSG00000108010"
            ],
            "expression_sum": {
                "class1": 3646.0,
                "class2": 1787.0
            },
            "ens_id": "ENSG00000067955",
            "uniprot_ids": [
                "Q13951",
                "H3BSC0",
                "J3KS23",
                "A0A494C0A9",
                "J3KTD8",
                "J3KRT0"
            ],
            "net_id": 0,
            "source_num": 2,
            "target_num": 23
        },
        "ENSG00000162599": {
            "id": "ENSG00000162599",
            "type": "TF",
            "symbol": "NFIA",
            "source": [
                "ENSG00000175745",
                "ENSG00000181449"
            ],
            "target": [
                "ENSG00000134138",
                "ENSG00000134245",
                "ENSG00000122545",
                "ENSG00000175745",
                "ENSG00000215386",
                "ENSG00000120885",
                "ENSG00000134308"
            ],
            "expression_sum": {
                "class1": 1610.0,
                "class2": 14292.0
            },
            "ens_id": "ENSG00000162599",
            "uniprot_ids": [
                "Q12857",
                "A0A590UJU2",
                "A0A590UK38",
                "B1AKN8",
                "A0A590UJ67",
                "S4R3W2",
                "S4R3K5",
                "S4R3K4",
                "S4R3W6",
                "A0A590UJA3",
                "A0A590UK09",
                "A0A590UJ82",
                "B1AKN6",
                "S4R308",
                "B1AKN5",
                "B1AKN7",
                "A0A590UK73"
            ],
            "net_id": 0,
            "source_num": 2,
            "target_num": 3
        },
        "ENSG00000168610": {
            "id": "ENSG00000168610",
            "type": "TF",
            "symbol": "STAT3",
            "source": [],
            "target": [
                "ENSG00000128564",
                "ENSG00000122012",
                "ENSG00000197959",
                "ENSG00000265972",
                "ENSG00000154556",
                "ENSG00000183742",
                "ENSG00000107438",
                "ENSG00000135406",
                "ENSG00000132698",
                "ENSG00000010278",
                "ENSG00000134138",
                "ENSG00000221866",
                "ENSG00000139970",
                "ENSG00000089356",
                "ENSG00000181885",
                "ENSG00000118816"
            ],
            "expression_sum": {
                "class1": 5863.0,
                "class2": 8873.0
            },
            "ens_id": "ENSG00000168610",
            "uniprot_ids": [
                "P40763",
                "A0A7I2V5N9",
                "A0A7I2V2T1",
                "A0A7I2V4C8",
                "A0A7I2V4R3",
                "A0A7I2V395",
                "A0A7I2V2G1",
                "A0A7I2V552",
                "G8JLH9",
                "A0A7I2YQD2",
                "A0A7I2V3V0",
                "A0A7I2YQR5",
                "A0A7I2V444",
                "A0A7I2V4F6",
                "A0A7I2YQI1",
                "A0A7I2V4R2"
            ],
            "net_id": 0,
            "source_num": 0,
            "target_num": 8
        },
        "ENSG00000114861": {
            "id": "ENSG00000114861",
            "type": "TF",
            "symbol": "FOXP1",
            "source": [
                "ENSG00000119547",
                "ENSG00000134138"
            ],
            "target": [
                "ENSG00000078668",
                "ENSG00000232119",
                "ENSG00000107223",
                "ENSG00000162704",
                "ENSG00000196531",
                "ENSG00000163634",
                "ENSG00000111716",
                "ENSG00000082153",
                "ENSG00000114784",
                "ENSG00000184752",
                "ENSG00000169567",
                "ENSG00000129355",
                "ENSG00000242485",
                "ENSG00000140941",
                "ENSG00000134248",
                "ENSG00000131143",
                "ENSG00000168653",
                "ENSG00000102144",
                "ENSG00000076554",
                "ENSG00000130770",
                "ENSG00000156467",
                "ENSG00000131507",
                "ENSG00000075415",
                "ENSG00000184076",
                "ENSG00000171314",
                "ENSG00000154556",
                "ENSG00000178741",
                "ENSG00000057757",
                "ENSG00000177889",
                "ENSG00000173267",
                "ENSG00000147684",
                "ENSG00000166913",
                "ENSG00000173812",
                "ENSG00000152256",
                "ENSG00000173915",
                "ENSG00000065911",
                "ENSG00000111229",
                "ENSG00000118816",
                "ENSG00000124570",
                "ENSG00000041357",
                "ENSG00000115310",
                "ENSG00000163875",
                "ENSG00000167136",
                "ENSG00000149547",
                "ENSG00000164924",
                "ENSG00000096092",
                "ENSG00000147403",
                "ENSG00000172115",
                "ENSG00000119547",
                "ENSG00000101558",
                "ENSG00000162545",
                "ENSG00000167283",
                "ENSG00000170035",
                "ENSG00000137726",
                "ENSG00000106615",
                "ENSG00000099341",
                "ENSG00000198886",
                "ENSG00000230989",
                "ENSG00000154518",
                "ENSG00000111640",
                "ENSG00000125868",
                "ENSG00000104904",
                "ENSG00000165637",
                "ENSG00000125977",
                "ENSG00000241685",
                "ENSG00000173726",
                "ENSG00000125971",
                "ENSG00000089693",
                "ENSG00000173674",
                "ENSG00000161970",
                "ENSG00000156976",
                "ENSG00000128245",
                "ENSG00000129625",
                "ENSG00000089220",
                "ENSG00000134138",
                "ENSG00000101460",
                "ENSG00000117758",
                "ENSG00000134717",
                "ENSG00000099624",
                "ENSG00000114023",
                "ENSG00000120265",
                "ENSG00000067225",
                "ENSG00000154723",
                "ENSG00000113712",
                "ENSG00000105438",
                "ENSG00000138069",
                "ENSG00000110321",
                "ENSG00000109654",
                "ENSG00000145495",
                "ENSG00000254999",
                "ENSG00000112335",
                "ENSG00000133872",
                "ENSG00000130558",
                "ENSG00000150459",
                "ENSG00000034713",
                "ENSG00000152234",
                "ENSG00000113387",
                "ENSG00000095380",
                "ENSG00000198668",
                "ENSG00000108107",
                "ENSG00000134308",
                "ENSG00000156508",
                "ENSG00000034510",
                "ENSG00000134684",
                "ENSG00000117410",
                "ENSG00000145362",
                "ENSG00000123144",
                "ENSG00000204525",
                "ENSG00000263001",
                "ENSG00000145741",
                "ENSG00000143256",
                "ENSG00000149084",
                "ENSG00000174437",
                "ENSG00000166619",
                "ENSG00000074800",
                "ENSG00000197111",
                "ENSG00000165502",
                "ENSG00000123562",
                "ENSG00000122034",
                "ENSG00000051108",
                "ENSG00000111669",
                "ENSG00000163041",
                "ENSG00000134825",
                "ENSG00000009307",
                "ENSG00000004779",
                "ENSG00000150991",
                "ENSG00000152795",
                "ENSG00000169139",
                "ENSG00000118363",
                "ENSG00000142669",
                "ENSG00000135940",
                "ENSG00000175756",
                "ENSG00000070831",
                "ENSG00000134333",
                "ENSG00000100568",
                "ENSG00000122705",
                "ENSG00000132432",
                "ENSG00000102317",
                "ENSG00000110700",
                "ENSG00000175416",
                "ENSG00000128609",
                "ENSG00000160014",
                "ENSG00000063177",
                "ENSG00000105649",
                "ENSG00000173660",
                "ENSG00000108953",
                "ENSG00000105220",
                "ENSG00000186184",
                "ENSG00000100804",
                "ENSG00000155368",
                "ENSG00000169564",
                "ENSG00000114125",
                "ENSG00000176871",
                "ENSG00000240972",
                "ENSG00000111775",
                "ENSG00000105379",
                "ENSG00000197102",
                "ENSG00000135390",
                "ENSG00000149806",
                "ENSG00000132824",
                "ENSG00000213585",
                "ENSG00000131508",
                "ENSG00000131437",
                "ENSG00000167658",
                "ENSG00000188612",
                "ENSG00000180817",
                "ENSG00000112695",
                "ENSG00000116288",
                "ENSG00000136810",
                "ENSG00000174444",
                "ENSG00000220205",
                "ENSG00000127184",
                "ENSG00000128524",
                "ENSG00000165678",
                "ENSG00000133318",
                "ENSG00000265241",
                "ENSG00000136238",
                "ENSG00000143933",
                "ENSG00000165629",
                "ENSG00000132963",
                "ENSG00000164096",
                "ENSG00000087460",
                "ENSG00000182670",
                "ENSG00000171530",
                "ENSG00000148450",
                "ENSG00000123349",
                "ENSG00000114115",
                "ENSG00000178127",
                "ENSG00000113558",
                "ENSG00000140990",
                "ENSG00000143153",
                "ENSG00000143198",
                "ENSG00000136628",
                "ENSG00000014641",
                "ENSG00000109180",
                "ENSG00000123983",
                "ENSG00000133639",
                "ENSG00000126267",
                "ENSG00000143742",
                "ENSG00000100129",
                "ENSG00000109472"
            ],
            "expression_sum": {
                "class1": 7147.0,
                "class2": 6582.0
            },
            "ens_id": "ENSG00000114861",
            "uniprot_ids": [
                "Q9H334",
                "G5E965",
                "A0A3B3IRW5",
                "A0A3B3ITZ4",
                "A0A3B3IT66",
                "A0A0B4J2F3",
                "A0A3B3IRS5",
                "A0A3B3IU08",
                "A0A3B3IST6",
                "A0A3B3IS87",
                "A0A3B3IT14",
                "A0A087WVT2",
                "A0A3B3IT26",
                "A0A3B3IRL7",
                "A0A3B3ISQ6",
                "A0A087X2G3",
                "A0A3B3IST0",
                "A0A3B3ISQ3",
                "C9IYY1",
                "A0A3B3IUB6",
                "C9J0F0",
                "C9J5T4",
                "A0A3B3ISQ7",
                "A0A3B3IRY1",
                "A0A3B3ISA4",
                "H0Y882",
                "A0A3B3IU15"
            ],
            "net_id": 0,
            "source_num": 2,
            "target_num": 62
        },
        "ENSG00000221869": {
            "id": "ENSG00000221869",
            "type": "TF",
            "symbol": "CEBPD",
            "source": [
                "ENSG00000163435",
                "ENSG00000160113",
                "ENSG00000164683",
                "ENSG00000130522",
                "ENSG00000177606"
            ],
            "target": [
                "ENSG00000206573",
                "ENSG00000106261",
                "ENSG00000115944",
                "ENSG00000162704",
                "ENSG00000196531",
                "ENSG00000126005",
                "ENSG00000171604",
                "ENSG00000116030",
                "ENSG00000099622",
                "ENSG00000180011",
                "ENSG00000129559",
                "ENSG00000184752",
                "ENSG00000169567",
                "ENSG00000125534",
                "ENSG00000177606",
                "ENSG00000160113",
                "ENSG00000174748",
                "ENSG00000131143",
                "ENSG00000168653",
                "ENSG00000241837",
                "ENSG00000156467",
                "ENSG00000142864",
                "ENSG00000161203",
                "ENSG00000075415",
                "ENSG00000166165",
                "ENSG00000141753",
                "ENSG00000181885",
                "ENSG00000184076",
                "ENSG00000171314",
                "ENSG00000266472",
                "ENSG00000150779",
                "ENSG00000154556",
                "ENSG00000178741",
                "ENSG00000163466",
                "ENSG00000198931",
                "ENSG00000166913",
                "ENSG00000173812",
                "ENSG00000141279",
                "ENSG00000099795",
                "ENSG00000111229",
                "ENSG00000151729",
                "ENSG00000118816",
                "ENSG00000088832",
                "ENSG00000162585",
                "ENSG00000051620",
                "ENSG00000122545",
                "ENSG00000149547",
                "ENSG00000159335",
                "ENSG00000164924",
                "ENSG00000189043",
                "ENSG00000150687",
                "ENSG00000142192",
                "ENSG00000277443",
                "ENSG00000101558",
                "ENSG00000162545",
                "ENSG00000101040",
                "ENSG00000128739",
                "ENSG00000109475",
                "ENSG00000154518",
                "ENSG00000196262",
                "ENSG00000111640",
                "ENSG00000104904",
                "ENSG00000165637",
                "ENSG00000183978",
                "ENSG00000241685",
                "ENSG00000173726",
                "ENSG00000091164",
                "ENSG00000172757",
                "ENSG00000125971",
                "ENSG00000130332",
                "ENSG00000116586",
                "ENSG00000188290",
                "ENSG00000156976",
                "ENSG00000173674",
                "ENSG00000147548",
                "ENSG00000089220",
                "ENSG00000134138",
                "ENSG00000119335",
                "ENSG00000115286",
                "ENSG00000134717",
                "ENSG00000119396",
                "ENSG00000136156",
                "ENSG00000177879",
                "ENSG00000105640",
                "ENSG00000099624",
                "ENSG00000171159",
                "ENSG00000067225",
                "ENSG00000154723",
                "ENSG00000166347",
                "ENSG00000213719",
                "ENSG00000113712",
                "ENSG00000110321",
                "ENSG00000233927",
                "ENSG00000254999",
                "ENSG00000004059",
                "ENSG00000152234",
                "ENSG00000137106",
                "ENSG00000177556",
                "ENSG00000034713",
                "ENSG00000113387",
                "ENSG00000100592",
                "ENSG00000198668",
                "ENSG00000134308",
                "ENSG00000034510",
                "ENSG00000123144",
                "ENSG00000115091",
                "ENSG00000149257",
                "ENSG00000143256",
                "ENSG00000149084",
                "ENSG00000126934",
                "ENSG00000154639",
                "ENSG00000168118",
                "ENSG00000106367",
                "ENSG00000197111",
                "ENSG00000214253",
                "ENSG00000101182",
                "ENSG00000162496",
                "ENSG00000142676",
                "ENSG00000171421",
                "ENSG00000129515",
                "ENSG00000100387",
                "ENSG00000140319",
                "ENSG00000206503",
                "ENSG00000122034",
                "ENSG00000187514",
                "ENSG00000111669",
                "ENSG00000163041",
                "ENSG00000118680",
                "ENSG00000243317",
                "ENSG00000175826",
                "ENSG00000184007",
                "ENSG00000009307",
                "ENSG00000178980",
                "ENSG00000107262",
                "ENSG00000004779",
                "ENSG00000182117",
                "ENSG00000124172",
                "ENSG00000182196",
                "ENSG00000169139",
                "ENSG00000168214",
                "ENSG00000118363",
                "ENSG00000162191",
                "ENSG00000100911",
                "ENSG00000070831",
                "ENSG00000162076",
                "ENSG00000107438",
                "ENSG00000122705",
                "ENSG00000166482",
                "ENSG00000110700",
                "ENSG00000184009",
                "ENSG00000075624",
                "ENSG00000095059",
                "ENSG00000173660",
                "ENSG00000103187",
                "ENSG00000108953",
                "ENSG00000186184",
                "ENSG00000100804",
                "ENSG00000155368",
                "ENSG00000272047",
                "ENSG00000114125",
                "ENSG00000185787",
                "ENSG00000163931",
                "ENSG00000164713",
                "ENSG00000130638",
                "ENSG00000240972",
                "ENSG00000111775",
                "ENSG00000105379",
                "ENSG00000123636",
                "ENSG00000135390",
                "ENSG00000149806",
                "ENSG00000239672",
                "ENSG00000213585",
                "ENSG00000167658",
                "ENSG00000005022",
                "ENSG00000188612",
                "ENSG00000128654",
                "ENSG00000112715",
                "ENSG00000180817",
                "ENSG00000112695",
                "ENSG00000116288",
                "ENSG00000090266",
                "ENSG00000158710",
                "ENSG00000205542",
                "ENSG00000160948",
                "ENSG00000136810",
                "ENSG00000174444",
                "ENSG00000169764",
                "ENSG00000168028",
                "ENSG00000127184",
                "ENSG00000136238",
                "ENSG00000104341",
                "ENSG00000130522",
                "ENSG00000224032",
                "ENSG00000167815",
                "ENSG00000122873",
                "ENSG00000126457",
                "ENSG00000176340",
                "ENSG00000167996",
                "ENSG00000132963",
                "ENSG00000065978",
                "ENSG00000087460",
                "ENSG00000170759",
                "ENSG00000171530",
                "ENSG00000148450",
                "ENSG00000177410",
                "ENSG00000178127",
                "ENSG00000113558",
                "ENSG00000101745",
                "ENSG00000140990",
                "ENSG00000182718",
                "ENSG00000105518",
                "ENSG00000163435",
                "ENSG00000133639",
                "ENSG00000126267",
                "ENSG00000100129",
                "ENSG00000170017",
                "ENSG00000092841",
                "ENSG00000145494"
            ],
            "expression_sum": {
                "class1": 1701.0,
                "class2": 15658.0
            },
            "ens_id": "ENSG00000221869",
            "uniprot_ids": [
                "P49716"
            ],
            "net_id": 0,
            "source_num": 5,
            "target_num": 72
        },
        "ENSG00000157557": {
            "id": "ENSG00000157557",
            "type": "TF",
            "symbol": "ETS2",
            "source": [
                "ENSG00000160113",
                "ENSG00000119950",
                "ENSG00000168214",
                "ENSG00000119547",
                "ENSG00000177606"
            ],
            "target": [
                "ENSG00000206573",
                "ENSG00000181061",
                "ENSG00000214530",
                "ENSG00000165655",
                "ENSG00000099622",
                "ENSG00000111640",
                "ENSG00000243317",
                "ENSG00000067836",
                "ENSG00000120071",
                "ENSG00000110955",
                "ENSG00000156976",
                "ENSG00000102144",
                "ENSG00000169432",
                "ENSG00000136710",
                "ENSG00000104112",
                "ENSG00000168214",
                "ENSG00000115738",
                "ENSG00000143933",
                "ENSG00000114023",
                "ENSG00000132424",
                "ENSG00000186352",
                "ENSG00000130338",
                "ENSG00000067225",
                "ENSG00000137500",
                "ENSG00000160714",
                "ENSG00000122873",
                "ENSG00000176340",
                "ENSG00000178741",
                "ENSG00000140416",
                "ENSG00000055609",
                "ENSG00000181722",
                "ENSG00000164830",
                "ENSG00000172115",
                "ENSG00000181315",
                "ENSG00000151729",
                "ENSG00000143158",
                "ENSG00000118816",
                "ENSG00000272398",
                "ENSG00000186472",
                "ENSG00000101745",
                "ENSG00000105649",
                "ENSG00000163875",
                "ENSG00000100320",
                "ENSG00000119547"
            ],
            "expression_sum": {
                "class1": 2470.0,
                "class2": 7721.0
            },
            "ens_id": "ENSG00000157557",
            "uniprot_ids": [
                "P15036",
                "A0A590UJP9",
                "C9JAG2",
                "A0A590UJR2",
                "A0A0C4DG47"
            ],
            "net_id": 0,
            "source_num": 5,
            "target_num": 15
        },
        "ENSG00000181722": {
            "id": "ENSG00000181722",
            "type": "TF",
            "symbol": "ZBTB20",
            "source": [
                "ENSG00000157557",
                "ENSG00000113580",
                "ENSG00000164330",
                "ENSG00000100219",
                "ENSG00000139352",
                "ENSG00000160113"
            ],
            "target": [
                "ENSG00000146278",
                "ENSG00000181061",
                "ENSG00000143321",
                "ENSG00000065613",
                "ENSG00000145675",
                "ENSG00000257727",
                "ENSG00000152558",
                "ENSG00000114902",
                "ENSG00000178467",
                "ENSG00000099622",
                "ENSG00000180011",
                "ENSG00000164093",
                "ENSG00000104964",
                "ENSG00000163399",
                "ENSG00000167397",
                "ENSG00000165637",
                "ENSG00000179981",
                "ENSG00000244187",
                "ENSG00000102804",
                "ENSG00000160113",
                "ENSG00000108176",
                "ENSG00000077782",
                "ENSG00000129625",
                "ENSG00000134248",
                "ENSG00000113732",
                "ENSG00000220205",
                "ENSG00000116209",
                "ENSG00000117155",
                "ENSG00000109472",
                "ENSG00000132424",
                "ENSG00000184076",
                "ENSG00000135968",
                "ENSG00000198125",
                "ENSG00000184164",
                "ENSG00000068697",
                "ENSG00000198168",
                "ENSG00000134333",
                "ENSG00000140939",
                "ENSG00000143013",
                "ENSG00000125851",
                "ENSG00000054356",
                "ENSG00000102007",
                "ENSG00000087460",
                "ENSG00000205981",
                "ENSG00000205922",
                "ENSG00000143774",
                "ENSG00000152128",
                "ENSG00000143870",
                "ENSG00000138085",
                "ENSG00000120742",
                "ENSG00000197261",
                "ENSG00000172663",
                "ENSG00000133639",
                "ENSG00000185624",
                "ENSG00000089063",
                "ENSG00000119888",
                "ENSG00000113580",
                "ENSG00000166228",
                "ENSG00000143545",
                "ENSG00000124155",
                "ENSG00000064601",
                "ENSG00000143543",
                "ENSG00000151552",
                "ENSG00000185650",
                "ENSG00000114115",
                "ENSG00000115828",
                "ENSG00000225783",
                "ENSG00000104870"
            ],
            "expression_sum": {
                "class1": 10960.0,
                "class2": 13009.0
            },
            "ens_id": "ENSG00000181722",
            "uniprot_ids": [
                "Q9HC78",
                "C9JUH0",
                "C9JCX0"
            ],
            "net_id": 0,
            "source_num": 6,
            "target_num": 11
        },
        "ENSG00000177606": {
            "id": "ENSG00000177606",
            "type": "TF",
            "symbol": "JUN",
            "source": [
                "ENSG00000100219",
                "ENSG00000221869",
                "ENSG00000139352"
            ],
            "target": [
                "ENSG00000176887",
                "ENSG00000198840",
                "ENSG00000132698",
                "ENSG00000185664",
                "ENSG00000164093",
                "ENSG00000122756",
                "ENSG00000165973",
                "ENSG00000112773",
                "ENSG00000169435",
                "ENSG00000115461",
                "ENSG00000148834",
                "ENSG00000010278",
                "ENSG00000104870",
                "ENSG00000182481",
                "ENSG00000189143",
                "ENSG00000166145",
                "ENSG00000198125",
                "ENSG00000181885",
                "ENSG00000132386",
                "ENSG00000166598",
                "ENSG00000117691",
                "ENSG00000135535",
                "ENSG00000164305",
                "ENSG00000107438",
                "ENSG00000167642",
                "ENSG00000161249",
                "ENSG00000187244",
                "ENSG00000198938",
                "ENSG00000198899",
                "ENSG00000167641",
                "ENSG00000015592",
                "ENSG00000143870",
                "ENSG00000135549",
                "ENSG00000213937",
                "ENSG00000157557",
                "ENSG00000119888",
                "ENSG00000100219",
                "ENSG00000221869",
                "ENSG00000185650",
                "ENSG00000108679",
                "ENSG00000172915",
                "ENSG00000179218"
            ],
            "expression_sum": {
                "class1": 17839.0,
                "class2": 10689.0
            },
            "ens_id": "ENSG00000177606",
            "uniprot_ids": [
                "P05412"
            ],
            "net_id": 0,
            "source_num": 3,
            "target_num": 11
        },
        "ENSG00000124766": {
            "id": "ENSG00000124766",
            "type": "TF",
            "symbol": "SOX4",
            "source": [],
            "target": [
                "ENSG00000132718",
                "ENSG00000149557",
                "ENSG00000148516",
                "ENSG00000139116",
                "ENSG00000132698",
                "ENSG00000109339",
                "ENSG00000074696",
                "ENSG00000164929",
                "ENSG00000185664",
                "ENSG00000050165",
                "ENSG00000182534",
                "ENSG00000080824",
                "ENSG00000154080",
                "ENSG00000166226",
                "ENSG00000166257",
                "ENSG00000112531",
                "ENSG00000130513",
                "ENSG00000147889",
                "ENSG00000171055",
                "ENSG00000172183",
                "ENSG00000104112",
                "ENSG00000090273",
                "ENSG00000182481",
                "ENSG00000123416",
                "ENSG00000108106",
                "ENSG00000181885",
                "ENSG00000113141",
                "ENSG00000138758",
                "ENSG00000092964",
                "ENSG00000166963",
                "ENSG00000172159",
                "ENSG00000105258",
                "ENSG00000173267",
                "ENSG00000152082",
                "ENSG00000175792",
                "ENSG00000120885",
                "ENSG00000198668",
                "ENSG00000054523",
                "ENSG00000146425",
                "ENSG00000188042",
                "ENSG00000163435",
                "ENSG00000143198",
                "ENSG00000135406",
                "ENSG00000119640",
                "ENSG00000122545",
                "ENSG00000143799",
                "ENSG00000141404",
                "ENSG00000169862",
                "ENSG00000205726",
                "ENSG00000185650",
                "ENSG00000163931",
                "ENSG00000140905"
            ],
            "expression_sum": {
                "class1": 103965.0,
                "class2": 44155.0
            },
            "ens_id": "ENSG00000124766",
            "uniprot_ids": [
                "Q06945"
            ],
            "net_id": 0,
            "source_num": 0,
            "target_num": 18
        },
        "ENSG00000116044": {
            "id": "ENSG00000116044",
            "type": "TF",
            "symbol": "NFE2L2",
            "source": [
                "ENSG00000164683",
                "ENSG00000163435"
            ],
            "target": [
                "ENSG00000130066",
                "ENSG00000182718",
                "ENSG00000163435",
                "ENSG00000143153",
                "ENSG00000109062",
                "ENSG00000169764",
                "ENSG00000131981",
                "ENSG00000164093",
                "ENSG00000118816",
                "ENSG00000135046"
            ],
            "expression_sum": {
                "class1": 4576.0,
                "class2": 10732.0
            },
            "ens_id": "ENSG00000116044",
            "uniprot_ids": [
                "Q16236",
                "K7ER33",
                "K7EIW5",
                "C9J1A8",
                "C9J3K4",
                "C9JBM0",
                "H7C498",
                "C9JFL6",
                "C9J0Y1"
            ],
            "net_id": 0,
            "source_num": 2,
            "target_num": 9
        },
        "ENSG00000128610": {
            "id": "ENSG00000128610",
            "type": "TF",
            "symbol": "FEZF1",
            "source": [
                "ENSG00000181449"
            ],
            "target": [
                "ENSG00000181449",
                "ENSG00000114353",
                "ENSG00000043355",
                "ENSG00000173404",
                "ENSG00000150687",
                "ENSG00000163931",
                "ENSG00000125462",
                "ENSG00000245526"
            ],
            "expression_sum": {
                "class1": 1699.0,
                "class2": 607.0
            },
            "ens_id": "ENSG00000128610",
            "uniprot_ids": [
                "A0PJY2",
                "C9J0C5"
            ],
            "net_id": 0,
            "source_num": 1,
            "target_num": 3
        },
        "ENSG00000130522": {
            "id": "ENSG00000130522",
            "type": "TF",
            "symbol": "JUND",
            "source": [
                "ENSG00000181449",
                "ENSG00000221869"
            ],
            "target": [
                "ENSG00000181449",
                "ENSG00000162496",
                "ENSG00000176887",
                "ENSG00000172020",
                "ENSG00000087085",
                "ENSG00000185664",
                "ENSG00000107105",
                "ENSG00000213190",
                "ENSG00000162188",
                "ENSG00000115461",
                "ENSG00000128564",
                "ENSG00000130303",
                "ENSG00000182481",
                "ENSG00000108106",
                "ENSG00000182916",
                "ENSG00000125354",
                "ENSG00000104435",
                "ENSG00000164305",
                "ENSG00000108515",
                "ENSG00000124194",
                "ENSG00000015592",
                "ENSG00000135046",
                "ENSG00000090097",
                "ENSG00000177675",
                "ENSG00000221869",
                "ENSG00000185650",
                "ENSG00000149926",
                "ENSG00000104870"
            ],
            "expression_sum": {
                "class1": 87153.0,
                "class2": 33345.0
            },
            "ens_id": "ENSG00000130522",
            "uniprot_ids": [
                "P17535",
                "U3KPR5"
            ],
            "net_id": 0,
            "source_num": 2,
            "target_num": 8
        },
        "ENSG00000119547": {
            "id": "ENSG00000119547",
            "type": "TF",
            "symbol": "ONECUT2",
            "source": [
                "ENSG00000185551",
                "ENSG00000157557",
                "ENSG00000134138",
                "ENSG00000167081",
                "ENSG00000113580",
                "ENSG00000148516",
                "ENSG00000103495",
                "ENSG00000108055",
                "ENSG00000114861",
                "ENSG00000139352"
            ],
            "target": [
                "ENSG00000090621",
                "ENSG00000135486",
                "ENSG00000162373",
                "ENSG00000058668",
                "ENSG00000173272",
                "ENSG00000162704",
                "ENSG00000135334",
                "ENSG00000196531",
                "ENSG00000156475",
                "ENSG00000111716",
                "ENSG00000175745",
                "ENSG00000164093",
                "ENSG00000184752",
                "ENSG00000196230",
                "ENSG00000117395",
                "ENSG00000182004",
                "ENSG00000179915",
                "ENSG00000065135",
                "ENSG00000104408",
                "ENSG00000103495",
                "ENSG00000171055",
                "ENSG00000149231",
                "ENSG00000147604",
                "ENSG00000182481",
                "ENSG00000123416",
                "ENSG00000142864",
                "ENSG00000100823",
                "ENSG00000198125",
                "ENSG00000108175",
                "ENSG00000140988",
                "ENSG00000197728",
                "ENSG00000102466",
                "ENSG00000028277",
                "ENSG00000077380",
                "ENSG00000167642",
                "ENSG00000163466",
                "ENSG00000096746",
                "ENSG00000138413",
                "ENSG00000134779",
                "ENSG00000071054",
                "ENSG00000109536",
                "ENSG00000111728",
                "ENSG00000069275",
                "ENSG00000118816",
                "ENSG00000041357",
                "ENSG00000119888",
                "ENSG00000122545",
                "ENSG00000121766",
                "ENSG00000153234",
                "ENSG00000189043",
                "ENSG00000123352",
                "ENSG00000180879",
                "ENSG00000277443",
                "ENSG00000183654",
                "ENSG00000233429",
                "ENSG00000171858",
                "ENSG00000115350",
                "ENSG00000138650",
                "ENSG00000148516",
                "ENSG00000187109",
                "ENSG00000106615",
                "ENSG00000156482",
                "ENSG00000196262",
                "ENSG00000091164",
                "ENSG00000182628",
                "ENSG00000167460",
                "ENSG00000172757",
                "ENSG00000171951",
                "ENSG00000060982",
                "ENSG00000188290",
                "ENSG00000100632",
                "ENSG00000161970",
                "ENSG00000141447",
                "ENSG00000134138",
                "ENSG00000164114",
                "ENSG00000119335",
                "ENSG00000168894",
                "ENSG00000108828",
                "ENSG00000176840",
                "ENSG00000077264",
                "ENSG00000177879",
                "ENSG00000215386",
                "ENSG00000101132",
                "ENSG00000188243",
                "ENSG00000125691",
                "ENSG00000143549",
                "ENSG00000138758",
                "ENSG00000133083",
                "ENSG00000143013",
                "ENSG00000233927",
                "ENSG00000117139",
                "ENSG00000137106",
                "ENSG00000157216",
                "ENSG00000168785",
                "ENSG00000108107",
                "ENSG00000138326",
                "ENSG00000134308",
                "ENSG00000122958",
                "ENSG00000022267",
                "ENSG00000034510",
                "ENSG00000133703",
                "ENSG00000137575",
                "ENSG00000135549",
                "ENSG00000154380",
                "ENSG00000151789",
                "ENSG00000064601",
                "ENSG00000104695",
                "ENSG00000113580",
                "ENSG00000075142",
                "ENSG00000106028",
                "ENSG00000175161",
                "ENSG00000115091",
                "ENSG00000115053",
                "ENSG00000117152",
                "ENSG00000127022",
                "ENSG00000213741",
                "ENSG00000074800",
                "ENSG00000145592",
                "ENSG00000069849",
                "ENSG00000133026",
                "ENSG00000147649",
                "ENSG00000123562",
                "ENSG00000163041",
                "ENSG00000130255",
                "ENSG00000184007",
                "ENSG00000143995",
                "ENSG00000174498",
                "ENSG00000185551",
                "ENSG00000197747",
                "ENSG00000105223",
                "ENSG00000164687",
                "ENSG00000106355",
                "ENSG00000057608",
                "ENSG00000174469",
                "ENSG00000065548",
                "ENSG00000130396",
                "ENSG00000118363",
                "ENSG00000108106",
                "ENSG00000005108",
                "ENSG00000117691",
                "ENSG00000112972",
                "ENSG00000006625",
                "ENSG00000116251",
                "ENSG00000071794",
                "ENSG00000122705",
                "ENSG00000110700",
                "ENSG00000114861",
                "ENSG00000196776",
                "ENSG00000125944",
                "ENSG00000054523",
                "ENSG00000063177",
                "ENSG00000100380",
                "ENSG00000115758",
                "ENSG00000113328",
                "ENSG00000129167",
                "ENSG00000155368",
                "ENSG00000117906",
                "ENSG00000105185",
                "ENSG00000105372",
                "ENSG00000157557",
                "ENSG00000094916",
                "ENSG00000205726",
                "ENSG00000128050",
                "ENSG00000043355",
                "ENSG00000163931",
                "ENSG00000164713",
                "ENSG00000176973",
                "ENSG00000105379",
                "ENSG00000186063",
                "ENSG00000107863",
                "ENSG00000136930",
                "ENSG00000164985",
                "ENSG00000239672",
                "ENSG00000144724",
                "ENSG00000146386",
                "ENSG00000213585",
                "ENSG00000131508",
                "ENSG00000127955",
                "ENSG00000180957",
                "ENSG00000131469",
                "ENSG00000188846",
                "ENSG00000137942",
                "ENSG00000080824",
                "ENSG00000166226",
                "ENSG00000108468",
                "ENSG00000181163",
                "ENSG00000136810",
                "ENSG00000142937",
                "ENSG00000078018",
                "ENSG00000196465",
                "ENSG00000112531",
                "ENSG00000174444",
                "ENSG00000104549",
                "ENSG00000136238",
                "ENSG00000117155",
                "ENSG00000103363",
                "ENSG00000079459",
                "ENSG00000138430",
                "ENSG00000047849",
                "ENSG00000168916",
                "ENSG00000132437",
                "ENSG00000167996",
                "ENSG00000065978",
                "ENSG00000170144",
                "ENSG00000152082",
                "ENSG00000100664",
                "ENSG00000189159",
                "ENSG00000242265",
                "ENSG00000143198",
                "ENSG00000122565",
                "ENSG00000054148",
                "ENSG00000134824",
                "ENSG00000165119"
            ],
            "expression_sum": {
                "class1": 33288.0,
                "class2": 3372.0
            },
            "ens_id": "ENSG00000119547",
            "uniprot_ids": [
                "O95948"
            ],
            "net_id": 0,
            "source_num": 10,
            "target_num": 53
        },
        "ENSG00000175745": {
            "id": "ENSG00000175745",
            "type": "TF",
            "symbol": "NR2F1",
            "source": [
                "ENSG00000162599",
                "ENSG00000113580",
                "ENSG00000122691",
                "ENSG00000164330",
                "ENSG00000108001",
                "ENSG00000184486",
                "ENSG00000119547",
                "ENSG00000143995"
            ],
            "target": [
                "ENSG00000147862",
                "ENSG00000140937",
                "ENSG00000109099",
                "ENSG00000144366",
                "ENSG00000185664",
                "ENSG00000198910",
                "ENSG00000143995",
                "ENSG00000084628",
                "ENSG00000164199",
                "ENSG00000105810",
                "ENSG00000070961",
                "ENSG00000134138",
                "ENSG00000110492",
                "ENSG00000197043",
                "ENSG00000185650",
                "ENSG00000130005",
                "ENSG00000104112",
                "ENSG00000133302",
                "ENSG00000122884",
                "ENSG00000185201",
                "ENSG00000162599",
                "ENSG00000196562",
                "ENSG00000181744",
                "ENSG00000132437",
                "ENSG00000213420",
                "ENSG00000187957",
                "ENSG00000026025",
                "ENSG00000131037",
                "ENSG00000149294",
                "ENSG00000147655",
                "ENSG00000123240",
                "ENSG00000185630",
                "ENSG00000196141",
                "ENSG00000113580",
                "ENSG00000159409",
                "ENSG00000123989",
                "ENSG00000047457",
                "ENSG00000018408",
                "ENSG00000164056",
                "ENSG00000129244"
            ],
            "expression_sum": {
                "class1": 32875.0,
                "class2": 29213.0
            },
            "ens_id": "ENSG00000175745",
            "uniprot_ids": [
                "P10589",
                "F1DAL7",
                "F1DAL9"
            ],
            "net_id": 0,
            "source_num": 7,
            "target_num": 10
        },
        "ENSG00000185551": {
            "id": "ENSG00000185551",
            "type": "TF",
            "symbol": "NR2F2",
            "source": [
                "ENSG00000119547"
            ],
            "target": [
                "ENSG00000174938",
                "ENSG00000198727",
                "ENSG00000198886",
                "ENSG00000104964",
                "ENSG00000118680",
                "ENSG00000156675",
                "ENSG00000198763",
                "ENSG00000245526",
                "ENSG00000153956",
                "ENSG00000179915",
                "ENSG00000104112",
                "ENSG00000198712",
                "ENSG00000167642",
                "ENSG00000198938",
                "ENSG00000145247",
                "ENSG00000198899",
                "ENSG00000242265",
                "ENSG00000251562",
                "ENSG00000148053",
                "ENSG00000127990",
                "ENSG00000102109",
                "ENSG00000215845",
                "ENSG00000119547"
            ],
            "expression_sum": {
                "class1": 14979.0,
                "class2": 8249.0
            },
            "ens_id": "ENSG00000185551",
            "uniprot_ids": [
                "P24468",
                "H3BTC2"
            ],
            "net_id": 0,
            "source_num": 1,
            "target_num": 4
        },
        "ENSG00000171634": {
            "id": "ENSG00000171634",
            "type": "TF",
            "symbol": "BPTF",
            "source": [
                "ENSG00000016082",
                "ENSG00000181449",
                "ENSG00000113580",
                "ENSG00000164330"
            ],
            "target": [
                "ENSG00000149557",
                "ENSG00000113580",
                "ENSG00000111728"
            ],
            "expression_sum": {
                "class1": 14482.0,
                "class2": 13915.0
            },
            "ens_id": "ENSG00000171634",
            "uniprot_ids": [
                "Q12830",
                "J3QQQ8",
                "A0A0A0MR81",
                "F5H176",
                "J3KSK9",
                "E7ETD6",
                "F5GXF5",
                "J3QQK4",
                "A0A2R8Y7Q1",
                "E9PE19",
                "J3QLW4",
                "J3KSY0"
            ],
            "net_id": 0,
            "source_num": 4,
            "target_num": 1
        },
        "ENSG00000113580": {
            "id": "ENSG00000113580",
            "type": "TF",
            "symbol": "NR3C1",
            "source": [
                "ENSG00000171634",
                "ENSG00000125968",
                "ENSG00000105698",
                "ENSG00000175745",
                "ENSG00000181722",
                "ENSG00000119950",
                "ENSG00000067955",
                "ENSG00000168214",
                "ENSG00000119547",
                "ENSG00000160113",
                "ENSG00000092199"
            ],
            "target": [
                "ENSG00000078668",
                "ENSG00000206573",
                "ENSG00000141741",
                "ENSG00000232119",
                "ENSG00000017260",
                "ENSG00000162704",
                "ENSG00000163634",
                "ENSG00000165655",
                "ENSG00000132698",
                "ENSG00000153130",
                "ENSG00000111674",
                "ENSG00000111716",
                "ENSG00000139644",
                "ENSG00000175745",
                "ENSG00000005810",
                "ENSG00000169612",
                "ENSG00000099622",
                "ENSG00000079950",
                "ENSG00000090060",
                "ENSG00000005020",
                "ENSG00000140941",
                "ENSG00000122641",
                "ENSG00000134248",
                "ENSG00000124785",
                "ENSG00000131143",
                "ENSG00000075945",
                "ENSG00000145817",
                "ENSG00000146247",
                "ENSG00000196419",
                "ENSG00000075415",
                "ENSG00000022840",
                "ENSG00000105993",
                "ENSG00000102172",
                "ENSG00000160714",
                "ENSG00000178567",
                "ENSG00000102466",
                "ENSG00000265972",
                "ENSG00000077380",
                "ENSG00000154556",
                "ENSG00000163466",
                "ENSG00000119421",
                "ENSG00000134283",
                "ENSG00000114439",
                "ENSG00000173812",
                "ENSG00000141279",
                "ENSG00000183091",
                "ENSG00000152256",
                "ENSG00000237190",
                "ENSG00000198899",
                "ENSG00000147416",
                "ENSG00000065911",
                "ENSG00000173915",
                "ENSG00000151729",
                "ENSG00000272398",
                "ENSG00000129636",
                "ENSG00000123240",
                "ENSG00000081181",
                "ENSG00000144320",
                "ENSG00000178913",
                "ENSG00000163875",
                "ENSG00000167136",
                "ENSG00000138083",
                "ENSG00000164924",
                "ENSG00000074319",
                "ENSG00000197930",
                "ENSG00000141759",
                "ENSG00000162545",
                "ENSG00000165264",
                "ENSG00000180773",
                "ENSG00000146007",
                "ENSG00000084764",
                "ENSG00000277791",
                "ENSG00000104964",
                "ENSG00000170242",
                "ENSG00000139910",
                "ENSG00000125868",
                "ENSG00000165637",
                "ENSG00000241685",
                "ENSG00000153815",
                "ENSG00000047249",
                "ENSG00000165672",
                "ENSG00000173674",
                "ENSG00000147548",
                "ENSG00000125968",
                "ENSG00000131236",
                "ENSG00000129625",
                "ENSG00000055163",
                "ENSG00000089220",
                "ENSG00000116898",
                "ENSG00000117758",
                "ENSG00000167733",
                "ENSG00000196072",
                "ENSG00000104112",
                "ENSG00000255302",
                "ENSG00000187164",
                "ENSG00000188243",
                "ENSG00000067225",
                "ENSG00000109390",
                "ENSG00000123933",
                "ENSG00000072401",
                "ENSG00000107771",
                "ENSG00000140939",
                "ENSG00000113712",
                "ENSG00000136143",
                "ENSG00000150459",
                "ENSG00000148444",
                "ENSG00000034713",
                "ENSG00000182247",
                "ENSG00000100592",
                "ENSG00000144746",
                "ENSG00000157168",
                "ENSG00000133703",
                "ENSG00000063854",
                "ENSG00000133606",
                "ENSG00000103769",
                "ENSG00000146425",
                "ENSG00000138071",
                "ENSG00000117410",
                "ENSG00000131100",
                "ENSG00000273841",
                "ENSG00000124802",
                "ENSG00000263001",
                "ENSG00000196405",
                "ENSG00000174437",
                "ENSG00000126934",
                "ENSG00000154639",
                "ENSG00000215845",
                "ENSG00000185808",
                "ENSG00000225783",
                "ENSG00000078140",
                "ENSG00000129515",
                "ENSG00000100519",
                "ENSG00000144366",
                "ENSG00000121741",
                "ENSG00000144635",
                "ENSG00000163041",
                "ENSG00000155115",
                "ENSG00000160194",
                "ENSG00000141027",
                "ENSG00000092199",
                "ENSG00000110955",
                "ENSG00000107262",
                "ENSG00000164346",
                "ENSG00000168137",
                "ENSG00000010256",
                "ENSG00000120053",
                "ENSG00000065518",
                "ENSG00000174953",
                "ENSG00000169100",
                "ENSG00000162191",
                "ENSG00000118363",
                "ENSG00000122417",
                "ENSG00000135940",
                "ENSG00000108654",
                "ENSG00000163888",
                "ENSG00000198836",
                "ENSG00000172336",
                "ENSG00000107581",
                "ENSG00000033800",
                "ENSG00000131966",
                "ENSG00000106052",
                "ENSG00000147677",
                "ENSG00000107438",
                "ENSG00000244754",
                "ENSG00000085224",
                "ENSG00000102317",
                "ENSG00000265491",
                "ENSG00000180370",
                "ENSG00000213465",
                "ENSG00000173404",
                "ENSG00000109332",
                "ENSG00000163170",
                "ENSG00000079246",
                "ENSG00000075785",
                "ENSG00000171222",
                "ENSG00000095059",
                "ENSG00000244879",
                "ENSG00000105185",
                "ENSG00000166974",
                "ENSG00000189241",
                "ENSG00000141562",
                "ENSG00000100320",
                "ENSG00000175166",
                "ENSG00000168303",
                "ENSG00000203879",
                "ENSG00000240972",
                "ENSG00000054267",
                "ENSG00000136783",
                "ENSG00000162852",
                "ENSG00000196655",
                "ENSG00000172915",
                "ENSG00000141030",
                "ENSG00000075292",
                "ENSG00000105419",
                "ENSG00000135390",
                "ENSG00000153310",
                "ENSG00000174938",
                "ENSG00000213585",
                "ENSG00000131508",
                "ENSG00000197223",
                "ENSG00000131437",
                "ENSG00000005022",
                "ENSG00000127616",
                "ENSG00000076043",
                "ENSG00000128654",
                "ENSG00000160271",
                "ENSG00000160613",
                "ENSG00000090266",
                "ENSG00000136810",
                "ENSG00000143862",
                "ENSG00000198015",
                "ENSG00000171634",
                "ENSG00000048649",
                "ENSG00000125656",
                "ENSG00000127184",
                "ENSG00000176731",
                "ENSG00000072832",
                "ENSG00000130312",
                "ENSG00000136950",
                "ENSG00000160551",
                "ENSG00000104388",
                "ENSG00000100095",
                "ENSG00000168159",
                "ENSG00000119314",
                "ENSG00000135472",
                "ENSG00000267100",
                "ENSG00000064607",
                "ENSG00000171530",
                "ENSG00000151914",
                "ENSG00000123349",
                "ENSG00000105583",
                "ENSG00000165943",
                "ENSG00000127914",
                "ENSG00000014641",
                "ENSG00000126267",
                "ENSG00000175582",
                "ENSG00000152377",
                "ENSG00000104765",
                "ENSG00000171130",
                "ENSG00000198612",
                "ENSG00000168300",
                "ENSG00000176171",
                "ENSG00000130725",
                "ENSG00000100129",
                "ENSG00000112305",
                "ENSG00000142507",
                "ENSG00000253738",
                "ENSG00000047932",
                "ENSG00000237765",
                "ENSG00000165119",
                "ENSG00000135250",
                "ENSG00000139163",
                "ENSG00000170871",
                "ENSG00000126005",
                "ENSG00000198840",
                "ENSG00000082153",
                "ENSG00000113657",
                "ENSG00000247095",
                "ENSG00000129559",
                "ENSG00000089737",
                "ENSG00000114784",
                "ENSG00000184752",
                "ENSG00000169567",
                "ENSG00000197302",
                "ENSG00000198258",
                "ENSG00000275052",
                "ENSG00000135002",
                "ENSG00000127445",
                "ENSG00000129355",
                "ENSG00000183172",
                "ENSG00000145545",
                "ENSG00000146282",
                "ENSG00000170027",
                "ENSG00000241837",
                "ENSG00000023287",
                "ENSG00000076554",
                "ENSG00000164190",
                "ENSG00000120509",
                "ENSG00000198125",
                "ENSG00000131495",
                "ENSG00000171314",
                "ENSG00000168291",
                "ENSG00000164975",
                "ENSG00000172159",
                "ENSG00000090238",
                "ENSG00000170315",
                "ENSG00000103266",
                "ENSG00000147684",
                "ENSG00000134779",
                "ENSG00000077254",
                "ENSG00000152932",
                "ENSG00000067715",
                "ENSG00000119950",
                "ENSG00000049759",
                "ENSG00000099795",
                "ENSG00000129084",
                "ENSG00000144747",
                "ENSG00000111229",
                "ENSG00000069345",
                "ENSG00000189060",
                "ENSG00000103145",
                "ENSG00000051620",
                "ENSG00000113742",
                "ENSG00000163939",
                "ENSG00000137941",
                "ENSG00000167283",
                "ENSG00000137100",
                "ENSG00000123106",
                "ENSG00000196199",
                "ENSG00000198886",
                "ENSG00000267313",
                "ENSG00000102934",
                "ENSG00000155506",
                "ENSG00000116661",
                "ENSG00000122643",
                "ENSG00000196262",
                "ENSG00000104904",
                "ENSG00000125971",
                "ENSG00000173726",
                "ENSG00000130332",
                "ENSG00000155959",
                "ENSG00000137692",
                "ENSG00000116586",
                "ENSG00000089693",
                "ENSG00000105254",
                "ENSG00000168092",
                "ENSG00000119335",
                "ENSG00000115286",
                "ENSG00000105698",
                "ENSG00000092051",
                "ENSG00000169432",
                "ENSG00000101132",
                "ENSG00000114023",
                "ENSG00000154723",
                "ENSG00000166347",
                "ENSG00000151327",
                "ENSG00000131174",
                "ENSG00000166405",
                "ENSG00000143013",
                "ENSG00000198843",
                "ENSG00000109654",
                "ENSG00000145495",
                "ENSG00000168288",
                "ENSG00000254999",
                "ENSG00000004059",
                "ENSG00000089289",
                "ENSG00000183779",
                "ENSG00000177556",
                "ENSG00000130558",
                "ENSG00000113387",
                "ENSG00000151893",
                "ENSG00000158864",
                "ENSG00000137409",
                "ENSG00000082701",
                "ENSG00000241468",
                "ENSG00000136888",
                "ENSG00000100442",
                "ENSG00000121653",
                "ENSG00000106028",
                "ENSG00000173917",
                "ENSG00000100697",
                "ENSG00000168118",
                "ENSG00000106367",
                "ENSG00000121022",
                "ENSG00000164615",
                "ENSG00000197111",
                "ENSG00000101182",
                "ENSG00000146729",
                "ENSG00000142949",
                "ENSG00000105701",
                "ENSG00000111669",
                "ENSG00000136100",
                "ENSG00000159720",
                "ENSG00000148798",
                "ENSG00000134825",
                "ENSG00000184007",
                "ENSG00000140307",
                "ENSG00000167863",
                "ENSG00000197747",
                "ENSG00000004779",
                "ENSG00000184277",
                "ENSG00000127995",
                "ENSG00000169139",
                "ENSG00000117318",
                "ENSG00000162368",
                "ENSG00000004478",
                "ENSG00000116459",
                "ENSG00000186868",
                "ENSG00000175756",
                "ENSG00000135968",
                "ENSG00000070831",
                "ENSG00000137500",
                "ENSG00000134333",
                "ENSG00000091640",
                "ENSG00000250479",
                "ENSG00000147123",
                "ENSG00000112320",
                "ENSG00000161960",
                "ENSG00000144674",
                "ENSG00000143575",
                "ENSG00000038219",
                "ENSG00000130560",
                "ENSG00000091844",
                "ENSG00000182050",
                "ENSG00000160014",
                "ENSG00000184203",
                "ENSG00000125818",
                "ENSG00000183617",
                "ENSG00000105649",
                "ENSG00000103187",
                "ENSG00000105220",
                "ENSG00000053254",
                "ENSG00000272047",
                "ENSG00000198791",
                "ENSG00000011295",
                "ENSG00000176871",
                "ENSG00000165152",
                "ENSG00000007168",
                "ENSG00000164713",
                "ENSG00000134049",
                "ENSG00000134744",
                "ENSG00000139146",
                "ENSG00000105379",
                "ENSG00000126067",
                "ENSG00000173692",
                "ENSG00000197457",
                "ENSG00000214530",
                "ENSG00000111237",
                "ENSG00000132670",
                "ENSG00000132824",
                "ENSG00000111911",
                "ENSG00000254615",
                "ENSG00000135070",
                "ENSG00000104886",
                "ENSG00000151366",
                "ENSG00000214413",
                "ENSG00000145782",
                "ENSG00000112695",
                "ENSG00000162188",
                "ENSG00000108176",
                "ENSG00000165416",
                "ENSG00000203667",
                "ENSG00000136710",
                "ENSG00000100354",
                "ENSG00000130414",
                "ENSG00000115738",
                "ENSG00000138430",
                "ENSG00000169738",
                "ENSG00000169213",
                "ENSG00000166780",
                "ENSG00000126214",
                "ENSG00000176340",
                "ENSG00000097033",
                "ENSG00000258289",
                "ENSG00000150753",
                "ENSG00000049618",
                "ENSG00000156587",
                "ENSG00000273611",
                "ENSG00000182670",
                "ENSG00000177459",
                "ENSG00000148943",
                "ENSG00000181315",
                "ENSG00000146701",
                "ENSG00000115514",
                "ENSG00000186472",
                "ENSG00000101745",
                "ENSG00000140990",
                "ENSG00000182718",
                "ENSG00000173436",
                "ENSG00000069956",
                "ENSG00000063046",
                "ENSG00000163541",
                "ENSG00000102109",
                "ENSG00000169439",
                "ENSG00000164258",
                "ENSG00000169925",
                "ENSG00000109472",
                "ENSG00000114956",
                "ENSG00000149926",
                "ENSG00000106261",
                "ENSG00000171604",
                "ENSG00000101210",
                "ENSG00000143155",
                "ENSG00000146066",
                "ENSG00000065135",
                "ENSG00000129235",
                "ENSG00000132639",
                "ENSG00000102144",
                "ENSG00000130770",
                "ENSG00000156467",
                "ENSG00000168280",
                "ENSG00000119013",
                "ENSG00000161203",
                "ENSG00000175324",
                "ENSG00000184076",
                "ENSG00000105887",
                "ENSG00000165434",
                "ENSG00000266472",
                "ENSG00000101782",
                "ENSG00000177889",
                "ENSG00000166913",
                "ENSG00000181722",
                "ENSG00000109046",
                "ENSG00000159199",
                "ENSG00000170873",
                "ENSG00000151743",
                "ENSG00000162434",
                "ENSG00000121851",
                "ENSG00000118816",
                "ENSG00000105290",
                "ENSG00000147669",
                "ENSG00000107679",
                "ENSG00000119888",
                "ENSG00000204442",
                "ENSG00000163468",
                "ENSG00000137312",
                "ENSG00000074071",
                "ENSG00000091039",
                "ENSG00000096092",
                "ENSG00000111371",
                "ENSG00000134056",
                "ENSG00000172115",
                "ENSG00000119547",
                "ENSG00000101558",
                "ENSG00000101040",
                "ENSG00000091542",
                "ENSG00000175130",
                "ENSG00000128789",
                "ENSG00000111640",
                "ENSG00000169750",
                "ENSG00000120071",
                "ENSG00000124596",
                "ENSG00000138078",
                "ENSG00000115241",
                "ENSG00000253719",
                "ENSG00000156976",
                "ENSG00000128245",
                "ENSG00000134138",
                "ENSG00000130294",
                "ENSG00000078369",
                "ENSG00000119396",
                "ENSG00000077264",
                "ENSG00000205531",
                "ENSG00000099624",
                "ENSG00000120265",
                "ENSG00000187231",
                "ENSG00000138495",
                "ENSG00000154001",
                "ENSG00000101489",
                "ENSG00000138069",
                "ENSG00000083099",
                "ENSG00000187257",
                "ENSG00000171033",
                "ENSG00000110321",
                "ENSG00000198420",
                "ENSG00000138107",
                "ENSG00000055609",
                "ENSG00000005483",
                "ENSG00000139726",
                "ENSG00000204568",
                "ENSG00000175203",
                "ENSG00000111642",
                "ENSG00000102409",
                "ENSG00000145362",
                "ENSG00000123144",
                "ENSG00000101557",
                "ENSG00000127125",
                "ENSG00000113575",
                "ENSG00000143256",
                "ENSG00000149084",
                "ENSG00000102931",
                "ENSG00000166619",
                "ENSG00000074800",
                "ENSG00000104863",
                "ENSG00000165502",
                "ENSG00000114302",
                "ENSG00000084733",
                "ENSG00000119705",
                "ENSG00000075151",
                "ENSG00000122034",
                "ENSG00000154582",
                "ENSG00000067955",
                "ENSG00000067836",
                "ENSG00000120533",
                "ENSG00000136854",
                "ENSG00000214113",
                "ENSG00000114209",
                "ENSG00000109107",
                "ENSG00000168214",
                "ENSG00000214517",
                "ENSG00000135535",
                "ENSG00000137996",
                "ENSG00000115524",
                "ENSG00000188994",
                "ENSG00000174775",
                "ENSG00000108946",
                "ENSG00000112237",
                "ENSG00000164830",
                "ENSG00000175416",
                "ENSG00000128609",
                "ENSG00000172586",
                "ENSG00000103316",
                "ENSG00000160953",
                "ENSG00000108953",
                "ENSG00000155368",
                "ENSG00000169564",
                "ENSG00000094916",
                "ENSG00000132305",
                "ENSG00000138663",
                "ENSG00000086061",
                "ENSG00000166922",
                "ENSG00000145675",
                "ENSG00000149806",
                "ENSG00000140386",
                "ENSG00000140740",
                "ENSG00000198727",
                "ENSG00000204272",
                "ENSG00000167658",
                "ENSG00000142168",
                "ENSG00000189180",
                "ENSG00000084070",
                "ENSG00000129946",
                "ENSG00000159176",
                "ENSG00000068745",
                "ENSG00000143761",
                "ENSG00000108848",
                "ENSG00000136193",
                "ENSG00000196586",
                "ENSG00000104427",
                "ENSG00000109919",
                "ENSG00000198786",
                "ENSG00000143933",
                "ENSG00000172172",
                "ENSG00000165629",
                "ENSG00000166503",
                "ENSG00000167815",
                "ENSG00000126457",
                "ENSG00000177971",
                "ENSG00000132963",
                "ENSG00000131089",
                "ENSG00000070087",
                "ENSG00000168522",
                "ENSG00000177311",
                "ENSG00000119048",
                "ENSG00000198408",
                "ENSG00000087470",
                "ENSG00000103254",
                "ENSG00000131370",
                "ENSG00000148450",
                "ENSG00000178127",
                "ENSG00000131876",
                "ENSG00000104381",
                "ENSG00000174456",
                "ENSG00000168701",
                "ENSG00000167699",
                "ENSG00000010244",
                "ENSG00000136628",
                "ENSG00000143198",
                "ENSG00000133639",
                "ENSG00000092148",
                "ENSG00000151552",
                "ENSG00000145494",
                "ENSG00000156411",
                "ENSG00000061676",
                "ENSG00000108960",
                "ENSG00000177200",
                "ENSG00000115944",
                "ENSG00000174939",
                "ENSG00000138443",
                "ENSG00000135334",
                "ENSG00000196531",
                "ENSG00000100138",
                "ENSG00000116030",
                "ENSG00000091592",
                "ENSG00000109133",
                "ENSG00000169435",
                "ENSG00000242485",
                "ENSG00000160113",
                "ENSG00000160075",
                "ENSG00000179915",
                "ENSG00000120992",
                "ENSG00000128872",
                "ENSG00000170043",
                "ENSG00000168653",
                "ENSG00000131507",
                "ENSG00000156261",
                "ENSG00000181885",
                "ENSG00000130723",
                "ENSG00000178741",
                "ENSG00000156515",
                "ENSG00000107758",
                "ENSG00000170606",
                "ENSG00000101266",
                "ENSG00000171206",
                "ENSG00000177885",
                "ENSG00000166595",
                "ENSG00000143774",
                "ENSG00000165209",
                "ENSG00000182512",
                "ENSG00000088832",
                "ENSG00000182463",
                "ENSG00000166136",
                "ENSG00000122545",
                "ENSG00000134330",
                "ENSG00000143612",
                "ENSG00000135720",
                "ENSG00000136485",
                "ENSG00000166848",
                "ENSG00000196683",
                "ENSG00000149547",
                "ENSG00000166206",
                "ENSG00000125870",
                "ENSG00000102189",
                "ENSG00000149600",
                "ENSG00000112514",
                "ENSG00000181061",
                "ENSG00000150787",
                "ENSG00000128739",
                "ENSG00000086666",
                "ENSG00000137726",
                "ENSG00000159023",
                "ENSG00000114902",
                "ENSG00000089818",
                "ENSG00000099341",
                "ENSG00000173273",
                "ENSG00000230989",
                "ENSG00000108309",
                "ENSG00000154518",
                "ENSG00000087302",
                "ENSG00000213064",
                "ENSG00000091164",
                "ENSG00000172757",
                "ENSG00000178449",
                "ENSG00000101460",
                "ENSG00000138175",
                "ENSG00000126858",
                "ENSG00000134717",
                "ENSG00000050426",
                "ENSG00000151778",
                "ENSG00000119782",
                "ENSG00000143252",
                "ENSG00000141367",
                "ENSG00000213719",
                "ENSG00000198168",
                "ENSG00000105438",
                "ENSG00000155099",
                "ENSG00000152234",
                "ENSG00000133872",
                "ENSG00000131094",
                "ENSG00000198668",
                "ENSG00000134308",
                "ENSG00000143158",
                "ENSG00000134684",
                "ENSG00000008324",
                "ENSG00000166228",
                "ENSG00000204525",
                "ENSG00000243449",
                "ENSG00000213619",
                "ENSG00000164919",
                "ENSG00000153774",
                "ENSG00000129197",
                "ENSG00000138757",
                "ENSG00000108561",
                "ENSG00000033627",
                "ENSG00000178982",
                "ENSG00000067900",
                "ENSG00000087448",
                "ENSG00000214253",
                "ENSG00000138382",
                "ENSG00000155876",
                "ENSG00000125821",
                "ENSG00000137547",
                "ENSG00000100387",
                "ENSG00000114573",
                "ENSG00000140319",
                "ENSG00000132466",
                "ENSG00000196233",
                "ENSG00000181704",
                "ENSG00000128923",
                "ENSG00000163399",
                "ENSG00000243317",
                "ENSG00000070669",
                "ENSG00000118680",
                "ENSG00000009307",
                "ENSG00000205937",
                "ENSG00000018189",
                "ENSG00000138138",
                "ENSG00000168958",
                "ENSG00000204856",
                "ENSG00000124172",
                "ENSG00000057608",
                "ENSG00000165169",
                "ENSG00000152795",
                "ENSG00000065548",
                "ENSG00000182195",
                "ENSG00000137876",
                "ENSG00000017797",
                "ENSG00000138398",
                "ENSG00000212907",
                "ENSG00000142669",
                "ENSG00000162076",
                "ENSG00000100568",
                "ENSG00000107281",
                "ENSG00000164323",
                "ENSG00000025293",
                "ENSG00000122705",
                "ENSG00000114631",
                "ENSG00000132432",
                "ENSG00000140374",
                "ENSG00000178104",
                "ENSG00000115306",
                "ENSG00000023734",
                "ENSG00000114405",
                "ENSG00000184009",
                "ENSG00000183726",
                "ENSG00000173660",
                "ENSG00000164292",
                "ENSG00000214941",
                "ENSG00000186184",
                "ENSG00000100804",
                "ENSG00000119640",
                "ENSG00000113648",
                "ENSG00000104643",
                "ENSG00000114125",
                "ENSG00000185787",
                "ENSG00000137947",
                "ENSG00000105568",
                "ENSG00000100554",
                "ENSG00000130638",
                "ENSG00000169045",
                "ENSG00000111775",
                "ENSG00000123636",
                "ENSG00000133142",
                "ENSG00000139684",
                "ENSG00000197102",
                "ENSG00000128989",
                "ENSG00000114120",
                "ENSG00000166562",
                "ENSG00000168724",
                "ENSG00000237651",
                "ENSG00000165704",
                "ENSG00000102003",
                "ENSG00000188612",
                "ENSG00000196781",
                "ENSG00000104763",
                "ENSG00000180817",
                "ENSG00000086065",
                "ENSG00000116288",
                "ENSG00000008018",
                "ENSG00000160948",
                "ENSG00000111530",
                "ENSG00000188322",
                "ENSG00000165487",
                "ENSG00000220205",
                "ENSG00000128524",
                "ENSG00000134313",
                "ENSG00000167232",
                "ENSG00000165678",
                "ENSG00000198712",
                "ENSG00000133302",
                "ENSG00000167113",
                "ENSG00000136238",
                "ENSG00000132424",
                "ENSG00000130338",
                "ENSG00000224032",
                "ENSG00000047849",
                "ENSG00000183527",
                "ENSG00000221914",
                "ENSG00000122873",
                "ENSG00000113851",
                "ENSG00000103415",
                "ENSG00000151923",
                "ENSG00000170759",
                "ENSG00000087460",
                "ENSG00000145247",
                "ENSG00000113558",
                "ENSG00000176978",
                "ENSG00000143153",
                "ENSG00000109180",
                "ENSG00000213937",
                "ENSG00000115233",
                "ENSG00000123983",
                "ENSG00000156052",
                "ENSG00000143742",
                "ENSG00000198648",
                "ENSG00000133169",
                "ENSG00000106244",
                "ENSG00000104419",
                "ENSG00000164167",
                "ENSG00000100201",
                "ENSG00000114115",
                "ENSG00000198888",
                "ENSG00000173163"
            ],
            "expression_sum": {
                "class1": 4215.0,
                "class2": 5563.0
            },
            "ens_id": "ENSG00000113580",
            "uniprot_ids": [
                "P04150",
                "Q3MSN4",
                "A0A494C0P1",
                "D6RDA9"
            ],
            "net_id": 0,
            "source_num": 11,
            "target_num": 114
        },
        "ENSG00000125968": {
            "id": "ENSG00000125968",
            "type": "TF",
            "symbol": "ID1",
            "source": [
                "ENSG00000113580",
                "ENSG00000139352"
            ],
            "target": [
                "ENSG00000113580"
            ],
            "expression_sum": {
                "class1": 45370.0,
                "class2": 24066.0
            },
            "ens_id": "ENSG00000125968",
            "uniprot_ids": [
                "P41134"
            ],
            "net_id": 0,
            "source_num": 2,
            "target_num": 1
        },
        "ENSG00000165462": {
            "id": "ENSG00000165462",
            "type": "TF",
            "symbol": "PHOX2A",
            "source": [
                "ENSG00000016082",
                "ENSG00000108055"
            ],
            "target": [
                "ENSG00000105419",
                "ENSG00000113448",
                "ENSG00000097021",
                "ENSG00000115297"
            ],
            "expression_sum": {
                "class1": 3582.0,
                "class2": 18.0
            },
            "ens_id": "ENSG00000165462",
            "uniprot_ids": [
                "O14813",
                "H0YGU5"
            ],
            "net_id": 0,
            "source_num": 2,
            "target_num": 1
        },
        "ENSG00000105419": {
            "id": "ENSG00000105419",
            "type": "TF",
            "symbol": "MEIS3",
            "source": [
                "ENSG00000167081",
                "ENSG00000113580",
                "ENSG00000103495",
                "ENSG00000177030",
                "ENSG00000196591",
                "ENSG00000165462"
            ],
            "target": [
                "ENSG00000117632",
                "ENSG00000162545",
                "ENSG00000197728",
                "ENSG00000137285"
            ],
            "expression_sum": {
                "class1": 17649.0,
                "class2": 4228.0
            },
            "ens_id": "ENSG00000105419",
            "uniprot_ids": [
                "Q99687",
                "H0YNA8",
                "H3BLU4",
                "H0YMV2",
                "U3KQV2",
                "U3KQJ9",
                "H0YLL0"
            ],
            "net_id": 0,
            "source_num": 6,
            "target_num": 2
        },
        "ENSG00000177030": {
            "id": "ENSG00000177030",
            "type": "TF",
            "symbol": "DEAF1",
            "source": [
                "ENSG00000167081",
                "ENSG00000164330"
            ],
            "target": [
                "ENSG00000105419",
                "ENSG00000005810",
                "ENSG00000113161",
                "ENSG00000169750",
                "ENSG00000114784",
                "ENSG00000136810",
                "ENSG00000117569",
                "ENSG00000170027",
                "ENSG00000133318",
                "ENSG00000136238",
                "ENSG00000115756",
                "ENSG00000079459",
                "ENSG00000165434",
                "ENSG00000143702",
                "ENSG00000112972",
                "ENSG00000171033",
                "ENSG00000152503",
                "ENSG00000052802",
                "ENSG00000189159",
                "ENSG00000272398",
                "ENSG00000105649",
                "ENSG00000134684",
                "ENSG00000273841",
                "ENSG00000100320",
                "ENSG00000162961"
            ],
            "expression_sum": {
                "class1": 7678.0,
                "class2": 4341.0
            },
            "ens_id": "ENSG00000177030",
            "uniprot_ids": [
                "O75398",
                "H0YCH1",
                "A0A804HIS1",
                "H0YCY2"
            ],
            "net_id": 0,
            "source_num": 2,
            "target_num": 7
        },
        "ENSG00000139352": {
            "id": "ENSG00000139352",
            "type": "TF",
            "symbol": "ASCL1",
            "source": [],
            "target": [
                "ENSG00000078668",
                "ENSG00000206573",
                "ENSG00000073803",
                "ENSG00000165655",
                "ENSG00000156475",
                "ENSG00000111674",
                "ENSG00000005810",
                "ENSG00000099622",
                "ENSG00000177606",
                "ENSG00000140941",
                "ENSG00000134248",
                "ENSG00000146247",
                "ENSG00000075415",
                "ENSG00000105993",
                "ENSG00000160714",
                "ENSG00000102466",
                "ENSG00000265972",
                "ENSG00000154556",
                "ENSG00000173812",
                "ENSG00000183091",
                "ENSG00000152256",
                "ENSG00000187244",
                "ENSG00000198899",
                "ENSG00000173915",
                "ENSG00000065911",
                "ENSG00000151729",
                "ENSG00000129636",
                "ENSG00000163875",
                "ENSG00000074319",
                "ENSG00000180879",
                "ENSG00000242173",
                "ENSG00000139767",
                "ENSG00000165264",
                "ENSG00000180773",
                "ENSG00000104964",
                "ENSG00000125868",
                "ENSG00000165637",
                "ENSG00000153815",
                "ENSG00000106565",
                "ENSG00000125356",
                "ENSG00000125968",
                "ENSG00000129625",
                "ENSG00000055163",
                "ENSG00000167733",
                "ENSG00000126778",
                "ENSG00000104112",
                "ENSG00000255302",
                "ENSG00000067225",
                "ENSG00000103657",
                "ENSG00000113712",
                "ENSG00000144485",
                "ENSG00000100592",
                "ENSG00000157168",
                "ENSG00000109113",
                "ENSG00000063854",
                "ENSG00000133606",
                "ENSG00000103769",
                "ENSG00000117410",
                "ENSG00000273841",
                "ENSG00000124802",
                "ENSG00000263001",
                "ENSG00000196405",
                "ENSG00000174437",
                "ENSG00000215845",
                "ENSG00000110680",
                "ENSG00000225783",
                "ENSG00000185808",
                "ENSG00000146242",
                "ENSG00000144366",
                "ENSG00000110955",
                "ENSG00000010256",
                "ENSG00000170634",
                "ENSG00000163888",
                "ENSG00000198836",
                "ENSG00000131966",
                "ENSG00000166959",
                "ENSG00000173404",
                "ENSG00000109332",
                "ENSG00000163170",
                "ENSG00000095059",
                "ENSG00000244879",
                "ENSG00000124155",
                "ENSG00000166974",
                "ENSG00000141562",
                "ENSG00000100320",
                "ENSG00000240972",
                "ENSG00000172915",
                "ENSG00000196655",
                "ENSG00000160050",
                "ENSG00000153310",
                "ENSG00000184672",
                "ENSG00000128654",
                "ENSG00000160613",
                "ENSG00000090266",
                "ENSG00000143862",
                "ENSG00000048649",
                "ENSG00000162374",
                "ENSG00000072832",
                "ENSG00000130312",
                "ENSG00000136950",
                "ENSG00000160551",
                "ENSG00000104388",
                "ENSG00000168159",
                "ENSG00000135472",
                "ENSG00000267100",
                "ENSG00000064607",
                "ENSG00000198938",
                "ENSG00000127914",
                "ENSG00000145730",
                "ENSG00000002933",
                "ENSG00000175582",
                "ENSG00000109103",
                "ENSG00000104765",
                "ENSG00000171130",
                "ENSG00000168300",
                "ENSG00000130725",
                "ENSG00000112305",
                "ENSG00000108679",
                "ENSG00000253738",
                "ENSG00000047932",
                "ENSG00000237765",
                "ENSG00000135250",
                "ENSG00000198840",
                "ENSG00000247095",
                "ENSG00000114784",
                "ENSG00000198258",
                "ENSG00000169567",
                "ENSG00000197302",
                "ENSG00000129355",
                "ENSG00000183172",
                "ENSG00000076554",
                "ENSG00000198125",
                "ENSG00000135821",
                "ENSG00000131495",
                "ENSG00000172159",
                "ENSG00000090238",
                "ENSG00000167642",
                "ENSG00000054356",
                "ENSG00000077254",
                "ENSG00000152932",
                "ENSG00000067715",
                "ENSG00000119950",
                "ENSG00000111229",
                "ENSG00000103145",
                "ENSG00000113742",
                "ENSG00000137941",
                "ENSG00000137100",
                "ENSG00000198886",
                "ENSG00000102934",
                "ENSG00000122643",
                "ENSG00000104904",
                "ENSG00000173726",
                "ENSG00000105254",
                "ENSG00000115286",
                "ENSG00000105698",
                "ENSG00000092051",
                "ENSG00000169432",
                "ENSG00000166347",
                "ENSG00000151327",
                "ENSG00000131174",
                "ENSG00000166405",
                "ENSG00000143013",
                "ENSG00000145495",
                "ENSG00000254999",
                "ENSG00000004059",
                "ENSG00000130558",
                "ENSG00000113387",
                "ENSG00000158864",
                "ENSG00000082701",
                "ENSG00000100442",
                "ENSG00000121653",
                "ENSG00000100697",
                "ENSG00000197111",
                "ENSG00000146729",
                "ENSG00000133134",
                "ENSG00000136100",
                "ENSG00000159720",
                "ENSG00000148798",
                "ENSG00000004779",
                "ENSG00000184277",
                "ENSG00000071553",
                "ENSG00000158604",
                "ENSG00000169139",
                "ENSG00000117318",
                "ENSG00000186868",
                "ENSG00000116459",
                "ENSG00000070831",
                "ENSG00000137500",
                "ENSG00000250479",
                "ENSG00000120885",
                "ENSG00000182050",
                "ENSG00000125818",
                "ENSG00000105220",
                "ENSG00000272047",
                "ENSG00000198791",
                "ENSG00000134049",
                "ENSG00000214530",
                "ENSG00000183742",
                "ENSG00000111237",
                "ENSG00000182985",
                "ENSG00000135070",
                "ENSG00000134769",
                "ENSG00000108176",
                "ENSG00000165416",
                "ENSG00000203667",
                "ENSG00000136710",
                "ENSG00000131149",
                "ENSG00000100354",
                "ENSG00000115738",
                "ENSG00000166780",
                "ENSG00000111664",
                "ENSG00000126214",
                "ENSG00000176340",
                "ENSG00000097033",
                "ENSG00000150753",
                "ENSG00000049618",
                "ENSG00000156587",
                "ENSG00000182670",
                "ENSG00000177459",
                "ENSG00000146701",
                "ENSG00000186472",
                "ENSG00000101745",
                "ENSG00000173436",
                "ENSG00000163541",
                "ENSG00000169925",
                "ENSG00000109472",
                "ENSG00000164898",
                "ENSG00000079102",
                "ENSG00000101210",
                "ENSG00000158321",
                "ENSG00000198830",
                "ENSG00000143155",
                "ENSG00000132639",
                "ENSG00000156467",
                "ENSG00000119013",
                "ENSG00000161203",
                "ENSG00000184076",
                "ENSG00000181722",
                "ENSG00000159199",
                "ENSG00000170873",
                "ENSG00000151743",
                "ENSG00000105290",
                "ENSG00000147669",
                "ENSG00000119888",
                "ENSG00000168003",
                "ENSG00000096092",
                "ENSG00000111371",
                "ENSG00000119547",
                "ENSG00000101558",
                "ENSG00000101040",
                "ENSG00000120071",
                "ENSG00000124596",
                "ENSG00000253719",
                "ENSG00000156976",
                "ENSG00000134138",
                "ENSG00000130294",
                "ENSG00000119396",
                "ENSG00000106484",
                "ENSG00000099624",
                "ENSG00000138495",
                "ENSG00000101489",
                "ENSG00000138069",
                "ENSG00000083099",
                "ENSG00000110321",
                "ENSG00000138814",
                "ENSG00000005483",
                "ENSG00000055609",
                "ENSG00000129158",
                "ENSG00000175203",
                "ENSG00000156642",
                "ENSG00000145362",
                "ENSG00000149084",
                "ENSG00000166619",
                "ENSG00000104863",
                "ENSG00000084733",
                "ENSG00000075151",
                "ENSG00000067836",
                "ENSG00000136854",
                "ENSG00000105223",
                "ENSG00000174469",
                "ENSG00000077279",
                "ENSG00000188994",
                "ENSG00000174775",
                "ENSG00000108946",
                "ENSG00000132510",
                "ENSG00000164830",
                "ENSG00000175416",
                "ENSG00000128609",
                "ENSG00000160953",
                "ENSG00000183775",
                "ENSG00000169564",
                "ENSG00000106415",
                "ENSG00000132305",
                "ENSG00000115828",
                "ENSG00000166922",
                "ENSG00000145675",
                "ENSG00000140386",
                "ENSG00000100243",
                "ENSG00000189180",
                "ENSG00000084070",
                "ENSG00000129946",
                "ENSG00000159176",
                "ENSG00000068745",
                "ENSG00000108848",
                "ENSG00000184545",
                "ENSG00000198786",
                "ENSG00000143933",
                "ENSG00000165629",
                "ENSG00000166503",
                "ENSG00000167815",
                "ENSG00000126457",
                "ENSG00000070087",
                "ENSG00000119048",
                "ENSG00000087470",
                "ENSG00000148450",
                "ENSG00000178127",
                "ENSG00000104381",
                "ENSG00000174456",
                "ENSG00000133639",
                "ENSG00000145494",
                "ENSG00000177200",
                "ENSG00000115944",
                "ENSG00000169435",
                "ENSG00000153956",
                "ENSG00000179915",
                "ENSG00000120992",
                "ENSG00000128872",
                "ENSG00000170043",
                "ENSG00000131507",
                "ENSG00000181885",
                "ENSG00000130723",
                "ENSG00000178741",
                "ENSG00000156515",
                "ENSG00000171206",
                "ENSG00000177885",
                "ENSG00000165209",
                "ENSG00000088832",
                "ENSG00000251562",
                "ENSG00000135720",
                "ENSG00000149547",
                "ENSG00000166206",
                "ENSG00000102189",
                "ENSG00000112514",
                "ENSG00000181061",
                "ENSG00000150787",
                "ENSG00000137726",
                "ENSG00000159023",
                "ENSG00000089818",
                "ENSG00000108309",
                "ENSG00000213064",
                "ENSG00000103460",
                "ENSG00000178449",
                "ENSG00000101460",
                "ENSG00000138175",
                "ENSG00000110492",
                "ENSG00000134717",
                "ENSG00000198168",
                "ENSG00000155099",
                "ENSG00000163331",
                "ENSG00000152234",
                "ENSG00000077327",
                "ENSG00000198668",
                "ENSG00000143158",
                "ENSG00000168772",
                "ENSG00000135549",
                "ENSG00000166228",
                "ENSG00000213619",
                "ENSG00000164919",
                "ENSG00000033627",
                "ENSG00000214253",
                "ENSG00000104870",
                "ENSG00000100387",
                "ENSG00000114573",
                "ENSG00000132466",
                "ENSG00000163399",
                "ENSG00000243317",
                "ENSG00000156675",
                "ENSG00000009307",
                "ENSG00000124172",
                "ENSG00000212907",
                "ENSG00000138398",
                "ENSG00000185088",
                "ENSG00000107281",
                "ENSG00000025293",
                "ENSG00000178104",
                "ENSG00000115306",
                "ENSG00000023734",
                "ENSG00000173660",
                "ENSG00000148925",
                "ENSG00000164292",
                "ENSG00000214941",
                "ENSG00000119640",
                "ENSG00000104643",
                "ENSG00000114125",
                "ENSG00000100554",
                "ENSG00000111775",
                "ENSG00000183741",
                "ENSG00000123636",
                "ENSG00000197102",
                "ENSG00000128989",
                "ENSG00000114120",
                "ENSG00000144834",
                "ENSG00000166562",
                "ENSG00000165704",
                "ENSG00000188612",
                "ENSG00000104763",
                "ENSG00000160948",
                "ENSG00000081019",
                "ENSG00000188322",
                "ENSG00000198712",
                "ENSG00000132424",
                "ENSG00000183527",
                "ENSG00000122873",
                "ENSG00000087460",
                "ENSG00000198774",
                "ENSG00000145247",
                "ENSG00000176978",
                "ENSG00000143153",
                "ENSG00000213937",
                "ENSG00000123983",
                "ENSG00000143742",
                "ENSG00000104419",
                "ENSG00000198888"
            ],
            "expression_sum": {
                "class1": 9481.0,
                "class2": 11679.0
            },
            "ens_id": "ENSG00000139352",
            "uniprot_ids": [
                "P50553"
            ],
            "net_id": 0,
            "source_num": 0,
            "target_num": 60
        },
        "ENSG00000004487": {
            "id": "ENSG00000004487",
            "type": "TF",
            "symbol": "KDM1A",
            "source": [
                "ENSG00000167081"
            ],
            "target": [
                "ENSG00000099864",
                "ENSG00000103034",
                "ENSG00000176788",
                "ENSG00000176887",
                "ENSG00000144339",
                "ENSG00000172020",
                "ENSG00000149557",
                "ENSG00000156011",
                "ENSG00000165410",
                "ENSG00000169856",
                "ENSG00000087085",
                "ENSG00000169783",
                "ENSG00000185664",
                "ENSG00000101210",
                "ENSG00000164093",
                "ENSG00000107105",
                "ENSG00000182534",
                "ENSG00000213190",
                "ENSG00000198910",
                "ENSG00000188290",
                "ENSG00000078018",
                "ENSG00000197959",
                "ENSG00000162374",
                "ENSG00000196361",
                "ENSG00000105409",
                "ENSG00000197747",
                "ENSG00000072832",
                "ENSG00000145491",
                "ENSG00000010270",
                "ENSG00000131711",
                "ENSG00000155980",
                "ENSG00000075618",
                "ENSG00000151320",
                "ENSG00000168280",
                "ENSG00000186868",
                "ENSG00000182481",
                "ENSG00000198125",
                "ENSG00000213760",
                "ENSG00000137285",
                "ENSG00000092964",
                "ENSG00000143702",
                "ENSG00000144218",
                "ENSG00000104435",
                "ENSG00000122012",
                "ENSG00000213420",
                "ENSG00000004838",
                "ENSG00000016082",
                "ENSG00000173267",
                "ENSG00000111897",
                "ENSG00000111728",
                "ENSG00000154917",
                "ENSG00000149294",
                "ENSG00000034510",
                "ENSG00000170091",
                "ENSG00000131016",
                "ENSG00000015592",
                "ENSG00000139970",
                "ENSG00000167081",
                "ENSG00000135406",
                "ENSG00000104833",
                "ENSG00000137267",
                "ENSG00000177675",
                "ENSG00000135447",
                "ENSG00000018236",
                "ENSG00000189043",
                "ENSG00000163931",
                "ENSG00000277586"
            ],
            "expression_sum": {
                "class1": 13324.0,
                "class2": 5734.0
            },
            "ens_id": "ENSG00000004487",
            "uniprot_ids": [
                "O60341",
                "R4GMP9",
                "R4GMQ1"
            ],
            "net_id": 0,
            "source_num": 1,
            "target_num": 31
        },
        "ENSG00000103495": {
            "id": "ENSG00000103495",
            "type": "TF",
            "symbol": "MAZ",
            "source": [
                "ENSG00000181449",
                "ENSG00000109132",
                "ENSG00000167081",
                "ENSG00000118263",
                "ENSG00000164330",
                "ENSG00000119547"
            ],
            "target": [
                "ENSG00000162373",
                "ENSG00000162704",
                "ENSG00000111674",
                "ENSG00000111716",
                "ENSG00000153130",
                "ENSG00000005810",
                "ENSG00000137955",
                "ENSG00000197959",
                "ENSG00000075945",
                "ENSG00000077380",
                "ENSG00000162490",
                "ENSG00000052802",
                "ENSG00000184697",
                "ENSG00000087258",
                "ENSG00000111728",
                "ENSG00000154917",
                "ENSG00000272398",
                "ENSG00000015592",
                "ENSG00000129636",
                "ENSG00000115540",
                "ENSG00000123240",
                "ENSG00000143545",
                "ENSG00000081181",
                "ENSG00000242173",
                "ENSG00000162545",
                "ENSG00000139112",
                "ENSG00000125868",
                "ENSG00000165637",
                "ENSG00000213190",
                "ENSG00000060982",
                "ENSG00000131236",
                "ENSG00000196072",
                "ENSG00000104112",
                "ENSG00000187164",
                "ENSG00000144642",
                "ENSG00000144218",
                "ENSG00000213420",
                "ENSG00000107130",
                "ENSG00000034713",
                "ENSG00000105696",
                "ENSG00000157168",
                "ENSG00000034510",
                "ENSG00000117410",
                "ENSG00000150722",
                "ENSG00000273841",
                "ENSG00000138698",
                "ENSG00000177519",
                "ENSG00000144635",
                "ENSG00000130540",
                "ENSG00000176749",
                "ENSG00000125744",
                "ENSG00000169314",
                "ENSG00000215217",
                "ENSG00000164687",
                "ENSG00000129682",
                "ENSG00000163888",
                "ENSG00000213760",
                "ENSG00000143702",
                "ENSG00000131966",
                "ENSG00000112972",
                "ENSG00000084731",
                "ENSG00000166974",
                "ENSG00000167552",
                "ENSG00000141505",
                "ENSG00000125814",
                "ENSG00000100320",
                "ENSG00000163931",
                "ENSG00000203879",
                "ENSG00000105419",
                "ENSG00000250303",
                "ENSG00000239672",
                "ENSG00000153310",
                "ENSG00000213585",
                "ENSG00000131437",
                "ENSG00000154277",
                "ENSG00000128654",
                "ENSG00000137942",
                "ENSG00000105784",
                "ENSG00000088727",
                "ENSG00000143862",
                "ENSG00000174021",
                "ENSG00000072832",
                "ENSG00000198513",
                "ENSG00000135205",
                "ENSG00000079459",
                "ENSG00000104388",
                "ENSG00000103154",
                "ENSG00000168159",
                "ENSG00000135472",
                "ENSG00000097021",
                "ENSG00000165943",
                "ENSG00000131016",
                "ENSG00000168824",
                "ENSG00000138587",
                "ENSG00000067064",
                "ENSG00000185668",
                "ENSG00000087085",
                "ENSG00000113657",
                "ENSG00000114784",
                "ENSG00000129355",
                "ENSG00000145545",
                "ENSG00000170027",
                "ENSG00000132872",
                "ENSG00000076554",
                "ENSG00000010270",
                "ENSG00000198125",
                "ENSG00000108175",
                "ENSG00000172159",
                "ENSG00000173267",
                "ENSG00000138413",
                "ENSG00000079462",
                "ENSG00000134202",
                "ENSG00000155959",
                "ENSG00000105254",
                "ENSG00000166257",
                "ENSG00000131711",
                "ENSG00000068366",
                "ENSG00000155980",
                "ENSG00000101132",
                "ENSG00000163539",
                "ENSG00000109654",
                "ENSG00000130558",
                "ENSG00000160188",
                "ENSG00000170091",
                "ENSG00000078328",
                "ENSG00000170445",
                "ENSG00000185650",
                "ENSG00000168118",
                "ENSG00000178234",
                "ENSG00000134986",
                "ENSG00000277586",
                "ENSG00000111275",
                "ENSG00000176788",
                "ENSG00000172020",
                "ENSG00000133134",
                "ENSG00000151640",
                "ENSG00000182093",
                "ENSG00000148798",
                "ENSG00000132821",
                "ENSG00000105409",
                "ENSG00000197747",
                "ENSG00000184277",
                "ENSG00000162368",
                "ENSG00000186868",
                "ENSG00000070831",
                "ENSG00000092964",
                "ENSG00000122012",
                "ENSG00000152503",
                "ENSG00000164221",
                "ENSG00000166736",
                "ENSG00000105649",
                "ENSG00000105220",
                "ENSG00000104833",
                "ENSG00000164330",
                "ENSG00000176871",
                "ENSG00000151150",
                "ENSG00000135447",
                "ENSG00000105379",
                "ENSG00000181449",
                "ENSG00000167508",
                "ENSG00000144339",
                "ENSG00000050165",
                "ENSG00000165443",
                "ENSG00000162188",
                "ENSG00000171224",
                "ENSG00000133318",
                "ENSG00000166780",
                "ENSG00000137285",
                "ENSG00000126214",
                "ENSG00000124194",
                "ENSG00000090097",
                "ENSG00000112290",
                "ENSG00000069956",
                "ENSG00000159409",
                "ENSG00000149926",
                "ENSG00000103034",
                "ENSG00000169856",
                "ENSG00000141698",
                "ENSG00000101210",
                "ENSG00000164093",
                "ENSG00000105255",
                "ENSG00000186106",
                "ENSG00000141526",
                "ENSG00000132639",
                "ENSG00000168280",
                "ENSG00000165434",
                "ENSG00000105088",
                "ENSG00000117632",
                "ENSG00000105290",
                "ENSG00000018236",
                "ENSG00000096092",
                "ENSG00000119547",
                "ENSG00000164287",
                "ENSG00000169783",
                "ENSG00000175130",
                "ENSG00000169750",
                "ENSG00000084628",
                "ENSG00000156976",
                "ENSG00000128245",
                "ENSG00000130294",
                "ENSG00000108828",
                "ENSG00000145491",
                "ENSG00000120265",
                "ENSG00000187231",
                "ENSG00000125354",
                "ENSG00000171033",
                "ENSG00000175203",
                "ENSG00000102409",
                "ENSG00000158164",
                "ENSG00000145362",
                "ENSG00000141404",
                "ENSG00000149084",
                "ENSG00000104863",
                "ENSG00000084733",
                "ENSG00000136717",
                "ENSG00000149557",
                "ENSG00000173207",
                "ENSG00000185664",
                "ENSG00000145335",
                "ENSG00000198910",
                "ENSG00000134198",
                "ENSG00000109107",
                "ENSG00000005108",
                "ENSG00000016082",
                "ENSG00000175416",
                "ENSG00000128609",
                "ENSG00000146267",
                "ENSG00000177875",
                "ENSG00000176973",
                "ENSG00000166922",
                "ENSG00000161082",
                "ENSG00000005249",
                "ENSG00000113161",
                "ENSG00000072041",
                "ENSG00000142549",
                "ENSG00000157851",
                "ENSG00000182287",
                "ENSG00000104427",
                "ENSG00000186469",
                "ENSG00000115756",
                "ENSG00000165629",
                "ENSG00000167996",
                "ENSG00000148154",
                "ENSG00000184117",
                "ENSG00000143198",
                "ENSG00000167081",
                "ENSG00000135406",
                "ENSG00000177675",
                "ENSG00000108960",
                "ENSG00000137343",
                "ENSG00000099864",
                "ENSG00000130643",
                "ENSG00000115944",
                "ENSG00000174939",
                "ENSG00000080822",
                "ENSG00000161682",
                "ENSG00000107105",
                "ENSG00000122756",
                "ENSG00000182534",
                "ENSG00000086300",
                "ENSG00000100167",
                "ENSG00000120992",
                "ENSG00000128872",
                "ENSG00000131507",
                "ENSG00000182481",
                "ENSG00000168589",
                "ENSG00000160753",
                "ENSG00000170312",
                "ENSG00000156515",
                "ENSG00000119865",
                "ENSG00000177885",
                "ENSG00000156298",
                "ENSG00000115665",
                "ENSG00000115310",
                "ENSG00000186973",
                "ENSG00000122545",
                "ENSG00000166848",
                "ENSG00000099194",
                "ENSG00000104722",
                "ENSG00000128739",
                "ENSG00000137726",
                "ENSG00000157514",
                "ENSG00000221946",
                "ENSG00000089818",
                "ENSG00000187714",
                "ENSG00000109084",
                "ENSG00000066557",
                "ENSG00000124302",
                "ENSG00000149313",
                "ENSG00000160752",
                "ENSG00000119782",
                "ENSG00000075618",
                "ENSG00000198961",
                "ENSG00000155099",
                "ENSG00000152669",
                "ENSG00000133872",
                "ENSG00000172687",
                "ENSG00000163879",
                "ENSG00000118263",
                "ENSG00000068615",
                "ENSG00000243449",
                "ENSG00000033627",
                "ENSG00000164176",
                "ENSG00000111726",
                "ENSG00000125821",
                "ENSG00000153789",
                "ENSG00000118680",
                "ENSG00000171551",
                "ENSG00000018189",
                "ENSG00000138138",
                "ENSG00000128564",
                "ENSG00000004838",
                "ENSG00000114631",
                "ENSG00000120437",
                "ENSG00000149294",
                "ENSG00000170899",
                "ENSG00000137267",
                "ENSG00000130638",
                "ENSG00000157005",
                "ENSG00000166562",
                "ENSG00000109881",
                "ENSG00000102003",
                "ENSG00000171724",
                "ENSG00000180817",
                "ENSG00000152578",
                "ENSG00000107643",
                "ENSG00000117569",
                "ENSG00000220205",
                "ENSG00000134313",
                "ENSG00000221866",
                "ENSG00000104549",
                "ENSG00000144369",
                "ENSG00000047849",
                "ENSG00000103415",
                "ENSG00000131037",
                "ENSG00000139970",
                "ENSG00000133169",
                "ENSG00000113810",
                "ENSG00000034239"
            ],
            "expression_sum": {
                "class1": 16130.0,
                "class2": 8065.0
            },
            "ens_id": "ENSG00000103495",
            "uniprot_ids": [
                "P56270",
                "I3L0M3",
                "H3BPU3",
                "H3BQI4",
                "H3BQS2",
                "A0A087WWR2",
                "I3L4D3",
                "I3L411",
                "H3BTS8",
                "H3BRC5",
                "I3L4Y2",
                "I3L2Z5"
            ],
            "net_id": 0,
            "source_num": 6,
            "target_num": 72
        },
        "ENSG00000108055": {
            "id": "ENSG00000108055",
            "type": "TF",
            "symbol": "SMC3",
            "source": [
                "ENSG00000181449",
                "ENSG00000164330"
            ],
            "target": [
                "ENSG00000103034",
                "ENSG00000169856",
                "ENSG00000087085",
                "ENSG00000005810",
                "ENSG00000113657",
                "ENSG00000101210",
                "ENSG00000198830",
                "ENSG00000164093",
                "ENSG00000107105",
                "ENSG00000182534",
                "ENSG00000129355",
                "ENSG00000137804",
                "ENSG00000186106",
                "ENSG00000187134",
                "ENSG00000197959",
                "ENSG00000170027",
                "ENSG00000132872",
                "ENSG00000010270",
                "ENSG00000168280",
                "ENSG00000148773",
                "ENSG00000165434",
                "ENSG00000108175",
                "ENSG00000170312",
                "ENSG00000066279",
                "ENSG00000173267",
                "ENSG00000138413",
                "ENSG00000184697",
                "ENSG00000087258",
                "ENSG00000175063",
                "ENSG00000105088",
                "ENSG00000111728",
                "ENSG00000154917",
                "ENSG00000272398",
                "ENSG00000176890",
                "ENSG00000134369",
                "ENSG00000123240",
                "ENSG00000105270",
                "ENSG00000099194",
                "ENSG00000119547",
                "ENSG00000104722",
                "ENSG00000162545",
                "ENSG00000176887",
                "ENSG00000169783",
                "ENSG00000157514",
                "ENSG00000221946",
                "ENSG00000167654",
                "ENSG00000204130",
                "ENSG00000260230",
                "ENSG00000166257",
                "ENSG00000196361",
                "ENSG00000131711",
                "ENSG00000068366",
                "ENSG00000155980",
                "ENSG00000144642",
                "ENSG00000144218",
                "ENSG00000163539",
                "ENSG00000125354",
                "ENSG00000155099",
                "ENSG00000107130",
                "ENSG00000157456",
                "ENSG00000034510",
                "ENSG00000170091",
                "ENSG00000118263",
                "ENSG00000134986",
                "ENSG00000277586",
                "ENSG00000117399",
                "ENSG00000176788",
                "ENSG00000172020",
                "ENSG00000114573",
                "ENSG00000173207",
                "ENSG00000185664",
                "ENSG00000145335",
                "ENSG00000198910",
                "ENSG00000099250",
                "ENSG00000197183",
                "ENSG00000134198",
                "ENSG00000018189",
                "ENSG00000128564",
                "ENSG00000105409",
                "ENSG00000197747",
                "ENSG00000164687",
                "ENSG00000151320",
                "ENSG00000186868",
                "ENSG00000005108",
                "ENSG00000213760",
                "ENSG00000092964",
                "ENSG00000143702",
                "ENSG00000122012",
                "ENSG00000016082",
                "ENSG00000142156",
                "ENSG00000152503",
                "ENSG00000149294",
                "ENSG00000105649",
                "ENSG00000084731",
                "ENSG00000166974",
                "ENSG00000104833",
                "ENSG00000174521",
                "ENSG00000164330",
                "ENSG00000141505",
                "ENSG00000137267",
                "ENSG00000177875",
                "ENSG00000151150",
                "ENSG00000125814",
                "ENSG00000100320",
                "ENSG00000163931",
                "ENSG00000135447",
                "ENSG00000162852",
                "ENSG00000181449",
                "ENSG00000250303",
                "ENSG00000188486",
                "ENSG00000161082",
                "ENSG00000005249",
                "ENSG00000131437",
                "ENSG00000165462",
                "ENSG00000080824",
                "ENSG00000166803",
                "ENSG00000105784",
                "ENSG00000117724",
                "ENSG00000174021",
                "ENSG00000162374",
                "ENSG00000134313",
                "ENSG00000104427",
                "ENSG00000221866",
                "ENSG00000197694",
                "ENSG00000186469",
                "ENSG00000115756",
                "ENSG00000047849",
                "ENSG00000104435",
                "ENSG00000126214",
                "ENSG00000167996",
                "ENSG00000124194",
                "ENSG00000148154",
                "ENSG00000131037",
                "ENSG00000131747",
                "ENSG00000111897",
                "ENSG00000177410",
                "ENSG00000131016",
                "ENSG00000139970",
                "ENSG00000143153",
                "ENSG00000135406",
                "ENSG00000168824",
                "ENSG00000159409",
                "ENSG00000175582",
                "ENSG00000113810",
                "ENSG00000169439",
                "ENSG00000177675"
            ],
            "expression_sum": {
                "class1": 16950.0,
                "class2": 11336.0
            },
            "ens_id": "ENSG00000108055",
            "uniprot_ids": [
                "Q9UQE7"
            ],
            "net_id": 0,
            "source_num": 2,
            "target_num": 52
        },
        "ENSG00000116560": {
            "id": "ENSG00000116560",
            "type": "TF",
            "symbol": "SFPQ",
            "source": [
                "ENSG00000181449",
                "ENSG00000176887",
                "ENSG00000109132",
                "ENSG00000167081",
                "ENSG00000164330"
            ],
            "target": [
                "ENSG00000103034",
                "ENSG00000174939",
                "ENSG00000185668",
                "ENSG00000005810",
                "ENSG00000113657",
                "ENSG00000198830",
                "ENSG00000164093",
                "ENSG00000182534",
                "ENSG00000129355",
                "ENSG00000137804",
                "ENSG00000128872",
                "ENSG00000186106",
                "ENSG00000187134",
                "ENSG00000170027",
                "ENSG00000132639",
                "ENSG00000075945",
                "ENSG00000168280",
                "ENSG00000148773",
                "ENSG00000164611",
                "ENSG00000108175",
                "ENSG00000172159",
                "ENSG00000170312",
                "ENSG00000066279",
                "ENSG00000138413",
                "ENSG00000067715",
                "ENSG00000118200",
                "ENSG00000087258",
                "ENSG00000175063",
                "ENSG00000111728",
                "ENSG00000272398",
                "ENSG00000134369",
                "ENSG00000123240",
                "ENSG00000115310",
                "ENSG00000166848",
                "ENSG00000105270",
                "ENSG00000018236",
                "ENSG00000096092",
                "ENSG00000099194",
                "ENSG00000162545",
                "ENSG00000176887",
                "ENSG00000169783",
                "ENSG00000157514",
                "ENSG00000204130",
                "ENSG00000125868",
                "ENSG00000084628",
                "ENSG00000130294",
                "ENSG00000100077",
                "ENSG00000104112",
                "ENSG00000131711",
                "ENSG00000187231",
                "ENSG00000144642",
                "ENSG00000163539",
                "ENSG00000123975",
                "ENSG00000213420",
                "ENSG00000171033",
                "ENSG00000155099",
                "ENSG00000130558",
                "ENSG00000157168",
                "ENSG00000170091",
                "ENSG00000206579",
                "ENSG00000145362",
                "ENSG00000170445",
                "ENSG00000141404",
                "ENSG00000118263",
                "ENSG00000149257",
                "ENSG00000185650",
                "ENSG00000164104",
                "ENSG00000134986",
                "ENSG00000176788",
                "ENSG00000172020",
                "ENSG00000133134",
                "ENSG00000149557",
                "ENSG00000145428",
                "ENSG00000114573",
                "ENSG00000173207",
                "ENSG00000130540",
                "ENSG00000198910",
                "ENSG00000018189",
                "ENSG00000138138",
                "ENSG00000128564",
                "ENSG00000105409",
                "ENSG00000151320",
                "ENSG00000005108",
                "ENSG00000213760",
                "ENSG00000092964",
                "ENSG00000143702",
                "ENSG00000142156",
                "ENSG00000152503",
                "ENSG00000149294",
                "ENSG00000171533",
                "ENSG00000166974",
                "ENSG00000174521",
                "ENSG00000164330",
                "ENSG00000137267",
                "ENSG00000151150",
                "ENSG00000100320",
                "ENSG00000181449",
                "ENSG00000118402",
                "ENSG00000188486",
                "ENSG00000131437",
                "ENSG00000113161",
                "ENSG00000050165",
                "ENSG00000082397",
                "ENSG00000117724",
                "ENSG00000171224",
                "ENSG00000143862",
                "ENSG00000162374",
                "ENSG00000134313",
                "ENSG00000104427",
                "ENSG00000221866",
                "ENSG00000198513",
                "ENSG00000197694",
                "ENSG00000115756",
                "ENSG00000126214",
                "ENSG00000131037",
                "ENSG00000131747",
                "ENSG00000131016",
                "ENSG00000139970",
                "ENSG00000135406",
                "ENSG00000159409",
                "ENSG00000113810",
                "ENSG00000169439",
                "ENSG00000177675",
                "ENSG00000137343"
            ],
            "expression_sum": {
                "class1": 30451.0,
                "class2": 15243.0
            },
            "ens_id": "ENSG00000116560",
            "uniprot_ids": [
                "P23246",
                "H0Y9K7",
                "H0Y9U2"
            ],
            "net_id": 0,
            "source_num": 5,
            "target_num": 39
        },
        "ENSG00000176887": {
            "id": "ENSG00000176887",
            "type": "TF",
            "symbol": "SOX11",
            "source": [
                "ENSG00000004487",
                "ENSG00000108055",
                "ENSG00000168214",
                "ENSG00000116560",
                "ENSG00000130522",
                "ENSG00000177606"
            ],
            "target": [
                "ENSG00000099901",
                "ENSG00000251562",
                "ENSG00000172757",
                "ENSG00000147274",
                "ENSG00000206503",
                "ENSG00000203667",
                "ENSG00000189403",
                "ENSG00000116560",
                "ENSG00000125691",
                "ENSG00000163939",
                "ENSG00000100823",
                "ENSG00000135387"
            ],
            "expression_sum": {
                "class1": 51157.0,
                "class2": 10806.0
            },
            "ens_id": "ENSG00000176887",
            "uniprot_ids": [
                "P35716"
            ],
            "net_id": 0,
            "source_num": 6,
            "target_num": 4
        },
        "ENSG00000162992": {
            "id": "ENSG00000162992",
            "type": "TF",
            "symbol": "NEUROD1",
            "source": [],
            "target": [
                "ENSG00000205542",
                "ENSG00000110492",
                "ENSG00000130558",
                "ENSG00000163191",
                "ENSG00000139289",
                "ENSG00000227716"
            ],
            "expression_sum": {
                "class1": 2538.0,
                "class2": 1057.0
            },
            "ens_id": "ENSG00000162992",
            "uniprot_ids": [
                "Q13562"
            ],
            "net_id": 0,
            "source_num": 0,
            "target_num": 3
        },
        "ENSG00000164754": {
            "id": "ENSG00000164754",
            "type": "TF",
            "symbol": "RAD21",
            "source": [
                "ENSG00000181449",
                "ENSG00000164330"
            ],
            "target": [
                "ENSG00000181449",
                "ENSG00000143401",
                "ENSG00000117399",
                "ENSG00000029993",
                "ENSG00000172020",
                "ENSG00000188486",
                "ENSG00000134057",
                "ENSG00000169856",
                "ENSG00000005249",
                "ENSG00000173207",
                "ENSG00000203760",
                "ENSG00000167654",
                "ENSG00000198830",
                "ENSG00000107105",
                "ENSG00000148798",
                "ENSG00000136108",
                "ENSG00000137804",
                "ENSG00000117724",
                "ENSG00000198910",
                "ENSG00000166257",
                "ENSG00000100167",
                "ENSG00000174021",
                "ENSG00000197959",
                "ENSG00000162374",
                "ENSG00000132872",
                "ENSG00000105409",
                "ENSG00000134313",
                "ENSG00000077152",
                "ENSG00000100526",
                "ENSG00000168393",
                "ENSG00000155980",
                "ENSG00000186868",
                "ENSG00000182481",
                "ENSG00000108106",
                "ENSG00000148773",
                "ENSG00000120802",
                "ENSG00000088325",
                "ENSG00000126950",
                "ENSG00000164611",
                "ENSG00000165434",
                "ENSG00000167553",
                "ENSG00000143702",
                "ENSG00000104435",
                "ENSG00000123975",
                "ENSG00000016082",
                "ENSG00000066279",
                "ENSG00000170312",
                "ENSG00000170540",
                "ENSG00000124194",
                "ENSG00000173267",
                "ENSG00000157456",
                "ENSG00000087258",
                "ENSG00000131747",
                "ENSG00000111897",
                "ENSG00000175063",
                "ENSG00000149294",
                "ENSG00000131016",
                "ENSG00000170091",
                "ENSG00000106268",
                "ENSG00000139970",
                "ENSG00000206579",
                "ENSG00000135406",
                "ENSG00000168824",
                "ENSG00000164330",
                "ENSG00000068615",
                "ENSG00000113810",
                "ENSG00000151150",
                "ENSG00000164104"
            ],
            "expression_sum": {
                "class1": 10941.0,
                "class2": 9238.0
            },
            "ens_id": "ENSG00000164754",
            "uniprot_ids": [
                "O60216",
                "E5RJK5",
                "E5RJW1",
                "E5RG18",
                "E5RFV8",
                "E5RI01",
                "E5RIN7",
                "E5RFZ5"
            ],
            "net_id": 0,
            "source_num": 2,
            "target_num": 31
        },
        "ENSG00000108001": {
            "id": "ENSG00000108001",
            "type": "TF",
            "symbol": "EBF3",
            "source": [],
            "target": [
                "ENSG00000067064",
                "ENSG00000135486",
                "ENSG00000162704",
                "ENSG00000138443",
                "ENSG00000111716",
                "ENSG00000171604",
                "ENSG00000139644",
                "ENSG00000175745",
                "ENSG00000182534",
                "ENSG00000065135",
                "ENSG00000134248",
                "ENSG00000089157",
                "ENSG00000168653",
                "ENSG00000131507",
                "ENSG00000189403",
                "ENSG00000142864",
                "ENSG00000075415",
                "ENSG00000184076",
                "ENSG00000144713",
                "ENSG00000197728",
                "ENSG00000163466",
                "ENSG00000099795",
                "ENSG00000167085",
                "ENSG00000272398",
                "ENSG00000135441",
                "ENSG00000122545",
                "ENSG00000189043",
                "ENSG00000099194",
                "ENSG00000277443",
                "ENSG00000101558",
                "ENSG00000148908",
                "ENSG00000172809",
                "ENSG00000074696",
                "ENSG00000154518",
                "ENSG00000196262",
                "ENSG00000172757",
                "ENSG00000160752",
                "ENSG00000143549",
                "ENSG00000138758",
                "ENSG00000213719",
                "ENSG00000133083",
                "ENSG00000113712",
                "ENSG00000126698",
                "ENSG00000198668",
                "ENSG00000108107",
                "ENSG00000146425",
                "ENSG00000241468",
                "ENSG00000153048",
                "ENSG00000104695",
                "ENSG00000075142",
                "ENSG00000106028",
                "ENSG00000115091",
                "ENSG00000100300",
                "ENSG00000149084",
                "ENSG00000127022",
                "ENSG00000108561",
                "ENSG00000197111",
                "ENSG00000101182",
                "ENSG00000140319",
                "ENSG00000163041",
                "ENSG00000243317",
                "ENSG00000178980",
                "ENSG00000167863",
                "ENSG00000197747",
                "ENSG00000182117",
                "ENSG00000065548",
                "ENSG00000108654",
                "ENSG00000162191",
                "ENSG00000070831",
                "ENSG00000122705",
                "ENSG00000143977",
                "ENSG00000184009",
                "ENSG00000075624",
                "ENSG00000183726",
                "ENSG00000108953",
                "ENSG00000094916",
                "ENSG00000227051",
                "ENSG00000164713",
                "ENSG00000163931",
                "ENSG00000111775",
                "ENSG00000136930",
                "ENSG00000131508",
                "ENSG00000188612",
                "ENSG00000196781",
                "ENSG00000080824",
                "ENSG00000116288",
                "ENSG00000153187",
                "ENSG00000142937",
                "ENSG00000115738",
                "ENSG00000104388",
                "ENSG00000047849",
                "ENSG00000167996",
                "ENSG00000150753",
                "ENSG00000132963",
                "ENSG00000170144",
                "ENSG00000042753",
                "ENSG00000138385",
                "ENSG00000151914",
                "ENSG00000182718",
                "ENSG00000143198",
                "ENSG00000122566",
                "ENSG00000165119"
            ],
            "expression_sum": {
                "class1": 4199.0,
                "class2": 731.0
            },
            "ens_id": "ENSG00000108001",
            "uniprot_ids": [
                "Q9H4W6",
                "H0Y3W9"
            ],
            "net_id": 0,
            "source_num": 0,
            "target_num": 30
        },
        "ENSG00000189403": {
            "id": "ENSG00000189403",
            "type": "TF",
            "symbol": "HMGB1",
            "source": [
                "ENSG00000181449",
                "ENSG00000176887",
                "ENSG00000109132",
                "ENSG00000167081",
                "ENSG00000118263",
                "ENSG00000164330",
                "ENSG00000108001"
            ],
            "target": [
                "ENSG00000164176",
                "ENSG00000162545",
                "ENSG00000099864",
                "ENSG00000161082",
                "ENSG00000087085",
                "ENSG00000161682",
                "ENSG00000107105",
                "ENSG00000050165",
                "ENSG00000130540",
                "ENSG00000086065",
                "ENSG00000160271",
                "ENSG00000099250",
                "ENSG00000106976",
                "ENSG00000084628",
                "ENSG00000171224",
                "ENSG00000128872",
                "ENSG00000124302",
                "ENSG00000100167",
                "ENSG00000188322",
                "ENSG00000108828",
                "ENSG00000221866",
                "ENSG00000168393",
                "ENSG00000198513",
                "ENSG00000186868",
                "ENSG00000115756",
                "ENSG00000160213",
                "ENSG00000143702",
                "ENSG00000103154",
                "ENSG00000171033",
                "ENSG00000105696",
                "ENSG00000118200",
                "ENSG00000164104",
                "ENSG00000087258",
                "ENSG00000072062",
                "ENSG00000272398",
                "ENSG00000206579",
                "ENSG00000167081",
                "ENSG00000141404",
                "ENSG00000113810",
                "ENSG00000137267",
                "ENSG00000100320",
                "ENSG00000203879",
                "ENSG00000134986"
            ],
            "expression_sum": {
                "class1": 62780.0,
                "class2": 78276.0
            },
            "ens_id": "ENSG00000189403",
            "uniprot_ids": [
                "P09429",
                "Q5T7C4"
            ],
            "net_id": 0,
            "source_num": 7,
            "target_num": 4
        },
        "ENSG00000137203": {
            "id": "ENSG00000137203",
            "type": "TF",
            "symbol": "TFAP2A",
            "source": [
                "ENSG00000028277",
                "ENSG00000134138"
            ],
            "target": [
                "ENSG00000086061",
                "ENSG00000067064",
                "ENSG00000167508",
                "ENSG00000115350",
                "ENSG00000128989",
                "ENSG00000172809",
                "ENSG00000135334",
                "ENSG00000213585",
                "ENSG00000177519",
                "ENSG00000184226",
                "ENSG00000187398",
                "ENSG00000099622",
                "ENSG00000167658",
                "ENSG00000163041",
                "ENSG00000109084",
                "ENSG00000165443",
                "ENSG00000101856",
                "ENSG00000070669",
                "ENSG00000114784",
                "ENSG00000165637",
                "ENSG00000118432",
                "ENSG00000156976",
                "ENSG00000134138",
                "ENSG00000168028",
                "ENSG00000140538",
                "ENSG00000131143",
                "ENSG00000134717",
                "ENSG00000102144",
                "ENSG00000182287",
                "ENSG00000072832",
                "ENSG00000220205",
                "ENSG00000104549",
                "ENSG00000160752",
                "ENSG00000254682",
                "ENSG00000115738",
                "ENSG00000079459",
                "ENSG00000224032",
                "ENSG00000105993",
                "ENSG00000165434",
                "ENSG00000186187",
                "ENSG00000028277",
                "ENSG00000182667",
                "ENSG00000162490",
                "ENSG00000171033",
                "ENSG00000164096",
                "ENSG00000182670",
                "ENSG00000052802",
                "ENSG00000130558",
                "ENSG00000133872",
                "ENSG00000166913",
                "ENSG00000091664",
                "ENSG00000105088",
                "ENSG00000134308",
                "ENSG00000105649",
                "ENSG00000118816",
                "ENSG00000185008",
                "ENSG00000152128",
                "ENSG00000170091",
                "ENSG00000183775",
                "ENSG00000078328",
                "ENSG00000101745",
                "ENSG00000105220",
                "ENSG00000123983",
                "ENSG00000164326",
                "ENSG00000170899",
                "ENSG00000169862",
                "ENSG00000137267",
                "ENSG00000104765",
                "ENSG00000100129",
                "ENSG00000149084",
                "ENSG00000128050",
                "ENSG00000152154",
                "ENSG00000149926",
                "ENSG00000139767"
            ],
            "expression_sum": {
                "class1": 8598.0,
                "class2": 91.0
            },
            "ens_id": "ENSG00000137203",
            "uniprot_ids": [
                "P05549",
                "A0A6E1XE14",
                "C1K3N0",
                "C9J6N8",
                "F8WDC8",
                "C9JXZ2",
                "H7C4N4",
                "H7C5E5",
                "F8WEX2"
            ],
            "net_id": 0,
            "source_num": 2,
            "target_num": 16
        },
        "ENSG00000028277": {
            "id": "ENSG00000028277",
            "type": "TF",
            "symbol": "POU2F2",
            "source": [
                "ENSG00000119547",
                "ENSG00000137203"
            ],
            "target": [
                "ENSG00000176788",
                "ENSG00000171033",
                "ENSG00000105409",
                "ENSG00000072832",
                "ENSG00000137267",
                "ENSG00000137203",
                "ENSG00000129355",
                "ENSG00000105649",
                "ENSG00000170091"
            ],
            "expression_sum": {
                "class1": 8686.0,
                "class2": 988.0
            },
            "ens_id": "ENSG00000028277",
            "uniprot_ids": [
                "P09086",
                "U3KQ96",
                "M0R1I6",
                "H0YNG1",
                "A0A1B0GW97",
                "U3KQN1",
                "H0YLL6",
                "H0YKT2",
                "A0A1B0GUT9",
                "A0A1B0GV81",
                "E9PR37",
                "H0YNW1",
                "A0A0D9SF23",
                "B5ME60"
            ],
            "net_id": 0,
            "source_num": 2,
            "target_num": 4
        },
        "ENSG00000006128": {
            "id": "ENSG00000006128",
            "type": "TF",
            "symbol": "TAC1",
            "source": [],
            "target": [
                "ENSG00000277586",
                "ENSG00000172209"
            ],
            "expression_sum": {
                "class1": 43069.0,
                "class2": 15659.0
            },
            "ens_id": "ENSG00000006128",
            "uniprot_ids": [
                "P20366"
            ],
            "net_id": 0,
            "source_num": 0,
            "target_num": 1
        },
        "ENSG00000095794": {
            "id": "ENSG00000095794",
            "type": "TF",
            "symbol": "CREM",
            "source": [],
            "target": [
                "ENSG00000109610",
                "ENSG00000145730",
                "ENSG00000128564",
                "ENSG00000182054",
                "ENSG00000166562",
                "ENSG00000137331",
                "ENSG00000135346",
                "ENSG00000169347",
                "ENSG00000111057"
            ],
            "expression_sum": {
                "class1": 5192.0,
                "class2": 4210.0
            },
            "ens_id": "ENSG00000095794",
            "uniprot_ids": [
                "Q03060",
                "E9PAR2",
                "C9IYM9",
                "A0A0A0MSU8",
                "C9J5A7",
                "E9PB41",
                "C9J2J0",
                "C9K092",
                "H7C4X0",
                "R4GN75",
                "F8WB03",
                "J3KR46",
                "F8WDF2",
                "G5E998"
            ],
            "net_id": 0,
            "source_num": 0,
            "target_num": 2
        },
        "ENSG00000118263": {
            "id": "ENSG00000118263",
            "type": "TF",
            "symbol": "KLF7",
            "source": [
                "ENSG00000100811",
                "ENSG00000106261",
                "ENSG00000016082",
                "ENSG00000148516",
                "ENSG00000103495",
                "ENSG00000164330",
                "ENSG00000108055",
                "ENSG00000067955",
                "ENSG00000116560",
                "ENSG00000196591",
                "ENSG00000128272"
            ],
            "target": [
                "ENSG00000078668",
                "ENSG00000173272",
                "ENSG00000139644",
                "ENSG00000197756",
                "ENSG00000185651",
                "ENSG00000182004",
                "ENSG00000065135",
                "ENSG00000134248",
                "ENSG00000103495",
                "ENSG00000010270",
                "ENSG00000189403",
                "ENSG00000108424",
                "ENSG00000156261",
                "ENSG00000101752",
                "ENSG00000197728",
                "ENSG00000166913",
                "ENSG00000079785",
                "ENSG00000173812",
                "ENSG00000099795",
                "ENSG00000118816",
                "ENSG00000135441",
                "ENSG00000104671",
                "ENSG00000215712",
                "ENSG00000115310",
                "ENSG00000113742",
                "ENSG00000159335",
                "ENSG00000108518",
                "ENSG00000145912",
                "ENSG00000278311",
                "ENSG00000172809",
                "ENSG00000155366",
                "ENSG00000172757",
                "ENSG00000168092",
                "ENSG00000131236",
                "ENSG00000115286",
                "ENSG00000077721",
                "ENSG00000119185",
                "ENSG00000141367",
                "ENSG00000144381",
                "ENSG00000115541",
                "ENSG00000133083",
                "ENSG00000162244",
                "ENSG00000133703",
                "ENSG00000135900",
                "ENSG00000034510",
                "ENSG00000146425",
                "ENSG00000103769",
                "ENSG00000241468",
                "ENSG00000104695",
                "ENSG00000141404",
                "ENSG00000273841",
                "ENSG00000106028",
                "ENSG00000113575",
                "ENSG00000126934",
                "ENSG00000145592",
                "ENSG00000121022",
                "ENSG00000175334",
                "ENSG00000092020",
                "ENSG00000117450",
                "ENSG00000184007",
                "ENSG00000184220",
                "ENSG00000178980",
                "ENSG00000164346",
                "ENSG00000167863",
                "ENSG00000150991",
                "ENSG00000169100",
                "ENSG00000108106",
                "ENSG00000070831",
                "ENSG00000092964",
                "ENSG00000171497",
                "ENSG00000110958",
                "ENSG00000116251",
                "ENSG00000025293",
                "ENSG00000175792",
                "ENSG00000102317",
                "ENSG00000184009",
                "ENSG00000124357",
                "ENSG00000125818",
                "ENSG00000100380",
                "ENSG00000104643",
                "ENSG00000167552",
                "ENSG00000137267",
                "ENSG00000132305",
                "ENSG00000100320",
                "ENSG00000112701",
                "ENSG00000099901",
                "ENSG00000164985",
                "ENSG00000128989",
                "ENSG00000239672",
                "ENSG00000131469",
                "ENSG00000164587",
                "ENSG00000142937",
                "ENSG00000158710",
                "ENSG00000136938",
                "ENSG00000108826",
                "ENSG00000044115",
                "ENSG00000136238",
                "ENSG00000197694",
                "ENSG00000143933",
                "ENSG00000182154",
                "ENSG00000221914",
                "ENSG00000126214",
                "ENSG00000172354",
                "ENSG00000168522",
                "ENSG00000152082",
                "ENSG00000177600",
                "ENSG00000151914",
                "ENSG00000174456",
                "ENSG00000166946",
                "ENSG00000221983",
                "ENSG00000135406",
                "ENSG00000174886",
                "ENSG00000137710",
                "ENSG00000106244",
                "ENSG00000092841",
                "ENSG00000165119"
            ],
            "expression_sum": {
                "class1": 7577.0,
                "class2": 1532.0
            },
            "ens_id": "ENSG00000118263",
            "uniprot_ids": [
                "O75840",
                "E7EQY2",
                "C9JX20",
                "E7EUU0",
                "E9PHC9"
            ],
            "net_id": 0,
            "source_num": 11,
            "target_num": 16
        },
        "ENSG00000164093": {
            "id": "ENSG00000164093",
            "type": "TF",
            "symbol": "PITX2",
            "source": [
                "ENSG00000100811",
                "ENSG00000004487",
                "ENSG00000103495",
                "ENSG00000181722",
                "ENSG00000108055",
                "ENSG00000116560",
                "ENSG00000168214",
                "ENSG00000196591",
                "ENSG00000116044",
                "ENSG00000119547",
                "ENSG00000177606"
            ],
            "target": [
                "ENSG00000244879",
                "ENSG00000132475",
                "ENSG00000256537",
                "ENSG00000090487",
                "ENSG00000177889",
                "ENSG00000254999",
                "ENSG00000170776",
                "ENSG00000162409",
                "ENSG00000119950",
                "ENSG00000175416",
                "ENSG00000185633",
                "ENSG00000134308",
                "ENSG00000112695",
                "ENSG00000164631"
            ],
            "expression_sum": {
                "class1": 4850.0,
                "class2": 13703.0
            },
            "ens_id": "ENSG00000164093",
            "uniprot_ids": [
                "Q99697",
                "D6RBG8",
                "U3KQ81",
                "D6RFI4"
            ],
            "net_id": 0,
            "source_num": 11,
            "target_num": 3
        },
        "ENSG00000164683": {
            "id": "ENSG00000164683",
            "type": "TF",
            "symbol": "HEY1",
            "source": [
                "ENSG00000181449"
            ],
            "target": [
                "ENSG00000104529",
                "ENSG00000137573",
                "ENSG00000118181",
                "ENSG00000152558",
                "ENSG00000154640",
                "ENSG00000166710",
                "ENSG00000129355",
                "ENSG00000102265",
                "ENSG00000105854",
                "ENSG00000084207",
                "ENSG00000126458",
                "ENSG00000166165",
                "ENSG00000135821",
                "ENSG00000245532",
                "ENSG00000142534",
                "ENSG00000265972",
                "ENSG00000103260",
                "ENSG00000117632",
                "ENSG00000130066",
                "ENSG00000229117",
                "ENSG00000143545",
                "ENSG00000135404",
                "ENSG00000221869",
                "ENSG00000150687",
                "ENSG00000147403",
                "ENSG00000187193",
                "ENSG00000144560",
                "ENSG00000100316",
                "ENSG00000155366",
                "ENSG00000133112",
                "ENSG00000131981",
                "ENSG00000125691",
                "ENSG00000122884",
                "ENSG00000213719",
                "ENSG00000181744",
                "ENSG00000233927",
                "ENSG00000129521",
                "ENSG00000108107",
                "ENSG00000138326",
                "ENSG00000156508",
                "ENSG00000116044",
                "ENSG00000153944",
                "ENSG00000185650",
                "ENSG00000110092",
                "ENSG00000145592",
                "ENSG00000162496",
                "ENSG00000142676",
                "ENSG00000119655",
                "ENSG00000148180",
                "ENSG00000177954",
                "ENSG00000186868",
                "ENSG00000185088",
                "ENSG00000163110",
                "ENSG00000051523",
                "ENSG00000107438",
                "ENSG00000003436",
                "ENSG00000063177",
                "ENSG00000135046",
                "ENSG00000120742",
                "ENSG00000105372",
                "ENSG00000137267",
                "ENSG00000170889",
                "ENSG00000143819",
                "ENSG00000008988",
                "ENSG00000135390",
                "ENSG00000008394",
                "ENSG00000154080",
                "ENSG00000129951",
                "ENSG00000137154",
                "ENSG00000060138",
                "ENSG00000158710",
                "ENSG00000174021",
                "ENSG00000168938",
                "ENSG00000164111",
                "ENSG00000109861",
                "ENSG00000158195",
                "ENSG00000140264",
                "ENSG00000177600",
                "ENSG00000166441",
                "ENSG00000134419",
                "ENSG00000149273",
                "ENSG00000152518",
                "ENSG00000182718",
                "ENSG00000221983",
                "ENSG00000133639",
                "ENSG00000109062",
                "ENSG00000185624",
                "ENSG00000138640",
                "ENSG00000018408",
                "ENSG00000114115",
                "ENSG00000089009"
            ],
            "expression_sum": {
                "class1": 2632.0,
                "class2": 5970.0
            },
            "ens_id": "ENSG00000164683",
            "uniprot_ids": [
                "Q9Y5J3",
                "A0A6I8PRS3",
                "B4DEI9",
                "A0A6I8PRC2",
                "E5RHK6",
                "A0A6I8PIY0",
                "A0A6I8PU82"
            ],
            "net_id": 0,
            "source_num": 1,
            "target_num": 20
        },
        "ENSG00000122691": {
            "id": "ENSG00000122691",
            "type": "TF",
            "symbol": "TWIST1",
            "source": [],
            "target": [
                "ENSG00000141756",
                "ENSG00000175745",
                "ENSG00000078018",
                "ENSG00000122641",
                "ENSG00000077782",
                "ENSG00000070961",
                "ENSG00000139329",
                "ENSG00000185420",
                "ENSG00000185650",
                "ENSG00000117122",
                "ENSG00000105854",
                "ENSG00000163191",
                "ENSG00000168542",
                "ENSG00000161203",
                "ENSG00000131116",
                "ENSG00000178531",
                "ENSG00000087460",
                "ENSG00000128606",
                "ENSG00000166482",
                "ENSG00000106819",
                "ENSG00000182809",
                "ENSG00000087245",
                "ENSG00000111341",
                "ENSG00000120708",
                "ENSG00000142089",
                "ENSG00000011465",
                "ENSG00000159335",
                "ENSG00000018408",
                "ENSG00000189043",
                "ENSG00000114115",
                "ENSG00000134986",
                "ENSG00000104870"
            ],
            "expression_sum": {
                "class1": 1346.0,
                "class2": 2882.0
            },
            "ens_id": "ENSG00000122691",
            "uniprot_ids": [
                "Q15672",
                "H7BY00",
                "H7C4D7"
            ],
            "net_id": 0,
            "source_num": 0,
            "target_num": 7
        },
        "ENSG00000169136": {
            "id": "ENSG00000169136",
            "type": "TF",
            "symbol": "ATF5",
            "source": [
                "ENSG00000150347",
                "ENSG00000164330"
            ],
            "target": [
                "ENSG00000115310",
                "ENSG00000075142"
            ],
            "expression_sum": {
                "class1": 18613.0,
                "class2": 7541.0
            },
            "ens_id": "ENSG00000169136",
            "uniprot_ids": [
                "Q9Y2D1",
                "M0R040",
                "M0R0D5",
                "M0QZD2",
                "A0A6Q8PGU9"
            ],
            "net_id": 0,
            "source_num": 2,
            "target_num": 2
        },
        "ENSG00000150347": {
            "id": "ENSG00000150347",
            "type": "TF",
            "symbol": "ARID5B",
            "source": [],
            "target": [
                "ENSG00000144713",
                "ENSG00000089157",
                "ENSG00000162244",
                "ENSG00000072110",
                "ENSG00000169136",
                "ENSG00000115268",
                "ENSG00000105193",
                "ENSG00000063177",
                "ENSG00000143933",
                "ENSG00000067225",
                "ENSG00000122026",
                "ENSG00000140941"
            ],
            "expression_sum": {
                "class1": 1993.0,
                "class2": 5771.0
            },
            "ens_id": "ENSG00000150347",
            "uniprot_ids": [
                "Q14865",
                "A0A2R8Y5F2",
                "A0A7P0TAD2"
            ],
            "net_id": 0,
            "source_num": 0,
            "target_num": 4
        }
    },
    "grps": {
        "ENSG00000160113_ENSG00000100219": {
            "id": "ENSG00000160113_ENSG00000100219",
            "type": "Standard",
            "score": 0.11149087516331603,
            "reversable": true,
            "correlations": {
                "class1": 0.2900114100966345,
                "class2": 0.6055192079666344
            },
            "regulatory_source": "ENSG00000160113",
            "regulatory_target": "ENSG00000100219",
            "net_id": 0
        },
        "ENSG00000167081_ENSG00000134138": {
            "id": "ENSG00000167081_ENSG00000134138",
            "type": "Mix",
            "score": 15.283601865989178,
            "reversable": true,
            "correlations": {
                "class1": 0.5296672475883527,
                "class2": 0.0777948803454681
            },
            "regulatory_source": "ENSG00000167081",
            "regulatory_target": "ENSG00000134138",
            "net_id": 0
        },
        "ENSG00000143995_ENSG00000134138": {
            "id": "ENSG00000143995_ENSG00000134138",
            "type": "Standard",
            "score": 1.49513127721154,
            "reversable": true,
            "correlations": {
                "class1": 0.563417452388825,
                "class2": 0.10004723602000329
            },
            "regulatory_source": "ENSG00000143995",
            "regulatory_target": "ENSG00000134138",
            "net_id": 0
        },
        "ENSG00000170365_ENSG00000016082": {
            "id": "ENSG00000170365_ENSG00000016082",
            "type": "Bridge",
            "score": 0,
            "reversable": true,
            "correlations": {
                "class1": 0.26309107059678555,
                "class2": 0.0
            },
            "regulatory_source": "ENSG00000170365",
            "regulatory_target": "ENSG00000016082",
            "net_id": 0
        },
        "ENSG00000100811_ENSG00000016082": {
            "id": "ENSG00000100811_ENSG00000016082",
            "type": "Bridge",
            "score": 0,
            "reversable": false,
            "correlations": {
                "class1": 0.2633993400240361,
                "class2": 0.0
            },
            "regulatory_source": "ENSG00000100811",
            "regulatory_target": "ENSG00000016082",
            "net_id": 0
        },
        "ENSG00000130726_ENSG00000016082": {
            "id": "ENSG00000130726_ENSG00000016082",
            "type": "Bridge",
            "score": 0,
            "reversable": false,
            "correlations": {
                "class1": 0.2146665643105613,
                "class2": 0.0
            },
            "regulatory_source": "ENSG00000130726",
            "regulatory_target": "ENSG00000016082",
            "net_id": 0
        },
        "ENSG00000109132_ENSG00000016082": {
            "id": "ENSG00000109132_ENSG00000016082",
            "type": "Bridge",
            "score": 0,
            "reversable": true,
            "correlations": {
                "class1": 0.5178363684232455,
                "class2": 0.0
            },
            "regulatory_source": "ENSG00000109132",
            "regulatory_target": "ENSG00000016082",
            "net_id": 0
        },
        "ENSG00000153147_ENSG00000016082": {
            "id": "ENSG00000153147_ENSG00000016082",
            "type": "Bridge",
            "score": 0,
            "reversable": false,
            "correlations": {
                "class1": 0.20518642367813436,
                "class2": 0.0
            },
            "regulatory_source": "ENSG00000153147",
            "regulatory_target": "ENSG00000016082",
            "net_id": 0
        },
        "ENSG00000198911_ENSG00000016082": {
            "id": "ENSG00000198911_ENSG00000016082",
            "type": "Bridge",
            "score": 0,
            "reversable": false,
            "correlations": {
                "class1": 0.2526636685700158,
                "class2": 0.0
            },
            "regulatory_source": "ENSG00000198911",
            "regulatory_target": "ENSG00000016082",
            "net_id": 0
        },
        "ENSG00000196591_ENSG00000016082": {
            "id": "ENSG00000196591_ENSG00000016082",
            "type": "Bridge",
            "score": 0,
            "reversable": false,
            "correlations": {
                "class1": 0.3387403904681899,
                "class2": 0.026042353626923153
            },
            "regulatory_source": "ENSG00000196591",
            "regulatory_target": "ENSG00000016082",
            "net_id": 0
        },
        "ENSG00000167081_ENSG00000105698": {
            "id": "ENSG00000167081_ENSG00000105698",
            "type": "Bridge",
            "score": 0,
            "reversable": true,
            "correlations": {
                "class1": 0.2553091410364716,
                "class2": 0.0
            },
            "regulatory_source": "ENSG00000167081",
            "regulatory_target": "ENSG00000105698",
            "net_id": 0
        },
        "ENSG00000213015_ENSG00000167081": {
            "id": "ENSG00000213015_ENSG00000167081",
            "type": "Bridge",
            "score": 0,
            "reversable": true,
            "correlations": {
                "class1": 0.2642361189755856,
                "class2": 0.021055737640306004
            },
            "regulatory_source": "ENSG00000167081",
            "regulatory_target": "ENSG00000213015",
            "net_id": 0
        },
        "ENSG00000167081_ENSG00000119950": {
            "id": "ENSG00000167081_ENSG00000119950",
            "type": "Bridge",
            "score": 0,
            "reversable": true,
            "correlations": {
                "class1": 0.2740979624559404,
                "class2": 0.03094223155896498
            },
            "regulatory_source": "ENSG00000167081",
            "regulatory_target": "ENSG00000119950",
            "net_id": 0
        },
        "ENSG00000167081_ENSG00000092199": {
            "id": "ENSG00000167081_ENSG00000092199",
            "type": "Bridge",
            "score": 0,
            "reversable": true,
            "correlations": {
                "class1": 0.3046820693507447,
                "class2": 0.0482197572190567
            },
            "regulatory_source": "ENSG00000167081",
            "regulatory_target": "ENSG00000092199",
            "net_id": 0
        },
        "ENSG00000164330_ENSG00000106261": {
            "id": "ENSG00000164330_ENSG00000106261",
            "type": "Bridge",
            "score": 0,
            "reversable": false,
            "correlations": {
                "class1": 0.29651699852419433,
                "class2": 0.03409191380708228
            },
            "regulatory_source": "ENSG00000106261",
            "regulatory_target": "ENSG00000164330",
            "net_id": 0
        },
        "ENSG00000164330_ENSG00000148516": {
            "id": "ENSG00000164330_ENSG00000148516",
            "type": "Bridge",
            "score": 0,
            "reversable": true,
            "correlations": {
                "class1": 0.2938579427554105,
                "class2": 0.07730901020342613
            },
            "regulatory_source": "ENSG00000164330",
            "regulatory_target": "ENSG00000148516",
            "net_id": 0
        },
        "ENSG00000164330_ENSG00000143799": {
            "id": "ENSG00000164330_ENSG00000143799",
            "type": "Bridge",
            "score": 0,
            "reversable": true,
            "correlations": {
                "class1": 0.3299685750903966,
                "class2": 0.0218934858198555
            },
            "regulatory_source": "ENSG00000143799",
            "regulatory_target": "ENSG00000164330",
            "net_id": 0
        },
        "ENSG00000164330_ENSG00000128272": {
            "id": "ENSG00000164330_ENSG00000128272",
            "type": "Bridge",
            "score": 0,
            "reversable": false,
            "correlations": {
                "class1": 0.31515569804592886,
                "class2": 0.04922327354615721
            },
            "regulatory_source": "ENSG00000128272",
            "regulatory_target": "ENSG00000164330",
            "net_id": 0
        },
        "ENSG00000164330_ENSG00000163848": {
            "id": "ENSG00000164330_ENSG00000163848",
            "type": "Bridge",
            "score": 0,
            "reversable": true,
            "correlations": {
                "class1": 0.26569635185838386,
                "class2": 0.05372488672004042
            },
            "regulatory_source": "ENSG00000163848",
            "regulatory_target": "ENSG00000164330",
            "net_id": 0
        },
        "ENSG00000165512_ENSG00000164330": {
            "id": "ENSG00000165512_ENSG00000164330",
            "type": "Bridge",
            "score": 0,
            "reversable": true,
            "correlations": {
                "class1": 0.2920799643181508,
                "class2": 0.0
            },
            "regulatory_source": "ENSG00000164330",
            "regulatory_target": "ENSG00000165512",
            "net_id": 0
        },
        "ENSG00000164330_ENSG00000067955": {
            "id": "ENSG00000164330_ENSG00000067955",
            "type": "Bridge",
            "score": 0,
            "reversable": true,
            "correlations": {
                "class1": 0.34600214119145356,
                "class2": 0.0
            },
            "regulatory_source": "ENSG00000164330",
            "regulatory_target": "ENSG00000067955",
            "net_id": 0
        },
        "ENSG00000162599_ENSG00000134138": {
            "id": "ENSG00000162599_ENSG00000134138",
            "type": "Bridge",
            "score": 0,
            "reversable": false,
            "correlations": {
                "class1": -0.05070187391377254,
                "class2": 0.19593784741325468
            },
            "regulatory_source": "ENSG00000162599",
            "regulatory_target": "ENSG00000134138",
            "net_id": 0
        },
        "ENSG00000168610_ENSG00000134138": {
            "id": "ENSG00000168610_ENSG00000134138",
            "type": "Bridge",
            "score": 0,
            "reversable": false,
            "correlations": {
                "class1": 0.05311907949346681,
                "class2": 0.2584405399581128
            },
            "regulatory_source": "ENSG00000168610",
            "regulatory_target": "ENSG00000134138",
            "net_id": 0
        },
        "ENSG00000134138_ENSG00000114861": {
            "id": "ENSG00000134138_ENSG00000114861",
            "type": "Bridge",
            "score": 0,
            "reversable": true,
            "correlations": {
                "class1": -0.05542300318111832,
                "class2": 0.28616522030659164
            },
            "regulatory_source": "ENSG00000114861",
            "regulatory_target": "ENSG00000134138",
            "net_id": 0
        },
        "ENSG00000221869_ENSG00000134138": {
            "id": "ENSG00000221869_ENSG00000134138",
            "type": "Bridge",
            "score": 0,
            "reversable": false,
            "correlations": {
                "class1": -0.06807576284944307,
                "class2": 0.19725863553053494
            },
            "regulatory_source": "ENSG00000221869",
            "regulatory_target": "ENSG00000134138",
            "net_id": 0
        },
        "ENSG00000168214_ENSG00000157557": {
            "id": "ENSG00000168214_ENSG00000157557",
            "type": "Bridge",
            "score": 0,
            "reversable": true,
            "correlations": {
                "class1": 0.1097965776503432,
                "class2": 0.36233005421537706
            },
            "regulatory_source": "ENSG00000168214",
            "regulatory_target": "ENSG00000157557",
            "net_id": 0
        },
        "ENSG00000181722_ENSG00000160113": {
            "id": "ENSG00000181722_ENSG00000160113",
            "type": "Bridge",
            "score": 0,
            "reversable": true,
            "correlations": {
                "class1": 0.15507753018920503,
                "class2": 0.35995214402359216
            },
            "regulatory_source": "ENSG00000181722",
            "regulatory_target": "ENSG00000160113",
            "net_id": 0
        },
        "ENSG00000177606_ENSG00000100219": {
            "id": "ENSG00000177606_ENSG00000100219",
            "type": "Bridge",
            "score": 0,
            "reversable": true,
            "correlations": {
                "class1": 0.05834601723672387,
                "class2": 0.278417872419843
            },
            "regulatory_source": "ENSG00000177606",
            "regulatory_target": "ENSG00000100219",
            "net_id": 0
        },
        "ENSG00000163435_ENSG00000124766": {
            "id": "ENSG00000163435_ENSG00000124766",
            "type": "Bridge",
            "score": 0,
            "reversable": false,
            "correlations": {
                "class1": 0.023234676717406602,
                "class2": 0.2379235065637544
            },
            "regulatory_source": "ENSG00000124766",
            "regulatory_target": "ENSG00000163435",
            "net_id": 0
        },
        "ENSG00000163435_ENSG00000116044": {
            "id": "ENSG00000163435_ENSG00000116044",
            "type": "Bridge",
            "score": 0,
            "reversable": true,
            "correlations": {
                "class1": 0.14293663351423422,
                "class2": 0.3469941623027341
            },
            "regulatory_source": "ENSG00000163435",
            "regulatory_target": "ENSG00000116044",
            "net_id": 0
        },
        "ENSG00000181449_ENSG00000128610": {
            "id": "ENSG00000181449_ENSG00000128610",
            "type": "Bridge",
            "score": 0,
            "reversable": true,
            "correlations": {
                "class1": 0.3506118280161127,
                "class2": 0.08656649786604453
            },
            "regulatory_source": "ENSG00000181449",
            "regulatory_target": "ENSG00000128610",
            "net_id": 0
        },
        "ENSG00000181449_ENSG00000130522": {
            "id": "ENSG00000181449_ENSG00000130522",
            "type": "Bridge",
            "score": 0,
            "reversable": true,
            "correlations": {
                "class1": 0.035396974547115524,
                "class2": 0.32243196764249643
            },
            "regulatory_source": "ENSG00000181449",
            "regulatory_target": "ENSG00000130522",
            "net_id": 0
        },
        "ENSG00000167081_ENSG00000119547": {
            "id": "ENSG00000167081_ENSG00000119547",
            "type": "Bridge",
            "score": 0,
            "reversable": false,
            "correlations": {
                "class1": 0.4106535765802696,
                "class2": 0.13800480004434365
            },
            "regulatory_source": "ENSG00000167081",
            "regulatory_target": "ENSG00000119547",
            "net_id": 0
        },
        "ENSG00000175745_ENSG00000164330": {
            "id": "ENSG00000175745_ENSG00000164330",
            "type": "Bridge",
            "score": 0,
            "reversable": false,
            "correlations": {
                "class1": 0.27751713944166506,
                "class2": 0.047820812197996825
            },
            "regulatory_source": "ENSG00000164330",
            "regulatory_target": "ENSG00000175745",
            "net_id": 0
        },
        "ENSG00000134138_ENSG00000119547": {
            "id": "ENSG00000134138_ENSG00000119547",
            "type": "Bridge",
            "score": 0,
            "reversable": true,
            "correlations": {
                "class1": 0.40846067566590705,
                "class2": 0.19250857819247655
            },
            "regulatory_source": "ENSG00000134138",
            "regulatory_target": "ENSG00000119547",
            "net_id": 0
        },
        "ENSG00000175745_ENSG00000134138": {
            "id": "ENSG00000175745_ENSG00000134138",
            "type": "Bridge",
            "score": 0,
            "reversable": false,
            "correlations": {
                "class1": 0.45380280691923197,
                "class2": 0.17095140842378917
            },
            "regulatory_source": "ENSG00000175745",
            "regulatory_target": "ENSG00000134138",
            "net_id": 0
        },
        "ENSG00000143995_ENSG00000119547": {
            "id": "ENSG00000143995_ENSG00000119547",
            "type": "Bridge",
            "score": 0,
            "reversable": false,
            "correlations": {
                "class1": 0.30442947516852986,
                "class2": 0.0
            },
            "regulatory_source": "ENSG00000119547",
            "regulatory_target": "ENSG00000143995",
            "net_id": 0
        },
        "ENSG00000175745_ENSG00000143995": {
            "id": "ENSG00000175745_ENSG00000143995",
            "type": "Bridge",
            "score": 0,
            "reversable": true,
            "correlations": {
                "class1": 0.36794476219426137,
                "class2": 0.1433253657393741
            },
            "regulatory_source": "ENSG00000143995",
            "regulatory_target": "ENSG00000175745",
            "net_id": 0
        },
        "ENSG00000148516_ENSG00000119547": {
            "id": "ENSG00000148516_ENSG00000119547",
            "type": "Bridge",
            "score": 0,
            "reversable": true,
            "correlations": {
                "class1": 0.25571824823061107,
                "class2": 0.0
            },
            "regulatory_source": "ENSG00000148516",
            "regulatory_target": "ENSG00000119547",
            "net_id": 0
        },
        "ENSG00000175745_ENSG00000162599": {
            "id": "ENSG00000175745_ENSG00000162599",
            "type": "Bridge",
            "score": 0,
            "reversable": true,
            "correlations": {
                "class1": 0.045407166417237285,
                "class2": 0.3503279791591227
            },
            "regulatory_source": "ENSG00000162599",
            "regulatory_target": "ENSG00000175745",
            "net_id": 0
        },
        "ENSG00000119547_ENSG00000114861": {
            "id": "ENSG00000119547_ENSG00000114861",
            "type": "Bridge",
            "score": 0,
            "reversable": true,
            "correlations": {
                "class1": 0.0,
                "class2": 0.20980690705336055
            },
            "regulatory_source": "ENSG00000114861",
            "regulatory_target": "ENSG00000119547",
            "net_id": 0
        },
        "ENSG00000157557_ENSG00000119547": {
            "id": "ENSG00000157557_ENSG00000119547",
            "type": "Bridge",
            "score": 0,
            "reversable": true,
            "correlations": {
                "class1": 0.0,
                "class2": 0.27287168558931296
            },
            "regulatory_source": "ENSG00000119547",
            "regulatory_target": "ENSG00000157557",
            "net_id": 0
        },
        "ENSG00000175745_ENSG00000119547": {
            "id": "ENSG00000175745_ENSG00000119547",
            "type": "Mix",
            "score": 16.074801420983203,
            "reversable": false,
            "correlations": {
                "class1": 0.3963438653120717,
                "class2": 0.0
            },
            "regulatory_source": "ENSG00000119547",
            "regulatory_target": "ENSG00000175745",
            "net_id": 0
        },
        "ENSG00000185551_ENSG00000119547": {
            "id": "ENSG00000185551_ENSG00000119547",
            "type": "Bridge",
            "score": 0,
            "reversable": true,
            "correlations": {
                "class1": 0.20974200724484765,
                "class2": 0.0
            },
            "regulatory_source": "ENSG00000185551",
            "regulatory_target": "ENSG00000119547",
            "net_id": 0
        },
        "ENSG00000171634_ENSG00000016082": {
            "id": "ENSG00000171634_ENSG00000016082",
            "type": "Bridge",
            "score": 0,
            "reversable": false,
            "correlations": {
                "class1": 0.21971541752452425,
                "class2": 0.0
            },
            "regulatory_source": "ENSG00000016082",
            "regulatory_target": "ENSG00000171634",
            "net_id": 0
        },
        "ENSG00000171634_ENSG00000164330": {
            "id": "ENSG00000171634_ENSG00000164330",
            "type": "Bridge",
            "score": 0,
            "reversable": false,
            "correlations": {
                "class1": 0.27775416469659275,
                "class2": 0.0566645617747423
            },
            "regulatory_source": "ENSG00000164330",
            "regulatory_target": "ENSG00000171634",
            "net_id": 0
        },
        "ENSG00000134138_ENSG00000113580": {
            "id": "ENSG00000134138_ENSG00000113580",
            "type": "Bridge",
            "score": 0,
```
