## Supplement 2 (DAN vs. Cocul) for "AGEAS: Automated Machine Learning based Genetic Regulatory Element Extraction System": meta_GRN.js

```
{
    "genes": {
        "ENSG00000004487": {
            "id": "ENSG00000004487",
            "type": "TF",
            "symbol": "KDM1A",
            "source": [
                "ENSG00000167081"
            ],
            "target": [
                "ENSG00000099864",
                "ENSG00000103034",
                "ENSG00000176788",
                "ENSG00000176887",
                "ENSG00000144339",
                "ENSG00000172020",
                "ENSG00000149557",
                "ENSG00000156011",
                "ENSG00000165410",
                "ENSG00000169856",
                "ENSG00000087085",
                "ENSG00000169783",
                "ENSG00000185664",
                "ENSG00000101210",
                "ENSG00000164093",
                "ENSG00000107105",
                "ENSG00000182534",
                "ENSG00000213190",
                "ENSG00000198910",
                "ENSG00000188290",
                "ENSG00000078018",
                "ENSG00000197959",
                "ENSG00000162374",
                "ENSG00000196361",
                "ENSG00000105409",
                "ENSG00000197747",
                "ENSG00000072832",
                "ENSG00000145491",
                "ENSG00000010270",
                "ENSG00000131711",
                "ENSG00000155980",
                "ENSG00000075618",
                "ENSG00000151320",
                "ENSG00000168280",
                "ENSG00000186868",
                "ENSG00000182481",
                "ENSG00000198125",
                "ENSG00000213760",
                "ENSG00000137285",
                "ENSG00000092964",
                "ENSG00000143702",
                "ENSG00000144218",
                "ENSG00000104435",
                "ENSG00000122012",
                "ENSG00000213420",
                "ENSG00000004838",
                "ENSG00000016082",
                "ENSG00000173267",
                "ENSG00000111897",
                "ENSG00000111728",
                "ENSG00000154917",
                "ENSG00000149294",
                "ENSG00000034510",
                "ENSG00000170091",
                "ENSG00000131016",
                "ENSG00000015592",
                "ENSG00000139970",
                "ENSG00000167081",
                "ENSG00000135406",
                "ENSG00000104833",
                "ENSG00000137267",
                "ENSG00000177675",
                "ENSG00000135447",
                "ENSG00000018236",
                "ENSG00000189043",
                "ENSG00000163931",
                "ENSG00000277586"
            ],
            "expression_sum": {
                "class1": 13324.0,
                "class2": 5734.0
            },
            "ens_id": "ENSG00000004487",
            "uniprot_ids": [
                "O60341",
                "R4GMP9",
                "R4GMQ1"
            ]
        },
        "ENSG00000188290": {
            "id": "ENSG00000188290",
            "type": "Gene",
            "symbol": "HES4",
            "source": [
                "ENSG00000181449",
                "ENSG00000152518",
                "ENSG00000114315",
                "ENSG00000134138",
                "ENSG00000167081",
                "ENSG00000004487",
                "ENSG00000164330",
                "ENSG00000221869",
                "ENSG00000196591",
                "ENSG00000151729",
                "ENSG00000119547"
            ],
            "target": [],
            "expression_sum": {
                "class1": 22971.0,
                "class2": 31592.0
            },
            "ens_id": "ENSG00000188290",
            "uniprot_ids": [
                "Q9HCC6",
                "E9PB28",
                "D6REB3"
            ]
        },
        "ENSG00000162374": {
            "id": "ENSG00000162374",
            "type": "Gene",
            "symbol": "ELAVL4",
            "source": [
                "ENSG00000100811",
                "ENSG00000130726",
                "ENSG00000109132",
                "ENSG00000016082",
                "ENSG00000167081",
                "ENSG00000148516",
                "ENSG00000004487",
                "ENSG00000108055",
                "ENSG00000139352",
                "ENSG00000164754",
                "ENSG00000116560",
                "ENSG00000196591",
                "ENSG00000143995"
            ],
            "target": [],
            "expression_sum": {
                "class1": 31005.0,
                "class2": 1862.0
            },
            "ens_id": "ENSG00000162374",
            "uniprot_ids": [
                "P26378",
                "B1APY9",
                "A0A0R4J2E6",
                "A0A494C0Q5",
                "A0A494C015",
                "A0A494BZW4",
                "A0A494C173",
                "A0A494C0M6",
                "A0A494C011",
                "A0A494C0F8",
                "A0A494C147",
                "A0A494C1D9"
            ]
        },
        "ENSG00000213190": {
            "id": "ENSG00000213190",
            "type": "Gene",
            "symbol": "MLLT11",
            "source": [
                "ENSG00000130726",
                "ENSG00000004487",
                "ENSG00000103495",
                "ENSG00000168214",
                "ENSG00000130522"
            ],
            "target": [],
            "expression_sum": {
                "class1": 108753.0,
                "class2": 11983.0
            },
            "ens_id": "ENSG00000213190",
            "uniprot_ids": [
                "Q13015"
            ]
        },
        "ENSG00000197747": {
            "id": "ENSG00000197747",
            "type": "Gene",
            "symbol": "S100A10",
            "source": [
                "ENSG00000163435",
                "ENSG00000167081",
                "ENSG00000113580",
                "ENSG00000148516",
                "ENSG00000004487",
                "ENSG00000103495",
                "ENSG00000164330",
                "ENSG00000198911",
                "ENSG00000108001",
                "ENSG00000108055",
                "ENSG00000213015",
                "ENSG00000196591",
                "ENSG00000119547"
            ],
            "target": [],
            "expression_sum": {
                "class1": 57918.0,
                "class2": 29600.0
            },
            "ens_id": "ENSG00000197747",
            "uniprot_ids": [
                "P60903"
            ]
        },
        "ENSG00000197959": {
            "id": "ENSG00000197959",
            "type": "Gene",
            "symbol": "DNM3",
            "source": [
                "ENSG00000147862",
                "ENSG00000100811",
                "ENSG00000109132",
                "ENSG00000016082",
                "ENSG00000143799",
                "ENSG00000004487",
                "ENSG00000103495",
                "ENSG00000168610",
                "ENSG00000108055",
                "ENSG00000213015",
                "ENSG00000164754",
                "ENSG00000196591",
                "ENSG00000129351",
                "ENSG00000092199"
            ],
            "target": [],
            "expression_sum": {
                "class1": 3569.0,
                "class2": 735.0
            },
            "ens_id": "ENSG00000197959",
            "uniprot_ids": [
                "Q9UQ16",
                "H0YBC6",
                "E5RIK2",
                "E5RHK8"
            ]
        },
        "ENSG00000143702": {
            "id": "ENSG00000143702",
            "type": "Gene",
            "symbol": "CEP170",
            "source": [
                "ENSG00000106261",
                "ENSG00000148516",
                "ENSG00000067955",
                "ENSG00000189180",
                "ENSG00000160113",
                "ENSG00000092199",
                "ENSG00000109132",
                "ENSG00000103495",
                "ENSG00000198911",
                "ENSG00000189403",
                "ENSG00000164754",
                "ENSG00000168214",
                "ENSG00000196591",
                "ENSG00000100811",
                "ENSG00000177030",
                "ENSG00000213015",
                "ENSG00000163848",
                "ENSG00000181315",
                "ENSG00000170365",
                "ENSG00000143799",
                "ENSG00000004487",
                "ENSG00000108055",
                "ENSG00000116560",
                "ENSG00000147140"
            ],
            "target": [],
            "expression_sum": {
                "class1": 19098.0,
                "class2": 4709.0
            },
            "ens_id": "ENSG00000143702",
            "uniprot_ids": [
                "Q5SW79",
                "E7EWM2",
                "H0Y4T4",
                "E5RIH6",
                "E7EMW0",
                "E5RJT5",
                "H0YB66",
                "E5RG47",
                "E5RGW7",
                "H0YB92",
                "H0Y2V6",
                "E5RFU8"
            ]
        },
        "ENSG00000176887": {
            "id": "ENSG00000176887",
            "type": "TF",
            "symbol": "SOX11",
            "source": [
                "ENSG00000004487",
                "ENSG00000108055",
                "ENSG00000168214",
                "ENSG00000116560",
                "ENSG00000130522",
                "ENSG00000177606"
            ],
            "target": [
                "ENSG00000099901",
                "ENSG00000251562",
                "ENSG00000172757",
                "ENSG00000147274",
                "ENSG00000206503",
                "ENSG00000203667",
                "ENSG00000189403",
                "ENSG00000116560",
                "ENSG00000125691",
                "ENSG00000163939",
                "ENSG00000100823",
                "ENSG00000135387"
            ],
            "expression_sum": {
                "class1": 51157.0,
                "class2": 10806.0
            },
            "ens_id": "ENSG00000176887",
            "uniprot_ids": [
                "P35716"
            ]
        },
        "ENSG00000034510": {
            "id": "ENSG00000034510",
            "type": "Gene",
            "symbol": "TMSB10",
            "source": [
                "ENSG00000181449",
                "ENSG00000114315",
                "ENSG00000167081",
                "ENSG00000118263",
                "ENSG00000004487",
                "ENSG00000164330",
                "ENSG00000103495",
                "ENSG00000198911",
                "ENSG00000108055",
                "ENSG00000114861",
                "ENSG00000221869",
                "ENSG00000067955",
                "ENSG00000196591",
                "ENSG00000119547",
                "ENSG00000140307"
            ],
            "target": [],
            "expression_sum": {
                "class1": 637074.0,
                "class2": 243924.0
            },
            "ens_id": "ENSG00000034510",
            "uniprot_ids": [
                "P63313"
            ]
        },
        "ENSG00000144218": {
            "id": "ENSG00000144218",
            "type": "Gene",
            "symbol": "AFF3",
            "source": [
                "ENSG00000100811",
                "ENSG00000109132",
                "ENSG00000016082",
                "ENSG00000004487",
                "ENSG00000103495",
                "ENSG00000164330",
                "ENSG00000198911",
                "ENSG00000108055",
                "ENSG00000067955",
                "ENSG00000168214",
                "ENSG00000196591"
            ],
            "target": [],
            "expression_sum": {
                "class1": 4010.0,
                "class2": 1071.0
            },
            "ens_id": "ENSG00000144218",
            "uniprot_ids": [
                "P51826",
                "A0A5F9ZHV9",
                "C9JC67",
                "A0A5F9ZHV2",
                "H7C4C2",
                "A0A5F9ZHZ1",
                "A0A5F9ZHU9",
                "C9JZ66",
                "C9JUC4",
                "C9J622",
                "A0A5F9ZHH7",
                "C9JIB1",
                "C9JUY1",
                "C9JMS1",
                "A0A5F9ZHE5",
                "H7C3W3",
                "C9J847"
            ]
        },
        "ENSG00000168280": {
            "id": "ENSG00000168280",
            "type": "Gene",
            "symbol": "KIF5C",
            "source": [
                "ENSG00000100811",
                "ENSG00000113580",
                "ENSG00000148516",
                "ENSG00000004487",
                "ENSG00000103495",
                "ENSG00000198911",
                "ENSG00000108055",
                "ENSG00000213015",
                "ENSG00000196591",
                "ENSG00000116560",
                "ENSG00000128272",
                "ENSG00000129351"
            ],
            "target": [],
            "expression_sum": {
                "class1": 66280.0,
                "class2": 8729.0
            },
            "ens_id": "ENSG00000168280",
            "uniprot_ids": [
                "O60282",
                "A0A7I2V352",
                "A0A7I2V3V4",
                "A0A7I2V492",
                "A0A7I2YQQ7",
                "A0A7I2V3N5",
                "A0A7I2YQB4",
                "A0A7I2V581",
                "A0A7I2V4K6"
            ]
        },
        "ENSG00000144339": {
            "id": "ENSG00000144339",
            "type": "Gene",
            "symbol": "TMEFF2",
            "source": [
                "ENSG00000170365",
                "ENSG00000016082",
                "ENSG00000004487",
                "ENSG00000103495",
                "ENSG00000198911",
                "ENSG00000196591",
                "ENSG00000160113"
            ],
            "target": [],
            "expression_sum": {
                "class1": 7361.0,
                "class2": 1097.0
            },
            "ens_id": "ENSG00000144339",
            "uniprot_ids": [
                "Q9UIK5"
            ]
        },
        "ENSG00000078018": {
            "id": "ENSG00000078018",
            "type": "Gene",
            "symbol": "MAP2",
            "source": [
                "ENSG00000109132",
                "ENSG00000134138",
                "ENSG00000016082",
                "ENSG00000122691",
                "ENSG00000004487",
                "ENSG00000196591",
                "ENSG00000119547"
            ],
            "target": [],
            "expression_sum": {
                "class1": 40450.0,
                "class2": 18433.0
            },
            "ens_id": "ENSG00000078018",
            "uniprot_ids": [
                "P11137",
                "A8MZ31",
                "A0A669KB77",
                "E7EV03",
                "H7BZB9"
            ]
        },
        "ENSG00000004838": {
            "id": "ENSG00000004838",
            "type": "Gene",
            "symbol": "ZMYND10",
            "source": [
                "ENSG00000004487",
                "ENSG00000103495"
            ],
            "target": [],
            "expression_sum": {
                "class1": 326.0,
                "class2": 2317.0
            },
            "ens_id": "ENSG00000004838",
            "uniprot_ids": [
                "O75800",
                "F2Z3M9",
                "C9JUQ8"
            ]
        },
        "ENSG00000163931": {
            "id": "ENSG00000163931",
            "type": "Gene",
            "symbol": "TKT",
            "source": [
                "ENSG00000167081",
                "ENSG00000004487",
                "ENSG00000103495",
                "ENSG00000108001",
                "ENSG00000108055",
                "ENSG00000221869",
                "ENSG00000196591",
                "ENSG00000124766",
                "ENSG00000128610",
                "ENSG00000119547",
                "ENSG00000160113"
            ],
            "target": [],
            "expression_sum": {
                "class1": 22858.0,
                "class2": 21397.0
            },
            "ens_id": "ENSG00000163931",
            "uniprot_ids": [
                "P29401",
                "F8W888",
                "F8WAX4",
                "A0A0B4J1R6",
                "E9PFF2"
            ]
        },
        "ENSG00000172020": {
            "id": "ENSG00000172020",
            "type": "Gene",
            "symbol": "GAP43",
            "source": [
                "ENSG00000100811",
                "ENSG00000130726",
                "ENSG00000100644",
                "ENSG00000109132",
                "ENSG00000004487",
                "ENSG00000103495",
                "ENSG00000108055",
                "ENSG00000164754",
                "ENSG00000116560",
                "ENSG00000196591",
                "ENSG00000128272",
                "ENSG00000130522"
            ],
            "target": [],
            "expression_sum": {
                "class1": 117923.0,
                "class2": 15761.0
            },
            "ens_id": "ENSG00000172020",
            "uniprot_ids": [
                "P17677"
            ]
        },
        "ENSG00000154917": {
            "id": "ENSG00000154917",
            "type": "Gene",
            "symbol": "RAB6B",
            "source": [
                "ENSG00000148516",
                "ENSG00000004487",
                "ENSG00000103495",
                "ENSG00000198911",
                "ENSG00000108055",
                "ENSG00000196591"
            ],
            "target": [],
            "expression_sum": {
                "class1": 8766.0,
                "class2": 2639.0
            },
            "ens_id": "ENSG00000154917",
            "uniprot_ids": [
                "Q9NRW1",
                "J3KR73",
                "C9JB90",
                "C9J0I2",
                "C9JU14"
            ]
        },
        "ENSG00000072832": {
            "id": "ENSG00000072832",
            "type": "Gene",
            "symbol": "CRMP1",
            "source": [
                "ENSG00000100811",
                "ENSG00000130726",
                "ENSG00000028277",
                "ENSG00000137203",
                "ENSG00000167081",
                "ENSG00000113580",
                "ENSG00000004487",
                "ENSG00000103495",
                "ENSG00000105698",
                "ENSG00000198911",
                "ENSG00000119950",
                "ENSG00000139352",
                "ENSG00000196591",
                "ENSG00000168214"
            ],
            "target": [],
            "expression_sum": {
                "class1": 49646.0,
                "class2": 6418.0
            },
            "ens_id": "ENSG00000072832",
            "uniprot_ids": [
                "Q14194",
                "E9PD68"
            ]
        },
        "ENSG00000164093": {
            "id": "ENSG00000164093",
            "type": "TF",
            "symbol": "PITX2",
            "source": [
                "ENSG00000100811",
                "ENSG00000004487",
                "ENSG00000103495",
                "ENSG00000181722",
                "ENSG00000108055",
                "ENSG00000116560",
                "ENSG00000168214",
                "ENSG00000196591",
                "ENSG00000116044",
                "ENSG00000119547",
                "ENSG00000177606"
            ],
            "target": [
                "ENSG00000244879",
                "ENSG00000132475",
                "ENSG00000256537",
                "ENSG00000090487",
                "ENSG00000177889",
                "ENSG00000254999",
                "ENSG00000170776",
                "ENSG00000162409",
                "ENSG00000119950",
                "ENSG00000175416",
                "ENSG00000185633",
                "ENSG00000134308",
                "ENSG00000112695",
                "ENSG00000164631"
            ],
            "expression_sum": {
                "class1": 4850.0,
                "class2": 13703.0
            },
            "ens_id": "ENSG00000164093",
            "uniprot_ids": [
                "Q99697",
                "D6RBG8",
                "U3KQ81",
                "D6RFI4"
            ]
        },
        "ENSG00000145491": {
            "id": "ENSG00000145491",
            "type": "Gene",
            "symbol": "ROPN1L",
            "source": [
                "ENSG00000004487",
                "ENSG00000103495"
            ],
            "target": [],
            "expression_sum": {
                "class1": 281.0,
                "class2": 2386.0
            },
            "ens_id": "ENSG00000145491",
            "uniprot_ids": [
                "Q96C74"
            ]
        },
        "ENSG00000176788": {
            "id": "ENSG00000176788",
            "type": "Gene",
            "symbol": "BASP1",
            "source": [
                "ENSG00000170365",
                "ENSG00000100811",
                "ENSG00000130726",
                "ENSG00000028277",
                "ENSG00000004487",
                "ENSG00000103495",
                "ENSG00000105698",
                "ENSG00000198911",
                "ENSG00000108055",
                "ENSG00000213015",
                "ENSG00000163848",
                "ENSG00000168214",
                "ENSG00000116560",
                "ENSG00000196591"
            ],
            "target": [],
            "expression_sum": {
                "class1": 191113.0,
                "class2": 35258.0
            },
            "ens_id": "ENSG00000176788",
            "uniprot_ids": [
                "P80723",
                "U3KQP0"
            ]
        },
        "ENSG00000016082": {
            "id": "ENSG00000016082",
            "type": "TF",
            "symbol": "ISL1",
            "source": [
                "ENSG00000170365",
                "ENSG00000100811",
                "ENSG00000130726",
                "ENSG00000109132",
                "ENSG00000153147",
                "ENSG00000004487",
                "ENSG00000103495",
                "ENSG00000198911",
                "ENSG00000108055",
                "ENSG00000164754",
                "ENSG00000168214",
                "ENSG00000196591"
            ],
            "target": [
                "ENSG00000017260",
                "ENSG00000162704",
                "ENSG00000111716",
                "ENSG00000005810",
                "ENSG00000140941",
                "ENSG00000197959",
                "ENSG00000145817",
                "ENSG00000075415",
                "ENSG00000077380",
                "ENSG00000154556",
                "ENSG00000118733",
                "ENSG00000052802",
                "ENSG00000087274",
                "ENSG00000165669",
                "ENSG00000065911",
                "ENSG00000106443",
                "ENSG00000129636",
                "ENSG00000197061",
                "ENSG00000242173",
                "ENSG00000139767",
                "ENSG00000174574",
                "ENSG00000139116",
                "ENSG00000075914",
                "ENSG00000047249",
                "ENSG00000060982",
                "ENSG00000141447",
                "ENSG00000175581",
                "ENSG00000067225",
                "ENSG00000144218",
                "ENSG00000117139",
                "ENSG00000140105",
                "ENSG00000159685",
                "ENSG00000205981",
                "ENSG00000157216",
                "ENSG00000144746",
                "ENSG00000157168",
                "ENSG00000133606",
                "ENSG00000125835",
                "ENSG00000075142",
                "ENSG00000150722",
                "ENSG00000263001",
                "ENSG00000213741",
                "ENSG00000086598",
                "ENSG00000129515",
                "ENSG00000076604",
                "ENSG00000142546",
                "ENSG00000110955",
                "ENSG00000175216",
                "ENSG00000168137",
                "ENSG00000149187",
                "ENSG00000213760",
                "ENSG00000107581",
                "ENSG00000112972",
                "ENSG00000147677",
                "ENSG00000231500",
                "ENSG00000204842",
                "ENSG00000167196",
                "ENSG00000180370",
                "ENSG00000079246",
                "ENSG00000168275",
                "ENSG00000057019",
                "ENSG00000111850",
                "ENSG00000105185",
                "ENSG00000166974",
                "ENSG00000138801",
                "ENSG00000100320",
                "ENSG00000114391",
                "ENSG00000164985",
                "ENSG00000135390",
                "ENSG00000239672",
                "ENSG00000141867",
                "ENSG00000153310",
                "ENSG00000256060",
                "ENSG00000197223",
                "ENSG00000154277",
                "ENSG00000131469",
                "ENSG00000128654",
                "ENSG00000090266",
                "ENSG00000171634",
                "ENSG00000162374",
                "ENSG00000169764",
                "ENSG00000166200",
                "ENSG00000104388",
                "ENSG00000204954",
                "ENSG00000143222",
                "ENSG00000151914",
                "ENSG00000161057",
                "ENSG00000177565",
                "ENSG00000125827",
                "ENSG00000171130",
                "ENSG00000073578",
                "ENSG00000081154",
                "ENSG00000122566",
                "ENSG00000143878",
                "ENSG00000135486",
                "ENSG00000109670",
                "ENSG00000275052",
                "ENSG00000128951",
                "ENSG00000100575",
                "ENSG00000172159",
                "ENSG00000167792",
                "ENSG00000138413",
                "ENSG00000070761",
                "ENSG00000163848",
                "ENSG00000069275",
                "ENSG00000119729",
                "ENSG00000113742",
                "ENSG00000170515",
                "ENSG00000144895",
                "ENSG00000115875",
                "ENSG00000198886",
                "ENSG00000164929",
                "ENSG00000196262",
                "ENSG00000115355",
                "ENSG00000214078",
                "ENSG00000173726",
                "ENSG00000134287",
                "ENSG00000089280",
                "ENSG00000139117",
                "ENSG00000101132",
                "ENSG00000122026",
                "ENSG00000151327",
                "ENSG00000106400",
                "ENSG00000140836",
                "ENSG00000113387",
                "ENSG00000156508",
                "ENSG00000196937",
                "ENSG00000082701",
                "ENSG00000234608",
                "ENSG00000170142",
                "ENSG00000173141",
                "ENSG00000084463",
                "ENSG00000185630",
                "ENSG00000100442",
                "ENSG00000115091",
                "ENSG00000204628",
                "ENSG00000178234",
                "ENSG00000121022",
                "ENSG00000197111",
                "ENSG00000163069",
                "ENSG00000134825",
                "ENSG00000184007",
                "ENSG00000099250",
                "ENSG00000197345",
                "ENSG00000116251",
                "ENSG00000038219",
                "ENSG00000053254",
                "ENSG00000164330",
                "ENSG00000156639",
                "ENSG00000175550",
                "ENSG00000144339",
                "ENSG00000156011",
                "ENSG00000135070",
                "ENSG00000214413",
                "ENSG00000145782",
                "ENSG00000153187",
                "ENSG00000174444",
                "ENSG00000110711",
                "ENSG00000106829",
                "ENSG00000138430",
                "ENSG00000126214",
                "ENSG00000097033",
                "ENSG00000273611",
                "ENSG00000164733",
                "ENSG00000179933",
                "ENSG00000148943",
                "ENSG00000189159",
                "ENSG00000177410",
                "ENSG00000186472",
                "ENSG00000170365",
                "ENSG00000182718",
                "ENSG00000089053",
                "ENSG00000122565",
                "ENSG00000163541",
                "ENSG00000164258",
                "ENSG00000147862",
                "ENSG00000006715",
                "ENSG00000079102",
                "ENSG00000197756",
                "ENSG00000065135",
                "ENSG00000008294",
                "ENSG00000104408",
                "ENSG00000105373",
                "ENSG00000265681",
                "ENSG00000084207",
                "ENSG00000058272",
                "ENSG00000165434",
                "ENSG00000122068",
                "ENSG00000170873",
                "ENSG00000162434",
                "ENSG00000118816",
                "ENSG00000182827",
                "ENSG00000204442",
                "ENSG00000163468",
                "ENSG00000172115",
                "ENSG00000118257",
                "ENSG00000156976",
                "ENSG00000078369",
                "ENSG00000171033",
                "ENSG00000138107",
                "ENSG00000149485",
                "ENSG00000071994",
                "ENSG00000157778",
                "ENSG00000154380",
                "ENSG00000182621",
                "ENSG00000172531",
                "ENSG00000148303",
                "ENSG00000054118",
                "ENSG00000023228",
                "ENSG00000136717",
                "ENSG00000168175",
                "ENSG00000136854",
                "ENSG00000196305",
                "ENSG00000117859",
                "ENSG00000113845",
                "ENSG00000197958",
                "ENSG00000077279",
                "ENSG00000188994",
                "ENSG00000108946",
                "ENSG00000071794",
                "ENSG00000172586",
                "ENSG00000063177",
                "ENSG00000094916",
                "ENSG00000103035",
                "ENSG00000138663",
                "ENSG00000140386",
                "ENSG00000072041",
                "ENSG00000113108",
                "ENSG00000165462",
                "ENSG00000128699",
                "ENSG00000108848",
                "ENSG00000108826",
                "ENSG00000156928",
                "ENSG00000186469",
                "ENSG00000166352",
                "ENSG00000126457",
                "ENSG00000070087",
                "ENSG00000174547",
                "ENSG00000131876",
                "ENSG00000148700",
                "ENSG00000167699",
                "ENSG00000092148",
                "ENSG00000135406",
                "ENSG00000134824",
                "ENSG00000177675",
                "ENSG00000151552",
                "ENSG00000061676",
                "ENSG00000080822",
                "ENSG00000168268",
                "ENSG00000138594",
                "ENSG00000179915",
                "ENSG00000159593",
                "ENSG00000180228",
                "ENSG00000139613",
                "ENSG00000131116",
                "ENSG00000197728",
                "ENSG00000104823",
                "ENSG00000130779",
                "ENSG00000229117",
                "ENSG00000166206",
                "ENSG00000126773",
                "ENSG00000115350",
                "ENSG00000086666",
                "ENSG00000154518",
                "ENSG00000087302",
                "ENSG00000182628",
                "ENSG00000090263",
                "ENSG00000105669",
                "ENSG00000160752",
                "ENSG00000136521",
                "ENSG00000204220",
                "ENSG00000164741",
                "ENSG00000155330",
                "ENSG00000108100",
                "ENSG00000122958",
                "ENSG00000111832",
                "ENSG00000118263",
                "ENSG00000068615",
                "ENSG00000153774",
                "ENSG00000165525",
                "ENSG00000083845",
                "ENSG00000132466",
                "ENSG00000128923",
                "ENSG00000184220",
                "ENSG00000009307",
                "ENSG00000101882",
                "ENSG00000018189",
                "ENSG00000107372",
                "ENSG00000109132",
                "ENSG00000204856",
                "ENSG00000182117",
                "ENSG00000150991",
                "ENSG00000212907",
                "ENSG00000007080",
                "ENSG00000104679",
                "ENSG00000142875",
                "ENSG00000173660",
                "ENSG00000214941",
                "ENSG00000186184",
                "ENSG00000170899",
                "ENSG00000123636",
                "ENSG00000162604",
                "ENSG00000163964",
                "ENSG00000102003",
                "ENSG00000010810",
                "ENSG00000196781",
                "ENSG00000188846",
                "ENSG00000078018",
                "ENSG00000111530",
                "ENSG00000070961",
                "ENSG00000103275",
                "ENSG00000167232",
                "ENSG00000134313",
                "ENSG00000198712",
                "ENSG00000113448",
                "ENSG00000186468",
                "ENSG00000183527",
                "ENSG00000100567",
                "ENSG00000144567",
                "ENSG00000123983",
                "ENSG00000172270",
                "ENSG00000198888",
                "ENSG00000089009"
            ],
            "expression_sum": {
                "class1": 6727.0,
                "class2": 11.0
            },
            "ens_id": "ENSG00000016082",
            "uniprot_ids": [
                "P61371",
                "D6RBJ1"
            ]
        },
        "ENSG00000131711": {
            "id": "ENSG00000131711",
            "type": "Gene",
            "symbol": "MAP1B",
            "source": [
                "ENSG00000134138",
                "ENSG00000004487",
                "ENSG00000103495",
                "ENSG00000108055",
                "ENSG00000168214",
                "ENSG00000116560",
                "ENSG00000196591",
                "ENSG00000128272",
                "ENSG00000092199"
            ],
            "target": [],
            "expression_sum": {
                "class1": 292659.0,
                "class2": 58925.0
            },
            "ens_id": "ENSG00000131711",
            "uniprot_ids": [
                "P46821",
                "D6RA32",
                "D6RA40",
                "D6RGJ3",
                "D6RCL2"
            ]
        },
        "ENSG00000122012": {
            "id": "ENSG00000122012",
            "type": "Gene",
            "symbol": "SV2C",
            "source": [
                "ENSG00000100811",
                "ENSG00000100644",
                "ENSG00000109132",
                "ENSG00000140836",
                "ENSG00000004487",
                "ENSG00000103495",
                "ENSG00000198911",
                "ENSG00000168610",
                "ENSG00000126778",
                "ENSG00000108055",
                "ENSG00000168214",
                "ENSG00000196591"
            ],
            "target": [],
            "expression_sum": {
                "class1": 2500.0,
                "class2": 356.0
            },
            "ens_id": "ENSG00000122012",
            "uniprot_ids": [
                "Q496J9",
                "B3KT41"
            ]
        },
        "ENSG00000170091": {
            "id": "ENSG00000170091",
            "type": "Gene",
            "symbol": "NSG2",
            "source": [
                "ENSG00000100811",
                "ENSG00000106261",
                "ENSG00000028277",
                "ENSG00000137203",
                "ENSG00000122482",
                "ENSG00000148516",
                "ENSG00000143799",
                "ENSG00000004487",
                "ENSG00000103495",
                "ENSG00000108055",
                "ENSG00000213015",
                "ENSG00000119950",
                "ENSG00000164754",
                "ENSG00000168214",
                "ENSG00000116560",
                "ENSG00000196591",
                "ENSG00000160113",
                "ENSG00000139613"
            ],
            "target": [],
            "expression_sum": {
                "class1": 14357.0,
                "class2": 2555.0
            },
            "ens_id": "ENSG00000170091",
            "uniprot_ids": [
                "Q9Y328",
                "G3XAM5",
                "E5RHS6",
                "E5RH73",
                "E5RGE8"
            ]
        },
        "ENSG00000137267": {
            "id": "ENSG00000137267",
            "type": "Gene",
            "symbol": "TUBB2A",
            "source": [
                "ENSG00000106261",
                "ENSG00000160113",
                "ENSG00000140307",
                "ENSG00000134138",
                "ENSG00000105698",
                "ENSG00000103495",
                "ENSG00000189403",
                "ENSG00000168214",
                "ENSG00000196591",
                "ENSG00000100811",
                "ENSG00000028277",
                "ENSG00000213015",
                "ENSG00000119950",
                "ENSG00000137203",
                "ENSG00000164683",
                "ENSG00000118263",
                "ENSG00000004487",
                "ENSG00000108055",
                "ENSG00000137947",
                "ENSG00000116560",
                "ENSG00000147140"
            ],
            "target": [],
            "expression_sum": {
                "class1": 101835.0,
                "class2": 18713.0
            },
            "ens_id": "ENSG00000137267",
            "uniprot_ids": [
                "Q13885"
            ]
        },
        "ENSG00000137285": {
            "id": "ENSG00000137285",
            "type": "Gene",
            "symbol": "TUBB2B",
            "source": [
                "ENSG00000105419",
                "ENSG00000181449",
                "ENSG00000100811",
                "ENSG00000109132",
                "ENSG00000134138",
                "ENSG00000122482",
                "ENSG00000004487",
                "ENSG00000103495",
                "ENSG00000213015",
                "ENSG00000168214",
                "ENSG00000196591",
                "ENSG00000147140",
                "ENSG00000128272"
            ],
            "target": [],
            "expression_sum": {
                "class1": 280173.0,
                "class2": 80355.0
            },
            "ens_id": "ENSG00000137285",
            "uniprot_ids": [
                "Q9BVA1"
            ]
        },
        "ENSG00000213760": {
            "id": "ENSG00000213760",
            "type": "Gene",
            "symbol": "ATP6V1G2",
            "source": [
                "ENSG00000170365",
                "ENSG00000100811",
                "ENSG00000109132",
                "ENSG00000175197",
                "ENSG00000016082",
                "ENSG00000004487",
                "ENSG00000103495",
                "ENSG00000128272",
                "ENSG00000108055",
                "ENSG00000213015",
                "ENSG00000168214",
                "ENSG00000116560",
                "ENSG00000196591",
                "ENSG00000147140",
                "ENSG00000160113"
            ],
            "target": [],
            "expression_sum": {
                "class1": 21218.0,
                "class2": 3011.0
            },
            "ens_id": "ENSG00000213760",
            "uniprot_ids": [
                "O95670",
                "H0Y474"
            ]
        },
        "ENSG00000111897": {
            "id": "ENSG00000111897",
            "type": "Gene",
            "symbol": "SERINC1",
            "source": [
                "ENSG00000109132",
                "ENSG00000004487",
                "ENSG00000108055",
                "ENSG00000164754",
                "ENSG00000196591"
            ],
            "target": [],
            "expression_sum": {
                "class1": 24585.0,
                "class2": 8207.0
            },
            "ens_id": "ENSG00000111897",
            "uniprot_ids": [
                "Q9NRX5"
            ]
        },
        "ENSG00000131016": {
            "id": "ENSG00000131016",
            "type": "Gene",
            "symbol": "AKAP12",
            "source": [
                "ENSG00000170365",
                "ENSG00000143799",
                "ENSG00000004487",
                "ENSG00000103495",
                "ENSG00000198911",
                "ENSG00000108055",
                "ENSG00000213015",
                "ENSG00000164754",
                "ENSG00000116560",
                "ENSG00000196591",
                "ENSG00000181315",
                "ENSG00000165512"
            ],
            "target": [],
            "expression_sum": {
                "class1": 12532.0,
                "class2": 7292.0
            },
            "ens_id": "ENSG00000131016",
            "uniprot_ids": [
                "Q02952"
            ]
        },
        "ENSG00000075618": {
            "id": "ENSG00000075618",
            "type": "Gene",
            "symbol": "FSCN1",
            "source": [
                "ENSG00000004487",
                "ENSG00000103495",
                "ENSG00000164330",
                "ENSG00000137947",
                "ENSG00000067955",
                "ENSG00000168214",
                "ENSG00000196591",
                "ENSG00000160113",
                "ENSG00000140307"
            ],
            "target": [],
            "expression_sum": {
                "class1": 19210.0,
                "class2": 6066.0
            },
            "ens_id": "ENSG00000075618",
            "uniprot_ids": [
                "Q16658",
                "A0A0A0MSB2",
                "C9JFC0",
                "C9JPH9"
            ]
        },
        "ENSG00000189043": {
            "id": "ENSG00000189043",
            "type": "Gene",
            "symbol": "NDUFA4",
            "source": [
                "ENSG00000181449",
                "ENSG00000114315",
                "ENSG00000167081",
                "ENSG00000004487",
                "ENSG00000122691",
                "ENSG00000164330",
                "ENSG00000108001",
                "ENSG00000137947",
                "ENSG00000221869",
                "ENSG00000196591",
                "ENSG00000067955",
                "ENSG00000119547"
            ],
            "target": [],
            "expression_sum": {
                "class1": 124471.0,
                "class2": 73711.0
            },
            "ens_id": "ENSG00000189043",
            "uniprot_ids": [
                "O00483"
            ]
        },
        "ENSG00000010270": {
            "id": "ENSG00000010270",
            "type": "Gene",
            "symbol": "STARD3NL",
            "source": [
                "ENSG00000109132",
                "ENSG00000167081",
                "ENSG00000118263",
                "ENSG00000004487",
                "ENSG00000164330",
                "ENSG00000103495",
                "ENSG00000108055",
                "ENSG00000213015",
                "ENSG00000196591"
            ],
            "target": [],
            "expression_sum": {
                "class1": 24483.0,
                "class2": 6035.0
            },
            "ens_id": "ENSG00000010270",
            "uniprot_ids": [
                "O95772",
                "C9J3F5",
                "C9JKL2",
                "C9JJ39",
                "C9JPX5"
            ]
        },
        "ENSG00000213420": {
            "id": "ENSG00000213420",
            "type": "Gene",
            "symbol": "GPC2",
            "source": [
                "ENSG00000004487",
                "ENSG00000103495",
                "ENSG00000175745",
                "ENSG00000213015",
                "ENSG00000196591",
                "ENSG00000147140",
                "ENSG00000116560",
                "ENSG00000092199"
            ],
            "target": [],
            "expression_sum": {
                "class1": 14071.0,
                "class2": 1872.0
            },
            "ens_id": "ENSG00000213420",
            "uniprot_ids": [
                "Q8N158",
                "A0A0J9YXG7"
            ]
        },
        "ENSG00000087085": {
            "id": "ENSG00000087085",
            "type": "Gene",
            "symbol": "ACHE",
            "source": [
                "ENSG00000100811",
                "ENSG00000130726",
                "ENSG00000148516",
                "ENSG00000143799",
                "ENSG00000004487",
                "ENSG00000103495",
                "ENSG00000108055",
                "ENSG00000189403",
                "ENSG00000147140",
                "ENSG00000196591",
                "ENSG00000130522",
                "ENSG00000160113",
                "ENSG00000092199"
            ],
            "target": [],
            "expression_sum": {
                "class1": 7846.0,
                "class2": 803.0
            },
            "ens_id": "ENSG00000087085",
            "uniprot_ids": [
                "P22303",
                "A0A498U6H7",
                "C9JD78",
                "F8WD68",
                "C9JZL6",
                "F8WAR7",
                "F8WD34",
                "C9J2S3"
            ]
        },
        "ENSG00000198910": {
            "id": "ENSG00000198910",
            "type": "Gene",
            "symbol": "L1CAM",
            "source": [
                "ENSG00000100811",
                "ENSG00000130726",
                "ENSG00000100644",
                "ENSG00000004487",
                "ENSG00000103495",
                "ENSG00000105698",
                "ENSG00000198911",
                "ENSG00000175745",
                "ENSG00000108055",
                "ENSG00000163848",
                "ENSG00000196591",
                "ENSG00000116560",
                "ENSG00000164754"
            ],
            "target": [],
            "expression_sum": {
                "class1": 11121.0,
                "class2": 788.0
            },
            "ens_id": "ENSG00000198910",
            "uniprot_ids": [
                "P32004",
                "E7EMY4",
                "E9PHJ4",
                "H0Y5C3",
                "E7EPI4",
                "E7EVM4",
                "H3BLW5"
            ]
        },
        "ENSG00000156011": {
            "id": "ENSG00000156011",
            "type": "Gene",
            "symbol": "PSD3",
            "source": [
                "ENSG00000016082",
                "ENSG00000004487"
            ],
            "target": [],
            "expression_sum": {
                "class1": 6126.0,
                "class2": 2704.0
            },
            "ens_id": "ENSG00000156011",
            "uniprot_ids": [
                "Q9NYI0",
                "B4DKF8",
                "E5RIH3",
                "E5RJ29",
                "H0YC20",
                "E5RJE4",
                "E5RH02",
                "A0A087X1Z0",
                "H0YAQ7",
                "A0A087WXD8",
                "E5RFG7"
            ]
        },
        "ENSG00000277586": {
            "id": "ENSG00000277586",
            "type": "Gene",
            "symbol": "NEFL",
            "source": [
                "ENSG00000100811",
                "ENSG00000130726",
                "ENSG00000004487",
                "ENSG00000103495",
                "ENSG00000198911",
                "ENSG00000108055",
                "ENSG00000196591",
                "ENSG00000006128"
            ],
            "target": [],
            "expression_sum": {
                "class1": 143369.0,
                "class2": 7977.0
            },
            "ens_id": "ENSG00000277586",
            "uniprot_ids": [
                "P07196"
            ]
        },
        "ENSG00000092964": {
            "id": "ENSG00000092964",
            "type": "Gene",
            "symbol": "DPYSL2",
            "source": [
                "ENSG00000100811",
                "ENSG00000129351",
                "ENSG00000106261",
                "ENSG00000118263",
                "ENSG00000004487",
                "ENSG00000103495",
                "ENSG00000105698",
                "ENSG00000198911",
                "ENSG00000108055",
                "ENSG00000168214",
                "ENSG00000116560",
                "ENSG00000124766",
                "ENSG00000196591",
                "ENSG00000147140",
                "ENSG00000092199"
            ],
            "target": [],
            "expression_sum": {
                "class1": 30945.0,
                "class2": 8602.0
            },
            "ens_id": "ENSG00000092964",
            "uniprot_ids": [
                "Q16555",
                "E5RFU4",
                "A0A1C7CYX9"
            ]
        },
        "ENSG00000015592": {
            "id": "ENSG00000015592",
            "type": "Gene",
            "symbol": "STMN4",
            "source": [
                "ENSG00000004487",
                "ENSG00000103495",
                "ENSG00000196591",
                "ENSG00000130522",
                "ENSG00000177606"
            ],
            "target": [],
            "expression_sum": {
                "class1": 63735.0,
                "class2": 6333.0
            },
            "ens_id": "ENSG00000015592",
            "uniprot_ids": [
                "Q9H169",
                "E5RIR6",
                "E7EVN3"
            ]
        },
        "ENSG00000104435": {
            "id": "ENSG00000104435",
            "type": "Gene",
            "symbol": "STMN2",
            "source": [
                "ENSG00000100811",
                "ENSG00000109132",
                "ENSG00000148516",
                "ENSG00000004487",
                "ENSG00000108055",
                "ENSG00000164754",
                "ENSG00000196591",
                "ENSG00000130522"
            ],
            "target": [],
            "expression_sum": {
                "class1": 348399.0,
                "class2": 24456.0
            },
            "ens_id": "ENSG00000104435",
            "uniprot_ids": [
                "Q93045",
                "E5RGX5"
            ]
        },
        "ENSG00000107105": {
            "id": "ENSG00000107105",
            "type": "Gene",
            "symbol": "ELAVL2",
            "source": [
                "ENSG00000100811",
                "ENSG00000130726",
                "ENSG00000109132",
                "ENSG00000153147",
                "ENSG00000148516",
                "ENSG00000004487",
                "ENSG00000103495",
                "ENSG00000105698",
                "ENSG00000198911",
                "ENSG00000108055",
                "ENSG00000189403",
                "ENSG00000213015",
                "ENSG00000164754",
                "ENSG00000168214",
                "ENSG00000196591",
                "ENSG00000130522"
            ],
            "target": [],
            "expression_sum": {
                "class1": 20427.0,
                "class2": 2682.0
            },
            "ens_id": "ENSG00000107105",
            "uniprot_ids": [
                "Q12926",
                "C9JB16",
                "B1AM48",
                "A0A0A0MRX1"
            ]
        },
        "ENSG00000167081": {
            "id": "ENSG00000167081",
            "type": "TF",
            "symbol": "PBX3",
            "source": [
                "ENSG00000100811",
                "ENSG00000134138",
                "ENSG00000004487",
                "ENSG00000103495",
                "ENSG00000105698",
                "ENSG00000213015",
                "ENSG00000189403",
                "ENSG00000119950",
                "ENSG00000196591",
                "ENSG00000168214",
                "ENSG00000092199"
            ],
            "target": [
                "ENSG00000078668",
                "ENSG00000090621",
                "ENSG00000206573",
                "ENSG00000112081",
                "ENSG00000232119",
                "ENSG00000107223",
                "ENSG00000162704",
                "ENSG00000163634",
                "ENSG00000165655",
                "ENSG00000153130",
                "ENSG00000111674",
                "ENSG00000111716",
                "ENSG00000139644",
                "ENSG00000005810",
                "ENSG00000169612",
                "ENSG00000099622",
                "ENSG00000137955",
                "ENSG00000185651",
                "ENSG00000090060",
                "ENSG00000005020",
                "ENSG00000106603",
                "ENSG00000140941",
                "ENSG00000161016",
                "ENSG00000134248",
                "ENSG00000127540",
                "ENSG00000075945",
                "ENSG00000125743",
                "ENSG00000099797",
                "ENSG00000119632",
                "ENSG00000189403",
                "ENSG00000159377",
                "ENSG00000136261",
                "ENSG00000075415",
                "ENSG00000147155",
                "ENSG00000077380",
                "ENSG00000154556",
                "ENSG00000052802",
                "ENSG00000134283",
                "ENSG00000198931",
                "ENSG00000173812",
                "ENSG00000197451",
                "ENSG00000152256",
                "ENSG00000237190",
                "ENSG00000173915",
                "ENSG00000065911",
                "ENSG00000151729",
                "ENSG00000172292",
                "ENSG00000170310",
                "ENSG00000129636",
                "ENSG00000115540",
                "ENSG00000123240",
                "ENSG00000041357",
                "ENSG00000102103",
                "ENSG00000141858",
                "ENSG00000178913",
                "ENSG00000163875",
                "ENSG00000163319",
                "ENSG00000167136",
                "ENSG00000164924",
                "ENSG00000074319",
                "ENSG00000149196",
                "ENSG00000141759",
                "ENSG00000174574",
                "ENSG00000146007",
                "ENSG00000106554",
                "ENSG00000084764",
                "ENSG00000277791",
                "ENSG00000125868",
                "ENSG00000153815",
                "ENSG00000047249",
                "ENSG00000147548",
                "ENSG00000171703",
                "ENSG00000196504",
                "ENSG00000131236",
                "ENSG00000089220",
                "ENSG00000055163",
                "ENSG00000141447",
                "ENSG00000116898",
                "ENSG00000175581",
                "ENSG00000196072",
                "ENSG00000179431",
                "ENSG00000187164",
                "ENSG00000188243",
                "ENSG00000067225",
                "ENSG00000109390",
                "ENSG00000113712",
                "ENSG00000126698",
                "ENSG00000136143",
                "ENSG00000233927",
                "ENSG00000117139",
                "ENSG00000148444",
                "ENSG00000140105",
                "ENSG00000159685",
                "ENSG00000034713",
                "ENSG00000130811",
                "ENSG00000144746",
                "ENSG00000095380",
                "ENSG00000133703",
                "ENSG00000063854",
                "ENSG00000103769",
                "ENSG00000034510",
                "ENSG00000146425",
                "ENSG00000005194",
                "ENSG00000138071",
                "ENSG00000203778",
                "ENSG00000117410",
                "ENSG00000131100",
                "ENSG00000075142",
                "ENSG00000273841",
                "ENSG00000124802",
                "ENSG00000263001",
                "ENSG00000142657",
                "ENSG00000196405",
                "ENSG00000183011",
                "ENSG00000174437",
                "ENSG00000114520",
                "ENSG00000213741",
                "ENSG00000126934",
                "ENSG00000145592",
                "ENSG00000187051",
                "ENSG00000069849",
                "ENSG00000240344",
                "ENSG00000129515",
                "ENSG00000100519",
                "ENSG00000144635",
                "ENSG00000187514",
                "ENSG00000163041",
                "ENSG00000155115",
                "ENSG00000160194",
                "ENSG00000141027",
                "ENSG00000092199",
                "ENSG00000110955",
                "ENSG00000178980",
                "ENSG00000204463",
                "ENSG00000164346",
                "ENSG00000150316",
                "ENSG00000010256",
                "ENSG00000120053",
                "ENSG00000174953",
                "ENSG00000169100",
                "ENSG00000162191",
                "ENSG00000118363",
                "ENSG00000108654",
                "ENSG00000135940",
                "ENSG00000170634",
                "ENSG00000108106",
                "ENSG00000198836",
                "ENSG00000107581",
                "ENSG00000033800",
                "ENSG00000105258",
                "ENSG00000106052",
                "ENSG00000147677",
                "ENSG00000231500",
                "ENSG00000108666",
                "ENSG00000153037",
                "ENSG00000175792",
                "ENSG00000102317",
                "ENSG00000265491",
                "ENSG00000213465",
                "ENSG00000109332",
                "ENSG00000075624",
                "ENSG00000072062",
                "ENSG00000168275",
                "ENSG00000075785",
                "ENSG00000184900",
                "ENSG00000095059",
                "ENSG00000110696",
                "ENSG00000111850",
                "ENSG00000102898",
                "ENSG00000107937",
                "ENSG00000166974",
                "ENSG00000063322",
                "ENSG00000232388",
                "ENSG00000141562",
                "ENSG00000114391",
                "ENSG00000163931",
                "ENSG00000175166",
                "ENSG00000168303",
                "ENSG00000167088",
                "ENSG00000100320",
                "ENSG00000162852",
                "ENSG00000105419",
                "ENSG00000141030",
                "ENSG00000075292",
                "ENSG00000164985",
                "ENSG00000239672",
                "ENSG00000105723",
                "ENSG00000153310",
                "ENSG00000256060",
                "ENSG00000131508",
                "ENSG00000197223",
                "ENSG00000005022",
                "ENSG00000180957",
                "ENSG00000131469",
                "ENSG00000080824",
                "ENSG00000160271",
                "ENSG00000090266",
                "ENSG00000136810",
                "ENSG00000142937",
                "ENSG00000048649",
                "ENSG00000162374",
                "ENSG00000169764",
                "ENSG00000106682",
                "ENSG00000116209",
                "ENSG00000072832",
                "ENSG00000130312",
                "ENSG00000136950",
                "ENSG00000079459",
                "ENSG00000104341",
                "ENSG00000197170",
                "ENSG00000104388",
                "ENSG00000083750",
                "ENSG00000168159",
                "ENSG00000267100",
                "ENSG00000170144",
                "ENSG00000097021",
                "ENSG00000152082",
                "ENSG00000087191",
                "ENSG00000257704",
                "ENSG00000168710",
                "ENSG00000127914",
                "ENSG00000124614",
                "ENSG00000014641",
                "ENSG00000126267",
                "ENSG00000175582",
                "ENSG00000104765",
                "ENSG00000171130",
                "ENSG00000198612",
                "ENSG00000120860",
                "ENSG00000081154",
                "ENSG00000122566",
                "ENSG00000131473",
                "ENSG00000112305",
                "ENSG00000108774",
                "ENSG00000269893",
                "ENSG00000047932",
                "ENSG00000091140",
                "ENSG00000253738",
                "ENSG00000135250",
                "ENSG00000165119",
                "ENSG00000067064",
                "ENSG00000163636",
                "ENSG00000135486",
                "ENSG00000115128",
                "ENSG00000126005",
                "ENSG00000137818",
                "ENSG00000082153",
                "ENSG00000241343",
                "ENSG00000100347",
                "ENSG00000105677",
                "ENSG00000089737",
                "ENSG00000114784",
                "ENSG00000184752",
                "ENSG00000197302",
                "ENSG00000129355",
                "ENSG00000188343",
                "ENSG00000275052",
                "ENSG00000135002",
                "ENSG00000100575",
                "ENSG00000127445",
                "ENSG00000183172",
                "ENSG00000114744",
                "ENSG00000146282",
                "ENSG00000170027",
                "ENSG00000147274",
                "ENSG00000171055",
                "ENSG00000078674",
                "ENSG00000076554",
                "ENSG00000010270",
                "ENSG00000103495",
                "ENSG00000147604",
                "ENSG00000204308",
                "ENSG00000214046",
                "ENSG00000123416",
                "ENSG00000171314",
                "ENSG00000168291",
                "ENSG00000090238",
                "ENSG00000130520",
                "ENSG00000170315",
                "ENSG00000167792",
                "ENSG00000138413",
                "ENSG00000147684",
                "ENSG00000096746",
                "ENSG00000103266",
                "ENSG00000079785",
                "ENSG00000134779",
                "ENSG00000119950",
                "ENSG00000099795",
                "ENSG00000129084",
                "ENSG00000069275",
                "ENSG00000138777",
                "ENSG00000072042",
                "ENSG00000126756",
                "ENSG00000162585",
                "ENSG00000103145",
                "ENSG00000163714",
                "ENSG00000051620",
                "ENSG00000028839",
                "ENSG00000114200",
                "ENSG00000189043",
                "ENSG00000163939",
                "ENSG00000117523",
                "ENSG00000152154",
                "ENSG00000137100",
                "ENSG00000123106",
                "ENSG00000100764",
                "ENSG00000109475",
                "ENSG00000196199",
                "ENSG00000187109",
                "ENSG00000198886",
                "ENSG00000147533",
                "ENSG00000155506",
                "ENSG00000122643",
                "ENSG00000196262",
                "ENSG00000104904",
                "ENSG00000100316",
                "ENSG00000125971",
                "ENSG00000173726",
                "ENSG00000134287",
                "ENSG00000155959",
                "ENSG00000137692",
                "ENSG00000116586",
                "ENSG00000089693",
                "ENSG00000105254",
                "ENSG00000168092",
                "ENSG00000115286",
                "ENSG00000105698",
                "ENSG00000120438",
                "ENSG00000084234",
                "ENSG00000105640",
                "ENSG00000114023",
                "ENSG00000125691",
                "ENSG00000154723",
                "ENSG00000122026",
                "ENSG00000166347",
                "ENSG00000149357",
                "ENSG00000198843",
                "ENSG00000125901",
                "ENSG00000109654",
                "ENSG00000168288",
                "ENSG00000130956",
                "ENSG00000130558",
                "ENSG00000234745",
                "ENSG00000113387",
                "ENSG00000158864",
                "ENSG00000137409",
                "ENSG00000006125",
                "ENSG00000156508",
                "ENSG00000196937",
                "ENSG00000135900",
                "ENSG00000082701",
                "ENSG00000110717",
                "ENSG00000120727",
                "ENSG00000100442",
                "ENSG00000143799",
                "ENSG00000106028",
                "ENSG00000173917",
                "ENSG00000115091",
                "ENSG00000204628",
                "ENSG00000115053",
                "ENSG00000178234",
                "ENSG00000134986",
                "ENSG00000121022",
                "ENSG00000164615",
                "ENSG00000113369",
                "ENSG00000197111",
                "ENSG00000163682",
                "ENSG00000175334",
                "ENSG00000146729",
                "ENSG00000171421",
                "ENSG00000256537",
                "ENSG00000124380",
                "ENSG00000123562",
                "ENSG00000105701",
                "ENSG00000166260",
                "ENSG00000111669",
                "ENSG00000136100",
                "ENSG00000159720",
                "ENSG00000184007",
                "ENSG00000140307",
                "ENSG00000132821",
                "ENSG00000102401",
                "ENSG00000167863",
                "ENSG00000105409",
                "ENSG00000197747",
                "ENSG00000004779",
                "ENSG00000184277",
                "ENSG00000106355",
                "ENSG00000090273",
                "ENSG00000162368",
                "ENSG00000254682",
                "ENSG00000004478",
                "ENSG00000175756",
                "ENSG00000134333",
                "ENSG00000197345",
                "ENSG00000116251",
                "ENSG00000091640",
                "ENSG00000147123",
                "ENSG00000161960",
                "ENSG00000143575",
                "ENSG00000038219",
                "ENSG00000130560",
                "ENSG00000159140",
                "ENSG00000160014",
                "ENSG00000125818",
                "ENSG00000105649",
                "ENSG00000146731",
                "ENSG00000103187",
                "ENSG00000105220",
                "ENSG00000105372",
                "ENSG00000053254",
                "ENSG00000111678",
                "ENSG00000011295",
                "ENSG00000176871",
                "ENSG00000007168",
                "ENSG00000169288",
                "ENSG00000111142",
                "ENSG00000164713",
                "ENSG00000132507",
                "ENSG00000176209",
                "ENSG00000105379",
                "ENSG00000175550",
                "ENSG00000099901",
                "ENSG00000167508",
                "ENSG00000173692",
                "ENSG00000111237",
                "ENSG00000132824",
                "ENSG00000112110",
                "ENSG00000023191",
                "ENSG00000111911",
                "ENSG00000135070",
                "ENSG00000151366",
                "ENSG00000108298",
                "ENSG00000145782",
                "ENSG00000166226",
                "ENSG00000112695",
                "ENSG00000174444",
                "ENSG00000165416",
                "ENSG00000203667",
                "ENSG00000133318",
                "ENSG00000130414",
                "ENSG00000115738",
                "ENSG00000103429",
                "ENSG00000138430",
                "ENSG00000169738",
                "ENSG00000169213",
                "ENSG00000166780",
                "ENSG00000176340",
                "ENSG00000172354",
                "ENSG00000273611",
                "ENSG00000182670",
                "ENSG00000177410",
                "ENSG00000115514",
                "ENSG00000101745",
                "ENSG00000145337",
                "ENSG00000113068",
                "ENSG00000182718",
                "ENSG00000221983",
                "ENSG00000173436",
                "ENSG00000175348",
                "ENSG00000069956",
                "ENSG00000063046",
                "ENSG00000122565",
                "ENSG00000163541",
                "ENSG00000146416",
                "ENSG00000164258",
                "ENSG00000116560",
                "ENSG00000106261",
                "ENSG00000198492",
                "ENSG00000141002",
                "ENSG00000171604",
                "ENSG00000179262",
                "ENSG00000183207",
                "ENSG00000073792",
                "ENSG00000143355",
                "ENSG00000178952",
                "ENSG00000196230",
                "ENSG00000225663",
                "ENSG00000186106",
                "ENSG00000198034",
                "ENSG00000129518",
                "ENSG00000104408",
                "ENSG00000105373",
                "ENSG00000129235",
                "ENSG00000173418",
                "ENSG00000102144",
                "ENSG00000167526",
                "ENSG00000130770",
                "ENSG00000265681",
                "ENSG00000142864",
                "ENSG00000119013",
                "ENSG00000161203",
                "ENSG00000100823",
                "ENSG00000142534",
                "ENSG00000151746",
                "ENSG00000101782",
                "ENSG00000177889",
                "ENSG00000142541",
                "ENSG00000109971",
                "ENSG00000109536",
                "ENSG00000156136",
                "ENSG00000118816",
                "ENSG00000121851",
                "ENSG00000151743",
                "ENSG00000135624",
                "ENSG00000104671",
                "ENSG00000147669",
                "ENSG00000004487",
                "ENSG00000163468",
                "ENSG00000111843",
                "ENSG00000137312",
                "ENSG00000091039",
                "ENSG00000167522",
                "ENSG00000099800",
                "ENSG00000147403",
                "ENSG00000134056",
                "ENSG00000167220",
                "ENSG00000277443",
                "ENSG00000172115",
                "ENSG00000119547",
                "ENSG00000145912",
                "ENSG00000233429",
                "ENSG00000006451",
                "ENSG00000177156",
                "ENSG00000145293",
                "ENSG00000167468",
                "ENSG00000187634",
                "ENSG00000111640",
                "ENSG00000125652",
                "ENSG00000135387",
                "ENSG00000124596",
                "ENSG00000084628",
                "ENSG00000115241",
                "ENSG00000156976",
                "ENSG00000180964",
                "ENSG00000111481",
                "ENSG00000253719",
                "ENSG00000134138",
                "ENSG00000161970",
                "ENSG00000160888",
                "ENSG00000128245",
                "ENSG00000077721",
                "ENSG00000177879",
                "ENSG00000205531",
                "ENSG00000154001",
                "ENSG00000138758",
                "ENSG00000115541",
                "ENSG00000069329",
                "ENSG00000138069",
                "ENSG00000162244",
                "ENSG00000083099",
                "ENSG00000144034",
                "ENSG00000171033",
                "ENSG00000071082",
                "ENSG00000225733",
                "ENSG00000071994",
                "ENSG00000005483",
                "ENSG00000115484",
                "ENSG00000177030",
                "ENSG00000138107",
                "ENSG00000126261",
                "ENSG00000130748",
                "ENSG00000138326",
                "ENSG00000204568",
                "ENSG00000157778",
                "ENSG00000175203",
                "ENSG00000111642",
                "ENSG00000154380",
                "ENSG00000104695",
                "ENSG00000123144",
                "ENSG00000100216",
                "ENSG00000101557",
                "ENSG00000172531",
                "ENSG00000113575",
                "ENSG00000149084",
                "ENSG00000127022",
                "ENSG00000074800",
                "ENSG00000162961",
                "ENSG00000148303",
                "ENSG00000104863",
                "ENSG00000225921",
                "ENSG00000165502",
                "ENSG00000114302",
                "ENSG00000084733",
                "ENSG00000142676",
                "ENSG00000119705",
                "ENSG00000165283",
                "ENSG00000075151",
                "ENSG00000122034",
                "ENSG00000185664",
                "ENSG00000100528",
                "ENSG00000154582",
                "ENSG00000130255",
                "ENSG00000067836",
                "ENSG00000214113",
                "ENSG00000096384",
                "ENSG00000109107",
                "ENSG00000147687",
                "ENSG00000168214",
                "ENSG00000197958",
                "ENSG00000100911",
                "ENSG00000160213",
                "ENSG00000171497",
                "ENSG00000188994",
                "ENSG00000174775",
                "ENSG00000143947",
                "ENSG00000164978",
                "ENSG00000112237",
                "ENSG00000164830",
                "ENSG00000213015",
                "ENSG00000175416",
                "ENSG00000128609",
                "ENSG00000090470",
                "ENSG00000170632",
                "ENSG00000063177",
                "ENSG00000155368",
                "ENSG00000198918",
                "ENSG00000169564",
                "ENSG00000094916",
                "ENSG00000103035",
                "ENSG00000132305",
                "ENSG00000138663",
                "ENSG00000086061",
                "ENSG00000204310",
                "ENSG00000070756",
                "ENSG00000136930",
                "ENSG00000166922",
                "ENSG00000149806",
                "ENSG00000140386",
                "ENSG00000140740",
                "ENSG00000146386",
                "ENSG00000204272",
                "ENSG00000166012",
                "ENSG00000167658",
                "ENSG00000113161",
                "ENSG00000197006",
                "ENSG00000185608",
                "ENSG00000142549",
                "ENSG00000084070",
                "ENSG00000174173",
                "ENSG00000068745",
                "ENSG00000196465",
                "ENSG00000168028",
                "ENSG00000108848",
                "ENSG00000136193",
                "ENSG00000156928",
                "ENSG00000104427",
                "ENSG00000124702",
                "ENSG00000111786",
                "ENSG00000109919",
                "ENSG00000143933",
                "ENSG00000166352",
                "ENSG00000165629",
                "ENSG00000136603",
                "ENSG00000167815",
                "ENSG00000177971",
                "ENSG00000126457",
                "ENSG00000167996",
                "ENSG00000132963",
                "ENSG00000042753",
                "ENSG00000175390",
                "ENSG00000168522",
                "ENSG00000148154",
                "ENSG00000119048",
                "ENSG00000198408",
                "ENSG00000177600",
                "ENSG00000148450",
                "ENSG00000134419",
                "ENSG00000178127",
                "ENSG00000174456",
                "ENSG00000184117",
                "ENSG00000167699",
                "ENSG00000166946",
                "ENSG00000092148",
                "ENSG00000181610",
                "ENSG00000054148",
                "ENSG00000270170",
                "ENSG00000143621",
                "ENSG00000156411",
                "ENSG00000108960",
                "ENSG00000137343",
                "ENSG00000177200",
                "ENSG00000115944",
                "ENSG00000080822",
                "ENSG00000111639",
                "ENSG00000196531",
                "ENSG00000100138",
                "ENSG00000174695",
                "ENSG00000116030",
                "ENSG00000129968",
                "ENSG00000115268",
                "ENSG00000182534",
                "ENSG00000093144",
                "ENSG00000125534",
                "ENSG00000160075",
                "ENSG00000170043",
                "ENSG00000089157",
                "ENSG00000149231",
                "ENSG00000131435",
                "ENSG00000130723",
                "ENSG00000198363",
                "ENSG00000140988",
                "ENSG00000168090",
                "ENSG00000122033",
                "ENSG00000197728",
                "ENSG00000178741",
                "ENSG00000156515",
                "ENSG00000115947",
                "ENSG00000101266",
                "ENSG00000177885",
                "ENSG00000233016",
                "ENSG00000143774",
                "ENSG00000182512",
                "ENSG00000088832",
                "ENSG00000179271",
                "ENSG00000122545",
                "ENSG00000134330",
                "ENSG00000143612",
                "ENSG00000135720",
                "ENSG00000007520",
                "ENSG00000166848",
                "ENSG00000149547",
                "ENSG00000125870",
                "ENSG00000159335",
                "ENSG00000120686",
                "ENSG00000102189",
                "ENSG00000123352",
                "ENSG00000108518",
                "ENSG00000181061",
                "ENSG00000128739",
                "ENSG00000086666",
                "ENSG00000278311",
                "ENSG00000172809",
                "ENSG00000145425",
                "ENSG00000114902",
                "ENSG00000157514",
                "ENSG00000106615",
                "ENSG00000107341",
                "ENSG00000089818",
                "ENSG00000099341",
                "ENSG00000173273",
                "ENSG00000172500",
                "ENSG00000230989",
                "ENSG00000087302",
                "ENSG00000091164",
                "ENSG00000136146",
                "ENSG00000066557",
                "ENSG00000090263",
                "ENSG00000172757",
                "ENSG00000105669",
                "ENSG00000188290",
                "ENSG00000149313",
                "ENSG00000063245",
                "ENSG00000136942",
                "ENSG00000126858",
                "ENSG00000185658",
                "ENSG00000050426",
                "ENSG00000119185",
                "ENSG00000160752",
                "ENSG00000143252",
                "ENSG00000141367",
                "ENSG00000144381",
                "ENSG00000136521",
                "ENSG00000100811",
                "ENSG00000183648",
                "ENSG00000159352",
                "ENSG00000213246",
                "ENSG00000155099",
                "ENSG00000172687",
                "ENSG00000133872",
                "ENSG00000137106",
                "ENSG00000179010",
                "ENSG00000198668",
                "ENSG00000108107",
                "ENSG00000134308",
                "ENSG00000111832",
                "ENSG00000061794",
                "ENSG00000134684",
                "ENSG00000008324",
                "ENSG00000166228",
                "ENSG00000213619",
                "ENSG00000153774",
                "ENSG00000129197",
                "ENSG00000138757",
                "ENSG00000108561",
                "ENSG00000033627",
                "ENSG00000178982",
                "ENSG00000108010",
                "ENSG00000083845",
                "ENSG00000138382",
                "ENSG00000111726",
                "ENSG00000125821",
                "ENSG00000137547",
                "ENSG00000100387",
                "ENSG00000140319",
                "ENSG00000132466",
                "ENSG00000149809",
                "ENSG00000181704",
                "ENSG00000172428",
                "ENSG00000118680",
                "ENSG00000243317",
                "ENSG00000070669",
                "ENSG00000165233",
                "ENSG00000117450",
                "ENSG00000184220",
                "ENSG00000103485",
                "ENSG00000009307",
                "ENSG00000143995",
                "ENSG00000138138",
                "ENSG00000204856",
                "ENSG00000124172",
                "ENSG00000182196",
                "ENSG00000057608",
                "ENSG00000150991",
                "ENSG00000065548",
                "ENSG00000152795",
                "ENSG00000212907",
                "ENSG00000196591",
                "ENSG00000137210",
                "ENSG00000166681",
                "ENSG00000132432",
                "ENSG00000184009",
                "ENSG00000023734",
                "ENSG00000120437",
                "ENSG00000125944",
                "ENSG00000054523",
                "ENSG00000183726",
                "ENSG00000148925",
                "ENSG00000128626",
                "ENSG00000214941",
                "ENSG00000113328",
                "ENSG00000186184",
                "ENSG00000166164",
                "ENSG00000119640",
                "ENSG00000113648",
                "ENSG00000170899",
                "ENSG00000185787",
                "ENSG00000170889",
                "ENSG00000137947",
                "ENSG00000100554",
                "ENSG00000130638",
                "ENSG00000169045",
                "ENSG00000111775",
                "ENSG00000123636",
                "ENSG00000087502",
                "ENSG00000139684",
                "ENSG00000197102",
                "ENSG00000114120",
                "ENSG00000166562",
                "ENSG00000168724",
                "ENSG00000126602",
                "ENSG00000180817",
                "ENSG00000086065",
                "ENSG00000164587",
                "ENSG00000116288",
                "ENSG00000181163",
                "ENSG00000008018",
                "ENSG00000160948",
                "ENSG00000117569",
                "ENSG00000198898",
                "ENSG00000115685",
                "ENSG00000215021",
                "ENSG00000165487",
                "ENSG00000103275",
                "ENSG00000134313",
                "ENSG00000101654",
                "ENSG00000104549",
                "ENSG00000044115",
                "ENSG00000265241",
                "ENSG00000136238",
                "ENSG00000091527",
                "ENSG00000123545",
                "ENSG00000224032",
                "ENSG00000047849",
                "ENSG00000182154",
                "ENSG00000221914",
                "ENSG00000122873",
                "ENSG00000103415",
                "ENSG00000065978",
                "ENSG00000087460",
                "ENSG00000100567",
                "ENSG00000151247",
                "ENSG00000113558",
                "ENSG00000143153",
                "ENSG00000109180",
                "ENSG00000123983",
                "ENSG00000116489",
                "ENSG00000143742",
                "ENSG00000133169",
                "ENSG00000205629",
                "ENSG00000174886",
                "ENSG00000164167",
                "ENSG00000092841",
                "ENSG00000109929",
                "ENSG00000089009",
                "ENSG00000173163"
            ],
            "expression_sum": {
                "class1": 19184.0,
                "class2": 2356.0
            },
            "ens_id": "ENSG00000167081",
            "uniprot_ids": [
                "P40426",
                "H3BLX0",
                "Q5JS98",
                "U3KQA2",
                "H0Y5D0"
            ]
        },
        "ENSG00000149294": {
            "id": "ENSG00000149294",
            "type": "Gene",
            "symbol": "NCAM1",
            "source": [
                "ENSG00000100811",
                "ENSG00000122482",
                "ENSG00000148516",
                "ENSG00000143799",
                "ENSG00000004487",
                "ENSG00000103495",
                "ENSG00000175745",
                "ENSG00000108055",
                "ENSG00000213015",
                "ENSG00000164754",
                "ENSG00000116560",
                "ENSG00000168214",
                "ENSG00000196591",
                "ENSG00000189180",
                "ENSG00000160113",
                "ENSG00000143995"
            ],
            "target": [],
            "expression_sum": {
                "class1": 30352.0,
                "class2": 5877.0
            },
            "ens_id": "ENSG00000149294",
            "uniprot_ids": [
                "P13591",
                "A0A0D9SF30",
                "A0A0C4DGS4",
                "A0A087WV75",
                "A0A087X1V2",
                "A0A087WTF6",
                "A0A087WWJ5",
                "A0A087WVU1",
                "A0A087WVD0",
                "A0A087WTR3",
                "A0A087WZS4",
                "A0A0D9SF98",
                "A0A087WWD4",
                "A0A087WX77",
                "H7BYX6",
                "A0A087WTE4"
            ]
        },
        "ENSG00000149557": {
            "id": "ENSG00000149557",
            "type": "Gene",
            "symbol": "FEZ1",
            "source": [
                "ENSG00000181449",
                "ENSG00000171634",
                "ENSG00000134138",
                "ENSG00000004487",
                "ENSG00000103495",
                "ENSG00000164330",
                "ENSG00000043355",
                "ENSG00000105698",
                "ENSG00000163848",
                "ENSG00000168214",
                "ENSG00000116560",
                "ENSG00000124766",
                "ENSG00000196591"
            ],
            "target": [],
            "expression_sum": {
                "class1": 23424.0,
                "class2": 9868.0
            },
            "ens_id": "ENSG00000149557",
            "uniprot_ids": [
                "Q99689",
                "J3KTR6",
                "E9PNH2"
            ]
        },
        "ENSG00000173267": {
            "id": "ENSG00000173267",
            "type": "Gene",
            "symbol": "SNCG",
            "source": [
                "ENSG00000100811",
                "ENSG00000148516",
                "ENSG00000143799",
                "ENSG00000004487",
                "ENSG00000103495",
                "ENSG00000164330",
                "ENSG00000108055",
                "ENSG00000114861",
                "ENSG00000164754",
                "ENSG00000196591",
                "ENSG00000124766",
                "ENSG00000128272",
                "ENSG00000160113",
                "ENSG00000139613"
            ],
            "target": [],
            "expression_sum": {
                "class1": 68268.0,
                "class2": 2638.0
            },
            "ens_id": "ENSG00000173267",
            "uniprot_ids": [
                "O76070",
                "F8W754"
            ]
        },
        "ENSG00000177675": {
            "id": "ENSG00000177675",
            "type": "Gene",
            "symbol": "CD163L1",
            "source": [
                "ENSG00000100811",
                "ENSG00000109132",
                "ENSG00000016082",
                "ENSG00000004487",
                "ENSG00000103495",
                "ENSG00000108055",
                "ENSG00000116560",
                "ENSG00000196591",
                "ENSG00000130522",
                "ENSG00000137309"
            ],
            "target": [],
            "expression_sum": {
                "class1": 2932.0,
                "class2": 45.0
            },
            "ens_id": "ENSG00000177675",
            "uniprot_ids": [
                "Q9NR16",
                "H0YFE6",
                "F5H7R7",
                "H0YGF0",
                "H0YFR4"
            ]
        },
        "ENSG00000111728": {
            "id": "ENSG00000111728",
            "type": "Gene",
            "symbol": "ST8SIA1",
            "source": [
                "ENSG00000100811",
                "ENSG00000171634",
                "ENSG00000122482",
                "ENSG00000143799",
                "ENSG00000004487",
                "ENSG00000103495",
                "ENSG00000164330",
                "ENSG00000105698",
                "ENSG00000198911",
                "ENSG00000108055",
                "ENSG00000168214",
                "ENSG00000116560",
                "ENSG00000196591",
                "ENSG00000119547"
            ],
            "target": [],
            "expression_sum": {
                "class1": 4857.0,
                "class2": 1397.0
            },
            "ens_id": "ENSG00000111728",
            "uniprot_ids": [
                "Q92185",
                "Q86X71",
                "H0YG41",
                "H0YGV9",
                "H0YFU1",
                "F8WCS8"
            ]
        },
        "ENSG00000018236": {
            "id": "ENSG00000018236",
            "type": "Gene",
            "symbol": "CNTN1",
            "source": [
                "ENSG00000143799",
                "ENSG00000004487",
                "ENSG00000103495",
                "ENSG00000196591",
                "ENSG00000116560"
            ],
            "target": [],
            "expression_sum": {
                "class1": 4784.0,
                "class2": 1757.0
            },
            "ens_id": "ENSG00000018236",
            "uniprot_ids": [
                "Q12860",
                "F8VUI9",
                "F8VQW3",
                "H0YIJ1",
                "F8VUI8",
                "F8VX96"
            ]
        },
        "ENSG00000135406": {
            "id": "ENSG00000135406",
            "type": "Gene",
            "symbol": "PRPH",
            "source": [
                "ENSG00000160113",
                "ENSG00000092199",
                "ENSG00000130726",
                "ENSG00000109132",
                "ENSG00000103495",
                "ENSG00000198911",
                "ENSG00000164754",
                "ENSG00000196591",
                "ENSG00000100811",
                "ENSG00000016082",
                "ENSG00000213015",
                "ENSG00000163848",
                "ENSG00000170365",
                "ENSG00000143799",
                "ENSG00000118263",
                "ENSG00000004487",
                "ENSG00000164330",
                "ENSG00000168610",
                "ENSG00000108055",
                "ENSG00000116560",
                "ENSG00000124766"
            ],
            "target": [],
            "expression_sum": {
                "class1": 45461.0,
                "class2": 258.0
            },
            "ens_id": "ENSG00000135406",
            "uniprot_ids": [
                "P41219",
                "H7C5W5",
                "F8W835"
            ]
        },
        "ENSG00000135447": {
            "id": "ENSG00000135447",
            "type": "Gene",
            "symbol": "PPP1R1A",
            "source": [
                "ENSG00000100811",
                "ENSG00000130726",
                "ENSG00000004487",
                "ENSG00000103495",
                "ENSG00000108055",
                "ENSG00000196591"
            ],
            "target": [],
            "expression_sum": {
                "class1": 21556.0,
                "class2": 1036.0
            },
            "ens_id": "ENSG00000135447",
            "uniprot_ids": [
                "Q13522",
                "H0YIU4",
                "F8VR76",
                "R4GMQ8"
            ]
        },
        "ENSG00000185664": {
            "id": "ENSG00000185664",
            "type": "Gene",
            "symbol": "PMEL",
            "source": [
                "ENSG00000134138",
                "ENSG00000167081",
                "ENSG00000148516",
                "ENSG00000143799",
                "ENSG00000004487",
                "ENSG00000103495",
                "ENSG00000164330",
                "ENSG00000175745",
                "ENSG00000108055",
                "ENSG00000137947",
                "ENSG00000196591",
                "ENSG00000147140",
                "ENSG00000124766",
                "ENSG00000130522",
                "ENSG00000177606",
                "ENSG00000092199"
            ],
            "target": [],
            "expression_sum": {
                "class1": 18655.0,
                "class2": 717.0
            },
            "ens_id": "ENSG00000185664",
            "uniprot_ids": [
                "P40967",
                "F8VXH8",
                "F8VYZ1",
                "F8VZC6",
                "H0YIJ2",
                "F8VZV2",
                "F8VUB1",
                "F8W1D1"
            ]
        },
        "ENSG00000155980": {
            "id": "ENSG00000155980",
            "type": "Gene",
            "symbol": "KIF5A",
            "source": [
                "ENSG00000100811",
                "ENSG00000130726",
                "ENSG00000106261",
                "ENSG00000148516",
                "ENSG00000143799",
                "ENSG00000004487",
                "ENSG00000103495",
                "ENSG00000108055",
                "ENSG00000164754",
                "ENSG00000147140",
                "ENSG00000196591"
            ],
            "target": [],
            "expression_sum": {
                "class1": 8142.0,
                "class2": 1345.0
            },
            "ens_id": "ENSG00000155980",
            "uniprot_ids": [
                "Q12840",
                "A0A6Q8PFR6",
                "A0A6Q8PGG7",
                "J3KNA1",
                "A0A6Q8PH74",
                "A0A6Q8PEZ8",
                "A0A6Q8PGJ3",
                "A0A6Q8PFV6",
                "A0A6Q8PH72",
                "A0A6Q8PFD6",
                "A0A6Q8PHA9",
                "A0A6Q8PFG5",
                "A0A6Q8PGT2",
                "A0A6Q8PGP0",
                "A0A6Q8PGP8",
                "A0A6Q8PH22",
                "A0A6Q8PFB8",
                "A0A6Q8PGN1",
                "A0A6Q8PFQ3"
            ]
        },
        "ENSG00000151320": {
            "id": "ENSG00000151320",
            "type": "Gene",
            "symbol": "AKAP6",
            "source": [
                "ENSG00000100811",
                "ENSG00000109132",
                "ENSG00000143799",
                "ENSG00000004487",
                "ENSG00000108055",
                "ENSG00000116560",
                "ENSG00000196591",
                "ENSG00000128272"
            ],
            "target": [],
            "expression_sum": {
                "class1": 6984.0,
                "class2": 1702.0
            },
            "ens_id": "ENSG00000151320",
            "uniprot_ids": [
                "Q13023",
                "G3V3H2",
                "G3V3B5",
                "G3V3H6",
                "H0YJT7",
                "G3V569",
                "G3V3H7"
            ]
        },
        "ENSG00000165410": {
            "id": "ENSG00000165410",
            "type": "Gene",
            "symbol": "CFL2",
            "source": [
                "ENSG00000004487",
                "ENSG00000164330"
            ],
            "target": [],
            "expression_sum": {
                "class1": 7323.0,
                "class2": 4085.0
            },
            "ens_id": "ENSG00000165410",
            "uniprot_ids": [
                "Q9Y281",
                "G3V2U0",
                "F8WDN3"
            ]
        },
        "ENSG00000139970": {
            "id": "ENSG00000139970",
            "type": "Gene",
            "symbol": "RTN1",
            "source": [
                "ENSG00000148516",
                "ENSG00000092199",
                "ENSG00000130726",
                "ENSG00000109132",
                "ENSG00000103495",
                "ENSG00000198911",
                "ENSG00000164754",
                "ENSG00000168214",
                "ENSG00000196591",
                "ENSG00000100811",
                "ENSG00000100644",
                "ENSG00000119950",
                "ENSG00000170365",
                "ENSG00000143799",
                "ENSG00000004487",
                "ENSG00000168610",
                "ENSG00000100219",
                "ENSG00000108055",
                "ENSG00000116560"
            ],
            "target": [],
            "expression_sum": {
                "class1": 122526.0,
                "class2": 7092.0
            },
            "ens_id": "ENSG00000139970",
            "uniprot_ids": [
                "Q16799",
                "Q2NKQ5",
                "A8MT72"
            ]
        },
        "ENSG00000169856": {
            "id": "ENSG00000169856",
            "type": "TF",
            "symbol": "ONECUT1",
            "source": [
                "ENSG00000100811",
                "ENSG00000122482",
                "ENSG00000004487",
                "ENSG00000103495",
                "ENSG00000108055",
                "ENSG00000213015",
                "ENSG00000164754",
                "ENSG00000196591",
                "ENSG00000147140",
                "ENSG00000168214",
                "ENSG00000160113"
            ],
            "target": [
                "ENSG00000145708"
            ],
            "expression_sum": {
                "class1": 4967.0,
                "class2": 133.0
            },
            "ens_id": "ENSG00000169856",
            "uniprot_ids": [
                "Q9UBC0",
                "U3KQR8"
            ]
        },
        "ENSG00000169783": {
            "id": "ENSG00000169783",
            "type": "Gene",
            "symbol": "LINGO1",
            "source": [
                "ENSG00000100811",
                "ENSG00000143799",
                "ENSG00000004487",
                "ENSG00000103495",
                "ENSG00000198911",
                "ENSG00000108055",
                "ENSG00000119950",
                "ENSG00000168214",
                "ENSG00000116560",
                "ENSG00000196591",
                "ENSG00000147140",
                "ENSG00000145734"
            ],
            "target": [],
            "expression_sum": {
                "class1": 9617.0,
                "class2": 1035.0
            },
            "ens_id": "ENSG00000169783",
            "uniprot_ids": [
                "Q96FE5",
                "H3BMN3",
                "H3BN48",
                "H0YMX3",
                "H3BQ49",
                "H0YNK7",
                "H3BS32",
                "H3BMW2",
                "H3BM59"
            ]
        },
        "ENSG00000103034": {
            "id": "ENSG00000103034",
            "type": "Gene",
            "symbol": "NDRG4",
            "source": [
                "ENSG00000100811",
                "ENSG00000106261",
                "ENSG00000148516",
                "ENSG00000004487",
                "ENSG00000103495",
                "ENSG00000108055",
                "ENSG00000196591",
                "ENSG00000116560",
                "ENSG00000092199"
            ],
            "target": [],
            "expression_sum": {
                "class1": 8687.0,
                "class2": 1931.0
            },
            "ens_id": "ENSG00000103034",
            "uniprot_ids": [
                "Q9ULP0",
                "B7Z9X4",
                "H3BQJ4",
                "H3BPT6",
                "H3BPY2",
                "H3BM44",
                "H3BM40",
                "H3BSC3",
                "H3BQ86",
                "H3BMG5",
                "H3BPH8",
                "H3BUH4",
                "H3BS80",
                "H3BST8",
                "H3BQ01",
                "H3BU25",
                "H3BNQ1",
                "H3BNB4",
                "H3BR92",
                "H3BN97",
                "H3BNX9",
                "H3BRT9",
                "H3BVF3",
                "H3BNE7",
                "H3BPQ3",
                "H3BMR6",
                "H3BUK1",
                "H3BVB0",
                "H3BU08",
                "H3BM83",
                "H3BN53",
                "H3BNK4"
            ]
        },
        "ENSG00000186868": {
            "id": "ENSG00000186868",
            "type": "Gene",
            "symbol": "MAPT",
            "source": [
                "ENSG00000100811",
                "ENSG00000115816",
                "ENSG00000130726",
                "ENSG00000109132",
                "ENSG00000164683",
                "ENSG00000113580",
                "ENSG00000004487",
                "ENSG00000103495",
                "ENSG00000198911",
                "ENSG00000108055",
                "ENSG00000189403",
                "ENSG00000139352",
                "ENSG00000164754",
                "ENSG00000196591",
                "ENSG00000147140",
                "ENSG00000213015",
                "ENSG00000129351",
                "ENSG00000160113"
            ],
            "target": [],
            "expression_sum": {
                "class1": 19355.0,
                "class2": 4424.0
            },
            "ens_id": "ENSG00000186868",
            "uniprot_ids": [
                "P10636",
                "I3L2Z2",
                "A0A7P0T936",
                "A0A7I2PLE3",
                "A0A7I2PJZ2"
            ]
        },
        "ENSG00000182481": {
            "id": "ENSG00000182481",
            "type": "Gene",
            "symbol": "KPNA2",
            "source": [
                "ENSG00000181449",
                "ENSG00000123933",
                "ENSG00000170365",
                "ENSG00000109132",
                "ENSG00000004487",
                "ENSG00000103495",
                "ENSG00000164330",
                "ENSG00000105698",
                "ENSG00000198911",
                "ENSG00000213015",
                "ENSG00000137947",
                "ENSG00000164754",
                "ENSG00000196591",
                "ENSG00000119950",
                "ENSG00000124766",
                "ENSG00000130522",
                "ENSG00000119547",
                "ENSG00000177606"
            ],
            "target": [],
            "expression_sum": {
                "class1": 21185.0,
                "class2": 6929.0
            },
            "ens_id": "ENSG00000182481",
            "uniprot_ids": [
                "P52292",
                "A0A7I2V487",
                "A0A7I2V351",
                "A0A7I2V3Z3",
                "A0A7I2YQE1",
                "J3KS65",
                "J3QLL0",
                "J3QL07"
            ]
        },
        "ENSG00000182534": {
            "id": "ENSG00000182534",
            "type": "Gene",
            "symbol": "MXRA7",
            "source": [
                "ENSG00000181449",
                "ENSG00000167081",
                "ENSG00000004487",
                "ENSG00000103495",
                "ENSG00000164330",
                "ENSG00000108001",
                "ENSG00000108055",
                "ENSG00000116560",
                "ENSG00000196591",
                "ENSG00000124766",
                "ENSG00000160113"
            ],
            "target": [],
            "expression_sum": {
                "class1": 18870.0,
                "class2": 6178.0
            },
            "ens_id": "ENSG00000182534",
            "uniprot_ids": [
                "P84157",
                "K7EQX8",
                "Q6ZR64",
                "K7EPS4"
            ]
        },
        "ENSG00000101210": {
            "id": "ENSG00000101210",
            "type": "Gene",
            "symbol": "EEF1A2",
            "source": [
                "ENSG00000113580",
                "ENSG00000004487",
                "ENSG00000103495",
                "ENSG00000164330",
                "ENSG00000108055",
                "ENSG00000139352",
                "ENSG00000196591"
            ],
            "target": [],
            "expression_sum": {
                "class1": 37906.0,
                "class2": 8677.0
            },
            "ens_id": "ENSG00000101210",
            "uniprot_ids": [
                "Q05639",
                "A0A2R8Y488",
                "A0A6Q8PFK6",
                "A0A2U3TZH3",
                "A0A2R8YDN5",
                "A0A2R8Y660"
            ]
        },
        "ENSG00000099864": {
            "id": "ENSG00000099864",
            "type": "Gene",
            "symbol": "PALM",
            "source": [
                "ENSG00000004487",
                "ENSG00000103495",
                "ENSG00000164330",
                "ENSG00000189403",
                "ENSG00000196591"
            ],
            "target": [],
            "expression_sum": {
                "class1": 10041.0,
                "class2": 3592.0
            },
            "ens_id": "ENSG00000099864",
            "uniprot_ids": [
                "O75781",
                "A0A087WWY4",
                "A0A087X1D6",
                "A0A087WTK8"
            ]
        },
        "ENSG00000104833": {
            "id": "ENSG00000104833",
            "type": "Gene",
            "symbol": "TUBB4A",
            "source": [
                "ENSG00000100811",
                "ENSG00000004487",
                "ENSG00000103495",
                "ENSG00000105698",
                "ENSG00000108055",
                "ENSG00000147140",
                "ENSG00000168214"
            ],
            "target": [],
            "expression_sum": {
                "class1": 5452.0,
                "class2": 789.0
            },
            "ens_id": "ENSG00000104833",
            "uniprot_ids": [
                "P04350",
                "M0QYM7",
                "M0R2D3",
                "M0R042",
                "M0R0M1",
                "M0R1I1",
                "M0QX14",
                "M0QY37",
                "M0QZL7",
                "M0R2T4",
                "M0QY85",
                "M0R278",
                "M0R0X0"
            ]
        },
        "ENSG00000196361": {
            "id": "ENSG00000196361",
            "type": "Gene",
            "symbol": "ELAVL3",
            "source": [
                "ENSG00000100811",
                "ENSG00000130726",
                "ENSG00000148516",
                "ENSG00000004487",
                "ENSG00000108055",
                "ENSG00000196591",
                "ENSG00000168214",
                "ENSG00000092199"
            ],
            "target": [],
            "expression_sum": {
                "class1": 23450.0,
                "class2": 4184.0
            },
            "ens_id": "ENSG00000196361",
            "uniprot_ids": [
                "Q14576",
                "K7EPB5"
            ]
        },
        "ENSG00000105409": {
            "id": "ENSG00000105409",
            "type": "Gene",
            "symbol": "ATP1A3",
            "source": [
                "ENSG00000147862",
                "ENSG00000100811",
                "ENSG00000130726",
                "ENSG00000028277",
                "ENSG00000167081",
                "ENSG00000143799",
                "ENSG00000004487",
                "ENSG00000120837",
                "ENSG00000103495",
                "ENSG00000198911",
                "ENSG00000108055",
                "ENSG00000119950",
                "ENSG00000164754",
                "ENSG00000196591",
                "ENSG00000116560",
                "ENSG00000092199"
            ],
            "target": [],
            "expression_sum": {
                "class1": 17624.0,
                "class2": 1727.0
            },
            "ens_id": "ENSG00000105409",
            "uniprot_ids": [
                "P13637",
                "A0A0A0MT26",
                "M0R116",
                "M0QXF2"
            ]
        },
        "ENSG00000198125": {
            "id": "ENSG00000198125",
            "type": "Gene",
            "symbol": "MB",
            "source": [
                "ENSG00000100811",
                "ENSG00000106261",
                "ENSG00000160113",
                "ENSG00000113580",
                "ENSG00000153147",
                "ENSG00000004487",
                "ENSG00000103495",
                "ENSG00000105698",
                "ENSG00000181722",
                "ENSG00000139352",
                "ENSG00000168214",
                "ENSG00000119547",
                "ENSG00000177606"
            ],
            "target": [],
            "expression_sum": {
                "class1": 427.0,
                "class2": 5139.0
            },
            "ens_id": "ENSG00000198125",
            "uniprot_ids": [
                "P02144",
                "F2Z2F1",
                "Q8WVH6",
                "B0QYF7",
                "F2Z337",
                "B0QYF8"
            ]
        },
        "ENSG00000116560": {
            "id": "ENSG00000116560",
            "type": "TF",
            "symbol": "SFPQ",
            "source": [
                "ENSG00000181449",
                "ENSG00000176887",
                "ENSG00000109132",
                "ENSG00000167081",
                "ENSG00000164330"
            ],
            "target": [
                "ENSG00000103034",
                "ENSG00000174939",
                "ENSG00000185668",
                "ENSG00000005810",
                "ENSG00000113657",
                "ENSG00000198830",
                "ENSG00000164093",
                "ENSG00000182534",
                "ENSG00000129355",
                "ENSG00000137804",
                "ENSG00000128872",
                "ENSG00000186106",
                "ENSG00000187134",
                "ENSG00000170027",
                "ENSG00000132639",
                "ENSG00000075945",
                "ENSG00000168280",
                "ENSG00000148773",
                "ENSG00000164611",
                "ENSG00000108175",
                "ENSG00000172159",
                "ENSG00000170312",
                "ENSG00000066279",
                "ENSG00000138413",
                "ENSG00000067715",
                "ENSG00000118200",
                "ENSG00000087258",
                "ENSG00000175063",
                "ENSG00000111728",
                "ENSG00000272398",
                "ENSG00000134369",
                "ENSG00000123240",
                "ENSG00000115310",
                "ENSG00000166848",
                "ENSG00000105270",
                "ENSG00000018236",
                "ENSG00000096092",
                "ENSG00000099194",
                "ENSG00000162545",
                "ENSG00000176887",
                "ENSG00000169783",
                "ENSG00000157514",
                "ENSG00000204130",
                "ENSG00000125868",
                "ENSG00000084628",
                "ENSG00000130294",
                "ENSG00000100077",
                "ENSG00000104112",
                "ENSG00000131711",
                "ENSG00000187231",
                "ENSG00000144642",
                "ENSG00000163539",
                "ENSG00000123975",
                "ENSG00000213420",
                "ENSG00000171033",
                "ENSG00000155099",
                "ENSG00000130558",
                "ENSG00000157168",
                "ENSG00000170091",
                "ENSG00000206579",
                "ENSG00000145362",
                "ENSG00000170445",
                "ENSG00000141404",
                "ENSG00000118263",
                "ENSG00000149257",
                "ENSG00000185650",
                "ENSG00000164104",
                "ENSG00000134986",
                "ENSG00000176788",
                "ENSG00000172020",
                "ENSG00000133134",
                "ENSG00000149557",
                "ENSG00000145428",
                "ENSG00000114573",
                "ENSG00000173207",
                "ENSG00000130540",
                "ENSG00000198910",
                "ENSG00000018189",
                "ENSG00000138138",
                "ENSG00000128564",
                "ENSG00000105409",
                "ENSG00000151320",
                "ENSG00000005108",
                "ENSG00000213760",
                "ENSG00000092964",
                "ENSG00000143702",
                "ENSG00000142156",
                "ENSG00000152503",
                "ENSG00000149294",
                "ENSG00000171533",
                "ENSG00000166974",
                "ENSG00000174521",
                "ENSG00000164330",
                "ENSG00000137267",
                "ENSG00000151150",
                "ENSG00000100320",
                "ENSG00000181449",
                "ENSG00000118402",
                "ENSG00000188486",
                "ENSG00000131437",
                "ENSG00000113161",
                "ENSG00000050165",
                "ENSG00000082397",
                "ENSG00000117724",
                "ENSG00000171224",
                "ENSG00000143862",
                "ENSG00000162374",
                "ENSG00000134313",
                "ENSG00000104427",
                "ENSG00000221866",
                "ENSG00000198513",
                "ENSG00000197694",
                "ENSG00000115756",
                "ENSG00000126214",
                "ENSG00000131037",
                "ENSG00000131747",
                "ENSG00000131016",
                "ENSG00000139970",
                "ENSG00000135406",
                "ENSG00000159409",
                "ENSG00000113810",
                "ENSG00000169439",
                "ENSG00000177675",
                "ENSG00000137343"
            ],
            "expression_sum": {
                "class1": 30451.0,
                "class2": 15243.0
            },
            "ens_id": "ENSG00000116560",
            "uniprot_ids": [
                "P23246",
                "H0Y9K7",
                "H0Y9U2"
            ]
        },
        "ENSG00000162545": {
            "id": "ENSG00000162545",
            "type": "Gene",
            "symbol": "CAMK2N1",
            "source": [
                "ENSG00000105419",
                "ENSG00000113580",
                "ENSG00000103495",
                "ENSG00000108055",
                "ENSG00000114861",
                "ENSG00000221869",
                "ENSG00000189403",
                "ENSG00000196591",
                "ENSG00000116560"
            ],
            "target": [],
            "expression_sum": {
                "class1": 27273.0,
                "class2": 7911.0
            },
            "ens_id": "ENSG00000162545",
            "uniprot_ids": [
                "Q7Z7J9"
            ]
        },
        "ENSG00000198830": {
            "id": "ENSG00000198830",
            "type": "Gene",
            "symbol": "HMGN2",
            "source": [
                "ENSG00000108055",
                "ENSG00000139352",
                "ENSG00000164754",
                "ENSG00000116560"
            ],
            "target": [],
            "expression_sum": {
                "class1": 45932.0,
                "class2": 79295.0
            },
            "ens_id": "ENSG00000198830",
            "uniprot_ids": [
                "P05204"
            ]
        },
        "ENSG00000084628": {
            "id": "ENSG00000084628",
            "type": "Gene",
            "symbol": "NKAIN1",
            "source": [
                "ENSG00000100811",
                "ENSG00000167081",
                "ENSG00000103495",
                "ENSG00000175745",
                "ENSG00000213015",
                "ENSG00000189403",
                "ENSG00000119950",
                "ENSG00000196591",
                "ENSG00000116560",
                "ENSG00000165512",
                "ENSG00000160113",
                "ENSG00000092199"
            ],
            "target": [],
            "expression_sum": {
                "class1": 7870.0,
                "class2": 558.0
            },
            "ens_id": "ENSG00000084628",
            "uniprot_ids": [
                "Q4KMZ8",
                "R4GNE2",
                "H0YDA0"
            ]
        },
        "ENSG00000185668": {
            "id": "ENSG00000185668",
            "type": "TF",
            "symbol": "POU3F1",
            "source": [
                "ENSG00000103495",
                "ENSG00000116560"
            ],
            "target": [
                "ENSG00000105379"
            ],
            "expression_sum": {
                "class1": 3944.0,
                "class2": 381.0
            },
            "ens_id": "ENSG00000185668",
            "uniprot_ids": [
                "Q03052"
            ]
        },
        "ENSG00000159409": {
            "id": "ENSG00000159409",
            "type": "Gene",
            "symbol": "CELF3",
            "source": [
                "ENSG00000100811",
                "ENSG00000103495",
                "ENSG00000175745",
                "ENSG00000108055",
                "ENSG00000196591",
                "ENSG00000116560"
            ],
            "target": [],
            "expression_sum": {
                "class1": 6727.0,
                "class2": 866.0
            },
            "ens_id": "ENSG00000159409",
            "uniprot_ids": [
                "Q5SZQ8",
                "H0Y623"
            ]
        },
        "ENSG00000173207": {
            "id": "ENSG00000173207",
            "type": "Gene",
            "symbol": "CKS1B",
            "source": [
                "ENSG00000108055",
                "ENSG00000103495",
                "ENSG00000164754",
                "ENSG00000116560"
            ],
            "target": [],
            "expression_sum": {
                "class1": 3247.0,
                "class2": 8182.0
            },
            "ens_id": "ENSG00000173207",
            "uniprot_ids": [
                "P61024",
                "Q9BZU3",
                "Q5T179"
            ]
        },
        "ENSG00000075945": {
            "id": "ENSG00000075945",
            "type": "Gene",
            "symbol": "KIFAP3",
            "source": [
                "ENSG00000167081",
                "ENSG00000113580",
                "ENSG00000148516",
                "ENSG00000103495",
                "ENSG00000164330",
                "ENSG00000196591",
                "ENSG00000116560"
            ],
            "target": [],
            "expression_sum": {
                "class1": 11304.0,
                "class2": 3987.0
            },
            "ens_id": "ENSG00000075945",
            "uniprot_ids": [
                "Q92845"
            ]
        },
        "ENSG00000066279": {
            "id": "ENSG00000066279",
            "type": "Gene",
            "symbol": "ASPM",
            "source": [
                "ENSG00000108055",
                "ENSG00000164754",
                "ENSG00000116560"
            ],
            "target": [],
            "expression_sum": {
                "class1": 50.0,
                "class2": 1619.0
            },
            "ens_id": "ENSG00000066279",
            "uniprot_ids": [
                "Q8IZT6",
                "A0A7P0Z4R7",
                "A0A7P0T8U9",
                "A0A7P0Z491"
            ]
        },
        "ENSG00000118200": {
            "id": "ENSG00000118200",
            "type": "Gene",
            "symbol": "CAMSAP2",
            "source": [
                "ENSG00000189403",
                "ENSG00000067955",
                "ENSG00000116560"
            ],
            "target": [],
            "expression_sum": {
                "class1": 5298.0,
                "class2": 2098.0
            },
            "ens_id": "ENSG00000118200",
            "uniprot_ids": [
                "Q08AD1",
                "H0Y4Z1",
                "H0YE13"
            ]
        },
        "ENSG00000134369": {
            "id": "ENSG00000134369",
            "type": "Gene",
            "symbol": "NAV1",
            "source": [
                "ENSG00000196591",
                "ENSG00000164330",
                "ENSG00000116560",
                "ENSG00000108055"
            ],
            "target": [],
            "expression_sum": {
                "class1": 6132.0,
                "class2": 2317.0
            },
            "ens_id": "ENSG00000134369",
            "uniprot_ids": [
                "Q8NEY1",
                "A0A0A0MRJ3",
                "H7BZD9",
                "H0YA94",
                "H0Y6F6"
            ]
        },
        "ENSG00000143862": {
            "id": "ENSG00000143862",
            "type": "Gene",
            "symbol": "ARL8A",
            "source": [
                "ENSG00000113580",
                "ENSG00000103495",
                "ENSG00000164330",
                "ENSG00000139352",
                "ENSG00000196591",
                "ENSG00000116560"
            ],
            "target": [],
            "expression_sum": {
                "class1": 14517.0,
                "class2": 4707.0
            },
            "ens_id": "ENSG00000143862",
            "uniprot_ids": [
                "Q96BM9",
                "A0A087X2J2"
            ]
        },
        "ENSG00000117724": {
            "id": "ENSG00000117724",
            "type": "Gene",
            "symbol": "CENPF",
            "source": [
                "ENSG00000108055",
                "ENSG00000164754",
                "ENSG00000116560"
            ],
            "target": [],
            "expression_sum": {
                "class1": 802.0,
                "class2": 8188.0
            },
            "ens_id": "ENSG00000117724",
            "uniprot_ids": [
                "P49454",
                "A0A087WTY4"
            ]
        },
        "ENSG00000134313": {
            "id": "ENSG00000134313",
            "type": "Gene",
            "symbol": "KIDINS220",
            "source": [
                "ENSG00000016082",
                "ENSG00000113580",
                "ENSG00000167081",
                "ENSG00000103495",
                "ENSG00000108055",
                "ENSG00000164754",
                "ENSG00000116560"
            ],
            "target": [],
            "expression_sum": {
                "class1": 18866.0,
                "class2": 6369.0
            },
            "ens_id": "ENSG00000134313",
            "uniprot_ids": [
                "Q9ULH0",
                "H0Y8E4",
                "F8WAY8",
                "A0A1W2PPY4",
                "A0A1W2PPB7",
                "E9PH70",
                "H7C584"
            ]
        },
        "ENSG00000115756": {
            "id": "ENSG00000115756",
            "type": "Gene",
            "symbol": "HPCAL1",
            "source": [
                "ENSG00000103495",
                "ENSG00000164330",
                "ENSG00000177030",
                "ENSG00000108055",
                "ENSG00000189403",
                "ENSG00000213015",
                "ENSG00000116560"
            ],
            "target": [],
            "expression_sum": {
                "class1": 7034.0,
                "class2": 781.0
            },
            "ens_id": "ENSG00000115756",
            "uniprot_ids": [
                "P37235",
                "H7BZC1",
                "E9PC71",
                "C9JW46"
            ]
        },
        "ENSG00000115310": {
            "id": "ENSG00000115310",
            "type": "Gene",
            "symbol": "RTN4",
            "source": [
                "ENSG00000181449",
                "ENSG00000115816",
                "ENSG00000169136",
                "ENSG00000148516",
                "ENSG00000118263",
                "ENSG00000103495",
                "ENSG00000164330",
                "ENSG00000114861",
                "ENSG00000067955",
                "ENSG00000116560",
                "ENSG00000196591",
                "ENSG00000181315",
                "ENSG00000128272",
                "ENSG00000160113"
            ],
            "target": [],
            "expression_sum": {
                "class1": 92423.0,
                "class2": 34405.0
            },
            "ens_id": "ENSG00000115310",
            "uniprot_ids": [
                "Q9NQC3",
                "C9J685",
                "A0A0U1RQR6",
                "F8W914",
                "H7C106"
            ]
        },
        "ENSG00000187231": {
            "id": "ENSG00000187231",
            "type": "Gene",
            "symbol": "SESTD1",
            "source": [
                "ENSG00000163435",
                "ENSG00000113580",
                "ENSG00000148516",
                "ENSG00000103495",
                "ENSG00000164330",
                "ENSG00000116560"
            ],
            "target": [],
            "expression_sum": {
                "class1": 13173.0,
                "class2": 4286.0
            },
            "ens_id": "ENSG00000187231",
            "uniprot_ids": [
                "Q86VW0",
                "F8WFB2",
                "C9J4X8",
                "H7BZR8",
                "C9JHW6",
                "H7BXT9",
                "H0Y774"
            ]
        },
        "ENSG00000118263": {
            "id": "ENSG00000118263",
            "type": "TF",
            "symbol": "KLF7",
            "source": [
                "ENSG00000100811",
                "ENSG00000106261",
                "ENSG00000016082",
                "ENSG00000148516",
                "ENSG00000103495",
                "ENSG00000164330",
                "ENSG00000108055",
                "ENSG00000067955",
                "ENSG00000116560",
                "ENSG00000196591",
                "ENSG00000128272"
            ],
            "target": [
                "ENSG00000078668",
                "ENSG00000173272",
                "ENSG00000139644",
                "ENSG00000197756",
                "ENSG00000185651",
                "ENSG00000182004",
                "ENSG00000065135",
                "ENSG00000134248",
                "ENSG00000103495",
                "ENSG00000010270",
                "ENSG00000189403",
                "ENSG00000108424",
                "ENSG00000156261",
                "ENSG00000101752",
                "ENSG00000197728",
                "ENSG00000166913",
                "ENSG00000079785",
                "ENSG00000173812",
                "ENSG00000099795",
                "ENSG00000118816",
                "ENSG00000135441",
                "ENSG00000104671",
                "ENSG00000215712",
                "ENSG00000115310",
                "ENSG00000113742",
                "ENSG00000159335",
                "ENSG00000108518",
                "ENSG00000145912",
                "ENSG00000278311",
                "ENSG00000172809",
                "ENSG00000155366",
                "ENSG00000172757",
                "ENSG00000168092",
                "ENSG00000131236",
                "ENSG00000115286",
                "ENSG00000077721",
                "ENSG00000119185",
                "ENSG00000141367",
                "ENSG00000144381",
                "ENSG00000115541",
                "ENSG00000133083",
                "ENSG00000162244",
                "ENSG00000133703",
                "ENSG00000135900",
                "ENSG00000034510",
                "ENSG00000146425",
                "ENSG00000103769",
                "ENSG00000241468",
                "ENSG00000104695",
                "ENSG00000141404",
                "ENSG00000273841",
                "ENSG00000106028",
                "ENSG00000113575",
                "ENSG00000126934",
                "ENSG00000145592",
                "ENSG00000121022",
                "ENSG00000175334",
                "ENSG00000092020",
                "ENSG00000117450",
                "ENSG00000184007",
                "ENSG00000184220",
                "ENSG00000178980",
                "ENSG00000164346",
                "ENSG00000167863",
                "ENSG00000150991",
                "ENSG00000169100",
                "ENSG00000108106",
                "ENSG00000070831",
                "ENSG00000092964",
                "ENSG00000171497",
                "ENSG00000110958",
                "ENSG00000116251",
                "ENSG00000025293",
                "ENSG00000175792",
                "ENSG00000102317",
                "ENSG00000184009",
                "ENSG00000124357",
                "ENSG00000125818",
                "ENSG00000100380",
                "ENSG00000104643",
                "ENSG00000167552",
                "ENSG00000137267",
                "ENSG00000132305",
                "ENSG00000100320",
                "ENSG00000112701",
                "ENSG00000099901",
                "ENSG00000164985",
                "ENSG00000128989",
                "ENSG00000239672",
                "ENSG00000131469",
                "ENSG00000164587",
                "ENSG00000142937",
                "ENSG00000158710",
                "ENSG00000136938",
                "ENSG00000108826",
                "ENSG00000044115",
                "ENSG00000136238",
                "ENSG00000197694",
                "ENSG00000143933",
                "ENSG00000182154",
                "ENSG00000221914",
                "ENSG00000126214",
                "ENSG00000172354",
                "ENSG00000168522",
                "ENSG00000152082",
                "ENSG00000177600",
                "ENSG00000151914",
                "ENSG00000174456",
                "ENSG00000166946",
                "ENSG00000221983",
                "ENSG00000135406",
                "ENSG00000174886",
                "ENSG00000137710",
                "ENSG00000106244",
                "ENSG00000092841",
                "ENSG00000165119"
            ],
            "expression_sum": {
                "class1": 7577.0,
                "class2": 1532.0
            },
            "ens_id": "ENSG00000118263",
            "uniprot_ids": [
                "O75840",
                "E7EQY2",
                "C9JX20",
                "E7EUU0",
                "E9PHC9"
            ]
        },
        "ENSG00000138413": {
            "id": "ENSG00000138413",
            "type": "Gene",
            "symbol": "IDH1",
            "source": [
                "ENSG00000016082",
                "ENSG00000167081",
                "ENSG00000148516",
                "ENSG00000103495",
                "ENSG00000164330",
                "ENSG00000108055",
                "ENSG00000067955",
                "ENSG00000116560",
                "ENSG00000196591",
                "ENSG00000128272",
                "ENSG00000119547"
            ],
            "target": [],
            "expression_sum": {
                "class1": 11773.0,
                "class2": 4323.0
            },
            "ens_id": "ENSG00000138413",
            "uniprot_ids": [
                "O75874",
                "C9J4N6",
                "C9JJE5",
                "C9JLU6"
            ]
        },
        "ENSG00000130294": {
            "id": "ENSG00000130294",
            "type": "Gene",
            "symbol": "KIF1A",
            "source": [
                "ENSG00000181449",
                "ENSG00000113580",
                "ENSG00000148516",
                "ENSG00000103495",
                "ENSG00000164330",
                "ENSG00000139352",
                "ENSG00000116560",
                "ENSG00000128272"
            ],
            "target": [],
            "expression_sum": {
                "class1": 18749.0,
                "class2": 11355.0
            },
            "ens_id": "ENSG00000130294",
            "uniprot_ids": [
                "Q12756",
                "A0A6Q8PFE9",
                "A0A3B3ITW6",
                "A0A3B3ITE5",
                "A0A6Q8PFR1",
                "H7C3Y8",
                "A0A3B3ITK3",
                "A0A3B3ITW4",
                "A0A6Q8PH56",
                "A0A3B3ITF7",
                "A0A3B3IU40",
                "A0A3B3IUA1",
                "C9JBH1",
                "A0A3F2YNW9",
                "A0A3B3IT28",
                "A0A6Q8PFJ8",
                "H7C0K6",
                "A0A6Q8PFL2",
                "A0A6Q8PGW8",
                "A0A6Q8PFR5",
                "A0A6Q8PGE4",
                "A0A6Q8PHQ5"
            ]
        },
        "ENSG00000144642": {
            "id": "ENSG00000144642",
            "type": "Gene",
            "symbol": "RBMS3",
            "source": [
                "ENSG00000170365",
                "ENSG00000109132",
                "ENSG00000143799",
                "ENSG00000103495",
                "ENSG00000164330",
                "ENSG00000198911",
                "ENSG00000108055",
                "ENSG00000116560",
                "ENSG00000196591",
                "ENSG00000181315",
                "ENSG00000137309"
            ],
            "target": [],
            "expression_sum": {
                "class1": 3101.0,
                "class2": 920.0
            },
            "ens_id": "ENSG00000144642",
            "uniprot_ids": [
                "Q6XE24",
                "A0A1B0GTI7",
                "C9JIJ9",
                "A0A1B0GUW5",
                "C9J9B2"
            ]
        },
        "ENSG00000163539": {
            "id": "ENSG00000163539",
            "type": "Gene",
            "symbol": "CLASP2",
            "source": [
                "ENSG00000148516",
                "ENSG00000103495",
                "ENSG00000198911",
                "ENSG00000108055",
                "ENSG00000147140",
                "ENSG00000116560"
            ],
            "target": [],
            "expression_sum": {
                "class1": 10088.0,
                "class2": 2698.0
            },
            "ens_id": "ENSG00000163539",
            "uniprot_ids": [
                "O75122",
                "B3KR06",
                "E7ERI8",
                "H7C4I5",
                "H7C5M8",
                "E3W994",
                "A0A804HJG7",
                "E7EW49",
                "H7C4X8",
                "D6RBU8",
                "J3KR49",
                "A0A0U1RQI6",
                "E7ENG2",
                "H7C4M5",
                "C9J668"
            ]
        },
        "ENSG00000114573": {
            "id": "ENSG00000114573",
            "type": "Gene",
            "symbol": "ATP6V1A",
            "source": [
                "ENSG00000163435",
                "ENSG00000113580",
                "ENSG00000148516",
                "ENSG00000108055",
                "ENSG00000139352",
                "ENSG00000116560"
            ],
            "target": [],
            "expression_sum": {
                "class1": 7620.0,
                "class2": 3687.0
            },
            "ens_id": "ENSG00000114573",
            "uniprot_ids": [
                "P38606",
                "F8WDJ3",
                "C9JVW8",
                "C9JA17"
            ]
        },
        "ENSG00000113810": {
            "id": "ENSG00000113810",
            "type": "Gene",
            "symbol": "SMC4",
            "source": [
                "ENSG00000103495",
                "ENSG00000108055",
                "ENSG00000189403",
                "ENSG00000164754",
                "ENSG00000116560"
            ],
            "target": [],
            "expression_sum": {
                "class1": 625.0,
                "class2": 3489.0
            },
            "ens_id": "ENSG00000113810",
            "uniprot_ids": [
                "Q9NTJ3",
                "C9IYK2",
                "E9PD53",
                "C9J578",
                "C9JR83",
                "C9J9E4",
                "C9JVD8",
                "C9JJ64",
                "C9JWF0",
                "F8WCB7"
            ]
        },
        "ENSG00000181449": {
            "id": "ENSG00000181449",
            "type": "TF",
            "symbol": "SOX2",
            "source": [
                "ENSG00000100811",
                "ENSG00000105698",
                "ENSG00000103495",
                "ENSG00000108055",
                "ENSG00000163848",
                "ENSG00000116560",
                "ENSG00000164754",
                "ENSG00000128610",
                "ENSG00000130522"
            ],
            "target": [
                "ENSG00000078668",
                "ENSG00000206573",
                "ENSG00000112081",
                "ENSG00000107223",
                "ENSG00000162704",
                "ENSG00000134884",
                "ENSG00000163634",
                "ENSG00000165655",
                "ENSG00000153130",
                "ENSG00000111716",
                "ENSG00000139644",
                "ENSG00000099622",
                "ENSG00000090060",
                "ENSG00000116161",
                "ENSG00000106603",
                "ENSG00000117395",
                "ENSG00000116350",
                "ENSG00000161016",
                "ENSG00000134248",
                "ENSG00000131143",
                "ENSG00000184983",
                "ENSG00000127540",
                "ENSG00000099797",
                "ENSG00000119632",
                "ENSG00000189403",
                "ENSG00000108424",
                "ENSG00000159377",
                "ENSG00000196419",
                "ENSG00000075415",
                "ENSG00000107874",
                "ENSG00000105993",
                "ENSG00000102172",
                "ENSG00000149212",
                "ENSG00000213722",
                "ENSG00000150779",
                "ENSG00000165898",
                "ENSG00000077380",
                "ENSG00000154556",
                "ENSG00000048740",
                "ENSG00000163466",
                "ENSG00000119421",
                "ENSG00000052802",
                "ENSG00000134283",
                "ENSG00000173812",
                "ENSG00000117592",
                "ENSG00000173915",
                "ENSG00000167085",
                "ENSG00000151729",
                "ENSG00000135441",
                "ENSG00000106443",
                "ENSG00000134453",
                "ENSG00000041357",
                "ENSG00000182220",
                "ENSG00000163875",
                "ENSG00000163319",
                "ENSG00000164924",
                "ENSG00000149196",
                "ENSG00000141759",
                "ENSG00000135069",
                "ENSG00000162664",
                "ENSG00000204392",
                "ENSG00000165264",
                "ENSG00000224877",
                "ENSG00000139116",
                "ENSG00000114354",
                "ENSG00000277791",
                "ENSG00000156482",
                "ENSG00000125868",
                "ENSG00000165637",
                "ENSG00000165672",
                "ENSG00000173674",
                "ENSG00000147548",
                "ENSG00000171703",
                "ENSG00000167775",
                "ENSG00000129625",
                "ENSG00000089220",
                "ENSG00000116898",
                "ENSG00000196072",
                "ENSG00000179431",
                "ENSG00000255302",
                "ENSG00000100526",
                "ENSG00000188243",
                "ENSG00000161542",
                "ENSG00000067225",
                "ENSG00000109390",
                "ENSG00000141552",
                "ENSG00000113712",
                "ENSG00000126698",
                "ENSG00000136143",
                "ENSG00000174851",
                "ENSG00000150459",
                "ENSG00000170540",
                "ENSG00000233927",
                "ENSG00000205581",
                "ENSG00000159685",
                "ENSG00000026025",
                "ENSG00000034713",
                "ENSG00000144746",
                "ENSG00000034510",
                "ENSG00000146425",
                "ENSG00000138071",
                "ENSG00000120963",
                "ENSG00000117410",
                "ENSG00000170584",
                "ENSG00000075142",
                "ENSG00000273841",
                "ENSG00000124802",
                "ENSG00000263001",
                "ENSG00000166275",
                "ENSG00000183011",
                "ENSG00000126934",
                "ENSG00000213741",
                "ENSG00000215845",
                "ENSG00000175701",
                "ENSG00000121774",
                "ENSG00000069849",
                "ENSG00000174917",
                "ENSG00000147649",
                "ENSG00000129515",
                "ENSG00000100519",
                "ENSG00000187514",
                "ENSG00000163041",
                "ENSG00000155115",
                "ENSG00000141027",
                "ENSG00000092199",
                "ENSG00000110955",
                "ENSG00000178980",
                "ENSG00000175216",
                "ENSG00000089335",
                "ENSG00000083807",
                "ENSG00000150316",
                "ENSG00000010256",
                "ENSG00000172239",
                "ENSG00000065518",
                "ENSG00000174953",
                "ENSG00000152767",
                "ENSG00000109775",
                "ENSG00000162191",
                "ENSG00000118363",
                "ENSG00000129562",
                "ENSG00000135940",
                "ENSG00000108654",
                "ENSG00000108106",
                "ENSG00000162599",
                "ENSG00000172336",
                "ENSG00000107581",
                "ENSG00000131966",
                "ENSG00000105258",
                "ENSG00000112972",
                "ENSG00000204842",
                "ENSG00000085224",
                "ENSG00000153037",
                "ENSG00000113013",
                "ENSG00000265491",
                "ENSG00000092621",
                "ENSG00000106245",
                "ENSG00000213465",
                "ENSG00000109332",
                "ENSG00000163170",
                "ENSG00000079246",
                "ENSG00000168275",
                "ENSG00000075785",
                "ENSG00000198937",
                "ENSG00000075624",
                "ENSG00000244879",
                "ENSG00000171222",
                "ENSG00000095059",
                "ENSG00000105185",
                "ENSG00000131051",
                "ENSG00000167552",
                "ENSG00000063322",
                "ENSG00000205726",
                "ENSG00000175166",
                "ENSG00000168303",
                "ENSG00000167088",
                "ENSG00000240972",
                "ENSG00000196655",
                "ENSG00000029993",
                "ENSG00000164985",
                "ENSG00000239672",
                "ENSG00000213585",
                "ENSG00000131508",
                "ENSG00000005022",
                "ENSG00000127616",
                "ENSG00000154277",
                "ENSG00000131469",
                "ENSG00000099817",
                "ENSG00000171603",
                "ENSG00000128654",
                "ENSG00000047410",
                "ENSG00000169714",
                "ENSG00000090266",
                "ENSG00000142937",
                "ENSG00000136810",
                "ENSG00000171634",
                "ENSG00000125656",
                "ENSG00000232112",
                "ENSG00000127184",
                "ENSG00000106682",
                "ENSG00000130312",
                "ENSG00000197694",
                "ENSG00000079459",
                "ENSG00000104388",
                "ENSG00000110917",
                "ENSG00000168159",
                "ENSG00000170144",
                "ENSG00000064607",
                "ENSG00000171530",
                "ENSG00000152082",
                "ENSG00000087191",
                "ENSG00000166734",
                "ENSG00000123349",
                "ENSG00000105583",
                "ENSG00000143870",
                "ENSG00000147655",
                "ENSG00000014641",
                "ENSG00000167614",
                "ENSG00000164683",
                "ENSG00000106803",
                "ENSG00000126267",
                "ENSG00000104765",
                "ENSG00000120860",
                "ENSG00000198612",
                "ENSG00000137710",
                "ENSG00000081154",
                "ENSG00000122566",
                "ENSG00000182199",
                "ENSG00000130725",
                "ENSG00000100129",
                "ENSG00000112305",
                "ENSG00000142507",
                "ENSG00000108774",
                "ENSG00000237765",
                "ENSG00000165119",
                "ENSG00000067064",
                "ENSG00000135486",
                "ENSG00000115128",
                "ENSG00000132718",
                "ENSG00000137818",
                "ENSG00000129559",
                "ENSG00000068796",
                "ENSG00000105677",
                "ENSG00000114784",
                "ENSG00000184752",
                "ENSG00000169567",
                "ENSG00000198258",
                "ENSG00000183172",
                "ENSG00000127445",
                "ENSG00000128951",
                "ENSG00000100575",
                "ENSG00000105810",
                "ENSG00000188186",
                "ENSG00000171735",
                "ENSG00000241837",
                "ENSG00000147274",
                "ENSG00000103495",
                "ENSG00000204308",
                "ENSG00000214046",
                "ENSG00000176101",
                "ENSG00000123416",
                "ENSG00000131495",
                "ENSG00000144713",
                "ENSG00000155957",
                "ENSG00000130520",
                "ENSG00000084623",
                "ENSG00000086758",
                "ENSG00000108671",
                "ENSG00000100852",
                "ENSG00000167792",
                "ENSG00000147684",
                "ENSG00000096746",
                "ENSG00000170315",
                "ENSG00000079785",
                "ENSG00000119950",
                "ENSG00000163848",
                "ENSG00000099795",
                "ENSG00000129084",
                "ENSG00000069275",
                "ENSG00000079462",
                "ENSG00000138777",
                "ENSG00000072042",
                "ENSG00000069345",
                "ENSG00000162585",
                "ENSG00000163714",
                "ENSG00000189043",
                "ENSG00000099246",
                "ENSG00000170515",
                "ENSG00000142192",
                "ENSG00000127838",
                "ENSG00000131828",
                "ENSG00000234545",
                "ENSG00000137100",
                "ENSG00000167283",
                "ENSG00000164405",
                "ENSG00000170035",
                "ENSG00000196199",
                "ENSG00000187109",
                "ENSG00000155506",
                "ENSG00000044574",
                "ENSG00000196262",
                "ENSG00000104904",
                "ENSG00000115355",
                "ENSG00000125977",
                "ENSG00000170348",
                "ENSG00000173726",
                "ENSG00000089280",
                "ENSG00000155959",
                "ENSG00000137692",
                "ENSG00000116586",
                "ENSG00000089693",
                "ENSG00000105254",
                "ENSG00000117419",
                "ENSG00000119335",
                "ENSG00000115286",
                "ENSG00000077152",
                "ENSG00000144028",
                "ENSG00000084234",
                "ENSG00000114023",
                "ENSG00000127774",
                "ENSG00000101367",
                "ENSG00000172893",
                "ENSG00000154723",
                "ENSG00000122026",
                "ENSG00000166598",
                "ENSG00000167978",
                "ENSG00000198729",
                "ENSG00000149357",
                "ENSG00000131174",
                "ENSG00000106400",
                "ENSG00000143013",
                "ENSG00000109654",
                "ENSG00000179091",
                "ENSG00000254999",
                "ENSG00000133935",
                "ENSG00000177556",
                "ENSG00000113387",
                "ENSG00000158864",
                "ENSG00000137409",
                "ENSG00000006125",
                "ENSG00000188690",
                "ENSG00000153944",
                "ENSG00000241468",
                "ENSG00000136888",
                "ENSG00000110717",
                "ENSG00000120727",
                "ENSG00000127884",
                "ENSG00000185630",
                "ENSG00000100442",
                "ENSG00000143799",
                "ENSG00000106028",
                "ENSG00000140612",
                "ENSG00000115091",
                "ENSG00000204628",
                "ENSG00000115053",
                "ENSG00000102241",
                "ENSG00000106367",
                "ENSG00000121022",
                "ENSG00000164615",
                "ENSG00000139168",
                "ENSG00000197111",
                "ENSG00000111275",
                "ENSG00000139343",
                "ENSG00000175334",
                "ENSG00000146729",
                "ENSG00000171421",
                "ENSG00000085231",
                "ENSG00000133134",
                "ENSG00000123562",
                "ENSG00000099783",
                "ENSG00000105701",
                "ENSG00000111669",
                "ENSG00000136100",
                "ENSG00000182093",
                "ENSG00000134825",
                "ENSG00000185745",
                "ENSG00000140307",
                "ENSG00000132463",
                "ENSG00000167863",
                "ENSG00000004779",
                "ENSG00000106355",
                "ENSG00000071553",
                "ENSG00000165280",
                "ENSG00000169139",
                "ENSG00000254682",
                "ENSG00000116459",
                "ENSG00000175756",
                "ENSG00000203950",
                "ENSG00000174684",
                "ENSG00000134333",
                "ENSG00000197345",
                "ENSG00000006625",
                "ENSG00000091640",
                "ENSG00000147123",
                "ENSG00000143575",
                "ENSG00000143977",
                "ENSG00000038219",
                "ENSG00000249915",
                "ENSG00000130560",
                "ENSG00000110700",
                "ENSG00000167004",
                "ENSG00000160014",
                "ENSG00000100201",
                "ENSG00000213639",
                "ENSG00000159140",
                "ENSG00000183617",
                "ENSG00000125818",
                "ENSG00000146540",
                "ENSG00000105220",
                "ENSG00000143353",
                "ENSG00000170892",
                "ENSG00000111678",
                "ENSG00000067560",
                "ENSG00000163001",
                "ENSG00000130724",
                "ENSG00000007168",
                "ENSG00000156639",
                "ENSG00000164713",
                "ENSG00000116649",
                "ENSG00000132507",
                "ENSG00000176209",
                "ENSG00000105379",
                "ENSG00000175550",
                "ENSG00000099901",
                "ENSG00000143727",
                "ENSG00000126067",
                "ENSG00000173692",
                "ENSG00000111237",
                "ENSG00000132670",
                "ENSG00000112110",
                "ENSG00000023191",
                "ENSG00000215301",
                "ENSG00000104886",
                "ENSG00000151366",
                "ENSG00000108298",
                "ENSG00000214413",
                "ENSG00000166226",
                "ENSG00000112695",
                "ENSG00000116747",
                "ENSG00000153187",
                "ENSG00000106628",
                "ENSG00000077782",
                "ENSG00000174444",
                "ENSG00000203667",
                "ENSG00000100354",
                "ENSG00000133318",
                "ENSG00000130414",
                "ENSG00000169738",
                "ENSG00000185236",
                "ENSG00000137285",
                "ENSG00000176340",
                "ENSG00000172354",
                "ENSG00000150753",
                "ENSG00000164096",
                "ENSG00000049618",
                "ENSG00000136824",
                "ENSG00000138385",
                "ENSG00000083457",
                "ENSG00000140264",
                "ENSG00000189159",
                "ENSG00000116754",
                "ENSG00000101745",
                "ENSG00000145337",
                "ENSG00000140990",
                "ENSG00000221983",
                "ENSG00000173436",
                "ENSG00000175348",
                "ENSG00000063046",
                "ENSG00000122565",
                "ENSG00000163541",
                "ENSG00000008952",
                "ENSG00000127483",
                "ENSG00000146416",
                "ENSG00000230124",
                "ENSG00000164258",
                "ENSG00000116560",
                "ENSG00000100941",
                "ENSG00000109472",
                "ENSG00000114956",
                "ENSG00000147862",
                "ENSG00000140391",
                "ENSG00000173272",
                "ENSG00000137815",
                "ENSG00000141002",
                "ENSG00000171604",
                "ENSG00000179262",
                "ENSG00000196976",
                "ENSG00000197756",
                "ENSG00000178952",
                "ENSG00000143155",
                "ENSG00000196230",
                "ENSG00000225663",
                "ENSG00000182004",
                "ENSG00000141526",
                "ENSG00000124733",
                "ENSG00000173418",
                "ENSG00000102144",
                "ENSG00000272325",
                "ENSG00000130770",
                "ENSG00000029363",
                "ENSG00000156467",
                "ENSG00000167526",
                "ENSG00000100028",
                "ENSG00000168259",
                "ENSG00000142864",
                "ENSG00000119013",
                "ENSG00000161203",
                "ENSG00000182944",
                "ENSG00000184076",
                "ENSG00000137288",
                "ENSG00000105887",
                "ENSG00000100836",
                "ENSG00000266472",
                "ENSG00000101782",
                "ENSG00000177889",
                "ENSG00000142541",
                "ENSG00000109971",
                "ENSG00000166913",
                "ENSG00000204435",
                "ENSG00000159199",
                "ENSG00000118816",
                "ENSG00000121851",
                "ENSG00000135624",
                "ENSG00000104671",
                "ENSG00000147669",
                "ENSG00000168003",
                "ENSG00000126524",
                "ENSG00000163468",
                "ENSG00000111843",
                "ENSG00000074071",
                "ENSG00000099800",
                "ENSG00000150687",
                "ENSG00000140905",
                "ENSG00000134056",
                "ENSG00000277443",
                "ENSG00000172115",
                "ENSG00000101558",
                "ENSG00000145912",
                "ENSG00000140350",
                "ENSG00000145293",
                "ENSG00000181924",
                "ENSG00000074696",
                "ENSG00000175130",
                "ENSG00000167468",
                "ENSG00000111640",
                "ENSG00000197746",
                "ENSG00000183978",
                "ENSG00000125652",
                "ENSG00000115241",
                "ENSG00000253719",
                "ENSG00000156976",
                "ENSG00000100632",
                "ENSG00000161970",
                "ENSG00000130294",
                "ENSG00000078369",
                "ENSG00000119396",
                "ENSG00000105404",
                "ENSG00000177879",
                "ENSG00000205531",
                "ENSG00000099624",
                "ENSG00000120265",
                "ENSG00000154001",
                "ENSG00000138758",
                "ENSG00000115541",
                "ENSG00000168066",
                "ENSG00000133083",
                "ENSG00000069329",
                "ENSG00000138069",
                "ENSG00000083099",
                "ENSG00000110321",
                "ENSG00000225733",
                "ENSG00000111875",
                "ENSG00000071994",
                "ENSG00000130748",
                "ENSG00000115484",
                "ENSG00000169689",
                "ENSG00000168785",
                "ENSG00000110651",
                "ENSG00000138326",
                "ENSG00000204568",
                "ENSG00000175203",
                "ENSG00000102409",
                "ENSG00000154380",
                "ENSG00000145833",
                "ENSG00000104695",
                "ENSG00000123144",
                "ENSG00000113575",
                "ENSG00000143256",
                "ENSG00000127022",
                "ENSG00000149084",
                "ENSG00000074800",
                "ENSG00000128610",
                "ENSG00000054118",
                "ENSG00000105193",
                "ENSG00000084733",
                "ENSG00000142676",
                "ENSG00000138668",
                "ENSG00000179950",
                "ENSG00000119705",
                "ENSG00000133026",
                "ENSG00000149557",
                "ENSG00000165283",
                "ENSG00000075151",
                "ENSG00000122034",
                "ENSG00000154582",
                "ENSG00000130255",
                "ENSG00000120533",
                "ENSG00000162734",
                "ENSG00000196305",
                "ENSG00000103024",
                "ENSG00000096384",
                "ENSG00000022277",
                "ENSG00000105223",
                "ENSG00000160310",
                "ENSG00000174469",
                "ENSG00000151665",
                "ENSG00000164902",
                "ENSG00000144224",
                "ENSG00000188994",
                "ENSG00000174775",
                "ENSG00000143947",
                "ENSG00000071794",
                "ENSG00000213015",
                "ENSG00000175416",
                "ENSG00000128609",
                "ENSG00000063177",
                "ENSG00000108953",
                "ENSG00000155368",
                "ENSG00000050393",
                "ENSG00000125611",
                "ENSG00000169564",
                "ENSG00000094916",
                "ENSG00000103035",
                "ENSG00000132305",
                "ENSG00000174721",
                "ENSG00000128050",
                "ENSG00000242372",
                "ENSG00000138663",
                "ENSG00000133706",
                "ENSG00000111011",
                "ENSG00000245694",
                "ENSG00000086061",
                "ENSG00000070756",
                "ENSG00000136930",
                "ENSG00000140740",
                "ENSG00000149806",
                "ENSG00000166012",
                "ENSG00000167658",
                "ENSG00000142168",
                "ENSG00000186577",
                "ENSG00000197006",
                "ENSG00000185608",
                "ENSG00000108468",
                "ENSG00000196465",
                "ENSG00000168028",
                "ENSG00000108826",
                "ENSG00000108848",
                "ENSG00000031698",
                "ENSG00000111786",
                "ENSG00000109919",
                "ENSG00000130731",
                "ENSG00000143933",
                "ENSG00000172172",
                "ENSG00000165629",
                "ENSG00000166503",
                "ENSG00000126457",
                "ENSG00000132963",
                "ENSG00000042753",
                "ENSG00000100664",
                "ENSG00000177600",
                "ENSG00000148450",
                "ENSG00000136522",
                "ENSG00000178127",
                "ENSG00000131876",
                "ENSG00000174456",
                "ENSG00000148700",
                "ENSG00000167699",
                "ENSG00000105518",
                "ENSG00000143198",
                "ENSG00000054148",
                "ENSG00000143621",
                "ENSG00000145494",
                "ENSG00000122378",
                "ENSG00000156411",
                "ENSG00000061676",
                "ENSG00000177200",
                "ENSG00000115944",
                "ENSG00000111639",
                "ENSG00000138443",
                "ENSG00000135334",
                "ENSG00000196531",
                "ENSG00000100138",
                "ENSG00000116030",
                "ENSG00000115268",
                "ENSG00000182534",
                "ENSG00000081138",
                "ENSG00000160075",
                "ENSG00000174748",
                "ENSG00000170043",
                "ENSG00000089157",
                "ENSG00000168653",
                "ENSG00000141504",
                "ENSG00000182481",
                "ENSG00000156261",
                "ENSG00000180228",
                "ENSG00000126432",
                "ENSG00000198363",
                "ENSG00000151465",
                "ENSG00000131116",
                "ENSG00000197728",
                "ENSG00000156515",
                "ENSG00000057757",
                "ENSG00000006015",
                "ENSG00000171206",
                "ENSG00000104823",
                "ENSG00000233016",
                "ENSG00000122406",
                "ENSG00000166595",
                "ENSG00000143774",
                "ENSG00000182512",
                "ENSG00000179271",
                "ENSG00000166136",
                "ENSG00000196141",
                "ENSG00000115310",
                "ENSG00000122545",
                "ENSG00000134330",
                "ENSG00000143612",
                "ENSG00000159884",
                "ENSG00000167969",
                "ENSG00000007520",
                "ENSG00000135720",
                "ENSG00000196683",
                "ENSG00000149547",
                "ENSG00000108055",
                "ENSG00000159335",
                "ENSG00000136485",
                "ENSG00000149600",
                "ENSG00000123352",
                "ENSG00000099194",
                "ENSG00000137133",
                "ENSG00000115350",
                "ENSG00000128739",
                "ENSG00000172809",
                "ENSG00000145425",
                "ENSG00000114902",
                "ENSG00000107341",
                "ENSG00000106615",
                "ENSG00000099341",
                "ENSG00000172500",
                "ENSG00000230989",
                "ENSG00000154518",
                "ENSG00000109084",
                "ENSG00000087302",
                "ENSG00000101856",
                "ENSG00000091164",
                "ENSG00000136146",
                "ENSG00000103342",
                "ENSG00000090263",
                "ENSG00000172757",
                "ENSG00000182628",
                "ENSG00000105669",
                "ENSG00000188290",
                "ENSG00000101460",
                "ENSG00000136942",
                "ENSG00000138175",
                "ENSG00000105176",
                "ENSG00000134717",
                "ENSG00000169599",
                "ENSG00000119185",
                "ENSG00000215386",
                "ENSG00000160752",
                "ENSG00000143252",
                "ENSG00000198961",
                "ENSG00000141367",
                "ENSG00000144381",
                "ENSG00000119318",
                "ENSG00000136521",
                "ENSG00000132688",
                "ENSG00000100811",
                "ENSG00000145354",
                "ENSG00000104177",
                "ENSG00000159352",
                "ENSG00000060762",
                "ENSG00000166794",
                "ENSG00000204220",
                "ENSG00000112335",
                "ENSG00000152234",
                "ENSG00000133872",
                "ENSG00000113811",
                "ENSG00000179010",
                "ENSG00000198668",
                "ENSG00000108107",
                "ENSG00000134308",
                "ENSG00000108100",
                "ENSG00000143158",
                "ENSG00000132341",
                "ENSG00000111832",
                "ENSG00000153774",
                "ENSG00000129197",
                "ENSG00000164919",
                "ENSG00000145741",
                "ENSG00000138757",
                "ENSG00000108561",
                "ENSG00000118418",
                "ENSG00000178982",
                "ENSG00000108010",
                "ENSG00000083845",
                "ENSG00000214253",
                "ENSG00000138382",
                "ENSG00000155876",
                "ENSG00000171867",
                "ENSG00000102316",
                "ENSG00000125821",
                "ENSG00000137547",
                "ENSG00000140319",
                "ENSG00000132466",
                "ENSG00000172428",
                "ENSG00000175826",
                "ENSG00000184220",
                "ENSG00000009307",
                "ENSG00000018625",
                "ENSG00000168958",
                "ENSG00000107372",
                "ENSG00000182117",
                "ENSG00000124172",
                "ENSG00000057608",
                "ENSG00000150991",
                "ENSG00000065548",
                "ENSG00000152795",
                "ENSG00000137876",
                "ENSG00000138398",
                "ENSG00000137210",
                "ENSG00000136527",
                "ENSG00000162076",
                "ENSG00000110958",
                "ENSG00000100568",
                "ENSG00000107281",
                "ENSG00000007080",
                "ENSG00000164323",
                "ENSG00000166681",
                "ENSG00000122705",
                "ENSG00000132432",
                "ENSG00000140374",
                "ENSG00000115306",
                "ENSG00000184009",
                "ENSG00000023734",
                "ENSG00000114405",
                "ENSG00000120437",
                "ENSG00000125944",
                "ENSG00000054523",
                "ENSG00000100380",
                "ENSG00000173660",
                "ENSG00000186184",
                "ENSG00000100804",
                "ENSG00000119640",
                "ENSG00000113648",
                "ENSG00000114125",
                "ENSG00000185787",
                "ENSG00000179222",
                "ENSG00000105568",
                "ENSG00000160799",
                "ENSG00000130638",
                "ENSG00000143333",
                "ENSG00000169045",
                "ENSG00000111775",
                "ENSG00000179218",
                "ENSG00000139684",
                "ENSG00000128989",
                "ENSG00000162604",
                "ENSG00000144834",
                "ENSG00000114120",
                "ENSG00000163964",
                "ENSG00000126602",
                "ENSG00000188612",
                "ENSG00000010810",
                "ENSG00000180817",
                "ENSG00000086065",
                "ENSG00000181163",
                "ENSG00000116288",
                "ENSG00000008018",
                "ENSG00000112531",
                "ENSG00000136938",
                "ENSG00000070961",
                "ENSG00000198898",
                "ENSG00000082898",
                "ENSG00000115685",
                "ENSG00000220205",
                "ENSG00000128524",
                "ENSG00000101654",
                "ENSG00000101940",
                "ENSG00000104549",
                "ENSG00000044115",
                "ENSG00000167113",
                "ENSG00000265241",
                "ENSG00000136238",
                "ENSG00000164754",
                "ENSG00000143183",
                "ENSG00000132424",
                "ENSG00000130522",
                "ENSG00000047849",
                "ENSG00000182154",
                "ENSG00000221914",
                "ENSG00000122873",
                "ENSG00000065978",
                "ENSG00000151923",
                "ENSG00000158195",
                "ENSG00000087460",
                "ENSG00000100567",
                "ENSG00000151247",
                "ENSG00000183513",
                "ENSG00000113558",
                "ENSG00000181817",
                "ENSG00000115457",
                "ENSG00000109180",
                "ENSG00000187735",
                "ENSG00000115233",
                "ENSG00000156052",
                "ENSG00000143742",
                "ENSG00000174886",
                "ENSG00000172270",
                "ENSG00000139637",
                "ENSG00000106244",
                "ENSG00000164167",
                "ENSG00000092841",
                "ENSG00000129351",
                "ENSG00000173163"
            ],
            "expression_sum": {
                "class1": 3253.0,
                "class2": 24950.0
            },
            "ens_id": "ENSG00000181449",
            "uniprot_ids": [
                "P48431"
            ]
        },
        "ENSG00000018189": {
            "id": "ENSG00000018189",
            "type": "Gene",
            "symbol": "RUFY3",
            "source": [
                "ENSG00000109132",
                "ENSG00000016082",
                "ENSG00000113580",
                "ENSG00000148516",
                "ENSG00000143799",
                "ENSG00000103495",
                "ENSG00000164330",
                "ENSG00000108055",
                "ENSG00000116560",
                "ENSG00000196591",
                "ENSG00000147140",
                "ENSG00000137309"
            ],
            "target": [],
            "expression_sum": {
                "class1": 26912.0,
                "class2": 9027.0
            },
            "ens_id": "ENSG00000018189",
            "uniprot_ids": [
                "Q7L099",
                "D6RCQ1",
                "H0Y8I0",
                "H0Y8G6",
                "D6RH24",
                "D6REM9"
            ]
        },
        "ENSG00000145362": {
            "id": "ENSG00000145362",
            "type": "Gene",
            "symbol": "ANK2",
            "source": [
                "ENSG00000109132",
                "ENSG00000113580",
                "ENSG00000148516",
                "ENSG00000143799",
                "ENSG00000103495",
                "ENSG00000164330",
                "ENSG00000114861",
                "ENSG00000139352",
                "ENSG00000116560",
                "ENSG00000128272"
            ],
            "target": [],
            "expression_sum": {
                "class1": 12281.0,
                "class2": 5101.0
            },
            "ens_id": "ENSG00000145362",
            "uniprot_ids": [
                "Q01484",
                "A0A5F9ZI15",
                "A0A5F9ZHD2",
                "B7Z651",
                "A0A5F9ZHN0",
                "I6L894",
                "A0A5F9ZHJ6",
                "A0A5F9ZHR2",
                "A0A5F9ZGX3",
                "A0A5F9ZI56",
                "A0A5F9ZH38",
                "A0A5F9ZHY6",
                "A0A5F9ZHV4",
                "A0A5F9ZGS5",
                "A0A5F9ZGS7",
                "A0A5F9ZH70",
                "A0A5F9ZH10",
                "A0A5F9ZI41",
                "A0A5K1VW73",
                "A0A5F9ZH99",
                "A0A5F9ZH34",
                "A0A5F9ZHJ4",
                "A0A5F9ZHN5",
                "A0A5F9ZHT4",
                "A0A5F9ZI45",
                "H0Y8Y2",
                "E9PHW9",
                "A0A5F9ZHQ5",
                "A0A5F9ZHL9",
                "A0A5F9ZGX8",
                "A0A5F9ZGZ5",
                "A0A5F9ZH19",
                "A0A087WZU3",
                "H0Y931",
                "A0A5F9ZHS1",
                "A0A5F9ZI16",
                "A0A5F9ZH39",
                "A0A5F9ZHL3",
                "A0A5F9ZGY1",
                "A0A5F9ZH35",
                "A0A5F9ZH18",
                "A0A5F9ZI40",
                "A0A5F9ZI18",
                "A0A669KBJ6",
                "A0A5F9ZH17",
                "A0A5F9ZI81",
                "A0A5F9ZHG3",
                "D6RHE1",
                "A0A5F9ZI69",
                "A0A5F9ZHT8",
                "A0A5F9ZGY3",
                "A0A5F9ZHQ3",
                "A0A0U1RQN6",
                "A0A5F9ZI30",
                "A0A5F9ZHE4",
                "A0A5F9ZI53",
                "A0A669KB19",
                "A0A5F9ZGZ1",
                "A0A5F9ZHE2",
                "A0A5F9ZHL5",
                "A0A5F9ZHF7",
                "A0A5F9ZHA1",
                "A0A5F9ZI65",
                "A0A5F9ZH03",
                "A0A5F9ZH30",
                "A0A5F9ZHC9",
                "A0A5F9ZI36",
                "A0A5F9ZHR8",
                "A0A5F9ZGY4",
                "A0A5F9ZI46",
                "H0Y8X8",
                "E9PCH6",
                "A0A5F9ZI73",
                "A0A5F9ZHM6",
                "D6RIY9",
                "A0A5F9ZI08",
                "H0Y933",
                "H0YAG3",
                "A0A5F9ZH58"
            ]
        },
        "ENSG00000145428": {
            "id": "ENSG00000145428",
            "type": "Gene",
            "symbol": "RNF175",
            "source": [
                "ENSG00000168214",
                "ENSG00000100811",
                "ENSG00000116560"
            ],
            "target": [],
            "expression_sum": {
                "class1": 4757.0,
                "class2": 336.0
            },
            "ens_id": "ENSG00000145428",
            "uniprot_ids": [
                "Q8N4F7",
                "D6RID2",
                "D6RAJ1",
                "D6RB06",
                "D6RB46"
            ]
        },
        "ENSG00000164104": {
            "id": "ENSG00000164104",
            "type": "Gene",
            "symbol": "HMGB2",
            "source": [
                "ENSG00000189403",
                "ENSG00000164754",
                "ENSG00000116560"
            ],
            "target": [],
            "expression_sum": {
                "class1": 15296.0,
                "class2": 20962.0
            },
            "ens_id": "ENSG00000164104",
            "uniprot_ids": [
                "P26583",
                "D6R9A6"
            ]
        },
        "ENSG00000113161": {
            "id": "ENSG00000113161",
            "type": "Gene",
            "symbol": "HMGCR",
            "source": [
                "ENSG00000109132",
                "ENSG00000167081",
                "ENSG00000103495",
                "ENSG00000164330",
                "ENSG00000177030",
                "ENSG00000198911",
                "ENSG00000116560",
                "ENSG00000196591",
                "ENSG00000128272"
            ],
            "target": [],
            "expression_sum": {
                "class1": 18056.0,
                "class2": 4548.0
            },
            "ens_id": "ENSG00000113161",
            "uniprot_ids": [
                "P04035",
                "C9JKX7",
                "H0Y8K6",
                "H0Y8F6",
                "A0A7P0Z481",
                "A0A7P0TBP1"
            ]
        },
        "ENSG00000134986": {
            "id": "ENSG00000134986",
            "type": "Gene",
            "symbol": "NREP",
            "source": [
                "ENSG00000109132",
                "ENSG00000167081",
                "ENSG00000122691",
                "ENSG00000103495",
                "ENSG00000164330",
                "ENSG00000108055",
                "ENSG00000189403",
                "ENSG00000116560",
                "ENSG00000196591"
            ],
            "target": [],
            "expression_sum": {
                "class1": 48616.0,
                "class2": 17909.0
            },
            "ens_id": "ENSG00000134986",
            "uniprot_ids": [
                "Q16612",
                "D6R982",
                "D6REJ5",
                "D6R9Y7",
                "D6RIC9"
            ]
        },
        "ENSG00000152503": {
            "id": "ENSG00000152503",
            "type": "Gene",
            "symbol": "TRIM36",
            "source": [
                "ENSG00000143799",
                "ENSG00000103495",
                "ENSG00000177030",
                "ENSG00000128272",
                "ENSG00000108055",
                "ENSG00000213015",
                "ENSG00000168214",
                "ENSG00000147140",
                "ENSG00000116560",
                "ENSG00000196591",
                "ENSG00000092199"
            ],
            "target": [],
            "expression_sum": {
                "class1": 4770.0,
                "class2": 966.0
            },
            "ens_id": "ENSG00000152503",
            "uniprot_ids": [
                "Q9NQ86",
                "E9PBG3",
                "D6RAU9"
            ]
        },
        "ENSG00000131437": {
            "id": "ENSG00000131437",
            "type": "Gene",
            "symbol": "KIF3A",
            "source": [
                "ENSG00000113580",
                "ENSG00000148516",
                "ENSG00000143799",
                "ENSG00000103495",
                "ENSG00000164330",
                "ENSG00000108055",
                "ENSG00000114861",
                "ENSG00000116560",
                "ENSG00000196591"
            ],
            "target": [],
            "expression_sum": {
                "class1": 19916.0,
                "class2": 5143.0
            },
            "ens_id": "ENSG00000131437",
            "uniprot_ids": [
                "Q9Y496",
                "A0A087X011",
                "J3KPF9",
                "C9J9X2",
                "H7C2F6",
                "E9PES4"
            ]
        },
        "ENSG00000170445": {
            "id": "ENSG00000170445",
            "type": "Gene",
            "symbol": "HARS",
            "source": [
                "ENSG00000109132",
                "ENSG00000164330",
                "ENSG00000103495",
                "ENSG00000116560"
            ],
            "target": [],
            "expression_sum": {
                "class1": 7557.0,
                "class2": 2878.0
            },
            "ens_id": "ENSG00000170445",
            "uniprot_ids": [
                "P12081",
                "A0A2R8YCI2",
                "B4DDD8",
                "B4E1C5",
                "A0A6Q8PHA5",
                "A0A6Q8PFN6",
                "A0A6Q8PHE3",
                "A0A2R8YFR1",
                "D6RF05",
                "A0A6Q8PF07",
                "A0A6Q8PFF2",
                "A0A6Q8PF47",
                "A0A6Q8PFG9",
                "A0A6Q8PHQ0",
                "E7ETE2"
            ]
        },
        "ENSG00000113657": {
            "id": "ENSG00000113657",
            "type": "Gene",
            "symbol": "DPYSL3",
            "source": [
                "ENSG00000113580",
                "ENSG00000103495",
                "ENSG00000198911",
                "ENSG00000108055",
                "ENSG00000196591",
                "ENSG00000116560"
            ],
            "target": [],
            "expression_sum": {
                "class1": 19968.0,
                "class2": 10015.0
            },
            "ens_id": "ENSG00000113657",
            "uniprot_ids": [
                "Q14195",
                "H0YB87",
                "H0YBT4",
                "D6RF19"
            ]
        },
        "ENSG00000164330": {
            "id": "ENSG00000164330",
            "type": "TF",
            "symbol": "EBF1",
            "source": [
                "ENSG00000100811",
                "ENSG00000106261",
                "ENSG00000016082",
                "ENSG00000148516",
                "ENSG00000143799",
                "ENSG00000105698",
                "ENSG00000103495",
                "ENSG00000128272",
                "ENSG00000108055",
                "ENSG00000119950",
                "ENSG00000163848",
                "ENSG00000168214",
                "ENSG00000116560",
                "ENSG00000196591",
                "ENSG00000164754",
                "ENSG00000165512",
                "ENSG00000067955"
            ],
            "target": [
                "ENSG00000206573",
                "ENSG00000005810",
                "ENSG00000117395",
                "ENSG00000134248",
                "ENSG00000189403",
                "ENSG00000022840",
                "ENSG00000164091",
                "ENSG00000144029",
                "ENSG00000102172",
                "ENSG00000150779",
                "ENSG00000077380",
                "ENSG00000154556",
                "ENSG00000134283",
                "ENSG00000173812",
                "ENSG00000117592",
                "ENSG00000106443",
                "ENSG00000123240",
                "ENSG00000102103",
                "ENSG00000178913",
                "ENSG00000074319",
                "ENSG00000146007",
                "ENSG00000106554",
                "ENSG00000157045",
                "ENSG00000163923",
                "ENSG00000125356",
                "ENSG00000255302",
                "ENSG00000144218",
                "ENSG00000113712",
                "ENSG00000153317",
                "ENSG00000026025",
                "ENSG00000157216",
                "ENSG00000133606",
                "ENSG00000022267",
                "ENSG00000103769",
                "ENSG00000170584",
                "ENSG00000075142",
                "ENSG00000273841",
                "ENSG00000124802",
                "ENSG00000142657",
                "ENSG00000196405",
                "ENSG00000126934",
                "ENSG00000151500",
                "ENSG00000086598",
                "ENSG00000121774",
                "ENSG00000149136",
                "ENSG00000127314",
                "ENSG00000142546",
                "ENSG00000185627",
                "ENSG00000092199",
                "ENSG00000107262",
                "ENSG00000175216",
                "ENSG00000174953",
                "ENSG00000118363",
                "ENSG00000122417",
                "ENSG00000112972",
                "ENSG00000231500",
                "ENSG00000265491",
                "ENSG00000113013",
                "ENSG00000109332",
                "ENSG00000079246",
                "ENSG00000057019",
                "ENSG00000095059",
                "ENSG00000232388",
                "ENSG00000114391",
                "ENSG00000168303",
                "ENSG00000112701",
                "ENSG00000162852",
                "ENSG00000135390",
                "ENSG00000131508",
                "ENSG00000154277",
                "ENSG00000198015",
                "ENSG00000048649",
                "ENSG00000169764",
                "ENSG00000120694",
                "ENSG00000127184",
                "ENSG00000160551",
                "ENSG00000103363",
                "ENSG00000064607",
                "ENSG00000171530",
                "ENSG00000149273",
                "ENSG00000165943",
                "ENSG00000137207",
                "ENSG00000108774",
                "ENSG00000115128",
                "ENSG00000134152",
                "ENSG00000138760",
                "ENSG00000183172",
                "ENSG00000169567",
                "ENSG00000275052",
                "ENSG00000114744",
                "ENSG00000147274",
                "ENSG00000123179",
                "ENSG00000101752",
                "ENSG00000108175",
                "ENSG00000084623",
                "ENSG00000096746",
                "ENSG00000119950",
                "ENSG00000138777",
                "ENSG00000069345",
                "ENSG00000162585",
                "ENSG00000075089",
                "ENSG00000028839",
                "ENSG00000189043",
                "ENSG00000163939",
                "ENSG00000075790",
                "ENSG00000099246",
                "ENSG00000115307",
                "ENSG00000116661",
                "ENSG00000134287",
                "ENSG00000105254",
                "ENSG00000119335",
                "ENSG00000120438",
                "ENSG00000127774",
                "ENSG00000172893",
                "ENSG00000106400",
                "ENSG00000122482",
                "ENSG00000125901",
                "ENSG00000113387",
                "ENSG00000149100",
                "ENSG00000158864",
                "ENSG00000137409",
                "ENSG00000006125",
                "ENSG00000136888",
                "ENSG00000170142",
                "ENSG00000127884",
                "ENSG00000121653",
                "ENSG00000140612",
                "ENSG00000163069",
                "ENSG00000111669",
                "ENSG00000124383",
                "ENSG00000197183",
                "ENSG00000102401",
                "ENSG00000132463",
                "ENSG00000124098",
                "ENSG00000197982",
                "ENSG00000165280",
                "ENSG00000116251",
                "ENSG00000091640",
                "ENSG00000147123",
                "ENSG00000143575",
                "ENSG00000176485",
                "ENSG00000184203",
                "ENSG00000183617",
                "ENSG00000105372",
                "ENSG00000143353",
                "ENSG00000111678",
                "ENSG00000169288",
                "ENSG00000130706",
                "ENSG00000161547",
                "ENSG00000135070",
                "ENSG00000108298",
                "ENSG00000174444",
                "ENSG00000130414",
                "ENSG00000169738",
                "ENSG00000172354",
                "ENSG00000150753",
                "ENSG00000138385",
                "ENSG00000164733",
                "ENSG00000115514",
                "ENSG00000113068",
                "ENSG00000221983",
                "ENSG00000069956",
                "ENSG00000122565",
                "ENSG00000164258",
                "ENSG00000137815",
                "ENSG00000141002",
                "ENSG00000171604",
                "ENSG00000170430",
                "ENSG00000166582",
                "ENSG00000225663",
                "ENSG00000065135",
                "ENSG00000105373",
                "ENSG00000124733",
                "ENSG00000272325",
                "ENSG00000100612",
                "ENSG00000166295",
                "ENSG00000168259",
                "ENSG00000119013",
                "ENSG00000161203",
                "ENSG00000025796",
                "ENSG00000151746",
                "ENSG00000101782",
                "ENSG00000099800",
                "ENSG00000147403",
                "ENSG00000140905",
                "ENSG00000101040",
                "ENSG00000277258",
                "ENSG00000115241",
                "ENSG00000161970",
                "ENSG00000100632",
                "ENSG00000129083",
                "ENSG00000078369",
                "ENSG00000133112",
                "ENSG00000138069",
                "ENSG00000110321",
                "ENSG00000115484",
                "ENSG00000110651",
                "ENSG00000157778",
                "ENSG00000145362",
                "ENSG00000182621",
                "ENSG00000123144",
                "ENSG00000100216",
                "ENSG00000143256",
                "ENSG00000149084",
                "ENSG00000102931",
                "ENSG00000165502",
                "ENSG00000114302",
                "ENSG00000104660",
                "ENSG00000185664",
                "ENSG00000067955",
                "ENSG00000130255",
                "ENSG00000214113",
                "ENSG00000022277",
                "ENSG00000147687",
                "ENSG00000160310",
                "ENSG00000158417",
                "ENSG00000197958",
                "ENSG00000171497",
                "ENSG00000198380",
                "ENSG00000108946",
                "ENSG00000213015",
                "ENSG00000175416",
                "ENSG00000170632",
                "ENSG00000172586",
                "ENSG00000063177",
                "ENSG00000160953",
                "ENSG00000155368",
                "ENSG00000169564",
                "ENSG00000103035",
                "ENSG00000128050",
                "ENSG00000165775",
                "ENSG00000086061",
                "ENSG00000167658",
                "ENSG00000113161",
                "ENSG00000189180",
                "ENSG00000196465",
                "ENSG00000128699",
                "ENSG00000143761",
                "ENSG00000111860",
                "ENSG00000031698",
                "ENSG00000109919",
                "ENSG00000167815",
                "ENSG00000126457",
                "ENSG00000177971",
                "ENSG00000070087",
                "ENSG00000177600",
                "ENSG00000148450",
                "ENSG00000136522",
                "ENSG00000166946",
                "ENSG00000136628",
                "ENSG00000092148",
                "ENSG00000054148",
                "ENSG00000070010",
                "ENSG00000181904",
                "ENSG00000104529",
                "ENSG00000118181",
                "ENSG00000080822",
                "ENSG00000186591",
                "ENSG00000168653",
                "ENSG00000159593",
                "ENSG00000156261",
                "ENSG00000198363",
                "ENSG00000151465",
                "ENSG00000140988",
                "ENSG00000122033",
                "ENSG00000163950",
                "ENSG00000143774",
                "ENSG00000165209",
                "ENSG00000182512",
                "ENSG00000110367",
                "ENSG00000100142",
                "ENSG00000115310",
                "ENSG00000122545",
                "ENSG00000175029",
                "ENSG00000108055",
                "ENSG00000099194",
                "ENSG00000086666",
                "ENSG00000188290",
                "ENSG00000167515",
                "ENSG00000134717",
                "ENSG00000169599",
                "ENSG00000141367",
                "ENSG00000213719",
                "ENSG00000105438",
                "ENSG00000112335",
                "ENSG00000204217",
                "ENSG00000132341",
                "ENSG00000124562",
                "ENSG00000100300",
                "ENSG00000145741",
                "ENSG00000087448",
                "ENSG00000128923",
                "ENSG00000175826",
                "ENSG00000009307",
                "ENSG00000138138",
                "ENSG00000168958",
                "ENSG00000057608",
                "ENSG00000165169",
                "ENSG00000065548",
                "ENSG00000150991",
                "ENSG00000017797",
                "ENSG00000107281",
                "ENSG00000132432",
                "ENSG00000184009",
                "ENSG00000120437",
                "ENSG00000054523",
                "ENSG00000146457",
                "ENSG00000100380",
                "ENSG00000128626",
                "ENSG00000166164",
                "ENSG00000119640",
                "ENSG00000104979",
                "ENSG00000114062",
                "ENSG00000128989",
                "ENSG00000213123",
                "ENSG00000168724",
                "ENSG00000126602",
                "ENSG00000036257",
                "ENSG00000164587",
                "ENSG00000008018",
                "ENSG00000160948",
                "ENSG00000112531",
                "ENSG00000082898",
                "ENSG00000115685",
                "ENSG00000101654",
                "ENSG00000167232",
                "ENSG00000144369",
                "ENSG00000091527",
                "ENSG00000182154",
                "ENSG00000181817",
                "ENSG00000143153",
                "ENSG00000187735",
                "ENSG00000087365",
                "ENSG00000174886",
                "ENSG00000017260",
                "ENSG00000111716",
                "ENSG00000169612",
                "ENSG00000099622",
                "ENSG00000114416",
                "ENSG00000131143",
                "ENSG00000184983",
                "ENSG00000127540",
                "ENSG00000180304",
                "ENSG00000108424",
                "ENSG00000147155",
                "ENSG00000051128",
                "ENSG00000250317",
                "ENSG00000141279",
                "ENSG00000237190",
                "ENSG00000135441",
                "ENSG00000172292",
                "ENSG00000129636",
                "ENSG00000134453",
                "ENSG00000182220",
                "ENSG00000141858",
                "ENSG00000167136",
                "ENSG00000149196",
                "ENSG00000204392",
                "ENSG00000114354",
                "ENSG00000153815",
                "ENSG00000196504",
                "ENSG00000055163",
                "ENSG00000037241",
                "ENSG00000109390",
                "ENSG00000170540",
                "ENSG00000144746",
                "ENSG00000133703",
                "ENSG00000146425",
                "ENSG00000120963",
                "ENSG00000131100",
                "ENSG00000176261",
                "ENSG00000114520",
                "ENSG00000215845",
                "ENSG00000069849",
                "ENSG00000240344",
                "ENSG00000123200",
                "ENSG00000147649",
                "ENSG00000144635",
                "ENSG00000187514",
                "ENSG00000076604",
                "ENSG00000160194",
                "ENSG00000178980",
                "ENSG00000083807",
                "ENSG00000168137",
                "ENSG00000172239",
                "ENSG00000129562",
                "ENSG00000108106",
                "ENSG00000116857",
                "ENSG00000256053",
                "ENSG00000105258",
                "ENSG00000263465",
                "ENSG00000169710",
                "ENSG00000163170",
                "ENSG00000184900",
                "ENSG00000110696",
                "ENSG00000088986",
                "ENSG00000107937",
                "ENSG00000131051",
                "ENSG00000166974",
                "ENSG00000029993",
                "ENSG00000239672",
                "ENSG00000005022",
                "ENSG00000131469",
                "ENSG00000047410",
                "ENSG00000136810",
                "ENSG00000142937",
                "ENSG00000171634",
                "ENSG00000232112",
                "ENSG00000106682",
                "ENSG00000079459",
                "ENSG00000178719",
                "ENSG00000119314",
                "ENSG00000170144",
                "ENSG00000143222",
                "ENSG00000127914",
                "ENSG00000177565",
                "ENSG00000106803",
                "ENSG00000175582",
                "ENSG00000171130",
                "ENSG00000198612",
                "ENSG00000130725",
                "ENSG00000122566",
                "ENSG00000142507",
                "ENSG00000047932",
                "ENSG00000237765",
                "ENSG00000165410",
                "ENSG00000089737",
                "ENSG00000130313",
                "ENSG00000129355",
                "ENSG00000188343",
                "ENSG00000128951",
                "ENSG00000100575",
                "ENSG00000120656",
                "ENSG00000241837",
                "ENSG00000010270",
                "ENSG00000056097",
                "ENSG00000131495",
                "ENSG00000173267",
                "ENSG00000147684",
                "ENSG00000072042",
                "ENSG00000126756",
                "ENSG00000103145",
                "ENSG00000137941",
                "ENSG00000167283",
                "ENSG00000187109",
                "ENSG00000115875",
                "ENSG00000164929",
                "ENSG00000204130",
                "ENSG00000196262",
                "ENSG00000125977",
                "ENSG00000089280",
                "ENSG00000130332",
                "ENSG00000168092",
                "ENSG00000115286",
                "ENSG00000105698",
                "ENSG00000105640",
                "ENSG00000114023",
                "ENSG00000166347",
                "ENSG00000131174",
                "ENSG00000106588",
                "ENSG00000196937",
                "ENSG00000188690",
                "ENSG00000135900",
                "ENSG00000170445",
                "ENSG00000143799",
                "ENSG00000204628",
                "ENSG00000102241",
                "ENSG00000134986",
                "ENSG00000121022",
                "ENSG00000113369",
                "ENSG00000163682",
                "ENSG00000171421",
                "ENSG00000085231",
                "ENSG00000124380",
                "ENSG00000105701",
                "ENSG00000134825",
                "ENSG00000090013",
                "ENSG00000167863",
                "ENSG00000125037",
                "ENSG00000004478",
                "ENSG00000105058",
                "ENSG00000175756",
                "ENSG00000145734",
                "ENSG00000197345",
                "ENSG00000005448",
                "ENSG00000250479",
                "ENSG00000160014",
                "ENSG00000146731",
                "ENSG00000130724",
                "ENSG00000164713",
                "ENSG00000132507",
                "ENSG00000175550",
                "ENSG00000173692",
                "ENSG00000132824",
                "ENSG00000164163",
                "ENSG00000134769",
                "ENSG00000166226",
                "ENSG00000103429",
                "ENSG00000102309",
                "ENSG00000167674",
                "ENSG00000258289",
                "ENSG00000164096",
                "ENSG00000156587",
                "ENSG00000116754",
                "ENSG00000175348",
                "ENSG00000141429",
                "ENSG00000100941",
                "ENSG00000164898",
                "ENSG00000152944",
                "ENSG00000141698",
                "ENSG00000162521",
                "ENSG00000196230",
                "ENSG00000100028",
                "ENSG00000029363",
                "ENSG00000100823",
                "ENSG00000105887",
                "ENSG00000169136",
                "ENSG00000160131",
                "ENSG00000109971",
                "ENSG00000159199",
                "ENSG00000118816",
                "ENSG00000172590",
                "ENSG00000151474",
                "ENSG00000137312",
                "ENSG00000167522",
                "ENSG00000111371",
                "ENSG00000172115",
                "ENSG00000101558",
                "ENSG00000125970",
                "ENSG00000006451",
                "ENSG00000167468",
                "ENSG00000111640",
                "ENSG00000183978",
                "ENSG00000125652",
                "ENSG00000124596",
                "ENSG00000169021",
                "ENSG00000130294",
                "ENSG00000077721",
                "ENSG00000099624",
                "ENSG00000120265",
                "ENSG00000138758",
                "ENSG00000150093",
                "ENSG00000162441",
                "ENSG00000130748",
                "ENSG00000111642",
                "ENSG00000137575",
                "ENSG00000145833",
                "ENSG00000103121",
                "ENSG00000104412",
                "ENSG00000113575",
                "ENSG00000225921",
                "ENSG00000105193",
                "ENSG00000009954",
                "ENSG00000084733",
                "ENSG00000136717",
                "ENSG00000149557",
                "ENSG00000206503",
                "ENSG00000100528",
                "ENSG00000120533",
                "ENSG00000162734",
                "ENSG00000168214",
                "ENSG00000071794",
                "ENSG00000112237",
                "ENSG00000106591",
                "ENSG00000130311",
                "ENSG00000130985",
                "ENSG00000094916",
                "ENSG00000138663",
                "ENSG00000186063",
                "ENSG00000140740",
                "ENSG00000140995",
                "ENSG00000146386",
                "ENSG00000197006",
                "ENSG00000185608",
                "ENSG00000108468",
                "ENSG00000205302",
                "ENSG00000104427",
                "ENSG00000115756",
                "ENSG00000113141",
                "ENSG00000166503",
                "ENSG00000132963",
                "ENSG00000175390",
                "ENSG00000168522",
                "ENSG00000148154",
                "ENSG00000119048",
                "ENSG00000172775",
                "ENSG00000134419",
                "ENSG00000174456",
                "ENSG00000135406",
                "ENSG00000143621",
                "ENSG00000177200",
                "ENSG00000135334",
                "ENSG00000100138",
                "ENSG00000168268",
                "ENSG00000107798",
                "ENSG00000125534",
                "ENSG00000131435",
                "ENSG00000126432",
                "ENSG00000130723",
                "ENSG00000124767",
                "ENSG00000178741",
                "ENSG00000156515",
                "ENSG00000101266",
                "ENSG00000233016",
                "ENSG00000166595",
                "ENSG00000134330",
                "ENSG00000184207",
                "ENSG00000167969",
                "ENSG00000007520",
                "ENSG00000159335",
                "ENSG00000137726",
                "ENSG00000141522",
                "ENSG00000114902",
                "ENSG00000099341",
                "ENSG00000116095",
                "ENSG00000103342",
                "ENSG00000172757",
                "ENSG00000105669",
                "ENSG00000050426",
                "ENSG00000136156",
                "ENSG00000119185",
                "ENSG00000169020",
                "ENSG00000171159",
                "ENSG00000144381",
                "ENSG00000136521",
                "ENSG00000213246",
                "ENSG00000155330",
                "ENSG00000112667",
                "ENSG00000133872",
                "ENSG00000198668",
                "ENSG00000134308",
                "ENSG00000134684",
                "ENSG00000089063",
                "ENSG00000204525",
                "ENSG00000168439",
                "ENSG00000138757",
                "ENSG00000067900",
                "ENSG00000214253",
                "ENSG00000138382",
                "ENSG00000118680",
                "ENSG00000184220",
                "ENSG00000205937",
                "ENSG00000115694",
                "ENSG00000065000",
                "ENSG00000125445",
                "ENSG00000165916",
                "ENSG00000182196",
                "ENSG00000152795",
                "ENSG00000165716",
                "ENSG00000072803",
                "ENSG00000100568",
                "ENSG00000176407",
                "ENSG00000125944",
                "ENSG00000183726",
                "ENSG00000214941",
                "ENSG00000186184",
                "ENSG00000100804",
                "ENSG00000179222",
                "ENSG00000163964",
                "ENSG00000166562",
                "ENSG00000104763",
                "ENSG00000181163",
                "ENSG00000158710",
                "ENSG00000215021",
                "ENSG00000122873",
                "ENSG00000113851",
                "ENSG00000145247",
                "ENSG00000137776",
                "ENSG00000139637",
                "ENSG00000106244",
                "ENSG00000078668",
                "ENSG00000090621",
                "ENSG00000100410",
                "ENSG00000116171",
                "ENSG00000162704",
                "ENSG00000139644",
                "ENSG00000175745",
                "ENSG00000182552",
                "ENSG00000090060",
                "ENSG00000116161",
                "ENSG00000160209",
                "ENSG00000161016",
                "ENSG00000125743",
                "ENSG00000105819",
                "ENSG00000179958",
                "ENSG00000075415",
                "ENSG00000107874",
                "ENSG00000152904",
                "ENSG00000163466",
                "ENSG00000198931",
                "ENSG00000197451",
                "ENSG00000152256",
                "ENSG00000173915",
                "ENSG00000111728",
                "ENSG00000167085",
                "ENSG00000170310",
                "ENSG00000115540",
                "ENSG00000041357",
                "ENSG00000163875",
                "ENSG00000163319",
                "ENSG00000164924",
                "ENSG00000141759",
                "ENSG00000165264",
                "ENSG00000174574",
                "ENSG00000110619",
                "ENSG00000139116",
                "ENSG00000277791",
                "ENSG00000149923",
                "ENSG00000118482",
                "ENSG00000116898",
                "ENSG00000179431",
                "ENSG00000188243",
                "ENSG00000100591",
                "ENSG00000140105",
                "ENSG00000034713",
                "ENSG00000130811",
                "ENSG00000005194",
                "ENSG00000153048",
                "ENSG00000125835",
                "ENSG00000183011",
                "ENSG00000174437",
                "ENSG00000187051",
                "ENSG00000129515",
                "ENSG00000155115",
                "ENSG00000204463",
                "ENSG00000010256",
                "ENSG00000120053",
                "ENSG00000135940",
                "ENSG00000172336",
                "ENSG00000131966",
                "ENSG00000107438",
                "ENSG00000153037",
                "ENSG00000106245",
                "ENSG00000075624",
                "ENSG00000072062",
                "ENSG00000168275",
                "ENSG00000102898",
                "ENSG00000152700",
                "ENSG00000167088",
                "ENSG00000240972",
                "ENSG00000196655",
                "ENSG00000141030",
                "ENSG00000164985",
                "ENSG00000250303",
                "ENSG00000141867",
                "ENSG00000165948",
                "ENSG00000197223",
                "ENSG00000180957",
                "ENSG00000099817",
                "ENSG00000143862",
                "ENSG00000176731",
                "ENSG00000161981",
                "ENSG00000162910",
                "ENSG00000104388",
                "ENSG00000097021",
                "ENSG00000087191",
                "ENSG00000166734",
                "ENSG00000257704",
                "ENSG00000130204",
                "ENSG00000014641",
                "ENSG00000121067",
                "ENSG00000176171",
                "ENSG00000131473",
                "ENSG00000112305",
                "ENSG00000253738",
                "ENSG00000165119",
                "ENSG00000126005",
                "ENSG00000241343",
                "ENSG00000184752",
                "ENSG00000198258",
                "ENSG00000170027",
                "ENSG00000023287",
                "ENSG00000111707",
                "ENSG00000176101",
                "ENSG00000123416",
                "ENSG00000171314",
                "ENSG00000144713",
                "ENSG00000067248",
                "ENSG00000164975",
                "ENSG00000130520",
                "ENSG00000170315",
                "ENSG00000167792",
                "ENSG00000138413",
                "ENSG00000163848",
                "ENSG00000099795",
                "ENSG00000079462",
                "ENSG00000166037",
                "ENSG00000163344",
                "ENSG00000051620",
                "ENSG00000119729",
                "ENSG00000135404",
                "ENSG00000113742",
                "ENSG00000142192",
                "ENSG00000131828",
                "ENSG00000171858",
                "ENSG00000144895",
                "ENSG00000123106",
                "ENSG00000147533",
                "ENSG00000155506",
                "ENSG00000100316",
                "ENSG00000170348",
                "ENSG00000140259",
                "ENSG00000143771",
                "ENSG00000089693",
                "ENSG00000169432",
                "ENSG00000144028",
                "ENSG00000164182",
                "ENSG00000122026",
                "ENSG00000198843",
                "ENSG00000109654",
                "ENSG00000145495",
                "ENSG00000254999",
                "ENSG00000130741",
                "ENSG00000177556",
                "ENSG00000186222",
                "ENSG00000156508",
                "ENSG00000082701",
                "ENSG00000120727",
                "ENSG00000101052",
                "ENSG00000106367",
                "ENSG00000101126",
                "ENSG00000197111",
                "ENSG00000164615",
                "ENSG00000169813",
                "ENSG00000175334",
                "ENSG00000099783",
                "ENSG00000159720",
                "ENSG00000182093",
                "ENSG00000184007",
                "ENSG00000140307",
                "ENSG00000106355",
                "ENSG00000169139",
                "ENSG00000109736",
                "ENSG00000203950",
                "ENSG00000070831",
                "ENSG00000161960",
                "ENSG00000167004",
                "ENSG00000249915",
                "ENSG00000110700",
                "ENSG00000159140",
                "ENSG00000170892",
                "ENSG00000067560",
                "ENSG00000176871",
                "ENSG00000156639",
                "ENSG00000143727",
                "ENSG00000167508",
                "ENSG00000111237",
                "ENSG00000182985",
                "ENSG00000023191",
                "ENSG00000151366",
                "ENSG00000214413",
                "ENSG00000145782",
                "ENSG00000116747",
                "ENSG00000153187",
                "ENSG00000106628",
                "ENSG00000100354",
                "ENSG00000131149",
                "ENSG00000183520",
                "ENSG00000138430",
                "ENSG00000176340",
                "ENSG00000273611",
                "ENSG00000166441",
                "ENSG00000101745",
                "ENSG00000145337",
                "ENSG00000170365",
                "ENSG00000182718",
                "ENSG00000140990",
                "ENSG00000163541",
                "ENSG00000146416",
                "ENSG00000028203",
                "ENSG00000112245",
                "ENSG00000092201",
                "ENSG00000141543",
                "ENSG00000173272",
                "ENSG00000179262",
                "ENSG00000101210",
                "ENSG00000197756",
                "ENSG00000178952",
                "ENSG00000260032",
                "ENSG00000129235",
                "ENSG00000103005",
                "ENSG00000117448",
                "ENSG00000142864",
                "ENSG00000137288",
                "ENSG00000142534",
                "ENSG00000211450",
                "ENSG00000106399",
                "ENSG00000142541",
                "ENSG00000122068",
                "ENSG00000181722",
                "ENSG00000109046",
                "ENSG00000143753",
                "ENSG00000135624",
                "ENSG00000163468",
                "ENSG00000155096",
                "ENSG00000091039",
                "ENSG00000117713",
                "ENSG00000177156",
                "ENSG00000140350",
                "ENSG00000128789",
                "ENSG00000197746",
                "ENSG00000135387",
                "ENSG00000160888",
                "ENSG00000205531",
                "ENSG00000143549",
                "ENSG00000187231",
                "ENSG00000225733",
                "ENSG00000177030",
                "ENSG00000171490",
                "ENSG00000154380",
                "ENSG00000127125",
                "ENSG00000127022",
                "ENSG00000074800",
                "ENSG00000162961",
                "ENSG00000148303",
                "ENSG00000133026",
                "ENSG00000122034",
                "ENSG00000138592",
                "ENSG00000154582",
                "ENSG00000067836",
                "ENSG00000114209",
                "ENSG00000186298",
                "ENSG00000164902",
                "ENSG00000144224",
                "ENSG00000137996",
                "ENSG00000188994",
                "ENSG00000164830",
                "ENSG00000133226",
                "ENSG00000135241",
                "ENSG00000108953",
                "ENSG00000105655",
                "ENSG00000132305",
                "ENSG00000204310",
                "ENSG00000136930",
                "ENSG00000149806",
                "ENSG00000132475",
                "ENSG00000108848",
                "ENSG00000156928",
                "ENSG00000111786",
                "ENSG00000124702",
                "ENSG00000143933",
                "ENSG00000167996",
                "ENSG00000131876",
                "ENSG00000134824",
                "ENSG00000156411",
                "ENSG00000099864",
                "ENSG00000138443",
                "ENSG00000196531",
                "ENSG00000115268",
                "ENSG00000182534",
                "ENSG00000242485",
                "ENSG00000086300",
                "ENSG00000170043",
                "ENSG00000149231",
                "ENSG00000182481",
                "ENSG00000188229",
                "ENSG00000118292",
                "ENSG00000180228",
                "ENSG00000126247",
                "ENSG00000136758",
                "ENSG00000006015",
                "ENSG00000173457",
                "ENSG00000088832",
                "ENSG00000135720",
                "ENSG00000136485",
                "ENSG00000013275",
                "ENSG00000125870",
                "ENSG00000123352",
                "ENSG00000112514",
                "ENSG00000137133",
                "ENSG00000278311",
                "ENSG00000172809",
                "ENSG00000157514",
                "ENSG00000107341",
                "ENSG00000172500",
                "ENSG00000091164",
                "ENSG00000136146",
                "ENSG00000155366",
                "ENSG00000160752",
                "ENSG00000075618",
                "ENSG00000143252",
                "ENSG00000134809",
                "ENSG00000198961",
                "ENSG00000159352",
                "ENSG00000155099",
                "ENSG00000137106",
                "ENSG00000008324",
                "ENSG00000166228",
                "ENSG00000118263",
                "ENSG00000213619",
                "ENSG00000129197",
                "ENSG00000164919",
                "ENSG00000165525",
                "ENSG00000033627",
                "ENSG00000140319",
                "ENSG00000132466",
                "ENSG00000149809",
                "ENSG00000243317",
                "ENSG00000165233",
                "ENSG00000107372",
                "ENSG00000085719",
                "ENSG00000137876",
                "ENSG00000138398",
                "ENSG00000136527",
                "ENSG00000136819",
                "ENSG00000110958",
                "ENSG00000101608",
                "ENSG00000164323",
                "ENSG00000122705",
                "ENSG00000178035",
                "ENSG00000140374",
                "ENSG00000173660",
                "ENSG00000113648",
                "ENSG00000104643",
                "ENSG00000227051",
                "ENSG00000162385",
                "ENSG00000169045",
                "ENSG00000165732",
                "ENSG00000008988",
                "ENSG00000196781",
                "ENSG00000180817",
                "ENSG00000086065",
                "ENSG00000136938",
                "ENSG00000220205",
                "ENSG00000136238",
                "ENSG00000164754",
                "ENSG00000143183",
                "ENSG00000123545",
                "ENSG00000224032",
                "ENSG00000105401",
                "ENSG00000221914",
                "ENSG00000103415",
                "ENSG00000065978",
                "ENSG00000087460",
                "ENSG00000151247",
                "ENSG00000170906",
                "ENSG00000109180",
                "ENSG00000123983",
                "ENSG00000143742",
                "ENSG00000205629",
                "ENSG00000172270",
                "ENSG00000177733",
                "ENSG00000092841",
                "ENSG00000129351",
                "ENSG00000089009",
                "ENSG00000141741",
                "ENSG00000134884",
                "ENSG00000163634",
                "ENSG00000095787",
                "ENSG00000169976",
                "ENSG00000106603",
                "ENSG00000140941",
                "ENSG00000184840",
                "ENSG00000075945",
                "ENSG00000099797",
                "ENSG00000196419",
                "ENSG00000105993",
                "ENSG00000213722",
                "ENSG00000088833",
                "ENSG00000119421",
                "ENSG00000114439",
                "ENSG00000087274",
                "ENSG00000165669",
                "ENSG00000184697",
                "ENSG00000065911",
                "ENSG00000151729",
                "ENSG00000272398",
                "ENSG00000197061",
                "ENSG00000125995",
                "ENSG00000135829",
                "ENSG00000135069",
                "ENSG00000162664",
                "ENSG00000112308",
                "ENSG00000148908",
                "ENSG00000139112",
                "ENSG00000156482",
                "ENSG00000165637",
                "ENSG00000241685",
                "ENSG00000060982",
                "ENSG00000147548",
                "ENSG00000171703",
                "ENSG00000131236",
                "ENSG00000129625",
                "ENSG00000196072",
                "ENSG00000100077",
                "ENSG00000067225",
                "ENSG00000144642",
                "ENSG00000141552",
                "ENSG00000136143",
                "ENSG00000205581",
                "ENSG00000117139",
                "ENSG00000104131",
                "ENSG00000095380",
                "ENSG00000034510",
                "ENSG00000138071",
                "ENSG00000263001",
                "ENSG00000213741",
                "ENSG00000145592",
                "ENSG00000134001",
                "ENSG00000138698",
                "ENSG00000123143",
                "ENSG00000110955",
                "ENSG00000164687",
                "ENSG00000169100",
                "ENSG00000162191",
                "ENSG00000147677",
                "ENSG00000108666",
                "ENSG00000070770",
                "ENSG00000175792",
                "ENSG00000167196",
                "ENSG00000180370",
                "ENSG00000171222",
                "ENSG00000181222",
                "ENSG00000105185",
                "ENSG00000153310",
                "ENSG00000100348",
                "ENSG00000213585",
                "ENSG00000145919",
                "ENSG00000146223",
                "ENSG00000131437",
                "ENSG00000143106",
                "ENSG00000127616",
                "ENSG00000174720",
                "ENSG00000128654",
                "ENSG00000080824",
                "ENSG00000160271",
                "ENSG00000166200",
                "ENSG00000130312",
                "ENSG00000136950",
                "ENSG00000197694",
                "ENSG00000115816",
                "ENSG00000168159",
                "ENSG00000152082",
                "ENSG00000115677",
                "ENSG00000151914",
                "ENSG00000168710",
                "ENSG00000124614",
                "ENSG00000092203",
                "ENSG00000104765",
                "ENSG00000073578",
                "ENSG00000182199",
                "ENSG00000269893",
                "ENSG00000067064",
                "ENSG00000135486",
                "ENSG00000163636",
                "ENSG00000109670",
                "ENSG00000137818",
                "ENSG00000082153",
                "ENSG00000002330",
                "ENSG00000129559",
                "ENSG00000134265",
                "ENSG00000105677",
                "ENSG00000135002",
                "ENSG00000171735",
                "ENSG00000146282",
                "ENSG00000078674",
                "ENSG00000103495",
                "ENSG00000126768",
                "ENSG00000147604",
                "ENSG00000204308",
                "ENSG00000165512",
                "ENSG00000086758",
                "ENSG00000134779",
                "ENSG00000079785",
                "ENSG00000070761",
                "ENSG00000185825",
                "ENSG00000134369",
                "ENSG00000163714",
                "ENSG00000173473",
                "ENSG00000196636",
                "ENSG00000170515",
                "ENSG00000164405",
                "ENSG00000196199",
                "ENSG00000104904",
                "ENSG00000134970",
                "ENSG00000173726",
                "ENSG00000167460",
                "ENSG00000155959",
                "ENSG00000116586",
                "ENSG00000186010",
                "ENSG00000084234",
                "ENSG00000125691",
                "ENSG00000151327",
                "ENSG00000167978",
                "ENSG00000186187",
                "ENSG00000089289",
                "ENSG00000168288",
                "ENSG00000004059",
                "ENSG00000144848",
                "ENSG00000095739",
                "ENSG00000234608",
                "ENSG00000241468",
                "ENSG00000110717",
                "ENSG00000115053",
                "ENSG00000111275",
                "ENSG00000142949",
                "ENSG00000136100",
                "ENSG00000099250",
                "ENSG00000197747",
                "ENSG00000162368",
                "ENSG00000254682",
                "ENSG00000100353",
                "ENSG00000116459",
                "ENSG00000134375",
                "ENSG00000134333",
                "ENSG00000038219",
                "ENSG00000086232",
                "ENSG00000125818",
                "ENSG00000103187",
                "ENSG00000115758",
                "ENSG00000105220",
                "ENSG00000053254",
                "ENSG00000272047",
                "ENSG00000011295",
                "ENSG00000119707",
                "ENSG00000007168",
                "ENSG00000176209",
                "ENSG00000112110",
                "ENSG00000067369",
                "ENSG00000104886",
                "ENSG00000097033",
                "ENSG00000029725",
                "ENSG00000177410",
                "ENSG00000146701",
                "ENSG00000116560",
                "ENSG00000114956",
                "ENSG00000198492",
                "ENSG00000183207",
                "ENSG00000073792",
                "ENSG00000182004",
                "ENSG00000198034",
                "ENSG00000141526",
                "ENSG00000104408",
                "ENSG00000173418",
                "ENSG00000160633",
                "ENSG00000102144",
                "ENSG00000167526",
                "ENSG00000084207",
                "ENSG00000058272",
                "ENSG00000184076",
                "ENSG00000177889",
                "ENSG00000166913",
                "ENSG00000182774",
                "ENSG00000111906",
                "ENSG00000204435",
                "ENSG00000109536",
                "ENSG00000162434",
                "ENSG00000090487",
                "ENSG00000126524",
                "ENSG00000081870",
                "ENSG00000111843",
                "ENSG00000074071",
                "ENSG00000109255",
                "ENSG00000134056",
                "ENSG00000145293",
                "ENSG00000181924",
                "ENSG00000148516",
                "ENSG00000074696",
                "ENSG00000169750",
                "ENSG00000253719",
                "ENSG00000156976",
                "ENSG00000169230",
                "ENSG00000105404",
                "ENSG00000177879",
                "ENSG00000154001",
                "ENSG00000115541",
                "ENSG00000168066",
                "ENSG00000162244",
                "ENSG00000083099",
                "ENSG00000198420",
                "ENSG00000149485",
                "ENSG00000071994",
                "ENSG00000109270",
                "ENSG00000104695",
                "ENSG00000141404",
                "ENSG00000172531",
                "ENSG00000054118",
                "ENSG00000179950",
                "ENSG00000142676",
                "ENSG00000165283",
                "ENSG00000054116",
                "ENSG00000196305",
                "ENSG00000096384",
                "ENSG00000102753",
                "ENSG00000109107",
                "ENSG00000113845",
                "ENSG00000124486",
                "ENSG00000198900",
                "ENSG00000100911",
                "ENSG00000214517",
                "ENSG00000115524",
                "ENSG00000174775",
                "ENSG00000143947",
                "ENSG00000128609",
                "ENSG00000182809",
                "ENSG00000198918",
                "ENSG00000147140",
                "ENSG00000133706",
                "ENSG00000205155",
                "ENSG00000161013",
                "ENSG00000088930",
                "ENSG00000168028",
                "ENSG00000042753",
                "ENSG00000198408",
                "ENSG00000178127",
                "ENSG00000184117",
                "ENSG00000167699",
                "ENSG00000105518",
                "ENSG00000143198",
                "ENSG00000170017",
                "ENSG00000145494",
                "ENSG00000122378",
                "ENSG00000115944",
                "ENSG00000111639",
                "ENSG00000093144",
                "ENSG00000160113",
                "ENSG00000138594",
                "ENSG00000160075",
                "ENSG00000120992",
                "ENSG00000089157",
                "ENSG00000166165",
                "ENSG00000131116",
                "ENSG00000168090",
                "ENSG00000197728",
                "ENSG00000104823",
                "ENSG00000177885",
                "ENSG00000130779",
                "ENSG00000122406",
                "ENSG00000215712",
                "ENSG00000179271",
                "ENSG00000196683",
                "ENSG00000149547",
                "ENSG00000120686",
                "ENSG00000102189",
                "ENSG00000149600",
                "ENSG00000108518",
                "ENSG00000106615",
                "ENSG00000087302",
                "ENSG00000136942",
                "ENSG00000105176",
                "ENSG00000119318",
                "ENSG00000166963",
                "ENSG00000104177",
                "ENSG00000152234",
                "ENSG00000108107",
                "ENSG00000143158",
                "ENSG00000111832",
                "ENSG00000061794",
                "ENSG00000153774",
                "ENSG00000108561",
                "ENSG00000178982",
                "ENSG00000108010",
                "ENSG00000083845",
                "ENSG00000155876",
                "ENSG00000171867",
                "ENSG00000111726",
                "ENSG00000125821",
                "ENSG00000070669",
                "ENSG00000117450",
                "ENSG00000018189",
                "ENSG00000101220",
                "ENSG00000182117",
                "ENSG00000124172",
                "ENSG00000196591",
                "ENSG00000142669",
                "ENSG00000100075",
                "ENSG00000025293",
                "ENSG00000178104",
                "ENSG00000115306",
                "ENSG00000114125",
                "ENSG00000185787",
                "ENSG00000105568",
                "ENSG00000100554",
                "ENSG00000111775",
                "ENSG00000133142",
                "ENSG00000114120",
                "ENSG00000163938",
                "ENSG00000188612",
                "ENSG00000137154",
                "ENSG00000116288",
                "ENSG00000205542",
                "ENSG00000103275",
                "ENSG00000128524",
                "ENSG00000137274",
                "ENSG00000044115",
                "ENSG00000047849",
                "ENSG00000130402",
                "ENSG00000198648",
                "ENSG00000085733",
                "ENSG00000114115",
                "ENSG00000181610",
                "ENSG00000173163"
            ],
            "expression_sum": {
                "class1": 14988.0,
                "class2": 1997.0
            },
            "ens_id": "ENSG00000164330",
            "uniprot_ids": [
                "Q9UH73",
                "E5RFQ1",
                "A0A087WTE5"
            ]
        },
        "ENSG00000164611": {
            "id": "ENSG00000164611",
            "type": "Gene",
            "symbol": "PTTG1",
            "source": [
                "ENSG00000164754",
                "ENSG00000116560"
            ],
            "target": [],
            "expression_sum": {
                "class1": 1824.0,
                "class2": 9438.0
            },
            "ens_id": "ENSG00000164611",
            "uniprot_ids": [
                "O95997",
                "E5RJR4"
            ]
        },
        "ENSG00000137343": {
            "id": "ENSG00000137343",
            "type": "Gene",
            "symbol": "ATAT1",
            "source": [
                "ENSG00000196591",
                "ENSG00000167081",
                "ENSG00000103495",
                "ENSG00000116560"
            ],
            "target": [],
            "expression_sum": {
                "class1": 8884.0,
                "class2": 2269.0
            },
            "ens_id": "ENSG00000137343",
            "uniprot_ids": [
                "Q5SQI0"
            ]
        },
        "ENSG00000096092": {
            "id": "ENSG00000096092",
            "type": "Gene",
            "symbol": "TMEM14A",
            "source": [
                "ENSG00000113580",
                "ENSG00000103495",
                "ENSG00000114861",
                "ENSG00000139352",
                "ENSG00000116560"
            ],
            "target": [],
            "expression_sum": {
                "class1": 13225.0,
                "class2": 4854.0
            },
            "ens_id": "ENSG00000096092",
            "uniprot_ids": [
                "Q9Y6G1"
            ]
        },
        "ENSG00000118402": {
            "id": "ENSG00000118402",
            "type": "Gene",
            "symbol": "ELOVL4",
            "source": [
                "ENSG00000116560"
            ],
            "target": [],
            "expression_sum": {
                "class1": 5190.0,
                "class2": 1437.0
            },
            "ens_id": "ENSG00000118402",
            "uniprot_ids": [
                "Q9GZR5"
            ]
        },
        "ENSG00000272398": {
            "id": "ENSG00000272398",
            "type": "Gene",
            "symbol": "CD24",
            "source": [
                "ENSG00000157557",
                "ENSG00000109132",
                "ENSG00000113580",
                "ENSG00000148516",
                "ENSG00000103495",
                "ENSG00000164330",
                "ENSG00000177030",
                "ENSG00000108001",
                "ENSG00000108055",
                "ENSG00000189403",
                "ENSG00000213015",
                "ENSG00000116560",
                "ENSG00000196591"
            ],
            "target": [],
            "expression_sum": {
                "class1": 89639.0,
                "class2": 41020.0
            },
            "ens_id": "ENSG00000272398",
            "uniprot_ids": [
                "P25063",
                "A0A087WW33",
                "A0A087WU21",
                "A0A087WU87"
            ]
        },
        "ENSG00000005108": {
            "id": "ENSG00000005108",
            "type": "Gene",
            "symbol": "THSD7A",
            "source": [
                "ENSG00000100811",
                "ENSG00000103495",
                "ENSG00000108055",
                "ENSG00000116560",
                "ENSG00000119547",
                "ENSG00000092199"
            ],
            "target": [],
            "expression_sum": {
                "class1": 13004.0,
                "class2": 2102.0
            },
            "ens_id": "ENSG00000005108",
            "uniprot_ids": [
                "Q9UPZ6",
                "A0A087WY08"
            ]
        },
        "ENSG00000170027": {
            "id": "ENSG00000170027",
            "type": "Gene",
            "symbol": "YWHAG",
            "source": [
                "ENSG00000109132",
                "ENSG00000167081",
                "ENSG00000113580",
                "ENSG00000148516",
                "ENSG00000103495",
                "ENSG00000164330",
                "ENSG00000177030",
                "ENSG00000108055",
                "ENSG00000116560",
                "ENSG00000196591",
                "ENSG00000129351"
            ],
            "target": [],
            "expression_sum": {
                "class1": 23627.0,
                "class2": 6746.0
            },
            "ens_id": "ENSG00000170027",
            "uniprot_ids": [
                "P61981"
            ]
        },
        "ENSG00000128564": {
            "id": "ENSG00000128564",
            "type": "Gene",
            "symbol": "VGF",
            "source": [
                "ENSG00000100811",
                "ENSG00000095794",
                "ENSG00000175197",
                "ENSG00000103495",
                "ENSG00000168610",
                "ENSG00000100219",
                "ENSG00000108055",
                "ENSG00000213015",
                "ENSG00000140307",
                "ENSG00000196591",
                "ENSG00000116560",
                "ENSG00000130522",
                "ENSG00000160113",
                "ENSG00000092199"
            ],
            "target": [],
            "expression_sum": {
                "class1": 15599.0,
                "class2": 4375.0
            },
            "ens_id": "ENSG00000128564",
            "uniprot_ids": [
                "O15240",
                "Q05BJ3"
            ]
        },
        "ENSG00000221866": {
            "id": "ENSG00000221866",
            "type": "Gene",
            "symbol": "PLXNA4",
            "source": [
                "ENSG00000170365",
                "ENSG00000100811",
                "ENSG00000130726",
                "ENSG00000100644",
                "ENSG00000143799",
                "ENSG00000103495",
                "ENSG00000168610",
                "ENSG00000108055",
                "ENSG00000189403",
                "ENSG00000168214",
                "ENSG00000116560",
                "ENSG00000196591",
                "ENSG00000092199"
            ],
            "target": [],
            "expression_sum": {
                "class1": 2209.0,
                "class2": 256.0
            },
            "ens_id": "ENSG00000221866",
            "uniprot_ids": [
                "Q9HCM2"
            ]
        },
        "ENSG00000133134": {
            "id": "ENSG00000133134",
            "type": "Gene",
            "symbol": "BEX2",
            "source": [
                "ENSG00000181449",
                "ENSG00000148516",
                "ENSG00000103495",
                "ENSG00000139352",
                "ENSG00000116560"
            ],
            "target": [],
            "expression_sum": {
                "class1": 58007.0,
                "class2": 27940.0
            },
            "ens_id": "ENSG00000133134",
            "uniprot_ids": [
                "Q9BXY8",
                "X6RDN0"
            ]
        },
        "ENSG00000157514": {
            "id": "ENSG00000157514",
            "type": "Gene",
            "symbol": "TSC22D3",
            "source": [
                "ENSG00000167081",
                "ENSG00000103495",
                "ENSG00000164330",
                "ENSG00000108055",
                "ENSG00000067955",
                "ENSG00000196591",
                "ENSG00000116560"
            ],
            "target": [],
            "expression_sum": {
                "class1": 6493.0,
                "class2": 2468.0
            },
            "ens_id": "ENSG00000157514",
            "uniprot_ids": [
                "Q99576",
                "D6RAX8",
                "Q5JRJ0",
                "D6R8Z7",
                "D6RBL1",
                "E7EWD5",
                "D6RCK9",
                "D6RD92",
                "D6RDZ7",
                "Q5JRJ2",
                "D6R9J1",
                "D6RD37",
                "D6RDZ9"
            ]
        },
        "ENSG00000157168": {
            "id": "ENSG00000157168",
            "type": "TF",
            "symbol": "NRG1",
            "source": [
                "ENSG00000109132",
                "ENSG00000016082",
                "ENSG00000113580",
                "ENSG00000103495",
                "ENSG00000139352",
                "ENSG00000116560"
            ],
            "target": [
                "ENSG00000070669",
                "ENSG00000134198",
                "ENSG00000101134"
            ],
            "expression_sum": {
                "class1": 5138.0,
                "class2": 954.0
            },
            "ens_id": "ENSG00000157168",
            "uniprot_ids": [
                "Q02297",
                "H0YBA3",
                "A0A494C114",
                "A0A494C0Q4",
                "A0A494C1B5",
                "E5RHQ1",
                "A0A087X0X9",
                "A0A5F9ZHA5",
                "E5RHP6",
                "A0A494C1F5",
                "A0A494C0K4",
                "A0A494BZT4",
                "E5RIG8",
                "A0A494C0L9",
                "H0YCP0",
                "A0A494C054",
                "A0A494C1F8",
                "H0YDC2",
                "A0A494C0T5",
                "A0A494C1G2",
                "F6RJN6",
                "A0A494C043"
            ]
        },
        "ENSG00000206579": {
            "id": "ENSG00000206579",
            "type": "Gene",
            "symbol": "XKR4",
            "source": [
                "ENSG00000189403",
                "ENSG00000164754",
                "ENSG00000196591",
                "ENSG00000116560",
                "ENSG00000145734"
            ],
            "target": [],
            "expression_sum": {
                "class1": 4620.0,
                "class2": 1180.0
            },
            "ens_id": "ENSG00000206579",
            "uniprot_ids": [
                "Q5GH76"
            ]
        },
        "ENSG00000171033": {
            "id": "ENSG00000171033",
            "type": "Gene",
            "symbol": "PKIA",
            "source": [
                "ENSG00000100811",
                "ENSG00000028277",
                "ENSG00000016082",
                "ENSG00000113580",
                "ENSG00000167081",
                "ENSG00000103495",
                "ENSG00000177030",
                "ENSG00000189403",
                "ENSG00000116560",
                "ENSG00000137203",
                "ENSG00000196591",
                "ENSG00000147140",
                "ENSG00000137309",
                "ENSG00000092199"
            ],
            "target": [],
            "expression_sum": {
                "class1": 23269.0,
                "class2": 4338.0
            },
            "ens_id": "ENSG00000171033",
            "uniprot_ids": [
                "P61925"
            ]
        },
        "ENSG00000104427": {
            "id": "ENSG00000104427",
            "type": "Gene",
            "symbol": "ZC2HC1A",
            "source": [
                "ENSG00000167081",
                "ENSG00000113580",
                "ENSG00000148516",
                "ENSG00000143799",
                "ENSG00000103495",
                "ENSG00000164330",
                "ENSG00000198911",
                "ENSG00000108055",
                "ENSG00000213015",
                "ENSG00000196591",
                "ENSG00000116560"
            ],
            "target": [],
            "expression_sum": {
                "class1": 16871.0,
                "class2": 5087.0
            },
            "ens_id": "ENSG00000104427",
            "uniprot_ids": [
                "Q96GY0",
                "H0YAP0"
            ]
        },
        "ENSG00000155099": {
            "id": "ENSG00000155099",
            "type": "Gene",
            "symbol": "PIP4P2",
            "source": [
                "ENSG00000167081",
                "ENSG00000113580",
                "ENSG00000103495",
                "ENSG00000164330",
                "ENSG00000108055",
                "ENSG00000139352",
                "ENSG00000196591",
                "ENSG00000116560"
            ],
            "target": [],
            "expression_sum": {
                "class1": 9934.0,
                "class2": 2663.0
            },
            "ens_id": "ENSG00000155099",
            "uniprot_ids": [
                "Q8N4L2",
                "E5RJC2",
                "E5RIP9",
                "E5RFT4",
                "E5RIY0"
            ]
        },
        "ENSG00000169439": {
            "id": "ENSG00000169439",
            "type": "Gene",
            "symbol": "SDC2",
            "source": [
                "ENSG00000108055",
                "ENSG00000113580",
                "ENSG00000148516",
                "ENSG00000116560"
            ],
            "target": [],
            "expression_sum": {
                "class1": 4325.0,
                "class2": 11216.0
            },
            "ens_id": "ENSG00000169439",
            "uniprot_ids": [
                "P34741",
                "E9PBI9",
                "E7ESK6",
                "E5RJB8",
                "E5RHU3"
            ]
        },
        "ENSG00000186106": {
            "id": "ENSG00000186106",
            "type": "Gene",
            "symbol": "ANKRD46",
            "source": [
                "ENSG00000167081",
                "ENSG00000148516",
                "ENSG00000143799",
                "ENSG00000103495",
                "ENSG00000108055",
                "ENSG00000116560",
                "ENSG00000137309"
            ],
            "target": [],
            "expression_sum": {
                "class1": 8822.0,
                "class2": 1889.0
            },
            "ens_id": "ENSG00000186106",
            "uniprot_ids": [
                "Q86W74",
                "J3KP26",
                "E5RHV2",
                "E5RI04",
                "E5RFM7",
                "E5RI20"
            ]
        },
        "ENSG00000172159": {
            "id": "ENSG00000172159",
            "type": "Gene",
            "symbol": "FRMD3",
            "source": [
                "ENSG00000109132",
                "ENSG00000016082",
                "ENSG00000113580",
                "ENSG00000148516",
                "ENSG00000103495",
                "ENSG00000139352",
                "ENSG00000116560",
                "ENSG00000124766",
                "ENSG00000137309"
            ],
            "target": [],
            "expression_sum": {
                "class1": 3118.0,
                "class2": 2014.0
            },
            "ens_id": "ENSG00000172159",
            "uniprot_ids": [
                "A2A2Y4",
                "Q5JV61"
            ]
        },
        "ENSG00000123975": {
            "id": "ENSG00000123975",
            "type": "Gene",
            "symbol": "CKS2",
            "source": [
                "ENSG00000164754",
                "ENSG00000116560"
            ],
            "target": [],
            "expression_sum": {
                "class1": 4350.0,
                "class2": 7742.0
            },
            "ens_id": "ENSG00000123975",
            "uniprot_ids": [
                "P33552"
            ]
        },
        "ENSG00000197694": {
            "id": "ENSG00000197694",
            "type": "Gene",
            "symbol": "SPTAN1",
            "source": [
                "ENSG00000181449",
                "ENSG00000143799",
                "ENSG00000118263",
                "ENSG00000164330",
                "ENSG00000198911",
                "ENSG00000108055",
                "ENSG00000067955",
                "ENSG00000116560",
                "ENSG00000196591",
                "ENSG00000137309"
            ],
            "target": [],
            "expression_sum": {
                "class1": 14828.0,
                "class2": 6300.0
            },
            "ens_id": "ENSG00000197694",
            "uniprot_ids": [
                "Q13813",
                "A0A1B0GWE2",
                "A0A0D9SFH4",
                "A0A1B0GV13",
                "A0A1B0GUN2",
                "A0A1B0GUH3",
                "A0A1B0GTB7",
                "A0A0D9SFF6",
                "A0A5F9ZHC3",
                "A0A1B0GTD1",
                "A0A0D9SF54",
                "A0A1B0GW19",
                "A0A0D9SGF6"
            ]
        },
        "ENSG00000130558": {
            "id": "ENSG00000130558",
            "type": "Gene",
            "symbol": "OLFM1",
            "source": [
                "ENSG00000162992",
                "ENSG00000167081",
                "ENSG00000113580",
                "ENSG00000103495",
                "ENSG00000114861",
                "ENSG00000139352",
                "ENSG00000116560",
                "ENSG00000137203"
            ],
            "target": [],
            "expression_sum": {
                "class1": 31680.0,
                "class2": 4589.0
            },
            "ens_id": "ENSG00000130558",
            "uniprot_ids": [
                "Q99784",
                "Q6IMJ6",
                "F5GZQ2",
                "F5H810",
                "Q5VZL8",
                "H0YFB9",
                "F8W870"
            ]
        },
        "ENSG00000050165": {
            "id": "ENSG00000050165",
            "type": "Gene",
            "symbol": "DKK3",
            "source": [
                "ENSG00000103495",
                "ENSG00000189403",
                "ENSG00000116560",
                "ENSG00000196591",
                "ENSG00000124766"
            ],
            "target": [],
            "expression_sum": {
                "class1": 11448.0,
                "class2": 4310.0
            },
            "ens_id": "ENSG00000050165",
            "uniprot_ids": [
                "Q9UBP4",
                "E9PHY3",
                "E9PKK9",
                "E9PKW6",
                "F6SYF8"
            ]
        },
        "ENSG00000149257": {
            "id": "ENSG00000149257",
            "type": "Gene",
            "symbol": "SERPINH1",
            "source": [
                "ENSG00000148516",
                "ENSG00000221869",
                "ENSG00000116560"
            ],
            "target": [],
            "expression_sum": {
                "class1": 2272.0,
                "class2": 8437.0
            },
            "ens_id": "ENSG00000149257",
            "uniprot_ids": [
                "P50454",
                "E9PPV6",
                "E9PR70",
                "E9PK86",
                "E9PRS3",
                "E9PLA6",
                "E9PNX1",
                "H0YEP8",
                "E9PMI5",
                "E9PKH2",
                "E9PIG2",
                "E9PJH8",
                "E9PQ34"
            ]
        },
        "ENSG00000171533": {
            "id": "ENSG00000171533",
            "type": "Gene",
            "symbol": "MAP6",
            "source": [
                "ENSG00000196591",
                "ENSG00000116560",
                "ENSG00000092199"
            ],
            "target": [],
            "expression_sum": {
                "class1": 22683.0,
                "class2": 5492.0
            },
            "ens_id": "ENSG00000171533",
            "uniprot_ids": [
                "Q96JE9"
            ]
        },
        "ENSG00000188486": {
            "id": "ENSG00000188486",
            "type": "Gene",
            "symbol": "H2AFX",
            "source": [
                "ENSG00000108055",
                "ENSG00000164754",
                "ENSG00000116560"
            ],
            "target": [],
            "expression_sum": {
                "class1": 1524.0,
                "class2": 3961.0
            },
            "ens_id": "ENSG00000188486",
            "uniprot_ids": [
                "P16104"
            ]
        },
        "ENSG00000187134": {
            "id": "ENSG00000187134",
            "type": "Gene",
            "symbol": "AKR1C1",
            "source": [
                "ENSG00000134138",
                "ENSG00000108055",
                "ENSG00000213015",
                "ENSG00000116560",
                "ENSG00000160113"
            ],
            "target": [],
            "expression_sum": {
                "class1": 5984.0,
                "class2": 433.0
            },
            "ens_id": "ENSG00000187134",
            "uniprot_ids": [
                "Q04828",
                "A6NHU4",
                "H0Y804"
            ]
        },
        "ENSG00000123240": {
            "id": "ENSG00000123240",
            "type": "Gene",
            "symbol": "OPTN",
            "source": [
                "ENSG00000163435",
                "ENSG00000109132",
                "ENSG00000167081",
                "ENSG00000113580",
                "ENSG00000148516",
                "ENSG00000143799",
                "ENSG00000103495",
                "ENSG00000164330",
                "ENSG00000175745",
                "ENSG00000108055",
                "ENSG00000116560"
            ],
            "target": [],
            "expression_sum": {
                "class1": 12771.0,
                "class2": 3711.0
            },
            "ens_id": "ENSG00000123240",
            "uniprot_ids": [
                "Q96CV9",
                "X6RKL2",
                "H7C1H4",
                "A0A087X2G2",
                "A0A087WY28"
            ]
        },
        "ENSG00000151150": {
            "id": "ENSG00000151150",
            "type": "Gene",
            "symbol": "ANK3",
            "source": [
                "ENSG00000092199",
                "ENSG00000134138",
                "ENSG00000148516",
                "ENSG00000143799",
                "ENSG00000103495",
                "ENSG00000108055",
                "ENSG00000164754",
                "ENSG00000116560",
                "ENSG00000143995"
            ],
            "target": [],
            "expression_sum": {
                "class1": 17550.0,
                "class2": 4091.0
            },
            "ens_id": "ENSG00000151150",
            "uniprot_ids": [
                "Q12955",
                "A0A087WVC2",
                "D6RBY7",
                "B1AQT1",
                "H0Y951",
                "A0A087WX55",
                "H0Y9E9",
                "H0Y8Z4",
                "A0A087WY90",
                "A0A087WTE8",
                "A0A087X0B4",
                "A0A087WZ26",
                "D6RHY3",
                "H0Y3A4",
                "D6RF31",
                "D6RFK6",
                "A0A087WZ65",
                "A0A087WWI5",
                "H0YA66",
                "A0A087WTF3",
                "H0YAH5",
                "A0A087WTP5",
                "A0A087X0L3"
            ]
        },
        "ENSG00000170312": {
            "id": "ENSG00000170312",
            "type": "Gene",
            "symbol": "CDK1",
            "source": [
                "ENSG00000103495",
                "ENSG00000108055",
                "ENSG00000164754",
                "ENSG00000147140",
                "ENSG00000116560"
            ],
            "target": [],
            "expression_sum": {
                "class1": 314.0,
                "class2": 3002.0
            },
            "ens_id": "ENSG00000170312",
            "uniprot_ids": [
                "P06493",
                "A0A024QZP7",
                "E5RIU6",
                "A0A087WZZ9"
            ]
        },
        "ENSG00000204130": {
            "id": "ENSG00000204130",
            "type": "Gene",
            "symbol": "RUFY2",
            "source": [
                "ENSG00000108055",
                "ENSG00000164330",
                "ENSG00000116560",
                "ENSG00000148516"
            ],
            "target": [],
            "expression_sum": {
                "class1": 9840.0,
                "class2": 2849.0
            },
            "ens_id": "ENSG00000204130",
            "uniprot_ids": [
                "Q8WXA3",
                "S4R3P2",
                "H0YD93"
            ]
        },
        "ENSG00000171224": {
            "id": "ENSG00000171224",
            "type": "Gene",
            "symbol": "FAM241B",
            "source": [
                "ENSG00000189403",
                "ENSG00000103495",
                "ENSG00000116560"
            ],
            "target": [],
            "expression_sum": {
                "class1": 11808.0,
                "class2": 2941.0
            },
            "ens_id": "ENSG00000171224",
            "uniprot_ids": [
                "Q96D05",
                "Q5VVH9"
            ]
        },
        "ENSG00000108175": {
            "id": "ENSG00000108175",
            "type": "Gene",
            "symbol": "ZMIZ1",
            "source": [
                "ENSG00000109132",
                "ENSG00000148516",
                "ENSG00000103495",
                "ENSG00000164330",
                "ENSG00000108055",
                "ENSG00000067955",
                "ENSG00000116560",
                "ENSG00000196591",
                "ENSG00000119547"
            ],
            "target": [],
            "expression_sum": {
                "class1": 7150.0,
                "class2": 2680.0
            },
            "ens_id": "ENSG00000108175",
            "uniprot_ids": [
                "Q9ULJ6",
                "A0A087WYE6"
            ]
        },
        "ENSG00000138138": {
            "id": "ENSG00000138138",
            "type": "Gene",
            "symbol": "ATAD1",
            "source": [
                "ENSG00000167081",
                "ENSG00000113580",
                "ENSG00000148516",
                "ENSG00000103495",
                "ENSG00000164330",
                "ENSG00000196591",
                "ENSG00000116560"
            ],
            "target": [],
            "expression_sum": {
                "class1": 13877.0,
                "class2": 4960.0
            },
            "ens_id": "ENSG00000138138",
            "uniprot_ids": [
                "Q8NBU5",
                "A0A7P0Z4J1",
                "A0A7P0T7Z9",
                "A0A7P0T9U2",
                "A0A7P0TAM5",
                "A0A7P0T9B4"
            ]
        },
        "ENSG00000099194": {
            "id": "ENSG00000099194",
            "type": "Gene",
            "symbol": "SCD",
            "source": [
                "ENSG00000181449",
                "ENSG00000109132",
                "ENSG00000103495",
                "ENSG00000164330",
                "ENSG00000108001",
                "ENSG00000108055",
                "ENSG00000168214",
                "ENSG00000116560",
                "ENSG00000196591",
                "ENSG00000160113"
            ],
            "target": [],
            "expression_sum": {
                "class1": 15199.0,
                "class2": 7009.0
            },
            "ens_id": "ENSG00000099194",
            "uniprot_ids": [
                "O00767"
            ]
        },
        "ENSG00000148773": {
            "id": "ENSG00000148773",
            "type": "Gene",
            "symbol": "MKI67",
            "source": [
                "ENSG00000108055",
                "ENSG00000164754",
                "ENSG00000116560"
            ],
            "target": [],
            "expression_sum": {
                "class1": 38.0,
                "class2": 2290.0
            },
            "ens_id": "ENSG00000148773",
            "uniprot_ids": [
                "P46013"
            ]
        },
        "ENSG00000067715": {
            "id": "ENSG00000067715",
            "type": "Gene",
            "symbol": "SYT1",
            "source": [
                "ENSG00000109132",
                "ENSG00000113580",
                "ENSG00000148516",
                "ENSG00000139352",
                "ENSG00000116560"
            ],
            "target": [],
            "expression_sum": {
                "class1": 41280.0,
                "class2": 10286.0
            },
            "ens_id": "ENSG00000067715",
            "uniprot_ids": [
                "P21579",
                "F8VZY3",
                "J3KQA0",
                "F8W1U9",
                "F8VXH0",
                "F8VYH8",
                "C9JX50"
            ]
        },
        "ENSG00000005810": {
            "id": "ENSG00000005810",
            "type": "Gene",
            "symbol": "MYCBP2",
            "source": [
                "ENSG00000016082",
                "ENSG00000113580",
                "ENSG00000167081",
                "ENSG00000148516",
                "ENSG00000103495",
                "ENSG00000164330",
                "ENSG00000177030",
                "ENSG00000108055",
                "ENSG00000139352",
                "ENSG00000067955",
                "ENSG00000116560",
                "ENSG00000128272"
            ],
            "target": [],
            "expression_sum": {
                "class1": 12827.0,
                "class2": 4708.0
            },
            "ens_id": "ENSG00000005810",
            "uniprot_ids": [
                "O75592",
                "A0A804HKQ1",
                "A0A499FJI4",
                "A0A804HJ25",
                "H7C3U4",
                "A0A804HIR9",
                "A0A804HL12"
            ]
        },
        "ENSG00000198513": {
            "id": "ENSG00000198513",
            "type": "Gene",
            "symbol": "ATL1",
            "source": [
                "ENSG00000148516",
                "ENSG00000143799",
                "ENSG00000103495",
                "ENSG00000213015",
                "ENSG00000189403",
                "ENSG00000116560",
                "ENSG00000128272"
            ],
            "target": [],
            "expression_sum": {
                "class1": 9902.0,
                "class2": 2161.0
            },
            "ens_id": "ENSG00000198513",
            "uniprot_ids": [
                "Q8WXF7",
                "A0A6I8PUC8",
                "A0A6I8PIS8",
                "H0YJA7",
                "H0YJ65",
                "A0A6I8PRS8",
                "G3V5T4",
                "G3V334",
                "G3V4Y8",
                "A0A804HLC1",
                "G3V321"
            ]
        },
        "ENSG00000185650": {
            "id": "ENSG00000185650",
            "type": "TF",
            "symbol": "ZFP36L1",
            "source": [
                "ENSG00000106261",
                "ENSG00000175745",
                "ENSG00000128272",
                "ENSG00000160113",
                "ENSG00000177606",
                "ENSG00000092199",
                "ENSG00000103495",
                "ENSG00000168214",
                "ENSG00000196591",
                "ENSG00000130522",
                "ENSG00000100811",
                "ENSG00000181722",
                "ENSG00000213015",
                "ENSG00000163435",
                "ENSG00000185630",
                "ENSG00000164683",
                "ENSG00000143799",
                "ENSG00000122691",
                "ENSG00000116560",
                "ENSG00000147140",
                "ENSG00000124766"
            ],
            "target": [
                "ENSG00000277791",
                "ENSG00000115286",
                "ENSG00000140941"
            ],
            "expression_sum": {
                "class1": 1965.0,
                "class2": 16300.0
            },
            "ens_id": "ENSG00000185650",
            "uniprot_ids": [
                "Q07352",
                "G3V2P5",
                "G3V515",
                "G3V2D5"
            ]
        },
        "ENSG00000126214": {
            "id": "ENSG00000126214",
            "type": "Gene",
            "symbol": "KLC1",
            "source": [
                "ENSG00000016082",
                "ENSG00000113580",
                "ENSG00000148516",
                "ENSG00000143799",
                "ENSG00000118263",
                "ENSG00000103495",
                "ENSG00000108055",
                "ENSG00000213015",
                "ENSG00000139352",
                "ENSG00000116560",
                "ENSG00000196591"
            ],
            "target": [],
            "expression_sum": {
                "class1": 25362.0,
                "class2": 5579.0
            },
            "ens_id": "ENSG00000126214",
            "uniprot_ids": [
                "Q07866",
                "G3V2P7",
                "G3V2E7",
                "H0YJL0",
                "H0YGB8",
                "H0YG16",
                "F8W6L3",
                "G3V3H3",
                "H0YJK1",
                "H0YJT3",
                "H0YJU9",
                "G5E9S8",
                "G3V5R9"
            ]
        },
        "ENSG00000137804": {
            "id": "ENSG00000137804",
            "type": "Gene",
            "symbol": "NUSAP1",
            "source": [
                "ENSG00000108055",
                "ENSG00000164754",
                "ENSG00000116560"
            ],
            "target": [],
            "expression_sum": {
                "class1": 184.0,
                "class2": 3565.0
            },
            "ens_id": "ENSG00000137804",
            "uniprot_ids": [
                "Q9BXS6",
                "H0YKA7",
                "H0YM18",
                "H0YMD2"
            ]
        },
        "ENSG00000104112": {
            "id": "ENSG00000104112",
            "type": "Gene",
            "symbol": "SCG3",
            "source": [
                "ENSG00000185551",
                "ENSG00000157557",
                "ENSG00000113580",
                "ENSG00000148516",
                "ENSG00000103495",
                "ENSG00000175745",
                "ENSG00000139352",
                "ENSG00000116560",
                "ENSG00000196591",
                "ENSG00000124766"
            ],
            "target": [],
            "expression_sum": {
                "class1": 31748.0,
                "class2": 18965.0
            },
            "ens_id": "ENSG00000104112",
            "uniprot_ids": [
                "Q8WXD2",
                "H0YKC2"
            ]
        },
        "ENSG00000128872": {
            "id": "ENSG00000128872",
            "type": "Gene",
            "symbol": "TMOD2",
            "source": [
                "ENSG00000113580",
                "ENSG00000148516",
                "ENSG00000103495",
                "ENSG00000189403",
                "ENSG00000139352",
                "ENSG00000116560"
            ],
            "target": [],
            "expression_sum": {
                "class1": 5274.0,
                "class2": 1728.0
            },
            "ens_id": "ENSG00000128872",
            "uniprot_ids": [
                "Q9NZR1",
                "G5EA42",
                "H0YMA2"
            ]
        },
        "ENSG00000174939": {
            "id": "ENSG00000174939",
            "type": "Gene",
            "symbol": "ASPHD1",
            "source": [
                "ENSG00000113580",
                "ENSG00000103495",
                "ENSG00000116560"
            ],
            "target": [],
            "expression_sum": {
                "class1": 6348.0,
                "class2": 2185.0
            },
            "ens_id": "ENSG00000174939",
            "uniprot_ids": [
                "Q5U4P2",
                "I3L2A5",
                "I3L2Y2"
            ]
        },
        "ENSG00000087258": {
            "id": "ENSG00000087258",
            "type": "Gene",
            "symbol": "GNAO1",
            "source": [
                "ENSG00000170365",
                "ENSG00000100811",
                "ENSG00000103495",
                "ENSG00000108055",
                "ENSG00000189403",
                "ENSG00000164754",
                "ENSG00000168214",
                "ENSG00000116560",
                "ENSG00000196591",
                "ENSG00000128272",
                "ENSG00000092199"
            ],
            "target": [],
            "expression_sum": {
                "class1": 9314.0,
                "class2": 1944.0
            },
            "ens_id": "ENSG00000087258",
            "uniprot_ids": [
                "P09471",
                "A0A1W2PRE1",
                "A0A6Q8PGR1",
                "H3BNR5",
                "A0A1W2PP38",
                "H3BTM2",
                "A0A1W2PRJ7",
                "A0A1W2PQK2",
                "A0A1W2PP87",
                "A0A1W2PS82",
                "A0A1W2PPG6",
                "A0A087WTB6",
                "A0A1W2PQ24"
            ]
        },
        "ENSG00000166848": {
            "id": "ENSG00000166848",
            "type": "Gene",
            "symbol": "TERF2IP",
            "source": [
                "ENSG00000109132",
                "ENSG00000167081",
                "ENSG00000113580",
                "ENSG00000148516",
                "ENSG00000103495",
                "ENSG00000116560"
            ],
            "target": [],
            "expression_sum": {
                "class1": 44037.0,
                "class2": 16237.0
            },
            "ens_id": "ENSG00000166848",
            "uniprot_ids": [
                "Q9NYB0",
                "H3BR63",
                "A0A590UJT3",
                "H3BMI8"
            ]
        },
        "ENSG00000131747": {
            "id": "ENSG00000131747",
            "type": "Gene",
            "symbol": "TOP2A",
            "source": [
                "ENSG00000108055",
                "ENSG00000164754",
                "ENSG00000116560"
            ],
            "target": [],
            "expression_sum": {
                "class1": 141.0,
                "class2": 4638.0
            },
            "ens_id": "ENSG00000131747",
            "uniprot_ids": [
                "P11388",
                "J3QR57",
                "J3KTB7"
            ]
        },
        "ENSG00000082397": {
            "id": "ENSG00000082397",
            "type": "Gene",
            "symbol": "EPB41L3",
            "source": [
                "ENSG00000109132",
                "ENSG00000128272",
                "ENSG00000148516",
                "ENSG00000116560"
            ],
            "target": [],
            "expression_sum": {
                "class1": 5388.0,
                "class2": 1550.0
            },
            "ens_id": "ENSG00000082397",
            "uniprot_ids": [
                "Q9Y2J2",
                "J3QS55",
                "J3QR33",
                "A8K968",
                "J3QKK4",
                "A0A0J9YWN0",
                "A0A0J9YWS4",
                "A0A0A0MRA8",
                "J3KT37",
                "A0A0J9YW31",
                "J3KRD1",
                "A0A0A0MSA4",
                "J3QKY2",
                "J3KS70",
                "A0A1B0GTF8",
                "A0A0J9YWE1",
                "A0A0J9YY18",
                "J3QS83",
                "J3QRQ6",
                "A0A0U1RR92",
                "A0A0J9YW26",
                "J3QLU5"
            ]
        },
        "ENSG00000141404": {
            "id": "ENSG00000141404",
            "type": "Gene",
            "symbol": "GNAL",
            "source": [
                "ENSG00000148516",
                "ENSG00000118263",
                "ENSG00000103495",
                "ENSG00000164330",
                "ENSG00000189403",
                "ENSG00000116560",
                "ENSG00000196591",
                "ENSG00000124766"
            ],
            "target": [],
            "expression_sum": {
                "class1": 4467.0,
                "class2": 1678.0
            },
            "ens_id": "ENSG00000141404",
            "uniprot_ids": [
                "P38405",
                "K7EMY6",
                "K7EQ80",
                "K7EPE2"
            ]
        },
        "ENSG00000166974": {
            "id": "ENSG00000166974",
            "type": "Gene",
            "symbol": "MAPRE2",
            "source": [
                "ENSG00000109132",
                "ENSG00000016082",
                "ENSG00000113580",
                "ENSG00000167081",
                "ENSG00000143799",
                "ENSG00000148516",
                "ENSG00000103495",
                "ENSG00000164330",
                "ENSG00000108055",
                "ENSG00000139352",
                "ENSG00000116560",
                "ENSG00000196591"
            ],
            "target": [],
            "expression_sum": {
                "class1": 8544.0,
                "class2": 2412.0
            },
            "ens_id": "ENSG00000166974",
            "uniprot_ids": [
                "Q15555",
                "M0QX52",
                "K7ERD8",
                "K7EL66",
                "K7ENB3"
            ]
        },
        "ENSG00000132639": {
            "id": "ENSG00000132639",
            "type": "Gene",
            "symbol": "SNAP25",
            "source": [
                "ENSG00000113580",
                "ENSG00000139352",
                "ENSG00000103495",
                "ENSG00000116560"
            ],
            "target": [],
            "expression_sum": {
                "class1": 17016.0,
                "class2": 3053.0
            },
            "ens_id": "ENSG00000132639",
            "uniprot_ids": [
                "P60880",
                "A0A0A0MSS0"
            ]
        },
        "ENSG00000125868": {
            "id": "ENSG00000125868",
            "type": "Gene",
            "symbol": "DSTN",
            "source": [
                "ENSG00000181449",
                "ENSG00000163435",
                "ENSG00000167081",
                "ENSG00000113580",
                "ENSG00000148516",
                "ENSG00000103495",
                "ENSG00000114861",
                "ENSG00000139352",
                "ENSG00000116560"
            ],
            "target": [],
            "expression_sum": {
                "class1": 55115.0,
                "class2": 31050.0
            },
            "ens_id": "ENSG00000125868",
            "uniprot_ids": [
                "P60981",
                "F6RFD5"
            ]
        },
        "ENSG00000175063": {
            "id": "ENSG00000175063",
            "type": "Gene",
            "symbol": "UBE2C",
            "source": [
                "ENSG00000108055",
                "ENSG00000164754",
                "ENSG00000116560"
            ],
            "target": [],
            "expression_sum": {
                "class1": 219.0,
                "class2": 5228.0
            },
            "ens_id": "ENSG00000175063",
            "uniprot_ids": [
                "O00762",
                "K4DI81",
                "A0A087WVK1",
                "A0A0A0MSE8"
            ]
        },
        "ENSG00000129355": {
            "id": "ENSG00000129355",
            "type": "Gene",
            "symbol": "CDKN2D",
            "source": [
                "ENSG00000028277",
                "ENSG00000164683",
                "ENSG00000113580",
                "ENSG00000167081",
                "ENSG00000103495",
                "ENSG00000164330",
                "ENSG00000198911",
                "ENSG00000108055",
                "ENSG00000114861",
                "ENSG00000139352",
                "ENSG00000213015",
                "ENSG00000196591",
                "ENSG00000116560",
                "ENSG00000147140",
                "ENSG00000160113"
            ],
            "target": [],
            "expression_sum": {
                "class1": 32811.0,
                "class2": 5558.0
            },
            "ens_id": "ENSG00000129355",
            "uniprot_ids": [
                "P55273"
            ]
        },
        "ENSG00000105270": {
            "id": "ENSG00000105270",
            "type": "Gene",
            "symbol": "CLIP3",
            "source": [
                "ENSG00000196591",
                "ENSG00000108055",
                "ENSG00000116560"
            ],
            "target": [],
            "expression_sum": {
                "class1": 5637.0,
                "class2": 1956.0
            },
            "ens_id": "ENSG00000105270",
            "uniprot_ids": [
                "Q96DZ5",
                "K7ESF3"
            ]
        },
        "ENSG00000174521": {
            "id": "ENSG00000174521",
            "type": "Gene",
            "symbol": "TTC9B",
            "source": [
                "ENSG00000100811",
                "ENSG00000105698",
                "ENSG00000108055",
                "ENSG00000196591",
                "ENSG00000116560",
                "ENSG00000092199"
            ],
            "target": [],
            "expression_sum": {
                "class1": 13312.0,
                "class2": 794.0
            },
            "ens_id": "ENSG00000174521",
            "uniprot_ids": [
                "Q8N6N2"
            ]
        },
        "ENSG00000131037": {
            "id": "ENSG00000131037",
            "type": "Gene",
            "symbol": "EPS8L1",
            "source": [
                "ENSG00000134138",
                "ENSG00000148516",
                "ENSG00000103495",
                "ENSG00000175745",
                "ENSG00000108055",
                "ENSG00000213015",
                "ENSG00000196591",
                "ENSG00000116560"
            ],
            "target": [],
            "expression_sum": {
                "class1": 6869.0,
                "class2": 704.0
            },
            "ens_id": "ENSG00000131037",
            "uniprot_ids": [
                "Q8TE68",
                "B4DKV7",
                "K7EN83",
                "K7ELW3",
                "K7EKX9"
            ]
        },
        "ENSG00000100077": {
            "id": "ENSG00000100077",
            "type": "Gene",
            "symbol": "GRK3",
            "source": [
                "ENSG00000164330",
                "ENSG00000116560",
                "ENSG00000148516"
            ],
            "target": [],
            "expression_sum": {
                "class1": 3683.0,
                "class2": 1294.0
            },
            "ens_id": "ENSG00000100077",
            "uniprot_ids": [
                "P35626",
                "Q8N433",
                "H7C099"
            ]
        },
        "ENSG00000100320": {
            "id": "ENSG00000100320",
            "type": "Gene",
            "symbol": "RBFOX2",
            "source": [
                "ENSG00000157557",
                "ENSG00000016082",
                "ENSG00000113580",
                "ENSG00000167081",
                "ENSG00000148516",
                "ENSG00000118263",
                "ENSG00000103495",
                "ENSG00000177030",
                "ENSG00000108055",
                "ENSG00000189403",
                "ENSG00000139352",
                "ENSG00000116560"
            ],
            "target": [],
            "expression_sum": {
                "class1": 19889.0,
                "class2": 6382.0
            },
            "ens_id": "ENSG00000100320",
            "uniprot_ids": [
                "O43251",
                "B0QYY4",
                "S4R469",
                "B0QYV1",
                "B0QYY6",
                "B0QYY7",
                "S4R3K7"
            ]
        },
        "ENSG00000130540": {
            "id": "ENSG00000130540",
            "type": "Gene",
            "symbol": "SULT4A1",
            "source": [
                "ENSG00000100811",
                "ENSG00000103495",
                "ENSG00000189403",
                "ENSG00000196591",
                "ENSG00000116560"
            ],
            "target": [],
            "expression_sum": {
                "class1": 5857.0,
                "class2": 706.0
            },
            "ens_id": "ENSG00000130540",
            "uniprot_ids": [
                "Q9BR01",
                "F8WE22"
            ]
        },
        "ENSG00000142156": {
            "id": "ENSG00000142156",
            "type": "Gene",
            "symbol": "COL6A1",
            "source": [
                "ENSG00000168214",
                "ENSG00000108055",
                "ENSG00000116560",
                "ENSG00000196591"
            ],
            "target": [],
            "expression_sum": {
                "class1": 10544.0,
                "class2": 5451.0
            },
            "ens_id": "ENSG00000142156",
            "uniprot_ids": [
                "P12109",
                "A0A087X0S5",
                "A0A804HJ28"
            ]
        },
        "ENSG00000065978": {
            "id": "ENSG00000065978",
            "type": "TF",
            "symbol": "YBX1",
            "source": [
                "ENSG00000181449",
                "ENSG00000109132",
                "ENSG00000167081",
                "ENSG00000164330",
                "ENSG00000221869",
                "ENSG00000137947",
                "ENSG00000067955",
                "ENSG00000119547"
            ],
            "target": [
                "ENSG00000111642",
                "ENSG00000103154",
                "ENSG00000153317",
                "ENSG00000007168"
            ],
            "expression_sum": {
                "class1": 134893.0,
                "class2": 88777.0
            },
            "ens_id": "ENSG00000065978",
            "uniprot_ids": [
                "P67809",
                "C9J5V9",
                "H0Y449"
            ]
        },
        "ENSG00000153317": {
            "id": "ENSG00000153317",
            "type": "Gene",
            "symbol": "ASAP1",
            "source": [
                "ENSG00000065978",
                "ENSG00000164330"
            ],
            "target": [],
            "expression_sum": {
                "class1": 6781.0,
                "class2": 2684.0
            },
            "ens_id": "ENSG00000153317",
            "uniprot_ids": [
                "Q9ULH1",
                "A0A0A0MRE5",
                "E5RHA9",
                "H0YBF7",
                "E5RFD9",
                "E5RHD7",
                "H0YBM4"
            ]
        },
        "ENSG00000111642": {
            "id": "ENSG00000111642",
            "type": "Gene",
            "symbol": "CHD4",
            "source": [
                "ENSG00000167081",
                "ENSG00000113580",
                "ENSG00000065978",
                "ENSG00000148516",
                "ENSG00000164330"
            ],
            "target": [],
            "expression_sum": {
                "class1": 20771.0,
                "class2": 10087.0
            },
            "ens_id": "ENSG00000111642",
            "uniprot_ids": [
                "Q14839",
                "A0A2R8YFD8",
                "A0A2R8YDW2",
                "A0A2R8YDJ9",
                "F5GWX5",
                "A0A2R8Y425",
                "A0A0C4DGG9",
                "A0A2R8Y8C1",
                "A0A2R8YD40",
                "A0A2R8Y6G9",
                "A0A2R8Y212",
                "A0A2R8Y795",
                "A0A2R8Y8B3",
                "A0A2R8Y521",
                "A0A2R8Y5J0",
                "A0A2R8Y4X2",
                "A0A2U3TZM0",
                "A0A2R8Y512",
                "A0A2R8Y5Z7",
                "A0A2R8YER1",
                "A0A2R8YE38",
                "A0A2R8Y539",
                "F5H6N4",
                "F5H596",
                "A0A2R8YFK9",
                "A0A2R8Y5M9",
                "A0A2R8Y7X1",
                "A0A2R8Y7I0",
                "A0A2R8Y685",
                "A0A2R8Y7M9"
            ]
        },
        "ENSG00000103154": {
            "id": "ENSG00000103154",
            "type": "Gene",
            "symbol": "NECAB2",
            "source": [
                "ENSG00000100811",
                "ENSG00000215021",
                "ENSG00000065978",
                "ENSG00000148516",
                "ENSG00000103495",
                "ENSG00000189403",
                "ENSG00000196591",
                "ENSG00000092199"
            ],
            "target": [],
            "expression_sum": {
                "class1": 5795.0,
                "class2": 691.0
            },
            "ens_id": "ENSG00000103154",
            "uniprot_ids": [
                "Q7Z6G3",
                "H3BPH6",
                "H3BQC7"
            ]
        },
        "ENSG00000007168": {
            "id": "ENSG00000007168",
            "type": "Gene",
            "symbol": "PAFAH1B1",
            "source": [
                "ENSG00000181449",
                "ENSG00000167081",
                "ENSG00000113580",
                "ENSG00000065978",
                "ENSG00000148516",
                "ENSG00000164330"
            ],
            "target": [],
            "expression_sum": {
                "class1": 17220.0,
                "class2": 8734.0
            },
            "ens_id": "ENSG00000007168",
            "uniprot_ids": [
                "P43034",
                "A0A6Q8PFT2",
                "I3L3N5",
                "I3L495",
                "I3L384",
                "A0A6Q8PFU3",
                "A0A6Q8PH33",
                "A0A6Q8PG63",
                "A0A6Q8PGF8"
            ]
        },
        "ENSG00000177606": {
            "id": "ENSG00000177606",
            "type": "TF",
            "symbol": "JUN",
            "source": [
                "ENSG00000100219",
                "ENSG00000221869",
                "ENSG00000139352"
            ],
            "target": [
                "ENSG00000176887",
                "ENSG00000198840",
                "ENSG00000132698",
                "ENSG00000185664",
                "ENSG00000164093",
                "ENSG00000122756",
                "ENSG00000165973",
                "ENSG00000112773",
                "ENSG00000169435",
                "ENSG00000115461",
                "ENSG00000148834",
                "ENSG00000010278",
                "ENSG00000104870",
                "ENSG00000182481",
                "ENSG00000189143",
                "ENSG00000166145",
                "ENSG00000198125",
                "ENSG00000181885",
                "ENSG00000132386",
                "ENSG00000166598",
                "ENSG00000117691",
                "ENSG00000135535",
                "ENSG00000164305",
                "ENSG00000107438",
                "ENSG00000167642",
                "ENSG00000161249",
                "ENSG00000187244",
                "ENSG00000198938",
                "ENSG00000198899",
                "ENSG00000167641",
                "ENSG00000015592",
                "ENSG00000143870",
                "ENSG00000135549",
                "ENSG00000213937",
                "ENSG00000157557",
                "ENSG00000119888",
                "ENSG00000100219",
                "ENSG00000221869",
                "ENSG00000185650",
                "ENSG00000108679",
                "ENSG00000172915",
                "ENSG00000179218"
            ],
            "expression_sum": {
                "class1": 17839.0,
                "class2": 10689.0
            },
            "ens_id": "ENSG00000177606",
            "uniprot_ids": [
                "P05412"
            ]
        },
        "ENSG00000132698": {
            "id": "ENSG00000132698",
            "type": "Gene",
            "symbol": "RAB25",
            "source": [
                "ENSG00000100811",
                "ENSG00000163435",
                "ENSG00000113580",
                "ENSG00000168610",
                "ENSG00000100219",
                "ENSG00000168214",
                "ENSG00000124766",
                "ENSG00000177606"
            ],
            "target": [],
            "expression_sum": {
                "class1": 427.0,
                "class2": 7248.0
            },
            "ens_id": "ENSG00000132698",
            "uniprot_ids": [
                "P57735"
            ]
        },
        "ENSG00000117691": {
            "id": "ENSG00000117691",
            "type": "Gene",
            "symbol": "NENF",
            "source": [
                "ENSG00000105698",
                "ENSG00000119950",
                "ENSG00000168214",
                "ENSG00000119547",
                "ENSG00000177606"
            ],
            "target": [],
            "expression_sum": {
                "class1": 10998.0,
                "class2": 28082.0
            },
            "ens_id": "ENSG00000117691",
            "uniprot_ids": [
                "Q9UMX5"
            ]
        },
        "ENSG00000143870": {
            "id": "ENSG00000143870",
            "type": "Gene",
            "symbol": "PDIA6",
            "source": [
                "ENSG00000181449",
                "ENSG00000181722",
                "ENSG00000177606"
            ],
            "target": [],
            "expression_sum": {
                "class1": 18509.0,
                "class2": 34028.0
            },
            "ens_id": "ENSG00000143870",
            "uniprot_ids": [
                "Q15084",
                "C9JNG5"
            ]
        },
        "ENSG00000119888": {
            "id": "ENSG00000119888",
            "type": "Gene",
            "symbol": "EPCAM",
            "source": [
                "ENSG00000163435",
                "ENSG00000160113",
                "ENSG00000113580",
                "ENSG00000105698",
                "ENSG00000181722",
                "ENSG00000139352",
                "ENSG00000173575",
                "ENSG00000168214",
                "ENSG00000119547",
                "ENSG00000177606",
                "ENSG00000092199"
            ],
            "target": [],
            "expression_sum": {
                "class1": 3257.0,
                "class2": 27181.0
            },
            "ens_id": "ENSG00000119888",
            "uniprot_ids": [
                "P16422",
                "B5MCA4",
                "C9JKY3"
            ]
        },
        "ENSG00000115461": {
            "id": "ENSG00000115461",
            "type": "Gene",
            "symbol": "IGFBP5",
            "source": [
                "ENSG00000126778",
                "ENSG00000163848",
                "ENSG00000168214",
                "ENSG00000130522",
                "ENSG00000177606"
            ],
            "target": [],
            "expression_sum": {
                "class1": 18010.0,
                "class2": 76740.0
            },
            "ens_id": "ENSG00000115461",
            "uniprot_ids": [
                "P24593",
                "C9JXX4"
            ]
        },
        "ENSG00000169435": {
            "id": "ENSG00000169435",
            "type": "Gene",
            "symbol": "RASSF6",
            "source": [
                "ENSG00000168214",
                "ENSG00000113580",
                "ENSG00000139352",
                "ENSG00000177606"
            ],
            "target": [],
            "expression_sum": {
                "class1": 219.0,
                "class2": 4609.0
            },
            "ens_id": "ENSG00000169435",
            "uniprot_ids": [
                "Q6ZTQ3"
            ]
        },
        "ENSG00000164305": {
            "id": "ENSG00000164305",
            "type": "Gene",
            "symbol": "CASP3",
            "source": [
                "ENSG00000130522",
                "ENSG00000177606"
            ],
            "target": [],
            "expression_sum": {
                "class1": 14692.0,
                "class2": 1984.0
            },
            "ens_id": "ENSG00000164305",
            "uniprot_ids": [
                "P42574",
                "A8MVM1",
                "C9JXR7"
            ]
        },
        "ENSG00000112773": {
            "id": "ENSG00000112773",
            "type": "Gene",
            "symbol": "TENT5A",
            "source": [
                "ENSG00000170365",
                "ENSG00000160113",
                "ENSG00000119950",
                "ENSG00000168214",
                "ENSG00000177606"
            ],
            "target": [],
            "expression_sum": {
                "class1": 3503.0,
                "class2": 6784.0
            },
            "ens_id": "ENSG00000112773",
            "uniprot_ids": [
                "Q96IP4",
                "Q5TF85",
                "H0Y6D7",
                "H0Y5Y3"
            ]
        },
        "ENSG00000135535": {
            "id": "ENSG00000135535",
            "type": "Gene",
            "symbol": "CD164",
            "source": [
                "ENSG00000106261",
                "ENSG00000160113",
                "ENSG00000113580",
                "ENSG00000168214",
                "ENSG00000177606"
            ],
            "target": [],
            "expression_sum": {
                "class1": 4307.0,
                "class2": 11073.0
            },
            "ens_id": "ENSG00000135535",
            "uniprot_ids": [
                "Q04900",
                "D6R9B4"
            ]
        },
        "ENSG00000135549": {
            "id": "ENSG00000135549",
            "type": "Gene",
            "symbol": "PKIB",
            "source": [
                "ENSG00000109132",
                "ENSG00000134138",
                "ENSG00000140262",
                "ENSG00000139352",
                "ENSG00000119547",
                "ENSG00000177606"
            ],
            "target": [],
            "expression_sum": {
                "class1": 21371.0,
                "class2": 13785.0
            },
            "ens_id": "ENSG00000135549",
            "uniprot_ids": [
                "Q9C010",
                "Q5T0Z6"
            ]
        },
        "ENSG00000189143": {
            "id": "ENSG00000189143",
            "type": "Gene",
            "symbol": "CLDN4",
            "source": [
                "ENSG00000168214",
                "ENSG00000100219",
                "ENSG00000163435",
                "ENSG00000177606"
            ],
            "target": [],
            "expression_sum": {
                "class1": 451.0,
                "class2": 7773.0
            },
            "ens_id": "ENSG00000189143",
            "uniprot_ids": [
                "O14493"
            ]
        },
        "ENSG00000221869": {
            "id": "ENSG00000221869",
            "type": "TF",
            "symbol": "CEBPD",
            "source": [
                "ENSG00000163435",
                "ENSG00000160113",
                "ENSG00000164683",
                "ENSG00000130522",
                "ENSG00000177606"
            ],
            "target": [
                "ENSG00000206573",
                "ENSG00000106261",
                "ENSG00000115944",
                "ENSG00000162704",
                "ENSG00000196531",
                "ENSG00000126005",
                "ENSG00000171604",
                "ENSG00000116030",
                "ENSG00000099622",
                "ENSG00000180011",
                "ENSG00000129559",
                "ENSG00000184752",
                "ENSG00000169567",
                "ENSG00000125534",
                "ENSG00000177606",
                "ENSG00000160113",
                "ENSG00000174748",
                "ENSG00000131143",
                "ENSG00000168653",
                "ENSG00000241837",
                "ENSG00000156467",
                "ENSG00000142864",
                "ENSG00000161203",
                "ENSG00000075415",
                "ENSG00000166165",
                "ENSG00000141753",
                "ENSG00000181885",
                "ENSG00000184076",
                "ENSG00000171314",
                "ENSG00000266472",
                "ENSG00000150779",
                "ENSG00000154556",
                "ENSG00000178741",
                "ENSG00000163466",
                "ENSG00000198931",
                "ENSG00000166913",
                "ENSG00000173812",
                "ENSG00000141279",
                "ENSG00000099795",
                "ENSG00000111229",
                "ENSG00000151729",
                "ENSG00000118816",
                "ENSG00000088832",
                "ENSG00000162585",
                "ENSG00000051620",
                "ENSG00000122545",
                "ENSG00000149547",
                "ENSG00000159335",
                "ENSG00000164924",
                "ENSG00000189043",
                "ENSG00000150687",
                "ENSG00000142192",
                "ENSG00000277443",
                "ENSG00000101558",
                "ENSG00000162545",
                "ENSG00000101040",
                "ENSG00000128739",
                "ENSG00000109475",
                "ENSG00000154518",
                "ENSG00000196262",
                "ENSG00000111640",
                "ENSG00000104904",
                "ENSG00000165637",
                "ENSG00000183978",
                "ENSG00000241685",
                "ENSG00000173726",
                "ENSG00000091164",
                "ENSG00000172757",
                "ENSG00000125971",
                "ENSG00000130332",
                "ENSG00000116586",
                "ENSG00000188290",
                "ENSG00000156976",
                "ENSG00000173674",
                "ENSG00000147548",
                "ENSG00000089220",
                "ENSG00000134138",
                "ENSG00000119335",
                "ENSG00000115286",
                "ENSG00000134717",
                "ENSG00000119396",
                "ENSG00000136156",
                "ENSG00000177879",
                "ENSG00000105640",
                "ENSG00000099624",
                "ENSG00000171159",
                "ENSG00000067225",
                "ENSG00000154723",
                "ENSG00000166347",
                "ENSG00000213719",
                "ENSG00000113712",
                "ENSG00000110321",
                "ENSG00000233927",
                "ENSG00000254999",
                "ENSG00000004059",
                "ENSG00000152234",
                "ENSG00000137106",
                "ENSG00000177556",
                "ENSG00000034713",
                "ENSG00000113387",
                "ENSG00000100592",
                "ENSG00000198668",
                "ENSG00000134308",
                "ENSG00000034510",
                "ENSG00000123144",
                "ENSG00000115091",
                "ENSG00000149257",
                "ENSG00000143256",
                "ENSG00000149084",
                "ENSG00000126934",
                "ENSG00000154639",
                "ENSG00000168118",
                "ENSG00000106367",
                "ENSG00000197111",
                "ENSG00000214253",
                "ENSG00000101182",
                "ENSG00000162496",
                "ENSG00000142676",
                "ENSG00000171421",
                "ENSG00000129515",
                "ENSG00000100387",
                "ENSG00000140319",
                "ENSG00000206503",
                "ENSG00000122034",
                "ENSG00000187514",
                "ENSG00000111669",
                "ENSG00000163041",
                "ENSG00000118680",
                "ENSG00000243317",
                "ENSG00000175826",
                "ENSG00000184007",
                "ENSG00000009307",
                "ENSG00000178980",
                "ENSG00000107262",
                "ENSG00000004779",
                "ENSG00000182117",
                "ENSG00000124172",
                "ENSG00000182196",
                "ENSG00000169139",
                "ENSG00000168214",
                "ENSG00000118363",
                "ENSG00000162191",
                "ENSG00000100911",
                "ENSG00000070831",
                "ENSG00000162076",
                "ENSG00000107438",
                "ENSG00000122705",
                "ENSG00000166482",
                "ENSG00000110700",
                "ENSG00000184009",
                "ENSG00000075624",
                "ENSG00000095059",
                "ENSG00000173660",
                "ENSG00000103187",
                "ENSG00000108953",
                "ENSG00000186184",
                "ENSG00000100804",
                "ENSG00000155368",
                "ENSG00000272047",
                "ENSG00000114125",
                "ENSG00000185787",
                "ENSG00000163931",
                "ENSG00000164713",
                "ENSG00000130638",
                "ENSG00000240972",
                "ENSG00000111775",
                "ENSG00000105379",
                "ENSG00000123636",
                "ENSG00000135390",
                "ENSG00000149806",
                "ENSG00000239672",
                "ENSG00000213585",
                "ENSG00000167658",
                "ENSG00000005022",
                "ENSG00000188612",
                "ENSG00000128654",
                "ENSG00000112715",
                "ENSG00000180817",
                "ENSG00000112695",
                "ENSG00000116288",
                "ENSG00000090266",
                "ENSG00000158710",
                "ENSG00000205542",
                "ENSG00000160948",
                "ENSG00000136810",
                "ENSG00000174444",
                "ENSG00000169764",
                "ENSG00000168028",
                "ENSG00000127184",
                "ENSG00000136238",
                "ENSG00000104341",
                "ENSG00000130522",
                "ENSG00000224032",
                "ENSG00000167815",
                "ENSG00000122873",
                "ENSG00000126457",
                "ENSG00000176340",
                "ENSG00000167996",
                "ENSG00000132963",
                "ENSG00000065978",
                "ENSG00000087460",
                "ENSG00000170759",
                "ENSG00000171530",
                "ENSG00000148450",
                "ENSG00000177410",
                "ENSG00000178127",
                "ENSG00000113558",
                "ENSG00000101745",
                "ENSG00000140990",
                "ENSG00000182718",
                "ENSG00000105518",
                "ENSG00000163435",
                "ENSG00000133639",
                "ENSG00000126267",
                "ENSG00000100129",
                "ENSG00000170017",
                "ENSG00000092841",
                "ENSG00000145494"
            ],
            "expression_sum": {
                "class1": 1701.0,
                "class2": 15658.0
            },
            "ens_id": "ENSG00000221869",
            "uniprot_ids": [
                "P49716"
            ]
        },
        "ENSG00000122756": {
            "id": "ENSG00000122756",
            "type": "Gene",
            "symbol": "CNTFR",
            "source": [
                "ENSG00000196591",
                "ENSG00000100811",
                "ENSG00000103495",
                "ENSG00000177606"
            ],
            "target": [],
            "expression_sum": {
                "class1": 10465.0,
                "class2": 1444.0
            },
            "ens_id": "ENSG00000122756",
            "uniprot_ids": [
                "P26992",
                "Q5T8H6"
            ]
        },
        "ENSG00000165973": {
            "id": "ENSG00000165973",
            "type": "Gene",
            "symbol": "NELL1",
            "source": [
                "ENSG00000177606"
            ],
            "target": [],
            "expression_sum": {
                "class1": 586.0,
                "class2": 3478.0
            },
            "ens_id": "ENSG00000165973",
            "uniprot_ids": [
                "Q92832",
                "J3KNC5",
                "F5H6I3"
            ]
        },
        "ENSG00000107438": {
            "id": "ENSG00000107438",
            "type": "Gene",
            "symbol": "PDLIM1",
            "source": [
                "ENSG00000163435",
                "ENSG00000160113",
                "ENSG00000164683",
                "ENSG00000113580",
                "ENSG00000164330",
                "ENSG00000168610",
                "ENSG00000221869",
                "ENSG00000168214",
                "ENSG00000177606",
                "ENSG00000092199"
            ],
            "target": [],
            "expression_sum": {
                "class1": 3744.0,
                "class2": 10802.0
            },
            "ens_id": "ENSG00000107438",
            "uniprot_ids": [
                "O00151"
            ]
        },
        "ENSG00000148834": {
            "id": "ENSG00000148834",
            "type": "Gene",
            "symbol": "GSTO1",
            "source": [
                "ENSG00000168214",
                "ENSG00000105698",
                "ENSG00000177606",
                "ENSG00000160113"
            ],
            "target": [],
            "expression_sum": {
                "class1": 3927.0,
                "class2": 14823.0
            },
            "ens_id": "ENSG00000148834",
            "uniprot_ids": [
                "P78417",
                "Q5TA02",
                "Q5TA01"
            ]
        },
        "ENSG00000010278": {
            "id": "ENSG00000010278",
            "type": "Gene",
            "symbol": "CD9",
            "source": [
                "ENSG00000177606",
                "ENSG00000168610"
            ],
            "target": [],
            "expression_sum": {
                "class1": 657.0,
                "class2": 36069.0
            },
            "ens_id": "ENSG00000010278",
            "uniprot_ids": [
                "P21926",
                "A6NNI4",
                "A0A7I2V3T7",
                "A0A087WU13",
                "F5GXT1",
                "A0A2R8Y478",
                "A0A7I2V3T0",
                "G8JLH6"
            ]
        },
        "ENSG00000166598": {
            "id": "ENSG00000166598",
            "type": "Gene",
            "symbol": "HSP90B1",
            "source": [
                "ENSG00000181449",
                "ENSG00000177606"
            ],
            "target": [],
            "expression_sum": {
                "class1": 26448.0,
                "class2": 45254.0
            },
            "ens_id": "ENSG00000166598",
            "uniprot_ids": [
                "P14625",
                "A0A7P0T917",
                "A0A7P0TAY2",
                "A0A087WT78",
                "F8W026",
                "A0A7P0TAG4",
                "A0A7P0TAE1",
                "A0A7P0Z405",
                "A0A7P0TAT8",
                "A0A7P0TBC2",
                "A0A7P0T8R3",
                "A0A7P0T823",
                "H0YIV0",
                "A0A7P0TAC2",
                "A0A7P0T885",
                "A0A7P0T8C8",
                "A0A7P0Z4B4"
            ]
        },
        "ENSG00000172915": {
            "id": "ENSG00000172915",
            "type": "Gene",
            "symbol": "NBEA",
            "source": [
                "ENSG00000163435",
                "ENSG00000113580",
                "ENSG00000100219",
                "ENSG00000139352",
                "ENSG00000168214",
                "ENSG00000177606"
            ],
            "target": [],
            "expression_sum": {
                "class1": 8107.0,
                "class2": 7514.0
            },
            "ens_id": "ENSG00000172915",
            "uniprot_ids": [
                "Q8NFP9",
                "A0A0D9SF28",
                "Q5T321"
            ]
        },
        "ENSG00000166145": {
            "id": "ENSG00000166145",
            "type": "Gene",
            "symbol": "SPINT1",
            "source": [
                "ENSG00000163435",
                "ENSG00000177606",
                "ENSG00000092199"
            ],
            "target": [],
            "expression_sum": {
                "class1": 330.0,
                "class2": 4243.0
            },
            "ens_id": "ENSG00000166145",
            "uniprot_ids": [
                "O43278",
                "H3BR01",
                "H3BSM2",
                "H3BVD9",
                "H3BNW5",
                "H3BTQ8",
                "H3BMB6"
            ]
        },
        "ENSG00000213937": {
            "id": "ENSG00000213937",
            "type": "Gene",
            "symbol": "CLDN9",
            "source": [
                "ENSG00000113580",
                "ENSG00000139352",
                "ENSG00000177606",
                "ENSG00000160113"
            ],
            "target": [],
            "expression_sum": {
                "class1": 167.0,
                "class2": 6832.0
            },
            "ens_id": "ENSG00000213937",
            "uniprot_ids": [
                "O95484"
            ]
        },
        "ENSG00000132386": {
            "id": "ENSG00000132386",
            "type": "Gene",
            "symbol": "SERPINF1",
            "source": [
                "ENSG00000177606"
            ],
            "target": [],
            "expression_sum": {
                "class1": 9904.0,
                "class2": 16597.0
            },
            "ens_id": "ENSG00000132386",
            "uniprot_ids": [
                "P36955",
                "I3L1U4",
                "I3L4N7",
                "I3L3Z3",
                "I3L4Z0",
                "I3L2R7",
                "I3L425",
                "I3L107",
                "I3L4F9"
            ]
        },
        "ENSG00000181885": {
            "id": "ENSG00000181885",
            "type": "Gene",
            "symbol": "CLDN7",
            "source": [
                "ENSG00000123933",
                "ENSG00000163435",
                "ENSG00000160113",
                "ENSG00000113580",
                "ENSG00000153147",
                "ENSG00000105698",
                "ENSG00000168610",
                "ENSG00000100219",
                "ENSG00000221869",
                "ENSG00000139352",
                "ENSG00000168214",
                "ENSG00000196591",
                "ENSG00000124766",
                "ENSG00000173575",
                "ENSG00000177606"
            ],
            "target": [],
            "expression_sum": {
                "class1": 425.0,
                "class2": 17786.0
            },
            "ens_id": "ENSG00000181885",
            "uniprot_ids": [
                "O95471",
                "I3L3X1",
                "I3L3L6",
                "K7EL87",
                "F5H496",
                "K7EP40"
            ]
        },
        "ENSG00000108679": {
            "id": "ENSG00000108679",
            "type": "Gene",
            "symbol": "LGALS3BP",
            "source": [
                "ENSG00000163435",
                "ENSG00000160113",
                "ENSG00000139352",
                "ENSG00000168214",
                "ENSG00000177606"
            ],
            "target": [],
            "expression_sum": {
                "class1": 9477.0,
                "class2": 21066.0
            },
            "ens_id": "ENSG00000108679",
            "uniprot_ids": [
                "Q08380",
                "K7EM15",
                "K7EJD3",
                "K7ER67",
                "K7ES75",
                "K7EP36",
                "K7ERZ6",
                "K7EKQ5",
                "K7ELA1",
                "K7EJY8",
                "K7EQT9",
                "K7ESM3",
                "K7EN99"
            ]
        },
        "ENSG00000179218": {
            "id": "ENSG00000179218",
            "type": "Gene",
            "symbol": "CALR",
            "source": [
                "ENSG00000168214",
                "ENSG00000181449",
                "ENSG00000177606"
            ],
            "target": [],
            "expression_sum": {
                "class1": 22508.0,
                "class2": 47691.0
            },
            "ens_id": "ENSG00000179218",
            "uniprot_ids": [
                "P27797",
                "K7EJB9",
                "K7ELE2",
                "K7EL50",
                "A0A7P0T861"
            ]
        },
        "ENSG00000161249": {
            "id": "ENSG00000161249",
            "type": "Gene",
            "symbol": "DMKN",
            "source": [
                "ENSG00000168214",
                "ENSG00000163435",
                "ENSG00000177606",
                "ENSG00000160113"
            ],
            "target": [],
            "expression_sum": {
                "class1": 785.0,
                "class2": 6238.0
            },
            "ens_id": "ENSG00000161249",
            "uniprot_ids": [
                "Q6E0U4",
                "M0QZU6",
                "C9JCN1",
                "A3EZ84",
                "C9J186",
                "E9PAQ7",
                "M0R0W2",
                "M0R265",
                "H7BZ95",
                "S4R427",
                "M0R0A5",
                "S4R2X7",
                "C9J3V3",
                "M0R1M0",
                "E7EMS7",
                "R4GMQ0",
                "S4R3L7",
                "M0R3F8",
                "C9IYI1"
            ]
        },
        "ENSG00000167642": {
            "id": "ENSG00000167642",
            "type": "Gene",
            "symbol": "SPINT2",
            "source": [
                "ENSG00000106261",
                "ENSG00000163435",
                "ENSG00000185551",
                "ENSG00000160113",
                "ENSG00000139352",
                "ENSG00000168214",
                "ENSG00000119547",
                "ENSG00000177606"
            ],
            "target": [],
            "expression_sum": {
                "class1": 29236.0,
                "class2": 49863.0
            },
            "ens_id": "ENSG00000167642",
            "uniprot_ids": [
                "O43291",
                "K7ESI5",
                "K7EKQ0",
                "K7ES27",
                "K7EKC8",
                "K7EM91",
                "K7EJS4"
            ]
        },
        "ENSG00000167641": {
            "id": "ENSG00000167641",
            "type": "Gene",
            "symbol": "PPP1R14A",
            "source": [
                "ENSG00000177606"
            ],
            "target": [],
            "expression_sum": {
                "class1": 3827.0,
                "class2": 310.0
            },
            "ens_id": "ENSG00000167641",
            "uniprot_ids": [
                "Q96A00",
                "A0A0B4J2A3",
                "K7EJB8",
                "K7EMN0"
            ]
        },
        "ENSG00000187244": {
            "id": "ENSG00000187244",
            "type": "Gene",
            "symbol": "BCAM",
            "source": [
                "ENSG00000168214",
                "ENSG00000139352",
                "ENSG00000177606"
            ],
            "target": [],
            "expression_sum": {
                "class1": 5040.0,
                "class2": 10800.0
            },
            "ens_id": "ENSG00000187244",
            "uniprot_ids": [
                "P50895",
                "A0A087WXM8",
                "K7ENU8",
                "K7ERB7"
            ]
        },
        "ENSG00000104870": {
            "id": "ENSG00000104870",
            "type": "Gene",
            "symbol": "FCGRT",
            "source": [
                "ENSG00000100811",
                "ENSG00000106261",
                "ENSG00000163435",
                "ENSG00000160113",
                "ENSG00000153147",
                "ENSG00000122691",
                "ENSG00000181722",
                "ENSG00000139352",
                "ENSG00000168214",
                "ENSG00000130522",
                "ENSG00000177606",
                "ENSG00000092199"
            ],
            "target": [],
            "expression_sum": {
                "class1": 1131.0,
                "class2": 9174.0
            },
            "ens_id": "ENSG00000104870",
            "uniprot_ids": [
                "P55899",
                "M0R2T3",
                "M0QYN9",
                "M0QYM4",
                "M0R0K7",
                "M0R0H0",
                "M0QYN1",
                "M0R266",
                "M0R220",
                "M0R0A9"
            ]
        },
        "ENSG00000100219": {
            "id": "ENSG00000100219",
            "type": "TF",
            "symbol": "XBP1",
            "source": [
                "ENSG00000168214",
                "ENSG00000106261",
                "ENSG00000177606",
                "ENSG00000160113"
            ],
            "target": [
                "ENSG00000106261",
                "ENSG00000152558",
                "ENSG00000126005",
                "ENSG00000166710",
                "ENSG00000198840",
                "ENSG00000132698",
                "ENSG00000171604",
                "ENSG00000160113",
                "ENSG00000177606",
                "ENSG00000076554",
                "ENSG00000163191",
                "ENSG00000189143",
                "ENSG00000184076",
                "ENSG00000181885",
                "ENSG00000245532",
                "ENSG00000265972",
                "ENSG00000154556",
                "ENSG00000114439",
                "ENSG00000181722",
                "ENSG00000141279",
                "ENSG00000111229",
                "ENSG00000051620",
                "ENSG00000150687",
                "ENSG00000197930",
                "ENSG00000135069",
                "ENSG00000101040",
                "ENSG00000198886",
                "ENSG00000144560",
                "ENSG00000134138",
                "ENSG00000166347",
                "ENSG00000213719",
                "ENSG00000121039",
                "ENSG00000143013",
                "ENSG00000148444",
                "ENSG00000026025",
                "ENSG00000129521",
                "ENSG00000263001",
                "ENSG00000109099",
                "ENSG00000163399",
                "ENSG00000138772",
                "ENSG00000156675",
                "ENSG00000118680",
                "ENSG00000128564",
                "ENSG00000168214",
                "ENSG00000124107",
                "ENSG00000070831",
                "ENSG00000144674",
                "ENSG00000106537",
                "ENSG00000114405",
                "ENSG00000091844",
                "ENSG00000172915",
                "ENSG00000030582",
                "ENSG00000002586",
                "ENSG00000123636",
                "ENSG00000214530",
                "ENSG00000174938",
                "ENSG00000198727",
                "ENSG00000147676",
                "ENSG00000104763",
                "ENSG00000160613",
                "ENSG00000060138",
                "ENSG00000169764",
                "ENSG00000196586",
                "ENSG00000198786",
                "ENSG00000146674",
                "ENSG00000118640",
                "ENSG00000170421",
                "ENSG00000164096",
                "ENSG00000049618",
                "ENSG00000139970",
                "ENSG00000182718",
                "ENSG00000163435",
                "ENSG00000133639",
                "ENSG00000109062",
                "ENSG00000138640",
                "ENSG00000104419",
                "ENSG00000198888"
            ],
            "expression_sum": {
                "class1": 15335.0,
                "class2": 32457.0
            },
            "ens_id": "ENSG00000100219",
            "uniprot_ids": [
                "P17861",
                "B1AHH2",
                "B1AHH1"
            ]
        },
        "ENSG00000157557": {
            "id": "ENSG00000157557",
            "type": "TF",
            "symbol": "ETS2",
            "source": [
                "ENSG00000160113",
                "ENSG00000119950",
                "ENSG00000168214",
                "ENSG00000119547",
                "ENSG00000177606"
            ],
            "target": [
                "ENSG00000206573",
                "ENSG00000181061",
                "ENSG00000214530",
                "ENSG00000165655",
                "ENSG00000099622",
                "ENSG00000111640",
                "ENSG00000243317",
                "ENSG00000067836",
                "ENSG00000120071",
                "ENSG00000110955",
                "ENSG00000156976",
                "ENSG00000102144",
                "ENSG00000169432",
                "ENSG00000136710",
                "ENSG00000104112",
                "ENSG00000168214",
                "ENSG00000115738",
                "ENSG00000143933",
                "ENSG00000114023",
                "ENSG00000132424",
                "ENSG00000186352",
                "ENSG00000130338",
                "ENSG00000067225",
                "ENSG00000137500",
                "ENSG00000160714",
                "ENSG00000122873",
                "ENSG00000176340",
                "ENSG00000178741",
                "ENSG00000140416",
                "ENSG00000055609",
                "ENSG00000181722",
                "ENSG00000164830",
                "ENSG00000172115",
                "ENSG00000181315",
                "ENSG00000151729",
                "ENSG00000143158",
                "ENSG00000118816",
                "ENSG00000272398",
                "ENSG00000186472",
                "ENSG00000101745",
                "ENSG00000105649",
                "ENSG00000163875",
                "ENSG00000100320",
                "ENSG00000119547"
            ],
            "expression_sum": {
                "class1": 2470.0,
                "class2": 7721.0
            },
            "ens_id": "ENSG00000157557",
            "uniprot_ids": [
                "P15036",
                "A0A590UJP9",
                "C9JAG2",
                "A0A590UJR2",
                "A0A0C4DG47"
            ]
        },
        "ENSG00000198899": {
            "id": "ENSG00000198899",
            "type": "Gene",
            "symbol": "MT-ATP6",
            "source": [
                "ENSG00000185551",
                "ENSG00000163435",
                "ENSG00000113580",
                "ENSG00000139352",
                "ENSG00000168214",
                "ENSG00000177606"
            ],
            "target": [],
            "expression_sum": {
                "class1": 576191.0,
                "class2": 1242074.0
            },
            "ens_id": "ENSG00000198899",
            "uniprot_ids": [
                "P00846"
            ]
        },
        "ENSG00000198938": {
            "id": "ENSG00000198938",
            "type": "Gene",
            "symbol": "MT-CO3",
            "source": [
                "ENSG00000185551",
                "ENSG00000163435",
                "ENSG00000139352",
                "ENSG00000168214",
                "ENSG00000177606"
            ],
            "target": [],
            "expression_sum": {
                "class1": 761150.0,
                "class2": 1643056.0
            },
            "ens_id": "ENSG00000198938",
            "uniprot_ids": [
                "P00414"
            ]
        },
        "ENSG00000198840": {
            "id": "ENSG00000198840",
            "type": "Gene",
            "symbol": "MT-ND3",
            "source": [
                "ENSG00000163435",
                "ENSG00000113580",
                "ENSG00000105698",
                "ENSG00000100219",
                "ENSG00000139352",
                "ENSG00000168214",
                "ENSG00000184486",
                "ENSG00000177606"
            ],
            "target": [],
            "expression_sum": {
                "class1": 368107.0,
                "class2": 655850.0
            },
            "ens_id": "ENSG00000198840",
            "uniprot_ids": [
                "P03897"
            ]
        },
        "ENSG00000162599": {
            "id": "ENSG00000162599",
            "type": "TF",
            "symbol": "NFIA",
            "source": [
                "ENSG00000175745",
                "ENSG00000181449"
            ],
            "target": [
                "ENSG00000134138",
                "ENSG00000134245",
                "ENSG00000122545",
                "ENSG00000175745",
                "ENSG00000215386",
                "ENSG00000120885",
                "ENSG00000134308"
            ],
            "expression_sum": {
                "class1": 1610.0,
                "class2": 14292.0
            },
            "ens_id": "ENSG00000162599",
            "uniprot_ids": [
                "Q12857",
                "A0A590UJU2",
                "A0A590UK38",
                "B1AKN8",
                "A0A590UJ67",
                "S4R3W2",
                "S4R3K5",
                "S4R3K4",
                "S4R3W6",
                "A0A590UJA3",
                "A0A590UK09",
                "A0A590UJ82",
                "B1AKN6",
                "S4R308",
                "B1AKN5",
                "B1AKN7",
                "A0A590UK73"
            ]
        },
        "ENSG00000134245": {
            "id": "ENSG00000134245",
            "type": "Gene",
            "symbol": "WNT2B",
            "source": [
                "ENSG00000162599"
            ],
            "target": [],
            "expression_sum": {
                "class1": 797.0,
                "class2": 4330.0
            },
            "ens_id": "ENSG00000134245",
            "uniprot_ids": [
                "Q93097",
                "Q5TEH8"
            ]
        },
        "ENSG00000134308": {
            "id": "ENSG00000134308",
            "type": "Gene",
            "symbol": "YWHAQ",
            "source": [
                "ENSG00000181449",
                "ENSG00000114315",
                "ENSG00000167081",
                "ENSG00000113580",
                "ENSG00000164330",
                "ENSG00000164093",
                "ENSG00000114861",
                "ENSG00000221869",
                "ENSG00000137203",
                "ENSG00000119547",
                "ENSG00000162599"
            ],
            "target": [],
            "expression_sum": {
                "class1": 79969.0,
                "class2": 29325.0
            },
            "ens_id": "ENSG00000134308",
            "uniprot_ids": [
                "P27348",
                "E9PG15"
            ]
        },
        "ENSG00000175745": {
            "id": "ENSG00000175745",
            "type": "TF",
            "symbol": "NR2F1",
            "source": [
                "ENSG00000162599",
                "ENSG00000113580",
                "ENSG00000122691",
                "ENSG00000164330",
                "ENSG00000108001",
                "ENSG00000184486",
                "ENSG00000119547",
                "ENSG00000143995"
            ],
            "target": [
                "ENSG00000147862",
                "ENSG00000140937",
                "ENSG00000109099",
                "ENSG00000144366",
                "ENSG00000185664",
                "ENSG00000198910",
                "ENSG00000143995",
                "ENSG00000084628",
                "ENSG00000164199",
                "ENSG00000105810",
                "ENSG00000070961",
                "ENSG00000134138",
                "ENSG00000110492",
                "ENSG00000197043",
                "ENSG00000185650",
                "ENSG00000130005",
                "ENSG00000104112",
                "ENSG00000133302",
                "ENSG00000122884",
                "ENSG00000185201",
                "ENSG00000162599",
                "ENSG00000196562",
                "ENSG00000181744",
                "ENSG00000132437",
                "ENSG00000213420",
                "ENSG00000187957",
                "ENSG00000026025",
                "ENSG00000131037",
                "ENSG00000149294",
                "ENSG00000147655",
                "ENSG00000123240",
                "ENSG00000185630",
                "ENSG00000196141",
                "ENSG00000113580",
                "ENSG00000159409",
                "ENSG00000123989",
                "ENSG00000047457",
                "ENSG00000018408",
                "ENSG00000164056",
                "ENSG00000129244"
            ],
            "expression_sum": {
                "class1": 32875.0,
                "class2": 29213.0
            },
            "ens_id": "ENSG00000175745",
            "uniprot_ids": [
                "P10589",
                "F1DAL7",
                "F1DAL9"
            ]
        },
        "ENSG00000122545": {
            "id": "ENSG00000122545",
            "type": "Gene",
            "symbol": "SEPT7",
            "source": [
                "ENSG00000181449",
                "ENSG00000167081",
                "ENSG00000113580",
                "ENSG00000103495",
                "ENSG00000164330",
                "ENSG00000108001",
                "ENSG00000221869",
                "ENSG00000196591",
                "ENSG00000124766",
                "ENSG00000119547",
                "ENSG00000162599"
            ],
            "target": [],
            "expression_sum": {
                "class1": 47839.0,
                "class2": 32964.0
            },
            "ens_id": "ENSG00000122545",
            "uniprot_ids": [
                "Q16181",
                "Q5JXL7",
                "A0A0U1RQW0",
                "A0A0U1RRD1",
                "A0A0U1RRM2",
                "G3V1Q4",
                "A0A0U1RRE1",
                "A0A5F9ZGZ7",
                "A0A0U1RRH9",
                "E7EPK1",
                "E7ES33"
            ]
        },
        "ENSG00000120885": {
            "id": "ENSG00000120885",
            "type": "Gene",
            "symbol": "CLU",
            "source": [
                "ENSG00000124766",
                "ENSG00000139352",
                "ENSG00000162599"
            ],
            "target": [],
            "expression_sum": {
                "class1": 27511.0,
                "class2": 28432.0
            },
            "ens_id": "ENSG00000120885",
            "uniprot_ids": [
                "P10909",
                "E5RG36",
                "E5RGB0",
                "H0YAS8",
                "E7ERK6",
                "E5RH61",
                "E5RJZ5",
                "H0YLK8",
                "H0YC35",
                "E5RJD6",
                "E7ETB4"
            ]
        },
        "ENSG00000134138": {
            "id": "ENSG00000134138",
            "type": "TF",
            "symbol": "MEIS2",
            "source": [
                "ENSG00000162599",
                "ENSG00000167081",
                "ENSG00000113580",
                "ENSG00000168610",
                "ENSG00000175745",
                "ENSG00000100219",
                "ENSG00000114861",
                "ENSG00000221869",
                "ENSG00000139352",
                "ENSG00000137203",
                "ENSG00000119547",
                "ENSG00000143995"
            ],
            "target": [
                "ENSG00000049130",
                "ENSG00000196230",
                "ENSG00000164199",
                "ENSG00000102265",
                "ENSG00000187134",
                "ENSG00000117984",
                "ENSG00000135506",
                "ENSG00000184486",
                "ENSG00000265972",
                "ENSG00000151746",
                "ENSG00000162490",
                "ENSG00000104805",
                "ENSG00000183091",
                "ENSG00000163956",
                "ENSG00000137203",
                "ENSG00000166250",
                "ENSG00000150593",
                "ENSG00000124570",
                "ENSG00000196141",
                "ENSG00000251562",
                "ENSG00000135404",
                "ENSG00000153234",
                "ENSG00000180879",
                "ENSG00000112514",
                "ENSG00000119547",
                "ENSG00000065613",
                "ENSG00000175130",
                "ENSG00000170266",
                "ENSG00000213064",
                "ENSG00000197746",
                "ENSG00000188290",
                "ENSG00000130005",
                "ENSG00000169432",
                "ENSG00000255302",
                "ENSG00000131711",
                "ENSG00000122884",
                "ENSG00000153113",
                "ENSG00000068697",
                "ENSG00000140939",
                "ENSG00000181744",
                "ENSG00000143013",
                "ENSG00000174851",
                "ENSG00000130511",
                "ENSG00000163382",
                "ENSG00000205981",
                "ENSG00000135549",
                "ENSG00000166228",
                "ENSG00000162496",
                "ENSG00000119655",
                "ENSG00000149557",
                "ENSG00000185664",
                "ENSG00000163399",
                "ENSG00000123131",
                "ENSG00000244187",
                "ENSG00000102804",
                "ENSG00000143995",
                "ENSG00000196220",
                "ENSG00000105223",
                "ENSG00000071553",
                "ENSG00000158604",
                "ENSG00000131773",
                "ENSG00000144674",
                "ENSG00000114861",
                "ENSG00000106537",
                "ENSG00000103316",
                "ENSG00000120742",
                "ENSG00000146540",
                "ENSG00000124155",
                "ENSG00000185115",
                "ENSG00000143543",
                "ENSG00000137267",
                "ENSG00000151150",
                "ENSG00000146278",
                "ENSG00000090932",
                "ENSG00000145675",
                "ENSG00000100345",
                "ENSG00000005249",
                "ENSG00000157851",
                "ENSG00000078018",
                "ENSG00000137880",
                "ENSG00000132475",
                "ENSG00000197043",
                "ENSG00000133318",
                "ENSG00000026297",
                "ENSG00000213614",
                "ENSG00000137285",
                "ENSG00000131037",
                "ENSG00000189159",
                "ENSG00000112290",
                "ENSG00000185624",
                "ENSG00000167081",
                "ENSG00000172270",
                "ENSG00000138640",
                "ENSG00000107872"
            ],
            "expression_sum": {
                "class1": 55941.0,
                "class2": 12368.0
            },
            "ens_id": "ENSG00000134138",
            "uniprot_ids": [
                "O14770",
                "H0YM65",
                "H0YMX5",
                "H0YKE5",
                "U3KQI2",
                "H0YKN2",
                "U3KQ95"
            ]
        },
        "ENSG00000215386": {
            "id": "ENSG00000215386",
            "type": "Gene",
            "symbol": "MIR99AHG",
            "source": [
                "ENSG00000181449",
                "ENSG00000119547",
                "ENSG00000162599"
            ],
            "target": [],
            "expression_sum": {
                "class1": 3924.0,
                "class2": 9201.0
            },
            "ens_id": "ENSG00000215386"
        },
        "ENSG00000137947": {
            "id": "ENSG00000137947",
            "type": "TF",
            "symbol": "GTF2B",
            "source": [
                "ENSG00000167081",
                "ENSG00000113580"
            ],
            "target": [
                "ENSG00000060982",
                "ENSG00000123416",
                "ENSG00000065978",
                "ENSG00000111716",
                "ENSG00000170540",
                "ENSG00000164032",
                "ENSG00000137267",
                "ENSG00000185664",
                "ENSG00000182481",
                "ENSG00000075618",
                "ENSG00000189159",
                "ENSG00000189043",
                "ENSG00000108106",
                "ENSG00000196230"
            ],
            "expression_sum": {
                "class1": 8365.0,
                "class2": 4489.0
            },
            "ens_id": "ENSG00000137947",
            "uniprot_ids": [
                "Q00403",
                "B1APE1",
                "B1APE2"
            ]
        },
        "ENSG00000164032": {
            "id": "ENSG00000164032",
            "type": "Gene",
            "symbol": "H2AFZ",
            "source": [
                "ENSG00000213015",
                "ENSG00000137947"
            ],
            "target": [],
            "expression_sum": {
                "class1": 58889.0,
                "class2": 54897.0
            },
            "ens_id": "ENSG00000164032",
            "uniprot_ids": [
                "P0C0S5",
                "A0A494C189"
            ]
        },
        "ENSG00000196230": {
            "id": "ENSG00000196230",
            "type": "Gene",
            "symbol": "TUBB",
            "source": [
                "ENSG00000181449",
                "ENSG00000106261",
                "ENSG00000134138",
                "ENSG00000167081",
                "ENSG00000164330",
                "ENSG00000137947",
                "ENSG00000119950",
                "ENSG00000196591",
                "ENSG00000067955",
                "ENSG00000119547",
                "ENSG00000160113"
            ],
            "target": [],
            "expression_sum": {
                "class1": 163962.0,
                "class2": 60361.0
            },
            "ens_id": "ENSG00000196230",
            "uniprot_ids": [
                "P07437",
                "Q5JP53",
                "Q5ST81"
            ]
        },
        "ENSG00000111716": {
            "id": "ENSG00000111716",
            "type": "Gene",
            "symbol": "LDHB",
            "source": [
                "ENSG00000181449",
                "ENSG00000109132",
                "ENSG00000016082",
                "ENSG00000113580",
                "ENSG00000167081",
                "ENSG00000103495",
                "ENSG00000164330",
                "ENSG00000108001",
                "ENSG00000114861",
                "ENSG00000137947",
                "ENSG00000119547",
                "ENSG00000196591",
                "ENSG00000181315",
                "ENSG00000137309"
            ],
            "target": [],
            "expression_sum": {
                "class1": 74907.0,
                "class2": 46249.0
            },
            "ens_id": "ENSG00000111716",
            "uniprot_ids": [
                "P07195",
                "F5H793",
                "A0A5F9ZHM4",
                "A8MW50",
                "C9J7H8"
            ]
        },
        "ENSG00000060982": {
            "id": "ENSG00000060982",
            "type": "Gene",
            "symbol": "BCAT1",
            "source": [
                "ENSG00000106261",
                "ENSG00000109132",
                "ENSG00000016082",
                "ENSG00000105698",
                "ENSG00000103495",
                "ENSG00000164330",
                "ENSG00000198911",
                "ENSG00000137947",
                "ENSG00000163848",
                "ENSG00000168214",
                "ENSG00000119547",
                "ENSG00000128272",
                "ENSG00000145734",
                "ENSG00000160113",
                "ENSG00000092199"
            ],
            "target": [],
            "expression_sum": {
                "class1": 6236.0,
                "class2": 2016.0
            },
            "ens_id": "ENSG00000060982",
            "uniprot_ids": [
                "P54687",
                "A0A087WYF2",
                "F5H2F2"
            ]
        },
        "ENSG00000123416": {
            "id": "ENSG00000123416",
            "type": "Gene",
            "symbol": "TUBA1B",
            "source": [
                "ENSG00000181449",
                "ENSG00000106261",
                "ENSG00000167081",
                "ENSG00000164330",
                "ENSG00000213015",
                "ENSG00000137947",
                "ENSG00000067955",
                "ENSG00000168214",
                "ENSG00000196591",
                "ENSG00000124766",
                "ENSG00000119547",
                "ENSG00000160113",
                "ENSG00000140307"
            ],
            "target": [],
            "expression_sum": {
                "class1": 103195.0,
                "class2": 85502.0
            },
            "ens_id": "ENSG00000123416",
            "uniprot_ids": [
                "P68363",
                "F8VWV9",
                "F8VVB9",
                "F8VX09",
                "F8VRK0",
                "F8VU66"
            ]
        },
        "ENSG00000170540": {
            "id": "ENSG00000170540",
            "type": "Gene",
            "symbol": "ARL6IP1",
            "source": [
                "ENSG00000181449",
                "ENSG00000164330",
                "ENSG00000137947",
                "ENSG00000164754",
                "ENSG00000160113"
            ],
            "target": [],
            "expression_sum": {
                "class1": 13668.0,
                "class2": 9485.0
            },
            "ens_id": "ENSG00000170540",
            "uniprot_ids": [
                "Q15041",
                "H3BTX6",
                "H3BPJ2",
                "H3BS91"
            ]
        },
        "ENSG00000189159": {
            "id": "ENSG00000189159",
            "type": "Gene",
            "symbol": "JPT1",
            "source": [
                "ENSG00000181449",
                "ENSG00000106261",
                "ENSG00000134138",
                "ENSG00000016082",
                "ENSG00000177030",
                "ENSG00000213015",
                "ENSG00000137947",
                "ENSG00000067955",
                "ENSG00000196591",
                "ENSG00000119547",
                "ENSG00000140307"
            ],
            "target": [],
            "expression_sum": {
                "class1": 128540.0,
                "class2": 39286.0
            },
            "ens_id": "ENSG00000189159",
            "uniprot_ids": [
                "Q9UK76",
                "J3KSH8",
                "J3KT51",
                "F2Z3M5"
            ]
        },
        "ENSG00000108106": {
            "id": "ENSG00000108106",
            "type": "Gene",
            "symbol": "UBE2S",
            "source": [
                "ENSG00000181449",
                "ENSG00000167081",
                "ENSG00000118263",
                "ENSG00000164330",
                "ENSG00000213015",
                "ENSG00000137947",
                "ENSG00000164754",
                "ENSG00000124766",
                "ENSG00000130522",
                "ENSG00000119547",
                "ENSG00000160113"
            ],
            "target": [],
            "expression_sum": {
                "class1": 28118.0,
                "class2": 14945.0
            },
            "ens_id": "ENSG00000108106",
            "uniprot_ids": [
                "Q16763",
                "K7EM75",
                "K7EPJ1"
            ]
        },
        "ENSG00000122482": {
            "id": "ENSG00000122482",
            "type": "TF",
            "symbol": "ZNF644",
            "source": [
                "ENSG00000164330"
            ],
            "target": [
                "ENSG00000169856",
                "ENSG00000111728",
                "ENSG00000149294",
                "ENSG00000170091",
                "ENSG00000137285"
            ],
            "expression_sum": {
                "class1": 8299.0,
                "class2": 6270.0
            },
            "ens_id": "ENSG00000122482",
            "uniprot_ids": [
                "Q9H582",
                "A0A087WZL9"
            ]
        },
        "ENSG00000185630": {
            "id": "ENSG00000185630",
            "type": "TF",
            "symbol": "PBX1",
            "source": [
                "ENSG00000016082",
                "ENSG00000181449",
                "ENSG00000175745"
            ],
            "target": [
                "ENSG00000162496",
                "ENSG00000185624",
                "ENSG00000185650",
                "ENSG00000123989",
                "ENSG00000166736",
                "ENSG00000118418"
            ],
            "expression_sum": {
                "class1": 26051.0,
                "class2": 19887.0
            },
            "ens_id": "ENSG00000185630",
            "uniprot_ids": [
                "P40424",
                "H0YLB0",
                "S4R377",
                "H0YLD4",
                "H0YLT4",
                "A0A0A0MTI0",
                "F8WA05",
                "H0YKH1",
                "A0A087X0P4",
                "H0YLF5",
                "H0YLM3"
            ]
        },
        "ENSG00000162496": {
            "id": "ENSG00000162496",
            "type": "Gene",
            "symbol": "DHRS3",
            "source": [
                "ENSG00000134138",
                "ENSG00000185630",
                "ENSG00000164683",
                "ENSG00000221869",
                "ENSG00000130522",
                "ENSG00000092199"
            ],
            "target": [],
            "expression_sum": {
                "class1": 267.0,
                "class2": 9837.0
            },
            "ens_id": "ENSG00000162496",
            "uniprot_ids": [
                "O75911",
                "Q5SUY4",
                "A0A087WTY3"
            ]
        },
        "ENSG00000123989": {
            "id": "ENSG00000123989",
            "type": "Gene",
            "symbol": "CHPF",
            "source": [
                "ENSG00000185630",
                "ENSG00000175745"
            ],
            "target": [],
            "expression_sum": {
                "class1": 4681.0,
                "class2": 8392.0
            },
            "ens_id": "ENSG00000123989",
            "uniprot_ids": [
                "Q8IZ52",
                "F8W6H2"
            ]
        },
        "ENSG00000118418": {
            "id": "ENSG00000118418",
            "type": "Gene",
            "symbol": "HMGN3",
            "source": [
                "ENSG00000185630",
                "ENSG00000181449",
                "ENSG00000105698",
                "ENSG00000092199"
            ],
            "target": [],
            "expression_sum": {
                "class1": 18067.0,
                "class2": 37547.0
            },
            "ens_id": "ENSG00000118418",
            "uniprot_ids": [
                "Q15651",
                "A0A087WZE9"
            ]
        },
        "ENSG00000166736": {
            "id": "ENSG00000166736",
            "type": "Gene",
            "symbol": "HTR3A",
            "source": [
                "ENSG00000147862",
                "ENSG00000185630",
                "ENSG00000103495",
                "ENSG00000126778",
                "ENSG00000196591"
            ],
            "target": [],
            "expression_sum": {
                "class1": 4339.0,
                "class2": 66.0
            },
            "ens_id": "ENSG00000166736",
            "uniprot_ids": [
                "P46098",
                "A0A0B4J205"
            ]
        },
        "ENSG00000185624": {
            "id": "ENSG00000185624",
            "type": "Gene",
            "symbol": "P4HB",
            "source": [
                "ENSG00000100811",
                "ENSG00000134138",
                "ENSG00000185630",
                "ENSG00000164683",
                "ENSG00000181722",
                "ENSG00000139613"
            ],
            "target": [],
            "expression_sum": {
                "class1": 15518.0,
                "class2": 19940.0
            },
            "ens_id": "ENSG00000185624",
            "uniprot_ids": [
                "P07237",
                "A0A7P0Z4J0",
                "B3KTQ9",
                "A0A7P0T9S9",
                "I3L3U6",
                "A0A7P0Z4B2",
                "I3L1Y5",
                "A0A7P0T8X1",
                "I3L514",
                "A0A7P0T9K9",
                "A0A7P0TBP8",
                "A0A7P0Z476",
                "A0A7P0Z4F8",
                "H0Y3Z3",
                "H7BZ94",
                "A0A7P0TBA3",
                "A0A7P0TA71",
                "A0A7P0T9U6",
                "I3NI03",
                "I3L0S0",
                "A0A7P0Z4S2",
                "A0A7P0TAX9",
                "A0A7P0TA35",
                "A0A7P0T940",
                "A0A7P0TB26",
                "I3L4M2",
                "A0A7P0TA97",
                "A0A7P0T8C0",
                "A0A7P0T8J3",
                "I3L3P5",
                "A0A7P0T9D6",
                "A0A7P0T9E3"
            ]
        },
        "ENSG00000143355": {
            "id": "ENSG00000143355",
            "type": "TF",
            "symbol": "LHX9",
            "source": [
                "ENSG00000167081",
                "ENSG00000148737"
            ],
            "target": [
                "ENSG00000148737",
                "ENSG00000187957",
                "ENSG00000184144"
            ],
            "expression_sum": {
                "class1": 9937.0,
                "class2": 2421.0
            },
            "ens_id": "ENSG00000143355",
            "uniprot_ids": [
                "Q9NQ69",
                "H0Y330",
                "A0A0C4DGY4",
                "H0YL54",
                "A0A087X083"
            ]
        },
        "ENSG00000184144": {
            "id": "ENSG00000184144",
            "type": "Gene",
            "symbol": "CNTN2",
            "source": [
                "ENSG00000143799",
                "ENSG00000143355"
            ],
            "target": [],
            "expression_sum": {
                "class1": 8158.0,
                "class2": 1943.0
            },
            "ens_id": "ENSG00000184144",
            "uniprot_ids": [
                "Q02246",
                "A0A1W2PPY1",
                "A0A1W2PR60",
                "A0A1W2PQJ4",
                "A0A1W2PPQ9",
                "A0A1B0GTL7",
                "A0A1W2PQ86",
                "A0A1W2PQ11"
            ]
        },
        "ENSG00000187957": {
            "id": "ENSG00000187957",
            "type": "Gene",
            "symbol": "DNER",
            "source": [
                "ENSG00000175745",
                "ENSG00000143355"
            ],
            "target": [],
            "expression_sum": {
                "class1": 15990.0,
                "class2": 4017.0
            },
            "ens_id": "ENSG00000187957",
            "uniprot_ids": [
                "Q8NFT8"
            ]
        },
        "ENSG00000148737": {
            "id": "ENSG00000148737",
            "type": "TF",
            "symbol": "TCF7L2",
            "source": [
                "ENSG00000143355"
            ],
            "target": [
                "ENSG00000143355"
            ],
            "expression_sum": {
                "class1": 1609.0,
                "class2": 3370.0
            },
            "ens_id": "ENSG00000148737",
            "uniprot_ids": [
                "Q9NQB0",
                "E2GH26",
                "C6ZRJ6",
                "C6ZRK5",
                "C6ZRJ7",
                "Q5VVR7",
                "Q5JRY5",
                "Q5JRY3",
                "A0A1B0GU84",
                "Q5VVR8",
                "A0A0D9SGH8",
                "A0A1B0GUV5",
                "Q5JRY4",
                "A0A1B0GVE1",
                "A0A0A0MTL7",
                "H0YEE5",
                "Q5VVR5"
            ]
        },
        "ENSG00000163435": {
            "id": "ENSG00000163435",
            "type": "TF",
            "symbol": "ELF3",
            "source": [
                "ENSG00000106261",
                "ENSG00000100219",
                "ENSG00000221869",
                "ENSG00000196591",
                "ENSG00000168214",
                "ENSG00000124766",
                "ENSG00000116044",
                "ENSG00000160113"
            ],
            "target": [
                "ENSG00000206573",
                "ENSG00000141741",
                "ENSG00000162704",
                "ENSG00000163634",
                "ENSG00000132698",
                "ENSG00000139644",
                "ENSG00000090060",
                "ENSG00000173467",
                "ENSG00000134248",
                "ENSG00000131143",
                "ENSG00000127540",
                "ENSG00000160932",
                "ENSG00000165215",
                "ENSG00000146247",
                "ENSG00000108424",
                "ENSG00000113282",
                "ENSG00000188483",
                "ENSG00000265972",
                "ENSG00000154556",
                "ENSG00000163466",
                "ENSG00000198931",
                "ENSG00000114439",
                "ENSG00000173812",
                "ENSG00000141279",
                "ENSG00000198899",
                "ENSG00000123240",
                "ENSG00000041357",
                "ENSG00000008517",
                "ENSG00000163875",
                "ENSG00000167136",
                "ENSG00000164924",
                "ENSG00000075213",
                "ENSG00000197930",
                "ENSG00000180879",
                "ENSG00000104964",
                "ENSG00000156482",
                "ENSG00000125868",
                "ENSG00000165637",
                "ENSG00000241685",
                "ENSG00000125356",
                "ENSG00000147548",
                "ENSG00000196504",
                "ENSG00000129625",
                "ENSG00000116898",
                "ENSG00000130513",
                "ENSG00000147889",
                "ENSG00000067225",
                "ENSG00000153113",
                "ENSG00000113712",
                "ENSG00000233927",
                "ENSG00000150459",
                "ENSG00000117139",
                "ENSG00000148444",
                "ENSG00000205581",
                "ENSG00000161249",
                "ENSG00000129521",
                "ENSG00000062716",
                "ENSG00000095380",
                "ENSG00000086548",
                "ENSG00000133703",
                "ENSG00000133606",
                "ENSG00000116044",
                "ENSG00000103769",
                "ENSG00000138071",
                "ENSG00000170293",
                "ENSG00000153048",
                "ENSG00000263001",
                "ENSG00000110092",
                "ENSG00000213741",
                "ENSG00000215845",
                "ENSG00000111907",
                "ENSG00000187514",
                "ENSG00000076604",
                "ENSG00000163041",
                "ENSG00000092199",
                "ENSG00000110955",
                "ENSG00000107262",
                "ENSG00000164346",
                "ENSG00000172239",
                "ENSG00000174953",
                "ENSG00000064651",
                "ENSG00000169100",
                "ENSG00000162191",
                "ENSG00000108654",
                "ENSG00000135940",
                "ENSG00000124107",
                "ENSG00000107581",
                "ENSG00000092010",
                "ENSG00000106052",
                "ENSG00000147677",
                "ENSG00000107438",
                "ENSG00000079246",
                "ENSG00000171222",
                "ENSG00000135046",
                "ENSG00000095059",
                "ENSG00000138801",
                "ENSG00000240972",
                "ENSG00000054267",
                "ENSG00000172915",
                "ENSG00000030582",
                "ENSG00000135390",
                "ENSG00000136770",
                "ENSG00000005022",
                "ENSG00000127616",
                "ENSG00000128654",
                "ENSG00000153250",
                "ENSG00000047410",
                "ENSG00000136810",
                "ENSG00000174021",
                "ENSG00000169764",
                "ENSG00000127184",
                "ENSG00000116209",
                "ENSG00000136950",
                "ENSG00000104388",
                "ENSG00000105835",
                "ENSG00000119314",
                "ENSG00000170144",
                "ENSG00000171530",
                "ENSG00000198938",
                "ENSG00000168710",
                "ENSG00000127914",
                "ENSG00000126267",
                "ENSG00000108679",
                "ENSG00000253738",
                "ENSG00000152558",
                "ENSG00000154640",
                "ENSG00000126005",
                "ENSG00000166710",
                "ENSG00000198840",
                "ENSG00000082153",
                "ENSG00000247095",
                "ENSG00000101443",
                "ENSG00000085871",
                "ENSG00000184752",
                "ENSG00000169567",
                "ENSG00000198258",
                "ENSG00000023287",
                "ENSG00000076554",
                "ENSG00000164190",
                "ENSG00000189143",
                "ENSG00000135821",
                "ENSG00000131495",
                "ENSG00000171314",
                "ENSG00000067248",
                "ENSG00000090238",
                "ENSG00000167642",
                "ENSG00000111229",
                "ENSG00000130066",
                "ENSG00000126756",
                "ENSG00000006007",
                "ENSG00000051620",
                "ENSG00000173473",
                "ENSG00000163939",
                "ENSG00000059804",
                "ENSG00000117523",
                "ENSG00000127838",
                "ENSG00000167283",
                "ENSG00000198886",
                "ENSG00000122643",
                "ENSG00000104904",
                "ENSG00000125977",
                "ENSG00000170348",
                "ENSG00000116586",
                "ENSG00000119335",
                "ENSG00000084234",
                "ENSG00000154723",
                "ENSG00000166347",
                "ENSG00000131174",
                "ENSG00000143013",
                "ENSG00000168288",
                "ENSG00000004059",
                "ENSG00000234745",
                "ENSG00000113387",
                "ENSG00000196937",
                "ENSG00000106028",
                "ENSG00000100697",
                "ENSG00000168118",
                "ENSG00000106367",
                "ENSG00000185650",
                "ENSG00000100815",
                "ENSG00000197111",
                "ENSG00000101182",
                "ENSG00000142949",
                "ENSG00000092820",
                "ENSG00000185043",
                "ENSG00000184007",
                "ENSG00000090013",
                "ENSG00000197747",
                "ENSG00000004779",
                "ENSG00000004478",
                "ENSG00000135968",
                "ENSG00000070831",
                "ENSG00000051523",
                "ENSG00000250479",
                "ENSG00000144674",
                "ENSG00000106537",
                "ENSG00000146731",
                "ENSG00000053254",
                "ENSG00000105048",
                "ENSG00000164713",
                "ENSG00000143819",
                "ENSG00000111237",
                "ENSG00000183742",
                "ENSG00000132824",
                "ENSG00000254505",
                "ENSG00000112695",
                "ENSG00000118640",
                "ENSG00000176340",
                "ENSG00000097033",
                "ENSG00000117335",
                "ENSG00000170421",
                "ENSG00000049618",
                "ENSG00000164733",
                "ENSG00000140264",
                "ENSG00000177410",
                "ENSG00000101745",
                "ENSG00000140990",
                "ENSG00000182718",
                "ENSG00000138640",
                "ENSG00000164258",
                "ENSG00000106261",
                "ENSG00000198492",
                "ENSG00000171604",
                "ENSG00000233098",
                "ENSG00000129235",
                "ENSG00000102144",
                "ENSG00000172183",
                "ENSG00000130770",
                "ENSG00000084207",
                "ENSG00000186395",
                "ENSG00000117448",
                "ENSG00000166295",
                "ENSG00000142864",
                "ENSG00000119013",
                "ENSG00000184076",
                "ENSG00000196154",
                "ENSG00000211450",
                "ENSG00000166337",
                "ENSG00000166913",
                "ENSG00000162434",
                "ENSG00000118816",
                "ENSG00000119888",
                "ENSG00000100906",
                "ENSG00000221869",
                "ENSG00000150687",
                "ENSG00000134056",
                "ENSG00000101040",
                "ENSG00000177156",
                "ENSG00000187193",
                "ENSG00000197956",
                "ENSG00000144560",
                "ENSG00000117525",
                "ENSG00000156976",
                "ENSG00000129083",
                "ENSG00000160888",
                "ENSG00000113732",
                "ENSG00000197448",
                "ENSG00000133112",
                "ENSG00000131981",
                "ENSG00000166145",
                "ENSG00000124762",
                "ENSG00000187231",
                "ENSG00000121039",
                "ENSG00000150093",
                "ENSG00000138069",
                "ENSG00000110321",
                "ENSG00000144034",
                "ENSG00000156642",
                "ENSG00000123144",
                "ENSG00000127125",
                "ENSG00000143256",
                "ENSG00000149084",
                "ENSG00000165502",
                "ENSG00000084733",
                "ENSG00000138061",
                "ENSG00000206503",
                "ENSG00000122034",
                "ENSG00000138592",
                "ENSG00000138772",
                "ENSG00000149932",
                "ENSG00000114209",
                "ENSG00000168214",
                "ENSG00000100911",
                "ENSG00000160213",
                "ENSG00000144224",
                "ENSG00000115524",
                "ENSG00000138764",
                "ENSG00000155368",
                "ENSG00000096696",
                "ENSG00000169564",
                "ENSG00000143543",
                "ENSG00000070756",
                "ENSG00000140740",
                "ENSG00000146386",
                "ENSG00000198727",
                "ENSG00000167658",
                "ENSG00000145632",
                "ENSG00000147676",
                "ENSG00000185608",
                "ENSG00000060138",
                "ENSG00000132475",
                "ENSG00000135480",
                "ENSG00000196586",
                "ENSG00000168938",
                "ENSG00000143933",
                "ENSG00000146674",
                "ENSG00000165629",
                "ENSG00000167996",
                "ENSG00000132963",
                "ENSG00000100664",
                "ENSG00000136522",
                "ENSG00000178127",
                "ENSG00000010244",
                "ENSG00000105518",
                "ENSG00000133639",
                "ENSG00000092148",
                "ENSG00000170017",
                "ENSG00000124214",
                "ENSG00000163683",
                "ENSG00000116030",
                "ENSG00000186591",
                "ENSG00000160113",
                "ENSG00000160075",
                "ENSG00000120992",
                "ENSG00000174748",
                "ENSG00000148346",
                "ENSG00000163191",
                "ENSG00000126432",
                "ENSG00000181885",
                "ENSG00000126247",
                "ENSG00000245532",
                "ENSG00000197728",
                "ENSG00000178741",
                "ENSG00000085063",
                "ENSG00000171206",
                "ENSG00000173457",
                "ENSG00000088832",
                "ENSG00000108518",
                "ENSG00000278311",
                "ENSG00000086666",
                "ENSG00000114902",
                "ENSG00000230989",
                "ENSG00000154518",
                "ENSG00000178449",
                "ENSG00000106541",
                "ENSG00000134717",
                "ENSG00000136156",
                "ENSG00000125844",
                "ENSG00000171159",
                "ENSG00000134809",
                "ENSG00000213719",
                "ENSG00000112335",
                "ENSG00000204525",
                "ENSG00000164919",
                "ENSG00000145741",
                "ENSG00000178982",
                "ENSG00000067900",
                "ENSG00000111057",
                "ENSG00000104870",
                "ENSG00000100387",
                "ENSG00000114573",
                "ENSG00000140319",
                "ENSG00000148180",
                "ENSG00000163399",
                "ENSG00000118680",
                "ENSG00000117450",
                "ENSG00000244187",
                "ENSG00000117472",
                "ENSG00000198763",
                "ENSG00000009307",
                "ENSG00000107372",
                "ENSG00000135679",
                "ENSG00000124172",
                "ENSG00000150991",
                "ENSG00000196591",
                "ENSG00000017797",
                "ENSG00000100568",
                "ENSG00000101608",
                "ENSG00000122705",
                "ENSG00000183726",
                "ENSG00000113648",
                "ENSG00000111775",
                "ENSG00000123636",
                "ENSG00000087502",
                "ENSG00000197102",
                "ENSG00000163938",
                "ENSG00000008394",
                "ENSG00000188612",
                "ENSG00000154080",
                "ENSG00000104763",
                "ENSG00000205542",
                "ENSG00000158710",
                "ENSG00000160948",
                "ENSG00000137880",
                "ENSG00000115685",
                "ENSG00000128524",
                "ENSG00000167232",
                "ENSG00000165678",
                "ENSG00000198712",
                "ENSG00000044115",
                "ENSG00000136238",
                "ENSG00000186352",
                "ENSG00000130402",
                "ENSG00000182154",
                "ENSG00000122873",
                "ENSG00000087460",
                "ENSG00000158195",
                "ENSG00000170759",
                "ENSG00000145247",
                "ENSG00000113558",
                "ENSG00000143153",
                "ENSG00000109062",
                "ENSG00000113407",
                "ENSG00000143742",
                "ENSG00000104419",
                "ENSG00000085733",
                "ENSG00000092841",
                "ENSG00000173230"
            ],
            "expression_sum": {
                "class1": 152.0,
                "class2": 9183.0
            },
            "ens_id": "ENSG00000163435",
            "uniprot_ids": [
                "P78545",
                "Q5SR36"
            ]
        },
        "ENSG00000160075": {
            "id": "ENSG00000160075",
            "type": "Gene",
            "symbol": "SSU72",
            "source": [
                "ENSG00000181449",
                "ENSG00000163435",
                "ENSG00000167081",
                "ENSG00000113580",
                "ENSG00000164330"
            ],
            "target": [],
            "expression_sum": {
                "class1": 18034.0,
                "class2": 12133.0
            },
            "ens_id": "ENSG00000160075",
            "uniprot_ids": [
                "Q9NP77"
            ]
        },
        "ENSG00000070831": {
            "id": "ENSG00000070831",
            "type": "Gene",
            "symbol": "CDC42",
            "source": [
                "ENSG00000163435",
                "ENSG00000113580",
                "ENSG00000118263",
                "ENSG00000103495",
                "ENSG00000164330",
                "ENSG00000108001",
                "ENSG00000100219",
                "ENSG00000114861",
                "ENSG00000221869",
                "ENSG00000139352"
            ],
            "target": [],
            "expression_sum": {
                "class1": 72396.0,
                "class2": 31017.0
            },
            "ens_id": "ENSG00000070831",
            "uniprot_ids": [
                "P60953",
                "Q5JYX0",
                "A0A590UJK8",
                "A0A3B3IRV0",
                "A0A494C1M1",
                "A0A494BZX6"
            ]
        },
        "ENSG00000183726": {
            "id": "ENSG00000183726",
            "type": "TF",
            "symbol": "TMEM50A",
            "source": [
                "ENSG00000163435",
                "ENSG00000167081",
                "ENSG00000113580",
                "ENSG00000164330",
                "ENSG00000108001"
            ],
            "target": [],
            "expression_sum": {
                "class1": 17781.0,
                "class2": 12945.0
            },
            "ens_id": "ENSG00000183726",
            "uniprot_ids": [
                "O95807"
            ]
        },
        "ENSG00000158195": {
            "id": "ENSG00000158195",
            "type": "Gene",
            "symbol": "WASF2",
            "source": [
                "ENSG00000164683",
                "ENSG00000181449",
                "ENSG00000163435"
            ],
            "target": [],
            "expression_sum": {
                "class1": 6235.0,
                "class2": 14299.0
            },
            "ens_id": "ENSG00000158195",
            "uniprot_ids": [
                "Q9Y6W5"
            ]
        },
        "ENSG00000130770": {
            "id": "ENSG00000130770",
            "type": "Gene",
            "symbol": "ATP5IF1",
            "source": [
                "ENSG00000181449",
                "ENSG00000163435",
                "ENSG00000167081",
                "ENSG00000113580",
                "ENSG00000114861"
            ],
            "target": [],
            "expression_sum": {
                "class1": 57363.0,
                "class2": 45854.0
            },
            "ens_id": "ENSG00000130770",
            "uniprot_ids": [
                "Q9UII2",
                "A0A0B4J230"
            ]
        },
        "ENSG00000198492": {
            "id": "ENSG00000198492",
            "type": "Gene",
            "symbol": "YTHDF2",
            "source": [
                "ENSG00000167081",
                "ENSG00000163435",
                "ENSG00000164330"
            ],
            "target": [],
            "expression_sum": {
                "class1": 14000.0,
                "class2": 7607.0
            },
            "ens_id": "ENSG00000198492",
            "uniprot_ids": [
                "Q9Y5A9",
                "S4R3J8",
                "S4R3V3"
            ]
        },
        "ENSG00000184007": {
            "id": "ENSG00000184007",
            "type": "Gene",
            "symbol": "PTP4A2",
            "source": [
                "ENSG00000163435",
                "ENSG00000109132",
                "ENSG00000016082",
                "ENSG00000113580",
                "ENSG00000167081",
                "ENSG00000118263",
                "ENSG00000164330",
                "ENSG00000221869",
                "ENSG00000119547"
            ],
            "target": [],
            "expression_sum": {
                "class1": 19169.0,
                "class2": 14000.0
            },
            "ens_id": "ENSG00000184007",
            "uniprot_ids": [
                "Q12974",
                "E9PRR9",
                "E9PML8",
                "E9PMY3",
                "E9PJC0",
                "E9PL34"
            ]
        },
        "ENSG00000116898": {
            "id": "ENSG00000116898",
            "type": "Gene",
            "symbol": "MRPS15",
            "source": [
                "ENSG00000181449",
                "ENSG00000163435",
                "ENSG00000167081",
                "ENSG00000113580",
                "ENSG00000164330"
            ],
            "target": [],
            "expression_sum": {
                "class1": 10800.0,
                "class2": 9370.0
            },
            "ens_id": "ENSG00000116898",
            "uniprot_ids": [
                "P82914"
            ]
        },
        "ENSG00000163875": {
            "id": "ENSG00000163875",
            "type": "Gene",
            "symbol": "MEAF6",
            "source": [
                "ENSG00000181449",
                "ENSG00000163435",
                "ENSG00000157557",
                "ENSG00000167081",
                "ENSG00000113580",
                "ENSG00000164330",
                "ENSG00000114861",
                "ENSG00000139352"
            ],
            "target": [],
            "expression_sum": {
                "class1": 25848.0,
                "class2": 11914.0
            },
            "ens_id": "ENSG00000163875",
            "uniprot_ids": [
                "Q9HAF1",
                "B1AK63"
            ]
        },
        "ENSG00000127125": {
            "id": "ENSG00000127125",
            "type": "Gene",
            "symbol": "PPCS",
            "source": [
                "ENSG00000113580",
                "ENSG00000163435",
                "ENSG00000164330"
            ],
            "target": [],
            "expression_sum": {
                "class1": 7313.0,
                "class2": 8125.0
            },
            "ens_id": "ENSG00000127125",
            "uniprot_ids": [
                "Q9HAB8",
                "Q5VVM3",
                "Q5VVM2"
            ]
        },
        "ENSG00000142949": {
            "id": "ENSG00000142949",
            "type": "Gene",
            "symbol": "PTPRF",
            "source": [
                "ENSG00000113580",
                "ENSG00000148516",
                "ENSG00000163435",
                "ENSG00000164330"
            ],
            "target": [],
            "expression_sum": {
                "class1": 6848.0,
                "class2": 5729.0
            },
            "ens_id": "ENSG00000142949",
            "uniprot_ids": [
                "P10586",
                "A2A437",
                "H0Y380",
                "H0Y4H1",
                "H0Y7Z9",
                "H0Y6Z7"
            ]
        },
        "ENSG00000117450": {
            "id": "ENSG00000117450",
            "type": "Gene",
            "symbol": "PRDX1",
            "source": [
                "ENSG00000167081",
                "ENSG00000118263",
                "ENSG00000163435",
                "ENSG00000164330"
            ],
            "target": [],
            "expression_sum": {
                "class1": 45889.0,
                "class2": 53053.0
            },
            "ens_id": "ENSG00000117450",
            "uniprot_ids": [
                "Q06830",
                "A0A0A0MSI0",
                "A0A0A0MRQ5"
            ]
        },
        "ENSG00000117448": {
            "id": "ENSG00000117448",
            "type": "Gene",
            "symbol": "AKR1A1",
            "source": [
                "ENSG00000163435",
                "ENSG00000164330"
            ],
            "target": [],
            "expression_sum": {
                "class1": 11441.0,
                "class2": 12946.0
            },
            "ens_id": "ENSG00000117448",
            "uniprot_ids": [
                "P14550",
                "Q5T621",
                "V9GYP9",
                "V9GYG2"
            ]
        },
        "ENSG00000117472": {
            "id": "ENSG00000117472",
            "type": "Gene",
            "symbol": "TSPAN1",
            "source": [
                "ENSG00000163435",
                "ENSG00000160113"
            ],
            "target": [],
            "expression_sum": {
                "class1": 158.0,
                "class2": 6952.0
            },
            "ens_id": "ENSG00000117472",
            "uniprot_ids": [
                "O60635"
            ]
        },
        "ENSG00000134717": {
            "id": "ENSG00000134717",
            "type": "Gene",
            "symbol": "BTF3L4",
            "source": [
                "ENSG00000181449",
                "ENSG00000163435",
                "ENSG00000113580",
                "ENSG00000164330",
                "ENSG00000114861",
                "ENSG00000221869",
                "ENSG00000139352",
                "ENSG00000137203"
            ],
            "target": [],
            "expression_sum": {
                "class1": 31391.0,
                "class2": 13620.0
            },
            "ens_id": "ENSG00000134717",
            "uniprot_ids": [
                "Q96K17",
                "E9PPC3",
                "E9PL10"
            ]
        },
        "ENSG00000116209": {
            "id": "ENSG00000116209",
            "type": "Gene",
            "symbol": "TMEM59",
            "source": [
                "ENSG00000167081",
                "ENSG00000181722",
                "ENSG00000163435"
            ],
            "target": [],
            "expression_sum": {
                "class1": 31027.0,
                "class2": 32573.0
            },
            "ens_id": "ENSG00000116209",
            "uniprot_ids": [
                "Q9BXS4",
                "Q5T706",
                "Q5T6Z8",
                "Q5T704",
                "Q5T703"
            ]
        },
        "ENSG00000162434": {
            "id": "ENSG00000162434",
            "type": "Gene",
            "symbol": "JAK1",
            "source": [
                "ENSG00000016082",
                "ENSG00000113580",
                "ENSG00000163435",
                "ENSG00000164330"
            ],
            "target": [],
            "expression_sum": {
                "class1": 5238.0,
                "class2": 5045.0
            },
            "ens_id": "ENSG00000162434",
            "uniprot_ids": [
                "P23458",
                "A0A5F9ZH73",
                "A0A5F9ZI39",
                "A0A5F9ZHI1",
                "A0A5F9ZHK2",
                "A0A5F9ZHN8",
                "A0A5F9ZHW0",
                "A0A5F9ZH07",
                "A0A5F9ZI01",
                "A0A5F9ZH32"
            ]
        },
        "ENSG00000142864": {
            "id": "ENSG00000142864",
            "type": "Gene",
            "symbol": "SERBP1",
            "source": [
                "ENSG00000181449",
                "ENSG00000163435",
                "ENSG00000167081",
                "ENSG00000164330",
                "ENSG00000108001",
                "ENSG00000221869",
                "ENSG00000119547"
            ],
            "target": [],
            "expression_sum": {
                "class1": 23798.0,
                "class2": 32363.0
            },
            "ens_id": "ENSG00000142864",
            "uniprot_ids": [
                "Q8NC51"
            ]
        },
        "ENSG00000174021": {
            "id": "ENSG00000174021",
            "type": "Gene",
            "symbol": "GNG5",
            "source": [
                "ENSG00000100811",
                "ENSG00000163435",
                "ENSG00000092199",
                "ENSG00000164683",
                "ENSG00000120837",
                "ENSG00000105698",
                "ENSG00000103495",
                "ENSG00000108055",
                "ENSG00000164754",
                "ENSG00000160113",
                "ENSG00000139613"
            ],
            "target": [],
            "expression_sum": {
                "class1": 20141.0,
                "class2": 40581.0
            },
            "ens_id": "ENSG00000174021",
            "uniprot_ids": [
                "P63218"
            ]
        },
        "ENSG00000097033": {
            "id": "ENSG00000097033",
            "type": "Gene",
            "symbol": "SH3GLB1",
            "source": [
                "ENSG00000163435",
                "ENSG00000016082",
                "ENSG00000113580",
                "ENSG00000148516",
                "ENSG00000164330",
                "ENSG00000139352"
            ],
            "target": [],
            "expression_sum": {
                "class1": 9484.0,
                "class2": 8054.0
            },
            "ens_id": "ENSG00000097033",
            "uniprot_ids": [
                "Q9Y371",
                "A0A087WW40"
            ]
        },
        "ENSG00000143013": {
            "id": "ENSG00000143013",
            "type": "Gene",
            "symbol": "LMO4",
            "source": [
                "ENSG00000181449",
                "ENSG00000163435",
                "ENSG00000134138",
                "ENSG00000113580",
                "ENSG00000140262",
                "ENSG00000181722",
                "ENSG00000100219",
                "ENSG00000139352",
                "ENSG00000119547"
            ],
            "target": [],
            "expression_sum": {
                "class1": 31120.0,
                "class2": 26163.0
            },
            "ens_id": "ENSG00000143013",
            "uniprot_ids": [
                "P61968"
            ]
        },
        "ENSG00000117525": {
            "id": "ENSG00000117525",
            "type": "Gene",
            "symbol": "F3",
            "source": [
                "ENSG00000163435",
                "ENSG00000168214"
            ],
            "target": [],
            "expression_sum": {
                "class1": 601.0,
                "class2": 9926.0
            },
            "ens_id": "ENSG00000117525",
            "uniprot_ids": [
                "P13726"
            ]
        },
        "ENSG00000168710": {
            "id": "ENSG00000168710",
            "type": "Gene",
            "symbol": "AHCYL1",
            "source": [
                "ENSG00000167081",
                "ENSG00000163435",
                "ENSG00000164330"
            ],
            "target": [],
            "expression_sum": {
                "class1": 8967.0,
                "class2": 8741.0
            },
            "ens_id": "ENSG00000168710",
            "uniprot_ids": [
                "O43865"
            ]
        },
        "ENSG00000134248": {
            "id": "ENSG00000134248",
            "type": "Gene",
            "symbol": "LAMTOR5",
            "source": [
                "ENSG00000181449",
                "ENSG00000163435",
                "ENSG00000167081",
                "ENSG00000113580",
                "ENSG00000118263",
                "ENSG00000164330",
                "ENSG00000108001",
                "ENSG00000181722",
                "ENSG00000114861",
                "ENSG00000139352"
            ],
            "target": [],
            "expression_sum": {
                "class1": 36322.0,
                "class2": 34170.0
            },
            "ens_id": "ENSG00000134248",
            "uniprot_ids": [
                "O43504",
                "E9PLX3",
                "A0A0C4DGV4",
                "R4GMU8"
            ]
        },
        "ENSG00000009307": {
            "id": "ENSG00000009307",
            "type": "Gene",
            "symbol": "CSDE1",
            "source": [
                "ENSG00000181449",
                "ENSG00000163435",
                "ENSG00000016082",
                "ENSG00000113580",
                "ENSG00000167081",
                "ENSG00000148516",
                "ENSG00000164330",
                "ENSG00000114861",
                "ENSG00000221869",
                "ENSG00000139352"
            ],
            "target": [],
            "expression_sum": {
                "class1": 33475.0,
                "class2": 20721.0
            },
            "ens_id": "ENSG00000009307",
            "uniprot_ids": [
                "O75534",
                "E9PLD4",
                "E9PJK7",
                "E9PLT0",
                "A0A1Y8EMC9",
                "E9PNG3",
                "E9PKN4"
            ]
        },
        "ENSG00000163399": {
            "id": "ENSG00000163399",
            "type": "Gene",
            "symbol": "ATP1A1",
            "source": [
                "ENSG00000106261",
                "ENSG00000163435",
                "ENSG00000134138",
                "ENSG00000113580",
                "ENSG00000008196",
                "ENSG00000100219",
                "ENSG00000181722",
                "ENSG00000139352",
                "ENSG00000168214",
                "ENSG00000160113"
            ],
            "target": [],
            "expression_sum": {
                "class1": 4983.0,
                "class2": 9911.0
            },
            "ens_id": "ENSG00000163399",
            "uniprot_ids": [
                "P05023",
                "Q5TC02",
                "Q5TC05",
                "Q5TC01"
            ]
        },
        "ENSG00000265972": {
            "id": "ENSG00000265972",
            "type": "Gene",
            "symbol": "TXNIP",
            "source": [
                "ENSG00000163435",
                "ENSG00000134138",
                "ENSG00000164683",
                "ENSG00000113580",
                "ENSG00000168610",
                "ENSG00000100219",
                "ENSG00000139352",
                "ENSG00000067955",
                "ENSG00000168214",
                "ENSG00000043355",
                "ENSG00000160113"
            ],
            "target": [],
            "expression_sum": {
                "class1": 48258.0,
                "class2": 49488.0
            },
            "ens_id": "ENSG00000265972",
            "uniprot_ids": [
                "Q9H3M7"
            ]
        },
        "ENSG00000163191": {
            "id": "ENSG00000163191",
            "type": "Gene",
            "symbol": "S100A11",
            "source": [
                "ENSG00000106261",
                "ENSG00000163435",
                "ENSG00000162992",
                "ENSG00000122691",
                "ENSG00000100219",
                "ENSG00000168214",
                "ENSG00000160113"
            ],
            "target": [],
            "expression_sum": {
                "class1": 16397.0,
                "class2": 61940.0
            },
            "ens_id": "ENSG00000163191",
            "uniprot_ids": [
                "P31949"
            ]
        },
        "ENSG00000197956": {
            "id": "ENSG00000197956",
            "type": "Gene",
            "symbol": "S100A6",
            "source": [
                "ENSG00000163435"
            ],
            "target": [],
            "expression_sum": {
                "class1": 6160.0,
                "class2": 13980.0
            },
            "ens_id": "ENSG00000197956",
            "uniprot_ids": [
                "P06703",
                "R4GN98"
            ]
        },
        "ENSG00000196154": {
            "id": "ENSG00000196154",
            "type": "Gene",
            "symbol": "S100A4",
            "source": [
                "ENSG00000163435"
            ],
            "target": [],
            "expression_sum": {
                "class1": 384.0,
                "class2": 3536.0
            },
            "ens_id": "ENSG00000196154",
            "uniprot_ids": [
                "P26447"
            ]
        },
        "ENSG00000143543": {
            "id": "ENSG00000143543",
            "type": "Gene",
            "symbol": "JTB",
            "source": [
                "ENSG00000181722",
                "ENSG00000163435",
                "ENSG00000134138"
            ],
            "target": [],
            "expression_sum": {
                "class1": 18636.0,
                "class2": 19530.0
            },
            "ens_id": "ENSG00000143543",
            "uniprot_ids": [
                "O76095"
            ]
        },
        "ENSG00000116586": {
            "id": "ENSG00000116586",
            "type": "Gene",
            "symbol": "LAMTOR2",
            "source": [
                "ENSG00000181449",
                "ENSG00000163435",
                "ENSG00000109132",
                "ENSG00000167081",
                "ENSG00000113580",
                "ENSG00000164330",
                "ENSG00000221869"
            ],
            "target": [],
            "expression_sum": {
                "class1": 11913.0,
                "class2": 13076.0
            },
            "ens_id": "ENSG00000116586",
            "uniprot_ids": [
                "Q9Y2Q5"
            ]
        },
        "ENSG00000158710": {
            "id": "ENSG00000158710",
            "type": "Gene",
            "symbol": "TAGLN2",
            "source": [
                "ENSG00000163435",
                "ENSG00000114315",
                "ENSG00000164683",
                "ENSG00000118263",
                "ENSG00000164330",
                "ENSG00000221869"
            ],
            "target": [],
            "expression_sum": {
                "class1": 26976.0,
                "class2": 16109.0
            },
            "ens_id": "ENSG00000158710",
            "uniprot_ids": [
                "P37802",
                "X6RJP6"
            ]
        },
        "ENSG00000215845": {
            "id": "ENSG00000215845",
            "type": "Gene",
            "symbol": "TSTD1",
            "source": [
                "ENSG00000181449",
                "ENSG00000185551",
                "ENSG00000163435",
                "ENSG00000113580",
                "ENSG00000164330",
                "ENSG00000139352",
                "ENSG00000168214"
            ],
            "target": [],
            "expression_sum": {
                "class1": 9023.0,
                "class2": 21409.0
            },
            "ens_id": "ENSG00000215845",
            "uniprot_ids": [
                "Q8NFU3",
                "R4GNF9"
            ]
        },
        "ENSG00000143256": {
            "id": "ENSG00000143256",
            "type": "Gene",
            "symbol": "PFDN2",
            "source": [
                "ENSG00000181449",
                "ENSG00000163435",
                "ENSG00000113580",
                "ENSG00000164330",
                "ENSG00000114861",
                "ENSG00000221869"
            ],
            "target": [],
            "expression_sum": {
                "class1": 46979.0,
                "class2": 22540.0
            },
            "ens_id": "ENSG00000143256",
            "uniprot_ids": [
                "Q9UHV9"
            ]
        },
        "ENSG00000143153": {
            "id": "ENSG00000143153",
            "type": "Gene",
            "symbol": "ATP1B1",
            "source": [
                "ENSG00000163435",
                "ENSG00000109132",
                "ENSG00000167081",
                "ENSG00000113580",
                "ENSG00000148516",
                "ENSG00000164330",
                "ENSG00000108055",
                "ENSG00000114861",
                "ENSG00000139352",
                "ENSG00000116044"
            ],
            "target": [],
            "expression_sum": {
                "class1": 31529.0,
                "class2": 34067.0
            },
            "ens_id": "ENSG00000143153",
            "uniprot_ids": [
                "P05026",
                "V9GYR2"
            ]
        },
        "ENSG00000117523": {
            "id": "ENSG00000117523",
            "type": "Gene",
            "symbol": "PRRC2C",
            "source": [
                "ENSG00000167081",
                "ENSG00000163435",
                "ENSG00000109132"
            ],
            "target": [],
            "expression_sum": {
                "class1": 22624.0,
                "class2": 18205.0
            },
            "ens_id": "ENSG00000117523",
            "uniprot_ids": [
                "Q9Y520",
                "E7EPN9",
                "A0A2R8Y424",
                "A0A2R8Y7F4",
                "H7C5N8",
                "A0A2R8YET2",
                "A0A0A0MS30"
            ]
        },
        "ENSG00000135821": {
            "id": "ENSG00000135821",
            "type": "Gene",
            "symbol": "GLUL",
            "source": [
                "ENSG00000164683",
                "ENSG00000163435",
                "ENSG00000139352"
            ],
            "target": [],
            "expression_sum": {
                "class1": 7046.0,
                "class2": 7437.0
            },
            "ens_id": "ENSG00000135821",
            "uniprot_ids": [
                "P15104",
                "A0A2R8YDT1"
            ]
        },
        "ENSG00000162704": {
            "id": "ENSG00000162704",
            "type": "Gene",
            "symbol": "ARPC5",
            "source": [
                "ENSG00000181449",
                "ENSG00000163435",
                "ENSG00000109132",
                "ENSG00000016082",
                "ENSG00000113580",
                "ENSG00000167081",
                "ENSG00000103495",
                "ENSG00000164330",
                "ENSG00000108001",
                "ENSG00000114861",
                "ENSG00000221869",
                "ENSG00000196591",
                "ENSG00000119547"
            ],
            "target": [],
            "expression_sum": {
                "class1": 34566.0,
                "class2": 18610.0
            },
            "ens_id": "ENSG00000162704",
            "uniprot_ids": [
                "O15511",
                "B1ALC0"
            ]
        },
        "ENSG00000047410": {
            "id": "ENSG00000047410",
            "type": "Gene",
            "symbol": "TPR",
            "source": [
                "ENSG00000181449",
                "ENSG00000163435",
                "ENSG00000164330"
            ],
            "target": [],
            "expression_sum": {
                "class1": 15160.0,
                "class2": 15130.0
            },
            "ens_id": "ENSG00000047410",
            "uniprot_ids": [
                "P12270",
                "Q5SWX9"
            ]
        },
        "ENSG00000117139": {
            "id": "ENSG00000117139",
            "type": "Gene",
            "symbol": "KDM5B",
            "source": [
                "ENSG00000163435",
                "ENSG00000016082",
                "ENSG00000167081",
                "ENSG00000148516",
                "ENSG00000164330",
                "ENSG00000119547"
            ],
            "target": [],
            "expression_sum": {
                "class1": 17261.0,
                "class2": 9723.0
            },
            "ens_id": "ENSG00000117139",
            "uniprot_ids": [
                "Q9UGL1",
                "B3KV94",
                "A0A3B3ITJ3",
                "A0A3B3ITA9",
                "A0A3B3IT25",
                "A0A3B3ITE7",
                "A0A3B3ISE6",
                "A0A3B3IS31",
                "A0A3B3IT85",
                "A0A3B3IT51",
                "A0A3B3ITJ7",
                "A0A3B3ITR1",
                "A0A3B3ITR3",
                "A0A3B3ITA8",
                "A0A3B3IS40",
                "A0A3B3ISK2",
                "R4GN45",
                "A0A3B3ITN3"
            ]
        },
        "ENSG00000117335": {
            "id": "ENSG00000117335",
            "type": "Gene",
            "symbol": "CD46",
            "source": [
                "ENSG00000163435"
            ],
            "target": [],
            "expression_sum": {
                "class1": 7135.0,
                "class2": 6505.0
            },
            "ens_id": "ENSG00000117335",
            "uniprot_ids": [
                "P15529",
                "R4GN18"
            ]
        },
        "ENSG00000143742": {
            "id": "ENSG00000143742",
            "type": "Gene",
            "symbol": "SRP9",
            "source": [
                "ENSG00000181449",
                "ENSG00000163435",
                "ENSG00000109132",
                "ENSG00000167081",
                "ENSG00000113580",
                "ENSG00000164330",
                "ENSG00000114861",
                "ENSG00000139352"
            ],
            "target": [],
            "expression_sum": {
                "class1": 75597.0,
                "class2": 44289.0
            },
            "ens_id": "ENSG00000143742",
            "uniprot_ids": [
                "P49458",
                "Q659G3",
                "E9PE20",
                "Q6P2S0"
            ]
        },
        "ENSG00000143819": {
            "id": "ENSG00000143819",
            "type": "Gene",
            "symbol": "EPHX1",
            "source": [
                "ENSG00000164683",
                "ENSG00000163435",
                "ENSG00000168214"
            ],
            "target": [],
            "expression_sum": {
                "class1": 2173.0,
                "class2": 6837.0
            },
            "ens_id": "ENSG00000143819",
            "uniprot_ids": [
                "P07099",
                "B1AQP8"
            ]
        },
        "ENSG00000163041": {
            "id": "ENSG00000163041",
            "type": "Gene",
            "symbol": "H3F3A",
            "source": [
                "ENSG00000181449",
                "ENSG00000163435",
                "ENSG00000109132",
                "ENSG00000114315",
                "ENSG00000167081",
                "ENSG00000113580",
                "ENSG00000148516",
                "ENSG00000108001",
                "ENSG00000114861",
                "ENSG00000221869",
                "ENSG00000137203",
                "ENSG00000119547"
            ],
            "target": [],
            "expression_sum": {
                "class1": 312781.0,
                "class2": 231113.0
            },
            "ens_id": "ENSG00000163041",
            "uniprot_ids": [
                "P84243",
                "B4DEB1",
                "A0A590UJJ6"
            ]
        },
        "ENSG00000168118": {
            "id": "ENSG00000168118",
            "type": "Gene",
            "symbol": "RAB4A",
            "source": [
                "ENSG00000163435",
                "ENSG00000113580",
                "ENSG00000103495",
                "ENSG00000221869",
                "ENSG00000196591"
            ],
            "target": [],
            "expression_sum": {
                "class1": 13309.0,
                "class2": 6564.0
            },
            "ens_id": "ENSG00000168118",
            "uniprot_ids": [
                "P20338",
                "A0A087WYT5"
            ]
        },
        "ENSG00000054267": {
            "id": "ENSG00000054267",
            "type": "Gene",
            "symbol": "ARID4B",
            "source": [
                "ENSG00000113580",
                "ENSG00000163435"
            ],
            "target": [],
            "expression_sum": {
                "class1": 11018.0,
                "class2": 9651.0
            },
            "ens_id": "ENSG00000054267",
            "uniprot_ids": [
                "Q4LE39",
                "H0Y7Y8",
                "H7C5R6",
                "X6RCR8"
            ]
        },
        "ENSG00000084733": {
            "id": "ENSG00000084733",
            "type": "Gene",
            "symbol": "RAB10",
            "source": [
                "ENSG00000181449",
                "ENSG00000163435",
                "ENSG00000167081",
                "ENSG00000113580",
                "ENSG00000148516",
                "ENSG00000103495",
                "ENSG00000164330",
                "ENSG00000139352"
            ],
            "target": [],
            "expression_sum": {
                "class1": 13359.0,
                "class2": 6614.0
            },
            "ens_id": "ENSG00000084733",
            "uniprot_ids": [
                "P61026"
            ]
        },
        "ENSG00000138061": {
            "id": "ENSG00000138061",
            "type": "Gene",
            "symbol": "CYP1B1",
            "source": [
                "ENSG00000163435"
            ],
            "target": [],
            "expression_sum": {
                "class1": 293.0,
                "class2": 6003.0
            },
            "ens_id": "ENSG00000138061",
            "uniprot_ids": [
                "Q16678",
                "A0A087WW26",
                "A0A087WUQ7"
            ]
        },
        "ENSG00000143933": {
            "id": "ENSG00000143933",
            "type": "Gene",
            "symbol": "CALM2",
            "source": [
                "ENSG00000181449",
                "ENSG00000163435",
                "ENSG00000157557",
                "ENSG00000109132",
                "ENSG00000167081",
                "ENSG00000113580",
                "ENSG00000148516",
                "ENSG00000118263",
                "ENSG00000164330",
                "ENSG00000140262",
                "ENSG00000150347",
                "ENSG00000114861",
                "ENSG00000139352"
            ],
            "target": [],
            "expression_sum": {
                "class1": 252716.0,
                "class2": 75750.0
            },
            "ens_id": "ENSG00000143933",
            "uniprot_ids": [
                "P0DP24",
                "M0QZ52",
                "Q96HY3",
                "E7EMB3",
                "F8WBR5",
                "A0A590UJE2",
                "A0A590UJI2",
                "A0A590UJC0"
            ]
        },
        "ENSG00000169764": {
            "id": "ENSG00000169764",
            "type": "Gene",
            "symbol": "UGP2",
            "source": [
                "ENSG00000163435",
                "ENSG00000016082",
                "ENSG00000167081",
                "ENSG00000164330",
                "ENSG00000100219",
                "ENSG00000221869",
                "ENSG00000116044"
            ],
            "target": [],
            "expression_sum": {
                "class1": 6011.0,
                "class2": 9350.0
            },
            "ens_id": "ENSG00000169764",
            "uniprot_ids": [
                "Q16851",
                "F2Z3P4",
                "F2Z3H1",
                "C9J3M0",
                "F8WC70",
                "A0A7I2V4T0",
                "A0A7I2YQ74",
                "C9JTZ5",
                "C9JWG0",
                "A0A087WYS1",
                "C9JUW1",
                "C9JNZ1",
                "F8WDA2",
                "A0A7I2V676",
                "C9JVG3",
                "A0A7I2V2Q3",
                "C9J6Q0",
                "E7EUC7",
                "H7C500",
                "C9JQU9"
            ]
        },
        "ENSG00000138069": {
            "id": "ENSG00000138069",
            "type": "Gene",
            "symbol": "RAB1A",
            "source": [
                "ENSG00000181449",
                "ENSG00000163435",
                "ENSG00000167081",
                "ENSG00000113580",
                "ENSG00000164330",
                "ENSG00000114861",
                "ENSG00000139352"
            ],
            "target": [],
            "expression_sum": {
                "class1": 18248.0,
                "class2": 9163.0
            },
            "ens_id": "ENSG00000138069",
            "uniprot_ids": [
                "P62820",
                "E7END7"
            ]
        },
        "ENSG00000138071": {
            "id": "ENSG00000138071",
            "type": "Gene",
            "symbol": "ACTR2",
            "source": [
                "ENSG00000181449",
                "ENSG00000163435",
                "ENSG00000167081",
                "ENSG00000113580",
                "ENSG00000164330"
            ],
            "target": [],
            "expression_sum": {
                "class1": 12763.0,
                "class2": 10019.0
            },
            "ens_id": "ENSG00000138071",
            "uniprot_ids": [
                "P61160",
                "F5H6T1",
                "A0A1B0GWD0",
```
