## Supplement 2 (DAN vs. Cocul) for "AGEAS: Automated Machine Learning based Genetic Regulatory Element Extraction System": outlier_grps.js

```
{
    "ENSG00000173726_ENSG00000016082": 48.95918030455923,
    "ENSG00000196262_ENSG00000016082": 40.84264476474827,
    "ENSG00000131469_ENSG00000016082": 34.867362904229275,
    "ENSG00000111716_ENSG00000016082": 30.834047627605567,
    "ENSG00000239672_ENSG00000016082": 27.550812915243025,
    "ENSG00000204628_ENSG00000016082": 25.588300071263834,
    "ENSG00000167081_ENSG00000135486": 21.999579882695006,
    "ENSG00000189043_ENSG00000167081": 14.473321944763978,
    "ENSG00000188846_ENSG00000016082": 11.943398410062617,
    "ENSG00000016082_ENSG00000009307": 11.88303917384639,
    "ENSG00000197111_ENSG00000016082": 9.157166835171061,
    "ENSG00000162704_ENSG00000016082": 8.865973056632688,
    "ENSG00000167081_ENSG00000089220": 7.827204557291017,
    "ENSG00000126214_ENSG00000016082": 7.7106038041430836,
    "ENSG00000105185_ENSG00000016082": 5.332046457326884,
    "ENSG00000070961_ENSG00000016082": 5.184013327836308,
    "ENSG00000182718_ENSG00000016082": 5.048787317095599,
    "ENSG00000178741_ENSG00000167081": 4.804690528492886,
    "ENSG00000196262_ENSG00000167081": 4.0843788208275225,
    "ENSG00000167658_ENSG00000167081": 4.062090891228483,
    "ENSG00000213760_ENSG00000016082": 3.6189224499649972,
    "ENSG00000277443_ENSG00000119547": 3.5913823859283536,
    "ENSG00000167081_ENSG00000135940": 3.5741172379828026,
    "ENSG00000167081_ENSG00000099622": 3.5161792616219416,
    "ENSG00000139970_ENSG00000103495": 3.4863639958381216,
    "ENSG00000181449_ENSG00000118363": 3.367299315437635,
    "ENSG00000164330_ENSG00000122545": 3.229488909166083,
    "ENSG00000164330_ENSG00000143198": 3.133590506044683,
    "ENSG00000166974_ENSG00000016082": 92.95914606548877,
    "ENSG00000172159_ENSG00000016082": 68.90485861799202,
    "ENSG00000163435_ENSG00000124107": 51.26246679742017
}
```
