## Supplement 2 (DAN vs. Cocul) for "AGEAS: Automated Machine Learning based Genetic Regulatory Element Extraction System": outlier_grps.js

```
{
    "ENSG00000083845_ENSG00000016082": 61.698793641956165,
    "ENSG00000128951_ENSG00000016082": 54.95828260364267,
    "ENSG00000111716_ENSG00000016082": 33.27442351984571,
    "ENSG00000277443_ENSG00000119547": 16.039046222843137,
    "ENSG00000186468_ENSG00000016082": 12.892312034957227,
    "ENSG00000196262_ENSG00000016082": 9.681328065623001,
    "ENSG00000065911_ENSG00000016082": 8.856292823361972,
    "ENSG00000163435_ENSG00000124107": 8.619732388935969,
    "ENSG00000164330_ENSG00000111716": 7.909290618330186,
    "ENSG00000089009_ENSG00000016082": 7.842633979272429,
    "ENSG00000131469_ENSG00000016082": 7.706995913371163,
    "ENSG00000126214_ENSG00000016082": 7.641529591195022,
    "ENSG00000139116_ENSG00000016082": 7.528091595192645,
    "ENSG00000239672_ENSG00000016082": 7.3078214026813715,
    "ENSG00000154277_ENSG00000016082": 5.499675962855292,
    "ENSG00000168824_ENSG00000103495": 5.461738964839991,
    "ENSG00000167081_ENSG00000140319": 4.982645300079471,
    "ENSG00000196262_ENSG00000164330": 4.827283896533073,
    "ENSG00000167081_ENSG00000112695": 4.397212207525605,
    "ENSG00000163435_ENSG00000154080": 4.384405129874934,
    "ENSG00000075142_ENSG00000016082": 4.196946922379371,
    "ENSG00000162704_ENSG00000016082": 4.193038268352663,
    "ENSG00000144339_ENSG00000016082": 4.179894386461948,
    "ENSG00000198910_ENSG00000103495": 3.7300226504026956,
    "ENSG00000164330_ENSG00000143198": 3.6509641158234682,
    "ENSG00000164330_ENSG00000162704": 3.635492808353958,
    "ENSG00000177410_ENSG00000163435": 3.6010966479719095,
    "ENSG00000167996_ENSG00000167081": 3.589851275063444,
    "ENSG00000139116_ENSG00000109132": 3.544733755701645,
    "ENSG00000167081_ENSG00000118816": 3.3718848311708696,
    "ENSG00000167081_ENSG00000163041": 3.1563895651157856,
    "ENSG00000164330_ENSG00000108953": 3.14981730958232,
    "ENSG00000143799_ENSG00000139970": 3.086223660088463,
    "ENSG00000171530_ENSG00000113580": 3.0471334518336572,
    "ENSG00000182718_ENSG00000016082": 92.94368907550789,
    "ENSG00000166974_ENSG00000016082": 88.94695293516254
}
```
