## Supplement 2 (DAN vs. Cocul) for "AGEAS: Automated Machine Learning based Genetic Regulatory Element Extraction System": outlier_grps.js

```
{
    "ENSG00000162704_ENSG00000016082": 63.38877026436225,
    "ENSG00000182718_ENSG00000016082": 41.1916863166532,
    "ENSG00000164330_ENSG00000122705": 33.750723958567235,
    "ENSG00000154518_ENSG00000016082": 23.600813524956315,
    "ENSG00000196531_ENSG00000167081": 18.922370675196078,
    "ENSG00000148303_ENSG00000016082": 16.976840235255324,
    "ENSG00000173726_ENSG00000016082": 16.038253289330257,
    "ENSG00000016082_ENSG00000009307": 11.050750360190644,
    "ENSG00000182718_ENSG00000163435": 9.319374081779717,
    "ENSG00000018189_ENSG00000016082": 8.980175193031863,
    "ENSG00000128951_ENSG00000016082": 8.750186375535304,
    "ENSG00000075142_ENSG00000016082": 8.314658665616204,
    "ENSG00000111716_ENSG00000016082": 8.197261949574827,
    "ENSG00000277443_ENSG00000119547": 6.927224285357188,
    "ENSG00000137285_ENSG00000105419": 5.607472155442066,
    "ENSG00000108055_ENSG00000104435": 5.363713689184388,
    "ENSG00000156976_ENSG00000016082": 4.965866316381353,
    "ENSG00000167081_ENSG00000103187": 4.868664062143716,
    "ENSG00000231500_ENSG00000016082": 4.826485629487524,
    "ENSG00000174444_ENSG00000016082": 4.758520634148669,
    "ENSG00000070961_ENSG00000016082": 4.753343962642168,
    "ENSG00000139352_ENSG00000101489": 4.319834261823555,
    "ENSG00000175745_ENSG00000119547": 3.8278903499438557,
    "ENSG00000164330_ENSG00000034510": 3.5303757357320342,
    "ENSG00000164330_ENSG00000075142": 3.488731461034531,
    "ENSG00000181449_ENSG00000141552": 3.4209798896559405,
    "ENSG00000173812_ENSG00000167081": 3.402341780480591,
    "ENSG00000167081_ENSG00000111716": 3.38429137647371,
    "ENSG00000243317_ENSG00000167081": 3.1383040895545933,
    "ENSG00000164330_ENSG00000163634": 3.102116986327041,
    "ENSG00000105409_ENSG00000103495": 3.0723814416104043,
    "ENSG00000167081_ENSG00000134138": 3.03669079494983,
    "ENSG00000239672_ENSG00000016082": 91.8048162924822,
    "ENSG00000166974_ENSG00000016082": 88.95487753140348
}
```
