## Supplement 2 (DAN vs. Cocul) for "AGEAS: Automated Machine Learning based Genetic Regulatory Element Extraction System": outlier_grps.js

```
{
    "ENSG00000166974_ENSG00000016082": 53.51904107672877,
    "ENSG00000173726_ENSG00000016082": 50.37755316572672,
    "ENSG00000162704_ENSG00000016082": 24.667569776493874,
    "ENSG00000188846_ENSG00000016082": 18.849809637929106,
    "ENSG00000167081_ENSG00000111775": 17.413128977062833,
    "ENSG00000231500_ENSG00000016082": 15.713501228767804,
    "ENSG00000113387_ENSG00000016082": 13.259358169292623,
    "ENSG00000171314_ENSG00000167081": 13.053502823112272,
    "ENSG00000186468_ENSG00000016082": 13.010028940301723,
    "ENSG00000128951_ENSG00000016082": 12.629671820599784,
    "ENSG00000144339_ENSG00000016082": 12.263969046430546,
    "ENSG00000070961_ENSG00000016082": 12.190859562545642,
    "ENSG00000154277_ENSG00000016082": 9.833858875832666,
    "ENSG00000177410_ENSG00000163435": 9.545969147583675,
    "ENSG00000167081_ENSG00000100387": 9.40149714403423,
    "ENSG00000163435_ENSG00000124107": 7.691006533651579,
    "ENSG00000140941_ENSG00000016082": 7.499447834522921,
    "ENSG00000016082_ENSG00000009307": 7.166814006405288,
    "ENSG00000156508_ENSG00000016082": 7.136920671969489,
    "ENSG00000164741_ENSG00000016082": 6.801370009252474,
    "ENSG00000171530_ENSG00000113580": 6.616887235827834,
    "ENSG00000167081_ENSG00000135486": 5.991991485574572,
    "ENSG00000172757_ENSG00000167081": 5.4608780779793005,
    "ENSG00000137285_ENSG00000105419": 5.267324460896663,
    "ENSG00000164330_ENSG00000162704": 5.180681640241476,
    "ENSG00000277443_ENSG00000119547": 5.14391641001233,
    "ENSG00000265972_ENSG00000100219": 5.143775026315463,
    "ENSG00000167081_ENSG00000149806": 5.063520492654456,
    "ENSG00000254999_ENSG00000164093": 4.959815744183508,
    "ENSG00000204628_ENSG00000016082": 4.9142935569651485,
    "ENSG00000182718_ENSG00000163435": 4.899729247500798,
    "ENSG00000167658_ENSG00000167081": 4.688026879580922,
    "ENSG00000196591_ENSG00000172020": 4.521667363042547,
    "ENSG00000163435_ENSG00000051620": 4.422966784675709,
    "ENSG00000167081_ENSG00000118816": 4.31474230891816,
    "ENSG00000182117_ENSG00000016082": 3.8927906481221655,
    "ENSG00000124107_ENSG00000100219": 3.8630708921766264,
    "ENSG00000089009_ENSG00000016082": 3.7938864819142353,
    "ENSG00000164326_ENSG00000137203": 3.73358937796099,
    "ENSG00000101132_ENSG00000016082": 3.7288504147065593,
    "ENSG00000213741_ENSG00000016082": 3.6695880439151316,
    "ENSG00000109670_ENSG00000016082": 3.6688003332924577,
    "ENSG00000156976_ENSG00000016082": 3.6427752489801484,
    "ENSG00000083845_ENSG00000016082": 3.637672189583531,
    "ENSG00000144642_ENSG00000109132": 3.6344358552507003,
    "ENSG00000126214_ENSG00000016082": 3.6103407801061267,
    "ENSG00000105185_ENSG00000016082": 3.6065450455845465,
    "ENSG00000067225_ENSG00000016082": 3.6023189221753547,
    "ENSG00000173660_ENSG00000016082": 3.5726269979115575,
    "ENSG00000213760_ENSG00000016082": 3.568224273336408,
    "ENSG00000075142_ENSG00000016082": 3.553197562833687,
    "ENSG00000175550_ENSG00000016082": 3.546631674195061,
    "ENSG00000182718_ENSG00000016082": 3.530035444801055,
    "ENSG00000167081_ENSG00000156976": 3.5135399622320262,
    "ENSG00000163435_ENSG00000160113": 3.494942486802738,
    "ENSG00000196262_ENSG00000016082": 3.4890900709849944,
    "ENSG00000168028_ENSG00000167081": 3.4888144447729887,
    "ENSG00000131469_ENSG00000016082": 3.488410700365316,
    "ENSG00000239672_ENSG00000016082": 3.4730732133169577,
    "ENSG00000164330_ENSG00000122705": 3.3698097857769835,
    "ENSG00000172159_ENSG00000016082": 3.310060496285341,
    "ENSG00000167081_ENSG00000163634": 3.2504625430869063,
    "ENSG00000167081_ENSG00000112695": 3.2178607577041007,
    "ENSG00000204272_ENSG00000167081": 3.050924864298701,
    "ENSG00000139352_ENSG00000104112": 3.033058210752633,
    "ENSG00000137203_ENSG00000091664": 3.01301548434259,
    "ENSG00000111716_ENSG00000016082": 92.95790102460248,
    "ENSG00000174444_ENSG00000016082": 80.91935494369413
}
```
