## Supplement 4 (CCl4 a6w HSC vs. PF) for "AGEAS: Automated Machine Learning based Genetic Regulatory Element Extraction System": full_atlas.js

```
{
    "network_0": {
        "genes": {
            "ENSMUSG00000029322": {
                "id": "ENSMUSG00000029322",
                "type": "TF",
                "symbol": "Plac8",
                "source": [
                    "ENSMUSG00000057098",
                    "ENSMUSG00000032035"
                ],
                "target": [
                    "ENSMUSG00000021377",
                    "ENSMUSG00000006360",
                    "ENSMUSG00000026087",
                    "ENSMUSG00000036309",
                    "ENSMUSG00000008348",
                    "ENSMUSG00000021606",
                    "ENSMUSG00000004665",
                    "ENSMUSG00000022216",
                    "ENSMUSG00000036438",
                    "ENSMUSG00000026238",
                    "ENSMUSG00000023505",
                    "ENSMUSG00000028797",
                    "ENSMUSG00000042485",
                    "ENSMUSG00000016252",
                    "ENSMUSG00000073616",
                    "ENSMUSG00000031311",
                    "ENSMUSG00000026421",
                    "ENSMUSG00000031198",
                    "ENSMUSG00000074676",
                    "ENSMUSG00000057137",
                    "ENSMUSG00000048126",
                    "ENSMUSG00000037894",
                    "ENSMUSG00000031119",
                    "ENSMUSG00000007872",
                    "ENSMUSG00000027170",
                    "ENSMUSG00000020738",
                    "ENSMUSG00000076432",
                    "ENSMUSG00000061353",
                    "ENSMUSG00000031059",
                    "ENSMUSG00000006304",
                    "ENSMUSG00000055302",
                    "ENSMUSG00000028648",
                    "ENSMUSG00000014294",
                    "ENSMUSG00000020053",
                    "ENSMUSG00000028063",
                    "ENSMUSG00000026207",
                    "ENSMUSG00000037999",
                    "ENSMUSG00000069601",
                    "ENSMUSG00000034353",
                    "ENSMUSG00000036533",
                    "ENSMUSG00000032562",
                    "ENSMUSG00000025511",
                    "ENSMUSG00000015092",
                    "ENSMUSG00000030122",
                    "ENSMUSG00000055322",
                    "ENSMUSG00000031246",
                    "ENSMUSG00000068245",
                    "ENSMUSG00000041481",
                    "ENSMUSG00000019804",
                    "ENSMUSG00000031783",
                    "ENSMUSG00000041741",
                    "ENSMUSG00000048332",
                    "ENSMUSG00000026768",
                    "ENSMUSG00000022587",
                    "ENSMUSG00000045767",
                    "ENSMUSG00000035268",
                    "ENSMUSG00000038633",
                    "ENSMUSG00000016382",
                    "ENSMUSG00000031818",
                    "ENSMUSG00000068798",
                    "ENSMUSG00000030220",
                    "ENSMUSG00000002778",
                    "ENSMUSG00000079523",
                    "ENSMUSG00000016494",
                    "ENSMUSG00000098274",
                    "ENSMUSG00000024290",
                    "ENSMUSG00000061518",
                    "ENSMUSG00000003746",
                    "ENSMUSG00000022665",
                    "ENSMUSG00000041577",
                    "ENSMUSG00000026032",
                    "ENSMUSG00000032135",
                    "ENSMUSG00000064358",
                    "ENSMUSG00000026895",
                    "ENSMUSG00000030272",
                    "ENSMUSG00000026104",
                    "ENSMUSG00000090272",
                    "ENSMUSG00000023904",
                    "ENSMUSG00000050628",
                    "ENSMUSG00000026568",
                    "ENSMUSG00000022450",
                    "ENSMUSG00000041571",
                    "ENSMUSG00000038467",
                    "ENSMUSG00000006299",
                    "ENSMUSG00000024576",
                    "ENSMUSG00000034708",
                    "ENSMUSG00000033306",
                    "ENSMUSG00000026728",
                    "ENSMUSG00000006356",
                    "ENSMUSG00000025204",
                    "ENSMUSG00000027907",
                    "ENSMUSG00000026278",
                    "ENSMUSG00000067818",
                    "ENSMUSG00000054808",
                    "ENSMUSG00000022108",
                    "ENSMUSG00000008136",
                    "ENSMUSG00000024248",
                    "ENSMUSG00000092341",
                    "ENSMUSG00000041736",
                    "ENSMUSG00000040681",
                    "ENSMUSG00000001289",
                    "ENSMUSG00000020585",
                    "ENSMUSG00000013593",
                    "ENSMUSG00000036256",
                    "ENSMUSG00000004285",
                    "ENSMUSG00000025993",
                    "ENSMUSG00000022234",
                    "ENSMUSG00000026078",
                    "ENSMUSG00000066551",
                    "ENSMUSG00000035929",
                    "ENSMUSG00000015932",
                    "ENSMUSG00000060802",
                    "ENSMUSG00000056515",
                    "ENSMUSG00000079017"
                ],
                "expression_sum": {
                    "class1": 85825.0,
                    "class2": 4703.0
                },
                "ens_id": "ENSMUSG00000029322",
                "uniprot_ids": [
                    "Q9JI48"
                ]
            },
            "ENSMUSG00000014294": {
                "id": "ENSMUSG00000014294",
                "type": "Gene",
                "symbol": "Ndufa2",
                "source": [
                    "ENSMUSG00000029322",
                    "ENSMUSG00000000247"
                ],
                "target": [],
                "expression_sum": {
                    "class1": 20804.0,
                    "class2": 15777.0
                },
                "ens_id": "ENSMUSG00000014294",
                "uniprot_ids": [
                    "Q9CQ75"
                ]
            },
            "ENSMUSG00000031818": {
                "id": "ENSMUSG00000031818",
                "type": "Gene",
                "symbol": "Cox4i1",
                "source": [
                    "ENSMUSG00000057098",
                    "ENSMUSG00000047638",
                    "ENSMUSG00000029322",
                    "ENSMUSG00000000247"
                ],
                "target": [],
                "expression_sum": {
                    "class1": 38815.0,
                    "class2": 30450.0
                },
                "ens_id": "ENSMUSG00000031818",
                "uniprot_ids": [
                    "P19783",
                    "M0QWX2",
                    "M0QWX7"
                ]
            },
            "ENSMUSG00000007872": {
                "id": "ENSMUSG00000007872",
                "type": "Gene",
                "symbol": "Id3",
                "source": [
                    "ENSMUSG00000041515",
                    "ENSMUSG00000056501",
                    "ENSMUSG00000052684",
                    "ENSMUSG00000037868",
                    "ENSMUSG00000052837",
                    "ENSMUSG00000047638",
                    "ENSMUSG00000030103",
                    "ENSMUSG00000000247",
                    "ENSMUSG00000038418",
                    "ENSMUSG00000053477",
                    "ENSMUSG00000021250",
                    "ENSMUSG00000029322",
                    "ENSMUSG00000003032",
                    "ENSMUSG00000005583",
                    "ENSMUSG00000018899",
                    "ENSMUSG00000071076"
                ],
                "target": [],
                "expression_sum": {
                    "class1": 106976.0,
                    "class2": 38464.0
                },
                "ens_id": "ENSMUSG00000007872",
                "uniprot_ids": [
                    "P41133"
                ]
            },
            "ENSMUSG00000000247": {
                "id": "ENSMUSG00000000247",
                "type": "TF",
                "symbol": "Lhx2",
                "source": [
                    "ENSMUSG00000032035",
                    "ENSMUSG00000056501",
                    "ENSMUSG00000052684",
                    "ENSMUSG00000023034",
                    "ENSMUSG00000052837",
                    "ENSMUSG00000038418",
                    "ENSMUSG00000021250",
                    "ENSMUSG00000038193",
                    "ENSMUSG00000071076"
                ],
                "target": [
                    "ENSMUSG00000019539",
                    "ENSMUSG00000019505",
                    "ENSMUSG00000024063",
                    "ENSMUSG00000059291",
                    "ENSMUSG00000014077",
                    "ENSMUSG00000058546",
                    "ENSMUSG00000028639",
                    "ENSMUSG00000031812",
                    "ENSMUSG00000021477",
                    "ENSMUSG00000026457",
                    "ENSMUSG00000022283",
                    "ENSMUSG00000068523",
                    "ENSMUSG00000040952",
                    "ENSMUSG00000048076",
                    "ENSMUSG00000030695",
                    "ENSMUSG00000041881",
                    "ENSMUSG00000053560",
                    "ENSMUSG00000025967",
                    "ENSMUSG00000026939",
                    "ENSMUSG00000030744",
                    "ENSMUSG00000036438",
                    "ENSMUSG00000026238",
                    "ENSMUSG00000000325",
                    "ENSMUSG00000060036",
                    "ENSMUSG00000008668",
                    "ENSMUSG00000061787",
                    "ENSMUSG00000038717",
                    "ENSMUSG00000020372",
                    "ENSMUSG00000041417",
                    "ENSMUSG00000008682",
                    "ENSMUSG00000023175",
                    "ENSMUSG00000021127",
                    "ENSMUSG00000026970",
                    "ENSMUSG00000031198",
                    "ENSMUSG00000023861",
                    "ENSMUSG00000061904",
                    "ENSMUSG00000034892",
                    "ENSMUSG00000022010",
                    "ENSMUSG00000005161",
                    "ENSMUSG00000020612",
                    "ENSMUSG00000032231",
                    "ENSMUSG00000071415",
                    "ENSMUSG00000021660",
                    "ENSMUSG00000061477",
                    "ENSMUSG00000060743",
                    "ENSMUSG00000007872",
                    "ENSMUSG00000056201",
                    "ENSMUSG00000020738",
                    "ENSMUSG00000022884",
                    "ENSMUSG00000061983",
                    "ENSMUSG00000004980",
                    "ENSMUSG00000037805",
                    "ENSMUSG00000027750",
                    "ENSMUSG00000006304",
                    "ENSMUSG00000028081",
                    "ENSMUSG00000063316",
                    "ENSMUSG00000055302",
                    "ENSMUSG00000017404",
                    "ENSMUSG00000070493",
                    "ENSMUSG00000021546",
                    "ENSMUSG00000046364",
                    "ENSMUSG00000004070",
                    "ENSMUSG00000022312",
                    "ENSMUSG00000060636",
                    "ENSMUSG00000055839",
                    "ENSMUSG00000014294",
                    "ENSMUSG00000064370",
                    "ENSMUSG00000068874",
                    "ENSMUSG00000037999",
                    "ENSMUSG00000069601",
                    "ENSMUSG00000028603",
                    "ENSMUSG00000042043",
                    "ENSMUSG00000037742",
                    "ENSMUSG00000025781",
                    "ENSMUSG00000022892",
                    "ENSMUSG00000032562",
                    "ENSMUSG00000025511",
                    "ENSMUSG00000029810",
                    "ENSMUSG00000038803",
                    "ENSMUSG00000052684",
                    "ENSMUSG00000012405",
                    "ENSMUSG00000000184",
                    "ENSMUSG00000020719",
                    "ENSMUSG00000036371",
                    "ENSMUSG00000090862",
                    "ENSMUSG00000021807",
                    "ENSMUSG00000022051",
                    "ENSMUSG00000030824",
                    "ENSMUSG00000064351",
                    "ENSMUSG00000027642",
                    "ENSMUSG00000003970",
                    "ENSMUSG00000003617",
                    "ENSMUSG00000021870",
                    "ENSMUSG00000018326",
                    "ENSMUSG00000020460",
                    "ENSMUSG00000035885",
                    "ENSMUSG00000027706",
                    "ENSMUSG00000055044",
                    "ENSMUSG00000029781",
                    "ENSMUSG00000042812",
                    "ENSMUSG00000025508",
                    "ENSMUSG00000063882",
                    "ENSMUSG00000025290",
                    "ENSMUSG00000039105",
                    "ENSMUSG00000037712",
                    "ENSMUSG00000009927",
                    "ENSMUSG00000033880",
                    "ENSMUSG00000031353",
                    "ENSMUSG00000045128",
                    "ENSMUSG00000022587",
                    "ENSMUSG00000025794",
                    "ENSMUSG00000023944",
                    "ENSMUSG00000024065",
                    "ENSMUSG00000023367",
                    "ENSMUSG00000024608",
                    "ENSMUSG00000001687",
                    "ENSMUSG00000029713",
                    "ENSMUSG00000039001",
                    "ENSMUSG00000036381",
                    "ENSMUSG00000006333",
                    "ENSMUSG00000058600",
                    "ENSMUSG00000062647",
                    "ENSMUSG00000010406",
                    "ENSMUSG00000030432",
                    "ENSMUSG00000046330",
                    "ENSMUSG00000074129",
                    "ENSMUSG00000090733",
                    "ENSMUSG00000031818",
                    "ENSMUSG00000020432",
                    "ENSMUSG00000010376",
                    "ENSMUSG00000002778",
                    "ENSMUSG00000032946",
                    "ENSMUSG00000098274",
                    "ENSMUSG00000027404",
                    "ENSMUSG00000016559",
                    "ENSMUSG00000060126",
                    "ENSMUSG00000003072",
                    "ENSMUSG00000061315",
                    "ENSMUSG00000057322",
                    "ENSMUSG00000027673",
                    "ENSMUSG00000012848",
                    "ENSMUSG00000060591",
                    "ENSMUSG00000023942",
                    "ENSMUSG00000049517",
                    "ENSMUSG00000025362",
                    "ENSMUSG00000019960",
                    "ENSMUSG00000044533",
                    "ENSMUSG00000029614",
                    "ENSMUSG00000022136",
                    "ENSMUSG00000025428",
                    "ENSMUSG00000073702",
                    "ENSMUSG00000003814",
                    "ENSMUSG00000062006",
                    "ENSMUSG00000062683",
                    "ENSMUSG00000040128",
                    "ENSMUSG00000032216",
                    "ENSMUSG00000079435",
                    "ENSMUSG00000047215",
                    "ENSMUSG00000062270",
                    "ENSMUSG00000022450",
                    "ENSMUSG00000041571",
                    "ENSMUSG00000053565",
                    "ENSMUSG00000032666",
                    "ENSMUSG00000004207",
                    "ENSMUSG00000031029",
                    "ENSMUSG00000028673",
                    "ENSMUSG00000024576",
                    "ENSMUSG00000035242",
                    "ENSMUSG00000050708",
                    "ENSMUSG00000014769",
                    "ENSMUSG00000015656",
                    "ENSMUSG00000067288",
                    "ENSMUSG00000058558",
                    "ENSMUSG00000029076",
                    "ENSMUSG00000056851",
                    "ENSMUSG00000022971",
                    "ENSMUSG00000052146",
                    "ENSMUSG00000060938",
                    "ENSMUSG00000047675",
                    "ENSMUSG00000025393",
                    "ENSMUSG00000034994",
                    "ENSMUSG00000041841",
                    "ENSMUSG00000067274",
                    "ENSMUSG00000054808",
                    "ENSMUSG00000024308",
                    "ENSMUSG00000071644",
                    "ENSMUSG00000057863",
                    "ENSMUSG00000026566",
                    "ENSMUSG00000022108",
                    "ENSMUSG00000032288",
                    "ENSMUSG00000014313",
                    "ENSMUSG00000024248",
                    "ENSMUSG00000028266",
                    "ENSMUSG00000024953",
                    "ENSMUSG00000030057",
                    "ENSMUSG00000041736",
                    "ENSMUSG00000045690",
                    "ENSMUSG00000041560",
                    "ENSMUSG00000005881",
                    "ENSMUSG00000049751",
                    "ENSMUSG00000001289",
                    "ENSMUSG00000024678",
                    "ENSMUSG00000036751",
                    "ENSMUSG00000032179",
                    "ENSMUSG00000013593",
                    "ENSMUSG00000008683",
                    "ENSMUSG00000055762",
                    "ENSMUSG00000043716",
                    "ENSMUSG00000021250",
                    "ENSMUSG00000017778",
                    "ENSMUSG00000055447",
                    "ENSMUSG00000022890",
                    "ENSMUSG00000000740",
                    "ENSMUSG00000007892",
                    "ENSMUSG00000027523",
                    "ENSMUSG00000024190",
                    "ENSMUSG00000028618",
                    "ENSMUSG00000057841",
                    "ENSMUSG00000079641",
                    "ENSMUSG00000027593",
                    "ENSMUSG00000035530",
                    "ENSMUSG00000028757",
                    "ENSMUSG00000059070",
                    "ENSMUSG00000031532",
                    "ENSMUSG00000027620",
                    "ENSMUSG00000047907",
                    "ENSMUSG00000052837",
                    "ENSMUSG00000048758",
                    "ENSMUSG00000029632",
                    "ENSMUSG00000022708",
                    "ENSMUSG00000028234",
                    "ENSMUSG00000093674",
                    "ENSMUSG00000028936",
                    "ENSMUSG00000042682",
                    "ENSMUSG00000044786",
                    "ENSMUSG00000028495",
                    "ENSMUSG00000059005",
                    "ENSMUSG00000028478",
                    "ENSMUSG00000071076",
                    "ENSMUSG00000041453",
                    "ENSMUSG00000003429",
                    "ENSMUSG00000018293",
                    "ENSMUSG00000029465",
                    "ENSMUSG00000038418",
                    "ENSMUSG00000030849",
                    "ENSMUSG00000021134",
                    "ENSMUSG00000000563",
                    "ENSMUSG00000031633",
                    "ENSMUSG00000037563",
                    "ENSMUSG00000032399"
                ],
                "expression_sum": {
                    "class1": 14582.0,
                    "class2": 111.0
                },
                "ens_id": "ENSMUSG00000000247",
                "uniprot_ids": [
                    "Q9Z0S2",
                    "F6Z9H5",
                    "A0A0A0MQK9",
                    "F6SM03",
                    "H3BKI3"
                ]
            },
            "ENSMUSG00000060636": {
                "id": "ENSMUSG00000060636",
                "type": "Gene",
                "symbol": "Rpl35a",
                "source": [
                    "ENSMUSG00000057098",
                    "ENSMUSG00000048387",
                    "ENSMUSG00000038193",
                    "ENSMUSG00000000247"
                ],
                "target": [],
                "expression_sum": {
                    "class1": 134254.0,
                    "class2": 76798.0
                },
                "ens_id": "ENSMUSG00000060636",
                "uniprot_ids": [
                    "O55142"
                ]
            },
            "ENSMUSG00000062647": {
                "id": "ENSMUSG00000062647",
                "type": "Gene",
                "symbol": "Rpl7a",
                "source": [
                    "ENSMUSG00000022528",
                    "ENSMUSG00000000247",
                    "ENSMUSG00000057098",
                    "ENSMUSG00000038193",
                    "ENSMUSG00000005583",
                    "ENSMUSG00000031965",
                    "ENSMUSG00000053007"
                ],
                "target": [],
                "expression_sum": {
                    "class1": 61371.0,
                    "class2": 28548.0
                },
                "ens_id": "ENSMUSG00000062647",
                "uniprot_ids": [
                    "P12970"
                ]
            },
            "ENSMUSG00000024608": {
                "id": "ENSMUSG00000024608",
                "type": "Gene",
                "symbol": "Rps14",
                "source": [
                    "ENSMUSG00000057098",
                    "ENSMUSG00000048387",
                    "ENSMUSG00000038193",
                    "ENSMUSG00000000247"
                ],
                "target": [],
                "expression_sum": {
                    "class1": 161001.0,
                    "class2": 83793.0
                },
                "ens_id": "ENSMUSG00000024608",
                "uniprot_ids": [
                    "P62264",
                    "A0A494BAS4",
                    "D3Z7I1",
                    "D3YVF4"
                ]
            },
            "ENSMUSG00000048387": {
                "id": "ENSMUSG00000048387",
                "type": "TF",
                "symbol": "Osr1",
                "source": [
                    "ENSMUSG00000022346",
                    "ENSMUSG00000028565"
                ],
                "target": [
                    "ENSMUSG00000019505",
                    "ENSMUSG00000031447",
                    "ENSMUSG00000058546",
                    "ENSMUSG00000007655",
                    "ENSMUSG00000028691",
                    "ENSMUSG00000022365",
                    "ENSMUSG00000025213",
                    "ENSMUSG00000040952",
                    "ENSMUSG00000031320",
                    "ENSMUSG00000027422",
                    "ENSMUSG00000073676",
                    "ENSMUSG00000027447",
                    "ENSMUSG00000056629",
                    "ENSMUSG00000024909",
                    "ENSMUSG00000021877",
                    "ENSMUSG00000034868",
                    "ENSMUSG00000060036",
                    "ENSMUSG00000008668",
                    "ENSMUSG00000036275",
                    "ENSMUSG00000008682",
                    "ENSMUSG00000030659",
                    "ENSMUSG00000018567",
                    "ENSMUSG00000034892",
                    "ENSMUSG00000029103",
                    "ENSMUSG00000071415",
                    "ENSMUSG00000053317",
                    "ENSMUSG00000061983",
                    "ENSMUSG00000029616",
                    "ENSMUSG00000020368",
                    "ENSMUSG00000063316",
                    "ENSMUSG00000060636",
                    "ENSMUSG00000028367",
                    "ENSMUSG00000037742",
                    "ENSMUSG00000036372",
                    "ENSMUSG00000037266",
                    "ENSMUSG00000038650",
                    "ENSMUSG00000090862",
                    "ENSMUSG00000090000",
                    "ENSMUSG00000030824",
                    "ENSMUSG00000003970",
                    "ENSMUSG00000009927",
                    "ENSMUSG00000025508",
                    "ENSMUSG00000042737",
                    "ENSMUSG00000023944",
                    "ENSMUSG00000078812",
                    "ENSMUSG00000024608",
                    "ENSMUSG00000029713",
                    "ENSMUSG00000039001",
                    "ENSMUSG00000037373",
                    "ENSMUSG00000006333",
                    "ENSMUSG00000058600",
                    "ENSMUSG00000046330",
                    "ENSMUSG00000074129",
                    "ENSMUSG00000090733",
                    "ENSMUSG00000028517",
                    "ENSMUSG00000002778",
                    "ENSMUSG00000079523",
                    "ENSMUSG00000098274",
                    "ENSMUSG00000026674",
                    "ENSMUSG00000060126",
                    "ENSMUSG00000026511",
                    "ENSMUSG00000058569",
                    "ENSMUSG00000012848",
                    "ENSMUSG00000025362",
                    "ENSMUSG00000027074",
                    "ENSMUSG00000029614",
                    "ENSMUSG00000021871",
                    "ENSMUSG00000038612",
                    "ENSMUSG00000073702",
                    "ENSMUSG00000044408",
                    "ENSMUSG00000062270",
                    "ENSMUSG00000045817",
                    "ENSMUSG00000024048",
                    "ENSMUSG00000028565",
                    "ENSMUSG00000038274",
                    "ENSMUSG00000037072",
                    "ENSMUSG00000020009",
                    "ENSMUSG00000028673",
                    "ENSMUSG00000021876",
                    "ENSMUSG00000020088",
                    "ENSMUSG00000034708",
                    "ENSMUSG00000096145",
                    "ENSMUSG00000062997",
                    "ENSMUSG00000021610",
                    "ENSMUSG00000046841",
                    "ENSMUSG00000025232",
                    "ENSMUSG00000058558",
                    "ENSMUSG00000003363",
                    "ENSMUSG00000006050",
                    "ENSMUSG00000023010",
                    "ENSMUSG00000029076",
                    "ENSMUSG00000041084",
                    "ENSMUSG00000008475",
                    "ENSMUSG00000041126",
                    "ENSMUSG00000060938",
                    "ENSMUSG00000052146",
                    "ENSMUSG00000067274",
                    "ENSMUSG00000057863",
                    "ENSMUSG00000022108",
                    "ENSMUSG00000074170",
                    "ENSMUSG00000024248",
                    "ENSMUSG00000030934",
                    "ENSMUSG00000020849",
                    "ENSMUSG00000018593",
                    "ENSMUSG00000002064",
                    "ENSMUSG00000045690",
                    "ENSMUSG00000027248",
                    "ENSMUSG00000040681",
                    "ENSMUSG00000062328",
                    "ENSMUSG00000020585",
                    "ENSMUSG00000022346",
                    "ENSMUSG00000055762",
                    "ENSMUSG00000043716",
                    "ENSMUSG00000071074",
                    "ENSMUSG00000024190",
                    "ENSMUSG00000020458",
                    "ENSMUSG00000027763",
                    "ENSMUSG00000006369",
                    "ENSMUSG00000059070",
                    "ENSMUSG00000066551",
                    "ENSMUSG00000048758",
                    "ENSMUSG00000078578",
                    "ENSMUSG00000028234",
                    "ENSMUSG00000035227",
                    "ENSMUSG00000093674",
                    "ENSMUSG00000044786",
                    "ENSMUSG00000028495",
                    "ENSMUSG00000025579",
                    "ENSMUSG00000079037",
                    "ENSMUSG00000041453",
                    "ENSMUSG00000019087",
                    "ENSMUSG00000029465",
                    "ENSMUSG00000029231",
                    "ENSMUSG00000032399"
                ],
                "expression_sum": {
                    "class1": 11.0,
                    "class2": 4732.0
                },
                "ens_id": "ENSMUSG00000048387",
                "uniprot_ids": [
                    "Q9WVG7",
                    "A0A1W2P7S2"
                ]
            },
            "ENSMUSG00000031447": {
                "id": "ENSMUSG00000031447",
                "type": "Gene",
                "symbol": "Lamp1",
                "source": [
                    "ENSMUSG00000032035",
                    "ENSMUSG00000022528",
                    "ENSMUSG00000057098",
                    "ENSMUSG00000038193",
                    "ENSMUSG00000003032",
                    "ENSMUSG00000005583",
                    "ENSMUSG00000048387"
                ],
                "target": [],
                "expression_sum": {
                    "class1": 37763.0,
                    "class2": 19814.0
                },
                "ens_id": "ENSMUSG00000031447",
                "uniprot_ids": [
                    "P11438"
                ]
            },
            "ENSMUSG00000012848": {
                "id": "ENSMUSG00000012848",
                "type": "Gene",
                "symbol": "Rps5",
                "source": [
                    "ENSMUSG00000023034",
                    "ENSMUSG00000000247",
                    "ENSMUSG00000057098",
                    "ENSMUSG00000038193",
                    "ENSMUSG00000005583",
                    "ENSMUSG00000048387"
                ],
                "target": [],
                "expression_sum": {
                    "class1": 125292.0,
                    "class2": 72878.0
                },
                "ens_id": "ENSMUSG00000012848",
                "uniprot_ids": [
                    "Q91V55",
                    "D3Z1S8",
                    "D3YYM6"
                ]
            },
            "ENSMUSG00000057098": {
                "id": "ENSMUSG00000057098",
                "type": "TF",
                "symbol": "Ebf1",
                "source": [
                    "ENSMUSG00000021250",
                    "ENSMUSG00000003032",
                    "ENSMUSG00000071076"
                ],
                "target": [
                    "ENSMUSG00000031447",
                    "ENSMUSG00000062456",
                    "ENSMUSG00000032518",
                    "ENSMUSG00000024011",
                    "ENSMUSG00000026878",
                    "ENSMUSG00000014599",
                    "ENSMUSG00000016256",
                    "ENSMUSG00000047187",
                    "ENSMUSG00000008668",
                    "ENSMUSG00000019877",
                    "ENSMUSG00000071415",
                    "ENSMUSG00000053317",
                    "ENSMUSG00000002900",
                    "ENSMUSG00000029616",
                    "ENSMUSG00000020368",
                    "ENSMUSG00000028081",
                    "ENSMUSG00000055302",
                    "ENSMUSG00000025781",
                    "ENSMUSG00000012405",
                    "ENSMUSG00000038650",
                    "ENSMUSG00000020737",
                    "ENSMUSG00000022820",
                    "ENSMUSG00000090862",
                    "ENSMUSG00000030824",
                    "ENSMUSG00000073639",
                    "ENSMUSG00000025508",
                    "ENSMUSG00000063882",
                    "ENSMUSG00000011752",
                    "ENSMUSG00000045128",
                    "ENSMUSG00000017466",
                    "ENSMUSG00000078812",
                    "ENSMUSG00000029455",
                    "ENSMUSG00000044080",
                    "ENSMUSG00000026536",
                    "ENSMUSG00000072235",
                    "ENSMUSG00000006333",
                    "ENSMUSG00000030432",
                    "ENSMUSG00000074129",
                    "ENSMUSG00000031818",
                    "ENSMUSG00000026674",
                    "ENSMUSG00000012848",
                    "ENSMUSG00000027074",
                    "ENSMUSG00000028964",
                    "ENSMUSG00000003032",
                    "ENSMUSG00000073702",
                    "ENSMUSG00000032216",
                    "ENSMUSG00000062270",
                    "ENSMUSG00000045817",
                    "ENSMUSG00000027712",
                    "ENSMUSG00000053565",
                    "ENSMUSG00000007891",
                    "ENSMUSG00000024048",
                    "ENSMUSG00000031029",
                    "ENSMUSG00000001891",
                    "ENSMUSG00000050708",
                    "ENSMUSG00000026478",
                    "ENSMUSG00000057278",
                    "ENSMUSG00000014846",
                    "ENSMUSG00000017493",
                    "ENSMUSG00000025351",
                    "ENSMUSG00000074884",
                    "ENSMUSG00000020849",
                    "ENSMUSG00000025066",
                    "ENSMUSG00000045690",
                    "ENSMUSG00000027248",
                    "ENSMUSG00000049751",
                    "ENSMUSG00000043716",
                    "ENSMUSG00000027523",
                    "ENSMUSG00000035530",
                    "ENSMUSG00000036781",
                    "ENSMUSG00000041801",
                    "ENSMUSG00000073411",
                    "ENSMUSG00000022174",
                    "ENSMUSG00000004610",
                    "ENSMUSG00000035227",
                    "ENSMUSG00000060802",
                    "ENSMUSG00000041453",
                    "ENSMUSG00000003429",
                    "ENSMUSG00000038418",
                    "ENSMUSG00000032399",
                    "ENSMUSG00000038900",
                    "ENSMUSG00000019505",
                    "ENSMUSG00000006360",
                    "ENSMUSG00000059291",
                    "ENSMUSG00000022841",
                    "ENSMUSG00000040824",
                    "ENSMUSG00000028691",
                    "ENSMUSG00000022283",
                    "ENSMUSG00000068523",
                    "ENSMUSG00000036199",
                    "ENSMUSG00000030695",
                    "ENSMUSG00000030744",
                    "ENSMUSG00000026238",
                    "ENSMUSG00000028822",
                    "ENSMUSG00000023175",
                    "ENSMUSG00000022336",
                    "ENSMUSG00000026970",
                    "ENSMUSG00000021702",
                    "ENSMUSG00000002107",
                    "ENSMUSG00000009549",
                    "ENSMUSG00000032231",
                    "ENSMUSG00000059796",
                    "ENSMUSG00000004455",
                    "ENSMUSG00000037805",
                    "ENSMUSG00000063316",
                    "ENSMUSG00000017404",
                    "ENSMUSG00000028648",
                    "ENSMUSG00000025132",
                    "ENSMUSG00000021917",
                    "ENSMUSG00000042043",
                    "ENSMUSG00000037266",
                    "ENSMUSG00000030790",
                    "ENSMUSG00000029642",
                    "ENSMUSG00000020719",
                    "ENSMUSG00000036371",
                    "ENSMUSG00000019804",
                    "ENSMUSG00000007039",
                    "ENSMUSG00000022090",
                    "ENSMUSG00000032014",
                    "ENSMUSG00000040249",
                    "ENSMUSG00000023092",
                    "ENSMUSG00000029070",
                    "ENSMUSG00000015839",
                    "ENSMUSG00000010406",
                    "ENSMUSG00000051695",
                    "ENSMUSG00000002778",
                    "ENSMUSG00000030654",
                    "ENSMUSG00000098274",
                    "ENSMUSG00000026511",
                    "ENSMUSG00000064080",
                    "ENSMUSG00000005973",
                    "ENSMUSG00000057322",
                    "ENSMUSG00000003161",
                    "ENSMUSG00000021025",
                    "ENSMUSG00000032301",
                    "ENSMUSG00000042079",
                    "ENSMUSG00000003814",
                    "ENSMUSG00000052040",
                    "ENSMUSG00000015671",
                    "ENSMUSG00000047215",
                    "ENSMUSG00000020205",
                    "ENSMUSG00000037072",
                    "ENSMUSG00000020009",
                    "ENSMUSG00000004207",
                    "ENSMUSG00000026728",
                    "ENSMUSG00000096145",
                    "ENSMUSG00000014769",
                    "ENSMUSG00000042312",
                    "ENSMUSG00000062997",
                    "ENSMUSG00000067288",
                    "ENSMUSG00000058558",
                    "ENSMUSG00000075254",
                    "ENSMUSG00000023010",
                    "ENSMUSG00000060938",
                    "ENSMUSG00000047675",
                    "ENSMUSG00000041841",
                    "ENSMUSG00000021242",
                    "ENSMUSG00000008892",
                    "ENSMUSG00000024248",
                    "ENSMUSG00000028837",
                    "ENSMUSG00000062328",
                    "ENSMUSG00000020585",
                    "ENSMUSG00000019818",
                    "ENSMUSG00000025492",
                    "ENSMUSG00000055762",
                    "ENSMUSG00000007041",
                    "ENSMUSG00000055447",
                    "ENSMUSG00000024190",
                    "ENSMUSG00000057841",
                    "ENSMUSG00000020458",
                    "ENSMUSG00000022234",
                    "ENSMUSG00000009291",
                    "ENSMUSG00000078578",
                    "ENSMUSG00000028234",
                    "ENSMUSG00000029322",
                    "ENSMUSG00000035585",
                    "ENSMUSG00000090137",
                    "ENSMUSG00000021131",
                    "ENSMUSG00000037062",
                    "ENSMUSG00000031765",
                    "ENSMUSG00000028639",
                    "ENSMUSG00000040952",
                    "ENSMUSG00000048076",
                    "ENSMUSG00000026021",
                    "ENSMUSG00000006699",
                    "ENSMUSG00000060036",
                    "ENSMUSG00000045962",
                    "ENSMUSG00000038717",
                    "ENSMUSG00000016252",
                    "ENSMUSG00000023004",
                    "ENSMUSG00000031198",
                    "ENSMUSG00000025130",
                    "ENSMUSG00000056737",
                    "ENSMUSG00000056201",
                    "ENSMUSG00000022884",
                    "ENSMUSG00000061983",
                    "ENSMUSG00000022032",
                    "ENSMUSG00000004980",
                    "ENSMUSG00000028967",
                    "ENSMUSG00000022400",
                    "ENSMUSG00000021546",
                    "ENSMUSG00000018585",
                    "ENSMUSG00000034165",
                    "ENSMUSG00000003380",
                    "ENSMUSG00000038803",
                    "ENSMUSG00000022051",
                    "ENSMUSG00000020460",
                    "ENSMUSG00000055044",
                    "ENSMUSG00000009927",
                    "ENSMUSG00000023944",
                    "ENSMUSG00000028416",
                    "ENSMUSG00000024608",
                    "ENSMUSG00000062647",
                    "ENSMUSG00000060591",
                    "ENSMUSG00000049517",
                    "ENSMUSG00000027940",
                    "ENSMUSG00000001794",
                    "ENSMUSG00000028692",
                    "ENSMUSG00000025780",
                    "ENSMUSG00000030795",
                    "ENSMUSG00000035242",
                    "ENSMUSG00000006356",
                    "ENSMUSG00000015656",
                    "ENSMUSG00000067274",
                    "ENSMUSG00000071644",
                    "ENSMUSG00000057863",
                    "ENSMUSG00000022108",
                    "ENSMUSG00000025510",
                    "ENSMUSG00000028249",
                    "ENSMUSG00000024953",
                    "ENSMUSG00000053012",
                    "ENSMUSG00000026670",
                    "ENSMUSG00000021250",
                    "ENSMUSG00000001833",
                    "ENSMUSG00000028618",
                    "ENSMUSG00000027620",
                    "ENSMUSG00000032966",
                    "ENSMUSG00000052837",
                    "ENSMUSG00000032324",
                    "ENSMUSG00000031708",
                    "ENSMUSG00000028936",
                    "ENSMUSG00000059005",
                    "ENSMUSG00000028478",
                    "ENSMUSG00000060073",
                    "ENSMUSG00000001847",
                    "ENSMUSG00000019428",
                    "ENSMUSG00000029465",
                    "ENSMUSG00000039208",
                    "ENSMUSG00000030057",
                    "ENSMUSG00000031633",
                    "ENSMUSG00000037563",
                    "ENSMUSG00000058135",
                    "ENSMUSG00000007815",
                    "ENSMUSG00000002602",
                    "ENSMUSG00000058546",
                    "ENSMUSG00000008348",
                    "ENSMUSG00000037822",
                    "ENSMUSG00000025967",
                    "ENSMUSG00000031556",
                    "ENSMUSG00000021877",
                    "ENSMUSG00000034868",
                    "ENSMUSG00000063480",
                    "ENSMUSG00000029649",
                    "ENSMUSG00000028797",
                    "ENSMUSG00000061787",
                    "ENSMUSG00000044627",
                    "ENSMUSG00000020372",
                    "ENSMUSG00000021939",
                    "ENSMUSG00000001240",
                    "ENSMUSG00000036880",
                    "ENSMUSG00000061904",
                    "ENSMUSG00000028369",
                    "ENSMUSG00000029621",
                    "ENSMUSG00000060743",
                    "ENSMUSG00000069744",
                    "ENSMUSG00000020738",
                    "ENSMUSG00000037499",
                    "ENSMUSG00000026547",
                    "ENSMUSG00000006304",
                    "ENSMUSG00000062825",
                    "ENSMUSG00000060636",
                    "ENSMUSG00000046364",
                    "ENSMUSG00000075706",
                    "ENSMUSG00000020440",
                    "ENSMUSG00000028367",
                    "ENSMUSG00000040385",
                    "ENSMUSG00000063457",
                    "ENSMUSG00000032314",
                    "ENSMUSG00000037742",
                    "ENSMUSG00000036372",
                    "ENSMUSG00000015092",
                    "ENSMUSG00000025016",
                    "ENSMUSG00000029767",
                    "ENSMUSG00000018326",
                    "ENSMUSG00000057113",
                    "ENSMUSG00000002233",
                    "ENSMUSG00000020267",
                    "ENSMUSG00000025794",
                    "ENSMUSG00000029713",
                    "ENSMUSG00000039001",
                    "ENSMUSG00000028452",
                    "ENSMUSG00000060019",
                    "ENSMUSG00000028530",
                    "ENSMUSG00000090733",
                    "ENSMUSG00000040212",
                    "ENSMUSG00000001123",
                    "ENSMUSG00000016559",
                    "ENSMUSG00000060126",
                    "ENSMUSG00000020149",
                    "ENSMUSG00000061315",
                    "ENSMUSG00000044533",
                    "ENSMUSG00000062006",
                    "ENSMUSG00000002015",
                    "ENSMUSG00000052727",
                    "ENSMUSG00000013622",
                    "ENSMUSG00000038274",
                    "ENSMUSG00000028108",
                    "ENSMUSG00000021996",
                    "ENSMUSG00000006519",
                    "ENSMUSG00000056851",
                    "ENSMUSG00000008475",
                    "ENSMUSG00000052146",
                    "ENSMUSG00000034994",
                    "ENSMUSG00000024308",
                    "ENSMUSG00000021218",
                    "ENSMUSG00000026872",
                    "ENSMUSG00000060860",
                    "ENSMUSG00000071866",
                    "ENSMUSG00000005881",
                    "ENSMUSG00000024914",
                    "ENSMUSG00000020048",
                    "ENSMUSG00000042747",
                    "ENSMUSG00000007892",
                    "ENSMUSG00000032383",
                    "ENSMUSG00000079641",
                    "ENSMUSG00000031422",
                    "ENSMUSG00000029622",
                    "ENSMUSG00000059070",
                    "ENSMUSG00000066551",
                    "ENSMUSG00000022708",
                    "ENSMUSG00000034118",
                    "ENSMUSG00000029659",
                    "ENSMUSG00000093674",
                    "ENSMUSG00000029430",
                    "ENSMUSG00000042682",
                    "ENSMUSG00000028495",
                    "ENSMUSG00000008540",
                    "ENSMUSG00000029484",
                    "ENSMUSG00000022548",
                    "ENSMUSG00000000563"
                ],
                "expression_sum": {
                    "class1": 205.0,
                    "class2": 16536.0
                },
                "ens_id": "ENSMUSG00000057098",
                "uniprot_ids": [
                    "Q07802",
                    "Q5SWK4",
                    "Q5SWK8"
                ]
            },
            "ENSMUSG00000063457": {
                "id": "ENSMUSG00000063457",
                "type": "Gene",
                "symbol": "Rps15",
                "source": [
                    "ENSMUSG00000031965",
                    "ENSMUSG00000038193",
                    "ENSMUSG00000047638",
                    "ENSMUSG00000057098"
                ],
                "target": [],
                "expression_sum": {
                    "class1": 120156.0,
                    "class2": 63027.0
                },
                "ens_id": "ENSMUSG00000063457",
                "uniprot_ids": [
                    "P62843",
                    "D3YTQ9"
                ]
            },
            "ENSMUSG00000008668": {
                "id": "ENSMUSG00000008668",
                "type": "Gene",
                "symbol": "Rps18",
                "source": [
                    "ENSMUSG00000023034",
                    "ENSMUSG00000000247",
                    "ENSMUSG00000057098",
                    "ENSMUSG00000038193",
                    "ENSMUSG00000005583",
                    "ENSMUSG00000048387"
                ],
                "target": [],
                "expression_sum": {
                    "class1": 133850.0,
                    "class2": 65639.0
                },
                "ens_id": "ENSMUSG00000008668",
                "uniprot_ids": [
                    "P62270",
                    "A0A3Q4EGP3",
                    "S4R1N6"
                ]
            },
            "ENSMUSG00000022108": {
                "id": "ENSMUSG00000022108",
                "type": "Gene",
                "symbol": "Itm2b",
                "source": [
                    "ENSMUSG00000032035",
                    "ENSMUSG00000047638",
                    "ENSMUSG00000000247",
                    "ENSMUSG00000057098",
                    "ENSMUSG00000038193",
                    "ENSMUSG00000029322",
                    "ENSMUSG00000048387",
                    "ENSMUSG00000071076"
                ],
                "target": [],
                "expression_sum": {
                    "class1": 233347.0,
                    "class2": 92392.0
                },
                "ens_id": "ENSMUSG00000022108",
                "uniprot_ids": [
                    "O89051",
                    "A0A2I3BQ30",
                    "A0A2I3BPW7"
                ]
            },
            "ENSMUSG00000003429": {
                "id": "ENSMUSG00000003429",
                "type": "Gene",
                "symbol": "Rps11",
                "source": [
                    "ENSMUSG00000057098",
                    "ENSMUSG00000038193",
                    "ENSMUSG00000000247"
                ],
                "target": [],
                "expression_sum": {
                    "class1": 98135.0,
                    "class2": 48915.0
                },
                "ens_id": "ENSMUSG00000003429",
                "uniprot_ids": [
                    "P62281",
                    "A0A1B0GSE8",
                    "A0A1B0GRR3"
                ]
            },
            "ENSMUSG00000032399": {
                "id": "ENSMUSG00000032399",
                "type": "Gene",
                "symbol": "Rpl4",
                "source": [
                    "ENSMUSG00000057098",
                    "ENSMUSG00000048387",
                    "ENSMUSG00000038193",
                    "ENSMUSG00000000247"
                ],
                "target": [],
                "expression_sum": {
                    "class1": 85485.0,
                    "class2": 44235.0
                },
                "ens_id": "ENSMUSG00000032399",
                "uniprot_ids": [
                    "Q9D8E6"
                ]
            },
            "ENSMUSG00000030744": {
                "id": "ENSMUSG00000030744",
                "type": "Gene",
                "symbol": "Rps3",
                "source": [
                    "ENSMUSG00000057098",
                    "ENSMUSG00000038193",
                    "ENSMUSG00000000247"
                ],
                "target": [],
                "expression_sum": {
                    "class1": 114154.0,
                    "class2": 53681.0
                },
                "ens_id": "ENSMUSG00000030744",
                "uniprot_ids": [
                    "P62908",
                    "D3YV43",
                    "A0A140LI77"
                ]
            },
            "ENSMUSG00000041841": {
                "id": "ENSMUSG00000041841",
                "type": "Gene",
                "symbol": "Rpl37",
                "source": [
                    "ENSMUSG00000057098",
                    "ENSMUSG00000038193",
                    "ENSMUSG00000000247"
                ],
                "target": [],
                "expression_sum": {
                    "class1": 163253.0,
                    "class2": 80407.0
                },
                "ens_id": "ENSMUSG00000041841",
                "uniprot_ids": [
                    "Q9D823"
                ]
            },
            "ENSMUSG00000037742": {
                "id": "ENSMUSG00000037742",
                "type": "Gene",
                "symbol": "Eef1a1",
                "source": [
                    "ENSMUSG00000056501",
                    "ENSMUSG00000023034",
                    "ENSMUSG00000022528",
                    "ENSMUSG00000047638",
                    "ENSMUSG00000000247",
                    "ENSMUSG00000057098",
                    "ENSMUSG00000038193",
                    "ENSMUSG00000048387"
                ],
                "target": [],
                "expression_sum": {
                    "class1": 331050.0,
                    "class2": 109025.0
                },
                "ens_id": "ENSMUSG00000037742",
                "uniprot_ids": [
                    "P10126",
                    "D3YZ68",
                    "D3Z3I8"
                ]
            },
            "ENSMUSG00000059070": {
                "id": "ENSMUSG00000059070",
                "type": "Gene",
                "symbol": "Rpl18",
                "source": [
                    "ENSMUSG00000057098",
                    "ENSMUSG00000048387",
                    "ENSMUSG00000000247"
                ],
                "target": [],
                "expression_sum": {
                    "class1": 68248.0,
                    "class2": 37919.0
                },
                "ens_id": "ENSMUSG00000059070",
                "uniprot_ids": [
                    "P35980",
                    "A0A1B0GSF7",
                    "A0A1B0GSS8",
                    "A0A1B0GS28",
                    "A0A1B0GSA8",
                    "A0A1B0GRZ3",
                    "A0A1B0GQU8"
                ]
            },
            "ENSMUSG00000003970": {
                "id": "ENSMUSG00000003970",
                "type": "Gene",
                "symbol": "Rpl8",
                "source": [
                    "ENSMUSG00000047638",
                    "ENSMUSG00000000247",
                    "ENSMUSG00000038193",
                    "ENSMUSG00000031965",
                    "ENSMUSG00000048387"
                ],
                "target": [],
                "expression_sum": {
                    "class1": 111387.0,
                    "class2": 56751.0
                },
                "ens_id": "ENSMUSG00000003970",
                "uniprot_ids": [
                    "P62918"
                ]
            },
            "ENSMUSG00000060938": {
                "id": "ENSMUSG00000060938",
                "type": "Gene",
                "symbol": "Rpl26",
                "source": [
                    "ENSMUSG00000023034",
                    "ENSMUSG00000000247",
                    "ENSMUSG00000057098",
                    "ENSMUSG00000038193",
                    "ENSMUSG00000076431",
                    "ENSMUSG00000031965",
                    "ENSMUSG00000048387"
                ],
                "target": [],
                "expression_sum": {
                    "class1": 124210.0,
                    "class2": 63185.0
                },
                "ens_id": "ENSMUSG00000060938",
                "uniprot_ids": [
                    "P61255",
                    "B1ARA3",
                    "B1ARA5"
                ]
            },
            "ENSMUSG00000074129": {
                "id": "ENSMUSG00000074129",
                "type": "Gene",
                "symbol": "Rpl13a",
                "source": [
                    "ENSMUSG00000023034",
                    "ENSMUSG00000047638",
                    "ENSMUSG00000000247",
                    "ENSMUSG00000057098",
                    "ENSMUSG00000038193",
                    "ENSMUSG00000048387"
                ],
                "target": [],
                "expression_sum": {
                    "class1": 181504.0,
                    "class2": 83293.0
                },
                "ens_id": "ENSMUSG00000074129",
                "uniprot_ids": [
                    "P19253",
                    "A0A1B0GSQ6",
                    "A0A1B0GTA1",
                    "A0A1B0GSC2",
                    "A0A1B0GRH1",
                    "A0A1B0GT00",
                    "A0A1B0GSL5",
                    "A0A1B0GQW6",
                    "A0A1B0GSF0",
                    "A0A1B0GSJ2",
                    "A0A1B0GSB2"
                ]
            },
            "ENSMUSG00000031320": {
                "id": "ENSMUSG00000031320",
                "type": "Gene",
                "symbol": "Rps4x",
                "source": [
                    "ENSMUSG00000056501",
                    "ENSMUSG00000023034",
                    "ENSMUSG00000038193",
                    "ENSMUSG00000005583",
                    "ENSMUSG00000048387"
                ],
                "target": [],
                "expression_sum": {
                    "class1": 147664.0,
                    "class2": 77702.0
                },
                "ens_id": "ENSMUSG00000031320",
                "uniprot_ids": [
                    "P62702"
                ]
            },
            "ENSMUSG00000090862": {
                "id": "ENSMUSG00000090862",
                "type": "Gene",
                "symbol": "Rps13",
                "source": [
                    "ENSMUSG00000022528",
                    "ENSMUSG00000000247",
                    "ENSMUSG00000057098",
                    "ENSMUSG00000005583",
                    "ENSMUSG00000048387"
                ],
                "target": [],
                "expression_sum": {
                    "class1": 84526.0,
                    "class2": 40637.0
                },
                "ens_id": "ENSMUSG00000090862",
                "uniprot_ids": [
                    "P62301",
                    "Q921R2",
                    "A0A0U1RQ71"
                ]
            },
            "ENSMUSG00000020585": {
                "id": "ENSMUSG00000020585",
                "type": "Gene",
                "symbol": "Laptm4a",
                "source": [
                    "ENSMUSG00000032035",
                    "ENSMUSG00000052684",
                    "ENSMUSG00000048387",
                    "ENSMUSG00000057098",
                    "ENSMUSG00000038193",
                    "ENSMUSG00000029322",
                    "ENSMUSG00000003032",
                    "ENSMUSG00000005583",
                    "ENSMUSG00000028565",
                    "ENSMUSG00000071076"
                ],
                "target": [],
                "expression_sum": {
                    "class1": 51795.0,
                    "class2": 34765.0
                },
                "ens_id": "ENSMUSG00000020585",
                "uniprot_ids": [
                    "E9QLR3"
                ]
            },
            "ENSMUSG00000005583": {
                "id": "ENSMUSG00000005583",
                "type": "TF",
                "symbol": "Mef2c",
                "source": [
                    "ENSMUSG00000055435",
                    "ENSMUSG00000032238",
                    "ENSMUSG00000022528",
                    "ENSMUSG00000071076"
                ],
                "target": [
                    "ENSMUSG00000031431",
                    "ENSMUSG00000040723",
                    "ENSMUSG00000062515",
                    "ENSMUSG00000031447",
                    "ENSMUSG00000025381",
                    "ENSMUSG00000032518",
                    "ENSMUSG00000007655",
                    "ENSMUSG00000025213",
                    "ENSMUSG00000053007",
                    "ENSMUSG00000026879",
                    "ENSMUSG00000032330",
                    "ENSMUSG00000014599",
                    "ENSMUSG00000016256",
                    "ENSMUSG00000027447",
                    "ENSMUSG00000064373",
                    "ENSMUSG00000025856",
                    "ENSMUSG00000015354",
                    "ENSMUSG00000042613",
                    "ENSMUSG00000000325",
                    "ENSMUSG00000008668",
                    "ENSMUSG00000078974",
                    "ENSMUSG00000018567",
                    "ENSMUSG00000028005",
                    "ENSMUSG00000031520",
                    "ENSMUSG00000071415",
                    "ENSMUSG00000030605",
                    "ENSMUSG00000008575",
                    "ENSMUSG00000046516",
                    "ENSMUSG00000053317",
                    "ENSMUSG00000075701",
                    "ENSMUSG00000022836",
                    "ENSMUSG00000031502",
                    "ENSMUSG00000020467",
                    "ENSMUSG00000048960",
                    "ENSMUSG00000021390",
                    "ENSMUSG00000002900",
                    "ENSMUSG00000029616",
                    "ENSMUSG00000036570",
                    "ENSMUSG00000026365",
                    "ENSMUSG00000036545",
                    "ENSMUSG00000024140",
                    "ENSMUSG00000017707",
                    "ENSMUSG00000034807",
                    "ENSMUSG00000019907",
                    "ENSMUSG00000014956",
                    "ENSMUSG00000000184",
                    "ENSMUSG00000020737",
                    "ENSMUSG00000090862",
                    "ENSMUSG00000030824",
                    "ENSMUSG00000027642",
                    "ENSMUSG00000036478",
                    "ENSMUSG00000031616",
                    "ENSMUSG00000003617",
                    "ENSMUSG00000032271",
                    "ENSMUSG00000033585",
                    "ENSMUSG00000027706",
                    "ENSMUSG00000025290",
                    "ENSMUSG00000027800",
                    "ENSMUSG00000031586",
                    "ENSMUSG00000041828",
                    "ENSMUSG00000022587",
                    "ENSMUSG00000045128",
                    "ENSMUSG00000017466",
                    "ENSMUSG00000026536",
                    "ENSMUSG00000072235",
                    "ENSMUSG00000024277",
                    "ENSMUSG00000006333",
                    "ENSMUSG00000038463",
                    "ENSMUSG00000002985",
                    "ENSMUSG00000068798",
                    "ENSMUSG00000028517",
                    "ENSMUSG00000016494",
                    "ENSMUSG00000020029",
                    "ENSMUSG00000062210",
                    "ENSMUSG00000024290",
                    "ENSMUSG00000026674",
                    "ENSMUSG00000075705",
                    "ENSMUSG00000012848",
                    "ENSMUSG00000015094",
                    "ENSMUSG00000063229",
                    "ENSMUSG00000022665",
                    "ENSMUSG00000027074",
                    "ENSMUSG00000021871",
                    "ENSMUSG00000024659",
                    "ENSMUSG00000024085",
                    "ENSMUSG00000024338",
                    "ENSMUSG00000027712",
                    "ENSMUSG00000007891",
                    "ENSMUSG00000024048",
                    "ENSMUSG00000028673",
                    "ENSMUSG00000034708",
                    "ENSMUSG00000001270",
                    "ENSMUSG00000050708",
                    "ENSMUSG00000026478",
                    "ENSMUSG00000019920",
                    "ENSMUSG00000017493",
                    "ENSMUSG00000025351",
                    "ENSMUSG00000074884",
                    "ENSMUSG00000055435",
                    "ENSMUSG00000001020",
                    "ENSMUSG00000034993",
                    "ENSMUSG00000021200",
                    "ENSMUSG00000027248",
                    "ENSMUSG00000052957",
                    "ENSMUSG00000051579",
                    "ENSMUSG00000004460",
                    "ENSMUSG00000026864",
                    "ENSMUSG00000031548",
                    "ENSMUSG00000028757",
                    "ENSMUSG00000073411",
                    "ENSMUSG00000024012",
                    "ENSMUSG00000009876",
                    "ENSMUSG00000029661",
                    "ENSMUSG00000060802",
                    "ENSMUSG00000040659",
                    "ENSMUSG00000017969",
                    "ENSMUSG00000004085",
                    "ENSMUSG00000018293",
                    "ENSMUSG00000035783",
                    "ENSMUSG00000014867",
                    "ENSMUSG00000038900",
                    "ENSMUSG00000006360",
                    "ENSMUSG00000059291",
                    "ENSMUSG00000031320",
                    "ENSMUSG00000009281",
                    "ENSMUSG00000022371",
                    "ENSMUSG00000036438",
                    "ENSMUSG00000029467",
                    "ENSMUSG00000023175",
                    "ENSMUSG00000018339",
                    "ENSMUSG00000035493",
                    "ENSMUSG00000002107",
                    "ENSMUSG00000073889",
                    "ENSMUSG00000022010",
                    "ENSMUSG00000032231",
                    "ENSMUSG00000007872",
                    "ENSMUSG00000027848",
                    "ENSMUSG00000073557",
                    "ENSMUSG00000041329",
                    "ENSMUSG00000029426",
                    "ENSMUSG00000029718",
                    "ENSMUSG00000057666",
                    "ENSMUSG00000028600",
                    "ENSMUSG00000047420",
                    "ENSMUSG00000020321",
                    "ENSMUSG00000025491",
                    "ENSMUSG00000021756",
                    "ENSMUSG00000024245",
                    "ENSMUSG00000036446",
                    "ENSMUSG00000031538",
                    "ENSMUSG00000019978",
                    "ENSMUSG00000022816",
                    "ENSMUSG00000026482",
                    "ENSMUSG00000031636",
                    "ENSMUSG00000002980",
                    "ENSMUSG00000035885",
                    "ENSMUSG00000030403",
                    "ENSMUSG00000007039",
                    "ENSMUSG00000029781",
                    "ENSMUSG00000026193",
                    "ENSMUSG00000022090",
                    "ENSMUSG00000027834",
                    "ENSMUSG00000032014",
                    "ENSMUSG00000038146",
                    "ENSMUSG00000068735",
                    "ENSMUSG00000029061",
                    "ENSMUSG00000040249",
                    "ENSMUSG00000032334",
                    "ENSMUSG00000023092",
                    "ENSMUSG00000029070",
                    "ENSMUSG00000023367",
                    "ENSMUSG00000022324",
                    "ENSMUSG00000023886",
                    "ENSMUSG00000027276",
                    "ENSMUSG00000005397",
                    "ENSMUSG00000030208",
                    "ENSMUSG00000022817",
                    "ENSMUSG00000030536",
                    "ENSMUSG00000063694",
                    "ENSMUSG00000003420",
                    "ENSMUSG00000020315",
                    "ENSMUSG00000058569",
                    "ENSMUSG00000018217",
                    "ENSMUSG00000023046",
                    "ENSMUSG00000058056",
                    "ENSMUSG00000064080",
                    "ENSMUSG00000005973",
                    "ENSMUSG00000042745",
                    "ENSMUSG00000001520",
                    "ENSMUSG00000016534",
                    "ENSMUSG00000003161",
                    "ENSMUSG00000032359",
                    "ENSMUSG00000041577",
                    "ENSMUSG00000001911",
                    "ENSMUSG00000022136",
                    "ENSMUSG00000018819",
                    "ENSMUSG00000003814",
                    "ENSMUSG00000061232",
                    "ENSMUSG00000027861",
                    "ENSMUSG00000002059",
                    "ENSMUSG00000020205",
                    "ENSMUSG00000037072",
                    "ENSMUSG00000020009",
                    "ENSMUSG00000004207",
                    "ENSMUSG00000020088",
                    "ENSMUSG00000026728",
                    "ENSMUSG00000096145",
                    "ENSMUSG00000042312",
                    "ENSMUSG00000062997",
                    "ENSMUSG00000032997",
                    "ENSMUSG00000025232",
                    "ENSMUSG00000067288",
                    "ENSMUSG00000075254",
                    "ENSMUSG00000031799",
                    "ENSMUSG00000055053",
                    "ENSMUSG00000041084",
                    "ENSMUSG00000022971",
                    "ENSMUSG00000025393",
                    "ENSMUSG00000021242",
                    "ENSMUSG00000074170",
                    "ENSMUSG00000002014",
                    "ENSMUSG00000014313",
                    "ENSMUSG00000032547",
                    "ENSMUSG00000092341",
                    "ENSMUSG00000026688",
                    "ENSMUSG00000087006",
                    "ENSMUSG00000032238",
                    "ENSMUSG00000021843",
                    "ENSMUSG00000062328",
                    "ENSMUSG00000006800",
                    "ENSMUSG00000020585",
                    "ENSMUSG00000032353",
                    "ENSMUSG00000025492",
                    "ENSMUSG00000046402",
                    "ENSMUSG00000055447",
                    "ENSMUSG00000057841",
                    "ENSMUSG00000046718",
                    "ENSMUSG00000027408",
                    "ENSMUSG00000020840",
                    "ENSMUSG00000028234",
                    "ENSMUSG00000037206",
                    "ENSMUSG00000015932",
                    "ENSMUSG00000060703",
                    "ENSMUSG00000029231",
                    "ENSMUSG00000073489",
                    "ENSMUSG00000020902",
                    "ENSMUSG00000035258",
                    "ENSMUSG00000001025",
                    "ENSMUSG00000019539",
                    "ENSMUSG00000032925",
                    "ENSMUSG00000031765",
                    "ENSMUSG00000019916",
                    "ENSMUSG00000045636",
                    "ENSMUSG00000040952",
                    "ENSMUSG00000022528",
                    "ENSMUSG00000032575",
                    "ENSMUSG00000079111",
                    "ENSMUSG00000042485",
                    "ENSMUSG00000030659",
                    "ENSMUSG00000012422",
                    "ENSMUSG00000025130",
                    "ENSMUSG00000020612",
                    "ENSMUSG00000038872",
                    "ENSMUSG00000056737",
                    "ENSMUSG00000020928",
                    "ENSMUSG00000061983",
                    "ENSMUSG00000022032",
                    "ENSMUSG00000076432",
                    "ENSMUSG00000038086",
                    "ENSMUSG00000048096",
                    "ENSMUSG00000028461",
                    "ENSMUSG00000059439",
                    "ENSMUSG00000034165",
                    "ENSMUSG00000043091",
                    "ENSMUSG00000032085",
                    "ENSMUSG00000022892",
                    "ENSMUSG00000031626",
                    "ENSMUSG00000029810",
                    "ENSMUSG00000070436",
                    "ENSMUSG00000015133",
                    "ENSMUSG00000027188",
                    "ENSMUSG00000027204",
                    "ENSMUSG00000020460",
                    "ENSMUSG00000028464",
                    "ENSMUSG00000028776",
                    "ENSMUSG00000037060",
                    "ENSMUSG00000021253",
                    "ENSMUSG00000053477",
                    "ENSMUSG00000016382",
                    "ENSMUSG00000062647",
                    "ENSMUSG00000046330",
                    "ENSMUSG00000079523",
                    "ENSMUSG00000059895",
                    "ENSMUSG00000073418",
                    "ENSMUSG00000046727",
                    "ENSMUSG00000016194",
                    "ENSMUSG00000038991",
                    "ENSMUSG00000060591",
                    "ENSMUSG00000023942",
                    "ENSMUSG00000049517",
                    "ENSMUSG00000038530",
                    "ENSMUSG00000024371",
                    "ENSMUSG00000069662",
                    "ENSMUSG00000021876",
                    "ENSMUSG00000024164",
                    "ENSMUSG00000006356",
                    "ENSMUSG00000020473",
                    "ENSMUSG00000001663",
                    "ENSMUSG00000023224",
                    "ENSMUSG00000026223",
                    "ENSMUSG00000049086",
                    "ENSMUSG00000027907",
                    "ENSMUSG00000067274",
                    "ENSMUSG00000049892",
                    "ENSMUSG00000069833",
                    "ENSMUSG00000033910",
                    "ENSMUSG00000039168",
                    "ENSMUSG00000020484",
                    "ENSMUSG00000015568",
                    "ENSMUSG00000040462",
                    "ENSMUSG00000066705",
                    "ENSMUSG00000036427",
                    "ENSMUSG00000028618",
                    "ENSMUSG00000026748",
                    "ENSMUSG00000047907",
                    "ENSMUSG00000023064",
                    "ENSMUSG00000028936",
                    "ENSMUSG00000036437",
                    "ENSMUSG00000022912",
                    "ENSMUSG00000025579",
                    "ENSMUSG00000079037",
                    "ENSMUSG00000079017",
                    "ENSMUSG00000052997",
                    "ENSMUSG00000000093",
                    "ENSMUSG00000058135",
                    "ENSMUSG00000040652",
                    "ENSMUSG00000029761",
                    "ENSMUSG00000028111",
                    "ENSMUSG00000021477",
                    "ENSMUSG00000028949",
                    "ENSMUSG00000097328",
                    "ENSMUSG00000027422",
                    "ENSMUSG00000020644",
                    "ENSMUSG00000056481",
                    "ENSMUSG00000025647",
                    "ENSMUSG00000029287",
                    "ENSMUSG00000031556",
                    "ENSMUSG00000021877",
                    "ENSMUSG00000034868",
                    "ENSMUSG00000026043",
                    "ENSMUSG00000021939",
                    "ENSMUSG00000041417",
                    "ENSMUSG00000008682",
                    "ENSMUSG00000020402",
                    "ENSMUSG00000001240",
                    "ENSMUSG00000014329",
                    "ENSMUSG00000040713",
                    "ENSMUSG00000034892",
                    "ENSMUSG00000028369",
                    "ENSMUSG00000060572",
                    "ENSMUSG00000012428",
                    "ENSMUSG00000029309",
                    "ENSMUSG00000061353",
                    "ENSMUSG00000031595",
                    "ENSMUSG00000021943",
                    "ENSMUSG00000009681",
                    "ENSMUSG00000060904",
                    "ENSMUSG00000064370",
                    "ENSMUSG00000020053",
                    "ENSMUSG00000040666",
                    "ENSMUSG00000026042",
                    "ENSMUSG00000067212",
                    "ENSMUSG00000020440",
                    "ENSMUSG00000039007",
                    "ENSMUSG00000032366",
                    "ENSMUSG00000001119",
                    "ENSMUSG00000075010",
                    "ENSMUSG00000029390",
                    "ENSMUSG00000059173",
                    "ENSMUSG00000029767",
                    "ENSMUSG00000031246",
                    "ENSMUSG00000090000",
                    "ENSMUSG00000021760",
                    "ENSMUSG00000032744",
                    "ENSMUSG00000048546",
                    "ENSMUSG00000021248",
                    "ENSMUSG00000001506",
                    "ENSMUSG00000021870",
                    "ENSMUSG00000041959",
                    "ENSMUSG00000017754",
                    "ENSMUSG00000004791",
                    "ENSMUSG00000020814",
                    "ENSMUSG00000053279",
                    "ENSMUSG00000033880",
                    "ENSMUSG00000025794",
                    "ENSMUSG00000038145",
                    "ENSMUSG00000015085",
                    "ENSMUSG00000020486",
                    "ENSMUSG00000090733",
                    "ENSMUSG00000040212",
                    "ENSMUSG00000050666",
                    "ENSMUSG00000032946",
                    "ENSMUSG00000001123",
                    "ENSMUSG00000037966",
                    "ENSMUSG00000016559",
                    "ENSMUSG00000055172",
                    "ENSMUSG00000060126",
                    "ENSMUSG00000026014",
                    "ENSMUSG00000037706",
                    "ENSMUSG00000001175",
                    "ENSMUSG00000032387",
                    "ENSMUSG00000040562",
                    "ENSMUSG00000032135",
                    "ENSMUSG00000019139",
                    "ENSMUSG00000030272",
                    "ENSMUSG00000025428",
                    "ENSMUSG00000062006",
                    "ENSMUSG00000052727",
                    "ENSMUSG00000078202",
                    "ENSMUSG00000027230",
                    "ENSMUSG00000020571",
                    "ENSMUSG00000002732",
                    "ENSMUSG00000025147",
                    "ENSMUSG00000006205",
                    "ENSMUSG00000051855",
                    "ENSMUSG00000006519",
                    "ENSMUSG00000045658",
                    "ENSMUSG00000029076",
                    "ENSMUSG00000030717",
                    "ENSMUSG00000049775",
                    "ENSMUSG00000039601",
                    "ENSMUSG00000024150",
                    "ENSMUSG00000029359",
                    "ENSMUSG00000067818",
                    "ENSMUSG00000024308",
                    "ENSMUSG00000046318",
                    "ENSMUSG00000024538",
                    "ENSMUSG00000090841",
                    "ENSMUSG00000041736",
                    "ENSMUSG00000005881",
                    "ENSMUSG00000016319",
                    "ENSMUSG00000004105",
                    "ENSMUSG00000038370",
                    "ENSMUSG00000020048",
                    "ENSMUSG00000042747",
                    "ENSMUSG00000041355",
                    "ENSMUSG00000000740",
                    "ENSMUSG00000007892",
                    "ENSMUSG00000029096",
                    "ENSMUSG00000021186",
                    "ENSMUSG00000026837",
                    "ENSMUSG00000032383",
                    "ENSMUSG00000079641",
                    "ENSMUSG00000022048",
                    "ENSMUSG00000006369",
                    "ENSMUSG00000031422",
                    "ENSMUSG00000029622",
                    "ENSMUSG00000001435",
                    "ENSMUSG00000039286",
                    "ENSMUSG00000004044",
                    "ENSMUSG00000022708",
                    "ENSMUSG00000034118",
                    "ENSMUSG00000070348",
                    "ENSMUSG00000019929",
                    "ENSMUSG00000029659",
                    "ENSMUSG00000029641",
                    "ENSMUSG00000026574",
                    "ENSMUSG00000018920",
                    "ENSMUSG00000016458",
                    "ENSMUSG00000008540",
                    "ENSMUSG00000021268",
                    "ENSMUSG00000018830",
                    "ENSMUSG00000090698",
                    "ENSMUSG00000075702",
                    "ENSMUSG00000004951",
                    "ENSMUSG00000057530"
                ],
                "expression_sum": {
                    "class1": 2579.0,
                    "class2": 9123.0
                },
                "ens_id": "ENSMUSG00000005583",
                "uniprot_ids": [
                    "Q8CFN5",
                    "A0A0G2JDK0",
                    "A0A0G2JEC2",
                    "A0A0G2JFL7",
                    "A0A0H2UH28",
                    "A0A0G2JFX2",
                    "A0A0G2JEC3",
                    "A0A0G2JFI7",
                    "A0A0H2UKB6",
                    "A0A0G2JFX1",
                    "A0A0G2JGK3",
                    "A0A0G2JE18",
                    "A0A0G2JGL3",
                    "A0A0G2JE38",
                    "A0A0G2JFM7",
                    "A0A0G2JFS4",
                    "A0A0G2JET3",
                    "A0A0G2JES4",
                    "A0A0G2JEM8",
                    "A0A0G2JDQ1",
                    "A0A0G2JDT0",
                    "A0A0G2JF80"
                ]
            },
            "ENSMUSG00000059895": {
                "id": "ENSMUSG00000059895",
                "type": "Gene",
                "symbol": "Ptp4a3",
                "source": [
                    "ENSMUSG00000023034",
                    "ENSMUSG00000058756",
                    "ENSMUSG00000022528",
                    "ENSMUSG00000005583",
                    "ENSMUSG00000020484"
                ],
                "target": [],
                "expression_sum": {
                    "class1": 1549.0,
                    "class2": 20820.0
                },
                "ens_id": "ENSMUSG00000059895",
                "uniprot_ids": [
                    "Q9D658",
                    "A0A2R8VHQ8",
                    "E9PXS4"
                ]
            },
            "ENSMUSG00000019907": {
                "id": "ENSMUSG00000019907",
                "type": "Gene",
                "symbol": "Ppp1r12a",
                "source": [
                    "ENSMUSG00000052684",
                    "ENSMUSG00000058756",
                    "ENSMUSG00000022346",
                    "ENSMUSG00000047638",
                    "ENSMUSG00000005583",
                    "ENSMUSG00000038418",
                    "ENSMUSG00000053477",
                    "ENSMUSG00000008575",
                    "ENSMUSG00000055435",
                    "ENSMUSG00000071076"
                ],
                "target": [],
                "expression_sum": {
                    "class1": 2417.0,
                    "class2": 7901.0
                },
                "ens_id": "ENSMUSG00000019907",
                "uniprot_ids": [
                    "Q9DBR7",
                    "A0A1W2P750"
                ]
            },
            "ENSMUSG00000002107": {
                "id": "ENSMUSG00000002107",
                "type": "Gene",
                "symbol": "Celf2",
                "source": [
                    "ENSMUSG00000032238",
                    "ENSMUSG00000052684",
                    "ENSMUSG00000022346",
                    "ENSMUSG00000057098",
                    "ENSMUSG00000038418",
                    "ENSMUSG00000053477",
                    "ENSMUSG00000003032",
                    "ENSMUSG00000005583",
                    "ENSMUSG00000028565",
                    "ENSMUSG00000071076"
                ],
                "target": [],
                "expression_sum": {
                    "class1": 1715.0,
                    "class2": 5708.0
                },
                "ens_id": "ENSMUSG00000002107",
                "uniprot_ids": [
                    "Q9Z0H4",
                    "A0A0R4J2B0",
                    "E9QA47",
                    "S4R2U7",
                    "S4R2U1",
                    "S4R2J2",
                    "V9GX43",
                    "A3KGT0",
                    "S4R2L5",
                    "S4R1S7",
                    "S4R2S7"
                ]
            },
            "ENSMUSG00000037868": {
                "id": "ENSMUSG00000037868",
                "type": "TF",
                "symbol": "Egr2",
                "source": [
                    "ENSMUSG00000056501",
                    "ENSMUSG00000052684",
                    "ENSMUSG00000023034",
                    "ENSMUSG00000052837",
                    "ENSMUSG00000026628",
                    "ENSMUSG00000038418",
                    "ENSMUSG00000021250",
                    "ENSMUSG00000003032",
                    "ENSMUSG00000071076",
                    "ENSMUSG00000018899"
                ],
                "target": [
                    "ENSMUSG00000023034",
                    "ENSMUSG00000053113",
                    "ENSMUSG00000003545",
                    "ENSMUSG00000033538",
                    "ENSMUSG00000052684",
                    "ENSMUSG00000021025",
                    "ENSMUSG00000015312",
                    "ENSMUSG00000008384",
                    "ENSMUSG00000021250",
                    "ENSMUSG00000003032",
                    "ENSMUSG00000055148",
                    "ENSMUSG00000021127",
                    "ENSMUSG00000015501",
                    "ENSMUSG00000040128",
                    "ENSMUSG00000035356",
                    "ENSMUSG00000052837",
                    "ENSMUSG00000026628",
                    "ENSMUSG00000020423",
                    "ENSMUSG00000097296",
                    "ENSMUSG00000092274",
                    "ENSMUSG00000007872",
                    "ENSMUSG00000044786",
                    "ENSMUSG00000018899",
                    "ENSMUSG00000027829",
                    "ENSMUSG00000006403",
                    "ENSMUSG00000028214",
                    "ENSMUSG00000032501",
                    "ENSMUSG00000035828",
                    "ENSMUSG00000038418",
                    "ENSMUSG00000009681"
                ],
                "expression_sum": {
                    "class1": 3930.0,
                    "class2": 381.0
                },
                "ens_id": "ENSMUSG00000037868",
                "uniprot_ids": [
                    "P08152",
                    "D3Z4G7",
                    "D3Z4X9",
                    "D3YYY2",
                    "D3Z3K7",
                    "D3YXS2"
                ]
            },
            "ENSMUSG00000021250": {
                "id": "ENSMUSG00000021250",
                "type": "TF",
                "symbol": "Fos",
                "source": [
                    "ENSMUSG00000041515",
                    "ENSMUSG00000056501",
                    "ENSMUSG00000052684",
                    "ENSMUSG00000037868",
                    "ENSMUSG00000023034",
                    "ENSMUSG00000052837",
                    "ENSMUSG00000030103",
                    "ENSMUSG00000000247",
                    "ENSMUSG00000042812",
                    "ENSMUSG00000057098",
                    "ENSMUSG00000003545",
                    "ENSMUSG00000003032",
                    "ENSMUSG00000018899",
                    "ENSMUSG00000071076"
                ],
                "target": [
                    "ENSMUSG00000019505",
                    "ENSMUSG00000006360",
                    "ENSMUSG00000031765",
                    "ENSMUSG00000024561",
                    "ENSMUSG00000030222",
                    "ENSMUSG00000028691",
                    "ENSMUSG00000003545",
                    "ENSMUSG00000036199",
                    "ENSMUSG00000025289",
                    "ENSMUSG00000032330",
                    "ENSMUSG00000030930",
                    "ENSMUSG00000004665",
                    "ENSMUSG00000053560",
                    "ENSMUSG00000036040",
                    "ENSMUSG00000037411",
                    "ENSMUSG00000036438",
                    "ENSMUSG00000006699",
                    "ENSMUSG00000008384",
                    "ENSMUSG00000055148",
                    "ENSMUSG00000018339",
                    "ENSMUSG00000035493",
                    "ENSMUSG00000021127",
                    "ENSMUSG00000023004",
                    "ENSMUSG00000029380",
                    "ENSMUSG00000002416",
                    "ENSMUSG00000007872",
                    "ENSMUSG00000009013",
                    "ENSMUSG00000036655",
                    "ENSMUSG00000009681",
                    "ENSMUSG00000036570",
                    "ENSMUSG00000055839",
                    "ENSMUSG00000075706",
                    "ENSMUSG00000020053",
                    "ENSMUSG00000037999",
                    "ENSMUSG00000069601",
                    "ENSMUSG00000032085",
                    "ENSMUSG00000053113",
                    "ENSMUSG00000037266",
                    "ENSMUSG00000033538",
                    "ENSMUSG00000084786",
                    "ENSMUSG00000021520",
                    "ENSMUSG00000052684",
                    "ENSMUSG00000030122",
                    "ENSMUSG00000055681",
                    "ENSMUSG00000031762",
                    "ENSMUSG00000003038",
                    "ENSMUSG00000015501",
                    "ENSMUSG00000038690",
                    "ENSMUSG00000023915",
                    "ENSMUSG00000030103",
                    "ENSMUSG00000042812",
                    "ENSMUSG00000028464",
                    "ENSMUSG00000032515",
                    "ENSMUSG00000074457",
                    "ENSMUSG00000027829",
                    "ENSMUSG00000037060",
                    "ENSMUSG00000028195",
                    "ENSMUSG00000031375",
                    "ENSMUSG00000021253",
                    "ENSMUSG00000005873",
                    "ENSMUSG00000038463",
                    "ENSMUSG00000002985",
                    "ENSMUSG00000035678",
                    "ENSMUSG00000059534",
                    "ENSMUSG00000072625",
                    "ENSMUSG00000015085",
                    "ENSMUSG00000068220",
                    "ENSMUSG00000023034",
                    "ENSMUSG00000030654",
                    "ENSMUSG00000054364",
                    "ENSMUSG00000016494",
                    "ENSMUSG00000056708",
                    "ENSMUSG00000038393",
                    "ENSMUSG00000037706",
                    "ENSMUSG00000046727",
                    "ENSMUSG00000001175",
                    "ENSMUSG00000032359",
                    "ENSMUSG00000061518",
                    "ENSMUSG00000019960",
                    "ENSMUSG00000015312",
                    "ENSMUSG00000003032",
                    "ENSMUSG00000037211",
                    "ENSMUSG00000030247",
                    "ENSMUSG00000020423",
                    "ENSMUSG00000024048",
                    "ENSMUSG00000023089",
                    "ENSMUSG00000021876",
                    "ENSMUSG00000037868",
                    "ENSMUSG00000022122",
                    "ENSMUSG00000006205",
                    "ENSMUSG00000035828",
                    "ENSMUSG00000057098",
                    "ENSMUSG00000064215",
                    "ENSMUSG00000018286",
                    "ENSMUSG00000017493",
                    "ENSMUSG00000039601",
                    "ENSMUSG00000067818",
                    "ENSMUSG00000056501",
                    "ENSMUSG00000037152",
                    "ENSMUSG00000024014",
                    "ENSMUSG00000000628",
                    "ENSMUSG00000090841",
                    "ENSMUSG00000000753",
                    "ENSMUSG00000071866",
                    "ENSMUSG00000028864",
                    "ENSMUSG00000036751",
                    "ENSMUSG00000038370",
                    "ENSMUSG00000001833",
                    "ENSMUSG00000026385",
                    "ENSMUSG00000024190",
                    "ENSMUSG00000041515",
                    "ENSMUSG00000007097",
                    "ENSMUSG00000035356",
                    "ENSMUSG00000052837",
                    "ENSMUSG00000047945",
                    "ENSMUSG00000073411",
                    "ENSMUSG00000029632",
                    "ENSMUSG00000030342",
                    "ENSMUSG00000022956",
                    "ENSMUSG00000044786",
                    "ENSMUSG00000008540",
                    "ENSMUSG00000018899",
                    "ENSMUSG00000075702",
                    "ENSMUSG00000027004",
                    "ENSMUSG00000006403",
                    "ENSMUSG00000018293",
                    "ENSMUSG00000000247",
                    "ENSMUSG00000038418",
                    "ENSMUSG00000035783",
                    "ENSMUSG00000030849",
                    "ENSMUSG00000058135"
                ],
                "expression_sum": {
                    "class1": 165566.0,
                    "class2": 132750.0
                },
                "ens_id": "ENSMUSG00000021250",
                "uniprot_ids": [
                    "P01101"
                ]
            },
            "ENSMUSG00000053560": {
                "id": "ENSMUSG00000053560",
                "type": "Gene",
                "symbol": "Ier2",
                "source": [
                    "ENSMUSG00000052684",
                    "ENSMUSG00000026628",
                    "ENSMUSG00000000247",
                    "ENSMUSG00000038193",
                    "ENSMUSG00000021250",
                    "ENSMUSG00000003545",
                    "ENSMUSG00000055148",
                    "ENSMUSG00000018899"
                ],
                "target": [],
                "expression_sum": {
                    "class1": 33337.0,
                    "class2": 23871.0
                },
                "ens_id": "ENSMUSG00000053560",
                "uniprot_ids": [
                    "P17950"
                ]
            },
            "ENSMUSG00000052684": {
                "id": "ENSMUSG00000052684",
                "type": "TF",
                "symbol": "Jun",
                "source": [
                    "ENSMUSG00000055435",
                    "ENSMUSG00000056501",
                    "ENSMUSG00000037868",
                    "ENSMUSG00000023034",
                    "ENSMUSG00000022528",
                    "ENSMUSG00000052837",
                    "ENSMUSG00000026628",
                    "ENSMUSG00000000247",
                    "ENSMUSG00000038418",
                    "ENSMUSG00000052534",
                    "ENSMUSG00000021250",
                    "ENSMUSG00000038193",
                    "ENSMUSG00000003545",
                    "ENSMUSG00000008575",
                    "ENSMUSG00000018899"
                ],
                "target": [
                    "ENSMUSG00000031431",
                    "ENSMUSG00000040723",
                    "ENSMUSG00000029761",
                    "ENSMUSG00000025381",
                    "ENSMUSG00000032925",
                    "ENSMUSG00000091345",
                    "ENSMUSG00000028111",
                    "ENSMUSG00000030222",
                    "ENSMUSG00000021477",
                    "ENSMUSG00000028691",
                    "ENSMUSG00000003545",
                    "ENSMUSG00000022365",
                    "ENSMUSG00000025213",
                    "ENSMUSG00000025289",
                    "ENSMUSG00000026879",
                    "ENSMUSG00000009281",
                    "ENSMUSG00000053560",
                    "ENSMUSG00000037411",
                    "ENSMUSG00000064373",
                    "ENSMUSG00000022528",
                    "ENSMUSG00000034868",
                    "ENSMUSG00000042613",
                    "ENSMUSG00000006699",
                    "ENSMUSG00000000325",
                    "ENSMUSG00000027239",
                    "ENSMUSG00000008384",
                    "ENSMUSG00000026043",
                    "ENSMUSG00000044627",
                    "ENSMUSG00000079111",
                    "ENSMUSG00000055148",
                    "ENSMUSG00000042485",
                    "ENSMUSG00000018339",
                    "ENSMUSG00000041417",
                    "ENSMUSG00000012422",
                    "ENSMUSG00000078974",
                    "ENSMUSG00000023004",
                    "ENSMUSG00000026421",
                    "ENSMUSG00000002107",
                    "ENSMUSG00000073889",
                    "ENSMUSG00000014329",
                    "ENSMUSG00000031007",
                    "ENSMUSG00000022010",
                    "ENSMUSG00000028369",
                    "ENSMUSG00000029103",
                    "ENSMUSG00000003541",
                    "ENSMUSG00000032231",
                    "ENSMUSG00000008575",
                    "ENSMUSG00000030218",
                    "ENSMUSG00000038872",
                    "ENSMUSG00000007872",
                    "ENSMUSG00000053317",
                    "ENSMUSG00000027848",
                    "ENSMUSG00000022836",
                    "ENSMUSG00000028597",
                    "ENSMUSG00000029309",
                    "ENSMUSG00000031845",
                    "ENSMUSG00000022032",
                    "ENSMUSG00000029718",
                    "ENSMUSG00000028967",
                    "ENSMUSG00000021390",
                    "ENSMUSG00000027828",
                    "ENSMUSG00000031595",
                    "ENSMUSG00000036655",
                    "ENSMUSG00000009681",
                    "ENSMUSG00000060904",
                    "ENSMUSG00000064370",
                    "ENSMUSG00000020053",
                    "ENSMUSG00000021917",
                    "ENSMUSG00000041120",
                    "ENSMUSG00000028339",
                    "ENSMUSG00000037999",
                    "ENSMUSG00000069601",
                    "ENSMUSG00000052534",
                    "ENSMUSG00000053113",
                    "ENSMUSG00000024140",
                    "ENSMUSG00000036372",
                    "ENSMUSG00000037266",
                    "ENSMUSG00000033538",
                    "ENSMUSG00000019907",
                    "ENSMUSG00000001119",
                    "ENSMUSG00000036446",
                    "ENSMUSG00000075010",
                    "ENSMUSG00000031451",
                    "ENSMUSG00000014956",
                    "ENSMUSG00000029390",
                    "ENSMUSG00000024076",
                    "ENSMUSG00000055681",
                    "ENSMUSG00000090000",
                    "ENSMUSG00000021760",
                    "ENSMUSG00000019278",
                    "ENSMUSG00000003477",
                    "ENSMUSG00000021248",
                    "ENSMUSG00000031616",
                    "ENSMUSG00000001506",
                    "ENSMUSG00000003038",
                    "ENSMUSG00000021870",
                    "ENSMUSG00000028871",
                    "ENSMUSG00000015501",
                    "ENSMUSG00000041959",
                    "ENSMUSG00000023915",
                    "ENSMUSG00000033585",
                    "ENSMUSG00000026628",
                    "ENSMUSG00000030103",
                    "ENSMUSG00000027706",
                    "ENSMUSG00000042812",
                    "ENSMUSG00000019579",
                    "ENSMUSG00000041828",
                    "ENSMUSG00000031586",
                    "ENSMUSG00000092274",
                    "ENSMUSG00000032515",
                    "ENSMUSG00000068735",
                    "ENSMUSG00000028776",
                    "ENSMUSG00000042737",
                    "ENSMUSG00000018672",
                    "ENSMUSG00000017466",
                    "ENSMUSG00000074457",
                    "ENSMUSG00000024065",
                    "ENSMUSG00000027829",
                    "ENSMUSG00000029061",
                    "ENSMUSG00000028195",
                    "ENSMUSG00000032334",
                    "ENSMUSG00000004631",
                    "ENSMUSG00000021253",
                    "ENSMUSG00000072235",
                    "ENSMUSG00000005873",
                    "ENSMUSG00000038463",
                    "ENSMUSG00000035678",
                    "ENSMUSG00000016024",
                    "ENSMUSG00000072625",
                    "ENSMUSG00000015085",
                    "ENSMUSG00000030208",
                    "ENSMUSG00000020486",
                    "ENSMUSG00000068220",
                    "ENSMUSG00000028517",
                    "ENSMUSG00000040212",
                    "ENSMUSG00000023034",
                    "ENSMUSG00000079523",
                    "ENSMUSG00000030654",
                    "ENSMUSG00000016494",
                    "ENSMUSG00000038393",
                    "ENSMUSG00000042453",
                    "ENSMUSG00000037706",
                    "ENSMUSG00000018217",
                    "ENSMUSG00000023046",
                    "ENSMUSG00000058056",
                    "ENSMUSG00000042745",
                    "ENSMUSG00000003161",
                    "ENSMUSG00000032359",
                    "ENSMUSG00000060591",
                    "ENSMUSG00000025784",
                    "ENSMUSG00000040562",
                    "ENSMUSG00000072812",
                    "ENSMUSG00000015312",
                    "ENSMUSG00000001911",
                    "ENSMUSG00000031661",
                    "ENSMUSG00000021871",
                    "ENSMUSG00000025880",
                    "ENSMUSG00000025724",
                    "ENSMUSG00000018819",
                    "ENSMUSG00000037211",
                    "ENSMUSG00000031239",
                    "ENSMUSG00000020423",
                    "ENSMUSG00000044408",
                    "ENSMUSG00000027712",
                    "ENSMUSG00000042254",
                    "ENSMUSG00000024048",
                    "ENSMUSG00000037072",
                    "ENSMUSG00000020009",
                    "ENSMUSG00000002732",
                    "ENSMUSG00000021876",
                    "ENSMUSG00000025780",
                    "ENSMUSG00000024164",
                    "ENSMUSG00000096145",
                    "ENSMUSG00000025147",
                    "ENSMUSG00000037868",
                    "ENSMUSG00000022122",
                    "ENSMUSG00000042312",
                    "ENSMUSG00000006205",
                    "ENSMUSG00000001663",
                    "ENSMUSG00000020473",
                    "ENSMUSG00000035828",
                    "ENSMUSG00000000594",
                    "ENSMUSG00000064215",
                    "ENSMUSG00000006050",
                    "ENSMUSG00000006519",
                    "ENSMUSG00000045658",
                    "ENSMUSG00000030717",
                    "ENSMUSG00000031799",
                    "ENSMUSG00000017493",
                    "ENSMUSG00000023010",
                    "ENSMUSG00000041084",
                    "ENSMUSG00000025351",
                    "ENSMUSG00000027907",
                    "ENSMUSG00000074884",
                    "ENSMUSG00000039601",
                    "ENSMUSG00000054808",
                    "ENSMUSG00000056501",
                    "ENSMUSG00000021242",
                    "ENSMUSG00000000628",
                    "ENSMUSG00000002014",
                    "ENSMUSG00000092341",
                    "ENSMUSG00000039899",
                    "ENSMUSG00000055435",
                    "ENSMUSG00000033910",
                    "ENSMUSG00000087006",
                    "ENSMUSG00000018593",
                    "ENSMUSG00000000753",
                    "ENSMUSG00000052957",
                    "ENSMUSG00000015568",
                    "ENSMUSG00000028864",
                    "ENSMUSG00000006800",
                    "ENSMUSG00000020585",
                    "ENSMUSG00000031380",
                    "ENSMUSG00000032353",
                    "ENSMUSG00000025492",
                    "ENSMUSG00000051579",
                    "ENSMUSG00000066705",
                    "ENSMUSG00000046402",
                    "ENSMUSG00000021250",
                    "ENSMUSG00000042747",
                    "ENSMUSG00000041355",
                    "ENSMUSG00000029096",
                    "ENSMUSG00000026837",
                    "ENSMUSG00000031548",
                    "ENSMUSG00000028618",
                    "ENSMUSG00000032383",
                    "ENSMUSG00000026748",
                    "ENSMUSG00000007097",
                    "ENSMUSG00000054203",
                    "ENSMUSG00000006369",
                    "ENSMUSG00000031422",
                    "ENSMUSG00000035356",
                    "ENSMUSG00000027859",
                    "ENSMUSG00000047945",
                    "ENSMUSG00000049107",
                    "ENSMUSG00000052837",
                    "ENSMUSG00000030342",
                    "ENSMUSG00000022174",
                    "ENSMUSG00000035227",
                    "ENSMUSG00000024012",
                    "ENSMUSG00000029659",
                    "ENSMUSG00000040170",
                    "ENSMUSG00000029641",
                    "ENSMUSG00000029661",
                    "ENSMUSG00000044786",
                    "ENSMUSG00000008540",
                    "ENSMUSG00000018899",
                    "ENSMUSG00000021268",
                    "ENSMUSG00000079037",
                    "ENSMUSG00000090698",
                    "ENSMUSG00000075702",
                    "ENSMUSG00000006403",
                    "ENSMUSG00000060703",
                    "ENSMUSG00000000247",
                    "ENSMUSG00000038418",
                    "ENSMUSG00000052997",
                    "ENSMUSG00000030849",
                    "ENSMUSG00000001025",
                    "ENSMUSG00000058135"
                ],
                "expression_sum": {
                    "class1": 84935.0,
                    "class2": 57932.0
                },
                "ens_id": "ENSMUSG00000052684",
                "uniprot_ids": [
                    "P05627"
                ]
            },
            "ENSMUSG00000038418": {
                "id": "ENSMUSG00000038418",
                "type": "TF",
                "symbol": "Egr1",
                "source": [
                    "ENSMUSG00000052684",
                    "ENSMUSG00000037868",
                    "ENSMUSG00000023034",
                    "ENSMUSG00000022528",
                    "ENSMUSG00000052837",
                    "ENSMUSG00000026628",
                    "ENSMUSG00000000247",
                    "ENSMUSG00000057098",
                    "ENSMUSG00000021250",
                    "ENSMUSG00000003545",
                    "ENSMUSG00000003032",
                    "ENSMUSG00000008575",
                    "ENSMUSG00000018899"
                ],
                "target": [
                    "ENSMUSG00000019505",
                    "ENSMUSG00000075706",
                    "ENSMUSG00000040723",
                    "ENSMUSG00000040212",
                    "ENSMUSG00000021917",
                    "ENSMUSG00000023034",
                    "ENSMUSG00000021242",
                    "ENSMUSG00000024014",
                    "ENSMUSG00000028367",
                    "ENSMUSG00000021477",
                    "ENSMUSG00000032314",
                    "ENSMUSG00000053113",
                    "ENSMUSG00000028249",
                    "ENSMUSG00000030934",
                    "ENSMUSG00000025213",
                    "ENSMUSG00000033910",
                    "ENSMUSG00000020315",
                    "ENSMUSG00000018217",
                    "ENSMUSG00000016194",
                    "ENSMUSG00000003161",
                    "ENSMUSG00000019907",
                    "ENSMUSG00000052684",
                    "ENSMUSG00000006800",
                    "ENSMUSG00000022528",
                    "ENSMUSG00000028822",
                    "ENSMUSG00000034868",
                    "ENSMUSG00000032353",
                    "ENSMUSG00000008384",
                    "ENSMUSG00000003032",
                    "ENSMUSG00000042747",
                    "ENSMUSG00000079111",
                    "ENSMUSG00000055148",
                    "ENSMUSG00000021248",
                    "ENSMUSG00000026837",
                    "ENSMUSG00000003038",
                    "ENSMUSG00000032383",
                    "ENSMUSG00000015501",
                    "ENSMUSG00000006369",
                    "ENSMUSG00000040128",
                    "ENSMUSG00000031422",
                    "ENSMUSG00000092274",
                    "ENSMUSG00000002107",
                    "ENSMUSG00000052837",
                    "ENSMUSG00000026628",
                    "ENSMUSG00000020423",
                    "ENSMUSG00000035356",
                    "ENSMUSG00000027706",
                    "ENSMUSG00000029103",
                    "ENSMUSG00000035227",
                    "ENSMUSG00000024048",
                    "ENSMUSG00000007891",
                    "ENSMUSG00000040170",
                    "ENSMUSG00000020571",
                    "ENSMUSG00000031119",
                    "ENSMUSG00000037072",
                    "ENSMUSG00000007872",
                    "ENSMUSG00000008540",
                    "ENSMUSG00000018899",
                    "ENSMUSG00000044786",
                    "ENSMUSG00000020009",
                    "ENSMUSG00000027829",
                    "ENSMUSG00000022400",
                    "ENSMUSG00000037868",
                    "ENSMUSG00000060703",
                    "ENSMUSG00000037499",
                    "ENSMUSG00000035828",
                    "ENSMUSG00000000247",
                    "ENSMUSG00000027828",
                    "ENSMUSG00000029465",
                    "ENSMUSG00000021765",
                    "ENSMUSG00000075254",
                    "ENSMUSG00000023010",
                    "ENSMUSG00000060904",
                    "ENSMUSG00000025351",
                    "ENSMUSG00000074884"
                ],
                "expression_sum": {
                    "class1": 38873.0,
                    "class2": 25496.0
                },
                "ens_id": "ENSMUSG00000038418",
                "uniprot_ids": [
                    "P08046"
                ]
            },
            "ENSMUSG00000052837": {
                "id": "ENSMUSG00000052837",
                "type": "TF",
                "symbol": "Junb",
                "source": [
                    "ENSMUSG00000041515",
                    "ENSMUSG00000052684",
                    "ENSMUSG00000037868",
                    "ENSMUSG00000023034",
                    "ENSMUSG00000026628",
                    "ENSMUSG00000030103",
                    "ENSMUSG00000000247",
                    "ENSMUSG00000038418",
                    "ENSMUSG00000057098",
                    "ENSMUSG00000021250",
                    "ENSMUSG00000003545",
                    "ENSMUSG00000003032",
                    "ENSMUSG00000018899",
                    "ENSMUSG00000071076"
                ],
                "target": [
                    "ENSMUSG00000019505",
                    "ENSMUSG00000020053",
                    "ENSMUSG00000097266",
                    "ENSMUSG00000023034",
                    "ENSMUSG00000037999",
                    "ENSMUSG00000000628",
                    "ENSMUSG00000069601",
                    "ENSMUSG00000024014",
                    "ENSMUSG00000054072",
                    "ENSMUSG00000028691",
                    "ENSMUSG00000056708",
                    "ENSMUSG00000053113",
                    "ENSMUSG00000003545",
                    "ENSMUSG00000037266",
                    "ENSMUSG00000042453",
                    "ENSMUSG00000033538",
                    "ENSMUSG00000030930",
                    "ENSMUSG00000037411",
                    "ENSMUSG00000028864",
                    "ENSMUSG00000052684",
                    "ENSMUSG00000017493",
                    "ENSMUSG00000031451",
                    "ENSMUSG00000008384",
                    "ENSMUSG00000019278",
                    "ENSMUSG00000021250",
                    "ENSMUSG00000068220",
                    "ENSMUSG00000003032",
                    "ENSMUSG00000055148",
                    "ENSMUSG00000003038",
                    "ENSMUSG00000021127",
                    "ENSMUSG00000037211",
                    "ENSMUSG00000041515",
                    "ENSMUSG00000024190",
                    "ENSMUSG00000007097",
                    "ENSMUSG00000015501",
                    "ENSMUSG00000040128",
                    "ENSMUSG00000023915",
                    "ENSMUSG00000029380",
                    "ENSMUSG00000047945",
                    "ENSMUSG00000026628",
                    "ENSMUSG00000030103",
                    "ENSMUSG00000020423",
                    "ENSMUSG00000002416",
                    "ENSMUSG00000035356",
                    "ENSMUSG00000097296",
                    "ENSMUSG00000042812",
                    "ENSMUSG00000040170",
                    "ENSMUSG00000032515",
                    "ENSMUSG00000007872",
                    "ENSMUSG00000044786",
                    "ENSMUSG00000022956",
                    "ENSMUSG00000008540",
                    "ENSMUSG00000071076",
                    "ENSMUSG00000018899",
                    "ENSMUSG00000021876",
                    "ENSMUSG00000027829",
                    "ENSMUSG00000049115",
                    "ENSMUSG00000037868",
                    "ENSMUSG00000022122",
                    "ENSMUSG00000006205",
                    "ENSMUSG00000028214",
                    "ENSMUSG00000032501",
                    "ENSMUSG00000035828",
                    "ENSMUSG00000000247",
                    "ENSMUSG00000064215",
                    "ENSMUSG00000034855",
                    "ENSMUSG00000039601",
                    "ENSMUSG00000015085",
                    "ENSMUSG00000030849",
                    "ENSMUSG00000038418",
                    "ENSMUSG00000045658",
                    "ENSMUSG00000090084",
                    "ENSMUSG00000035678",
                    "ENSMUSG00000009681",
                    "ENSMUSG00000001025",
                    "ENSMUSG00000058135"
                ],
                "expression_sum": {
                    "class1": 117091.0,
                    "class2": 64489.0
                },
                "ens_id": "ENSMUSG00000052837",
                "uniprot_ids": [
                    "P09450"
                ]
            },
            "ENSMUSG00000008540": {
                "id": "ENSMUSG00000008540",
                "type": "Gene",
                "symbol": "Mgst1",
                "source": [
                    "ENSMUSG00000032238",
                    "ENSMUSG00000052684",
                    "ENSMUSG00000052837",
                    "ENSMUSG00000022346",
                    "ENSMUSG00000005583",
                    "ENSMUSG00000057098",
                    "ENSMUSG00000038418",
                    "ENSMUSG00000053477",
                    "ENSMUSG00000021250",
                    "ENSMUSG00000008575",
                    "ENSMUSG00000028565",
                    "ENSMUSG00000020484",
                    "ENSMUSG00000071076"
                ],
                "target": [],
                "expression_sum": {
                    "class1": 1719.0,
                    "class2": 11272.0
                },
                "ens_id": "ENSMUSG00000008540",
                "uniprot_ids": [
                    "Q91VS7",
                    "E9QJW0",
                    "A0A0N4SVZ0",
                    "D3YVR3",
                    "A0A0N4SV17",
                    "D3YU60"
                ]
            },
            "ENSMUSG00000071076": {
                "id": "ENSMUSG00000071076",
                "type": "TF",
                "symbol": "Jund",
                "source": [
                    "ENSMUSG00000056501",
                    "ENSMUSG00000052837",
                    "ENSMUSG00000026628",
                    "ENSMUSG00000000247",
                    "ENSMUSG00000008575"
                ],
                "target": [
                    "ENSMUSG00000031431",
                    "ENSMUSG00000040723",
                    "ENSMUSG00000025381",
                    "ENSMUSG00000019188",
                    "ENSMUSG00000022365",
                    "ENSMUSG00000024011",
                    "ENSMUSG00000025213",
                    "ENSMUSG00000053007",
                    "ENSMUSG00000026879",
                    "ENSMUSG00000016256",
                    "ENSMUSG00000047187",
                    "ENSMUSG00000030298",
                    "ENSMUSG00000064373",
                    "ENSMUSG00000056629",
                    "ENSMUSG00000042613",
                    "ENSMUSG00000044600",
                    "ENSMUSG00000000325",
                    "ENSMUSG00000067847",
                    "ENSMUSG00000027239",
                    "ENSMUSG00000078713",
                    "ENSMUSG00000078974",
                    "ENSMUSG00000018567",
                    "ENSMUSG00000058672",
                    "ENSMUSG00000031007",
                    "ENSMUSG00000008575",
                    "ENSMUSG00000053317",
                    "ENSMUSG00000075701",
                    "ENSMUSG00000022836",
                    "ENSMUSG00000021390",
                    "ENSMUSG00000020368",
                    "ENSMUSG00000031825",
                    "ENSMUSG00000084786",
                    "ENSMUSG00000019907",
                    "ENSMUSG00000031565",
                    "ENSMUSG00000020737",
                    "ENSMUSG00000030824",
                    "ENSMUSG00000037379",
                    "ENSMUSG00000020099",
                    "ENSMUSG00000027706",
                    "ENSMUSG00000031586",
                    "ENSMUSG00000032515",
                    "ENSMUSG00000017466",
                    "ENSMUSG00000072235",
                    "ENSMUSG00000005873",
                    "ENSMUSG00000038463",
                    "ENSMUSG00000028517",
                    "ENSMUSG00000016494",
                    "ENSMUSG00000020029",
                    "ENSMUSG00000024290",
                    "ENSMUSG00000038393",
                    "ENSMUSG00000015094",
                    "ENSMUSG00000022665",
                    "ENSMUSG00000015312",
                    "ENSMUSG00000028964",
                    "ENSMUSG00000021871",
                    "ENSMUSG00000024659",
                    "ENSMUSG00000025724",
                    "ENSMUSG00000062270",
                    "ENSMUSG00000027712",
                    "ENSMUSG00000007891",
                    "ENSMUSG00000024048",
                    "ENSMUSG00000028673",
                    "ENSMUSG00000001891",
                    "ENSMUSG00000050708",
                    "ENSMUSG00000026478",
                    "ENSMUSG00000037868",
                    "ENSMUSG00000022122",
                    "ENSMUSG00000000594",
                    "ENSMUSG00000019920",
                    "ENSMUSG00000057098",
                    "ENSMUSG00000014846",
                    "ENSMUSG00000017493",
                    "ENSMUSG00000025351",
                    "ENSMUSG00000074884",
                    "ENSMUSG00000024014",
                    "ENSMUSG00000055435",
                    "ENSMUSG00000020849",
                    "ENSMUSG00000034993",
                    "ENSMUSG00000000753",
                    "ENSMUSG00000051579",
                    "ENSMUSG00000004460",
                    "ENSMUSG00000028757",
                    "ENSMUSG00000035356",
                    "ENSMUSG00000027859",
                    "ENSMUSG00000022174",
                    "ENSMUSG00000029048",
                    "ENSMUSG00000035227",
                    "ENSMUSG00000024012",
                    "ENSMUSG00000060802",
                    "ENSMUSG00000014867",
                    "ENSMUSG00000019505",
                    "ENSMUSG00000059291",
                    "ENSMUSG00000030222",
                    "ENSMUSG00000028691",
                    "ENSMUSG00000003545",
                    "ENSMUSG00000030930",
                    "ENSMUSG00000009281",
                    "ENSMUSG00000028822",
                    "ENSMUSG00000008384",
                    "ENSMUSG00000023175",
                    "ENSMUSG00000026421",
                    "ENSMUSG00000031839",
                    "ENSMUSG00000002107",
                    "ENSMUSG00000032231",
                    "ENSMUSG00000007872",
                    "ENSMUSG00000027848",
                    "ENSMUSG00000052419",
                    "ENSMUSG00000029718",
                    "ENSMUSG00000028600",
                    "ENSMUSG00000025132",
                    "ENSMUSG00000021917",
                    "ENSMUSG00000028339",
                    "ENSMUSG00000037266",
                    "ENSMUSG00000001525",
                    "ENSMUSG00000031538",
                    "ENSMUSG00000019978",
                    "ENSMUSG00000022816",
                    "ENSMUSG00000007039",
                    "ENSMUSG00000029781",
                    "ENSMUSG00000022090",
                    "ENSMUSG00000019579",
                    "ENSMUSG00000068735",
                    "ENSMUSG00000074457",
                    "ENSMUSG00000040249",
                    "ENSMUSG00000032334",
                    "ENSMUSG00000026335",
                    "ENSMUSG00000029070",
                    "ENSMUSG00000035678",
                    "ENSMUSG00000016024",
                    "ENSMUSG00000030208",
                    "ENSMUSG00000023034",
                    "ENSMUSG00000030654",
                    "ENSMUSG00000003420",
                    "ENSMUSG00000020315",
                    "ENSMUSG00000058569",
                    "ENSMUSG00000018217",
                    "ENSMUSG00000023046",
                    "ENSMUSG00000058056",
                    "ENSMUSG00000005973",
                    "ENSMUSG00000042745",
                    "ENSMUSG00000016534",
                    "ENSMUSG00000003161",
                    "ENSMUSG00000032359",
                    "ENSMUSG00000001911",
                    "ENSMUSG00000005583",
                    "ENSMUSG00000022136",
                    "ENSMUSG00000018819",
                    "ENSMUSG00000032301",
                    "ENSMUSG00000020709",
                    "ENSMUSG00000044408",
                    "ENSMUSG00000002059",
                    "ENSMUSG00000019362",
                    "ENSMUSG00000041571",
                    "ENSMUSG00000037072",
                    "ENSMUSG00000020009",
                    "ENSMUSG00000004207",
                    "ENSMUSG00000020088",
                    "ENSMUSG00000026728",
                    "ENSMUSG00000096145",
                    "ENSMUSG00000036752",
                    "ENSMUSG00000042312",
                    "ENSMUSG00000035828",
                    "ENSMUSG00000075254",
                    "ENSMUSG00000023010",
                    "ENSMUSG00000031799",
                    "ENSMUSG00000041084",
                    "ENSMUSG00000041126",
                    "ENSMUSG00000054808",
                    "ENSMUSG00000021242",
                    "ENSMUSG00000002014",
                    "ENSMUSG00000092341",
                    "ENSMUSG00000039899",
                    "ENSMUSG00000030934",
                    "ENSMUSG00000011114",
                    "ENSMUSG00000028864",
                    "ENSMUSG00000006800",
                    "ENSMUSG00000020585",
                    "ENSMUSG00000015575",
                    "ENSMUSG00000032353",
                    "ENSMUSG00000025492",
                    "ENSMUSG00000007041",
                    "ENSMUSG00000055447",
                    "ENSMUSG00000020458",
                    "ENSMUSG00000032959",
                    "ENSMUSG00000009291",
                    "ENSMUSG00000037206",
                    "ENSMUSG00000060703",
                    "ENSMUSG00000030849",
                    "ENSMUSG00000029231",
                    "ENSMUSG00000021131",
                    "ENSMUSG00000035258",
                    "ENSMUSG00000001025",
                    "ENSMUSG00000045636",
                    "ENSMUSG00000048076",
                    "ENSMUSG00000042712",
                    "ENSMUSG00000026021",
                    "ENSMUSG00000022528",
                    "ENSMUSG00000006699",
                    "ENSMUSG00000032575",
                    "ENSMUSG00000079111",
                    "ENSMUSG00000030659",
                    "ENSMUSG00000012422",
                    "ENSMUSG00000023004",
                    "ENSMUSG00000031198",
                    "ENSMUSG00000025130",
                    "ENSMUSG00000003541",
                    "ENSMUSG00000029103",
                    "ENSMUSG00000038872",
                    "ENSMUSG00000022452",
                    "ENSMUSG00000056737",
                    "ENSMUSG00000022032",
                    "ENSMUSG00000027828",
                    "ENSMUSG00000022400",
                    "ENSMUSG00000055839",
                    "ENSMUSG00000034165",
                    "ENSMUSG00000032085",
                    "ENSMUSG00000053113",
                    "ENSMUSG00000052681",
                    "ENSMUSG00000003380",
                    "ENSMUSG00000038803",
                    "ENSMUSG00000046768",
                    "ENSMUSG00000026628",
                    "ENSMUSG00000042737",
                    "ENSMUSG00000021253",
                    "ENSMUSG00000053477",
                    "ENSMUSG00000079523",
                    "ENSMUSG00000038991",
                    "ENSMUSG00000060591",
                    "ENSMUSG00000001794",
                    "ENSMUSG00000028843",
                    "ENSMUSG00000020423",
                    "ENSMUSG00000069662",
                    "ENSMUSG00000021876",
                    "ENSMUSG00000025780",
                    "ENSMUSG00000024164",
                    "ENSMUSG00000001663",
                    "ENSMUSG00000064215",
                    "ENSMUSG00000006050",
                    "ENSMUSG00000004897",
                    "ENSMUSG00000027907",
                    "ENSMUSG00000022108",
                    "ENSMUSG00000000628",
                    "ENSMUSG00000020219",
                    "ENSMUSG00000028249",
                    "ENSMUSG00000069833",
                    "ENSMUSG00000033910",
                    "ENSMUSG00000039168",
                    "ENSMUSG00000018593",
                    "ENSMUSG00000013698",
                    "ENSMUSG00000001418",
                    "ENSMUSG00000015568",
                    "ENSMUSG00000021250",
                    "ENSMUSG00000028618",
                    "ENSMUSG00000054203",
                    "ENSMUSG00000052837",
                    "ENSMUSG00000023272",
                    "ENSMUSG00000021576",
                    "ENSMUSG00000079037",
                    "ENSMUSG00000079017",
                    "ENSMUSG00000006403",
                    "ENSMUSG00000000247",
                    "ENSMUSG00000019087",
                    "ENSMUSG00000029465",
                    "ENSMUSG00000052997",
                    "ENSMUSG00000005054",
                    "ENSMUSG00000058135",
                    "ENSMUSG00000007815",
                    "ENSMUSG00000029761",
                    "ENSMUSG00000002602",
                    "ENSMUSG00000028111",
                    "ENSMUSG00000031875",
                    "ENSMUSG00000021477",
                    "ENSMUSG00000037822",
                    "ENSMUSG00000027422",
                    "ENSMUSG00000097328",
                    "ENSMUSG00000020803",
                    "ENSMUSG00000025647",
                    "ENSMUSG00000044337",
                    "ENSMUSG00000031556",
                    "ENSMUSG00000021877",
                    "ENSMUSG00000068196",
                    "ENSMUSG00000034868",
                    "ENSMUSG00000044627",
                    "ENSMUSG00000021939",
                    "ENSMUSG00000055148",
                    "ENSMUSG00000001240",
                    "ENSMUSG00000014329",
                    "ENSMUSG00000040713",
                    "ENSMUSG00000028369",
                    "ENSMUSG00000022769",
                    "ENSMUSG00000029309",
                    "ENSMUSG00000026547",
                    "ENSMUSG00000009013",
                    "ENSMUSG00000027808",
                    "ENSMUSG00000031595",
                    "ENSMUSG00000009681",
                    "ENSMUSG00000024327",
                    "ENSMUSG00000060904",
                    "ENSMUSG00000075706",
                    "ENSMUSG00000020053",
                    "ENSMUSG00000026042",
                    "ENSMUSG00000037999",
                    "ENSMUSG00000020440",
                    "ENSMUSG00000028367",
                    "ENSMUSG00000032314",
                    "ENSMUSG00000032366",
                    "ENSMUSG00000036372",
                    "ENSMUSG00000033538",
                    "ENSMUSG00000020810",
                    "ENSMUSG00000025016",
                    "ENSMUSG00000001119",
                    "ENSMUSG00000021759",
                    "ENSMUSG00000075010",
                    "ENSMUSG00000029390",
                    "ENSMUSG00000029767",
                    "ENSMUSG00000024076",
                    "ENSMUSG00000055681",
                    "ENSMUSG00000090000",
                    "ENSMUSG00000021760",
                    "ENSMUSG00000021248",
                    "ENSMUSG00000001506",
                    "ENSMUSG00000003038",
                    "ENSMUSG00000021870",
                    "ENSMUSG00000041959",
                    "ENSMUSG00000018672",
                    "ENSMUSG00000028195",
                    "ENSMUSG00000037373",
                    "ENSMUSG00000068220",
                    "ENSMUSG00000020486",
                    "ENSMUSG00000040212",
                    "ENSMUSG00000001123",
                    "ENSMUSG00000020149",
                    "ENSMUSG00000037706",
                    "ENSMUSG00000025784",
                    "ENSMUSG00000070394",
                    "ENSMUSG00000031965",
                    "ENSMUSG00000002015",
                    "ENSMUSG00000028128",
                    "ENSMUSG00000013622",
                    "ENSMUSG00000020571",
                    "ENSMUSG00000028108",
                    "ENSMUSG00000002732",
                    "ENSMUSG00000006205",
                    "ENSMUSG00000006519",
                    "ENSMUSG00000045658",
                    "ENSMUSG00000029076",
                    "ENSMUSG00000030717",
                    "ENSMUSG00000039601",
                    "ENSMUSG00000024308",
                    "ENSMUSG00000056501",
                    "ENSMUSG00000024038",
                    "ENSMUSG00000041736",
                    "ENSMUSG00000002064",
                    "ENSMUSG00000020048",
                    "ENSMUSG00000042747",
                    "ENSMUSG00000041355",
                    "ENSMUSG00000029096",
                    "ENSMUSG00000026837",
                    "ENSMUSG00000032383",
                    "ENSMUSG00000022048",
                    "ENSMUSG00000031422",
                    "ENSMUSG00000029622",
                    "ENSMUSG00000047945",
                    "ENSMUSG00000066551",
                    "ENSMUSG00000039286",
                    "ENSMUSG00000030342",
                    "ENSMUSG00000022708",
                    "ENSMUSG00000019929",
                    "ENSMUSG00000029659",
                    "ENSMUSG00000071984",
                    "ENSMUSG00000029641",
                    "ENSMUSG00000044786",
                    "ENSMUSG00000008540",
                    "ENSMUSG00000018899",
                    "ENSMUSG00000021268",
                    "ENSMUSG00000090698",
                    "ENSMUSG00000075702",
                    "ENSMUSG00000014769"
                ],
                "expression_sum": {
                    "class1": 45888.0,
                    "class2": 59450.0
                },
                "ens_id": "ENSMUSG00000071076",
                "uniprot_ids": [
                    "P15066"
                ]
            },
            "ENSMUSG00000021390": {
                "id": "ENSMUSG00000021390",
                "type": "Gene",
                "symbol": "Ogn",
                "source": [
                    "ENSMUSG00000052684",
                    "ENSMUSG00000003032",
                    "ENSMUSG00000005583",
                    "ENSMUSG00000028565",
                    "ENSMUSG00000071076"
                ],
                "target": [],
                "expression_sum": {
                    "class1": 1146.0,
                    "class2": 16441.0
                },
                "ens_id": "ENSMUSG00000021390",
                "uniprot_ids": [
                    "Q62000"
                ]
            },
            "ENSMUSG00000029096": {
                "id": "ENSMUSG00000029096",
                "type": "Gene",
                "symbol": "Htra3",
                "source": [
                    "ENSMUSG00000032238",
                    "ENSMUSG00000052684",
                    "ENSMUSG00000022346",
                    "ENSMUSG00000005583",
                    "ENSMUSG00000053477",
                    "ENSMUSG00000003032",
                    "ENSMUSG00000008575",
                    "ENSMUSG00000028565",
                    "ENSMUSG00000071076"
                ],
                "target": [],
                "expression_sum": {
                    "class1": 7593.0,
                    "class2": 27781.0
                },
                "ens_id": "ENSMUSG00000029096",
                "uniprot_ids": [
                    "Q9D236",
                    "A0A0J9YV92",
                    "A0A0J9YVC4",
                    "A0A0J9YU80"
                ]
            },
            "ENSMUSG00000058135": {
                "id": "ENSMUSG00000058135",
                "type": "Gene",
                "symbol": "Gstm1",
                "source": [
                    "ENSMUSG00000055435",
                    "ENSMUSG00000032238",
                    "ENSMUSG00000032035",
                    "ENSMUSG00000052684",
                    "ENSMUSG00000052837",
                    "ENSMUSG00000005583",
                    "ENSMUSG00000057098",
                    "ENSMUSG00000021250",
                    "ENSMUSG00000003032",
                    "ENSMUSG00000008575",
                    "ENSMUSG00000028565",
                    "ENSMUSG00000071076"
                ],
                "target": [],
                "expression_sum": {
                    "class1": 18288.0,
                    "class2": 43266.0
                },
                "ens_id": "ENSMUSG00000058135",
                "uniprot_ids": [
                    "P10649",
                    "F6WHQ7",
                    "A2AE89"
                ]
            },
            "ENSMUSG00000075702": {
                "id": "ENSMUSG00000075702",
                "type": "Gene",
                "symbol": "Selm",
                "source": [
                    "ENSMUSG00000052684",
                    "ENSMUSG00000021250",
                    "ENSMUSG00000003032",
                    "ENSMUSG00000005583",
                    "ENSMUSG00000071076"
                ],
                "target": [],
                "expression_sum": {
                    "class1": 17418.0,
                    "class2": 20444.0
                },
                "ens_id": "ENSMUSG00000075702",
                "uniprot_ids": [
                    "Q8VHC3"
                ]
            },
            "ENSMUSG00000032231": {
                "id": "ENSMUSG00000032231",
                "type": "Gene",
                "symbol": "Anxa2",
                "source": [
                    "ENSMUSG00000032035",
                    "ENSMUSG00000052684",
                    "ENSMUSG00000005583",
                    "ENSMUSG00000000247",
                    "ENSMUSG00000057098",
                    "ENSMUSG00000003032",
                    "ENSMUSG00000055435",
                    "ENSMUSG00000028565",
                    "ENSMUSG00000071076"
                ],
                "target": [],
                "expression_sum": {
                    "class1": 27881.0,
                    "class2": 12495.0
                },
                "ens_id": "ENSMUSG00000032231",
                "uniprot_ids": [
                    "P07356",
                    "B0V2N7",
                    "B0V2N5",
                    "B0V2N8"
                ]
            },
            "ENSMUSG00000021876": {
                "id": "ENSMUSG00000021876",
                "type": "Gene",
                "symbol": "Rnase4",
                "source": [
                    "ENSMUSG00000055435",
                    "ENSMUSG00000052684",
                    "ENSMUSG00000021877",
                    "ENSMUSG00000052837",
                    "ENSMUSG00000022346",
                    "ENSMUSG00000005583",
                    "ENSMUSG00000048387",
                    "ENSMUSG00000021250",
                    "ENSMUSG00000003032",
                    "ENSMUSG00000008575",
                    "ENSMUSG00000028565",
                    "ENSMUSG00000020484",
                    "ENSMUSG00000071076"
                ],
                "target": [],
                "expression_sum": {
                    "class1": 7086.0,
                    "class2": 18776.0
                },
                "ens_id": "ENSMUSG00000021876",
                "uniprot_ids": [
                    "Q8C7E4"
                ]
            },
            "ENSMUSG00000029659": {
                "id": "ENSMUSG00000029659",
                "type": "Gene",
                "symbol": "Medag",
                "source": [
                    "ENSMUSG00000052684",
                    "ENSMUSG00000005583",
                    "ENSMUSG00000057098",
                    "ENSMUSG00000003032",
                    "ENSMUSG00000008575",
                    "ENSMUSG00000028565",
                    "ENSMUSG00000071076"
                ],
                "target": [],
                "expression_sum": {
                    "class1": 404.0,
                    "class2": 3598.0
                },
                "ens_id": "ENSMUSG00000029659",
                "uniprot_ids": [
                    "Q14BA6"
                ]
            },
            "ENSMUSG00000000753": {
                "id": "ENSMUSG00000000753",
                "type": "Gene",
                "symbol": "Serpinf1",
                "source": [
                    "ENSMUSG00000032238",
                    "ENSMUSG00000052684",
                    "ENSMUSG00000053477",
                    "ENSMUSG00000021250",
                    "ENSMUSG00000071076"
                ],
                "target": [],
                "expression_sum": {
                    "class1": 620.0,
                    "class2": 37372.0
                },
                "ens_id": "ENSMUSG00000000753",
                "uniprot_ids": [
                    "P97298",
                    "E9PWS2",
                    "Q5ND37",
                    "E9PZX2",
                    "E9PXR9",
                    "B7ZC25",
                    "F6S1M4"
                ]
            },
            "ENSMUSG00000020053": {
                "id": "ENSMUSG00000020053",
                "type": "Gene",
                "symbol": "Igf1",
                "source": [
                    "ENSMUSG00000032238",
                    "ENSMUSG00000052684",
                    "ENSMUSG00000052837",
                    "ENSMUSG00000022346",
                    "ENSMUSG00000053477",
                    "ENSMUSG00000021250",
                    "ENSMUSG00000029322",
                    "ENSMUSG00000003032",
                    "ENSMUSG00000005583",
                    "ENSMUSG00000020484",
                    "ENSMUSG00000071076"
                ],
                "target": [],
                "expression_sum": {
                    "class1": 1900.0,
                    "class2": 12864.0
                },
                "ens_id": "ENSMUSG00000020053",
                "uniprot_ids": [
                    "P05017",
                    "Q8CAR0",
                    "E9Q138",
                    "Q4VJB9",
                    "D3Z7M4",
                    "E9PU89"
                ]
            },
            "ENSMUSG00000053113": {
                "id": "ENSMUSG00000053113",
                "type": "Gene",
                "symbol": "Socs3",
                "source": [
                    "ENSMUSG00000041515",
                    "ENSMUSG00000056501",
                    "ENSMUSG00000052684",
                    "ENSMUSG00000037868",
                    "ENSMUSG00000042406",
                    "ENSMUSG00000023034",
                    "ENSMUSG00000052837",
                    "ENSMUSG00000026628",
                    "ENSMUSG00000030103",
                    "ENSMUSG00000022528",
                    "ENSMUSG00000038418",
                    "ENSMUSG00000038193",
                    "ENSMUSG00000021250",
                    "ENSMUSG00000055148",
                    "ENSMUSG00000003032",
                    "ENSMUSG00000055435",
                    "ENSMUSG00000018899",
                    "ENSMUSG00000071076"
                ],
                "target": [],
                "expression_sum": {
                    "class1": 31900.0,
                    "class2": 9658.0
                },
                "ens_id": "ENSMUSG00000053113",
                "uniprot_ids": [
                    "O35718"
                ]
            },
            "ENSMUSG00000090698": {
                "id": "ENSMUSG00000090698",
                "type": "Gene",
                "symbol": "Apold1",
                "source": [
                    "ENSMUSG00000000078",
                    "ENSMUSG00000052684",
                    "ENSMUSG00000026628",
                    "ENSMUSG00000030103",
                    "ENSMUSG00000003032",
                    "ENSMUSG00000005583",
                    "ENSMUSG00000055148",
                    "ENSMUSG00000021127",
                    "ENSMUSG00000071076"
                ],
                "target": [],
                "expression_sum": {
                    "class1": 8080.0,
                    "class2": 1314.0
                },
                "ens_id": "ENSMUSG00000090698",
                "uniprot_ids": [
                    "E9Q0X2"
                ]
            },
            "ENSMUSG00000055148": {
                "id": "ENSMUSG00000055148",
                "type": "TF",
                "symbol": "Klf2",
                "source": [
                    "ENSMUSG00000041515",
                    "ENSMUSG00000056501",
                    "ENSMUSG00000037868",
                    "ENSMUSG00000052684",
                    "ENSMUSG00000023034",
                    "ENSMUSG00000052837",
                    "ENSMUSG00000022346",
                    "ENSMUSG00000030103",
                    "ENSMUSG00000038418",
                    "ENSMUSG00000021250",
                    "ENSMUSG00000003032",
                    "ENSMUSG00000071076"
                ],
                "target": [
                    "ENSMUSG00000067818",
                    "ENSMUSG00000006360",
                    "ENSMUSG00000031765",
                    "ENSMUSG00000000628",
                    "ENSMUSG00000053113",
                    "ENSMUSG00000056708",
                    "ENSMUSG00000003545",
                    "ENSMUSG00000053560",
                    "ENSMUSG00000037411",
                    "ENSMUSG00000028864",
                    "ENSMUSG00000037405",
                    "ENSMUSG00000017009",
                    "ENSMUSG00000027074",
                    "ENSMUSG00000008384",
                    "ENSMUSG00000031762",
                    "ENSMUSG00000007877",
                    "ENSMUSG00000042622",
                    "ENSMUSG00000021127",
                    "ENSMUSG00000024190",
                    "ENSMUSG00000037211",
                    "ENSMUSG00000073411",
                    "ENSMUSG00000020423",
                    "ENSMUSG00000023064",
                    "ENSMUSG00000022010",
                    "ENSMUSG00000024371",
                    "ENSMUSG00000026255",
                    "ENSMUSG00000024232",
                    "ENSMUSG00000044786",
                    "ENSMUSG00000018899",
                    "ENSMUSG00000090698",
                    "ENSMUSG00000061353",
                    "ENSMUSG00000057666",
                    "ENSMUSG00000034855",
                    "ENSMUSG00000023224",
                    "ENSMUSG00000009681"
                ],
                "expression_sum": {
                    "class1": 44466.0,
                    "class2": 38463.0
                },
                "ens_id": "ENSMUSG00000055148",
                "uniprot_ids": [
                    "Q60843"
                ]
            },
            "ENSMUSG00000003545": {
                "id": "ENSMUSG00000003545",
                "type": "TF",
                "symbol": "Fosb",
                "source": [
                    "ENSMUSG00000056501",
                    "ENSMUSG00000052684",
                    "ENSMUSG00000023034",
                    "ENSMUSG00000037868",
                    "ENSMUSG00000052837",
                    "ENSMUSG00000026628",
                    "ENSMUSG00000030103",
                    "ENSMUSG00000042812",
                    "ENSMUSG00000021250",
                    "ENSMUSG00000055148",
                    "ENSMUSG00000003032",
                    "ENSMUSG00000018899",
                    "ENSMUSG00000071076"
                ],
                "target": [
                    "ENSMUSG00000023034",
                    "ENSMUSG00000069601",
                    "ENSMUSG00000054364",
                    "ENSMUSG00000024014",
                    "ENSMUSG00000028691",
                    "ENSMUSG00000056708",
                    "ENSMUSG00000037266",
                    "ENSMUSG00000087006",
                    "ENSMUSG00000053560",
                    "ENSMUSG00000052684",
                    "ENSMUSG00000022528",
                    "ENSMUSG00000015312",
                    "ENSMUSG00000008384",
                    "ENSMUSG00000021250",
                    "ENSMUSG00000003032",
                    "ENSMUSG00000021127",
                    "ENSMUSG00000024190",
                    "ENSMUSG00000037211",
                    "ENSMUSG00000015501",
                    "ENSMUSG00000040128",
                    "ENSMUSG00000047945",
                    "ENSMUSG00000026628",
                    "ENSMUSG00000020423",
                    "ENSMUSG00000052837",
                    "ENSMUSG00000032501",
                    "ENSMUSG00000028967",
                    "ENSMUSG00000038418",
                    "ENSMUSG00000039601"
                ],
                "expression_sum": {
                    "class1": 20507.0,
                    "class2": 9028.0
                },
                "ens_id": "ENSMUSG00000003545",
                "uniprot_ids": [
                    "P13346",
                    "A0A140LIT4",
                    "A0A140LHS3",
                    "A0A140LJH0",
                    "A0A140LIE4",
                    "A0A140LHW0"
                ]
            },
            "ENSMUSG00000015312": {
                "id": "ENSMUSG00000015312",
                "type": "Gene",
                "symbol": "Gadd45b",
                "source": [
                    "ENSMUSG00000037868",
                    "ENSMUSG00000052684",
                    "ENSMUSG00000030103",
                    "ENSMUSG00000021250",
                    "ENSMUSG00000003545",
                    "ENSMUSG00000003032",
                    "ENSMUSG00000018899",
                    "ENSMUSG00000071076"
                ],
                "target": [],
                "expression_sum": {
                    "class1": 5736.0,
                    "class2": 6541.0
                },
                "ens_id": "ENSMUSG00000015312",
                "uniprot_ids": [
                    "P22339",
                    "A0A1W2P6W4"
                ]
            },
            "ENSMUSG00000031451": {
                "id": "ENSMUSG00000031451",
                "type": "Gene",
                "symbol": "Gas6",
                "source": [
                    "ENSMUSG00000022346",
                    "ENSMUSG00000052837",
                    "ENSMUSG00000028565",
                    "ENSMUSG00000052684"
                ],
                "target": [],
                "expression_sum": {
                    "class1": 2284.0,
                    "class2": 24732.0
                },
                "ens_id": "ENSMUSG00000031451",
                "uniprot_ids": [
                    "Q61592"
                ]
            },
            "ENSMUSG00000026628": {
                "id": "ENSMUSG00000026628",
                "type": "TF",
                "symbol": "Atf3",
                "source": [
                    "ENSMUSG00000041515",
                    "ENSMUSG00000037868",
                    "ENSMUSG00000052684",
                    "ENSMUSG00000042406",
                    "ENSMUSG00000052837",
                    "ENSMUSG00000030103",
                    "ENSMUSG00000038418",
                    "ENSMUSG00000003545",
                    "ENSMUSG00000003032",
                    "ENSMUSG00000018899",
                    "ENSMUSG00000071076"
                ],
                "target": [
                    "ENSMUSG00000006360",
                    "ENSMUSG00000031765",
                    "ENSMUSG00000024014",
                    "ENSMUSG00000000628",
                    "ENSMUSG00000053113",
                    "ENSMUSG00000056708",
                    "ENSMUSG00000090841",
                    "ENSMUSG00000003545",
                    "ENSMUSG00000033538",
                    "ENSMUSG00000001627",
                    "ENSMUSG00000053560",
                    "ENSMUSG00000037411",
                    "ENSMUSG00000052684",
                    "ENSMUSG00000037405",
                    "ENSMUSG00000021025",
                    "ENSMUSG00000038370",
                    "ENSMUSG00000042406",
                    "ENSMUSG00000036438",
                    "ENSMUSG00000019960",
                    "ENSMUSG00000008384",
                    "ENSMUSG00000031762",
                    "ENSMUSG00000003032",
                    "ENSMUSG00000026864",
                    "ENSMUSG00000021127",
                    "ENSMUSG00000024190",
                    "ENSMUSG00000037211",
                    "ENSMUSG00000041515",
                    "ENSMUSG00000015501",
                    "ENSMUSG00000023904",
                    "ENSMUSG00000029380",
                    "ENSMUSG00000047945",
                    "ENSMUSG00000052837",
                    "ENSMUSG00000020423",
                    "ENSMUSG00000030103",
                    "ENSMUSG00000035356",
                    "ENSMUSG00000073411",
                    "ENSMUSG00000097296",
                    "ENSMUSG00000032515",
                    "ENSMUSG00000020205",
                    "ENSMUSG00000044786",
                    "ENSMUSG00000071076",
                    "ENSMUSG00000018899",
                    "ENSMUSG00000027829",
                    "ENSMUSG00000090698",
                    "ENSMUSG00000037868",
                    "ENSMUSG00000006403",
                    "ENSMUSG00000032501",
                    "ENSMUSG00000038418",
                    "ENSMUSG00000034855",
                    "ENSMUSG00000035678",
                    "ENSMUSG00000073489",
                    "ENSMUSG00000009681",
                    "ENSMUSG00000039601"
                ],
                "expression_sum": {
                    "class1": 16194.0,
                    "class2": 10101.0
                },
                "ens_id": "ENSMUSG00000026628",
                "uniprot_ids": [
                    "Q60765"
                ]
            },
            "ENSMUSG00000032359": {
                "id": "ENSMUSG00000032359",
                "type": "Gene",
                "symbol": "Ctsh",
                "source": [
                    "ENSMUSG00000052684",
                    "ENSMUSG00000022346",
                    "ENSMUSG00000021250",
                    "ENSMUSG00000008575",
                    "ENSMUSG00000005583",
                    "ENSMUSG00000071076"
                ],
                "target": [],
                "expression_sum": {
                    "class1": 271.0,
                    "class2": 7940.0
                },
                "ens_id": "ENSMUSG00000032359",
                "uniprot_ids": [
                    "P49935",
                    "A0A087WR20",
                    "D7UEQ9",
                    "D3Z437",
                    "D3YVI3"
                ]
            },
            "ENSMUSG00000017493": {
                "id": "ENSMUSG00000017493",
                "type": "Gene",
                "symbol": "Igfbp4",
                "source": [
                    "ENSMUSG00000032238",
                    "ENSMUSG00000052684",
                    "ENSMUSG00000052837",
                    "ENSMUSG00000022346",
                    "ENSMUSG00000005583",
                    "ENSMUSG00000057098",
                    "ENSMUSG00000021250",
                    "ENSMUSG00000003032",
                    "ENSMUSG00000008575",
                    "ENSMUSG00000031965",
                    "ENSMUSG00000028565",
                    "ENSMUSG00000020484",
                    "ENSMUSG00000071076"
                ],
                "target": [],
                "expression_sum": {
                    "class1": 20106.0,
                    "class2": 39416.0
                },
                "ens_id": "ENSMUSG00000017493",
                "uniprot_ids": [
                    "P47879",
                    "H3BJT2",
                    "H3BLB7"
                ]
            },
            "ENSMUSG00000006369": {
                "id": "ENSMUSG00000006369",
                "type": "Gene",
                "symbol": "Fbln1",
                "source": [
                    "ENSMUSG00000052684",
                    "ENSMUSG00000022346",
                    "ENSMUSG00000038418",
                    "ENSMUSG00000053477",
                    "ENSMUSG00000008575",
                    "ENSMUSG00000005583",
                    "ENSMUSG00000048387"
                ],
                "target": [],
                "expression_sum": {
                    "class1": 1519.0,
                    "class2": 35920.0
                },
                "ens_id": "ENSMUSG00000006369",
                "uniprot_ids": [
                    "Q08879"
                ]
            },
            "ENSMUSG00000008575": {
                "id": "ENSMUSG00000008575",
                "type": "TF",
                "symbol": "Nfib",
                "source": [
                    "ENSMUSG00000038193",
                    "ENSMUSG00000005583",
                    "ENSMUSG00000052684",
                    "ENSMUSG00000071076"
                ],
                "target": [
                    "ENSMUSG00000031431",
                    "ENSMUSG00000024063",
                    "ENSMUSG00000029761",
                    "ENSMUSG00000020439",
                    "ENSMUSG00000031765",
                    "ENSMUSG00000021477",
                    "ENSMUSG00000022365",
                    "ENSMUSG00000025213",
                    "ENSMUSG00000026879",
                    "ENSMUSG00000028413",
                    "ENSMUSG00000036438",
                    "ENSMUSG00000021877",
                    "ENSMUSG00000000325",
                    "ENSMUSG00000030659",
                    "ENSMUSG00000026421",
                    "ENSMUSG00000028003",
                    "ENSMUSG00000073889",
                    "ENSMUSG00000030605",
                    "ENSMUSG00000030218",
                    "ENSMUSG00000004473",
                    "ENSMUSG00000031740",
                    "ENSMUSG00000031595",
                    "ENSMUSG00000028600",
                    "ENSMUSG00000041120",
                    "ENSMUSG00000037999",
                    "ENSMUSG00000069601",
                    "ENSMUSG00000032085",
                    "ENSMUSG00000034353",
                    "ENSMUSG00000024140",
                    "ENSMUSG00000017707",
                    "ENSMUSG00000032366",
                    "ENSMUSG00000027215",
                    "ENSMUSG00000019907",
                    "ENSMUSG00000052684",
                    "ENSMUSG00000021759",
                    "ENSMUSG00000014956",
                    "ENSMUSG00000029390",
                    "ENSMUSG00000031246",
                    "ENSMUSG00000001506",
                    "ENSMUSG00000021870",
                    "ENSMUSG00000028871",
                    "ENSMUSG00000002980",
                    "ENSMUSG00000035885",
                    "ENSMUSG00000029781",
                    "ENSMUSG00000026768",
                    "ENSMUSG00000027834",
                    "ENSMUSG00000068735",
                    "ENSMUSG00000074457",
                    "ENSMUSG00000032334",
                    "ENSMUSG00000026335",
                    "ENSMUSG00000021253",
                    "ENSMUSG00000072235",
                    "ENSMUSG00000021765",
                    "ENSMUSG00000038463",
                    "ENSMUSG00000016024",
                    "ENSMUSG00000037166",
                    "ENSMUSG00000068220",
                    "ENSMUSG00000030208",
                    "ENSMUSG00000002289",
                    "ENSMUSG00000028517",
                    "ENSMUSG00000052310",
                    "ENSMUSG00000016494",
                    "ENSMUSG00000028465",
                    "ENSMUSG00000020029",
                    "ENSMUSG00000024290",
                    "ENSMUSG00000042453",
                    "ENSMUSG00000020315",
                    "ENSMUSG00000018217",
                    "ENSMUSG00000058056",
                    "ENSMUSG00000032359",
                    "ENSMUSG00000038991",
                    "ENSMUSG00000001911",
                    "ENSMUSG00000032135",
                    "ENSMUSG00000019139",
                    "ENSMUSG00000018819",
                    "ENSMUSG00000024048",
                    "ENSMUSG00000020205",
                    "ENSMUSG00000020009",
                    "ENSMUSG00000021876",
                    "ENSMUSG00000024164",
                    "ENSMUSG00000050708",
                    "ENSMUSG00000025147",
                    "ENSMUSG00000006356",
                    "ENSMUSG00000006205",
                    "ENSMUSG00000000594",
                    "ENSMUSG00000019920",
                    "ENSMUSG00000064215",
                    "ENSMUSG00000014846",
                    "ENSMUSG00000030717",
                    "ENSMUSG00000017493",
                    "ENSMUSG00000025393",
                    "ENSMUSG00000067818",
                    "ENSMUSG00000054808",
                    "ENSMUSG00000021242",
                    "ENSMUSG00000014313",
                    "ENSMUSG00000002014",
                    "ENSMUSG00000090841",
                    "ENSMUSG00000092341",
                    "ENSMUSG00000026688",
                    "ENSMUSG00000039168",
                    "ENSMUSG00000052957",
                    "ENSMUSG00000031380",
                    "ENSMUSG00000032353",
                    "ENSMUSG00000004460",
                    "ENSMUSG00000020048",
                    "ENSMUSG00000042747",
                    "ENSMUSG00000029096",
                    "ENSMUSG00000024190",
                    "ENSMUSG00000006369",
                    "ENSMUSG00000028757",
                    "ENSMUSG00000023272",
                    "ENSMUSG00000004044",
                    "ENSMUSG00000020840",
                    "ENSMUSG00000015932",
                    "ENSMUSG00000024012",
                    "ENSMUSG00000029659",
                    "ENSMUSG00000040170",
                    "ENSMUSG00000008540",
                    "ENSMUSG00000071076",
                    "ENSMUSG00000079037",
                    "ENSMUSG00000004085",
                    "ENSMUSG00000060703",
                    "ENSMUSG00000038418",
                    "ENSMUSG00000030849",
                    "ENSMUSG00000042284",
                    "ENSMUSG00000001025",
                    "ENSMUSG00000058135"
                ],
                "expression_sum": {
                    "class1": 17518.0,
                    "class2": 11173.0
                },
                "ens_id": "ENSMUSG00000008575",
                "uniprot_ids": [
                    "P97863",
                    "I7HIP8",
                    "A2BG75",
                    "A2BG76",
                    "A2BG77",
                    "A2ADI3"
                ]
            },
            "ENSMUSG00000035356": {
                "id": "ENSMUSG00000035356",
                "type": "Gene",
                "symbol": "Nfkbiz",
                "source": [
                    "ENSMUSG00000041515",
                    "ENSMUSG00000056501",
                    "ENSMUSG00000052684",
                    "ENSMUSG00000037868",
                    "ENSMUSG00000042406",
                    "ENSMUSG00000023034",
                    "ENSMUSG00000052837",
                    "ENSMUSG00000026628",
                    "ENSMUSG00000030103",
                    "ENSMUSG00000038418",
                    "ENSMUSG00000021250",
                    "ENSMUSG00000003032",
                    "ENSMUSG00000018899",
                    "ENSMUSG00000071076"
                ],
                "target": [],
                "expression_sum": {
                    "class1": 7633.0,
                    "class2": 2441.0
                },
                "ens_id": "ENSMUSG00000035356",
                "uniprot_ids": [
                    "Q9EST8"
                ]
            },
            "ENSMUSG00000060703": {
                "id": "ENSMUSG00000060703",
                "type": "Gene",
                "symbol": "Cd302",
                "source": [
                    "ENSMUSG00000052684",
                    "ENSMUSG00000038418",
                    "ENSMUSG00000003032",
                    "ENSMUSG00000005583",
                    "ENSMUSG00000008575",
                    "ENSMUSG00000071076"
                ],
                "target": [],
                "expression_sum": {
                    "class1": 30299.0,
                    "class2": 10955.0
                },
                "ens_id": "ENSMUSG00000060703",
                "uniprot_ids": [
                    "Q9DCG2"
                ]
            },
            "ENSMUSG00000064373": {
                "id": "ENSMUSG00000064373",
                "type": "Gene",
                "symbol": "Sepp1",
                "source": [
                    "ENSMUSG00000032238",
                    "ENSMUSG00000052684",
                    "ENSMUSG00000022346",
                    "ENSMUSG00000038193",
                    "ENSMUSG00000003032",
                    "ENSMUSG00000005583",
                    "ENSMUSG00000028565",
                    "ENSMUSG00000071076"
                ],
                "target": [],
                "expression_sum": {
                    "class1": 37388.0,
                    "class2": 28480.0
                },
                "ens_id": "ENSMUSG00000064373",
                "uniprot_ids": [
                    "P70274",
                    "E0CX80",
                    "E0CZ76",
                    "E0CY03"
                ]
            },
            "ENSMUSG00000057666": {
                "id": "ENSMUSG00000057666",
                "type": "Gene",
                "symbol": "Gapdh",
                "source": [
                    "ENSMUSG00000055148",
                    "ENSMUSG00000020484",
                    "ENSMUSG00000005583"
                ],
                "target": [],
                "expression_sum": {
                    "class1": 28269.0,
                    "class2": 24340.0
                },
                "ens_id": "ENSMUSG00000057666",
                "uniprot_ids": [
                    "P16858",
                    "S4R1W8",
                    "S4R1N5",
                    "A0A0A0MQF6",
                    "S4R257"
                ]
            },
            "ENSMUSG00000002980": {
                "id": "ENSMUSG00000002980",
                "type": "Gene",
                "symbol": "Bcam",
                "source": [
                    "ENSMUSG00000023034",
                    "ENSMUSG00000058756",
                    "ENSMUSG00000038193",
                    "ENSMUSG00000008575",
                    "ENSMUSG00000005583"
                ],
                "target": [],
                "expression_sum": {
                    "class1": 445.0,
                    "class2": 15153.0
                },
                "ens_id": "ENSMUSG00000002980",
                "uniprot_ids": [
                    "Q9R069",
                    "D6RE44",
                    "D3YTK7"
                ]
            },
            "ENSMUSG00000009876": {
                "id": "ENSMUSG00000009876",
                "type": "Gene",
                "symbol": "Cox4i2",
                "source": [
                    "ENSMUSG00000038193",
                    "ENSMUSG00000005583"
                ],
                "target": [],
                "expression_sum": {
                    "class1": 1164.0,
                    "class2": 9694.0
                },
                "ens_id": "ENSMUSG00000009876",
                "uniprot_ids": [
                    "Q91W29"
                ]
            },
            "ENSMUSG00000006360": {
                "id": "ENSMUSG00000006360",
                "type": "Gene",
                "symbol": "Crip1",
                "source": [
                    "ENSMUSG00000023034",
                    "ENSMUSG00000026628",
                    "ENSMUSG00000057098",
                    "ENSMUSG00000021250",
                    "ENSMUSG00000029322",
                    "ENSMUSG00000005583",
                    "ENSMUSG00000055148"
                ],
                "target": [],
                "expression_sum": {
                    "class1": 12842.0,
                    "class2": 203119.0
                },
                "ens_id": "ENSMUSG00000006360",
                "uniprot_ids": [
                    "P63254",
                    "A0A0G2JEK2"
                ]
            },
            "ENSMUSG00000079523": {
                "id": "ENSMUSG00000079523",
                "type": "Gene",
                "symbol": "Tmsb10",
                "source": [
                    "ENSMUSG00000032035",
                    "ENSMUSG00000052684",
                    "ENSMUSG00000022528",
                    "ENSMUSG00000005583",
                    "ENSMUSG00000038193",
                    "ENSMUSG00000029322",
                    "ENSMUSG00000003032",
                    "ENSMUSG00000055435",
                    "ENSMUSG00000048387",
                    "ENSMUSG00000071076"
                ],
                "target": [],
                "expression_sum": {
                    "class1": 239963.0,
                    "class2": 44526.0
                },
                "ens_id": "ENSMUSG00000079523",
                "uniprot_ids": [
                    "Q6ZWY8",
                    "A0A0N4SVF0"
                ]
            },
            "ENSMUSG00000021477": {
                "id": "ENSMUSG00000021477",
                "type": "Gene",
                "symbol": "Ctsl",
                "source": [
                    "ENSMUSG00000055435",
                    "ENSMUSG00000032035",
                    "ENSMUSG00000052684",
                    "ENSMUSG00000058756",
                    "ENSMUSG00000005583",
                    "ENSMUSG00000000247",
                    "ENSMUSG00000038418",
                    "ENSMUSG00000038193",
                    "ENSMUSG00000003032",
                    "ENSMUSG00000008575",
                    "ENSMUSG00000028565",
                    "ENSMUSG00000071076"
                ],
                "target": [],
                "expression_sum": {
                    "class1": 37920.0,
                    "class2": 20428.0
                },
                "ens_id": "ENSMUSG00000021477",
                "uniprot_ids": [
                    "P06797",
                    "A0A1Y7VP49",
                    "A0A1Y7VNM3",
                    "A0A1Y7VNQ4",
                    "A0A1Y7VJC1"
                ]
            },
            "ENSMUSG00000029309": {
                "id": "ENSMUSG00000029309",
                "type": "Gene",
                "symbol": "Sparcl1",
                "source": [
                    "ENSMUSG00000052534",
                    "ENSMUSG00000005583",
                    "ENSMUSG00000052684",
                    "ENSMUSG00000071076"
                ],
                "target": [],
                "expression_sum": {
                    "class1": 20109.0,
                    "class2": 38424.0
                },
                "ens_id": "ENSMUSG00000029309",
                "uniprot_ids": [
                    "P70663",
                    "A0A0G2JFH7"
                ]
            },
            "ENSMUSG00000042747": {
                "id": "ENSMUSG00000042747",
                "type": "Gene",
                "symbol": "Krtcap2",
                "source": [
                    "ENSMUSG00000052684",
                    "ENSMUSG00000005583",
                    "ENSMUSG00000057098",
                    "ENSMUSG00000038418",
                    "ENSMUSG00000003032",
                    "ENSMUSG00000008575",
                    "ENSMUSG00000071076"
                ],
                "target": [],
                "expression_sum": {
                    "class1": 13423.0,
                    "class2": 10239.0
                },
                "ens_id": "ENSMUSG00000042747",
                "uniprot_ids": [
                    "Q5RL79",
                    "A0A0A0MQN1"
                ]
            },
            "ENSMUSG00000016319": {
                "id": "ENSMUSG00000016319",
                "type": "Gene",
                "symbol": "Slc25a5",
                "source": [
                    "ENSMUSG00000020484",
                    "ENSMUSG00000005583"
                ],
                "target": [],
                "expression_sum": {
                    "class1": 10940.0,
                    "class2": 10803.0
                },
                "ens_id": "ENSMUSG00000016319",
                "uniprot_ids": [
                    "P51881"
                ]
            },
            "ENSMUSG00000020928": {
                "id": "ENSMUSG00000020928",
                "type": "Gene",
                "symbol": "Higd1b",
                "source": [
                    "ENSMUSG00000005583"
                ],
                "target": [],
                "expression_sum": {
                    "class1": 21.0,
                    "class2": 5472.0
                },
                "ens_id": "ENSMUSG00000020928",
                "uniprot_ids": [
                    "Q99JY6"
                ]
            },
            "ENSMUSG00000040170": {
                "id": "ENSMUSG00000040170",
                "type": "Gene",
                "symbol": "Fmo2",
                "source": [
                    "ENSMUSG00000032238",
                    "ENSMUSG00000052684",
                    "ENSMUSG00000052837",
                    "ENSMUSG00000038418",
                    "ENSMUSG00000008575",
                    "ENSMUSG00000020484"
                ],
                "target": [],
                "expression_sum": {
                    "class1": 109.0,
                    "class2": 11806.0
                },
                "ens_id": "ENSMUSG00000040170",
                "uniprot_ids": [
                    "Q8K2I3",
                    "D3YVR1"
                ]
            },
            "ENSMUSG00000016494": {
                "id": "ENSMUSG00000016494",
                "type": "Gene",
                "symbol": "Cd34",
                "source": [
                    "ENSMUSG00000052684",
                    "ENSMUSG00000022346",
                    "ENSMUSG00000005583",
                    "ENSMUSG00000021250",
                    "ENSMUSG00000029322",
                    "ENSMUSG00000003032",
                    "ENSMUSG00000008575",
                    "ENSMUSG00000028565",
                    "ENSMUSG00000071076"
                ],
                "target": [],
                "expression_sum": {
                    "class1": 613.0,
                    "class2": 14659.0
                },
                "ens_id": "ENSMUSG00000016494",
                "uniprot_ids": [
                    "Q64314",
                    "A0A0A6YX84"
                ]
            },
            "ENSMUSG00000037266": {
                "id": "ENSMUSG00000037266",
                "type": "Gene",
                "symbol": "Rsrp1",
                "source": [
                    "ENSMUSG00000052684",
                    "ENSMUSG00000052837",
                    "ENSMUSG00000057098",
                    "ENSMUSG00000021250",
                    "ENSMUSG00000003545",
                    "ENSMUSG00000003032",
                    "ENSMUSG00000048387",
                    "ENSMUSG00000071076"
                ],
                "target": [],
                "expression_sum": {
                    "class1": 28028.0,
                    "class2": 18827.0
                },
                "ens_id": "ENSMUSG00000037266",
                "uniprot_ids": [
                    "Q3UC65"
                ]
            },
            "ENSMUSG00000018899": {
                "id": "ENSMUSG00000018899",
                "type": "TF",
                "symbol": "Irf1",
                "source": [
                    "ENSMUSG00000041515",
                    "ENSMUSG00000056501",
                    "ENSMUSG00000052684",
                    "ENSMUSG00000037868",
                    "ENSMUSG00000023034",
                    "ENSMUSG00000042406",
                    "ENSMUSG00000052837",
                    "ENSMUSG00000026628",
                    "ENSMUSG00000030103",
                    "ENSMUSG00000022346",
                    "ENSMUSG00000038418",
                    "ENSMUSG00000021250",
                    "ENSMUSG00000003032",
                    "ENSMUSG00000055148",
                    "ENSMUSG00000071076"
                ],
                "target": [
                    "ENSMUSG00000056501",
                    "ENSMUSG00000023034",
                    "ENSMUSG00000031765",
                    "ENSMUSG00000024014",
                    "ENSMUSG00000000628",
                    "ENSMUSG00000054072",
                    "ENSMUSG00000053113",
                    "ENSMUSG00000056708",
                    "ENSMUSG00000003545",
                    "ENSMUSG00000033538",
                    "ENSMUSG00000001627",
                    "ENSMUSG00000053560",
                    "ENSMUSG00000037411",
                    "ENSMUSG00000052684",
                    "ENSMUSG00000037405",
                    "ENSMUSG00000021025",
                    "ENSMUSG00000022346",
                    "ENSMUSG00000015312",
                    "ENSMUSG00000008384",
                    "ENSMUSG00000031762",
                    "ENSMUSG00000021250",
                    "ENSMUSG00000003032",
                    "ENSMUSG00000035493",
                    "ENSMUSG00000021127",
                    "ENSMUSG00000024190",
                    "ENSMUSG00000041515",
                    "ENSMUSG00000015501",
                    "ENSMUSG00000040128",
                    "ENSMUSG00000035356",
                    "ENSMUSG00000052837",
                    "ENSMUSG00000026628",
                    "ENSMUSG00000030103",
                    "ENSMUSG00000020423",
                    "ENSMUSG00000097296",
                    "ENSMUSG00000032515",
                    "ENSMUSG00000007872",
                    "ENSMUSG00000044786",
                    "ENSMUSG00000020205",
                    "ENSMUSG00000027829",
                    "ENSMUSG00000037868",
                    "ENSMUSG00000006403",
                    "ENSMUSG00000032501",
                    "ENSMUSG00000035828",
                    "ENSMUSG00000038418",
                    "ENSMUSG00000034855",
                    "ENSMUSG00000030849",
                    "ENSMUSG00000035678",
                    "ENSMUSG00000073489"
                ],
                "expression_sum": {
                    "class1": 9123.0,
                    "class2": 1422.0
                },
                "ens_id": "ENSMUSG00000018899",
                "uniprot_ids": [
                    "P15314",
                    "Q3U5M1",
                    "A8Y5B9",
                    "Q5SX15",
                    "Q5SX14",
                    "D6RH71",
                    "Q5SX16"
                ]
            },
            "ENSMUSG00000033538": {
                "id": "ENSMUSG00000033538",
                "type": "Gene",
                "symbol": "Casp4",
                "source": [
                    "ENSMUSG00000056501",
                    "ENSMUSG00000037868",
                    "ENSMUSG00000052684",
                    "ENSMUSG00000023034",
                    "ENSMUSG00000052837",
                    "ENSMUSG00000026628",
                    "ENSMUSG00000021250",
                    "ENSMUSG00000018899",
                    "ENSMUSG00000071076"
                ],
                "target": [],
                "expression_sum": {
                    "class1": 3929.0,
                    "class2": 647.0
                },
                "ens_id": "ENSMUSG00000033538",
                "uniprot_ids": [
                    "P70343",
                    "F6SUE5",
                    "E0CY92"
                ]
            },
            "ENSMUSG00000035885": {
                "id": "ENSMUSG00000035885",
                "type": "Gene",
                "symbol": "Cox8a",
                "source": [
                    "ENSMUSG00000000247",
                    "ENSMUSG00000053477",
                    "ENSMUSG00000008575",
                    "ENSMUSG00000005583",
                    "ENSMUSG00000020484"
                ],
                "target": [],
                "expression_sum": {
                    "class1": 33978.0,
                    "class2": 36956.0
                },
                "ens_id": "ENSMUSG00000035885",
                "uniprot_ids": [
                    "Q64445",
                    "A0A494B971"
                ]
            },
            "ENSMUSG00000042453": {
                "id": "ENSMUSG00000042453",
                "type": "Gene",
                "symbol": "Reln",
                "source": [
                    "ENSMUSG00000032035",
                    "ENSMUSG00000056501",
                    "ENSMUSG00000052684",
                    "ENSMUSG00000052837",
                    "ENSMUSG00000038193",
                    "ENSMUSG00000015846",
                    "ENSMUSG00000076431",
                    "ENSMUSG00000008575",
                    "ENSMUSG00000020484"
                ],
                "target": [],
                "expression_sum": {
                    "class1": 66813.0,
                    "class2": 37.0
                },
                "ens_id": "ENSMUSG00000042453",
                "uniprot_ids": [
                    "Q60841",
                    "F6U2B2"
                ]
            },
            "ENSMUSG00000038193": {
                "id": "ENSMUSG00000038193",
                "type": "TF",
                "symbol": "Hand2",
                "source": [],
                "target": [
                    "ENSMUSG00000019539",
                    "ENSMUSG00000059291",
                    "ENSMUSG00000031447",
                    "ENSMUSG00000025381",
                    "ENSMUSG00000058546",
                    "ENSMUSG00000019916",
                    "ENSMUSG00000032518",
                    "ENSMUSG00000007655",
                    "ENSMUSG00000021477",
                    "ENSMUSG00000040952",
                    "ENSMUSG00000031320",
                    "ENSMUSG00000030930",
                    "ENSMUSG00000009281",
                    "ENSMUSG00000016256",
                    "ENSMUSG00000027447",
                    "ENSMUSG00000030744",
                    "ENSMUSG00000053560",
                    "ENSMUSG00000022371",
                    "ENSMUSG00000021877",
                    "ENSMUSG00000064373",
                    "ENSMUSG00000034868",
                    "ENSMUSG00000008668",
                    "ENSMUSG00000020372",
                    "ENSMUSG00000021939",
                    "ENSMUSG00000041417",
                    "ENSMUSG00000008682",
                    "ENSMUSG00000035493",
                    "ENSMUSG00000001240",
                    "ENSMUSG00000026970",
                    "ENSMUSG00000073889",
                    "ENSMUSG00000034892",
                    "ENSMUSG00000071415",
                    "ENSMUSG00000008575",
                    "ENSMUSG00000061477",
                    "ENSMUSG00000031119",
                    "ENSMUSG00000027848",
                    "ENSMUSG00000050212",
                    "ENSMUSG00000061983",
                    "ENSMUSG00000029426",
                    "ENSMUSG00000050953",
                    "ENSMUSG00000029718",
                    "ENSMUSG00000031740",
                    "ENSMUSG00000036655",
                    "ENSMUSG00000002900",
                    "ENSMUSG00000048096",
                    "ENSMUSG00000028081",
                    "ENSMUSG00000060636",
                    "ENSMUSG00000026365",
                    "ENSMUSG00000036545",
                    "ENSMUSG00000022312",
                    "ENSMUSG00000069601",
                    "ENSMUSG00000063457",
                    "ENSMUSG00000037742",
                    "ENSMUSG00000053113",
                    "ENSMUSG00000039007",
                    "ENSMUSG00000032366",
                    "ENSMUSG00000029810",
                    "ENSMUSG00000020810",
                    "ENSMUSG00000052684",
                    "ENSMUSG00000070436",
                    "ENSMUSG00000001119",
                    "ENSMUSG00000036446",
                    "ENSMUSG00000012405",
                    "ENSMUSG00000030824",
                    "ENSMUSG00000027642",
                    "ENSMUSG00000003970",
                    "ENSMUSG00000026482",
                    "ENSMUSG00000002980",
                    "ENSMUSG00000020460",
                    "ENSMUSG00000026814",
                    "ENSMUSG00000030249",
                    "ENSMUSG00000056091",
                    "ENSMUSG00000029781",
                    "ENSMUSG00000025508",
                    "ENSMUSG00000025290",
                    "ENSMUSG00000028464",
                    "ENSMUSG00000032014",
                    "ENSMUSG00000033880",
                    "ENSMUSG00000022587",
                    "ENSMUSG00000025794",
                    "ENSMUSG00000040249",
                    "ENSMUSG00000037060",
                    "ENSMUSG00000023367",
                    "ENSMUSG00000029070",
                    "ENSMUSG00000024608",
                    "ENSMUSG00000001687",
                    "ENSMUSG00000053477",
                    "ENSMUSG00000039001",
                    "ENSMUSG00000006333",
                    "ENSMUSG00000002985",
                    "ENSMUSG00000058600",
                    "ENSMUSG00000062647",
                    "ENSMUSG00000022894",
                    "ENSMUSG00000046330",
                    "ENSMUSG00000074129",
                    "ENSMUSG00000090733",
                    "ENSMUSG00000020432",
                    "ENSMUSG00000002778",
                    "ENSMUSG00000079523",
                    "ENSMUSG00000003420",
                    "ENSMUSG00000060126",
                    "ENSMUSG00000042453",
                    "ENSMUSG00000058569",
                    "ENSMUSG00000026014",
                    "ENSMUSG00000046727",
                    "ENSMUSG00000005973",
                    "ENSMUSG00000012848",
                    "ENSMUSG00000016534",
                    "ENSMUSG00000060591",
                    "ENSMUSG00000023942",
                    "ENSMUSG00000049517",
                    "ENSMUSG00000019960",
                    "ENSMUSG00000022665",
                    "ENSMUSG00000027074",
                    "ENSMUSG00000029614",
                    "ENSMUSG00000022136",
                    "ENSMUSG00000003814",
                    "ENSMUSG00000062006",
                    "ENSMUSG00000061232",
                    "ENSMUSG00000032216",
                    "ENSMUSG00000079435",
                    "ENSMUSG00000047215",
                    "ENSMUSG00000094724",
                    "ENSMUSG00000027712",
                    "ENSMUSG00000024371",
                    "ENSMUSG00000069662",
                    "ENSMUSG00000032666",
                    "ENSMUSG00000028108",
                    "ENSMUSG00000004207",
                    "ENSMUSG00000031029",
                    "ENSMUSG00000028673",
                    "ENSMUSG00000024232",
                    "ENSMUSG00000002732",
                    "ENSMUSG00000034708",
                    "ENSMUSG00000050708",
                    "ENSMUSG00000019970",
                    "ENSMUSG00000022122",
                    "ENSMUSG00000062997",
                    "ENSMUSG00000028214",
                    "ENSMUSG00000028173",
                    "ENSMUSG00000058558",
                    "ENSMUSG00000067288",
                    "ENSMUSG00000029076",
                    "ENSMUSG00000023224",
                    "ENSMUSG00000026223",
                    "ENSMUSG00000049086",
                    "ENSMUSG00000022971",
                    "ENSMUSG00000025351",
                    "ENSMUSG00000060938",
                    "ENSMUSG00000041841",
                    "ENSMUSG00000052146",
                    "ENSMUSG00000039601",
                    "ENSMUSG00000067818",
                    "ENSMUSG00000067274",
                    "ENSMUSG00000057863",
                    "ENSMUSG00000026566",
                    "ENSMUSG00000022108",
                    "ENSMUSG00000024538",
                    "ENSMUSG00000032547",
                    "ENSMUSG00000024248",
                    "ENSMUSG00000020484",
                    "ENSMUSG00000060550",
                    "ENSMUSG00000027248",
                    "ENSMUSG00000005881",
                    "ENSMUSG00000062328",
                    "ENSMUSG00000046491",
                    "ENSMUSG00000020585",
                    "ENSMUSG00000004105",
                    "ENSMUSG00000032353",
                    "ENSMUSG00000019055",
                    "ENSMUSG00000008683",
                    "ENSMUSG00000043716",
                    "ENSMUSG00000055447",
                    "ENSMUSG00000000740",
                    "ENSMUSG00000021186",
                    "ENSMUSG00000028618",
                    "ENSMUSG00000057841",
                    "ENSMUSG00000079641",
                    "ENSMUSG00000022048",
                    "ENSMUSG00000032383",
                    "ENSMUSG00000025993",
                    "ENSMUSG00000047907",
                    "ENSMUSG00000048758",
                    "ENSMUSG00000022708",
                    "ENSMUSG00000028234",
                    "ENSMUSG00000037206",
                    "ENSMUSG00000019929",
                    "ENSMUSG00000009876",
                    "ENSMUSG00000028936",
                    "ENSMUSG00000028495",
                    "ENSMUSG00000018830",
                    "ENSMUSG00000027004",
                    "ENSMUSG00000003429",
                    "ENSMUSG00000000247",
                    "ENSMUSG00000090137",
                    "ENSMUSG00000035783",
                    "ENSMUSG00000030849",
                    "ENSMUSG00000029231",
                    "ENSMUSG00000020902",
                    "ENSMUSG00000037563",
                    "ENSMUSG00000032399",
                    "ENSMUSG00000031253"
                ],
                "expression_sum": {
                    "class1": 22064.0,
                    "class2": 3667.0
                },
                "ens_id": "ENSMUSG00000038193",
                "uniprot_ids": [
                    "Q61039"
                ]
            },
            "ENSMUSG00000032334": {
                "id": "ENSMUSG00000032334",
                "type": "Gene",
                "symbol": "Loxl1",
                "source": [
                    "ENSMUSG00000052684",
                    "ENSMUSG00000005583",
                    "ENSMUSG00000053477",
                    "ENSMUSG00000003032",
                    "ENSMUSG00000008575",
                    "ENSMUSG00000028565",
                    "ENSMUSG00000071076"
                ],
                "target": [],
                "expression_sum": {
                    "class1": 6154.0,
                    "class2": 11476.0
                },
                "ens_id": "ENSMUSG00000032334",
                "uniprot_ids": [
                    "A0A0R4J0Q4"
                ]
            },
            "ENSMUSG00000018339": {
                "id": "ENSMUSG00000018339",
                "type": "Gene",
                "symbol": "Gpx3",
                "source": [
                    "ENSMUSG00000022346",
                    "ENSMUSG00000021250",
                    "ENSMUSG00000052684",
                    "ENSMUSG00000005583"
                ],
                "target": [],
                "expression_sum": {
                    "class1": 18458.0,
                    "class2": 58950.0
                },
                "ens_id": "ENSMUSG00000018339",
                "uniprot_ids": [
                    "P46412",
                    "A0A1C7ZMZ5",
                    "A0A1C7ZMZ7"
                ]
            },
            "ENSMUSG00000068798": {
                "id": "ENSMUSG00000068798",
                "type": "Gene",
                "symbol": "Rap1a",
                "source": [
                    "ENSMUSG00000029322",
                    "ENSMUSG00000005583"
                ],
                "target": [],
                "expression_sum": {
                    "class1": 7485.0,
                    "class2": 10382.0
                },
                "ens_id": "ENSMUSG00000068798",
                "uniprot_ids": [
                    "P62835",
                    "A0A0G2JDL9",
                    "A0A0G2JED9",
                    "A0A0G2JE52"
                ]
            },
            "ENSMUSG00000024164": {
                "id": "ENSMUSG00000024164",
                "type": "Gene",
                "symbol": "C3",
                "source": [
                    "ENSMUSG00000052684",
                    "ENSMUSG00000022346",
                    "ENSMUSG00000005583",
                    "ENSMUSG00000008575",
                    "ENSMUSG00000028565",
                    "ENSMUSG00000020484",
                    "ENSMUSG00000071076"
                ],
                "target": [],
                "expression_sum": {
                    "class1": 2405.0,
                    "class2": 19434.0
                },
                "ens_id": "ENSMUSG00000024164",
                "uniprot_ids": [
                    "P01027",
                    "H3BL60",
                    "H3BKW9"
                ]
            },
            "ENSMUSG00000047945": {
                "id": "ENSMUSG00000047945",
                "type": "Gene",
                "symbol": "Marcksl1",
                "source": [
                    "ENSMUSG00000056501",
                    "ENSMUSG00000052684",
                    "ENSMUSG00000023034",
                    "ENSMUSG00000052837",
                    "ENSMUSG00000026628",
                    "ENSMUSG00000030103",
                    "ENSMUSG00000022346",
                    "ENSMUSG00000021250",
                    "ENSMUSG00000003545",
                    "ENSMUSG00000003032",
                    "ENSMUSG00000071076"
                ],
                "target": [],
                "expression_sum": {
                    "class1": 40746.0,
                    "class2": 794.0
                },
                "ens_id": "ENSMUSG00000047945",
                "uniprot_ids": [
                    "P28667"
                ]
            },
            "ENSMUSG00000020484": {
                "id": "ENSMUSG00000020484",
                "type": "TF",
                "symbol": "Xbp1",
                "source": [
                    "ENSMUSG00000038193",
                    "ENSMUSG00000023034",
                    "ENSMUSG00000005583"
                ],
                "target": [
                    "ENSMUSG00000025393",
                    "ENSMUSG00000031431",
                    "ENSMUSG00000020053",
                    "ENSMUSG00000024063",
                    "ENSMUSG00000054808",
                    "ENSMUSG00000064370",
                    "ENSMUSG00000028339",
                    "ENSMUSG00000069601",
                    "ENSMUSG00000021242",
                    "ENSMUSG00000002014",
                    "ENSMUSG00000062210",
                    "ENSMUSG00000059895",
                    "ENSMUSG00000090841",
                    "ENSMUSG00000045636",
                    "ENSMUSG00000017707",
                    "ENSMUSG00000032366",
                    "ENSMUSG00000042453",
                    "ENSMUSG00000025213",
                    "ENSMUSG00000058056",
                    "ENSMUSG00000027215",
                    "ENSMUSG00000016319",
                    "ENSMUSG00000038991",
                    "ENSMUSG00000006800",
                    "ENSMUSG00000031565",
                    "ENSMUSG00000031556",
                    "ENSMUSG00000022528",
                    "ENSMUSG00000036438",
                    "ENSMUSG00000042613",
                    "ENSMUSG00000014956",
                    "ENSMUSG00000032353",
                    "ENSMUSG00000074227",
                    "ENSMUSG00000003477",
                    "ENSMUSG00000025809",
                    "ENSMUSG00000030659",
                    "ENSMUSG00000021870",
                    "ENSMUSG00000064354",
                    "ENSMUSG00000026421",
                    "ENSMUSG00000035885",
                    "ENSMUSG00000030403",
                    "ENSMUSG00000023272",
                    "ENSMUSG00000028128",
                    "ENSMUSG00000020423",
                    "ENSMUSG00000037712",
                    "ENSMUSG00000022010",
                    "ENSMUSG00000015932",
                    "ENSMUSG00000028464",
                    "ENSMUSG00000020612",
                    "ENSMUSG00000024012",
                    "ENSMUSG00000040170",
                    "ENSMUSG00000008540",
                    "ENSMUSG00000006342",
                    "ENSMUSG00000074457",
                    "ENSMUSG00000021876",
                    "ENSMUSG00000021268",
                    "ENSMUSG00000004085",
                    "ENSMUSG00000033306",
                    "ENSMUSG00000096145",
                    "ENSMUSG00000025147",
                    "ENSMUSG00000024164",
                    "ENSMUSG00000006205",
                    "ENSMUSG00000057666",
                    "ENSMUSG00000064215",
                    "ENSMUSG00000014846",
                    "ENSMUSG00000016024",
                    "ENSMUSG00000017493",
                    "ENSMUSG00000041084",
                    "ENSMUSG00000025351"
                ],
                "expression_sum": {
                    "class1": 12449.0,
                    "class2": 6214.0
                },
                "ens_id": "ENSMUSG00000020484",
                "uniprot_ids": [
                    "O35426",
                    "G3UYH6",
                    "H3BLF6",
                    "A0A5F8MPD3"
                ]
            },
            "ENSMUSG00000032515": {
                "id": "ENSMUSG00000032515",
                "type": "Gene",
                "symbol": "Csrnp1",
                "source": [
                    "ENSMUSG00000041515",
                    "ENSMUSG00000056501",
                    "ENSMUSG00000052684",
                    "ENSMUSG00000023034",
                    "ENSMUSG00000052837",
                    "ENSMUSG00000026628",
                    "ENSMUSG00000030103",
                    "ENSMUSG00000021250",
                    "ENSMUSG00000003032",
                    "ENSMUSG00000018899",
                    "ENSMUSG00000071076"
                ],
                "target": [],
                "expression_sum": {
                    "class1": 2983.0,
                    "class2": 1546.0
                },
                "ens_id": "ENSMUSG00000032515",
                "uniprot_ids": [
                    "P59054",
                    "A0A1L1SRY6",
                    "A0A1L1SRU4"
                ]
            },
            "ENSMUSG00000020423": {
                "id": "ENSMUSG00000020423",
                "type": "Gene",
                "symbol": "Btg2",
                "source": [
                    "ENSMUSG00000041515",
                    "ENSMUSG00000056501",
                    "ENSMUSG00000052684",
                    "ENSMUSG00000037868",
                    "ENSMUSG00000052837",
                    "ENSMUSG00000026628",
                    "ENSMUSG00000030103",
                    "ENSMUSG00000022346",
                    "ENSMUSG00000038418",
                    "ENSMUSG00000021250",
                    "ENSMUSG00000055148",
                    "ENSMUSG00000003545",
                    "ENSMUSG00000003032",
                    "ENSMUSG00000018899",
                    "ENSMUSG00000020484",
                    "ENSMUSG00000071076"
                ],
                "target": [],
                "expression_sum": {
                    "class1": 26463.0,
                    "class2": 18373.0
                },
                "ens_id": "ENSMUSG00000020423",
                "uniprot_ids": [
                    "Q04211"
                ]
            },
            "ENSMUSG00000046402": {
                "id": "ENSMUSG00000046402",
                "type": "Gene",
                "symbol": "Rbp1",
                "source": [
                    "ENSMUSG00000032035",
                    "ENSMUSG00000052684",
                    "ENSMUSG00000005583"
                ],
                "target": [],
                "expression_sum": {
                    "class1": 108824.0,
                    "class2": 14592.0
                },
                "ens_id": "ENSMUSG00000046402",
                "uniprot_ids": [
                    "Q00915"
                ]
            },
            "ENSMUSG00000003032": {
                "id": "ENSMUSG00000003032",
                "type": "TF",
                "symbol": "Klf4",
                "source": [
                    "ENSMUSG00000037868",
                    "ENSMUSG00000023034",
                    "ENSMUSG00000052837",
                    "ENSMUSG00000026628",
                    "ENSMUSG00000057098",
                    "ENSMUSG00000038418",
                    "ENSMUSG00000021250",
                    "ENSMUSG00000003545",
                    "ENSMUSG00000018899"
                ],
                "target": [
                    "ENSMUSG00000030116",
                    "ENSMUSG00000007815",
                    "ENSMUSG00000005338",
                    "ENSMUSG00000019505",
                    "ENSMUSG00000059291",
                    "ENSMUSG00000031447",
                    "ENSMUSG00000002602",
                    "ENSMUSG00000031765",
                    "ENSMUSG00000028111",
                    "ENSMUSG00000031875",
                    "ENSMUSG00000007655",
                    "ENSMUSG00000021477",
                    "ENSMUSG00000028691",
                    "ENSMUSG00000003545",
                    "ENSMUSG00000024011",
                    "ENSMUSG00000053007",
                    "ENSMUSG00000026879",
                    "ENSMUSG00000037822",
                    "ENSMUSG00000048076",
                    "ENSMUSG00000056481",
                    "ENSMUSG00000037411",
                    "ENSMUSG00000022371",
                    "ENSMUSG00000044337",
                    "ENSMUSG00000064373",
                    "ENSMUSG00000028822",
                    "ENSMUSG00000015354",
                    "ENSMUSG00000034868",
                    "ENSMUSG00000006699",
                    "ENSMUSG00000044600",
                    "ENSMUSG00000029649",
                    "ENSMUSG00000008384",
                    "ENSMUSG00000055148",
                    "ENSMUSG00000001240",
                    "ENSMUSG00000012422",
                    "ENSMUSG00000023004",
                    "ENSMUSG00000039405",
                    "ENSMUSG00000002107",
                    "ENSMUSG00000069763",
                    "ENSMUSG00000003541",
                    "ENSMUSG00000060572",
                    "ENSMUSG00000032231",
                    "ENSMUSG00000007872",
                    "ENSMUSG00000056737",
                    "ENSMUSG00000020467",
                    "ENSMUSG00000035212",
                    "ENSMUSG00000022032",
                    "ENSMUSG00000037499",
                    "ENSMUSG00000021390",
                    "ENSMUSG00000009013",
                    "ENSMUSG00000009681",
                    "ENSMUSG00000018585",
                    "ENSMUSG00000020053",
                    "ENSMUSG00000025132",
                    "ENSMUSG00000020440",
                    "ENSMUSG00000028367",
                    "ENSMUSG00000053113",
                    "ENSMUSG00000037266",
                    "ENSMUSG00000084786",
                    "ENSMUSG00000031626",
                    "ENSMUSG00000036446",
                    "ENSMUSG00000020737",
                    "ENSMUSG00000029767",
                    "ENSMUSG00000055681",
                    "ENSMUSG00000031538",
                    "ENSMUSG00000019978",
                    "ENSMUSG00000019278",
                    "ENSMUSG00000021760",
                    "ENSMUSG00000090000",
                    "ENSMUSG00000036478",
                    "ENSMUSG00000022816",
                    "ENSMUSG00000027204",
                    "ENSMUSG00000041959",
                    "ENSMUSG00000033585",
                    "ENSMUSG00000026628",
                    "ENSMUSG00000007039",
                    "ENSMUSG00000027706",
                    "ENSMUSG00000074743",
                    "ENSMUSG00000022090",
                    "ENSMUSG00000041828",
                    "ENSMUSG00000032014",
                    "ENSMUSG00000032515",
                    "ENSMUSG00000092274",
                    "ENSMUSG00000017466",
                    "ENSMUSG00000074457",
                    "ENSMUSG00000027829",
                    "ENSMUSG00000032334",
                    "ENSMUSG00000023092",
                    "ENSMUSG00000072235",
                    "ENSMUSG00000038463",
                    "ENSMUSG00000005397",
                    "ENSMUSG00000022894",
                    "ENSMUSG00000015085",
                    "ENSMUSG00000030208",
                    "ENSMUSG00000054452",
                    "ENSMUSG00000068220",
                    "ENSMUSG00000028517",
                    "ENSMUSG00000040212",
                    "ENSMUSG00000023034",
                    "ENSMUSG00000079523",
                    "ENSMUSG00000030654",
                    "ENSMUSG00000016494",
                    "ENSMUSG00000001123",
                    "ENSMUSG00000026674",
                    "ENSMUSG00000020315",
                    "ENSMUSG00000037706",
                    "ENSMUSG00000018217",
                    "ENSMUSG00000017002",
                    "ENSMUSG00000003161",
                    "ENSMUSG00000060591",
                    "ENSMUSG00000025784",
                    "ENSMUSG00000040562",
                    "ENSMUSG00000072812",
                    "ENSMUSG00000015312",
                    "ENSMUSG00000040280",
                    "ENSMUSG00000024659",
                    "ENSMUSG00000018819",
                    "ENSMUSG00000032301",
                    "ENSMUSG00000020423",
                    "ENSMUSG00000027712",
                    "ENSMUSG00000024048",
                    "ENSMUSG00000069662",
                    "ENSMUSG00000028108",
                    "ENSMUSG00000037072",
                    "ENSMUSG00000002732",
                    "ENSMUSG00000021876",
                    "ENSMUSG00000001891",
                    "ENSMUSG00000026478",
                    "ENSMUSG00000037868",
                    "ENSMUSG00000042312",
                    "ENSMUSG00000057098",
                    "ENSMUSG00000031722",
                    "ENSMUSG00000075254",
                    "ENSMUSG00000023010",
                    "ENSMUSG00000045658",
                    "ENSMUSG00000031799",
                    "ENSMUSG00000014846",
                    "ENSMUSG00000017493",
                    "ENSMUSG00000041084",
                    "ENSMUSG00000039601",
                    "ENSMUSG00000027907",
                    "ENSMUSG00000074884",
                    "ENSMUSG00000024308",
                    "ENSMUSG00000024014",
                    "ENSMUSG00000000628",
                    "ENSMUSG00000028249",
                    "ENSMUSG00000095385",
                    "ENSMUSG00000020849",
                    "ENSMUSG00000069833",
                    "ENSMUSG00000041736",
                    "ENSMUSG00000087006",
                    "ENSMUSG00000034993",
                    "ENSMUSG00000026670",
                    "ENSMUSG00000075602",
                    "ENSMUSG00000020585",
                    "ENSMUSG00000025492",
                    "ENSMUSG00000007041",
                    "ENSMUSG00000021250",
                    "ENSMUSG00000042747",
                    "ENSMUSG00000029096",
                    "ENSMUSG00000026837",
                    "ENSMUSG00000028618",
                    "ENSMUSG00000032383",
                    "ENSMUSG00000026748",
                    "ENSMUSG00000031422",
                    "ENSMUSG00000047907",
                    "ENSMUSG00000047945",
                    "ENSMUSG00000052837",
                    "ENSMUSG00000035356",
                    "ENSMUSG00000037206",
                    "ENSMUSG00000024012",
                    "ENSMUSG00000029659",
                    "ENSMUSG00000060802",
                    "ENSMUSG00000026574",
                    "ENSMUSG00000044786",
                    "ENSMUSG00000018899",
                    "ENSMUSG00000079037",
                    "ENSMUSG00000090698",
                    "ENSMUSG00000075702",
                    "ENSMUSG00000060703",
                    "ENSMUSG00000019087",
                    "ENSMUSG00000022261",
                    "ENSMUSG00000038418",
                    "ENSMUSG00000029484",
                    "ENSMUSG00000040026",
                    "ENSMUSG00000001025",
                    "ENSMUSG00000058135"
                ],
                "expression_sum": {
                    "class1": 8719.0,
                    "class2": 10220.0
                },
                "ens_id": "ENSMUSG00000003032",
                "uniprot_ids": [
                    "Q60793",
                    "B7ZCH2",
                    "B7ZCH1"
                ]
            },
            "ENSMUSG00000004791": {
                "id": "ENSMUSG00000004791",
                "type": "Gene",
                "symbol": "Pgf",
                "source": [
                    "ENSMUSG00000005583"
                ],
                "target": [],
                "expression_sum": {
                    "class1": 37.0,
                    "class2": 3853.0
                },
                "ens_id": "ENSMUSG00000004791",
                "uniprot_ids": [
                    "P49764",
                    "A0A1Y7VIW3"
                ]
            },
            "ENSMUSG00000042485": {
                "id": "ENSMUSG00000042485",
                "type": "Gene",
                "symbol": "Mustn1",
                "source": [
                    "ENSMUSG00000052684",
                    "ENSMUSG00000053477",
                    "ENSMUSG00000029322",
                    "ENSMUSG00000005583",
                    "ENSMUSG00000028565"
                ],
                "target": [],
                "expression_sum": {
                    "class1": 47208.0,
                    "class2": 55732.0
                },
                "ens_id": "ENSMUSG00000042485",
                "uniprot_ids": [
                    "Q99JI1"
                ]
            },
            "ENSMUSG00000068735": {
                "id": "ENSMUSG00000068735",
                "type": "Gene",
                "symbol": "Trp53i11",
                "source": [
                    "ENSMUSG00000052684",
                    "ENSMUSG00000023034",
                    "ENSMUSG00000047638",
                    "ENSMUSG00000008575",
                    "ENSMUSG00000005583",
                    "ENSMUSG00000071076"
                ],
                "target": [],
                "expression_sum": {
                    "class1": 7085.0,
                    "class2": 4141.0
                },
                "ens_id": "ENSMUSG00000068735",
                "uniprot_ids": [
                    "Q4QQM4",
                    "A2AGS6"
                ]
            },
            "ENSMUSG00000001520": {
                "id": "ENSMUSG00000001520",
                "type": "Gene",
                "symbol": "Nrip2",
                "source": [
                    "ENSMUSG00000022346",
                    "ENSMUSG00000023034",
                    "ENSMUSG00000025408",
                    "ENSMUSG00000005583"
                ],
                "target": [],
                "expression_sum": {
                    "class1": 6.0,
                    "class2": 7268.0
                },
                "ens_id": "ENSMUSG00000001520",
                "uniprot_ids": [
                    "Q9JHR9",
                    "A0A0N4SUN5",
                    "F7BIJ5",
                    "E9QAD1"
                ]
            },
            "ENSMUSG00000025784": {
                "id": "ENSMUSG00000025784",
                "type": "Gene",
                "symbol": "Clec3b",
                "source": [
                    "ENSMUSG00000052684",
                    "ENSMUSG00000022346",
                    "ENSMUSG00000052534",
                    "ENSMUSG00000003032",
                    "ENSMUSG00000028565",
                    "ENSMUSG00000071076"
                ],
                "target": [],
                "expression_sum": {
                    "class1": 197.0,
                    "class2": 48804.0
                },
                "ens_id": "ENSMUSG00000025784",
                "uniprot_ids": [
                    "Q8CFZ6"
                ]
            },
            "ENSMUSG00000024290": {
                "id": "ENSMUSG00000024290",
                "type": "Gene",
                "symbol": "Rock1",
                "source": [
                    "ENSMUSG00000053477",
                    "ENSMUSG00000029322",
                    "ENSMUSG00000008575",
                    "ENSMUSG00000005583",
                    "ENSMUSG00000020644",
                    "ENSMUSG00000071076"
                ],
                "target": [],
                "expression_sum": {
                    "class1": 3778.0,
                    "class2": 7101.0
                },
                "ens_id": "ENSMUSG00000024290",
                "uniprot_ids": [
                    "P70335",
                    "A0A3Q4L2X0",
                    "A0A3Q4EGR2"
                ]
            },
            "ENSMUSG00000006205": {
                "id": "ENSMUSG00000006205",
                "type": "Gene",
                "symbol": "Htra1",
                "source": [
                    "ENSMUSG00000052684",
                    "ENSMUSG00000052837",
                    "ENSMUSG00000022346",
                    "ENSMUSG00000005583",
                    "ENSMUSG00000053477",
                    "ENSMUSG00000021250",
                    "ENSMUSG00000008575",
                    "ENSMUSG00000020484",
                    "ENSMUSG00000071076"
                ],
                "target": [],
                "expression_sum": {
                    "class1": 1974.0,
                    "class2": 10796.0
                },
                "ens_id": "ENSMUSG00000006205",
                "uniprot_ids": [
                    "Q9R118"
                ]
            },
            "ENSMUSG00000072235": {
                "id": "ENSMUSG00000072235",
                "type": "Gene",
                "symbol": "Tuba1a",
                "source": [
                    "ENSMUSG00000055435",
                    "ENSMUSG00000032035",
                    "ENSMUSG00000052684",
                    "ENSMUSG00000005583",
                    "ENSMUSG00000057098",
                    "ENSMUSG00000052534",
                    "ENSMUSG00000003032",
                    "ENSMUSG00000008575",
                    "ENSMUSG00000071076"
                ],
                "target": [],
                "expression_sum": {
                    "class1": 27755.0,
                    "class2": 25732.0
                },
                "ens_id": "ENSMUSG00000072235",
                "uniprot_ids": [
                    "P68369"
                ]
            },
            "ENSMUSG00000023034": {
                "id": "ENSMUSG00000023034",
                "type": "TF",
                "symbol": "Nr4a1",
                "source": [
                    "ENSMUSG00000041515",
                    "ENSMUSG00000037868",
                    "ENSMUSG00000052684",
                    "ENSMUSG00000052837",
                    "ENSMUSG00000030103",
                    "ENSMUSG00000038418",
                    "ENSMUSG00000021250",
                    "ENSMUSG00000003545",
                    "ENSMUSG00000003032",
                    "ENSMUSG00000018899",
                    "ENSMUSG00000071076"
                ],
                "target": [
                    "ENSMUSG00000006360",
                    "ENSMUSG00000029761",
                    "ENSMUSG00000031765",
                    "ENSMUSG00000028949",
                    "ENSMUSG00000003545",
                    "ENSMUSG00000022283",
                    "ENSMUSG00000036854",
                    "ENSMUSG00000040952",
                    "ENSMUSG00000026879",
                    "ENSMUSG00000031320",
                    "ENSMUSG00000032330",
                    "ENSMUSG00000037405",
                    "ENSMUSG00000036438",
                    "ENSMUSG00000008384",
                    "ENSMUSG00000008668",
                    "ENSMUSG00000020372",
                    "ENSMUSG00000021939",
                    "ENSMUSG00000008682",
                    "ENSMUSG00000055148",
                    "ENSMUSG00000035493",
                    "ENSMUSG00000021127",
                    "ENSMUSG00000097296",
                    "ENSMUSG00000034892",
                    "ENSMUSG00000071415",
                    "ENSMUSG00000041329",
                    "ENSMUSG00000061983",
                    "ENSMUSG00000061353",
                    "ENSMUSG00000031740",
                    "ENSMUSG00000048096",
                    "ENSMUSG00000036545",
                    "ENSMUSG00000040666",
                    "ENSMUSG00000032085",
                    "ENSMUSG00000037742",
                    "ENSMUSG00000053113",
                    "ENSMUSG00000032366",
                    "ENSMUSG00000021520",
                    "ENSMUSG00000033538",
                    "ENSMUSG00000018171",
                    "ENSMUSG00000020810",
                    "ENSMUSG00000052684",
                    "ENSMUSG00000001119",
                    "ENSMUSG00000014956",
                    "ENSMUSG00000031762",
                    "ENSMUSG00000032744",
                    "ENSMUSG00000027188",
                    "ENSMUSG00000002980",
                    "ENSMUSG00000030103",
                    "ENSMUSG00000025290",
                    "ENSMUSG00000028464",
                    "ENSMUSG00000038146",
                    "ENSMUSG00000032515",
                    "ENSMUSG00000068735",
                    "ENSMUSG00000028776",
                    "ENSMUSG00000045128",
                    "ENSMUSG00000022587",
                    "ENSMUSG00000027829",
                    "ENSMUSG00000002985",
                    "ENSMUSG00000031328",
                    "ENSMUSG00000030432",
                    "ENSMUSG00000074129",
                    "ENSMUSG00000090733",
                    "ENSMUSG00000022817",
                    "ENSMUSG00000059895",
                    "ENSMUSG00000056708",
                    "ENSMUSG00000060126",
                    "ENSMUSG00000046727",
                    "ENSMUSG00000001175",
                    "ENSMUSG00000001520",
                    "ENSMUSG00000012848",
                    "ENSMUSG00000032387",
                    "ENSMUSG00000021025",
                    "ENSMUSG00000049517",
                    "ENSMUSG00000027074",
                    "ENSMUSG00000029614",
                    "ENSMUSG00000003032",
                    "ENSMUSG00000062006",
                    "ENSMUSG00000040128",
                    "ENSMUSG00000079435",
                    "ENSMUSG00000078202",
                    "ENSMUSG00000020205",
                    "ENSMUSG00000024232",
                    "ENSMUSG00000031029",
                    "ENSMUSG00000023089",
                    "ENSMUSG00000001270",
                    "ENSMUSG00000050708",
                    "ENSMUSG00000037868",
                    "ENSMUSG00000028214",
                    "ENSMUSG00000035828",
                    "ENSMUSG00000067288",
                    "ENSMUSG00000023224",
                    "ENSMUSG00000052146",
                    "ENSMUSG00000039601",
                    "ENSMUSG00000060938",
                    "ENSMUSG00000067818",
                    "ENSMUSG00000067274",
                    "ENSMUSG00000024014",
                    "ENSMUSG00000000628",
                    "ENSMUSG00000074170",
                    "ENSMUSG00000090841",
                    "ENSMUSG00000028266",
                    "ENSMUSG00000020484",
                    "ENSMUSG00000071866",
                    "ENSMUSG00000028864",
                    "ENSMUSG00000008683",
                    "ENSMUSG00000021250",
                    "ENSMUSG00000007877",
                    "ENSMUSG00000000740",
                    "ENSMUSG00000007892",
                    "ENSMUSG00000021186",
                    "ENSMUSG00000024190",
                    "ENSMUSG00000057841",
                    "ENSMUSG00000041515",
                    "ENSMUSG00000035356",
                    "ENSMUSG00000052837",
                    "ENSMUSG00000047945",
                    "ENSMUSG00000022708",
                    "ENSMUSG00000028234",
                    "ENSMUSG00000015932",
                    "ENSMUSG00000028936",
                    "ENSMUSG00000074578",
                    "ENSMUSG00000018899",
                    "ENSMUSG00000000247",
                    "ENSMUSG00000038418",
                    "ENSMUSG00000035783",
                    "ENSMUSG00000037563"
                ],
                "expression_sum": {
                    "class1": 11016.0,
                    "class2": 10485.0
                },
                "ens_id": "ENSMUSG00000023034",
                "uniprot_ids": [
                    "P12813",
                    "A0A2R8VJZ1"
                ]
            },
            "ENSMUSG00000023064": {
                "id": "ENSMUSG00000023064",
                "type": "Gene",
                "symbol": "Sncg",
                "source": [
                    "ENSMUSG00000055148",
                    "ENSMUSG00000005583"
                ],
                "target": [],
                "expression_sum": {
                    "class1": 1.0,
                    "class2": 15774.0
                },
                "ens_id": "ENSMUSG00000023064",
                "uniprot_ids": [
                    "Q9Z0F7"
                ]
            },
            "ENSMUSG00000032387": {
                "id": "ENSMUSG00000032387",
                "type": "Gene",
                "symbol": "Rbpms2",
                "source": [
                    "ENSMUSG00000058756",
                    "ENSMUSG00000020644",
                    "ENSMUSG00000023034",
                    "ENSMUSG00000005583"
                ],
                "target": [],
                "expression_sum": {
                    "class1": 485.0,
                    "class2": 4556.0
                },
                "ens_id": "ENSMUSG00000032387",
                "uniprot_ids": [
                    "Q8VC52",
                    "Q9CWY6",
                    "A0A1L1SVH5",
                    "A0A1L1SRM3"
                ]
            },
            "ENSMUSG00000026879": {
                "id": "ENSMUSG00000026879",
                "type": "Gene",
                "symbol": "Gsn",
                "source": [
                    "ENSMUSG00000052684",
                    "ENSMUSG00000023034",
                    "ENSMUSG00000022528",
                    "ENSMUSG00000022346",
                    "ENSMUSG00000005583",
                    "ENSMUSG00000003032",
                    "ENSMUSG00000008575",
                    "ENSMUSG00000028565",
                    "ENSMUSG00000071076"
                ],
                "target": [],
                "expression_sum": {
                    "class1": 20316.0,
                    "class2": 411941.0
                },
                "ens_id": "ENSMUSG00000026879",
                "uniprot_ids": [
                    "P13020",
                    "A6PWS5",
                    "A0A0J9YUQ8",
                    "A0A0J9YUJ8"
                ]
            },
            "ENSMUSG00000026574": {
                "id": "ENSMUSG00000026574",
                "type": "Gene",
                "symbol": "Dpt",
                "source": [
                    "ENSMUSG00000022346",
                    "ENSMUSG00000066551",
                    "ENSMUSG00000003032",
                    "ENSMUSG00000005583",
                    "ENSMUSG00000028565",
                    "ENSMUSG00000021127"
                ],
                "target": [],
                "expression_sum": {
                    "class1": 8606.0,
                    "class2": 48325.0
                },
                "ens_id": "ENSMUSG00000026574",
                "uniprot_ids": [
                    "Q9QZZ6"
                ]
            },
            "ENSMUSG00000044786": {
                "id": "ENSMUSG00000044786",
                "type": "Gene",
                "symbol": "Zfp36",
                "source": [
                    "ENSMUSG00000041515",
                    "ENSMUSG00000056501",
                    "ENSMUSG00000052684",
                    "ENSMUSG00000037868",
                    "ENSMUSG00000052837",
                    "ENSMUSG00000026628",
                    "ENSMUSG00000030103",
                    "ENSMUSG00000048387",
                    "ENSMUSG00000000247",
                    "ENSMUSG00000038418",
                    "ENSMUSG00000021250",
                    "ENSMUSG00000055148",
                    "ENSMUSG00000003032",
                    "ENSMUSG00000018899",
                    "ENSMUSG00000071076"
                ],
                "target": [],
                "expression_sum": {
                    "class1": 49942.0,
                    "class2": 16051.0
                },
                "ens_id": "ENSMUSG00000044786",
                "uniprot_ids": [
                    "P22893",
                    "Q9DBE5"
                ]
            },
            "ENSMUSG00000052997": {
                "id": "ENSMUSG00000052997",
                "type": "Gene",
                "symbol": "Uba2",
                "source": [
                    "ENSMUSG00000052684",
                    "ENSMUSG00000047638",
                    "ENSMUSG00000053477",
                    "ENSMUSG00000005583",
                    "ENSMUSG00000071076"
                ],
                "target": [],
                "expression_sum": {
                    "class1": 2483.0,
                    "class2": 7173.0
                },
                "ens_id": "ENSMUSG00000052997",
                "uniprot_ids": [
                    "Q9Z1F9",
                    "H3BLM2",
                    "A2BH29",
                    "H3BJQ2",
                    "H3BLR3"
                ]
            },
            "ENSMUSG00000087006": {
                "id": "ENSMUSG00000087006",
                "type": "Gene",
                "symbol": "Gm13889",
                "source": [
                    "ENSMUSG00000041515",
                    "ENSMUSG00000052684",
                    "ENSMUSG00000022346",
                    "ENSMUSG00000003545",
                    "ENSMUSG00000003032",
                    "ENSMUSG00000005583"
                ],
                "target": [],
                "expression_sum": {
                    "class1": 20589.0,
                    "class2": 22077.0
                },
                "ens_id": "ENSMUSG00000087006",
                "uniprot_ids": [
                    "A0A087WSH6",
                    "A0A087WRN5"
                ]
            },
            "ENSMUSG00000025351": {
                "id": "ENSMUSG00000025351",
                "type": "Gene",
                "symbol": "Cd63",
                "source": [
                    "ENSMUSG00000052684",
                    "ENSMUSG00000057098",
                    "ENSMUSG00000038418",
                    "ENSMUSG00000038193",
                    "ENSMUSG00000005583",
                    "ENSMUSG00000020484",
                    "ENSMUSG00000071076"
                ],
                "target": [],
                "expression_sum": {
                    "class1": 51939.0,
                    "class2": 32064.0
                },
                "ens_id": "ENSMUSG00000025351",
                "uniprot_ids": [
                    "P41731",
                    "A0A1W2P8C6"
                ]
            },
            "ENSMUSG00000078202": {
                "id": "ENSMUSG00000078202",
                "type": "Gene",
                "symbol": "Nrarp",
                "source": [
                    "ENSMUSG00000023034",
                    "ENSMUSG00000005583"
                ],
                "target": [],
                "expression_sum": {
                    "class1": 3.0,
                    "class2": 6494.0
                },
                "ens_id": "ENSMUSG00000078202",
                "uniprot_ids": [
                    "Q91ZA8"
                ]
            },
            "ENSMUSG00000041959": {
                "id": "ENSMUSG00000041959",
                "type": "Gene",
                "symbol": "S100a10",
                "source": [
                    "ENSMUSG00000052684",
                    "ENSMUSG00000005583",
                    "ENSMUSG00000003032",
                    "ENSMUSG00000055435",
                    "ENSMUSG00000028565",
                    "ENSMUSG00000071076"
                ],
                "target": [],
                "expression_sum": {
                    "class1": 19414.0,
                    "class2": 23795.0
                },
                "ens_id": "ENSMUSG00000041959",
                "uniprot_ids": [
                    "P08207"
                ]
            },
            "ENSMUSG00000042312": {
                "id": "ENSMUSG00000042312",
                "type": "Gene",
                "symbol": "S100a13",
                "source": [
                    "ENSMUSG00000052684",
                    "ENSMUSG00000057098",
                    "ENSMUSG00000003032",
                    "ENSMUSG00000005583",
                    "ENSMUSG00000071076"
                ],
                "target": [],
                "expression_sum": {
                    "class1": 4095.0,
                    "class2": 6574.0
                },
                "ens_id": "ENSMUSG00000042312",
                "uniprot_ids": [
                    "P97352",
                    "A0A0A0MQ90"
                ]
            },
            "ENSMUSG00000019929": {
                "id": "ENSMUSG00000019929",
                "type": "Gene",
                "symbol": "Dcn",
                "source": [
                    "ENSMUSG00000032035",
                    "ENSMUSG00000056501",
                    "ENSMUSG00000038193",
                    "ENSMUSG00000005583",
                    "ENSMUSG00000028565",
                    "ENSMUSG00000071076"
                ],
                "target": [],
                "expression_sum": {
                    "class1": 1401173.0,
                    "class2": 310161.0
                },
                "ens_id": "ENSMUSG00000019929",
                "uniprot_ids": [
                    "P28654",
                    "A0A1W2P6I8"
                ]
            },
            "ENSMUSG00000025147": {
                "id": "ENSMUSG00000025147",
                "type": "Gene",
                "symbol": "Mob2",
                "source": [
                    "ENSMUSG00000052684",
                    "ENSMUSG00000047638",
                    "ENSMUSG00000008575",
                    "ENSMUSG00000005583",
                    "ENSMUSG00000020484"
                ],
                "target": [],
                "expression_sum": {
                    "class1": 1886.0,
                    "class2": 6267.0
                },
                "ens_id": "ENSMUSG00000025147",
                "uniprot_ids": [
                    "Q8VI63",
                    "A0A1B0GT55",
                    "A0A1B0GR18",
                    "A0A1B0GRX6"
                ]
            },
            "ENSMUSG00000029359": {
                "id": "ENSMUSG00000029359",
                "type": "Gene",
                "symbol": "Tesc",
                "source": [
                    "ENSMUSG00000005583"
                ],
                "target": [],
                "expression_sum": {
                    "class1": 2.0,
                    "class2": 3837.0
                },
                "ens_id": "ENSMUSG00000029359",
                "uniprot_ids": [
                    "Q9JKL5",
                    "D0EM46"
                ]
            },
            "ENSMUSG00000039601": {
                "id": "ENSMUSG00000039601",
                "type": "Gene",
                "symbol": "Rcan2",
                "source": [
                    "ENSMUSG00000056501",
                    "ENSMUSG00000052684",
                    "ENSMUSG00000023034",
                    "ENSMUSG00000052837",
                    "ENSMUSG00000026628",
                    "ENSMUSG00000015839",
                    "ENSMUSG00000052534",
                    "ENSMUSG00000038193",
                    "ENSMUSG00000021250",
                    "ENSMUSG00000003545",
                    "ENSMUSG00000003032",
                    "ENSMUSG00000005583",
                    "ENSMUSG00000071076"
                ],
                "target": [],
                "expression_sum": {
                    "class1": 138.0,
                    "class2": 39503.0
                },
                "ens_id": "ENSMUSG00000039601",
                "uniprot_ids": [
                    "Q9JHG2",
                    "Q8VIP4",
                    "A0A2R8W6E8",
                    "A0A2R8VKI0",
                    "A0A2R8W6X3"
                ]
            },
            "ENSMUSG00000042745": {
                "id": "ENSMUSG00000042745",
                "type": "Gene",
                "symbol": "Id1",
                "source": [
                    "ENSMUSG00000005583",
                    "ENSMUSG00000052684",
                    "ENSMUSG00000071076"
                ],
                "target": [],
                "expression_sum": {
                    "class1": 1865.0,
                    "class2": 10252.0
                },
                "ens_id": "ENSMUSG00000042745",
                "uniprot_ids": [
                    "P20067",
                    "A2AHY3"
                ]
            },
            "ENSMUSG00000068220": {
                "id": "ENSMUSG00000068220",
                "type": "Gene",
                "symbol": "Lgals1",
                "source": [
                    "ENSMUSG00000032238",
                    "ENSMUSG00000000078",
                    "ENSMUSG00000052684",
                    "ENSMUSG00000052837",
                    "ENSMUSG00000021250",
                    "ENSMUSG00000003032",
                    "ENSMUSG00000008575",
                    "ENSMUSG00000071076"
                ],
                "target": [],
                "expression_sum": {
                    "class1": 36183.0,
                    "class2": 64109.0
                },
                "ens_id": "ENSMUSG00000068220",
                "uniprot_ids": [
                    "P16045",
                    "A0A2R8VHJ0"
                ]
            },
            "ENSMUSG00000024190": {
                "id": "ENSMUSG00000024190",
                "type": "Gene",
                "symbol": "Dusp1",
                "source": [
                    "ENSMUSG00000023034",
                    "ENSMUSG00000022528",
                    "ENSMUSG00000052837",
                    "ENSMUSG00000026628",
                    "ENSMUSG00000000247",
                    "ENSMUSG00000057098",
                    "ENSMUSG00000021250",
                    "ENSMUSG00000055148",
                    "ENSMUSG00000003545",
                    "ENSMUSG00000008575",
                    "ENSMUSG00000048387",
                    "ENSMUSG00000018899"
                ],
                "target": [],
                "expression_sum": {
                    "class1": 59919.0,
                    "class2": 46620.0
                },
                "ens_id": "ENSMUSG00000024190",
                "uniprot_ids": [
                    "P28563",
                    "A0A494B924",
                    "A0A494B9K9"
                ]
            },
            "ENSMUSG00000022032": {
                "id": "ENSMUSG00000022032",
                "type": "Gene",
                "symbol": "Scara5",
                "source": [
                    "ENSMUSG00000052684",
                    "ENSMUSG00000022346",
                    "ENSMUSG00000057098",
                    "ENSMUSG00000028639",
                    "ENSMUSG00000003032",
                    "ENSMUSG00000005583",
                    "ENSMUSG00000071076"
                ],
                "target": [],
                "expression_sum": {
                    "class1": 23.0,
                    "class2": 3706.0
                },
                "ens_id": "ENSMUSG00000022032",
                "uniprot_ids": [
                    "Q8K299"
                ]
            },
            "ENSMUSG00000002732": {
                "id": "ENSMUSG00000002732",
                "type": "Gene",
                "symbol": "Fkbp7",
                "source": [
                    "ENSMUSG00000052684",
                    "ENSMUSG00000038193",
                    "ENSMUSG00000003032",
                    "ENSMUSG00000005583",
                    "ENSMUSG00000071076"
                ],
                "target": [],
                "expression_sum": {
                    "class1": 9183.0,
                    "class2": 5862.0
                },
                "ens_id": "ENSMUSG00000002732",
                "uniprot_ids": [
                    "O54998",
                    "H3BJI5"
                ]
            },
            "ENSMUSG00000027712": {
                "id": "ENSMUSG00000027712",
                "type": "Gene",
                "symbol": "Anxa5",
                "source": [
                    "ENSMUSG00000032238",
                    "ENSMUSG00000032035",
                    "ENSMUSG00000052684",
                    "ENSMUSG00000057098",
                    "ENSMUSG00000038193",
                    "ENSMUSG00000003032",
                    "ENSMUSG00000005583",
                    "ENSMUSG00000028565",
                    "ENSMUSG00000071076"
                ],
                "target": [],
                "expression_sum": {
                    "class1": 26423.0,
                    "class2": 18213.0
                },
                "ens_id": "ENSMUSG00000027712",
                "uniprot_ids": [
                    "P48036",
                    "A0A0G2JGQ0"
                ]
            },
            "ENSMUSG00000007655": {
                "id": "ENSMUSG00000007655",
                "type": "Gene",
                "symbol": "Cav1",
                "source": [
                    "ENSMUSG00000048387",
                    "ENSMUSG00000038193",
                    "ENSMUSG00000003032",
                    "ENSMUSG00000005583"
                ],
                "target": [],
                "expression_sum": {
                    "class1": 18.0,
                    "class2": 7289.0
                },
                "ens_id": "ENSMUSG00000007655",
                "uniprot_ids": [
                    "P49817",
                    "D3Z0J2",
                    "D3Z148",
                    "H3BLQ4",
                    "H3BKG0"
                ]
            },
            "ENSMUSG00000029231": {
                "id": "ENSMUSG00000029231",
                "type": "Gene",
                "symbol": "Pdgfra",
                "source": [
                    "ENSMUSG00000048387",
                    "ENSMUSG00000038193",
                    "ENSMUSG00000005583",
                    "ENSMUSG00000071076"
                ],
                "target": [],
                "expression_sum": {
                    "class1": 12310.0,
                    "class2": 6070.0
                },
                "ens_id": "ENSMUSG00000029231",
                "uniprot_ids": [
                    "P26618",
                    "A0A0J9YV49",
                    "A0A0J9YUD9",
                    "A0A0J9YVF3",
                    "A0A0J9YV87"
                ]
            },
            "ENSMUSG00000026688": {
                "id": "ENSMUSG00000026688",
                "type": "Gene",
                "symbol": "Mgst3",
                "source": [
                    "ENSMUSG00000005583",
                    "ENSMUSG00000008575"
                ],
                "target": [],
                "expression_sum": {
                    "class1": 1458.0,
                    "class2": 5685.0
                },
                "ens_id": "ENSMUSG00000026688",
                "uniprot_ids": [
                    "Q9CPU4"
                ]
            },
            "ENSMUSG00000031586": {
                "id": "ENSMUSG00000031586",
                "type": "Gene",
                "symbol": "Rbpms",
                "source": [
                    "ENSMUSG00000032238",
                    "ENSMUSG00000052684",
                    "ENSMUSG00000022346",
                    "ENSMUSG00000047638",
                    "ENSMUSG00000005583",
                    "ENSMUSG00000071076"
                ],
                "target": [],
                "expression_sum": {
                    "class1": 14836.0,
                    "class2": 11417.0
                },
                "ens_id": "ENSMUSG00000031586",
                "uniprot_ids": [
                    "Q9WVB0",
                    "T1ECW4",
                    "S4R1Y2",
                    "S4R1A1",
                    "S4R2P3",
                    "S4R280",
                    "S4R1T1"
                ]
            },
            "ENSMUSG00000061477": {
                "id": "ENSMUSG00000061477",
                "type": "Gene",
                "symbol": "Rps7",
                "source": [
                    "ENSMUSG00000031965",
                    "ENSMUSG00000038193",
                    "ENSMUSG00000000247"
                ],
                "target": [],
                "expression_sum": {
                    "class1": 155375.0,
                    "class2": 51594.0
                },
                "ens_id": "ENSMUSG00000061477",
                "uniprot_ids": [
                    "P62082"
                ]
            },
            "ENSMUSG00000028691": {
                "id": "ENSMUSG00000028691",
                "type": "Gene",
                "symbol": "Prdx1",
                "source": [
                    "ENSMUSG00000052684",
                    "ENSMUSG00000052837",
                    "ENSMUSG00000057098",
                    "ENSMUSG00000021250",
                    "ENSMUSG00000003545",
                    "ENSMUSG00000003032",
                    "ENSMUSG00000048387",
                    "ENSMUSG00000071076"
                ],
                "target": [],
                "expression_sum": {
                    "class1": 33551.0,
                    "class2": 23914.0
                },
                "ens_id": "ENSMUSG00000028691",
                "uniprot_ids": [
                    "P35700",
                    "B1AXW6",
                    "B1AXW5",
                    "B1AXW4"
                ]
            },
            "ENSMUSG00000024076": {
                "id": "ENSMUSG00000024076",
                "type": "Gene",
                "symbol": "Vit",
                "source": [
                    "ENSMUSG00000022346",
                    "ENSMUSG00000052684",
                    "ENSMUSG00000071076"
                ],
                "target": [],
                "expression_sum": {
                    "class1": 4.0,
                    "class2": 4194.0
                },
                "ens_id": "ENSMUSG00000024076",
                "uniprot_ids": [
                    "Q8VHI5",
                    "A0A3B2WCI9"
                ]
            },
            "ENSMUSG00000014956": {
                "id": "ENSMUSG00000014956",
                "type": "Gene",
                "symbol": "Ppp1cb",
                "source": [
                    "ENSMUSG00000032238",
                    "ENSMUSG00000052684",
                    "ENSMUSG00000023034",
                    "ENSMUSG00000058756",
                    "ENSMUSG00000022528",
                    "ENSMUSG00000005583",
                    "ENSMUSG00000008575",
                    "ENSMUSG00000020644",
                    "ENSMUSG00000020484"
                ],
                "target": [],
                "expression_sum": {
                    "class1": 5143.0,
                    "class2": 14510.0
                },
                "ens_id": "ENSMUSG00000014956",
                "uniprot_ids": [
                    "P62141",
                    "A0A0J9YUG2",
                    "A0A0J9YUU8"
                ]
            },
            "ENSMUSG00000056501": {
                "id": "ENSMUSG00000056501",
                "type": "TF",
                "symbol": "Cebpb",
                "source": [
                    "ENSMUSG00000041515",
                    "ENSMUSG00000052684",
                    "ENSMUSG00000030103",
                    "ENSMUSG00000021250",
                    "ENSMUSG00000071076",
                    "ENSMUSG00000018899"
                ],
                "target": [
                    "ENSMUSG00000031765",
                    "ENSMUSG00000003545",
                    "ENSMUSG00000031320",
                    "ENSMUSG00000030930",
                    "ENSMUSG00000037411",
                    "ENSMUSG00000037405",
                    "ENSMUSG00000028268",
                    "ENSMUSG00000008384",
                    "ENSMUSG00000055148",
                    "ENSMUSG00000035493",
                    "ENSMUSG00000021127",
                    "ENSMUSG00000028003",
                    "ENSMUSG00000029380",
                    "ENSMUSG00000031119",
                    "ENSMUSG00000007872",
                    "ENSMUSG00000049115",
                    "ENSMUSG00000031845",
                    "ENSMUSG00000061983",
                    "ENSMUSG00000050953",
                    "ENSMUSG00000061353",
                    "ENSMUSG00000036655",
                    "ENSMUSG00000009681",
                    "ENSMUSG00000037999",
                    "ENSMUSG00000037742",
                    "ENSMUSG00000053113",
                    "ENSMUSG00000033538",
                    "ENSMUSG00000052684",
                    "ENSMUSG00000070436",
                    "ENSMUSG00000031762",
                    "ENSMUSG00000027188",
                    "ENSMUSG00000015501",
                    "ENSMUSG00000030103",
                    "ENSMUSG00000029163",
                    "ENSMUSG00000022587",
                    "ENSMUSG00000032515",
                    "ENSMUSG00000045128",
                    "ENSMUSG00000033880",
                    "ENSMUSG00000023944",
                    "ENSMUSG00000027829",
                    "ENSMUSG00000031375",
                    "ENSMUSG00000039377",
                    "ENSMUSG00000034855",
                    "ENSMUSG00000035678",
                    "ENSMUSG00000002985",
                    "ENSMUSG00000072625",
                    "ENSMUSG00000054072",
                    "ENSMUSG00000056708",
                    "ENSMUSG00000060126",
                    "ENSMUSG00000042453",
                    "ENSMUSG00000021025",
                    "ENSMUSG00000022665",
                    "ENSMUSG00000025498",
                    "ENSMUSG00000040128",
                    "ENSMUSG00000035373",
                    "ENSMUSG00000020423",
                    "ENSMUSG00000024371",
                    "ENSMUSG00000020205",
                    "ENSMUSG00000037868",
                    "ENSMUSG00000028214",
                    "ENSMUSG00000023224",
                    "ENSMUSG00000039601",
                    "ENSMUSG00000024014",
                    "ENSMUSG00000000628",
                    "ENSMUSG00000060550",
                    "ENSMUSG00000028864",
                    "ENSMUSG00000062328",
                    "ENSMUSG00000032179",
                    "ENSMUSG00000021250",
                    "ENSMUSG00000007892",
                    "ENSMUSG00000035385",
                    "ENSMUSG00000041515",
                    "ENSMUSG00000035356",
                    "ENSMUSG00000047945",
                    "ENSMUSG00000073411",
                    "ENSMUSG00000019929",
                    "ENSMUSG00000044786",
                    "ENSMUSG00000071076",
                    "ENSMUSG00000018899",
                    "ENSMUSG00000027004",
                    "ENSMUSG00000006403",
                    "ENSMUSG00000032501",
                    "ENSMUSG00000000247",
                    "ENSMUSG00000030849",
                    "ENSMUSG00000073489"
                ],
                "expression_sum": {
                    "class1": 56111.0,
                    "class2": 21063.0
                },
                "ens_id": "ENSMUSG00000056501",
                "uniprot_ids": [
                    "P28033"
                ]
            },
            "ENSMUSG00000040212": {
                "id": "ENSMUSG00000040212",
                "type": "Gene",
                "symbol": "Emp3",
                "source": [
                    "ENSMUSG00000052684",
                    "ENSMUSG00000057098",
                    "ENSMUSG00000038418",
                    "ENSMUSG00000003032",
                    "ENSMUSG00000005583",
                    "ENSMUSG00000071076"
                ],
                "target": [],
                "expression_sum": {
                    "class1": 10517.0,
                    "class2": 12826.0
                },
                "ens_id": "ENSMUSG00000040212",
                "uniprot_ids": [
                    "O35912",
                    "A0A1B0GSZ1"
                ]
            },
            "ENSMUSG00000018217": {
                "id": "ENSMUSG00000018217",
                "type": "Gene",
                "symbol": "Pmp22",
                "source": [
                    "ENSMUSG00000052684",
                    "ENSMUSG00000005583",
                    "ENSMUSG00000038418",
                    "ENSMUSG00000003032",
                    "ENSMUSG00000008575",
                    "ENSMUSG00000028565",
                    "ENSMUSG00000071076"
                ],
                "target": [],
                "expression_sum": {
                    "class1": 7272.0,
                    "class2": 13323.0
                },
                "ens_id": "ENSMUSG00000018217",
                "uniprot_ids": [
                    "Q5SXS3"
                ]
            },
            "ENSMUSG00000038370": {
                "id": "ENSMUSG00000038370",
                "type": "Gene",
                "symbol": "Pcp4l1",
                "source": [
                    "ENSMUSG00000058756",
                    "ENSMUSG00000026628",
                    "ENSMUSG00000021250",
                    "ENSMUSG00000005583"
                ],
                "target": [],
                "expression_sum": {
                    "class1": 16.0,
                    "class2": 15855.0
                },
                "ens_id": "ENSMUSG00000038370",
                "uniprot_ids": [
                    "Q6W8Q3"
                ]
            },
            "ENSMUSG00000045658": {
                "id": "ENSMUSG00000045658",
                "type": "Gene",
                "symbol": "Pid1",
                "source": [
                    "ENSMUSG00000032238",
                    "ENSMUSG00000052684",
                    "ENSMUSG00000052837",
                    "ENSMUSG00000022346",
                    "ENSMUSG00000003032",
                    "ENSMUSG00000005583",
                    "ENSMUSG00000071076"
                ],
                "target": [],
                "expression_sum": {
                    "class1": 543.0,
                    "class2": 7123.0
                },
                "ens_id": "ENSMUSG00000045658",
                "uniprot_ids": [
                    "Q3UBG2",
                    "H3BKK5",
                    "H3BJT8",
                    "H3BJF2",
                    "H3BJG6"
                ]
            },
            "ENSMUSG00000021200": {
                "id": "ENSMUSG00000021200",
                "type": "Gene",
                "symbol": "Asb2",
                "source": [
                    "ENSMUSG00000058756",
                    "ENSMUSG00000005583"
                ],
                "target": [],
                "expression_sum": {
                    "class1": 10.0,
                    "class2": 3325.0
                },
                "ens_id": "ENSMUSG00000021200",
                "uniprot_ids": [
                    "Q8K0L0"
                ]
            },
            "ENSMUSG00000029761": {
                "id": "ENSMUSG00000029761",
                "type": "Gene",
                "symbol": "Cald1",
                "source": [
                    "ENSMUSG00000032238",
                    "ENSMUSG00000052684",
                    "ENSMUSG00000023034",
                    "ENSMUSG00000022346",
                    "ENSMUSG00000047638",
                    "ENSMUSG00000005583",
                    "ENSMUSG00000053477",
                    "ENSMUSG00000008575",
                    "ENSMUSG00000020644",
                    "ENSMUSG00000071076"
                ],
                "target": [],
                "expression_sum": {
                    "class1": 30167.0,
                    "class2": 39391.0
                },
                "ens_id": "ENSMUSG00000029761",
                "uniprot_ids": [
                    "Q8VCQ8",
                    "E9Q9F3",
                    "F6QLP8",
                    "F6ZAW1",
                    "D3Z6I7",
                    "E9Q0M9",
                    "F6T2Z7",
                    "F6RGN9",
                    "E9QA16",
                    "E9QA15",
                    "S4R1T7"
                ]
            },
            "ENSMUSG00000001025": {
                "id": "ENSMUSG00000001025",
                "type": "Gene",
                "symbol": "S100a6",
                "source": [
                    "ENSMUSG00000052684",
                    "ENSMUSG00000052837",
                    "ENSMUSG00000022346",
                    "ENSMUSG00000005583",
                    "ENSMUSG00000003032",
                    "ENSMUSG00000008575",
                    "ENSMUSG00000071076"
                ],
                "target": [],
                "expression_sum": {
                    "class1": 2262.0,
                    "class2": 63154.0
                },
                "ens_id": "ENSMUSG00000001025",
                "uniprot_ids": [
                    "P14069"
                ]
            },
            "ENSMUSG00000049517": {
                "id": "ENSMUSG00000049517",
                "type": "Gene",
                "symbol": "Rps23",
                "source": [
                    "ENSMUSG00000032035",
                    "ENSMUSG00000023034",
                    "ENSMUSG00000000247",
                    "ENSMUSG00000057098",
                    "ENSMUSG00000038193",
                    "ENSMUSG00000005583"
                ],
                "target": [],
                "expression_sum": {
                    "class1": 163131.0,
                    "class2": 90799.0
                },
                "ens_id": "ENSMUSG00000049517",
                "uniprot_ids": [
                    "P62267"
                ]
            },
            "ENSMUSG00000041329": {
                "id": "ENSMUSG00000041329",
                "type": "Gene",
                "symbol": "Atp1b2",
                "source": [
                    "ENSMUSG00000023034",
                    "ENSMUSG00000005583"
                ],
                "target": [],
                "expression_sum": {
                    "class1": 717.0,
                    "class2": 4844.0
                },
                "ens_id": "ENSMUSG00000041329",
                "uniprot_ids": [
                    "P14231",
                    "I7HIF3"
                ]
            },
            "ENSMUSG00000025492": {
                "id": "ENSMUSG00000025492",
                "type": "Gene",
                "symbol": "Ifitm3",
                "source": [
                    "ENSMUSG00000032035",
                    "ENSMUSG00000052684",
                    "ENSMUSG00000057098",
                    "ENSMUSG00000003032",
                    "ENSMUSG00000005583",
                    "ENSMUSG00000071076"
                ],
                "target": [],
                "expression_sum": {
                    "class1": 297680.0,
                    "class2": 49594.0
                },
                "ens_id": "ENSMUSG00000025492",
                "uniprot_ids": [
                    "Q9CQW9"
                ]
            },
            "ENSMUSG00000027800": {
                "id": "ENSMUSG00000027800",
                "type": "Gene",
                "symbol": "Tm4sf1",
                "source": [
                    "ENSMUSG00000005583"
                ],
                "target": [],
                "expression_sum": {
                    "class1": 1837.0,
                    "class2": 14406.0
                },
                "ens_id": "ENSMUSG00000027800",
                "uniprot_ids": [
                    "Q64302",
                    "A0A0G2JGU1"
                ]
            },
            "ENSMUSG00000017466": {
                "id": "ENSMUSG00000017466",
                "type": "Gene",
                "symbol": "Timp2",
                "source": [
                    "ENSMUSG00000052684",
                    "ENSMUSG00000022528",
                    "ENSMUSG00000057098",
                    "ENSMUSG00000003032",
                    "ENSMUSG00000005583",
                    "ENSMUSG00000071076"
                ],
                "target": [],
                "expression_sum": {
                    "class1": 26369.0,
                    "class2": 29345.0
                },
                "ens_id": "ENSMUSG00000017466",
                "uniprot_ids": [
                    "Q6PI17",
                    "B1AQJ3"
                ]
            },
            "ENSMUSG00000023224": {
                "id": "ENSMUSG00000023224",
                "type": "Gene",
                "symbol": "Serping1",
                "source": [
                    "ENSMUSG00000056501",
                    "ENSMUSG00000023034",
                    "ENSMUSG00000058756",
                    "ENSMUSG00000038193",
                    "ENSMUSG00000005583",
                    "ENSMUSG00000055148"
                ],
                "target": [],
                "expression_sum": {
                    "class1": 139210.0,
                    "class2": 54316.0
                },
                "ens_id": "ENSMUSG00000023224",
                "uniprot_ids": [
                    "P97290",
                    "A2ATR8"
                ]
            },
            "ENSMUSG00000021268": {
                "id": "ENSMUSG00000021268",
                "type": "Gene",
                "symbol": "Meg3",
                "source": [
                    "ENSMUSG00000052684",
                    "ENSMUSG00000022346",
                    "ENSMUSG00000005583",
                    "ENSMUSG00000020484",
                    "ENSMUSG00000071076"
                ],
                "target": [],
                "expression_sum": {
                    "class1": 2358.0,
                    "class2": 18502.0
                },
                "ens_id": "ENSMUSG00000021268"
            },
            "ENSMUSG00000036256": {
                "id": "ENSMUSG00000036256",
                "type": "Gene",
                "symbol": "Igfbp7",
                "source": [
                    "ENSMUSG00000029322"
                ],
                "target": [],
                "expression_sum": {
                    "class1": 953413.0,
                    "class2": 162752.0
                },
                "ens_id": "ENSMUSG00000036256",
                "uniprot_ids": [
                    "F8WH23",
                    "E9Q5D9"
                ]
            },
            "ENSMUSG00000064358": {
                "id": "ENSMUSG00000064358",
                "type": "Gene",
                "symbol": "mt-Co3",
                "source": [
                    "ENSMUSG00000029322",
                    "ENSMUSG00000022708"
                ],
                "target": [],
                "expression_sum": {
                    "class1": 176112.0,
                    "class2": 212702.0
                },
                "ens_id": "ENSMUSG00000064358",
                "uniprot_ids": [
                    "P00416"
                ]
            },
            "ENSMUSG00000030122": {
                "id": "ENSMUSG00000030122",
                "type": "Gene",
                "symbol": "Ptms",
                "source": [
                    "ENSMUSG00000021250",
                    "ENSMUSG00000029322"
                ],
                "target": [],
                "expression_sum": {
                    "class1": 22835.0,
                    "class2": 18753.0
                },
                "ens_id": "ENSMUSG00000030122",
                "uniprot_ids": [
                    "Q9D0J8"
                ]
            },
            "ENSMUSG00000026238": {
                "id": "ENSMUSG00000026238",
                "type": "Gene",
                "symbol": "Ptma",
                "source": [
                    "ENSMUSG00000025498",
                    "ENSMUSG00000047638",
                    "ENSMUSG00000000247",
                    "ENSMUSG00000057098",
                    "ENSMUSG00000029322"
                ],
                "target": [],
                "expression_sum": {
                    "class1": 99702.0,
                    "class2": 63901.0
                },
                "ens_id": "ENSMUSG00000026238",
                "uniprot_ids": [
                    "P26350",
                    "A0A087WQN2",
                    "A0A087WP98",
                    "A0A087WPN6"
                ]
            },
            "ENSMUSG00000006356": {
                "id": "ENSMUSG00000006356",
                "type": "Gene",
                "symbol": "Crip2",
                "source": [
                    "ENSMUSG00000047638",
                    "ENSMUSG00000057098",
                    "ENSMUSG00000029322",
                    "ENSMUSG00000015846",
                    "ENSMUSG00000008575",
                    "ENSMUSG00000005583"
                ],
                "target": [],
                "expression_sum": {
                    "class1": 63331.0,
                    "class2": 24982.0
                },
                "ens_id": "ENSMUSG00000006356",
                "uniprot_ids": [
                    "Q9DCT8",
                    "A0A0G2JF37"
                ]
            },
            "ENSMUSG00000031246": {
                "id": "ENSMUSG00000031246",
                "type": "Gene",
                "symbol": "Sh3bgrl",
                "source": [
                    "ENSMUSG00000029322",
                    "ENSMUSG00000005583",
                    "ENSMUSG00000008575"
                ],
                "target": [],
                "expression_sum": {
                    "class1": 11989.0,
                    "class2": 11409.0
                },
                "ens_id": "ENSMUSG00000031246",
                "uniprot_ids": [
                    "Q9JJU8"
                ]
            },
            "ENSMUSG00000036438": {
                "id": "ENSMUSG00000036438",
                "type": "Gene",
                "symbol": "Calm2",
                "source": [
                    "ENSMUSG00000023034",
                    "ENSMUSG00000047638",
                    "ENSMUSG00000026628",
                    "ENSMUSG00000005583",
                    "ENSMUSG00000000247",
                    "ENSMUSG00000021250",
                    "ENSMUSG00000029322",
                    "ENSMUSG00000008575",
                    "ENSMUSG00000020484"
                ],
                "target": [],
                "expression_sum": {
                    "class1": 28520.0,
                    "class2": 38596.0
                },
                "ens_id": "ENSMUSG00000036438",
                "uniprot_ids": [
                    "P0DP26",
                    "P0DP27",
                    "P0DP28",
                    "A0A3Q4EHJ0",
                    "A0A571BE58"
                ]
            },
            "ENSMUSG00000061787": {
                "id": "ENSMUSG00000061787",
                "type": "Gene",
                "symbol": "Rps17",
                "source": [
                    "ENSMUSG00000057098",
                    "ENSMUSG00000000247"
                ],
                "target": [],
                "expression_sum": {
                    "class1": 90787.0,
                    "class2": 46553.0
                },
                "ens_id": "ENSMUSG00000061787",
                "uniprot_ids": [
                    "P63276"
                ]
            },
            "ENSMUSG00000055435": {
                "id": "ENSMUSG00000055435",
                "type": "TF",
                "symbol": "Maf",
                "source": [
                    "ENSMUSG00000005583",
                    "ENSMUSG00000052684",
                    "ENSMUSG00000071076"
                ],
                "target": [
                    "ENSMUSG00000024308",
                    "ENSMUSG00000024063",
                    "ENSMUSG00000034165",
                    "ENSMUSG00000079523",
                    "ENSMUSG00000021477",
                    "ENSMUSG00000053113",
                    "ENSMUSG00000056708",
                    "ENSMUSG00000028249",
                    "ENSMUSG00000055172",
                    "ENSMUSG00000019907",
                    "ENSMUSG00000052684",
                    "ENSMUSG00000021877",
                    "ENSMUSG00000075010",
                    "ENSMUSG00000020737",
                    "ENSMUSG00000029767",
                    "ENSMUSG00000026249",
                    "ENSMUSG00000055447",
                    "ENSMUSG00000020048",
                    "ENSMUSG00000005583",
                    "ENSMUSG00000041959",
                    "ENSMUSG00000022010",
                    "ENSMUSG00000074743",
                    "ENSMUSG00000032231",
                    "ENSMUSG00000024048",
                    "ENSMUSG00000021876",
                    "ENSMUSG00000025780",
                    "ENSMUSG00000026478",
                    "ENSMUSG00000079017",
                    "ENSMUSG00000001847",
                    "ENSMUSG00000026536",
                    "ENSMUSG00000064215",
                    "ENSMUSG00000062825",
                    "ENSMUSG00000025283",
                    "ENSMUSG00000073489",
                    "ENSMUSG00000072235",
                    "ENSMUSG00000075254",
                    "ENSMUSG00000031799",
                    "ENSMUSG00000058135"
                ],
                "expression_sum": {
                    "class1": 5180.0,
                    "class2": 2486.0
                },
                "ens_id": "ENSMUSG00000055435",
                "uniprot_ids": [
                    "P54843"
                ]
            },
            "ENSMUSG00000022528": {
                "id": "ENSMUSG00000022528",
                "type": "TF",
                "symbol": "Hes1",
                "source": [
                    "ENSMUSG00000052684",
                    "ENSMUSG00000047638",
                    "ENSMUSG00000038418",
                    "ENSMUSG00000003545",
                    "ENSMUSG00000005583",
                    "ENSMUSG00000020484",
                    "ENSMUSG00000071076"
                ],
                "target": [
                    "ENSMUSG00000026365",
                    "ENSMUSG00000036545",
                    "ENSMUSG00000031447",
                    "ENSMUSG00000031765",
                    "ENSMUSG00000079523",
                    "ENSMUSG00000032518",
                    "ENSMUSG00000028949",
                    "ENSMUSG00000037742",
                    "ENSMUSG00000053113",
                    "ENSMUSG00000060126",
                    "ENSMUSG00000032366",
                    "ENSMUSG00000059895",
                    "ENSMUSG00000022892",
                    "ENSMUSG00000040952",
                    "ENSMUSG00000026879",
                    "ENSMUSG00000046727",
                    "ENSMUSG00000021756",
                    "ENSMUSG00000027248",
                    "ENSMUSG00000052684",
                    "ENSMUSG00000012405",
                    "ENSMUSG00000021877",
                    "ENSMUSG00000014956",
                    "ENSMUSG00000060036",
                    "ENSMUSG00000090862",
                    "ENSMUSG00000031762",
                    "ENSMUSG00000043716",
                    "ENSMUSG00000055447",
                    "ENSMUSG00000020048",
                    "ENSMUSG00000005583",
                    "ENSMUSG00000021939",
                    "ENSMUSG00000027642",
                    "ENSMUSG00000022136",
                    "ENSMUSG00000021127",
                    "ENSMUSG00000024190",
                    "ENSMUSG00000032383",
                    "ENSMUSG00000032216",
                    "ENSMUSG00000022708",
                    "ENSMUSG00000029781",
                    "ENSMUSG00000028234",
                    "ENSMUSG00000025508",
                    "ENSMUSG00000024371",
                    "ENSMUSG00000028936",
                    "ENSMUSG00000020205",
                    "ENSMUSG00000028495",
                    "ENSMUSG00000017466",
                    "ENSMUSG00000040249",
                    "ENSMUSG00000019970",
                    "ENSMUSG00000061983",
                    "ENSMUSG00000037805",
                    "ENSMUSG00000039377",
                    "ENSMUSG00000038418",
                    "ENSMUSG00000039001",
                    "ENSMUSG00000020902",
                    "ENSMUSG00000029076",
                    "ENSMUSG00000062647",
                    "ENSMUSG00000022971"
                ],
                "expression_sum": {
                    "class1": 22719.0,
                    "class2": 11710.0
                },
                "ens_id": "ENSMUSG00000022528",
                "uniprot_ids": [
                    "P35428"
                ]
            },
            "ENSMUSG00000052534": {
                "id": "ENSMUSG00000052534",
                "type": "TF",
                "symbol": "Pbx1",
                "source": [
                    "ENSMUSG00000052684"
                ],
                "target": [
                    "ENSMUSG00000024063",
                    "ENSMUSG00000029309",
                    "ENSMUSG00000052684",
                    "ENSMUSG00000026335",
                    "ENSMUSG00000025784",
                    "ENSMUSG00000026566",
                    "ENSMUSG00000072235",
                    "ENSMUSG00000024048",
                    "ENSMUSG00000035258",
                    "ENSMUSG00000016194",
                    "ENSMUSG00000022636",
                    "ENSMUSG00000039601"
                ],
                "expression_sum": {
                    "class1": 9798.0,
                    "class2": 8386.0
                },
                "ens_id": "ENSMUSG00000052534",
                "uniprot_ids": [
                    "P41778",
                    "D9J2V6"
                ]
            },
            "ENSMUSG00000041515": {
                "id": "ENSMUSG00000041515",
                "type": "TF",
                "symbol": "Irf8",
                "source": [
                    "ENSMUSG00000056501",
                    "ENSMUSG00000023034",
                    "ENSMUSG00000052837",
                    "ENSMUSG00000026628",
                    "ENSMUSG00000021250",
                    "ENSMUSG00000018899"
                ],
                "target": [
                    "ENSMUSG00000056501",
                    "ENSMUSG00000023034",
                    "ENSMUSG00000032515",
                    "ENSMUSG00000052837",
                    "ENSMUSG00000026628",
                    "ENSMUSG00000020423",
                    "ENSMUSG00000028967",
                    "ENSMUSG00000044786",
                    "ENSMUSG00000035356",
                    "ENSMUSG00000008384",
                    "ENSMUSG00000053113",
                    "ENSMUSG00000021250",
                    "ENSMUSG00000055148",
                    "ENSMUSG00000007872",
                    "ENSMUSG00000087006",
                    "ENSMUSG00000016194",
                    "ENSMUSG00000018899"
                ],
                "expression_sum": {
                    "class1": 4392.0,
                    "class2": 265.0
                },
                "ens_id": "ENSMUSG00000041515",
                "uniprot_ids": [
                    "P23611",
                    "F7C1A3",
                    "E0CZE4"
                ]
            },
            "ENSMUSG00000030103": {
                "id": "ENSMUSG00000030103",
                "type": "TF",
                "symbol": "Bhlhe40",
                "source": [
                    "ENSMUSG00000056501",
                    "ENSMUSG00000052684",
                    "ENSMUSG00000023034",
                    "ENSMUSG00000052837",
                    "ENSMUSG00000026628",
                    "ENSMUSG00000021250",
                    "ENSMUSG00000018899"
                ],
                "target": [
                    "ENSMUSG00000056501",
                    "ENSMUSG00000023034",
                    "ENSMUSG00000053113",
                    "ENSMUSG00000003545",
                    "ENSMUSG00000020644",
                    "ENSMUSG00000015312",
                    "ENSMUSG00000021250",
                    "ENSMUSG00000055148",
                    "ENSMUSG00000040128",
                    "ENSMUSG00000035356",
                    "ENSMUSG00000047945",
                    "ENSMUSG00000026628",
                    "ENSMUSG00000020423",
                    "ENSMUSG00000052837",
                    "ENSMUSG00000032515",
                    "ENSMUSG00000007872",
                    "ENSMUSG00000044786",
                    "ENSMUSG00000018899",
                    "ENSMUSG00000090698",
                    "ENSMUSG00000028967",
                    "ENSMUSG00000035828",
                    "ENSMUSG00000009681"
                ],
                "expression_sum": {
                    "class1": 3879.0,
                    "class2": 961.0
                },
                "ens_id": "ENSMUSG00000030103",
                "uniprot_ids": [
                    "O35185",
                    "E9Q2C3"
                ]
            },
            "ENSMUSG00000022346": {
                "id": "ENSMUSG00000022346",
                "type": "TF",
                "symbol": "Myc",
                "source": [
                    "ENSMUSG00000048387",
                    "ENSMUSG00000018899"
                ],
                "target": [
                    "ENSMUSG00000020053",
                    "ENSMUSG00000054808",
                    "ENSMUSG00000029761",
                    "ENSMUSG00000016494",
                    "ENSMUSG00000025408",
                    "ENSMUSG00000032366",
                    "ENSMUSG00000033910",
                    "ENSMUSG00000026879",
                    "ENSMUSG00000087006",
                    "ENSMUSG00000001520",
                    "ENSMUSG00000032359",
                    "ENSMUSG00000019907",
                    "ENSMUSG00000025784",
                    "ENSMUSG00000064373",
                    "ENSMUSG00000031451",
                    "ENSMUSG00000015354",
                    "ENSMUSG00000024076",
                    "ENSMUSG00000019278",
                    "ENSMUSG00000003477",
                    "ENSMUSG00000055148",
                    "ENSMUSG00000029096",
                    "ENSMUSG00000018339",
                    "ENSMUSG00000054203",
                    "ENSMUSG00000026421",
                    "ENSMUSG00000006369",
                    "ENSMUSG00000031239",
                    "ENSMUSG00000002107",
                    "ENSMUSG00000047945",
                    "ENSMUSG00000020423",
                    "ENSMUSG00000028005",
                    "ENSMUSG00000060572",
                    "ENSMUSG00000031586",
                    "ENSMUSG00000026574",
                    "ENSMUSG00000028776",
                    "ENSMUSG00000008540",
                    "ENSMUSG00000018899",
                    "ENSMUSG00000074457",
                    "ENSMUSG00000021876",
                    "ENSMUSG00000021268",
                    "ENSMUSG00000022836",
                    "ENSMUSG00000024164",
                    "ENSMUSG00000020473",
                    "ENSMUSG00000006205",
                    "ENSMUSG00000022032",
                    "ENSMUSG00000045658",
                    "ENSMUSG00000038463",
                    "ENSMUSG00000017493",
                    "ENSMUSG00000048387",
                    "ENSMUSG00000001025"
                ],
                "expression_sum": {
                    "class1": 5651.0,
                    "class2": 1793.0
                },
                "ens_id": "ENSMUSG00000022346",
                "uniprot_ids": [
                    "P01108",
                    "F6PX41",
                    "F8WID3",
                    "E0CZD1"
                ]
            },
            "ENSMUSG00000042812": {
                "id": "ENSMUSG00000042812",
                "type": "TF",
                "symbol": "Foxf1",
                "source": [
                    "ENSMUSG00000052837",
                    "ENSMUSG00000021250",
                    "ENSMUSG00000052684",
                    "ENSMUSG00000000247"
                ],
                "target": [
                    "ENSMUSG00000031375",
                    "ENSMUSG00000021250",
                    "ENSMUSG00000030849",
                    "ENSMUSG00000003545",
                    "ENSMUSG00000031119",
                    "ENSMUSG00000035493"
                ],
                "expression_sum": {
                    "class1": 8363.0,
                    "class2": 1724.0
                },
                "ens_id": "ENSMUSG00000042812",
                "uniprot_ids": [
                    "Q61080"
                ]
            },
            "ENSMUSG00000042406": {
                "id": "ENSMUSG00000042406",
                "type": "TF",
                "symbol": "Atf4",
                "source": [
                    "ENSMUSG00000026628"
                ],
                "target": [
                    "ENSMUSG00000022836",
                    "ENSMUSG00000035356",
                    "ENSMUSG00000026628",
                    "ENSMUSG00000053113",
                    "ENSMUSG00000018899"
                ],
                "expression_sum": {
                    "class1": 8359.0,
                    "class2": 3788.0
                },
                "ens_id": "ENSMUSG00000042406",
                "uniprot_ids": [
                    "Q06507",
                    "A0A2R8VI82"
                ]
            },
            "ENSMUSG00000032238": {
                "id": "ENSMUSG00000032238",
                "type": "TF",
                "symbol": "Rora",
                "source": [
                    "ENSMUSG00000005583"
                ],
                "target": [
                    "ENSMUSG00000064370",
                    "ENSMUSG00000020053",
                    "ENSMUSG00000064363",
                    "ENSMUSG00000029761",
                    "ENSMUSG00000028339",
                    "ENSMUSG00000014313",
                    "ENSMUSG00000090841",
                    "ENSMUSG00000024140",
                    "ENSMUSG00000058056",
                    "ENSMUSG00000000753",
                    "ENSMUSG00000052957",
                    "ENSMUSG00000064373",
                    "ENSMUSG00000014956",
                    "ENSMUSG00000068220",
                    "ENSMUSG00000005583",
                    "ENSMUSG00000029096",
                    "ENSMUSG00000021870",
                    "ENSMUSG00000026421",
                    "ENSMUSG00000002107",
                    "ENSMUSG00000024012",
                    "ENSMUSG00000031586",
                    "ENSMUSG00000027712",
                    "ENSMUSG00000024048",
                    "ENSMUSG00000040170",
                    "ENSMUSG00000020009",
                    "ENSMUSG00000008540",
                    "ENSMUSG00000045658",
                    "ENSMUSG00000016024",
                    "ENSMUSG00000017493",
                    "ENSMUSG00000058135"
                ],
                "expression_sum": {
                    "class1": 4893.0,
                    "class2": 2920.0
                },
                "ens_id": "ENSMUSG00000032238",
                "uniprot_ids": [
                    "P51448",
                    "G3UZ02"
                ]
            },
            "ENSMUSG00000028565": {
                "id": "ENSMUSG00000028565",
                "type": "TF",
                "symbol": "Nfia",
                "source": [
                    "ENSMUSG00000048387"
                ],
                "target": [
                    "ENSMUSG00000028517",
                    "ENSMUSG00000016494",
                    "ENSMUSG00000021477",
                    "ENSMUSG00000024140",
                    "ENSMUSG00000026879",
                    "ENSMUSG00000018217",
                    "ENSMUSG00000020585",
                    "ENSMUSG00000025784",
                    "ENSMUSG00000040562",
                    "ENSMUSG00000064373",
                    "ENSMUSG00000031451",
                    "ENSMUSG00000042485",
                    "ENSMUSG00000029096",
                    "ENSMUSG00000041959",
                    "ENSMUSG00000002107",
                    "ENSMUSG00000028369",
                    "ENSMUSG00000019929",
                    "ENSMUSG00000032231",
                    "ENSMUSG00000029659",
                    "ENSMUSG00000027712",
                    "ENSMUSG00000024048",
                    "ENSMUSG00000028108",
                    "ENSMUSG00000020009",
                    "ENSMUSG00000004207",
                    "ENSMUSG00000026574",
                    "ENSMUSG00000008540",
                    "ENSMUSG00000020205",
                    "ENSMUSG00000021876",
                    "ENSMUSG00000079037",
                    "ENSMUSG00000022836",
                    "ENSMUSG00000024164",
                    "ENSMUSG00000032334",
                    "ENSMUSG00000000594",
                    "ENSMUSG00000021390",
                    "ENSMUSG00000015085",
                    "ENSMUSG00000017493",
                    "ENSMUSG00000048387",
                    "ENSMUSG00000058135"
                ],
                "expression_sum": {
                    "class1": 12541.0,
                    "class2": 12297.0
                },
                "ens_id": "ENSMUSG00000028565",
                "uniprot_ids": [
                    "Q02780",
                    "B1AUB9",
                    "B1AUC0",
                    "F7CDR2",
                    "B1AUB8",
                    "B1AUB6"
                ]
            },
            "ENSMUSG00000032035": {
                "id": "ENSMUSG00000032035",
                "type": "TF",
                "symbol": "Ets1",
                "source": [
                    "ENSMUSG00000015846"
                ],
                "target": [
                    "ENSMUSG00000019505",
                    "ENSMUSG00000005338",
                    "ENSMUSG00000064363",
                    "ENSMUSG00000031447",
                    "ENSMUSG00000021477",
                    "ENSMUSG00000027447",
                    "ENSMUSG00000024909",
                    "ENSMUSG00000034868",
                    "ENSMUSG00000023175",
                    "ENSMUSG00000064354",
                    "ENSMUSG00000026970",
                    "ENSMUSG00000059588",
                    "ENSMUSG00000025130",
                    "ENSMUSG00000025790",
                    "ENSMUSG00000034892",
                    "ENSMUSG00000032231",
                    "ENSMUSG00000023336",
                    "ENSMUSG00000031021",
                    "ENSMUSG00000029426",
                    "ENSMUSG00000029718",
                    "ENSMUSG00000036655",
                    "ENSMUSG00000062825",
                    "ENSMUSG00000020368",
                    "ENSMUSG00000024678",
                    "ENSMUSG00000069601",
                    "ENSMUSG00000034353",
                    "ENSMUSG00000028603",
                    "ENSMUSG00000034845",
                    "ENSMUSG00000022892",
                    "ENSMUSG00000032562",
                    "ENSMUSG00000056025",
                    "ENSMUSG00000021756",
                    "ENSMUSG00000038803",
                    "ENSMUSG00000070436",
                    "ENSMUSG00000029767",
                    "ENSMUSG00000030824",
                    "ENSMUSG00000064351",
                    "ENSMUSG00000021248",
                    "ENSMUSG00000027642",
                    "ENSMUSG00000030249",
                    "ENSMUSG00000029163",
                    "ENSMUSG00000026193",
                    "ENSMUSG00000033880",
                    "ENSMUSG00000022587",
                    "ENSMUSG00000023944",
                    "ENSMUSG00000024065",
                    "ENSMUSG00000029455",
                    "ENSMUSG00000031375",
                    "ENSMUSG00000072235",
                    "ENSMUSG00000036381",
                    "ENSMUSG00000026860",
                    "ENSMUSG00000046330",
                    "ENSMUSG00000020432",
                    "ENSMUSG00000079523",
                    "ENSMUSG00000001123",
                    "ENSMUSG00000060126",
                    "ENSMUSG00000042453",
                    "ENSMUSG00000058569",
                    "ENSMUSG00000026014",
                    "ENSMUSG00000005973",
                    "ENSMUSG00000016534",
                    "ENSMUSG00000060591",
                    "ENSMUSG00000032492",
                    "ENSMUSG00000034707",
                    "ENSMUSG00000049517",
                    "ENSMUSG00000022665",
                    "ENSMUSG00000031661",
                    "ENSMUSG00000061232",
                    "ENSMUSG00000027712",
                    "ENSMUSG00000024048",
                    "ENSMUSG00000069662",
                    "ENSMUSG00000032666",
                    "ENSMUSG00000028108",
                    "ENSMUSG00000004207",
                    "ENSMUSG00000020205",
                    "ENSMUSG00000035242",
                    "ENSMUSG00000034708",
                    "ENSMUSG00000026478",
                    "ENSMUSG00000032528",
                    "ENSMUSG00000070407",
                    "ENSMUSG00000020689",
                    "ENSMUSG00000026223",
                    "ENSMUSG00000029076",
                    "ENSMUSG00000022971",
                    "ENSMUSG00000074884",
                    "ENSMUSG00000054808",
                    "ENSMUSG00000022108",
                    "ENSMUSG00000041736",
                    "ENSMUSG00000028132",
                    "ENSMUSG00000027248",
                    "ENSMUSG00000020585",
                    "ENSMUSG00000032179",
                    "ENSMUSG00000025492",
                    "ENSMUSG00000046402",
                    "ENSMUSG00000034158",
                    "ENSMUSG00000055447",
                    "ENSMUSG00000000740",
                    "ENSMUSG00000007892",
                    "ENSMUSG00000027523",
                    "ENSMUSG00000028618",
                    "ENSMUSG00000032383",
                    "ENSMUSG00000035530",
                    "ENSMUSG00000047907",
                    "ENSMUSG00000073411",
                    "ENSMUSG00000048758",
                    "ENSMUSG00000022708",
                    "ENSMUSG00000029322",
                    "ENSMUSG00000019929",
                    "ENSMUSG00000060802",
                    "ENSMUSG00000001847",
                    "ENSMUSG00000000247",
                    "ENSMUSG00000030849",
                    "ENSMUSG00000020902",
                    "ENSMUSG00000073489",
                    "ENSMUSG00000024109",
                    "ENSMUSG00000058135"
                ],
                "expression_sum": {
                    "class1": 11559.0,
                    "class2": 1092.0
                },
                "ens_id": "ENSMUSG00000032035",
                "uniprot_ids": [
                    "P27577",
                    "V9GXE2",
                    "E9PWI8",
                    "V9GX28",
                    "A0A5F8MPR7"
                ]
            },
            "ENSMUSG00000076432": {
                "id": "ENSMUSG00000076432",
                "type": "Gene",
                "symbol": "Ywhaq",
                "source": [
                    "ENSMUSG00000053477",
                    "ENSMUSG00000047638",
                    "ENSMUSG00000029322",
                    "ENSMUSG00000005583"
                ],
                "target": [],
                "expression_sum": {
                    "class1": 12373.0,
                    "class2": 15117.0
                },
                "ens_id": "ENSMUSG00000076432",
                "uniprot_ids": [
                    "P68254"
                ]
            },
            "ENSMUSG00000015846": {
                "id": "ENSMUSG00000015846",
                "type": "TF",
                "symbol": "Rxra",
                "source": [],
                "target": [
                    "ENSMUSG00000020486",
                    "ENSMUSG00000032035",
                    "ENSMUSG00000026678",
                    "ENSMUSG00000032946",
                    "ENSMUSG00000034353",
                    "ENSMUSG00000024140",
                    "ENSMUSG00000025491",
                    "ENSMUSG00000034845",
                    "ENSMUSG00000001552",
                    "ENSMUSG00000042453",
                    "ENSMUSG00000028132",
                    "ENSMUSG00000021756",
                    "ENSMUSG00000022371",
                    "ENSMUSG00000032179",
                    "ENSMUSG00000000184",
                    "ENSMUSG00000038591",
                    "ENSMUSG00000041577",
                    "ENSMUSG00000000325",
                    "ENSMUSG00000038742",
                    "ENSMUSG00000059588",
                    "ENSMUSG00000028003",
                    "ENSMUSG00000012428",
                    "ENSMUSG00000023336",
                    "ENSMUSG00000069662",
                    "ENSMUSG00000032666",
                    "ENSMUSG00000024065",
                    "ENSMUSG00000006356",
                    "ENSMUSG00000032528",
                    "ENSMUSG00000070407",
                    "ENSMUSG00000020689",
                    "ENSMUSG00000036655",
                    "ENSMUSG00000072625"
                ],
                "expression_sum": {
                    "class1": 4376.0,
                    "class2": 460.0
                },
                "ens_id": "ENSMUSG00000015846",
                "uniprot_ids": [
                    "P28700",
                    "Q6LC96",
                    "A2AJP2",
                    "A0A5F8MP91",
                    "A0A5F8MQ23"
                ]
            },
            "ENSMUSG00000074457": {
                "id": "ENSMUSG00000074457",
                "type": "Gene",
                "symbol": "S100a16",
                "source": [
                    "ENSMUSG00000052684",
                    "ENSMUSG00000022346",
                    "ENSMUSG00000021250",
                    "ENSMUSG00000003032",
                    "ENSMUSG00000008575",
                    "ENSMUSG00000020484",
                    "ENSMUSG00000071076"
                ],
                "target": [],
                "expression_sum": {
                    "class1": 407.0,
                    "class2": 6586.0
                },
                "ens_id": "ENSMUSG00000074457",
                "uniprot_ids": [
                    "Q9D708",
                    "D3Z2Y6"
                ]
            },
            "ENSMUSG00000020737": {
                "id": "ENSMUSG00000020737",
                "type": "Gene",
                "symbol": "Hn1",
                "source": [
                    "ENSMUSG00000005583",
                    "ENSMUSG00000057098",
                    "ENSMUSG00000003032",
                    "ENSMUSG00000055435",
                    "ENSMUSG00000071076"
                ],
                "target": [],
                "expression_sum": {
                    "class1": 5625.0,
                    "class2": 4800.0
                },
                "ens_id": "ENSMUSG00000020737",
                "uniprot_ids": [
                    "P97825"
                ]
            },
            "ENSMUSG00000028005": {
                "id": "ENSMUSG00000028005",
                "type": "Gene",
                "symbol": "Gucy1b3",
                "source": [
                    "ENSMUSG00000022346",
                    "ENSMUSG00000005583"
                ],
                "target": [],
                "expression_sum": {
                    "class1": 11360.0,
                    "class2": 5707.0
                },
                "ens_id": "ENSMUSG00000028005",
                "uniprot_ids": [
                    "O54865",
                    "Q80YP4"
                ]
            },
            "ENSMUSG00000022371": {
                "id": "ENSMUSG00000022371",
                "type": "Gene",
                "symbol": "Col14a1",
                "source": [
                    "ENSMUSG00000026872",
                    "ENSMUSG00000038193",
                    "ENSMUSG00000015846",
                    "ENSMUSG00000003032",
                    "ENSMUSG00000005583"
                ],
                "target": [],
                "expression_sum": {
                    "class1": 81209.0,
                    "class2": 13551.0
                },
                "ens_id": "ENSMUSG00000022371",
                "uniprot_ids": [
                    "Q80X19",
                    "B7ZNH7",
                    "K3W4R4",
                    "F7D5Y4"
                ]
            },
            "ENSMUSG00000031765": {
                "id": "ENSMUSG00000031765",
                "type": "Gene",
                "symbol": "Mt1",
                "source": [
                    "ENSMUSG00000056501",
                    "ENSMUSG00000023034",
                    "ENSMUSG00000022528",
                    "ENSMUSG00000026628",
                    "ENSMUSG00000005583",
                    "ENSMUSG00000057098",
                    "ENSMUSG00000021250",
                    "ENSMUSG00000003032",
                    "ENSMUSG00000008575",
                    "ENSMUSG00000055148",
                    "ENSMUSG00000018899"
                ],
                "target": [],
                "expression_sum": {
                    "class1": 115205.0,
                    "class2": 41751.0
                },
                "ens_id": "ENSMUSG00000031765",
                "uniprot_ids": [
                    "P02802",
                    "A0A1D5RLN7",
                    "A0A1D5RMG1"
                ]
            },
            "ENSMUSG00000009681": {
                "id": "ENSMUSG00000009681",
                "type": "Gene",
                "symbol": "Bcr",
                "source": [
                    "ENSMUSG00000056501",
                    "ENSMUSG00000052684",
                    "ENSMUSG00000037868",
                    "ENSMUSG00000052837",
                    "ENSMUSG00000026628",
                    "ENSMUSG00000030103",
                    "ENSMUSG00000021250",
                    "ENSMUSG00000003032",
                    "ENSMUSG00000005583",
                    "ENSMUSG00000055148",
                    "ENSMUSG00000071076"
                ],
                "target": [],
                "expression_sum": {
                    "class1": 7907.0,
                    "class2": 839.0
                },
                "ens_id": "ENSMUSG00000009681",
                "uniprot_ids": [
                    "Q6PAJ1",
                    "A0A1W2P6J3",
                    "A0A1W2P6I7",
                    "A0A571BEM4"
                ]
            },
            "ENSMUSG00000090733": {
                "id": "ENSMUSG00000090733",
                "type": "Gene",
                "symbol": "Rps27",
                "source": [
                    "ENSMUSG00000023034",
                    "ENSMUSG00000000247",
                    "ENSMUSG00000057098",
                    "ENSMUSG00000038193",
                    "ENSMUSG00000005583",
                    "ENSMUSG00000048387"
                ],
                "target": [],
                "expression_sum": {
                    "class1": 229761.0,
                    "class2": 107039.0
                },
                "ens_id": "ENSMUSG00000090733",
                "uniprot_ids": [
                    "Q6ZWU9",
                    "A0A0G2JDW7",
                    "A0A0G2JG29",
                    "A0A0G2JEX7"
                ]
            },
            "ENSMUSG00000040952": {
                "id": "ENSMUSG00000040952",
                "type": "Gene",
                "symbol": "Rps19",
                "source": [
                    "ENSMUSG00000023034",
                    "ENSMUSG00000022528",
                    "ENSMUSG00000000247",
                    "ENSMUSG00000057098",
                    "ENSMUSG00000038193",
                    "ENSMUSG00000005583",
                    "ENSMUSG00000031965",
                    "ENSMUSG00000048387"
                ],
                "target": [],
                "expression_sum": {
                    "class1": 156248.0,
                    "class2": 83886.0
                },
                "ens_id": "ENSMUSG00000040952",
                "uniprot_ids": [
                    "Q9CZX8",
                    "D3YUG3",
                    "D3YUT3",
                    "S4R223",
                    "D3Z722",
                    "D3Z5R8"
                ]
            },
            "ENSMUSG00000058600": {
                "id": "ENSMUSG00000058600",
                "type": "Gene",
                "symbol": "Rpl30",
                "source": [
                    "ENSMUSG00000048387",
                    "ENSMUSG00000038193",
                    "ENSMUSG00000000247"
                ],
                "target": [],
                "expression_sum": {
                    "class1": 67497.0,
                    "class2": 32572.0
                },
                "ens_id": "ENSMUSG00000058600",
                "uniprot_ids": [
                    "P62889",
                    "A0A2I3BQF4"
                ]
            },
            "ENSMUSG00000047215": {
                "id": "ENSMUSG00000047215",
                "type": "Gene",
                "symbol": "Rpl9",
                "source": [
                    "ENSMUSG00000057098",
                    "ENSMUSG00000038193",
                    "ENSMUSG00000000247"
                ],
                "target": [],
                "expression_sum": {
                    "class1": 127172.0,
                    "class2": 60716.0
                },
                "ens_id": "ENSMUSG00000047215",
                "uniprot_ids": [
                    "P51410",
                    "D3Z629",
                    "A0A0G2JFQ3",
                    "A0A0G2JES3",
                    "D3YZT0"
                ]
            },
            "ENSMUSG00000041453": {
                "id": "ENSMUSG00000041453",
                "type": "Gene",
                "symbol": "Rpl21",
                "source": [
                    "ENSMUSG00000057098",
                    "ENSMUSG00000048387",
                    "ENSMUSG00000047638",
                    "ENSMUSG00000000247"
                ],
                "target": [],
                "expression_sum": {
                    "class1": 140870.0,
                    "class2": 81337.0
                },
                "ens_id": "ENSMUSG00000041453",
                "uniprot_ids": [
                    "Q9CQM8"
                ]
            },
            "ENSMUSG00000028495": {
                "id": "ENSMUSG00000028495",
                "type": "Gene",
                "symbol": "Rps6",
                "source": [
                    "ENSMUSG00000022528",
                    "ENSMUSG00000047638",
                    "ENSMUSG00000000247",
                    "ENSMUSG00000057098",
                    "ENSMUSG00000038193",
                    "ENSMUSG00000031965",
                    "ENSMUSG00000048387"
                ],
                "target": [],
                "expression_sum": {
                    "class1": 117047.0,
                    "class2": 57082.0
                },
                "ens_id": "ENSMUSG00000028495",
                "uniprot_ids": [
                    "P62754"
                ]
            },
            "ENSMUSG00000019278": {
                "id": "ENSMUSG00000019278",
                "type": "Gene",
                "symbol": "Dpep1",
                "source": [
                    "ENSMUSG00000022346",
                    "ENSMUSG00000052837",
                    "ENSMUSG00000052684",
                    "ENSMUSG00000003032"
                ],
                "target": [],
                "expression_sum": {
                    "class1": 353.0,
                    "class2": 12028.0
                },
                "ens_id": "ENSMUSG00000019278",
                "uniprot_ids": [
                    "P31428",
                    "A0A1D5RML1",
                    "A0A1D5RM32"
                ]
            },
            "ENSMUSG00000015085": {
                "id": "ENSMUSG00000015085",
                "type": "Gene",
                "symbol": "Entpd2",
                "source": [
                    "ENSMUSG00000052684",
                    "ENSMUSG00000052837",
                    "ENSMUSG00000021250",
                    "ENSMUSG00000003032",
                    "ENSMUSG00000005583",
                    "ENSMUSG00000028565"
                ],
                "target": [],
                "expression_sum": {
                    "class1": 548.0,
                    "class2": 7057.0
                },
                "ens_id": "ENSMUSG00000015085",
                "uniprot_ids": [
                    "O55026"
                ]
            },
            "ENSMUSG00000029718": {
                "id": "ENSMUSG00000029718",
                "type": "Gene",
                "symbol": "Pcolce",
                "source": [
                    "ENSMUSG00000032035",
                    "ENSMUSG00000052684",
                    "ENSMUSG00000038193",
                    "ENSMUSG00000005583",
                    "ENSMUSG00000071076"
                ],
                "target": [],
                "expression_sum": {
                    "class1": 41666.0,
                    "class2": 26645.0
                },
                "ens_id": "ENSMUSG00000029718",
                "uniprot_ids": [
                    "Q61398",
                    "D3YWH0",
                    "D3YZE3",
                    "A0A0G2JE29",
                    "D3YUE2"
                ]
            },
            "ENSMUSG00000035783": {
                "id": "ENSMUSG00000035783",
                "type": "Gene",
                "symbol": "Acta2",
                "source": [
                    "ENSMUSG00000023034",
                    "ENSMUSG00000058756",
                    "ENSMUSG00000038193",
                    "ENSMUSG00000021250",
                    "ENSMUSG00000005583"
                ],
                "target": [],
                "expression_sum": {
                    "class1": 8125.0,
                    "class2": 217388.0
                },
                "ens_id": "ENSMUSG00000035783",
                "uniprot_ids": [
                    "P62737",
                    "A0A494B9T3"
                ]
            },
            "ENSMUSG00000064215": {
                "id": "ENSMUSG00000064215",
                "type": "Gene",
                "symbol": "Ifi27",
                "source": [
                    "ENSMUSG00000052684",
                    "ENSMUSG00000052837",
                    "ENSMUSG00000021250",
                    "ENSMUSG00000008575",
                    "ENSMUSG00000055435",
                    "ENSMUSG00000020484",
                    "ENSMUSG00000071076"
                ],
                "target": [],
                "expression_sum": {
                    "class1": 6199.0,
                    "class2": 10432.0
                },
                "ens_id": "ENSMUSG00000064215",
                "uniprot_ids": [
                    "Q8C6G0",
                    "Q70LN0",
                    "Q70LM9",
                    "Q9D1E1",
                    "D3Z4L8",
                    "E9Q1C5"
                ]
            },
            "ENSMUSG00000020486": {
                "id": "ENSMUSG00000020486",
                "type": "Gene",
                "symbol": "Sept4",
                "source": [
                    "ENSMUSG00000052684",
                    "ENSMUSG00000028639",
                    "ENSMUSG00000015846",
                    "ENSMUSG00000005583",
                    "ENSMUSG00000071076"
                ],
                "target": [],
                "expression_sum": {
                    "class1": 68998.0,
                    "class2": 5830.0
                },
                "ens_id": "ENSMUSG00000020486",
                "uniprot_ids": [
                    "Q5ND19",
                    "P28661",
                    "A0A5F8MP96",
                    "Q5ND13",
                    "Q5ND09"
                ]
            },
            "ENSMUSG00000028639": {
                "id": "ENSMUSG00000028639",
                "type": "TF",
                "symbol": "Ybx1",
                "source": [
                    "ENSMUSG00000057098",
                    "ENSMUSG00000000247"
                ],
                "target": [
                    "ENSMUSG00000020486",
                    "ENSMUSG00000023367",
                    "ENSMUSG00000022032",
                    "ENSMUSG00000041577",
                    "ENSMUSG00000068522",
                    "ENSMUSG00000018417"
                ],
                "expression_sum": {
                    "class1": 29858.0,
                    "class2": 27229.0
                },
                "ens_id": "ENSMUSG00000028639",
                "uniprot_ids": [
                    "P62960",
                    "A2BGG7"
                ]
            },
            "ENSMUSG00000016024": {
                "id": "ENSMUSG00000016024",
                "type": "Gene",
                "symbol": "Lbp",
                "source": [
                    "ENSMUSG00000032238",
                    "ENSMUSG00000052684",
                    "ENSMUSG00000008575",
                    "ENSMUSG00000020484",
                    "ENSMUSG00000071076"
                ],
                "target": [],
                "expression_sum": {
                    "class1": 129.0,
                    "class2": 2661.0
                },
                "ens_id": "ENSMUSG00000016024",
                "uniprot_ids": [
                    "Q61805",
                    "A2AC65",
                    "F6XKX9"
                ]
            },
            "ENSMUSG00000024048": {
                "id": "ENSMUSG00000024048",
                "type": "Gene",
                "symbol": "Myl12a",
                "source": [
                    "ENSMUSG00000055435",
                    "ENSMUSG00000032238",
                    "ENSMUSG00000032035",
                    "ENSMUSG00000052684",
                    "ENSMUSG00000005583",
                    "ENSMUSG00000048387",
                    "ENSMUSG00000057098",
                    "ENSMUSG00000038418",
                    "ENSMUSG00000052534",
                    "ENSMUSG00000021250",
                    "ENSMUSG00000003032",
                    "ENSMUSG00000008575",
                    "ENSMUSG00000028565",
                    "ENSMUSG00000071076"
                ],
                "target": [],
                "expression_sum": {
                    "class1": 52081.0,
                    "class2": 15595.0
                },
                "ens_id": "ENSMUSG00000024048",
                "uniprot_ids": [
                    "Q6ZWQ9",
                    "D3Z249",
                    "D3YV37"
                ]
            },
            "ENSMUSG00000020029": {
                "id": "ENSMUSG00000020029",
                "type": "Gene",
                "symbol": "Nudt4",
                "source": [
                    "ENSMUSG00000053477",
                    "ENSMUSG00000071076",
                    "ENSMUSG00000005583",
                    "ENSMUSG00000008575"
                ],
                "target": [],
                "expression_sum": {
                    "class1": 2459.0,
                    "class2": 8855.0
                },
                "ens_id": "ENSMUSG00000020029",
                "uniprot_ids": [
                    "Q8R2U6"
                ]
            },
            "ENSMUSG00000024140": {
                "id": "ENSMUSG00000024140",
                "type": "TF",
                "symbol": "Epas1",
                "source": [
                    "ENSMUSG00000032238",
                    "ENSMUSG00000052684",
                    "ENSMUSG00000005583",
                    "ENSMUSG00000015846",
                    "ENSMUSG00000008575",
                    "ENSMUSG00000028565"
                ],
                "target": [
                    "ENSMUSG00000054452",
                    "ENSMUSG00000032420",
                    "ENSMUSG00000046330",
                    "ENSMUSG00000049439"
                ],
                "expression_sum": {
                    "class1": 14142.0,
                    "class2": 5104.0
                },
                "ens_id": "ENSMUSG00000024140",
                "uniprot_ids": [
                    "P97481"
                ]
            },
            "ENSMUSG00000021917": {
                "id": "ENSMUSG00000021917",
                "type": "Gene",
                "symbol": "Spcs1",
                "source": [
                    "ENSMUSG00000057098",
                    "ENSMUSG00000038418",
                    "ENSMUSG00000052684",
                    "ENSMUSG00000071076"
                ],
                "target": [],
                "expression_sum": {
                    "class1": 16694.0,
                    "class2": 7303.0
                },
                "ens_id": "ENSMUSG00000021917",
                "uniprot_ids": [
                    "Q9D958",
                    "A0A2I3BQG3",
                    "A0A2I3BQF0",
                    "A0A2I3BRW0"
                ]
            },
            "ENSMUSG00000027834": {
                "id": "ENSMUSG00000027834",
                "type": "Gene",
                "symbol": "Serpini1",
                "source": [
                    "ENSMUSG00000005583",
                    "ENSMUSG00000008575"
                ],
                "target": [],
                "expression_sum": {
                    "class1": 1521.0,
                    "class2": 4668.0
                },
                "ens_id": "ENSMUSG00000027834",
                "uniprot_ids": [
                    "O35684",
                    "E0CYW0"
                ]
            },
            "ENSMUSG00000009927": {
                "id": "ENSMUSG00000009927",
                "type": "Gene",
                "symbol": "Rps25",
                "source": [
                    "ENSMUSG00000057098",
                    "ENSMUSG00000048387",
                    "ENSMUSG00000000247"
                ],
                "target": [],
                "expression_sum": {
                    "class1": 76284.0,
                    "class2": 47742.0
                },
                "ens_id": "ENSMUSG00000009927",
                "uniprot_ids": [
                    "P62852",
                    "A0A1L1SQA8"
                ]
            },
            "ENSMUSG00000046364": {
                "id": "ENSMUSG00000046364",
                "type": "Gene",
                "symbol": "Rpl27a",
                "source": [
                    "ENSMUSG00000031965",
                    "ENSMUSG00000057098",
                    "ENSMUSG00000000247"
                ],
                "target": [],
                "expression_sum": {
                    "class1": 88935.0,
                    "class2": 55155.0
                },
                "ens_id": "ENSMUSG00000046364",
                "uniprot_ids": [
                    "P14115",
                    "G3UZV5"
                ]
            },
            "ENSMUSG00000023367": {
                "id": "ENSMUSG00000023367",
                "type": "Gene",
                "symbol": "Tmem176a",
                "source": [
                    "ENSMUSG00000028639",
                    "ENSMUSG00000038193",
                    "ENSMUSG00000005583",
                    "ENSMUSG00000000247"
                ],
                "target": [],
                "expression_sum": {
                    "class1": 90134.0,
                    "class2": 10306.0
                },
                "ens_id": "ENSMUSG00000023367",
                "uniprot_ids": [
                    "Q9DCS1",
                    "A0A0N4SUL5",
                    "A0A0N4SVW3",
                    "A0A0N4SVU8"
                ]
            },
            "ENSMUSG00000037072": {
                "id": "ENSMUSG00000037072",
                "type": "Gene",
                "symbol": "Sep15",
                "source": [
                    "ENSMUSG00000052684",
                    "ENSMUSG00000057098",
                    "ENSMUSG00000038418",
                    "ENSMUSG00000003032",
                    "ENSMUSG00000005583",
                    "ENSMUSG00000048387",
                    "ENSMUSG00000071076"
                ],
                "target": [],
                "expression_sum": {
                    "class1": 31733.0,
                    "class2": 18092.0
                },
                "ens_id": "ENSMUSG00000037072",
                "uniprot_ids": [
                    "A0A0R4J0K1",
                    "A0A1C7ZMY4",
                    "A0A0G2JEF3"
                ]
            },
            "ENSMUSG00000031422": {
                "id": "ENSMUSG00000031422",
                "type": "Gene",
                "symbol": "Morf4l2",
                "source": [
                    "ENSMUSG00000052684",
                    "ENSMUSG00000057098",
                    "ENSMUSG00000038418",
                    "ENSMUSG00000003032",
                    "ENSMUSG00000005583",
                    "ENSMUSG00000071076"
                ],
                "target": [],
                "expression_sum": {
                    "class1": 10829.0,
                    "class2": 7400.0
                },
                "ens_id": "ENSMUSG00000031422",
                "uniprot_ids": [
                    "Q9R0Q4",
                    "A2AEB6",
                    "A2AEB7",
                    "A2AEB5",
                    "A2AEB8",
                    "A2AEB9",
                    "A2AEB3",
                    "A2AEB4"
                ]
            },
            "ENSMUSG00000060591": {
                "id": "ENSMUSG00000060591",
                "type": "Gene",
                "symbol": "Ifitm2",
                "source": [
                    "ENSMUSG00000032035",
                    "ENSMUSG00000052684",
                    "ENSMUSG00000000247",
                    "ENSMUSG00000057098",
                    "ENSMUSG00000038193",
                    "ENSMUSG00000003032",
                    "ENSMUSG00000005583",
                    "ENSMUSG00000071076"
                ],
                "target": [],
                "expression_sum": {
                    "class1": 107145.0,
                    "class2": 33004.0
                },
                "ens_id": "ENSMUSG00000060591",
                "uniprot_ids": [
                    "Q99J93"
                ]
            },
            "ENSMUSG00000004207": {
                "id": "ENSMUSG00000004207",
                "type": "Gene",
                "symbol": "Psap",
                "source": [
                    "ENSMUSG00000032035",
                    "ENSMUSG00000000247",
                    "ENSMUSG00000057098",
                    "ENSMUSG00000038193",
                    "ENSMUSG00000005583",
                    "ENSMUSG00000028565",
                    "ENSMUSG00000071076"
                ],
                "target": [],
                "expression_sum": {
                    "class1": 45580.0,
                    "class2": 24159.0
                },
                "ens_id": "ENSMUSG00000004207",
                "uniprot_ids": [
                    "Q61207",
                    "Q8BFQ1",
                    "J3QPG5",
                    "E9PZ00",
                    "K3W4L3"
                ]
            },
            "ENSMUSG00000020612": {
                "id": "ENSMUSG00000020612",
                "type": "Gene",
                "symbol": "Prkar1a",
                "source": [
                    "ENSMUSG00000047638",
                    "ENSMUSG00000020484",
                    "ENSMUSG00000005583",
                    "ENSMUSG00000000247"
                ],
                "target": [],
                "expression_sum": {
                    "class1": 12222.0,
                    "class2": 11236.0
                },
                "ens_id": "ENSMUSG00000020612",
                "uniprot_ids": [
                    "Q9DBC7",
                    "A2AI69"
                ]
            },
            "ENSMUSG00000029614": {
                "id": "ENSMUSG00000029614",
                "type": "Gene",
                "symbol": "Rpl6",
                "source": [
                    "ENSMUSG00000048387",
                    "ENSMUSG00000038193",
                    "ENSMUSG00000023034",
                    "ENSMUSG00000000247"
                ],
                "target": [],
                "expression_sum": {
                    "class1": 114722.0,
                    "class2": 53657.0
                },
                "ens_id": "ENSMUSG00000029614",
                "uniprot_ids": [
                    "P47911",
                    "A0A0J9YU32"
                ]
            },
            "ENSMUSG00000031375": {
                "id": "ENSMUSG00000031375",
                "type": "Gene",
                "symbol": "Bgn",
                "source": [
                    "ENSMUSG00000042812",
                    "ENSMUSG00000032035",
                    "ENSMUSG00000021250",
                    "ENSMUSG00000056501"
                ],
                "target": [],
                "expression_sum": {
                    "class1": 393016.0,
                    "class2": 87690.0
                },
                "ens_id": "ENSMUSG00000031375",
                "uniprot_ids": [
                    "P28653"
                ]
            },
            "ENSMUSG00000036446": {
                "id": "ENSMUSG00000036446",
                "type": "Gene",
                "symbol": "Lum",
                "source": [
                    "ENSMUSG00000052684",
                    "ENSMUSG00000053477",
                    "ENSMUSG00000038193",
                    "ENSMUSG00000003032",
                    "ENSMUSG00000005583"
                ],
                "target": [],
                "expression_sum": {
                    "class1": 88603.0,
                    "class2": 46293.0
                },
                "ens_id": "ENSMUSG00000036446",
                "uniprot_ids": [
                    "P51885"
                ]
            },
            "ENSMUSG00000024014": {
                "id": "ENSMUSG00000024014",
                "type": "Gene",
                "symbol": "Pim1",
                "source": [
                    "ENSMUSG00000056501",
                    "ENSMUSG00000023034",
                    "ENSMUSG00000052837",
                    "ENSMUSG00000026628",
                    "ENSMUSG00000038418",
                    "ENSMUSG00000021250",
                    "ENSMUSG00000003545",
                    "ENSMUSG00000003032",
                    "ENSMUSG00000018899",
                    "ENSMUSG00000071076"
                ],
                "target": [],
                "expression_sum": {
                    "class1": 7590.0,
                    "class2": 2414.0
                },
                "ens_id": "ENSMUSG00000024014",
                "uniprot_ids": [
                    "P06803",
                    "A0A3Q4EGX3",
                    "A0A3Q4EGB5"
                ]
            },
            "ENSMUSG00000001506": {
                "id": "ENSMUSG00000001506",
                "type": "Gene",
                "symbol": "Col1a1",
                "source": [
                    "ENSMUSG00000071076",
                    "ENSMUSG00000005583",
                    "ENSMUSG00000052684",
                    "ENSMUSG00000008575"
                ],
                "target": [],
                "expression_sum": {
                    "class1": 33962.0,
                    "class2": 34306.0
                },
                "ens_id": "ENSMUSG00000001506",
                "uniprot_ids": [
                    "P11087"
                ]
            },
            "ENSMUSG00000001911": {
                "id": "ENSMUSG00000001911",
                "type": "TF",
                "symbol": "Nfix",
                "source": [
                    "ENSMUSG00000071076",
                    "ENSMUSG00000005583",
                    "ENSMUSG00000052684",
                    "ENSMUSG00000008575"
                ],
                "target": [
                    "ENSMUSG00000027931"
                ],
                "expression_sum": {
                    "class1": 3870.0,
                    "class2": 5667.0
                },
                "ens_id": "ENSMUSG00000001911",
                "uniprot_ids": [
                    "P70257",
                    "Q3TYK3",
                    "D3YZ00",
                    "E9PUH7"
                ]
            },
            "ENSMUSG00000041084": {
                "id": "ENSMUSG00000041084",
                "type": "Gene",
                "symbol": "Ostc",
                "source": [
                    "ENSMUSG00000052684",
                    "ENSMUSG00000003032",
                    "ENSMUSG00000005583",
                    "ENSMUSG00000048387",
                    "ENSMUSG00000020484",
                    "ENSMUSG00000071076"
                ],
                "target": [],
                "expression_sum": {
                    "class1": 8276.0,
                    "class2": 5532.0
                },
                "ens_id": "ENSMUSG00000041084",
                "uniprot_ids": [
                    "Q78XF5"
                ]
            },
            "ENSMUSG00000035493": {
                "id": "ENSMUSG00000035493",
                "type": "Gene",
                "symbol": "Tgfbi",
                "source": [
                    "ENSMUSG00000056501",
                    "ENSMUSG00000023034",
                    "ENSMUSG00000042812",
                    "ENSMUSG00000038193",
                    "ENSMUSG00000021250",
                    "ENSMUSG00000005583",
                    "ENSMUSG00000018899"
                ],
                "target": [],
                "expression_sum": {
                    "class1": 102105.0,
                    "class2": 9504.0
                },
                "ens_id": "ENSMUSG00000035493",
                "uniprot_ids": [
                    "P82198",
                    "Q3UXJ2"
                ]
            },
            "ENSMUSG00000020644": {
                "id": "ENSMUSG00000020644",
                "type": "TF",
                "symbol": "Id2",
                "source": [
                    "ENSMUSG00000030103",
                    "ENSMUSG00000005583"
                ],
                "target": [
                    "ENSMUSG00000032387",
                    "ENSMUSG00000029761",
                    "ENSMUSG00000001175",
                    "ENSMUSG00000034566",
                    "ENSMUSG00000014956",
                    "ENSMUSG00000018293",
                    "ENSMUSG00000037712",
                    "ENSMUSG00000022010",
                    "ENSMUSG00000024290",
                    "ENSMUSG00000029580",
                    "ENSMUSG00000032330",
                    "ENSMUSG00000058056"
                ],
                "expression_sum": {
                    "class1": 7989.0,
                    "class2": 16476.0
                },
                "ens_id": "ENSMUSG00000020644",
                "uniprot_ids": [
                    "P41136",
                    "A0A1Y7VMT3"
                ]
            },
            "ENSMUSG00000028517": {
                "id": "ENSMUSG00000028517",
                "type": "Gene",
                "symbol": "Plpp3",
                "source": [
                    "ENSMUSG00000052684",
                    "ENSMUSG00000005583",
                    "ENSMUSG00000048387",
                    "ENSMUSG00000003032",
                    "ENSMUSG00000008575",
                    "ENSMUSG00000028565",
                    "ENSMUSG00000071076"
                ],
                "target": [],
                "expression_sum": {
                    "class1": 1847.0,
                    "class2": 13633.0
                },
                "ens_id": "ENSMUSG00000028517",
                "uniprot_ids": [
                    "Q99JY8"
                ]
            }
        },
        "grps": {
            "ENSMUSG00000029322_ENSMUSG00000014294": {
                "id": "ENSMUSG00000029322_ENSMUSG00000014294",
                "type": "Mix",
                "score": 5.534167394363044,
                "reversable": false,
                "correlations": {
                    "class1": 0.3983076592094462,
                    "class2": 0.0786880997602173
                },
                "regulatory_source": "ENSMUSG00000029322",
                "regulatory_target": "ENSMUSG00000014294"
            },
            "ENSMUSG00000031818_ENSMUSG00000029322": {
                "id": "ENSMUSG00000031818_ENSMUSG00000029322",
                "type": "Standard",
                "score": 0.775817831734018,
                "reversable": false,
                "correlations": {
                    "class1": 0.5149750262543713,
                    "class2": 0.13024749915687156
                },
                "regulatory_source": "ENSMUSG00000029322",
                "regulatory_target": "ENSMUSG00000031818"
            },
            "ENSMUSG00000029322_ENSMUSG00000007872": {
                "id": "ENSMUSG00000029322_ENSMUSG00000007872",
                "type": "Mix",
                "score": 5.728981334502581,
                "reversable": false,
                "correlations": {
                    "class1": 0.3569527385510402,
                    "class2": 0.0
                },
                "regulatory_source": "ENSMUSG00000029322",
                "regulatory_target": "ENSMUSG00000007872"
            },
            "ENSMUSG00000060636_ENSMUSG00000000247": {
                "id": "ENSMUSG00000060636_ENSMUSG00000000247",
                "type": "Mix",
                "score": 6.59820306700032,
                "reversable": false,
                "correlations": {
                    "class1": 0.3512008182192743,
                    "class2": 0.03545341342514245
                },
                "regulatory_source": "ENSMUSG00000000247",
                "regulatory_target": "ENSMUSG00000060636"
            },
            "ENSMUSG00000062647_ENSMUSG00000000247": {
                "id": "ENSMUSG00000062647_ENSMUSG00000000247",
                "type": "Standard",
                "score": 0.9410486516609298,
                "reversable": false,
                "correlations": {
                    "class1": 0.36044003265263236,
                    "class2": 0.05657419631453429
                },
                "regulatory_source": "ENSMUSG00000000247",
                "regulatory_target": "ENSMUSG00000062647"
            },
            "ENSMUSG00000024608_ENSMUSG00000000247": {
                "id": "ENSMUSG00000024608_ENSMUSG00000000247",
                "type": "Standard",
                "score": 0.5350961130932429,
                "reversable": false,
                "correlations": {
                    "class1": 0.35937470713243075,
                    "class2": 0.038573822364074556
                },
                "regulatory_source": "ENSMUSG00000000247",
                "regulatory_target": "ENSMUSG00000024608"
            },
            "ENSMUSG00000048387_ENSMUSG00000031447": {
                "id": "ENSMUSG00000048387_ENSMUSG00000031447",
                "type": "Standard",
                "score": 0.6313244860731053,
                "reversable": false,
                "correlations": {
                    "class1": 0.0,
                    "class2": 0.37087825069910707
                },
                "regulatory_source": "ENSMUSG00000048387",
                "regulatory_target": "ENSMUSG00000031447"
            },
            "ENSMUSG00000048387_ENSMUSG00000012848": {
                "id": "ENSMUSG00000048387_ENSMUSG00000012848",
                "type": "Standard",
                "score": 0.5672083826211916,
                "reversable": false,
                "correlations": {
                    "class1": 0.060909072316776794,
                    "class2": 0.4123206325393834
                },
                "regulatory_source": "ENSMUSG00000048387",
                "regulatory_target": "ENSMUSG00000012848"
            },
            "ENSMUSG00000048387_ENSMUSG00000024608": {
                "id": "ENSMUSG00000048387_ENSMUSG00000024608",
                "type": "Standard",
                "score": 0.45847352038978145,
                "reversable": false,
                "correlations": {
                    "class1": 0.06463152398609764,
                    "class2": 0.38478821223534954
                },
                "regulatory_source": "ENSMUSG00000048387",
                "regulatory_target": "ENSMUSG00000024608"
            },
            "ENSMUSG00000057098_ENSMUSG00000024608": {
                "id": "ENSMUSG00000057098_ENSMUSG00000024608",
                "type": "Mix",
                "score": 14.919033599086804,
                "reversable": false,
                "correlations": {
                    "class1": 0.0,
                    "class2": 0.38928289704040947
                },
                "regulatory_source": "ENSMUSG00000057098",
                "regulatory_target": "ENSMUSG00000024608"
            },
            "ENSMUSG00000063457_ENSMUSG00000057098": {
                "id": "ENSMUSG00000063457_ENSMUSG00000057098",
                "type": "Mix",
                "score": 5.61747930722422,
                "reversable": false,
                "correlations": {
                    "class1": 0.03762571400967034,
                    "class2": 0.3834251904053063
                },
                "regulatory_source": "ENSMUSG00000057098",
                "regulatory_target": "ENSMUSG00000063457"
            },
            "ENSMUSG00000057098_ENSMUSG00000008668": {
                "id": "ENSMUSG00000057098_ENSMUSG00000008668",
                "type": "Mix",
                "score": 5.463778209982271,
                "reversable": false,
                "correlations": {
                    "class1": 0.0,
                    "class2": 0.3790949261202038
                },
                "regulatory_source": "ENSMUSG00000057098",
                "regulatory_target": "ENSMUSG00000008668"
            },
            "ENSMUSG00000057098_ENSMUSG00000022108": {
                "id": "ENSMUSG00000057098_ENSMUSG00000022108",
                "type": "Standard",
                "score": 0.6441225137860037,
                "reversable": false,
                "correlations": {
                    "class1": 0.0,
                    "class2": 0.35107894513456317
                },
                "regulatory_source": "ENSMUSG00000057098",
                "regulatory_target": "ENSMUSG00000022108"
            },
            "ENSMUSG00000057098_ENSMUSG00000003429": {
                "id": "ENSMUSG00000057098_ENSMUSG00000003429",
                "type": "Standard",
                "score": 0.5602482230211044,
                "reversable": false,
                "correlations": {
                    "class1": 0.0,
                    "class2": 0.4041494920189355
                },
                "regulatory_source": "ENSMUSG00000057098",
                "regulatory_target": "ENSMUSG00000003429"
            },
            "ENSMUSG00000057098_ENSMUSG00000032399": {
                "id": "ENSMUSG00000057098_ENSMUSG00000032399",
                "type": "Mix",
                "score": 6.26147944068973,
                "reversable": false,
                "correlations": {
                    "class1": 0.0,
                    "class2": 0.375627370573335
                },
                "regulatory_source": "ENSMUSG00000057098",
                "regulatory_target": "ENSMUSG00000032399"
            },
            "ENSMUSG00000057098_ENSMUSG00000030744": {
                "id": "ENSMUSG00000057098_ENSMUSG00000030744",
                "type": "Mix",
                "score": 6.468213841313619,
                "reversable": false,
                "correlations": {
                    "class1": 0.027503184427258777,
                    "class2": 0.3798724584603384
                },
                "regulatory_source": "ENSMUSG00000057098",
                "regulatory_target": "ENSMUSG00000030744"
            },
            "ENSMUSG00000057098_ENSMUSG00000041841": {
                "id": "ENSMUSG00000057098_ENSMUSG00000041841",
                "type": "Standard",
                "score": 0.557027923471079,
                "reversable": false,
                "correlations": {
                    "class1": 0.029309955387579387,
                    "class2": 0.3618323689585674
                },
                "regulatory_source": "ENSMUSG00000057098",
                "regulatory_target": "ENSMUSG00000041841"
            },
            "ENSMUSG00000057098_ENSMUSG00000037742": {
                "id": "ENSMUSG00000057098_ENSMUSG00000037742",
                "type": "Mix",
                "score": 5.416335591381316,
                "reversable": false,
                "correlations": {
                    "class1": 0.0,
                    "class2": 0.41647513900100464
                },
                "regulatory_source": "ENSMUSG00000057098",
                "regulatory_target": "ENSMUSG00000037742"
            },
            "ENSMUSG00000037742_ENSMUSG00000000247": {
                "id": "ENSMUSG00000037742_ENSMUSG00000000247",
                "type": "Mix",
                "score": 7.74574153122967,
                "reversable": false,
                "correlations": {
                    "class1": 0.43565129936792324,
                    "class2": 0.04224571805120184
                },
                "regulatory_source": "ENSMUSG00000000247",
                "regulatory_target": "ENSMUSG00000037742"
            },
            "ENSMUSG00000059070_ENSMUSG00000057098": {
                "id": "ENSMUSG00000059070_ENSMUSG00000057098",
                "type": "Standard",
                "score": 0.49233206664839974,
                "reversable": false,
                "correlations": {
                    "class1": 0.030377671107013424,
                    "class2": 0.35596307825009754
                },
                "regulatory_source": "ENSMUSG00000057098",
                "regulatory_target": "ENSMUSG00000059070"
            },
            "ENSMUSG00000030744_ENSMUSG00000000247": {
                "id": "ENSMUSG00000030744_ENSMUSG00000000247",
                "type": "Standard",
                "score": 0.35757688589844205,
                "reversable": false,
                "correlations": {
                    "class1": 0.34541220237470344,
                    "class2": 0.03603023742333218
                },
                "regulatory_source": "ENSMUSG00000000247",
                "regulatory_target": "ENSMUSG00000030744"
            },
            "ENSMUSG00000003970_ENSMUSG00000000247": {
                "id": "ENSMUSG00000003970_ENSMUSG00000000247",
                "type": "Mix",
                "score": 9.946048104893595,
                "reversable": false,
                "correlations": {
                    "class1": 0.36865573845727817,
                    "class2": 0.05158566508268787
                },
                "regulatory_source": "ENSMUSG00000000247",
                "regulatory_target": "ENSMUSG00000003970"
            },
            "ENSMUSG00000060938_ENSMUSG00000048387": {
                "id": "ENSMUSG00000060938_ENSMUSG00000048387",
                "type": "Standard",
                "score": 0.5772235966249396,
                "reversable": false,
                "correlations": {
                    "class1": 0.0638261443489527,
                    "class2": 0.38777538608716544
                },
                "regulatory_source": "ENSMUSG00000048387",
                "regulatory_target": "ENSMUSG00000060938"
            },
            "ENSMUSG00000074129_ENSMUSG00000057098": {
                "id": "ENSMUSG00000074129_ENSMUSG00000057098",
                "type": "Standard",
                "score": 0.3336860290978623,
                "reversable": false,
                "correlations": {
                    "class1": 0.0,
                    "class2": 0.3549428215877327
                },
                "regulatory_source": "ENSMUSG00000057098",
                "regulatory_target": "ENSMUSG00000074129"
            },
            "ENSMUSG00000041841_ENSMUSG00000000247": {
                "id": "ENSMUSG00000041841_ENSMUSG00000000247",
                "type": "Mix",
                "score": 5.926969215515621,
                "reversable": false,
                "correlations": {
                    "class1": 0.35937680034982683,
                    "class2": 0.03729373291632408
                },
                "regulatory_source": "ENSMUSG00000000247",
                "regulatory_target": "ENSMUSG00000041841"
            },
            "ENSMUSG00000048387_ENSMUSG00000031320": {
                "id": "ENSMUSG00000048387_ENSMUSG00000031320",
                "type": "Standard",
                "score": 0.9554206441146518,
                "reversable": false,
                "correlations": {
                    "class1": 0.07759264388901752,
                    "class2": 0.43302239656213515
                },
                "regulatory_source": "ENSMUSG00000048387",
                "regulatory_target": "ENSMUSG00000031320"
            },
            "ENSMUSG00000090862_ENSMUSG00000000247": {
                "id": "ENSMUSG00000090862_ENSMUSG00000000247",
                "type": "Mix",
                "score": 8.975555162646321,
                "reversable": false,
                "correlations": {
                    "class1": 0.34113743823225656,
                    "class2": 0.04928848624182726
                },
                "regulatory_source": "ENSMUSG00000000247",
                "regulatory_target": "ENSMUSG00000090862"
            },
            "ENSMUSG00000048387_ENSMUSG00000020585": {
                "id": "ENSMUSG00000048387_ENSMUSG00000020585",
                "type": "Mix",
                "score": 6.238709422012988,
                "reversable": false,
                "correlations": {
                    "class1": 0.0,
                    "class2": 0.3926534984761467
                },
                "regulatory_source": "ENSMUSG00000048387",
                "regulatory_target": "ENSMUSG00000020585"
            },
            "ENSMUSG00000059895_ENSMUSG00000005583": {
                "id": "ENSMUSG00000059895_ENSMUSG00000005583",
                "type": "Standard",
                "score": 1.3785905140656636,
                "reversable": false,
                "correlations": {
                    "class1": 0.08054707703459893,
                    "class2": 0.43772367480509605
                },
                "regulatory_source": "ENSMUSG00000005583",
                "regulatory_target": "ENSMUSG00000059895"
            },
            "ENSMUSG00000019907_ENSMUSG00000005583": {
                "id": "ENSMUSG00000019907_ENSMUSG00000005583",
                "type": "Mix",
                "score": 8.992227808541104,
                "reversable": false,
                "correlations": {
                    "class1": 0.04371462553078975,
                    "class2": 0.4901687962295429
                },
                "regulatory_source": "ENSMUSG00000005583",
                "regulatory_target": "ENSMUSG00000019907"
            },
            "ENSMUSG00000005583_ENSMUSG00000002107": {
                "id": "ENSMUSG00000005583_ENSMUSG00000002107",
                "type": "Mix",
                "score": 10.947855760501891,
                "reversable": false,
                "correlations": {
                    "class1": 0.0,
                    "class2": -0.30269577300963824
                },
                "regulatory_source": "ENSMUSG00000005583",
                "regulatory_target": "ENSMUSG00000002107"
            },
            "ENSMUSG00000037868_ENSMUSG00000021250": {
                "id": "ENSMUSG00000037868_ENSMUSG00000021250",
                "type": "Mix",
                "score": 5.611695544374111,
                "reversable": true,
                "correlations": {
                    "class1": 0.511602981350903,
                    "class2": 0.10137532326055317
                },
                "regulatory_source": "ENSMUSG00000037868",
                "regulatory_target": "ENSMUSG00000021250"
            },
            "ENSMUSG00000053560_ENSMUSG00000021250": {
                "id": "ENSMUSG00000053560_ENSMUSG00000021250",
                "type": "Mix",
                "score": 8.171255069000967,
                "reversable": false,
                "correlations": {
                    "class1": 0.6938031413597321,
                    "class2": 0.4234083942937145
                },
                "regulatory_source": "ENSMUSG00000021250",
                "regulatory_target": "ENSMUSG00000053560"
            },
            "ENSMUSG00000052684_ENSMUSG00000038418": {
                "id": "ENSMUSG00000052684_ENSMUSG00000038418",
                "type": "Mix",
                "score": 11.947891079741218,
                "reversable": true,
                "correlations": {
                    "class1": 0.7664433043909842,
                    "class2": 0.486543404345432
                },
                "regulatory_source": "ENSMUSG00000052684",
                "regulatory_target": "ENSMUSG00000038418"
            },
            "ENSMUSG00000052837_ENSMUSG00000038418": {
                "id": "ENSMUSG00000052837_ENSMUSG00000038418",
                "type": "Mix",
                "score": 13.674555392069976,
                "reversable": true,
                "correlations": {
                    "class1": 0.816953701569463,
                    "class2": 0.5696919058794143
                },
                "regulatory_source": "ENSMUSG00000052837",
                "regulatory_target": "ENSMUSG00000038418"
            },
            "ENSMUSG00000052684_ENSMUSG00000008540": {
                "id": "ENSMUSG00000052684_ENSMUSG00000008540",
                "type": "Mix",
                "score": 9.414890690797678,
                "reversable": false,
                "correlations": {
                    "class1": 0.0,
                    "class2": 0.4708711183763723
                },
                "regulatory_source": "ENSMUSG00000052684",
                "regulatory_target": "ENSMUSG00000008540"
            },
            "ENSMUSG00000071076_ENSMUSG00000021390": {
                "id": "ENSMUSG00000071076_ENSMUSG00000021390",
                "type": "Mix",
                "score": 5.526325055742347,
                "reversable": false,
                "correlations": {
                    "class1": 0.05504122209584657,
                    "class2": 0.4507139948726272
                },
                "regulatory_source": "ENSMUSG00000071076",
                "regulatory_target": "ENSMUSG00000021390"
            },
            "ENSMUSG00000071076_ENSMUSG00000029096": {
                "id": "ENSMUSG00000071076_ENSMUSG00000029096",
                "type": "Mix",
                "score": 5.699395184995124,
                "reversable": false,
                "correlations": {
                    "class1": 0.16270941377798304,
                    "class2": 0.5689505604443278
                },
                "regulatory_source": "ENSMUSG00000071076",
                "regulatory_target": "ENSMUSG00000029096"
            },
            "ENSMUSG00000071076_ENSMUSG00000058135": {
                "id": "ENSMUSG00000071076_ENSMUSG00000058135",
                "type": "Standard",
                "score": 1.1318553391128547,
                "reversable": false,
                "correlations": {
                    "class1": 0.10173575170565055,
                    "class2": 0.4992405792689435
                },
                "regulatory_source": "ENSMUSG00000071076",
                "regulatory_target": "ENSMUSG00000058135"
            },
            "ENSMUSG00000075702_ENSMUSG00000071076": {
                "id": "ENSMUSG00000075702_ENSMUSG00000071076",
                "type": "Standard",
                "score": 1.033885260717099,
                "reversable": false,
                "correlations": {
                    "class1": 0.14441541564946575,
                    "class2": 0.5057471001944507
                },
                "regulatory_source": "ENSMUSG00000071076",
                "regulatory_target": "ENSMUSG00000075702"
            },
            "ENSMUSG00000071076_ENSMUSG00000032231": {
                "id": "ENSMUSG00000071076_ENSMUSG00000032231",
                "type": "Mix",
                "score": 8.589637426747593,
                "reversable": false,
                "correlations": {
                    "class1": 0.09131678291065735,
                    "class2": 0.47673079157771725
                },
                "regulatory_source": "ENSMUSG00000071076",
                "regulatory_target": "ENSMUSG00000032231"
            },
            "ENSMUSG00000071076_ENSMUSG00000021876": {
                "id": "ENSMUSG00000071076_ENSMUSG00000021876",
                "type": "Mix",
                "score": 12.116067695806787,
                "reversable": false,
                "correlations": {
                    "class1": 0.09691444896673937,
                    "class2": 0.5914264320590543
                },
                "regulatory_source": "ENSMUSG00000071076",
                "regulatory_target": "ENSMUSG00000021876"
            },
            "ENSMUSG00000071076_ENSMUSG00000029659": {
                "id": "ENSMUSG00000071076_ENSMUSG00000029659",
                "type": "Standard",
                "score": 0.8822730468055532,
                "reversable": false,
                "correlations": {
                    "class1": 0.08759311064378093,
                    "class2": 0.4262426461615084
                },
                "regulatory_source": "ENSMUSG00000071076",
                "regulatory_target": "ENSMUSG00000029659"
            },
            "ENSMUSG00000071076_ENSMUSG00000008540": {
                "id": "ENSMUSG00000071076_ENSMUSG00000008540",
                "type": "Mix",
                "score": 12.117623450641636,
                "reversable": false,
                "correlations": {
                    "class1": 0.08176840400100349,
                    "class2": 0.5245678510596467
                },
                "regulatory_source": "ENSMUSG00000071076",
                "regulatory_target": "ENSMUSG00000008540"
            },
            "ENSMUSG00000052684_ENSMUSG00000000753": {
                "id": "ENSMUSG00000052684_ENSMUSG00000000753",
                "type": "Mix",
                "score": 14.977130481998238,
                "reversable": false,
                "correlations": {
                    "class1": 0.03437712335857798,
                    "class2": 0.43356757378057553
                },
                "regulatory_source": "ENSMUSG00000052684",
                "regulatory_target": "ENSMUSG00000000753"
            },
            "ENSMUSG00000052684_ENSMUSG00000020053": {
                "id": "ENSMUSG00000052684_ENSMUSG00000020053",
                "type": "Mix",
                "score": 18.850769681477235,
                "reversable": false,
                "correlations": {
                    "class1": -0.07502879935792303,
                    "class2": 0.40516312762048773
                },
                "regulatory_source": "ENSMUSG00000052684",
                "regulatory_target": "ENSMUSG00000020053"
            },
            "ENSMUSG00000071076_ENSMUSG00000020053": {
                "id": "ENSMUSG00000071076_ENSMUSG00000020053",
                "type": "Mix",
                "score": 13.332142974899995,
                "reversable": false,
                "correlations": {
                    "class1": -0.028028671985073837,
                    "class2": 0.4461161908027037
                },
                "regulatory_source": "ENSMUSG00000071076",
                "regulatory_target": "ENSMUSG00000020053"
            },
            "ENSMUSG00000053113_ENSMUSG00000052684": {
                "id": "ENSMUSG00000053113_ENSMUSG00000052684",
                "type": "Mix",
                "score": 7.03491541026321,
                "reversable": false,
                "correlations": {
                    "class1": 0.7260928050980276,
                    "class2": 0.4200604160119371
                },
                "regulatory_source": "ENSMUSG00000052684",
                "regulatory_target": "ENSMUSG00000053113"
            },
            "ENSMUSG00000090698_ENSMUSG00000052684": {
                "id": "ENSMUSG00000090698_ENSMUSG00000052684",
                "type": "Mix",
                "score": 8.110685410599022,
                "reversable": false,
                "correlations": {
                    "class1": 0.35912839603911373,
                    "class2": -0.07170878293356608
                },
                "regulatory_source": "ENSMUSG00000052684",
                "regulatory_target": "ENSMUSG00000090698"
            },
            "ENSMUSG00000071076_ENSMUSG00000055148": {
                "id": "ENSMUSG00000071076_ENSMUSG00000055148",
                "type": "Standard",
                "score": 0.8925988455170332,
                "reversable": false,
                "correlations": {
                    "class1": 0.5650328779947075,
                    "class2": 0.24800406666903702
                },
                "regulatory_source": "ENSMUSG00000071076",
                "regulatory_target": "ENSMUSG00000055148"
            },
            "ENSMUSG00000055148_ENSMUSG00000003545": {
                "id": "ENSMUSG00000055148_ENSMUSG00000003545",
                "type": "Mix",
                "score": 14.881027125911217,
                "reversable": false,
                "correlations": {
                    "class1": 0.5666618661792494,
                    "class2": 0.19337881173911073
                },
                "regulatory_source": "ENSMUSG00000055148",
                "regulatory_target": "ENSMUSG00000003545"
            },
            "ENSMUSG00000015312_ENSMUSG00000003545": {
                "id": "ENSMUSG00000015312_ENSMUSG00000003545",
                "type": "Standard",
                "score": 1.0762441534764344,
                "reversable": false,
                "correlations": {
                    "class1": 0.44338913725403223,
                    "class2": 0.11452195516190655
                },
                "regulatory_source": "ENSMUSG00000003545",
                "regulatory_target": "ENSMUSG00000015312"
            },
            "ENSMUSG00000053560_ENSMUSG00000003545": {
                "id": "ENSMUSG00000053560_ENSMUSG00000003545",
                "type": "Standard",
                "score": 1.021304356332249,
                "reversable": false,
                "correlations": {
                    "class1": 0.589485191505976,
                    "class2": 0.22349812022466964
                },
                "regulatory_source": "ENSMUSG00000003545",
                "regulatory_target": "ENSMUSG00000053560"
            },
            "ENSMUSG00000071076_ENSMUSG00000052837": {
                "id": "ENSMUSG00000071076_ENSMUSG00000052837",
                "type": "Mix",
                "score": 6.036426962904938,
                "reversable": true,
                "correlations": {
                    "class1": 0.7238904426938447,
                    "class2": 0.4376071659527366
                },
                "regulatory_source": "ENSMUSG00000071076",
                "regulatory_target": "ENSMUSG00000052837"
            },
            "ENSMUSG00000055148_ENSMUSG00000038418": {
                "id": "ENSMUSG00000055148_ENSMUSG00000038418",
                "type": "Mix",
                "score": 8.189327875064823,
                "reversable": false,
                "correlations": {
                    "class1": 0.5798601054470557,
                    "class2": 0.22569929951045264
                },
                "regulatory_source": "ENSMUSG00000038418",
                "regulatory_target": "ENSMUSG00000055148"
            },
            "ENSMUSG00000052684_ENSMUSG00000031451": {
                "id": "ENSMUSG00000052684_ENSMUSG00000031451",
                "type": "Standard",
                "score": 0.6199535942467939,
                "reversable": false,
                "correlations": {
                    "class1": -0.05539840876026362,
                    "class2": 0.3508439446337546
                },
                "regulatory_source": "ENSMUSG00000052684",
                "regulatory_target": "ENSMUSG00000031451"
            },
            "ENSMUSG00000055148_ENSMUSG00000053560": {
                "id": "ENSMUSG00000055148_ENSMUSG00000053560",
                "type": "Standard",
                "score": 0.6092912499842723,
                "reversable": false,
                "correlations": {
                    "class1": 0.5155082130652904,
                    "class2": 0.19806272926237303
                },
                "regulatory_source": "ENSMUSG00000055148",
                "regulatory_target": "ENSMUSG00000053560"
            },
            "ENSMUSG00000038418_ENSMUSG00000026628": {
                "id": "ENSMUSG00000038418_ENSMUSG00000026628",
                "type": "Mix",
                "score": 7.6345749976519315,
                "reversable": true,
                "correlations": {
                    "class1": 0.5998243543116571,
                    "class2": 0.2822248149600215
                },
                "regulatory_source": "ENSMUSG00000026628",
                "regulatory_target": "ENSMUSG00000038418"
            },
            "ENSMUSG00000052684_ENSMUSG00000032359": {
                "id": "ENSMUSG00000052684_ENSMUSG00000032359",
                "type": "Mix",
                "score": 17.792270397119857,
                "reversable": false,
                "correlations": {
                    "class1": 0.03656817221589649,
                    "class2": 0.3604616066524656
                },
                "regulatory_source": "ENSMUSG00000052684",
                "regulatory_target": "ENSMUSG00000032359"
            },
            "ENSMUSG00000052684_ENSMUSG00000017493": {
                "id": "ENSMUSG00000052684_ENSMUSG00000017493",
                "type": "Mix",
                "score": 14.669761823148434,
                "reversable": false,
                "correlations": {
                    "class1": -0.028759260948858063,
                    "class2": 0.41717336202319866
                },
                "regulatory_source": "ENSMUSG00000052684",
                "regulatory_target": "ENSMUSG00000017493"
            },
            "ENSMUSG00000038418_ENSMUSG00000006369": {
                "id": "ENSMUSG00000038418_ENSMUSG00000006369",
                "type": "Mix",
                "score": 9.121181332573904,
                "reversable": false,
                "correlations": {
                    "class1": 0.08274986227505732,
                    "class2": 0.4682086503522818
                },
                "regulatory_source": "ENSMUSG00000038418",
                "regulatory_target": "ENSMUSG00000006369"
            },
            "ENSMUSG00000008575_ENSMUSG00000006369": {
                "id": "ENSMUSG00000008575_ENSMUSG00000006369",
                "type": "Mix",
                "score": 12.836378577517912,
                "reversable": false,
                "correlations": {
                    "class1": 0.06607959523021795,
                    "class2": 0.48604526390772135
                },
                "regulatory_source": "ENSMUSG00000008575",
                "regulatory_target": "ENSMUSG00000006369"
            },
            "ENSMUSG00000052837_ENSMUSG00000035356": {
                "id": "ENSMUSG00000052837_ENSMUSG00000035356",
                "type": "Mix",
                "score": 6.123969228370881,
                "reversable": false,
                "correlations": {
                    "class1": 0.6240309810899616,
                    "class2": 0.2866735418432276
                },
                "regulatory_source": "ENSMUSG00000052837",
                "regulatory_target": "ENSMUSG00000035356"
            },
            "ENSMUSG00000071076_ENSMUSG00000002107": {
                "id": "ENSMUSG00000071076_ENSMUSG00000002107",
                "type": "Standard",
                "score": 0.8200708206264424,
                "reversable": false,
                "correlations": {
                    "class1": 0.11274356872521135,
                    "class2": 0.49457513636281797
                },
                "regulatory_source": "ENSMUSG00000071076",
                "regulatory_target": "ENSMUSG00000002107"
            },
            "ENSMUSG00000060703_ENSMUSG00000005583": {
                "id": "ENSMUSG00000060703_ENSMUSG00000005583",
                "type": "Mix",
                "score": 13.309318177830036,
                "reversable": false,
                "correlations": {
                    "class1": 0.10028355228674272,
                    "class2": -0.30992647395571093
                },
                "regulatory_source": "ENSMUSG00000005583",
                "regulatory_target": "ENSMUSG00000060703"
            },
            "ENSMUSG00000064373_ENSMUSG00000005583": {
                "id": "ENSMUSG00000064373_ENSMUSG00000005583",
                "type": "Mix",
                "score": 9.688359935588611,
                "reversable": false,
                "correlations": {
                    "class1": 0.12065204024706741,
                    "class2": -0.3187467272755233
                },
                "regulatory_source": "ENSMUSG00000005583",
                "regulatory_target": "ENSMUSG00000064373"
            },
            "ENSMUSG00000057666_ENSMUSG00000005583": {
                "id": "ENSMUSG00000057666_ENSMUSG00000005583",
                "type": "Mix",
                "score": 5.576216841527998,
                "reversable": false,
                "correlations": {
                    "class1": 0.0389966067346583,
                    "class2": 0.4532066274242551
                },
                "regulatory_source": "ENSMUSG00000005583",
                "regulatory_target": "ENSMUSG00000057666"
            },
            "ENSMUSG00000005583_ENSMUSG00000002980": {
                "id": "ENSMUSG00000005583_ENSMUSG00000002980",
                "type": "Mix",
                "score": 19.582772869822723,
                "reversable": false,
                "correlations": {
                    "class1": 0.07128599961773534,
                    "class2": 0.5916496739167013
                },
                "regulatory_source": "ENSMUSG00000005583",
                "regulatory_target": "ENSMUSG00000002980"
            },
            "ENSMUSG00000009876_ENSMUSG00000005583": {
                "id": "ENSMUSG00000009876_ENSMUSG00000005583",
                "type": "Standard",
                "score": 0.6883529301622776,
                "reversable": false,
                "correlations": {
                    "class1": 0.08897760158211071,
                    "class2": 0.5515706069326294
                },
                "regulatory_source": "ENSMUSG00000005583",
                "regulatory_target": "ENSMUSG00000009876"
            },
            "ENSMUSG00000006360_ENSMUSG00000005583": {
                "id": "ENSMUSG00000006360_ENSMUSG00000005583",
                "type": "Standard",
                "score": 0.5543217373113999,
                "reversable": false,
                "correlations": {
                    "class1": 0.13566777206356434,
                    "class2": 0.48791491617849947
                },
                "regulatory_source": "ENSMUSG00000005583",
                "regulatory_target": "ENSMUSG00000006360"
            },
            "ENSMUSG00000079523_ENSMUSG00000005583": {
                "id": "ENSMUSG00000079523_ENSMUSG00000005583",
                "type": "Mix",
                "score": 13.12986647593557,
                "reversable": false,
                "correlations": {
                    "class1": 0.1688006938384951,
                    "class2": -0.3155909069100486
                },
                "regulatory_source": "ENSMUSG00000005583",
                "regulatory_target": "ENSMUSG00000079523"
            },
            "ENSMUSG00000021477_ENSMUSG00000005583": {
                "id": "ENSMUSG00000021477_ENSMUSG00000005583",
                "type": "Standard",
                "score": 0.5341637004951337,
                "reversable": false,
                "correlations": {
                    "class1": 0.09596746575262713,
                    "class2": -0.2913924755817878
                },
                "regulatory_source": "ENSMUSG00000005583",
                "regulatory_target": "ENSMUSG00000021477"
            },
            "ENSMUSG00000029309_ENSMUSG00000005583": {
                "id": "ENSMUSG00000029309_ENSMUSG00000005583",
                "type": "Standard",
                "score": 0.4914154364073667,
                "reversable": false,
                "correlations": {
                    "class1": 0.08542986811277582,
                    "class2": 0.5034291261286029
                },
                "regulatory_source": "ENSMUSG00000005583",
                "regulatory_target": "ENSMUSG00000029309"
            },
            "ENSMUSG00000071076_ENSMUSG00000042747": {
                "id": "ENSMUSG00000071076_ENSMUSG00000042747",
                "type": "Mix",
                "score": 11.660320438610958,
                "reversable": false,
                "correlations": {
                    "class1": 0.19432085940056046,
                    "class2": 0.5209955423970228
                },
                "regulatory_source": "ENSMUSG00000071076",
                "regulatory_target": "ENSMUSG00000042747"
            },
            "ENSMUSG00000016319_ENSMUSG00000005583": {
                "id": "ENSMUSG00000016319_ENSMUSG00000005583",
                "type": "Mix",
                "score": 5.3686921771469756,
                "reversable": false,
                "correlations": {
                    "class1": 0.09077840274461077,
                    "class2": 0.4889666584961277
                },
                "regulatory_source": "ENSMUSG00000005583",
                "regulatory_target": "ENSMUSG00000016319"
            },
            "ENSMUSG00000020928_ENSMUSG00000005583": {
                "id": "ENSMUSG00000020928_ENSMUSG00000005583",
                "type": "Mix",
                "score": 8.869349544035082,
                "reversable": false,
                "correlations": {
                    "class1": 0.0,
                    "class2": 0.428114002130053
                },
                "regulatory_source": "ENSMUSG00000005583",
                "regulatory_target": "ENSMUSG00000020928"
            },
            "ENSMUSG00000040170_ENSMUSG00000008575": {
                "id": "ENSMUSG00000040170_ENSMUSG00000008575",
                "type": "Standard",
                "score": 0.7861759909249663,
                "reversable": false,
                "correlations": {
                    "class1": 0.0,
                    "class2": 0.48702439902241246
                },
                "regulatory_source": "ENSMUSG00000008575",
                "regulatory_target": "ENSMUSG00000040170"
            },
            "ENSMUSG00000052684_ENSMUSG00000016494": {
                "id": "ENSMUSG00000052684_ENSMUSG00000016494",
                "type": "Standard",
                "score": 0.4422319731263898,
                "reversable": false,
                "correlations": {
                    "class1": -0.03051117643725209,
                    "class2": 0.3768979668301148
                },
                "regulatory_source": "ENSMUSG00000052684",
                "regulatory_target": "ENSMUSG00000016494"
            },
            "ENSMUSG00000071076_ENSMUSG00000037266": {
                "id": "ENSMUSG00000071076_ENSMUSG00000037266",
                "type": "Standard",
                "score": 0.43867169540620726,
                "reversable": false,
                "correlations": {
                    "class1": 0.09126887782331845,
                    "class2": 0.4498613888409563
                },
                "regulatory_source": "ENSMUSG00000071076",
                "regulatory_target": "ENSMUSG00000037266"
            },
            "ENSMUSG00000018899_ENSMUSG00000003545": {
                "id": "ENSMUSG00000018899_ENSMUSG00000003545",
                "type": "Standard",
                "score": 0.43237237902334835,
                "reversable": false,
                "correlations": {
                    "class1": 0.4622492265489637,
                    "class2": 0.0906129101328391
                },
                "regulatory_source": "ENSMUSG00000018899",
                "regulatory_target": "ENSMUSG00000003545"
            },
            "ENSMUSG00000033538_ENSMUSG00000021250": {
                "id": "ENSMUSG00000033538_ENSMUSG00000021250",
                "type": "Mix",
                "score": 9.654077367939992,
                "reversable": false,
                "correlations": {
                    "class1": 0.46092618703248966,
                    "class2": 0.08938745631576397
                },
                "regulatory_source": "ENSMUSG00000021250",
                "regulatory_target": "ENSMUSG00000033538"
            },
            "ENSMUSG00000035885_ENSMUSG00000005583": {
                "id": "ENSMUSG00000035885_ENSMUSG00000005583",
                "type": "Standard",
                "score": 0.43333898940403154,
                "reversable": false,
                "correlations": {
                    "class1": 0.14935009248041906,
                    "class2": 0.48780643010466096
                },
                "regulatory_source": "ENSMUSG00000005583",
                "regulatory_target": "ENSMUSG00000035885"
            },
            "ENSMUSG00000071076_ENSMUSG00000032359": {
                "id": "ENSMUSG00000071076_ENSMUSG00000032359",
                "type": "Standard",
                "score": 0.7783690007661582,
                "reversable": false,
                "correlations": {
                    "class1": 0.07611838004584123,
                    "class2": 0.4450517696652375
                },
                "regulatory_source": "ENSMUSG00000071076",
                "regulatory_target": "ENSMUSG00000032359"
            },
            "ENSMUSG00000042453_ENSMUSG00000008575": {
                "id": "ENSMUSG00000042453_ENSMUSG00000008575",
                "type": "Standard",
                "score": 0.4539445287259852,
                "reversable": false,
                "correlations": {
                    "class1": 0.3703202968197738,
                    "class2": 0.0
                },
                "regulatory_source": "ENSMUSG00000008575",
                "regulatory_target": "ENSMUSG00000042453"
            },
            "ENSMUSG00000042453_ENSMUSG00000038193": {
                "id": "ENSMUSG00000042453_ENSMUSG00000038193",
                "type": "Standard",
                "score": 0.581072168527888,
                "reversable": false,
                "correlations": {
                    "class1": 0.36568445216382395,
                    "class2": 0.0
                },
                "regulatory_source": "ENSMUSG00000038193",
                "regulatory_target": "ENSMUSG00000042453"
            },
            "ENSMUSG00000071076_ENSMUSG00000032334": {
                "id": "ENSMUSG00000071076_ENSMUSG00000032334",
                "type": "Standard",
                "score": 0.41575496245270654,
                "reversable": false,
                "correlations": {
                    "class1": 0.1664556373257322,
                    "class2": 0.5521150212847061
                },
                "regulatory_source": "ENSMUSG00000071076",
                "regulatory_target": "ENSMUSG00000032334"
            },
            "ENSMUSG00000052684_ENSMUSG00000018339": {
                "id": "ENSMUSG00000052684_ENSMUSG00000018339",
                "type": "Mix",
                "score": 15.545402058622585,
                "reversable": false,
                "correlations": {
                    "class1": 0.0,
                    "class2": 0.4401536111660592
                },
                "regulatory_source": "ENSMUSG00000052684",
                "regulatory_target": "ENSMUSG00000018339"
            },
            "ENSMUSG00000068798_ENSMUSG00000005583": {
                "id": "ENSMUSG00000068798_ENSMUSG00000005583",
                "type": "Standard",
                "score": 0.385762479532573,
                "reversable": false,
                "correlations": {
                    "class1": 0.08114385446843378,
                    "class2": 0.3743316071649902
                },
                "regulatory_source": "ENSMUSG00000005583",
                "regulatory_target": "ENSMUSG00000068798"
            },
            "ENSMUSG00000024164_ENSMUSG00000005583": {
                "id": "ENSMUSG00000024164_ENSMUSG00000005583",
                "type": "Standard",
                "score": 0.37359583969792554,
                "reversable": false,
                "correlations": {
                    "class1": 0.0,
                    "class2": -0.2641466861949991
                },
                "regulatory_source": "ENSMUSG00000005583",
                "regulatory_target": "ENSMUSG00000024164"
            },
            "ENSMUSG00000037868_ENSMUSG00000003545": {
                "id": "ENSMUSG00000037868_ENSMUSG00000003545",
                "type": "Standard",
                "score": 0.35682246903337206,
                "reversable": false,
                "correlations": {
                    "class1": 0.4463381561221811,
                    "class2": 0.06207258592573403
                },
                "regulatory_source": "ENSMUSG00000037868",
                "regulatory_target": "ENSMUSG00000003545"
            },
            "ENSMUSG00000052837_ENSMUSG00000047945": {
                "id": "ENSMUSG00000052837_ENSMUSG00000047945",
                "type": "Standard",
                "score": 0.4157737869530733,
                "reversable": false,
                "correlations": {
                    "class1": 0.42439188525229576,
                    "class2": 0.04393349606684859
                },
                "regulatory_source": "ENSMUSG00000052837",
                "regulatory_target": "ENSMUSG00000047945"
            },
            "ENSMUSG00000020484_ENSMUSG00000008540": {
                "id": "ENSMUSG00000020484_ENSMUSG00000008540",
                "type": "Mix",
                "score": 7.254722911487969,
                "reversable": false,
                "correlations": {
                    "class1": 0.07893916610386924,
                    "class2": 0.4268194806832527
                },
                "regulatory_source": "ENSMUSG00000020484",
                "regulatory_target": "ENSMUSG00000008540"
            },
            "ENSMUSG00000071076_ENSMUSG00000032515": {
                "id": "ENSMUSG00000071076_ENSMUSG00000032515",
                "type": "Standard",
                "score": 0.5212189605800789,
                "reversable": false,
                "correlations": {
                    "class1": 0.43457228694507316,
                    "class2": 0.05067465123963916
                },
                "regulatory_source": "ENSMUSG00000071076",
                "regulatory_target": "ENSMUSG00000032515"
            },
            "ENSMUSG00000037868_ENSMUSG00000020423": {
                "id": "ENSMUSG00000037868_ENSMUSG00000020423",
                "type": "Standard",
                "score": 0.5168013553832186,
                "reversable": false,
                "correlations": {
                    "class1": 0.42012187176186133,
                    "class2": 0.058471460237173714
                },
                "regulatory_source": "ENSMUSG00000037868",
                "regulatory_target": "ENSMUSG00000020423"
            },
            "ENSMUSG00000052837_ENSMUSG00000037868": {
                "id": "ENSMUSG00000052837_ENSMUSG00000037868",
                "type": "Standard",
                "score": 0.33804457166687746,
                "reversable": true,
                "correlations": {
                    "class1": 0.5311628853922231,
                    "class2": 0.15277836594022196
                },
                "regulatory_source": "ENSMUSG00000037868",
                "regulatory_target": "ENSMUSG00000052837"
            },
            "ENSMUSG00000053113_ENSMUSG00000038418": {
                "id": "ENSMUSG00000053113_ENSMUSG00000038418",
                "type": "Standard",
                "score": 0.33559133607951197,
                "reversable": false,
                "correlations": {
                    "class1": 0.7122110598603091,
                    "class2": 0.396598414344347
                },
                "regulatory_source": "ENSMUSG00000038418",
                "regulatory_target": "ENSMUSG00000053113"
            },
            "ENSMUSG00000046402_ENSMUSG00000005583": {
                "id": "ENSMUSG00000046402_ENSMUSG00000005583",
                "type": "Standard",
                "score": 0.3227102727493203,
                "reversable": false,
                "correlations": {
                    "class1": 0.11994206687215267,
                    "class2": -0.28114918778287634
                },
                "regulatory_source": "ENSMUSG00000005583",
                "regulatory_target": "ENSMUSG00000046402"
            },
            "ENSMUSG00000071076_ENSMUSG00000035356": {
                "id": "ENSMUSG00000071076_ENSMUSG00000035356",
                "type": "Standard",
                "score": 0.6667131795224761,
                "reversable": false,
                "correlations": {
                    "class1": 0.5024347221954248,
                    "class2": 0.156026469607008
                },
                "regulatory_source": "ENSMUSG00000071076",
                "regulatory_target": "ENSMUSG00000035356"
            },
            "ENSMUSG00000035356_ENSMUSG00000018899": {
                "id": "ENSMUSG00000035356_ENSMUSG00000018899",
                "type": "Standard",
                "score": 0.47788519356480486,
                "reversable": false,
                "correlations": {
                    "class1": 0.47968902708672667,
                    "class2": 0.09737655523455271
                },
                "regulatory_source": "ENSMUSG00000018899",
                "regulatory_target": "ENSMUSG00000035356"
            },
            "ENSMUSG00000020423_ENSMUSG00000003032": {
                "id": "ENSMUSG00000020423_ENSMUSG00000003032",
                "type": "Standard",
                "score": 0.334914396256949,
                "reversable": false,
                "correlations": {
                    "class1": 0.48722033614385385,
                    "class2": 0.057622128849269656
                },
                "regulatory_source": "ENSMUSG00000003032",
                "regulatory_target": "ENSMUSG00000020423"
            },
            "ENSMUSG00000005583_ENSMUSG00000004791": {
                "id": "ENSMUSG00000005583_ENSMUSG00000004791",
                "type": "Standard",
                "score": 0.312979628034223,
                "reversable": false,
                "correlations": {
                    "class1": 0.0,
                    "class2": 0.36629146622203956
                },
                "regulatory_source": "ENSMUSG00000005583",
                "regulatory_target": "ENSMUSG00000004791"
            },
            "ENSMUSG00000042485_ENSMUSG00000005583": {
                "id": "ENSMUSG00000042485_ENSMUSG00000005583",
                "type": "Outer",
                "score": 39.79276204826006,
                "reversable": false,
                "correlations": {
                    "class1": 0.04674400822977806,
                    "class2": 0.47841374701896716
                },
                "regulatory_source": "ENSMUSG00000005583",
                "regulatory_target": "ENSMUSG00000042485"
            },
            "ENSMUSG00000068735_ENSMUSG00000005583": {
                "id": "ENSMUSG00000068735_ENSMUSG00000005583",
                "type": "Outer",
                "score": 23.147887403472026,
                "reversable": false,
                "correlations": {
                    "class1": 0.07342333918553179,
                    "class2": 0.4965447454803944
                },
                "regulatory_source": "ENSMUSG00000005583",
                "regulatory_target": "ENSMUSG00000068735"
            },
            "ENSMUSG00000005583_ENSMUSG00000001520": {
                "id": "ENSMUSG00000005583_ENSMUSG00000001520",
                "type": "Outer",
                "score": 21.92534176296431,
                "reversable": false,
                "correlations": {
                    "class1": 0.0,
                    "class2": 0.5838057376776824
                },
                "regulatory_source": "ENSMUSG00000005583",
                "regulatory_target": "ENSMUSG00000001520"
            },
            "ENSMUSG00000071076_ENSMUSG00000025784": {
                "id": "ENSMUSG00000071076_ENSMUSG00000025784",
                "type": "Mix",
                "score": 17.395885082093162,
                "reversable": false,
                "correlations": {
                    "class1": 0.0,
                    "class2": 0.5469995439617341
                },
                "regulatory_source": "ENSMUSG00000071076",
                "regulatory_target": "ENSMUSG00000025784"
            },
            "ENSMUSG00000024290_ENSMUSG00000005583": {
                "id": "ENSMUSG00000024290_ENSMUSG00000005583",
                "type": "Mix",
                "score": 16.609385933221724,
                "reversable": false,
                "correlations": {
                    "class1": 0.052725090567346006,
                    "class2": 0.4558572196103253
                },
                "regulatory_source": "ENSMUSG00000005583",
                "regulatory_target": "ENSMUSG00000024290"
            },
            "ENSMUSG00000052684_ENSMUSG00000006205": {
                "id": "ENSMUSG00000052684_ENSMUSG00000006205",
                "type": "Outer",
                "score": 13.638128841156307,
                "reversable": false,
                "correlations": {
                    "class1": -0.06983883310238564,
                    "class2": 0.4112271023768541
                },
                "regulatory_source": "ENSMUSG00000052684",
                "regulatory_target": "ENSMUSG00000006205"
            },
            "ENSMUSG00000072235_ENSMUSG00000071076": {
                "id": "ENSMUSG00000072235_ENSMUSG00000071076",
                "type": "Outer",
                "score": 12.762260145171464,
                "reversable": false,
                "correlations": {
                    "class1": 0.1924934197667652,
                    "class2": 0.5615620571673006
                },
                "regulatory_source": "ENSMUSG00000071076",
                "regulatory_target": "ENSMUSG00000072235"
            },
            "ENSMUSG00000071076_ENSMUSG00000060703": {
                "id": "ENSMUSG00000071076_ENSMUSG00000060703",
                "type": "Mix",
                "score": 12.552019431565961,
                "reversable": false,
                "correlations": {
                    "class1": 0.17294606861296122,
                    "class2": 0.5472400500692081
                },
                "regulatory_source": "ENSMUSG00000071076",
                "regulatory_target": "ENSMUSG00000060703"
            },
            "ENSMUSG00000052684_ENSMUSG00000021250": {
                "id": "ENSMUSG00000052684_ENSMUSG00000021250",
                "type": "Outer",
                "score": 19.024522880672137,
                "reversable": true,
                "correlations": {
                    "class1": 0.8363165091507709,
                    "class2": 0.5472563359064866
                },
                "regulatory_source": "ENSMUSG00000052684",
                "regulatory_target": "ENSMUSG00000021250"
            },
            "ENSMUSG00000038418_ENSMUSG00000021250": {
                "id": "ENSMUSG00000038418_ENSMUSG00000021250",
                "type": "Outer",
                "score": 11.591868893707016,
                "reversable": false,
                "correlations": {
                    "class1": 0.8121305836268662,
                    "class2": 0.4677018533225259
                },
                "regulatory_source": "ENSMUSG00000021250",
                "regulatory_target": "ENSMUSG00000038418"
            },
            "ENSMUSG00000052684_ENSMUSG00000023034": {
                "id": "ENSMUSG00000052684_ENSMUSG00000023034",
                "type": "Outer",
                "score": 13.017814959839429,
                "reversable": true,
                "correlations": {
                    "class1": 0.538127822132192,
                    "class2": 0.09376026005645308
                },
                "regulatory_source": "ENSMUSG00000052684",
                "regulatory_target": "ENSMUSG00000023034"
            },
            "ENSMUSG00000035356_ENSMUSG00000021250": {
                "id": "ENSMUSG00000035356_ENSMUSG00000021250",
                "type": "Mix",
                "score": 10.044555998517025,
                "reversable": false,
                "correlations": {
                    "class1": 0.5610132718339899,
                    "class2": 0.23612559109598347
                },
                "regulatory_source": "ENSMUSG00000021250",
                "regulatory_target": "ENSMUSG00000035356"
            },
            "ENSMUSG00000023064_ENSMUSG00000005583": {
                "id": "ENSMUSG00000023064_ENSMUSG00000005583",
                "type": "Outer",
                "score": 17.11153510951664,
                "reversable": false,
                "correlations": {
                    "class1": 0.0,
                    "class2": 0.46819730889432354
                },
                "regulatory_source": "ENSMUSG00000005583",
                "regulatory_target": "ENSMUSG00000023064"
            },
            "ENSMUSG00000032387_ENSMUSG00000005583": {
                "id": "ENSMUSG00000032387_ENSMUSG00000005583",
                "type": "Outer",
                "score": 20.667882740951086,
                "reversable": false,
                "correlations": {
                    "class1": 0.0,
                    "class2": 0.44998655648149855
                },
                "regulatory_source": "ENSMUSG00000005583",
                "regulatory_target": "ENSMUSG00000032387"
            },
            "ENSMUSG00000023034_ENSMUSG00000003545": {
                "id": "ENSMUSG00000023034_ENSMUSG00000003545",
                "type": "Outer",
                "score": 15.563020345950088,
                "reversable": true,
                "correlations": {
                    "class1": 0.5335436705169669,
                    "class2": 0.08802224186371972
                },
                "regulatory_source": "ENSMUSG00000003545",
                "regulatory_target": "ENSMUSG00000023034"
            },
            "ENSMUSG00000021250_ENSMUSG00000020423": {
                "id": "ENSMUSG00000021250_ENSMUSG00000020423",
                "type": "Outer",
                "score": 8.99526902203732,
                "reversable": false,
                "correlations": {
                    "class1": 0.68506172172299,
                    "class2": 0.32264805129399154
                },
                "regulatory_source": "ENSMUSG00000021250",
                "regulatory_target": "ENSMUSG00000020423"
            },
            "ENSMUSG00000026879_ENSMUSG00000005583": {
                "id": "ENSMUSG00000026879_ENSMUSG00000005583",
                "type": "Outer",
                "score": 16.782767456502782,
                "reversable": false,
                "correlations": {
                    "class1": 0.0745273843753412,
                    "class2": -0.3116592680084563
                },
                "regulatory_source": "ENSMUSG00000005583",
                "regulatory_target": "ENSMUSG00000026879"
            },
            "ENSMUSG00000071076_ENSMUSG00000026628": {
                "id": "ENSMUSG00000071076_ENSMUSG00000026628",
                "type": "Mix",
                "score": 8.903213017748817,
                "reversable": true,
                "correlations": {
                    "class1": 0.5223556542852718,
                    "class2": 0.22161000063545727
                },
                "regulatory_source": "ENSMUSG00000026628",
                "regulatory_target": "ENSMUSG00000071076"
            },
            "ENSMUSG00000026574_ENSMUSG00000005583": {
                "id": "ENSMUSG00000026574_ENSMUSG00000005583",
                "type": "Outer",
                "score": 19.014921882516084,
                "reversable": false,
                "correlations": {
                    "class1": 0.1037367807758741,
                    "class2": -0.3269058640999696
                },
                "regulatory_source": "ENSMUSG00000005583",
                "regulatory_target": "ENSMUSG00000026574"
            },
            "ENSMUSG00000038418_ENSMUSG00000020423": {
                "id": "ENSMUSG00000038418_ENSMUSG00000020423",
                "type": "Outer",
                "score": 13.037064430136912,
                "reversable": false,
                "correlations": {
                    "class1": 0.6678098415232274,
                    "class2": 0.29041195585192636
                },
                "regulatory_source": "ENSMUSG00000038418",
                "regulatory_target": "ENSMUSG00000020423"
            },
            "ENSMUSG00000052684_ENSMUSG00000044786": {
                "id": "ENSMUSG00000052684_ENSMUSG00000044786",
                "type": "Outer",
                "score": 14.559563245868608,
                "reversable": false,
                "correlations": {
                    "class1": 0.7612085498989385,
                    "class2": 0.4382832636744016
                },
                "regulatory_source": "ENSMUSG00000052684",
                "regulatory_target": "ENSMUSG00000044786"
            },
            "ENSMUSG00000052997_ENSMUSG00000005583": {
                "id": "ENSMUSG00000052997_ENSMUSG00000005583",
                "type": "Outer",
                "score": 24.826864386543935,
                "reversable": false,
                "correlations": {
                    "class1": 0.05896005715408981,
                    "class2": 0.5237165720942244
                },
                "regulatory_source": "ENSMUSG00000005583",
                "regulatory_target": "ENSMUSG00000052997"
            },
            "ENSMUSG00000087006_ENSMUSG00000005583": {
                "id": "ENSMUSG00000087006_ENSMUSG00000005583",
                "type": "Mix",
                "score": 15.880160868019923,
                "reversable": false,
                "correlations": {
                    "class1": 0.03788482209910931,
                    "class2": 0.5272971227385967
                },
                "regulatory_source": "ENSMUSG00000005583",
                "regulatory_target": "ENSMUSG00000087006"
            },
            "ENSMUSG00000071076_ENSMUSG00000025351": {
                "id": "ENSMUSG00000071076_ENSMUSG00000025351",
                "type": "Mix",
                "score": 7.673515326734044,
                "reversable": false,
                "correlations": {
                    "class1": 0.2334496297307922,
                    "class2": 0.556720222381856
                },
                "regulatory_source": "ENSMUSG00000071076",
                "regulatory_target": "ENSMUSG00000025351"
            },
            "ENSMUSG00000071076_ENSMUSG00000021477": {
                "id": "ENSMUSG00000071076_ENSMUSG00000021477",
                "type": "Mix",
                "score": 10.401309379224493,
                "reversable": false,
                "correlations": {
                    "class1": 0.14018575174157172,
                    "class2": 0.5556585216797687
                },
                "regulatory_source": "ENSMUSG00000071076",
                "regulatory_target": "ENSMUSG00000021477"
            },
            "ENSMUSG00000078202_ENSMUSG00000005583": {
                "id": "ENSMUSG00000078202_ENSMUSG00000005583",
                "type": "Mix",
                "score": 7.899451580762552,
                "reversable": false,
                "correlations": {
                    "class1": 0.0,
                    "class2": 0.5337774538866751
                },
                "regulatory_source": "ENSMUSG00000005583",
                "regulatory_target": "ENSMUSG00000078202"
            },
            "ENSMUSG00000053113_ENSMUSG00000052837": {
                "id": "ENSMUSG00000053113_ENSMUSG00000052837",
                "type": "Mix",
                "score": 14.325234638170969,
                "reversable": false,
                "correlations": {
                    "class1": 0.7774853678239327,
                    "class2": 0.40688012760558817
                },
                "regulatory_source": "ENSMUSG00000052837",
                "regulatory_target": "ENSMUSG00000053113"
            },
            "ENSMUSG00000071076_ENSMUSG00000041959": {
                "id": "ENSMUSG00000071076_ENSMUSG00000041959",
                "type": "Outer",
                "score": 19.922980703349563,
                "reversable": false,
                "correlations": {
                    "class1": 0.10150407086500257,
                    "class2": 0.5516425029111981
                },
                "regulatory_source": "ENSMUSG00000071076",
                "regulatory_target": "ENSMUSG00000041959"
            },
            "ENSMUSG00000053113_ENSMUSG00000023034": {
                "id": "ENSMUSG00000053113_ENSMUSG00000023034",
                "type": "Mix",
                "score": 7.680315682061678,
                "reversable": false,
                "correlations": {
                    "class1": 0.5764425800870968,
                    "class2": 0.1418538091861824
                },
                "regulatory_source": "ENSMUSG00000023034",
                "regulatory_target": "ENSMUSG00000053113"
            },
            "ENSMUSG00000071076_ENSMUSG00000042312": {
                "id": "ENSMUSG00000071076_ENSMUSG00000042312",
                "type": "Outer",
                "score": 12.301936025276312,
                "reversable": false,
                "correlations": {
                    "class1": 0.09332613347447885,
                    "class2": 0.4107535082427858
                },
                "regulatory_source": "ENSMUSG00000071076",
                "regulatory_target": "ENSMUSG00000042312"
            },
            "ENSMUSG00000038193_ENSMUSG00000019929": {
                "id": "ENSMUSG00000038193_ENSMUSG00000019929",
                "type": "Outer",
                "score": 19.8126364152465,
                "reversable": false,
                "correlations": {
                    "class1": 0.4115732693143844,
                    "class2": -0.060129979674140986
                },
                "regulatory_source": "ENSMUSG00000038193",
                "regulatory_target": "ENSMUSG00000019929"
            },
            "ENSMUSG00000053113_ENSMUSG00000026628": {
                "id": "ENSMUSG00000053113_ENSMUSG00000026628",
                "type": "Mix",
                "score": 7.612415598515531,
                "reversable": false,
                "correlations": {
                    "class1": 0.5549277034403501,
                    "class2": 0.20625975930029494
                },
                "regulatory_source": "ENSMUSG00000026628",
                "regulatory_target": "ENSMUSG00000053113"
            },
            "ENSMUSG00000025147_ENSMUSG00000005583": {
                "id": "ENSMUSG00000025147_ENSMUSG00000005583",
                "type": "Mix",
                "score": 14.66252352901201,
                "reversable": false,
                "correlations": {
                    "class1": 0.02976097936901441,
                    "class2": 0.43224178133904945
                },
                "regulatory_source": "ENSMUSG00000005583",
                "regulatory_target": "ENSMUSG00000025147"
            },
            "ENSMUSG00000071076_ENSMUSG00000044786": {
                "id": "ENSMUSG00000071076_ENSMUSG00000044786",
                "type": "Outer",
                "score": 7.261824384906225,
                "reversable": false,
                "correlations": {
                    "class1": 0.7195629129491783,
                    "class2": 0.43462143903306705
                },
                "regulatory_source": "ENSMUSG00000071076",
                "regulatory_target": "ENSMUSG00000044786"
            },
            "ENSMUSG00000071076_ENSMUSG00000020423": {
                "id": "ENSMUSG00000071076_ENSMUSG00000020423",
                "type": "Outer",
                "score": 29.699639484170422,
                "reversable": false,
                "correlations": {
                    "class1": 0.6305303250748671,
                    "class2": 0.1360759856265647
                },
                "regulatory_source": "ENSMUSG00000071076",
                "regulatory_target": "ENSMUSG00000020423"
            },
            "ENSMUSG00000020423_ENSMUSG00000003545": {
                "id": "ENSMUSG00000020423_ENSMUSG00000003545",
                "type": "Mix",
                "score": 24.477951223720282,
                "reversable": false,
                "correlations": {
                    "class1": 0.6150586419877898,
                    "class2": 0.14771988980339001
                },
                "regulatory_source": "ENSMUSG00000003545",
                "regulatory_target": "ENSMUSG00000020423"
            },
            "ENSMUSG00000052837_ENSMUSG00000044786": {
                "id": "ENSMUSG00000052837_ENSMUSG00000044786",
                "type": "Mix",
                "score": 20.110871403882523,
                "reversable": false,
                "correlations": {
                    "class1": 0.8013166393989419,
                    "class2": 0.5358748557334677
                },
                "regulatory_source": "ENSMUSG00000052837",
                "regulatory_target": "ENSMUSG00000044786"
            },
            "ENSMUSG00000029359_ENSMUSG00000005583": {
                "id": "ENSMUSG00000029359_ENSMUSG00000005583",
                "type": "Mix",
                "score": 16.636466942280986,
                "reversable": false,
                "correlations": {
                    "class1": 0.0,
                    "class2": 0.3972766992095762
                },
                "regulatory_source": "ENSMUSG00000005583",
                "regulatory_target": "ENSMUSG00000029359"
            },
            "ENSMUSG00000044786_ENSMUSG00000021250": {
                "id": "ENSMUSG00000044786_ENSMUSG00000021250",
                "type": "Outer",
                "score": 16.077484406992387,
                "reversable": false,
                "correlations": {
                    "class1": 0.8084256567644267,
                    "class2": 0.45534947164997497
                },
                "regulatory_source": "ENSMUSG00000021250",
                "regulatory_target": "ENSMUSG00000044786"
            },
            "ENSMUSG00000071076_ENSMUSG00000006205": {
                "id": "ENSMUSG00000071076_ENSMUSG00000006205",
                "type": "Outer",
                "score": 15.962924439330877,
                "reversable": false,
                "correlations": {
                    "class1": 0.0,
                    "class2": 0.4612242450129689
                },
                "regulatory_source": "ENSMUSG00000071076",
                "regulatory_target": "ENSMUSG00000006205"
            },
            "ENSMUSG00000039601_ENSMUSG00000021250": {
                "id": "ENSMUSG00000039601_ENSMUSG00000021250",
                "type": "Mix",
                "score": 17.380240849864325,
                "reversable": false,
                "correlations": {
                    "class1": 0.0,
                    "class2": 0.4197913609072639
                },
                "regulatory_source": "ENSMUSG00000021250",
                "regulatory_target": "ENSMUSG00000039601"
            },
            "ENSMUSG00000052684_ENSMUSG00000003545": {
                "id": "ENSMUSG00000052684_ENSMUSG00000003545",
                "type": "Mix",
                "score": 18.44593690735485,
                "reversable": true,
                "correlations": {
                    "class1": 0.7252397828162958,
                    "class2": 0.4389445314638871
                },
                "regulatory_source": "ENSMUSG00000052684",
                "regulatory_target": "ENSMUSG00000003545"
            },
            "ENSMUSG00000052837_ENSMUSG00000052684": {
                "id": "ENSMUSG00000052837_ENSMUSG00000052684",
                "type": "Mix",
                "score": 14.306139403707094,
                "reversable": true,
                "correlations": {
                    "class1": 0.8218419773024086,
                    "class2": 0.4292453022171103
                },
                "regulatory_source": "ENSMUSG00000052684",
                "regulatory_target": "ENSMUSG00000052837"
            },
            "ENSMUSG00000042745_ENSMUSG00000005583": {
                "id": "ENSMUSG00000042745_ENSMUSG00000005583",
                "type": "Outer",
                "score": 15.308536504766225,
                "reversable": false,
                "correlations": {
                    "class1": 0.05273125917423713,
                    "class2": 0.5192564767975391
                },
                "regulatory_source": "ENSMUSG00000005583",
                "regulatory_target": "ENSMUSG00000042745"
            },
            "ENSMUSG00000052684_ENSMUSG00000020423": {
                "id": "ENSMUSG00000052684_ENSMUSG00000020423",
                "type": "Outer",
                "score": 25.55111497611776,
                "reversable": false,
                "correlations": {
                    "class1": 0.6598968913082331,
                    "class2": 0.13890513210590136
                },
                "regulatory_source": "ENSMUSG00000052684",
                "regulatory_target": "ENSMUSG00000020423"
            },
            "ENSMUSG00000019929_ENSMUSG00000005583": {
                "id": "ENSMUSG00000019929_ENSMUSG00000005583",
                "type": "Mix",
                "score": 20.154316197746116,
                "reversable": false,
                "correlations": {
                    "class1": 0.07132537501486069,
                    "class2": -0.34957069155628173
                },
                "regulatory_source": "ENSMUSG00000005583",
                "regulatory_target": "ENSMUSG00000019929"
            },
            "ENSMUSG00000052684_ENSMUSG00000021876": {
                "id": "ENSMUSG00000052684_ENSMUSG00000021876",
                "type": "Mix",
                "score": 10.39251078177558,
                "reversable": false,
                "correlations": {
                    "class1": 0.030802171968629964,
                    "class2": 0.45499805220100015
                },
                "regulatory_source": "ENSMUSG00000052684",
                "regulatory_target": "ENSMUSG00000021876"
            },
            "ENSMUSG00000038418_ENSMUSG00000003545": {
                "id": "ENSMUSG00000038418_ENSMUSG00000003545",
                "type": "Outer",
                "score": 10.460085965718816,
                "reversable": false,
                "correlations": {
                    "class1": 0.7383230112763797,
                    "class2": 0.293287548256507
                },
                "regulatory_source": "ENSMUSG00000003545",
                "regulatory_target": "ENSMUSG00000038418"
            },
            "ENSMUSG00000071076_ENSMUSG00000023034": {
                "id": "ENSMUSG00000071076_ENSMUSG00000023034",
                "type": "Mix",
                "score": 9.004490883220216,
                "reversable": false,
                "correlations": {
                    "class1": 0.5195877483764404,
                    "class2": 0.11148371947566481
                },
                "regulatory_source": "ENSMUSG00000071076",
                "regulatory_target": "ENSMUSG00000023034"
            },
            "ENSMUSG00000071076_ENSMUSG00000068220": {
                "id": "ENSMUSG00000071076_ENSMUSG00000068220",
                "type": "Mix",
                "score": 10.071252936449493,
                "reversable": false,
                "correlations": {
```
