## Supplement 4 (CCl4 a6w HSC vs. PF) for "AGEAS: Automated Machine Learning based Genetic Regulatory Element Extraction System": key_atlas.js

```
{
    "genes": {
        "ENSMUSG00000029322": {
            "id": "ENSMUSG00000029322",
            "type": "TF",
            "symbol": "Plac8",
            "source": [
                "ENSMUSG00000057098",
                "ENSMUSG00000032035"
            ],
            "target": [
                "ENSMUSG00000021377",
                "ENSMUSG00000006360",
                "ENSMUSG00000026087",
                "ENSMUSG00000036309",
                "ENSMUSG00000008348",
                "ENSMUSG00000021606",
                "ENSMUSG00000004665",
                "ENSMUSG00000022216",
                "ENSMUSG00000036438",
                "ENSMUSG00000026238",
                "ENSMUSG00000023505",
                "ENSMUSG00000028797",
                "ENSMUSG00000042485",
                "ENSMUSG00000016252",
                "ENSMUSG00000073616",
                "ENSMUSG00000031311",
                "ENSMUSG00000026421",
                "ENSMUSG00000031198",
                "ENSMUSG00000074676",
                "ENSMUSG00000057137",
                "ENSMUSG00000048126",
                "ENSMUSG00000037894",
                "ENSMUSG00000031119",
                "ENSMUSG00000007872",
                "ENSMUSG00000027170",
                "ENSMUSG00000020738",
                "ENSMUSG00000076432",
                "ENSMUSG00000061353",
                "ENSMUSG00000031059",
                "ENSMUSG00000006304",
                "ENSMUSG00000055302",
                "ENSMUSG00000028648",
                "ENSMUSG00000014294",
                "ENSMUSG00000020053",
                "ENSMUSG00000028063",
                "ENSMUSG00000026207",
                "ENSMUSG00000037999",
                "ENSMUSG00000069601",
                "ENSMUSG00000034353",
                "ENSMUSG00000036533",
                "ENSMUSG00000032562",
                "ENSMUSG00000025511",
                "ENSMUSG00000015092",
                "ENSMUSG00000030122",
                "ENSMUSG00000055322",
                "ENSMUSG00000031246",
                "ENSMUSG00000068245",
                "ENSMUSG00000041481",
                "ENSMUSG00000019804",
                "ENSMUSG00000031783",
                "ENSMUSG00000041741",
                "ENSMUSG00000048332",
                "ENSMUSG00000026768",
                "ENSMUSG00000022587",
                "ENSMUSG00000045767",
                "ENSMUSG00000035268",
                "ENSMUSG00000038633",
                "ENSMUSG00000016382",
                "ENSMUSG00000031818",
                "ENSMUSG00000068798",
                "ENSMUSG00000030220",
                "ENSMUSG00000002778",
                "ENSMUSG00000079523",
                "ENSMUSG00000016494",
                "ENSMUSG00000098274",
                "ENSMUSG00000024290",
                "ENSMUSG00000061518",
                "ENSMUSG00000003746",
                "ENSMUSG00000022665",
                "ENSMUSG00000041577",
                "ENSMUSG00000026032",
                "ENSMUSG00000032135",
                "ENSMUSG00000064358",
                "ENSMUSG00000026895",
                "ENSMUSG00000030272",
                "ENSMUSG00000026104",
                "ENSMUSG00000090272",
                "ENSMUSG00000023904",
                "ENSMUSG00000050628",
                "ENSMUSG00000026568",
                "ENSMUSG00000022450",
                "ENSMUSG00000041571",
                "ENSMUSG00000038467",
                "ENSMUSG00000006299",
                "ENSMUSG00000024576",
                "ENSMUSG00000034708",
                "ENSMUSG00000033306",
                "ENSMUSG00000026728",
                "ENSMUSG00000006356",
                "ENSMUSG00000025204",
                "ENSMUSG00000027907",
                "ENSMUSG00000026278",
                "ENSMUSG00000067818",
                "ENSMUSG00000054808",
                "ENSMUSG00000022108",
                "ENSMUSG00000008136",
                "ENSMUSG00000024248",
                "ENSMUSG00000092341",
                "ENSMUSG00000041736",
                "ENSMUSG00000040681",
                "ENSMUSG00000001289",
                "ENSMUSG00000020585",
                "ENSMUSG00000013593",
                "ENSMUSG00000036256",
                "ENSMUSG00000004285",
                "ENSMUSG00000025993",
                "ENSMUSG00000022234",
                "ENSMUSG00000026078",
                "ENSMUSG00000066551",
                "ENSMUSG00000035929",
                "ENSMUSG00000015932",
                "ENSMUSG00000060802",
                "ENSMUSG00000056515",
                "ENSMUSG00000079017"
            ],
            "expression_sum": {
                "class1": 85825.0,
                "class2": 4703.0
            },
            "ens_id": "ENSMUSG00000029322",
            "uniprot_ids": [
                "Q9JI48"
            ],
            "net_id": 0,
            "source_num": 2,
            "target_num": 20
        },
        "ENSMUSG00000000247": {
            "id": "ENSMUSG00000000247",
            "type": "TF",
            "symbol": "Lhx2",
            "source": [
                "ENSMUSG00000032035",
                "ENSMUSG00000056501",
                "ENSMUSG00000052684",
                "ENSMUSG00000023034",
                "ENSMUSG00000052837",
                "ENSMUSG00000038418",
                "ENSMUSG00000021250",
                "ENSMUSG00000038193",
                "ENSMUSG00000071076"
            ],
            "target": [
                "ENSMUSG00000019539",
                "ENSMUSG00000019505",
                "ENSMUSG00000024063",
                "ENSMUSG00000059291",
                "ENSMUSG00000014077",
                "ENSMUSG00000058546",
                "ENSMUSG00000028639",
                "ENSMUSG00000031812",
                "ENSMUSG00000021477",
                "ENSMUSG00000026457",
                "ENSMUSG00000022283",
                "ENSMUSG00000068523",
                "ENSMUSG00000040952",
                "ENSMUSG00000048076",
                "ENSMUSG00000030695",
                "ENSMUSG00000041881",
                "ENSMUSG00000053560",
                "ENSMUSG00000025967",
                "ENSMUSG00000026939",
                "ENSMUSG00000030744",
                "ENSMUSG00000036438",
                "ENSMUSG00000026238",
                "ENSMUSG00000000325",
                "ENSMUSG00000060036",
                "ENSMUSG00000008668",
                "ENSMUSG00000061787",
                "ENSMUSG00000038717",
                "ENSMUSG00000020372",
                "ENSMUSG00000041417",
                "ENSMUSG00000008682",
                "ENSMUSG00000023175",
                "ENSMUSG00000021127",
                "ENSMUSG00000026970",
                "ENSMUSG00000031198",
                "ENSMUSG00000023861",
                "ENSMUSG00000061904",
                "ENSMUSG00000034892",
                "ENSMUSG00000022010",
                "ENSMUSG00000005161",
                "ENSMUSG00000020612",
                "ENSMUSG00000032231",
                "ENSMUSG00000071415",
                "ENSMUSG00000021660",
                "ENSMUSG00000061477",
                "ENSMUSG00000060743",
                "ENSMUSG00000007872",
                "ENSMUSG00000056201",
                "ENSMUSG00000020738",
                "ENSMUSG00000022884",
                "ENSMUSG00000061983",
                "ENSMUSG00000004980",
                "ENSMUSG00000037805",
                "ENSMUSG00000027750",
                "ENSMUSG00000006304",
                "ENSMUSG00000028081",
                "ENSMUSG00000063316",
                "ENSMUSG00000055302",
                "ENSMUSG00000017404",
                "ENSMUSG00000070493",
                "ENSMUSG00000021546",
                "ENSMUSG00000046364",
                "ENSMUSG00000004070",
                "ENSMUSG00000022312",
                "ENSMUSG00000060636",
                "ENSMUSG00000055839",
                "ENSMUSG00000014294",
                "ENSMUSG00000064370",
                "ENSMUSG00000068874",
                "ENSMUSG00000037999",
                "ENSMUSG00000069601",
                "ENSMUSG00000028603",
                "ENSMUSG00000042043",
                "ENSMUSG00000037742",
                "ENSMUSG00000025781",
                "ENSMUSG00000022892",
                "ENSMUSG00000032562",
                "ENSMUSG00000025511",
                "ENSMUSG00000029810",
                "ENSMUSG00000038803",
                "ENSMUSG00000052684",
                "ENSMUSG00000012405",
                "ENSMUSG00000000184",
                "ENSMUSG00000020719",
                "ENSMUSG00000036371",
                "ENSMUSG00000090862",
                "ENSMUSG00000021807",
                "ENSMUSG00000022051",
                "ENSMUSG00000030824",
                "ENSMUSG00000064351",
                "ENSMUSG00000027642",
                "ENSMUSG00000003970",
                "ENSMUSG00000003617",
                "ENSMUSG00000021870",
                "ENSMUSG00000018326",
                "ENSMUSG00000020460",
                "ENSMUSG00000035885",
                "ENSMUSG00000027706",
                "ENSMUSG00000055044",
                "ENSMUSG00000029781",
                "ENSMUSG00000042812",
                "ENSMUSG00000025508",
                "ENSMUSG00000063882",
                "ENSMUSG00000025290",
                "ENSMUSG00000039105",
                "ENSMUSG00000037712",
                "ENSMUSG00000009927",
                "ENSMUSG00000033880",
                "ENSMUSG00000031353",
                "ENSMUSG00000045128",
                "ENSMUSG00000022587",
                "ENSMUSG00000025794",
                "ENSMUSG00000023944",
                "ENSMUSG00000024065",
                "ENSMUSG00000023367",
                "ENSMUSG00000024608",
                "ENSMUSG00000001687",
                "ENSMUSG00000029713",
                "ENSMUSG00000039001",
                "ENSMUSG00000036381",
                "ENSMUSG00000006333",
                "ENSMUSG00000058600",
                "ENSMUSG00000062647",
                "ENSMUSG00000010406",
                "ENSMUSG00000030432",
                "ENSMUSG00000046330",
                "ENSMUSG00000074129",
                "ENSMUSG00000090733",
                "ENSMUSG00000031818",
                "ENSMUSG00000020432",
                "ENSMUSG00000010376",
                "ENSMUSG00000002778",
                "ENSMUSG00000032946",
                "ENSMUSG00000098274",
                "ENSMUSG00000027404",
                "ENSMUSG00000016559",
                "ENSMUSG00000060126",
                "ENSMUSG00000003072",
                "ENSMUSG00000061315",
                "ENSMUSG00000057322",
                "ENSMUSG00000027673",
                "ENSMUSG00000012848",
                "ENSMUSG00000060591",
                "ENSMUSG00000023942",
                "ENSMUSG00000049517",
                "ENSMUSG00000025362",
                "ENSMUSG00000019960",
                "ENSMUSG00000044533",
                "ENSMUSG00000029614",
                "ENSMUSG00000022136",
                "ENSMUSG00000025428",
                "ENSMUSG00000073702",
                "ENSMUSG00000003814",
                "ENSMUSG00000062006",
                "ENSMUSG00000062683",
                "ENSMUSG00000040128",
                "ENSMUSG00000032216",
                "ENSMUSG00000079435",
                "ENSMUSG00000047215",
                "ENSMUSG00000062270",
                "ENSMUSG00000022450",
                "ENSMUSG00000041571",
                "ENSMUSG00000053565",
                "ENSMUSG00000032666",
                "ENSMUSG00000004207",
                "ENSMUSG00000031029",
                "ENSMUSG00000028673",
                "ENSMUSG00000024576",
                "ENSMUSG00000035242",
                "ENSMUSG00000050708",
                "ENSMUSG00000014769",
                "ENSMUSG00000015656",
                "ENSMUSG00000067288",
                "ENSMUSG00000058558",
                "ENSMUSG00000029076",
                "ENSMUSG00000056851",
                "ENSMUSG00000022971",
                "ENSMUSG00000052146",
                "ENSMUSG00000060938",
                "ENSMUSG00000047675",
                "ENSMUSG00000025393",
                "ENSMUSG00000034994",
                "ENSMUSG00000041841",
                "ENSMUSG00000067274",
                "ENSMUSG00000054808",
                "ENSMUSG00000024308",
                "ENSMUSG00000071644",
                "ENSMUSG00000057863",
                "ENSMUSG00000026566",
                "ENSMUSG00000022108",
                "ENSMUSG00000032288",
                "ENSMUSG00000014313",
                "ENSMUSG00000024248",
                "ENSMUSG00000028266",
                "ENSMUSG00000024953",
                "ENSMUSG00000030057",
                "ENSMUSG00000041736",
                "ENSMUSG00000045690",
                "ENSMUSG00000041560",
                "ENSMUSG00000005881",
                "ENSMUSG00000049751",
                "ENSMUSG00000001289",
                "ENSMUSG00000024678",
                "ENSMUSG00000036751",
                "ENSMUSG00000032179",
                "ENSMUSG00000013593",
                "ENSMUSG00000008683",
                "ENSMUSG00000055762",
                "ENSMUSG00000043716",
                "ENSMUSG00000021250",
                "ENSMUSG00000017778",
                "ENSMUSG00000055447",
                "ENSMUSG00000022890",
                "ENSMUSG00000000740",
                "ENSMUSG00000007892",
                "ENSMUSG00000027523",
                "ENSMUSG00000024190",
                "ENSMUSG00000028618",
                "ENSMUSG00000057841",
                "ENSMUSG00000079641",
                "ENSMUSG00000027593",
                "ENSMUSG00000035530",
                "ENSMUSG00000028757",
                "ENSMUSG00000059070",
                "ENSMUSG00000031532",
                "ENSMUSG00000027620",
                "ENSMUSG00000047907",
                "ENSMUSG00000052837",
                "ENSMUSG00000048758",
                "ENSMUSG00000029632",
                "ENSMUSG00000022708",
                "ENSMUSG00000028234",
                "ENSMUSG00000093674",
                "ENSMUSG00000028936",
                "ENSMUSG00000042682",
                "ENSMUSG00000044786",
                "ENSMUSG00000028495",
                "ENSMUSG00000059005",
                "ENSMUSG00000028478",
                "ENSMUSG00000071076",
                "ENSMUSG00000041453",
                "ENSMUSG00000003429",
                "ENSMUSG00000018293",
                "ENSMUSG00000029465",
                "ENSMUSG00000038418",
                "ENSMUSG00000030849",
                "ENSMUSG00000021134",
                "ENSMUSG00000000563",
                "ENSMUSG00000031633",
                "ENSMUSG00000037563",
                "ENSMUSG00000032399"
            ],
            "expression_sum": {
                "class1": 14582.0,
                "class2": 111.0
            },
            "ens_id": "ENSMUSG00000000247",
            "uniprot_ids": [
                "Q9Z0S2",
                "F6Z9H5",
                "A0A0A0MQK9",
                "F6SM03",
                "H3BKI3"
            ],
            "net_id": 0,
            "source_num": 9,
            "target_num": 50
        },
        "ENSMUSG00000048387": {
            "id": "ENSMUSG00000048387",
            "type": "TF",
            "symbol": "Osr1",
            "source": [
                "ENSMUSG00000022346",
                "ENSMUSG00000028565"
            ],
            "target": [
                "ENSMUSG00000019505",
                "ENSMUSG00000031447",
                "ENSMUSG00000058546",
                "ENSMUSG00000007655",
                "ENSMUSG00000028691",
                "ENSMUSG00000022365",
                "ENSMUSG00000025213",
                "ENSMUSG00000040952",
                "ENSMUSG00000031320",
                "ENSMUSG00000027422",
                "ENSMUSG00000073676",
                "ENSMUSG00000027447",
                "ENSMUSG00000056629",
                "ENSMUSG00000024909",
                "ENSMUSG00000021877",
                "ENSMUSG00000034868",
                "ENSMUSG00000060036",
                "ENSMUSG00000008668",
                "ENSMUSG00000036275",
                "ENSMUSG00000008682",
                "ENSMUSG00000030659",
                "ENSMUSG00000018567",
                "ENSMUSG00000034892",
                "ENSMUSG00000029103",
                "ENSMUSG00000071415",
                "ENSMUSG00000053317",
                "ENSMUSG00000061983",
                "ENSMUSG00000029616",
                "ENSMUSG00000020368",
                "ENSMUSG00000063316",
                "ENSMUSG00000060636",
                "ENSMUSG00000028367",
                "ENSMUSG00000037742",
                "ENSMUSG00000036372",
                "ENSMUSG00000037266",
                "ENSMUSG00000038650",
                "ENSMUSG00000090862",
                "ENSMUSG00000090000",
                "ENSMUSG00000030824",
                "ENSMUSG00000003970",
                "ENSMUSG00000009927",
                "ENSMUSG00000025508",
                "ENSMUSG00000042737",
                "ENSMUSG00000023944",
                "ENSMUSG00000078812",
                "ENSMUSG00000024608",
                "ENSMUSG00000029713",
                "ENSMUSG00000039001",
                "ENSMUSG00000037373",
                "ENSMUSG00000006333",
                "ENSMUSG00000058600",
                "ENSMUSG00000046330",
                "ENSMUSG00000074129",
                "ENSMUSG00000090733",
                "ENSMUSG00000028517",
                "ENSMUSG00000002778",
                "ENSMUSG00000079523",
                "ENSMUSG00000098274",
                "ENSMUSG00000026674",
                "ENSMUSG00000060126",
                "ENSMUSG00000026511",
                "ENSMUSG00000058569",
                "ENSMUSG00000012848",
                "ENSMUSG00000025362",
                "ENSMUSG00000027074",
                "ENSMUSG00000029614",
                "ENSMUSG00000021871",
                "ENSMUSG00000038612",
                "ENSMUSG00000073702",
                "ENSMUSG00000044408",
                "ENSMUSG00000062270",
                "ENSMUSG00000045817",
                "ENSMUSG00000024048",
                "ENSMUSG00000028565",
                "ENSMUSG00000038274",
                "ENSMUSG00000037072",
                "ENSMUSG00000020009",
                "ENSMUSG00000028673",
                "ENSMUSG00000021876",
                "ENSMUSG00000020088",
                "ENSMUSG00000034708",
                "ENSMUSG00000096145",
                "ENSMUSG00000062997",
                "ENSMUSG00000021610",
                "ENSMUSG00000046841",
                "ENSMUSG00000025232",
                "ENSMUSG00000058558",
                "ENSMUSG00000003363",
                "ENSMUSG00000006050",
                "ENSMUSG00000023010",
                "ENSMUSG00000029076",
                "ENSMUSG00000041084",
                "ENSMUSG00000008475",
                "ENSMUSG00000041126",
                "ENSMUSG00000060938",
                "ENSMUSG00000052146",
                "ENSMUSG00000067274",
                "ENSMUSG00000057863",
                "ENSMUSG00000022108",
                "ENSMUSG00000074170",
                "ENSMUSG00000024248",
                "ENSMUSG00000030934",
                "ENSMUSG00000020849",
                "ENSMUSG00000018593",
                "ENSMUSG00000002064",
                "ENSMUSG00000045690",
                "ENSMUSG00000027248",
                "ENSMUSG00000040681",
                "ENSMUSG00000062328",
                "ENSMUSG00000020585",
                "ENSMUSG00000022346",
                "ENSMUSG00000055762",
                "ENSMUSG00000043716",
                "ENSMUSG00000071074",
                "ENSMUSG00000024190",
                "ENSMUSG00000020458",
                "ENSMUSG00000027763",
                "ENSMUSG00000006369",
                "ENSMUSG00000059070",
                "ENSMUSG00000066551",
                "ENSMUSG00000048758",
                "ENSMUSG00000078578",
                "ENSMUSG00000028234",
                "ENSMUSG00000035227",
                "ENSMUSG00000093674",
                "ENSMUSG00000044786",
                "ENSMUSG00000028495",
                "ENSMUSG00000025579",
                "ENSMUSG00000079037",
                "ENSMUSG00000041453",
                "ENSMUSG00000019087",
                "ENSMUSG00000029465",
                "ENSMUSG00000029231",
                "ENSMUSG00000032399"
            ],
            "expression_sum": {
                "class1": 11.0,
                "class2": 4732.0
            },
            "ens_id": "ENSMUSG00000048387",
            "uniprot_ids": [
                "Q9WVG7",
                "A0A1W2P7S2"
            ],
            "net_id": 0,
            "source_num": 2,
            "target_num": 37
        },
        "ENSMUSG00000057098": {
            "id": "ENSMUSG00000057098",
            "type": "TF",
            "symbol": "Ebf1",
            "source": [
                "ENSMUSG00000021250",
                "ENSMUSG00000003032",
                "ENSMUSG00000071076"
            ],
            "target": [
                "ENSMUSG00000031447",
                "ENSMUSG00000062456",
                "ENSMUSG00000032518",
                "ENSMUSG00000024011",
                "ENSMUSG00000026878",
                "ENSMUSG00000014599",
                "ENSMUSG00000016256",
                "ENSMUSG00000047187",
                "ENSMUSG00000008668",
                "ENSMUSG00000019877",
                "ENSMUSG00000071415",
                "ENSMUSG00000053317",
                "ENSMUSG00000002900",
                "ENSMUSG00000029616",
                "ENSMUSG00000020368",
                "ENSMUSG00000028081",
                "ENSMUSG00000055302",
                "ENSMUSG00000025781",
                "ENSMUSG00000012405",
                "ENSMUSG00000038650",
                "ENSMUSG00000020737",
                "ENSMUSG00000022820",
                "ENSMUSG00000090862",
                "ENSMUSG00000030824",
                "ENSMUSG00000073639",
                "ENSMUSG00000025508",
                "ENSMUSG00000063882",
                "ENSMUSG00000011752",
                "ENSMUSG00000045128",
                "ENSMUSG00000017466",
                "ENSMUSG00000078812",
                "ENSMUSG00000029455",
                "ENSMUSG00000044080",
                "ENSMUSG00000026536",
                "ENSMUSG00000072235",
                "ENSMUSG00000006333",
                "ENSMUSG00000030432",
                "ENSMUSG00000074129",
                "ENSMUSG00000031818",
                "ENSMUSG00000026674",
                "ENSMUSG00000012848",
                "ENSMUSG00000027074",
                "ENSMUSG00000028964",
                "ENSMUSG00000003032",
                "ENSMUSG00000073702",
                "ENSMUSG00000032216",
                "ENSMUSG00000062270",
                "ENSMUSG00000045817",
                "ENSMUSG00000027712",
                "ENSMUSG00000053565",
                "ENSMUSG00000007891",
                "ENSMUSG00000024048",
                "ENSMUSG00000031029",
                "ENSMUSG00000001891",
                "ENSMUSG00000050708",
                "ENSMUSG00000026478",
                "ENSMUSG00000057278",
                "ENSMUSG00000014846",
                "ENSMUSG00000017493",
                "ENSMUSG00000025351",
                "ENSMUSG00000074884",
                "ENSMUSG00000020849",
                "ENSMUSG00000025066",
                "ENSMUSG00000045690",
                "ENSMUSG00000027248",
                "ENSMUSG00000049751",
                "ENSMUSG00000043716",
                "ENSMUSG00000027523",
                "ENSMUSG00000035530",
                "ENSMUSG00000036781",
                "ENSMUSG00000041801",
                "ENSMUSG00000073411",
                "ENSMUSG00000022174",
                "ENSMUSG00000004610",
                "ENSMUSG00000035227",
                "ENSMUSG00000060802",
                "ENSMUSG00000041453",
                "ENSMUSG00000003429",
                "ENSMUSG00000038418",
                "ENSMUSG00000032399",
                "ENSMUSG00000038900",
                "ENSMUSG00000019505",
                "ENSMUSG00000006360",
                "ENSMUSG00000059291",
                "ENSMUSG00000022841",
                "ENSMUSG00000040824",
                "ENSMUSG00000028691",
                "ENSMUSG00000022283",
                "ENSMUSG00000068523",
                "ENSMUSG00000036199",
                "ENSMUSG00000030695",
                "ENSMUSG00000030744",
                "ENSMUSG00000026238",
                "ENSMUSG00000028822",
                "ENSMUSG00000023175",
                "ENSMUSG00000022336",
                "ENSMUSG00000026970",
                "ENSMUSG00000021702",
                "ENSMUSG00000002107",
                "ENSMUSG00000009549",
                "ENSMUSG00000032231",
                "ENSMUSG00000059796",
                "ENSMUSG00000004455",
                "ENSMUSG00000037805",
                "ENSMUSG00000063316",
                "ENSMUSG00000017404",
                "ENSMUSG00000028648",
                "ENSMUSG00000025132",
                "ENSMUSG00000021917",
                "ENSMUSG00000042043",
                "ENSMUSG00000037266",
                "ENSMUSG00000030790",
                "ENSMUSG00000029642",
                "ENSMUSG00000020719",
                "ENSMUSG00000036371",
                "ENSMUSG00000019804",
                "ENSMUSG00000007039",
                "ENSMUSG00000022090",
                "ENSMUSG00000032014",
                "ENSMUSG00000040249",
                "ENSMUSG00000023092",
                "ENSMUSG00000029070",
                "ENSMUSG00000015839",
                "ENSMUSG00000010406",
                "ENSMUSG00000051695",
                "ENSMUSG00000002778",
                "ENSMUSG00000030654",
                "ENSMUSG00000098274",
                "ENSMUSG00000026511",
                "ENSMUSG00000064080",
                "ENSMUSG00000005973",
                "ENSMUSG00000057322",
                "ENSMUSG00000003161",
                "ENSMUSG00000021025",
                "ENSMUSG00000032301",
                "ENSMUSG00000042079",
                "ENSMUSG00000003814",
                "ENSMUSG00000052040",
                "ENSMUSG00000015671",
                "ENSMUSG00000047215",
                "ENSMUSG00000020205",
                "ENSMUSG00000037072",
                "ENSMUSG00000020009",
                "ENSMUSG00000004207",
                "ENSMUSG00000026728",
                "ENSMUSG00000096145",
                "ENSMUSG00000014769",
                "ENSMUSG00000042312",
                "ENSMUSG00000062997",
                "ENSMUSG00000067288",
                "ENSMUSG00000058558",
                "ENSMUSG00000075254",
                "ENSMUSG00000023010",
                "ENSMUSG00000060938",
                "ENSMUSG00000047675",
                "ENSMUSG00000041841",
                "ENSMUSG00000021242",
                "ENSMUSG00000008892",
                "ENSMUSG00000024248",
                "ENSMUSG00000028837",
                "ENSMUSG00000062328",
                "ENSMUSG00000020585",
                "ENSMUSG00000019818",
                "ENSMUSG00000025492",
                "ENSMUSG00000055762",
                "ENSMUSG00000007041",
                "ENSMUSG00000055447",
                "ENSMUSG00000024190",
                "ENSMUSG00000057841",
                "ENSMUSG00000020458",
                "ENSMUSG00000022234",
                "ENSMUSG00000009291",
                "ENSMUSG00000078578",
                "ENSMUSG00000028234",
                "ENSMUSG00000029322",
                "ENSMUSG00000035585",
                "ENSMUSG00000090137",
                "ENSMUSG00000021131",
                "ENSMUSG00000037062",
                "ENSMUSG00000031765",
                "ENSMUSG00000028639",
                "ENSMUSG00000040952",
                "ENSMUSG00000048076",
                "ENSMUSG00000026021",
                "ENSMUSG00000006699",
                "ENSMUSG00000060036",
                "ENSMUSG00000045962",
                "ENSMUSG00000038717",
                "ENSMUSG00000016252",
                "ENSMUSG00000023004",
                "ENSMUSG00000031198",
                "ENSMUSG00000025130",
                "ENSMUSG00000056737",
                "ENSMUSG00000056201",
                "ENSMUSG00000022884",
                "ENSMUSG00000061983",
                "ENSMUSG00000022032",
                "ENSMUSG00000004980",
                "ENSMUSG00000028967",
                "ENSMUSG00000022400",
                "ENSMUSG00000021546",
                "ENSMUSG00000018585",
                "ENSMUSG00000034165",
                "ENSMUSG00000003380",
                "ENSMUSG00000038803",
                "ENSMUSG00000022051",
                "ENSMUSG00000020460",
                "ENSMUSG00000055044",
                "ENSMUSG00000009927",
                "ENSMUSG00000023944",
                "ENSMUSG00000028416",
                "ENSMUSG00000024608",
                "ENSMUSG00000062647",
                "ENSMUSG00000060591",
                "ENSMUSG00000049517",
                "ENSMUSG00000027940",
                "ENSMUSG00000001794",
                "ENSMUSG00000028692",
                "ENSMUSG00000025780",
                "ENSMUSG00000030795",
                "ENSMUSG00000035242",
                "ENSMUSG00000006356",
                "ENSMUSG00000015656",
                "ENSMUSG00000067274",
                "ENSMUSG00000071644",
                "ENSMUSG00000057863",
                "ENSMUSG00000022108",
                "ENSMUSG00000025510",
                "ENSMUSG00000028249",
                "ENSMUSG00000024953",
                "ENSMUSG00000053012",
                "ENSMUSG00000026670",
                "ENSMUSG00000021250",
                "ENSMUSG00000001833",
                "ENSMUSG00000028618",
                "ENSMUSG00000027620",
                "ENSMUSG00000032966",
                "ENSMUSG00000052837",
                "ENSMUSG00000032324",
                "ENSMUSG00000031708",
                "ENSMUSG00000028936",
                "ENSMUSG00000059005",
                "ENSMUSG00000028478",
                "ENSMUSG00000060073",
                "ENSMUSG00000001847",
                "ENSMUSG00000019428",
                "ENSMUSG00000029465",
                "ENSMUSG00000039208",
                "ENSMUSG00000030057",
                "ENSMUSG00000031633",
                "ENSMUSG00000037563",
                "ENSMUSG00000058135",
                "ENSMUSG00000007815",
                "ENSMUSG00000002602",
                "ENSMUSG00000058546",
                "ENSMUSG00000008348",
                "ENSMUSG00000037822",
                "ENSMUSG00000025967",
                "ENSMUSG00000031556",
                "ENSMUSG00000021877",
                "ENSMUSG00000034868",
                "ENSMUSG00000063480",
                "ENSMUSG00000029649",
                "ENSMUSG00000028797",
                "ENSMUSG00000061787",
                "ENSMUSG00000044627",
                "ENSMUSG00000020372",
                "ENSMUSG00000021939",
                "ENSMUSG00000001240",
                "ENSMUSG00000036880",
                "ENSMUSG00000061904",
                "ENSMUSG00000028369",
                "ENSMUSG00000029621",
                "ENSMUSG00000060743",
                "ENSMUSG00000069744",
                "ENSMUSG00000020738",
                "ENSMUSG00000037499",
                "ENSMUSG00000026547",
                "ENSMUSG00000006304",
                "ENSMUSG00000062825",
                "ENSMUSG00000060636",
                "ENSMUSG00000046364",
                "ENSMUSG00000075706",
                "ENSMUSG00000020440",
                "ENSMUSG00000028367",
                "ENSMUSG00000040385",
                "ENSMUSG00000063457",
                "ENSMUSG00000032314",
                "ENSMUSG00000037742",
                "ENSMUSG00000036372",
                "ENSMUSG00000015092",
                "ENSMUSG00000025016",
                "ENSMUSG00000029767",
                "ENSMUSG00000018326",
                "ENSMUSG00000057113",
                "ENSMUSG00000002233",
                "ENSMUSG00000020267",
                "ENSMUSG00000025794",
                "ENSMUSG00000029713",
                "ENSMUSG00000039001",
                "ENSMUSG00000028452",
                "ENSMUSG00000060019",
                "ENSMUSG00000028530",
                "ENSMUSG00000090733",
                "ENSMUSG00000040212",
                "ENSMUSG00000001123",
                "ENSMUSG00000016559",
                "ENSMUSG00000060126",
                "ENSMUSG00000020149",
                "ENSMUSG00000061315",
                "ENSMUSG00000044533",
                "ENSMUSG00000062006",
                "ENSMUSG00000002015",
                "ENSMUSG00000052727",
                "ENSMUSG00000013622",
                "ENSMUSG00000038274",
                "ENSMUSG00000028108",
                "ENSMUSG00000021996",
                "ENSMUSG00000006519",
                "ENSMUSG00000056851",
                "ENSMUSG00000008475",
                "ENSMUSG00000052146",
                "ENSMUSG00000034994",
                "ENSMUSG00000024308",
                "ENSMUSG00000021218",
                "ENSMUSG00000026872",
                "ENSMUSG00000060860",
                "ENSMUSG00000071866",
                "ENSMUSG00000005881",
                "ENSMUSG00000024914",
                "ENSMUSG00000020048",
                "ENSMUSG00000042747",
                "ENSMUSG00000007892",
                "ENSMUSG00000032383",
                "ENSMUSG00000079641",
                "ENSMUSG00000031422",
                "ENSMUSG00000029622",
                "ENSMUSG00000059070",
                "ENSMUSG00000066551",
                "ENSMUSG00000022708",
                "ENSMUSG00000034118",
                "ENSMUSG00000029659",
                "ENSMUSG00000093674",
                "ENSMUSG00000029430",
                "ENSMUSG00000042682",
                "ENSMUSG00000028495",
                "ENSMUSG00000008540",
                "ENSMUSG00000029484",
                "ENSMUSG00000022548",
                "ENSMUSG00000000563"
            ],
            "expression_sum": {
                "class1": 205.0,
                "class2": 16536.0
            },
            "ens_id": "ENSMUSG00000057098",
            "uniprot_ids": [
                "Q07802",
                "Q5SWK4",
                "Q5SWK8"
            ],
            "net_id": 0,
            "source_num": 3,
            "target_num": 63
        },
        "ENSMUSG00000005583": {
            "id": "ENSMUSG00000005583",
            "type": "TF",
            "symbol": "Mef2c",
            "source": [
                "ENSMUSG00000055435",
                "ENSMUSG00000032238",
                "ENSMUSG00000022528",
                "ENSMUSG00000071076"
            ],
            "target": [
                "ENSMUSG00000031431",
                "ENSMUSG00000040723",
                "ENSMUSG00000062515",
                "ENSMUSG00000031447",
                "ENSMUSG00000025381",
                "ENSMUSG00000032518",
                "ENSMUSG00000007655",
                "ENSMUSG00000025213",
                "ENSMUSG00000053007",
                "ENSMUSG00000026879",
                "ENSMUSG00000032330",
                "ENSMUSG00000014599",
                "ENSMUSG00000016256",
                "ENSMUSG00000027447",
                "ENSMUSG00000064373",
                "ENSMUSG00000025856",
                "ENSMUSG00000015354",
                "ENSMUSG00000042613",
                "ENSMUSG00000000325",
                "ENSMUSG00000008668",
                "ENSMUSG00000078974",
                "ENSMUSG00000018567",
                "ENSMUSG00000028005",
                "ENSMUSG00000031520",
                "ENSMUSG00000071415",
                "ENSMUSG00000030605",
                "ENSMUSG00000008575",
                "ENSMUSG00000046516",
                "ENSMUSG00000053317",
                "ENSMUSG00000075701",
                "ENSMUSG00000022836",
                "ENSMUSG00000031502",
                "ENSMUSG00000020467",
                "ENSMUSG00000048960",
                "ENSMUSG00000021390",
                "ENSMUSG00000002900",
                "ENSMUSG00000029616",
                "ENSMUSG00000036570",
                "ENSMUSG00000026365",
                "ENSMUSG00000036545",
                "ENSMUSG00000024140",
                "ENSMUSG00000017707",
                "ENSMUSG00000034807",
                "ENSMUSG00000019907",
                "ENSMUSG00000014956",
                "ENSMUSG00000000184",
                "ENSMUSG00000020737",
                "ENSMUSG00000090862",
                "ENSMUSG00000030824",
                "ENSMUSG00000027642",
                "ENSMUSG00000036478",
                "ENSMUSG00000031616",
                "ENSMUSG00000003617",
                "ENSMUSG00000032271",
                "ENSMUSG00000033585",
                "ENSMUSG00000027706",
                "ENSMUSG00000025290",
                "ENSMUSG00000027800",
                "ENSMUSG00000031586",
                "ENSMUSG00000041828",
                "ENSMUSG00000022587",
                "ENSMUSG00000045128",
                "ENSMUSG00000017466",
                "ENSMUSG00000026536",
                "ENSMUSG00000072235",
                "ENSMUSG00000024277",
                "ENSMUSG00000006333",
                "ENSMUSG00000038463",
                "ENSMUSG00000002985",
                "ENSMUSG00000068798",
                "ENSMUSG00000028517",
                "ENSMUSG00000016494",
                "ENSMUSG00000020029",
                "ENSMUSG00000062210",
                "ENSMUSG00000024290",
                "ENSMUSG00000026674",
                "ENSMUSG00000075705",
                "ENSMUSG00000012848",
                "ENSMUSG00000015094",
                "ENSMUSG00000063229",
                "ENSMUSG00000022665",
                "ENSMUSG00000027074",
                "ENSMUSG00000021871",
                "ENSMUSG00000024659",
                "ENSMUSG00000024085",
                "ENSMUSG00000024338",
                "ENSMUSG00000027712",
                "ENSMUSG00000007891",
                "ENSMUSG00000024048",
                "ENSMUSG00000028673",
                "ENSMUSG00000034708",
                "ENSMUSG00000001270",
                "ENSMUSG00000050708",
                "ENSMUSG00000026478",
                "ENSMUSG00000019920",
                "ENSMUSG00000017493",
                "ENSMUSG00000025351",
                "ENSMUSG00000074884",
                "ENSMUSG00000055435",
                "ENSMUSG00000001020",
                "ENSMUSG00000034993",
                "ENSMUSG00000021200",
                "ENSMUSG00000027248",
                "ENSMUSG00000052957",
                "ENSMUSG00000051579",
                "ENSMUSG00000004460",
                "ENSMUSG00000026864",
                "ENSMUSG00000031548",
                "ENSMUSG00000028757",
                "ENSMUSG00000073411",
                "ENSMUSG00000024012",
                "ENSMUSG00000009876",
                "ENSMUSG00000029661",
                "ENSMUSG00000060802",
                "ENSMUSG00000040659",
                "ENSMUSG00000017969",
                "ENSMUSG00000004085",
                "ENSMUSG00000018293",
                "ENSMUSG00000035783",
                "ENSMUSG00000014867",
                "ENSMUSG00000038900",
                "ENSMUSG00000006360",
                "ENSMUSG00000059291",
                "ENSMUSG00000031320",
                "ENSMUSG00000009281",
                "ENSMUSG00000022371",
                "ENSMUSG00000036438",
                "ENSMUSG00000029467",
                "ENSMUSG00000023175",
                "ENSMUSG00000018339",
                "ENSMUSG00000035493",
                "ENSMUSG00000002107",
                "ENSMUSG00000073889",
                "ENSMUSG00000022010",
                "ENSMUSG00000032231",
                "ENSMUSG00000007872",
                "ENSMUSG00000027848",
                "ENSMUSG00000073557",
                "ENSMUSG00000041329",
                "ENSMUSG00000029426",
                "ENSMUSG00000029718",
                "ENSMUSG00000057666",
                "ENSMUSG00000028600",
                "ENSMUSG00000047420",
                "ENSMUSG00000020321",
                "ENSMUSG00000025491",
                "ENSMUSG00000021756",
                "ENSMUSG00000024245",
                "ENSMUSG00000036446",
                "ENSMUSG00000031538",
                "ENSMUSG00000019978",
                "ENSMUSG00000022816",
                "ENSMUSG00000026482",
                "ENSMUSG00000031636",
                "ENSMUSG00000002980",
                "ENSMUSG00000035885",
                "ENSMUSG00000030403",
                "ENSMUSG00000007039",
                "ENSMUSG00000029781",
                "ENSMUSG00000026193",
                "ENSMUSG00000022090",
                "ENSMUSG00000027834",
                "ENSMUSG00000032014",
                "ENSMUSG00000038146",
                "ENSMUSG00000068735",
                "ENSMUSG00000029061",
                "ENSMUSG00000040249",
                "ENSMUSG00000032334",
                "ENSMUSG00000023092",
                "ENSMUSG00000029070",
                "ENSMUSG00000023367",
                "ENSMUSG00000022324",
                "ENSMUSG00000023886",
                "ENSMUSG00000027276",
                "ENSMUSG00000005397",
                "ENSMUSG00000030208",
                "ENSMUSG00000022817",
                "ENSMUSG00000030536",
                "ENSMUSG00000063694",
                "ENSMUSG00000003420",
                "ENSMUSG00000020315",
                "ENSMUSG00000058569",
                "ENSMUSG00000018217",
                "ENSMUSG00000023046",
                "ENSMUSG00000058056",
                "ENSMUSG00000064080",
                "ENSMUSG00000005973",
                "ENSMUSG00000042745",
                "ENSMUSG00000001520",
                "ENSMUSG00000016534",
                "ENSMUSG00000003161",
                "ENSMUSG00000032359",
                "ENSMUSG00000041577",
                "ENSMUSG00000001911",
                "ENSMUSG00000022136",
                "ENSMUSG00000018819",
                "ENSMUSG00000003814",
                "ENSMUSG00000061232",
                "ENSMUSG00000027861",
                "ENSMUSG00000002059",
                "ENSMUSG00000020205",
                "ENSMUSG00000037072",
                "ENSMUSG00000020009",
                "ENSMUSG00000004207",
                "ENSMUSG00000020088",
                "ENSMUSG00000026728",
                "ENSMUSG00000096145",
                "ENSMUSG00000042312",
                "ENSMUSG00000062997",
                "ENSMUSG00000032997",
                "ENSMUSG00000025232",
                "ENSMUSG00000067288",
                "ENSMUSG00000075254",
                "ENSMUSG00000031799",
                "ENSMUSG00000055053",
                "ENSMUSG00000041084",
                "ENSMUSG00000022971",
                "ENSMUSG00000025393",
                "ENSMUSG00000021242",
                "ENSMUSG00000074170",
                "ENSMUSG00000002014",
                "ENSMUSG00000014313",
                "ENSMUSG00000032547",
                "ENSMUSG00000092341",
                "ENSMUSG00000026688",
                "ENSMUSG00000087006",
                "ENSMUSG00000032238",
                "ENSMUSG00000021843",
                "ENSMUSG00000062328",
                "ENSMUSG00000006800",
                "ENSMUSG00000020585",
                "ENSMUSG00000032353",
                "ENSMUSG00000025492",
                "ENSMUSG00000046402",
                "ENSMUSG00000055447",
                "ENSMUSG00000057841",
                "ENSMUSG00000046718",
                "ENSMUSG00000027408",
                "ENSMUSG00000020840",
                "ENSMUSG00000028234",
                "ENSMUSG00000037206",
                "ENSMUSG00000015932",
                "ENSMUSG00000060703",
                "ENSMUSG00000029231",
                "ENSMUSG00000073489",
                "ENSMUSG00000020902",
                "ENSMUSG00000035258",
                "ENSMUSG00000001025",
                "ENSMUSG00000019539",
                "ENSMUSG00000032925",
                "ENSMUSG00000031765",
                "ENSMUSG00000019916",
                "ENSMUSG00000045636",
                "ENSMUSG00000040952",
                "ENSMUSG00000022528",
                "ENSMUSG00000032575",
                "ENSMUSG00000079111",
                "ENSMUSG00000042485",
                "ENSMUSG00000030659",
                "ENSMUSG00000012422",
                "ENSMUSG00000025130",
                "ENSMUSG00000020612",
                "ENSMUSG00000038872",
                "ENSMUSG00000056737",
                "ENSMUSG00000020928",
                "ENSMUSG00000061983",
                "ENSMUSG00000022032",
                "ENSMUSG00000076432",
                "ENSMUSG00000038086",
                "ENSMUSG00000048096",
                "ENSMUSG00000028461",
                "ENSMUSG00000059439",
                "ENSMUSG00000034165",
                "ENSMUSG00000043091",
                "ENSMUSG00000032085",
                "ENSMUSG00000022892",
                "ENSMUSG00000031626",
                "ENSMUSG00000029810",
                "ENSMUSG00000070436",
                "ENSMUSG00000015133",
                "ENSMUSG00000027188",
                "ENSMUSG00000027204",
                "ENSMUSG00000020460",
                "ENSMUSG00000028464",
                "ENSMUSG00000028776",
                "ENSMUSG00000037060",
                "ENSMUSG00000021253",
                "ENSMUSG00000053477",
                "ENSMUSG00000016382",
                "ENSMUSG00000062647",
                "ENSMUSG00000046330",
                "ENSMUSG00000079523",
                "ENSMUSG00000059895",
                "ENSMUSG00000073418",
                "ENSMUSG00000046727",
                "ENSMUSG00000016194",
                "ENSMUSG00000038991",
                "ENSMUSG00000060591",
                "ENSMUSG00000023942",
                "ENSMUSG00000049517",
                "ENSMUSG00000038530",
                "ENSMUSG00000024371",
                "ENSMUSG00000069662",
                "ENSMUSG00000021876",
                "ENSMUSG00000024164",
                "ENSMUSG00000006356",
                "ENSMUSG00000020473",
                "ENSMUSG00000001663",
                "ENSMUSG00000023224",
                "ENSMUSG00000026223",
                "ENSMUSG00000049086",
                "ENSMUSG00000027907",
                "ENSMUSG00000067274",
                "ENSMUSG00000049892",
                "ENSMUSG00000069833",
                "ENSMUSG00000033910",
                "ENSMUSG00000039168",
                "ENSMUSG00000020484",
                "ENSMUSG00000015568",
                "ENSMUSG00000040462",
                "ENSMUSG00000066705",
                "ENSMUSG00000036427",
                "ENSMUSG00000028618",
                "ENSMUSG00000026748",
                "ENSMUSG00000047907",
                "ENSMUSG00000023064",
                "ENSMUSG00000028936",
                "ENSMUSG00000036437",
                "ENSMUSG00000022912",
                "ENSMUSG00000025579",
                "ENSMUSG00000079037",
                "ENSMUSG00000079017",
                "ENSMUSG00000052997",
                "ENSMUSG00000000093",
                "ENSMUSG00000058135",
                "ENSMUSG00000040652",
                "ENSMUSG00000029761",
                "ENSMUSG00000028111",
                "ENSMUSG00000021477",
                "ENSMUSG00000028949",
                "ENSMUSG00000097328",
                "ENSMUSG00000027422",
                "ENSMUSG00000020644",
                "ENSMUSG00000056481",
                "ENSMUSG00000025647",
                "ENSMUSG00000029287",
                "ENSMUSG00000031556",
                "ENSMUSG00000021877",
                "ENSMUSG00000034868",
                "ENSMUSG00000026043",
                "ENSMUSG00000021939",
                "ENSMUSG00000041417",
                "ENSMUSG00000008682",
                "ENSMUSG00000020402",
                "ENSMUSG00000001240",
                "ENSMUSG00000014329",
                "ENSMUSG00000040713",
                "ENSMUSG00000034892",
                "ENSMUSG00000028369",
                "ENSMUSG00000060572",
                "ENSMUSG00000012428",
                "ENSMUSG00000029309",
                "ENSMUSG00000061353",
                "ENSMUSG00000031595",
                "ENSMUSG00000021943",
                "ENSMUSG00000009681",
                "ENSMUSG00000060904",
                "ENSMUSG00000064370",
                "ENSMUSG00000020053",
                "ENSMUSG00000040666",
                "ENSMUSG00000026042",
                "ENSMUSG00000067212",
                "ENSMUSG00000020440",
                "ENSMUSG00000039007",
                "ENSMUSG00000032366",
                "ENSMUSG00000001119",
                "ENSMUSG00000075010",
                "ENSMUSG00000029390",
                "ENSMUSG00000059173",
                "ENSMUSG00000029767",
                "ENSMUSG00000031246",
                "ENSMUSG00000090000",
                "ENSMUSG00000021760",
                "ENSMUSG00000032744",
                "ENSMUSG00000048546",
                "ENSMUSG00000021248",
                "ENSMUSG00000001506",
                "ENSMUSG00000021870",
                "ENSMUSG00000041959",
                "ENSMUSG00000017754",
                "ENSMUSG00000004791",
                "ENSMUSG00000020814",
                "ENSMUSG00000053279",
                "ENSMUSG00000033880",
                "ENSMUSG00000025794",
                "ENSMUSG00000038145",
                "ENSMUSG00000015085",
                "ENSMUSG00000020486",
                "ENSMUSG00000090733",
                "ENSMUSG00000040212",
                "ENSMUSG00000050666",
                "ENSMUSG00000032946",
                "ENSMUSG00000001123",
                "ENSMUSG00000037966",
                "ENSMUSG00000016559",
                "ENSMUSG00000055172",
                "ENSMUSG00000060126",
                "ENSMUSG00000026014",
                "ENSMUSG00000037706",
                "ENSMUSG00000001175",
                "ENSMUSG00000032387",
                "ENSMUSG00000040562",
                "ENSMUSG00000032135",
                "ENSMUSG00000019139",
                "ENSMUSG00000030272",
                "ENSMUSG00000025428",
                "ENSMUSG00000062006",
                "ENSMUSG00000052727",
                "ENSMUSG00000078202",
                "ENSMUSG00000027230",
                "ENSMUSG00000020571",
                "ENSMUSG00000002732",
                "ENSMUSG00000025147",
                "ENSMUSG00000006205",
                "ENSMUSG00000051855",
                "ENSMUSG00000006519",
                "ENSMUSG00000045658",
                "ENSMUSG00000029076",
                "ENSMUSG00000030717",
                "ENSMUSG00000049775",
                "ENSMUSG00000039601",
                "ENSMUSG00000024150",
                "ENSMUSG00000029359",
                "ENSMUSG00000067818",
                "ENSMUSG00000024308",
                "ENSMUSG00000046318",
                "ENSMUSG00000024538",
                "ENSMUSG00000090841",
                "ENSMUSG00000041736",
                "ENSMUSG00000005881",
                "ENSMUSG00000016319",
                "ENSMUSG00000004105",
                "ENSMUSG00000038370",
                "ENSMUSG00000020048",
                "ENSMUSG00000042747",
                "ENSMUSG00000041355",
                "ENSMUSG00000000740",
                "ENSMUSG00000007892",
                "ENSMUSG00000029096",
                "ENSMUSG00000021186",
                "ENSMUSG00000026837",
                "ENSMUSG00000032383",
                "ENSMUSG00000079641",
                "ENSMUSG00000022048",
                "ENSMUSG00000006369",
                "ENSMUSG00000031422",
                "ENSMUSG00000029622",
                "ENSMUSG00000001435",
                "ENSMUSG00000039286",
                "ENSMUSG00000004044",
                "ENSMUSG00000022708",
                "ENSMUSG00000034118",
                "ENSMUSG00000070348",
                "ENSMUSG00000019929",
                "ENSMUSG00000029659",
                "ENSMUSG00000029641",
                "ENSMUSG00000026574",
                "ENSMUSG00000018920",
                "ENSMUSG00000016458",
                "ENSMUSG00000008540",
                "ENSMUSG00000021268",
                "ENSMUSG00000018830",
                "ENSMUSG00000090698",
                "ENSMUSG00000075702",
                "ENSMUSG00000004951",
                "ENSMUSG00000057530"
            ],
            "expression_sum": {
                "class1": 2579.0,
                "class2": 9123.0
            },
            "ens_id": "ENSMUSG00000005583",
            "uniprot_ids": [
                "Q8CFN5",
                "A0A0G2JDK0",
                "A0A0G2JEC2",
                "A0A0G2JFL7",
                "A0A0H2UH28",
                "A0A0G2JFX2",
                "A0A0G2JEC3",
                "A0A0G2JFI7",
                "A0A0H2UKB6",
                "A0A0G2JFX1",
                "A0A0G2JGK3",
                "A0A0G2JE18",
                "A0A0G2JGL3",
                "A0A0G2JE38",
                "A0A0G2JFM7",
                "A0A0G2JFS4",
                "A0A0G2JET3",
                "A0A0G2JES4",
                "A0A0G2JEM8",
                "A0A0G2JDQ1",
                "A0A0G2JDT0",
                "A0A0G2JF80"
            ],
            "net_id": 0,
            "source_num": 4,
            "target_num": 124
        },
        "ENSMUSG00000037868": {
            "id": "ENSMUSG00000037868",
            "type": "TF",
            "symbol": "Egr2",
            "source": [
                "ENSMUSG00000056501",
                "ENSMUSG00000052684",
                "ENSMUSG00000023034",
                "ENSMUSG00000052837",
                "ENSMUSG00000026628",
                "ENSMUSG00000038418",
                "ENSMUSG00000021250",
                "ENSMUSG00000003032",
                "ENSMUSG00000071076",
                "ENSMUSG00000018899"
            ],
            "target": [
                "ENSMUSG00000023034",
                "ENSMUSG00000053113",
                "ENSMUSG00000003545",
                "ENSMUSG00000033538",
                "ENSMUSG00000052684",
                "ENSMUSG00000021025",
                "ENSMUSG00000015312",
                "ENSMUSG00000008384",
                "ENSMUSG00000021250",
                "ENSMUSG00000003032",
                "ENSMUSG00000055148",
                "ENSMUSG00000021127",
                "ENSMUSG00000015501",
                "ENSMUSG00000040128",
                "ENSMUSG00000035356",
                "ENSMUSG00000052837",
                "ENSMUSG00000026628",
                "ENSMUSG00000020423",
                "ENSMUSG00000097296",
                "ENSMUSG00000092274",
                "ENSMUSG00000007872",
                "ENSMUSG00000044786",
                "ENSMUSG00000018899",
                "ENSMUSG00000027829",
                "ENSMUSG00000006403",
                "ENSMUSG00000028214",
                "ENSMUSG00000032501",
                "ENSMUSG00000035828",
                "ENSMUSG00000038418",
                "ENSMUSG00000009681"
            ],
            "expression_sum": {
                "class1": 3930.0,
                "class2": 381.0
            },
            "ens_id": "ENSMUSG00000037868",
            "uniprot_ids": [
                "P08152",
                "D3Z4G7",
                "D3Z4X9",
                "D3YYY2",
                "D3Z3K7",
                "D3YXS2"
            ],
            "net_id": 0,
            "source_num": 10,
            "target_num": 18
        },
        "ENSMUSG00000021250": {
            "id": "ENSMUSG00000021250",
            "type": "TF",
            "symbol": "Fos",
            "source": [
                "ENSMUSG00000041515",
                "ENSMUSG00000056501",
                "ENSMUSG00000052684",
                "ENSMUSG00000037868",
                "ENSMUSG00000023034",
                "ENSMUSG00000052837",
                "ENSMUSG00000030103",
                "ENSMUSG00000000247",
                "ENSMUSG00000042812",
                "ENSMUSG00000057098",
                "ENSMUSG00000003545",
                "ENSMUSG00000003032",
                "ENSMUSG00000018899",
                "ENSMUSG00000071076"
            ],
            "target": [
                "ENSMUSG00000019505",
                "ENSMUSG00000006360",
                "ENSMUSG00000031765",
                "ENSMUSG00000024561",
                "ENSMUSG00000030222",
                "ENSMUSG00000028691",
                "ENSMUSG00000003545",
                "ENSMUSG00000036199",
                "ENSMUSG00000025289",
                "ENSMUSG00000032330",
                "ENSMUSG00000030930",
                "ENSMUSG00000004665",
                "ENSMUSG00000053560",
                "ENSMUSG00000036040",
                "ENSMUSG00000037411",
                "ENSMUSG00000036438",
                "ENSMUSG00000006699",
                "ENSMUSG00000008384",
                "ENSMUSG00000055148",
                "ENSMUSG00000018339",
                "ENSMUSG00000035493",
                "ENSMUSG00000021127",
                "ENSMUSG00000023004",
                "ENSMUSG00000029380",
                "ENSMUSG00000002416",
                "ENSMUSG00000007872",
                "ENSMUSG00000009013",
                "ENSMUSG00000036655",
                "ENSMUSG00000009681",
                "ENSMUSG00000036570",
                "ENSMUSG00000055839",
                "ENSMUSG00000075706",
                "ENSMUSG00000020053",
                "ENSMUSG00000037999",
                "ENSMUSG00000069601",
                "ENSMUSG00000032085",
                "ENSMUSG00000053113",
                "ENSMUSG00000037266",
                "ENSMUSG00000033538",
                "ENSMUSG00000084786",
                "ENSMUSG00000021520",
                "ENSMUSG00000052684",
                "ENSMUSG00000030122",
                "ENSMUSG00000055681",
                "ENSMUSG00000031762",
                "ENSMUSG00000003038",
                "ENSMUSG00000015501",
                "ENSMUSG00000038690",
                "ENSMUSG00000023915",
                "ENSMUSG00000030103",
                "ENSMUSG00000042812",
                "ENSMUSG00000028464",
                "ENSMUSG00000032515",
                "ENSMUSG00000074457",
                "ENSMUSG00000027829",
                "ENSMUSG00000037060",
                "ENSMUSG00000028195",
                "ENSMUSG00000031375",
                "ENSMUSG00000021253",
                "ENSMUSG00000005873",
                "ENSMUSG00000038463",
                "ENSMUSG00000002985",
                "ENSMUSG00000035678",
                "ENSMUSG00000059534",
                "ENSMUSG00000072625",
                "ENSMUSG00000015085",
                "ENSMUSG00000068220",
                "ENSMUSG00000023034",
                "ENSMUSG00000030654",
                "ENSMUSG00000054364",
                "ENSMUSG00000016494",
                "ENSMUSG00000056708",
                "ENSMUSG00000038393",
                "ENSMUSG00000037706",
                "ENSMUSG00000046727",
                "ENSMUSG00000001175",
                "ENSMUSG00000032359",
                "ENSMUSG00000061518",
                "ENSMUSG00000019960",
                "ENSMUSG00000015312",
                "ENSMUSG00000003032",
                "ENSMUSG00000037211",
                "ENSMUSG00000030247",
                "ENSMUSG00000020423",
                "ENSMUSG00000024048",
                "ENSMUSG00000023089",
                "ENSMUSG00000021876",
                "ENSMUSG00000037868",
                "ENSMUSG00000022122",
                "ENSMUSG00000006205",
                "ENSMUSG00000035828",
                "ENSMUSG00000057098",
                "ENSMUSG00000064215",
                "ENSMUSG00000018286",
                "ENSMUSG00000017493",
                "ENSMUSG00000039601",
                "ENSMUSG00000067818",
                "ENSMUSG00000056501",
                "ENSMUSG00000037152",
                "ENSMUSG00000024014",
                "ENSMUSG00000000628",
                "ENSMUSG00000090841",
                "ENSMUSG00000000753",
                "ENSMUSG00000071866",
                "ENSMUSG00000028864",
                "ENSMUSG00000036751",
                "ENSMUSG00000038370",
                "ENSMUSG00000001833",
                "ENSMUSG00000026385",
                "ENSMUSG00000024190",
                "ENSMUSG00000041515",
                "ENSMUSG00000007097",
                "ENSMUSG00000035356",
                "ENSMUSG00000052837",
                "ENSMUSG00000047945",
                "ENSMUSG00000073411",
                "ENSMUSG00000029632",
                "ENSMUSG00000030342",
                "ENSMUSG00000022956",
                "ENSMUSG00000044786",
                "ENSMUSG00000008540",
                "ENSMUSG00000018899",
                "ENSMUSG00000075702",
                "ENSMUSG00000027004",
                "ENSMUSG00000006403",
                "ENSMUSG00000018293",
                "ENSMUSG00000000247",
                "ENSMUSG00000038418",
                "ENSMUSG00000035783",
                "ENSMUSG00000030849",
                "ENSMUSG00000058135"
            ],
            "expression_sum": {
                "class1": 165566.0,
                "class2": 132750.0
            },
            "ens_id": "ENSMUSG00000021250",
            "uniprot_ids": [
                "P01101"
            ],
            "net_id": 0,
            "source_num": 14,
            "target_num": 55
        },
        "ENSMUSG00000052684": {
            "id": "ENSMUSG00000052684",
            "type": "TF",
            "symbol": "Jun",
            "source": [
                "ENSMUSG00000055435",
                "ENSMUSG00000056501",
                "ENSMUSG00000037868",
                "ENSMUSG00000023034",
                "ENSMUSG00000022528",
                "ENSMUSG00000052837",
                "ENSMUSG00000026628",
                "ENSMUSG00000000247",
                "ENSMUSG00000038418",
                "ENSMUSG00000052534",
                "ENSMUSG00000021250",
                "ENSMUSG00000038193",
                "ENSMUSG00000003545",
                "ENSMUSG00000008575",
                "ENSMUSG00000018899"
            ],
            "target": [
                "ENSMUSG00000031431",
                "ENSMUSG00000040723",
                "ENSMUSG00000029761",
                "ENSMUSG00000025381",
                "ENSMUSG00000032925",
                "ENSMUSG00000091345",
                "ENSMUSG00000028111",
                "ENSMUSG00000030222",
                "ENSMUSG00000021477",
                "ENSMUSG00000028691",
                "ENSMUSG00000003545",
                "ENSMUSG00000022365",
                "ENSMUSG00000025213",
                "ENSMUSG00000025289",
                "ENSMUSG00000026879",
                "ENSMUSG00000009281",
                "ENSMUSG00000053560",
                "ENSMUSG00000037411",
                "ENSMUSG00000064373",
                "ENSMUSG00000022528",
                "ENSMUSG00000034868",
                "ENSMUSG00000042613",
                "ENSMUSG00000006699",
                "ENSMUSG00000000325",
                "ENSMUSG00000027239",
                "ENSMUSG00000008384",
                "ENSMUSG00000026043",
                "ENSMUSG00000044627",
                "ENSMUSG00000079111",
                "ENSMUSG00000055148",
                "ENSMUSG00000042485",
                "ENSMUSG00000018339",
                "ENSMUSG00000041417",
                "ENSMUSG00000012422",
                "ENSMUSG00000078974",
                "ENSMUSG00000023004",
                "ENSMUSG00000026421",
                "ENSMUSG00000002107",
                "ENSMUSG00000073889",
                "ENSMUSG00000014329",
                "ENSMUSG00000031007",
                "ENSMUSG00000022010",
                "ENSMUSG00000028369",
                "ENSMUSG00000029103",
                "ENSMUSG00000003541",
                "ENSMUSG00000032231",
                "ENSMUSG00000008575",
                "ENSMUSG00000030218",
                "ENSMUSG00000038872",
                "ENSMUSG00000007872",
                "ENSMUSG00000053317",
                "ENSMUSG00000027848",
                "ENSMUSG00000022836",
                "ENSMUSG00000028597",
                "ENSMUSG00000029309",
                "ENSMUSG00000031845",
                "ENSMUSG00000022032",
                "ENSMUSG00000029718",
                "ENSMUSG00000028967",
                "ENSMUSG00000021390",
                "ENSMUSG00000027828",
                "ENSMUSG00000031595",
                "ENSMUSG00000036655",
                "ENSMUSG00000009681",
                "ENSMUSG00000060904",
                "ENSMUSG00000064370",
                "ENSMUSG00000020053",
                "ENSMUSG00000021917",
                "ENSMUSG00000041120",
                "ENSMUSG00000028339",
                "ENSMUSG00000037999",
                "ENSMUSG00000069601",
                "ENSMUSG00000052534",
                "ENSMUSG00000053113",
                "ENSMUSG00000024140",
                "ENSMUSG00000036372",
                "ENSMUSG00000037266",
                "ENSMUSG00000033538",
                "ENSMUSG00000019907",
                "ENSMUSG00000001119",
                "ENSMUSG00000036446",
                "ENSMUSG00000075010",
                "ENSMUSG00000031451",
                "ENSMUSG00000014956",
                "ENSMUSG00000029390",
                "ENSMUSG00000024076",
                "ENSMUSG00000055681",
                "ENSMUSG00000090000",
                "ENSMUSG00000021760",
                "ENSMUSG00000019278",
                "ENSMUSG00000003477",
                "ENSMUSG00000021248",
                "ENSMUSG00000031616",
                "ENSMUSG00000001506",
                "ENSMUSG00000003038",
                "ENSMUSG00000021870",
                "ENSMUSG00000028871",
                "ENSMUSG00000015501",
                "ENSMUSG00000041959",
                "ENSMUSG00000023915",
                "ENSMUSG00000033585",
                "ENSMUSG00000026628",
                "ENSMUSG00000030103",
                "ENSMUSG00000027706",
                "ENSMUSG00000042812",
                "ENSMUSG00000019579",
                "ENSMUSG00000041828",
                "ENSMUSG00000031586",
                "ENSMUSG00000092274",
                "ENSMUSG00000032515",
                "ENSMUSG00000068735",
                "ENSMUSG00000028776",
                "ENSMUSG00000042737",
                "ENSMUSG00000018672",
                "ENSMUSG00000017466",
                "ENSMUSG00000074457",
                "ENSMUSG00000024065",
                "ENSMUSG00000027829",
                "ENSMUSG00000029061",
                "ENSMUSG00000028195",
                "ENSMUSG00000032334",
                "ENSMUSG00000004631",
                "ENSMUSG00000021253",
                "ENSMUSG00000072235",
                "ENSMUSG00000005873",
                "ENSMUSG00000038463",
                "ENSMUSG00000035678",
                "ENSMUSG00000016024",
                "ENSMUSG00000072625",
                "ENSMUSG00000015085",
                "ENSMUSG00000030208",
                "ENSMUSG00000020486",
                "ENSMUSG00000068220",
                "ENSMUSG00000028517",
                "ENSMUSG00000040212",
                "ENSMUSG00000023034",
                "ENSMUSG00000079523",
                "ENSMUSG00000030654",
                "ENSMUSG00000016494",
                "ENSMUSG00000038393",
                "ENSMUSG00000042453",
                "ENSMUSG00000037706",
                "ENSMUSG00000018217",
                "ENSMUSG00000023046",
                "ENSMUSG00000058056",
                "ENSMUSG00000042745",
                "ENSMUSG00000003161",
                "ENSMUSG00000032359",
                "ENSMUSG00000060591",
                "ENSMUSG00000025784",
                "ENSMUSG00000040562",
                "ENSMUSG00000072812",
                "ENSMUSG00000015312",
                "ENSMUSG00000001911",
                "ENSMUSG00000031661",
                "ENSMUSG00000021871",
                "ENSMUSG00000025880",
                "ENSMUSG00000025724",
                "ENSMUSG00000018819",
                "ENSMUSG00000037211",
                "ENSMUSG00000031239",
                "ENSMUSG00000020423",
                "ENSMUSG00000044408",
                "ENSMUSG00000027712",
                "ENSMUSG00000042254",
                "ENSMUSG00000024048",
                "ENSMUSG00000037072",
                "ENSMUSG00000020009",
                "ENSMUSG00000002732",
                "ENSMUSG00000021876",
                "ENSMUSG00000025780",
                "ENSMUSG00000024164",
                "ENSMUSG00000096145",
                "ENSMUSG00000025147",
                "ENSMUSG00000037868",
                "ENSMUSG00000022122",
                "ENSMUSG00000042312",
                "ENSMUSG00000006205",
                "ENSMUSG00000001663",
                "ENSMUSG00000020473",
                "ENSMUSG00000035828",
                "ENSMUSG00000000594",
                "ENSMUSG00000064215",
                "ENSMUSG00000006050",
                "ENSMUSG00000006519",
                "ENSMUSG00000045658",
                "ENSMUSG00000030717",
                "ENSMUSG00000031799",
                "ENSMUSG00000017493",
                "ENSMUSG00000023010",
                "ENSMUSG00000041084",
                "ENSMUSG00000025351",
                "ENSMUSG00000027907",
                "ENSMUSG00000074884",
                "ENSMUSG00000039601",
                "ENSMUSG00000054808",
                "ENSMUSG00000056501",
                "ENSMUSG00000021242",
                "ENSMUSG00000000628",
                "ENSMUSG00000002014",
                "ENSMUSG00000092341",
                "ENSMUSG00000039899",
                "ENSMUSG00000055435",
                "ENSMUSG00000033910",
                "ENSMUSG00000087006",
                "ENSMUSG00000018593",
                "ENSMUSG00000000753",
                "ENSMUSG00000052957",
                "ENSMUSG00000015568",
                "ENSMUSG00000028864",
                "ENSMUSG00000006800",
                "ENSMUSG00000020585",
                "ENSMUSG00000031380",
                "ENSMUSG00000032353",
                "ENSMUSG00000025492",
                "ENSMUSG00000051579",
                "ENSMUSG00000066705",
                "ENSMUSG00000046402",
                "ENSMUSG00000021250",
                "ENSMUSG00000042747",
                "ENSMUSG00000041355",
                "ENSMUSG00000029096",
                "ENSMUSG00000026837",
                "ENSMUSG00000031548",
                "ENSMUSG00000028618",
                "ENSMUSG00000032383",
                "ENSMUSG00000026748",
                "ENSMUSG00000007097",
                "ENSMUSG00000054203",
                "ENSMUSG00000006369",
                "ENSMUSG00000031422",
                "ENSMUSG00000035356",
                "ENSMUSG00000027859",
                "ENSMUSG00000047945",
                "ENSMUSG00000049107",
                "ENSMUSG00000052837",
                "ENSMUSG00000030342",
                "ENSMUSG00000022174",
                "ENSMUSG00000035227",
                "ENSMUSG00000024012",
                "ENSMUSG00000029659",
                "ENSMUSG00000040170",
                "ENSMUSG00000029641",
                "ENSMUSG00000029661",
                "ENSMUSG00000044786",
                "ENSMUSG00000008540",
                "ENSMUSG00000018899",
                "ENSMUSG00000021268",
                "ENSMUSG00000079037",
                "ENSMUSG00000090698",
                "ENSMUSG00000075702",
                "ENSMUSG00000006403",
                "ENSMUSG00000060703",
                "ENSMUSG00000000247",
                "ENSMUSG00000038418",
                "ENSMUSG00000052997",
                "ENSMUSG00000030849",
                "ENSMUSG00000001025",
                "ENSMUSG00000058135"
            ],
            "expression_sum": {
                "class1": 84935.0,
                "class2": 57932.0
            },
            "ens_id": "ENSMUSG00000052684",
            "uniprot_ids": [
                "P05627"
            ],
            "net_id": 0,
            "source_num": 15,
            "target_num": 108
        },
        "ENSMUSG00000038418": {
            "id": "ENSMUSG00000038418",
            "type": "TF",
            "symbol": "Egr1",
            "source": [
                "ENSMUSG00000052684",
                "ENSMUSG00000037868",
                "ENSMUSG00000023034",
                "ENSMUSG00000022528",
                "ENSMUSG00000052837",
                "ENSMUSG00000026628",
                "ENSMUSG00000000247",
                "ENSMUSG00000057098",
                "ENSMUSG00000021250",
                "ENSMUSG00000003545",
                "ENSMUSG00000003032",
                "ENSMUSG00000008575",
                "ENSMUSG00000018899"
            ],
            "target": [
                "ENSMUSG00000019505",
                "ENSMUSG00000075706",
                "ENSMUSG00000040723",
                "ENSMUSG00000040212",
                "ENSMUSG00000021917",
                "ENSMUSG00000023034",
                "ENSMUSG00000021242",
                "ENSMUSG00000024014",
                "ENSMUSG00000028367",
                "ENSMUSG00000021477",
                "ENSMUSG00000032314",
                "ENSMUSG00000053113",
                "ENSMUSG00000028249",
                "ENSMUSG00000030934",
                "ENSMUSG00000025213",
                "ENSMUSG00000033910",
                "ENSMUSG00000020315",
                "ENSMUSG00000018217",
                "ENSMUSG00000016194",
                "ENSMUSG00000003161",
                "ENSMUSG00000019907",
                "ENSMUSG00000052684",
                "ENSMUSG00000006800",
                "ENSMUSG00000022528",
                "ENSMUSG00000028822",
                "ENSMUSG00000034868",
                "ENSMUSG00000032353",
                "ENSMUSG00000008384",
                "ENSMUSG00000003032",
                "ENSMUSG00000042747",
                "ENSMUSG00000079111",
                "ENSMUSG00000055148",
                "ENSMUSG00000021248",
                "ENSMUSG00000026837",
                "ENSMUSG00000003038",
                "ENSMUSG00000032383",
                "ENSMUSG00000015501",
                "ENSMUSG00000006369",
                "ENSMUSG00000040128",
                "ENSMUSG00000031422",
                "ENSMUSG00000092274",
                "ENSMUSG00000002107",
                "ENSMUSG00000052837",
                "ENSMUSG00000026628",
                "ENSMUSG00000020423",
                "ENSMUSG00000035356",
                "ENSMUSG00000027706",
                "ENSMUSG00000029103",
                "ENSMUSG00000035227",
                "ENSMUSG00000024048",
                "ENSMUSG00000007891",
                "ENSMUSG00000040170",
                "ENSMUSG00000020571",
                "ENSMUSG00000031119",
                "ENSMUSG00000037072",
                "ENSMUSG00000007872",
                "ENSMUSG00000008540",
                "ENSMUSG00000018899",
                "ENSMUSG00000044786",
                "ENSMUSG00000020009",
                "ENSMUSG00000027829",
                "ENSMUSG00000022400",
                "ENSMUSG00000037868",
                "ENSMUSG00000060703",
                "ENSMUSG00000037499",
                "ENSMUSG00000035828",
                "ENSMUSG00000000247",
                "ENSMUSG00000027828",
                "ENSMUSG00000029465",
                "ENSMUSG00000021765",
                "ENSMUSG00000075254",
                "ENSMUSG00000023010",
                "ENSMUSG00000060904",
                "ENSMUSG00000025351",
                "ENSMUSG00000074884"
            ],
            "expression_sum": {
                "class1": 38873.0,
                "class2": 25496.0
            },
            "ens_id": "ENSMUSG00000038418",
            "uniprot_ids": [
                "P08046"
            ],
            "net_id": 0,
            "source_num": 13,
            "target_num": 31
        },
        "ENSMUSG00000052837": {
            "id": "ENSMUSG00000052837",
            "type": "TF",
            "symbol": "Junb",
            "source": [
                "ENSMUSG00000041515",
                "ENSMUSG00000052684",
                "ENSMUSG00000037868",
                "ENSMUSG00000023034",
                "ENSMUSG00000026628",
                "ENSMUSG00000030103",
                "ENSMUSG00000000247",
                "ENSMUSG00000038418",
                "ENSMUSG00000057098",
                "ENSMUSG00000021250",
                "ENSMUSG00000003545",
                "ENSMUSG00000003032",
                "ENSMUSG00000018899",
                "ENSMUSG00000071076"
            ],
            "target": [
                "ENSMUSG00000019505",
                "ENSMUSG00000020053",
                "ENSMUSG00000097266",
                "ENSMUSG00000023034",
                "ENSMUSG00000037999",
                "ENSMUSG00000000628",
                "ENSMUSG00000069601",
                "ENSMUSG00000024014",
                "ENSMUSG00000054072",
                "ENSMUSG00000028691",
                "ENSMUSG00000056708",
                "ENSMUSG00000053113",
                "ENSMUSG00000003545",
                "ENSMUSG00000037266",
                "ENSMUSG00000042453",
                "ENSMUSG00000033538",
                "ENSMUSG00000030930",
                "ENSMUSG00000037411",
                "ENSMUSG00000028864",
                "ENSMUSG00000052684",
                "ENSMUSG00000017493",
                "ENSMUSG00000031451",
                "ENSMUSG00000008384",
                "ENSMUSG00000019278",
                "ENSMUSG00000021250",
                "ENSMUSG00000068220",
                "ENSMUSG00000003032",
                "ENSMUSG00000055148",
                "ENSMUSG00000003038",
                "ENSMUSG00000021127",
                "ENSMUSG00000037211",
                "ENSMUSG00000041515",
                "ENSMUSG00000024190",
                "ENSMUSG00000007097",
                "ENSMUSG00000015501",
                "ENSMUSG00000040128",
                "ENSMUSG00000023915",
                "ENSMUSG00000029380",
                "ENSMUSG00000047945",
                "ENSMUSG00000026628",
                "ENSMUSG00000030103",
                "ENSMUSG00000020423",
                "ENSMUSG00000002416",
                "ENSMUSG00000035356",
                "ENSMUSG00000097296",
                "ENSMUSG00000042812",
                "ENSMUSG00000040170",
                "ENSMUSG00000032515",
                "ENSMUSG00000007872",
                "ENSMUSG00000044786",
                "ENSMUSG00000022956",
                "ENSMUSG00000008540",
                "ENSMUSG00000071076",
                "ENSMUSG00000018899",
                "ENSMUSG00000021876",
                "ENSMUSG00000027829",
                "ENSMUSG00000049115",
                "ENSMUSG00000037868",
                "ENSMUSG00000022122",
                "ENSMUSG00000006205",
                "ENSMUSG00000028214",
                "ENSMUSG00000032501",
                "ENSMUSG00000035828",
                "ENSMUSG00000000247",
                "ENSMUSG00000064215",
                "ENSMUSG00000034855",
                "ENSMUSG00000039601",
                "ENSMUSG00000015085",
                "ENSMUSG00000030849",
                "ENSMUSG00000038418",
                "ENSMUSG00000045658",
                "ENSMUSG00000090084",
                "ENSMUSG00000035678",
                "ENSMUSG00000009681",
                "ENSMUSG00000001025",
                "ENSMUSG00000058135"
            ],
            "expression_sum": {
                "class1": 117091.0,
                "class2": 64489.0
            },
            "ens_id": "ENSMUSG00000052837",
            "uniprot_ids": [
                "P09450"
            ],
            "net_id": 0,
            "source_num": 14,
            "target_num": 44
        },
        "ENSMUSG00000071076": {
            "id": "ENSMUSG00000071076",
            "type": "TF",
            "symbol": "Jund",
            "source": [
                "ENSMUSG00000056501",
                "ENSMUSG00000052837",
                "ENSMUSG00000026628",
                "ENSMUSG00000000247",
                "ENSMUSG00000008575"
            ],
            "target": [
                "ENSMUSG00000031431",
                "ENSMUSG00000040723",
                "ENSMUSG00000025381",
                "ENSMUSG00000019188",
                "ENSMUSG00000022365",
                "ENSMUSG00000024011",
                "ENSMUSG00000025213",
                "ENSMUSG00000053007",
                "ENSMUSG00000026879",
                "ENSMUSG00000016256",
                "ENSMUSG00000047187",
                "ENSMUSG00000030298",
                "ENSMUSG00000064373",
                "ENSMUSG00000056629",
                "ENSMUSG00000042613",
                "ENSMUSG00000044600",
                "ENSMUSG00000000325",
                "ENSMUSG00000067847",
                "ENSMUSG00000027239",
                "ENSMUSG00000078713",
                "ENSMUSG00000078974",
                "ENSMUSG00000018567",
                "ENSMUSG00000058672",
                "ENSMUSG00000031007",
                "ENSMUSG00000008575",
                "ENSMUSG00000053317",
                "ENSMUSG00000075701",
                "ENSMUSG00000022836",
                "ENSMUSG00000021390",
                "ENSMUSG00000020368",
                "ENSMUSG00000031825",
                "ENSMUSG00000084786",
                "ENSMUSG00000019907",
                "ENSMUSG00000031565",
                "ENSMUSG00000020737",
                "ENSMUSG00000030824",
                "ENSMUSG00000037379",
                "ENSMUSG00000020099",
                "ENSMUSG00000027706",
                "ENSMUSG00000031586",
                "ENSMUSG00000032515",
                "ENSMUSG00000017466",
                "ENSMUSG00000072235",
                "ENSMUSG00000005873",
                "ENSMUSG00000038463",
                "ENSMUSG00000028517",
                "ENSMUSG00000016494",
                "ENSMUSG00000020029",
                "ENSMUSG00000024290",
                "ENSMUSG00000038393",
                "ENSMUSG00000015094",
                "ENSMUSG00000022665",
                "ENSMUSG00000015312",
                "ENSMUSG00000028964",
                "ENSMUSG00000021871",
                "ENSMUSG00000024659",
                "ENSMUSG00000025724",
                "ENSMUSG00000062270",
                "ENSMUSG00000027712",
                "ENSMUSG00000007891",
                "ENSMUSG00000024048",
                "ENSMUSG00000028673",
                "ENSMUSG00000001891",
                "ENSMUSG00000050708",
                "ENSMUSG00000026478",
                "ENSMUSG00000037868",
                "ENSMUSG00000022122",
                "ENSMUSG00000000594",
                "ENSMUSG00000019920",
                "ENSMUSG00000057098",
                "ENSMUSG00000014846",
                "ENSMUSG00000017493",
                "ENSMUSG00000025351",
                "ENSMUSG00000074884",
                "ENSMUSG00000024014",
                "ENSMUSG00000055435",
                "ENSMUSG00000020849",
                "ENSMUSG00000034993",
                "ENSMUSG00000000753",
                "ENSMUSG00000051579",
                "ENSMUSG00000004460",
                "ENSMUSG00000028757",
                "ENSMUSG00000035356",
                "ENSMUSG00000027859",
                "ENSMUSG00000022174",
                "ENSMUSG00000029048",
                "ENSMUSG00000035227",
                "ENSMUSG00000024012",
                "ENSMUSG00000060802",
                "ENSMUSG00000014867",
                "ENSMUSG00000019505",
                "ENSMUSG00000059291",
                "ENSMUSG00000030222",
                "ENSMUSG00000028691",
                "ENSMUSG00000003545",
                "ENSMUSG00000030930",
                "ENSMUSG00000009281",
                "ENSMUSG00000028822",
                "ENSMUSG00000008384",
                "ENSMUSG00000023175",
                "ENSMUSG00000026421",
                "ENSMUSG00000031839",
                "ENSMUSG00000002107",
                "ENSMUSG00000032231",
                "ENSMUSG00000007872",
                "ENSMUSG00000027848",
                "ENSMUSG00000052419",
                "ENSMUSG00000029718",
                "ENSMUSG00000028600",
                "ENSMUSG00000025132",
                "ENSMUSG00000021917",
                "ENSMUSG00000028339",
                "ENSMUSG00000037266",
                "ENSMUSG00000001525",
                "ENSMUSG00000031538",
                "ENSMUSG00000019978",
                "ENSMUSG00000022816",
                "ENSMUSG00000007039",
                "ENSMUSG00000029781",
                "ENSMUSG00000022090",
                "ENSMUSG00000019579",
                "ENSMUSG00000068735",
                "ENSMUSG00000074457",
                "ENSMUSG00000040249",
                "ENSMUSG00000032334",
                "ENSMUSG00000026335",
                "ENSMUSG00000029070",
                "ENSMUSG00000035678",
                "ENSMUSG00000016024",
                "ENSMUSG00000030208",
                "ENSMUSG00000023034",
                "ENSMUSG00000030654",
                "ENSMUSG00000003420",
                "ENSMUSG00000020315",
                "ENSMUSG00000058569",
                "ENSMUSG00000018217",
                "ENSMUSG00000023046",
                "ENSMUSG00000058056",
                "ENSMUSG00000005973",
                "ENSMUSG00000042745",
                "ENSMUSG00000016534",
                "ENSMUSG00000003161",
                "ENSMUSG00000032359",
                "ENSMUSG00000001911",
                "ENSMUSG00000005583",
                "ENSMUSG00000022136",
                "ENSMUSG00000018819",
                "ENSMUSG00000032301",
                "ENSMUSG00000020709",
                "ENSMUSG00000044408",
                "ENSMUSG00000002059",
                "ENSMUSG00000019362",
                "ENSMUSG00000041571",
                "ENSMUSG00000037072",
                "ENSMUSG00000020009",
                "ENSMUSG00000004207",
                "ENSMUSG00000020088",
                "ENSMUSG00000026728",
                "ENSMUSG00000096145",
                "ENSMUSG00000036752",
                "ENSMUSG00000042312",
                "ENSMUSG00000035828",
                "ENSMUSG00000075254",
                "ENSMUSG00000023010",
                "ENSMUSG00000031799",
                "ENSMUSG00000041084",
                "ENSMUSG00000041126",
                "ENSMUSG00000054808",
                "ENSMUSG00000021242",
                "ENSMUSG00000002014",
                "ENSMUSG00000092341",
                "ENSMUSG00000039899",
                "ENSMUSG00000030934",
                "ENSMUSG00000011114",
                "ENSMUSG00000028864",
                "ENSMUSG00000006800",
                "ENSMUSG00000020585",
                "ENSMUSG00000015575",
                "ENSMUSG00000032353",
                "ENSMUSG00000025492",
                "ENSMUSG00000007041",
                "ENSMUSG00000055447",
                "ENSMUSG00000020458",
                "ENSMUSG00000032959",
                "ENSMUSG00000009291",
                "ENSMUSG00000037206",
                "ENSMUSG00000060703",
                "ENSMUSG00000030849",
                "ENSMUSG00000029231",
                "ENSMUSG00000021131",
                "ENSMUSG00000035258",
                "ENSMUSG00000001025",
                "ENSMUSG00000045636",
                "ENSMUSG00000048076",
                "ENSMUSG00000042712",
                "ENSMUSG00000026021",
                "ENSMUSG00000022528",
                "ENSMUSG00000006699",
                "ENSMUSG00000032575",
                "ENSMUSG00000079111",
                "ENSMUSG00000030659",
                "ENSMUSG00000012422",
                "ENSMUSG00000023004",
                "ENSMUSG00000031198",
                "ENSMUSG00000025130",
                "ENSMUSG00000003541",
                "ENSMUSG00000029103",
                "ENSMUSG00000038872",
                "ENSMUSG00000022452",
                "ENSMUSG00000056737",
                "ENSMUSG00000022032",
                "ENSMUSG00000027828",
                "ENSMUSG00000022400",
                "ENSMUSG00000055839",
                "ENSMUSG00000034165",
                "ENSMUSG00000032085",
                "ENSMUSG00000053113",
                "ENSMUSG00000052681",
                "ENSMUSG00000003380",
                "ENSMUSG00000038803",
                "ENSMUSG00000046768",
                "ENSMUSG00000026628",
                "ENSMUSG00000042737",
                "ENSMUSG00000021253",
                "ENSMUSG00000053477",
                "ENSMUSG00000079523",
                "ENSMUSG00000038991",
                "ENSMUSG00000060591",
                "ENSMUSG00000001794",
                "ENSMUSG00000028843",
                "ENSMUSG00000020423",
                "ENSMUSG00000069662",
                "ENSMUSG00000021876",
                "ENSMUSG00000025780",
                "ENSMUSG00000024164",
                "ENSMUSG00000001663",
                "ENSMUSG00000064215",
                "ENSMUSG00000006050",
                "ENSMUSG00000004897",
                "ENSMUSG00000027907",
                "ENSMUSG00000022108",
                "ENSMUSG00000000628",
                "ENSMUSG00000020219",
                "ENSMUSG00000028249",
                "ENSMUSG00000069833",
                "ENSMUSG00000033910",
                "ENSMUSG00000039168",
                "ENSMUSG00000018593",
                "ENSMUSG00000013698",
                "ENSMUSG00000001418",
                "ENSMUSG00000015568",
                "ENSMUSG00000021250",
                "ENSMUSG00000028618",
                "ENSMUSG00000054203",
                "ENSMUSG00000052837",
                "ENSMUSG00000023272",
                "ENSMUSG00000021576",
                "ENSMUSG00000079037",
                "ENSMUSG00000079017",
                "ENSMUSG00000006403",
                "ENSMUSG00000000247",
                "ENSMUSG00000019087",
                "ENSMUSG00000029465",
                "ENSMUSG00000052997",
                "ENSMUSG00000005054",
                "ENSMUSG00000058135",
                "ENSMUSG00000007815",
                "ENSMUSG00000029761",
                "ENSMUSG00000002602",
                "ENSMUSG00000028111",
                "ENSMUSG00000031875",
                "ENSMUSG00000021477",
                "ENSMUSG00000037822",
                "ENSMUSG00000027422",
                "ENSMUSG00000097328",
                "ENSMUSG00000020803",
                "ENSMUSG00000025647",
                "ENSMUSG00000044337",
                "ENSMUSG00000031556",
                "ENSMUSG00000021877",
                "ENSMUSG00000068196",
                "ENSMUSG00000034868",
                "ENSMUSG00000044627",
                "ENSMUSG00000021939",
                "ENSMUSG00000055148",
                "ENSMUSG00000001240",
                "ENSMUSG00000014329",
                "ENSMUSG00000040713",
                "ENSMUSG00000028369",
                "ENSMUSG00000022769",
                "ENSMUSG00000029309",
                "ENSMUSG00000026547",
                "ENSMUSG00000009013",
                "ENSMUSG00000027808",
                "ENSMUSG00000031595",
                "ENSMUSG00000009681",
                "ENSMUSG00000024327",
                "ENSMUSG00000060904",
                "ENSMUSG00000075706",
                "ENSMUSG00000020053",
                "ENSMUSG00000026042",
                "ENSMUSG00000037999",
                "ENSMUSG00000020440",
                "ENSMUSG00000028367",
                "ENSMUSG00000032314",
                "ENSMUSG00000032366",
                "ENSMUSG00000036372",
                "ENSMUSG00000033538",
                "ENSMUSG00000020810",
                "ENSMUSG00000025016",
                "ENSMUSG00000001119",
                "ENSMUSG00000021759",
                "ENSMUSG00000075010",
                "ENSMUSG00000029390",
                "ENSMUSG00000029767",
                "ENSMUSG00000024076",
                "ENSMUSG00000055681",
                "ENSMUSG00000090000",
                "ENSMUSG00000021760",
                "ENSMUSG00000021248",
                "ENSMUSG00000001506",
                "ENSMUSG00000003038",
                "ENSMUSG00000021870",
                "ENSMUSG00000041959",
                "ENSMUSG00000018672",
                "ENSMUSG00000028195",
                "ENSMUSG00000037373",
                "ENSMUSG00000068220",
                "ENSMUSG00000020486",
                "ENSMUSG00000040212",
                "ENSMUSG00000001123",
                "ENSMUSG00000020149",
                "ENSMUSG00000037706",
                "ENSMUSG00000025784",
                "ENSMUSG00000070394",
                "ENSMUSG00000031965",
                "ENSMUSG00000002015",
                "ENSMUSG00000028128",
                "ENSMUSG00000013622",
                "ENSMUSG00000020571",
                "ENSMUSG00000028108",
                "ENSMUSG00000002732",
                "ENSMUSG00000006205",
                "ENSMUSG00000006519",
                "ENSMUSG00000045658",
                "ENSMUSG00000029076",
                "ENSMUSG00000030717",
                "ENSMUSG00000039601",
                "ENSMUSG00000024308",
                "ENSMUSG00000056501",
                "ENSMUSG00000024038",
                "ENSMUSG00000041736",
                "ENSMUSG00000002064",
                "ENSMUSG00000020048",
                "ENSMUSG00000042747",
                "ENSMUSG00000041355",
                "ENSMUSG00000029096",
                "ENSMUSG00000026837",
                "ENSMUSG00000032383",
                "ENSMUSG00000022048",
                "ENSMUSG00000031422",
                "ENSMUSG00000029622",
                "ENSMUSG00000047945",
                "ENSMUSG00000066551",
                "ENSMUSG00000039286",
                "ENSMUSG00000030342",
                "ENSMUSG00000022708",
                "ENSMUSG00000019929",
                "ENSMUSG00000029659",
                "ENSMUSG00000071984",
                "ENSMUSG00000029641",
                "ENSMUSG00000044786",
                "ENSMUSG00000008540",
                "ENSMUSG00000018899",
                "ENSMUSG00000021268",
                "ENSMUSG00000090698",
                "ENSMUSG00000075702",
                "ENSMUSG00000014769"
            ],
            "expression_sum": {
                "class1": 45888.0,
                "class2": 59450.0
            },
            "ens_id": "ENSMUSG00000071076",
            "uniprot_ids": [
                "P15066"
            ],
            "net_id": 0,
            "source_num": 5,
            "target_num": 99
        },
        "ENSMUSG00000055148": {
            "id": "ENSMUSG00000055148",
            "type": "TF",
            "symbol": "Klf2",
            "source": [
                "ENSMUSG00000041515",
                "ENSMUSG00000056501",
                "ENSMUSG00000037868",
                "ENSMUSG00000052684",
                "ENSMUSG00000023034",
                "ENSMUSG00000052837",
                "ENSMUSG00000022346",
                "ENSMUSG00000030103",
                "ENSMUSG00000038418",
                "ENSMUSG00000021250",
                "ENSMUSG00000003032",
                "ENSMUSG00000071076"
            ],
            "target": [
                "ENSMUSG00000067818",
                "ENSMUSG00000006360",
                "ENSMUSG00000031765",
                "ENSMUSG00000000628",
                "ENSMUSG00000053113",
                "ENSMUSG00000056708",
                "ENSMUSG00000003545",
                "ENSMUSG00000053560",
                "ENSMUSG00000037411",
                "ENSMUSG00000028864",
                "ENSMUSG00000037405",
                "ENSMUSG00000017009",
                "ENSMUSG00000027074",
                "ENSMUSG00000008384",
                "ENSMUSG00000031762",
                "ENSMUSG00000007877",
                "ENSMUSG00000042622",
                "ENSMUSG00000021127",
                "ENSMUSG00000024190",
                "ENSMUSG00000037211",
                "ENSMUSG00000073411",
                "ENSMUSG00000020423",
                "ENSMUSG00000023064",
                "ENSMUSG00000022010",
                "ENSMUSG00000024371",
                "ENSMUSG00000026255",
                "ENSMUSG00000024232",
                "ENSMUSG00000044786",
                "ENSMUSG00000018899",
                "ENSMUSG00000090698",
                "ENSMUSG00000061353",
                "ENSMUSG00000057666",
                "ENSMUSG00000034855",
                "ENSMUSG00000023224",
                "ENSMUSG00000009681"
            ],
            "expression_sum": {
                "class1": 44466.0,
                "class2": 38463.0
            },
            "ens_id": "ENSMUSG00000055148",
            "uniprot_ids": [
                "Q60843"
            ],
            "net_id": 0,
            "source_num": 12,
            "target_num": 14
        },
        "ENSMUSG00000003545": {
            "id": "ENSMUSG00000003545",
            "type": "TF",
            "symbol": "Fosb",
            "source": [
                "ENSMUSG00000056501",
                "ENSMUSG00000052684",
                "ENSMUSG00000023034",
                "ENSMUSG00000037868",
                "ENSMUSG00000052837",
                "ENSMUSG00000026628",
                "ENSMUSG00000030103",
                "ENSMUSG00000042812",
                "ENSMUSG00000021250",
                "ENSMUSG00000055148",
                "ENSMUSG00000003032",
                "ENSMUSG00000018899",
                "ENSMUSG00000071076"
            ],
            "target": [
                "ENSMUSG00000023034",
                "ENSMUSG00000069601",
                "ENSMUSG00000054364",
                "ENSMUSG00000024014",
                "ENSMUSG00000028691",
                "ENSMUSG00000056708",
                "ENSMUSG00000037266",
                "ENSMUSG00000087006",
                "ENSMUSG00000053560",
                "ENSMUSG00000052684",
                "ENSMUSG00000022528",
                "ENSMUSG00000015312",
                "ENSMUSG00000008384",
                "ENSMUSG00000021250",
                "ENSMUSG00000003032",
                "ENSMUSG00000021127",
                "ENSMUSG00000024190",
                "ENSMUSG00000037211",
                "ENSMUSG00000015501",
                "ENSMUSG00000040128",
                "ENSMUSG00000047945",
                "ENSMUSG00000026628",
                "ENSMUSG00000020423",
                "ENSMUSG00000052837",
                "ENSMUSG00000032501",
                "ENSMUSG00000028967",
                "ENSMUSG00000038418",
                "ENSMUSG00000039601"
            ],
            "expression_sum": {
                "class1": 20507.0,
                "class2": 9028.0
            },
            "ens_id": "ENSMUSG00000003545",
            "uniprot_ids": [
                "P13346",
                "A0A140LIT4",
                "A0A140LHS3",
                "A0A140LJH0",
                "A0A140LIE4",
                "A0A140LHW0"
            ],
            "net_id": 0,
            "source_num": 13,
            "target_num": 18
        },
        "ENSMUSG00000026628": {
            "id": "ENSMUSG00000026628",
            "type": "TF",
            "symbol": "Atf3",
            "source": [
                "ENSMUSG00000041515",
                "ENSMUSG00000037868",
                "ENSMUSG00000052684",
                "ENSMUSG00000042406",
                "ENSMUSG00000052837",
                "ENSMUSG00000030103",
                "ENSMUSG00000038418",
                "ENSMUSG00000003545",
                "ENSMUSG00000003032",
                "ENSMUSG00000018899",
                "ENSMUSG00000071076"
            ],
            "target": [
                "ENSMUSG00000006360",
                "ENSMUSG00000031765",
                "ENSMUSG00000024014",
                "ENSMUSG00000000628",
                "ENSMUSG00000053113",
                "ENSMUSG00000056708",
                "ENSMUSG00000090841",
                "ENSMUSG00000003545",
                "ENSMUSG00000033538",
                "ENSMUSG00000001627",
                "ENSMUSG00000053560",
                "ENSMUSG00000037411",
                "ENSMUSG00000052684",
                "ENSMUSG00000037405",
                "ENSMUSG00000021025",
                "ENSMUSG00000038370",
                "ENSMUSG00000042406",
                "ENSMUSG00000036438",
                "ENSMUSG00000019960",
                "ENSMUSG00000008384",
                "ENSMUSG00000031762",
                "ENSMUSG00000003032",
                "ENSMUSG00000026864",
                "ENSMUSG00000021127",
                "ENSMUSG00000024190",
                "ENSMUSG00000037211",
                "ENSMUSG00000041515",
                "ENSMUSG00000015501",
                "ENSMUSG00000023904",
                "ENSMUSG00000029380",
                "ENSMUSG00000047945",
                "ENSMUSG00000052837",
                "ENSMUSG00000020423",
                "ENSMUSG00000030103",
                "ENSMUSG00000035356",
                "ENSMUSG00000073411",
                "ENSMUSG00000097296",
                "ENSMUSG00000032515",
                "ENSMUSG00000020205",
                "ENSMUSG00000044786",
                "ENSMUSG00000071076",
                "ENSMUSG00000018899",
                "ENSMUSG00000027829",
                "ENSMUSG00000090698",
                "ENSMUSG00000037868",
                "ENSMUSG00000006403",
                "ENSMUSG00000032501",
                "ENSMUSG00000038418",
                "ENSMUSG00000034855",
                "ENSMUSG00000035678",
                "ENSMUSG00000073489",
                "ENSMUSG00000009681",
                "ENSMUSG00000039601"
            ],
            "expression_sum": {
                "class1": 16194.0,
                "class2": 10101.0
            },
            "ens_id": "ENSMUSG00000026628",
            "uniprot_ids": [
                "Q60765"
            ],
            "net_id": 0,
            "source_num": 11,
            "target_num": 28
        },
        "ENSMUSG00000008575": {
            "id": "ENSMUSG00000008575",
            "type": "TF",
            "symbol": "Nfib",
            "source": [
                "ENSMUSG00000038193",
                "ENSMUSG00000005583",
                "ENSMUSG00000052684",
                "ENSMUSG00000071076"
            ],
            "target": [
                "ENSMUSG00000031431",
                "ENSMUSG00000024063",
                "ENSMUSG00000029761",
                "ENSMUSG00000020439",
                "ENSMUSG00000031765",
                "ENSMUSG00000021477",
                "ENSMUSG00000022365",
                "ENSMUSG00000025213",
                "ENSMUSG00000026879",
                "ENSMUSG00000028413",
                "ENSMUSG00000036438",
                "ENSMUSG00000021877",
                "ENSMUSG00000000325",
                "ENSMUSG00000030659",
                "ENSMUSG00000026421",
                "ENSMUSG00000028003",
                "ENSMUSG00000073889",
                "ENSMUSG00000030605",
                "ENSMUSG00000030218",
                "ENSMUSG00000004473",
                "ENSMUSG00000031740",
                "ENSMUSG00000031595",
                "ENSMUSG00000028600",
                "ENSMUSG00000041120",
                "ENSMUSG00000037999",
                "ENSMUSG00000069601",
                "ENSMUSG00000032085",
                "ENSMUSG00000034353",
                "ENSMUSG00000024140",
                "ENSMUSG00000017707",
                "ENSMUSG00000032366",
                "ENSMUSG00000027215",
                "ENSMUSG00000019907",
                "ENSMUSG00000052684",
                "ENSMUSG00000021759",
                "ENSMUSG00000014956",
                "ENSMUSG00000029390",
                "ENSMUSG00000031246",
                "ENSMUSG00000001506",
                "ENSMUSG00000021870",
                "ENSMUSG00000028871",
                "ENSMUSG00000002980",
                "ENSMUSG00000035885",
                "ENSMUSG00000029781",
                "ENSMUSG00000026768",
                "ENSMUSG00000027834",
                "ENSMUSG00000068735",
                "ENSMUSG00000074457",
                "ENSMUSG00000032334",
                "ENSMUSG00000026335",
                "ENSMUSG00000021253",
                "ENSMUSG00000072235",
                "ENSMUSG00000021765",
                "ENSMUSG00000038463",
                "ENSMUSG00000016024",
                "ENSMUSG00000037166",
                "ENSMUSG00000068220",
                "ENSMUSG00000030208",
                "ENSMUSG00000002289",
                "ENSMUSG00000028517",
                "ENSMUSG00000052310",
                "ENSMUSG00000016494",
                "ENSMUSG00000028465",
                "ENSMUSG00000020029",
                "ENSMUSG00000024290",
                "ENSMUSG00000042453",
                "ENSMUSG00000020315",
                "ENSMUSG00000018217",
                "ENSMUSG00000058056",
                "ENSMUSG00000032359",
                "ENSMUSG00000038991",
                "ENSMUSG00000001911",
                "ENSMUSG00000032135",
                "ENSMUSG00000019139",
                "ENSMUSG00000018819",
                "ENSMUSG00000024048",
                "ENSMUSG00000020205",
                "ENSMUSG00000020009",
                "ENSMUSG00000021876",
                "ENSMUSG00000024164",
                "ENSMUSG00000050708",
                "ENSMUSG00000025147",
                "ENSMUSG00000006356",
                "ENSMUSG00000006205",
                "ENSMUSG00000000594",
                "ENSMUSG00000019920",
                "ENSMUSG00000064215",
                "ENSMUSG00000014846",
                "ENSMUSG00000030717",
                "ENSMUSG00000017493",
                "ENSMUSG00000025393",
                "ENSMUSG00000067818",
                "ENSMUSG00000054808",
                "ENSMUSG00000021242",
                "ENSMUSG00000014313",
                "ENSMUSG00000002014",
                "ENSMUSG00000090841",
                "ENSMUSG00000092341",
                "ENSMUSG00000026688",
                "ENSMUSG00000039168",
                "ENSMUSG00000052957",
                "ENSMUSG00000031380",
                "ENSMUSG00000032353",
                "ENSMUSG00000004460",
                "ENSMUSG00000020048",
                "ENSMUSG00000042747",
                "ENSMUSG00000029096",
                "ENSMUSG00000024190",
                "ENSMUSG00000006369",
                "ENSMUSG00000028757",
                "ENSMUSG00000023272",
                "ENSMUSG00000004044",
                "ENSMUSG00000020840",
                "ENSMUSG00000015932",
                "ENSMUSG00000024012",
                "ENSMUSG00000029659",
                "ENSMUSG00000040170",
                "ENSMUSG00000008540",
                "ENSMUSG00000071076",
                "ENSMUSG00000079037",
                "ENSMUSG00000004085",
                "ENSMUSG00000060703",
                "ENSMUSG00000038418",
                "ENSMUSG00000030849",
                "ENSMUSG00000042284",
                "ENSMUSG00000001025",
                "ENSMUSG00000058135"
            ],
            "expression_sum": {
                "class1": 17518.0,
                "class2": 11173.0
            },
            "ens_id": "ENSMUSG00000008575",
            "uniprot_ids": [
                "P97863",
                "I7HIP8",
                "A2BG75",
                "A2BG76",
                "A2BG77",
                "A2ADI3"
            ],
            "net_id": 0,
            "source_num": 4,
            "target_num": 49
        },
        "ENSMUSG00000018899": {
            "id": "ENSMUSG00000018899",
            "type": "TF",
            "symbol": "Irf1",
            "source": [
                "ENSMUSG00000041515",
                "ENSMUSG00000056501",
                "ENSMUSG00000052684",
                "ENSMUSG00000037868",
                "ENSMUSG00000023034",
                "ENSMUSG00000042406",
                "ENSMUSG00000052837",
                "ENSMUSG00000026628",
                "ENSMUSG00000030103",
                "ENSMUSG00000022346",
                "ENSMUSG00000038418",
                "ENSMUSG00000021250",
                "ENSMUSG00000003032",
                "ENSMUSG00000055148",
                "ENSMUSG00000071076"
            ],
            "target": [
                "ENSMUSG00000056501",
                "ENSMUSG00000023034",
                "ENSMUSG00000031765",
                "ENSMUSG00000024014",
                "ENSMUSG00000000628",
                "ENSMUSG00000054072",
                "ENSMUSG00000053113",
                "ENSMUSG00000056708",
                "ENSMUSG00000003545",
                "ENSMUSG00000033538",
                "ENSMUSG00000001627",
                "ENSMUSG00000053560",
                "ENSMUSG00000037411",
                "ENSMUSG00000052684",
                "ENSMUSG00000037405",
                "ENSMUSG00000021025",
                "ENSMUSG00000022346",
                "ENSMUSG00000015312",
                "ENSMUSG00000008384",
                "ENSMUSG00000031762",
                "ENSMUSG00000021250",
                "ENSMUSG00000003032",
                "ENSMUSG00000035493",
                "ENSMUSG00000021127",
                "ENSMUSG00000024190",
                "ENSMUSG00000041515",
                "ENSMUSG00000015501",
                "ENSMUSG00000040128",
                "ENSMUSG00000035356",
                "ENSMUSG00000052837",
                "ENSMUSG00000026628",
                "ENSMUSG00000030103",
                "ENSMUSG00000020423",
                "ENSMUSG00000097296",
                "ENSMUSG00000032515",
                "ENSMUSG00000007872",
                "ENSMUSG00000044786",
                "ENSMUSG00000020205",
                "ENSMUSG00000027829",
                "ENSMUSG00000037868",
                "ENSMUSG00000006403",
                "ENSMUSG00000032501",
                "ENSMUSG00000035828",
                "ENSMUSG00000038418",
                "ENSMUSG00000034855",
                "ENSMUSG00000030849",
                "ENSMUSG00000035678",
                "ENSMUSG00000073489"
            ],
            "expression_sum": {
                "class1": 9123.0,
                "class2": 1422.0
            },
            "ens_id": "ENSMUSG00000018899",
            "uniprot_ids": [
                "P15314",
                "Q3U5M1",
                "A8Y5B9",
                "Q5SX15",
                "Q5SX14",
                "D6RH71",
                "Q5SX16"
            ],
            "net_id": 0,
            "source_num": 15,
            "target_num": 26
        },
        "ENSMUSG00000038193": {
            "id": "ENSMUSG00000038193",
            "type": "TF",
            "symbol": "Hand2",
            "source": [],
            "target": [
                "ENSMUSG00000019539",
                "ENSMUSG00000059291",
                "ENSMUSG00000031447",
                "ENSMUSG00000025381",
                "ENSMUSG00000058546",
                "ENSMUSG00000019916",
                "ENSMUSG00000032518",
                "ENSMUSG00000007655",
                "ENSMUSG00000021477",
                "ENSMUSG00000040952",
                "ENSMUSG00000031320",
                "ENSMUSG00000030930",
                "ENSMUSG00000009281",
                "ENSMUSG00000016256",
                "ENSMUSG00000027447",
                "ENSMUSG00000030744",
                "ENSMUSG00000053560",
                "ENSMUSG00000022371",
                "ENSMUSG00000021877",
                "ENSMUSG00000064373",
                "ENSMUSG00000034868",
                "ENSMUSG00000008668",
                "ENSMUSG00000020372",
                "ENSMUSG00000021939",
                "ENSMUSG00000041417",
                "ENSMUSG00000008682",
                "ENSMUSG00000035493",
                "ENSMUSG00000001240",
                "ENSMUSG00000026970",
                "ENSMUSG00000073889",
                "ENSMUSG00000034892",
                "ENSMUSG00000071415",
                "ENSMUSG00000008575",
                "ENSMUSG00000061477",
                "ENSMUSG00000031119",
                "ENSMUSG00000027848",
                "ENSMUSG00000050212",
                "ENSMUSG00000061983",
                "ENSMUSG00000029426",
                "ENSMUSG00000050953",
                "ENSMUSG00000029718",
                "ENSMUSG00000031740",
                "ENSMUSG00000036655",
                "ENSMUSG00000002900",
                "ENSMUSG00000048096",
                "ENSMUSG00000028081",
                "ENSMUSG00000060636",
                "ENSMUSG00000026365",
                "ENSMUSG00000036545",
                "ENSMUSG00000022312",
                "ENSMUSG00000069601",
                "ENSMUSG00000063457",
                "ENSMUSG00000037742",
                "ENSMUSG00000053113",
                "ENSMUSG00000039007",
                "ENSMUSG00000032366",
                "ENSMUSG00000029810",
                "ENSMUSG00000020810",
                "ENSMUSG00000052684",
                "ENSMUSG00000070436",
                "ENSMUSG00000001119",
                "ENSMUSG00000036446",
                "ENSMUSG00000012405",
                "ENSMUSG00000030824",
                "ENSMUSG00000027642",
                "ENSMUSG00000003970",
                "ENSMUSG00000026482",
                "ENSMUSG00000002980",
                "ENSMUSG00000020460",
                "ENSMUSG00000026814",
                "ENSMUSG00000030249",
                "ENSMUSG00000056091",
                "ENSMUSG00000029781",
                "ENSMUSG00000025508",
                "ENSMUSG00000025290",
                "ENSMUSG00000028464",
                "ENSMUSG00000032014",
                "ENSMUSG00000033880",
                "ENSMUSG00000022587",
                "ENSMUSG00000025794",
                "ENSMUSG00000040249",
                "ENSMUSG00000037060",
                "ENSMUSG00000023367",
                "ENSMUSG00000029070",
                "ENSMUSG00000024608",
                "ENSMUSG00000001687",
                "ENSMUSG00000053477",
                "ENSMUSG00000039001",
                "ENSMUSG00000006333",
                "ENSMUSG00000002985",
                "ENSMUSG00000058600",
                "ENSMUSG00000062647",
                "ENSMUSG00000022894",
                "ENSMUSG00000046330",
                "ENSMUSG00000074129",
                "ENSMUSG00000090733",
                "ENSMUSG00000020432",
                "ENSMUSG00000002778",
                "ENSMUSG00000079523",
                "ENSMUSG00000003420",
                "ENSMUSG00000060126",
                "ENSMUSG00000042453",
                "ENSMUSG00000058569",
                "ENSMUSG00000026014",
                "ENSMUSG00000046727",
                "ENSMUSG00000005973",
                "ENSMUSG00000012848",
                "ENSMUSG00000016534",
                "ENSMUSG00000060591",
                "ENSMUSG00000023942",
                "ENSMUSG00000049517",
                "ENSMUSG00000019960",
                "ENSMUSG00000022665",
                "ENSMUSG00000027074",
                "ENSMUSG00000029614",
                "ENSMUSG00000022136",
                "ENSMUSG00000003814",
                "ENSMUSG00000062006",
                "ENSMUSG00000061232",
                "ENSMUSG00000032216",
                "ENSMUSG00000079435",
                "ENSMUSG00000047215",
                "ENSMUSG00000094724",
                "ENSMUSG00000027712",
                "ENSMUSG00000024371",
                "ENSMUSG00000069662",
                "ENSMUSG00000032666",
                "ENSMUSG00000028108",
                "ENSMUSG00000004207",
                "ENSMUSG00000031029",
                "ENSMUSG00000028673",
                "ENSMUSG00000024232",
                "ENSMUSG00000002732",
                "ENSMUSG00000034708",
                "ENSMUSG00000050708",
                "ENSMUSG00000019970",
                "ENSMUSG00000022122",
                "ENSMUSG00000062997",
                "ENSMUSG00000028214",
                "ENSMUSG00000028173",
                "ENSMUSG00000058558",
                "ENSMUSG00000067288",
                "ENSMUSG00000029076",
                "ENSMUSG00000023224",
                "ENSMUSG00000026223",
                "ENSMUSG00000049086",
                "ENSMUSG00000022971",
                "ENSMUSG00000025351",
                "ENSMUSG00000060938",
                "ENSMUSG00000041841",
                "ENSMUSG00000052146",
                "ENSMUSG00000039601",
                "ENSMUSG00000067818",
                "ENSMUSG00000067274",
                "ENSMUSG00000057863",
                "ENSMUSG00000026566",
                "ENSMUSG00000022108",
                "ENSMUSG00000024538",
                "ENSMUSG00000032547",
                "ENSMUSG00000024248",
                "ENSMUSG00000020484",
                "ENSMUSG00000060550",
                "ENSMUSG00000027248",
                "ENSMUSG00000005881",
                "ENSMUSG00000062328",
                "ENSMUSG00000046491",
                "ENSMUSG00000020585",
                "ENSMUSG00000004105",
                "ENSMUSG00000032353",
                "ENSMUSG00000019055",
                "ENSMUSG00000008683",
                "ENSMUSG00000043716",
                "ENSMUSG00000055447",
                "ENSMUSG00000000740",
                "ENSMUSG00000021186",
                "ENSMUSG00000028618",
                "ENSMUSG00000057841",
                "ENSMUSG00000079641",
                "ENSMUSG00000022048",
                "ENSMUSG00000032383",
                "ENSMUSG00000025993",
                "ENSMUSG00000047907",
                "ENSMUSG00000048758",
                "ENSMUSG00000022708",
                "ENSMUSG00000028234",
                "ENSMUSG00000037206",
                "ENSMUSG00000019929",
                "ENSMUSG00000009876",
                "ENSMUSG00000028936",
                "ENSMUSG00000028495",
                "ENSMUSG00000018830",
                "ENSMUSG00000027004",
                "ENSMUSG00000003429",
                "ENSMUSG00000000247",
                "ENSMUSG00000090137",
                "ENSMUSG00000035783",
                "ENSMUSG00000030849",
                "ENSMUSG00000029231",
                "ENSMUSG00000020902",
                "ENSMUSG00000037563",
                "ENSMUSG00000032399",
                "ENSMUSG00000031253"
            ],
            "expression_sum": {
                "class1": 22064.0,
                "class2": 3667.0
            },
            "ens_id": "ENSMUSG00000038193",
            "uniprot_ids": [
                "Q61039"
            ],
            "net_id": 0,
            "source_num": 0,
            "target_num": 54
        },
        "ENSMUSG00000020484": {
            "id": "ENSMUSG00000020484",
            "type": "TF",
            "symbol": "Xbp1",
            "source": [
                "ENSMUSG00000038193",
                "ENSMUSG00000023034",
                "ENSMUSG00000005583"
            ],
            "target": [
                "ENSMUSG00000025393",
                "ENSMUSG00000031431",
                "ENSMUSG00000020053",
                "ENSMUSG00000024063",
                "ENSMUSG00000054808",
                "ENSMUSG00000064370",
                "ENSMUSG00000028339",
                "ENSMUSG00000069601",
                "ENSMUSG00000021242",
                "ENSMUSG00000002014",
                "ENSMUSG00000062210",
                "ENSMUSG00000059895",
                "ENSMUSG00000090841",
                "ENSMUSG00000045636",
                "ENSMUSG00000017707",
                "ENSMUSG00000032366",
                "ENSMUSG00000042453",
                "ENSMUSG00000025213",
                "ENSMUSG00000058056",
                "ENSMUSG00000027215",
                "ENSMUSG00000016319",
                "ENSMUSG00000038991",
                "ENSMUSG00000006800",
                "ENSMUSG00000031565",
                "ENSMUSG00000031556",
                "ENSMUSG00000022528",
                "ENSMUSG00000036438",
                "ENSMUSG00000042613",
                "ENSMUSG00000014956",
                "ENSMUSG00000032353",
                "ENSMUSG00000074227",
                "ENSMUSG00000003477",
                "ENSMUSG00000025809",
                "ENSMUSG00000030659",
                "ENSMUSG00000021870",
                "ENSMUSG00000064354",
                "ENSMUSG00000026421",
                "ENSMUSG00000035885",
                "ENSMUSG00000030403",
                "ENSMUSG00000023272",
                "ENSMUSG00000028128",
                "ENSMUSG00000020423",
                "ENSMUSG00000037712",
                "ENSMUSG00000022010",
                "ENSMUSG00000015932",
                "ENSMUSG00000028464",
                "ENSMUSG00000020612",
                "ENSMUSG00000024012",
                "ENSMUSG00000040170",
                "ENSMUSG00000008540",
                "ENSMUSG00000006342",
                "ENSMUSG00000074457",
                "ENSMUSG00000021876",
                "ENSMUSG00000021268",
                "ENSMUSG00000004085",
                "ENSMUSG00000033306",
                "ENSMUSG00000096145",
                "ENSMUSG00000025147",
                "ENSMUSG00000024164",
                "ENSMUSG00000006205",
                "ENSMUSG00000057666",
                "ENSMUSG00000064215",
                "ENSMUSG00000014846",
                "ENSMUSG00000016024",
                "ENSMUSG00000017493",
                "ENSMUSG00000041084",
                "ENSMUSG00000025351"
            ],
            "expression_sum": {
                "class1": 12449.0,
                "class2": 6214.0
            },
            "ens_id": "ENSMUSG00000020484",
            "uniprot_ids": [
                "O35426",
                "G3UYH6",
                "H3BLF6",
                "A0A5F8MPD3"
            ],
            "net_id": 0,
            "source_num": 3,
            "target_num": 24
        },
        "ENSMUSG00000003032": {
            "id": "ENSMUSG00000003032",
            "type": "TF",
            "symbol": "Klf4",
            "source": [
                "ENSMUSG00000037868",
                "ENSMUSG00000023034",
                "ENSMUSG00000052837",
                "ENSMUSG00000026628",
                "ENSMUSG00000057098",
                "ENSMUSG00000038418",
                "ENSMUSG00000021250",
                "ENSMUSG00000003545",
                "ENSMUSG00000018899"
            ],
            "target": [
                "ENSMUSG00000030116",
                "ENSMUSG00000007815",
                "ENSMUSG00000005338",
                "ENSMUSG00000019505",
                "ENSMUSG00000059291",
                "ENSMUSG00000031447",
                "ENSMUSG00000002602",
                "ENSMUSG00000031765",
                "ENSMUSG00000028111",
                "ENSMUSG00000031875",
                "ENSMUSG00000007655",
                "ENSMUSG00000021477",
                "ENSMUSG00000028691",
                "ENSMUSG00000003545",
                "ENSMUSG00000024011",
                "ENSMUSG00000053007",
                "ENSMUSG00000026879",
                "ENSMUSG00000037822",
                "ENSMUSG00000048076",
                "ENSMUSG00000056481",
                "ENSMUSG00000037411",
                "ENSMUSG00000022371",
                "ENSMUSG00000044337",
                "ENSMUSG00000064373",
                "ENSMUSG00000028822",
                "ENSMUSG00000015354",
                "ENSMUSG00000034868",
                "ENSMUSG00000006699",
                "ENSMUSG00000044600",
                "ENSMUSG00000029649",
                "ENSMUSG00000008384",
                "ENSMUSG00000055148",
                "ENSMUSG00000001240",
                "ENSMUSG00000012422",
                "ENSMUSG00000023004",
                "ENSMUSG00000039405",
                "ENSMUSG00000002107",
                "ENSMUSG00000069763",
                "ENSMUSG00000003541",
                "ENSMUSG00000060572",
                "ENSMUSG00000032231",
                "ENSMUSG00000007872",
                "ENSMUSG00000056737",
                "ENSMUSG00000020467",
                "ENSMUSG00000035212",
                "ENSMUSG00000022032",
                "ENSMUSG00000037499",
                "ENSMUSG00000021390",
                "ENSMUSG00000009013",
                "ENSMUSG00000009681",
                "ENSMUSG00000018585",
                "ENSMUSG00000020053",
                "ENSMUSG00000025132",
                "ENSMUSG00000020440",
                "ENSMUSG00000028367",
                "ENSMUSG00000053113",
                "ENSMUSG00000037266",
                "ENSMUSG00000084786",
                "ENSMUSG00000031626",
                "ENSMUSG00000036446",
                "ENSMUSG00000020737",
                "ENSMUSG00000029767",
                "ENSMUSG00000055681",
                "ENSMUSG00000031538",
                "ENSMUSG00000019978",
                "ENSMUSG00000019278",
                "ENSMUSG00000021760",
                "ENSMUSG00000090000",
                "ENSMUSG00000036478",
                "ENSMUSG00000022816",
                "ENSMUSG00000027204",
                "ENSMUSG00000041959",
                "ENSMUSG00000033585",
                "ENSMUSG00000026628",
                "ENSMUSG00000007039",
                "ENSMUSG00000027706",
                "ENSMUSG00000074743",
                "ENSMUSG00000022090",
                "ENSMUSG00000041828",
                "ENSMUSG00000032014",
                "ENSMUSG00000032515",
                "ENSMUSG00000092274",
                "ENSMUSG00000017466",
                "ENSMUSG00000074457",
                "ENSMUSG00000027829",
                "ENSMUSG00000032334",
                "ENSMUSG00000023092",
                "ENSMUSG00000072235",
                "ENSMUSG00000038463",
                "ENSMUSG00000005397",
                "ENSMUSG00000022894",
                "ENSMUSG00000015085",
                "ENSMUSG00000030208",
                "ENSMUSG00000054452",
                "ENSMUSG00000068220",
                "ENSMUSG00000028517",
                "ENSMUSG00000040212",
                "ENSMUSG00000023034",
                "ENSMUSG00000079523",
                "ENSMUSG00000030654",
                "ENSMUSG00000016494",
                "ENSMUSG00000001123",
                "ENSMUSG00000026674",
                "ENSMUSG00000020315",
                "ENSMUSG00000037706",
                "ENSMUSG00000018217",
                "ENSMUSG00000017002",
                "ENSMUSG00000003161",
                "ENSMUSG00000060591",
                "ENSMUSG00000025784",
                "ENSMUSG00000040562",
                "ENSMUSG00000072812",
                "ENSMUSG00000015312",
                "ENSMUSG00000040280",
                "ENSMUSG00000024659",
                "ENSMUSG00000018819",
                "ENSMUSG00000032301",
                "ENSMUSG00000020423",
                "ENSMUSG00000027712",
                "ENSMUSG00000024048",
                "ENSMUSG00000069662",
                "ENSMUSG00000028108",
                "ENSMUSG00000037072",
                "ENSMUSG00000002732",
                "ENSMUSG00000021876",
                "ENSMUSG00000001891",
                "ENSMUSG00000026478",
                "ENSMUSG00000037868",
                "ENSMUSG00000042312",
                "ENSMUSG00000057098",
                "ENSMUSG00000031722",
                "ENSMUSG00000075254",
                "ENSMUSG00000023010",
                "ENSMUSG00000045658",
                "ENSMUSG00000031799",
                "ENSMUSG00000014846",
                "ENSMUSG00000017493",
                "ENSMUSG00000041084",
                "ENSMUSG00000039601",
                "ENSMUSG00000027907",
                "ENSMUSG00000074884",
                "ENSMUSG00000024308",
                "ENSMUSG00000024014",
                "ENSMUSG00000000628",
                "ENSMUSG00000028249",
                "ENSMUSG00000095385",
                "ENSMUSG00000020849",
                "ENSMUSG00000069833",
                "ENSMUSG00000041736",
                "ENSMUSG00000087006",
                "ENSMUSG00000034993",
                "ENSMUSG00000026670",
                "ENSMUSG00000075602",
                "ENSMUSG00000020585",
                "ENSMUSG00000025492",
                "ENSMUSG00000007041",
                "ENSMUSG00000021250",
                "ENSMUSG00000042747",
                "ENSMUSG00000029096",
                "ENSMUSG00000026837",
                "ENSMUSG00000028618",
                "ENSMUSG00000032383",
                "ENSMUSG00000026748",
                "ENSMUSG00000031422",
                "ENSMUSG00000047907",
                "ENSMUSG00000047945",
                "ENSMUSG00000052837",
                "ENSMUSG00000035356",
                "ENSMUSG00000037206",
                "ENSMUSG00000024012",
                "ENSMUSG00000029659",
                "ENSMUSG00000060802",
                "ENSMUSG00000026574",
                "ENSMUSG00000044786",
                "ENSMUSG00000018899",
                "ENSMUSG00000079037",
                "ENSMUSG00000090698",
                "ENSMUSG00000075702",
                "ENSMUSG00000060703",
                "ENSMUSG00000019087",
                "ENSMUSG00000022261",
                "ENSMUSG00000038418",
                "ENSMUSG00000029484",
                "ENSMUSG00000040026",
                "ENSMUSG00000001025",
                "ENSMUSG00000058135"
            ],
            "expression_sum": {
                "class1": 8719.0,
                "class2": 10220.0
            },
            "ens_id": "ENSMUSG00000003032",
            "uniprot_ids": [
                "Q60793",
                "B7ZCH2",
                "B7ZCH1"
            ],
            "net_id": 0,
            "source_num": 9,
            "target_num": 74
        },
        "ENSMUSG00000023034": {
            "id": "ENSMUSG00000023034",
            "type": "TF",
            "symbol": "Nr4a1",
            "source": [
                "ENSMUSG00000041515",
                "ENSMUSG00000037868",
                "ENSMUSG00000052684",
                "ENSMUSG00000052837",
                "ENSMUSG00000030103",
                "ENSMUSG00000038418",
                "ENSMUSG00000021250",
                "ENSMUSG00000003545",
                "ENSMUSG00000003032",
                "ENSMUSG00000018899",
                "ENSMUSG00000071076"
            ],
            "target": [
                "ENSMUSG00000006360",
                "ENSMUSG00000029761",
                "ENSMUSG00000031765",
                "ENSMUSG00000028949",
                "ENSMUSG00000003545",
                "ENSMUSG00000022283",
                "ENSMUSG00000036854",
                "ENSMUSG00000040952",
                "ENSMUSG00000026879",
                "ENSMUSG00000031320",
                "ENSMUSG00000032330",
                "ENSMUSG00000037405",
                "ENSMUSG00000036438",
                "ENSMUSG00000008384",
                "ENSMUSG00000008668",
                "ENSMUSG00000020372",
                "ENSMUSG00000021939",
                "ENSMUSG00000008682",
                "ENSMUSG00000055148",
                "ENSMUSG00000035493",
                "ENSMUSG00000021127",
                "ENSMUSG00000097296",
                "ENSMUSG00000034892",
                "ENSMUSG00000071415",
                "ENSMUSG00000041329",
                "ENSMUSG00000061983",
                "ENSMUSG00000061353",
                "ENSMUSG00000031740",
                "ENSMUSG00000048096",
                "ENSMUSG00000036545",
                "ENSMUSG00000040666",
                "ENSMUSG00000032085",
                "ENSMUSG00000037742",
                "ENSMUSG00000053113",
                "ENSMUSG00000032366",
                "ENSMUSG00000021520",
                "ENSMUSG00000033538",
                "ENSMUSG00000018171",
                "ENSMUSG00000020810",
                "ENSMUSG00000052684",
                "ENSMUSG00000001119",
                "ENSMUSG00000014956",
                "ENSMUSG00000031762",
                "ENSMUSG00000032744",
                "ENSMUSG00000027188",
                "ENSMUSG00000002980",
                "ENSMUSG00000030103",
                "ENSMUSG00000025290",
                "ENSMUSG00000028464",
                "ENSMUSG00000038146",
                "ENSMUSG00000032515",
                "ENSMUSG00000068735",
                "ENSMUSG00000028776",
                "ENSMUSG00000045128",
                "ENSMUSG00000022587",
                "ENSMUSG00000027829",
                "ENSMUSG00000002985",
                "ENSMUSG00000031328",
                "ENSMUSG00000030432",
                "ENSMUSG00000074129",
                "ENSMUSG00000090733",
                "ENSMUSG00000022817",
                "ENSMUSG00000059895",
                "ENSMUSG00000056708",
                "ENSMUSG00000060126",
                "ENSMUSG00000046727",
                "ENSMUSG00000001175",
                "ENSMUSG00000001520",
                "ENSMUSG00000012848",
                "ENSMUSG00000032387",
                "ENSMUSG00000021025",
                "ENSMUSG00000049517",
                "ENSMUSG00000027074",
                "ENSMUSG00000029614",
                "ENSMUSG00000003032",
                "ENSMUSG00000062006",
                "ENSMUSG00000040128",
                "ENSMUSG00000079435",
                "ENSMUSG00000078202",
                "ENSMUSG00000020205",
                "ENSMUSG00000024232",
                "ENSMUSG00000031029",
                "ENSMUSG00000023089",
                "ENSMUSG00000001270",
                "ENSMUSG00000050708",
                "ENSMUSG00000037868",
                "ENSMUSG00000028214",
                "ENSMUSG00000035828",
                "ENSMUSG00000067288",
                "ENSMUSG00000023224",
                "ENSMUSG00000052146",
                "ENSMUSG00000039601",
                "ENSMUSG00000060938",
                "ENSMUSG00000067818",
                "ENSMUSG00000067274",
                "ENSMUSG00000024014",
                "ENSMUSG00000000628",
                "ENSMUSG00000074170",
                "ENSMUSG00000090841",
                "ENSMUSG00000028266",
                "ENSMUSG00000020484",
                "ENSMUSG00000071866",
                "ENSMUSG00000028864",
                "ENSMUSG00000008683",
                "ENSMUSG00000021250",
                "ENSMUSG00000007877",
                "ENSMUSG00000000740",
                "ENSMUSG00000007892",
                "ENSMUSG00000021186",
                "ENSMUSG00000024190",
                "ENSMUSG00000057841",
                "ENSMUSG00000041515",
                "ENSMUSG00000035356",
                "ENSMUSG00000052837",
                "ENSMUSG00000047945",
                "ENSMUSG00000022708",
                "ENSMUSG00000028234",
                "ENSMUSG00000015932",
                "ENSMUSG00000028936",
                "ENSMUSG00000074578",
                "ENSMUSG00000018899",
                "ENSMUSG00000000247",
                "ENSMUSG00000038418",
                "ENSMUSG00000035783",
                "ENSMUSG00000037563"
            ],
            "expression_sum": {
                "class1": 11016.0,
                "class2": 10485.0
            },
            "ens_id": "ENSMUSG00000023034",
            "uniprot_ids": [
                "P12813",
                "A0A2R8VJZ1"
            ],
            "net_id": 0,
            "source_num": 11,
            "target_num": 47
        },
        "ENSMUSG00000056501": {
            "id": "ENSMUSG00000056501",
            "type": "TF",
            "symbol": "Cebpb",
            "source": [
                "ENSMUSG00000041515",
                "ENSMUSG00000052684",
                "ENSMUSG00000030103",
                "ENSMUSG00000021250",
                "ENSMUSG00000071076",
                "ENSMUSG00000018899"
            ],
            "target": [
                "ENSMUSG00000031765",
                "ENSMUSG00000003545",
                "ENSMUSG00000031320",
                "ENSMUSG00000030930",
                "ENSMUSG00000037411",
                "ENSMUSG00000037405",
                "ENSMUSG00000028268",
                "ENSMUSG00000008384",
                "ENSMUSG00000055148",
                "ENSMUSG00000035493",
                "ENSMUSG00000021127",
                "ENSMUSG00000028003",
                "ENSMUSG00000029380",
                "ENSMUSG00000031119",
                "ENSMUSG00000007872",
                "ENSMUSG00000049115",
                "ENSMUSG00000031845",
                "ENSMUSG00000061983",
                "ENSMUSG00000050953",
                "ENSMUSG00000061353",
                "ENSMUSG00000036655",
                "ENSMUSG00000009681",
                "ENSMUSG00000037999",
                "ENSMUSG00000037742",
                "ENSMUSG00000053113",
                "ENSMUSG00000033538",
                "ENSMUSG00000052684",
                "ENSMUSG00000070436",
                "ENSMUSG00000031762",
                "ENSMUSG00000027188",
                "ENSMUSG00000015501",
                "ENSMUSG00000030103",
                "ENSMUSG00000029163",
                "ENSMUSG00000022587",
                "ENSMUSG00000032515",
                "ENSMUSG00000045128",
                "ENSMUSG00000033880",
                "ENSMUSG00000023944",
                "ENSMUSG00000027829",
                "ENSMUSG00000031375",
                "ENSMUSG00000039377",
                "ENSMUSG00000034855",
                "ENSMUSG00000035678",
                "ENSMUSG00000002985",
                "ENSMUSG00000072625",
                "ENSMUSG00000054072",
                "ENSMUSG00000056708",
                "ENSMUSG00000060126",
                "ENSMUSG00000042453",
                "ENSMUSG00000021025",
                "ENSMUSG00000022665",
                "ENSMUSG00000025498",
                "ENSMUSG00000040128",
                "ENSMUSG00000035373",
                "ENSMUSG00000020423",
                "ENSMUSG00000024371",
                "ENSMUSG00000020205",
                "ENSMUSG00000037868",
                "ENSMUSG00000028214",
                "ENSMUSG00000023224",
                "ENSMUSG00000039601",
                "ENSMUSG00000024014",
                "ENSMUSG00000000628",
                "ENSMUSG00000060550",
                "ENSMUSG00000028864",
                "ENSMUSG00000062328",
                "ENSMUSG00000032179",
                "ENSMUSG00000021250",
                "ENSMUSG00000007892",
                "ENSMUSG00000035385",
                "ENSMUSG00000041515",
                "ENSMUSG00000035356",
                "ENSMUSG00000047945",
                "ENSMUSG00000073411",
                "ENSMUSG00000019929",
                "ENSMUSG00000044786",
                "ENSMUSG00000071076",
                "ENSMUSG00000018899",
                "ENSMUSG00000027004",
                "ENSMUSG00000006403",
                "ENSMUSG00000032501",
                "ENSMUSG00000000247",
                "ENSMUSG00000030849",
                "ENSMUSG00000073489"
            ],
            "expression_sum": {
                "class1": 56111.0,
                "class2": 21063.0
            },
            "ens_id": "ENSMUSG00000056501",
            "uniprot_ids": [
                "P28033"
            ],
            "net_id": 0,
            "source_num": 6,
            "target_num": 29
        },
        "ENSMUSG00000055435": {
            "id": "ENSMUSG00000055435",
            "type": "TF",
            "symbol": "Maf",
            "source": [
                "ENSMUSG00000005583",
                "ENSMUSG00000052684",
                "ENSMUSG00000071076"
            ],
            "target": [
                "ENSMUSG00000024308",
                "ENSMUSG00000024063",
                "ENSMUSG00000034165",
                "ENSMUSG00000079523",
                "ENSMUSG00000021477",
                "ENSMUSG00000053113",
                "ENSMUSG00000056708",
                "ENSMUSG00000028249",
                "ENSMUSG00000055172",
                "ENSMUSG00000019907",
                "ENSMUSG00000052684",
                "ENSMUSG00000021877",
                "ENSMUSG00000075010",
                "ENSMUSG00000020737",
                "ENSMUSG00000029767",
                "ENSMUSG00000026249",
                "ENSMUSG00000055447",
                "ENSMUSG00000020048",
                "ENSMUSG00000005583",
                "ENSMUSG00000041959",
                "ENSMUSG00000022010",
                "ENSMUSG00000074743",
                "ENSMUSG00000032231",
                "ENSMUSG00000024048",
                "ENSMUSG00000021876",
                "ENSMUSG00000025780",
                "ENSMUSG00000026478",
                "ENSMUSG00000079017",
                "ENSMUSG00000001847",
                "ENSMUSG00000026536",
                "ENSMUSG00000064215",
                "ENSMUSG00000062825",
                "ENSMUSG00000025283",
                "ENSMUSG00000073489",
                "ENSMUSG00000072235",
                "ENSMUSG00000075254",
                "ENSMUSG00000031799",
                "ENSMUSG00000058135"
            ],
            "expression_sum": {
                "class1": 5180.0,
                "class2": 2486.0
            },
            "ens_id": "ENSMUSG00000055435",
            "uniprot_ids": [
                "P54843"
            ],
            "net_id": 0,
            "source_num": 3,
            "target_num": 14
        },
        "ENSMUSG00000022528": {
            "id": "ENSMUSG00000022528",
            "type": "TF",
            "symbol": "Hes1",
            "source": [
                "ENSMUSG00000052684",
                "ENSMUSG00000047638",
                "ENSMUSG00000038418",
                "ENSMUSG00000003545",
                "ENSMUSG00000005583",
                "ENSMUSG00000020484",
                "ENSMUSG00000071076"
            ],
            "target": [
                "ENSMUSG00000026365",
                "ENSMUSG00000036545",
                "ENSMUSG00000031447",
                "ENSMUSG00000031765",
                "ENSMUSG00000079523",
                "ENSMUSG00000032518",
                "ENSMUSG00000028949",
                "ENSMUSG00000037742",
                "ENSMUSG00000053113",
                "ENSMUSG00000060126",
                "ENSMUSG00000032366",
                "ENSMUSG00000059895",
                "ENSMUSG00000022892",
                "ENSMUSG00000040952",
                "ENSMUSG00000026879",
                "ENSMUSG00000046727",
                "ENSMUSG00000021756",
                "ENSMUSG00000027248",
                "ENSMUSG00000052684",
                "ENSMUSG00000012405",
                "ENSMUSG00000021877",
                "ENSMUSG00000014956",
                "ENSMUSG00000060036",
                "ENSMUSG00000090862",
                "ENSMUSG00000031762",
                "ENSMUSG00000043716",
                "ENSMUSG00000055447",
                "ENSMUSG00000020048",
                "ENSMUSG00000005583",
                "ENSMUSG00000021939",
                "ENSMUSG00000027642",
                "ENSMUSG00000022136",
                "ENSMUSG00000021127",
                "ENSMUSG00000024190",
                "ENSMUSG00000032383",
                "ENSMUSG00000032216",
                "ENSMUSG00000022708",
                "ENSMUSG00000029781",
                "ENSMUSG00000028234",
                "ENSMUSG00000025508",
                "ENSMUSG00000024371",
                "ENSMUSG00000028936",
                "ENSMUSG00000020205",
                "ENSMUSG00000028495",
                "ENSMUSG00000017466",
                "ENSMUSG00000040249",
                "ENSMUSG00000019970",
                "ENSMUSG00000061983",
                "ENSMUSG00000037805",
                "ENSMUSG00000039377",
                "ENSMUSG00000038418",
                "ENSMUSG00000039001",
                "ENSMUSG00000020902",
                "ENSMUSG00000029076",
                "ENSMUSG00000062647",
                "ENSMUSG00000022971"
            ],
            "expression_sum": {
                "class1": 22719.0,
                "class2": 11710.0
            },
            "ens_id": "ENSMUSG00000022528",
            "uniprot_ids": [
                "P35428"
            ],
            "net_id": 0,
            "source_num": 6,
            "target_num": 17
        },
        "ENSMUSG00000052534": {
            "id": "ENSMUSG00000052534",
            "type": "TF",
            "symbol": "Pbx1",
            "source": [
                "ENSMUSG00000052684"
            ],
            "target": [
                "ENSMUSG00000024063",
                "ENSMUSG00000029309",
                "ENSMUSG00000052684",
                "ENSMUSG00000026335",
                "ENSMUSG00000025784",
                "ENSMUSG00000026566",
                "ENSMUSG00000072235",
                "ENSMUSG00000024048",
                "ENSMUSG00000035258",
                "ENSMUSG00000016194",
                "ENSMUSG00000022636",
                "ENSMUSG00000039601"
            ],
            "expression_sum": {
                "class1": 9798.0,
                "class2": 8386.0
            },
            "ens_id": "ENSMUSG00000052534",
            "uniprot_ids": [
                "P41778",
                "D9J2V6"
            ],
            "net_id": 0,
            "source_num": 1,
            "target_num": 6
        },
        "ENSMUSG00000041515": {
            "id": "ENSMUSG00000041515",
            "type": "TF",
            "symbol": "Irf8",
            "source": [
                "ENSMUSG00000056501",
                "ENSMUSG00000023034",
                "ENSMUSG00000052837",
                "ENSMUSG00000026628",
                "ENSMUSG00000021250",
                "ENSMUSG00000018899"
            ],
            "target": [
                "ENSMUSG00000056501",
                "ENSMUSG00000023034",
                "ENSMUSG00000032515",
                "ENSMUSG00000052837",
                "ENSMUSG00000026628",
                "ENSMUSG00000020423",
                "ENSMUSG00000028967",
                "ENSMUSG00000044786",
                "ENSMUSG00000035356",
                "ENSMUSG00000008384",
                "ENSMUSG00000053113",
                "ENSMUSG00000021250",
                "ENSMUSG00000055148",
                "ENSMUSG00000007872",
                "ENSMUSG00000087006",
                "ENSMUSG00000016194",
                "ENSMUSG00000018899"
            ],
            "expression_sum": {
                "class1": 4392.0,
                "class2": 265.0
            },
            "ens_id": "ENSMUSG00000041515",
            "uniprot_ids": [
                "P23611",
                "F7C1A3",
                "E0CZE4"
            ],
            "net_id": 0,
            "source_num": 6,
            "target_num": 14
        },
        "ENSMUSG00000030103": {
            "id": "ENSMUSG00000030103",
            "type": "TF",
            "symbol": "Bhlhe40",
            "source": [
                "ENSMUSG00000056501",
                "ENSMUSG00000052684",
                "ENSMUSG00000023034",
                "ENSMUSG00000052837",
                "ENSMUSG00000026628",
                "ENSMUSG00000021250",
                "ENSMUSG00000018899"
            ],
            "target": [
                "ENSMUSG00000056501",
                "ENSMUSG00000023034",
                "ENSMUSG00000053113",
                "ENSMUSG00000003545",
                "ENSMUSG00000020644",
                "ENSMUSG00000015312",
                "ENSMUSG00000021250",
                "ENSMUSG00000055148",
                "ENSMUSG00000040128",
                "ENSMUSG00000035356",
                "ENSMUSG00000047945",
                "ENSMUSG00000026628",
                "ENSMUSG00000020423",
                "ENSMUSG00000052837",
                "ENSMUSG00000032515",
                "ENSMUSG00000007872",
                "ENSMUSG00000044786",
                "ENSMUSG00000018899",
                "ENSMUSG00000090698",
                "ENSMUSG00000028967",
                "ENSMUSG00000035828",
                "ENSMUSG00000009681"
            ],
            "expression_sum": {
                "class1": 3879.0,
                "class2": 961.0
            },
            "ens_id": "ENSMUSG00000030103",
            "uniprot_ids": [
                "O35185",
                "E9Q2C3"
            ],
            "net_id": 0,
            "source_num": 7,
            "target_num": 19
        },
        "ENSMUSG00000022346": {
            "id": "ENSMUSG00000022346",
            "type": "TF",
            "symbol": "Myc",
            "source": [
                "ENSMUSG00000048387",
                "ENSMUSG00000018899"
            ],
            "target": [
                "ENSMUSG00000020053",
                "ENSMUSG00000054808",
                "ENSMUSG00000029761",
                "ENSMUSG00000016494",
                "ENSMUSG00000025408",
                "ENSMUSG00000032366",
                "ENSMUSG00000033910",
                "ENSMUSG00000026879",
                "ENSMUSG00000087006",
                "ENSMUSG00000001520",
                "ENSMUSG00000032359",
                "ENSMUSG00000019907",
                "ENSMUSG00000025784",
                "ENSMUSG00000064373",
                "ENSMUSG00000031451",
                "ENSMUSG00000015354",
                "ENSMUSG00000024076",
                "ENSMUSG00000019278",
                "ENSMUSG00000003477",
                "ENSMUSG00000055148",
                "ENSMUSG00000029096",
                "ENSMUSG00000018339",
                "ENSMUSG00000054203",
                "ENSMUSG00000026421",
                "ENSMUSG00000006369",
                "ENSMUSG00000031239",
                "ENSMUSG00000002107",
                "ENSMUSG00000047945",
                "ENSMUSG00000020423",
                "ENSMUSG00000028005",
                "ENSMUSG00000060572",
                "ENSMUSG00000031586",
                "ENSMUSG00000026574",
                "ENSMUSG00000028776",
                "ENSMUSG00000008540",
                "ENSMUSG00000018899",
                "ENSMUSG00000074457",
                "ENSMUSG00000021876",
                "ENSMUSG00000021268",
                "ENSMUSG00000022836",
                "ENSMUSG00000024164",
                "ENSMUSG00000020473",
                "ENSMUSG00000006205",
                "ENSMUSG00000022032",
                "ENSMUSG00000045658",
                "ENSMUSG00000038463",
                "ENSMUSG00000017493",
                "ENSMUSG00000048387",
                "ENSMUSG00000001025"
            ],
            "expression_sum": {
                "class1": 5651.0,
                "class2": 1793.0
            },
            "ens_id": "ENSMUSG00000022346",
            "uniprot_ids": [
                "P01108",
                "F6PX41",
                "F8WID3",
                "E0CZD1"
            ],
            "net_id": 0,
            "source_num": 2,
            "target_num": 35
        },
        "ENSMUSG00000042812": {
            "id": "ENSMUSG00000042812",
            "type": "TF",
            "symbol": "Foxf1",
            "source": [
                "ENSMUSG00000052837",
                "ENSMUSG00000021250",
                "ENSMUSG00000052684",
                "ENSMUSG00000000247"
            ],
            "target": [
                "ENSMUSG00000031375",
                "ENSMUSG00000021250",
                "ENSMUSG00000030849",
                "ENSMUSG00000003545",
                "ENSMUSG00000031119",
                "ENSMUSG00000035493"
            ],
            "expression_sum": {
                "class1": 8363.0,
                "class2": 1724.0
            },
            "ens_id": "ENSMUSG00000042812",
            "uniprot_ids": [
                "Q61080"
            ],
            "net_id": 0,
            "source_num": 4,
            "target_num": 4
        },
        "ENSMUSG00000042406": {
            "id": "ENSMUSG00000042406",
            "type": "TF",
            "symbol": "Atf4",
            "source": [
                "ENSMUSG00000026628"
            ],
            "target": [
                "ENSMUSG00000022836",
                "ENSMUSG00000035356",
                "ENSMUSG00000026628",
                "ENSMUSG00000053113",
                "ENSMUSG00000018899"
            ],
            "expression_sum": {
                "class1": 8359.0,
                "class2": 3788.0
            },
            "ens_id": "ENSMUSG00000042406",
            "uniprot_ids": [
                "Q06507",
                "A0A2R8VI82"
            ],
            "net_id": 0,
            "source_num": 1,
            "target_num": 4
        },
        "ENSMUSG00000032238": {
            "id": "ENSMUSG00000032238",
            "type": "TF",
            "symbol": "Rora",
            "source": [
                "ENSMUSG00000005583"
            ],
            "target": [
                "ENSMUSG00000064370",
                "ENSMUSG00000020053",
                "ENSMUSG00000064363",
                "ENSMUSG00000029761",
                "ENSMUSG00000028339",
                "ENSMUSG00000014313",
                "ENSMUSG00000090841",
                "ENSMUSG00000024140",
                "ENSMUSG00000058056",
                "ENSMUSG00000000753",
                "ENSMUSG00000052957",
                "ENSMUSG00000064373",
                "ENSMUSG00000014956",
                "ENSMUSG00000068220",
                "ENSMUSG00000005583",
                "ENSMUSG00000029096",
                "ENSMUSG00000021870",
                "ENSMUSG00000026421",
                "ENSMUSG00000002107",
                "ENSMUSG00000024012",
                "ENSMUSG00000031586",
                "ENSMUSG00000027712",
                "ENSMUSG00000024048",
                "ENSMUSG00000040170",
                "ENSMUSG00000020009",
                "ENSMUSG00000008540",
                "ENSMUSG00000045658",
                "ENSMUSG00000016024",
                "ENSMUSG00000017493",
                "ENSMUSG00000058135"
            ],
            "expression_sum": {
                "class1": 4893.0,
                "class2": 2920.0
            },
            "ens_id": "ENSMUSG00000032238",
            "uniprot_ids": [
                "P51448",
                "G3UZ02"
            ],
            "net_id": 0,
            "source_num": 1,
            "target_num": 19
        },
        "ENSMUSG00000028565": {
            "id": "ENSMUSG00000028565",
            "type": "TF",
            "symbol": "Nfia",
            "source": [
                "ENSMUSG00000048387"
            ],
            "target": [
                "ENSMUSG00000028517",
                "ENSMUSG00000016494",
                "ENSMUSG00000021477",
                "ENSMUSG00000024140",
                "ENSMUSG00000026879",
                "ENSMUSG00000018217",
                "ENSMUSG00000020585",
                "ENSMUSG00000025784",
                "ENSMUSG00000040562",
                "ENSMUSG00000064373",
                "ENSMUSG00000031451",
                "ENSMUSG00000042485",
                "ENSMUSG00000029096",
                "ENSMUSG00000041959",
                "ENSMUSG00000002107",
                "ENSMUSG00000028369",
                "ENSMUSG00000019929",
                "ENSMUSG00000032231",
                "ENSMUSG00000029659",
                "ENSMUSG00000027712",
                "ENSMUSG00000024048",
                "ENSMUSG00000028108",
                "ENSMUSG00000020009",
                "ENSMUSG00000004207",
                "ENSMUSG00000026574",
                "ENSMUSG00000008540",
                "ENSMUSG00000020205",
                "ENSMUSG00000021876",
                "ENSMUSG00000079037",
                "ENSMUSG00000022836",
                "ENSMUSG00000024164",
                "ENSMUSG00000032334",
                "ENSMUSG00000000594",
                "ENSMUSG00000021390",
                "ENSMUSG00000015085",
                "ENSMUSG00000017493",
                "ENSMUSG00000048387",
                "ENSMUSG00000058135"
            ],
            "expression_sum": {
                "class1": 12541.0,
                "class2": 12297.0
            },
            "ens_id": "ENSMUSG00000028565",
            "uniprot_ids": [
                "Q02780",
                "B1AUB9",
                "B1AUC0",
                "F7CDR2",
                "B1AUB8",
                "B1AUB6"
            ],
            "net_id": 0,
            "source_num": 1,
            "target_num": 30
        },
        "ENSMUSG00000032035": {
            "id": "ENSMUSG00000032035",
            "type": "TF",
            "symbol": "Ets1",
            "source": [
                "ENSMUSG00000015846"
            ],
            "target": [
                "ENSMUSG00000019505",
                "ENSMUSG00000005338",
                "ENSMUSG00000064363",
                "ENSMUSG00000031447",
                "ENSMUSG00000021477",
                "ENSMUSG00000027447",
                "ENSMUSG00000024909",
                "ENSMUSG00000034868",
                "ENSMUSG00000023175",
                "ENSMUSG00000064354",
                "ENSMUSG00000026970",
                "ENSMUSG00000059588",
                "ENSMUSG00000025130",
                "ENSMUSG00000025790",
                "ENSMUSG00000034892",
                "ENSMUSG00000032231",
                "ENSMUSG00000023336",
                "ENSMUSG00000031021",
                "ENSMUSG00000029426",
                "ENSMUSG00000029718",
                "ENSMUSG00000036655",
                "ENSMUSG00000062825",
                "ENSMUSG00000020368",
                "ENSMUSG00000024678",
                "ENSMUSG00000069601",
                "ENSMUSG00000034353",
                "ENSMUSG00000028603",
                "ENSMUSG00000034845",
                "ENSMUSG00000022892",
                "ENSMUSG00000032562",
                "ENSMUSG00000056025",
                "ENSMUSG00000021756",
                "ENSMUSG00000038803",
                "ENSMUSG00000070436",
                "ENSMUSG00000029767",
                "ENSMUSG00000030824",
                "ENSMUSG00000064351",
                "ENSMUSG00000021248",
                "ENSMUSG00000027642",
                "ENSMUSG00000030249",
                "ENSMUSG00000029163",
                "ENSMUSG00000026193",
                "ENSMUSG00000033880",
                "ENSMUSG00000022587",
                "ENSMUSG00000023944",
                "ENSMUSG00000024065",
                "ENSMUSG00000029455",
                "ENSMUSG00000031375",
                "ENSMUSG00000072235",
                "ENSMUSG00000036381",
                "ENSMUSG00000026860",
                "ENSMUSG00000046330",
                "ENSMUSG00000020432",
                "ENSMUSG00000079523",
                "ENSMUSG00000001123",
                "ENSMUSG00000060126",
                "ENSMUSG00000042453",
                "ENSMUSG00000058569",
                "ENSMUSG00000026014",
                "ENSMUSG00000005973",
                "ENSMUSG00000016534",
                "ENSMUSG00000060591",
                "ENSMUSG00000032492",
                "ENSMUSG00000034707",
                "ENSMUSG00000049517",
                "ENSMUSG00000022665",
                "ENSMUSG00000031661",
                "ENSMUSG00000061232",
                "ENSMUSG00000027712",
                "ENSMUSG00000024048",
                "ENSMUSG00000069662",
                "ENSMUSG00000032666",
                "ENSMUSG00000028108",
                "ENSMUSG00000004207",
                "ENSMUSG00000020205",
                "ENSMUSG00000035242",
                "ENSMUSG00000034708",
                "ENSMUSG00000026478",
                "ENSMUSG00000032528",
                "ENSMUSG00000070407",
                "ENSMUSG00000020689",
                "ENSMUSG00000026223",
                "ENSMUSG00000029076",
                "ENSMUSG00000022971",
                "ENSMUSG00000074884",
                "ENSMUSG00000054808",
                "ENSMUSG00000022108",
                "ENSMUSG00000041736",
                "ENSMUSG00000028132",
                "ENSMUSG00000027248",
                "ENSMUSG00000020585",
                "ENSMUSG00000032179",
                "ENSMUSG00000025492",
                "ENSMUSG00000046402",
                "ENSMUSG00000034158",
                "ENSMUSG00000055447",
                "ENSMUSG00000000740",
                "ENSMUSG00000007892",
                "ENSMUSG00000027523",
                "ENSMUSG00000028618",
                "ENSMUSG00000032383",
                "ENSMUSG00000035530",
                "ENSMUSG00000047907",
                "ENSMUSG00000073411",
                "ENSMUSG00000048758",
                "ENSMUSG00000022708",
                "ENSMUSG00000029322",
                "ENSMUSG00000019929",
                "ENSMUSG00000060802",
                "ENSMUSG00000001847",
                "ENSMUSG00000000247",
                "ENSMUSG00000030849",
                "ENSMUSG00000020902",
                "ENSMUSG00000073489",
                "ENSMUSG00000024109",
                "ENSMUSG00000058135"
            ],
            "expression_sum": {
                "class1": 11559.0,
                "class2": 1092.0
            },
            "ens_id": "ENSMUSG00000032035",
            "uniprot_ids": [
                "P27577",
                "V9GXE2",
                "E9PWI8",
                "V9GX28",
                "A0A5F8MPR7"
            ],
            "net_id": 0,
            "source_num": 1,
            "target_num": 21
        },
        "ENSMUSG00000015846": {
            "id": "ENSMUSG00000015846",
            "type": "TF",
            "symbol": "Rxra",
            "source": [],
            "target": [
                "ENSMUSG00000020486",
                "ENSMUSG00000032035",
                "ENSMUSG00000026678",
                "ENSMUSG00000032946",
                "ENSMUSG00000034353",
                "ENSMUSG00000024140",
                "ENSMUSG00000025491",
                "ENSMUSG00000034845",
                "ENSMUSG00000001552",
                "ENSMUSG00000042453",
                "ENSMUSG00000028132",
                "ENSMUSG00000021756",
                "ENSMUSG00000022371",
                "ENSMUSG00000032179",
                "ENSMUSG00000000184",
                "ENSMUSG00000038591",
                "ENSMUSG00000041577",
                "ENSMUSG00000000325",
                "ENSMUSG00000038742",
                "ENSMUSG00000059588",
                "ENSMUSG00000028003",
                "ENSMUSG00000012428",
                "ENSMUSG00000023336",
                "ENSMUSG00000069662",
                "ENSMUSG00000032666",
                "ENSMUSG00000024065",
                "ENSMUSG00000006356",
                "ENSMUSG00000032528",
                "ENSMUSG00000070407",
                "ENSMUSG00000020689",
                "ENSMUSG00000036655",
                "ENSMUSG00000072625"
            ],
            "expression_sum": {
                "class1": 4376.0,
                "class2": 460.0
            },
            "ens_id": "ENSMUSG00000015846",
            "uniprot_ids": [
                "P28700",
                "Q6LC96",
                "A2AJP2",
                "A0A5F8MP91",
                "A0A5F8MQ23"
            ],
            "net_id": 0,
            "source_num": 0,
            "target_num": 6
        },
        "ENSMUSG00000028639": {
            "id": "ENSMUSG00000028639",
            "type": "TF",
            "symbol": "Ybx1",
            "source": [
                "ENSMUSG00000057098",
                "ENSMUSG00000000247"
            ],
            "target": [
                "ENSMUSG00000020486",
                "ENSMUSG00000023367",
                "ENSMUSG00000022032",
                "ENSMUSG00000041577",
                "ENSMUSG00000068522",
                "ENSMUSG00000018417"
            ],
            "expression_sum": {
                "class1": 29858.0,
                "class2": 27229.0
            },
            "ens_id": "ENSMUSG00000028639",
            "uniprot_ids": [
                "P62960",
                "A2BGG7"
            ],
            "net_id": 0,
            "source_num": 2,
            "target_num": 3
        },
        "ENSMUSG00000020644": {
            "id": "ENSMUSG00000020644",
            "type": "TF",
            "symbol": "Id2",
            "source": [
                "ENSMUSG00000030103",
                "ENSMUSG00000005583"
            ],
            "target": [
                "ENSMUSG00000032387",
                "ENSMUSG00000029761",
                "ENSMUSG00000001175",
                "ENSMUSG00000034566",
                "ENSMUSG00000014956",
                "ENSMUSG00000018293",
                "ENSMUSG00000037712",
                "ENSMUSG00000022010",
                "ENSMUSG00000024290",
                "ENSMUSG00000029580",
                "ENSMUSG00000032330",
                "ENSMUSG00000058056"
            ],
            "expression_sum": {
                "class1": 7989.0,
                "class2": 16476.0
            },
            "ens_id": "ENSMUSG00000020644",
            "uniprot_ids": [
                "P41136",
                "A0A1Y7VMT3"
            ],
            "net_id": 0,
            "source_num": 2,
            "target_num": 4
        }
    },
    "grps": {
        "ENSMUSG00000037868_ENSMUSG00000021250": {
            "id": "ENSMUSG00000037868_ENSMUSG00000021250",
            "type": "Mix",
            "score": 5.611695544374111,
            "reversable": true,
            "correlations": {
                "class1": 0.511602981350903,
                "class2": 0.10137532326055317
            },
            "regulatory_source": "ENSMUSG00000037868",
            "regulatory_target": "ENSMUSG00000021250",
            "net_id": 0
        },
        "ENSMUSG00000052684_ENSMUSG00000038418": {
            "id": "ENSMUSG00000052684_ENSMUSG00000038418",
            "type": "Mix",
            "score": 11.947891079741218,
            "reversable": true,
            "correlations": {
                "class1": 0.7664433043909842,
                "class2": 0.486543404345432
            },
            "regulatory_source": "ENSMUSG00000052684",
            "regulatory_target": "ENSMUSG00000038418",
            "net_id": 0
        },
        "ENSMUSG00000052837_ENSMUSG00000038418": {
            "id": "ENSMUSG00000052837_ENSMUSG00000038418",
            "type": "Mix",
            "score": 13.674555392069976,
            "reversable": true,
            "correlations": {
                "class1": 0.816953701569463,
                "class2": 0.5696919058794143
            },
            "regulatory_source": "ENSMUSG00000052837",
            "regulatory_target": "ENSMUSG00000038418",
            "net_id": 0
        },
        "ENSMUSG00000071076_ENSMUSG00000055148": {
            "id": "ENSMUSG00000071076_ENSMUSG00000055148",
            "type": "Standard",
            "score": 0.8925988455170332,
            "reversable": false,
            "correlations": {
                "class1": 0.5650328779947075,
                "class2": 0.24800406666903702
            },
            "regulatory_source": "ENSMUSG00000071076",
            "regulatory_target": "ENSMUSG00000055148",
            "net_id": 0
        },
        "ENSMUSG00000055148_ENSMUSG00000003545": {
            "id": "ENSMUSG00000055148_ENSMUSG00000003545",
            "type": "Mix",
            "score": 14.881027125911217,
            "reversable": false,
            "correlations": {
                "class1": 0.5666618661792494,
                "class2": 0.19337881173911073
            },
            "regulatory_source": "ENSMUSG00000055148",
            "regulatory_target": "ENSMUSG00000003545",
            "net_id": 0
        },
        "ENSMUSG00000071076_ENSMUSG00000052837": {
            "id": "ENSMUSG00000071076_ENSMUSG00000052837",
            "type": "Mix",
            "score": 6.036426962904938,
            "reversable": true,
            "correlations": {
                "class1": 0.7238904426938447,
                "class2": 0.4376071659527366
            },
            "regulatory_source": "ENSMUSG00000071076",
            "regulatory_target": "ENSMUSG00000052837",
            "net_id": 0
        },
        "ENSMUSG00000055148_ENSMUSG00000038418": {
            "id": "ENSMUSG00000055148_ENSMUSG00000038418",
            "type": "Mix",
            "score": 8.189327875064823,
            "reversable": false,
            "correlations": {
                "class1": 0.5798601054470557,
                "class2": 0.22569929951045264
            },
            "regulatory_source": "ENSMUSG00000038418",
            "regulatory_target": "ENSMUSG00000055148",
            "net_id": 0
        },
        "ENSMUSG00000038418_ENSMUSG00000026628": {
            "id": "ENSMUSG00000038418_ENSMUSG00000026628",
            "type": "Mix",
            "score": 7.6345749976519315,
            "reversable": true,
            "correlations": {
                "class1": 0.5998243543116571,
                "class2": 0.2822248149600215
            },
            "regulatory_source": "ENSMUSG00000026628",
            "regulatory_target": "ENSMUSG00000038418",
            "net_id": 0
        },
        "ENSMUSG00000018899_ENSMUSG00000003545": {
            "id": "ENSMUSG00000018899_ENSMUSG00000003545",
            "type": "Standard",
            "score": 0.43237237902334835,
            "reversable": false,
            "correlations": {
                "class1": 0.4622492265489637,
                "class2": 0.0906129101328391
            },
            "regulatory_source": "ENSMUSG00000018899",
            "regulatory_target": "ENSMUSG00000003545",
            "net_id": 0
        },
        "ENSMUSG00000037868_ENSMUSG00000003545": {
            "id": "ENSMUSG00000037868_ENSMUSG00000003545",
            "type": "Standard",
            "score": 0.35682246903337206,
            "reversable": false,
            "correlations": {
                "class1": 0.4463381561221811,
                "class2": 0.06207258592573403
            },
            "regulatory_source": "ENSMUSG00000037868",
            "regulatory_target": "ENSMUSG00000003545",
            "net_id": 0
        },
        "ENSMUSG00000052837_ENSMUSG00000037868": {
            "id": "ENSMUSG00000052837_ENSMUSG00000037868",
            "type": "Standard",
            "score": 0.33804457166687746,
            "reversable": true,
            "correlations": {
                "class1": 0.5311628853922231,
                "class2": 0.15277836594022196
            },
            "regulatory_source": "ENSMUSG00000037868",
            "regulatory_target": "ENSMUSG00000052837",
            "net_id": 0
        },
        "ENSMUSG00000052684_ENSMUSG00000021250": {
            "id": "ENSMUSG00000052684_ENSMUSG00000021250",
            "type": "Outer",
            "score": 19.024522880672137,
            "reversable": true,
            "correlations": {
                "class1": 0.8363165091507709,
                "class2": 0.5472563359064866
            },
            "regulatory_source": "ENSMUSG00000052684",
            "regulatory_target": "ENSMUSG00000021250",
            "net_id": 0
        },
        "ENSMUSG00000038418_ENSMUSG00000021250": {
            "id": "ENSMUSG00000038418_ENSMUSG00000021250",
            "type": "Outer",
            "score": 11.591868893707016,
            "reversable": false,
            "correlations": {
                "class1": 0.8121305836268662,
                "class2": 0.4677018533225259
            },
            "regulatory_source": "ENSMUSG00000021250",
            "regulatory_target": "ENSMUSG00000038418",
            "net_id": 0
        },
        "ENSMUSG00000052684_ENSMUSG00000023034": {
            "id": "ENSMUSG00000052684_ENSMUSG00000023034",
            "type": "Outer",
            "score": 13.017814959839429,
            "reversable": true,
            "correlations": {
                "class1": 0.538127822132192,
                "class2": 0.09376026005645308
            },
            "regulatory_source": "ENSMUSG00000052684",
            "regulatory_target": "ENSMUSG00000023034",
            "net_id": 0
        },
        "ENSMUSG00000023034_ENSMUSG00000003545": {
            "id": "ENSMUSG00000023034_ENSMUSG00000003545",
            "type": "Outer",
            "score": 15.563020345950088,
            "reversable": true,
            "correlations": {
                "class1": 0.5335436705169669,
                "class2": 0.08802224186371972
            },
            "regulatory_source": "ENSMUSG00000003545",
            "regulatory_target": "ENSMUSG00000023034",
            "net_id": 0
        },
        "ENSMUSG00000071076_ENSMUSG00000026628": {
            "id": "ENSMUSG00000071076_ENSMUSG00000026628",
            "type": "Mix",
            "score": 8.903213017748817,
            "reversable": true,
            "correlations": {
                "class1": 0.5223556542852718,
                "class2": 0.22161000063545727
            },
            "regulatory_source": "ENSMUSG00000026628",
            "regulatory_target": "ENSMUSG00000071076",
            "net_id": 0
        },
        "ENSMUSG00000052684_ENSMUSG00000003545": {
            "id": "ENSMUSG00000052684_ENSMUSG00000003545",
            "type": "Mix",
            "score": 18.44593690735485,
            "reversable": true,
            "correlations": {
                "class1": 0.7252397828162958,
                "class2": 0.4389445314638871
            },
            "regulatory_source": "ENSMUSG00000052684",
            "regulatory_target": "ENSMUSG00000003545",
            "net_id": 0
        },
        "ENSMUSG00000052837_ENSMUSG00000052684": {
            "id": "ENSMUSG00000052837_ENSMUSG00000052684",
            "type": "Mix",
            "score": 14.306139403707094,
            "reversable": true,
            "correlations": {
                "class1": 0.8218419773024086,
                "class2": 0.4292453022171103
            },
            "regulatory_source": "ENSMUSG00000052684",
            "regulatory_target": "ENSMUSG00000052837",
            "net_id": 0
        },
        "ENSMUSG00000038418_ENSMUSG00000003545": {
            "id": "ENSMUSG00000038418_ENSMUSG00000003545",
            "type": "Outer",
            "score": 10.460085965718816,
            "reversable": false,
            "correlations": {
                "class1": 0.7383230112763797,
                "class2": 0.293287548256507
            },
            "regulatory_source": "ENSMUSG00000003545",
            "regulatory_target": "ENSMUSG00000038418",
            "net_id": 0
        },
        "ENSMUSG00000071076_ENSMUSG00000023034": {
            "id": "ENSMUSG00000071076_ENSMUSG00000023034",
            "type": "Mix",
            "score": 9.004490883220216,
            "reversable": false,
            "correlations": {
                "class1": 0.5195877483764404,
                "class2": 0.11148371947566481
            },
            "regulatory_source": "ENSMUSG00000071076",
            "regulatory_target": "ENSMUSG00000023034",
            "net_id": 0
        },
        "ENSMUSG00000055148_ENSMUSG00000052684": {
            "id": "ENSMUSG00000055148_ENSMUSG00000052684",
            "type": "Outer",
            "score": 10.379065907956804,
            "reversable": false,
            "correlations": {
                "class1": 0.608857883048902,
                "class2": 0.21972937889656577
            },
            "regulatory_source": "ENSMUSG00000052684",
            "regulatory_target": "ENSMUSG00000055148",
            "net_id": 0
        },
        "ENSMUSG00000023034_ENSMUSG00000021250": {
            "id": "ENSMUSG00000023034_ENSMUSG00000021250",
            "type": "Mix",
            "score": 7.128264578854652,
            "reversable": true,
            "correlations": {
                "class1": 0.6106406075093944,
                "class2": 0.31267852734765933
            },
            "regulatory_source": "ENSMUSG00000021250",
            "regulatory_target": "ENSMUSG00000023034",
            "net_id": 0
        },
        "ENSMUSG00000052837_ENSMUSG00000003545": {
            "id": "ENSMUSG00000052837_ENSMUSG00000003545",
            "type": "Outer",
            "score": 25.321612504269716,
            "reversable": true,
            "correlations": {
                "class1": 0.7419223691421334,
                "class2": 0.2884698241138656
            },
            "regulatory_source": "ENSMUSG00000003545",
            "regulatory_target": "ENSMUSG00000052837",
            "net_id": 0
        },
        "ENSMUSG00000056501_ENSMUSG00000021250": {
            "id": "ENSMUSG00000056501_ENSMUSG00000021250",
            "type": "Outer",
            "score": 14.445440473045448,
            "reversable": true,
            "correlations": {
                "class1": 0.5800351968689449,
                "class2": 0.3157183980906766
            },
            "regulatory_source": "ENSMUSG00000056501",
            "regulatory_target": "ENSMUSG00000021250",
            "net_id": 0
        },
        "ENSMUSG00000021250_ENSMUSG00000003545": {
            "id": "ENSMUSG00000021250_ENSMUSG00000003545",
            "type": "Outer",
            "score": 9.483693398907551,
            "reversable": true,
            "correlations": {
                "class1": 0.7599432458376313,
                "class2": 0.4565047111958447
            },
            "regulatory_source": "ENSMUSG00000003545",
            "regulatory_target": "ENSMUSG00000021250",
            "net_id": 0
        },
        "ENSMUSG00000021250_ENSMUSG00000018899": {
            "id": "ENSMUSG00000021250_ENSMUSG00000018899",
            "type": "Mix",
            "score": 8.52131077363052,
            "reversable": true,
            "correlations": {
                "class1": 0.4884707833501266,
                "class2": 0.13285732004398826
            },
            "regulatory_source": "ENSMUSG00000018899",
            "regulatory_target": "ENSMUSG00000021250",
            "net_id": 0
        },
        "ENSMUSG00000055148_ENSMUSG00000052837": {
            "id": "ENSMUSG00000055148_ENSMUSG00000052837",
            "type": "Mix",
            "score": 7.188994776785808,
            "reversable": false,
            "correlations": {
                "class1": 0.6546782421362538,
                "class2": 0.3989626630420214
            },
            "regulatory_source": "ENSMUSG00000052837",
            "regulatory_target": "ENSMUSG00000055148",
            "net_id": 0
        },
        "ENSMUSG00000052684_ENSMUSG00000026628": {
            "id": "ENSMUSG00000052684_ENSMUSG00000026628",
            "type": "Mix",
            "score": 6.969643652888925,
            "reversable": true,
            "correlations": {
                "class1": 0.5864999242817666,
                "class2": 0.2287600775760728
            },
            "regulatory_source": "ENSMUSG00000026628",
            "regulatory_target": "ENSMUSG00000052684",
            "net_id": 0
        },
        "ENSMUSG00000052837_ENSMUSG00000021250": {
            "id": "ENSMUSG00000052837_ENSMUSG00000021250",
            "type": "Outer",
            "score": 24.793569765305428,
            "reversable": true,
            "correlations": {
                "class1": 0.8634234826228351,
                "class2": 0.5452250516649075
            },
            "regulatory_source": "ENSMUSG00000052837",
            "regulatory_target": "ENSMUSG00000021250",
            "net_id": 0
        },
        "ENSMUSG00000055435_ENSMUSG00000052684": {
            "id": "ENSMUSG00000055435_ENSMUSG00000052684",
            "type": "Bridge",
            "score": 0,
            "reversable": true,
            "correlations": {
                "class1": 0.04897615132528646,
                "class2": 0.2631439322421368
            },
            "regulatory_source": "ENSMUSG00000052684",
            "regulatory_target": "ENSMUSG00000055435",
            "net_id": 0
        },
        "ENSMUSG00000052684_ENSMUSG00000022528": {
            "id": "ENSMUSG00000052684_ENSMUSG00000022528",
            "type": "Bridge",
            "score": 0,
            "reversable": true,
            "correlations": {
                "class1": 0.30944264922087644,
                "class2": 0.0
            },
            "regulatory_source": "ENSMUSG00000052684",
            "regulatory_target": "ENSMUSG00000022528",
            "net_id": 0
        },
        "ENSMUSG00000052684_ENSMUSG00000052534": {
            "id": "ENSMUSG00000052684_ENSMUSG00000052534",
            "type": "Bridge",
            "score": 0,
            "reversable": true,
            "correlations": {
                "class1": 0.08638584424929044,
                "class2": 0.2876643227993388
            },
            "regulatory_source": "ENSMUSG00000052534",
            "regulatory_target": "ENSMUSG00000052684",
            "net_id": 0
        },
        "ENSMUSG00000052837_ENSMUSG00000041515": {
            "id": "ENSMUSG00000052837_ENSMUSG00000041515",
            "type": "Bridge",
            "score": 0,
            "reversable": true,
            "correlations": {
                "class1": 0.2894967982165958,
                "class2": 0.0704198584676184
            },
            "regulatory_source": "ENSMUSG00000052837",
            "regulatory_target": "ENSMUSG00000041515",
            "net_id": 0
        },
        "ENSMUSG00000052837_ENSMUSG00000030103": {
            "id": "ENSMUSG00000052837_ENSMUSG00000030103",
            "type": "Bridge",
            "score": 0,
            "reversable": true,
            "correlations": {
                "class1": 0.4305144860301364,
                "class2": 0.18192329163268683
            },
            "regulatory_source": "ENSMUSG00000030103",
            "regulatory_target": "ENSMUSG00000052837",
            "net_id": 0
        },
        "ENSMUSG00000055148_ENSMUSG00000022346": {
            "id": "ENSMUSG00000055148_ENSMUSG00000022346",
            "type": "Bridge",
            "score": 0,
            "reversable": false,
            "correlations": {
                "class1": 0.2823366557987358,
                "class2": 0.07164639304568834
            },
            "regulatory_source": "ENSMUSG00000022346",
            "regulatory_target": "ENSMUSG00000055148",
            "net_id": 0
        },
        "ENSMUSG00000042812_ENSMUSG00000003545": {
            "id": "ENSMUSG00000042812_ENSMUSG00000003545",
            "type": "Bridge",
            "score": 0,
            "reversable": false,
            "correlations": {
                "class1": 0.3163113394205811,
                "class2": 0.0930664232693993
            },
            "regulatory_source": "ENSMUSG00000042812",
            "regulatory_target": "ENSMUSG00000003545",
            "net_id": 0
        },
        "ENSMUSG00000042406_ENSMUSG00000026628": {
            "id": "ENSMUSG00000042406_ENSMUSG00000026628",
            "type": "Bridge",
            "score": 0,
            "reversable": true,
            "correlations": {
                "class1": 0.3155121108408707,
                "class2": 0.11190693132970479
            },
            "regulatory_source": "ENSMUSG00000026628",
            "regulatory_target": "ENSMUSG00000042406",
            "net_id": 0
        },
        "ENSMUSG00000032238_ENSMUSG00000005583": {
            "id": "ENSMUSG00000032238_ENSMUSG00000005583",
            "type": "Bridge",
            "score": 0,
            "reversable": true,
            "correlations": {
                "class1": 0.0757326703396924,
                "class2": -0.1335361777349255
            },
            "regulatory_source": "ENSMUSG00000032238",
            "regulatory_target": "ENSMUSG00000005583",
            "net_id": 0
        },
        "ENSMUSG00000048387_ENSMUSG00000028565": {
            "id": "ENSMUSG00000048387_ENSMUSG00000028565",
            "type": "Bridge",
            "score": 0,
            "reversable": true,
            "correlations": {
                "class1": 0.0,
                "class2": 0.2074994862931572
            },
            "regulatory_source": "ENSMUSG00000028565",
            "regulatory_target": "ENSMUSG00000048387",
            "net_id": 0
        },
        "ENSMUSG00000032035_ENSMUSG00000000247": {
            "id": "ENSMUSG00000032035_ENSMUSG00000000247",
            "type": "Bridge",
            "score": 0,
            "reversable": false,
            "correlations": {
                "class1": 0.20187640931490597,
                "class2": 0.0
            },
            "regulatory_source": "ENSMUSG00000032035",
            "regulatory_target": "ENSMUSG00000000247",
            "net_id": 0
        },
        "ENSMUSG00000071076_ENSMUSG00000003545": {
            "id": "ENSMUSG00000071076_ENSMUSG00000003545",
            "type": "Bridge",
            "score": 0,
            "reversable": false,
            "correlations": {
                "class1": 0.6661142511739162,
                "class2": 0.3854597276981423
            },
            "regulatory_source": "ENSMUSG00000071076",
            "regulatory_target": "ENSMUSG00000003545",
            "net_id": 0
        },
        "ENSMUSG00000071076_ENSMUSG00000037868": {
            "id": "ENSMUSG00000071076_ENSMUSG00000037868",
            "type": "Bridge",
            "score": 0,
            "reversable": false,
            "correlations": {
                "class1": 0.40457905909603975,
                "class2": 0.1114806443693004
            },
            "regulatory_source": "ENSMUSG00000071076",
            "regulatory_target": "ENSMUSG00000037868",
            "net_id": 0
        },
        "ENSMUSG00000071076_ENSMUSG00000021250": {
            "id": "ENSMUSG00000071076_ENSMUSG00000021250",
            "type": "Bridge",
            "score": 0,
            "reversable": false,
            "correlations": {
                "class1": 0.7417102274436516,
                "class2": 0.5358630159734392
            },
            "regulatory_source": "ENSMUSG00000071076",
            "regulatory_target": "ENSMUSG00000021250",
            "net_id": 0
        },
        "ENSMUSG00000071076_ENSMUSG00000008575": {
            "id": "ENSMUSG00000071076_ENSMUSG00000008575",
            "type": "Bridge",
            "score": 0,
            "reversable": true,
            "correlations": {
                "class1": 0.17571564133909143,
                "class2": 0.451832909910978
            },
            "regulatory_source": "ENSMUSG00000008575",
            "regulatory_target": "ENSMUSG00000071076",
            "net_id": 0
        },
        "ENSMUSG00000071076_ENSMUSG00000005583": {
            "id": "ENSMUSG00000071076_ENSMUSG00000005583",
            "type": "Bridge",
            "score": 0,
            "reversable": false,
            "correlations": {
                "class1": 0.07056412739683127,
                "class2": -0.14959169375504733
            },
            "regulatory_source": "ENSMUSG00000071076",
            "regulatory_target": "ENSMUSG00000005583",
            "net_id": 0
        },
        "ENSMUSG00000071076_ENSMUSG00000018899": {
            "id": "ENSMUSG00000071076_ENSMUSG00000018899",
            "type": "Bridge",
            "score": 0,
            "reversable": false,
            "correlations": {
                "class1": 0.41611333855067467,
                "class2": 0.13519004196566423
            },
            "regulatory_source": "ENSMUSG00000071076",
            "regulatory_target": "ENSMUSG00000018899",
            "net_id": 0
        },
        "ENSMUSG00000071076_ENSMUSG00000000247": {
            "id": "ENSMUSG00000071076_ENSMUSG00000000247",
            "type": "Bridge",
            "score": 0,
            "reversable": true,
            "correlations": {
                "class1": 0.22573770863076817,
                "class2": 0.0
            },
            "regulatory_source": "ENSMUSG00000000247",
            "regulatory_target": "ENSMUSG00000071076",
            "net_id": 0
        },
        "ENSMUSG00000071076_ENSMUSG00000057098": {
            "id": "ENSMUSG00000071076_ENSMUSG00000057098",
            "type": "Bridge",
            "score": 0,
            "reversable": false,
            "correlations": {
                "class1": 0.0,
                "class2": 0.26284909883450763
            },
            "regulatory_source": "ENSMUSG00000071076",
            "regulatory_target": "ENSMUSG00000057098",
            "net_id": 0
        },
        "ENSMUSG00000071076_ENSMUSG00000056501": {
            "id": "ENSMUSG00000071076_ENSMUSG00000056501",
            "type": "Bridge",
            "score": 0,
            "reversable": true,
            "correlations": {
                "class1": 0.5340783081084048,
                "class2": 0.2531676777172529
            },
            "regulatory_source": "ENSMUSG00000056501",
            "regulatory_target": "ENSMUSG00000071076",
            "net_id": 0
        },
        "ENSMUSG00000071076_ENSMUSG00000055435": {
            "id": "ENSMUSG00000071076_ENSMUSG00000055435",
            "type": "Bridge",
            "score": 0,
            "reversable": false,
            "correlations": {
                "class1": 0.055712117939097215,
                "class2": 0.28418246416275755
            },
            "regulatory_source": "ENSMUSG00000071076",
            "regulatory_target": "ENSMUSG00000055435",
            "net_id": 0
        },
        "ENSMUSG00000071076_ENSMUSG00000022528": {
            "id": "ENSMUSG00000071076_ENSMUSG00000022528",
            "type": "Bridge",
            "score": 0,
            "reversable": false,
            "correlations": {
                "class1": 0.2935903767142557,
                "class2": 0.0
            },
            "regulatory_source": "ENSMUSG00000071076",
            "regulatory_target": "ENSMUSG00000022528",
            "net_id": 0
        },
        "ENSMUSG00000052684_ENSMUSG00000037868": {
            "id": "ENSMUSG00000052684_ENSMUSG00000037868",
            "type": "Standard",
            "score": 0.33361658147132583,
            "reversable": true,
            "correlations": {
                "class1": 0.47283157082885174,
                "class2": 0.09844575066028119
            },
            "regulatory_source": "ENSMUSG00000052684",
            "regulatory_target": "ENSMUSG00000037868",
            "net_id": 0
        },
        "ENSMUSG00000052684_ENSMUSG00000008575": {
            "id": "ENSMUSG00000052684_ENSMUSG00000008575",
            "type": "Bridge",
            "score": 0,
            "reversable": true,
            "correlations": {
                "class1": 0.1639864502448749,
                "class2": 0.40382237752143146
            },
            "regulatory_source": "ENSMUSG00000008575",
            "regulatory_target": "ENSMUSG00000052684",
            "net_id": 0
        },
        "ENSMUSG00000052684_ENSMUSG00000018899": {
            "id": "ENSMUSG00000052684_ENSMUSG00000018899",
            "type": "Bridge",
            "score": 0,
            "reversable": true,
            "correlations": {
                "class1": 0.5059442115248347,
                "class2": 0.1539983259065759
            },
            "regulatory_source": "ENSMUSG00000052684",
            "regulatory_target": "ENSMUSG00000018899",
            "net_id": 0
        },
        "ENSMUSG00000052684_ENSMUSG00000038193": {
            "id": "ENSMUSG00000052684_ENSMUSG00000038193",
            "type": "Bridge",
            "score": 0,
            "reversable": false,
            "correlations": {
                "class1": 0.28694760886044174,
                "class2": 0.039183204276180605
            },
            "regulatory_source": "ENSMUSG00000038193",
            "regulatory_target": "ENSMUSG00000052684",
            "net_id": 0
        },
        "ENSMUSG00000052684_ENSMUSG00000000247": {
            "id": "ENSMUSG00000052684_ENSMUSG00000000247",
            "type": "Bridge",
            "score": 0,
            "reversable": true,
            "correlations": {
                "class1": 0.2691844387631659,
                "class2": 0.0
            },
            "regulatory_source": "ENSMUSG00000000247",
            "regulatory_target": "ENSMUSG00000052684",
            "net_id": 0
        },
        "ENSMUSG00000056501_ENSMUSG00000052684": {
            "id": "ENSMUSG00000056501_ENSMUSG00000052684",
            "type": "Bridge",
            "score": 0,
            "reversable": true,
            "correlations": {
                "class1": 0.5391570469555416,
                "class2": 0.23910625170370356
            },
            "regulatory_source": "ENSMUSG00000056501",
            "regulatory_target": "ENSMUSG00000052684",
            "net_id": 0
        },
        "ENSMUSG00000052684_ENSMUSG00000030103": {
            "id": "ENSMUSG00000052684_ENSMUSG00000030103",
            "type": "Bridge",
            "score": 0,
            "reversable": false,
            "correlations": {
                "class1": 0.3691058460572213,
                "class2": 0.14041804830162619
            },
            "regulatory_source": "ENSMUSG00000052684",
            "regulatory_target": "ENSMUSG00000030103",
            "net_id": 0
        },
        "ENSMUSG00000052684_ENSMUSG00000042812": {
            "id": "ENSMUSG00000052684_ENSMUSG00000042812",
            "type": "Bridge",
            "score": 0,
            "reversable": false,
            "correlations": {
                "class1": 0.3455794346895246,
                "class2": 0.12044690658610233
            },
            "regulatory_source": "ENSMUSG00000052684",
            "regulatory_target": "ENSMUSG00000042812",
            "net_id": 0
        },
        "ENSMUSG00000038418_ENSMUSG00000037868": {
            "id": "ENSMUSG00000038418_ENSMUSG00000037868",
            "type": "Bridge",
            "score": 0,
            "reversable": true,
            "correlations": {
                "class1": 0.5268507931116221,
                "class2": 0.2126703622134834
            },
            "regulatory_source": "ENSMUSG00000037868",
            "regulatory_target": "ENSMUSG00000038418",
            "net_id": 0
        },
        "ENSMUSG00000038418_ENSMUSG00000008575": {
            "id": "ENSMUSG00000038418_ENSMUSG00000008575",
            "type": "Bridge",
            "score": 0,
            "reversable": false,
            "correlations": {
                "class1": 0.1656629347288697,
                "class2": 0.3972304066329651
            },
            "regulatory_source": "ENSMUSG00000008575",
            "regulatory_target": "ENSMUSG00000038418",
            "net_id": 0
        },
        "ENSMUSG00000038418_ENSMUSG00000018899": {
            "id": "ENSMUSG00000038418_ENSMUSG00000018899",
            "type": "Bridge",
            "score": 0,
            "reversable": true,
            "correlations": {
                "class1": 0.5190921168740867,
                "class2": 0.188554731581964
            },
            "regulatory_source": "ENSMUSG00000018899",
            "regulatory_target": "ENSMUSG00000038418",
            "net_id": 0
        },
        "ENSMUSG00000038418_ENSMUSG00000003032": {
            "id": "ENSMUSG00000038418_ENSMUSG00000003032",
            "type": "Bridge",
            "score": 0,
            "reversable": true,
            "correlations": {
                "class1": 0.5097088086655946,
                "class2": 0.2978186122803058
            },
            "regulatory_source": "ENSMUSG00000003032",
            "regulatory_target": "ENSMUSG00000038418",
            "net_id": 0
        },
        "ENSMUSG00000038418_ENSMUSG00000023034": {
            "id": "ENSMUSG00000038418_ENSMUSG00000023034",
            "type": "Bridge",
            "score": 0,
            "reversable": true,
            "correlations": {
                "class1": 0.5962212316012241,
                "class2": 0.3012418939572574
            },
            "regulatory_source": "ENSMUSG00000023034",
            "regulatory_target": "ENSMUSG00000038418",
            "net_id": 0
        },
        "ENSMUSG00000038418_ENSMUSG00000000247": {
            "id": "ENSMUSG00000038418_ENSMUSG00000000247",
            "type": "Bridge",
            "score": 0,
            "reversable": true,
            "correlations": {
                "class1": 0.2744649111709719,
                "class2": 0.0
            },
            "regulatory_source": "ENSMUSG00000000247",
            "regulatory_target": "ENSMUSG00000038418",
            "net_id": 0
        },
        "ENSMUSG00000057098_ENSMUSG00000038418": {
            "id": "ENSMUSG00000057098_ENSMUSG00000038418",
            "type": "Bridge",
            "score": 0,
            "reversable": false,
            "correlations": {
                "class1": 0.0,
                "class2": 0.24044655137730508
            },
            "regulatory_source": "ENSMUSG00000057098",
            "regulatory_target": "ENSMUSG00000038418",
            "net_id": 0
        },
        "ENSMUSG00000038418_ENSMUSG00000022528": {
            "id": "ENSMUSG00000038418_ENSMUSG00000022528",
            "type": "Bridge",
            "score": 0,
            "reversable": true,
            "correlations": {
                "class1": 0.3117544236129782,
                "class2": 0.09095658077051604
            },
            "regulatory_source": "ENSMUSG00000022528",
            "regulatory_target": "ENSMUSG00000038418",
            "net_id": 0
        },
        "ENSMUSG00000052837_ENSMUSG00000026628": {
            "id": "ENSMUSG00000052837_ENSMUSG00000026628",
            "type": "Bridge",
            "score": 0,
            "reversable": true,
            "correlations": {
                "class1": 0.6139476303232696,
                "class2": 0.36601273301829085
            },
            "regulatory_source": "ENSMUSG00000026628",
            "regulatory_target": "ENSMUSG00000052837",
            "net_id": 0
        },
        "ENSMUSG00000052837_ENSMUSG00000018899": {
            "id": "ENSMUSG00000052837_ENSMUSG00000018899",
            "type": "Standard",
            "score": 0.5838490115109743,
            "reversable": true,
            "correlations": {
                "class1": 0.5657972494086193,
                "class2": 0.20554838595203948
            },
            "regulatory_source": "ENSMUSG00000052837",
            "regulatory_target": "ENSMUSG00000018899",
            "net_id": 0
        },
        "ENSMUSG00000052837_ENSMUSG00000003032": {
            "id": "ENSMUSG00000052837_ENSMUSG00000003032",
            "type": "Bridge",
            "score": 0,
            "reversable": true,
            "correlations": {
                "class1": 0.5289166145781354,
                "class2": 0.2798282568860291
            },
            "regulatory_source": "ENSMUSG00000003032",
            "regulatory_target": "ENSMUSG00000052837",
            "net_id": 0
        },
        "ENSMUSG00000052837_ENSMUSG00000023034": {
            "id": "ENSMUSG00000052837_ENSMUSG00000023034",
            "type": "Bridge",
            "score": 0,
            "reversable": true,
            "correlations": {
                "class1": 0.6227927315028381,
                "class2": 0.42023806508960526
            },
            "regulatory_source": "ENSMUSG00000052837",
            "regulatory_target": "ENSMUSG00000023034",
            "net_id": 0
        },
        "ENSMUSG00000052837_ENSMUSG00000000247": {
            "id": "ENSMUSG00000052837_ENSMUSG00000000247",
            "type": "Bridge",
            "score": 0,
            "reversable": true,
            "correlations": {
                "class1": 0.2644143897050098,
                "class2": -0.030137184023743904
            },
            "regulatory_source": "ENSMUSG00000000247",
            "regulatory_target": "ENSMUSG00000052837",
            "net_id": 0
        },
        "ENSMUSG00000057098_ENSMUSG00000052837": {
            "id": "ENSMUSG00000057098_ENSMUSG00000052837",
            "type": "Bridge",
            "score": 0,
            "reversable": false,
            "correlations": {
                "class1": 0.0,
                "class2": 0.23290360794216175
            },
            "regulatory_source": "ENSMUSG00000057098",
            "regulatory_target": "ENSMUSG00000052837",
            "net_id": 0
        },
        "ENSMUSG00000052837_ENSMUSG00000042812": {
            "id": "ENSMUSG00000052837_ENSMUSG00000042812",
            "type": "Bridge",
            "score": 0,
            "reversable": false,
            "correlations": {
                "class1": 0.3821532041350245,
                "class2": 0.17255097629798793
            },
            "regulatory_source": "ENSMUSG00000052837",
            "regulatory_target": "ENSMUSG00000042812",
            "net_id": 0
        },
        "ENSMUSG00000055148_ENSMUSG00000037868": {
            "id": "ENSMUSG00000055148_ENSMUSG00000037868",
            "type": "Bridge",
            "score": 0,
            "reversable": false,
            "correlations": {
                "class1": 0.37702090997462606,
                "class2": 0.029652061929184424
            },
            "regulatory_source": "ENSMUSG00000037868",
            "regulatory_target": "ENSMUSG00000055148",
            "net_id": 0
        },
        "ENSMUSG00000055148_ENSMUSG00000021250": {
            "id": "ENSMUSG00000055148_ENSMUSG00000021250",
            "type": "Bridge",
            "score": 0,
            "reversable": false,
            "correlations": {
                "class1": 0.6488783854802571,
                "class2": 0.42165602274726977
            },
            "regulatory_source": "ENSMUSG00000021250",
            "regulatory_target": "ENSMUSG00000055148",
            "net_id": 0
        },
        "ENSMUSG00000055148_ENSMUSG00000018899": {
            "id": "ENSMUSG00000055148_ENSMUSG00000018899",
            "type": "Bridge",
            "score": 0,
            "reversable": false,
            "correlations": {
                "class1": 0.4427015205484326,
                "class2": 0.10387789749163544
            },
            "regulatory_source": "ENSMUSG00000055148",
            "regulatory_target": "ENSMUSG00000018899",
            "net_id": 0
        },
        "ENSMUSG00000055148_ENSMUSG00000003032": {
            "id": "ENSMUSG00000055148_ENSMUSG00000003032",
            "type": "Bridge",
            "score": 0,
            "reversable": false,
            "correlations": {
                "class1": 0.5083398327191408,
                "class2": 0.2753652941655167
            },
            "regulatory_source": "ENSMUSG00000003032",
            "regulatory_target": "ENSMUSG00000055148",
            "net_id": 0
        },
        "ENSMUSG00000055148_ENSMUSG00000023034": {
            "id": "ENSMUSG00000055148_ENSMUSG00000023034",
            "type": "Bridge",
            "score": 0,
            "reversable": false,
            "correlations": {
                "class1": 0.5327611550037461,
                "class2": 0.3272088935794265
            },
            "regulatory_source": "ENSMUSG00000023034",
            "regulatory_target": "ENSMUSG00000055148",
            "net_id": 0
        },
        "ENSMUSG00000056501_ENSMUSG00000055148": {
            "id": "ENSMUSG00000056501_ENSMUSG00000055148",
            "type": "Bridge",
            "score": 0,
            "reversable": false,
            "correlations": {
                "class1": 0.49520793805896324,
                "class2": 0.279784580569881
            },
            "regulatory_source": "ENSMUSG00000056501",
            "regulatory_target": "ENSMUSG00000055148",
            "net_id": 0
        },
        "ENSMUSG00000055148_ENSMUSG00000041515": {
            "id": "ENSMUSG00000055148_ENSMUSG00000041515",
            "type": "Bridge",
            "score": 0,
            "reversable": false,
            "correlations": {
                "class1": 0.2551153842175999,
                "class2": 0.0
            },
            "regulatory_source": "ENSMUSG00000041515",
            "regulatory_target": "ENSMUSG00000055148",
            "net_id": 0
        },
        "ENSMUSG00000055148_ENSMUSG00000030103": {
            "id": "ENSMUSG00000055148_ENSMUSG00000030103",
            "type": "Bridge",
            "score": 0,
            "reversable": false,
            "correlations": {
                "class1": 0.3553506771966051,
                "class2": 0.0985258890333429
            },
            "regulatory_source": "ENSMUSG00000030103",
            "regulatory_target": "ENSMUSG00000055148",
            "net_id": 0
        },
        "ENSMUSG00000026628_ENSMUSG00000003545": {
            "id": "ENSMUSG00000026628_ENSMUSG00000003545",
            "type": "Bridge",
            "score": 0,
            "reversable": true,
            "correlations": {
                "class1": 0.5416606293768768,
                "class2": 0.21769179459708643
            },
            "regulatory_source": "ENSMUSG00000026628",
            "regulatory_target": "ENSMUSG00000003545",
            "net_id": 0
        },
        "ENSMUSG00000003545_ENSMUSG00000003032": {
            "id": "ENSMUSG00000003545_ENSMUSG00000003032",
            "type": "Bridge",
            "score": 0,
            "reversable": true,
            "correlations": {
                "class1": 0.493082770080834,
                "class2": 0.19603815570936658
            },
            "regulatory_source": "ENSMUSG00000003032",
            "regulatory_target": "ENSMUSG00000003545",
            "net_id": 0
        },
        "ENSMUSG00000056501_ENSMUSG00000003545": {
            "id": "ENSMUSG00000056501_ENSMUSG00000003545",
            "type": "Bridge",
            "score": 0,
            "reversable": false,
            "correlations": {
                "class1": 0.48625587041643803,
                "class2": 0.14725017963761106
            },
            "regulatory_source": "ENSMUSG00000056501",
            "regulatory_target": "ENSMUSG00000003545",
            "net_id": 0
        },
        "ENSMUSG00000022528_ENSMUSG00000003545": {
            "id": "ENSMUSG00000022528_ENSMUSG00000003545",
            "type": "Bridge",
            "score": 0,
            "reversable": false,
            "correlations": {
                "class1": 0.27395899398120127,
                "class2": 0.0716529618935336
            },
            "regulatory_source": "ENSMUSG00000003545",
            "regulatory_target": "ENSMUSG00000022528",
            "net_id": 0
        },
        "ENSMUSG00000030103_ENSMUSG00000003545": {
            "id": "ENSMUSG00000030103_ENSMUSG00000003545",
            "type": "Bridge",
            "score": 0,
            "reversable": false,
            "correlations": {
                "class1": 0.3588044369869844,
                "class2": 0.08915076999811186
            },
            "regulatory_source": "ENSMUSG00000030103",
            "regulatory_target": "ENSMUSG00000003545",
            "net_id": 0
        },
        "ENSMUSG00000037868_ENSMUSG00000026628": {
            "id": "ENSMUSG00000037868_ENSMUSG00000026628",
            "type": "Bridge",
            "score": 0,
            "reversable": true,
            "correlations": {
                "class1": 0.4063797448880915,
                "class2": 0.0
            },
            "regulatory_source": "ENSMUSG00000026628",
            "regulatory_target": "ENSMUSG00000037868",
            "net_id": 0
        },
        "ENSMUSG00000037868_ENSMUSG00000018899": {
            "id": "ENSMUSG00000037868_ENSMUSG00000018899",
            "type": "Bridge",
            "score": 0,
            "reversable": true,
            "correlations": {
                "class1": 0.3524583540848284,
                "class2": 0.04617866736520871
            },
            "regulatory_source": "ENSMUSG00000037868",
            "regulatory_target": "ENSMUSG00000018899",
            "net_id": 0
        },
        "ENSMUSG00000037868_ENSMUSG00000003032": {
            "id": "ENSMUSG00000037868_ENSMUSG00000003032",
            "type": "Bridge",
            "score": 0,
            "reversable": true,
            "correlations": {
                "class1": 0.3220763055992768,
                "class2": 0.08888618214494533
            },
            "regulatory_source": "ENSMUSG00000003032",
            "regulatory_target": "ENSMUSG00000037868",
            "net_id": 0
        },
        "ENSMUSG00000037868_ENSMUSG00000023034": {
            "id": "ENSMUSG00000037868_ENSMUSG00000023034",
            "type": "Bridge",
            "score": 0,
            "reversable": true,
            "correlations": {
                "class1": 0.4154982566003549,
                "class2": 0.10177517111288847
            },
            "regulatory_source": "ENSMUSG00000037868",
            "regulatory_target": "ENSMUSG00000023034",
            "net_id": 0
        },
        "ENSMUSG00000056501_ENSMUSG00000037868": {
            "id": "ENSMUSG00000056501_ENSMUSG00000037868",
            "type": "Bridge",
            "score": 0,
            "reversable": false,
            "correlations": {
                "class1": 0.34456796733082223,
                "class2": 0.12587320524582973
            },
            "regulatory_source": "ENSMUSG00000056501",
            "regulatory_target": "ENSMUSG00000037868",
            "net_id": 0
        },
        "ENSMUSG00000021250_ENSMUSG00000003032": {
            "id": "ENSMUSG00000021250_ENSMUSG00000003032",
            "type": "Bridge",
            "score": 0,
            "reversable": true,
            "correlations": {
                "class1": 0.5037288969641436,
                "class2": 0.2762462140233046
            },
            "regulatory_source": "ENSMUSG00000003032",
            "regulatory_target": "ENSMUSG00000021250",
            "net_id": 0
        },
        "ENSMUSG00000021250_ENSMUSG00000000247": {
            "id": "ENSMUSG00000021250_ENSMUSG00000000247",
            "type": "Bridge",
            "score": 0,
            "reversable": true,
            "correlations": {
                "class1": 0.28651730790481883,
                "class2": 0.0
            },
            "regulatory_source": "ENSMUSG00000000247",
            "regulatory_target": "ENSMUSG00000021250",
            "net_id": 0
        },
        "ENSMUSG00000057098_ENSMUSG00000021250": {
            "id": "ENSMUSG00000057098_ENSMUSG00000021250",
            "type": "Bridge",
            "score": 0,
            "reversable": true,
            "correlations": {
                "class1": 0.0,
                "class2": 0.21223431697525263
            },
            "regulatory_source": "ENSMUSG00000057098",
            "regulatory_target": "ENSMUSG00000021250",
            "net_id": 0
        },
        "ENSMUSG00000041515_ENSMUSG00000021250": {
            "id": "ENSMUSG00000041515_ENSMUSG00000021250",
            "type": "Bridge",
            "score": 0,
            "reversable": true,
            "correlations": {
                "class1": 0.2891361459930216,
                "class2": 0.0
            },
            "regulatory_source": "ENSMUSG00000041515",
            "regulatory_target": "ENSMUSG00000021250",
            "net_id": 0
        },
        "ENSMUSG00000030103_ENSMUSG00000021250": {
            "id": "ENSMUSG00000030103_ENSMUSG00000021250",
            "type": "Bridge",
            "score": 0,
            "reversable": true,
            "correlations": {
                "class1": 0.4017224713299765,
                "class2": 0.11741532314537408
            },
            "regulatory_source": "ENSMUSG00000030103",
            "regulatory_target": "ENSMUSG00000021250",
            "net_id": 0
        },
        "ENSMUSG00000042812_ENSMUSG00000021250": {
            "id": "ENSMUSG00000042812_ENSMUSG00000021250",
            "type": "Bridge",
            "score": 0,
            "reversable": true,
            "correlations": {
                "class1": 0.37890289877717953,
                "class2": 0.13150746269099306
            },
            "regulatory_source": "ENSMUSG00000042812",
            "regulatory_target": "ENSMUSG00000021250",
            "net_id": 0
        },
        "ENSMUSG00000026628_ENSMUSG00000018899": {
            "id": "ENSMUSG00000026628_ENSMUSG00000018899",
            "type": "Bridge",
            "score": 0,
            "reversable": true,
            "correlations": {
                "class1": 0.4620167973110394,
                "class2": 0.11032155798095769
            },
            "regulatory_source": "ENSMUSG00000026628",
            "regulatory_target": "ENSMUSG00000018899",
            "net_id": 0
        },
        "ENSMUSG00000026628_ENSMUSG00000003032": {
            "id": "ENSMUSG00000026628_ENSMUSG00000003032",
            "type": "Bridge",
            "score": 0,
            "reversable": true,
            "correlations": {
                "class1": 0.4361333631428031,
                "class2": 0.15993480357109063
            },
            "regulatory_source": "ENSMUSG00000026628",
            "regulatory_target": "ENSMUSG00000003032",
            "net_id": 0
        },
        "ENSMUSG00000041515_ENSMUSG00000026628": {
            "id": "ENSMUSG00000041515_ENSMUSG00000026628",
            "type": "Bridge",
            "score": 0,
            "reversable": true,
            "correlations": {
                "class1": 0.20585304568274204,
                "class2": -0.0345276717374778
            },
            "regulatory_source": "ENSMUSG00000026628",
            "regulatory_target": "ENSMUSG00000041515",
            "net_id": 0
        },
        "ENSMUSG00000030103_ENSMUSG00000026628": {
            "id": "ENSMUSG00000030103_ENSMUSG00000026628",
            "type": "Bridge",
            "score": 0,
            "reversable": true,
            "correlations": {
                "class1": 0.27058256684972004,
                "class2": 0.06191142393687492
            },
            "regulatory_source": "ENSMUSG00000026628",
            "regulatory_target": "ENSMUSG00000030103",
            "net_id": 0
        },
        "ENSMUSG00000008575_ENSMUSG00000005583": {
            "id": "ENSMUSG00000008575_ENSMUSG00000005583",
            "type": "Bridge",
            "score": 0,
            "reversable": false,
            "correlations": {
                "class1": 0.09446259624146577,
                "class2": -0.17653270521043482
            },
            "regulatory_source": "ENSMUSG00000005583",
            "regulatory_target": "ENSMUSG00000008575",
            "net_id": 0
        },
        "ENSMUSG00000038193_ENSMUSG00000008575": {
            "id": "ENSMUSG00000038193_ENSMUSG00000008575",
            "type": "Bridge",
            "score": 0,
            "reversable": false,
            "correlations": {
                "class1": 0.2559180024093437,
                "class2": 0.0
            },
            "regulatory_source": "ENSMUSG00000038193",
            "regulatory_target": "ENSMUSG00000008575",
            "net_id": 0
        },
        "ENSMUSG00000020484_ENSMUSG00000005583": {
            "id": "ENSMUSG00000020484_ENSMUSG00000005583",
            "type": "Bridge",
            "score": 0,
            "reversable": false,
            "correlations": {
                "class1": 0.07277737973626353,
                "class2": -0.15550150609476054
            },
            "regulatory_source": "ENSMUSG00000005583",
            "regulatory_target": "ENSMUSG00000020484",
            "net_id": 0
        },
        "ENSMUSG00000055435_ENSMUSG00000005583": {
            "id": "ENSMUSG00000055435_ENSMUSG00000005583",
            "type": "Bridge",
            "score": 0,
            "reversable": true,
            "correlations": {
                "class1": 0.03570969930100298,
                "class2": -0.19157702259293735
            },
            "regulatory_source": "ENSMUSG00000055435",
            "regulatory_target": "ENSMUSG00000005583",
            "net_id": 0
        },
        "ENSMUSG00000022528_ENSMUSG00000005583": {
            "id": "ENSMUSG00000022528_ENSMUSG00000005583",
            "type": "Bridge",
            "score": 0,
            "reversable": true,
            "correlations": {
                "class1": 0.06754484462471692,
                "class2": 0.3000855748305624
            },
            "regulatory_source": "ENSMUSG00000005583",
            "regulatory_target": "ENSMUSG00000022528",
            "net_id": 0
        },
        "ENSMUSG00000018899_ENSMUSG00000003032": {
            "id": "ENSMUSG00000018899_ENSMUSG00000003032",
            "type": "Bridge",
            "score": 0,
            "reversable": true,
            "correlations": {
                "class1": 0.3416211914026175,
                "class2": 0.07031236702558624
            },
            "regulatory_source": "ENSMUSG00000003032",
            "regulatory_target": "ENSMUSG00000018899",
            "net_id": 0
        },
        "ENSMUSG00000023034_ENSMUSG00000018899": {
            "id": "ENSMUSG00000023034_ENSMUSG00000018899",
            "type": "Bridge",
            "score": 0,
            "reversable": true,
            "correlations": {
                "class1": 0.42659015486606466,
                "class2": 0.09712207370873793
            },
            "regulatory_source": "ENSMUSG00000018899",
            "regulatory_target": "ENSMUSG00000023034",
            "net_id": 0
        },
        "ENSMUSG00000056501_ENSMUSG00000018899": {
            "id": "ENSMUSG00000056501_ENSMUSG00000018899",
            "type": "Bridge",
            "score": 0,
            "reversable": true,
            "correlations": {
                "class1": 0.44764444607399667,
                "class2": 0.13564226130234508
            },
            "regulatory_source": "ENSMUSG00000056501",
            "regulatory_target": "ENSMUSG00000018899",
            "net_id": 0
        },
        "ENSMUSG00000041515_ENSMUSG00000018899": {
            "id": "ENSMUSG00000041515_ENSMUSG00000018899",
            "type": "Bridge",
            "score": 0,
            "reversable": true,
            "correlations": {
                "class1": 0.2987049056662975,
                "class2": 0.07379600005850777
            },
            "regulatory_source": "ENSMUSG00000041515",
            "regulatory_target": "ENSMUSG00000018899",
            "net_id": 0
        },
        "ENSMUSG00000030103_ENSMUSG00000018899": {
            "id": "ENSMUSG00000030103_ENSMUSG00000018899",
            "type": "Bridge",
            "score": 0,
            "reversable": true,
            "correlations": {
                "class1": 0.28967740822917215,
                "class2": 0.06609765089652861
            },
            "regulatory_source": "ENSMUSG00000030103",
            "regulatory_target": "ENSMUSG00000018899",
            "net_id": 0
        },
        "ENSMUSG00000022346_ENSMUSG00000018899": {
            "id": "ENSMUSG00000022346_ENSMUSG00000018899",
            "type": "Bridge",
            "score": 0,
            "reversable": true,
            "correlations": {
                "class1": 0.31162484317659334,
                "class2": 0.07878125796866468
            },
            "regulatory_source": "ENSMUSG00000018899",
            "regulatory_target": "ENSMUSG00000022346",
            "net_id": 0
        },
        "ENSMUSG00000042406_ENSMUSG00000018899": {
            "id": "ENSMUSG00000042406_ENSMUSG00000018899",
            "type": "Bridge",
            "score": 0,
            "reversable": false,
            "correlations": {
                "class1": 0.2782115321349435,
                "class2": 0.06516281782394828
            },
            "regulatory_source": "ENSMUSG00000042406",
            "regulatory_target": "ENSMUSG00000018899",
            "net_id": 0
        },
        "ENSMUSG00000038193_ENSMUSG00000020484": {
            "id": "ENSMUSG00000038193_ENSMUSG00000020484",
            "type": "Bridge",
            "score": 0,
            "reversable": false,
            "correlations": {
                "class1": 0.22006528532540173,
                "class2": 0.0
            },
            "regulatory_source": "ENSMUSG00000038193",
            "regulatory_target": "ENSMUSG00000020484",
            "net_id": 0
        },
        "ENSMUSG00000038193_ENSMUSG00000000247": {
            "id": "ENSMUSG00000038193_ENSMUSG00000000247",
            "type": "Bridge",
            "score": 0,
            "reversable": false,
            "correlations": {
                "class1": 0.320483784973289,
                "class2": 0.027428079856191428
            },
            "regulatory_source": "ENSMUSG00000038193",
            "regulatory_target": "ENSMUSG00000000247",
            "net_id": 0
        },
        "ENSMUSG00000023034_ENSMUSG00000020484": {
            "id": "ENSMUSG00000023034_ENSMUSG00000020484",
            "type": "Bridge",
            "score": 0,
            "reversable": false,
            "correlations": {
                "class1": 0.20325637917960476,
                "class2": 0.0
            },
            "regulatory_source": "ENSMUSG00000023034",
            "regulatory_target": "ENSMUSG00000020484",
            "net_id": 0
        },
        "ENSMUSG00000022528_ENSMUSG00000020484": {
            "id": "ENSMUSG00000022528_ENSMUSG00000020484",
            "type": "Bridge",
            "score": 0,
            "reversable": false,
            "correlations": {
                "class1": 0.19037658522941078,
                "class2": -0.04185243092052634
            },
            "regulatory_source": "ENSMUSG00000020484",
            "regulatory_target": "ENSMUSG00000022528",
            "net_id": 0
        },
        "ENSMUSG00000023034_ENSMUSG00000003032": {
            "id": "ENSMUSG00000023034_ENSMUSG00000003032",
            "type": "Bridge",
            "score": 0,
            "reversable": true,
            "correlations": {
                "class1": 0.42701224797172904,
                "class2": 0.11349242150073609
            },
            "regulatory_source": "ENSMUSG00000003032",
            "regulatory_target": "ENSMUSG00000023034",
            "net_id": 0
        },
        "ENSMUSG00000057098_ENSMUSG00000003032": {
            "id": "ENSMUSG00000057098_ENSMUSG00000003032",
            "type": "Bridge",
            "score": 0,
            "reversable": true,
            "correlations": {
                "class1": 0.07841903401280102,
                "class2": 0.32375805075567277
            },
            "regulatory_source": "ENSMUSG00000003032",
            "regulatory_target": "ENSMUSG00000057098",
            "net_id": 0
        },
        "ENSMUSG00000023034_ENSMUSG00000000247": {
            "id": "ENSMUSG00000023034_ENSMUSG00000000247",
            "type": "Bridge",
            "score": 0,
            "reversable": false,
            "correlations": {
                "class1": 0.18810663843122466,
                "class2": -0.041273943748537295
            },
            "regulatory_source": "ENSMUSG00000023034",
            "regulatory_target": "ENSMUSG00000000247",
            "net_id": 0
        },
        "ENSMUSG00000041515_ENSMUSG00000023034": {
            "id": "ENSMUSG00000041515_ENSMUSG00000023034",
            "type": "Bridge",
            "score": 0,
            "reversable": true,
            "correlations": {
                "class1": 0.22416182379663774,
                "class2": -0.044447214579342406
            },
            "regulatory_source": "ENSMUSG00000041515",
            "regulatory_target": "ENSMUSG00000023034",
            "net_id": 0
        },
        "ENSMUSG00000030103_ENSMUSG00000023034": {
            "id": "ENSMUSG00000030103_ENSMUSG00000023034",
            "type": "Bridge",
            "score": 0,
            "reversable": true,
            "correlations": {
                "class1": 0.37953707754189986,
                "class2": 0.08455151647419093
            },
            "regulatory_source": "ENSMUSG00000030103",
            "regulatory_target": "ENSMUSG00000023034",
            "net_id": 0
        },
        "ENSMUSG00000048387_ENSMUSG00000022346": {
            "id": "ENSMUSG00000048387_ENSMUSG00000022346",
            "type": "Bridge",
            "score": 0,
            "reversable": true,
            "correlations": {
                "class1": 0.0,
                "class2": 0.2001949148118965
            },
            "regulatory_source": "ENSMUSG00000048387",
            "regulatory_target": "ENSMUSG00000022346",
            "net_id": 0
        },
        "ENSMUSG00000056501_ENSMUSG00000000247": {
            "id": "ENSMUSG00000056501_ENSMUSG00000000247",
            "type": "Bridge",
            "score": 0,
            "reversable": false,
            "correlations": {
                "class1": 0.20750755324609726,
                "class2": 0.0
            },
            "regulatory_source": "ENSMUSG00000056501",
            "regulatory_target": "ENSMUSG00000000247",
            "net_id": 0
        },
        "ENSMUSG00000042812_ENSMUSG00000000247": {
            "id": "ENSMUSG00000042812_ENSMUSG00000000247",
            "type": "Bridge",
            "score": 0,
            "reversable": false,
            "correlations": {
                "class1": 0.2519458084343931,
                "class2": 0.03279768643123066
            },
            "regulatory_source": "ENSMUSG00000000247",
            "regulatory_target": "ENSMUSG00000042812",
            "net_id": 0
        },
        "ENSMUSG00000057098_ENSMUSG00000029322": {
            "id": "ENSMUSG00000057098_ENSMUSG00000029322",
            "type": "Bridge",
            "score": 0,
            "reversable": false,
            "correlations": {
                "class1": 0.0,
                "class2": 0.23115942033960685
            },
            "regulatory_source": "ENSMUSG00000057098",
            "regulatory_target": "ENSMUSG00000029322",
            "net_id": 0
        },
        "ENSMUSG00000056501_ENSMUSG00000041515": {
            "id": "ENSMUSG00000056501_ENSMUSG00000041515",
            "type": "Bridge",
            "score": 0,
            "reversable": true,
            "correlations": {
                "class1": 0.2729074590935114,
                "class2": 0.042441543877284924
            },
            "regulatory_source": "ENSMUSG00000056501",
            "regulatory_target": "ENSMUSG00000041515",
            "net_id": 0
        },
        "ENSMUSG00000056501_ENSMUSG00000030103": {
            "id": "ENSMUSG00000056501_ENSMUSG00000030103",
            "type": "Bridge",
            "score": 0,
            "reversable": true,
            "correlations": {
                "class1": 0.3820252477406918,
                "class2": 0.11188027728042743
            },
            "regulatory_source": "ENSMUSG00000056501",
            "regulatory_target": "ENSMUSG00000030103",
            "net_id": 0
        },
        "ENSMUSG00000032035_ENSMUSG00000029322": {
            "id": "ENSMUSG00000032035_ENSMUSG00000029322",
            "type": "Bridge",
            "score": 0,
            "reversable": false,
            "correlations": {
                "class1": 0.18458180501786528,
                "class2": -0.03625181858454962
            },
            "regulatory_source": "ENSMUSG00000032035",
            "regulatory_target": "ENSMUSG00000029322",
            "net_id": 0
        },
        "ENSMUSG00000032035_ENSMUSG00000015846": {
            "id": "ENSMUSG00000032035_ENSMUSG00000015846",
            "type": "Bridge",
            "score": 0,
            "reversable": false,
            "correlations": {
                "class1": 0.2132775107072164,
                "class2": 0.0
            },
            "regulatory_source": "ENSMUSG00000015846",
            "regulatory_target": "ENSMUSG00000032035",
            "net_id": 0
        },
        "ENSMUSG00000028639_ENSMUSG00000000247": {
            "id": "ENSMUSG00000028639_ENSMUSG00000000247",
            "type": "Bridge",
            "score": 0,
            "reversable": false,
            "correlations": {
                "class1": 0.24771885582010925,
                "class2": 0.0
            },
            "regulatory_source": "ENSMUSG00000000247",
            "regulatory_target": "ENSMUSG00000028639",
            "net_id": 0
        },
        "ENSMUSG00000057098_ENSMUSG00000028639": {
            "id": "ENSMUSG00000057098_ENSMUSG00000028639",
            "type": "Bridge",
            "score": 0,
            "reversable": false,
            "correlations": {
                "class1": 0.0475896700096964,
                "class2": 0.30769158319752227
            },
            "regulatory_source": "ENSMUSG00000057098",
            "regulatory_target": "ENSMUSG00000028639",
            "net_id": 0
        },
        "ENSMUSG00000020644_ENSMUSG00000005583": {
            "id": "ENSMUSG00000020644_ENSMUSG00000005583",
            "type": "Bridge",
            "score": 0,
            "reversable": false,
            "correlations": {
                "class1": 0.0,
                "class2": 0.23437047544831524
            },
            "regulatory_source": "ENSMUSG00000005583",
            "regulatory_target": "ENSMUSG00000020644",
            "net_id": 0
        },
        "ENSMUSG00000030103_ENSMUSG00000020644": {
            "id": "ENSMUSG00000030103_ENSMUSG00000020644",
            "type": "Bridge",
            "score": 0,
            "reversable": false,
            "correlations": {
                "class1": 0.21201816555561567,
                "class2": 0.0
            },
            "regulatory_source": "ENSMUSG00000030103",
            "regulatory_target": "ENSMUSG00000020644",
            "net_id": 0
        }
    }
}
```
