## Supplement 4 (CCl4 a6w HSC vs. PF) for "AGEAS: Automated Machine Learning based Genetic Regulatory Element Extraction System": meta_GRN.js

```
{
    "genes": {
        "ENSMUSG00000052534": {
            "id": "ENSMUSG00000052534",
            "type": "TF",
            "symbol": "Pbx1",
            "source": [
                "ENSMUSG00000052684"
            ],
            "target": [
                "ENSMUSG00000024063",
                "ENSMUSG00000029309",
                "ENSMUSG00000052684",
                "ENSMUSG00000026335",
                "ENSMUSG00000025784",
                "ENSMUSG00000026566",
                "ENSMUSG00000072235",
                "ENSMUSG00000024048",
                "ENSMUSG00000035258",
                "ENSMUSG00000016194",
                "ENSMUSG00000022636",
                "ENSMUSG00000039601"
            ],
            "expression_sum": {
                "class1": 9798.0,
                "class2": 8386.0
            },
            "ens_id": "ENSMUSG00000052534",
            "uniprot_ids": [
                "P41778",
                "D9J2V6"
            ]
        },
        "ENSMUSG00000026335": {
            "id": "ENSMUSG00000026335",
            "type": "Gene",
            "symbol": "Pam",
            "source": [
                "ENSMUSG00000052534",
                "ENSMUSG00000005836",
                "ENSMUSG00000071076",
                "ENSMUSG00000008575"
            ],
            "target": [],
            "expression_sum": {
                "class1": 64954.0,
                "class2": 17233.0
            },
            "ens_id": "ENSMUSG00000026335",
            "uniprot_ids": [
                "P97467",
                "E9Q704",
                "F6TGI4",
                "F8VQA4",
                "F7D4K4",
                "F6T0D3"
            ]
        },
        "ENSMUSG00000026566": {
            "id": "ENSMUSG00000026566",
            "type": "Gene",
            "symbol": "Mpzl1",
            "source": [
                "ENSMUSG00000052534",
                "ENSMUSG00000038193",
                "ENSMUSG00000000247"
            ],
            "target": [],
            "expression_sum": {
                "class1": 7509.0,
                "class2": 1360.0
            },
            "ens_id": "ENSMUSG00000026566",
            "uniprot_ids": [
                "Q3TEW6",
                "A0A0A6YWA2",
                "A0A0A6YXL8",
                "A0A0A6YXX8",
                "A0A0A6YX14",
                "A0A0A6YW22",
                "A0A0A6YXW1",
                "A0A0R4J0V2"
            ]
        },
        "ENSMUSG00000016194": {
            "id": "ENSMUSG00000016194",
            "type": "Gene",
            "symbol": "Hsd11b1",
            "source": [
                "ENSMUSG00000038418",
                "ENSMUSG00000041515",
                "ENSMUSG00000052534",
                "ENSMUSG00000005583"
            ],
            "target": [],
            "expression_sum": {
                "class1": 58577.0,
                "class2": 5077.0
            },
            "ens_id": "ENSMUSG00000016194",
            "uniprot_ids": [
                "P50172",
                "Q3TJI8",
                "F6TSI8",
                "F2Z3U6"
            ]
        },
        "ENSMUSG00000052684": {
            "id": "ENSMUSG00000052684",
            "type": "TF",
            "symbol": "Jun",
            "source": [
                "ENSMUSG00000055435",
                "ENSMUSG00000056501",
                "ENSMUSG00000037868",
                "ENSMUSG00000023034",
                "ENSMUSG00000022528",
                "ENSMUSG00000052837",
                "ENSMUSG00000026628",
                "ENSMUSG00000000247",
                "ENSMUSG00000038418",
                "ENSMUSG00000052534",
                "ENSMUSG00000021250",
                "ENSMUSG00000038193",
                "ENSMUSG00000003545",
                "ENSMUSG00000008575",
                "ENSMUSG00000018899"
            ],
            "target": [
                "ENSMUSG00000031431",
                "ENSMUSG00000040723",
                "ENSMUSG00000029761",
                "ENSMUSG00000025381",
                "ENSMUSG00000032925",
                "ENSMUSG00000091345",
                "ENSMUSG00000028111",
                "ENSMUSG00000030222",
                "ENSMUSG00000021477",
                "ENSMUSG00000028691",
                "ENSMUSG00000003545",
                "ENSMUSG00000022365",
                "ENSMUSG00000025213",
                "ENSMUSG00000025289",
                "ENSMUSG00000026879",
                "ENSMUSG00000009281",
                "ENSMUSG00000053560",
                "ENSMUSG00000037411",
                "ENSMUSG00000064373",
                "ENSMUSG00000022528",
                "ENSMUSG00000034868",
                "ENSMUSG00000042613",
                "ENSMUSG00000006699",
                "ENSMUSG00000000325",
                "ENSMUSG00000027239",
                "ENSMUSG00000008384",
                "ENSMUSG00000026043",
                "ENSMUSG00000044627",
                "ENSMUSG00000079111",
                "ENSMUSG00000055148",
                "ENSMUSG00000042485",
                "ENSMUSG00000018339",
                "ENSMUSG00000041417",
                "ENSMUSG00000012422",
                "ENSMUSG00000078974",
                "ENSMUSG00000023004",
                "ENSMUSG00000026421",
                "ENSMUSG00000002107",
                "ENSMUSG00000073889",
                "ENSMUSG00000014329",
                "ENSMUSG00000031007",
                "ENSMUSG00000022010",
                "ENSMUSG00000028369",
                "ENSMUSG00000029103",
                "ENSMUSG00000003541",
                "ENSMUSG00000032231",
                "ENSMUSG00000008575",
                "ENSMUSG00000030218",
                "ENSMUSG00000038872",
                "ENSMUSG00000007872",
                "ENSMUSG00000053317",
                "ENSMUSG00000027848",
                "ENSMUSG00000022836",
                "ENSMUSG00000028597",
                "ENSMUSG00000029309",
                "ENSMUSG00000031845",
                "ENSMUSG00000022032",
                "ENSMUSG00000029718",
                "ENSMUSG00000028967",
                "ENSMUSG00000021390",
                "ENSMUSG00000027828",
                "ENSMUSG00000031595",
                "ENSMUSG00000036655",
                "ENSMUSG00000009681",
                "ENSMUSG00000060904",
                "ENSMUSG00000064370",
                "ENSMUSG00000020053",
                "ENSMUSG00000021917",
                "ENSMUSG00000041120",
                "ENSMUSG00000028339",
                "ENSMUSG00000037999",
                "ENSMUSG00000069601",
                "ENSMUSG00000052534",
                "ENSMUSG00000053113",
                "ENSMUSG00000024140",
                "ENSMUSG00000036372",
                "ENSMUSG00000037266",
                "ENSMUSG00000033538",
                "ENSMUSG00000019907",
                "ENSMUSG00000001119",
                "ENSMUSG00000036446",
                "ENSMUSG00000075010",
                "ENSMUSG00000031451",
                "ENSMUSG00000014956",
                "ENSMUSG00000029390",
                "ENSMUSG00000024076",
                "ENSMUSG00000055681",
                "ENSMUSG00000090000",
                "ENSMUSG00000021760",
                "ENSMUSG00000019278",
                "ENSMUSG00000003477",
                "ENSMUSG00000021248",
                "ENSMUSG00000031616",
                "ENSMUSG00000001506",
                "ENSMUSG00000003038",
                "ENSMUSG00000021870",
                "ENSMUSG00000028871",
                "ENSMUSG00000015501",
                "ENSMUSG00000041959",
                "ENSMUSG00000023915",
                "ENSMUSG00000033585",
                "ENSMUSG00000026628",
                "ENSMUSG00000030103",
                "ENSMUSG00000027706",
                "ENSMUSG00000042812",
                "ENSMUSG00000019579",
                "ENSMUSG00000041828",
                "ENSMUSG00000031586",
                "ENSMUSG00000092274",
                "ENSMUSG00000032515",
                "ENSMUSG00000068735",
                "ENSMUSG00000028776",
                "ENSMUSG00000042737",
                "ENSMUSG00000018672",
                "ENSMUSG00000017466",
                "ENSMUSG00000074457",
                "ENSMUSG00000024065",
                "ENSMUSG00000027829",
                "ENSMUSG00000029061",
                "ENSMUSG00000028195",
                "ENSMUSG00000032334",
                "ENSMUSG00000004631",
                "ENSMUSG00000021253",
                "ENSMUSG00000072235",
                "ENSMUSG00000005873",
                "ENSMUSG00000038463",
                "ENSMUSG00000035678",
                "ENSMUSG00000016024",
                "ENSMUSG00000072625",
                "ENSMUSG00000015085",
                "ENSMUSG00000030208",
                "ENSMUSG00000020486",
                "ENSMUSG00000068220",
                "ENSMUSG00000028517",
                "ENSMUSG00000040212",
                "ENSMUSG00000023034",
                "ENSMUSG00000079523",
                "ENSMUSG00000030654",
                "ENSMUSG00000016494",
                "ENSMUSG00000038393",
                "ENSMUSG00000042453",
                "ENSMUSG00000037706",
                "ENSMUSG00000018217",
                "ENSMUSG00000023046",
                "ENSMUSG00000058056",
                "ENSMUSG00000042745",
                "ENSMUSG00000003161",
                "ENSMUSG00000032359",
                "ENSMUSG00000060591",
                "ENSMUSG00000025784",
                "ENSMUSG00000040562",
                "ENSMUSG00000072812",
                "ENSMUSG00000015312",
                "ENSMUSG00000001911",
                "ENSMUSG00000031661",
                "ENSMUSG00000021871",
                "ENSMUSG00000025880",
                "ENSMUSG00000025724",
                "ENSMUSG00000018819",
                "ENSMUSG00000037211",
                "ENSMUSG00000031239",
                "ENSMUSG00000020423",
                "ENSMUSG00000044408",
                "ENSMUSG00000027712",
                "ENSMUSG00000042254",
                "ENSMUSG00000024048",
                "ENSMUSG00000037072",
                "ENSMUSG00000020009",
                "ENSMUSG00000002732",
                "ENSMUSG00000021876",
                "ENSMUSG00000025780",
                "ENSMUSG00000024164",
                "ENSMUSG00000096145",
                "ENSMUSG00000025147",
                "ENSMUSG00000037868",
                "ENSMUSG00000022122",
                "ENSMUSG00000042312",
                "ENSMUSG00000006205",
                "ENSMUSG00000001663",
                "ENSMUSG00000020473",
                "ENSMUSG00000035828",
                "ENSMUSG00000000594",
                "ENSMUSG00000064215",
                "ENSMUSG00000006050",
                "ENSMUSG00000006519",
                "ENSMUSG00000045658",
                "ENSMUSG00000030717",
                "ENSMUSG00000031799",
                "ENSMUSG00000017493",
                "ENSMUSG00000023010",
                "ENSMUSG00000041084",
                "ENSMUSG00000025351",
                "ENSMUSG00000027907",
                "ENSMUSG00000074884",
                "ENSMUSG00000039601",
                "ENSMUSG00000054808",
                "ENSMUSG00000056501",
                "ENSMUSG00000021242",
                "ENSMUSG00000000628",
                "ENSMUSG00000002014",
                "ENSMUSG00000092341",
                "ENSMUSG00000039899",
                "ENSMUSG00000055435",
                "ENSMUSG00000033910",
                "ENSMUSG00000087006",
                "ENSMUSG00000018593",
                "ENSMUSG00000000753",
                "ENSMUSG00000052957",
                "ENSMUSG00000015568",
                "ENSMUSG00000028864",
                "ENSMUSG00000006800",
                "ENSMUSG00000020585",
                "ENSMUSG00000031380",
                "ENSMUSG00000032353",
                "ENSMUSG00000025492",
                "ENSMUSG00000051579",
                "ENSMUSG00000066705",
                "ENSMUSG00000046402",
                "ENSMUSG00000021250",
                "ENSMUSG00000042747",
                "ENSMUSG00000041355",
                "ENSMUSG00000029096",
                "ENSMUSG00000026837",
                "ENSMUSG00000031548",
                "ENSMUSG00000028618",
                "ENSMUSG00000032383",
                "ENSMUSG00000026748",
                "ENSMUSG00000007097",
                "ENSMUSG00000054203",
                "ENSMUSG00000006369",
                "ENSMUSG00000031422",
                "ENSMUSG00000035356",
                "ENSMUSG00000027859",
                "ENSMUSG00000047945",
                "ENSMUSG00000049107",
                "ENSMUSG00000052837",
                "ENSMUSG00000030342",
                "ENSMUSG00000022174",
                "ENSMUSG00000035227",
                "ENSMUSG00000024012",
                "ENSMUSG00000029659",
                "ENSMUSG00000040170",
                "ENSMUSG00000029641",
                "ENSMUSG00000029661",
                "ENSMUSG00000044786",
                "ENSMUSG00000008540",
                "ENSMUSG00000018899",
                "ENSMUSG00000021268",
                "ENSMUSG00000079037",
                "ENSMUSG00000090698",
                "ENSMUSG00000075702",
                "ENSMUSG00000006403",
                "ENSMUSG00000060703",
                "ENSMUSG00000000247",
                "ENSMUSG00000038418",
                "ENSMUSG00000052997",
                "ENSMUSG00000030849",
                "ENSMUSG00000001025",
                "ENSMUSG00000058135"
            ],
            "expression_sum": {
                "class1": 84935.0,
                "class2": 57932.0
            },
            "ens_id": "ENSMUSG00000052684",
            "uniprot_ids": [
                "P05627"
            ]
        },
        "ENSMUSG00000029309": {
            "id": "ENSMUSG00000029309",
            "type": "Gene",
            "symbol": "Sparcl1",
            "source": [
                "ENSMUSG00000052534",
                "ENSMUSG00000005583",
                "ENSMUSG00000052684",
                "ENSMUSG00000071076"
            ],
            "target": [],
            "expression_sum": {
                "class1": 20109.0,
                "class2": 38424.0
            },
            "ens_id": "ENSMUSG00000029309",
            "uniprot_ids": [
                "P70663",
                "A0A0G2JFH7"
            ]
        },
        "ENSMUSG00000025784": {
            "id": "ENSMUSG00000025784",
            "type": "Gene",
            "symbol": "Clec3b",
            "source": [
                "ENSMUSG00000052684",
                "ENSMUSG00000022346",
                "ENSMUSG00000052534",
                "ENSMUSG00000003032",
                "ENSMUSG00000028565",
                "ENSMUSG00000071076"
            ],
            "target": [],
            "expression_sum": {
                "class1": 197.0,
                "class2": 48804.0
            },
            "ens_id": "ENSMUSG00000025784",
            "uniprot_ids": [
                "Q8CFZ6"
            ]
        },
        "ENSMUSG00000072235": {
            "id": "ENSMUSG00000072235",
            "type": "Gene",
            "symbol": "Tuba1a",
            "source": [
                "ENSMUSG00000055435",
                "ENSMUSG00000032035",
                "ENSMUSG00000052684",
                "ENSMUSG00000005583",
                "ENSMUSG00000057098",
                "ENSMUSG00000052534",
                "ENSMUSG00000003032",
                "ENSMUSG00000008575",
                "ENSMUSG00000071076"
            ],
            "target": [],
            "expression_sum": {
                "class1": 27755.0,
                "class2": 25732.0
            },
            "ens_id": "ENSMUSG00000072235",
            "uniprot_ids": [
                "P68369"
            ]
        },
        "ENSMUSG00000022636": {
            "id": "ENSMUSG00000022636",
            "type": "Gene",
            "symbol": "Alcam",
            "source": [
                "ENSMUSG00000052534"
            ],
            "target": [],
            "expression_sum": {
                "class1": 10520.0,
                "class2": 341.0
            },
            "ens_id": "ENSMUSG00000022636",
            "uniprot_ids": [
                "Q61490",
                "Q5MPX5",
                "E9Q4G8",
                "F6QH25",
                "E9Q3Q6"
            ]
        },
        "ENSMUSG00000035258": {
            "id": "ENSMUSG00000035258",
            "type": "Gene",
            "symbol": "Abi3bp",
            "source": [
                "ENSMUSG00000052534",
                "ENSMUSG00000005583",
                "ENSMUSG00000071076"
            ],
            "target": [],
            "expression_sum": {
                "class1": 11775.0,
                "class2": 5297.0
            },
            "ens_id": "ENSMUSG00000035258",
            "uniprot_ids": [
                "Q59IW9",
                "Q59IW7",
                "Q59IW6",
                "Q59IW5",
                "A0A338P6S8",
                "A0A338P6W9",
                "E9Q8E3"
            ]
        },
        "ENSMUSG00000039601": {
            "id": "ENSMUSG00000039601",
            "type": "Gene",
            "symbol": "Rcan2",
            "source": [
                "ENSMUSG00000056501",
                "ENSMUSG00000052684",
                "ENSMUSG00000023034",
                "ENSMUSG00000052837",
                "ENSMUSG00000026628",
                "ENSMUSG00000015839",
                "ENSMUSG00000052534",
                "ENSMUSG00000038193",
                "ENSMUSG00000021250",
                "ENSMUSG00000003545",
                "ENSMUSG00000003032",
                "ENSMUSG00000005583",
                "ENSMUSG00000071076"
            ],
            "target": [],
            "expression_sum": {
                "class1": 138.0,
                "class2": 39503.0
            },
            "ens_id": "ENSMUSG00000039601",
            "uniprot_ids": [
                "Q9JHG2",
                "Q8VIP4",
                "A0A2R8W6E8",
                "A0A2R8VKI0",
                "A0A2R8W6X3"
            ]
        },
        "ENSMUSG00000024048": {
            "id": "ENSMUSG00000024048",
            "type": "Gene",
            "symbol": "Myl12a",
            "source": [
                "ENSMUSG00000055435",
                "ENSMUSG00000032238",
                "ENSMUSG00000032035",
                "ENSMUSG00000052684",
                "ENSMUSG00000005583",
                "ENSMUSG00000048387",
                "ENSMUSG00000057098",
                "ENSMUSG00000038418",
                "ENSMUSG00000052534",
                "ENSMUSG00000021250",
                "ENSMUSG00000003032",
                "ENSMUSG00000008575",
                "ENSMUSG00000028565",
                "ENSMUSG00000071076"
            ],
            "target": [],
            "expression_sum": {
                "class1": 52081.0,
                "class2": 15595.0
            },
            "ens_id": "ENSMUSG00000024048",
            "uniprot_ids": [
                "Q6ZWQ9",
                "D3Z249",
                "D3YV37"
            ]
        },
        "ENSMUSG00000024063": {
            "id": "ENSMUSG00000024063",
            "type": "Gene",
            "symbol": "Lbh",
            "source": [
                "ENSMUSG00000000247",
                "ENSMUSG00000053477",
                "ENSMUSG00000052534",
                "ENSMUSG00000008575",
                "ENSMUSG00000055435",
                "ENSMUSG00000020484"
            ],
            "target": [],
            "expression_sum": {
                "class1": 23121.0,
                "class2": 10505.0
            },
            "ens_id": "ENSMUSG00000024063",
            "uniprot_ids": [
                "Q9CX60",
                "F6UVG6"
            ]
        },
        "ENSMUSG00000026628": {
            "id": "ENSMUSG00000026628",
            "type": "TF",
            "symbol": "Atf3",
            "source": [
                "ENSMUSG00000041515",
                "ENSMUSG00000037868",
                "ENSMUSG00000052684",
                "ENSMUSG00000042406",
                "ENSMUSG00000052837",
                "ENSMUSG00000030103",
                "ENSMUSG00000038418",
                "ENSMUSG00000003545",
                "ENSMUSG00000003032",
                "ENSMUSG00000018899",
                "ENSMUSG00000071076"
            ],
            "target": [
                "ENSMUSG00000006360",
                "ENSMUSG00000031765",
                "ENSMUSG00000024014",
                "ENSMUSG00000000628",
                "ENSMUSG00000053113",
                "ENSMUSG00000056708",
                "ENSMUSG00000090841",
                "ENSMUSG00000003545",
                "ENSMUSG00000033538",
                "ENSMUSG00000001627",
                "ENSMUSG00000053560",
                "ENSMUSG00000037411",
                "ENSMUSG00000052684",
                "ENSMUSG00000037405",
                "ENSMUSG00000021025",
                "ENSMUSG00000038370",
                "ENSMUSG00000042406",
                "ENSMUSG00000036438",
                "ENSMUSG00000019960",
                "ENSMUSG00000008384",
                "ENSMUSG00000031762",
                "ENSMUSG00000003032",
                "ENSMUSG00000026864",
                "ENSMUSG00000021127",
                "ENSMUSG00000024190",
                "ENSMUSG00000037211",
                "ENSMUSG00000041515",
                "ENSMUSG00000015501",
                "ENSMUSG00000023904",
                "ENSMUSG00000029380",
                "ENSMUSG00000047945",
                "ENSMUSG00000052837",
                "ENSMUSG00000020423",
                "ENSMUSG00000030103",
                "ENSMUSG00000035356",
                "ENSMUSG00000073411",
                "ENSMUSG00000097296",
                "ENSMUSG00000032515",
                "ENSMUSG00000020205",
                "ENSMUSG00000044786",
                "ENSMUSG00000071076",
                "ENSMUSG00000018899",
                "ENSMUSG00000027829",
                "ENSMUSG00000090698",
                "ENSMUSG00000037868",
                "ENSMUSG00000006403",
                "ENSMUSG00000032501",
                "ENSMUSG00000038418",
                "ENSMUSG00000034855",
                "ENSMUSG00000035678",
                "ENSMUSG00000073489",
                "ENSMUSG00000009681",
                "ENSMUSG00000039601"
            ],
            "expression_sum": {
                "class1": 16194.0,
                "class2": 10101.0
            },
            "ens_id": "ENSMUSG00000026628",
            "uniprot_ids": [
                "Q60765"
            ]
        },
        "ENSMUSG00000020423": {
            "id": "ENSMUSG00000020423",
            "type": "Gene",
            "symbol": "Btg2",
            "source": [
                "ENSMUSG00000041515",
                "ENSMUSG00000056501",
                "ENSMUSG00000052684",
                "ENSMUSG00000037868",
                "ENSMUSG00000052837",
                "ENSMUSG00000026628",
                "ENSMUSG00000030103",
                "ENSMUSG00000022346",
                "ENSMUSG00000038418",
                "ENSMUSG00000021250",
                "ENSMUSG00000055148",
                "ENSMUSG00000003545",
                "ENSMUSG00000003032",
                "ENSMUSG00000018899",
                "ENSMUSG00000020484",
                "ENSMUSG00000071076"
            ],
            "target": [],
            "expression_sum": {
                "class1": 26463.0,
                "class2": 18373.0
            },
            "ens_id": "ENSMUSG00000020423",
            "uniprot_ids": [
                "Q04211"
            ]
        },
        "ENSMUSG00000056708": {
            "id": "ENSMUSG00000056708",
            "type": "Gene",
            "symbol": "Ier5",
            "source": [
                "ENSMUSG00000056501",
                "ENSMUSG00000023034",
                "ENSMUSG00000052837",
                "ENSMUSG00000026628",
                "ENSMUSG00000021250",
                "ENSMUSG00000003545",
                "ENSMUSG00000055435",
                "ENSMUSG00000055148",
                "ENSMUSG00000018899"
            ],
            "target": [],
            "expression_sum": {
                "class1": 9593.0,
                "class2": 4353.0
            },
            "ens_id": "ENSMUSG00000056708",
            "uniprot_ids": [
                "O89113"
            ]
        },
        "ENSMUSG00000038370": {
            "id": "ENSMUSG00000038370",
            "type": "Gene",
            "symbol": "Pcp4l1",
            "source": [
                "ENSMUSG00000058756",
                "ENSMUSG00000026628",
                "ENSMUSG00000021250",
                "ENSMUSG00000005583"
            ],
            "target": [],
            "expression_sum": {
                "class1": 16.0,
                "class2": 15855.0
            },
            "ens_id": "ENSMUSG00000038370",
            "uniprot_ids": [
                "Q6W8Q3"
            ]
        },
        "ENSMUSG00000006403": {
            "id": "ENSMUSG00000006403",
            "type": "Gene",
            "symbol": "Adamts4",
            "source": [
                "ENSMUSG00000056501",
                "ENSMUSG00000037868",
                "ENSMUSG00000052684",
                "ENSMUSG00000026628",
                "ENSMUSG00000021250",
                "ENSMUSG00000018899",
                "ENSMUSG00000071076"
            ],
            "target": [],
            "expression_sum": {
                "class1": 2530.0,
                "class2": 700.0
            },
            "ens_id": "ENSMUSG00000006403",
            "uniprot_ids": [
                "Q8BNJ2",
                "Q3TNX8",
                "E9PV91"
            ]
        },
        "ENSMUSG00000073489": {
            "id": "ENSMUSG00000073489",
            "type": "Gene",
            "symbol": "Ifi204",
            "source": [
                "ENSMUSG00000032035",
                "ENSMUSG00000056501",
                "ENSMUSG00000026628",
                "ENSMUSG00000005583",
                "ENSMUSG00000055435",
                "ENSMUSG00000018899"
            ],
            "target": [],
            "expression_sum": {
                "class1": 11148.0,
                "class2": 3014.0
            },
            "ens_id": "ENSMUSG00000073489",
            "uniprot_ids": [
                "P0DOV2"
            ]
        },
        "ENSMUSG00000026864": {
            "id": "ENSMUSG00000026864",
            "type": "Gene",
            "symbol": "Hspa5",
            "source": [
                "ENSMUSG00000026628",
                "ENSMUSG00000005583"
            ],
            "target": [],
            "expression_sum": {
                "class1": 40128.0,
                "class2": 24709.0
            },
            "ens_id": "ENSMUSG00000026864",
            "uniprot_ids": [
                "P20029",
                "A0A0A6YXF5"
            ]
        },
        "ENSMUSG00000037211": {
            "id": "ENSMUSG00000037211",
            "type": "Gene",
            "symbol": "Spry1",
            "source": [
                "ENSMUSG00000052684",
                "ENSMUSG00000052837",
                "ENSMUSG00000026628",
                "ENSMUSG00000021250",
                "ENSMUSG00000003545",
                "ENSMUSG00000055148"
            ],
            "target": [],
            "expression_sum": {
                "class1": 12964.0,
                "class2": 1769.0
            },
            "ens_id": "ENSMUSG00000037211",
            "uniprot_ids": [
                "Q9QXV9"
            ]
        },
        "ENSMUSG00000027829": {
            "id": "ENSMUSG00000027829",
            "type": "Gene",
            "symbol": "Ccnl1",
            "source": [
                "ENSMUSG00000056501",
                "ENSMUSG00000037868",
                "ENSMUSG00000052684",
                "ENSMUSG00000023034",
                "ENSMUSG00000052837",
                "ENSMUSG00000026628",
                "ENSMUSG00000038418",
                "ENSMUSG00000021250",
                "ENSMUSG00000003032",
                "ENSMUSG00000018899"
            ],
            "target": [],
            "expression_sum": {
                "class1": 5751.0,
                "class2": 3266.0
            },
            "ens_id": "ENSMUSG00000027829",
            "uniprot_ids": [
                "Q52KE7",
                "F6WYR6",
                "F6WST0",
                "F7BI63",
                "F7BER3"
            ]
        },
        "ENSMUSG00000003032": {
            "id": "ENSMUSG00000003032",
            "type": "TF",
            "symbol": "Klf4",
            "source": [
                "ENSMUSG00000037868",
                "ENSMUSG00000023034",
                "ENSMUSG00000052837",
                "ENSMUSG00000026628",
                "ENSMUSG00000057098",
                "ENSMUSG00000038418",
                "ENSMUSG00000021250",
                "ENSMUSG00000003545",
                "ENSMUSG00000018899"
            ],
            "target": [
                "ENSMUSG00000030116",
                "ENSMUSG00000007815",
                "ENSMUSG00000005338",
                "ENSMUSG00000019505",
                "ENSMUSG00000059291",
                "ENSMUSG00000031447",
                "ENSMUSG00000002602",
                "ENSMUSG00000031765",
                "ENSMUSG00000028111",
                "ENSMUSG00000031875",
                "ENSMUSG00000007655",
                "ENSMUSG00000021477",
                "ENSMUSG00000028691",
                "ENSMUSG00000003545",
                "ENSMUSG00000024011",
                "ENSMUSG00000053007",
                "ENSMUSG00000026879",
                "ENSMUSG00000037822",
                "ENSMUSG00000048076",
                "ENSMUSG00000056481",
                "ENSMUSG00000037411",
                "ENSMUSG00000022371",
                "ENSMUSG00000044337",
                "ENSMUSG00000064373",
                "ENSMUSG00000028822",
                "ENSMUSG00000015354",
                "ENSMUSG00000034868",
                "ENSMUSG00000006699",
                "ENSMUSG00000044600",
                "ENSMUSG00000029649",
                "ENSMUSG00000008384",
                "ENSMUSG00000055148",
                "ENSMUSG00000001240",
                "ENSMUSG00000012422",
                "ENSMUSG00000023004",
                "ENSMUSG00000039405",
                "ENSMUSG00000002107",
                "ENSMUSG00000069763",
                "ENSMUSG00000003541",
                "ENSMUSG00000060572",
                "ENSMUSG00000032231",
                "ENSMUSG00000007872",
                "ENSMUSG00000056737",
                "ENSMUSG00000020467",
                "ENSMUSG00000035212",
                "ENSMUSG00000022032",
                "ENSMUSG00000037499",
                "ENSMUSG00000021390",
                "ENSMUSG00000009013",
                "ENSMUSG00000009681",
                "ENSMUSG00000018585",
                "ENSMUSG00000020053",
                "ENSMUSG00000025132",
                "ENSMUSG00000020440",
                "ENSMUSG00000028367",
                "ENSMUSG00000053113",
                "ENSMUSG00000037266",
                "ENSMUSG00000084786",
                "ENSMUSG00000031626",
                "ENSMUSG00000036446",
                "ENSMUSG00000020737",
                "ENSMUSG00000029767",
                "ENSMUSG00000055681",
                "ENSMUSG00000031538",
                "ENSMUSG00000019978",
                "ENSMUSG00000019278",
                "ENSMUSG00000021760",
                "ENSMUSG00000090000",
                "ENSMUSG00000036478",
                "ENSMUSG00000022816",
                "ENSMUSG00000027204",
                "ENSMUSG00000041959",
                "ENSMUSG00000033585",
                "ENSMUSG00000026628",
                "ENSMUSG00000007039",
                "ENSMUSG00000027706",
                "ENSMUSG00000074743",
                "ENSMUSG00000022090",
                "ENSMUSG00000041828",
                "ENSMUSG00000032014",
                "ENSMUSG00000032515",
                "ENSMUSG00000092274",
                "ENSMUSG00000017466",
                "ENSMUSG00000074457",
                "ENSMUSG00000027829",
                "ENSMUSG00000032334",
                "ENSMUSG00000023092",
                "ENSMUSG00000072235",
                "ENSMUSG00000038463",
                "ENSMUSG00000005397",
                "ENSMUSG00000022894",
                "ENSMUSG00000015085",
                "ENSMUSG00000030208",
                "ENSMUSG00000054452",
                "ENSMUSG00000068220",
                "ENSMUSG00000028517",
                "ENSMUSG00000040212",
                "ENSMUSG00000023034",
                "ENSMUSG00000079523",
                "ENSMUSG00000030654",
                "ENSMUSG00000016494",
                "ENSMUSG00000001123",
                "ENSMUSG00000026674",
                "ENSMUSG00000020315",
                "ENSMUSG00000037706",
                "ENSMUSG00000018217",
                "ENSMUSG00000017002",
                "ENSMUSG00000003161",
                "ENSMUSG00000060591",
                "ENSMUSG00000025784",
                "ENSMUSG00000040562",
                "ENSMUSG00000072812",
                "ENSMUSG00000015312",
                "ENSMUSG00000040280",
                "ENSMUSG00000024659",
                "ENSMUSG00000018819",
                "ENSMUSG00000032301",
                "ENSMUSG00000020423",
                "ENSMUSG00000027712",
                "ENSMUSG00000024048",
                "ENSMUSG00000069662",
                "ENSMUSG00000028108",
                "ENSMUSG00000037072",
                "ENSMUSG00000002732",
                "ENSMUSG00000021876",
                "ENSMUSG00000001891",
                "ENSMUSG00000026478",
                "ENSMUSG00000037868",
                "ENSMUSG00000042312",
                "ENSMUSG00000057098",
                "ENSMUSG00000031722",
                "ENSMUSG00000075254",
                "ENSMUSG00000023010",
                "ENSMUSG00000045658",
                "ENSMUSG00000031799",
                "ENSMUSG00000014846",
                "ENSMUSG00000017493",
                "ENSMUSG00000041084",
                "ENSMUSG00000039601",
                "ENSMUSG00000027907",
                "ENSMUSG00000074884",
                "ENSMUSG00000024308",
                "ENSMUSG00000024014",
                "ENSMUSG00000000628",
                "ENSMUSG00000028249",
                "ENSMUSG00000095385",
                "ENSMUSG00000020849",
                "ENSMUSG00000069833",
                "ENSMUSG00000041736",
                "ENSMUSG00000087006",
                "ENSMUSG00000034993",
                "ENSMUSG00000026670",
                "ENSMUSG00000075602",
                "ENSMUSG00000020585",
                "ENSMUSG00000025492",
                "ENSMUSG00000007041",
                "ENSMUSG00000021250",
                "ENSMUSG00000042747",
                "ENSMUSG00000029096",
                "ENSMUSG00000026837",
                "ENSMUSG00000028618",
                "ENSMUSG00000032383",
                "ENSMUSG00000026748",
                "ENSMUSG00000031422",
                "ENSMUSG00000047907",
                "ENSMUSG00000047945",
                "ENSMUSG00000052837",
                "ENSMUSG00000035356",
                "ENSMUSG00000037206",
                "ENSMUSG00000024012",
                "ENSMUSG00000029659",
                "ENSMUSG00000060802",
                "ENSMUSG00000026574",
                "ENSMUSG00000044786",
                "ENSMUSG00000018899",
                "ENSMUSG00000079037",
                "ENSMUSG00000090698",
                "ENSMUSG00000075702",
                "ENSMUSG00000060703",
                "ENSMUSG00000019087",
                "ENSMUSG00000022261",
                "ENSMUSG00000038418",
                "ENSMUSG00000029484",
                "ENSMUSG00000040026",
                "ENSMUSG00000001025",
                "ENSMUSG00000058135"
            ],
            "expression_sum": {
                "class1": 8719.0,
                "class2": 10220.0
            },
            "ens_id": "ENSMUSG00000003032",
            "uniprot_ids": [
                "Q60793",
                "B7ZCH2",
                "B7ZCH1"
            ]
        },
        "ENSMUSG00000047945": {
            "id": "ENSMUSG00000047945",
            "type": "Gene",
            "symbol": "Marcksl1",
            "source": [
                "ENSMUSG00000056501",
                "ENSMUSG00000052684",
                "ENSMUSG00000023034",
                "ENSMUSG00000052837",
                "ENSMUSG00000026628",
                "ENSMUSG00000030103",
                "ENSMUSG00000022346",
                "ENSMUSG00000021250",
                "ENSMUSG00000003545",
                "ENSMUSG00000003032",
                "ENSMUSG00000071076"
            ],
            "target": [],
            "expression_sum": {
                "class1": 40746.0,
                "class2": 794.0
            },
            "ens_id": "ENSMUSG00000047945",
            "uniprot_ids": [
                "P28667"
            ]
        },
        "ENSMUSG00000029380": {
            "id": "ENSMUSG00000029380",
            "type": "Gene",
            "symbol": "Cxcl1",
            "source": [
                "ENSMUSG00000052837",
                "ENSMUSG00000026628",
                "ENSMUSG00000056501",
                "ENSMUSG00000021250"
            ],
            "target": [],
            "expression_sum": {
                "class1": 16842.0,
                "class2": 4497.0
            },
            "ens_id": "ENSMUSG00000029380",
            "uniprot_ids": [
                "P12850"
            ]
        },
        "ENSMUSG00000034855": {
            "id": "ENSMUSG00000034855",
            "type": "Gene",
            "symbol": "Cxcl10",
            "source": [
                "ENSMUSG00000056501",
                "ENSMUSG00000052837",
                "ENSMUSG00000026628",
                "ENSMUSG00000055148",
                "ENSMUSG00000018899"
            ],
            "target": [],
            "expression_sum": {
                "class1": 17741.0,
                "class2": 188.0
            },
            "ens_id": "ENSMUSG00000034855",
            "uniprot_ids": [
                "P17515",
                "D3YW23"
            ]
        },
        "ENSMUSG00000037411": {
            "id": "ENSMUSG00000037411",
            "type": "Gene",
            "symbol": "Serpine1",
            "source": [
                "ENSMUSG00000056501",
                "ENSMUSG00000052684",
                "ENSMUSG00000052837",
                "ENSMUSG00000026628",
                "ENSMUSG00000021250",
                "ENSMUSG00000003032",
                "ENSMUSG00000055148",
                "ENSMUSG00000018899"
            ],
            "target": [],
            "expression_sum": {
                "class1": 11759.0,
                "class2": 3356.0
            },
            "ens_id": "ENSMUSG00000037411",
            "uniprot_ids": [
                "G5E899"
            ]
        },
        "ENSMUSG00000000628": {
            "id": "ENSMUSG00000000628",
            "type": "Gene",
            "symbol": "Hk2",
            "source": [
                "ENSMUSG00000056501",
                "ENSMUSG00000052684",
                "ENSMUSG00000023034",
                "ENSMUSG00000052837",
                "ENSMUSG00000026628",
                "ENSMUSG00000021250",
                "ENSMUSG00000055148",
                "ENSMUSG00000003032",
                "ENSMUSG00000018899",
                "ENSMUSG00000071076"
            ],
            "target": [],
            "expression_sum": {
                "class1": 4645.0,
                "class2": 638.0
            },
            "ens_id": "ENSMUSG00000000628",
            "uniprot_ids": [
                "O08528",
                "E9Q9M6",
                "E9Q5B5"
            ]
        },
        "ENSMUSG00000030103": {
            "id": "ENSMUSG00000030103",
            "type": "TF",
            "symbol": "Bhlhe40",
            "source": [
                "ENSMUSG00000056501",
                "ENSMUSG00000052684",
                "ENSMUSG00000023034",
                "ENSMUSG00000052837",
                "ENSMUSG00000026628",
                "ENSMUSG00000021250",
                "ENSMUSG00000018899"
            ],
            "target": [
                "ENSMUSG00000056501",
                "ENSMUSG00000023034",
                "ENSMUSG00000053113",
                "ENSMUSG00000003545",
                "ENSMUSG00000020644",
                "ENSMUSG00000015312",
                "ENSMUSG00000021250",
                "ENSMUSG00000055148",
                "ENSMUSG00000040128",
                "ENSMUSG00000035356",
                "ENSMUSG00000047945",
                "ENSMUSG00000026628",
                "ENSMUSG00000020423",
                "ENSMUSG00000052837",
                "ENSMUSG00000032515",
                "ENSMUSG00000007872",
                "ENSMUSG00000044786",
                "ENSMUSG00000018899",
                "ENSMUSG00000090698",
                "ENSMUSG00000028967",
                "ENSMUSG00000035828",
                "ENSMUSG00000009681"
            ],
            "expression_sum": {
                "class1": 3879.0,
                "class2": 961.0
            },
            "ens_id": "ENSMUSG00000030103",
            "uniprot_ids": [
                "O35185",
                "E9Q2C3"
            ]
        },
        "ENSMUSG00000090698": {
            "id": "ENSMUSG00000090698",
            "type": "Gene",
            "symbol": "Apold1",
            "source": [
                "ENSMUSG00000000078",
                "ENSMUSG00000052684",
                "ENSMUSG00000026628",
                "ENSMUSG00000030103",
                "ENSMUSG00000003032",
                "ENSMUSG00000005583",
                "ENSMUSG00000055148",
                "ENSMUSG00000021127",
                "ENSMUSG00000071076"
            ],
            "target": [],
            "expression_sum": {
                "class1": 8080.0,
                "class2": 1314.0
            },
            "ens_id": "ENSMUSG00000090698",
            "uniprot_ids": [
                "E9Q0X2"
            ]
        },
        "ENSMUSG00000003545": {
            "id": "ENSMUSG00000003545",
            "type": "TF",
            "symbol": "Fosb",
            "source": [
                "ENSMUSG00000056501",
                "ENSMUSG00000052684",
                "ENSMUSG00000023034",
                "ENSMUSG00000037868",
                "ENSMUSG00000052837",
                "ENSMUSG00000026628",
                "ENSMUSG00000030103",
                "ENSMUSG00000042812",
                "ENSMUSG00000021250",
                "ENSMUSG00000055148",
                "ENSMUSG00000003032",
                "ENSMUSG00000018899",
                "ENSMUSG00000071076"
            ],
            "target": [
                "ENSMUSG00000023034",
                "ENSMUSG00000069601",
                "ENSMUSG00000054364",
                "ENSMUSG00000024014",
                "ENSMUSG00000028691",
                "ENSMUSG00000056708",
                "ENSMUSG00000037266",
                "ENSMUSG00000087006",
                "ENSMUSG00000053560",
                "ENSMUSG00000052684",
                "ENSMUSG00000022528",
                "ENSMUSG00000015312",
                "ENSMUSG00000008384",
                "ENSMUSG00000021250",
                "ENSMUSG00000003032",
                "ENSMUSG00000021127",
                "ENSMUSG00000024190",
                "ENSMUSG00000037211",
                "ENSMUSG00000015501",
                "ENSMUSG00000040128",
                "ENSMUSG00000047945",
                "ENSMUSG00000026628",
                "ENSMUSG00000020423",
                "ENSMUSG00000052837",
                "ENSMUSG00000032501",
                "ENSMUSG00000028967",
                "ENSMUSG00000038418",
                "ENSMUSG00000039601"
            ],
            "expression_sum": {
                "class1": 20507.0,
                "class2": 9028.0
            },
            "ens_id": "ENSMUSG00000003545",
            "uniprot_ids": [
                "P13346",
                "A0A140LIT4",
                "A0A140LHS3",
                "A0A140LJH0",
                "A0A140LIE4",
                "A0A140LHW0"
            ]
        },
        "ENSMUSG00000008384": {
            "id": "ENSMUSG00000008384",
            "type": "Gene",
            "symbol": "Sertad1",
            "source": [
                "ENSMUSG00000041515",
                "ENSMUSG00000056501",
                "ENSMUSG00000052684",
                "ENSMUSG00000037868",
                "ENSMUSG00000023034",
                "ENSMUSG00000052837",
                "ENSMUSG00000026628",
                "ENSMUSG00000038418",
                "ENSMUSG00000021250",
                "ENSMUSG00000055148",
                "ENSMUSG00000003545",
                "ENSMUSG00000003032",
                "ENSMUSG00000018899",
                "ENSMUSG00000071076"
            ],
            "target": [],
            "expression_sum": {
                "class1": 4146.0,
                "class2": 2027.0
            },
            "ens_id": "ENSMUSG00000008384",
            "uniprot_ids": [
                "Q9JL10",
                "D3Z2X7"
            ]
        },
        "ENSMUSG00000044786": {
            "id": "ENSMUSG00000044786",
            "type": "Gene",
            "symbol": "Zfp36",
            "source": [
                "ENSMUSG00000041515",
                "ENSMUSG00000056501",
                "ENSMUSG00000052684",
                "ENSMUSG00000037868",
                "ENSMUSG00000052837",
                "ENSMUSG00000026628",
                "ENSMUSG00000030103",
                "ENSMUSG00000048387",
                "ENSMUSG00000000247",
                "ENSMUSG00000038418",
                "ENSMUSG00000021250",
                "ENSMUSG00000055148",
                "ENSMUSG00000003032",
                "ENSMUSG00000018899",
                "ENSMUSG00000071076"
            ],
            "target": [],
            "expression_sum": {
                "class1": 49942.0,
                "class2": 16051.0
            },
            "ens_id": "ENSMUSG00000044786",
            "uniprot_ids": [
                "P22893",
                "Q9DBE5"
            ]
        },
        "ENSMUSG00000015501": {
            "id": "ENSMUSG00000015501",
            "type": "TF",
            "symbol": "Hivep2",
            "source": [
                "ENSMUSG00000056501",
                "ENSMUSG00000052684",
                "ENSMUSG00000037868",
                "ENSMUSG00000052837",
                "ENSMUSG00000026628",
                "ENSMUSG00000038418",
                "ENSMUSG00000021250",
                "ENSMUSG00000003545",
                "ENSMUSG00000018899"
            ],
            "target": [],
            "expression_sum": {
                "class1": 4910.0,
                "class2": 1760.0
            },
            "ens_id": "ENSMUSG00000015501",
            "uniprot_ids": [
                "Q3UHF7",
                "A0A087WQF9"
            ]
        },
        "ENSMUSG00000037868": {
            "id": "ENSMUSG00000037868",
            "type": "TF",
            "symbol": "Egr2",
            "source": [
                "ENSMUSG00000056501",
                "ENSMUSG00000052684",
                "ENSMUSG00000023034",
                "ENSMUSG00000052837",
                "ENSMUSG00000026628",
                "ENSMUSG00000038418",
                "ENSMUSG00000021250",
                "ENSMUSG00000003032",
                "ENSMUSG00000071076",
                "ENSMUSG00000018899"
            ],
            "target": [
                "ENSMUSG00000023034",
                "ENSMUSG00000053113",
                "ENSMUSG00000003545",
                "ENSMUSG00000033538",
                "ENSMUSG00000052684",
                "ENSMUSG00000021025",
                "ENSMUSG00000015312",
                "ENSMUSG00000008384",
                "ENSMUSG00000021250",
                "ENSMUSG00000003032",
                "ENSMUSG00000055148",
                "ENSMUSG00000021127",
                "ENSMUSG00000015501",
                "ENSMUSG00000040128",
                "ENSMUSG00000035356",
                "ENSMUSG00000052837",
                "ENSMUSG00000026628",
                "ENSMUSG00000020423",
                "ENSMUSG00000097296",
                "ENSMUSG00000092274",
                "ENSMUSG00000007872",
                "ENSMUSG00000044786",
                "ENSMUSG00000018899",
                "ENSMUSG00000027829",
                "ENSMUSG00000006403",
                "ENSMUSG00000028214",
                "ENSMUSG00000032501",
                "ENSMUSG00000035828",
                "ENSMUSG00000038418",
                "ENSMUSG00000009681"
            ],
            "expression_sum": {
                "class1": 3930.0,
                "class2": 381.0
            },
            "ens_id": "ENSMUSG00000037868",
            "uniprot_ids": [
                "P08152",
                "D3Z4G7",
                "D3Z4X9",
                "D3YYY2",
                "D3Z3K7",
                "D3YXS2"
            ]
        },
        "ENSMUSG00000009681": {
            "id": "ENSMUSG00000009681",
            "type": "Gene",
            "symbol": "Bcr",
            "source": [
                "ENSMUSG00000056501",
                "ENSMUSG00000052684",
                "ENSMUSG00000037868",
                "ENSMUSG00000052837",
                "ENSMUSG00000026628",
                "ENSMUSG00000030103",
                "ENSMUSG00000021250",
                "ENSMUSG00000003032",
                "ENSMUSG00000005583",
                "ENSMUSG00000055148",
                "ENSMUSG00000071076"
            ],
            "target": [],
            "expression_sum": {
                "class1": 7907.0,
                "class2": 839.0
            },
            "ens_id": "ENSMUSG00000009681",
            "uniprot_ids": [
                "Q6PAJ1",
                "A0A1W2P6J3",
                "A0A1W2P6I7",
                "A0A571BEM4"
            ]
        },
        "ENSMUSG00000019960": {
            "id": "ENSMUSG00000019960",
            "type": "Gene",
            "symbol": "Dusp6",
            "source": [
                "ENSMUSG00000038193",
                "ENSMUSG00000026628",
                "ENSMUSG00000021250",
                "ENSMUSG00000000247"
            ],
            "target": [],
            "expression_sum": {
                "class1": 10239.0,
                "class2": 1639.0
            },
            "ens_id": "ENSMUSG00000019960",
            "uniprot_ids": [
                "Q9DBB1",
                "A0A1W2P715"
            ]
        },
        "ENSMUSG00000020205": {
            "id": "ENSMUSG00000020205",
            "type": "Gene",
            "symbol": "Phlda1",
            "source": [
                "ENSMUSG00000032035",
                "ENSMUSG00000056501",
                "ENSMUSG00000023034",
                "ENSMUSG00000022528",
                "ENSMUSG00000026628",
                "ENSMUSG00000005583",
                "ENSMUSG00000057098",
                "ENSMUSG00000008575",
                "ENSMUSG00000028565",
                "ENSMUSG00000018899"
            ],
            "target": [],
            "expression_sum": {
                "class1": 51213.0,
                "class2": 22408.0
            },
            "ens_id": "ENSMUSG00000020205",
            "uniprot_ids": [
                "Q62392"
            ]
        },
        "ENSMUSG00000090841": {
            "id": "ENSMUSG00000090841",
            "type": "Gene",
            "symbol": "Myl6",
            "source": [
                "ENSMUSG00000032238",
                "ENSMUSG00000023034",
                "ENSMUSG00000026628",
                "ENSMUSG00000021250",
                "ENSMUSG00000008575",
                "ENSMUSG00000005583",
                "ENSMUSG00000020484"
            ],
            "target": [],
            "expression_sum": {
                "class1": 45020.0,
                "class2": 125993.0
            },
            "ens_id": "ENSMUSG00000090841",
            "uniprot_ids": [
                "Q60605",
                "A0A1W2P7Q9",
                "A0A1W2P6G5",
                "A0A1W2P6F6",
                "A0A1W2P8F0",
                "A0A1W2P888"
            ]
        },
        "ENSMUSG00000071076": {
            "id": "ENSMUSG00000071076",
            "type": "TF",
            "symbol": "Jund",
            "source": [
                "ENSMUSG00000056501",
                "ENSMUSG00000052837",
                "ENSMUSG00000026628",
                "ENSMUSG00000000247",
                "ENSMUSG00000008575"
            ],
            "target": [
                "ENSMUSG00000031431",
                "ENSMUSG00000040723",
                "ENSMUSG00000025381",
                "ENSMUSG00000019188",
                "ENSMUSG00000022365",
                "ENSMUSG00000024011",
                "ENSMUSG00000025213",
                "ENSMUSG00000053007",
                "ENSMUSG00000026879",
                "ENSMUSG00000016256",
                "ENSMUSG00000047187",
                "ENSMUSG00000030298",
                "ENSMUSG00000064373",
                "ENSMUSG00000056629",
                "ENSMUSG00000042613",
                "ENSMUSG00000044600",
                "ENSMUSG00000000325",
                "ENSMUSG00000067847",
                "ENSMUSG00000027239",
                "ENSMUSG00000078713",
                "ENSMUSG00000078974",
                "ENSMUSG00000018567",
                "ENSMUSG00000058672",
                "ENSMUSG00000031007",
                "ENSMUSG00000008575",
                "ENSMUSG00000053317",
                "ENSMUSG00000075701",
                "ENSMUSG00000022836",
                "ENSMUSG00000021390",
                "ENSMUSG00000020368",
                "ENSMUSG00000031825",
                "ENSMUSG00000084786",
                "ENSMUSG00000019907",
                "ENSMUSG00000031565",
                "ENSMUSG00000020737",
                "ENSMUSG00000030824",
                "ENSMUSG00000037379",
                "ENSMUSG00000020099",
                "ENSMUSG00000027706",
                "ENSMUSG00000031586",
                "ENSMUSG00000032515",
                "ENSMUSG00000017466",
                "ENSMUSG00000072235",
                "ENSMUSG00000005873",
                "ENSMUSG00000038463",
                "ENSMUSG00000028517",
                "ENSMUSG00000016494",
                "ENSMUSG00000020029",
                "ENSMUSG00000024290",
                "ENSMUSG00000038393",
                "ENSMUSG00000015094",
                "ENSMUSG00000022665",
                "ENSMUSG00000015312",
                "ENSMUSG00000028964",
                "ENSMUSG00000021871",
                "ENSMUSG00000024659",
                "ENSMUSG00000025724",
                "ENSMUSG00000062270",
                "ENSMUSG00000027712",
                "ENSMUSG00000007891",
                "ENSMUSG00000024048",
                "ENSMUSG00000028673",
                "ENSMUSG00000001891",
                "ENSMUSG00000050708",
                "ENSMUSG00000026478",
                "ENSMUSG00000037868",
                "ENSMUSG00000022122",
                "ENSMUSG00000000594",
                "ENSMUSG00000019920",
                "ENSMUSG00000057098",
                "ENSMUSG00000014846",
                "ENSMUSG00000017493",
                "ENSMUSG00000025351",
                "ENSMUSG00000074884",
                "ENSMUSG00000024014",
                "ENSMUSG00000055435",
                "ENSMUSG00000020849",
                "ENSMUSG00000034993",
                "ENSMUSG00000000753",
                "ENSMUSG00000051579",
                "ENSMUSG00000004460",
                "ENSMUSG00000028757",
                "ENSMUSG00000035356",
                "ENSMUSG00000027859",
                "ENSMUSG00000022174",
                "ENSMUSG00000029048",
                "ENSMUSG00000035227",
                "ENSMUSG00000024012",
                "ENSMUSG00000060802",
                "ENSMUSG00000014867",
                "ENSMUSG00000019505",
                "ENSMUSG00000059291",
                "ENSMUSG00000030222",
                "ENSMUSG00000028691",
                "ENSMUSG00000003545",
                "ENSMUSG00000030930",
                "ENSMUSG00000009281",
                "ENSMUSG00000028822",
                "ENSMUSG00000008384",
                "ENSMUSG00000023175",
                "ENSMUSG00000026421",
                "ENSMUSG00000031839",
                "ENSMUSG00000002107",
                "ENSMUSG00000032231",
                "ENSMUSG00000007872",
                "ENSMUSG00000027848",
                "ENSMUSG00000052419",
                "ENSMUSG00000029718",
                "ENSMUSG00000028600",
                "ENSMUSG00000025132",
                "ENSMUSG00000021917",
                "ENSMUSG00000028339",
                "ENSMUSG00000037266",
                "ENSMUSG00000001525",
                "ENSMUSG00000031538",
                "ENSMUSG00000019978",
                "ENSMUSG00000022816",
                "ENSMUSG00000007039",
                "ENSMUSG00000029781",
                "ENSMUSG00000022090",
                "ENSMUSG00000019579",
                "ENSMUSG00000068735",
                "ENSMUSG00000074457",
                "ENSMUSG00000040249",
                "ENSMUSG00000032334",
                "ENSMUSG00000026335",
                "ENSMUSG00000029070",
                "ENSMUSG00000035678",
                "ENSMUSG00000016024",
                "ENSMUSG00000030208",
                "ENSMUSG00000023034",
                "ENSMUSG00000030654",
                "ENSMUSG00000003420",
                "ENSMUSG00000020315",
                "ENSMUSG00000058569",
                "ENSMUSG00000018217",
                "ENSMUSG00000023046",
                "ENSMUSG00000058056",
                "ENSMUSG00000005973",
                "ENSMUSG00000042745",
                "ENSMUSG00000016534",
                "ENSMUSG00000003161",
                "ENSMUSG00000032359",
                "ENSMUSG00000001911",
                "ENSMUSG00000005583",
                "ENSMUSG00000022136",
                "ENSMUSG00000018819",
                "ENSMUSG00000032301",
                "ENSMUSG00000020709",
                "ENSMUSG00000044408",
                "ENSMUSG00000002059",
                "ENSMUSG00000019362",
                "ENSMUSG00000041571",
                "ENSMUSG00000037072",
                "ENSMUSG00000020009",
                "ENSMUSG00000004207",
                "ENSMUSG00000020088",
                "ENSMUSG00000026728",
                "ENSMUSG00000096145",
                "ENSMUSG00000036752",
                "ENSMUSG00000042312",
                "ENSMUSG00000035828",
                "ENSMUSG00000075254",
                "ENSMUSG00000023010",
                "ENSMUSG00000031799",
                "ENSMUSG00000041084",
                "ENSMUSG00000041126",
                "ENSMUSG00000054808",
                "ENSMUSG00000021242",
                "ENSMUSG00000002014",
                "ENSMUSG00000092341",
                "ENSMUSG00000039899",
                "ENSMUSG00000030934",
                "ENSMUSG00000011114",
                "ENSMUSG00000028864",
                "ENSMUSG00000006800",
                "ENSMUSG00000020585",
                "ENSMUSG00000015575",
                "ENSMUSG00000032353",
                "ENSMUSG00000025492",
                "ENSMUSG00000007041",
                "ENSMUSG00000055447",
                "ENSMUSG00000020458",
                "ENSMUSG00000032959",
                "ENSMUSG00000009291",
                "ENSMUSG00000037206",
                "ENSMUSG00000060703",
                "ENSMUSG00000030849",
                "ENSMUSG00000029231",
                "ENSMUSG00000021131",
                "ENSMUSG00000035258",
                "ENSMUSG00000001025",
                "ENSMUSG00000045636",
                "ENSMUSG00000048076",
                "ENSMUSG00000042712",
                "ENSMUSG00000026021",
                "ENSMUSG00000022528",
                "ENSMUSG00000006699",
                "ENSMUSG00000032575",
                "ENSMUSG00000079111",
                "ENSMUSG00000030659",
                "ENSMUSG00000012422",
                "ENSMUSG00000023004",
                "ENSMUSG00000031198",
                "ENSMUSG00000025130",
                "ENSMUSG00000003541",
                "ENSMUSG00000029103",
                "ENSMUSG00000038872",
                "ENSMUSG00000022452",
                "ENSMUSG00000056737",
                "ENSMUSG00000022032",
                "ENSMUSG00000027828",
                "ENSMUSG00000022400",
                "ENSMUSG00000055839",
                "ENSMUSG00000034165",
                "ENSMUSG00000032085",
                "ENSMUSG00000053113",
                "ENSMUSG00000052681",
                "ENSMUSG00000003380",
                "ENSMUSG00000038803",
                "ENSMUSG00000046768",
                "ENSMUSG00000026628",
                "ENSMUSG00000042737",
                "ENSMUSG00000021253",
                "ENSMUSG00000053477",
                "ENSMUSG00000079523",
                "ENSMUSG00000038991",
                "ENSMUSG00000060591",
                "ENSMUSG00000001794",
                "ENSMUSG00000028843",
                "ENSMUSG00000020423",
                "ENSMUSG00000069662",
                "ENSMUSG00000021876",
                "ENSMUSG00000025780",
                "ENSMUSG00000024164",
                "ENSMUSG00000001663",
                "ENSMUSG00000064215",
                "ENSMUSG00000006050",
                "ENSMUSG00000004897",
                "ENSMUSG00000027907",
                "ENSMUSG00000022108",
                "ENSMUSG00000000628",
                "ENSMUSG00000020219",
                "ENSMUSG00000028249",
                "ENSMUSG00000069833",
                "ENSMUSG00000033910",
                "ENSMUSG00000039168",
                "ENSMUSG00000018593",
                "ENSMUSG00000013698",
                "ENSMUSG00000001418",
                "ENSMUSG00000015568",
                "ENSMUSG00000021250",
                "ENSMUSG00000028618",
                "ENSMUSG00000054203",
                "ENSMUSG00000052837",
                "ENSMUSG00000023272",
                "ENSMUSG00000021576",
                "ENSMUSG00000079037",
                "ENSMUSG00000079017",
                "ENSMUSG00000006403",
                "ENSMUSG00000000247",
                "ENSMUSG00000019087",
                "ENSMUSG00000029465",
                "ENSMUSG00000052997",
                "ENSMUSG00000005054",
                "ENSMUSG00000058135",
                "ENSMUSG00000007815",
                "ENSMUSG00000029761",
                "ENSMUSG00000002602",
                "ENSMUSG00000028111",
                "ENSMUSG00000031875",
                "ENSMUSG00000021477",
                "ENSMUSG00000037822",
                "ENSMUSG00000027422",
                "ENSMUSG00000097328",
                "ENSMUSG00000020803",
                "ENSMUSG00000025647",
                "ENSMUSG00000044337",
                "ENSMUSG00000031556",
                "ENSMUSG00000021877",
                "ENSMUSG00000068196",
                "ENSMUSG00000034868",
                "ENSMUSG00000044627",
                "ENSMUSG00000021939",
                "ENSMUSG00000055148",
                "ENSMUSG00000001240",
                "ENSMUSG00000014329",
                "ENSMUSG00000040713",
                "ENSMUSG00000028369",
                "ENSMUSG00000022769",
                "ENSMUSG00000029309",
                "ENSMUSG00000026547",
                "ENSMUSG00000009013",
                "ENSMUSG00000027808",
                "ENSMUSG00000031595",
                "ENSMUSG00000009681",
                "ENSMUSG00000024327",
                "ENSMUSG00000060904",
                "ENSMUSG00000075706",
                "ENSMUSG00000020053",
                "ENSMUSG00000026042",
                "ENSMUSG00000037999",
                "ENSMUSG00000020440",
                "ENSMUSG00000028367",
                "ENSMUSG00000032314",
                "ENSMUSG00000032366",
                "ENSMUSG00000036372",
                "ENSMUSG00000033538",
                "ENSMUSG00000020810",
                "ENSMUSG00000025016",
                "ENSMUSG00000001119",
                "ENSMUSG00000021759",
                "ENSMUSG00000075010",
                "ENSMUSG00000029390",
                "ENSMUSG00000029767",
                "ENSMUSG00000024076",
                "ENSMUSG00000055681",
                "ENSMUSG00000090000",
                "ENSMUSG00000021760",
                "ENSMUSG00000021248",
                "ENSMUSG00000001506",
                "ENSMUSG00000003038",
                "ENSMUSG00000021870",
                "ENSMUSG00000041959",
                "ENSMUSG00000018672",
                "ENSMUSG00000028195",
                "ENSMUSG00000037373",
                "ENSMUSG00000068220",
                "ENSMUSG00000020486",
                "ENSMUSG00000040212",
                "ENSMUSG00000001123",
                "ENSMUSG00000020149",
                "ENSMUSG00000037706",
                "ENSMUSG00000025784",
                "ENSMUSG00000070394",
                "ENSMUSG00000031965",
                "ENSMUSG00000002015",
                "ENSMUSG00000028128",
                "ENSMUSG00000013622",
                "ENSMUSG00000020571",
                "ENSMUSG00000028108",
                "ENSMUSG00000002732",
                "ENSMUSG00000006205",
                "ENSMUSG00000006519",
                "ENSMUSG00000045658",
                "ENSMUSG00000029076",
                "ENSMUSG00000030717",
                "ENSMUSG00000039601",
                "ENSMUSG00000024308",
                "ENSMUSG00000056501",
                "ENSMUSG00000024038",
                "ENSMUSG00000041736",
                "ENSMUSG00000002064",
                "ENSMUSG00000020048",
                "ENSMUSG00000042747",
                "ENSMUSG00000041355",
                "ENSMUSG00000029096",
                "ENSMUSG00000026837",
                "ENSMUSG00000032383",
                "ENSMUSG00000022048",
                "ENSMUSG00000031422",
                "ENSMUSG00000029622",
                "ENSMUSG00000047945",
                "ENSMUSG00000066551",
                "ENSMUSG00000039286",
                "ENSMUSG00000030342",
                "ENSMUSG00000022708",
                "ENSMUSG00000019929",
                "ENSMUSG00000029659",
                "ENSMUSG00000071984",
                "ENSMUSG00000029641",
                "ENSMUSG00000044786",
                "ENSMUSG00000008540",
                "ENSMUSG00000018899",
                "ENSMUSG00000021268",
                "ENSMUSG00000090698",
                "ENSMUSG00000075702",
                "ENSMUSG00000014769"
            ],
            "expression_sum": {
                "class1": 45888.0,
                "class2": 59450.0
            },
            "ens_id": "ENSMUSG00000071076",
            "uniprot_ids": [
                "P15066"
            ]
        },
        "ENSMUSG00000097296": {
            "id": "ENSMUSG00000097296",
            "type": "Gene",
            "symbol": "Gm26532",
            "source": [
                "ENSMUSG00000037868",
                "ENSMUSG00000023034",
                "ENSMUSG00000052837",
                "ENSMUSG00000026628",
                "ENSMUSG00000018899"
            ],
            "target": [],
            "expression_sum": {
                "class1": 3192.0,
                "class2": 2757.0
            },
            "ens_id": "ENSMUSG00000097296"
        },
        "ENSMUSG00000053560": {
            "id": "ENSMUSG00000053560",
            "type": "Gene",
            "symbol": "Ier2",
            "source": [
                "ENSMUSG00000052684",
                "ENSMUSG00000026628",
                "ENSMUSG00000000247",
                "ENSMUSG00000038193",
                "ENSMUSG00000021250",
                "ENSMUSG00000003545",
                "ENSMUSG00000055148",
                "ENSMUSG00000018899"
            ],
            "target": [],
            "expression_sum": {
                "class1": 33337.0,
                "class2": 23871.0
            },
            "ens_id": "ENSMUSG00000053560",
            "uniprot_ids": [
                "P17950"
            ]
        },
        "ENSMUSG00000052837": {
            "id": "ENSMUSG00000052837",
            "type": "TF",
            "symbol": "Junb",
            "source": [
                "ENSMUSG00000041515",
                "ENSMUSG00000052684",
                "ENSMUSG00000037868",
                "ENSMUSG00000023034",
                "ENSMUSG00000026628",
                "ENSMUSG00000030103",
                "ENSMUSG00000000247",
                "ENSMUSG00000038418",
                "ENSMUSG00000057098",
                "ENSMUSG00000021250",
                "ENSMUSG00000003545",
                "ENSMUSG00000003032",
                "ENSMUSG00000018899",
                "ENSMUSG00000071076"
            ],
            "target": [
                "ENSMUSG00000019505",
                "ENSMUSG00000020053",
                "ENSMUSG00000097266",
                "ENSMUSG00000023034",
                "ENSMUSG00000037999",
                "ENSMUSG00000000628",
                "ENSMUSG00000069601",
                "ENSMUSG00000024014",
                "ENSMUSG00000054072",
                "ENSMUSG00000028691",
                "ENSMUSG00000056708",
                "ENSMUSG00000053113",
                "ENSMUSG00000003545",
                "ENSMUSG00000037266",
                "ENSMUSG00000042453",
                "ENSMUSG00000033538",
                "ENSMUSG00000030930",
                "ENSMUSG00000037411",
                "ENSMUSG00000028864",
                "ENSMUSG00000052684",
                "ENSMUSG00000017493",
                "ENSMUSG00000031451",
                "ENSMUSG00000008384",
                "ENSMUSG00000019278",
                "ENSMUSG00000021250",
                "ENSMUSG00000068220",
                "ENSMUSG00000003032",
                "ENSMUSG00000055148",
                "ENSMUSG00000003038",
                "ENSMUSG00000021127",
                "ENSMUSG00000037211",
                "ENSMUSG00000041515",
                "ENSMUSG00000024190",
                "ENSMUSG00000007097",
                "ENSMUSG00000015501",
                "ENSMUSG00000040128",
                "ENSMUSG00000023915",
                "ENSMUSG00000029380",
                "ENSMUSG00000047945",
                "ENSMUSG00000026628",
                "ENSMUSG00000030103",
                "ENSMUSG00000020423",
                "ENSMUSG00000002416",
                "ENSMUSG00000035356",
                "ENSMUSG00000097296",
                "ENSMUSG00000042812",
                "ENSMUSG00000040170",
                "ENSMUSG00000032515",
                "ENSMUSG00000007872",
                "ENSMUSG00000044786",
                "ENSMUSG00000022956",
                "ENSMUSG00000008540",
                "ENSMUSG00000071076",
                "ENSMUSG00000018899",
                "ENSMUSG00000021876",
                "ENSMUSG00000027829",
                "ENSMUSG00000049115",
                "ENSMUSG00000037868",
                "ENSMUSG00000022122",
                "ENSMUSG00000006205",
                "ENSMUSG00000028214",
                "ENSMUSG00000032501",
                "ENSMUSG00000035828",
                "ENSMUSG00000000247",
                "ENSMUSG00000064215",
                "ENSMUSG00000034855",
                "ENSMUSG00000039601",
                "ENSMUSG00000015085",
                "ENSMUSG00000030849",
                "ENSMUSG00000038418",
                "ENSMUSG00000045658",
                "ENSMUSG00000090084",
                "ENSMUSG00000035678",
                "ENSMUSG00000009681",
                "ENSMUSG00000001025",
                "ENSMUSG00000058135"
            ],
            "expression_sum": {
                "class1": 117091.0,
                "class2": 64489.0
            },
            "ens_id": "ENSMUSG00000052837",
            "uniprot_ids": [
                "P09450"
            ]
        },
        "ENSMUSG00000031762": {
            "id": "ENSMUSG00000031762",
            "type": "Gene",
            "symbol": "Mt2",
            "source": [
                "ENSMUSG00000056501",
                "ENSMUSG00000023034",
                "ENSMUSG00000022528",
                "ENSMUSG00000026628",
                "ENSMUSG00000021250",
                "ENSMUSG00000055148",
                "ENSMUSG00000018899"
            ],
            "target": [],
            "expression_sum": {
                "class1": 53885.0,
                "class2": 16534.0
            },
            "ens_id": "ENSMUSG00000031762",
            "uniprot_ids": [
                "P02798",
                "A0A1D5RLC3"
            ]
        },
        "ENSMUSG00000031765": {
            "id": "ENSMUSG00000031765",
            "type": "Gene",
            "symbol": "Mt1",
            "source": [
                "ENSMUSG00000056501",
                "ENSMUSG00000023034",
                "ENSMUSG00000022528",
                "ENSMUSG00000026628",
                "ENSMUSG00000005583",
                "ENSMUSG00000057098",
                "ENSMUSG00000021250",
                "ENSMUSG00000003032",
                "ENSMUSG00000008575",
                "ENSMUSG00000055148",
                "ENSMUSG00000018899"
            ],
            "target": [],
            "expression_sum": {
                "class1": 115205.0,
                "class2": 41751.0
            },
            "ens_id": "ENSMUSG00000031765",
            "uniprot_ids": [
                "P02802",
                "A0A1D5RLN7",
                "A0A1D5RMG1"
            ]
        },
        "ENSMUSG00000041515": {
            "id": "ENSMUSG00000041515",
            "type": "TF",
            "symbol": "Irf8",
            "source": [
                "ENSMUSG00000056501",
                "ENSMUSG00000023034",
                "ENSMUSG00000052837",
                "ENSMUSG00000026628",
                "ENSMUSG00000021250",
                "ENSMUSG00000018899"
            ],
            "target": [
                "ENSMUSG00000056501",
                "ENSMUSG00000023034",
                "ENSMUSG00000032515",
                "ENSMUSG00000052837",
                "ENSMUSG00000026628",
                "ENSMUSG00000020423",
                "ENSMUSG00000028967",
                "ENSMUSG00000044786",
                "ENSMUSG00000035356",
                "ENSMUSG00000008384",
                "ENSMUSG00000053113",
                "ENSMUSG00000021250",
                "ENSMUSG00000055148",
                "ENSMUSG00000007872",
                "ENSMUSG00000087006",
                "ENSMUSG00000016194",
                "ENSMUSG00000018899"
            ],
            "expression_sum": {
                "class1": 4392.0,
                "class2": 265.0
            },
            "ens_id": "ENSMUSG00000041515",
            "uniprot_ids": [
                "P23611",
                "F7C1A3",
                "E0CZE4"
            ]
        },
        "ENSMUSG00000033538": {
            "id": "ENSMUSG00000033538",
            "type": "Gene",
            "symbol": "Casp4",
            "source": [
                "ENSMUSG00000056501",
                "ENSMUSG00000037868",
                "ENSMUSG00000052684",
                "ENSMUSG00000023034",
                "ENSMUSG00000052837",
                "ENSMUSG00000026628",
                "ENSMUSG00000021250",
                "ENSMUSG00000018899",
                "ENSMUSG00000071076"
            ],
            "target": [],
            "expression_sum": {
                "class1": 3929.0,
                "class2": 647.0
            },
            "ens_id": "ENSMUSG00000033538",
            "uniprot_ids": [
                "P70343",
                "F6SUE5",
                "E0CY92"
            ]
        },
        "ENSMUSG00000037405": {
            "id": "ENSMUSG00000037405",
            "type": "Gene",
            "symbol": "Icam1",
            "source": [
                "ENSMUSG00000056501",
                "ENSMUSG00000023034",
                "ENSMUSG00000026628",
                "ENSMUSG00000055148",
                "ENSMUSG00000018899"
            ],
            "target": [],
            "expression_sum": {
                "class1": 5406.0,
                "class2": 943.0
            },
            "ens_id": "ENSMUSG00000037405",
            "uniprot_ids": [
                "P13597"
            ]
        },
        "ENSMUSG00000032515": {
            "id": "ENSMUSG00000032515",
            "type": "Gene",
            "symbol": "Csrnp1",
            "source": [
                "ENSMUSG00000041515",
                "ENSMUSG00000056501",
                "ENSMUSG00000052684",
                "ENSMUSG00000023034",
                "ENSMUSG00000052837",
                "ENSMUSG00000026628",
                "ENSMUSG00000030103",
                "ENSMUSG00000021250",
                "ENSMUSG00000003032",
                "ENSMUSG00000018899",
                "ENSMUSG00000071076"
            ],
            "target": [],
            "expression_sum": {
                "class1": 2983.0,
                "class2": 1546.0
            },
            "ens_id": "ENSMUSG00000032515",
            "uniprot_ids": [
                "P59054",
                "A0A1L1SRY6",
                "A0A1L1SRU4"
            ]
        },
        "ENSMUSG00000018899": {
            "id": "ENSMUSG00000018899",
            "type": "TF",
            "symbol": "Irf1",
            "source": [
                "ENSMUSG00000041515",
                "ENSMUSG00000056501",
                "ENSMUSG00000052684",
                "ENSMUSG00000037868",
                "ENSMUSG00000023034",
                "ENSMUSG00000042406",
                "ENSMUSG00000052837",
                "ENSMUSG00000026628",
                "ENSMUSG00000030103",
                "ENSMUSG00000022346",
                "ENSMUSG00000038418",
                "ENSMUSG00000021250",
                "ENSMUSG00000003032",
                "ENSMUSG00000055148",
                "ENSMUSG00000071076"
            ],
            "target": [
                "ENSMUSG00000056501",
                "ENSMUSG00000023034",
                "ENSMUSG00000031765",
                "ENSMUSG00000024014",
                "ENSMUSG00000000628",
                "ENSMUSG00000054072",
                "ENSMUSG00000053113",
                "ENSMUSG00000056708",
                "ENSMUSG00000003545",
                "ENSMUSG00000033538",
                "ENSMUSG00000001627",
                "ENSMUSG00000053560",
                "ENSMUSG00000037411",
                "ENSMUSG00000052684",
                "ENSMUSG00000037405",
                "ENSMUSG00000021025",
                "ENSMUSG00000022346",
                "ENSMUSG00000015312",
                "ENSMUSG00000008384",
                "ENSMUSG00000031762",
                "ENSMUSG00000021250",
                "ENSMUSG00000003032",
                "ENSMUSG00000035493",
                "ENSMUSG00000021127",
                "ENSMUSG00000024190",
                "ENSMUSG00000041515",
                "ENSMUSG00000015501",
                "ENSMUSG00000040128",
                "ENSMUSG00000035356",
                "ENSMUSG00000052837",
                "ENSMUSG00000026628",
                "ENSMUSG00000030103",
                "ENSMUSG00000020423",
                "ENSMUSG00000097296",
                "ENSMUSG00000032515",
                "ENSMUSG00000007872",
                "ENSMUSG00000044786",
                "ENSMUSG00000020205",
                "ENSMUSG00000027829",
                "ENSMUSG00000037868",
                "ENSMUSG00000006403",
                "ENSMUSG00000032501",
                "ENSMUSG00000035828",
                "ENSMUSG00000038418",
                "ENSMUSG00000034855",
                "ENSMUSG00000030849",
                "ENSMUSG00000035678",
                "ENSMUSG00000073489"
            ],
            "expression_sum": {
                "class1": 9123.0,
                "class2": 1422.0
            },
            "ens_id": "ENSMUSG00000018899",
            "uniprot_ids": [
                "P15314",
                "Q3U5M1",
                "A8Y5B9",
                "Q5SX15",
                "Q5SX14",
                "D6RH71",
                "Q5SX16"
            ]
        },
        "ENSMUSG00000053113": {
            "id": "ENSMUSG00000053113",
            "type": "Gene",
            "symbol": "Socs3",
            "source": [
                "ENSMUSG00000041515",
                "ENSMUSG00000056501",
                "ENSMUSG00000052684",
                "ENSMUSG00000037868",
                "ENSMUSG00000042406",
                "ENSMUSG00000023034",
                "ENSMUSG00000052837",
                "ENSMUSG00000026628",
                "ENSMUSG00000030103",
                "ENSMUSG00000022528",
                "ENSMUSG00000038418",
                "ENSMUSG00000038193",
                "ENSMUSG00000021250",
                "ENSMUSG00000055148",
                "ENSMUSG00000003032",
                "ENSMUSG00000055435",
                "ENSMUSG00000018899",
                "ENSMUSG00000071076"
            ],
            "target": [],
            "expression_sum": {
                "class1": 31900.0,
                "class2": 9658.0
            },
            "ens_id": "ENSMUSG00000053113",
            "uniprot_ids": [
                "O35718"
            ]
        },
        "ENSMUSG00000001627": {
            "id": "ENSMUSG00000001627",
            "type": "Gene",
            "symbol": "Ifrd1",
            "source": [
                "ENSMUSG00000026628",
                "ENSMUSG00000018899"
            ],
            "target": [],
            "expression_sum": {
                "class1": 3732.0,
                "class2": 2466.0
            },
            "ens_id": "ENSMUSG00000001627",
            "uniprot_ids": [
                "P19182",
                "E9Q949",
                "E9Q1E6",
                "E9PVS0",
                "F6TBT6",
                "E9QA38",
                "E9QA50"
            ]
        },
        "ENSMUSG00000021025": {
            "id": "ENSMUSG00000021025",
            "type": "Gene",
            "symbol": "Nfkbia",
            "source": [
                "ENSMUSG00000056501",
                "ENSMUSG00000037868",
                "ENSMUSG00000023034",
                "ENSMUSG00000026628",
                "ENSMUSG00000057098",
                "ENSMUSG00000018899"
            ],
            "target": [],
            "expression_sum": {
                "class1": 26787.0,
                "class2": 10367.0
            },
            "ens_id": "ENSMUSG00000021025",
            "uniprot_ids": [
                "Q9Z1E3"
            ]
        },
        "ENSMUSG00000021127": {
            "id": "ENSMUSG00000021127",
            "type": "TF",
            "symbol": "Zfp36l1",
            "source": [
                "ENSMUSG00000056501",
                "ENSMUSG00000037868",
                "ENSMUSG00000023034",
                "ENSMUSG00000022528",
                "ENSMUSG00000052837",
                "ENSMUSG00000026628",
                "ENSMUSG00000000247",
                "ENSMUSG00000021250",
                "ENSMUSG00000003545",
                "ENSMUSG00000055148",
                "ENSMUSG00000018899"
            ],
            "target": [
                "ENSMUSG00000090698",
                "ENSMUSG00000026421",
                "ENSMUSG00000030849",
                "ENSMUSG00000038742",
                "ENSMUSG00000026574"
            ],
            "expression_sum": {
                "class1": 32513.0,
                "class2": 13232.0
            },
            "ens_id": "ENSMUSG00000021127",
            "uniprot_ids": [
                "P23950",
                "A0A1W2P7G3",
                "A0A1W2P6I1"
            ]
        },
        "ENSMUSG00000006360": {
            "id": "ENSMUSG00000006360",
            "type": "Gene",
            "symbol": "Crip1",
            "source": [
                "ENSMUSG00000023034",
                "ENSMUSG00000026628",
                "ENSMUSG00000057098",
                "ENSMUSG00000021250",
                "ENSMUSG00000029322",
                "ENSMUSG00000005583",
                "ENSMUSG00000055148"
            ],
            "target": [],
            "expression_sum": {
                "class1": 12842.0,
                "class2": 203119.0
            },
            "ens_id": "ENSMUSG00000006360",
            "uniprot_ids": [
                "P63254",
                "A0A0G2JEK2"
            ]
        },
        "ENSMUSG00000032501": {
            "id": "ENSMUSG00000032501",
            "type": "Gene",
            "symbol": "Trib1",
            "source": [
                "ENSMUSG00000056501",
                "ENSMUSG00000037868",
                "ENSMUSG00000052837",
                "ENSMUSG00000026628",
                "ENSMUSG00000003545",
                "ENSMUSG00000018899"
            ],
            "target": [],
            "expression_sum": {
                "class1": 2868.0,
                "class2": 1055.0
            },
            "ens_id": "ENSMUSG00000032501",
            "uniprot_ids": [
                "Q8K4K4"
            ]
        },
        "ENSMUSG00000042406": {
            "id": "ENSMUSG00000042406",
            "type": "TF",
            "symbol": "Atf4",
            "source": [
                "ENSMUSG00000026628"
            ],
            "target": [
                "ENSMUSG00000022836",
                "ENSMUSG00000035356",
                "ENSMUSG00000026628",
                "ENSMUSG00000053113",
                "ENSMUSG00000018899"
            ],
            "expression_sum": {
                "class1": 8359.0,
                "class2": 3788.0
            },
            "ens_id": "ENSMUSG00000042406",
            "uniprot_ids": [
                "Q06507",
                "A0A2R8VI82"
            ]
        },
        "ENSMUSG00000035356": {
            "id": "ENSMUSG00000035356",
            "type": "Gene",
            "symbol": "Nfkbiz",
            "source": [
                "ENSMUSG00000041515",
                "ENSMUSG00000056501",
                "ENSMUSG00000052684",
                "ENSMUSG00000037868",
                "ENSMUSG00000042406",
                "ENSMUSG00000023034",
                "ENSMUSG00000052837",
                "ENSMUSG00000026628",
                "ENSMUSG00000030103",
                "ENSMUSG00000038418",
                "ENSMUSG00000021250",
                "ENSMUSG00000003032",
                "ENSMUSG00000018899",
                "ENSMUSG00000071076"
            ],
            "target": [],
            "expression_sum": {
                "class1": 7633.0,
                "class2": 2441.0
            },
            "ens_id": "ENSMUSG00000035356",
            "uniprot_ids": [
                "Q9EST8"
            ]
        },
        "ENSMUSG00000023904": {
            "id": "ENSMUSG00000023904",
            "type": "Gene",
            "symbol": "Hcfc1r1",
            "source": [
                "ENSMUSG00000026628",
                "ENSMUSG00000029322"
            ],
            "target": [],
            "expression_sum": {
                "class1": 7601.0,
                "class2": 12054.0
            },
            "ens_id": "ENSMUSG00000023904",
            "uniprot_ids": [
                "Q9CYQ5",
                "J3QPD4"
            ]
        },
        "ENSMUSG00000024190": {
            "id": "ENSMUSG00000024190",
            "type": "Gene",
            "symbol": "Dusp1",
            "source": [
                "ENSMUSG00000023034",
                "ENSMUSG00000022528",
                "ENSMUSG00000052837",
                "ENSMUSG00000026628",
                "ENSMUSG00000000247",
                "ENSMUSG00000057098",
                "ENSMUSG00000021250",
                "ENSMUSG00000055148",
                "ENSMUSG00000003545",
                "ENSMUSG00000008575",
                "ENSMUSG00000048387",
                "ENSMUSG00000018899"
            ],
            "target": [],
            "expression_sum": {
                "class1": 59919.0,
                "class2": 46620.0
            },
            "ens_id": "ENSMUSG00000024190",
            "uniprot_ids": [
                "P28563",
                "A0A494B924",
                "A0A494B9K9"
            ]
        },
        "ENSMUSG00000024014": {
            "id": "ENSMUSG00000024014",
            "type": "Gene",
            "symbol": "Pim1",
            "source": [
                "ENSMUSG00000056501",
                "ENSMUSG00000023034",
                "ENSMUSG00000052837",
                "ENSMUSG00000026628",
                "ENSMUSG00000038418",
                "ENSMUSG00000021250",
                "ENSMUSG00000003545",
                "ENSMUSG00000003032",
                "ENSMUSG00000018899",
                "ENSMUSG00000071076"
            ],
            "target": [],
            "expression_sum": {
                "class1": 7590.0,
                "class2": 2414.0
            },
            "ens_id": "ENSMUSG00000024014",
            "uniprot_ids": [
                "P06803",
                "A0A3Q4EGX3",
                "A0A3Q4EGB5"
            ]
        },
        "ENSMUSG00000073411": {
            "id": "ENSMUSG00000073411",
            "type": "Gene",
            "symbol": "H2-D1",
            "source": [
                "ENSMUSG00000032035",
                "ENSMUSG00000056501",
                "ENSMUSG00000026628",
                "ENSMUSG00000057098",
                "ENSMUSG00000021250",
                "ENSMUSG00000005583",
                "ENSMUSG00000055148"
            ],
            "target": [],
            "expression_sum": {
                "class1": 261015.0,
                "class2": 20567.0
            },
            "ens_id": "ENSMUSG00000073411",
            "uniprot_ids": [
                "P01899",
                "G3UZP7"
            ]
        },
        "ENSMUSG00000035678": {
            "id": "ENSMUSG00000035678",
            "type": "Gene",
            "symbol": "Tnfsf9",
            "source": [
                "ENSMUSG00000056501",
                "ENSMUSG00000052684",
                "ENSMUSG00000052837",
                "ENSMUSG00000026628",
                "ENSMUSG00000021250",
                "ENSMUSG00000018899",
                "ENSMUSG00000071076"
            ],
            "target": [],
            "expression_sum": {
                "class1": 2277.0,
                "class2": 304.0
            },
            "ens_id": "ENSMUSG00000035678",
            "uniprot_ids": [
                "P41274"
            ]
        },
        "ENSMUSG00000036438": {
            "id": "ENSMUSG00000036438",
            "type": "Gene",
            "symbol": "Calm2",
            "source": [
                "ENSMUSG00000023034",
                "ENSMUSG00000047638",
                "ENSMUSG00000026628",
                "ENSMUSG00000005583",
                "ENSMUSG00000000247",
                "ENSMUSG00000021250",
                "ENSMUSG00000029322",
                "ENSMUSG00000008575",
                "ENSMUSG00000020484"
            ],
            "target": [],
            "expression_sum": {
                "class1": 28520.0,
                "class2": 38596.0
            },
            "ens_id": "ENSMUSG00000036438",
            "uniprot_ids": [
                "P0DP26",
                "P0DP27",
                "P0DP28",
                "A0A3Q4EHJ0",
                "A0A571BE58"
            ]
        },
        "ENSMUSG00000038418": {
            "id": "ENSMUSG00000038418",
            "type": "TF",
            "symbol": "Egr1",
            "source": [
                "ENSMUSG00000052684",
                "ENSMUSG00000037868",
                "ENSMUSG00000023034",
                "ENSMUSG00000022528",
                "ENSMUSG00000052837",
                "ENSMUSG00000026628",
                "ENSMUSG00000000247",
                "ENSMUSG00000057098",
                "ENSMUSG00000021250",
                "ENSMUSG00000003545",
                "ENSMUSG00000003032",
                "ENSMUSG00000008575",
                "ENSMUSG00000018899"
            ],
            "target": [
                "ENSMUSG00000019505",
                "ENSMUSG00000075706",
                "ENSMUSG00000040723",
                "ENSMUSG00000040212",
                "ENSMUSG00000021917",
                "ENSMUSG00000023034",
                "ENSMUSG00000021242",
                "ENSMUSG00000024014",
                "ENSMUSG00000028367",
                "ENSMUSG00000021477",
                "ENSMUSG00000032314",
                "ENSMUSG00000053113",
                "ENSMUSG00000028249",
                "ENSMUSG00000030934",
                "ENSMUSG00000025213",
                "ENSMUSG00000033910",
                "ENSMUSG00000020315",
                "ENSMUSG00000018217",
                "ENSMUSG00000016194",
                "ENSMUSG00000003161",
                "ENSMUSG00000019907",
                "ENSMUSG00000052684",
                "ENSMUSG00000006800",
                "ENSMUSG00000022528",
                "ENSMUSG00000028822",
                "ENSMUSG00000034868",
                "ENSMUSG00000032353",
                "ENSMUSG00000008384",
                "ENSMUSG00000003032",
                "ENSMUSG00000042747",
                "ENSMUSG00000079111",
                "ENSMUSG00000055148",
                "ENSMUSG00000021248",
                "ENSMUSG00000026837",
                "ENSMUSG00000003038",
                "ENSMUSG00000032383",
                "ENSMUSG00000015501",
                "ENSMUSG00000006369",
                "ENSMUSG00000040128",
                "ENSMUSG00000031422",
                "ENSMUSG00000092274",
                "ENSMUSG00000002107",
                "ENSMUSG00000052837",
                "ENSMUSG00000026628",
                "ENSMUSG00000020423",
                "ENSMUSG00000035356",
                "ENSMUSG00000027706",
                "ENSMUSG00000029103",
                "ENSMUSG00000035227",
                "ENSMUSG00000024048",
                "ENSMUSG00000007891",
                "ENSMUSG00000040170",
                "ENSMUSG00000020571",
                "ENSMUSG00000031119",
                "ENSMUSG00000037072",
                "ENSMUSG00000007872",
                "ENSMUSG00000008540",
                "ENSMUSG00000018899",
                "ENSMUSG00000044786",
                "ENSMUSG00000020009",
                "ENSMUSG00000027829",
                "ENSMUSG00000022400",
                "ENSMUSG00000037868",
                "ENSMUSG00000060703",
                "ENSMUSG00000037499",
                "ENSMUSG00000035828",
                "ENSMUSG00000000247",
                "ENSMUSG00000027828",
                "ENSMUSG00000029465",
                "ENSMUSG00000021765",
                "ENSMUSG00000075254",
                "ENSMUSG00000023010",
                "ENSMUSG00000060904",
                "ENSMUSG00000025351",
                "ENSMUSG00000074884"
            ],
            "expression_sum": {
                "class1": 38873.0,
                "class2": 25496.0
            },
            "ens_id": "ENSMUSG00000038418",
            "uniprot_ids": [
                "P08046"
            ]
        },
        "ENSMUSG00000015846": {
            "id": "ENSMUSG00000015846",
            "type": "TF",
            "symbol": "Rxra",
            "source": [],
            "target": [
                "ENSMUSG00000020486",
                "ENSMUSG00000032035",
                "ENSMUSG00000026678",
                "ENSMUSG00000032946",
                "ENSMUSG00000034353",
                "ENSMUSG00000024140",
                "ENSMUSG00000025491",
                "ENSMUSG00000034845",
                "ENSMUSG00000001552",
                "ENSMUSG00000042453",
                "ENSMUSG00000028132",
                "ENSMUSG00000021756",
                "ENSMUSG00000022371",
                "ENSMUSG00000032179",
                "ENSMUSG00000000184",
                "ENSMUSG00000038591",
                "ENSMUSG00000041577",
                "ENSMUSG00000000325",
                "ENSMUSG00000038742",
                "ENSMUSG00000059588",
                "ENSMUSG00000028003",
                "ENSMUSG00000012428",
                "ENSMUSG00000023336",
                "ENSMUSG00000069662",
                "ENSMUSG00000032666",
                "ENSMUSG00000024065",
                "ENSMUSG00000006356",
                "ENSMUSG00000032528",
                "ENSMUSG00000070407",
                "ENSMUSG00000020689",
                "ENSMUSG00000036655",
                "ENSMUSG00000072625"
            ],
            "expression_sum": {
                "class1": 4376.0,
                "class2": 460.0
            },
            "ens_id": "ENSMUSG00000015846",
            "uniprot_ids": [
                "P28700",
                "Q6LC96",
                "A2AJP2",
                "A0A5F8MP91",
                "A0A5F8MQ23"
            ]
        },
        "ENSMUSG00000034353": {
            "id": "ENSMUSG00000034353",
            "type": "Gene",
            "symbol": "Ramp1",
            "source": [
                "ENSMUSG00000032035",
                "ENSMUSG00000047638",
                "ENSMUSG00000029322",
                "ENSMUSG00000015846",
                "ENSMUSG00000008575"
            ],
            "target": [],
            "expression_sum": {
                "class1": 54268.0,
                "class2": 7630.0
            },
            "ens_id": "ENSMUSG00000034353",
            "uniprot_ids": [
                "Q9WTJ5",
                "A0A087WPC6",
                "E9Q915"
            ]
        },
        "ENSMUSG00000041577": {
            "id": "ENSMUSG00000041577",
            "type": "Gene",
            "symbol": "Prelp",
            "source": [
                "ENSMUSG00000028639",
                "ENSMUSG00000029322",
                "ENSMUSG00000015846",
                "ENSMUSG00000005583"
            ],
            "target": [],
            "expression_sum": {
                "class1": 110026.0,
                "class2": 9770.0
            },
            "ens_id": "ENSMUSG00000041577",
            "uniprot_ids": [
                "Q9JK53"
            ]
        },
        "ENSMUSG00000032666": {
            "id": "ENSMUSG00000032666",
            "type": "Gene",
            "symbol": "1700025G04Rik",
            "source": [
                "ENSMUSG00000032035",
                "ENSMUSG00000038193",
                "ENSMUSG00000015846",
                "ENSMUSG00000000247"
            ],
            "target": [],
            "expression_sum": {
                "class1": 14602.0,
                "class2": 1573.0
            },
            "ens_id": "ENSMUSG00000032666",
            "uniprot_ids": [
                "Q8K207",
                "A0A0A0MQF7"
            ]
        },
        "ENSMUSG00000026678": {
            "id": "ENSMUSG00000026678",
            "type": "Gene",
            "symbol": "Rgs5",
            "source": [
                "ENSMUSG00000015846"
            ],
            "target": [],
            "expression_sum": {
                "class1": 120678.0,
                "class2": 58286.0
            },
            "ens_id": "ENSMUSG00000026678",
            "uniprot_ids": [
                "O08850",
                "A0A0A6YWD0"
            ]
        },
        "ENSMUSG00000059588": {
            "id": "ENSMUSG00000059588",
            "type": "Gene",
            "symbol": "Calcrl",
            "source": [
                "ENSMUSG00000032035",
                "ENSMUSG00000015846"
            ],
            "target": [],
            "expression_sum": {
                "class1": 9701.0,
                "class2": 268.0
            },
            "ens_id": "ENSMUSG00000059588",
            "uniprot_ids": [
                "Q9R1W5"
            ]
        },
        "ENSMUSG00000028003": {
            "id": "ENSMUSG00000028003",
            "type": "Gene",
            "symbol": "Lrat",
            "source": [
                "ENSMUSG00000015846",
                "ENSMUSG00000056501",
                "ENSMUSG00000008575"
            ],
            "target": [],
            "expression_sum": {
                "class1": 79466.0,
                "class2": 231.0
            },
            "ens_id": "ENSMUSG00000028003",
            "uniprot_ids": [
                "Q9JI60"
            ]
        },
        "ENSMUSG00000028132": {
            "id": "ENSMUSG00000028132",
            "type": "Gene",
            "symbol": "Tmem56",
            "source": [
                "ENSMUSG00000032035",
                "ENSMUSG00000015846"
            ],
            "target": [],
            "expression_sum": {
                "class1": 44781.0,
                "class2": 28.0
            },
            "ens_id": "ENSMUSG00000028132",
            "uniprot_ids": [
                "Q8CGF5",
                "D3Z3E6"
            ]
        },
        "ENSMUSG00000012428": {
            "id": "ENSMUSG00000012428",
            "type": "Gene",
            "symbol": "Steap4",
            "source": [
                "ENSMUSG00000015846",
                "ENSMUSG00000005583"
            ],
            "target": [],
            "expression_sum": {
                "class1": 59378.0,
                "class2": 2952.0
            },
            "ens_id": "ENSMUSG00000012428",
            "uniprot_ids": [
                "Q923B6"
            ]
        },
        "ENSMUSG00000042453": {
            "id": "ENSMUSG00000042453",
            "type": "Gene",
            "symbol": "Reln",
            "source": [
                "ENSMUSG00000032035",
                "ENSMUSG00000056501",
                "ENSMUSG00000052684",
                "ENSMUSG00000052837",
                "ENSMUSG00000038193",
                "ENSMUSG00000015846",
                "ENSMUSG00000076431",
                "ENSMUSG00000008575",
                "ENSMUSG00000020484"
            ],
            "target": [],
            "expression_sum": {
                "class1": 66813.0,
                "class2": 37.0
            },
            "ens_id": "ENSMUSG00000042453",
            "uniprot_ids": [
                "Q60841",
                "F6U2B2"
            ]
        },
        "ENSMUSG00000000184": {
            "id": "ENSMUSG00000000184",
            "type": "Gene",
            "symbol": "Ccnd2",
            "source": [
                "ENSMUSG00000005583",
                "ENSMUSG00000015846",
                "ENSMUSG00000000247"
            ],
            "target": [],
            "expression_sum": {
                "class1": 24456.0,
                "class2": 11087.0
            },
            "ens_id": "ENSMUSG00000000184",
            "uniprot_ids": [
                "P30280",
                "Q9D8L9",
                "A0A0J9YU51"
            ]
        },
        "ENSMUSG00000025491": {
            "id": "ENSMUSG00000025491",
            "type": "Gene",
            "symbol": "Ifitm1",
            "source": [
                "ENSMUSG00000015846",
                "ENSMUSG00000005583"
            ],
            "target": [],
            "expression_sum": {
                "class1": 130845.0,
                "class2": 6646.0
            },
            "ens_id": "ENSMUSG00000025491",
            "uniprot_ids": [
                "Q9D103"
            ]
        },
        "ENSMUSG00000069662": {
            "id": "ENSMUSG00000069662",
            "type": "Gene",
            "symbol": "Marcks",
            "source": [
                "ENSMUSG00000032035",
                "ENSMUSG00000038193",
                "ENSMUSG00000015846",
                "ENSMUSG00000003032",
                "ENSMUSG00000005583",
                "ENSMUSG00000071076"
            ],
            "target": [],
            "expression_sum": {
                "class1": 27744.0,
                "class2": 3686.0
            },
            "ens_id": "ENSMUSG00000069662",
            "uniprot_ids": [
                "P26645"
            ]
        },
        "ENSMUSG00000034845": {
            "id": "ENSMUSG00000034845",
            "type": "Gene",
            "symbol": "Plvap",
            "source": [
                "ENSMUSG00000032035",
                "ENSMUSG00000015846"
            ],
            "target": [],
            "expression_sum": {
                "class1": 211610.0,
                "class2": 718.0
            },
            "ens_id": "ENSMUSG00000034845",
            "uniprot_ids": [
                "G3X924"
            ]
        },
        "ENSMUSG00000023336": {
            "id": "ENSMUSG00000023336",
            "type": "Gene",
            "symbol": "Wfdc1",
            "source": [
                "ENSMUSG00000032035",
                "ENSMUSG00000015846"
            ],
            "target": [],
            "expression_sum": {
                "class1": 23695.0,
                "class2": 4922.0
            },
            "ens_id": "ENSMUSG00000023336",
            "uniprot_ids": [
                "Q9ESH5",
                "Q9CQZ8"
            ]
        },
        "ENSMUSG00000072625": {
            "id": "ENSMUSG00000072625",
            "type": "Gene",
            "symbol": "Gdf2",
            "source": [
                "ENSMUSG00000021250",
                "ENSMUSG00000052684",
                "ENSMUSG00000015846",
                "ENSMUSG00000056501"
            ],
            "target": [],
            "expression_sum": {
                "class1": 35762.0,
                "class2": 48.0
            },
            "ens_id": "ENSMUSG00000072625",
            "uniprot_ids": [
                "Q9WV56"
            ]
        },
        "ENSMUSG00000038742": {
            "id": "ENSMUSG00000038742",
            "type": "Gene",
            "symbol": "Angptl6",
            "source": [
                "ENSMUSG00000015846",
                "ENSMUSG00000021127"
            ],
            "target": [],
            "expression_sum": {
                "class1": 61922.0,
                "class2": 271.0
            },
            "ens_id": "ENSMUSG00000038742",
            "uniprot_ids": [
                "Q8R0Z6"
            ]
        },
        "ENSMUSG00000032035": {
            "id": "ENSMUSG00000032035",
            "type": "TF",
            "symbol": "Ets1",
            "source": [
                "ENSMUSG00000015846"
            ],
            "target": [
                "ENSMUSG00000019505",
                "ENSMUSG00000005338",
                "ENSMUSG00000064363",
                "ENSMUSG00000031447",
                "ENSMUSG00000021477",
                "ENSMUSG00000027447",
                "ENSMUSG00000024909",
                "ENSMUSG00000034868",
                "ENSMUSG00000023175",
                "ENSMUSG00000064354",
                "ENSMUSG00000026970",
                "ENSMUSG00000059588",
                "ENSMUSG00000025130",
                "ENSMUSG00000025790",
                "ENSMUSG00000034892",
                "ENSMUSG00000032231",
                "ENSMUSG00000023336",
                "ENSMUSG00000031021",
                "ENSMUSG00000029426",
                "ENSMUSG00000029718",
                "ENSMUSG00000036655",
                "ENSMUSG00000062825",
                "ENSMUSG00000020368",
                "ENSMUSG00000024678",
                "ENSMUSG00000069601",
                "ENSMUSG00000034353",
                "ENSMUSG00000028603",
                "ENSMUSG00000034845",
                "ENSMUSG00000022892",
                "ENSMUSG00000032562",
                "ENSMUSG00000056025",
                "ENSMUSG00000021756",
                "ENSMUSG00000038803",
                "ENSMUSG00000070436",
                "ENSMUSG00000029767",
                "ENSMUSG00000030824",
                "ENSMUSG00000064351",
                "ENSMUSG00000021248",
                "ENSMUSG00000027642",
                "ENSMUSG00000030249",
                "ENSMUSG00000029163",
                "ENSMUSG00000026193",
                "ENSMUSG00000033880",
                "ENSMUSG00000022587",
                "ENSMUSG00000023944",
                "ENSMUSG00000024065",
                "ENSMUSG00000029455",
                "ENSMUSG00000031375",
                "ENSMUSG00000072235",
                "ENSMUSG00000036381",
                "ENSMUSG00000026860",
                "ENSMUSG00000046330",
                "ENSMUSG00000020432",
                "ENSMUSG00000079523",
                "ENSMUSG00000001123",
                "ENSMUSG00000060126",
                "ENSMUSG00000042453",
                "ENSMUSG00000058569",
                "ENSMUSG00000026014",
                "ENSMUSG00000005973",
                "ENSMUSG00000016534",
                "ENSMUSG00000060591",
                "ENSMUSG00000032492",
                "ENSMUSG00000034707",
                "ENSMUSG00000049517",
                "ENSMUSG00000022665",
                "ENSMUSG00000031661",
                "ENSMUSG00000061232",
                "ENSMUSG00000027712",
                "ENSMUSG00000024048",
                "ENSMUSG00000069662",
                "ENSMUSG00000032666",
                "ENSMUSG00000028108",
                "ENSMUSG00000004207",
                "ENSMUSG00000020205",
                "ENSMUSG00000035242",
                "ENSMUSG00000034708",
                "ENSMUSG00000026478",
                "ENSMUSG00000032528",
                "ENSMUSG00000070407",
                "ENSMUSG00000020689",
                "ENSMUSG00000026223",
                "ENSMUSG00000029076",
                "ENSMUSG00000022971",
                "ENSMUSG00000074884",
                "ENSMUSG00000054808",
                "ENSMUSG00000022108",
                "ENSMUSG00000041736",
                "ENSMUSG00000028132",
                "ENSMUSG00000027248",
                "ENSMUSG00000020585",
                "ENSMUSG00000032179",
                "ENSMUSG00000025492",
                "ENSMUSG00000046402",
                "ENSMUSG00000034158",
                "ENSMUSG00000055447",
                "ENSMUSG00000000740",
                "ENSMUSG00000007892",
                "ENSMUSG00000027523",
                "ENSMUSG00000028618",
                "ENSMUSG00000032383",
                "ENSMUSG00000035530",
                "ENSMUSG00000047907",
                "ENSMUSG00000073411",
                "ENSMUSG00000048758",
                "ENSMUSG00000022708",
                "ENSMUSG00000029322",
                "ENSMUSG00000019929",
                "ENSMUSG00000060802",
                "ENSMUSG00000001847",
                "ENSMUSG00000000247",
                "ENSMUSG00000030849",
                "ENSMUSG00000020902",
                "ENSMUSG00000073489",
                "ENSMUSG00000024109",
                "ENSMUSG00000058135"
            ],
            "expression_sum": {
                "class1": 11559.0,
                "class2": 1092.0
            },
            "ens_id": "ENSMUSG00000032035",
            "uniprot_ids": [
                "P27577",
                "V9GXE2",
                "E9PWI8",
                "V9GX28",
                "A0A5F8MPR7"
            ]
        },
        "ENSMUSG00000032179": {
            "id": "ENSMUSG00000032179",
            "type": "Gene",
            "symbol": "Bmp5",
            "source": [
                "ENSMUSG00000032035",
                "ENSMUSG00000015846",
                "ENSMUSG00000056501",
                "ENSMUSG00000000247"
            ],
            "target": [],
            "expression_sum": {
                "class1": 25665.0,
                "class2": 557.0
            },
            "ens_id": "ENSMUSG00000032179",
            "uniprot_ids": [
                "P49003"
            ]
        },
        "ENSMUSG00000032528": {
            "id": "ENSMUSG00000032528",
            "type": "Gene",
            "symbol": "Vipr1",
            "source": [
                "ENSMUSG00000032035",
                "ENSMUSG00000015846"
            ],
            "target": [],
            "expression_sum": {
                "class1": 26571.0,
                "class2": 67.0
            },
            "ens_id": "ENSMUSG00000032528",
            "uniprot_ids": [
                "P97751"
            ]
        },
        "ENSMUSG00000070407": {
            "id": "ENSMUSG00000070407",
            "type": "Gene",
            "symbol": "Hs3st3b1",
            "source": [
                "ENSMUSG00000032035",
                "ENSMUSG00000015846"
            ],
            "target": [],
            "expression_sum": {
                "class1": 15166.0,
                "class2": 72.0
            },
            "ens_id": "ENSMUSG00000070407",
            "uniprot_ids": [
                "Q9QZS6"
            ]
        },
        "ENSMUSG00000020486": {
            "id": "ENSMUSG00000020486",
            "type": "Gene",
            "symbol": "Sept4",
            "source": [
                "ENSMUSG00000052684",
                "ENSMUSG00000028639",
                "ENSMUSG00000015846",
                "ENSMUSG00000005583",
                "ENSMUSG00000071076"
            ],
            "target": [],
            "expression_sum": {
                "class1": 68998.0,
                "class2": 5830.0
            },
            "ens_id": "ENSMUSG00000020486",
            "uniprot_ids": [
                "Q5ND19",
                "P28661",
                "A0A5F8MP96",
                "Q5ND13",
                "Q5ND09"
            ]
        },
        "ENSMUSG00000001552": {
            "id": "ENSMUSG00000001552",
            "type": "Gene",
            "symbol": "Jup",
            "source": [
                "ENSMUSG00000047638",
                "ENSMUSG00000015846"
            ],
            "target": [],
            "expression_sum": {
                "class1": 10105.0,
                "class2": 1090.0
            },
            "ens_id": "ENSMUSG00000001552",
            "uniprot_ids": [
                "Q02257"
            ]
        },
        "ENSMUSG00000020689": {
            "id": "ENSMUSG00000020689",
            "type": "Gene",
            "symbol": "Itgb3",
            "source": [
                "ENSMUSG00000032035",
                "ENSMUSG00000015846"
            ],
            "target": [],
            "expression_sum": {
                "class1": 5792.0,
                "class2": 313.0
            },
            "ens_id": "ENSMUSG00000020689",
            "uniprot_ids": [
                "O54890"
            ]
        },
        "ENSMUSG00000021756": {
            "id": "ENSMUSG00000021756",
            "type": "Gene",
            "symbol": "Il6st",
            "source": [
                "ENSMUSG00000022528",
                "ENSMUSG00000032035",
                "ENSMUSG00000015846",
                "ENSMUSG00000005583"
            ],
            "target": [],
            "expression_sum": {
                "class1": 20193.0,
                "class2": 2185.0
            },
            "ens_id": "ENSMUSG00000021756",
            "uniprot_ids": [
                "Q00560",
                "Q3TDT5",
                "V9GX52",
                "V9GXA2",
                "V9GXJ5",
                "V9GX00",
                "V9GX73",
                "V9GXT4"
            ]
        },
        "ENSMUSG00000036655": {
            "id": "ENSMUSG00000036655",
            "type": "Gene",
            "symbol": "Colec11",
            "source": [
                "ENSMUSG00000032035",
                "ENSMUSG00000056501",
                "ENSMUSG00000052684",
                "ENSMUSG00000038193",
                "ENSMUSG00000021250",
                "ENSMUSG00000015846"
            ],
            "target": [],
            "expression_sum": {
                "class1": 276981.0,
                "class2": 2345.0
            },
            "ens_id": "ENSMUSG00000036655",
            "uniprot_ids": [
                "A0A0R4J0M6",
                "A0A1Y7VKG6",
                "A0A1Y7VLI6"
            ]
        },
        "ENSMUSG00000006356": {
            "id": "ENSMUSG00000006356",
            "type": "Gene",
            "symbol": "Crip2",
            "source": [
                "ENSMUSG00000047638",
                "ENSMUSG00000057098",
                "ENSMUSG00000029322",
                "ENSMUSG00000015846",
                "ENSMUSG00000008575",
                "ENSMUSG00000005583"
            ],
            "target": [],
            "expression_sum": {
                "class1": 63331.0,
                "class2": 24982.0
            },
            "ens_id": "ENSMUSG00000006356",
            "uniprot_ids": [
                "Q9DCT8",
                "A0A0G2JF37"
            ]
        },
        "ENSMUSG00000038591": {
            "id": "ENSMUSG00000038591",
            "type": "Gene",
            "symbol": "Colec10",
            "source": [
                "ENSMUSG00000015846"
            ],
            "target": [],
            "expression_sum": {
                "class1": 42714.0,
                "class2": 6.0
            },
            "ens_id": "ENSMUSG00000038591",
            "uniprot_ids": [
                "Q8CF98"
            ]
        },
        "ENSMUSG00000022371": {
            "id": "ENSMUSG00000022371",
            "type": "Gene",
            "symbol": "Col14a1",
            "source": [
                "ENSMUSG00000026872",
                "ENSMUSG00000038193",
                "ENSMUSG00000015846",
                "ENSMUSG00000003032",
                "ENSMUSG00000005583"
            ],
            "target": [],
            "expression_sum": {
                "class1": 81209.0,
                "class2": 13551.0
            },
            "ens_id": "ENSMUSG00000022371",
            "uniprot_ids": [
                "Q80X19",
                "B7ZNH7",
                "K3W4R4",
                "F7D5Y4"
            ]
        },
        "ENSMUSG00000000325": {
            "id": "ENSMUSG00000000325",
            "type": "Gene",
            "symbol": "Arvcf",
            "source": [
                "ENSMUSG00000052684",
                "ENSMUSG00000005583",
                "ENSMUSG00000000247",
                "ENSMUSG00000015846",
                "ENSMUSG00000008575",
                "ENSMUSG00000071076"
            ],
            "target": [],
            "expression_sum": {
                "class1": 12601.0,
                "class2": 1652.0
            },
            "ens_id": "ENSMUSG00000000325"
        },
        "ENSMUSG00000024065": {
            "id": "ENSMUSG00000024065",
            "type": "Gene",
            "symbol": "Ehd3",
            "source": [
                "ENSMUSG00000032035",
                "ENSMUSG00000015846",
                "ENSMUSG00000052684",
                "ENSMUSG00000000247"
            ],
            "target": [],
            "expression_sum": {
                "class1": 38270.0,
                "class2": 820.0
            },
            "ens_id": "ENSMUSG00000024065",
            "uniprot_ids": [
                "Q9QXY6"
            ]
        },
        "ENSMUSG00000024140": {
            "id": "ENSMUSG00000024140",
            "type": "TF",
            "symbol": "Epas1",
            "source": [
                "ENSMUSG00000032238",
                "ENSMUSG00000052684",
                "ENSMUSG00000005583",
                "ENSMUSG00000015846",
                "ENSMUSG00000008575",
                "ENSMUSG00000028565"
            ],
            "target": [
                "ENSMUSG00000054452",
                "ENSMUSG00000032420",
                "ENSMUSG00000046330",
                "ENSMUSG00000049439"
            ],
            "expression_sum": {
                "class1": 14142.0,
                "class2": 5104.0
            },
            "ens_id": "ENSMUSG00000024140",
            "uniprot_ids": [
                "P97481"
            ]
        },
        "ENSMUSG00000032946": {
            "id": "ENSMUSG00000032946",
            "type": "Gene",
            "symbol": "Rasgrp2",
            "source": [
                "ENSMUSG00000047638",
                "ENSMUSG00000000247",
                "ENSMUSG00000053477",
                "ENSMUSG00000015846",
                "ENSMUSG00000005583"
            ],
            "target": [],
            "expression_sum": {
                "class1": 15073.0,
                "class2": 3231.0
            },
            "ens_id": "ENSMUSG00000032946",
            "uniprot_ids": [
                "Q9QUG9",
                "D3YZP1",
                "D3YZP2",
                "D3YZX2",
                "D3Z2F9",
                "D3Z3N3",
                "D3YXD6",
                "D6RDD3",
                "D3YZP4"
            ]
        },
        "ENSMUSG00000000247": {
            "id": "ENSMUSG00000000247",
            "type": "TF",
            "symbol": "Lhx2",
            "source": [
                "ENSMUSG00000032035",
                "ENSMUSG00000056501",
                "ENSMUSG00000052684",
                "ENSMUSG00000023034",
                "ENSMUSG00000052837",
                "ENSMUSG00000038418",
                "ENSMUSG00000021250",
                "ENSMUSG00000038193",
                "ENSMUSG00000071076"
            ],
            "target": [
                "ENSMUSG00000019539",
                "ENSMUSG00000019505",
                "ENSMUSG00000024063",
                "ENSMUSG00000059291",
                "ENSMUSG00000014077",
                "ENSMUSG00000058546",
                "ENSMUSG00000028639",
                "ENSMUSG00000031812",
                "ENSMUSG00000021477",
                "ENSMUSG00000026457",
                "ENSMUSG00000022283",
                "ENSMUSG00000068523",
                "ENSMUSG00000040952",
                "ENSMUSG00000048076",
                "ENSMUSG00000030695",
                "ENSMUSG00000041881",
                "ENSMUSG00000053560",
                "ENSMUSG00000025967",
                "ENSMUSG00000026939",
                "ENSMUSG00000030744",
                "ENSMUSG00000036438",
                "ENSMUSG00000026238",
                "ENSMUSG00000000325",
                "ENSMUSG00000060036",
                "ENSMUSG00000008668",
                "ENSMUSG00000061787",
                "ENSMUSG00000038717",
                "ENSMUSG00000020372",
                "ENSMUSG00000041417",
                "ENSMUSG00000008682",
                "ENSMUSG00000023175",
                "ENSMUSG00000021127",
                "ENSMUSG00000026970",
                "ENSMUSG00000031198",
                "ENSMUSG00000023861",
                "ENSMUSG00000061904",
                "ENSMUSG00000034892",
                "ENSMUSG00000022010",
                "ENSMUSG00000005161",
                "ENSMUSG00000020612",
                "ENSMUSG00000032231",
                "ENSMUSG00000071415",
                "ENSMUSG00000021660",
                "ENSMUSG00000061477",
                "ENSMUSG00000060743",
                "ENSMUSG00000007872",
                "ENSMUSG00000056201",
                "ENSMUSG00000020738",
                "ENSMUSG00000022884",
                "ENSMUSG00000061983",
                "ENSMUSG00000004980",
                "ENSMUSG00000037805",
                "ENSMUSG00000027750",
                "ENSMUSG00000006304",
                "ENSMUSG00000028081",
                "ENSMUSG00000063316",
                "ENSMUSG00000055302",
                "ENSMUSG00000017404",
                "ENSMUSG00000070493",
                "ENSMUSG00000021546",
                "ENSMUSG00000046364",
                "ENSMUSG00000004070",
                "ENSMUSG00000022312",
                "ENSMUSG00000060636",
                "ENSMUSG00000055839",
                "ENSMUSG00000014294",
                "ENSMUSG00000064370",
                "ENSMUSG00000068874",
                "ENSMUSG00000037999",
                "ENSMUSG00000069601",
                "ENSMUSG00000028603",
                "ENSMUSG00000042043",
                "ENSMUSG00000037742",
                "ENSMUSG00000025781",
                "ENSMUSG00000022892",
                "ENSMUSG00000032562",
                "ENSMUSG00000025511",
                "ENSMUSG00000029810",
                "ENSMUSG00000038803",
                "ENSMUSG00000052684",
                "ENSMUSG00000012405",
                "ENSMUSG00000000184",
                "ENSMUSG00000020719",
                "ENSMUSG00000036371",
                "ENSMUSG00000090862",
                "ENSMUSG00000021807",
                "ENSMUSG00000022051",
                "ENSMUSG00000030824",
                "ENSMUSG00000064351",
                "ENSMUSG00000027642",
                "ENSMUSG00000003970",
                "ENSMUSG00000003617",
                "ENSMUSG00000021870",
                "ENSMUSG00000018326",
                "ENSMUSG00000020460",
                "ENSMUSG00000035885",
                "ENSMUSG00000027706",
                "ENSMUSG00000055044",
                "ENSMUSG00000029781",
                "ENSMUSG00000042812",
                "ENSMUSG00000025508",
                "ENSMUSG00000063882",
                "ENSMUSG00000025290",
                "ENSMUSG00000039105",
                "ENSMUSG00000037712",
                "ENSMUSG00000009927",
                "ENSMUSG00000033880",
                "ENSMUSG00000031353",
                "ENSMUSG00000045128",
                "ENSMUSG00000022587",
                "ENSMUSG00000025794",
                "ENSMUSG00000023944",
                "ENSMUSG00000024065",
                "ENSMUSG00000023367",
                "ENSMUSG00000024608",
                "ENSMUSG00000001687",
                "ENSMUSG00000029713",
                "ENSMUSG00000039001",
                "ENSMUSG00000036381",
                "ENSMUSG00000006333",
                "ENSMUSG00000058600",
                "ENSMUSG00000062647",
                "ENSMUSG00000010406",
                "ENSMUSG00000030432",
                "ENSMUSG00000046330",
                "ENSMUSG00000074129",
                "ENSMUSG00000090733",
                "ENSMUSG00000031818",
                "ENSMUSG00000020432",
                "ENSMUSG00000010376",
                "ENSMUSG00000002778",
                "ENSMUSG00000032946",
                "ENSMUSG00000098274",
                "ENSMUSG00000027404",
                "ENSMUSG00000016559",
                "ENSMUSG00000060126",
                "ENSMUSG00000003072",
                "ENSMUSG00000061315",
                "ENSMUSG00000057322",
                "ENSMUSG00000027673",
                "ENSMUSG00000012848",
                "ENSMUSG00000060591",
                "ENSMUSG00000023942",
                "ENSMUSG00000049517",
                "ENSMUSG00000025362",
                "ENSMUSG00000019960",
                "ENSMUSG00000044533",
                "ENSMUSG00000029614",
                "ENSMUSG00000022136",
                "ENSMUSG00000025428",
                "ENSMUSG00000073702",
                "ENSMUSG00000003814",
                "ENSMUSG00000062006",
                "ENSMUSG00000062683",
                "ENSMUSG00000040128",
                "ENSMUSG00000032216",
                "ENSMUSG00000079435",
                "ENSMUSG00000047215",
                "ENSMUSG00000062270",
                "ENSMUSG00000022450",
                "ENSMUSG00000041571",
                "ENSMUSG00000053565",
                "ENSMUSG00000032666",
                "ENSMUSG00000004207",
                "ENSMUSG00000031029",
                "ENSMUSG00000028673",
                "ENSMUSG00000024576",
                "ENSMUSG00000035242",
                "ENSMUSG00000050708",
                "ENSMUSG00000014769",
                "ENSMUSG00000015656",
                "ENSMUSG00000067288",
                "ENSMUSG00000058558",
                "ENSMUSG00000029076",
                "ENSMUSG00000056851",
                "ENSMUSG00000022971",
                "ENSMUSG00000052146",
                "ENSMUSG00000060938",
                "ENSMUSG00000047675",
                "ENSMUSG00000025393",
                "ENSMUSG00000034994",
                "ENSMUSG00000041841",
                "ENSMUSG00000067274",
                "ENSMUSG00000054808",
                "ENSMUSG00000024308",
                "ENSMUSG00000071644",
                "ENSMUSG00000057863",
                "ENSMUSG00000026566",
                "ENSMUSG00000022108",
                "ENSMUSG00000032288",
                "ENSMUSG00000014313",
                "ENSMUSG00000024248",
                "ENSMUSG00000028266",
                "ENSMUSG00000024953",
                "ENSMUSG00000030057",
                "ENSMUSG00000041736",
                "ENSMUSG00000045690",
                "ENSMUSG00000041560",
                "ENSMUSG00000005881",
                "ENSMUSG00000049751",
                "ENSMUSG00000001289",
                "ENSMUSG00000024678",
                "ENSMUSG00000036751",
                "ENSMUSG00000032179",
                "ENSMUSG00000013593",
                "ENSMUSG00000008683",
                "ENSMUSG00000055762",
                "ENSMUSG00000043716",
                "ENSMUSG00000021250",
                "ENSMUSG00000017778",
                "ENSMUSG00000055447",
                "ENSMUSG00000022890",
                "ENSMUSG00000000740",
                "ENSMUSG00000007892",
                "ENSMUSG00000027523",
                "ENSMUSG00000024190",
                "ENSMUSG00000028618",
                "ENSMUSG00000057841",
                "ENSMUSG00000079641",
                "ENSMUSG00000027593",
                "ENSMUSG00000035530",
                "ENSMUSG00000028757",
                "ENSMUSG00000059070",
                "ENSMUSG00000031532",
                "ENSMUSG00000027620",
                "ENSMUSG00000047907",
                "ENSMUSG00000052837",
                "ENSMUSG00000048758",
                "ENSMUSG00000029632",
                "ENSMUSG00000022708",
                "ENSMUSG00000028234",
                "ENSMUSG00000093674",
                "ENSMUSG00000028936",
                "ENSMUSG00000042682",
                "ENSMUSG00000044786",
                "ENSMUSG00000028495",
                "ENSMUSG00000059005",
                "ENSMUSG00000028478",
                "ENSMUSG00000071076",
                "ENSMUSG00000041453",
                "ENSMUSG00000003429",
                "ENSMUSG00000018293",
                "ENSMUSG00000029465",
                "ENSMUSG00000038418",
                "ENSMUSG00000030849",
                "ENSMUSG00000021134",
                "ENSMUSG00000000563",
                "ENSMUSG00000031633",
                "ENSMUSG00000037563",
                "ENSMUSG00000032399"
            ],
            "expression_sum": {
                "class1": 14582.0,
                "class2": 111.0
            },
            "ens_id": "ENSMUSG00000000247",
            "uniprot_ids": [
                "Q9Z0S2",
                "F6Z9H5",
                "A0A0A0MQK9",
                "F6SM03",
                "H3BKI3"
            ]
        },
        "ENSMUSG00000043716": {
            "id": "ENSMUSG00000043716",
            "type": "Gene",
            "symbol": "Rpl7",
            "source": [
                "ENSMUSG00000022528",
                "ENSMUSG00000000247",
                "ENSMUSG00000057098",
                "ENSMUSG00000038193",
                "ENSMUSG00000048387"
            ],
            "target": [],
            "expression_sum": {
                "class1": 89824.0,
                "class2": 43999.0
            },
            "ens_id": "ENSMUSG00000043716",
            "uniprot_ids": [
                "P14148",
                "F6XI62"
            ]
        },
        "ENSMUSG00000073702": {
            "id": "ENSMUSG00000073702",
            "type": "Gene",
            "symbol": "Rpl31",
            "source": [
                "ENSMUSG00000057098",
                "ENSMUSG00000048387",
                "ENSMUSG00000000247"
            ],
            "target": [],
            "expression_sum": {
                "class1": 96594.0,
                "class2": 49398.0
            },
            "ens_id": "ENSMUSG00000073702",
            "uniprot_ids": [
                "P62900",
                "A0A0A6YXL3",
                "A0A0A6YX26"
            ]
        },
        "ENSMUSG00000025967": {
            "id": "ENSMUSG00000025967",
            "type": "Gene",
            "symbol": "Eef1b2",
            "source": [
                "ENSMUSG00000057098",
                "ENSMUSG00000000247"
            ],
            "target": [],
            "expression_sum": {
                "class1": 23925.0,
                "class2": 11243.0
            },
            "ens_id": "ENSMUSG00000025967",
            "uniprot_ids": [
                "O70251",
                "A0A087WS46",
                "M0QW60",
                "M0QWK5",
                "G3UZ47",
                "G3UX43",
                "M0QWH8"
            ]
        },
        "ENSMUSG00000046330": {
            "id": "ENSMUSG00000046330",
            "type": "Gene",
            "symbol": "Rpl37a",
            "source": [
                "ENSMUSG00000032035",
                "ENSMUSG00000000247",
                "ENSMUSG00000038193",
                "ENSMUSG00000024140",
                "ENSMUSG00000005583",
                "ENSMUSG00000048387"
            ],
            "target": [],
            "expression_sum": {
                "class1": 185640.0,
                "class2": 85588.0
            },
            "ens_id": "ENSMUSG00000046330",
            "uniprot_ids": [
                "P61514",
                "A0A1D5RL86"
            ]
        },
        "ENSMUSG00000006304": {
            "id": "ENSMUSG00000006304",
            "type": "Gene",
            "symbol": "Arpc2",
            "source": [
                "ENSMUSG00000057098",
                "ENSMUSG00000029322",
                "ENSMUSG00000000247"
            ],
            "target": [],
            "expression_sum": {
                "class1": 21631.0,
                "class2": 13022.0
            },
            "ens_id": "ENSMUSG00000006304",
            "uniprot_ids": [
                "Q9CVB6",
                "D3YXG6",
                "A0A087WRT2"
            ]
        },
        "ENSMUSG00000026238": {
            "id": "ENSMUSG00000026238",
            "type": "Gene",
            "symbol": "Ptma",
            "source": [
                "ENSMUSG00000025498",
                "ENSMUSG00000047638",
                "ENSMUSG00000000247",
                "ENSMUSG00000057098",
                "ENSMUSG00000029322"
            ],
            "target": [],
            "expression_sum": {
                "class1": 99702.0,
                "class2": 63901.0
            },
            "ens_id": "ENSMUSG00000026238",
            "uniprot_ids": [
                "P26350",
                "A0A087WQN2",
                "A0A087WP98",
                "A0A087WPN6"
            ]
        },
        "ENSMUSG00000026457": {
            "id": "ENSMUSG00000026457",
            "type": "Gene",
            "symbol": "Adipor1",
            "source": [
                "ENSMUSG00000000247"
            ],
            "target": [],
            "expression_sum": {
                "class1": 5762.0,
                "class2": 2768.0
            },
            "ens_id": "ENSMUSG00000026457",
            "uniprot_ids": [
                "Q91VH1"
            ]
        },
        "ENSMUSG00000013593": {
            "id": "ENSMUSG00000013593",
            "type": "Gene",
            "symbol": "Ndufs2",
            "source": [
                "ENSMUSG00000029322",
                "ENSMUSG00000000247"
            ],
            "target": [],
            "expression_sum": {
                "class1": 8460.0,
                "class2": 5508.0
            },
            "ens_id": "ENSMUSG00000013593",
            "uniprot_ids": [
                "Q91WD5",
                "A0A0A6YXD3",
                "A0A0A6YW30",
                "D3YXT0"
            ]
        },
        "ENSMUSG00000060743": {
            "id": "ENSMUSG00000060743",
            "type": "Gene",
            "symbol": "H3f3a",
            "source": [
                "ENSMUSG00000057098",
                "ENSMUSG00000047638",
                "ENSMUSG00000000247"
            ],
            "target": [],
            "expression_sum": {
                "class1": 40847.0,
                "class2": 23481.0
            },
            "ens_id": "ENSMUSG00000060743",
            "uniprot_ids": [
                "P84244",
                "E0CYN1",
                "E0CYR7",
                "E0CZ27",
                "F8WI35"
            ]
        },
        "ENSMUSG00000025781": {
            "id": "ENSMUSG00000025781",
            "type": "Gene",
            "symbol": "Atp5c1",
            "source": [
                "ENSMUSG00000057098",
                "ENSMUSG00000000247"
            ],
            "target": [],
            "expression_sum": {
                "class1": 12268.0,
                "class2": 9241.0
            },
            "ens_id": "ENSMUSG00000025781",
            "uniprot_ids": [
                "Q91VR2",
                "Q8C2Q8",
                "A2AKV2",
                "A2AKV3",
                "A2AKV1",
                "A2AKV0",
                "A2AKU9"
            ]
        },
        "ENSMUSG00000026939": {
            "id": "ENSMUSG00000026939",
            "type": "Gene",
            "symbol": "Tmem141",
            "source": [
                "ENSMUSG00000000247"
            ],
            "target": [],
            "expression_sum": {
                "class1": 9467.0,
                "class2": 616.0
            },
            "ens_id": "ENSMUSG00000026939",
            "uniprot_ids": [
                "A2AJB2",
                "F8WGV9"
            ]
        },
        "ENSMUSG00000062647": {
            "id": "ENSMUSG00000062647",
            "type": "Gene",
            "symbol": "Rpl7a",
            "source": [
                "ENSMUSG00000022528",
                "ENSMUSG00000000247",
                "ENSMUSG00000057098",
                "ENSMUSG00000038193",
                "ENSMUSG00000005583",
                "ENSMUSG00000031965",
                "ENSMUSG00000053007"
            ],
            "target": [],
            "expression_sum": {
                "class1": 61371.0,
                "class2": 28548.0
            },
            "ens_id": "ENSMUSG00000062647",
            "uniprot_ids": [
                "P12970"
            ]
        },
        "ENSMUSG00000026970": {
            "id": "ENSMUSG00000026970",
            "type": "Gene",
            "symbol": "Rbms1",
            "source": [
                "ENSMUSG00000057098",
                "ENSMUSG00000032035",
                "ENSMUSG00000038193",
                "ENSMUSG00000000247"
            ],
            "target": [],
            "expression_sum": {
                "class1": 18702.0,
                "class2": 5125.0
            },
            "ens_id": "ENSMUSG00000026970",
            "uniprot_ids": [
                "Q91W59",
                "E9PZ21"
            ]
        },
        "ENSMUSG00000059005": {
            "id": "ENSMUSG00000059005",
            "type": "Gene",
            "symbol": "Hnrnpa3",
            "source": [
                "ENSMUSG00000057098",
                "ENSMUSG00000000247"
            ],
            "target": [],
            "expression_sum": {
                "class1": 14355.0,
                "class2": 9765.0
            },
            "ens_id": "ENSMUSG00000059005",
            "uniprot_ids": [
                "Q8BG05",
                "A2AL12",
                "A2AL13"
            ]
        },
        "ENSMUSG00000014077": {
            "id": "ENSMUSG00000014077",
            "type": "Gene",
            "symbol": "Chp1",
            "source": [
                "ENSMUSG00000000247"
            ],
            "target": [],
            "expression_sum": {
                "class1": 6626.0,
                "class2": 2771.0
            },
            "ens_id": "ENSMUSG00000014077",
            "uniprot_ids": [
                "P61022",
                "B0R091",
                "B0R092"
            ]
        },
        "ENSMUSG00000027404": {
            "id": "ENSMUSG00000027404",
            "type": "Gene",
            "symbol": "Snrpb",
            "source": [
                "ENSMUSG00000000247"
            ],
            "target": [],
            "expression_sum": {
                "class1": 14832.0,
                "class2": 8727.0
            },
            "ens_id": "ENSMUSG00000027404",
            "uniprot_ids": [
                "P27048",
                "A0A0G2JGN4"
            ]
        },
        "ENSMUSG00000027593": {
            "id": "ENSMUSG00000027593",
            "type": "Gene",
            "symbol": "Raly",
            "source": [
                "ENSMUSG00000000247"
            ],
            "target": [],
            "expression_sum": {
                "class1": 11537.0,
                "class2": 4801.0
            },
            "ens_id": "ENSMUSG00000027593",
            "uniprot_ids": [
                "Q64012",
                "A2AU61",
                "A2AU60",
                "A2AU62"
            ]
        },
        "ENSMUSG00000005881": {
            "id": "ENSMUSG00000005881",
            "type": "Gene",
            "symbol": "Ergic3",
            "source": [
                "ENSMUSG00000057098",
                "ENSMUSG00000038193",
                "ENSMUSG00000005583",
                "ENSMUSG00000000247"
            ],
            "target": [],
            "expression_sum": {
                "class1": 17991.0,
                "class2": 6149.0
            },
            "ens_id": "ENSMUSG00000005881",
            "uniprot_ids": [
                "Q9CQE7",
                "F6UIS1",
                "F6RK81"
            ]
        },
        "ENSMUSG00000027620": {
            "id": "ENSMUSG00000027620",
            "type": "Gene",
            "symbol": "Rbm39",
            "source": [
                "ENSMUSG00000057098",
                "ENSMUSG00000000247"
            ],
            "target": [],
            "expression_sum": {
                "class1": 26232.0,
                "class2": 12490.0
            },
            "ens_id": "ENSMUSG00000027620",
            "uniprot_ids": [
                "Q8VH51",
                "D6RCZ8",
                "B7ZD63",
                "B7ZD64",
                "E9Q8F0",
                "E9Q999",
                "D6RJ85",
                "E9PZF9",
                "B7ZD61",
                "F7AA45"
            ]
        },
        "ENSMUSG00000027642": {
            "id": "ENSMUSG00000027642",
            "type": "Gene",
            "symbol": "Rpn2",
            "source": [
                "ENSMUSG00000032035",
                "ENSMUSG00000022528",
                "ENSMUSG00000000247",
                "ENSMUSG00000038193",
                "ENSMUSG00000005583"
            ],
            "target": [],
            "expression_sum": {
                "class1": 17341.0,
                "class2": 7428.0
            },
            "ens_id": "ENSMUSG00000027642",
            "uniprot_ids": [
                "Q9DBG6",
                "A2ACG7"
            ]
        },
        "ENSMUSG00000018326": {
            "id": "ENSMUSG00000018326",
            "type": "Gene",
            "symbol": "Ywhab",
            "source": [
                "ENSMUSG00000057098",
                "ENSMUSG00000000247"
            ],
            "target": [],
            "expression_sum": {
                "class1": 11705.0,
                "class2": 6040.0
            },
            "ens_id": "ENSMUSG00000018326",
            "uniprot_ids": [
                "Q9CQV8",
                "A2A5N1"
            ]
        },
        "ENSMUSG00000047907": {
            "id": "ENSMUSG00000047907",
            "type": "Gene",
            "symbol": "Tshz2",
            "source": [
                "ENSMUSG00000032035",
                "ENSMUSG00000000247",
                "ENSMUSG00000038193",
                "ENSMUSG00000003032",
                "ENSMUSG00000005583",
                "ENSMUSG00000030551"
            ],
            "target": [],
            "expression_sum": {
                "class1": 28389.0,
                "class2": 12971.0
            },
            "ens_id": "ENSMUSG00000047907",
            "uniprot_ids": [
                "Q68FE9",
                "Q8K350",
                "A0A0A0MQI3",
                "A0A087WS38"
            ]
        },
        "ENSMUSG00000027523": {
            "id": "ENSMUSG00000027523",
            "type": "Gene",
            "symbol": "Gnas",
            "source": [
                "ENSMUSG00000057098",
                "ENSMUSG00000032035",
                "ENSMUSG00000047638",
                "ENSMUSG00000000247"
            ],
            "target": [],
            "expression_sum": {
                "class1": 72560.0,
                "class2": 44043.0
            },
            "ens_id": "ENSMUSG00000027523",
            "uniprot_ids": [
                "Q6R0H6",
                "Q6R0H7",
                "P63094",
                "Q9Z0F1",
                "Z4YKV1",
                "A6PW71",
                "A6PW70",
                "A0A571BGH5",
                "A0A571BDL4",
                "A0A571BEM2",
                "A0A571BE25",
                "A0A571BE38",
                "A0A571BEG2",
                "A0A571BEI3",
                "A2A610"
            ]
        },
        "ENSMUSG00000039001": {
            "id": "ENSMUSG00000039001",
            "type": "Gene",
            "symbol": "Rps21",
            "source": [
                "ENSMUSG00000022528",
                "ENSMUSG00000000247",
                "ENSMUSG00000057098",
                "ENSMUSG00000038193",
                "ENSMUSG00000048387"
            ],
            "target": [],
            "expression_sum": {
                "class1": 73873.0,
                "class2": 42184.0
            },
            "ens_id": "ENSMUSG00000039001",
            "uniprot_ids": [
                "Q9CQR2"
            ]
        },
        "ENSMUSG00000079641": {
            "id": "ENSMUSG00000079641",
            "type": "Gene",
            "symbol": "Rpl39",
            "source": [
                "ENSMUSG00000057098",
                "ENSMUSG00000038193",
                "ENSMUSG00000005583",
                "ENSMUSG00000000247"
            ],
            "target": [],
            "expression_sum": {
                "class1": 143212.0,
                "class2": 71972.0
            },
            "ens_id": "ENSMUSG00000079641",
            "uniprot_ids": [
                "P62892"
            ]
        },
        "ENSMUSG00000008682": {
            "id": "ENSMUSG00000008682",
            "type": "Gene",
            "symbol": "Rpl10",
            "source": [
                "ENSMUSG00000023034",
                "ENSMUSG00000000247",
                "ENSMUSG00000038193",
                "ENSMUSG00000005583",
                "ENSMUSG00000048387"
            ],
            "target": [],
            "expression_sum": {
                "class1": 170301.0,
                "class2": 89736.0
            },
            "ens_id": "ENSMUSG00000008682",
            "uniprot_ids": [
                "Q6ZWV3",
                "I7HLV2",
                "A0A1B0GXC3"
            ]
        },
        "ENSMUSG00000031198": {
            "id": "ENSMUSG00000031198",
            "type": "Gene",
            "symbol": "Fundc2",
            "source": [
                "ENSMUSG00000057098",
                "ENSMUSG00000071076",
                "ENSMUSG00000029322",
                "ENSMUSG00000000247"
            ],
            "target": [],
            "expression_sum": {
                "class1": 13035.0,
                "class2": 4847.0
            },
            "ens_id": "ENSMUSG00000031198",
            "uniprot_ids": [
                "Q9D6K8"
            ]
        },
        "ENSMUSG00000079435": {
            "id": "ENSMUSG00000079435",
            "type": "Gene",
            "symbol": "Rpl36a",
            "source": [
                "ENSMUSG00000038193",
                "ENSMUSG00000023034",
                "ENSMUSG00000000247"
            ],
            "target": [],
            "expression_sum": {
                "class1": 72731.0,
                "class2": 39449.0
            },
            "ens_id": "ENSMUSG00000079435",
            "uniprot_ids": [
                "P83882",
                "E9Q131"
            ]
        },
        "ENSMUSG00000031353": {
            "id": "ENSMUSG00000031353",
            "type": "Gene",
            "symbol": "Rbbp7",
            "source": [
                "ENSMUSG00000000247"
            ],
            "target": [],
            "expression_sum": {
                "class1": 9398.0,
                "class2": 4601.0
            },
            "ens_id": "ENSMUSG00000031353",
            "uniprot_ids": [
                "Q60973",
                "A2AFJ1",
                "F6ZLC6",
                "A2AFI9",
                "F6U539"
            ]
        },
        "ENSMUSG00000003617": {
            "id": "ENSMUSG00000003617",
            "type": "Gene",
            "symbol": "Cp",
            "source": [
                "ENSMUSG00000005583",
                "ENSMUSG00000000247"
            ],
            "target": [],
            "expression_sum": {
                "class1": 15477.0,
                "class2": 4385.0
            },
            "ens_id": "ENSMUSG00000003617",
            "uniprot_ids": [
                "Q61147",
                "G3X8Q5",
                "E9PZD8",
                "G3UXD2",
                "G3UXG1",
                "G3UWP5",
                "G3X9T8",
                "G3UZ53"
            ]
        },
        "ENSMUSG00000027706": {
            "id": "ENSMUSG00000027706",
            "type": "Gene",
            "symbol": "Sec62",
            "source": [
                "ENSMUSG00000052684",
                "ENSMUSG00000000247",
                "ENSMUSG00000038418",
                "ENSMUSG00000003032",
                "ENSMUSG00000005583",
                "ENSMUSG00000071076"
            ],
            "target": [],
            "expression_sum": {
                "class1": 19914.0,
                "class2": 9499.0
            },
            "ens_id": "ENSMUSG00000027706",
            "uniprot_ids": [
                "Q8BU14"
            ]
        },
        "ENSMUSG00000027673": {
            "id": "ENSMUSG00000027673",
            "type": "Gene",
            "symbol": "Ndufb5",
            "source": [
                "ENSMUSG00000000247"
            ],
            "target": [],
            "expression_sum": {
                "class1": 10211.0,
                "class2": 5729.0
            },
            "ens_id": "ENSMUSG00000027673",
            "uniprot_ids": [
                "Q9CQH3",
                "D3Z568",
                "D3YX99",
                "F6Y6V5",
                "D3Z6W9",
                "F8WI84",
                "F6VXF9"
            ]
        },
        "ENSMUSG00000027750": {
            "id": "ENSMUSG00000027750",
            "type": "Gene",
            "symbol": "Postn",
            "source": [
                "ENSMUSG00000000247"
            ],
            "target": [],
            "expression_sum": {
                "class1": 20859.0,
                "class2": 2016.0
            },
            "ens_id": "ENSMUSG00000027750",
            "uniprot_ids": [
                "Q62009",
                "F7C9H0"
            ]
        },
        "ENSMUSG00000036381": {
            "id": "ENSMUSG00000036381",
            "type": "Gene",
            "symbol": "P2ry14",
            "source": [
                "ENSMUSG00000032035",
                "ENSMUSG00000000247"
            ],
            "target": [],
            "expression_sum": {
                "class1": 9888.0,
                "class2": 195.0
            },
            "ens_id": "ENSMUSG00000036381",
            "uniprot_ids": [
                "Q9ESG6",
                "A0A0G2JGB4",
                "A0A0G2JEW9",
                "A0A0G2JE58"
            ]
        },
        "ENSMUSG00000028081": {
            "id": "ENSMUSG00000028081",
            "type": "Gene",
            "symbol": "Rps3a1",
            "source": [
                "ENSMUSG00000031965",
                "ENSMUSG00000038193",
                "ENSMUSG00000057098",
                "ENSMUSG00000000247"
            ],
            "target": [],
            "expression_sum": {
                "class1": 130652.0,
                "class2": 62008.0
            },
            "ens_id": "ENSMUSG00000028081",
            "uniprot_ids": [
                "P97351"
            ]
        },
        "ENSMUSG00000090733": {
            "id": "ENSMUSG00000090733",
            "type": "Gene",
            "symbol": "Rps27",
            "source": [
                "ENSMUSG00000023034",
                "ENSMUSG00000000247",
                "ENSMUSG00000057098",
                "ENSMUSG00000038193",
                "ENSMUSG00000005583",
                "ENSMUSG00000048387"
            ],
            "target": [],
            "expression_sum": {
                "class1": 229761.0,
                "class2": 107039.0
            },
            "ens_id": "ENSMUSG00000090733",
            "uniprot_ids": [
                "Q6ZWU9",
                "A0A0G2JDW7",
                "A0A0G2JG29",
                "A0A0G2JEX7"
            ]
        },
        "ENSMUSG00000068874": {
            "id": "ENSMUSG00000068874",
            "type": "Gene",
            "symbol": "Selenbp1",
            "source": [
                "ENSMUSG00000000247"
            ],
            "target": [],
            "expression_sum": {
                "class1": 13950.0,
                "class2": 5345.0
            },
            "ens_id": "ENSMUSG00000068874",
            "uniprot_ids": [
                "P17563",
                "D6RHN2"
            ]
        },
        "ENSMUSG00000000563": {
            "id": "ENSMUSG00000000563",
            "type": "Gene",
            "symbol": "Atp5f1",
            "source": [
                "ENSMUSG00000057098",
                "ENSMUSG00000000247"
            ],
            "target": [],
            "expression_sum": {
                "class1": 10012.0,
                "class2": 8175.0
            },
            "ens_id": "ENSMUSG00000000563",
            "uniprot_ids": [
                "Q9CQQ7",
                "A0A0G2JGX3",
                "D6RGL7"
            ]
        },
        "ENSMUSG00000062006": {
            "id": "ENSMUSG00000062006",
            "type": "Gene",
            "symbol": "Rpl34",
            "source": [
                "ENSMUSG00000023034",
                "ENSMUSG00000000247",
                "ENSMUSG00000057098",
                "ENSMUSG00000038193",
                "ENSMUSG00000005583"
            ],
            "target": [],
            "expression_sum": {
                "class1": 110034.0,
                "class2": 50976.0
            },
            "ens_id": "ENSMUSG00000062006",
            "uniprot_ids": [
                "Q9D1R9",
                "A0A0G2JEY6",
                "A0A0G2JGY8"
            ]
        },
        "ENSMUSG00000028266": {
            "id": "ENSMUSG00000028266",
            "type": "Gene",
            "symbol": "Lmo4",
            "source": [
                "ENSMUSG00000023034",
                "ENSMUSG00000000247"
            ],
            "target": [],
            "expression_sum": {
                "class1": 9209.0,
                "class2": 2872.0
            },
            "ens_id": "ENSMUSG00000028266",
            "uniprot_ids": [
                "P61969",
                "A0A0G2JDX9",
                "A0A0G2JF61"
            ]
        },
        "ENSMUSG00000068523": {
            "id": "ENSMUSG00000068523",
            "type": "Gene",
            "symbol": "Gng5",
            "source": [
                "ENSMUSG00000057098",
                "ENSMUSG00000000247"
            ],
            "target": [],
            "expression_sum": {
                "class1": 30544.0,
                "class2": 14457.0
            },
            "ens_id": "ENSMUSG00000068523",
            "uniprot_ids": [
                "Q80SZ7",
                "D3Z7Q3"
            ]
        },
        "ENSMUSG00000028234": {
            "id": "ENSMUSG00000028234",
            "type": "Gene",
            "symbol": "Rps20",
            "source": [
                "ENSMUSG00000023034",
                "ENSMUSG00000022528",
                "ENSMUSG00000000247",
                "ENSMUSG00000057098",
                "ENSMUSG00000038193",
                "ENSMUSG00000005583",
                "ENSMUSG00000048387"
            ],
            "target": [],
            "expression_sum": {
                "class1": 82795.0,
                "class2": 44093.0
            },
            "ens_id": "ENSMUSG00000028234",
            "uniprot_ids": [
                "P60867"
            ]
        },
        "ENSMUSG00000040128": {
            "id": "ENSMUSG00000040128",
            "type": "Gene",
            "symbol": "Pnrc1",
            "source": [
                "ENSMUSG00000056501",
                "ENSMUSG00000037868",
                "ENSMUSG00000023034",
                "ENSMUSG00000052837",
                "ENSMUSG00000030103",
                "ENSMUSG00000000247",
                "ENSMUSG00000038418",
                "ENSMUSG00000003545",
                "ENSMUSG00000018899"
            ],
            "target": [],
            "expression_sum": {
                "class1": 15614.0,
                "class2": 8919.0
            },
            "ens_id": "ENSMUSG00000040128",
            "uniprot_ids": [
                "Q3TWH3"
            ]
        },
        "ENSMUSG00000028478": {
            "id": "ENSMUSG00000028478",
            "type": "Gene",
            "symbol": "Clta",
            "source": [
                "ENSMUSG00000057098",
                "ENSMUSG00000000247"
            ],
            "target": [],
            "expression_sum": {
                "class1": 19712.0,
                "class2": 8314.0
            },
            "ens_id": "ENSMUSG00000028478",
            "uniprot_ids": [
                "Q6PFA2",
                "B1AWE1",
                "B1AWD8",
                "B1AWD9",
                "B1AWE0"
            ]
        },
        "ENSMUSG00000039105": {
            "id": "ENSMUSG00000039105",
            "type": "Gene",
            "symbol": "Atp6v1g1",
            "source": [
                "ENSMUSG00000000247"
            ],
            "target": [],
            "expression_sum": {
                "class1": 11706.0,
                "class2": 7998.0
            },
            "ens_id": "ENSMUSG00000039105",
            "uniprot_ids": [
                "Q9CR51"
            ]
        },
        "ENSMUSG00000028495": {
            "id": "ENSMUSG00000028495",
            "type": "Gene",
            "symbol": "Rps6",
            "source": [
                "ENSMUSG00000022528",
                "ENSMUSG00000047638",
                "ENSMUSG00000000247",
                "ENSMUSG00000057098",
                "ENSMUSG00000038193",
                "ENSMUSG00000031965",
                "ENSMUSG00000048387"
            ],
            "target": [],
            "expression_sum": {
                "class1": 117047.0,
                "class2": 57082.0
            },
            "ens_id": "ENSMUSG00000028495",
            "uniprot_ids": [
                "P62754"
            ]
        },
        "ENSMUSG00000028618": {
            "id": "ENSMUSG00000028618",
            "type": "Gene",
            "symbol": "Tmem59",
            "source": [
                "ENSMUSG00000032035",
                "ENSMUSG00000052684",
                "ENSMUSG00000000247",
                "ENSMUSG00000057098",
                "ENSMUSG00000038193",
                "ENSMUSG00000003032",
                "ENSMUSG00000005583",
                "ENSMUSG00000071076"
            ],
            "target": [],
            "expression_sum": {
                "class1": 29466.0,
                "class2": 12479.0
            },
            "ens_id": "ENSMUSG00000028618",
            "uniprot_ids": [
                "Q9QY73",
                "D3Z0H8",
                "D3YVM2",
                "D6RI96"
            ]
        },
        "ENSMUSG00000028603": {
            "id": "ENSMUSG00000028603",
            "type": "Gene",
            "symbol": "Scp2",
            "source": [
                "ENSMUSG00000032035",
                "ENSMUSG00000000247"
            ],
            "target": [],
            "expression_sum": {
                "class1": 12508.0,
                "class2": 6926.0
            },
            "ens_id": "ENSMUSG00000028603",
            "uniprot_ids": [
                "P32020",
                "A2APS5",
                "A2APS4"
            ]
        },
        "ENSMUSG00000063882": {
            "id": "ENSMUSG00000063882",
            "type": "Gene",
            "symbol": "Uqcrh",
            "source": [
                "ENSMUSG00000025498",
                "ENSMUSG00000022708",
                "ENSMUSG00000000247",
                "ENSMUSG00000057098",
                "ENSMUSG00000053477"
            ],
            "target": [],
            "expression_sum": {
                "class1": 27135.0,
                "class2": 19213.0
            },
            "ens_id": "ENSMUSG00000063882",
            "uniprot_ids": [
                "P99028",
                "Q9D2Q7"
            ]
        },
        "ENSMUSG00000047675": {
            "id": "ENSMUSG00000047675",
            "type": "Gene",
            "symbol": "Rps8",
            "source": [
                "ENSMUSG00000057098",
                "ENSMUSG00000000247"
            ],
            "target": [],
            "expression_sum": {
                "class1": 147875.0,
                "class2": 98782.0
            },
            "ens_id": "ENSMUSG00000047675",
            "uniprot_ids": [
                "P62242"
            ]
        },
        "ENSMUSG00000028639": {
            "id": "ENSMUSG00000028639",
            "type": "TF",
            "symbol": "Ybx1",
            "source": [
                "ENSMUSG00000057098",
                "ENSMUSG00000000247"
            ],
            "target": [
                "ENSMUSG00000020486",
                "ENSMUSG00000023367",
                "ENSMUSG00000022032",
                "ENSMUSG00000041577",
                "ENSMUSG00000068522",
                "ENSMUSG00000018417"
            ],
            "expression_sum": {
                "class1": 29858.0,
                "class2": 27229.0
            },
            "ens_id": "ENSMUSG00000028639",
            "uniprot_ids": [
                "P62960",
                "A2BGG7"
            ]
        },
        "ENSMUSG00000028673": {
            "id": "ENSMUSG00000028673",
            "type": "Gene",
            "symbol": "Fuca1",
            "source": [
                "ENSMUSG00000000247",
                "ENSMUSG00000038193",
                "ENSMUSG00000005583",
                "ENSMUSG00000048387",
                "ENSMUSG00000071076"
            ],
            "target": [],
            "expression_sum": {
                "class1": 10874.0,
                "class2": 5021.0
            },
            "ens_id": "ENSMUSG00000028673",
            "uniprot_ids": [
                "Q99LJ1"
            ]
        },
        "ENSMUSG00000059291": {
            "id": "ENSMUSG00000059291",
            "type": "Gene",
            "symbol": "Rpl11",
            "source": [
                "ENSMUSG00000000247",
                "ENSMUSG00000057098",
                "ENSMUSG00000038193",
                "ENSMUSG00000003032",
                "ENSMUSG00000005583",
                "ENSMUSG00000071076"
            ],
            "target": [],
            "expression_sum": {
                "class1": 93116.0,
                "class2": 44256.0
            },
            "ens_id": "ENSMUSG00000059291",
            "uniprot_ids": [
                "Q9CXW4",
                "A2BH06"
            ]
        },
        "ENSMUSG00000007872": {
            "id": "ENSMUSG00000007872",
            "type": "Gene",
            "symbol": "Id3",
            "source": [
                "ENSMUSG00000041515",
                "ENSMUSG00000056501",
                "ENSMUSG00000052684",
                "ENSMUSG00000037868",
                "ENSMUSG00000052837",
                "ENSMUSG00000047638",
                "ENSMUSG00000030103",
                "ENSMUSG00000000247",
                "ENSMUSG00000038418",
                "ENSMUSG00000053477",
                "ENSMUSG00000021250",
                "ENSMUSG00000029322",
                "ENSMUSG00000003032",
                "ENSMUSG00000005583",
                "ENSMUSG00000018899",
                "ENSMUSG00000071076"
            ],
            "target": [],
            "expression_sum": {
                "class1": 106976.0,
                "class2": 38464.0
            },
            "ens_id": "ENSMUSG00000007872",
            "uniprot_ids": [
                "P41133"
            ]
        },
        "ENSMUSG00000028757": {
            "id": "ENSMUSG00000028757",
            "type": "Gene",
            "symbol": "Ddost",
            "source": [
                "ENSMUSG00000071076",
                "ENSMUSG00000005583",
                "ENSMUSG00000008575",
                "ENSMUSG00000000247"
            ],
            "target": [],
            "expression_sum": {
                "class1": 13007.0,
                "class2": 6960.0
            },
            "ens_id": "ENSMUSG00000028757",
            "uniprot_ids": [
                "O54734"
            ]
        },
        "ENSMUSG00000028936": {
            "id": "ENSMUSG00000028936",
            "type": "Gene",
            "symbol": "Rpl22",
            "source": [
                "ENSMUSG00000023034",
                "ENSMUSG00000022528",
                "ENSMUSG00000000247",
                "ENSMUSG00000057098",
                "ENSMUSG00000038193",
                "ENSMUSG00000005583"
            ],
            "target": [],
            "expression_sum": {
                "class1": 49974.0,
                "class2": 21647.0
            },
            "ens_id": "ENSMUSG00000028936",
            "uniprot_ids": [
                "P67984"
            ]
        },
        "ENSMUSG00000029076": {
            "id": "ENSMUSG00000029076",
            "type": "Gene",
            "symbol": "Sdf4",
            "source": [
                "ENSMUSG00000032035",
                "ENSMUSG00000022528",
                "ENSMUSG00000000247",
                "ENSMUSG00000038193",
                "ENSMUSG00000005583",
                "ENSMUSG00000048387",
                "ENSMUSG00000071076"
            ],
            "target": [],
            "expression_sum": {
                "class1": 20455.0,
                "class2": 6489.0
            },
            "ens_id": "ENSMUSG00000029076",
            "uniprot_ids": [
                "Q61112"
            ]
        },
        "ENSMUSG00000038803": {
            "id": "ENSMUSG00000038803",
            "type": "Gene",
            "symbol": "Ost4",
            "source": [
                "ENSMUSG00000057098",
                "ENSMUSG00000071076",
                "ENSMUSG00000032035",
                "ENSMUSG00000000247"
            ],
            "target": [],
            "expression_sum": {
                "class1": 25364.0,
                "class2": 9331.0
            },
            "ens_id": "ENSMUSG00000038803",
            "uniprot_ids": [
                "Q99LX8"
            ]
        },
        "ENSMUSG00000055302": {
            "id": "ENSMUSG00000055302",
            "type": "Gene",
            "symbol": "Mrfap1",
            "source": [
                "ENSMUSG00000057098",
                "ENSMUSG00000029322",
                "ENSMUSG00000000247"
            ],
            "target": [],
            "expression_sum": {
                "class1": 24289.0,
                "class2": 13316.0
            },
            "ens_id": "ENSMUSG00000055302",
            "uniprot_ids": [
                "Q9CQL7"
            ]
        },
        "ENSMUSG00000037999": {
            "id": "ENSMUSG00000037999",
            "type": "Gene",
            "symbol": "Arap2",
            "source": [
                "ENSMUSG00000056501",
                "ENSMUSG00000052684",
                "ENSMUSG00000052837",
                "ENSMUSG00000000247",
                "ENSMUSG00000021250",
                "ENSMUSG00000029322",
                "ENSMUSG00000008575",
                "ENSMUSG00000071076"
            ],
            "target": [],
            "expression_sum": {
                "class1": 10851.0,
                "class2": 599.0
            },
            "ens_id": "ENSMUSG00000037999",
            "uniprot_ids": [
                "E9QP44",
                "E0CXD0"
            ]
        },
        "ENSMUSG00000047215": {
            "id": "ENSMUSG00000047215",
            "type": "Gene",
            "symbol": "Rpl9",
            "source": [
                "ENSMUSG00000057098",
                "ENSMUSG00000038193",
                "ENSMUSG00000000247"
            ],
            "target": [],
            "expression_sum": {
                "class1": 127172.0,
                "class2": 60716.0
            },
            "ens_id": "ENSMUSG00000047215",
            "uniprot_ids": [
                "P51410",
                "D3Z629",
                "A0A0G2JFQ3",
                "A0A0G2JES3",
                "D3YZT0"
            ]
        },
        "ENSMUSG00000058558": {
            "id": "ENSMUSG00000058558",
            "type": "Gene",
            "symbol": "Rpl5",
            "source": [
                "ENSMUSG00000057098",
                "ENSMUSG00000048387",
                "ENSMUSG00000038193",
                "ENSMUSG00000000247"
            ],
            "target": [],
            "expression_sum": {
                "class1": 50219.0,
                "class2": 21787.0
            },
            "ens_id": "ENSMUSG00000058558",
            "uniprot_ids": [
                "P47962",
                "D3YYV8"
            ]
        },
        "ENSMUSG00000067274": {
            "id": "ENSMUSG00000067274",
            "type": "Gene",
            "symbol": "Rplp0",
            "source": [
                "ENSMUSG00000023034",
                "ENSMUSG00000000247",
                "ENSMUSG00000057098",
                "ENSMUSG00000038193",
                "ENSMUSG00000005583",
                "ENSMUSG00000048387"
            ],
            "target": [],
            "expression_sum": {
                "class1": 123999.0,
                "class2": 66138.0
            },
            "ens_id": "ENSMUSG00000067274",
            "uniprot_ids": [
                "P14869",
                "S4R1N1",
                "D3YVM5"
            ]
        },
        "ENSMUSG00000029614": {
            "id": "ENSMUSG00000029614",
            "type": "Gene",
            "symbol": "Rpl6",
            "source": [
                "ENSMUSG00000048387",
                "ENSMUSG00000038193",
                "ENSMUSG00000023034",
                "ENSMUSG00000000247"
            ],
            "target": [],
            "expression_sum": {
                "class1": 114722.0,
                "class2": 53657.0
            },
            "ens_id": "ENSMUSG00000029614",
            "uniprot_ids": [
                "P47911",
                "A0A0J9YU32"
            ]
        },
        "ENSMUSG00000029465": {
            "id": "ENSMUSG00000029465",
            "type": "Gene",
            "symbol": "Arpc3",
            "source": [
                "ENSMUSG00000000247",
                "ENSMUSG00000057098",
                "ENSMUSG00000038418",
                "ENSMUSG00000048387",
                "ENSMUSG00000071076"
            ],
            "target": [],
            "expression_sum": {
                "class1": 16637.0,
                "class2": 6564.0
            },
            "ens_id": "ENSMUSG00000029465",
            "uniprot_ids": [
                "Q9JM76",
                "D3Z2F7",
                "H7BWZ3",
                "A0A0G2JFK7",
                "D3Z2F8"
            ]
        },
        "ENSMUSG00000070493": {
            "id": "ENSMUSG00000070493",
            "type": "Gene",
            "symbol": "Chchd2",
            "source": [
                "ENSMUSG00000025498",
                "ENSMUSG00000000247"
            ],
            "target": [],
            "expression_sum": {
                "class1": 37033.0,
                "class2": 25804.0
            },
            "ens_id": "ENSMUSG00000070493",
            "uniprot_ids": [
                "Q9D1L0"
            ]
        },
        "ENSMUSG00000029713": {
            "id": "ENSMUSG00000029713",
            "type": "Gene",
            "symbol": "Gnb2",
            "source": [
                "ENSMUSG00000057098",
                "ENSMUSG00000048387",
                "ENSMUSG00000000247"
            ],
            "target": [],
            "expression_sum": {
                "class1": 20371.0,
                "class2": 12083.0
            },
            "ens_id": "ENSMUSG00000029713",
            "uniprot_ids": [
                "P62880",
                "D3YZX3",
                "E9PWM7",
                "D3Z1M1",
                "D3Z1T4",
                "E9QKR0"
            ]
        },
        "ENSMUSG00000041453": {
            "id": "ENSMUSG00000041453",
            "type": "Gene",
            "symbol": "Rpl21",
            "source": [
                "ENSMUSG00000057098",
                "ENSMUSG00000048387",
                "ENSMUSG00000047638",
                "ENSMUSG00000000247"
            ],
            "target": [],
            "expression_sum": {
                "class1": 140870.0,
                "class2": 81337.0
            },
            "ens_id": "ENSMUSG00000041453",
            "uniprot_ids": [
                "Q9CQM8"
            ]
        },
        "ENSMUSG00000001687": {
            "id": "ENSMUSG00000001687",
            "type": "Gene",
            "symbol": "Ubl3",
            "source": [
                "ENSMUSG00000038193",
                "ENSMUSG00000000247"
            ],
            "target": [],
            "expression_sum": {
                "class1": 9218.0,
                "class2": 2452.0
            },
            "ens_id": "ENSMUSG00000001687",
            "uniprot_ids": [
                "Q9Z2M6"
            ]
        },
        "ENSMUSG00000029632": {
            "id": "ENSMUSG00000029632",
            "type": "Gene",
            "symbol": "Ndufa4",
            "source": [
                "ENSMUSG00000047638",
                "ENSMUSG00000021250",
                "ENSMUSG00000000247"
            ],
            "target": [],
            "expression_sum": {
                "class1": 20402.0,
                "class2": 14036.0
            },
            "ens_id": "ENSMUSG00000029632",
            "uniprot_ids": [
                "Q62425",
                "A0A0N4SVQ1"
            ]
        },
        "ENSMUSG00000029810": {
            "id": "ENSMUSG00000029810",
            "type": "Gene",
            "symbol": "Tmem176b",
            "source": [
                "ENSMUSG00000038193",
                "ENSMUSG00000005583",
                "ENSMUSG00000000247"
            ],
            "target": [],
            "expression_sum": {
                "class1": 87719.0,
                "class2": 14659.0
            },
            "ens_id": "ENSMUSG00000029810",
            "uniprot_ids": [
                "Q9R1Q6",
                "A0A0N4SVT4",
                "A0A0N4SUS9",
                "A0A0N4SV46",
                "A0A0N4SUY1",
                "A0A0N4SUM7"
            ]
        },
        "ENSMUSG00000023367": {
            "id": "ENSMUSG00000023367",
            "type": "Gene",
            "symbol": "Tmem176a",
            "source": [
                "ENSMUSG00000028639",
                "ENSMUSG00000038193",
                "ENSMUSG00000005583",
                "ENSMUSG00000000247"
            ],
            "target": [],
            "expression_sum": {
                "class1": 90134.0,
                "class2": 10306.0
            },
            "ens_id": "ENSMUSG00000023367",
            "uniprot_ids": [
                "Q9DCS1",
                "A0A0N4SUL5",
                "A0A0N4SVW3",
                "A0A0N4SVU8"
            ]
        },
        "ENSMUSG00000004980": {
            "id": "ENSMUSG00000004980",
            "type": "Gene",
            "symbol": "Hnrnpa2b1",
            "source": [
                "ENSMUSG00000057098",
                "ENSMUSG00000000247"
            ],
            "target": [],
            "expression_sum": {
                "class1": 16714.0,
                "class2": 11709.0
            },
            "ens_id": "ENSMUSG00000004980",
            "uniprot_ids": [
                "O88569",
                "A0A0N4SUM2"
            ]
        },
        "ENSMUSG00000029781": {
            "id": "ENSMUSG00000029781",
            "type": "Gene",
            "symbol": "Fkbp9",
            "source": [
                "ENSMUSG00000022528",
                "ENSMUSG00000000247",
                "ENSMUSG00000038193",
                "ENSMUSG00000008575",
                "ENSMUSG00000005583",
                "ENSMUSG00000071076"
            ],
            "target": [],
            "expression_sum": {
                "class1": 13346.0,
                "class2": 5792.0
            },
            "ens_id": "ENSMUSG00000029781",
            "uniprot_ids": [
                "Q9Z247"
            ]
        },
        "ENSMUSG00000036371": {
            "id": "ENSMUSG00000036371",
            "type": "Gene",
            "symbol": "Serbp1",
            "source": [
                "ENSMUSG00000057098",
                "ENSMUSG00000000247"
            ],
            "target": [],
            "expression_sum": {
                "class1": 14959.0,
                "class2": 9422.0
            },
            "ens_id": "ENSMUSG00000036371",
            "uniprot_ids": [
                "Q9CY58",
                "Q3UMP4",
                "A0A0N4SV32",
                "A0A0N4SVK5",
                "A0A0N4SUQ1",
                "A0A0N4SV40",
                "A0A0N4SUN8",
                "A0A0N4SWH2"
            ]
        },
        "ENSMUSG00000030057": {
            "id": "ENSMUSG00000030057",
            "type": "Gene",
            "symbol": "Cnbp",
            "source": [
                "ENSMUSG00000057098",
                "ENSMUSG00000000247"
            ],
            "target": [],
            "expression_sum": {
                "class1": 19531.0,
                "class2": 12390.0
            },
            "ens_id": "ENSMUSG00000030057",
            "uniprot_ids": [
                "P53996",
                "A0A0N4SVS6"
            ]
        },
        "ENSMUSG00000057841": {
            "id": "ENSMUSG00000057841",
            "type": "Gene",
            "symbol": "Rpl32",
            "source": [
                "ENSMUSG00000023034",
                "ENSMUSG00000000247",
                "ENSMUSG00000057098",
                "ENSMUSG00000038193",
                "ENSMUSG00000005583"
            ],
            "target": [],
            "expression_sum": {
                "class1": 134863.0,
                "class2": 63059.0
            },
            "ens_id": "ENSMUSG00000057841",
            "uniprot_ids": [
                "P62911"
            ]
        },
        "ENSMUSG00000006333": {
            "id": "ENSMUSG00000006333",
            "type": "Gene",
            "symbol": "Rps9",
            "source": [
                "ENSMUSG00000000247",
                "ENSMUSG00000057098",
                "ENSMUSG00000038193",
                "ENSMUSG00000005583",
                "ENSMUSG00000048387"
            ],
            "target": [],
            "expression_sum": {
                "class1": 96715.0,
                "class2": 49515.0
            },
            "ens_id": "ENSMUSG00000006333",
            "uniprot_ids": [
                "Q6ZWN5",
                "Q9CXW7",
                "D3YWH9",
                "D3YUV6",
                "F7CJS8",
                "D3Z673"
            ]
        },
        "ENSMUSG00000030432": {
            "id": "ENSMUSG00000030432",
            "type": "Gene",
            "symbol": "Rpl28",
            "source": [
                "ENSMUSG00000057098",
                "ENSMUSG00000023034",
                "ENSMUSG00000000247"
            ],
            "target": [],
            "expression_sum": {
                "class1": 82928.0,
                "class2": 42524.0
            },
            "ens_id": "ENSMUSG00000030432",
            "uniprot_ids": [
                "P41105",
                "F6Z0X0"
            ]
        },
        "ENSMUSG00000012848": {
            "id": "ENSMUSG00000012848",
            "type": "Gene",
            "symbol": "Rps5",
            "source": [
                "ENSMUSG00000023034",
                "ENSMUSG00000000247",
                "ENSMUSG00000057098",
                "ENSMUSG00000038193",
                "ENSMUSG00000005583",
                "ENSMUSG00000048387"
            ],
            "target": [],
            "expression_sum": {
                "class1": 125292.0,
                "class2": 72878.0
            },
            "ens_id": "ENSMUSG00000012848",
            "uniprot_ids": [
                "Q91V55",
                "D3Z1S8",
                "D3YYM6"
            ]
        },
        "ENSMUSG00000041571": {
            "id": "ENSMUSG00000041571",
            "type": "Gene",
            "symbol": "Sepw1",
            "source": [
                "ENSMUSG00000071076",
                "ENSMUSG00000029322",
                "ENSMUSG00000000247"
            ],
            "target": [],
            "expression_sum": {
                "class1": 39230.0,
                "class2": 32611.0
            },
            "ens_id": "ENSMUSG00000041571",
            "uniprot_ids": [
                "P63300"
            ]
        },
        "ENSMUSG00000041560": {
            "id": "ENSMUSG00000041560",
            "type": "Gene",
            "symbol": "Gltscr2",
            "source": [
                "ENSMUSG00000000247"
            ],
            "target": [],
            "expression_sum": {
                "class1": 19708.0,
                "class2": 8229.0
            },
            "ens_id": "ENSMUSG00000041560",
            "uniprot_ids": [
                "Q8BK35"
            ]
        },
        "ENSMUSG00000040952": {
            "id": "ENSMUSG00000040952",
            "type": "Gene",
            "symbol": "Rps19",
            "source": [
                "ENSMUSG00000023034",
                "ENSMUSG00000022528",
                "ENSMUSG00000000247",
                "ENSMUSG00000057098",
                "ENSMUSG00000038193",
                "ENSMUSG00000005583",
                "ENSMUSG00000031965",
                "ENSMUSG00000048387"
            ],
            "target": [],
            "expression_sum": {
                "class1": 156248.0,
                "class2": 83886.0
            },
            "ens_id": "ENSMUSG00000040952",
            "uniprot_ids": [
                "Q9CZX8",
                "D3YUG3",
                "D3YUT3",
                "S4R223",
                "D3Z722",
                "D3Z5R8"
            ]
        },
        "ENSMUSG00000037563": {
            "id": "ENSMUSG00000037563",
            "type": "Gene",
            "symbol": "Rps16",
            "source": [
                "ENSMUSG00000057098",
                "ENSMUSG00000038193",
                "ENSMUSG00000023034",
                "ENSMUSG00000000247"
            ],
            "target": [],
            "expression_sum": {
                "class1": 180448.0,
                "class2": 70623.0
            },
            "ens_id": "ENSMUSG00000037563",
            "uniprot_ids": [
                "P14131"
            ]
        },
        "ENSMUSG00000054808": {
            "id": "ENSMUSG00000054808",
            "type": "Gene",
            "symbol": "Actn4",
            "source": [
                "ENSMUSG00000032035",
                "ENSMUSG00000052684",
                "ENSMUSG00000022346",
                "ENSMUSG00000047638",
                "ENSMUSG00000000247",
                "ENSMUSG00000053477",
                "ENSMUSG00000029322",
                "ENSMUSG00000008575",
                "ENSMUSG00000020484",
                "ENSMUSG00000071076"
            ],
            "target": [],
            "expression_sum": {
                "class1": 29837.0,
                "class2": 6022.0
            },
            "ens_id": "ENSMUSG00000054808",
            "uniprot_ids": [
                "P57780",
                "E9Q2W9",
                "D3Z761",
                "D3Z0L8",
                "A0A1L1SVJ6",
                "A0A1L1SV25"
            ]
        },
        "ENSMUSG00000053565": {
            "id": "ENSMUSG00000053565",
            "type": "Gene",
            "symbol": "Eif3k",
            "source": [
                "ENSMUSG00000057098",
                "ENSMUSG00000000247"
            ],
            "target": [],
            "expression_sum": {
                "class1": 17038.0,
                "class2": 8827.0
            },
            "ens_id": "ENSMUSG00000053565",
            "uniprot_ids": [
                "Q9DBZ5",
                "Q3TY56",
                "A0A140LJ59"
            ]
        },
        "ENSMUSG00000036751": {
            "id": "ENSMUSG00000036751",
            "type": "Gene",
            "symbol": "Cox6b1",
            "source": [
                "ENSMUSG00000021250",
                "ENSMUSG00000000247"
            ],
            "target": [],
            "expression_sum": {
                "class1": 20299.0,
                "class2": 18243.0
            },
            "ens_id": "ENSMUSG00000036751",
            "uniprot_ids": [
                "P56391",
                "A0A140LIU3"
            ]
        },
        "ENSMUSG00000019539": {
            "id": "ENSMUSG00000019539",
            "type": "Gene",
            "symbol": "Rcn3",
            "source": [
                "ENSMUSG00000038193",
                "ENSMUSG00000005583",
                "ENSMUSG00000000247"
            ],
            "target": [],
            "expression_sum": {
                "class1": 28157.0,
                "class2": 14971.0
            },
            "ens_id": "ENSMUSG00000019539",
            "uniprot_ids": [
                "Q8BH97",
                "A0A1B0GS22",
                "A0A1B0GRI0",
                "A0A1B0GSK5",
                "A0A1B0GR19",
                "A0A1B0GR86"
            ]
        },
        "ENSMUSG00000003429": {
            "id": "ENSMUSG00000003429",
            "type": "Gene",
            "symbol": "Rps11",
            "source": [
                "ENSMUSG00000057098",
                "ENSMUSG00000038193",
                "ENSMUSG00000000247"
            ],
            "target": [],
            "expression_sum": {
                "class1": 98135.0,
                "class2": 48915.0
            },
            "ens_id": "ENSMUSG00000003429",
            "uniprot_ids": [
                "P62281",
                "A0A1B0GSE8",
                "A0A1B0GRR3"
            ]
        },
        "ENSMUSG00000074129": {
            "id": "ENSMUSG00000074129",
            "type": "Gene",
            "symbol": "Rpl13a",
            "source": [
                "ENSMUSG00000023034",
                "ENSMUSG00000047638",
                "ENSMUSG00000000247",
                "ENSMUSG00000057098",
                "ENSMUSG00000038193",
                "ENSMUSG00000048387"
            ],
            "target": [],
            "expression_sum": {
                "class1": 181504.0,
                "class2": 83293.0
            },
            "ens_id": "ENSMUSG00000074129",
            "uniprot_ids": [
                "P19253",
                "A0A1B0GSQ6",
                "A0A1B0GTA1",
                "A0A1B0GSC2",
                "A0A1B0GRH1",
                "A0A1B0GT00",
                "A0A1B0GSL5",
                "A0A1B0GQW6",
                "A0A1B0GSF0",
                "A0A1B0GSJ2",
                "A0A1B0GSB2"
            ]
        },
        "ENSMUSG00000050708": {
            "id": "ENSMUSG00000050708",
            "type": "Gene",
            "symbol": "Ftl1",
            "source": [
                "ENSMUSG00000023034",
                "ENSMUSG00000005583",
                "ENSMUSG00000000247",
                "ENSMUSG00000057098",
                "ENSMUSG00000038193",
                "ENSMUSG00000008575",
                "ENSMUSG00000071076"
            ],
            "target": [],
            "expression_sum": {
                "class1": 189303.0,
                "class2": 94835.0
            },
            "ens_id": "ENSMUSG00000050708",
            "uniprot_ids": [
                "Q9CPX4",
                "A0A1B0GRH4",
                "A0A1B0GR60"
            ]
        },
        "ENSMUSG00000030824": {
            "id": "ENSMUSG00000030824",
            "type": "Gene",
            "symbol": "Nucb1",
            "source": [
                "ENSMUSG00000032035",
                "ENSMUSG00000000247",
                "ENSMUSG00000057098",
                "ENSMUSG00000038193",
                "ENSMUSG00000005583",
                "ENSMUSG00000048387",
                "ENSMUSG00000071076"
            ],
            "target": [],
            "expression_sum": {
                "class1": 16302.0,
                "class2": 4504.0
            },
            "ens_id": "ENSMUSG00000030824",
            "uniprot_ids": [
                "Q02819",
                "A0A1B0GR92",
                "A0A1B0GR41",
                "A0A1C7CYU3",
                "A0A1B0GT83",
                "A0A1B0GS16"
            ]
        },
        "ENSMUSG00000059070": {
            "id": "ENSMUSG00000059070",
            "type": "Gene",
            "symbol": "Rpl18",
            "source": [
                "ENSMUSG00000057098",
                "ENSMUSG00000048387",
                "ENSMUSG00000000247"
            ],
            "target": [],
            "expression_sum": {
                "class1": 68248.0,
                "class2": 37919.0
            },
            "ens_id": "ENSMUSG00000059070",
            "uniprot_ids": [
                "P35980",
                "A0A1B0GSF7",
                "A0A1B0GSS8",
                "A0A1B0GS28",
                "A0A1B0GSA8",
                "A0A1B0GRZ3",
                "A0A1B0GQU8"
            ]
        },
        "ENSMUSG00000002778": {
            "id": "ENSMUSG00000002778",
            "type": "Gene",
            "symbol": "Kdelr1",
            "source": [
                "ENSMUSG00000000247",
                "ENSMUSG00000057098",
                "ENSMUSG00000038193",
                "ENSMUSG00000029322",
                "ENSMUSG00000048387"
            ],
            "target": [],
            "expression_sum": {
                "class1": 14034.0,
                "class2": 5711.0
            },
            "ens_id": "ENSMUSG00000002778",
            "uniprot_ids": [
                "Q99JH8",
                "A0A1B0GT97",
                "A0A1B0GQW0"
            ]
        },
        "ENSMUSG00000061787": {
            "id": "ENSMUSG00000061787",
            "type": "Gene",
            "symbol": "Rps17",
            "source": [
                "ENSMUSG00000057098",
                "ENSMUSG00000000247"
            ],
            "target": [],
            "expression_sum": {
                "class1": 90787.0,
                "class2": 46553.0
            },
            "ens_id": "ENSMUSG00000061787",
            "uniprot_ids": [
                "P63276"
            ]
        },
        "ENSMUSG00000030744": {
            "id": "ENSMUSG00000030744",
            "type": "Gene",
            "symbol": "Rps3",
            "source": [
                "ENSMUSG00000057098",
                "ENSMUSG00000038193",
                "ENSMUSG00000000247"
            ],
            "target": [],
            "expression_sum": {
                "class1": 114154.0,
                "class2": 53681.0
            },
            "ens_id": "ENSMUSG00000030744",
            "uniprot_ids": [
                "P62908",
                "D3YV43",
                "A0A140LI77"
            ]
        },
        "ENSMUSG00000031029": {
            "id": "ENSMUSG00000031029",
            "type": "Gene",
            "symbol": "Eif3f",
            "source": [
                "ENSMUSG00000057098",
                "ENSMUSG00000038193",
                "ENSMUSG00000023034",
                "ENSMUSG00000000247"
            ],
            "target": [],
            "expression_sum": {
                "class1": 36747.0,
                "class2": 13031.0
            },
            "ens_id": "ENSMUSG00000031029",
            "uniprot_ids": [
                "Q9DCH4"
            ]
        },
        "ENSMUSG00000046364": {
            "id": "ENSMUSG00000046364",
            "type": "Gene",
            "symbol": "Rpl27a",
            "source": [
                "ENSMUSG00000031965",
                "ENSMUSG00000057098",
                "ENSMUSG00000000247"
            ],
            "target": [],
            "expression_sum": {
                "class1": 88935.0,
                "class2": 55155.0
            },
            "ens_id": "ENSMUSG00000046364",
            "uniprot_ids": [
                "P14115",
                "G3UZV5"
            ]
        },
        "ENSMUSG00000090862": {
            "id": "ENSMUSG00000090862",
            "type": "Gene",
            "symbol": "Rps13",
            "source": [
                "ENSMUSG00000022528",
                "ENSMUSG00000000247",
                "ENSMUSG00000057098",
                "ENSMUSG00000005583",
                "ENSMUSG00000048387"
            ],
            "target": [],
            "expression_sum": {
                "class1": 84526.0,
                "class2": 40637.0
            },
            "ens_id": "ENSMUSG00000090862",
            "uniprot_ids": [
                "P62301",
                "Q921R2",
                "A0A0U1RQ71"
            ]
        },
        "ENSMUSG00000008683": {
            "id": "ENSMUSG00000008683",
            "type": "Gene",
            "symbol": "Rps15a",
            "source": [
                "ENSMUSG00000031965",
                "ENSMUSG00000038193",
                "ENSMUSG00000023034",
                "ENSMUSG00000000247"
            ],
            "target": [],
            "expression_sum": {
                "class1": 127990.0,
                "class2": 57587.0
            },
            "ens_id": "ENSMUSG00000008683",
            "uniprot_ids": [
                "P62245",
                "D3Z712",
                "F8WJ41",
                "D3YVB4"
            ]
        },
        "ENSMUSG00000030695": {
            "id": "ENSMUSG00000030695",
            "type": "Gene",
            "symbol": "Aldoa",
            "source": [
                "ENSMUSG00000057098",
                "ENSMUSG00000000247"
            ],
            "target": [],
            "expression_sum": {
                "class1": 30651.0,
                "class2": 24631.0
            },
            "ens_id": "ENSMUSG00000030695",
            "uniprot_ids": [
                "P05064",
                "A6ZI44",
                "D3YWI1",
                "A0A0U1RPT5",
                "D3Z510",
                "A0A0U1RPN8",
                "D3YV98"
            ]
        },
        "ENSMUSG00000030849": {
            "id": "ENSMUSG00000030849",
            "type": "Gene",
            "symbol": "Fgfr2",
            "source": [
                "ENSMUSG00000032035",
                "ENSMUSG00000056501",
                "ENSMUSG00000052684",
                "ENSMUSG00000052837",
                "ENSMUSG00000047638",
                "ENSMUSG00000000247",
                "ENSMUSG00000042812",
                "ENSMUSG00000053477",
                "ENSMUSG00000038193",
                "ENSMUSG00000021250",
                "ENSMUSG00000076431",
                "ENSMUSG00000008575",
                "ENSMUSG00000018899",
                "ENSMUSG00000021127",
                "ENSMUSG00000071076"
            ],
            "target": [],
            "expression_sum": {
                "class1": 21406.0,
                "class2": 159.0
            },
            "ens_id": "ENSMUSG00000030849",
            "uniprot_ids": [
                "E9Q7T0",
                "F2Z480",
                "E9Q700",
                "F7BDU9",
                "E9Q7E6",
                "E9Q7E8",
                "E9Q5C3",
                "E9Q5C2",
                "E9QK53",
                "E9PXV8",
                "E9PX60",
                "D6RJK5",
                "E9PX88",
                "D3Z5M8",
                "E9PX67",
                "E9PX90",
                "E9Q7C7",
                "E9PVU6",
                "E9Q708"
            ]
        },
        "ENSMUSG00000060591": {
            "id": "ENSMUSG00000060591",
            "type": "Gene",
            "symbol": "Ifitm2",
            "source": [
                "ENSMUSG00000032035",
                "ENSMUSG00000052684",
                "ENSMUSG00000000247",
                "ENSMUSG00000057098",
                "ENSMUSG00000038193",
                "ENSMUSG00000003032",
                "ENSMUSG00000005583",
                "ENSMUSG00000071076"
            ],
            "target": [],
            "expression_sum": {
                "class1": 107145.0,
                "class2": 33004.0
            },
            "ens_id": "ENSMUSG00000060591",
            "uniprot_ids": [
                "Q99J93"
            ]
        },
        "ENSMUSG00000025508": {
            "id": "ENSMUSG00000025508",
            "type": "Gene",
            "symbol": "Rplp2",
            "source": [
                "ENSMUSG00000022528",
                "ENSMUSG00000000247",
                "ENSMUSG00000057098",
                "ENSMUSG00000038193",
                "ENSMUSG00000048387"
            ],
            "target": [],
            "expression_sum": {
                "class1": 104423.0,
                "class2": 55541.0
            },
            "ens_id": "ENSMUSG00000025508",
            "uniprot_ids": [
                "P99027"
            ]
        },
        "ENSMUSG00000025511": {
            "id": "ENSMUSG00000025511",
            "type": "Gene",
            "symbol": "Tspan4",
            "source": [
                "ENSMUSG00000029322",
                "ENSMUSG00000000247"
            ],
            "target": [],
            "expression_sum": {
                "class1": 22837.0,
                "class2": 3474.0
            },
            "ens_id": "ENSMUSG00000025511",
            "uniprot_ids": [
                "Q9DCK3",
                "D3YXS4",
                "A0A668KLD1",
                "D3Z0T3",
                "D3Z0B7"
            ]
        },
        "ENSMUSG00000061983": {
            "id": "ENSMUSG00000061983",
            "type": "Gene",
            "symbol": "Rps12",
            "source": [
                "ENSMUSG00000056501",
                "ENSMUSG00000023034",
                "ENSMUSG00000022528",
                "ENSMUSG00000000247",
                "ENSMUSG00000057098",
                "ENSMUSG00000038193",
                "ENSMUSG00000005583",
                "ENSMUSG00000048387"
            ],
            "target": [],
            "expression_sum": {
                "class1": 91446.0,
                "class2": 42911.0
            },
            "ens_id": "ENSMUSG00000061983",
            "uniprot_ids": [
                "Q6ZWZ6",
                "A0A1W2P810",
                "F7AEH4",
                "A0A1W2P7A1"
            ]
        },
        "ENSMUSG00000004207": {
            "id": "ENSMUSG00000004207",
            "type": "Gene",
            "symbol": "Psap",
            "source": [
                "ENSMUSG00000032035",
                "ENSMUSG00000000247",
                "ENSMUSG00000057098",
                "ENSMUSG00000038193",
                "ENSMUSG00000005583",
                "ENSMUSG00000028565",
                "ENSMUSG00000071076"
            ],
            "target": [],
            "expression_sum": {
                "class1": 45580.0,
                "class2": 24159.0
            },
            "ens_id": "ENSMUSG00000004207",
            "uniprot_ids": [
                "Q61207",
                "Q8BFQ1",
                "J3QPG5",
                "E9PZ00",
                "K3W4L3"
            ]
        },
        "ENSMUSG00000069601": {
            "id": "ENSMUSG00000069601",
            "type": "Gene",
            "symbol": "Ank3",
            "source": [
                "ENSMUSG00000032035",
                "ENSMUSG00000052684",
                "ENSMUSG00000026872",
                "ENSMUSG00000052837",
                "ENSMUSG00000000247",
                "ENSMUSG00000053477",
                "ENSMUSG00000038193",
                "ENSMUSG00000021250",
                "ENSMUSG00000003545",
                "ENSMUSG00000029322",
                "ENSMUSG00000008575",
                "ENSMUSG00000020484"
            ],
            "target": [],
            "expression_sum": {
                "class1": 13983.0,
                "class2": 90.0
            },
            "ens_id": "ENSMUSG00000069601",
            "uniprot_ids": [
                "G5E8K5",
                "Q3UVY0",
                "G3X971",
                "S4R162",
                "S4R208",
                "S4R2F5",
                "W4VSQ0",
                "S4R165",
                "S4R278",
                "S4R2J6",
                "S4R1S2",
                "S4R229",
                "G5E8K3",
                "A0A087WRU2",
                "S4R2K9",
                "S4R1X7",
                "S4R1U4",
                "A0A087WRP9",
                "A0A087WNU5",
                "G5E8K2",
                "S4R236",
                "S4R2S8",
                "A0A1W2P812",
                "S4R187",
                "S4R2C1"
            ]
        },
        "ENSMUSG00000023175": {
            "id": "ENSMUSG00000023175",
            "type": "Gene",
            "symbol": "Bsg",
            "source": [
                "ENSMUSG00000032035",
                "ENSMUSG00000000247",
                "ENSMUSG00000057098",
                "ENSMUSG00000005583",
                "ENSMUSG00000071076"
            ],
            "target": [],
            "expression_sum": {
                "class1": 35669.0,
                "class2": 16989.0
            },
            "ens_id": "ENSMUSG00000023175",
            "uniprot_ids": [
                "P18572",
                "J3QP71",
                "K3W4Q8"
            ]
        },
        "ENSMUSG00000003072": {
            "id": "ENSMUSG00000003072",
            "type": "Gene",
            "symbol": "Atp5d",
            "source": [
                "ENSMUSG00000000247"
            ],
            "target": [],
            "expression_sum": {
                "class1": 15252.0,
                "class2": 13679.0
            },
            "ens_id": "ENSMUSG00000003072",
            "uniprot_ids": [
                "Q9D3D9",
                "D3Z7S4"
            ]
        },
        "ENSMUSG00000035242": {
            "id": "ENSMUSG00000035242",
            "type": "Gene",
            "symbol": "Oaz1",
            "source": [
                "ENSMUSG00000057098",
                "ENSMUSG00000032035",
                "ENSMUSG00000000247"
            ],
            "target": [],
            "expression_sum": {
                "class1": 40203.0,
                "class2": 31071.0
            },
            "ens_id": "ENSMUSG00000035242",
            "uniprot_ids": [
                "P54369",
                "J3QM96",
                "F8WI29",
                "J3QMV2",
                "J3QM60",
                "J3QME8"
            ]
        },
        "ENSMUSG00000034994": {
            "id": "ENSMUSG00000034994",
            "type": "Gene",
            "symbol": "Eef2",
            "source": [
                "ENSMUSG00000057098",
                "ENSMUSG00000076431",
                "ENSMUSG00000000247"
            ],
            "target": [],
            "expression_sum": {
                "class1": 78381.0,
                "class2": 35842.0
            },
            "ens_id": "ENSMUSG00000034994",
            "uniprot_ids": [
                "P58252"
            ]
        },
        "ENSMUSG00000061904": {
            "id": "ENSMUSG00000061904",
            "type": "Gene",
            "symbol": "Slc25a3",
            "source": [
                "ENSMUSG00000057098",
                "ENSMUSG00000000247"
            ],
            "target": [],
            "expression_sum": {
                "class1": 17186.0,
                "class2": 11680.0
            },
            "ens_id": "ENSMUSG00000061904",
            "uniprot_ids": [
                "Q8VEM8",
                "G5E902",
                "E9PX16"
            ]
        },
        "ENSMUSG00000061315": {
            "id": "ENSMUSG00000061315",
            "type": "Gene",
            "symbol": "Naca",
            "source": [
                "ENSMUSG00000057098",
                "ENSMUSG00000000247"
            ],
            "target": [],
            "expression_sum": {
                "class1": 52251.0,
                "class2": 25707.0
            },
            "ens_id": "ENSMUSG00000061315",
            "uniprot_ids": [
                "P70670",
                "Q60817"
            ]
        },
        "ENSMUSG00000025393": {
            "id": "ENSMUSG00000025393",
            "type": "Gene",
            "symbol": "Atp5b",
            "source": [
                "ENSMUSG00000025498",
                "ENSMUSG00000047638",
                "ENSMUSG00000005583",
                "ENSMUSG00000000247",
                "ENSMUSG00000053477",
                "ENSMUSG00000008575",
                "ENSMUSG00000020484"
            ],
            "target": [],
            "expression_sum": {
                "class1": 23132.0,
                "class2": 24589.0
            },
            "ens_id": "ENSMUSG00000025393",
            "uniprot_ids": [
                "P56480"
            ]
        },
        "ENSMUSG00000093674": {
            "id": "ENSMUSG00000093674",
            "type": "Gene",
            "symbol": "Rpl41",
            "source": [
                "ENSMUSG00000057098",
                "ENSMUSG00000048387",
                "ENSMUSG00000000247"
            ],
            "target": [],
            "expression_sum": {
                "class1": 196686.0,
                "class2": 128718.0
            },
            "ens_id": "ENSMUSG00000093674",
            "uniprot_ids": [
                "P62947",
                "H3BK91"
            ]
        },
        "ENSMUSG00000025362": {
            "id": "ENSMUSG00000025362",
            "type": "Gene",
            "symbol": "Rps26",
            "source": [
                "ENSMUSG00000048387",
                "ENSMUSG00000000247"
            ],
            "target": [],
            "expression_sum": {
                "class1": 61330.0,
                "class2": 34531.0
            },
            "ens_id": "ENSMUSG00000025362",
            "uniprot_ids": [
                "P62855"
            ]
        },
        "ENSMUSG00000031532": {
            "id": "ENSMUSG00000031532",
            "type": "Gene",
            "symbol": "Saraf",
            "source": [
                "ENSMUSG00000000247"
            ],
            "target": [],
            "expression_sum": {
                "class1": 14070.0,
                "class2": 5568.0
            },
            "ens_id": "ENSMUSG00000031532",
            "uniprot_ids": [
                "Q8R3Q0",
                "A0A0R4J0D1"
            ]
        },
        "ENSMUSG00000031633": {
            "id": "ENSMUSG00000031633",
            "type": "Gene",
            "symbol": "Slc25a4",
            "source": [
                "ENSMUSG00000057098",
                "ENSMUSG00000000247"
            ],
            "target": [],
            "expression_sum": {
                "class1": 25055.0,
                "class2": 20589.0
            },
            "ens_id": "ENSMUSG00000031633",
            "uniprot_ids": [
                "P48962"
            ]
        },
        "ENSMUSG00000045128": {
            "id": "ENSMUSG00000045128",
            "type": "Gene",
            "symbol": "Rpl18a",
            "source": [
                "ENSMUSG00000056501",
                "ENSMUSG00000023034",
                "ENSMUSG00000000247",
                "ENSMUSG00000057098",
                "ENSMUSG00000005583"
            ],
            "target": [],
            "expression_sum": {
                "class1": 182128.0,
                "class2": 79882.0
            },
            "ens_id": "ENSMUSG00000045128",
            "uniprot_ids": [
                "P62717",
                "A0A1D5RMC7",
                "A0A1D5RM79",
                "A0A1D5RLW5",
                "A0A1D5RM85",
                "A0A1D5RLA0",
                "A0A1D5RME4"
            ]
        },
        "ENSMUSG00000003814": {
            "id": "ENSMUSG00000003814",
            "type": "Gene",
            "symbol": "Calr",
            "source": [
                "ENSMUSG00000057098",
                "ENSMUSG00000038193",
                "ENSMUSG00000005583",
                "ENSMUSG00000000247"
            ],
            "target": [],
            "expression_sum": {
                "class1": 46743.0,
                "class2": 17355.0
            },
            "ens_id": "ENSMUSG00000003814",
            "uniprot_ids": [
                "P14211"
            ]
        },
        "ENSMUSG00000005161": {
            "id": "ENSMUSG00000005161",
            "type": "Gene",
            "symbol": "Prdx2",
            "source": [
                "ENSMUSG00000000247"
            ],
            "target": [],
            "expression_sum": {
                "class1": 12992.0,
                "class2": 7267.0
            },
            "ens_id": "ENSMUSG00000005161",
            "uniprot_ids": [
                "Q61171",
                "D3Z4A4"
            ]
        },
        "ENSMUSG00000031818": {
            "id": "ENSMUSG00000031818",
            "type": "Gene",
            "symbol": "Cox4i1",
            "source": [
                "ENSMUSG00000057098",
                "ENSMUSG00000047638",
                "ENSMUSG00000029322",
                "ENSMUSG00000000247"
            ],
            "target": [],
            "expression_sum": {
                "class1": 38815.0,
                "class2": 30450.0
            },
            "ens_id": "ENSMUSG00000031818",
            "uniprot_ids": [
                "P19783",
                "M0QWX2",
                "M0QWX7"
            ]
        },
        "ENSMUSG00000042812": {
            "id": "ENSMUSG00000042812",
            "type": "TF",
            "symbol": "Foxf1",
            "source": [
                "ENSMUSG00000052837",
                "ENSMUSG00000021250",
                "ENSMUSG00000052684",
                "ENSMUSG00000000247"
            ],
            "target": [
                "ENSMUSG00000031375",
                "ENSMUSG00000021250",
                "ENSMUSG00000030849",
                "ENSMUSG00000003545",
                "ENSMUSG00000031119",
                "ENSMUSG00000035493"
            ],
            "expression_sum": {
                "class1": 8363.0,
                "class2": 1724.0
            },
            "ens_id": "ENSMUSG00000042812",
            "uniprot_ids": [
                "Q61080"
            ]
        },
        "ENSMUSG00000031812": {
            "id": "ENSMUSG00000031812",
            "type": "Gene",
            "symbol": "Map1lc3b",
            "source": [
                "ENSMUSG00000000247"
            ],
            "target": [],
            "expression_sum": {
                "class1": 19863.0,
                "class2": 14560.0
            },
            "ens_id": "ENSMUSG00000031812",
            "uniprot_ids": [
                "Q9CQV6",
                "M0QWC2",
                "A0A1D5RLG5",
                "M0QWT8"
            ]
        },
        "ENSMUSG00000000740": {
            "id": "ENSMUSG00000000740",
            "type": "Gene",
            "symbol": "Rpl13",
            "source": [
                "ENSMUSG00000032035",
                "ENSMUSG00000023034",
                "ENSMUSG00000000247",
                "ENSMUSG00000038193",
                "ENSMUSG00000005583"
            ],
            "target": [],
            "expression_sum": {
                "class1": 199675.0,
                "class2": 112601.0
            },
            "ens_id": "ENSMUSG00000000740",
            "uniprot_ids": [
                "P47963"
            ]
        },
        "ENSMUSG00000012405": {
            "id": "ENSMUSG00000012405",
            "type": "Gene",
            "symbol": "Rpl15",
            "source": [
                "ENSMUSG00000057098",
                "ENSMUSG00000022528",
                "ENSMUSG00000038193",
                "ENSMUSG00000000247"
            ],
            "target": [],
            "expression_sum": {
                "class1": 55020.0,
                "class2": 25838.0
            },
            "ens_id": "ENSMUSG00000012405",
            "uniprot_ids": [
                "Q9CZM2",
                "B8JKK2"
            ]
        },
        "ENSMUSG00000021807": {
            "id": "ENSMUSG00000021807",
            "type": "Gene",
            "symbol": "2700060E02Rik",
            "source": [
                "ENSMUSG00000000247"
            ],
            "target": [],
            "expression_sum": {
                "class1": 15633.0,
                "class2": 6922.0
            },
            "ens_id": "ENSMUSG00000021807",
            "uniprot_ids": [
                "Q9CQE8"
            ]
        },
        "ENSMUSG00000025290": {
            "id": "ENSMUSG00000025290",
            "type": "Gene",
            "symbol": "Rps24",
            "source": [
                "ENSMUSG00000038193",
                "ENSMUSG00000023034",
                "ENSMUSG00000005583",
                "ENSMUSG00000000247"
            ],
            "target": [],
            "expression_sum": {
                "class1": 119120.0,
                "class2": 64607.0
            },
            "ens_id": "ENSMUSG00000025290",
            "uniprot_ids": [
                "P62849",
                "A0A286YEB7"
            ]
        },
        "ENSMUSG00000021870": {
            "id": "ENSMUSG00000021870",
            "type": "Gene",
            "symbol": "Slmap",
            "source": [
                "ENSMUSG00000025498",
                "ENSMUSG00000032238",
                "ENSMUSG00000052684",
                "ENSMUSG00000047638",
                "ENSMUSG00000005583",
                "ENSMUSG00000000247",
                "ENSMUSG00000053477",
                "ENSMUSG00000008575",
                "ENSMUSG00000020484",
                "ENSMUSG00000071076"
            ],
            "target": [],
            "expression_sum": {
                "class1": 13277.0,
                "class2": 4050.0
            },
            "ens_id": "ENSMUSG00000021870",
            "uniprot_ids": [
                "Q3URD3",
                "F6WY34",
                "F6UV57",
                "F8WIH0",
                "F6YCM8",
                "D3Z7V3",
                "H7BX64"
            ]
        },
        "ENSMUSG00000042682": {
            "id": "ENSMUSG00000042682",
            "type": "Gene",
            "symbol": "Selk",
            "source": [
                "ENSMUSG00000057098",
                "ENSMUSG00000000247"
            ],
            "target": [],
            "expression_sum": {
                "class1": 27791.0,
                "class2": 14848.0
            },
            "ens_id": "ENSMUSG00000042682",
            "uniprot_ids": [
                "Q9JLJ1",
                "A0A286YD19"
            ]
        },
        "ENSMUSG00000037712": {
            "id": "ENSMUSG00000037712",
            "type": "Gene",
            "symbol": "Fermt2",
            "source": [
                "ENSMUSG00000047638",
                "ENSMUSG00000000247",
                "ENSMUSG00000053477",
                "ENSMUSG00000020644",
                "ENSMUSG00000020484"
            ],
            "target": [],
            "expression_sum": {
                "class1": 33032.0,
                "class2": 5882.0
            },
            "ens_id": "ENSMUSG00000037712",
            "uniprot_ids": [
                "Q8CIB5",
                "A6X941",
                "A6X940",
                "A0A2I3BPQ7"
            ]
        },
        "ENSMUSG00000010406": {
            "id": "ENSMUSG00000010406",
            "type": "Gene",
            "symbol": "Mrpl52",
            "source": [
                "ENSMUSG00000057098",
                "ENSMUSG00000000247"
            ],
            "target": [],
            "expression_sum": {
                "class1": 13280.0,
                "class2": 6416.0
            },
            "ens_id": "ENSMUSG00000010406",
            "uniprot_ids": [
                "Q9D0Y8",
                "A0A0G2JE60"
            ]
        },
        "ENSMUSG00000010376": {
            "id": "ENSMUSG00000010376",
            "type": "Gene",
            "symbol": "Nedd8",
            "source": [
                "ENSMUSG00000000247"
            ],
            "target": [],
            "expression_sum": {
                "class1": 14814.0,
                "class2": 9559.0
            },
            "ens_id": "ENSMUSG00000010376",
            "uniprot_ids": [
                "P29595",
                "A0A2I3BRG0"
            ]
        },
        "ENSMUSG00000022051": {
            "id": "ENSMUSG00000022051",
            "type": "Gene",
            "symbol": "Bnip3l",
            "source": [
                "ENSMUSG00000057098",
                "ENSMUSG00000000247"
            ],
            "target": [],
            "expression_sum": {
                "class1": 9544.0,
                "class2": 4928.0
            },
            "ens_id": "ENSMUSG00000022051",
            "uniprot_ids": [
                "Q9Z2F7",
                "Q91Z78"
            ]
        },
        "ENSMUSG00000022108": {
            "id": "ENSMUSG00000022108",
            "type": "Gene",
            "symbol": "Itm2b",
            "source": [
                "ENSMUSG00000032035",
                "ENSMUSG00000047638",
                "ENSMUSG00000000247",
                "ENSMUSG00000057098",
                "ENSMUSG00000038193",
                "ENSMUSG00000029322",
                "ENSMUSG00000048387",
                "ENSMUSG00000071076"
            ],
            "target": [],
            "expression_sum": {
                "class1": 233347.0,
                "class2": 92392.0
            },
            "ens_id": "ENSMUSG00000022108",
            "uniprot_ids": [
                "O89051",
                "A0A2I3BQ30",
                "A0A2I3BPW7"
            ]
        },
        "ENSMUSG00000060126": {
            "id": "ENSMUSG00000060126",
            "type": "Gene",
            "symbol": "Tpt1",
            "source": [
                "ENSMUSG00000032035",
                "ENSMUSG00000056501",
                "ENSMUSG00000023034",
                "ENSMUSG00000022528",
                "ENSMUSG00000000247",
                "ENSMUSG00000057098",
                "ENSMUSG00000038193",
                "ENSMUSG00000005583",
                "ENSMUSG00000031965",
                "ENSMUSG00000048387"
            ],
            "target": [],
            "expression_sum": {
                "class1": 130237.0,
                "class2": 69022.0
            },
            "ens_id": "ENSMUSG00000060126",
            "uniprot_ids": [
                "P63028",
                "D3YU75"
            ]
        },
        "ENSMUSG00000022010": {
            "id": "ENSMUSG00000022010",
            "type": "Gene",
            "symbol": "Tsc22d1",
            "source": [
                "ENSMUSG00000052684",
                "ENSMUSG00000047638",
                "ENSMUSG00000005583",
                "ENSMUSG00000000247",
                "ENSMUSG00000053477",
                "ENSMUSG00000055148",
                "ENSMUSG00000055435",
                "ENSMUSG00000020644",
                "ENSMUSG00000020484"
            ],
            "target": [],
            "expression_sum": {
                "class1": 18550.0,
                "class2": 24021.0
            },
            "ens_id": "ENSMUSG00000022010",
            "uniprot_ids": [
                "P62500",
                "H3BLI9",
                "H3BK98",
                "H3BJ00",
                "D3Z0V7",
                "H3BLH4",
                "E9QLZ1",
                "D3YW86",
                "H3BJV9",
                "H3BK07"
            ]
        },
        "ENSMUSG00000022136": {
            "id": "ENSMUSG00000022136",
            "type": "Gene",
            "symbol": "Dnajc3",
            "source": [
                "ENSMUSG00000022528",
                "ENSMUSG00000000247",
                "ENSMUSG00000038193",
                "ENSMUSG00000005583",
                "ENSMUSG00000071076"
            ],
            "target": [],
            "expression_sum": {
                "class1": 14905.0,
                "class2": 6703.0
            },
            "ens_id": "ENSMUSG00000022136",
            "uniprot_ids": [
                "Q91YW3"
            ]
        },
        "ENSMUSG00000015656": {
            "id": "ENSMUSG00000015656",
            "type": "Gene",
            "symbol": "Hspa8",
            "source": [
                "ENSMUSG00000025498",
                "ENSMUSG00000047638",
                "ENSMUSG00000057098",
                "ENSMUSG00000000247"
            ],
            "target": [],
            "expression_sum": {
                "class1": 74964.0,
                "class2": 37393.0
            },
            "ens_id": "ENSMUSG00000015656",
            "uniprot_ids": [
                "P63017",
                "Q504P4",
                "D3Z5E2"
            ]
        },
        "ENSMUSG00000009927": {
            "id": "ENSMUSG00000009927",
            "type": "Gene",
            "symbol": "Rps25",
            "source": [
                "ENSMUSG00000057098",
                "ENSMUSG00000048387",
                "ENSMUSG00000000247"
            ],
            "target": [],
            "expression_sum": {
                "class1": 76284.0,
                "class2": 47742.0
            },
            "ens_id": "ENSMUSG00000009927",
            "uniprot_ids": [
                "P62852",
                "A0A1L1SQA8"
            ]
        },
        "ENSMUSG00000038717": {
            "id": "ENSMUSG00000038717",
            "type": "Gene",
            "symbol": "Atp5l",
            "source": [
                "ENSMUSG00000057098",
                "ENSMUSG00000000247"
            ],
            "target": [],
            "expression_sum": {
                "class1": 26274.0,
                "class2": 20749.0
            },
            "ens_id": "ENSMUSG00000038717",
            "uniprot_ids": [
                "Q9CPQ8"
            ]
        },
        "ENSMUSG00000032288": {
            "id": "ENSMUSG00000032288",
            "type": "Gene",
            "symbol": "Imp3",
            "source": [
                "ENSMUSG00000000247"
            ],
            "target": [],
            "expression_sum": {
                "class1": 10003.0,
                "class2": 3061.0
            },
            "ens_id": "ENSMUSG00000032288",
            "uniprot_ids": [
                "Q921Y2"
            ]
        },
        "ENSMUSG00000007892": {
            "id": "ENSMUSG00000007892",
            "type": "Gene",
            "symbol": "Rplp1",
            "source": [
                "ENSMUSG00000032035",
                "ENSMUSG00000056501",
                "ENSMUSG00000023034",
                "ENSMUSG00000000247",
                "ENSMUSG00000057098",
                "ENSMUSG00000005583"
            ],
            "target": [],
            "expression_sum": {
                "class1": 155837.0,
                "class2": 82969.0
            },
            "ens_id": "ENSMUSG00000007892",
            "uniprot_ids": [
                "P47955"
            ]
        },
        "ENSMUSG00000032399": {
            "id": "ENSMUSG00000032399",
            "type": "Gene",
            "symbol": "Rpl4",
            "source": [
                "ENSMUSG00000057098",
                "ENSMUSG00000048387",
                "ENSMUSG00000038193",
                "ENSMUSG00000000247"
            ],
            "target": [],
            "expression_sum": {
                "class1": 85485.0,
                "class2": 44235.0
            },
            "ens_id": "ENSMUSG00000032399",
            "uniprot_ids": [
                "Q9D8E6"
            ]
        },
        "ENSMUSG00000032231": {
            "id": "ENSMUSG00000032231",
            "type": "Gene",
            "symbol": "Anxa2",
            "source": [
                "ENSMUSG00000032035",
                "ENSMUSG00000052684",
                "ENSMUSG00000005583",
                "ENSMUSG00000000247",
                "ENSMUSG00000057098",
                "ENSMUSG00000003032",
                "ENSMUSG00000055435",
                "ENSMUSG00000028565",
                "ENSMUSG00000071076"
            ],
            "target": [],
            "expression_sum": {
                "class1": 27881.0,
                "class2": 12495.0
            },
            "ens_id": "ENSMUSG00000032231",
            "uniprot_ids": [
                "P07356",
                "B0V2N7",
                "B0V2N5",
                "B0V2N8"
            ]
        },
        "ENSMUSG00000032216": {
            "id": "ENSMUSG00000032216",
            "type": "Gene",
            "symbol": "Nedd4",
            "source": [
                "ENSMUSG00000057098",
                "ENSMUSG00000022528",
                "ENSMUSG00000038193",
                "ENSMUSG00000000247"
            ],
            "target": [],
            "expression_sum": {
                "class1": 22705.0,
                "class2": 13086.0
            },
            "ens_id": "ENSMUSG00000032216",
            "uniprot_ids": [
                "P46935",
                "Q3V335",
                "V9GXS6",
                "A0A571BEP7",
                "V9GWV8",
                "V9GXK3",
                "A0A571BDN7"
            ]
        },
        "ENSMUSG00000037742": {
            "id": "ENSMUSG00000037742",
            "type": "Gene",
            "symbol": "Eef1a1",
            "source": [
                "ENSMUSG00000056501",
                "ENSMUSG00000023034",
                "ENSMUSG00000022528",
                "ENSMUSG00000047638",
                "ENSMUSG00000000247",
                "ENSMUSG00000057098",
                "ENSMUSG00000038193",
                "ENSMUSG00000048387"
            ],
            "target": [],
            "expression_sum": {
                "class1": 331050.0,
                "class2": 109025.0
            },
            "ens_id": "ENSMUSG00000037742",
            "uniprot_ids": [
                "P10126",
                "D3YZ68",
                "D3Z3I8"
            ]
        },
        "ENSMUSG00000062270": {
            "id": "ENSMUSG00000062270",
            "type": "Gene",
            "symbol": "Morf4l1",
            "source": [
                "ENSMUSG00000057098",
                "ENSMUSG00000071076",
                "ENSMUSG00000048387",
                "ENSMUSG00000000247"
            ],
            "target": [],
            "expression_sum": {
                "class1": 28312.0,
                "class2": 24558.0
            },
            "ens_id": "ENSMUSG00000062270",
            "uniprot_ids": [
                "P60762",
                "A0A087WQ34",
                "A0A087WSL1",
                "A0A087WQB0"
            ]
        },
        "ENSMUSG00000048758": {
            "id": "ENSMUSG00000048758",
            "type": "Gene",
            "symbol": "Rpl29",
            "source": [
                "ENSMUSG00000032035",
                "ENSMUSG00000038193",
                "ENSMUSG00000048387",
                "ENSMUSG00000000247"
            ],
            "target": [],
            "expression_sum": {
                "class1": 75718.0,
                "class2": 29184.0
            },
            "ens_id": "ENSMUSG00000048758",
            "uniprot_ids": [
                "P47915",
                "A0A1L1SUN1",
                "A0A1L1SS27",
                "A0A1L1STJ3"
            ]
        },
        "ENSMUSG00000032562": {
            "id": "ENSMUSG00000032562",
            "type": "Gene",
            "symbol": "Gnai2",
            "source": [
                "ENSMUSG00000032035",
                "ENSMUSG00000029322",
                "ENSMUSG00000000247"
            ],
            "target": [],
            "expression_sum": {
                "class1": 31735.0,
                "class2": 9536.0
            },
            "ens_id": "ENSMUSG00000032562",
            "uniprot_ids": [
                "P08752",
                "A0A0A6YXC2",
                "A0A0A6YXN0",
                "A0A0A6YWA9"
            ]
        },
        "ENSMUSG00000025794": {
            "id": "ENSMUSG00000025794",
            "type": "Gene",
            "symbol": "Rpl14",
            "source": [
                "ENSMUSG00000057098",
                "ENSMUSG00000038193",
                "ENSMUSG00000005583",
                "ENSMUSG00000000247"
            ],
            "target": [],
            "expression_sum": {
                "class1": 94055.0,
                "class2": 60922.0
            },
            "ens_id": "ENSMUSG00000025794",
            "uniprot_ids": [
                "Q9CR57",
                "A0A1L1SUF6"
            ]
        },
        "ENSMUSG00000020432": {
            "id": "ENSMUSG00000020432",
            "type": "Gene",
            "symbol": "Tcn2",
            "source": [
                "ENSMUSG00000032035",
                "ENSMUSG00000038193",
                "ENSMUSG00000000247"
            ],
            "target": [],
            "expression_sum": {
                "class1": 15094.0,
                "class2": 2545.0
            },
            "ens_id": "ENSMUSG00000020432",
            "uniprot_ids": [
                "O88968"
            ]
        },
        "ENSMUSG00000020460": {
            "id": "ENSMUSG00000020460",
            "type": "Gene",
            "symbol": "Rps27a",
            "source": [
                "ENSMUSG00000057098",
                "ENSMUSG00000038193",
                "ENSMUSG00000005583",
                "ENSMUSG00000000247"
            ],
            "target": [],
            "expression_sum": {
                "class1": 117810.0,
                "class2": 61478.0
            },
            "ens_id": "ENSMUSG00000020460",
            "uniprot_ids": [
                "P62983"
            ]
        },
        "ENSMUSG00000020372": {
            "id": "ENSMUSG00000020372",
            "type": "Gene",
            "symbol": "Gnb2l1",
            "source": [
                "ENSMUSG00000057098",
                "ENSMUSG00000038193",
                "ENSMUSG00000023034",
                "ENSMUSG00000000247"
            ],
            "target": [],
            "expression_sum": {
                "class1": 47929.0,
                "class2": 17452.0
            },
            "ens_id": "ENSMUSG00000020372",
            "uniprot_ids": [
                "P68040"
            ]
        },
        "ENSMUSG00000048076": {
            "id": "ENSMUSG00000048076",
            "type": "Gene",
            "symbol": "Arf1",
            "source": [
                "ENSMUSG00000057098",
                "ENSMUSG00000071076",
                "ENSMUSG00000003032",
                "ENSMUSG00000000247"
            ],
            "target": [],
            "expression_sum": {
                "class1": 17528.0,
                "class2": 11805.0
            },
            "ens_id": "ENSMUSG00000048076",
            "uniprot_ids": [
                "P84078"
            ]
        },
        "ENSMUSG00000019505": {
            "id": "ENSMUSG00000019505",
            "type": "Gene",
            "symbol": "Ubb",
            "source": [
                "ENSMUSG00000025498",
                "ENSMUSG00000032035",
                "ENSMUSG00000052837",
                "ENSMUSG00000047638",
                "ENSMUSG00000000247",
                "ENSMUSG00000057098",
                "ENSMUSG00000038418",
                "ENSMUSG00000021250",
                "ENSMUSG00000003032",
                "ENSMUSG00000048387",
                "ENSMUSG00000071076"
            ],
            "target": [],
            "expression_sum": {
                "class1": 174128.0,
                "class2": 87692.0
            },
            "ens_id": "ENSMUSG00000019505",
            "uniprot_ids": [
                "P0CG49",
                "Q5SX22"
            ]
        },
        "ENSMUSG00000060938": {
            "id": "ENSMUSG00000060938",
            "type": "Gene",
            "symbol": "Rpl26",
            "source": [
                "ENSMUSG00000023034",
                "ENSMUSG00000000247",
                "ENSMUSG00000057098",
                "ENSMUSG00000038193",
                "ENSMUSG00000076431",
                "ENSMUSG00000031965",
                "ENSMUSG00000048387"
            ],
            "target": [],
            "expression_sum": {
                "class1": 124210.0,
                "class2": 63185.0
            },
            "ens_id": "ENSMUSG00000060938",
            "uniprot_ids": [
                "P61255",
                "B1ARA3",
                "B1ARA5"
            ]
        },
        "ENSMUSG00000018293": {
            "id": "ENSMUSG00000018293",
            "type": "Gene",
            "symbol": "Pfn1",
            "source": [
                "ENSMUSG00000047638",
                "ENSMUSG00000000247",
                "ENSMUSG00000053477",
                "ENSMUSG00000021250",
                "ENSMUSG00000005583",
                "ENSMUSG00000020644"
            ],
            "target": [],
            "expression_sum": {
                "class1": 43639.0,
                "class2": 32834.0
            },
            "ens_id": "ENSMUSG00000018293",
            "uniprot_ids": [
                "P62962",
                "Q5SX49",
                "J3QMC2"
            ]
        },
        "ENSMUSG00000058546": {
            "id": "ENSMUSG00000058546",
            "type": "Gene",
            "symbol": "Rpl23a",
            "source": [
                "ENSMUSG00000057098",
                "ENSMUSG00000048387",
                "ENSMUSG00000038193",
                "ENSMUSG00000000247"
            ],
            "target": [],
            "expression_sum": {
                "class1": 119621.0,
                "class2": 78343.0
            },
            "ens_id": "ENSMUSG00000058546",
            "uniprot_ids": [
                "P62751"
            ]
        },
        "ENSMUSG00000071415": {
            "id": "ENSMUSG00000071415",
            "type": "Gene",
            "symbol": "Rpl23",
            "source": [
                "ENSMUSG00000023034",
                "ENSMUSG00000000247",
                "ENSMUSG00000057098",
                "ENSMUSG00000038193",
                "ENSMUSG00000005583",
                "ENSMUSG00000048387"
            ],
            "target": [],
            "expression_sum": {
                "class1": 130606.0,
                "class2": 60285.0
            },
            "ens_id": "ENSMUSG00000071415",
            "uniprot_ids": [
                "P62830",
                "A2A6F8"
            ]
        },
        "ENSMUSG00000017404": {
            "id": "ENSMUSG00000017404",
            "type": "Gene",
            "symbol": "Rpl19",
            "source": [
                "ENSMUSG00000057098",
                "ENSMUSG00000000247"
            ],
            "target": [],
            "expression_sum": {
                "class1": 105247.0,
                "class2": 55383.0
            },
            "ens_id": "ENSMUSG00000017404",
            "uniprot_ids": [
                "P84099",
                "A2A547"
            ]
        },
        "ENSMUSG00000035530": {
            "id": "ENSMUSG00000035530",
            "type": "Gene",
            "symbol": "Eif1",
            "source": [
                "ENSMUSG00000057098",
                "ENSMUSG00000032035",
                "ENSMUSG00000000247"
            ],
            "target": [],
            "expression_sum": {
                "class1": 87157.0,
                "class2": 60158.0
            },
            "ens_id": "ENSMUSG00000035530",
            "uniprot_ids": [
                "P48024",
                "J3QN87",
                "J3QPZ8"
            ]
        },
        "ENSMUSG00000063316": {
            "id": "ENSMUSG00000063316",
            "type": "Gene",
            "symbol": "Rpl27",
            "source": [
                "ENSMUSG00000031965",
                "ENSMUSG00000048387",
                "ENSMUSG00000057098",
                "ENSMUSG00000000247"
            ],
            "target": [],
            "expression_sum": {
                "class1": 59130.0,
                "class2": 28442.0
            },
            "ens_id": "ENSMUSG00000063316",
            "uniprot_ids": [
                "P61358",
                "A2A4Q0"
            ]
        },
        "ENSMUSG00000020719": {
            "id": "ENSMUSG00000020719",
            "type": "Gene",
            "symbol": "Ddx5",
            "source": [
                "ENSMUSG00000057098",
                "ENSMUSG00000047638",
                "ENSMUSG00000000247"
            ],
            "target": [],
            "expression_sum": {
                "class1": 21092.0,
                "class2": 19905.0
            },
            "ens_id": "ENSMUSG00000020719",
            "uniprot_ids": [
                "Q8BTS0",
                "B1ARC0",
                "B1ARB9",
                "S4R1I6",
                "S4R1E3"
            ]
        },
        "ENSMUSG00000020612": {
            "id": "ENSMUSG00000020612",
            "type": "Gene",
            "symbol": "Prkar1a",
            "source": [
                "ENSMUSG00000047638",
                "ENSMUSG00000020484",
                "ENSMUSG00000005583",
                "ENSMUSG00000000247"
            ],
            "target": [],
            "expression_sum": {
                "class1": 12222.0,
                "class2": 11236.0
            },
            "ens_id": "ENSMUSG00000020612",
            "uniprot_ids": [
                "Q9DBC7",
                "A2AI69"
            ]
        },
        "ENSMUSG00000057322": {
            "id": "ENSMUSG00000057322",
            "type": "Gene",
            "symbol": "Rpl38",
            "source": [
                "ENSMUSG00000057098",
                "ENSMUSG00000000247"
            ],
            "target": [],
            "expression_sum": {
                "class1": 95716.0,
                "class2": 55698.0
            },
            "ens_id": "ENSMUSG00000057322",
            "uniprot_ids": [
                "Q9JJI8"
            ]
        },
        "ENSMUSG00000020738": {
            "id": "ENSMUSG00000020738",
            "type": "Gene",
            "symbol": "Sumo2",
            "source": [
                "ENSMUSG00000057098",
                "ENSMUSG00000029322",
                "ENSMUSG00000000247"
            ],
            "target": [],
            "expression_sum": {
                "class1": 22911.0,
                "class2": 17041.0
            },
            "ens_id": "ENSMUSG00000020738",
            "uniprot_ids": [
                "P61957",
                "D3YW94",
                "H7BWX9",
                "D3Z794"
            ]
        },
        "ENSMUSG00000016559": {
            "id": "ENSMUSG00000016559",
            "type": "Gene",
            "symbol": "H3f3b",
            "source": [
                "ENSMUSG00000057098",
                "ENSMUSG00000005583",
                "ENSMUSG00000000247"
            ],
            "target": [],
            "expression_sum": {
                "class1": 48143.0,
                "class2": 40325.0
            },
            "ens_id": "ENSMUSG00000016559",
            "uniprot_ids": [
                "P84244"
            ]
        },
        "ENSMUSG00000033880": {
            "id": "ENSMUSG00000033880",
            "type": "Gene",
            "symbol": "Lgals3bp",
            "source": [
                "ENSMUSG00000025498",
                "ENSMUSG00000032035",
                "ENSMUSG00000056501",
                "ENSMUSG00000047638",
                "ENSMUSG00000000247",
                "ENSMUSG00000038193",
                "ENSMUSG00000005583"
            ],
            "target": [],
            "expression_sum": {
                "class1": 60217.0,
                "class2": 2141.0
            },
            "ens_id": "ENSMUSG00000033880",
            "uniprot_ids": [
                "Q07797",
                "F6VRP8",
                "E9Q5X5"
            ]
        },
        "ENSMUSG00000021546": {
            "id": "ENSMUSG00000021546",
            "type": "Gene",
            "symbol": "Hnrnpk",
            "source": [
                "ENSMUSG00000057098",
                "ENSMUSG00000000247"
            ],
            "target": [],
            "expression_sum": {
                "class1": 19438.0,
                "class2": 14296.0
            },
            "ens_id": "ENSMUSG00000021546",
            "uniprot_ids": [
                "P61979",
                "H3BJ43",
                "Q8BT23",
                "B2M1R6",
                "H3BLP7",
                "H3BKD0",
                "A0A286YDM3",
                "A0A286YE41",
                "H3BJS9",
                "A0A286YCM2",
                "A0A286YDH1",
                "H3BLL4",
                "H3BKI8",
                "H3BK96",
                "A0A286YEC4"
            ]
        },
        "ENSMUSG00000021477": {
            "id": "ENSMUSG00000021477",
            "type": "Gene",
            "symbol": "Ctsl",
            "source": [
                "ENSMUSG00000055435",
                "ENSMUSG00000032035",
                "ENSMUSG00000052684",
                "ENSMUSG00000058756",
                "ENSMUSG00000005583",
                "ENSMUSG00000000247",
                "ENSMUSG00000038418",
                "ENSMUSG00000038193",
                "ENSMUSG00000003032",
                "ENSMUSG00000008575",
                "ENSMUSG00000028565",
                "ENSMUSG00000071076"
            ],
            "target": [],
            "expression_sum": {
                "class1": 37920.0,
                "class2": 20428.0
            },
            "ens_id": "ENSMUSG00000021477",
            "uniprot_ids": [
                "P06797",
                "A0A1Y7VP49",
                "A0A1Y7VNM3",
                "A0A1Y7VNQ4",
                "A0A1Y7VJC1"
            ]
        },
        "ENSMUSG00000017778": {
            "id": "ENSMUSG00000017778",
            "type": "Gene",
            "symbol": "Cox7c",
            "source": [
                "ENSMUSG00000000247"
            ],
            "target": [],
            "expression_sum": {
                "class1": 28143.0,
                "class2": 20935.0
            },
            "ens_id": "ENSMUSG00000017778",
            "uniprot_ids": [
                "P17665",
                "B8JJA9",
                "B8JJB0"
            ]
        },
        "ENSMUSG00000049517": {
            "id": "ENSMUSG00000049517",
            "type": "Gene",
            "symbol": "Rps23",
            "source": [
                "ENSMUSG00000032035",
                "ENSMUSG00000023034",
                "ENSMUSG00000000247",
                "ENSMUSG00000057098",
                "ENSMUSG00000038193",
                "ENSMUSG00000005583"
            ],
            "target": [],
            "expression_sum": {
                "class1": 163131.0,
                "class2": 90799.0
            },
            "ens_id": "ENSMUSG00000049517",
            "uniprot_ids": [
                "P62267"
            ]
        },
        "ENSMUSG00000042043": {
            "id": "ENSMUSG00000042043",
            "type": "Gene",
            "symbol": "Tbca",
            "source": [
                "ENSMUSG00000057098",
                "ENSMUSG00000000247"
            ],
            "target": [],
            "expression_sum": {
                "class1": 11502.0,
                "class2": 6979.0
            },
            "ens_id": "ENSMUSG00000042043",
            "uniprot_ids": [
                "P48428",
                "A0A1Y7VLW1",
                "A0A1Y7VJ48"
            ]
        },
        "ENSMUSG00000021660": {
            "id": "ENSMUSG00000021660",
            "type": "Gene",
            "symbol": "Btf3",
            "source": [
                "ENSMUSG00000000247"
            ],
            "target": [],
            "expression_sum": {
                "class1": 29518.0,
                "class2": 14563.0
            },
            "ens_id": "ENSMUSG00000021660",
            "uniprot_ids": [
                "Q64152"
            ]
        },
        "ENSMUSG00000041417": {
            "id": "ENSMUSG00000041417",
            "type": "Gene",
            "symbol": "Pik3r1",
            "source": [
                "ENSMUSG00000038193",
                "ENSMUSG00000005583",
                "ENSMUSG00000052684",
                "ENSMUSG00000000247"
            ],
            "target": [],
            "expression_sum": {
                "class1": 10688.0,
                "class2": 3342.0
            },
            "ens_id": "ENSMUSG00000041417",
            "uniprot_ids": [
                "P26450",
                "Q80UI5",
                "P70305",
                "A0A087WQM2"
            ]
        },
        "ENSMUSG00000061477": {
            "id": "ENSMUSG00000061477",
            "type": "Gene",
            "symbol": "Rps7",
            "source": [
                "ENSMUSG00000031965",
                "ENSMUSG00000038193",
                "ENSMUSG00000000247"
            ],
            "target": [],
            "expression_sum": {
                "class1": 155375.0,
                "class2": 51594.0
            },
            "ens_id": "ENSMUSG00000061477",
            "uniprot_ids": [
                "P62082"
            ]
        },
        "ENSMUSG00000034892": {
            "id": "ENSMUSG00000034892",
            "type": "Gene",
            "symbol": "Rps29",
            "source": [
                "ENSMUSG00000032035",
                "ENSMUSG00000023034",
                "ENSMUSG00000000247",
                "ENSMUSG00000038193",
                "ENSMUSG00000005583",
                "ENSMUSG00000048387"
            ],
            "target": [],
            "expression_sum": {
                "class1": 209500.0,
                "class2": 131464.0
            },
            "ens_id": "ENSMUSG00000034892",
            "uniprot_ids": [
                "P62274"
            ]
        },
        "ENSMUSG00000049751": {
            "id": "ENSMUSG00000049751",
            "type": "Gene",
            "symbol": "Rpl36al",
            "source": [
                "ENSMUSG00000057098",
                "ENSMUSG00000000247"
            ],
            "target": [],
            "expression_sum": {
                "class1": 23995.0,
                "class2": 14118.0
            },
            "ens_id": "ENSMUSG00000049751",
            "uniprot_ids": [
                "P83882"
            ]
        },
        "ENSMUSG00000045690": {
            "id": "ENSMUSG00000045690",
            "type": "Gene",
            "symbol": "Wdr89",
            "source": [
                "ENSMUSG00000057098",
                "ENSMUSG00000048387",
                "ENSMUSG00000000247"
            ],
            "target": [],
            "expression_sum": {
                "class1": 20604.0,
                "class2": 11574.0
            },
            "ens_id": "ENSMUSG00000045690",
            "uniprot_ids": [
                "Q9D0R9"
            ]
        },
        "ENSMUSG00000021134": {
            "id": "ENSMUSG00000021134",
            "type": "Gene",
            "symbol": "Srsf5",
            "source": [
                "ENSMUSG00000000247"
            ],
            "target": [],
            "expression_sum": {
                "class1": 9204.0,
                "class2": 8472.0
            },
            "ens_id": "ENSMUSG00000021134",
            "uniprot_ids": [
                "O35326",
                "Q9D8S5",
                "S4R2G3"
            ]
        },
        "ENSMUSG00000021250": {
            "id": "ENSMUSG00000021250",
            "type": "TF",
            "symbol": "Fos",
            "source": [
                "ENSMUSG00000041515",
                "ENSMUSG00000056501",
                "ENSMUSG00000052684",
                "ENSMUSG00000037868",
                "ENSMUSG00000023034",
                "ENSMUSG00000052837",
                "ENSMUSG00000030103",
                "ENSMUSG00000000247",
                "ENSMUSG00000042812",
                "ENSMUSG00000057098",
                "ENSMUSG00000003545",
                "ENSMUSG00000003032",
                "ENSMUSG00000018899",
                "ENSMUSG00000071076"
            ],
            "target": [
                "ENSMUSG00000019505",
                "ENSMUSG00000006360",
                "ENSMUSG00000031765",
                "ENSMUSG00000024561",
                "ENSMUSG00000030222",
                "ENSMUSG00000028691",
                "ENSMUSG00000003545",
                "ENSMUSG00000036199",
                "ENSMUSG00000025289",
                "ENSMUSG00000032330",
                "ENSMUSG00000030930",
                "ENSMUSG00000004665",
                "ENSMUSG00000053560",
                "ENSMUSG00000036040",
                "ENSMUSG00000037411",
                "ENSMUSG00000036438",
                "ENSMUSG00000006699",
                "ENSMUSG00000008384",
                "ENSMUSG00000055148",
                "ENSMUSG00000018339",
                "ENSMUSG00000035493",
                "ENSMUSG00000021127",
                "ENSMUSG00000023004",
                "ENSMUSG00000029380",
                "ENSMUSG00000002416",
                "ENSMUSG00000007872",
                "ENSMUSG00000009013",
                "ENSMUSG00000036655",
                "ENSMUSG00000009681",
                "ENSMUSG00000036570",
                "ENSMUSG00000055839",
                "ENSMUSG00000075706",
                "ENSMUSG00000020053",
                "ENSMUSG00000037999",
                "ENSMUSG00000069601",
                "ENSMUSG00000032085",
                "ENSMUSG00000053113",
                "ENSMUSG00000037266",
                "ENSMUSG00000033538",
                "ENSMUSG00000084786",
                "ENSMUSG00000021520",
                "ENSMUSG00000052684",
                "ENSMUSG00000030122",
                "ENSMUSG00000055681",
                "ENSMUSG00000031762",
                "ENSMUSG00000003038",
                "ENSMUSG00000015501",
                "ENSMUSG00000038690",
                "ENSMUSG00000023915",
                "ENSMUSG00000030103",
                "ENSMUSG00000042812",
                "ENSMUSG00000028464",
                "ENSMUSG00000032515",
                "ENSMUSG00000074457",
                "ENSMUSG00000027829",
                "ENSMUSG00000037060",
                "ENSMUSG00000028195",
                "ENSMUSG00000031375",
                "ENSMUSG00000021253",
                "ENSMUSG00000005873",
                "ENSMUSG00000038463",
                "ENSMUSG00000002985",
                "ENSMUSG00000035678",
                "ENSMUSG00000059534",
                "ENSMUSG00000072625",
                "ENSMUSG00000015085",
                "ENSMUSG00000068220",
                "ENSMUSG00000023034",
                "ENSMUSG00000030654",
                "ENSMUSG00000054364",
                "ENSMUSG00000016494",
                "ENSMUSG00000056708",
                "ENSMUSG00000038393",
                "ENSMUSG00000037706",
                "ENSMUSG00000046727",
                "ENSMUSG00000001175",
                "ENSMUSG00000032359",
                "ENSMUSG00000061518",
                "ENSMUSG00000019960",
                "ENSMUSG00000015312",
                "ENSMUSG00000003032",
                "ENSMUSG00000037211",
                "ENSMUSG00000030247",
                "ENSMUSG00000020423",
                "ENSMUSG00000024048",
                "ENSMUSG00000023089",
                "ENSMUSG00000021876",
                "ENSMUSG00000037868",
                "ENSMUSG00000022122",
                "ENSMUSG00000006205",
                "ENSMUSG00000035828",
                "ENSMUSG00000057098",
                "ENSMUSG00000064215",
                "ENSMUSG00000018286",
                "ENSMUSG00000017493",
                "ENSMUSG00000039601",
                "ENSMUSG00000067818",
                "ENSMUSG00000056501",
                "ENSMUSG00000037152",
                "ENSMUSG00000024014",
                "ENSMUSG00000000628",
                "ENSMUSG00000090841",
                "ENSMUSG00000000753",
                "ENSMUSG00000071866",
                "ENSMUSG00000028864",
                "ENSMUSG00000036751",
                "ENSMUSG00000038370",
                "ENSMUSG00000001833",
                "ENSMUSG00000026385",
                "ENSMUSG00000024190",
                "ENSMUSG00000041515",
                "ENSMUSG00000007097",
                "ENSMUSG00000035356",
                "ENSMUSG00000052837",
                "ENSMUSG00000047945",
                "ENSMUSG00000073411",
                "ENSMUSG00000029632",
                "ENSMUSG00000030342",
                "ENSMUSG00000022956",
                "ENSMUSG00000044786",
                "ENSMUSG00000008540",
                "ENSMUSG00000018899",
                "ENSMUSG00000075702",
                "ENSMUSG00000027004",
                "ENSMUSG00000006403",
                "ENSMUSG00000018293",
                "ENSMUSG00000000247",
                "ENSMUSG00000038418",
                "ENSMUSG00000035783",
                "ENSMUSG00000030849",
                "ENSMUSG00000058135"
            ],
            "expression_sum": {
                "class1": 165566.0,
                "class2": 132750.0
            },
            "ens_id": "ENSMUSG00000021250",
            "uniprot_ids": [
                "P01101"
            ]
        },
        "ENSMUSG00000041841": {
            "id": "ENSMUSG00000041841",
            "type": "Gene",
            "symbol": "Rpl37",
            "source": [
                "ENSMUSG00000057098",
                "ENSMUSG00000038193",
                "ENSMUSG00000000247"
            ],
            "target": [],
            "expression_sum": {
                "class1": 163253.0,
                "class2": 80407.0
            },
            "ens_id": "ENSMUSG00000041841",
            "uniprot_ids": [
                "Q9D823"
            ]
        },
        "ENSMUSG00000058600": {
            "id": "ENSMUSG00000058600",
            "type": "Gene",
            "symbol": "Rpl30",
            "source": [
                "ENSMUSG00000048387",
                "ENSMUSG00000038193",
                "ENSMUSG00000000247"
            ],
            "target": [],
            "expression_sum": {
                "class1": 67497.0,
                "class2": 32572.0
            },
            "ens_id": "ENSMUSG00000058600",
            "uniprot_ids": [
                "P62889",
                "A0A2I3BQF4"
            ]
        },
        "ENSMUSG00000014313": {
            "id": "ENSMUSG00000014313",
            "type": "Gene",
            "symbol": "Cox6c",
            "source": [
                "ENSMUSG00000025498",
                "ENSMUSG00000032238",
                "ENSMUSG00000047638",
                "ENSMUSG00000005583",
                "ENSMUSG00000022708",
                "ENSMUSG00000000247",
                "ENSMUSG00000008575"
            ],
            "target": [],
            "expression_sum": {
                "class1": 46812.0,
                "class2": 33150.0
            },
            "ens_id": "ENSMUSG00000014313",
            "uniprot_ids": [
                "Q9CPQ1",
                "A0A0G2JDA6"
            ]
        },
        "ENSMUSG00000022283": {
            "id": "ENSMUSG00000022283",
            "type": "Gene",
            "symbol": "Pabpc1",
            "source": [
                "ENSMUSG00000057098",
                "ENSMUSG00000023034",
                "ENSMUSG00000000247"
            ],
            "target": [],
            "expression_sum": {
                "class1": 24894.0,
                "class2": 11628.0
            },
            "ens_id": "ENSMUSG00000022283",
            "uniprot_ids": [
                "P29341",
                "F6ZAX1",
                "A0A2I3BR37"
            ]
        },
        "ENSMUSG00000022312": {
            "id": "ENSMUSG00000022312",
            "type": "Gene",
            "symbol": "Eif3h",
            "source": [
                "ENSMUSG00000038193",
                "ENSMUSG00000000247"
            ],
            "target": [],
            "expression_sum": {
                "class1": 23378.0,
                "class2": 9687.0
            },
            "ens_id": "ENSMUSG00000022312",
            "uniprot_ids": [
                "Q91WK2"
            ]
        },
        "ENSMUSG00000022587": {
            "id": "ENSMUSG00000022587",
            "type": "Gene",
            "symbol": "Ly6e",
            "source": [
                "ENSMUSG00000032035",
                "ENSMUSG00000056501",
                "ENSMUSG00000023034",
                "ENSMUSG00000058756",
                "ENSMUSG00000000247",
                "ENSMUSG00000038193",
                "ENSMUSG00000029322",
                "ENSMUSG00000005583"
            ],
            "target": [],
            "expression_sum": {
                "class1": 220229.0,
                "class2": 12348.0
            },
            "ens_id": "ENSMUSG00000022587",
            "uniprot_ids": [
                "Q64253",
                "A0A087WPY4",
                "A0A087WNT2",
                "A0A087WQ65",
                "A0A087WRZ2",
                "A0A087WNS9"
            ]
        },
        "ENSMUSG00000055762": {
            "id": "ENSMUSG00000055762",
            "type": "Gene",
            "symbol": "Eef1d",
            "source": [
                "ENSMUSG00000057098",
                "ENSMUSG00000048387",
                "ENSMUSG00000000247"
            ],
            "target": [],
            "expression_sum": {
                "class1": 21520.0,
                "class2": 13050.0
            },
            "ens_id": "ENSMUSG00000055762",
            "uniprot_ids": [
                "A0A0R4J1L2",
                "Q9CVV2",
                "Q91VK2",
                "A0A0R4J1E2",
                "Q80T06",
                "D3YY68",
                "F6ZFU0",
                "D3Z7N2",
                "D3YZT9",
                "D3Z356",
                "D3YUQ9",
                "E9QN08",
                "E9PUM6"
            ]
        },
        "ENSMUSG00000003970": {
            "id": "ENSMUSG00000003970",
            "type": "Gene",
            "symbol": "Rpl8",
            "source": [
                "ENSMUSG00000047638",
                "ENSMUSG00000000247",
                "ENSMUSG00000038193",
                "ENSMUSG00000031965",
                "ENSMUSG00000048387"
            ],
            "target": [],
            "expression_sum": {
                "class1": 111387.0,
                "class2": 56751.0
            },
            "ens_id": "ENSMUSG00000003970",
            "uniprot_ids": [
                "P62918"
            ]
        },
        "ENSMUSG00000060036": {
            "id": "ENSMUSG00000060036",
            "type": "Gene",
            "symbol": "Rpl3",
            "source": [
                "ENSMUSG00000057098",
                "ENSMUSG00000022528",
                "ENSMUSG00000048387",
                "ENSMUSG00000000247"
            ],
            "target": [],
            "expression_sum": {
                "class1": 75012.0,
                "class2": 42977.0
            },
            "ens_id": "ENSMUSG00000060036",
            "uniprot_ids": [
                "P27659",
                "A0A2R8VHN4",
                "A0A087WQK0",
                "A0A087WNS0"
            ]
        },
        "ENSMUSG00000022450": {
            "id": "ENSMUSG00000022450",
            "type": "Gene",
            "symbol": "Ndufa6",
            "source": [
                "ENSMUSG00000047638",
                "ENSMUSG00000029322",
                "ENSMUSG00000000247"
            ],
            "target": [],
            "expression_sum": {
                "class1": 16151.0,
                "class2": 11374.0
            },
            "ens_id": "ENSMUSG00000022450",
            "uniprot_ids": [
                "Q9CQZ5",
                "A0A2R8W6V1",
                "A0A2R8VHX1",
                "A0A2R8VI90"
            ]
        },
        "ENSMUSG00000041736": {
            "id": "ENSMUSG00000041736",
            "type": "Gene",
            "symbol": "Tspo",
            "source": [
                "ENSMUSG00000032035",
                "ENSMUSG00000047638",
                "ENSMUSG00000000247",
                "ENSMUSG00000029322",
                "ENSMUSG00000003032",
                "ENSMUSG00000005583",
                "ENSMUSG00000071076"
            ],
            "target": [],
            "expression_sum": {
                "class1": 24678.0,
                "class2": 9465.0
            },
            "ens_id": "ENSMUSG00000041736",
            "uniprot_ids": [
                "P50637",
                "A0A140LIU9"
            ]
        },
        "ENSMUSG00000001289": {
            "id": "ENSMUSG00000001289",
            "type": "Gene",
            "symbol": "Pfdn5",
            "source": [
                "ENSMUSG00000029322",
                "ENSMUSG00000000247"
            ],
            "target": [],
            "expression_sum": {
                "class1": 37320.0,
                "class2": 18586.0
            },
            "ens_id": "ENSMUSG00000001289",
            "uniprot_ids": [
                "Q9WU28",
                "E9PVG3",
                "E9Q093",
                "H7BWX1",
                "E9PZ62",
                "E9Q5Q8"
            ]
        },
        "ENSMUSG00000056851": {
            "id": "ENSMUSG00000056851",
            "type": "Gene",
            "symbol": "Pcbp2",
            "source": [
                "ENSMUSG00000057098",
                "ENSMUSG00000000247"
            ],
            "target": [],
            "expression_sum": {
                "class1": 18132.0,
                "class2": 9385.0
            },
            "ens_id": "ENSMUSG00000056851",
            "uniprot_ids": [
                "Q61990",
                "B2M1R7",
                "A0A2R8VHP9",
                "Q3TT81",
                "A0A2R8W6H3",
                "A0A2R8VHG2",
                "A0A2R8VI12",
                "A0A2R8VI92",
                "A0A2R8VHZ0",
                "A0A2R8VKF0",
                "A0A2R8VHL8",
                "A0A2R8W6L5",
                "A0A2R8VI15",
                "A0A2R8VHI3",
                "A0A2R8VHY9",
                "A0A2R8VJY1",
                "A0A2R8VI71",
                "A0A2R8VI73",
                "A0A2R8VI25",
                "A0A2R8VKN0",
                "A0A2R8W6U6",
                "A0A2R8VHN3"
            ]
        },
        "ENSMUSG00000062683": {
            "id": "ENSMUSG00000062683",
            "type": "Gene",
            "symbol": "Atp5g2",
            "source": [
                "ENSMUSG00000000247"
            ],
            "target": [],
            "expression_sum": {
                "class1": 19624.0,
                "class2": 10947.0
            },
            "ens_id": "ENSMUSG00000062683",
            "uniprot_ids": [
                "P56383",
                "A0A087WQS5",
                "A0A087WSE2"
            ]
        },
        "ENSMUSG00000004070": {
            "id": "ENSMUSG00000004070",
            "type": "Gene",
            "symbol": "Hmox2",
            "source": [
                "ENSMUSG00000000247"
            ],
            "target": [],
            "expression_sum": {
                "class1": 10046.0,
                "class2": 2277.0
            },
            "ens_id": "ENSMUSG00000004070",
            "uniprot_ids": [
                "O70252",
                "D3Z4A2",
                "D3YX62",
                "D3YXN4"
            ]
        },
        "ENSMUSG00000022884": {
            "id": "ENSMUSG00000022884",
            "type": "Gene",
            "symbol": "Eif4a2",
            "source": [
                "ENSMUSG00000057098",
                "ENSMUSG00000047638",
                "ENSMUSG00000000247"
            ],
            "target": [],
            "expression_sum": {
                "class1": 26256.0,
                "class2": 12521.0
            },
            "ens_id": "ENSMUSG00000022884",
            "uniprot_ids": [
                "P10630",
                "Q8BTU6",
                "D6RJ11",
                "D6RJ60",
                "E9Q561",
                "A0A338P6X5"
            ]
        },
        "ENSMUSG00000060636": {
            "id": "ENSMUSG00000060636",
            "type": "Gene",
            "symbol": "Rpl35a",
            "source": [
                "ENSMUSG00000057098",
                "ENSMUSG00000048387",
                "ENSMUSG00000038193",
                "ENSMUSG00000000247"
            ],
            "target": [],
            "expression_sum": {
                "class1": 134254.0,
                "class2": 76798.0
            },
            "ens_id": "ENSMUSG00000060636",
            "uniprot_ids": [
                "O55142"
            ]
        },
        "ENSMUSG00000022708": {
            "id": "ENSMUSG00000022708",
            "type": "TF",
            "symbol": "Zbtb20",
            "source": [
                "ENSMUSG00000032035",
                "ENSMUSG00000023034",
                "ENSMUSG00000022528",
                "ENSMUSG00000000247",
                "ENSMUSG00000057098",
                "ENSMUSG00000038193",
                "ENSMUSG00000005583",
                "ENSMUSG00000071076"
            ],
            "target": [
                "ENSMUSG00000064354",
                "ENSMUSG00000002289",
                "ENSMUSG00000033585",
                "ENSMUSG00000014313",
                "ENSMUSG00000059173",
                "ENSMUSG00000063882",
                "ENSMUSG00000027931",
                "ENSMUSG00000064358",
                "ENSMUSG00000030930"
            ],
            "expression_sum": {
                "class1": 32148.0,
                "class2": 17872.0
            },
            "ens_id": "ENSMUSG00000022708",
            "uniprot_ids": [
                "Q8K0L9",
                "E0CX28",
                "E0CXK1",
                "E0CYH3",
                "E0CYJ9",
                "E0CXT9"
            ]
        },
        "ENSMUSG00000055447": {
            "id": "ENSMUSG00000055447",
            "type": "Gene",
            "symbol": "Cd47",
            "source": [
                "ENSMUSG00000032035",
                "ENSMUSG00000022528",
                "ENSMUSG00000005583",
                "ENSMUSG00000000247",
                "ENSMUSG00000057098",
                "ENSMUSG00000038193",
                "ENSMUSG00000055435",
                "ENSMUSG00000071076"
            ],
            "target": [],
            "expression_sum": {
                "class1": 22139.0,
                "class2": 9890.0
            },
            "ens_id": "ENSMUSG00000055447",
            "uniprot_ids": [
                "Q61735",
                "A0A2R8VJU9",
                "D3Z187",
                "A0A2R8VK70",
                "A0A2R8VI94",
                "A0A2R8W6P0",
                "A0A2R8VI30"
            ]
        },
        "ENSMUSG00000098274": {
            "id": "ENSMUSG00000098274",
            "type": "Gene",
            "symbol": "Rpl24",
            "source": [
                "ENSMUSG00000057098",
                "ENSMUSG00000048387",
                "ENSMUSG00000029322",
                "ENSMUSG00000000247"
            ],
            "target": [],
            "expression_sum": {
                "class1": 93017.0,
                "class2": 48216.0
            },
            "ens_id": "ENSMUSG00000098274",
            "uniprot_ids": [
                "Q8BP67"
            ]
        },
        "ENSMUSG00000022890": {
            "id": "ENSMUSG00000022890",
            "type": "Gene",
            "symbol": "Atp5j",
            "source": [
                "ENSMUSG00000000247"
            ],
            "target": [],
            "expression_sum": {
                "class1": 21353.0,
                "class2": 17339.0
            },
            "ens_id": "ENSMUSG00000022890",
            "uniprot_ids": [
                "P97450",
                "E9QAD6"
            ]
        },
        "ENSMUSG00000022892": {
            "id": "ENSMUSG00000022892",
            "type": "Gene",
            "symbol": "App",
            "source": [
                "ENSMUSG00000022528",
                "ENSMUSG00000032035",
                "ENSMUSG00000005583",
                "ENSMUSG00000000247"
            ],
            "target": [],
            "expression_sum": {
                "class1": 36897.0,
                "class2": 10799.0
            },
            "ens_id": "ENSMUSG00000022892",
            "uniprot_ids": [
                "P12023",
                "Q3TWF3",
                "A0A2I3BR03",
                "A0A2I3BQZ9",
                "A0A2I3BPT1"
            ]
        },
        "ENSMUSG00000022971": {
            "id": "ENSMUSG00000022971",
            "type": "Gene",
            "symbol": "Ifnar2",
            "source": [
                "ENSMUSG00000032035",
                "ENSMUSG00000022528",
                "ENSMUSG00000000247",
                "ENSMUSG00000038193",
                "ENSMUSG00000005583"
            ],
            "target": [],
            "expression_sum": {
                "class1": 24166.0,
                "class2": 2693.0
            },
            "ens_id": "ENSMUSG00000022971",
            "uniprot_ids": [
                "O35664",
                "H3BJ04",
                "H3BKB3"
            ]
        },
        "ENSMUSG00000023861": {
            "id": "ENSMUSG00000023861",
            "type": "Gene",
            "symbol": "Mpc1",
            "source": [
                "ENSMUSG00000025498",
                "ENSMUSG00000047638",
                "ENSMUSG00000000247"
            ],
            "target": [],
            "expression_sum": {
                "class1": 8887.0,
                "class2": 4917.0
            },
            "ens_id": "ENSMUSG00000023861",
            "uniprot_ids": [
                "P63030",
                "D3YWY6",
                "E9Q0V4",
                "D3Z5S0",
                "D3Z786"
            ]
        },
        "ENSMUSG00000014769": {
            "id": "ENSMUSG00000014769",
            "type": "Gene",
            "symbol": "Psmb1",
            "source": [
                "ENSMUSG00000057098",
                "ENSMUSG00000071076",
                "ENSMUSG00000000247"
            ],
            "target": [],
            "expression_sum": {
                "class1": 13539.0,
                "class2": 7755.0
            },
            "ens_id": "ENSMUSG00000014769",
            "uniprot_ids": [
                "O09061",
                "A0A338P7C0",
                "A0A338P7F1"
            ]
        },
        "ENSMUSG00000055839": {
            "id": "ENSMUSG00000055839",
            "type": "Gene",
            "symbol": "Tceb2",
            "source": [
                "ENSMUSG00000071076",
                "ENSMUSG00000021250",
                "ENSMUSG00000000247"
            ],
            "target": [],
            "expression_sum": {
                "class1": 33734.0,
                "class2": 19455.0
            },
            "ens_id": "ENSMUSG00000055839",
            "uniprot_ids": [
                "P62869",
                "A0A3B2WBM3"
            ]
        },
        "ENSMUSG00000044533": {
            "id": "ENSMUSG00000044533",
            "type": "Gene",
            "symbol": "Rps2",
            "source": [
                "ENSMUSG00000031965",
                "ENSMUSG00000057098",
                "ENSMUSG00000000247"
            ],
            "target": [],
            "expression_sum": {
                "class1": 97750.0,
                "class2": 62387.0
            },
            "ens_id": "ENSMUSG00000044533",
            "uniprot_ids": [
                "P25444",
                "D3YWJ3",
                "D3YVC1"
            ]
        },
        "ENSMUSG00000052146": {
            "id": "ENSMUSG00000052146",
            "type": "Gene",
            "symbol": "Rps10",
            "source": [
                "ENSMUSG00000023034",
                "ENSMUSG00000000247",
                "ENSMUSG00000057098",
                "ENSMUSG00000038193",
                "ENSMUSG00000048387"
            ],
            "target": [],
            "expression_sum": {
                "class1": 73238.0,
                "class2": 34552.0
            },
            "ens_id": "ENSMUSG00000052146",
            "uniprot_ids": [
                "P63325",
                "Q3UW83",
                "A0A338P731",
                "A0A338P7K4"
            ]
        },
        "ENSMUSG00000037805": {
            "id": "ENSMUSG00000037805",
            "type": "Gene",
            "symbol": "Rpl10a",
            "source": [
                "ENSMUSG00000022528",
                "ENSMUSG00000000247",
                "ENSMUSG00000057098",
                "ENSMUSG00000031965",
                "ENSMUSG00000053007"
            ],
            "target": [],
            "expression_sum": {
                "class1": 65589.0,
                "class2": 36794.0
            },
            "ens_id": "ENSMUSG00000037805",
            "uniprot_ids": [
                "Q5XJF6",
                "D6RE43",
                "A0A3B2WCA5",
                "A0A3B2WDD2",
                "A0A3B2W820",
                "A0A3B2WBL1",
                "A0A3B2W824"
            ]
        },
        "ENSMUSG00000067288": {
            "id": "ENSMUSG00000067288",
            "type": "Gene",
            "symbol": "Rps28",
            "source": [
                "ENSMUSG00000023034",
                "ENSMUSG00000000247",
                "ENSMUSG00000057098",
                "ENSMUSG00000038193",
                "ENSMUSG00000005583",
                "ENSMUSG00000031965"
            ],
            "target": [],
            "expression_sum": {
                "class1": 121311.0,
                "class2": 76057.0
            },
            "ens_id": "ENSMUSG00000067288",
            "uniprot_ids": [
                "P62858",
                "G3UYV7"
            ]
        },
        "ENSMUSG00000041881": {
            "id": "ENSMUSG00000041881",
            "type": "Gene",
            "symbol": "Ndufa7",
            "source": [
                "ENSMUSG00000000247"
            ],
            "target": [],
            "expression_sum": {
                "class1": 17440.0,
                "class2": 11894.0
            },
            "ens_id": "ENSMUSG00000041881",
            "uniprot_ids": [
                "Q9Z1P6",
                "G3UXZ4"
            ]
        },
        "ENSMUSG00000024308": {
            "id": "ENSMUSG00000024308",
            "type": "Gene",
            "symbol": "Tapbp",
            "source": [
                "ENSMUSG00000005583",
                "ENSMUSG00000000247",
                "ENSMUSG00000057098",
                "ENSMUSG00000003032",
                "ENSMUSG00000055435",
                "ENSMUSG00000071076"
            ],
            "target": [],
            "expression_sum": {
                "class1": 14455.0,
                "class2": 5011.0
            },
            "ens_id": "ENSMUSG00000024308",
            "uniprot_ids": [
                "Q9R233",
                "Q3TCU5",
                "G3UZZ2",
                "A0A3Q4L2T5",
                "A0A3Q4EGL0"
            ]
        },
        "ENSMUSG00000008668": {
            "id": "ENSMUSG00000008668",
            "type": "Gene",
            "symbol": "Rps18",
            "source": [
                "ENSMUSG00000023034",
                "ENSMUSG00000000247",
                "ENSMUSG00000057098",
                "ENSMUSG00000038193",
                "ENSMUSG00000005583",
                "ENSMUSG00000048387"
            ],
            "target": [],
            "expression_sum": {
                "class1": 133850.0,
                "class2": 65639.0
            },
            "ens_id": "ENSMUSG00000008668",
            "uniprot_ids": [
                "P62270",
                "A0A3Q4EGP3",
                "S4R1N6"
            ]
        },
        "ENSMUSG00000023944": {
            "id": "ENSMUSG00000023944",
            "type": "Gene",
            "symbol": "Hsp90ab1",
            "source": [
                "ENSMUSG00000025498",
                "ENSMUSG00000032035",
                "ENSMUSG00000056501",
                "ENSMUSG00000047638",
                "ENSMUSG00000000247",
                "ENSMUSG00000057098",
                "ENSMUSG00000048387"
            ],
            "target": [],
            "expression_sum": {
                "class1": 80261.0,
                "class2": 31355.0
            },
            "ens_id": "ENSMUSG00000023944",
            "uniprot_ids": [
                "P11499",
                "E9PX27",
                "E9Q0C3",
                "E9Q3D6",
                "D3Z1R1"
            ]
        },
        "ENSMUSG00000023942": {
            "id": "ENSMUSG00000023942",
            "type": "Gene",
            "symbol": "Slc29a1",
            "source": [
                "ENSMUSG00000038193",
                "ENSMUSG00000005583",
                "ENSMUSG00000000247"
            ],
            "target": [],
            "expression_sum": {
                "class1": 14253.0,
                "class2": 6240.0
            },
            "ens_id": "ENSMUSG00000023942",
            "uniprot_ids": [
                "Q9JIM1",
                "E9PWY7",
                "E9Q0Q5",
                "E9Q503",
                "E9Q221",
                "E9PVH9",
                "E9PXQ6",
                "E9PXM6",
                "E9Q7R8",
                "E9Q5E9",
                "E9PWD6",
                "E9PZV0",
                "E9Q3D1"
            ]
        },
        "ENSMUSG00000057863": {
            "id": "ENSMUSG00000057863",
            "type": "Gene",
            "symbol": "Rpl36",
            "source": [
                "ENSMUSG00000057098",
                "ENSMUSG00000048387",
                "ENSMUSG00000038193",
                "ENSMUSG00000000247"
            ],
            "target": [],
            "expression_sum": {
                "class1": 99641.0,
                "class2": 51865.0
            },
            "ens_id": "ENSMUSG00000057863",
            "uniprot_ids": [
                "Q6ZWZ4"
            ]
        },
        "ENSMUSG00000024248": {
            "id": "ENSMUSG00000024248",
            "type": "Gene",
            "symbol": "Cox7a2l",
            "source": [
                "ENSMUSG00000000247",
                "ENSMUSG00000057098",
                "ENSMUSG00000038193",
                "ENSMUSG00000029322",
                "ENSMUSG00000048387"
            ],
            "target": [],
            "expression_sum": {
                "class1": 32809.0,
                "class2": 11956.0
            },
            "ens_id": "ENSMUSG00000024248",
            "uniprot_ids": [
                "Q61387",
                "E9PZS8"
            ]
        },
        "ENSMUSG00000014294": {
            "id": "ENSMUSG00000014294",
            "type": "Gene",
            "symbol": "Ndufa2",
            "source": [
                "ENSMUSG00000029322",
                "ENSMUSG00000000247"
            ],
            "target": [],
            "expression_sum": {
                "class1": 20804.0,
                "class2": 15777.0
            },
            "ens_id": "ENSMUSG00000014294",
            "uniprot_ids": [
                "Q9CQ75"
            ]
        },
        "ENSMUSG00000024608": {
            "id": "ENSMUSG00000024608",
            "type": "Gene",
            "symbol": "Rps14",
            "source": [
                "ENSMUSG00000057098",
                "ENSMUSG00000048387",
                "ENSMUSG00000038193",
                "ENSMUSG00000000247"
            ],
            "target": [],
            "expression_sum": {
                "class1": 161001.0,
                "class2": 83793.0
            },
            "ens_id": "ENSMUSG00000024608",
            "uniprot_ids": [
                "P62264",
                "A0A494BAS4",
                "D3Z7I1",
                "D3YVF4"
            ]
        },
        "ENSMUSG00000024576": {
            "id": "ENSMUSG00000024576",
            "type": "Gene",
            "symbol": "Csnk1a1",
            "source": [
                "ENSMUSG00000029322",
                "ENSMUSG00000000247"
            ],
            "target": [],
            "expression_sum": {
                "class1": 15248.0,
                "class2": 7129.0
            },
            "ens_id": "ENSMUSG00000024576",
            "uniprot_ids": [
                "Q8BK63",
                "F6YBC9",
                "E9Q3W1",
                "H7BXB1",
                "E9Q2U6",
                "E9Q4G7",
                "E9PWB2"
            ]
        },
        "ENSMUSG00000025428": {
            "id": "ENSMUSG00000025428",
            "type": "Gene",
            "symbol": "Atp5a1",
            "source": [
                "ENSMUSG00000047638",
                "ENSMUSG00000005583",
                "ENSMUSG00000000247"
            ],
            "target": [],
            "expression_sum": {
                "class1": 12569.0,
                "class2": 11946.0
            },
            "ens_id": "ENSMUSG00000025428",
            "uniprot_ids": [
                "Q03265",
                "D6RJ16",
                "D3Z6F5"
            ]
        },
        "ENSMUSG00000056201": {
            "id": "ENSMUSG00000056201",
            "type": "Gene",
            "symbol": "Cfl1",
            "source": [
                "ENSMUSG00000057098",
                "ENSMUSG00000000247"
            ],
            "target": [],
            "expression_sum": {
                "class1": 43733.0,
                "class2": 22035.0
            },
            "ens_id": "ENSMUSG00000056201",
            "uniprot_ids": [
                "P18760",
                "A0A494B9A7",
                "F8WGL3"
            ]
        },
        "ENSMUSG00000024953": {
            "id": "ENSMUSG00000024953",
            "type": "Gene",
            "symbol": "Prdx5",
            "source": [
                "ENSMUSG00000057098",
                "ENSMUSG00000000247"
            ],
            "target": [],
            "expression_sum": {
                "class1": 14444.0,
                "class2": 8439.0
            },
            "ens_id": "ENSMUSG00000024953",
            "uniprot_ids": [
                "P99029",
                "A0A494BAZ4",
                "H3BJQ7",
                "G3UZJ4"
            ]
        },
        "ENSMUSG00000035885": {
            "id": "ENSMUSG00000035885",
            "type": "Gene",
            "symbol": "Cox8a",
            "source": [
                "ENSMUSG00000000247",
                "ENSMUSG00000053477",
                "ENSMUSG00000008575",
                "ENSMUSG00000005583",
                "ENSMUSG00000020484"
            ],
            "target": [],
            "expression_sum": {
                "class1": 33978.0,
                "class2": 36956.0
            },
            "ens_id": "ENSMUSG00000035885",
            "uniprot_ids": [
                "Q64445",
                "A0A494B971"
            ]
        },
        "ENSMUSG00000071644": {
            "id": "ENSMUSG00000071644",
            "type": "Gene",
            "symbol": "Eef1g",
            "source": [
                "ENSMUSG00000057098",
                "ENSMUSG00000000247"
            ],
            "target": [],
            "expression_sum": {
                "class1": 39745.0,
                "class2": 18974.0
            },
            "ens_id": "ENSMUSG00000071644",
            "uniprot_ids": [
                "Q9D8N0"
            ]
        },
        "ENSMUSG00000024678": {
            "id": "ENSMUSG00000024678",
            "type": "Gene",
            "symbol": "Ms4a4d",
            "source": [
                "ENSMUSG00000032035",
                "ENSMUSG00000000247"
            ],
            "target": [],
            "expression_sum": {
                "class1": 25630.0,
                "class2": 5742.0
            },
            "ens_id": "ENSMUSG00000024678",
            "uniprot_ids": [
                "Q99N05"
            ]
        },
        "ENSMUSG00000055044": {
            "id": "ENSMUSG00000055044",
            "type": "Gene",
            "symbol": "Pdlim1",
            "source": [
                "ENSMUSG00000057098",
                "ENSMUSG00000000247"
            ],
            "target": [],
            "expression_sum": {
                "class1": 23558.0,
                "class2": 2266.0
            },
            "ens_id": "ENSMUSG00000055044",
            "uniprot_ids": [
                "O70400",
                "S4R1V0"
            ]
        },
        "ENSMUSG00000064351": {
            "id": "ENSMUSG00000064351",
            "type": "Gene",
            "symbol": "mt-Co1",
            "source": [
                "ENSMUSG00000032035",
                "ENSMUSG00000047638",
                "ENSMUSG00000000247"
            ],
            "target": [],
            "expression_sum": {
                "class1": 169485.0,
                "class2": 153327.0
            },
            "ens_id": "ENSMUSG00000064351",
            "uniprot_ids": [
                "P00397"
            ]
        },
        "ENSMUSG00000064370": {
            "id": "ENSMUSG00000064370",
            "type": "Gene",
            "symbol": "mt-Cytb",
            "source": [
                "ENSMUSG00000032238",
                "ENSMUSG00000052684",
                "ENSMUSG00000000247",
                "ENSMUSG00000005583",
                "ENSMUSG00000020484"
            ],
            "target": [],
            "expression_sum": {
                "class1": 90043.0,
                "class2": 129761.0
            },
            "ens_id": "ENSMUSG00000064370",
            "uniprot_ids": [
                "P00158"
            ]
        },
        "ENSMUSG00000015839": {
            "id": "ENSMUSG00000015839",
            "type": "TF",
            "symbol": "Nfe2l2",
            "source": [
                "ENSMUSG00000057098"
            ],
            "target": [
                "ENSMUSG00000015354",
                "ENSMUSG00000014846",
                "ENSMUSG00000038463",
                "ENSMUSG00000053007",
                "ENSMUSG00000039601"
            ],
            "expression_sum": {
                "class1": 5415.0,
                "class2": 3955.0
            },
            "ens_id": "ENSMUSG00000015839",
            "uniprot_ids": [
                "Q60795"
            ]
        },
        "ENSMUSG00000038463": {
            "id": "ENSMUSG00000038463",
            "type": "Gene",
            "symbol": "Olfml2b",
            "source": [
                "ENSMUSG00000052684",
                "ENSMUSG00000022346",
                "ENSMUSG00000005583",
                "ENSMUSG00000015839",
                "ENSMUSG00000021250",
                "ENSMUSG00000003032",
                "ENSMUSG00000008575",
                "ENSMUSG00000071076"
            ],
            "target": [],
            "expression_sum": {
                "class1": 566.0,
                "class2": 4694.0
            },
            "ens_id": "ENSMUSG00000038463",
            "uniprot_ids": [
                "Q3V1G4"
            ]
        },
        "ENSMUSG00000053007": {
            "id": "ENSMUSG00000053007",
            "type": "TF",
            "symbol": "Creb5",
            "source": [
                "ENSMUSG00000015839",
                "ENSMUSG00000005583",
                "ENSMUSG00000003032",
                "ENSMUSG00000071076"
            ],
            "target": [
                "ENSMUSG00000038991",
                "ENSMUSG00000027204",
                "ENSMUSG00000044337",
                "ENSMUSG00000037805",
                "ENSMUSG00000026193",
                "ENSMUSG00000004460",
                "ENSMUSG00000062647"
            ],
            "expression_sum": {
                "class1": 272.0,
                "class2": 2430.0
            },
            "ens_id": "ENSMUSG00000053007",
            "uniprot_ids": [
                "A0A0N4SV71",
                "A0A0N4SV26",
                "A0A0N4SUK5",
                "A0A0N4SUW5"
            ]
        },
        "ENSMUSG00000014846": {
            "id": "ENSMUSG00000014846",
            "type": "Gene",
            "symbol": "Tppp3",
            "source": [
                "ENSMUSG00000015839",
                "ENSMUSG00000057098",
                "ENSMUSG00000003032",
                "ENSMUSG00000008575",
                "ENSMUSG00000020484",
                "ENSMUSG00000071076"
            ],
            "target": [],
            "expression_sum": {
                "class1": 163.0,
                "class2": 20143.0
            },
            "ens_id": "ENSMUSG00000014846",
            "uniprot_ids": [
                "Q9CRB6"
            ]
        },
        "ENSMUSG00000015354": {
            "id": "ENSMUSG00000015354",
            "type": "Gene",
            "symbol": "Pcolce2",
            "source": [
                "ENSMUSG00000015839",
                "ENSMUSG00000022346",
                "ENSMUSG00000003032",
                "ENSMUSG00000005583"
            ],
            "target": [],
            "expression_sum": {
                "class1": 29.0,
                "class2": 7559.0
            },
            "ens_id": "ENSMUSG00000015354",
            "uniprot_ids": [
                "Q8R4W6"
            ]
        },
        "ENSMUSG00000056501": {
            "id": "ENSMUSG00000056501",
            "type": "TF",
            "symbol": "Cebpb",
            "source": [
                "ENSMUSG00000041515",
                "ENSMUSG00000052684",
                "ENSMUSG00000030103",
                "ENSMUSG00000021250",
                "ENSMUSG00000071076",
                "ENSMUSG00000018899"
            ],
            "target": [
                "ENSMUSG00000031765",
                "ENSMUSG00000003545",
                "ENSMUSG00000031320",
                "ENSMUSG00000030930",
                "ENSMUSG00000037411",
                "ENSMUSG00000037405",
                "ENSMUSG00000028268",
                "ENSMUSG00000008384",
                "ENSMUSG00000055148",
                "ENSMUSG00000035493",
                "ENSMUSG00000021127",
                "ENSMUSG00000028003",
                "ENSMUSG00000029380",
                "ENSMUSG00000031119",
                "ENSMUSG00000007872",
                "ENSMUSG00000049115",
                "ENSMUSG00000031845",
                "ENSMUSG00000061983",
                "ENSMUSG00000050953",
                "ENSMUSG00000061353",
                "ENSMUSG00000036655",
                "ENSMUSG00000009681",
                "ENSMUSG00000037999",
                "ENSMUSG00000037742",
                "ENSMUSG00000053113",
                "ENSMUSG00000033538",
                "ENSMUSG00000052684",
                "ENSMUSG00000070436",
                "ENSMUSG00000031762",
                "ENSMUSG00000027188",
                "ENSMUSG00000015501",
                "ENSMUSG00000030103",
                "ENSMUSG00000029163",
                "ENSMUSG00000022587",
                "ENSMUSG00000032515",
                "ENSMUSG00000045128",
                "ENSMUSG00000033880",
                "ENSMUSG00000023944",
                "ENSMUSG00000027829",
                "ENSMUSG00000031375",
                "ENSMUSG00000039377",
                "ENSMUSG00000034855",
                "ENSMUSG00000035678",
                "ENSMUSG00000002985",
                "ENSMUSG00000072625",
                "ENSMUSG00000054072",
                "ENSMUSG00000056708",
                "ENSMUSG00000060126",
                "ENSMUSG00000042453",
                "ENSMUSG00000021025",
                "ENSMUSG00000022665",
                "ENSMUSG00000025498",
                "ENSMUSG00000040128",
                "ENSMUSG00000035373",
                "ENSMUSG00000020423",
                "ENSMUSG00000024371",
                "ENSMUSG00000020205",
                "ENSMUSG00000037868",
                "ENSMUSG00000028214",
                "ENSMUSG00000023224",
                "ENSMUSG00000039601",
                "ENSMUSG00000024014",
                "ENSMUSG00000000628",
                "ENSMUSG00000060550",
                "ENSMUSG00000028864",
                "ENSMUSG00000062328",
                "ENSMUSG00000032179",
                "ENSMUSG00000021250",
                "ENSMUSG00000007892",
                "ENSMUSG00000035385",
                "ENSMUSG00000041515",
                "ENSMUSG00000035356",
                "ENSMUSG00000047945",
                "ENSMUSG00000073411",
                "ENSMUSG00000019929",
                "ENSMUSG00000044786",
                "ENSMUSG00000071076",
                "ENSMUSG00000018899",
                "ENSMUSG00000027004",
                "ENSMUSG00000006403",
                "ENSMUSG00000032501",
                "ENSMUSG00000000247",
                "ENSMUSG00000030849",
                "ENSMUSG00000073489"
            ],
            "expression_sum": {
                "class1": 56111.0,
                "class2": 21063.0
            },
            "ens_id": "ENSMUSG00000056501",
            "uniprot_ids": [
                "P28033"
            ]
        },
        "ENSMUSG00000039377": {
            "id": "ENSMUSG00000039377",
            "type": "TF",
            "symbol": "Hlx",
            "source": [
                "ENSMUSG00000022528",
                "ENSMUSG00000056501"
            ],
            "target": [],
            "expression_sum": {
                "class1": 12466.0,
                "class2": 2119.0
            },
            "ens_id": "ENSMUSG00000039377",
            "uniprot_ids": [
                "Q61670",
                "G3V024"
            ]
        },
        "ENSMUSG00000027004": {
            "id": "ENSMUSG00000027004",
            "type": "Gene",
            "symbol": "Frzb",
            "source": [
                "ENSMUSG00000038193",
                "ENSMUSG00000056501",
                "ENSMUSG00000021250"
            ],
            "target": [],
            "expression_sum": {
                "class1": 10303.0,
                "class2": 120.0
            },
            "ens_id": "ENSMUSG00000027004",
            "uniprot_ids": [
                "P97401"
            ]
        },
        "ENSMUSG00000023224": {
            "id": "ENSMUSG00000023224",
            "type": "Gene",
            "symbol": "Serping1",
            "source": [
                "ENSMUSG00000056501",
                "ENSMUSG00000023034",
                "ENSMUSG00000058756",
                "ENSMUSG00000038193",
                "ENSMUSG00000005583",
                "ENSMUSG00000055148"
            ],
            "target": [],
            "expression_sum": {
                "class1": 139210.0,
                "class2": 54316.0
            },
            "ens_id": "ENSMUSG00000023224",
            "uniprot_ids": [
                "P97290",
                "A2ATR8"
            ]
        },
        "ENSMUSG00000027188": {
            "id": "ENSMUSG00000027188",
            "type": "Gene",
            "symbol": "Pamr1",
            "source": [
                "ENSMUSG00000023034",
                "ENSMUSG00000056501",
                "ENSMUSG00000005583"
            ],
            "target": [],
            "expression_sum": {
                "class1": 8160.0,
                "class2": 1252.0
            },
            "ens_id": "ENSMUSG00000027188",
            "uniprot_ids": [
                "Q8BU25"
            ]
        },
        "ENSMUSG00000031119": {
            "id": "ENSMUSG00000031119",
            "type": "Gene",
            "symbol": "Gpc4",
            "source": [
                "ENSMUSG00000056501",
                "ENSMUSG00000042812",
                "ENSMUSG00000038418",
                "ENSMUSG00000038193",
                "ENSMUSG00000029322"
            ],
            "target": [],
            "expression_sum": {
                "class1": 19021.0,
                "class2": 1236.0
            },
            "ens_id": "ENSMUSG00000031119",
            "uniprot_ids": [
                "P51655"
            ]
        },
        "ENSMUSG00000031375": {
            "id": "ENSMUSG00000031375",
            "type": "Gene",
            "symbol": "Bgn",
            "source": [
                "ENSMUSG00000042812",
                "ENSMUSG00000032035",
                "ENSMUSG00000021250",
                "ENSMUSG00000056501"
            ],
            "target": [],
            "expression_sum": {
                "class1": 393016.0,
                "class2": 87690.0
            },
            "ens_id": "ENSMUSG00000031375",
            "uniprot_ids": [
                "P28653"
            ]
        },
        "ENSMUSG00000031320": {
            "id": "ENSMUSG00000031320",
            "type": "Gene",
            "symbol": "Rps4x",
            "source": [
                "ENSMUSG00000056501",
                "ENSMUSG00000023034",
                "ENSMUSG00000038193",
                "ENSMUSG00000005583",
                "ENSMUSG00000048387"
            ],
            "target": [],
            "expression_sum": {
                "class1": 147664.0,
                "class2": 77702.0
            },
            "ens_id": "ENSMUSG00000031320",
            "uniprot_ids": [
                "P62702"
            ]
        },
        "ENSMUSG00000028268": {
            "id": "ENSMUSG00000028268",
            "type": "Gene",
            "symbol": "Gbp3",
            "source": [
                "ENSMUSG00000056501"
            ],
            "target": [],
            "expression_sum": {
                "class1": 7171.0,
                "class2": 361.0
            },
            "ens_id": "ENSMUSG00000028268",
            "uniprot_ids": [
                "Q61107",
                "D3YVW6",
                "D3YUH5",
                "A0A0G2JDL1"
            ]
        },
        "ENSMUSG00000028214": {
            "id": "ENSMUSG00000028214",
            "type": "Gene",
            "symbol": "Gem",
            "source": [
                "ENSMUSG00000025498",
                "ENSMUSG00000056501",
                "ENSMUSG00000037868",
                "ENSMUSG00000023034",
                "ENSMUSG00000052837",
                "ENSMUSG00000038193"
            ],
            "target": [],
            "expression_sum": {
                "class1": 23169.0,
                "class2": 7430.0
            },
            "ens_id": "ENSMUSG00000028214",
            "uniprot_ids": [
                "P55041"
            ]
        },
        "ENSMUSG00000028864": {
            "id": "ENSMUSG00000028864",
            "type": "Gene",
            "symbol": "Hgf",
            "source": [
                "ENSMUSG00000056501",
                "ENSMUSG00000052684",
                "ENSMUSG00000023034",
                "ENSMUSG00000052837",
                "ENSMUSG00000021250",
                "ENSMUSG00000055148",
                "ENSMUSG00000071076"
            ],
            "target": [],
            "expression_sum": {
                "class1": 19906.0,
                "class2": 466.0
            },
            "ens_id": "ENSMUSG00000028864",
            "uniprot_ids": [
                "Q08048"
            ]
        },
        "ENSMUSG00000029163": {
            "id": "ENSMUSG00000029163",
            "type": "Gene",
            "symbol": "Emilin1",
            "source": [
                "ENSMUSG00000032035",
                "ENSMUSG00000056501"
            ],
            "target": [],
            "expression_sum": {
                "class1": 38332.0,
                "class2": 6016.0
            },
            "ens_id": "ENSMUSG00000029163",
            "uniprot_ids": [
                "Q99K41"
            ]
        },
        "ENSMUSG00000061353": {
            "id": "ENSMUSG00000061353",
            "type": "Gene",
            "symbol": "Cxcl12",
            "source": [
                "ENSMUSG00000056501",
                "ENSMUSG00000023034",
                "ENSMUSG00000029322",
                "ENSMUSG00000005583",
                "ENSMUSG00000055148"
            ],
            "target": [],
            "expression_sum": {
                "class1": 297139.0,
                "class2": 20671.0
            },
            "ens_id": "ENSMUSG00000061353",
            "uniprot_ids": [
                "P40224",
                "H7BX38"
            ]
        },
        "ENSMUSG00000002985": {
            "id": "ENSMUSG00000002985",
            "type": "Gene",
            "symbol": "Apoe",
            "source": [
                "ENSMUSG00000056501",
                "ENSMUSG00000023034",
                "ENSMUSG00000021250",
                "ENSMUSG00000038193",
                "ENSMUSG00000076431",
                "ENSMUSG00000005583"
            ],
            "target": [],
            "expression_sum": {
                "class1": 406049.0,
                "class2": 100210.0
            },
            "ens_id": "ENSMUSG00000002985",
            "uniprot_ids": [
                "P08226",
                "G3UWW2",
                "G3UZM8",
                "A0A1B0GX15",
                "G3UWN5"
            ]
        },
        "ENSMUSG00000070436": {
            "id": "ENSMUSG00000070436",
            "type": "Gene",
            "symbol": "Serpinh1",
            "source": [
                "ENSMUSG00000032035",
                "ENSMUSG00000056501",
                "ENSMUSG00000058756",
                "ENSMUSG00000038193",
                "ENSMUSG00000005583"
            ],
            "target": [],
            "expression_sum": {
                "class1": 113657.0,
                "class2": 49248.0
            },
            "ens_id": "ENSMUSG00000070436",
            "uniprot_ids": [
                "P19324",
                "A0A140LHR4",
                "A0A140LHD6",
                "A0A140LHK0"
            ]
        },
        "ENSMUSG00000030930": {
            "id": "ENSMUSG00000030930",
            "type": "Gene",
            "symbol": "Chst15",
            "source": [
                "ENSMUSG00000056501",
                "ENSMUSG00000052837",
                "ENSMUSG00000022708",
                "ENSMUSG00000021250",
                "ENSMUSG00000038193",
                "ENSMUSG00000071076"
            ],
            "target": [],
            "expression_sum": {
                "class1": 13454.0,
                "class2": 1073.0
            },
            "ens_id": "ENSMUSG00000030930",
            "uniprot_ids": [
                "Q91XQ5"
            ]
        },
        "ENSMUSG00000025498": {
            "id": "ENSMUSG00000025498",
            "type": "TF",
            "symbol": "Irf7",
            "source": [
                "ENSMUSG00000056501"
            ],
            "target": [
                "ENSMUSG00000025393",
                "ENSMUSG00000019505",
                "ENSMUSG00000067212",
                "ENSMUSG00000014313",
                "ENSMUSG00000008348",
                "ENSMUSG00000022285",
                "ENSMUSG00000028410",
                "ENSMUSG00000027215",
                "ENSMUSG00000025647",
                "ENSMUSG00000014226",
                "ENSMUSG00000021270",
                "ENSMUSG00000026238",
                "ENSMUSG00000024966",
                "ENSMUSG00000021870",
                "ENSMUSG00000061232",
                "ENSMUSG00000026104",
                "ENSMUSG00000023861",
                "ENSMUSG00000063882",
                "ENSMUSG00000033880",
                "ENSMUSG00000023944",
                "ENSMUSG00000034708",
                "ENSMUSG00000015656",
                "ENSMUSG00000028214",
                "ENSMUSG00000031167",
                "ENSMUSG00000056116",
                "ENSMUSG00000070493"
            ],
            "expression_sum": {
                "class1": 21500.0,
                "class2": 520.0
            },
            "ens_id": "ENSMUSG00000025498",
            "uniprot_ids": [
                "P70434",
                "Q3TW14",
                "D3Z4U9",
                "F6TQ89",
                "A0A1B0GRH7"
            ]
        },
        "ENSMUSG00000050953": {
            "id": "ENSMUSG00000050953",
            "type": "Gene",
            "symbol": "Gja1",
            "source": [
                "ENSMUSG00000038193",
                "ENSMUSG00000056501"
            ],
            "target": [],
            "expression_sum": {
                "class1": 6186.0,
                "class2": 3105.0
            },
            "ens_id": "ENSMUSG00000050953",
            "uniprot_ids": [
                "P23242",
                "A0A1W2P793",
                "A0A1W2P8A3"
            ]
        },
        "ENSMUSG00000019929": {
            "id": "ENSMUSG00000019929",
            "type": "Gene",
            "symbol": "Dcn",
            "source": [
                "ENSMUSG00000032035",
                "ENSMUSG00000056501",
                "ENSMUSG00000038193",
                "ENSMUSG00000005583",
                "ENSMUSG00000028565",
                "ENSMUSG00000071076"
            ],
            "target": [],
            "expression_sum": {
                "class1": 1401173.0,
                "class2": 310161.0
            },
            "ens_id": "ENSMUSG00000019929",
            "uniprot_ids": [
                "P28654",
                "A0A1W2P6I8"
            ]
        },
        "ENSMUSG00000055148": {
            "id": "ENSMUSG00000055148",
            "type": "TF",
            "symbol": "Klf2",
            "source": [
                "ENSMUSG00000041515",
                "ENSMUSG00000056501",
                "ENSMUSG00000037868",
                "ENSMUSG00000052684",
                "ENSMUSG00000023034",
                "ENSMUSG00000052837",
                "ENSMUSG00000022346",
                "ENSMUSG00000030103",
                "ENSMUSG00000038418",
                "ENSMUSG00000021250",
                "ENSMUSG00000003032",
                "ENSMUSG00000071076"
            ],
            "target": [
                "ENSMUSG00000067818",
                "ENSMUSG00000006360",
                "ENSMUSG00000031765",
                "ENSMUSG00000000628",
                "ENSMUSG00000053113",
                "ENSMUSG00000056708",
                "ENSMUSG00000003545",
                "ENSMUSG00000053560",
                "ENSMUSG00000037411",
                "ENSMUSG00000028864",
                "ENSMUSG00000037405",
                "ENSMUSG00000017009",
                "ENSMUSG00000027074",
                "ENSMUSG00000008384",
                "ENSMUSG00000031762",
                "ENSMUSG00000007877",
                "ENSMUSG00000042622",
                "ENSMUSG00000021127",
                "ENSMUSG00000024190",
                "ENSMUSG00000037211",
                "ENSMUSG00000073411",
                "ENSMUSG00000020423",
                "ENSMUSG00000023064",
                "ENSMUSG00000022010",
                "ENSMUSG00000024371",
                "ENSMUSG00000026255",
                "ENSMUSG00000024232",
                "ENSMUSG00000044786",
                "ENSMUSG00000018899",
                "ENSMUSG00000090698",
                "ENSMUSG00000061353",
                "ENSMUSG00000057666",
                "ENSMUSG00000034855",
                "ENSMUSG00000023224",
                "ENSMUSG00000009681"
            ],
            "expression_sum": {
                "class1": 44466.0,
                "class2": 38463.0
            },
            "ens_id": "ENSMUSG00000055148",
            "uniprot_ids": [
                "Q60843"
            ]
        },
        "ENSMUSG00000031845": {
            "id": "ENSMUSG00000031845",
            "type": "Gene",
            "symbol": "Bco1",
            "source": [
                "ENSMUSG00000056501",
                "ENSMUSG00000052684"
            ],
            "target": [],
            "expression_sum": {
                "class1": 10153.0,
                "class2": 11.0
            },
            "ens_id": "ENSMUSG00000031845",
            "uniprot_ids": [
                "Q9JJS6",
                "E9Q321"
            ]
        },
        "ENSMUSG00000035385": {
            "id": "ENSMUSG00000035385",
            "type": "Gene",
            "symbol": "Ccl2",
            "source": [
                "ENSMUSG00000056501"
            ],
            "target": [],
            "expression_sum": {
                "class1": 21475.0,
                "class2": 503.0
            },
            "ens_id": "ENSMUSG00000035385",
            "uniprot_ids": [
                "P10148"
            ]
        },
        "ENSMUSG00000035373": {
            "id": "ENSMUSG00000035373",
            "type": "Gene",
            "symbol": "Ccl7",
            "source": [
                "ENSMUSG00000056501"
            ],
            "target": [],
            "expression_sum": {
                "class1": 3576.0,
                "class2": 760.0
            },
            "ens_id": "ENSMUSG00000035373",
            "uniprot_ids": [
                "Q03366"
            ]
        },
        "ENSMUSG00000049115": {
            "id": "ENSMUSG00000049115",
            "type": "Gene",
            "symbol": "Agtr1a",
            "source": [
                "ENSMUSG00000052837",
                "ENSMUSG00000056501"
            ],
            "target": [],
            "expression_sum": {
                "class1": 37643.0,
                "class2": 775.0
            },
            "ens_id": "ENSMUSG00000049115",
            "uniprot_ids": [
                "P29754",
                "A0A1Y7VIT7",
                "A0A1Y7VMR7",
                "A0A1Y7VL16",
                "A0A1Y7VJY2"
            ]
        },
        "ENSMUSG00000035493": {
            "id": "ENSMUSG00000035493",
            "type": "Gene",
            "symbol": "Tgfbi",
            "source": [
                "ENSMUSG00000056501",
                "ENSMUSG00000023034",
                "ENSMUSG00000042812",
                "ENSMUSG00000038193",
                "ENSMUSG00000021250",
                "ENSMUSG00000005583",
                "ENSMUSG00000018899"
            ],
            "target": [],
            "expression_sum": {
                "class1": 102105.0,
                "class2": 9504.0
            },
            "ens_id": "ENSMUSG00000035493",
            "uniprot_ids": [
                "P82198",
                "Q3UXJ2"
            ]
        },
        "ENSMUSG00000022665": {
            "id": "ENSMUSG00000022665",
            "type": "Gene",
            "symbol": "Ccdc80",
            "source": [
                "ENSMUSG00000032035",
                "ENSMUSG00000056501",
                "ENSMUSG00000038193",
                "ENSMUSG00000029322",
                "ENSMUSG00000005583",
                "ENSMUSG00000071076"
            ],
            "target": [],
            "expression_sum": {
                "class1": 69127.0,
                "class2": 34631.0
            },
            "ens_id": "ENSMUSG00000022665",
            "uniprot_ids": [
                "Q8R2G6"
            ]
        },
        "ENSMUSG00000024371": {
            "id": "ENSMUSG00000024371",
            "type": "Gene",
            "symbol": "C2",
            "source": [
                "ENSMUSG00000056501",
                "ENSMUSG00000022528",
                "ENSMUSG00000038193",
                "ENSMUSG00000005583",
                "ENSMUSG00000055148"
            ],
            "target": [],
            "expression_sum": {
                "class1": 9726.0,
                "class2": 1545.0
            },
            "ens_id": "ENSMUSG00000024371",
            "uniprot_ids": [
                "P21180",
                "B8JJM9",
                "B8JJN2",
                "F6ZCY7"
            ]
        },
        "ENSMUSG00000060550": {
            "id": "ENSMUSG00000060550",
            "type": "Gene",
            "symbol": "H2-Q7",
            "source": [
                "ENSMUSG00000038193",
                "ENSMUSG00000056501"
            ],
            "target": [],
            "expression_sum": {
                "class1": 21652.0,
                "class2": 218.0
            },
            "ens_id": "ENSMUSG00000060550",
            "uniprot_ids": [
                "P14429",
                "E9PWT4",
                "L7N260",
                "E9QJR9"
            ]
        },
        "ENSMUSG00000054072": {
            "id": "ENSMUSG00000054072",
            "type": "Gene",
            "symbol": "Iigp1",
            "source": [
                "ENSMUSG00000052837",
                "ENSMUSG00000056501",
                "ENSMUSG00000018899"
            ],
            "target": [],
            "expression_sum": {
                "class1": 24156.0,
                "class2": 1008.0
            },
            "ens_id": "ENSMUSG00000054072",
            "uniprot_ids": [
                "Q9QZ85"
            ]
        },
        "ENSMUSG00000062328": {
            "id": "ENSMUSG00000062328",
            "type": "Gene",
            "symbol": "Rpl17",
            "source": [
                "ENSMUSG00000056501",
                "ENSMUSG00000057098",
                "ENSMUSG00000038193",
                "ENSMUSG00000005583",
                "ENSMUSG00000031965",
                "ENSMUSG00000048387"
            ],
            "target": [],
            "expression_sum": {
                "class1": 131255.0,
                "class2": 62755.0
            },
            "ens_id": "ENSMUSG00000062328",
            "uniprot_ids": [
                "Q6ZWZ7"
            ]
        },
        "ENSMUSG00000045658": {
            "id": "ENSMUSG00000045658",
            "type": "Gene",
            "symbol": "Pid1",
            "source": [
                "ENSMUSG00000032238",
                "ENSMUSG00000052684",
                "ENSMUSG00000052837",
                "ENSMUSG00000022346",
                "ENSMUSG00000003032",
                "ENSMUSG00000005583",
                "ENSMUSG00000071076"
            ],
            "target": [],
            "expression_sum": {
                "class1": 543.0,
                "class2": 7123.0
            },
            "ens_id": "ENSMUSG00000045658",
            "uniprot_ids": [
                "Q3UBG2",
                "H3BKK5",
                "H3BJT8",
                "H3BJF2",
                "H3BJG6"
            ]
        },
        "ENSMUSG00000044337": {
            "id": "ENSMUSG00000044337",
            "type": "Gene",
            "symbol": "Ackr3",
            "source": [
                "ENSMUSG00000053007",
                "ENSMUSG00000003032",
                "ENSMUSG00000071076"
            ],
            "target": [],
            "expression_sum": {
                "class1": 171.0,
                "class2": 3549.0
            },
            "ens_id": "ENSMUSG00000044337",
            "uniprot_ids": [
                "P56485"
            ]
        },
        "ENSMUSG00000026478": {
            "id": "ENSMUSG00000026478",
            "type": "Gene",
            "symbol": "Lamc1",
            "source": [
                "ENSMUSG00000032035",
                "ENSMUSG00000005583",
                "ENSMUSG00000057098",
                "ENSMUSG00000003032",
                "ENSMUSG00000055435",
                "ENSMUSG00000071076"
            ],
            "target": [],
            "expression_sum": {
                "class1": 10652.0,
                "class2": 3771.0
            },
            "ens_id": "ENSMUSG00000026478",
            "uniprot_ids": [
                "F8VQJ3",
                "F6TLW1",
                "F6SFQ3"
            ]
        },
        "ENSMUSG00000026574": {
            "id": "ENSMUSG00000026574",
            "type": "Gene",
            "symbol": "Dpt",
            "source": [
                "ENSMUSG00000022346",
                "ENSMUSG00000066551",
                "ENSMUSG00000003032",
                "ENSMUSG00000005583",
                "ENSMUSG00000028565",
                "ENSMUSG00000021127"
            ],
            "target": [],
            "expression_sum": {
                "class1": 8606.0,
                "class2": 48325.0
            },
            "ens_id": "ENSMUSG00000026574",
            "uniprot_ids": [
                "Q9QZZ6"
            ]
        },
        "ENSMUSG00000026674": {
            "id": "ENSMUSG00000026674",
            "type": "Gene",
            "symbol": "Ddr2",
            "source": [
                "ENSMUSG00000057098",
                "ENSMUSG00000048387",
                "ENSMUSG00000003032",
                "ENSMUSG00000005583"
            ],
            "target": [],
            "expression_sum": {
                "class1": 7577.0,
                "class2": 3016.0
            },
            "ens_id": "ENSMUSG00000026674",
            "uniprot_ids": [
                "Q62371",
                "A0A0A6YXY2"
            ]
        },
        "ENSMUSG00000026670": {
            "id": "ENSMUSG00000026670",
            "type": "Gene",
            "symbol": "Uap1",
            "source": [
                "ENSMUSG00000057098",
                "ENSMUSG00000003032"
            ],
            "target": [],
            "expression_sum": {
                "class1": 1709.0,
                "class2": 3502.0
            },
            "ens_id": "ENSMUSG00000026670",
            "uniprot_ids": [
                "Q3UHZ7",
                "A0A0R4J1F6",
                "A0A0R4J085"
            ]
        },
        "ENSMUSG00000005338": {
            "id": "ENSMUSG00000005338",
            "type": "Gene",
            "symbol": "Cadm3",
            "source": [
                "ENSMUSG00000032035",
                "ENSMUSG00000003032"
            ],
            "target": [],
            "expression_sum": {
                "class1": 11519.0,
                "class2": 1102.0
            },
            "ens_id": "ENSMUSG00000005338",
            "uniprot_ids": [
                "Q99N28",
                "K4DI58"
            ]
        },
        "ENSMUSG00000037499": {
            "id": "ENSMUSG00000037499",
            "type": "Gene",
            "symbol": "Nenf",
            "source": [
                "ENSMUSG00000057098",
                "ENSMUSG00000038418",
                "ENSMUSG00000003032"
            ],
            "target": [],
            "expression_sum": {
                "class1": 14893.0,
                "class2": 16790.0
            },
            "ens_id": "ENSMUSG00000037499",
            "uniprot_ids": [
                "Q9CQ45"
            ]
        },
        "ENSMUSG00000016494": {
            "id": "ENSMUSG00000016494",
            "type": "Gene",
            "symbol": "Cd34",
            "source": [
                "ENSMUSG00000052684",
                "ENSMUSG00000022346",
                "ENSMUSG00000005583",
                "ENSMUSG00000021250",
                "ENSMUSG00000029322",
                "ENSMUSG00000003032",
                "ENSMUSG00000008575",
                "ENSMUSG00000028565",
                "ENSMUSG00000071076"
            ],
            "target": [],
            "expression_sum": {
                "class1": 613.0,
                "class2": 14659.0
            },
            "ens_id": "ENSMUSG00000016494",
            "uniprot_ids": [
                "Q64314",
                "A0A0A6YX84"
            ]
        },
        "ENSMUSG00000002107": {
            "id": "ENSMUSG00000002107",
            "type": "Gene",
            "symbol": "Celf2",
            "source": [
                "ENSMUSG00000032238",
                "ENSMUSG00000052684",
                "ENSMUSG00000022346",
                "ENSMUSG00000057098",
                "ENSMUSG00000038418",
                "ENSMUSG00000053477",
                "ENSMUSG00000003032",
                "ENSMUSG00000005583",
                "ENSMUSG00000028565",
                "ENSMUSG00000071076"
            ],
            "target": [],
            "expression_sum": {
                "class1": 1715.0,
                "class2": 5708.0
            },
            "ens_id": "ENSMUSG00000002107",
            "uniprot_ids": [
                "Q9Z0H4",
                "A0A0R4J2B0",
                "E9QA47",
                "S4R2U7",
                "S4R2U1",
                "S4R2J2",
                "V9GX43",
                "A3KGT0",
                "S4R2L5",
                "S4R1S7",
                "S4R2S7"
            ]
        },
        "ENSMUSG00000026748": {
            "id": "ENSMUSG00000026748",
            "type": "Gene",
            "symbol": "Plxdc2",
            "source": [
                "ENSMUSG00000052684",
                "ENSMUSG00000003032",
                "ENSMUSG00000005583"
            ],
            "target": [],
            "expression_sum": {
                "class1": 1809.0,
                "class2": 5378.0
            },
            "ens_id": "ENSMUSG00000026748",
            "uniprot_ids": [
                "Q9DC11",
                "B1AY85",
                "B1AY86"
            ]
        },
        "ENSMUSG00000015085": {
            "id": "ENSMUSG00000015085",
            "type": "Gene",
            "symbol": "Entpd2",
            "source": [
                "ENSMUSG00000052684",
                "ENSMUSG00000052837",
                "ENSMUSG00000021250",
                "ENSMUSG00000003032",
                "ENSMUSG00000005583",
                "ENSMUSG00000028565"
            ],
            "target": [],
            "expression_sum": {
                "class1": 548.0,
                "class2": 7057.0
            },
            "ens_id": "ENSMUSG00000015085",
            "uniprot_ids": [
                "O55026"
            ]
        },
        "ENSMUSG00000026837": {
            "id": "ENSMUSG00000026837",
            "type": "Gene",
            "symbol": "Col5a1",
            "source": [
                "ENSMUSG00000052684",
                "ENSMUSG00000038418",
                "ENSMUSG00000003032",
                "ENSMUSG00000005583",
                "ENSMUSG00000071076"
            ],
            "target": [],
            "expression_sum": {
                "class1": 9557.0,
                "class2": 2313.0
            },
            "ens_id": "ENSMUSG00000026837",
            "uniprot_ids": [
                "O88207",
                "A3KGE7",
                "A0A5F8MQ70"
            ]
        },
        "ENSMUSG00000026879": {
            "id": "ENSMUSG00000026879",
            "type": "Gene",
            "symbol": "Gsn",
            "source": [
                "ENSMUSG00000052684",
                "ENSMUSG00000023034",
                "ENSMUSG00000022528",
                "ENSMUSG00000022346",
                "ENSMUSG00000005583",
                "ENSMUSG00000003032",
                "ENSMUSG00000008575",
                "ENSMUSG00000028565",
                "ENSMUSG00000071076"
            ],
            "target": [],
            "expression_sum": {
                "class1": 20316.0,
                "class2": 411941.0
            },
            "ens_id": "ENSMUSG00000026879",
            "uniprot_ids": [
                "P13020",
                "A6PWS5",
                "A0A0J9YUQ8",
                "A0A0J9YUJ8"
            ]
        },
        "ENSMUSG00000060703": {
            "id": "ENSMUSG00000060703",
            "type": "Gene",
            "symbol": "Cd302",
            "source": [
                "ENSMUSG00000052684",
                "ENSMUSG00000038418",
                "ENSMUSG00000003032",
                "ENSMUSG00000005583",
                "ENSMUSG00000008575",
                "ENSMUSG00000071076"
            ],
            "target": [],
            "expression_sum": {
                "class1": 30299.0,
                "class2": 10955.0
            },
            "ens_id": "ENSMUSG00000060703",
            "uniprot_ids": [
                "Q9DCG2"
            ]
        },
        "ENSMUSG00000002732": {
            "id": "ENSMUSG00000002732",
            "type": "Gene",
            "symbol": "Fkbp7",
            "source": [
                "ENSMUSG00000052684",
                "ENSMUSG00000038193",
                "ENSMUSG00000003032",
                "ENSMUSG00000005583",
                "ENSMUSG00000071076"
            ],
            "target": [],
            "expression_sum": {
                "class1": 9183.0,
                "class2": 5862.0
            },
            "ens_id": "ENSMUSG00000002732",
            "uniprot_ids": [
                "O54998",
                "H3BJI5"
            ]
        },
        "ENSMUSG00000087006": {
            "id": "ENSMUSG00000087006",
            "type": "Gene",
            "symbol": "Gm13889",
            "source": [
                "ENSMUSG00000041515",
                "ENSMUSG00000052684",
                "ENSMUSG00000022346",
                "ENSMUSG00000003545",
                "ENSMUSG00000003032",
                "ENSMUSG00000005583"
            ],
            "target": [],
            "expression_sum": {
                "class1": 20589.0,
                "class2": 22077.0
            },
            "ens_id": "ENSMUSG00000087006",
            "uniprot_ids": [
                "A0A087WSH6",
                "A0A087WRN5"
            ]
        },
        "ENSMUSG00000074884": {
            "id": "ENSMUSG00000074884",
            "type": "Gene",
            "symbol": "Serf2",
            "source": [
                "ENSMUSG00000032035",
                "ENSMUSG00000052684",
                "ENSMUSG00000057098",
                "ENSMUSG00000038418",
                "ENSMUSG00000003032",
                "ENSMUSG00000005583",
                "ENSMUSG00000071076"
            ],
            "target": [],
            "expression_sum": {
                "class1": 66250.0,
                "class2": 30981.0
            },
            "ens_id": "ENSMUSG00000074884",
            "uniprot_ids": [
                "P84102",
                "A2ARQ4",
                "D6RCY2",
                "D6RIP7",
                "A2ARQ2"
            ]
        },
        "ENSMUSG00000060802": {
            "id": "ENSMUSG00000060802",
            "type": "Gene",
            "symbol": "B2m",
            "source": [
                "ENSMUSG00000032035",
                "ENSMUSG00000047638",
                "ENSMUSG00000057098",
                "ENSMUSG00000029322",
                "ENSMUSG00000003032",
                "ENSMUSG00000005583",
                "ENSMUSG00000071076"
            ],
            "target": [],
            "expression_sum": {
                "class1": 569087.0,
                "class2": 43809.0
            },
            "ens_id": "ENSMUSG00000060802",
            "uniprot_ids": [
                "P01887"
            ]
        },
        "ENSMUSG00000027204": {
            "id": "ENSMUSG00000027204",
            "type": "Gene",
            "symbol": "Fbn1",
            "source": [
                "ENSMUSG00000053007",
                "ENSMUSG00000003032",
                "ENSMUSG00000005583"
            ],
            "target": [],
            "expression_sum": {
                "class1": 5789.0,
                "class2": 5318.0
            },
            "ens_id": "ENSMUSG00000027204",
            "uniprot_ids": [
                "Q61554"
            ]
        },
        "ENSMUSG00000079037": {
            "id": "ENSMUSG00000079037",
            "type": "Gene",
            "symbol": "Prnp",
            "source": [
                "ENSMUSG00000052684",
                "ENSMUSG00000005583",
                "ENSMUSG00000048387",
                "ENSMUSG00000003032",
                "ENSMUSG00000008575",
                "ENSMUSG00000028565",
                "ENSMUSG00000071076"
            ],
            "target": [],
            "expression_sum": {
                "class1": 13438.0,
                "class2": 8029.0
            },
            "ens_id": "ENSMUSG00000079037",
            "uniprot_ids": [
                "P04925"
            ]
        },
        "ENSMUSG00000074743": {
            "id": "ENSMUSG00000074743",
            "type": "Gene",
            "symbol": "Thbd",
            "source": [
                "ENSMUSG00000003032",
                "ENSMUSG00000055435"
            ],
            "target": [],
            "expression_sum": {
                "class1": 952.0,
                "class2": 2621.0
            },
            "ens_id": "ENSMUSG00000074743",
            "uniprot_ids": [
                "P15306"
            ]
        },
        "ENSMUSG00000017002": {
            "id": "ENSMUSG00000017002",
            "type": "Gene",
            "symbol": "Slpi",
            "source": [
                "ENSMUSG00000003032"
            ],
            "target": [],
            "expression_sum": {
                "class1": 412.0,
                "class2": 123.0
            },
            "ens_id": "ENSMUSG00000017002",
            "uniprot_ids": [
                "P97430",
                "E9Q748",
                "E9Q4A3"
            ]
        },
        "ENSMUSG00000023092": {
            "id": "ENSMUSG00000023092",
            "type": "Gene",
            "symbol": "Fhl1",
            "source": [
                "ENSMUSG00000057098",
                "ENSMUSG00000003032",
                "ENSMUSG00000005583"
            ],
            "target": [],
            "expression_sum": {
                "class1": 20379.0,
                "class2": 7952.0
            },
            "ens_id": "ENSMUSG00000023092",
            "uniprot_ids": [
                "P97447",
                "A2AEX7",
                "A2AEY0",
                "A2AEY2",
                "A2AEX8",
                "A2AEX6",
                "A2AEX9",
                "A2AEY1"
            ]
        },
        "ENSMUSG00000019087": {
            "id": "ENSMUSG00000019087",
            "type": "Gene",
            "symbol": "Atp6ap1",
            "source": [
                "ENSMUSG00000048387",
                "ENSMUSG00000003032",
                "ENSMUSG00000071076"
            ],
            "target": [],
            "expression_sum": {
                "class1": 7337.0,
                "class2": 4770.0
            },
            "ens_id": "ENSMUSG00000019087",
            "uniprot_ids": [
                "Q9R1Q9",
                "B7FAU7",
                "B7FAU3",
                "Q3TKX1",
                "F6ZE56",
                "F6X9J0",
                "F7AYP0"
            ]
        },
        "ENSMUSG00000031422": {
            "id": "ENSMUSG00000031422",
            "type": "Gene",
            "symbol": "Morf4l2",
            "source": [
                "ENSMUSG00000052684",
                "ENSMUSG00000057098",
                "ENSMUSG00000038418",
                "ENSMUSG00000003032",
                "ENSMUSG00000005583",
                "ENSMUSG00000071076"
            ],
            "target": [],
            "expression_sum": {
                "class1": 10829.0,
                "class2": 7400.0
            },
            "ens_id": "ENSMUSG00000031422",
            "uniprot_ids": [
                "Q9R0Q4",
                "A2AEB6",
                "A2AEB7",
                "A2AEB5",
                "A2AEB8",
                "A2AEB9",
                "A2AEB3",
                "A2AEB4"
            ]
        },
        "ENSMUSG00000027712": {
            "id": "ENSMUSG00000027712",
            "type": "Gene",
            "symbol": "Anxa5",
            "source": [
                "ENSMUSG00000032238",
                "ENSMUSG00000032035",
                "ENSMUSG00000052684",
                "ENSMUSG00000057098",
                "ENSMUSG00000038193",
                "ENSMUSG00000003032",
                "ENSMUSG00000005583",
                "ENSMUSG00000028565",
                "ENSMUSG00000071076"
            ],
            "target": [],
            "expression_sum": {
                "class1": 26423.0,
                "class2": 18213.0
            },
            "ens_id": "ENSMUSG00000027712",
            "uniprot_ids": [
                "P48036",
                "A0A0G2JGQ0"
            ]
        },
        "ENSMUSG00000042747": {
            "id": "ENSMUSG00000042747",
            "type": "Gene",
            "symbol": "Krtcap2",
            "source": [
                "ENSMUSG00000052684",
                "ENSMUSG00000005583",
                "ENSMUSG00000057098",
                "ENSMUSG00000038418",
                "ENSMUSG00000003032",
                "ENSMUSG00000008575",
                "ENSMUSG00000071076"
            ],
            "target": [],
            "expression_sum": {
                "class1": 13423.0,
                "class2": 10239.0
            },
            "ens_id": "ENSMUSG00000042747",
            "uniprot_ids": [
                "Q5RL79",
                "A0A0A0MQN1"
            ]
        },
        "ENSMUSG00000042312": {
            "id": "ENSMUSG00000042312",
            "type": "Gene",
            "symbol": "S100a13",
            "source": [
                "ENSMUSG00000052684",
                "ENSMUSG00000057098",
                "ENSMUSG00000003032",
                "ENSMUSG00000005583",
                "ENSMUSG00000071076"
            ],
            "target": [],
            "expression_sum": {
                "class1": 4095.0,
                "class2": 6574.0
            },
            "ens_id": "ENSMUSG00000042312",
            "uniprot_ids": [
                "P97352",
                "A0A0A0MQ90"
            ]
        },
        "ENSMUSG00000074457": {
            "id": "ENSMUSG00000074457",
            "type": "Gene",
            "symbol": "S100a16",
            "source": [
                "ENSMUSG00000052684",
                "ENSMUSG00000022346",
                "ENSMUSG00000021250",
                "ENSMUSG00000003032",
                "ENSMUSG00000008575",
                "ENSMUSG00000020484",
                "ENSMUSG00000071076"
            ],
            "target": [],
            "expression_sum": {
                "class1": 407.0,
                "class2": 6586.0
            },
            "ens_id": "ENSMUSG00000074457",
            "uniprot_ids": [
                "Q9D708",
                "D3Z2Y6"
            ]
        },
        "ENSMUSG00000001025": {
            "id": "ENSMUSG00000001025",
            "type": "Gene",
            "symbol": "S100a6",
            "source": [
                "ENSMUSG00000052684",
                "ENSMUSG00000052837",
                "ENSMUSG00000022346",
                "ENSMUSG00000005583",
                "ENSMUSG00000003032",
                "ENSMUSG00000008575",
                "ENSMUSG00000071076"
            ],
            "target": [],
            "expression_sum": {
                "class1": 2262.0,
                "class2": 63154.0
            },
            "ens_id": "ENSMUSG00000001025",
            "uniprot_ids": [
                "P14069"
            ]
        },
        "ENSMUSG00000027907": {
            "id": "ENSMUSG00000027907",
            "type": "Gene",
            "symbol": "S100a11",
            "source": [
                "ENSMUSG00000052684",
                "ENSMUSG00000029322",
                "ENSMUSG00000003032",
                "ENSMUSG00000005583",
                "ENSMUSG00000071076"
            ],
            "target": [],
            "expression_sum": {
                "class1": 57022.0,
                "class2": 35522.0
            },
            "ens_id": "ENSMUSG00000027907",
            "uniprot_ids": [
                "P50543"
            ]
        },
        "ENSMUSG00000041959": {
            "id": "ENSMUSG00000041959",
            "type": "Gene",
            "symbol": "S100a10",
            "source": [
                "ENSMUSG00000052684",
                "ENSMUSG00000005583",
                "ENSMUSG00000003032",
                "ENSMUSG00000055435",
                "ENSMUSG00000028565",
                "ENSMUSG00000071076"
            ],
            "target": [],
            "expression_sum": {
                "class1": 19414.0,
                "class2": 23795.0
            },
            "ens_id": "ENSMUSG00000041959",
            "uniprot_ids": [
                "P08207"
            ]
        },
        "ENSMUSG00000028111": {
            "id": "ENSMUSG00000028111",
            "type": "Gene",
            "symbol": "Ctsk",
            "source": [
                "ENSMUSG00000052684",
                "ENSMUSG00000005583",
                "ENSMUSG00000003032",
                "ENSMUSG00000071076"
            ],
            "target": [],
            "expression_sum": {
                "class1": 4439.0,
                "class2": 1936.0
            },
            "ens_id": "ENSMUSG00000028111",
            "uniprot_ids": [
                "P55097"
            ]
        },
        "ENSMUSG00000028108": {
            "id": "ENSMUSG00000028108",
            "type": "Gene",
            "symbol": "Ecm1",
            "source": [
                "ENSMUSG00000032035",
                "ENSMUSG00000057098",
                "ENSMUSG00000038193",
                "ENSMUSG00000003032",
                "ENSMUSG00000028565",
                "ENSMUSG00000071076"
            ],
            "target": [],
            "expression_sum": {
                "class1": 274737.0,
                "class2": 16809.0
            },
            "ens_id": "ENSMUSG00000028108",
            "uniprot_ids": [
                "Q61508",
                "D3Z038",
                "F8WI14",
                "D3Z772",
                "D3Z0U9",
                "D3YV22",
                "D3YV64",
                "D3YZV0"
            ]
        },
        "ENSMUSG00000040562": {
            "id": "ENSMUSG00000040562",
            "type": "Gene",
            "symbol": "Gstm2",
            "source": [
                "ENSMUSG00000028565",
                "ENSMUSG00000052684",
                "ENSMUSG00000003032",
                "ENSMUSG00000005583"
            ],
            "target": [],
            "expression_sum": {
                "class1": 9933.0,
                "class2": 8169.0
            },
            "ens_id": "ENSMUSG00000040562",
            "uniprot_ids": [
                "P15626",
                "D3YX76"
            ]
        },
        "ENSMUSG00000058135": {
            "id": "ENSMUSG00000058135",
            "type": "Gene",
            "symbol": "Gstm1",
            "source": [
                "ENSMUSG00000055435",
                "ENSMUSG00000032238",
                "ENSMUSG00000032035",
                "ENSMUSG00000052684",
                "ENSMUSG00000052837",
                "ENSMUSG00000005583",
                "ENSMUSG00000057098",
                "ENSMUSG00000021250",
                "ENSMUSG00000003032",
                "ENSMUSG00000008575",
                "ENSMUSG00000028565",
                "ENSMUSG00000071076"
            ],
            "target": [],
            "expression_sum": {
                "class1": 18288.0,
                "class2": 43266.0
            },
            "ens_id": "ENSMUSG00000058135",
            "uniprot_ids": [
                "P10649",
                "F6WHQ7",
                "A2AE89"
            ]
        },
        "ENSMUSG00000041084": {
            "id": "ENSMUSG00000041084",
            "type": "Gene",
            "symbol": "Ostc",
            "source": [
                "ENSMUSG00000052684",
                "ENSMUSG00000003032",
                "ENSMUSG00000005583",
                "ENSMUSG00000048387",
                "ENSMUSG00000020484",
                "ENSMUSG00000071076"
            ],
            "target": [],
            "expression_sum": {
                "class1": 8276.0,
                "class2": 5532.0
            },
            "ens_id": "ENSMUSG00000041084",
            "uniprot_ids": [
                "Q78XF5"
            ]
        },
        "ENSMUSG00000037072": {
            "id": "ENSMUSG00000037072",
            "type": "Gene",
            "symbol": "Sep15",
            "source": [
                "ENSMUSG00000052684",
                "ENSMUSG00000057098",
                "ENSMUSG00000038418",
                "ENSMUSG00000003032",
                "ENSMUSG00000005583",
                "ENSMUSG00000048387",
                "ENSMUSG00000071076"
            ],
            "target": [],
            "expression_sum": {
                "class1": 31733.0,
                "class2": 18092.0
            },
            "ens_id": "ENSMUSG00000037072",
            "uniprot_ids": [
                "A0A0R4J0K1",
                "A0A1C7ZMY4",
                "A0A0G2JEF3"
            ]
        },
        "ENSMUSG00000028249": {
            "id": "ENSMUSG00000028249",
            "type": "Gene",
            "symbol": "Sdcbp",
            "source": [
                "ENSMUSG00000057098",
                "ENSMUSG00000038418",
                "ENSMUSG00000003032",
                "ENSMUSG00000055435",
                "ENSMUSG00000071076"
            ],
            "target": [],
            "expression_sum": {
                "class1": 11327.0,
                "class2": 9125.0
            },
            "ens_id": "ENSMUSG00000028249",
            "uniprot_ids": [
                "O08992",
                "Q3TMX0",
                "H3BLG5",
                "A2AKJ6",
                "A2AKJ9",
                "A2AKJ5"
            ]
        },
        "ENSMUSG00000028367": {
            "id": "ENSMUSG00000028367",
            "type": "Gene",
            "symbol": "Txn1",
            "source": [
                "ENSMUSG00000057098",
                "ENSMUSG00000038418",
                "ENSMUSG00000003032",
                "ENSMUSG00000048387",
                "ENSMUSG00000071076"
            ],
            "target": [],
            "expression_sum": {
                "class1": 14610.0,
                "class2": 14066.0
            },
            "ens_id": "ENSMUSG00000028367",
            "uniprot_ids": [
                "P10639"
            ]
        },
        "ENSMUSG00000035212": {
            "id": "ENSMUSG00000035212",
            "type": "Gene",
            "symbol": "Leprot",
            "source": [
                "ENSMUSG00000003032"
            ],
            "target": [],
            "expression_sum": {
                "class1": 6488.0,
                "class2": 3075.0
            },
            "ens_id": "ENSMUSG00000035212",
            "uniprot_ids": [
                "O89013"
            ]
        },
        "ENSMUSG00000028517": {
            "id": "ENSMUSG00000028517",
            "type": "Gene",
            "symbol": "Plpp3",
            "source": [
                "ENSMUSG00000052684",
                "ENSMUSG00000005583",
                "ENSMUSG00000048387",
                "ENSMUSG00000003032",
                "ENSMUSG00000008575",
                "ENSMUSG00000028565",
                "ENSMUSG00000071076"
            ],
            "target": [],
            "expression_sum": {
                "class1": 1847.0,
                "class2": 13633.0
            },
            "ens_id": "ENSMUSG00000028517",
            "uniprot_ids": [
                "Q99JY8"
            ]
        },
        "ENSMUSG00000028691": {
            "id": "ENSMUSG00000028691",
            "type": "Gene",
            "symbol": "Prdx1",
            "source": [
                "ENSMUSG00000052684",
                "ENSMUSG00000052837",
                "ENSMUSG00000057098",
                "ENSMUSG00000021250",
                "ENSMUSG00000003545",
                "ENSMUSG00000003032",
                "ENSMUSG00000048387",
                "ENSMUSG00000071076"
            ],
            "target": [],
            "expression_sum": {
                "class1": 33551.0,
                "class2": 23914.0
            },
            "ens_id": "ENSMUSG00000028691",
            "uniprot_ids": [
                "P35700",
                "B1AXW6",
                "B1AXW5",
                "B1AXW4"
            ]
        },
        "ENSMUSG00000028822": {
            "id": "ENSMUSG00000028822",
            "type": "Gene",
            "symbol": "Tmem50a",
            "source": [
                "ENSMUSG00000057098",
                "ENSMUSG00000038418",
                "ENSMUSG00000003032",
                "ENSMUSG00000071076"
            ],
            "target": [],
            "expression_sum": {
                "class1": 17865.0,
                "class2": 8285.0
            },
            "ens_id": "ENSMUSG00000028822",
            "uniprot_ids": [
                "Q9CXL1"
            ]
        },
        "ENSMUSG00000037266": {
            "id": "ENSMUSG00000037266",
            "type": "Gene",
            "symbol": "Rsrp1",
            "source": [
                "ENSMUSG00000052684",
                "ENSMUSG00000052837",
                "ENSMUSG00000057098",
                "ENSMUSG00000021250",
                "ENSMUSG00000003545",
                "ENSMUSG00000003032",
                "ENSMUSG00000048387",
                "ENSMUSG00000071076"
            ],
            "target": [],
            "expression_sum": {
                "class1": 28028.0,
                "class2": 18827.0
            },
            "ens_id": "ENSMUSG00000037266",
            "uniprot_ids": [
                "Q3UC65"
            ]
        },
        "ENSMUSG00000006699": {
            "id": "ENSMUSG00000006699",
            "type": "Gene",
            "symbol": "Cdc42",
            "source": [
                "ENSMUSG00000052684",
                "ENSMUSG00000057098",
                "ENSMUSG00000021250",
                "ENSMUSG00000003032",
                "ENSMUSG00000071076"
            ],
            "target": [],
            "expression_sum": {
                "class1": 19323.0,
                "class2": 13218.0
            },
            "ens_id": "ENSMUSG00000006699",
            "uniprot_ids": [
                "P60766"
            ]
        },
        "ENSMUSG00000060572": {
            "id": "ENSMUSG00000060572",
            "type": "Gene",
            "symbol": "Mfap2",
            "source": [
                "ENSMUSG00000022346",
                "ENSMUSG00000003032",
                "ENSMUSG00000005583"
            ],
            "target": [],
            "expression_sum": {
                "class1": 2684.0,
                "class2": 4741.0
            },
            "ens_id": "ENSMUSG00000060572",
            "uniprot_ids": [
                "Q99PM0"
            ]
        },
        "ENSMUSG00000003161": {
            "id": "ENSMUSG00000003161",
            "type": "Gene",
            "symbol": "Sri",
            "source": [
                "ENSMUSG00000052684",
                "ENSMUSG00000057098",
                "ENSMUSG00000038418",
                "ENSMUSG00000003032",
                "ENSMUSG00000005583",
                "ENSMUSG00000071076"
            ],
            "target": [],
            "expression_sum": {
                "class1": 9625.0,
                "class2": 5701.0
            },
            "ens_id": "ENSMUSG00000003161",
            "uniprot_ids": [
                "Q6P069"
            ]
        },
        "ENSMUSG00000029096": {
            "id": "ENSMUSG00000029096",
            "type": "Gene",
            "symbol": "Htra3",
            "source": [
                "ENSMUSG00000032238",
                "ENSMUSG00000052684",
                "ENSMUSG00000022346",
                "ENSMUSG00000005583",
                "ENSMUSG00000053477",
                "ENSMUSG00000003032",
                "ENSMUSG00000008575",
                "ENSMUSG00000028565",
                "ENSMUSG00000071076"
            ],
            "target": [],
            "expression_sum": {
                "class1": 7593.0,
                "class2": 27781.0
            },
            "ens_id": "ENSMUSG00000029096",
            "uniprot_ids": [
                "Q9D236",
                "A0A0J9YV92",
                "A0A0J9YVC4",
                "A0A0J9YU80"
            ]
        },
        "ENSMUSG00000037822": {
            "id": "ENSMUSG00000037822",
            "type": "Gene",
            "symbol": "Smim14",
            "source": [
                "ENSMUSG00000057098",
                "ENSMUSG00000003032",
                "ENSMUSG00000071076"
            ],
            "target": [],
            "expression_sum": {
                "class1": 8590.0,
                "class2": 10062.0
            },
            "ens_id": "ENSMUSG00000037822",
            "uniprot_ids": [
                "Q91VT8",
                "F8WHW8",
                "D3YYP6"
            ]
        },
        "ENSMUSG00000029484": {
            "id": "ENSMUSG00000029484",
            "type": "Gene",
            "symbol": "Anxa3",
            "source": [
                "ENSMUSG00000057098",
                "ENSMUSG00000003032"
            ],
            "target": [],
            "expression_sum": {
                "class1": 3483.0,
                "class2": 3190.0
            },
            "ens_id": "ENSMUSG00000029484",
            "uniprot_ids": [
                "O35639",
                "Q3TET3",
                "A0A0G2JGL7",
                "A0A0G2JDV9",
                "A0A0G2JFB3"
            ]
        },
        "ENSMUSG00000009013": {
            "id": "ENSMUSG00000009013",
            "type": "Gene",
            "symbol": "Dynll1",
            "source": [
                "ENSMUSG00000021250",
                "ENSMUSG00000003032",
                "ENSMUSG00000071076"
            ],
            "target": [],
            "expression_sum": {
                "class1": 28050.0,
                "class2": 25859.0
            },
            "ens_id": "ENSMUSG00000009013",
            "uniprot_ids": [
                "P63168"
            ]
        },
        "ENSMUSG00000029649": {
            "id": "ENSMUSG00000029649",
            "type": "Gene",
            "symbol": "Pomp",
            "source": [
                "ENSMUSG00000057098",
                "ENSMUSG00000003032"
            ],
            "target": [],
            "expression_sum": {
                "class1": 9881.0,
                "class2": 6311.0
            },
            "ens_id": "ENSMUSG00000029649",
            "uniprot_ids": [
                "Q9CQT5",
                "A0A0J9YVJ2",
                "A0A0J9YUI3"
            ]
        },
        "ENSMUSG00000029659": {
            "id": "ENSMUSG00000029659",
            "type": "Gene",
            "symbol": "Medag",
            "source": [
                "ENSMUSG00000052684",
                "ENSMUSG00000005583",
                "ENSMUSG00000057098",
                "ENSMUSG00000003032",
                "ENSMUSG00000008575",
                "ENSMUSG00000028565",
                "ENSMUSG00000071076"
            ],
            "target": [],
            "expression_sum": {
                "class1": 404.0,
                "class2": 3598.0
            },
            "ens_id": "ENSMUSG00000029659",
            "uniprot_ids": [
                "Q14BA6"
            ]
        },
        "ENSMUSG00000007655": {
            "id": "ENSMUSG00000007655",
            "type": "Gene",
            "symbol": "Cav1",
            "source": [
                "ENSMUSG00000048387",
                "ENSMUSG00000038193",
                "ENSMUSG00000003032",
                "ENSMUSG00000005583"
            ],
            "target": [],
            "expression_sum": {
                "class1": 18.0,
                "class2": 7289.0
            },
            "ens_id": "ENSMUSG00000007655",
            "uniprot_ids": [
                "P49817",
                "D3Z0J2",
                "D3Z148",
                "H3BLQ4",
                "H3BKG0"
            ]
        },
        "ENSMUSG00000020440": {
            "id": "ENSMUSG00000020440",
            "type": "Gene",
            "symbol": "Arf5",
            "source": [
                "ENSMUSG00000057098",
                "ENSMUSG00000005583",
                "ENSMUSG00000003032",
                "ENSMUSG00000071076"
            ],
            "target": [],
            "expression_sum": {
                "class1": 15221.0,
                "class2": 8233.0
            },
            "ens_id": "ENSMUSG00000020440",
            "uniprot_ids": [
                "P84084"
            ]
        },
        "ENSMUSG00000029767": {
            "id": "ENSMUSG00000029767",
            "type": "Gene",
            "symbol": "Calu",
            "source": [
                "ENSMUSG00000032035",
                "ENSMUSG00000005583",
                "ENSMUSG00000057098",
                "ENSMUSG00000003032",
                "ENSMUSG00000055435",
                "ENSMUSG00000071076"
            ],
            "target": [],
            "expression_sum": {
                "class1": 9548.0,
                "class2": 5449.0
            },
            "ens_id": "ENSMUSG00000029767",
            "uniprot_ids": [
                "O35887",
                "Q6XLQ8",
                "G3UWR0",
                "G3UWV3",
                "G3UXA3",
                "G3UXA8",
                "G3V004",
                "G3UY49"
            ]
        },
        "ENSMUSG00000056737": {
            "id": "ENSMUSG00000056737",
            "type": "Gene",
            "symbol": "Capg",
            "source": [
                "ENSMUSG00000057098",
                "ENSMUSG00000005583",
                "ENSMUSG00000003032",
                "ENSMUSG00000071076"
            ],
            "target": [],
            "expression_sum": {
                "class1": 4867.0,
                "class2": 3153.0
            },
            "ens_id": "ENSMUSG00000056737",
            "uniprot_ids": [
                "Q99LB4",
                "D3Z014",
                "D3Z4K5",
                "D3YZN3",
                "D3YU77",
                "D3YTL5"
            ]
        },
        "ENSMUSG00000079523": {
            "id": "ENSMUSG00000079523",
            "type": "Gene",
            "symbol": "Tmsb10",
            "source": [
                "ENSMUSG00000032035",
                "ENSMUSG00000052684",
                "ENSMUSG00000022528",
                "ENSMUSG00000005583",
                "ENSMUSG00000038193",
                "ENSMUSG00000029322",
                "ENSMUSG00000003032",
                "ENSMUSG00000055435",
                "ENSMUSG00000048387",
                "ENSMUSG00000071076"
            ],
            "target": [],
            "expression_sum": {
                "class1": 239963.0,
                "class2": 44526.0
            },
            "ens_id": "ENSMUSG00000079523",
            "uniprot_ids": [
                "Q6ZWY8",
                "A0A0N4SVF0"
            ]
        },
        "ENSMUSG00000030116": {
            "id": "ENSMUSG00000030116",
            "type": "Gene",
            "symbol": "Mfap5",
            "source": [
                "ENSMUSG00000003032"
            ],
            "target": [],
            "expression_sum": {
                "class1": 344.0,
                "class2": 15837.0
            },
            "ens_id": "ENSMUSG00000030116",
            "uniprot_ids": [
                "Q9QZJ6",
                "F8WJ99",
                "D3Z0U1",
                "D3Z4D7",
                "D3YW93",
                "D3Z3L7"
            ]
        },
        "ENSMUSG00000030208": {
            "id": "ENSMUSG00000030208",
            "type": "Gene",
            "symbol": "Emp1",
            "source": [
                "ENSMUSG00000052684",
                "ENSMUSG00000053477",
                "ENSMUSG00000003032",
                "ENSMUSG00000005583",
                "ENSMUSG00000008575",
                "ENSMUSG00000071076"
            ],
            "target": [],
            "expression_sum": {
                "class1": 4935.0,
                "class2": 6051.0
            },
            "ens_id": "ENSMUSG00000030208",
            "uniprot_ids": [
                "P47801",
                "S4R2U4"
            ]
        },
        "ENSMUSG00000002602": {
            "id": "ENSMUSG00000002602",
            "type": "Gene",
            "symbol": "Axl",
            "source": [
                "ENSMUSG00000057098",
                "ENSMUSG00000003032",
                "ENSMUSG00000071076"
            ],
            "target": [],
            "expression_sum": {
                "class1": 4235.0,
                "class2": 8968.0
            },
            "ens_id": "ENSMUSG00000002602",
            "uniprot_ids": [
                "Q00993",
                "Q6PE80",
                "F6YPR4"
            ]
        },
        "ENSMUSG00000040212": {
            "id": "ENSMUSG00000040212",
            "type": "Gene",
            "symbol": "Emp3",
            "source": [
                "ENSMUSG00000052684",
                "ENSMUSG00000057098",
                "ENSMUSG00000038418",
                "ENSMUSG00000003032",
                "ENSMUSG00000005583",
                "ENSMUSG00000071076"
            ],
            "target": [],
            "expression_sum": {
                "class1": 10517.0,
                "class2": 12826.0
            },
            "ens_id": "ENSMUSG00000040212",
            "uniprot_ids": [
                "O35912",
                "A0A1B0GSZ1"
            ]
        },
        "ENSMUSG00000040026": {
            "id": "ENSMUSG00000040026",
            "type": "Gene",
            "symbol": "Saa3",
            "source": [
                "ENSMUSG00000003032"
            ],
            "target": [],
            "expression_sum": {
                "class1": 23891.0,
                "class2": 404.0
            },
            "ens_id": "ENSMUSG00000040026",
            "uniprot_ids": [
                "P04918",
                "A0A1B0GT10"
            ]
        },
        "ENSMUSG00000033585": {
            "id": "ENSMUSG00000033585",
            "type": "Gene",
            "symbol": "Ndn",
            "source": [
                "ENSMUSG00000052684",
                "ENSMUSG00000022708",
                "ENSMUSG00000003032",
                "ENSMUSG00000005583"
            ],
            "target": [],
            "expression_sum": {
                "class1": 1898.0,
                "class2": 3737.0
            },
            "ens_id": "ENSMUSG00000033585",
            "uniprot_ids": [
                "P25233"
            ]
        },
        "ENSMUSG00000039405": {
            "id": "ENSMUSG00000039405",
            "type": "Gene",
            "symbol": "Prss23",
            "source": [
                "ENSMUSG00000003032"
            ],
            "target": [],
            "expression_sum": {
                "class1": 18315.0,
                "class2": 13836.0
            },
            "ens_id": "ENSMUSG00000039405",
            "uniprot_ids": [
                "Q9D6X6",
                "A0A140LHE7",
                "A0A140LIL0",
                "A0A140LHF1",
                "A0A140LJH8",
                "A0A140LIH7"
            ]
        },
        "ENSMUSG00000030654": {
            "id": "ENSMUSG00000030654",
            "type": "Gene",
            "symbol": "Arl6ip1",
            "source": [
                "ENSMUSG00000052684",
                "ENSMUSG00000057098",
                "ENSMUSG00000021250",
                "ENSMUSG00000003032",
                "ENSMUSG00000071076"
            ],
            "target": [],
            "expression_sum": {
                "class1": 9288.0,
                "class2": 4039.0
            },
            "ens_id": "ENSMUSG00000030654",
            "uniprot_ids": [
                "Q9JKW0",
                "A0A0U1RPY6",
                "A0A0N4SW86",
                "A0A0N4SV01"
            ]
        },
        "ENSMUSG00000025492": {
            "id": "ENSMUSG00000025492",
            "type": "Gene",
            "symbol": "Ifitm3",
            "source": [
                "ENSMUSG00000032035",
                "ENSMUSG00000052684",
                "ENSMUSG00000057098",
                "ENSMUSG00000003032",
                "ENSMUSG00000005583",
                "ENSMUSG00000071076"
            ],
            "target": [],
            "expression_sum": {
                "class1": 297680.0,
                "class2": 49594.0
            },
            "ens_id": "ENSMUSG00000025492",
            "uniprot_ids": [
                "Q9CQW9"
            ]
        },
        "ENSMUSG00000018819": {
            "id": "ENSMUSG00000018819",
            "type": "Gene",
            "symbol": "Lsp1",
            "source": [
                "ENSMUSG00000052684",
                "ENSMUSG00000053477",
                "ENSMUSG00000003032",
                "ENSMUSG00000005583",
                "ENSMUSG00000008575",
                "ENSMUSG00000071076"
            ],
            "target": [],
            "expression_sum": {
                "class1": 159.0,
                "class2": 3538.0
            },
            "ens_id": "ENSMUSG00000018819",
            "uniprot_ids": [
                "P19973",
                "A0A1B0GRF5",
                "A0A1B0GSH0",
                "A2A6J7",
                "A2A6J4"
            ]
        },
        "ENSMUSG00000037706": {
            "id": "ENSMUSG00000037706",
            "type": "Gene",
            "symbol": "Cd81",
            "source": [
                "ENSMUSG00000052684",
                "ENSMUSG00000021250",
                "ENSMUSG00000003032",
                "ENSMUSG00000005583",
                "ENSMUSG00000033863",
                "ENSMUSG00000071076"
            ],
            "target": [],
            "expression_sum": {
                "class1": 52771.0,
                "class2": 42729.0
            },
            "ens_id": "ENSMUSG00000037706",
            "uniprot_ids": [
                "P35762",
                "A0A140LJL0",
                "A0A140LHT1"
            ]
        },
        "ENSMUSG00000019978": {
            "id": "ENSMUSG00000019978",
            "type": "Gene",
            "symbol": "Epb41l2",
            "source": [
                "ENSMUSG00000005583",
                "ENSMUSG00000003032",
                "ENSMUSG00000071076"
            ],
            "target": [],
            "expression_sum": {
                "class1": 5219.0,
                "class2": 2324.0
            },
            "ens_id": "ENSMUSG00000019978",
            "uniprot_ids": [
                "O70318",
                "Q80UE4",
                "Q80UE5",
                "A0A1W2P6I5",
                "Q8C928",
                "A0A1W2P6H2",
                "A0A1W2P896",
                "A0A1W2P8C0",
                "A0A1W2P7I4",
                "A0A1W2P7I2",
                "A0A1W2P7H7"
            ]
        },
        "ENSMUSG00000015312": {
            "id": "ENSMUSG00000015312",
            "type": "Gene",
            "symbol": "Gadd45b",
            "source": [
                "ENSMUSG00000037868",
                "ENSMUSG00000052684",
                "ENSMUSG00000030103",
                "ENSMUSG00000021250",
                "ENSMUSG00000003545",
                "ENSMUSG00000003032",
                "ENSMUSG00000018899",
                "ENSMUSG00000071076"
            ],
            "target": [],
            "expression_sum": {
                "class1": 5736.0,
                "class2": 6541.0
            },
            "ens_id": "ENSMUSG00000015312",
            "uniprot_ids": [
                "P22339",
                "A0A1W2P6W4"
            ]
        },
        "ENSMUSG00000054452": {
            "id": "ENSMUSG00000054452",
            "type": "Gene",
            "symbol": "Aes",
            "source": [
                "ENSMUSG00000024140",
                "ENSMUSG00000003032"
            ],
            "target": [],
            "expression_sum": {
                "class1": 14020.0,
                "class2": 9206.0
            },
            "ens_id": "ENSMUSG00000054452",
            "uniprot_ids": [
                "P63002",
                "A0A1W2P757"
            ]
        },
        "ENSMUSG00000020053": {
            "id": "ENSMUSG00000020053",
            "type": "Gene",
            "symbol": "Igf1",
            "source": [
                "ENSMUSG00000032238",
                "ENSMUSG00000052684",
                "ENSMUSG00000052837",
                "ENSMUSG00000022346",
                "ENSMUSG00000053477",
                "ENSMUSG00000021250",
                "ENSMUSG00000029322",
                "ENSMUSG00000003032",
                "ENSMUSG00000005583",
                "ENSMUSG00000020484",
                "ENSMUSG00000071076"
            ],
            "target": [],
            "expression_sum": {
                "class1": 1900.0,
                "class2": 12864.0
            },
            "ens_id": "ENSMUSG00000020053",
            "uniprot_ids": [
                "P05017",
                "Q8CAR0",
                "E9Q138",
                "Q4VJB9",
                "D3Z7M4",
                "E9PU89"
            ]
        },
        "ENSMUSG00000036478": {
            "id": "ENSMUSG00000036478",
            "type": "Gene",
            "symbol": "Btg1",
            "source": [
                "ENSMUSG00000003032",
                "ENSMUSG00000005583"
            ],
            "target": [],
            "expression_sum": {
                "class1": 5559.0,
                "class2": 10662.0
            },
            "ens_id": "ENSMUSG00000036478",
            "uniprot_ids": [
                "P62325",
                "A0A1W2P7F9"
            ]
        },
        "ENSMUSG00000036446": {
            "id": "ENSMUSG00000036446",
            "type": "Gene",
            "symbol": "Lum",
            "source": [
                "ENSMUSG00000052684",
                "ENSMUSG00000053477",
                "ENSMUSG00000038193",
                "ENSMUSG00000003032",
                "ENSMUSG00000005583"
            ],
            "target": [],
            "expression_sum": {
                "class1": 88603.0,
                "class2": 46293.0
            },
            "ens_id": "ENSMUSG00000036446",
            "uniprot_ids": [
                "P51885"
            ]
        },
        "ENSMUSG00000040280": {
            "id": "ENSMUSG00000040280",
            "type": "Gene",
            "symbol": "Ndufa4l2",
            "source": [
                "ENSMUSG00000003032"
            ],
            "target": [],
            "expression_sum": {
                "class1": 7296.0,
                "class2": 13026.0
            },
            "ens_id": "ENSMUSG00000040280",
            "uniprot_ids": [
                "Q4FZG9"
            ]
        },
        "ENSMUSG00000031447": {
            "id": "ENSMUSG00000031447",
            "type": "Gene",
            "symbol": "Lamp1",
            "source": [
                "ENSMUSG00000032035",
                "ENSMUSG00000022528",
                "ENSMUSG00000057098",
                "ENSMUSG00000038193",
                "ENSMUSG00000003032",
                "ENSMUSG00000005583",
                "ENSMUSG00000048387"
            ],
            "target": [],
            "expression_sum": {
                "class1": 37763.0,
                "class2": 19814.0
            },
            "ens_id": "ENSMUSG00000031447",
            "uniprot_ids": [
                "P11438"
            ]
        },
        "ENSMUSG00000031538": {
            "id": "ENSMUSG00000031538",
            "type": "Gene",
            "symbol": "Plat",
            "source": [
                "ENSMUSG00000005583",
                "ENSMUSG00000003032",
                "ENSMUSG00000071076"
            ],
            "target": [],
            "expression_sum": {
                "class1": 3375.0,
                "class2": 4396.0
            },
            "ens_id": "ENSMUSG00000031538",
            "uniprot_ids": [
                "P11214"
            ]
        },
        "ENSMUSG00000031626": {
            "id": "ENSMUSG00000031626",
            "type": "Gene",
            "symbol": "Sorbs2",
            "source": [
                "ENSMUSG00000003032",
                "ENSMUSG00000005583"
            ],
            "target": [],
            "expression_sum": {
                "class1": 15.0,
                "class2": 5421.0
            },
            "ens_id": "ENSMUSG00000031626",
            "uniprot_ids": [
                "Q3UTJ2",
                "B2RXQ9",
                "B7ZWM6",
                "B9EKP8",
                "F7BEU1",
                "F6V513",
                "A0A1B0GRL6",
                "D3Z0N2",
                "D3Z080",
                "A0A1B0GRC3",
                "D3YUT0",
                "Z4YJR7",
                "F6ZWT0",
                "D3Z203",
                "D3Z2C7",
                "A0A0R4J1W0",
                "D3Z4W6",
                "D3Z0E9",
                "A0A0R4J1X5",
                "D3YY43",
                "A0A1B0GSU3",
                "A0A0R4J1T1",
                "A0A5F8MPQ4",
                "D3YTN2"
            ]
        },
        "ENSMUSG00000055681": {
            "id": "ENSMUSG00000055681",
            "type": "Gene",
            "symbol": "Cope",
            "source": [
                "ENSMUSG00000021250",
                "ENSMUSG00000052684",
                "ENSMUSG00000003032",
                "ENSMUSG00000071076"
            ],
            "target": [],
            "expression_sum": {
                "class1": 8299.0,
                "class2": 6573.0
            },
            "ens_id": "ENSMUSG00000055681",
            "uniprot_ids": [
                "O89079",
                "F6XIG5",
                "D3Z315",
                "F6YFR7",
                "E9Q6I5"
            ]
        },
        "ENSMUSG00000031799": {
            "id": "ENSMUSG00000031799",
            "type": "Gene",
            "symbol": "Tpm4",
            "source": [
                "ENSMUSG00000052684",
                "ENSMUSG00000005583",
                "ENSMUSG00000003032",
                "ENSMUSG00000055435",
                "ENSMUSG00000071076"
            ],
            "target": [],
            "expression_sum": {
                "class1": 19180.0,
                "class2": 7540.0
            },
            "ens_id": "ENSMUSG00000031799",
            "uniprot_ids": [
                "Q6IRU2",
                "A0A571BEU1"
            ]
        },
        "ENSMUSG00000044600": {
            "id": "ENSMUSG00000044600",
            "type": "Gene",
            "symbol": "Smim7",
            "source": [
                "ENSMUSG00000003032",
                "ENSMUSG00000071076"
            ],
            "target": [],
            "expression_sum": {
                "class1": 4854.0,
                "class2": 3804.0
            },
            "ens_id": "ENSMUSG00000044600",
            "uniprot_ids": [
                "Q5RKS2",
                "F8WIU9"
            ]
        },
        "ENSMUSG00000031875": {
            "id": "ENSMUSG00000031875",
            "type": "Gene",
            "symbol": "Cmtm3",
            "source": [
                "ENSMUSG00000003032",
                "ENSMUSG00000071076"
            ],
            "target": [],
            "expression_sum": {
                "class1": 6874.0,
                "class2": 2079.0
            },
            "ens_id": "ENSMUSG00000031875",
            "uniprot_ids": [
                "Q99LJ5",
                "A0A1D5RLE8",
                "A0A1D5RM50",
                "A0A1D5RLU9"
            ]
        },
        "ENSMUSG00000031722": {
            "id": "ENSMUSG00000031722",
            "type": "Gene",
            "symbol": "Hp",
            "source": [
                "ENSMUSG00000003032"
            ],
            "target": [],
            "expression_sum": {
                "class1": 1572.0,
                "class2": 190.0
            },
            "ens_id": "ENSMUSG00000031722",
            "uniprot_ids": [
                "Q61646"
            ]
        },
        "ENSMUSG00000019278": {
            "id": "ENSMUSG00000019278",
            "type": "Gene",
            "symbol": "Dpep1",
            "source": [
                "ENSMUSG00000022346",
                "ENSMUSG00000052837",
                "ENSMUSG00000052684",
                "ENSMUSG00000003032"
            ],
            "target": [],
            "expression_sum": {
                "class1": 353.0,
                "class2": 12028.0
            },
            "ens_id": "ENSMUSG00000019278",
            "uniprot_ids": [
                "P31428",
                "A0A1D5RML1",
                "A0A1D5RM32"
            ]
        },
        "ENSMUSG00000021876": {
            "id": "ENSMUSG00000021876",
            "type": "Gene",
            "symbol": "Rnase4",
            "source": [
                "ENSMUSG00000055435",
                "ENSMUSG00000052684",
                "ENSMUSG00000021877",
                "ENSMUSG00000052837",
                "ENSMUSG00000022346",
                "ENSMUSG00000005583",
                "ENSMUSG00000048387",
                "ENSMUSG00000021250",
                "ENSMUSG00000003032",
                "ENSMUSG00000008575",
                "ENSMUSG00000028565",
                "ENSMUSG00000020484",
                "ENSMUSG00000071076"
            ],
            "target": [],
            "expression_sum": {
                "class1": 7086.0,
                "class2": 18776.0
            },
            "ens_id": "ENSMUSG00000021876",
            "uniprot_ids": [
                "Q8C7E4"
            ]
        },
        "ENSMUSG00000022032": {
            "id": "ENSMUSG00000022032",
            "type": "Gene",
            "symbol": "Scara5",
            "source": [
                "ENSMUSG00000052684",
                "ENSMUSG00000022346",
                "ENSMUSG00000057098",
                "ENSMUSG00000028639",
                "ENSMUSG00000003032",
                "ENSMUSG00000005583",
                "ENSMUSG00000071076"
            ],
            "target": [],
            "expression_sum": {
                "class1": 23.0,
                "class2": 3706.0
            },
            "ens_id": "ENSMUSG00000022032",
            "uniprot_ids": [
                "Q8K299"
            ]
        },
        "ENSMUSG00000022090": {
            "id": "ENSMUSG00000022090",
            "type": "Gene",
            "symbol": "Pdlim2",
            "source": [
                "ENSMUSG00000057098",
                "ENSMUSG00000005583",
                "ENSMUSG00000003032",
                "ENSMUSG00000071076"
            ],
            "target": [],
            "expression_sum": {
                "class1": 11244.0,
                "class2": 5118.0
            },
            "ens_id": "ENSMUSG00000022090",
            "uniprot_ids": [
                "Q8R1G6",
                "A0A087WPL1",
                "E9Q996",
                "F7C957"
            ]
        },
        "ENSMUSG00000084786": {
            "id": "ENSMUSG00000084786",
            "type": "Gene",
            "symbol": "Ubl5",
            "source": [
                "ENSMUSG00000021250",
                "ENSMUSG00000003032",
                "ENSMUSG00000071076"
            ],
            "target": [],
            "expression_sum": {
                "class1": 10889.0,
                "class2": 8519.0
            },
            "ens_id": "ENSMUSG00000084786",
            "uniprot_ids": [
                "Q9EPV8",
                "E0CYW1",
                "E0CY38",
                "E0CY91",
                "E0CZB3"
            ]
        },
        "ENSMUSG00000032014": {
            "id": "ENSMUSG00000032014",
            "type": "Gene",
            "symbol": "Oaf",
            "source": [
                "ENSMUSG00000057098",
                "ENSMUSG00000038193",
                "ENSMUSG00000003032",
                "ENSMUSG00000005583"
            ],
            "target": [],
            "expression_sum": {
                "class1": 17282.0,
                "class2": 3948.0
            },
            "ens_id": "ENSMUSG00000032014",
            "uniprot_ids": [
                "Q8QZR4",
                "A0A1L1SUK4"
            ]
        },
        "ENSMUSG00000095385": {
            "id": "ENSMUSG00000095385",
            "type": "Gene",
            "symbol": "D630033O11Rik",
            "source": [
                "ENSMUSG00000003032"
            ],
            "target": [],
            "expression_sum": {
                "class1": 5361.0,
                "class2": 295.0
            },
            "ens_id": "ENSMUSG00000095385"
        },
        "ENSMUSG00000032301": {
            "id": "ENSMUSG00000032301",
            "type": "Gene",
            "symbol": "Psma4",
            "source": [
                "ENSMUSG00000057098",
                "ENSMUSG00000003032",
                "ENSMUSG00000071076"
            ],
            "target": [],
            "expression_sum": {
                "class1": 7508.0,
                "class2": 4407.0
            },
            "ens_id": "ENSMUSG00000032301",
            "uniprot_ids": [
                "Q9R1P0",
                "E9PW69",
                "E9Q0X0"
            ]
        },
        "ENSMUSG00000037206": {
            "id": "ENSMUSG00000037206",
            "type": "Gene",
            "symbol": "Islr",
            "source": [
                "ENSMUSG00000038193",
                "ENSMUSG00000005583",
                "ENSMUSG00000003032",
                "ENSMUSG00000071076"
            ],
            "target": [],
            "expression_sum": {
                "class1": 13630.0,
                "class2": 11700.0
            },
            "ens_id": "ENSMUSG00000037206",
            "uniprot_ids": [
                "Q6GU68",
                "A0A1L1STF4"
            ]
        },
        "ENSMUSG00000032334": {
            "id": "ENSMUSG00000032334",
            "type": "Gene",
            "symbol": "Loxl1",
            "source": [
                "ENSMUSG00000052684",
                "ENSMUSG00000005583",
                "ENSMUSG00000053477",
                "ENSMUSG00000003032",
                "ENSMUSG00000008575",
                "ENSMUSG00000028565",
                "ENSMUSG00000071076"
            ],
            "target": [],
            "expression_sum": {
                "class1": 6154.0,
                "class2": 11476.0
            },
            "ens_id": "ENSMUSG00000032334",
            "uniprot_ids": [
                "A0A0R4J0Q4"
            ]
        },
        "ENSMUSG00000032383": {
            "id": "ENSMUSG00000032383",
            "type": "Gene",
            "symbol": "Ppib",
            "source": [
                "ENSMUSG00000032035",
                "ENSMUSG00000052684",
                "ENSMUSG00000022528",
                "ENSMUSG00000057098",
                "ENSMUSG00000038418",
                "ENSMUSG00000038193",
                "ENSMUSG00000003032",
                "ENSMUSG00000005583",
                "ENSMUSG00000071076"
            ],
            "target": [],
            "expression_sum": {
                "class1": 33381.0,
                "class2": 15590.0
            },
            "ens_id": "ENSMUSG00000032383",
            "uniprot_ids": [
                "P24369"
            ]
        },
        "ENSMUSG00000007815": {
            "id": "ENSMUSG00000007815",
            "type": "Gene",
            "symbol": "Rhoa",
            "source": [
                "ENSMUSG00000057098",
                "ENSMUSG00000003032",
                "ENSMUSG00000071076"
            ],
            "target": [],
            "expression_sum": {
                "class1": 20164.0,
                "class2": 19742.0
            },
            "ens_id": "ENSMUSG00000007815",
            "uniprot_ids": [
                "Q9QUI0",
                "A0A0A6YWJ1",
                "A0A0A6YXF6"
            ]
        },
        "ENSMUSG00000075702": {
            "id": "ENSMUSG00000075702",
            "type": "Gene",
            "symbol": "Selm",
            "source": [
                "ENSMUSG00000052684",
                "ENSMUSG00000021250",
                "ENSMUSG00000003032",
                "ENSMUSG00000005583",
                "ENSMUSG00000071076"
            ],
            "target": [],
            "expression_sum": {
                "class1": 17418.0,
                "class2": 20444.0
            },
            "ens_id": "ENSMUSG00000075702",
            "uniprot_ids": [
                "Q8VHC3"
            ]
        },
        "ENSMUSG00000001891": {
            "id": "ENSMUSG00000001891",
            "type": "Gene",
            "symbol": "Ugp2",
            "source": [
                "ENSMUSG00000057098",
                "ENSMUSG00000003032",
                "ENSMUSG00000071076"
            ],
            "target": [],
            "expression_sum": {
                "class1": 2386.0,
                "class2": 4475.0
            },
            "ens_id": "ENSMUSG00000001891",
            "uniprot_ids": [
                "Q91ZJ5"
            ]
        },
        "ENSMUSG00000020467": {
            "id": "ENSMUSG00000020467",
            "type": "Gene",
            "symbol": "Efemp1",
            "source": [
                "ENSMUSG00000003032",
                "ENSMUSG00000005583"
            ],
            "target": [],
            "expression_sum": {
                "class1": 15374.0,
                "class2": 5286.0
            },
            "ens_id": "ENSMUSG00000020467",
            "uniprot_ids": [
                "Q8BPB5",
                "F6ZFS0"
            ]
        },
        "ENSMUSG00000020315": {
            "id": "ENSMUSG00000020315",
            "type": "Gene",
            "symbol": "Sptbn1",
            "source": [
                "ENSMUSG00000038418",
                "ENSMUSG00000003032",
                "ENSMUSG00000005583",
                "ENSMUSG00000008575",
                "ENSMUSG00000071076"
            ],
            "target": [],
            "expression_sum": {
                "class1": 10746.0,
                "class2": 9430.0
            },
            "ens_id": "ENSMUSG00000020315",
            "uniprot_ids": [
                "Q62261",
                "A0A0A0MQG2"
            ]
        },
        "ENSMUSG00000057098": {
            "id": "ENSMUSG00000057098",
            "type": "TF",
            "symbol": "Ebf1",
            "source": [
                "ENSMUSG00000021250",
                "ENSMUSG00000003032",
                "ENSMUSG00000071076"
            ],
            "target": [
                "ENSMUSG00000031447",
                "ENSMUSG00000062456",
                "ENSMUSG00000032518",
                "ENSMUSG00000024011",
                "ENSMUSG00000026878",
                "ENSMUSG00000014599",
                "ENSMUSG00000016256",
                "ENSMUSG00000047187",
                "ENSMUSG00000008668",
                "ENSMUSG00000019877",
                "ENSMUSG00000071415",
                "ENSMUSG00000053317",
                "ENSMUSG00000002900",
                "ENSMUSG00000029616",
                "ENSMUSG00000020368",
                "ENSMUSG00000028081",
                "ENSMUSG00000055302",
                "ENSMUSG00000025781",
                "ENSMUSG00000012405",
                "ENSMUSG00000038650",
                "ENSMUSG00000020737",
                "ENSMUSG00000022820",
                "ENSMUSG00000090862",
                "ENSMUSG00000030824",
                "ENSMUSG00000073639",
                "ENSMUSG00000025508",
                "ENSMUSG00000063882",
                "ENSMUSG00000011752",
                "ENSMUSG00000045128",
                "ENSMUSG00000017466",
                "ENSMUSG00000078812",
                "ENSMUSG00000029455",
                "ENSMUSG00000044080",
                "ENSMUSG00000026536",
                "ENSMUSG00000072235",
                "ENSMUSG00000006333",
                "ENSMUSG00000030432",
                "ENSMUSG00000074129",
                "ENSMUSG00000031818",
                "ENSMUSG00000026674",
                "ENSMUSG00000012848",
                "ENSMUSG00000027074",
                "ENSMUSG00000028964",
                "ENSMUSG00000003032",
                "ENSMUSG00000073702",
                "ENSMUSG00000032216",
                "ENSMUSG00000062270",
                "ENSMUSG00000045817",
                "ENSMUSG00000027712",
                "ENSMUSG00000053565",
                "ENSMUSG00000007891",
                "ENSMUSG00000024048",
                "ENSMUSG00000031029",
                "ENSMUSG00000001891",
                "ENSMUSG00000050708",
                "ENSMUSG00000026478",
                "ENSMUSG00000057278",
                "ENSMUSG00000014846",
                "ENSMUSG00000017493",
                "ENSMUSG00000025351",
                "ENSMUSG00000074884",
                "ENSMUSG00000020849",
                "ENSMUSG00000025066",
                "ENSMUSG00000045690",
                "ENSMUSG00000027248",
                "ENSMUSG00000049751",
                "ENSMUSG00000043716",
                "ENSMUSG00000027523",
                "ENSMUSG00000035530",
                "ENSMUSG00000036781",
                "ENSMUSG00000041801",
                "ENSMUSG00000073411",
                "ENSMUSG00000022174",
                "ENSMUSG00000004610",
                "ENSMUSG00000035227",
                "ENSMUSG00000060802",
                "ENSMUSG00000041453",
                "ENSMUSG00000003429",
                "ENSMUSG00000038418",
                "ENSMUSG00000032399",
                "ENSMUSG00000038900",
                "ENSMUSG00000019505",
                "ENSMUSG00000006360",
                "ENSMUSG00000059291",
                "ENSMUSG00000022841",
                "ENSMUSG00000040824",
                "ENSMUSG00000028691",
                "ENSMUSG00000022283",
                "ENSMUSG00000068523",
                "ENSMUSG00000036199",
                "ENSMUSG00000030695",
                "ENSMUSG00000030744",
                "ENSMUSG00000026238",
                "ENSMUSG00000028822",
                "ENSMUSG00000023175",
                "ENSMUSG00000022336",
                "ENSMUSG00000026970",
                "ENSMUSG00000021702",
                "ENSMUSG00000002107",
                "ENSMUSG00000009549",
                "ENSMUSG00000032231",
                "ENSMUSG00000059796",
                "ENSMUSG00000004455",
                "ENSMUSG00000037805",
                "ENSMUSG00000063316",
                "ENSMUSG00000017404",
                "ENSMUSG00000028648",
                "ENSMUSG00000025132",
                "ENSMUSG00000021917",
                "ENSMUSG00000042043",
                "ENSMUSG00000037266",
                "ENSMUSG00000030790",
                "ENSMUSG00000029642",
                "ENSMUSG00000020719",
                "ENSMUSG00000036371",
                "ENSMUSG00000019804",
                "ENSMUSG00000007039",
                "ENSMUSG00000022090",
                "ENSMUSG00000032014",
                "ENSMUSG00000040249",
                "ENSMUSG00000023092",
                "ENSMUSG00000029070",
                "ENSMUSG00000015839",
                "ENSMUSG00000010406",
                "ENSMUSG00000051695",
                "ENSMUSG00000002778",
                "ENSMUSG00000030654",
                "ENSMUSG00000098274",
                "ENSMUSG00000026511",
                "ENSMUSG00000064080",
                "ENSMUSG00000005973",
                "ENSMUSG00000057322",
                "ENSMUSG00000003161",
                "ENSMUSG00000021025",
                "ENSMUSG00000032301",
                "ENSMUSG00000042079",
                "ENSMUSG00000003814",
                "ENSMUSG00000052040",
                "ENSMUSG00000015671",
                "ENSMUSG00000047215",
                "ENSMUSG00000020205",
                "ENSMUSG00000037072",
                "ENSMUSG00000020009",
                "ENSMUSG00000004207",
                "ENSMUSG00000026728",
                "ENSMUSG00000096145",
                "ENSMUSG00000014769",
                "ENSMUSG00000042312",
                "ENSMUSG00000062997",
                "ENSMUSG00000067288",
                "ENSMUSG00000058558",
                "ENSMUSG00000075254",
                "ENSMUSG00000023010",
                "ENSMUSG00000060938",
                "ENSMUSG00000047675",
                "ENSMUSG00000041841",
                "ENSMUSG00000021242",
                "ENSMUSG00000008892",
                "ENSMUSG00000024248",
                "ENSMUSG00000028837",
                "ENSMUSG00000062328",
                "ENSMUSG00000020585",
                "ENSMUSG00000019818",
                "ENSMUSG00000025492",
                "ENSMUSG00000055762",
                "ENSMUSG00000007041",
                "ENSMUSG00000055447",
                "ENSMUSG00000024190",
                "ENSMUSG00000057841",
                "ENSMUSG00000020458",
                "ENSMUSG00000022234",
                "ENSMUSG00000009291",
                "ENSMUSG00000078578",
                "ENSMUSG00000028234",
                "ENSMUSG00000029322",
                "ENSMUSG00000035585",
                "ENSMUSG00000090137",
                "ENSMUSG00000021131",
                "ENSMUSG00000037062",
                "ENSMUSG00000031765",
                "ENSMUSG00000028639",
                "ENSMUSG00000040952",
                "ENSMUSG00000048076",
                "ENSMUSG00000026021",
                "ENSMUSG00000006699",
                "ENSMUSG00000060036",
                "ENSMUSG00000045962",
                "ENSMUSG00000038717",
                "ENSMUSG00000016252",
                "ENSMUSG00000023004",
                "ENSMUSG00000031198",
                "ENSMUSG00000025130",
                "ENSMUSG00000056737",
                "ENSMUSG00000056201",
                "ENSMUSG00000022884",
                "ENSMUSG00000061983",
                "ENSMUSG00000022032",
                "ENSMUSG00000004980",
                "ENSMUSG00000028967",
                "ENSMUSG00000022400",
                "ENSMUSG00000021546",
                "ENSMUSG00000018585",
                "ENSMUSG00000034165",
                "ENSMUSG00000003380",
                "ENSMUSG00000038803",
                "ENSMUSG00000022051",
                "ENSMUSG00000020460",
                "ENSMUSG00000055044",
                "ENSMUSG00000009927",
                "ENSMUSG00000023944",
                "ENSMUSG00000028416",
                "ENSMUSG00000024608",
                "ENSMUSG00000062647",
                "ENSMUSG00000060591",
                "ENSMUSG00000049517",
                "ENSMUSG00000027940",
                "ENSMUSG00000001794",
                "ENSMUSG00000028692",
                "ENSMUSG00000025780",
                "ENSMUSG00000030795",
                "ENSMUSG00000035242",
                "ENSMUSG00000006356",
                "ENSMUSG00000015656",
                "ENSMUSG00000067274",
                "ENSMUSG00000071644",
                "ENSMUSG00000057863",
                "ENSMUSG00000022108",
                "ENSMUSG00000025510",
                "ENSMUSG00000028249",
                "ENSMUSG00000024953",
                "ENSMUSG00000053012",
                "ENSMUSG00000026670",
                "ENSMUSG00000021250",
                "ENSMUSG00000001833",
                "ENSMUSG00000028618",
                "ENSMUSG00000027620",
                "ENSMUSG00000032966",
                "ENSMUSG00000052837",
                "ENSMUSG00000032324",
                "ENSMUSG00000031708",
                "ENSMUSG00000028936",
                "ENSMUSG00000059005",
                "ENSMUSG00000028478",
                "ENSMUSG00000060073",
                "ENSMUSG00000001847",
                "ENSMUSG00000019428",
                "ENSMUSG00000029465",
                "ENSMUSG00000039208",
                "ENSMUSG00000030057",
                "ENSMUSG00000031633",
                "ENSMUSG00000037563",
                "ENSMUSG00000058135",
                "ENSMUSG00000007815",
                "ENSMUSG00000002602",
                "ENSMUSG00000058546",
                "ENSMUSG00000008348",
                "ENSMUSG00000037822",
                "ENSMUSG00000025967",
                "ENSMUSG00000031556",
                "ENSMUSG00000021877",
                "ENSMUSG00000034868",
                "ENSMUSG00000063480",
                "ENSMUSG00000029649",
                "ENSMUSG00000028797",
                "ENSMUSG00000061787",
                "ENSMUSG00000044627",
                "ENSMUSG00000020372",
                "ENSMUSG00000021939",
                "ENSMUSG00000001240",
                "ENSMUSG00000036880",
                "ENSMUSG00000061904",
                "ENSMUSG00000028369",
                "ENSMUSG00000029621",
                "ENSMUSG00000060743",
                "ENSMUSG00000069744",
                "ENSMUSG00000020738",
                "ENSMUSG00000037499",
                "ENSMUSG00000026547",
                "ENSMUSG00000006304",
                "ENSMUSG00000062825",
                "ENSMUSG00000060636",
                "ENSMUSG00000046364",
                "ENSMUSG00000075706",
                "ENSMUSG00000020440",
                "ENSMUSG00000028367",
                "ENSMUSG00000040385",
                "ENSMUSG00000063457",
                "ENSMUSG00000032314",
                "ENSMUSG00000037742",
                "ENSMUSG00000036372",
                "ENSMUSG00000015092",
                "ENSMUSG00000025016",
                "ENSMUSG00000029767",
                "ENSMUSG00000018326",
                "ENSMUSG00000057113",
                "ENSMUSG00000002233",
                "ENSMUSG00000020267",
                "ENSMUSG00000025794",
                "ENSMUSG00000029713",
                "ENSMUSG00000039001",
                "ENSMUSG00000028452",
                "ENSMUSG00000060019",
                "ENSMUSG00000028530",
                "ENSMUSG00000090733",
                "ENSMUSG00000040212",
                "ENSMUSG00000001123",
                "ENSMUSG00000016559",
                "ENSMUSG00000060126",
                "ENSMUSG00000020149",
                "ENSMUSG00000061315",
                "ENSMUSG00000044533",
                "ENSMUSG00000062006",
                "ENSMUSG00000002015",
                "ENSMUSG00000052727",
                "ENSMUSG00000013622",
                "ENSMUSG00000038274",
                "ENSMUSG00000028108",
                "ENSMUSG00000021996",
                "ENSMUSG00000006519",
                "ENSMUSG00000056851",
                "ENSMUSG00000008475",
                "ENSMUSG00000052146",
                "ENSMUSG00000034994",
                "ENSMUSG00000024308",
                "ENSMUSG00000021218",
                "ENSMUSG00000026872",
                "ENSMUSG00000060860",
                "ENSMUSG00000071866",
                "ENSMUSG00000005881",
                "ENSMUSG00000024914",
                "ENSMUSG00000020048",
                "ENSMUSG00000042747",
                "ENSMUSG00000007892",
                "ENSMUSG00000032383",
                "ENSMUSG00000079641",
                "ENSMUSG00000031422",
                "ENSMUSG00000029622",
                "ENSMUSG00000059070",
                "ENSMUSG00000066551",
                "ENSMUSG00000022708",
                "ENSMUSG00000034118",
                "ENSMUSG00000029659",
                "ENSMUSG00000093674",
                "ENSMUSG00000029430",
                "ENSMUSG00000042682",
                "ENSMUSG00000028495",
                "ENSMUSG00000008540",
                "ENSMUSG00000029484",
                "ENSMUSG00000022548",
                "ENSMUSG00000000563"
            ],
            "expression_sum": {
                "class1": 205.0,
                "class2": 16536.0
            },
```
