## Supplement 4 (CCl4 a6w HSC vs. PF) for "AGEAS: Automated Machine Learning based Genetic Regulatory Element Extraction System": outlier_grps.js

```
{
    "ENSMUSG00000042485_ENSMUSG00000005583": 36.57886552330235,
    "ENSMUSG00000068735_ENSMUSG00000005583": 19.933990878514315,
    "ENSMUSG00000005583_ENSMUSG00000001520": 18.7114452380066,
    "ENSMUSG00000071076_ENSMUSG00000025784": 14.18198855713545,
    "ENSMUSG00000024290_ENSMUSG00000005583": 13.39548940826401,
    "ENSMUSG00000052684_ENSMUSG00000006205": 10.424232316198596,
    "ENSMUSG00000072235_ENSMUSG00000071076": 9.548363620213753,
    "ENSMUSG00000071076_ENSMUSG00000060703": 9.33812290660825,
    "ENSMUSG00000052684_ENSMUSG00000021250": 8.404562951697514,
    "ENSMUSG00000038418_ENSMUSG00000021250": 8.377972368749305,
    "ENSMUSG00000052684_ENSMUSG00000023034": 8.05586498608715,
    "ENSMUSG00000035356_ENSMUSG00000021250": 6.830659473559313,
    "ENSMUSG00000023064_ENSMUSG00000005583": 6.0019777072277405,
    "ENSMUSG00000032387_ENSMUSG00000005583": 5.967966519910633,
    "ENSMUSG00000023034_ENSMUSG00000003545": 5.944988865060106,
    "ENSMUSG00000021250_ENSMUSG00000020423": 5.781372497079608,
    "ENSMUSG00000090862_ENSMUSG00000000247": 5.761658637688609,
    "ENSMUSG00000026879_ENSMUSG00000005583": 5.694593766594694,
    "ENSMUSG00000071076_ENSMUSG00000026628": 5.6893164927911055,
    "ENSMUSG00000026574_ENSMUSG00000005583": 5.68499041999284,
    "ENSMUSG00000038418_ENSMUSG00000020423": 5.683283841086697,
    "ENSMUSG00000052684_ENSMUSG00000044786": 5.576345925925922,
    "ENSMUSG00000052997_ENSMUSG00000005583": 4.665289987835879,
    "ENSMUSG00000064358_ENSMUSG00000029322": 4.62009352928469,
    "ENSMUSG00000087006_ENSMUSG00000005583": 4.522679591915329,
    "ENSMUSG00000071076_ENSMUSG00000025351": 4.459618801776332,
    "ENSMUSG00000071076_ENSMUSG00000021477": 4.415425890813217,
    "ENSMUSG00000078202_ENSMUSG00000005583": 4.31181833067954,
    "ENSMUSG00000053113_ENSMUSG00000052837": 4.186090169334792,
    "ENSMUSG00000071076_ENSMUSG00000041959": 3.378000023956146,
    "ENSMUSG00000053113_ENSMUSG00000023034": 3.3414330568102044,
    "ENSMUSG00000071076_ENSMUSG00000042312": 3.3223915579839565,
    "ENSMUSG00000038193_ENSMUSG00000019929": 3.2872461358979104,
    "ENSMUSG00000030122_ENSMUSG00000029322": 3.210557778800001,
    "ENSMUSG00000053113_ENSMUSG00000026628": 3.14438148225427,
    "ENSMUSG00000025147_ENSMUSG00000005583": 3.104851323450022,
    "ENSMUSG00000071076_ENSMUSG00000044786": 3.0821218868888884,
    "ENSMUSG00000048387_ENSMUSG00000020585": 3.024812897055277,
    "ENSMUSG00000071076_ENSMUSG00000020423": 26.48574295921271,
    "ENSMUSG00000020423_ENSMUSG00000003545": 21.26405469876257,
    "ENSMUSG00000052837_ENSMUSG00000044786": 16.896974878924812,
    "ENSMUSG00000029359_ENSMUSG00000005583": 13.422570417323275,
    "ENSMUSG00000044786_ENSMUSG00000021250": 12.863587882034674,
    "ENSMUSG00000071076_ENSMUSG00000006205": 12.749027914373166,
    "ENSMUSG00000039601_ENSMUSG00000021250": 11.368429143134037,
    "ENSMUSG00000052684_ENSMUSG00000003545": 11.368214571731489,
    "ENSMUSG00000052837_ENSMUSG00000052684": 11.092242878749383,
    "ENSMUSG00000042745_ENSMUSG00000005583": 7.701540689728936,
    "ENSMUSG00000052684_ENSMUSG00000020423": 7.454527339129341,
    "ENSMUSG00000019929_ENSMUSG00000005583": 7.4521147393135765,
    "ENSMUSG00000052684_ENSMUSG00000021876": 7.17861425681787,
    "ENSMUSG00000038418_ENSMUSG00000003545": 5.818469451229697,
    "ENSMUSG00000071076_ENSMUSG00000023034": 5.790594358262504,
    "ENSMUSG00000071076_ENSMUSG00000068220": 5.741269926207475,
    "ENSMUSG00000024190_ENSMUSG00000003545": 5.64759953527284,
    "ENSMUSG00000071076_ENSMUSG00000022032": 5.592389946757354,
    "ENSMUSG00000052837_ENSMUSG00000039601": 5.553114482750674,
    "ENSMUSG00000071076_ENSMUSG00000002732": 5.48333175656681,
    "ENSMUSG00000029322_ENSMUSG00000026238": 5.465158443841482,
    "ENSMUSG00000029096_ENSMUSG00000005583": 5.323408072135334,
    "ENSMUSG00000071076_ENSMUSG00000020585": 5.317727532706944,
    "ENSMUSG00000053113_ENSMUSG00000021250": 5.182487366642451,
    "ENSMUSG00000052684_ENSMUSG00000021477": 4.054247486398131,
    "ENSMUSG00000071076_ENSMUSG00000027712": 4.031893030478303,
    "ENSMUSG00000007655_ENSMUSG00000005583": 3.9968177945989076,
    "ENSMUSG00000029231_ENSMUSG00000005583": 3.959638310605863,
    "ENSMUSG00000055148_ENSMUSG00000052684": 3.9461710586530705,
    "ENSMUSG00000023034_ENSMUSG00000021250": 3.9143680538969403,
    "ENSMUSG00000052684_ENSMUSG00000006369": 3.861487643719066,
    "ENSMUSG00000026688_ENSMUSG00000005583": 3.8594784126373773,
    "ENSMUSG00000029322_ENSMUSG00000006356": 3.8366731401884695,
    "ENSMUSG00000031586_ENSMUSG00000005583": 3.7583648322637857,
    "ENSMUSG00000061477_ENSMUSG00000000247": 3.683740246114078,
    "ENSMUSG00000071076_ENSMUSG00000028691": 3.5236534707048843,
    "ENSMUSG00000071076_ENSMUSG00000000753": 3.428707324573982,
    "ENSMUSG00000052837_ENSMUSG00000003545": 22.107715979312005,
    "ENSMUSG00000071076_ENSMUSG00000024076": 18.0599771582601,
    "ENSMUSG00000014956_ENSMUSG00000005583": 13.295769586429365,
    "ENSMUSG00000016494_ENSMUSG00000005583": 13.07476851013246,
    "ENSMUSG00000055148_ENSMUSG00000044786": 12.90658312737477,
    "ENSMUSG00000071076_ENSMUSG00000064373": 11.991584276624076,
    "ENSMUSG00000056501_ENSMUSG00000021250": 11.231543948087737,
    "ENSMUSG00000052684_ENSMUSG00000025784": 10.902758504427938,
    "ENSMUSG00000071076_ENSMUSG00000040212": 9.915676376540516,
    "ENSMUSG00000052684_ENSMUSG00000032515": 9.779323260316838,
    "ENSMUSG00000031246_ENSMUSG00000029322": 7.841633085959625,
    "ENSMUSG00000021876_ENSMUSG00000005583": 7.781144869808331,
    "ENSMUSG00000055148_ENSMUSG00000053113": 7.596098710876831,
    "ENSMUSG00000071076_ENSMUSG00000016494": 7.534985005837032,
    "ENSMUSG00000071076_ENSMUSG00000018217": 7.441133152794328,
    "ENSMUSG00000038370_ENSMUSG00000005583": 7.359406183726755,
    "ENSMUSG00000052684_ENSMUSG00000045658": 7.189011630327358,
    "ENSMUSG00000021250_ENSMUSG00000003545": 6.269796873949839,
    "ENSMUSG00000021200_ENSMUSG00000005583": 6.163689914617984,
    "ENSMUSG00000064373_ENSMUSG00000052684": 5.53866828300545,
    "ENSMUSG00000029761_ENSMUSG00000005583": 5.484093906837928,
    "ENSMUSG00000071076_ENSMUSG00000001025": 5.415200209092442,
    "ENSMUSG00000052684_ENSMUSG00000035356": 5.408381245346356,
    "ENSMUSG00000049517_ENSMUSG00000000247": 5.339596107050707,
    "ENSMUSG00000021250_ENSMUSG00000018899": 5.307414248672807,
    "ENSMUSG00000041329_ENSMUSG00000005583": 5.072552986807257,
    "ENSMUSG00000036438_ENSMUSG00000029322": 4.259127514782113,
    "ENSMUSG00000052684_ENSMUSG00000025492": 4.00467841638815,
    "ENSMUSG00000027800_ENSMUSG00000005583": 3.988410038640975,
    "ENSMUSG00000055148_ENSMUSG00000052837": 3.975098251828096,
    "ENSMUSG00000008540_ENSMUSG00000005583": 3.9249957387553995,
    "ENSMUSG00000071076_ENSMUSG00000026879": 3.8869825620229657,
    "ENSMUSG00000017466_ENSMUSG00000005583": 3.8092819737903296,
    "ENSMUSG00000023224_ENSMUSG00000005583": 3.773977927482205,
    "ENSMUSG00000016494_ENSMUSG00000008575": 3.761959563259081,
    "ENSMUSG00000052684_ENSMUSG00000026628": 3.7557471279312136,
    "ENSMUSG00000052684_ENSMUSG00000021268": 3.7464592836248247,
    "ENSMUSG00000052837_ENSMUSG00000021250": 3.7082288407786397,
    "ENSMUSG00000071076_ENSMUSG00000045658": 3.6304982818435048,
    "ENSMUSG00000052684_ENSMUSG00000007872": 3.5665492641059044,
    "ENSMUSG00000061787_ENSMUSG00000057098": 3.074135623125704
}
```
