## Supplement 4 (CCl4 a6w HSC vs. PF) for "AGEAS: Automated Machine Learning based Genetic Regulatory Element Extraction System": outlier_grps.js

```
{
    "ENSMUSG00000032387_ENSMUSG00000005583": 18.456987015093837,
    "ENSMUSG00000064358_ENSMUSG00000029322": 15.809282157915314,
    "ENSMUSG00000039601_ENSMUSG00000021250": 15.169345124007076,
    "ENSMUSG00000026879_ENSMUSG00000005583": 14.571871730645535,
    "ENSMUSG00000071076_ENSMUSG00000002732": 13.908603443972678,
    "ENSMUSG00000042745_ENSMUSG00000005583": 13.097640778908977,
    "ENSMUSG00000042485_ENSMUSG00000005583": 12.895754744347677,
    "ENSMUSG00000055148_ENSMUSG00000003545": 12.670131400053968,
    "ENSMUSG00000071076_ENSMUSG00000018217": 12.114414119762618,
    "ENSMUSG00000053113_ENSMUSG00000052837": 12.11433891231372,
    "ENSMUSG00000038418_ENSMUSG00000020423": 10.826168704279663,
    "ENSMUSG00000071076_ENSMUSG00000006205": 8.706147952762011,
    "ENSMUSG00000052684_ENSMUSG00000006205": 8.317692098019089,
    "ENSMUSG00000038418_ENSMUSG00000003545": 8.249190239861568,
    "ENSMUSG00000041329_ENSMUSG00000005583": 8.172538305982982,
    "ENSMUSG00000052684_ENSMUSG00000021250": 8.164147986134926,
    "ENSMUSG00000052837_ENSMUSG00000003545": 8.048402855817065,
    "ENSMUSG00000071076_ENSMUSG00000020053": 8.042421551096114,
    "ENSMUSG00000021200_ENSMUSG00000005583": 7.919701759967408,
    "ENSMUSG00000071076_ENSMUSG00000025784": 6.422079727526764,
    "ENSMUSG00000071076_ENSMUSG00000032231": 6.3787417008903455,
    "ENSMUSG00000071076_ENSMUSG00000041959": 6.3401245157804,
    "ENSMUSG00000052684_ENSMUSG00000021477": 6.3096726807765675,
    "ENSMUSG00000026688_ENSMUSG00000005583": 6.300347582030746,
    "ENSMUSG00000052837_ENSMUSG00000039601": 6.26535067304768,
    "ENSMUSG00000005583_ENSMUSG00000001520": 6.229160685207106,
    "ENSMUSG00000029322_ENSMUSG00000026238": 6.17901428621611,
    "ENSMUSG00000020423_ENSMUSG00000003545": 6.0529182505595855,
    "ENSMUSG00000055148_ENSMUSG00000038418": 5.978432149207574,
    "ENSMUSG00000021250_ENSMUSG00000020423": 5.638402327251855,
    "ENSMUSG00000052684_ENSMUSG00000009681": 4.775961411003325,
    "ENSMUSG00000071076_ENSMUSG00000016494": 4.649881317280981,
    "ENSMUSG00000032035_ENSMUSG00000019929": 4.607304893920046,
    "ENSMUSG00000071076_ENSMUSG00000027712": 4.47163701306768,
    "ENSMUSG00000071076_ENSMUSG00000020585": 4.407664090238277,
    "ENSMUSG00000052684_ENSMUSG00000025784": 4.2886262784818285,
    "ENSMUSG00000029231_ENSMUSG00000005583": 4.251718755346755,
    "ENSMUSG00000061477_ENSMUSG00000000247": 4.232426412556465,
    "ENSMUSG00000044786_ENSMUSG00000021250": 4.168131338408985,
    "ENSMUSG00000049517_ENSMUSG00000000247": 3.6923994292181272,
    "ENSMUSG00000052837_ENSMUSG00000020423": 3.489220911219769,
    "ENSMUSG00000055148_ENSMUSG00000053113": 3.2870275488875755,
    "ENSMUSG00000053560_ENSMUSG00000052684": 3.0129464188505604,
    "ENSMUSG00000052837_ENSMUSG00000021250": 22.58267403944818,
    "ENSMUSG00000026574_ENSMUSG00000005583": 16.804026156658836,
    "ENSMUSG00000007655_ENSMUSG00000005583": 16.281257742770386,
    "ENSMUSG00000052684_ENSMUSG00000020423": 13.663048374608257,
    "ENSMUSG00000023034_ENSMUSG00000003545": 13.35212462009284,
    "ENSMUSG00000055148_ENSMUSG00000044786": 12.712839177864476,
    "ENSMUSG00000038193_ENSMUSG00000019929": 12.315127206685565,
    "ENSMUSG00000071076_ENSMUSG00000020423": 11.343220757047531,
    "ENSMUSG00000053113_ENSMUSG00000021250": 11.284898564024248,
    "ENSMUSG00000052837_ENSMUSG00000052684": 10.67878393158208,
    "ENSMUSG00000071076_ENSMUSG00000021876": 9.905171969949539,
    "ENSMUSG00000071076_ENSMUSG00000021477": 8.190413653367244,
    "ENSMUSG00000064373_ENSMUSG00000005583": 7.477464209731362,
    "ENSMUSG00000038418_ENSMUSG00000021250": 7.453913842917469,
    "ENSMUSG00000052684_ENSMUSG00000044786": 7.234762094570343,
    "ENSMUSG00000052684_ENSMUSG00000008540": 7.2039949649404305,
    "ENSMUSG00000071076_ENSMUSG00000064373": 6.064032073974086,
    "ENSMUSG00000053560_ENSMUSG00000021250": 5.960359343143718,
    "ENSMUSG00000071076_ENSMUSG00000001025": 5.906693509177,
    "ENSMUSG00000071076_ENSMUSG00000026879": 5.882964504968961,
    "ENSMUSG00000071076_ENSMUSG00000040212": 5.863168377080311,
    "ENSMUSG00000052684_ENSMUSG00000019278": 5.721292516269784,
    "ENSMUSG00000019929_ENSMUSG00000005583": 5.55582806228042,
    "ENSMUSG00000071076_ENSMUSG00000017493": 5.468204200523547,
    "ENSMUSG00000038418_ENSMUSG00000026628": 5.423679271794684,
    "ENSMUSG00000029322_ENSMUSG00000006356": 5.393288669360508,
    "ENSMUSG00000008540_ENSMUSG00000005583": 5.373827224696968,
    "ENSMUSG00000052684_ENSMUSG00000015085": 5.321736474643411,
    "ENSMUSG00000072235_ENSMUSG00000071076": 4.263237588088282,
    "ENSMUSG00000008575_ENSMUSG00000006369": 4.203404004845539,
    "ENSMUSG00000024290_ENSMUSG00000005583": 4.020835577478923,
    "ENSMUSG00000021876_ENSMUSG00000005583": 4.013725921980883,
    "ENSMUSG00000055148_ENSMUSG00000052684": 3.999171530518951,
    "ENSMUSG00000071076_ENSMUSG00000052837": 3.8255312370476893,
    "ENSMUSG00000071076_ENSMUSG00000026628": 3.7992899649205536,
    "ENSMUSG00000021250_ENSMUSG00000003545": 3.7963808809599477,
    "ENSMUSG00000052684_ENSMUSG00000035356": 3.7912465320568685,
    "ENSMUSG00000071076_ENSMUSG00000029718": 3.7819067151786654,
    "ENSMUSG00000041841_ENSMUSG00000000247": 3.716073489658373,
    "ENSMUSG00000057098_ENSMUSG00000008668": 3.2528824841250223,
    "ENSMUSG00000071076_ENSMUSG00000000753": 3.2240585133424795,
    "ENSMUSG00000052684_ENSMUSG00000006369": 18.33412658094532,
    "ENSMUSG00000014956_ENSMUSG00000005583": 17.74545603022185,
    "ENSMUSG00000052684_ENSMUSG00000020053": 16.639873955619986,
    "ENSMUSG00000052684_ENSMUSG00000003545": 16.2350411814976,
    "ENSMUSG00000023064_ENSMUSG00000005583": 14.900639383659394,
    "ENSMUSG00000087006_ENSMUSG00000005583": 13.669265142162674,
    "ENSMUSG00000052684_ENSMUSG00000000753": 12.332202167086725,
    "ENSMUSG00000052684_ENSMUSG00000023034": 10.80691923398218,
    "ENSMUSG00000030122_ENSMUSG00000029322": 9.832312969455119,
    "ENSMUSG00000071076_ENSMUSG00000060703": 7.893378058192474,
    "ENSMUSG00000071076_ENSMUSG00000068220": 7.860357210592245,
    "ENSMUSG00000033538_ENSMUSG00000021250": 7.443181642082744,
    "ENSMUSG00000016494_ENSMUSG00000008575": 7.3780846610970094,
    "ENSMUSG00000090698_ENSMUSG00000052684": 5.899789684741774,
    "ENSMUSG00000052997_ENSMUSG00000005583": 5.8523814614777505,
    "ENSMUSG00000078202_ENSMUSG00000005583": 5.688555854905304,
    "ENSMUSG00000035783_ENSMUSG00000005583": 5.671209863306862,
    "ENSMUSG00000027800_ENSMUSG00000005583": 5.667503894556684,
    "ENSMUSG00000071076_ENSMUSG00000064215": 5.456827927445072,
    "ENSMUSG00000038370_ENSMUSG00000005583": 5.419465269765831,
    "ENSMUSG00000053113_ENSMUSG00000026628": 5.401519872658282,
    "ENSMUSG00000029096_ENSMUSG00000005583": 5.295256936615337,
    "ENSMUSG00000028639_ENSMUSG00000020486": 5.264383759287212,
    "ENSMUSG00000071076_ENSMUSG00000044786": 5.050928659048977,
    "ENSMUSG00000020484_ENSMUSG00000008540": 5.043827185630721,
    "ENSMUSG00000053113_ENSMUSG00000052684": 4.824019684405961,
    "ENSMUSG00000052684_ENSMUSG00000045658": 4.394989334731063,
    "ENSMUSG00000053113_ENSMUSG00000023034": 4.204034080099018,
    "ENSMUSG00000068735_ENSMUSG00000005583": 4.108810951014345,
    "ENSMUSG00000031586_ENSMUSG00000005583": 3.96298158026521,
    "ENSMUSG00000052837_ENSMUSG00000035356": 3.9130735025136327,
    "ENSMUSG00000071076_ENSMUSG00000028691": 3.8788555733828995,
    "ENSMUSG00000052684_ENSMUSG00000017493": 3.7388793087459575,
    "ENSMUSG00000023224_ENSMUSG00000005583": 3.591641476894881,
    "ENSMUSG00000071076_ENSMUSG00000029096": 3.4884994591378757,
    "ENSMUSG00000057098_ENSMUSG00000037742": 3.2054398655240677,
    "ENSMUSG00000016319_ENSMUSG00000005583": 3.157796451289727
}
```
