## Supplement 4 (CCl4 a6w HSC vs. PF) for "AGEAS: Automated Machine Learning based Genetic Regulatory Element Extraction System": outlier_grps.js

```
{
    "ENSMUSG00000052684_ENSMUSG00000006369": 25.440573156837726,
    "ENSMUSG00000038370_ENSMUSG00000005583": 14.494861310097262,
    "ENSMUSG00000005583_ENSMUSG00000001520": 13.363067507860599,
    "ENSMUSG00000042485_ENSMUSG00000005583": 13.053887094665848,
    "ENSMUSG00000052684_ENSMUSG00000025784": 12.782107907222452,
    "ENSMUSG00000071076_ENSMUSG00000006205": 12.4131810717328,
    "ENSMUSG00000023064_ENSMUSG00000005583": 12.139592601673822,
    "ENSMUSG00000057098_ENSMUSG00000024608": 11.974181889397727,
    "ENSMUSG00000025147_ENSMUSG00000005583": 11.717671819322932,
    "ENSMUSG00000008575_ENSMUSG00000006369": 9.891526867828835,
    "ENSMUSG00000052684_ENSMUSG00000006205": 9.240365815331876,
    "ENSMUSG00000042745_ENSMUSG00000005583": 9.067997730207097,
    "ENSMUSG00000071076_ENSMUSG00000041959": 9.013417380481028,
    "ENSMUSG00000071076_ENSMUSG00000016494": 8.878593231415797,
    "ENSMUSG00000071076_ENSMUSG00000018217": 8.853005905261298,
    "ENSMUSG00000052684_ENSMUSG00000020423": 8.485156292156734,
    "ENSMUSG00000053113_ENSMUSG00000021250": 8.25881832324793,
    "ENSMUSG00000052837_ENSMUSG00000003545": 8.185295588750874,
    "ENSMUSG00000071076_ENSMUSG00000021876": 6.64872135706455,
    "ENSMUSG00000014956_ENSMUSG00000005583": 6.546738180292915,
    "ENSMUSG00000071076_ENSMUSG00000001025": 6.520166554983235,
    "ENSMUSG00000029322_ENSMUSG00000026238": 6.372880438357425,
    "ENSMUSG00000023034_ENSMUSG00000003545": 6.357283309218541,
    "ENSMUSG00000071076_ENSMUSG00000025784": 6.344969501686503,
    "ENSMUSG00000038418_ENSMUSG00000003545": 6.3375126884202455,
    "ENSMUSG00000021250_ENSMUSG00000003545": 6.103910890314635,
    "ENSMUSG00000026879_ENSMUSG00000005583": 6.058404476759627,
    "ENSMUSG00000052837_ENSMUSG00000044786": 5.732816470405035,
    "ENSMUSG00000047215_ENSMUSG00000000247": 5.129017810821576,
    "ENSMUSG00000026688_ENSMUSG00000005583": 4.9534407363320785,
    "ENSMUSG00000071076_ENSMUSG00000020423": 4.872559399397797,
    "ENSMUSG00000021200_ENSMUSG00000005583": 4.758222960571351,
    "ENSMUSG00000053113_ENSMUSG00000023034": 4.7354639723726,
    "ENSMUSG00000038418_ENSMUSG00000020423": 4.437373393196543,
    "ENSMUSG00000024290_ENSMUSG00000005583": 4.0785447496362774,
    "ENSMUSG00000052684_ENSMUSG00000008540": 3.5806589921857035,
    "ENSMUSG00000057098_ENSMUSG00000030744": 3.523362131624542,
    "ENSMUSG00000071076_ENSMUSG00000000753": 3.3400506961961773,
    "ENSMUSG00000007655_ENSMUSG00000005583": 3.2550933179187718,
    "ENSMUSG00000017466_ENSMUSG00000005583": 3.1678181370245206,
    "ENSMUSG00000038193_ENSMUSG00000019929": 16.867784705557426,
    "ENSMUSG00000052684_ENSMUSG00000021250": 16.07967117098306,
    "ENSMUSG00000052684_ENSMUSG00000032359": 14.847418687430782,
    "ENSMUSG00000029322_ENSMUSG00000006356": 13.146029003922,
    "ENSMUSG00000039601_ENSMUSG00000021250": 12.602085244314035,
    "ENSMUSG00000052684_ENSMUSG00000018339": 12.600550348933508,
    "ENSMUSG00000052684_ENSMUSG00000007872": 10.418710977779241,
    "ENSMUSG00000079523_ENSMUSG00000005583": 10.185014766246493,
    "ENSMUSG00000071076_ENSMUSG00000064373": 9.41322001378614,
    "ENSMUSG00000071076_ENSMUSG00000042312": 9.357084315587235,
    "ENSMUSG00000064358_ENSMUSG00000029322": 9.224394187888361,
    "ENSMUSG00000071076_ENSMUSG00000008540": 9.172771740952559,
    "ENSMUSG00000068735_ENSMUSG00000005583": 9.167742375355983,
    "ENSMUSG00000055148_ENSMUSG00000044786": 8.870100847101963,
    "ENSMUSG00000052837_ENSMUSG00000021250": 8.582465142041922,
    "ENSMUSG00000072235_ENSMUSG00000071076": 6.912632883352934,
    "ENSMUSG00000052684_ENSMUSG00000020053": 6.5452087495353455,
    "ENSMUSG00000064373_ENSMUSG00000005583": 6.535456276275448,
    "ENSMUSG00000052684_ENSMUSG00000023034": 6.485829674972452,
    "ENSMUSG00000008540_ENSMUSG00000005583": 6.455647315139727,
    "ENSMUSG00000038418_ENSMUSG00000021250": 6.290905292275546,
    "ENSMUSG00000052684_ENSMUSG00000044786": 6.249230882576476,
    "ENSMUSG00000044786_ENSMUSG00000021250": 6.23401524955718,
    "ENSMUSG00000052837_ENSMUSG00000038418": 6.201053773580283,
    "ENSMUSG00000038418_ENSMUSG00000006369": 6.176329622884828,
    "ENSMUSG00000052684_ENSMUSG00000021477": 6.095519706638691,
    "ENSMUSG00000019907_ENSMUSG00000005583": 6.047376098852027,
    "ENSMUSG00000020928_ENSMUSG00000005583": 5.924497834346004,
    "ENSMUSG00000071076_ENSMUSG00000002732": 5.599943104892119,
    "ENSMUSG00000021250_ENSMUSG00000020423": 5.540516065124768,
    "ENSMUSG00000031586_ENSMUSG00000005583": 5.347000925457579,
    "ENSMUSG00000030122_ENSMUSG00000029322": 5.015549986038612,
    "ENSMUSG00000052997_ENSMUSG00000005583": 4.92101099397032,
    "ENSMUSG00000036438_ENSMUSG00000029322": 4.899599419395903,
    "ENSMUSG00000071076_ENSMUSG00000020585": 4.896673737150206,
    "ENSMUSG00000032035_ENSMUSG00000019929": 4.86882341986349,
    "ENSMUSG00000037742_ENSMUSG00000000247": 4.800889821540593,
    "ENSMUSG00000029359_ENSMUSG00000005583": 4.760579123683676,
    "ENSMUSG00000026574_ENSMUSG00000005583": 4.666454434503261,
    "ENSMUSG00000032387_ENSMUSG00000005583": 4.66572234853218,
    "ENSMUSG00000041329_ENSMUSG00000005583": 4.63132925256935,
    "ENSMUSG00000029096_ENSMUSG00000005583": 4.62748887705168,
    "ENSMUSG00000055148_ENSMUSG00000052684": 4.465520865453047,
    "ENSMUSG00000024190_ENSMUSG00000003545": 3.8464405231790404,
    "ENSMUSG00000052684_ENSMUSG00000017493": 3.1345802871235806,
    "ENSMUSG00000071076_ENSMUSG00000027712": 5.334259134174607
}
```
