## Supplement 4 (CCl4 a6w HSC vs. PF) for "AGEAS: Automated Machine Learning based Genetic Regulatory Element Extraction System": outlier_grps.js

```
{
    "ENSMUSG00000052684_ENSMUSG00000020423": 23.10384939058331,
    "ENSMUSG00000068735_ENSMUSG00000005583": 20.10095666947771,
    "ENSMUSG00000071076_ENSMUSG00000027712": 15.349380469857302,
    "ENSMUSG00000020423_ENSMUSG00000003545": 14.382186164951225,
    "ENSMUSG00000041329_ENSMUSG00000005583": 12.693487365575894,
    "ENSMUSG00000071076_ENSMUSG00000020423": 12.179766323654846,
    "ENSMUSG00000052684_ENSMUSG00000044786": 12.112297660334159,
    "ENSMUSG00000052837_ENSMUSG00000021250": 11.956408531083175,
    "ENSMUSG00000016494_ENSMUSG00000005583": 11.845197830047823,
    "ENSMUSG00000052837_ENSMUSG00000038418": 11.227289806535527,
    "ENSMUSG00000052684_ENSMUSG00000006369": 10.948231835339529,
    "ENSMUSG00000025147_ENSMUSG00000005583": 9.603688120608805,
    "ENSMUSG00000064373_ENSMUSG00000052684": 8.865976791050782,
    "ENSMUSG00000052684_ENSMUSG00000020053": 8.081525968715717,
    "ENSMUSG00000017466_ENSMUSG00000005583": 8.076980141395138,
    "ENSMUSG00000072235_ENSMUSG00000071076": 7.946563575263189,
    "ENSMUSG00000071076_ENSMUSG00000060703": 7.8283125384274515,
    "ENSMUSG00000029096_ENSMUSG00000005583": 7.798793891207554,
    "ENSMUSG00000052684_ENSMUSG00000025784": 7.749284163085138,
    "ENSMUSG00000042485_ENSMUSG00000005583": 6.076745937747976,
    "ENSMUSG00000038418_ENSMUSG00000003545": 5.786821051122658,
    "ENSMUSG00000016024_ENSMUSG00000008575": 5.67479674925595,
    "ENSMUSG00000052837_ENSMUSG00000003545": 5.67251057249319,
    "ENSMUSG00000038418_ENSMUSG00000020423": 5.601315158927894,
    "ENSMUSG00000071076_ENSMUSG00000021876": 4.907280974043525,
    "ENSMUSG00000030122_ENSMUSG00000029322": 4.719339309452852,
    "ENSMUSG00000064358_ENSMUSG00000029322": 4.652935195028962,
    "ENSMUSG00000071076_ENSMUSG00000040212": 4.489039329362933,
    "ENSMUSG00000052684_ENSMUSG00000008540": 4.340394109107228,
    "ENSMUSG00000008540_ENSMUSG00000005583": 4.26163034317282,
    "ENSMUSG00000060636_ENSMUSG00000000247": 4.1509374814658715,
    "ENSMUSG00000038418_ENSMUSG00000021250": 4.061610076217153,
    "ENSMUSG00000053113_ENSMUSG00000021250": 4.035642498271137,
    "ENSMUSG00000071076_ENSMUSG00000026879": 3.989113491553725,
    "ENSMUSG00000023064_ENSMUSG00000005583": 3.7944098208319867,
    "ENSMUSG00000071076_ENSMUSG00000007872": 3.6159198658763176,
    "ENSMUSG00000052837_ENSMUSG00000052684": 3.4909962869466216,
    "ENSMUSG00000071076_ENSMUSG00000029718": 3.27598454876941,
    "ENSMUSG00000052684_ENSMUSG00000015312": 3.1294039982400537,
    "ENSMUSG00000029322_ENSMUSG00000026238": 22.833059721286237,
    "ENSMUSG00000052997_ENSMUSG00000005583": 22.379598801009486,
    "ENSMUSG00000071076_ENSMUSG00000041959": 17.475715117815113,
    "ENSMUSG00000026688_ENSMUSG00000005583": 14.296269844523774,
    "ENSMUSG00000071076_ENSMUSG00000016494": 13.969724418149967,
    "ENSMUSG00000052684_ENSMUSG00000000753": 12.529864896463788,
    "ENSMUSG00000071076_ENSMUSG00000018217": 11.959613961538071,
    "ENSMUSG00000071076_ENSMUSG00000020053": 10.884877389365545,
    "ENSMUSG00000060703_ENSMUSG00000005583": 10.862052592295587,
    "ENSMUSG00000042745_ENSMUSG00000005583": 7.631769760507675,
    "ENSMUSG00000057098_ENSMUSG00000024608": 7.613261276253695,
    "ENSMUSG00000057098_ENSMUSG00000046364": 7.549497918596536,
    "ENSMUSG00000055148_ENSMUSG00000003545": 7.403093904160759,
    "ENSMUSG00000029322_ENSMUSG00000006356": 7.1981661792758835,
    "ENSMUSG00000044786_ENSMUSG00000021250": 7.166452003200196,
    "ENSMUSG00000052684_ENSMUSG00000021250": 7.100289311127596,
    "ENSMUSG00000071076_ENSMUSG00000001025": 5.8533937695473455,
    "ENSMUSG00000052684_ENSMUSG00000023034": 5.399203942833648,
    "ENSMUSG00000029231_ENSMUSG00000005583": 5.2959265064051415,
    "ENSMUSG00000005583_ENSMUSG00000001520": 5.262496594216042,
    "ENSMUSG00000052684_ENSMUSG00000006205": 5.254691284214657,
    "ENSMUSG00000071076_ENSMUSG00000023034": 5.120603181686382,
    "ENSMUSG00000019907_ENSMUSG00000005583": 5.101851058330509,
    "ENSMUSG00000021250_ENSMUSG00000020423": 5.074562702243464,
    "ENSMUSG00000071076_ENSMUSG00000006205": 5.07080838494897,
    "ENSMUSG00000055148_ENSMUSG00000053113": 4.795406254604542,
    "ENSMUSG00000038193_ENSMUSG00000019929": 4.1881101664933205,
    "ENSMUSG00000014956_ENSMUSG00000005583": 4.1464286243221435,
    "ENSMUSG00000071076_ENSMUSG00000064373": 4.145619055550113,
    "ENSMUSG00000026574_ENSMUSG00000005583": 3.9060952548378767,
    "ENSMUSG00000021250_ENSMUSG00000003545": 3.8580756741382056,
    "ENSMUSG00000057098_ENSMUSG00000032399": 3.8142138551552804,
    "ENSMUSG00000038193_ENSMUSG00000035493": 3.8120050015950104,
    "ENSMUSG00000057098_ENSMUSG00000028495": 3.806230286298036,
    "ENSMUSG00000026879_ENSMUSG00000005583": 3.798984407953283,
    "ENSMUSG00000071076_ENSMUSG00000021477": 3.727722593507311,
    "ENSMUSG00000027834_ENSMUSG00000005583": 3.0250118134150084,
    "ENSMUSG00000021876_ENSMUSG00000005583": 32.535866844256105,
    "ENSMUSG00000019929_ENSMUSG00000005583": 17.707050612211667,
    "ENSMUSG00000005583_ENSMUSG00000002980": 17.135507284288273,
    "ENSMUSG00000052684_ENSMUSG00000017493": 12.222496237613985,
    "ENSMUSG00000021200_ENSMUSG00000005583": 11.829814695806805,
    "ENSMUSG00000052684_ENSMUSG00000038418": 9.50062549420677,
    "ENSMUSG00000071076_ENSMUSG00000042747": 9.213054853076509,
    "ENSMUSG00000005583_ENSMUSG00000002107": 8.500590174967442,
    "ENSMUSG00000023034_ENSMUSG00000003545": 8.409025734204434,
    "ENSMUSG00000071076_ENSMUSG00000002732": 8.173118898587465,
    "ENSMUSG00000052837_ENSMUSG00000044786": 7.997950871612651,
    "ENSMUSG00000055148_ENSMUSG00000052684": 7.9318003224223546,
    "ENSMUSG00000003970_ENSMUSG00000000247": 7.498782519359147,
    "ENSMUSG00000020644_ENSMUSG00000014956": 3.7703077717603053,
    "ENSMUSG00000087006_ENSMUSG00000005583": 3.2931798244763506,
    "ENSMUSG00000029322_ENSMUSG00000007872": 3.281715748968132,
    "ENSMUSG00000071076_ENSMUSG00000032231": 3.254615312793969,
    "ENSMUSG00000071076_ENSMUSG00000029096": 3.2352020958761907,
    "ENSMUSG00000090698_ENSMUSG00000052684": 3.221098882000108,
    "ENSMUSG00000026879_ENSMUSG00000008575": 3.1850814341401765,
    "ENSMUSG00000032387_ENSMUSG00000005583": 3.1801451322725645,
    "ENSMUSG00000063457_ENSMUSG00000057098": 3.170213721689771,
    "ENSMUSG00000037868_ENSMUSG00000021250": 3.164429958839663,
    "ENSMUSG00000052684_ENSMUSG00000021477": 3.1571615230610806,
    "ENSMUSG00000057666_ENSMUSG00000005583": 3.128951255993549,
    "ENSMUSG00000064373_ENSMUSG00000005583": 3.116242428311167,
    "ENSMUSG00000029322_ENSMUSG00000014294": 3.086901808828595,
    "ENSMUSG00000048387_ENSMUSG00000029614": 3.0807392528529736,
    "ENSMUSG00000071076_ENSMUSG00000021390": 3.079059470207898,
    "ENSMUSG00000029359_ENSMUSG00000005583": 3.071949721255059,
    "ENSMUSG00000023224_ENSMUSG00000005583": 3.0685879804800926,
    "ENSMUSG00000016494_ENSMUSG00000008575": 3.065128394120254,
    "ENSMUSG00000071076_ENSMUSG00000004207": 3.059449734440589,
    "ENSMUSG00000052684_ENSMUSG00000035356": 3.051277916122737,
    "ENSMUSG00000071076_ENSMUSG00000028517": 3.033951522613752,
    "ENSMUSG00000053113_ENSMUSG00000018899": 3.0317494761744506,
    "ENSMUSG00000071076_ENSMUSG00000024076": 3.029068711047727,
    "ENSMUSG00000056501_ENSMUSG00000021250": 3.024593010204168
}
```
